# Identification of druggable host targets needed for SARS-CoV-2 infection by combined pharmacological evaluation and cellular network directed prioritization both in vitro and in vivo

**DOI:** 10.1101/2021.04.20.440626

**Authors:** J.J. Patten, Patrick T. Keiser, Deisy Gysi, Giulia Menichetti, Hiroyuki Mori, Callie J. Donahue, Xiao Gan, Italo do Valle, Kathleen Geoghegan-Barek, Manu Anantpadma, RuthMabel Boytz, Jacob L. Berrigan, Sarah Hulsey-Stubbs, Tess Ayazika, Colin O’Leary, Sallieu Jalloh, Florence Wagner, Seyoum Ayehunie, Stephen J. Elledge, Deborah Anderson, Joseph Loscalzo, Marinka Zitnik, Suryaram Gummuluru, Mark N. Namchuk, Albert-László Barabási, Robert A. Davey

## Abstract

Identification of host factors contributing to replication of viruses and resulting disease progression remains a promising approach for development of new therapeutics. Here, we evaluated 6710 clinical and preclinical compounds targeting 2183 host proteins by immunocytofluorescence-based screening to identify SARS-CoV-2 infection inhibitors. Computationally integrating relationships between small molecule structure, dose-response antiviral activity, host target and cell interactome networking produced cellular networks important for infection. This analysis revealed 389 small molecules, >12 scaffold classes and 813 host targets with micromolar to low nanomolar activities. From these classes, representatives were extensively evaluated for mechanism of action in stable and primary human cell models, and additionally against Beta and Delta SARS-CoV-2 variants and MERS-CoV. One promising candidate, obatoclax, significantly reduced SARS-CoV-2 viral lung load in mice. Ultimately, this work establishes a rigorous approach for future pharmacological and computational identification of novel host factor dependencies and treatments for viral diseases.

## Introduction

In December 2019, an unidentified pneumonia was reported in Wuhan, China, and by early January 2020, the causative agent had been identified as a novel coronavirus now called SARS-CoV-2. SARS-CoV-2, of the family *Coronaviridae,* is a single-stranded, positive-sense RNA virus with a genome of approximately 29kb ^1, 2^. SARS-CoV-2 is the third coronavirus in recent history to produce an epidemic, after SARS-CoV and MERS-CoV ^3, 4^. The virus has rapidly spread across the world causing over 290 million cases globally of COVID-19, as of January 2022, resulting in more than 5.4 million deaths ^5^ and causing economic contraction, mass unemployment, disruption of education, and increasing poverty levels ^6^. Fortunately, effective vaccines have become quickly available and are being distributed on an emergency basis. However, given the emergence of more contagious and potentially pathogenic variants, and the need for more effective therapies for patients, the search for new therapeutics remains a priority ^7, 8^. These efforts were launched by a series of repurposing screens of varying sizes. The largest was a screen of a 12,000 compound library, termed ReFRAME ^9^, and a recently reported screen of a 3000 compound library by Ditmar *et al* ^10^. Both studies identified PIKfyve inhibitors, numerous protease inhibitors (for example MG-132), several classes of kinase inhibitors and cyclosporin and analogues. While it was encouraging to see overlap between these studies, each also highlighted the interassay variability seen in evaluation of small molecule inhibitors of SARS-CoV-2 replication to date and the need for additional and more detailed studies.

Clinical successes to date have largely been confined to repurposed antivirals ^11^, highlighting the need for further efforts to identify compounds with novel mechanisms of action and better define pathways and targets of interest for future SARS-CoV2 therapeutics. The current study focused on screening of the Drug Repurposing Hub (DRH) library, a collection of 6710 compounds highly enriched with molecules which have been FDA approved, entered clinical trials (4.8% Phase 1, 2.0% Phase 2, 7.5% Phase 3, 4.4% Phase 4), or have been extensively pre-clinically characterized ^12^. The library comprises clusters of structurally related molecules that, with a well annotated database for host drug targets, provides a unique opportunity to capture preliminary structure-activity relationships of active molecules as well as identification of host targets of action that could be used to better design novel, specific therapeutics against SARS-CoV-2. The primary screen was the first to be conducted using multiple treatment doses at this scale, providing a data set that grades all compounds by activity from strongest to least active. These data were used to expand our previous computational studies in establishing SARS CoV-2 associated protein networks that may be enriched in host targets for future drug discovery efforts ^13^. The most promising hits from the screening effort were assessed for efficacy in a series of orthogonal assays conducted in human cell lines (Huh-7 and A549 cells), and human primary cell-based tissue models. Our studies confirmed the activity of several previously identified (inhibitors of PIKfyve, cathepsins, protein synthesis) as well as novel compound classes. The most promising small molecule treatment to emerge from the screen, obatoclax, demonstrated consistent activity across all cell-based assays and virus strains tested, including against MERS-CoV and reduced SARS-CoV-2 titers by up to 10-fold in a mouse infection model using clinically achievable compound exposures.

## Results

### Immunofluorescence-based screening of small compound library identifies potent inhibitors of SARS-CoV-2 infection of cells

The DRH compound library (Fig. 1A) was initially evaluated using an immunofluorescence-based assay to detect SARS-CoV-2 N protein expression. The screen was followed by computational analysis to prioritize the most potent for follow up in mechanistic assays evaluating impact on cell entry, genome replication and egress of progeny viruses as well as evaluation in primary human cell models (Fig. 1B). In order to capture multiple rounds of viral replication ^14^ and obtain strong N protein staining, the primary screen was conducted for 36-48 hours. While suitability of other cell types was evaluated for screening, VeroE6 cells, derived from African green monkey kidney cells, supported the most robust and consistent infection compatible with high-throughput screening at high biocontainment. Using automated analysis of stained cells (Fig. 1C, upper) the assay gave an average >80-fold difference between infection of untreated cells and those treated with a previously reported inhibitor of SARS-CoV-2 replication, the protease inhibitor, E64d ^15^ and yielded an assay Z’ of 0.6, indicating suitability for a high throughput screen ^16^ (Fig. 1C, lower).

**Figure 1.**
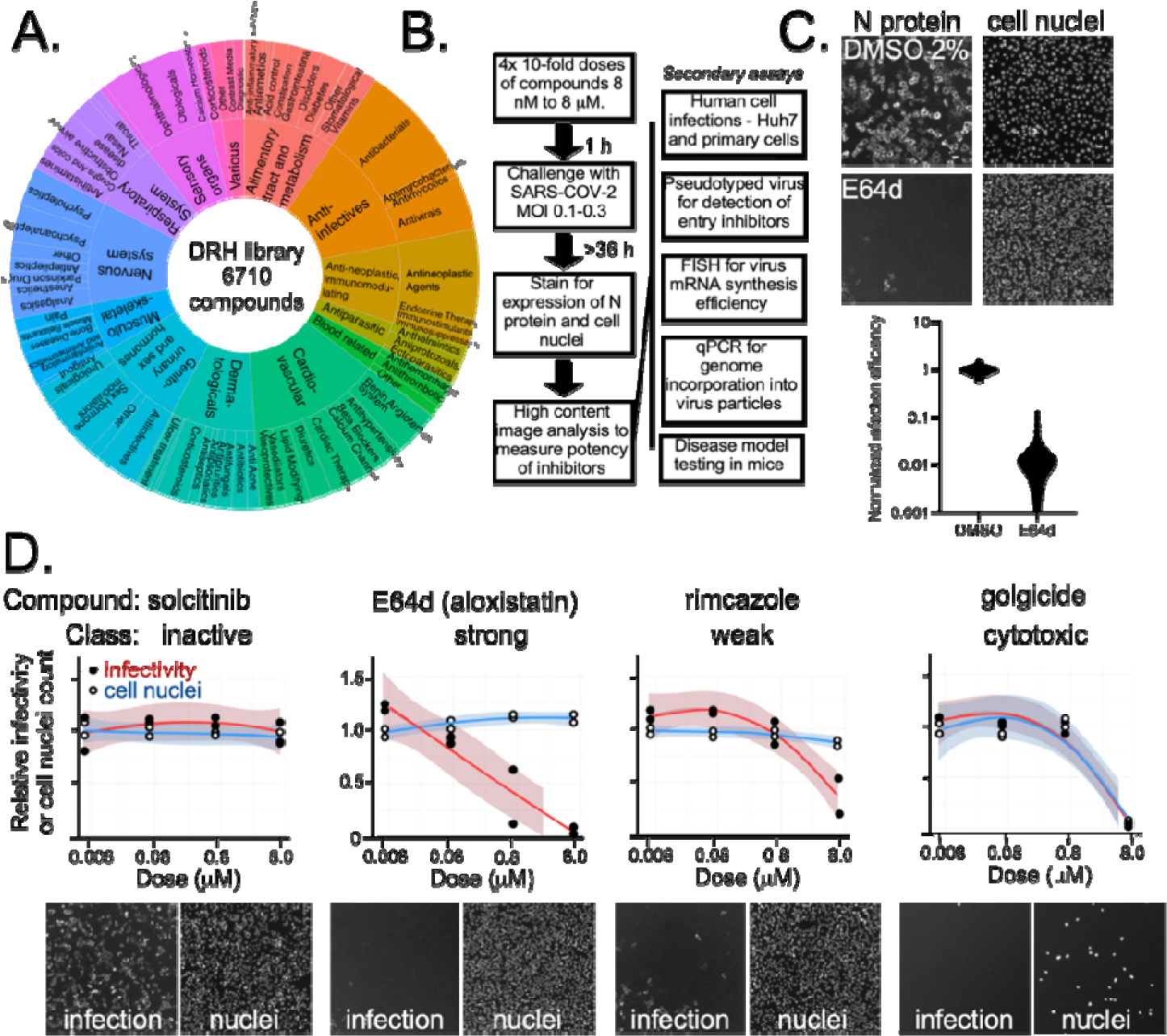
HTS screen for inhibitors of SARS-CoV-2. A) Composition of the Drug Repurposing Hub library based on ATC classifications for compounds (4,277 of the 6710). B) Workflow of immunofluorescence-based infection focus assay developed for screening of SARS-CoV-2 inhibitors and secondary mechanistic assays performed on active compounds. C) Dynamic range of the HTS assay evaluated using 5 µM E64d (Aloxistatin) versus 2% DMSO as vehicle. Upper panel: Examples of microscope images of infected cells stained with SARS- COV-2 N specific antibody and cell nuclei stained with Hoeschst 33342 for DMSO or E64d- treated cells. Lower panel: Outcomes for 190 wells of each treatment are shown. The assay gave a Z-factor of 0.6 in 384 well plates and, thus, was suitable for HTS. D) Examples of concentration response curve classes seen in the screen and cell images with N protein staining at left and cell nuclei at right. The indicated compounds show: no effect, strong activity with no cell loss (E64d), weak reduction in infectivity and no cell loss (rimcazole), reduction in infection that parallels cell loss that is likely due to cytotoxicity (golgicide).

To identify compounds with concentration responses and to grade each for potency, four doses of each compound were tested, ranging from 8 nM to 8 μM in 10-fold increments. Compounds were initially grouped using a log-logistic regression for viral load (Wald test; padj < 0.01; Bonferroni correction) to classify compounds (Table S1) on potency as strong (80% viral reduction, or Z score > 2.5), weak (50-80% viral reduction, Z score 1.5-2.5) inactive (padj >= 0.01), or cytotoxic (>60% cell loss by nuclei count and Wald test on a log-logistic regression for cell count; padj < 0.01; Bonferroni correction).We identified 172 (2.56%) strongly active, 217 (3.23%) weakly active and 1.6% as cytotoxic (examples shown in Fig. 1D). The data is deposited in the DRH database: https://www.broadinstitute.org/drug-repurposing-hub with active compounds summarized in Fig. 2A.

**Figure 2.**
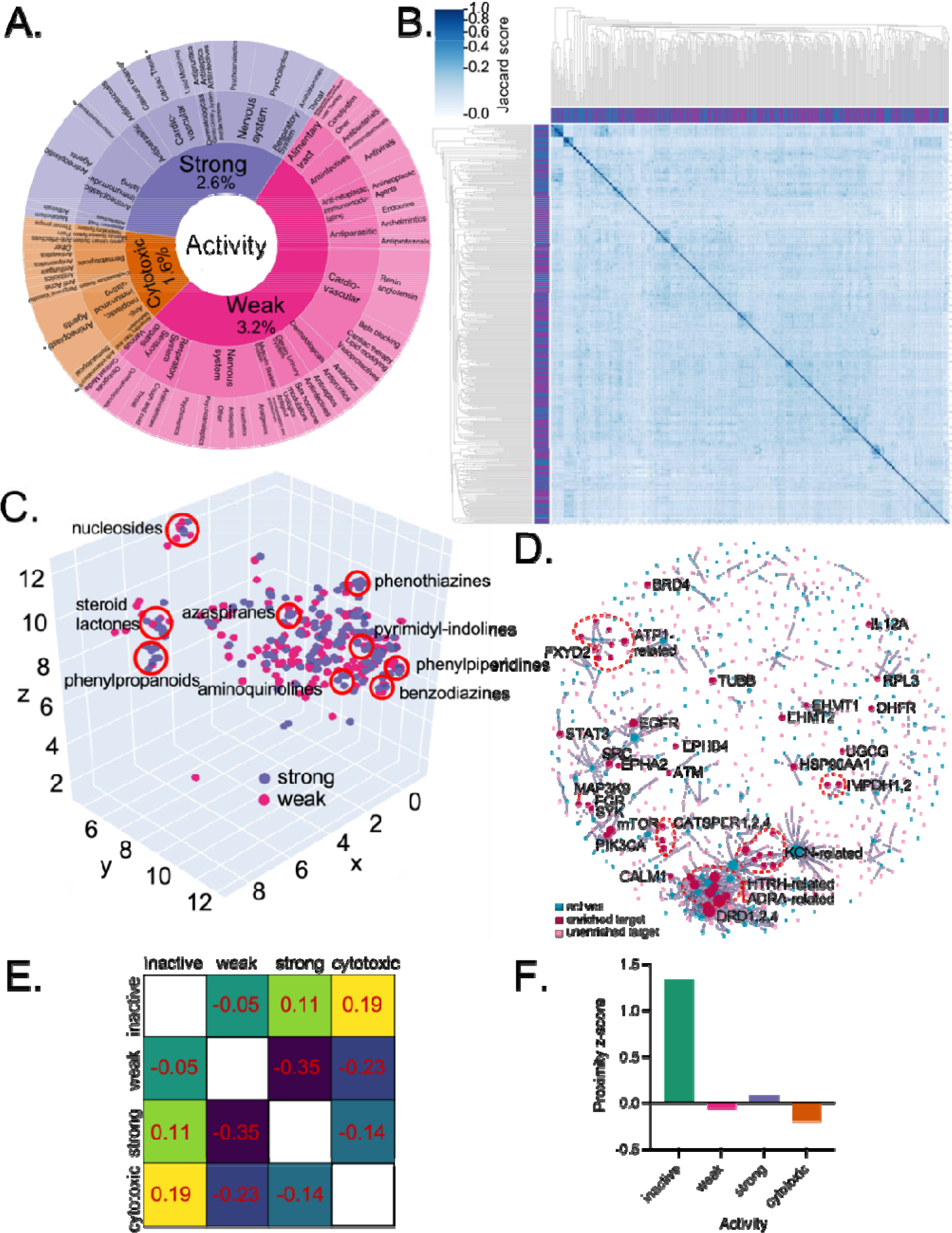
Computational analysis of primary screen outcomes. A) Chart showing the active compounds by ATC classifiers for strongly active, weakly active and cytotoxic compounds. The fraction of compounds in each category is shown as a percentage of all compounds in the library. B) Pairwise comparison of active compounds based on computed structural similarity using Morgan fingerprints and Jaccard analysis. C) 3D representation of structural similarity of active molecules based on reduced dimensionality of the molecular bit vectors. Localized clusters of major structural classes are indicated by red circles and are approximate locations in the plot. Please refer to Fig. S3 for an interactive version with higher detail. D) Relationship of enriched protein drug targets (red circles) or unenriched targets (pink circles) to active compounds (blue circles) for compounds with annotated targets in the DRH database. E) Separation heatmap between the targets of different activity classes within the human protein-protein interaction network. Negative network separation values reflect overlapping neighborhoods. F) Proximity z-score between the drug targets of each category and the SARS-CoV-2 protein-binding host factors in the human protein-protein interaction network. Drugs with treatment activity have Z-scores close to zero, much lower than inactive drugs, indicating they hit targets in network proximity of SARS-CoV-2 binding proteins.

### Structure-activity analysis reveals compound classes effective against SARS-CoV-2 replication

To reveal the structural characteristics that defined active compounds, structures were encoded using Morgan Fingerprints (around molecular heavy atoms up to radius 3, and hashed each molecular fingerprint to a binary vector of 8,192 bits, Table S2) and similarity searches performed. While the library showed diverse structural characteristics, active molecules showed a higher median of active bits (Fig. S1), with strongly active compounds showing significant enrichment for 89 bits. Hierarchical clustering of the Jaccard similarity matrix showed no clear connections to experimental classes for the entire library (Fig. S2) as would be expected from a diverse library of this size but actives showed small clusters of structurally related molecules appearing close to the diagonal (Fig. 2B). Reducing the dimensionality of the bit vector representation to 3D with UMAP ^17^ revealed the presence of local clusters (Figure 2C and in interactive Fig. S3) rather than by global structural properties characterizing each experimental class. Of these, some have been previously reported, such as aminoquinolines (amodiaquine- related), nucleosides (Remdesivir-like) and phenothiazines (chlorpromazine-like, antipsychotic drugs) but others have not been previously discussed in detail (pyrimidyl-indolines, similar to GSK2606414), azaspiranes, phenylpiperidines, benzodiazines, phenylpropanoids and steroid lactones (cardiac glycosides).

We next evaluated if the drugs that presented structural similarity were also targeting the same or homologous proteins. For that, we selected all pairwise molecule structures with Jaccard similarity higher than 0.5 for which both had at least one protein target described in the Broad Library. We next calculated the gene overlap of those drugs, and found that out of the 32 pairwise similar drugs, 29 compound pairs shared at least one protein target (Fig. S4). For the remaining 3 pairs, we compared the protein target sequences and found each were related. Only one pair (sertraline and indraline) share structural similarity without sharing protein target similarity. This analysis suggests that for these related pairs of molecules, activity may correspond to homologous or similar protein targets.

### Computational analysis reveals enriched drug targets and pathways that are involved in SARS-CoV-2 replication

Of the 6710 compounds, 4,277 had attributable host targets annotated in the DRH database. GO enrichment showed a similar profile with active drugs enriched for processes related to rRNA, microtubules, collagen and proteoglycan binding, and GPCR receptor activity, GTP metabolism, viral transcription processes, ion channel function and protein folding (Table S3). It is interesting that angiotensin-related drugs were identified as SARS-CoV-2 uses angiotensin converting enzyme 2 (ACE2) as a receptor. While a topic of interest early in the pandemic, patients already taking such drugs have not shown a beneficial outcome ^18^. Furthermore, effort was made to prevent oversampling of promiscuous compounds and over-represented targets but GPCR-targeting compounds are one of the most abundant drug classes and, thus, must be interpreted with caution. Fisher’s exact test was used to identify if active compounds were enriched for known drug-targets. Of the 389 active compounds, 813 targets were identified and only 51 targets appeared enriched (Fisher’s exact test; p < 0.05). These were related to sodium ion export, membrane repolarization, regularization of cardiac conduction (Table S4). The drug-target network analysis for these compounds revealed two large modules of drugs sharing its targets, comprising 61 drugs, and 119 targets, and 43 drugs and 66 targets. We also identified 105 additional smaller modules suggesting compounds targeted a discrete group of host targets (Fig. 2D).

To understand where these drug-targets are found in a human protein-protein interaction network, we first calculated the largest connected component (LCC) targets in each outcome class and the significance of the module size using a degree-preserving approach ^19^, preventing the same high degree nodes being selected repeatedly. We find that the LCC of the drug targets are statistically significantly larger than random for strong (149; 126.02 ± 10.91; Z-Score 2.11) and weak (211; 173.84 ± 15.18; Z-Score 2.45) classes. The combined class was also significant (311; 277 ± 18.69; Z-Score 1.8) (Fig. S5). Taken together, the analysis suggests clustering of drug targets within the interaction network.

To better understand the relationships of active compounds the network separation between targets of each category was computed (Fig. 2E). Similar to our previous study ^20^, targets of active compounds have a negative network separation (7._?,-N\_= -0.35), indicating that each targets the same neighborhood in the human protein-protein interaction network. In contrast, inactive compounds have close-to-zero or positive separation from the active compounds (7._V\.-N\_= -0.05, 7._V\.-?,_= 0.11), indicating inactive compounds target a different network neighborhood. Interestingly, when we asked if targets were related to identified SARS-CoV-2 host protein interactions ^21^ we found, similar to our previous study ^20^, that the relative network proximity of each target module to the COVID-binding proteins is predictive of efficacy. Active compounds have z-scores close to zero (Fig. 2F) whereas, inactive compounds have a much stronger positive proximity z-score, indicating their targets are far from COVID-binding proteins than random expectation. Taken together, these data show that active and inactive compounds target distinct network neighborhoods in the human protein-protein interaction network, and their network proximity to the COVID-binding proteins is predictive of drug efficacy.

In addition to analyzing gene targets, 6,150 of the compounds were annotated for mechanism of action in the DRH ^12^. Using this information, we checked for over- and under-represented MOA that could be driving the drug response using a ’}^S^-test (p < 0.05, FDR-BH). Here, inhibitors of mTOR, antimalarial agents, ATPase, HSP, bromodomain, and tubulin polymerization were significantly enriched. Furthermore, mapping of compounds to the Anatomical Therapeutic Chemical Classification (ATC) drug categories ^22^ and chemical taxonomy identified compound classes that were over-represented in the hits (Table 1). These included previously identified classes of compounds active against SARS-CoV-2 *in vitro*, including phenothiazines, benzothiazines, and other antipsychotic agents, as well as new classes such as PERK inhibitors and the cardiac glycosides. Each of these classes were also found clustered in the structural analysis (Fig. 2C) supporting the relationship between drug class and host targets.

**Table 1.** Relationship of active compounds based on the Anatomical Therapeutic Chemical (ATC) Classification and drug categories. Drug category and therapeutic indication were evaluated for overrepresentation using a one-sided binomial exact test and Benjamini-Hochberg correction for multiple hypothesis testing to calculate the p-value.

|  | Classification | Unadjusted *p*-value |
| --- | --- | --- |
| Drug category | Phenothiazines | 0.0004 |
|  | Antipsychotic Agents (First Generation [Typical]) | 0.0394 |
| Level 2 ATC Code | Antimigraine Preparations | 0.0483 |
|  | Cardiac Glycosides | 0.0082 |
|  | Antipsychotics | 0.0006 |
|  | Antidepressants | 0.0321 |
|  | Agents Against Amoebiasis And Other Protozoal Diseases | 0.0303 |
|  | Angiotensin Ii Receptor Blockers (Arbs), Combinations | 0.0483 |
|  | Plant Alkaloids And Other Natural Products | 0.0235 |
|  | Angiotensin Ii Receptor Blockers (Arbs), Plain | 0.0235 |

### Rescreening and orthogonal validation of hits with human cell lines

Based on performance in the primary screen and compound classification using the structural similarity clustering and ATC classifications, 40 representative compounds were chosen for detailed potency determination, and mechanistic analysis. The compounds were initially rescreened in VeroE6 cells to confirm activity before evaluation in human cell lines and timing was set to capture limited virus replication. In general, most compounds yielded EC_50_ values similar to that expected from the primary screen potency determination (Table 2, third column, Figure S6 and examples in Fig. 3A). For some compounds, such as methotrexate, a complex inhibition pattern was observed upon dose titration, being inhibitory at low doses but reaching a plateau above the 50% infection level and so an EC_50_ was not calculated.

**Table 2.** Effects of small molecule treatment on virus infection. Shown are the annotated mechanism of action of each compound, the EC_50_ values obtained for infection inhibition in VeroE6 cells and subsequent testing in Huh7 cells for virus mRNA production and virus- associated gRNA release into culture supernatant. For the VeroE6 cell tests, 10 concentrations from 2 nM up to 4 µM were used to construct dose response curves (Fig. S6) and EC_50_ values calculated. For the Huh7 cells, treatments started at this EC_50_ value ± 4-fold and tested again at reduced concentrations if higher than expected potency was seen. Measurements were performed in triplicate and concentrations giving the indicated reduction in signal are shown.

| Compound | DRH mechanism of action target | EC<sub>50</sub> (µM) in VeroE6 cells | gRNA release inhibitory concentration >50% (µM) | Virus mRNA synthesis inhibitory concentration >50% (µM) | mRNA:gRNA fold difference | <sup>b</sup>Indicated site of action<sup>a</sup> |
| --- | --- | --- | --- | --- | --- | --- |
| Narasin | Antiprotozoal agent | 0.01 | 0.15 | 0.05 | 0.4 |  |
| K-strophanthidin | ATPase | 0.75 | 0.15 | 0.45 | 3 |  |
| Ouabain | ATPase | 0.045 | 0.15 | 0.15 | 1 |  |
| VE-822 | ATR kinase | 1.33 | 0.45 | 1.30 | 2.9 |  |
| Nanchangmycin | Autophagy | 0.033 | 0.15 | 0.15 | 1 |  |
| Obatoclax | BCL | 0.048 | 0.02 | 0.05 | 3 |  |
| BET-BAY-002 | BET | >4 | 0.15 | >10 | >10 | egress |
| BMS-986158 | BET | >4 | >10 | >10 | - |  |
| CPI-0610 | BET | >4 | >10 | >10 | - |  |
| Mivebresib | BET | >4 | 0.15 | >10 | >10 | egress |
| Calpeptin | Calpain protease | 2 | 1.30 | 4.00 | 3.1 |  |
| Proscillaridin | Na<sup>+</sup>/K<sup>+</sup> channel | 0.001 | 0.01 | 0.05 | 3.9 |  |
| VBY-825 | Cathepsin protease | 1.8 | 1.30 | 1.30 | 1 |  |
| Talniflumate | Cyclooxygenase | >4 | 1.30 | >10 | >10 | egress |
| Methotrexate | DHFR | <sup>a</sup>- | >10 | >10 | - |  |
| Pralatrexate | DHFR | >4 | 0.15 | >10 | >10 | egress |
| BAY-2402234 | DHODH | 0.006 | 0.02 | 0.01 | 0.4 |  |
| Sangivamycin | DNA synthesis | 0.045 | 0.02 | 0.05 | 3 |  |
| Anisomycin | DNA synthesis | 0.049 | >10 | >10 | - |  |
| Eliglustat | Glycosyl transferase | >4 | 4.00 | 12.00 | 3 |  |
| A-485 | Histone acetyltransferase | 0.65 | 1.00 | 4.00 | 4 |  |
| BIX-01294 | Histone lysine methyltransferase | 0.82 | 0.45 | >10 | >10 | egress |
| AT13387 | Heat shock protein | >4 | 0.15 | 0.15 | 1 |  |
| Ganetespib | Heat shock protein | >4 | >10 | 0.45 | <0.1 | replication |
| NVP-HSP990 | Heat shock protein | >4 | >10 | >10 | - |  |
| SNX-2112 | Heat shock protein | >4 | >10 | 0.45 | <0.1 | replication |
| Apilimod (STA-5326) | Interleukin synthesis | 0.1 | 0.45 | >10 | >10 | egress |
| Omipalisib (GSK2126458) | mTOR/PI 3K | >4 | 0.05 | 0.15 | 3 |  |
| Deslanoside | Na/K-ATPase | 0.165 | 0.45 | 0.45 | 1 |  |
| Mepacrine | NFkB pathway | 1.33 | 0.05 | 0.45 | 9 | egress |
| APY0201 | Phosphoinositide dependent kinase | 0.001 | 1.30 | 1.30 | 1 |  |
| MG-132 | Proteasome | 0.15 | 0.45 | 1.30 | 2.9 |  |
| Bruceantin | Protein synthesis | 0.11 | 0.15 | 0.45 | 3 |  |
| Emetine | Protein synthesis | 0.125 | 0.05 | 0.05 | 1 |  |
| Harringtonine | Protein synthesis inhibitor | 0.12 | 0.01 | 0.05 | 5 | egress |
| Homoharringtonine | Protein synthesis | 0.43 | 0.02 | 0.15 | 8.9 | egress |
| Tioguanine | Purine antagonist | 1.2 | 0.45 | 4.00 | 8.9 | egress |
| Niclosamide | STAT/DNA replication | >4 | 1.30 | 4.00 | 3.1 |  |
| Bafilomycin A1 | V-ATPase | 0.12 | 0.05 | 0.05 | 1 |  |
| GSK-2606414 | PERK<br>inhibitor | >0.36 | ND | ND | - |  |

<sup>a</sup> Gave a dose response curve but did not cross 50% infection threshold and so EC<sub>50</sub> was not calculated
<sup>b</sup> Site of action is based on the ratio of potencies for inhibition of virus mRNA production in cells and gRNA release into culture supernatants. Potent inhibition of mRNA indicates inhibition of RNA replication, whereas a lack of replication inhibition but more potent inhibition of gRNA release indicates selective inhibition of egress.
ND – Not determined.

**Figure 3.**
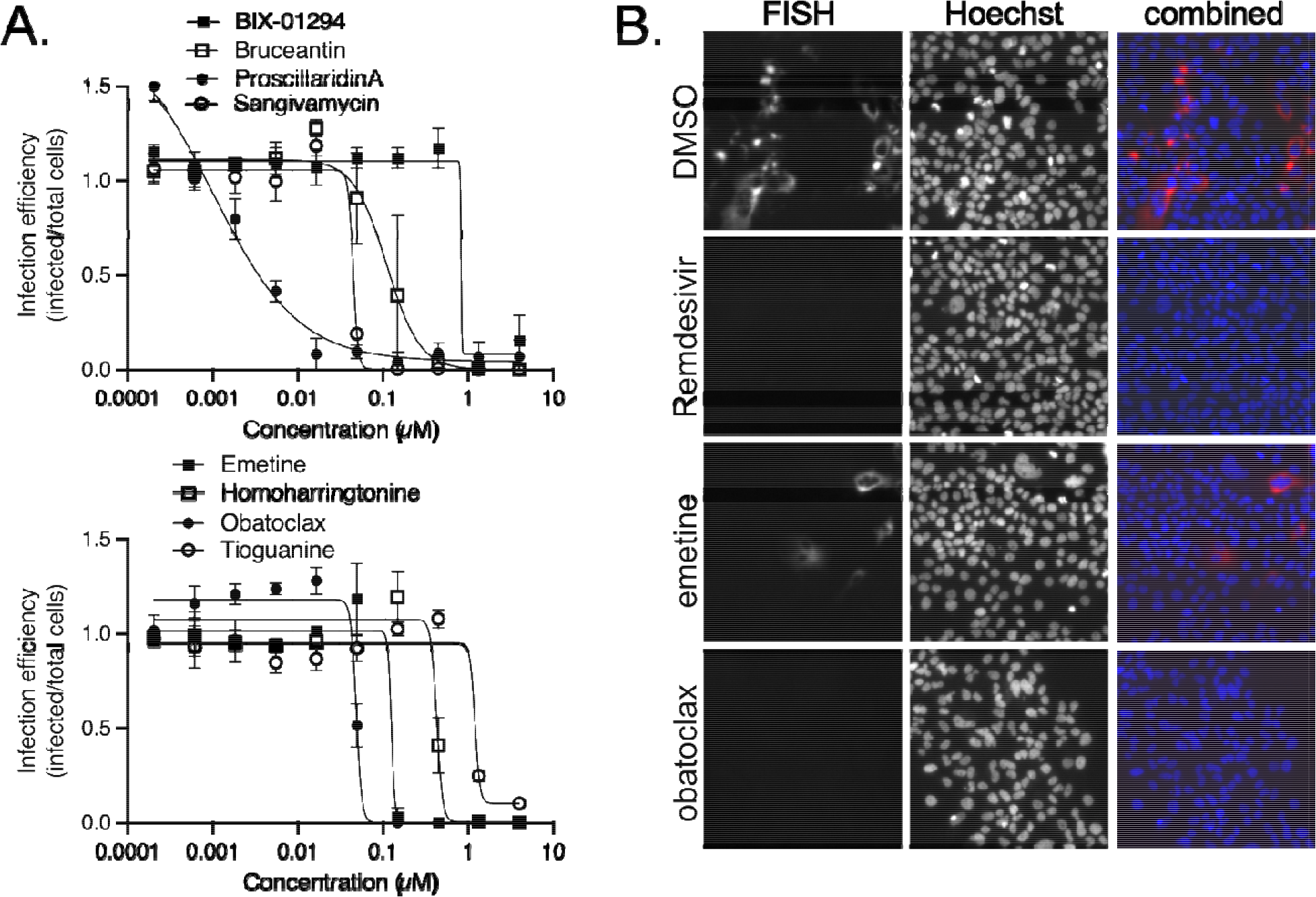
Concentration response curves for active compounds and measurement of small molecule effect on mRNA production from the virus subgenomic promoter by FISH assay in Huh7 cells. A) Examples of response curves are shown for the indicated active compounds using treatment concentrations ranging from 4 uM down to 0.2 nM. Each concentration was repeated in triplicate and average and standard deviations shown and is normalized to vehicle (DMSO) treated controls. B) Virus mRNA production was measured by in situ oligonucleotide- based detection and is summarized in Table 3. Examples of images are shown for the indicated compounds. Cells were stained for production of virus mRNA encoding the N protein by the smiFISH method using virus gene specific oligonucleotides and a Cy5 labeled oligonucleotide that bound to a shared complementary region on each (red). Cell nuclei were stained with Hoechst 33342 (blue).

Compounds were then evaluated using a human cell line, Huh7, that is intrinsically susceptible to SARS-CoV-2 infection. Orthogonal assays measured cell-associated virus RNA levels by a FISH assay and qPCR. The FISH assays was used on fixed cells and the qPCR was used on virus containing cell supernatants to measure replication and then release respectively. Since the qPCR and FISH assays have low throughput, initial concentrations of compound were limited to the EC_90_ values from VeroE6 cell tests +/- 4-fold. For the PCR based assay, strong and statistically significant effects of harringtonine, homoharringtonine, proscillaridin, BAY-2402234, obatoclax, and sangivamycin were seen at <20 nM producing 74 to 93% (P<0.01) decreases in gRNA in the culture supernatant (Table 2, fourth column). While most of the compounds showed potencies similar to those in VeroE6 cells, the mTOR inhibitor, omipalisib, showed inhibition below the lowest concentration tested indicating >10-fold stronger potency than seen in VeroE6 cells. Differences were also observed for two each of the HSP90 and BET protein inhibitors that were classed as weak inhibitors in Vero cells. For the HSP90 inhibitor AT13387, activity at the highest dose tested (150 nM) was observed. BET inhibitors, BET-BAY-002 and mivebresib, were also active. Overall, such changes in activity suggest differences in dependency on BET and HSP90 proteins or drug action in Vero versus human Huh7 cells.

The impact of compounds on virus mRNA production was measured using a FISH assay targeting RNA production from the viral subgenomic promoter that controls expression of structural proteins (Fig. 3B) and image analysis to score replication efficiency. In general, activity in this assay reflected that seen in the qPCR-based assay with potency typically being within 3-fold of each other (Table 2). The DHODH inhibitor, BAY-2402234, was the most potent, inhibiting infection by 50% at 10 nM. The protein synthesis inhibitors, emetine, harringtonine, and homoharringtonine, were equally active, blocking >50% RNA production at 50 nM (Fig. 3B). Other strongly active compounds were the nucleoside analog, sangivamycin, as well the mTOR and BCL inhibitors, omipalisib and obatoclax, respectively. Cardiac glycosides, proscillaridin and ouabain, were also active at 50 and 150 nM, respectively. Of note was the large difference in activity of harringtonine and homoharringtonine being over 5 to 8-fold more potent in the virus genome release assay over the RNA production assay, which would be consistent with disruption of virus assembly and may reflect the need for balanced stoichiometry of structural proteins easily disrupted by protein synthesis inhibition. Similarly, differences were noted for compounds that inhibited genome release from cells in the qPCR assay but had negligible impact on the viral mRNA signal. These compounds were the DHFR inhibitor, pralatrexate, the COX inhibitor, talniflumate, the BET inhibitors, BET-BAY-002 and mivebresib, and the methyltransferase inhibitor, BIX-01294, suggesting that each also worked to disrupt packaging of viral genomes into new virus particles and/or their release into the culture medium. A recent report indicated that methotrexate, a paralog of pralatrexate, can block release of SARS-CoV-2 genomic RNA into the culture medium ^23^. For the COX inhibitor, talniflumate, COX activity has also been shown to be important for packaging of pseudorabies genomes into capsids ^24^. Furthermore, BET proteins were identified as interacting with SARS-CoV-2 E, a protein important for maturation and release of virions from cells ^21^ and were identified as a COVID-network associated drug target class l analysis in our computational analysis (Fig. 2E, F). To our knowledge, a role for methyltransferases in production and packaging of virus genomes has not been reported and will require additional work to understand its role.

### Pseudotype assay to test entry inhibition

We next sought to evaluate the impact of compounds that displayed the most potent effects on virus replication on SARS-CoV-2 spike glycoprotein (S)-mediated entry. SARS-CoV-2 S protein mediates virus entry upon binding to the angiotensin-converting enzyme 2 (ACE2) receptor and subsequent proteolytic activation by either cell surface expressed serine proteases, such as TMPRSS2 or endosomal cathepsins, CatL or CatS, to trigger fusion of viral and host membranes ^15^. Eleven compounds were tested at the Vero E6 EC_90_ concentrations in A549 cells expressing recombinant ACE2 (Fig. S7) and challenged with SARS-CoV-2 S pseudotyped lentiviral vectors (LV). Of the compounds evaluated, omipalisib and BAY-2402234 most potently inhibited SARS-CoV-2 S pseudotyped LV entry by >95%, P<0.05 (Fig. 4), though similar inhibition was also observed against vesicular stomatitis virus glycoprotein (VSV-G) pseudotyped LV, suggesting these molecules likely targeted a post-entry step. Nanchangmycin also similarly affected both SARS-CoV-2 S and VSV-G pseudotype LV entry (75% inhibition, P<0.05). In contrast, Narasin displayed selective inhibition (∼80%) of SARS-CoV-2 S pseudotypes without significantly affecting VSV-G pseudotype LV entry. Salinomycin, a methylated analogue of narasin, has been shown to alter lysosome function ^25^, which is suggestive of an effect of narasin on endosomal uptake of the virus. Similarly, proscillaridin also appeared to specifically block entry of SARS-CoV-2 S pseudotype LV. Proscillaridin was shown to reduce receptor availability for hepatitis B virus ^26^ and thus may affect a membrane function important for SARS-CoV-2 infection. Obatoclax displayed partial inhibition of SARS-CoV-2 S pseudotype LV entry. Despite being characterized as a BCL inhibitor, obatoclax has been reported to interfere with endosomal acidification pathways needed for entry of alpha and flaviviruses into cells ^27^. The partial inhibition of SARS-CoV-2 S-mediated entry by proscillaridin and obatoclax compared to the almost complete inhibition of virus replication (robust decrease in N protein expression and viral RNA copy number in cell supernatants) suggests each drug may act at an entry as well as othervirus replication steps. The methyltransferase inhibitor BIX-01294, the PERK inhibitor GSK2606414, the CREBBP/p300 inhibitor A485, the nucleoside analog sangivamycin, and the protein synthesis inhibitor emetine, were each ineffective in blocking SARS-CoV-2 S-mediated entry, and based on activity in the replication assays, must act at steps after virus entry into cells.

**Figure 4.**
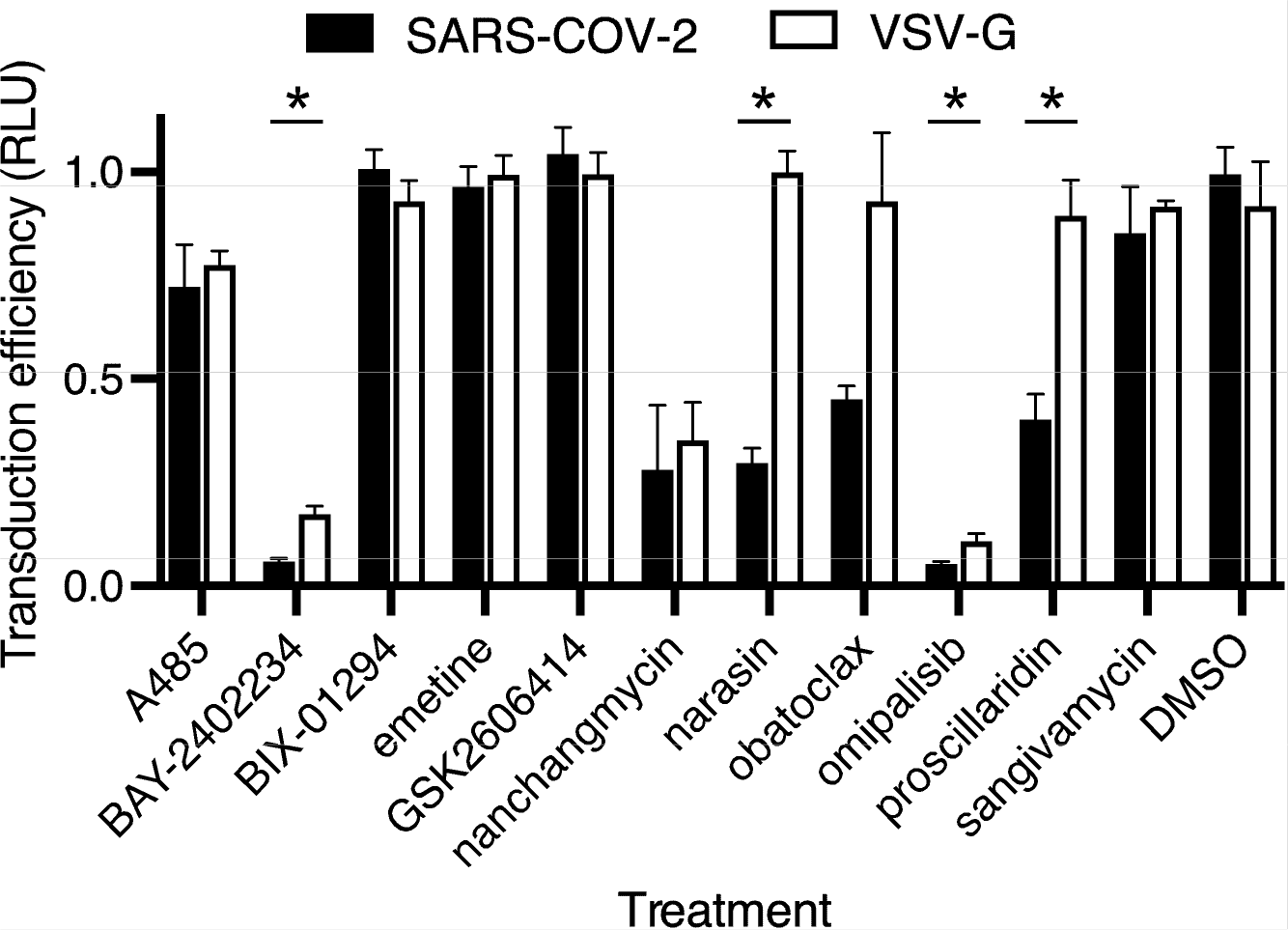
Effect of compounds on infection by virus glycoprotein pseudotyped viruses. A549 cells expressing recombinant ACE2 protein were pretreated with each indicated compound at the EC_90_ concentration from assays using wild type virus on VeroE6 cells. Lentivirus pseudotypes, encoding firefly luciferase, as a marker of infection, and bearing the SARS-CoV-2 or VSV glycoproteins (VSV-G) were used to transduce cells at 20 ng of p24 capsid equivalent per infection. Transduction efficiency was measured by luciferase activity as relative light units (RLU) and normalized to DMSO-treated controls. Measurements used 3 replicates with the average and standard deviation shown. Multiple t-tests compensating for false discovery (Q=2) were used to identify significant differences (P<0.05) between the VSV-G and SARS-CoV-2 pseudotype infection efficiencies indicated by *.

### Prioritized compounds do not show phospholipidosis

Recently, it was reported that many compounds from repurposing libraries showing antiviral activity act by by inducing phospholipidosis (PLD) and disrupting cell lipid metabolism ^28, 29^. Since, some of the compounds of interest had potentially protonatable amines, a hallmark of PLD inducers, we evaluated each by NBD-PE staining, but found that none induced PLD (Fig. S8) even when tested at >100 times higher than their anti-viral active concentrations. We conclude that the compounds act through non-PLD mechanisms.

### Confirmation of SARS-CoV-2 inhibition in a physiologically relevant 3D tissue model

Four of the most active treatments were evaluated using primary lung and gastrointestinal multicellular models. While COVID19 causes severe respiratory distress, several studies have reported over half of patients also suffer from gastrointestinal symptoms, including diarrhea, abdominal pain, and vomiting. The gut is also a prominent site for virus replication ^30^. Human donor-derived stratified epithelial cell models representing the airway and gut were obtained commercially ^31^. Similar to previous reports ^32^, infection in the lung model was variable between batches and not easily used for compound evaluation. However, the gastrointestinal model cells expressed ACE2 mRNA and protein (Fig. S9) and gave robust, uniform infection (Fig. 5A). By morphology, the infected cells appeared to be epithelial. The compounds tested in this model, including Remdesivir as an active control ^33^ were used at the VeroE6 EC_90_ concentration. Emetine, GSK2606414 and obatoclax were as effective as Remdesivir, preventing >80% of cell infection. Proscillaridin was less effective, showing inhibition ranging from 50 to 80%. These observed differences helped further refine compound activities, with the most promising showing consistent activity across cell lines and human tissue models.

**Figure 5.**
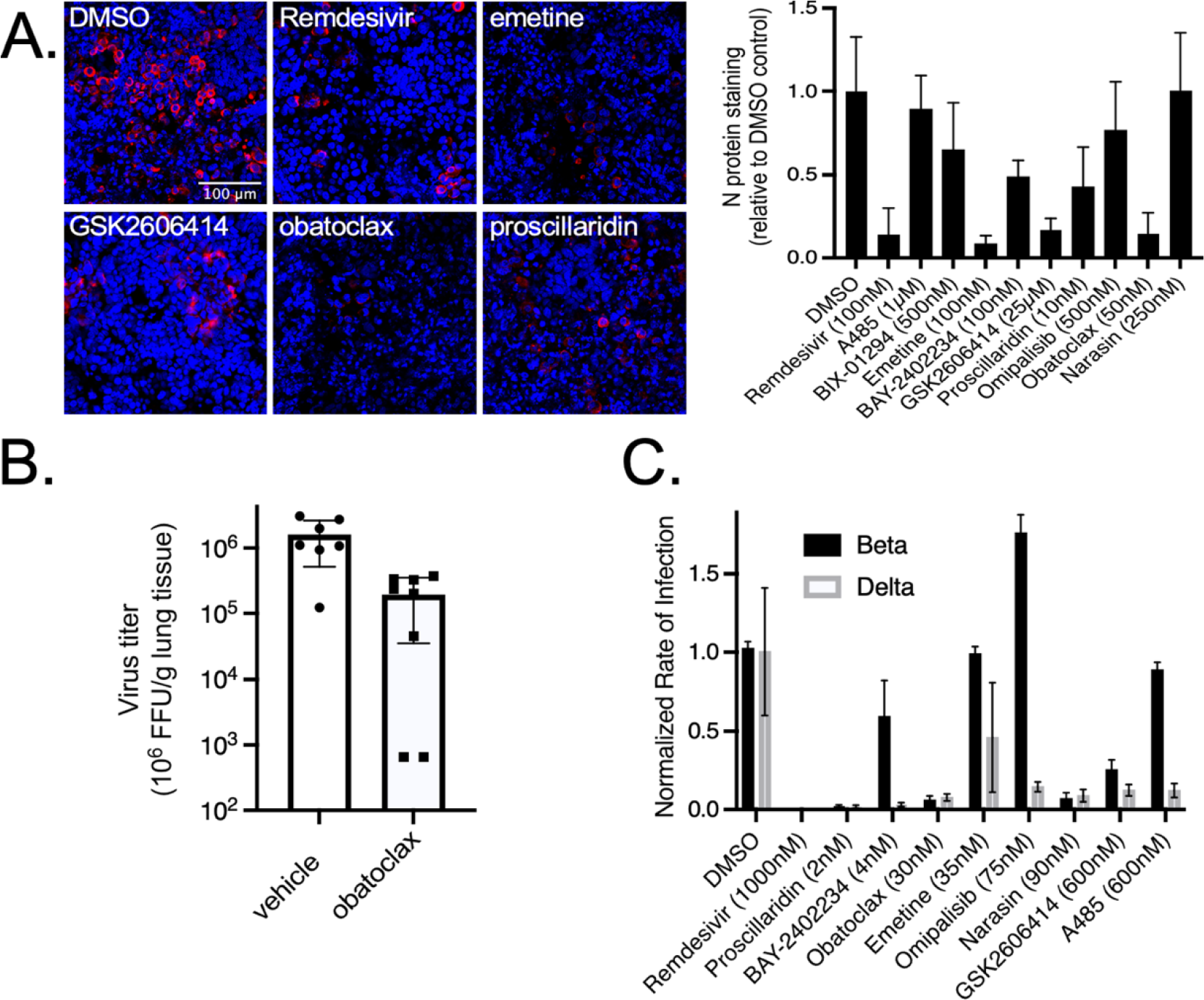
Inhibition of infection in primary human cell model and efficacy testing in the K18 ACE2 mouse model of disease. A) Primary intestinal epithelial cell cultures were challenged with SARS-CoV-2 and after 3 days fixed in formalin. Cells were stained for N protein using a specific antibody (red) and cell nuclei using Hoechst 33342 (blue) and examples of active compounds shown (left panel). Infection efficiency was measured by total fluorescence relative to DMSO-treated controls and normalized to the cell nuclei signal (right panel). Each test was performed in triplicate with average and standard deviations shown. B) Mice were challenged by the intranasal route with SARS-CoV-2 and starting 6 hours later, daily treatment with obatoclax at 3 mg/kg. Virus load in the lungs (FFU/g tissue) was measured 4 days post-challenge. For each mouse tissue sample, two measurements were made and averaged. Two female treated mice gave virus loads below and at the the limit of detection for the assay respectively. Each were plotted at 50% the LOD. C) Beta and delta virus strains were tested for susceptibility to small molecule treatments tested at the indicated doses (close to EC90) for that seen in the Washington strain using A549-ACE2 cells. Each was performed in triplicate and averages with standard deviations are shown.

### In vivo evaluation in the mouse disease model

Obatoclaxdemonstrated potent and consistent antiviral activity across al in vitro assessments, and has been reported to have favorable pharmacokinetics and tolerability in mice ^34^. We used K18- hACE2 transgenic mice, which express recombinant human ACE2 (hACE2) protein under control of a K18 cytokeratin promoter to evaluate each compound as a treatment for disease. Others have shown that challenge by the intranasal route results in virus replication in the lungs, heart, brain, kidneys and gut ^35, 36^ with titers increasing over 2-3 days and then subsiding after day 5 in most tissues, but with titers in the brain continuing to rise to >10^8^ pfu/g tissue. The latter is thought to result in severe symptoms, which may not be relevant to human disease outcomes. To focus on treatments effective in the respiratory system, animals were treated daily for 4 days after virus challenge and virus load in the lungs was measured. Vehicle control animals reached titers consistent with previously reported levels of 1-3 x 10^6^ FFU/g lung tissue. At a dose of 3 mg/kg, obatoclax gave a consistent and significant reduction in average virus load by >88% in both male and female mice (p=0.0028). For two of the female treated mice, lung virus load fell below the limits of detection of the assay, although one did have trace amounts of detectable virus (Fig. 5B). This suggests that obatoclax may be useful as a COVID-19 treatment candidate.

Lastly, we tested efficacy of each of the potent hits against Beta and Delta variants of SARS-CoV-2 (Fig. 5C) and the distantly related MERS-CoV virus (Fig. S10). Surprisingly, the Beta variant showed several differences in susceptibility to drug treatment. Of note, Omipalisib appeared to elevate infection, while A485 and BAY-2402234, showed little activity. In contrast, the Delta variant showed a similar profile to that seen for the Washington strain used for the initial screening. This suggests that the Beta variant may have a different dependency profile on cell processes disrupted by the compounds.

## Discussion

While multiple vaccine candidates are now available, alternative therapeutic avenues will be needed to control COVID19 disease. Here, we evaluated 6,710 small molecules for inhibition of SARS-CoV-2 infection using *in vitro* and *in vivo* disease models. Of the approximately 200 active compounds, multiple candidates had potencies that are clinically achievable. We performed detailed mechanistic analysis for 40 inhibitors of SARS-CoV-2 replication with differing ascribed mechanisms of action, including proscillaridin, emetine, obatoclax, sangivamycin, omipalisib, GSK2606414, and BAY2402234. Each demonstrated *in vitro* potency that may be clinically useful given further evaluation in the context of disease. Structural analysis revealed distinct clusters of related molecules with conserved molecular motifs that could be further developed into additional active compounds. Further analysis of reported mechanism of action and associated gene targets suggested discrete families of host factors involved in SARS-CoV-2 replication, related to microtubule function, mTOR, ER kinases, protein synthesis and folding and ion channel function. Mechanistic evaluation of the antiviral effects of the lead candidates revealed activities impacting each step of virus replication from cell entry through to egress and may provide new tools to better understand the infection cycle of this virus. These results highlight the host dependency factors necessary for SARS-CoV-2 replication and potential sites of virus vulnerability for further development of antiviral therapeutics.

Obatoclax consistently showed potent antiviral activity across all *in vitro* systems, as well efficacy in the mouse disease model. The 10-fold decrease in lung titers is promising given that a similar reduction of virus load has been associated with decreased mortality in COVID-19 patients ^37^. Obatoclax was originally developed as a broadly acting BCL-2 homology domain 3 inhibitor for cancer treatment ^38^ with activities including inducing apoptosis and elevating autophagy ^38, 39^, each potentially virucidal outcomes. However, the 7 other BCL-2 inhibitors that we tested were weak or inactive, indicating that BCL-2 inhibition is less likely to be a specific mechanistic target. A previous report showed inhibition of endosomal acidification by obatoclax but not by other BCL-2 inhibitors and explained inhibition of infection by alphaviruses through preventing pH-dependent endosomal escape ^27^. Consistent with this mechanism, the pseudotype infection assay (Fig. 4), showed obatoclax partially blocking infection for the SARS-CoV-2 pseudotype. However, another report showed that influenza virus, which also requires endosomal acidification to infect cells, was unaffected by obatoclax ^40^. Taken together, while SARS-CoV-2 requires acidified endosomes for cell entry and entry is partly blocked by obatoclax, an additional entry-independent modality may be involved in infection inhibition and will require more detailed characterization.

One of the most potent classes of inhibitors identified were the cardiac glycosides, such as proscillaridin, that have Na+/K+ channel blocking activity. These steroid-like molecules affect cardiac function ^41^, have been reported to alter cell membrane fluidity ^42^, affect receptor function for a range of ligands, and can induce apoptosis ^43^ of cancer cells. Based on the potent cardiac effects and narrow therapeutic windows, they are unlikely to be viable for repurposing but may provide information on virus infection mechanism. The pseudotype entry assay suggested that proscillaridin, like obatoclax, partially interfered with the virus entry step. Cardiac glycosides were reported to inhibit hepatitis B entry into cells by interfering with binding to its receptor, NTCPA3, as well as a post-entry replication step ^26^. Hepatitis B virus showed a similar spectrum of cardiac glycoside type and potency infection inhibition to that seen here for SARS-CoV-2, suggesting a similar mechanism may aid SARS-CoV-2 infection. In contrast, other cardiac glycosides, including ouabain, also a SARS-CoV-2 inhibitor, were found active against HIV-1 gene transcription by altering RNA processing ^44^ with disruption of MEK/ERK1/2 signaling being responsible for infection inhibition. Another action of cardiac glycosides is to inhibit protein synthesis ^45^. Indeed, other known protein synthesis inhibitors, such as emetine, harringtonine, and homoharringtonine, rivalled the cardiac glycosides as potent inhibitors, in some cases showing efficacy at low nanomolar concentrations. These agents have previously been well characterized as broad-spectrum antivirals ^46^ and likely reflect the importance of balanced protein production needed during virus replication and assembly.

Another class of compounds that have not been reported to affect SARS-CoV-2 infection are the BET inhibitors. These compounds, like mivebresib, affected SARS-CoV-2 infection in both cell types tested and were identified in our protein network analysis (Fig. 2). Previous work showed two of the four BET proteins, BRD2 and BRD4, were bound by virus E protein ^21^. The E protein is involved in assembly and budding of newly formed coronaviruses from the cell. Our finding that the BET inhibitors block viral egress (Table 4) is consistent with this mechanism of action. Being able to identify such egress inhibitors also serves to demonstrate the utility of the secondary orthogonal assays to detect the effects of candidate compounds on virus at different points in its lifecycle.

In addition to identifying individual compounds that can inhibit SARS-CoV-2, we have demonstrated that the data from the high-throughput screen can be used to identify the potential importance of cellular pathways required for virus replication that in turn may lead to new drug targets not part of the screen. Recent publications have demonstrated a convergence of data obtained from genetic screens, such as genome wide CRISPR and small molecule screens. For example, both genetic and compound screens have repeatedly suggested endosomal trafficking as an important pathway for viral entry and replication, including for coronaviruses ^47–49^. This view is bolstered by the reproducible activity of apilimod in our study and others ^50^, and the positive activity observed in our screen with APY0201, a second PIKfyve inhibitor. However, we also found distinct differences between closely genetically related virus strains, suggesting that even subtle changes in the virus genome can induce complicated changes in host factor dependency and replication behavior. This finding emphasizes the need to evaluate multiple virus strains for susceptibility to small molecule inhibitors.

The two largest compound screens reported to date are this study and the screen of the 12,000 compound ReFRAME library ^9^. Significant overlap was observed in the classes of molecules observed to have activity, but new active compounds were also observed. The main submicromolar potency leads identified in the ReFRAME study included apilimod, VBY-825, ONO 5334, Z LVG CHN2, and MDL 28170. Of these, apilimod and VBY-825 were present in the DRH library giving similar activities in both studies. In the present study, we also identified additional active drug classes including the BET inhibitors. The main difference in the assay design of the two screens was use of a single treatment dose (5 µM) and measuring infection by virus-induced cell death for the REFRAME screen versus multiple concentrations and virus protein expression used here and likely accounts for differences in outcomes. A recent report indicated that many drug repurposing efforts for antivirals, may enrich for compounds that induce phospholipidosis (PLD) by disrupting cell lipid metabolism. Most of these compounds have micromolar potencies against virus that parallel effects on PLD. By examining only the most potent compounds in our screen, we likely avoided such effects, as indeed none of the compounds showed PLD even when tested at over 100 times that needed to inhibit virus infection.

In summary, using a high-throughput drug screen, we have elucidated both potent antivirals and host factors strongly entwined in the SARS-CoV-2 lifecycle. We aimed here to provide both a more detailed understanding of how the virus hijacks host machinery to replicate, as well as methods to interrupt these dependencies. By making all the data publicly available through the DRH we anticipate that both the active and inactive compounds will inform future studies. Due to the pressing need for antivirals to combat COVID-19, our findings will aid scientists and clinicians with identifying, prioritizing, and testing novel therapeutics, and help alleviate the burden this pandemic has placed on our society. While these findings are encouraging, and suggest drug-repurposing as a productive approach for rapidly identifying treatments, it is important to be mindful that novel drug-disease interactions may occur when used for a different purpose, necessitating prospective clinical trial assessments.

## Materials and Methods

### Wet Lab methods

#### Cell and virus cultivation

Vero E6 cells were obtained from ATCC (Manassas, VA, USA) and maintained in DMEM supplemented with 10% fetal bovine serum (FBS) at 37°C in a humidified CO_2_ incubator. The SARS-CoV-2 strain USA-WA1/2020, was isolated from a traveler returning to Washington State, USA from Wuhan, China, and was obtained from BEI resources (Manassas, VA, USA). The virus stock was passaged twice on Vero E6 cells by challenging the cells at an MOI of less than 0.01 and incubating until cytopathology was seen (typically 3 days after inoculation). A sample of the culture supernatant was sequenced by NGS and was consistent with the original isolate without evidence of other viral or bacterial contaminants. The virus stock was stored at - 80°C. SARS-CoV-2 B.1.351, designated Beta (hCoV-19/South Africa/KRISP-K005325/2020), and MERS-CoV (EMC/2012) were also obtained from BEI and grown as above. We thank Jacquelyn Turcinovic, Scott Seitz, and John H. Connor, NEIDL, Boston University, for isolation and sequence analysis of the Delta variant, SARS-CoV-2 B.1.617 designated as hCoV-19/USA/MA-NEIDL-01399/2021. The virus was cultivated by Devin Kenney and provided by Dr. Florian Douam, NEIDL, Boston University.

Cells used for testing SARS-CoV-2 Beta and Delta variants were made by transduction of A549 cells with pHAGE-EF1-ACE2. ACE2 expression measured by FACS after staining with ACE2 specific antibody and appropriate secondary fluorescently labeled antibody. The cell pool was sorted by FACS, without prior selection, into four pools based on ACE2 surface expression levels. Cells were then passaged and ACE2 expression was confirmed. The lowest ACE2 expressing cells were used for testing efficacy of small molecules for inhibition of SARS-CoV-2 variants.

A549 cells stably expressing DPP4 were generated by transducing A549 cells (ATCC CCL-185) with DPP4 lentivirus. To generate DPP4 lentivirus, 293FT cells were transfected with pLEX307- DPP4-puro (Addgene plasmid #158451), pLP1, pLP2, and pVSV-G using calcium phosphate. 16 hours post transfection, supernatant was discarded and replaced with fresh media. 48 hours after transfection, cell supernatant was collected, clarified by centrifugation, and used to inoculate A549 cells. Two days after transduction, cells were put under 1 µg/ml puromycin selection. For FACS, 5 x10^6^ transduced cells were stained with 1.25µg DPP4 antibody (R&D Systems AF1180) for one hour at room temperature. After washing cells 2X with PBS, Chicken anti-goat-FITC secondary antibody (Invitrogen A21467) was applied for 45 mins and then washed 2X with PBS before sorting. Cells were sorted into four bins based on DPP4 expression.

#### Primary high throughput drug screening against SARS-CoV-2

A total of 6,710 compounds from the Broad Institute DRH (Cambridge, MA, USA) were Echo plated at 4 doses in 384 well plates by Broad Institute staff. The night prior to screening, 7x10^3^ Vero E6 cells were seeded into each well of a 384 well plate. For evaluation of small molecule efficacy against infection with wild type SARS-CoV-2 virus, compounds were first dissolved in DMSO and then diluted into culture medium before addition to cells (final concentration of DMSO <0.5%). The cells were incubated for a minimum of 1 hour, moved to the biocontainment laboratory, and challenged with virus at an MOI between 0.1 and 0.3. Dosing was at a final concentration of 8, 0.8, 0.08, and 0.008 µM. As a positive control, 5 µM E-64d was used as it was previously reported to inhibit SARS-CoV-2 infection ^15^. Negative controls were treated with DMSO at 0.5%. After 36 to 48 hours, cells were submerged in 10% neutral buffered formalin for at least 6 hours, removed from the containment lab, and washed in PBS. Cells were permeabilized in 0.1% (v/v) Triton X-100 for 15 minutes and blocked in 3.5% BSA for 1 hour. Virus antigen was stained with SARS-CoV-2 specific antibody MM05 (Sino Biologicals, Beijing, China) overnight at 4°C. Alexa Fluor 488-labeled goat anti-mouse antibody from ThermoFisher (Waltham, MA, USA) was added to cells for 2 hours, cells were washed in PBS, and Hoechst 33342 dye was added to stain cell nuclei. Plates were imaged on a Biotek Cytation 1 automated imager and CellProfiler software (Broad Institute, MA, USA) was used for image analysis incorporating a customized processing pipeline (available on https://github.com/RDaveyLab/COVID19/). Infection efficiency was calculated as the ratio of infected cells to total cell nuclei. Reduction of nuclei was used to flag treatments as indicative of potential cytotoxicity. The assay was performed in duplicate. Results from the assay will be available at https://repurposing.broadinstitute.org/AssayResultsViewer/Results.

#### Reconfirmation of top hits

To verify the results of the initial screen, small molecules were chosen for full dose-response evaluation based on initial potency and drug class. The night before the screen, 2x10^4^ Vero E6 cells were seeded in each well of a 96 well plate. As before, compounds were diluted in culture medium and incubated on cells for a minimum of 1 hour, with a range of final concentration from 4 µM to 0.2 nM in a three-fold dilution series. Infection, fixation, staining, and analysis were performed as described above. The assay was performed in triplicate.

#### Validation of inhibitors in human cells

Compounds that yielded consistent dose-responses were further evaluated using qPCR to detect virus genomes and in situ virus mRNA detection as orthogonal assays. They were also evaluated in the intrinsically infectable human cell line, Huh7.

#### RT-qPCR detection of viral genomes from cell supernatants

To measure virus assembly and release, qPCR was used to detect viral genomic RNA in the cell culture supernatant using a method modified from Suzuki *et al.* ^51^. The cell supernatant was collected at 48 h post infection, and then mixed with 2x virus lysis buffer (0.25% Triton X-100, 50 mM KCl, 100 mM Tris-HCl pH 7.4, 40% glycerol) at an equal volume for 10 min at room temperature. Five microliters of the mixture was added to Luna Universal Probe One-Step RT-qPCR mixture (NEB, MA, USA) to a final volume of 20 µl. PCR amplification was detected and validated by CFX96 Touch Real-Time PCR Detection System (Bio-Rad, CA, USA). The cycling protocol used was 55°C for 10 min, 95°C for 1 min, followed by 40 cycles of 95°C for 10 sec, and 60°C for 30 sec. The primer and probe sets were from 2019-nCoV RUO Kit (IDT, IA, USA) and nCoV_N2 forward and reverse primers. The primer sequences were forward primer 2019- nCoV_N2-F 5’-GAC CCC AAA ATC AGC GAA AT-3’, reverse primer 2019-nCoV_N2-R 5’- TCT GGT TAC TGC CAG TTG AAT CTG-3’ and probe 2019-nCoV_N2-P 5’-FAM-ACCCCG CAT TAC GTT TGG TGG ACC-BHQ1-3’.

As a positive control for RT-qPCR of SARS-CoV2 RNA, an RNA fragment was synthesized. In brief, 2019-nCoV_N_Positive Control (IDT) DNA was amplified using T7 promoter-containing forward primer 5’-TAATACGACTCACTATAGGGTAAAGGCCAACAACAACAAG-3’ and reverse primer 5’-GAGTCAGCACTGCTCATGGATTG-3’ from GENEWIZ (MA, USA). After electrophoresis and gel extraction by Monarch DNA Gel Extraction Kit (NEB), the PCR product was transcribed using HiScribe T7 High Yield RNA Synthesis Kit (NEB) according to the manufacturer’s instructions, followed by DNA template removal by DNase I (NEB) treatment and purification using Monarch Cleanup Kit (NEB). RNA copy number was calculated from its molecular weight and absorbance measured by a NanoDrop 1000 (Thermo Fisher Scientific, MA, USA). A standard curve was generated using dilutions of RNA to relate genome copy number to qPCR cycle threshold (Ct). For each qPCR reaction set, 4 of the standards were included to ensure assay performance.

#### smiFISH detection of viral mRNA

RNA fluorescence in situ hybridization (FISH) was used to measure the amount of virus subgenomic mRNA being produced and its localization in cells following a method adapted from Tsanov et al. ^52^ and optimized for SARS-CoV-2. Thirty-one oligonucleotide probes were designed to hybridize to ORF3a and S subgenomic mRNA using Oligostan software (see table below). Each had a common tail that bound to a complementary sequence on an oligonucleotide conjugated to a Cy5 fluor. Cell nuclei were stained with Hoechst 33342 dye and samples were imaged on a Cytation 1 microscope (Biotek, VT, USA). Images were quantified using CellProfiler ^53^, (using pipelines available on https://github.com/RDaveyLab/COVID19/) and infection efficiencies calculated from the percentage of RNAFISH positive cells in each sample.

#### Pseudotype evaluation of treatment effects on cell entry

Single round SARS-CoV-2 S or VSV-G pseudotyped luciferase-expressing lentivectors were generated via calcium phosphate mediated transient co-transfection of HEK293T cells with HIVΔenv/luc and SARS-CoV-2 S/gp41 plasmids ^54^. The SARS-CoV2 S/gp41 expression plasmid was a gracious gift of Dr. Nir Hacohen (Broad Institute), that expresses a codon-optimized version of CoV-2 S and modified to include the eight most membrane-proximal residues of the HIV-1 envelope glycoprotein cytoplasmic domain after residue 1246 of the S protein. The VSV-G expression plasmid (HCMV-G) has been described previously ^54^. Virus-containing cell supernatants were harvested 2 days post transfection and filtered using 0.45µm syringe filters, aliquoted and stored at -80°C until further use. The p24^gag^ content of the virus stocks was quantified using a p24^gag^ ELISA, as described before ^55^. ACE2-expressing lentivectors were generated via transient co-transfection of HEK293T cells with RRL.sin.cPPT.SFFV/Ace2.IRES-puro (Addgene Plasmid #145839), psPAX2 and VSV-G. A549 cells were transduced with ACE2 lentivectors, selected for ACE2 expression by culturing in puromycin-containing media, and cell surface expression of ACE2 was confirmed by FACS. For entry inhibition assays, A549/ACE2 cells (1x10^4^ cells per well of 96 well flat bottom plate) were pretreated with the compounds at the indicated concentrations for 1 h prior to spinoculation with 20 ng of p24^gag^ equivalent SARS-CoV-2 S or VSV-G pseudotyped lentivirus particles. Cells were lysed 48 h post infection, and cell lysates processed for measurement of luciferase activity.

#### Phospholipidosis assay

A549-ACE2 cells were plated in 96-well plates in RPMI supplemented with 10% FBS and allowed to attach overnight. NBD-PE dissolved in DMSO was added to cells for a final concentration of 17µM, and cells were incubated with reagent for 2 hours at 37LC. After 2 hours, compound dose curves, beginning at 10µM and following a 9-point 5-fold dilution series, were added to cells and cells were incubated overnight at 37L positive control. After 22 to 23 hours post compound addition, cells were fixed in 10% neutral buffered formalin for 30 minutes at room temperature, then stained with Hoechst at 1:10,000 in PBS. Plates were imaged at 4x on a Cytation Cell Imaging Multimode Reader using Gen 3.08 acquisition software. Phospholipidosis (PLD) was quantified using a CellProfiler pipeline. Briefly, Hoechst-stained nuclei were expanded to encompass approximate cell area and PLD, defined as GFP signal above background, was classified as an object. PLD falling within the cell area was attached to each cell to define a new object, and the total intensity of PLD-positive cells in each image was measured. PLD was normalized to fold change over DMSO treatment and compared to amodiaquine (a known PLD inducer) to determine PLD activity.

#### Reconstructed in vitro 3D tissue model of small intestine

Human small intestine epithelial cells were obtained and used to produce a reconstructed tissue model as described previously ^31^. Briefly, cryoprotected intestinal epithelial cells were seeded onto collagen coated cell culture inserts (MatTek Corporation, 0.6 cm2) in medium (SMI-100- MM, MatTek Corporation, Ashland, MA). Cells were cultivated submerged for 24 hours, followed by cultivation for 13 days at the air-liquid interface (ALI) at 37°C, with 5% CO2 and 98% relative humidity. During the ALI culture period, tissues (designated SMI-200 or EpiIntestinal) were fed basolaterally through the membrane of the cell culture inserts of the 24- well plate. During this culture period, epithelial cells and fibroblasts self-assemble in the correct orientation. Under this culture condition, the organotypic tissues stratify, differentiate, and form a distinct apical-basolateral polarity. The polarized organotypic small intestinal full thickness tissues form “villi-like” tissue structure with an apical epithelial architecture on top of a fibroblast substrate. To complete cellular differentiation and stratification, cells were cultured for a total of 14 days prior to their shipment for the SARS CoV-2 infection studies.

Tissues were supplied growing on permeable membranes in transwell cultures. The tissues were challenged on the apical and basolateral sides at an MOI of 1 (estimated from virus titer on Vero E6 cells and 10^6^ cells in the tissue model). After 1 hour, the inoculum was washed off and the culture incubated for an additional 3 days. Tissues were then fixed in formalin, and stained for SARS-CoV-2 N protein and Hoechst 33342 as above. The tissue was mounted on a glass slide and imaged using a Zeiss LSM 700 laser scanning confocal microscope.

#### Drug testing against variants of concern and MERS-CoV

The evening prior to infection, 1.5x10^4^ A549 cells expressing ACE2 or DPP4 were seeded into 96 well plates. Compounds were dosed out at three concentrations- their respective IC90s, plus 5-fold above and below. After a one hour incubation, virus was added to the pretreated cells (approximately 100 FFUs/well of SARS-CoV-2 Beta, 200 FFUs/well of SARS-CoV-2 Delta, and 200 PFUs/well of MERS-CoV). The SARS-CoV-2 infected cells were fixed after 36-48 hours, immunostained for viral infection, and quantified as described above. The MERS-CoV infected cells were allowed to incubate for five days prior to fixation. Cell nuclei were stained with Hoechst and counted. Treated/untreated and infected wells were normalized to untreated and uninfected wells to determine if the addition of drug rescued cell counts.

#### Mouse model treatment and infection focus assay detection of virus load in lungs

*B6.Cg-Tg(K18-ACE2)2Prlmn/J (*K18-hACE2) transgenic mice were purchased from Jackson laboratories (Farmington, CT). Mice were challenged at 10-15 weeks of age in groups of 4 or 5 by the intranasal route with 10^5^ FFU SARS-CoV-2 per nare. Starting at 6 hours after virus challenge and once each day, animals were dosed by the intraperitoneal route with the indicated treatment. Four days after challenge, mice were euthanized and lungs removed at necropsy. A biopsy punch (4 mm) was used to collect 2 samples of lung tissue and was stored frozen in PBS. For evaluation of virus load, the lung tissue was homogenized using a TissueLyser II (Qiagen, MD, USA) for two cycles at 30 Hz for 2 min. Debris was pelleted for 10 min at 16,800 xg in a centrifuge (Eppendorf, CT, USA). The supernatant was titrated onto Vero E6 cells and after 1 hour overlaid with a 5% solution of methylcellulose in DMEM and incubated overnight. Cells were fixed in 10% formalin and stained with the N specific antibody used for screening. Foci of infected cells were counted and titers calculated.

#### Computational methods. Drug-Response Classifier

In order to classify the drug response by potency, the infection efficiency and total cell nuclei count were normalized per plate to the average of in-plate untreated control wells. Infection efficiency was calculated as the ratio of total infected cells divided by the total cell nuclei count per well. Wells with an infection efficiency greater than one were removed from further analysis as these indicated a fault in the imaging. Treatments that resulted in >60% loss of cell nuclei compared to untreated controls were flagged as being potentially cytotoxic and were deprioritized. Treatments resulting in >80% cell loss were not evaluated as the virus MOI would be significantly altered together with a loss in calculation accuracy.

The classification of the drug-response outcomes was performed using a drug response curve (DRC) model. We used the R package drc ^56^ to calculate the DRCs using a log-logistic model that estimates four parameters (Hill slope, IC50, min, and max). Each drug-response was classified by inspecting cell nuclei count, and then evaluating the drug effect on the inhibition of viral infection efficiency. Each drug-response was classified in two steps: first inspecting toxicity by nuclei count and then evaluating the drug effect on the inhibition of viral proliferation using the model given by Eq. 1.

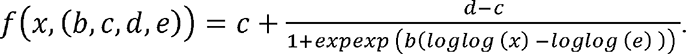

Eq. 1 DRC log-logistic Model with four parameters, where b is hill, c is the min value, d is the max value and e is the EC_20_

To inspect the cell nuclei count for each drug, we first estimated the model parameters using as response variable the normalized nuclei count in the treated cells. We tested the dose-response effect for all drugs using a ’J^s^ test for goodness of fit, and treatments with p < 0.01 (FDR- Bonferroni correction) were defined as potentially cytotoxic causing substantial nuclei loss. If a drug had a nuclei count reduction less than 40% of untreated, or if toxicity was observed only at the highest (8 μM) concentration it was not considered cytotoxic. To evaluate inhibition of viral replication, we used as response for the DRC model the normalized number of infected cells in the treated well. A drug was considered to have a dose-response effect by using a ’J^s^ test for goodness of fit (P < 0.01, FDR-Bonferroni correction), and the significant drugs were defined as strong (Z score >2.5 corresponding to >80% inhibition), weak (Z-score 1.5-2.5 with 50-80% inhibition), or not effective (Z score <1.5 and <50% inhibition) over the range of concentrations. Z-scores were calculated based on the standard deviation and mean signals of vehicle treated cells from all plates.

Treatments that showed cell nuclei loss greater than infection inhibition for at least half of doses tested were classified as being potentially cytotoxic and were de-prioritized for follow-up studies.

#### Accuracy

All outcomes were manually inspected in order to validate the two step-model and annotation compared to the computational model derived outcome.

#### GO enrichment

Using all targets within each drug category, we performed a GO enrichment analysis in the biological process and molecular function mode using the enrichR tool.

#### Gene-target network analysis

Using genes that were statistically significantly enriched in each category, we built a bipartite network, from drugs and targets for each category.

#### Network effect

In order to understand where the drug-targets were placed in a human protein-protein interaction network, we first calculated the largest connected component (LCC) of each drug category and calculated the significance of the module size using a degree-preserving approach ^19^, preventing the same high degree nodes being selected repeatedly by choosing 100 bins in 1,000 simulations. For that purpose, the R package NetSci was used.

#### Target enrichment

To understand if there was an enrichment of any gene as a drug target in each category, we used a Fisher’s exact test, and considered a gene to be enriched in any category if its p-value is lower than 0.05 (complete results can be found in Table S3). For each set of genes, we performed a biological function enrichment using Enrichr ^57^.

#### Pathway analysis

We aggregated the targets of the compounds classified in the different outcome categories (e.g., strong, weak) and performed pathway enrichment analysis (Reactome) using the R package ReactomePA ^58^.

#### Mechanism of action

We retrieved mechanism of action annotations for 6,150 drugs from the Drug Repurposing Hub^.12^. For each mechanism of action, we checked over- or under-representation of drugs in the different drug outcomes (e.g., strong, weak) by using a Chi-Square test (p < 0.05, FDR-BH).

#### Chemical structural relationship analysis

To evaluate structure similarity between molecules, similarity searches and clustering were performed using molecular fingerprints (FPs). The python package RDKit ^59^ was used to standardize SMILES and InChIKeys associated with each drug tested in the experiments, and generate Morgan FPs (also known as extended-connectivity fingerprints). All molecular substructures up to radius 3 were assigned to unique identifiers and hashed to vectors of 8,192 bits in order to capture fragments of bigger size and reduce the potential bit collision. For all molecules with experimental outcome, structural similarity was quantified by Jaccard (or Tanimoto) similarity. We further leveraged the Jaccard metric to reduce the dimensionality of the bit vector representation to 3D with UMAP ^17^. In the bit representation of Morgan FPs each bit is mapped to multiple structural fragments appearing in molecules. We evaluated the significance of each bit for ligand binding by calculating the average number of non-hydrogen atoms in the associated fragments, finding that in the overall database the median size of the expected chemical fragment associated with each bit is 9. The enrichment of each experimental class in active bits (hypergeometric test, BH multiple testing correction with u: 0 ) was then calculated. As shown in Figure S1, the significant bits for strong drugs appear to be moderately bigger and potentially more meaningful for binding, with 22 bits in the Strong Class with average size greater than the database median, compared to only 1 bit for ‘Cyto’, cytotoxic class.

#### Statistics

Aside from the tests described above and within the text, statistical significance between treatments and across experiments was calculated using Student’s T test or one-way ANOVA with Tukey’s post-test for comparison of multiple groups. For the animal experiments, nested ANOVA was used to compare multiple replicates for different samples within a single experiment. Dose-response curves were generated and IC_50_ values calculated by fitting a four parameter [Inhibitor] versus response with variable slope curve to the data. Each test and curve fitting were performed using GraphPad Prism version 8.0.0 (GraphPad Software, GraphPad Software, San Diego, California USA).

## Acknowledgements

JJP, CJD, HM, PK, KGB, RB, SHS and RAD were supported by NIH grants P01AI120943, 5R01AI128364-02, 5R01AI125453-05, UC7AI095321 and by a grant from the Massachusetts Consortium on Pathogen Readiness. JLB, SJ and SG were supported by NIH grants 3RO1AI064099-15. ALB is supported by NIH grant 1P01HL132825, American Heart Association grant 151708, and ERC grant 810115-DYNASET. MZ is supported, in part, by NSF grants IIS-2030459 and IIS-2033384, and by Harvard Data Science Initiative. JL was supported, in part, by NIH grants HG007690, HL108630, and HL119145, and by American Heart Association grants D700382 and CV-19; MN was supported by a grant from the Massachusetts Consortium on Pathogen Readiness. SJE is an investigator with the Howard Hughes Medical Institute. We acknowledge the hard work and dedication of the NEIDL veterinary services team who performed disease treatment testing in the high biological containment laboratory under institutionally approved IACUC protocols to RAD. \We thank the Center for the Development of Therapeutics and Repurposing Hub at the Broad Institute for providing the compound library. We thank Tim Mitichson for critical reading of the manuscript. We thank John H. Connor, Jacquelyn Turcinovic, Scott Seitz, Florian Douam, and Devin Kenney for isolation and production of SARS-CoV-2 Delta (B.1.617), and Anthony Griffiths and Anna Honko for production of MERS-CoV.

## Data and materials availability

Supporting data for this study are shown or available in the Supplementary Information. The primary screening data is also available through the Drug Repurposing Hub portal: https://www.broadinstitute.org/drug-repurposing-hub or https://clue.io/repurposing-app. Computer coding scripts for image and data analysis are available on https://github.com/RDaveyLab/COVID19/.

## Supplementary figures and tables

### Supplementary figures

**Figure S1.**
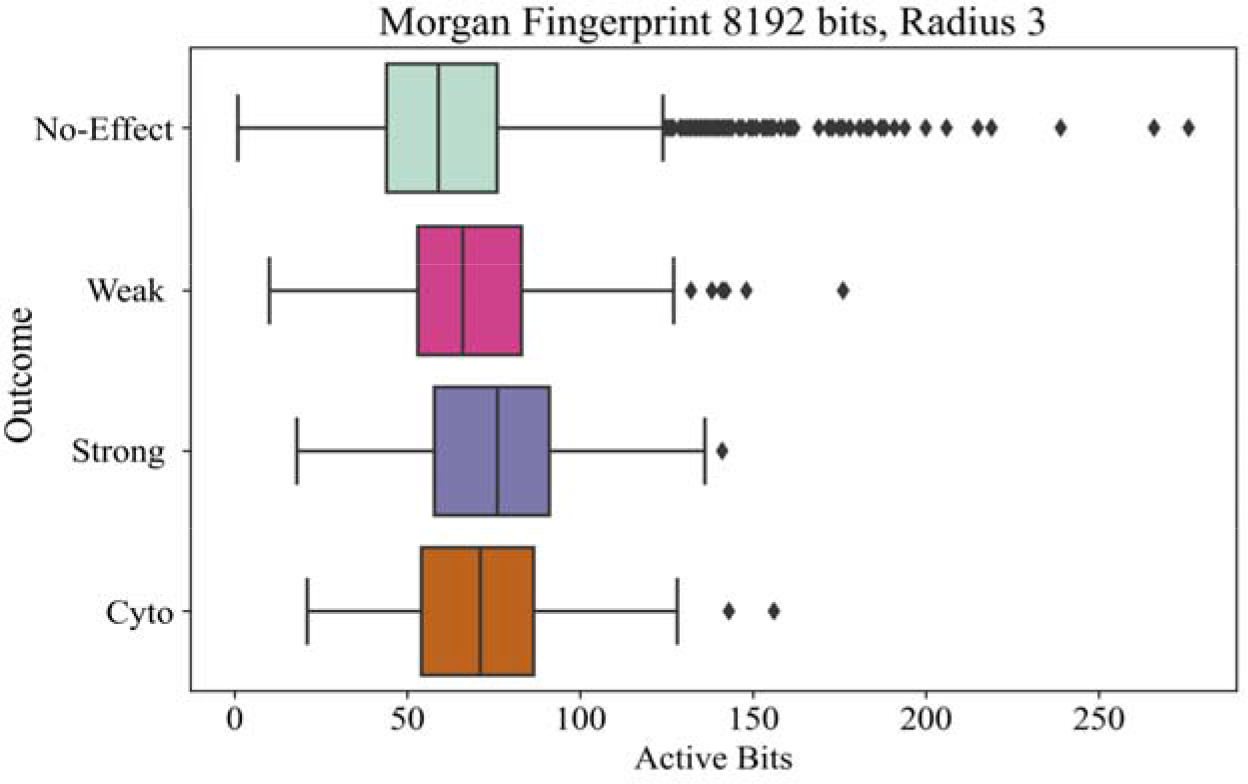
Active bit enrichment for compounds. Structures of small molecules were encoded using Morgan Fingerprints with atoms up to radius 3, and hashed to a binary vector of 8,192 bits. Active small molecules showed higher medians of active bits with strongly active compounds showing significant enrichment for 93 bits, suggesting clusters of structurally similar molecules.

**Figure S2.**
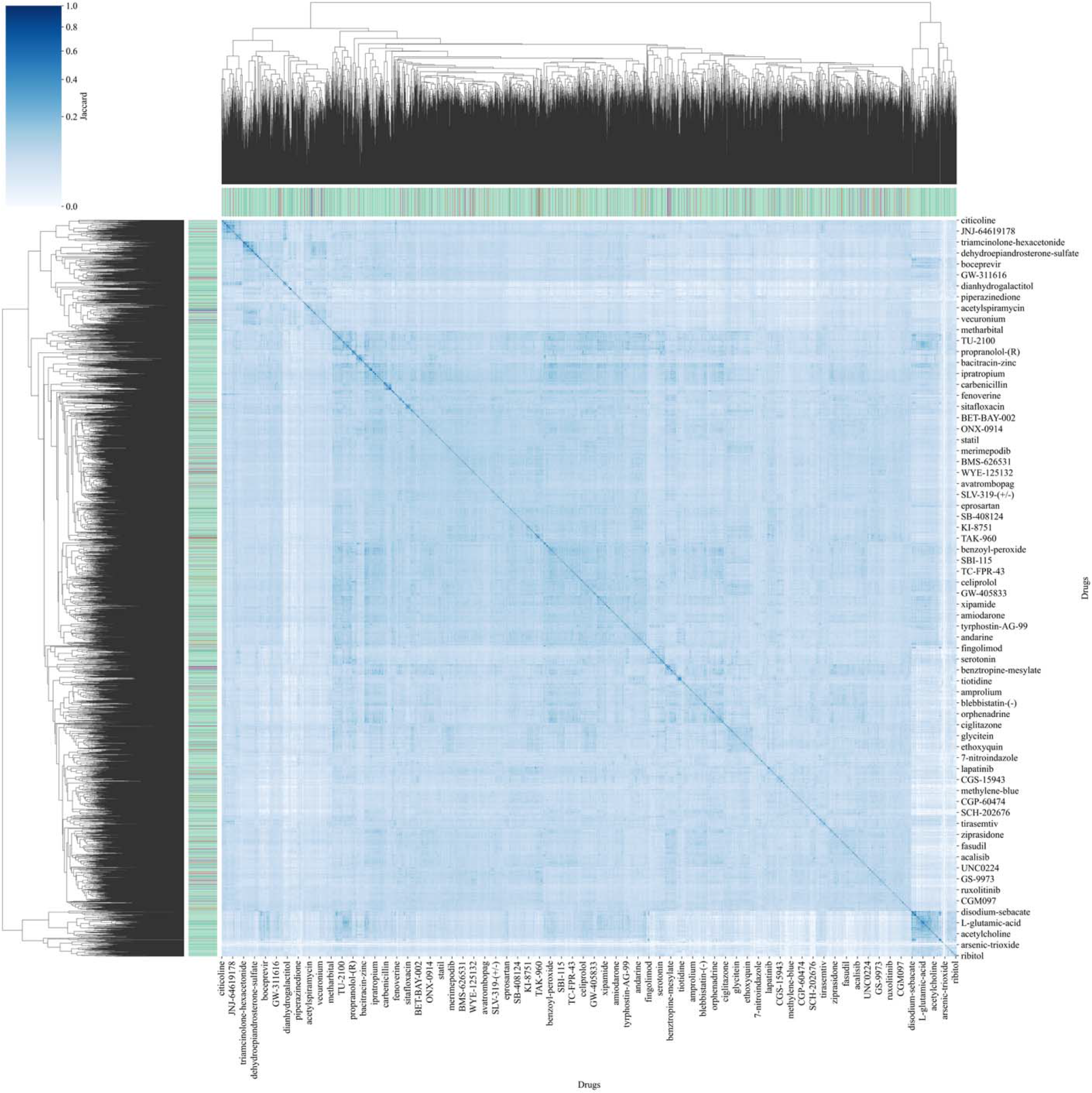
Jaccard similarity matrix for the entire DRH library. The figure shows the pairwise comparison of all compounds in the small molecule library used for the screen based on computed structural similarity using Morgan fingerprints and Jaccard analysis.

**Figure S3.**
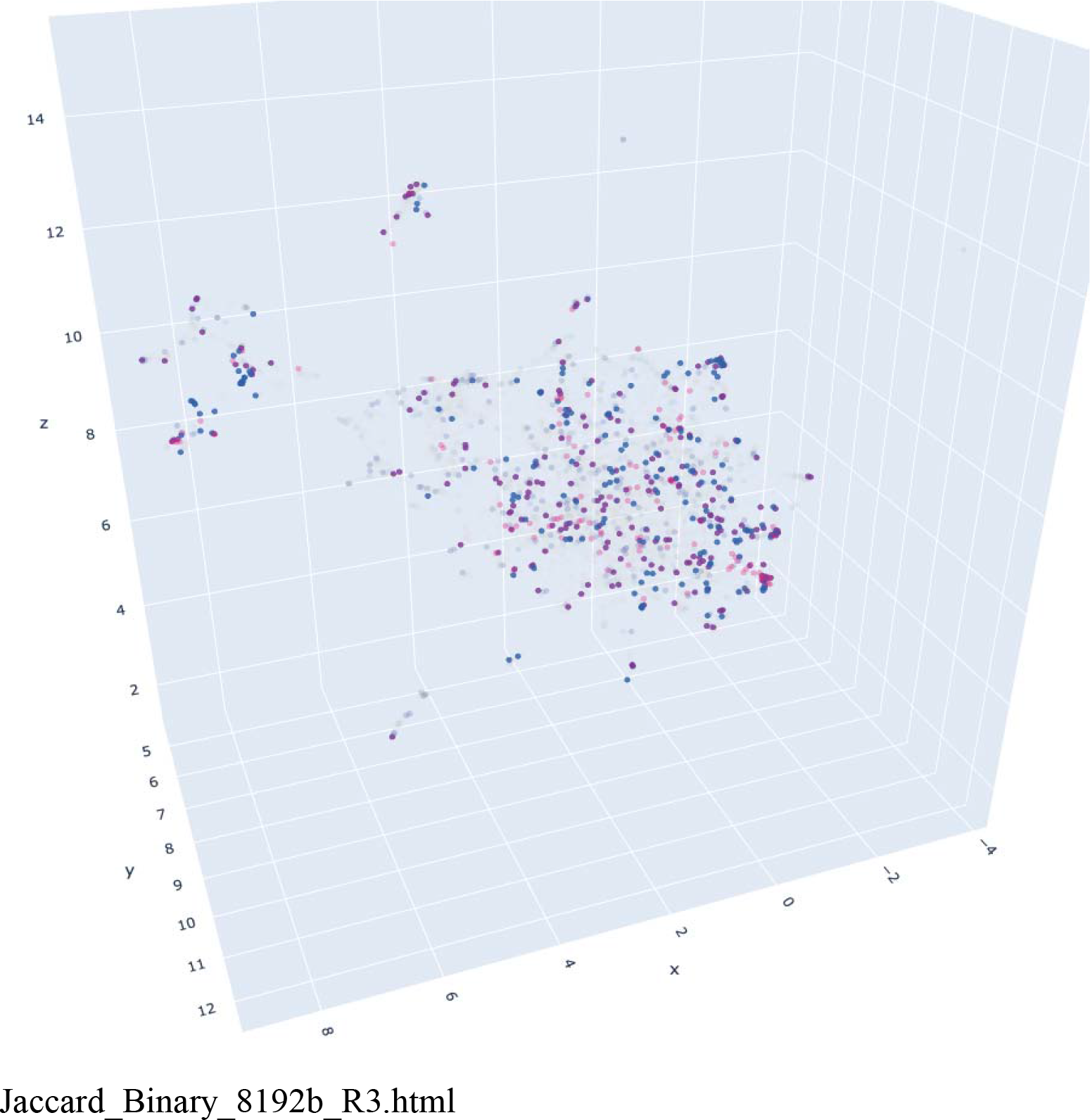
Interactive 3D representation of structural similarity of active molecules. The plot is based on the alignment of all molecules with outcomes in the primary screen and on the reduced dimensionality of the molecular bit vectors encoding each molecule. Classes of active molecules can be selected by clicking the legend. The graph can be reorientated by click- dragging on the plot using a mouse. Only the strong and weak activity classes were considered for most data analysis. The very weak and low classes (gave 35% and 25% inhibition respectively at the highest concentrations tested) are included for completeness but have Z-scores <1.5 from controls.

**Fig. S4.**
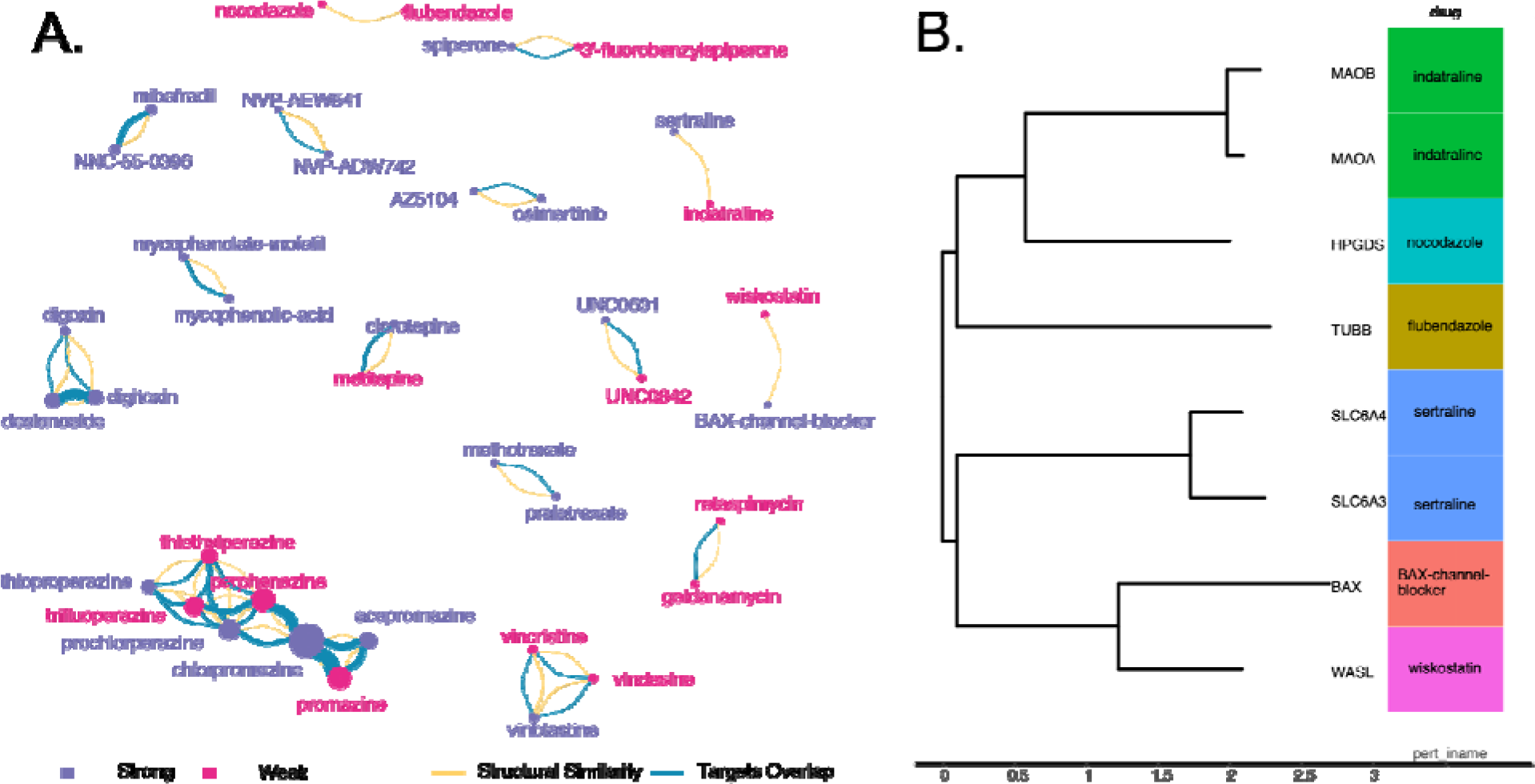
Analysis of structurally related pairs of active molecules and relationship to protein drug targets. Similar small molecules are connected by yellow links if they share structural similarity, and by a blue link if they share the same host protein target. B) For 3 pairs of small molecules where no identical protein target was shared were analyzed for the relatedness of the known target. 2 of the pairs (wiskostatin and BAX) and (nocadalozone and flubendazole) share related protein targets.

**Figure S5.**
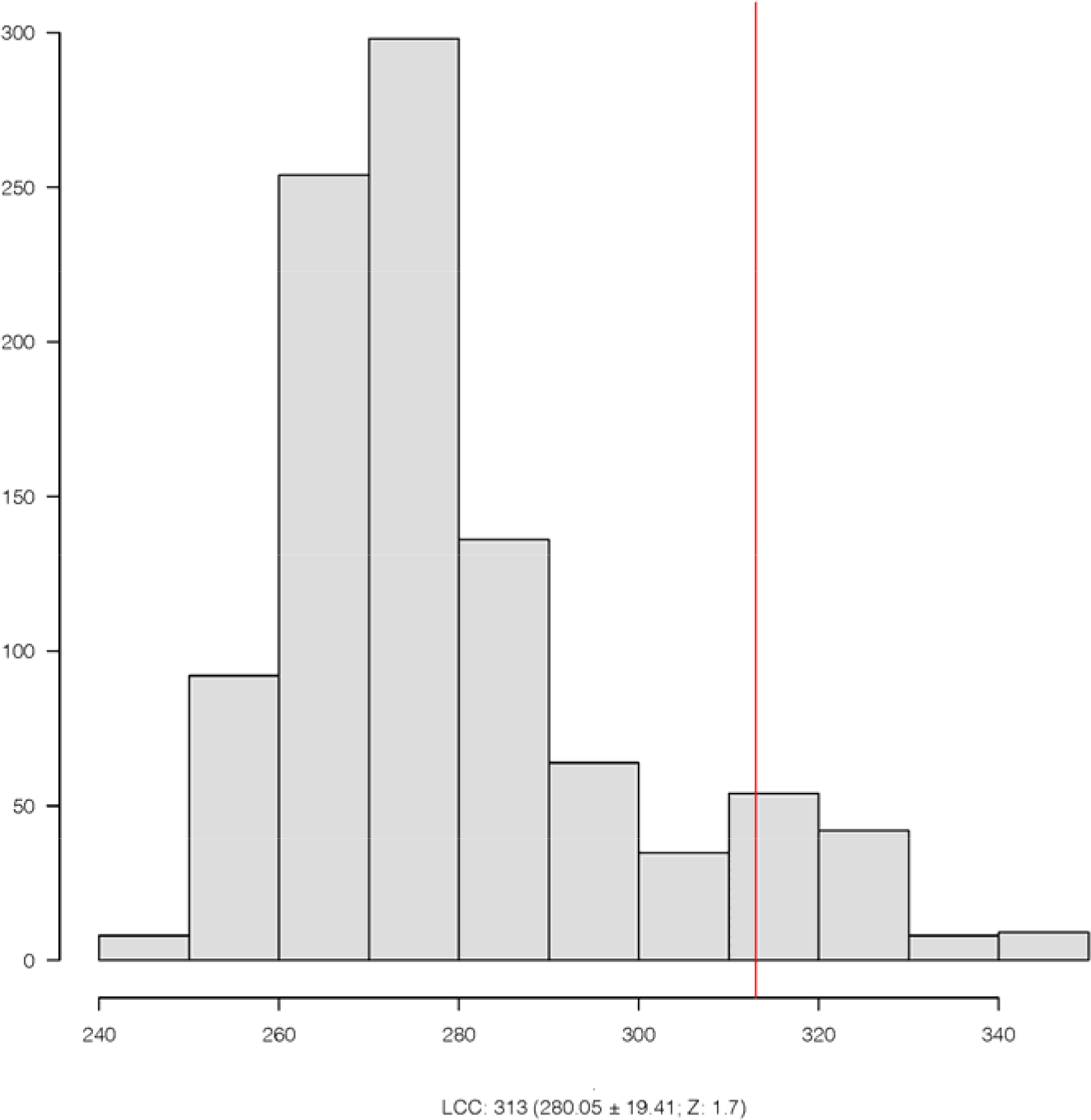
Largest Connected Component significance of targets from each drug category. Active small molecules have a larger component by chance (Z-score > 1.5), suggesting that the compound protein targets responsible for inhibition of virus infection reside in a related PPI neighborhood.

**Figure S6.**
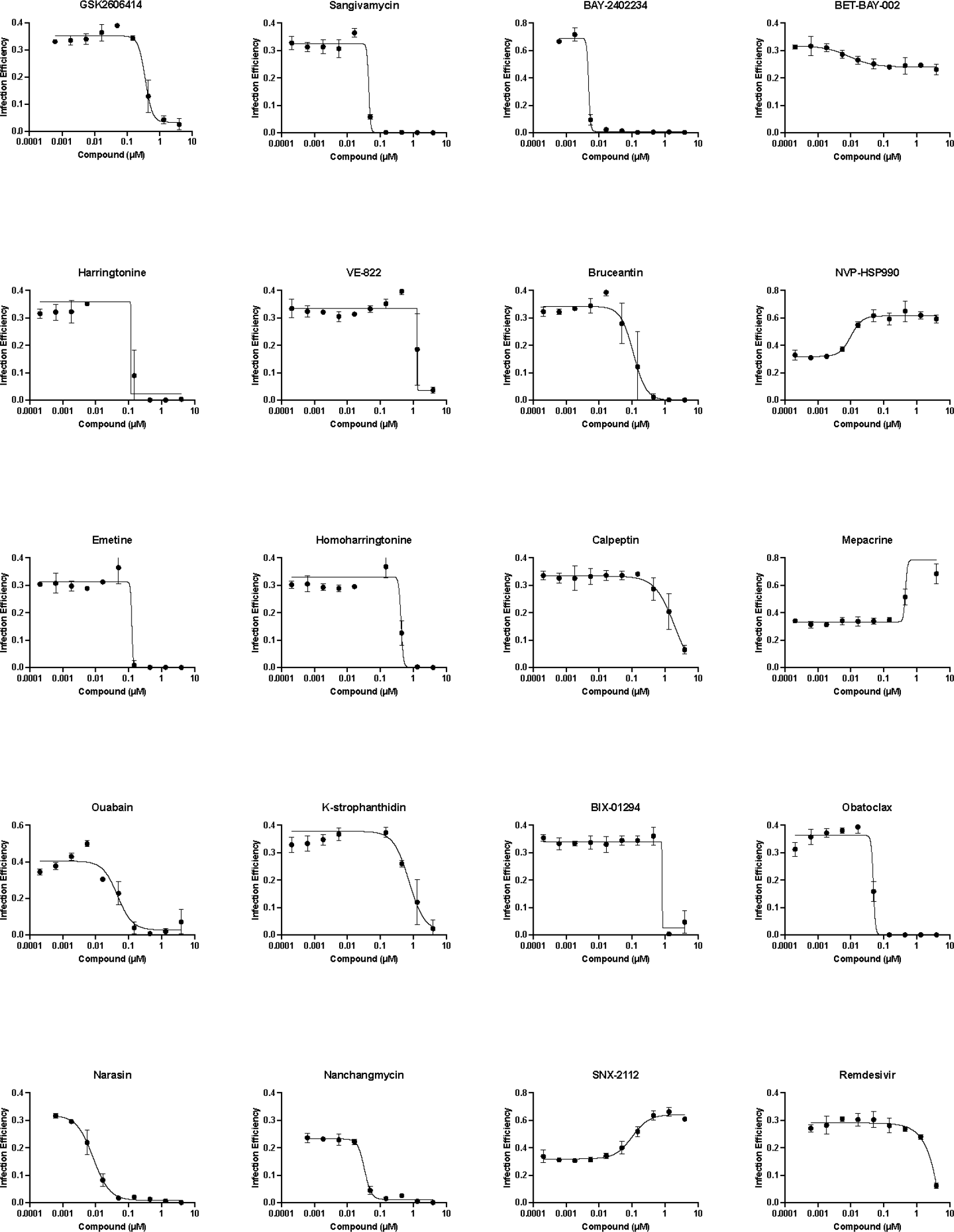
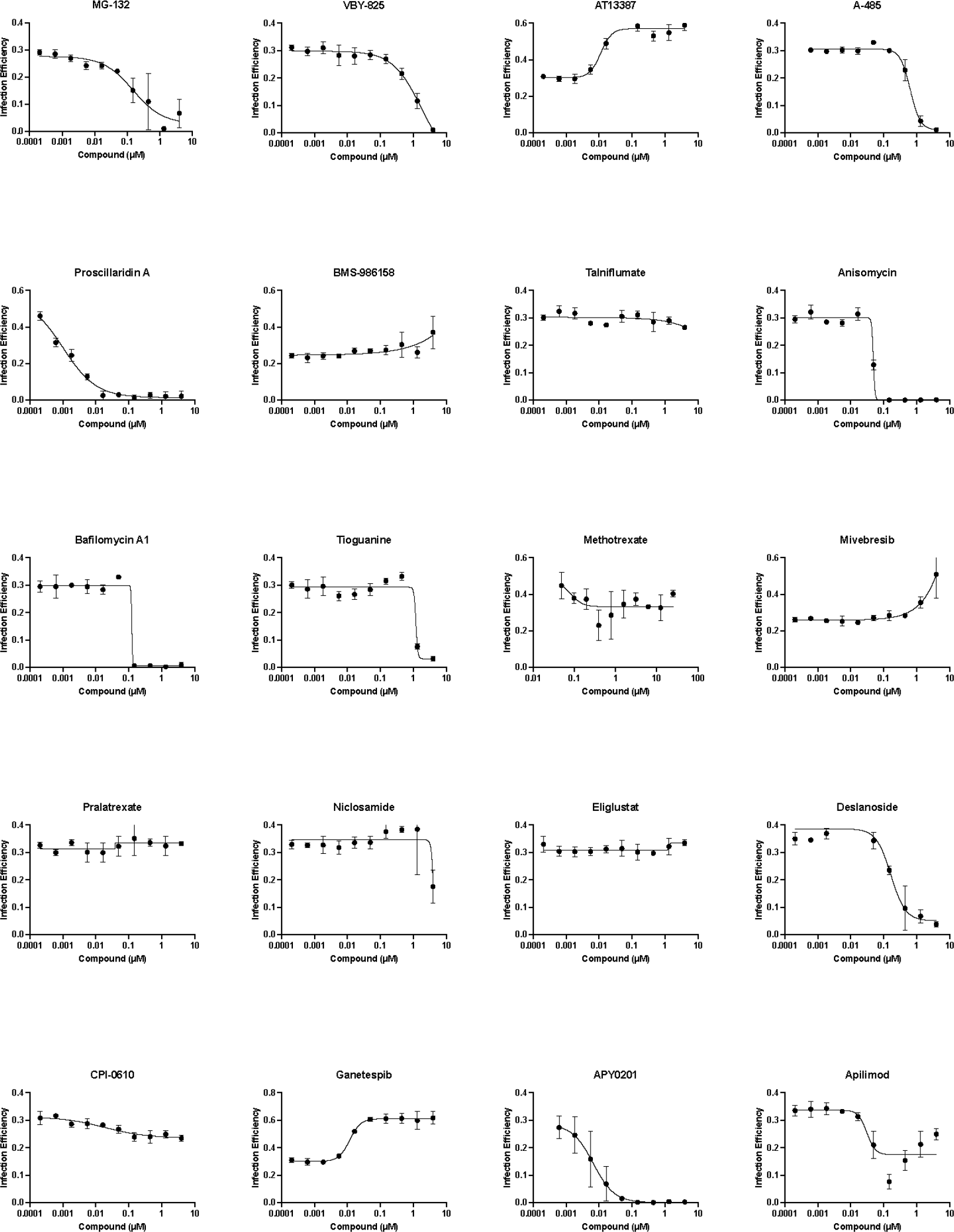
Concentration response curves. The activity of the indicated small molecules was evaluated in VeroE6 cells at the indicated concentrations (μM). Infection efficiency was measured by dividing the number of infected cells by the total cell nuclei present. Each concentration was performed in triplicate with averages and standard deviations indicated. Curves were fitted using Graphpad Prism software using the [Inhibitor] vs. response with variable slope, four parameter equation. EC50 values were calculated from the curve fits.

**Figure S7.**
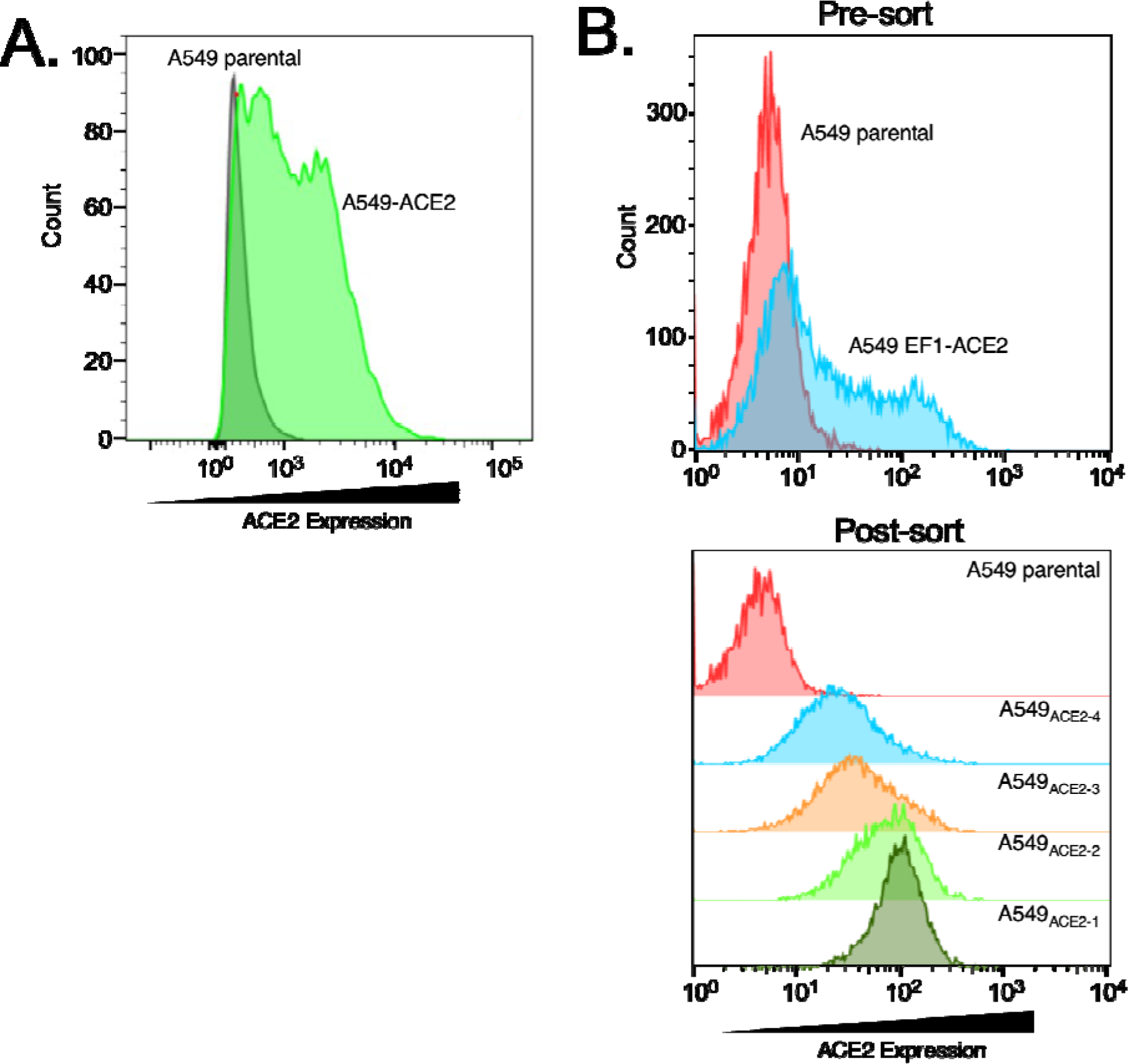
Characterization of recombinant A549 cells expressing human ACE2. Two types of A549-ACE2 were developed. A. Cells used for the pseudotyped virus work were a pool of cells transduced with a lentivirus encoding a recombinant human ACE2 protein. The cells were selected by a resistance marker on the lentivirus vector and then analyzed by FACS using an ACE2 specific antibody. B. Cells used for testing SARS-CoV-2 beta and delta variants were made by transduction of A549 cells with pHAGE-EF1-ACE2. Upper panel shows ACE2 expression measured by FACS after staining with ACE2 specific antibody. The cell pool was sorted by FACS, without prior selection, into four bins (A549_ACE2-1_ – A549_ACE2-4_) based on ACE2 surface expression levels. Cells were then passaged and ACE2 expression in each bin was assessed via flow cytometry. A549_ACE2-4_ cells were used for testing efficacy of small molecules for inhibition of SARS-CoV-2 variants.

**Figure S8.**
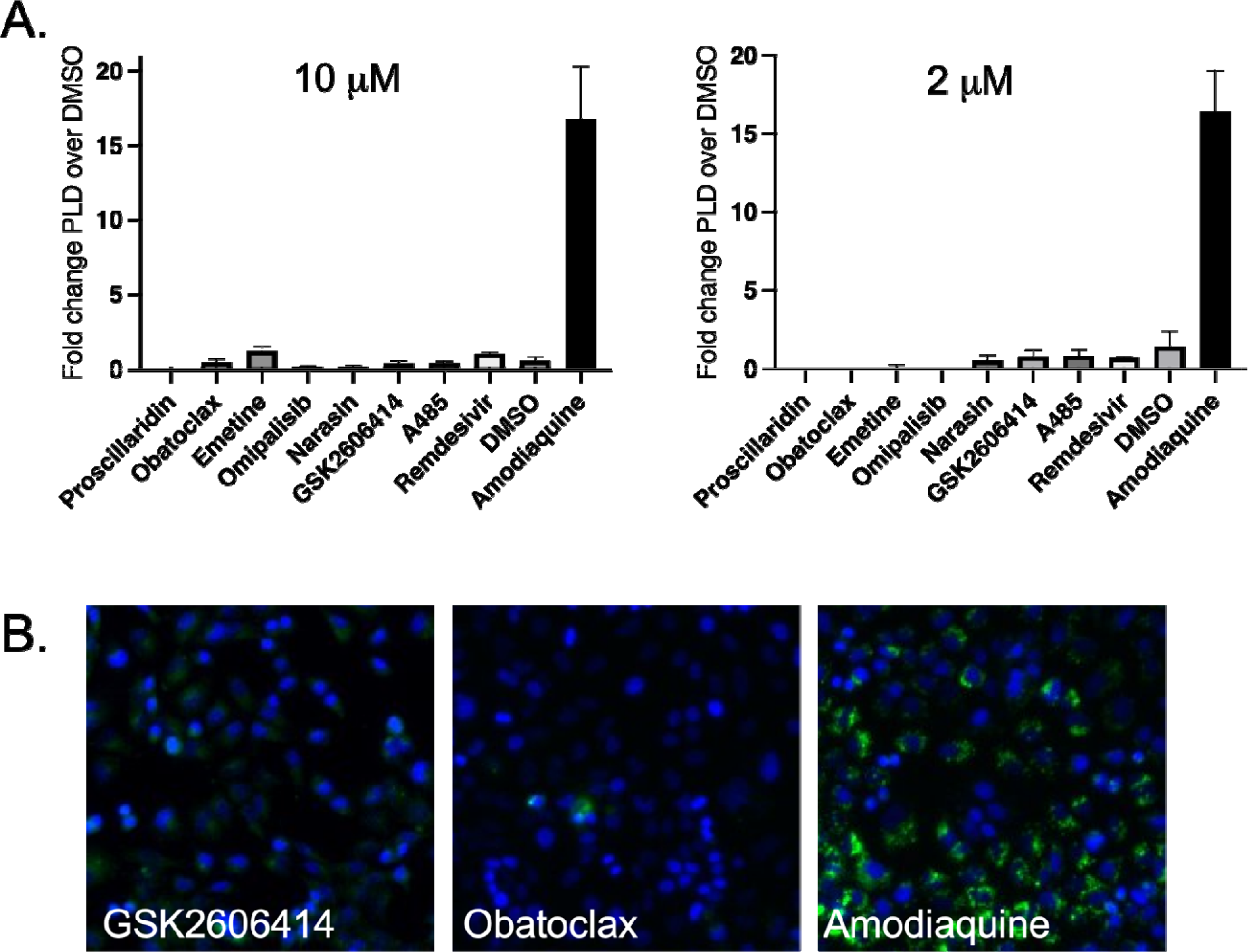
Phospholipidosis induction by compounds. A549-ACE2 cells were incubated in the presence of NBD-PE and the amount of retained NBD-PE measured by fluorescence. A. Quantitive analysis of 10 and 2 μM levels for each drug measured in triplicate. C. Representative images of NBD staiing in cells (green) and cell nuclei stained with Hoeschst 33342 (blue).

**Figure S9.**
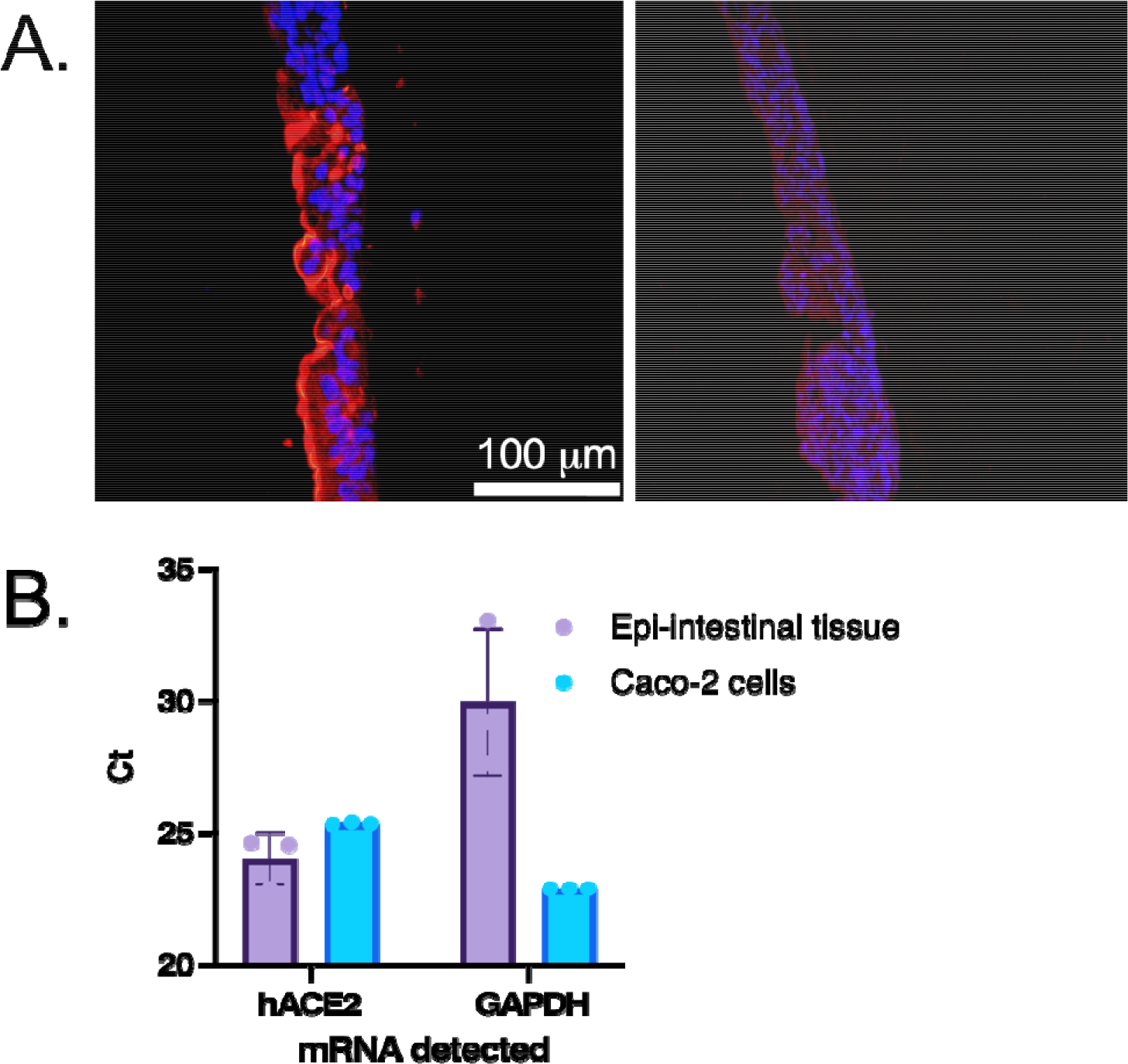
ACE2 expression in Mattek epiintestinal tissue models. A. Epi-intestinal tissue models were stained with hACE2 specific antibody (red) and cell nuclei identified by DAPI (blue). Left panel shows staining with specific antibody, Right panel is signal without primary antibody. B. qPCR was perfiormed on cell lysates from epi-intestinal tissues or Caco-2 cells, a known expressor of hACE2. Both hACE2 and GAPDH signals were measured and Ct’s shown without normalization. Despite a higher GAPDH Ct, therefore a smaller amount of cellular mRNA in the sample, the tissue showed similar levels of hACE2 mRNA,

**Figure S10.**
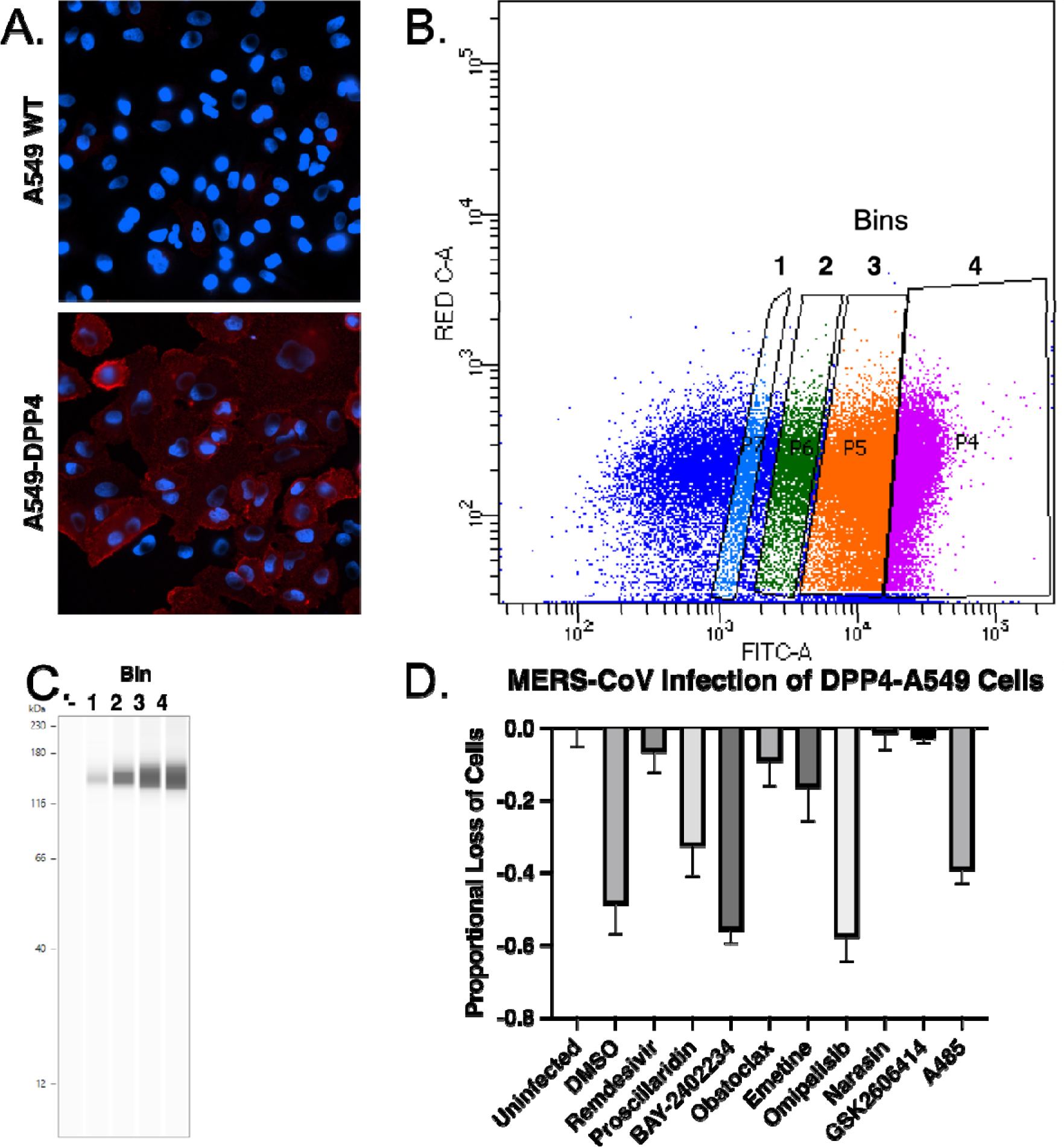
Selection of A549 cells expressing DPP4 (MERS-CoV receptor) and testing of activity of compounds. Cells were transfected with plasmid expression vectors expression encoding DPP4. A. Expression levels measured by immunofluorecence staining with specific antibody (red) and cell nuclei stained with Hoeschst 33342 (blue). B. After selection in puromycin, the cells were sorted by FACS into 4 bins based on level of DPP4 expression. C. Expression was verified by immunoblot. D, The indicated compounds were tested for inhibition of MERS-CoV in Bin 1 DPP4-A549 cells using reduction of virus induced cytopathy to measure changes in virus infection. Each compound was used at the EC90 seen for SARS-COV-2 Washington strain.

### Supplementary tables

Table S1. **All data from the primary screen. (separate file).** Shown is the name of compound tested, abbreviated Drug Repurposing Hub identifier, SMILES formula, InChIKey formula, outcome in the screening assay. In this classification curves were graded on a scale of strong, weak, very weak, low, no-effect or cytotoxic. Only the strong and weak categories are considered statistically significant based on a Z-score >1.5. Cytotoxic compounds are those that reduced cell nuclei counts by >60% relative to the vehicle treated controls. In other work, vehicle treated cells (DMSO at 0.2%) showed no significant difference to untreated cells for nuclei counts. A file showing the structures and dose response curves is available on our Github link.

Table S2. **Morgan FP data. (separate file)** This file contains the encoding used to construct the Morgan Fingerprints with atoms up to radius 3, as hashed to a binary vector of 8,192 bits used for the structural alignment of the small molecules in the library.

Table S3 **Enrichment drug targets by genes and GO terms.** Fisher’s exact test was used to identify enriched gene categories and biological function enrichment was then performed using Enrichr. Enriched genes are shown at the bottom of the table.

**Table S4.**
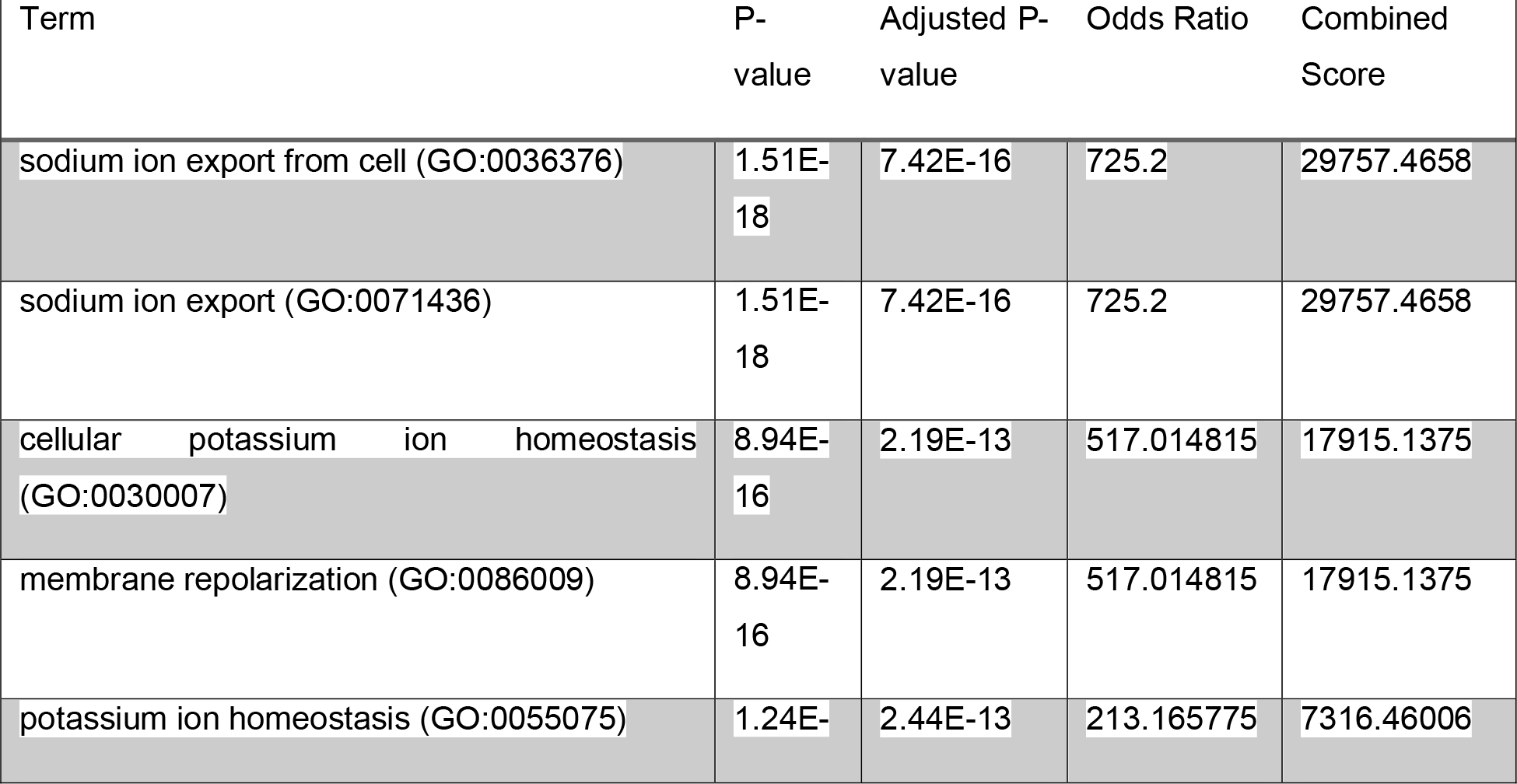
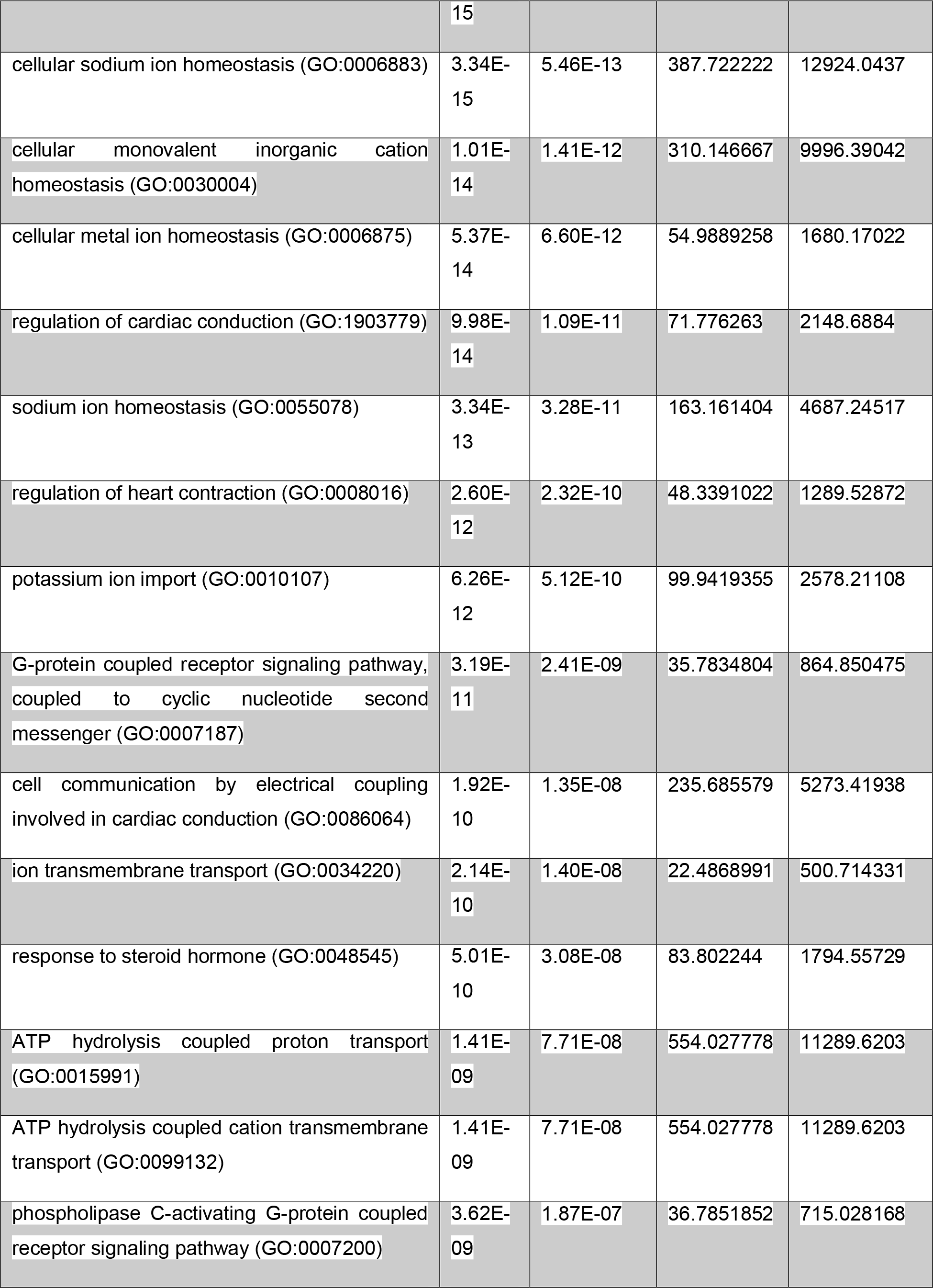
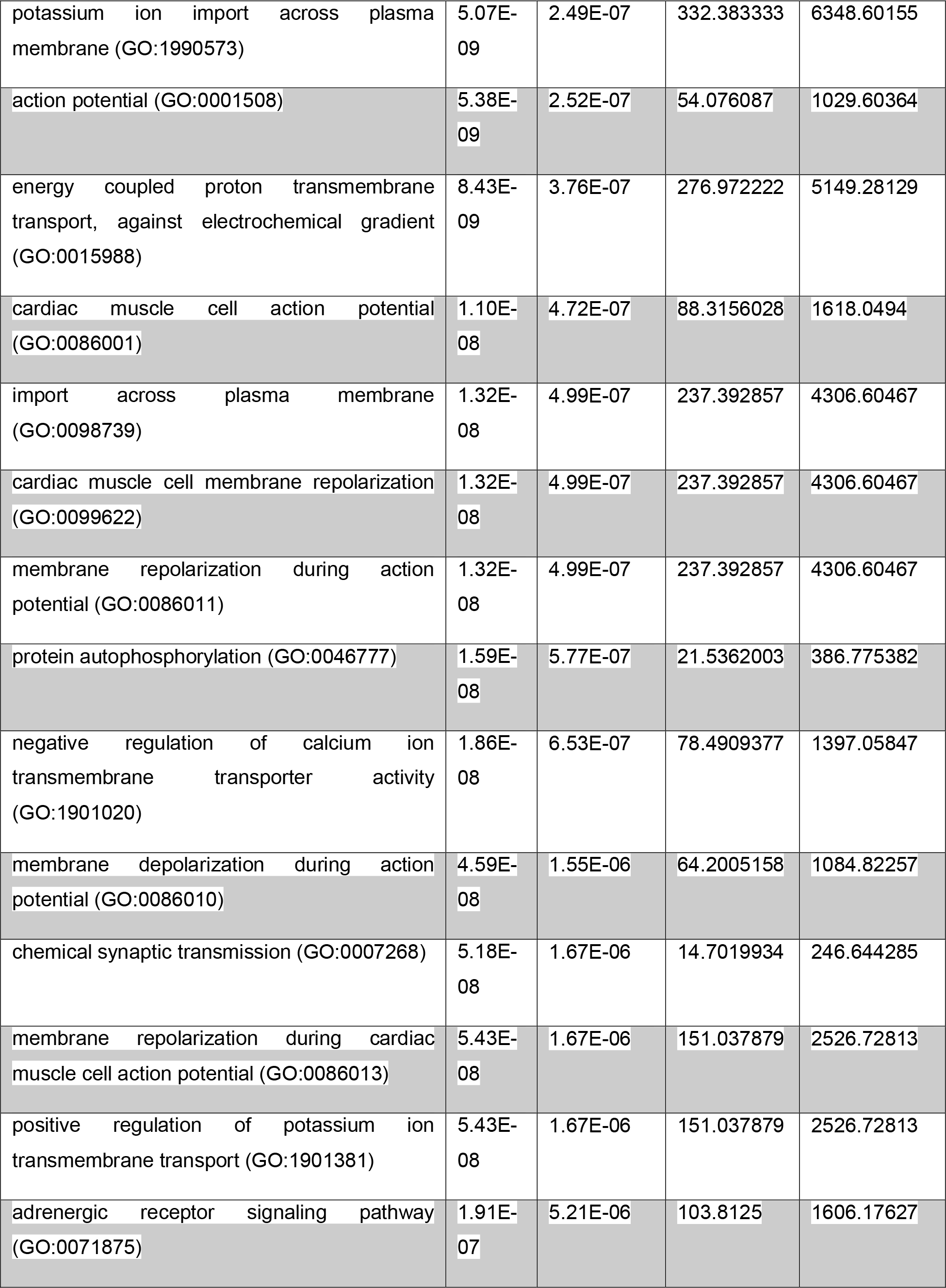
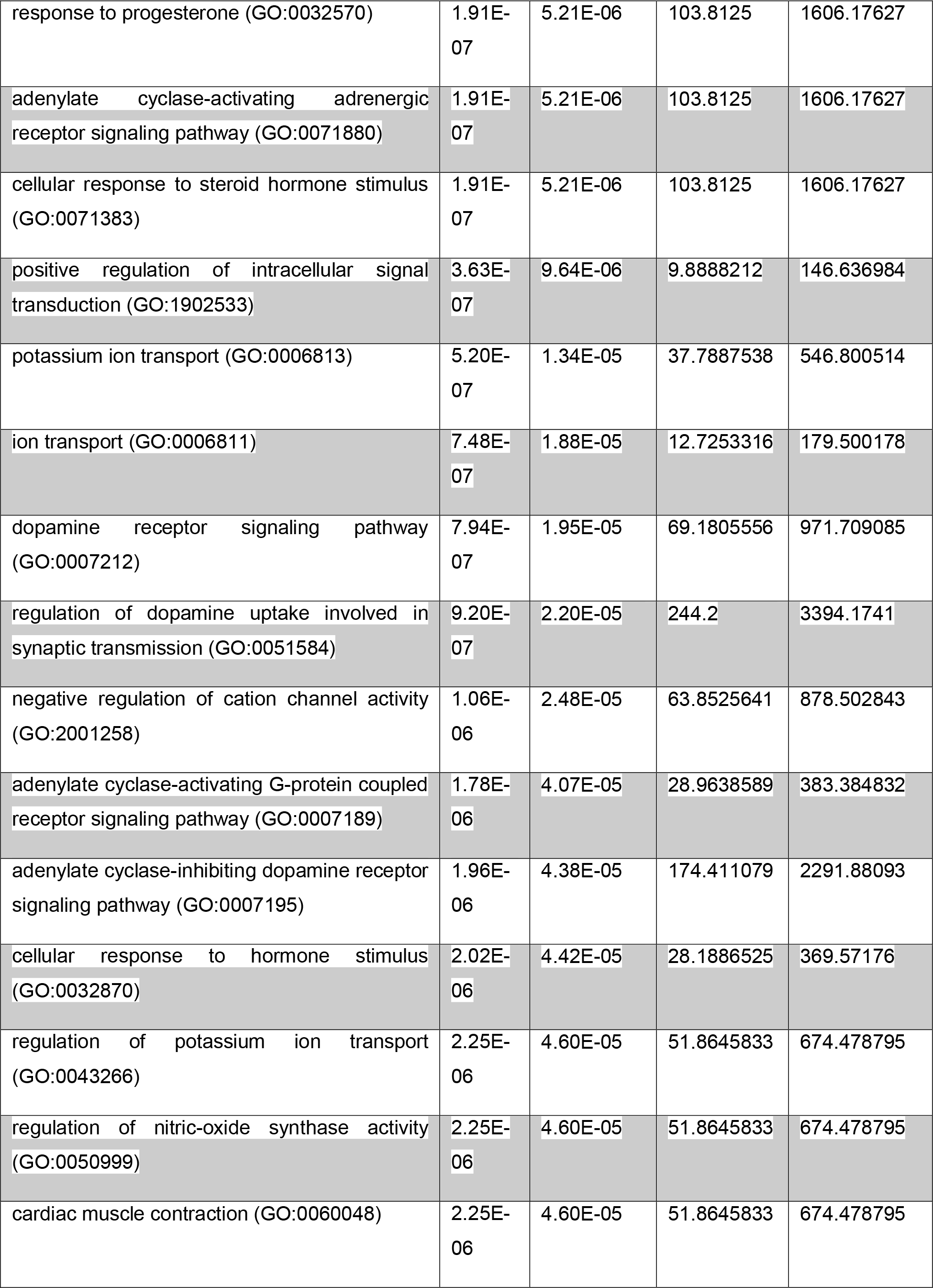
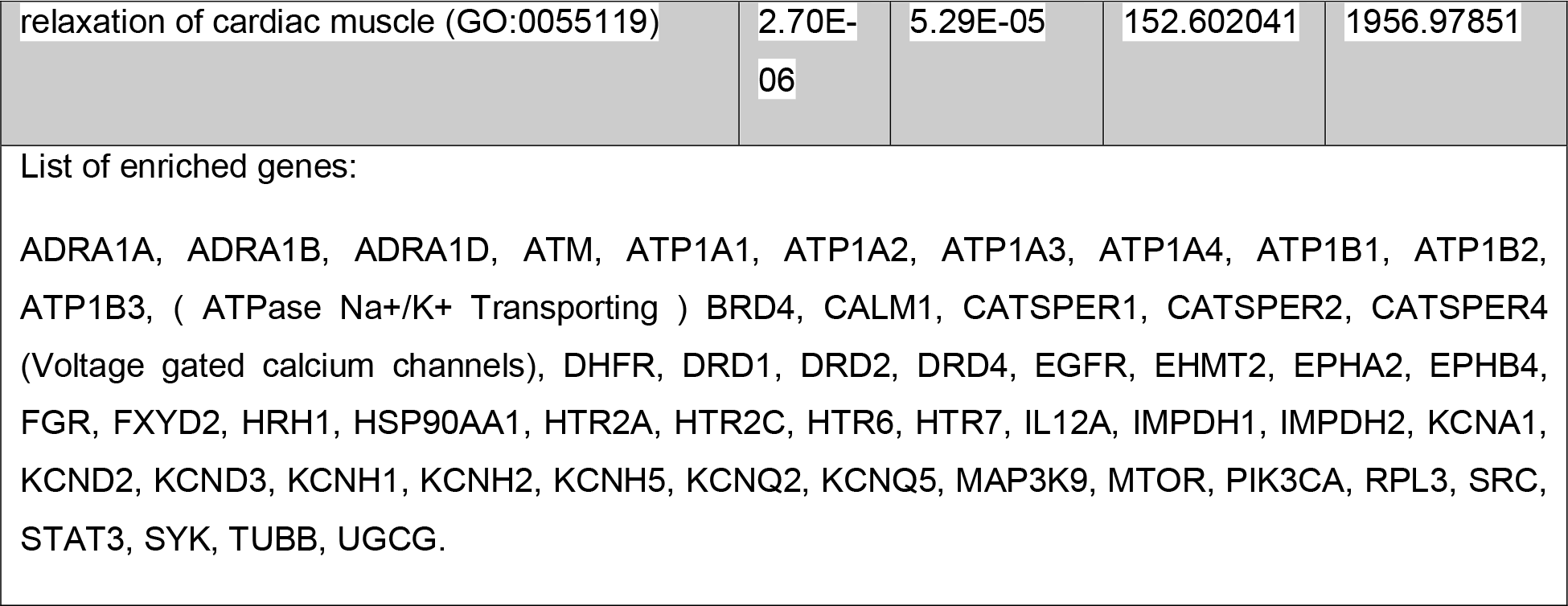
Enrichment of known drug-targets. (separate file) For the 389 active compounds (strong and weak classes), protein targets were assigned as indicated.

