## Supplemental Table 2 for "Identification of druggable host targets needed for SARS-CoV-2 infection by combined pharmacological evaluation and cellular network directed prioritization both in vitro and in vivo"

| PUBCHEM_SUBSTANCE_SYNONYM | DRH ID code | PUBCHEM_EXT_DATASOURCE_SMILES | InChIKey | Outcome | |
|---|---|---|---|---|---|
| 7-hydroxystaurosporine | BRD-K45293975 | CN[C@@H]1C[C@H]2O[C@@](C)([C@@H]1OC)n1c3ccccc3c3c4c(c5c6ccccc6n2c5c31)C(=O)N[C@H]4O | PBCZSGKMGDDXIJ-KRUBCLEUSA-N | Cyto | inactive |
| A-674563 | BRD-K78177893 | Cc1[nH]nc2ccc(-c3cncc(OC[C@@H](N)Cc4ccccc4)c3)cc12 | BPNUQXPIQBZCMR-IBGZPJMESA-N | Cyto | inactive |
| aclarubicin | BRD-K31965398 | CC[C@@]1(O)C[C@H](O[C@H]2C[C@H](N(C)C)[C@H](O[C@H]3C[C@H](O)[C@H](O[C@H]4CCC(=O)[C@H](C)O4)[C@H](C)O3)[C@H](C)O2)c2c(cc3c(c2O)C(=O)c2c(O)cccc2C3=O)[C@H]1C(=O)OC | USZYSDMBJDPRIF-SVEJIMAYSA-N | Cyto | inactive |
| adapalene | BRD-K33127281 | COc1ccc(-c2ccc3cc(C(=O)O)ccc3c2)cc1C12CC3CC(CC(C3)C1)C2 | LZCDAPDGXCYOEH-UHFFFAOYSA-N | Cyto | inactive |
| alaproclate | BRD-K01826783 | C[C@@H](N)C(=O)OC(C)(C)Cc1ccc(Cl)cc1 | FZSPJBYOKQPKCD-SECBINFHSA-N | Cyto | inactive |
| alvespimycin | BRD-K83988098 | CO[C@H]1/C=C\C=C(/C)C(=O)NC2=CC(=O)C(NCCN(C)C)=C(C[C@@H](C)C[C@H](OC)[C@H](O)[C@@H](C)/C=C(\C)[C@@H]1OC(N)=O)C2=O | KUFRQPKVAWMTJO-LMZWQJSESA-N | Cyto | inactive |
| alvocidib | BRD-K87909389 | CN1CC[C@H](c2c(O)cc(O)c3c(=O)cc(-c4ccccc4Cl)oc23)[C@H](O)C1 | BIIVYFLTOXDAOV-YVEFUNNKSA-N | Cyto | inactive |
| anisomycin | BRD-K91370081 | COc1ccc(C[C@H]2NC[C@H](O)[C@H]2OC(C)=O)cc1 | YKJYKKNCCRKFSL-RDBSUJKOSA-N | Cyto | inactive |
| ansamitocin-p-3 | BRD-K00003196 | COc1cc2cc(c1Cl)N(C)C(=O)C[C@H](OC(=O)C(C)C)[C@]1(C)O[C@H]1[C@H](C)[C@@H]1C[C@@](O)(NC(=O)O1)[C@H](OC)/C=C/C=C(\C)C2 | OPQNCARIZFLNLF-JBHFWYGFSA-N | Cyto | inactive |
| anthralin | BRD-K04725460 | O=C1c2c(O)cccc2Cc2cccc(O)c21 | NUZWLKWWNNJHPT-UHFFFAOYSA-N | Cyto | inactive |
| API-1 | BRD-K97840621 | NC(=O)c1cn([C@@H]2O[C@H](CO)[C@@H](O)[C@H]2O)c2ncnc(N)c2c1=O | SPBWHPXCWJLQRU-FITJORAGSA-N | Cyto | inactive |
| AR-42 | BRD-K13167095 | CC(C)[C@H](C(=O)Nc1ccc(C(=O)NO)cc1)c1ccccc1 | LAMIXXKAWNLXOC-INIZCTEOSA-N | Cyto | inactive |
| avitinib | BRD-K00003294 | C=CC(=O)Nc1cccc(Oc2nc(Nc3ccc(N4CCN(C)CC4)c(F)c3)nc3[nH]ccc23)c1 | UOFYSRZSLXWIQB-UHFFFAOYSA-N | Cyto | inactive |
| AZD7762 | BRD-K86525559 | NC(=O)Nc1cc(-c2cccc(F)c2)sc1C(=O)N[C@H]1CCCNC1 | IAYGCINLNONXHY-LBPRGKRZSA-N | Cyto | inactive |
| BAY-1251152 | BRD-K00003533 | COc1cc(F)ccc1-c1cc(Nc2cc(C[S@](C)(=N)=O)ccn2)ncc1F | YZCUMZWULWOUMD-MUUNZHRXSA-N | Cyto | inactive |
| BEBT-908 | BRD-K00003291 | CNc1ccc(-c2nc(N3CCOCC3)c3sc(CN(C)c4ncc(C(=O)NO)cn4)cc3n2)cn1 | TWJZFXHSPBBPNI-UHFFFAOYSA-N | Cyto | inactive |
| belinostat | BRD-K17743125 | O=C(/C=C/c1cccc(S(=O)(=O)Nc2ccccc2)c1)NO | NCNRHFGMJRPRSK-MDZDMXLPSA-N | Cyto | inactive |
| BI-2536 | BRD-K64890080 | CC[C@@H]1C(=O)N(C)c2cnc(Nc3ccc(C(=O)NC4CCN(C)CC4)cc3OC)nc2N1C1CCCC1 | XQVVPGYIWAGRNI-JOCHJYFZSA-N | Cyto | inactive |
| BI-D1870 | BRD-A81370665 | CC(C)CCN1c2nc(Nc3cc(F)c(O)c(F)c3)ncc2N(C)C(=O)[C@@H]1C | DTEKTGDVSARYDS-NSHDSACASA-N | Cyto | inactive |
| bortezomib | BRD-K88510285 | CC(C)C[C@H](NC(=O)[C@H](Cc1ccccc1)NC(=O)c1cnccn1)B(O)O | GXJABQQUPOEUTA-RDJZCZTQSA-N | Cyto | inactive |
| brefeldin-A | BRD-K77841042 | C[C@H]1CCC/C=C/[C@@H]2C[C@H](O)C[C@H]2[C@H](O)/C=C/C(=O)O1 | KQNZDYYTLMIZCT-KQPMLPITSA-N | Cyto | inactive |
| brilliant-green | BRD-K60348325 | CCN(CC)c1ccc([C+](c2ccccc2)c2ccc(N(CC)CC)cc2)cc1 | HXCILVUBKWANLN-UHFFFAOYSA-N | Cyto | inactive |
| brolitene | BRD-K76021681 | O=C(/C=C/c1ccccc1)NCCO | OSCTXCOERRNGLW-VOTSOKGWSA-N | Cyto | inactive |
| buparlisib | BRD-K42191735 | Nc1cc(C(F)(F)F)c(-c2cc(N3CCOCC3)nc(N3CCOCC3)n2)cn1 | CWHUFRVAEUJCEF-UHFFFAOYSA-N | Cyto | inactive |
| caffeic-acid-phenethyl-ester | BRD-K96188950 | O=C(/C=C/c1ccc(O)c(O)c1)OCCc1ccccc1 | SWUARLUWKZWEBQ-VQHVLOKHSA-N | Cyto | inactive |
| carmofur | BRD-K11630072 | CCCCCCNC(=O)n1cc(F)c(=O)[nH]c1=O | AOCCBINRVIKJHY-UHFFFAOYSA-N | Cyto | inactive |
| cbl0137 | BRD-K00003273 | CC(=O)c1ccc2c(c1)c1cc(C(C)=O)ccc1n2CCNC(C)C | JKCSODVERGVDLT-UHFFFAOYSA-N | Cyto | inactive |
| cetrimonium | BRD-K47079459 | CCCCCCCCCCCCCCCC[N+](C)(C)C | RLGQACBPNDBWTB-UHFFFAOYSA-N | Cyto | inactive |
| CGP-60474 | BRD-K79090631 | OCCCNc1cc(-c2ccnc(Nc3cccc(Cl)c3)n2)ccn1 | IYNDTACKOAXKBJ-UHFFFAOYSA-N | Cyto | inactive |
| ciclopirox | BRD-K13044802 | Cc1cc(C2CCCCC2)n(O)c(=O)c1 | SCKYRAXSEDYPSA-UHFFFAOYSA-N | Cyto | inactive |
| cinaciguat | BRD-K73440166 | O=C(O)CCCCN(CCc1ccccc1OCc1ccc(CCc2ccccc2)cc1)Cc1ccc(C(=O)O)cc1 | WPYWMXNXEZFMAK-UHFFFAOYSA-N | Cyto | inactive |
| cinromide | BRD-K66241279 | CCNC(=O)/C=C/c1cccc(Br)c1 | LDCXGZCEMNMWIL-VOTSOKGWSA-N | Cyto | inactive |
| colchicine | BRD-K00259736 | COc1cc2c(c(OC)c1OC)-c1ccc(OC)c(=O)cc1[C@@H](NC(C)=O)CC2 | IAKHMKGGTNLKSZ-INIZCTEOSA-N | Cyto | inactive |
| crolibulin | BRD-K00003162 | COc1cc([C@H]2C(C#N)=C(N)Oc3c2ccc(N)c3N)cc(Br)c1OC | JXONINOYTKKXQQ-CQSZACIVSA-N | Cyto | inactive |
| CT7001 | BRD-K00003453 | CC(C)c1cnn2c(NCc3ccccc3)cc(NC[C@H]3CCNC[C@@H]3O)nc12 | YCVGLKWJKIKVBI-MJGOQNOKSA-N | Cyto | inactive |
| CUDC-907 | BRD-K77638923 | COc1ccc(-c2nc(N3CCOCC3)c3sc(CN(C)c4ncc(C(=O)NO)cn4)cc3n2)cn1 | JOWXJLIFIIOYMS-UHFFFAOYSA-N | Cyto | inactive |
| CYC065 | BRD-K00003404 | CC[C@H](Nc1nc(NCc2cnc(C)cc2C)c2ncn(C(C)C)c2n1)[C@@H](C)O | DLPIYBKBHMZCJI-WBVHZDCISA-N | Cyto | inactive |
| CYT-997 | BRD-K23363278 | CCC[C@H](Nc1nc(-c2ccc(NC(=O)NCC)c(OC)c2)ncc1C)c1cccnc1 | MTJHLONVHHPNSI-IBGZPJMESA-N | Cyto | inactive |
| D-64131 | BRD-K26997899 | COc1ccc2[nH]c(C(=O)c3ccccc3)cc2c1 | ICMIJSRDISNKOC-UHFFFAOYSA-N | Cyto | inactive |
| dacinostat | BRD-K56957086 | O=C(/C=C/c1ccc(CN(CCO)CCc2c[nH]c3ccccc23)cc1)NO | BWDQBBCUWLSASG-MDZDMXLPSA-N | Cyto | inactive |
| dactinomycin | BRD-K70578146 | Cc1c2oc3c(C)ccc(C(=O)N[C@@H]4C(=O)N[C@H](C(C)C)C(=O)N5CCC[C@H]5C(=O)N(C)CC(=O)N(C)[C@@H](C(C)C)C(=O)O[C@@H]4C)c3nc-2c(C(=O)N[C@@H]2C(=O)N[C@H](C(C)C)C(=O)N3CCC[C@H]3C(=O)N(C)CC(=O)N(C)[C@@H](C(C)C)C(=O)O[C@@H]2C)c(N)c1=O | RJURFGZVJUQBHK-IIXSONLDSA-N | Cyto | inactive |
| daunorubicin | BRD-K43389675 | COc1cccc2c1C(=O)c1c(O)c3c(c(O)c1C2=O)C[C@@](O)(C(C)=O)C[C@@H]3O[C@H]1C[C@H](N)[C@H](O)[C@H](C)O1 | STQGQHZAVUOBTE-VGBVRHCVSA-N | Cyto | inactive |
| delanzomib | BRD-K59325863 | CC(C)C[C@H](NC(=O)[C@@H](NC(=O)c1cccc(-c2ccccc2)n1)[C@@H](C)O)B(O)O | SJFBTAPEPRWNKH-CCKFTAQKSA-N | Cyto | inactive |
| dinaciclib | BRD-K13662825 | CCc1cnn2c(NCc3ccc[n+]([O-])c3)cc(N3CCCC[C@H]3CCO)nc12 | PIMQWRZWLQKKBJ-SFHVURJKSA-N | Cyto | inactive |
| domiphen | BRD-K30431096 | CCCCCCCCCCCC[N+](C)(C)CCOc1ccccc1 | YXUPZGKORWTXID-UHFFFAOYSA-N | Cyto | inactive |
| ecamsule-triethanolamine | BRD-A60635434 | CC1(C)[C@H]2CC[C@]1(CS(=O)(=O)O)C(=O)C2=Cc1ccc(C=C2C(=O)[C@@]3(CS(=O)(=O)O)CC[C@@H]2C3(C)C)cc1 | HEAHZSUCFKFERC-MPPVQRIUSA-N | Cyto | inactive |
| emamectin | BRD-K00003516 | CN[C@@H]1[C@@H](C)O[C@@H](O[C@H]2[C@H](C)O[C@@H](O[C@@H]3/C(C)=C/C[C@@H]4C[C@@H](C[C@]5(C=C[C@H](C)[C@@H](C(C)C)O5)O4)OC(=O)[C@@H]4C=C(C)[C@@H](O)[C@H]5OC/C(=C\C=C\[C@@H]3C)[C@@]45O)C[C@@H]2OC)C[C@H]1OC | DXIOOXFZLKCVHK-HGIWSLISSA-N | Cyto | inactive |
| entasobulin | BRD-K00004610 | O=C(Nc1ccc2ncccc2c1)C(=O)c1cn(Cc2ccc(Cl)cc2)c2ccccc12 | LJIUXAHLYIPHOK-UHFFFAOYSA-N | Cyto | inactive |
| epiandrosterone | BRD-K03972660 | C[C@]12CC[C@H]3[C@@H](CC[C@H]4C[C@@H](O)CC[C@@]43C)[C@@H]1CCC2=O | QGXBDMJGAMFCBF-LUJOEAJASA-N | Cyto | inactive |
| ER-27319 | BRD-K11927976 | Cc1ccc2c(=O)c3ccccc3n(CCCN)c2c1C | CFPDEQBPNROZDC-UHFFFAOYSA-N | Cyto | inactive |
| fedratinib | BRD-K12502280 | Cc1cnc(Nc2ccc(OCCN3CCCC3)cc2)nc1Nc1cccc(S(=O)(=O)NC(C)(C)C)c1 | JOOXLOJCABQBSG-UHFFFAOYSA-N | Cyto | inactive |
| fidaxomicin | BRD-K00003591 | CCc1c(Cl)c(O)c(Cl)c(O)c1C(=O)O[C@H]1[C@H](O)[C@H](OC)[C@H](OC/C2=C\C=C\C[C@H](O)/C(C)=C/[C@H](CC)[C@@H](O[C@@H]3OC(C)(C)[C@@H](OC(=O)C(C)C)[C@H](O)[C@@H]3O)/C(C)=C/C(C)=C/C[C@@H]([C@@H](C)O)OC2=O)O[C@@H]1C | ZVGNESXIJDCBKN-UUEYKCAUSA-N | Cyto | inactive |
| FRAX486 | BRD-K57243773 | CCn1c(=O)c(-c2ccc(Cl)cc2Cl)cc2cnc(Nc3ccc(N4CCNCC4)c(F)c3)nc21 | DHKFOIHIUYFSOF-UHFFFAOYSA-N | Cyto | inactive |

| | | | | | |
|---|---|---|---|---|---|
| gilteritinib | BRD-K72280606 | CCc1nc(C(N)=O)c(Nc2ccc(N3CCC(N4CCN(C)CC4)CC3)c(OC)c2)nc1NC1CCOCC1 | GYQYAJJFPNQOOW-UHFFFAOYSA-N | Cyto | inactive |
| givinostat | BRD-K13810148 | CCN(CC)Cc1ccc2cc(COC(=O)Nc3ccc(C(=O)NO)cc3)ccc2c1 | YALNUENQHAQXEA-UHFFFAOYSA-N | Cyto | inactive |
| GSK2334470 | BRD-K70642949 | CNc1nc(-c2ccc3c(N)[nH]nc3c2)cc(N2C[C@@H](C(=O)NC3CCCCC3)CC[C@H]2C)n1 | QLPHOXTXAKOFMU-WBVHZDCISA-N | Cyto | inactive |
| GSK3117391 | BRD-K00004614 | O=C(CCc1ccc(CN[C@H](C(=O)OC2CCCC2)C2CCCCC2)cn1)NO | AFDPFLDWOXXHQM-NRFANRHFSA-N | Cyto | inactive |
| ID-8 | BRD-K39166528 | COc1ccc(-n2c(C)c([N+](=O)[O-])c3ccc(O)cc32)cc1 | VVZNWYXIOADGSW-UHFFFAOYSA-N | Cyto | inactive |
| ixazomib-citrate | BRD-A66419424 | CC(C)C[C@H](NC(=O)CNC(=O)c1cc(Cl)ccc1Cl)B1OC(=O)C[C@](CC(=O)O)(C(=O)O)O1 | YTXSYWAKVMZICI-VBKZILBWSA-N | Cyto | inactive |
| JNJ-26481585 | BRD-K83837640 | Cn1cc(CNCC2CCN(c3ncc(C(=O)NO)cn3)CC2)c2ccccc21 | PAWIYAYFNXQGAP-UHFFFAOYSA-N | Cyto | inactive |
| JNJ-7706621 | BRD-K11636097 | Nc1nc(Nc2ccc(S(N)(=O)=O)cc2)nn1C(=O)c1c(F)cccc1F | KDKUVYLMPJIGKA-UHFFFAOYSA-N | Cyto | inactive |
| KF-38789 | BRD-A54927599 | COc1ccc([C@H]2CC(c3c(O)cc(C)oc3=O)=NCCS2)c(OC)c1 | NHFIAAAMCYVRIW-QGZVFWFLSA-N | Cyto | inactive |
| LMK-235 | BRD-K49921286 | Cc1cc(C)cc(C(=O)NOCCCCCC(=O)NO)c1 | VRYZCEONIWEUAV-UHFFFAOYSA-N | Cyto | inactive |
| M-344 | BRD-K45528773 | CN(C)c1ccc(C(=O)NCCCCCCC(=O)NO)cc1 | MXWDSZWTBOCWBK-UHFFFAOYSA-N | Cyto | inactive |
| maytansinol-isobutyrate | BRD-K15898725 | COc1cc2cc(c1Cl)N(C)C(=O)C[C@H](OC(=O)C(C)C)[C@]1(C)O[C@H]1[C@H](C)[C@@H]1C[C@@](O)(NC(=O)O1)[C@H](OC)/C=C/C=C(\C)C2 | OPQNCARIZFLNLF-JBHFWYGFSA-N | Cyto | inactive |
| mepacrine | BRD-A45889380 | CCN(CC)CCC[C@@H](C)Nc1c2ccc(Cl)cc2nc2ccc(OC)cc12 | GPKJTRJOBQGKQK-MRXNPFEDSA-N | Cyto | inactive |
| MG-132 | BRD-K60230970 | CC(C)C[C@@H](C=O)NC(=O)[C@H](CC(C)C)NC(=O)[C@H](CC(C)C)NC(=O)OCc1ccccc1 | TZYWCYJVHRLUCT-VABKMULXSA-N | Cyto | inactive |
| mivobulin | BRD-K39576764 | CCOC(=O)Nc1cc2c(c(N)n1)NC(c1ccccc1)C(C)=N2 | XOJFFEJRKAIVCH-UHFFFAOYSA-N | Cyto | inactive |
| NMS-873 | BRD-K26253904 | Cc1cc(OCc2nnc(SC3CCCC3)n2-c2cccnc2)ccc1-c1ccc(S(C)(=O)=O)cc1 | UJGTUKMAJVCBIS-UHFFFAOYSA-N | Cyto | inactive |
| NSC-632839 | BRD-K81197548 | Cc1ccc(/C=C2\CNC/C(=C\c3ccc(C)cc3)C2=O)cc1 | PKCYYPHSCUSQDK-AYKLPDECSA-N | Cyto | inactive |
| NVP-BEZ235 | BRD-K12184916 | Cn1c(=O)n(-c2ccc(C(C)(C)C#N)cc2)c2c3cc(-c4cnc5ccccc5c4)ccc3ncc21 | JOGKUKXHTYWRGZ-UHFFFAOYSA-N | Cyto | inactive |
| omtriptolide | BRD-K00003212 | CC(C)[C@]12O[C@H]1[C@@H]1O[C@@]13[C@@]1(C)CCC4=C(COC4=O)[C@@H]1C[C@@H]1O[C@@]13[C@@H]2OC(=O)CCC(=O)O | HROMYAWHLUOUPY-AHCCQAQQSA-N | Cyto | inactive |
| oprozomib | BRD-K91438851 | COC[C@H](NC(=O)c1cnc(C)s1)C(=O)N[C@@H](COC)C(=O)N[C@@H](Cc1ccccc1)C(=O)[C@@]1(C)CO1 | SWZXEVABPLUDIO-WSZYKNRRSA-N | Cyto | inactive |
| OTS514 | BRD-K38018935 | Cc1cc(O)c(-c2ccc([C@@H](C)CN)cc2)c2c1[nH]c(=O)c1sccc12 | OETLNMOJNONWOY-LBPRGKRZSA-N | Cyto | inactive |
| OTX015 | BRD-K43471985 | Cc1sc2c(c1C)C(c1ccc(Cl)cc1)=N[C@@H](CC(=O)Nc1ccc(O)cc1)c1nnc(C)n1-2 | GNMUEVRJHCWKTO-FQEVSTJZSA-N | Cyto | inactive |
| oxaliplatin | BRD-K78960041 | O=C1O[Pt]2(N[C@@H]3CCCC[C@H]3N2)OC1=O | DWAFYCQODLXJNR-BNTLRKBRSA-L | Cyto | inactive |
| P276-00 | BRD-K51791723 | CN1CC[C@@H](c2c(O)cc(O)c3c(=O)cc(-c4ccccc4Cl)oc23)[C@@H]1CO | QLUYMIVVAYRECT-OCCSQVGLSA-N | Cyto | inactive |
| palmitoylcarnitine | BRD-K01826784 | CCCCCCCCCCCCCCCC(=O)O[C@@H](CC(=O)O)C[N+](C)(C)C | XOMRRQXKHMYMOC-NRFANRHFSA-O | Cyto | inactive |
| panobinostat | BRD-K02130563 | Cc1[nH]c2ccccc2c1CCNCc1ccc(/C=C/C(=O)NO)cc1 | FPOHNWQLNRZRFC-ZHACJKMWSA-N | Cyto | inactive |
| PD-166285 | BRD-K93217218 | CCN(CC)CCOc1ccc(Nc2ncc3cc(-c4c(Cl)cccc4Cl)c(=O)n(C)c3n2)cc1 | IFPPYSWJNWHOLQ-UHFFFAOYSA-N | Cyto | inactive |
| PF-431396 | BRD-K93788137 | CN(c1ccccc1CNc1nc(Nc2ccc3c(c2)CC(=O)N3)ncc1C(F)(F)F)S(C)(=O)=O | POJZIZBONPAWIV-UHFFFAOYSA-N | Cyto | inactive |
| PHA-848125 | BRD-K60997853 | CNC(=O)c1nn(C)c2c1C(C)(C)Cc1cnc(Nc3ccc(N4CCN(C)CC4)cc3)nc1-2 | RXZMYLDMFYNEIM-UHFFFAOYSA-N | Cyto | inactive |
| podophyllotoxin | BRD-K47869605 | COc1cc([C@@H]2c3cc4c(cc3[C@H](O)[C@H]3COC(=O)[C@H]23)OCO4)cc(OC)c1OC | YJGVMLPVUAXIQN-XVVDYKMHSA-N | Cyto | inactive |
| propranolol-(R) | BRD-K92830582 | CC(C)NC[C@@H](O)COc1cccc2ccccc12 | AQHHHDLHHXJYJD-CQSZACIVSA-N | Cyto | inactive |
| PSNCBAM-1 | BRD-K33502629 | O=C(Nc1ccc(Cl)cc1)Nc1cccc(-c2cccc(N3CCCC3)n2)c1 | HDAYFSFWIPRJSO-UHFFFAOYSA-N | Cyto | inactive |
| R306465 | BRD-K86687746 | O=C(NO)c1cnc(N2CCN(S(=O)(=O)c3ccc4ccccc4c3)CC2)nc1 | MUTBJZVSRNUIHA-UHFFFAOYSA-N | Cyto | inactive |
| R547 | BRD-K52233191 | COc1ccc(F)c(F)c1C(=O)c1cnc(NC2CCN(S(C)(=O)=O)CC2)nc1N | JRNJNYBQQYBCLE-UHFFFAOYSA-N | Cyto | inactive |
| rigosertib | BRD-K55187425 | COc1cc(OC)c(/C=C/S(=O)(=O)Cc2ccc(OC)c(NCC(=O)O)c2)c(OC)c1 | OWBFCJROIKNMGD-BQYQJAHWSA-N | Cyto | inactive |
| ripazepam | BRD-K57427145 | CCn1nc(C)c2c1C(c1ccccc1)=NCC(=O)N2 | YFHYNLHGFKXAIQ-UHFFFAOYSA-N | Cyto | inactive |
| RN-1747 | BRD-K45519571 | O=[N+]([O-])c1cc(Cl)ccc1S(=O)(=O)N1CCN(Cc2ccccc2)CC1 | ZNLVYSJQUMALEO-UHFFFAOYSA-N | Cyto | inactive |
| Ro-1138452 | BRD-K77220725 | CC(C)Oc1ccc(Cc2ccc(NC3=NCCN3)cc2)cc1 | GYYRMJMXXLJZAB-UHFFFAOYSA-N | Cyto | inactive |
| romidepsin | BRD-K61397605 | C/C=C1\NC(=O)[C@H]2CSSCC/C=C/[C@H](CC(=O)N[C@H](C(C)C)C(=O)N2)OC(=O)[C@H](C(C)C)NC1=O | OHRURASPPZQGQM-GCCNXGTGSA-N | Cyto | inactive |
| saccharin | BRD-K46493214 | O=C1NS(=O)(=O)c2ccccc21 | CVHZOJJKTDOEJC-UHFFFAOYSA-N | Cyto | inactive |
| SB-218078 | BRD-A06352508 | O=C1NC(=O)c2c1c1c3ccccc3n3c1c1c2c2ccccc2n1[C@H]1CC[C@H]3O1 | OTPNDVKVEAIXTI-HZPDHXFCSA-N | Cyto | inactive |
| SB-939 | BRD-K86797399 | CCCCc1nc2cc(/C=C/C(=O)NO)ccc2n1CCN(CC)CC | JHDKZFFAIZKUCU-ZRDIBKRKSA-N | Cyto | inactive |
| STF-62247 | BRD-K85479953 | Cc1cccc(Nc2nc(-c3ccncc3)cs2)c1 | KATNUHQNJGNLPW-UHFFFAOYSA-N | Cyto | inactive |
| TAK-960 | BRD-K48059512 | COc1cc(C(=O)NC2CCN(C)CC2)c(F)cc1Nc1ncc2c(n1)N(C1CCCC1)CC(F)(F)C(=O)N2C | GWRSATNRNFYMDI-UHFFFAOYSA-N | Cyto | inactive |
| TEN-010 | BRD-K00005242 | Cc1sc2c(c1C)C(c1ccc(Cl)cc1)=N[C@@H](CC(=O)NCCCN1CCN(C)CC1)c1nnc(C)n1-2 | PKQXLRYFPSZKDU-QFIPXVFZSA-N | Cyto | inactive |
| teniposide | BRD-K10362825 | COc1cc([C@@H]2c3cc4c(cc3[C@@H](O[C@@H]3O[C@@H]5CO[C@@H](c6cccs6)O[C@H]5[C@H](O)[C@H]3O)[C@H]3COC(=O)[C@H]23)OCO4)cc(OC)c1O | NRUKOCRGYNPUPR-QBPJDGROSA-N | Cyto | inactive |
| TG-02 | BRD-K14560436 | CN1C/C=C/CCOc2cccc(c2)-c2ccnc(n2)Nc2cccc(c2)C1 | VXBAJLGYBMTJCY-NSCUHMNNSA-N | Cyto | inactive |
| TG-101209 | BRD-K39841531 | Cc1cnc(Nc2ccc(N3CCN(C)CC3)cc2)nc1Nc1cccc(S(=O)(=O)NC(C)(C)C)c1 | JVDOKQYTTYUYDV-UHFFFAOYSA-N | Cyto | inactive |
| torin-2 | BRD-K68174511 | Nc1ccc(-c2ccc3ncc4ccc(=O)n(-c5cccc(C(F)(F)F)c5)c4c3c2)cn1 | GUXXEUUYCAYESJ-UHFFFAOYSA-N | Cyto | inactive |
| TP-0903 | BRD-K84034737 | CN1CCN(Cc2ccc(Nc3ncc(Cl)c(Nc4ccccc4S(=O)(=O)N(C)C)n3)cc2)CC1 | YUAALFPUEOYPNX-UHFFFAOYSA-N | Cyto | inactive |
| trichostatin-a | BRD-K68202742 | CC(/C=C/C(=O)NO)=C\[C@@H](C)C(=O)c1ccc(N(C)C)cc1 | RTKIYFITIVXBLE-QEQCGCAPSA-N | Cyto | inactive |
| TW-37 | BRD-K28360340 | CC(C)c1ccccc1Cc1cc(C(=O)Nc2ccc(S(=O)(=O)c3ccccc3C(C)(C)C)cc2)c(O)c(O)c1O | PQAPVTKIEGUPRN-UHFFFAOYSA-N | Cyto | inactive |
| vinburnine | BRD-K40227168 | CC[C@]12CCCN3CCc4c(n(c5ccccc45)C(=O)C1)[C@@H]32 | WYJAPUKIYAZSEM-MOPGFXCFSA-N | Cyto | inactive |
| volasertib | BRD-K69776681 | CC[C@@H]1C(=O)N(C)c2cnc(Nc3ccc(C(=O)N[C@H]4CC[C@H](N5CCN(CC6CC6)CC5)CC4)cc3OC)nc2N1C(C)C | SXNJFOWDRLKDSF-STROYTFGSA-N | Cyto | inactive |
| VUF10166 | BRD-K11046126 | CN1CCN(c2nc3ccccc3nc2Cl)CC1 | FFXVTQDGTKEXHF-UHFFFAOYSA-N | Cyto | inactive |
| WZ-4002 | BRD-K72420232 | C=CC(=O)Nc1cccc(Oc2nc(Nc3ccc(N4CCN(C)CC4)cc3OC)ncc2Cl)c1 | ITTRLTNMFYIYPA-UHFFFAOYSA-N | Cyto | inactive |
| ziritaxestat | BRD-K00003431 | CCc1nc2c(C)cc(N3CCN(CC(=O)N4CC(O)C4)CC3)cn2c1N(C)c1nc(-c2ccc(F)cc2)c(C#N)s1 | REQQVBGILUTQNN-UHFFFAOYSA-N | Cyto | inactive |
| 12-O-tetradecanoylphorbol-13-acetate | BRD-K68552125 | CCCCCCCCCCCCCC(=O)O[C@@H]1[C@@H](C)[C@@]2(O)[C@@H](C=C(CO)C[C@]3(O)C(=O)C(C)=C[C@@H]23)[C@@H]2C(C)(C)[C@]12OC(C)=O | PHEDXBVPIONUQT-RGYGYFBISA-N | Low | inactive |

| | | | | | |
|---|---|---|---|---|---|
| 2-(3-mercaptopropyl)pentanedioic-acid | BRD-A66788520 | O=C(O)CC[C@@H](CCCS)C(=O)O | FNLNSQHJKVQCBP-ZCFIWIBFSA-N | Low | inactive |
| 2-chloroadenosine | BRD-K79511609 | Nc1nc(Cl)nc2c1ncn2[C@@H]1O[C@H](CO)[C@@H](O)[C@H]1O | BIXYYZIIJIXVFW-UUOKFMHZSA-N | Low | inactive |
| 2-TEDC | BRD-K89072800 | N#C/C(=C\c1ccc(O)c(O)c1)C(=O)OCCc1cccs1 | KCGLTUBCAGZLHP-XYOKQWHBSA-N | Low | inactive |
| ABT-751 | BRD-K91623615 | COc1ccc(S(=O)(=O)Nc2cccnc2Nc2ccc(O)cc2)cc1 | URCVCIZFVQDVPM-UHFFFAOYSA-N | Low | inactive |
| AC-55649 | BRD-K93176058 | CCCCCCCCc1ccc(-c2ccc(C(=O)O)cc2)cc1 | HXBKPYIEQLLNBK-UHFFFAOYSA-N | Low | inactive |
| acarbose | BRD-K44276885 | C[C@H]1O[C@H](O[C@@H]2[C@@H](CO)O[C@H](O[C@@H]3[C@@H](CO)O[C@@H](O)[C@H](O)[C@H]3O)[C@H](O)[C@H]2O)[C@H](O)[C@@H](O)[C@@H]1N[C@H]1C=C(CO)[C@@H](O)[C@H](O)[C@H]1O | XUFXOAAUWZOOIT-SXARVLRPSA-N | Low | inactive |
| adelmidrol | BRD-K78097265 | O=C(CCCCCCCC(=O)NCCO)NCCO | PAHZPHDAJQIETD-UHFFFAOYSA-N | Low | inactive |
| afamelanotide | BRD-K00003330 | CCCC[C@H](NC(=O)[C@H](CO)NC(=O)[C@H](Cc1ccc(O)cc1)NC(=O)[C@H](CO)NC(C)=O)C(=O)N[C@@H](CCC(=O)O)C(=O)N[C@@H](Cc1cnc[nH]1)C(=O)N[C@H](Cc1ccccc1)C(=O)N[C@@H](CCCNC(=N)N)C(=O)N[C@@H](Cc1c[nH]c2ccccc12)C(=O)NCC(=O)N[C@@H](CCCCN)C(=O)N1CCC[C@H]1C(=O)N[C@H](C(N)=O)C(C)C | UAHFGYDRQSXQEB-LEBBXHLNSA-N | Low | inactive |
| alfadolone-acetate | BRD-K51751936 | CC(=O)OCC(=O)[C@H]1CC[C@H]2[C@@H]3CC[C@H]4C[C@H](O)CC[C@]4(C)[C@H]3C(=O)C[C@]12C | QRJOQYLXZPQQMX-FWROMSNXSA-N | Low | inactive |
| aliskiren-hemifumarate | BRD-K30020243 | COCCCOc1cc(C[C@@H](C[C@H](N)[C@@H](O)C[C@H](C(=O)NCC(C)(C)C(N)=O)C(C)C)C(C)C)ccc1OC | UXOWGYHJODZGMF-QORCZRPOSA-N | Low | inactive |
| altretamine | BRD-K67043667 | CN(C)c1nc(N(C)C)nc(N(C)C)n1 | UUVWYPNAQBNQJQ-UHFFFAOYSA-N | Low | inactive |
| amcinonide | BRD-A36010170 | CC(=O)OCC(=O)[C@@]12OC3(CCCC3)O[C@@H]1C[C@H]1[C@H]3CCC4=CC(=O)C=C[C@]4(C)[C@@]3(F)[C@@H](O)C[C@@]12C | ILKJAFIWWBXGDU-KNPIAJNZSA-N | Low | inactive |
| AMG-208 | BRD-K32285926 | COc1ccc2c(OCc3nnc4ccc(-c5ccccc5)nn34)ccnc2c1 | HEAIZQNMNCHNFD-UHFFFAOYSA-N | Low | inactive |
| aminosalicylate | BRD-K80267133 | Nc1ccc(C(=O)O)c(O)c1 | WUBBRNOQWQTFEX-UHFFFAOYSA-N | Low | inactive |
| aminothiazole | BRD-K13421763 | Nc1nccs1 | RAIPHJJURHTUIC-UHFFFAOYSA-N | Low | inactive |
| amphotericin-b | BRD-K27352494 | C[C@@H]1OC(=O)C[C@H](O)C[C@H](O)CC[C@@H](O)[C@H](O)C[C@H](O)C[C@]2(O)C[C@H](O)[C@@H](C(=O)O)[C@H](C[C@@H](O[C@@H]3O[C@H](C)[C@@H](O)[C@H](N)[C@@H]3O)/C=C/C=C/C=C/C=C/C=C/C=C/C=C/[C@H](C)[C@@H](O)[C@H]1C)O2 | APKFDSVGJQXUKY-INPOYWNPSA-N | Low | inactive |
| ampiroxicam | BRD-A39172021 | CCOC(=O)O[C@@H](C)OC1=C(C(=O)Nc2ccccn2)N(C)S(=O)(=O)c2ccccc21 | LSNWBKACGXCGAJ-ZDUSSCGKSA-N | Low | inactive |
| anisodamine | BRD-A29623586 | CN1[C@@H]2C[C@H](OC(=O)[C@H](CO)c3ccccc3)C[C@@H]1[C@@H](O)C2 | WTQYWNWRJNXDEG-JKJDWNRSSA-N | Low | inactive |
| antimony-potassium | BRD-K00003675 | O=C1O[SbH]23OC(=O)[C@H](O2)[C@H]2O[SbH]4(OC(=O)[C@H](O4)[C@H]1O3)OC2=O | ZCIXFSVENQDMCK-SOMOIXMJSA-J | Low | inactive |
| AQ-RA741 | BRD-K81729199 | CCN(CC)CCCCC1CCN(CC(=O)N2c3ccccc3C(=O)Nc3cccnc32)CC1 | BCUGCHZRMKTPMU-UHFFFAOYSA-N | Low | inactive |
| arachidonic-acid | BRD-K03070961 | CCCCC/C=C\C/C=C\C/C=C\C/C=C\CCCC(=O)O | YZXBAPSDXZZRGB-DOFZRALJSA-N | Low | inactive |
| aripiprazole-lauroxil | BRD-K00004574 | CCCCCCCCCCCC(=O)OCN1C(=O)CCc2ccc(OCCCCN3CCN(c4cccc(Cl)c4Cl)CC3)cc21 | DDINXHAORAAYAD-UHFFFAOYSA-N | Low | inactive |
| artemotil | BRD-K96006160 | CCO[C@H]1O[C@@H]2O[C@@]3(C)CC[C@H]4[C@H](C)CC[C@@H]([C@H]1C)[C@]42OO3 | NLYNIRQVMRLPIQ-XQLAAWPRSA-N | Low | inactive |
| atosiban | BRD-A85261566 | CCOc1ccc(C[C@H]2NC(=O)CCSSC[C@@H](C(=O)N3CCC[C@H]3C(=O)N[C@@H](CCCN)C(=O)NCC(N)=O)NC(=O)[C@H](CC(N)=O)NC(=O)[C@H]([C@@H](C)O)NC(=O)[C@H]([C@@H](C)CC)NC2=O)cc1 | VWXRQYYUEIYXCZ-OBIMUBPZSA-N | Low | inactive |
| atuveciclib | BRD-K00003144 | COc1cc(F)ccc1-c1ncnc(Nc2cccc(C[S@](C)(=N)=O)c2)n1 | ACWKGTGIJRCOOM-HHHXNRCGSA-N | Low | inactive |
| AZD5991 | BRD-K00005264 | Cc1c2c(nn1C)CSCc1cc(n(C)n1)CSc1cc(c3ccccc3c1)OCCCc1c(C(=O)O)n(C)c3c-2c(Cl)ccc13 | KBQCEQAXHPIRTF-UHFFFAOYSA-N | Low | inactive |
| azelnidipine | BRD-A01098288 | CC1=C(C(=O)OC(C)C)[C@@H](c2cccc([N+](=O)[O-])c2)C(C(=O)OC2CN(C(c3ccccc3)c3ccccc3)C2)=C(N)N1 | ZKFQEACEUNWPMT-MUUNZHRXSA-N | Low | inactive |
| azosemide | BRD-K68770713 | NS(=O)(=O)c1cc(-c2nn[nH]n2)c(NCc2cccs2)cc1Cl | HMEDEBAJARCKCT-UHFFFAOYSA-N | Low | inactive |
| baloxavir-marboxil | BRD-K00003495 | COC(=O)OCOc1c2n(ccc1=O)N([C@@H]1c3ccccc3SCc3c1ccc(F)c3F)[C@@H]1COCCN1C2=O | RZVPBGBYGMDSBG-GGAORHGYSA-N | Low | inactive |
| benzo[d]thiazole-2(3H)-thione | BRD-K55160477 | S=c1[nH]c2ccccc2s1 | YXIWHUQXZSMYRE-UHFFFAOYSA-N | Low | inactive |
| benztropine-mesylate | BRD-K68804560 | CN1[C@H]2CC[C@@H]1C[C@H](OC(c1ccccc1)c1ccccc1)C2 | GIJXKZJWITVLHI-PMOLBWCYSA-N | Low | inactive |
| bepotastine | BRD-K98624455 | O=C(O)CCCN1CCC(O[C@@H](c2ccc(Cl)cc2)c2ccccn2)CC1 | YWGDOWXRIALTES-NRFANRHFSA-N | Low | inactive |
| betamethasone-butyrate-propionate | BRD-K00004736 | CCCC(=O)O[C@]1(C(=O)COC(=O)CC)[C@@H](C)C[C@H]2[C@@H]3CCC4=CC(=O)C=C[C@]4(C)[C@@]3(F)[C@@H](O)C[C@@]21C | VXOWJCTXWVWLLC-REGDIAEZSA-N | Low | inactive |
| betamethasone-dipropionate | BRD-K58148589 | CCC(=O)OCC(=O)[C@@]1(OC(=O)CC)[C@@H](C)C[C@H]2[C@@H]3CCC4=CC(=O)C=C[C@]4(C)[C@@]3(F)[C@@H](O)C[C@@]21C | CIWBQSYVNNPZIQ-XYWKZLDCSA-N | Low | inactive |
| bethanechol | BRD-K00003637 | C[C@@H](C[N+](C)(C)C)OC(N)=O | NZUPCNDJBJXXRF-LURJTMIESA-O | Low | inactive |
| BGT226 | BRD-K29415052 | COc1ccc(-c2ccc3ncc4c(c3c2)n(-c2ccc(N3CCNCC3)c(C(F)(F)F)c2)c(=O)n4C)cn1 | BMMXYEBLEBULND-UHFFFAOYSA-N | Low | inactive |
| BIBR-1532 | BRD-K04623885 | C/C(=C\C(=O)Nc1ccccc1C(=O)O)c1ccc2ccccc2c1 | PGFQXGLPJUCTOI-WYMLVPIESA-N | Low | inactive |
| bicalutamide | BRD-A29485665 | CC(O)(CS(=O)(=O)c1ccc(F)cc1)C(=O)Nc1ccc(C#N)c(C(F)(F)F)c1 | LKJPYSCBVHEWIU-UHFFFAOYSA-N | Low | inactive |
| BIMU-8 | BRD-K00004300 | CC(C)n1c(=O)n(C(=O)N[C@@H]2C[C@@H]3CC[C@H](C2)N3C)c2ccccc21 | AFOFVIBWSLOHFR-QDMKHBRRSA-N | Low | inactive |
| blebbistatin-(+/-) | BRD-A75817871 | Cc1ccc2c(c1)C(=O)[C@]1(O)CCN(c3ccccc3)C1=N2 | LZAXPYOBKSJSEX-GOSISDBHSA-N | Low | inactive |
| BMS-470539 | BRD-K22000702 | CCCC(=O)C1(c2ccccc2)CCN(C(=O)[C@@H](Cc2ccc(OC)cc2)NC(=O)[C@@H](N)Cc2cncn2C)CC1 | GULVVTHHUIIMRL-WUFINQPMSA-N | Low | inactive |
| BU226 | BRD-K56115039 | c1ccc2cc(C3=NCCN3)ncc2c1 | ZXRSVNHYTSSFMN-UHFFFAOYSA-N | Low | inactive |
| camicinal | BRD-K23248433 | C[C@H]1CN(Cc2ccc(CC(=O)N3CCC(Nc4cccc(F)c4)CC3)cc2)CCN1 | RZKDEGZIFSJVNA-IBGZPJMESA-N | Low | inactive |
| candicidin | BRD-A37348498 | C[C@@H](C[C@@H](C)[C@H]1OC(=O)CC(=O)CCCC(=O)C[C@@H](O)C[C@@H](O)C[C@@H](O)CC(=O)C[C@@H](O)[C@H](C(=O)O)[C@@H](O)C[C@@H](O[C@@H]2O[C@H](C)[C@H](O)[C@@H](O)[C@@H]2N)/C=C/C=C/C=C/C=C/C=C/C=C/C=C/[C@@H]1C)[C@@H](O)CC(=O)c1ccc(N)cc1 | HYIZCDMXFDRDFE-GCCXFADDSA-N | Low | inactive |
| canertinib | BRD-K50168500 | C=CC(=O)Nc1cc2c(Nc3ccc(F)c(Cl)c3)ncnc2cc1OCCCN1CCOCC1 | OMZCMEYTWSXEPZ-UHFFFAOYSA-N | Low | inactive |
| capreomycin | BRD-K00005140 | C[C@@H]1NC(=O)[C@@H](N)CNC(=O)[C@H]([C@H]2CCN=C(N)N2)NC(=O)/C(=C\NC(N)=O)NC(=O)[C@H](CNC(=O)C[C@@H](N)CCCN)NC1=O | FRXNXDHFQYZYNA-OJBVOROPSA-N | Low | inactive |
| caprylic-acid | BRD-K35170555 | CCCCCCCC(=O)O | WWZKQHOCKIZLMA-UHFFFAOYSA-N | Low | inactive |

| | | | | | |
|---|---|---|---|---|---|
| carbenicillin | BRD-A13078313 | CC1(C)S[C@@H]2[C@H](NC(=O)[C@@H](C(=O)O)c3ccccc3)C(=O)N2[C@H]1C(=O)O | FPPNZSSZRUTDAP-BBGACYKPSA-N | Low | inactive |
| carbinoxamine | BRD-K01826745 | CN(C)CCO[C@@H](c1ccc(Cl)cc1)c1ccccn1 | OJFSXZCBGQGRNV-INIZCTEOSA-N | Low | inactive |
| carprofen | BRD-A17411484 | C[C@@H](C(=O)O)c1ccc2c(c1)[nH]c1ccc(Cl)cc12 | PUXBGTOOZJQSKH-MRVPVSSYSA-N | Low | inactive |
| CE3F4 | BRD-A47940219 | C[C@H]1CCc2c(cc(Br)c(F)c2Br)N1C=O | ZZLQPWXVZCPUGC-LURJTMIESA-N | Low | inactive |
| cefonicid | BRD-K34058848 | O=C(O)C1=C(CSc2nnnn2CS(=O)(=O)O)CS[C@@H]2[C@H](NC(=O)[C@H](O)c3ccccc3)C(=O)N12 | DYAIAHUQIPBDIP-AXAPSJFSSA-N | Low | inactive |
| cefoselis | BRD-K72339976 | CO/N=C(\C(=O)N[C@@H]1C(=O)N2C(C(=O)O)=C(Cn3ccc(=N)n3CCO)CS[C@H]12)c1csc(N)n1 | ZINFAXPQMLDEEJ-GFVOIPPFSA-N | Low | inactive |
| cefpirome | BRD-K56542719 | CO/N=C(/C(=O)N[C@@H]1C(=O)N2C(C(=O)O)=C(C[n+]3cccc4c3CCC4)CS[C@H]12)c1csc(N)n1 | DKOQGJHPHLTOJR-PUIFZAHNSA-O | Low | inactive |
| cetirizine | BRD-K01826520 | O=C(O)COCCN1CCN([C@@H](c2ccccc2)c2ccc(Cl)cc2)CC1 | ZKLPARSLTMPFCP-NRFANRHFSA-N | Low | inactive |
| CFI-402257 | BRD-K00003409 | Cc1cc(-c2cnn3c(NC[C@H]4C[C@@](C)(O)C4)cc(Oc4cccnc4)nc23)ccc1C(=O)NC1CC1 | PMQUGSPFUBGJCZ-CHOKWEPUSA-N | Low | inactive |
| CGP-37849 | BRD-A90272038 | C/C(=C\[C@@H](N)C(=O)O)CP(=O)(O)O | BDYHNCZIGYIOGJ-XWCPEMDWSA-N | Low | inactive |
| CHIR-124 | BRD-K02241333 | O=c1[nH]c2ccc(Cl)cc2c(N[C@@H]2CN3CCC2CC3)c1-c1nc2ccccc2[nH]1 | MOVBBVMDHIRCTG-LJQANCHMSA-N | Low | inactive |
| CI-923 | BRD-A50016482 | CCN[C@@H](C)CN1CCc2c(c(=O)oc3ccccc23)C1 | MJRICTKAEYTJAV-LBPRGKRZSA-N | Low | inactive |
| ciaftalan-zinc | BRD-K01423020 | c1ccc2c(c1)C1=N/C2=N\c2c3ccccc3c3n2[Zn]n2/c(c4ccccc4/c2=N/C2=N/C(=N\3)c3ccccc32)=N\1 | PODBBOVVOGJETB-UHFFFAOYSA-N | Low | inactive |
| cipemastat | BRD-K95362002 | CN1C(=O)N(C[C@H](C(=O)NO)[C@@H](CC2CCCC2)C(=O)N2CCCCC2)C(=O)C1(C)C | GFUITADOEPNRML-SJORKVTESA-N | Low | inactive |
| ciproxifan | BRD-K01767299 | O=C(c1ccc(OCCCc2cnc[nH]2)cc1)C1CC1 | ACQBHJXEAYTHCY-UHFFFAOYSA-N | Low | inactive |
| clofedanol | BRD-A15687940 | CN(C)CC[C@@](O)(c1ccccc1)c1ccccc1Cl | WRCHFMBCVFFYEQ-QGZVFWFLSA-N | Low | inactive |
| clofoctol | BRD-K02900412 | CC(C)(C)CC(C)(C)c1ccc(O)c(Cc2ccc(Cl)cc2Cl)c1 | HQVZOORKDNCGCK-UHFFFAOYSA-N | Low | inactive |
| cloperastine | BRD-A80908310 | Clc1ccc([C@@H](OCCN2CCCCC2)c2ccccc2)cc1 | FLNXBVJLPJNOSI-FQEVSTJZSA-N | Low | inactive |
| copanlisib | BRD-K24666289 | COc1c(OCCCN2CCOCC2)ccc2c1N=C(NC(=O)c1cnc(N)nc1)N1CCN=C21 | PZBCKZWLPGJMAO-UHFFFAOYSA-N | Low | inactive |
| curcumol | BRD-K41933240 | C=C1C[C@@]2(O)O[C@@]3(C[C@H]2C(C)C)[C@@H](C)CC[C@@H]13 | QRMPRVXWPCLVNI-YYFQZIEXSA-N | Low | inactive |
| cyclobenzaprine | BRD-K42348709 | CN(C)CCC=C1c2ccccc2C=Cc2ccccc21 | JURKNVYFZMSNLP-UHFFFAOYSA-N | Low | inactive |
| D-7193 | BRD-K32372024 | CC(=O)N[C@@H](CSSC[C@H](NC(C)=O)C(=O)O)C(=O)O | YTPQSLLEROSACP-YUMQZZPRSA-N | Low | inactive |
| D-delta-tocopherol | BRD-K04657451 | Cc1cc(O)cc2c1O[C@](C)(CCC[C@H](C)CCC[C@H](C)CCCC(C)C)CC2 | GZIFEOYASATJEH-VHFRWLAGSA-N | Low | inactive |
| danofloxacin | BRD-K45916615 | CN1C[C@@H]2C[C@H]1CN2c1cc2c(cc1F)c(=O)c(C(=O)O)cn2C1CC1 | QMLVECGLEOSESV-RYUDHWBXSA-N | Low | inactive |
| dapoxetine | BRD-K51579475 | CN(C)[C@@H](CCOc1cccc2ccccc12)c1ccccc1 | USRHYDPUVLEVMC-FQEVSTJZSA-N | Low | inactive |
| delivert | BRD-K45503887 | CC[C@H](C)[C@H](NC(=O)[C@H](Cc1ccc(O)cc1)NC(=O)[C@@H](NC(=O)[C@@H](CCCNC(=N)N)NC(=O)[C@@H](N)CC(=O)O)C(C)C)C(=O)N[C@@H](Cc1cnc[nH]1)C(=O)N1CCC[C@@H]1C(=O)N[C@H](Cc1ccccc1)C(=O)O | CZGUSIXMZVURDU-FCWOTDAASA-N | Low | inactive |
| dexmedetomidine | BRD-K18423410 | Cc1cccc([C@H](C)c2cnc[nH]2)c1C | CUHVIMMYOGQXCV-NSHDSACASA-N | Low | inactive |
| dextrose | BRD-K51431759 | O=C[C@H](O)[C@@H](O)[C@H](O)[C@H](O)CO | GZCGUPFRVQAUEE-SLPGGIOYSA-N | Low | inactive |
| dibenzothiophene | BRD-K46863940 | c1ccc2c(c1)sc1ccccc12 | IYYZUPMFVPLQIF-UHFFFAOYSA-N | Low | inactive |
| dienogest | BRD-K50853363 | C[C@]12CCC3=C4CCC(=O)C=C4CC[C@H]3[C@@H]1CC[C@@]2(O)CC#N | AZFLJNIPTRTECV-FUMNGEBKSA-N | Low | inactive |
| difluprednate | BRD-K72992801 | CCCC(=O)O[C@]1(C(=O)COC(C)=O)CC[C@H]2[C@@H]3C[C@H](F)C4=CC(=O)C=C[C@]4(C)[C@@]3(F)[C@@H](O)C[C@@]21C | WYQPLTPSGFELIB-JTQPXKBDSA-N | Low | inactive |
| dilmapimod | BRD-K00003459 | Cc1cc(F)ccc1-c1nc(NC(CO)CO)nc2c1ccc(=O)n2-c1c(F)cccc1F | ORVNHOYNEHYKJG-UHFFFAOYSA-N | Low | inactive |
| dipyridamole | BRD-K86301799 | OCCN(CCO)c1nc(N2CCCCC2)c2nc(N(CCO)CCO)nc(N3CCCCC3)c2n1 | IZEKFCXSFNUWAM-UHFFFAOYSA-N | Low | inactive |
| disopyramide | BRD-A29734509 | CC(C)N(CCC(C(N)=O)(c1ccccc1)c1ccccn1)C(C)C | UVTNFZQICZKOEM-UHFFFAOYSA-N | Low | inactive |
| DMP-777 | BRD-K54936858 | CCC[C@@H](NC(=O)N1C(=O)C(CC)(CC)[C@@H]1Oc1ccc(C(=O)N2CCN(C)CC2)cc1)c1ccc2c(c1)OCO2 | ZSDCIRYNTCVTMF-GIGWZHCTSA-N | Low | inactive |
| dodecyl-sulfate | BRD-K05680146 | CCCCCCCCCCCCOS(=O)(=O)O | MOTZDAYCYVMXPC-UHFFFAOYSA-N | Low | inactive |
| doxercalciferol | BRD-K82928892 | C=C1/C(=C/C=C2\CCC[C@@]3(C)[C@H]2CC[C@@H]3[C@H](C)/C=C/[C@@H](C)C(C)C)C[C@@H](O)C[C@@H]1O | HKXBNHCUPKIYDM-ZKVKAWQFSA-N | Low | inactive |
| drotaverine | BRD-K00633793 | CCOc1ccc(CC2=NCCc3cc(OCC)c(OCC)cc32)cc1OCC | NCVXEYAKRVJMTQ-UHFFFAOYSA-N | Low | inactive |
| DS2-(806622) | BRD-K09619578 | O=C(Nc1c(-c2cccs2)nc2ccccn12)c1ccc(Cl)cc1 | AZKMWHRDICVYEI-UHFFFAOYSA-N | Low | inactive |
| duvelisib | BRD-K93468883 | C[C@H](Nc1ncnc2nc[nH]c12)c1cc2cccc(Cl)c2c(=O)n1-c1ccccc1 | SJVQHLPISAIATJ-ZDUSSCGKSA-N | Low | inactive |
| ebrotidine | BRD-K22421059 | N=C(N)Nc1nc(CSCCN/C=N/S(=O)(=O)c2ccc(Br)cc2)cs1 | ZQHFZHPUZXNPMF-UHFFFAOYSA-N | Low | inactive |
| ebselen | BRD-K29359156 | O=c1c2ccccc2[se]n1-c1ccccc1 | DYEFUKCXAQOFHX-UHFFFAOYSA-N | Low | inactive |
| EC-144 | BRD-K41705163 | COc1c(C)cnc(Cn2cc(C#CCC(C)(C)O)c3c(Cl)nc(N)nc32)c1C | VOASEWXFCTZRDF-UHFFFAOYSA-N | Low | inactive |
| emiglitate | BRD-K00003206 | CCOC(=O)c1ccc(OCCN2C[C@H](O)[C@@H](O)[C@H](O)[C@H]2CO)cc1 | NWWORXYTJRPSMC-QKPAOTATSA-N | Low | inactive |
| enalapril | BRD-K57545991 | CCOC(=O)[C@H](CCc1ccccc1)N[C@@H](C)C(=O)N1CCC[C@H]1C(=O)O | GBXSMTUPTTWBMN-XIRDDKMYSA-N | Low | inactive |
| eprobemide | BRD-K17979676 | O=C(NCCCN1CCOCC1)c1ccc(Cl)cc1 | YYFGRAGNYHYWEZ-UHFFFAOYSA-N | Low | inactive |
| esomeprazole | BRD-K61443506 | COc1ccc2nc([S@@](=O)Cc3ncc(C)c(OC)c3C)[nH]c2c1 | SUBDBMMJDZJVOS-DEOSSOPVSA-N | Low | inactive |
| estradiol-benzoate | BRD-K44412991 | C[C@]12CC[C@@H]3c4ccc(OC(=O)c5ccccc5)cc4CC[C@H]3[C@@H]1CC[C@@H]2O | UYIFTLBWAOGQBI-BZDYCCQFSA-N | Low | inactive |
| ethimizol | BRD-K19672466 | CCn1cnc(C(=O)NC)c1C(=O)NC | RYRFAMRQBZNEPX-UHFFFAOYSA-N | Low | inactive |
| ethotoin | BRD-A30205217 | CCN1C(=O)N[C@H](c2ccccc2)C1=O | SZQIFWWUIBRPBZ-SECBINFHSA-N | Low | inactive |
| ethoxzolamide | BRD-K18131774 | CCOc1ccc2nc(S(N)(=O)=O)sc2c1 | OUZWUKMCLIBBOG-UHFFFAOYSA-N | Low | inactive |
| ethyl-2-(carbamoyloxy)benzoate | BRD-K21892374 | CCOC(=O)c1ccccc1OC(N)=O | JFMIIMPCFQKAAD-UHFFFAOYSA-N | Low | inactive |
| eticlopride | BRD-K50417881 | CCc1cc(Cl)c(OC)c(C(=O)NC[C@@H]2CCCN2CC)c1O | AADCDMQTJNYOSS-LBPRGKRZSA-N | Low | inactive |
| ETP-45658 | BRD-K15858023 | Cn1ncc2c(N3CCOCC3)nc(-c3cccc(O)c3)nc21 | BJVRNXSHJLDZJR-UHFFFAOYSA-N | Low | inactive |
| ETP-46464 | BRD-K24556407 | CC(C)(C#N)c1ccc(N2C(=O)OCc3cnc4ccc(-c5cnc6ccccc6c5)cc4c32)cc1 | DPLMXAYKJZOTKO-UHFFFAOYSA-N | Low | inactive |
| falecalcitriol | BRD-K08251020 | C=C1/C(=C\C=C2/CCC[C@@]3(C)[C@H]2CC[C@@H]3[C@H](C)CCCC(O)(C(F)(F)F)C(F)(F)F)C[C@@H](O)C[C@@H]1O | XPYGGHVSFMUHLH-UUSULHAXSA-N | Low | inactive |
| fantofarone | BRD-K00003465 | COc1ccc(CCN(C)CCCOc2ccc(S(=O)(=O)c3c(C(C)C)cn4ccccc34)cc2)cc1OC | ITAMRBIZWGDOHW-UHFFFAOYSA-N | Low | inactive |
| faropenem-medoxomil | BRD-K59045783 | Cc1oc(=O)oc1COC(=O)C1=C([C@H]2CCCO2)S[C@@H]2[C@@H]([C@@H](C)O)C(=O)N12 | JQBKWZPHJOEQAO-DVPVEWDBSA-N | Low | inactive |
| fasudil | BRD-K76617868 | O=S(=O)(c1cccc2cnccc12)N1CCCNCC1 | NGOGFTYYXHNFQH-UHFFFAOYSA-N | Low | inactive |
| felodipine | BRD-A30815329 | CCOC(=O)C1=C(C)NC(C)=C(C(=O)OC)[C@@H]1c1cccc(Cl)c1Cl | RZTAMFZIAATZDJ-HNNXBMFYSA-N | Low | inactive |

| | | | | | |
|---|---|---|---|---|---|
| fenbufen | BRD-K12513978 | O=C(O)CCC(=O)c1ccc(-c2ccccc2)cc1 | ZPAKPRAICRBAOD-UHFFFAOYSA-N | Low | inactive |
| fenoldopam | BRD-K01826790 | Oc1ccc([C@H]2CNCCc3c2cc(O)c(O)c3Cl)cc1 | TVURRHSHRRELCG-CYBMUJFWSA-N | Low | inactive |
| fexofenadine | BRD-A73368467 | CC(C)(C(=O)O)c1ccc([C@@H](O)CCCN2CCC(C(O)(c3ccccc3)c3ccccc3)CC2)cc1 | RWTNPBWLLIMQHL-LJAQVGFWSA-N | Low | inactive |
| FK-866 | BRD-K58550667 | O=C(/C=C/c1cccnc1)NCCCCC1CCN(C(=O)c2ccccc2)CC1 | KPBNHDGDUADAGP-VAWYXSNFSA-N | Low | inactive |
| flufenamic-acid | BRD-K44067360 | O=C(O)c1ccccc1Nc1cccc(C(F)(F)F)c1 | LPEPZBJOKDYZAD-UHFFFAOYSA-N | Low | inactive |
| flumethasone | BRD-K61496577 | C[C@@H]1C[C@H]2[C@@H]3C[C@H](F)C4=CC(=O)C=C[C@]4(C)[C@@]3(F)[C@@H](O)C[C@]2(C)[C@@]1(O)C(=O)CO | WXURHACBFYSXBI-GQKYHHCASA-N | Low | inactive |
| fluoxetine | BRD-A31159102 | CNCCC(Oc1ccc(C(F)(F)F)cc1)c1ccccc1 | RTHCYVBBDHJXIQ-UHFFFAOYSA-N | Low | inactive |
| flupentixol | BRD-K70487031 | OCCN1CCN(CC/C=C2/c3ccccc3Sc3ccc(C(F)(F)F)cc32)CC1 | NJMYODHXAKYRHW-DVZOWYKESA-N | Low | inactive |
| fluphenazine | BRD-K55127134 | OCCN1CCN(CCCN2c3ccccc3Sc3ccc(C(F)(F)F)cc32)CC1 | PLDUPXSUYLZYBN-UHFFFAOYSA-N | Low | inactive |
| fluticasone-propionate | BRD-K62310379 | CCC(=O)O[C@]1(C(=O)SCF)[C@H](C)C[C@H]2[C@@H]3C[C@H](F)C4=CC(=O)C=C[C@]4(C)[C@@]3(F)[C@@H](O)C[C@@]21C | WMWTYOKRWGGJOA-CENSZEJFSA-N | Low | inactive |
| flutrimazole | BRD-A00218260 | Fc1ccc(C(c2ccccc2)(c2ccccc2F)n2ccnc2)cc1 | QHMWCHQXCUNUAK-UHFFFAOYSA-N | Low | inactive |
| folic-acid | BRD-K18673820 | Nc1nc(=O)c2nc(CNc3ccc(C(=O)N[C@@H](CCC(=O)O)C(=O)O)cc3)cnc2[nH]1 | OVBPIULPVIDEAO-LBPRGKRZSA-N | Low | inactive |
| fosaprepitant-dimeglumine | BRD-K23848629 | C[C@@H](O[C@H]1OCCN(Cc2nn(P(=O)(O)O)c(=O)[nH]2)[C@H]1c1ccc(F)cc1)c1cc(C(F)(F)F)cc(C(F)(F)F)c1 | BARDROPHSZEBKC-OITMNORJSA-N | Low | inactive |
| fosfomycin | BRD-K81101512 | C[C@@H]1O[C@@H]1P(=O)(O)O | YMDXZJFXQJVXBF-STHAYSLISA-N | Low | inactive |
| furagin | BRD-K00003314 | O=C1CN(/N=C/C=C/c2ccc([N+](=O)[O-])o2)C(=O)N1 | DECBQELQORZLLP-UAIOPKHMSA-N | Low | inactive |
| furegrelate | BRD-K55529781 | O=C(O)c1cc2cc(Cc3cccnc3)ccc2o1 | VHWFITPGPFLBGT-UHFFFAOYSA-N | Low | inactive |
| gadobutrol | BRD-K00003579 | O=C1CN2CCN3CCN([C@@H](CO)[C@@H](O)CO)CCN(CC2)CC(=O)O[Gd](O1)OC(=O)C3 | ZPDFIIGFYAHNSK-YYLIZZNMSA-K | Low | inactive |
| gadodiamide | BRD-K38140108 | CNC(=O)CN1CCN2CCN(CC(=O)NC)CC(=O)O[Gd](OC(=O)C2)OC(=O)C1 | HZHFFEYYPYZMNU-UHFFFAOYSA-K | Low | inactive |
| GDC-0810 | BRD-K61923022 | CC/C(=C(/c1ccc(/C=C/C(=O)O)cc1)c1ccc2n[nH]cc2c1)c1ccc(F)cc1Cl | BURHGPHDEVGCEZ-KJGLQBJMSA-N | Low | inactive |
| gemcitabine | BRD-K15108141 | Nc1ccn([C@@H]2O[C@H](CO)[C@@H](O)C2(F)F)c(=O)n1 | SDUQYLNIPVEERB-QPPQHZFASA-N | Low | inactive |
| geniposidic-acid | BRD-K73771095 | O=C(O)C1=CO[C@@H](O[C@@H]2O[C@H](CO)[C@@H](O)[C@H](O)[C@H]2O)[C@@H]2C(CO)=CC[C@H]12 | ZJDOESGVOWAULF-OGJQONSISA-N | Low | inactive |
| gepefrine | BRD-A04447196 | C[C@@H](N)Cc1cccc(O)c1 | WTDGMHYYGNJEKQ-SSDOTTSWSA-N | Low | inactive |
| glycopyrrolate | BRD-A81233518 | C[N+]1(C)CC[C@H](OC(=O)[C@](O)(c2ccccc2)C2CCCC2)C1 | ANGKOCUUWGHLCE-HKUYNNGSSA-N | Low | inactive |
| GSK2110183 | BRD-K25412176 | Cn1ncc(Cl)c1-c1cc(C(=O)N[C@H](CN)Cc2cccc(F)c2)sc1Cl | AFJRDFWMXUECEW-LBPRGKRZSA-N | Low | inactive |
| GSK269962 | BRD-K89464729 | CCn1c(-c2nonc2N)nc2cnc(Oc3cccc(NC(=O)c4ccc(OCCN5CCOCC5)cc4)c3)cc21 | YOVNFNXUCOWYSG-UHFFFAOYSA-N | Low | inactive |
| GSK650394 | BRD-K14991967 | O=C(O)c1ccc(-c2c[nH]c3ncc(-c4ccccc4)cc23)cc1C1CCCC1 | WVSBGSNVCDAMCF-UHFFFAOYSA-N | Low | inactive |
| guancydine | BRD-K00005334 | CCC(C)(C)NC(=N)NC#N | VZVGEDRCVUKSEL-UHFFFAOYSA-N | Low | inactive |
| GW-9662 | BRD-K93258693 | O=C(Nc1ccccc1)c1cc([N+](=O)[O-])ccc1Cl | DNTSIBUQMRRYIU-UHFFFAOYSA-N | Low | inactive |
| halothane | BRD-A20550616 | FC(F)(F)[C@@H](Cl)Br | BCQZXOMGPXTTIC-PVQJCKRUSA-N | Low | inactive |
| heclin | BRD-K49788188 | CCc1ccc(/C=C/C(=O)Nc2ccc(C(C)=O)cc2)o1 | SPTWXRJNCFIDRQ-ZHACJKMWSA-N | Low | inactive |
| hematoporphyrin | BRD-K01825010 | CC1=C([C@@H](C)O)c2cc3nc(cc4[nH]c(cc5[nH]c(cc1n2)c(C)c5CCC(=O)O)c(CCC(=O)O)c4C)C([C@@H](C)O)=C3C | DWZDGMDDUHAIQA-WOJBJXKFSA-N | Low | inactive |
| hesperetin | BRD-K30553453 | COc1ccc([C@@H]2CC(=O)c3c(O)cc(O)cc3O2)cc1O | AIONOLUJZLIMTK-AWEZNQCLSA-N | Low | inactive |
| HQK-1001 | BRD-K44963273 | CCC(C)(C)C(=O)O | VUAXHMVRKOTJKP-UHFFFAOYSA-N | Low | inactive |
| hydrocortisone-phosphate | BRD-K01824977 | C[C@@]12CCC(=O)C=C1CC[C@@H]1[C@@H]2[C@@H](O)C[C@]2(C)[C@H]1CC[C@]2(O)C(=O)COP(=O)(O)O | BGSOJVFOEQLVMH-SBXZOHCASA-N | Low | inactive |
| ibuproxam | BRD-A40940854 | CC(C)Cc1ccc([C@@H](C)C(=O)NO)cc1 | BYPIURIATSUHDW-SNVBAGLBSA-N | Low | inactive |
| ICA-069673 | BRD-K59839632 | O=C(Nc1cnc(Cl)nc1)c1ccc(F)c(F)c1 | IIBSHMFXVWTQSJ-UHFFFAOYSA-N | Low | inactive |
| ICI-199441 | BRD-K73290745 | CN(C(=O)Cc1ccc(Cl)c(Cl)c1)[C@H](CN1CCCC1)c1ccccc1 | AEJOEPSMZCEYJN-HXUWFJFHSA-N | Low | inactive |
| idazoxan | BRD-A18696154 | c1ccc2c(c1)OC[C@H](C1=NCCN1)O2 | HPMRFMKYPGXPEP-SNVBAGLBSA-N | Low | inactive |
| imidapril | BRD-K11466259 | CCOC(=O)[C@H](CCc1ccccc1)N[C@@H](C)C(=O)N1C(=O)N(C)C[C@H]1C(=O)O | KLZWOWYOHUKJIG-BPUTZDHNSA-N | Low | inactive |
| imidurea | BRD-K00003682 | O=C(NCNC(=O)N[C@H]1C(=O)NC(=O)N1CO)N[C@H]1C(=O)NC(=O)N1CO | ZCTXEAQXZGPWFG-RFZPGFLSSA-N | Low | inactive |
| INCB-3284 | BRD-K67628093 | COc1ccc([C@]2(O)CC[C@H](N3CC[C@@H](NC(=O)CNC(=O)c4cccc(C(F)(F)F)c4)C3)CC2)cn1 | NXZNYBUBXWWKCP-DNRQZRRGSA-N | Low | inactive |
| indirubin | BRD-K17894950 | O=C1Nc2ccccc2/C1=C1/Nc2ccccc2C1=O | CRDNMYFJWFXOCH-YPKPFQOOSA-N | Low | inactive |
| INX-08189 | BRD-K00004752 | COc1nc(N)nc2c1ncn2[C@@H]1O[C@H](CO[P@@](=O)(N[C@@H](C)C(=O)OCC(C)(C)C)Oc2cccc3ccccc23)[C@@H](O)[C@@]1(C)O | YFXGICNMLCGLHJ-KGSOWIGMSA-N | Low | inactive |
| ipragliflozin-L-proline | BRD-K51247865 | OC[C@H]1O[C@@H](c2ccc(F)c(Cc3cc4ccccc4s3)c2)[C@H](O)[C@@H](O)[C@@H]1O | AHFWIQIYAXSLBA-RQXATKFSSA-N | Low | inactive |
| irsogladine | BRD-K74195153 | Nc1nc(N)nc(-c2cc(Cl)ccc2Cl)n1 | ATCGGEJZONJOCL-UHFFFAOYSA-N | Low | inactive |
| isoleucine | BRD-A02232681 | CC[C@@H](C)[C@@H](N)C(=O)O | AGPKZVBTJJNPAG-RFZPGFLSSA-N | Low | inactive |
| isotretinoin | BRD-K76723084 | CC1=C(/C=C/C(C)=C/C=C/C(C)=C\C(=O)O)C(C)(C)CCC1 | SHGAZHPCJJPHSC-XFYACQKRSA-N | Low | inactive |
| istradefylline | BRD-K87774949 | CCn1c(=O)c2c(nc(/C=C/c3ccc(OC)c(OC)c3)n2C)n(CC)c1=O | IQVRBWUUXZMOPW-PKNBQFBNSA-N | Low | inactive |
| JK-184 | BRD-K38860038 | CCOc1ccc(Nc2nc(-c3c(C)nc4ccccn34)cs2)cc1 | ROYXIPOUVGDTAO-UHFFFAOYSA-N | Low | inactive |
| JLK-6 | BRD-K22010301 | COc1oc(=O)c2cc(N)ccc2c1Cl | AMDGKLWVCUXONP-UHFFFAOYSA-N | Low | inactive |
| JNJ-63533054 | BRD-K19482962 | C[C@H](NC(=O)CNC(=O)c1cccc(Cl)c1)c1ccccc1 | MWDVCHRYCKXEBY-LBPRGKRZSA-N | Low | inactive |
| kanamycin | BRD-K08327065 | NC[C@H]1O[C@H](O[C@@H]2[C@@H](N)C[C@@H](N)[C@H](O[C@H]3O[C@H](CO)[C@@H](O)[C@H](N)[C@H]3O)[C@H]2O)[C@H](O)[C@@H](O)[C@@H]1O | SBUJHOSQTJFQJX-NOAMYHISSA-N | Low | inactive |
| kasugamycin | BRD-K91644281 | C[C@H]1O[C@H](O[C@@H]2[C@@H](O)[C@@H](O)[C@H](O)[C@H](O)[C@H]2O)[C@@H](N)C[C@@H]1NC(=N)C(=O)O | PVTHJAPFENJVNC-MHRBZPPQSA-N | Low | inactive |
| KU-55933 | BRD-K25311561 | O=c1cc(-c2cccc3c2Sc2ccccc2S3)oc(N2CCOCC2)c1 | XRKYMMUGXMWDAO-UHFFFAOYSA-N | Low | inactive |
| L-741742 | BRD-K13211965 | Cc1c(C2CCN(CCc3ccccc3)CC2)noc1-c1ccc(Cl)cc1 | QDMAKDIDFJTXEL-UHFFFAOYSA-N | Low | inactive |
| L-ascorbyl-6-palmitate | BRD-A45419686 | CCCCCCCCCCCCCCCC(=O)OC[C@H](O)[C@H]1OC(=O)C(=O)[C@@H]1O | CYJPUZLKFAUFHF-HFSMHLIXSA-N | Low | inactive |
| lafutidine | BRD-K00003615 | O=C(C[S@@](=O)Cc1ccco1)NC/C=C\COc1cc(CN2CCCCC2)ccn1 | KMZQAVXSMUKBPD-FCRIZLRQSA-N | Low | inactive |
| larotrectinib | BRD-K36574127 | O=C(Nc1cnn2ccc(N3CCC[C@@H]3c3cc(F)ccc3F)nc12)N1CC[C@H](O)C1 | NYNZQNWKBKUAII-KBXCAEBGSA-N | Low | inactive |
| lifirafenib | BRD-K00003275 | O=C1CCc2c(Oc3ccc4c(c3)[C@@H]3[C@H](O4)[C@H]3c3nc4ccc(C(F)(F)F)cc4[nH]3)ccnc2N1 | NGFFVZQXSRKHBM-FKBYEOEOSA-N | Low | inactive |
| lincomycin | BRD-K49384696 | CCC[C@@H]1C[C@@H](C(=O)N[C@H]([C@@H](C)O)[C@H]2O[C@H](SC)[C@H](O)[C@@H](O)[C@H]2O)N(C)C1 | OJMMVQQUTAEWLP-KIDUDLJLSA-N | Low | inactive |
| linezolid | BRD-K66876909 | CC(=O)NC[C@H]1CN(c2ccc(N3CCOCC3)c(F)c2)C(=O)O1 | TYZROVQLWOKYKF-ZDUSSCGKSA-N | Low | inactive |
| linifanib | BRD-K99749624 | Cc1ccc(F)c(NC(=O)Nc2ccc(-c3cccc4[nH]nc(N)c34)cc2)c1 | MPVGZUGXCQEXTM-UHFFFAOYSA-N | Low | inactive |
| lisuride | BRD-K67943528 | CCN(CC)C(=O)N[C@H]1C=C2c3cccc4[nH]cc(c34)C[C@H]2N(C)C1 | BKRGVLQUQGGVSM-KBXCAEBGSA-N | Low | inactive |

| | | | | | |
|---|---|---|---|---|---|
| LTX-315 | BRD-K00003300 | NCCCC[C@H](N)C(=O)N[C@@H](CCCCN)C(=O)N[C@@H](Cc1c[nH]c2ccccc12)C(=O)N[C@@H](Cc1c[nH]c2ccccc12)C(=O)N[C@@H](CCCCN)C(=O)N[C@@H](CCCCN)C(=O)N[C@@H](Cc1c[nH]c2ccccc12)C(=O)N[C@H](C(=O)N[C@@H](CCCCN)C(N)=O)C(c1ccccc1)c1ccccc1 | GGAKLYWEFZCVIT-TVEKFXMRSA-N | Low | inactive |
| LY255283 | BRD-K92492521 | CCc1cc(C(C)=O)c(O)cc1OCCCCCC(C)(C)c1nn[nH]n1 | WCGXJPFHTHQNJL-UHFFFAOYSA-N | Low | inactive |
| LY2979165 | BRD-K68220839 | C[C@H](N)C(=O)N[C@@]1(C(=O)O)C[C@@H](Sc2nnc[nH]2)[C@H]2[C@H](C(=O)O)[C@H]21 | BBGHHIUQOKQCBW-LDZWZCGGSA-N | Low | inactive |
| LY411575 | BRD-K20013301 | C[C@H](NC(=O)[C@@H](O)c1cc(F)cc(F)c1)C(=O)N[C@@H]1C(=O)N(C)c2ccccc2-c2ccccc21 | ULSSJYNJIZWPSB-CVRXJBIPSA-N | Low | inactive |
| lypressin | BRD-K93331255 | NCCCC[C@H](NC(=O)[C@@H]1CCCN1C(=O)[C@@H]1CSSC[C@H](N)C(=O)N[C@@H](Cc2ccc(O)cc2)C(=O)N[C@@H](Cc2ccccc2)C(=O)N[C@@H](CCC(N)=O)C(=O)N[C@@H](CC(N)=O)C(=O)N1)C(=O)NCC(N)=O | BJFIDCADFRDPIO-DZCXQCEKSA-N | Low | inactive |
| maprotiline | BRD-K03319035 | CNCCC[C@]12CC[C@H](c3ccccc31)c1ccccc12 | QSLMDECMDJKHMQ-GSXCWMCISA-N | Low | inactive |
| meclofenamic-acid | BRD-K50398167 | Cc1ccc(Cl)c(Nc2ccccc2C(=O)O)c1Cl | SBDNJUWAMKYJOX-UHFFFAOYSA-N | Low | inactive |
| meclofenoxate | BRD-K10314788 | CN(C)CCOC(=O)COc1ccc(Cl)cc1 | XZTYGFHCIAKPGJ-UHFFFAOYSA-N | Low | inactive |
| medrysone | BRD-K56515112 | CC(=O)[C@H]1CC[C@H]2[C@@H]3C[C@H](C)C4=CC(=O)CC[C@]4(C)[C@H]3[C@@H](O)C[C@]12C | GZENKSODFLBBHQ-ILSZZQPISA-N | Low | inactive |
| megestrol | BRD-K00004691 | CC(=O)[C@@]1(O)CC[C@H]2[C@@H]3C=C(C)C4=CC(=O)CC[C@]4(C)[C@H]3CC[C@@]21C | VXIMPSPISRVBPZ-NWUMPJBXSA-N | Low | inactive |
| mesalazine | BRD-K28849549 | Nc1ccc(O)c(C(=O)O)c1 | KBOPZPXVLCULAV-UHFFFAOYSA-N | Low | inactive |
| metergoline | BRD-K56699285 | CN1C[C@H](CNC(=O)OCc2ccccc2)C[C@@H]2c3cccc4c3c(cn4C)C[C@H]21 | WZHJKEUHNJHDLS-QTGUNEKASA-N | Low | inactive |
| methocarbamol | BRD-A31521121 | COc1ccccc1OC[C@@H](O)COC(N)=O | GNXFOGHNGIVQEH-MRVPVSSYSA-N | Low | inactive |
| micronomicin | BRD-K18552138 | CNC[C@@H]1CC[C@@H](N)[C@@H](O[C@@H]2[C@@H](N)C[C@@H](N)[C@H](O[C@H]3OC[C@](C)(O)[C@H](NC)[C@H]3O)[C@H]2O)O1 | DNYGXMICFMACRA-XHEDQWPISA-N | Low | inactive |
| mifepristone | BRD-K37270826 | CC#C[C@]1(O)CC[C@H]2[C@@H]3CCC4=CC(=O)CCC4=C3[C@@H](c3ccc(N(C)C)cc3)C[C@@]21C | VKHAHZOOUSRJNA-GCNJZUOMSA-N | Low | inactive |
| miglitol | BRD-K44779798 | OCCN1C[C@H](O)[C@@H](O)[C@H](O)[C@H]1CO | IBAQFPQHRJAVAV-ULAWRXDQSA-N | Low | inactive |
| mivacurium | BRD-A48465967 | COc1cc2c(cc1OC)[C@@H](Cc1cc(OC)c(OC)c(OC)c1)[N@+]1(C)CCCOC(=O)CC/C=C/CCC(=O)OCCC[N@@+]1(C)CCc3cc(OC)c(OC)cc3[C@H]1Cc1cc(OC)c(OC)c(OC)c1)CC2 | ILVYCEVXHALBSC-GZICNWBJSA-N | Low | inactive |
| MK-6096 | BRD-K03894385 | Cc1ccc(-c2ncccn2)c(C(=O)N2C[C@H](COc3ccc(F)cn3)CC[C@H]2C)c1 | NPFDWHQSDBWQLH-QZTJIDSGSA-N | Low | inactive |
| MN-64 | BRD-K16183658 | CC(C)c1ccc(-c2cc(=O)c3ccccc3o2)cc1 | PYTOHIUBXSJKQH-UHFFFAOYSA-N | Low | inactive |
| modaline | BRD-K25317177 | Cc1nccnc1N1CCCCC1 | BJHCGMHPEKLROM-UHFFFAOYSA-N | Low | inactive |
| montelukast | BRD-K69470111 | CC(C)(O)c1ccccc1CC[C@@H](SCC1(CC(=O)O)CC1)c1cccc(/C=C/c2ccc3ccc(Cl)cc3n2)c1 | UCHDWCPVSPXUMX-TZIWLTJVSA-N | Low | inactive |
| moroxydine | BRD-K88786763 | N=C(N)/N=C(\N)N1CCOCC1 | KJHOZAZQWVKILO-UHFFFAOYSA-N | Low | inactive |
| N-acetyl-D-glucosamine | BRD-K80653534 | CC(=O)N[C@@H](C=O)[C@@H](O)[C@H](O)[C@H](O)CO | MBLBDJOUHNCFQT-LXGUWJNJSA-N | Low | inactive |
| naftopidil | BRD-K01826561 | COc1ccccc1N1CCN(C[C@@H](O)COc2cccc3ccccc23)CC1 | HRRBJVNMSRJFHQ-HXUWFJFHSA-N | Low | inactive |
| nebivolol | BRD-K00003617 | O[C@H](CNC[C@@H](O)[C@H]1CCc2cc(F)ccc2O1)[C@@H]1CCc2cc(F)ccc2O1 | KOHIRBRYDXPAMZ-YHBROIRLSA-N | Low | inactive |
| neomycin | BRD-K71013094 | NC[C@@H]1O[C@H](O[C@@H]2[C@@H](CO)O[C@@H](O[C@@H]3[C@@H](O)[C@H](N)C[C@H](N)[C@H]3O[C@H]3O[C@H](CN)[C@@H](O)[C@H](O)[C@H]3N)[C@@H]2O)[C@H](N)[C@@H](O)[C@@H]1O | PGBHMTALBVVCIT-VCIWKGPPSA-N | Low | inactive |
| niceritrol | BRD-K11256763 | O=C(OCC(COC(=O)c1cccnc1)(COC(=O)c1cccnc1)COC(=O)c1cccnc1)c1cccnc1 | KUEUWHJGRZKESU-UHFFFAOYSA-N | Low | inactive |
| nifursol | BRD-K85757157 | O=C(N/N=C/c1ccc([N+](=O)[O-])o1)c1cc([N+](=O)[O-])cc([N+](=O)[O-])c1O | XXUXXCZCUGIGPP-WLRTZDKTSA-N | Low | inactive |
| NN-DNJ | BRD-K00004640 | CCCCCCCCCN1C[C@H](O)[C@@H](O)[C@H](O)[C@H]1CO | FTSCEGKYKXESFF-LXTVHRRPSA-N | Low | inactive |
| nomegestrol-acetate | BRD-K27351809 | CC(=O)O[C@]1(C(C)=O)CC[C@H]2[C@@H]3C=C(C)C4=CC(=O)CC[C@@H]4[C@H]3CC[C@@]21C | IIVBFTNIGYRNQY-YQLZSBIMSA-N | Low | inactive |
| nonoxynol-9 | BRD-K88625236 | CCCCCCCCCc1ccc(OCCOCCOCCOCCOCCOCCOCCOCCOCCO)cc1 | FBWNMEQMRUMQSO-UHFFFAOYSA-N | Low | inactive |
| nortriptyline | BRD-K91263825 | CNCCC=C1c2ccccc2CCc2ccccc21 | PHVGLTMQBUFIQQ-UHFFFAOYSA-N | Low | inactive |
| NU-6027 | BRD-K03294269 | Nc1nc(N)c(N=O)c(OCC2CCCCC2)n1 | DGWXOLHKVGDQLN-UHFFFAOYSA-N | Low | inactive |
| OC000459 | BRD-K36463073 | Cc1c(Cc2ccc3ccccc3n2)c2cc(F)ccc2n1CC(=O)O | FATGTHLOZSXOBC-UHFFFAOYSA-N | Low | inactive |
| oxytetracycline | BRD-K01826559 | CN(C)[C@@H]1C(=O)C(C(N)=O)=C(O)[C@@]2(O)C(O)=C3C(=O)c4c(O)cccc4[C@@](C)(O)[C@H]3[C@H](O)[C@@H]12 | KTUXFHGUHYRDNF-PXOLEDIWSA-N | Low | inactive |
| pantethine | BRD-K68764924 | CC(C)(CO)[C@@H](O)C(=O)NCCC(=O)NCCSSCCNC(=O)CCNC(=O)[C@H](O)C(C)(C)CO | DJWYOLJPSHDSAL-ROUUACIJSA-N | Low | inactive |
| PD-102807 | BRD-A89337244 | CCOC(=O)c1c(C)[nH]c2ccc3c(c12)CN1CCc2cc(OC)ccc2[C@H]1O3 | VDDUJINYXKGZLV-JOCHJYFZSA-N | Low | inactive |
| PD-118057 | BRD-K34657753 | O=C(O)c1ccccc1Nc1ccc(CCc2ccc(Cl)c(Cl)c2)cc1 | ZCQOSCDABPVAFB-UHFFFAOYSA-N | Low | inactive |
| PD-173074 | BRD-K97764662 | CCN(CC)CCCCNc1ncc2cc(-c3cc(OC)cc(OC)c3)c(NC(=O)NC(C)(C)C)nc2n1 | DXCUKNQANPLTEJ-UHFFFAOYSA-N | Low | inactive |
| PDE10-IN-1 | BRD-K38145871 | Cc1ncc(C)n2nc([C@@H]3C[C@H]3c3nc4c5cccnc5ccn4c3C)nc12 | ZLHNYULRKIEGAY-HZPDHXFCSA-N | Low | inactive |
| pefloxacin | BRD-K55034111 | CCn1cc(C(=O)O)c(=O)c2cc(F)c(N3CCN(C)CC3)cc21 | FHFYDNQZQSQIAI-UHFFFAOYSA-N | Low | inactive |
| pempidine | BRD-K71075093 | CN1C(C)(C)CCCC1(C)C | XULIXFLCVXWHRF-UHFFFAOYSA-N | Low | inactive |
| pentamidine | BRD-K13183738 | N=C(N)c1ccc(OCCCCCOc2ccc(C(=N)N)cc2)cc1 | XDRYMKDFEDOLFX-UHFFFAOYSA-N | Low | inactive |
| perfluamine | BRD-K16758059 | FC(F)(F)C(F)(F)C(F)(F)N(C(F)(F)C(F)(F)C(F)(F)F)C(F)(F)C(F)(F)C(F)(F)F | JAJLKEVKNDUJBG-UHFFFAOYSA-N | Low | inactive |
| perflubron | BRD-K92988025 | FC(F)(F)C(F)(F)C(F)(F)C(F)(F)C(F)(F)C(F)(F)C(F)(F)C(F)(F)Br | WTWWXOGTJWMJHI-UHFFFAOYSA-N | Low | inactive |
| PF-02545920 | BRD-K31519811 | Cn1cc(-c2ccncc2)c(-c2ccc(OCc3ccc4ccccc4n3)cc2)n1 | AZEXWHKOMMASPA-UHFFFAOYSA-N | Low | inactive |
| PF-06409577 | BRD-K00003731 | O=C(O)c1c[nH]c2cc(Cl)c(-c3ccc(C4(O)CCC4)cc3)cc12 | FHQXLWCFSUSXBF-UHFFFAOYSA-N | Low | inactive |
| PFK-158 | BRD-K28010364 | O=C(/C=C/c1ccc2ccc(C(F)(F)F)cc2n1)c1ccncc1 | IAJOMYABKVAZCN-AATRIKPKSA-N | Low | inactive |
| phenindione | BRD-K70592963 | O=C1c2ccccc2C(=O)C1c1ccccc1 | NFBAXHOPROOJAW-UHFFFAOYSA-N | Low | inactive |
| PI3Kd-IN-2 | BRD-K00004592 | COc1ncc(-c2nc(N3CCOCC3)nc3c(CN4CCC(C(C)(C)O)CC4)cc(F)cc23)cc1NS(C)(=O)=O | NVWKNQGHVMMAJW-UHFFFAOYSA-N | Low | inactive |
| pikamilone | BRD-K56691760 | O=C(O)CCCNC(=O)c1cccnc1 | NAJVRARAUNYNDX-UHFFFAOYSA-N | Low | inactive |
| pipotiazine | BRD-K00004560 | CN(C)S(=O)(=O)c1ccc2c(c1)N(CCCN1CCC(CCO)CC1)c1ccccc1S2 | JOMHSQGEWSNUKU-UHFFFAOYSA-N | Low | inactive |
| piribedil | BRD-K47936004 | c1cnc(N2CCN(Cc3ccc4c(c3)OCO4)CC2)nc1 | OQDPVLVUJFGPGQ-UHFFFAOYSA-N | Low | inactive |
| pomalidomide | BRD-A12994259 | Nc1cccc2c1C(=O)N([C@H]1CCC(=O)NC1=O)C2=O | UVSMNLNDYGZFPF-QMMMGPOBSA-N | Low | inactive |
| potassium-iodide | BRD-M55321172 | I.[K] | FJWLWIRHZOHPIY-UHFFFAOYSA-N | Low | inactive |
| PRE-084 | BRD-K57304726 | O=C(OCCN1CCOCC1)C1(c2ccccc2)CCCCC1 | RQHKZUBCUZVZEF-UHFFFAOYSA-N | Low | inactive |

| | | | | | |
|---|---|---|---|---|---|
| prilocaine | BRD-K01825595 | CCCN[C@@H](C)C(=O)Nc1ccccc1C | MVFGUOIZUNYYSO-NSHDSACASA-N | Low | inactive |
| propyl-benzoate | BRD-K15329706 | CCCOC(=O)c1ccccc1 | UDEWPOVQBGFNGE-UHFFFAOYSA-N | Low | inactive |
| proquazone | BRD-K54339150 | Cc1ccc2c(-c3ccccc3)nc(=O)n(C(C)C)c2c1 | JTIGKVIOEQASGT-UHFFFAOYSA-N | Low | inactive |
| pyridoxine | BRD-K14349461 | Cc1ncc(CO)c(CO)c1O | LXNHXLLTXMVWPM-UHFFFAOYSA-N | Low | inactive |
| pyrintegrin | BRD-K94659840 | O=S(=O)(NCC1CC1)c1ccc(Nc2nccc(N3CCCc4cc(O)ccc43)n2)cc1 | QRJTZIJWDLJKQO-UHFFFAOYSA-N | Low | inactive |
| ramatroban | BRD-K08586861 | O=C(O)CCn1c2c(c3ccccc31)C[C@H](NS(=O)(=O)c1ccc(F)cc1)CC2 | LDXDSHIEDAPSSA-OAHLLOKOSA-N | Low | inactive |
| ramelteon | BRD-K28761891 | CCC(=O)NCC[C@@H]1CCc2ccc3c(c21)CCO3 | YLXDSYKOBKBWJQ-LBPRGKRZSA-N | Low | inactive |
| repaglinide | BRD-K82846253 | CCOc1cc(CC(=O)N[C@@H](CC(C)C)c2ccccc2N2CCCCC2)ccc1C(=O)O | FAEKWTJYAYMJKF-QHCPKHFHSA-N | Low | inactive |
| repsox | BRD-K80480517 | Cc1cccc(-c2n[nH]cc2-c2ccc3ncccc3n2)n1 | LBPKYPYHDKKRFS-UHFFFAOYSA-N | Low | inactive |
| resibufogenin | BRD-K25970317 | C[C@]12CC[C@H](O)C[C@H]1CC[C@@H]1[C@@H]2CC[C@]2(C)[C@@H](c3ccc(=O)oc3)C[C@H]3O[C@@]312 | ATLJNLYIJOCWJE-CWMZOUAVSA-N | Low | inactive |
| rilmenidine | BRD-K01638814 | C1COC(NC(C2CC2)C2CC2)=N1 | CQXADFVORZEARL-UHFFFAOYSA-N | Low | inactive |
| rilpivirine | BRD-K06240250 | Cc1cc(/C=C/C#N)cc(C)c1Nc1ccnc(Nc2ccc(C#N)cc2)n1 | YIBOMRUWOWDFLG-ONEGZZNKSA-N | Low | inactive |
| Ro-10-5824 | BRD-K79684402 | Cc1ncc(CN2CC=C(c3ccccc3)CC2)c(N)n1 | KABDATZAOUSYES-UHFFFAOYSA-N | Low | inactive |
| ronidazole | BRD-K94266545 | Cn1c([N+](=O)[O-])cnc1COC(N)=O | PQFRTXSWDXZRRS-UHFFFAOYSA-N | Low | inactive |
| rosuvastatin | BRD-K82941592 | CC(C)c1nc(N(C)S(C)(=O)=O)nc(-c2ccc(F)cc2)c1/C=C/[C@@H](O)C[C@@H](O)CC(=O)O | BPRHUIZQVSMCRT-VEUZHWNKSA-N | Low | inactive |
| RTA-408 | BRD-K56211775 | CC1(C)CC[C@]2(NC(=O)C(C)(F)F)CC[C@]3(C)[C@H](C(=O)C=C4[C@@]3(C)CC[C@H]3C(C)(C)C(=O)C(C#N)=C[C@]43C)[C@@H]2C1 | RJCWBNBKOKFWNY-IDPLTSGASA-N | Low | inactive |
| ruboxistaurin | BRD-K27677357 | CN(C)C[C@@H]1CCn2cc(c3ccccc32)C2=C(C(=O)NC2=O)c2cn(c3ccccc23)CCO1 | ZCBUQCWBWNUWSU-SFHVURJKSA-N | Low | inactive |
| salicylic-acid | BRD-K93632104 | O=C(O)c1ccccc1O | YGSDEFSMJLZEOE-UHFFFAOYSA-N | Low | inactive |
| sanguinarium-chloride | BRD-K66898851 | C[n+]1cc2c3c(ccc2c2ccc4cc5c(cc4c21)OCO5)OCO3 | INVGWHRKADIJHF-UHFFFAOYSA-N | Low | inactive |
| SCH-28080 | BRD-K55748775 | Cc1nc2c(OCc3ccccc3)cccn2c1CC#N | PYKJFEPAUKAXNN-UHFFFAOYSA-N | Low | inactive |
| SD-2590 | BRD-K51102532 | COCCN1CCC(C(=O)NO)(S(=O)(=O)c2ccc(Oc3ccc(OC(F)(F)F)cc3)cc2)CC1 | NXUHVWMBPOFLMA-UHFFFAOYSA-N | Low | inactive |
| SDZ-SER-082 | BRD-K31339597 | CN1CC[C@@H]2CN3CCc4cccc(c43)[C@@H]2C1 | YASBOGFWAMXINH-TZMCWYRMSA-N | Low | inactive |
| Se-methylselenocysteine | BRD-K74227499 | C[Se]C[C@H](N)C(=O)O | XDSSPSLGNGIIHP-VKHMYHEASA-N | Low | inactive |
| segesterone-acetate | BRD-K00003150 | C=C1C[C@H]2[C@@H]3CCC4=CC(=O)CC[C@@H]4[C@H]3CC[C@]2(C)[C@@]1(OC(C)=O)C(C)=O | CKFBRGLGTWAVLG-GOMYTPFNSA-N | Low | inactive |
| sertindole | BRD-K69116396 | O=C1NCCN1CCN1CCC(c2cn(-c3ccc(F)cc3)c3ccc(Cl)cc23)CC1 | GZKLJWGUPQBVJQ-UHFFFAOYSA-N | Low | inactive |
| simeprevir | BRD-K00003545 | COc1ccc2c(O[C@@H]3C[C@H]4C(=O)N[C@]5(C(=O)NS(=O)(=O)C6CC6)C[C@H]5/C=C\CCCCN(C)C(=O)[C@@H]4C3)cc(-c3nc(C(C)C)cs3)nc2c1C | JTZZSQYMACOLNN-VDWJNHBNSA-N | Low | inactive |
| SirReal-2 | BRD-K16472653 | Cc1cc(C)nc(SCC(=O)Nc2ncc(Cc3cccc4ccccc34)s2)n1 | MENNDDDTIIZDDN-UHFFFAOYSA-N | Low | inactive |
| sitafloxacin | BRD-K10836013 | N[C@@H]1CN(c2c(F)cc3c(=O)c(C(=O)O)cn([C@@H]4C[C@@H]4F)c3c2Cl)CC12CC2 | PNUZDKCDAWUEGK-CYZMBNFOSA-N | Low | inactive |
| sitravatinib | BRD-K00003238 | COCCNCc1ccc(-c2cc3nccc(Oc4ccc(NC(=O)C5(C(=O)Nc6ccc(F)cc6)CC5)cc4F)c3s2)nc1 | WLAVZAAODLTUSW-UHFFFAOYSA-N | Low | inactive |
| sivelestat | BRD-K02152879 | CC(C)(C)C(=O)Oc1ccc(S(=O)(=O)Nc2ccccc2C(=O)NCC(=O)O)cc1 | BTGNGJJLZOIYID-UHFFFAOYSA-N | Low | inactive |
| SLV-320 | BRD-K34445261 | O[C@H]1CC[C@H](Nc2nc(-c3ccccc3)nc3[nH]ccc23)CC1 | RBZNJGHIKXAKQE-HDJSIYSDSA-N | Low | inactive |
| SM-108 | BRD-K70053378 | NC(=O)C1N=CNC1=O | PXQCVBRDIZZRGM-UHFFFAOYSA-N | Low | inactive |
| SNAP | BRD-A67006024 | CC(=O)N[C@@H](C(=O)O)C(C)(C)S[NH2+][O-] | MLXCQEWOIZBBDM-YFKPBYRVSA-N | Low | inactive |
| SP-141 | BRD-K28571773 | COc1ccc2[nH]c3c(-c4cccc5ccccc45)nccc3c2c1 | AABFWJDLCCDJJN-UHFFFAOYSA-N | Low | inactive |
| sparteine-(+) | BRD-K38203080 | C1CCN2C[C@H]3C[C@H](CN4CCCC[C@H]34)[C@@H]2C1 | SLRCCWJSBJZJBV-TUVASFSCSA-N | Low | inactive |
| SQ-109 | BRD-A08772153 | CC(C)=CCC/C(C)=C/CNCCNC1[C@H]2C[C@@H]3C[C@@H](C[C@H]1C3)C2 | JFIBVDBTCDTBRH-UNRAVDGJSA-N | Low | inactive |
| SR-1078 | BRD-K26501486 | O=C(Nc1ccc(C(O)(C(F)(F)F)C(F)(F)F)cc1)c1ccc(C(F)(F)F)cc1 | DUXWIYXHHGNUJU-UHFFFAOYSA-N | Low | inactive |
| sucrose-octaacetate | BRD-K01294196 | CC(=O)OC[C@H]1O[C@H](O[C@]2(COC(C)=O)O[C@H](COC(C)=O)[C@@H](OC(C)=O)[C@@H]2OC(C)=O)[C@H](OC(C)=O)[C@@H](OC(C)=O)[C@@H]1OC(C)=O | ZIJKGAXBCRWEOL-SAXBRCJISA-N | Low | inactive |
| sulfabenzamide | BRD-K59983611 | Nc1ccc(S(=O)(=O)NC(=O)c2ccccc2)cc1 | PBCZLFBEBARBBI-UHFFFAOYSA-N | Low | inactive |
| sulfacetamide | BRD-K21520694 | CC(=O)NS(=O)(=O)c1ccc(N)cc1 | SKIVFJLNDNKQPD-UHFFFAOYSA-N | Low | inactive |
| suramin | BRD-K70327167 | Cc1ccc(C(=O)Nc2ccc(S(=O)(=O)O)c3cc(S(=O)(=O)O)cc(S(=O)(=O)O)c23)cc1NC(=O)c1cccc(NC(=O)Nc2cccc(C(=O)Nc3cc(C(=O)Nc4ccc(S(=O)(=O)O)c5cc(S(=O)(=O)O)cc(S(=O)(=O)O)c45)ccc3C)c2)c1 | FIAFUQMPZJWCLV-UHFFFAOYSA-N | Low | inactive |
| tafluprost | BRD-K65059905 | CC(C)OC(=O)CCC/C=C\C[C@H]1[C@@H](O)C[C@@H](O)[C@@H]1/C=C/C(F)(F)COc1ccccc1 | WSNODXPBBALQOF-VEJSHDCNSA-N | Low | inactive |
| talaporfin | BRD-K00004965 | C=Cc1c(C)c2cc3nc(c(CC(=O)N[C@@H](CC(=O)O)C(=O)O)c4nc(cc5[nH]c(cc1[nH]2)c(C)c5CC)C(C)=C4C(=O)O)[C@@H](CCC(=O)O)[C@@H]3C | SIEXFRDYNDREBM-BRXYURPTSA-N | Low | inactive |
| tamsulosin | BRD-K74339692 | CCOc1ccccc1OCCN[C@H](C)Cc1ccc(OC)c(S(N)(=O)=O)c1 | DRHKJLXJIQTDTD-OAHLLOKOSA-N | Low | inactive |
| tannic-acid | BRD-K01584643 | O=C(OC[C@H]1O[C@@H](OC(=O)c2cc(O)c(O)c(OC(=O)c3cc(O)c(O)c(O)c3)c2)[C@H](OC(=O)c2cc(O)c(O)c(OC(=O)c3cc(O)c(O)c(O)c3)c2)[C@@H](OC(=O)c2cc(O)c(O)c(OC(=O)c3cc(O)c(O)c(O)c3)c2)[C@@H]1OC(=O)c1cc(O)c(O)c(OC(=O)c2cc(O)c(O)c(O)c2)c1)c1cc(O)c(O)c(OC(=O)c2cc(O)c(O)c(O)c2)c1 | LRBQNJMCXXYXIU-PPKXGCFTSA-N | Low | inactive |
| tazarotene | BRD-K76841105 | CCOC(=O)c1ccc(C#Cc2ccc3c(c2)C(C)(C)CCS3)nc1 | OGQICQVSFDPSEI-UHFFFAOYSA-N | Low | inactive |
| TC-O-9311 | BRD-K54297077 | COc1cc(OC)cc(C(=O)NNC(=O)Nc2cccc3ccccc23)c1 | KPTMSQHTGZMEFU-UHFFFAOYSA-N | Low | inactive |
| TC-S-7005 | BRD-K89584803 | C[C@H](Nc1cc2c(-c3ccc4c(c3)OCO4)noc2cn1)c1ccccc1 | DAAGOCSHXCDWDY-ZDUSSCGKSA-N | Low | inactive |
| TCS-HDAC6-20b | BRD-K68358632 | CC(C)C(=O)SCCCCC[C@H](NC(=O)OC(C)(C)C)C(=O)NC12CC3CC(CC(C3)C1)C2 | DQZMLPMCOMRJOR-MOVGKWKRSA-N | Low | inactive |
| telmesteine | BRD-A05523972 | CCOC(=O)N1CSC[C@@H]1C(=O)O | XBJWOGLKABXFJE-RXMQYKEDSA-N | Low | inactive |
| tesaglitazar | BRD-K15507868 | CCO[C@@H](Cc1ccc(OCCc2ccc(OS(C)(=O)=O)cc2)cc1)C(=O)O | CXGTZJYQWSUFET-IBGZPJMESA-N | Low | inactive |
| thiamet-g | BRD-K38380126 | CCNC1=N[C@H]2[C@H](O[C@H](CO)[C@@H](O)[C@@H]2O)S1 | PPAIMZHKIXDJRN-FMDGEEDCSA-N | Low | inactive |
| thiocolchicoside | BRD-K26619122 | COc1c(O[C@@H]2O[C@H](CO)[C@@H](O)[C@H](O)[C@H]2O)cc2c(c1OC)-c1ccc(SC)c(=O)cc1[C@@H](NC(C)=O)CC2 | LEQAKWQJCITZNK-AXHKHJLKSA-N | Low | inactive |
| THK5351 | BRD-K00003401 | CNc1ccc(-c2ccc3cc(OC[C@H](O)CF)ccc3n2)cn1 | DLVXFZWSPCOWSN-CQSZACIVSA-N | Low | inactive |
| thymol | BRD-K17458574 | Cc1ccc(C(C)C)c(O)c1 | MGSRCZKZVOBKFT-UHFFFAOYSA-N | Low | inactive |
| tiaramide | BRD-K27293958 | O=C(Cn1c(=O)sc2ccc(Cl)cc21)N1CCN(CCO)CC1 | HTJXMOGUGMSZOG-UHFFFAOYSA-N | Low | inactive |
| ticagrelor | BRD-K25838957 | CCCSc1nc(N[C@@H]2C[C@H]2c2ccc(F)c(F)c2)c2nnn([C@@H]3C[C@H](OCCO)[C@@H](O)[C@H]3O)c2n1 | OEKWJQXRCDYSHL-FNOIDJSQSA-N | Low | inactive |
| tinoridine | BRD-K84313779 | CCOC(=O)c1c(N)sc2c1CCN(Cc1ccccc1)C2 | PFENFDGYVLAFBR-UHFFFAOYSA-N | Low | inactive |

| | | | | | |
|---|---|---|---|---|---|
| TMC647055 | BRD-K49659468 | COc1ccc2c(c1)C=C1Cn3c-2c(C2CCCCC2)c2ccc(cc23)C(=O)NS(=O)(=O)N(C)CCOCCN(C)C1=O | UOBYJVFBFSLCTQ-UHFFFAOYSA-N | Low | inactive |
| tofisopam | BRD-K94830213 | CCC1=C(C)NN=C(c2ccc(OC)c(OC)c2)c2cc(OC)c(OC)cc21 | LOUZDJSQOFXLAP-UHFFFAOYSA-N | Low | inactive |
| tolazoline | BRD-K46211610 | c1ccc(CC2=NCCN2)cc1 | JIVZKJJQOZQXQB-UHFFFAOYSA-N | Low | inactive |
| topiramate | BRD-K29653726 | CC1(C)O[C@@H]2CO[C@@]3(COS(N)(=O)=O)OC(C)(C)O[C@H]3[C@@H]2O1 | KJADKKWYZYXHBB-XBWDGYHZSA-N | Low | inactive |
| tricaprylin | BRD-K24790130 | CCCCCCCC(=O)OCC(COC(=O)CCCCCCC)OC(=O)CCCCCCC | VLPFTAMPNXLGLX-UHFFFAOYSA-N | Low | inactive |
| trifluridine | BRD-K03243820 | O=c1[nH]c(=O)n([C@H]2C[C@H](O)[C@@H](CO)O2)cc1C(F)(F)F | VSQQQLOSPVPRAZ-RRKCRQDMSA-N | Low | inactive |
| TRIM | BRD-K36851334 | FC(F)(F)c1ccccc1-n1ccnc1 | WZBWBNCQUTXYEL-UHFFFAOYSA-N | Low | inactive |
| trimethadione | BRD-K16606819 | CN1C(=O)OC(C)(C)C1=O | IRYJRGCIQBGHIV-UHFFFAOYSA-N | Low | inactive |
| trimipramine | BRD-K01825162 | C[C@@H](CN(C)C)CN1c2ccccc2CCc2ccccc21 | ZSCDBOWYZJWBIY-INIZCTEOSA-N | Low | inactive |
| tripelennamine | BRD-K57033106 | CN(C)CCN(Cc1ccccc1)c1ccccn1 | UFLGIAIHIAPJJC-UHFFFAOYSA-N | Low | inactive |
| tryptophan | BRD-K83776863 | N[C@@H](Cc1c[nH]c2ccccc12)C(=O)O | QIVBCDIJIAJPQS-VIFPVBQESA-N | Low | inactive |
| valbenazine | BRD-K00003233 | COc1cc2c(cc1OC)[C@H]1C[C@@H](OC(=O)[C@@H](N)C(C)C)[C@H](CC(C)C)CN1CC2 | GEJDGVNQKABXKG-CFKGEZKQSA-N | Low | inactive |
| vanitiolide | BRD-K26502378 | COc1cc(C(=S)N2CCOCC2)ccc1O | WQYRHRAZNNRDIA-UHFFFAOYSA-N | Low | inactive |
| vincamine | BRD-K40902647 | CC[C@]12CCCN3CCc4c(n(c5ccccc45)[C@@](O)(C(=O)OC)C1)[C@@H]32 | RXPRRQLKFXBCSJ-GIVPXCGWSA-N | Low | inactive |
| VP-20629 | BRD-K21071223 | O=C(O)CCc1c[nH]c2ccccc12 | GOLXRNDWAUTYKT-UHFFFAOYSA-N | Low | inactive |
| WAY-316606 | BRD-K61871885 | O=S(=O)(NC1CCNCC1)c1cc(S(=O)(=O)c2ccccc2)ccc1C(F)(F)F | ITBGJNVZJBVPLJ-UHFFFAOYSA-N | Low | inactive |
| WZ8040 | BRD-K81728688 | C=CC(=O)Nc1cccc(Sc2nc(Nc3ccc(N4CCN(C)CC4)cc3)ncc2Cl)c1 | KIISCIGBPUVZBF-UHFFFAOYSA-N | Low | inactive |
| Y-26763 | BRD-K45117373 | CC(=O)N(O)[C@@H]1c2cc(C#N)ccc2OC(C)(C)[C@H]1O | IWTCFIIOUXJOOV-OLZOCXBDSA-N | Low | inactive |
| YM-58483 | BRD-K42563464 | Cc1nnsc1C(=O)Nc1ccc(-n2nc(C(F)(F)F)cc2C(F)(F)F)cc1 | XPRZIORDEVHURQ-UHFFFAOYSA-N | Low | inactive |
| YM-976 | BRD-K12932420 | CCc1ccc2c(-c3cccc(Cl)c3)nc(=O)n(CC)c2n1 | MNHXYNNKDDXKNP-UHFFFAOYSA-N | Low | inactive |
| ZK-93426 | BRD-K68392338 | CCOC(=O)c1ncc2[nH]c3cccc(OC(C)C)c3c2c1C | VMDUABMKBUKKPG-UHFFFAOYSA-N | Low | inactive |
| zomepirac | BRD-K81326768 | Cc1cc(CC(=O)O)n(C)c1C(=O)c1ccc(Cl)cc1 | ZXVNMYWKKDOREA-UHFFFAOYSA-N | Low | inactive |
| zotepine | BRD-K64557645 | CN(C)CCOC1=Cc2ccccc2Sc2ccc(Cl)cc21 | HDOZVRUNCMBHFH-UHFFFAOYSA-N | Low | inactive |
| (R)-(-)-apomorphine | BRD-K76022557 | CN1CCc2cccc3c2[C@H]1Cc1ccc(O)c(O)c1-3 | VMWNQDUVQKEIOC-CYBMUJFWSA-N | No-Effect | inactive |
| (R)-(-)-rolipram | BRD-K75516118 | COc1ccc([C@@H]2CNC(=O)C2)cc1OC1CCCC1 | HJORMJIFDVBMOB-LBPRGKRZSA-N | No-Effect | inactive |
| (R)-baclofen | BRD-K62353271 | NC[C@H](CC(=O)O)c1ccc(Cl)cc1 | KPYSYYIEGFHWSV-QMMMGPOBSA-N | No-Effect | inactive |
| (S)-(+)-rolipram | BRD-K65856711 | COc1ccc([C@H]2CNC(=O)C2)cc1OC1CCCC1 | HJORMJIFDVBMOB-GFCCVEGCSA-N | No-Effect | inactive |
| [sar9,met(o2)11]-substance-p | BRD-K89787693 | CC(C)C[C@H](NC(=O)CN(C)C(=O)[C@H](Cc1ccccc1)NC(=O)[C@H](Cc1ccccc1)NC(=O)[C@H](CCC(N)=O)NC(=O)[C@H](CCC(N)=O)NC(=O)[C@@H]1CCCN1C(=O)[C@H](CCCCN)NC(=O)[C@@H]1CCCN1C(=O)[C@@H](N)CCCNC(=N)N)C(=O)N[C@@H](CCS(C)(=O)=O)C(N)=O | OUPXSLGGCPUZJJ-SARDKLJWSA-N | No-Effect | inactive |
| 1-((Z)-3-chloroallyl)-1,3,5,7-tetraazaadamantan-1-ium | BRD-K88956297 | Cl/C=C\C[N+]12CN3CN(CN(C3)C1)C2 | LDLCEGCJYSDJLX-UPHRSURJSA-N | No-Effect | inactive |
| 1-(1,2-diphenylethyl)piperidine-(+/-) | BRD-K01825681 | c1ccc(C[C@H](c2ccccc2)N2CCCCC2)cc1 | JQWJJJYHVHNXJH-LJQANCHMSA-N | No-Effect | inactive |
| 1-(2-chloro-5-methylphenoxy)-3-(isopropylamino)-2-propanol | BRD-A95802703 | Cc1ccc(Cl)c(OC[C@H](O)CNC(C)C)c1 | NJEIOWSBPCZKTL-LLVKDONJSA-N | No-Effect | inactive |
| 1-acetyl-4-methylpiperazine | BRD-K96055017 | CC(=O)N1CCN(C)CC1 | YSDBJKNOEWSFGA-UHFFFAOYSA-N | No-Effect | inactive |
| 1-azakenpaullone | BRD-K03273112 | O=C1Cc2c([nH]c3ccc(Br)cc23)-c2ncccc2N1 | NTSBZVCEIVPKBJ-UHFFFAOYSA-N | No-Effect | inactive |
| 1-deoxymannojirimycin | BRD-K89665618 | OC[C@H]1NC[C@@H](O)[C@@H](O)[C@@H]1O | LXBIFEVIBLOUGU-KVTDHHQDSA-N | No-Effect | inactive |
| 1-EBIO | BRD-K70586315 | CCn1c(=O)[nH]c2ccccc21 | CXUCKELNYMZTRT-UHFFFAOYSA-N | No-Effect | inactive |
| 1-ethyl-2-pyrrolidone | BRD-K08288352 | CCN1CCCC1=O | ZFPGARUNNKGOBB-UHFFFAOYSA-N | No-Effect | inactive |
| 1-hexadecanal | BRD-K35114608 | CCCCCCCCCCCCCCCC=O | NIOYUNMRJMEDGI-UHFFFAOYSA-N | No-Effect | inactive |
| 1-hexadecanol | BRD-K37203094 | CCCCCCCCCCCCCCCCO | BXWNKGSJHAJOGX-UHFFFAOYSA-N | No-Effect | inactive |
| 1-naphthyl-PP1 | BRD-K29542628 | CC(C)(C)n1nc(-c2cccc3ccccc23)c2c(N)ncnc21 | XSHQBIXMLULFEV-UHFFFAOYSA-N | No-Effect | inactive |
| 1-octacosanol | BRD-K03083257 | CCCCCCCCCCCCCCCCCCCCCCCCCCCCO | CNNRPFQICPFDPO-UHFFFAOYSA-N | No-Effect | inactive |
| 1-octanol | BRD-K09338665 | CCCCCCCCO | KBPLFHHGFOOTCA-UHFFFAOYSA-N | No-Effect | inactive |
| 1-phenylbiguanide | BRD-K31491153 | N=C(N=C(N)N)Nc1ccccc1 | CUQCMXFWIMOWRP-UHFFFAOYSA-N | No-Effect | inactive |
| 1,2,3,4,5,6-hexabromocyclohexane | BRD-K06817181 | BrC1C(Br)C(Br)C(Br)C(Br)C1Br | QFQZKISCBJKVHI-UHFFFAOYSA-N | No-Effect | inactive |
| 1,3-dipropyl-8-phenylxanthine | BRD-K41853443 | CCCn1c(=O)c2[nH]c(-c3ccccc3)nc2n(CCC)c1=O | CLIGSMOZKDCDRZ-UHFFFAOYSA-N | No-Effect | inactive |
| 1,4-butanediol | BRD-K22346679 | OCCCCO | WERYXYBDKMZEQL-UHFFFAOYSA-N | No-Effect | inactive |
| 1,5-dicaffeoylquinic-acid | BRD-K53083666 | O=C(/C=C/c1ccc(O)c(O)c1)O[C@@H]1C[C@](OC(=O)/C=C/c2ccc(O)c(O)c2)(C(=O)O)C[C@@H](O)[C@H]1O | YDDUMTOHNYZQPO-PSEXTPKNSA-N | No-Effect | inactive |
| 10-deacetylbaccatin | BRD-K96631475 | CC(=O)O[C@@]12CO[C@@H]1C[C@H](O)[C@@]1(C)C(=O)[C@H](O)C3=C(C)[C@@H](O)C[C@@](O)([C@@H](OC(=O)c4ccccc4)[C@@H]12)C3(C)C | YWLXLRUDGLRYDR-ZHPRIASZSA-N | No-Effect | inactive |
| 10-hydroxycamptothecin | BRD-K63784565 | CC[C@@]1(O)C(=O)OCc2c1cc1n(c2=O)Cc2cc3cc(O)ccc3nc2-1 | HAWSQZCWOQZXHI-FQEVSTJZSA-N | No-Effect | inactive |
| 10058-F4 | BRD-K49460120 | CCc1ccc(/C=C2/SC(=S)NC2=O)cc1 | SVXDHPADAXBMFB-JXMROGBWSA-N | No-Effect | inactive |
| 1400W | BRD-K30237152 | CC(=N)NCc1cccc(CN)c1 | RODUKNYOEVZQPR-UHFFFAOYSA-N | No-Effect | inactive |
| 16,16-dimethylprostaglandin-e2 | BRD-K92301463 | CCCCC(C)(C)[C@H](O)/C=C/[C@H]1[C@H](O)CC(=O)[C@@H]1C/C=C\CCCC(=O)O | QAOBBBBDJSWHMU-WMBBNPMCSA-N | No-Effect | inactive |
| 17-hydroxyprogesterone-caproate | BRD-K70264088 | CCCCCC(=O)O[C@]1(C(C)=O)CC[C@H]2[C@@H]3CCC4=CC(=O)CC[C@]4(C)[C@H]3CC[C@@]21C | DOMWKUIIPQCAJU-LJHIYBGHSA-N | No-Effect | inactive |
| 17-PA | BRD-K07395323 | C[C@]12CC[C@H]3[C@@H](CC[C@H]4C[C@H](O)CC[C@@]43C)[C@@H]1CC=C2c1ccccc1 | SINAMTXBCYKFDL-WBJZGETLSA-N | No-Effect | inactive |
| 1E-1-(2-hydroxy-5-methylphenyl)-1-dodecanone-oxime | BRD-A37752546 | CCCCCCCCCCC[C@@H](N=O)c1cc(C)ccc1O | NFONIVRMILHYLH-GOSISDBHSA-N | No-Effect | inactive |

| | | | | | |
|---|---|---|---|---|---|
| 1S,2R-phenylpropanolamine | BRD-K27853494 | C[C@@H](N)[C@@H](O)c1ccccc1 | DLNKOYKMWOXYQA-VXNVDRBHSA-N | No-Effect | inactive |
| 2-(1-adamantylamino)benzoic-acid | BRD-K75167327 | O=C(O)c1ccccc1NC(=O)C12CC3CC(CC(C3)C1)C2 | XKZLSYWCDJZTMW-UHFFFAOYSA-N | No-Effect | inactive |
| 2-(chloromethyl)-5,6,7,8-tetrahydro[1]benzothieno[2,3-d]pyrimidin-4(3H)-one | BRD-K46989997 | O=C1NC(=CCl)Nc2sc3c(c21)CCCC3 | UTWSZXBGTLNPOV-UHFFFAOYSA-N | No-Effect | inactive |
| 2-[1-(4-piperonyl)piperazinyl]benzothiazole | BRD-K91868854 | c1ccc2sc(N3CCN(Cc4ccc5c(c4)OCO5)CC3)nc2c1 | BYHKGNWKJMGHGE-UHFFFAOYSA-N | No-Effect | inactive |
| 2-aminobenzenesulfonamide | BRD-K32681104 | Nc1ccccc1S(N)(=O)=O | YAZSBRQTAHVVGE-UHFFFAOYSA-N | No-Effect | inactive |
| 2-APB | BRD-K13346085 | NCCOB(c1ccccc1)c1ccccc1 | BLZVCIGGICSWIG-UHFFFAOYSA-N | No-Effect | inactive |
| 2-BFI | BRD-K22745370 | c1ccc2oc(C3=NCCN3)cc2c1 | YTJOHEUHUVBKSB-UHFFFAOYSA-N | No-Effect | inactive |
| 2-chloro-N6-cyclopentyladenosine | BRD-K70088346 | OC[C@H]1O[C@@H](n2cnc3c(NC4CCCC4)nc(Cl)nc32)[C@H](O)[C@@H]1O | XSMYYYQVWPZWIZ-IDTAVKCVSA-N | No-Effect | inactive |
| 2-chloropyrazine | BRD-K84075000 | Clc1cnccn1 | GELVZYOEQVJIRR-UHFFFAOYSA-N | No-Effect | inactive |
| 2-CMDO | BRD-K92588747 | CN1CCN(C2=Cc3ccccc3Oc3ccc(Cl)cc32)CC1 | OTKUTGIVEREJLL-UHFFFAOYSA-N | No-Effect | inactive |
| 2-deoxy-2-([methyl(nitroso)carbamoyl]amino)hexose | BRD-A55902763 | CN([NH2+][O-])C(=O)N[C@@H](C=O)[C@@H](O)[C@@H](O)[C@@H](O)CO | VUVHIMLGTHRGQZ-YTLHQDLWSA-N | No-Effect | inactive |
| 2-deoxyglucose | BRD-K97808269 | OC[C@H]1O[C@H](O)C[C@@H](O)[C@@H]1O | PMMURAAUARKVCB-ZXXMMSQZSA-N | No-Effect | inactive |
| 2-ethoxybenzoic-acid | BRD-K04843302 | CCOc1ccccc1C(=O)O | XDZMPRGFOOFSBL-UHFFFAOYSA-N | No-Effect | inactive |
| 2-ethyl-1,3-hexanediol | BRD-A55455283 | CCC[C@@H](O)[C@@H](CC)CO | RWLALWYNXFYRGW-JGVFFNPUSA-N | No-Effect | inactive |
| 2-fluoro-2-deoxy-D-galactose | BRD-A53037300 | OC[C@H]1O[C@@H](O)[C@@H](F)[C@@H](O)[C@@H]1O | ZCXUVYAZINUVJD-AIECOIEWSA-N | No-Effect | inactive |
| 2-fluorofucose | BRD-K00004677 | C[C@@H]1O[C@H](O)[C@@H](F)[C@H](O)[C@@H]1O | IRKXGKIPOMIQOD-QYESYBIKSA-N | No-Effect | inactive |
| 2-hydroxy-4-((E)-3-(4-hydroxyphenyl)acryloyl)-2-((2R,3R,4S,5S,6R)-3,4,5-trihydroxy-6-(hydroxymethyl)tetrahydro-2H-pyran-2-yl)-6-((2S,3R,4R,5S,6R)-3,4,5-trihydroxy-6-(hydroxymethyl)tetrahydro-2H-pyran-2-yl)cyclohexane-1,3,5-trione | BRD-A52169337 | O=C1C(=C(O)/C=C/c2ccc(O)cc2)C(=O)[C@@](O)([C@@H]2O[C@H](CO)[C@@H](O)[C@H](O)[C@H]2O)C(=O)[C@H]1[C@@H]1O[C@H](CO)[C@@H](O)[C@H](O)[C@H]1O | BWKBPVRNWJFLSC-FROSJQSHSA-N | No-Effect | inactive |
| 2-hydroxyethyl-salicylate | BRD-K41866823 | O=C(OCCO)c1ccccc1O | LVYLCBNXHHHPSB-UHFFFAOYSA-N | No-Effect | inactive |
| 2-hydroxyflutamide | BRD-K82027074 | CC(C)(O)C(=O)Nc1ccc([N+](=O)[O-])c(C(F)(F)F)c1 | YPQLFJODEKMJEF-UHFFFAOYSA-N | No-Effect | inactive |
| 2-hydroxysaclofen | BRD-A27924917 | NC[C@@](O)(CS(=O)(=O)O)c1ccc(Cl)cc1 | WBSMZVIMANOCNX-SECBINFHSA-N | No-Effect | inactive |
| 2-iminobiotin | BRD-K07954936 | NC1=N[C@H]2CS[C@@H](CCCCC(=O)O)[C@H]2N1 | WWVANQJRLPIHNS-ZKWXMUAHSA-N | No-Effect | inactive |
| 2-iodohippuric-acid | BRD-K56634222 | O=C(O)CNC(=O)c1ccccc1I | CORFWQGVBFFZHF-UHFFFAOYSA-N | No-Effect | inactive |
| 2-iodomelatonin | BRD-K25649279 | COc1ccc2[nH]c(I)c(CCNC(C)=O)c2c1 | FJDDSMSDZHURBJ-UHFFFAOYSA-N | No-Effect | inactive |
| 2-methoxyestradiol | BRD-K44408410 | COc1cc2c(cc1O)CC[C@@H]1[C@@H]2CC[C@]2(C)[C@@H](O)CC[C@@H]12 | CQOQDQWUFQDJMK-SSTWWWIQSA-N | No-Effect | inactive |
| 2-methyl-5-hydroxytryptamine | BRD-K64137799 | Cc1[nH]c2ccc(O)cc2c1CCN | WYWNEDARFVJQSG-UHFFFAOYSA-N | No-Effect | inactive |
| 2-methyl-5-nitrophenol | BRD-K22327730 | Cc1ccc([N+](=O)[O-])cc1O | UMFDLIXUUJMPSI-UHFFFAOYSA-N | No-Effect | inactive |
| 2-methylimidazole | BRD-K68534611 | Cc1ncc[nH]1 | LXBGSDVWAMZHDD-UHFFFAOYSA-N | No-Effect | inactive |
| 2-octyldodecan-1-ol | BRD-A51407134 | CCCCCCCCCC[C@@H](CO)CCCCCCCC | LEACJMVNYZDSKR-FQEVSTJZSA-N | No-Effect | inactive |
| 2-oleoylglycerol | BRD-K77087341 | CCCCCCCC/C=C\CCCCCCCC(=O)OC(CO)CO | UPWGQKDVAURUGE-KTKRTIGZSA-N | No-Effect | inactive |
| 2-oxoglutaric-acid | BRD-K89712525 | O=C(O)CCC(=O)C(=O)O | KPGXRSRHYNQIFN-UHFFFAOYSA-N | No-Effect | inactive |
| 2-oxopropanoate | BRD-K86587225 | C=C(O)C(=O)O | FEWFXBUNENSNBQ-UHFFFAOYSA-N | No-Effect | inactive |
| 2-phenylmelatonin | BRD-K41869275 | COc1ccc2[nH]c(-c3ccccc3)c(CCNC(C)=O)c2c1 | OFCLARYYBGKCHN-UHFFFAOYSA-N | No-Effect | inactive |
| 2-PMDQ | BRD-A04661934 | O=C1Nc2ccccc2C2=N[C@@H](CN3CCN(c4ccccc4)CC3)CN12 | ONSOQRPYFKDOEB-INIZCTEOSA-N | No-Effect | inactive |
| 2-pyridylethylamine | BRD-K17874705 | NCCc1ccccn1 | XPQIPUZPSLAZDV-UHFFFAOYSA-N | No-Effect | inactive |
| 2-thiouracil | BRD-K58819470 | O=c1cc[nH]c(=S)[nH]1 | ZEMGGZBWXRYJHK-UHFFFAOYSA-N | No-Effect | inactive |
| 2,3-cis/exo-camphanediol | BRD-K44602126 | CC1(C)[C@@H]2CC[C@@]1(C)[C@H](O)[C@@H]2O | AYEOSGBMQHXVER-DQUBFYRCSA-N | No-Effect | inactive |
| 2,3-cis/exo-pinanediol | BRD-A75263867 | CC1(C)C2CC1[C@](C)(O)[C@H](O)C2 | MOILFCKRQFQVFS-VUMZSGCYSA-N | No-Effect | inactive |
| 2,3-DCPE | BRD-K61734568 | OCCNCCCOc1cccc(Cl)c1Cl | QVEIRCZEBQRCTR-UHFFFAOYSA-N | No-Effect | inactive |
| 2,4-dinitrochlorobenzene | BRD-K97833335 | O=[N+]([O-])c1ccc(Cl)c([N+](=O)[O-])c1 | VYZAHLCBVHPDDF-UHFFFAOYSA-N | No-Effect | inactive |
| 2,4-dinitrophenol | BRD-K21910317 | O=[N+]([O-])c1ccc(O)c([N+](=O)[O-])c1 | UFBJCMHMOXMLKC-UHFFFAOYSA-N | No-Effect | inactive |
| 2,5-furandimethanol | BRD-K29791386 | OCc1ccc(CO)o1 | DSLRVRBSNLHVBH-UHFFFAOYSA-N | No-Effect | inactive |
| 2,6-dimethylpiperidine | BRD-A21276153 | C[C@H]1CCC[C@@H](C)N1 | SDGKUVSVPIIUCF-KNVOCYPGSA-N | No-Effect | inactive |
| 2'-C-methylguanosine | BRD-K78092779 | C[C@@]1(O)[C@H](O)[C@@H](CO)O[C@H]1n1cnc2c(=O)nc(N)[nH]c21 | NVKAMPJSWMHVDK-GITKWUPZSA-N | No-Effect | inactive |
| 2'-MeCCPA | BRD-K63932022 | C[C@@]1(O)[C@H](O)[C@@H](CO)O[C@H]1n1cnc2c(NC3CCCC3)nc(Cl)nc21 | MMPAUXMIDJWGFO-ROMFRFKVSA-N | No-Effect | inactive |
| 20-HETE | BRD-K00005345 | O=C(O)CCC/C=C\C/C=C\C/C=C\C/C=C\CCCCCO | NNDIXBJHNLFJJP-DTLRTWKJSA-N | No-Effect | inactive |
| 20-hydroxyecdysone | BRD-K28343777 | CC(C)(O)CC[C@@H](O)[C@](C)(O)[C@@H]1CC[C@@]2(O)C3=CC(=O)[C@@H]4C[C@@H](O)[C@@H](O)C[C@]4(C)[C@@H]3CC[C@]12C | NKDFYOWSKOHCCO-HMIIXLFTSA-N | No-Effect | inactive |

| | | | | | |
|---|---|---|---|---|---|
| 3-(4-methylbenzylidene)camphor | BRD-A55464252 | Cc1ccc(/C=C2\C(=O)[C@]3(C)CC[C@H]2C3(C)C)cc1 | HEOCBCNFKCOKBX-LBONJMQPSA-N | No-Effect | inactive |
| 3-alpha-bis-(4-fluorophenyl)-methoxytropane | BRD-K00004218 | CN1[C@H]2CC[C@@H]1C[C@H](OC(c1ccc(F)cc1)c1ccc(F)cc1)C2 | MHNSOBBJZCWUGS-PMOLBWCYSA-N | No-Effect | inactive |
| 3-alpha-hydroxy-5-beta-androstan-17-one | BRD-A43598550 | C[C@]12CC[C@H]3[C@@H](CC[C@@H]4C[C@H](O)CC[C@]34C)[C@@H]1CCC2=O | QGXBDMJGAMFCBF-BNSUEQOYSA-N | No-Effect | inactive |
| 3-amino-benzamide | BRD-K08703257 | NC(=O)c1cccc(N)c1 | GSCPDZHWVNUUFI-UHFFFAOYSA-N | No-Effect | inactive |
| 3-anilinopropan-1-ol | BRD-K49721826 | OCCCNc1ccccc1 | GZYBSURBYCLNSM-UHFFFAOYSA-N | No-Effect | inactive |
| 3-AQC | BRD-K38370968 | C=CCN1CCN(c2nc3ccccc3nc2C#N)CC1 | IDOXKFDPDOIUMG-UHFFFAOYSA-N | No-Effect | inactive |
| 3-bromo-7-nitroindazole | BRD-K24689407 | O=[N+]([O-])c1cccc2c(Br)[nH]nc12 | NFSTZPMYAZRZPC-UHFFFAOYSA-N | No-Effect | inactive |
| 3-bromocamphor | BRD-K00004535 | CC1(C)[C@H]2CC[C@]1(C)C(=O)[C@H]2Br | NJQADTYRAYFBJN-BYULHYEWSA-N | No-Effect | inactive |
| 3-bromopyruvate | BRD-K92980438 | O=C(O)C(O)=CBr | CHNJCELMWMIDSC-UHFFFAOYSA-N | No-Effect | inactive |
| 3-carboxy-4-hydroxyphenylglycine-(R) | BRD-K76592088 | N[C@@H](C(=O)O)c1ccc(O)c(C(=O)O)c1 | CHZBCZTXSTWCIG-SSDOTTSWSA-N | No-Effect | inactive |
| 3-carboxy-4-hydroxyphenylglycine-(S) | BRD-K61585876 | N[C@H](C(=O)O)c1ccc(O)c(C(=O)O)c1 | CHZBCZTXSTWCIG-ZETCQYMHSA-N | No-Effect | inactive |
| 3-CPMT | BRD-K00004217 | CN1[C@H]2CC[C@@H]1C[C@H](O[C@H](c1ccccc1)c1ccc(Cl)cc1)C2 | OCAXFDULERPAJM-IVAOSVALSA-N | No-Effect | inactive |
| 3-deazaadenosine | BRD-A79431551 | Nc1nccc2c1ncn2[C@@H]1O[C@H](CO)[C@@H](O)[C@@H]1O | DBZQFUNLCALWDY-GPUHXXMPSA-N | No-Effect | inactive |
| 3-deazaneplanocin-A | BRD-K77791657 | Nc1nccc2c1ncn2[C@@H]1C=C(CO)[C@@H](O)[C@H]1O | OMKHWTRUYNAGFG-IEBDPFPHSA-N | No-Effect | inactive |
| 3-deazauridine | BRD-A26105646 | O=c1cc(O)ccn1[C@H]1O[C@H](CO)[C@@H](O)[C@@H]1O | CBOKZNLSFMZJJA-PRQTWWFRSA-N | No-Effect | inactive |
| 3-hydroxy-3-phenylpentanamide | BRD-A66143912 | CC[C@@](O)(CC(N)=O)c1ccccc1 | MOHYRCCDARWQRM-LLVKDONJSA-N | No-Effect | inactive |
| 3-indolebutyric-acid | BRD-K43187796 | O=C(O)CCCc1c[nH]c2ccccc12 | JTEDVYBZBROSJT-UHFFFAOYSA-N | No-Effect | inactive |
| 3-MATIDA | BRD-A87125127 | Cc1cc(C(=O)O)sc1[C@@H](N)C(=O)O | KOMWRBFEDDEWEP-RXMQYKEDSA-N | No-Effect | inactive |
| 3-methyl-GABA | BRD-K01825682 | C[C@@H](CN)CC(=O)O | CZGLBWZXGIAIBU-SCSAIBSYSA-N | No-Effect | inactive |
| 3-methyladenine | BRD-K81647657 | Cn1cnc(=N)c2[nH]cnc21 | ZPBYVFQJHWLTFB-UHFFFAOYSA-N | No-Effect | inactive |
| 3-MPPI | BRD-K62581435 | COc1ccccc1N1CCN(CCn2c(=O)[nH]c3c([nH]c4ccccc43)c2=O)CC1 | AQASGOHUMGAWJJ-UHFFFAOYSA-N | No-Effect | inactive |
| 3,3'-dichlorobenzaldazine | BRD-K82903013 | Clc1cccc(/C=N/N=C/c2cccc(Cl)c2)c1 | XMOVWXSCYLINBJ-BEQMOXJMSA-N | No-Effect | inactive |
| 3,3'-diindolylmethane | BRD-K37846922 | c1ccc2c(Cc3c[nH]c4ccccc34)c[nH]c2c1 | VFTRKSBEFQDZKX-UHFFFAOYSA-N | No-Effect | inactive |
| 3,4-DCPG-(+/-) | BRD-A49947412 | N[C@@H](C(=O)O)c1ccc(C(=O)O)c(C(=O)O)c1 | IJVMOGKBEVRBPP-SSDOTTSWSA-N | No-Effect | inactive |
| 3,4-DCPG-(R) | BRD-K06204668 | N[C@@H](C(=O)O)c1ccc(C(=O)O)c(C(=O)O)c1 | IJVMOGKBEVRBPP-SSDOTTSWSA-N | No-Effect | inactive |
| 3,4-DCPG-(S) | BRD-K86305586 | N[C@H](C(=O)O)c1ccc(C(=O)O)c(C(=O)O)c1 | IJVMOGKBEVRBPP-ZETCQYMHSA-N | No-Effect | inactive |
| 3,5-DHPG-(S) | BRD-K57796080 | N[C@H](C(=O)O)c1cc(O)cc(O)c1 | HOOWCUZPEFNHDT-ZETCQYMHSA-N | No-Effect | inactive |
| 3PO | BRD-K58413118 | O=C(/C=C/c1cccnc1)c1ccncc1 | UOWGYMNWMDNSTL-ONEGZZNKSA-N | No-Effect | inactive |
| 4-(4-fluorobenzoyl)-1-(4-phenylbutyl)-piperidine | BRD-K47337578 | O=C(c1ccc(F)cc1)C1CCN(CCCCc2ccccc2)CC1 | FTJXZACAQRDRNT-UHFFFAOYSA-N | No-Effect | inactive |
| 4-(aminomethyl)benzoic-acid | BRD-K00003517 | NCc1ccc(C(=O)O)cc1 | QCTBMLYLENLHLA-UHFFFAOYSA-N | No-Effect | inactive |
| 4-acetyl-1,1-dimethylpiperazinium | BRD-K97019106 | CC(=O)N1CC[N+](C)(C)CC1 | MSBLMBWXUVQCDY-UHFFFAOYSA-N | No-Effect | inactive |
| 4-aminohippuric-acid | BRD-K65544384 | Nc1ccc(C(=O)NCC(=O)O)cc1 | HSMNQINEKMPTIC-UHFFFAOYSA-N | No-Effect | inactive |
| 4-carboxy-3-hydroxyphenylglycine-(RS) | BRD-A80383043 | N[C@@H](C(=O)O)c1ccc(C(=O)O)c(O)c1 | GXZSAQLJWLCLOX-SSDOTTSWSA-N | No-Effect | inactive |
| 4-carboxy-3-hydroxyphenylglycine-(S) | BRD-K74133873 | N[C@H](C(=O)O)c1ccc(C(=O)O)c(O)c1 | GXZSAQLJWLCLOX-ZETCQYMHSA-N | No-Effect | inactive |
| 4-chlorophenylguanidine | BRD-K39160765 | N=C(N)Nc1ccc(Cl)cc1 | ZSJNJAJDBNFVCA-UHFFFAOYSA-N | No-Effect | inactive |
| 4-CMTB | BRD-A54139254 | CC(C)[C@@H](C(=O)Nc1nccs1)c1ccc(Cl)cc1 | AZYDQCGCBDQYFSE-GFCCVEGCSA-N | No-Effect | inactive |
| 4-DAMP | BRD-K29313239 | C[N+]1(C)CCC(OC(=O)C(c2ccccc2)c2ccccc2)CC1 | HYJRTXSYDAFGJK-UHFFFAOYSA-N | No-Effect | inactive |
| 4-galactosyllactose | BRD-A32796301 | OC[C@H]1O[C@@H](O)[C@H](O)[C@@H](O)[C@@H]1O[C@@H]1O[C@H](CO)[C@H](O[C@@H]2O[C@H](CO)[C@H](O)[C@H](O)[C@H]2O)[C@H](O)[C@H]1O | FYGDTMLNYKFZSV-QGAOIPDPSA-N | No-Effect | inactive |
| 4-HQN | BRD-K91758890 | O=c1nc[nH]c2ccccc12 | QMNUDYFKZYBWQX-UHFFFAOYSA-N | No-Effect | inactive |
| 4-hydroxy-phenazone | BRD-K49759007 | Cc1c(O)c(=O)n(-c2ccccc2)n1C | SKVPTPMWXJSBTF-UHFFFAOYSA-N | No-Effect | inactive |
| 4-IBP | BRD-K62537556 | O=C(NC1CCN(Cc2ccccc2)CC1)c1ccc(I)cc1 | HELCSESNNDZLFM-UHFFFAOYSA-N | No-Effect | inactive |
| 4-iodo-6-phenylpyrimidine | BRD-K25690923 | Ic1cc(-c2ccccc2)ncn1 | ZTCJXHNJVLUUMR-UHFFFAOYSA-N | No-Effect | inactive |
| 4-iodo-L-phenylalanine | BRD-K66664625 | N[C@H](Cc1ccc(I)cc1)C(=O)O | PZNQZSRPDOEBMS-MRVPVSSYSA-N | No-Effect | inactive |
| 4-methylgenistein | BRD-K73303757 | COc1ccc(-c2coc3cc(O)cc(O)c3c2=O)cc1 | WUADCCWRTIWANL-UHFFFAOYSA-N | No-Effect | inactive |
| 4-methylhistamine | BRD-K70821460 | Cc1nc[nH]c1CCN | UGYXPZQILZRKJJ-UHFFFAOYSA-N | No-Effect | inactive |
| 4-mu-8C | BRD-K81316007 | Cc1cc(=O)oc2c(C=O)c(O)ccc12 | RTHHSXOVIJWFQP-UHFFFAOYSA-N | No-Effect | inactive |
| 4-P-PDOT | BRD-K00004224 | CCC(=O)N[C@@H]1Cc2ccccc2[C@H](c2ccccc2)C1 | RCYLUNPFECYGDW-AEFFLSMTSA-N | No-Effect | inactive |
| 4-phenolsulfonic-acid | BRD-K41024817 | O=S(=O)(O)c1ccc(O)cc1 | FEPBITJSIHRMRT-UHFFFAOYSA-N | No-Effect | inactive |
| 4-phenyl-1,2,3,4-tetrahydroisoquinoline | BRD-A40504327 | c1ccc([C@@H]2CNCc3ccccc32)cc1 | OSZMNJRKIPAVOS-HNNXBMFYSA-N | No-Effect | inactive |
| 4-PPBP | BRD-K78061844 | c1ccc(CCCCN2CCC(c3ccccc3)CC2)cc1 | HQGDPZPNAXRCSA-UHFFFAOYSA-N | No-Effect | inactive |
| 4-propylbenzoic-acid | BRD-K42108215 | CCCc1ccc(C(=O)O)cc1 | ATZHGRNFEFVDDJ-UHFFFAOYSA-N | No-Effect | inactive |
| 4-pyrimidinecarbonitrile | BRD-K04552268 | N#Cc1ccncn1 | ZIEWSZYVEDTXGH-UHFFFAOYSA-N | No-Effect | inactive |
| 4-tert-butylphenol | BRD-K06338106 | CC(C)(C)c1ccc(O)cc1 | QHPQWRBYOIRBIT-UHFFFAOYSA-N | No-Effect | inactive |

| | | | | | |
|---|---|---|---|---|---|
| 4,4-pentamethylenepiperidine | BRD-K49545903 | C1CCC2(CC1)CCNCC2 | LIZKZVQBLDHKCY-UHFFFAOYSA-N | No-Effect | inactive |
| 4,4'-DDT | BRD-K51799616 | Clc1ccc(C(c2ccc(Cl)cc2)C(Cl)(Cl)Cl)cc1 | YVGGHNCTFXOJCH-UHFFFAOYSA-N | No-Effect | inactive |
| 4,5,6,7-tetrabromobenzotriazole | BRD-K97118047 | Brc1c(Br)c(Br)c2n[nH]nc2c1Br | OMZYUVOATZSGJY-UHFFFAOYSA-N | No-Effect | inactive |
| 4BP-TQS | BRD-A18598741 | NS(=O)(=O)c1ccc2c(c1)[C@H]1C=CC[C@@H]1[C@@H](c1ccc(Br)cc1)N2 | YNCXHXYZTLIZTO-RLFYNMQTSA-N | No-Effect | inactive |
| 4E1RCat | BRD-K67173685 | O=C(O)c1ccc(N2C(=O)/C(=C/c3ccc(-c4ccc([N+](=O)[O-])cc4)o3)C=C2c2ccccc2)cc1 | BBQRBOIMSKMFFO-LTGZKZEYSA-N | No-Effect | inactive |
| 4EGI-1 | BRD-K00004308 | O=C(O)C(Cc1ccccc1[N+](=O)[O-])=NNc1nc(-c2ccc(Cl)c(Cl)c2)cs1 | KFRKRECSIYXARE-UHFFFAOYSA-N | No-Effect | inactive |
| 4SC-202 | BRD-K11773281 | Cn1cc(-c2ccc(S(=O)(=O)n3ccc(/C=C/C(=O)Nc4ccccc4N)c3)cc2)cn1 | PRXXYMVLYKJITB-IZZDOVSWSA-N | No-Effect | inactive |
| 5-(4chlorophenyl)-4-ethyl-2,4-dihydro-3H-1,2,4-triazol-3-one | BRD-K58146797 | CCn1c(-c2ccc(Cl)cc2)n[nH]c1=O | PXINDYJACXPZIN-UHFFFAOYSA-N | No-Effect | inactive |
| 5-amino-3-D-ribofuranosylthiazolo[4,5-d]pyrimidin-2,7(3H,6H)-dione | BRD-K87535339 | Nc1nc(=O)c2sc(=O)n([C@@H]3O[C@H](CO)[C@@H](O)[C@H]3O)c2[nH]1 | TZYVRXZQAWPIAB-FCLHUMLKSA-N | No-Effect | inactive |
| 5-aminolevulinic-acid | BRD-K57631554 | NCC(=O)CCC(=O)O | ZGXJTSGNIOSYLO-UHFFFAOYSA-N | No-Effect | inactive |
| 5-BDBD | BRD-K33690734 | O=C1CN=C(c2cccc(Br)c2)c2oc3ccccc3c2N1 | NKYMVQPXXTZHSF-UHFFFAOYSA-N | No-Effect | inactive |
| 5-carboxamidotryptamine | BRD-K92382976 | NCCc1c[nH]c2ccc(C(N)=O)cc12 | WKZLNEWVIAGNAW-UHFFFAOYSA-N | No-Effect | inactive |
| 5-fluoro-3-pyridyl-methanol | BRD-K73381198 | OCc1cncc(F)c1 | GGGJYJXAFSEWNM-UHFFFAOYSA-N | No-Effect | inactive |
| 5-fluoropyrimidine | BRD-K00004679 | Fc1cncnc1 | KSPDSMOWMQFPBL-UHFFFAOYSA-N | No-Effect | inactive |
| 5-fluorouracil | BRD-K24844714 | O=c1[nH]cc(F)c(=O)[nH]1 | GHASVSINZRGABV-UHFFFAOYSA-N | No-Effect | inactive |
| 5-FP | BRD-K34437622 | O=c1ncc(F)c[nH]1 | HPABFFGQPLJKBP-UHFFFAOYSA-N | No-Effect | inactive |
| 5-HMF | BRD-K86799112 | O=Cc1ccc(CO)o1 | NOEGNKMFWQHSLB-UHFFFAOYSA-N | No-Effect | inactive |
| 5-hydroxydecanoic-acid | BRD-K01666923 | CCCCC[C@@H](O)CCCC(=O)O | LMHJFKYQYDSOQO-SECBINFHSA-N | No-Effect | inactive |
| 5-hydroxyectoine | BRD-K03871161 | CC1=NC[C@H](O)[C@@H](C(=O)O)N1 | KIIBBJKLKFTNQO-WHFBIAKZSA-N | No-Effect | inactive |
| 5-hydroxymethyl-tolterodine | BRD-K24603946 | CC(C)N(CC[C@H](c1ccccc1)c1cc(CO)ccc1O)C(C)C | DUXZAXCGJSBGDW-HXUWFJFHSA-N | No-Effect | inactive |
| 5-hydroxytryptophan | BRD-A73930134 | N[C@@H](Cc1c[nH]c2ccc(O)cc12)C(=O)O | LDCYZAJDBXYCGN-VIFPVBQESA-N | No-Effect | inactive |
| 5-iodo-A-85380 | BRD-K90065682 | Ic1cncc(OC[C@@H]2CCN2)c1 | RKVRGRXOYDTUEY-QMMMGPOBSA-N | No-Effect | inactive |
| 5-methylfurmethiodide | BRD-K45437867 | Cc1ccc(C[N+](C)(C)C)o1 | KOWVJDFMEZKDDT-UHFFFAOYSA-N | No-Effect | inactive |
| 5-methylhydantoin-(D) | BRD-K06050544 | C[C@H]1NC(=O)NC1=O | VMAQYKGITHDWKL-UWTATZPHSA-N | No-Effect | inactive |
| 5-methylhydantoin-(L) | BRD-K56033645 | C[C@@H]1NC(=O)NC1=O | VMAQYKGITHDWKL-REOHCLBHSA-N | No-Effect | inactive |
| 5-octanoylsalicylic-acid | BRD-K00004681 | CCCCCCCC(=O)c1ccc(O)c(C(=O)O)c1 | IXIGWKNBFPKCCD-UHFFFAOYSA-N | No-Effect | inactive |
| 5,7-dichlorokynurenic-acid | BRD-K60287130 | O=C(O)c1cc(=O)c2c(Cl)cc(Cl)cc2[nH]1 | BGKFPRIGXAVYNX-UHFFFAOYSA-N | No-Effect | inactive |
| 5'-chloro-5'-deoxy-ENBA-(+/-) | BRD-K00004289 | O[C@@H]1[C@@H](CCl)O[C@@H](n2cnc3c(N[C@H]4C[C@@H]5CC[C@H]4C5)ncnc32)[C@@H]1O | PVJGDYDNVNCGBT-CYDCOVMPSA-N | No-Effect | inactive |
| 6-aminochrysene | BRD-K00656370 | Nc1cc2c3ccccc3ccc2c2ccccc12 | KIVUHCNVDWYUNP-UHFFFAOYSA-N | No-Effect | inactive |
| 6-aminopenicillanic-acid | BRD-K30284967 | CC1(C)S[C@@H]2[C@H](N)C(=O)N2[C@H]1C(=O)O | NGHVIOIJCVXTGV-ALEPSDHESA-N | No-Effect | inactive |
| 6-benzylaminopurine | BRD-K62929068 | c1ccc(CNc2ncnc3nc[nH]c23)cc1 | NWBJYWHLCVSVIJ-UHFFFAOYSA-N | No-Effect | inactive |
| 6-chloromelatonin | BRD-K34663752 | COc1cc2c(CCNC(C)=O)c[nH]c2cc1Cl | LUINDDOUWHRIPW-UHFFFAOYSA-N | No-Effect | inactive |
| 6-diazo-5-oxo-L-norleucine | BRD-K84924563 | N#[N+]CC(=O)CC[C@H](N)C(=O)O | UUTYCZUIECRVAL-YFKPBYRVSA-O | No-Effect | inactive |
| 6-iodo-nordihydrocapsaicin | BRD-K49236613 | CCCCCCCCC(=O)NCc1cc(OC)c(O)cc1I | AAORACFZMYMFCG-UHFFFAOYSA-N | No-Effect | inactive |
| 7-aminocephalosporanic-acid | BRD-K65377893 | CC(=O)OCC1=C(C(=O)O)N2C(=O)[C@@H](N)[C@H]2SC1 | HSHGZXNAXBPPDL-HZGVNTEJSA-N | No-Effect | inactive |
| 7-chlorokynurenic-acid | BRD-K84214706 | O=C(O)c1cc(=O)c2ccc(Cl)cc2[nH]1 | UAWVRVFHMOSAPU-UHFFFAOYSA-N | No-Effect | inactive |
| 7-hydroxy-DPAT | BRD-K01826780 | CCCN(CCC)[C@H]1CCc2ccc(O)cc2C1 | BLYMJBIZMIGWFK-HNNXBMFYSA-N | No-Effect | inactive |
| 7-hydroxy-PIPAT | BRD-K01825667 | CCCN(C/C=C/I)[C@H]1CCc2ccc(O)cc2C1 | RTMIJLQPWFKAFE-GZTQLTBSSA-N | No-Effect | inactive |
| 7-keto-DHEA | BRD-K07785030 | CC(=O)O[C@H]1CC[C@@]2(C)C(=CC(=O)[C@H]3[C@@H]4CCC(=O)[C@@]4(C)CC[C@@H]32)C1 | VVSMJVQHDZUPIL-XFKPDKBWSA-N | No-Effect | inactive |
| 7-methoxytacrine | BRD-K53156626 | COc1ccc2nc3c(c(N)c2c1)CCCC3 | APQPVVOYBLOJDY-UHFFFAOYSA-N | No-Effect | inactive |
| 7-methylxanthine | BRD-K41592905 | Cn1cnc2[nH]c(=O)[nH]c(=O)c21 | PFWLFWPASULGAN-UHFFFAOYSA-N | No-Effect | inactive |
| 7-nitroindazole | BRD-K04430056 | O=[N+]([O-])c1cccc2c[nH]nc12 | PQCAUHUKTBHUSA-UHFFFAOYSA-N | No-Effect | inactive |
| 7,8,9,10-tetrahydroazepino[2,1-b]quinazolin-12(6H)-one | BRD-K77756533 | O=C1c2ccccc2NC2=CCCCCN12 | HBEJFHWHFIAMAI-UHFFFAOYSA-N | No-Effect | inactive |
| 8-bromo-cAMP | BRD-A09554849 | Nc1ncnc2c1nc(Br)n2[C@@H]1O[C@@H]2CO[P@@](=O)(O)O[C@H]2[C@H]1O | DVKQVRZMKBDMDH-UUOKFMHZSA-N | No-Effect | inactive |
| 8-bromo-cGMP | BRD-A00077618 | Nc1nc(=O)c2nc(Br)n([C@@H]3O[C@@H]4CO[P@@](=O)(O)O[C@H]4[C@H]3O)c2[nH]1 | YUFCOOWNNHGGOD-UMMCILCDSA-N | No-Effect | inactive |
| 8-hydroxy-DPAT | BRD-K01826732 | CCCN(CCC)[C@H]1CCc2cccc(O)c2C1 | ASXGJMSKWNBENU-AWEZNQCLSA-N | No-Effect | inactive |
| 8-hydroxy-PIPAT | BRD-A60594020 | CCCN(C/C=C/I)[C@H]1CCc2cccc(O)c2C1 | QBXHUZJZYDSLRH-MDDUGHTDSA-N | No-Effect | inactive |
| 8-M-PDOT | BRD-A07232941 | CCC(=O)N[C@H]1CCc2cccc(OC)c2C1 | RVIGBTUDFAGRTQ-NSHDSACASA-N | No-Effect | inactive |
| 80841-78-7 | BRD-K03044000 | Cc1oc(=O)oc1CCl | QCLFSYYUWPUWQR-UHFFFAOYSA-N | No-Effect | inactive |
| 9-aminoacridine | BRD-K00535541 | Nc1c2ccccc2nc2ccccc12 | XJGFWWJLMVZSIG-UHFFFAOYSA-N | No-Effect | inactive |
| 9-aminocamptothecin | BRD-K09291936 | CC[C@@]1(O)C(=O)OCc2c1cc1n(c2=O)Cc2cc3c(N)cccc3nc2-1 | FUXVKZWTXQUGMW-FQEVSTJZSA-N | No-Effect | inactive |
| 9-anthracenecarboxylic-acid | BRD-K62607075 | O=C(O)c1c2ccccc2cc2ccccc12 | XGWFJBFNAQHLEF-UHFFFAOYSA-N | No-Effect | inactive |

| | | | | | |
|---|---|---|---|---|---|
| A-1070722 | BRD-K70329400 | COc1ccc2c(NC(=O)Nc3cccc(C(F)(F)F)n3)ccnc2c1 | VQPBIJGXSXEOCU-UHFFFAOYSA-N | No-Effect | inactive |
| A-1120 | BRD-K59060513 | O=C(O)c1ccccc1NC(=O)N1CCC(c2ccccc2C(F)(F)F)CC1 | MEAQCLPMSVEOQF-UHFFFAOYSA-N | No-Effect | inactive |
| A-317491 | BRD-K38019854 | O=C(O)c1cc(C(=O)O)c(C(=O)N(Cc2cccc(Oc3ccccc3)c2)[C@H]2CCCc3ccccc32)cc1C(=O)O | VQGBOYBIENNKMI-LJAQVGFWSA-N | No-Effect | inactive |
| A-33903 | BRD-A40302156 | CC(=O)N[C@@H]1N=C(c2ccccc2)c2ccccc2NC1=O | MMXSDWJOQSEPSM-MRXNPFEDSA-N | No-Effect | inactive |
| A-366 | BRD-K06182768 | COc1cc2c(cc1OCCCN1CCCC1)N=C(N)C21CCC1 | BKCDJTRMYWSXMC-UHFFFAOYSA-N | No-Effect | inactive |
| A-381393 | BRD-K00005339 | Cc1ccc(N2CCN(Cc3nc4ccccc4[nH]3)CC2)cc1C | SAQMCVDGOIRQTC-UHFFFAOYSA-N | No-Effect | inactive |
| A-412997 | BRD-K62066682 | Cc1cccc(NC(=O)CN2CCC(c3ccccn3)CC2)c1 | JFCDMGGMCUKHST-UHFFFAOYSA-N | No-Effect | inactive |
| A-438079 | BRD-K13223831 | Clc1cccc(-c2nnnn2Cc2cccnc2)c1Cl | MMPAULQSJLVKHP-UHFFFAOYSA-N | No-Effect | inactive |
| A-582941 | BRD-K79743788 | CN1C[C@H]2CN(c3ccc(-c4ccccc4)nn3)C[C@H]2C1 | GTMRUYCIJSNXGB-GASCZTMLSA-N | No-Effect | inactive |
| A-61603 | BRD-K00004225 | CS(=O)(=O)Nc1c(O)ccc2c1CCC[C@@H]2C1=NCCN1 | OQFCXJDXHCDLHX-NSHDSACASA-N | No-Effect | inactive |
| A-7 | BRD-K03301001 | NCCCCCCCCCCNS(=O)(=O)c1cccc2c(Cl)cccc12 | JLUUJHIQVIXTGT-UHFFFAOYSA-N | No-Effect | inactive |
| A-769662 | BRD-K41918892 | N#Cc1c(O)c2c(-c3ccc(-c4ccccc4O)cc3)csc2[nH]c1=O | CTESJDQKVOEUOY-UHFFFAOYSA-N | No-Effect | inactive |
| A-784168 | BRD-K87665527 | O=C(Nc1ccc(S(=O)(=O)C(F)(F)F)cc1)C1=CCN(c2ncccc2C(F)(F)F)CC1 | SDUAWRFBHRAFBM-UHFFFAOYSA-N | No-Effect | inactive |
| A-803467 | BRD-K28570407 | COc1cc(NC(=O)c2ccc(-c3ccc(Cl)cc3)o2)cc(OC)c1 | VHKBTPQDHDSBSP-UHFFFAOYSA-N | No-Effect | inactive |
| A-804598 | BRD-K94313941 | C[C@H](N/C(=N\c1cccc2ncccc12)NC#N)c1ccccc1 | PQYCRDPLPKGSME-AWEZNQCLSA-N | No-Effect | inactive |
| A-839977 | BRD-K60427831 | Clc1cccc(-n2nnnc2NCc2ccccc2Oc2ccccn2)c1Cl | GMVNBKZQJFRFAR-UHFFFAOYSA-N | No-Effect | inactive |
| A-867744 | BRD-K33059059 | CCC(=O)c1cc(-c2ccc(Cl)cc2)n(-c2ccc(S(N)(=O)=O)cc2)c1C | ABACVOXFUHDKNZ-UHFFFAOYSA-N | No-Effect | inactive |
| A-887826 | BRD-K20239555 | CCCCOc1ccc(-c2cncc(C(=O)NCc3cccnc3N3CCOCC3)c2)cc1Cl | JPJGFWKHSMUKFO-UHFFFAOYSA-N | No-Effect | inactive |
| A-922500 | BRD-K81614372 | O=C(Nc1ccccc1)Nc1ccc(-c2ccc(C(=O)[C@@H]3CCC[C@H]3C(=O)O)cc2)cc1 | BOZRFEQDOFSZBV-DHIUTWEWSA-N | No-Effect | inactive |
| A-939572 | BRD-K99308908 | CNC(=O)c1cccc(NC(=O)N2CCC(Oc3ccccc3Cl)CC2)c1 | DPYTYQFYDLYWHZ-UHFFFAOYSA-N | No-Effect | inactive |
| A-967079 | BRD-K70744942 | CCC(=N\O)/C(C)=C/c1ccc(F)cc1 | HKROEBDHHKMNBZ-CHBKHGQFSA-N | No-Effect | inactive |
| A-987306 | BRD-K68982262 | Nc1nc2c(c(N3CCNCC3)n1)CCC1=C2[C@@H]2CCCC[C@@H]2O1 | DJKJVWJQAVGLHJ-YPMHNXCESA-N | No-Effect | inactive |
| A0001 | BRD-K57865272 | CC1=C(C)C(=O)C(CC[C@](C)(O)CCC[C@H](C)CCC[C@H](C)CCCC(C)C)=C(C)C1=O | LTVDFSLWFKLJDQ-IEOSBIPESA-N | No-Effect | inactive |
| A12B4C3 | BRD-A30000356 | CCCCCCCCCC[C@@H](O)[C@H]1C=C[C@@H]2C(=O)N(c3ccccc3)C(=O)[C@@H]2N1Nc1ccc([N+](=O)[O-])cc1 | KVHGJAKTBPFFNV-KUXCXQDQSA-N | No-Effect | inactive |
| A205804 | BRD-K77627880 | Cc1ccc(Sc2cncc3sc(C(N)=O)cc23)cc1 | QQGWEXFLMJGCAL-UHFFFAOYSA-N | No-Effect | inactive |
| A61603 | BRD-A83650191 | CS(=O)(=O)Nc1c(O)ccc2c1CCC[C@H]2C1=NCCN1 | OQFCXJDXHCDLHX-LLVKDONJSA-N | No-Effect | inactive |
| A66 | BRD-K49055432 | Cc1nc(NC(=O)N2CCC[C@H]2C(N)=O)sc1-c1csc(C(C)(C)C)n1 | HBPXWEPKNBHKAX-NSHDSACASA-N | No-Effect | inactive |
| A740003 | BRD-A78952587 | COc1ccc(CC(=O)N[C@@H](N/C(=N/C#N)Nc2cccc3ncccc23)C(C)(C)C)cc1OC | PUHSRMSFDASMAE-DEOSSOPVSA-N | No-Effect | inactive |
| A77636 | BRD-K72226571 | NC[C@@H]1O[C@H](C23CC4CC(CC(C4)C2)C3)Cc2c1ccc(O)c2O | QLJOSZATCBCBDR-DFKUFRTHSA-N | No-Effect | inactive |
| A922500 | BRD-K28861668 | O=C(Nc1ccccc1)Nc1ccc(-c2ccc(C(=O)[C@@H]3CCC[C@@H]3C(=O)O)cc2)cc1 | BOZRFEQDOFSZBV-PKTZIBPZSA-N | No-Effect | inactive |
| AA-29504 | BRD-K13944729 | CCOC(=O)Nc1ccc(NCc2c(C)cc(C)cc2C)cc1N | SCOOTUMXCXKEAD-UHFFFAOYSA-N | No-Effect | inactive |
| AAI101 | BRD-K00003450 | C[n+]1ccn(C[C@@]2(C)[C@H](C(=O)O)N3C(=O)C[C@H]3S2(=O)=O)n1 | HFZITXBUTWITPT-YWVKMMECSA-O | No-Effect | inactive |
| AB-423 | BRD-K00003109 | CC[C@@H](C)NS(=O)(=O)c1ccc(F)c(C(=O)Nc2ccc(F)c(F)c2)c1 | BBLXLHYPDOMJMO-SNVBAGLBSA-N | No-Effect | inactive |
| abacavir | BRD-K17443395 | Nc1nc(NC2CC2)c2ncn([C@H]3C=C[C@@H](CO)C3)c2n1 | MCGSCOLBFJQGHM-SCZZXKLOSA-N | No-Effect | inactive |
| abafungin | BRD-K77432251 | Cc1ccc(Oc2ccccc2-c2csc(NC3=NCCCN3)n2)c(C)c1 | TYBHXIFFPVFXQW-UHFFFAOYSA-N | No-Effect | inactive |
| abamectin | BRD-K00004970 | CO[C@H]1C[C@H](O[C@H]2[C@H](C)O[C@@H](O[C@@H]3/C(C)=C/C[C@@H]4C[C@@H](C[C@]5(C=C[C@H](C)[C@@H](C(C)C)O5)O4)OC(=O)[C@@H]4C=C(C)[C@@H](O)[C@H]5OC/C(=C\C=C\[C@@H]3C)[C@@]45O)C[C@@H]2OC)O[C@@H](C)[C@@H]1O | ZFUKERYTFURFGA-PVVXTEPVSA-N | No-Effect | inactive |
| abametapir | BRD-K65498798 | Cc1ccc(-c2ccc(C)cn2)nc1 | PTRATZCAGVBFIQ-UHFFFAOYSA-N | No-Effect | inactive |
| abarelix | BRD-K00003158 | CC(=O)N[C@H](Cc1ccc2ccccc2c1)C(=O)N[C@H](Cc1ccc(Cl)cc1)C(=O)N[C@H](Cc1cccnc1)C(=O)N[C@@H](CO)C(=O)N(C)[C@@H](Cc1ccc(O)cc1)C(=O)N[C@H](CC(N)=O)C(=O)N[C@@H](CC(C)C)C(=O)N[C@@H](CCCCNC(C)C)C(=O)N1CCC[C@H]1C(=O)N[C@H](C)C(N)=O | AIWRTTMUVOZGPW-HSPKUQOVSA-N | No-Effect | inactive |
| ABBV-744 | BRD-K00003108 | CCNC(=O)c1cc2c(-c3cc(C(C)(C)O)ccc3Oc3c(C)cc(F)cc3C)cn(C)c(O)c-2n1 | LQPFCYWRBQRRGE-UHFFFAOYSA-N | No-Effect | inactive |
| ABC-294640 | BRD-K01825984 | O=C(NCc1ccncc1)[C@@]12C[C@H]3C[C@@H](C1)C[C@](c1ccc(Cl)cc1)(C3)C2 | CAOTVXGYTWCKQE-BRNYJPRKSA-N | No-Effect | inactive |
| abemaciclib | BRD-K33622447 | CCN1CCN(Cc2ccc(Nc3ncc(F)c(-c4cc(F)c5nc(C)n(C(C)C)c5c4)n3)nc2)CC1 | UZWDCWONPYILKI-UHFFFAOYSA-N | No-Effect | inactive |
| abiraterone | BRD-K50071428 | C[C@]12CC[C@H]3[C@@H](CC=C4C[C@@H](O)CC[C@@]43C)[C@@H]1CC=C2c1cccnc1 | GZOSMCIZMLWJML-VJLLXTKPSA-N | No-Effect | inactive |
| abiraterone-acetate | BRD-K24048528 | CC(=O)O[C@H]1CC[C@@]2(C)C(=CC[C@H]3[C@@H]4CC=C(c5cccnc5)[C@@]4(C)CC[C@@H]32)C1 | UVIQSJCZCSLXRZ-UBUQANBQSA-N | No-Effect | inactive |
| ABT-072 | BRD-K00004730 | COc1c(/C=C/c2ccc(NS(C)(=O)=O)cc2)cc(-n2ccc(=O)[nH]c2=O)cc1C(C)(C)C | XMZSTQYSBYEENY-RMKNXTFCSA-N | No-Effect | inactive |
| ABT-202 | BRD-A29844814 | N[C@H]1CCN(c2cccnc2)C1 | LVGMMVAWLISWJD-QMMMGPOBSA-N | No-Effect | inactive |
| ABT-239 | BRD-K22749967 | C[C@@H]1CCCN1CCc1cc2cc(-c3ccc(C#N)cc3)ccc2o1 | KFHYZKCRXNRKRC-MRXNPFEDSA-N | No-Effect | inactive |
| ABT-491 | BRD-K97233161 | C#Cc1cccc2c1c(C(=O)c1ccc(Cn3c(C)nc4cnccc43)c(F)c1)cn2C(=O)N(C)C | GDLNHSUSOZEAOR-UHFFFAOYSA-N | No-Effect | inactive |
| ABT-639 | BRD-K11792081 | O=C(c1cc(S(=O)(=O)Nc2ccccc2F)c(F)cc1Cl)N1CCN2CCC[C@@H]2C1 | AGPIHNZOZNKRGT-CYBMUJFWSA-N | No-Effect | inactive |
| ABT-702 | BRD-K36021395 | Nc1ncnc2nc(-c3ccc(N4CCOCC4)nc3)cc(-c3cccc(Br)c3)c12 | RQCXKDWOCUJWQZ-UHFFFAOYSA-N | No-Effect | inactive |
| ABT-724 | BRD-K37714784 | c1ccc(N2CCN(Cc3nc4ccccc4[nH]3)CC2)nc1 | FRPJGTNLZNXQEX-UHFFFAOYSA-N | No-Effect | inactive |
| ABT-737 | BRD-K56301217 | CN(C)CC[C@H](CSc1ccccc1)Nc1ccc(S(=O)(=O)NC(=O)c2ccc(N3CCN(Cc4ccccc4-c4ccc(Cl)cc4)CC3)cc2)cc1[N+](=O)[O-] | HPLNQCPCUACXLM-PGUFJCEWSA-N | No-Effect | inactive |
| ABX-1431 | BRD-K00004603 | O=C(OC(C(F)(F)F)C(F)(F)F)N1CCN(Cc2ccc(C(F)(F)F)cc2N2CCCC2)CC1 | SQZJGTOZFRNWCX-UHFFFAOYSA-N | No-Effect | inactive |
| ABX464 | BRD-K00003394 | FC(F)(F)Oc1ccc(Nc2ccc3cccc(Cl)c3n2)cc1 | OZOGDCZJYVSUBR-UHFFFAOYSA-N | No-Effect | inactive |
| AC-186 | BRD-K53788420 | Oc1ccc(C2(c3ccccc3F)CCC(F)(F)CC2)cc1 | HSMUKSLNDJOZQG-UHFFFAOYSA-N | No-Effect | inactive |
| AC-261066 | BRD-K31144453 | CCCCOCCOc1nc(-c2ccc(C(=O)O)c(F)c2)sc1C | HSAOETBFVAWNRP-UHFFFAOYSA-N | No-Effect | inactive |
| AC-55541 | BRD-A27911617 | C/C(=N\NC(=O)[C@@H](NC(=O)c1ccccc1)c1n[nH]c(=O)c2ccccc12)c1cccc(Br)c1 | UCUHFWIFSHROPY-SQEMJAKJSA-N | No-Effect | inactive |
| AC-7954-(+/-) | BRD-A49544621 | CN(C)CC[C@@]1(c2ccc(Cl)cc2)Cc2ccccc2C(=O)O1 | HIVBATDUVFEJFZ-IBGZPJMESA-N | No-Effect | inactive |
| AC1NDSS5 | BRD-K10226125 | CCCCCCCCCCCCCCCC[S+](CCC)c1ccccc1C(=O)OCC | VZXULFIUJKSVJY-UHFFFAOYSA-N | No-Effect | inactive |
| AC260584 | BRD-K00004583 | CCCCC1CCN(CCCN2C(=O)COc3cc(F)ccc32)CC1 | BWAKMLKESHKOHK-UHFFFAOYSA-N | No-Effect | inactive |
| acadesine | BRD-K53448858 | NC(=O)c1ncn([C@@H]2O[C@H](CO)[C@@H](O)[C@H]2O)c1N | RTRQQBHATOEIAF-UUOKFMHZSA-N | No-Effect | inactive |
| acalabrutinib | BRD-K64034691 | CC#CC(=O)N1CCC[C@H]1c1nc(-c2ccc(C(=O)Nc3ccccn3)cc2)c2c(N)nccn12 | WDENQIQQYWYTPO-IBGZPJMESA-N | No-Effect | inactive |
| acalisib | BRD-K92517771 | C[C@H](Nc1ncnc2nc[nH]c12)c1nc2ccc(F)cc2c(=O)n1-c1ccccc1 | DOCINCLJNAXZQF-LBPRGKRZSA-N | No-Effect | inactive |
| acamprosate | BRD-K26262077 | CC(=O)NCCCS(=O)(=O)O | AFCGFAGUEYAMAO-UHFFFAOYSA-N | No-Effect | inactive |

| | | | | | |
|---|---|---|---|---|---|
| ACDPP | BRD-K68633617 | CN(C)c1nc(N)c(C(=O)Nc2cccnc2)nc1Cl | RNGRUHWJJPTYNR-UHFFFAOYSA-N | No-Effect | inactive |
| acebilustat | BRD-K00003255 | O=C(O)c1ccc(CN2C[C@@H]3C[C@H]2CN3Cc2ccc(Oc3ccc(-c4ncco4)cc3)cc2)cc1 | GERJIEKMNDGSCS-DQEYMECFSA-N | No-Effect | inactive |
| acebutolol | BRD-A29260609 | CCCC(=O)Nc1ccc(OCC(O)CNC(C)C)c(C(C)=O)c1 | GOEMGAFJFRBGGG-UHFFFAOYSA-N | No-Effect | inactive |
| acecainide | BRD-K07753030 | CCN(CC)CCNC(=O)c1ccc(NC(C)=O)cc1 | KEECCEWTUVWFCV-UHFFFAOYSA-N | No-Effect | inactive |
| aceclidine | BRD-K01825000 | CC(=O)O[C@H]1CN2CCC1CC2 | WRJPSSPFHGNBMG-VIFPVBQESA-N | No-Effect | inactive |
| aceclofenac | BRD-K68538666 | O=C(O)COC(=O)Cc1ccccc1Nc1c(Cl)cccc1Cl | MNIPYSSQXLZQLJ-UHFFFAOYSA-N | No-Effect | inactive |
| acedapsone | BRD-K97417064 | CC(=O)Nc1ccc(S(=O)(=O)c2ccc(NC(C)=O)cc2)cc1 | AMTPYFGPPVFBBI-UHFFFAOYSA-N | No-Effect | inactive |
| acedoben | BRD-K47195926 | CC(=O)Nc1ccc(C(=O)O)cc1 | QCXJEYYXVJIFCE-UHFFFAOYSA-N | No-Effect | inactive |
| acefylline | BRD-K44004064 | Cn1c(=O)c2c(ncn2CC(=O)O)n(C)c1=O | HCYFGRCYSCXKNQ-UHFFFAOYSA-N | No-Effect | inactive |
| aceglutamide | BRD-K13619941 | CC(=O)N[C@@H](CCC(N)=O)C(=O)O | KSMRODHGGIIXDV-YFKPBYRVSA-N | No-Effect | inactive |
| acelarin | BRD-K00003531 | C[C@H](N[P@@](=O)(OC[C@H]1O[C@@H](n2ccc(N)nc2=O)C(F)(F)[C@@H]1O)Oc1ccccc1)C(=O)OCc1ccccc1 | NHTKGYOMICWFQZ-LHFSRKHSSA-N | No-Effect | inactive |
| aceneuramic-acid | BRD-K36262466 | CC(=O)N[C@@H]1[C@@H](O)C[C@](O)(C(=O)O)O[C@H]1[C@H](O)[C@H](O)CO | SQVRNKJHWKZAKO-PFQGKNLYSA-N | No-Effect | inactive |
| acenocoumarol | BRD-A65051990 | CC(=O)C[C@@H](c1ccc([N+](=O)[O-])cc1)c1c(O)c2ccccc2oc1=O | VABCILAOYCMVPS-HNNXBMFYSA-N | No-Effect | inactive |
| acesulfame-potassium | BRD-K50850824 | CC1=CC(=O)NS(=O)(=O)O1 | YGCFIWIQZPHFLU-UHFFFAOYSA-N | No-Effect | inactive |
| acetanilide | BRD-K11094367 | CC(=O)Nc1ccccc1 | FZERHIULMFGESH-UHFFFAOYSA-N | No-Effect | inactive |
| acetarsol | BRD-K99048103 | CC(=O)Nc1cc([As](=O)(O)O)ccc1O | ODFJOVXVLFUVNQ-UHFFFAOYSA-N | No-Effect | inactive |
| acetazolamide | BRD-K43457670 | CC(=O)Nc1nnc(S(N)(=O)=O)s1 | BZKPWHYZMXOIDC-UHFFFAOYSA-N | No-Effect | inactive |
| acetohexamide | BRD-K52960356 | CC(=O)c1ccc(S(=O)(=O)NC(=O)NC2CCCCC2)cc1 | VGZSUPCWNCWDAN-UHFFFAOYSA-N | No-Effect | inactive |
| acetohydroxamic-acid | BRD-K21615095 | CC(=O)NO | RRUDCFGSUDOHDG-UHFFFAOYSA-N | No-Effect | inactive |
| acetophenazine | BRD-K42938903 | CC(=O)c1ccc2c(c1)N(CCCN1CCN(CCO)CC1)c1ccccc1S2 | WNTYBHLDCKXEOT-UHFFFAOYSA-N | No-Effect | inactive |
| acetriazoic-acid | BRD-K46416130 | CC(=O)Nc1c(I)cc(I)c(C(=O)O)c1I | GNOGSFBXBWBTIG-UHFFFAOYSA-N | No-Effect | inactive |
| acetyl-11-keto-beta-boswellic-acid | BRD-A44696962 | CC(=O)O[C@@H]1CC[C@]2(C)[C@@H]3C(=O)C=C4[C@@H]5[C@@H](C)[C@H](C)CC[C@]5(C)CC[C@@]4(C)[C@]3(C)CC[C@H]2[C@@]1(C)C(=O)O | HMMGKOVEOFBCAU-OQJDUFEHSA-N | No-Effect | inactive |
| acetyl-farnesyl-cysteine | BRD-K79437791 | CC(=O)N[C@@H](CSC/C=C(\C)CC/C=C(\C)CCC=C(C)C)C(=O)O | XTURYZYJYQRJDO-BNAHBJSTSA-N | No-Effect | inactive |
| acetyl-l-leucine | BRD-K27406233 | CC(=O)N[C@@H](CC(C)C)C(=O)O | WXNXCEHXYPACJF-ZETCQYMHSA-N | No-Effect | inactive |
| acetylcholine | BRD-K82381502 | CC(=O)OCC[N+](C)(C)C | OIPILFWXSMYKGL-UHFFFAOYSA-N | No-Effect | inactive |
| acetylcysteine | BRD-K59058747 | CC(=O)N[C@@H](CS)C(=O)O | PWKSKIMOESPYIA-BYPYZUCNSA-N | No-Effect | inactive |
| acetylspiramycin | BRD-K00004968 | CO[C@@H]1[C@@H](O[C@@H]2O[C@H](C)[C@@H](O[C@H]3C[C@@](C)(O)[C@@H](O)[C@H](C)O3)[C@H](N(C)C)[C@H]2O)[C@@H](CC=O)C[C@@H](C)[C@@H](O[C@H]2CC[C@H](N(C)C)[C@@H](C)O2)/C=C/C=C/C[C@@H](C)OC(=O)C[C@H]1OC(C)=O | ZPCCSZFPOXBNDL-ZSTSFXQOSA-N | No-Effect | inactive |
| acexamic-acid | BRD-K72215350 | CC(=O)NCCCCCC(=O)O | WDSCBUNMANHPFH-UHFFFAOYSA-N | No-Effect | inactive |
| acifran | BRD-A29520968 | C[C@@]1(c2ccccc2)OC(C(=O)O)=CC1=O | DFDGRKNOFOJBAJ-LBPRGKRZSA-N | No-Effect | inactive |
| acipimox | BRD-K63736853 | Cc1cnc(C(=O)O)c[n+]1[O-] | DJQOOSBJCLSSEY-UHFFFAOYSA-N | No-Effect | inactive |
| acitazanolast | BRD-K74236708 | O=C(O)C(=O)Nc1cccc(-c2nn[nH]n2)c1 | VWQZJJZGISNFOE-UHFFFAOYSA-N | No-Effect | inactive |
| acitretin | BRD-K62012036 | COc1cc(C)c(/C=C/C(C)=C/C=C/C(C)=C/C(=O)O)c(C)c1C | IHUNBGSDBOWDMA-AQFIFDHZSA-N | No-Effect | inactive |
| acivicin | BRD-K13050541 | N[C@@H](C(=O)O)[C@@H]1CC(Cl)=NO1 | QAWIHIJWNYOLBE-ZAFYKAAXSA-N | No-Effect | inactive |
| aclidinium | BRD-K74363950 | O=C(O[C@H]1C[N+]2(CCCOc3ccccc3)CCC1CC2)C(O)(c1cccs1)c1cccs1 | ASMXXROZKSBQIH-VITNCHFBSA-N | No-Effect | inactive |
| acoziborole | BRD-K00003303 | CC1(C)OB(O)c2cc(NC(=O)c3ccc(F)cc3C(F)(F)F)ccc21 | PTYGDEXEGLDNAZ-UHFFFAOYSA-N | No-Effect | inactive |
| ACPC | BRD-K66465025 | NC1(C(=O)O)CC1 | PAJPWUMXBYXFCZ-UHFFFAOYSA-N | No-Effect | inactive |
| acriflavine | BRD-M87138257 | C[n+]1c2cc(N)ccc2cc2ccc(N)cc21.Nc1ccc2cc3ccc(N)cc3nc2c1 | BCBIBQWDFMQRSJ-UHFFFAOYSA-O | No-Effect | inactive |
| acriflavinium | BRD-K97874492 | C[n+]1c2cc(N)ccc2cc2ccc(N)cc21 | XSIOKTWDEOJMGG-UHFFFAOYSA-O | No-Effect | inactive |
| acrisorcin | BRD-M49658242 | CCCCCCc1ccc(O)cc1O.Nc1c2ccccc2nc2ccccc12 | YZODJQFXMFEJRM-UHFFFAOYSA-N | No-Effect | inactive |
| acrivastine | BRD-K50163129 | Cc1ccc(/C(=C\CN2CCCC2)c2cccc(/C=C/C(=O)O)n2)cc1 | PWACSDKDOHSSQD-IUTFFREVSA-N | No-Effect | inactive |
| acrylate | BRD-M00184283 | C=CC(=O)O.[Na] | USNWAMPROKAEIT-UHFFFAOYSA-N | No-Effect | inactive |
| ACT-132577 | BRD-K34843150 | NS(=O)(=O)Nc1ncnc(OCCOc2ncc(Br)cn2)c1-c1ccc(Br)cc1 | DKULOVKANLVDEA-UHFFFAOYSA-N | No-Effect | inactive |
| ACT-462206 | BRD-K19781342 | COc1ccc(S(=O)(=O)N2CCC[C@H]2C(=O)Nc2cc(C)cc(C)c2)cc1 | NHPQGZOBHSVTAQ-IBGZPJMESA-N | No-Effect | inactive |
| actarit | BRD-K33483813 | CC(=O)Nc1ccc(CC(=O)O)cc1 | MROJXXOCABQVEF-UHFFFAOYSA-N | No-Effect | inactive |
| ACTB-1003 | BRD-K35936936 | COCc1c(-c2ccc(NC(=O)Nc3cc(C(F)(F)F)ccc3F)c(F)c2)c2c(N)ncnn2c1CN1CCOCC1 | GZPJCJKUZPUFAL-UHFFFAOYSA-N | No-Effect | inactive |
| actinoquinol | BRD-K08837089 | CCOc1ccc(S(=O)(=O)O)c2cccnc12 | YAMVZYRZAMBCED-UHFFFAOYSA-N | No-Effect | inactive |
| acumapimod | BRD-K00003220 | Cc1ccc(C(=O)NC2CC2)cc1-n1ncc(C(=O)c2cccc(C#N)c2)c1N | VGUSQKZDZHAAEE-UHFFFAOYSA-N | No-Effect | inactive |
| ACY-1215 | BRD-K82928847 | O=C(CCCCCCNC(=O)c1cnc(N(c2ccccc2)c2ccccc2)nc1)NO | QGZYDVAGYRLSKP-UHFFFAOYSA-N | No-Effect | inactive |
| AD-5467 | BRD-A80280426 | CC(C)c1cccc2c1O[C@H](C(C)C)C(=S)N2CC(=O)O | CLDJCRWXLDLJLO-CQSZACIVSA-N | No-Effect | inactive |
| adaprev | BRD-A73427433 | O=P(O)(O)OC[C@H]1O[C@H](O)[C@H](O)[C@@H](O)[C@@H]1O | NBSCHQHZLSJFNQ-DVKNGEFBSA-N | No-Effect | inactive |
| adaptavir | BRD-K37151368 | C[C@@H](N)C(=O)N[C@@H](CO)C(=O)N[C@H](C(=O)N[C@H](C(=O)N[C@H](C(=O)N[C@@H](CC(N)=O)C(=O)N[C@@H](Cc1ccc(O)cc1)C(=O)N[C@H](C(N)=O)[C@@H](C)O)[C@@H](C)O)[C@@H](C)O)[C@@H](C)O | AKWRNBWMGFUAMF-ZESMOPTKSA-N | No-Effect | inactive |
| adarotene | BRD-K00152668 | O=C(O)/C=C/c1ccc(-c2ccc(O)c(C34CC5CC(CC(C5)C3)C4)c2)cc1 | QAWBIEIZDDIEMW-FPYGCLRLSA-N | No-Effect | inactive |
| adatanserin | BRD-K67789209 | O=C(NCCN1CCN(c2ncccn2)CC1)C12CC3CC(CC(C3)C1)C2 | HPFLVTSWRFCPCV-UHFFFAOYSA-N | No-Effect | inactive |
| adavivint | BRD-K00003499 | CC(C)CC(=O)Nc1cncc(-c2ccc3[nH]nc(-c4nc5cncc(-c6cccc(F)c6)c5[nH]4)c3c2)c1 | AQDWDWAYVBQMAM-UHFFFAOYSA-N | No-Effect | inactive |
| ADD-233089 | BRD-K43055376 | O=c1[nH]c(N2CCOCC2)cn1-c1ccc(Cl)cc1 | OKVOKULYZXODKU-UHFFFAOYSA-N | No-Effect | inactive |
| adefovir | BRD-K77484724 | Nc1ncnc2c1ncn2CCOCP(=O)(O)O | SUPKOOSCJHTBAH-UHFFFAOYSA-N | No-Effect | inactive |
| adefovir-dipivoxil | BRD-K96159438 | CC(C)(C)C(=O)OCOP(=O)(COCCn1cnc2c(N)ncnc21)OCOC(=O)C(C)(C)C | WOZSCQDILHKSGG-UHFFFAOYSA-N | No-Effect | inactive |
| ademetionine | BRD-K78185697 | C[S+](CC[C@H](N)C(=O)O)C[C@H]1O[C@@H](n2cnc3c(N)ncnc32)[C@H](O)[C@@H]1O | MEFKEPWMEQBLKI-AIRLBKTGSA-O | No-Effect | inactive |
| adenine | BRD-K26060080 | Nc1ncnc2nc[nH]c12 | GFFGJBXGBJISGV-UHFFFAOYSA-N | No-Effect | inactive |
| adenosine | BRD-K01577834 | Nc1ncnc2c1ncn2[C@@H]1O[C@H](CO)[C@@H](O)[C@H]1O | OIRDTQYFTABQOQ-KQYNXXCUSA-N | No-Effect | inactive |
| adenosine-phosphate | BRD-K85767870 | Nc1ncnc2c1ncn2[C@@H]1O[C@H](COP(=O)(O)O)[C@@H](O)[C@H]1O | UDMBCSSLTHHNCD-KQYNXXCUSA-N | No-Effect | inactive |
| adenosine-triphosphate | BRD-K00003709 | Nc1ncnc2c1ncn2[C@@H]1O[C@H](COP(=O)(O)OP(=O)(O)OP(=O)(O)O)[C@@H](O)[C@H]1O | ZKHQWZAMYRWXGA-KQYNXXCUSA-N | No-Effect | inactive |
| adiphenine | BRD-K60907894 | CCN(CC)CCOC(=O)C(c1ccccc1)c1ccccc1 | JGOAIQNSOGZNBX-UHFFFAOYSA-N | No-Effect | inactive |
| adipic-acid | BRD-K10656395 | O=C(O)CCCCC(=O)O | WNLRTRBMVRJNCN-UHFFFAOYSA-N | No-Effect | inactive |
| adiporon | BRD-K68810443 | O=C(COc1ccc(C(=O)c2ccccc2)cc1)NC1CCN(Cc2ccccc2)CC1 | SHHUPGSHGSNPDB-UHFFFAOYSA-N | No-Effect | inactive |
| ADL5859 | BRD-K39624409 | CCN(CC)C(=O)c1ccc(C2=CC3(CCNCC3)Oc3cccc(O)c32)cc1 | OPIKUXLJQFYMSC-UHFFFAOYSA-N | No-Effect | inactive |
| adomeglivant | BRD-K00003302 | Cc1cc(O[C@@H](CCC(F)(F)F)c2ccc(C(=O)NCCC(=O)O)cc2)cc(C)c1-c1ccc(C(C)(C)C)cc1 | FASLTMSUPQDLIB-MHZLTWQESA-N | No-Effect | inactive |

| | | | | | |
|---|---|---|---|---|---|
| adoprazine | BRD-K34870043 | Fc1ccc(-c2cncc(CN3CCN(c4cccc5c4OCCO5)CC3)c2)cc1 | IUVSEUFHPNITEQ-UHFFFAOYSA-N | No-Effect | inactive |
| adrafinil | BRD-A90926615 | O=C(C[S@@](=O)C(c1ccccc1)c1ccccc1)NO | CGNMLOKEMNBUAI-HXUWFJFHSA-N | No-Effect | inactive |
| adrenalone | BRD-K82343404 | CNCC(=O)c1ccc(O)c(O)c1 | PZMVOUYYNKPMSI-UHFFFAOYSA-N | No-Effect | inactive |
| adrenosterone | BRD-K17368287 | C[C@]12CC(=O)[C@H]3[C@@H](CCC4=CC(=O)CC[C@@]43C)[C@@H]1CCC2=O | RZRPTBIGEANTGU-IRIMSJTPSA-N | No-Effect | inactive |
| ADX-10059 | BRD-K77911775 | Cc1cc(C)c(N)c(C#Cc2cccc(F)c2)n1 | MEDCLNYIYBERKO-UHFFFAOYSA-N | No-Effect | inactive |
| ADX-47273 | BRD-K56543881 | O=C(c1ccc(F)cc1)N1CCC[C@H](c2nc(-c3ccc(F)cc3)no2)C1 | VXQCCZHCFBHTTD-HNNXBMFYSA-N | No-Effect | inactive |
| AEE788 | BRD-K40718343 | CCN1CCN(Cc2ccc(-c3cc4c(N[C@H](C)c5ccccc5)ncnc4[nH]3)cc2)CC1 | OONFNUWBHFSNBT-HXUWFJFHSA-N | No-Effect | inactive |
| AF38469 | BRD-K47816470 | Cc1cccc(NC(=O)c2ccc(C(F)(F)F)cc2C(=O)O)n1 | JWCUSQCZMQIBMR-UHFFFAOYSA-N | No-Effect | inactive |
| afalanine | BRD-K53205568 | CC(=O)N[C@@H](Cc1ccccc1)C(=O)O | CBQJSKKFNMDLON-JTQLQIEISA-N | No-Effect | inactive |
| afimoxifene | BRD-K04210847 | CC/C(=C(\c1ccc(O)cc1)c1ccc(OCCN(C)C)cc1)c1ccccc1 | TXUZVZSFRXZGTL-QPLCGJKRSA-N | No-Effect | inactive |
| afloqualone | BRD-K69247067 | Cc1ccccc1-n1c(CF)nc2ccc(N)cc2c1=O | VDOSWXIDETXFET-UHFFFAOYSA-N | No-Effect | inactive |
| AFN-1252 | BRD-K23791635 | Cc1c(CN(C)C(=O)/C=C/c2cnc3c(c2)CCC(=O)N3)oc2ccccc12 | QXTWSUQCXCWEHF-JXMROGBWSA-N | No-Effect | inactive |
| afobazole | BRD-K37793595 | CCOc1ccc2nc(SCCN3CCOCC3)[nH]c2c1 | WWNUCVSRRUDYPP-UHFFFAOYSA-N | No-Effect | inactive |
| AG-1024 | BRD-K08310154 | CC(C)(C)c1cc(C=C(C#N)C#N)cc(Br)c1O | ABBADGFSRBWENF-UHFFFAOYSA-N | No-Effect | inactive |
| AG-14361 | BRD-K00615600 | CN(C)Cc1ccc(-c2nc3cccc4c3n2CCNC4=O)cc1 | SEKJSSBJKFLZIT-UHFFFAOYSA-N | No-Effect | inactive |
| AG-490 | BRD-K47105409 | N#C/C(=C\c1ccc(O)c(O)c1)C(=O)NCc1ccccc1 | TUCIOBMMDDOEMM-RIYZIHGNSA-N | No-Effect | inactive |
| AG-555 | BRD-K34428485 | N#C/C(=C\c1ccc(O)c(O)c1)C(=O)NCCCc1ccccc1 | GSQOBTOAOGXIFL-LFIBNONCSA-N | No-Effect | inactive |
| AG-556 | BRD-K81048131 | N#C/C(=C\c1ccc(O)c(O)c1)C(=O)NCCCCc1ccccc1 | GWCNJMUSWLTSCW-SFQUDFHCSA-N | No-Effect | inactive |
| AGI-5198 | BRD-A56371469 | Cc1ccccc1[C@H](C(=O)NC1CCCCC1)N(C(=O)Cn1ccnc1C)c1cccc(F)c1 | FNYGWXSATBUBER-AREMUKBSSA-N | No-Effect | inactive |
| AGI-6780 | BRD-K04691817 | O=C(Nc1cccc(C(F)(F)F)c1)Nc1cc(S(=O)(=O)NC2CC2)ccc1-c1ccsc1 | CCAWRGNYALGPQH-UHFFFAOYSA-N | No-Effect | inactive |
| AGM-1470 | BRD-A78111609 | CO[C@@H]1[C@H](OC(=O)NC(=O)CCl)CC[C@]2(CO2)[C@@H]1[C@@]1(C)O[C@@H]1CC=C(C)C | MSHZHSPISPJWHW-FRDQJGBZSA-N | No-Effect | inactive |
| agmatine | BRD-K49738308 | N=C(N)NCCCCN | QYPPJABKJHAVHS-UHFFFAOYSA-N | No-Effect | inactive |
| AGN-192403 | BRD-A18074780 | CC(C)[C@@H]1[C@H]2CC[C@@H](C2)[C@H]1N | SHXVJSFOJHMEEQ-AXTSPUMRSA-N | No-Effect | inactive |
| AGN-194310 | BRD-K70262971 | CCc1ccc(C2=CC(C)(C)Sc3ccc(C#Cc4ccc(C(=O)O)cc4)cc32)cc1 | LHUPKWKWYWOMSK-UHFFFAOYSA-N | No-Effect | inactive |
| agomelatine | BRD-K84698278 | COc1ccc2cccc(CCNC(C)=O)c2c1 | YJYPHIXNFHFHND-UHFFFAOYSA-N | No-Effect | inactive |
| AGP-103 | BRD-K91495480 | CCCCCCCCCCCCCCCCCCCCC(=O)O | UKMSUNONTOPOIO-UHFFFAOYSA-N | No-Effect | inactive |
| AH-7614 | BRD-K10852420 | Cc1ccc(S(=O)(=O)NC2c3ccccc3Oc3ccccc32)cc1 | OZCQEUZTOAAWDK-UHFFFAOYSA-N | No-Effect | inactive |
| AH11110 | BRD-A64808605 | N=C(C[C@@H](O)COc1ccccc1-c1ccccc1)N1CCCCC1 | HSVZQQYIMKLNLN-GOSISDBHSA-N | No-Effect | inactive |
| AH6809 | BRD-K59339270 | CC(C)Oc1ccc2c(=O)c3cc(C(=O)O)ccc3oc2c1 | AQFFXPQJLZFABJ-UHFFFAOYSA-N | No-Effect | inactive |
| AI-10-49 | BRD-K89899948 | FC(F)(F)Oc1ccc2nc(-c3ccc(OCCOCCOc4ccc(-c5nc6ccc(OC(F)(F)F)cc6[nH]5)nc4)cn3)[nH]c2c1 | WJBSSBFGPKTMQQ-UHFFFAOYSA-N | No-Effect | inactive |
| AIM-100 | BRD-K85139301 | c1ccc(-c2oc3ncnc(NC[C@@H]4CCCO4)c3c2-c2ccccc2)cc1 | XNFHHOXCDUAYSR-SFHVURJKSA-N | No-Effect | inactive |
| AJ76-(+) | BRD-K76677554 | CCCN[C@@H]1CCc2c(OC)cccc2[C@@H]1C | YGHLYBIUVOLKCV-SMDDNHRTSA-N | No-Effect | inactive |
| ajmaline | BRD-K00005347 | CC[C@H]1[C@H]2C[C@H]3[C@@H]4N(C)c5ccccc5[C@]45C[C@@H]([C@H]2[C@H]5O)N3[C@@H]1O | CJDRUOGAGYHKKD-BDXZSHBPSA-N | No-Effect | inactive |
| AK-7 | BRD-K09143580 | O=C(Nc1cccc(Br)c1)c1cccc(S(=O)(=O)N2CCCCCC2)c1 | IYAYHZZWYNXHEQ-UHFFFAOYSA-N | No-Effect | inactive |
| AKBA | BRD-K54907283 | CC(=O)O[C@@H]1CC[C@@]2(C)[C@@H](CC[C@]3(C)[C@@H]2C(=O)C=C2[C@@H]4[C@@H](C)[C@H](C)CC[C@]4(C)CC[C@]23C)[C@@]1(C)C(=O)O | HMMGKOVEOFBCAU-BCDBGHSCSA-N | No-Effect | inactive |
| aklomide | BRD-K26807903 | NC(=O)c1ccc([N+](=O)[O-])cc1Cl | GFGSZUNNBQXGMK-UHFFFAOYSA-N | No-Effect | inactive |
| AL-8697 | BRD-K44731351 | Cc1c(F)cc(C(=O)NC2CC2)cc1-c1c(F)cn2c(C(C)(C)C)nnc2c1F | ZVBTZTQYHOXIBC-UHFFFAOYSA-N | No-Effect | inactive |
| alacepril | BRD-K51784806 | CC(=O)SC[C@@H](C)C(=O)N1CCC[C@H]1C(=O)N[C@@H](Cc1ccccc1)C(=O)O | FHHHOYXPRDYHEZ-COXVUDFISA-N | No-Effect | inactive |
| alafosfalin | BRD-K80403280 | C[C@H](N)C(=O)N[C@@H](C)P(=O)(O)O | BHAYDBSYOBONRV-IUYQGCFVSA-N | No-Effect | inactive |
| alagebrium | BRD-K96029467 | Cc1sc[n+](CC(=O)c2ccccc2)c1C | LYLFQLCLUXOFOL-UHFFFAOYSA-N | No-Effect | inactive |
| alanine | BRD-A33317514 | C[C@@H](N)C(=O)O | QNAYBMKLOCPYGJ-UWTATZPHSA-N | No-Effect | inactive |
| alanosine | BRD-K00004718 | N[C@@H](CN(O)[N+][O-])C(=O)O | MLFKVJCWGUZWNV-REOHCLBHSA-N | No-Effect | inactive |
| alantolactone | BRD-K53731810 | C=C1C(=O)O[C@@H]2C[C@@]3(C)CCC[C@H](C)C3=C[C@H]12 | PXOYOCNNSUAQNS-AGNJHWRGSA-N | No-Effect | inactive |
| alarelin | BRD-K42140276 | CCNC(=O)[C@@H]1CCCN1C(=O)[C@H](CCCNC(=N)N)NC(=O)[C@H](CC(C)C)NC(=O)[C@@H](C)NC(=O)[C@H](Cc1ccc(O)cc1)NC(=O)[C@H](CO)NC(=O)[C@H](Cc1c[nH]c2ccccc12)NC(=O)[C@H](Cc1cnc[nH]1)NC(=O)[C@@H]1CCC(=O)N1 | QLGKMLRGKPKGKI-QWQCQAQZSA-N | No-Effect | inactive |
| albendazole | BRD-K79131256 | CCCSc1ccc2nc(NC(=O)OC)[nH]c2c1 | HXHWSAZORRCQMX-UHFFFAOYSA-N | No-Effect | inactive |
| albendazole-oxide | BRD-A67060370 | CCC[S@@](=O)c1ccc2nc(NC(=O)OC)[nH]c2c1 | VXTGHWHFYNYFFV-LJQANCHMSA-N | No-Effect | inactive |
| albuterol | BRD-K01825118 | CC(C)(C)NC[C@@H](O)c1ccc(O)c(CO)c1 | NDAUXUAQIAJITI-GFCCVEGCSA-N | No-Effect | inactive |
| alcaftadine | BRD-K55991142 | CN1CCC(=C2c3ccccc3CCn3c(C=O)cnc32)CC1 | MWTBKTRZPHJQLH-UHFFFAOYSA-N | No-Effect | inactive |
| alclofenac | BRD-K47492008 | C=CCOc1ccc(CC(=O)O)cc1Cl | ARHWPKZXBHOEEE-UHFFFAOYSA-N | No-Effect | inactive |
| alclometasone-dipropionate | BRD-A17407635 | CCC(=O)OCC(=O)[C@@]1(OC(=O)CC)[C@H](C)C[C@H]2[C@@H]3[C@H]([C@@H](O)C[C@@]21C)[C@@]1(C)C=CC(=O)C=C1C[C@H]3Cl | DJHCCTTVDRAMEH-FQEREKBASA-N | No-Effect | inactive |
| alcuronium | BRD-A84907376 | C=CC[N@@+]12CC[C@@]34c5ccccc5N5/C=C6/[C@H]7C[C@H]8[C@@]9(CC[N@@+]8(CC=C)C/C7=C\CO)c7ccccc7N(/C=C(/[C@@H](C[C@@H]31)/C(=C/CO)C2)[C@@H]54)[C@H]69 | MUQUYTSLDVKIOF-JGNNFNAWSA-N | No-Effect | inactive |
| alda-1 | BRD-K06890060 | O=C(NCc1ccc2c(c1)OCO2)c1c(Cl)cccc1Cl | NMKJFZCBCIUYHI-UHFFFAOYSA-N | No-Effect | inactive |
| aldoxorubicin | BRD-K68928792 | COc1cccc2c1C(=O)c1c(O)c3c(c(O)c1C2=O)C[C@@](O)(/C(CO)=N/NC(=O)CCCCCN1C(=O)C=CC1=O)C[C@@H]3O[C@H]1C[C@H](N)[C@@H](O)[C@H](C)O1 | OBMJQRLIQQTJLR-USGQOSEYSA-N | No-Effect | inactive |
| alendronate | BRD-K75527158 | NCCCC(O)(P(=O)(O)O)P(=O)(O)O | OGSPWJRAVKPPFI-UHFFFAOYSA-N | No-Effect | inactive |
| alfacalcidol | BRD-K49522529 | C=C1/C(=C\C=C2/CCC[C@@]3(C)[C@H]2CC[C@@H]3[C@H](C)CCCC(C)C)C[C@@H](O)C[C@@H]1O | OFHCOWSQAMBJIW-AVJTYSNKSA-N | No-Effect | inactive |
| algestone-acetophenide | BRD-K00005348 | CC(=O)[C@@]12O[C@](C)(c3ccccc3)O[C@@H]1C[C@H]1[C@@H]3CCC4=CC(=O)CC[C@]4(C)[C@H]3CC[C@@]12C | AHBKIEXBQNRDNL-FVCOMRFXSA-N | No-Effect | inactive |
| alibendol | BRD-K60102752 | C=CCc1cc(OC)c(O)c(C(=O)NCCO)c1 | UMJHTFHIQDEGKB-UHFFFAOYSA-N | No-Effect | inactive |
| alimemazine | BRD-K01827280 | C[C@@H](CN(C)C)CN1c2ccccc2Sc2ccccc21 | ZZHLYYDVIOPZBE-AWEZNQCLSA-N | No-Effect | inactive |
| alinidine | BRD-K00004709 | C=CCN(C1=NCCN1)c1c(Cl)cccc1Cl | OXTYVEUAQHPPMV-UHFFFAOYSA-N | No-Effect | inactive |
| alisertib | BRD-K75295174 | COc1cc(Nc2ncc3c(n2)-c2ccc(Cl)cc2C(c2c(F)cccc2OC)=NC3)ccc1C(=O)O | ZLHFILGSQDJULK-UHFFFAOYSA-N | No-Effect | inactive |
| alisporivir | BRD-K00003135 | C/C=C/C[C@@H](C)[C@@H](O)[C@H]1C(=O)N[C@@H](CC)C(=O)N(C)[C@H](C)C(=O)N(CC)[C@@H](C(C)C)C(=O)N[C@@H](C(C)C)C(=O)N(C)[C@@H](CC(C)C)C(=O)N[C@@H](C)C(=O)N[C@H](C)C(=O)N(C)[C@@H](CC(C)C)C(=O)N(C)[C@@H](CC(C)C)C(=O)N(C)[C@@H](C(C)C)C(=O)N1C | OLROWHGDTNFZBH-XEMWPYQTSA-N | No-Effect | inactive |
| alizapride | BRD-K00003607 | C=CCN1CCC[C@@H]1CNC(=O)c1cc2nn[nH]c2cc1OC | KSEYRUGYKHXGFW-LLVKDONJSA-N | No-Effect | inactive |
| alizarin | BRD-K73191876 | O=C1c2ccccc2C(=O)c2c1ccc(O)c2O | RGCKGOZRHPZPFP-UHFFFAOYSA-N | No-Effect | inactive |
| allantoin | BRD-A96407378 | NC(=O)N[C@H]1NC(=O)NC1=O | POJWUDADGALRAB-SFOWXEAESA-N | No-Effect | inactive |
| allicin | BRD-A60402836 | C=CCS[S@@](=O)CC=C | JDLKFOPOAOFWQN-SECBINFHSA-N | No-Effect | inactive |

| | | | | | |
|---|---|---|---|---|---|
| allopurinol | BRD-K86307448 | O=c1nc[nH]c2n[nH]cc12 | OFCNXPDARWKPPY-UHFFFAOYSA-N | No-Effect | inactive |
| allopurinol-riboside | BRD-A93407926-K000 | OC[C@H]1O[C@@H](n2ncc3c(O)ncnc32)[C@H](O)[C@@H]1O.O=c1nc[nH]c2c1cnn2[C@@H]1O[C@H](CO)[C@H](O)[C@@H]1O | DLMWPDAKCTTXOJ-IKZRVYOSSA-N | No-Effect | inactive |
| allylestrenol | BRD-K12872829 | C=CC[C@]1(O)CC[C@H]2[C@@H]3CCC4=CCCC[C@@H]4[C@H]3CC[C@@]21C | ATXHVCQZZJYMCF-XUDSTZEESA-N | No-Effect | inactive |
| allylisothiocyanate | BRD-K33219225 | C=CCN=C=S | ZOJBYZNEUISWFT-UHFFFAOYSA-N | No-Effect | inactive |
| allylthiourea | BRD-K89634775 | C=CCNC(N)=S | HTKFORQRBXIQHD-UHFFFAOYSA-N | No-Effect | inactive |
| almitrine | BRD-K82677201 | C=CCNc1nc(NCC=C)nc(N2CCN(C(c3ccc(F)cc3)c3ccc(F)cc3)CC2)n1 | OBDOVFRMEYHSQB-UHFFFAOYSA-N | No-Effect | inactive |
| almorexant | BRD-K84810405 | CNC(=O)[C@@H](c1ccccc1)N1CCc2cc(OC)c(OC)cc2[C@@H]1CCc1ccc(C(F)(F)F)cc1 | DKMACHNQISHMDN-RPLLCQBOSA-N | No-Effect | inactive |
| almotriptan | BRD-K67601717 | CN(C)CCc1c[nH]c2ccc(CS(=O)(=O)N3CCCC3)cc12 | WKEMJKQOLOHJLZ-UHFFFAOYSA-N | No-Effect | inactive |
| aloe-emodin | BRD-K04851519 | O=C1c2cccc(O)c2C(=O)c2c(O)cc(CO)cc21 | YDQWDHRMZQUTBA-UHFFFAOYSA-N | No-Effect | inactive |
| alogliptin | BRD-K83003151 | Cn1c(=O)cc(N2CCC[C@@H](N)C2)n(Cc2ccccc2C#N)c1=O | ZSBOMTDTBDDKMP-OAHLLOKOSA-N | No-Effect | inactive |
| aloperine | BRD-K96391852 | C1=C2CCCN[C@H]2[C@@H]2C[C@H]1[C@H]1CCCCN1C2 | SKOLRLSBMUGVOY-GBJTYRQASA-N | No-Effect | inactive |
| alosetron | BRD-K46742498 | Cc1nc[nH]c1CN1CCc2c(c3ccccc3n2C)C1=O | JSWZEAMFRNKZNL-UHFFFAOYSA-N | No-Effect | inactive |
| alovudine | BRD-K00004697 | Cc1cn([C@H]2C[C@H](F)[C@@H](CO)O2)c(=O)[nH]c1=O | UXCAQJAQSWSNPQ-XLPZGREQSA-N | No-Effect | inactive |
| alpelisib | BRD-K54997624 | Cc1nc(NC(=O)N2CCC[C@H]2C(N)=O)sc1-c1ccnc(C(C)(C)C(F)(F)F)c1 | STUWGJZDJHPWGZ-LBPRGKRZSA-N | No-Effect | inactive |
| alpha-asarone | BRD-K72536715 | C/C=C/c1cc(OC)c(OC)cc1OC | RKFAZBXYICVSKP-AATRIKPKSA-N | No-Effect | inactive |
| alpha-D-glucopyranose | BRD-K87424134 | OC[C@H]1O[C@H](O)[C@H](O)[C@@H](O)[C@@H]1O | WQZGKKKJIJFFOK-DVKNGEFBSA-N | No-Effect | inactive |
| alpha-glucosyl-hesperidin | BRD-K00005238 | COc1ccc([C@@H]2CC(=O)c3c(O)cc(O[C@@H]4O[C@H](CO[C@@H]5O[C@@H](C)[C@H](O)[C@@H](O)[C@H]5O)[C@@H](O[C@H]5O[C@H](CO)[C@@H](O)[C@H](O)[C@H]5O)[C@H](O)[C@H]4O)cc3O2)cc1O | IMMBLRJLSYJQIZ-RBWYFKFWSA-N | No-Effect | inactive |
| alpha-linolenic-acid | BRD-K33396764 | CC/C=C\C/C=C\C/C=C\CCCCCCCC(=O)O | DTOSIQBPPRVQHS-PDBXOOCHSA-N | No-Effect | inactive |
| alpha-methylhistamine-dihydrobromide-(R)-(-) | BRD-K80028314 | C[C@@H](N)Cc1cnc[nH]1 | XNQIOISZPFVUFG-RXMQYKEDSA-N | No-Effect | inactive |
| alpha-methylhistamine-dihydrobromide-(S)-(+) | BRD-K28308851 | C[C@H](N)Cc1cnc[nH]1 | XNQIOISZPFVUFG-YFKPBYRVSA-N | No-Effect | inactive |
| alpha-methylserotonin | BRD-K01826806 | C[C@@H](N)Cc1c[nH]c2ccc(O)cc12 | LYPCGXKCQDYTFV-SSDOTTSWSA-N | No-Effect | inactive |
| alpha-tochopherol | BRD-A82892199 | Cc1c(C)c2c(c(C)c1O)CC[C@](C)(CCC[C@H](C)CCC[C@H](C)CCCC(C)C)O2 | GVJHHUAWPYXKBD-QLVXXPONSA-N | No-Effect | inactive |
| alpha-tochopheryl-acetate | BRD-A61850409 | CC(=O)Oc1c(C)c(C)c2c(c1C)CC[C@](C)(CCC[C@H](C)CCC[C@H](C)CCCC(C)C)O2 | ZAKOWWREFLAJOT-PKIAKLFDSA-N | No-Effect | inactive |
| alpidem | BRD-K66896751 | CCCN(CCC)C(=O)Cc1c(-c2ccc(Cl)cc2)nc2ccc(Cl)cn12 | JRTIDHTUMYMPRU-UHFFFAOYSA-N | No-Effect | inactive |
| alprenolol | BRD-K01825817 | C=CCc1ccccc1OC[C@H](O)CNC(C)C | PAZJSJFMUHDSTF-CQSZACIVSA-N | No-Effect | inactive |
| alprostadil | BRD-K52459643 | CCCCC[C@H](O)/C=C/[C@H]1[C@H](O)CC(=O)[C@@H]1CCCCCCC(=O)O | GMVPRGQOIOIIMI-DWKJAMRDSA-N | No-Effect | inactive |
| alrestatin | BRD-K35498378 | O=C(O)CN1C(=O)c2cccc3cccc(c23)C1=O | GCUCIFQCGJIRNT-UHFFFAOYSA-N | No-Effect | inactive |
| ALS-8112 | BRD-K21415491 | Nc1ccn([C@@H]2O[C@@](CO)(CCl)[C@@H](O)[C@H]2F)c(=O)n1 | AWSRKKBIPSQHOJ-IBCQBUCCSA-N | No-Effect | inactive |
| ALS-8176 | BRD-K00004605 | CC(C)C(=O)OC[C@@]1(CCl)O[C@@H](n2ccc(N)nc2=O)[C@H](F)[C@@H]1OC(=O)C(C)C | MJVKYGMNSQJLIN-KYZVSKTDSA-N | No-Effect | inactive |
| altanserin | BRD-K00610438 | O=C(c1ccc(F)cc1)C1CCN(CCn2c(=S)[nH]c3ccccc3c2=O)CC1 | SMYALUSCZJXWHG-UHFFFAOYSA-N | No-Effect | inactive |
| althiazide | BRD-A56675431 | C=CCSC[C@@H]1Nc2cc(Cl)c(S(N)(=O)=O)cc2S(=O)(=O)N1 | VGLGVJVUHYTIIU-LLVKDONJSA-N | No-Effect | inactive |
| altinicline | BRD-K52681925 | C#Cc1cncc([C@@H]2CCCN2C)c1 | NUPUDYKEEJNZRG-LBPRGKRZSA-N | No-Effect | inactive |
| altiratinib | BRD-K78627070 | O=C(Nc1cc(Oc2cc(F)c(NC(=O)C3(C(=O)Nc4ccc(F)cc4)CC3)cc2F)ccn1)C1CC1 | GNNDEPIMDAZHRQ-UHFFFAOYSA-N | No-Effect | inactive |
| altrenogest | BRD-K50276048 | C=CC[C@]1(O)CC[C@H]2[C@@H]3CCC4=CC(=O)CCC4=C3C=C[C@@]21C | VWAUPFMBXBWEQY-ANULTFPQSA-N | No-Effect | inactive |
| alverine | BRD-K89055274 | CCN(CCCc1ccccc1)CCCc1ccccc1 | ZPFXAOWNKLFJDN-UHFFFAOYSA-N | No-Effect | inactive |
| alvimopan | BRD-K07857022 | C[C@H]1CN(C[C@H](Cc2ccccc2)C(=O)NCC(=O)O)CC[C@@]1(C)c1cccc(O)c1 | UPNUIXSCZBYVBB-JVFUWBCBSA-N | No-Effect | inactive |
| ALX-40-4C | BRD-K00003337 | CC(=O)N[C@H](CCCNC(=N)N)C(=O)N[C@H](CCCNC(=N)N)C(=O)N[C@H](CCCNC(=N)N)C(=O)N[C@H](CCCNC(=N)N)C(=O)N[C@H](CCCNC(=N)N)C(=O)N[C@H](CCCNC(=N)N)C(=O)N[C@H](CCCNC(=N)N)C(=O)N[C@H](CCCNC(=N)N)C(=O)N[C@H](CCCNC(=N)N)C(N)=O | LGLVVVCSQBZONM-HCCLCSBVSA-N | No-Effect | inactive |
| ALX-5407 | BRD-K53979406 | CN(CC[C@@H](Oc1ccc(-c2ccccc2)cc1)c1ccc(F)cc1)CC(=O)O | FDORQEIHOKEJNX-HSZRJFAPSA-N | No-Effect | inactive |
| AM-24 | BRD-K59013864 | Oc1c(I)cc(I)cc1I | VAPDZNUFNKUROY-UHFFFAOYSA-N | No-Effect | inactive |
| AM-251 | BRD-K92000912 | Cc1c(C(=O)NN2CCCCC2)nn(-c2ccc(Cl)cc2Cl)c1-c1ccc(I)cc1 | BUZAJRPLUGXRAB-UHFFFAOYSA-N | No-Effect | inactive |
| AM-404 | BRD-K55462201 | CCCCC/C=C\C/C=C\C/C=C\C/C=C\CCCC(=O)Nc1ccc(O)cc1 | IJBZOOZRAXHERC-DOFZRALJSA-N | No-Effect | inactive |
| AM-580 | BRD-K06854232 | CC1(C)CCC(C)(C)c2cc(C(=O)Nc3ccc(C(=O)O)cc3)ccc21 | SZWKGOZKRMMLAJ-UHFFFAOYSA-N | No-Effect | inactive |
| AM-630 | BRD-K63533170 | COc1ccc(C(=O)c2c(C)n(CCN3CCOCC3)c3cc(I)ccc23)cc1 | JHOTYHDSLIUKCJ-UHFFFAOYSA-N | No-Effect | inactive |
| AM-92016 | BRD-K01826811 | CN(CCc1ccc(Cl)c(Cl)c1)C[C@@H](O)COc1ccc(NS(C)(=O)=O)cc1 | BQBSHJUQVIWEFM-MRXNPFEDSA-N | No-Effect | inactive |
| AM095 | BRD-K23952234 | Cc1noc(-c2ccc(-c3ccc(CC(=O)O)cc3)cc2)c1NC(=O)O[C@H](C)c1ccccc1 | LNDDRUPAICPXIN-GOSISDBHSA-N | No-Effect | inactive |
| AM211 | BRD-K00003152 | CCN(Cc1cc(C(F)(F)F)ccc1-c1cc(CC(=O)O)ccc1OC)C(=O)NCc1ccccc1 | OPXIRFWNLBDKQB-UHFFFAOYSA-N | No-Effect | inactive |
| AM679 | BRD-K88443722 | COc1cnc(-c2ccc(Cn3c(CC(C)(C)C(=O)O)c(SC(C)(C)C)c4cc(OC[C@@H]5Cc6ccccc6N5C(C)=O)ccc43)cc2)nc1 | VYXWHVDEWWHDLH-LJAQVGFWSA-N | No-Effect | inactive |
| amantadine | BRD-K70330367 | NC12CC3CC(CC(C3)C1)C2 | DKNWSYNQZKUICI-UHFFFAOYSA-N | No-Effect | inactive |
| ambazone | BRD-K11244467 | N=C(N)N=Nc1ccc(NNC(N)=S)cc1 | HQDSAVMTLKVTGJ-UHFFFAOYSA-N | No-Effect | inactive |
| ambenonium | BRD-K11292736 | CC[N+](CC)(CCNC(=O)C(=O)NCC[N+](CC)(CC)Cc1ccccc1Cl)Cc1ccccc1Cl | OMHBPUNFVFNHJK-UHFFFAOYSA-P | No-Effect | inactive |
| ambrisentan | BRD-K41185545 | COC(c1ccccc1)(c1ccccc1)[C@H](Oc1nc(C)cc(C)n1)C(=O)O | OUJTZYPIHDYQMC-LJQANCHMSA-N | No-Effect | inactive |
| ambroxol | BRD-K56558538 | Nc1c(Br)cc(Br)cc1CN[C@H]1CC[C@H](O)CC1 | JBDGDEWWOUBZPM-XYPYZODXSA-N | No-Effect | inactive |
| amcasertib | BRD-K00003244 | CCN(CC)CCNC(=O)c1c(C)[nH]c(/C=C2/C(=O)Nc3ccc(-c4csc(-c5ccccc5)n4)cc32)c1C | QDWKGEFGLQMDAM-JJIBRWJFSA-N | No-Effect | inactive |
| AMD-3465 | BRD-K03019184 | c1ccc(CNCc2ccc(CN3CCCNCCNCCCNCC3)cc2)nc1 | CWJJHESJXJQCJA-UHFFFAOYSA-N | No-Effect | inactive |
| AMD11070 | BRD-K42948882 | NCCCCN(Cc1nc2ccccc2[nH]1)[C@H]1CCCc2cccnc21 | WVLHHLRVNDMIAR-IBGZPJMESA-N | No-Effect | inactive |
| amdinocillin | BRD-K41051431 | CC1(C)S[C@@H]2[C@H](/N=C/N3CCCCCC3)C(=O)N2[C@H]1C(=O)O | BWWVAEOLVKTZFQ-ISVUSNJMSA-N | No-Effect | inactive |

| | | | | | |
|---|---|---|---|---|---|
| ameltolide | BRD-K48173020 | Cc1cccc(C)c1NC(=O)c1ccc(N)cc1 | HZIWGOAXOBPQGY-UHFFFAOYSA-N | No-Effect | inactive |
| amenamevir | BRD-K00004966 | Cc1cccc(C)c1N(CC(=O)Nc1ccc(-c2ncon2)cc1)C(=O)C1CCS(=O)(=O)CC1 | MNHNIVNAFBSLLX-UHFFFAOYSA-N | No-Effect | inactive |
| ametantrone | BRD-K00004726 | O=C1c2ccccc2C(=O)c2c(NCCNCCO)ccc(NCCNCCO)c21 | FFGSXKJJVBXWCY-UHFFFAOYSA-N | No-Effect | inactive |
| amezinium | BRD-K51387999 | COc1cc(N)cn[n+]1-c1ccccc1 | VXROHTDSRBRJLN-UHFFFAOYSA-O | No-Effect | inactive |
| amfenac | BRD-K82122102 | Nc1c(CC(=O)O)cccc1C(=O)c1ccccc1 | SOYCMDCMZDHQFP-UHFFFAOYSA-N | No-Effect | inactive |
| amflutizole | BRD-K50813454 | Nc1c(-c2cccc(C(F)(F)F)c2)nsc1C(=O)O | KVMCEGAWQYTFKC-UHFFFAOYSA-N | No-Effect | inactive |
| AMG-176 | BRD-K43900408 | CO[C@H]1/C=C/C[C@H](C)[C@@H](C)S(=O)(=O)NC(=O)c2ccc3c(c2)N(C[C@@H]2CC[C@H]21)C[C@@]1(CCCc2cc(Cl)ccc21)CO3 | JQNINBDKGLWYMU-GEAQBIRJSA-N | No-Effect | inactive |
| AMG-232 | BRD-K64925568 | CC(C)[C@@H](CS(=O)(=O)C(C)C)N1C(=O)[C@@](C)(CC(=O)O)C[C@H](c2cccc(Cl)c2)[C@H]1c1ccc(Cl)cc1 | DRLCSJFKKILATL-YWCVFVGNSA-N | No-Effect | inactive |
| AMG-319 | BRD-K21077415 | C[C@H](Nc1ncnc2nc[nH]c12)c1cc2ccc(F)cc2nc1-c1ccccn1 | KWRYMZHCQIOOEB-LBPRGKRZSA-N | No-Effect | inactive |
| AMG-337 | BRD-K27061362 | COCCOc1cnc2ccn([C@H](C)c3nnc4c(F)cc(-c5cnn(C)c5)cn34)c(=O)c2c1 | DWHXUGDWKAIASB-CQSZACIVSA-N | No-Effect | inactive |
| AMG-487 | BRD-K90239174 | CCOc1ccc(-n2c([C@@H](C)N(Cc3cccnc3)C(=O)Cc3ccc(OC(F)(F)F)cc3)nc3ncccc3c2=O)cc1 | WQTKNBPCJKRYPA-OAQYLSRUSA-N | No-Effect | inactive |
| AMG-487-(+/-) | BRD-A01307728 | CCOc1ccc(-n2c([C@@H](C)N(Cc3cccnc3)C(=O)Cc3ccc(OC(F)(F)F)cc3)nc3ncccc3c2=O)cc1 | WQTKNBPCJKRYPA-OAQYLSRUSA-N | No-Effect | inactive |
| AMG-510 | BRD-K00010385 | C=CC(=O)N1CCN(c2nc(=O)n(-c3c(C)ccnc3C(C)C)c3nc(-c4c(O)cccc4F)c(F)cc23)[C@@H](C)C1 | NXQKSXLFSAEQCZ-SFHVURJKSA-N | No-Effect | inactive |
| AMG-517 | BRD-K34185671 | CC(=O)Nc1nc2c(Oc3cc(-c4ccc(C(F)(F)F)cc4)ncn3)cccc2s1 | YUTIXVXZQIQWGY-UHFFFAOYSA-N | No-Effect | inactive |
| AMG-548 | BRD-K86673452 | Cn1c(NC[C@@H](N)Cc2ccccc2)nc(-c2ccncc2)c(-c2ccc3ccccc3c2)c1=O | RQVKVJIRFKVPBF-VWLOTQADSA-N | No-Effect | inactive |
| AMG-837 | BRD-K43802723 | CC#C[C@@H](CC(=O)O)c1ccc(OCc2cccc(-c3ccc(C(F)(F)F)cc3)c2)cc1 | ZOPNBMMVVZRSGH-NRFANRHFSA-N | No-Effect | inactive |
| AMG-925 | BRD-K97452254 | C[C@H]1CC[C@H](n2c3cnccc3c3cnc(Nc4ccc5c(n4)CCN(C(=O)CO)C5)nc32)CC1 | BBUVDDPUURMFOX-SAABIXHNSA-N | No-Effect | inactive |
| AMG-9810 | BRD-K01995783 | CC(C)(C)c1ccc(/C=C/C(=O)Nc2ccc3c(c2)OCCO3)cc1 | GZTFUVZVLYUPRG-IZZDOVSWSA-N | No-Effect | inactive |
| AMG319 | BRD-A58947127 | C[C@@H](Nc1ncnc2nc[nH]c12)c1cc2ccc(F)cc2nc1-c1ccccn1 | KWRYMZHCQIOOEB-GFCCVEGCSA-N | No-Effect | inactive |
| AMG458 | BRD-K21782625 | COc1ccc2c(Oc3ccc(NC(=O)c4c(C)n(CC(C)(C)O)n(-c5ccccc5)c4=O)nc3)ccnc2c1 | GLBZSOQDAOLMGC-UHFFFAOYSA-N | No-Effect | inactive |
| AMG900 | BRD-K21728777 | Cc1csc(-c2nnc(Nc3ccc(Oc4ncccc4-c4ccnc(N)n4)cc3)c3ccccc23)c1 | IVUGFMLRJOCGAS-UHFFFAOYSA-N | No-Effect | inactive |
| AMI-1 | BRD-K75114721 | O=C(Nc1ccc2c(O)cc(S(=O)(=O)O)cc2c1)Nc1ccc2c(O)cc(S(=O)(=O)O)cc2c1 | PCGISRHGYLRXSR-UHFFFAOYSA-N | No-Effect | inactive |
| amibegron | BRD-K24610819 | CCOC(=O)COc1ccc2c(c1)C[C@@H](NC[C@H](O)c1cccc(Cl)c1)CC2 | RDJQCOBTKKAQAH-FPOVZHCZSA-N | No-Effect | inactive |
| amidopyrine | BRD-K12568846 | Cc1c(N(C)C)c(=O)n(-c2ccccc2)n1C | RMMXTBMQSGEXHJ-UHFFFAOYSA-N | No-Effect | inactive |
| amifampridine | BRD-K80262108 | Nc1ccncc1N | OYTKINVCDFNREN-UHFFFAOYSA-N | No-Effect | inactive |
| amiflamine | BRD-A19401019 | Cc1cc(N(C)C)ccc1C[C@@H](C)N | HFQMYSHATTXRTC-SNVBAGLBSA-N | No-Effect | inactive |
| amifostine | BRD-K73947551 | NCCCNCCSP(=O)(O)O | JKOQGQFVAUAYPM-UHFFFAOYSA-N | No-Effect | inactive |
| amikacin | BRD-K88043978 | NCC[C@H](O)C(=O)N[C@@H]1C[C@H](N)[C@@H](O[C@H]2O[C@H](CN)[C@@H](O)[C@H](O)[C@H]2O)[C@H](O)[C@H]1O[C@H]1O[C@H](CO)[C@@H](O)[C@H](N)[C@H]1O | LKCWBDHBTVXHDL-RMDFUYIESA-N | No-Effect | inactive |
| amiloride | BRD-K97181089 | N=C(N)NC(=O)c1nc(Cl)c(N)nc1N | XSDQTOBWRPYKKA-UHFFFAOYSA-N | No-Effect | inactive |
| aminocaproic-acid | BRD-K64931368 | NCCCCCC(=O)O | SLXKOJJOQWFEFD-UHFFFAOYSA-N | No-Effect | inactive |
| aminoglutethimide | BRD-A25234499 | CC[C@@]1(c2ccc(N)cc2)CCC(=O)NC1=O | ROBVIMPUHSLWNV-ZDUSSCGKSA-N | No-Effect | inactive |
| aminoguanidine | BRD-K25114078 | NN=C(N)N | HAMNKKUPIHEESI-UHFFFAOYSA-N | No-Effect | inactive |
| aminohydroxybutyric-acid | BRD-A40777960 | NC[C@@H](O)CC(=O)O | YQGDEPYYFWUPGO-VKHMYHEASA-N | No-Effect | inactive |
| aminolevulinic-acid-benzyl-ester | BRD-K59998349 | NCC(=O)CCC(=O)OCc1ccccc1 | AFFZCQDLQIEQGB-UHFFFAOYSA-N | No-Effect | inactive |
| aminomethyltransferase | BRD-A28318179 | C[C@H]1CCN=C(N)S1 | BATVOUKHGLKDGQ-BYPYZUCNSA-N | No-Effect | inactive |
| aminophylline | BRD-K97799481 | Cn1c(=O)c2[nH]cnc2n(C)c1=O | ZFXYFBGIUFBOJW-UHFFFAOYSA-N | No-Effect | inactive |
| aminopterin | BRD-K87544030 | Nc1nc(N)c2nc(CNc3ccc(C(=O)N[C@@H](CCC(=O)O)C(=O)O)cc3)cnc2n1 | TVZGACDUOSZQKY-LBPRGKRZSA-N | No-Effect | inactive |
| aminopurvalanol-a | BRD-K07762753 | CC(C)[C@H](CO)Nc1nc(Nc2cc(N)cc(Cl)c2)c2ncn(C(C)C)c2n1 | RAMROQQYRRQPDL-HNNXBMFYSA-N | No-Effect | inactive |
| aminothiadiazole | BRD-K94837153 | Nc1nncs1 | QUKGLNCXGVWCJX-UHFFFAOYSA-N | No-Effect | inactive |
| amiprilose | BRD-K97643133 | CN(C)CCCO[C@H]1[C@@H]([C@H](O)CO)O[C@@H]2OC(C)(C)O[C@H]12 | YXBQLONCIPUQKO-UJPOAAIJSA-N | No-Effect | inactive |
| amiselimod | BRD-K00003223 | CCCCCCCOc1ccc(CCC(N)(CO)CO)cc1C(F)(F)F | JVCPIJKPAKAIIP-UHFFFAOYSA-N | No-Effect | inactive |
| amisulpride | BRD-A60197193 | CCN1CCC[C@@H]1CNC(=O)c1cc(S(=O)(=O)CC)c(N)cc1OC | NTJOBXMMWNYJFB-GFCCVEGCSA-N | No-Effect | inactive |
| amitifadine | BRD-A68942014 | Clc1ccc([C@]23CNC[C@H]2C3)cc1Cl | BSMNRYCSBFHEMQ-KCJUWKMLSA-N | No-Effect | inactive |
| amitraz | BRD-K33335792 | Cc1ccc(/N=C/N(C)/C=N/c2ccc(C)cc2C)c(C)c1 | QXAITBQSYVNQDR-ZIOPAAQOSA-N | No-Effect | inactive |
| amlexanox | BRD-K29530284 | CC(C)c1ccc2oc3nc(N)c(C(=O)O)cc3c(=O)c2c1 | SGRYPYWGNKJSDL-UHFFFAOYSA-N | No-Effect | inactive |
| amlodipine | BRD-K01825117 | CCOC(=O)C1=C(COCCN)NC(C)=C(C(=O)OC)[C@H]1c1ccccc1Cl | HTIQEAQVCYTUBX-QGZVFWFLSA-N | No-Effect | inactive |
| ammonium-lactate | BRD-M29182745 | C[C@@H](O)C(=O)O.N | RZOBLYBZQXQGFY-HSHFZTNMSA-N | No-Effect | inactive |
| ammonium-perfluorocaprylate | BRD-K32531356 | O=C(O)C(F)(F)C(F)(F)C(F)(F)C(F)(F)C(F)(F)C(F)(F)C(F)(F)F | SNGREZUHAYWORS-UHFFFAOYSA-N | No-Effect | inactive |
| AMN-082 | BRD-K23335153 | c1ccc(C(NCCNC(c2ccccc2)c2ccccc2)c2ccccc2)cc1 | DTZDSNQYNPNCPK-UHFFFAOYSA-N | No-Effect | inactive |
| amonafide | BRD-K56334280 | CN(C)CCN1C(=O)c2cccc3cc(N)cc(c23)C1=O | UPALIKSFLSVKIS-UHFFFAOYSA-N | No-Effect | inactive |
| amorolfine | BRD-A37390343 | CCC(C)(C)c1ccc(C[C@@H](C)CN2C[C@H](C)O[C@H](C)C2)cc1 | MQHLMHIZUIDKOO-KURKYZTESA-N | No-Effect | inactive |
| amoxapine | BRD-K02265150 | Clc1ccc2c(c1)C(N1CCNCC1)=Nc1ccccc1O2 | QWGDMFLQWFTERH-UHFFFAOYSA-N | No-Effect | inactive |
| amoxicillin | BRD-K55044200 | CC1(C)S[C@@H]2[C@H](NC(=O)[C@H](N)c3ccc(O)cc3)C(=O)N2[C@H]1C(=O)O | LSQZJLSUYDQPKJ-NJBDSQKTSA-N | No-Effect | inactive |
| AMPA-(RS) | BRD-K01826817 | Cc1o[nH]c(=O)c1C[C@@H](N)C(=O)O | UUDAMDVQRQNNHZ-RXMQYKEDSA-N | No-Effect | inactive |
| AMPA-(S) | BRD-K92493880 | Cc1o[nH]c(=O)c1C[C@H](N)C(=O)O | UUDAMDVQRQNNHZ-YFKPBYRVSA-N | No-Effect | inactive |
| ampalex | BRD-K24771047 | O=C(c1ccc2nccnc2c1)N1CCCCC1 | ANDGGVOPIJEHOF-UHFFFAOYSA-N | No-Effect | inactive |
| amperozide | BRD-K52397688 | CCNC(=O)N1CCN(CCCC(c2ccc(F)cc2)c2ccc(F)cc2)CC1 | NNAIYOXJNVGUOM-UHFFFAOYSA-N | No-Effect | inactive |
| amprenavir | BRD-K47827687 | CC(C)CN(C[C@@H](O)[C@H](Cc1ccccc1)NC(=O)O[C@H]1CCOC1)S(=O)(=O)c1ccc(N)cc1 | YMARZQAQMVYCKC-OEMFJLHTSA-N | No-Effect | inactive |
| amprolium | BRD-K90027121 | CCCc1ncc(C[n+]2ccccc2C)c(N)n1 | IPZFPROOBOUEIG-UHFFFAOYSA-N | No-Effect | inactive |
| ampyrone | BRD-K20484910 | Cc1c(N)c(=O)n(-c2ccccc2)n1C | RLFWWDJHLFCNIJ-UHFFFAOYSA-N | No-Effect | inactive |
| amrubicin | BRD-K13843533 | CC(=O)[C@]1(N)Cc2c(O)c3c(c(O)c2[C@@H](O[C@H]2C[C@H](O)[C@@H](O)CO2)C1)C(=O)c1ccccc1C3=O | VJZITPJGSQKZMX-XDPRQOKASA-N | No-Effect | inactive |
| amsacrine | BRD-K68346641 | COc1cc(NS(C)(=O)=O)ccc1Nc1c2ccccc2nc2ccccc12 | XCPGHVQEEXUHNC-UHFFFAOYSA-N | No-Effect | inactive |
| amthamine | BRD-K39670393 | Cc1nc(N)sc1CCN | LHVRFUVVRXGZPV-UHFFFAOYSA-N | No-Effect | inactive |
| amtolmetin-guacil | BRD-K70924353 | COc1ccccc1OC(=O)CNC(=O)Cc1ccc(C(=O)c2ccc(C)cc2)n1C | CWJNMKKMGIAGDK-UHFFFAOYSA-N | No-Effect | inactive |

| | | | | | |
|---|---|---|---|---|---|
| amuvatinib | BRD-K66538826 | S=C(NCc1ccc2c(c1)OCO2)N1CCN(c2ncnc3c2oc2ccccc23)CC1 | FOFDIMHVKGYHRU-UHFFFAOYSA-N | No-Effect | inactive |
| amygdalin | BRD-K03504028 | N#C[C@H](O[C@@H]1O[C@H](CO[C@@H]2O[C@H](CO)[C@@H](O)[C@H](O)[C@H]2O)[C@@H](O)[C@H](O)[C@H]1O)c1ccccc1 | XUCIJNAGGSZNQT-JHSLDZJXSA-N | No-Effect | inactive |
| amyl-nitrite | BRD-K03326591 | CCCCCO[NH2+][O-] | GVGUCRAJNQYOSE-UHFFFAOYSA-N | No-Effect | inactive |
| amyleine | BRD-A09062839 | CC[C@@](C)(CN(C)C)OC(=O)c1ccccc1 | FDMBBCOBEAVDAO-AWEZNQCLSA-N | No-Effect | inactive |
| amylene-hydrate | BRD-K93502141 | CCC(C)(C)O | MSXVEPNJUHWQHW-UHFFFAOYSA-N | No-Effect | inactive |
| AMZ30 | BRD-K00256256 | N#C/C(=C\c1cccn1S(=O)(=O)c1cccc([N+](=O)[O-])c1)S(=O)(=O)c1ccc(F)cc1 | GUCUORSUHTZMBW-YBFXNURJSA-N | No-Effect | inactive |
| AN2718 | BRD-K21920947 | OB1OCc2cc(Cl)ccc21 | HMAFTPZYFJFEHK-UHFFFAOYSA-N | No-Effect | inactive |
| ANA-12 | BRD-A19609491 | O=C(Nc1ccccc1C(=O)N[C@H]1CCCCNC1=O)c1cc2ccccc2s1 | TUSCYCAIGRVBMD-KRWDZBQOSA-N | No-Effect | inactive |
| anabasine | BRD-K01826509 | c1cncc([C@H]2CCCCN2)c1 | MTXSIJUGVMTTMU-SNVBAGLBSA-N | No-Effect | inactive |
| anacetrapib | BRD-K77118600 | COc1cc(F)c(C(C)C)cc1-c1ccc(C(F)(F)F)cc1CN1C(=O)O[C@H](c2cc(C(F)(F)F)cc(C(F)(F)F)c2)[C@@H]1C | MZZLGJHLQGUVPN-HAWMADMCSA-N | No-Effect | inactive |
| anagliptin | BRD-K05355328 | Cc1cc2ncc(C(=O)NCC(C)(C)NCC(=O)N3CCC[C@H]3C#N)cn2n1 | LDXYBEHACFJIEL-HNNXBMFYSA-N | No-Effect | inactive |
| anagrelide | BRD-K62200014 | O=C1CN2Cc3c(ccc(Cl)c3Cl)N=C2N1 | OTBXOEAOVRKTNQ-UHFFFAOYSA-N | No-Effect | inactive |
| anamorelin | BRD-K01174345 | CN(C)N(C)C(=O)[C@@]1(Cc2ccccc2)CCCN(C(=O)[C@@H](Cc2c[nH]c3ccccc23)NC(=O)C(C)(C)N)C1 | VQPFSIRUEPQQPP-MXBOTTGLSA-N | No-Effect | inactive |
| anandamide | BRD-K42352790 | CCCCC/C=C\C/C=C\C/C=C\C/C=C\CCCC(=O)NCCO | LGEQQWMQCRIYKG-DOFZRALJSA-N | No-Effect | inactive |
| anastrozole | BRD-K52172416 | CC(C)(C#N)c1cc(Cn2cncn2)cc(C(C)(C)C#N)c1 | YBBLVLTVTVSKRW-UHFFFAOYSA-N | No-Effect | inactive |
| andarine | BRD-K82164249 | CC(=O)Nc1ccc(OC[C@](C)(O)C(=O)Nc2ccc([N+](=O)[O-])c(C(F)(F)F)c2)cc1 | YVXVTLGIDOACBJ-SFHVURJKSA-N | No-Effect | inactive |
| andrographolide | BRD-K89282837 | C=C1CC[C@H]2[C@@](C)(CC[C@@H](O)[C@@]2(C)CO)[C@@H]1C/C=C1/C(=O)OC[C@H]1O | BOJKULTULYSRAS-OTESTREVSA-N | No-Effect | inactive |
| androstenone | BRD-A64011898 | C[C@]12C=CC[C@H]1[C@@H]1CC[C@H]3CC(=O)CC[C@@]3(C)[C@H]1CC2 | HFVMLYAGWXSTQI-UGRKVPRCSA-N | No-Effect | inactive |
| androsterone | BRD-K12329651 | C[C@]12CC[C@H]3[C@@H](CC[C@H]4C[C@H](O)CC[C@@]43C)[C@@H]1CCC2=O | QGXBDMJGAMFCBF-HLUDHZFRSA-N | No-Effect | inactive |
| anecortave-acetate | BRD-K00005346 | CC(=O)OCC(=O)[C@@]1(O)CC[C@H]2[C@@H]3CCC4=CC(=O)CC[C@]4(C)C3=CC[C@@]21C | YUWPMEXLKGOSBF-GACAOOTBSA-N | No-Effect | inactive |
| anethole | BRD-K49060658 | C/C=C/c1ccc(OC)cc1 | RUVINXPYWBROJD-ONEGZZNKSA-N | No-Effect | inactive |
| anethole-trithione | BRD-K92490248 | COc1ccc(-c2cc(=S)ss2)cc1 | KYLIZBIRMBGUOP-UHFFFAOYSA-N | No-Effect | inactive |
| anidulafungin | BRD-K00003560 | CCCCCOc1ccc(-c2ccc(-c3ccc(C(=O)N[C@H]4C[C@@H](O)[C@@H](O)NC(=O)[C@@H]5[C@@H](O)[C@@H](C)CN5C(=O)[C@H]([C@@H](C)O)NC(=O)[C@H]([C@H](O)[C@@H](O)c5ccc(O)cc5)NC(=O)[C@@H]5C[C@@H](O)CN5C(=O)[C@H]([C@@H](C)O)NC4=O)cc3)cc2)cc1 | JHVAMHSQVVQIOT-MFAJLEFUSA-N | No-Effect | inactive |
| aniracetam | BRD-K88611939 | COc1ccc(C(=O)N2CCCC2=O)cc1 | ZXNRTKGTQJPIJK-UHFFFAOYSA-N | No-Effect | inactive |
| anisindione | BRD-K47458712 | COc1ccc(C2C(=O)c3ccccc3C2=O)cc1 | XRCFXMGQEVUZFC-UHFFFAOYSA-N | No-Effect | inactive |
| anisotropine | BRD-K71125570 | CCCC(CCC)C(=O)O[C@@H]1C[C@@H]2CC[C@H](C1)[N+]2(C)C | XGGHHHBGPSNXFE-ZSHCYNCHSA-N | No-Effect | inactive |
| anlotinib | BRD-K36949735 | COc1cc2c(Oc3ccc4[nH]c(C)cc4c3F)ccnc2cc1OCC1(N)CC1 | KSMZEXLVHXZPEF-UHFFFAOYSA-N | No-Effect | inactive |
| anpirtoline | BRD-K55424922 | Clc1cccc(SC2CCNCC2)n1 | GGALEXMXDMUMDM-UHFFFAOYSA-N | No-Effect | inactive |
| ANR-94 | BRD-K31170746 | CCOc1nc2c(N)ncnc2n1CC | QUGDTMONBLMLLD-UHFFFAOYSA-N | No-Effect | inactive |
| antagonist-g | BRD-K25863791 | CSCC[C@H](NC(=O)[C@H](CC(C)C)NC(=O)[C@@H](Cc1c[nH]c2ccccc12)NC(=O)[C@H](Cc1ccccc1)N(C)C(=O)[C@@H](Cc1c[nH]c2ccccc12)NC(=O)[C@@H](N)CCCNC(=N)N)C(N)=O | CUCSSYAUKKIDJV-FAXBSAIASA-N | No-Effect | inactive |
| antalarmin | BRD-K01363385 | CCCCN(CC)c1nc(C)nc2c1c(C)c(C)n2-c1c(C)cc(C)cc1C | IXPROWGEHNVJOY-UHFFFAOYSA-N | No-Effect | inactive |
| antazoline | BRD-K99300445 | c1ccc(CN(CC2=NCCN2)c2ccccc2)cc1 | REYFJDPCWQRWAA-UHFFFAOYSA-N | No-Effect | inactive |
| anthraquinone | BRD-K15957397 | O=C1c2ccccc2C(=O)c2ccccc21 | RZVHIXYEVGDQDX-UHFFFAOYSA-N | No-Effect | inactive |
| antimonyl | BRD-A74328360 | O=C1O[SbH]23OC(=O)C(O2)C2O[SbH]4(OC(=O)C(O4)C1O3)OC2=O | ZCIXFSVENQDMCK-UHFFFAOYSA-J | No-Effect | inactive |
| antimycin-A | BRD-K90152265 | CCCCCC[C@H]1C(=O)O[C@H](C)[C@H](NC(=O)c2cccc(NC=O)c2O)C(=O)O[C@@H](C)[C@@H]1OC(=O)CC(C)C | UIFFUZWRFRDZJC-SBOOETFBSA-N | No-Effect | inactive |
| antioxine | BRD-K15541734 | Cc1ccc(C(C)C)cc1O | RECUKUPTGUEGMW-UHFFFAOYSA-N | No-Effect | inactive |
| AP-18 | BRD-K32143970 | CC(=C\c1ccc(Cl)cc1)/C(C)=N/O | MHTJEUOFLVQMCL-NJHPPEEMSA-N | No-Effect | inactive |
| AP1903 | BRD-K28383148 | CC[C@H](C(=O)N1CCCC[C@H]1C(=O)O[C@H](CCc1ccc(OC)c(OC)c1)c1cccc(OCC(=O)NCCNC(=O)COc2cccc([C@@H](CCc3ccc(OC)c(OC)c3)OC(=O)[C@@H]3CCCCN3C(=O)[C@@H](CC)c3cc(OC)c(OC)c(OC)c3)c2)c1)c1cc(OC)c(OC)c(OC)c1 | GQLCLPLEEOUJQC-ZTQDTCGGSA-N | No-Effect | inactive |
| AP26113 | BRD-K56032964 | COc1cc(N2CCC(N(C)C)CC2)ccc1Nc1ncc(Cl)c(Nc2ccccc2P(C)(C)=O)n1 | OVDSPTSBIQCAIN-UHFFFAOYSA-N | No-Effect | inactive |
| apabetalone | BRD-K20001883 | COc1cc(OC)c2c(=O)nc(-c3cc(C)c(OCCO)c(C)c3)[nH]c2c1 | NETXMUIMUZJUTB-UHFFFAOYSA-N | No-Effect | inactive |
| apafant | BRD-K28115081 | Cc1nnc2n1-c1sc(CCC(=O)N3CCOCC3)cc1C(c1ccccc1Cl)=NC2 | JGPJQFOROWSRRS-UHFFFAOYSA-N | No-Effect | inactive |
| apalutamide | BRD-K91308639 | CNC(=O)c1ccc(N2C(=S)N(c3cnc(C#N)c(C(F)(F)F)c3)C(=O)C23CCC3)cc1F | HJBWBFZLDZWPHF-UHFFFAOYSA-N | No-Effect | inactive |
| apararenone | BRD-K00003492 | CC1(C)Oc2cc(NS(C)(=O)=O)ccc2N(c2ccc(F)cc2)C1=O | AZNHWXAFPBYFGH-UHFFFAOYSA-N | No-Effect | inactive |
| apatinib | BRD-K25244359 | N#CC1(c2ccc(NC(=O)c3cccnc3NCc3ccncc3)cc2)CCCC1 | WPEWQEMJFLWMLV-UHFFFAOYSA-N | No-Effect | inactive |
| APC-100 | BRD-K07449869 | Cc1c(C)c2c(c(C)c1O)CCC(C)(C)O2 | SEBPXHSZHLFWRL-UHFFFAOYSA-N | No-Effect | inactive |
| APcK-110 | BRD-K72748448 | COc1cc(OC)cc(-c2ccc3c(-c4ccc(F)cc4)[nH]nc3n2)c1 | CAMGZGMEPKFVMF-UHFFFAOYSA-N | No-Effect | inactive |
| APD597 | BRD-K86226872 | COc1c(Nc2ccc(S(C)(=O)=O)nc2C)ncnc1OC1CCN(C(=O)OC(C)C)CC1 | WPDCHTSXOPUOII-UHFFFAOYSA-N | No-Effect | inactive |
| APD668 | BRD-K10859802 | CC(C)OC(=O)N1CCC(Oc2ncnc3c2cnn3-c2ccc(S(C)(=O)=O)cc2F)CC1 | XTRUQJBVQBUKSQ-UHFFFAOYSA-N | No-Effect | inactive |
| aphidicolin | BRD-K85140930 | C[C@]12CC[C@@H](O)[C@@](C)(CO)[C@@H]1CC[C@H]1C[C@@H]3C[C@@]12CC[C@]3(O)CO | NOFOAYPPHIUXJR-APNQCZIXSA-N | No-Effect | inactive |
| API-001 | BRD-K26325692 | Cc1cc(OCCN2CCOCC2)nn1-c1ccc2ccccc2c1 | DGPGXHRHNRYVDH-UHFFFAOYSA-N | No-Effect | inactive |
| apixaban | BRD-K67121414 | COc1ccc(-n2nc(C(N)=O)c3c2C(=O)N(c2ccc(N4CCCCC4=O)cc2)CC3)cc1 | QNZCBYKSOIHPEH-UHFFFAOYSA-N | No-Effect | inactive |
| apocynin | BRD-K64990520 | COc1cc(C(C)=O)ccc1O | DFYRUELUNQRZTB-UHFFFAOYSA-N | No-Effect | inactive |
| apoptosis-activator-II | BRD-K15164005 | O=C1C(=O)N(Cc2ccc(Cl)c(Cl)c2)c2ccccc21 | KGRJPLRFGLMQMV-UHFFFAOYSA-N | No-Effect | inactive |
| APR-246 | BRD-A09004963 | COC[C@]1(CO)C(=O)C2CCN1CC2 | BGBNULCRKBVAKL-SNVBAGLBSA-N | No-Effect | inactive |
| apramycin | BRD-K96737756 | CN[C@@H]1[C@@H](O[C@H]2O[C@H](CO)[C@@H](N)[C@H](O)[C@H]2O)O[C@H]2C[C@@H](N)[C@@H](O[C@@H]3[C@@H](N)C[C@@H](N)[C@H](O)[C@H]3O)O[C@@H]2[C@@H]1O | XZNUGFQTQHRASN-XQENGBIVSA-N | No-Effect | inactive |
| apratastat | BRD-K06883221 | CC1(C)SCCN(S(=O)(=O)c2ccc(OCC#CCO)cc2)[C@H]1C(=O)NO | MAVDNGWEBZTACC-HNNXBMFYSA-N | No-Effect | inactive |
| apremilast | BRD-K93208532 | CCOc1cc([C@@H](CS(C)(=O)=O)N2C(=O)c3cccc(NC(C)=O)c3C2=O)ccc1OC | IMOZEMNVLZVGJZ-QGZVFWFLSA-N | No-Effect | inactive |
| aprepitant | BRD-K52827117 | C[C@@H](O[C@H]1OCCN(Cc2n[nH]c(=O)[nH]2)[C@H]1c1ccc(F)cc1)c1cc(C(F)(F)F)cc(C(F)(F)F)c1 | ATALOFNDEOCMKK-OITMNORJSA-N | No-Effect | inactive |
| apricitabine | BRD-K55172746 | Nc1ccn([C@H]2CO[C@@H](CO)S2)c(=O)n1 | RYMCFYKJDVMSIR-RNFRBKRXSA-N | No-Effect | inactive |

| | | | | | |
|---|---|---|---|---|---|
| aprindine | BRD-K53236343 | CCN(CC)CCCN(c1ccccc1)C1Cc2ccccc2C1 | NZLBHDRPUJLHCE-UHFFFAOYSA-N | No-Effect | inactive |
| apronal | BRD-K00003524 | C=CC[C@H](C(=O)NC(N)=O)C(C)C | KSUUMAWCGDNLFK-ZETCQYMHSA-N | No-Effect | inactive |
| aptiganel | BRD-K63919159 | CCc1cccc(N(C)C(=N)Nc2cccc3ccccc23)c1 | BFNCJMURTMZBTE-UHFFFAOYSA-N | No-Effect | inactive |
| APTO-253 | BRD-K68003372 | Cc1[nH]c2ccc(F)cc2c1-c1nc2c3cccnc3c3ncccc3c2[nH]1 | NIRXBXIPHUTNNI-UHFFFAOYSA-N | No-Effect | inactive |
| APY-29 | BRD-K17511755 | c1cc(Nc2cc(C3CC3)n[nH]2)nc(Nc2ccc3nc[nH]c3c2)n1 | WJNBSTLIALIIEW-UHFFFAOYSA-N | No-Effect | inactive |
| apyramide | BRD-K00004738 | COc1ccc2c(c1)c(CC(=O)Oc1ccc(NC(C)=O)cc1)c(C)n2C(=O)c1ccc(Cl)cc1 | KWUFTKVMXUYTBF-UHFFFAOYSA-N | No-Effect | inactive |
| AQW-051 | BRD-K00004705 | Cc1ccc(-c2ccc(O[C@H]3CN4CCC3CC4)cn2)cc1 | NPDLTEZXGWRMLQ-IBGZPJMESA-N | No-Effect | inactive |
| AR-12 | BRD-K90497590 | NCC(=O)Nc1ccc(-n2nc(C(F)(F)F)cc2-c2ccc3c(ccc4ccccc43)c2)cc1 | YULUCECVQOCQFQ-UHFFFAOYSA-N | No-Effect | inactive |
| AR-A014418 | BRD-K67860401 | COc1ccc(CNC(=O)Nc2ncc([N+](=O)[O-])s2)cc1 | YAEMHJKFIIIULI-UHFFFAOYSA-N | No-Effect | inactive |
| AR-C155858 | BRD-K42325600 | Cc1n[nH]c(C)c1Cc1sc2c(c1C(=O)N1C[C@H](O)CO1)c(=O)n(C)c(=O)n2CC(C)C | ISIVOJWVBJIOFM-ZDUSSCGKSA-N | No-Effect | inactive |
| aramchol | BRD-K00004646 | CCCCCCCCCCCCCCCCCCC(=O)N[C@H]1CC[C@@]2(C)[C@@H](C1)C[C@@H](O)[C@@H]1[C@@H]2C[C@H](O)[C@@]2(C)[C@H]1CC[C@@H]2[C@H](C)CCC(=O)O | SHKXZIQNFMOPBS-OOMQYRRCSA-N | No-Effect | inactive |
| arbutin | BRD-K08037784 | OC[C@H]1O[C@@H](Oc2ccc(O)cc2)[C@H](O)[C@@H](O)[C@@H]1O | BJRNKVDFDLYUGJ-RMPHRYRLSA-N | No-Effect | inactive |
| ARC-239 | BRD-K93188295 | COc1ccccc1N1CCN(CCN2C(=O)c3ccccc3C(C)(C)C2=O)CC1 | JFNKXGOEOQCXDM-UHFFFAOYSA-N | No-Effect | inactive |
| arctigenin | BRD-K53523901 | COc1cc(C[C@H]2C(=O)OC[C@@H]2Cc2ccc(OC)c(OC)c2)ccc1O | NQWVSMVXKMHKTF-JKSUJKDBSA-N | No-Effect | inactive |
| arcyriaflavin-a | BRD-K72726508 | O=C1NC(=O)c2c1c1c3ccccc3[nH]c1c1[nH]c3ccccc3c21 | KAJXOWFGKYKMMZ-UHFFFAOYSA-N | No-Effect | inactive |
| ardeparin | BRD-K00003590 | CO[C@H]1[C@H](O)[C@@H](OS(=O)(=O)O)[C@H](O[C@@H]2[C@@H](COS(=O)(=O)O)O[C@H](OC)[C@H](NS(=O)(=O)O)[C@H]2O)O[C@H]1C(=O)O | HXSDFQWQRCUQHF-IDOMHHBZSA-N | No-Effect | inactive |
| arecaidine-but-2-ynyl-ester | BRD-K63792901 | CC#CCOC(=O)C1=CCCN(C)C1 | LBFWTPRSXIRKMZ-UHFFFAOYSA-N | No-Effect | inactive |
| arecaidine-propargyl-ester | BRD-K23922020 | C#CCOC(=O)C1=CCCN(C)C1 | SPHRJZBOFYIKMC-UHFFFAOYSA-N | No-Effect | inactive |
| arecoline | BRD-K88646909 | COC(=O)C1=CCCN(C)C1 | HJJPJSXJAXAIPN-UHFFFAOYSA-N | No-Effect | inactive |
| argatroban | BRD-A48261811 | C[C@H]1CNc2c(cccc2S(=O)(=O)N[C@@H](CCCNC(=N)N)C(=O)N2CC[C@@H](C)C[C@@H]2C(=O)O)C1 | KXNPVXPOPUZYGB-XYVMCAHJSA-N | No-Effect | inactive |
| argireline | BRD-K00003332 | CSCC[C@H](NC(=O)[C@H](CCC(=O)O)NC(=O)[C@H](CCC(=O)O)NC(C)=O)C(=O)N[C@@H](CCC(N)=O)C(=O)N[C@@H](CCCNC(=N)N)C(=O)N[C@@H](CCCNC(=N)N)C(N)=O | RJZNPROJTJSYLC-LLINQDLYSA-N | No-Effect | inactive |
| arglabin | BRD-K62734952 | C=C1C(=O)O[C@H]2[C@H]1CC[C@]1(C)O[C@@]13CC=C(C)[C@H]23 | UVJYAKBJSGRTHA-CUZKYEQNSA-N | No-Effect | inactive |
| aripiprazole | BRD-K70358946 | O=C1CCc2ccc(OCCCCN3CCN(c4cccc(Cl)c4Cl)CC3)cc2N1 | CEUORZQYGODEFX-UHFFFAOYSA-N | No-Effect | inactive |
| arofylline | BRD-K33852358 | CCCn1c(=O)c2[nH]cnc2n(-c2ccc(Cl)cc2)c1=O | GVTLDPJNRVMCAL-UHFFFAOYSA-N | No-Effect | inactive |
| arotinolol | BRD-A28044053 | CC(C)(C)NC[C@@H](O)CSc1nc(-c2ccc(C(N)=O)s2)cs1 | BHIAIPWSVYSKJS-SECBINFHSA-N | No-Effect | inactive |
| ARQ-092 | BRD-K60171656 | Nc1ncccc1-c1nc2ccc(-c3ccccc3)nc2n1-c1ccc(C2(N)CCC2)cc1 | HNFMVVHMKGFCMB-UHFFFAOYSA-N | No-Effect | inactive |
| ARQ-621 | BRD-K43436647 | C#CC[C@H](c1nc2cc(Cl)ccc2c(=O)n1Nc1ccccc1)N(CCCN)C(=O)c1ccc(Cl)c1F | UPJSUQWHUVLLNW-XMMPIXPASA-N | No-Effect | inactive |
| ARRY-334543 | BRD-K46386702 | C[C@@H]1COC(Nc2ccc3ncnc(Nc4ccc(OCc5nccs5)c(Cl)c4)c3c2)=N1 | UWXSAYUXVSFDBQ-CYBMUJFWSA-N | No-Effect | inactive |
| arsanilic-acid | BRD-K24179883 | Nc1ccc([As](=O)(O)O)cc1 | XKNKHVGWJDPIRJ-UHFFFAOYSA-N | No-Effect | inactive |
| artemether | BRD-K39621635 | CO[C@H]1O[C@@H]2O[C@@]3(C)CC[C@H]4[C@H](C)CC[C@@H]([C@H]1C)[C@]42OO3 | SXYIRMFQILZOAM-HVNFFKDJSA-N | No-Effect | inactive |
| artemisinin | BRD-K13112821 | C[C@@H]1CC[C@H]2[C@@H](C)C(=O)O[C@@H]3O[C@@]4(C)CC[C@@H]1[C@]32OO4 | BLUAFEHZUWYNDE-NNWCWBAJSA-N | No-Effect | inactive |
| artesunate | BRD-K54634444 | C[C@@H]1CC[C@H]2[C@@H](C)[C@H](OC(=O)CCC(=O)O)O[C@@H]3O[C@@]4(C)CC[C@@H]1[C@]23OO4 | FIHJKUPKCHIPAT-AHIGJZGOSA-N | No-Effect | inactive |
| articaine | BRD-A70514680 | CCCN[C@@H](C)C(=O)Nc1c(C)csc1C(=O)OC | QTGIAADRBBLJGA-VIFPVBQESA-N | No-Effect | inactive |
| arundic-acid | BRD-K49223707 | CCCCCC[C@@H](CCC)C(=O)O | YCYMCMYLORLIJX-SNVBAGLBSA-N | No-Effect | inactive |
| ARV-825 | BRD-A00047421 | Cc1sc2c(c1C)C(c1ccc(Cl)cc1)=N[C@@H](CC(=O)Nc1ccc(OCCOCCOCCOCCNc3cccc4c3C(=O)N([C@H]3CCC(=O)NC3=O)C4=O)cc1)c1nnc(C)n1-2 | RWLOGRLTDKDANT-ZPGRZCPFSA-N | No-Effect | inactive |
| AS-1269574 | BRD-K07656463 | Cc1cc(NCCO)nc(-c2ccc(Br)cc2)n1 | DUKPGOOUJNUIOI-UHFFFAOYSA-N | No-Effect | inactive |
| AS-1892802 | BRD-K27051047 | O=C(Nc1ccc(-c2ccncc2)cc1)N[C@H](CO)c1ccccc1 | WDTFYYZHMRBVHK-LJQANCHMSA-N | No-Effect | inactive |
| AS-1949490 | BRD-K61936403 | C[C@H](NC(=O)c1sccc1OCc1ccc(Cl)cc1)c1ccccc1 | RFZPGNRLOKVZJY-AWEZNQCLSA-N | No-Effect | inactive |
| AS-2034178 | BRD-K75254015 | O=C(O)CCc1ccc(NCc2cccc3c2CCCN3CCOc2ccccc2)cc1F | VENOXIKBBUVHRY-UHFFFAOYSA-N | No-Effect | inactive |
| AS-2444697 | BRD-K73565918 | Cc1cc(-c2nc(C(=O)Nc3cn(C4CCOCC4)nc3C(N)=O)co2)ccn1 | JANBIXNDYNRYQP-UHFFFAOYSA-N | No-Effect | inactive |
| AS-252424 | BRD-K14988953 | O=C1NC(=O)/C(=C/c2ccc(-c3ccc(F)cc3O)o2)S1 | OYYVWNDMOQPMGE-SDQBBNPISA-N | No-Effect | inactive |
| AS-604850 | BRD-K63915849 | O=C1NC(=O)/C(=C/c2ccc3c(c2)OC(F)(F)O3)S1 | SRLVNYDXMUGOFI-YWEYNIOJSA-N | No-Effect | inactive |
| AS-703026 | BRD-K89014967 | O=C(NC[C@H](O)CO)c1ccncc1Nc1ccc(I)cc1F | VIUAUNHCRHHYNE-JTQLQIEISA-N | No-Effect | inactive |
| AS-77 | BRD-K62427771 | O=c1ccc2c(OCCCCOc3ccccc3)c3ccoc3cc2o1 | KINMYBBFQRSVLL-UHFFFAOYSA-N | No-Effect | inactive |
| ASA-404 | BRD-K27938825 | Cc1ccc2c(=O)c3cccc(CC(=O)O)c3oc2c1C | XGOYIMQSIKSOBS-UHFFFAOYSA-N | No-Effect | inactive |
| asapiprant | BRD-K00003230 | CC(C)Oc1ccc(S(=O)(=O)N2CCN(c3ccc(-c4ncco4)c(OCC(=O)O)c3)CC2)cc1 | ZMZNWNTZRWXTJU-UHFFFAOYSA-N | No-Effect | inactive |
| asaraldehyde | BRD-K88219015 | COc1cc(OC)c(OC)cc1C=O | IAJBQAYHSQIQRE-UHFFFAOYSA-N | No-Effect | inactive |
| ASC-J9 | BRD-K30331124 | COc1ccc(/C=C/C(=O)C=C(O)/C=C/c2ccc(OC)c(OC)c2)cc1OC | ZMGUKFHHNQMKJI-URFDALOKSA-N | No-Effect | inactive |
| asciminib | BRD-K00003456 | O=C(Nc1ccc(OC(F)(F)Cl)cc1)c1cnc(N2CC[C@@H](O)C2)c(-c2cc[nH]n2)c1 | VOVZXURTCKPRDQ-CQSZACIVSA-N | No-Effect | inactive |
| ascomycin | BRD-K88882192 | CC[C@@H]1/C=C(\C)C[C@H](C)C[C@H](OC)[C@H]2O[C@@](O)(C(=O)C(=O)N3CCCC[C@H]3C(=O)O[C@H](/C(C)=C/[C@@H]3CC[C@@H](O)[C@H](OC)C3)[C@H](C)[C@@H](O)CC1=O)[C@H](C)C[C@@H]2OC | ZDQSOHOQTUFQEM-NURRSENYSA-N | No-Effect | inactive |
| ascorbic-acid | BRD-A85242401 | O=C1O[C@H]([C@@H](O)CO)[C@@H](O)C1=O | KRGQEOSDQHTZMX-TXQLHFEISA-N | No-Effect | inactive |
| asimadoline | BRD-K00003477 | CN(C(=O)C(c1ccccc1)c1ccccc1)[C@H](CN1CC[C@H](O)C1)c1ccccc1 | JHLHNYVMZCADTC-LOSJGSFVSA-N | No-Effect | inactive |
| asivatrep | BRD-K00003138 | CCCc1nc(C(F)(F)F)ccc1/C=C/C(=O)N[C@H](C)c1cc(F)c(NS(C)(=O)=O)c(F)c1 | UKGJZDSUJSPAJL-YPUOHESYSA-N | No-Effect | inactive |
| ASP-2535 | BRD-K07150325 | CC(C)c1nnc(-c2ccc(-c3ccccc3)nc2)n1-c1cccc2nonc12 | FQGLDGKVKDPVLO-UHFFFAOYSA-N | No-Effect | inactive |
| ASP-6537 | BRD-K82239282 | COc1ccc(-c2nc(OC)nn2-c2ccc(OC)cc2)cc1 | OBAICMYWONZOSZ-UHFFFAOYSA-N | No-Effect | inactive |
| ASP3026 | BRD-K06542892 | COc1cc(N2CCC(N3CCN(C)CC3)CC2)ccc1Nc1ncnc(Nc2ccccc2S(=O)(=O)C(C)C)n1 | MGGBYMDAPCCKCT-UHFFFAOYSA-N | No-Effect | inactive |
| aspartame | BRD-K78841970 | COC(=O)[C@H](Cc1ccccc1)NC(=O)[C@@H](N)CC(=O)O | IAOZJIPTCAWIRG-QWRGUYRKSA-N | No-Effect | inactive |
| aspirin | BRD-K11433652 | CC(=O)Oc1ccccc1C(=O)O | BSYNRYMUTXBXSQ-UHFFFAOYSA-N | No-Effect | inactive |
| AST-1306 | BRD-K26838195 | C=CC(=O)Nc1ccc2ncnc(Nc3ccc(OCc4cccc(F)c4)c(Cl)c3)c2c1 | MVZGYPSXNDCANY-UHFFFAOYSA-N | No-Effect | inactive |
| astaxanthin | BRD-K51832961 | CC1=C(/C=C/C(C)=C/C=C/C(C)=C/C=C/C=C(C)/C=C/C=C(C)/C=C/C2=C(C)C(=O)[C@@H](O)CC2(C)C)C(C)(C)C[C@H](O)C1=O | MQZIGYBFDRPAKN-UWFIBFSHSA-N | No-Effect | inactive |

| | | | | | |
|---|---|---|---|---|---|
| astilbin | BRD-K44845912 | C[C@@H]1O[C@@H](O[C@H]2C(=O)c3c(O)cc(O)cc3O[C@@H]2c2ccc(O)c(O)c2)[C@H](O)[C@H](O)[C@H]1O | ZROGCCBNZBKLEL-MPRHSVQHSA-N | No-Effect | inactive |
| astragaloside-a | BRD-K08188887 | CC(C)(O)[C@H]1CC[C@@](C)([C@H]2[C@@H](O)C[C@@]3(C)[C@@H]4C[C@H](O[C@@H]5O[C@H](CO)[C@@H](O)[C@H](O)[C@H]5O)[C@H]5C(C)(C)[C@@H](O[C@@H]6OC[C@@H](O)[C@H](O)[C@H]6O)CC[C@@]56C[C@@]46CC[C@]23C)O1 | QMNWISYXSJWHRY-AUJDEUPOSA-N | No-Effect | inactive |
| ASTX660 | BRD-K00003508 | C[C@@H]1CN(CC(=O)N2CC(C)(C)c3nc(CO)c(Cc4ccc(F)cc4)cc32)[C@@H](CN2CCOC[C@H]2C)CN1 | YCXOHEXZVKOGEV-DNRQZRRGSA-N | No-Effect | inactive |
| asunaprevir | BRD-K87028197 | C=C[C@@H]1C[C@]1(NC(=O)[C@@H]1C[C@@H](Oc2ncc(OC)c3ccc(Cl)cc23)CN1C(=O)[C@@H](NC(=O)OC(C)(C)C)C(C)(C)C)C(=O)NS(=O)(=O)C1CC1 | XRWSZZJLZRKHHD-WVWIJVSJSA-N | No-Effect | inactive |
| asymmetrical-dimethylarginine | BRD-K36269323 | CN(C)C(=N)NCCC[C@H](N)C(=O)O | YDGMGEXADBMOMJ-LURJTMIESA-N | No-Effect | inactive |
| AT-1015 | BRD-K06496777 | O=CN1CCC(C(=O)NCCN2CCC(=C3c4ccccc4C=Cc4ccccc43)CC2)CC1 | BSPMYKLEFGUTSI-UHFFFAOYSA-N | No-Effect | inactive |
| AT-7519 | BRD-K13390322 | O=C(NC1CCNCC1)c1[nH]ncc1NC(=O)c1c(Cl)cccc1Cl | OVPNQJVDAFNBDN-UHFFFAOYSA-N | No-Effect | inactive |
| AT-9283 | BRD-K24576554 | O=C(Nc1c[nH]nc1-c1nc2ccc(CN3CCOCC3)cc2[nH]1)NC1CC1 | LOLPPWBBNUVNQZ-UHFFFAOYSA-N | No-Effect | inactive |
| AT13148 | BRD-K32127727 | NC[C@@](O)(c1ccc(Cl)cc1)c1ccc(-c2cn[nH]c2)cc1 | IIRWNGPLJQXWFJ-KRWDZBQOSA-N | No-Effect | inactive |
| AT7867 | BRD-K12040459 | Clc1ccc(C2(c3ccc(-c4cn[nH]c4)cc3)CCNCC2)cc1 | LZMOSYUFVYJEPY-UHFFFAOYSA-N | No-Effect | inactive |
| ataluren | BRD-K94830329 | O=C(O)c1cccc(-c2noc(-c3ccccc3F)n2)c1 | OOUGLTULBSNHNF-UHFFFAOYSA-N | No-Effect | inactive |
| atazanavir | BRD-K09757388 | COC(=O)N[C@H](C(=O)N[C@@H](Cc1ccccc1)[C@@H](O)CN(Cc1ccc(-c2ccccn2)cc1)NC(=O)[C@@H](NC(=O)OC)C(C)(C)C)C(C)(C)C | AXRYRYVKAWYZBR-GASGPIRDSA-N | No-Effect | inactive |
| ATB-346 | BRD-A52551912 | COc1ccc2cc([C@@H](C)C(=O)Oc3ccc(C(N)=S)cc3)ccc2c1 | YCNMAPLPQYQJFC-CYBMUJFWSA-N | No-Effect | inactive |
| atenolol-(-) | BRD-K44993696 | CC(C)NC[C@H](O)COc1ccc(CC(N)=O)cc1 | METKIMKYRPQLGS-LBPRGKRZSA-N | No-Effect | inactive |
| atenolol-(+) | BRD-K68075732 | CC(C)NC[C@@H](O)COc1ccc(CC(N)=O)cc1 | METKIMKYRPQLGS-GFCCVEGCSA-N | No-Effect | inactive |
| atenolol-(+/-) | BRD-A20239487 | CC(C)NC[C@@H](O)COc1ccc(CC(N)=O)cc1 | METKIMKYRPQLGS-GFCCVEGCSA-N | No-Effect | inactive |
| atglistatin | BRD-K43363084 | CN(C)C(=O)Nc1cccc(-c2ccc(N(C)C)cc2)c1 | AWOPBSAJHCUSAS-UHFFFAOYSA-N | No-Effect | inactive |
| aticaprant | BRD-K00003342 | Cc1cc(C)cc([C@@H]2CCCN2Cc2ccc(Oc3ccc(C(N)=O)cc3F)cc2)c1 | ZHPMYDSXGRRERG-DEOSSOPVSA-N | No-Effect | inactive |
| atipamezole | BRD-K39252998 | CCC1(c2cnc[nH]2)Cc2ccccc2C1 | HSWPZIDYAHLZDD-UHFFFAOYSA-N | No-Effect | inactive |
| atizoram | BRD-K76614248 | COc1ccc(C2CNC(=O)NC2)cc1O[C@H]1C[C@@H]2CC[C@H]1C2 | LITNEAPWQHVPOK-FFSVYQOJSA-N | No-Effect | inactive |
| ATN-161 | BRD-K00521308 | CC(=O)N1CCC[C@H]1C(=O)N[C@@H](Cc1cnc[nH]1)C(=O)N[C@@H](CO)C(=O)N[C@@H](CS)C(=O)N[C@@H](CC(N)=O)C(N)=O | MMHDBUJXLOFTLC-WOYTXXSLSA-N | No-Effect | inactive |
| atomoxetine | BRD-K20141153 | CNCC[C@@H](Oc1ccccc1C)c1ccccc1 | VHGCDTVCOLNTBX-QGZVFWFLSA-N | No-Effect | inactive |
| atorvastatin | BRD-K69726342 | CC(C)c1c(C(=O)Nc2ccccc2)c(-c2ccccc2)c(-c2ccc(F)cc2)n1CC[C@@H](O)C[C@@H](O)CC(=O)O | XUKUURHRXDUEBC-KAYWLYCHSA-N | No-Effect | inactive |
| atovaquone | BRD-A19795905 | O=C1C(=O)[C@H]([C@H]2CC[C@H](c3ccc(Cl)cc3)CC2)C(=O)c2ccccc21 | UYFLKEIEOREOFC-ZXYWRSMDSA-N | No-Effect | inactive |
| ATPA | BRD-A65145453 | CC(C)(C)c1o[nH]c(=O)c1C[C@@H](N)C(=O)O | PIXJURSCCVBKRF-ZCFIWIBFSA-N | No-Effect | inactive |
| atractylenolide-i | BRD-K00003732 | C=C1CCC[C@]2(C)C=C3OC(=O)C(C)=C3C[C@@H]12 | ZTVSGQPHMUYCRS-SWLSCSKDSA-N | No-Effect | inactive |
| atrasentan | BRD-K16295392 | CCCCN(CCCC)C(=O)CN1C[C@H](c2ccc3c(c2)OCO3)[C@@H](C(=O)O)[C@@H]1c1ccc(OC)cc1 | MOTJMGVDPWRKOC-QPVYNBJUSA-N | No-Effect | inactive |
| atropine | BRD-K01825666 | CN1[C@H]2CC[C@@H]1C[C@H](OC(=O)[C@@H](CO)c1ccccc1)C2 | RKUNBYITZUJHSG-JJXSEGSLSA-N | No-Effect | inactive |
| atropine-oxide | BRD-K00004526 | C[N@@+]1([O-])[C@H]2CC[C@@H]1C[C@H](OC(=O)[C@H](CO)c1ccccc1)C2 | HGWPFSBHDACWNL-JFBPSJKJSA-N | No-Effect | inactive |
| auranofin | BRD-K45995181 | CCP(CC)CC)=[Au]S[C@@H]1O[C@H](COC(C)=O)[C@@H](OC(C)=O)[C@H](OC(C)=O)[C@H]1OC(C)=O | AUJRCFUBUPVWSZ-XTZHGVARSA-M | No-Effect | inactive |
| aurora-a-inhibitor-i | BRD-K53561341 | CCN1CCN(C(=O)Cc2ccc(Nc3ncc(F)c(Nc4ccc(C(=O)Nc5ccccc5Cl)cc4)n3)cc2)CC1 | AKSIZPIFQAYJGF-UHFFFAOYSA-N | No-Effect | inactive |
| aurothioglucose | BRD-K00003601 | OC[C@H]1O[C@H](S[Au])[C@H](O)[C@@H](O)[C@@H]1O | XHVAWZZCDCWGBK-WYRLRVFGSA-M | No-Effect | inactive |
| AUT-1 | BRD-K00005330 | CC[C@H]1NC(=O)N(c2ccc(Oc3ccc(C)c(OC)c3)nc2)C1=O | AMAOXEGBJHLCSF-CQSZACIVSA-N | No-Effect | inactive |
| autotaxin-modulator-1 | BRD-A94640958 | C[C@H](c1ccc2ccc(O[C@H]3CC[C@@H](C(F)(F)F)CC3)c(C(F)(F)F)c2c1)N1[C@H]2CC[C@@H]1C[C@H](C(=O)O)C2 | PZASAAIJIFDWSB-PQGZPOPSSA-N | No-Effect | inactive |
| AV-412 | BRD-K23190681 | C=CC(=O)Nc1cc2c(Nc3ccc(F)c(Cl)c3)ncnc2cc1C#CC(C)(C)N1CCN(C)CC1 | ZAJXXUDARPGGOC-UHFFFAOYSA-N | No-Effect | inactive |
| AV-608 | BRD-K96615647 | O=C(N[C@H]1CCN(C(=O)c2cc(C(F)(F)F)cc(C(F)(F)F)c2)[C@H](Cc2ccc(Cl)cc2)C1)c1ccnc2ccccc12 | NXLUTEDAEFXMQR-BJKOFHAPSA-N | No-Effect | inactive |
| avacopan | BRD-K00003256 | Cc1cccc(NC(=O)[C@H]2CCCN(C(=O)c3c(C)cccc3F)[C@H]2c2ccc(NC3CCCC3)cc2)cc1C(F)(F)F | PUKBOVABABRILL-YZNIXAGQSA-N | No-Effect | inactive |
| avadomide | BRD-A88644406 | Cc1nc2cccc(N)c2c(=O)n1[C@H]1CCC(=O)NC1=O | RSNPAKAFCAAMBH-JTQLQIEISA-N | No-Effect | inactive |
| avagacestat | BRD-K83720053 | NC(=O)[C@@H](CCC(F)(F)F)N(Cc1ccc(-c2ncon2)cc1F)S(=O)(=O)c1ccc(Cl)cc1 | XEAOPVUAMONVLA-QGZVFWFLSA-N | No-Effect | inactive |
| avanafil | BRD-K65781196 | COc1ccc(CNc2nc(N3CCC[C@H]3CO)ncc2C(=O)NCc2ncccn2)cc1Cl | WEAJZXNPAWBCOA-INIZCTEOSA-N | No-Effect | inactive |
| avapritinib | BRD-K00003419 | Cn1cc(-c2cc3c(N4CCN(c5ncc([C@@](C)(N)c6ccc(F)cc6)cn5)CC4)ncnn3c2)cn1 | DWYRIWUZIJHQKQ-SANMLTNESA-N | No-Effect | inactive |
| avasimibe | BRD-K67901620 | CC(C)c1cc(C(C)C)c(CC(=O)NS(=O)(=O)Oc2c(C(C)C)cccc2C(C)C)c(C(C)C)c1 | PTQXTEKSNBVPQJ-UHFFFAOYSA-N | No-Effect | inactive |
| avatrombopag | BRD-K68544925 | O=C(Nc1nc(-c2cc(Cl)cs2)c(N2CCN(C3CCCCC3)CC2)s1)c1cnc(N2CCC(C(=O)O)CC2)c(Cl)c1 | OFZJKCQENFPZBH-UHFFFAOYSA-N | No-Effect | inactive |
| AVE-0991 | BRD-K36154108 | CCNC(=O)NS(=O)(=O)c1sc(CC(C)C)cc1-c1ccc(Cn2c(-c3ccccc3)nc(OC)c2C=O)cc1 | QTOZBSNPDCWHPV-UHFFFAOYSA-N | No-Effect | inactive |
| AVE-3085 | BRD-K00004734 | O=C(NC1Cc2ccccc2C1)c1ccc2c(c1)OC(F)(F)O2 | OKCJNSDIVCXYAL-UHFFFAOYSA-N | No-Effect | inactive |
| AVex-73 | BRD-K00003435 | CN(C)C[C@H]1CCOC1(c1ccccc1)c1ccccc1 | BOTHKNZTGGXFEQ-GOSISDBHSA-N | No-Effect | inactive |
| avibactam | BRD-K85882401 | NC(=O)[C@@H]1CC[C@@H]2CN1C(=O)N2OS(=O)(=O)O | NDCUAPJVLWFHHB-UHNVWZDZSA-N | No-Effect | inactive |
| AVL-292 | BRD-K87782578 | C=CC(=O)Nc1cccc(Nc2nc(Nc3ccc(OCCOC)cc3)ncc2F)c1 | KXBDTLQSDKGAEB-UHFFFAOYSA-N | No-Effect | inactive |
| AVN-492 | BRD-K00003436 | CNc1nn2c(C)c(N(C)C)c(C)nc2c1S(=O)(=O)c1ccccc1 | SNPPEHMSSOEYDH-UHFFFAOYSA-N | No-Effect | inactive |
| AVN-944 | BRD-K62196610 | CC[C@H](CC#N)OC(=O)N[C@@H](C)c1cccc(NC(=O)Nc2ccc(-c3cnco3)c(OC)c2)c1 | GYCPCOJTCINIFZ-OXJNMPFZSA-N | No-Effect | inactive |
| avobenzone | BRD-K14401183 | COc1ccc(C(=O)C=C(O)c2ccc(C(C)(C)C)cc2)cc1 | LKIIIESDXWZZRE-UHFFFAOYSA-N | No-Effect | inactive |
| avoralstat | BRD-K00003224 | C=Cc1cc(C(=O)Nc2ccc(C(=N)N)cc2)c(-c2ccc(C(=O)NCC3CC3)nc2C(=O)O)cc1OC | TUWMKPVJGGWGNL-UHFFFAOYSA-N | No-Effect | inactive |
| avridine | BRD-K26665484 | CCCCCCCCCCCCCCCCCCN(CCCCCCCCCCCCCCCCCC)CCCN(CCO)CCO | WXNRAKRZUCLRBP-UHFFFAOYSA-N | No-Effect | inactive |
| AX-024 | BRD-K00003479 | COc1ccc2c(c1)C(c1ccc(F)cc1)=C(CN1CCCC1)CO2 | VMMKGVRPILDZML-UHFFFAOYSA-N | No-Effect | inactive |
| axitinib | BRD-K29905972 | CNC(=O)c1ccccc1Sc1ccc2c(/C=C/c3ccccn3)[nH]nc2c1 | RITAVMQDGBJQJZ-FMIVXFBMSA-N | No-Effect | inactive |
| AZ-10417808 | BRD-K36258877 | C=CCNC(=O)c1cc([N+](=O)[O-])cc2c(=O)nc(Nc3ccc(Cl)c(Cl)c3)[nH]c12 | VUBILPOZTRGZJK-UHFFFAOYSA-N | No-Effect | inactive |

| | | | | | |
|---|---|---|---|---|---|
| AZ-10606120 | BRD-K09907507 | O=C(CC12CC3CC(CC(C3)C1)C2)Nc1cccc2nc(NCCNCCO)ccc12 | FQMZXMVHHKXGTM-UHFFFAOYSA-N | No-Effect | inactive |
| AZ-12080282 | BRD-K05973499 | Cc1ccc(-c2ncc[nH]2)cc1NC(=O)c1ccc(OCc2ccccn2)cc1 | KMSKKYBHDQYHMC-UHFFFAOYSA-N | No-Effect | inactive |
| AZ-628 | BRD-K05804044 | Cc1ccc(NC(=O)c2cccc(C(C)(C)C#N)c2)cc1Nc1ccc2ncn(C)c(=O)c2c1 | ZGBGPEDJXCYQPH-UHFFFAOYSA-N | No-Effect | inactive |
| AZ-7371 | BRD-K14661019 | COc1ncnc(Cn2cc(C(=O)NCCO)c3ncc(C)cc32)c1C | VDRYGTNDKXIPSK-UHFFFAOYSA-N | No-Effect | inactive |
| AZ191 | BRD-K80935598 | COc1cc(N2CCN(C)CC2)ccc1Nc1nccc(-c2cn(C)c3cnccc23)n1 | ZYVXTMKTGDARKR-UHFFFAOYSA-N | No-Effect | inactive |
| AZ20 | BRD-K42436189 | C[C@@H]1COCCN1c1cc(C2(S(C)(=O)=O)CC2)nc(-c2cccc3[nH]ccc23)n1 | SCGCBAAYLFTIJU-CQSZACIVSA-N | No-Effect | inactive |
| AZ505 | BRD-K76244476 | O=C1COc2c(CCNCCN(C(=O)CCNCCc3ccc(Cl)c(Cl)c3)C3CCCCC3)ccc(O)c2N1 | LIBVHXXKHSODII-UHFFFAOYSA-N | No-Effect | inactive |
| AZ960 | BRD-K64888243 | Cc1cc(Nc2nc(N[C@@H](C)c3ccc(F)cc3)c(C#N)cc2F)[nH]n1 | SUNXHXDJOIXABJ-NSHDSACASA-N | No-Effect | inactive |
| azacitidine | BRD-K03406345 | Nc1ncn([C@@H]2O[C@H](CO)[C@@H](O)[C@H]2O)c(=O)n1 | NMUSYJAQQFHJEW-KVTDHHQDSA-N | No-Effect | inactive |
| azacyclonol | BRD-K97061094 | OC(c1ccccc1)(c1ccccc1)C1CCNCC1 | ZMISODWVFHHWNR-UHFFFAOYSA-N | No-Effect | inactive |
| azaguanine-8 | BRD-K53114651 | Nc1nc2nn[nH]c2c(=O)[nH]1 | LPXQRXLUHJKZIE-UHFFFAOYSA-N | No-Effect | inactive |
| azalomycin-B | BRD-K38292725 | CC[C@@H]1[C@@H](C)O[C@@](O)([C@@H](C)[C@H](O)[C@H](C)[C@H]2OC(=O)/C=C/C=C/[C@H](C)[C@@H]([C@@H](C)[C@@H](O)[C@H](C)[C@@]3(O)C[C@@H](O[C@H]4C[C@H](O)[C@H](O)[C@H](C)O4)[C@H](CC)[C@@H](C)O3)OC(=O)/C=C/C=C/[C@@H]2C)C[C@H]1O[C@H]1C[C@H](O)[C@H](O)[C@H](C)O1 | OSERMIPXNLXAPD-MJMYBOKFSA-N | No-Effect | inactive |
| azaperone | BRD-K45861246 | O=C(CCCN1CCN(c2ccccn2)CC1)c1ccc(F)cc1 | XTKDAFGWCDAMPY-UHFFFAOYSA-N | No-Effect | inactive |
| azapropazone | BRD-A70182876 | CCC[C@H]1C(=O)N2C(N(C)C)=Nc3ccc(C)cc3N2C1=O | MPHPHYZQRGLTBO-LLVKDONJSA-N | No-Effect | inactive |
| azaserine | BRD-K47464228 | N#[N+]CC(=O)OC[C@H](N)C(=O)O | YXEASJLRXGZRHS-VKHMYHEASA-O | No-Effect | inactive |
| azasetron | BRD-K01826751 | CN1C(=O)COc2c(C(=O)N[C@H]3CN4CCC3CC4)cc(Cl)cc21 | WUKZPHOXUVCQOR-ZDUSSCGKSA-N | No-Effect | inactive |
| azatadine | BRD-K77821588 | CN1CCC(=C2c3ccccc3CCc3cccnc32)CC1 | SEBMTIQKRHYNIT-UHFFFAOYSA-N | No-Effect | inactive |
| azathioprine | BRD-K32821942 | Cn1cnc([N+](=O)[O-])c1Sc1ncnc2nc[nH]c12 | LMEKQMALGUDUQG-UHFFFAOYSA-N | No-Effect | inactive |
| AZD1080 | BRD-K61041538 | N#Cc1ccc2c(c1)C(c1ccc(CN3CCOCC3)cn1)C(=O)N2 | RDXUCTYBYOIJGA-UHFFFAOYSA-N | No-Effect | inactive |
| AZD1208 | BRD-K21908111 | N[C@@H]1CCCN(c2c(/C=C3\SC(=O)NC3=O)cccc2-c2ccccc2)C1 | MCUJKPPARUPFJM-UWCCDQBKSA-N | No-Effect | inactive |
| AZD1283 | BRD-K77060810 | CCOC(=O)c1cc(C#N)c(N2CCC(C(=O)NS(=O)(=O)Cc3ccccc3)CC2)nc1C | NEMHKCNXXRQYRF-UHFFFAOYSA-N | No-Effect | inactive |
| AZD1446 | BRD-A44188509 | O=C(c1ccc(Cl)o1)N1C[C@H]2CNC[C@@H]2C1 | GTUIQNHJSXQMKW-HTQZYQBOSA-N | No-Effect | inactive |
| AZD1480 | BRD-K65928735 | Cc1cc(Nc2nc(N[C@@H](C)c3ncc(F)cn3)ncc2Cl)[nH]n1 | PDOQBOJDRPLBQU-QMMMGPOBSA-N | No-Effect | inactive |
| AZD1981 | BRD-K83186168 | CC(=O)Nc1cccc2c1c(Sc1ccc(Cl)cc1)c(C)n2CC(=O)O | JWYIGNODXSRKGP-UHFFFAOYSA-N | No-Effect | inactive |
| AZD2461 | BRD-K85014146 | COC1CCN(C(=O)c2cc(Cc3n[nH]c(=O)c4ccccc34)ccc2F)CC1 | HYNBNUYQTQIHJK-UHFFFAOYSA-N | No-Effect | inactive |
| AZD3264 | BRD-K14287049 | Cc1noc(C)c1-c1ccc(-c2cc(C(N)=O)c(NC(N)=O)s2)c(O[C@H]2CCNC2)c1 | NMFIXEFNQPDISY-ZDUSSCGKSA-N | No-Effect | inactive |
| AZD3514 | BRD-K84996356 | CC(=O)N1CCN(CCOc2ccc(C3CCN(C4=Nn5c(nnc5C(F)(F)F)CC4)CC3)cc2)CC1 | JMEYDSHPKCSIJC-UHFFFAOYSA-N | No-Effect | inactive |
| AZD3759 | BRD-K41626799 | COc1cc2ncnc(Nc3cccc(Cl)c3F)c2cc1OC(=O)N1CCN(C)C[C@H]1C | MXDSJQHFFDGFDK-CYBMUJFWSA-N | No-Effect | inactive |
| AZD3839 | BRD-K00004606 | NC1=N[C@@](c2cccc(-c3cncnc3)c2)(c2ccnc(C(F)F)c2)c2cccc(F)c21 | MRXBCEQZNKUUIP-DEOSSOPVSA-N | No-Effect | inactive |
| AZD3965 | BRD-K25210703 | Cc1[nH]nc(C(F)(F)F)c1Cc1sc2c(c1C(=O)N1C[C@](C)(O)CO1)c(=O)n(C)c(=O)n2C(C)C | PRNXOFBDXNTIFG-FQEVSTJZSA-N | No-Effect | inactive |
| AZD3988 | BRD-K73984371 | O=C(O)C[C@H]1CC[C@H](c2ccc(NC(=O)c3nnc(Nc4ccc(F)c(F)c4)o3)cc2)CC1 | NGEBYTLALFOQKI-HDJSIYSDSA-N | No-Effect | inactive |
| AZD4282 | BRD-K72369578 | NCC(=O)O | DHMQDGOQFOQNFH-UHFFFAOYSA-N | No-Effect | inactive |
| AZD4547 | BRD-K28392481 | COc1cc(CCc2cc(NC(=O)c3ccc(N4C[C@H](C)N[C@H](C)C4)cc3)[nH]n2)cc(OC)c1 | VRQMAABPASPXMW-HDICACEKSA-N | No-Effect | inactive |
| AZD4635 | BRD-K00003443 | Cc1cc(-c2nnc(N)nc2-c2ccc(F)cc2)cc(Cl)n1 | NCWQLHHDGDXIJN-UHFFFAOYSA-N | No-Effect | inactive |
| AZD5069 | BRD-K00003297 | C[C@@H](Oc1cc(NS(=O)(=O)N2CCC2)nc(SCc2cccc(F)c2F)n1)[C@@H](O)CO | QZECRCLSIGFCIO-RISCZKNCSA-N | No-Effect | inactive |
| AZD5363 | BRD-K99023089 | NC1(C(=O)N[C@@H](CCO)c2ccc(Cl)cc2)CCN(c2ncnc3[nH]ccc23)CC1 | JDUBGYFRJFOXQC-KRWDZBQOSA-N | No-Effect | inactive |
| AZD5582 | BRD-K88767641 | CN[C@@H](C)C(=O)N[C@H](C(=O)N1CCC[C@H]1C(=O)N[C@H]1c2ccccc2C[C@H]1OCC#CC#CCO[C@@H]1Cc2ccccc2[C@@H]1NC(=O)[C@@H]1CCCN1C(=O)[C@@H](NC(=O)[C@H](C)NC)C1CCCCC1)C1CCCCC1 | WLMCRYCCYXHPQF-ZVMUOSSASA-N | No-Effect | inactive |
| AZD6482 | BRD-K58772419 | Cc1cc([C@@H](C)Nc2ccccc2C(=O)O)c2nc(N3CCOCC3)cc(=O)n2c1 | IRTDIKMSKMREGO-OAHLLOKOSA-N | No-Effect | inactive |
| AZD6738 | BRD-A36013532 | C[C@@H]1COCCN1c1cc(C2([S@@](C)(=N)=O)CC2)nc(-c2ccnc3[nH]ccc23)n1 | OHUHVTCQTUDPIJ-MUWSIPGASA-N | No-Effect | inactive |
| AZD6765 | BRD-A94033181 | N[C@@H](Cc1ccccn1)c1ccccc1 | FWUQWDCOOWEXRY-ZDUSSCGKSA-N | No-Effect | inactive |
| AZD7545 | BRD-K52836380 | CN(C)C(=O)c1ccc(S(=O)(=O)c2ccc(NC(=O)[C@@](C)(O)C(F)(F)F)c(Cl)c2)cc1 | DTDZLJHKVNTQGZ-GOSISDBHSA-N | No-Effect | inactive |
| AZD7594 | BRD-K00003100 | C[C@H](NC(=O)C(C)(F)F)[C@H](Oc1ccc2c(cnn2-c2cccc(C(=O)N[C@@H]3CCOC3)c2)c1)c1ccc2c(c1)OCCO2 | ZZWJKLGCDHYVMB-BWGXUDETSA-N | No-Effect | inactive |
| AZD7687 | BRD-K31495718 | Cc1nc(C)c(-c2ccc([C@H]3CC[C@H](CC(=O)O)CC3)cc2)nc1C(N)=O | YXFNPRHZMOGREC-SHTZXODSSA-N | No-Effect | inactive |
| AZD7986 | BRD-K00003398 | Cn1c(=O)oc2ccc(-c3ccc(C[C@@H](C#N)NC(=O)[C@@H]4CNCCCO4)cc3)cc21 | AEXFXNFMSAAELR-RXVVDRJESA-N | No-Effect | inactive |
| AZD8055 | BRD-K69932463 | COc1ccc(-c2ccc3c(N4CCOC[C@@H]4C)nc(N4CCOC[C@@H]4C)nc3n2)cc1CO | KVLFRAWTRWDEDF-IRXDYDNUSA-N | No-Effect | inactive |
| AZD8186 | BRD-K47289408 | C[C@@H](Nc1cc(F)cc(F)c1)c1cc(C(=O)N(C)C)cc2c(=O)cc(N3CCOCC3)oc12 | LMJFJIDLEAWOQJ-CQSZACIVSA-N | No-Effect | inactive |
| AZD8330 | BRD-K37687095 | Cc1cc(C(=O)NOCCO)c(Nc2ccc(I)cc2F)n(C)c1=O | RWEVIPRMPFNTLO-UHFFFAOYSA-N | No-Effect | inactive |
| AZD8797 | BRD-K03336321 | CC(C)C[C@H](CO)Nc1nc(S[C@@H](C)c2ccccc2)nc2nc(N)sc12 | ZMQSLMZOWVGBSM-GXTWGEPZSA-N | No-Effect | inactive |
| AZD8835 | BRD-K82894950 | CCn1nc(C2CCN(C(=O)CCO)CC2)nc1-c1cnc(N)c(-c2nnc(C(C)(C)C)o2)n1 | ZGRDYKFVDCFJCZ-UHFFFAOYSA-N | No-Effect | inactive |
| AZD8931 | BRD-K98572433 | CNC(=O)CN1CCC(Oc2cc3c(Nc4cccc(Cl)c4F)ncnc3cc2OC)CC1 | DFJSJLGUIXFDJP-UHFFFAOYSA-N | No-Effect | inactive |
| AZD9496 | BRD-K73235735 | C[C@@H]1Cc2c([nH]c3ccccc23)[C@@H](c2c(F)cc(/C=C/C(=O)O)cc2F)N1CC(C)(C)F | DFBDRVGWBHBJNR-BBNFHIFMSA-N | No-Effect | inactive |
| AZD9668 | BRD-K31476763 | Cc1c(-c2ccnn2C)cc(C(=O)NCc2ccc(S(C)(=O)=O)cn2)c(=O)n1-c1cccc(C(F)(F)F)c1 | QNQZWEGMKJBHEM-UHFFFAOYSA-N | No-Effect | inactive |
| azelaic-acid | BRD-K08976401 | O=C(O)CCCCCCCC(=O)O | BDJRBEYXGGNYIS-UHFFFAOYSA-N | No-Effect | inactive |
| azelastine | BRD-A68888262 | CN1CCC[C@H](n2nc(Cc3ccc(Cl)cc3)c3ccccc3c2=O)CC1 | MBUVEWMHONZEQD-SFHVURJKSA-N | No-Effect | inactive |
| azeliragon | BRD-K51899933 | CCCCc1nc(-c2ccc(OCCCN(CC)CC)cc2)cn1-c1ccc(Oc2ccc(Cl)cc2)cc1 | KJNNWYBAOPXVJY-UHFFFAOYSA-N | No-Effect | inactive |
| azithromycin | BRD-K74501079 | CC[C@H]1OC(=O)[C@H](C)[C@@H](O[C@H]2C[C@@](C)(OC)[C@@H](O)[C@H](C)O2)[C@H](C)[C@@H](O[C@@H]2O[C@H](C)C[C@H](N(C)C)[C@H]2O)[C@](C)(O)C[C@@H](C)CN(C)[C@H](C)[C@@H](O)[C@]1(C)O | MQTOSJVFKKJCRP-BICOPXKESA-N | No-Effect | inactive |

| | | | | | |
|---|---|---|---|---|---|
| azlocillin | BRD-K60663764 | CC1(C)S[C@@H]2[C@H](NC(=O)[C@H](NC(=O)N3CCNC3=O)c3ccccc3)C(=O)N2[C@H]1C(=O)O | JTWOMNBEOCYFNV-NFFDBFGFSA-N | No-Effect | inactive |
| AZM-475271 | BRD-K51329597 | COc1ccc(Cl)c(Nc2ncnc3cc(OCC4CCN(C)CC4)c(OC)cc23)c1 | WPOXAFXHRJYEIC-UHFFFAOYSA-N | No-Effect | inactive |
| azodicarbonamide | BRD-K87036601 | NC(=O)/N=N/C(N)=O | XOZUGNYVDXMRKW-AATRIKPKSA-N | No-Effect | inactive |
| azomycin-(2-nitroimidazole) | BRD-K26824830 | O=[N+]([O-])c1ncc[nH]1 | YZEUHQHUFTYLPH-UHFFFAOYSA-N | No-Effect | inactive |
| aztreonam | BRD-K62607865 | C[C@H]1[C@H](NC(=O)/C(=N\OC(C)(C)C(=O)O)c2csc(N)n2)C(=O)N1S(=O)(=O)O | WZPBZJONDBGPKJ-VEHQQRBSSA-N | No-Effect | inactive |
| bacampicillin | BRD-A41698174 | CCOC(=O)O[C@@H](C)OC(=O)[C@@H]1N2C(=O)[C@@H](NC(=O)[C@H](N)c3ccccc3)[C@H]2SC1(C)C | PFOLLRNADZZWEX-XIJZRVNUSA-N | No-Effect | inactive |
| bacitracin | BRD-K14079729 | CC[C@H](C)[C@H](N)C1=N[C@H](C(=O)N[C@@H](CC(C)C)C(=O)N[C@H](CCC(=O)O)C(=O)N[C@H](C(=O)N[C@H]2CCCCNC(=O)[C@H](CC(N)=O)NC(=O)[C@@H](CC(=O)O)NC(=O)[C@H](Cc3cnc[nH]3)NC(=O)[C@@H](Cc3ccccc3)NC(=O)[C@H]([C@@H](C)CC)NC(=O)[C@@H](CCCN)NC2=O)[C@@H](C)CC)CS1 | CLKOFPXJLQSYAH-ABRJDSQDSA-N | No-Effect | inactive |
| bacitracin-zinc | BRD-K00003625 | CC[C@H](C)[C@H](N)C1=N[C@H](C(=O)N[C@@H](CC(C)C)C(=O)N[C@@H]2CCC(=O)O[Zn]OC(=O)C[C@H]3NC(=O)[C@H](Cc4c[nH]cn4)NC(=O)[C@@H](Cc4ccccc4)NC(=O)[C@H]([C@@H](C)CC)NC(=O)[C@@H](CCCN)NC(=O)[C@H](CCCCNC(=O)[C@H](CC(N)=O)NC3=O)NC(=O)[C@H]([C@@H](C)CC)NC2=O)CS1 | UCRLQOPRDMGYOA-DFTDUNEMSA-L | No-Effect | inactive |
| baclofen | BRD-K01826724 | NC[C@@H](CC(=O)O)c1ccc(Cl)cc1 | KPYSYYIEGFHWSV-MRVPVSSYSA-N | No-Effect | inactive |
| baicalein | BRD-K72327355 | O=c1c(O)c(O)cc2oc(-c3ccccc3)cc(O)c1-2 | WTNAEULBGCSUSK-UHFFFAOYSA-N | No-Effect | inactive |
| bakuchiol | BRD-K09785578 | C=C[C@@](C)(/C=C/c1ccc(O)cc1)CCC=C(C)C | LFYJSSARVMHQJB-QIXNEVBVSA-N | No-Effect | inactive |
| balaglitazone | BRD-A00147595 | Cn1c(COc2ccc(C[C@H]3SC(=O)NC3=O)cc2)nc2ccccc2c1=O | IETKPTYAGKZLKY-MRXNPFEDSA-N | No-Effect | inactive |
| balapiravir | BRD-K47976015 | CC(C)C(=O)OC[C@@]1(N=[N+]=[N-])O[C@@H](n2ccc(N)nc2=O)[C@H](OC(=O)C(C)C)[C@@H]1OC(=O)C(C)C | VKXWOLCNTHXCLF-DXEZIKHYSA-N | No-Effect | inactive |
| balofloxacin | BRD-A47790386 | CN[C@H]1CCCN(c2c(F)cc3c(=O)c(C(=O)O)cn(C4CC4)c3c2OC)C1 | MGQLHRYJBWGORO-NSHDSACASA-N | No-Effect | inactive |
| baloxavir | BRD-K00003496 | O=C1c2c(O)c(=O)ccn2N([C@@H]2c3ccccc3SCc3c2ccc(F)c3F)[C@@H]2COCCN12 | FIDLLEYNNRGVFR-CTNGQTDRSA-N | No-Effect | inactive |
| balsalazide | BRD-K00003656 | O=C(O)CCNC(=O)c1ccc(/N=N/c2ccc(O)c(C(=O)O)c2)cc1 | IPOKCKJONYRRHP-FMQUCBEESA-N | No-Effect | inactive |
| BAM7 | BRD-K07442505 | CCOc1ccccc1N/N=C1/C(=O)N(c2nc(-c3ccccc3)cs2)N=C1C | KOCVKGYKBLJEPK-LYBHJNIJSA-N | No-Effect | inactive |
| bamaquimast | BRD-K00003411 | CCCn1c(=O)c(CCCOC(=O)NC)nc2ccccc21 | HNQSZPBGDAJIJO-UHFFFAOYSA-N | No-Effect | inactive |
| bambuterol | BRD-A17462676 | CN(C)C(=O)Oc1cc(OC(=O)N(C)C)cc(C(O)CNC(C)(C)C)c1 | ANZXOIAKUNOVQU-UHFFFAOYSA-N | No-Effect | inactive |
| BAN-ORL-24 | BRD-K47049295 | O=C(NCCCN1CCC2(CC1)OCc1ccccc12)[C@H]1CCCN1Cc1ccccc1 | MAKMQGKJURAJEN-RUZDIDTESA-N | No-Effect | inactive |
| banoxantrone | BRD-K00003159 | C[N+](C)([O-])CCNc1ccc(NCC[N+](C)(C)[O-])c2c1C(=O)c1c(O)ccc(O)c1C2=O | YZBAXVICWUUHGG-UHFFFAOYSA-N | No-Effect | inactive |
| barasertib | BRD-K31928526 | CCN(CCCOc1ccc2c(Nc3cc(CC(=O)Nc4cccc(F)c4)n[nH]3)ncnc2c1)CCOP(=O)(O)O | GBJVVSCPOBPEIT-UHFFFAOYSA-N | No-Effect | inactive |
| barasertib-HQPA | BRD-K63923597 | CCN(CCO)CCCOc1ccc2c(Nc3cc(CC(=O)Nc4cccc(F)c4)n[nH]3)ncnc2c1 | QYZOGCMHVIGURT-UHFFFAOYSA-N | No-Effect | inactive |
| bardoxolone-methyl | BRD-K69694239 | COC(=O)[C@]12CCC(C)(C)C[C@H]1[C@H]1C(=O)C=C3[C@@](C)(CC[C@H]4C(C)(C)C(=O)C(C#N)=C[C@]34C)[C@]1(C)CC2 | WPTTVJLTNAWYAO-KPOXMGGZSA-N | No-Effect | inactive |
| baricitinib | BRD-K53581288 | CCS(=O)(=O)N1CC(CC#N)(n2cc(-c3ncnc4[nH]ccc34)cn2)C1 | XUZMWHLSFXCVMG-UHFFFAOYSA-N | No-Effect | inactive |
| barium-6-O-phosphonato-D-glucose | BRD-K67760597 | O=C[C@H](O)[C@@H](O)[C@H](O)[C@H](O)COP(=O)(O)O | VFRROHXSMXFLSN-SLPGGIOYSA-N | No-Effect | inactive |
| barnidipine | BRD-K78781788 | COC(=O)C1=C(C)NC(C)=C(C(=O)O[C@H]2CCN(Cc3ccccc3)C2)[C@H]1c1cccc([N+](=O)[O-])c1 | VXMOONUMYLCFJD-DHLKQENFSA-N | No-Effect | inactive |
| basic-fuchsin | BRD-K26362911 | CC1=CC(=C(c2ccc(N)cc2)c2ccc(N)cc2)C=CC1=N | YDCMWLQSPFTWCS-UHFFFAOYSA-N | No-Effect | inactive |
| basimglurant | BRD-K62387885 | Cc1nc(C#Cc2ccnc(Cl)c2)c(C)n1-c1ccc(F)cc1 | UPZWINBEAHDTLA-UHFFFAOYSA-N | No-Effect | inactive |
| basmisanil | BRD-K00003221 | Cc1onc(-c2ccc(F)cc2)c1COc1ccc(C(=O)N2CCS(=O)(=O)CC2)cn1 | VCGRFBXVSFAGGA-UHFFFAOYSA-N | No-Effect | inactive |
| batefenterol | BRD-K00003148 | COc1cc(NC(=O)CCN2CCC(OC(=O)Nc3ccccc3-c3ccccc3)CC2)c(Cl)cc1CNC[C@H](O)c1ccc(O)c2nc(O)ccc12 | URWYQGVSPQJGGB-DHUJRADRSA-N | No-Effect | inactive |
| batimastat | BRD-K82818427 | CNC(=O)[C@H](Cc1ccccc1)NC(=O)[C@H](CC(C)C)[C@H](CSc1cccs1)C(=O)NO | XFILPEOLDIKJHX-QYZOEREBSA-N | No-Effect | inactive |
| bavisant | BRD-K79595931 | O=C(c1ccc(CN2CCOCC2)cc1)N1CCN(C2CC2)CC1 | BGBVSGSIXIIREO-UHFFFAOYSA-N | No-Effect | inactive |
| BAY-11-7082 | BRD-K15025317 | Cc1ccc(S(=O)(=O)/C=C/C#N)cc1 | DOEWDSDBFRHVAP-KRXBUXKQSA-N | No-Effect | inactive |
| BAY-11-7085 | BRD-K48684885 | CC(C)(C)c1ccc(S(=O)(=O)/C=C/C#N)cc1 | VHKZGNPOHPFPER-ONNFQVAWSA-N | No-Effect | inactive |
| BAY-1436032 | BRD-K00003359 | C[C@H]1C[C@@H](n2c(Nc3ccc(OC(F)(F)F)cc3)nc3cc(CCC(=O)O)ccc32)CC(C)(C)C1 | RNMAUIMMNAHKQR-QFBILLFUSA-N | No-Effect | inactive |
| BAY-41-2272 | BRD-K32033120 | Nc1nc(-c2nn(Cc3ccccc3F)c3ncccc23)ncc1C1CC1 | ATOAHNRJAXSBOR-UHFFFAOYSA-N | No-Effect | inactive |
| BAY-60-6583 | BRD-K32952627 | N#Cc1c(N)nc(SCC(N)=O)c(C#N)c1-c1ccc(OCC2CC2)cc1 | ZTYHZMAZUWOXNC-UHFFFAOYSA-N | No-Effect | inactive |
| BAY-60-7550 | BRD-K86932920 | COc1ccc(Cc2nc(=O)c3c(C)nc([C@@H](CCCc4ccccc4)[C@@H](C)O)n3[nH]2)cc1OC | MYTWFJKBZGMYCS-NQIIRXRSSA-N | No-Effect | inactive |
| BAY-61-3606 | BRD-K53281329 | COc1ccc(-c2cc3nccn3c(Nc3ncccc3C(N)=O)n2)cc1OC | JWQOJVOKBAAAAR-UHFFFAOYSA-N | No-Effect | inactive |
| BAY-8002 | BRD-K00005331 | O=C(Nc1ccccc1C(=O)O)c1cc(S(=O)(=O)c2ccccc2)ccc1Cl | CLAUJSRBKSRTGQ-UHFFFAOYSA-N | No-Effect | inactive |
| BAY-85-8050 | BRD-A11731518 | N[C@@H](C[C@@H](F)C(=O)O)C(=O)O | JPSHPWJJSVEEAX-GBXIJSLDSA-N | No-Effect | inactive |
| BAY-87-2243 | BRD-K70463136 | Cc1cc(-c2nc(-c3ccc(OC(F)(F)F)cc3)no2)nn1Cc1ccnc(N2CCN(C3CC3)CC2)c1 | CDJNNOJINJAXPV-UHFFFAOYSA-N | No-Effect | inactive |
| BAY-K-8644-(+/-) | BRD-A05457250 | COC(=O)C1=C(C)NC(C)=C([N+](=O)[O-])[C@@H]1c1ccccc1C(F)(F)F | ZFLWDHHVRRZMEI-CYBMUJFWSA-N | No-Effect | inactive |
| BAY-K-8644-(s)-(-) | BRD-K14329163 | COC(=O)C1=C(C)NC(C)=C([N+](=O)[O-])[C@H]1c1ccccc1C(F)(F)F | ZFLWDHHVRRZMEI-ZDUSSCGKSA-N | No-Effect | inactive |
| BAY-W-9798 | BRD-K44471774 | COC(=O)C1=C(C)N(C)C(C)=C(C(=O)OC)C1c1ccc(C(F)(F)F)cc1 | AVFYPHPSIBAAME-UHFFFAOYSA-N | No-Effect | inactive |
| BAY1217389 | BRD-K00003143 | COc1ccc(Oc2cc(NCCC(F)(F)F)c3ncc(-c4ccc(C(=O)NC5CC5)c(C)c4)n3n2)c(F)c1F | WNEILUNVMHVMPH-UHFFFAOYSA-N | No-Effect | inactive |
| BAZ2-ICR | BRD-K66830666 | Cn1cc(CCn2cnc(-c3cnn(C)c3)c2-c2ccc(C#N)cc2)cn1 | RRZVGDGTWNQAPW-UHFFFAOYSA-N | No-Effect | inactive |
| bazedoxifene | BRD-K90195324 | Cc1c(-c2ccc(O)cc2)n(Cc2ccc(OCCN3CCCCCC3)cc2)c2ccc(O)cc12 | UCJGJABZCDBEDK-UHFFFAOYSA-N | No-Effect | inactive |
| BC-11 | BRD-K14646328 | N=C(N)SCc1ccc(B(O)O)cc1 | KDQVMGQHPAKJAN-UHFFFAOYSA-N | No-Effect | inactive |
| BCI-540 | BRD-K23672206 | Cc1oc2nc3c(c(NC(=O)CN4CCCC4=O)c2c1C)CCCC3 | PSPGQHXMUKWNDI-UHFFFAOYSA-N | No-Effect | inactive |
| BCTC | BRD-K66541497 | CC(C)(C)c1ccc(NC(=O)N2CCN(c3ncccc3Cl)CC2)cc1 | ROGUAPYLUCHQGK-UHFFFAOYSA-N | No-Effect | inactive |
| BCX-1470 | BRD-K90472149 | N=C(N)c1cc2ccc(OC(=O)c3cccs3)cc2s1 | OTGQTQBPQCRNRG-UHFFFAOYSA-N | No-Effect | inactive |
| BD-1008 | BRD-K17008822 | CN(CCc1ccc(Cl)c(Cl)c1)CCN1CCCC1 | ASGIQUHBAVIOTI-UHFFFAOYSA-N | No-Effect | inactive |
| BD-1047 | BRD-K36864847 | CN(C)CCN(C)CCc1ccc(Cl)c(Cl)c1 | MGVRNMUKTZOQOW-UHFFFAOYSA-N | No-Effect | inactive |
| BD-1063 | BRD-K29668683 | CN1CCN(CCc2ccc(Cl)c(Cl)c2)CC1 | SUIZRDJCBVPASY-UHFFFAOYSA-N | No-Effect | inactive |
| BE-2254 | BRD-A24429032 | O=C1c2ccccc2CC[C@@H]1CNCCc1ccc(O)cc1 | PZZOEXPDTYIBPI-MRXNPFEDSA-N | No-Effect | inactive |
| beclabuvir | BRD-K00003128 | COc1ccc2c(c1)[C@@H]1C[C@]1(C(=O)N1[C@H]3CC[C@@H]1CN(C)C3)Cn1c-2c(C2CCCCC2)c2ccc(C(=O)NS(=O)(=O)N(C)C)cc21 | ZTTKEBYSXUCBSE-QDFUAKMASA-N | No-Effect | inactive |
| beclamide | BRD-K44824027 | O=C(CCCl)NCc1ccccc1 | JPYQFYIEOUVJDU-UHFFFAOYSA-N | No-Effect | inactive |

| | | | | | |
|---|---|---|---|---|---|
| beclomethasone | BRD-A73385064 | C[C@H]1C[C@H]2[C@H]3CCC4=CC(=O)C=C[C@@]4(C)[C@@]3(Cl)[C@@H](O)C[C@@]2(C)[C@@]1(O)C(=O)CO | NBMKJKDGKREAPL-DGLOSGAVSA-N | No-Effect | inactive |
| beclomethasone-dipropionate | BRD-K97810537 | CCC(=O)OCC(=O)[C@@]1(OC(=O)CC)[C@@H](C)C[C@H]2[C@@H]3CCC4=CC(=O)C=C[C@]4(C)[C@@]3(Cl)[C@@H](O)C[C@@]21C | KUVIULQEHSCUHY-XYWKZLDCSA-N | No-Effect | inactive |
| bedaquiline | BRD-K90817094 | COc1nc2ccc(Br)cc2cc1[C@@H](c1ccccc1)[C@@](O)(CCN(C)C)c1cccc2ccccc12 | QUIJNHUBAXPXFS-XLJNKUFUSA-N | No-Effect | inactive |
| befuraline | BRD-K99077012 | O=C(c1cc2ccccc2o1)N1CCN(Cc2ccccc2)CC1 | SRIJFPBZWUFLFD-UHFFFAOYSA-N | No-Effect | inactive |
| begacestat | BRD-K58319700 | O=S(=O)(N[C@H](CO)C(C(F)(F)F)C(F)(F)F)c1ccc(Cl)s1 | PSXOKXJMVRSARX-SCSAIBSYSA-N | No-Effect | inactive |
| bekanamycin | BRD-K73425385 | NC[C@H]1O[C@H](O[C@@H]2[C@@H](N)C[C@@H](N)[C@H](O[C@H]3O[C@H](CO)[C@@H](O)[C@H](N)[C@H]3O)[C@H]2O)[C@H](N)[C@@H](O)[C@@H]1O | SKKLOUVUUNMCJE-FQSMHNGLSA-N | No-Effect | inactive |
| belotecan | BRD-K00003160 | CC[C@@]1(O)C(=O)OCc2c1cc1n(c2=O)Cc2c-1nc1ccccc1c2CCNC(C)C | LNHWXBUNXOXMRL-VWLOTQADSA-N | No-Effect | inactive |
| bemegride | BRD-K14993104 | CCC1(C)CC(=O)NC(=O)C1 | ORRZGUBHBVWWOP-UHFFFAOYSA-N | No-Effect | inactive |
| bemesetron | BRD-K00004211 | CN1[C@H]2CC[C@@H]1C[C@H](OC(=O)c1cc(Cl)cc(Cl)c1)C2 | MNJNPLVXBISNSX-WDNDVIMCSA-N | No-Effect | inactive |
| bemotrizinol | BRD-A92974210 | CCCC[C@H](CC)COc1ccc(-c2nc(-c3ccc(OC)cc3)nc(-c3ccc(OC[C@H](CC)CCCC)cc3O)n2)c(O)c1 | XVAMCHGMPYWHNL-MKPDMIMOSA-N | No-Effect | inactive |
| benactyzine | BRD-K85030058 | CCN(CC)CCOC(=O)C(O)(c1ccccc1)c1ccccc1 | IVQOFBKHQCTVQV-UHFFFAOYSA-N | No-Effect | inactive |
| benaxibine | BRD-K00004531 | O=C(O)c1ccc(N[C@@H]2O[C@H](CO)[C@@H](O)[C@H](O)[C@H]2O)cc1 | VXMVXDITZWJXJN-RMPHRYRLSA-N | No-Effect | inactive |
| benazepril | BRD-K49807096 | CCOC(=O)[C@H](CCc1ccccc1)N[C@H]1CCc2ccccc2N(CC(=O)O)C1=O | XPCFTKFZXHTYIP-PMACEKPBSA-N | No-Effect | inactive |
| bendamustine | BRD-K17068645 | Cn1c(CCCC(=O)O)nc2cc(N(CCCl)CCCl)ccc21 | YTKUWDBFDASYHO-UHFFFAOYSA-N | No-Effect | inactive |
| bendazac | BRD-K19515365 | O=C(O)COc1nn(Cc2ccccc2)c2ccccc12 | BYFMCKSPFYVMOU-UHFFFAOYSA-N | No-Effect | inactive |
| bendazol | BRD-K60379054 | c1ccc(Cc2nc3ccccc3[nH]2)cc1 | YTLQFZVCLXFFRK-UHFFFAOYSA-N | No-Effect | inactive |
| bendroflumethiazide | BRD-A80017228 | NS(=O)(=O)c1cc2c(cc1C(F)(F)F)N[C@@H](Cc1ccccc1)NS2(=O)=O | HDWIHXWEUNVBIY-CQSZACIVSA-N | No-Effect | inactive |
| benethamine | BRD-K13473780 | c1ccc(CCNCc2ccccc2)cc1 | UPABQMWFWCMOFV-UHFFFAOYSA-N | No-Effect | inactive |
| benfluorex | BRD-K00003672 | C[C@@H](Cc1cccc(C(F)(F)F)c1)NCCOC(=O)c1ccccc1 | CJAVTWRYCDNHSM-AWEZNQCLSA-N | No-Effect | inactive |
| benfluralin | BRD-K33903083 | CCCCN(CC)c1c([N+](=O)[O-])cc(C(F)(F)F)cc1[N+](=O)[O-] | SMDHCQAYESWHAE-UHFFFAOYSA-N | No-Effect | inactive |
| benfotiamine | BRD-K05319475 | C/C(=C(\CCOP(=O)(O)O)SC(=O)c1ccccc1)N(C=O)Cc1cnc(C)nc1N | BTNNPSLJPBRMLZ-LGMDPLHJSA-N | No-Effect | inactive |
| benidipine | BRD-K97181463 | COC(=O)C1=C(C)NC(C)=C(C(=O)O[C@@H]2CCCN(Cc3ccccc3)C2)[C@@H]1c1cccc([N+](=O)[O-])c1 | QZVNQOLPLYWLHQ-ZEQKJWHPSA-N | No-Effect | inactive |
| benperidol | BRD-K79425933 | O=C(CCCN1CCC(n2c(=O)[nH]c3ccccc32)CC1)c1ccc(F)cc1 | FEBOTPHFXYHVPL-UHFFFAOYSA-N | No-Effect | inactive |
| benproperine | BRD-K01826024 | C[C@@H](COc1ccccc1Cc1ccccc1)N1CCCCC1 | JTUQXGZRVLWBCR-SFHVURJKSA-N | No-Effect | inactive |
| benserazide | BRD-A61194565 | NC(CO)C(=O)NNCc1ccc(O)c(O)c1O | BNQDCRGUHNALGH-UHFFFAOYSA-N | No-Effect | inactive |
| benurestat | BRD-K88553858 | O=C(CNC(=O)c1ccc(Cl)cc1)NO | JFZGBMJPJZDNNT-UHFFFAOYSA-N | No-Effect | inactive |
| benzalkonium | BRD-K34234316 | CCCCCCCCCCCC[N+](C)(C)Cc1ccccc1 | VVZBFOKBSDGVGZ-UHFFFAOYSA-N | No-Effect | inactive |
| benzamil | BRD-K87158025 | NC(=NC(=O)c1nc(Cl)c(N)nc1N)NCc1ccccc1 | KXDROGADUISDGY-UHFFFAOYSA-N | No-Effect | inactive |
| benzbromarone | BRD-K11717138 | CCc1oc2ccccc2c1C(=O)c1cc(Br)c(O)c(Br)c1 | WHQCHUCQKNIQEC-UHFFFAOYSA-N | No-Effect | inactive |
| benzenemethanol,-2,5-dimethyl-[[(1-methylethyl)amino]methyl] | BRD-A98083480 | Cc1ccc(C)c([C@H](O)CNC(C)C)c1 | PHFWPIKQCFQPJY-CYBMUJFWSA-N | No-Effect | inactive |
| benzethonium | BRD-K72723676 | CC(C)(C)CC(C)(C)c1ccc(OCCOCC[N+](C)(C)Cc2ccccc2)cc1 | SIYLLGKDQZGJHK-UHFFFAOYSA-N | No-Effect | inactive |
| benznidazole | BRD-K56156805 | O=C(Cn1ccnc1[N+](=O)[O-])NCc1ccccc1 | CULUWZNBISUWAS-UHFFFAOYSA-N | No-Effect | inactive |
| benzoclidine | BRD-A45929034 | O=C(O[C@H]1CN2CCC1CC2)c1ccccc1 | AHKAOMZZTQULDS-ZDUSSCGKSA-N | No-Effect | inactive |
| benzofenac | BRD-K26556907 | O=C(O)Cc1ccc(OCc2ccccc2)c(Cl)c1 | DFHBBMJTBBLQSA-UHFFFAOYSA-N | No-Effect | inactive |
| benzoic-acid | BRD-K53397409 | O=C(O)c1ccccc1 | WPYMKLBDIGXBTP-UHFFFAOYSA-N | No-Effect | inactive |
| benzoin | BRD-A55748233 | O=C(c1ccccc1)[C@H](O)c1ccccc1 | ISAOCJYIOMOJEB-CYBMUJFWSA-N | No-Effect | inactive |
| benzonatate | BRD-K91699951 | CCCCNc1ccc(C(=O)OCCOCCOCCOCCOCCOCCOCCOCCOC)cc1 | MAFMQEKGGFWBAB-UHFFFAOYSA-N | No-Effect | inactive |
| benzoquinonium-dibromide | BRD-K18772613 | CC[N+](CC)(CCCNC1=CC(=O)C(NCCC[N+](CC)(CC)Cc2ccccc2)=CC1=O)Cc1ccccc1 | YERABYSOHUZTPQ-UHFFFAOYSA-P | No-Effect | inactive |
| benzotript | BRD-K33219230 | O=C(N[C@@H](Cc1c[nH]c2ccccc12)C(=O)O)c1ccc(Cl)cc1 | QJERBBQXOMUURJ-INIZCTEOSA-N | No-Effect | inactive |
| benzoxiquine | BRD-K70327191 | O=C(Oc1cccc2cccnc12)c1ccccc1 | BHKPHCKISVSDGV-UHFFFAOYSA-N | No-Effect | inactive |
| benzoyl-peroxide | BRD-K59986511 | O=C(OOC(=O)c1ccccc1)c1ccccc1 | OMPJBNCRMGITSC-UHFFFAOYSA-N | No-Effect | inactive |
| benzoylpas | BRD-K20431737 | O=C(Nc1ccc(C(=O)O)c(O)c1)c1ccccc1 | GNZCRYWFWKFIDM-UHFFFAOYSA-N | No-Effect | inactive |
| benzthiazide | BRD-K21450440 | NS(=O)(=O)c1cc2c(cc1Cl)NC(=CSCc1ccccc1)NS2(=O)=O | MQMCZGVMQZPYQA-UHFFFAOYSA-N | No-Effect | inactive |
| benzydamine | BRD-K28542495 | CN(C)CCCOc1nn(Cc2ccccc2)c2ccccc12 | CNBGNNVCVSKAQZ-UHFFFAOYSA-N | No-Effect | inactive |
| benzyl-alcohol | BRD-K98494274 | OCc1ccccc1 | WVDDGKGOMKODPV-UHFFFAOYSA-N | No-Effect | inactive |
| benzyl-benzoate | BRD-K52072429 | O=C(OCc1ccccc1)c1ccccc1 | SESFRYSPDFLNCH-UHFFFAOYSA-N | No-Effect | inactive |
| benzyl-isothiocyanate | BRD-K80253852 | S=C=NCc1ccccc1 | MDKCFLQDBWCQCV-UHFFFAOYSA-N | No-Effect | inactive |
| benzyldimethylhexadecylammonium | BRD-K23403298 | CCCCCCCCCCCCCCCC[N+](C)(C)Cc1ccccc1 | QDYLMAYUEZBUFO-UHFFFAOYSA-N | No-Effect | inactive |
| benzyldimethyloctylammonium | BRD-K11576103 | CCCCCCCC[N+](C)(C)Cc1ccccc1 | SHFLYPPECXRCFO-UHFFFAOYSA-N | No-Effect | inactive |
| benzylpenicillin | BRD-K55191674 | CC1(C)S[C@@H]2[C@H](NC(=O)Cc3ccccc3)C(=O)N2[C@H]1C(=O)O | JGSARLDLIJGVTE-MBNYWOFBSA-N | No-Effect | inactive |
| bephenium-hydroxynaphthoate | BRD-K47878171 | C[N+](C)(CCOc1ccccc1)Cc1ccccc1 | AVWWVJUMXRXPNF-UHFFFAOYSA-N | No-Effect | inactive |
| bepridil | BRD-A91008255 | CC(C)COCC(CN(Cc1ccccc1)c1ccccc1)N1CCCC1 | UIEATEWHFDRYRU-UHFFFAOYSA-N | No-Effect | inactive |
| berberine | BRD-K14796088 | COc1ccc2cc3[n+](cc2c1OC)CCc1cc2c(cc1-3)OCO2 | YBHILYKTIRIUTE-UHFFFAOYSA-N | No-Effect | inactive |
| bergapten | BRD-K12968785 | COc1c2ccoc2cc2oc(=O)ccc12 | BGEBZHIAGXMEMV-UHFFFAOYSA-N | No-Effect | inactive |
| bergenin | BRD-K04176066 | COc1c(O)cc2c(c1O)[C@@H]1O[C@H](CO)[C@@H](O)[C@H](O)[C@H]1OC2=O | YWJXCIXBAKGUKZ-HJJNZUOJSA-N | No-Effect | inactive |
| besifloxacin | BRD-K10196357 | N[C@@H]1CCCCN(c2c(F)cc3c(=O)c(C(=O)O)cn(C4CC4)c3c2Cl)C1 | QFFGVLORLPOAEC-SNVBAGLBSA-N | No-Effect | inactive |
| beta-amyloid-synthesis-inhibitor | BRD-K38687824 | Oc1cccc(Nc2ccc(Nc3cccc(O)c3)nn2)c1 | GYNDFOLSPPBBIK-UHFFFAOYSA-N | No-Effect | inactive |
| beta-carotene | BRD-K74225533 | CC1=C(/C=C/C(C)=C/C=C/C(C)=C/C=C/C=C(C)/C=C/C=C(C)/C=C/C2=C(C)CCCC2(C)C)C(C)(C)CCC1 | OENHQHLEOONYIE-JLTXGRSLSA-N | No-Effect | inactive |
| beta-CCB | BRD-K87349602 | CCCCOC(=O)c1cc2c(cn1)[nH]c1ccccc12 | WGNGIELOOKACSB-UHFFFAOYSA-N | No-Effect | inactive |
| beta-elemene | BRD-K21045639 | C=C[C@]1(C)CC[C@@H](C(=C)C)C[C@H]1C(=C)C | OPFTUNCRGUEPRZ-QLFBSQMISA-N | No-Effect | inactive |
| beta-funaltrexamine | BRD-K95110148 | COC(=O)/C=C/C(=O)N[C@@H]1CC[C@@]2(O)[C@H]3Cc4ccc(O)c5c4[C@@]2(CCN3CC2CC2)[C@H]1O5 | PQKHESYTSKMWFP-WZJCLRDWSA-N | No-Effect | inactive |
| beta-glycerophosphoric-acid | BRD-K39061174 | O=P(O)(O)OC(CO)CO | DHCLVCXQIBBOPH-UHFFFAOYSA-N | No-Effect | inactive |

| | | | | | |
|---|---|---|---|---|---|
| beta-hydroxy-beta-methylbutyrate | BRD-M02899238 | CC(C)(O)CC(=O)O.CC(C)(O)CC(=O)O | KORPMXSVHOHPMS-UHFFFAOYSA-N | No-Effect | inactive |
| beta-hydroxybutyrate | BRD-A41387824 | CC(O)CC(=O)O | WHBMMWSBFZVSSR-UHFFFAOYSA-N | No-Effect | inactive |
| beta-lapachone | BRD-K28061410 | CC1(C)CCC2=C(O1)c1ccccc1C(=O)C2=O | QZPQTZZNNJUOLS-UHFFFAOYSA-N | No-Effect | inactive |
| beta-naphthol | BRD-K21164796 | Oc1ccc2ccccc2c1 | JWAZRIHNYRIHIV-UHFFFAOYSA-N | No-Effect | inactive |
| betaine | BRD-K12014493 | C[N+](C)(C)CC(=O)O | KWIUHFFTVRNATP-UHFFFAOYSA-O | No-Effect | inactive |
| betamethasone | BRD-K39188321 | C[C@H]1C[C@H]2[C@@H]3CCC4=CC(=O)C=C[C@]4(C)[C@@]3(F)[C@@H](O)C[C@]2(C)[C@@]1(O)C(=O)CO | UREBDLICKHMUKA-DVTGEIKXSA-N | No-Effect | inactive |
| betamethasone-acetate | BRD-K00025439 | CC(=O)OCC(=O)[C@@]1(O)[C@@H](C)C[C@H]2[C@@H]3CCC4=CC(=O)C=C[C@]4(C)[C@@]3(F)[C@@H](O)C[C@@]21C | AKUJBENLRBOFTD-QZIXMDIESA-N | No-Effect | inactive |
| betamethasone-valerate | BRD-K34032314 | CCCCC(=O)O[C@]1(C(=O)CO)[C@@H](C)C[C@H]2[C@@H]3CCC4=CC(=O)C=C[C@]4(C)[C@@]3(F)[C@@H](O)C[C@@]21C | SNHRLVCMMWUAJD-SUYDQAKGSA-N | No-Effect | inactive |
| betamipron | BRD-K82142815 | O=C(O)CCNC(=O)c1ccccc1 | CWXYHOHYCJXYFQ-UHFFFAOYSA-N | No-Effect | inactive |
| betaxolol | BRD-K01825592 | CC(C)NC[C@@H](O)COc1ccc(CCOCC2CC2)cc1 | NWIUTZDMDHAVTP-QGZVFWFLSA-N | No-Effect | inactive |
| betazole | BRD-K20075662 | NCCc1ccn[nH]1 | JXDFEQONERDKSS-UHFFFAOYSA-N | No-Effect | inactive |
| BETP | BRD-K00004305 | CC[S@](=O)c1nc(-c2cccc(OCc3ccccc3)c2)cc(C(F)(F)F)n1 | NTDFYGSSDDMNHI-NDEPHWFRSA-N | No-Effect | inactive |
| betrixaban | BRD-K45275534 | COc1ccc(NC(=O)c2ccc(C(=N)N(C)C)cc2)c(C(=O)Nc2ccc(Cl)cn2)c1 | XHOLNRLADUSQLD-UHFFFAOYSA-N | No-Effect | inactive |
| betulinic-acid | BRD-K45401373 | C=C(C)[C@@H]1CC[C@]2(C(=O)O)CC[C@]3(C)[C@H](CC[C@@H]4[C@@]5(C)CC[C@H](O)C(C)(C)[C@@H]5CC[C@]43C)[C@@H]12 | QGJZLNKBHJESQX-FZFNOLFKSA-N | No-Effect | inactive |
| bevantolol | BRD-K01825622 | COc1ccc(CCNC[C@@H](O)COc2cccc(C)c2)cc1OC | HXLAFSUPPDYFEO-QGZVFWFLSA-N | No-Effect | inactive |
| bevenopran | BRD-K00004680 | COc1cc(CNCCC2CCOCC2)ccc1Oc1cnc(C(N)=O)cn1 | ZGCYVRNZWGUXNQ-UHFFFAOYSA-N | No-Effect | inactive |
| bevirimat-dimeglumine | BRD-K16898998 | C=C(C)[C@@H]1CC[C@]2(C(=O)O)CC[C@]3(C)[C@H](CC[C@@H]4[C@@]5(C)CC[C@H](OC(=O)CC(C)(C)C(=O)O)C(C)(C)[C@@H]5CC[C@]43C)[C@@H]12 | YJEJKUQEXFSVCJ-WRFMNRASSA-N | No-Effect | inactive |
| bexagliflozin | BRD-K00003246 | OC[C@H]1O[C@@H](c2ccc(Cl)c(Cc3ccc(OCCOC4CC4)cc3)c2)[C@H](O)[C@@H](O)[C@@H]1O | BTCRKOKVYTVOLU-SJSRKZJXSA-N | No-Effect | inactive |
| bexarotene | BRD-K92441787 | C=C(c1ccc(C(=O)O)cc1)c1cc2c(cc1C)C(C)(C)CCC2(C)C | NAVMQTYZDKMPEU-UHFFFAOYSA-N | No-Effect | inactive |
| bezafibrate | BRD-K46018455 | CC(C)(Oc1ccc(CCNC(=O)c2ccc(Cl)cc2)cc1)C(=O)O | IIBYAHWJQTYFKB-UHFFFAOYSA-N | No-Effect | inactive |
| BFH772 | BRD-K00003368 | O=C(Nc1cccc(C(F)(F)F)c1)c1cccc2cc(Oc3cc(CO)ncn3)ccc12 | JNLSTLQFDDAULK-UHFFFAOYSA-N | No-Effect | inactive |
| BGP-15 | BRD-K00003393 | N=C(NOC[C@@H](O)CN1CCCCC1)c1cccnc1 | MVLOQULXIYSERZ-ZDUSSCGKSA-N | No-Effect | inactive |
| BHQ | BRD-K95603879 | CC(C)(C)c1cc(O)c(C(C)(C)C)cc1O | JZODKRWQWUWGCD-UHFFFAOYSA-N | No-Effect | inactive |
| BI-224436 | BRD-K53422148 | Cc1nc2ccccc2c(-c2ccc3c4c(ccnc24)CCO3)c1[C@H](OC(C)(C)C)C(=O)O | MIXIIJCBELCMCZ-VWLOTQADSA-N | No-Effect | inactive |
| BI-409306 | BRD-K00003120 | Oc1nc(Cc2ccccn2)nc2c1cnn2C1CCOCC1 | BZTIJCSHNVZMES-UHFFFAOYSA-N | No-Effect | inactive |
| BI-78D3 | BRD-K73982490 | O=c1[nH]nc(Sc2ncc([N+](=O)[O-])s2)n1-c1ccc2c(c1)OCCO2 | QFRLDZGQEZCCJZ-UHFFFAOYSA-N | No-Effect | inactive |
| BI-847325 | BRD-K37492701 | CCNC(=O)C#Cc1ccc2c(c1)NC(=O)/C2=C(/Nc1ccc(CN(C)C)cc1)c1ccccc1 | FLBNLJLONKAPLR-BYYHNAKLSA-N | No-Effect | inactive |
| BIA-10-2474 | BRD-K00004735 | CN(C(=O)n1cnc(-c2ccc[n+]([O-])c2)c1)C1CCCCC1 | DOWVMJFBDGWVML-UHFFFAOYSA-N | No-Effect | inactive |
| biapenem | BRD-A80321291 | C[C@@H](O)[C@H]1C(=O)N2C(C(=O)O)=C(SC3Cn4cnc[n+]4C3)[C@H](C)[C@H]12 | MRMBZHPJVKCOMA-YJFSRANCSA-O | No-Effect | inactive |
| BIBN4096 | BRD-K43927145 | NCCCC[C@H](NC(=O)[C@@H](Cc1cc(Br)c(O)c(Br)c1)NC(=O)N1CCC(N2Cc3ccccc3NC2=O)CC1)C(=O)N1CCN(c2ccncc2)CC1 | ITIXDWVDFFXNEG-JHOUSYSJSA-N | No-Effect | inactive |
| bicifadine | BRD-K00004299 | Cc1ccc([C@@]23CNC[C@@H]2C3)cc1 | OFYVIGTWSQPCLF-NWDGAFQWSA-N | No-Effect | inactive |
| bictegravir | BRD-K00003247 | O=C(NCc1c(F)cc(F)cc1F)c1cn2c(c(O)c1=O)C(=O)N1[C@H]3CC[C@H](C3)O[C@@H]1C2 | SOLUWJRYJLAZCX-LYOVBCGYSA-N | No-Effect | inactive |
| bicuculline-(+) | BRD-K84363966 | CN1CCc2cc3c(cc2[C@H]1[C@@H]1OC(=O)c2c1ccc1c2OCO1)OCO3 | IYGYMKDQCDOMRE-ZWKOTPCHSA-N | No-Effect | inactive |
| bicuculline-methochloride-(-) | BRD-K19094907 | C[N+]1(C)CCc2cc3c(cc2[C@H]1[C@@H]1OC(=O)c2c1ccc1c2OCO1)OCO3 | WDIQXKYUSINZME-RBUKOAKNSA-N | No-Effect | inactive |
| bicyclol | BRD-K16546375 | COC(=O)c1cc(OC)c2c(c1-c1c(CO)cc(OC)c3c1OCO3)OCO2 | KXMTXZACPVCDMH-UHFFFAOYSA-N | No-Effect | inactive |
| bifemelane | BRD-K18779551 | CNCCCCOc1ccccc1Cc1ccccc1 | QSQQPMHPCBLLGX-UHFFFAOYSA-N | No-Effect | inactive |
| bifendate | BRD-K26581543 | COC(=O)c1cc(OC)c2c(c1-c1c(C(=O)OC)cc(OC)c3c1OCO3)OCO2 | JMZOMFYRADAWOG-UHFFFAOYSA-N | No-Effect | inactive |
| bifonazole | BRD-A94543220 | c1ccc(-c2ccc([C@H](c3ccccc3)n3ccnc3)cc2)cc1 | OCAPBUJLXMYKEJ-QFIPXVFZSA-N | No-Effect | inactive |
| bilastine | BRD-K51937257 | CCOCCn1c(C2CCN(CCc3ccc(C(C)(C)C(=O)O)cc3)CC2)nc2ccccc21 | ACCMWZWAEFYUGZ-UHFFFAOYSA-N | No-Effect | inactive |
| bilobalide | BRD-K32039267 | CC(C)(C)[C@]1(O)C[C@@H]2OC(=O)C[C@@]23C(=O)O[C@@H]2OC(=O)[C@H](O)[C@@]231 | MOLPUWBMSBJXER-YDGSQGCISA-N | No-Effect | inactive |
| bimatoprost | BRD-K00003616 | CCNC(=O)CCC/C=C\C[C@H]1[C@@H](O)C[C@@H](O)[C@@H]1/C=C/[C@@H](O)CCc1ccccc1 | AQOKCDNYWBIDND-FTOWTWDKSA-N | No-Effect | inactive |
| BINA | BRD-K00004296 | Cc1c(OCc2cccc(-c3ccc(C(=O)O)cc3)c2)cc2c(c1C)C(=O)[C@@H](C1CCCC1)C2 | KMKBEESNZAPKMP-AREMUKBSSA-N | No-Effect | inactive |
| bindarit | BRD-K36467523 | CC(C)(OCc1nn(Cc2ccccc2)c2ccccc12)C(=O)O | MTHORRSSURHQPZ-UHFFFAOYSA-N | No-Effect | inactive |
| binimetinib | BRD-K82244583 | Cn1cnc2c(F)c(Nc3ccc(Br)cc3F)c(C(=O)NOCCO)cc21 | ACWZRVQXLIRSDF-UHFFFAOYSA-N | No-Effect | inactive |
| BIO-1211 | BRD-K03399463 | Cc1ccccc1NC(=O)Nc1ccc(CC(=O)N[C@@H](CC(C)C)C(=O)N[C@@H](CC(=O)O)C(=O)N[C@H](C(=O)N2CCC[C@H]2C(=O)O)C(C)C)cc1 | NVVGCQABIHSJSQ-KFZSMJGVSA-N | No-Effect | inactive |
| BIO-5192 | BRD-K21318844 | Cc1ccccc1NC(=O)Nc1ccc(CC(=O)N(C)[C@@H](CC(C)C)C(=O)NCC[C@H](NC(=O)[C@@H]2CCCN2S(=O)(=O)c2cc(Cl)cc(Cl)c2)C(=O)O)cc1 | MNQBPRHHZPXCKZ-ZDCRTTOTSA-N | No-Effect | inactive |
| biotin | BRD-K89210380 | O=C(O)CCCC[C@@H]1SC[C@@H]2NC(=O)N[C@H]12 | YBJHBAHKTGYVGT-ZKWXMUAHSA-N | No-Effect | inactive |
| bipenamol | BRD-K47690382 | NCc1ccccc1Sc1ccccc1CO | HLAFSNJRKZLMPT-UHFFFAOYSA-N | No-Effect | inactive |
| birinapant | BRD-K71221037 | CC[C@H](NC(=O)[C@H](C)NC)C(=O)N1C[C@@H](O)C[C@H]1Cc1c(-c2[nH]c3cc(F)ccc3c2C[C@@H]2C[C@H](O)CN2C(=O)[C@H](CC)NC(=O)[C@H](C)NC)[nH]c2cc(F)ccc12 | PKWRMUKBEYJEIX-DXXQBUJASA-N | No-Effect | inactive |
| BIRT-377 | BRD-K64880136 | CN1C(=O)N(c2cc(Cl)cc(Cl)c2)C(=O)[C@@]1(C)Cc1ccc(Br)cc1 | FJNJHZQMQRVZEE-GOSISDBHSA-N | No-Effect | inactive |
| bis(maltolato)oxovanadium(IV) | BRD-K16180792 | Cc1occc(=O)c1O[V](=O)Oc1c(C)occc1=O | XUOLEICXAPEOSI-UHFFFAOYSA-L | No-Effect | inactive |
| bisacodyl | BRD-K39987650 | CC(=O)Oc1ccc(C(c2ccc(OC(C)=O)cc2)c2ccccn2)cc1 | KHOITXIGCFIULA-UHFFFAOYSA-N | No-Effect | inactive |
| bisantrene | BRD-K00003395 | C(=N/NC1=NCCN1)\c1c2ccccc2c(/C=N/NC2=NCCN2)c2ccccc12 | NJSMWLQOCQIOPE-OCHFTUDZSA-N | No-Effect | inactive |
| bisindolylmaleimide-IX | BRD-K06543683 | Cn1cc(C2=C(c3cn(CCCSC(=N)N)c4ccccc34)C(=O)NC2=O)c2ccccc21 | DSXXEELGXBCYNQ-UHFFFAOYSA-N | No-Effect | inactive |
| bismuth-oxalate | BRD-M02179497 | O=C(O)C(=O)O.O=C(O)C(=O)O.O=C(O)C(=O)O.[Bi].[Bi] | CQNPQVYSAJWZIP-UHFFFAOYSA-N | No-Effect | inactive |
| bismuth-subcitrate-potassium | BRD-K01666936 | O=C(O)C[C@](O)(CC(=O)O[Bi]1OC(=O)C[C@@](O)(CC(=O)O)C(=O)O1)C(=O)O | KNOSZBQOKCZXTI-UHFFFAOYSA-K | No-Effect | inactive |
| bismuth-subgallate | BRD-K65978409 | O=C(O)c1cc(O)c2c(c1)O[Bi](O)O2 | JAONZGLTYYUPCT-UHFFFAOYSA-K | No-Effect | inactive |
| bismuth(III)-trifluoromethanesulfonate | BRD-M56276249 | O=S(=O)(O)C(F)(F)F.O=S(=O)(O)C(F)(F)F.O=S(=O)(O)C(F)(F)F.[Bi] | HUXHLVOKNDXSRQ-UHFFFAOYSA-N | No-Effect | inactive |

| | | | | | |
|---|---|---|---|---|---|
| bisoctrizole | BRD-K00228584 | CC(C)(C)CC(C)(C)c1cc(Cc2cc(C(C)(C)CC(C)(C)C)cc(-n3nc4ccccc4n3)c2O)c(O)c(-n2nc3ccccc3n2)c1 | FQUNFJULCYSSOP-UHFFFAOYSA-N | No-Effect | inactive |
| bisoprolol | BRD-A89175223 | CC(C)NC[C@@H](O)COc1ccc(COCCOC(C)C)cc1 | VHYCDWMUTMEGQY-QGZVFWFLSA-N | No-Effect | inactive |
| bisphenol-A | BRD-K41494493 | CC(C)(c1ccc(O)cc1)c1ccc(O)cc1 | IISBACLAFKSPIT-UHFFFAOYSA-N | No-Effect | inactive |
| bitopertin | BRD-K13756951 | C[C@H](Oc1ccc(S(C)(=O)=O)cc1C(=O)N1CCN(c2ncc(C(F)(F)F)cc2F)CC1)C(F)(F)F | YUUGYIUSCYNSQR-LBPRGKRZSA-N | No-Effect | inactive |
| bitoscanate | BRD-K77548206 | S=C=Nc1ccc(N=C=S)cc1 | OMWQUXGVXQELIX-UHFFFAOYSA-N | No-Effect | inactive |
| bivalirudin | BRD-K73504359 | CC[C@H](C)[C@H](NC(=O)[C@H](CCC(=O)O)NC(=O)[C@H](CCC(=O)O)NC(=O)[C@H](Cc1ccccc1)NC(=O)[C@H](CC(=O)O)NC(=O)CNC(=O)[C@H](CC(N)=O)NC(=O)CNC(=O)CNC(=O)CNC(=O)CNC(=O)[C@@H]1CCCN1C(=O)[C@H](CCCNC(=N)N)NC(=O)[C@@H]1CCCN1C(=O)[C@H](N)Cc1ccccc1)C(=O)N1CCC[C@H]1C(=O)N[C@@H](CCC(=O)O)C(=O)N[C@@H](CCC(=O)O)C(=O)N[C@@H](Cc1ccc(O)cc1)C(=O)N[C@@H](CC(C)C)C(=O)O | OIRCOABEOLEUMC-GEJPAHFPSA-N | No-Effect | inactive |
| BIX-01294 | BRD-K26818574 | COc1cc2nc(N3CCCN(C)CC3)nc(NC3CCN(Cc4ccccc4)CC3)c2cc1OC | OSXFATOLZGZLSK-UHFFFAOYSA-N | No-Effect | inactive |
| BIX-02188 | BRD-K20290250 | CN(C)Cc1cccc(N/C(=C2\C(=O)Nc3cc(C(N)=O)ccc32)c2ccccc2)c1 | FSZPIAXLCCQFCM-FCQUAONHSA-N | No-Effect | inactive |
| BIX-02189 | BRD-K73368362 | CN(C)Cc1cccc(N/C(=C2\C(=O)Nc3cc(C(=O)N(C)C)ccc32)c2ccccc2)c1 | HOMJAAIVTDVQJA-IZHYLOQSSA-N | No-Effect | inactive |
| BL-5583 | BRD-K75282704 | O=C(O)c1cc2ccccc2s1 | DYSJMQABFPKAQM-UHFFFAOYSA-N | No-Effect | inactive |
| blebbistatin-(-) | BRD-K61295575 | Cc1ccc2c(c1)C(=O)[C@]1(O)CCN(c3ccccc3)C1=N2 | LZAXPYOBKSJSEX-GOSISDBHSA-N | No-Effect | inactive |
| bleomycetin | BRD-K00004533 | Cc1c(N)nc([C@H](CC(N)=O)NC[C@H](N)C(N)=O)nc1C(=O)N[C@H](C(=O)N[C@H](C)[C@@H](O)[C@H](C)C(=O)N[C@H](C(=O)NCCc1nc(-c2nc(C(=O)NCCCNCCCCN)cs2)cs1)[C@@H](C)O)[C@@H](O[C@@H]1O[C@@H](CO)[C@@H](O)[C@H](O)[C@@H]1O[C@@H]1O[C@H](CO)[C@@H](O)[C@H](OC(N)=O)[C@@H]1O)c1c[nH]cn1 | QYOAUOAXCQAEMW-DDZPZTJXSA-N | No-Effect | inactive |
| bleomycin | BRD-K00003588 | Cc1c(N)nc([C@H](CC(N)=O)NC[C@@H](N)C(N)=O)nc1C(=O)N[C@H](C(=O)N[C@@H](C)[C@H](O)[C@@H](C)C(=O)N[C@@H](C(=O)NCCc1nc(-c2nc(C(=O)NCCC[S+](C)C)cs2)cs1)[C@@H](C)O)[C@@H](O[C@H]1O[C@H](CO)[C@@H](O)[C@@H](O)[C@@H]1O[C@H]1O[C@H](CO)[C@@H](O)[C@H](OC(N)=O)[C@@H]1O)c1c[nH]cn1 | OYVAGSVQBOHSSS-WMBAVFRVSA-O | No-Effect | inactive |
| blonanserin | BRD-K25943794 | CCN1CCN(c2cc(-c3ccc(F)cc3)c3c(n2)CCCCCC3)CC1 | XVGOZDAJGBALKS-UHFFFAOYSA-N | No-Effect | inactive |
| BLU9931 | BRD-K72898849 | C=CC(=O)Nc1cccc(C)c1Nc1ncc2cc(-c3c(Cl)c(OC)cc(OC)c3Cl)ccc2n1 | TXEBNKKOLVBTFK-UHFFFAOYSA-N | No-Effect | inactive |
| BLZ945 | BRD-K83157152 | CNC(=O)c1cc(Oc2ccc3nc(N[C@@H]4CCCC[C@H]4O)sc3c2)ccn1 | ADZBMFGQQWPHMJ-RHSMWYFYSA-N | No-Effect | inactive |
| BML-190 | BRD-K94379058 | COc1ccc2c(c1)c(CC(=O)N1CCOCC1)c(C)n2C(=O)c1ccc(Cl)cc1 | BJSDNVVWJYDOLK-UHFFFAOYSA-N | No-Effect | inactive |
| BML-284 | BRD-K42624714 | COc1cccc(-c2cc(NCc3ccc4c(c3)OCO4)nc(N)n2)c1 | FABQUVYDAXWUQP-UHFFFAOYSA-N | No-Effect | inactive |
| BMS-182874 | BRD-K56509348 | Cc1noc(NS(=O)(=O)c2cccc3c(N(C)C)cccc23)c1C | MJRGSRRZKSJHOE-UHFFFAOYSA-N | No-Effect | inactive |
| BMS-191011 | BRD-K95609758 | O=c1oc(-c2ccc(C(F)(F)F)cc2)nn1Cc1cc(Cl)ccc1O | QKOWACXSXTXRKA-UHFFFAOYSA-N | No-Effect | inactive |
| BMS-191095 | BRD-K01231377 | CC1(C)Oc2ccc(C#N)cc2[C@H](N(Cc2ncc[nH]2)c2ccc(Cl)cc2)[C@H]1O | SMIKIPXIDLITMP-LEWJYISDSA-N | No-Effect | inactive |
| BMS-207940 | BRD-K00004634 | Cc1onc(NS(=O)(=O)c2ccccc2-c2ccc(-c3ncco3)cc2CN(C)C(=O)CC(C)(C)C)c1C | ORJRYNKVKJAJPY-UHFFFAOYSA-N | No-Effect | inactive |
| BMS-214662 | BRD-K00003203 | N#Cc1ccc2c(c1)CN(S(=O)(=O)c1cccs1)[C@H](Cc1ccccc1)CN2Cc1cnc[nH]1 | OLCWFLWEHWLBTO-HSZRJFAPSA-N | No-Effect | inactive |
| BMS-265246 | BRD-K62814476 | CCCCOc1c(C(=O)c2c(F)cc(C)cc2F)cnc2n[nH]cc12 | SCFMWQIQBVZOQR-UHFFFAOYSA-N | No-Effect | inactive |
| BMS-299897 | BRD-K02950022 | C[C@H](c1ccc(F)cc1CCCC(=O)O)N(c1cc(F)ccc1F)S(=O)(=O)c1ccc(Cl)cc1 | IZAOBRWCUGOKNH-OAHLLOKOSA-N | No-Effect | inactive |
| BMS-309403 | BRD-K87956243 | CCc1c(-c2ccccc2)c(-c2ccccc2)nn1-c1ccccc1-c1cccc(OCC(=O)O)c1 | SJRVJRYZAQYCEE-UHFFFAOYSA-N | No-Effect | inactive |
| BMS-345541 | BRD-K13566078 | Cc1ccc2nc(NCCN)c3ncc(C)n3c2c1 | PSPFQEBFYXJZEV-UHFFFAOYSA-N | No-Effect | inactive |
| BMS-387032 | BRD-K43389698 | CC(C)(C)c1cnc(CSc2cnc(NC(=O)C3CCNCC3)s2)o1 | OUSFTKFNBAZUKL-UHFFFAOYSA-N | No-Effect | inactive |
| BMS-536924 | BRD-K34581968 | Cc1cc(N2CCOCC2)cc2[nH]c(-c3c(NC[C@@H](O)c4cccc(Cl)c4)cc[nH]c3=O)nc12 | ZWVZORIKUNOTCS-OAQYLSRUSA-N | No-Effect | inactive |
| BMS-582949 | BRD-K00003168 | CCCNC(=O)c1cn2ncnc(Nc3cc(C(=O)NC4CC4)ccc3C)c2c1C | GDTQLZHHDRRBEB-UHFFFAOYSA-N | No-Effect | inactive |
| BMS-587101 | BRD-K20093108 | CN1C(=O)N(c2cc(Cl)cc(Cl)c2)C(=O)[C@]12CN(Cc1cc(C(=O)O)cs1)C[C@H]2c1ccc(C#N)cc1 | NXNKJLOEGWSJGI-BKMJKUGQSA-N | No-Effect | inactive |
| BMS-599626 | BRD-K07106112 | Cc1c(NC(=O)OC[C@@H]2COCCN2)cn2ncnc(Nc3ccc4c(cnn4Cc4cccc(F)c4)c3)c12 | LUJZZYWHBDHDQX-QFIPXVFZSA-N | No-Effect | inactive |
| BMS-626529 | BRD-K97883416 | COc1cnc(-n2cnc(C)n2)c2[nH]cc(C(=O)C(=O)N3CCN(C(=O)c4ccccc4)CC3)c12 | QRPZBKAMSFHVRW-UHFFFAOYSA-N | No-Effect | inactive |
| BMS-626531 | BRD-K00004550 | Cc1ccc(C(=O)NC2CC2)cc1NC(=O)c1cnn(-c2ncccc2C(F)(F)F)c1C | ZZZJRULMCGMYRH-UHFFFAOYSA-N | No-Effect | inactive |
| BMS-649 | BRD-K25991633 | CC1(C)CCC(C)(C)c2cc(C3(c4ccc(C(=O)O)cc4)OCCO3)ccc21 | ZZUKALQMHNSWTK-UHFFFAOYSA-N | No-Effect | inactive |
| BMS-688521 | BRD-K85649978 | CN1C(=O)N(c2cc(Cl)cc(Cl)c2)C(=O)[C@]12CN(c1ccc(C(=O)O)cn1)C[C@H]2c1ccc(C#N)cc1 | LILGMDXLRPEBNH-HFZDXXHNSA-N | No-Effect | inactive |
| BMS-690514 | BRD-K76239644 | COc1cccc(Nc2ncnn3ccc(CN4CC[C@@H](N)[C@H](O)C4)c23)c1 | CSGQVNMSRKWUSH-IAGOWNOFSA-N | No-Effect | inactive |
| BMS-707035 | BRD-K49555808 | Cn1c(N2CCCCS2(=O)=O)nc(C(=O)NCc2ccc(F)cc2)c(O)c1=O | VNIWZCGZPBJWBI-UHFFFAOYSA-N | No-Effect | inactive |
| BMS-754807 | BRD-K13049116 | C[C@@]1(C(=O)Nc2ccc(F)nc2)CCCN1c1nc(Nc2cc(C3CC3)n[nH]2)c2cccn2n1 | LQVXSNNAFNGRAH-QHCPKHFHSA-N | No-Effect | inactive |
| BMS-770767 | BRD-K00004655 | O[C@]12CC[C@](c3nnc4c(Oc5ccccc5Cl)cccn43)(CC1)C2 | HNXGIFYHJKEXNA-KDURUIRLSA-N | No-Effect | inactive |
| BMS-777607 | BRD-K40738845 | CCOc1ccn(-c2ccc(F)cc2)c(=O)c1C(=O)Nc1ccc(Oc2ccnc(N)c2Cl)c(F)c1 | VNBRGSXVFBYQNN-UHFFFAOYSA-N | No-Effect | inactive |
| BMS-779788 | BRD-K00003305 | CC(C)(O)c1cn(-c2ccc(-c3cccc(S(C)(=O)=O)c3)cc2)c(C(C)(C)c2ccccc2Cl)n1 | JLPURTXCSILYLW-UHFFFAOYSA-N | No-Effect | inactive |
| BMS-806 | BRD-K55773932 | COc1ccnc2[nH]cc(C(=O)C(=O)N3CCN(C(=O)c4ccccc4)C[C@H]3C)c12 | OKGPFTLYBPQBIX-CQSZACIVSA-N | No-Effect | inactive |
| BMS-813160 | BRD-K00003512 | CC(=O)N[C@@H]1C[C@H](NC(C)(C)C)CC[C@@H]1N1CC[C@H](Nc2ncnc3cc(C(C)(C)C)nn23)C1=O | CMVHFGNTABZQJU-HCXYKTFWSA-N | No-Effect | inactive |
| BMS-817378 | BRD-K81026553 | Nc1nccc(Oc2ccc(NC(=O)c3cn(COP(=O)(O)O)cc(-c4ccc(F)cc4)c3=O)cc2F)c1Cl | KXDZWUPUSDCGDD-UHFFFAOYSA-N | No-Effect | inactive |
| BMS-863233 | BRD-K12511835 | O=c1[nH]c([C@@H]2CCCN2)nc2c1oc1ccc(Cl)cc12 | JJWLXRKVUJDJKG-VIFPVBQESA-N | No-Effect | inactive |
| BMS-906024 | BRD-K29343434 | CN1C(=O)[C@@H](NC(=O)[C@H](CCC(F)(F)F)[C@H](CCC(F)(F)F)C(N)=O)N=C(c2ccccc2)c2ccccc21 | AYOUDDAETNMCBW-GSHUGGBRSA-N | No-Effect | inactive |
| BMS-911543 | BRD-K00003189 | CCn1c(C(=O)N(C2CC2)C2CC2)cc2c3c(ncn3C)c(Nc3cc(C)n(C)n3)nc21 | JCINBYQJBYJGDM-UHFFFAOYSA-N | No-Effect | inactive |
| BMS-935177 | BRD-K00004721 | Cc1c(-c2ccc(C(N)=O)c3[nH]c4cc(C(C)(C)O)ccc4c23)cccc1-n1cnc2ccccc2c1=O | TVJRDCQUZMGBAB-UHFFFAOYSA-N | No-Effect | inactive |
| BMS-983970 | BRD-K22316123 | NC(=O)[C@@H](CC1CC1)[C@@H](CCC(F)(F)F)C(=O)N[C@H]1N=C(c2ccccc2)c2cccc(F)c2NC1=O | RANYIWAZSQWERN-WLENULPWSA-N | No-Effect | inactive |

| | | | | | |
|---|---|---|---|---|---|
| BMS-986020 | BRD-K00004586 | Cc1noc(-c2ccc(-c3ccc(C4(C(=O)O)CC4)cc3)cc2)c1NC(=O)O[C@H](C)c1ccccc1 | GQBRZBHEPUQRPL-LJQANCHMSA-N | No-Effect | inactive |
| BMS-986142 | BRD-K00004591 | Cc1c(-c2c(F)cc(C(N)=O)c3[nH]c4c(c23)CC[C@H](C(C)(C)O)C4)cccc1-n1c(=O)c2cccc(F)c2n(C)c1=O | ZRYMMWAJAFUANM-INIZCTEOSA-N | No-Effect | inactive |
| BMS-986195 | BRD-K00003110 | CC#CC(=O)N[C@H]1CCCN(c2c(F)cc(C(N)=O)c3[nH]c(C)c(C)c23)C1 | VJPPLCNBDLZIFG-ZDUSSCGKSA-N | No-Effect | inactive |
| BMS-986205 | BRD-K00003418 | C[C@@H](C(=O)Nc1ccc(Cl)cc1)[C@H]1CC[C@@H](c2ccnc3ccc(F)cc32)CC1 | KRTIYQIPSAGSBP-ZACQAIPSSA-N | No-Effect | inactive |
| BMS-CCR2-22 | BRD-K15119094 | CSc1ccc(C(=O)N[C@@H]2CCCC[C@@H]2NC(=O)CNC(=O)c2cc(C(F)(F)F)ccc2NC(=O)NC(C)C)cc1 | IBPXYDUJQWENPM-XZOQPEGZSA-N | No-Effect | inactive |
| BMY-14802 | BRD-A15435692 | O[C@@H](CCCN1CCN(c2ncc(F)cn2)CC1)c1ccc(F)cc1 | ZXUYYZPJUGQHLQ-KRWDZBQOSA-N | No-Effect | inactive |
| BMY-45778 | BRD-K84895041 | O=C(O)COc1cccc(-c2ocnc2-c2nc(-c3ccccc3)c(-c3ccccc3)o2)c1 | DSRSEEYZGWTODH-UHFFFAOYSA-N | No-Effect | inactive |
| BMY-7378 | BRD-K31054881 | COc1ccccc1N1CCN(CCN2C(=O)CC3(CCCC3)CC2=O)CC1 | AYYCFGDXLUPJAQ-UHFFFAOYSA-N | No-Effect | inactive |
| BNTX | BRD-K18722736 | O=C1/C(=C/c2ccccc2)C[C@@]2(O)[C@H]3Cc4ccc(O)c5c4[C@@]2(CCN3CC2CC2)[C@H]1O5 | WXOUFNFMPVMGFZ-BDQAUFNLSA-N | No-Effect | inactive |
| boceprevir | BRD-A09735281 | CC(C)(C)NC(=O)N[C@H](C(=O)N1C[C@H]2[C@@H]([C@H]1C(=O)N[C@@H](CC1CCC1)C(=O)C(N)=O)C2(C)C)C(C)(C)C | LHHCSNFAOIFYRV-ZYNAIFEFSA-N | No-Effect | inactive |
| bonaphthone | BRD-K34149328 | O=C1C=Cc2cc(Br)ccc2C1=O | MXWZRRPNVLCHMY-UHFFFAOYSA-N | No-Effect | inactive |
| bopindolol | BRD-K01825685 | Cc1cc2c(OC[C@@H](CNC(C)(C)C)OC(=O)c3ccccc3)cccc2[nH]1 | UUOJIACWOAYWEZ-GOSISDBHSA-N | No-Effect | inactive |
| bornyl-acetate | BRD-A41385909 | CC(=O)O[C@H]1C[C@H]2CC[C@@]1(C)C2(C)C | KGEKLUUHTZCSIP-JFGNBEQYSA-N | No-Effect | inactive |
| boronophenylalanine | BRD-K50122911 | N[C@@H](Cc1ccc(B(O)O)cc1)C(=O)O | NFIVJOSXJDORSP-QMMMGPOBSA-N | No-Effect | inactive |
| BOS-172722 | BRD-K00003111 | CCOc1cc(-c2nncn2C)ccc1Nc1ncc2cc(C)nc(NCC(C)(C)C)c2n1 | SGWLRDAOCLITOM-UHFFFAOYSA-N | No-Effect | inactive |
| bosentan | BRD-K74763371 | COc1ccccc1Oc1c(NS(=O)(=O)c2ccc(C(C)(C)C)cc2)nc(-c2ncccn2)nc1OCCO | GJPICJJJRGTNOD-UHFFFAOYSA-N | No-Effect | inactive |
| bosutinib | BRD-K99964838 | COc1cc(Nc2c(C#N)cnc3cc(OCCCN4CCN(C)CC4)c(OC)cc23)c(Cl)cc1Cl | UBPYILGKFZZVDX-UHFFFAOYSA-N | No-Effect | inactive |
| BP-554 | BRD-K45479396 | c1ccc(N2CCN(CCCOc3ccc4c(c3)OCO4)CC2)cc1 | VUSNBQDZGKZEDL-UHFFFAOYSA-N | No-Effect | inactive |
| BP-897 | BRD-K17378184 | COc1ccccc1N1CCN(CCCCNC(=O)c2ccc3ccccc3c2)CC1 | MNHDKMDLOJSCGN-UHFFFAOYSA-N | No-Effect | inactive |
| BQ-123 | BRD-K00003554 | CC(C)C[C@@H]1NC(=O)[C@@H](C(C)C)NC(=O)[C@@H]2CCCN2C(=O)[C@@H](CC(=O)O)NC(=O)[C@@H](Cc2c[nH]c3ccccc23)NC1=O | VYCMAAOURFJIHD-PJNXIOHISA-N | No-Effect | inactive |
| BQ-788 | BRD-K02490038 | CCCC[C@@H](NC(=O)[C@@H](Cc1cn(C(=O)OC)c2ccccc12)NC(=O)[C@H](CC(C)(C)C)NC(=O)N1[C@@H](C)CCC[C@H]1C)C(=O)O | LPAHKJMGDSJDRG-DJYQTOCQSA-N | No-Effect | inactive |
| BQU57 | BRD-A40257118 | Cc1nn(C)c2c1[C@@H](c1ccc(C(F)(F)F)cc1)C(C#N)C(=N)O2 | QFUFJOJDXRWBNA-YUZLPWPTSA-N | No-Effect | inactive |
| BRD4770 | BRD-K72264770 | COC(=O)c1ccc2c(c1)nc(NC(=O)c1ccccc1)n2CCCc1ccccc1 | UCGWYCMPZXDHNR-UHFFFAOYSA-N | No-Effect | inactive |
| BRD7389 | BRD-K15107389 | O=C1c2ccccc2-c2c(NCCc3ccccc3)c(=O)[nH]c3cccc1c23 | XASCINRGTHLHGM-UHFFFAOYSA-N | No-Effect | inactive |
| BRD9876 | BRD-K89329876 | CC(C)(C)c1ccc2cc(C#N)c(C#N)cc2c1 | MKILROQBJOOZKC-UHFFFAOYSA-N | No-Effect | inactive |
| bremelanotide | BRD-K06328344 | CCCC[C@H](NC(C)=O)C(=O)N[C@H]1CC(=O)NCCCC[C@@H](C(=O)O)NC(=O)[C@H](Cc2c[nH]c3ccccc23)NC(=O)[C@H](CCCNC(=N)N)NC(=O)[C@@H](Cc2ccccc2)NC(=O)[C@H](Cc2cnc[nH]2)NC1=O | FFHBJDQSGDNCIV-MFVUMRCOSA-N | No-Effect | inactive |
| bretylium | BRD-K49669424 | CC[N+](C)(C)Cc1ccccc1Br | AAQOQKQBGPPFNS-UHFFFAOYSA-N | No-Effect | inactive |
| brexpiprazole | BRD-K00911143 | O=c1ccc2ccc(OCCCCN3CCN(c4cccc5sccc45)CC3)cc2[nH]1 | ZKIAIYBUSXZPLP-UHFFFAOYSA-N | No-Effect | inactive |
| brimonidine | BRD-K68264559 | Brc1c(NC2=NCCN2)ccc2nccnc12 | XYLJNLCSTIOKRM-UHFFFAOYSA-N | No-Effect | inactive |
| brincidofovir | BRD-K00004672 | CCCCCCCCCCCCCCCCOCCCOP(=O)(O)CO[C@H](CO)Cn1ccc(N)nc1=O | WXJFKKQWPMNTIM-VWLOTQADSA-N | No-Effect | inactive |
| brinzolamide | BRD-K74913225 | CCN[C@H]1CN(CCCOC)S(=O)(=O)c2sc(S(N)(=O)=O)cc21 | HCRKCZRJWPKOAR-JTQLQIEISA-N | No-Effect | inactive |
| brivanib | BRD-K28428262 | Cc1cc2c(F)c(Oc3ncnn4cc(OC[C@@H](C)O)c(C)c34)ccc2[nH]1 | WCWUXEGQKLTGDX-LLVKDONJSA-N | No-Effect | inactive |
| brivanib-alaninate | BRD-K57187269 | Cc1cc2c(F)c(Oc3ncnn4cc(OC[C@@H](C)OC(=O)[C@H](C)N)c(C)c34)ccc2[nH]1 | LTEJRLHKIYCEOX-OCCSQVGLSA-N | No-Effect | inactive |
| BRL-15572 | BRD-K01826777 | O[C@@H](CN1CCN(c2cccc(Cl)c2)CC1)C(c1ccccc1)c1ccccc1 | QJHCTHPYUOXOGM-DEOSSOPVSA-N | No-Effect | inactive |
| BRL-26314 | BRD-A57806482 | O=C(O)[C@@H](Cc1ccccc1)NCc1ccc(Cl)cc1 | SAOHCSVJQIAGDD-OAHLLOKOSA-N | No-Effect | inactive |
| BRL-37344 | BRD-K07507905 | C[C@H](Cc1ccc(OCC(=O)O)cc1)NC[C@H](O)c1cccc(Cl)c1 | ZGGNJJJYUVRADP-ACJLOTCBSA-N | No-Effect | inactive |
| BRL-44408 | BRD-K01825677 | C[C@@H]1c2ccccc2CN1CC1=NCCN1 | SGOFAUSEYBZKDQ-SNVBAGLBSA-N | No-Effect | inactive |
| BRL-52537 | BRD-A37347161 | O=C(Cc1ccc(Cl)c(Cl)c1)N1CCCC[C@@H]1CN1CCCC1 | GHCCBWMZKJQGLS-OAHLLOKOSA-N | No-Effect | inactive |
| BRL-54443 | BRD-K17868609 | CN1CCC(c2c[nH]c3ccc(O)cc23)CC1 | WKNFADCGOAHBPG-UHFFFAOYSA-N | No-Effect | inactive |
| bromantan | BRD-K86358349 | Brc1ccc(NC2C3CC4CC(C3)CC2C4)cc1 | LWJALJDRFBXHKX-UHFFFAOYSA-N | No-Effect | inactive |
| bromebric-acid | BRD-K17874929 | COc1ccc(C(=O)/C(Br)=C/C(=O)O)cc1 | UPZFHUODAYGHDZ-TWGQIWQCSA-N | No-Effect | inactive |
| bromfenac | BRD-K47679368 | Nc1c(CC(=O)O)cccc1C(=O)c1ccc(Br)cc1 | ZBPLOVFIXSTCRZ-UHFFFAOYSA-N | No-Effect | inactive |
| bromhexine | BRD-K47631482 | CN(Cc1cc(Br)cc(Br)c1N)C1CCCCC1 | OJGDCBLYJGHCIH-UHFFFAOYSA-N | No-Effect | inactive |
| bromindione | BRD-K50490982 | O=C1c2ccccc2C(=O)C1c1ccc(Br)cc1 | NPUZIGSOEWMFKK-UHFFFAOYSA-N | No-Effect | inactive |
| bromisoval | BRD-K00003522 | CC(C)[C@@H](Br)C(=O)NC(N)=O | CMCCHHWTTBEZNM-SCSAIBSYSA-N | No-Effect | inactive |
| bromocriptine | BRD-K14496212 | CC(C)C[C@H]1C(=O)N2CCC[C@H]2[C@]2(O)O[C@](NC(=O)[C@@H]3C=C4c5cccc6[nH]c(Br)c(c56)C[C@H]4N(C)C3)(C(C)C)C(=O)N12 | OZVBMTJYIDMWIL-AYFBDAFISA-N | No-Effect | inactive |
| bromopride | BRD-K73642618 | CCN(CC)CCNC(=O)c1cc(Br)c(N)cc1OC | GIYAQDDTCWHPPL-UHFFFAOYSA-N | No-Effect | inactive |
| bromperidol | BRD-K78643075 | O=C(CCCN1CCC(O)(c2ccc(Br)cc2)CC1)c1ccc(F)cc1 | RKLNONIVDFXQRX-UHFFFAOYSA-N | No-Effect | inactive |
| brompheniramine | BRD-K01825159 | CN(C)CC[C@@H](c1ccc(Br)cc1)c1ccccn1 | ZDIGNSYAACHWNL-HNNXBMFYSA-N | No-Effect | inactive |
| bronopol | BRD-K33457401 | O=[N+]([O-])C(Br)(CO)CO | LVDKZNITIUWNER-UHFFFAOYSA-N | No-Effect | inactive |
| bropirimine | BRD-K17500877 | Nc1nc(=O)c(Br)c(-c2ccccc2)[nH]1 | CIUUIPMOFZIWIZ-UHFFFAOYSA-N | No-Effect | inactive |
| broxaldine | BRD-K36222532 | Cc1ccc2c(Br)cc(Br)c(OC(=O)c3ccccc3)c2n1 | IJTPLVAAROHGGB-UHFFFAOYSA-N | No-Effect | inactive |
| broxaterol | BRD-A08316590 | CC(C)(C)NC[C@@H](O)c1cc(Br)no1 | JBRBWHCVRGURBA-ZCFIWIBFSA-N | No-Effect | inactive |
| broxuridine | BRD-K79095980 | O=c1[nH]c(=O)n([C@H]2C[C@H](O)[C@@H](CO)O2)cc1Br | WOVKYSAHUYNSMH-RRKCRQDMSA-N | No-Effect | inactive |
| broxyquinoline | BRD-K23301018 | Oc1c(Br)cc(Br)c2cccnc12 | ZDASUJMDVPTNTF-UHFFFAOYSA-N | No-Effect | inactive |
| BS-181 | BRD-K31313613 | CC(C)c1cnn2c(NCc3ccccc3)cc(NCCCCCCN)nc12 | DNYBIOICMDTDAP-UHFFFAOYSA-N | No-Effect | inactive |
| BT-11 | BRD-K00003444 | O=C(c1cccc(-c2nc3ccccc3[nH]2)n1)N1CCN(C(=O)c2cccc(-c3nc4ccccc4[nH]3)n2)CC1 | MVHWZNBAQIGPOQ-UHFFFAOYSA-N | No-Effect | inactive |
| BTB06584 | BRD-K37933329 | O=C(Oc1cc(S(=O)(=O)c2ccccc2)ccc1[N+](=O)[O-])c1ccc(Cl)cc1 | WNDWKKPBLAKXMI-UHFFFAOYSA-N | No-Effect | inactive |
| BTB1 | BRD-K89000287 | O=[N+]([O-])c1cc(Cl)ccc1S(=O)(=O)c1ccccc1 | VZDUQPHKUBZMLW-UHFFFAOYSA-N | No-Effect | inactive |
| BTS | BRD-K64178227 | Cc1ccc(S(=O)(=O)NCc2ccccc2)cc1 | WTHKAJZQYNKTCJ-UHFFFAOYSA-N | No-Effect | inactive |
| BTS-54505 | BRD-A68304895 | CC(C)C[C@@H](N)C1(c2ccc(Cl)cc2)CCC1 | WQSACWZKKZPCHN-CQSZACIVSA-N | No-Effect | inactive |
| BTS-71321 | BRD-K00010938 | CC(C)(NCCCn1ccnc1)c1ccc(Cl)cc1 | GRKYRJREJPZXIU-UHFFFAOYSA-N | No-Effect | inactive |
| BTS-72664 | BRD-A65793993 | C[C@@H](Oc1ccc(Cl)cc1)c1ccnc2ncnn12 | FCEPUTGEUCVRRI-SECBINFHSA-N | No-Effect | inactive |
| BTT-3033 | BRD-K01492390 | CN(c1ccc(NC(=O)Nc2ccccc2)cc1)S(=O)(=O)c1cnn(-c2ccc(F)cc2)c1 | NSLIQOPYDUKWTA-UHFFFAOYSA-N | No-Effect | inactive |
| BTZ043-racemate | BRD-A53037217 | C[C@H]1COC2(CCN(c3nc(=O)c4cc(C(F)(F)F)cc([N+](=O)[O-])c4s3)CC2)O1 | GTUIRORNXIOHQR-VIFPVBQESA-N | No-Effect | inactive |
| BU-224 | BRD-K55344148 | c1ccc2nc(C3=NCCN3)ccc2c1 | OSUVWJGCKQEMHK-UHFFFAOYSA-N | No-Effect | inactive |
| BU-239 | BRD-K11696279 | c1ccc2nc(C3=NCCN3)cnc2c1 | MQSSFHXQKUSFIW-UHFFFAOYSA-N | No-Effect | inactive |
| bucetin | BRD-A88684804 | CCOc1ccc(NC(=O)C[C@@H](C)O)cc1 | LIAWQASKBFCRNR-SECBINFHSA-N | No-Effect | inactive |
| bucillamine | BRD-K24697665 | CC(C)(S)C(=O)N[C@@H](CS)C(=O)O | VUAFHZCUKUDDBC-BYPYZUCNSA-N | No-Effect | inactive |

| | | | | | |
|---|---|---|---|---|---|
| bucladesine | BRD-A94624445 | CCCC(=O)Nc1ncnc2c1ncn2[C@@H]1O[C@@H]2CO[P@@](=O)(O)O[C@H]2[C@H]1OC(=O)CCC | CJGYSWNGNKCJSB-YVLZZHOMSA-N | No-Effect | inactive |
| buclizine | BRD-A91444184 | CC(C)(C)c1ccc(CN2CCN([C@@H](c3ccccc3)c3ccc(Cl)cc3)CC2)cc1 | MOYGZHXDRJNJEP-MHZLTWQESA-N | No-Effect | inactive |
| budesonide | BRD-A34299591 | CCC[C@H]1O[C@@H]2C[C@H]3[C@@H]4CCC5=CC(=O)C=C[C@]5(C)[C@H]4[C@@H](O)C[C@]3(C)[C@]2(C(=O)CO)O1 | VOVIALXJUBGFJZ-AMTWPJRWSA-N | No-Effect | inactive |
| budipine | BRD-K18324993 | CC(C)(C)N1CCC(c2ccccc2)(c2ccccc2)CC1 | QIHLUZAFSSMXHQ-UHFFFAOYSA-N | No-Effect | inactive |
| bufexamac | BRD-K36660044 | CCCCOc1ccc(CC(=O)NO)cc1 | MXJWRABVEGLYDG-UHFFFAOYSA-N | No-Effect | inactive |
| buflomedil | BRD-K19462402 | COc1cc(OC)c(C(=O)CCCN2CCCC2)c(OC)c1 | OWYLAEYXIQKAOL-UHFFFAOYSA-N | No-Effect | inactive |
| bumetanide | BRD-K38197229 | CCCCNc1cc(C(=O)O)cc(S(N)(=O)=O)c1Oc1ccccc1 | MAEIEVLCKWDQJH-UHFFFAOYSA-N | No-Effect | inactive |
| bunazosin | BRD-K35034482 | CCCC(=O)N1CCCN(c2nc(N)c3cc(OC)c(OC)cc3n2)CC1 | RHLJLALHBZGAFM-UHFFFAOYSA-N | No-Effect | inactive |
| buphenine | BRD-A36267905 | CC(CCc1ccccc1)NC(C)C(O)c1ccc(O)cc1 | PTGXAUBQBSGPKF-UHFFFAOYSA-N | No-Effect | inactive |
| bupicomide | BRD-K51700582 | CCCCc1ccc(C(N)=O)nc1 | VKSPIPWLHGKJQO-UHFFFAOYSA-N | No-Effect | inactive |
| bupivacaine | BRD-K00003636 | CCCCN1CCCC[C@H]1C(=O)Nc1c(C)cccc1C | LEBVLXFERQHONN-INIZCTEOSA-N | No-Effect | inactive |
| bupranolol | BRD-A48420483 | Cc1ccc(Cl)c(OC[C@@H](O)CNC(C)(C)C)c1 | HQIRNZOQPUAHHV-NSHDSACASA-N | No-Effect | inactive |
| bupropion | BRD-K01826729 | C[C@@H](NC(C)(C)C)C(=O)c1cccc(Cl)c1 | SNPPWIUOZRMYNY-SECBINFHSA-N | No-Effect | inactive |
| buramate | BRD-K30360804 | O=C(NCc1ccccc1)OCCO | YFLRYAVDPKONNX-UHFFFAOYSA-N | No-Effect | inactive |
| buspirone | BRD-K93461745 | O=C1CC2(CCCC2)CC(=O)N1CCCCN1CCN(c2ncccn2)CC1 | QWCRAEMEVRGPNT-UHFFFAOYSA-N | No-Effect | inactive |
| busulfan | BRD-K23204545 | CS(=O)(=O)OCCCCOS(C)(=O)=O | COVZYZSDYWQREU-UHFFFAOYSA-N | No-Effect | inactive |
| butabindide | BRD-K40581315 | CCCCNC(=O)[C@@H]1Cc2ccccc2N1C(=O)[C@@H](N)CC | HAQIRIMIVLFYIX-ZFWWWQNUSA-N | No-Effect | inactive |
| butacaine | BRD-K99622919 | CCCCN(CCCC)CCCOC(=O)c1ccc(N)cc1 | HQFWVSGBVLEQGA-UHFFFAOYSA-N | No-Effect | inactive |
| butaclamol | BRD-K20620780 | CC(C)(C)[C@]1(O)CCN2C[C@H]3c4ccccc4CCc4cccc(c43)[C@@H]2C1 | ZZJYIKPMDIWRSN-HWBMXIPRSA-N | No-Effect | inactive |
| butaprost | BRD-K00004715 | CCCC1([C@H](O)C/C=C/[C@H]2[C@H](O)CC(=O)[C@@H]2CCCCCC(=O)OC)CCC1 | XRISENIKJUKIHD-LHQZMKCDSA-N | No-Effect | inactive |
| butenafine | BRD-K90630139 | CN(Cc1ccc(C(C)(C)C)cc1)Cc1cccc2ccccc12 | ABJKWBDEJIDSJZ-UHFFFAOYSA-N | No-Effect | inactive |
| buthionine-sulfoximine | BRD-A47706533 | CCCCS(=N)(=O)CC[C@H](N)C(=O)O | KJQFBVYMGADDTQ-CVSPRKDYSA-N | No-Effect | inactive |
| butibufen | BRD-A46136514 | CC[C@@H](C(=O)O)c1ccc(CC(C)C)cc1 | UULSXYSSHHRCQK-CYBMUJFWSA-N | No-Effect | inactive |
| butoconazole | BRD-K01824994 | Clc1ccc(CC[C@@H](Cn2ccnc2)Sc2c(Cl)cccc2Cl)cc1 | SWLMUYACZKCSHZ-INIZCTEOSA-N | No-Effect | inactive |
| butofilolol | BRD-A27274504 | CCCC(=O)c1cc(F)ccc1OC[C@@H](O)CNC(C)(C)C | NMBNQRJDEPOXCP-ZDUSSCGKSA-N | No-Effect | inactive |
| butyl-paraben | BRD-K08287586 | CCCCOC(=O)c1ccc(O)cc1 | QFOHBWFCKVYLES-UHFFFAOYSA-N | No-Effect | inactive |
| butylated-hydroxyanisole | BRD-K93035859 | COc1ccc(O)c(C(C)(C)C)c1 | MRBKEAMVRSLQPH-UHFFFAOYSA-N | No-Effect | inactive |
| butylated-hydroxytoluene | BRD-K53153417 | Cc1cc(C(C)(C)C)c(O)c(C(C)(C)C)c1 | NLZUEZXRPGMBCV-UHFFFAOYSA-N | No-Effect | inactive |
| butylidenephthalide | BRD-K02260986 | CCC/C=C1\OC(=O)c2ccccc21 | WMBOCUXXNSOQHM-FLIBITNWSA-N | No-Effect | inactive |
| butylscopolamine-bromide | BRD-K01825841 | CCCC[N@@+]1(C)[C@H]2C[C@H](OC(=O)[C@H](CO)c3ccccc3)C[C@@H]1[C@H]1O[C@@H]21 | YBCNXCRZPWQOBR-FRNBUXIHSA-N | No-Effect | inactive |
| BVD-523 | BRD-K28965160 | CC(C)Nc1cc(-c2c[nH]c(C(=O)N[C@H](CO)c3cccc(Cl)c3)c2)c(Cl)cn1 | KSERXGMCDHOLSS-LJQANCHMSA-N | No-Effect | inactive |
| BVT-2733 | BRD-K01742705 | Cc1c(Cl)cccc1S(=O)(=O)Nc1nc(CC(=O)N2CCN(C)CC2)cs1 | YDPRNGAPPNPYQQ-UHFFFAOYSA-N | No-Effect | inactive |
| BVT-948 | BRD-K68567222 | CC1(C)C(=O)NC2=C1C(=O)C(=O)c1ccccc12 | LLPBUXODFQZPFH-UHFFFAOYSA-N | No-Effect | inactive |
| BW-180C | BRD-K00003548 | CC(C)C[C@@H](NC(=O)[C@H](Cc1ccccc1)NC(=O)CNC(=O)[C@@H](C)NC(=O)[C@@H](N)Cc1ccc(O)cc1)C(=O)O | ZHUJMSMQIPIPTF-IBURTVSXSA-N | No-Effect | inactive |
| BW-348U87 | BRD-K00004593 | C/C(=N\NC(=S)NNC(=S)Nc1ccccc1Cl)c1ccccn1 | YYLKOELWSMRYHV-VXLYETTFSA-N | No-Effect | inactive |
| BW-373U86 | BRD-K74990253 | C=CCN1C[C@H](C)N([C@H](c2ccc(C(=O)N(CC)CC)cc2)c2cccc(O)c2)C[C@H]1C | LBLDMHBSVIVJPM-YZIHRLCOSA-N | No-Effect | inactive |
| BW-616U | BRD-K07061232 | CNC(=O)c1ccc2c(c1)S(=O)(=O)c1ccccc1C2=O | MLPSDBJCWIYSEA-UHFFFAOYSA-N | No-Effect | inactive |
| BW-A4C | BRD-K80194255 | CC(=O)N(O)C/C=C/c1cccc(Oc2ccccc2)c1 | CEUDWZXMLMKPNN-SOFGYWHQSA-N | No-Effect | inactive |
| BW-B70C | BRD-A55946879 | C[C@@H](/C=C/c1cccc(Oc2ccc(F)cc2)c1)N(O)C(N)=O | UAIYNMRLUHHRMF-FYJFLYSWSA-N | No-Effect | inactive |
| BX-795 | BRD-K47983010 | O=C(NCCCNc1nc(Nc2cccc(NC(=O)N3CCCC3)c2)ncc1I)c1cccs1 | VAVXGGRQQJZYBL-UHFFFAOYSA-N | No-Effect | inactive |
| BX-912 | BRD-K49669041 | O=C(Nc1cccc(Nc2ncc(Br)c(NCCc3cnc[nH]3)n2)c1)N1CCCC1 | DMMILYKXNCVKOJ-UHFFFAOYSA-N | No-Effect | inactive |
| BYK-204165 | BRD-K61665160 | Cn1cccc1/C=C1\C(=O)NC(=O)c2ccccc21 | BTYSIDSTHDDAJW-LCYFTJDESA-N | No-Effect | inactive |
| C-1 | BRD-K72462751 | O=S(=O)(c1cccc2cnccc12)N1CCNCC1 | UPTYCYWTFGTCCG-UHFFFAOYSA-N | No-Effect | inactive |
| C-751 | BRD-K15337272 | CCOC(=O)c1cnc[nH]1 | KLWYPRNPRNPORS-UHFFFAOYSA-N | No-Effect | inactive |
| C106 | BRD-M12328004 | CN1CCC[C@@H]1c1cccnc1.Cc1ccc(C(=O)O[C@H](C(=O)O)[C@@H](OC(=O)c2ccc(C)cc2)C(=O)O)cc1 | GEGPCGHEPLKDLY-DRHSTVBOSA-N | No-Effect | inactive |
| C11-Acetate | BRD-K83914624 | [Y+3] | GRTBAGCGDOYUBE-UHFFFAOYSA-N | No-Effect | inactive |
| C188-9 | BRD-K00003114 | COc1ccc(S(=O)(=O)Nc2cc(-c3c(O)ccc4ccccc34)c(O)c3ccccc23)cc1 | QDCJDYWGYVPBDO-UHFFFAOYSA-N | No-Effect | inactive |
| C34 | BRD-K43167535 | CC(=O)N[C@H]1[C@@H](OC(C)C)O[C@H](COC(C)=O)[C@@H](OC(C)=O)[C@@H]1OC(C)=O | KMIQMFHPUJUDMC-HHARLNAUSA-N | No-Effect | inactive |
| C646 | BRD-K73383190 | CC1=NN(c2ccc(C(=O)O)cc2)C(=O)C1=Cc1ccc(-c2cc(C)c(C)cc2[N+](=O)[O-])o1 | HEKJYZZSCQBJGB-UHFFFAOYSA-N | No-Effect | inactive |
| CA-4948 | BRD-K00003511 | Nc1ccc(-c2cccc(C(=O)Nc3cnc4nc(N5CCOCC5)oc4c3)n2)cn1 | RWIMETUXCNDSLE-UHFFFAOYSA-N | No-Effect | inactive |
| cabagin | BRD-K41504985 | C[S+](C)CC[C@H](N)C(=O)O | YDBYJHTYSHBBAU-YFKPBYRVSA-O | No-Effect | inactive |
| cabaletta | BRD-K54028654 | OC[C@H]1O[C@H](O[C@H]2O[C@H](CO)[C@@H](O)[C@H](O)[C@H]2O)[C@H](O)[C@@H](O)[C@@H]1O | HDTRYLNUVZCQOY-LIZSDCNHSA-N | | inactive |
| cabazitaxel | BRD-K10384176 | CO[C@H]1C[C@H]2OC[C@@]2(OC(C)=O)[C@H]2[C@H](OC(=O)c3ccccc3)[C@]3(O)C[C@H](OC(=O)[C@H](O)[C@@H](NC(=O)OC(C)(C)C)c4ccccc4)C(C)=C([C@@H](OC)C(=O)[C@]12C)C3(C)C | BMQGVNUXMIRLCK-OAGWZNDDSA-N | No-Effect | inactive |
| cabergoline | BRD-K86882815 | C=CCN1C[C@H](C(=O)N(CCCN(C)C)C(=O)NCC)C[C@@H]2c3cccc4[nH]cc(c34)C[C@H]21 | KORNTPPJEAJQIU-KJXAQDMKSA-N | No-Effect | inactive |
| cabotegravir | BRD-K76819217 | C[C@H]1CO[C@@H]2Cn3cc(C(=O)NCc4ccc(F)cc4F)c(=O)c(O)c3C(=O)N12 | WCWSTNLSLKSJPK-LKFCYVNXSA-N | No-Effect | inactive |
| cabozantinib | BRD-K51544265 | COc1cc2nccc(Oc3ccc(NC(=O)C4(C(=O)Nc5ccc(F)cc5)CC4)cc3)c2cc1OC | ONIQOQHATWINJY-UHFFFAOYSA-N | No-Effect | inactive |
| CaCCinh-A01 | BRD-A90751343 | CC(C)(C)[C@H]1CCc2c(sc(NC(=O)c3ccco3)c2C(=O)O)C1 | ACLUEOBQFRYTQS-JTQLQIEISA-N | No-Effect | inactive |
| cadazolid | BRD-K00003370 | O=C(O)c1cn(C2CC2)c2cc(N3CCC(O)(COc4ccc(N5C[C@H](CO)OC5=O)cc4F)CC3)c(F)cc2c1=O | XWFCFMXQTBGXQW-GOSISDBHSA-N | No-Effect | inactive |
| cadralazine | BRD-A78134170 | CCOC(=O)NNc1ccc(N(CC)C[C@@H](C)O)nn1 | QLTVVOATEHFXLT-SECBINFHSA-N | No-Effect | inactive |
| cafestol | BRD-K46741531 | C[C@@]12CCc3occc3[C@H]1CC[C@@]13C[C@@H](CC[C@H]12)[C@@](O)(CO)C3 | DNJVYWXIDISQRD-HWUKTEKMSA-N | No-Effect | inactive |
| caffeic-acid | BRD-K09900591 | O=C(O)/C=C/c1ccc(O)c(O)c1 | QAIPRVGONGVQAS-DUXPYHPUSA-N | No-Effect | inactive |
| caffeine | BRD-K02404261 | Cn1c(=O)c2c(ncn2C)n(C)c1=O | RYYVLZVUVIJVGH-UHFFFAOYSA-N | No-Effect | inactive |
| calcifediol | BRD-K77175907 | C=C1CC[C@H](O)C/C1=C/C=C1\CCC[C@@]2(C)[C@H]1CC[C@@H]2[C@H](C)CCCC(C)(C)O | JWUBBDSIWDLEOM-DTOXIADCSA-N | No-Effect | inactive |
| calcipotriol | BRD-K56429665 | C=C1/C(=C\C=C2/CCC[C@@]3(C)[C@H]2CC[C@@H]3[C@H](C)/C=C/[C@@H](O)C2CC2)C[C@@H](O)C[C@@H]1O | LWQQLNNNIPYSNX-UROSTWAQSA-N | No-Effect | inactive |

| | | | | | |
|---|---|---|---|---|---|
| calcitonin | BRD-K00005332 | CC(C)C[C@H](NC(=O)[C@H](CCCCN)NC(=O)CNC(=O)[C@H](CC(C)C)NC(=O)[C@@H](NC(=O)[C@@H]1CSSC[C@H](N)C(=O)N[C@@H](CO)C(=O)N[C@@H](CC(N)=O)C(=O)N[C@@H](CC(C)C)C(=O)N[C@@H](CO)C(=O)N[C@@H]([C@@H](C)O)C(=O)N1)C(C)C)C(=O)N[C@@H](CO)C(=O)N[C@@H](CCC(N)=O)C(=O)N[C@@H](CCC(=O)O)C(=O)N[C@@H](CC(C)C)C(=O)N[C@@H](Cc1c[nH]cn1)C(=O)N[C@@H](CCCCN)C(=O)N[C@@H](CC(C)C)C(=O)N[C@@H](CCC(N)=O)C(=O)N[C@H](C(=O)N[C@@H](Cc1ccc(O)cc1)C(=O)N1CCC[C@H]1C(=O)N[C@@H](CCCNC(=N)N)C(=O)N[C@H](C(=O)N[C@@H](CC(=O)O)C(=O)N[C@H](C(=O)NCC(=O)N[C@@H](C)C(=O)NCC(=O)N[C@H](C(=O)N1CCC[C@H]1C(N)=O)[C@@H](C)O)C(C)C)[C@@H](C)O)[C@@H](C)O | JDJALSWDQPEHEJ-LMVCGNDWSA-N | No-Effect | inactive |
| calcium-gluceptate | BRD-K35952844 | O=C(O)[C@H](O)[C@H](O)[C@@H](O)[C@H](O)[C@H](O)CO | KWMLJOLKUYYJFJ-VFUOTHLCSA-N | No-Effect | inactive |
| calcium-levofolinate | BRD-K97009491 | Nc1nc(=O)c2c([nH]1)NC[C@H](CNc1ccc(C(=O)N[C@@H](CCC(=O)O)C(=O)O)cc1)N2C=O | VVIAGPKUTFNRDU-STQMWFEESA-N | No-Effect | inactive |
| Calhex-231 | BRD-K19811997 | C[C@@H](N[C@H]1CCCC[C@@H]1NC(=O)c1ccc(Cl)cc1)c1cccc2ccccc12 | YTFUQWWKTIWYEY-CQLNOVPUSA-N | No-Effect | inactive |
| CaMKII-IN-1 | BRD-K35772771 | Cc1c(Cl)cccc1S(=O)(=O)Nc1nc(NCCc2ccccc2)nc2c1CN(Cc1ccccc1)CC2 | LGUUETDTMQTHML-UHFFFAOYSA-N | No-Effect | inactive |
| camobucol | BRD-K00004732 | CC(C)(Sc1cc(C(C)(C)C)c(O)c(C(C)(C)C)c1)Sc1cc(C(C)(C)C)c(OCC(=O)O)c(C(C)(C)C)c1 | FGBGXESDYFKUFX-UHFFFAOYSA-N | No-Effect | inactive |
| camostat-mesilate | BRD-K43806473 | CN(C)C(=O)COC(=O)Cc1ccc(OC(=O)c2ccc(NC(=N)N)cc2)cc1 | XASIMHXSUQUHLV-UHFFFAOYSA-N | No-Effect | inactive |
| camphor-(+) | BRD-K10034334 | CC1(C)[C@@H]2CC[C@@]1(C)C(=O)C2 | DSSYKIVIOFKYAU-XCBNKYQSSA-N | No-Effect | inactive |
| camphor-(1R) | BRD-K15714472 | CC1(C)[C@@H]2CC[C@]1(C)C(=O)C2 | DSSYKIVIOFKYAU-GMSGAONNSA-N | No-Effect | inactive |
| camptothecin | BRD-K37890730 | CC[C@@]1(O)C(=O)OCc2c1cc1n(c2=O)Cc2cc3ccccc3nc2-1 | VSJKWCGYPAHWDS-FQEVSTJZSA-N | No-Effect | inactive |
| canagliflozin | BRD-K90868879 | Cc1ccc([C@@H]2O[C@H](CO)[C@@H](O)[C@H](O)[C@H]2O)cc1Cc1ccc(-c2ccc(F)cc2)s1 | XTNGUQKDFGDXSJ-ZXGKGEBGSA-N | No-Effect | inactive |
| candesartan-cilexetil | BRD-A65671304 | CCOc1nc2cccc(C(=O)O[C@@H](C)OC(=O)OC3CCCCC3)c2n1Cc1ccc(-c2ccccc2-c2nn[nH]n2)cc1 | GHOSNRCGJFBJIB-OAQYLSRUSA-N | No-Effect | inactive |
| cangrelor | BRD-K00004669 | CSCCNc1nc(SCCC(F)(F)F)nc2c1ncn2[C@@H]1O[C@H](COP(=O)(O)OP(=O)(O)C(Cl)(Cl)P(=O)(O)O)[C@@H](O)[C@H]1O | PAEBIVWUMLRPSK-IDTAVKCVSA-N | No-Effect | inactive |
| canrenone | BRD-K81644801 | C[C@]12CC[C@H]3[C@@H](C=CC4=CC(=O)CC[C@@]43C)[C@@H]1CC[C@@]21CCC(=O)O1 | UJVLDDZCTMKXJK-WNHSNXHDSA-N | No-Effect | inactive |
| cantharidin | BRD-K80684056 | C[C@]12C(=O)OC(=O)[C@@]1(C)[C@@H]1CC[C@H]2O1 | DHZBEENLJMYSHQ-XCVPVQRUSA-N | No-Effect | inactive |
| capadenoson | BRD-K97025174 | N#Cc1c(N)nc(SCc2csc(-c3ccc(Cl)cc3)n2)c(C#N)c1-c1ccc(OCCO)cc1 | CITWCLNVRIKQAF-UHFFFAOYSA-N | No-Effect | inactive |
| capecitabine | BRD-K61192372 | CCCCCOC(=O)Nc1nc(=O)n([C@@H]2O[C@H](C)[C@@H](O)[C@H]2O)cc1F | GAGWJHPBXLXJQN-UORFTKCHSA-N | No-Effect | inactive |
| capobenic-acid | BRD-K04282412 | COc1cc(C(=O)NCCCCCC(=O)O)cc(OC)c1OC | MPTXLVRHYGBOQY-UHFFFAOYSA-N | No-Effect | inactive |
| capromorelin | BRD-K00003188 | CN1N=C2CCN(C(=O)[C@@H](COCc3ccccc3)NC(=O)C(C)(C)N)C[C@@]2(Cc2ccccc2)C1=O | KVLLHLWBPNCVNR-SKCUWOTOSA-N | No-Effect | inactive |
| capsaicin | BRD-K50590187 | COc1cc(CNC(=O)CCCC/C=C/C(C)C)ccc1O | YKPUWZUDDOIDPM-SOFGYWHQSA-N | No-Effect | inactive |
| capsazepine | BRD-K44849676 | Oc1cc2c(cc1O)CN(C(=S)NCCc1ccc(Cl)cc1)CCC2 | DRCMAZOSEIMCHM-UHFFFAOYSA-N | No-Effect | inactive |
| captamine | BRD-K81645907 | CN(C)CCS | DENMGZODXQRYAR-UHFFFAOYSA-N | No-Effect | inactive |
| captan | BRD-A79857384 | O=C1[C@H]2CC=CC[C@@H]2C(=O)N1SC(Cl)(Cl)Cl | LDVVMCZRFWMZSG-WDSKDSINSA-N | No-Effect | inactive |
| captopril | BRD-K54529596 | C[C@H](CS)C(=O)N1CCC[C@H]1C(=O)O | FAKRSMQSSFJEIM-RQJHMYQMSA-N | No-Effect | inactive |
| caracemide | BRD-K62790656 | CNC(=O)ON(C(C)=O)C(=O)NC | JURAJLFHWXNPHG-UHFFFAOYSA-N | No-Effect | inactive |
| caramiphen | BRD-K59895502 | CCN(CC)CCOC(=O)C1(c2ccccc2)CCCC1 | OFAIGZWCDGNZGT-UHFFFAOYSA-N | No-Effect | inactive |
| carazolol | BRD-A50675490 | CC(C)NC[C@@H](O)COc1cccc2[nH]c3ccccc3c12 | BQXQGZPYHWWCEB-CYBMUJFWSA-N | No-Effect | inactive |
| carbachol | BRD-K75037734 | C[N+](C)(C)CCOC(N)=O | VPJXQGSRWJZDOB-UHFFFAOYSA-O | No-Effect | inactive |
| carbadox | BRD-K56735750 | COC(=O)N/N=C/c1c[n+]([O-])c2ccccc2[n+]1[O-] | OVGGLBAWFMIPPY-WUXMJOGZSA-N | No-Effect | inactive |
| carbamazepine | BRD-K71799949 | NC(=O)N1c2ccccc2C=Cc2ccccc21 | FFGPTBGBLSHEPO-UHFFFAOYSA-N | No-Effect | inactive |
| carbaril | BRD-K54208592 | CNC(=O)Oc1cccc2ccccc12 | CVXBEEMKQHEXEN-UHFFFAOYSA-N | No-Effect | inactive |
| carbarsone | BRD-K94848716 | NC(=O)Nc1ccc([As](=O)(O)O)cc1 | WWXBHTZSYYGCSG-UHFFFAOYSA-N | No-Effect | inactive |
| carbazochrome | BRD-A11070335 | CN1C2=CC(=O)/C(=N\NC(N)=O)C=C2C[C@H]1S(=O)(=O)O | OZCACMPSTYQSMM-DKBJHAQFSA-N | No-Effect | inactive |
| carbendazim | BRD-K12906202 | COC(=O)Nc1nc2ccccc2[nH]1 | TWFZGCMQGLPBSX-UHFFFAOYSA-N | No-Effect | inactive |
| carbenoxolone | BRD-K12765235 | CC1(C)[C@@H](OC(=O)CCC(=O)O)CC[C@@]2(C)[C@H]1CC[C@]1(C)[C@@H]2C(=O)C=C2[C@@H]3C[C@@](C)(C(=O)O)CC[C@]3(C)CC[C@]21C | OBZHEBDUNPOCJG-WBXJDKIVSA-N | No-Effect | inactive |
| carbetapentane | BRD-K06181161 | CCN(CC)CCOCCOC(=O)C1(c2ccccc2)CCCC1 | CFJMRBQWBDQYMK-UHFFFAOYSA-N | No-Effect | inactive |
| carbetocin | BRD-K56107812 | CC[C@H](C)[C@@H]1NC(=O)[C@H](Cc2ccc(OC)cc2)NC(=O)CCCSC[C@@H](C(=O)N2CCC[C@H]2C(=O)N[C@@H](CC(C)C)C(=O)NCC(N)=O)NC(=O)[C@H](CC(N)=O)NC(=O)[C@H](CCC(N)=O)NC1=O | NSTRIRCPWQHTIA-DTRKZRJBSA-N | No-Effect | inactive |
| carboplatin | BRD-K90947825 | N[Pt]1(N)OC(=O)C2(CCC2)C(=O)O1 | YAYRGNWWLMLWJE-UHFFFAOYSA-L | No-Effect | inactive |
| carboxyamidotriazole | BRD-K68408772 | NC(=O)c1nnn(Cc2cc(Cl)c(C(=O)c3ccc(Cl)cc3)c(Cl)c2)c1N | WNRZHQBJSXRYJK-UHFFFAOYSA-N | No-Effect | inactive |
| carboxylosartan | BRD-K23533833 | CCCCc1nc(Cl)c(C(=O)O)n1Cc1ccc(-c2ccccc2-c2nn[nH]n2)cc1 | ZEUXAIYYDDCIRX-UHFFFAOYSA-N | No-Effect | inactive |
| carboxypyridine-disulfide | BRD-K26530649 | O=C(O)c1ccc(SSc2ccc(C(=O)O)cn2)nc1 | GSASOFRDSIKDSN-UHFFFAOYSA-N | No-Effect | inactive |
| cardiogenol-C | BRD-K99174507 | COc1ccc(Nc2nccc(NCCO)n2)cc1 | LKBSMPFEKIBRGC-UHFFFAOYSA-N | No-Effect | inactive |
| cardionogen-1 | BRD-K50905564 | c1coc(-c2nnc3sc(C4CCCCC4)nn23)c1 | SWTUPSNBTSPBIH-UHFFFAOYSA-N | No-Effect | inactive |
| carebastine | BRD-K49554218 | CC(C)(C(=O)O)c1ccc(C(=O)CCCN2CCC(OC(c3ccccc3)c3ccccc3)CC2)cc1 | XGHOVGYJHWQGCC-UHFFFAOYSA-N | No-Effect | inactive |
| carglumic-acid | BRD-K60240222 | NC(=O)N[C@@H](CCC(=O)O)C(=O)O | LCQLHJZYVOQKHU-VKHMYHEASA-N | No-Effect | inactive |
| caricotamide | BRD-K90215590 | NC(=O)CN1C=CCC(C(N)=O)=C1 | WEJRSYKFMCUCRQ-UHFFFAOYSA-N | No-Effect | inactive |
| cariporide | BRD-K21670884 | CC(C)c1ccc(C(=O)N=C(N)N)cc1S(C)(=O)=O | IWXNYAIICFKCTM-UHFFFAOYSA-N | No-Effect | inactive |
| cariprazine | BRD-K57764956 | CN(C)C(=O)N[C@H]1CC[C@H](CCN2CCN(c3cccc(Cl)c3Cl)CC2)CC1 | KPWSJANDNDDRMB-QAQDUYKDSA-N | No-Effect | inactive |
| carmoterol | BRD-K25095405 | COc1ccc(C[C@@H](C)NC[C@H](O)c2ccc(O)c3[nH]c(=O)ccc23)cc1 | IHOXNOQMRZISPV-YJYMSZOUSA-N | No-Effect | inactive |
| carmoxirole | BRD-K82484965 | O=C(O)c1ccc2[nH]cc(CCCCN3CC=C(c4ccccc4)CC3)c2c1 | AFSOIHMEOKEZJF-UHFFFAOYSA-N | No-Effect | inactive |
| carmustine | BRD-K36234266 | O=C(NCCCl)N(CCCl)[NH2+][O-] | IIJOAQBKDPUMEO-UHFFFAOYSA-N | No-Effect | inactive |
| carnitine-(D/L) | BRD-A89594340 | C[N+](C)(C)C[C@@H](O)CC(=O)O | PHIQHXFUZVPYII-LURJTMIESA-O | No-Effect | inactive |
| carnosine | BRD-A96051074 | NCCC(=O)N[C@@H](Cc1cnc[nH]1)C(=O)O | CQOVPNPJLQNMDC-ZETCQYMHSA-N | No-Effect | inactive |
| caroverin | BRD-K70161083 | CCN(CC)CCn1c(=O)c(Cc2ccc(OC)cc2)nc2ccccc21 | MSPRUJDUTKRMLM-UHFFFAOYSA-N | No-Effect | inactive |
| caroxazone | BRD-K92129045 | NC(=O)CN1Cc2ccccc2OC1=O | KYCBWEZLKCTALM-UHFFFAOYSA-N | No-Effect | inactive |
| carsalam | BRD-K13238618 | O=c1[nH]c(=O)c2ccccc2o1 | OAYRYNVEFFWSHK-UHFFFAOYSA-N | No-Effect | inactive |
| carteolol | BRD-A42167015 | CC(C)(C)NCC(O)COc1cccc2c1CCC(=O)N2 | LWAFSWPYPHEXKX-UHFFFAOYSA-N | No-Effect | inactive |
| carumonam | BRD-K71026158 | NC(=O)OC[C@@H]1[C@H](NC(=O)/C(=N\OCC(=O)O)c2csc(N)n2)C(=O)N1S(=O)(=O)O | UIMOJFJSJSIGLV-JNHMLNOCSA-N | No-Effect | inactive |
| carvedilol | BRD-K01826535 | COc1ccccc1OCCNC[C@@H](O)COc1cccc2[nH]c3ccccc3c12 | OGHNVEJMJSYVRP-QGZVFWFLSA-N | No-Effect | inactive |
| carzenide | BRD-K09295674 | NS(=O)(=O)c1ccc(C(=O)O)cc1 | UCAGLBKTLXCODC-UHFFFAOYSA-N | No-Effect | inactive |

| | | | | | |
|---|---|---|---|---|---|
| casanthranol-variant | BRD-K12413266 | OCc1cc(O)c2c(O)c3c(O)cccc3c(O[C@@H]3O[C@H](CO)[C@H](O)[C@@H](O)[C@H]3O)c2c1 | XMCJWIFCIIXRGI-BAEVKLAGSA-N | No-Effect | inactive |
| caspofungin | BRD-K00002625 | CC[C@H](C)C[C@H](C)CCCCCCCCC(=O)N[C@H]1C[C@@H](O)[C@@H](NCCN)NC(=O)[C@@H]2[C@@H](O)CCN2C(=O)[C@H]([C@H](O)CCN)NC(=O)[C@H]([C@H](O)[C@@H](O)c2ccc(O)cc2)NC(=O)[C@@H]2C[C@@H](O)CN2C(=O)[C@H]([C@@H](C)O)NC1=O | JYIKNQVWKBUSNH-WVDDFWQHSA-N | No-Effect | inactive |
| castanospermine | BRD-K20897876 | O[C@H]1CN2CC[C@H](O)[C@@H]2[C@@H](O)[C@@H]1O | JDVVGAQPNNXQDW-TVNFTVLESA-N | No-Effect | inactive |
| catharanthine | BRD-K92377961 | CCC1=C[C@@H]2CN3CCc4c([nH]c5ccccc45)[C@@](C(=O)OC)(C2)[C@@H]13 | CMKFQVZJOWHHDV-NQZBTDCJSA-N | No-Effect | inactive |
| CB-03-01 | BRD-K41779183 | CCC(=O)O[C@]1(C(=O)CO)CC[C@H]2[C@@H]3CCC4=CC(=O)CC[C@]4(C)[C@H]3CC[C@@]21C | GPNHMOZDMYNCPO-PDUMRIMRSA-N | No-Effect | inactive |
| CB-10-277 | BRD-K59929863 | CN(C)/N=N/c1ccc(C(=O)O)cc1 | GUAZPUYTLMUTMA-ZHACJKMWSA-N | No-Effect | inactive |
| CB-5083 | BRD-K49048664 | Cc1cc2c(C(N)=O)cccc2n1-c1nc2c(c(NCc3ccccc3)n1)COCC2 | RDALZZCKQFLGJP-UHFFFAOYSA-N | No-Effect | inactive |
| CB-839 | BRD-K58070273 | O=C(Cc1cccc(OC(F)(F)F)c1)Nc1ccc(CCCCc2nnc(NC(=O)Cc3ccccn3)s2)nn1 | PRAAPINBUWJLGA-UHFFFAOYSA-N | No-Effect | inactive |
| CBiPES | BRD-K34675210 | CCS(=O)(=O)N(Cc1cccnc1)c1cccc(-c2ccc(C#N)cc2)c1 | HDVYXILCBYGKGU-UHFFFAOYSA-N | No-Effect | inactive |
| CC-223 | BRD-K88737571 | CO[C@H]1CC[C@H](N2C(=O)CNc3ncc(-c4ccc(C(C)(C)O)nc4)nc32)CC1 | UFKLYTOEMRFKAD-SHTZXODSSA-N | No-Effect | inactive |
| CC-401 | BRD-K00004320 | c1cc(OCCN2CCCCC2)cc(-c2n[nH]c3ccc(-c4nnc[nH]4)cc23)c1 | XDJCLCLBSGGNKS-UHFFFAOYSA-N | No-Effect | inactive |
| CC-90003 | BRD-K00003116 | C=CC(=O)Nc1cc(C)ccc1Nc1nc(Nc2cc(OC)ncc2C)ncc1C(F)(F)F | ILUKRINUNLAVMH-UHFFFAOYSA-N | No-Effect | inactive |
| CC-930 | BRD-K48213016 | O[C@H]1CC[C@H](Nc2ncc3nc(Nc4c(F)cc(F)cc4F)n([C@H]4CCOC4)c3n2)CC1 | IBGLGMOPHJQDJB-IHRRRGAJSA-N | No-Effect | inactive |
| CC4 | BRD-K17751100 | O=c1cccc2n1C[C@H]1C[C@@H]2CN(CCN2C[C@@H]3C[C@H](C2)c2cccc(=O)n2C3)C1 | LEXDAVFCJPDCNA-VNTMZGSJSA-N | No-Effect | inactive |
| CCG-1423 | BRD-A79726147 | C[C@@H](ONC(=O)c1cc(C(F)(F)F)cc(C(F)(F)F)c1)C(=O)Nc1ccc(Cl)cc1 | DSMXVSGJIDFLKP-SECBINFHSA-N | No-Effect | inactive |
| CCG-50014 | BRD-K08893438 | Cc1ccc(-n2sc(=O)n(Cc3ccc(F)cc3)c2=O)cc1 | QUIIIYITNGOFEI-UHFFFAOYSA-N | No-Effect | inactive |
| CCG-63802 | BRD-K26970495 | Cc1cccc(Oc2nc3c(C)cccn3c(=O)c2/C=C(\C#N)c2nc3ccccc3s2)c1 | VFSVKVQMZDJFQX-NBVRZTHBSA-N | No-Effect | inactive |
| CCG-63808 | BRD-K68474505 | Cc1cccn2c(=O)c(/C=C(\C#N)c3nc4ccccc4s3)c(Oc3ccc(F)cc3)nc12 | IPZHFKHGSYRBNT-DTQAZKPQSA-N | No-Effect | inactive |
| CCMI | BRD-K16338837 | Cc1cc(C(=CNc2ccc(Cl)cc2)C(=O)Nc2ccc(Cl)cc2)on1 | VMAKIACTLSBBIY-UHFFFAOYSA-N | No-Effect | inactive |
| CCMQ | BRD-K06159959 | O=C(O)Cc1cc2ccccc2nc1C(=O)O | IFXJDEHFADWURH-UHFFFAOYSA-N | No-Effect | inactive |
| CCT018159 | BRD-K65503129 | CCc1cc(-c2n[nH]c(C)c2-c2ccc3c(c2)OCCO3)c(O)cc1O | OWPMENVYXDJDOW-UHFFFAOYSA-N | No-Effect | inactive |
| CCT128930 | BRD-K76406695 | NC1(Cc2ccc(Cl)cc2)CCN(c2ncnc3[nH]ccc23)CC1 | RZIDZIGAXXNODG-UHFFFAOYSA-N | No-Effect | inactive |
| CCT129202 | BRD-K57136142 | CN(C)c1ccc(-c2nc3ncc(Cl)c(N4CCN(CC(=O)Nc5nccs5)CC4)c3[nH]2)cc1 | QYKHWEFPFAGNEV-UHFFFAOYSA-N | No-Effect | inactive |
| CCT137690 | BRD-K03289018 | Cc1cc(CN2CCN(c3c(Br)cnc4nc(-c5ccc(N6CCN(C)CC6)cc5)[nH]c34)CC2)no1 | GFLQCBTXTRCREJ-UHFFFAOYSA-N | No-Effect | inactive |
| CCT196969 | BRD-K00005244 | CC(C)(C)c1cc(NC(=O)Nc2ccc(Oc3ccnc4nc(O)cnc34)cc2F)n(-c2ccccc2)n1 | KYYKGSDLXXKQCR-UHFFFAOYSA-N | No-Effect | inactive |
| CCT245737 | BRD-K00003276 | N#Cc1cnc(Nc2cc(NC[C@H]3CNCCO3)c(C(F)(F)F)cn2)cn1 | YBYYWUUUGCNAHQ-LLVKDONJSA-N | No-Effect | inactive |
| CD-437 | BRD-K28907958 | O=C(O)c1ccc2cc(-c3ccc(O)c(C45CC6CC(CC(C6)C4)C5)c3)ccc2c1 | LDGIHZJOIQSHPB-UHFFFAOYSA-N | No-Effect | inactive |
| CDBA | BRD-A08079565 | OC[C@H]1O[C@H]2O[C@@H]3[C@@H](O)[C@H](O)[C@@H](O[C@@H]3CO)O[C@@H]3[C@@H](O)[C@H](O)[C@@H](O[C@@H]3CO)O[C@@H]3[C@@H](O)[C@H](O)[C@@H](O[C@@H]3CO)O[C@@H]3[C@@H](O)[C@H](O)[C@@H](O[C@@H]3CO)O[C@@H]3[C@@H](O)[C@H](O)[C@@H](O[C@@H]3CO)O[C@@H]3[C@@H](O)[C@H](O)[C@@H](O[C@@H]3CO)O[C@@H]1[C@@H](O)[C@@H]2O | WHGYBXFWUBPSRW-GUJBRCDDSA-N | No-Effect | inactive |
| CDK1-5-inhibitor | BRD-K87932577 | Nc1[nH]nc2nc3ccccc3nc12 | DWHVZCLBMTZRQM-UHFFFAOYSA-N | No-Effect | inactive |
| CDK9-IN-6 | BRD-K43914553 | COC[C@H](C)N[C@H]1CC[C@H](Nc2cc(-c3cccc(NCC4(C#N)CCOCC4)n3)c(Cl)cn2)CC1 | XWQVQSXLXAXOPJ-ACRUOGEOSA-N | No-Effect | inactive |
| CDPPB | BRD-K02213468 | N#Cc1cccc(C(=O)Nc2cc(-c3ccccc3)nn2-c2ccccc2)c1 | BKUIZWILNWHFHD-UHFFFAOYSA-N | No-Effect | inactive |
| cebranopadol | BRD-K48141473 | CN(C)[C@]1(c2ccccc2)CC[C@]2(CC1)OCCc1c3cc(F)ccc3[nH]c12 | CSMVOZKEWSOFER-RQNOJGIXSA-N | No-Effect | inactive |
| cediranib | BRD-K86930074 | COc1cc2c(Oc3ccc4[nH]c(C)cc4c3F)ncnc2cc1OCCCN1CCCC1 | XXJWYDDUDKYVKI-UHFFFAOYSA-N | No-Effect | inactive |
| cedrol | BRD-A61188230 | C[C@H]1CC[C@H]2C(C)(C)[C@H]3C[C@@]12CC[C@@]3(C)O | SVURIXNDRWRAFU-OBORUHMCSA-N | No-Effect | inactive |
| cedrol-(+) | BRD-K96362839 | C[C@@H]1CC[C@H]2C(C)(C)[C@H]3C[C@]21CC[C@@]3(C)O | SVURIXNDRWRAFU-OGMFBOKVSA-N | No-Effect | inactive |
| cefaclor | BRD-K20338176 | N[C@@H](C(=O)N[C@@H]1C(=O)N2C(C(=O)O)=C(Cl)CS[C@H]12)c1ccccc1 | QYIYFLOTGYLRGG-GPCCPHFNSA-N | No-Effect | inactive |
| cefadroxil | BRD-K63242551 | CC1=C(C(=O)O)N2C(=O)[C@@H](NC(=O)[C@H](N)c3ccc(O)cc3)[C@H]2SC1 | BOEGTKLJZSQCCD-UEKVPHQBSA-N | No-Effect | inactive |
| cefalonium | BRD-K15514357 | NC(=O)c1cc[n+](CC2=C(C(=O)O)N3C(=O)[C@@H](NC(=O)Cc4cccs4)[C@H]3SC2)cc1 | FMZXNVLFJHCSAF-DNVCBOLYSA-O | No-Effect | inactive |
| cefamandole | BRD-K27130738 | Cn1nnnc1SCC1=C(C(=O)O)N2C(=O)[C@@H](NC(=O)[C@H](O)c3ccccc3)[C@H]2SC1 | OLVCFLKTBJRLHI-AXAPSJFSSA-N | No-Effect | inactive |
| cefamandole-nafate | BRD-K17903788 | Cn1nnnc1SCC1=C(C(=O)O)N2C(=O)[C@@H](NC(=O)[C@H](OC=O)c3ccccc3)[C@H]2SC1 | RRJHESVQVSRQEX-SUYBPPKGSA-N | No-Effect | inactive |
| cefathiamidine | BRD-K63835048 | CC(=O)OCC1=C(C(=O)O)N2C(=O)[C@@H](NC(=O)CS/C(=N/C(C)C)NC(C)C)[C@H]2SC1 | JYXACOFERDBGGQ-RHSMWYFYSA-N | No-Effect | inactive |
| cefazolin | BRD-K99750640 | Cc1nnc(SCC2=C(C(=O)O)N3C(=O)[C@@H](NC(=O)Cn4cnnn4)[C@H]3SC2)s1 | MLYYVTUWGNIJIB-BXKDBHETSA-N | No-Effect | inactive |
| cefcapene-pivoxil | BRD-K27515606 | CC/C=C(\C(=O)N[C@@H]1C(=O)N2C(C(=O)OCOC(=O)C(C)(C)C)=C(COC(N)=O)CS[C@H]12)c1csc(N)n1 | WVPAABNYMHNFJG-QDVBXLKVSA-N | No-Effect | inactive |
| cefdinir | BRD-K15766189 | C=CC1=C(C(=O)O)N2C(=O)[C@@H](NC(=O)/C(=N\O)c3csc(N)n3)[C@H]2SC1 | RTXOFQZKPXMALH-GHXIOONMSA-N | No-Effect | inactive |
| cefditoren-pivoxil | BRD-K72167406 | CO/N=C(\C(=O)N[C@@H]1C(=O)N2C(C(=O)OCOC(=O)C(C)(C)C)=C(/C=C\c3scnc3C)CS[C@H]12)c1csc(N)n1 | AFZFFLVORLEPPO-UVYJNCLZSA-N | No-Effect | inactive |
| cefepime | BRD-A12077521 | CO/N=C(/C(=O)N[C@@H]1C(=O)N2C(C(=O)O)=C(C[N+]3(C)CCCC3)CS[C@H]12)c1csc(N)n1 | HVFLCNVBZFFHBT-FKULVZFESA-O | No-Effect | inactive |
| cefetamet | BRD-K27798832 | CO/N=C(\C(=O)N[C@@H]1C(=O)N2C(C(=O)O)=C(C)CS[C@H]12)c1csc(N)n1 | MQLRYUCJDNBWMV-GHXIOONMSA-N | No-Effect | inactive |
| cefetamet-pivoxil | BRD-K21372554 | CO/N=C(\C(=O)N[C@@H]1C(=O)N2C(C(=O)OCOC(=O)C(C)(C)C)=C(C)CS[C@H]12)c1csc(N)n1 | DASYMCLQENWCJG-XUKDPADISA-N | No-Effect | inactive |
| cefiderocol | BRD-K00003257 | CC(C)(O/N=C(\C(=O)N[C@@H]1C(=O)N2C(C(=O)O)=C(C[N+]3(CCNC(=O)c4ccc(O)c(O)c4Cl)CCCC3)CS[C@H]12)c1csc(N)n1)C(=O)O | DBPPRLRVDVJOCL-FQRUVTKNSA-O | No-Effect | inactive |
| cefixime | BRD-K71059170 | C=CC1=C(C(=O)O)N2C(=O)[C@@H](NC(=O)/C(=N\OCC(=O)O)c3csc(N)n3)[C@H]2SC1 | OKBVVJOGVLARMR-QSWIMTSFSA-N | No-Effect | inactive |
| cefmenoxime | BRD-K00003661 | CO/N=C(\C(=O)N[C@@H]1C(=O)N2C(C(=O)O)=C(CSc3nnnn3C)CS[C@H]12)c1csc(N)n1 | HJJDBAOLQAWBMH-YCRCPZNHSA-N | No-Effect | inactive |
| cefmetazole | BRD-K48470486 | CO[C@@]1(NC(=O)CSCC#N)C(=O)N2C(C(=O)O)=C(CSc3nnnn3C)CS[C@@H]21 | SNBUBQHDYVFSQF-HIFRSBDPSA-N | No-Effect | inactive |

| | | | | | |
|---|---|---|---|---|---|
| cefminox | BRD-K77073709 | CO[C@@]1(NC(=O)CSC[C@@H](N)C(=O)O)C(=O)N2C(C(=O)O)=C(CSc3nnnn3C)CS[C@@H]21 | JSDXOWVAHXDYCU-VXSYNFHWSA-N | No-Effect | inactive |
| cefoperazone | BRD-K02292852 | CCN1CCN(C(=O)N[C@@H](C(=O)N[C@@H]2C(=O)N3C(C(=O)O)=C(CSc4nnnn4C)CS[C@H]23)c2ccc(O)cc2)C(=O)C1=O | GCFBRXLSHGKWDP-XCGNWRKASA-N | No-Effect | inactive |
| cefonicide | BRD-K37848908 | NCc1ccccc1CC(=O)N[C@@H]1C(=O)N2C(C(=O)O)=C(CSc3nnnn3CC(=O)O)CS[C@H]12 | SLAYUXIURFNXPG-CRAIPNDOSA-N | No-Effect | inactive |
| cefotaxime | BRD-K78364995 | CO/N=C(\C(=O)N[C@@H]1C(=O)N2C(C(=O)O)=C(COC(C)=O)CS[C@H]12)c1csc(N)n1 | GPRBEKHLDVQUJE-QSWIMTSFSA-N | No-Effect | inactive |
| cefotetan | BRD-A78723049 | CO[C@@]1(NC(=O)C2SC(=C(C(N)=O)C(=O)O)S2)C(=O)N2C(C(=O)O)=C(CSc3nnnn3C)CS[C@@H]21 | SRZNHPXWXCNNDU-RHBCBLIFSA-N | No-Effect | inactive |
| cefotiam | BRD-K02275692 | CN(C)CCn1nnnc1SCC1=C(C(=O)O)N2C(=O)[C@@H](NC(=O)Cc3csc(N)n3)[C@H]2SC1 | QYQDKDWGWDOFFU-IUODEOHRSA-N | No-Effect | inactive |
| cefotiam-cilexetil | BRD-A76661185 | C[C@@H](OC(=O)OC1CCCCC1)OC(=O)C1=C(CSc2nnnn2CCN(C)C)CS[C@H]2[C@@H](NC(=O)Cc3csc(N)n3)C(=O)N12 | VVFDMWZLBPUKTD-XLHILCRZSA-N | No-Effect | inactive |
| cefoxitin | BRD-K70976396 | CO[C@@]1(NC(=O)Cc2cccs2)C(=O)N2C(C(=O)O)=C(COC(N)=O)CS[C@@H]21 | WZOZEZRFJCJXNZ-ZBFHGGJFSA-N | No-Effect | inactive |
| cefpiramide | BRD-K92872987 | Cc1cc(=O)c(C(=O)N[C@@H](C(=O)N[C@@H]2C(=O)N3C(C(=O)O)=C(CSc4nnnn4C)CS[C@H]23)c2ccc(O)cc2)c[nH]1 | PWAUCHMQEXVFJR-PMAPCBKXSA-N | No-Effect | inactive |
| cefpodoxime | BRD-A34205213 | COCC1=C(C(=O)O[C@@H](C)OC(=O)OC(C)C)N2C(=O)[C@@H](NC(=O)/C(=N\OC)c3csc(N)n3)[C@H]2SC1 | LTINZAODLRIQIX-OLEGMKQCSA-N | No-Effect | inactive |
| cefradine | BRD-K13797099 | CC1=C(C(=O)O)N2C(=O)[C@@H](NC(=O)[C@H](N)C3=CCC=CC3)[C@H]2SC1 | RDLPVSKMFDYCOR-UEKVPHQBSA-N | No-Effect | inactive |
| cefsulodin | BRD-K03907181 | NC(=O)c1cc[n+](CC2=C(C(=O)O)N3C(=O)[C@@H](NC(=O)[C@@H](c4ccccc4)S(=O)(=O)O)[C@H]3SC2)cc1 | SYLKGLMBLAAGSC-QLVMHMETSA-O | No-Effect | inactive |
| ceftaroline-fosamil | BRD-K37252028 | CCO/N=C(/C(=O)N[C@@H]1C(=O)N2C(C(=O)O)=C(Sc3nc(-c4cc[n+](C)cc4)cs3)CS[C@H]12)c1nc(=NP(=O)(O)O)s[nH]1 | ZCCUWMICIWSJIX-XHNDKCDBSA-O | No-Effect | inactive |
| ceftazidime | BRD-K52117081 | CC(C)(O/N=C(\C(=O)N[C@@H]1C(=O)N2C(C(=O)O)=C(C[n+]3ccccc3)CS[C@H]12)c1csc(N)n1)C(=O)O | ORFOPKXBNMVMKC-DWVKKRMSSA-O | No-Effect | inactive |
| ceftiofur | BRD-K82960980 | CO/N=C(\C(=O)N[C@@H]1C(=O)N2C(C(=O)O)=C(CSC(=O)c3ccco3)CS[C@H]12)c1csc(N)n1 | ZBHXIWJRIFEVQY-IHMPYVIRSA-N | No-Effect | inactive |
| ceftizoxim | BRD-K04301924 | CO/N=C(\C(=O)N[C@@H]1C(=O)N2C(C(=O)O)=CCS[C@H]12)c1csc(N)n1 | NNULBSISHYWZJU-LLKWHZGFSA-N | No-Effect | inactive |
| ceftobiprole | BRD-K00003117 | Nc1nc(/C(=N/O)C(=O)N[C@@H]2C(=O)N3C(C(=O)O)=C(/C=C4\CCN([C@@H]5CCNC5)C4=O)CS[C@H]23)ns1 | VOAZJEPQLGBXGO-SDAWRPRTSA-N | No-Effect | inactive |
| ceftriaxone | BRD-K96734425 | CO/N=C(\C(=O)N[C@@H]1C(=O)N2C(C(=O)O)=C(CSc3nc(=O)c(O)nn3C)CS[C@H]12)c1csc(N)n1 | VAAUVRVFOQPIGI-SPQHTLEESA-N | No-Effect | inactive |
| cefuroxime | BRD-K02733959 | CO/N=C(\C(=O)N[C@@H]1C(=O)N2C(C(=O)O)=C(COC(N)=O)CS[C@H]12)c1ccco1 | JFPVXVDWJQMJEE-IZRZKJBUSA-N | No-Effect | inactive |
| cefuroxime-axetil | BRD-A22303829 | CO/N=C(\C(=O)N[C@@H]1C(=O)N2C(C(=O)O[C@@H](C)OC(C)=O)=C(COC(N)=O)CS[C@H]12)c1ccco1 | KEJCWVGMRLCZQQ-QFDKGRJDSA-N | No-Effect | inactive |
| celecoxib | BRD-K02637541 | Cc1ccc(-c2cc(C(F)(F)F)nn2-c2ccc(S(N)(=O)=O)cc2)cc1 | RZEKVGVHFLEQIL-UHFFFAOYSA-N | No-Effect | inactive |
| celiprolol | BRD-A95602221 | CCN(CC)C(=O)Nc1ccc(OC[C@@H](O)CNC(C)(C)C)c(C(C)=O)c1 | JOATXPAWOHTVSZ-INIZCTEOSA-N | No-Effect | inactive |
| cenerimod | BRD-K00003248 | CCc1cc(-c2noc(-c3cc(OC)nc(C4CCCC4)c3)n2)cc(C)c1OC[C@@H](O)CO | KJKKMMMRWISKRF-FQEVSTJZSA-N | No-Effect | inactive |
| cenicriviroc | BRD-K00003181 | CCCCOCCOc1ccc(-c2ccc3c(c2)/C=C(/C(=O)Nc2ccc([S@@](=O)Cc4cncn4CCC)cc2)CCCN3CC(C)C)cc1 | PNDKCRDVVKJPKG-WHERJAGFSA-N | No-Effect | inactive |
| cenobamate | BRD-K00003249 | NC(=O)O[C@@H](Cn1ncnn1)c1ccccc1Cl | GFHAXPJGXSQLPT-VIFPVBQESA-N | No-Effect | inactive |
| centanafadine | BRD-K00004588 | c1ccc2cc([C@]34CNC[C@H]3C4)ccc2c1 | HKHCSWPSUSWGLI-CABCVRRESA-N | No-Effect | inactive |
| centazolone | BRD-K31329721 | Nn1cnc2cc3ccccc3cc2c1=O | CZXUQBVMAUWSBX-UHFFFAOYSA-N | No-Effect | inactive |
| centhaquin | BRD-K00004619 | Cc1cccc(N2CCN(CCc3ccc4ccccc4n3)CC2)c1 | UJNWGFBJUHIJKK-UHFFFAOYSA-N | No-Effect | inactive |
| centpropazine | BRD-A28958257 | CCC(=O)c1ccc(OC[C@@H](O)CN2CCN(c3ccccc3)CC2)cc1 | ZQPXSRTZFYHSFB-FQEVSTJZSA-N | No-Effect | inactive |
| CEP-32496 | BRD-K20722021 | COc1cc2ncnc(Oc3cccc(NC(=O)Nc4cc(C(C)(C)C(F)(F)F)on4)c3)c2cc1OC | DKNUPRMJNUQNHR-UHFFFAOYSA-N | No-Effect | inactive |
| CEP-33779 | BRD-K37237504 | CN1CCN(c2cccc(Nc3nc4c(-c5ccc(S(C)(=O)=O)cc5)cccn4n3)c2)CC1 | RFZKSQIFOZZIAQ-UHFFFAOYSA-N | No-Effect | inactive |
| CEP-40783 | BRD-K00003396 | COc1cc2nccc(Oc3ccc(NC(=O)c4cn(C(C)C)c(=O)n(-c5ccc(F)cc5)c4=O)cc3F)c2cc1OC | FKCWHHYUMFGOPY-UHFFFAOYSA-N | No-Effect | inactive |
| cephalexin | BRD-K90733503 | CC1=C(C(=O)O)N2C(=O)[C@@H](NC(=O)[C@H](N)c3ccccc3)[C@H]2SC1 | ZAIPMKNFIOOWCQ-UEKVPHQBSA-N | No-Effect | inactive |
| cephalomannine | BRD-K68904758 | C/C=C(\C)C(=O)N[C@@H](c1ccccc1)[C@@H](O)C(=O)O[C@H]1C[C@@]2(O)[C@@H](OC(=O)c3ccccc3)[C@H]3[C@](C)(C(=O)[C@H](OC(C)=O)C(=C1C)C2(C)C)[C@@H](O)C[C@H]1OC[C@]13OC(C)=O | DBXFAPJCZABTDR-WBYYIXQISA-N | No-Effect | inactive |
| cephalosporin-c-zn | BRD-K00025564 | CC(=O)OCC1=C2C(=O)O[Zn]OC(=O)[C@@H](N)CCCC(=O)N[C@@H]3C(=O)N2[C@@H]3SC1 | TVMXJQZBKOMFQK-CYSXAYAHSA-L | No-Effect | inactive |
| cephalothin | BRD-K28210218 | CC(=O)OCC1=C(C(=O)O)N2C(=O)[C@@H](NC(=O)Cc3cccs3)[C@H]2SC1 | XIURVHNZVLADCM-IUODEOHRSA-N | No-Effect | inactive |
| cephapirin | BRD-K49880111 | CC(=O)OCC1=C(C(=O)O)N2C(=O)[C@@H](NC(=O)CSc3ccncc3)[C@H]2SC1 | UQLLWWBDSUHNEB-CZUORRHYSA-N | No-Effect | inactive |
| cepharanthine | BRD-K96194081 | COc1ccc2cc1Oc1ccc(cc1)C[C@H]1c3c(cc4c(c3Oc3cc5c(cc3OC)CCN(C)[C@@H]5C2)OCO4)CCN1C | YVPXVXANRNDGTA-WDYNHAJCSA-N | No-Effect | inactive |
| cephradine | BRD-A90807417 | CC1=C(C(=O)O)N2C(=O)[C@@H](NC(=O)[C@H](N)C3=CCC=CC3)[C@@H]2SC1 | RDLPVSKMFDYCOR-HFAKWTLXSA-N | No-Effect | inactive |
| ceramide | BRD-K41707108 | CCCCCCCCCCCCC/C=C/[C@@H](O)[C@H](CO)NC(C)=O | BLTCBVOJNNKFKC-QUDYQQOWSA-N | No-Effect | inactive |
| ceritinib | BRD-K22149900 | Cc1cc(Nc2ncc(Cl)c(Nc3ccccc3S(=O)(=O)C(C)C)n2)c(OC(C)C)cc1C1CCNCC1 | VERWOWGGCGHDQE-UHFFFAOYSA-N | No-Effect | inactive |
| cerulenin | BRD-K52075040 | C/C=C/C/C=C/CCC(=O)[C@H]1O[C@H]1C(N)=O | GVEZIHKRYBHEFX-NQQPLRFYSA-N | No-Effect | inactive |
| ceruletide | BRD-K00003320 | CSCC[C@H](NC(=O)[C@H](Cc1c[nH]c2ccccc12)NC(=O)CNC(=O)[C@@H](NC(=O)[C@H](Cc1ccc(OS(=O)(=O)O)cc1)NC(=O)[C@H](CC(=O)O)NC(=O)[C@H](CCC(N)=O)NC(=O)[C@@H]1CCC(=O)N1)[C@@H](C)O)C(=O)N[C@@H](CC(=O)O)C(=O)N[C@@H](Cc1ccccc1)C(N)=O | YRALAIOMGQZKOW-HYAOXDFASA-N | No-Effect | inactive |
| cetilistat | BRD-K94863486 | CCCCCCCCCCCCCCCCOc1nc2ccc(C)cc2c(=O)o1 | MVCQKIKWYUURMU-UHFFFAOYSA-N | No-Effect | inactive |
| cetrorelix | BRD-K76315403 | CC(=O)N[C@H](Cc1ccc2ccccc2c1)C(=O)N[C@H](Cc1ccc(Cl)cc1)C(=O)N[C@H](Cc1cccnc1)C(=O)N[C@@H](CO)C(=O)N[C@@H](Cc1ccc(O)cc1)C(=O)N[C@H](CCCNC(N)=O)C(=O)N[C@@H](CC(C)C)C(=O)N[C@@H](CCCNC(=N)N)C(=O)N1CCC[C@H]1C(=O)N[C@H](C)C(N)=O | SBNPWPIBESPSIF-MHWMIDJBSA-N | No-Effect | inactive |
| cevimeline | BRD-K00004291 | C[C@H]1O[C@]2(CS1)CN1CCC2CC1 | WUTYZMFRCNBCHQ-WPRPVWTQSA-N | No-Effect | inactive |
| cevipabulin | BRD-K00003570 | CNCCCOc1cc(F)c(-c2c(Cl)nc3ncnn3c2N[C@@H](C)C(F)(F)F)c(F)c1 | ZUZPCOQWSYNWLU-VIFPVBQESA-N | No-Effect | inactive |

| | | | | | |
|---|---|---|---|---|---|
| CF102 | BRD-K52396582 | CNC(=O)[C@H]1O[C@@H](n2cnc3c(NCc4cccc(I)c4)nc(Cl)nc32)[C@H](O)[C@@H]1O | IPSYPUKKXMNCNQ-PFHKOEEOSA-N | No-Effect | inactive |
| CFI-400945 | BRD-K27040396 | COc1ccc2c(c1)[C@]1(C[C@H]1c1ccc3c(/C=C/c4ccc(CN5C[C@H](C)O[C@H](C)C5)cc4)[nH]nc3c1)C(=O)N2 | DADASRPKWOGKCU-FVTQAUBDSA-N | No-Effect | inactive |
| CFM-1571 | BRD-K64341947 | COc1ccc(NC(=O)c2cc(OCCCN(C)C)nn2Cc2ccccc2)cc1 | YZCSJBGQLATPMH-UHFFFAOYSA-N | No-Effect | inactive |
| CFM-2 | BRD-K42859542 | COc1cc2c(cc1OC)C(c1ccc(N)cc1)=NNC(=O)C2 | MJKADKZSYQWGLL-UHFFFAOYSA-N | No-Effect | inactive |
| CFTRinh-172 | BRD-K72004355 | O=C(O)c1ccc(/C=C2/SC(=S)N(c3cccc(C(F)(F)F)c3)C2=O)cc1 | JIMHYXZZCWVCMI-RIYZIHGNSA-N | No-Effect | inactive |
| CG-400549 | BRD-K48888725 | Cc1c(N)cccc1Cn1ccc(OCCc2cccs2)cc1=O | YCLREGRRHGLOAK-UHFFFAOYSA-N | No-Effect | inactive |
| CGH2466 | BRD-K82028950 | Nc1nc(-c2ccc(Cl)c(Cl)c2)c(-c2ccncc2)s1 | BEPGKLOHQTXUHX-UHFFFAOYSA-N | No-Effect | inactive |
| CGK-733 | BRD-A50737080 | O=C(N[C@H](NC(=S)Nc1ccc(F)c([N+](=O)[O-])c1)C(Cl)(Cl)Cl)C(c1ccccc1)c1ccccc1 | HLCDNLNLQNYZTK-OAQYLSRUSA-N | No-Effect | inactive |
| CGM097 | BRD-K79584249 | COc1cc2c(cc1OC(C)C)[C@H](c1ccc(Cl)cc1)N(c1ccc(N(C)C[C@H]3CC[C@H](N4CCN(C)C(=O)C4)CC3)cc1)C(=O)C2 | CLRSLRWKONPSRQ-IIPSPAQQSA-N | No-Effect | inactive |
| CGP-12177 | BRD-A40084411 | CC(C)(C)NC[C@@H](O)COc1cccc2[nH]c(=O)[nH]c12 | UMQUQWCJKFOUGV-SECBINFHSA-N | No-Effect | inactive |
| CGP-13501 | BRD-K95851186 | CC(C)(C=O)Cc1cc(C(C)(C)C)c(O)c(C(C)(C)C)c1 | XGWATTXMMMANFJ-UHFFFAOYSA-N | No-Effect | inactive |
| CGP-20712A | BRD-K01827291 | CS(=O)(=O)O.Cn1cc(C(F)(F)F)nc1-c1ccc(OC[C@H](O)CNCCOc2ccc(O)c(C(N)=O)c2)cc1 | VFPOVCXWKBYDNF-XFULWGLBSA-N | No-Effect | inactive |
| CGP-25454A | BRD-K00003372 | CCN(CC)CCNC(=O)c1cc(C#N)c(Cl)cc1OC | JEMKCYHUYUPGFN-UHFFFAOYSA-N | No-Effect | inactive |
| CGP-37157 | BRD-A35623999 | O=C1CS[C@@H](c2ccccc2Cl)c2cc(Cl)ccc2N1 | KQEPIRKXSUIUTH-HNNXBMFYSA-N | No-Effect | inactive |
| CGP-39551 | BRD-K00004626 | CCOC(=O)[C@H](N)/C=C(\C)CP(=O)(O)O | OKDOWCKDTWNRCB-PTYLAXBQSA-N | No-Effect | inactive |
| CGP-52411 | BRD-K13087974 | O=C1NC(=O)c2cc(Nc3ccccc3)c(Nc3ccccc3)cc21 | AAALVYBICLMAMA-UHFFFAOYSA-N | No-Effect | inactive |
| CGP-53353 | BRD-K32292990 | O=C1NC(=O)c2cc(Nc3ccc(F)cc3)c(Nc3ccc(F)cc3)cc21 | RONQPWQYDRPRGG-UHFFFAOYSA-N | No-Effect | inactive |
| CGP-54626 | BRD-A74208568 | C[C@H](NC[C@H](O)CP(=O)(O)CC1CCCCC1)c1ccc(Cl)c(Cl)c1 | JGGVBBYJRQOPPA-BBRMVZONSA-N | No-Effect | inactive |
| CGP-55845 | BRD-K00004232 | C[C@H](NC[C@H](O)CP(=O)(O)Cc1ccccc1)c1ccc(Cl)c(Cl)c1 | ZODSPDOOCZZEIM-BBRMVZONSA-N | No-Effect | inactive |
| CGP-57380 | BRD-K42500029 | Nc1ncnc2n[nH]c(Nc3ccc(F)cc3)c12 | UQPMANVRZYYQMD-UHFFFAOYSA-N | No-Effect | inactive |
| CGP-71683 | BRD-K34321528 | Nc1nc(NC[C@H]2CC[C@H](CNS(=O)(=O)c3cccc4ccccc34)CC2)nc2ccccc12 | UULIGRNKXHCLQN-WGSAOQKQSA-N | No-Effect | inactive |
| CGP-74514 | BRD-K56239177 | CCn1cnc2c(Nc3cccc(Cl)c3)nc(N[C@@H]3CCCC[C@@H]3N)nc21 | UTBSBSOBZHXMHI-LSDHHAIUSA-N | No-Effect | inactive |
| CGP-78608 | BRD-K60946979 | C[C@@H](NCc1cc(Br)cc2[nH]c(=O)c(=O)[nH]c12)P(=O)(O)O | DPFHVUSPVHRVFL-YFKPBYRVSA-N | No-Effect | inactive |
| CGP-7930 | BRD-K65786282 | CC(C)(CO)Cc1cc(C(C)(C)C)c(O)c(C(C)(C)C)c1 | XLWJPQQFJNGUPA-UHFFFAOYSA-N | No-Effect | inactive |
| CGS-15943 | BRD-K49049886 | Nc1nc2ccc(Cl)cc2c2nc(-c3ccco3)nn12 | MSJODEOZODDVGW-UHFFFAOYSA-N | No-Effect | inactive |
| CGS-20625 | BRD-K68103045 | COc1ccc(-n2[nH]c3c4c(ncc3c2=O)CCCC4)cc1 | UBLXQFIFWUEVGJ-UHFFFAOYSA-N | No-Effect | inactive |
| CGS-21680 | BRD-K27176996 | CCNC(=O)[C@H]1O[C@@H](n2cnc3c(N)nc(NCCc4ccc(CCC(=O)O)cc4)nc32)[C@H](O)[C@@H]1O | PAOANWZGLPPROA-RQXXJAGISA-N | No-Effect | inactive |
| CGS-9896 | BRD-K54093161 | O=c1c2cnc3ccccc3c2[nH]n1-c1ccc(Cl)cc1 | QCBUAKLOWCOUCR-UHFFFAOYSA-N | No-Effect | inactive |
| CH-170 | BRD-K24820488 | Cc1nc(Cn2cnc3c2c(=O)n(C)c(=O)n3C)no1 | JBXSBZBZASARMY-UHFFFAOYSA-N | No-Effect | inactive |
| CH-5183284 | BRD-K11683436 | Cc1nc2ccc(-n3ncc(C(=O)c4cc5ccccc5[nH]4)c3N)cc2[nH]1 | BEMNJULZEQTDJY-UHFFFAOYSA-N | No-Effect | inactive |
| CH223191 | BRD-K22314899 | Cc1ccccc1/N=N/c1ccc(NC(=O)c2ccnn2C)c(C)c1 | LKTNEXPODAWWFM-GHVJWSGMSA-N | No-Effect | inactive |
| CH5132799 | BRD-K28352084 | CS(=O)(=O)N1CCc2c(-c3cnc(N)nc3)nc(N3CCOCC3)nc21 | JEGHXKRHKHPBJD-UHFFFAOYSA-N | No-Effect | inactive |
| CH55 | BRD-K71487808 | CC(C)(C)c1cc(C(=O)/C=C/c2ccc(C(=O)O)cc2)cc(C(C)(C)C)c1 | FOUVTBKPJRMLPE-FMIVXFBMSA-N | No-Effect | inactive |
| CHC | BRD-K60302405 | N#C/C(=C\c1ccc(O)cc1)C(=O)O | AFVLVVWMAFSXCK-VMPITWQZSA-N | No-Effect | inactive |
| chenodeoxycholic-acid | BRD-K18135438 | C[C@H](CCC(=O)O)[C@H]1CC[C@H]2[C@H]3[C@H](CC[C@@]21C)[C@@]1(C)CC[C@@H](O)C[C@H]1C[C@H]3O | RUDATBOHQWOJDD-BSWAIDMHSA-N | No-Effect | inactive |
| CHF5074 | BRD-K34508702 | O=C(O)C1(c2ccc(-c3ccc(Cl)c(Cl)c3)c(F)c2)CC1 | LIYLTQQDABRNRX-UHFFFAOYSA-N | No-Effect | inactive |
| chicago-sky-blue-6b | BRD-K56483981 | COc1cc(-c2ccc(NN=C3C=Cc4c(S(=O)(=O)O)cc(S(=O)(=O)O)c(N)c4C3=O)c(OC)c2)ccc1NN=C1C=Cc2c(S(=O)(=O)O)cc(S(=O)(=O)O)c(N)c2C1=O | BGULJNMSWJQIEK-UHFFFAOYSA-N | No-Effect | inactive |
| chidamide | BRD-K28537285 | Nc1ccc(F)cc1NC(=O)c1ccc(CNC(=O)/C=C/c2cccnc2)cc1 | WXHHICFWKXDFOW-BJMVGYQFSA-N | No-Effect | inactive |
| chiniofon | BRD-K52284881 | O=S(=O)(O)c1cc(I)c(O)c2ncccc12 | ZBJWWKFMHOAPNS-UHFFFAOYSA-N | No-Effect | inactive |
| CHIR-99021 | BRD-K16189898 | Cc1c[nH]c(-c2cnc(NCCNc3ccc(C#N)cn3)nc2-c2ccc(Cl)cc2Cl)n1 | AQGNHMOJWBZFQQ-UHFFFAOYSA-N | No-Effect | inactive |
| chloralose | BRD-K00004971 | OC[C@@H](O)[C@H]1O[C@@H]2O[C@H](C(Cl)(Cl)Cl)O[C@@H]2[C@H]1O | OJYGBLRPYBAHRT-IPQSZEQASA-N | No-Effect | inactive |
| chlorambucil | BRD-K29458283 | O=C(O)CCCc1ccc(N(CCCl)CCCl)cc1 | JCKYGMPEJWAADB-UHFFFAOYSA-N | No-Effect | inactive |
| chloramine-T | BRD-K93546328 | Cc1ccc(S(=O)(=O)NCl)cc1 | NXTVQNIVUKXOIL-UHFFFAOYSA-N | No-Effect | inactive |
| chloramphenicol | BRD-K08111712 | O=C(N[C@H](CO)[C@H](O)c1ccc([N+](=O)[O-])cc1)C(Cl)Cl | WIIZWVCIJKGZOK-RKDXNWHRSA-N | No-Effect | inactive |
| chloramphenicol-sodium-succinate | BRD-K25209357 | O=C(O)CCC(=O)OC[C@@H](NC(=O)C(Cl)Cl)[C@H](O)c1ccc([N+](=O)[O-])cc1 | LIRCDOVJWUGTMW-ZWNOBZJWSA-N | No-Effect | inactive |
| chlorazanil | BRD-K21452876 | Nc1ncnc(Nc2ccc(Cl)cc2)n1 | YRZQHIVOIFJEEE-UHFFFAOYSA-N | No-Effect | inactive |
| chlorfenson | BRD-K89409436 | O=S(=O)(Oc1ccc(Cl)cc1)c1ccc(Cl)cc1 | RZXLPPRPEOUENN-UHFFFAOYSA-N | No-Effect | inactive |
| chlorhexidine | BRD-K52256627 | N=C(N=C(N)Nc1ccc(Cl)cc1)NCCCCCCNC(=N)N=C(N)Nc1ccc(Cl)cc1 | GHXZTYHSJHQHIJ-UHFFFAOYSA-N | No-Effect | inactive |
| chlorindanol | BRD-K01653460 | Oc1ccc(Cl)c2c1CCC2 | ATAJVFBUUIBIEO-UHFFFAOYSA-N | No-Effect | inactive |
| chlorindione | BRD-K61802864 | O=C1c2ccccc2C(=O)C1c1ccc(Cl)cc1 | NJDUWAXIURWWLN-UHFFFAOYSA-N | No-Effect | inactive |
| chlorisondamine-diiodide | BRD-K18030938 | C[N+](C)(C)CC[N+]1(C)Cc2c(Cl)c(Cl)c(Cl)c(Cl)c2C1 | IXWDUZLHWJKVPX-UHFFFAOYSA-N | No-Effect | inactive |
| chlormadinone-acetate | BRD-K38704569 | CC(=O)O[C@]1(C(C)=O)CC[C@H]2[C@@H]3C=C(Cl)C4=CC(=O)CC[C@]4(C)[C@H]3CC[C@@]21C | QMBJSIBWORFWQT-DFXBJWIESA-N | No-Effect | inactive |
| chlormezanone | BRD-A20348246 | CN1C(=O)CCS(=O)(=O)[C@@H]1c1ccc(Cl)cc1 | WEQAYVWKMWHEJO-LLVKDONJSA-N | No-Effect | inactive |
| chlormidazole | BRD-K60443845 | Cc1nc2ccccc2n1Cc1ccc(Cl)cc1 | WNAQOLSMVPFGTE-UHFFFAOYSA-N | No-Effect | inactive |
| chlorocresol | BRD-K89056082 | Cc1cc(O)ccc1Cl | CFKMVGJGLGKFKI-UHFFFAOYSA-N | No-Effect | inactive |
| chlorogenic-acid | BRD-K47114202 | O=C(/C=C/c1ccc(O)c(O)c1)O[C@@H]1C[C@](O)(C(=O)O)C[C@@H](O)[C@H]1O | CWVRJTMFETXNAD-JUHZACGLSA-N | No-Effect | inactive |
| chlorophyllin-copper | BRD-A92960111 | C=Cc1c(C)c2n3c1/C=C1\N=C(/C=c4/c(C)c(C(=O)O)/c(n4[Cu]3)=C(\CC(=O)O)C3=N/C(=C\2)[C@@H](C)[C@@H]3CCC(=O)O)C(CC)=C1C | ZDOYGJNADZJRFB-PVMVIUQGSA-L | No-Effect | inactive |
| chloroprocaine | BRD-A86864073 | CCN(CC)CCOC(=O)c1ccc(N)cc1Cl | VDANGULDQQJODZ-UHFFFAOYSA-N | No-Effect | inactive |
| chloropyramine | BRD-K83257731 | CN(C)CCN(Cc1ccc(Cl)cc1)c1ccccn1 | ICKFFNBDFNZJSX-UHFFFAOYSA-N | No-Effect | inactive |
| chlorothiazide | BRD-K88682005 | NS(=O)(=O)c1cc2c(cc1Cl)N=CNS2(=O)=O | JBMKAUGHUNFTOL-UHFFFAOYSA-N | No-Effect | inactive |
| chlorotrianisene | BRD-K27062708 | COc1ccc(C(Cl)=C(c2ccc(OC)cc2)c2ccc(OC)cc2)cc1 | BFPSDSIWYFKGBC-UHFFFAOYSA-N | No-Effect | inactive |
| chloroxine | BRD-K17075857 | Oc1c(Cl)cc(Cl)c2cccnc12 | WDFKMLRRRCGAKS-UHFFFAOYSA-N | No-Effect | inactive |
| chloroxoquinoline | BRD-K83335125 | O=c1cc[nH]c2cc(Cl)ccc12 | XMFXTXKSWIDMER-UHFFFAOYSA-N | No-Effect | inactive |
| chloroxylenol | BRD-K17223896 | Cc1cc(O)cc(C)c1Cl | OSDLLIBGSJNGJE-UHFFFAOYSA-N | No-Effect | inactive |
| chlorphenamine | BRD-K01826567 | CN(C)CC[C@@H](c1ccc(Cl)cc1)c1ccccn1 | SOYKEARSMXGVTM-HNNXBMFYSA-N | No-Effect | inactive |
| chlorphenesin | BRD-A92262698 | OC[C@@H](O)COc1ccc(Cl)cc1 | MXOAEAUPQDYUQM-MRVPVSSYSA-N | No-Effect | inactive |
| chlorphenoxamine | BRD-K00003537 | CN(C)CCO[C@@](C)(c1ccccc1)c1ccc(Cl)cc1 | KKHPNPMTPORSQE-SFHVURJKSA-N | No-Effect | inactive |
| chlorphensin-carbamate | BRD-A39230911 | NC(=O)OC[C@@H](O)COc1ccc(Cl)cc1 | SKPLBLUECSEIFO-QMMMGPOBSA-N | No-Effect | inactive |
| chlorpropamide | BRD-K97746869 | CCCNC(=O)NS(=O)(=O)c1ccc(Cl)cc1 | RKWGIWYCVPQPMF-UHFFFAOYSA-N | No-Effect | inactive |
| chlorpropham | BRD-K59590127 | CC(C)OC(=O)Nc1cccc(Cl)c1 | CWJSHJJYOPWUGX-UHFFFAOYSA-N | No-Effect | inactive |
| chlorprothixene | BRD-K36207157 | CN(C)CC/C=C1/c2ccccc2Sc2ccc(Cl)cc21 | WSPOMRSOLSGNFJ-AUWJEWJLSA-N | No-Effect | inactive |
| chlorpyrifos | BRD-K08303368 | CCOP(=S)(OCC)Oc1nc(Cl)c(Cl)cc1Cl | SBPBAQFWLVIOKP-UHFFFAOYSA-N | No-Effect | inactive |

| | | | | | |
|---|---|---|---|---|---|
| chlortetracycline | BRD-K00003353 | CN(C)[C@@H]1C(=O)[C@@H](C(N)=O)C(=O)[C@@]2(O)C(=O)[C@H]3C(=O)c4c(O)ccc(Cl)c4[C@@](C)(O)[C@H]3C[C@@H]12 | RNPLSXLZRGPQCU-YPOKJENWSA-N | No-Effect | inactive |
| chlorthalidone | BRD-A26384407 | NS(=O)(=O)c1cc([C@@]2(O)NC(=O)c3ccccc32)ccc1Cl | JIVPVXMEBJLZRO-AWEZNQCLSA-N | No-Effect | inactive |
| chlorzoxazone | BRD-K98174813 | O=c1[nH]c2cc(Cl)ccc2o1 | TZFWDZFKRBELIQ-UHFFFAOYSA-N | No-Effect | inactive |
| cholecalciferol | BRD-K62779383 | C=C1CC[C@H](O)C/C1=C/C=C1\CCC[C@@]2(C)[C@H]1CC[C@@H]2[C@H](C)CCCC(C)C | QYSXJUFSXHHAJI-YRZJJWOYSA-N | No-Effect | inactive |
| cholesterol | BRD-K19338361 | CC(C)CCC[C@@H](C)[C@H]1CC[C@H]2[C@@H]3CC=C4C[C@@H](O)CC[C@]4(C)[C@H]3CC[C@@]21C | HVYWMOMLDIMFJA-DPAQBDIFSA-N | No-Effect | inactive |
| choline | BRD-K77300776 | C[N+](C)(C)CCO | OEYIOHPDSNJKLS-UHFFFAOYSA-N | No-Effect | inactive |
| choline-alfoscerate | BRD-K00003587 | C[N+](C)(C)CCOP(=O)(O)OC[C@H](O)CO | SUHOQUVVVLNYQR-MRVPVSSYSA-O | No-Effect | inactive |
| CHPG | BRD-A19381518 | N[C@@H](C(=O)O)c1cc(O)ccc1Cl | UNIDAFCQFPGYJJ-SSDOTTSWSA-N | No-Effect | inactive |
| chromanol-(+/-) | BRD-M02075109 | CCS(=O)(=O)N(C)[C@@H]1c2cc(C#N)ccc2OC(C)(C)[C@H]1O.CCS(=O)(=O)N(C)[C@H]1c2cc(C#N)ccc2OC(C)(C)[C@@H]1O | VCIPBXUEKWCJAU-DZYJZKBLSA-N | No-Effect | inactive |
| chromium-picolinate | BRD-K01029106 | O=C(O[Cr](OC(=O)c1ccccn1)OC(=O)c1ccccn1)c1ccccn1 | CBDQOLKNTOMMTL-UHFFFAOYSA-K | No-Effect | inactive |
| chromocarb | BRD-K94720315 | O=C(O)c1cc(=O)c2ccccc2o1 | RVMGXWBCQGAWBR-UHFFFAOYSA-N | No-Effect | inactive |
| chrysin | BRD-K22861715 | O=c1cc(-c2ccccc2)oc2cc(O)cc(O)c12 | RTIXKCRFFJGDFG-UHFFFAOYSA-N | No-Effect | inactive |
| chuanxiongzine | BRD-K25691735 | Cc1nc(C)c(C)nc1C | FINHMKGKINIASC-UHFFFAOYSA-N | No-Effect | inactive |
| CI-844 | BRD-K83022601 | c1ccc(Oc2cccnc2)cc1 | KDGUIKPOZGYLQJ-UHFFFAOYSA-N | No-Effect | inactive |
| CI-953 | BRD-K00004748 | Cc1cccc(Cl)c1NC(=O)Nc1ccncc1 | ZSBWDKCIIZYQIR-UHFFFAOYSA-N | No-Effect | inactive |
| CI-966 | BRD-K67680372 | O=C(O)C1=CCCN(CCOC(c2ccc(C(F)(F)F)cc2)c2ccc(C(F)(F)F)cc2)C1 | CMHQDSBIBSKHFP-UHFFFAOYSA-N | No-Effect | inactive |
| CI-976 | BRD-K88544581 | CCCCCCCCCCCC(C)(C)C(=O)Nc1c(OC)cc(OC)cc1OC | WAFNZAURAWBNDZ-UHFFFAOYSA-N | No-Effect | inactive |
| cianidanol | BRD-K58736316 | Oc1cc(O)c2c(c1)O[C@H](c1ccc(O)c(O)c1)[C@@H](O)C2 | PFTAWBLQPZVEMU-DZGCQCFKSA-N | No-Effect | inactive |
| cibenzoline | BRD-A20742938 | c1ccc(C2(c3ccccc3)C[C@H]2C2=NCCN2)cc1 | IPOBOOXFSRWSHL-INIZCTEOSA-N | No-Effect | inactive |
| ciclesonide | BRD-K35245662 | CC(C)C(=O)OCC(=O)[C@@]12O[C@H](C3CCCCC3)O[C@@H]1C[C@H]1[C@@H]3CCC4=CC(=O)C=C[C@]4(C)[C@H]3[C@@H](O)C[C@@]12C | LUKZNWIVRBCLON-GXOBDPJESA-N | No-Effect | inactive |
| cicloprofen | BRD-A94965935 | C[C@@H](C(=O)O)c1ccc2c(c1)Cc1ccccc1-2 | LRXFKKPEBXIPMW-SNVBAGLBSA-N | No-Effect | inactive |
| CID-16020046 | BRD-K00004311 | Cc1ccc(-c2n[nH]c3c2[C@H](c2cccc(O)c2)N(c2ccc(C(=O)O)cc2)C3=O)cc1 | VGUQVYZXABOXCX-QHCPKHFHSA-N | No-Effect | inactive |
| CID-2011756 | BRD-K92781136 | O=C(Nc1ccc(CN2CCOCC2)cc1)c1ccc(-c2cccc(Cl)c2)o1 | XQJWTJLJEYIUDZ-UHFFFAOYSA-N | No-Effect | inactive |
| CID-2745687 | BRD-K06405316 | COC(=O)c1cnn(-c2ccc(F)cc2F)c1/C=N/NC(=S)NC(C)(C)C | CYNLZIBKERMMOA-AWQFTUOYSA-N | No-Effect | inactive |
| CID-5458317 | BRD-A08840375 | OC1=c2ccccc2=C(NC2CCCCC2)/C1=C\c1ccccc1 | BJJVCNWHNQAAGB-HMMYKYKNSA-N | No-Effect | inactive |
| cidofovir | BRD-K96153583 | Nc1ccn(C[C@@H](CO)OCP(=O)(O)O)c(=O)n1 | VWFCHDSQECPREK-LURJTMIESA-N | No-Effect | inactive |
| ciglitazone | BRD-A93000692 | CC1(COc2ccc(C[C@H]3SC(=O)NC3=O)cc2)CCCCC1 | YZFWTZACSRHJQD-OAHLLOKOSA-N | No-Effect | inactive |
| cilastatin | BRD-K85518562 | CC1(C)C[C@@H]1C(=O)N/C(=C\CCCCSC[C@H](N)C(=O)O)C(=O)O | DHSUYTOATWAVLW-WFVMDLQDSA-N | No-Effect | inactive |
| cilazapril | BRD-K96177243 | CCOC(=O)[C@H](CCc1ccccc1)N[C@H]1CCCN2CCC[C@@H](C(=O)O)N2C1=O | HHHKFGXWKKUNCY-FHWLQOOXSA-N | No-Effect | inactive |
| cilengitide | BRD-K57252450 | CC(C)[C@H]1C(=O)N[C@@H](CCCNC(=N)N)C(=O)NCC(=O)N[C@@H](CC(=O)O)C(=O)N[C@H](Cc2ccccc2)C(=O)N1C | AMLYAMJWYAIXIA-VWNVYAMZSA-N | No-Effect | inactive |
| cilnidipine | BRD-K00004281 | COCCOC(=O)C1=C(C)NC(C)=C(C(=O)OC/C=C/c2ccccc2)[C@@H]1c1cccc([N+](=O)[O-])c1 | KJEBULYHNRNJTE-HGJFNOKFSA-N | No-Effect | inactive |
| cilofexor | BRD-K00004595 | O=C(O)c1ccnc(N2CC(O)(c3ccc(OCc4c(-c5c(Cl)cccc5Cl)noc4C4CC4)cc3Cl)C2)c1 | KZSKGLFYQAYZCO-UHFFFAOYSA-N | No-Effect | inactive |
| cilomilast | BRD-K00003550 | COc1ccc([C@]2(C#N)CC[C@@H](C(=O)O)CC2)cc1OC1CCCC1 | CFBUZOUXXHZCFB-OYOVHJISSA-N | No-Effect | inactive |
| cilostamide | BRD-K28578425 | CN(C(=O)CCCOc1ccc2[nH]c(=O)ccc2c1)C1CCCCC1 | UIAYVIIHMORPSJ-UHFFFAOYSA-N | No-Effect | inactive |
| cilostazol | BRD-K67017579 | O=C1CCc2cc(OCCCCc3nnnn3C3CCCCC3)ccc2N1 | RRGUKTPIGVIEKM-UHFFFAOYSA-N | No-Effect | inactive |
| ciluprevir | BRD-K00004667 | COc1ccc2c(O[C@@H]3C[C@H]4C(=O)N[C@]5(C(=O)O)C[C@H]5/C=C\CCCCC[C@H](NC(=O)OC5CCCC5)C(=O)N4C3)cc(-c3csc(NC(C)C)n3)nc2c1 | PJZPDFUUXKKDNB-KNINVFKUSA-N | No-Effect | inactive |
| cimaterol | BRD-A65440446 | CC(C)NC[C@@H](O)c1ccc(N)c(C#N)c1 | BUXRLJCGHZZYNE-GFCCVEGCSA-N | No-Effect | inactive |
| cimetidine | BRD-K34157611 | CN/C(=N\C#N)NCCSCc1[nH]cnc1C | AQIXAKUUQRKLND-UHFFFAOYSA-N | No-Effect | inactive |
| cimetropium | BRD-A99899486 | C[N@@+]1(CC2CC2)[C@H]2C[C@H](OC(=O)[C@H](CO)c3ccccc3)C[C@@H]1[C@H]1O[C@@H]21 | QVVOZYKELHAIPX-IGDPVYPXSA-N | No-Effect | inactive |
| cinacalcet | BRD-K73838513 | C[C@@H](NCCCc1cccc(C(F)(F)F)c1)c1cccc2ccccc12 | VDHAWDNDOKGFTD-MRXNPFEDSA-N | No-Effect | inactive |
| cinalukast | BRD-K53720352 | CCC(CC)(CC(=O)Nc1cccc(/C=C/c2nc(C3CCC3)cs2)c1)C(=O)O | BZMKNPGKXJAIDV-VAWYXSNFSA-N | No-Effect | inactive |
| cinanserin | BRD-K40901640 | CN(C)CCCSc1ccccc1NC(=O)/C=C/c1ccccc1 | RSUVYMGADVXGOU-BUHFOSPRSA-N | No-Effect | inactive |
| cinchonidine | BRD-K39079086 | C=C[C@H]1CN2CC[C@H]1C[C@H]2[C@H](O)c1ccnc2ccccc12 | KMPWYEUPVWOPIM-KODHJQJWSA-N | No-Effect | inactive |
| cinchonine | BRD-K36551084 | C=C[C@H]1CN2CC[C@H]1C[C@@H]2[C@@H](O)c1ccnc2ccccc12 | KMPWYEUPVWOPIM-QAMTZSDWSA-N | No-Effect | inactive |
| cinepazet | BRD-K00531707 | CCOC(=O)CN1CCN(C(=O)/C=C/c2cc(OC)c(OC)c(OC)c2)CC1 | XDUOTWNXVDBCDY-VOTSOKGWSA-N | No-Effect | inactive |
| cinepazide | BRD-K46839850 | COc1cc(/C=C/C(=O)N2CCN(CC(=O)N3CCCC3)CC2)cc(OC)c1OC | RCUDFXMNPQNBDU-VOTSOKGWSA-N | No-Effect | inactive |
| cinflumide | BRD-K25154476 | O=C(/C=C/c1cccc(F)c1)NC1CC1 | NCOOUEIQXVWKTO-QPJJXVBHSA-N | No-Effect | inactive |
| cinnamaldehyde | BRD-K26875402 | O=C/C=C/c1ccccc1 | KJPRLNWUNMBNBZ-QPJJXVBHSA-N | No-Effect | inactive |
| cinnarazine | BRD-K07220430 | C(=C/c1ccccc1)\CN1CCN(C(c2ccccc2)c2ccccc2)CC1 | DERZBLKQOCDDDZ-JLHYYAGUSA-N | No-Effect | inactive |
| cinoctramide | BRD-K08699567 | COc1cc(/C=C/C(=O)N2CCCCCCC2)cc(OC)c1OC | MDGVCMGGLSOVIQ-MDZDMXLPSA-N | No-Effect | inactive |
| cinoxacin | BRD-K14704277 | CCn1nc(C(=O)O)c(=O)c2cc3c(cc21)OCO3 | VDUWPHTZYNWKRN-UHFFFAOYSA-N | No-Effect | inactive |
| cinoxate | BRD-K17828029 | CCOCCOC(=O)/C=C/c1ccc(OC)cc1 | CMDKPGRTAQVGFQ-RMKNXTFCSA-N | No-Effect | inactive |
| CINPA-1 | BRD-K24570280 | CCOC(=O)Nc1ccc2c(c1)N(C(=O)CN(CC)CC)c1ccccc1CC2 | AYQBYSPEGRYKFA-UHFFFAOYSA-N | No-Effect | inactive |
| cintirorgon | BRD-K00003458 | CC(C)(C[C@H]1CN(S(=O)(=O)c2cccc(C(F)(F)F)c2)c2cc(-c3cc(F)cc(OC(F)F)c3)ccc2O1)C(=O)O | GULSIMHVQYBADX-FQEVSTJZSA-N | No-Effect | inactive |
| cintriamide | BRD-K24507548 | COc1cc(/C=C/C(N)=O)cc(OC)c1OC | LRLKZVMLJBNNPE-SNAWJCMRSA-N | No-Effect | inactive |
| cipamfylline | BRD-K00004625 | Nc1nc2c([nH]1)c(=O)n(CC1CC1)c(=O)n2CC1CC1 | KSPYMJJKQMWWNB-UHFFFAOYSA-N | No-Effect | inactive |
| cipargamin | BRD-K27811785 | C[C@H]1Cc2c([nH]c3cc(Cl)c(F)cc23)[C@@]2(N1)C(=O)Nc1ccc(Cl)cc12 | CKLPLPZSUQEDRT-WPCRTTGESA-N | No-Effect | inactive |
| ciprofibrate | BRD-A49358627 | CC(C)(Oc1ccc([C@H]2CC2(Cl)Cl)cc1)C(=O)O | KPSRODZRAIWAKH-SNVBAGLBSA-N | No-Effect | inactive |
| ciprofloxacin | BRD-K04804440 | O=C(O)c1cn(C2CC2)c2cc(N3CCNCC3)c(F)cc2c1=O | MYSWGUAQZAJSOK-UHFFFAOYSA-N | No-Effect | inactive |
| CIQ | BRD-A35425966 | COc1ccc(OC[C@@H]2c3cc(OC)c(OC)cc3CCN2C(=O)c2cccc(Cl)c2)cc1 | VYMILMYEENZHAR-HSZRJFAPSA-N | No-Effect | inactive |
| cirazoline | BRD-K54142781 | c1ccc(C2CC2)c(OCC2=NCCN2)c1 | YAORIDZYZDUZCM-UHFFFAOYSA-N | No-Effect | inactive |
| cis-9,trans-11-conjugated-linoleic-acid | BRD-K80025777 | CCCCCC/C=C/C=C\CCCCCCCC(=O)O | JBYXPOFIGCOSSB-GOJKSUSPSA-N | No-Effect | inactive |
| cis-aconitic-acid | BRD-K71756122 | O=C(O)/C=C(/CC(=O)O)C(=O)O | GTZCVFVGUGFEME-IWQZZHSRSA-N | No-Effect | inactive |
| cis-ACPD | BRD-K00004205 | N[C@]1(C(=O)O)CC[C@@H](C(=O)O)C1 | YFYNOWXBIBKGHB-CLZZGJSISA-N | No-Effect | inactive |
| cis-exo-camphanediol-2,3 | BRD-K41962502 | CC1(C)[C@H]2CC[C@]1(C)[C@@H](O)[C@H]2O | AYEOSGBMQHXVER-AZQAYCESSA-N | No-Effect | inactive |
| cis-urocanic-acid | BRD-K52670952-K251 | O=C(O)/C=C/c1cnc[nH]1.O=C(O)/C=C\c1cnc[nH]1 | JHBGDMJSZRFFDD-KDTZGSNLSA-N | No-Effect | inactive |

| | | | | | |
|---|---|---|---|---|---|
| cisapride | BRD-K06895174 | COc1cc(N)c(Cl)cc1C(=O)N[C@@H]1CCN(CCCOc2ccc(F)cc2)C[C@@H]1OC | DCSUBABJRXZOMT-IRLDBZIGSA-N | No-Effect | inactive |
| cisatracurium | BRD-K17498618 | COc1ccc(C[C@@H]2c3cc(OC)c(OC)cc3CC[N@+]2(C)CCC(=O)OCCCCCOC(=O)CC[N@@+]2(C)CCc3cc(OC)c(OC)cc3[C@H]2Cc2ccc(OC)c(OC)c2)cc1OC | YXSLJKQTIDHPOT-LJCJQEJUSA-N | No-Effect | inactive |
| cisplatin | BRD-K69172251 | N[Pt](N)(Cl)Cl | DQLATGHUWYMOKM-UHFFFAOYSA-L | No-Effect | inactive |
| citalopram | BRD-A47598013 | CN(C)CCC[C@@]1(c2ccc(F)cc2)OCc2cc(C#N)ccc21 | WSEQXVZVJXJVFP-FQEVSTJZSA-N | No-Effect | inactive |
| citarinostat | BRD-K00003202 | O=C(CCCCCCNC(=O)c1cnc(N(c2ccccc2)c2ccccc2Cl)nc1)NO | VLIUIBXPEDFJRF-UHFFFAOYSA-N | No-Effect | inactive |
| CITCO | BRD-K53263234 | Clc1ccc(-c2nc3sccn3c2/C=N/OCc2ccc(Cl)c(Cl)c2)cc1 | ZQWBOKJVVYNKTL-AUEPDCJTSA-N | No-Effect | inactive |
| citicoline | BRD-K01827268 | C[N+](C)(C)CCOP(=O)(O)OP(=O)(O)OC[C@H]1O[C@@H](n2ccc(N)nc2=O)[C@H](O)[C@@H]1O | RZZPDXZPRHQOCG-OJAKKHQRSA-O | No-Effect | inactive |
| citiolone | BRD-A57957888 | CC(=O)N[C@H]1CCSC1=O | NRFJZTXWLKPZAV-YFKPBYRVSA-N | No-Effect | inactive |
| citric-acid | BRD-K76792669 | O=C(O)CC(O)(CC(=O)O)C(=O)O | KRKNYBCHXYNGOX-UHFFFAOYSA-N | No-Effect | inactive |
| CJ-033466 | BRD-K21306330 | Cc1cn2c(N)c(Cl)cc(C(=O)NCC3CCN(CC(C)C)CC3)c2n1 | ISKHMDNIWXPUGR-UHFFFAOYSA-N | No-Effect | inactive |
| CJ-13610 | BRD-K00004624 | Cc1nccn1-c1ccc(Sc2cccc(C3(C(N)=O)CCOCC3)c2)cc1 | VPTONMHDLLMOOV-UHFFFAOYSA-N | No-Effect | inactive |
| CK-101 | BRD-K00005246 | C=CC(=O)Nc1cccc(-c2cccc3cnc(Nc4ccc(N5CCN(CCO)CC5)c(F)c4F)nc23)c1 | IDRGFNPZDVBSSE-UHFFFAOYSA-N | No-Effect | inactive |
| CK-636 | BRD-K81572616 | Cc1[nH]c2ccccc2c1CCNC(=O)c1cccs1 | ACAKNPKRLPMONU-UHFFFAOYSA-N | No-Effect | inactive |
| CKD-712 | BRD-A26458246 | Oc1cc2c(cc1O)[C@H](Cc1cccc3ccccc13)NCC2 | YGCQFKVNIBDJFW-SFHVURJKSA-N | No-Effect | inactive |
| CKI-7 | BRD-K34303607 | NCCNS(=O)(=O)c1ccc(Cl)c2ccncc12 | OGKYMFFYOWUTKV-UHFFFAOYSA-N | No-Effect | inactive |
| CL-218872 | BRD-K00662280 | Cc1nnc2ccc(-c3cccc(C(F)(F)F)c3)nn12 | GUOQUXNJZHGPQF-UHFFFAOYSA-N | No-Effect | inactive |
| CL-225385 | BRD-K87573013 | N#CCC(=O)c1ccccc1 | ZJRCIQAMTAINCB-UHFFFAOYSA-N | No-Effect | inactive |
| CL316243 | BRD-K06467130 | C[C@H](Cc1ccc2c(c1)OC(C(=O)O)(C(=O)O)O2)NC[C@H](O)c1cccc(Cl)c1 | JEDJMKTVUPSHFW-ABAIWWIYSA-N | No-Effect | inactive |
| cladribine | BRD-K93034159 | Nc1nc(Cl)nc2c1ncn2[C@H]1C[C@H](O)[C@@H](CO)O1 | PTOAARAWEBMLNO-KVQBGUIXSA-N | No-Effect | inactive |
| clamikalant | BRD-K00004576 | CNC(=S)NS(=O)(=O)c1cc(CCNC(=O)c2cc(Cl)ccc2OC)ccc1OC | VXTKXGKPBOLHRY-UHFFFAOYSA-N | No-Effect | inactive |
| clanfenur | BRD-K62378877 | CN(C)c1cccc(F)c1C(=O)NC(=O)Nc1ccc(Cl)cc1 | SRLPZQAEBMZCIJ-UHFFFAOYSA-N | No-Effect | inactive |
| clarithromycin | BRD-K49668410 | CC[C@H]1OC(=O)[C@H](C)[C@@H](O[C@H]2C[C@@](C)(OC)[C@@H](O)[C@H](C)O2)[C@H](C)[C@@H](O[C@@H]2O[C@H](C)C[C@H](N(C)C)[C@H]2O)[C@](C)(OC)C[C@@H](C)C(=O)[C@H](C)[C@@H](O)[C@]1(C)O | AGOYDEPGAOXOCK-KCBOHYOISA-N | No-Effect | inactive |
| clavulanate | BRD-K94250879 | O=C(O)[C@H]1/C(=C/CO)O[C@@H]2CC(=O)N21 | HZZVJAQRINQKSD-PBFISZAISA-N | No-Effect | inactive |
| clebopride | BRD-K17294426 | COc1cc(N)c(Cl)cc1C(=O)NC1CCN(Cc2ccccc2)CC1 | BVPWJMCABCPUQY-UHFFFAOYSA-N | No-Effect | inactive |
| clemizole | BRD-K04704168 | Clc1ccc(Cn2c(CN3CCCC3)nc3ccccc32)cc1 | CJXAEXPPLWQRFR-UHFFFAOYSA-N | No-Effect | inactive |
| cleviprex | BRD-A01563671 | CCCC(=O)OCOC(=O)C1=C(C)NC(C)=C(C(=O)OC)[C@@H]1c1cccc(Cl)c1Cl | KPBZROQVTHLCDU-SFHVURJKSA-N | No-Effect | inactive |
| clevudine | BRD-K00003562 | Cc1cn([C@H]2O[C@@H](CO)[C@H](O)[C@H]2F)c(=O)[nH]c1=O | GBBJCSTXCAQSSJ-XQXXSGGOSA-N | No-Effect | inactive |
| clidinium | BRD-K01827265 | C[N+]12CCC(CC1)[C@H](OC(=O)C(O)(c1ccccc1)c1ccccc1)C2 | HOOSGZJRQIVJSZ-WEQDKIBNSA-N | No-Effect | inactive |
| climbazole | BRD-A61676498 | CC(C)(C)C(=O)[C@@H](Oc1ccc(Cl)cc1)n1ccnc1 | OWEGWHBOCFMBLP-CQSZACIVSA-N | No-Effect | inactive |
| clinafloxacin | BRD-K01825499 | N[C@H]1CCN(c2c(F)cc3c(=O)c(C(=O)O)cn(C4CC4)c3c2Cl)C1 | QGPKADBNRMWEQR-QMMMGPOBSA-N | No-Effect | inactive |
| clindamycin | BRD-K23586533 | CCC[C@@H]1C[C@@H](C(=O)N[C@H]([C@H](C)Cl)[C@H]2O[C@H](SC)[C@H](O)[C@@H](O)[C@H]2O)N(C)C1 | KDLRVYVGXIQJDK-AWPVFWJPSA-N | No-Effect | inactive |
| clindamycin-palmitate | BRD-K10759270 | CCCCCCCCCCCCCCCC(=O)O[C@@H]1[C@@H](O)[C@@H](O)[C@@H]([C@H](NC(=O)[C@@H]2C[C@@H](CCC)CN2C)[C@H](C)Cl)O[C@@H]1SC | OYSKUZDIHNKWLV-PRUAPSLNSA-N | No-Effect | inactive |
| clindamycin-phosphate | BRD-K27771035 | CCC[C@@H]1C[C@@H](C(=O)N[C@H]([C@H](C)Cl)[C@H]2O[C@H](SC)[C@H](OP(=O)(O)O)[C@@H](O)[C@H]2O)N(C)C1 | UFUVLHLTWXBHGZ-MGZQPHGTSA-N | No-Effect | inactive |
| clinofibrate | BRD-A90625545 | CCC(C)(Oc1ccc(C2(c3ccc(O[C@](C)(CC)C(=O)O)cc3)CCCCC2)cc1)C(=O)O | BMOVQUBVGICXQN-AVJYQCBHSA-N | No-Effect | inactive |
| clioquinol | BRD-K09255212 | Oc1c(I)cc(Cl)c2cccnc12 | QCDFBFJGMNKBDO-UHFFFAOYSA-N | No-Effect | inactive |
| clobenpropit | BRD-K71430621 | N=C(NCc1ccc(Cl)cc1)SCCCc1cnc[nH]1 | UCAIEVHKDLMIFL-UHFFFAOYSA-N | No-Effect | inactive |
| clobetasone-butyrate | BRD-K58810291 | CCCC(=O)O[C@]1(C(=O)CCl)[C@@H](C)C[C@H]2[C@@H]3CCC4=CC(=O)C=C[C@]4(C)[C@@]3(F)C(=O)C[C@@]21C | FBRAWBYQGRLCEK-AVVSTMBFSA-N | No-Effect | inactive |
| clobutinol | BRD-A43809092 | C[C@@H](CN(C)C)C(C)(O)Cc1ccc(Cl)cc1 | KVHHQGIIZCJATJ-ZSOXZCCMSA-N | No-Effect | inactive |
| clocortolone-pivalate | BRD-K38003476 | C[C@@H]1C[C@H]2[C@@H]3C[C@H](F)C4=CC(=O)C=C[C@]4(C)[C@@]3(Cl)[C@@H](O)C[C@]2(C)[C@H]1C(=O)COC(=O)C(C)(C)C | SXYZQZLHAIHKKY-GSTUPEFVSA-N | No-Effect | inactive |
| clodronic-acid | BRD-K72542090 | O=P(O)(O)C(Cl)(Cl)P(=O)(O)O | ACSIXWWBWUQEHA-UHFFFAOYSA-N | No-Effect | inactive |
| clofarabine | BRD-K34022604 | Nc1nc(Cl)nc2c1ncn2[C@@H]1O[C@H](CO)[C@@H](O)[C@@H]1F | WDDPHFBMKLOVOX-AYQXTPAHSA-N | No-Effect | inactive |
| clofibrate | BRD-K45252063 | CCOC(=O)C(C)(C)Oc1ccc(Cl)cc1 | KNHUKKLJHYUCFP-UHFFFAOYSA-N | No-Effect | inactive |
| clofibric-acid | BRD-K19111024 | CC(C)(Oc1ccc(Cl)cc1)C(=O)O | TXCGAZHTZHNUAI-UHFFFAOYSA-N | No-Effect | inactive |
| clofilium | BRD-K02680176 | CCCCCCC[N+](CC)(CC)CCCCc1ccc(Cl)cc1 | WPSYTTKBGAZSCX-UHFFFAOYSA-N | No-Effect | inactive |
| clomesone | BRD-K39230570 | CS(=O)(=O)CS(=O)(=O)OCCCl | SEHSPJCWCBQHPF-UHFFFAOYSA-N | No-Effect | inactive |
| clomethiazole | BRD-K50422030 | Cc1ncsc1CCCl | PCLITLDOTJTVDJ-UHFFFAOYSA-N | No-Effect | inactive |
| clonidine | BRD-K98530306 | Clc1cccc(Cl)c1NC1=NCCN1 | GJSURZIOUXUGAL-UHFFFAOYSA-N | No-Effect | inactive |
| clonixin | BRD-K18399440 | Cc1c(Cl)cccc1Nc1ncccc1C(=O)O | CLOMYZFHNHFSIQ-UHFFFAOYSA-N | No-Effect | inactive |
| clonixin-lysinate | BRD-M63173034 | Cc1c(Cl)cccc1Nc1ncccc1C(=O)O.NCCCC[C@H](N)C(=O)O | CVNFYQCHAWFYQI-ZSCHJXSPSA-N | No-Effect | inactive |
| clopamide | BRD-K15567837 | C[C@H]1CCC[C@@H](C)N1NC(=O)c1ccc(Cl)c(S(N)(=O)=O)c1 | LBXHRAWDUMTPSE-AOOOYVTPSA-N | No-Effect | inactive |
| cloperastine-fendizoate | BRD-K00003529 | Clc1ccc(C(OCCN2CCCCC2)c2ccccc2)cc1 | FLNXBVJLPJNOSI-UHFFFAOYSA-N | No-Effect | inactive |
| clopidogrel | BRD-K27721098 | COC(=O)[C@H](c1ccccc1Cl)N1CCc2sccc2C1 | GKTWGGQPFAXNFI-HNNXBMFYSA-N | No-Effect | inactive |
| clopidol | BRD-K80036624 | Cc1[nH]c(C)c(Cl)c(=O)c1Cl | ZDPIZLCVJAAHHR-UHFFFAOYSA-N | No-Effect | inactive |
| cloprostenol-(+/-) | BRD-K17850764 | O=C(O)CCC/C=C\C[C@H]1[C@@H](O)C[C@@H](O)[C@@H]1/C=C/[C@@H](O)COc1cccc(Cl)c1 | VJGGHXVGBSZVMZ-QIZQQNKQSA-N | No-Effect | inactive |
| cloranolol | BRD-A04971881 | CC(C)(C)NC[C@@H](O)COc1cc(Cl)ccc1Cl | XYCMOTOFHFTUIU-SNVBAGLBSA-N | No-Effect | inactive |
| clorgiline | BRD-K73251053 | C#CCN(C)CCCOc1ccc(Cl)cc1Cl | BTFHLQRNAMSNLC-UHFFFAOYSA-N | No-Effect | inactive |
| clorprenaline | BRD-K01825579 | CC(C)NC[C@@H](O)c1ccccc1Cl | SSMSBSWKLKKXGG-LLVKDONJSA-N | No-Effect | inactive |
| clorsulon | BRD-K97521363 | Nc1cc(C(Cl)=C(Cl)Cl)c(S(N)(=O)=O)cc1S(N)(=O)=O | QOVTVIYTBRHADL-UHFFFAOYSA-N | No-Effect | inactive |
| closantel | BRD-K01825680 | Cc1cc([C@@H](C#N)c2ccc(Cl)cc2)c(Cl)cc1NC(=O)c1cc(I)cc(I)c1O | JMPFSEBWVLAJKM-KRWDZBQOSA-N | No-Effect | inactive |
| clotiapine | BRD-K10990317 | CN1CCN(C2=Nc3ccccc3Sc3ccc(Cl)cc32)CC1 | KAAZGXDPUNNEFN-UHFFFAOYSA-N | No-Effect | inactive |
| clotrimazole | BRD-K15916496 | Clc1ccccc1C(c1ccccc1)(c1ccccc1)n1ccnc1 | VNFPBHJOKIVQEB-UHFFFAOYSA-N | No-Effect | inactive |
| cloxacillin | BRD-K01244426 | Cc1onc(-c2ccccc2Cl)c1C(=O)N[C@@H]1C(=O)N2[C@@H](C(=O)O)C(C)(C)S[C@H]12 | LQOLIRLGBULYKD-JKIFEVAISA-N | No-Effect | inactive |
| cloxyquin | BRD-K46982791 | Oc1ccc(Cl)c2cccnc12 | CTQMJYWDVABFRZ-UHFFFAOYSA-N | No-Effect | inactive |
| CM-4620 | BRD-K00003438 | Cc1cccc(F)c1C(=O)Nc1cnc(-c2cc3c(cc2Cl)OC(F)(F)O3)cn1 | QQMKTHUGOQDEIL-UHFFFAOYSA-N | No-Effect | inactive |
| CMPD-1 | BRD-K54095730 | O=C(CCCc1ccc(-c2ccccc2F)cc1)Nc1ccc(O)cc1 | ODYAQBDIXCVKAE-UHFFFAOYSA-N | No-Effect | inactive |
| CNQX | BRD-K19438463 | N#Cc1cc2[nH]c(=O)c(=O)[nH]c2cc1[N+](=O)[O-] | RPXVIAFEQBNEAX-UHFFFAOYSA-N | No-Effect | inactive |
| CNX-2006 | BRD-K24869513 | C=CC(=O)Nc1cccc(Nc2nc(Nc3ccc(NC4CN(CCF)C4)cc3OC)ncc2C(F)(F)F)c1 | BFSRTTWIPACGMI-UHFFFAOYSA-N | No-Effect | inactive |

| | | | | | |
|---|---|---|---|---|---|
| CO-101244 | BRD-K55677650 | Cc1ccc(CC2(O)CCN(CCOc3ccc(O)cc3)CC2)cc1 | VKMFDKYCIKEDMR-UHFFFAOYSA-N | No-Effect | inactive |
| CO-102862 | BRD-K70912147 | NC(=O)N/N=C/c1ccc(Oc2ccc(F)cc2)cc1 | MHUUDVZSPFRUSK-RQZCQDPDSA-N | No-Effect | inactive |
| cobicistat | BRD-K51040301 | CC(C)c1nc(CN(C)C(=O)N[C@@H](CCN2CCOCC2)C(=O)N[C@H](CC[C@H](Cc2ccccc2)NC(=O)OCc2cncs2)Cc2ccccc2)cs1 | ZCIGNRJZKPOIKD-CQXVEOKZSA-N | No-Effect | inactive |
| coenzyme-I | BRD-K01825730 | NC(=O)C1=CN([C@@H]2O[C@H](COP(=O)(O)OP(=O)(O)OC[C@H]3O[C@@H](n4cnc5c(N)ncnc54)[C@H](O)[C@@H]3O)[C@@H](O)[C@H]2O)C=CC1 | BOPGDPNILDQYTO-NNYOXOHSSA-N | No-Effect | inactive |
| coenzyme-Q10 | BRD-K75043328 | COC1=C(OC)C(=O)C(C/C=C(\C)CC/C=C(\C)CC/C=C(\C)CC/C=C(\C)CC/C=C(\C)CC/C=C(\C)CC/C=C(\C)CC/C=C(\C)CC/C=C(\C)CCC=C(C)C)=C(C)C1=O | ACTIUHUUMQJHFO-UPTCCGCDSA-N | No-Effect | inactive |
| colforsin | BRD-A55416093 | C=C[C@@]1(C)CC(=O)[C@@]2(O)[C@](C)(O1)[C@@H](OC(C)=O)[C@@H](O)[C@H]1C(C)(C)CC[C@H](O)[C@@]12C | OHCQJHSOBUTRHG-KGGHGJDLSA-N | No-Effect | inactive |
| colforsin-daproate | BRD-K19388745 | C=C[C@@]1(C)CC(=O)[C@@]2(O)[C@](C)(O1)[C@@H](OC(C)=O)[C@@H](OC(=O)CCN(C)C)[C@H]1C(C)(C)CC[C@H](O)[C@@]12C | RSOZZQTUMVBTMR-XGUNBQNXSA-N | No-Effect | inactive |
| colfosceril-palmitate | BRD-K01825113 | CCCCCCCCCCCCCCCC(=O)OC[C@H](CO[P@@](=O)(O)OCC[N+](C)(C)C)OC(=O)CCCCCCCCCCCCCCC | KILNVBDSWZSGLL-KXQOOQHDSA-O | No-Effect | inactive |
| colistin | BRD-K01666935 | CC[C@@H](C)CCCC(=O)N[C@@H](CCN)C(=O)N[C@H](C(=O)N[C@@H](CCN)C(=O)N[C@H]1CCNC(=O)[C@H]([C@@H](C)O)NC(=O)[C@H](CCN)NC(=O)[C@H](CCN)NC(=O)[C@H](CC(C)C)NC(=O)[C@@H](CC(C)C)NC(=O)[C@H](CCN)NC1=O)[C@@H](C)O | YKQOSKADJPQZHB-XIVCCLANSA-N | No-Effect | inactive |
| compound-401 | BRD-K19655798 | O=c1cc(N2CCOCC2)nc2c3ccccc3ccn12 | BVRDQVRQVGRNHG-UHFFFAOYSA-N | No-Effect | inactive |
| compound-58112 | BRD-K76139886 | COc1cccc(CN2C[Si](C)(C)O[Si](C)(C)C2)c1 | CUYLJVJMQPPTOS-UHFFFAOYSA-N | No-Effect | inactive |
| compound-w | BRD-K98143539 | O=C(O)c1cc(Oc2ccc([N+](=O)[O-])cc2)cc(Oc2ccc([N+](=O)[O-])cc2)c1 | JOSXKPZXMVHRKU-UHFFFAOYSA-N | No-Effect | inactive |
| conivaptan | BRD-K37385792 | Cc1nc2c([nH]1)CCN(C(=O)c1ccc(NC(=O)c3ccccc3-c3ccccc3)cc1)c1ccccc1-2 | IKENVDNFQMCRTR-UHFFFAOYSA-N | No-Effect | inactive |
| copper-histidine | BRD-K36449075 | N[C@H](Cc1cnc[nH]1)C(=O)O | HNDVDQJCIGZPNO-RXMQYKEDSA-N | No-Effect | inactive |
| COR-170 | BRD-K74965187 | CCCCCn1cc(C(=O)NC23CC4CC(CC(C4)C2)C3)c(=O)c2cc(-c3ccccc3)ccc21 | MSJISJDTJJYBFT-UHFFFAOYSA-N | No-Effect | inactive |
| cordycepin | BRD-K65428967 | Nc1ncnc2c1ncn2[C@@H]1O[C@H](CO)C[C@H]1O | OFEZSBMBBKLLBJ-BAJZRUMYSA-N | No-Effect | inactive |
| corosolic-acid | BRD-K27885593 | C[C@@H]1[C@H]2C3=CC[C@@H]4[C@@]5(C)C[C@@H](O)[C@H](O)C(C)(C)[C@@H]5CC[C@@]4(C)[C@]3(C)CC[C@@]2(C(=O)O)CC[C@H]1C | HFGSQOYIOKBQOW-ZSDYHTTISA-N | No-Effect | inactive |
| CORT-108297 | BRD-K00004700 | CCOC[C@]12Cc3cnn(-c4ccc(F)cc4)c3C=C1CCN(S(=O)(=O)c1ccc(C(F)(F)F)cc1)C2 | SLKURXRZHJOZOD-RUZDIDTESA-N | No-Effect | inactive |
| corticosterone | BRD-K73589401 | C[C@]12C[C@H](O)[C@H]3[C@@H](CCC4=CC(=O)CC[C@@]43C)[C@@H]1CC[C@@H]2C(=O)CO | OMFXVFTZEKFJBZ-HJTSIMOOSA-N | No-Effect | inactive |
| cortisone | BRD-K00025572 | C[C@]12CC(=O)[C@H]3[C@@H](CCC4=CC(=O)CC[C@@]43C)[C@@H]1CC[C@]2(O)C(=O)CO | MFYSYFVPBJMHGN-ZPOLXVRWSA-N | No-Effect | inactive |
| cortisone-acetate | BRD-K86161929 | CC(=O)OCC(=O)[C@@]1(O)CC[C@H]2[C@@H]3CCC4=CC(=O)CC[C@]4(C)[C@H]3C(=O)C[C@@]21C | ITRJWOMZKQRYTA-RFZYENFJSA-N | No-Effect | inactive |
| cortodoxone | BRD-K32891457 | C[C@]12CC[C@H]3[C@@H](CCC4=CC(=O)CC[C@@]43C)[C@@H]1CC[C@]2(O)C(=O)CO | WHBHBVVOGNECLV-OBQKJFGGSA-N | No-Effect | inactive |
| costunolide | BRD-K26833429 | C=C1C(=O)O[C@@H]2/C=C(\C)CC/C=C(\C)CC[C@@H]12 | HRYLQFBHBWLLLL-AHNJNIBGSA-N | No-Effect | inactive |
| cot-inhibitor-1 | BRD-K98710228 | N#Cc1cnc2c(Cl)cc(NCc3cn(CCN4CCCCCC4)nn3)cc2c1Nc1ccc(F)c(Cl)c1 | ZWFKJFUFYPOTKL-UHFFFAOYSA-N | No-Effect | inactive |
| cot-inhibitor-2 | BRD-K49468759 | CCN1CCC(n2cc(CNc3cc(Cl)c4ncc(C#N)c(Nc5ccc(F)c(Cl)c5)c4c3)nn2)CC1 | PHNZIIMWDVXPGG-UHFFFAOYSA-N | No-Effect | inactive |
| COTI-2 | BRD-K00003348 | S=C(N/N=C1\CCCc2cccnc21)N1CCN(c2ccccn2)CC1 | UTDAKQMBNSHJJB-CJLVFECKSA-N | No-Effect | inactive |
| cotinine | BRD-K94144010 | CN1C(=O)CC[C@H]1c1cccnc1 | UIKROCXWUNQSPJ-VIFPVBQESA-N | No-Effect | inactive |
| CP-105,696 | BRD-K00004654 | O=C(O)C1(c2ccc3c(c2)OC[C@H](Cc2ccc(-c4ccccc4)cc2)[C@H]3O)CCCC1 | KMNLXCBYBKHKSK-BKMJKUGQSA-N | No-Effect | inactive |
| CP-316819 | BRD-K53913008 | CON(C)C(=O)[C@H](O)[C@H](Cc1ccccc1)NC(=O)c1cc2cc(Cl)ccc2[nH]1 | WEQLRDLTRCEUHG-PKOBYXMFSA-N | No-Effect | inactive |
| CP-339818 | BRD-K00004233 | CCCCC/N=c1\ccn(Cc2ccccc2)c2ccccc12 | MMGAVKCAGQCFHS-LSDHQDQOSA-N | No-Effect | inactive |
| CP-376395 | BRD-K31394475 | CCC(CC)Nc1cc(C)nc(Oc2c(C)cc(C)cc2C)c1C | VIZBSVDBNLAVAW-UHFFFAOYSA-N | No-Effect | inactive |
| CP-471474 | BRD-K33234937 | CC(C)(NS(=O)(=O)c1ccc(Oc2ccc(F)cc2)cc1)C(=O)NO | QCOQJYRPDUMCNP-UHFFFAOYSA-N | No-Effect | inactive |
| CP-532623 | BRD-K36611520 | CC[C@@H]1C[C@H](N(Cc2cc(C(F)(F)F)cc(C(F)(F)F)c2)C(=O)c2cc(C(F)(F)F)ccc2N1C(=O)OC(C)C | TUPKOWFPVAXQFP-OFNKIYASSA-N | No-Effect | inactive |
| CP-547632 | BRD-K00004719 | NC(=O)c1c(OCc2c(F)cc(Br)cc2F)nsc1NC(=O)NCCCCN1CCCC1 | HXHAJRMTJXHJJZ-UHFFFAOYSA-N | No-Effect | inactive |
| CP-673451 | BRD-K43621685 | COCCOc1ccc2c(c1)ncn2-c1ccc2cccc(N3CCC(N)CC3)c2n1 | DEEOXSOLTLIWMG-UHFFFAOYSA-N | No-Effect | inactive |
| CP-724714 | BRD-K76908866 | COCC(=O)NC/C=C/c1ccc2ncnc(Nc3ccc(Oc4ccc(C)nc4)c(C)c3)c2c1 | LLVZBTWPGQVVLW-SNAWJCMRSA-N | No-Effect | inactive |
| CP-93129 | BRD-K81876028 | O=c1ccc2[nH]cc(C3=CCNCC3)c2[nH]1 | PJYVGMRFPFNZCT-UHFFFAOYSA-N | No-Effect | inactive |
| CP-94253 | BRD-K33860217 | CCCOc1ccc2[nH]cc(C3=CCNCC3)c2n1 | KWQWBZIGHIOKIO-UHFFFAOYSA-N | No-Effect | inactive |
| CP-945,598 | BRD-K09090949 | CCNC1(C(N)=O)CCN(c2ncnc3c2nc(-c2ccccc2Cl)n3-c2ccc(Cl)cc2)CC1 | UNAZAADNBYXMIV-UHFFFAOYSA-N | No-Effect | inactive |
| CP-99994 | BRD-K09946180 | COc1ccccc1CN[C@H]1CCCN[C@H]1c1ccccc1 | DTQNEFOKTXXQKV-HKUYNNGSSA-N | No-Effect | inactive |
| CPA-inhibitor | BRD-A69126730 | O=C(O)[C@H](Cc1ccccc1)CN(O)C(=O)Cc1ccccc1 | PONANXDRJJIGPG-MRXNPFEDSA-N | No-Effect | inactive |
| CPCCOEt | BRD-K00004223 | CCOC(=O)[C@]12C[C@H]1/C(=N\O)c1ccccc1O2 | FXCTZFMSAHZQTR-PFJVYOFESA-N | No-Effect | inactive |
| CPI-1189 | BRD-K11073688 | CC(=O)Nc1ccc(C(=O)NC(C)(C)C)cc1 | DJKNRCWSXSZACF-UHFFFAOYSA-N | No-Effect | inactive |
| CPI-1205 | BRD-K00003360 | COc1cc(C)[nH]c(=O)c1CNC(=O)c1c(C)n([C@H](C)C2CCN(CC(F)(F)F)CC2)c2ccccc12 | HPODOLXTMDHLLC-QGZVFWFLSA-N | No-Effect | inactive |
| CPI-169 | BRD-K97181972 | CCS(=O)(=O)N1CCC([C@@H](C)n2c(C)c(C(=O)NCc3c(OC)cc(C)[nH]c3=O)c3ccccc32)CC1 | LHGUZCKPFXXVPV-GOSISDBHSA-N | No-Effect | inactive |
| CPI-203 | BRD-K15811707 | Cc1sc2c(c1C)C(c1ccc(Cl)cc1)=N[C@@H](CC(N)=O)c1nnc(C)n1-2 | QECMENZMDBOLDR-AWEZNQCLSA-N | No-Effect | inactive |
| CPI-360 | BRD-K18319422 | COc1cc(C)[nH]c(=O)c1CNC(=O)c1c(C)n([C@H](C)C2CCOCC2)c2ccccc12 | PFPSFENQCNMITC-MRXNPFEDSA-N | No-Effect | inactive |
| CPI-444 | BRD-K00003442 | Cc1ccc(-c2nc(N)nc3c2nnn3Cc2cccc(CO[C@H]3CCOC3)n2)o1 | KURQKNMKCGYWRJ-HNNXBMFYSA-N | No-Effect | inactive |
| CPI-613 | BRD-A06916187 | O=C(O)CCCC[C@@H](CCSCc1ccccc1)SCc1ccccc1 | ZYRLHJIMTROTBO-NRFANRHFSA-N | No-Effect | inactive |
| CPP | BRD-K66094457 | O=C(O)[C@H]1CN(CCCP(=O)(O)O)CCN1 | CUVGUPIVTLGRGI-SSDOTTSWSA-N | No-Effect | inactive |
| CPSI-1306-(+/-) | BRD-A81603255 | O=C(C[C@H]1CC(c2ccc(F)cc2F)=NO1)N1CCOCC1 | GUWOSEVHCSNYFK-LLVKDONJSA-N | No-Effect | inactive |
| CQS | BRD-K56839624 | Nc1ccc(S(=O)(=O)Nc2cnc3c(Cl)cccc3n2)cc1 | CTSNHMQGVWXIEG-UHFFFAOYSA-N | No-Effect | inactive |
| CR8-(R) | BRD-K40331046 | CC[C@H](CO)Nc1nc(NCc2ccc(-c3ccccn3)cc2)c2ncn(C(C)C)c2n1 | HOCBJBNQIQQQGT-LJQANCHMSA-N | No-Effect | inactive |
| creatine | BRD-K19615002 | CN(CC(=O)O)C(=N)N | CVSVTCORWBXHQV-UHFFFAOYSA-N | No-Effect | inactive |
| creatinine | BRD-K13495773 | CN1CC(=O)NC1=N | DDRJAANPRJIHGJ-UHFFFAOYSA-N | No-Effect | inactive |
| creatinol-phosphate | BRD-K42991124 | CN(CCOP(=O)(O)O)C(=N)N | FOIPWTMKYXWFGC-UHFFFAOYSA-N | No-Effect | inactive |
| crenigacestat | BRD-K00003129 | C[C@H](NC(=O)CCC(F)(F)F)C(=O)N[C@@H]1C(=O)N(CCO)c2ncccc2-c2ccccc21 | YCBAQKQAINQRFW-UGSOOPFHSA-N | No-Effect | inactive |
| crenolanib | BRD-K22127577 | CC1(COc2ccc3c(c2)ncn3-c2ccc3cccc(N4CCC(N)CC4)c3n2)COC1 | DYNHJHQFHQTFTP-UHFFFAOYSA-N | No-Effect | inactive |
| cresol | BRD-K56940914 | Cc1ccc(O)cc1 | IWDCLRJOBJJRNH-UHFFFAOYSA-N | No-Effect | inactive |

| | | | | | |
|---|---|---|---|---|---|
| cresopirine | BRD-K58648841 | CC(=O)Oc1c(C)cccc1C(=O)O | XRBMKGUDDJPAMH-UHFFFAOYSA-N | No-Effect | inactive |
| cridanimod | BRD-K97939847 | O=C(O)Cn1c2ccccc2c(=O)c2ccccc21 | UOMKBIIXHQIERR-UHFFFAOYSA-N | No-Effect | inactive |
| crisaborole | BRD-K43230258 | N#Cc1ccc(Oc2ccc3c(c2)COB3O)cc1 | USZAGAREISWJDP-UHFFFAOYSA-N | No-Effect | inactive |
| crisdesalazine | BRD-K00004545 | O=C(O)c1cc(NCCc2ccc(C(F)(F)F)cc2)ccc1O | UTMVACIBQLDZLP-UHFFFAOYSA-N | No-Effect | inactive |
| crizotinib | BRD-K78431006 | C[C@@H](Oc1cc(-c2cnn(C3CCNCC3)c2)cnc1N)c1c(Cl)ccc(F)c1Cl | KTEIFNKAUNYNJU-GFCCVEGCSA-N | No-Effect | inactive |
| croconazole | BRD-K73701464 | C=C(c1ccccc1OCc1cccc(Cl)c1)n1ccnc1 | WHPAGCJNPTUGGD-UHFFFAOYSA-N | No-Effect | inactive |
| cromakalim | BRD-M89827113 | CC1(C)Oc2ccc(C#N)cc2[C@@H](N2CCCC2=O)[C@@H]1O.CC1(C)Oc2ccc(C#N)cc2[C@H](N2CCCC2=O)[C@H]1O | BVMDRSUIADWFLV-PNSAJKJISA-N | No-Effect | inactive |
| crotamiton | BRD-K57179821 | C/C=C/C(=O)N(CC)c1ccccc1C | DNTGGZPQPQTDQF-XBXARRHUSA-N | No-Effect | inactive |
| CRT0044876 | BRD-K02816415 | O=C(O)c1cc2cccc([N+](=O)[O-])c2[nH]1 | BIUCOFQROHIAEO-UHFFFAOYSA-N | No-Effect | inactive |
| cryptotanshinone | BRD-K33336844 | C[C@H]1COC2=C1C(=O)C(=O)c1c2ccc2c1CCCC2(C)C | GVKKJJOMQCNPGB-JTQLQIEISA-N | No-Effect | inactive |
| crystal-violet | BRD-K00003324 | CN(C)c1ccc(C(=C2C=CC(=[N+](C)C)C=C2)c2ccc(N(C)C)cc2)cc1 | LGLFFNDHMLKUMI-UHFFFAOYSA-N | No-Effect | inactive |
| CS-110266 | BRD-K15519488 | c1ccc2sc(C3(N4CCCCC4)CCCCC3)cc2c1 | RGSVXQJPSWZXOP-UHFFFAOYSA-N | No-Effect | inactive |
| CS-917 | BRD-K35601925 | CC(C)Cc1sc(N)nc1-c1ccc(P(=O)(O)O)o1 | XJMYIJPPDSZOPN-UHFFFAOYSA-N | No-Effect | inactive |
| CT-7758 | BRD-K40870905 | O=C(O)[C@H](Cc1ccc(Nc2nccc3ccncc23)cc1)NC1=C(Br)C(=O)C12CCCCC2 | KYHVWHYLKOHLKA-FQEVSTJZSA-N | No-Effect | inactive |
| CTEP | BRD-K30572193 | Cc1nc(C#Cc2ccnc(Cl)c2)c(C)n1-c1ccc(OC(F)(F)F)cc1 | GOHCTCOGYKAJLZ-UHFFFAOYSA-N | No-Effect | inactive |
| CTS-1027 | BRD-K53508936 | O=C(NO)C1(CS(=O)(=O)c2ccc(Oc3ccc(Cl)cc3)cc2)CCOCC1 | ROSNVSQTEGHUKU-UHFFFAOYSA-N | No-Effect | inactive |
| CU-CPT-4a | BRD-K58234744 | O=C(N[C@H](Cc1ccccc1)C(=O)O)c1sc2cc(F)ccc2c1Cl | IAASQMCXDRISAV-CYBMUJFWSA-N | No-Effect | inactive |
| CU-T12-9 | BRD-K89462779 | CNc1ccc([N+](=O)[O-])cc1-n1cnc(-c2ccc(C(F)(F)F)cc2)c1 | LITXVDAFEYLWQE-UHFFFAOYSA-N | No-Effect | inactive |
| CUDC-101 | BRD-K41278234 | C#Cc1cccc(Nc2ncnc3cc(OC)c(OCCCCCCC(=O)NO)cc23)c1 | PLIVFNIUGLLCEK-UHFFFAOYSA-N | No-Effect | inactive |
| CUDC-427 | BRD-K69351586 | CN[C@@H](C)C(=O)N[C@H](C(=O)N1CCC[C@H]1C(=O)Nc1sc(-c2ncco2)nc1-c1ccccc1)C1CCCCC1 | HSHPBORBOJIXSQ-HARLFGEKSA-N | No-Effect | inactive |
| CUR-61414 | BRD-K69624312 | COc1cccc(CN(C(=O)CC(C)(C)C)[C@H]2C[C@@H](C(=O)N3CCNCC3)N(Cc3ccc4c(c3)OCO4)C2)c1 | WPHXYKUPFJRJDK-AHWVRZQESA-N | No-Effect | inactive |
| curcumin | BRD-K07572174 | COc1cc(/C=C/C(=O)C=C(O)/C=C/c2ccc(O)c(OC)c2)ccc1O | ZIUSSTSXXLLKKK-SMSFQYLMSA-N | No-Effect | inactive |
| cutamesine | BRD-K90777969 | COc1ccc(CCN2CCN(CCCc3ccccc3)CC2)cc1OC | UVSWWUWQVAQPJR-UHFFFAOYSA-N | No-Effect | inactive |
| CV-1808 | BRD-K49197823 | Nc1nc(Nc2ccccc2)nc2c1ncn2[C@@H]1O[C@H](CO)[C@@H](O)[C@H]1O | SCNILGOVBBRMBK-SDBHATRESA-N | No-Effect | inactive |
| CVT-10216 | BRD-K60659193 | CS(=O)(=O)Nc1ccc(-c2coc3cc(OCc4cccc(C(=O)O)c4)ccc3c2=O)cc1 | YYOOFJZTRCPVFD-UHFFFAOYSA-N | No-Effect | inactive |
| CW-008 | BRD-K84068331 | COc1ccn2ncc(-c3nc4n[nH]c(NC(=O)c5ccc(F)cc5)c4cc3F)c2c1 | KCIMKTNQCUDTFI-UHFFFAOYSA-N | No-Effect | inactive |
| CX-516 | BRD-K25140590 | O=C(c1ccc2c(c1)OCO2)N1CCCCC1 | BXBNADAPIHHXJQ-UHFFFAOYSA-N | No-Effect | inactive |
| CX-5461 | BRD-K12787259 | Cc1cnc(CNC(=O)c2c(=O)c3ccc(N4CCCN(C)CC4)nc3n3c2sc2ccccc23)cn1 | XGPBJCHFROADCK-UHFFFAOYSA-N | No-Effect | inactive |
| CXD101 | BRD-K00003388 | Cc1nn(C)cc1CN1CCC(c2ccc(C(=O)Nc3ccccc3N)cc2)CC1 | JHDZMASHNBKTPS-UHFFFAOYSA-N | No-Effect | inactive |
| CY208-243 | BRD-K79353516 | CN1Cc2ccccc2[C@@H]2c3cccc4[nH]cc(c34)C[C@H]21 | WRNKIDLXXXIELU-IEBWSBKVSA-N | No-Effect | inactive |
| cyacetacide | BRD-K39371216 | N#CCC(=O)NN | HPHBOJANXDKUQD-UHFFFAOYSA-N | No-Effect | inactive |
| cyamemazine | BRD-A42881122 | C[C@@H](CN(C)C)CN1c2ccccc2Sc2ccc(C#N)cc21 | SLFGIOIONGJGRT-AWEZNQCLSA-N | No-Effect | inactive |
| cyanocobalamin | BRD-A18627414 | C/C1=C2/N=C(/C=C3\N=C(/C(C)=C4/[C@@H](CCC(N)=O)[C@](C)(CC(N)=O)[C@](C)([C@@H]5N=C1[C@](C)(CCC(=O)NC[C@@H](C)O[P@](=O)(O)O[C@@H]1[C@@H](CO)O[C@H](n6cnc7cc(C)c(C)cc76)[C@@H]1O)[C@H]5CC(N)=O)N4[Co+]C#N)[C@@](C)(CC(N)=O)[C@@H]3CCC(N)=O)C(C)(C)[C@@H]2CCC(N)=O | RMRCNWBMXRMIRW-WZHZPDAFSA-M | No-Effect | inactive |
| cyanopindolol | BRD-K01825675 | CC(C)(C)NC[C@@H](O)COc1cccc2[nH]c(C#N)cc12 | QXIUMMLTJVHILT-GFCCVEGCSA-N | No-Effect | inactive |
| CYC116 | BRD-K64866502 | Cc1nc(N)sc1-c1ccnc(Nc2ccc(N3CCOCC3)cc2)n1 | GPSZYOIFQZPWEJ-UHFFFAOYSA-N | No-Effect | inactive |
| cyclamic-acid | BRD-K50785021 | O=S(=O)(O)NC1CCCCC1 | HCAJEUSONLESMK-UHFFFAOYSA-N | No-Effect | inactive |
| cyclandelate | BRD-A27370296 | C[C@H]1C[C@H](OC(=O)[C@@H](O)c2ccccc2)CC(C)(C)C1 | WZHCOOQXZCIUNC-QEJZJMRPSA-N | No-Effect | inactive |
| cyclic-AMP | BRD-A41840684 | Nc1ncnc2c1ncn2[C@H]1O[C@H]2CO[P@@](=O)(O)O[C@@H]2[C@@H]1O | IVOMOUWHDPKRLL-DEGSGYPDSA-N | No-Effect | inactive |
| cyclocreatine | BRD-K14379200 | NC1=NCCN1CC(=O)O | AMHZIUVRYRVYBA-UHFFFAOYSA-N | No-Effect | inactive |
| cyclocytidine | BRD-K67536197 | N=c1ccn2c(n1)O[C@H]1[C@H](O)[C@@H](CO)O[C@H]12 | BBDAGFIXKZCXAH-CCXZUQQUSA-N | No-Effect | inactive |
| cyclopamine | BRD-K58938839 | CC1=C2C[C@H]3[C@@H](CC=C4C[C@@H](O)CC[C@@]43C)[C@@H]2CC[C@]12O[C@@H]1C[C@H](C)CN[C@H]1[C@H]2C | QASFUMOKHFSJGL-LAFRSMQTSA-N | No-Effect | inactive |
| cyclopenthiazide | BRD-A41250306 | NS(=O)(=O)c1cc2c(cc1Cl)N[C@@H](CC1CCCC1)NS2(=O)=O | BKYKPTRYDKTTJY-CYBMUJFWSA-N | No-Effect | inactive |
| cyclopentolate | BRD-A77291778 | CN(C)CCOC(=O)[C@@H](c1ccccc1)C1(O)CCCC1 | SKYSRIRYMSLOIN-OAHLLOKOSA-N | No-Effect | inactive |
| cyclophosphamide | BRD-A09722536 | O=[P@]1(N(CCCl)CCCl)NCCCO1 | CMSMOCZEIVJLDB-CQSZACIVSA-N | No-Effect | inactive |
| cyclopiazonic-acid | BRD-A10420615 | CC(=O)C1=C(O)[C@@H]2[C@H]3c4c[nH]c5cccc(c45)C[C@H]3C(C)(C)N2C1=O | SZINUGQCTHLQAZ-DQYPLSBCSA-N | No-Effect | inactive |
| cycloserine-(D) | BRD-K87226815 | N[C@@H]1CONC1=O | DYDCUQKUCUHJBH-UWTATZPHSA-N | No-Effect | inactive |
| cyclothiazide | BRD-A38675539 | NS(=O)(=O)c1cc2c(cc1Cl)N[C@H]([C@H]1C[C@@H]3C=C[C@H]1C3)NS2(=O)=O | BOCUKUHCLICSIY-JWUFEOHASA-N | No-Effect | inactive |
| CycLuc1 | BRD-K00005335 | O=C(O)[C@H]1CSC(c2nc3cc4c(cc3s2)NCC4)=N1 | HBZBAMXERPYTFS-SECBINFHSA-N | No-Effect | inactive |
| CYM-50260 | BRD-K69877120 | FCc1ccc(OCCOc2ccc(Cl)cc2Cl)c(Cl)n1 | FHPOTBQOUBMMCI-UHFFFAOYSA-N | No-Effect | inactive |
| CYM-50769 | BRD-K27732927 | COc1ccc(Oc2c(Cl)cnn(C3c4ccccc4-c4ccccc43)c2=O)cc1 | QHVSQUYCVUHYKT-UHFFFAOYSA-N | No-Effect | inactive |
| CYM-5520 | BRD-K29151923 | Cc1cc(C(=O)Cn2cc(C#N)ccc2=O)c(C)n1Cc1ccccc1 | FMYGNANMYYHBSU-UHFFFAOYSA-N | No-Effect | inactive |
| CYM-5541 | BRD-K75940276 | O=C(c1cc(C2CC2)on1)N(C1CCCCC1)C1CCCCC1 | NDKGACIWVAOUQH-UHFFFAOYSA-N | No-Effect | inactive |
| cypermethrin | BRD-A62798249 | CC1(C)[C@@H](C(=O)O[C@@H](C#N)c2cccc(Oc3ccccc3)c2)[C@@H]1C=C(Cl)Cl | KAATUXNTWXVJKI-CMKODMSKSA-N | No-Effect | inactive |
| cyproheptadine | BRD-K28143534 | CN1CCC(=C2c3ccccc3C=Cc3ccccc32)CC1 | JJCFRYNCJDLXIK-UHFFFAOYSA-N | No-Effect | inactive |
| cyproterone-acetate | BRD-K41141507 | CC(=O)O[C@]1(C(C)=O)CC[C@H]2[C@@H]3C=C(Cl)C4=CC(=O)[C@@H]5C[C@@H]5[C@]4(C)[C@H]3CC[C@@]21C | UWFYSQMTEOIJJG-FDTZYFLXSA-N | No-Effect | inactive |
| cyromazine | BRD-K36711084 | Nc1nc(N)nc(NC2CC2)n1 | LVQDKIWDGQRHTE-UHFFFAOYSA-N | No-Effect | inactive |
| cysteamine | BRD-K92611362 | NCCS | UFULAYFCSOUIOV-UHFFFAOYSA-N | No-Effect | inactive |
| cyt387 | BRD-K87737963 | N#CCNC(=O)c1ccc(-c2ccnc(Nc3ccc(N4CCOCC4)cc3)n2)cc1 | ZVHNDZWQTBEVRY-UHFFFAOYSA-N | No-Effect | inactive |
| cytarabine | BRD-K33106058 | Nc1ccn([C@@H]2O[C@H](CO)[C@@H](O)[C@@H]2O)c(=O)n1 | UHDGCWIWMRVCDJ-CCXZUQQUSA-N | No-Effect | inactive |
| cytidine | BRD-K71847383 | Nc1ccn([C@@H]2O[C@H](CO)[C@@H](O)[C@H]2O)c(=O)n1 | UHDGCWIWMRVCDJ-XVFCMESISA-N | No-Effect | inactive |
| cytisine | BRD-K74186897 | O=c1cccc2n1C[C@@H]1CNC[C@H]2C1 | ANJTVLIZGCUXLD-DTWKUNHWSA-N | No-Effect | inactive |
| cytochalasin-B | BRD-K26664453 | C=C1[C@@H](C)[C@H]2[C@H](Cc3ccccc3)NC(=O)[C@]23OC(=O)/C=C/[C@H](O)CCC[C@@H](C)C/C=C/[C@H]3[C@@H]1O | GBOGMAARMMDZGR-TYHYBEHESA-N | No-Effect | inactive |
| cytochlor | BRD-K48988407 | Nc1nc(=O)n([C@H]2C[C@@H](O)[C@@H](CO)O2)cc1Cl | NGGRGTWYSXYVDK-FSDSQADBSA-N | No-Effect | inactive |
| CZC-54252 | BRD-K87021635 | COc1cc(N2CCOCC2)ccc1Nc1ncc(Cl)c(Nc2ccccc2NS(C)(=O)=O)n1 | CLGWUCNXOBLWFM-UHFFFAOYSA-N | No-Effect | inactive |
| CZC24832 | BRD-K01377300 | CC(C)(C)NS(=O)(=O)c1cncc(-c2cc(F)c3nc(N)nn3c2)c1 | RXRZPHQBTHQXSV-UHFFFAOYSA-N | No-Effect | inactive |
| D-4476 | BRD-K09132007 | NC(=O)c1ccc(-c2nc(-c3ccccn3)c(-c3ccc4c(c3)OCCO4)[nH]2)cc1 | DPDZHVCKYBCJHW-UHFFFAOYSA-N | No-Effect | inactive |
| D-alpha-tocopheryl-succinate | BRD-K64636206 | Cc1c(C)c2c(c(C)c1OC(=O)CCC(=O)O)CC[C@@](C)(CCC[C@H](C)CCC[C@H](C)CCCC(C)C)O2 | IELOKBJPULMYRW-NJQVLOCASA-N | No-Effect | inactive |
| D-limonene | BRD-K20879694 | C=C(C)[C@H]1CC=C(C)CC1 | XMGQYMWWDOXHJM-JTQLQIEISA-N | No-Effect | inactive |
| D-phenylalanine | BRD-K00911807 | N[C@H](Cc1ccccc1)C(=O)O | COLNVLDHVKWLRT-MRVPVSSYSA-N | No-Effect | inactive |
| D-serine | BRD-K71822263 | N[C@H](CO)C(=O)O | MTCFGRXMJLQNBG-UWTATZPHSA-N | No-Effect | inactive |
| D,L-mevalonic-acid-lactone | BRD-A05674712 | C[C@@]1(O)CCOC(=O)C1 | JYVXNLLUYHCIIH-ZCFIWIBFSA-N | No-Effect | inactive |

| | | | | | |
|---|---|---|---|---|---|
| D609 | BRD-A77118605 | S=C(S)OC1CC2CC1C1CCCC21 | PWLXJBSAUKKGCM-UHFFFAOYSA-N | No-Effect | inactive |
| DAA-1106 | BRD-K01080633 | COc1ccc(OC)c(CN(C(C)=O)c2cc(F)ccc2Oc2ccccc2)c1 | DCRZYADKQRHHSF-UHFFFAOYSA-N | No-Effect | inactive |
| dabigatran | BRD-K38168441 | Cn1c(CNc2ccc(C(=N)N)cc2)nc2cc(C(=O)N(CCC(=O)O)c3ccccn3)ccc21 | YBSJFWOBGCMAKL-UHFFFAOYSA-N | No-Effect | inactive |
| dabigatran-etexilate | BRD-K57041787 | CCCCCCOC(=O)/N=C(/N)c1ccc(NCc2nc3cc(C(=O)N(CCC(=O)OCC)c4ccccn4)ccc3n2C)cc1 | KSGXQBZTULBEEQ-UHFFFAOYSA-N | No-Effect | inactive |
| dacarbazine | BRD-K35520305 | CN(C)/N=N/c1nc[nH]c1C(N)=O | FDKXTQMXEQVLRF-ZHACJKMWSA-N | No-Effect | inactive |
| daclatasvir | BRD-K82861010 | COC(=O)N[C@H](C(=O)N1CCC[C@H]1c1nc(-c2ccc(-c3ccc(-c4c[nH]c([C@@H]5CCCN5C(=O)[C@@H](NC(=O)OC)C(C)C)n4)cc3)cc2)c[nH]1)C(C)C | FKRSSPOQAMALKA-CUPIEXAXSA-N | No-Effect | inactive |
| dacomitinib | BRD-K57169635 | COc1cc2ncnc(Nc3ccc(F)c(Cl)c3)c2cc1NC(=O)/C=C/CN1CCCCC1 | LVXJQMNHJWSHET-AATRIKPKSA-N | No-Effect | inactive |
| daidzein | BRD-K42095107 | O=c1c(-c2ccc(O)cc2)coc2cc(O)ccc12 | ZQSIJRDFPHDXIC-UHFFFAOYSA-N | No-Effect | inactive |
| daidzin | BRD-K12979078 | O=c1c(-c2ccc(O)cc2)coc2cc(O[C@@H]3O[C@H](CO)[C@@H](O)[C@H](O)[C@H]3O)ccc12 | KYQZWONCHDNPDP-QNDFHXLGSA-N | No-Effect | inactive |
| dalargin | BRD-A89196962 | CC(C)C[C@@H](NC(=O)[C@@H](Cc1ccccc1)NC(=O)CNC(=O)[C@H](C)NC(=O)[C@@H](N)Cc1ccc(O)cc1)C(=O)N[C@H](CCCNC(=N)N)C(=O)O | GDPHPXYFLPDZGH-ABDYMBPTSA-N | No-Effect | inactive |
| dalbavancin | BRD-K30177383 | CN[C@H]1C(=O)N[C@@H]2Cc3ccc(cc3)Oc3cc4cc(c3O[C@@H]3O[C@H](C(=O)O)[C@@H](O)[C@H](O)[C@H]3NC(=O)CCCCCCCCC(C)C)Oc3ccc(cc3Cl)[C@@H](O)[C@@H]3NC(=O)[C@H](NC(=O)[C@@H]4NC(=O)[C@@H](NC2=O)c2cc(cc(O)c2Cl)Oc2cc1ccc2O)c1ccc(O)c(c1)-c1c(O[C@H]2O[C@H](CO)[C@@H](O)[C@H](O)[C@@H]2O)cc(O)cc1[C@@H](C(=O)NCCCN(C)C)NC3=O | KGPGQDLTDHGEGT-SZUNQUCBSA-N | No-Effect | inactive |
| dalcetrapib | BRD-K18849474 | CCC(CC)CC1(C(=O)Nc2ccccc2SC(=O)C(C)C)CCCCC1 | YZQLWPMZQVHJED-UHFFFAOYSA-N | No-Effect | inactive |
| dalfopristin | BRD-K00003321 | CCN(CC)CCS(=O)(=O)[C@@H]1CCN2C(=O)c3coc(n3)CC(=O)C[C@H](O)/C=C(C)/C=C\CNC(=O)/C=C/[C@@H](C)[C@H](C(C)C)OC(=O)[C@@H]12 | SUYRLXYYZQTJHF-FENKCRFESA-N | No-Effect | inactive |
| daltroban | BRD-K26813177 | O=C(O)Cc1ccc(CCNS(=O)(=O)c2ccc(Cl)cc2)cc1 | IULOBWFWYDMECP-UHFFFAOYSA-N | No-Effect | inactive |
| daminozide | BRD-K83452553 | CN(C)NC(=O)CCC(=O)O | NOQGZXFMHARMLW-UHFFFAOYSA-N | No-Effect | inactive |
| danazol | BRD-K48970916 | C#C[C@]1(O)CC[C@H]2[C@@H]3CCC4=Cc5oncc5C[C@]4(C)[C@H]3CC[C@@]21C | POZRVZJJTULAOH-LHZXLZLDSA-N | No-Effect | inactive |
| danegaptide | BRD-K35589454 | NCC(=O)N1C[C@H](NC(=O)c2ccccc2)C[C@H]1C(=O)O | BIZKIHUJGMSVFD-MNOVXSKESA-N | No-Effect | inactive |
| danirixin | BRD-K13769342 | Cc1c(F)cccc1NC(=O)Nc1ccc(Cl)c(S(=O)(=O)[C@H]2CCCNC2)c1O | NGYNBSHYFOFVLS-LBPRGKRZSA-N | No-Effect | inactive |
| danoprevir | BRD-K08111240 | CC(C)(C)OC(=O)N[C@H]1CCCCC/C=C\[C@@H]2C[C@@]2(C(=O)NS(=O)(=O)C2CC2)NC(=O)[C@@H]2C[C@@H](OC(=O)N3Cc4cccc(F)c4C3)CN2C1=O | ZVTDLPBHTSMEJZ-JSZLBQEHSA-N | No-Effect | inactive |
| dantrolene | BRD-K81272440 | O=C1CN(/N=C/c2ccc(-c3ccc([N+](=O)[O-])cc3)o2)C(=O)N1 | OZOMQRBLCMDCEG-VIZOYTHASA-N | No-Effect | inactive |
| danusertib | BRD-K07881437 | CO[C@@H](C(=O)N1Cc2n[nH]c(NC(=O)c3ccc(N4CCN(C)CC4)cc3)c2C1)c1ccccc1 | XKFTZKGMDDZMJI-HSZRJFAPSA-N | No-Effect | inactive |
| dapansutrile | BRD-K00004659 | CS(=O)(=O)CCC#N | LQFRYKBDZNPJSW-UHFFFAOYSA-N | No-Effect | inactive |
| daphnetin | BRD-K61269089 | O=c1ccc2ccc(O)c(O)c2o1 | ATEFPOUAMCWAQS-UHFFFAOYSA-N | No-Effect | inactive |
| dapiprazole | BRD-K33133047 | Cc1ccccc1N1CCN(CCc2nnc3n2CCCC3)CC1 | RFWZESUMWJKKRN-UHFFFAOYSA-N | No-Effect | inactive |
| dapivirine | BRD-K12885236 | Cc1cc(C)c(Nc2ccnc(Nc3ccc(C#N)cc3)n2)c(C)c1 | ILAYIAGXTHKHNT-UHFFFAOYSA-N | No-Effect | inactive |
| daprodustat | BRD-K06455935 | O=C(O)CNC(=O)c1c(O)n(C2CCCCC2)c(=O)n(C2CCCCC2)c1=O | NVTKJBXOBFRPLQ-UHFFFAOYSA-N | No-Effect | inactive |
| dapsone | BRD-K62363391 | Nc1ccc(S(=O)(=O)c2ccc(N)cc2)cc1 | MQJKPEGWNLWLTK-UHFFFAOYSA-N | No-Effect | inactive |
| DAPT | BRD-K47177710 | C[C@H](NC(=O)Cc1cc(F)cc(F)c1)C(=O)N[C@H](C(=O)OC(C)(C)C)c1ccccc1 | DWJXYEABWRJFSP-XOBRGWDASA-N | No-Effect | inactive |
| daptomycin | BRD-K00003611 | CCCCCCCCCC(=O)N[C@@H](Cc1c[nH]c2ccccc12)C(=O)N[C@H](CC(N)=O)C(=O)N[C@@H](CC(=O)O)C(=O)N[C@@H]1C(=O)NCC(=O)N[C@@H](CCCN)C(=O)N[C@@H](CC(=O)O)C(=O)N[C@H](C)C(=O)N[C@@H](CC(=O)O)C(=O)NCC(=O)N[C@H](CO)C(=O)N[C@@H]([C@H](C)CC(=O)O)C(=O)N[C@@H](CC(=O)c2ccccc2N)C(=O)O[C@@H]1C | DOAKLVKFURWEDJ-QCMAZARJSA-N | No-Effect | inactive |
| darapladib | BRD-K19085898 | CCN(CC)CCN(Cc1ccc(-c2ccc(C(F)(F)F)cc2)cc1)C(=O)Cn1c(SCc2ccc(F)cc2)nc(=O)c2c1CCC2 | WDPFJWLDPVQCAJ-UHFFFAOYSA-N | No-Effect | inactive |
| darglitazone | BRD-K82908348 | Cc1oc(-c2ccccc2)nc1CCC(=O)c1ccc(CC2SC(=O)NC2=O)cc1 | QQKNSPHAFATFNQ-UHFFFAOYSA-N | No-Effect | inactive |
| darifenacin | BRD-K95773607 | NC(=O)C(c1ccccc1)(c1ccccc1)[C@@H]1CCN(CCc2ccc3c(c2)CCO3)C1 | HXGBXQDTNZMWGS-RUZDIDTESA-N | No-Effect | inactive |
| darolutamide | BRD-A44971162 | C[C@@H](Cn1ccc(-c2ccc(C#N)c(Cl)c2)n1)NC(=O)c1cc([C@@H](C)O)n[nH]1 | BLIJXOOIHRSQRB-NWDGAFQWSA-N | No-Effect | inactive |
| darunavir | BRD-K84986517 | CC(C)CN(C[C@@H](O)[C@H](Cc1ccccc1)NC(=O)O[C@H]1CO[C@H]2OCC[C@@H]12)S(=O)(=O)c1ccc(N)cc1 | CJBJHOAVZSMMDJ-HEXNFIEUSA-N | No-Effect | inactive |
| darusentan | BRD-K54863166 | COc1cc(OC)nc(O[C@H](C(=O)O)C(OC)(c2ccccc2)c2ccccc2)n1 | FEJVSJIALLTFRP-LJQANCHMSA-N | No-Effect | inactive |
| dasabuvir | BRD-K02943813 | COc1c(-c2ccc3cc(NS(C)(=O)=O)ccc3c2)cc(-n2ccc(=O)[nH]c2=O)cc1C(C)(C)C | NBRBXGKOEOGLOI-UHFFFAOYSA-N | No-Effect | inactive |
| dasatinib | BRD-K49328571 | Cc1nc(Nc2ncc(C(=O)Nc3c(C)cccc3Cl)s2)cc(N2CCN(CCO)CC2)n1 | ZBNZXTGUTAYRHI-UHFFFAOYSA-N | No-Effect | inactive |
| DAU-5884 | BRD-K31633810 | CN1[C@H]2CC[C@@H]1C[C@H](OC(=O)N1Cc3ccccc3NC1=O)C2 | NWXCDQHBVVPPTL-WDNDVIMCSA-N | No-Effect | inactive |
| daucosterol | BRD-K14276241 | CC[C@H](CC[C@@H](C)[C@H]1CC[C@H]2[C@@H]3CC=C4C[C@@H](O[C@@H]5O[C@H](CO)[C@@H](O)[C@H](O)[C@H]5O)CC[C@]4(C)[C@H]3CC[C@@]21C)C(C)C | NPJICTMALKLTFW-OFUAXYCQSA-N | No-Effect | inactive |
| dazmegrel | BRD-K20738689 | Cc1c(Cn2ccnc2)c2ccccc2n1CCC(=O)O | DEQLGSOHGTZKFB-UHFFFAOYSA-N | No-Effect | inactive |
| dazoxiben | BRD-K82767007 | O=C(O)c1ccc(OCCn2ccnc2)cc1 | XQGZSYKGWHUSDH-UHFFFAOYSA-N | No-Effect | inactive |
| DBeQ | BRD-K59204667 | c1ccc(CNc2nc(NCc3ccccc3)c3ccccc3n2)cc1 | QAIMUUJJAJBPCL-UHFFFAOYSA-N | No-Effect | inactive |
| DBPR-211 | BRD-K00004704 | O=C(NN1CCCCC1)c1nn(-c2ccc(Cl)cc2Cl)c(-c2ccc(C#Cc3ccc(C(F)(F)F)cc3)s2)c1CNS(=O)(=O)N1CCCC1 | KAIYFJUWIBTYPK-UHFFFAOYSA-N | No-Effect | inactive |
| DBPR108 | BRD-K83020231 | CC(C)(CC(=O)N1CCCC1)NCC(=O)N1C[C@@H](F)C[C@H]1C#N | VQKSCYBKUIDZEI-STQMWFEESA-N | No-Effect | inactive |
| DC-260126 | BRD-K10596918 | CCCCc1ccc(NS(=O)(=O)c2ccc(F)cc2)cc1 | CNGHPXKWPGIDSK-UHFFFAOYSA-N | No-Effect | inactive |
| DCC-2618 | BRD-K49246350 | Cn1cc(-c2cc(Oc3cc(F)c(NC(=O)C4(C(=O)Nc5ccccc5)CC4)cc3F)ccn2)cn1 | WWOXKWLDMLMYQY-UHFFFAOYSA-N | No-Effect | inactive |
| DCEBIO | BRD-K89299012 | CCn1c(=O)[nH]c2cc(Cl)c(Cl)cc21 | LKHRMULASXZCLG-UHFFFAOYSA-N | No-Effect | inactive |
| DCP-LA | BRD-K00005344 | CCCCC[C@H]1C[C@H]1C[C@H]1C[C@H]1CCCCCCCC(=O)O | CONYTTFKIUJZOF-ZSYWTGECSA-N | No-Effect | inactive |
| DCPIB | BRD-A39255369 | CCCC[C@@]1(C2CCCC2)Cc2cc(OCCCC(=O)O)c(Cl)c(Cl)c2C1=O | KHKGTPJPBOQECW-QFIPXVFZSA-N | No-Effect | inactive |
| DDR1-IN-1 | BRD-K61845293 | CCN1CCN(Cc2ccc(C(=O)Nc3ccc(C)c(Oc4ccc5c(c4)CC(=O)N5)c3)cc2C(F)(F)F)CC1 | AOZPVMOOEJAZGK-UHFFFAOYSA-N | No-Effect | inactive |
| debrisoquin | BRD-K80419150 | N=C(N)N1CCc2ccccc2C1 | JWPGJSVJDAJRLW-UHFFFAOYSA-N | No-Effect | inactive |
| decamethonium | BRD-K49783248 | C[N+](C)(C)CCCCCCCCCC[N+](C)(C)C | MTCUAOILFDZKCO-UHFFFAOYSA-N | No-Effect | inactive |

| | | | | | |
|---|---|---|---|---|---|
| decamethoxine | BRD-A56320380 | CC(C)[C@H]1CC[C@H](C)C[C@@H]1OC(=O)C[N+](C)(C)CCCCCCCCCC[N+](C)(C)CC(=O)O[C@H]1C[C@H](C)CC[C@@H]1C(C)C | ZADGGFKUEYIBJC-BPBUSIJASA-N | No-Effect | inactive |
| decernotinib | BRD-A95300432 | CC[C@@](C)(Nc1ccnc(-c2c[nH]c3ncccc23)n1)C(=O)NCC(F)(F)F | ASUGUQWIHMTFJL-QGZVFWFLSA-N | No-Effect | inactive |
| decitabine | BRD-K79254416 | Nc1ncn([C@H]2C[C@H](O)[C@@H](CO)O2)c(=O)n1 | XAUDJQYHKZQPEU-KVQBGUIXSA-N | No-Effect | inactive |
| declopramide | BRD-K00004661 | CCN(CC)CCNC(=O)c1ccc(N)c(Cl)c1 | YEYAKZXEBSVURO-UHFFFAOYSA-N | No-Effect | inactive |
| decloxizine | BRD-K00003318 | OCCOCCN1CCN(C(c2ccccc2)c2ccccc2)CC1 | OXBBIHZWNDPBMQ-UHFFFAOYSA-N | No-Effect | inactive |
| decoquinate | BRD-K17641316 | CCCCCCCCCCOc1cc2c(=O)c(C(=O)OCC)c[nH]c2cc1OCC | JHAYEQICABJSTP-UHFFFAOYSA-N | No-Effect | inactive |
| defactinib | BRD-K43578482 | CNC(=O)c1ccc(Nc2ncc(C(F)(F)F)c(NCc3nccnc3N(C)S(C)(=O)=O)n2)cc1 | FWLMVFUGMHIOAA-UHFFFAOYSA-N | No-Effect | inactive |
| deferasirox | BRD-K97010173 | O=C(O)c1ccc(-n2nc(-c3ccccc3O)nc2-c2ccccc2O)cc1 | BOFQWVMAQOTZIW-UHFFFAOYSA-N | No-Effect | inactive |
| deferasirox-iron-chelate | BRD-K00003219 | O=C(O)c1ccc(-n2nc3nc2-c2ccccc2O[Fe]Oc2ccccc2-3)cc1 | XLHBYAVMDSNIEM-UHFFFAOYSA-L | No-Effect | inactive |
| deferiprone | BRD-K06878038 | Cc1c(O)c(=O)ccn1C | TZXKOCQBRNJULO-UHFFFAOYSA-N | No-Effect | inactive |
| deferoxamine-mesylate | BRD-K09821361 | CC(=O)N(O)CCCCCNC(=O)CCC(=O)N(O)CCCCCNC(=O)CCC(=O)N(O)CCCCCN | UBQYURCVBFRUQT-UHFFFAOYSA-N | No-Effect | inactive |
| deflazacort | BRD-K30468934 | CC(=O)OCC(=O)[C@@]12N=C(C)O[C@@H]1C[C@H]1[C@@H]3CCC4=CC(=O)C=C[C@]4(C)[C@H]3[C@@H](O)C[C@@]12C | FBHSPRKOSMHSIF-GRMWVWQJSA-N | No-Effect | inactive |
| deforolimus | BRD-K62304154 | CO[C@H]1C[C@@H]2CC[C@@H](C)[C@@](O)(O2)C(=O)C(=O)N2CCCC[C@H]2C(=O)O[C@H]([C@H](C)C[C@@H]2CC[C@@H](OP(C)(C)=O)[C@H](OC)C2)CC(=O)[C@H](C)/C=C(\C)[C@@H](O)[C@@H](OC)C(=O)[C@H](C)C[C@H](C)/C=C/C=C/C=C/1C | BUROJSBIWGDYCN-GAUTUEMISA-N | No-Effect | inactive |
| degarelix | BRD-K00003578 | CC(=O)N[C@H](Cc1ccc2ccccc2c1)C(=O)N[C@H](Cc1ccc(Cl)cc1)C(=O)N[C@H](Cc1cccnc1)C(=O)N[C@@H](CO)C(=O)N[C@@H](Cc1ccc(NC(=O)[C@@H]2CC(=O)NC(=O)N2)cc1)C(=O)N[C@H](Cc1ccc(NC(N)=O)cc1)C(=O)N[C@@H](CC(C)C)C(=O)N[C@@H](CCCCNC(C)C)C(=O)N1CCC[C@H]1C(=O)N[C@H](C)C(N)=O | MEUCPCLKGZSHTA-XYAYPHGZSA-N | No-Effect | inactive |
| dehydroacetic-acid | BRD-K00025573 | CC(=O)c1c(O)cc(C)oc1=O | PKLPQOJFHFGVBS-UHFFFAOYSA-N | No-Effect | inactive |
| dehydrocholate | BRD-A64125466 | C[C@H](CCC(=O)O)[C@H]1CC[C@H]2[C@@H]3C(=O)C[C@H]4CC(=O)CC[C@]4(C)[C@@H]3CC(=O)[C@@]21C | OHXPGWPVLFPUSM-IVFZZBRKSA-N | No-Effect | inactive |
| dehydrocorydaline | BRD-K67186445 | COc1cc2c(cc1OC)-c1c(C)c3ccc(OC)c(OC)c3c[n+]1CC2 | RFKQJTRWODZPHF-UHFFFAOYSA-N | No-Effect | inactive |
| dehydroepiandrosterone | BRD-K41256143 | C[C@]12CC[C@H]3[C@@H](CC=C4C[C@@H](O)CC[C@@]43C)[C@@H]1CCC2=O | FMGSKLZLMKYGDP-USOAJAOKSA-N | No-Effect | inactive |
| dehydroepiandrosterone-sulfate | BRD-K15360631 | C[C@]12CC[C@H]3[C@@H](CC=C4C[C@@H](OS(=O)(=O)O)CC[C@@]43C)[C@@H]1CCC2=O | CZWCKYRVOZZJNM-USOAJAOKSA-N | No-Effect | inactive |
| delafloxacin | BRD-K90991629 | Nc1nc(-n2cc(C(=O)O)c(=O)c3cc(F)c(N4CC(O)C4)c(Cl)c32)c(F)cc1F | DYDCPNMLZGFQTM-UHFFFAOYSA-N | No-Effect | inactive |
| delamanid | BRD-K17480742 | C[C@]1(COc2ccc(N3CCC(Oc4ccc(OC(F)(F)F)cc4)CC3)cc2)Cn2cc([N+](=O)[O-])nc2O1 | XDAOLTSRNUSPPH-XMMPIXPASA-N | No-Effect | inactive |
| delapril | BRD-K00573984 | CCOC(=O)[C@H](CCc1ccccc1)N[C@@H](C)C(=O)N(CC(=O)O)C1Cc2ccccc2C1 | WOUOLAUOZXOLJQ-MBSDFSHPSA-N | No-Effect | inactive |
| delavirdine | BRD-K34068325 | CC(C)Nc1cccnc1N1CCN(C(=O)c2cc3cc(NS(C)(=O)=O)ccc3[nH]2)CC1 | WHBIGIKBNXZKFE-UHFFFAOYSA-N | No-Effect | inactive |
| delcasertib | BRD-K00003468 | CC(C)C[C@H](NC(=O)[C@H](CO)NC(=O)CNC(=O)[C@H](CC(C)C)NC(=O)[C@H](CCC(=O)O)NC(=O)[C@H](Cc1ccc(O)cc1)NC(=O)[C@H](CO)NC(=O)[C@H](CC(N)=O)NC(=O)[C@H](Cc1ccccc1)NC(=O)[C@H](CO)NC(=O)[C@@H](N)CSSC[C@H](N)C(=O)N[C@@H](Cc1ccc(O)cc1)C(=O)NCC(=O)N[C@@H](CCCNC(=N)N)C(=O)N[C@@H](CCCCN)C(=O)N[C@@H](CCCCN)C(=O)N[C@@H](CCCNC(=N)N)C(=O)N[C@@H](CCCNC(=N)N)C(=O)N[C@@H](CCC(N)=O)C(=O)N[C@@H](CCCNC(=N)N)C(=O)N[C@@H](CCCNC(=N)N)C(=O)N[C@@H](CCCNC(=N)N)C(=O)O)C(=O)O | XPXZTVWPRKJTAA-PJKOMPQUSA-N | No-Effect | inactive |
| delgocitinib | BRD-K00003500 | C[C@H]1CN(C(=O)CC#N)[C@]12CCN(c1ncnc3[nH]ccc13)C2 | LOWWYYZBZNSPDT-ZBEGNZNMSA-N | No-Effect | inactive |
| delpazolid | BRD-K00003363 | CN1CCN(c2ccc(N3C[C@H](CO)OC3=O)cc2F)C=N1 | QLUWQAFDTNAYPN-LLVKDONJSA-N | No-Effect | inactive |
| delphinidin | BRD-K79107392 | Oc1cc(O)c2cc(O)c(-c3cc(O)c(O)c(O)c3)[o+]c2c1 | JKHRCGUTYDNCLE-UHFFFAOYSA-O | No-Effect | inactive |
| delta-tocotrienol | BRD-K85776940 | CC(C)=CCC/C(C)=C/CC/C(C)=C/CC[C@]1(C)CCc2cc(O)cc(C)c2O1 | ODADKLYLWWCHNB-LDYBVBFYSA-N | No-Effect | inactive |
| demecarium | BRD-K51471001 | CN(CCCCCCCCCCN(C)C(=O)Oc1cccc([N+](C)(C)C)c1)C(=O)Oc1cccc([N+](C)(C)C)c1 | RWZVPVOZTJJMNU-UHFFFAOYSA-N | No-Effect | inactive |
| demeclocycline | BRD-K00003347 | CN(C)[C@@H]1C(=O)[C@@H](C(N)=O)C(=O)[C@@]2(O)C(=O)[C@H]3C(=O)c4c(O)ccc(Cl)c4[C@@H](O)[C@H]3C[C@@H]12 | JHOHXJYRVSLQCB-AWLLFJHESA-N | No-Effect | inactive |
| denotivir | BRD-K01773083 | Cc1nsc(NC(=O)c2ccccc2)c1C(=O)Nc1ccc(Cl)cc1 | ZPBLNADJHWHOEP-UHFFFAOYSA-N | No-Effect | inactive |
| deoxyarbutin | BRD-A62562325 | Oc1ccc(O[C@H]2CCCCO2)cc1 | GFBCWCDNXDKFRH-NSHDSACASA-N | No-Effect | inactive |
| deoxycholic-acid | BRD-K41170226 | C[C@H](CCC(=O)O)[C@H]1CC[C@H]2[C@@H]3CC[C@@H]4C[C@H](O)CC[C@]4(C)[C@H]3C[C@H](O)[C@@]21C | KXGVEGMKQFWNSR-LLQZFEROSA-N | No-Effect | inactive |
| deoxyepinephrine | BRD-K33151569 | CNCCc1ccc(O)c(O)c1 | NGKZFDYBISXGGS-UHFFFAOYSA-N | No-Effect | inactive |
| dequalinium | BRD-K77685957 | Cc1cc(N)c2ccccc2[n+]1CCCCCCCCCC[n+]1c(C)cc(N)c2ccccc21 | PCSWXVJAIHCTMO-UHFFFAOYSA-P | No-Effect | inactive |
| deracoxib | BRD-K68558722 | COc1ccc(-c2cc(C(F)F)nn2-c2ccc(S(N)(=O)=O)cc2)cc1F | WAZQAZKAZLXFMK-UHFFFAOYSA-N | No-Effect | inactive |
| derazantinib | BRD-K00003310 | COCCNCCc1cccc(Nc2ncc3c(n2)-c2ccccc2[C@H](c2ccccc2F)C3)c1 | KPJDVVCDVBFRMU-AREMUKBSSA-N | No-Effect | inactive |
| desidustat | BRD-K00003452 | O=C(O)CNC(=O)c1c(O)c2ccccc2n(OCC2CC2)c1=O | IKRKQQLJYBAPQT-UHFFFAOYSA-N | No-Effect | inactive |
| desipramine | BRD-K60762818 | CNCCCN1c2ccccc2CCc2ccccc21 | HCYAFALTSJYZDH-UHFFFAOYSA-N | No-Effect | inactive |
| desisobutyryl-ciclesonide | BRD-K00003103 | C[C@]12C[C@H](O)[C@H]3[C@@H](CCC4=CC(=O)C=C[C@@]43C)[C@@H]1C[C@@H]1O[C@@H](C3CCCCC3)O[C@]12C(=O)CO | OXPLANUPKBHPMS-FYSAVPLMSA-N | No-Effect | inactive |
| desloratadine | BRD-K82357231 | Clc1ccc2c(c1)CCc1cccnc1C2=C1CCNCC1 | JAUOIFJMECXRGI-UHFFFAOYSA-N | No-Effect | inactive |
| deslorelin | BRD-K00003134 | CCNC(=O)[C@@H]1CCCN1C(=O)[C@H](CCCN=C(N)N)NC(=O)[C@H](CC(C)C)NC(=O)[C@@H](Cc1c[nH]c2ccccc12)NC(=O)[C@H](Cc1ccc(O)cc1)NC(=O)[C@H](CO)NC(=O)[C@H](Cc1c[nH]c2ccccc12)NC(=O)[C@H](Cc1c[nH]cn1)NC(=O)[C@@H]1CCC(=O)N1 | GJKXGJCSJWBJEZ-XRSSZCMZSA-N | No-Effect | inactive |
| desmethylclozapine | BRD-K10042277 | Clc1ccc2c(c1)N=C(N1CCNCC1)c1ccccc1N2 | JNNOSTQEZICQQP-UHFFFAOYSA-N | No-Effect | inactive |
| desmopressin-acetate | BRD-K00003582 | N=C(N)NCCC[C@@H](NC(=O)[C@@H]1CCCN1C(=O)[C@H]1CSSCCC(=O)N[C@@H](Cc2ccc(O)cc2)C(=O)N[C@@H](Cc2ccccc2)C(=O)N[C@@H](CCC(N)=O)C(=O)N[C@@H](CC(N)=O)C(=O)N1)C(=O)NCC(N)=O | NFLWUMRGJYTJIN-ZYWOLCKZSA-N | No-Effect | inactive |
| desogestrel | BRD-K46061689 | C#C[C@]1(O)CC[C@H]2[C@@H]3CCC4=CCCC[C@@H]4[C@H]3C(=C)C[C@@]21CC | RPLCPCMSCLEKRS-BPIQYHPVSA-N | No-Effect | inactive |
| desonide | BRD-K21528677 | CC1(C)O[C@@H]2C[C@H]3[C@@H]4CCC5=CC(=O)C=C[C@]5(C)[C@H]4[C@@H](O)C[C@]3(C)[C@]2(C(=O)CO)O1 | WBGKWQHBNHJJPZ-LECWWXJVSA-N | No-Effect | inactive |
| desoximetasone | BRD-K30697463 | C[C@@H]1C[C@H]2[C@@H]3CCC4=CC(=O)C=C[C@]4(C)[C@@]3(F)[C@@H](O)C[C@]2(C)[C@H]1C(=O)CO | VWVSBHGCDBMOOT-IIEHVVJPSA-N | No-Effect | inactive |
| desoxycorticosterone-pivalate | BRD-K26024145 | CC(C)(C)C(=O)OCC(=O)[C@H]1CC[C@H]2[C@@H]3CCC4=CC(=O)CC[C@]4(C)[C@H]3CC[C@]12C | VVOIQBFMTVCINR-WWMZEODYSA-N | No-Effect | inactive |
| desoxycortone | BRD-K98521173 | C[C@]12CC[C@H]3[C@@H](CCC4=CC(=O)CC[C@@]43C)[C@@H]1CC[C@@H]2C(=O)CO | ZESRJSPZRDMNHY-YFWFAHHUSA-N | No-Effect | inactive |

| | | | | | |
|---|---|---|---|---|---|
| desoxypeganine | BRD-K13819402 | c1ccc2c(c1)CN1CCCC1=N2 | WUFQLZTXIWKION-UHFFFAOYSA-N | No-Effect | inactive |
| desvenlafaxine | BRD-K01825598 | CN(C)C[C@@H](c1ccc(O)cc1)C1(O)CCCCC1 | KYYIDSXMWOZKMP-HNNXBMFYSA-N | No-Effect | inactive |
| detomidine | BRD-K77597856 | Cc1cccc(Cc2cnc[nH]2)c1C | RHDJRPPFURBGLQ-UHFFFAOYSA-N | No-Effect | inactive |
| devazepide | BRD-K31238592 | CN1C(=O)[C@@H](NC(=O)c2cc3ccccc3[nH]2)N=C(c2ccccc2)c2ccccc21 | NFHRQQKPEBFUJK-HSZRJFAPSA-N | No-Effect | inactive |
| dexamethasone | BRD-K38775274 | C[C@@H]1C[C@H]2[C@@H]3CCC4=CC(=O)C=C[C@]4(C)[C@@]3(F)[C@@H](O)C[C@]2(C)[C@@]1(O)C(=O)CO | UREBDLICKHMUKA-CXSFZGCWSA-N | No-Effect | inactive |
| dexamethasone-acetate | BRD-K47635719 | CC(=O)OCC(=O)[C@@]1(O)[C@H](C)C[C@H]2[C@@H]3CCC4=CC(=O)C=C[C@]4(C)[C@@]3(F)[C@@H](O)C[C@@]21C | AKUJBENLRBOFTD-RPRRAYFGSA-N | No-Effect | inactive |
| dexchlorpheniramine | BRD-K39462424 | CN(C)CC[C@@H](c1ccc(Cl)cc1)c1ccccn1 | SOYKEARSMXGVTM-HNNXBMFYSA-N | No-Effect | inactive |
| dexfosfoserine | BRD-K30019776 | N[C@@H](COP(=O)(O)O)C(=O)O | BZQFBWGGLXLEPQ-REOHCLBHSA-N | No-Effect | inactive |
| dexlansoprazole | BRD-K05900209 | Cc1c(OCC(F)(F)F)ccnc1C[S@@](=O)c1nc2ccccc2[nH]1 | MJIHNNLFOKEZEW-RUZDIDTESA-N | No-Effect | inactive |
| dexloxiglumide | BRD-K07609981 | CCCCCN(CCCOC)C(=O)[C@@H](CCC(=O)O)NC(=O)c1ccc(Cl)c(Cl)c1 | QNQZBKQEIFTHFZ-GOSISDBHSA-N | No-Effect | inactive |
| dexniguldipine | BRD-K44309363 | COC(=O)C1=C(C)NC(C)=C(C(=O)OCCCN2CCC(c3ccccc3)(c3ccccc3)CC2)[C@@H]1c1cccc([N+](=O)[O-])c1 | SVJMLYUFVDMUHP-MGBGTMOVSA-N | No-Effect | inactive |
| dexpanthenol | BRD-K71210479 | CC(C)(CO)[C@@H](O)C(=O)NCCCO | SNPLKNRPJHDVJA-ZETCQYMHSA-N | No-Effect | inactive |
| dexrazoxane | BRD-K07265709 | C[C@@H](CN1CC(=O)NC(=O)C1)N1CC(=O)NC(=O)C1 | BMKDZUISNHGIBY-ZETCQYMHSA-N | No-Effect | inactive |
| dextran | BRD-K67117832 | O=C[C@H](O)[C@@H](O)[C@H](O)[C@H](O)CO[C@H]1O[C@H](CO[C@H]2O[C@H](CO)[C@@H](O)[C@H](O)[C@H]2O)[C@@H](O)[C@H](O)[C@H]1O | FZWBNHMXJMCXLU-BLAUPYHCSA-N | No-Effect | inactive |
| dextromethorphan | BRD-K33211335 | COc1ccc2c(c1)[C@]13CCCC[C@@H]1[C@H](C2)N(C)CC3 | MKXZASYAUGDDCJ-NJAFHUGGSA-N | No-Effect | inactive |
| dextrorotation-nimorazole-phosphate-ester | BRD-K00003272 | Cc1ncc([N+](=O)[O-])n1C[C@@H](CN1CCOCC1)OP(=O)(O)O | CCDHHPOGWLYHAX-SNVBAGLBSA-N | No-Effect | inactive |
| DFB | BRD-K82002057 | Fc1cccc(/C=N/N=C/c2cccc(F)c2)c1 | YYMCVDNIIFNDJK-BEQMOXJMSA-N | No-Effect | inactive |
| DG-172 | BRD-K75748943 | CN1CCN(c2ccc(/C=C(\C#N)c3ccccc3Br)cc2)CC1 | ZJMDTBBCNMGFMS-SAPNQHFASA-N | No-Effect | inactive |
| DG051 | BRD-K00003486 | O=C(O)CCCN1CCC[C@H]1COc1ccc(Oc2ccc(Cl)cc2)cc1 | PVCTYSQBVIGZRU-KRWDZBQOSA-N | No-Effect | inactive |
| DH-97 | BRD-K06014311 | CCCCC(=O)NCCc1c(Cc2ccccc2)[nH]c2ccccc12 | HDOIPCLEKCEANF-UHFFFAOYSA-N | No-Effect | inactive |
| DHBP | BRD-K41262162 | CCCCCCC[n+]1ccc(-c2cc[n+](CCCCCCC)cc2)cc1 | XJGZGUSMZSXHJI-UHFFFAOYSA-N | No-Effect | inactive |
| diacerein | BRD-K69122748 | CC(=O)Oc1cccc2c1C(=O)c1c(OC(C)=O)cc(C(=O)O)cc1C2=O | TYNLGDBUJLVSMA-UHFFFAOYSA-N | No-Effect | inactive |
| diacetamate | BRD-K93611241 | CC(=O)Nc1ccc(OC(C)=O)cc1 | UJAOSPFULOFZRR-UHFFFAOYSA-N | No-Effect | inactive |
| diadenosine-tetraphosphate | BRD-A52394261 | Nc1ncnc2c1ncn2[C@@H]1O[C@H](COP(=O)(O)OP(=O)(O)OP(=O)(O)OP(=O)(O)OC[C@H]2O[C@@H](n3cnc4c(N)ncnc43)[C@H](O)[C@@H]2O)[C@@H](O)[C@H]1O | YOAHKNVSNCMZGQ-XPWFQUROSA-N | No-Effect | inactive |
| dianhydrogalactitol | BRD-K68502831 | O[C@H]([C@@H]1CO1)[C@@H](O)[C@H]1CO1 | AAFJXZWCNVJTMK-GUCUJZIJSA-N | No-Effect | inactive |
| diarylpropionitrile | BRD-A27143604 | N#C[C@H](Cc1ccc(O)cc1)c1ccc(O)cc1 | GHZHWDWADLAOIQ-ZDUSSCGKSA-N | No-Effect | inactive |
| diatrizoic-acid | BRD-K48247567 | CC(=O)Nc1c(I)c(NC(C)=O)c(I)c(C(=O)O)c1I | YVPYQUNUQOZFHG-UHFFFAOYSA-N | No-Effect | inactive |
| diaveridine | BRD-K97464279 | COc1ccc(Cc2cnc(N)nc2N)cc1OC | LDBTVAXGKYIFHO-UHFFFAOYSA-N | No-Effect | inactive |
| diazoxide | BRD-K73109821 | CC1=Nc2ccc(Cl)cc2S(=O)(=O)N1 | GDLBFKVLRPITMI-UHFFFAOYSA-N | No-Effect | inactive |
| dibekacin | BRD-K48427910 | NC[C@@H]1CC[C@@H](N)[C@@H](O[C@@H]2[C@@H](N)C[C@@H](N)[C@H](O[C@H]3O[C@H](CO)[C@@H](O)[C@H](N)[C@H]3O)[C@@H]2O)O1 | JJCQSGDBDPYCEO-XVZSLQNASA-N | No-Effect | inactive |
| dibenzepine | BRD-K79145749 | CN(C)CCN1C(=O)c2ccccc2N(C)c2ccccc21 | QPGGEKPRGVJKQB-UHFFFAOYSA-N | No-Effect | inactive |
| dibutyl-phthalate | BRD-K73477617 | CCCCOC(=O)c1ccccc1C(=O)OCCCC | DOIRQSBPFJWKBE-UHFFFAOYSA-N | No-Effect | inactive |
| dichlorisone-acetate | BRD-K54187111 | CC(=O)OCC(=O)[C@@]1(O)CC[C@H]2[C@@H]3CCC4=CC(=O)C=C[C@]4(C)[C@@]3(Cl)[C@@H](Cl)C[C@@]21C | YNNURTVKPVJVEI-GSLJADNHSA-N | No-Effect | inactive |
| dichloroacetate | BRD-M97302542 | O=C(O)C(Cl)Cl.[Na] | ZOXHFYPCZQICQD-UHFFFAOYSA-N | No-Effect | inactive |
| dichlorophen | BRD-K62326629 | Oc1ccc(Cl)cc1Cc1cc(Cl)ccc1O | MDNWOSOZYLHTCG-UHFFFAOYSA-N | No-Effect | inactive |
| diclazuril | BRD-A06426627 | N#C[C@H](c1ccc(Cl)cc1)c1c(Cl)cc(-n2ncc(=O)[nH]c2=O)cc1Cl | ZSZFUDFOPOMEET-GFCCVEGCSA-N | No-Effect | inactive |
| diclofenac | BRD-K08252256 | O=C(O)Cc1ccccc1Nc1c(Cl)cccc1Cl | DCOPUUMXTXDBNB-UHFFFAOYSA-N | No-Effect | inactive |
| diclofenamide | BRD-K71499074 | NS(=O)(=O)c1cc(Cl)c(Cl)c(S(N)(=O)=O)c1 | GJQPMPFPNINLKP-UHFFFAOYSA-N | No-Effect | inactive |
| diclofensine | BRD-A51078674 | COc1ccc2c(c1)CN(C)C[C@H]2c1ccc(Cl)c(Cl)c1 | ZJDCGVDEEHWEIG-HNNXBMFYSA-N | No-Effect | inactive |
| dicloralurea | BRD-A41657926 | O=C(N[C@@H](O)C(Cl)(Cl)Cl)N[C@H](O)C(Cl)(Cl)Cl | PPJXIHLNYDVTDI-XIXRPRMCSA-N | No-Effect | inactive |
| dicloxacillin | BRD-K05673000 | Cc1onc(-c2c(Cl)cccc2Cl)c1C(=O)N[C@@H]1C(=O)N2[C@@H](C(=O)O)C(C)(C)S[C@H]12 | YFAGHNZHGGCZAX-JKIFEVAISA-N | No-Effect | inactive |
| dicoumarol | BRD-K82236179 | O=c1oc2ccccc2c(O)c1Cc1c(O)c2ccccc2oc1=O | DOBMPNYZJYQDGZ-UHFFFAOYSA-N | No-Effect | inactive |
| didanosine | BRD-K99063460 | O=c1nc[nH]c2c1ncn2[C@H]1CC[C@@H](CO)O1 | BXZVVICBKDXVGW-NKWVEPMBSA-N | No-Effect | inactive |
| dideoxyadenosine | BRD-K84794093 | Nc1ncnc2c1ncn2[C@H]1CC[C@@H](CO)O1 | WVXRAFOPTSTNLL-NKWVEPMBSA-N | No-Effect | inactive |
| didox | BRD-K94146583 | O=C(NO)c1ccc(O)c(O)c1 | QJMCKEPOKRERLN-UHFFFAOYSA-N | No-Effect | inactive |
| dienestrol | BRD-K95309561 | C/C=C(C(=C/C)/c1ccc(O)cc1)\c1ccc(O)cc1 | NFDFQCUYFHCNBW-SCGPFSFSSA-N | No-Effect | inactive |
| diethylcarbamazine | BRD-K45542189 | CCN(CC)C(=O)N1CCN(C)CC1 | RCKMWOKWVGPNJF-UHFFFAOYSA-N | No-Effect | inactive |
| diethylstilbestrol | BRD-K45330754 | CC/C(=C(/CC)c1ccc(O)cc1)c1ccc(O)cc1 | RGLYKWWBQGJZGM-ISLYRVAYSA-N | No-Effect | inactive |
| diethyltoluamide | BRD-K94080537 | CCN(CC)C(=O)c1cccc(C)c1 | MMOXZBCLCQITDF-UHFFFAOYSA-N | No-Effect | inactive |
| diflorasone-diacetate | BRD-A66869276 | CC(=O)OCC(=O)[C@@]1(OC(C)=O)[C@@H](C)C[C@H]2[C@H]3C[C@H](F)C4=CC(=O)C=C[C@]4(C)[C@@]3(F)[C@@H](O)C[C@@]21C | BOBLHFUVNSFZPJ-OAEHMYPASA-N | No-Effect | inactive |
| difloxacin | BRD-K73923631 | CN1CCN(c2cc3c(cc2F)c(=O)c(C(=O)O)cn3-c2ccc(F)cc2)CC1 | NOCJXYPHIIZEHN-UHFFFAOYSA-N | No-Effect | inactive |
| diflunisal | BRD-K22031190 | O=C(O)c1cc(-c2ccc(F)cc2F)ccc1O | HUPFGZXOMWLGNK-UHFFFAOYSA-N | No-Effect | inactive |
| diftalone | BRD-K80076813 | O=C1c2ccccc2CN2C(=O)c3ccccc3CN12 | CXSJGNHRBWJXEA-UHFFFAOYSA-N | No-Effect | inactive |
| dihomo-gamma-linolenic-acid | BRD-K20152659 | CCCCC/C=C\C/C=C\C/C=C\CCCCCCC(=O)O | HOBAELRKJCKHQD-QNEBEIHSSA-N | No-Effect | inactive |
| dihydrexidine | BRD-K31471398 | Oc1cc2c(cc1O)[C@H]1c3ccccc3CN[C@@H]1CC2 | BGOQGUHWXBGXJW-RHSMWYFYSA-N | No-Effect | inactive |
| dihydroergocristine | BRD-K97440753 | CC(C)[C@@]1(NC(=O)[C@@H]2C[C@@H]3c4cccc5[nH]cc(c45)C[C@H]3N(C)C2)O[C@@]2(O)[C@@H]3CCCN3C(=O)[C@H](Cc3ccccc3)N2C1=O | DEQITUUQPICUMR-HJPBWRTMSA-N | No-Effect | inactive |
| dihydroergotamine | BRD-K72166146 | CN1C[C@H](C(=O)N[C@]2(C)O[C@@]3(O)[C@@H]4CCCN4C(=O)[C@H](Cc4ccccc4)N3C2=O)C[C@@H]2c3cccc4[nH]cc(c34)C[C@H]21 | LUZRJRNZXALNLM-JGRZULCMSA-N | No-Effect | inactive |
| dihydromyricetin | BRD-K01614093 | O=C1c2c(O)cc(O)cc2O[C@H](c2cc(O)c(O)c(O)c2)[C@H]1O | KJXSIXMJHKAJOD-LSDHHAIUSA-N | No-Effect | inactive |
| dihydrostreptomycin | BRD-M15413738 | CN[C@@H]1[C@H](O[C@H]2[C@H](O[C@@H]3[C@@H](O)[C@H](O)[C@@H](NC(=N)N)[C@H](O)[C@H]3NC(=N)N)O[C@@H](C)[C@]2(O)CO)O[C@@H](CO)[C@H](O)[C@H]1O.CN[C@@H]1[C@H](O[C@H]2[C@H](O[C@@H]3[C@@H](O)[C@H](O)[C@@H](NC(=N)N)[C@H](O)[C@H]3NC(=N)N)O[C@@H](C)[C@]2(O)CO)O[C@@H](CO)[C@H](O)[C@H]1O | DLNWBHHJUACHNB-SNGKZNBCSA-N | No-Effect | inactive |
| dihydrotachysterol | BRD-A34455052 | CC(C)[C@@H](C)/C=C/[C@@H](C)[C@H]1CC[C@@H]2/C(=C/C=C3\C[C@@H](O)CC[C@@H]3C)CCC[C@]21C | ILYCWAKSDCYMBB-CEQNRCNESA-N | No-Effect | inactive |
| dihydroxyacetone | BRD-K18637542 | O=C(CO)CO | RXKJFZQQPQGTFL-UHFFFAOYSA-N | No-Effect | inactive |

| | | | | | |
|---|---|---|---|---|---|
| dihydroxyphenylglycine | BRD-A20589515 | N[C@@H](C(=O)O)c1cc(O)cc(O)c1 | HOOWCUZPEFNHDT-SSDOTTSWSA-N | No-Effect | inactive |
| diiodothyropropionic-acid | BRD-K36802417 | O=C(O)CCc1cc(I)c(Oc2ccc(O)cc2)c(I)c1 | WONYMNWUJVKVII-UHFFFAOYSA-N | No-Effect | inactive |
| dilazep | BRD-K48722258 | COc1cc(C(=O)OCCCN2CCCN(CCCOC(=O)c3cc(OC)c(OC)c(OC)c3)CC2)cc(OC)c1OC | QVZCXCJXTMIDME-UHFFFAOYSA-N | No-Effect | inactive |
| diloxanide | BRD-K10974103 | CN(C(=O)C(Cl)Cl)c1ccc(OC(=O)c2ccco2)cc1 | BDYYDXJSHYEDGB-UHFFFAOYSA-N | No-Effect | inactive |
| diltiazem | BRD-K24023109 | COc1ccc([C@@H]2Sc3ccccc3N(CCN(C)C)C(=O)[C@@H]2OC(C)=O)cc1 | HSUGRBWQSSZJOP-RTWAWAEBSA-N | No-Effect | inactive |
| DIM-C-pPhCO2Me | BRD-K00005337 | COC(=O)c1ccc(C(c2c[nH]c3ccccc23)c2c[nH]c3ccccc23)cc1 | BBAOKSZCULLDIW-UHFFFAOYSA-N | No-Effect | inactive |
| dimaprit | BRD-K23541596 | CN(C)CCCSC(=N)N | OLHQOJYVQUNWPL-UHFFFAOYSA-N | No-Effect | inactive |
| dimemorfan | BRD-K29812331 | Cc1ccc2c(c1)[C@]13CCCC[C@@H]1[C@H](C2)N(C)CC3 | KBEZZLAAKIIPFK-NJAFHUGGSA-N | No-Effect | inactive |
| dimenhydrinate | BRD-M98649031 | CN(C)CCOC(c1ccccc1)c1ccccc1.Cn1c(=O)c2[nH]c(Cl)nc2n(C)c1=O | NFLLKCVHYJRNRH-UHFFFAOYSA-N | No-Effect | inactive |
| dimercaprol | BRD-A56152450 | OC[C@@H](S)CS | WQABCVAJNWAXTE-GSVOUGTGSA-N | No-Effect | inactive |
| dimercaptosuccinic-acid | BRD-K88701661 | O=C(O)[C@H](S)[C@H](S)C(=O)O | ACTRVOBWPAIOHC-XIXRPRMCSA-N | No-Effect | inactive |
| dimesna | BRD-K30126819 | O=S(=O)(O)CCSSCCS(=O)(=O)O | BYUKOOOZTSTOOH-UHFFFAOYSA-N | No-Effect | inactive |
| dimethadione | BRD-K43687498 | CC1(C)OC(=O)NC1=O | JYJFNDQBESEHJQ-UHFFFAOYSA-N | No-Effect | inactive |
| dimethicone | BRD-K26674746 | C[Si](C)(C)O[Si](C)(C)C | UQEAIHBTYFGYIE-UHFFFAOYSA-N | No-Effect | inactive |
| dimethindene-(S)-(+) | BRD-K83192048 | C[C@@H](C1=C(CCN(C)C)Cc2ccccc21)c1ccccn1 | MVMQESMQSYOVGV-OAHLLOKOSA-N | No-Effect | inactive |
| dimethyl-fumarate | BRD-K31111078 | COC(=O)/C=C/C(=O)OC | LDCRTTXIJACKKU-ONEGZZNKSA-N | No-Effect | inactive |
| dimethyl-isosorbide | BRD-A17224697 | CO[C@H]1CO[C@@H]2[C@H]1OC[C@H]2OC | MEJYDZQQVZJMPP-YWIQKCBGSA-N | No-Effect | inactive |
| dimetindene | BRD-K01826026 | C[C@@H](C1=C(CCN(C)C)Cc2ccccc21)c1ccccn1 | MVMQESMQSYOVGV-OAHLLOKOSA-N | No-Effect | inactive |
| dimetridazole | BRD-K72728560 | Cc1ncc([N+](=O)[O-])n1C | IBXPYPUJPLLOIN-UHFFFAOYSA-N | No-Effect | inactive |
| diminazene-aceturate | BRD-K92848252 | N=C(N)c1ccc(/N=N/Nc2ccc(C(=N)N)cc2)cc1 | XNYZHCFCZNMTFY-UHFFFAOYSA-N | No-Effect | inactive |
| dimpylate | BRD-K60567437 | CCOP(=S)(OCC)Oc1cc(C)nc(C(C)C)n1 | FHIVAFMUCKRCQO-UHFFFAOYSA-N | No-Effect | inactive |
| dinoprost | BRD-K81672972 | CCCCC[C@H](O)/C=C/[C@H]1[C@H](O)C[C@H](O)[C@@H]1C/C=C\CCCC(=O)O | PXGPLTODNUVGFL-YNNPMVKQSA-N | No-Effect | inactive |
| dinoprostone | BRD-K26521938 | CCCCC[C@H](O)/C=C/[C@H]1[C@H](O)CC(=O)[C@@H]1C/C=C\CCCC(=O)O | XEYBRNLFEZDVAW-ARSRFYASSA-N | No-Effect | inactive |
| dioscin | BRD-K67340225 | C[C@@H]1O[C@@H](O[C@H]2[C@H](O)[C@@H](O[C@@H]3O[C@@H](C)[C@H](O)[C@@H](O)[C@H]3O)[C@H](O[C@H]3CC[C@@]4(C)C(=CC[C@H]5[C@@H]6C[C@@H]7O[C@]8(CC[C@@H](C)CO8)[C@@H](C)[C@@H]7[C@@]6(C)CC[C@@H]54)C3)O[C@@H]2CO)[C@H](O)[C@H](O)[C@H]1O | VNONINPVFQTJOC-ZGXDEBHDSA-N | No-Effect | inactive |
| diosgenin | BRD-K28690501 | C[C@H]1[C@H]2[C@H](C[C@H]3[C@@H]4CC=C5C[C@@H](O)CC[C@]5(C)[C@H]4CC[C@@]32C)O[C@]12CC[C@@H](C)CO2 | WQLVFSAGQJTQCK-VKROHFNGSA-N | No-Effect | inactive |
| diosmetin | BRD-K26862302 | COc1ccc(-c2cc(=O)c3c(O)cc(O)cc3o2)cc1O | MBNGWHIJMBWFHU-UHFFFAOYSA-N | No-Effect | inactive |
| diosmin | BRD-K94436377 | COc1ccc(-c2cc(=O)c3c(O)cc(O[C@@H]4O[C@H](CO[C@@H]5O[C@@H](C)[C@@H](O)[C@@H](O)[C@H]5O)[C@@H](O)[C@H](O)[C@H]4O)cc3o2)cc1O | GZSOSUNBTXMUFQ-YFAPSIMESA-N | No-Effect | inactive |
| dioxybenzone | BRD-K22193694 | COc1ccc(C(=O)c2ccccc2O)c(O)c1 | MEZZCSHVIGVWFI-UHFFFAOYSA-N | No-Effect | inactive |
| dipeptamin | BRD-K11595419 | C[C@H](N)C(=O)N[C@@H](CCC(N)=O)C(=O)O | HJCMDXDYPOUFDY-WHFBIAKZSA-N | No-Effect | inactive |
| diphemanil | BRD-K93441486 | C[N+]1(C)CCC(=C(c2ccccc2)c2ccccc2)CC1 | LCTZPQRFOZKZNK-UHFFFAOYSA-N | No-Effect | inactive |
| diphencyprone | BRD-K51730347 | O=c1c(-c2ccccc2)c1-c1ccccc1 | HCIBTBXNLVOFER-UHFFFAOYSA-N | No-Effect | inactive |
| diphenhydramine | BRD-K47278471 | CN(C)CCOC(c1ccccc1)c1ccccc1 | ZZVUWRFHKOJYTH-UHFFFAOYSA-N | No-Effect | inactive |
| diphenidol | BRD-K01663662 | OC(CCCN1CCCCC1)(c1ccccc1)c1ccccc1 | OGAKLTJNUQRZJU-UHFFFAOYSA-N | No-Effect | inactive |
| diphenyleneiodonium | BRD-K65814004 | c1ccc2c(c1)[I]c1ccccc1-2 | ZSZPLBNCHKXQRY-UHFFFAOYSA-N | No-Effect | inactive |
| diphenylguanidine | BRD-K12362304 | N=C(Nc1ccccc1)Nc1ccccc1 | OWRCNXZUPFZXOS-UHFFFAOYSA-N | No-Effect | inactive |
| diphenylpyraline | BRD-K22936972 | CN1CCC(OC(c2ccccc2)c2ccccc2)CC1 | OWQUZNMMYNAXSL-UHFFFAOYSA-N | No-Effect | inactive |
| dipivefrine | BRD-A47494775 | CNC[C@@H](O)c1ccc(OC(=O)C(C)(C)C)c(OC(=O)C(C)(C)C)c1 | OCUJLLGVOUDECM-CYBMUJFWSA-N | No-Effect | inactive |
| dipraglurant | BRD-K64838454 | Fc1ccc2nc(CCC#Cc3ccccn3)cn2c1 | LZXMUJCJAWVHPZ-UHFFFAOYSA-N | No-Effect | inactive |
| DIPT | BRD-K75385383 | CC(C)N(CCc1c[nH]c2ccccc12)C(C)C | ZRVAAGAZUWXRIP-UHFFFAOYSA-N | No-Effect | inactive |
| dipyrocetyl | BRD-K47608922 | CC(=O)Oc1cccc(C(=O)O)c1OC(C)=O | NYIZXMGNIUSNKL-UHFFFAOYSA-N | No-Effect | inactive |
| diquafosol | BRD-K00003514 | O=c1ccn([C@@H]2O[C@H](COP(=O)(O)OP(=O)(O)OP(=O)(O)OP(=O)(O)OC[C@H]3O[C@@H](n4ccc(=O)[nH]c4=O)[C@H](O)[C@@H]3O)[C@@H](O)[C@H]2O)c(=O)[nH]1 | NMLMACJWHPHKGR-NCOIDOBVSA-N | No-Effect | inactive |
| dirithromycin | BRD-K52756523 | CC[C@H]1OC(=O)[C@H](C)[C@@H](O[C@H]2C[C@@](C)(OC)[C@@H](O)[C@H](C)O2)[C@H](C)[C@@H](O[C@@H]2O[C@H](C)C[C@H](N(C)C)[C@H]2O)[C@](C)(O)C[C@@H](C)[C@@H]2N[C@@H](COCCOC)O[C@H]([C@H]2C)[C@]1(C)O | WLOHNSSYAXHWNR-DWIOZXRMSA-N | No-Effect | inactive |
| diroximel-fumarate | BRD-K87386013 | COC(=O)/C=C/C(=O)OCCN1C(=O)CCC1=O | YIMYDTCOUQIDMT-SNAWJCMRSA-N | No-Effect | inactive |
| disodium-sebacate | BRD-K66672047 | O=C(O)CCCCCCCCC(=O)O | CXMXRPHRNRROMY-UHFFFAOYSA-N | No-Effect | inactive |
| disulfiram | BRD-K32744045 | CCN(CC)C(=S)SSC(=S)N(CC)CC | AUZONCFQVSMFAP-UHFFFAOYSA-N | No-Effect | inactive |
| dithranol | BRD-K51807679 | Oc1cccc2cc3cccc(O)c3c(O)c12 | YUTJCNNFTOIOGT-UHFFFAOYSA-N | No-Effect | inactive |
| ditiocarb-sodium-trihydrate | BRD-K40654626 | CCN(CC)C(=S)S | LMBWSYZSUOEYSN-UHFFFAOYSA-N | No-Effect | inactive |
| ditolylguanidine | BRD-K33459542 | Cc1ccccc1NC(=N)Nc1ccccc1C | OPNUROKCUBTKLF-UHFFFAOYSA-N | No-Effect | inactive |
| dixanthogen | BRD-K05919151 | CCOC(=S)SSC(=S)OCC | FVIGODVHAVLZOO-UHFFFAOYSA-N | No-Effect | inactive |
| dixyrazine | BRD-K00003339 | C[C@H](CN1CCN(CCOCCO)CC1)CN1c2ccccc2Sc2ccccc21 | MSYUMPGNGDNTIQ-HXUWFJFHSA-N | No-Effect | inactive |
| dizocilpine-(-) | BRD-K23163214 | C[C@@]12N[C@@H](Cc3ccccc31)c1ccccc12 | LBOJYSIDWZQNJS-JKSUJKDBSA-N | No-Effect | inactive |
| dizocilpine-(+) | BRD-K58930050 | C[C@]12N[C@H](Cc3ccccc31)c1ccccc12 | LBOJYSIDWZQNJS-CVEARBPZSA-N | No-Effect | inactive |
| DMAB-anabaseine | BRD-K26573499 | CN(C)c1ccc(/C=C2\CCCN=C2c2cccnc2)cc1 | KHUUTHZSRXABAC-DTQAZKPQSA-N | No-Effect | inactive |
| DMeOB | BRD-K36775483 | COc1cccc(/C=N/N=C/c2cccc(OC)c2)c1 | FBNPHFBYHYNMHC-JYFOCSDGSA-N | No-Effect | inactive |
| DMH1 | BRD-K39381259 | CC(C)Oc1ccc(-c2cnc3c(-c4ccnc5ccccc45)cnn3c2)cc1 | JMIFGARJSWXZSH-UHFFFAOYSA-N | No-Effect | inactive |
| DMH4 | BRD-K40575944 | c1ccc(-c2cnn3cc(-c4ccc(OCCN5CCOCC5)cc4)cnc23)cc1 | SKZQZGSPYYHTQG-UHFFFAOYSA-N | No-Effect | inactive |
| DMNB | BRD-K03564313 | COc1cc(C=O)c([N+](=O)[O-])cc1OC | YWSPWKXREVSQCA-UHFFFAOYSA-N | No-Effect | inactive |
| DMP-543 | BRD-K89274813 | O=C1c2ccccc2C(Cc2ccnc(F)c2)(Cc2ccnc(F)c2)c2ccccc21 | MUJBUUDUXGDXLW-UHFFFAOYSA-N | No-Effect | inactive |
| DMPS | BRD-A22971209 | O=S(=O)(O)C[C@@H](S)CS | JLVSRWOIZZXQAD-VKHMYHEASA-N | No-Effect | inactive |
| DMSO | BRD-M56212252 | CS(C)=O.CS(C)=O | CETRZFQIITUQQL-UHFFFAOYSA-N | No-Effect | inactive |
| DNQX | BRD-K64400208 | O=c1[nH]c2cc([N+](=O)[O-])c([N+](=O)[O-])cc2[nH]c1=O | RWVIMCIPOAXUDG-UHFFFAOYSA-N | No-Effect | inactive |
| dobutamine | BRD-A78322124 | CC(CCc1ccc(O)cc1)NCCc1ccc(O)c(O)c1 | JRWZLRBJNMZMFE-UHFFFAOYSA-N | No-Effect | inactive |
| docebenone | BRD-K08290224 | CC1=C(C)C(=O)C(CCCCC#CCCCC#CCO)=C(C)C1=O | WDEABJKSGGRCQA-UHFFFAOYSA-N | No-Effect | inactive |
| docetaxel | BRD-K30577245 | CC(=O)O[C@@]12CO[C@@H]1C[C@H](O)[C@@]1(C)C(=O)[C@H](O)C3=C(C)[C@@H](OC(=O)[C@H](O)[C@@H](NC(=O)OC(C)(C)C)c4ccccc4)C[C@@](O)([C@@H](OC(=O)c4ccccc4)[C@H]12)C3(C)C | ZDZOTLJHXYCWBA-VCVYQWHSSA-N | No-Effect | inactive |

| | | | | | |
|---|---|---|---|---|---|
| doconexent | BRD-K39965020 | CC/C=C\C/C=C\C/C=C\C/C=C\C/C=C\C/C=C\CCC(=O)O | MBMBGCFOFBJSGT-KUBAVDMBSA-N | No-Effect | inactive |
| docosanol | BRD-K50050533 | CCCCCCCCCCCCCCCCCCCCCCO | NOPFSRXAKWQILS-UHFFFAOYSA-N | No-Effect | inactive |
| docusate | BRD-A00092689 | CCCC[C@H](CC)COC(=O)[C@H](CC(=O)OC[C@H](CC)CCCC)S(=O)(=O)O | HNSDLXPSAYFUHK-SQNIBIBYSA-N | No-Effect | inactive |
| dofequidar | BRD-A14941520 | O=C(C(c1ccccc1)c1ccccc1)N1CCN(CC(O)COc2cccc3ncccc23)CC1 | KLWUUPVJTLHYIM-UHFFFAOYSA-N | No-Effect | inactive |
| dofetilide | BRD-K86887724 | CN(CCOc1ccc(NS(C)(=O)=O)cc1)CCc1ccc(NS(C)(=O)=O)cc1 | IXTMWRCNAAVVAI-UHFFFAOYSA-N | No-Effect | inactive |
| dolasetron | BRD-K28029915 | O=C(O[C@@H]1C[C@@H]2C[C@@H]3C[C@H](C1)N2CC3=O)c1c[nH]c2ccccc12 | UKTAZPQNNNJVKR-YXSUXZIUSA-N | No-Effect | inactive |
| dolutegravir | BRD-K67463151 | C[C@@H]1CCO[C@H]2Cn3cc(C(=O)NCc4ccc(F)cc4F)c(=O)c(O)c3C(=O)N21 | RHWKPHLQXYSBKR-BMIGLBTASA-N | No-Effect | inactive |
| domperidone | BRD-K38305202 | O=c1[nH]c2ccccc2n1CCCN1CCC(n2c(=O)[nH]c3cc(Cl)ccc32)CC1 | FGXWKSZFVQUSTL-UHFFFAOYSA-N | No-Effect | inactive |
| donepezil | BRD-A49160188 | COc1cc2c(cc1OC)C(=O)[C@H](CC1CCN(Cc3ccccc3)CC1)C2 | ADEBPBSSDYVVLD-HXUWFJFHSA-N | No-Effect | inactive |
| donitriptan | BRD-K31092604 | N#Cc1ccc(N2CCN(C(=O)COc3ccc4[nH]cc(CCN)c4c3)CC2)cc1 | SOHCKWZVTCTQBG-UHFFFAOYSA-N | No-Effect | inactive |
| dopamine | BRD-K43887077 | NCCc1ccc(O)c(O)c1 | VYFYYTLLBUKUHU-UHFFFAOYSA-N | No-Effect | inactive |
| doramapimod | BRD-K74065929 | Cc1ccc(-n2nc(C(C)(C)C)cc2NC(=O)Nc2ccc(OCCN3CCOCC3)c3ccccc23)cc1 | MVCOAUNKQVWQHZ-UHFFFAOYSA-N | No-Effect | inactive |
| doramectin | BRD-K00003581 | CO[C@H]1C[C@H](O[C@H]2[C@H](C)O[C@@H](O[C@@H]3/C(C)=C/C[C@@H]4C[C@@H](C[C@]5(C=C[C@H](C)[C@@H](C6CCCCC6)O5)O4)OC(=O)[C@@H]4C=C(C)[C@@H](O)[C@H]5OC/C(=C\C=C\[C@@H]3C)[C@@]45O)C[C@@H]2OC)O[C@@H](C)[C@@H]1O | QLFZZSKTJWDQOS-YDBLARSUSA-N | No-Effect | inactive |
| doravirine | BRD-K00003231 | Cn1c(Cn2ccc(C(F)(F)F)c(Oc3cc(Cl)cc(C#N)c3)c2=O)n[nH]c1=O | ZIAOVIPSKUPPQW-UHFFFAOYSA-N | No-Effect | inactive |
| doripenem | BRD-K45319714 | C[C@@H](O)[C@H]1C(=O)N2C(C(=O)O)=C(S[C@@H]3CN[C@H](CNS(N)(=O)=O)C3)[C@H](C)[C@H]12 | AVAACINZEOAHHE-VFZPANTDSA-N | No-Effect | inactive |
| dorzolamide | BRD-K48617017 | CCN[C@H]1C[C@H](C)S(=O)(=O)c2sc(S(N)(=O)=O)cc21 | IAVUPMFITXYVAF-XPUUQOCRSA-N | No-Effect | inactive |
| dosulepin | BRD-K54759182 | CN(C)CC/C=C1\c2ccccc2CSc2ccccc21 | PHTUQLWOUWZIMZ-GZTJUZNOSA-N | No-Effect | inactive |
| DOTMP | BRD-K00004678 | O=P(O)(O)CN1CCN(CP(=O)(O)O)CCN(CP(=O)(O)O)CCN(CP(=O)(O)O)CC1 | RCXMQNIDOFXYDO-UHFFFAOYSA-N | No-Effect | inactive |
| dovitinib | BRD-K85402309 | CN1CCN(c2ccc3nc(-c4c(N)c5c(F)cccc5[nH]c4=O)[nH]c3c2)CC1 | PIQCTGMSNWUMAF-UHFFFAOYSA-N | No-Effect | inactive |
| doxapram | BRD-K01825594 | CCN1C[C@@H](CCN2CCOCC2)C(c2ccccc2)(c2ccccc2)C1=O | XFDJYSQDBULQSI-JOCHJYFZSA-N | No-Effect | inactive |
| doxazosin | BRD-K01825125 | COc1cc2nc(N3CCN(C(=O)[C@H]4COc5ccccc5O4)CC3)nc(N)c2cc1OC | RUZYUOTYCVRMRZ-HXUWFJFHSA-N | No-Effect | inactive |
| doxepin | BRD-K37694030 | CN(C)CC/C=C1/c2ccccc2COc2ccccc21 | ODQWQRRAPPTVAG-BOPFTXTBSA-N | No-Effect | inactive |
| doxifluridine | BRD-K58262659 | C[C@H]1O[C@@H](n2cc(F)c(=O)[nH]c2=O)[C@H](O)[C@@H]1O | ZWAOHEXOSAUJHY-ZIYNGMLESA-N | No-Effect | inactive |
| doxofylline | BRD-K70507123 | Cn1c(=O)c2c(ncn2CC2OCCO2)n(C)c1=O | HWXIGFIVGWUZAO-UHFFFAOYSA-N | No-Effect | inactive |
| doxorubicin | BRD-K92093830 | COc1cccc2c1C(=O)c1c(O)c3c(c(O)c1C2=O)C[C@@](O)(C(=O)CO)C[C@@H]3O[C@H]1C[C@H](N)[C@H](O)[C@H](C)O1 | AOJJSUZBOXZQNB-TZSSRYMLSA-N | No-Effect | inactive |
| doxycycline | BRD-K01825679 | C[C@H]1c2cccc(O)c2C(=O)C2=C(O)[C@]3(O)C(=O)C(C(N)=O)=C(O)[C@@H](N(C)C)[C@@H]3[C@@H](O)[C@@H]21 | JBIWCJUYHHGXTC-AKNGSSGZSA-N | No-Effect | inactive |
| doxylamine | BRD-A44008656 | CN(C)CCOC(C)(c1ccccc1)c1ccccn1 | HCFDWZZGGLSKEP-UHFFFAOYSA-N | No-Effect | inactive |
| DpC | BRD-K00004636 | CN(C(=S)NN=C(c1ccccn1)c1ccccn1)C1CCCCC1 | GNLZNQJBZNOUBM-UHFFFAOYSA-N | No-Effect | inactive |
| DPC-681 | BRD-K37087535 | CC(C)CN(C[C@@H](O)[C@H](Cc1ccccc1)NC(=O)[C@@H](NC(=O)CNCc1cccc(F)c1)C(C)(C)C)S(=O)(=O)c1cccc(N)c1 | PNIFFZXGBAYVMQ-RKKDRKJOSA-N | No-Effect | inactive |
| DPCPX | BRD-K01824921 | CCCn1c(=O)c2[nH]c(C3CCCC3)nc2n(CCC)c1=O | FFBDFADSZUINTG-UHFFFAOYSA-N | No-Effect | inactive |
| DPI-201106 | BRD-A64553394 | N#Cc1cc2c(OC[C@H](O)CN3CCN(C(c4ccccc4)c4ccccc4)CC3)cccc2[nH]1 | BYBYHCOEAFHGJL-RUZDIDTESA-N | No-Effect | inactive |
| DPO-1 | BRD-K99922388 | CC(C)[C@@H]1CC[C@@H](C)C[C@@H]1P(=O)(c1ccccc1)c1ccccc1 | BPCNGVCAHAIZEE-COPCDDAFSA-N | No-Effect | inactive |
| DPPE | BRD-K80315159 | CCN(CC)CCOc1ccc(Cc2ccccc2)cc1 | NFIXBCVWIPOYCD-UHFFFAOYSA-N | No-Effect | inactive |
| DQP-1105 | BRD-A67976398 | Cc1ccc2[nH]c(=O)c(C3=NN(C(=O)CCC(=O)O)[C@H](c4ccc(Br)cc4)C3)c(-c3ccccc3)c2c1 | MVYNQMRZRULRIE-DEOSSOPVSA-N | No-Effect | inactive |
| DR-2313 | BRD-K94920105 | Cc1nc(=O)c2c([nH]1)CCSC2 | HRYKZAKEAVZGJD-UHFFFAOYSA-N | No-Effect | inactive |
| DR-4485 | BRD-A39794701 | O=C1Nc2ccc(Cl)c3c2[C@@]1(CCCCN1CC=C(c2ccc(Cl)cc2)CC1)CCC3 | SBTRHJHOLCAPFT-AREMUKBSSA-N | No-Effect | inactive |
| DRF053-(R) | BRD-K25486274 | CC[C@H](CO)Nc1nc(Nc2cccc(-c3ccccn3)c2)c2ncn(C(C)C)c2n1 | LWANFAFTTOKZAX-QGZVFWFLSA-N | No-Effect | inactive |
| droloxifene | BRD-K38477637 | CC/C(=C(/c1ccc(OCCN(C)C)cc1)c1cccc(O)c1)c1ccccc1 | ZQZFYGIXNQKOAV-OCEACIFDSA-N | No-Effect | inactive |
| dromostanolone-propionate | BRD-A38706605 | CCC(=O)O[C@H]1CC[C@H]2[C@H]3CC[C@H]4CC(=O)[C@@H](C)C[C@@]4(C)[C@@H]3CC[C@@]12C | NOTIQUSPUUHHEH-QOMQUSCHSA-N | No-Effect | inactive |
| dronedarone | BRD-K05524748 | CCCCc1oc2ccc(NS(C)(=O)=O)cc2c1C(=O)c1ccc(OCCCN(CCCC)CCCC)cc1 | ZQTNQVWKHCQYLQ-UHFFFAOYSA-N | No-Effect | inactive |
| droperidol | BRD-K97158071 | O=C(CCCN1CC=C(n2c(=O)[nH]c3ccccc32)CC1)c1ccc(F)cc1 | RMEDXOLNCUSCGS-UHFFFAOYSA-N | No-Effect | inactive |
| dropropizine | BRD-A29349577 | OC[C@@H](O)CN1CCN(c2ccccc2)CC1 | PTVWPYVOOKLBCG-ZDUSSCGKSA-N | No-Effect | inactive |
| drospirenone | BRD-K04394237 | C[C@]12CC[C@H]3[C@H]([C@@H]1[C@@H]1C[C@@H]1[C@@]21CCC(=O)O1)[C@H]1C[C@H]1C1=CC(=O)CC[C@@]13C | METQSPRSQINEEU-HXCATZOESA-N | No-Effect | inactive |
| droxicam | BRD-K17219082 | CN1c2c(oc(=O)n(-c3ccccn3)c2=O)-c2ccccc2S1(=O)=O | OEHFRZLKGRKFAS-UHFFFAOYSA-N | No-Effect | inactive |
| droxidopa | BRD-K20372197 | N[C@H](C(=O)O)[C@H](O)c1ccc(O)c(O)c1 | QXWYKJLNLSIPIN-JGVFFNPUSA-N | No-Effect | inactive |
| droxinostat | BRD-K11558771 | Cc1cc(Cl)ccc1OCCCC(=O)NO | JHSXDAWGLCZYSM-UHFFFAOYSA-N | No-Effect | inactive |
| DSM265 | BRD-K35498911 | Cc1cc(Nc2ccc(S(F)(F)(F)(F)F)cc2)n2nc(C(C)(F)F)nc2n1 | OIZSVTOIBNSVOS-UHFFFAOYSA-N | No-Effect | inactive |
| DSPC | BRD-K00004570 | CCCCCCCCCCCCCCCCCC(=O)OC[C@@H](COP(=O)(O)OCC[N+](C)(C)C)OC(=O)CCCCCCCCCCCCCCCCC | NRJAVPSFFCBXDT-WBCKFURZSA-O | No-Effect | inactive |
| DSR-6434 | BRD-K46384285 | CCCCNc1nc(N)c2[nH]c(=O)n(Cc3ccc(OCCN(C)C)nc3)c2n1 | SSZHESNDOMBSRV-UHFFFAOYSA-N | No-Effect | inactive |
| DU-728 | BRD-K08859203 | N=C(N)NCCC[C@H](N)C(=O)NCC(=O)N[C@@H](CC(=O)O)C(=O)N[C@@H](CO)C(=O)O | NNRFRJQMBSBXGO-CIUDSAMLSA-N | No-Effect | inactive |
| DUP-697 | BRD-K06221026 | CS(=O)(=O)c1ccc(-c2cc(Br)sc2-c2ccc(F)cc2)cc1 | AJFTZWGGHJXZOB-UHFFFAOYSA-N | No-Effect | inactive |
| dutasteride | BRD-K30373883 | C[C@]12CC[C@H]3[C@@H](CC[C@H]4NC(=O)C=C[C@]34C)[C@@H]1CC[C@@H]2C(=O)Nc1cc(C(F)(F)F)ccc1C(F)(F)F | JWJOTENAMICLJG-QWBYCMEYSA-N | No-Effect | inactive |
| DVD-111 | BRD-K22520627 | CCOC(=O)CNC(=O)[C@@H]1CCCN1C(=O)Cc1ccccc1 | PJNSMUBMSNAEEN-AWEZNQCLSA-N | No-Effect | inactive |
| DWK-1339 | BRD-K00003171 | COCCCc1ccc2oc(-c3ccc(OC)c(OC)c3)cc2c1 | CUEPJIGXQLIOIK-UHFFFAOYSA-N | No-Effect | inactive |
| dynasore | BRD-K87718015 | O=C(N/N=C/c1ccc(O)c(O)c1)c1cc2ccccc2cc1O | SYNDQCRDGGCQRZ-VXLYETTFSA-N | No-Effect | inactive |
| dynole-34-2 | BRD-K23479488 | CCCCCCCCNC(=O)/C(C#N)=C/c1cn(CCCN(C)C)c2ccccc12 | MYIMRONQBLZJFI-DYTRJAOYSA-N | No-Effect | inactive |
| dyphylline | BRD-A00827783 | Cn1c(=O)c2c(ncn2C[C@@H](O)CO)n(C)c1=O | KSCFJBIXMNOVSH-ZCFIWIBFSA-N | No-Effect | inactive |
| E-2001 | BRD-K00004561 | O=C(CN1CCC(C(=O)c2ccc(F)cc2)CC1)c1ccc2ccccc2c1 | WPFPDSUMDURRRW-UHFFFAOYSA-N | No-Effect | inactive |
| E-2012 | BRD-K50564936 | COc1cc(/C=C2\CCCN([C@@H](C)c3ccc(F)cc3)C2=O)ccc1-n1cnc(C)c1 | PUOAETJYKQITMO-LANLRWRYSA-N | No-Effect | inactive |
| E-4031 | BRD-K41713976 | Cc1cccc(CCN2CCC(C(=O)c3ccc(NS(C)(=O)=O)cc3)CC2)n1 | SRUISGSHWFJION-UHFFFAOYSA-N | No-Effect | inactive |
| E-64 | BRD-K00004314 | CC(C)C[C@H](NC(=O)[C@H]1O[C@@H]1C(=O)O)C(=O)NCCCCNC(=N)N | LTLYEAJONXGNFG-DCAQKATOSA-N | No-Effect | inactive |
| E3330 | BRD-K00004649 | CCCCCCCCC/C(=C\C1=C(C)C(=O)C(OC)=C(OC)C1=O)C(=O)O | AALSSIXXBDPENJ-FYWRMAATSA-N | No-Effect | inactive |
| E7046 | BRD-K00003449 | C[C@H](NC(=O)c1c(C(F)F)nn(C)c1Oc1cccc(C(F)(F)F)c1)c1ccc(C(=O)O)cc1 | MKLKAQMPKHNQPR-NSHDSACASA-N | No-Effect | inactive |

| | | | | | |
|---|---|---|---|---|---|
| E7449 | BRD-K05444225 | O=C1NNc2nc(CN3Cc4ccccc4C3)nc3cccc1c23 | JLFSBHQQXIAQEC-UHFFFAOYSA-N | No-Effect | inactive |
| E7820 | BRD-K00003172 | Cc1ccc(NS(=O)(=O)c2cccc(C#N)c2)c2[nH]cc(C#N)c12 | LWGUASZLXHYWIV-UHFFFAOYSA-N | No-Effect | inactive |
| eact | BRD-K53030960 | COCCN(C(=O)c1cc(OC)c(OC)c(OC)c1)c1nc(-c2ccccc2)cs1 | ZUXNHFFVQWADJL-UHFFFAOYSA-N | No-Effect | inactive |
| EB-47 | BRD-K66998413 | Nc1ncnc2c1ncn2[C@H]1O[C@@H](C(=O)N2CCN(CC(=O)Nc3cccc4c3CNC4=O)CC2)[C@H](O)[C@@H]1O | DDFLFKTXUWPNMV-LFALDJFMSA-N | No-Effect | inactive |
| ebastine | BRD-K13314656 | CC(C)(C)c1ccc(C(=O)CCCN2CCC(OC(c3ccccc3)c3ccccc3)CC2)cc1 | MJJALKDDGIKVBE-UHFFFAOYSA-N | No-Effect | inactive |
| EBPC | BRD-K00004209 | CCOC(=O)[C@@H]1CN(Cc2ccccc2)C(=O)C1=O | FQMSSTZJKSWSMP-LLVKDONJSA-N | No-Effect | inactive |
| EC-23 | BRD-K81004638 | CC1(C)CCC(C)(C)c2cc(C#Cc3ccc(C(=O)O)cc3)ccc21 | OQVLOWLEEHYBJH-UHFFFAOYSA-N | No-Effect | inactive |
| ecabapide | BRD-K47371940 | CNC(=O)c1cccc(NCC(=O)NCCc2ccc(OC)c(OC)c2)c1 | JTAGHJPZEDNHHA-UHFFFAOYSA-N | No-Effect | inactive |
| ecabet | BRD-K03353589 | CC(C)c1cc2c(cc1S(=O)(=O)O)[C@@]1(C)CCC[C@@](C)(C(=O)O)[C@@H]1CC2 | IWCWQNVIUXZOMJ-MISYRCLQSA-N | No-Effect | inactive |
| econazole | BRD-A51820102 | Clc1ccc(CO[C@@H](Cn2ccnc2)c2ccc(Cl)cc2Cl)cc1 | LEZWWPYKPKIXLL-SFHVURJKSA-N | No-Effect | inactive |
| ectoine | BRD-K34566599 | CC1=NCC[C@@H](C(=O)O)N1 | WQXNXVUDBPYKBA-YFKPBYRVSA-N | No-Effect | inactive |
| edaglitazone | BRD-K00004306 | Cc1oc(-c2ccccc2)nc1CCOc1ccc(C[C@@H]2SC(=O)NC2=O)c2sccc12 | HAAXAFNSRADSMK-FQEVSTJZSA-N | No-Effect | inactive |
| edaravone | BRD-K35458079 | Cc1cc(=O)n(-c2ccccc2)[nH]1 | KZQYIMCESJLPQH-UHFFFAOYSA-N | No-Effect | inactive |
| edasalonexent | BRD-K00003258 | CC/C=C\C/C=C\C/C=C\C/C=C\C/C=C\C/C=C\CCC(=O)NCCNC(=O)c1ccccc1O | JQLBBYLGWHUHRW-KUBAVDMBSA-N | No-Effect | inactive |
| edelfosine | BRD-K01825687 | CCCCCCCCCCCCCCCCCCOC[C@H](COP(=O)(O)OCC[N+](C)(C)C)OC | MHFRGQHAERHWKZ-HHHXNRCGSA-O | No-Effect | inactive |
| editol | BRD-A78588743 | C[C@@H](O)CN(CCN(C[C@@H](C)O)C[C@@H](C)O)C[C@@H](C)O | NSOXQYCFHDMMGV-AAVRWANBSA-N | No-Effect | inactive |
| EDO-S101 | BRD-K00003432 | Cn1c(CCCCCCC(=O)NO)nc2cc(N(CCCl)CCCl)ccc21 | GISXTRIGVCKQBX-UHFFFAOYSA-N | No-Effect | inactive |
| edonerpic-maleate | BRD-K00003259 | OC1CN(CCCOCCc2ccc3sccc3c2)C1 | HQNACSFBDBYLJP-UHFFFAOYSA-N | No-Effect | inactive |
| edoxaban | BRD-K97058489 | CN1CCc2nc(C(=O)N[C@@H]3C[C@@H](C(=O)N(C)C)CC[C@@H]3NC(=O)C(=O)Nc3ccc(Cl)cn3)sc2C1 | HGVDHZBSSITLCT-JLJPHGGASA-N | No-Effect | inactive |
| edoxudine | BRD-K86892782 | CCc1cn([C@H]2C[C@H](O)[C@@H](CO)O2)c(=O)[nH]c1=O | XACKNLSZYYIACO-DJLDLDEBSA-N | No-Effect | inactive |
| edrophonium | BRD-K81128206 | CC[N+](C)(C)c1cccc(O)c1 | VWLHWLSRQJQWRG-UHFFFAOYSA-O | No-Effect | inactive |
| EED226 | BRD-K19002914 | CS(=O)(=O)c1ccc(-c2cnc(NCc3ccco3)n3cnnc23)cc1 | DYIRSNMPIZZNBK-UHFFFAOYSA-N | No-Effect | inactive |
| EF5 | BRD-K00004643 | O=C(Cn1ccnc1[N+](=O)[O-])NCC(F)(F)C(F)(F)F | JGGDSDPOPRWSCX-UHFFFAOYSA-N | No-Effect | inactive |
| efaproxiral | BRD-K37355418 | Cc1cc(C)cc(NC(=O)Cc2ccc(OC(C)(C)C(=O)O)cc2)c1 | BNFRJXLZYUTIII-UHFFFAOYSA-N | No-Effect | inactive |
| efaroxan | BRD-A10739734 | CCC1(C2=NCCN2)Cc2ccccc2O1 | RATZLMXRALDSJW-UHFFFAOYSA-N | No-Effect | inactive |
| efatutazone | BRD-A24560335 | Cc1cc(Oc2ccc3nc(COc4ccc(C[C@H]5SC(=O)NC5=O)cc4)n(C)c3c2)cc(C)c1N | JCYNMRJCUYVDBC-HSZRJFAPSA-N | No-Effect | inactive |
| efavirenz | BRD-K40758068 | O=C1Nc2ccc(Cl)cc2[C@@](C#CC2CC2)(C(F)(F)F)O1 | XPOQHMRABVBWPR-ZDUSSCGKSA-N | No-Effect | inactive |
| efinaconazole | BRD-A30774155 | C=C1CCN([C@H](C)[C@@](O)(Cn2cncn2)c2ccc(F)cc2F)CC1 | NFEZZTICAUWDHU-KDOFPFPSSA-N | No-Effect | inactive |
| efletirizine | BRD-K00004551 | O=C(O)COCCN1CCN(C(c2ccc(F)cc2)c2ccc(F)cc2)CC1 | BAWMMJAUVBLLEE-UHFFFAOYSA-N | No-Effect | inactive |
| eflornithine | BRD-A96292436 | NCCCC(N)(C(=O)O)C(F)F | VLCYCQAOQCDTCN-UHFFFAOYSA-N | No-Effect | inactive |
| efloxate | BRD-K55166090 | CCOC(=O)COc1ccc2c(=O)cc(-c3ccccc3)oc2c1 | ZVXBAHLOGZCFTP-UHFFFAOYSA-N | No-Effect | inactive |
| efonidipine | BRD-K00004292 | CC1=C(C(=O)OCCN(Cc2ccccc2)c2ccccc2)[C@H](c2cccc([N+](=O)[O-])c2)C(P2(=O)OCC(C)(C)CO2)=C(C)N1 | NSVFSAJIGAJDMR-HKBQPEDESA-N | No-Effect | inactive |
| EG00229 | BRD-K50653584 | N=C(N)NCCC[C@H](NC(=O)c1sccc1NS(=O)(=O)c1cccc2nsnc12)C(=O)O | ZWWMEDURALZMEV-NSHDSACASA-N | No-Effect | inactive |
| eglumetad | BRD-K65359424 | N[C@@]1(C(=O)O)CC[C@H]2[C@H](C(=O)O)[C@H]21 | VTAARTQTOOYTES-RGDLXGNYSA-N | No-Effect | inactive |
| EHop-016 | BRD-K32289541 | CCn1c2ccccc2c2cc(Nc3ccnc(NCCCN4CCOCC4)n3)ccc21 | AFTZZRFCMOAFCR-UHFFFAOYSA-N | No-Effect | inactive |
| EI1 | BRD-K91535048 | CCC(CC)n1ccc2c(C(=O)NCc3c(C)cc(C)[nH]c3=O)cc(C#N)cc21 | PFHDWRIVDDIFRP-UHFFFAOYSA-N | No-Effect | inactive |
| EIPA | BRD-K88120876 | CCN(c1nc(N)c(C(=O)N=C(N)N)nc1Cl)C(C)C | QDERNBXNXJCIQK-UHFFFAOYSA-N | No-Effect | inactive |
| EIT-hydrobromide | BRD-K81623406 | CCSC(=N)N | VFIZBHJTOHUOEK-UHFFFAOYSA-N | No-Effect | inactive |
| elacridar | BRD-K88807631 | COc1cc2c(cc1OC)CN(CCc1ccc(NC(=O)c3cccc4c(=O)c5cccc(OC)c5[nH]c34)cc1)CC2 | OSFCMRGOZNQUSW-UHFFFAOYSA-N | No-Effect | inactive |
| elacytarabine | BRD-K00004609 | CCCCCCCC/C=C/CCCCCCCC(=O)OC[C@H]1O[C@@H](n2ccc(N)nc2=O)[C@@H](O)[C@@H]1O | FLFGNMFWNBOBGE-FNNZEKJRSA-N | No-Effect | inactive |
| elafibranor | BRD-K03165895 | CSc1ccc(C(=O)/C=C/c2cc(C)c(OC(C)(C)C(=O)O)c(C)c2)cc1 | AFLFKFHDSCQHOL-IZZDOVSWSA-N | No-Effect | inactive |
| elagolix | BRD-K55808645 | COc1cccc(-c2c(C)n(Cc3c(F)cccc3C(F)(F)F)c(=O)n(C[C@H](NCCCC(=O)O)c3ccccc3)c2=O)c1F | HEAUOKZIVMZVQL-VWLOTQADSA-N | No-Effect | inactive |
| eldecalcitol | BRD-K00003316 | C=C1/C(=C\C=C2/CCC[C@@]3(C)[C@H]2CC[C@@H]3[C@H](C)CCCC(C)(C)O)C[C@@H](O)[C@@H](OCCCO)[C@@H]1O | FZEXGDDBXLBRTD-AYIMTCTASA-N | No-Effect | inactive |
| eleclazine | BRD-K00003225 | O=C1c2cc(-c3ccc(OC(F)(F)F)cc3)ccc2OCCN1Cc1ncccn1 | YNUAEEJQYHYLMS-UHFFFAOYSA-N | No-Effect | inactive |
| elemene | BRD-K79565391 | C=C[C@]1(C)CC[C@@H](C(=C)C)C[C@@H]1C(=C)C | OPFTUNCRGUEPRZ-RBSFLKMASA-N | No-Effect | inactive |
| elesclomol | BRD-K82135108 | CN(NC(=O)CC(=O)NN(C)C(=S)c1ccccc1)C(=S)c1ccccc1 | BKJIXTWSNXCKJH-UHFFFAOYSA-N | No-Effect | inactive |
| elinogrel | BRD-K59271229 | CNc1cc2[nH]c(=O)n(-c3ccc(NC(=O)NS(=O)(=O)c4ccc(Cl)s4)cc3)c(=O)c2cc1F | LGSDFTPAICUONK-UHFFFAOYSA-N | No-Effect | inactive |
| eliprodil | BRD-A61392169 | O[C@@H](CN1CCC(Cc2ccc(F)cc2)CC1)c1ccc(Cl)cc1 | GGUSQTSTQSHJAH-FQEVSTJZSA-N | No-Effect | inactive |
| ellagic-acid | BRD-K30466858 | O=c1oc2c(O)c(O)cc3c(=O)oc4c(O)c(O)cc1c4c23 | AFSDNFLWKVMVRB-UHFFFAOYSA-N | No-Effect | inactive |
| eltanexor | BRD-K00003369 | NC(=O)/C(=C\n1cnc(-c2cc(C(F)(F)F)cc(C(F)(F)F)c2)n1)c1cncnc1 | JFBAVWVBLRIWHM-MLPAPPSSSA-N | No-Effect | inactive |
| eltoprazine | BRD-K86171477 | c1cc2c(c(N3CCNCC3)c1)OCCO2 | WVLHGCRWEHCIOT-UHFFFAOYSA-N | No-Effect | inactive |
| eltrombopag | BRD-K00003572 | CC1=NN(c2ccc(C)c(C)c2)C(=O)/C1=N\Nc1cccc(-c2cccc(C(=O)O)c2)c1O | XDXWLKQMMKQXPV-QYQHSDTDSA-N | No-Effect | inactive |
| eluxadoline | BRD-K39560679 | COc1ccc(CN(C(=O)[C@@H](N)Cc2c(C)cc(C(N)=O)cc2C)[C@@H](C)c2nc(-c3ccccc3)c[nH]2)cc1C(=O)O | QFNHIDANIVGXPE-FNZWTVRRSA-N | No-Effect | inactive |
| elvitegravir | BRD-K54472332 | COc1cc2c(cc1Cc1cccc(Cl)c1F)c(=O)c(C(=O)O)cn2[C@H](CO)C(C)C | JUZYLCPPVHEVSV-LJQANCHMSA-N | No-Effect | inactive |
| embelin | BRD-K86727142 | CCCCCCCCCCCC1C(=O)C(O)=C(O)C1=O | HQWGBIUMYSOPFP-UHFFFAOYSA-N | No-Effect | inactive |
| EMD-1214063 | BRD-K97963946 | CN1CCC(COc2cnc(-c3cccc(Cn4nc(-c5cccc(C#N)c5)ccc4=O)c3)nc2)CC1 | AHYMHWXQRWRBKT-UHFFFAOYSA-N | No-Effect | inactive |
| EMD-386088 | BRD-K47659338 | Cc1[nH]c2ccc(Cl)cc2c1C1=CCNCC1 | BPPGPYJBCVXILI-UHFFFAOYSA-N | No-Effect | inactive |
| EMD-53998 | BRD-A65248799 | COc1ccc(C(=O)N2CCCc3cc(C4=NNC(=O)S[C@@H]4C)ccc32)cc1OC | IZLRMTJLQCLMKF-CYBMUJFWSA-N | No-Effect | inactive |
| EMD-66684 | BRD-K88759641 | CCCCc1nc2ccn(CC(=O)N(C)C)c(=O)c2n1Cc1ccc(-c2ccccc2-c2nn[nH]n2)cc1 | CQVJTTOZTTVVBK-UHFFFAOYSA-N | No-Effect | inactive |
| EMD534085 | BRD-K00003185 | CN(C)CCNC(=O)NC[C@H]1CC[C@@H]2[C@H](O1)c1cc(C(F)(F)F)ccc1N[C@H]2c1ccccc1 | MARIUIDCPUZLKZ-FUKQBSRTSA-N | No-Effect | inactive |
| emedastine | BRD-K15010214 | CCOCCn1c(N2CCCN(C)CC2)nc2ccccc21 | KBUZBQVCBVDWKX-UHFFFAOYSA-N | No-Effect | inactive |
| emeramide | BRD-K00003226 | O=C(NCCS)c1cccc(C(=O)NCCS)c1 | JUTBAVRYDAKVGQ-UHFFFAOYSA-N | No-Effect | inactive |
| emixustat | BRD-K00003290 | NCC[C@@H](O)c1cccc(OCC2CCCCC2)c1 | WJIGGYYSZBWCGC-MRXNPFEDSA-N | No-Effect | inactive |
| emodepside | BRD-K00003414 | CC(C)C[C@H]1C(=O)O[C@H](Cc2ccc(N3CCOCC3)cc2)C(=O)N(C)[C@H](CC(C)C)C(=O)O[C@H](C)C(=O)N(C)[C@@H](CC(C)C)C(=O)O[C@H](Cc2ccc(N3CCOCC3)cc2)C(=O)N(C)[C@@H](CC(C)C)C(=O)O[C@H](C)C(=O)N1C | ZMQMTKVVAMWKNY-CZOLZNPPSA-N | No-Effect | inactive |
| emodin | BRD-K58685305 | Cc1cc(O)c2c(c1)C(=O)c1cc(O)cc(O)c1C2=O | RHMXXJGYXNZAPX-UHFFFAOYSA-N | No-Effect | inactive |

| | | | | | |
|---|---|---|---|---|---|
| emorfazone | BRD-K90629600 | CCOc1c(N2CCOCC2)cnn(C)c1=O | URJQOOISAKEBKW-UHFFFAOYSA-N | No-Effect | inactive |
| emoxipin | BRD-K28982731 | CCc1nc(C)ccc1O | JPGDYIGSCHWQCC-UHFFFAOYSA-N | No-Effect | inactive |
| empagliflozin | BRD-K53765467 | OC[C@H]1O[C@@H](c2ccc(Cl)c(Cc3ccc(O[C@H]4CCOC4)cc3)c2)[C@H](O)[C@@H](O)[C@@H]1O | OBWASQILIWPZMG-QZMOQZSNSA-N | No-Effect | inactive |
| emricasan | BRD-K00023644 | C[C@H](NC(=O)C(=O)Nc1ccccc1C(C)(C)C)C(=O)N[C@@H](CC(=O)O)C(=O)COc1c(F)c(F)cc(F)c1F | SCVHJVCATBPIHN-SJCJKPOMSA-N | No-Effect | inactive |
| emtricitabine | BRD-K61452026 | Nc1nc(=O)n([C@@H]2CS[C@H](CO)O2)cc1F | XQSPYNMVSIKCOC-NTSWFWBYSA-N | No-Effect | inactive |
| EN460 | BRD-K86063171 | O=C(O)c1cc(N2N=C(C(F)(F)F)/C(=C/c3ccc(-c4ccccc4)o3)C2=O)ccc1Cl | RSLFQCNAOMQAIH-WJDWOHSUSA-N | No-Effect | inactive |
| enalaprilat | BRD-K99447003 | C[C@H](N[C@@H](CCc1ccccc1)C(=O)O)C(=O)N1CCC[C@H]1C(=O)O | LZFZMUMEGBBDTC-QEJZJMRPSA-N | No-Effect | inactive |
| enbucrilate | BRD-K66918327 | C=C(C#N)C(=O)OCCCC | JJJFUHOGVZWXNQ-UHFFFAOYSA-N | No-Effect | inactive |
| enciprazine | BRD-A62197763 | COc1ccccc1N1CCN(C[C@@H](O)COc2cc(OC)c(OC)c(OC)c2)CC1 | KSQCNASWXSCJTD-QGZVFWFLSA-N | No-Effect | inactive |
| encorafenib | BRD-K00003576 | COC(=O)N[C@@H](C)CNc1nccc(-c2cn(C(C)C)nc2-c2cc(Cl)cc(NS(C)(=O)=O)c2F)n1 | CMJCXYNUCSMDBY-ZDUSSCGKSA-N | No-Effect | inactive |
| endo-IWR-1 | BRD-K61314889 | O=C(Nc1cccc2cccnc12)c1ccc(N2C(=O)[C@H]3[C@H]4C=C[C@H](C4)[C@H]3C2=O)cc1 | ZGSXEXBYLJIOGF-ALFLXDJESA-N | No-Effect | inactive |
| enflurane | BRD-A51831848 | FC(F)OC(F)(F)[C@@H](F)Cl | JPGQOUSTVILISH-PVQJCKRUSA-N | No-Effect | inactive |
| enilconazole | BRD-K01825690 | C=CCO[C@H](Cn1ccnc1)c1ccc(Cl)cc1Cl | PZBPKYOVPCNPJY-CQSZACIVSA-N | No-Effect | inactive |
| eniluracil | BRD-K26425819 | C#Cc1c[nH]c(=O)[nH]c1=O | JOZGNYDSEBIJDH-UHFFFAOYSA-N | No-Effect | inactive |
| eniporide | BRD-K00004568 | Cc1cc(-n2cccc2)c(S(C)(=O)=O)cc1C(=O)NC(=N)N | UADMBZFZZOBWBB-UHFFFAOYSA-N | No-Effect | inactive |
| enisamium-iodide | BRD-K10029415 | C[n+]1ccc(C(=O)NCc2ccccc2)cc1 | QOEJDHLJOKRPJG-UHFFFAOYSA-O | No-Effect | inactive |
| ENMD-2076 | BRD-K68488863 | Cc1cc(Nc2cc(N3CCN(C)CC3)nc(/C=C/c3ccccc3)n2)[nH]n1 | BLQYVHBZHAISJM-CMDGGOBGSA-N | No-Effect | inactive |
| enocitabine | BRD-K32007074 | CCCCCCCCCCCCCCCCCCCCCC(=O)Nc1ccn([C@@H]2O[C@H](CO)[C@@H](O)[C@@H]2O)c(=O)n1 | SAMRUMKYXPVKPA-VFKOLLTISA-N | No-Effect | inactive |
| enoxacin | BRD-K78113049 | CCn1cc(C(=O)O)c(=O)c2cc(F)c(N3CCNCC3)nc21 | IDYZIJYBMGIQMJ-UHFFFAOYSA-N | No-Effect | inactive |
| enoximone | BRD-K21152241 | CSc1ccc(C(=O)c2[nH]c(=O)[nH]c2C)cc1 | ZJKNESGOIKRXQY-UHFFFAOYSA-N | No-Effect | inactive |
| enoxolone | BRD-K96137854 | CC1(C)[C@@H](O)CC[C@@]2(C)[C@H]1CC[C@]1(C)[C@@H]2C(=O)C=C2[C@@H]3C[C@@](C)(C(=O)O)CC[C@]3(C)CC[C@]21C | MPDGHEJMBKOTSU-YKLVYJNSSA-N | No-Effect | inactive |
| enprofylline | BRD-K35007173 | CCCn1c(=O)[nH]c(=O)c2[nH]cnc21 | SIQPXVQCUCHWDI-UHFFFAOYSA-N | No-Effect | inactive |
| enrofloxacin | BRD-K76534306 | CCN1CCN(c2cc3c(cc2F)c(=O)c(C(=O)O)cn3C2CC2)CC1 | SPFYMRJSYKOXGV-UHFFFAOYSA-N | No-Effect | inactive |
| entacapone | BRD-K83636919 | CCN(CC)C(=O)/C(C#N)=C/c1cc(O)c(O)c([N+](=O)[O-])c1 | JRURYQJSLYLRLN-BJMVGYQFSA-N | No-Effect | inactive |
| entecavir | BRD-K08206212 | C=C1[C@@H](n2cnc3c(=O)nc(N)[nH]c32)C[C@H](O)[C@H]1CO | QDGZDCVAUDNJFG-FXQIFTODSA-N | No-Effect | inactive |
| entinostat | BRD-K77908580 | Nc1ccccc1NC(=O)c1ccc(CNC(=O)OCc2cccnc2)cc1 | INVTYAOGFAGBOE-UHFFFAOYSA-N | No-Effect | inactive |
| entrectinib | BRD-K39009484 | CN1CCN(c2ccc(C(=O)Nc3[nH]nc4ccc(Cc5cc(F)cc(F)c5)cc34)c(NC3CCOCC3)c2)CC1 | HAYYBYPASCDWEQ-UHFFFAOYSA-N | No-Effect | inactive |
| enzalutamide | BRD-K56851771 | CNC(=O)c1ccc(N2C(=S)N(c3ccc(C#N)c(C(F)(F)F)c3)C(=O)C2(C)C)cc1F | WXCXUHSOUPDCQV-UHFFFAOYSA-N | No-Effect | inactive |
| enzastaurin | BRD-K79404599 | Cn1cc(C2=C(c3cn(C4CCN(Cc5ccccn5)CC4)c4ccccc34)C(=O)NC2=O)c2ccccc21 | AXRCEOKUDYDWLF-UHFFFAOYSA-N | No-Effect | inactive |
| EO-1428 | BRD-K42452249 | Cc1ccccc1C(=O)c1ccc(Nc2ccc(Br)cc2N)cc1Cl | HDCLCHNAEZNGNV-UHFFFAOYSA-N | No-Effect | inactive |
| epacadostat | BRD-K79450420 | NS(=O)(=O)NCCNc1nonc1C(=Nc1ccc(F)c(Br)c1)NO | FBKMWOJEPMPVTQ-UHFFFAOYSA-N | No-Effect | inactive |
| epalrestat | BRD-K98054437 | CC(/C=C1\SC(=S)N(CC(=O)O)C1=O)=C\c1ccccc1 | CHNUOJQWGUIOLD-NFZZJPOKSA-N | No-Effect | inactive |
| eperezolid | BRD-K47554101 | CC(=O)NC[C@H]1CN(c2ccc(N3CCN(C(=O)CO)CC3)c(F)c2)C(=O)O1 | SIMWTRCFFSTNMG-AWEZNQCLSA-N | No-Effect | inactive |
| epetraborole | BRD-K00003130 | NC[C@H]1OB(O)c2c(OCCCO)cccc21 | FXQIIDINBDJDKL-SNVBAGLBSA-N | No-Effect | inactive |
| ephedrine | BRD-K45014108 | CN[C@@H](C)[C@H](O)c1ccccc1 | KWGRBVOPPLSCSI-WPRPVWTQSA-N | No-Effect | inactive |
| ephedrine-(racemic) | BRD-A54236247 | CN[C@@H](C)[C@@H](O)c1ccccc1 | KWGRBVOPPLSCSI-WCBMZHEXSA-N | No-Effect | inactive |
| epicatechin-gallate-(-) | BRD-K50660797 | O=C(O[C@@H]1Cc2c(O)cc(O)cc2O[C@@H]1c1ccc(O)c(O)c1)c1cc(O)c(O)c(O)c1 | LSHVYAFMTMFKBA-TZIWHRDSSA-N | No-Effect | inactive |
| epiestriol | BRD-K00005352 | C[C@]12CC[C@@H]3c4ccc(O)cc4CC[C@H]3[C@@H]1C[C@H](O)[C@@H]2O | PROQIPRRNZUXQM-ZMSHIADSSA-N | No-Effect | inactive |
| epigallocatechin-(-) | BRD-K27529556 | Oc1cc(O)c2c(c1)O[C@H](c1cc(O)c(O)c(O)c1)[C@H](O)C2 | XMOCLSLCDHWDHP-IUODEOHRSA-N | No-Effect | inactive |
| epigallocatechin-gallate-(-) | BRD-K55591206 | O=C(O[C@@H]1Cc2c(O)cc(O)cc2O[C@@H]1c1cc(O)c(O)c(O)c1)c1cc(O)c(O)c(O)c1 | WMBWREPUVVBILR-WIYYLYMNSA-N | No-Effect | inactive |
| epinastine | BRD-A98990573 | NC1=NC[C@@H]2c3ccccc3Cc3ccccc3N12 | WHWZLSFABNNENI-OAHLLOKOSA-N | No-Effect | inactive |
| epinephrine | BRD-K63126190 | CNC[C@H](O)c1ccc(O)c(O)c1 | UCTWMZQNUQWSLP-VIFPVBQESA-N | No-Effect | inactive |
| epirizole | BRD-K39339537 | COc1cc(C)nc(-n2nc(C)cc2OC)n1 | RHAXSHUQNIEUEY-UHFFFAOYSA-N | No-Effect | inactive |
| epirubicin | BRD-K04548931 | COc1cccc2c1C(=O)c1c(O)c3c(c(O)c1C2=O)C[C@@](O)(C(=O)CO)C[C@@H]3O[C@H]1C[C@H](N)[C@@H](O)[C@H](C)O1 | AOJJSUZBOXZQNB-VTZDEGQISA-N | No-Effect | inactive |
| epitiostanol | BRD-K02314383 | C[C@]12CC[C@H]3[C@@H](CC[C@H]4C[C@@H]5S[C@@H]5C[C@@]43C)[C@@H]1CC[C@@H]2O | OBMLHUPNRURLOK-XGRAFVIBSA-N | No-Effect | inactive |
| eplerenone | BRD-K19761926 | COC(=O)[C@@H]1CC2=CC(=O)CC[C@]2(C)[C@@]23O[C@@H]2C[C@@]2(C)[C@@H](CC[C@@]24CCC(=O)O4)[C@H]13 | JUKPWJGBANNWMW-VWBFHTRKSA-N | No-Effect | inactive |
| epomediol | BRD-A59329714 | CC1(C)OC2(C)[C@@H](O)CC1C[C@@H]2O | JSNQSLSBBZFGBM-VHSUDJNHSA-N | No-Effect | inactive |
| epoprostenol | BRD-K64054020 | CCCCC[C@H](O)/C=C/[C@H]1[C@H](O)C[C@@H]2O/C(=C\CCCC(=O)O)C[C@@H]21 | KAQKFAOMNZTLHT-OZUDYXHBSA-N | No-Effect | inactive |
| epothilone-A | BRD-K71823332 | C/C(=C\c1csc(C)n1)[C@@H]1C[C@@H]2O[C@@H]2CCC[C@H](C)[C@H](O)[C@@H](C)C(=O)C(C)(C)[C@@H](O)CC(=O)O1 | HESCAJZNRMSMJG-KKQRBIROSA-N | No-Effect | inactive |
| epothilone-B | BRD-K48935217 | C/C(=C\c1csc(C)n1)[C@@H]1C[C@@H]2O[C@]2(C)CCC[C@H](C)[C@H](O)[C@@H](C)C(=O)C(C)(C)[C@@H](O)CC(=O)O1 | QXRSDHAAWVKZLJ-PVYNADRNSA-N | No-Effect | inactive |
| epothilone-D | BRD-K52762805 | C/C1=C/C[C@@H](/C(C)=C/c2csc(C)n2)OC(=O)C[C@H](O)C(C)(C)C(=O)[C@H](C)[C@@H](O)[C@@H](C)CCC1 | XOZIUKBZLSUILX-GIQCAXHBSA-N | No-Effect | inactive |
| EPPTB | BRD-K42536037 | CCOc1cccc(NC(=O)c2ccc(N3CCCC3)c(C(F)(F)F)c2)c1 | KLFVWQCQUXXLOU-UHFFFAOYSA-N | No-Effect | inactive |
| eprazinone | BRD-A72212290 | CCO[C@@H](CN1CCN(C[C@@H](C)C(=O)c2ccccc2)CC1)c1ccccc1 | BSHWLCACYCVCJE-OFNKIYASSA-N | No-Effect | inactive |
| eprinomectin | BRD-K00003555 | CC[C@H](C)[C@H]1O[C@]2(C=C[C@@H]1C)C[C@@H]1C[C@@H](C/C=C(\C)[C@@H](O[C@H]3C[C@H](OC)[C@@H](O[C@H]4C[C@H](OC)[C@@H](NC(C)=O)[C@H](C)O4)[C@H](C)O3)[C@@H](C)/C=C/C=C3\CO[C@@H]4[C@H](O)C(C)=C[C@@H](C(=O)O1)[C@]34O)O2 | ZKWQQXZUCOBISE-NQMKODOQSA-N | No-Effect | inactive |
| epristeride | BRD-K00003478 | CC(C)(C)NC(=O)[C@H]1CC[C@H]2[C@@H]3CC=C4C=C(C(=O)O)CC[C@]4(C)[C@H]3CC[C@]12C | VAPSMQAHNAZRKC-PQWRYPMOSA-N | No-Effect | inactive |
| eprodisate | BRD-K11463116 | O=S(=O)(O)CCCS(=O)(=O)O | MGNVWUDMMXZUDI-UHFFFAOYSA-N | No-Effect | inactive |
| eprosartan | BRD-K67977190 | CCCCc1ncc(/C=C(\Cc2cccs2)C(=O)O)n1Cc1ccc(C(=O)O)cc1 | OROAFUQRIXKEMV-LDADJPATSA-N | No-Effect | inactive |
| eptifibatide | BRD-K68830460 | N=C(N)NCCCC[C@@H]1NC(=O)CCSSC[C@@H](C(N)=O)NC(=O)[C@@H]2CCCN2C(=O)[C@H](Cc2c[nH]c3ccccc23)NC(=O)[C@H](CC(=O)O)NC(=O)CNC1=O | CZKPOZZJODAYPZ-LROMGURASA-N | No-Effect | inactive |
| EPZ-5676 | BRD-K26531771 | CC(C)N(C[C@H]1O[C@@H](n2cnc3c(N)ncnc32)[C@H](O)[C@@H]1O)[C@H]1C[C@@H](CCc2nc3ccc(C(C)(C)C)cc3[nH]2)C1 | LXFOLMYKSYSZQS-LURJZOHASA-N | No-Effect | inactive |

| | | | | | |
|---|---|---|---|---|---|
| EPZ004777 | BRD-K69070623 | CC(C)N(CCCNC(=O)Nc1ccc(C(C)(C)C)cc1)C[C@H]1O[C@@H](n2ccc3c(N)ncnc32)[C@H](O)[C@@H]1O | WXRGFPHDRFQODR-ICLZECGLSA-N | No-Effect | inactive |
| EPZ005687 | BRD-K57754230 | Cc1cc(C)c(CNC(=O)c2cc(-c3ccc(CN4CCOCC4)cc3)cc3c2cnn3C2CCCC2)c(=O)[nH]1 | ZOIBZSZLMJDVDQ-UHFFFAOYSA-N | No-Effect | inactive |
| EPZ011989 | BRD-K87468701 | CCN(c1cc(C#CCN2CCOCC2)cc(C(=O)NCc2c(C)cc(C)[nH]c2=O)c1C)[C@H]1CC[C@H](N(C)CCOC)CC1 | XQFINGFCBFHOPE-WXUXXXNLSA-N | No-Effect | inactive |
| EPZ015666 | BRD-K70446295 | O=C(NC[C@H](O)CN1CCc2ccccc2C1)c1cc(NC2COC2)ncn1 | ZKXZLIFRWWKZRY-KRWDZBQOSA-N | No-Effect | inactive |
| EPZ020411 | BRD-K33586289 | CNCCN(C)Cc1c[nH]nc1-c1ccc(O[C@H]2C[C@H](OCCC3CCOCC3)C2)cc1 | QMDKVNSQXPVCRD-RQNOJGIXSA-N | No-Effect | inactive |
| equilin | BRD-A88080608 | C[C@@]12CC[C@H]3C(=CCc4cc(O)ccc43)[C@@H]1CCC2=O | WKRLQDKEXYKHJB-UWWQBHOKSA-N | No-Effect | inactive |
| equol | BRD-K54847683 | Oc1ccc([C@H]2COc3cc(O)ccc3C2)cc1 | ADFCQWZHKCXPAJ-GFCCVEGCSA-N | No-Effect | inactive |
| erastin | BRD-A25004090 | CCOc1ccccc1-n1c([C@@H](C)N2CCN(C(=O)COc3ccc(Cl)cc3)CC2)nc2ccccc2c1=O | BKQFRNYHFIQEKN-OAQYLSRUSA-N | No-Effect | inactive |
| eravacycline | BRD-K00003240 | CN(C)[C@@H]1C(=O)[C@@H](C(N)=O)C(=O)[C@@]2(O)C(=O)[C@@H]3C(=O)c4c(O)c(NC(=O)CN5CCCC5)cc(F)c4C[C@H]3C[C@@H]12 | LSZZJAUTRJAFCO-KKCFVTAMSA-N | No-Effect | inactive |
| ercalcitriol | BRD-K84163249 | C=C1/C(=C\C=C2/CCC[C@@]3(C)[C@H]2CC[C@@H]3[C@H](C)/C=C/[C@H](C)C(C)(C)O)C[C@@H](O)C[C@@H]1O | ZGLHBRQAEXKACO-XJRQOBMKSA-N | No-Effect | inactive |
| erdafitinib | BRD-K84868168 | COc1cc(OC)cc(N(CCNC(C)C)c2ccc3ncc(-c4cnn(C)c4)nc3c2)c1 | OLAHOMJCDNXHFI-UHFFFAOYSA-N | No-Effect | inactive |
| erdosteine | BRD-A05515753 | O=C(O)CSCC(=O)N[C@H]1CCSC1=O | QGFORSXNKQLDNO-YFKPBYRVSA-N | No-Effect | inactive |
| ergocalciferol | BRD-K56155962 | C=C1CC[C@H](O)C/C1=C/C=C1\CCC[C@@]2(C)[C@H]1CC[C@@H]2[C@H](C)/C=C/[C@H](C)C(C)C | MECHNRXZTMCUDQ-RKHKHRCZSA-N | No-Effect | inactive |
| ergonovine | BRD-K17311468 | C[C@@H](CO)NC(=O)[C@@H]1C=C2c3cccc4[nH]cc(c34)C[C@H]2N(C)C1 | WVVSZNPYNCNODU-XTQGRXLLSA-N | No-Effect | inactive |
| ergotamine | BRD-K15380626 | CN1C[C@H](C(=O)N[C@]2(C)O[C@@]3(O)[C@@H]4CCCN4C(=O)[C@H](Cc4ccccc4)N3C2=O)C=C2c3cccc4[nH]cc(c34)C[C@H]21 | XCGSFFUVFURLIX-VFGNJEKYSA-N | No-Effect | inactive |
| eribulin | BRD-K00003156 | C=C1C[C@@H]2CC[C@@]34C[C@H]5O[C@H]6[C@@H](O3)[C@H]3O[C@H](CC[C@@H]3O[C@H]6[C@H]5O4)CC(=O)C[C@H]3[C@H](C[C@H]4O[C@@H](CC[C@@H]1O2)C[C@@H](C)C4=C)O[C@H](C[C@H](O)CN)[C@@H]3OC | UFNVPOGXISZXJD-JBQZKEIOSA-N | No-Effect | inactive |
| erlotinib | BRD-K70401845 | C#Cc1cccc(Nc2ncnc3cc(OCCOC)c(OCCOC)cc23)c1 | AAKJLRGGTJKAMG-UHFFFAOYSA-N | No-Effect | inactive |
| ertapenem | BRD-A40472231 | C[C@H]1C(S[C@@H]2CN[C@H](C(=O)Nc3cccc(C(=O)O)c3)C2)=C(C(=O)O)N2C(=O)[C@@H]([C@@H](C)O)[C@@H]12 | JUZNIMUFDBIJCM-FKTXGUMASA-N | No-Effect | inactive |
| erteberel | BRD-K80608265 | Oc1ccc([C@@H]2Oc3ccc(O)cc3[C@@H]3CCC[C@@H]32)cc1 | XIESSJVMWNJCGZ-VKJFTORMSA-N | No-Effect | inactive |
| ertugliflozin | BRD-K00003574 | CCOc1ccc(Cc2cc([C@]34OC[C@](CO)(O3)[C@@H](O)[C@H](O)[C@H]4O)ccc2Cl)cc1 | MCIACXAZCBVDEE-CUUWFGFTSA-N | No-Effect | inactive |
| erythritol | BRD-K79315489 | OC[C@H](O)[C@H](O)CO | UNXHWFMMPAWVPI-ZXZARUISSA-N | No-Effect | inactive |
| erythromycin | BRD-K63550407 | CC[C@H]1OC(=O)[C@H](C)[C@@H](O[C@H]2C[C@@](C)(OC)[C@@H](O)[C@H](C)O2)[C@H](C)[C@@H](O[C@@H]2O[C@H](C)C[C@H](N(C)C)[C@H]2O)[C@](C)(O)C[C@@H](C)C(=O)[C@H](C)[C@@H](O)[C@]1(C)O | ULGZDMOVFRHVEP-RWJQBGPGSA-N | No-Effect | inactive |
| erythromycin-estolate | BRD-K39746403 | CCC(=O)O[C@H]1[C@H](O[C@@H]2[C@@H](C)[C@H](O[C@H]3C[C@@](C)(OC)[C@@H](O)[C@H](C)O3)[C@@H](C)C(=O)O[C@H](CC)[C@@](C)(O)[C@H](O)[C@@H](C)C(=O)[C@H](C)C[C@@]2(C)O)O[C@H](C)C[C@@H]1N(C)C | TYQXKHPOXXXCTP-CSLYCKPJSA-N | No-Effect | inactive |
| erythromycin-ethylsuccinate | BRD-K51677086 | CCOC(=O)CCC(=O)O[C@H]1[C@H](O[C@@H]2[C@@H](C)[C@H](O[C@H]3C[C@@](C)(OC)[C@@H](O)[C@H](C)O3)[C@@H](C)C(=O)O[C@H](CC)[C@@](C)(O)[C@H](O)[C@@H](C)C(=O)[C@H](C)C[C@@]2(C)O)O[C@H](C)C[C@@H]1N(C)C | NSYZCCDSJNWWJL-YXOIYICCSA-N | No-Effect | inactive |
| erythrosine | BRD-K07549832 | O=C1OC2(c3ccccc31)c1cc(I)c(O)c(I)c1Oc1c2cc(I)c(O)c1I | OALHHIHQOFIMEF-UHFFFAOYSA-N | No-Effect | inactive |
| esaprazole | BRD-K92426617 | O=C(CN1CCNCC1)NC1CCCCC1 | RTFADALJKSFJDZ-UHFFFAOYSA-N | No-Effect | inactive |
| esaxerenone | BRD-K00003374 | Cc1c(C(=O)Nc2ccc(S(C)(=O)=O)cc2)cn(CCO)c1-c1ccccc1C(F)(F)F | NOSNHVJANRODGR-UHFFFAOYSA-N | No-Effect | inactive |
| escin | BRD-K00010936 | CC=C(C)C(=O)O[C@H]1[C@H](OC(C)=O)[C@@]2(CO)[C@@H](CC1(C)C)C1=CC[C@@H]3[C@@]4(C)CC[C@H](O[C@@H]5O[C@H](C(=O)O)[C@@H](O[C@@H]6O[C@H](CO)[C@@H](O)[C@H](O)[C@H]6O)[C@H](O)[C@H]5O[C@@H]5O[C@H](CO)[C@@H](O)[C@H](O)[C@@H]5O)[C@](C)(CO)[C@@H]4CC[C@]3(C)[C@]1(C)C[C@H]2O | AXNVHPCVMSNXNP-FDGZCRRKSA-N | No-Effect | inactive |
| escitalopram | BRD-K70301876 | CN(C)CCC[C@]1(c2ccc(F)cc2)OCc2cc(C#N)ccc21 | WSEQXVZVJXJVFP-FQEVSTJZSA-N | No-Effect | inactive |
| esculin | BRD-K51742987 | O=c1ccc2cc(O[C@@H]3O[C@H](CO)[C@@H](O)[C@H](O)[C@H]3O)c(O)cc2o1 | XHCADAYNFIFUHF-TVKJYDDYSA-N | No-Effect | inactive |
| eseroline-(-) | BRD-K13629382 | CN1CC[C@@]2(C)c3cc(O)ccc3N(C)[C@@H]12 | HKGWQUVGHPDEBZ-OLZOCXBDSA-N | No-Effect | inactive |
| ESI-09 | BRD-K77139516 | CC(C)(C)c1cc(C(=O)/C(C#N)=N/Nc2cccc(Cl)c2)no1 | DXEATJQGQHDURZ-DEDYPNTBSA-N | No-Effect | inactive |
| eslicarbazepine-acetate | BRD-K61237927 | CC(=O)O[C@H]1Cc2ccccc2N(C(N)=O)c2ccccc21 | QIALRBLEEWJACW-INIZCTEOSA-N | No-Effect | inactive |
| esmolol | BRD-K01825842 | COC(=O)CCc1ccc(OC[C@@H](O)CNC(C)C)cc1 | AQNDDEOPVVGCPG-AWEZNQCLSA-N | No-Effect | inactive |
| esonarimod | BRD-K00003282 | CC(=O)SC[C@H](CC(=O)c1ccc(C)cc1)C(=O)O | YRSSFEUQNAXQMX-LBPRGKRZSA-N | No-Effect | inactive |
| estetrol | BRD-K75445017 | C[C@]12CC[C@@H]3c4ccc(O)cc4CC[C@H]3[C@@H]1[C@@H](O)[C@@H](O)[C@@H]2O | AJIPIJNNOJSSQC-NYLIRDPKSA-N | No-Effect | inactive |
| estradiol | BRD-K18910433 | C[C@]12CC[C@@H]3c4ccc(O)cc4CC[C@H]3[C@@H]1CC[C@@H]2O | VOXZDWNPVJITMN-ZBRFXRBCSA-N | No-Effect | inactive |
| estradiol-3-O-sulfamate | BRD-K00004741 | C[C@]12CC[C@@H]3c4ccc(OS(N)(=O)=O)cc4CC[C@H]3[C@@H]1CC[C@@H]2O | YXYXCSOJKUAPJI-ZBRFXRBCSA-N | No-Effect | inactive |
| estradiol-acetate | BRD-K53319039 | CC(=O)Oc1ccc2c(c1)CC[C@@H]1[C@@H]2CC[C@]2(C)[C@@H](O)CC[C@@H]12 | FHXBMXJMKMWVRG-SLHNCBLASA-N | No-Effect | inactive |
| estradiol-cypionate | BRD-K64984393 | C[C@]12CC[C@@H]3c4ccc(O)cc4CC[C@H]3[C@@H]1CC[C@@H]2OC(=O)CCC1CCCC1 | UOACKFBJUYNSLK-XRKIENNPSA-N | No-Effect | inactive |
| estradiol-valerate | BRD-K66766661 | CCCCC(=O)O[C@H]1CC[C@H]2[C@@H]3CCc4cc(O)ccc4[C@H]3CC[C@]12C | RSEPBGGWRJCQGY-RBRWEJTLSA-N | No-Effect | inactive |
| estramustine | BRD-A70858459 | C[C@]12CC[C@@H]3c4ccc(OC(=O)N(CCCl)CCCl)cc4CC[C@H]3[C@@H]1CC[C@@H]2O | FRPJXPJMRWBBIH-RBRWEJTLSA-N | No-Effect | inactive |
| estramustine-phosphate | BRD-K00077635 | C[C@]12CC[C@@H]3c4ccc(OC(=O)N(CCCl)CCCl)cc4CC[C@H]3[C@@H]1CC[C@@H]2OP(=O)(O)O | ADFOJJHRTBFFOF-RBRWEJTLSA-N | No-Effect | inactive |
| estriol | BRD-K17016787 | C[C@]12CC[C@@H]3c4ccc(O)cc4CC[C@H]3[C@@H]1C[C@@H](O)[C@@H]2O | PROQIPRRNZUXQM-ZXXIGWHRSA-N | No-Effect | inactive |
| estropipate | BRD-K51095933 | C[C@]12CC[C@@H]3c4ccc(OS(=O)(=O)O)cc4CC[C@H]3[C@@H]1CCC2=O | JKKFKPJIXZFSSB-CBZIJGRNSA-N | No-Effect | inactive |
| etacrynic-acid | BRD-K63630713 | C=C(CC)C(=O)c1ccc(OCC(=O)O)c(Cl)c1Cl | AVOLMBLBETYQHX-UHFFFAOYSA-N | No-Effect | inactive |
| etamsylate | BRD-K71055333 | O=S(=O)(O)c1cc(O)ccc1O | IKQCSJBQLWJEPU-UHFFFAOYSA-N | No-Effect | inactive |

| | | | | | |
|---|---|---|---|---|---|
| etanidazole | BRD-K96042922 | O=C(Cn1ccnc1[N+](=O)[O-])NCCO | WCDWBPCFGJXFJZ-UHFFFAOYSA-N | No-Effect | inactive |
| etazolate | BRD-K64755930 | CCOC(=O)c1cnc2c(cnn2CC)c1NN=C(C)C | OPQRBXUBWHDHPQ-UHFFFAOYSA-N | No-Effect | inactive |
| ETC-159 | BRD-K54264768 | Cn1c(=O)c2c(ncn2CC(=O)Nc2ccc(-c3ccccc3)nn2)n(C)c1=O | QTRXIFVSTWXRJJ-UHFFFAOYSA-N | No-Effect | inactive |
| ETC-206 | BRD-K00003115 | N#Cc1ccc(-c2cnc3ccc(-c4ccc(C(=O)N5CCOCC5)cc4)cn23)cc1 | FWRFPHJSGLYXTD-UHFFFAOYSA-N | No-Effect | inactive |
| etebenecid | BRD-K24443618 | CCN(CC)S(=O)(=O)c1ccc(C(=O)O)cc1 | UACOQEQOBAQRDQ-UHFFFAOYSA-N | No-Effect | inactive |
| eterobarb | BRD-K64834129 | CCC1(c2ccccc2)C(=O)N(COC)C(=O)N(COC)C1=O | DACOQFZGGLCXMA-UHFFFAOYSA-N | No-Effect | inactive |
| ethacizin | BRD-K10667240 | CCOC(=O)Nc1ccc2c(c1)N(C(=O)CCN(CC)CC)c1ccccc1S2 | PQXGNJKJMFUPPM-UHFFFAOYSA-N | No-Effect | inactive |
| ethambutol | BRD-K93231391 | CC[C@@H](CO)NCCN[C@@H](CC)CO | AEUTYOVWOVBAKS-UWVGGRQHSA-N | No-Effect | inactive |
| ethamivan | BRD-K38055836 | CCN(CC)C(=O)c1ccc(O)c(OC)c1 | BQJODPIMMWWMFC-UHFFFAOYSA-N | No-Effect | inactive |
| ethanolamine-oleate | BRD-M75061779 | CCCCCCCC/C=C\CCCCCCCC(=O)O.NCCO | KGWDUNBJIMUFAP-KVVVOXFISA-N | No-Effect | inactive |
| ethaverine | BRD-K36756879 | CCOc1ccc(Cc2nccc3cc(OCC)c(OCC)cc23)cc1OCC | ZOWYFYXTIWQBEP-UHFFFAOYSA-N | No-Effect | inactive |
| ethenzamide | BRD-K88308881 | CCOc1ccccc1C(N)=O | SBNKFTQSBPKMBZ-UHFFFAOYSA-N | No-Effect | inactive |
| ethinyl-estradiol | BRD-K48195008 | C#C[C@]1(O)CC[C@H]2[C@@H]3CCc4cc(O)ccc4[C@H]3CC[C@@]21C | BFPYWIDHMRZLRN-SLHNCBLASA-N | No-Effect | inactive |
| ethopabate | BRD-K02475039 | CCOc1cc(NC(C)=O)ccc1C(=O)OC | GOVWOKSKFSBNGD-UHFFFAOYSA-N | No-Effect | inactive |
| ethosuximide | BRD-A99633051 | CC[C@@]1(C)CC(=O)NC1=O | HAPOVYFOVVWLRS-ZETCQYMHSA-N | No-Effect | inactive |
| ethoxyquin | BRD-K56792340 | CCOc1ccc2c(c1)C(C)=CC(C)(C)N2 | DECIPOUIJURFOJ-UHFFFAOYSA-N | No-Effect | inactive |
| ethyl-paraben | BRD-K02464583 | CCOC(=O)c1ccc(O)cc1 | NUVBSKCKDOMJSU-UHFFFAOYSA-N | No-Effect | inactive |
| ethyl-pyruvate | BRD-K47738867 | C=C(O)C(=O)OCC | PSYUBHDPOIZRST-UHFFFAOYSA-N | No-Effect | inactive |
| ethyl-vanillin | BRD-K82104468 | CCOc1cc(C=O)ccc1O | CBOQJANXLMLOSS-UHFFFAOYSA-N | No-Effect | inactive |
| ethylenediaminetetraacetic-acid | BRD-K82603084 | O=C(O)CN(CCN(CC(=O)O)CC(=O)O)CC(=O)O | KCXVZYZYPLLWCC-UHFFFAOYSA-N | No-Effect | inactive |
| ethylnorepinephrine | BRD-A68509429 | CC[C@@H](N)[C@@H](O)c1ccc(O)c(O)c1 | LENNRXOJHWNHSD-XCBNKYQSSA-N | No-Effect | inactive |
| ethynodiol-diacetate | BRD-K46331928 | C#C[C@]1(OC(C)=O)CC[C@H]2[C@@H]3CCC4=C[C@@H](OC(C)=O)CC[C@@H]4[C@H]3CC[C@@]21C | ONKUMRGIYFNPJW-KIEAKMPYSA-N | No-Effect | inactive |
| ethynylcytidine | BRD-K00003207 | C#C[C@@]1(O)[C@@H](CO)O[C@@H](n2ccc(N)nc2=O)[C@@H]1O | JFIWEPHGRUDAJN-DYUFWOLASA-N | No-Effect | inactive |
| etidronic-acid | BRD-K37949327 | CC(O)(P(=O)(O)O)P(=O)(O)O | DBVJJBKOTRCVKF-UHFFFAOYSA-N | No-Effect | inactive |
| etifenin | BRD-K63979671 | CCc1cccc(CC)c1NC(=O)CN(CC(=O)O)CC(=O)O | WNIDXAKKFOKNEF-UHFFFAOYSA-N | No-Effect | inactive |
| etilefrine | BRD-A09925278 | CCNCC(O)c1cccc(O)c1 | SQVIAVUSQAWMKL-UHFFFAOYSA-N | No-Effect | inactive |
| etilevodopa | BRD-K90982969 | CCOC(=O)[C@@H](N)Cc1ccc(O)c(O)c1 | NULMGOSOSZBEQL-QMMMGPOBSA-N | No-Effect | inactive |
| etizolam | BRD-K68232413 | CCc1cc2c(s1)-n1c(C)nnc1CN=C2c1ccccc1Cl | VMZUTJCNQWMAGF-UHFFFAOYSA-N | No-Effect | inactive |
| etodolac | BRD-A74667430 | CCc1cccc2c3c([nH]c12)[C@](CC)(CC(=O)O)OCC3 | NNYBQONXHNTVIJ-KRWDZBQOSA-N | No-Effect | inactive |
| etofenamate | BRD-K73541271 | O=C(OCCOCCO)c1ccccc1Nc1cccc(C(F)(F)F)c1 | XILVEPYQJIOVNB-UHFFFAOYSA-N | No-Effect | inactive |
| etofibrate | BRD-K49338325 | CC(C)(Oc1ccc(Cl)cc1)C(=O)OCCOC(=O)c1cccnc1 | XXRVYAFBUDSLJX-UHFFFAOYSA-N | No-Effect | inactive |
| etofylline | BRD-K83064458 | Cn1c(=O)c2c(ncn2CCO)n(C)c1=O | NWPRCRWQMGIBOT-UHFFFAOYSA-N | No-Effect | inactive |
| etofylline-clofibrate | BRD-K49840922 | Cn1c(=O)c2c(ncn2CCOC(=O)C(C)(C)Oc2ccc(Cl)cc2)n(C)c1=O | KYAKGJDISSNVPZ-UHFFFAOYSA-N | No-Effect | inactive |
| etomidate | BRD-K55055802 | CCOC(=O)c1cncn1[C@H](C)c1ccccc1 | NPUKDXXFDDZOKR-LLVKDONJSA-N | No-Effect | inactive |
| etomoxir | BRD-K32405725 | CCOC(=O)[C@@]1(CCCCCCOc2ccc(Cl)cc2)CO1 | DZLOHEOHWICNIL-QGZVFWFLSA-N | No-Effect | inactive |
| etonogestrel | BRD-K00003655 | C#C[C@]1(O)CC[C@H]2[C@@H]3CCC4=CC(=O)CC[C@@H]4[C@H]3C(=C)C[C@@]21CC | GCKFUYQCUCGESZ-BPIQYHPVSA-N | No-Effect | inactive |
| etoposide | BRD-K37798499 | COc1cc([C@@H]2c3cc4c(cc3[C@@H](O[C@@H]3O[C@@H]5CO[C@@H](C)O[C@H]5[C@H](O)[C@H]3O)[C@H]3COC(=O)[C@H]23)OCO4)cc(OC)c1O | VJJPUSNTGOMMGY-MRVIYFEKSA-N | No-Effect | inactive |
| etoricoxib | BRD-K54770957 | Cc1ccc(-c2ncc(Cl)cc2-c2ccc(S(C)(=O)=O)cc2)cn1 | MNJVRJDLRVPLFE-UHFFFAOYSA-N | No-Effect | inactive |
| etosalamide | BRD-K60914966 | CCOCCOc1ccccc1C(N)=O | XBNRCMFFZFCWMH-UHFFFAOYSA-N | No-Effect | inactive |
| etoxybamide | BRD-K51364817 | O=C(CCCO)NCCO | MFHPPMMWHSHDSI-UHFFFAOYSA-N | No-Effect | inactive |
| etrabamine | BRD-K00004658 | CN[C@H]1CCc2ncsc2C1 | YDSVAKPJAOSZJA-LURJTMIESA-N | No-Effect | inactive |
| etrasimod | BRD-K00003139 | O=C(O)C[C@H]1CCc2c1[nH]c1ccc(OCc3ccc(C4CCCC4)c(C(F)(F)F)c3)cc21 | MVGWUTBTXDYMND-QGZVFWFLSA-N | No-Effect | inactive |
| etravirine | BRD-K74717603 | Cc1cc(C#N)cc(C)c1Oc1nc(Nc2ccc(C#N)cc2)nc(N)c1Br | PYGWGZALEOIKDF-UHFFFAOYSA-N | No-Effect | inactive |
| etretinate | BRD-K49076555 | CCOC(=O)/C=C(C)/C=C/C=C(C)/C=C/c1c(C)cc(OC)c(C)c1C | HQMNCQVAMBCHCO-DJRRULDNSA-N | No-Effect | inactive |
| eucalyptol | BRD-K29996876 | CC12CCC(CC1)C(C)(C)O2 | WEEGYLXZBRQIMU-UHFFFAOYSA-N | No-Effect | inactive |
| eucatropine | BRD-A37942872 | C[C@H]1C[C@H](OC(=O)[C@@H](O)c2ccccc2)CC(C)(C)N1C | QSAVEGSLJISCDF-QEJZJMRPSA-N | No-Effect | inactive |
| eugenol | BRD-K32977963 | C=CCc1ccc(O)c(OC)c1 | RRAFCDWBNXTKKO-UHFFFAOYSA-N | No-Effect | inactive |
| EUK-134 | BRD-K19741031 | COc1cccc2c1O[Mn]13(Cl)Oc4c(cccc4OC)C=[N+]1CC[N+]3=C2 | QCAWBJDECIOVBU-UHFFFAOYSA-K | No-Effect | inactive |
| eupatilin | BRD-K47031502 | COc1ccc(-c2cc(=O)c3c(O)c(OC)c(O)cc3o2)cc1OC | DRRWBCNQOKKKOL-UHFFFAOYSA-N | No-Effect | inactive |
| evacetrapib | BRD-K16551357 | Cc1cc(C)c2c(c1)[C@@H](N(Cc1cc(C(F)(F)F)cc(C(F)(F)F)c1)c1nnn(C)n1)CCCN2C[C@H]1CC[C@H](C(=O)O)CC1 | IHIUGIVXARLYHP-YBXDKENTSA-N | No-Effect | inactive |
| evans-blue | BRD-K24548560 | Cc1cc(-c2ccc(NN=C3C=Cc4c(S(=O)(=O)O)cc(S(=O)(=O)O)c(N)c4C3=O)c(C)c2)ccc1NN=C1C=Cc2c(S(=O)(=O)O)cc(S(=O)(=O)O)c(N)c2C1=O | LIRUWPZKVZZTHC-UHFFFAOYSA-N | No-Effect | inactive |
| evatanepag | BRD-K11452913 | CC(C)(C)c1ccc(CN(Cc2cccc(OCC(=O)O)c2)S(=O)(=O)c2cccnc2)cc1 | WOHRHWDYFNWPNG-UHFFFAOYSA-N | No-Effect | inactive |
| evenamide | BRD-K00004688 | CCCCOc1cccc(CCNCC(=O)N(C)C)c1 | GRHBODILPPXVKN-UHFFFAOYSA-N | No-Effect | inactive |
| everolimus | BRD-K13514097 | CO[C@H]1C[C@@H]2CC[C@@H](C)[C@@](O)(O2)C(=O)C(=O)N2CCCC[C@H]2C(=O)O[C@H]([C@H](C)C[C@@H]2CC[C@@H](OCCO)[C@H](OC)C2)CC(=O)[C@H](C)/C=C(\C)[C@@H](O)[C@@H](OC)C(=O)[C@H](C)C[C@H](C)/C=C/C=C/C=C/1C | HKVAMNSJSFKALM-GKUWKFKPSA-N | No-Effect | inactive |
| evobrutinib | BRD-K00003405 | C=CC(=O)N1CCC(CNc2ncnc(N)c2-c2ccc(Oc3ccccc3)cc2)CC1 | QUIWHXQETADMGN-UHFFFAOYSA-N | No-Effect | inactive |
| evocalcet | BRD-K00003250 | C[C@@H](N[C@H]1CCN(c2ccc(CC(=O)O)cc2)C1)c1cccc2ccccc12 | RZNUIYPHQFXBAN-XLIONFOSSA-N | No-Effect | inactive |
| evodiamine | BRD-A68631409 | CN1c2ccccc2C(=O)N2CCc3c([nH]c4ccccc34)[C@H]21 | TXDUTHBFYKGSAH-SFHVURJKSA-N | No-Effect | inactive |
| EVP-6124 | BRD-K71603915 | O=C(N[C@H]1CN2CCC1CC2)c1cc2cccc(Cl)c2s1 | SSRDSYXGYPJKRR-ZDUSSCGKSA-N | No-Effect | inactive |
| EVP4593 | BRD-K68395654 | Nc1ccc2ncnc(NCCc3ccc(Oc4ccccc4)cc3)c2c1 | IBAKVEUZKHOWNG-UHFFFAOYSA-N | No-Effect | inactive |
| EW-7197 | BRD-K74396648 | Cc1cccc(-c2nc(CNc3ccccc3F)[nH]c2-c2ccc3ncnn3c2)n1 | FJCDSQATIJKQKA-UHFFFAOYSA-N | No-Effect | inactive |
| EX-527 | BRD-A73908300 | NC(=O)[C@H]1CCCc2c1[nH]c1ccc(Cl)cc21 | FUZYTVDVLBBXDL-VIFPVBQESA-N | No-Effect | inactive |
| exalamide | BRD-K12609457 | CCCCCCOc1ccccc1C(N)=O | CKSJXOVLXUMMFF-UHFFFAOYSA-N | No-Effect | inactive |
| examorelin | BRD-K35304636 | Cc1[nH]c2ccccc2c1C[C@@H](NC(=O)[C@@H](N)Cc1cnc[nH]1)C(=O)N[C@@H](C)C(=O)N[C@@H](Cc1c[nH]c2ccccc12)C(=O)N[C@H](Cc1ccccc1)C(=O)N[C@@H](CCCCN)C(N)=O | RVWNMGKSNGWLOL-GIIHNPQRSA-N | No-Effect | inactive |
| exatecan-mesylate | BRD-K92150287 | CC[C@@]1(O)C(=O)OCc2c1cc1n(c2=O)Cc2c-1nc1cc(F)c(C)c3c1c2[C@@H](N)CC3 | ZVYVPGLRVWUPMP-FYSMJZIKSA-N | No-Effect | inactive |
| exemestane | BRD-K33425534 | C=C1C[C@@H]2[C@H](CC[C@]3(C)C(=O)CC[C@@H]23)[C@@]2(C)C=CC(=O)C=C12 | BFYIZQONLCFLEV-DAELLWKTSA-N | No-Effect | inactive |

| | | | | | |
|---|---|---|---|---|---|
| exenatide | BRD-K00003559 | CC[C@H](C)[C@H](NC(=O)[C@H](Cc1ccccc1)NC(=O)[C@H](CC(C)C)NC(=O)[C@H](CCCNC(=N)N)NC(=O)[C@@H](NC(=O)[C@H](C)NC(=O)[C@H](CCC(=O)O)NC(=O)[C@H](CCC(=O)O)NC(=O)[C@H](CCC(=O)O)NC(=O)[C@H](CCSC)NC(=O)[C@H](CCC(N)=O)NC(=O)[C@H](CCCCN)NC(=O)[C@H](CO)NC(=O)[C@H](CC(C)C)NC(=O)[C@H](CC(=O)O)NC(=O)[C@H](CO)NC(=O)[C@@H](NC(=O)[C@H](Cc1ccccc1)NC(=O)[C@@H](NC(=O)CNC(=O)[C@H](CCC(=O)O)NC(=O)CNC(=O)[C@@H](N)Cc1c[nH]cn1)[C@@H](C)O)[C@@H](C)O)C(C)C)C(=O)N[C@@H](CCC(=O)O)C(=O)N[C@@H](Cc1c[nH]c2ccccc12)C(=O)N[C@@H](CC(C)C)C(=O)N[C@@H](CCCCN)C(=O)N[C@@H](CC(N)=O)C(=O)NCC(=O)NCC(=O)N1CCC[C@H]1C(=O)N[C@@H](CO)C(=O)N[C@@H](CO)C(=O)NCC(=O)N[C@@H](C)C(=O)N1CCC[C@H]1C(=O)N1CCC[C@H]1C(=O)N1CCC[C@H]1C(=O)N[C@@H](CO)C(N)=O | HTQBXNHDCUEHJF-XWLPCZSASA-N | No-Effect | inactive |
| exherin | BRD-K81795818 | CC(=O)N[C@H]1CSSC[C@@H](C(N)=O)NC(=O)[C@H](C(C)C)NC(=O)[C@H](C)NC(=O)[C@H](Cc2cnc[nH]2)NC1=O | FQVLRGLGWNWPSS-BXBUPLCLSA-N | No-Effect | inactive |
| exifone | BRD-K65716359 | O=C(c1cc(O)c(O)c(O)c1)c1ccc(O)c(O)c1O | XEDWWPGWIXPVRQ-UHFFFAOYSA-N | No-Effect | inactive |
| exisulind | BRD-K37708699 | CC1=C(CC(=O)O)c2cc(F)ccc2/C1=C\c1ccc(S(C)(=O)=O)cc1 | MVGSNCBCUWPVDA-MFOYZWKCSA-N | No-Effect | inactive |
| EXO-1 | BRD-K78633253 | COC(=O)c1ccccc1NC(=O)c1ccc(F)cc1 | KIAPWMKFHIKQOZ-UHFFFAOYSA-N | No-Effect | inactive |
| exo-IWR-1 | BRD-K96155658 | O=C(Nc1cccc2cccnc12)c1ccc(N2C(=O)[C@H]3[C@@H]4C=C[C@@H](C4)[C@H]3C2=O)cc1 | ZGSXEXBYLJIOGF-BTYSMDAFSA-N | No-Effect | inactive |
| ezatiostat | BRD-K98963219 | CCOC(=O)[C@@H](N)CCC(=O)N[C@@H](CSCc1ccccc1)C(=O)N[C@@H](C(=O)OCC)c1ccccc1 | GWEJFLVSOGNLSS-WPFOTENUSA-N | No-Effect | inactive |
| ezetimibe | BRD-K42260897 | O=C1[C@H](CC[C@H](O)c2ccc(F)cc2)[C@@H](c2ccc(O)cc2)N1c1ccc(F)cc1 | OLNTVTPDXPETLC-XPWALMASSA-N | No-Effect | inactive |
| ezutromid | BRD-K25570267 | CCN(CC)c1ccc(-n2nc3cc(C)c(N)cc3n2)cc1 | WRIPKYOZKAJHRL-UHFFFAOYSA-N | No-Effect | inactive |
| F-11440 | BRD-K87948281 | Cn1c(=O)cnn(CCCCN2CCN(c3ncccn3)CC2)c1=O | NMYAHEULKSYAPP-UHFFFAOYSA-N | No-Effect | inactive |
| F-16915 | BRD-K00003299 | CC/C=C\C/C=C\C/C=C\C/C=C\C/C=C\C/C=C\CCC(=O)OCc1cccnc1 | BUWVEXMRBAWBPJ-KUBAVDMBSA-N | No-Effect | inactive |
| F351 | BRD-K41267922 | Cc1ccc(=O)n(-c2ccc(O)cc2)c1 | NETTXQJYJRFTFS-UHFFFAOYSA-N | No-Effect | inactive |
| fadrozole | BRD-A25619068 | N#Cc1ccc([C@H]2CCCc3cncn32)cc1 | CLPFFLWZZBQMAO-CQSZACIVSA-N | No-Effect | inactive |
| fagomine | BRD-K61536264 | OC[C@H]1NCC[C@@H](O)[C@@H]1O | YZNNBIPIQWYLDM-HSUXUTPPSA-N | No-Effect | inactive |
| famciclovir | BRD-K45033733 | CC(=O)OCC(CCn1cnc2cnc(N)nc21)COC(C)=O | GGXKWVWZWMLJEH-UHFFFAOYSA-N | No-Effect | inactive |
| famotidine | BRD-K00673382 | N=C(N)Nc1nc(CSCC/C(N)=N\S(N)(=O)=O)cs1 | XUFQPHANEAPEMJ-UHFFFAOYSA-N | No-Effect | inactive |
| fananserin | BRD-K08998509 | O=S1(=O)c2cccc3cccc(c23)N1CCCN1CCN(c2ccc(F)cc2)CC1 | VGIGHGMPMUCLIQ-UHFFFAOYSA-N | No-Effect | inactive |
| fanetizole | BRD-K60798049 | c1ccc(CCNc2nc(-c3ccccc3)cs2)cc1 | WEEYMMXMBFJUAI-UHFFFAOYSA-N | No-Effect | inactive |
| farampator | BRD-K00003507 | O=C(c1ccc2nonc2c1)N1CCCCC1 | XFVRBYKKGGDPAJ-UHFFFAOYSA-N | No-Effect | inactive |
| farnesyl-thiosalicylic-acid-amide | BRD-K14590126 | CC(C)=CCC/C(C)=C/CC/C(C)=C/CSc1ccccc1C(N)=O | GZTMFRUGZMZCRD-CFBAGHHKSA-N | No-Effect | inactive |
| faropenem | BRD-A49622760 | C[C@@H](O)[C@@H]1C(=O)N2C(C(=O)O)=C([C@H]3CCCO3)S[C@H]12 | HGGAKXAHAYOLDJ-OJKLQORTSA-N | No-Effect | inactive |
| fasoracetam | BRD-K00004571 | O=C1CC[C@H](C(=O)N2CCCCC2)N1 | GOWRRBABHQUJMX-MRVPVSSYSA-N | No-Effect | inactive |
| favipiravir | BRD-K04264130 | NC(=O)c1nc(F)c[nH]c1=O | ZCGNOVWYSGBHAU-UHFFFAOYSA-N | No-Effect | inactive |
| FCE-22250 | BRD-K00004537 | CO[C@H]1/C=C/O[C@@]2(C)Oc3c(C)c(O)c4c(O)c(c(/C=N/N=C/N5CCCCC5)c(O)c4c3C2=O)NC(=O)/C(C)=C\C=C\[C@H](C)[C@H](O)[C@@H](C)[C@@H](O)[C@@H](C)[C@H](OC(C)=O)[C@@H]1C | JEGGQYICWGMXOZ-YNTMCLQCSA-N | No-Effect | inactive |
| fdcyd | BRD-K18157228 | Nc1nc(=O)n([C@H]2C[C@H](O)[C@@H](CO)O2)cc1F | IDYKCXHJJGMAEV-RRKCRQDMSA-N | No-Effect | inactive |
| febuprol | BRD-K00004566 | CCCCOC[C@@H](O)COc1ccccc1 | WBLXZPHICYCDGN-GFCCVEGCSA-N | No-Effect | inactive |
| febuxostat | BRD-K48367671 | Cc1nc(-c2ccc(OCC(C)C)c(C#N)c2)sc1C(=O)O | BQSJTQLCZDPROO-UHFFFAOYSA-N | No-Effect | inactive |
| felbamate | BRD-K99107520 | NC(=O)OCC(COC(N)=O)c1ccccc1 | WKGXYQFOCVYPAC-UHFFFAOYSA-N | No-Effect | inactive |
| felbinac | BRD-K91740057 | O=C(O)Cc1ccc(-c2ccccc2)cc1 | QRZAKQDHEVVFRX-UHFFFAOYSA-N | No-Effect | inactive |
| felbinac-ethyl | BRD-K26404903 | CCOC(=O)Cc1ccc(-c2ccccc2)cc1 | NPPJLSILDPVHCM-UHFFFAOYSA-N | No-Effect | inactive |
| felypressin | BRD-K00003319 | NCCCC[C@H](NC(=O)[C@@H]1CCCN1C(=O)[C@@H]1CSSC[C@H](N)C(=O)N[C@@H](Cc2ccccc2)C(=O)N[C@@H](Cc2ccccc2)C(=O)N[C@@H](CCC(N)=O)C(=O)N[C@@H](CC(N)=O)C(=O)N1)C(=O)NCC(N)=O | SFKQVVDKFKYTNA-DZCXQCEKSA-N | No-Effect | inactive |
| fenaclon | BRD-K08791496 | O=C(CCCl)NCCc1ccccc1 | BITWVVLBVUMFHU-UHFFFAOYSA-N | No-Effect | inactive |
| fenbendazole | BRD-K51318897 | COC(=O)Nc1nc2ccc(Sc3ccccc3)cc2[nH]1 | HDDSHPAODJUKPD-UHFFFAOYSA-N | No-Effect | inactive |
| fenclonine-(+/-) | BRD-K01825694 | N[C@@H](Cc1ccc(Cl)cc1)C(=O)O | NIGWMJHCCYYCSF-QMMMGPOBSA-N | No-Effect | inactive |
| fenebrutinib | BRD-K25290930 | C[C@H]1CN(C2COC2)CCN1c1ccc(Nc2cc(-c3ccnc(N4CCn5c(cc6c5CC(C)(C)C6)C4=O)c3CO)cn(C)c2=O)nc1 | WNEODWDFDXWOLU-QHCPKHFHSA-N | No-Effect | inactive |
| fenigam | BRD-A66920069 | NC[C@@H](CC(=O)O)c1ccccc1 | DAFOCGYVTAOKAJ-SECBINFHSA-N | No-Effect | inactive |
| fenipentol | BRD-A66284731 | CCCC[C@@H](O)c1ccccc1 | OVGORFFCBUIFIA-LLVKDONJSA-N | No-Effect | inactive |
| fenobam | BRD-K41160163 | CN1CC(=O)N=C1NC(=O)Nc1cccc(Cl)c1 | DWPQODZAOSWNHB-UHFFFAOYSA-N | No-Effect | inactive |
| fenofibrate | BRD-K50388907 | CC(C)OC(=O)C(C)(C)Oc1ccc(C(=O)c2ccc(Cl)cc2)cc1 | YMTINGFKWWXKFG-UHFFFAOYSA-N | No-Effect | inactive |
| fenofibric-acid | BRD-K30213273 | CC(C)(Oc1ccc(C(=O)c2ccc(Cl)cc2)cc1)C(=O)O | MQOBSOSZFYZQOK-UHFFFAOYSA-N | No-Effect | inactive |
| fenoprofen | BRD-K01827286 | C[C@@H](C(=O)O)c1cccc(Oc2ccccc2)c1 | RDJGLLICXDHJDY-LLVKDONJSA-N | No-Effect | inactive |
| fenoterol | BRD-K00003668 | C[C@@H](Cc1ccc(O)cc1)NC[C@@H](O)c1cc(O)cc(O)c1 | LSLYOANBFKQKPT-APPDUMDISA-N | No-Effect | inactive |
| fenoverine | BRD-K77564118 | O=C(CN1CCN(Cc2ccc3c(c2)OCO3)CC1)N1c2ccccc2Sc2ccccc21 | UBAJTZKNDCEGKL-UHFFFAOYSA-N | No-Effect | inactive |
| fenoxaprop-p-ethyl | BRD-K44777569 | CCOC(=O)[C@@H](C)Oc1ccc(Oc2nc3ccc(Cl)cc3o2)cc1 | PQKBPHSEKWERTG-LLVKDONJSA-N | No-Effect | inactive |
| fenretinide | BRD-K89085489 | CC1=C(/C=C/C(C)=C/C=C/C(C)=C/C(=O)Nc2ccc(O)cc2)C(C)(C)CCC1 | AKJHMTWEGVYYSE-FXILSDISSA-N | No-Effect | inactive |
| fenspiride | BRD-K26739552 | O=C1NCC2(CCN(CCc3ccccc3)CC2)O1 | FVNFBBAOMBJTST-UHFFFAOYSA-N | No-Effect | inactive |
| fenthion | BRD-K67217586 | COP(=S)(OC)Oc1ccc(SC)c(C)c1 | PNVJTZOFSHSLTO-UHFFFAOYSA-N | No-Effect | inactive |
| fentiazac | BRD-K16077845 | O=C(O)Cc1sc(-c2ccccc2)nc1-c1ccc(Cl)cc1 | JIEKMACRVQTPRC-UHFFFAOYSA-N | No-Effect | inactive |
| fenticonazole | BRD-A33697453 | Clc1ccc([C@@H](Cn2ccnc2)OCc2ccc(Sc3ccccc3)cc2)c(Cl)c1 | ZCJYUTQZBAIHBS-XMMPIXPASA-N | No-Effect | inactive |
| FERb-033 | BRD-K61642990 | O/N=C/c1ccc(-c2ccc(O)c(F)c2)c(Cl)c1O | LRRMQNGSYOUANY-OMCISZLKSA-N | No-Effect | inactive |
| ferrostatin-1 | BRD-K97375133 | CCOC(=O)c1ccc(NC2CCCCC2)c(N)c1 | UJHBVMHOBZBWMX-UHFFFAOYSA-N | No-Effect | inactive |
| ferulic-acid | BRD-K66295376 | COc1cc(/C=C/C(=O)O)ccc1O | KSEBMYQBYZTDHS-HWKANZROSA-N | No-Effect | inactive |
| fesoterodine | BRD-K46731503 | CC(C)C(=O)Oc1ccc(CO)cc1[C@H](CCN(C(C)C)C(C)C)c1ccccc1 | DCCSDBARQIPTGU-HSZRJFAPSA-N | No-Effect | inactive |
| fevipiprant | BRD-K00003232 | Cc1c(CC(=O)O)c2cccnc2n1Cc1ccc(S(C)(=O)=O)cc1C(F)(F)F | GFPPXZDRVCSVNR-UHFFFAOYSA-N | No-Effect | inactive |
| fexaramine | BRD-K08864679 | COC(=O)/C=C/c1cccc(N(Cc2ccc(-c3ccc(N(C)C)cc3)cc2)C(=O)C2CCCCC2)c1 | VLQTUNDJHLEFEQ-KGENOOAVSA-N | No-Effect | inactive |
| fexinidazole | BRD-K37069697 | CSc1ccc(OCc2ncc([N+](=O)[O-])n2C)cc1 | MIWWSGDADVMLTG-UHFFFAOYSA-N | No-Effect | inactive |
| fezolinetant | BRD-K00003287 | Cc1nsc(-c2nnc3n2CCN(C(=O)c2ccc(F)cc2)[C@@H]3C)n1 | PPSNFPASKFYPMN-SECBINFHSA-N | No-Effect | inactive |
| FG-2216 | BRD-K56807945 | O=C(O)CNC(=O)c1nc(Cl)c2ccccc2c1O | OUQVKRKGTAUJQA-UHFFFAOYSA-N | No-Effect | inactive |
| FG-4592 | BRD-K14385366 | Cc1nc(C(=O)NCC(=O)O)c(O)c2ccc(Oc3ccccc3)cc12 | YOZBGTLTNGAVFU-UHFFFAOYSA-N | No-Effect | inactive |
| FG-7142 | BRD-K61951118 | CNC(=O)c1cc2c(cn1)[nH]c1ccccc12 | QMCOPDWHWYSJSA-UHFFFAOYSA-N | No-Effect | inactive |
| FGIN-1-27 | BRD-K09778810 | CCCCCCN(CCCCCC)C(=O)Cc1c(-c2ccc(F)cc2)[nH]c2ccccc12 | VUWXAQFLTSBUDB-UHFFFAOYSA-N | No-Effect | inactive |
| FGIN-1-43 | BRD-K66093087 | CCCCCCN(CCCCCC)C(=O)Cc1c(-c2ccc(Cl)cc2)[nH]c2ccc(Cl)cc12 | XTZUPNNVXIMWAR-UHFFFAOYSA-N | No-Effect | inactive |
| FH-535 | BRD-K48112880 | Cc1cc([N+](=O)[O-])ccc1NS(=O)(=O)c1cc(Cl)ccc1Cl | AXNUEXXEQGQWPA-UHFFFAOYSA-N | No-Effect | inactive |

| | | | | | |
|---|---|---|---|---|---|
| FH1 | BRD-K44777625 | CC(=O)Nc1ccc(Cc2ccc(NC(C)=O)cc2)cc1 | OEXMNSOPAKOPEF-UHFFFAOYSA-N | No-Effect | inactive |
| fiacitabine | BRD-K07160047 | Nc1nc(=O)n([C@@H]2O[C@H](CO)[C@@H](O)[C@@H]2F)cc1I | GIMSJJHKKXRFGV-BYPJNBLXSA-N | No-Effect | inactive |
| fidarestat | BRD-A63752151 | NC(=O)[C@H]1C[C@]2(NC(=O)NC2=O)c2cc(F)ccc2O1 | WAAPEIZFCHNLKK-PELKAZGASA-N | No-Effect | inactive |
| filanesib | BRD-K37379014 | CON(C)C(=O)N1N=C(c2cc(F)ccc2F)S[C@@]1(CCCN)c1ccccc1 | LLXISKGBWFTGEI-FQEVSTJZSA-N | No-Effect | inactive |
| filgotinib | BRD-K16803204 | O=C(Nc1nc2cccc(-c3ccc(CN4CCS(=O)(=O)CC4)cc3)n2n1)C1CC1 | RIJLVEAXPNLDTC-UHFFFAOYSA-N | No-Effect | inactive |
| fimasartan | BRD-K21586122 | CCCCc1nc(C)c(CC(=S)N(C)C)c(=O)n1Cc1ccc(-c2ccccc2-c2nn[nH]n2)cc1 | AMEROGPZOLAFBN-UHFFFAOYSA-N | No-Effect | inactive |
| finafloxacin | BRD-K98384930 | N#Cc1c(N2C[C@@H]3NCCO[C@H]3C2)c(F)cc2c(=O)c(C(=O)O)cn(C3CC3)c12 | FYMHQCNFKNMJAV-HOTGVXAUSA-N | No-Effect | inactive |
| finasteride | BRD-K01095011 | CC(C)(C)NC(=O)[C@H]1CC[C@H]2[C@@H]3CC[C@H]4NC(=O)C=C[C@]4(C)[C@H]3CC[C@]12C | DBEPLOCGEIEOCV-WSBQPABSSA-N | No-Effect | inactive |
| fingolimod | BRD-K88025533 | CCCCCCCCc1ccc(CCC(N)(CO)CO)cc1 | KKGQTZUTZRNORY-UHFFFAOYSA-N | No-Effect | inactive |
| fipexide | BRD-K37688416 | O=C(COc1ccc(Cl)cc1)N1CCN(Cc2ccc3c(c2)OCO3)CC1 | BFUJHVVEMMWLHC-UHFFFAOYSA-N | No-Effect | inactive |
| FIPI | BRD-K61560897 | O=C(NCCN1CCC(n2c(=O)[nH]c3ccccc32)CC1)c1cc2cc(F)ccc2[nH]1 | LHABRXRGDLASIH-UHFFFAOYSA-N | No-Effect | inactive |
| fipronil | BRD-A50675702 | N#Cc1nn(-c2c(Cl)cc(C(F)(F)F)cc2Cl)c(N)c1[S@@](=O)C(F)(F)F | ZOCSXAVNDGMNBV-AREMUKBSSA-N | No-Effect | inactive |
| firategrast | BRD-K17513304 | CCOCc1cc(OC)c(-c2ccc(C[C@H](NC(=O)c3c(F)cccc3F)C(=O)O)cc2)c(OC)c1 | YLFZHHDVRSYTKT-NRFANRHFSA-N | No-Effect | inactive |
| firocoxib | BRD-K09090523 | CC1(C)OC(=O)C(OCC2CC2)=C1c1ccc(S(C)(=O)=O)cc1 | FULAPETWGIGNMT-UHFFFAOYSA-N | No-Effect | inactive |
| fisetin | BRD-K31836715 | O=c1c(O)c(-c2ccc(O)c(O)c2)oc2cc(O)ccc12 | XHEFDIBZLJXQHF-UHFFFAOYSA-N | No-Effect | inactive |
| fisogatinib | BRD-K00003377 | C=CC(=O)N[C@H]1CCOC[C@H]1Nc1ncc2cc(-c3c(Cl)c(OC)cc(OC)c3Cl)ccc2n1 | MGZKYOAQVGSSGC-DLBZAZTESA-N | No-Effect | inactive |
| FIT | BRD-K17896185 | CCC(=O)N(c1ccccc1)C1CCN(CCc2ccc(N=C=S)cc2)CC1 | VDKBIFPJULZUPU-UHFFFAOYSA-N | No-Effect | inactive |
| FK-33-824 | BRD-A01907367 | C[C@H](NC(=O)[C@@H](N)Cc1ccc(O)cc1)C(=O)NCC(=O)N(C)[C@@H](Cc1ccccc1)C(=O)N[C@H](CO)CC[S@@](C)=O | HYZHONGSQNXMPH-YJRDZJJMSA-N | No-Effect | inactive |
| FK-3311 | BRD-K38350380 | CC(=O)c1ccc(NS(C)(=O)=O)c(Oc2ccc(F)cc2F)c1 | DIIYLGZNZGPXRR-UHFFFAOYSA-N | No-Effect | inactive |
| FK-409 | BRD-K00005243 | CC/C(=C\C(=N\O)C(N)=O)[C@@H](C)[N+](=O)[O-] | MZAGXDHQGXUDDX-FGCOBAPNSA-N | No-Effect | inactive |
| FK-888 | BRD-K83508485 | CN(Cc1ccccc1)C(=O)[C@H](Cc1ccc2ccccc2c1)NC(=O)[C@@H]1C[C@@H](O)CN1C(=O)c1cn(C)c2ccccc12 | BFNKQTIJVFGCKQ-PDJGWCFMSA-N | No-Effect | inactive |
| FK-962 | BRD-K80607346 | CC(=O)N1CCC(NC(=O)c2ccc(F)cc2)CC1 | VBHVOHJOTMCSBQ-UHFFFAOYSA-N | No-Effect | inactive |
| flavin-adenine-dinucleotide | BRD-K01666925 | Cc1cc2nc3c(=O)[nH]c(=O)nc-3n(C[C@H](O)[C@H](O)[C@H](O)CO[P@](=O)(O)O[P@](=O)(O)OC[C@H]3O[C@@H](n4cnc5c(N)ncnc54)[C@H](O)[C@@H]3O)c2cc1C | VWWQXMAJTJZDQX-UYBVJOGSSA-N | No-Effect | inactive |
| flavoxate | BRD-K47639036 | Cc1c(-c2ccccc2)oc2c(C(=O)OCCN3CCCCC3)cccc2c1=O | SPIUTQOUKAMGCX-UHFFFAOYSA-N | No-Effect | inactive |
| flecainide | BRD-A09472452 | O=C(NCC1CCCCN1)c1cc(OCC(F)(F)F)ccc1OCC(F)(F)F | DJBNUMBKLMJRSA-UHFFFAOYSA-N | No-Effect | inactive |
| fleroxacin | BRD-K61717546 | CN1CCN(c2c(F)cc3c(=O)c(C(=O)O)cn(CCF)c3c2F)CC1 | XBJBPGROQZJDOJ-UHFFFAOYSA-N | No-Effect | inactive |
| FLI-06 | BRD-A44500574 | CC1=NC2=C(C(=O)CC(C)(C)C2)[C@H](c2ccc([N+](=O)[O-])cc2)C1C(=O)OC1CCCCC1 | DUGBYNJMBYPWGS-FOIFJWKZSA-N | No-Effect | inactive |
| flibanserin | BRD-K33637378 | O=c1[nH]c2ccccc2n1CCN1CCN(c2cccc(C(F)(F)F)c2)CC1 | PPRRDFIXUUSXRA-UHFFFAOYSA-N | No-Effect | inactive |
| flindokalner | BRD-K56740996 | COc1ccc(Cl)cc1[C@]1(F)C(=O)Nc2ccc(C(F)(F)F)cc21 | ULYONBAOIMCNEH-HNNXBMFYSA-N | No-Effect | inactive |
| floctafenine | BRD-A23418262 | O=C(OC[C@H](O)CO)c1ccccc1Nc1ccnc2c(C(F)(F)F)cccc12 | APQPGQGAWABJLN-GFCCVEGCSA-N | No-Effect | inactive |
| flopropione | BRD-K43383936 | CCC(=O)c1c(O)cc(O)cc1O | PTHLEKANMPKYDB-UHFFFAOYSA-N | No-Effect | inactive |
| florfenicol | BRD-K11298197 | CS(=O)(=O)c1ccc([C@@H](O)[C@@H](CF)NC(=O)C(Cl)Cl)cc1 | AYIRNRDRBQJXIF-NXEZZACHSA-N | No-Effect | inactive |
| florifenine | BRD-K00004567 | O=C(OCCN1CCCC1)c1ccccc1Nc1ccnc2cc(C(F)(F)F)ccc12 | DQOYUDHAZMAVLD-UHFFFAOYSA-N | No-Effect | inactive |
| flosequinan | BRD-A79643124 | Cn1cc([S@@](C)=O)c(=O)c2cc(F)cc21 | UYGONJYYUKVHDD-MRXNPFEDSA-N | No-Effect | inactive |
| floxuridine | BRD-K47832606 | O=c1[nH]c(=O)n([C@H]2C[C@H](O)[C@@H](CO)O2)cc1F | ODKNJVUHOIMIIZ-RRKCRQDMSA-N | No-Effect | inactive |
| flucloxacillin | BRD-K27871792 | Cc1onc(-c2c(F)cccc2Cl)c1C(=O)N[C@@H]1C(=O)N2[C@@H](C(=O)O)C(C)(C)S[C@H]12 | UIOFUWFRIANQPC-JKIFEVAISA-N | No-Effect | inactive |
| fluconazole | BRD-K05977355 | OC(Cn1cncn1)(Cn1cncn1)c1ccc(F)cc1F | RFHAOTPXVQNOHP-UHFFFAOYSA-N | No-Effect | inactive |
| flucytosine | BRD-K82143716 | Nc1nc(=O)[nH]cc1F | XRECTZIEBJDKEO-UHFFFAOYSA-N | No-Effect | inactive |
| fludarabine-phosphate | BRD-K71106091 | Nc1nc(F)nc2c1ncn2[C@@H]1O[C@H](COP(=O)(O)O)[C@@H](O)[C@@H]1O | GIUYCYHIANZCFB-FJFJXFQQSA-N | No-Effect | inactive |
| fludroxycortide | BRD-K00824317 | CC1(C)O[C@@H]2C[C@H]3[C@@H]4C[C@H](F)C5=CC(=O)CC[C@]5(C)[C@H]4[C@@H](O)C[C@]3(C)[C@]2(C(=O)CO)O1 | POPFMWWJOGLOIF-XWCQMRHXSA-N | No-Effect | inactive |
| flumazenil | BRD-K98769987 | CCOC(=O)c1ncn2c1CN(C)C(=O)c1cc(F)ccc1-2 | OFBIFZUFASYYRE-UHFFFAOYSA-N | No-Effect | inactive |
| flumequine | BRD-A69777949 | C[C@H]1CCc2cc(F)cc3c(=O)c(C(=O)O)cn1c23 | DPSPPJIUMHPXMA-ZETCQYMHSA-N | No-Effect | inactive |
| flumethasone-pivalate | BRD-K00025599 | C[C@@H]1C[C@H]2[C@@H]3C[C@H](F)C4=CC(=O)C=C[C@]4(C)[C@@]3(F)[C@@H](O)C[C@]2(C)[C@@]1(O)C(=O)COC(=O)C(C)(C)C | JWRMHDSINXPDHB-OJAGFMMFSA-N | No-Effect | inactive |
| flumexadol | BRD-A43518428 | FC(F)(F)c1cccc([C@H]2CNCCO2)c1 | GXPYCYWPUGKQIJ-SNVBAGLBSA-N | No-Effect | inactive |
| flunarizine | BRD-K29582677 | Fc1ccc(C(c2ccc(F)cc2)N2CCN(C/C=C/c3ccccc3)CC2)cc1 | SMANXXCATUTDDT-QPJJXVBHSA-N | No-Effect | inactive |
| flunisolide | BRD-K49577446 | CC1(C)O[C@@H]2C[C@H]3[C@@H]4C[C@H](F)C5=CC(=O)C=C[C@]5(C)[C@H]4[C@@H](O)C[C@]3(C)[C@]2(C(=O)CO)O1 | XSFJVAJPIHIPKU-XWCQMRHXSA-N | No-Effect | inactive |
| flunixin-meglumin | BRD-K99984802 | Cc1c(Nc2ncccc2C(=O)O)cccc1C(F)(F)F | NOOCSNJCXJYGPE-UHFFFAOYSA-N | No-Effect | inactive |
| fluocinolone-acetonide | BRD-K94353609 | CC1(C)O[C@@H]2C[C@H]3[C@@H]4C[C@H](F)C5=CC(=O)C=C[C@]5(C)[C@@]4(F)[C@@H](O)C[C@]3(C)[C@]2(C(=O)CO)O1 | FEBLZLNTKCEFIT-VSXGLTOVSA-N | No-Effect | inactive |
| fluocinonide | BRD-K57886322 | CC(=O)OCC(=O)[C@@]12OC(C)(C)O[C@@H]1C[C@H]1[C@@H]3C[C@H](F)C4=CC(=O)C=C[C@]4(C)[C@@]3(F)[C@@H](O)C[C@@]12C | WJOHZNCJWYWUJD-IUGZLZTKSA-N | No-Effect | inactive |
| fluorescein | BRD-K21913543 | O=C1OC2(c3ccc(O)cc3Oc3cc(O)ccc32)c2ccccc21 | GNBHRKFJIUUOQI-UHFFFAOYSA-N | No-Effect | inactive |
| fluorometholone | BRD-K64862097 | CC(=O)[C@@]1(O)CC[C@H]2[C@@H]3C[C@H](C)C4=CC(=O)C=C[C@]4(C)[C@@]3(F)[C@@H](O)C[C@@]21C | FAOZLTXFLGPHNG-KNAQIMQKSA-N | No-Effect | inactive |
| fluorometholone-acetate | BRD-K02528732 | CC(=O)O[C@]1(C(C)=O)CC[C@H]2[C@@H]3C[C@H](C)C4=CC(=O)C=C[C@]4(C)[C@@]3(F)[C@@H](O)C[C@@]21C | YRFXGQHBPBMFHW-SBTZIJSASA-N | No-Effect | inactive |
| fluoromethylcholine | BRD-K51199656 | C[N+](C)(CF)CCO | PBVFROWIWWGIFK-UHFFFAOYSA-N | No-Effect | inactive |
| fluphenazine-decanoate | BRD-K27898013 | CCCCCCCCCC(=O)OCCN1CCN(CCCN2c3ccccc3Sc3ccc(C(F)(F)F)cc32)CC1 | VIQCGTZFEYDQMR-UHFFFAOYSA-N | No-Effect | inactive |
| flupirtine | BRD-K50720187 | CCOC(=O)Nc1ccc(NCc2ccc(F)cc2)nc1N | JUUFBMODXQKSTD-UHFFFAOYSA-N | No-Effect | inactive |
| fluprazine | BRD-K44091134 | NC(=O)NCCN1CCN(c2cccc(C(F)(F)F)c2)CC1 | CHQKHXJPWDSTPY-UHFFFAOYSA-N | No-Effect | inactive |
| fluralaner | BRD-K00003239 | Cc1cc(C2=NO[C@](c3cc(Cl)cc(Cl)c3)(C(F)(F)F)C2)ccc1C(=O)NCC(=O)NCC(F)(F)F | MLBZKOGAMRTSKP-HXUWFJFHSA-N | No-Effect | inactive |
| flurbiprofen-(+/-) | BRD-A86044036 | C[C@@H](C(=O)O)c1ccc(-c2ccccc2)c(F)c1 | SYTBZMRGLBWNTM-SNVBAGLBSA-N | No-Effect | inactive |
| flurbiprofen-(S)-(+) | BRD-K69907333 | C[C@H](C(=O)O)c1ccc(-c2ccccc2)c(F)c1 | SYTBZMRGLBWNTM-JTQLQIEISA-N | No-Effect | inactive |
| flurbiprofen-axetil | BRD-K00004724 | CC(=O)O[C@@H](C)OC(=O)[C@H](C)c1ccc(-c2ccccc2)c(F)c1 | ALIVXCSEERJYHU-TZMCWYRMSA-N | No-Effect | inactive |
| flurizan | BRD-K75410565 | C[C@@H](C(=O)O)c1ccc(-c2ccccc2)c(F)c1 | SYTBZMRGLBWNTM-SNVBAGLBSA-N | No-Effect | inactive |
| flurofamide | BRD-K67445247 | NP(N)(=O)NC(=O)c1ccc(F)cc1 | QWZFVMCWPLMLTL-UHFFFAOYSA-N | No-Effect | inactive |
| flurothyl | BRD-K14763542 | FC(F)(F)COCC(F)(F)F | KGPPDNUWZNWPSI-UHFFFAOYSA-N | No-Effect | inactive |
| fluroxene | BRD-K56908511 | C=COCC(F)(F)F | DLEGDLSLRSOURQ-UHFFFAOYSA-N | No-Effect | inactive |

| | | | | | |
|---|---|---|---|---|---|
| fluspirilene | BRD-K77947974 | O=C1NCN(c2ccccc2)C12CCN(CCCC(c1ccc(F)cc1)c1ccc(F)cc1)CC2 | QOYHHIBFXOOADH-UHFFFAOYSA-N | No-Effect | inactive |
| flutamide | BRD-K28307902 | CC(C)C(=O)Nc1ccc([N+](=O)[O-])c(C(F)(F)F)c1 | MKXKFYHWDHIYRV-UHFFFAOYSA-N | No-Effect | inactive |
| fluvastatin | BRD-K66296774 | CC(C)n1c(/C=C/[C@@H](O)C[C@@H](O)CC(=O)O)c(-c2ccc(F)cc2)c2ccccc21 | FJLGEFLZQAZZCD-MCBHFWOFSA-N | No-Effect | inactive |
| fluvoxamine | BRD-K53517854 | COCCCC/C(=N\OCCN)c1ccc(C(F)(F)F)cc1 | CJOFXWAVKWHTFT-XSFVSMFZSA-N | No-Effect | inactive |
| fmoc-L-leucine | BRD-K15327298 | CC(C)C[C@H](NC(=O)OCC1c2ccccc2-c2ccccc21)C(=O)O | CBPJQFCAFFNICX-IBGZPJMESA-N | No-Effect | inactive |
| FN-1501 | BRD-K00003096 | CN1CCN(Cc2ccc(NC(=O)c3[nH]ncc3Nc3ncnc4[nH]ccc34)cc2)CC1 | VXLAKHWYGRKCGI-UHFFFAOYSA-N | No-Effect | inactive |
| fomepizole | BRD-K56810756 | Cc1cn[nH]c1 | RIKMMFOAQPJVMX-UHFFFAOYSA-N | No-Effect | inactive |
| fomocaine | BRD-K44942188 | c1ccc(OCc2ccc(CCCN3CCOCC3)cc2)cc1 | CVHGCWVMTZWGAY-UHFFFAOYSA-N | No-Effect | inactive |
| fondaparinux | BRD-K00003513 | CO[C@H]1O[C@H](COS(=O)(=O)O)[C@@H](O[C@@H]2O[C@@H](C(=O)O)[C@@H](O[C@H]3O[C@H](COS(=O)(=O)O)[C@@H](O[C@@H]4O[C@H](C(=O)O)[C@@H](O[C@H]5O[C@H](COS(=O)(=O)O)[C@@H](O)[C@H](O)[C@H]5NS(=O)(=O)O)[C@H](O)[C@H]4O)[C@H](OS(=O)(=O)O)[C@H]3NS(=O)(=O)O)[C@H](O)[C@H]2OS(=O)(=O)O)[C@H](O)[C@H]1NS(=O)(=O)O | KANJSNBRCNMZMV-ABRZTLGGSA-N | No-Effect | inactive |
| foretinib | BRD-K03449891 | COc1cc2c(Oc3ccc(NC(=O)C4(C(=O)Nc5ccc(F)cc5)CC4)cc3F)ccnc2cc1OCCCN1CCOCC1 | CXQHYVUVSFXTMY-UHFFFAOYSA-N | No-Effect | inactive |
| formestane | BRD-A26503646 | C[C@]12CC[C@H]3[C@@H](CC[C@H]4C(=O)C(=O)CC[C@]34C)[C@@H]1CCC2=O | ZMEULISFKSVRLQ-LZBOESDWSA-N | No-Effect | inactive |
| formononetin | BRD-K55567017 | COc1ccc(-c2coc3cc(O)ccc3c2=O)cc1 | HKQYGTCOTHHOMP-UHFFFAOYSA-N | No-Effect | inactive |
| formoterol | BRD-K54006094 | COc1ccc(C[C@H](C)NC[C@@H](O)c2ccc(O)c(NC=O)c2)cc1 | BPZSYCZIITTYBL-ORAYPTAESA-N | No-Effect | inactive |
| forodesine | BRD-K16505263 | O=c1nc[nH]c2c([C@@H]3N[C@H](CO)[C@@H](O)[C@H]3O)c[nH]c12 | IWKXDMQDITUYRK-KUBHLMPHSA-N | No-Effect | inactive |
| forskolin | BRD-K09602097 | C=C[C@@]1(C)CC(=O)[C@@]2(O)[C@](C)(O1)[C@@H](OC(C)=O)[C@@H](O)[C@H]1C(C)(C)CC[C@H](O)[C@@]12C | OHCQJHSOBUTRHG-KGGHGJDLSA-N | No-Effect | inactive |
| fosamprenavir | BRD-K59110290 | CC(C)CN(C[C@@H](OP(=O)(O)O)[C@H](Cc1ccccc1)NC(=O)O[C@H]1CCOC1)S(=O)(=O)c1ccc(N)cc1 | MLBVMOWEQCZNCC-OEMFJLHTSA-N | No-Effect | inactive |
| foscarnet | BRD-K85146014 | O=C(O)P(=O)(O)O | ZJAOAACCNHFJAH-UHFFFAOYSA-N | No-Effect | inactive |
| fosfestrol | BRD-K20801936 | CC/C(=C(/CC)c1ccc(OP(=O)(O)O)cc1)c1ccc(OP(=O)(O)O)cc1 | NLORYLAYLIXTID-ISLYRVAYSA-N | No-Effect | inactive |
| fosfluconazole | BRD-K00003384 | O=P(O)(O)OC(Cn1cncn1)(Cn1cncn1)c1ccc(F)cc1F | GHJWNRRCRIGGIO-UHFFFAOYSA-N | No-Effect | inactive |
| fosfosal | BRD-K28687144 | O=C(O)c1ccccc1OP(=O)(O)O | FFKUDWZICMJVPA-UHFFFAOYSA-N | No-Effect | inactive |
| fosfructose | BRD-K27963513 | O=C(COP(=O)(O)O)[C@@H](O)[C@H](O)[C@H](O)COP(=O)(O)O | XPYBSIWDXQFNMH-UYFOZJQFSA-N | No-Effect | inactive |
| fosinopril | BRD-A51382177 | CCC(=O)O[C@@H](OP(=O)(CCCCc1ccccc1)CC(=O)N1C[C@H](C2CCCCC2)C[C@H]1C(=O)O)C(C)C | BIDNLKIUORFRQP-AJTMIRJTSA-N | No-Effect | inactive |
| fosinoprilat | BRD-A33506612 | O=C(O)[C@@H]1C[C@@H](C2CCCCC2)CN1C(=O)C[P@@](=O)(O)CCCCc1ccccc1 | WOIWWYDXDVSWAZ-RTWAWAEBSA-N | No-Effect | inactive |
| fosphenytoin | BRD-K53809807 | O=C1NC(c2ccccc2)(c2ccccc2)C(=O)N1COP(=O)(O)O | XWLUWCNOOVRFPX-UHFFFAOYSA-N | No-Effect | inactive |
| fostamatinib | BRD-K85178109 | COc1cc(Nc2ncc(F)c(Nc3ccc4c(n3)N(COP(=O)(O)O)C(=O)C(C)(C)O4)n2)cc(OC)c1OC | GKDRMWXFWHEQQT-UHFFFAOYSA-N | No-Effect | inactive |
| fostemsavir | BRD-K00003191 | COc1cnc(-n2cnc(C)n2)c2c1c(C(=O)C(=O)N1CCN(C(=O)c3ccccc3)CC1)cn2COP(=O)(O)O | SWMDAPWAQQTBOG-UHFFFAOYSA-N | No-Effect | inactive |
| fotemustine | BRD-A62893078 | CCOP(=O)(OCC)[C@@H](C)NC(=O)N(CCCl)[NH2+][O-] | UETIXAMLWBFWOE-QMMMGPOBSA-N | No-Effect | inactive |
| foxy-5 | BRD-K13005598 | CSCC[C@H](NC=O)C(=O)N[C@@H](CC(=O)O)C(=O)NCC(=O)N[C@@H](CS)C(=O)N[C@@H](CCC(=O)O)C(=O)N[C@@H](CC(C)C)C(=O)O | WFZPJYYCTSHDJI-ATIWLJMLSA-N | No-Effect | inactive |
| FPA-124 | BRD-M17926088 | Cl[Cu]Cl.NC(=S)N/N=C/c1coc2ccccc2c1=O | HZRXQEVFXXQYFS-VBPDOGLUSA-L | No-Effect | inactive |
| FPL-12495 | BRD-A09368919 | CC(N)(Cc1ccccc1)c1ccccc1 | JINQHBRSXWQJAZ-UHFFFAOYSA-N | No-Effect | inactive |
| FPL-55712 | BRD-A96456596 | CCCc1c(OC[C@@H](O)COc2ccc3c(=O)cc(C(=O)O)oc3c2CCC)ccc(C(C)=O)c1O | LMQBMWHHGVZWMR-MRXNPFEDSA-N | No-Effect | inactive |
| FPL-62064 | BRD-K59875992 | COc1ccc(Nc2ccn(-c3ccccc3)n2)cc1 | WKLGNFJHVJIZPK-UHFFFAOYSA-N | No-Effect | inactive |
| FPL-64176 | BRD-K78959463 | COC(=O)c1c(C)[nH]c(C)c1C(=O)c1ccccc1Cc1ccccc1 | MDMWHKZANMNXTF-UHFFFAOYSA-N | No-Effect | inactive |
| FPS-ZM1 | BRD-K87691185 | O=C(c1ccc(Cl)cc1)N(Cc1ccccc1)C1CCCCC1 | XDFKWGIBQMHSOH-UHFFFAOYSA-N | No-Effect | inactive |
| FR-122047 | BRD-K30990140 | COc1ccc(-c2nc(C(=O)N3CCN(C)CC3)sc2-c2ccc(OC)cc2)cc1 | UBHKJRYGKOSQDJ-UHFFFAOYSA-N | No-Effect | inactive |
| FR-139317 | BRD-K19318328 | CC(C)C[C@H](NC(=O)N1CCCCCC1)C(=O)N[C@H](Cc1cn(C)c2ccccc12)C(=O)N[C@H](Cc1ccccn1)C(=O)O | LIOKMIQQPDDTNO-UPRLRBBYSA-N | No-Effect | inactive |
| FR-180204 | BRD-K68756823 | Nc1[nH]nc2nnc(-c3c(-c4ccccc4)nn4ccccc34)cc12 | XVECMUKVOMUNLE-UHFFFAOYSA-N | No-Effect | inactive |
| frentizole | BRD-K07202345 | COc1ccc2nc(NC(=O)Nc3ccccc3)sc2c1 | JHBWYQRKOUBPCA-UHFFFAOYSA-N | No-Effect | inactive |
| freselestat | BRD-K00004668 | CC(C)[C@@H](NC(=O)Cn1c(-c2ccccc2)ncc(N)c1=O)[C@@H](O)c1nnc(C(C)(C)C)o1 | HMPQTEPEMQZWQH-QZTJIDSGSA-N | No-Effect | inactive |
| fruquintinib | BRD-K70504253 | CNC(=O)c1c(C)oc2cc(Oc3ncnc4cc(OC)c(OC)cc34)ccc12 | BALLNEJQLSTPIO-UHFFFAOYSA-N | No-Effect | inactive |
| FT011 | BRD-K00003378 | C#CCOc1ccc(/C=C/C(=O)Nc2ccccc2C(=O)O)cc1OC | UIWZIDIJCUEOMT-PKNBQFBNSA-N | No-Effect | inactive |
| ftaxilide | BRD-K05724540 | Cc1cccc(C)c1NC(=O)c1ccccc1C(=O)O | LLECMGGNFBKPRH-UHFFFAOYSA-N | No-Effect | inactive |
| ftorafur | BRD-K99383816 | O=c1[nH]c(=O)n([C@H]2CCCO2)cc1F | WFWLQNSHRPWKFK-ZCFIWIBFSA-N | No-Effect | inactive |
| fucoxanthin | BRD-K00003355 | CC(=O)O[C@H]1CC(C)(C)C(=C=C/C(C)=C/C=C/C(C)=C/C=C/C=C(C)/C=C/C=C(\C)C(=O)C[C@]23O[C@@]2(C)C[C@H](O)CC3(C)C)[C@](C)(O)C1 | SJWWTRQNNRNTPU-WEFWQODPSA-N | No-Effect | inactive |
| fudosteine | BRD-K54838207 | N[C@@H](CSCCCO)C(=O)O | KINWYTAUPKOPCQ-YFKPBYRVSA-N | No-Effect | inactive |
| fulvestrant | BRD-A85667082 | C[C@]12CC[C@@H]3c4ccc(O)cc4C[C@@H](CCCCCCCCC[S@@](=O)CCCC(F)(F)C(F)(F)F)[C@H]3[C@@H]1CC[C@@H]2O | VWUXBMIQPBEWFH-CIAKRVSBSA-N | No-Effect | inactive |
| fumagillin | BRD-A99367815 | CO[C@@H]1[C@H](OC(=O)/C=C/C=C/C=C/C=C/C(=O)O)CC[C@]2(CO2)[C@@H]1[C@@]1(C)O[C@@H]1CC=C(C)C | NGGMYCMLYOUNGM-FKSJYSFPSA-N | No-Effect | inactive |
| fumonisin-B1 | BRD-K61915943 | CCCC[C@@H](C)[C@@H](OC(=O)C[C@@H](CC(=O)O)C(=O)O)[C@H](C[C@@H](C)C[C@H](O)CCCC[C@@H](O)C[C@H](O)[C@@H](C)N)OC(=O)C[C@@H](CC(=O)O)C(=O)O | UVBUBMSSQKOIBE-DSLOAKGESA-N | No-Effect | inactive |
| funapide | BRD-K00004702 | O=C1N(Cc2ccc(C(F)(F)F)o2)c2ccccc2[C@]12COc1cc3c(cc12)OCO3 | NEBUOXBYNAHKFV-NRFANRHFSA-N | No-Effect | inactive |
| furaltadone | BRD-K01824979 | O=C1O[C@H](CN2CCOCC2)CN1/N=C/c1ccc([N+](=O)[O-])o1 | YVQVOQKFMFRVGR-LLUUXILJSA-N | No-Effect | inactive |
| furamidine | BRD-K77758223 | N=C(N)c1ccc(-c2ccc(-c3ccc(C(=N)N)cc3)o2)cc1 | ZJHZBDRZEZEDGB-UHFFFAOYSA-N | No-Effect | inactive |
| furazolidone | BRD-K11756522 | O=C1OCCN1/N=C/c1ccc([N+](=O)[O-])o1 | PLHJDBGFXBMTGZ-WEVVVXLNSA-N | No-Effect | inactive |
| furosemide | BRD-K78010432 | NS(=O)(=O)c1cc(C(=O)O)c(NCc2ccco2)cc1Cl | ZZUFCTLCJUWOSV-UHFFFAOYSA-N | No-Effect | inactive |
| fursultiamine | BRD-A71157293 | C/C(=C(/CCO)SSC[C@H]1CCCO1)N(C=O)Cc1cnc(C)nc1N | JTLXCMOFVBXEKD-XYGUXCPDSA-N | No-Effect | inactive |
| furvina | BRD-K00004693 | O=[N+]([O-])/C(Br)=C\c1ccc(Br)o1 | MJPPGVVIDGQOQT-HYXAFXHYSA-N | No-Effect | inactive |
| G-1 | BRD-K62696033 | CC(=O)c1ccc2c(c1)[C@@H]1C=CC[C@@H]1[C@H](c1cc3c(cc1Br)OCO3)N2 | VHSVKVWHYFBIFJ-HKZYLEAXSA-N | No-Effect | inactive |
| G-15 | BRD-K00004290 | Brc1cc2c(cc1[C@H]1Nc3ccccc3[C@H]3C=CC[C@H]31)OCO2 | YOLTZIVRJAPVPH-MVBJNABHSA-N | No-Effect | inactive |
| g-caryophyllene | BRD-K26878113 | C=C1CC/C=C(\C)CC[C@@H]2[C@@H]1CC2(C)C | NPNUFJAVOOONJE-GFUGXAQUSA-N | No-Effect | inactive |
| G007-LK | BRD-K12198961 | CS(=O)(=O)c1ccc(-c2nnc(/C=C/c3nnc(-c4ccc(C#N)cc4)o3)n2-c2ccccc2Cl)nc1 | HIWVLHPKZNBSBE-OUKQBFOZSA-N | No-Effect | inactive |
| GABA-linoleamide | BRD-K46254452 | CCCCC/C=C\C/C=C\CCCCCCCC(=O)NCCCC(=O)O | YGFYZZQGVSUXJE-HZJYTTRNSA-N | No-Effect | inactive |

| | | | | | |
|---|---|---|---|---|---|
| gabapentin | BRD-K62737565 | NCC1(CC(=O)O)CCCCC1 | UGJMXCAKCUNAIE-UHFFFAOYSA-N | No-Effect | inactive |
| gabapentin-enacarbil | BRD-A86665761 | CC(C)C(=O)O[C@@H](C)OC(=O)NCC1(CC(=O)O)CCCCC1 | TZDUHAJSIBHXDL-GFCCVEGCSA-N | No-Effect | inactive |
| gabazine | BRD-K79905821 | COc1ccc(-c2ccc(=N)n(CCCC(=O)O)n2)cc1 | ACVGNKYJVGNLIL-UHFFFAOYSA-N | No-Effect | inactive |
| gaboxadol | BRD-K89104321 | O=c1[nH]oc2c1CCNC2 | ZXRVKCBLGJOCEE-UHFFFAOYSA-N | No-Effect | inactive |
| gadopentetic-acid | BRD-K25106841 | O=C(O)CN1CCN2CCN(CC(=O)O)CC(=O)O[Gd](OC(=O)C2)OC(=O)C1 | IZOOGPBRAOKZFK-UHFFFAOYSA-K | No-Effect | inactive |
| gadoterate-meglumine | BRD-K11099631 | O=C(O)CN1CCN(CC(=O)O)CCN(CC(=O)O)CCN(CC(=O)O)CC1 | WDLRUFUQRNWCPK-UHFFFAOYSA-N | No-Effect | inactive |
| gadoteridol | BRD-A72401848 | C[C@@H](O)CN1CCN2CCN3CCN(CC1)CC(=O)O[Gd](OC(=O)C3)OC(=O)C2 | DPNNNPAKRZOSMO-PFEQFJNWSA-K | No-Effect | inactive |
| GAL-021 | BRD-K00003410 | CCCNc1nc(NCCC)nc(N(C)OC)n1 | FJNLCHNQVJVCPY-UHFFFAOYSA-N | No-Effect | inactive |
| galantamine | BRD-K49481516 | COc1ccc2c3c1O[C@H]1C[C@@H](O)C=C[C@@]31CCN(C)C2 | ASUTZQLVASHGKV-JDFRZJQESA-N | No-Effect | inactive |
| galeterone | BRD-K00004319 | C[C@]12CC[C@H]3[C@@H](CC=C4C[C@@H](O)CC[C@@]43C)[C@@H]1CC=C2n1cnc2ccccc21 | PAFKTGFSEFKSQG-PAASFTFBSA-N | No-Effect | inactive |
| gallamine-triethiodide | BRD-K43106192 | CC[N+](CC)(CC)CCOc1cccc(OCC[N+](CC)(CC)CC)c1OCC[N+](CC)(CC)CC | OZLPUNFFCJDMJD-UHFFFAOYSA-N | No-Effect | inactive |
| gallic-acid | BRD-K77345217 | O=C(O)c1cc(O)c(O)c(O)c1 | LNTHITQWFMADLM-UHFFFAOYSA-N | No-Effect | inactive |
| gallium-triquinolin-8-olate | BRD-K46648676 | c1cnc2c(O[Ga](Oc3cccc4cccnc34)Oc3cccc4cccnc34)cccc2c1 | KWQNQSDKCINQQP-UHFFFAOYSA-K | No-Effect | inactive |
| gallopamil | BRD-A52922642 | COc1ccc(CCN(C)CCCC(C#N)(c2cc(OC)c(OC)c(OC)c2)C(C)C)cc1OC | XQLWNAFCTODIRK-UHFFFAOYSA-N | No-Effect | inactive |
| gambogic-acid | BRD-K08542803 | CC(C)=CCC[C@]1(C)C=Cc2c(O)c3c(c(CC=C(C)C)c2O1)O[C@]12C(=C[C@@H]4C[C@H]1C(C)(C)O[C@@]2(C/C=C(/C)C(=O)O)C4=O)C3=O | GEZHEQNLKAOMCA-RRZNCOCZSA-N | No-Effect | inactive |
| gamma-aminobutyric-acid | BRD-K77245796 | NCCCC(=O)O | BTCSSZJGUNDROE-UHFFFAOYSA-N | No-Effect | inactive |
| gamma-linolenic-acid | BRD-K18059238 | CCCCC/C=C\C/C=C\C/C=C\CCCCC(=O)O | VZCCETWTMQHEPK-QNEBEIHSSA-N | No-Effect | inactive |
| ganaxolone | BRD-K28217197 | CC(=O)[C@H]1CC[C@H]2[C@@H]3CC[C@H]4C[C@](C)(O)CC[C@]4(C)[C@H]3CC[C@]12C | PGTVWKLGGCQMBR-FLBATMFCSA-N | No-Effect | inactive |
| ganoderic-acid-a | BRD-K00003733 | C[C@H](CC(=O)C[C@@H](C)[C@H]1C[C@H](O)[C@@]2(C)C3=C(C(=O)C[C@]12C)[C@@]1(C)CCC(=O)C(C)(C)[C@@H]1C[C@@H]3O)C(=O)O | DYOKDAQBNHPJFD-JNTBEZBXSA-N | No-Effect | inactive |
| GANT-58 | BRD-K64451768 | c1cc(-c2sc(-c3ccncc3)c(-c3ccncc3)c2-c2ccncc2)ccn1 | USWLOKMMUTWFMD-UHFFFAOYSA-N | No-Effect | inactive |
| GANT-61 | BRD-K09485525 | CN(C)c1ccccc1CN1CCCN(Cc2ccccc2N(C)C)C1c1ccncc1 | KVQOGDQTWWCZFX-UHFFFAOYSA-N | No-Effect | inactive |
| garenoxacin | BRD-K77038618 | C[C@H]1NCc2cc(-c3ccc4c(=O)c(C(=O)O)cn(C5CC5)c4c3OC(F)F)ccc21 | NJDRXTDGYFKORP-LLVKDONJSA-N | No-Effect | inactive |
| gastrodin | BRD-K15447284 | OCc1ccc(O[C@@H]2O[C@H](CO)[C@@H](O)[C@H](O)[C@H]2O)cc1 | PUQSUZTXKPLAPR-UJPOAAIJSA-N | No-Effect | inactive |
| gavestinel | BRD-K81981783 | O=C(/C=C/c1c(C(=O)O)[nH]c2cc(Cl)cc(Cl)c12)Nc1ccccc1 | WZBNEZWCNKUOSM-VOTSOKGWSA-N | No-Effect | inactive |
| GBR-12783 | BRD-K92015269 | C(=C/c1ccccc1)\CN1CCN(CCOC(c2ccccc2)c2ccccc2)CC1 | XFBDGHFDKJITGC-JLHYYAGUSA-N | No-Effect | inactive |
| GBR-12935 | BRD-K50135270 | c1ccc(CCCN2CCN(CCOC(c3ccccc3)c3ccccc3)CC2)cc1 | RAQPOZGWANIDQT-UHFFFAOYSA-N | No-Effect | inactive |
| GBR-13069 | BRD-K92577649 | Fc1ccc(C(OCCN2CCN(C/C=C/c3ccccc3)CC2)c2ccc(F)cc2)cc1 | DASHZBBQOARCMQ-QPJJXVBHSA-N | No-Effect | inactive |
| GDC-0068 | BRD-K43002773 | CC(C)NC[C@@H](C(=O)N1CCN(c2ncnc3c2[C@H](C)C[C@H]3O)CC1)c1ccc(Cl)cc1 | GRZXWCHAXNAUHY-NSISKUIASA-N | No-Effect | inactive |
| GDC-0077 | BRD-K00003420 | C[C@H](Nc1ccc2c(c1)OCCn1cc(N3C(=O)OC[C@H]3C(F)F)nc1-2)C(N)=O | SGEUNORSOZVTOL-CABZTGNLSA-N | No-Effect | inactive |
| GDC-0152 | BRD-K68938568 | CN[C@@H](C)C(=O)N[C@H](C(=O)N1CCC[C@H]1C(=O)Nc1snnc1-c1ccccc1)C1CCCCC1 | WZRFLSDVFPIXOV-LRQRDZAKSA-N | No-Effect | inactive |
| GDC-0349 | BRD-K43187018 | CCNC(=O)Nc1ccc(-c2nc3c(c(N4CCOC[C@@H]4C)n2)CCN(C2COC2)C3)cc1 | RGJOJUGRHPQXGF-INIZCTEOSA-N | No-Effect | inactive |
| GDC-0575 | BRD-K00003112 | N[C@@H]1CCCN(c2c(Br)cnc3[nH]cc(NC(=O)C4CC4)c23)C1 | BAZRWWGASYWYGB-SNVBAGLBSA-N | No-Effect | inactive |
| GDC-0623 | BRD-K46908090 | O=C(NOCCO)c1ccc2cncn2c1Nc1ccc(I)cc1F | RFWVETIZUQEJEF-UHFFFAOYSA-N | No-Effect | inactive |
| GDC-0834 | BRD-A30984645 | Cc1c(NC(=O)c2cc3c(s2)CCCC3)cccc1-c1cn(C)c(=O)c(Nc2ccc([C@H]3C(=O)N(C)CCN3C)cc2)n1 | CDOOFZZILLRUQH-LJAQVGFWSA-N | No-Effect | inactive |
| GDC-0879 | BRD-K67578145 | OCCn1cc(-c2ccc3c(c2)CC/C3=N\O)c(-c2ccncc2)n1 | DEZZLWQELQORIU-RELWKKBWSA-N | No-Effect | inactive |
| GDC-0927 | BRD-K00003101 | CC1=C(c2cccc(O)c2)[C@H](c2ccc(OCCN3CC(CF)C3)cc2)Oc2ccc(O)cc21 | KJAAPZIFCQQQKX-NDEPHWFRSA-N | No-Effect | inactive |
| GDC-0994 | BRD-A20830788 | Cn1nccc1Nc1nccc(-c2ccn([C@@H](CO)c3ccc(Cl)c(F)c3)c(=O)c2)n1 | RZUOCXOYPYGSKL-SFHVURJKSA-N | No-Effect | inactive |
| gedunin | BRD-K02703275 | CC(=O)O[C@@H]1C[C@H]2C(C)(C)C(=O)C=C[C@]2(C)[C@H]2CC[C@@]3(C)[C@H](c4ccoc4)OC(=O)[C@H]4O[C@]43[C@]12C | YJXDGWUNRYLINJ-BHAPSIHVSA-N | No-Effect | inactive |
| gefapixant | BRD-K00004589 | COc1cc(C(C)C)c(Oc2cnc(N)nc2N)cc1S(N)(=O)=O | HLWURFKMDLAKOD-UHFFFAOYSA-N | No-Effect | inactive |
| gefitinib | BRD-K64052750 | COc1cc2ncnc(Nc3ccc(F)c(Cl)c3)c2cc1OCCCN1CCOCC1 | XGALLCVXEZPNRQ-UHFFFAOYSA-N | No-Effect | inactive |
| gemcadiol | BRD-K43890836 | CC(C)(CO)CCCCCCC(C)(C)CO | DJKWDKJYFYUEBQ-UHFFFAOYSA-N | No-Effect | inactive |
| gemcitabine-elaidate | BRD-K09456915 | CCCCCCCC/C=C/CCCCCCCC(=O)OC[C@H]1O[C@@H](n2ccc(N)nc2=O)C(F)(F)[C@@H]1O | HESSNRGIEVBPRB-QDDPNBLJSA-N | No-Effect | inactive |
| gemfibrozil | BRD-K11129031 | Cc1ccc(C)c(OCCCC(C)(C)C(=O)O)c1 | HEMJJKBWTPKOJG-UHFFFAOYSA-N | No-Effect | inactive |
| gemifloxacin | BRD-A40787240 | CO/N=C1\CN(c2nc3c(cc2F)c(=O)c(C(=O)O)cn3C2CC2)C[C@@H]1CN | ZRCVYEYHRGVLOC-NOTZLVJDSA-N | No-Effect | inactive |
| gemigliptin | BRD-K00003182 | N[C@@H](CC(=O)N1CCc2c(nc(C(F)(F)F)nc2C(F)(F)F)C1)CN1CC(F)(F)CCC1=O | ZWPRRQZNBDYKLH-VIFPVBQESA-N | No-Effect | inactive |
| genipin | BRD-K28824103 | COC(=O)C1=CO[C@@H](O)[C@@H]2C(CO)=CC[C@H]12 | AZKVWQKMDGGDSV-BCMRRPTOSA-N | No-Effect | inactive |
| geniposide | BRD-K57275767 | COC(=O)C1=CO[C@@H](O[C@@H]2O[C@H](CO)[C@@H](O)[C@H](O)[C@H]2O)[C@@H]2C(CO)=CC[C@H]12 | IBFYXTRXDNAPMM-BVTMAQQCSA-N | No-Effect | inactive |
| gentamycin | BRD-K97891829 | CN[C@H](C)[C@@H]1CC[C@@H](N)[C@@H](O[C@@H]2[C@@H](N)C[C@@H](N)[C@H](O[C@H]3OC[C@](C)(O)[C@H](NC)[C@H]3O)[C@H]2O)O1 | CEAZRRDELHUEMR-CAMVTXANSA-N | No-Effect | inactive |
| gentiopicrin | BRD-K33131085 | C=C[C@@H]1C2=CCOC(=O)C2=CO[C@H]1O[C@@H]1O[C@H](CO)[C@@H](O)[C@H](O)[C@H]1O | DUAGQYUORDTXOR-GPQRQXLASA-N | No-Effect | inactive |
| genz-123346 | BRD-K70693421 | CCCCCCCCC(=O)N[C@H](CN1CCCC1)[C@H](O)c1ccc2c(c1)OCCO2 | JMNXWOFCUJJYEO-HYBUGGRVSA-N | No-Effect | inactive |
| genz-644282 | BRD-K99475619 | CNCCn1c(=O)c2cc(OC)c(OC)cc2c2cnc3cc4c(cc3c21)OCO4 | BAORCAMWLWRZQG-UHFFFAOYSA-N | No-Effect | inactive |
| gepon | BRD-A56754222 | CC(C)[C@@H](NC(=O)[C@@H](NC(=O)[C@@H](CCC(=O)O)NC(=O)[C@H](CCCNC(=N)N)NC(=O)[C@@H](CCCNC(=N)N)NC(=O)[C@@H](CCCCN)NC(=O)[C@@H](CCCCN)NC(=O)[C@@H](CCC(=O)O)NC(=O)[C@H](N)[C@@H](C)O)[C@@H](C)O)C(=O)N[C@H](CCC(=O)O)C(=O)N[C@@H](CCCNC(=N)N)C(=O)N[C@@H](CCC(=O)O)C(=O)N[C@@H](CCCCN)C(=O)N[C@@H](CCC(=O)O)C(=O)O | HTQONPPEZULJKY-MUPFFGQUSA-N | No-Effect | inactive |
| gepotidacin | BRD-K33499285 | O=c1ccc2ncc(=O)n3c2n1C[C@H]3CN1CCC(NCc2cc3c(cn2)OCCC3)CC1 | PZFAZQUREQIODZ-LJQANCHMSA-N | No-Effect | inactive |

| | | | | | |
|---|---|---|---|---|---|
| geranyl-farnesylacetate | BRD-K02780233 | CC(C)=CCC/C(C)=C/CC/C(C)=C/CCC(=O)OC/C=C(\C)CCC=C(C)C | ZPACYDRSPFRDHO-ROBAGEODSA-N | No-Effect | inactive |
| gestodene | BRD-K97197005 | C#C[C@]1(O)C=C[C@H]2[C@@H]3CCC4=CC(=O)CC[C@@H]4[C@H]3CC[C@@]21CC | SIGSPDASOTUPFS-XUDSTZEESA-N | No-Effect | inactive |
| gestrinone | BRD-K97472745 | C#C[C@]1(O)CC[C@H]2[C@@H]3CCC4=CC(=O)CCC4=C3C=C[C@@]21CC | BJJXHLWLUDYTGC-ANULTFPQSA-N | No-Effect | inactive |
| GF109203X | BRD-K31342827 | CN(C)CCCn1cc(C2=C(c3c[nH]c4ccccc34)C(=O)NC2=O)c2ccccc21 | QMGUOJYZJKLOLH-UHFFFAOYSA-N | No-Effect | inactive |
| GGsTop | BRD-A25625670 | COP(=O)(CC[C@@H](N)C(=O)O)Oc1cccc(CC(=O)O)c1 | NTFPDEDRMYYPAC-FAYKFVSFSA-N | No-Effect | inactive |
| gimeracil | BRD-K35687421 | O=c1cc(O)c(Cl)c[nH]1 | ZPLQIPFOCGIIHV-UHFFFAOYSA-N | No-Effect | inactive |
| ginkgolide-B | BRD-K93816012 | C[C@@H]1C(=O)O[C@H]2[C@H](O)[C@@]34[C@H]5C[C@@H](C(C)(C)C)[C@]36[C@@H](OC(=O)[C@@H]6O)O[C@@]4(C(=O)O5)[C@@]12O | SQOJOAFXDQDRGF-MMQTXUMRSA-N | No-Effect | inactive |
| ginsenoside-C-K | BRD-K01942833 | CC(C)=CCC[C@](C)(O[C@@H]1O[C@H](CO)[C@@H](O)[C@H](O)[C@H]1O)[C@H]1CC[C@]2(C)[C@@H]1[C@H](O)C[C@@H]1[C@@]3(C)CC[C@H](O)C(C)(C)[C@@H]3CC[C@]12C | FVIZARNDLVOMSU-IRFFNABBSA-N | No-Effect | inactive |
| ginsenoside-RD | BRD-A84810646 | CC(C)=CCC[C@@](C)(O[C@@H]1O[C@H](CO)[C@@H](O)[C@H](O)[C@H]1O)[C@H]1CC[C@]2(C)[C@@H]1[C@H](O)C[C@@H]1[C@@]3(C)CC[C@H](O[C@@H]4O[C@H](CO)[C@@H](O)[C@H](O)[C@H]4O[C@@H]4O[C@H](CO)[C@@H](O)[C@H](O)[C@H]4O)C(C)(C)[C@@H]3CC[C@]12C | RLDVZILFNVRJTL-ZMHPOSLNSA-N | No-Effect | inactive |
| ginsenoside-RE | BRD-K00003326 | CC(C)=CCC[C@](C)(O[C@@H]1O[C@H](CO)[C@@H](O)[C@H](O)[C@H]1O)[C@H]1CC[C@]2(C)[C@@H]1[C@H](O)C[C@@H]1[C@@]3(C)CC[C@H](O)C(C)(C)[C@@H]3[C@@H](O[C@@H]3O[C@H](CO)[C@@H](O)[C@H](O)[C@H]3O[C@@H]3O[C@@H](C)[C@H](O)[C@@H](O)[C@H]3O)C[C@]12C | PWAOOJDMFUQOKB-WCZZMFLVSA-N | No-Effect | inactive |
| GK921 | BRD-K38272302 | C(#Cc1nc2ncccc2nc1OCCN1CCCC1)c1ccccc1 | MNYJJHBAEYKXEG-UHFFFAOYSA-N | No-Effect | inactive |
| GKA-50 | BRD-K03827870 | COC[C@H](C)Oc1cc(O[C@@H](C)Cc2ccccc2)cc(C(=O)Nc2ccc(C(=O)O)cn2)c1 | OCBMECSFDVUYQN-ROUUACIJSA-N | No-Effect | inactive |
| glafenine | BRD-K01826515 | O=C(OC[C@H](O)CO)c1ccccc1Nc1ccnc2cc(Cl)ccc12 | GWOFUCIGLDBNKM-CYBMUJFWSA-N | No-Effect | inactive |
| glasdegib | BRD-K34341643 | CN1CC[C@@H](NC(=O)Nc2ccc(C#N)cc2)C[C@@H]1c1nc2ccccc2[nH]1 | SFNSLLSYNZWZQG-VQIMIIECSA-N | No-Effect | inactive |
| glecaprevir | BRD-K00003260 | CC(C)(C)[C@@H]1NC(=O)O[C@@H]2CCC[C@H]2OC/C=C\C(F)(F)c2nc3ccccc3nc2O[C@@H]2C[C@@H](C(=O)N[C@]3(C(=O)NS(=O)(=O)C4(C)CC4)C[C@H]3C(F)F)N(C2)C1=O | MLSQGNCUYAMAHD-ZSXAAYSXSA-N | No-Effect | inactive |
| glesatinib | BRD-K00003288 | COCCNCc1ccc(-c2cc3nccc(Oc4ccc(NC(=S)NC(=O)Cc5ccc(F)cc5)cc4F)c3s2)nc1 | YRCHYHRCBXNYNU-UHFFFAOYSA-N | No-Effect | inactive |
| gliclazide | BRD-A61154809 | Cc1ccc(S(=O)(=O)NC(=O)NN2C[C@H]3CCC[C@@H]3C2)cc1 | BOVGTQGAOIONJV-CHWSQXEVSA-N | No-Effect | inactive |
| glimepiride | BRD-K34776109 | CCC1=C(C)CN(C(=O)NCCc2ccc(S(=O)(=O)NC(=O)N[C@H]3CC[C@H](C)CC3)cc2)C1=O | WIGIZIANZCJQQY-RUCARUNLSA-N | No-Effect | inactive |
| glipizide | BRD-K12219985 | Cc1cnc(C(=O)NCCc2ccc(S(=O)(=O)NC(=O)NC3CCCCC3)cc2)cn1 | ZJJXGWJIGJFDTL-UHFFFAOYSA-N | No-Effect | inactive |
| gliquidone | BRD-K80396088 | COc1ccc2c(c1)C(=O)N(CCc1ccc(S(=O)(=O)NC(=O)NC3CCCCC3)cc1)C(=O)C2(C)C | LLJFMFZYVVLQKT-UHFFFAOYSA-N | No-Effect | inactive |
| Gln-1062 | BRD-K00004731 | COc1ccc2c3c1O[C@H]1C[C@@H](OC(=O)c4ccccc4)C=C[C@@]31CCN(C)C2 | JKVNJTYHRABHIY-WXVUKLJWSA-N | No-Effect | inactive |
| GLPG0187 | BRD-K00003380 | COc1ccc(S(=O)(=O)N[C@@H](CNc2nc(C)nc(N3CCC(c4ccc5c(n4)NCCC5)CC3)c2C)C(=O)O)cc1 | CXHCNOMGODVIKB-VWLOTQADSA-N | No-Effect | inactive |
| GLPG0492 | BRD-K15893360 | CN1C(=O)N(c2ccc(C#N)c(C(F)(F)F)c2)C(=O)[C@@]1(CO)c1ccccc1 | VAJGULUVTFDTAS-GOSISDBHSA-N | No-Effect | inactive |
| GLPG0492-R-enantiomer | BRD-K04643927 | CN1C(=O)N(c2ccc(C#N)c(C(F)(F)F)c2)C(=O)[C@]1(CO)c1ccccc1 | VAJGULUVTFDTAS-SFHVURJKSA-N | No-Effect | inactive |
| GLPG1837 | BRD-K00003095 | CC1(C)Cc2c(sc(NC(=O)c3cc[nH]n3)c2C(N)=O)C(C)(C)O1 | GHTGYZMBQPXTCQ-UHFFFAOYSA-N | No-Effect | inactive |
| glucaric-acid | BRD-K60150658 | O=C(O)[C@@H](O)[C@@H](O)[C@H](O)[C@@H](O)C(=O)O | DSLZVSRJTYRBFB-LLEIAEIESA-N | No-Effect | inactive |
| gluconolactone | BRD-K04288422 | O=C1O[C@H](CO)[C@H](O)[C@@H](O)[C@@H]1O | PHOQVHQSTUBQQK-KLVWXMOXSA-N | No-Effect | inactive |
| glucosamine | BRD-K29918010 | N[C@H]1[C@H](O)O[C@H](CO)[C@@H](O)[C@@H]1O | MSWZFWKMSRAUBD-QZABAPFNSA-N | No-Effect | inactive |
| glutathione | BRD-K69346723 | N[C@@H](CCC(=O)N[C@@H](CS)C(=O)NCC(=O)O)C(=O)O | RWSXRVCMGQZWBV-WDSKDSINSA-N | No-Effect | inactive |
| glutathione-monoisopropyl-ester | BRD-K58937086 | CC(C)OC(=O)CNC(=O)[C@H](CS)NC(=O)CC[C@H](N)C(=O)O | MVNCPACIPXNJIW-IUCAKERBSA-N | No-Effect | inactive |
| gly-gln | BRD-K85066592 | NCC(=O)N[C@@H](CCC(N)=O)C(=O)O | PNMUAGGSDZXTHX-BYPYZUCNSA-N | No-Effect | inactive |
| glyburide | BRD-K36927236 | COc1ccc(Cl)cc1C(=O)NCCc1ccc(S(=O)(=O)NC(=O)NC2CCCCC2)cc1 | ZNNLBTZKUZBEKO-UHFFFAOYSA-N | No-Effect | inactive |
| glycerol-monolaurate | BRD-A10773072 | CCCCCCCCCCCC(=O)OC[C@@H](O)CO | ARIWANIATODDMH-AWEZNQCLSA-N | No-Effect | inactive |
| glycerol-phenylbutyrate | BRD-K00003520 | O=C(CCCc1ccccc1)OCC(COC(=O)CCCc1ccccc1)OC(=O)CCCc1ccccc1 | ZSDBFLMJVAGKOU-UHFFFAOYSA-N | No-Effect | inactive |
| glycitein | BRD-K78303961 | COc1cc2c(=O)c(-c3ccc(O)cc3)coc2cc1O | DXYUAIFZCFRPTH-UHFFFAOYSA-N | No-Effect | inactive |
| GM-1485 | BRD-K00004613 | CCC(C)(C)C(=O)C(=O)N1CCC[C@H]1C(=O)O | FOPALECPEUVCTL-QMMMGPOBSA-N | No-Effect | inactive |
| GMX1778 | BRD-K69832040 | N#CN/C(=N\c1ccncc1)NCCCCCCOc1ccc(Cl)cc1 | BOIPLTNGIAPDBY-UHFFFAOYSA-N | No-Effect | inactive |
| GNE-7915 | BRD-K33963447 | CCNc1nc(Nc2cc(F)c(C(=O)N3CCOCC3)cc2OC)ncc1C(F)(F)F | XCFLWTZSJYBCPF-UHFFFAOYSA-N | No-Effect | inactive |
| GNF-2 | BRD-K97056771 | NC(=O)c1cccc(-c2cc(Nc3ccc(OC(F)(F)F)cc3)ncn2)c1 | WEVYNIUIFUYDGI-UHFFFAOYSA-N | No-Effect | inactive |
| GNF-5 | BRD-K87632963 | O=C(NCCO)c1cccc(-c2cc(Nc3ccc(OC(F)(F)F)cc3)ncn2)c1 | IIQUYGWWHIHOCF-UHFFFAOYSA-N | No-Effect | inactive |
| GNF-5837 | BRD-K19975102 | Cc1ccc(NC(=O)Nc2cc(C(F)(F)F)ccc2F)cc1Nc1ccc2c(c1)NC(=O)/C2=C\c1ccc[nH]1 | YYDUWLSETXNJJT-MTJSOVHGSA-N | No-Effect | inactive |
| GNF-7 | BRD-K69075588 | Cc1ccc(Nc2ncc3c(n2)N(C)C(=O)N(c2cc(NC(=O)c4cccc(C(F)(F)F)c4)ccc2C)C3)cn1 | SZNYUUZOQHNEKB-UHFFFAOYSA-N | No-Effect | inactive |
| GNTI | BRD-K64748124 | N=C(N)Nc1ccc2[nH]c3c(c2c1)C[C@@]1(O)[C@H]2Cc4ccc(O)c5c4[C@@]1(CCN2CC1CC1)[C@H]3O5 | VLNHDKDBGWXJEE-GYHUNEDQSA-N | No-Effect | inactive |
| go-6983 | BRD-K36984403 | COc1ccc2c(c1)c(C1=C(c3c[nH]c4ccccc34)C(=O)NC1=O)cn2CCCN(C)C | LLJJDLHGZUOMQP-UHFFFAOYSA-N | No-Effect | inactive |
| golgicide-A | BRD-A57886255 | Fc1cc(F)c2c(c1)[C@H]1C=CC[C@H]1[C@@H](c1cccnc1)N2 | NJZHEQOUHLZCOX-WOSRLPQWSA-N | No-Effect | inactive |
| gonadorelin | BRD-K96874295 | CC(C)C[C@H](NC(=O)CNC(=O)[C@H](Cc1ccc(O)cc1)NC(=O)[C@H](CO)NC(=O)[C@H](Cc1c[nH]c2ccccc12)NC(=O)[C@H](Cc1cnc[nH]1)NC(=O)[C@@H]1CCC(=O)N1)C(=O)N[C@@H](CCCNC(=N)N)C(=O)N1CCC[C@H]1C(=O)NCC(N)=O | XLXSAKCOAKORKW-AQJXLSMYSA-N | No-Effect | inactive |
| goserelin-acetate | BRD-K99504665 | CC(C)C[C@H](NC(=O)[C@@H](COC(C)(C)C)NC(=O)[C@H](Cc1ccc(O)cc1)NC(=O)[C@H](CO)NC(=O)[C@H](Cc1c[nH]c2ccccc12)NC(=O)[C@H](Cc1cnc[nH]1)NC(=O)[C@@H]1CCC(=O)N1)C(=O)N[C@@H](CCCNC(=N)N)C(=O)N1CCC[C@H]1C(=O)NNC(N)=O | BLCLNMBMMGCOAS-URPVMXJPSA-N | No-Effect | inactive |
| gossypol | BRD-K19295594 | Cc1cc2c(C(C)C)c(O)c(O)c(C=O)c2c(O)c1-c1c(C)cc2c(C(C)C)c(O)c(O)c(C=O)c2c1O | QBKSWRVVCFFDOT-UHFFFAOYSA-N | No-Effect | inactive |
| GP1a | BRD-K65285700 | Cc1ccc2c(c1)Cc1c(C(=O)NN3CCCCC3)nn(-c3ccc(Cl)cc3Cl)c1-2 | FNOMTMVRTBHRET-UHFFFAOYSA-N | No-Effect | inactive |
| GP2a | BRD-K34608650 | Cc1ccc2c(c1)Cc1c(C(=O)NC3CCCCC3)nn(-c3ccc(Cl)cc3Cl)c1-2 | SPEFJYZGXZENAF-UHFFFAOYSA-N | No-Effect | inactive |
| GPBAR-A | BRD-K67397687 | O=C1COc2ncc(-c3ccccc3F)c2CN1Cc1cc(C(F)(F)F)cc(C(F)(F)F)c1 | ZIXNJVGTAXRKAP-UHFFFAOYSA-N | No-Effect | inactive |

| | | | | | |
|---|---|---|---|---|---|
| GPI-1046 | BRD-K44869961 | CCC(C)(C)C(=O)C(=O)N1CCC[C@H]1C(=O)OCCCc1cccnc1 | OQAHHWOPVDDWHD-INIZCTEOSA-N | No-Effect | inactive |
| GPi-688 | BRD-A90201073 | O=C(N[C@@H]1Cc2ccccc2N(C[C@@H](O)CO)C1=O)c1cc2cc(Cl)sc2[nH]1 | UICNBXVDHCBKCE-TZMCWYRMSA-N | No-Effect | inactive |
| GPR120-modulator-1 | BRD-K19409553 | Cc1cc(OCc2nc(-c3ccccc3Cl)cs2)ccc1OCC(=O)O | FONSSTYHKIFEBM-UHFFFAOYSA-N | No-Effect | inactive |
| GR-103691 | BRD-K50891186 | COc1ccccc1N1CCN(CCCCNC(=O)c2ccc(-c3ccc(C(C)=O)cc3)cc2)CC1 | JARNORYOPMINDY-UHFFFAOYSA-N | No-Effect | inactive |
| GR-113808 | BRD-K49945136 | Cn1cc(C(=O)OCC2CCN(CCNS(C)(=O)=O)CC2)c2ccccc21 | MOZPSIXKYJUTKI-UHFFFAOYSA-N | No-Effect | inactive |
| GR-135531 | BRD-K83023055 | COC(=O)Nc1ccc2[nH]cc(CCNC(C)=O)c2c1 | MPZVHKLZCUEJFO-UHFFFAOYSA-N | No-Effect | inactive |
| GR-144053 | BRD-K12120659 | N=C(N)c1ccc(N2CCN(C3CCN(CC(=O)O)CC3)CC2)cc1 | QGEGSJUSWLLZLH-UHFFFAOYSA-N | No-Effect | inactive |
| GR-159897 | BRD-K52394958 | COC1(C[S@@](=O)c2ccccc2)CCN(CCc2c[nH]c3ccc(F)cc23)CC1 | BANYJBHWTOJQDU-GDLZYMKVSA-N | No-Effect | inactive |
| GR-235 | BRD-K00004242 | CC=C1C(=O)C[C@H]2[C@@H]3CCC4=CC(=O)CC[C@]4(C)[C@H]3CC[C@]12C | WDXRGPWQVHZTQJ-NRJJLHBYSA-N | No-Effect | inactive |
| GR-79236 | BRD-K40578143 | OC[C@H]1O[C@@H](n2cnc3c(N[C@H]4CCC[C@@H]4O)ncnc32)[C@H](O)[C@@H]1O | GYWXTRVEUURNEW-TVDBPQCTSA-N | No-Effect | inactive |
| GR125487 | BRD-K62594868 | COc1[nH]c2ccc(F)cc2c1C(=O)OCC1CCN(CCNS(C)(=O)=O)CC1 | SSPCCAYAIDNNJX-UHFFFAOYSA-N | No-Effect | inactive |
| GR46611 | BRD-K98155362 | COc1ccc(CNC(=O)/C=C/c2ccc3[nH]cc(CCN(C)C)c3c2)cc1 | LBVZWEWTNUDWNS-YRNVUSSQSA-N | No-Effect | inactive |
| gramicidin | BRD-K45170529 | CC(C)C[C@@H](NC(=O)[C@H](C)NC(=O)CNC(=O)[C@@H](NC=O)C(C)C)C(=O)N[C@@H](C)C(=O)N[C@@H](C(=O)N[C@H](C(=O)N[C@@H](C(=O)N[C@@H](Cc1c[nH]c2ccccc12)C(=O)N[C@H](CC(C)C)C(=O)N[C@@H](Cc1c[nH]c2ccccc12)C(=O)N[C@H](CC(C)C)C(=O)N[C@@H](Cc1c[nH]c2ccccc12)C(=O)N[C@H](CC(C)C)C(=O)N[C@@H](Cc1c[nH]c2ccccc12)C(=O)NCCO)C(C)C)C(C)C)C(C)C | ZWCXYZRRTRDGQE-LUPIJMBPSA-N | No-Effect | inactive |
| gramine | BRD-K26005076 | CN(C)Cc1c[nH]c2ccccc12 | OCDGBSUVYYVKQZ-UHFFFAOYSA-N | No-Effect | inactive |
| granisetron | BRD-A10967948 | CN1[C@H]2CCC[C@@H]1C[C@H](NC(=O)c1nn(C)c3ccccc13)C2 | MFWNKCLOYSRHCJ-BTTYYORXSA-N | No-Effect | inactive |
| grapiprant | BRD-K62050508 | CCc1nc2c(C)nc(C)cc2n1-c1ccc(CCNC(=O)NS(=O)(=O)c2ccc(C)cc2)cc1 | HZVLFTCYCLXTGV-UHFFFAOYSA-N | No-Effect | inactive |
| grazoprevir | BRD-K24943235 | C=C[C@@H]1C[C@]1(NC(=O)[C@@H]1C[C@@H]2CN1C(=O)[C@H](C(C)(C)C)NC(=O)O[C@@H]1C[C@H]1CCCCCc1nc3ccc(OC)cc3nc1O2)C(=O)NS(=O)(=O)C1CC1 | OBMNJSNZOWALQB-NCQNOWPTSA-N | No-Effect | inactive |
| griseofulvin | BRD-K08273968 | COC1=CC(=O)C[C@@H](C)[C@]12Oc1c(Cl)c(OC)cc(OC)c1C2=O | DDUHZTYCFQRHIY-RBHXEPJQSA-N | No-Effect | inactive |
| GS-143 | BRD-K99916169 | O=C(O)c1ccc(N2N=C(Cc3ccccc3)/C(=C\c3ccc(-c4ccccc4F)o3)C2=O)cc1 | IZPMWFSVTDOCDI-HAVVHWLPSA-N | No-Effect | inactive |
| GS-39783 | BRD-K75478907 | CSc1nc(NC2CCCC2)c([N+](=O)[O-])c(NC2CCCC2)n1 | GSGVDKOCBKBMGG-UHFFFAOYSA-N | No-Effect | inactive |
| GS-6201 | BRD-K68453341 | CCCn1c(=O)c2[nH]c(-c3cnn(Cc4cccc(C(F)(F)F)c4)c3)nc2n(CC)c1=O | KOYXXLLNCXWUNF-UHFFFAOYSA-N | No-Effect | inactive |
| GS-9620 | BRD-K73585091 | CCCCOc1nc(N)c2c(n1)N(Cc1cccc(CN3CCCC3)c1)CC(=O)N2 | VFOKSTCIRGDTBR-UHFFFAOYSA-N | No-Effect | inactive |
| GS-967 | BRD-K26526496 | FC(F)(F)Oc1ccc(-c2ccc3nnc(C(F)(F)F)n3c2)cc1 | FEVBKJITJDHASC-UHFFFAOYSA-N | No-Effect | inactive |
| GS-9973 | BRD-K76719364 | c1cn2cc(-c3ccc4c[nH]nc4c3)nc(Nc3ccc(N4CCOCC4)cc3)c2n1 | XSMSNFMDVXXHGJ-UHFFFAOYSA-N | No-Effect | inactive |
| GSK-3-inhibitor-IX | BRD-K04923131 | O=C1Nc2cc(Br)ccc2/C1=C1/Nc2ccccc2/C1=N/O | DDLZLOKCJHBUHD-UJZLGWIISA-N | No-Effect | inactive |
| GSK-J1 | BRD-K32847255 | O=C(O)CCNc1cc(N2CCc3ccccc3CC2)nc(-c2ccccn2)n1 | AVZCPICCWKMZDT-UHFFFAOYSA-N | No-Effect | inactive |
| GSK-J2 | BRD-K82484347 | O=C(O)CCNc1cc(N2CCc3ccccc3CC2)nc(-c2cccnc2)n1 | LJIFOCRGDDQFJF-UHFFFAOYSA-N | No-Effect | inactive |
| GSK-J4 | BRD-K98203492 | CCOC(=O)CCNc1cc(N2CCc3ccccc3CC2)nc(-c2ccccn2)n1 | WBKCKEHGXNWYMO-UHFFFAOYSA-N | No-Effect | inactive |
| GSK-LSD-1 | BRD-K33521504 | c1ccc([C@@H]2C[C@H]2NC2CCNCC2)cc1 | BASFYRLYJAZPPL-UONOGXRCSA-N | No-Effect | inactive |
| GSK0660 | BRD-K98684188 | COC(=O)c1sccc1S(=O)(=O)Nc1ccc(Nc2ccccc2)cc1OC | NDFKBGWLUHKMFY-UHFFFAOYSA-N | No-Effect | inactive |
| GSK1059615 | BRD-K06750613 | O=C1NC(=O)/C(=C/c2ccc3nccc(-c4ccncc4)c3c2)S1 | QDITZBLZQQZVEE-YBEGLDIGSA-N | No-Effect | inactive |
| GSK1070916 | BRD-K36740062 | CCn1cc(-c2ccnc3[nH]c(-c4cccc(CN(C)C)c4)cc23)c(-c2ccc(NC(=O)N(C)C)cc2)n1 | QTBWCSQGBMPECM-UHFFFAOYSA-N | No-Effect | inactive |
| GSK1292263 | BRD-K60341624 | CC(C)c1noc(N2CCC(COc3ccc(-c4ccc(S(C)(=O)=O)cc4)nc3)CC2)n1 | AYJRTVVIBJSSKN-UHFFFAOYSA-N | No-Effect | inactive |
| GSK163090 | BRD-K69669039 | Cc1ccc2c(N3CCN(CCc4cccc(N5CCNC5=O)c4)CC3)cccc2n1 | ANGUXJDGJCHGOG-UHFFFAOYSA-N | No-Effect | inactive |
| GSK1838705A | BRD-K06749501 | CNC(=O)c1c(F)cccc1Nc1nc(Nc2cc3c(cc2OC)CCN3C(=O)CN(C)C)nc2[nH]ccc12 | HZTYDQRUAWIZRE-UHFFFAOYSA-N | No-Effect | inactive |
| GSK189254 | BRD-K00003166 | CNC(=O)c1ccc(Oc2ccc3c(c2)CCN(C2CCC2)CC3)nc1 | WROHEWWOCPRMIA-UHFFFAOYSA-N | No-Effect | inactive |
| GSK1904529A | BRD-K04833372 | CCc1cc(Nc2nccc(-c3c(-c4ccc(OC)c(C(=O)Nc5c(F)cccc5F)c4)nc4ccccn34)n2)c(OC)cc1N1CCC(N2CCN(S(C)(=O)=O)CC2)CC1 | MOSKATHMXWSZTQ-UHFFFAOYSA-N | No-Effect | inactive |
| GSK2190915 | BRD-K15691453 | CCOc1ccc(-c2ccc(Cn3c(CC(C)(C)C(=O)O)c(SC(C)(C)C)c4cc(OCc5ccc(C)cn5)ccc43)cc2)cn1 | DFQGDHBGRSTTHX-UHFFFAOYSA-N | No-Effect | inactive |
| GSK2193874 | BRD-K90737788 | O=C(NC1(c2ccccc2)CC1)c1c(CN2CCC(N3CCCCC3)CC2)c(-c2cccc(C(F)(F)F)c2)nc2cc(Br)ccc12 | UIVOZBSCHXCGPS-UHFFFAOYSA-N | No-Effect | inactive |
| GSK2239633A | BRD-K00003364 | COc1cccc2c1c(NS(=O)(=O)c1ccc(Cl)s1)nn2Cc1cccc(CNC(=O)C(C)(C)O)c1 | YTEVTHHGQMUPHC-UHFFFAOYSA-N | No-Effect | inactive |
| GSK2256098 | BRD-K00003379 | CONC(=O)c1ccccc1Nc1cc(Nc2cc(C)nn2C(C)C)ncc1Cl | BVAHPPKGOOJSPU-UHFFFAOYSA-N | No-Effect | inactive |
| GSK2256294A | BRD-K62738477 | CNc1nc(C)nc(N[C@H]2CCC[C@@H](C(=O)NCc3ccc(C#N)cc3C(F)(F)F)C2)n1 | LQHDJQIMETZMPH-ZBFHGGJFSA-N | No-Effect | inactive |
| GSK2330672 | BRD-K05858796 | CCCC[C@]1(CC)CS(=O)(=O)c2cc(CNC(CC(=O)O)CC(=O)O)c(OC)cc2[C@@H](c2ccccc2)N1 | CZGVOBIGEBDYTP-VSGBNLITSA-N | No-Effect | inactive |
| GSK239512 | BRD-K00004689 | O=C1CCCN1c1ccc(Oc2ccc3c(c2)CCN(C2CCC2)CC3)nc1 | YFRBKEVUUCQYOW-UHFFFAOYSA-N | No-Effect | inactive |
| GSK256066 | BRD-K37590257 | COc1cccc(Nc2c(C(N)=O)cnc3c(C)cc(S(=O)(=O)c4cccc(C(=O)N(C)C)c4)cc23)c1 | JFHROPTYMMSOLG-UHFFFAOYSA-N | No-Effect | inactive |
| GSK2801 | BRD-K48062199 | CCCOc1ccn2c(C(C)=O)cc(-c3ccccc3S(C)(=O)=O)c2c1 | KHWCPNJRJCNVRI-UHFFFAOYSA-N | No-Effect | inactive |
| GSK2816126 | BRD-K21867462 | CC[C@H](C)n1cc(C)c2c(C(=O)NCc3c(C)cc(C)[nH]c3=O)cc(-c3ccc(N4CCNCC4)nc3)cc21 | FKSFKBQGSFSOSM-QFIPXVFZSA-N | No-Effect | inactive |
| GSK2830371 | BRD-K17472821 | Cc1ncc(Cl)cc1NCc1ccc(C(=O)N[C@@H](CC2CCCC2)C(=O)NC2CC2)s1 | IVDUVEGCMXCMSO-FQEVSTJZSA-N | No-Effect | inactive |
| GSK2837808A | BRD-K50679358 | COc1ncc(-c2ccc3c(Nc4cc(Oc5cc(F)cc(F)c5)cc(C(=O)O)c4)c(S(=O)(=O)NC4CC4)cnc3c2)c(OC)n1 | RZBCPMYJIARMGV-UHFFFAOYSA-N | No-Effect | inactive |
| GSK2838232 | BRD-K08398903 | CC(C)C1=C2[C@H]3CC[C@@H]4[C@@]5(C)CC[C@H](OC(=O)CC(C)(C)C(=O)O)C(C)(C)[C@@H]5CC[C@@]4(C)[C@]3(C)CC[C@@]2([C@@H](O)CN(CCN(C)C)Cc2ccc(Cl)cc2)CC1=O | MEOWKUXNVNJAMY-PZFKGGKESA-N | No-Effect | inactive |
| GSK2879552 | BRD-K09223495 | O=C(O)c1ccc(CN2CCC(CN[C@@H]3C[C@H]3c3ccccc3)CC2)cc1 | LRULVYSBRWUVGR-FCHUYYIVSA-N | No-Effect | inactive |
| GSK2881078 | BRD-K97656802 | C[C@H](CS(C)(=O)=O)n1ccc2c(C(F)(F)F)c(C#N)ccc21 | SKDVMPZQJMZEAC-SECBINFHSA-N | No-Effect | inactive |
| GSK2981278 | BRD-K00003292 | CCc1ccc(N(CC(C)C)S(=O)(=O)c2ccc(OCC3CCOCC3)c(CO)c2)cc1 | LZLBRISQTJVZNP-UHFFFAOYSA-N | No-Effect | inactive |
| GSK2982772 | BRD-K00003429 | CN1C(=O)[C@@H](NC(=O)c2nc(Cc3ccccc3)n[nH]2)COc2ccccc21 | LYPAFUINURXJSG-AWEZNQCLSA-N | No-Effect | inactive |
| GSK3179106 | BRD-K00003373 | CCOc1cc(=O)[nH]cc1-c1ccc(CC(=O)Nc2cc(C(C)(C)C(F)(F)F)on2)c(F)c1 | IDXKJSSOUXWLDB-UHFFFAOYSA-N | No-Effect | inactive |
| GSK3326595 | BRD-K00003421 | CC(=O)N1CCC(Nc2cc(C(=O)NC[C@H](O)CN3CCc4ccccc4C3)ncn2)CC1 | JLCCNYVTIWRPIZ-NRFANRHFSA-N | No-Effect | inactive |

| | | | | | |
|---|---|---|---|---|---|
| GSK356278 | BRD-K00004711 | CCn1ncc2c(NC3CCOCC3)c(-c3nnc(Cc4sc(C)nc4C)o3)cnc21 | AWDJJMXJUOHGLC-UHFFFAOYSA-N | No-Effect | inactive |
| GSK37647 | BRD-K26514105 | COc1ccc(S(=O)(=O)Nc2c(C)cc(C)cc2C)cc1 | FQUAFMNPXPXOJE-UHFFFAOYSA-N | No-Effect | inactive |
| GSK3787 | BRD-K03155172 | O=C(NCCS(=O)(=O)c1ccc(C(F)(F)F)cn1)c1ccc(Cl)cc1 | JFUIMTGOQCQTPF-UHFFFAOYSA-N | No-Effect | inactive |
| GSK429286A | BRD-A10523515 | CC1=C(C(=O)Nc2cc3c[nH]nc3cc2F)[C@H](c2ccc(C(F)(F)F)cc2)CC(=O)N1 | OLIIUAHHAZEXEX-AWEZNQCLSA-N | No-Effect | inactive |
| GSK503 | BRD-K72293620 | Cc1cc(C)c(CNC(=O)c2cc(-c3ccc(N4CCN(C)CC4)nc3)cc3c2c(C)cn3C(C)C)c(=O)[nH]1 | HRDQQHUKUIKFHT-UHFFFAOYSA-N | No-Effect | inactive |
| GSK656 | BRD-K00003483 | NC[C@H]1OB(O)c2c(OCCO)ccc(Cl)c21 | DJUOWOXTPXUHDQ-MRVPVSSYSA-N | No-Effect | inactive |
| GSK690693 | BRD-K25325018 | CCn1c(-c2nonc2N)nc2c(C#CC(C)(C)O)ncc(OC[C@H]3CCCNC3)c21 | KGPGFQWBCSZGEL-ZDUSSCGKSA-N | No-Effect | inactive |
| GSK9027 | BRD-K29328421 | O=S(=O)(Cc1ccccc1)Nc1ccc(-c2ccc3c(cnn3-c3ccc(F)cc3)c2)c(C(F)(F)F)c1 | DXBJGDVBQPEMOB-UHFFFAOYSA-N | No-Effect | inactive |
| GSK923295 | BRD-K89015388 | CC(C)Oc1ccc(C(=O)N[C@H](CNC(=O)CN(C)C)Cc2ccc(-c3cn4cccc([C@H](C)O)c4n3)cc2)cc1Cl | WHMXDBPHBVLYRC-OFVILXPXSA-N | No-Effect | inactive |
| GTS21 | BRD-K14235254 | COc1ccc(/C=C2\CCCN=C2c2cccnc2)c(OC)c1 | RPYWXZCFYPVCNQ-RVDMUPIBSA-N | No-Effect | inactive |
| guacetisal | BRD-K00739965 | COc1ccccc1OC(=O)c1ccccc1OC(C)=O | HSJFYRYGGKLQBT-UHFFFAOYSA-N | No-Effect | inactive |
| guadecitabine | BRD-K00003187 | Nc1ncn([C@H]2C[C@H](OP(=O)(O)OC[C@H]3O[C@@H](n4cnc5c(=O)nc(N)[nH]c54)C[C@@H]3O)[C@@H](CO)O2)c(=O)n1 | GUWXKKAWLCENJA-WGWHJZDNSA-N | No-Effect | inactive |
| guaiacol | BRD-K96282009 | COc1ccccc1O | LHGVFZTZFXWLCP-UHFFFAOYSA-N | No-Effect | inactive |
| guaifenesin | BRD-A90515964 | COc1ccccc1OC[C@@H](O)CO | HSRJKNPTNIJEKV-QMMMGPOBSA-N | No-Effect | inactive |
| guanaben-acetate | BRD-K62736196 | N=C(N)N/N=C/c1c(Cl)cccc1Cl | WDZVGELJXXEGPV-YIXHJXPBSA-N | No-Effect | inactive |
| guanethidine | BRD-K62676709 | N=C(N)NCCN1CCCCCCC1 | ACGDKVXYNVEAGU-UHFFFAOYSA-N | No-Effect | inactive |
| guanfacine | BRD-K32830106 | N=C(N)NC(=O)Cc1c(Cl)cccc1Cl | INJOMKTZOLKMBF-UHFFFAOYSA-N | No-Effect | inactive |
| guanidine | BRD-K76037122 | N=C(N)N | ZRALSGWEFCBTJO-UHFFFAOYSA-N | No-Effect | inactive |
| guanidinoethyldisulfide-bicarbonate | BRD-K69191882 | N=C(N)NCCSSCCNC(=N)N | IRMTXMJNHRISQH-UHFFFAOYSA-N | No-Effect | inactive |
| guanidinopropionic-acid | BRD-K50781608 | N=C(N)NCCC(=O)O | KMXXSJLYVJEBHI-UHFFFAOYSA-N | No-Effect | inactive |
| Gue-1654 | BRD-K18745919 | CSc1nc2ccc3nc(NC(=O)C(c4ccccc4)c4ccccc4)sc3c2s1 | UFOBDFMYJABXGK-UHFFFAOYSA-N | No-Effect | inactive |
| guvacine | BRD-K01980392 | O=C(O)C1=CCCNC1 | QTDZOWFRBNTPQR-UHFFFAOYSA-N | No-Effect | inactive |
| GW-0742 | BRD-K82983861 | Cc1cc(SCc2sc(-c3ccc(C(F)(F)F)c(F)c3)nc2C)ccc1OCC(=O)O | HWVNEWGKWRGSRK-UHFFFAOYSA-N | No-Effect | inactive |
| GW-1100 | BRD-K23614267 | CCOC(=O)c1ccc(-n2cc(Cc3cnc(OCC)nc3)c(=O)nc2SCc2ccc(F)cc2)cc1 | PTPNCCWOTBBVJR-UHFFFAOYSA-N | No-Effect | inactive |
| GW-1929 | BRD-K00004238 | CN(CCOc1ccc(C[C@H](Nc2ccccc2C(=O)c2ccccc2)C(=O)O)cc1)c1ccccn1 | QTQMRBZOBKYXCG-MHZLTWQESA-N | No-Effect | inactive |
| GW-2580 | BRD-K24242599 | COc1ccc(COc2ccc(Cc3cnc(N)nc3N)cc2OC)cc1 | MYQAUKPBNJWPIE-UHFFFAOYSA-N | No-Effect | inactive |
| GW-311616 | BRD-K37864785 | CC(C)[C@@H]1C(=O)N(S(C)(=O)=O)[C@@H]2CCN(C(=O)/C=C/CN3CCCCC3)[C@@H]12 | NDNKNUMSTIMSHQ-URZKGLGPSA-N | No-Effect | inactive |
| GW-405833 | BRD-K10705233 | COc1ccc2c(c1)c(CCN1CCOCC1)c(C)n2C(=O)c1cccc(Cl)c1Cl | FSFZRNZSZYDVLI-UHFFFAOYSA-N | No-Effect | inactive |
| GW-4064 | BRD-K88186167 | CC(C)c1onc(-c2c(Cl)cccc2Cl)c1COc1ccc(/C=C/c2cccc(C(=O)O)c2)c(Cl)c1 | BYTNEISLBIENSA-MDZDMXLPSA-N | No-Effect | inactive |
| GW-438014A | BRD-K13039700 | O=C(Nc1nc2ccccc2n1CCc1ccccc1)c1ccccc1 | CDUDZZGLFOFINU-UHFFFAOYSA-N | No-Effect | inactive |
| GW-441756 | BRD-K41312087 | Cn1cc(/C=C2\C(=O)Nc3cccnc32)c2ccccc21 | NXNQLECPAXXYTR-LCYFTJDESA-N | No-Effect | inactive |
| GW-501516 | BRD-K14880289 | Cc1cc(SCc2sc(-c3ccc(C(F)(F)F)cc3)nc2C)ccc1OCC(=O)O | YDBLKRPLXZNVNB-UHFFFAOYSA-N | No-Effect | inactive |
| GW-5074 | BRD-K39520573 | O=C1Nc2ccc(I)cc2C1=Cc1cc(Br)c(O)c(Br)c1 | LMXYVLFTZRPNRV-UHFFFAOYSA-N | No-Effect | inactive |
| GW-542573X | BRD-K36485341 | COc1ccccc1NC(=O)OCC1CCN(C(=O)OC(C)(C)C)CC1 | SAXGSDIZIYFNKD-UHFFFAOYSA-N | No-Effect | inactive |
| GW-583340 | BRD-K79930101 | CS(=O)(=O)CCNCc1nc(-c2ccc3ncnc(Nc4ccc(OCc5cccc(F)c5)c(Cl)c4)c3c2)cs1 | MNNXRVSQOABRRP-UHFFFAOYSA-N | No-Effect | inactive |
| GW-6471 | BRD-K20828910 | CCC(=O)NC[C@H](Cc1ccc(OCCc2nc(-c3ccccc3)oc2C)cc1)N/C(C)=C\C(=O)c1ccc(C(F)(F)F)cc1 | TYEFSRMOUXWTDN-DYQICHDWSA-N | No-Effect | inactive |
| GW-7647 | BRD-K96263742 | CC(C)(Sc1ccc(CCN(CCCCC2CCCCC2)C(=O)NC2CCCCC2)cc1)C(=O)O | PKNYXWMTHFMHKD-UHFFFAOYSA-N | No-Effect | inactive |
| GW-788388 | BRD-K78086729 | O=C(NC1CCOCC1)c1ccc(-c2cc(-c3c[nH]nc3-c3ccccn3)ccn2)cc1 | SAGZIBJAQGBRQA-UHFFFAOYSA-N | No-Effect | inactive |
| GW-803430 | BRD-K65576800 | COc1cc(-n2cnc3cc(-c4ccc(Cl)cc4)sc3c2=O)ccc1OCCN1CCCC1 | MWULMTACIBZPGN-UHFFFAOYSA-N | No-Effect | inactive |
| GW-842166 | BRD-K26823213 | O=C(NCC1CCOCC1)c1cnc(Nc2ccc(Cl)cc2Cl)nc1C(F)(F)F | TWQYWUXBZHPIIV-UHFFFAOYSA-N | No-Effect | inactive |
| GW-843682X | BRD-K90382497 | COc1cc2ncn(-c3cc(OCc4ccccc4C(F)(F)F)c(C(N)=O)s3)c2cc1OC | JSKUWFIZUALZLX-UHFFFAOYSA-N | No-Effect | inactive |
| GW-9508 | BRD-K71534238 | O=C(O)CCc1ccc(NCc2cccc(Oc3ccccc3)c2)cc1 | DGENZVKCTGIDRZ-UHFFFAOYSA-N | No-Effect | inactive |
| GW274150 | BRD-K65840970 | CC(=N)NCCSCC[C@H](N)C(=O)O | MOLOJNHYNHBPCW-ZETCQYMHSA-N | No-Effect | inactive |
| GYKI-52466 | BRD-K24240364 | CC1=NN=C(c2ccc(N)cc2)c2cc3c(cc2C1)OCO3 | LFBZZHVSGAHQPP-UHFFFAOYSA-N | No-Effect | inactive |
| gynostemma-extract | BRD-K58775955 | CC(C)=CCC[C@](C)(O[C@@H]1O[C@H](CO[C@@H]2OC[C@@H](O)[C@H](O)[C@H]2O)[C@@H](O)[C@H](O)[C@H]1O)[C@H]1CC[C@]2(C)[C@@H]1[C@H](O)C[C@@H]1[C@@]3(C)CC[C@H](O[C@@H]4O[C@H](CO)[C@@H](O)[C@H](O)[C@H]4O)C(C)(C)[C@@H]3CC[C@]12C | ZTQSADJAYQOCDD-HUGMCNGHSA-N | No-Effect | inactive |
| H-151 | BRD-K00005333 | CCc1ccc(NC(=O)Nc2c[nH]c3ccccc23)cc1 | UJZDIKVQFMCLBE-UHFFFAOYSA-N | No-Effect | inactive |
| H-89 | BRD-K27737647 | O=S(=O)(NCCNC/C=C/c1ccc(Br)cc1)c1cccc2cnccc12 | ZKZXNDJNWUTGDK-NSCUHMNNSA-N | No-Effect | inactive |
| H2L-5765834 | BRD-K26439554 | O=C(O)c1ccc2c(c1)C(=O)N(c1cccc(Oc3ccc([N+](=O)[O-])cc3)c1)C2=O | HFYPTENHTPNXGP-UHFFFAOYSA-N | No-Effect | inactive |
| HA-1004 | BRD-K05434375 | N=C(N)NCCNS(=O)(=O)c1cccc2cnccc12 | MZNDNBFMSVMUCX-UHFFFAOYSA-N | No-Effect | inactive |
| HA-130 | BRD-K64452997 | O=C1S/C(=C\c2ccc(OCc3cccc(B(O)O)c3)cc2)C(=O)N1Cc1ccc(F)cc1 | VTNKMYWFWQTEHE-XKZIYDEJSA-N | No-Effect | inactive |
| HA-966-(R)-(+) | BRD-K58046914 | N[C@@H]1CCN(O)C1=O | HCKUBNLZMKAEIN-GSVOUGTGSA-N | No-Effect | inactive |
| HA-966-(S)-(-) | BRD-K41304947 | N[C@H]1CCN(O)C1=O | HCKUBNLZMKAEIN-VKHMYHEASA-N | No-Effect | inactive |
| HA14-1 | BRD-K18163608 | CCOC(=O)C1=C(N)Oc2ccc(Br)cc2[C@H]1[C@H](C#N)C(=O)OCC | SXJDCULZDFWMJC-AAEUAGOBSA-N | No-Effect | inactive |
| halazone | BRD-K68091065 | O=C(O)c1ccc(S(=O)(=O)N(Cl)Cl)cc1 | XPDVQPODLRGWPL-UHFFFAOYSA-N | No-Effect | inactive |
| halcinonide | BRD-K81709173 | CC1(C)O[C@@H]2C[C@H]3[C@@H]4CCC5=CC(=O)CC[C@]5(C)[C@@]4(F)[C@@H](O)C[C@]3(C)[C@]2(C(=O)CCl)O1 | MUQNGPZZQDCDFT-JNQJZLCISA-N | No-Effect | inactive |
| halofantrine | BRD-A51964809 | CCCCN(CCCC)CC[C@@H](O)c1cc2c(Cl)cc(Cl)cc2c2cc(C(F)(F)F)ccc12 | FOHHNHSLJDZUGQ-RUZDIDTESA-N | No-Effect | inactive |
| halopemide | BRD-K50055394 | O=C(NCCN1CCC(n2c(=O)[nH]c3cc(Cl)ccc32)CC1)c1ccc(F)cc1 | NBHPRWLFLUBAIE-UHFFFAOYSA-N | No-Effect | inactive |
| haloperidol | BRD-K67783091 | O=C(CCCN1CCC(O)(c2ccc(Cl)cc2)CC1)c1ccc(F)cc1 | LNEPOXFFQSENCJ-UHFFFAOYSA-N | No-Effect | inactive |
| haloperidol-decanoate | BRD-K80859331 | CCCCCCCCCC(=O)OC1(c2ccc(Cl)cc2)CCN(CCCC(=O)c2ccc(F)cc2)CC1 | GUTXTARXLVFHDK-UHFFFAOYSA-N | No-Effect | inactive |
| haloprogin | BRD-K13238168 | Clc1cc(Cl)c(OCC#CI)cc1Cl | CTETYYAZBPJBHE-UHFFFAOYSA-N | No-Effect | inactive |
| harmane | BRD-K12630395 | Cc1nccc2c1[nH]c1ccccc12 | PSFDQSOCUJVVGF-UHFFFAOYSA-N | No-Effect | inactive |
| HBED | BRD-K00004555 | O=C(O)CN(CCN(CC(=O)O)Cc1ccccc1O)Cc1ccccc1O | GRUVVLWKPGIYEG-UHFFFAOYSA-N | No-Effect | inactive |
| HC-030031 | BRD-K73815512 | CC(C)c1ccc(NC(=O)Cn2cnc3c2c(=O)n(C)c(=O)n3C)cc1 | HEQDZPHDVAOBLN-UHFFFAOYSA-N | No-Effect | inactive |
| HC-067047 | BRD-K03391209 | Cc1c(C(=O)Nc2cccc(C(F)(F)F)c2)cc(-c2ccccc2)n1CCCN1CCOCC1 | NCZYSQOTAYFTNM-UHFFFAOYSA-N | No-Effect | inactive |
| HC-070 | BRD-K00004601 | Cn1c(=O)n(CCCO)c(=O)c2c1nc(Oc1cccc(Cl)c1)n2Cc1ccc(Cl)cc1 | VYJPVPHNGWEIBT-UHFFFAOYSA-N | No-Effect | inactive |

| | | | | | |
|---|---|---|---|---|---|
| HEMADO | BRD-K34672903 | CCCCC#Cc1nc(NC)c2ncn([C@@H]3O[C@H](CO)[C@@H](O)[C@H]3O)c2n1 | KOCIMZNSNPOGOP-IWCJZZDYSA-N | No-Effect | inactive |
| hemin | BRD-K74057164 | C=CC1=C(C)C2=N/C1=C\c1c(C)c(CCC(=O)O)c3n1[Fe](O)n1/c(c(C)c(C=C)/c1=C/2)=C\C1=N/C(=C\3)C(CCC(=O)O)=C1C | BMUDPLZKKRQECS-KKXYAJQKSA-K | No-Effect | inactive |
| hemomex-s | BRD-K15231341 | CC(C)=CCC[C@](C)(O)[C@H]1CC[C@]2(C)[C@@H]1[C@H](O)C[C@@H]1[C@@]3(C)CC[C@H](O)C(C)(C)[C@@H]3[C@@H](O)C[C@]12C | SHCBCKBYTHZQGZ-CJPZEJHVSA-N | No-Effect | inactive |
| hepsulfam | BRD-K00004739 | NS(=O)(=O)OCCCCCCCOS(N)(=O)=O | GOJJWDOZNKBUSR-UHFFFAOYSA-N | No-Effect | inactive |
| heptaminol | BRD-A32988684 | C[C@@H](N)CCCC(C)(C)O | LREQLEBVOXIEOM-SSDOTTSWSA-N | No-Effect | inactive |
| HER2-Inhibitor-1 | BRD-K89709433 | Cc1cc(Nc2ncnc3ccc(-c4ccc(CNCCS(C)(=O)=O)o4)cc23)ccc1Oc1ccn2ncnc2c1 | QVMNYGOVNWWFKF-UHFFFAOYSA-N | No-Effect | inactive |
| hesperadin | BRD-K58122457 | CCS(=O)(=O)Nc1ccc2c(c1)/C(=C(/Nc1ccc(CN3CCCCC3)cc1)c1ccccc1)C(=O)N2 | GLDSKRNGVVYJAB-DQSJHHFOSA-N | No-Effect | inactive |
| hesperidin | BRD-K38903228 | COc1ccc([C@@H]2CC(=O)c3c(O)cc(O[C@@H]4O[C@H](CO[C@@H]5O[C@@H](C)[C@H](O)[C@@H](O)[C@H]5O)[C@@H](O)[C@H](O)[C@H]4O)cc3O2)cc1O | QUQPHWDTPGMPEX-QJBIFVCTSA-N | No-Effect | inactive |
| hetacillin | BRD-K00005353 | CC1(C)S[C@@H]2[C@H](N3C(=O)[C@@H](c4ccccc4)NC3(C)C)C(=O)N2[C@H]1C(=O)O | DXVUYOAEDJXBPY-NFFDBFGFSA-N | No-Effect | inactive |
| hexamethylenebisacetamide | BRD-K04710043 | CC(=O)NCCCCCCNC(C)=O | BNQSTAOJRULKNX-UHFFFAOYSA-N | No-Effect | inactive |
| hexaminolevulinate | BRD-K73753850 | CCCCCCOC(=O)CCC(=O)CN | RYQOILLJDKPETL-UHFFFAOYSA-N | No-Effect | inactive |
| hexasodium-phytate | BRD-K70007857 | O=P(O)(O)O[C@H]1[C@H](OP(=O)(O)O)[C@@H](OP(=O)(O)O)[C@H](OP(=O)(O)O)[C@@H](OP(=O)(O)O)[C@H]1OP(=O)(O)O | IMQLKJBTEOYOSI-GPIVLXJGSA-N | No-Effect | inactive |
| hexestrol | BRD-K69253806 | CC[C@H](c1ccc(O)cc1)[C@@H](CC)c1ccc(O)cc1 | PBBGSZCBWVPOOL-HDICACEKSA-N | No-Effect | inactive |
| hexetidine | BRD-A18605525 | CCCC[C@H](CC)CN1CN(C[C@H](CC)CCCC)C[C@](C)(N)C1 | DTOUUUZOYKYHEP-AERCQKQUSA-N | No-Effect | inactive |
| hexonic-acid | BRD-A63267145 | O=C(O)[C@H](O)[C@H](O)[C@H](O)[C@H](O)CO | RGHNJXZEOKUKBD-TXICZTDVSA-N | No-Effect | inactive |
| hexoprenaline | BRD-A33894669 | Oc1ccc([C@H](O)CNCCCCCCNC[C@H](O)c2ccc(O)c(O)c2)cc1O | OXLZNBCNGJWPRV-SZPZYZBQSA-N | No-Effect | inactive |
| hexylcaine | BRD-A97730597 | C[C@@H](CNC1CCCCC1)OC(=O)c1ccccc1 | DKLKMKYDWHYZTD-ZDUSSCGKSA-N | No-Effect | inactive |
| hexylene-glycol | BRD-A14234430 | C[C@@H](O)CC(C)(C)O | SVTBMSDMJJWYQN-RXMQYKEDSA-N | No-Effect | inactive |
| higenamine | BRD-A55204514 | Oc1ccc(C[C@H]2NCCc3cc(O)c(O)cc32)cc1 | WZRCQWQRFZITDX-CQSZACIVSA-N | No-Effect | inactive |
| hippuric-acid | BRD-K62914667 | O=C(O)CNC(=O)c1ccccc1 | QIAFMBKCNZACKA-UHFFFAOYSA-N | No-Effect | inactive |
| histamine | BRD-K01674964 | NCCc1cnc[nH]1 | NTYJJOPFIAHURM-UHFFFAOYSA-N | No-Effect | inactive |
| HJC-0350 | BRD-K34411045 | Cc1cc(C)c(S(=O)(=O)n2cc(C)cc2C)c(C)c1 | AFZWZVLPIMHLSE-UHFFFAOYSA-N | No-Effect | inactive |
| HKI-357 | BRD-K58811567 | CCOc1cc2ncc(C#N)c(Nc3ccc(OCc4cccc(F)c4)c(Cl)c3)c2cc1NC(=O)/C=C/CN(C)C | NERXPXBELDBEPZ-RMKNXTFCSA-N | No-Effect | inactive |
| HLCL-61 | BRD-K73746696 | CCn1c2ccccc2c2cc(CNCc3ccccc3OC)ccc21 | FIOIJUCPVZEPFX-UHFFFAOYSA-N | No-Effect | inactive |
| HM-30181 | BRD-K74319597 | COc1cc2c(cc1OC)CN(CCc1ccc(-n3nnc(-c4cc(OC)c(OC)cc4NC(=O)c4cc(=O)c5ccccc5o4)n3)cc1)CC2 | AHJUHHDDCJQACA-UHFFFAOYSA-N | No-Effect | inactive |
| HMN-214 | BRD-K70511574 | COc1ccc(S(=O)(=O)N(C(C)=O)c2ccccc2/C=C/c2cc[n+]([O-])cc2)cc1 | OCKHRKSTDPOHEN-BQYQJAHWSA-N | No-Effect | inactive |
| HOKU-81 | BRD-K00003598 | CC(C)(C)NC[C@H](O)c1ccc(O)cc1Cl | LIXBJWRFCNRAPA-NSHDSACASA-N | No-Effect | inactive |
| homatropine | BRD-K01826529 | CN1[C@H]2CC[C@@H]1C[C@H](OC(=O)[C@@H](O)c1ccccc1)C2 | ZTVIKZXZYLEVOL-YJNKXOJESA-N | No-Effect | inactive |
| homatropine-methylbromide | BRD-A54234250 | C[N+]1(C)[C@H]2CC[C@@H]1C[C@H](OC(=O)[C@@H](O)c1ccccc1)C2 | WKWOJBUWWZTUQV-JJXSEGSLSA-N | No-Effect | inactive |
| homidium | BRD-K28354016 | CC[n+]1c(-c2ccccc2)c2cc(N)ccc2c2ccc(N)cc21 | QTANTQQOYSUMLC-UHFFFAOYSA-O | No-Effect | inactive |
| homochlorcyclizine | BRD-K01826562 | CN1CCCN([C@@H](c2ccccc2)c2ccc(Cl)cc2)CC1 | WEUCDJCFJHYFRL-IBGZPJMESA-N | No-Effect | inactive |
| homoquinolinic-acid | BRD-K30519779 | O=C(O)Cc1cccnc1C(=O)O | HQPMJFFEXJELOQ-UHFFFAOYSA-N | No-Effect | inactive |
| homosalate | BRD-A34751532 | C[C@H]1C[C@H](OC(=O)c2ccccc2O)CC(C)(C)C1 | WSSJONWNBBTCMG-RYUDHWBXSA-N | No-Effect | inactive |
| homoveratrylamine | BRD-K59570838 | COc1ccc(CCN)cc1OC | ANOUKFYBOAKOIR-UHFFFAOYSA-N | No-Effect | inactive |
| honokiol | BRD-K98493452 | C=CCc1ccc(O)c(-c2ccc(O)c(CC=C)c2)c1 | FVYXIJYOAGAUQK-UHFFFAOYSA-N | No-Effect | inactive |
| hordenine | BRD-K93631643 | CN(C)CCc1ccc(O)cc1 | KUBCEEMXQZUPDQ-UHFFFAOYSA-N | No-Effect | inactive |
| HP-228 | BRD-K00004723 | CCCC[C@H](NC(C)=O)C(=O)N[C@@H](CCC(N)=O)C(=O)N[C@@H](Cc1c[nH]cn1)C(=O)N[C@H](Cc1ccccc1)C(=O)N[C@@H](CCCNC(=N)N)C(=O)N[C@H](Cc1c[nH]c2ccccc12)C(=O)NCC(N)=O | UHKZNDVUBSEJJO-XAYNKGEHSA-N | No-Effect | inactive |
| hPGDS-IN-1 | BRD-K00003140 | CC(C)(O)c1nc(-c2cccc(CNC(=O)c3cnc(-c4ccccn4)nc3)c2)no1 | VJCLAPUACUQZOV-UHFFFAOYSA-N | No-Effect | inactive |
| HPPH | BRD-K00003730 | CCCCCCO[C@H](C)c1c(C)c2cc3nc(c4c5[nH]c(cc6nc(cc1[nH]2)C(C)=C6CC)c(C)c5C(=O)C4)[C@@H](CCC(=O)O)[C@@H]3C | PVXGCBZIVFCMJK-NZJKTDFXSA-N | No-Effect | inactive |
| HSR6071 | BRD-K00004745 | O=C(Nc1nnn[nH]1)c1cncc(N2CCCC2)n1 | LUJDHCXCWJFNOJ-UHFFFAOYSA-N | No-Effect | inactive |
| HT-2157 | BRD-K00003386 | O=C1/C(=N\c2cccc(C(F)(F)F)c2)c2ccccc2N1c1ccccc1 | TXCGMRVPXUBHAL-PLRJNAJWSA-N | No-Effect | inactive |
| HTH-01-015 | BRD-K58726989 | Cc1nc(Nc2cnn(C3CCNCC3)c2)nc2c1N(C)C(=O)c1cc3ccccc3cc1N2C | CHSDJDLAKKAWCI-UHFFFAOYSA-N | No-Effect | inactive |
| HTHQ | BRD-K00003389 | CCCCCCOc1cc(C)c(O)c(C)c1C | ATMNQRRJNBCQJO-UHFFFAOYSA-N | No-Effect | inactive |
| HTMT | BRD-K01825665 | C[C@@H](CCCCC(=O)Nc1ccc(C(F)(F)F)cc1)NCCc1cnc[nH]1 | PMKJGGBYYNEYPA-AWEZNQCLSA-N | No-Effect | inactive |
| huperzine-A | BRD-K62240499 | C/C=C1\[C@H]2C=C(C)C[C@]1(N)c1ccc(=O)[nH]c1C2 | ZRJBHWIHUMBLCN-YQEJDHNASA-N | No-Effect | inactive |
| hyaluronic-acid | BRD-K59788572 | CC(=O)N[C@@H]1C[C@H](O)[C@@H](CO)O[C@H]1O[C@@H]1[C@@H](C(=O)O)O[C@@H](O)[C@H](O)[C@H]1O | WCDDVEOXEIYWFB-VXORFPGASA-N | No-Effect | inactive |
| hycanthone | BRD-K50406511 | CCN(CC)CCNc1ccc(CO)c2sc3ccccc3c(=O)c12 | MFZWMTSUNYWVBU-UHFFFAOYSA-N | No-Effect | inactive |
| hydralazine | BRD-K82103381 | NNc1nncc2ccccc12 | RPTUSVTUFVMDQK-UHFFFAOYSA-N | No-Effect | inactive |
| hydrastinine | BRD-K92726801 | C[N+]1=Cc2cc3c(cc2CC1)OCO3 | LGDCROILDUSHQX-UHFFFAOYSA-N | No-Effect | inactive |
| hydrochlorothiazide | BRD-K13078532 | NS(=O)(=O)c1cc2c(cc1Cl)NCNS2(=O)=O | JZUFKLXOESDKRF-UHFFFAOYSA-N | No-Effect | inactive |
| hydrocortisone-acetate | BRD-K11612998 | CC(=O)OCC(=O)[C@@]1(O)CC[C@H]2[C@@H]3CCC4=CC(=O)CC[C@]4(C)[C@H]3[C@@H](O)C[C@@]21C | ALEXXDVDDISNDU-JZYPGELDSA-N | No-Effect | inactive |
| hydrocortisone-butyrate | BRD-K99199077 | CCCC(=O)O[C@]1(C(=O)CO)CC[C@H]2[C@@H]3CCC4=CC(=O)CC[C@]4(C)[C@H]3[C@@H](O)C[C@@]21C | BMCQMVFGOVHVNG-TUFAYURCSA-N | No-Effect | inactive |
| hydrocortisone-hemisuccinate | BRD-K00025601 | C[C@]12C[C@H](O)[C@H]3[C@@H](CCC4=CC(=O)CC[C@@]43C)[C@@H]1CC[C@]2(O)C(=O)COC(=O)CCC(=O)O | VWQWXZAWFPZJDA-CGVGKPPMSA-N | No-Effect | inactive |
| hydrocortisone-valerate | BRD-K09968782 | CCCCC(=O)O[C@]1(C(=O)CO)CC[C@H]2[C@@H]3CCC4=CC(=O)CC[C@]4(C)[C@H]3[C@@H](O)C[C@@]21C | FZCHYNWYXKICIO-FZNHGJLXSA-N | No-Effect | inactive |
| hydroflumethiazide | BRD-K36862742 | NS(=O)(=O)c1cc2c(cc1C(F)(F)F)NCNS2(=O)=O | DMDGGSIALPNSEE-UHFFFAOYSA-N | No-Effect | inactive |
| hydroquinidine | BRD-K65866076 | CC[C@H]1CN2CC[C@H]1C[C@@H]2[C@@H](O)c1ccnc2ccc(OC)cc12 | LJOQGZACKSYWCH-LHHVKLHASA-N | No-Effect | inactive |
| hydroquinone | BRD-K11227669 | Oc1ccc(O)cc1 | QIGBRXMKCJKVMJ-UHFFFAOYSA-N | No-Effect | inactive |
| hydroxocobalamin | BRD-K00003525 | C/C1=C2/N=C(/C=C3/N=C(/C(C)=C4\N([Co+]O)[C@H]([C@H](CC(N)=O)[C@@]4(C)CCC(=O)NC[C@@H](C)OP(=O)(O)O[C@@H]4[C@@H](CO)O[C@H](n5cnc6cc(C)c(C)cc65)[C@@H]4O)[C@]4(C)N=C1[C@@H](CCC(N)=O)[C@]4(C)CC(N)=O)[C@@H](CCC(N)=O)C3(C)C)[C@@H](CCC(N)=O)[C@]2(C)CC(N)=O | YOZNUFWCRFCGIH-WZHZPDAFSA-L | No-Effect | inactive |
| hydroxyamphetamine | BRD-A80871782 | C[C@@H](N)Cc1ccc(O)cc1 | GIKNHHRFLCDOEU-SSDOTTSWSA-N | No-Effect | inactive |
| hydroxychloroquine | BRD-A99117172 | CCN(CCO)CCCC(C)Nc1ccnc2cc(Cl)ccc12 | XXSMGPRMXLTPCZ-UHFFFAOYSA-N | No-Effect | inactive |
| hydroxyfasudil | BRD-K77793136 | O=c1[nH]ccc2c(S(=O)(=O)N3CCCNCC3)cccc12 | ZAVGJDAFCZAWSZ-UHFFFAOYSA-N | No-Effect | inactive |

| | | | | | |
|---|---|---|---|---|---|
| hydroxyprogesterone | BRD-K44932450 | CC(=O)[C@@]1(O)CC[C@H]2[C@@H]3CCC4=CC(=O)CC[C@]4(C)[C@H]3CC[C@@]21C | DBPWSSGDRRHUNT-CEGNMAFCSA-N | No-Effect | inactive |
| hydroxyprogesterone-acetate | BRD-K92902018 | CC(=O)O[C@]1(C(C)=O)CC[C@H]2[C@@H]3CCC4=CC(=O)CC[C@]4(C)[C@H]3CC[C@@]21C | VTHUYJIXSMGYOQ-KOORYGTMSA-N | No-Effect | inactive |
| hydroxypropyl-beta-cyclodextrin | BRD-A01412266 | C[C@@H](O)COC[C@H]1O[C@@H]2O[C@@H]3[C@@H](COC[C@@H](C)O)O[C@H](O[C@@H]4[C@@H](COC[C@@H](C)O)O[C@H](O[C@@H]5[C@@H](COC[C@@H](C)O)O[C@H](O[C@@H]6[C@@H](COC[C@@H](C)O)O[C@H](O[C@@H]7[C@@H](COC[C@@H](C)O)O[C@H](O[C@@H]8[C@@H](COC[C@@H](C)O)O[C@H](O[C@H]1[C@H](O)[C@H]2O)[C@H](O)[C@H]8O)[C@H](O)[C@H]7O)[C@H](O)[C@H]6O)[C@H](O)[C@H]5O)[C@H](O)[C@H]4O)[C@H](O)[C@H]3O | ODLHGICHYURWBS-PGFAAJFJSA-N | No-Effect | inactive |
| hydroxysafflor-yellow-A | BRD-K00003327 | O=C(/C=C/c1ccc(O)cc1)[C@H]1C(=O)[C@H]([C@@H]2O[C@H](CO)[C@@H](O)[C@H](O)[C@H]2O)C(=O)[C@@](O)([C@@H]2O[C@H](CO)[C@@H](O)[C@H](O)[C@H]2O)C1=O | DKEZDPGUJMMUCC-GJVNFNJISA-N | No-Effect | inactive |
| hydroxystilbamidine | BRD-K62186992 | N=C(N)c1ccc(/C=C/c2ccc(C(=N)N)cc2O)cc1 | TUESWZZJYCLFNL-DAFODLJHSA-N | No-Effect | inactive |
| hydroxytacrine-maleate-(R,S) | BRD-K01825165 | Nc1c2c(nc3ccccc13)CCC[C@@H]2O | HLVVITIHAZBPKB-NSHDSACASA-N | No-Effect | inactive |
| hydroxytoluic-acid | BRD-K75608666 | Cc1cccc(C(=O)O)c1O | WHSXTWFYRGOBGO-UHFFFAOYSA-N | No-Effect | inactive |
| hydroxyurea | BRD-K51747290 | NC(=O)NO | VSNHCAURESNICA-UHFFFAOYSA-N | No-Effect | inactive |
| hydroxyzine | BRD-K01826510 | OCCOCCN1CCN([C@@H](c2ccccc2)c2ccc(Cl)cc2)CC1 | ZQDWXGKKHFNSQK-NRFANRHFSA-N | No-Effect | inactive |
| hygromycin-B | BRD-A24252652 | CN[C@H]1C[C@@H](N)[C@H](O)[C@@H](O[C@@H]2O[C@H](CO)[C@H](O)[C@H]3O[C@@]4(O[C@H]23)O[C@H]([C@@H](N)CO)[C@@H](O)[C@H](O)[C@H]4O)[C@@H]1O | GRRNUXAQVGOGFE-SIGCQSELSA-N | No-Effect | inactive |
| hymecromone | BRD-K46424862 | Cc1cc(=O)oc2cc(O)ccc12 | HSHNITRMYYLLCV-UHFFFAOYSA-N | No-Effect | inactive |
| hyodeoxycholic-acid | BRD-K44009692 | C[C@H](CCC(=O)O)[C@H]1CC[C@H]2[C@@H]3C[C@H](O)[C@@H]4C[C@H](O)CC[C@]4(C)[C@H]3CC[C@@]21C | DGABKXLVXPYZII-SIBKNCMHSA-N | No-Effect | inactive |
| hyoscyamine | BRD-K40530731 | CN1[C@H]2CC[C@@H]1C[C@H](OC(=O)[C@H](CO)c1ccccc1)C2 | RKUNBYITZUJHSG-FXUDXRNXSA-N | No-Effect | inactive |
| hypericin | BRD-K29673530 | Cc1cc(O)c2c(=O)c3c(O)cc(O)c4c5c(O)cc(O)c6c(=O)c7c(O)cc(C)c8c1c2c(c34)c(c78)c65 | BTXNYTINYBABQR-UHFFFAOYSA-N | No-Effect | inactive |
| hyperin | BRD-A63780829 | O=c1c(O[C@@H]2O[C@H](CO)[C@@H](O)[C@H](O)[C@@H]2O)c(-c2ccc(O)c(O)c2)oc2cc(O)cc(O)c12 | OVSQVDMCBVZWGM-FHYQVQSVSA-N | No-Effect | inactive |
| hypoxanthine | BRD-K03739921 | O=c1nc[nH]c2nc[nH]c12 | FDGQSTZJBFJUBT-UHFFFAOYSA-N | No-Effect | inactive |
| I-BET151 | BRD-K36363294 | COc1cc2c(cc1-c1c(C)noc1C)ncc1[nH]c(=O)n([C@H](C)c3ccccn3)c12 | VUVUVNZRUGEAHB-CYBMUJFWSA-N | No-Effect | inactive |
| I-BRD9 | BRD-K04972057 | CCn1cc(-c2cccc(C(F)(F)F)c2)c2sc(C(=N)NC3CCS(=O)(=O)CC3)cc2c1=O | WRUWGLUCNBMGPS-UHFFFAOYSA-N | No-Effect | inactive |
| I-BZA | BRD-K14762862 | CCN(CC)CCNC(=O)c1ccc(I)cc1 | HRVWCNMOEFNEON-UHFFFAOYSA-N | No-Effect | inactive |
| I-CBP-112 | BRD-K57090026 | CCC(=O)N1CCOc2c(cc(-c3ccc(OC)c(OC)c3)cc2OC[C@H]2CCCN(C)C2)C1 | YKNAKDFZAWQEEO-IBGZPJMESA-N | No-Effect | inactive |
| IACS-10759 | BRD-K00003105 | Cc1nc(-c2nc(-c3ccc(OC(F)(F)F)cc3)no2)nn1Cc1cccc(N2CCC(S(C)(=O)=O)CC2)c1 | HWJWNWZJUYCGKV-UHFFFAOYSA-N | No-Effect | inactive |
| IB-MECA | BRD-K31135544 | CNC(=O)[C@H]1O[C@@H](n2cnc3c(NCc4cccc(I)c4)ncnc32)[C@H](O)[C@@H]1O | HUJXGQILHAUCCV-MOROJQBDSA-N | No-Effect | inactive |
| ibandronate | BRD-K08681769 | CCCCCN(C)CCC(O)(P(=O)(O)O)P(=O)(O)O | MPBVHIBUJCELCL-UHFFFAOYSA-N | No-Effect | inactive |
| IBC-293 | BRD-K15196155 | CC(C)n1nnc2cc(C(=O)O)ccc21 | RUTVRAJKELSHCC-UHFFFAOYSA-N | No-Effect | inactive |
| iberdomide | BRD-K00003408 | O=C1CC[C@H](N2Cc3c(OCc4ccc(CN5CCOCC5)cc4)cccc3C2=O)C(=O)N1 | IXZOHGPZAQLIBH-NRFANRHFSA-N | No-Effect | inactive |
| ibiglustat | BRD-K14063023 | CC(C)(NC(=O)O[C@@H]1CN2CCC1CC2)c1csc(-c2ccc(F)cc2)n1 | YFHRCLAKZBDRHN-MRXNPFEDSA-N | No-Effect | inactive |
| IBMX | BRD-K94979336 | CC(C)Cn1c(=O)n(C)c(=O)c2[nH]cnc21 | APIXJSLKIYYUKG-UHFFFAOYSA-N | No-Effect | inactive |
| ibotenic-acid | BRD-A02971356 | N[C@@H](C(=O)O)c1cc(=O)[nH]o1 | IRJCBFDCFXCWGO-SCSAIBSYSA-N | No-Effect | inactive |
| ibrolipim | BRD-K62490786 | CCOP(=O)(Cc1ccc(C(=O)Nc2ccc(Br)cc2C#N)cc1)OCC | KPRTURMJVWXURQ-UHFFFAOYSA-N | No-Effect | inactive |
| ibrutinib | BRD-K70301465 | C=CC(=O)N1CCC[C@@H](n2nc(-c3ccc(Oc4ccccc4)cc3)c3c(N)ncnc32)C1 | XYFPWWZEPKGCCK-GOSISDBHSA-N | No-Effect | inactive |
| ibudilast | BRD-K16444452 | CC(C)C(=O)c1c(C(C)C)nn2ccccc12 | ZJVFLBOZORBYFE-UHFFFAOYSA-N | No-Effect | inactive |
| ibuprofen-(S) | BRD-K14965640 | CC(C)Cc1ccc([C@H](C)C(=O)O)cc1 | HEFNNWSXXWATRW-JTQLQIEISA-N | No-Effect | inactive |
| ibuprofen-lysine | BRD-M64062803 | CC(C)Cc1ccc([C@@H](C)C(=O)O)cc1.NCCCC[C@H](N)C(=O)O | IHHXIUAEPKVVII-APFIOPMWSA-N | No-Effect | inactive |
| ibuprofen-piconol | BRD-A75224450 | CC(C)Cc1ccc([C@@H](C)C(=O)OCc2ccccn2)cc1 | ACEWLPOYLGNNHV-OAHLLOKOSA-N | No-Effect | inactive |
| ibutamoren | BRD-K44771174 | CC(C)(N)C(=O)N[C@H](COCc1ccccc1)C(=O)N1CCC2(CC1)CN(S(C)(=O)=O)c1ccccc12 | UMUPQWIGCOZEOY-JOCHJYFZSA-N | No-Effect | inactive |
| ibutilide | BRD-A74391928 | CCCCCCCN(CC)CCC[C@@H](O)c1ccc(NS(C)(=O)=O)cc1 | ALOBUEHUHMBRLE-HXUWFJFHSA-N | No-Effect | inactive |
| IC261 | BRD-K09638361 | COc1cc(OC)c(/C=C2/C(=O)Nc3ccccc32)c(OC)c1 | JBJYTZXCZDNOJW-JLHYYAGUSA-N | No-Effect | inactive |
| ICA-110381 | BRD-K71686520 | O=C(Nc1ccc(Cl)nc1)c1ccc(Cl)cc1 | FOYOGIUAMYSVIT-UHFFFAOYSA-N | No-Effect | inactive |
| ICA-121431 | BRD-K03149601 | O=C(Nc1ccc(S(=O)(=O)Nc2nccs2)cc1)C(c1ccccc1)c1ccccc1 | URSQNPPONHUJDL-UHFFFAOYSA-N | No-Effect | inactive |
| icariin | BRD-K65639003 | COc1ccc(-c2oc3c(CC=C(C)C)c(O[C@@H]4O[C@H](CO)[C@@H](O)[C@H](O)[C@H]4O)cc(O)c3c(=O)c2O[C@@H]2O[C@@H](C)[C@H](O)[C@@H](O)[C@H]2O)cc1 | TZJALUIVHRYQQB-XLRXWWTNSA-N | No-Effect | inactive |
| icaritin | BRD-K24476327 | COc1ccc(-c2oc3c(CC=C(C)C)c(O)cc(O)c3c(=O)c2O)cc1 | TUUXBSASAQJECY-UHFFFAOYSA-N | No-Effect | inactive |
| icatibant-acetate | BRD-K63772874 | N=C(N)NCCC[C@H](NC(=O)[C@@H]1C[C@@H]2CCCC[C@@H]2N1C(=O)[C@H]1Cc2ccccc2CN1C(=O)[C@H](CO)NC(=O)[C@H](Cc1cccs1)NC(=O)CNC(=O)[C@@H]1C[C@@H](O)CN1C(=O)[C@@H]1CCCN1C(=O)[C@H](CCCNC(=N)N)NC(=O)[C@H](N)CCCNC(=N)N)C(=O)O | QURWXBZNHXJZBE-SKXRKSCCSA-N | No-Effect | inactive |
| ICG-001 | BRD-K73261812 | O=C1[C@H](Cc2ccc(O)cc2)N2C(=O)CCN(C(=O)NCc3ccccc3)[C@H]2CN1Cc1cccc2ccccc12 | HQWTUOLCGKIECB-XZWHSSHBSA-N | No-Effect | inactive |
| ICI-118,551 | BRD-K74603273 | Cc1ccc(OC[C@@H](O)[C@H](C)NC(C)C)c2c1CCC2 | VFIDUCMKNJIJTO-XJKSGUPXSA-N | No-Effect | inactive |
| ICI-162846 | BRD-K66782112 | N=C(NCC(F)(F)F)Nc1ccn(CCCCC(N)=O)n1 | ALCSGJCIESECFD-UHFFFAOYSA-N | No-Effect | inactive |
| ICI-169369 | BRD-K00004569 | CN(C)CCSc1nc2ccccc2cc1-c1ccccc1 | HYOLQGVNMQNERE-UHFFFAOYSA-N | No-Effect | inactive |
| ICI-185,282 | BRD-K29133151 | O=C(O)CCC/C=C\C[C@@H]1CO[C@H](C(F)(F)F)O[C@@H]1c1ccccc1O | ZWAVGFSZMACJHA-PMNBYGLBSA-N | No-Effect | inactive |
| ICI-192605 | BRD-K59433843 | O=C(O)CC/C=C\C[C@@H]1CO[C@H](c2ccccc2Cl)O[C@@H]1c1ccccc1O | WHUIENZXNGAHQI-GKZDNZBASA-N | No-Effect | inactive |
| ICI-215,001 | BRD-K52487250 | O=C(O)COc1ccc(OCCNC[C@H](O)COc2ccccc2)cc1 | MXWICZOXPUWAHY-HNNXBMFYSA-N | No-Effect | inactive |
| ICI-63197 | BRD-K52219182 | CCCn1c(=O)c(C)cn2nc(N)nc12 | UQDVRVNMIJAGRK-UHFFFAOYSA-N | No-Effect | inactive |
| ICI-89406 | BRD-A03359064 | N#Cc1ccccc1OC[C@@H](O)CNCCNC(=O)Nc1ccccc1 | HTLWRKRZKFAAAH-KRWDZBQOSA-N | No-Effect | inactive |
| icilin | BRD-K48869804 | O=C1NC(c2cccc([N+](=O)[O-])c2)=CCN1c1ccccc1O | RCEFMOGVOYEGJN-UHFFFAOYSA-N | No-Effect | inactive |
| iclaprim | BRD-K00003415 | COc1cc(Cc2cnc(N)nc2N)c2c(c1OC)O[C@H](C1CC1)C=C2 | HWJPWWYTGBZDEG-AWEZNQCLSA-N | No-Effect | inactive |
| icomucret | BRD-K41234004 | CCCCC[C@H](O)/C=C/C=C\C/C=C\C/C=C\CCCC(=O)O | JSFATNQSLKRBCI-VAEKSGALSA-N | No-Effect | inactive |
| icosapent | BRD-K47192521 | CC/C=C\C/C=C\C/C=C\C/C=C\C/C=C\CCCC(=O)O | JAZBEHYOTPTENJ-JLNKQSITSA-N | No-Effect | inactive |

| | | | | | |
|---|---|---|---|---|---|
| iCRT-14 | BRD-K18634484 | Cc1cc(/C=C2\SC(=O)N(c3ccccc3)C2=O)c(C)n1-c1cccnc1 | NCSHZXNGQYSKLR-UNOMPAQXSA-N | No-Effect | inactive |
| ID-1101 | BRD-K90305467 | C[C@H](O)[C@H](C)[C@H](N)C(=O)O | OSCCDBFHNMXNME-YUPRTTJUSA-N | No-Effect | inactive |
| idalopirdine | BRD-K79102359 | Fc1ccc2c(CCNCc3cccc(OCC(F)(F)C(F)F)c3)c[nH]c2c1 | YBAWYTYNMZWMMJ-UHFFFAOYSA-N | No-Effect | inactive |
| idasanutlin | BRD-K62627508 | COc1cc(C(=O)O)ccc1NC(=O)[C@@H]1N[C@@H](CC(C)(C)C)[C@](C#N)(c2ccc(Cl)cc2F)[C@H]1c1cccc(Cl)c1F | TVTXCJFHQKSQQM-LJQIRTBHSA-N | No-Effect | inactive |
| idebenone | BRD-K37516142 | COC1=C(OC)C(=O)C(CCCCCCCCCCO)=C(C)C1=O | JGPMMRGNQUBGND-UHFFFAOYSA-N | No-Effect | inactive |
| idelalisib | BRD-K60866521 | CC[C@H](Nc1ncnc2nc[nH]c12)c1nc2cccc(F)c2c(=O)n1-c1ccccc1 | IFSDAJWBUCMOAH-HNNXBMFYSA-N | No-Effect | inactive |
| IDF-11774 | BRD-K00003098 | CN1CCN(C(=O)COc2ccc(C34CC5CC(CC(C5)C3)C4)cc2)CC1 | QGBBBLPWBSWERZ-UHFFFAOYSA-N | No-Effect | inactive |
| IDH-305 | BRD-K00003457 | Cc1cc([C@H](C)Nc2nccc(N3C(=O)OC[C@@H]3[C@H](C)F)n2)ncc1-c1ccnc(C(F)(F)F)c1 | DCGDPJCUIKLTDU-SUNYJGFJSA-N | No-Effect | inactive |
| IDO5L | BRD-K22356203 | Nc1nonc1/C(=N\c1ccc(F)c(Cl)c1)NO | HGXSLPIXNPASGZ-UHFFFAOYSA-N | No-Effect | inactive |
| idoxuridine | BRD-K76634210 | O=c1[nH]c(=O)n([C@H]2C[C@H](O)[C@@H](CO)O2)cc1I | XQFRJNBWHJMXHO-RRKCRQDMSA-N | No-Effect | inactive |
| IDRA-21 | BRD-A14344385 | C[C@H]1Nc2ccc(Cl)cc2S(=O)(=O)N1 | VZRNTCHTJRLTMU-YFKPBYRVSA-N | No-Effect | inactive |
| idramantone | BRD-K00003671 | O=C1[C@H]2C[C@H]3C[C@@H]1C[C@](O)(C3)C2 | TZBDEVBNMSLVKT-XYYXLIQBSA-N | No-Effect | inactive |
| idronoxil | BRD-K74671368 | Oc1ccc(C2=Cc3ccc(O)cc3OC2)cc1 | ZZUBHVMHNVYXRR-UHFFFAOYSA-N | No-Effect | inactive |
| IEM1460 | BRD-K13829090 | C[N+](C)(C)CCCCCNCC12CC3CC(CC(C3)C1)C2 | IWYZMMIPAJDJBN-UHFFFAOYSA-N | No-Effect | inactive |
| IEM1754 | BRD-K80060353 | NCCCCCNCC12CC3CC(CC(C3)C1)C2 | UWWMQHVWAYFCDT-UHFFFAOYSA-N | No-Effect | inactive |
| ifenprodil | BRD-K00004210 | C[C@@H]([C@H](O)c1ccc(O)cc1)N1CCC(Cc2ccccc2)CC1 | UYNVMODNBIQBMV-KKSFZXQISA-N | No-Effect | inactive |
| ifosfamide | BRD-A67097164 | O=[P@]1(NCCCl)OCCCN1CCCl | HOMGKSMUEGBAAB-CQSZACIVSA-N | No-Effect | inactive |
| igmesine | BRD-A48376327 | CCC(C/C=C/c1ccccc1)(c1ccccc1)N(C)CC1CC1 | VCZSWYIFCKGTJI-JLHYYAGUSA-N | No-Effect | inactive |
| iguratimod | BRD-K15426076 | CS(=O)(=O)Nc1cc2occ(NC=O)c(=O)c2cc1Oc1ccccc1 | ANMATWQYLIFGOK-UHFFFAOYSA-N | No-Effect | inactive |
| IKK-16 | BRD-K14618467 | O=C(c1ccc(Nc2nccc(-c3cc4ccccc4s3)n2)cc1)N1CCC(N2CCCC2)CC1 | BWZJBXAPRCVCKQ-UHFFFAOYSA-N | No-Effect | inactive |
| IKK-2-inhibitor | BRD-K26373640 | NC(=O)c1sc(-c2ccsc2)cc1N | BMUACLADCKCNKZ-UHFFFAOYSA-N | No-Effect | inactive |
| ilepcimide | BRD-K60480335 | O=C(/C=C/c1ccc2c(c1)OCO2)N1CCCCC1 | BLPUOQGPBJPXRL-FNORWQNLSA-N | No-Effect | inactive |
| ilomastat | BRD-K51662849 | CNC(=O)[C@H](Cc1c[nH]c2ccccc12)NC(=O)[C@@H](CC(=O)NO)CC(C)C | NITYDPDXAAFEIT-DYVFJYSZSA-N | No-Effect | inactive |
| iloperidone | BRD-K48722833 | COc1cc(C(C)=O)ccc1OCCCN1CCC(c2noc3cc(F)ccc23)CC1 | XMXHEBAFVSFQEX-UHFFFAOYSA-N | No-Effect | inactive |
| iloprost | BRD-A45664787 | CC#CC[C@@H](C)[C@H](O)/C=C/[C@@H]1[C@H]2C/C(=C/CCCC(=O)O)C[C@H]2C[C@H]1O | HIFJCPQKFCZDDL-GBSCXWAGSA-N | No-Effect | inactive |
| Imazodan | BRD-K96298462 | O=C1CCC(c2ccc(-n3ccnc3)cc2)=NN1 | VXMYWVMXSWJFCV-UHFFFAOYSA-N | No-Effect | inactive |
| imeglimin | BRD-K55822903 | C[C@@H]1N=C(N)N=C(N(C)C)N1 | GFICWFZTBXUVIG-SCSAIBSYSA-N | No-Effect | inactive |
| imetit | BRD-K33183200 | N=C(N)SCCc1cnc[nH]1 | PEHSVUKQDJULKE-UHFFFAOYSA-N | No-Effect | inactive |
| imexon | BRD-A78195072 | NC1=NC(=O)N2C[C@@H]12 | BIXBBIPTYBJTRY-VBMBUWLQSA-N | No-Effect | inactive |
| imidacloprid | BRD-K76609936 | O=[N+]([O-])N=C1NCCN1Cc1ccc(Cl)nc1 | YWTYJOPNNQFBPC-UHFFFAOYSA-N | No-Effect | inactive |
| imidafenacin | BRD-K29626940 | Cc1nccn1CCC(C(N)=O)(c1ccccc1)c1ccccc1 | SQKXYSGRELMAAU-UHFFFAOYSA-N | No-Effect | inactive |
| imidazoacridinone | BRD-K00004617 | CCN(CC)CCNc1ccc2ncn3c4ccc(O)cc4c(=O)c1c23 | CUNDRHORZHFPLY-UHFFFAOYSA-N | No-Effect | inactive |
| imiloxan | BRD-A10903566 | CCn1ccnc1C[C@H]1COc2ccccc2O1 | UXABARREKCJULM-NSHDSACASA-N | No-Effect | inactive |
| imiquimod | BRD-K26657438 | CC(C)Cn1cnc2c(N)nc3ccccc3c21 | DOUYETYNHWVLEO-UHFFFAOYSA-N | No-Effect | inactive |
| imirestat | BRD-K00003208 | O=C1NC(=O)C2(N1)c1cc(F)ccc1-c1ccc(F)cc12 | QCCHBHSAIQIQGO-UHFFFAOYSA-N | No-Effect | inactive |
| imisopasem-manganese | BRD-K00003154 | Cl[Mn]123(Cl)N4CCN1[C@@H]1CCCC[C@H]1N2Cc1cccc(n1)CN3[C@@H]1CCCC[C@H]14 | UWYSKLMRWPKZES-XNPJUPKFSA-L | No-Effect | inactive |
| immepip | BRD-K25906698 | c1ncc(CC2CCNCC2)[nH]1 | MCNGUYXRBCIGOV-UHFFFAOYSA-N | No-Effect | inactive |
| immethridine | BRD-K49519092 | c1cc(Cc2cnc[nH]2)ccn1 | DFVSGZHJSIEEQQ-UHFFFAOYSA-N | No-Effect | inactive |
| impentamine | BRD-K20049318 | NCCCCCc1cnc[nH]1 | MZCJWLAXZRFUPI-UHFFFAOYSA-N | No-Effect | inactive |
| IMREG-1 | BRD-K00926464 | N[C@@H](Cc1ccc(O)cc1)C(=O)NCC(=O)NCC(=O)O | HIINQLBHPIQYHN-JTQLQIEISA-N | No-Effect | inactive |
| inarigivir | BRD-K00003532 | CO[C@@H]1[C@H](O[P@@](=O)(S)OC[C@H]2O[C@@H](n3cnc4c(N)ncnc43)C[C@@H]2O)[C@@H](CO)O[C@H]1n1ccc(=O)[nH]c1=O | LYMICVBGNUEHGE-NLKZJYKHSA-N | No-Effect | inactive |
| inarigivir-soproxil | BRD-K00004751 | CO[C@@H]1[C@H](O[P@](=O)(OC[C@H]2O[C@@H](n3cnc4c(N)ncnc43)C[C@@H]2O)SCOC(=O)OC(C)C)[C@@H](CO)O[C@H]1n1ccc(=O)[nH]c1=O | CJCYTUJOSMYXLE-NHMFLIHISA-N | No-Effect | inactive |
| INC-280 | BRD-K81016934 | CNC(=O)c1ccc(-c2cnc3ncc(Cc4ccc5ncccc5c4)n3n2)cc1F | LIOLIMKSCNQPLV-UHFFFAOYSA-N | No-Effect | inactive |
| INCA-6 | BRD-K33308633 | O=C1C=CC(=O)C2=C1C1c3ccccc3C2c2ccccc21 | GCHPUOHXXCNSQL-UHFFFAOYSA-N | No-Effect | inactive |
| INCB-003284 | BRD-K28059174 | COc1ccc(C2(O)CCC(N3CC[C@@H](NC(=O)CNC(=O)c4cccc(C(F)(F)F)c4)C3)CC2)cn1 | NXZNYBUBXWWKCP-JMOWIOHXSA-N | No-Effect | inactive |
| incyclinide | BRD-K00003344 | NC(=O)[C@@H]1C(=O)C[C@@H]2C[C@@H]3Cc4cccc(O)c4C(=O)[C@@H]3C(=O)[C@]2(O)C1=O | MDYVZYDNBUAVCN-WCAKBIDQSA-N | No-Effect | inactive |
| indacaterol | BRD-K89208535 | CCc1cc2c(cc1CC)CC(NC[C@H](O)c1ccc(O)c3[nH]c(=O)ccc13)C2 | QZZUEBNBZAPZLX-QFIPXVFZSA-N | No-Effect | inactive |
| indalpine | BRD-K64973364 | c1ccc2c(CCC3CCNCC3)c[nH]c2c1 | SADQVAVFGNTEOD-UHFFFAOYSA-N | No-Effect | inactive |
| indapamide | BRD-A95869247 | C[C@H]1Cc2ccccc2N1NC(=O)c1ccc(Cl)c(S(N)(=O)=O)c1 | NDDAHWYSQHTHNT-JTQLQIEISA-N | No-Effect | inactive |
| indeglitazar | BRD-K00003177 | COc1ccc(S(=O)(=O)n2cc(CCC(=O)O)c3cc(OC)ccc32)cc1 | YMPALHOKRBVHOJ-UHFFFAOYSA-N | No-Effect | inactive |
| indeloxazine | BRD-A59747053 | C1=Cc2cccc(OC[C@H]3CNCCO3)c2C1 | MADRVGBADLFHMO-GFCCVEGCSA-N | No-Effect | inactive |
| indibulin | BRD-K74057757 | O=C(Nc1ccncc1)C(=O)c1cn(Cc2ccc(Cl)cc2)c2ccccc12 | SOLIIYNRSAWTSQ-UHFFFAOYSA-N | No-Effect | inactive |
| indiplon | BRD-K90948141 | CC(=O)N(C)c1cccc(-c2ccnc3c(C(=O)c4cccs4)cnn23)c1 | CBIAWPMZSFFRGN-UHFFFAOYSA-N | No-Effect | inactive |
| indirubin-3-monoxime | BRD-K19136521 | O=C1Nc2ccccc2/C1=C1/Nc2ccccc2/C1=N\O | HBDSHCUSXQATPO-BGBJRWHRSA-N | No-Effect | inactive |
| indisulam | BRD-K17610631 | NS(=O)(=O)c1ccc(S(=O)(=O)Nc2cccc3c(Cl)c[nH]c23)cc1 | SETFNECMODOHTO-UHFFFAOYSA-N | No-Effect | inactive |
| indium-tri(2-propanolate) | BRD-K42332878 | CC(C)O[In](OC(C)C)OC(C)C | OVZUSPADPSOQQN-UHFFFAOYSA-N | No-Effect | inactive |
| indium(III)-isopropoxide | BRD-M08274809 | CC(C)O.CC(C)O.CC(C)O.[In] | ZLAWRQNZHDPZMF-UHFFFAOYSA-N | No-Effect | inactive |
| indobufen | BRD-A16296925 | CC[C@@H](C(=O)O)c1ccc(N2Cc3ccccc3C2=O)cc1 | AYDXAULLCROVIT-OAHLLOKOSA-N | No-Effect | inactive |
| indole-3-carbinol | BRD-K01815685 | OCc1c[nH]c2ccccc12 | IVYPNXXAYMYVSP-UHFFFAOYSA-N | No-Effect | inactive |
| indole-3-pyrubate | BRD-K09109825 | O=C(O)C(O)=Cc1c[nH]c2ccccc12 | KDCSRTPNDBPWQZ-UHFFFAOYSA-N | No-Effect | inactive |
| indomethacin | BRD-K57222227 | COc1ccc2c(c1)c(CC(=O)O)c(C)n2C(=O)c1ccc(Cl)cc1 | CGIGDMFJXJATDK-UHFFFAOYSA-N | No-Effect | inactive |
| indoprofen | BRD-A44090213 | C[C@@H](C(=O)O)c1ccc(N2Cc3ccccc3C2=O)cc1 | RJMIEHBSYVWVIN-LLVKDONJSA-N | No-Effect | inactive |
| indoramin | BRD-K76953762 | O=C(NC1CCN(CCc2c[nH]c3ccccc23)CC1)c1ccccc1 | JXZZEXZZKAWDSP-UHFFFAOYSA-N | No-Effect | inactive |
| indoximod | BRD-K93255255 | Cn1cc(C[C@@H](N)C(=O)O)c2ccccc21 | ZADWXFSZEAPBJS-SNVBAGLBSA-N | No-Effect | inactive |
| ingenol | BRD-K00003329 | CC1=C[C@]23C(=O)[C@@H](C=C(CO)[C@@H](O)[C@]2(O)[C@H]1O)[C@H]1[C@@H](C[C@H]3C)C1(C)C | VEBVPUXQAPLADL-POYOOMFHSA-N | No-Effect | inactive |
| INH1 | BRD-K70177501 | Cc1ccc(-c2csc(NC(=O)c3ccccc3)n2)c(C)c1 | JPMOKRWIYQGMJL-UHFFFAOYSA-N | No-Effect | inactive |
| inimur | BRD-A71407503 | CSC[C@H]1CN(/N=C/c2ccc([N+](=O)[O-])o2)C(=O)O1 | SRQKTCXJCCHINN-ORMNHXAVSA-N | No-Effect | inactive |
| iniparib | BRD-K11153516 | NC(=O)c1ccc(I)c([N+](=O)[O-])c1 | MDOJTZQKHMAPBK-UHFFFAOYSA-N | No-Effect | inactive |
| inosine | BRD-K79612754 | O=c1nc[nH]c2c1ncn2[C@@H]1O[C@H](CO)[C@@H](O)[C@H]1O | UGQMRVRMYYASKQ-KQYNXXCUSA-N | No-Effect | inactive |
| inositol-hexanicotinate | BRD-K82800842 | O=C(O[C@H]1[C@H](OC(=O)c2cccnc2)[C@@H](OC(=O)c2cccnc2)[C@H](OC(=O)c2cccnc2)[C@@H](OC(=O)c2cccnc2)[C@H]1OC(=O)c1cccnc1)c1cccnc1 | MFZCIDXOLLEMOO-GYSGTQPESA-N | No-Effect | inactive |
| INS316 | BRD-A29125379 | O=c1ccn([C@@H]2O[C@H](COP(=O)(O)OP(=O)(O)OP(=O)(O)O)[C@@H](O)[C@H]2O)c(=O)[nH]1 | PGAVKCOVUIYSFO-PXBUCIJWSA-N | No-Effect | inactive |

| | | | | | |
|---|---|---|---|---|---|
| INT-747 | BRD-K30968378 | CC[C@@H]1[C@@H]2C[C@H](O)CC[C@]2(C)[C@H]2CC[C@@]3(C)[C@@H](CC[C@@H]3[C@H](C)CCC(=O)O)[C@@H]2[C@@H]1O | ZXERDUOLZKYMJM-ZWECCWDJSA-N | No-Effect | inactive |
| INT-767 | BRD-K33067152 | CC[C@@H]1[C@@H]2C[C@H](O)CC[C@]2(C)[C@H]2CC[C@@]3(C)[C@@H](CC[C@@H]3[C@H](C)CCOS(=O)(=O)O)[C@@H]2[C@@H]1O | XGIYOABXZNJOHV-APIYUPOTSA-N | No-Effect | inactive |
| integrin-antagonist-1 | BRD-K88165431 | Cc1cc(C)n(-c2cccc([C@H](CC(=O)O)CN3CC[C@@H](CCc4ccc5c(n4)NCCC5)C3)c2)n1 | ZMXBIIQMSGOIRZ-RCZVLFRGSA-N | No-Effect | inactive |
| iobenguane | BRD-K43860855 | N=C(N)NCc1cccc(I)c1 | PDWUPXJEEYOOTR-UHFFFAOYSA-N | No-Effect | inactive |
| iocetamic-acid | BRD-A37492983 | CC(=O)N(C[C@H](C)C(=O)O)c1c(I)cc(I)c(N)c1I | GSVQIUGOUKJHRC-YFKPBYRVSA-N | No-Effect | inactive |
| iodixanol | BRD-A08660406 | CC(=O)N(CC(O)CN(C(C)=O)c1c(I)c(C(=O)NC[C@@H](O)CO)c(I)c(C(=O)NC[C@@H](O)CO)c1I)c1c(I)c(C(=O)NC[C@@H](O)CO)c(I)c(C(=O)NC[C@@H](O)CO)c1I | NBQNWMBBSKPBAY-BRSBDYLESA-N | No-Effect | inactive |
| iodoantipyrine | BRD-K69216798 | Cc1c(I)c(=O)n(-c2ccccc2)n1C | ZZOBLCHCPLOXCE-UHFFFAOYSA-N | No-Effect | inactive |
| iododexetimide | BRD-A46813902 | O=C1CC[C@](c2ccccc2)(C2CCN(Cc3ccc(I)cc3)CC2)C(=O)N1 | WJLRTFJPHDSXAF-QHCPKHFHSA-N | No-Effect | inactive |
| iodophenpropit | BRD-K51918615 | N=C(NCCc1ccc(I)cc1)SCCCc1cnc[nH]1 | UBHYDQAARZKHEZ-UHFFFAOYSA-N | No-Effect | inactive |
| iodoquinol | BRD-K75855670 | Oc1c(I)cc(I)c2cccnc12 | UXZFQZANDVDGMM-UHFFFAOYSA-N | No-Effect | inactive |
| iohexol | BRD-A60217728 | CC(=O)N(C[C@@H](O)CO)c1c(I)c(C(=O)NC[C@@H](O)CO)c(I)c(C(=O)NC[C@@H](O)CO)c1I | NTHXOOBQLCIOLC-GMTAPVOTSA-N | No-Effect | inactive |
| iopamidol | BRD-K75868704 | C[C@H](O)C(=O)Nc1c(I)c(C(=O)NC(CO)CO)c(I)c(C(=O)NC(CO)CO)c1I | XQZXYNRDCRIARQ-LURJTMIESA-N | No-Effect | inactive |
| iopanic-acid | BRD-A42628519 | CC[C@@H](Cc1c(I)cc(I)c(N)c1I)C(=O)O | OIRFJRBSRORBCM-YFKPBYRVSA-N | No-Effect | inactive |
| iopodic-acid | BRD-K51956333 | CN(C)/C=N/c1c(I)cc(I)c(CCC(=O)O)c1I | YQNFBOJPTAXAKV-OMCISZLKSA-N | No-Effect | inactive |
| iothalamic-acid-d3 | BRD-K98250023 | CNC(=O)c1c(I)c(NC(C)=O)c(I)c(C(=O)O)c1I | UXIGWFXRQKWHHA-UHFFFAOYSA-N | No-Effect | inactive |
| ioversol | BRD-A65818372 | O=C(NC[C@@H](O)CO)c1c(I)c(C(=O)NC[C@@H](O)CO)c(I)c(N(CCO)C(=O)CO)c1I | AMDBBAQNWSUWGN-RKDXNWHRSA-N | No-Effect | inactive |
| IOWH032 | BRD-K84892605 | O=C(NCc1ccc(Oc2ccccc2)cc1)c1nc(-c2cc(Br)c(O)c(Br)c2)no1 | DSFNLJXHXBIKDS-UHFFFAOYSA-N | No-Effect | inactive |
| IOX1 | BRD-K70751730 | O=C(O)c1ccc(O)c2ncccc12 | JGRPKOGHYBAVMW-UHFFFAOYSA-N | No-Effect | inactive |
| IOX2 | BRD-K98251413 | O=C(O)CNC(=O)c1c(O)c2ccccc2n(Cc2ccccc2)c1=O | CAOSCCRYLYQBES-UHFFFAOYSA-N | No-Effect | inactive |
| ioxaglic-acid | BRD-K79124250 | CNC(=O)c1c(I)c(C(=O)NCC(=O)Nc2c(I)c(C(=O)O)c(I)c(C(=O)NCCO)c2I)c(I)c(N(C)C(C)=O)c1I | TYYBFXNZMFNZJT-UHFFFAOYSA-N | No-Effect | inactive |
| ioxilan | BRD-A09370961 | CC(=O)N(C[C@@H](O)CO)c1c(I)c(C(=O)NCCO)c(I)c(C(=O)NC[C@@H](O)CO)c1I | UUMLTINZBQPNGF-NXEZZACHSA-N | No-Effect | inactive |
| IPA-3 | BRD-K38727704 | Oc1ccc2ccccc2c1SSc1c(O)ccc2ccccc12 | RFAXLXKIAKIUDT-UHFFFAOYSA-N | No-Effect | inactive |
| IPI549 | BRD-K00003385 | C[C@H](NC(=O)c1c(N)nn2cccnc12)c1cc2cccc(C#Cc3cnn(C)c3)c2c(=O)n1-c1ccccc1 | XUMALORDVCFWKV-IBGZPJMESA-N | No-Effect | inactive |
| ipidacrine | BRD-K66896231 | Nc1c2c(nc3c1CCC3)CCCC2 | YLUSMKAJIQOXPV-UHFFFAOYSA-N | No-Effect | inactive |
| ipratropium | BRD-K01826778 | CC(C)[N@@+]1(C)[C@H]2CC[C@@H]1C[C@H](OC(=O)[C@@H](CO)c1ccccc1)C2 | OEXHQOGQTVQTAT-KKKDIUQISA-N | No-Effect | inactive |
| ipriflavone | BRD-K93618743 | CC(C)Oc1ccc2c(=O)c(-c3ccccc3)coc2c1 | SFBODOKJTYAUCM-UHFFFAOYSA-N | No-Effect | inactive |
| iproniazid | BRD-K88568253 | CC(C)NNC(=O)c1ccncc1 | NYMGNSNKLVNMIA-UHFFFAOYSA-N | No-Effect | inactive |
| ipsapirone | BRD-K90574421 | O=C1c2ccccc2S(=O)(=O)N1CCCCN1CCN(c2ncccn2)CC1 | TZJUVVIWVWFLCD-UHFFFAOYSA-N | No-Effect | inactive |
| irbesartan | BRD-K60038276 | CCCCC1=NC2(CCCC2)C(=O)N1Cc1ccc(-c2ccccc2-c2nn[nH]n2)cc1 | YOSHYTLCDANDAN-UHFFFAOYSA-N | No-Effect | inactive |
| irganox-1010 | BRD-K00004549 | CC(C)(C)c1cc(CCC(=O)OCC(COC(=O)CCc2cc(C(C)(C)C)c(O)c(C(C)(C)C)c2)(COC(=O)CCc2cc(C(C)(C)C)c(O)c(C(C)(C)C)c2)COC(=O)CCc2cc(C(C)(C)C)c(O)c(C(C)(C)C)c2)cc(C(C)(C)C)c1O | BGYHLZZASRKEJE-UHFFFAOYSA-N | No-Effect | inactive |
| iRGD-peptide | BRD-K00003335 | N=C(N)NCCC[C@@H]1NC(=O)[C@@H](N)CSSC[C@@H](C(=O)O)NC(=O)[C@H](CC(=O)O)NC(=O)[C@@H]2CCCN2C(=O)CNC(=O)[C@H](CCCCN)NC(=O)[C@H](CC(=O)O)NC(=O)CNC1=O | YHTTWXCDIRTOQX-FQJIPJFPSA-N | No-Effect | inactive |
| irinotecan | BRD-K08547377 | CCc1c2c(nc3ccc(OC(=O)N4CCC(N5CCCCC5)CC4)cc13)-c1cc3c(c(=O)n1C2)COC(=O)[C@]3(O)CC | UWKQSNNFCGGAFS-XIFFEERXSA-N | No-Effect | inactive |
| IRL-2500 | BRD-K98357249 | Cc1cc(C)cc(C(=O)N(C)[C@H](Cc2ccc(-c3ccccc3)cc2)C(=O)N[C@@H](Cc2c[nH]c3ccccc23)C(=O)O)c1 | UZDORQWMYRRLQV-JHOUSYSJSA-N | No-Effect | inactive |
| irosustat | BRD-K87777313 | NS(=O)(=O)Oc1ccc2c3c(c(=O)oc2c1)CCCCC3 | DSLPMJSGSBLWRE-UHFFFAOYSA-N | No-Effect | inactive |
| isamoltane | BRD-K01825673 | CC(C)NC[C@@H](O)COc1ccccc1-n1cccc1 | XVTVPGKWYHWYAD-CQSZACIVSA-N | No-Effect | inactive |
| isavuconazole | BRD-A17009129 | C[C@@H](c1nc(-c2ccc(C#N)cc2)cs1)[C@](O)(Cn1cncn1)c1cc(F)ccc1F | DDFOUSQFMYRUQK-FPTDNZKUSA-N | No-Effect | inactive |
| isaxonine | BRD-K95957366 | CC(C)Nc1ncccn1 | FTCYIGBVOHNHCD-UHFFFAOYSA-N | No-Effect | inactive |
| isbufylline | BRD-K06762493 | CC(C)Cn1cnc2c1c(=O)n(C)c(=O)n2C | WHUWQSQEVISUMC-UHFFFAOYSA-N | No-Effect | inactive |
| isepamicin | BRD-K74759733 | CN[C@@H]1[C@@H](O)[C@@H](O[C@@H]2[C@@H](O)[C@H](O[C@H]3O[C@H](CN)[C@@H](O)[C@H](O)[C@H]3O)[C@@H](N)C[C@H]2NC(=O)[C@@H](O)CN)OC[C@]1(C)O | UDIIBEDMEYAVNG-ZKFPOVNWSA-N | No-Effect | inactive |
| ISO-1 | BRD-A94055480 | COC(=O)C[C@H]1CC(c2ccc(O)cc2)=NO1 | AIXMJTYHQHQJLU-SNVBAGLBSA-N | No-Effect | inactive |
| isobutamben | BRD-K96714749 | CC(C)COC(=O)c1ccc(N)cc1 | PUYOAVGNCWPANW-UHFFFAOYSA-N | No-Effect | inactive |
| isobutyramide | BRD-K54057105 | CC(C)C(N)=O | WFKAJVHLWXSISD-UHFFFAOYSA-N | No-Effect | inactive |
| isocarboxazid | BRD-K93332168 | Cc1cc(C(=O)NNCc2ccccc2)no1 | XKFPYPQQHFEXRZ-UHFFFAOYSA-N | No-Effect | inactive |
| isoconazole | BRD-K01825590 | Clc1ccc([C@@H](Cn2ccnc2)OCc2c(Cl)cccc2Cl)c(Cl)c1 | MPIPASJGOJYODL-GOSISDBHSA-N | No-Effect | inactive |
| isodibut | BRD-K60241851 | O=C(O)CCCN1C(=O)c2cccc3cccc(c23)C1=O | ZHXRDXTYPCPBTI-UHFFFAOYSA-N | No-Effect | inactive |
| isoetharine | BRD-A24587114 | CC[C@@H](NC(C)C)[C@@H](O)c1ccc(O)c(O)c1 | HUYWAWARQUIQLE-MFKMUULPSA-N | No-Effect | inactive |
| isofloxythepin | BRD-A07704283 | CC(C)c1ccc2c(c1)[C@H](N1CCN(CCO)CC1)Cc1ccc(F)cc1S2 | XDBMBHVYXPBOPL-OAQYLSRUSA-N | No-Effect | inactive |
| isoflupredone-acetate | BRD-K29173907 | CC(=O)OCC(=O)[C@@]1(O)CC[C@H]2[C@@H]3CCC4=CC(=O)C=C[C@]4(C)[C@@]3(F)[C@@H](O)C[C@@]21C | ZOCUOMKMBMEYQV-GSLJADNHSA-N | No-Effect | inactive |
| isoflurane | BRD-A31564021 | FC(F)O[C@@H](Cl)C(F)(F)F | PIWKPBJCKXDKJR-PVQJCKRUSA-N | No-Effect | inactive |
| isoflurophate | BRD-K56575995 | CC(C)OP(=O)(F)OC(C)C | MUCZHBLJLSDCSD-UHFFFAOYSA-N | No-Effect | inactive |
| isoguvacine | BRD-K66480997 | O=C(O)C1=CCNCC1 | KRVDMABBKYMBHG-UHFFFAOYSA-N | No-Effect | inactive |
| isoliquiritigenin | BRD-K33583600 | O=C(/C=C/c1ccc(O)cc1)c1ccc(O)cc1O | DXDRHHKMWQZJHT-FPYGCLRLSA-N | No-Effect | inactive |
| isoniazid | BRD-K87202646 | NNC(=O)c1ccncc1 | QRXWMOHMRWLFEY-UHFFFAOYSA-N | No-Effect | inactive |
| isopentyl-4-methoxycinnamate | BRD-K37851352 | COc1ccc(/C=C/C(=O)OCCC(C)C)cc1 | UBNYRXMKIIGMKK-RMKNXTFCSA-N | No-Effect | inactive |
| isoprenaline | BRD-A04322457 | CC(C)NC[C@@H](O)c1ccc(O)c(O)c1 | JWZZKOKVBUJMES-LLVKDONJSA-N | No-Effect | inactive |
| isopropamide-iodide | BRD-K14127446 | CC(C)[N+](C)(CCC(C(N)=O)(c1ccccc1)c1ccccc1)C(C)C | JTPUMZTWMWIVPA-UHFFFAOYSA-O | No-Effect | inactive |
| isopropyl-myristate | BRD-K13409143 | CCCCCCCCCCCCCC(=O)OC(C)C | AXISYYRBXTVTFY-UHFFFAOYSA-N | No-Effect | inactive |
| isopropyl-palmitate | BRD-K30758549 | CCCCCCCCCCCCCCCC(=O)OC(C)C | XUGNVMKQXJXZCD-UHFFFAOYSA-N | No-Effect | inactive |
| isoquercitrin | BRD-K73991644 | O=c1c(O[C@@H]2O[C@H](CO)[C@@H](O)[C@H](O)[C@H]2O)c(-c2ccc(O)c(O)c2)oc2cc(O)cc(O)c12 | OVSQVDMCBVZWGM-QSOFNFLRSA-N | No-Effect | inactive |
| Isoquinoline,-6,7-dimethoxy-1-methyl-1,2,3,4-tetrahydro, | BRD-K01826521 | COc1cc2c(cc1OC)[C@H](C)NCC2 | HMYJLVDKPJHJCF-QMMMGPOBSA-N | No-Effect | inactive |
| isosorbide | BRD-K47178855 | O[C@H]1CO[C@@H]2[C@H](O)CO[C@H]12 | KLDXJTOLSGUMSJ-JGWLITMVSA-N | No-Effect | inactive |

| | | | | | |
|---|---|---|---|---|---|
| isosorbide-dinitrate | BRD-K55853390 | O=[N+]([O-])O[C@H]1CO[C@H]2[C@@H]1OC[C@H]2O[N+](=O)[O-] | MOYKHGMNXAOIAT-JGWLITMVSA-N | No-Effect | inactive |
| isosorbide-mononitrate | BRD-K82225283 | O=[N+]([O-])O[C@@H]1CO[C@@H]2[C@@H](O)CO[C@H]12 | YWXYYJSYQOXTPL-SLPGGIOYSA-N | No-Effect | inactive |
| isosteviol | BRD-A96529864 | C[C@@]12CC[C@H]3[C@]4(C)CCC[C@@](C)(C(=O)O)[C@H]4CC[C@]3(CC1=O)C2 | KFVUFODCZDRVSS-XGBBNYNSSA-N | No-Effect | inactive |
| isotiquimide | BRD-A02933148 | Cc1ccnc2c1CCC[C@H]2C(N)=S | PWPWEYFVLITGNU-SECBINFHSA-N | No-Effect | inactive |
| isovaleramide | BRD-K90069354 | CC(C)CC(N)=O | SANOUVWGPVYVAV-UHFFFAOYSA-N | No-Effect | inactive |
| isoxepac | BRD-K92116564 | O=C(O)Cc1ccc2c(c1)C(=O)c1ccccc1CO2 | QFGMXJOBTNZHEL-UHFFFAOYSA-N | No-Effect | inactive |
| isoxicam | BRD-A75552914 | Cc1cc(NC(=O)C2=C(O)c3ccccc3S(=O)(=O)N2C)no1 | YYUAYBYLJSNDCX-UHFFFAOYSA-N | No-Effect | inactive |
| isoxsuprine | BRD-K00003690 | C[C@@H](COc1ccccc1)N[C@H](C)[C@@H](O)c1ccc(O)cc1 | BMUKKTUHUDJSNZ-PMUMKWKESA-N | No-Effect | inactive |
| ispinesib | BRD-K64881305 | Cc1ccc(C(=O)N(CCCN)[C@@H](c2nc3cc(Cl)ccc3c(=O)n2Cc2ccccc2)C(C)C)cc1 | QJZRFPJCWMNVAV-HHHXNRCGSA-N | No-Effect | inactive |
| isradipine | BRD-A90799790 | COC(=O)C1=C(C)NC(C)=C(C(=O)OC(C)C)[C@@H]1c1cccc2nonc12 | HMJIYCCIJYRONP-MRXNPFEDSA-N | No-Effect | inactive |
| ISRIB | BRD-K12826175 | O=C(COc1ccc(Cl)cc1)N[C@H]1CC[C@H](NC(=O)COc2ccc(Cl)cc2)CC1 | HJGMCDHQPXTGAV-IYARVYRRSA-N | No-Effect | inactive |
| ISX-9 | BRD-K53903639 | O=C(NC1CC1)c1cc(-c2cccs2)on1 | SYENTKHGMVKGAQ-UHFFFAOYSA-N | No-Effect | inactive |
| itacitinib | BRD-K67482377 | N#CCC1(n2cc(-c3ncnc4[nH]ccc34)cn2)CN(C2CCN(C(=O)c3ccnc(C(F)(F)F)c3F)CC2)C1 | KTBSXLIQKWEBRB-UHFFFAOYSA-N | No-Effect | inactive |
| ITE | BRD-K60298136 | COC(=O)c1csc(C(=O)c2c[nH]c3ccccc23)n1 | KDDXOGDIPZSCTM-UHFFFAOYSA-N | No-Effect | inactive |
| ITI214 | BRD-K77987496 | CN1C(=O)c2c(nn(Cc3ccc(-c4cccc(F)n4)cc3)c2Nc2ccccc2)N2C1=N[C@@H]1CCC[C@@H]12 | BBIPVJCGIASXJB-PKTZIBPZSA-N | No-Effect | inactive |
| itopride | BRD-K96740444 | COc1ccc(C(=O)NCc2ccc(OCCN(C)C)cc2)cc1OC | QQQIECGTIMUVDS-UHFFFAOYSA-N | No-Effect | inactive |
| itraconazole | BRD-K00004318 | CC[C@H](C)n1ncn(-c2ccc(N3CCN(c4ccc(OC[C@H]5CO[C@](Cn6cncn6)(c6ccc(Cl)cc6Cl)O5)cc4)CC3)cc2)c1=O | VHVPQPYKVGDNFY-TUJWMRSMSA-N | No-Effect | inactive |
| ITX3 | BRD-K36515744 | Cc1cc(/C=c2/sc3nc4ccccc4n3c2=O)c(C)n1-c1ccccc1 | SJMYMKPBODEZSH-DEDYPNTBSA-N | No-Effect | inactive |
| ITX5061 | BRD-K76911699 | COc1c(NC(=O)C(=O)c2ccc(OCCN3CCOCC3)c3ccccc23)cc(C(C)(C)C)cc1NS(C)(=O)=O | UUROSJLZNDSXRF-UHFFFAOYSA-N | No-Effect | inactive |
| IU1 | BRD-K45841694 | Cc1cc(C(=O)CN2CCCC2)c(C)n1-c1ccc(F)cc1 | JUWDSDKJBMFLHE-UHFFFAOYSA-N | No-Effect | inactive |
| ivabradine | BRD-K33972682 | COc1cc2c(cc1OC)CC(=O)N(CCCN(C)C[C@H]1Cc3cc(OC)c(OC)cc31)CC2 | ACRHBAYQBXXRTO-OAQYLSRUSA-N | No-Effect | inactive |
| ivacaftor | BRD-K52818472 | CC(C)(C)c1cc(C(C)(C)C)c(NC(=O)c2c[nH]c3ccccc3c2=O)cc1O | PURKAOJPTOLRMP-UHFFFAOYSA-N | No-Effect | inactive |
| ivermectin | BRD-K85554912 | CC[C@H](C)[C@H]1O[C@]2(CC[C@@H]1C)C[C@@H]1C[C@@H](C/C=C(\C)[C@@H](O[C@H]3C[C@H](OC)[C@@H](O[C@H]4C[C@H](OC)[C@@H](O)[C@H](C)O4)[C@H](C)O3)[C@@H](C)/C=C/C=C3\CO[C@@H]4[C@H](O)C(C)=C[C@@H](C(=O)O1)[C@]34O)O2 | AZSNMRSAGSSBNP-XPNPUAGNSA-N | No-Effect | inactive |
| ivosidenib | BRD-K93073858 | N#Cc1ccnc(N2C(=O)CC[C@H]2C(=O)N(c2cncc(F)c2)[C@H](C(=O)NC2CC(F)(F)C2)c2ccccc2Cl)c1 | WIJZXSAJMHAVGX-DHLKQENFSA-N | No-Effect | inactive |
| IWP-L6 | BRD-K53234107 | O=C(CSc1nc2c(c(=O)n1-c1ccccc1)SCC2)Nc1ccc(-c2ccccc2)cn1 | QESQGTFWEQMCMH-UHFFFAOYSA-N | No-Effect | inactive |
| ixabepilone | BRD-K03601870 | C/C(=C\c1csc(C)n1)[C@@H]1C[C@@H]2O[C@]2(C)CCC[C@H](C)[C@H](O)[C@@H](C)C(=O)C(C)(C)[C@@H](O)CC(=O)N1 | FABUFPQFXZVHFB-PVYNADRNSA-N | No-Effect | inactive |
| J147 | BRD-K00004699 | COc1cccc(/C=N/N(C(=O)C(F)(F)F)c2ccc(C)cc2C)c1 | HYMZAYGFKNNHDN-SSDVNMTOSA-N | No-Effect | inactive |
| JAK3-inhibitor-V | BRD-K95676198 | C=CC(=O)c1ccc2ccccc2c1 | FOTCGZPFSUWZBN-UHFFFAOYSA-N | No-Effect | inactive |
| JD-5037 | BRD-K00003271 | CC(C)[C@H](/N=C(/NS(=O)(=O)c1ccc(Cl)cc1)N1C[C@H](c2ccccc2)C(c2ccc(Cl)cc2)=N1)C(N)=O | GTCSIQFTNPTSLO-RPWUZVMVSA-N | No-Effect | inactive |
| JDTic | BRD-K95837862 | CC(C)[C@@H](CN1CC[C@@](C)(c2cccc(O)c2)[C@@H](C)C1)NC(=O)[C@H]1Cc2cc(O)ccc2CN1 | ZLVXBBHTMQJRSX-VMGNSXQWSA-N | No-Effect | inactive |
| JI-101 | BRD-K00003209 | COc1ccc(Br)cc1NC(=O)Nc1cccc2c1ccn2Cc1ccnc(N)c1 | ZXBFYBLSJMEBEP-UHFFFAOYSA-N | No-Effect | inactive |
| JIB04 | BRD-K76180386 | Clc1ccc(N/N=C(\c2ccccc2)c2ccccn2)nc1 | YHHFKWKMXWRVTJ-OQKWZONESA-N | No-Effect | inactive |
| JNJ-10191584 | BRD-K15086322 | CN1CCN(C(=O)c2nc3cc(Cl)cc3[nH]2)CC1 | MOIWSUQWIOVGRH-UHFFFAOYSA-N | No-Effect | inactive |
| JNJ-10397049 | BRD-K61721596 | CC1(C)OC[C@H](NC(=O)Nc2ccc(Br)cc2Br)[C@H](c2ccccc2)O1 | RBKIJGLHFFQHBE-IRXDYDNUSA-N | No-Effect | inactive |
| JNJ-16259685 | BRD-K25596805 | CO[C@H]1CC[C@@H](C(=O)c2ccc3nc4c(cc3c2)CCCO4)CC1 | QOTAQTRFJWLFCR-XFHMXUHZSA-N | No-Effect | inactive |
| JNJ-1661010 | BRD-K47376733 | O=C(Nc1ccccc1)N1CCN(c2nc(-c3ccccc3)ns2)CC1 | BHBOSTKQCZEAJM-UHFFFAOYSA-N | No-Effect | inactive |
| JNJ-17203212 | BRD-K51770398 | O=C(Nc1ccc(C(F)(F)F)cn1)N1CCN(c2ncccc2C(F)(F)F)CC1 | JFRYYGVYCWYIDQ-UHFFFAOYSA-N | No-Effect | inactive |
| JNJ-26990990 | BRD-K24864897 | NS(=O)(=O)NCc1csc2ccccc12 | AWSKBQNOSRREEY-UHFFFAOYSA-N | No-Effect | inactive |
| JNJ-27141491 | BRD-K41895652 | CC[C@@H](c1ccc(F)c(F)c1)n1c(C(=O)OC)c(-c2ccno2)[nH]c1=S | SYARXICKXVXWNA-LBPRGKRZSA-N | No-Effect | inactive |
| JNJ-37822681 | BRD-K31802667 | Fc1ccc(CN2CCC(Nc3ccc(C(F)(F)F)nn3)CC2)cc1F | UVUYWJWYRLJHEN-UHFFFAOYSA-N | No-Effect | inactive |
| JNJ-38877605 | BRD-K02123250 | Cn1cc(-c2ccc3nnc(C(F)(F)c4ccc5ncccc5c4)n3n2)cn1 | JRWCBEOAFGHNNU-UHFFFAOYSA-N | No-Effect | inactive |
| JNJ-38877618 | BRD-K00003094 | FC(F)(c1ccc2ncccc2c1)c1nnc2ccc(-c3ccncc3)nn12 | KOAWAWHSMVKCON-UHFFFAOYSA-N | No-Effect | inactive |
| JNJ-39758979 | BRD-K00003403 | CC(C)c1cc(N2CC[C@@H](N)C2)nc(N)n1 | COOGVHJHSCBOQT-MRVPVSSYSA-N | No-Effect | inactive |
| JNJ-40411813 | BRD-K55029398 | CCCCn1ccc(N2CCC(c3ccccc3)CC2)c(Cl)c1=O | HYOGJHCDLQSAHX-UHFFFAOYSA-N | No-Effect | inactive |
| JNJ-42041935 | BRD-K01036220 | O=C(O)c1cnn(-c2nc3cc(Cl)c(OC(F)(F)F)cc3[nH]2)c1 | FXHHASJVTYRJHH-UHFFFAOYSA-N | No-Effect | inactive |
| JNJ-42165279 | BRD-K70330032 | O=C(Nc1cnccc1Cl)N1CCN(Cc2ccc3c(c2)OC(F)(F)O3)CC1 | YWGYNGCRVZLMCS-UHFFFAOYSA-N | No-Effect | inactive |
| JNJ-54175446 | BRD-K00004575 | C[C@@H]1c2nnn(-c3ncc(F)cn3)c2CCN1C(=O)c1cccc(C(F)(F)F)c1Cl | CWFVVQFVGMFTBD-SECBINFHSA-N | No-Effect | inactive |
| JNJ-64619178 | BRD-K30421675 | Nc1nc2cc(CC[C@H]3C[C@@H](n4ccc5c(N)ncnc54)[C@H](O)[C@@H]3O)ccc2cc1Br | DBSMLQTUDJVICQ-CJODITQLSA-N | No-Effect | inactive |
| JNJ-678 | BRD-K00004600 | CS(=O)(=O)CCCn1c(Cn2c(=O)n(CC(F)(F)F)c3ccncc32)cc2cc(Cl)ccc21 | GTQTUABHRCWVLL-UHFFFAOYSA-N | No-Effect | inactive |
| JNJ-7777120 | BRD-K17774839 | CN1CCN(C(=O)c2cc3cc(Cl)ccc3[nH]2)CC1 | HUQJRYMLJBBEDO-UHFFFAOYSA-N | No-Effect | inactive |
| josamycin | BRD-K00003705 | CO[C@@H]1[C@@H](O[C@@H]2O[C@H](C)[C@@H](O[C@H]3C[C@@](C)(O)[C@@H](OC(=O)CC(C)C)[C@H](C)O3)[C@H](N(C)C)[C@H]2O)[C@@H](CC=O)C[C@@H](C)[C@@H](O)/C=C/C=C/C[C@@H](C)OC(=O)C[C@H]1OC(C)=O | XJSFLOJWULLJQS-NGVXBBESSA-N | No-Effect | inactive |
| JPH203 | BRD-K00003340 | Nc1cc(COc2c(Cl)cc(C[C@H](N)C(=O)O)cc2Cl)c2oc(-c3ccccc3)nc2c1 | XNRZJPQTMQZBCE-SFHVURJKSA-N | No-Effect | inactive |
| JQ1-(+) | BRD-K54606188 | Cc1sc2c(c1C)C(c1ccc(Cl)cc1)=N[C@@H](CC(=O)OC(C)(C)C)c1nnc(C)n1-2 | DNVXATUJJDPFDM-KRWDZBQOSA-N | No-Effect | inactive |
| JTE-013 | BRD-K43330982 | Cc1nn(C)c2nc(NNC(=O)Nc3cc(Cl)nc(Cl)c3)cc(C(C)C)c12 | RNSLRQNDXRSASX-UHFFFAOYSA-N | No-Effect | inactive |
| JTE-607 | BRD-K81405859 | CCOC(=O)[C@H](Cc1ccccc1)NC(=O)c1cc(Cl)c(OCCN2CCN(C)CC2)c(Cl)c1O | IPSSXIMJJXSJQB-FQEVSTJZSA-N | No-Effect | inactive |
| JTE-907 | BRD-K63150726 | CCCCCOc1c(OC)ccc2cc(C(=O)NCc3ccc4c(c3)OCO4)c(=O)[nH]c12 | GRAJFFFXJYFVOC-UHFFFAOYSA-N | No-Effect | inactive |
| JW-55 | BRD-K53328210 | COc1ccc(C2(CNC(=O)c3ccc(NC(=O)c4ccco4)cc3)CCOCC2)cc1 | ZJZWZIXSGNFWQQ-UHFFFAOYSA-N | No-Effect | inactive |
| JW-642 | BRD-K48424871 | O=C(OC(C(F)(F)F)C(F)(F)F)N1CCN(Cc2cccc(Oc3ccccc3)c2)CC1 | AVSCNEOUWSVZEY-UHFFFAOYSA-N | No-Effect | inactive |
| JW-67 | BRD-K24554037 | O=C1Nc2ccccc2C12OCC1(CO2)COC2(OC1)C(=O)Nc1ccccc12 | BTXRSHKJNDFHGA-UHFFFAOYSA-N | No-Effect | inactive |
| JW-74 | BRD-K75664313 | COc1ccc(-n2c(SCc3nc(-c4ccc(C)cc4)no3)nnc2-c2ccncc2)cc1 | KRIKILRRJCIWNB-UHFFFAOYSA-N | No-Effect | inactive |

| | | | | | |
|---|---|---|---|---|---|
| JX-401 | BRD-K05464208 | COc1cc(SC)ccc1C(=O)N1CCC(Cc2ccccc2)CC1 | OMGLGPKQUFSRNN-UHFFFAOYSA-N | No-Effect | inactive |
| JZL-184 | BRD-K45446451 | O=C(Oc1ccc([N+](=O)[O-])cc1)N1CCC(C(O)(c2ccc3c(c2)OCO3)c2ccc3c(c2)OCO3)CC1 | SEGYOKHGGFKMCX-UHFFFAOYSA-N | No-Effect | inactive |
| JZL-195 | BRD-K38875233 | O=C(Oc1ccc([N+](=O)[O-])cc1)N1CCN(Cc2cccc(Oc3ccccc3)c2)CC1 | QNYRAEKLMNDRFY-UHFFFAOYSA-N | No-Effect | inactive |
| K-247 | BRD-A74648924 | O=C(O)c1ccc(N[C@@H]2O[C@H](CO)[C@@H](O)[C@@H]2O)cc1 | QNUBRHYSSHJWDH-CHWFTXMASA-N | No-Effect | inactive |
| K-858 | BRD-A67748489 | CC(=O)NC1=NN(C(C)=O)[C@](C)(c2ccccc2)S1 | JEFVYQYZCAVNTP-ZDUSSCGKSA-N | No-Effect | inactive |
| K-MAP | BRD-A45906415 | O=C(O)c1ccc(N[C@@H]2O[C@H](CO)[C@@H](O)[C@H](O)[C@@H]2O)cc1 | VXMVXDITZWJXJN-LDMBFOFVSA-N | No-Effect | inactive |
| K-Ras(G12C)-inhibitor-12 | BRD-K90756851 | C=CC(=O)N1CCN(C(=O)CNc2cc(I)c(Cl)cc2O)CC1 | JFIFBWVNHLXJFY-UHFFFAOYSA-N | No-Effect | inactive |
| K145 | BRD-K12106308 | CCCCOc1ccc(CC/C=C2\SC(=O)N(CCN)C2=O)cc1 | MPZXLTZVPUSTFY-SOFYXZRVSA-N | No-Effect | inactive |
| kaempferol | BRD-K12807006 | O=c1c(O)c(-c2ccc(O)cc2)oc2cc(O)cc(O)c12 | IYRMWMYZSQPJKC-UHFFFAOYSA-N | No-Effect | inactive |
| KAF-156 | BRD-K00004578 | CC1(C)c2nc(-c3ccc(F)cc3)c(Nc3ccc(F)cc3)n2CCN1C(=O)CN | BUPRVECGWBHCQV-UHFFFAOYSA-N | No-Effect | inactive |
| kainic-acid | BRD-K52459018 | C=C(C)[C@H]1CN[C@H](C(=O)O)[C@H]1CC(=O)O | VLSMHEGGTFMBBZ-OOZYFLPDSA-N | No-Effect | inactive |
| kakonein | BRD-A07815743 | O=c1c(-c2ccc(O)cc2)coc2c([C@H]3O[C@H](CO)[C@@H](O)[C@H](O)[C@@H]3O)c(O)ccc12 | HKEAFJYKMMKDOR-YAJUCGGBSA-N | No-Effect | inactive |
| karenitecin | BRD-K00003176 | CC[C@@]1(O)C(=O)OCc2c1cc1n(c2=O)Cc2c-1nc1ccccc1c2CC[Si](C)(C)C | POADTFBBIXOWFJ-VWLOTQADSA-N | No-Effect | inactive |
| kartogenin | BRD-K27584835 | O=C(O)c1ccccc1C(=O)Nc1ccc(-c2ccccc2)cc1 | SLUINPGXGFUMLL-UHFFFAOYSA-N | No-Effect | inactive |
| KB-SRC-4 | BRD-K88154110 | Nc1ncnc2c1c(-c1ccc(Cl)cc1)nn2-c1cccc(-c2cnnn2Cc2cccc(-c3ccccc3)c2)c1 | UHIZYQVRKSWIFO-UHFFFAOYSA-N | No-Effect | inactive |
| KBG | BRD-A00376169 | O[C@H]1O[C@@H]2CO[C@H](c3ccccc3)O[C@H]2[C@H](O)[C@H]1O | FOLRUCXBTYDAQK-PTEOKISQSA-N | No-Effect | inactive |
| KD-023 | BRD-K71164191 | CCCCCCCCCCCCCCOc1ccc(C(=O)O)o1 | CZRCFAOMWRAFIC-UHFFFAOYSA-N | No-Effect | inactive |
| KD025 | BRD-K88506063 | CC(C)NC(=O)COc1cccc(-c2nc(Nc3ccc4n[nH]cc4c3)c3ccccc3n2)c1 | GKHIVNAUVKXIIY-UHFFFAOYSA-N | No-Effect | inactive |
| kenpaullone | BRD-K37312348 | O=C1Cc2c([nH]c3ccc(Br)cc23)-c2ccccc2N1 | QQUXFYAWXPMDOE-UHFFFAOYSA-N | No-Effect | inactive |
| ketanserin | BRD-K49671696 | O=C(c1ccc(F)cc1)C1CCN(CCn2c(=O)[nH]c3ccccc3c2=O)CC1 | FPCCSQOGAWCVBH-UHFFFAOYSA-N | No-Effect | inactive |
| ketoconazole | BRD-K29113274 | CC(=O)N1CCN(c2ccc(OC[C@@H]3CO[C@@](Cn4ccnc4)(c4ccc(Cl)cc4Cl)O3)cc2)CC1 | XMAYWYJOQHXEEK-ZEQKJWHPSA-N | No-Effect | inactive |
| ketohexokinase-inhibitor-1 | BRD-K00003341 | C[C@H]1CCN1c1nc(N2C[C@H]3[C@H](CC(=O)O)[C@H]3C2)cc(C(F)(F)F)n1 | MDUYWDNWFXSMJJ-XWLWVQCSSA-N | No-Effect | inactive |
| ketoprofen | BRD-A97739905 | C[C@@H](C(=O)O)c1cccc(C(=O)c2ccccc2)c1 | DKYWVDODHFEZIM-LLVKDONJSA-N | No-Effect | inactive |
| ketorolac | BRD-K01825750 | O=C(c1ccccc1)c1ccc2n1CC[C@@H]2C(=O)O | OZWKMVRBQXNZKK-NSHDSACASA-N | No-Effect | inactive |
| ketotifen | BRD-K28936863 | CN1CCC(=C2c3ccccc3C=C(O)c3sccc32)CC1 | OVJRNFMIYGGLDL-UHFFFAOYSA-N | No-Effect | inactive |
| kevetrin | BRD-K61339991 | N#CCCCSC(=N)N | PRDJGNVQBVXXEO-UHFFFAOYSA-N | No-Effect | inactive |
| KG-5 | BRD-K90747162 | CSc1nc(N)cc(Oc2ccc(-c3nnc(Nc4cccc(C(F)(F)F)c4)[nH]3)cc2)n1 | CMYHZFCJPORPHY-UHFFFAOYSA-N | No-Effect | inactive |
| KH-CB19 | BRD-K66430217 | CCOC(=O)c1c(C(C#N)C=N)c2ccc(Cl)c(Cl)c2n1C | DXZRBHUCOHBAHP-UHFFFAOYSA-N | No-Effect | inactive |
| khellin | BRD-K80353807 | COc1c2occc2c(OC)c2c(=O)cc(C)oc12 | HSMPDPBYAYSOBC-UHFFFAOYSA-N | No-Effect | inactive |
| KHK-IN-1 | BRD-K84605263 | CSc1ccccc1Nc1nc(N2CCNCC2)nc2c(NCC3CC3)ncnc12 | HFLMLZKGLUEWBU-UHFFFAOYSA-N | No-Effect | inactive |
| KHS-101 | BRD-K78978711 | CC(C)CNc1ccnc(NCc2csc(-c3ccccc3)n2)n1 | DGRJOOOHPBSAHD-UHFFFAOYSA-N | No-Effect | inactive |
| KI-16425 | BRD-A25569250 | Cc1noc(-c2ccc(CSCCC(=O)O)cc2)c1NC(=O)O[C@H](C)c1ccccc1Cl | LLIFMNUXGDHTRO-OAHLLOKOSA-N | No-Effect | inactive |
| KI-20227 | BRD-A88787317 | COc1cc(Oc2ccnc3cc(OC)c(OC)cc23)ccc1NC(=O)N[C@@H](C)c1nccs1 | SHPFDGWALWEPGS-AWEZNQCLSA-N | No-Effect | inactive |
| KI-8751 | BRD-K47150025 | COc1cc2nccc(Oc3ccc(NC(=O)Nc4ccc(F)cc4F)c(F)c3)c2cc1OC | LFKQSJNCVRGFCC-UHFFFAOYSA-N | No-Effect | inactive |
| kinetin | BRD-K65667145 | c1coc(CNc2ncnc3nc[nH]c23)c1 | QANMHLXAZMSUEX-UHFFFAOYSA-N | No-Effect | inactive |
| KL-001 | BRD-A94580299 | CS(=O)(=O)N(Cc1ccco1)C[C@@H](O)Cn1c2ccccc2c2ccccc21 | OQAFDLPAPSSOHY-MRXNPFEDSA-N | No-Effect | inactive |
| KM-0118 | BRD-K00005245 | CN(C)CCNC(=O)COc1c2cccc1Cc1cccc(c1OCC(=O)NCCN(C)C)Cc1cccc(c1OCC(=O)NCCN(C)C)Cc1cccc(c1OCC(=O)NCCN(C)C)C2 | CQVAQQNDZCZBSU-UHFFFAOYSA-N | No-Effect | inactive |
| KME-4 | BRD-K00004747 | CC(C)(C)c1cc(/C=C2\CCOC2=O)cc(C(C)(C)C)c1O | DFPYHQJPGCODSB-UKTHLTGXSA-N | No-Effect | inactive |
| KML29 | BRD-K46734292 | O=C(OC(C(F)(F)F)C(F)(F)F)N1CCC(C(O)(c2ccc3c(c2)OCO3)c2ccc3c(c2)OCO3)CC1 | SXHQLPHDBLTFPM-UHFFFAOYSA-N | No-Effect | inactive |
| KN-62 | BRD-A81177136 | CN([C@@H](Cc1ccc(OS(=O)(=O)c2cccc3cnccc23)cc1)C(=O)N1CCN(c2ccccc2)CC1)S(=O)(=O)c1cccc2cnccc12 | RJVLFQBBRSMWHX-DHUJRADRSA-N | No-Effect | inactive |
| KN-93 | BRD-K93480852 | COc1ccc(S(=O)(=O)N(CCO)c2ccccc2CN(C)C/C=C/c2ccc(Cl)cc2)cc1 | LLLQTDSSHZREGW-AATRIKPKSA-N | No-Effect | inactive |
| Ko143 | BRD-K64642496 | COc1ccc2c3c([nH]c2c1)[C@H](CC(C)C)N1C(=O)[C@H](CCC(=O)OC(C)(C)C)NC(=O)[C@@H]1C3 | NXNRAECHCJZNRF-JBACZVJFSA-N | No-Effect | inactive |
| KP-1212 | BRD-K46690591 | NC1=NC(=O)N([C@H]2C[C@@H](O)[C@@H](CO)O2)CN1 | LAOLDMMWVYDDID-HSUXUTPPSA-N | No-Effect | inactive |
| KPT-185 | BRD-K71467466 | COc1cc(-c2ncn(/C=C\C(=O)OC(C)C)n2)cc(C(F)(F)F)c1 | NLNGWFLRRRYNIL-PLNGDYQASA-N | No-Effect | inactive |
| KPT-276 | BRD-K75352575 | O=C(/C=C\n1cnc(-c2cc(C(F)(F)F)cc(C(F)(F)F)c2)n1)N1CCC(F)(F)C1 | JCHAWRDHMUSLMM-UPHRSURJSA-N | No-Effect | inactive |
| KR-33494 | BRD-K00004587 | O=C(CSc1ccc(Br)cc1)Nc1cn(CCc2ccccc2)nc1C(=O)O | ZRLJEHIUGYTTSZ-UHFFFAOYSA-N | No-Effect | inactive |
| KRCA-0008 | BRD-K40604024 | COc1cc(N2CCN(C(C)=O)CC2)ccc1Nc1ncc(Cl)c(Nc2ccc(N3CCN(C(C)=O)CC3)cc2OC)n1 | TXDIRJCYNAWBOS-UHFFFAOYSA-N | No-Effect | inactive |
| KRN-633 | BRD-K90993449 | CCCNC(=O)Nc1ccc(Oc2ncnc3cc(OC)c(OC)cc23)cc1Cl | VPBYZLCHOKSGRX-UHFFFAOYSA-N | No-Effect | inactive |
| KS-176 | BRD-K81053848 | O=C(Nc1ccccc1C(=O)Nc1ccc(CCO)cc1)c1ccc([N+](=O)[O-])cc1 | LTWQQWSXYYXVGA-UHFFFAOYSA-N | No-Effect | inactive |
| KT-433 | BRD-K66437909 | CCc1oc2ccccc2c1C(=O)c1ccc(OCC(=O)O)cc1 | XSDFWTIKVXKOGJ-UHFFFAOYSA-N | No-Effect | inactive |
| KU-0063794 | BRD-K67566344 | COc1ccc(-c2ccc3c(N4CCOCC4)nc(N4C[C@H](C)O[C@H](C)C4)nc3n2)cc1CO | RFSMUFRPPYDYRD-CALCHBBNSA-N | No-Effect | inactive |
| KU14R | BRD-A38747044 | CC[C@@]1(c2ncc[nH]2)Cc2ccccc2O1 | JCWVNNMJXQJVNC-ZDUSSCGKSA-N | No-Effect | inactive |
| kuromanin | BRD-K51337109 | OC[C@H]1O[C@@H](Oc2cc3c(O)cc(O)cc3[o+]c2-c2ccc(O)c(O)c2)[C@H](O)[C@@H](O)[C@@H]1O | RKWHWFONKJEUEF-GQUPQBGVSA-O | No-Effect | inactive |
| KW-2449 | BRD-K21718444 | O=C(c1ccc(/C=C/c2[nH]nc3ccccc23)cc1)N1CCNCC1 | YYLKKYCXAOBSRM-JXMROGBWSA-N | No-Effect | inactive |
| KW-2478 | BRD-K41213548 | CCc1c(O)cc(O)c(C(=O)c2ccc(OCCN3CCOCC3)c(OC)c2)c1CC(=O)N(COC)CCOC | VFUXSYAXEKYYMB-UHFFFAOYSA-N | No-Effect | inactive |
| KY02111 | BRD-K13302470 | COc1ccc(CCC(=O)Nc2nc3ccc(Cl)cc3s2)cc1OC | LXFKEVQQSKQXPR-UHFFFAOYSA-N | No-Effect | inactive |
| kynurenic-acid | BRD-K85872723 | O=C(O)c1cc(=O)c2ccccc2[nH]1 | HCZHHEIFKROPDY-UHFFFAOYSA-N | No-Effect | inactive |
| L-(+)-rhamnose-monohydrate | BRD-K00003714 | C[C@@H]1O[C@@H](O)[C@H](O)[C@H](O)[C@H]1O | SHZGCJCMOBCMKK-HGVZOGFYSA-N | No-Effect | inactive |
| L-152804 | BRD-K01827100 | CC1(C)CC(=O)C([C@@H]2C3=C(CC(C)(C)CC3=O)Oc3ccccc32)C(=O)C1 | XQDXIHYKAGHZRX-LJQANCHMSA-N | No-Effect | inactive |
| L-161982 | BRD-K39733634 | CCCCc1nn(-c2ccccc2C(F)(F)F)c(=O)n1Cc1ccc(-c2ccccc2S(=O)(=O)NC(=O)c2sccc2C)cc1 | MMDNKTXNUZFVKD-UHFFFAOYSA-N | No-Effect | inactive |
| L-165041 | BRD-K98372770 | CCCc1c(OCCCOc2ccc(OCC(=O)O)cc2)ccc(C(C)=O)c1O | HBBVCKCCQCQCTJ-UHFFFAOYSA-N | No-Effect | inactive |
| L-168049 | BRD-K18036262 | CCCOc1ccc(Br)cc1-c1cc(-c2ccc(Cl)cc2)[nH]c1-c1ccncc1 | HHBOWXZOLYQFNY-UHFFFAOYSA-N | No-Effect | inactive |
| L-365260 | BRD-K34277641 | Cc1cccc(NC(=O)N[C@@H]2N=C(c3ccccc3)c3ccccc3N(C)C2=O)c1 | KDFQABSFVYLGPM-QFIPXVFZSA-N | No-Effect | inactive |
| L-368899 | BRD-K00004282 | Cc1ccccc1N1CCN(S(=O)(=O)C[C@]23CC[C@H](C[C@@H]2NC(=O)[C@@H](N)CCS(C)(=O)=O)C3(C)C)CC1 | MWIASLNTAGRGGA-ZJPWWDJASA-N | No-Effect | inactive |
| L-655,708 | BRD-K13474508 | CCOC(=O)c1ncn2c1[C@@H]1CCCN1C(=O)c1cc(OC)ccc1-2 | YKYOQIXTECBVBB-AWEZNQCLSA-N | No-Effect | inactive |
| L-655240 | BRD-K89402695 | Cc1c(CC(C)(C)C(=O)O)n(Cc2ccc(Cl)cc2)c2ccc(F)cc12 | JLPYLHLUHJOPNL-UHFFFAOYSA-N | No-Effect | inactive |

| | | | | | |
|---|---|---|---|---|---|
| L-670596 | BRD-A85472596 | CS(=O)(=O)c1ccc(Cn2c3c(c4cc(F)cc(F)c42)CCC[C@@H]3CC(=O)O)cc1 | GMJWAIMJHBTYPD-CQSZACIVSA-N | No-Effect | inactive |
| L-689560 | BRD-K74284736 | O=C(Nc1ccccc1)N[C@H]1C[C@H](C(=O)O)Nc2cc(Cl)cc(Cl)c21 | UCKHICKHGAOGAP-UONOGXRCSA-N | No-Effect | inactive |
| L-690330 | BRD-K28470988 | CC(Oc1ccc(O)cc1)(P(=O)(O)O)P(=O)(O)O | JKOCAAWWDVHWKB-UHFFFAOYSA-N | No-Effect | inactive |
| L-693403 | BRD-K83322645 | c1ccc(CN2CCC3(CCc4ccccc43)CC2)cc1 | GSXJSGMDWKPVBY-UHFFFAOYSA-N | No-Effect | inactive |
| L-694247 | BRD-K18816859 | CS(=O)(=O)Nc1ccc(Cc2noc(-c3ccc4[nH]cc(CCN)c4c3)n2)cc1 | HKXMQLISPYELRD-UHFFFAOYSA-N | No-Effect | inactive |
| L-701252 | BRD-K10176267 | O=C(c1c(O)c2ccc(Cl)cc2[nH]c1=O)C1CC1 | MXEFWCFPCLDOOG-UHFFFAOYSA-N | No-Effect | inactive |
| L-701324 | BRD-K08109516 | O=c1[nH]c2cc(Cl)ccc2c(O)c1-c1cccc(Oc2ccccc2)c1 | FLVRDMUHUXVRET-UHFFFAOYSA-N | No-Effect | inactive |
| L-732,138 | BRD-K37952347 | CC(=O)N[C@H](Cc1c[nH]c2ccccc12)C(=O)OCc1cc(C(F)(F)F)cc(C(F)(F)F)c1 | BYYQYXVAWXAYQC-LJQANCHMSA-N | No-Effect | inactive |
| L-733060 | BRD-K15791587 | FC(F)(F)c1cc(CO[C@H]2CCCN[C@H]2c2ccccc2)cc(C(F)(F)F)c1 | FCDRFVCGMLUYPG-ROUUACIJSA-N | No-Effect | inactive |
| L-745870 | BRD-K05528470 | Clc1ccc(N2CCN(Cc3c[nH]c4ncccc34)CC2)cc1 | OGJGQVFWEPNYSB-UHFFFAOYSA-N | No-Effect | inactive |
| L-755507 | BRD-K39359968 | CCCCCCNC(=O)Nc1ccc(S(=O)(=O)Nc2ccc(CCNC[C@H](O)COc3ccc(O)cc3)cc2)cc1 | NYYJKMXNVNFOFQ-MHZLTWQESA-N | No-Effect | inactive |
| L-760735 | BRD-K30425481 | C[C@@H](O[C@H]1OCCN(Cc2n[nH]nc2CN(C)C)[C@H]1c1ccc(F)cc1)c1cc(C(F)(F)F)cc(C(F)(F)F)c1 | USRYEHHMJIRICK-ZNZBMKLDSA-N | No-Effect | inactive |
| L-778123 | BRD-K10958118 | N#Cc1ccc(Cn2cncc2CN2CCN(c3cccc(Cl)c3)C(=O)C2)cc1 | JNUGFGAVPBYSHF-UHFFFAOYSA-N | No-Effect | inactive |
| L-798106 | BRD-K43905307 | COc1ccc(Br)cc1S(=O)(=O)NC(=O)/C=C/c1ccccc1Cc1ccc2ccccc2c1 | ODTKFNUPVBULRJ-NTCAYCPXSA-N | No-Effect | inactive |
| L-838417 | BRD-K46647427 | Cn1ncnc1COc1nn2c(-c3cc(F)ccc3F)nnc2cc1C(C)(C)C | BQDUNOMMYOKHEP-UHFFFAOYSA-N | No-Effect | inactive |
| L-arginine | BRD-K06033833 | N=C(N)NCCC[C@H](N)C(=O)O | ODKSFYDXXFIFQN-BYPYZUCNSA-N | No-Effect | inactive |
| L-asparagine | BRD-K06665116 | NC(=O)C[C@H](N)C(=O)O | DCXYFEDJOCDNAF-REOHCLBHSA-N | No-Effect | inactive |
| L-asparagine-N-hydroxy | BRD-K82178523 | N[C@@H](CC(=O)NO)C(=O)O | ZBYVTTSIVDYQSO-REOHCLBHSA-N | No-Effect | inactive |
| L-citrulline | BRD-A12237696 | NC(=O)NCCC[C@H](N)C(=O)O | RHGKLRLOHDJJDR-BYPYZUCNSA-N | No-Effect | inactive |
| L-cysteine | BRD-K00795468 | N[C@@H](CS)C(=O)O | XUJNEKJLAYXESH-REOHCLBHSA-N | No-Effect | inactive |
| L-ergothioneine | BRD-K26341917 | C[N+](C)(C)[C@@H](Cc1c[nH]c(=S)[nH]1)C(=O)O | SSISHJJTAXXQAX-ZETCQYMHSA-O | No-Effect | inactive |
| L-glutamine | BRD-K83896451 | NC(=O)CC[C@H](N)C(=O)O | ZDXPYRJPNDTMRX-VKHMYHEASA-N | No-Effect | inactive |
| L-hydroxyproline | BRD-A12454076 | O=C(O)[C@@H]1C[C@@H](O)CN1 | PMMYEEVYMWASQN-DMTCNVIQSA-N | No-Effect | inactive |
| L-kynurenine | BRD-K20497775 | Nc1ccccc1C(=O)C[C@H](N)C(=O)O | YGPSJZOEDVAXAB-QMMMGPOBSA-N | No-Effect | inactive |
| L-leucine | BRD-K12853841 | CC(C)C[C@H](N)C(=O)O | ROHFNLRQFUQHCH-YFKPBYRVSA-N | No-Effect | inactive |
| L-lysine | BRD-K86016185 | NCCCC[C@H](N)C(=O)O | KDXKERNSBIXSRK-YFKPBYRVSA-N | No-Effect | inactive |
| L-methionine | BRD-K06502269 | CSCC[C@H](N)C(=O)O | FFEARJCKVFRZRR-BYPYZUCNSA-N | No-Effect | inactive |
| L-mimosine | BRD-K00004529 | N[C@@H](CN1C=CC(=O)C(=O)C1)C(=O)O | BLYDGIISUNWHLF-YFKPBYRVSA-N | No-Effect | inactive |
| L-monomethylarginine | BRD-K20946707 | CN[C@@H](CCCNC(=N)N)C(=O)O | NTWVQPHTOUKMDI-YFKPBYRVSA-N | No-Effect | inactive |
| L-NAME | BRD-K33141550 | COC(=O)[C@@H](N)CCCNC(N)=N[N+](=O)[O-] | KCWZGJVSDFYRIX-YFKPBYRVSA-N | No-Effect | inactive |
| L-NIL | BRD-K60036103 | CC(=N)NCCCC[C@H](N)C(=O)O | ONYFNWIHJBLQKE-ZETCQYMHSA-N | No-Effect | inactive |
| L-phenylisopropyladenosine | BRD-K25521961 | C[C@H](Cc1ccccc1)Nc1ncnc2c1ncn2[C@@H]1O[C@H](CO)[C@@H](O)[C@H]1O | RIRGCFBBHQEQQH-SSFGXONLSA-N | No-Effect | inactive |
| L-proline | BRD-K01666412 | O=C(O)[C@@H]1CCCN1 | ONIBWKKTOPOVIA-BYPYZUCNSA-N | No-Effect | inactive |
| L-quisqualic-acid | BRD-K92404805 | N[C@@H](Cn1oc(=O)[nH]c1=O)C(=O)O | ASNFTDCKZKHJSW-REOHCLBHSA-N | No-Effect | inactive |
| L-threonine | BRD-K75508643 | C[C@@H](O)[C@H](N)C(=O)O | AYFVYJQAPQTCCC-GBXIJSLDSA-N | No-Effect | inactive |
| L-valine | BRD-K59424860 | CC(C)[C@@H](N)C(=O)O | KZSNJWFQEVHDMF-SCSAIBSYSA-N | No-Effect | inactive |
| labetalol | BRD-K01825098 | C[C@@H](CCc1ccccc1)NC[C@@H](O)c1ccc(O)c(C(N)=O)c1 | SGUAFYQXFOLMHL-SCLBCKFNSA-N | No-Effect | inactive |
| lacidipine | BRD-K43586850 | CCOC(=O)C1=C(C)NC(C)=C(C(=O)OCC)C1c1ccccc1/C=C/C(=O)OC(C)(C)C | GKQPCPXONLDCMU-CCEZHUSRSA-N | No-Effect | inactive |
| lacitol | BRD-K40787673 | OC[C@H](O)[C@@H](O)[C@H](O[C@@H]1O[C@H](CO)[C@H](O)[C@H](O)[C@H]1O)[C@H](O)CO | VQHSOMBJVWLPSR-JVCRWLNRSA-N | No-Effect | inactive |
| lactulose | BRD-K77258318 | OC[C@H]1O[C@](O)(CO)[C@@H](O)[C@@H]1O[C@@H]1O[C@H](CO)[C@H](O)[C@H](O)[C@H]1O | JCQLYHFGKNRPGE-FCVZTGTOSA-N | No-Effect | inactive |
| lamivudine | BRD-K02992638 | Nc1ccn([C@@H]2CS[C@H](CO)O2)c(=O)n1 | JTEGQNOMFQHVDC-NKWVEPMBSA-N | No-Effect | inactive |
| lamotrigine | BRD-K93460210 | Nc1nnc(-c2cccc(Cl)c2Cl)c(N)n1 | PYZRQGJRPPTADH-UHFFFAOYSA-N | No-Effect | inactive |
| lanabecestat | BRD-K00003387 | CC#Cc1cncc(-c2ccc3c(c2)[C@@]2(N=C(C)C(N)=N2)[C@]2(CC[C@H](OC)CC2)C3)c1 | WKDNQONLGXOZRG-HRNNMHKYSA-N | No-Effect | inactive |
| landiolol | BRD-K86436199 | CC1(C)OC[C@@H](COC(=O)CCc2ccc(OC[C@@H](O)CNCCNC(=O)N3CCOCC3)cc2)O1 | WMDSZGFJQKSLLH-RBBKRZOGSA-N | No-Effect | inactive |
| lanifibranor | BRD-K00003462 | O=C(O)CCCc1cc2cc(Cl)ccc2n1S(=O)(=O)c1ccc2ncsc2c1 | OQDQIFQRNZIEEJ-UHFFFAOYSA-N | No-Effect | inactive |
| lanoconazole | BRD-A54275602 | N#C/C(=C1/SC[C@H](c2ccccc2Cl)S1)n1ccnc1 | ZRTQSJFIDWNVJW-MNWMYKRDSA-N | No-Effect | inactive |
| lansoprazole | BRD-K00004280 | Cc1c(OCC(F)(F)F)ccnc1C[S@@](=O)c1nc2ccccc2[nH]1 | MJIHNNLFOKEZEW-RUZDIDTESA-N | No-Effect | inactive |
| lapatinib | BRD-K19687926 | CS(=O)(=O)CCNCc1ccc(-c2ccc3ncnc(Nc4ccc(OCc5cccc(F)c5)c(Cl)c4)c3c2)o1 | BCFGMOOMADDAQU-UHFFFAOYSA-N | No-Effect | inactive |
| lappaconite | BRD-K00025602 | CCN1C[C@]2(OC(=O)c3ccccc3NC(C)=O)CC[C@H](OC)[C@@]34[C@@H]2C[C@@H]([C@@H]13)[C@@]1(O)C[C@H](OC)[C@H]2C[C@@H]4[C@]1(O)[C@H]2OC | NWBWCXBPKTTZNQ-QOQRDJBUSA-N | No-Effect | inactive |
| laquinimod | BRD-K60918217 | CCN(C(=O)c1c(O)c2c(Cl)cccc2n(C)c1=O)c1ccccc1 | GKWPCEFFIHSJOE-UHFFFAOYSA-N | No-Effect | inactive |
| larazotide-acetate | BRD-K00003469 | CC(C)C[C@H](NC(=O)[C@@H](NC(=O)CNC(=O)CN)C(C)C)C(=O)N[C@H](C(=O)N[C@@H](CCC(N)=O)C(=O)N1CCC[C@H]1C(=O)NCC(=O)O)C(C)C | ORFLZNAGUTZRLQ-ZMBVWFSWSA-N | No-Effect | inactive |
| laropiprant | BRD-K48894757 | CS(=O)(=O)c1cc(F)cc2c3c(n(Cc4ccc(Cl)cc4)c12)[C@@H](CC(=O)O)CC3 | NXFFJDQHYLNEJK-CYBMUJFWSA-N | No-Effect | inactive |
| lasmiditan | BRD-K00003178 | CN1CCC(C(=O)c2cccc(NC(=O)c3c(F)cc(F)cc3F)n2)CC1 | XEDHVZKDSYZQBF-UHFFFAOYSA-N | No-Effect | inactive |
| latrepirdine | BRD-K55703048 | Cc1ccc2c(c1)c1c(n2CCc2ccc(C)nc2)CCN(C)C1 | JNODQFNWMXFMEV-UHFFFAOYSA-N | No-Effect | inactive |
| lauric-acid | BRD-K67375056 | CCCCCCCCCCCC(=O)O | POULHZVOKOAJMA-UHFFFAOYSA-N | No-Effect | inactive |
| lauric-diethanolamide | BRD-K11548689 | CCCCCCCCCCCC(=O)N(CCO)CCO | AOMUHOFOVNGZAN-UHFFFAOYSA-N | No-Effect | inactive |
| laurocapram | BRD-K35698076 | CCCCCCCCCCCCN1CCCCCC1=O | AXTGDCSMTYGJND-UHFFFAOYSA-N | No-Effect | inactive |
| lazabemide | BRD-K51486818 | NCCNC(=O)c1ccc(Cl)cn1 | JZXRLKWWVNUZRB-UHFFFAOYSA-N | No-Effect | inactive |
| lazertinib | BRD-K00003501 | C=CC(=O)Nc1cc(Nc2nccc(-n3cc(CN(C)C)c(-c4ccccc4)n3)n2)c(OC)cc1N1CCOCC1 | RRMJMHOQSALEJJ-UHFFFAOYSA-N | No-Effect | inactive |
| LB-100 | BRD-K00025603 | CN1CCN(C(=O)[C@@H]2[C@H]3CC[C@H](O3)[C@@H]2C(=O)O)CC1 | JUQMLSGOTNKJKI-YTWAJWBKSA-N | No-Effect | inactive |
| LB42708 | BRD-K18834913 | O=C(c1cn(Cc2cncn2Cc2ccc(Br)cc2)cc1-c1cccc2ccccc12)N1CCOCC1 | GUUIRIMAQGOLHT-UHFFFAOYSA-N | No-Effect | inactive |
| LCL-161 | BRD-K08248804 | CN[C@@H](C)C(=O)N[C@H](C(=O)N1CCC[C@H]1c1nc(C(=O)c2ccc(F)cc2)cs1)C1CCCCC1 | UFPFGVNKHCLJJO-SSKFGXFMSA-N | No-Effect | inactive |
| LCQ908 | BRD-K20008181 | O=C(O)C[C@H]1CC[C@H](c2ccc(-c3ccc(Nc4ccc(C(F)(F)F)nc4)cn3)cc2)CC1 | GXALXAKNHIROPE-QAQDUYKDSA-N | No-Effect | inactive |

| | | | | | |
|---|---|---|---|---|---|
| LCZ696 | BRD-M31057929 | CCCCC(=O)N(Cc1ccc(-c2ccccc2-c2nn[nH]n2)cc1)[C@H](C(=O)O)C(C)C.CCOC(=O)[C@H](C)C[C@@H](Cc1ccc(-c2ccccc2)cc1)NC(=O)CCC(=O)O | XTKIDERFOUYBDE-QEPKIHTBSA-N | No-Effect | inactive |
| LDC1267 | BRD-K44715996 | CCOc1cn(-c2ccc(F)cc2C)nc1C(=O)Nc1ccc(Oc2ccnc3cc(OC)c(OC)cc23)c(F)c1 | ISPBCAXOSOLFME-UHFFFAOYSA-N | No-Effect | inactive |
| LDN-209929 | BRD-K61795638 | COc1ccc2nc3ccc(Cl)cc3c(SCCCN)c2c1 | XMMAOXMYUGQGJD-UHFFFAOYSA-N | No-Effect | inactive |
| LDN-212854 | BRD-K59831625 | c1cc(-c2cnn3cc(-c4ccc(N5CCNCC5)cc4)cnc23)c2cccnc2c1 | BBDGBGOVJPEFBT-UHFFFAOYSA-N | No-Effect | inactive |
| LDN-57444 | BRD-K94485812 | CC(=O)O/N=C1/C(=O)N(Cc2cc(Cl)ccc2Cl)c2ccc(Cl)cc21 | OPQRFPHLZZPCCH-LTGZKZEYSA-N | No-Effect | inactive |
| LDN193189 | BRD-K04853698 | c1ccc2c(-c3cnn4cc(-c5ccc(N6CCNCC6)cc5)cnc34)ccnc2c1 | CDOVNWNANFFLFJ-UHFFFAOYSA-N | No-Effect | inactive |
| LE-135 | BRD-K06593056 | CN1c2cc3c(cc2N=C(c2ccc(C(=O)O)cc2)c2ccccc21)C(C)(C)CCC3(C)C | YZZAIQOVMHVWBS-UHFFFAOYSA-N | No-Effect | inactive |
| ledipasvir | BRD-K05446112 | COC(=O)N[C@H](C(=O)N1CC2(CC2)C[C@H]1c1nc(-c2ccc3c(c2)C(F)(F)c2cc(-c4ccc5nc([C@@H]6[C@H]7CC[C@H](C7)N6C(=O)[C@@H](NC(=O)OC)C(C)C)[nH]c5c4)ccc2-3)c[nH]1)C(C)C | VRTWBAAJJOHBQU-KMWAZVGDSA-N | No-Effect | inactive |
| leflunomide | BRD-K78692225 | Cc1oncc1C(=O)Nc1ccc(C(F)(F)F)cc1 | VHOGYURTWQBHIL-UHFFFAOYSA-N | No-Effect | inactive |
| lemborexant | BRD-K78666663 | Cc1ncc(OC[C@@]2(c3cccc(F)c3)C[C@H]2C(=O)Nc2ccc(F)cn2)c(C)n1 | MUGXRYIUWFITCP-PGRDOPGGSA-N | No-Effect | inactive |
| leniolisib | BRD-K00003261 | CCC(=O)N1CC[C@H](Nc2ncnc3c2CN(c2cnc(OC)c(C(F)(F)F)c2)CC3)C1 | MWKYMZXCGYXLPL-ZDUSSCGKSA-N | No-Effect | inactive |
| lenvatinib | BRD-K39974922 | COc1cc2nccc(Oc3ccc(NC(=O)NC4CC4)c(Cl)c3)c2cc1C(N)=O | WOSKHXYHFSIKNG-UHFFFAOYSA-N | No-Effect | inactive |
| leonurine | BRD-K00004696 | COc1cc(C(=O)OCCCCNC(=N)N)cc(OC)c1O | WNGSUWLDMZFYNZ-UHFFFAOYSA-N | No-Effect | inactive |
| LEQ506 | BRD-K00003269 | Cc1c(Cc2ccccc2)nnc(N2CCN(c3cnc(C(C)(C)O)cn3)[C@H](C)C2)c1C | POERAARDVFVDLO-QGZVFWFLSA-N | No-Effect | inactive |
| lercanidipine | BRD-A18992208 | COC(=O)C1=C(C)NC(C)=C(C(=O)OC(C)(C)CN(C)CCC(c2ccccc2)c2ccccc2)[C@@H]1c1cccc([N+](=O)[O-])c1 | ZDXUKAKRHYTAKV-MGBGTMOVSA-N | No-Effect | inactive |
| lerisetron | BRD-K62398570 | c1ccc(Cn2c(N3CCNCC3)nc3ccccc32)cc1 | PWWDCRQZITYKDV-UHFFFAOYSA-N | No-Effect | inactive |
| lersivirine | BRD-K03289534 | CCc1nn(CCO)c(CC)c1Oc1cc(C#N)cc(C#N)c1 | MCPUZZJBAHRIPO-UHFFFAOYSA-N | No-Effect | inactive |
| lesinurad | BRD-K75024268 | O=C(O)CSc1nnc(Br)n1-c1ccc(C2CC2)c2ccccc12 | FGQFOYHRJSUHMR-UHFFFAOYSA-N | No-Effect | inactive |
| lestaurtinib | BRD-A27376179 | C[C@]12O[C@@H](C[C@]1(O)CO)n1c3ccccc3c3c4c(c5c6ccccc6n2c5c31)CNC4=O | UIARLYUEJFELEN-ATANMQQVSA-N | No-Effect | inactive |
| leteprinim | BRD-K21612934 | O=C(CCn1cnc2c(=O)nc[nH]c21)Nc1ccc(C(=O)O)cc1 | JMPOIZCOJJMTHI-UHFFFAOYSA-N | No-Effect | inactive |
| letermovir | BRD-K94288301 | COc1cccc(N2CCN(C3=Nc4c(F)cccc4[C@H](CC(=O)O)N3c3cc(C(F)(F)F)ccc3OC)CC2)c1 | FWYSMLBETOMXAG-QHCPKHFHSA-N | No-Effect | inactive |
| letosteine | BRD-A32597168 | CCOC(=O)CSCC[C@H]1N[C@H](C(=O)O)CS1 | IKOCLISPVJZJEA-YUMQZZPRSA-N | No-Effect | inactive |
| letrazuril | BRD-K00003474 | N#C[C@H](c1ccc(F)cc1)c1c(Cl)cc(-n2ncc(=O)[nH]c2=O)cc1Cl | XQKYUBTUOHHNDV-GFCCVEGCSA-N | No-Effect | inactive |
| letrozole | BRD-K88789588 | N#Cc1ccc(C(c2ccc(C#N)cc2)n2cncn2)cc1 | HPJKCIUCZWXJDR-UHFFFAOYSA-N | No-Effect | inactive |
| leucovorin | BRD-K01825581 | Nc1nc(=O)c2c([nH]1)NC[C@H](CNc1ccc(C(=O)N[C@@H](CCC(=O)O)C(=O)O)cc1)N2C=O | VVIAGPKUTFNRDU-STQMWFEESA-N | No-Effect | inactive |
| leucylleucine-methyl-ester | BRD-K81136890 | COC(=O)[C@H](CC(C)C)NC(=O)[C@@H](N)CC(C)C | AJMOLNFDYWTVQW-QWRGUYRKSA-N | No-Effect | inactive |
| leuprolide | BRD-K45300898 | CCNC(=O)[C@@H]1CCCN1C(=O)[C@H](CCCNC(=N)N)NC(=O)[C@H](CC(C)C)NC(=O)[C@@H](CC(C)C)NC(=O)[C@H](Cc1ccc(O)cc1)NC(=O)[C@H](CO)NC(=O)[C@H](Cc1c[nH]c2ccccc12)NC(=O)[C@H](Cc1cnc[nH]1)NC(=O)[C@@H]1CCC(=O)N1 | GFIJNRVAKGFPGQ-LIJARHBVSA-N | No-Effect | inactive |
| levalbuterol | BRD-K34469523 | CC(C)(C)NC[C@H](O)c1ccc(O)c(CO)c1 | NDAUXUAQIAJITI-LBPRGKRZSA-N | No-Effect | inactive |
| levcromakalim | BRD-K24526313 | CC1(C)Oc2ccc(C#N)cc2[C@@H](N2CCCC2=O)[C@@H]1O | TVZCRIROJQEVOT-CABCVRRESA-N | No-Effect | inactive |
| levo-phencynonate | BRD-A64698388 | CN1C[C@@H]2CCC[C@H](C1)[C@H]2OC(=O)[C@](O)(c1ccccc1)C1CCCC1 | ROZOEEGFKDFEFP-AXRVMRSRSA-N | No-Effect | inactive |
| levobetaxolol | BRD-K43883465 | CC(C)NC[C@H](O)COc1ccc(CCOCC2CC2)cc1 | NWIUTZDMDHAVTP-KRWDZBQOSA-N | No-Effect | inactive |
| levobunolol | BRD-K31812033 | CC(C)(C)NC[C@H](O)COc1cccc2c1CCCC2=O | IXHBTMCLRNMKHZ-LBPRGKRZSA-N | No-Effect | inactive |
| levobunolol-(+) | BRD-K73323637 | CC(C)(C)NC[C@@H](O)COc1cccc2c1CCCC2=O | IXHBTMCLRNMKHZ-GFCCVEGCSA-N | No-Effect | inactive |
| levobunolol-(+/-) | BRD-A46876182 | CC(C)(C)NC[C@@H](O)COc1cccc2c1CCCC2=O | IXHBTMCLRNMKHZ-GFCCVEGCSA-N | No-Effect | inactive |
| levocabastine | BRD-K33453211 | C[C@@H]1CN([C@H]2CC[C@](C#N)(c3ccc(F)cc3)CC2)CC[C@]1(C(=O)O)c1ccccc1 | ZCGOMHNNNFPNMX-KYTRFIICSA-N | No-Effect | inactive |
| levocarnitine | BRD-K59647213 | C[N+](C)(C)C[C@H](O)CC(=O)O | PHIQHXFUZVPYII-ZCFIWIBFSA-O | No-Effect | inactive |
| levocarnitine-propionate | BRD-K84366268 | CCC(=O)O[C@H](CC(=O)O)C[N+](C)(C)C | UFAHZIUFPNSHSL-MRVPVSSYSA-O | No-Effect | inactive |
| levocetirizine | BRD-K27184429 | O=C(O)COCCN1CCN([C@H](c2ccccc2)c2ccc(Cl)cc2)CC1 | ZKLPARSLTMPFCP-OAQYLSRUSA-N | No-Effect | inactive |
| levodopa | BRD-K34730807 | N[C@@H](Cc1ccc(O)c(O)c1)C(=O)O | WTDRDQBEARUVNC-LURJTMIESA-N | No-Effect | inactive |
| levodropropizine | BRD-K83551379 | OC[C@@H](O)CN1CCN(c2ccccc2)CC1 | PTVWPYVOOKLBCG-ZDUSSCGKSA-N | No-Effect | inactive |
| levofloxacin | BRD-K09471561 | C[C@H]1COc2c(N3CCN(C)CC3)c(F)cc3c(=O)c(C(=O)O)cn1c23 | GSDSWSVVBLHKDQ-JTQLQIEISA-N | No-Effect | inactive |
| levomenol | BRD-K62751771 | CC(C)=CCC[C@](C)(O)[C@@H]1CC=C(C)CC1 | RGZSQWQPBWRIAQ-CABCVRRESA-N | No-Effect | inactive |
| levomenthol | BRD-K50329370 | CC(C)[C@@H]1CC[C@@H](C)C[C@H]1O | NOOLISFMXDJSKH-KXUCPTDWSA-N | No-Effect | inactive |
| levonordefrin | BRD-K02976377 | C[C@H](N)[C@H](O)c1ccc(O)c(O)c1 | GEFQWZLICWMTKF-CDUCUWFYSA-N | No-Effect | inactive |
| levonorgestrel | BRD-K35189033 | C#C[C@]1(O)CC[C@H]2[C@@H]3CCC4=CC(=O)CC[C@@H]4[C@H]3CC[C@@]21CC | WWYNJERNGUHSAO-XUDSTZEESA-N | No-Effect | inactive |
| levopropoxyphene | BRD-A54635441 | CCC(=O)OC(Cc1ccccc1)(c1ccccc1)[C@@H](C)CN(C)C | XLMALTXPSGQGBX-HXBUSHRASA-N | No-Effect | inactive |
| levosimendan | BRD-K03095347 | C[C@@H]1CC(=O)NN=C1c1ccc(NN=C(C#N)C#N)cc1 | WHXMKTBCFHIYNQ-SECBINFHSA-N | No-Effect | inactive |
| LFM-A13 | BRD-A30655177 | CC(=O)[C@@H](C#N)C(=O)Nc1cc(Br)ccc1Br | BQLDGDMKDYAGNE-MRVPVSSYSA-N | No-Effect | inactive |
| LGD-6972 | BRD-K00003132 | Cc1cc(C)c(-c2ccc(NC(=O)[C@H](Cc3ccc(C(=O)NCCS(=O)(=O)O)cc3)c3ccc(-c4ccc(C(C)(C)C)cc4)cc3)cc2)c(C)c1 | HKJMCBYPVCGZFB-LDLOPFEMSA-N | No-Effect | inactive |
| LGK-974 | BRD-K16810791 | Cc1cc(-c2ncc(CC(=O)Nc3ccc(-c4cnccn4)cn3)cc2C)ccn1 | XXYGTCZJJLTAGH-UHFFFAOYSA-N | No-Effect | inactive |
| LH846 | BRD-K47409078 | Cc1cc2sc(NC(=O)Cc3ccccc3)nc2cc1Cl | DYHAMRNAHTWYKY-UHFFFAOYSA-N | No-Effect | inactive |
| liarozole | BRD-K00004284 | Clc1cccc([C@H](c2ccc3nc[nH]c3c2)n2ccnc2)c1 | UGFHIPBXIWJXNA-QGZVFWFLSA-N | No-Effect | inactive |
| licarbazepine | BRD-A22642447 | NC(=O)N1c2ccccc2C[C@@H](O)c2ccccc21 | BMPDWHIDQYTSHX-CQSZACIVSA-N | No-Effect | inactive |
| licochalcone-A | BRD-K99667445 | C=CC(C)(C)c1cc(/C=C/C(=O)c2ccc(O)cc2)c(OC)cc1O | KAZSKMJFUPEHHW-DHZHZOJOSA-N | No-Effect | inactive |
| licofelone | BRD-K08439945 | CC1(C)Cc2c(-c3ccccc3)c(-c3ccc(Cl)cc3)c(CC(=O)O)n2C1 | UAWXGRJVZSAUSZ-UHFFFAOYSA-N | No-Effect | inactive |
| licogliflozin | BRD-K00003504 | CCc1ccc([C@@H]2O[C@H](CO)[C@@H](O)[C@H](O)[C@H]2O)cc1Cc1ccc2c(c1)OCCO2 | XFJAMQQAAMJFGB-ZQGJOIPISA-N | No-Effect | inactive |
| licostinel | BRD-K81881550 | O=c1[nH]c2cc(Cl)c(Cl)c([N+](=O)[O-])c2[nH]c1=O | CHFSOFHQIZKQCR-UHFFFAOYSA-N | No-Effect | inactive |
| lidamidine | BRD-K68693535 | CNC(=N)NC(=O)Nc1c(C)cccc1C | RRHJHSBDJDZUGL-UHFFFAOYSA-N | No-Effect | inactive |
| lidocaine | BRD-K52662033 | CCN(CC)CC(=O)Nc1c(C)cccc1C | NNJVILVZKWQKPM-UHFFFAOYSA-N | No-Effect | inactive |
| lidoflazine | BRD-K62996583 | Cc1cccc(C)c1NC(=O)CN1CCN(CCCC(c2ccc(F)cc2)c2ccc(F)cc2)CC1 | ZBIAKUMOEKILTF-UHFFFAOYSA-N | No-Effect | inactive |
| lifitegrast | BRD-K60265345 | CS(=O)(=O)c1cccc(C[C@H](NC(=O)c2c(Cl)cc3c(c2Cl)CCN(C(=O)c2cc4ccoc4c2)C3)C(=O)O)c1 | JFOZKMSJYSPYLN-QHCPKHFHSA-N | No-Effect | inactive |
| ligustilide | BRD-K48875051 | CCC/C=C1\OC(=O)C2=C1CCC=C2 | IQVQXVFMNOFTMU-FLIBITNWSA-N | No-Effect | inactive |
| limaprost-alfadex | BRD-K25015158 | CCCC[C@H](C)C[C@H](O)/C=C/[C@H]1[C@H](O)CC(=O)[C@@H]1CCCC/C=C/C(=O)O | OJZYRQPMEIEQFC-UAWLTFRCSA-N | No-Effect | inactive |

| | | | | | |
|---|---|---|---|---|---|
| LIMKi-3 | BRD-K75373031 | CC(C)C(=O)Nc1ncc(-c2cc(C(F)F)nn2-c2c(Cl)cccc2Cl)s1 | IVUGBSGLHRJSSP-UHFFFAOYSA-N | No-Effect | inactive |
| limonin | BRD-K05906022 | CC1(C)O[C@H]2CC(=O)OC[C@@]23[C@H]1CC(=O)[C@]1(C)[C@@H]3CC[C@@]2(C)[C@H](c3ccoc3)OC(=O)[C@H]3O[C@]321 | KBDSLGBFQAGHBE-MSGMIQHVSA-N | No-Effect | inactive |
| linagliptin | BRD-K96550715 | CC#CCn1c(N2CCC[C@@H](N)C2)nc2c1c(=O)n(Cc1nc(C)c3ccccc3n1)c(=O)n2C | LTXREWYXXSTFRX-QGZVFWFLSA-N | No-Effect | inactive |
| linalool | BRD-K01827427 | C=C[C@](C)(O)CCC=C(C)C | CDOSHBSSFJOMGT-JTQLQIEISA-N | No-Effect | inactive |
| linaprazan | BRD-K00003366 | Cc1cccc(C)c1CNc1cc(C(=O)NCCO)cn2c(C)c(C)nc12 | GHVIMBCFLRTFHI-UHFFFAOYSA-N | No-Effect | inactive |
| lindane | BRD-K93814253 | Cl[C@H]1[C@H](Cl)[C@@H](Cl)[C@@H](Cl)[C@H](Cl)[C@H]1Cl | JLYXXMFPNIAWKQ-GNIYUCBRSA-N | No-Effect | inactive |
| linoleic-acid | BRD-K08973992 | CCCCC/C=C\C/C=C\CCCCCCCC(=O)O | OYHQOLUKZRVURQ-HZJYTTRNSA-N | No-Effect | inactive |
| linopirdine | BRD-K59332007 | O=C1N(c2ccccc2)c2ccccc2C1(Cc1ccncc1)Cc1ccncc1 | YEJCDKJIEMIWRQ-UHFFFAOYSA-N | No-Effect | inactive |
| linsidomine | BRD-K72922393 | Nc1c[n+](N2CCOCC2)no1 | XZYRDNUZENBYFI-UHFFFAOYSA-N | No-Effect | inactive |
| linsitinib | BRD-K86118762 | C[C@]1(O)C[C@@H](c2nc(-c3ccc4ccc(-c5ccccc5)nc4c3)c3c(N)nccn32)C1 | PKCDDUHJAFVJJB-VLZXCDOPSA-N | No-Effect | inactive |
| lintopride | BRD-K00004744 | CCN1CCN=C1CNC(=O)c1cc(Cl)c(N)cc1OC | MJCKISCWHDATBN-UHFFFAOYSA-N | No-Effect | inactive |
| liothyronine | BRD-K89152108 | N[C@@H](Cc1cc(I)c(Oc2ccc(O)c(I)c2)c(I)c1)C(=O)O | AUYYCJSJGJYCDS-LBPRGKRZSA-N | No-Effect | inactive |
| liothyronine-(isomer) | BRD-K46428823 | N[C@H](Cc1cc(I)c(Oc2ccc(O)c(I)c2)c(I)c1)C(=O)O | AUYYCJSJGJYCDS-GFCCVEGCSA-N | No-Effect | inactive |
| lipoamide | BRD-A14440173 | NC(=O)CCCC[C@H]1CCSS1 | FCCDDURTIIUXBY-ZETCQYMHSA-N | No-Effect | inactive |
| liquiritin | BRD-K39084158 | O=C1C[C@@H](c2ccc(O[C@@H]3O[C@H](CO)[C@@H](O)[C@H](O)[C@H]3O)cc2)Oc2cc(O)ccc21 | DEMKZLAVQYISIA-ZRWXNEIDSA-N | No-Effect | inactive |
| liranaftate | BRD-K35775715 | COc1cccc(N(C)C(=S)Oc2ccc3c(c2)CCCC3)n1 | VPHPQNGOVQYUMG-UHFFFAOYSA-N | No-Effect | inactive |
| lirimilast | BRD-K31339321 | CS(=O)(=O)Oc1ccc2c(NC(N)=O)c(C(=O)c3ccc(Cl)cc3Cl)oc2c1 | YPFLFUJKZDAXRA-UHFFFAOYSA-N | No-Effect | inactive |
| lisadimate | BRD-A76533773 | Nc1ccc(C(=O)OC[C@@H](O)CO)cc1 | WHQOKFZWSDOTQP-VIFPVBQESA-N | No-Effect | inactive |
| lisinopril | BRD-K67966701 | NCCCC[C@H](N[C@@H](CCc1ccccc1)C(=O)O)C(=O)N1CCC[C@H]1C(=O)O | RLAWWYSOJDYHDC-BZSNNMDCSA-N | No-Effect | inactive |
| lisofylline | BRD-A05334458 | C[C@@H](O)CCCCn1c(=O)c2c(ncn2C)n(C)c1=O | NSMXQKNUPPXBRG-SECBINFHSA-N | No-Effect | inactive |
| lithium-acetoacetate | BRD-K73234506 | CC(=O)CC(=O)O | WDJHALXBUFZDSR-UHFFFAOYSA-N | No-Effect | inactive |
| lithium-citrate | BRD-M41245765 | O=C(O)CC(O)(CC(=O)O)C(=O)O.[Li].[Li].[Li] | SODVLVKGOJVWCQ-UHFFFAOYSA-N | No-Effect | inactive |
| lithocholic-acid | BRD-K20986251 | C[C@H](CCC(=O)O)[C@H]1CC[C@H]2[C@@H]3CC[C@@H]4C[C@H](O)CC[C@]4(C)[C@H]3CC[C@@]21C | SMEROWZSTRWXGI-HVATVPOCSA-N | No-Effect | inactive |
| litronesib | BRD-K83029223 | CCNCCS(=O)(=O)NC[C@@]1(c2ccccc2)SC(NC(=O)C(C)(C)C)=NN1C(=O)C(C)(C)C | YVAFBXLHPINSIK-QHCPKHFHSA-N | No-Effect | inactive |
| lixivaptan | BRD-K59391348 | Cc1ccc(F)cc1C(=O)Nc1ccc(C(=O)N2Cc3cccn3Cc3ccccc32)c(Cl)c1 | PPHTXRNHTVLQED-UHFFFAOYSA-N | No-Effect | inactive |
| LM-22A4 | BRD-K70404782 | O=C(NCCO)c1cc(C(=O)NCCO)cc(C(=O)NCCO)c1 | RGWJKANXFYJKHN-UHFFFAOYSA-N | No-Effect | inactive |
| LM11A-31 | BRD-A04649065 | CC[C@@H](C)[C@@H](N)C(=O)NCCN1CCOCC1 | YNMUTYLWSRFTPX-GHMZBOCLSA-N | No-Effect | inactive |
| LMI070 | BRD-K08532119 | CC1(C)CC(Oc2ccc(-c3ccc(-c4cn[nH]c4)cc3O)nn2)CC(C)(C)N1 | STWTUEAWRAIWJG-UHFFFAOYSA-N | No-Effect | inactive |
| lobeline | BRD-K66206289 | CN1[C@H](C[C@H](O)c2ccccc2)CCC[C@@H]1CC(=O)c1ccccc1 | MXYUKLILVYORSK-HBMCJLEFSA-N | No-Effect | inactive |
| lobendazole | BRD-K15834839 | CCOC(=O)Nc1nc2ccccc2[nH]1 | OKOVSTKGUBOSTB-UHFFFAOYSA-N | No-Effect | inactive |
| lobenzarit | BRD-K55847762 | O=C(O)c1ccccc1Nc1cc(Cl)ccc1C(=O)O | UGDPYGKWIHHBMB-UHFFFAOYSA-N | No-Effect | inactive |
| lodenosine | BRD-K00004554 | Nc1ncnc2c1ncn2[C@@H]1O[C@H](CO)C[C@@H]1F | KBEMFSMODRNJHE-JFWOZONXSA-N | No-Effect | inactive |
| lodoxamide | BRD-K66937583 | N#Cc1cc(NC(=O)C(=O)O)c(Cl)c(NC(=O)C(=O)O)c1 | RVGLGHVJXCETIO-UHFFFAOYSA-N | No-Effect | inactive |
| lofemizole | BRD-K06388517 | Cc1[nH]cnc1-c1ccc(Cl)cc1 | MBKWNJVQSFBLQI-UHFFFAOYSA-N | No-Effect | inactive |
| lofepramine | BRD-K82147103 | CN(CCCN1c2ccccc2CCc2ccccc21)CC(=O)c1ccc(Cl)cc1 | SAPNXPWPAUFAJU-UHFFFAOYSA-N | No-Effect | inactive |
| lofexidine | BRD-A02990301 | C[C@@H](Oc1c(Cl)cccc1Cl)C1=NCCN1 | KSMAGQUYOIHWFS-SSDOTTSWSA-N | No-Effect | inactive |
| lomefloxacin | BRD-K01825701 | CCn1cc(C(=O)O)c(=O)c2cc(F)c(N3CCN[C@@H](C)C3)c(F)c21 | ZEKZLJVOYLTDKK-VIFPVBQESA-N | No-Effect | inactive |
| lomeguatrib | BRD-K27955832 | Nc1nc(OCc2cc(Br)cs2)c2[nH]cnc2n1 | JUJPKFNFCWJBCX-UHFFFAOYSA-N | No-Effect | inactive |
| lomerizine | BRD-K62858456 | COc1ccc(CN2CCN(C(c3ccc(F)cc3)c3ccc(F)cc3)CC2)c(OC)c1OC | JQSAYKKFZOSZGJ-UHFFFAOYSA-N | No-Effect | inactive |
| lomitapide | BRD-K92213669 | O=C(NC1CCN(CCCCC2(C(=O)NCC(F)(F)F)c3ccccc3-c3ccccc32)CC1)c1ccccc1-c1ccc(C(F)(F)F)cc1 | MBBCVAKAJPKAKM-UHFFFAOYSA-N | No-Effect | inactive |
| lomustine | BRD-K51768138 | O=C(NC1CCCCC1)N(CCCl)[NH2+][O-] | NEXPKUGJJKVILF-UHFFFAOYSA-N | No-Effect | inactive |
| lonafarnib | BRD-K41599323 | NC(=O)N1CCC(CC(=O)N2CCC([C@H]3c4ncc(Br)cc4CCc4cc(Cl)cc(Br)c43)CC2)CC1 | DHMTURDWPRKSOA-RUZDIDTESA-N | No-Effect | inactive |
| lonidamine | BRD-K78513633 | O=C(O)c1nn(Cc2ccc(Cl)cc2Cl)c2ccccc12 | WDRYRZXSPDWGEB-UHFFFAOYSA-N | No-Effect | inactive |
| loperamide | BRD-K61250553 | CN(C)C(=O)C(CCN1CCC(O)(c2ccc(Cl)cc2)CC1)(c1ccccc1)c1ccccc1 | RDOIQAHITMMDAJ-UHFFFAOYSA-N | No-Effect | inactive |
| lopinavir | BRD-K99451608 | Cc1cccc(C)c1OCC(=O)N[C@@H](Cc1ccccc1)[C@@H](O)C[C@H](Cc1ccccc1)NC(=O)[C@H](C(C)C)N1CCCNC1=O | KJHKTHWMRKYKJE-SUGCFTRWSA-N | No-Effect | inactive |
| loracarbef | BRD-K28935038 | N[C@@H](C(=O)N[C@@H]1C(=O)N2C(C(=O)O)=C(Cl)CC[C@H]12)c1ccccc1 | JAPHQRWPEGVNBT-UTUOFQBUSA-N | No-Effect | inactive |
| loratadine | BRD-K82795137 | CCOC(=O)N1CCC(=C2c3ccc(Cl)cc3CCc3cccnc32)CC1 | JCCNYMKQOSZNPW-UHFFFAOYSA-N | No-Effect | inactive |
| loreclezole | BRD-K09659835 | Cl/C(=C\n1cncn1)c1ccc(Cl)cc1Cl | XGLHZTBDUXXHOM-WMZJFQQLSA-N | No-Effect | inactive |
| lorediplon | BRD-K42066335 | CC(=O)N(C)c1cc(-c2ccnc3c(C(=O)c4cccs4)cnn23)ccc1F | NQPOCLFSADOXBR-UHFFFAOYSA-N | No-Effect | inactive |
| lorglumide | BRD-K01825818 | CCCCCN(CCCCC)C(=O)[C@@H](CCC(=O)O)NC(=O)c1ccc(Cl)c(Cl)c1 | IEKOTSCYBBDIJC-LJQANCHMSA-N | No-Effect | inactive |
| lorlatinib | BRD-K99879819 | C[C@H]1Oc2cc(cnc2N)-c2c(nn(C)c2C#N)CN(C)C(=O)c2ccc(F)cc21 | IIXWYSCJSQVBQM-LLVKDONJSA-N | No-Effect | inactive |
| lornoxicam | BRD-A90551241 | CN1C(C(=O)Nc2ccccn2)=C(O)c2sc(Cl)cc2S1(=O)=O | WLHQHAUOOXYABV-UHFFFAOYSA-N | No-Effect | inactive |
| losartan | BRD-K76205745 | CCCCc1nc(Cl)c(CO)n1Cc1ccc(-c2ccccc2-c2nn[nH]n2)cc1 | PSIFNNKUMBGKDQ-UHFFFAOYSA-N | No-Effect | inactive |
| losmapimod | BRD-K56195681 | Cc1c(F)cc(C(=O)NC2CC2)cc1-c1ccc(C(=O)NCC(C)(C)C)cn1 | KKYABQBFGDZVNQ-UHFFFAOYSA-N | No-Effect | inactive |
| lotamilast | BRD-K00003137 | CNc1nc(-c2cccc(NC(=O)c3ccc(C(=O)OC)cc3)c2)cc2cc(OC)c(OC)cc2n1 | BBTFKAOFCSOZMB-UHFFFAOYSA-N | No-Effect | inactive |
| loteprednol | BRD-K39983086 | CCOC(=O)O[C@]1(C(=O)OCCl)CC[C@H]2[C@@H]3CCC4=CC(=O)C=C[C@]4(C)[C@H]3[C@@H](O)C[C@@]21C | DMKSVUSAATWOCU-HROMYWEYSA-N | No-Effect | inactive |
| lovastatin | BRD-K09416995 | CC[C@H](C)C(=O)O[C@H]1C[C@@H](C)C=C2C=C[C@H](C)[C@H](CC[C@@H]3C[C@@H](O)CC(=O)O3)[C@H]21 | PCZOHLXUXFIOCF-BXMDZJJMSA-N | No-Effect | inactive |
| loxapine | BRD-K39915878 | CN1CCN(C2=Nc3ccccc3Oc3ccc(Cl)cc32)CC1 | XJGVXQDUIWGIRW-UHFFFAOYSA-N | No-Effect | inactive |
| loxistatin-acid | BRD-K21937671 | CC(C)CCNC(=O)[C@H](CC(C)C)NC(=O)[C@H]1O[C@@H]1C(=O)O | SCMSYZJDIQPSDI-SRVKXCTJSA-N | No-Effect | inactive |
| loxoprofen | BRD-K01825691 | C[C@@H](C(=O)O)c1ccc(C[C@H]2CCCC2=O)cc1 | YMBXTVYHTMGZDW-ZWNOBZJWSA-N | No-Effect | inactive |
| LP-533401 | BRD-K00004673 | Nc1nc(O[C@@H](c2ccc(-c3cccc(F)c3)cc2)C(F)(F)F)cc(-c2ccc(C[C@H](N)C(=O)O)cc2)n1 | JZWUKILTKYJLCN-URXFXBBRSA-N | No-Effect | inactive |
| LPA2-antagonist-1 | BRD-K32322923 | Cc1csc2c(N[C@@H](C)CN3CCN(S(=O)(=O)c4ccc(Cl)c(Cl)c4)CC3)ncnc12 | BPRNMVDTWIHULJ-AWEZNQCLSA-N | No-Effect | inactive |
| LRRK2-IN-1 | BRD-K01436366 | COc1cc(C(=O)N2CCC(N3CCN(C)CC3)CC2)ccc1Nc1ncc2c(n1)N(C)c1ccccc1C(=O)N2C | IWMCPJZTADUIFX-UHFFFAOYSA-N | No-Effect | inactive |
| LSN-2463359 | BRD-K79337272 | CC(C)NC(=O)c1ccc(C#Cc2ccncc2)cn1 | STAFRSGTRKNXHF-UHFFFAOYSA-N | No-Effect | inactive |
| LSZ-102 | BRD-K00004675 | CC(F)(F)c1cc(F)ccc1-c1sc2cc(O)ccc2c1Oc1ccc(/C=C/C(=O)O)cc1 | SJXNPGGVGZXKKI-NYYWCZLTSA-N | No-Effect | inactive |
| LTA | BRD-K52914072 | N=C(Nc1ccccc1)c1cccs1 | RCORNHOUCUDCAC-UHFFFAOYSA-N | No-Effect | inactive |
| LTB4 | BRD-K53124401 | CCCCC/C=C\C[C@@H](O)/C=C/C=C/C=C\[C@@H](O)CCCC(=O)O | VNYSSYRCGWBHLG-AMOLWHMGSA-N | No-Effect | inactive |
| Lu-AA-47070 | BRD-K83970940 | CC(C)(C)CC(=O)Nc1c(F)cc(C(=O)/N=c2\sccn2COP(=O)(O)O)cc1F | MSWIQSFUBYCFJE-PGMHBOJBSA-N | No-Effect | inactive |
| lubiprostone | BRD-K33814251 | CCCCC(F)(F)C(=O)CC[C@H]1[C@H](O)CC(=O)[C@@H]1CCCCCCC(=O)O | DBVFKLAGQHYVGQ-BFYDXBDKSA-N | No-Effect | inactive |
| lucanthone | BRD-K00003521 | CCN(CC)CCNc1ccc(C)c2sc3ccccc3c(=O)c12 | FBQPGGIHOFZRGH-UHFFFAOYSA-N | No-Effect | inactive |
| lucitanib | BRD-K31309378 | CNC(=O)c1cccc2cc(Oc3ccnc4cc(OCC5(N)CC5)c(OC)cc34)ccc12 | CUDVHEFYRIWYQD-UHFFFAOYSA-N | No-Effect | inactive |

| | | | | | |
|---|---|---|---|---|---|
| lufenuron | BRD-A71774530 | O=C(NC(=O)c1c(F)cccc1F)Nc1cc(Cl)c(OC(F)(F)[C@@H](F)C(F)(F)F)cc1Cl | PWPJGUXAGUPAHP-AWEZNQCLSA-N | No-Effect | inactive |
| luliconazole | BRD-K71058253 | N#C/C(=C1/SC[C@@H](c2ccc(Cl)cc2Cl)S1)n1ccnc1 | YTAOBBFIOAEMLL-REQDGWNSSA-N | No-Effect | inactive |
| lumacaftor | BRD-K69236721 | Cc1ccc(NC(=O)C2(c3ccc4c(c3)OC(F)(F)O4)CC2)nc1-c1cccc(C(=O)O)c1 | UFSKUSARDNFIRC-UHFFFAOYSA-N | No-Effect | inactive |
| lumefantrine | BRD-A05926236 | CCCCN(CCCC)C[C@@H](O)c1cc(Cl)cc2c1-c1ccc(Cl)cc1/C2=C/c1ccc(Cl)cc1 | DYLGFOYVTXJFJP-OXTLUBNBSA-N | No-Effect | inactive |
| lumiracoxib | BRD-K26901728 | Cc1ccc(Nc2c(F)cccc2Cl)c(CC(=O)O)c1 | KHPKQFYUPIUARC-UHFFFAOYSA-N | No-Effect | inactive |
| lupanine | BRD-K00004532 | O=C1CCC[C@@H]2[C@H]3C[C@@H](CN12)[C@@H]1CCCCN1C3 | JYIJIIVLEOETIQ-XDQVBPFNSA-N | No-Effect | inactive |
| lurasidone | BRD-K51143828 | O=C1[C@H]2[C@@H]3CC[C@@H](C3)[C@H]2C(=O)N1C[C@@H]1CCCC[C@H]1CN1CCN(c2nsc3ccccc23)CC1 | PQXKDMSYBGKCJA-CVTJIBDQSA-N | No-Effect | inactive |
| lusaperidone | BRD-K00004742 | Cc1nc2ccccn2c(=O)c1CCN1CCc2oc3ccccc3c2C1 | ZYXHQIPQIKTEDI-UHFFFAOYSA-N | No-Effect | inactive |
| lusutrombopag | BRD-K93228461 | CCCCCCO[C@@H](C)c1cccc(-c2csc(NC(=O)c3cc(Cl)c(/C=C(\C)C(=O)O)c(Cl)c3)n2)c1OC | NOZIJMHMKORZBA-KJCUYJGMSA-N | No-Effect | inactive |
| lutein | BRD-K16742848 | CC1=C[C@H](O)CC(C)(C)[C@H]1/C=C/C(C)=C/C=C/C(C)=C/C=C/C=C(C)/C=C/C=C(C)/C=C/C1=C(C)C[C@@H](O)CC1(C)C | KBPHJBAIARWVSC-RGZFRNHPSA-N | No-Effect | inactive |
| luteolin | BRD-K05236810 | O=c1cc(-c2ccc(O)c(O)c2)oc2cc(O)cc(O)c12 | IQPNAANSBPBGFQ-UHFFFAOYSA-N | No-Effect | inactive |
| LX1031 | BRD-K32101625 | COc1cccc(-c2ccc([C@@H](Oc3cc(-c4ccc(C[C@H](N)C(=O)O)cc4)nc(N)n3)C(F)(F)F)cc2)c1 | XNMUICFMGGQSMZ-WIOPSUGQSA-N | No-Effect | inactive |
| LX7101 | BRD-K21038058 | Cc1c[nH]c2ncnc(N3CCC(CN)(C(=O)Nc4cccc(OC(=O)N(C)C)c4)CC3)c12 | PWPNYABQEOGNNC-UHFFFAOYSA-N | No-Effect | inactive |
| LXR-623 | BRD-K54520417 | Fc1ccc(-c2c3cccc(C(F)(F)F)c3nn2Cc2ccc(F)cc2Cl)cc1 | KYWWJENKIMRJBI-UHFFFAOYSA-N | No-Effect | inactive |
| LY2090314 | BRD-K78055238 | O=C1NC(=O)C(c2cnc3ccccn23)=C1c1cn2c3c(cc(F)cc13)CN(C(=O)N1CCCCC1)CC2 | HRJWTAWVFDCTGO-UHFFFAOYSA-N | No-Effect | inactive |
| LY2109761 | BRD-K47557313 | c1ccc(-c2nn3c(c2-c2ccnc4cc(OCCN5CCOCC5)ccc24)CCC3)nc1 | IHLVSLOZUHKNMQ-UHFFFAOYSA-N | No-Effect | inactive |
| LY2140023 | BRD-K49519144 | N[C@@]1(C(=O)O)CS(=O)(=O)[C@H]2[C@H](C(=O)O)[C@H]21 | AVDUGNCTZRCAHH-MDASVERJSA-N | No-Effect | inactive |
| LY215490 | BRD-K34313798 | O=C(O)[C@@H]1C[C@H]2C[C@@H](CCc3nn[nH]n3)CC[C@H]2CN1 | ZXFRFPSZAKNPQQ-YTWAJWBKSA-N | No-Effect | inactive |
| LY2157299 | BRD-K78118466 | Cc1cccc(-c2nn3c(c2-c2ccnc4ccc(C(N)=O)cc24)CCC3)n1 | IVRXNBXKWIJUQB-UHFFFAOYSA-N | No-Effect | inactive |
| LY2183240 | BRD-K37865504 | CN(C)C(=O)n1nnnc1Cc1ccc(-c2ccccc2)cc1 | GZNIYOXWFCDBBJ-UHFFFAOYSA-N | No-Effect | inactive |
| LY223982 | BRD-K35367061 | COc1ccc(/C=C/CCCCOc2ccc(C(=O)c3cccc(C(=O)O)c3)cc2CCC(=O)O)cc1 | SYZSSLLFRVDRHL-QPJJXVBHSA-N | No-Effect | inactive |
| LY225910 | BRD-K33864865 | CC(C)Oc1cccc(-n2c(CCc3c[nH]c4ccc(Br)cc34)nc3ccccc3c2=O)c1 | KUECXUACQOYKNB-UHFFFAOYSA-N | No-Effect | inactive |
| LY231617 | BRD-K85804405 | CCNCc1cc(C(C)(C)C)c(O)c(C(C)(C)C)c1 | WQNYIWJBDGLKEB-UHFFFAOYSA-N | No-Effect | inactive |
| LY2334737 | BRD-K04993719 | CCCC(CCC)C(=O)Nc1ccn([C@@H]2O[C@H](CO)[C@@H](O)C2(F)F)c(=O)n1 | MEOYFIHNRBNEPI-UXIGCNINSA-N | No-Effect | inactive |
| LY2365109 | BRD-K13888115 | CN(CCOc1ccc(-c2ccc3c(c2)OCO3)cc1C(C)(C)C)CC(=O)O | FKPLJWGRBCQLTL-UHFFFAOYSA-N | No-Effect | inactive |
| LY2452473 | BRD-K90171280 | CC(C)OC(=O)N[C@H]1Cc2c(n(Cc3ccccn3)c3ccc(C#N)cc23)C1 | IHIWYQYVBNODSV-KRWDZBQOSA-N | No-Effect | inactive |
| LY2510924 | BRD-K00003131 | CC(C)NCCCC[C@H](NC(=O)[C@H]1CCC(=O)N[C@@H](Cc2ccccc2)C(=O)N[C@H](Cc2ccc(O)cc2)C(=O)N[C@@H](CCCCNC(C)C)C(=O)N[C@H](CCCN=C(N)N)C(=O)N[C@@H](Cc2ccc3ccccc3c2)C(=O)NCC(=O)N1)C(N)=O | IJHWVENTEFSNBC-OFPUNPKKSA-N | No-Effect | inactive |
| LY2584702 | BRD-K21833260 | Cn1cc(-c2ccc(F)c(C(F)(F)F)c2)nc1C1CCN(c2ncnc3n[nH]cc23)CC1 | FYXRSVDHGLUMHB-UHFFFAOYSA-N | No-Effect | inactive |
| LY2603618 | BRD-K14109347 | Cc1cnc(NC(=O)Nc2cc(Br)c(C)cc2OC[C@@H]2CNCCO2)cn1 | SYYBDNPGDKKJDU-ZDUSSCGKSA-N | No-Effect | inactive |
| LY2606368 | BRD-K20463415 | COc1cccc(OCCCN)c1-c1cc(Nc2cnc(C#N)cn2)[nH]n1 | DOTGPNHGTYJDEP-UHFFFAOYSA-N | No-Effect | inactive |
| LY2608204 | BRD-K12068470 | O=C(Nc1ncc(SCCN2CCCC2)s1)[C@]1(c2ccc(S(=O)(=O)C3CC3)cc2)C[C@H]1C1CCCCC1 | QIIVJLHCZUTGSD-CUBQBAPOSA-N | No-Effect | inactive |
| LY266097 | BRD-A64004759 | COc1ccc(C[C@H]2NCCc3c2[nH]c2ccc(C)cc32)c(Cl)c1OC | NJLHHCITDFZZSE-QGZVFWFLSA-N | No-Effect | inactive |
| LY2784544 | BRD-K25630527 | Cc1cc(Nc2cc(CN3CCOCC3)c3nc(Cc4ccc(Cl)cc4F)n3n2)[nH]n1 | SQSZANZGUXWJEA-UHFFFAOYSA-N | No-Effect | inactive |
| LY2795050 | BRD-K00003195 | NC(=O)c1ccc(Oc2ccc(CN3CCC[C@H]3c3cccnc3)cc2)c(Cl)c1 | LOOCZNLSXJHWTG-NRFANRHFSA-N | No-Effect | inactive |
| LY2801653 | BRD-K18961567 | Cc1ccc(C(=O)Nc2ccc(Oc3cc4cnn(C)c4cc3-c3cn[nH]c3)c(F)c2)c(=O)n1-c1ccc(F)cc1 | QHADVLVFMKEIIP-UHFFFAOYSA-N | No-Effect | inactive |
| LY2811376 | BRD-K94534639 | C[C@@]1(c2cc(-c3cncnc3)c(F)cc2F)CCSC(N)=N1 | MJQMRGWYPNIERM-HNNXBMFYSA-N | No-Effect | inactive |
| LY2828360 | BRD-K00005249 | Cc1nc(N2CCN(C)CC2)c2nc(-c3ccccc3Cl)n(C3CCOCC3)c2n1 | UCMNDPDJRSEZPL-UHFFFAOYSA-N | No-Effect | inactive |
| LY2874455 | BRD-K17203476 | C[C@@H](Oc1ccc2n[nH]c(/C=C/c3cnn(CCO)c3)c2c1)c1c(Cl)cncc1Cl | GKJCVYLDJWTWQU-CXLRFSCWSA-N | No-Effect | inactive |
| LY288513 | BRD-K24675965 | O=C1NN(C(=O)Nc2ccc(Br)cc2)[C@@H](c2ccccc2)[C@@H]1c1ccccc1 | LMUQHXHWJWQXSD-PMACEKPBSA-N | No-Effect | inactive |
| LY2886721 | BRD-K51258098 | NC1=N[C@@]2(c3cc(NC(=O)c4ccc(F)cn4)ccc3F)COC[C@H]2CS1 | NIDRNVHMMDAAIK-YPMLDQLKSA-N | No-Effect | inactive |
| LY294002 | BRD-K27305650 | O=c1cc(N2CCOCC2)oc2c(-c3ccccc3)cccc12 | CZQHHVNHHHRRDU-UHFFFAOYSA-N | No-Effect | inactive |
| LY3000328 | BRD-K21070437 | CNC(=O)O[C@H]1COc2ccc(N3CCN(C4COC4)CC3)cc2[C@@H]1NC(=O)c1ccc(F)cc1 | NDEBZCZEAVMSQF-GOTSBHOMSA-N | No-Effect | inactive |
| LY3009120 | BRD-K38385169 | CNc1ncc2cc(-c3cc(NC(=O)NCCC(C)(C)C)c(F)cc3C)c(C)nc2n1 | HHCBMISMPSAZBF-UHFFFAOYSA-N | No-Effect | inactive |
| LY303511 | BRD-K22385716 | O=c1cc(N2CCNCC2)oc2c(-c3ccccc3)cccc12 | NGAGMBNBKCDCDJ-UHFFFAOYSA-N | No-Effect | inactive |
| LY310762 | BRD-K41194171 | CC1(C)C(=O)N(CCN2CCC(C(=O)c3ccc(F)cc3)CC2)c2ccccc21 | KDXISMANFPJVJY-UHFFFAOYSA-N | No-Effect | inactive |
| LY311727 | BRD-K00004565 | CCc1c(CC(N)=O)c2cc(OCCCP(=O)(O)O)ccc2n1Cc1ccccc1 | OPWQYOUZRHDKBR-UHFFFAOYSA-N | No-Effect | inactive |
| LY3200882 | BRD-K00003446 | CC(C)(O)c1cc(Nc2cc(Oc3cn(C4CC4)nc3C3CCOCC3)ccn2)ccn1 | PNPFMWIDAKQFPY-UHFFFAOYSA-N | No-Effect | inactive |
| LY320135 | BRD-K78637815 | COc1ccc(-c2oc3cc(OC)ccc3c2C(=O)c2ccc(C#N)cc2)cc1 | RYNSGDFWBJWWSZ-UHFFFAOYSA-N | No-Effect | inactive |
| LY3214996 | BRD-K00003416 | Cn1nccc1Nc1nccc(-c2cc3c(s2)C(C)(C)N(CCN2CCOCC2)C3=O)n1 | JNPRPMBJODOFEC-UHFFFAOYSA-N | No-Effect | inactive |
| LY3295668 | BRD-K00004722 | Cc1cc(Nc2cc(F)c(C[C@@]3(C(=O)O)CCN(Cc4cccc(Cl)c4F)[C@H](C)C3)n2)n[nH]1 | YQQZZYYQTCPEAS-OYLFLEFRSA-N | No-Effect | inactive |
| LY344864 | BRD-K95899059 | CN(C)[C@@H]1CCc2[nH]c3ccc(NC(=O)c4ccc(F)cc4)cc3c2C1 | GKWHICIUSVVNGX-MRXNPFEDSA-N | No-Effect | inactive |
| LY364947 | BRD-K06234293 | c1ccc(-c2n[nH]cc2-c2ccnc3ccccc23)nc1 | IBCXZJCWDGCXQT-UHFFFAOYSA-N | No-Effect | inactive |
| LY393558 | BRD-K08534264 | CC(C)N1Cc2cc(S(C)(=O)=O)ccc2N(CCN2CC=C(c3c[nH]c4cc(F)ccc34)CC2)S1(=O)=O | QUSLYAPLTMMCFE-UHFFFAOYSA-N | No-Effect | inactive |
| LY404187 | BRD-A50062753 | CC(C)S(=O)(=O)NC[C@H](C)c1ccc(-c2ccc(C#N)cc2)cc1 | HOQAVGZLYRYHSO-HNNXBMFYSA-N | No-Effect | inactive |
| LY450108 | BRD-K54538117 | CC(C)S(=O)(=O)NC[C@H](C)c1ccc(NC(=O)c2cc(F)cc(F)c2)cc1 | ACOXQYLJOQAHST-ZDUSSCGKSA-N | No-Effect | inactive |
| LY456236 | BRD-K87696786 | COc1ccc(Nc2ncnc3ccc(OC)cc23)cc1 | XPJOMFBGKIABQW-UHFFFAOYSA-N | No-Effect | inactive |
| lycopene | BRD-K37083340 | CC(C)=CCC/C(C)=C/C=C/C(C)=C/C=C/C(C)=C/C=C/C=C(C)/C=C/C=C(C)/C=C/C=C(\C)CCC=C(C)C | OAIJSZIZWZSQBC-GYZMGTAESA-N | No-Effect | inactive |
| lymecycline | BRD-K00004536 | CN(C)[C@@H]1C(=O)[C@@H](C(=O)NCNCCCC[C@H](N)C(=O)O)C(=O)[C@@]2(O)C(=O)[C@@H]3C(=O)c4c(O)cccc4[C@@](C)(O)[C@H]3C[C@@H]12 | DZTKAWPWXGIREW-KKSKXDPKSA-N | No-Effect | inactive |
| lynestrenol | BRD-K39707084 | C#C[C@]1(O)CC[C@H]2[C@@H]3CCC4=CCCC[C@@H]4[C@H]3CC[C@@]21C | YNVGQYHLRCDXFQ-XGXHKTLJSA-N | No-Effect | inactive |
| M-14157 | BRD-A96977263 | CCCCCC(O)CCCC(=O)O | LMHJFKYQYDSOQO-UHFFFAOYSA-N | No-Effect | inactive |
| M-25 | BRD-K51265828 | CCCC(CCC)C(=O)NCc1ccc2c(cnn2-c2ccccc2OC)c1 | WATNXVHKRRTUFK-UHFFFAOYSA-N | No-Effect | inactive |
| m-3M3FBS | BRD-K09635314 | Cc1cc(C)c(S(=O)(=O)Nc2cccc(C(F)(F)F)c2)c(C)c1 | ZIIUUSVHCHPIQD-UHFFFAOYSA-N | No-Effect | inactive |

| | | | | | |
|---|---|---|---|---|---|
| m-chlorophenylbiguanide | BRD-K36965586 | N=C(N=C(N)N)Nc1cccc(Cl)c1 | DIHXJZHAIHGSAW-UHFFFAOYSA-N | No-Effect | inactive |
| m-THP | BRD-K91147684 | Oc1cccc(-c2c3nc(c(-c4cccc(O)c4)c4ccc([nH]4)c(-c4cccc(O)c4)c4cc([nH]4)c(-c4cccc(O)c4)c4nc2C=C4)C=C3)c1 | XAEPFBXVLMACNN-UHFFFAOYSA-N | No-Effect | inactive |
| M8-B | BRD-K90410034 | COc1cc(CN(CCN)C(=O)c2cccs2)ccc1OCc1ccccc1 | UOGGWHBYFVBUIY-UHFFFAOYSA-N | No-Effect | inactive |
| mabuprofen | BRD-A93683007 | CC(C)Cc1ccc([C@@H](C)C(=O)NCCO)cc1 | JVGUNCHERKJFCM-GFCCVEGCSA-N | No-Effect | inactive |
| mabuterol | BRD-A13116749 | CC(C)(C)NC[C@@H](O)c1cc(Cl)c(N)c(C(F)(F)F)c1 | JSJCTEKTBOKRST-SNVBAGLBSA-N | No-Effect | inactive |
| macelignan | BRD-K94948151 | COc1cc(C[C@H](C)[C@H](C)Cc2ccc3c(c2)OCO3)ccc1O | QDDILOVMGWUNGD-UONOGXRCSA-N | No-Effect | inactive |
| macitentan | BRD-K48812570 | CCCNS(=O)(=O)Nc1ncnc(OCCOc2ncc(Br)cn2)c1-c1ccc(Br)cc1 | JGCMEBMXRHSZKX-UHFFFAOYSA-N | No-Effect | inactive |
| macitentan-n-butyl-analogue | BRD-K00003167 | CCCCS(=O)(=O)Nc1ncnc(OCCOc2ncc(Br)cn2)c1-c1ccc(Br)cc1 | OZBDTKLWBSATSN-UHFFFAOYSA-N | No-Effect | inactive |
| macozinone | BRD-K16343284 | O=c1nc(N2CCN(CC3CCCCC3)CC2)sc2c([N+](=O)[O-])cc(C(F)(F)F)cc12 | BJDZBXGJNBMCAV-UHFFFAOYSA-N | No-Effect | inactive |
| madecassoside | BRD-K00003328 | C[C@@H]1[C@H]2C3=CC[C@@H]4[C@@]5(C)C[C@@H](O)[C@H](O)[C@@](C)(CO)[C@@H]5[C@H](O)C[C@@]4(C)[C@]3(C)CC[C@@]2(C(=O)O[C@@H]2O[C@H](CO[C@@H]3O[C@H](CO)[C@@H](O[C@@H]4O[C@@H](C)[C@H](O)[C@@H](O)[C@H]4O)[C@H](O)[C@H]3O)[C@@H](O)[C@H](O)[C@H]2O)CC[C@H]1C | BNMGUJRJUUDLHW-HCZMHFOYSA-N | No-Effect | inactive |
| madrasin | BRD-K48905682 | COc1ccc2c(C)nc(Nc3nc(=O)c(C)c(C)[nH]3)nc2c1 | QQJIYKXTEMDJFM-UHFFFAOYSA-N | No-Effect | inactive |
| mafenide | BRD-K30649484 | NCc1ccc(S(N)(=O)=O)cc1 | TYMRLRRVMHJFTF-UHFFFAOYSA-N | No-Effect | inactive |
| magnolol | BRD-K26168087 | C=CCc1ccc(O)c(-c2cc(CC=C)ccc2O)c1 | VVOAZFWZEDHOOU-UHFFFAOYSA-N | No-Effect | inactive |
| malathion | BRD-A01976766 | CCOC(=O)C[C@@H](SP(=S)(OC)OC)C(=O)OCC | JXSJBGJIGXNWCI-MRVPVSSYSA-N | No-Effect | inactive |
| malic-acid | BRD-A18396475 | O=C(O)C[C@H](O)C(=O)O | BJEPYKJPYRNKOW-REOHCLBHSA-N | No-Effect | inactive |
| malotilate | BRD-K97884852 | CC(C)OC(=O)C(C(=O)OC(C)C)=C1SC=CS1 | YPIQVCUJEKAZCP-UHFFFAOYSA-N | No-Effect | inactive |
| maltobionic-acid | BRD-K01624185 | O=C(O)[C@H](O)[C@@H](O)[C@H](O[C@@H]1O[C@H](CO)[C@H](O)[C@H](O)[C@H]1O)[C@H](O)CO | JYTUSYBCFIZPBE-AMTLMPIISA-N | No-Effect | inactive |
| mandelic-acid | BRD-A91448670 | O=C(O)[C@H](O)c1ccccc1 | IWYDHOAUDWTVEP-SSDOTTSWSA-N | No-Effect | inactive |
| mangafodipir | BRD-K57365211 | Cc1nc(COP(=O)(O)O)cc2c1O[Mn]Oc1c(cc(COP(=O)(O)O)nc1C)CN(CC(=O)O)CCN(CC(=O)O)C2 | CXFKOLCMCRBYPL-UHFFFAOYSA-L | No-Effect | inactive |
| manidipine | BRD-K01825587 | COC(=O)C1=C(C)NC(C)=C(C(=O)OCCN2CCN(C(c3ccccc3)c3ccccc3)CC2)[C@@H]1c1cccc([N+](=O)[O-])c1 | ANEBWFXPVPTEET-JGCGQSQUSA-N | No-Effect | inactive |
| manitimus | BRD-K00004729 | C#CCCC(=O)[C@@H](C#N)C(=O)Nc1ccc(C(F)(F)F)cc1 | MYMWDZLPEPMSCN-GFCCVEGCSA-N | No-Effect | inactive |
| mannitol-D | BRD-K30374899 | OC[C@@H](O)[C@@H](O)[C@H](O)[C@H](O)CO | FBPFZTCFMRRESA-KVTDHHQDSA-N | No-Effect | inactive |
| maraviroc | BRD-K00004293 | Cc1nnc(C(C)C)n1[C@H]1C[C@@H]2CC[C@H](C1)N2CC[C@H](NC(=O)C1CCC(F)(F)CC1)c1ccccc1 | GSNHKUDZZFZSJB-QYOOZWMWSA-N | No-Effect | inactive |
| maribavir | BRD-K59524082 | CC(C)Nc1nc2cc(Cl)c(Cl)cc2n1[C@H]1O[C@@H](CO)[C@H](O)[C@@H]1O | KJFBVJALEQWJBS-XUXIUFHCSA-N | No-Effect | inactive |
| marimastat | BRD-K21396683 | CNC(=O)[C@@H](NC(=O)[C@H](CC(C)C)[C@H](O)C(=O)NO)C(C)(C)C | OCSMOTCMPXTDND-OUAUKWLOSA-N | No-Effect | inactive |
| masoprocol | BRD-A87130939 | C[C@@H](Cc1ccc(O)c(O)c1)[C@@H](C)Cc1ccc(O)c(O)c1 | HCZKYJDFEPMADG-RYUDHWBXSA-N | No-Effect | inactive |
| matrine | BRD-K08623988 | O=C1CCC[C@@H]2[C@H]3CCCN4CCC[C@@H](CN12)[C@@H]34 | ZSBXGIUJOOQZMP-JLNYLFASSA-N | No-Effect | inactive |
| mavacamten | BRD-K00003497 | CC(C)n1c(=O)cc(N[C@@H](C)c2ccccc2)[nH]c1=O | RLCLASQCAPXVLM-NSHDSACASA-N | No-Effect | inactive |
| mavatrep | BRD-K00003237 | CC(C)(O)c1ccccc1-c1ccc2nc(/C=C/c3ccc(C(F)(F)F)cc3)[nH]c2c1 | ORDHXXHTBUZRCN-NTEUORMPSA-N | No-Effect | inactive |
| mavoglurant | BRD-K07790550 | COC(=O)N1CC[C@@H]2[C@H]1CCC[C@@]2(O)C#Cc1cccc(C)c1 | ZFPZEYHRWGMJCV-ZHALLVOQSA-N | No-Effect | inactive |
| maxacalcitol | BRD-K47318833 | C=C1/C(=C\C=C2/CCC[C@]3(C)[C@@H]([C@H](C)OCCC(C)(C)O)CC[C@@H]23)C[C@@H](O)C[C@@H]1O | DTXXSJZBSTYZKE-ZDQKKZTESA-N | No-Effect | inactive |
| MBX-2982 | BRD-K32218650 | CCc1cnc(N2CCC(c3nc(COc4ccc(-n5cnnn5)cc4)cs3)CC2)nc1 | NFTMKHWBOINJGM-UHFFFAOYSA-N | No-Effect | inactive |
| MC-1 | BRD-K50712474 | Cc1ncc(COP(=O)(O)O)c(C=O)c1O | NGVDGCNFYWLIFO-UHFFFAOYSA-N | No-Effect | inactive |
| MC1568 | BRD-K24031866 | Cn1c(/C=C/C(=O)NO)ccc1/C=C/C(=O)c1cccc(F)c1 | QQDIFLSJMFDTCQ-FIFLTTCUSA-N | No-Effect | inactive |
| MCC950 | BRD-K55418700 | CC(C)(O)c1coc(S(=O)(=O)NC(=O)Nc2c3c(cc4c2CCC4)CCC3)c1 | HUUSXLKCTQDPGL-UHFFFAOYSA-N | No-Effect | inactive |
| mCPP | BRD-K75844781 | Clc1cccc(N2CCNCC2)c1 | VHFVKMTVMIZMIK-UHFFFAOYSA-N | No-Effect | inactive |
| MD-920 | BRD-K92758126 | C=C1C[C@]23C[C@@]1(O)CC[C@H]2[C@@]12C=C[C@H](O)[C@@](C)(C(=O)O1)[C@H]2[C@@H]3C(=O)O | IXORZMNAPKEEDV-OBDJNFEBSA-N | No-Effect | inactive |
| mdivi-1 | BRD-K61128465 | COc1cc(-n2c(=S)[nH]c3ccccc3c2=O)c(Cl)cc1Cl | NZJKEVWTYMOYOR-UHFFFAOYSA-N | No-Effect | inactive |
| MDL-11939 | BRD-A62057054 | O[C@H](c1ccccc1)C1CCN(CCc2ccccc2)CC1 | AXNGJCOYCMDPQG-HXUWFJFHSA-N | No-Effect | inactive |
| MDL-27531 | BRD-K50888640 | Cn1c(-c2ccccc2)nnc1S(C)(=O)=O | DPGDRICHXKQXEN-UHFFFAOYSA-N | No-Effect | inactive |
| MDL-29951 | BRD-K59753853 | O=C(O)CCc1c(C(=O)O)[nH]c2cc(Cl)cc(Cl)c12 | KNBSYZNKEAWABY-UHFFFAOYSA-N | No-Effect | inactive |
| MDL-72832 | BRD-K01825683 | O=C1CC2(CCCC2)CC(=O)N1CCCCNC[C@H]1COc2ccccc2O1 | HCVRCHQFYYZPHT-KRWDZBQOSA-N | No-Effect | inactive |
| MDL-73005EF | BRD-K01826803 | O=C1CC2(CCCC2)CC(=O)N1CCNC[C@H]1COc2ccccc2O1 | BVMYCHKQPGEOSI-HNNXBMFYSA-N | No-Effect | inactive |
| MDMS | BRD-K70840370 | CS(=O)(=O)OCOS(C)(=O)=O | IQLZWWDXNXZGPK-UHFFFAOYSA-N | No-Effect | inactive |
| ME-0328 | BRD-K24283677 | C[C@H](NC(=O)CCc1nc(=O)c2ccccc2[nH]1)c1ccccc1 | QIHBWVVVRYYYRO-ZDUSSCGKSA-N | No-Effect | inactive |
| ME0328 | BRD-A63444178 | C[C@@H](NC(=O)CCc1nc(=O)c2ccccc2[nH]1)c1ccccc1 | QIHBWVVVRYYYRO-CYBMUJFWSA-N | No-Effect | inactive |
| mebendazole | BRD-K77987382 | COC(=O)Nc1nc2ccc(C(=O)c3ccccc3)cc2[nH]1 | OPXLLQIJSORQAM-UHFFFAOYSA-N | No-Effect | inactive |
| mebeverine | BRD-A09467419 | CCN(CCCCOC(=O)c1ccc(OC)c(OC)c1)[C@@H](C)Cc1ccc(OC)cc1 | VYVKHNNGDFVQGA-IBGZPJMESA-N | No-Effect | inactive |
| mebhydrolin | BRD-K29713308 | CN1CCc2c(c3ccccc3n2Cc2ccccc2)C1 | FQQIIPAOSKSOJM-UHFFFAOYSA-N | No-Effect | inactive |
| mebicar | BRD-K76606751 | CN1C(=O)N(C)C2C1N(C)C(=O)N2C | XIUUSFJTJXFNGH-UHFFFAOYSA-N | No-Effect | inactive |
| mebrofenin | BRD-K96691275 | Cc1cc(C)c(NC(=O)CN(CC(=O)O)CC(=O)O)c(C)c1Br | MHPZZZZLAQGTHT-UHFFFAOYSA-N | No-Effect | inactive |
| mecamylamine | BRD-A20119038 | CNC1(C)C2CCC(C2)C1(C)C | IMYZQPCYWPFTAG-UHFFFAOYSA-N | No-Effect | inactive |
| mecarbinate | BRD-K28405228 | CCOC(=O)c1c(C)n(C)c2ccc(O)cc12 | YTBNTDMBGXAOCG-UHFFFAOYSA-N | No-Effect | inactive |
| meclinertant | BRD-K25075681 | COc1cccc(OC)c1-c1cc(C(=O)NC2(C(=O)O)C3CC4CC(C3)CC2C4)nn1-c1ccnc2cc(Cl)ccc12 | DYLJVOXRWLXDIG-UHFFFAOYSA-N | No-Effect | inactive |
| meclocycline-sulfosalicylate | BRD-A07600638 | C=C1c2c(Cl)ccc(O)c2C(=O)C2=C(O)[C@]3(O)C(O)=C(C(N)=O)C(=O)[C@@H](N(C)C)[C@@H]3[C@@H](O)[C@H]12 | MEFKOOXNVFALTA-IWVLMIASSA-N | No-Effect | inactive |
| mecysteine | BRD-K81993098 | COC(=O)[C@@H](N)CS | MCYHPZGUONZRGO-VKHMYHEASA-N | No-Effect | inactive |
| medetomidine | BRD-A66563878 | Cc1cccc([C@@H](C)c2cnc[nH]2)c1C | CUHVIMMYOGQXCV-LLVKDONJSA-N | No-Effect | inactive |
| medica-16 | BRD-K92908289 | CC(C)(CCCCCCCCCCC(C)(C)CC(=O)O)CC(=O)O | HYSMCRNFENOHJH-UHFFFAOYSA-N | No-Effect | inactive |
| medorinone | BRD-K40715924 | Cc1nccc2[nH]c(=O)ccc12 | OCCZJXAHSUCJSA-UHFFFAOYSA-N | No-Effect | inactive |
| medroxyprogesterone | BRD-K99615199 | CC(=O)[C@@]1(O)CC[C@H]2[C@@H]3C[C@H](C)C4=CC(=O)CC[C@]4(C)[C@H]3CC[C@@]21C | FRQMUZJSZHZSGN-HBNHAYAOSA-N | No-Effect | inactive |
| medroxyprogesterone-acetate | BRD-K82216340 | CC(=O)O[C@]1(C(C)=O)CC[C@H]2[C@@H]3C[C@H](C)C4=CC(=O)CC[C@]4(C)[C@H]3CC[C@@]21C | PSGAAPLEWMOORI-PEINSRQWSA-N | No-Effect | inactive |
| mefenamic-acid | BRD-K92778217 | Cc1cccc(Nc2ccccc2C(=O)O)c1C | HYYBABOKPJLUIN-UHFFFAOYSA-N | No-Effect | inactive |
| mefexamide | BRD-K20655524 | CCN(CC)CCNC(=O)COc1ccc(OC)cc1 | HUNIPYLVUPMFCZ-UHFFFAOYSA-N | No-Effect | inactive |
| megestrol-acetate | BRD-K19507340 | CC(=O)O[C@]1(C(C)=O)CC[C@H]2[C@@H]3C=C(C)C4=CC(=O)CC[C@]4(C)[C@H]3CC[C@@]21C | RQZAXGRLVPAYTJ-GQFGMJRRSA-N | No-Effect | inactive |
| meglitinide | BRD-K51324732 | COc1ccc(Cl)cc1C(=O)NCCc1ccc(C(=O)O)cc1 | SWLAMJPTOQZTAE-UHFFFAOYSA-N | No-Effect | inactive |
| meglumine | BRD-K28595280 | CNC[C@H](O)[C@@H](O)[C@H](O)[C@H](O)CO | MBBZMMPHUWSWHV-BDVNFPICSA-N | No-Effect | inactive |

| | | | | | |
|---|---|---|---|---|---|
| meisoindigo | BRD-K14529030 | CN1C(=O)/C(=C2/C(=O)Nc3ccccc32)c2ccccc21 | QNOCRUSVMMAKSC-CCEZHUSRSA-N | No-Effect | inactive |
| MEK1-2-inhibitor | BRD-K12244279 | N#CC(C(=N)Sc1ccc(N)cc1)c1ccccc1C(F)(F)F | XQRVCODIGHFXKQ-UHFFFAOYSA-N | No-Effect | inactive |
| melanotan-ii | BRD-K75989555 | CCCC[C@H](NC(C)=O)C(=O)N[C@H]1CC(=O)NCCCC[C@@H](C(N)=O)NC(=O)[C@H](Cc2c[nH]c3ccccc23)NC(=O)[C@H](CCCNC(=N)N)NC(=O)[C@@H](Cc2ccccc2)NC(=O)[C@H](Cc2cnc[nH]2)NC1=O | JDKLPDJLXHXHNV-MFVUMRCOSA-N | No-Effect | inactive |
| melatonin | BRD-K97530723 | COc1ccc2[nH]cc(CCNC(C)=O)c2c1 | DRLFMBDRBRZALE-UHFFFAOYSA-N | No-Effect | inactive |
| meldonium | BRD-K79392608 | C[N+](C)(C)NCCC(=O)O | PVBQYTCFVWZSJK-UHFFFAOYSA-O | No-Effect | inactive |
| melengestrol-acetate | BRD-A42746051 | C=C1C[C@H]2[C@H]3C=C(C)C4=CC(=O)CC[C@]4(C)[C@@H]3CC[C@]2(C)[C@@]1(OC(C)=O)C(C)=O | UDKABVSQKJNZBH-AKZSSJSASA-N | No-Effect | inactive |
| melevodopa | BRD-K18749874 | COC(=O)[C@@H](N)Cc1ccc(O)c(O)c1 | XBBDACCLCFWBSI-ZETCQYMHSA-N | No-Effect | inactive |
| meloxicam | BRD-K01826742 | Cc1cnc(NC(=O)C2=C(O)c3ccccc3S(=O)(=O)N2C)s1 | ZRVUJXDFFKFLMG-UHFFFAOYSA-N | No-Effect | inactive |
| melperone | BRD-K92984783 | CC1CCN(CCCC(=O)c2ccc(F)cc2)CC1 | DKMFBWQBDIGMHM-UHFFFAOYSA-N | No-Effect | inactive |
| melphalan | BRD-K87827419 | N[C@@H](Cc1ccc(N(CCCl)CCCl)cc1)C(=O)O | SGDBTWWWUNNDEQ-LBPRGKRZSA-N | No-Effect | inactive |
| melphalan-n-oxide | BRD-K36394566 | N[C@@H](Cc1ccc([N+]([O-])(CCCl)CCCl)cc1)C(=O)O | GSKQMLGAUOTSKT-LBPRGKRZSA-N | No-Effect | inactive |
| meluadrine | BRD-A07164067 | CC(C)(C)NC[C@@H](O)c1ccc(O)cc1Cl | LIXBJWRFCNRAPA-LLVKDONJSA-N | No-Effect | inactive |
| memantine | BRD-K91938660 | C[C@]12C[C@@H]3C[C@](C)(C1)C[C@](N)(C3)C2 | BUGYDGFZZOZRHP-CDECOKDKSA-N | No-Effect | inactive |
| menadione | BRD-K78126613 | CC1=CC(=O)c2ccccc2C1=O | MJVAVZPDRWSRRC-UHFFFAOYSA-N | No-Effect | inactive |
| menadione-bisulfite | BRD-K01826025 | C[C@@]1(S(=O)(=O)O)CC(=O)c2ccccc2C1=O | WIXFIQKTHUVFDI-LLVKDONJSA-N | No-Effect | inactive |
| menatetrenone | BRD-K75864342 | CC(C)=CCC/C(C)=C/CC/C(C)=C/CC/C(C)=C/CC1=C(C)C(=O)c2ccccc2C1=O | DKHGMERMDICWDU-GHDNBGIDSA-N | No-Effect | inactive |
| menbutone | BRD-K53852914 | COc1ccc(C(=O)CCC(=O)O)c2ccccc12 | FHGJSJFIQNQBCK-UHFFFAOYSA-N | No-Effect | inactive |
| meparfylon | BRD-A11662333 | C#CC(C)(O)CC | QXLPXWSKPNOQLE-UHFFFAOYSA-N | No-Effect | inactive |
| mepazine | BRD-K01825575 | CN1CCC[C@@H](CN2c3ccccc3Sc3ccccc32)C1 | CBHCDHNUZWWAPP-OAHLLOKOSA-N | No-Effect | inactive |
| mepenzolate | BRD-A62421304 | C[N+]1(C)CCC[C@H](OC(=O)C(O)(c2ccccc2)c2ccccc2)C1 | GKNPSSNBBWDAGH-IBGZPJMESA-N | No-Effect | inactive |
| mephenesin | BRD-A55469827 | Cc1ccccc1OC[C@@H](O)CO | JWDYCNIAQWPBHD-VIFPVBQESA-N | No-Effect | inactive |
| mephentermine | BRD-K18194590 | CNC(C)(C)Cc1ccccc1 | RXQCGGRTAILOIN-UHFFFAOYSA-N | No-Effect | inactive |
| mephenytoin | BRD-A83937277 | CC[C@@]1(c2ccccc2)NC(=O)N(C)C1=O | GMHKMTDVRCWUDX-LBPRGKRZSA-N | No-Effect | inactive |
| mepiroxol | BRD-K78299798 | [O-][n+]1cccc(CO)c1 | LQYJAVWWKTWWKN-UHFFFAOYSA-N | No-Effect | inactive |
| mepivacaine | BRD-A03216249 | Cc1cccc(C)c1NC(=O)[C@H]1CCCCN1C | INWLQCZOYSRPNW-CYBMUJFWSA-N | No-Effect | inactive |
| meprednisone | BRD-K18207669 | C[C@H]1C[C@H]2[C@@H]3CCC4=CC(=O)C=C[C@]4(C)[C@H]3C(=O)C[C@]2(C)[C@@]1(O)C(=O)CO | PIDANAQULIKBQS-RNUIGHNZSA-N | No-Effect | inactive |
| meprylcaine | BRD-K65417056 | CCCNC(C)(C)COC(=O)c1ccccc1 | VXJABHHJLXLNMP-UHFFFAOYSA-N | No-Effect | inactive |
| meptazinol | BRD-A02710418 | CC[C@@]1(c2cccc(O)c2)CCCCN(C)C1 | JLICHNCFTLFZJN-OAHLLOKOSA-N | No-Effect | inactive |
| mepyramine | BRD-K97564742 | COc1ccc(CN(CCN(C)C)c2ccccn2)cc1 | YECBIJXISLIIDS-UHFFFAOYSA-N | No-Effect | inactive |
| mequinol | BRD-K45216060 | COc1ccc(O)cc1 | NWVVVBRKAWDGAB-UHFFFAOYSA-N | No-Effect | inactive |
| merbarone | BRD-K45152786 | O=C(Nc1ccccc1)c1c(O)[nH]c(=S)[nH]c1=O | GFYRZTLCYQQVHZ-UHFFFAOYSA-N | No-Effect | inactive |
| merbromin | BRD-K21771665 | O=C(O)c1ccccc1-c1c2cc(Br)c(=O)cc-2oc2c([Hg]O)c(O)c(Br)cc12 | OHCFGBLQZMYRTR-UHFFFAOYSA-M | No-Effect | inactive |
| mercaptopurine | BRD-K91601245 | S=c1nc[nH]c2nc[nH]c12 | GLVAUDGFNGKCSF-UHFFFAOYSA-N | No-Effect | inactive |
| mercaptosuccinic-acid | BRD-A43435802 | O=C(O)C[C@@H](S)C(=O)O | NJRXVEJTAYWCQJ-UWTATZPHSA-N | No-Effect | inactive |
| Merck60 | BRD-K04466929 | CC(=O)Nc1ccc(C(=O)Nc2cc(-c3cccs3)ccc2N)cc1 | ABZSPJVXTTUFAA-UHFFFAOYSA-N | No-Effect | inactive |
| mericitabine | BRD-K69726595 | CC(C)C(=O)OC[C@H]1O[C@@H](n2ccc(N)nc2=O)[C@](C)(F)[C@@H]1OC(=O)C(C)C | MLESJYFEMSJZLZ-MAAOGQSESA-N | No-Effect | inactive |
| merimepodib | BRD-K09372874 | COc1cc(NC(=O)Nc2cccc(CNC(=O)O[C@H]3CCOC3)c2)ccc1-c1cnco1 | JBPUGFODGPKTDW-SFHVURJKSA-N | No-Effect | inactive |
| meropenem | BRD-K01612348 | C[C@@H](O)[C@H]1C(=O)N2C(C(=O)O)=C(S[C@@H]3CN[C@H](C(=O)N(C)C)C3)[C@H](C)[C@H]12 | DMJNNHOOLUXYBV-PQTSNVLCSA-N | No-Effect | inactive |
| mesna | BRD-K49954789 | O=S(=O)(O)CCS | ZNEWHQLOPFWXOF-UHFFFAOYSA-N | No-Effect | inactive |
| mesoridazine | BRD-K01827307 | CN1CCCC[C@@H]1CCN1c2ccccc2Sc2ccc([S@@](C)=O)cc21 | SLVMESMUVMCQIY-AKJBCIBTSA-N | No-Effect | inactive |
| mestinon | BRD-K69688083 | CN(C)C(=O)Oc1ccc[n+](C)c1 | RVOLLAQWKVFTGE-UHFFFAOYSA-N | No-Effect | inactive |
| mestranol | BRD-K31920458 | C#C[C@]1(O)CC[C@H]2[C@@H]3CCc4cc(OC)ccc4[C@H]3CC[C@@]21C | IMSSROKUHAOUJS-MJCUULBUSA-N | No-Effect | inactive |
| mesulergine | BRD-K91336023 | CN1C[C@@H](NS(=O)(=O)N(C)C)C[C@@H]2c3cccc4c3c(cn4C)C[C@H]21 | JLVHTNZNKOSCNB-YSVLISHTSA-N | No-Effect | inactive |
| meta-cresyl-acetate | BRD-K33442414 | CC(=O)Oc1cccc(C)c1 | OTGAHJPFNKQGAE-UHFFFAOYSA-N | No-Effect | inactive |
| metacetamol | BRD-K16474819 | CC(=O)Nc1cccc(O)c1 | QLNWXBAGRTUKKI-UHFFFAOYSA-N | No-Effect | inactive |
| metafolin | BRD-K28580965 | CN1c2c([nH]c(N)nc2=O)NC[C@@H]1CNc1ccc(C(=O)N[C@@H](CCC(=O)O)C(=O)O)cc1 | ZNOVTXRBGFNYRX-STQMWFEESA-N | No-Effect | inactive |
| metamizole | BRD-K76812510 | Cc1c(N(C)CS(=O)(=O)O)c(=O)n(-c2ccccc2)n1C | LVWZTYCIRDMTEY-UHFFFAOYSA-N | No-Effect | inactive |
| metaproterenol | BRD-K01826731 | CC(C)NC[C@@H](O)c1cc(O)cc(O)c1 | LMOINURANNBYCM-LLVKDONJSA-N | No-Effect | inactive |
| metaraminol | BRD-K96253961 | C[C@H](N)[C@H](O)c1cccc(O)c1 | WXFIGDLSSYIKKV-RCOVLWMOSA-N | No-Effect | inactive |
| metatinib | BRD-K27182532 | Nc1nccc(Oc2ccc(NC(=O)c3c[nH]cc(-c4ccc(F)cc4)c3=O)cc2F)c1Cl | PDYXPCKITKHFOZ-UHFFFAOYSA-N | No-Effect | inactive |
| metaxalone | BRD-A94709349 | Cc1cc(C)cc(OC[C@H]2CNC(=O)O2)c1 | IMWZZHHPURKASS-LLVKDONJSA-N | No-Effect | inactive |
| metazosin | BRD-A78277353 | COc1cc2nc(N3CCN(C(=O)[C@H](C)OC)CC3)nc(N)c2cc1OC | YEOTYALSMRNXLJ-NSHDSACASA-N | No-Effect | inactive |
| metenkephalin | BRD-A38218502 | CSCC[C@@H](NC(=O)[C@@H](Cc1ccccc1)NC(=O)CNC(=O)CNC(=O)[C@@H](N)Cc1ccc(O)cc1)C(=O)O | YFGBQHOOROIVKG-BHDDXSALSA-N | No-Effect | inactive |
| metformin | BRD-K79602928 | CN(C)C(=N)N=C(N)N | XZWYZXLIPXDOLR-UHFFFAOYSA-N | No-Effect | inactive |
| methacholine | BRD-A18056626 | CC(=O)O[C@@H](C)C[N+](C)(C)C | NZWOPGCLSHLLPA-ZETCQYMHSA-N | No-Effect | inactive |
| methacycline | BRD-K00003351 | C=C1c2cccc(O)c2C(O)=C2C(=O)[C@@H]2C(=O)[C@]3(O)C(=O)[C@@H](C(N)=O)C(=O)[C@@H](N(C)C)[C@@H]3[C@@H](O)[C@H]12 | QNJKSUYWBCNWFI-IKHBEOKZSA-N | No-Effect | inactive |
| methanesulfonyl-fluoride | BRD-K53896891 | CS(=O)(=O)F | KNWQLFOXPQZGPX-UHFFFAOYSA-N | No-Effect | inactive |
| methantheline | BRD-K09859624 | CC[N+](C)(CC)CCOC(=O)C1c2ccccc2Oc2ccccc21 | GZHFODJQISUKAY-UHFFFAOYSA-N | No-Effect | inactive |
| methapyrilene | BRD-K47323024 | CN(C)CCN(Cc1cccs1)c1ccccn1 | HNJJXZKZRAWDPF-UHFFFAOYSA-N | No-Effect | inactive |
| metharbital | BRD-K88143662 | CCC1(CC)C(=O)NC(=O)N(C)C1=O | FWJKNZONDWOGMI-UHFFFAOYSA-N | No-Effect | inactive |
| methazolamide | BRD-K00002562 | CC(=O)N=c1sc(S(N)(=O)=O)nn1C | FLOSMHQXBMRNHR-UHFFFAOYSA-N | No-Effect | inactive |
| methenamine | BRD-K30114692 | C1N2CN3CN1CN(C2)C3 | VKYKSIONXSXAKP-UHFFFAOYSA-N | No-Effect | inactive |
| methiazole | BRD-K02764365 | COC(=O)Nc1nc2ccc(SC(C)C)cc2[nH]1 | SXJIZQPZESTWLD-UHFFFAOYSA-N | No-Effect | inactive |
| methicillin | BRD-K34388247 | COc1cccc(OC)c1C(=O)N[C@@H]1C(=O)N2[C@@H](C(=O)O)C(C)(C)S[C@H]12 | RJQXTJLFIWVMTO-TYNCELHUSA-N | No-Effect | inactive |
| methimazole | BRD-K54416256 | Cn1cc[nH]c1=S | PMRYVIKBURPHAH-UHFFFAOYSA-N | No-Effect | inactive |
| methionine | BRD-A39515206 | CSCC[C@@H](N)C(=O)O | FFEARJCKVFRZRR-SCSAIBSYSA-N | No-Effect | inactive |
| methiopril | BRD-K00004541 | CC(=O)SCCC(=O)N1[C@H](C)CCC[C@@H]1C(=O)O | JHFOAISYWJRTIM-PSASIEDQSA-N | No-Effect | inactive |
| methoprene-(s) | BRD-K34591242 | COC(C)(C)CCC[C@H](C)C/C=C/C(C)=C/C(=O)OC(C)C | NFGXHKASABOEEW-GYMWBFJFSA-N | No-Effect | inactive |
| methoxamine | BRD-A39189014 | COc1ccc(OC)c(C(O)C(C)N)c1 | WJAJPNHVVFWKKL-UHFFFAOYSA-N | No-Effect | inactive |
| methoxsalen | BRD-K63430059 | COc1c2occc2cc2ccc(=O)oc12 | QXKHYNVANLEOEG-UHFFFAOYSA-N | No-Effect | inactive |
| methoxyamine | BRD-K79277568 | CON | GMPKIPWJBDOURN-UHFFFAOYSA-N | No-Effect | inactive |
| methoxyflurane | BRD-K06152156 | COC(F)(F)C(Cl)Cl | RFKMCNOHBTXSMU-UHFFFAOYSA-N | No-Effect | inactive |
| methoxyphenamine | BRD-A21474126 | CN[C@@H](C)Cc1ccccc1OC | OEHAYUOVELTAPG-VIFPVBQESA-N | No-Effect | inactive |

| | | | | | |
|---|---|---|---|---|---|
| methscopolamine | BRD-K48619321 | C[N+]1(C)[C@H]2C[C@H](OC(=O)[C@H](CO)c3ccccc3)C[C@@H]1[C@H]1O[C@@H]21 | LZCOQTDXKCNBEE-IKIFYQGPSA-N | No-Effect | inactive |
| methsuximide | BRD-A57592049 | CN1C(=O)C[C@](C)(c2ccccc2)C1=O | AJXPJJZHWIXJCJ-GFCCVEGCSA-N | No-Effect | inactive |
| methyclothiazide | BRD-A95340155 | CN1[C@@H](CCl)Nc2cc(Cl)c(S(N)(=O)=O)cc2S1(=O)=O | CESYKOGBSMNBPD-VIFPVBQESA-N | No-Effect | inactive |
| methyl-5-hydroxytryptamine | BRD-K01827294 | C[C@@H](N)Cc1c[nH]c2ccc(O)cc12 | LYPCGXKCQDYTFV-SSDOTTSWSA-N | No-Effect | inactive |
| methyl-aminolevulinate | BRD-K01292611 | COC(=O)CCC(=O)CN | YUUAYBAIHCDHHD-UHFFFAOYSA-N | No-Effect | inactive |
| methyl-nicotinate | BRD-K10132944 | COC(=O)c1cccnc1 | YNBADRVTZLEFNH-UHFFFAOYSA-N | No-Effect | inactive |
| methyl-salicylate | BRD-K10103010 | COC(=O)c1ccccc1O | OSWPMRLSEDHDFF-UHFFFAOYSA-N | No-Effect | inactive |
| methylatropine-nitrate | BRD-K00004543 | C[N+]1(C)[C@H]2CC[C@@H]1C[C@H](OC(=O)[C@@H](CO)c1ccccc1)C2 | PIPAJLPNWZMYQA-HZMVEIRTSA-N | No-Effect | inactive |
| methylbenactyzium-bromide | BRD-K00003518 | CC[N+](C)(CC)CCOC(=O)C(O)(c1ccccc1)c1ccccc1 | HDAMOICMOAXFLJ-UHFFFAOYSA-N | No-Effect | inactive |
| methylbenzethonium-chloride | BRD-K46531780 | Cc1cc(C(C)(C)CC(C)(C)C)ccc1OCCOCC[N+](C)(C)Cc1ccccc1 | UQOOPIMXBSPVJI-UHFFFAOYSA-N | No-Effect | inactive |
| methylcobalamin | BRD-K00003534 | C[Co+]N1/C2=C(/C)C3=N/C(=C/C4=N/C(=C(/C)C5=N[C@@](C)([C@@H]1[C@H](CC(N)=O)[C@@]2(C)CCC(=O)NC[C@@H](C)OP(=O)(O)O[C@@H]1[C@@H](CO)O[C@H](n2cnc6cc(C)c(C)cc62)[C@@H]1O)[C@@](C)(CC(N)=O)[C@@H]5CCC(N)=O)[C@@](C)(CC(N)=O)[C@@H]4CCC(N)=O)C(C)(C)[C@@H]3CCC(N)=O | JEWJRMKHSMTXPP-WDJKBINPSA-M | No-Effect | inactive |
| methyldopa | BRD-K48578705 | C[C@](N)(Cc1ccc(O)c(O)c1)C(=O)O | CJCSPKMFHVPWAR-JTQLQIEISA-N | No-Effect | inactive |
| methyldopate | BRD-K82259334 | CCOC(=O)[C@@](C)(N)Cc1ccc(O)c(O)c1 | SVEBYYWCXTVYCR-LBPRGKRZSA-N | No-Effect | inactive |
| methylergometrine | BRD-K34685430 | CC[C@@H](CO)NC(=O)[C@@H]1C=C2c3cccc4[nH]cc(c34)C[C@H]2N(C)C1 | UNBRKDKAWYKMIV-QWQRMKEZSA-N | No-Effect | inactive |
| methyllycaconitine | BRD-K43192255 | CCN1C[C@]2(COC(=O)c3ccccc3N3C(=O)C[C@H](C)C3=O)CC[C@H](OC)[C@]34[C@H]1[C@](O)([C@@H](OC)[C@H]23)[C@@]1(O)C[C@H](OC)[C@H]2C[C@@H]4[C@@H]1[C@H]2OC | XLTANAWLDBYGFU-VTLKBQQISA-N | No-Effect | inactive |
| methylnaltrexone | BRD-K00003599 | C[N@+]1(CC2CC2)CC[C@]23c4c5ccc(O)c4O[C@H]2C(=O)CC[C@@]3(O)[C@H]1C5 | JVLBPIPGETUEET-WIXLDOGYSA-O | No-Effect | inactive |
| methylprednisolone | BRD-K35240538 | C[C@H]1C[C@H]2[C@@H]3CC[C@](O)(C(=O)CO)[C@@]3(C)C[C@H](O)[C@@H]2[C@@]2(C)C=CC(=O)C=C12 | VHRSUDSXCMQTMA-PJHHCJLFSA-N | No-Effect | inactive |
| methylprednisolone-aceponate | BRD-K24452555 | CCC(=O)O[C@]1(C(=O)COC(C)=O)CC[C@H]2[C@@H]3C[C@H](C)C4=CC(=O)C=C[C@]4(C)[C@H]3[C@@H](O)C[C@@]21C | DALKLAYLIPSCQL-YPYQNWSCSA-N | No-Effect | inactive |
| methylprednisolone-sodium-succinate | BRD-K00003673 | C[C@H]1C[C@H]2[C@@H]3CC[C@](O)(C(=O)COC(=O)CCC(=O)O)[C@@]3(C)C[C@H](O)[C@@H]2[C@@]2(C)C=CC(=O)C=C12 | IMBXEJJVJRTNOW-XYMSELFBSA-N | No-Effect | inactive |
| methylthiouracil | BRD-K11900042 | Cc1cc(=O)[nH]c(=S)[nH]1 | HWGBHCRJGXAGEU-UHFFFAOYSA-N | No-Effect | inactive |
| methysergide | BRD-K35941380 | CC[C@@H](CO)NC(=O)[C@@H]1C=C2c3cccc4c3c(cn4C)C[C@H]2N(C)C1 | KPJZHOPZRAFDTN-ZRGWGRIASA-N | No-Effect | inactive |
| meticrane | BRD-K58265391 | Cc1cc2c(cc1S(N)(=O)=O)S(=O)(=O)CCC2 | FNQQBFNIYODEMB-UHFFFAOYSA-N | No-Effect | inactive |
| metipranolol | BRD-A99388803 | CC(=O)Oc1c(C)cc(OC[C@H](O)CNC(C)C)c(C)c1C | BQIPXWYNLPYNHW-OAHLLOKOSA-N | No-Effect | inactive |
| metirosine | BRD-K83766205 | C[C@](N)(Cc1ccc(O)cc1)C(=O)O | NHTGHBARYWONDQ-JTQLQIEISA-N | No-Effect | inactive |
| metoclopramide | BRD-K75641298 | CCN(CC)CCNC(=O)c1cc(Cl)c(N)cc1OC | TTWJBBZEZQICBI-UHFFFAOYSA-N | No-Effect | inactive |
| metocurine | BRD-K16484091 | COc1ccc2cc1Oc1cc3c(cc1OC)CC[N+](C)(C)[C@H]3Cc1ccc(cc1)Oc1c(OC)c(OC)cc3c1[C@@H](C2)[N+](C)(C)CC3 | JFXBEKISTKFVAB-AJQTZOPKSA-N | No-Effect | inactive |
| metolazone | BRD-A61793559 | Cc1ccccc1N1C(=O)c2cc(S(N)(=O)=O)c(Cl)cc2N[C@@H]1C | AQCHWTWZEMGIFD-JTQLQIEISA-N | No-Effect | inactive |
| metoprolol | BRD-K01826568 | COCCc1ccc(OC[C@@H](O)CNC(C)C)cc1 | IUBSYMUCCVWXPE-AWEZNQCLSA-N | No-Effect | inactive |
| metoxibutropate | BRD-A71725768 | COc1ccccc1OC(=O)[C@@H](C)c1ccc(CC(C)C)cc1 | FVRWRGBTEHAPDI-HNNXBMFYSA-N | No-Effect | inactive |
| metrifonate | BRD-A22101059 | COP(=O)(OC)[C@@H](O)C(Cl)(Cl)Cl | NFACJZMKEDPNKN-GSVOUGTGSA-N | No-Effect | inactive |
| metrizamide | BRD-A45543382 | CC(=O)Nc1c(I)c(C(=O)N[C@H]2[C@@H](O)O[C@H](CO)[C@@H](O)[C@@H]2O)c(I)c(N(C)C(C)=O)c1I | BAQCROVBDNBEEB-KSDLJXBFSA-N | No-Effect | inactive |
| metronidazole | BRD-K52020312 | Cc1ncc([N+](=O)[O-])n1CCO | VAOCPAMSLUNLGC-UHFFFAOYSA-N | No-Effect | inactive |
| metyrapone | BRD-K46862739 | CC(C)(C(=O)c1cccnc1)c1cccnc1 | FJLBFSROUSIWMA-UHFFFAOYSA-N | No-Effect | inactive |
| mevastatin | BRD-K94441233 | CC[C@H](C)C(=O)O[C@H]1CCC=C2C=C[C@H](C)[C@H](CC[C@@H]3C[C@@H](O)CC(=O)O3)[C@H]21 | AJLFOPYRIVGYMJ-INTXDZFKSA-N | No-Effect | inactive |
| mexeneone | BRD-K18613660 | COc1ccc(C(=O)c2ccc(C)cc2)c(O)c1 | MJVGBKJNTFCUJM-UHFFFAOYSA-N | No-Effect | inactive |
| mexiletine | BRD-A64092382 | Cc1cccc(C)c1OC[C@@H](C)N | VLPIATFUUWWMKC-SNVBAGLBSA-N | No-Effect | inactive |
| mezlocillin | BRD-K45114938 | CC1(C)S[C@H]2[C@@H](NC(=O)[C@@H](NC(=O)N3CCN(S(C)(=O)=O)C3=O)c3ccccc3)C(=O)N2[C@@H]1C(=O)O | YPBATNHYBCGSSN-ZJOBFFGXSA-N | No-Effect | inactive |
| MF-101 | BRD-K34328244 | O=C1C[C@@H](c2ccc(O)cc2)Oc2cc(O)ccc21 | FURUXTVZLHCCNA-AWEZNQCLSA-N | No-Effect | inactive |
| MG-624 | BRD-K03128577 | CC[N+](CC)(CC)CCOc1ccc(/C=C/c2ccccc2)cc1 | YIMFXBPXUPTIEJ-OUKQBFOZSA-N | No-Effect | inactive |
| MGCD-265 | BRD-K56277358 | Cn1cnc(-c2cc3nccc(Oc4ccc(NC(=S)NC(=O)Cc5ccccc5)cc4F)c3s2)c1 | UFICVEHDQUKCEA-UHFFFAOYSA-N | No-Effect | inactive |
| mibampator | BRD-K42981054 | CC(C)S(=O)(=O)NC[C@H](C)c1ccc(-c2ccc(CCNS(C)(=O)=O)cc2)cc1 | ULRDYYKSPCRXAJ-KRWDZBQOSA-N | No-Effect | inactive |
| micafungin | BRD-K40407248 | CCCCCOc1ccc(-c2cc(-c3ccc(C(=O)N[C@H]4C[C@@H](O)[C@@H](O)NC(=O)[C@@H]5[C@@H](O)[C@@H](C)CN5C(=O)[C@H]([C@H](O)CC(N)=O)NC(=O)[C@H]([C@H](O)[C@@H](O)c5ccc(O)c(OS(=O)(=O)O)c5)NC(=O)[C@@H]5C[C@@H](O)CN5C(=O)[C@H]([C@@H](C)O)NC4=O)cc3)no2)cc1 | PIEUQSKUWLMALL-YABMTYFHSA-N | No-Effect | inactive |
| miconazole | BRD-K01826511 | Clc1ccc(CO[C@@H](Cn2ccnc2)c2ccc(Cl)cc2Cl)c(Cl)c1 | BYBLEWFAAKGYCD-SFHVURJKSA-N | No-Effect | inactive |
| midafotel | BRD-K45163704 | O=C(O)[C@H]1CN(C/C=C/P(=O)(O)O)CCN1 | VZXMZMJSGLFKQI-ABVWVHJUSA-N | No-Effect | inactive |
| midecamycin | BRD-K68822425 | CCC(=O)O[C@H]1[C@H](C)O[C@@H](O[C@@H]2[C@@H](C)O[C@@H](O[C@@H]3[C@@H](OC)[C@H](OC(=O)CC)CC(=O)O[C@H](C)C/C=C/C=C/[C@H](O)[C@H](C)C[C@@H]3CC=O)[C@H](O)[C@H]2N(C)C)C[C@@]1(C)O | DMUAPQTXSSNEDD-QALJCMCCSA-N | No-Effect | inactive |
| midodrine | BRD-A79981887 | COc1ccc(OC)c([C@@H](O)CNC(=O)CN)c1 | PTKSEFOSCHHMPD-JTQLQIEISA-N | No-Effect | inactive |
| midostaurin | BRD-K13646352 | CO[C@@H]1[C@H](N(C)C(=O)c2ccccc2)C[C@H]2O[C@]1(C)n1c3ccccc3c3c4c(c5c6ccccc6n2c5c31)C(=O)NC4 | BMGQWWVMWDBQGC-IIFHNQTCSA-N | No-Effect | inactive |
| mifobate | BRD-K00004244 | COP(=O)(OC)O[C@@H](c1ccc(Cl)cc1)P(=O)(OC)OC | VQHUQHAPWMNBLP-LLVKDONJSA-N | No-Effect | inactive |
| milacemide | BRD-K64338723 | CCCCCNCC(N)=O | GJNNXIYZWIZFRH-UHFFFAOYSA-N | No-Effect | inactive |
| milademetan | BRD-K00003406 | CC1(C)CCC2(CC1)N[C@@H](C(=O)N[C@@H]1CC[C@@H](C(N)=O)OC1)[C@H](c1ccnc(Cl)c1F)[C@]21C(=O)Nc2cc(Cl)ccc21 | RYAYYVTWKAOAJF-QISPRATLSA-N | No-Effect | inactive |
| milnacipran | BRD-K00004287 | CCN(CC)C(=O)[C@@]1(c2ccccc2)C[C@H]1CN | GJJFMKBJSRMPLA-DZGCQCFKSA-N | No-Effect | inactive |
| milrinone | BRD-K67080878 | Cc1[nH]c(=O)c(C#N)cc1-c1ccncc1 | PZRHRDRVRGEVNW-UHFFFAOYSA-N | No-Effect | inactive |
| miltefosine | BRD-K01824991 | CCCCCCCCCCCCCCCCOP(=O)(O)OCC[N+](C)(C)C | PQLXHQMOHUQAKB-UHFFFAOYSA-O | No-Effect | inactive |
| mimosine | BRD-K67642028 | N[C@@H](Cn1ccc(=O)c(O)c1)C(=O)O | WZNJWVWKTVETCG-YFKPBYRVSA-N | No-Effect | inactive |
| minaprine | BRD-K02867583 | Cc1cc(-c2ccccc2)nnc1NCCN1CCOCC1 | LDMWSLGGVTVJPG-UHFFFAOYSA-N | No-Effect | inactive |
| minerval | BRD-A74347210 | CCCCCCCC/C=C\CCCCCC[C@@H](O)C(=O)O | JBSOOFITVPOOSY-DOOKAGJSSA-N | No-Effect | inactive |

| | | | | | |
|---|---|---|---|---|---|
| minocycline | BRD-K00003346 | CN(C)c1ccc(O)c2c1C[C@H]1C[C@H]3[C@H](N(C)C)C(=O)[C@@H](C(N)=O)C(=O)[C@@]3(O)C(=O)[C@H]1C2=O | ZZZRUAITSXLWBH-NOHUXHDNSA-N | No-Effect | inactive |
| minodronic-acid | BRD-K89337853 | O=P(O)(O)C(O)(Cc1cnc2ccccn12)P(=O)(O)O | VMMKGHQPQIEGSQ-UHFFFAOYSA-N | No-Effect | inactive |
| minoxidil | BRD-K06902185 | Nc1cc(N2CCCCC2)nc(N)[n+]1[O-] | ZFMITUMMTDLWHR-UHFFFAOYSA-N | No-Effect | inactive |
| miocamycin | BRD-K30764557 | CCC(=O)O[C@H]1[C@H](C)O[C@@H](O[C@@H]2[C@@H](C)O[C@@H](O[C@@H]3[C@@H](OC)[C@H](OC(=O)CC)CC(=O)O[C@H](C)C/C=C/C=C/[C@H](OC(C)=O)[C@H](C)C[C@@H]3CC=O)[C@H](O)[C@@H]2N(C)C)C[C@@]1(C)OC(C)=O | GQNZGCARKRHPOH-RQIKCTSVSA-N | No-Effect | inactive |
| MIRA-1 | BRD-K39706510 | CCC(=O)OCN1C(=O)C=CC1=O | YXEWPGYLMHXLPS-UHFFFAOYSA-N | No-Effect | inactive |
| mirabegron | BRD-K47949423 | Nc1nc(CC(=O)Nc2ccc(CCNC[C@H](O)c3ccccc3)cc2)cs1 | PBAPPPCECJKMCM-IBGZPJMESA-N | No-Effect | inactive |
| miridesap | BRD-K00003434 | O=C(O)[C@H]1CCCN1C(=O)CCCCC(=O)N1CCC[C@@H]1C(=O)O | HZLAWYIBLZNRFZ-VXGBXAGGSA-N | No-Effect | inactive |
| mirin | BRD-K55420858 | NC1=NC(=O)/C(=C/c2ccc(O)cc2)S1 | YBHQCJILTOVLHD-YVMONPNESA-N | No-Effect | inactive |
| miriplatin | BRD-M78068755 | CCCCCCCCCCCCCC(=O)O.CCCCCCCCCCCCCC(=O)O.N[C@H]1CCCC[C@@H]1N.[Pt+2] | BGIHRZPJIYJKAZ-JPLLOCSESA-N | No-Effect | inactive |
| mirogabalin-besylate | BRD-K00003484 | CCC1=C[C@@H]2[C@H](C1)C[C@]2(CN)CC(=O)O | FTBQORVNHOIASH-CKYFFXLPSA-N | No-Effect | inactive |
| mirtazapine | BRD-A64977602 | CN1CCN2c3ncccc3Cc3ccccc3[C@@H]2C1 | RONZAEMNMFQXRA-INIZCTEOSA-N | No-Effect | inactive |
| misonidazole | BRD-A16263897 | COC[C@@H](O)Cn1ccnc1[N+](=O)[O-] | OBBCSXFCDPPXOL-LURJTMIESA-N | No-Effect | inactive |
| misoprostol | BRD-A72351335 | CCCCC(C)(O)C/C=C/[C@H]1[C@H](O)CC(=O)[C@@H]1CCCCCCC(=O)OC | OJLOPKGSLYJEMD-URPKTTJQSA-N | No-Effect | inactive |
| mitapivat | BRD-K00003136 | O=C(c1ccc(NS(=O)(=O)c2cccc3cccnc23)cc1)N1CCN(CC2CC2)CC1 | XAYGBKHKBBXDAK-UHFFFAOYSA-N | No-Effect | inactive |
| mitiglinide | BRD-K52959329 | O=C(O)[C@H](CC(=O)N1C[C@H]2CCCC[C@H]2C1)Cc1ccccc1 | WPGGHFDDFPHPOB-BBWFWOEESA-N | No-Effect | inactive |
| mitoflaxone | BRD-K35882976 | O=C(O)Cc1cccc2c(=O)cc(-c3ccccc3)oc12 | TZZNWMJZDWYJAZ-UHFFFAOYSA-N | No-Effect | inactive |
| mitomycin-C | BRD-K59670716 | CO[C@@]12[C@H](COC(N)=O)C3=C(C(=O)C(C)=C(N)C3=O)N1C[C@@H]1N[C@@H]12 | NWIBSHFKIJFRCO-WUDYKRTCSA-N | No-Effect | inactive |
| mitoquinone | BRD-K00003362 | COC1=C(OC)C(=O)C(CCCCCCCCCC[P+](c2ccccc2)(c2ccccc2)c2ccccc2)=C(C)C1=O | OIIMUKXVVLRCAF-UHFFFAOYSA-N | No-Effect | inactive |
| mitoxantrone | BRD-K21680192 | O=C1c2c(O)ccc(O)c2C(=O)c2c(NCCNCCO)ccc(NCCNCCO)c21 | KKZJGLLVHKMTCM-UHFFFAOYSA-N | No-Effect | inactive |
| MIW-815 | BRD-K00003146 | Nc1ncnc2c1ncn2[C@@H]1O[C@@H]2CO[P@@](=O)(S)O[C@@H]3[C@H](O)[C@@H](CO[P@@](=O)(S)O[C@H]2[C@H]1O)O[C@H]3n1cnc2c(N)ncnc21 | IZJJFUQKKZFVLH-YBVMPXGUSA-N | No-Effect | inactive |
| mizolastine | BRD-K21884483 | CN(c1nc(=O)cc[nH]1)C1CCN(c2nc3ccccc3n2Cc2ccc(F)cc2)CC1 | PVLJETXTTWAYEW-UHFFFAOYSA-N | No-Effect | inactive |
| mizoribine | BRD-K88061624 | NC(=O)C1N=CN([C@@H]2O[C@H](CO)[C@@H](O)[C@H]2O)C1=O | OPMVDWXPLVKNQN-AAXBYFMASA-N | No-Effect | inactive |
| MJ-15 | BRD-K71315565 | Cc1c(C(=O)NCc2cccnc2)nn(-c2ccc(Cl)cc2Cl)c1-c1ccc(Cl)cc1 | PJEIKJFNEFJFLA-UHFFFAOYSA-N | No-Effect | inactive |
| MK-0354 | BRD-K91405252 | C1Cc2[nH]nc(-c3nn[nH]n3)c2C1 | LTQYSJKGRPGMPO-UHFFFAOYSA-N | No-Effect | inactive |
| MK-0557 | BRD-K00003190 | O=C1O[C@]2(CC[C@@H](C(=O)Nc3ccn(-c4ccccc4F)n3)CC2)c2cnccc21 | RMYZIRFUCOMQRH-XLOAEROZSA-N | No-Effect | inactive |
| MK-0608 | BRD-K00004557 | C[C@@]1(O)[C@H](O)[C@@H](CO)O[C@H]1n1ccc2c(N)ncnc21 | IRZRJANZDIOOIF-GAJNKVMBSA-N | No-Effect | inactive |
| MK-0752 | BRD-K18850819 | O=C(O)CC[C@H]1CC[C@@](c2cc(F)ccc2F)(S(=O)(=O)c2ccc(Cl)cc2)CC1 | XCGJIFAKUZNNOR-QCKZDCLWSA-N | No-Effect | inactive |
| MK-0773 | BRD-K38868394 | CN1C(=O)C(F)=C[C@]2(C)[C@H]3CC[C@]4(C)[C@@H](C(=O)NCc5nc6ncccc6[nH]5)CC[C@H]4[C@@H]3CC[C@@H]12 | GBEUKTWTUSPHEE-JWJWXJQQSA-N | No-Effect | inactive |
| MK-0812 | BRD-K14175878 | CO[C@@H]1COCC[C@@H]1N[C@@H]1CC[C@@](C(=O)N2CCc3ncc(C(F)(F)F)cc3C2)(C(C)C)C1 | MTMDXAIUENDNDL-RJSMDTJLSA-N | No-Effect | inactive |
| MK-1064 | BRD-K00003304 | COc1ccc(CNC(=O)c2cc(-c3cncc(Cl)c3)cnc2-c2ccccn2)nc1OC | CKTWQGHVNRYNCM-UHFFFAOYSA-N | No-Effect | inactive |
| MK-1775 | BRD-K54256913 | C=CCn1c(=O)c2cnc(Nc3ccc(N4CCN(C)CC4)cc3)nc2n1-c1cccc(C(C)(C)O)n1 | BKWJAKQVGHWELA-UHFFFAOYSA-N | No-Effect | inactive |
| MK-1903 | BRD-K00004660 | O=C(O)c1n[nH]c2c1C[C@H]1C[C@@H]21 | CUTZNERBKDMLAP-QWWZWVQMSA-N | No-Effect | inactive |
| MK-2295 | BRD-K43316319 | FC(F)(F)c1ccc(Nc2ncnc3cc(-c4ncccc4C(F)(F)F)ccc23)cc1 | VTANGSDRFFLTSQ-UHFFFAOYSA-N | No-Effect | inactive |
| MK-2461 | BRD-K56405753 | CN(C[C@@H]1COCCO1)S(=O)(=O)Nc1ccc2ccc3ncc(-c4cnn(C)c4)cc3c(=O)c2c1 | JGEBLDKNWBUGRZ-HXUWFJFHSA-N | No-Effect | inactive |
| MK-2894 | BRD-K27983781 | Cc1sc(C)c(C(=O)NC2(c3ccc(C(=O)O)cc3)CC2)c1Cc1ccc(C(F)(F)F)cc1 | QJZQFVRFJCGDKF-UHFFFAOYSA-N | No-Effect | inactive |
| MK-3207 | BRD-K66911501 | O=C(CN1C(=O)C2(CCCC2)NC[C@H]1c1cc(F)cc(F)c1)Nc1ccc2c(c1)C[C@@]1(C2)C(=O)Nc2ncccc21 | AZAANWYREOQRFB-SETSBSEESA-N | No-Effect | inactive |
| MK-3697 | BRD-K95828359 | COc1ccc(CNC(=O)c2cc(-c3cncc(C)c3)ncc2-c2nccs2)nc1OC | VSOUDUXMPUHJEU-UHFFFAOYSA-N | No-Effect | inactive |
| MK-5046 | BRD-K03886848 | O[C@@](Cc1ncc(CC2(C(F)(F)F)CC2)[nH]1)(c1ccc(-n2cccn2)cc1)C(F)(F)F | UJINBEQCDMOAHM-SFHVURJKSA-N | No-Effect | inactive |
| MK-5108 | BRD-K53665955 | O=C(O)[C@]1(Cc2cccc(Nc3nccs3)n2)CC[C@@H](Oc2cccc(Cl)c2F)CC1 | LCVIRAZGMYMNNT-VVONHTQRSA-N | No-Effect | inactive |
| MK-571 | BRD-A73679382 | CN(C)C(=O)CCS[C@@H](SCCC(=O)O)c1cccc(/C=C/c2ccc3ccc(Cl)cc3n2)c1 | AXUZQJFHDNNPFG-JORPFSLRSA-N | No-Effect | inactive |
| MK-7246 | BRD-K00003199 | CN([C@@H]1CCc2c(CC(=O)O)c3ccccc3n2C1)S(=O)(=O)c1ccc(F)cc1 | JTCAGRAKUAAYDY-OAHLLOKOSA-N | No-Effect | inactive |
| MK-7622 | BRD-K00003194 | Cc1ccc(Cc2cc3c(=O)n([C@H]4CCCC[C@@H]4O)cnc3c3ccccc23)cn1 | JUVQLZBJFOGEEO-GOTSBHOMSA-N | No-Effect | inactive |
| MK-8033 | BRD-K55512740 | Cn1cc(-c2cnc3ccc4ccc(CS(=O)(=O)NCc5ccccn5)cc4c(=O)c3c2)cn1 | VMJFTOSOFDEKTM-UHFFFAOYSA-N | No-Effect | inactive |
| MK-8245 | BRD-K29656036 | O=C(O)Cn1nnc(-c2cc(N3CCC(Oc4cc(F)ccc4Br)CC3)no2)n1 | UJEAABFSXKCSGI-UHFFFAOYSA-N | No-Effect | inactive |
| MK-8617 | BRD-K00003397 | COc1ccc(C(NC(=O)c2cnc(-c3cccnn3)nc2O)c2ccc(OC)cc2)cc1 | WXLPERVDMILVIF-UHFFFAOYSA-N | No-Effect | inactive |
| MK-8745 | BRD-K25279140 | O=C(c1cccc(Cl)c1F)N1CCN(Cc2cccc(Nc3nccs3)n2)CC1 | YCRFPWKUUNKNDN-UHFFFAOYSA-N | No-Effect | inactive |
| MK-886 | BRD-K20168484 | CC(C)c1ccc2c(c1)c(SC(C)(C)C)c(CC(C)(C)C(=O)O)n2Cc1ccc(Cl)cc1 | QAOAOVKBIIKRNL-UHFFFAOYSA-N | No-Effect | inactive |
| MK-8998 | BRD-K86677390 | CC(C)c1ccc(CC(=O)N[C@H](C)c2ccc(OCC(F)(F)F)cn2)cc1 | IQIKXZMPPBEWAD-CQSZACIVSA-N | No-Effect | inactive |
| MK2-IN-1 | BRD-K90316462 | O=C(c1ccc(-c2ccc(Cl)cc2)o1)N(Cc1ccccn1)c1ccc(N2CCNCC2)cc1 | LIPNXDIBECKOMJ-UHFFFAOYSA-N | No-Effect | inactive |
| MKT-077 | BRD-K61695967 | CCn1c(=O)/c(=C2\Sc3ccccc3N2C)s/c1=C/c1cccc[n+]1CC | QLYHOWIZHHZGPV-XUTLUUPISA-N | No-Effect | inactive |
| ML10302 | BRD-K69343602 | COc1cc(N)c(Cl)cc1C(=O)OCCN1CCCCC1 | RVFIAQAAZUEPPE-UHFFFAOYSA-N | No-Effect | inactive |
| ML130 | BRD-K89306950 | Cc1ccc(S(=O)(=O)n2c(N)nc3ccccc32)cc1 | SRFABRWQVPCPRG-UHFFFAOYSA-N | No-Effect | inactive |
| ML133 | BRD-K33743361 | COc1ccc(CNCc2cccc3ccccc23)cc1 | GIFYKGIODMXYNT-UHFFFAOYSA-N | No-Effect | inactive |
| ML141 | BRD-A28422330 | COc1ccc([C@H]2CC(c3ccccc3)=NN2c2ccc(S(N)(=O)=O)cc2)cc1 | QBNZBMVRFYREHK-JOCHJYFZSA-N | No-Effect | inactive |
| ML161 | BRD-K59036917 | CCCC(=O)Nc1cccc(NC(=O)c2ccccc2Br)c1 | DFOVLSMXPWPCFH-UHFFFAOYSA-N | No-Effect | inactive |
| ML167 | BRD-K64835161 | Cc1ccc(CNc2ncnc3ccc(-c4ccc(CO)o4)cc23)o1 | ROCFOIBAEVAOLQ-UHFFFAOYSA-N | No-Effect | inactive |
| ML179 | BRD-K77547509 | O=c1cc(N2CCN(c3cccc(C(F)(F)F)c3)CC2)[nH]c(=O)n1C1CCCCC1 | MFABFKUNLDYKGH-UHFFFAOYSA-N | No-Effect | inactive |
| ML193 | BRD-K27539181 | Cc1cc(C)c2nc(-c3ccccn3)cc(C(=O)Nc3ccc(S(=O)(=O)Nc4onc(C)c4C)cc3)c2c1 | HTSLEZOTMYUPLU-UHFFFAOYSA-N | No-Effect | inactive |
| ML204 | BRD-K80021413 | Cc1cc(N2CCCCC2)nc2ccccc12 | USYRQXDHKXGTCK-UHFFFAOYSA-N | No-Effect | inactive |
| ML213 | BRD-K00004301 | Cc1cc(C)c(NC(=O)[C@@H]2C[C@H]3CC[C@@H]2C3)c(C)c1 | SIQGKPGBLYKQBB-RRFJBIMHSA-N | No-Effect | inactive |
| ML218 | BRD-K92995792 | CC(C)(C)CCN1C[C@H]2[C@H](CNC(=O)c3cc(Cl)cc(Cl)c3)[C@H]2C1 | GSJIGYLGKSBYBC-OSYLJGHBSA-N | No-Effect | inactive |
| ML228 | BRD-K44432556 | c1ccc(-c2ccc(CNc3nc(-c4ccccn4)nnc3-c3ccccc3)cc2)cc1 | QNRODODTMXCRKU-UHFFFAOYSA-N | No-Effect | inactive |
| ML277 | BRD-K60020910 | COc1ccc(-c2csc(NC(=O)[C@H]3CCCN3S(=O)(=O)c3ccc(C)cc3)n2)cc1 | OXQNLLVUVDAEHC-OAQYLSRUSA-N | No-Effect | inactive |

| | | | | | |
|---|---|---|---|---|---|
| ML281 | BRD-K48188942 | CC(C)c1ccc(NC(=O)c2cccs2)c(-c2nc3ccccc3[nH]c2=O)c1 | HWOYIOLMBQSTQS-UHFFFAOYSA-N | No-Effect | inactive |
| ML297 | BRD-K51137250 | Cc1cc(NC(=O)Nc2ccc(F)c(F)c2)n(-c2ccccc2)n1 | IEKSMUSSYJUQMY-UHFFFAOYSA-N | No-Effect | inactive |
| ML298 | BRD-K24600306 | O=C(NCCN1CCC2(CC1)C(=O)NCN2c1cccc(F)c1)c1ccc(F)c(F)c1 | XAPVAKKLQGLNOY-UHFFFAOYSA-N | No-Effect | inactive |
| ML314 | BRD-K92606079 | COc1cc2nc(C3CC3)nc(N3CCN(c4ccccc4OC)CC3)c2cc1OC | SWEOAXMICIJCQC-UHFFFAOYSA-N | No-Effect | inactive |
| ML323 | BRD-K93942811 | Cc1cnc(-c2ccccc2C(C)C)nc1NCc1ccc(-n2ccnn2)cc1 | VUIRVWPJNKZOSS-UHFFFAOYSA-N | No-Effect | inactive |
| ML324 | BRD-K25611237 | CN(C)CCCNC(=O)c1ccc(-c2cc(O)c3ncccc3c2)cc1 | QDBVSOZTVKXUES-UHFFFAOYSA-N | No-Effect | inactive |
| ML347 | BRD-K33892651 | COc1ccc(-c2cnc3c(-c4cccc5ncccc45)cnn3c2)cc1 | FVRYPYDPKSZGNS-UHFFFAOYSA-N | No-Effect | inactive |
| ML348 | BRD-K81044214 | O=C(CN1CCN(C(=O)c2ccco2)CC1)Nc1cc(C(F)(F)F)ccc1Cl | OXKNHBBDOIMFFQ-UHFFFAOYSA-N | No-Effect | inactive |
| ML365 | BRD-K02160913 | COc1ccccc1C(=O)Nc1cccc(NC(=O)c2cccc(C)c2)c1 | UTAJHKSGYJSZBR-UHFFFAOYSA-N | No-Effect | inactive |
| ML9 | BRD-K68402494 | O=S(=O)(c1cccc2c(Cl)cccc12)N1CCCNCC1 | OZSMSRIUUDGTEP-UHFFFAOYSA-N | No-Effect | inactive |
| MLN0128 | BRD-K59317601 | CC(C)n1nc(-c2ccc3oc(N)nc3c2)c2c(N)ncnc21 | GYLDXIAOMVERTK-UHFFFAOYSA-N | No-Effect | inactive |
| MLN1117 | BRD-K73877023 | Nc1nc2cc(-c3ccc4ncc(C(=O)N5CCOCC5)n4c3)ccc2o1 | BLGWHBSBBJNKJO-UHFFFAOYSA-N | No-Effect | inactive |
| MLN2480 | BRD-K43410529 | C[C@@H](NC(=O)c1ncnc(N)c1Cl)c1ncc(C(=O)Nc2cc(C(F)(F)F)c(Cl)cn2)s1 | VWMJHAFYPMOMGF-ZCFIWIBFSA-N | No-Effect | inactive |
| MLN8054 | BRD-K83963101 | O=C(O)c1ccc(Nc2ncc3c(n2)-c2ccc(Cl)cc2C(c2c(F)cccc2F)=NC3)cc1 | HHFBDROWDBDFBR-UHFFFAOYSA-N | No-Effect | inactive |
| MLN9708 | BRD-K15203492 | CC(C)C[C@H](NC(=O)CNC(=O)c1cc(Cl)ccc1Cl)B1OC(=O)C(CC(=O)O)(CC(=O)O)O1 | MBOMYENWWXQSNW-AWEZNQCLSA-N | No-Effect | inactive |
| MM-102 | BRD-K68474068 | CCC(CC)(NC(=O)C(C)C)C(=O)N[C@@H](CCCNC(=N)N)C(=O)NC1(C(=O)NC(c2ccc(F)cc2)c2ccc(F)cc2)CCCC1 | RZKSQRIPRKWVBU-MHZLTWQESA-N | No-Effect | inactive |
| MM-11253 | BRD-K98353094 | CC1(C)CCC(C)(C)c2cc(C3(c4ccc5cc(C(=O)O)ccc5c4)SCCS3)ccc21 | DPVGXUNWCVEIGT-UHFFFAOYSA-N | No-Effect | inactive |
| MM-206 | BRD-K00004622 | O=S(=O)(Nc1cc(Sc2ccccc2)c(O)c2ccccc12)c1c(F)c(F)c(F)c(F)c1F | ZXJHEANDHXHGMV-UHFFFAOYSA-N | No-Effect | inactive |
| MM77 | BRD-K50293352 | COc1ccccc1N1CCN(CCCCN2C(=O)CCC2=O)CC1 | MQHOGPLXAXSQAJ-UHFFFAOYSA-N | No-Effect | inactive |
| MMPX | BRD-K56077740 | COCc1nc2c([nH]1)c(=O)n(C)c(=O)n2CC(C)C | NBLBCGUCPBXKOV-UHFFFAOYSA-N | No-Effect | inactive |
| MMV-390048 | BRD-K78973789 | CS(=O)(=O)c1ccc(-c2cnc(N)c(-c3ccc(C(F)(F)F)nc3)c2)cc1 | RTJQABCNNLMCJF-UHFFFAOYSA-N | No-Effect | inactive |
| MNITMT | BRD-K63151507 | Cn1cnnc1Sc1c([N+](=O)[O-])ncn1C | NMBDIMPYACLUDE-UHFFFAOYSA-N | No-Effect | inactive |
| mocetinostat | BRD-K16485616 | Nc1ccccc1NC(=O)c1ccc(CNc2nccc(-c3cccnc3)n2)cc1 | HRNLUBSXIHFDHP-UHFFFAOYSA-N | No-Effect | inactive |
| moclobemide | BRD-K07237224 | O=C(NCCN1CCOCC1)c1ccc(Cl)cc1 | YHXISWVBGDMDLQ-UHFFFAOYSA-N | No-Effect | inactive |
| moexipril | BRD-K34441861 | CCOC(=O)[C@H](CCc1ccccc1)N[C@@H](C)C(=O)N1Cc2cc(OC)c(OC)cc2C[C@H]1C(=O)O | UWWDHYUMIORJTA-HSQYWUDLSA-N | No-Effect | inactive |
| mofegiline | BRD-K52598503 | NC/C(=C/F)CCc1ccc(F)cc1 | VXLBSYHAEKDUSU-JXMROGBWSA-N | No-Effect | inactive |
| mofezolac | BRD-K49372556 | COc1ccc(-c2noc(CC(=O)O)c2-c2ccc(OC)cc2)cc1 | DJEIHHYCDCTAAH-UHFFFAOYSA-N | No-Effect | inactive |
| moguisteine | BRD-A87673115 | CCOC(=O)CC(=O)N1CCS[C@@H]1COc1ccccc1OC | WSYVIAQNTFPTBI-OAHLLOKOSA-N | No-Effect | inactive |
| molidustat | BRD-K01291782 | O=c1c(-n2ccnn2)c[nH]n1-c1cc(N2CCOCC2)ncn1 | IJMBOKOTALXLKS-UHFFFAOYSA-N | No-Effect | inactive |
| molindone | BRD-A65280694 | CCc1c(C)[nH]c2c1C(=O)[C@H](CN1CCOCC1)CC2 | KLPWJLBORRMFGK-LBPRGKRZSA-N | No-Effect | inactive |
| molsidomine | BRD-K63861289 | CCOC(=O)Nc1c[n+](N2CCOCC2)no1 | XLFWDASMENKTKL-UHFFFAOYSA-O | No-Effect | inactive |
| mometasone | BRD-K50265942 | C[C@@H]1C[C@H]2[C@@H]3CCC4=CC(=O)C=C[C@]4(C)[C@@]3(Cl)[C@@H](O)C[C@]2(C)[C@@]1(O)C(=O)CCl | QLIIKPVHVRXHRI-CXSFZGCWSA-N | No-Effect | inactive |
| mometasone-furoate | BRD-K60640630 | C[C@@H]1C[C@H]2[C@@H]3CCC4=CC(=O)C=C[C@]4(C)[C@@]3(Cl)[C@@H](O)C[C@]2(C)[C@@]1(OC(=O)c1ccco1)C(=O)CCl | WOFMFGQZHJDGCX-ZULDAHANSA-N | No-Effect | inactive |
| monastrol | BRD-A78377521 | CCOC(=O)C1=C(C)NC(=S)N[C@@H]1c1cccc(O)c1 | LOBCDGHHHHGHFA-GFCCVEGCSA-N | No-Effect | inactive |
| monepantel | BRD-K00004607 | C[C@](C#N)(COc1cc(C#N)ccc1C(F)(F)F)NC(=O)c1ccc(SC(F)(F)F)cc1 | WTERNLDOAPYGJD-SFHVURJKSA-N | No-Effect | inactive |
| monobenzone | BRD-K54262262 | Oc1ccc(OCc2ccccc2)cc1 | VYQNWZOUAUKGHI-UHFFFAOYSA-N | No-Effect | inactive |
| monocrotaline | BRD-K65508953 | C[C@H]1C(=O)O[C@@H]2CCN3CC=C(COC(=O)[C@](C)(O)[C@]1(C)O)[C@H]23 | QVCMHGGNRFRMAD-XFGHUUIASA-N | No-Effect | inactive |
| monoctanoin | BRD-A48938994 | CCCCCCCC(=O)OC[C@@H](O)CO | GHBFNMLVSPCDGN-JTQLQIEISA-N | No-Effect | inactive |
| monomethyl-fumarate | BRD-K00004695 | COC(=O)/C=C/C(=O)O | NKHAVTQWNUWKEO-NSCUHMNNSA-N | No-Effect | inactive |
| monosodium-alpha-luminol | BRD-K79926543 | Nc1cccc2c(=O)[nH][nH]c(=O)c12 | HWYHZTIRURJOHG-UHFFFAOYSA-N | No-Effect | inactive |
| monostearin | BRD-A31454688 | CCCCCCCCCCCCCCCCCC(=O)OC[C@@H](O)CO | VBICKXHEKHSIBG-FQEVSTJZSA-N | No-Effect | inactive |
| moprolol | BRD-A51747092 | COc1ccccc1OC[C@@H](O)CNC(C)C | LFTFGCDECFPSQD-NSHDSACASA-N | No-Effect | inactive |
| moracizine | BRD-K21548250 | CCOC(=O)Nc1ccc2c(c1)N(C(=O)CCN1CCOCC1)c1ccccc1S2 | FUBVWMNBEHXPSU-UHFFFAOYSA-N | No-Effect | inactive |
| morin | BRD-K11590034 | O=c1c(O)c(-c2ccc(O)cc2O)oc2cc(O)cc(O)c12 | YXOLAZRVSSWPPT-UHFFFAOYSA-N | No-Effect | inactive |
| morinidazole | BRD-A40546373 | Cc1ncc([N+](=O)[O-])n1C[C@@H](O)CN1CCOCC1 | GAZGHCHCYRSPIV-JTQLQIEISA-N | No-Effect | inactive |
| morniflumate | BRD-K38938675 | O=C(OCCN1CCOCC1)c1cccnc1Nc1cccc(C(F)(F)F)c1 | LDXSPUSKBDTEKA-UHFFFAOYSA-N | No-Effect | inactive |
| morphothiadin | BRD-K00003491 | CCOC(=O)C1=C(CN2CCOCC2)NC(c2nccs2)=N[C@@H]1c1ccc(F)cc1Br | SQGRDKSRFFUBBU-GOSISDBHSA-N | No-Effect | inactive |
| mosapride | BRD-A39052811 | CCOc1cc(N)c(Cl)cc1C(=O)NC[C@H]1CN(Cc2ccc(F)cc2)CCO1 | YPELFRMCRYSPKZ-INIZCTEOSA-N | No-Effect | inactive |
| motapizone | BRD-K00004657 | CC1=C(c2cc(-n3ccnc3)cs2)NNC(=O)C1 | IQHAKYLSMONFMT-UHFFFAOYSA-N | No-Effect | inactive |
| motesanib | BRD-K99616396 | CC1(C)CNc2cc(NC(=O)c3cccnc3NCc3ccncc3)ccc21 | RAHBGWKEPAQNFF-UHFFFAOYSA-N | No-Effect | inactive |
| motolimod | BRD-K11785681 | CCCN(CCC)C(=O)C1=Cc2ccc(-c3ccc(C(=O)N4CCCC4)cc3)cc2N=C(N)C1 | QSPOQCXMGPDIHI-UHFFFAOYSA-N | No-Effect | inactive |
| moxalactam | BRD-A08899266 | CO[C@@]1(NC(=O)[C@@H](C(=O)O)c2ccc(O)cc2)C(=O)N2C(C(=O)O)=C(CSc3nnnn3C)CO[C@@H]21 | JWCSIUVGFCSJCK-UOXRKKOCSA-N | No-Effect | inactive |
| moxaverine | BRD-K12742203 | CCc1cc2cc(OC)c(OC)cc2c(Cc2ccccc2)n1 | MYCMTMIGRXJNSO-UHFFFAOYSA-N | No-Effect | inactive |
| moxisylyte | BRD-K81144366 | CC(=O)Oc1cc(C(C)C)c(OCCN(C)C)cc1C | VRYMTAVOXVTQEF-UHFFFAOYSA-N | No-Effect | inactive |
| moxonidine | BRD-K77771411 | COc1nc(C)nc(Cl)c1NC1=NCCN1 | WPNJAUFVNXKLIM-UHFFFAOYSA-N | No-Effect | inactive |
| mozavaptan | BRD-A31095847 | Cc1ccccc1C(=O)Nc1ccc(C(=O)N2CCC[C@H](N(C)C)c3ccccc32)cc1 | WRNXUQJJCIZICJ-DEOSSOPVSA-N | No-Effect | inactive |
| MPEP | BRD-K60690191 | Cc1cccc(C#Cc2ccccc2)n1 | NEWKHUASLBMWRE-UHFFFAOYSA-N | No-Effect | inactive |
| MPI-0479605 | BRD-K45551181 | Cc1cc(N2CCOCC2)ccc1Nc1nc(NC2CCCCC2)c2[nH]cnc2n1 | OVJBNYKNHXJGSA-UHFFFAOYSA-N | No-Effect | inactive |
| Mps-BAY-2a | BRD-K18530250 | CC(C)CNc1nc(-c2cccc3ncccc23)cn2c(-c3ccc(C(=O)NC4CC4)cc3)cnc12 | MDYKTGNHXNTATG-UHFFFAOYSA-N | No-Effect | inactive |
| Mps1-IN-1 | BRD-K94853001 | COc1cc(N2CCC(O)CC2)ccc1Nc1ncc(Nc2ccccc2S(=O)(=O)C(C)C)c2cc[nH]c2n1 | NMJMRSQTDLRCRQ-UHFFFAOYSA-N | No-Effect | inactive |
| Mps1-IN-5 | BRD-K43812936 | COc1cc(S(C)(=O)=O)ccc1Nc1nc2ccc(-c3ccc(NC(=O)[C@H](C)c4ccc(F)cc4)cc3)cn2n1 | NRJKIOCCERLIDG-GOSISDBHSA-N | No-Effect | inactive |
| MR-16728 | BRD-A30590053 | O=C(NCCN1CCCCCC1)[C@H](c1ccccc1)C1CCCCC1 | AGKAAJMNNVUNDK-JOCHJYFZSA-N | No-Effect | inactive |
| MR-948 | BRD-K84883208 | OCCc1ccc(OCCn2ccnc2)cc1 | LQQIZTXSZOYPPS-UHFFFAOYSA-N | No-Effect | inactive |
| MRK-016 | BRD-K44022883 | Cc1cc(-c2nncc3c(C(C)(C)C)c(OCc4ncnn4C)nn23)no1 | QYSYOGCIDRANAR-UHFFFAOYSA-N | No-Effect | inactive |
| MRK-409 | BRD-K57416838 | Cn1ncnc1COc1nn2c(-c3c(F)cccc3F)nnc2cc1C1CCC1 | GOIFCXRIFSYPFG-UHFFFAOYSA-N | No-Effect | inactive |
| MRK-560 | BRD-K68795758 | O=S(=O)(N[C@H]1CC[C@@](c2cc(F)ccc2F)(S(=O)(=O)c2ccc(Cl)cc2)CC1)C(F)(F)F | WDZVWDXOIGQJIO-UJKQEGAGSA-N | No-Effect | inactive |
| MRS-1220 | BRD-K37618799 | O=C(Cc1ccccc1)Nc1nc2ccc(Cl)cc2c2nc(-c3ccco3)nn12 | TWWFAXQOKNBUCR-UHFFFAOYSA-N | No-Effect | inactive |
| MRS-1334 | BRD-A06784547 | CCOC(=O)C1=C(C)NC(c2ccccc2)=C(C(=O)OCc2ccc([N+](=O)[O-])cc2)[C@@H]1C#Cc1ccccc1 | QFLOJAMZLQXHFS-AREMUKBSSA-N | No-Effect | inactive |
| MRS-1845 | BRD-A78093359 | C#CCN1C(C)=C(C(=O)OC)[C@H](c2cccc([N+](=O)[O-])c2)C(C(=O)OCC)=C1C | BITHABUTZRAUGT-IBGZPJMESA-N | No-Effect | inactive |
| MRS-2578 | BRD-K91135178 | S=C=Nc1cccc(NC(=S)NCCCCNC(=S)Nc2cccc(N=C=S)c2)c1 | QOHNRGHTJPFMSL-UHFFFAOYSA-N | No-Effect | inactive |
| MRS-3777 | BRD-K10575656 | c1ccc(Oc2nc(NC3CCCCC3)c3[nH]cnc3n2)cc1 | OGCQBCHLFMBCCE-UHFFFAOYSA-N | No-Effect | inactive |

| | | | | | |
|---|---|---|---|---|---|
| MS-424 | BRD-K00004683 | C[C@@H]1C=CNN1C(=O)c1cccnc1 | AEHLCVKEOAFFJC-MRVPVSSYSA-N | No-Effect | inactive |
| MSDC-0160 | BRD-K00003381 | CCc1ccc(C(=O)COc2ccc(C[C@@H]3SC(=O)NC3=O)cc2)nc1 | IRNJSRAGRIZIHD-KRWDZBQOSA-N | No-Effect | inactive |
| MSDC-0602K | BRD-K00004703 | COc1cccc(C(=O)COc2ccc(Cc3sc(O)nc3O)cc2)c1 | JIJXYXYLGRAIGV-UHFFFAOYSA-N | No-Effect | inactive |
| MTPG | BRD-A28329383 | C[C@@](N)(C(=O)O)c1ccc(-c2nn[nH]n2)cc1 | OZBFBAYESADVDX-JTQLQIEISA-N | No-Effect | inactive |
| mubritinib | BRD-K19061412 | FC(F)(F)c1ccc(/C=C/c2nc(COc3ccc(CCCCn4ccnn4)cc3)co2)cc1 | ZTFBIUXIQYRUNT-MDWZMJQESA-N | No-Effect | inactive |
| mupirocin | BRD-K15262564 | C/C(=C\C(=O)OCCCCCCCCC(=O)O)C[C@@H]1OC[C@H](C[C@@H]2O[C@H]2[C@@H](C)[C@H](C)O)[C@@H](O)[C@H]1O | MINDHVHHQZYEEK-HBBNESRFSA-N | No-Effect | inactive |
| muscimol | BRD-K60441002 | NCc1cc(=O)[nH]o1 | ZJQHPWUVQPJPQT-UHFFFAOYSA-N | No-Effect | inactive |
| MUT056399 | BRD-K72078047 | CCc1cc(O)c(Oc2ccc(C(N)=O)cc2F)cc1F | QUHARGDBJJUOEB-UHFFFAOYSA-N | No-Effect | inactive |
| MY-5445 | BRD-K90524085 | Clc1cccc(Nc2nnc(-c3ccccc3)c3ccccc23)c1 | CEHQLKSLMFIHBF-UHFFFAOYSA-N | No-Effect | inactive |
| myricetin | BRD-K43149758 | O=c1c(O)c(-c2cc(O)c(O)c(O)c2)oc2cc(O)cc(O)c12 | IKMDFBPHZNJCSN-UHFFFAOYSA-N | No-Effect | inactive |
| myristyl-nicotinate | BRD-K88153212 | CCCCCCCCCCCCCCOC(=O)c1cccnc1 | TVGLGJWCZSCAEM-UHFFFAOYSA-N | No-Effect | inactive |
| N-(2-chlorophenyl)-2-({(2E)-2-[1-(2-pyridinyl)ethylidene]hydrazino}carbothioyl)hydrazinecarbothioamide | BRD-K57027742 | CC(=NNC(=S)NNC(=S)Nc1ccccc1Cl)c1ccccn1 | YYLKOELWSMRYHV-UHFFFAOYSA-N | No-Effect | inactive |
| N-(p-amylcinnamoyl)-anthranilic-acid | BRD-K77817104 | CCCCCc1ccc(/C=C/C(=O)Nc2ccccc2C(=O)O)cc1 | GAMRBCZMOOMBSQ-CCEZHUSRSA-N | No-Effect | inactive |
| N-[2-(piperidinylamino)ethyl]-4-iodobenzamide | BRD-K78844995 | O=C(NCCN1CCCCC1)c1ccc(I)cc1 | WCMWVYQHPUQKHW-UHFFFAOYSA-N | No-Effect | inactive |
| N-acetyl-tyrosine | BRD-A24857382 | CC(=O)N[C@@H](Cc1ccc(O)cc1)C(=O)O | CAHKINHBCWCHCF-JTQLQIEISA-N | No-Effect | inactive |
| N-acetylcarnosine | BRD-K52523550 | CC(=O)NCCC(=O)N[C@@H](Cc1cnc[nH]1)C(=O)O | BKAYIFDRRZZKNF-VIFPVBQESA-N | No-Effect | inactive |
| N-acetylglycyl-D-glutamic-acid | BRD-K28121004 | CC(=O)NCC(=O)N[C@H](CCC(=O)O)C(=O)O | CQTYJLQESPLSOP-ZCFIWIBFSA-N | No-Effect | inactive |
| N-acetylmannosamine | BRD-A92872453 | CC(=O)N[C@H]1[C@@H](O)O[C@@H](CO)[C@@H](O)[C@H]1O | OVRNDRQMDRJTHS-SHGPDSBTSA-N | No-Effect | inactive |
| N-acetyltryptamine | BRD-K73700643 | CC(=O)NCCc1c[nH]c2ccccc12 | NVUGEQAEQJTCIX-UHFFFAOYSA-N | No-Effect | inactive |
| N-alpha-methylhistamine-dihydrochloride | BRD-K75390981 | CNCCc1cnc[nH]1 | PHSPJQZRQAJPPF-UHFFFAOYSA-N | No-Effect | inactive |
| N-benzylnaltrindole | BRD-K00004214 | Oc1ccc2c3c1O[C@H]1c4c(c5ccccc5n4Cc4ccccc4)C[C@@]4(O)[C@@H](C2)N(CC2CC2)CC[C@]314 | PSPANKCQVWSVHM-OUDZLNJNSA-N | No-Effect | inactive |
| N-cyclohexyl-2-benzothiazolesulfenamide | BRD-K64191834 | c1ccc2sc(SNC3CCCCC3)nc2c1 | DEQZTKGFXNUBJL-UHFFFAOYSA-N | No-Effect | inactive |
| N-hydroxynicotinamide | BRD-K28030264 | O=C(NO)c1cccnc1 | IYCHDNQCHLMLJZ-UHFFFAOYSA-N | No-Effect | inactive |
| N-methyl-(-)ephedrine-[1R,2S] | BRD-K82236108 | C[C@@H]([C@H](O)c1ccccc1)N(C)C | FMCGSUUBYTWNDP-ONGXEEELSA-N | No-Effect | inactive |
| N-methylformamide | BRD-K33445938 | CNC=O | ATHHXGZTWNVVOU-UHFFFAOYSA-N | No-Effect | inactive |
| N-methyllidocaine-iodide | BRD-K60060639 | CC[N+](C)(CC)CC(=O)Nc1c(C)cccc1C | HFRFLLYLVAKRMZ-UHFFFAOYSA-O | No-Effect | inactive |
| N-methylpyrrolidone | BRD-K52732207 | CN1CCCC1=O | SECXISVLQFMRJM-UHFFFAOYSA-N | No-Effect | inactive |
| N-methylquipazine | BRD-K62056274 | CN1CCN(c2ccc3ccccc3n2)CC1 | HOMWNUXPSJQSSU-UHFFFAOYSA-N | No-Effect | inactive |
| N-MPPP | BRD-K93806173 | CN(C(=O)Cc1ccccc1)[C@H](CN1CCCC1)c1ccccc1 | OVZWUDZIAAHNFG-HXUWFJFHSA-N | No-Effect | inactive |
| N-oxydiethylenebenzothiazole-2-sulfenamide | BRD-K97360717 | c1ccc2sc(SN3CCOCC3)nc2c1 | MHKLKWCYGIBEQF-UHFFFAOYSA-N | No-Effect | inactive |
| N-salicoylaminophenol | BRD-K39693230 | O=C(Nc1ccc(O)cc1)c1ccccc1O | LGCMKPRGGJRYGM-UHFFFAOYSA-N | No-Effect | inactive |
| N20C | BRD-K41143549 | NC(=O)CNCCC(c1ccccc1)c1ccccc1 | BYGXICRJQZKUPA-UHFFFAOYSA-N | No-Effect | inactive |
| N6-cyclopentyladenosine | BRD-K60150510 | OC[C@H]1O[C@@H](n2cnc3c(NC4CCCC4)ncnc32)[C@H](O)[C@@H]1O | SQMWSBKSHWARHU-SDBHATRESA-N | No-Effect | inactive |
| N6-methyladenosine | BRD-K81807412 | CNc1ncnc2c1ncn2[C@@H]1O[C@H](CO)[C@@H](O)[C@H]1O | VQAYFKKCNSOZKM-IOSLPCCCSA-N | No-Effect | inactive |
| N6022 | BRD-K82522873 | Cc1cc(C(N)=O)ccc1-n1c(CCC(=O)O)ccc1-c1ccc(-n2ccnc2)cc1 | YVPGZQLRPAGKLA-UHFFFAOYSA-N | No-Effect | inactive |
| NAB-2 | BRD-K36804890 | Cc1ccc(C)c(-n2cnc3cc(C(=O)NCc4ccccc4Cl)ccc32)c1 | CZSLEMCYYGEGKP-UHFFFAOYSA-N | No-Effect | inactive |
| nabumetone | BRD-K65146499 | COc1ccc2cc(CCC(C)=O)ccc2c1 | BLXXJMDCKKHMKV-UHFFFAOYSA-N | No-Effect | inactive |
| nadide | BRD-K01824972 | NC(=O)c1ccc[n+]([C@@H]2O[C@H](COP(=O)(O)OP(=O)(O)OC[C@H]3O[C@@H](n4cnc5c(N)ncnc54)[C@H](O)[C@@H]3O)[C@@H](O)[C@H]2O)c1 | BAWFJGJZGIEFAR-NNYOXOHSSA-O | No-Effect | inactive |
| nadifloxacin | BRD-A24397426 | C[C@H]1CCc2c(N3CCC(O)CC3)c(F)cc3c(=O)c(C(=O)O)cn1c23 | JYJTVFIEFKZWCJ-JTQLQIEISA-N | No-Effect | inactive |
| nadolol | BRD-A87606379 | CC(C)(C)NC[C@@H](O)COc1cccc2c1C[C@H](O)[C@H](O)C2 | VWPOSFSPZNDTMJ-YUELXQCFSA-N | No-Effect | inactive |
| NADPH | BRD-M06409173 | NC(=O)C1=CN([C@@H]2O[C@H](COP(=O)(O)OP(=O)(O)OC[C@H]3O[C@@H](n4cnc5c(N)ncnc54)[C@H](OP(=O)(O)O)[C@@H]3O)[C@@H](O)[C@H]2O)C=CC1.NC1CCCCC1.NC1CCCCC1.NC1CCCCC1.NC1CCCCC1 | PTKRUDMLGIIORX-ITGWJZMWSA-N | No-Effect | inactive |
| nafadotride | BRD-A71262238 | CCCCN1CCC[C@@H]1CNC(=O)c1cc(C#N)c2ccccc2c1OC | IDZASIQMRGPBCQ-QGZVFWFLSA-N | No-Effect | inactive |
| nafamostat | BRD-K06674495 | N=C(N)Nc1ccc(C(=O)Oc2ccc3cc(C(=N)N)ccc3c2)cc1 | MQQNFDZXWVTQEH-UHFFFAOYSA-N | No-Effect | inactive |
| nafcillin | BRD-K18574842 | CCOc1ccc2ccccc2c1C(=O)N[C@@H]1C(=O)N2[C@@H](C(=O)O)C(C)(C)S[C@H]12 | GPXLMGHLHQJAGZ-JTDSTZFVSA-N | No-Effect | inactive |
| naftidrofuryl | BRD-A67862938 | CCN(CC)CCOC(=O)[C@H](Cc1cccc2ccccc12)C[C@H]1CCCO1 | KBAFPSLPKGSANY-FGZHOGPDSA-N | No-Effect | inactive |
| naftifine | BRD-K74141488 | CN(C/C=C/c1ccccc1)Cc1cccc2ccccc12 | OZGNYLLQHRPOBR-DHZHZOJOSA-N | No-Effect | inactive |
| nalbuphine | BRD-K66404838 | Oc1ccc2c3c1O[C@H]1[C@@H](O)CC[C@@]4(O)[C@@H](C2)N(CC2CCC2)CC[C@]314 | NETZHAKZCGBWSS-CEDHKZHLSA-N | No-Effect | inactive |
| nalfurafine | BRD-K89272869 | CN(C(=O)/C=C/c1ccoc1)[C@@H]1CC[C@@]2(O)[C@H]3Cc4ccc(O)c5c4[C@@]2(CCN3CC2CC2)[C@H]1O5 | XGZZHZMWIXFATA-UEZBDDGYSA-N | No-Effect | inactive |
| nalidixic-acid | BRD-K47886988 | CCn1cc(C(=O)O)c(=O)c2ccc(C)nc21 | MHWLWQUZZRMNGJ-UHFFFAOYSA-N | No-Effect | inactive |
| nalmefene | BRD-K68407436 | C=C1CC[C@@]2(O)[C@H]3Cc4ccc(O)c5c4[C@@]2(CCN3CC2CC2)[C@H]1O5 | WJBLNOPPDWQMCH-MBPVOVBZSA-N | No-Effect | inactive |
| naloxegol | BRD-K41458421 | C=CCN1CC[C@]23c4c5ccc(O)c4O[C@H]2[C@@H](OCCOCCOCCOCCOCCOCCOCCOC)CC[C@@]3(O)[C@H]1C5 | XNKCCCKFOQNXKV-ZRSCBOBOSA-N | No-Effect | inactive |
| naloxone | BRD-K67511046 | C=CCN1CC[C@]23c4c5ccc(O)c4O[C@H]2C(=O)CC[C@@]3(O)[C@H]1C5 | UZHSEJADLWPNLE-GRGSLBFTSA-N | No-Effect | inactive |
| naloxone-benzoylhydrazone | BRD-K70898774 | C=CCN1CC[C@]23c4c5ccc(O)c4O[C@H]2/C(=N/NC(=O)c2ccccc2)CC[C@@]3(O)[C@H]1C5 | AKXCFAYOTIEFOH-XTNAHFASSA-N | No-Effect | inactive |

| | | | | | |
|---|---|---|---|---|---|
| naltrexone | BRD-K88172511 | O=C1CC[C@@]2(O)[C@H]3Cc4ccc(O)c5c4[C@@]2(CCN3CC2CC2)[C@H]1O5 | DQCKKXVULJGBQN-XFWGSAIBSA-N | No-Effect | inactive |
| naltriben | BRD-K81275354 | Oc1ccc2c3c1O[C@H]1c4oc5ccccc5c4C[C@@]4(O)[C@@H](C2)N(CC2CC2)CC[C@]314 | ZHVWWEYETMPAMX-IFKAHUTRSA-N | No-Effect | inactive |
| naltrindole | BRD-K34088084 | Oc1ccc2c3c1O[C@H]1c4[nH]c5ccccc5c4C[C@@]4(O)[C@@H](C2)N(CC2CC2)CC[C@]314 | WIYUZYBFCWCCQJ-IFKAHUTRSA-N | No-Effect | inactive |
| Nampt-IN-1 | BRD-K18280217 | CC(C)(O)CS(=O)(=O)Nc1ccc2c(c1)CCN(C(=O)COc1cccnc1)C2 | QHHSCLARESIWBH-UHFFFAOYSA-N | No-Effect | inactive |
| NAN-190 | BRD-K69195780 | COc1ccccc1N1CCN(CCCCN2C(=O)c3ccccc3C2=O)CC1 | SJDOMIRMMUGQQK-UHFFFAOYSA-N | No-Effect | inactive |
| napamezole | BRD-K00004563 | C1=C(CC2=NCCN2)CCc2ccccc21 | WETRBJOSGIDJHQ-UHFFFAOYSA-N | No-Effect | inactive |
| naphazoline | BRD-K77641333 | c1ccc2c(CC3=NCCN3)cccc2c1 | CNIIGCLFLJGOGP-UHFFFAOYSA-N | No-Effect | inactive |
| naphthoquine-phosphate | BRD-K13664493 | CC(C)(C)NCc1cc(Nc2ccnc3cc(Cl)ccc23)c2c(c1O)CCCC2 | VEVMYTDOWUQLGI-UHFFFAOYSA-N | No-Effect | inactive |
| napirimus | BRD-K91234829 | Cn1cc(C(=O)c2cccc3ccccc23)cc1C(=O)O | CNUPQURTHORUGH-UHFFFAOYSA-N | No-Effect | inactive |
| naproxen | BRD-K59197931 | COc1ccc2cc([C@H](C)C(=O)O)ccc2c1 | CMWTZPSULFXXJA-VIFPVBQESA-N | No-Effect | inactive |
| naratriptan | BRD-K61337602 | CNS(=O)(=O)CCc1ccc2[nH]cc(C3CCN(C)CC3)c2c1 | AMKVXSZCKVJAGH-UHFFFAOYSA-N | No-Effect | inactive |
| naringenin | BRD-K08832567 | O=C1C[C@@H](c2ccc(O)cc2)Oc2cc(O)cc(O)c21 | FTVWIRXFELQLPI-ZDUSSCGKSA-N | No-Effect | inactive |
| naringeninic-acid | BRD-K87122641 | O=C(O)/C=C/c1ccc(O)cc1 | NGSWKAQJJWESNS-ZZXKWVIFSA-N | No-Effect | inactive |
| naringin | BRD-K02953697 | C[C@@H]1O[C@@H](O[C@H]2[C@H](Oc3cc(O)c4c(c3)O[C@H](c3ccc(O)cc3)CC4=O)O[C@H](CO)[C@@H](O)[C@@H]2O)[C@H](O)[C@H](O)[C@H]1O | DFPMSGMNTNDNHN-ZPHOTFPESA-N | No-Effect | inactive |
| naringin-dihydrochalcone | BRD-K72522627 | C[C@@H]1O[C@@H](O[C@H]2[C@H](Oc3cc(O)c(C(=O)CCc4ccc(O)cc4)c(O)c3)O[C@H](CO)[C@@H](O)[C@@H]2O)[C@H](O)[C@H](O)[C@H]1O | CWBZAESOUBENAP-QVNVHUMTSA-N | No-Effect | inactive |
| narlaprevir | BRD-K38287497 | CCCC[C@H](NC(=O)[C@@H]1[C@@H]2[C@H](CN1C(=O)[C@@H](NC(=O)NC1(CS(=O)(=O)C(C)(C)C)CCCCC1)C(C)(C)C)C2(C)C)C(=O)C(=O)NC1CC1 | RICZEKWVNZFTNZ-LFGITCQGSA-N | No-Effect | inactive |
| NAS-181 | BRD-K00004235 | C1=C(CN2CCOCC2)COc2c1cccc2OC[C@H]1CNCCO1 | RTKDBEOSPDFLGD-QGZVFWFLSA-N | No-Effect | inactive |
| nastorazepide | BRD-K17295893 | CC(C)(C)C(=O)CN1C(=O)[C@H](NC(=O)Nc2cccc(C(=O)O)c2)CN(C2CCCC2)c2ccccc21 | VIJCCFFEBCOOIE-JOCHJYFZSA-N | No-Effect | inactive |
| natamycin | BRD-K90563805 | C[C@@H]1C/C=C/C=C/C=C/C=C/[C@H](O[C@@H]2O[C@H](C)[C@@H](O)[C@H](N)[C@@H]2O)C[C@@H]2O[C@](O)(C[C@@H](O)C[C@H]3O[C@@H]3/C=C/C(=O)O1)C[C@H](O)[C@H]2C(=O)O | NCXMLFZGDNKEPB-FFPOYIOWSA-N | No-Effect | inactive |
| nateglinide | BRD-K44353683 | CC(C)[C@H]1CC[C@H](C(=O)N[C@H](Cc2ccccc2)C(=O)O)CC1 | OELFLUMRDSZNSF-BRWVUGGUSA-N | No-Effect | inactive |
| NAV-26 | BRD-A84202322 | O=C(NCc1ccc(OC(F)(F)F)cc1)[C@@H]1c2ccccc2C(=O)N1C1CCOCC1 | ICGMZCVSHDKQTE-IBGZPJMESA-N | No-Effect | inactive |
| navarixin | BRD-K82850175 | CC[C@@H](Nc1c(Nc2cccc(C(=O)N(C)C)c2O)c(=O)c1=O)c1ccc(C)o1 | RXIUEIPPLAFSDF-CYBMUJFWSA-N | No-Effect | inactive |
| naxagolide | BRD-K00004670 | CCCN1CCO[C@@H]2c3cc(O)ccc3CC[C@H]21 | JCSREICEMHWFAY-HUUCEWRRSA-N | No-Effect | inactive |
| NB-001 | BRD-K00004647 | Nc1ncnc2c1ncn2CCNCCCCCO | CVPTTZZCRDVGSU-UHFFFAOYSA-N | No-Effect | inactive |
| NBI-27914 | BRD-K61177364 | CCCN(CC1CC1)c1nc(C)nc(Nc2c(Cl)cc(Cl)cc2Cl)c1Cl | KNADXBVKFAUMCR-UHFFFAOYSA-N | No-Effect | inactive |
| NBI-98782 | BRD-K15241725 | COc1cc2c(cc1OC)[C@H]1C[C@@H](O)[C@H](CC(C)C)CN1CC2 | WEQLWGNDNRARGE-DJIMGWMZSA-N | No-Effect | inactive |
| NBQX | BRD-K11796549 | NS(=O)(=O)c1cccc2c1c([N+](=O)[O-])cc1[nH]c(=O)c(=O)[nH]c12 | UQNAFPHGVPVTAL-UHFFFAOYSA-N | No-Effect | inactive |
| NCS-382 | BRD-K01826782 | O=C(O)/C=C1\CCCc2ccccc2[C@H]1O | UADPGHINQMWEAG-FROQITRMSA-N | No-Effect | inactive |
| NCX-1015 | BRD-K00004749 | C[C@]12C[C@H](O)[C@H]3[C@@H](CCC4=CC(=O)C=C[C@@]43C)[C@@H]1CC[C@]2(O)C(=O)COC(=O)c1ccc(CO[N+](=O)[O-])cc1 | MJHYBJOMJPGNMM-KGWLDMEJSA-N | No-Effect | inactive |
| NDD-094 | BRD-K00004664 | Cc1nccn1Cc1ccccn1 | CTUICISGLKIJLG-UHFFFAOYSA-N | No-Effect | inactive |
| NDT-9513727 | BRD-K17539220 | CCCCn1c(-c2ccccc2)nc(-c2ccccc2)c1CN(Cc1ccc2c(c1)OCO2)Cc1ccc2c(c1)OCO2 | ITACCRHKSPSKKL-UHFFFAOYSA-N | No-Effect | inactive |
| necrostatin-1 | BRD-K01825019 | CN1C(=O)[C@H](Cc2c[nH]c3ccccc23)NC1=S | TXUWMXQFNYDOEZ-NSHDSACASA-N | No-Effect | inactive |
| necrostatin-2 | BRD-K68851986 | CN1C(=O)N[C@H](Cc2c[nH]c3c(Cl)cccc23)C1=O | WIKGAEMMNQTUGL-SNVBAGLBSA-N | No-Effect | inactive |
| nefazodone | BRD-K90789829 | CCc1nn(CCCN2CCN(c3cccc(Cl)c3)CC2)c(=O)n1CCOc1ccccc1 | VRBKIVRKKCLPHA-UHFFFAOYSA-N | No-Effect | inactive |
| nefiracetam | BRD-K65678996 | Cc1cccc(C)c1NC(=O)CN1CCCC1=O | NGHTXZCKLWZPGK-UHFFFAOYSA-N | No-Effect | inactive |
| nefopam | BRD-K01826512 | CN1CCO[C@@H](c2ccccc2)c2ccccc2C1 | RGPDEAGGEXEMMM-KRWDZBQOSA-N | No-Effect | inactive |
| nelarabine | BRD-K84466663 | COc1nc(N)nc2c1ncn2[C@@H]1O[C@H](CO)[C@@H](O)[C@@H]1O | IXOXBSCIXZEQEQ-UHTZMRCNSA-N | No-Effect | inactive |
| nelociguat | BRD-K00003315 | COC(=O)Nc1c(N)nc(-c2nn(Cc3ccccc3F)c3ncccc23)nc1N | FTQHGWIXJSSWOY-UHFFFAOYSA-N | No-Effect | inactive |
| nelotanserin | BRD-K00003466 | COc1ccc(NC(=O)Nc2ccc(F)cc2F)cc1-c1c(Br)cnn1C | COSPVUFTLGQDQL-UHFFFAOYSA-N | No-Effect | inactive |
| nemazoline | BRD-K00004662 | Nc1c(Cl)cc(C=C2NCCN2)cc1Cl | OEOYJZUHFXEJEI-UHFFFAOYSA-N | No-Effect | inactive |
| nemonapride | BRD-K47289124 | CNc1cc(OC)c(C(=O)N[C@@H]2CCN(Cc3ccccc3)[C@@H]2C)cc1Cl | KRVOJOCLBAAKSJ-RDTXWAMCSA-N | No-Effect | inactive |
| nemorexant | BRD-K00003505 | COc1ccc(-n2nccn2)c(C(=O)N2CCC[C@@]2(C)c2nc3ccc(Cl)c(C)c3[nH]2)c1 | NBGABHGMJVIVBW-QHCPKHFHSA-N | No-Effect | inactive |
| neo-gilurytmal | BRD-A53539450 | CCC[N@@+]12[C@H]3C[C@@]45c6ccccc6N(C)[C@H]4[C@H]1C[C@@H]([C@H](CC)[C@H]2O)[C@H]3[C@@H]5O | UAUHEPXILIZYCU-JSQPETPRSA-N | No-Effect | inactive |
| neohesperidin | BRD-K00003712 | COc1ccc([C@@H]2CC(=O)c3c(O)cc(O[C@@H]4O[C@H](CO)[C@@H](O)[C@H](O)[C@H]4O[C@@H]4O[C@@H](C)[C@H](O)[C@@H](O)[C@H]4O)cc3O2)cc1O | ARGKVCXINMKCAZ-UZRWAPQLSA-N | No-Effect | inactive |
| neohesperidin-dihydrochalcone | BRD-K61032563 | COc1ccc(CCC(=O)c2c(O)cc(O[C@@H]3O[C@H](CO)[C@@H](O)[C@H](O)[C@H]3O[C@@H]3O[C@@H](C)[C@H](O)[C@@H](O)[C@H]3O)cc2O)cc1O | ITVGXXMINPYUHD-CUVHLRMHSA-N | No-Effect | inactive |
| nepicastat | BRD-K09126848 | NCc1c[nH]c(=S)n1[C@H]1CCc2c(F)cc(F)cc2C1 | YZZVIKDAOTXDEB-JTQLQIEISA-N | No-Effect | inactive |
| neratinib | BRD-K85606544 | CCOc1cc2ncc(C#N)c(Nc3ccc(OCc4ccccn4)c(Cl)c3)c2cc1NC(=O)/C=C/CN(C)C | JWNPDZNEKVCWMY-VQHVLOKHSA-N | No-Effect | inactive |
| nerbacadol | BRD-K43647122 | Cc1oncc1C(=O)N1CCCCC1 | ZWNGSOLJLMBZFZ-UHFFFAOYSA-N | No-Effect | inactive |
| neridronic-acid | BRD-K60212747 | NCCCCCC(O)(P(=O)(O)O)P(=O)(O)O | PUUSSSIBPPTKTP-UHFFFAOYSA-N | No-Effect | inactive |
| nesbuvir | BRD-K79989959 | CNC(=O)c1c(-c2ccc(F)cc2)oc2cc(N(CCO)S(C)(=O)=O)c(C3CC3)cc12 | WTDWVLJJJOTABN-UHFFFAOYSA-N | No-Effect | inactive |
| netilmicin | BRD-K00003604 | CCN[C@@H]1C[C@H](N)[C@@H](O[C@H]2OC(CN)=CC[C@H]2N)[C@H](O)[C@H]1O[C@H]1OC[C@](C)(O)[C@H](NC)[C@H]1O | CIDUJQMULVCIBT-MQDUPKMGSA-N | No-Effect | inactive |
| netupitant | BRD-K97428065 | Cc1ccccc1-c1cc(N2CCN(C)CC2)ncc1N(C)C(=O)C(C)(C)c1cc(C(F)(F)F)cc(C(F)(F)F)c1 | WAXQNWCZJDTGBU-UHFFFAOYSA-N | No-Effect | inactive |
| Neu2000 | BRD-K00003470 | O=C(O)c1cc(NCc2c(F)c(F)c(C(F)(F)F)c(F)c2F)ccc1O | HABROHXUHNHQMY-UHFFFAOYSA-N | No-Effect | inactive |
| neurodazine | BRD-K22631935 | COc1ccc(-c2nc(-c3ccc(-c4cccc(Cl)c4)o3)[nH]c2-c2ccc(OC)cc2)cc1 | FEEOFPAEDSMOTO-UHFFFAOYSA-N | No-Effect | inactive |
| neuropathiazol | BRD-K28567778 | CCOC(=O)c1ccc(N(C)c2csc(-c3ccccc3)n2)cc1 | MGKYEWWFHSESJN-UHFFFAOYSA-N | No-Effect | inactive |
| nevirapine | BRD-K15502390 | Cc1ccnc2c1NC(=O)c1cccnc1N2C1CC1 | NQDJXKOVJZTUJA-UHFFFAOYSA-N | No-Effect | inactive |
| nexturastat-A | BRD-K70402238 | CCCCN(Cc1ccc(C(=O)NO)cc1)C(=O)Nc1ccccc1 | JZWXMCPARMXZQV-UHFFFAOYSA-N | No-Effect | inactive |
| NFKB-activation-inhibitor-II | BRD-K83354763 | Cc1ccc(NCCCc2ccccc2)c(N)c1 | YMFNPBSZFWXMAD-UHFFFAOYSA-N | No-Effect | inactive |
| NGB-2904 | BRD-K05181084 | O=C(NCCCCN1CCN(c2cccc(Cl)c2Cl)CC1)c1ccc2c(c1)Cc1ccccc1-2 | YJPWVCIGSHWNON-UHFFFAOYSA-N | No-Effect | inactive |
| NGD-94-1 | BRD-K28639661 | c1ccc(-c2ncc(CN3CCN(c4ncccn4)CC3)[nH]2)cc1 | OTVQCHUIPJYASM-UHFFFAOYSA-N | No-Effect | inactive |

| | | | | | |
|---|---|---|---|---|---|
| NGD-98-2 | BRD-K81779590 | CCC(CC)Nc1nc(C)c(-c2ccc(OC(F)(F)F)cc2OC)nc1OC | PSUFQBRCUSJTDO-UHFFFAOYSA-N | No-Effect | inactive |
| NGP555 | BRD-K00003490 | CCc1cc(Nc2nc(-c3ccc(-n4cnc(C)c4)c(F)c3)cs2)c(C)cc1C | WDEKUGNKKOGFOA-UHFFFAOYSA-N | No-Effect | inactive |
| NH125 | BRD-K31086665 | CCCCCCCCCCCCCCCC[n+]1ccn(Cc2ccccc2)c1C | ZZXUNAFAKNKXFW-UHFFFAOYSA-N | No-Effect | inactive |
| NHI-2 | BRD-K05653025 | COC(=O)c1cc2c(C(F)(F)F)cc(-c3ccccc3)cc2n1O | YPPFWRWCZNXINO-UHFFFAOYSA-N | No-Effect | inactive |
| NI-57 | BRD-K03042257 | COc1cc(C#N)ccc1S(=O)(=O)Nc1ccc2c(c1)cc(C)c(=O)n2C | UEMQPCYDWCSVCU-UHFFFAOYSA-N | No-Effect | inactive |
| niacin | BRD-K60237333 | O=C(O)c1cccnc1 | PVNIIMVLHYAWGP-UHFFFAOYSA-N | No-Effect | inactive |
| nialamide | BRD-K12102668 | O=C(CCNNC(=O)c1ccncc1)NCc1ccccc1 | NOIIUHRQUVNIDD-UHFFFAOYSA-N | No-Effect | inactive |
| nibentan | BRD-A85876464 | CCN(CC)CCCC[C@@H](NC(=O)c1ccc([N+](=O)[O-])cc1)c1ccccc1 | YRFYIORBFFGPPZ-OAQYLSRUSA-N | No-Effect | inactive |
| nicainoprol | BRD-K00003382 | CC(C)NC[C@H](O)COc1cccc2c1N(C(=O)c1cccnc1)CCC2 | AUIHHZBJBKRDIE-SFHVURJKSA-N | No-Effect | inactive |
| nicaraven | BRD-A86871940 | C[C@@H](CNC(=O)c1cccnc1)NC(=O)c1cccnc1 | KTXBOOWDLPUROC-NSHDSACASA-N | No-Effect | inactive |
| nicardipine | BRD-A26711594 | COC(=O)C1=C(C)NC(C)=C(C(=O)OCCN(C)Cc2ccccc2)[C@@H]1c1cccc([N+](=O)[O-])c1 | ZBBHBTPTTSWHBA-XMMPIXPASA-N | No-Effect | inactive |
| nicergoline | BRD-K76810206 | CO[C@]12C[C@@H](COC(=O)c3cncc(Br)c3)CN(C)[C@@H]1Cc1cn(C)c3cccc2c13 | YSEXMKHXIOCEJA-FVFQAYNVSA-N | No-Effect | inactive |
| nicodicosapent | BRD-K00003262 | CC/C=C\C/C=C\C/C=C\C/C=C\C/C=C\CCCC(=O)NCCNC(=O)c1cccnc1 | YYQALOWPCKMFDQ-JLNKQSITSA-N | No-Effect | inactive |
| nicorandil | BRD-K97752965 | O=C(NCCO[N+](=O)[O-])c1cccnc1 | LBHIOVVIQHSOQN-UHFFFAOYSA-N | No-Effect | inactive |
| nicotinamide | BRD-K28912512 | NC(=O)c1cccnc1 | DFPAKSUCGFBDDF-UHFFFAOYSA-N | No-Effect | inactive |
| nicotine | BRD-K05395900 | CN1CCC[C@H]1c1cccnc1 | SNICXCGAKADSCV-JTQLQIEISA-N | No-Effect | inactive |
| nicotinyl-alcohol-tartrate | BRD-K76600340 | OCc1cccnc1 | MVQVNTPHUGQQHK-UHFFFAOYSA-N | No-Effect | inactive |
| NIDA-41020 | BRD-K17617556 | COc1ccc(-c2c(C)c(C(=O)NN3CCCCC3)nn2-c2ccc(Cl)cc2Cl)cc1 | KWDBQJRWPWTGPF-UHFFFAOYSA-N | No-Effect | inactive |
| nifedipine | BRD-K96354014 | COC(=O)C1=C(C)NC(C)=C(C(=O)OC)C1c1ccccc1[N+](=O)[O-] | HYIMSNHJOBLJNT-UHFFFAOYSA-N | No-Effect | inactive |
| nifekalant | BRD-K32311154 | Cn1c(NCCN(CCO)CCCc2ccc([N+](=O)[O-])cc2)cc(=O)n(C)c1=O | OEBPANQZQGQPHF-UHFFFAOYSA-N | No-Effect | inactive |
| nifenalol | BRD-A21280314 | CC(C)NC[C@@H](O)c1ccc([N+](=O)[O-])cc1 | UAORFCGRZIGNCI-LLVKDONJSA-N | No-Effect | inactive |
| nifenazone | BRD-K47407372 | Cc1c(NC(=O)c2cccnc2)c(=O)n(-c2ccccc2)n1C | BRZANEXCSZCZCI-UHFFFAOYSA-N | No-Effect | inactive |
| niflumic-acid | BRD-K98763141 | O=C(O)c1cccnc1Nc1cccc(C(F)(F)F)c1 | JZFPYUNJRRFVQU-UHFFFAOYSA-N | No-Effect | inactive |
| nifuroxazide | BRD-K68188368 | O=C(N/N=C/c1ccc([N+](=O)[O-])o1)c1ccc(O)cc1 | YCWSUKQGVSGXJO-NTUHNPAUSA-N | No-Effect | inactive |
| nifurtimox | BRD-A84020532 | C[C@H]1CS(=O)(=O)CCN1/N=C/c1ccc([N+](=O)[O-])o1 | ARFHIAQFJWUCFH-KIFNTFBYSA-N | No-Effect | inactive |
| niguldipine-(R)-(-) | BRD-K01825503 | COC(=O)C1=C(C)NC(C)=C(C(=O)OCCCN2CCC(c3ccccc3)(c3ccccc3)CC2)[C@@H]1c1cccc([N+](=O)[O-])c1 | SVJMLYUFVDMUHP-MGBGTMOVSA-N | No-Effect | inactive |
| niguldipine-(S)-(+) | BRD-K59333713 | COC(=O)C1=C(C)NC(C)=C(C(=O)OCCCN2CCC(c3ccccc3)(c3ccccc3)CC2)[C@H]1c1cccc([N+](=O)[O-])c1 | SVJMLYUFVDMUHP-XIFFEERXSA-N | No-Effect | inactive |
| nikethamide | BRD-K91544578 | CCN(CC)C(=O)c1cccnc1 | NCYVXEGFNDZQCU-UHFFFAOYSA-N | No-Effect | inactive |
| nilotinib | BRD-K81528515 | Cc1cn(-c2cc(NC(=O)c3ccc(C)c(Nc4nccc(-c5cccnc5)n4)c3)cc(C(F)(F)F)c2)cn1 | HHZIURLSWUIHRB-UHFFFAOYSA-N | No-Effect | inactive |
| nilutamide | BRD-K23566484 | CC1(C)NC(=O)N(c2ccc([N+](=O)[O-])c(C(F)(F)F)c2)C1=O | XWXYUMMDTVBTOU-UHFFFAOYSA-N | No-Effect | inactive |
| nilvadipine | BRD-A84687895 | COC(=O)C1=C(C#N)NC(C)=C(C(=O)OC(C)C)[C@@H]1c1cccc([N+](=O)[O-])c1 | FAIIFDPAEUKBEP-INIZCTEOSA-N | No-Effect | inactive |
| NIM811 | BRD-K00003331 | C/C=C/C[C@@H](C)[C@@H](O)[C@H]1C(=O)N[C@@H](CC)C(=O)N(C)CC(=O)N(C)[C@@H]([C@@H](C)CC)C(=O)N[C@@H](C(C)C)C(=O)N(C)[C@@H](CC(C)C)C(=O)N[C@@H](C)C(=O)N[C@H](C)C(=O)N(C)[C@@H](CC(C)C)C(=O)N(C)[C@@H](CC(C)C)C(=O)N(C)[C@@H](C(C)C)C(=O)N1C | RPJPZDVUUKWPGT-FOIHOXPVSA-N | No-Effect | inactive |
| nimesulide | BRD-K76775527 | CS(=O)(=O)Nc1ccc([N+](=O)[O-])cc1Oc1ccccc1 | HYWYRSMBCFDLJT-UHFFFAOYSA-N | No-Effect | inactive |
| nimodipine | BRD-A58048407 | COCCOC(=O)C1=C(C)NC(C)=C(C(=O)OC(C)C)[C@@H]1c1cccc([N+](=O)[O-])c1 | UIAGMCDKSXEBJQ-LJQANCHMSA-N | No-Effect | inactive |
| nimorazole | BRD-K79145628 | O=[N+]([O-])c1cncn1CCN1CCOCC1 | MDJFHRLTPRPZLY-UHFFFAOYSA-N | No-Effect | inactive |
| nimustine | BRD-K13240081 | Cc1ncc(CNC(=O)N(CCCl)[NH2+][O-])c(N)n1 | QMAZPCHGABINAB-UHFFFAOYSA-N | No-Effect | inactive |
| ningetinib-tosylate | BRD-K00003476 | Cc1c(C(=O)Nc2ccc(Oc3ccnc4cc(OCC(C)(C)O)ccc34)c(F)c2)c(=O)n(-c2ccccc2)n1C | VQYYQSZNRVQLIS-UHFFFAOYSA-N | No-Effect | inactive |
| nintedanib | BRD-K49075727 | COC(=O)c1ccc2c(c1)NC(=O)/C2=C(\Nc1ccc(N(C)C(=O)CN2CCN(C)CC2)cc1)c1ccccc1 | XZXHXSATPCNXJR-ZIADKAODSA-N | No-Effect | inactive |
| niraparib | BRD-K54955827 | NC(=O)c1cccc2cn(-c3ccc([C@@H]4CCCNC4)cc3)nc12 | PCHKPVIQAHNQLW-CQSZACIVSA-N | No-Effect | inactive |
| niridazole | BRD-K53123955 | O=C1NCCN1c1ncc([N+](=O)[O-])s1 | RDXLYGJSWZYTFJ-UHFFFAOYSA-N | No-Effect | inactive |
| nisoldipine | BRD-A84465106 | COC(=O)C1=C(C)NC(C)=C(C(=O)OCC(C)C)[C@@H]1c1ccccc1[N+](=O)[O-] | VKQFCGNPDRICFG-GOSISDBHSA-N | No-Effect | inactive |
| nisoxetine | BRD-K01825513 | CNCC[C@@H](Oc1ccccc1OC)c1ccccc1 | ITJNARMNRKSWTA-OAHLLOKOSA-N | No-Effect | inactive |
| nitarsone | BRD-K56707426 | O=[N+]([O-])c1ccc([As](=O)(O)O)cc1 | FUUFQLXAIUOWML-UHFFFAOYSA-N | No-Effect | inactive |
| nitecapone | BRD-K75551531 | CC(=O)C(=Cc1cc(O)c(O)c([N+](=O)[O-])c1)C(C)=O | UPMRZALMHVUCIN-UHFFFAOYSA-N | No-Effect | inactive |
| nitenpyram | BRD-K47887731 | CCN(Cc1ccc(Cl)nc1)/C(=C/[N+](=O)[O-])NC | CFRPSFYHXJZSBI-DHZHZOJOSA-N | No-Effect | inactive |
| nithiamide | BRD-K39467894 | CC(=O)Nc1ncc([N+](=O)[O-])s1 | UJRRDDHEMZLWFI-UHFFFAOYSA-N | No-Effect | inactive |
| nitisinone | BRD-K01109406 | O=C1CCCC(=O)C1=C(O)c1ccc(C(F)(F)F)cc1[N+](=O)[O-] | PMHVFNYNPNKNRO-UHFFFAOYSA-N | No-Effect | inactive |
| nitrendipine | BRD-A02006392 | CCOC(=O)C1=C(C)NC(C)=C(C(=O)OC)[C@@H]1c1cccc([N+](=O)[O-])c1 | PVHUJELLJLJGLN-INIZCTEOSA-N | No-Effect | inactive |
| nitrocaramiphen | BRD-K28453807 | CCN(CC)CCOC(=O)C1(c2ccc([N+](=O)[O-])cc2)CCCC1 | NSHMVLOWOAQFTP-UHFFFAOYSA-N | No-Effect | inactive |
| nitroflurbiprofen | BRD-K00004737 | C[C@@H](C(=O)OCCCCO[N+](=O)[O-])c1ccc(-c2ccccc2)c(F)c1 | DLWSRGHNJVLJAH-CQSZACIVSA-N | No-Effect | inactive |
| nitrofurantoin | BRD-K76927775 | O=C1CN(/N=C/c2ccc([N+](=O)[O-])o2)C(=O)N1 | NXFQHRVNIOXGAQ-YCRREMRBSA-N | No-Effect | inactive |
| nitrofurazone | BRD-K79092138 | NC(=O)N/N=C/c1ccc([N+](=O)[O-])o1 | IAIWVQXQOWNYOU-FPYGCLRLSA-N | No-Effect | inactive |
| nitromide | BRD-K76381435 | NC(=O)c1cc([N+](=O)[O-])cc([N+](=O)[O-])c1 | UUKWKUSGGZNXGA-UHFFFAOYSA-N | No-Effect | inactive |
| nitrosodimethylurea | BRD-K08927448 | CNC(=O)N(C)[NH2+][O-] | RCUSTCPNMVCLLN-UHFFFAOYSA-N | No-Effect | inactive |
| nitroxoline | BRD-K97534525 | O=[N+]([O-])c1ccc(O)c2ncccc12 | RJIWZDNTCBHXAL-UHFFFAOYSA-N | No-Effect | inactive |
| nizatidine | BRD-K73589491 | CN/C(=C\[N+](=O)[O-])NCCSCc1csc(CN(C)C)n1 | SGXXNSQHWDMGGP-IZZDOVSWSA-N | No-Effect | inactive |
| nizofenone | BRD-K84011460 | CCN(CC)Cc1nccn1-c1ccc([N+](=O)[O-])cc1C(=O)c1ccccc1Cl | WZGBZLHGOVJDET-UHFFFAOYSA-N | No-Effect | inactive |
| NK-252 | BRD-K47441935 | O=C(NCc1ccccn1)Nc1nnc(-c2ccco2)o1 | FNSCFQXZZNCDAI-UHFFFAOYSA-N | No-Effect | inactive |
| NKP-1339 | BRD-K74796838 | Cl[Ru+3](Cl)(Cl)(Cl)([n+]1cc2ccccc2[nH]1)[n+]1cc2ccccc2[nH]1 | KNLORLKZMHLRBH-UHFFFAOYSA-J | No-Effect | inactive |
| NKY-80 | BRD-A75015151 | Nc1ncc2c(n1)C[C@@H](c1ccco1)CC2=O | SOJUSNIBPPMLCC-SSDOTTSWSA-N | No-Effect | inactive |
| NLG919 | BRD-A62133859 | O[C@@H](C[C@H]1c2ccccc2-c2cncn21)C1CCCCC1 | YTRRAUACYORZLX-WMZOPIPTSA-N | No-Effect | inactive |
| NM107 | BRD-K79424231 | C[C@@]1(O)[C@H](O)[C@@H](CO)O[C@H]1n1ccc(N)nc1=O | PPUDLEUZKVJXSZ-VPCXQMTMSA-N | No-Effect | inactive |
| NMDA | BRD-K62414032 | CN[C@H](CC(=O)O)C(=O)O | HOKKHZGPKSLGJE-GSVOUGTGSA-N | No-Effect | inactive |
| NMS-1286937 | BRD-K83699324 | CN1CCN(c2ccc(OC(F)(F)F)c(Nc3ncc4c(n3)-c3c(c(C(N)=O)nn3CCO)CC4)c2)CC1 | QHLVBNKYJGBCQJ-UHFFFAOYSA-N | No-Effect | inactive |
| NMS-P715 | BRD-K54170555 | CCc1cccc(CC)c1NC(=O)c1nn(C)c2c1CCc1cnc(Nc3ccc(C(=O)NC4CCN(C)CC4)cc3OC(F)(F)F)nc1-2 | JFOAJUGFHDCBJJ-UHFFFAOYSA-N | No-Effect | inactive |
| NNC-63-0532 | BRD-K72904818 | COC(=O)CN1CN(c2ccccc2)C2(CCN(Cc3cccc4ccccc34)CC2)C1=O | AQMPIDSGLFVVPL-UHFFFAOYSA-N | No-Effect | inactive |
| NNC-711 | BRD-K42221274 | O=C(O)C1=CCCN(CCON=C(c2ccccc2)c2ccccc2)C1 | NGNALWDRPKNJGR-UHFFFAOYSA-N | No-Effect | inactive |
| NO-ASA | BRD-K08807999 | CC(=O)Oc1ccccc1C(=O)Oc1cccc(CO[N+](=O)[O-])c1 | IOJUJUOXKXMJNF-UHFFFAOYSA-N | No-Effect | inactive |
| nobiletin | BRD-K06753942 | COc1ccc(-c2cc(=O)c3c(OC)c(OC)c(OC)c(OC)c3o2)cc1OC | MRIAQLRQZPPODS-UHFFFAOYSA-N | No-Effect | inactive |
| nolatrexed | BRD-K36636143 | Cc1ccc2[nH]c(N)nc(=O)c2c1Sc1ccncc1 | XHWRWCSCBDLOLM-UHFFFAOYSA-N | No-Effect | inactive |
| nomifensine | BRD-A29644307 | CN1Cc2c(N)cccc2[C@H](c2ccccc2)C1 | XXPANQJNYNUNES-AWEZNQCLSA-N | No-Effect | inactive |

| | | | | | |
|---|---|---|---|---|---|
| nonivamide | BRD-K05907307 | CCCCCCCCC(=O)NCc1ccc(O)c(OC)c1 | RGOVYLWUIBMPGK-UHFFFAOYSA-N | No-Effect | inactive |
| nor-binaltorphimine | BRD-K36381762 | Oc1ccc2c3c1O[C@H]1c4[nH]c5c(c4C[C@@]4(O)[C@@H](C2)N(CC2CC2)CC[C@]314)C[C@@]1(O)[C@H]2Cc3ccc(O)c4c3[C@@]1(CCN2CC1CC1)[C@H]5O4 | APSUXPSYBJVPPS-YAUKWVCOSA-N | No-Effect | inactive |
| norcyclobenzaprine | BRD-K63165456 | CNCCC=C1c2ccccc2C=Cc2ccccc21 | XECQQDXTQRYYBH-UHFFFAOYSA-N | No-Effect | inactive |
| norepinephrine | BRD-K87349682 | NC[C@H](O)c1ccc(O)c(O)c1 | SFLSHLFXELFNJZ-QMMMGPOBSA-N | No-Effect | inactive |
| norethindrone | BRD-K92073408 | C#C[C@]1(O)CC[C@H]2[C@@H]3CCC4=CC(=O)CC[C@@H]4[C@H]3CC[C@@]21C | VIKNJXKGJWUCNN-XGXHKTLJSA-N | No-Effect | inactive |
| norethindrone-acetate | BRD-K96037667 | C#C[C@]1(OC(C)=O)CC[C@H]2[C@@H]3CCC4=CC(=O)CC[C@@H]4[C@H]3CC[C@@]21C | IMONTRJLAWHYGT-ZCPXKWAGSA-N | No-Effect | inactive |
| noretynodrel | BRD-A00758722 | C#C[C@]1(O)CC[C@H]2[C@@H]3CCC4=C(CCC(=O)C4)[C@H]3CC[C@@]21C | ICTXHFFSOAJUMG-SLHNCBLASA-N | No-Effect | inactive |
| norfloxacin | BRD-K11196887 | CCn1cc(C(=O)O)c(=O)c2cc(F)c(N3CCNCC3)cc21 | OGJPXUAPXNRGGI-UHFFFAOYSA-N | No-Effect | inactive |
| norgestimate | BRD-K38041788 | C#C[C@]1(OC(C)=O)CC[C@H]2[C@@H]3CCC4=C/C(=N/O)CC[C@@H]4[C@H]3CC[C@@]21CC | KIQQMECNKUGGKA-NMYWJIRASA-N | No-Effect | inactive |
| nornicotine | BRD-K00003718 | c1cncc([C@H]2CCCN2)c1 | MYKUKUCHPMASKF-SECBINFHSA-N | No-Effect | inactive |
| norvancomycin | BRD-K00003706 | CC(C)C[C@@H](N)C(=O)N[C@H]1C(=O)N[C@@H](CC(N)=O)C(=O)N[C@H]2C(=O)N[C@H]3C(=O)N[C@H](C(=O)N[C@H](C(=O)O)c4cc(O)cc(O)c4-c4cc3ccc4O)[C@H](O)c3ccc(c(Cl)c3)Oc3cc2cc(c3O[C@@H]2O[C@H](CO)[C@@H](O)[C@H](O)[C@H]2O[C@H]2C[C@](C)(N)[C@H](O)[C@@H](C)O2)Oc2ccc(cc2Cl)[C@H]1O | NUAQIRUAZSJTAI-YRPZDAAMSA-N | No-Effect | inactive |
| noscapine | BRD-K89237706 | COc1ccc2c(c1OC)C(=O)O[C@@H]2[C@H]1c2c(cc3c(c2OC)OCO3)CCN1C | AKNNEGZIBPJZJG-MSOLQXFVSA-N | No-Effect | inactive |
| NOV-002 | BRD-K11411329 | N[C@@H](CCC(=O)N[C@@H](CSSC[C@H](NC(=O)CC[C@@H](N)C(=O)O)C(=O)NCC(=O)O)C(=O)NCC(=O)O)C(=O)O | YPZRWBKMTBYPTK-USZNOCQGSA-N | No-Effect | inactive |
| novobiocin | BRD-K85307935 | CO[C@@H]1[C@@H](OC(N)=O)[C@@H](O)[C@H](Oc2ccc3c(O)c(NC(=O)c4ccc(O)c(CC=C(C)C)c4)c(=O)oc3c2C)OC1(C)C | YJQPYGGHQPGBLI-KGSXXDOSSA-N | No-Effect | inactive |
| NPC-01 | BRD-K58277159 | C[C@]12C[C@H](O)[C@H]3[C@@H](CCC4=CC(=O)CC[C@@]43C)[C@@H]1CCC2=O | WSCUHXPGYUMQEX-KCZNZURUSA-N | No-Effect | inactive |
| NPC-15199 | BRD-K65275554 | CC(C)C[C@@H](NC(=O)OCC1c2ccccc2-c2ccccc21)C(=O)O | CBPJQFCAFFNICX-LJQANCHMSA-N | No-Effect | inactive |
| NPPB | BRD-K89272762 | O=C(O)c1cc([N+](=O)[O-])ccc1NCCCc1ccccc1 | WBSMIPAMAXNXFS-UHFFFAOYSA-N | No-Effect | inactive |
| NPS-2407 | BRD-K00004686 | CC(C)(Cc1ccccc1)NC(=O)c1cnc2ccccc2n1 | DQEWVIJXZPUPQA-UHFFFAOYSA-N | No-Effect | inactive |
| NPY-5RA972 | BRD-K95600043 | Cc1c(NC(=O)N2CCOCC2)ccc2c1c1ccccc1n2C(C)C | ZNFMHLDBJPAFEQ-UHFFFAOYSA-N | No-Effect | inactive |
| NQDI-1 | BRD-K61918008 | CCOC(=O)c1c2c3c(cccc3[nH]c1=O)C(=O)c1ccccc1-2 | UFJGFNHRMPMALC-UHFFFAOYSA-N | No-Effect | inactive |
| NS-018 | BRD-K21865955 | C[C@H](Nc1cc(-c2cnn(C)c2)cc(Nc2cnccn2)n1)c1ccc(F)cc1 | UQTPDWDAYHAZNT-AWEZNQCLSA-N | No-Effect | inactive |
| NS-11394 | BRD-K00004548 | CC(C)(O)c1ccc2c(c1)ncn2-c1cccc(-c2ccccc2C#N)c1 | HLKYSQGBIIIQJN-UHFFFAOYSA-N | No-Effect | inactive |
| NS-1643 | BRD-K73738482 | O=C(Nc1cc(C(F)(F)F)ccc1O)Nc1cc(C(F)(F)F)ccc1O | NJFVQMRYJZHGME-UHFFFAOYSA-N | No-Effect | inactive |
| NS-19504 | BRD-K03808106 | Nc1ncc(Cc2ccc(Br)cc2)s1 | HGWLTZOMQZIUBW-UHFFFAOYSA-N | No-Effect | inactive |
| NS-309 | BRD-K00004295 | O=C1Nc2c(ccc(Cl)c2Cl)/C1=N/O | CVOUSAVHMDXCKG-UHFFFAOYSA-N | No-Effect | inactive |
| NS-3623 | BRD-K24434292 | O=C(Nc1cccc(C(F)(F)F)c1)Nc1ccc(Br)cc1-c1nn[nH]n1 | JXPULDIATMTIIN-UHFFFAOYSA-N | No-Effect | inactive |
| NS-3861 | BRD-K00004309 | CN1[C@@H]2C=C(c3sccc3Br)C[C@H]1CC2 | ZJCNRVHZJADROC-VHSXEESVSA-N | No-Effect | inactive |
| NS-5806 | BRD-K93314017 | O=C(Nc1cc(C(F)(F)F)cc(C(F)(F)F)c1)Nc1c(Br)cc(Br)cc1-c1nn[nH]n1 | UZWJWROOLOOCPQ-UHFFFAOYSA-N | No-Effect | inactive |
| NS-8 | BRD-K22667752 | Cc1c(-c2ccccc2F)[nH]c(N)c1C#N | KVDWOQGDMAEJLM-UHFFFAOYSA-N | No-Effect | inactive |
| NS-8593 | BRD-K34624869 | c1ccc2c(c1)CCC[C@H]2Nc1nc2ccccc2[nH]1 | XZIZUQSOFMLIIR-CQSZACIVSA-N | No-Effect | inactive |
| NS-9283 | BRD-K40176866 | N#Cc1cccc(-c2nc(-c3cccnc3)no2)c1 | HGFXDSQLRSWUBO-UHFFFAOYSA-N | No-Effect | inactive |
| NSC-23766 | BRD-K01826800 | CCN(CC)CCC[C@@H](C)Nc1nc(C)cc(Nc2ccc3nc(C)cc(N)c3c2)n1 | DEFBCZWQLILOJF-MRXNPFEDSA-N | No-Effect | inactive |
| NSC-3852 | BRD-K13169950 | O=Nc1ccc(O)c2ncccc12 | RZWRYPGAUIOOMK-UHFFFAOYSA-N | No-Effect | inactive |
| NSC-405020 | BRD-A67975844 | CCC[C@@H](C)NC(=O)c1ccc(Cl)c(Cl)c1 | ARDYECYBETXQFD-MRVPVSSYSA-N | No-Effect | inactive |
| NSC-4644 | BRD-K64874225 | C=C(C)[C@@H]1CC[C@]2(CO)CC[C@]3(C)[C@H](CC[C@@H]4[C@@]5(C)CC[C@H](O)C(C)(C)[C@@H]5CC[C@]43C)[C@@H]12 | FVWJYYTZTCVBKE-ROUWMTJPSA-N | No-Effect | inactive |
| NSC-5844 | BRD-K73799155 | Clc1ccc2c(NCCNc3ccnc4cc(Cl)ccc34)ccnc2c1 | SSXYXSMMVMVYEV-UHFFFAOYSA-N | No-Effect | inactive |
| NSC-625987 | BRD-K62949423 | COc1ccc(OC)c2c(=S)c3ccccc3[nH]c12 | KFAKESMKRPNZTM-UHFFFAOYSA-N | No-Effect | inactive |
| NSC-636819 | BRD-K61120504 | O=[N+]([O-])c1cc([N+](=O)[O-])c(/C=C/c2ccc(Cl)c(Cl)c2)cc1/C=C/c1ccc(Cl)c(Cl)c1 | YYZPXBXVBQSBDG-IJIVKGSJSA-N | No-Effect | inactive |
| NSC-663284 | BRD-K03109492 | O=C1C(Cl)=C(NCCN2CCOCC2)C(=O)c2ncccc21 | BMKPVDQDJQWBPD-UHFFFAOYSA-N | No-Effect | inactive |
| NSC-697923 | BRD-K20605374 | Cc1ccc(S(=O)(=O)c2ccc([N+](=O)[O-])o2)cc1 | GAUHIPWCDXOLCZ-UHFFFAOYSA-N | No-Effect | inactive |
| NSC-95397 | BRD-K68143200 | O=C1C(SCCO)=C(SCCO)C(=O)c2ccccc21 | MAASHDQFQDDECQ-UHFFFAOYSA-N | No-Effect | inactive |
| NSC-9965 | BRD-K49811169 | CO[C@H]1CC=C2CCN3CCC4=C(CC(=O)OC4)[C@]23C1 | ALSKYCOJJPXPFS-BBRMVZONSA-N | No-Effect | inactive |
| NSI-189 | BRD-K74371986 | CC(C)CCNc1ncccc1C(=O)N1CCN(Cc2ccccc2)CC1 | DYTOQURYRYYNOR-UHFFFAOYSA-N | No-Effect | inactive |
| NSP-805 | BRD-K00004733 | CC1=C(c2ccc(NC3=C(C)C(=O)CC3)cc2)NNC(=O)C1 | VSEWBFSTIMFCDL-UHFFFAOYSA-N | No-Effect | inactive |
| NT157 | BRD-K54098378 | Oc1cc(CNC(=S)/C=C/c2cc(O)c(O)c(Br)c2)cc(O)c1O | NIPUPOUEGOSAAO-OWOJBTEDSA-N | No-Effect | inactive |
| NTRC-824 | BRD-K07788774 | Cc1ccc(S(=O)(=O)Nc2ccc3c(c2)c(C(=O)C(F)(F)F)cn3CC(=O)N[C@@H](CC(C)C)C(=O)O)cc1 | SOOLLEBJNGCDOZ-FQEVSTJZSA-N | No-Effect | inactive |
| nTZDpa | BRD-K54708045 | O=C(O)c1c(Sc2ccccc2)c2cc(Cl)ccc2n1Cc1ccc(Cl)cc1 | VUPOTURDKDMIGQ-UHFFFAOYSA-N | No-Effect | inactive |
| NU-1025 | BRD-K48692744 | Cc1nc(=O)c2cccc(O)c2[nH]1 | YJDAOHJWLUNFLX-UHFFFAOYSA-N | No-Effect | inactive |
| NU-2058 | BRD-K20976221 | Nc1nc(OCC2CCCCC2)c2[nH]cnc2n1 | MWGXGTJJAOZBNW-UHFFFAOYSA-N | No-Effect | inactive |
| NU-7026 | BRD-K09537769 | O=c1cc(N2CCOCC2)oc2c1ccc1ccccc12 | KKTZALUTXUZPSN-UHFFFAOYSA-N | No-Effect | inactive |
| NU-7441 | BRD-K00337317 | O=c1cc(N2CCOCC2)oc2c(-c3cccc4c3sc3ccccc34)cccc12 | JAMULYFATHSZJM-UHFFFAOYSA-N | No-Effect | inactive |
| nuclomedone | BRD-A96466199 | O=C1N=C2SCCN2C(=O)[C@H]1Cc1ccc(Cl)cc1 | QSKVYHYMQQQAFS-JTQLQIEISA-N | No-Effect | inactive |
| nutlin-3 | BRD-K00002882 | COc1ccc(C2=N[C@@H](c3ccc(Cl)cc3)[C@@H](c3ccc(Cl)cc3)N2C(=O)N2CCNC(=O)C2)c(OC(C)C)c1 | BDUHCSBCVGXTJM-WUFINQPMSA-N | No-Effect | inactive |
| nuvenzepine | BRD-K00004743 | CN1CCC(C(=O)N2c3ccccc3NC(=O)c3cccnc32)CC1 | HPKYRXAEGNUARA-UHFFFAOYSA-N | No-Effect | inactive |
| NVP-231 | BRD-K01573278 | O=C(Nc1nc2ccc(NC(=O)C34CC5CC(CC(C5)C3)C4)cc2s1)c1ccccc1 | MVSSJPGNLQPWSW-UHFFFAOYSA-N | No-Effect | inactive |
| NVP-BGJ398 | BRD-K42728290 | CCN1CCN(c2ccc(Nc3cc(N(C)C(=O)Nc4c(Cl)c(OC)cc(OC)c4Cl)ncn3)cc2)CC1 | QADPYRIHXKWUSV-UHFFFAOYSA-N | No-Effect | inactive |
| NVP-BSK805 | BRD-K02017404 | Fc1cc(-c2cccc3ncc(-c4cnn(C5CCNCC5)c4)nc23)cc(F)c1CN1CCOCC1 | IBPVXAOOVUAOKJ-UHFFFAOYSA-N | No-Effect | inactive |
| NVP-BVU972 | BRD-K77943374 | Cn1cc(-c2ccc3ncc(Cc4ccc5ncccc5c4)n3n2)cn1 | RNCNPRCUHHDYPC-UHFFFAOYSA-N | No-Effect | inactive |
| NVP-DPP728 | BRD-K77396579 | N#Cc1ccc(NCCNCC(=O)N2CCC[C@H]2C#N)nc1 | VFFZWMWTUSXDCB-ZDUSSCGKSA-N | No-Effect | inactive |
| NVP-TAE226 | BRD-K21025364 | CNC(=O)c1ccccc1Nc1nc(Nc2ccc(N3CCOCC3)cc2OC)ncc1Cl | UYJNQQDJUOUFQJ-UHFFFAOYSA-N | No-Effect | inactive |
| NVP-TAE684 | BRD-K50140147 | COc1cc(N2CCC(N3CCN(C)CC3)CC2)ccc1Nc1ncc(Cl)c(Nc2ccccc2S(=O)(=O)C(C)C)n1 | QQWUGDVOUVUTOY-UHFFFAOYSA-N | No-Effect | inactive |
| NVP-TNKS656 | BRD-K87454346 | COc1ccc(C(=O)C2CCN(CC(=O)N(Cc3nc(=O)c4c([nH]3)CCOC4)CC3CC3)CC2)cc1 | DYGBNAYFDZEYBA-UHFFFAOYSA-N | No-Effect | inactive |
| NVS-PAK1-1 | BRD-K28132190 | CC(C)NC(=O)N1CC[C@H](NC2=Nc3cc(F)ccc3N(CC(F)F)c3ccc(Cl)cc32)C1 | OINGHOPGNMYCAB-INIZCTEOSA-N | No-Effect | inactive |
| NXY-059 | BRD-K46133855 | CC(C)(C)/[N+]([O-])=C/c1ccc(S(=O)(=O)O)cc1S(=O)(=O)O | OVTCHWSLKGENKP-GHXNOFRVSA-N | No-Effect | inactive |
| O-1918 | BRD-K18316707 | C=C(C)[C@@H]1CCC(C)=C[C@H]1c1c(OC)cc(C)cc1OC | ICHJMVMWPKLUKT-JKSUJKDBSA-N | No-Effect | inactive |
| O-304 | BRD-K00003113 | O=C(Nc1nc(=O)n(Cc2ccc(Cl)cc2)s1)c1ccc(Cl)cc1 | WEDWLYRQKUTOAX-UHFFFAOYSA-N | No-Effect | inactive |

| | | | | | |
|---|---|---|---|---|---|
| o-3M3FBS | BRD-K46384212 | Cc1cc(C)c(S(=O)(=O)Nc2ccccc2C(F)(F)F)c(C)c1 | SKJJIFRWCCSXGL-UHFFFAOYSA-N | No-Effect | inactive |
| O-acetyl-L-serine | BRD-K63247785 | CC(=O)OC[C@H](N)C(=O)O | VZXPDPZARILFQX-BYPYZUCNSA-N | No-Effect | inactive |
| o-mercapto-benzoic-acid | BRD-K36998189 | O=C(O)c1ccccc1S | NBOMNTLFRHMDEZ-UHFFFAOYSA-N | No-Effect | inactive |
| O4I1 | BRD-K53849769 | COc1ccc(COc2ccc(CC#N)cc2)cc1 | QJEJLARVLKHYFW-UHFFFAOYSA-N | No-Effect | inactive |
| O6-benzylguanine | BRD-K92041145 | Nc1nc(OCc2ccccc2)c2[nH]cnc2n1 | KRWMERLEINMZFT-UHFFFAOYSA-N | No-Effect | inactive |
| OAC1 | BRD-K10878147 | O=C(Nc1cc2cc[nH]c2cn1)c1ccccc1 | HWJRIFZDXJKJJN-UHFFFAOYSA-N | No-Effect | inactive |
| OAC2 | BRD-K18305454 | O=C(Nc1ccc2[nH]ccc2c1)c1ccccc1 | JCAFGYWSIWYMOX-UHFFFAOYSA-N | No-Effect | inactive |
| obatoclax | BRD-K15600710 | COC1=CC(c2cc3ccccc3[nH]2)=N/C1=C\c1[nH]c(C)cc1C | RFTSSZJZXOSICM-GRSHGNNSSA-N | Strong | active |
| obeticholic-acid | BRD-A75838038 | CC[C@@H]1[C@@H]2C[C@H](O)CC[C@]2(C)[C@H]2CC[C@]3(C)[C@H]([C@@H](C)CCC(=O)O)CC[C@@H]3[C@@H]2[C@@H]1O | ZXERDUOLZKYMJM-XNNZWSTBSA-N | No-Effect | inactive |
| obidoxime | BRD-K33588785 | O/N=C/c1cc[n+](COC[n+]2ccc(/C=N/O)cc2)cc1 | HIGRLDNHDGYWQJ-UHFFFAOYSA-P | No-Effect | inactive |
| ocaperidone | BRD-K00003399 | Cc1nc2c(C)cccn2c(=O)c1CCN1CCC(c2noc3cc(F)ccc23)CC1 | ZZQNEJILGNNOEP-UHFFFAOYSA-N | No-Effect | inactive |
| ochromycinone | BRD-K92524032 | C[C@@H]1CC(=O)c2c(ccc3c2C(=O)c2cccc(O)c2C3=O)C1 | ZAWXOCUFQSQDJS-VIFPVBQESA-N | No-Effect | inactive |
| ocinaplon | BRD-K92581468 | O=C(c1ccccn1)c1cnn2c(-c3ccncc3)ccnc12 | OQJFBUOFGHPMSR-UHFFFAOYSA-N | No-Effect | inactive |
| OCO-1112 | BRD-K22097830 | CC(C)c1ccc(C(=O)NCCOc2ccc(C(=O)O)cc2)cc1 | ODLDHXPJRGNTPC-UHFFFAOYSA-N | No-Effect | inactive |
| octadecan-1-ol | BRD-K20501889 | CCCCCCCCCCCCCCCCCCO | GLDOVTGHNKAZLK-UHFFFAOYSA-N | No-Effect | inactive |
| octamethyltrisiloxane | BRD-K73201113 | C[Si](C)(C)O[Si](C)(C)O[Si](C)(C)C | CXQXSVUQTKDNFP-UHFFFAOYSA-N | No-Effect | inactive |
| octenidine | BRD-K04568635 | CCCCCCCCN=c1ccn(CCCCCCCCCCn2ccc(=NCCCCCCCC)cc2)cc1 | ZVXNYZWXUADSRV-UHFFFAOYSA-N | No-Effect | inactive |
| octinoxate | BRD-K00003688 | CCCC[C@@H](CC)COC(=O)/C=C/c1ccc(OC)cc1 | YBGZDTIWKVFICR-NRMKIYEFSA-N | No-Effect | inactive |
| octisalate | BRD-K00003664 | CCCC[C@@H](CC)COC(=O)c1ccccc1O | FMRHJJZUHUTGKE-GFCCVEGCSA-N | No-Effect | inactive |
| octodrine | BRD-A37873789 | CC(C)CCC[C@@H](C)N | QNIVIMYXGGFTAK-MRVPVSSYSA-N | No-Effect | inactive |
| octopamine | BRD-A14240607 | NCC(O)c1ccc(O)cc1 | QHGUCRYDKWKLMG-UHFFFAOYSA-N | No-Effect | inactive |
| octreotide | BRD-A64173453 | C[C@@H](O)[C@@H]1NC(=O)[C@H](CCCCN)NC(=O)[C@H](Cc2c[nH]c3ccccc23)NC(=O)[C@H](Cc2ccccc2)NC(=O)[C@H](NC(=O)[C@@H](N)Cc2ccccc2)CSSC[C@H](C(=O)N[C@H](CO)[C@@H](C)O)NC1=O | DEQANNDTNATYII-XZNMVFOCSA-N | No-Effect | inactive |
| odanacatib | BRD-K22822991 | CC(C)(F)C[C@H](N[C@@H](c1ccc(-c2ccc(S(C)(=O)=O)cc2)cc1)C(F)(F)F)C(=O)NC1(C#N)CC1 | FWIVDMJALNEADT-SFTDATJTSA-N | No-Effect | inactive |
| ODQ | BRD-K26015241 | O=c1onc2cnc3ccccc3n12 | LZMHWZHOZLVYDL-UHFFFAOYSA-N | No-Effect | inactive |
| OF-1 | BRD-K92658264 | COc1cc2c(cc1NS(=O)(=O)c1ccc(Br)cc1C)n(C)c(=O)n2C | YUNQZQREIHWDQT-UHFFFAOYSA-N | No-Effect | inactive |
| ofloxacin | BRD-K01826479 | C[C@H]1COc2c(N3CCN(C)CC3)c(F)cc3c(=O)c(C(=O)O)cn1c23 | GSDSWSVVBLHKDQ-JTQLQIEISA-N | No-Effect | inactive |
| ogerin | BRD-K00005341 | Nc1nc(NCc2ccccc2)nc(-c2ccccc2CO)n1 | MDGIEDNDSFMSLP-UHFFFAOYSA-N | No-Effect | inactive |
| oglufanide | BRD-K00003164 | N[C@@H](CCC(=O)O)C(=O)N[C@@H](Cc1c[nH]c2ccccc12)C(=O)O | LLEUXCDZPQOJMY-AAEUAGOBSA-N | No-Effect | inactive |
| OICR-9429 | BRD-K94083718 | CN1CCN(c2ccc(-c3cccc(CN4CCOCC4)c3)cc2NC(=O)c2c[nH]c(=O)cc2C(F)(F)F)CC1 | DJOVLOYCGXNVPI-UHFFFAOYSA-N | No-Effect | inactive |
| olanzapine | BRD-K18895904 | Cc1cc2c(s1)Nc1ccccc1N=C2N1CCN(C)CC1 | KVWDHTXUZHCGIO-UHFFFAOYSA-N | No-Effect | inactive |
| OLDA | BRD-K95523387 | CCCCCCCC/C=C\CCCCCCCC(=O)NCCc1ccc(O)c(O)c1 | QQBPLXNESPTPNU-KTKRTIGZSA-N | No-Effect | inactive |
| oleamide | BRD-K06644925 | CCCCCCCC/C=C\CCCCCCCC(N)=O | FATBGEAMYMYZAF-KTKRTIGZSA-N | No-Effect | inactive |
| oleanolic-acid | BRD-K90647725 | CC1(C)CC[C@]2(C(=O)O)CC[C@]3(C)C(=CC[C@@H]4[C@@]5(C)CC[C@H](O)C(C)(C)[C@@H]5CC[C@]43C)[C@@H]2C1 | MIJYXULNPSFWEK-GTOFXWBISA-N | No-Effect | inactive |
| oleoylethanolamide | BRD-K66956375 | CCCCCCCC/C=C\CCCCCCCC(=O)NCCO | BOWVQLFMWHZBEF-KTKRTIGZSA-N | No-Effect | inactive |
| olesoxime | BRD-K18210059 | CC(C)CCC[C@@H](C)[C@H]1CC[C@H]2[C@@H]3CCC4=C/C(=N/O)CC[C@]4(C)[C@H]3CC[C@@]21C | QNTASHOAVRSLMD-SIWSWZRQSA-N | No-Effect | inactive |
| oleuropein | BRD-A00842753 | C/C=C1\[C@H](O[C@H]2O[C@@H](CO)[C@@H](O)[C@@H](O)[C@@H]2O)OC=C(C(=O)OC)[C@H]1CC(=O)OCCc1ccc(O)c(O)c1 | RFWGABANNQMHMZ-JIBQERFGSA-N | No-Effect | inactive |
| oligomycin-A | BRD-K00003580 | CC[C@H]1/C=C/C=C/C[C@@H](C)[C@H](O)[C@@](C)(O)C(=O)[C@@H](C)[C@H](O)[C@@H](C)C(=O)[C@@H](C)[C@H](O)[C@@H](C)/C=C/C(=O)O[C@H]2[C@H](C)[C@@]3(CC[C@H](C)[C@H](C[C@@H](C)O)O3)O[C@@H](CC1)[C@H]2C | MNULEGDCPYONBU-AWJDAWNUSA-N | No-Effect | inactive |
| olmesartan | BRD-K83144676 | CCCc1nc(C(C)(C)O)c(C(=O)O)n1Cc1ccc(-c2ccccc2-c2nn[nH]n2)cc1 | VTRAEEWXHOVJFV-UHFFFAOYSA-N | No-Effect | inactive |
| olmesartan-medoxomil | BRD-K78485176 | CCCc1nc(C(C)(C)O)c(C(=O)OCc2oc(=O)oc2C)n1Cc1ccc(-c2ccccc2-c2nn[nH]n2)cc1 | UQGKUQLKSCSZGY-UHFFFAOYSA-N | No-Effect | inactive |
| olodanrigan | BRD-K00003151 | COc1ccc2c(c1OCc1ccccc1)C[C@@H](C(=O)O)N(C(=O)C(c1ccccc1)c1ccccc1)C2 | GHBCIXGRCZIPNQ-MHZLTWQESA-N | No-Effect | inactive |
| olomoucine | BRD-K82731415 | Cn1cnc2c(NCc3ccccc3)nc(NCCO)nc21 | GTVPOLSIJWJJNY-UHFFFAOYSA-N | No-Effect | inactive |
| olopatadine | BRD-K67637637 | CN(C)CC/C=C1/c2ccccc2COc2ccc(CC(=O)O)cc21 | JBIMVDZLSHOPLA-LSCVHKIXSA-N | No-Effect | inactive |
| olprinone | BRD-K20745393 | Cc1[nH]c(=O)c(C#N)cc1-c1ccc2nccn2c1 | JPAWFIIYTJQOKW-UHFFFAOYSA-N | No-Effect | inactive |
| olsalazine | BRD-K57853941 | O=C(O)c1cc(/N=N/c2ccc(O)c(C(=O)O)c2)ccc1O | QQBDLJCYGRGAKP-FOCLMDBBSA-N | No-Effect | inactive |
| oltipraz | BRD-K86972824 | Cc1c(-c2cnccn2)ssc1=S | CKNAQFVBEHDJQV-UHFFFAOYSA-N | No-Effect | inactive |
| olvanil | BRD-K31560319 | CCCCCCCC/C=C\CCCCCCCC(=O)NCc1ccc(O)c(OC)c1 | OPZKBPQVWDSATI-KHPPLWFESA-N | No-Effect | inactive |
| omadacycline | BRD-K00003569 | CN(C)c1cc(CNCC(C)(C)C)c(O)c2c1C[C@H]1C[C@H]3[C@H](N(C)C)C(=O)[C@H](C(N)=O)C(=O)[C@@]3(O)C(=O)[C@@H]1C2=O | OVIWLBQFYMOVOF-NXSZOOMBSA-N | No-Effect | inactive |
| omapatrilat | BRD-K00003265 | O=C(N[C@H]1CCS[C@H]2CCC[C@@H](C(=O)O)N2C1=O)[C@@H](S)Cc1ccccc1 | LVRLSYPNFFBYCZ-VGWMRTNUSA-N | No-Effect | inactive |
| ombitasvir | BRD-K11072003 | COC(=O)N[C@H](C(=O)N1CCC[C@H]1C(=O)Nc1ccc([C@@H]2CC[C@@H](c3ccc(NC(=O)[C@@H]4CCCN4C(=O)[C@@H](NC(=O)OC)C(C)C)cc3)N2c2ccc(C(C)(C)C)cc2)cc1)C(C)C | PIDFDZJZLOTZTM-KHVQSSSXSA-N | No-Effect | inactive |
| OMDM-2 | BRD-K48830578 | CCCCCCCC/C=C\CCCCCCCC(=O)N[C@@H](CO)Cc1ccc(O)cc1 | ICDMLAQPOAVWNH-HAAQQRBASA-N | No-Effect | inactive |
| omecamtiv-mecarbil | BRD-K36680103 | COC(=O)N1CCN(Cc2cccc(NC(=O)Nc3ccc(C)nc3)c2F)CC1 | RFUBTTPMWSKEIW-UHFFFAOYSA-N | No-Effect | inactive |
| omega-(4-iodophenyl)pentadecanoic-acid | BRD-K53156781 | O=C(O)CCCCCCCCCCCCCCc1ccc(I)cc1 | YAONEUNUMVOKNQ-UHFFFAOYSA-N | No-Effect | inactive |
| omega-3-acid-esters | BRD-K31139048 | CC/C=C\C/C=C\C/C=C\C/C=C\C/C=C\CCCC(=O)OCC | SSQPWTVBQMWLSZ-AAQCHOMXSA-N | No-Effect | inactive |
| omeprazole | BRD-A55962179 | COc1ccc2nc([S@@](=O)Cc3ncc(C)c(OC)c3C)[nH]c2c1 | SUBDBMMJDZJVOS-DEOSSOPVSA-N | No-Effect | inactive |
| omeprazole-magnesium | BRD-A56892734 | COc1ccc2c(c1)nc([S@@](=O)Cc1ncc(C)c(OC)c1C)n2[Mg]n1c([S@@](=O)Cc2ncc(C)c(OC)c2C)nc2cc(OC)ccc21 | KWORUUGOSLYAGD-YPPDDXJESA-N | No-Effect | inactive |
| omeprazole-sulfide | BRD-K69035426 | COc1ccc2nc(SCc3ncc(C)c(OC)c3C)[nH]c2c1 | XURCIPRUUASYLR-UHFFFAOYSA-N | No-Effect | inactive |
| ON-01500 | BRD-K55790047 | COc1cc(OC)c(/C=C/S(=O)(=O)Cc2ccc(OC)c(N)c2)c(OC)c1 | KUTAVMCJCVADGZ-BQYQJAHWSA-N | No-Effect | inactive |
| ON123300 | BRD-K88554883 | CN1CCN(c2ccc(Nc3ncc4cc(C#N)c(=O)n(C5CCCC5)c4n3)cc2)CC1 | VADOZMZXXRBXNY-UHFFFAOYSA-N | No-Effect | inactive |
| ONC201 | BRD-K58529924 | Cc1ccccc1CN1C(=O)C2=C(CCN(Cc3ccccc3)C2)N2CCN=C12 | VLULRUCCHYVXOH-UHFFFAOYSA-N | No-Effect | inactive |
| oncrasin-1 | BRD-K43449605 | O=Cc1cn(Cc2ccc(Cl)cc2)c2ccccc12 | ZDRQMXCSSAPUMM-UHFFFAOYSA-N | No-Effect | inactive |
| ondansetron | BRD-K01826546 | Cc1nccn1C[C@H]1CCc2c(c3ccccc3n2C)C1=O | FELGMEQIXOGIFQ-CYBMUJFWSA-N | No-Effect | inactive |
| ONO-4059 | BRD-K00003197 | CC#CC(=O)N1CC[C@@H](n2c(=O)n(-c3ccc(Oc4ccccc4)cc3)c3c(N)ncnc32)C1 | SEJLPXCPMNSRAM-GOSISDBHSA-N | No-Effect | inactive |
| ONO-4817 | BRD-K58894520 | CCOCOC[C@H](C[C@H](C)C(=O)NO)NC(=O)c1ccc(Oc2ccccc2)cc1 | HDWWQELUBWGQGA-WMZOPIPTSA-N | No-Effect | inactive |
| ONO-8130 | BRD-K76725781 | Cc1csc(S(=O)(=O)N(CC(C)C)c2cc3c(cc2OCc2ccc(C(=O)O)cc2)CCC3)n1 | IOXMDBQCUCDLAF-UHFFFAOYSA-N | No-Effect | inactive |

| | | | | | |
|---|---|---|---|---|---|
| ONO-AE3-208 | BRD-A80567330 | C[C@@H](C(=O)Nc1cc(C#N)ccc1CCCC(=O)O)c1ccc(F)c2ccccc12 | MTDIMKNAJUQTIO-OAHLLOKOSA-N | No-Effect | inactive |
| ONX-0914 | BRD-K00003557 | COc1ccc(C[C@H](NC(=O)[C@H](C)NC(=O)CN2CCOCC2)C(=O)N[C@@H](Cc2ccccc2)C(=O)[C@@]2(C)CO2)cc1 | WQAVPPWWLLVGFK-VTNASVEKSA-N | No-Effect | inactive |
| OP-0595 | BRD-K00004708 | NCCONC(=O)[C@@H]1CC[C@@H]2CN1C(=O)N2OS(=O)(=O)O | RSBPYSTVZQAADE-RQJHMYQMSA-N | No-Effect | inactive |
| OPC-21268 | BRD-K19426040 | CC(=O)NCCCOc1ccc(C(=O)N2CCC(N3C(=O)CCc4ccccc43)CC2)cc1 | KSNUCNRMDYJBKT-UHFFFAOYSA-N | No-Effect | inactive |
| opicapone | BRD-K66192203 | Cc1c(Cl)c(C)[n+]([O-])c(Cl)c1-c1noc(-c2cc(O)c(O)c([N+](=O)[O-])c2)n1 | ASOADIZOVZTJSR-UHFFFAOYSA-N | No-Effect | inactive |
| opipramol | BRD-K43786866 | OCCN1CCN(CCCN2c3ccccc3C=Cc3ccccc32)CC1 | YNZFUWZUGRBMHL-UHFFFAOYSA-N | No-Effect | inactive |
| OR-486 | BRD-K18602506 | O=[N+]([O-])c1cc(O)c(O)c([N+](=O)[O-])c1 | VDCDWNDTNSWDFJ-UHFFFAOYSA-N | No-Effect | inactive |
| orantinib | BRD-K55966568 | Cc1[nH]c(/C=C2\C(=O)Nc3ccccc32)c(C)c1CCC(=O)O | NHFDRBXTEDBWCZ-ZROIWOOFSA-N | No-Effect | inactive |
| orbifloxacin | BRD-K63001556 | C[C@H]1CN(c2c(F)c(F)c3c(=O)c(C(=O)O)cn(C4CC4)c3c2F)C[C@@H](C)N1 | QIPQASLPWJVQMH-DTORHVGOSA-N | No-Effect | inactive |
| ORE1001 | BRD-K67068943 | CC(C)C[C@H](N[C@@H](Cc1cncn1Cc1cc(Cl)cc(Cl)c1)C(=O)O)C(=O)O | NTCCRGGIJNDEAB-IRXDYDNUSA-N | No-Effect | inactive |
| ORG-12962 | BRD-K07409305 | FC(F)(F)c1ccc(N2CCNCC2)nc1Cl | QZYYPQAYSFBKPW-UHFFFAOYSA-N | No-Effect | inactive |
| ORG-25543 | BRD-K32853922 | COc1cc(C(=O)NCC2(N(C)C)CCCC2)cc(OC)c1OCc1ccccc1 | ZCYOBMASPXQBSZ-UHFFFAOYSA-N | No-Effect | inactive |
| ORG-26576 | BRD-K40002998 | O=C1c2cccnc2OC[C@@H]2CCCN12 | FIKUEZUFASUKAH-QMMMGPOBSA-N | No-Effect | inactive |
| ORG-27569 | BRD-K57597776 | CCc1c(C(=O)NCCc2ccc(N3CCCCC3)cc2)[nH]c2ccc(Cl)cc12 | AHFZDNYNXFMRFQ-UHFFFAOYSA-N | No-Effect | inactive |
| ORG-9768 | BRD-K41434598 | CC1(CN)Cc2ccccc2C1 | JOLKQWPKJFRPOU-UHFFFAOYSA-N | No-Effect | inactive |
| orlistat | BRD-K63343048 | CCCCCCCCCCC[C@@H](C[C@@H]1OC(=O)[C@H]1CCCCCC)OC(=O)[C@H](CC(C)C)NC=O | AHLBNYSZXLDEJQ-FWEHEUNISA-N | No-Effect | inactive |
| ornidazole | BRD-A42759514 | Cc1ncc([N+](=O)[O-])n1C[C@@H](O)CCl | IPWKIXLWTCNBKN-LURJTMIESA-N | No-Effect | inactive |
| ornipressin | BRD-K00003333 | NCCC[C@H](NC(=O)[C@@H]1CCCN1C(=O)[C@@H]1CSSC[C@H](N)C(=O)N[C@@H](Cc2ccc(O)cc2)C(=O)N[C@@H](Cc2ccccc2)C(=O)N[C@@H](CCC(N)=O)C(=O)N[C@@H](CC(N)=O)C(=O)N1)C(=O)NCC(N)=O | MUNMIGOEDGHVLE-LGYYRGKSSA-N | No-Effect | inactive |
| ornithine | BRD-K01825132 | Cl.NCCC[C@@H](N)C(=O)O | GGTYBZJRPHEQDG-PGMHMLKASA-N | No-Effect | inactive |
| orotic-acid | BRD-K32572029 | O=C(O)c1cc(=O)[nH]c(=O)[nH]1 | PXQPEWDEAKTCGB-UHFFFAOYSA-N | No-Effect | inactive |
| orphanin-fq | BRD-K05717154 | CC(C)C[C@H](NC(=O)[C@H](CCCCN)NC(=O)[C@H](CCCNC(=N)N)NC(=O)[C@H](C)NC(=O)[C@H](CO)NC(=O)[C@H](CCCCN)NC(=O)[C@H](CCCNC(=N)N)NC(=O)[C@H](C)NC(=O)CNC(=O)[C@@H](NC(=O)[C@H](Cc1ccccc1)NC(=O)CNC(=O)CNC(=O)[C@@H](N)Cc1ccccc1)[C@@H](C)O)C(=O)N[C@@H](C)C(=O)N[C@@H](CC(N)=O)C(=O)N[C@@H](CCC(N)=O)C(=O)O | PULGYDLMFSFVBL-SMFNREODSA-N | No-Effect | inactive |
| orphenadrine | BRD-A53576514 | Cc1ccccc1[C@@H](OCCN(C)C)c1ccccc1 | QVYRGXJJSLMXQH-SFHVURJKSA-N | No-Effect | inactive |
| orteronel | BRD-K00674894 | CNC(=O)c1ccc2cc([C@]3(O)CCn4cncc43)ccc2c1 | OZPFIJIOIVJZMN-GOSISDBHSA-N | No-Effect | inactive |
| ORY-1001 | BRD-K59711701 | NC1CCC(N[C@@H]2C[C@H]2c2ccccc2)CC1 | ALHBJBCQLJZYON-PFSRBDOWSA-N | No-Effect | inactive |
| oseltamivir-carboxylate | BRD-K93767137 | CCC(CC)O[C@@H]1C=C(C(=O)O)C[C@H](N)[C@H]1NC(C)=O | NENPYTRHICXVCS-YNEHKIRRSA-N | No-Effect | inactive |
| oseltamivir-phosphate | BRD-K76011241 | CCOC(=O)C1=C[C@@H](OC(CC)CC)[C@H](NC(C)=O)[C@@H](N)C1 | VSZGPKBBMSAYNT-RRFJBIMHSA-N | No-Effect | inactive |
| osemozotan | BRD-K60371362 | c1ccc2c(c1)OC[C@H](CNCCCOc1ccc3c(c1)OCO3)O2 | MEEQBDCQPIZMLY-HNNXBMFYSA-N | No-Effect | inactive |
| OSI-420 | BRD-K73309154 | C#Cc1cccc(Nc2ncnc3cc(OCCOC)c(OCCO)cc23)c1 | KOQIAZNBAWFSQM-UHFFFAOYSA-N | No-Effect | inactive |
| OSI-930 | BRD-K25340465 | O=C(Nc1ccc(OC(F)(F)F)cc1)c1sccc1NCc1ccnc2ccccc12 | FGTCROZDHDSNIO-UHFFFAOYSA-N | No-Effect | inactive |
| osilodrostat | BRD-K00003210 | N#Cc1ccc([C@H]2CCc3cncn32)c(F)c1 | USUZGMWDZDXMDG-CYBMUJFWSA-N | No-Effect | inactive |
| ospemifene | BRD-K44184170 | OCCOc1ccc(/C(=C(/CCCl)c2ccccc2)c2ccccc2)cc1 | LUMKNAVTFCDUIE-VHXPQNKSSA-N | No-Effect | inactive |
| ostarine | BRD-K29735307 | C[C@](O)(COc1ccc(C#N)cc1)C(=O)Nc1ccc(C#N)c(C(F)(F)F)c1 | JNGVJMBLXIUVRD-SFHVURJKSA-N | No-Effect | inactive |
| osthol | BRD-K78294846 | COc1ccc2ccc(=O)oc2c1CC=C(C)C | MBRLOUHOWLUMFF-UHFFFAOYSA-N | No-Effect | inactive |
| OT-R-antagonist-1 | BRD-K42009284 | CO/N=C1/C[C@@H](C(=O)NC[C@@H](O)c2ccccc2)N(C(=O)c2ccc(-c3ccccc3C)cc2)C1 | IBXGJPAYWMFXSF-UEEONYLUSA-N | No-Effect | inactive |
| otamixaban | BRD-K80082640 | COC(=O)[C@H](Cc1cccc(C(=N)N)c1)[C@@H](C)NC(=O)c1ccc(-c2cc[n+]([O-])cc2)cc1 | PFGVNLZDWRZPJW-OPAMFIHVSA-N | No-Effect | inactive |
| otenzepad | BRD-A00520476 | CCN(CC)C[C@H]1CCCCN1CC(=O)N1c2ccccc2C(=O)Nc2cccnc21 | UBRKDAVQCKZSPO-GOSISDBHSA-N | No-Effect | inactive |
| otilonium | BRD-K85337334 | CCCCCCCCOc1ccccc1C(=O)Nc1ccc(C(=O)OCC[N+](C)(CC)CC)cc1 | NQHNLNLJPDMBFN-UHFFFAOYSA-O | No-Effect | inactive |
| OXA-06 | BRD-K98736329 | Fc1ccccc1CNCc1ccc(-c2ccnc3[nH]ccc23)cc1 | XUIUXXQVHPEQNM-UHFFFAOYSA-N | No-Effect | inactive |
| oxaceprol | BRD-K65230684 | CC(=O)N1C[C@H](O)C[C@H]1C(=O)O | BAPRUDZDYCKSOQ-RITPCOANSA-N | No-Effect | inactive |
| oxacillin | BRD-K96786677 | Cc1onc(-c2ccccc2)c1C(=O)N[C@@H]1C(=O)N2[C@@H](C(=O)O)C(C)(C)S[C@H]12 | UWYHMGVUTGAWSP-JKIFEVAISA-N | No-Effect | inactive |
| oxacyclohexadecan-2-one | BRD-K36982194 | O=C1CCCCCCCCCCCCCCO1 | FKUPPRZPSYCDRS-UHFFFAOYSA-N | No-Effect | inactive |
| oxaloacetate | BRD-K61279411 | O=C(O)C=C(O)C(=O)O | UWYVPFMHMJIBHE-UHFFFAOYSA-N | No-Effect | inactive |
| oxantel | BRD-K66019333 | CN1CCCN=C1/C=C/c1cccc(O)c1 | VRYKTHBAWRESFI-VOTSOKGWSA-N | No-Effect | inactive |
| oxaprozin | BRD-K25394294 | O=C(O)CCc1nc(-c2ccccc2)c(-c2ccccc2)o1 | OFPXSFXSNFPTHF-UHFFFAOYSA-N | No-Effect | inactive |
| oxatomide | BRD-K22556255 | O=c1[nH]c2ccccc2n1CCCN1CCN(C(c2ccccc2)c2ccccc2)CC1 | BAINIUMDFURPJM-UHFFFAOYSA-N | No-Effect | inactive |
| oxcarbazepine | BRD-K04196797 | NC(=O)N1c2ccccc2CC(=O)c2ccccc21 | CTRLABGOLIVAIY-UHFFFAOYSA-N | No-Effect | inactive |
| oxethazaine | BRD-K56940463 | CN(C(=O)CN(CCO)CC(=O)N(C)C(C)(C)Cc1ccccc1)C(C)(C)Cc1ccccc1 | FTLDJPRFCGDUFH-UHFFFAOYSA-N | No-Effect | inactive |
| oxfendazole | BRD-A33447119 | COC(=O)Nc1nc2ccc([S@@](=O)c3ccccc3)cc2[nH]1 | BEZZFPOZAYTVHN-QFIPXVFZSA-N | No-Effect | inactive |
| oxfenicine | BRD-K30373577 | N[C@@H](C(=O)O)c1ccc(O)cc1 | LJCWONGJFPCTTL-SSDOTTSWSA-N | No-Effect | inactive |
| oxibendazole | BRD-K52075715 | CCCOc1ccc2nc(NC(=O)OC)[nH]c2c1 | RAOCRURYZCVHMG-UHFFFAOYSA-N | No-Effect | inactive |
| oxiconazole | BRD-K56104152 | Clc1ccc(CO/N=C(\Cn2ccnc2)c2ccc(Cl)cc2Cl)c(Cl)c1 | QRJJEGAJXVEBNE-HKOYGPOVSA-N | No-Effect | inactive |
| oxidopamine | BRD-K41809988 | NCCc1cc(O)c(O)cc1O | DIVDFFZHCJEHGG-UHFFFAOYSA-N | No-Effect | inactive |
| oxiniacic-acid | BRD-K20417253 | O=C(O)c1ccc[n+]([O-])c1 | FJCFFCXMEXZEIM-UHFFFAOYSA-N | No-Effect | inactive |
| oxiperomide | BRD-K78096648 | O=c1[nH]c2ccccc2n1C1CCN(CCOc2ccccc2)CC1 | NVDBBGBUTKLRSN-UHFFFAOYSA-N | No-Effect | inactive |
| oxiracetam | BRD-K50677762 | NC(=O)CN1C[C@H](O)CC1=O | IHLAQQPQKRMGSS-SCSAIBSYSA-N | No-Effect | inactive |
| oxolamine | BRD-K42596464 | CCN(CC)CCc1nc(-c2ccccc2)no1 | IDCHQQSVJAAUQQ-UHFFFAOYSA-N | No-Effect | inactive |
| oxolinic-acid | BRD-K73394555 | CCn1cc(C(=O)O)c(=O)c2cc3c(cc21)OCO3 | KYGZCKSPAKDVKC-UHFFFAOYSA-N | No-Effect | inactive |
| oxonic-acid | BRD-K31468676 | O=C(O)c1nc(=O)[nH]c(=O)[nH]1 | RYYCJUAHISIHTL-UHFFFAOYSA-N | No-Effect | inactive |
| oxotremorine-M | BRD-K01942991 | C[N+](C)(C)CC#CCN1CCCC1=O | CANZROMYQDHYHR-UHFFFAOYSA-N | No-Effect | inactive |
| oxotremorine-sesquifumarate | BRD-K21936341 | O=C1CCCN1CC#CCN1CCCC1 | RSDOPYMFZBJHRL-UHFFFAOYSA-N | No-Effect | inactive |
| oxprenolol | BRD-A43671941 | C=CCOc1ccccc1OC[C@H](O)CNC(C)C | CEMAWMOMDPGJMB-CYBMUJFWSA-N | No-Effect | inactive |
| oxybutynin | BRD-K01825580 | CCN(CC)CC#CCOC(=O)[C@](O)(c1ccccc1)C1CCCCC1 | XIQVNETUBQGFHX-QFIPXVFZSA-N | No-Effect | inactive |
| oxyclozanide | BRD-K07283971 | O=C(Nc1cc(Cl)cc(Cl)c1O)c1c(O)c(Cl)cc(Cl)c1Cl | JYWIYHUXVMAGLG-UHFFFAOYSA-N | No-Effect | inactive |
| oxyfedrine | BRD-K73027814 | COc1cccc(C(=O)CCN[C@@H](C)[C@H](O)c2ccccc2)c1 | GDYUVHBMFVMBAF-LIRRHRJNSA-N | No-Effect | inactive |
| oxymatrine | BRD-K00003713 | O=C1CCC[C@H]2[C@H]3CCC[N@+]4([O-])CCC[C@@H](CN12)[C@@H]34 | XVPBINOPNYFXID-JARXUMMXSA-N | No-Effect | inactive |
| oxymetazoline | BRD-K16195444 | Cc1cc(C(C)(C)C)c(O)c(C)c1CC1=NCCN1 | WYWIFABBXFUGLM-UHFFFAOYSA-N | No-Effect | inactive |

| | | | | | |
|---|---|---|---|---|---|
| oxyphencyclimine | BRD-A58193911 | CN1CCCN=C1COC(=O)[C@](O)(c1ccccc1)C1CCCCC1 | DUDKAZCAISNGQN-FQEVSTJZSA-N | No-Effect | inactive |
| oxyphenonium | BRD-A14208071 | CC[N+](C)(CC)CCOC(=O)C(O)(c1ccccc1)C1CCCCC1 | GFRUPHOKLBPHTQ-UHFFFAOYSA-N | No-Effect | inactive |
| oxypurinol | BRD-K31841515 | O=c1nc2[nH][nH]cc-2c(=O)[nH]1 | HXNFUBHNUDHIGC-UHFFFAOYSA-N | No-Effect | inactive |
| oxyquinoline | BRD-K66808046 | Oc1cccc2cccnc12 | MCJGNVYPOGVAJF-UHFFFAOYSA-N | No-Effect | inactive |
| oxytocin | BRD-K25243230 | CC[C@H](C)[C@@H]1NC(=O)[C@H](Cc2ccc(O)cc2)NC(=O)[C@@H](N)CSSC[C@@H](C(=O)N2CCC[C@H]2C(=O)N[C@@H](CC(C)C)C(=O)NCC(N)=O)NC(=O)[C@H](CC(N)=O)NC(=O)[C@H](CCC(N)=O)NC1=O | XNOPRXBHLZRZKH-DSZYJQQASA-N | No-Effect | inactive |
| ozagrel | BRD-K53061490 | O=C(O)/C=C/c1ccc(Cn2ccnc2)cc1 | SHZKQBHERIJWAO-AATRIKPKSA-N | No-Effect | inactive |
| ozanimod | BRD-K46625559 | CC(C)Oc1ccc(-c2nc(-c3cccc4c3CC[C@@H]4NCCO)no2)cc1C#N | XRVDGNKRPOAQTN-FQEVSTJZSA-N | No-Effect | inactive |
| ozenoxacin | BRD-K00003184 | CNc1ncc(-c2ccc3c(=O)c(C(=O)O)cn(C4CC4)c3c2C)cc1C | XPIJWUTXQAGSLK-UHFFFAOYSA-N | No-Effect | inactive |
| p-dimethylinamyl-benzoate | BRD-K42336459 | CCCCCOC(=O)c1ccc(N(C)C)cc1 | LXTZRIBXKVRLOA-UHFFFAOYSA-N | No-Effect | inactive |
| P22077 | BRD-K97714174 | CC(=O)c1cc([N+](=O)[O-])c(Sc2ccc(F)cc2F)s1 | RMAMGGNACJHXHO-UHFFFAOYSA-N | No-Effect | inactive |
| P5091 | BRD-K13606314 | CC(=O)c1cc([N+](=O)[O-])c(Sc2cccc(Cl)c2Cl)s1 | LKZLGMAAKNEGCH-UHFFFAOYSA-N | No-Effect | inactive |
| P7C3 | BRD-A09487507 | O[C@@H](CNc1ccccc1)Cn1c2ccc(Br)cc2c2cc(Br)ccc21 | FZHHRERIIVOATI-KRWDZBQOSA-N | No-Effect | inactive |
| PAC-1 | BRD-K92991072 | C=CCc1cccc(/C=N/NC(=O)CN2CCN(Cc3ccccc3)CC2)c1O | YQNRVGJCPCNMKT-LFVJCYFKSA-N | No-Effect | inactive |
| paclitaxel | BRD-K62008436 | CC(=O)O[C@H]1C(=O)[C@]2(C)[C@@H](O)C[C@H]3OC[C@@]3(OC(C)=O)[C@H]2[C@H](OC(=O)c2ccccc2)[C@]2(O)C[C@H](OC(=O)[C@H](O)[C@@H](NC(=O)c3ccccc3)c3ccccc3)C(C)=C1C2(C)C | RCINICONZNJXQF-MZXODVADSA-N | No-Effect | inactive |
| PACOCF3 | BRD-K05677083 | CCCCCCCCCCCCCCCC(=O)C(F)(F)F | MAHYXYTYTLCTQD-UHFFFAOYSA-N | No-Effect | inactive |
| padsevonil | BRD-K00003493 | COCc1nn2c(CN3C[C@@H](CC(F)(F)Cl)CC3=O)c(C(F)(F)F)nc2s1 | DCXFIOLWWRXEQH-SSDOTTSWSA-N | No-Effect | inactive |
| paeonol | BRD-K94239562 | COc1ccc(C(C)=O)c(O)c1 | UILPJVPSNHJFIK-UHFFFAOYSA-N | No-Effect | inactive |
| pafuramidine | BRD-K68805147 | CON=C(N)c1ccc(-c2ccc(-c3ccc(C(N)=NOC)cc3)o2)cc1 | UKOQVLAXCBRRGH-UHFFFAOYSA-N | No-Effect | inactive |
| pagoclone | BRD-K00003428 | CC(C)CCC(=O)C[C@H]1c2ccccc2C(=O)N1c1ccc2ccc(Cl)nc2n1 | HIUPRQPBWVEQJJ-IBGZPJMESA-N | No-Effect | inactive |
| palbociclib | BRD-K51313569 | CC(=O)c1c(C)c2cnc(Nc3ccc(N4CCNCC4)cn3)nc2n(C2CCCC2)c1=O | AHJRHEGDXFFMBM-UHFFFAOYSA-N | No-Effect | inactive |
| palifosfamide | BRD-K12047401 | O=P(O)(NCCCl)NCCCl | BKCJZNIZRWYHBN-UHFFFAOYSA-N | No-Effect | inactive |
| paliperidone | BRD-A99888680 | Cc1nc2n(c(=O)c1CCN1CCC(c3noc4cc(F)ccc34)CC1)CCC[C@@H]2O | PMXMIIMHBWHSKN-IBGZPJMESA-N | No-Effect | inactive |
| palmatine-chloride | BRD-K58466253 | COc1cc2c(cc1OC)-c1cc3ccc(OC)c(OC)c3c[n+]1CC2 | QUCQEUCGKKTEBI-UHFFFAOYSA-N | No-Effect | inactive |
| palmitoleic-acid | BRD-K91356052 | CCCCCC/C=C\CCCCCCCC(=O)O | SECPZKHBENQXJG-FPLPWBNLSA-N | No-Effect | inactive |
| palmitoylethanolamide | BRD-K68095457 | CCCCCCCCCCCCCCCC(=O)NCCO | HXYVTAGFYLMHSO-UHFFFAOYSA-N | No-Effect | inactive |
| palomid-529 | BRD-A50998626 | COc1ccc(COc2ccc3oc(=O)c4cc([C@@H](C)O)ccc4c3cc2OC)cc1 | YEAHTLOYHVWAKW-CQSZACIVSA-N | No-Effect | inactive |
| palonosetron | BRD-K08924299 | O=C1c2cccc3c2[C@H](CCC3)CN1[C@@H]1CN2CCC1CC2 | CPZBLNMUGSZIPR-NVXWUHKLSA-N | No-Effect | inactive |
| palosuran | BRD-K07682953 | Cc1cc(NC(=O)NCCN2CCC(O)(Cc3ccccc3)CC2)c2ccccc2n1 | WYJCYXOCHXWTHG-UHFFFAOYSA-N | No-Effect | inactive |
| palovarotene | BRD-K83794243 | CC1(C)CCC(C)(C)c2cc(Cn3cccn3)c(/C=C/c3ccc(C(=O)O)cc3)cc21 | YTFHCXIPDIHOIA-DHZHZOJOSA-N | No-Effect | inactive |
| pamabrom | BRD-M67659203 | CC(C)(N)CO.Cn1c(=O)c2[nH]c(Br)nc2n(C)c1=O | ATOTUUBRFJHZQG-UHFFFAOYSA-N | No-Effect | inactive |
| pamidronate | BRD-K58513245 | NCCC(O)(P(=O)(O)O)P(=O)(O)O | WRUUGTRCQOWXEG-UHFFFAOYSA-N | No-Effect | inactive |
| pamiparib | BRD-K00003461 | C[C@]12CCCN1CC1=NNC(=O)c3cc(F)cc4[nH]c2c1c34 | DENYZIUJOTUUNY-MRXNPFEDSA-N | No-Effect | inactive |
| pancuronium | BRD-K50817623 | CC(=O)O[C@H]1C[C@@H]2CC[C@@H]3[C@H](CC[C@@]4(C)[C@H]3C[C@H]([N+]3(C)CCCCC3)[C@@H]4OC(C)=O)[C@@]2(C)C[C@@H]1[N+]1(C)CCCCC1 | GVEAYVLWDAFXET-XGHATYIMSA-N | No-Effect | inactive |
| panipenem | BRD-K00004671 | C=C(N)N1CC[C@H](SC2=C(C(=O)O)N3C(=O)[C@H]([C@@H](C)O)[C@H]3C2)C1 | UIWSCWBHZRPIKR-DLYFRVTGSA-N | No-Effect | inactive |
| pantothenic-acid | BRD-K29392254 | CC(C)(CO)[C@@H](O)C(=O)NCCC(=O)O | GHOKWGTUZJEAQD-ZETCQYMHSA-N | No-Effect | inactive |
| PAOPA | BRD-K18573821 | NC(=O)CN1CC[C@@H](NC(=O)[C@@H]2CCCN2)C1=O | QXHVGEXNEZRSGG-JGVFFNPUSA-N | No-Effect | inactive |
| paquinimod | BRD-K00003371 | CCc1cccc2c1c(O)c(C(=O)N(CC)c1ccccc1)c(=O)n2C | DIKSYHCCYVYKRO-UHFFFAOYSA-N | No-Effect | inactive |
| para-toluenesulfonamide | BRD-K62412084 | Cc1ccc(S(N)(=O)=O)cc1 | LMYRWZFENFIFIT-UHFFFAOYSA-N | No-Effect | inactive |
| paracetamol | BRD-K41524689 | CC(=O)Nc1ccc(O)cc1 | RZVAJINKPMORJF-UHFFFAOYSA-N | No-Effect | inactive |
| parachlorophenol | BRD-K40992116 | Oc1ccc(Cl)cc1 | WXNZTHHGJRFXKQ-UHFFFAOYSA-N | No-Effect | inactive |
| paramethadione | BRD-A22128695 | CC[C@@]1(C)OC(=O)N(C)C1=O | VQASKUSHBVDKGU-SSDOTTSWSA-N | No-Effect | inactive |
| paraxanthine | BRD-K24084088 | Cn1c(=O)[nH]c2ncn(C)c2c1=O | QUNWUDVFRNGTCO-UHFFFAOYSA-N | No-Effect | inactive |
| parbendazole | BRD-K02407574 | CCCCc1ccc2nc(NC(=O)OC)[nH]c2c1 | YRWLZFXJFBZBEY-UHFFFAOYSA-N | No-Effect | inactive |
| parcetasal | BRD-K00004740 | CC(=O)Nc1ccc(O[C@]2(C)OC(=O)c3ccccc3O2)cc1 | ZAPRLADYRFPQSH-KRWDZBQOSA-N | No-Effect | inactive |
| pardoprunox | BRD-K11851476 | CN1CCN(c2cccc3[nH]c(=O)oc23)CC1 | YVPUUUDAZYFFQT-UHFFFAOYSA-N | No-Effect | inactive |
| parecoxib | BRD-K13800121 | CCC(=O)NS(=O)(=O)c1ccc(-c2c(-c3ccccc3)noc2C)cc1 | TZRHLKRLEZJVIJ-UHFFFAOYSA-N | No-Effect | inactive |
| parethoxycaine | BRD-K54799382 | CCOc1ccc(C(=O)OCCN(CC)CC)cc1 | OWWVHQUOYSPNNE-UHFFFAOYSA-N | No-Effect | inactive |
| pargyline | BRD-K83597974 | C#CCN(C)Cc1ccccc1 | DPWPWRLQFGFJFI-UHFFFAOYSA-N | No-Effect | inactive |
| paricalcitol | BRD-K60585088 | C[C@@H](/C=C/[C@@H](C)[C@H]1CC[C@H]2/C(=C/C=C3C[C@@H](O)C[C@H](O)C3)CCC[C@@]21C)C(C)(C)O | BPKAHTKRCLCHEA-UBFJEZKGSA-N | No-Effect | inactive |
| paritaprevir | BRD-K89920762 | Cc1cnc(C(=O)N[C@H]2CCCCC/C=C\[C@@H]3C[C@@]3(C(=O)NS(=O)(=O)C3CC3)NC(=O)[C@@H]3C[C@@H](Oc4nc5ccccc5c5ccccc45)CN3C2=O)cn1 | UAUIUKWPKRJZJV-QPLHLKROSA-N | No-Effect | inactive |
| paromomycin | BRD-K36524108 | NC[C@@H]1O[C@H](O[C@@H]2[C@@H](CO)O[C@@H](O[C@@H]3[C@@H](O)[C@H](N)C[C@H](N)[C@H]3O[C@H]3O[C@H](CO)[C@@H](O)[C@H](O)[C@H]3N)[C@@H]2O)[C@H](N)[C@@H](O)[C@@H]1O | UOZODPSAJZTQNH-LSWIJEOBSA-N | No-Effect | inactive |
| paroxetine | BRD-K37991163 | Fc1ccc([C@@H]2CCNC[C@H]2COc2ccc3c(c2)OCO3)cc1 | AHOUBRCZNHFOSL-YOEHRIQHSA-N | No-Effect | inactive |
| paroxypropione | BRD-K07441213 | CCC(=O)c1ccc(O)cc1 | RARSHUDCJQSEFJ-UHFFFAOYSA-N | No-Effect | inactive |
| parsaclisib | BRD-K00004594 | CCOc1c([C@H](C)n2nc(C)c3c(N)ncnc32)cc(Cl)c(F)c1[C@@H]1CNC(=O)C1 | ZQPDJCIXJHUERQ-QWRGUYRKSA-N | No-Effect | inactive |
| parthenolide | BRD-K98548675 | C=C1C(=O)O[C@H]2[C@H]1CC/C(C)=C/CC[C@@]1(C)O[C@@H]21 | KTEXNACQROZXEV-SLXBATTESA-N | No-Effect | inactive |
| parthenolide-(-) | BRD-K34851558 | C=C1C(=O)O[C@H]2[C@H]1CC/C(C)=C\CC[C@@]1(C)O[C@@H]21 | KTEXNACQROZXEV-QLIGOWBFSA-N | No-Effect | inactive |
| parthenolide-(alternate-stereo) | BRD-K28120222 | C=C1C(=O)O[C@H]2[C@H]1CC/C(C)=C/CC[C@@]1(C)O[C@H]21 | KTEXNACQROZXEV-PVLRGYAZSA-N | No-Effect | inactive |
| pasireotide | BRD-K59715032 | NCCCC[C@@H]1NC(=O)[C@@H](Cc2c[nH]c3ccccc23)NC(=O)[C@H](c2ccccc2)NC(=O)[C@@H]2C[C@@H](OC(=O)NCCN)CN2C(=O)[C@H](Cc2ccccc2)NC(=O)[C@H](Cc2ccc(OCc3ccccc3)cc2)NC1=O | VMZMNAABQBOLAK-DBILLSOUSA-N | No-Effect | inactive |
| PAT-1251 | BRD-K00003481 | NCc1cc(Oc2cccc(C(=O)N3C[C@@H](O)[C@H](F)C3)c2)nc(C(F)(F)F)c1 | ODGXXYXJORZPHE-ZIAGYGMSSA-N | No-Effect | inactive |
| pavinetant | BRD-K00003170 | CC[C@H](NC(=O)c1c(NS(C)(=O)=O)c(-c2ccccc2)nc2ccccc12)c1ccccc1 | QYTBBBAHNIWFOD-NRFANRHFSA-N | No-Effect | inactive |
| pazopanib | BRD-K74514084 | Cc1ccc(Nc2nccc(N(C)c3ccc4c(C)n(C)nc4c3)n2)cc1S(N)(=O)=O | CUIHSIWYWATEQL-UHFFFAOYSA-N | No-Effect | inactive |
| pazufloxacin | BRD-K66412701 | C[C@H]1COc2c(C3(N)CC3)c(F)cc3c(=O)c(C(=O)O)cn1c23 | XAGMUUZPGZWTRP-ZETCQYMHSA-N | No-Effect | inactive |
| PBI-4050 | BRD-K00003391 | CCCCCc1cccc(CC(=O)O)c1 | PEGQOIGYZLJMIB-UHFFFAOYSA-N | No-Effect | inactive |

| | | | | | |
|---|---|---|---|---|---|
| PBT-1033 | BRD-K00004615 | CN(C)Cc1ccc2c(Cl)cc(Cl)c(O)c2n1 | YZPOQCQXOSEMAZ-UHFFFAOYSA-N | No-Effect | inactive |
| PCA-4248 | BRD-A29289453 | COC(=O)C1=C(C)NC(C)=C(C(=O)OCCSc2ccccc2)[C@@H]1C | DHCNAWNKZMNTIS-GFCCVEGCSA-N | No-Effect | inactive |
| PCI-24781 | BRD-K12867552 | CN(C)Cc1c(C(=O)NCCOc2ccc(C(=O)NO)cc2)oc2ccccc12 | MAUCONCHVWBMHK-UHFFFAOYSA-N | No-Effect | inactive |
| PCI-27483 | BRD-K00003213 | N=C(N)c1ccc2nc(-c3cc(CC(=O)N[C@@H](CC(=O)O)C(=O)O)cc(-c4cc(S(N)(=O)=O)ccc4O)c3O)[nH]c2c1 | WDJHHCAKBRKCLW-IBGZPJMESA-N | No-Effect | inactive |
| PCI-29732 | BRD-K71223042 | Nc1ncnc2c1c(-c1ccc(Oc3ccccc3)cc1)nn2C1CCCC1 | GMJUPMONHWAZCP-UHFFFAOYSA-N | No-Effect | inactive |
| PCI-34051 | BRD-K88742110 | COc1ccc(Cn2ccc3ccc(C(=O)NO)cc32)cc1 | AJRGHIGYPXNABY-UHFFFAOYSA-N | No-Effect | inactive |
| PCO-400 | BRD-K52721684 | CC1(C)Oc2ccc(C#N)cc2[C@@H](OC2=CC(=O)CC2)[C@@H]1O | WJBHHTPFTVKZCV-CVEARBPZSA-N | No-Effect | inactive |
| PD-0325901 | BRD-K49865102 | O=C(NOC[C@H](O)CO)c1ccc(F)c(F)c1Nc1ccc(I)cc1F | SUDAHWBOROXANE-SECBINFHSA-N | No-Effect | inactive |
| PD-123319 | BRD-K34713073 | Cc1cc(Cn2cnc3c2C[C@@H](C(=O)O)N(C(=O)C(c2ccccc2)c2ccccc2)C3)ccc1N(C)C | YSTVFDAKLDMYCR-NDEPHWFRSA-N | No-Effect | inactive |
| PD-128907 | BRD-K62456038 | CCCN1CCO[C@@H]2c3cc(O)ccc3OC[C@H]21 | YOILXOMTHPUMRG-TZMCWYRMSA-N | No-Effect | inactive |
| PD-156707 | BRD-K13826399 | COc1ccc(C(=O)/C(Cc2cc(OC)c(OC)c(OC)c2)=C(\C(=O)O)c2ccc3c(c2)OCO3)cc1 | OFCTYBMKKTVKCR-QQTULTPQSA-N | No-Effect | inactive |
| PD-158780 | BRD-K48735772 | CNc1cc2c(Nc3cccc(Br)c3)ncnc2cn1 | KFHMLBXBRCITHF-UHFFFAOYSA-N | No-Effect | inactive |
| PD-160170 | BRD-K12079898 | CC(C)c1ccccc1S(=O)(=O)c1cc(N)c2ncccc2c1[N+](=O)[O-] | YUVNGBZROXQYQH-UHFFFAOYSA-N | No-Effect | inactive |
| PD-166793 | BRD-K60984070 | CC(C)[C@H](NS(=O)(=O)c1ccc(-c2ccc(Br)cc2)cc1)C(=O)O | GJOCABIDMCKCEG-INIZCTEOSA-N | No-Effect | inactive |
| PD-168077 | BRD-K47761761 | Cc1cccc(C(=O)NCN2CCN(c3ccccc3C#N)CC2)c1 | DNULYRGWTFLJQL-UHFFFAOYSA-N | No-Effect | inactive |
| PD-168568 | BRD-A82013137 | Cc1ccc(N2CCN(CC[C@H]3NC(=O)c4ccccc43)CC2)cc1C | WQEPZBNLBWDIRZ-OAQYLSRUSA-N | No-Effect | inactive |
| PD-173212 | BRD-K04426574 | CC(C)C[C@@H](C(=O)N[C@@H](Cc1ccc(OCc2ccccc2)cc1)C(=O)NC(C)(C)C)N(C)Cc1ccc(C(C)(C)C)cc1 | GCDHMGROXQUFNR-HEVIKAOCSA-N | No-Effect | inactive |
| PD-184352 | BRD-K05104363 | O=C(NOCC1CC1)c1ccc(F)c(F)c1Nc1ccc(I)cc1Cl | GFMMXOIFOQCCGU-UHFFFAOYSA-N | No-Effect | inactive |
| PD-198306 | BRD-K88677950 | Cc1cc(I)ccc1Nc1c(C(=O)NOCC2CC2)cc(F)c(F)c1F | UHAXDAKQGVISBZ-UHFFFAOYSA-N | No-Effect | inactive |
| PD-318088 | BRD-A57798112 | O=C(NOC[C@H](O)CO)c1cc(Br)c(F)c(F)c1Nc1ccc(I)cc1F | XXSSGBYXSKOLAM-MRVPVSSYSA-N | No-Effect | inactive |
| PD-407824 | BRD-K82823804 | O=C1NC(=O)c2c1c(-c1ccccc1)cc1[nH]c3ccc(O)cc3c21 | IAUZTOZLTFSMIE-UHFFFAOYSA-N | No-Effect | inactive |
| PD-81723 | BRD-K77951111 | Cc1sc(N)c(C(=O)c2cccc(C(F)(F)F)c2)c1C | KKDKAWKYGCUOGR-UHFFFAOYSA-N | No-Effect | inactive |
| PD-98059 | BRD-K62810658 | COc1cccc(-c2cc(=O)c3ccccc3o2)c1N | QFWCYNPOPKQOKV-UHFFFAOYSA-N | No-Effect | inactive |
| PD1-PDL-inhibitor-1 | BRD-K04032780 | COc1cc(OCc2cccc(-c3ccccc3)c2C)cc(OC)c1CN1CCCC[C@H]1C(=O)O | ZBOYJODMIAUJHH-SANMLTNESA-N | No-Effect | inactive |
| PDD-00017273 | BRD-K00021095 | Cc1cc(Cn2c(=O)n(Cc3cnc(C)s3)c(=O)c3cc(S(=O)(=O)NC4(C)CC4)ccc32)n(C)n1 | IFWUBRBMMNTBRZ-UHFFFAOYSA-N | No-Effect | inactive |
| PDP-EA | BRD-K26358347 | CCCCCCCCCCCCCCCc1cccc(OCC(=O)NCCO)c1 | BIOVSNWTCKRADD-UHFFFAOYSA-N | No-Effect | inactive |
| peficitinib | BRD-A14577621 | NC(=O)c1cnc2[nH]ccc2c1N[C@H]1[C@H]2C[C@H]3C[C@@H]1C[C@](O)(C3)C2 | DREIJXJRTLTGJC-ZKVNVPQCSA-N | No-Effect | inactive |
| pelanserin | BRD-K96436774 | O=c1[nH]c2ccccc2c(=O)n1CCCN1CCN(c2ccccc2)CC1 | WPKPLSFHHBBLRY-UHFFFAOYSA-N | No-Effect | inactive |
| pelitinib | BRD-K08799216 | CCOc1cc2ncc(C#N)c(Nc3ccc(F)c(Cl)c3)c2cc1NC(=O)/C=C/CN(C)C | WVUNYSQLFKLYNI-AATRIKPKSA-N | No-Effect | inactive |
| pelrinone | BRD-K00004687 | Cc1nc(=O)c(C#N)c(NCc2cccnc2)[nH]1 | NZXHFDXOCVIYLO-UHFFFAOYSA-N | No-Effect | inactive |
| pemafibrate | BRD-K00003251 | CC[C@@H](Oc1cccc(CN(CCCOc2ccc(OC)cc2)c2nc3ccccc3o2)c1)C(=O)O | ZHKNLJLMDFQVHJ-RUZDIDTESA-N | No-Effect | inactive |
| pemirolast | BRD-K31731454 | Cc1cccn2c(=O)c(-c3nn[nH]n3)cnc12 | HIANJWSAHKJQTH-UHFFFAOYSA-N | No-Effect | inactive |
| penbutolol | BRD-K55705469 | CC(C)(C)NC[C@H](O)COc1ccccc1C1CCCC1 | KQXKVJAGOJTNJS-HNNXBMFYSA-N | No-Effect | inactive |
| penciclovir | BRD-K47780086 | Nc1nc(=O)c2ncn(CCC(CO)CO)c2[nH]1 | JNTOCHDNEULJHD-UHFFFAOYSA-N | No-Effect | inactive |
| penfluridol | BRD-K15409150 | OC1(c2ccc(Cl)c(C(F)(F)F)c2)CCN(CCCC(c2ccc(F)cc2)c2ccc(F)cc2)CC1 | MDLAAYDRRZXJIF-UHFFFAOYSA-N | No-Effect | inactive |
| penicillamine-(D) | BRD-K17518951 | CC(C)(S)[C@@H](N)C(=O)O | VVNCNSJFMMFHPL-VKHMYHEASA-N | No-Effect | inactive |
| penicillamine-(racemic) | BRD-K01825697 | CC(C)(S)[C@@H](N)C(=O)O | VVNCNSJFMMFHPL-VKHMYHEASA-N | No-Effect | inactive |
| penicillin-v-potassium | BRD-K43966364 | CC1(C)S[C@@H]2[C@H](NC(=O)COc3ccccc3)C(=O)N2[C@H]1C(=O)O | BPLBGHOLXOTWMN-MBNYWOFBSA-N | No-Effect | inactive |
| pentacosanoic-acid | BRD-K48996094 | CCCCCCCCCCCCCCCCCCCCCCCC(=O)O | MWMPEAHGUXCSMY-UHFFFAOYSA-N | No-Effect | inactive |
| pentagastrin | BRD-K78789465 | CSCC[C@H](NC(=O)[C@H](Cc1c[nH]c2ccccc12)NC(=O)CCNC(=O)OC(C)(C)C)C(=O)N[C@@H](CC(=O)O)C(=O)N[C@@H](Cc1ccccc1)C(N)=O | NEYNJQRKHLUJRU-DZUOILHNSA-N | No-Effect | inactive |
| pentetic-acid | BRD-K40621224 | O=C(O)CN(CCN(CC(=O)O)CC(=O)O)CCN(CC(=O)O)CC(=O)O | QPCDCPDFJACHGM-UHFFFAOYSA-N | No-Effect | inactive |
| pentolinium | BRD-K44942604 | C[N+]1(CCCCC[N+]2(C)CCCC2)CCCC1 | XSBSKEQEUFOSDD-UHFFFAOYSA-N | No-Effect | inactive |
| pentostatin | BRD-K91543828 | OC[C@H]1O[C@@H](n2cnc3c2NC=NC[C@H]3O)C[C@@H]1O | FPVKHBSQESCIEP-JQCXWYLXSA-N | No-Effect | inactive |
| pentoxifylline | BRD-K57569181 | CC(=O)CCCCn1c(=O)c2c(ncn2C)n(C)c1=O | BYPFEZZEUUWMEJ-UHFFFAOYSA-N | No-Effect | inactive |
| pentylenetetrazol | BRD-K57718010 | C1CCc2nnnn2CC1 | CWRVKFFCRWGWCS-UHFFFAOYSA-N | No-Effect | inactive |
| peposertib | BRD-K00003426 | COc1ccc([C@@H](O)c2cc(-c3ncnc4cc(N5CCOCC5)ccc34)c(F)cc2Cl)nn1 | MOWXJLUYGFNTAL-DEOSSOPVSA-N | No-Effect | inactive |
| pepstatin | BRD-K13571841 | CC(C)CC(=O)N[C@H](C(=O)N[C@H](C(=O)N[C@@H](CC(C)C)[C@@H](O)CC(=O)N[C@@H](C)C(=O)N[C@@H](CC(C)C)[C@@H](O)CC(=O)O)C(C)C)C(C)C | FAXGPCHRFPCXOO-LXTPJMTPSA-N | No-Effect | inactive |
| peramivir | BRD-K00519864 | CCC(CC)[C@H](NC(C)=O)[C@@H]1[C@H](O)[C@@H](C(=O)O)C[C@H]1NC(=N)N | XRQDFNLINLXZLB-CKIKVBCHSA-N | No-Effect | inactive |
| perchlozone | BRD-K00004530 | NC(=S)N/N=C/c1ccncc1 | QTSJJDIDTYFLFC-BJMVGYQFSA-N | No-Effect | inactive |
| peretinoin | BRD-K99431849 | CC(C)=CCC/C(C)=C/CC/C(C)=C/C=C/C(C)=C/C(=O)O | UUBHZHZSIKRVIV-KCXSXWJSSA-N | No-Effect | inactive |
| perfluorodecalin | BRD-K55401595 | FC1(F)C(F)(F)C(F)(F)C2(F)C(F)(F)C(F)(F)C(F)(F)C(F)(F)C2(F)C1(F)F | UWEYRJFJVCLAGH-UHFFFAOYSA-N | No-Effect | inactive |
| perfluorohexyloctane | BRD-K00004546 | CCCCCCCCC(F)(F)C(F)(F)C(F)(F)C(F)(F)C(F)(F)C(F)(F)F | WRYIIOKOQSICTB-UHFFFAOYSA-N | No-Effect | inactive |
| pergolide | BRD-K60770992 | CCCN1C[C@H](CSC)C[C@@H]2c3cccc4[nH]cc(c34)C[C@H]21 | YEHCICAEULNIGD-MZMPZRCHSA-N | No-Effect | inactive |
| perifosine | BRD-A89082344 | CCCCCCCCCCCCCCCCCCOP(=O)(O)OC1CC[N+](C)(C)CC1 | SZFPYBIJACMNJV-UHFFFAOYSA-O | No-Effect | inactive |
| perillyl-alcohol | BRD-A13323580 | C=C(C)[C@@H]1CC=C(CO)CC1 | NDTYTMIUWGWIMO-SNVBAGLBSA-N | No-Effect | inactive |
| perindopril | BRD-K92731339 | CCC[C@H](N[C@@H](C)C(=O)N1[C@H](C(=O)O)C[C@@H]2CCCC[C@@H]21)C(=O)OCC | IPVQLZZIHOAWMC-QXKUPLGCSA-N | No-Effect | inactive |
| perindoprilat | BRD-K00768353 | CCC[C@H](N[C@@H](C)C(=O)N1[C@H](C(=O)O)C[C@@H]2CCCC[C@@H]21)C(=O)O | ODAIHABQVKJNIY-PEDHHIEDSA-N | No-Effect | inactive |
| permethrin | BRD-A25366096 | CC1(C)[C@@H](C(=O)OCc2cccc(Oc3ccccc3)c2)[C@@H]1C=C(Cl)Cl | RLLPVAHGXHCWKJ-PKOBYXMFSA-N | No-Effect | inactive |
| perospirone | BRD-K85503079 | O=C1[C@H]2CCCC[C@H]2C(=O)N1CCCCN1CCN(c2nsc3ccccc23)CC1 | FBVFZWUMDDXLLG-HDICACEKSA-N | No-Effect | inactive |
| perzinfotel | BRD-K04643877 | O=c1c2c(c1=O)N(CCP(=O)(O)O)CCCN2 | BDABGOLMYNHHTR-UHFFFAOYSA-N | No-Effect | inactive |
| PETCM | BRD-A93477898 | O[C@@H](Cc1ccncc1)C(Cl)(Cl)Cl | NGTDJJKTGRNNAU-ZETCQYMHSA-N | No-Effect | inactive |
| pevonedistat | BRD-K67844266 | NS(=O)(=O)OC[C@@H]1C[C@@H](n2ccc3c(N[C@H]4CCc5ccccc54)ncnc32)C[C@@H]1O | MPUQHZXIXSTTDU-QXGSTGNESA-N | No-Effect | inactive |
| pexidartinib | BRD-K54519451 | FC(F)(F)c1ccc(CNc2ccc(Cc3c[nH]c4ncc(Cl)cc34)cn2)cn1 | JGWRKYUXBBNENE-UHFFFAOYSA-N | No-Effect | inactive |
| pexmetinib | BRD-K53405000 | Cc1ccc(-n2nc(C(C)(C)C)cc2NC(=O)NCc2cc(F)ccc2Oc2ccc3c(cnn3CCO)c2)cc1 | LNMRSSIMGCDUTP-UHFFFAOYSA-N | No-Effect | inactive |
| PF-02413873 | BRD-K00004632 | Cc1cc(Oc2c(C3CC3)nn(CS(C)(=O)=O)c2C)cc(C)c1C#N | QSFGZNVRVZHUGV-UHFFFAOYSA-N | No-Effect | inactive |
| PF-03049423 | BRD-K07070922 | CCCOCCn1c(=O)c(N2CCN(CCO)CC2)nc2cnc(-c3ccc(OC)nc3)cc21 | MNLNAGRCHNMKKJ-UHFFFAOYSA-N | No-Effect | inactive |

| | | | | | |
|---|---|---|---|---|---|
| PF-03084014 | BRD-K61691541 | CCC[C@H](N[C@H]1CCc2cc(F)cc(F)c2C1)C(=O)Nc1cn(C(C)(C)CNCC(C)(C)C)cn1 | VFCRKLWBYMDAED-REWPJTCUSA-N | No-Effect | inactive |
| PF-03758309 | BRD-K37764012 | Cc1nc(Nc2[nH]nc3c2CN(C(=O)N[C@H](CN(C)C)c2ccccc2)C3(C)C)c2sccc2n1 | AYCPARAPKDAOEN-LJQANCHMSA-N | No-Effect | inactive |
| PF-03814735 | BRD-K68747584 | CC(=O)NCC(=O)N1[C@H]2CC[C@@H]1c1cc(Nc3ncc(C(F)(F)F)c(NC4CCC4)n3)ccc12 | RYYNGWLOYLRZLK-RBUKOAKNSA-N | No-Effect | inactive |
| PF-03882845 | BRD-K51041191 | N#Cc1ccc(N2N=C3c4ccc(C(=O)O)cc4CCC3[C@@H]2C2CCCC2)cc1Cl | XNULRSOGWPFPBL-AKRCKQFNSA-N | No-Effect | inactive |
| PF-04136309 | BRD-K94420399 | O=C(NCC(=O)N1CC[C@H](N[C@H]2CC[C@@](O)(c3ccc(-c4ncccn4)cn3)CC2)C1)c1cccc(C(F)(F)F)c1 | ZNSVOHSYDRPBGI-LXWOLXCRSA-N | No-Effect | inactive |
| PF-04217903 | BRD-K73319509 | OCCn1cc(-c2cnc3nnn(Cc4ccc5ncccc5c4)c3n2)cn1 | PDMUGYOXRHVNMO-UHFFFAOYSA-N | No-Effect | inactive |
| PF-04418948 | BRD-K54871792 | COc1ccc2cc(OCC3(C(=O)O)CN(C(=O)c4ccc(F)cc4)C3)ccc2c1 | LWJGMYMNSNVCEM-UHFFFAOYSA-N | No-Effect | inactive |
| PF-04447943 | BRD-K50031829 | C[C@@H]1CN(Cc2ncccn2)C[C@H]1c1nc(=O)c2cnn(C3CCOCC3)c2[nH]1 | IWXUVYOOUMLUTQ-CZUORRHYSA-N | No-Effect | inactive |
| PF-04457845 | BRD-K86816506 | O=C(Nc1cccnn1)N1CCC(=Cc2cccc(Oc3ccc(C(F)(F)F)cn3)c2)CC1 | BATCTBJIJJEPHM-UHFFFAOYSA-N | No-Effect | inactive |
| PF-04620110 | BRD-K30550578 | Nc1ncnc2c1C(=O)N(c1ccc([C@H]3CC[C@H](CC(=O)O)CC3)cc1)CCO2 | GEVVQZHMFVFGLN-HDJSIYSDSA-N | No-Effect | inactive |
| PF-04885614 | BRD-K92435527 | CC(C)(N)c1nc(-c2ccc(OC(F)(F)F)cc2)c[nH]1 | AGORGFNWYAUYSU-UHFFFAOYSA-N | No-Effect | inactive |
| PF-04937319 | BRD-K27755542 | Cc1cnc(NC(=O)c2cc(Oc3cnc(C(=O)N(C)C)nc3)c3cc(C)oc3c2)cn1 | MASKQITXHVYVFL-UHFFFAOYSA-N | No-Effect | inactive |
| PF-04995274 | BRD-K00004635 | OC1(CN2CCC(COc3noc4cccc(O[C@@H]5CCOC5)c34)CC2)CCOCC1 | WLLOFQROROXOMO-GOSISDBHSA-N | No-Effect | inactive |
| PF-05089771 | BRD-K00003145 | Nc1[nH]ncc1-c1cc(Cl)ccc1Oc1cc(F)c(S(=O)(=O)Nc2cscn2)cc1Cl | ZYSCOUXLBXGGIM-UHFFFAOYSA-N | No-Effect | inactive |
| PF-05180999 | BRD-K00003097 | Cc1nc(-c2cnn(C)c2-c2ccc(C(F)(F)F)cn2)c2c(N3CCC3)ncnn12 | CLGCHUKGBICQTE-UHFFFAOYSA-N | No-Effect | inactive |
| PF-05190457 | BRD-K02554563 | Cc1cc(-c2ccc3c(c2)CC[C@H]3N2CC3(CCN(C(=O)Cc4cn5cc(C)sc5n4)CC3)C2)ncn1 | ZIUDADZJCKGWKR-AREMUKBSSA-N | No-Effect | inactive |
| PF-05212384 | BRD-K07955840 | CN(C)C1CCN(C(=O)c2ccc(NC(=O)Nc3ccc(-c4nc(N5CCOCC5)nc(N5CCOCC5)n4)cc3)cc2)CC1 | DWZAEMINVBZMHQ-UHFFFAOYSA-N | No-Effect | inactive |
| PF-06260414 | BRD-K00004644 | C[C@@H]1CNS(=O)(=O)N(c2ccc3c(C#N)cccc3c2)C1 | ZKAVFOXYJCREBQ-SNVBAGLBSA-N | No-Effect | inactive |
| PF-06273340 | BRD-K00004638 | CC(C)(CO)n1cc(C(=O)c2cncc(NC(=O)Cc3ccc(Cl)cn3)c2)c2cnc(N)nc21 | BPIWZDNVMQQBQX-UHFFFAOYSA-N | No-Effect | inactive |
| PF-06282999 | BRD-K00003279 | COc1ccc(Cl)cc1-c1cc(=O)[nH]c(=S)n1CC(N)=O | ICYNYWFGIDGBRD-UHFFFAOYSA-N | No-Effect | inactive |
| PF-06305591 | BRD-K00004712 | C[C@@H](C(N)=O)[C@H](N)c1nc2cc(C(C)(C)C)ccc2[nH]1 | APWZIFIAVVFPNT-PELKAZGASA-N | No-Effect | inactive |
| PF-06412562 | BRD-K00004652 | Cc1cc(Oc2nccc3[nH]ncc23)ccc1-c1c(C)ncnc1C | IDIUJOHYYBNCPC-UHFFFAOYSA-N | No-Effect | inactive |
| PF-06447475 | BRD-K83114366 | N#Cc1cccc(-c2c[nH]c3ncnc(N4CCOCC4)c23)c1 | BHTWDJBVZQBRKP-UHFFFAOYSA-N | No-Effect | inactive |
| PF-06459988 | BRD-K00003311 | C=CC(=O)N1C[C@H](COc2nc(Nc3cnn(C)c3)nc3[nH]cc(Cl)c23)[C@@H](OC)C1 | ODMXWZROLKITMS-RISCZKNCSA-N | No-Effect | inactive |
| PF-06463922 | BRD-A23124853 | C[C@H]1Oc2cc(cnc2N)-c2cc(nn(C)c2C#N)CN(C)C(=O)c2ccc(F)cc21 | IIXWYSCJSQVBQM-LLVKDONJSA-N | No-Effect | inactive |
| PF-06650833 | BRD-K00003296 | CC[C@H]1[C@@H](COc2nccc3cc(C(N)=O)c(OC)cc23)NC(=O)[C@H]1F | JKDGKIBAOAFRPJ-ZBINZKHDSA-N | No-Effect | inactive |
| PF-06651600 | BRD-K47084954 | C=CC(=O)N1C[C@H](Nc2ncnc3[nH]ccc23)CC[C@@H]1C | CBRJPFGIXUFMTM-WDEREUQCSA-N | No-Effect | inactive |
| PF-06747775 | BRD-K00004648 | C=CC(=O)N[C@@H]1CN(c2nc(Nc3cn(C)nc3OC)c3ncn(C)c3n2)C[C@H]1F | JYIUNVOCEFIUIU-GHMZBOCLSA-N | No-Effect | inactive |
| PF-06751979 | BRD-K00004598 | C[C@H]1C[C@H]2CSC(N)=N[C@@]2(c2nc(NC(=O)c3ccc(OC(F)F)cn3)cs2)CO1 | ZLZUHACSRMOLLV-RAALSFIWSA-N | No-Effect | inactive |
| PF-06840003 | BRD-K00003400 | O=C1C[C@H](c2c[nH]c3ccc(F)cc23)C(=O)N1 | MXKLDYKORJEOPR-MRVPVSSYSA-N | No-Effect | inactive |
| PF-06873600 | BRD-K00004750 | C[C@@]1(O)CCC[C@H]1n1c(=O)c(C(F)F)cc2cnc(NC3CCN(S(C)(=O)=O)CC3)nc21 | QIEKHLDZKRQLLN-FOIQADDNSA-N | No-Effect | inactive |
| PF-06928215 | BRD-K00005326 | O=C(N[C@H]1CCCC[C@H]1C(=O)O)c1cnn2c(O)cc(-c3ccccc3)nc12 | PHGTYMRDHDEHLI-HIFRSBDPSA-N | No-Effect | inactive |
| PF-3274167 | BRD-K31417912 | COCc1nnc(N2CC(Oc3ccc(F)cc3Cl)C2)n1-c1ccc(OC)nc1 | HNIFCPBQMKPRCX-UHFFFAOYSA-N | No-Effect | inactive |
| PF-3845 | BRD-K68997413 | O=C(Nc1cccnc1)N1CCC(Cc2cccc(Oc3ccc(C(F)(F)F)cn3)c2)CC1 | NBOJHRYUGLRASX-UHFFFAOYSA-N | No-Effect | inactive |
| PF-4191834 | BRD-K00004633 | Cn1nccc1-c1ccc(Sc2cccc(C3(C(N)=O)CCOCC3)c2)cc1 | DVNQWYLVSNPCJZ-UHFFFAOYSA-N | No-Effect | inactive |
| PF-4708671 | BRD-K14711741 | CCc1cncnc1N1CCN(Cc2nc3cc(C(F)(F)F)cc3[nH]2)CC1 | FBLPQCAQRNSVHB-UHFFFAOYSA-N | No-Effect | inactive |
| PF-477736 | BRD-K03063480 | Cn1cc(-c2[nH]c3cc(NC(=O)[C@H](N)C4CCCCC4)cc4c3c2C=NNC4=O)cn1 | NDEXUOWTGYUVGA-LJQANCHMSA-N | No-Effect | inactive |
| PF-4800567 | BRD-K43428468 | Nc1ncnc2c1c(COc1cccc(Cl)c1)nn2C1CCOCC1 | AUMDBEHGJRZSOO-UHFFFAOYSA-N | No-Effect | inactive |
| PF-4981517 | BRD-K78318619 | Cc1ccc(-c2c(-c3nn(C)c4ncnc(N5CC[C@H](N6CCCCC6)C5)c34)cnn2C)cc1 | WDWIMDKOXZZYHH-FQEVSTJZSA-N | No-Effect | inactive |
| PF-514273 | BRD-K00004642 | CC(F)(F)CN1CCOc2c(nn(-c3ccccc3Cl)c2-c2ccc(Cl)cc2)C1=O | FJMQJSUOOGOWBD-UHFFFAOYSA-N | No-Effect | inactive |
| PF-5274857 | BRD-K40302533 | Cc1cnc(-c2cc(N3CCN(C(=O)CCS(C)(=O)=O)CC3)ncc2Cl)c(C)c1 | BBVNTTZIOTWDSV-UHFFFAOYSA-N | No-Effect | inactive |
| PF-562271 | BRD-K99545815 | CN(c1ncccc1CNc1nc(Nc2ccc3c(c2)CC(=O)N3)ncc1C(F)(F)F)S(C)(=O)=O | MZDKLVOWGIOKTN-UHFFFAOYSA-N | No-Effect | inactive |
| PF-573228 | BRD-K79239947 | CS(=O)(=O)c1cccc(CNc2nc(Nc3ccc4c(c3)CCC(=O)N4)ncc2C(F)(F)F)c1 | HESLKTSGTIBHJU-UHFFFAOYSA-N | No-Effect | inactive |
| PF-670462 | BRD-K96295391 | Nc1nccc(-c2c(-c3ccc(F)cc3)ncn2C2CCCCC2)n1 | WUDBUIUHVNECTM-UHFFFAOYSA-N | No-Effect | inactive |
| PF-750 | BRD-K83213911 | O=C(Nc1ccccc1)N1CCC(Cc2cnc3ccccc3c2)CC1 | BIODYGOZWZNCAG-UHFFFAOYSA-N | No-Effect | inactive |
| PF-8380 | BRD-K50865540 | O=C(CCN1CCN(C(=O)OCc2cc(Cl)cc(Cl)c2)CC1)c1ccc2[nH]c(=O)oc2c1 | JMSUDQYHPSNBSN-UHFFFAOYSA-N | No-Effect | inactive |
| PF-915275 | BRD-K18289362 | N#Cc1ccc(-c2ccc(S(=O)(=O)Nc3cccc(N)n3)cc2)cc1 | ZESFDAKNYJQYKO-UHFFFAOYSA-N | No-Effect | inactive |
| PFI-3 | BRD-K61524559 | O=C(/C=C/N1C[C@H]2C[C@@H]1CN2c1ccccn1)c1ccccc1O | INAICWLVUAKEPB-QSTFCLMHSA-N | No-Effect | inactive |
| PFI-4 | BRD-K13106865 | COc1ccccc1C(=O)Nc1cc2c(cc1N1CCCC1)n(C)c(=O)n2C | QCIJLRJBZDBVDB-UHFFFAOYSA-N | No-Effect | inactive |
| PFK-015 | BRD-K95573441 | O=C(/C=C/c1ccc2ccccc2n1)c1ccncc1 | UJJUKZPBUMCSJZ-BQYQJAHWSA-N | No-Effect | inactive |
| PG-9 | BRD-K00004212 | C[C@@H](C(=O)O[C@@H]1C[C@@H]2CC[C@H](C1)N2C)c1ccc(Br)cc1 | GHDWIFTUSXQYGK-FWYOQMDTSA-N | No-Effect | inactive |
| PGL5001 | BRD-A53813595 | N#C[C@H](c1ccnc(OCc2ccc(CN3CCOCC3)cc2)n1)c1nc2ccccc2s1 | XCPPIJCBCWUBNT-HXUWFJFHSA-N | No-Effect | inactive |
| PH-797804 | BRD-K95880107 | CNC(=O)c1ccc(C)c(-n2c(C)cc(OCc3ccc(F)cc3F)c(Br)c2=O)c1 | KCAJXIDMCNPGHZ-UHFFFAOYSA-N | No-Effect | inactive |
| PHA-543613 | BRD-K00004641 | O=C(N[C@H]1CN2CCC1CC2)c1cc2cocc2cn1 | IPKZCLGGYKRDES-ZDUSSCGKSA-N | No-Effect | inactive |
| PHA-568487 | BRD-K48053228 | O=C(N[C@H]1CN2CCC1CC2)c1ccc2c(c1)OCCO2 | LUVXHMJTVXZFPD-ZDUSSCGKSA-N | No-Effect | inactive |
| PHA-665752 | BRD-K95435023 | Cc1[nH]c(/C=C2\C(=O)Nc3ccc(S(=O)(=O)Cc4c(Cl)cccc4Cl)cc32)c(C)c1C(=O)N1CCC[C@@H]1CN1CCCC1 | OYONTEXKYJZFHA-SSHUPFPWSA-N | No-Effect | inactive |
| PHA-680632 | BRD-K09443272 | CCc1cccc(CC)c1NC(=O)N1Cc2n[nH]c(NC(=O)c3ccc(N4CCN(C)CC4)cc3)c2C1 | OBWNXGOQPLDDPS-UHFFFAOYSA-N | No-Effect | inactive |
| PHA-767491 | BRD-K50000283 | O=C1NCCc2[nH]c(-c3ccncc3)cc21 | DKXHSOUZPMHNIZ-UHFFFAOYSA-N | No-Effect | inactive |
| PHA-793887 | BRD-K64800655 | CC(C)CC(=O)Nc1[nH]nc2c1CN(C(=O)C1CCN(C)CC1)C2(C)C | HUXYBQXJVXOMKX-UHFFFAOYSA-N | No-Effect | inactive |
| PHCCC | BRD-K00004222 | O=C(Nc1ccccc1)[C@]12C[C@H]1/C(=N\O)c1ccccc1O2 | FPXPIEZPAXSELW-RGNAGUPPSA-N | No-Effect | inactive |
| phenacaine | BRD-K81092066 | CCOc1ccc(N=C(C)Nc2ccc(OCC)cc2)cc1 | QXDAEKSDNVPFJG-UHFFFAOYSA-N | No-Effect | inactive |
| phenacemide | BRD-K40905133 | NC(=O)NC(=O)Cc1ccccc1 | XPFRXWCVYUEORT-UHFFFAOYSA-N | No-Effect | inactive |
| phenacetin | BRD-K38323065 | CCOc1ccc(NC(C)=O)cc1 | CPJSUEIXXCENMM-UHFFFAOYSA-N | No-Effect | inactive |
| phenazone | BRD-K46937689 | Cc1cc(=O)n(-c2ccccc2)n1C | VEQOALNAAJBPNY-UHFFFAOYSA-N | No-Effect | inactive |

| | | | | | |
|---|---|---|---|---|---|
| phenazopyridine | BRD-K76304753 | Nc1ccc(/N=N/c2ccccc2)c(N)n1 | QPFYXYFORQJZEC-FOCLMDBBSA-N | No-Effect | inactive |
| phenelzine | BRD-K87024524 | NNCCc1ccccc1 | RMUCZJUITONUFY-UHFFFAOYSA-N | No-Effect | inactive |
| phenethicillin | BRD-A66025870 | C[C@@H](Oc1ccccc1)C(=O)N[C@@H]1C(=O)N2[C@@H](C(=O)O)C(C)(C)S[C@H]12 | NONJJLVGHLVQQM-ADGXKJENSA-N | No-Effect | inactive |
| phenethyl-isothiocyanate | BRD-K56700933 | S=C=NCCc1ccccc1 | IZJDOKYDEWTZSO-UHFFFAOYSA-N | No-Effect | inactive |
| phenformin | BRD-K11399644 | N=C(N=C(N)N)NCCc1ccccc1 | ICFJFFQQTFMIBG-UHFFFAOYSA-N | No-Effect | inactive |
| phenindamine | BRD-A33178345 | CN1CCC2=C(C1)[C@H](c1ccccc1)c1ccccc12 | ISFHAYSTHMVOJR-LJQANCHMSA-N | No-Effect | inactive |
| pheniramine | BRD-A23072235 | CN(C)CC[C@@H](c1ccccc1)c1ccccn1 | IJHNSHDBIRRJRN-HNNXBMFYSA-N | No-Effect | inactive |
| phenol | BRD-K76549667 | Oc1ccccc1 | ISWSIDIOOBJBQZ-UHFFFAOYSA-N | No-Effect | inactive |
| phenolphthalein | BRD-K19227686 | O=C1OC(c2ccc(O)cc2)(c2ccc(O)cc2)c2ccccc21 | KJFMBFZCATUALV-UHFFFAOYSA-N | No-Effect | inactive |
| phenolsulfonphthalein | BRD-K79892937 | O=S1(=O)OC(c2ccc(O)cc2)(c2ccc(O)cc2)c2ccccc21 | BELBBZDIHDAJOR-UHFFFAOYSA-N | No-Effect | inactive |
| phenothiazine | BRD-K59597909 | c1ccc2c(c1)Nc1ccccc1S2 | WJFKNYWRSNBZNX-UHFFFAOYSA-N | No-Effect | inactive |
| phenoxybenzamine | BRD-A67799922 | CC(COc1ccccc1)N(CCCl)Cc1ccccc1 | QZVCTJOXCFMACW-UHFFFAOYSA-N | No-Effect | inactive |
| phenprobamate | BRD-K22009844 | NC(=O)OCCCc1ccccc1 | CAMYKONBWHRPDD-UHFFFAOYSA-N | No-Effect | inactive |
| phenprocoumon | BRD-A49838158 | CC[C@@H](c1ccccc1)c1c(O)c2ccccc2oc1=O | DQDAYGNAKTZFIW-ZDUSSCGKSA-N | No-Effect | inactive |
| phenserine | BRD-K46580984 | CN1CC[C@@]2(C)c3cc(OC(=O)Nc4ccccc4)ccc3N(C)[C@@H]12 | PBHFNBQPZCRWQP-QUCCMNQESA-N | No-Effect | inactive |
| phensuximide | BRD-A18043272 | CN1C(=O)C[C@H](c2ccccc2)C1=O | WLWFNJKHKGIJNW-SECBINFHSA-N | No-Effect | inactive |
| phentolamine | BRD-K90333595 | Cc1ccc(N(CC2=NCCN2)c2cccc(O)c2)cc1 | MRBDMNSDAVCSSF-UHFFFAOYSA-N | No-Effect | inactive |
| phenyl-aminosalicylate | BRD-K73157543 | Nc1ccc(C(=O)Oc2ccccc2)c(O)c1 | DNVVZWSVACQWJE-UHFFFAOYSA-N | No-Effect | inactive |
| phenyl-salicylate | BRD-K44091963 | O=C(Oc1ccccc1)c1ccccc1O | ZQBAKBUEJOMQEX-UHFFFAOYSA-N | No-Effect | inactive |
| phenylacetylglutamine | BRD-K70592483 | NC(=O)CC[C@H](NC(=O)Cc1ccccc1)C(=O)O | JFLIEFSWGNOPJJ-JTQLQIEISA-N | No-Effect | inactive |
| phenylbenzimidazole-sulfonic-acid | BRD-K02729857 | O=S(=O)(O)c1ccc2nc(-c3ccccc3)[nH]c2c1 | UVCJGUGAGLDPAA-UHFFFAOYSA-N | No-Effect | inactive |
| phenylbutazone | BRD-K10843433 | CCCCC1C(=O)N(c2ccccc2)N(c2ccccc2)C1=O | VYMDGNCVAMGZFE-UHFFFAOYSA-N | No-Effect | inactive |
| phenylbutyrate | BRD-K67102207 | O=C(O)CCCc1ccccc1 | OBKXEAXTFZPCHS-UHFFFAOYSA-N | No-Effect | inactive |
| phenylethyl-alcohol | BRD-K42371134 | OCCc1ccccc1 | WRMNZCZEMHIOCP-UHFFFAOYSA-N | No-Effect | inactive |
| phenylmercuric-acetate | BRD-K74193284 | CC(=O)O[Hg]c1ccccc1 | XEBWQGVWTUSTLN-UHFFFAOYSA-M | No-Effect | inactive |
| phenylmethylsulfonyl-fluoride | BRD-K33422401 | O=S(=O)(F)Cc1ccccc1 | YBYRMVIVWMBXKQ-UHFFFAOYSA-N | No-Effect | inactive |
| phenylpiracetam | BRD-A65683612 | NC(=O)CN1C[C@H](c2ccccc2)CC1=O | LYONXVJRBWWGQO-SNVBAGLBSA-N | No-Effect | inactive |
| phenytoin | BRD-K55930204 | O=C1NC(=O)C(c2ccccc2)(c2ccccc2)N1 | CXOFVDLJLONNDW-UHFFFAOYSA-N | No-Effect | inactive |
| PhiKan-083 | BRD-K44400149 | CCn1c2ccccc2c2cc(CNC)ccc21 | LBPNOEAFWYTTEB-UHFFFAOYSA-N | No-Effect | inactive |
| phloretin | BRD-K15563106 | O=C(CCc1ccc(O)cc1)c1c(O)cc(O)cc1O | VGEREEWJJVICBM-UHFFFAOYSA-N | No-Effect | inactive |
| phlorizin | BRD-K73756878 | O=C(CCc1ccc(O)cc1)c1c(O)cc(O)cc1O[C@@H]1O[C@H](CO)[C@@H](O)[C@H](O)[C@H]1O | IOUVKUPGCMBWBT-QNDFHXLGSA-N | No-Effect | inactive |
| phloroglucin | BRD-K00405906 | Oc1cc(O)cc(O)c1 | QCDYQQDYXPDABM-UHFFFAOYSA-N | No-Effect | inactive |
| phortress | BRD-K74558903 | Cc1cc(-c2nc3cc(F)ccc3s2)ccc1NC(=O)[C@@H](N)CCCCN | LFDOEFHQLNGBQQ-HNNXBMFYSA-N | No-Effect | inactive |
| phosphatidylcholine | BRD-A10131665 | CCCCC/C=C\C/C=C\CCCCCCCC(=O)O[C@H](COC(=O)CCCCCCCCCCCCCCC)CO[P@@](=O)(O)OCC[N+](C)(C)C | JLPULHDHAOZNQI-ZTIMHPMXSA-O | No-Effect | inactive |
| PHP-501 | BRD-K07460388 | On1ncc(C2CCNCC2)c1-c1cccc(-c2ccccc2)c1 | UHEYIOPHTBWBQN-UHFFFAOYSA-N | No-Effect | inactive |
| phthalylsulfacetamide | BRD-K77115421 | CC(=O)NS(=O)(=O)c1ccc(NC(=O)c2ccccc2C(=O)O)cc1 | SNWQKAWITMVCQW-UHFFFAOYSA-N | No-Effect | inactive |
| PHTPP | BRD-K76568384 | Oc1ccc(-c2c(-c3ccccc3)nn3c(C(F)(F)F)cc(C(F)(F)F)nc23)cc1 | AEZPAUSGTAHLOQ-UHFFFAOYSA-N | No-Effect | inactive |
| physostigmine | BRD-K25650355 | CNC(=O)Oc1ccc2c(c1)[C@]1(C)CCN(C)[C@@H]1N2C | PIJVFDBKTWXHHD-HIFRSBDPSA-N | No-Effect | inactive |
| phytonadione | BRD-K76661572 | CC1=C(C/C=C(\C)CCC[C@H](C)CCC[C@H](C)CCCC(C)C)C(=O)c2ccccc2C1=O | MBWXNTAXLNYFJB-NKFFZRIASA-N | No-Effect | inactive |
| phytosphingosine | BRD-K05048137 | CCCCCCCCCCCCCC[C@@H](O)[C@@H](O)[C@@H](N)CO | AERBNCYCJBRYDG-KSZLIROESA-N | No-Effect | inactive |
| PI3K-IN-2 | BRD-K62374002 | Nc1cc(C(F)(F)F)c(-c2nc(N3CCOCC3)nc(N3CCOCC3)n2)cn1 | ADGGYDAFIHSYFI-UHFFFAOYSA-N | No-Effect | inactive |
| PI4KIII-beta-inhibitor-1 | BRD-K75528696 | COc1ccc(-c2sc(NC(=O)C(C)(C)C)nc2C)cc1S(=O)(=O)Nc1ccc(O)cc1 | PLUYFBRIGUAKBR-UHFFFAOYSA-N | No-Effect | inactive |
| pibenzimol | BRD-K17026858 | CN1CCN(c2ccc3nc(-c4ccc5nc(-c6ccc(O)cc6)[nH]c5c4)[nH]c3c2)CC1 | INAAIJLSXJJHOZ-UHFFFAOYSA-N | No-Effect | inactive |
| piboserod | BRD-K02750403 | CCCCN1CCC(CNC(=O)c2c3n(c4ccccc24)CCCO3)CC1 | KVCSJPATKXABRQ-UHFFFAOYSA-N | No-Effect | inactive |
| picartamide | BRD-A71765060 | CNC(=S)[C@@]1(c2ccccn2)CCCS1 | ITNLONMDUMHEOK-NSHDSACASA-N | No-Effect | inactive |
| piceatannol | BRD-K91509126 | Oc1cc(O)cc(/C=C/c2ccc(O)c(O)c2)c1 | CDRPUGZCRXZLFL-OWOJBTEDSA-N | No-Effect | inactive |
| piclamilast | BRD-K47642186 | COc1ccc(C(=O)Nc2c(Cl)cncc2Cl)cc1OC1CCCC1 | RRRUXBQSQLKHEL-UHFFFAOYSA-N | No-Effect | inactive |
| picolamine | BRD-K68653965 | NCc1cccnc1 | HDOUGSFASVGDCS-UHFFFAOYSA-N | No-Effect | inactive |
| picolinic-acid | BRD-K60612034 | O=C(O)c1ccccn1 | SIOXPEMLGUPBBT-UHFFFAOYSA-N | No-Effect | inactive |
| piconol | BRD-K28492574 | OCc1ccccn1 | SHNUBALDGXWUJI-UHFFFAOYSA-N | No-Effect | inactive |
| picotamide | BRD-K67277431 | COc1ccc(C(=O)NCc2cccnc2)cc1C(=O)NCc1cccnc1 | KYWCWBXGRWWINE-UHFFFAOYSA-N | No-Effect | inactive |
| picrotin | BRD-A91822151 | CC(C)(O)[C@@H]1[C@@H]2C(=O)O[C@H]1[C@@H]1OC(=O)[C@@]34O[C@@H]3C[C@@]2(O)[C@]14C | RYEFFICCPKWYML-XSVQNFTRSA-N | No-Effect | inactive |
| picrotoxinin | BRD-K36461289 | C=C(C)[C@@H]1[C@H]2OC(=O)[C@H]1[C@]1(O)C[C@H]3O[C@]34C(=O)O[C@H]2[C@@]41C | PIMZUZSSNYHVCU-KBLUICEQSA-N | No-Effect | inactive |
| pidolic-acid | BRD-K57313110 | O=C1CC[C@@H](C(=O)O)N1 | ODHCTXKNWHHXJC-VKHMYHEASA-N | No-Effect | inactive |
| pifithrin-alpha | BRD-K66874953 | Cc1ccc(C(=O)Cn2c3c(sc2=N)CCCC3)cc1 | OLTZHXYLLRJLST-UHFFFAOYSA-N | No-Effect | inactive |
| pifithrin-cyclic | BRD-K31792052 | Cc1ccc(-c2cn3c4c(sc3n2)CCCC4)cc1 | IMUKUMUNZJILCG-UHFFFAOYSA-N | No-Effect | inactive |
| PIK-293 | BRD-K30977212 | Cc1ccccc1-n1c(Cn2ncc3c(N)ncnc32)nc2cccc(C)c2c1=O | KQDBVHKNIYROHU-UHFFFAOYSA-N | No-Effect | inactive |
| PIK-294 | BRD-K50631926 | Cc1ccccc1-n1c(Cn2nc(-c3cccc(O)c3)c3c(N)ncnc32)nc2cccc(C)c2c1=O | WFSLJOPRIJSOJR-UHFFFAOYSA-N | No-Effect | inactive |
| PIK-75 | BRD-K49371609 | Cc1ccc([N+](=O)[O-])cc1S(=O)(=O)N(C)/N=C/c1cnc2ccc(Br)cn12 | QTHCAAFKVUWAFI-DJKKODMXSA-N | No-Effect | inactive |
| piketoprofen | BRD-A85548292 | Cc1ccnc(NC(=O)[C@H](C)c2cccc(C(=O)c3ccccc3)c2)c1 | ASFKKFRSMGBFRO-MRXNPFEDSA-N | No-Effect | inactive |
| pilaralisib | BRD-K49456274 | COc1ccc(Cl)c(Nc2nc3ccccc3nc2NS(=O)(=O)c2cccc(NC(=O)C(C)(C)N)c2)c1 | QINPEPAQOBZPOF-UHFFFAOYSA-N | No-Effect | inactive |
| pilocarpine | BRD-K85090592 | CC[C@@H]1C(=O)OC[C@@H]1Cc1cncn1C | QCHFTSOMWOSFHM-WPRPVWTQSA-N | No-Effect | inactive |
| pilsicainide | BRD-K83245588 | Cc1cccc(C)c1NC(=O)CC12CCCN1CCC2 | BCQTVJKBTWGHCX-UHFFFAOYSA-N | No-Effect | inactive |
| PIM-1-Inhibitor-2 | BRD-K48832421 | Nc1ncc(-c2ccc3noc(-c4ccc(Cl)cc4)c3c2)n1 | AQVFETGXIRKVAQ-UHFFFAOYSA-N | No-Effect | inactive |
| PIM-447 | BRD-K00003280 | C[C@@H]1C[C@H](N)C[C@H](c2ccncc2NC(=O)c2ccc(F)c(-c3c(F)cccc3F)n2)C1 | VRQXRVAKPDCRCI-ZNMIVQPWSA-N | No-Effect | inactive |
| pimecrolimus | BRD-K92107055 | CC[C@@H]1/C=C(\C)C[C@H](C)C[C@H](OC)[C@H]2O[C@@](O)(C(=O)C(=O)N3CCCC[C@H]3C(=O)O[C@H](/C(C)=C/[C@@H]3CC[C@H](Cl)[C@H](OC)C3)[C@H](C)[C@@H](O)CC1=O)[C@H](C)C[C@@H]2OC | KASDHRXLYQOAKZ-XDSKOBMDSA-N | No-Effect | inactive |
| pimobendan | BRD-K01826269 | COc1ccc(-c2nc3ccc(C4=NNC(=O)C[C@H]4C)cc3[nH]2)cc1 | GLBJJMFZWDBELO-LLVKDONJSA-N | No-Effect | inactive |

| | | | | | |
|---|---|---|---|---|---|
| pimodivir | BRD-K00003127 | O=C(O)[C@H]1C2CCC(CC2)[C@@H]1Nc1nc(-c2c[nH]c3ncc(F)cc23)ncc1F | JGPXDNKSIXAZEQ-SBBZOCNPSA-N | No-Effect | inactive |
| pimonidazole | BRD-A67521319 | O=[N+]([O-])c1nccn1C[C@@H](O)CN1CCCCC1 | WVWOOAYQYLJEFD-JTQLQIEISA-N | No-Effect | inactive |
| pimozide | BRD-K01292756 | O=c1[nH]c2ccccc2n1C1CCN(CCCC(c2ccc(F)cc2)c2ccc(F)cc2)CC1 | YVUQSNJEYSNKRX-UHFFFAOYSA-N | No-Effect | inactive |
| pinacidil | BRD-A43882281 | C[C@@H](NC(=NC#N)Nc1ccncc1)C(C)(C)C | IVVNZDGDKPTYHK-SNVBAGLBSA-N | No-Effect | inactive |
| pinanediol | BRD-K32485706 | CC1(C)[C@H]2C[C@@H]1[C@](C)(O)[C@H](O)C2 | MOILFCKRQFQVFS-OORONAJNSA-N | No-Effect | inactive |
| pinaverium | BRD-A13175897 | COc1cc(Br)c(C[N+]2(CCOCC[C@H]3CC[C@@H]4C[C@H]3C4(C)C)CCOCC2)cc1OC | DDHUTBKXLWCZCO-CEMLEFRQSA-N | No-Effect | inactive |
| pincainide | BRD-K90727284 | Cc1cccc(C)c1NC(=O)CN1CCCCCC1 | RJOUHGWLHPOQSA-UHFFFAOYSA-N | No-Effect | inactive |
| pindolol | BRD-A97701745 | CC(C)NC[C@@H](O)COc1cccc2[nH]ccc12 | JZQKKSLKJUAGIC-LLVKDONJSA-N | No-Effect | inactive |
| pindolol-(-) | BRD-K95598440 | CC(C)NC[C@H](O)COc1cccc2[nH]ccc12 | JZQKKSLKJUAGIC-NSHDSACASA-N | No-Effect | inactive |
| pinocembrin | BRD-K94689771 | O=C1C[C@@H](c2ccccc2)Oc2cc(O)cc(O)c21 | URFCJEUYXNAHFI-ZDUSSCGKSA-N | No-Effect | inactive |
| pioglitazone | BRD-K01826904 | CCc1ccc(CCOc2ccc(C[C@H]3SC(=O)NC3=O)cc2)nc1 | HYAFETHFCAUJAY-QGZVFWFLSA-N | No-Effect | inactive |
| pipamperone | BRD-K26801045 | NC(=O)C1(N2CCCCC2)CCN(CCCC(=O)c2ccc(F)cc2)CC1 | AXKPFOAXAHJUAG-UHFFFAOYSA-N | No-Effect | inactive |
| pipecuronium | BRD-A21544762 | CC(=O)O[C@H]1C[C@@H]2CC[C@@H]3[C@H](CC[C@@]4(C)[C@H]3C[C@H](N3CC[N+](C)(C)CC3)[C@@H]4OC(C)=O)[C@@]2(C)C[C@@H]1N1CC[N+](C)(C)CC1 | OWWLUIWOFHMHOQ-XGHATYIMSA-N | No-Effect | inactive |
| pipemidic-acid | BRD-K08999871 | CCn1cc(C(=O)O)c(=O)c2cnc(N3CCNCC3)nc21 | JOHZPMXAZQZXHR-UHFFFAOYSA-N | No-Effect | inactive |
| pipenzolate | BRD-A53561827 | CC[N@@+]1(C)CCC[C@H](OC(=O)C(O)(c2ccccc2)c2ccccc2)C1 | WPUKUEMZZRVAKZ-REWPJTCUSA-N | No-Effect | inactive |
| piperacetazine | BRD-K16277217 | CC(=O)c1ccc2c(c1)N(CCCN1CCC(CCO)CC1)c1ccccc1S2 | BTFMCMVEUCGQDX-UHFFFAOYSA-N | No-Effect | inactive |
| piperacillin | BRD-K86873305 | CCN1CCN(C(=O)N[C@@H](C(=O)N[C@@H]2C(=O)N3[C@@H](C(=O)O)C(C)(C)S[C@H]23)c2ccccc2)C(=O)C1=O | IVBHGBMCVLDMKU-GXNBUGAJSA-N | No-Effect | inactive |
| piperaquine-phosphate | BRD-K27382019 | Clc1ccc2c(N3CCN(CCCN4CCN(c5ccnc6cc(Cl)ccc56)CC4)CC3)ccnc2c1 | UCRHFBCYFMIWHC-UHFFFAOYSA-N | No-Effect | inactive |
| piperazine | BRD-K13249881 | C1CNCCN1 | GLUUGHFHXGJENI-UHFFFAOYSA-N | No-Effect | inactive |
| piperazinedione | BRD-K95941975 | O=C1CNC(=O)CN1 | BXRNXXXXHLBUKK-UHFFFAOYSA-N | No-Effect | inactive |
| piperidolate | BRD-A97479839 | CCN1CCC[C@H](OC(=O)C(c2ccccc2)c2ccccc2)C1 | KTHVBAZBLKXIHZ-IBGZPJMESA-N | No-Effect | inactive |
| piperine | BRD-K59522102 | O=C(/C=C/C=C/c1ccc2c(c1)OCO2)N1CCCCC1 | MXXWOMGUGJBKIW-YPCIICBESA-N | No-Effect | inactive |
| piperonyl-butoxide | BRD-K76452401 | CCCCOCCOCCOCc1cc2c(cc1CCC)OCO2 | FIPWRIJSWJWJAI-UHFFFAOYSA-N | No-Effect | inactive |
| pipobroman | BRD-K15656090 | O=C(CCBr)N1CCN(C(=O)CCBr)CC1 | NJBFOOCLYDNZJN-UHFFFAOYSA-N | No-Effect | inactive |
| pipofezine | BRD-K00003317 | CN1CCN(c2cc3c(nn2)Oc2ccccc2N3C)CC1 | SDYYIRPAZHJOLM-UHFFFAOYSA-N | No-Effect | inactive |
| pipotiazine-palmitate | BRD-K48429278 | CCCCCCCCCCCCCCCC(=O)OCCC1CCN(CCCN2c3ccccc3Sc3ccc(S(=O)(=O)N(C)C)cc32)CC1 | KTOYYUONFQWSMW-UHFFFAOYSA-N | No-Effect | inactive |
| piracetam | BRD-K19456237 | NC(=O)CN1CCCC1=O | GMZVRMREEHBGGF-UHFFFAOYSA-N | No-Effect | inactive |
| pirarubicin | BRD-K83794624 | COc1cccc2c1C(=O)c1c(O)c3c(c(O)c1C2=O)C[C@@](O)(C(=O)CO)C[C@@H]3O[C@H]1C[C@H](N)[C@H](O[C@@H]2CCCCO2)[C@H](C)O1 | KMSKQZKKOZQFFG-YXRRJAAWSA-N | No-Effect | inactive |
| pirenoxine | BRD-K65992024 | O=C(O)c1cc(=O)c2c3nc4ccccc4oc-3cc(=O)c2[nH]1 | OKPNYGAWTYOBFZ-UHFFFAOYSA-N | No-Effect | inactive |
| pirenzepine | BRD-K89375097 | CN1CCN(CC(=O)N2c3ccccc3C(=O)Nc3cccnc32)CC1 | RMHMFHUVIITRHF-UHFFFAOYSA-N | No-Effect | inactive |
| piretanide | BRD-K87990216 | NS(=O)(=O)c1cc(C(=O)O)cc(N2CCCC2)c1Oc1ccccc1 | UJEWTUDSLQGTOA-UHFFFAOYSA-N | No-Effect | inactive |
| pirfenidone | BRD-K96862998 | Cc1ccc(=O)n(-c2ccccc2)c1 | ISWRGOKTTBVCFA-UHFFFAOYSA-N | No-Effect | inactive |
| piricapiron | BRD-K00004544 | CC1(C)[C@@H]2CC[C@@]1(C)C(=O)N(CCCCN1CCN(c3ncccn3)CC1)C2=O | NCMIPRCQUMYAEC-JPYJTQIMSA-N | No-Effect | inactive |
| pirinixic-acid | BRD-K01902415 | Cc1cccc(Nc2cc(Cl)nc(SCC(=O)O)n2)c1C | SZRPDCCEHVWOJX-UHFFFAOYSA-N | No-Effect | inactive |
| pirlindole | BRD-K01826702 | Cc1ccc2c(c1)c1c3n2CCN[C@H]3CCC1 | IWVRVEIKCBFZNF-ZDUSSCGKSA-N | No-Effect | inactive |
| pirmenol | BRD-K00003530 | C[C@H]1CCC[C@@H](C)N1CCC[C@@](O)(c1ccccc1)c1ccccn1 | APUDBKTWDCXQJA-XQBPLPMBSA-N | No-Effect | inactive |
| piroctone-olamine | BRD-A96510685 | Cc1cc(C[C@H](C)CC(C)(C)C)n(O)c(=O)c1 | OIQJEQLSYJSNDS-NSHDSACASA-N | No-Effect | inactive |
| pirodavir | BRD-K22261391 | CCOC(=O)c1ccc(OCCC2CCN(c3ccc(C)nn3)CC2)cc1 | KCHIOGFOPPOUJC-UHFFFAOYSA-N | No-Effect | inactive |
| piromidic-acid | BRD-K37682401 | CCn1cc(C(=O)O)c(=O)c2cnc(N3CCCC3)nc21 | RCIMBBZXSXFZBV-UHFFFAOYSA-N | No-Effect | inactive |
| piroxicam | BRD-K01826746 | CN1C(C(=O)Nc2ccccn2)=C(O)c2ccccc2S1(=O)=O | QYSPLQLAKJAUJT-UHFFFAOYSA-N | No-Effect | inactive |
| piroximone | BRD-K71977764 | CCc1[nH]c(=O)[nH]c1C(=O)c1ccncc1 | OQGWJZOWLHWFME-UHFFFAOYSA-N | No-Effect | inactive |
| pirozadil | BRD-K00004581 | COc1cc(C(=O)OCc2cccc(COC(=O)c3cc(OC)c(OC)c(OC)c3)n2)cc(OC)c1OC | DIIBXMIIOQXTHW-UHFFFAOYSA-N | No-Effect | inactive |
| pirquinozol | BRD-K63651590 | O=c1[nH]c2ccccc2c2cc(CO)nn12 | XLUKOGNIEDDIMV-UHFFFAOYSA-N | No-Effect | inactive |
| PIT | BRD-K43978949 | O=C1C(c2ccccn2)=[N+]([O-])c2ccccc21 | LXEZPPZLNVOLDI-UHFFFAOYSA-N | No-Effect | inactive |
| pivagabine | BRD-K62264492 | CC(C)(C)C(=O)NCCCC(=O)O | SRPNQDXRVRCTNK-UHFFFAOYSA-N | No-Effect | inactive |
| pivampicillin | BRD-K32791151 | CC(C)(C)C(=O)OCOC(=O)[C@@H]1N2C(=O)[C@@H](NC(=O)[C@H](N)c3ccccc3)[C@H]2SC1(C)C | ZEMIJUDPLILVNQ-ZXFNITATSA-N | No-Effect | inactive |
| pivanex | BRD-K20504455 | CCCC(=O)OCOC(=O)C(C)(C)C | GYKLFBYWXZYSOW-UHFFFAOYSA-N | No-Effect | inactive |
| pivmecillinam | BRD-K67100011 | CC(C)(C)C(=O)OCOC(=O)[C@@H]1N2C(=O)[C@@H](/N=C/N3CCCCCC3)[C@H]2SC1(C)C | NPGNOVNWUSPMDP-UTEPHESZSA-N | No-Effect | inactive |
| pixantrone | BRD-K48304345 | NCCNc1ccc(NCCN)c2c1C(=O)c1ccncc1C2=O | PEZPMAYDXJQYRV-UHFFFAOYSA-N | No-Effect | inactive |
| pizotifen | BRD-K75958195 | CN1CCC(=C2c3ccccc3CCc3sccc32)CC1 | FIADGNVRKBPQEU-UHFFFAOYSA-N | No-Effect | inactive |
| PJ-34 | BRD-K11853856 | CN(C)CC(=O)Nc1ccc2[nH]c(=O)c3ccccc3c2c1 | UYJZZVDLGDDTCL-UHFFFAOYSA-N | No-Effect | inactive |
| PK-11195 | BRD-A41451487 | CC[C@@H](C)N(C)C(=O)c1cc2ccccc2c(-c2ccccc2Cl)n1 | RAVIZVQZGXBOQO-CQSZACIVSA-N | No-Effect | inactive |
| PK-44 | BRD-K04035716 | N[C@@H](CC(=O)N1CCn2c(nnc2C(F)(F)F)C1)Cc1[nH]nc2c(F)c(F)ccc12 | LIPWZZYFIVYJSI-MRVPVSSYSA-N | No-Effect | inactive |
| PK-THPP | BRD-K44043890 | CCCC(=O)C1CCN(c2ncnc3c2CN(C(=O)c2ccc(-c4ccccc4)cc2)CC3)CC1 | CJZGRIRZVHNUSM-UHFFFAOYSA-N | No-Effect | inactive |
| PKI-166 | BRD-K10115470 | C[C@@H](Nc1ncnc2[nH]c(-c3ccc(O)cc3)cc12)c1ccccc1 | XRYJULCDUUATMC-CYBMUJFWSA-N | No-Effect | inactive |
| PKI-179 | BRD-A28467416 | O=C(Nc1ccncc1)Nc1ccc(-c2nc(N3CCOCC3)nc(N3[C@H]4CC[C@@H]3COC4)n2)cc1 | WXUUCRLKXQMWRY-OYRHEFFESA-N | No-Effect | inactive |
| pleconaril | BRD-K27625831 | Cc1cc(CCCOc2c(C)cc(-c3noc(C(F)(F)F)n3)cc2C)on1 | KQOXLKOJHVFTRN-UHFFFAOYSA-N | No-Effect | inactive |
| plerixafor | BRD-K33240821 | c1cc(CN2CCCNCCNCCCNCC2)ccc1CN1CCCNCCNCCCNCC1 | YIQPUIGJQJDJOS-UHFFFAOYSA-N | No-Effect | inactive |
| plovamer-acetate | BRD-A20781140 | N[C@@H](Cc1ccc(O)cc1)C(=O)O | OUYCCCASQSFEME-QMMMGPOBSA-N | No-Effect | inactive |
| plumbagin | BRD-K36137799 | CC1=CC(=O)c2c(O)cccc2C1=O | VCMMXZQDRFWYSE-UHFFFAOYSA-N | No-Effect | inactive |
| plurisin-1 | BRD-K99149715 | O=C(NNc1ccccc1)c1ccncc1 | HUDWXDLBWRHCKO-UHFFFAOYSA-N | No-Effect | inactive |
| PLX4720 | BRD-K16478699 | CCCS(=O)(=O)Nc1ccc(F)c(C(=O)c2c[nH]c3ncc(Cl)cc23)c1F | YZDJQTHVDDOVHR-UHFFFAOYSA-N | No-Effect | inactive |
| PLX647 | BRD-K79294107 | FC(F)(F)c1ccc(CNc2ccc(Cc3c[nH]c4ncccc34)cn2)cc1 | NODCQQSEMCESEC-UHFFFAOYSA-N | No-Effect | inactive |
| PLX8394 | BRD-K53723268 | O=C(c1c(F)ccc(NS(=O)(=O)N2CC[C@@H](F)C2)c1F)c1c[nH]c2ncc(-c3cnc(C4CC4)nc3)cc12 | YYACLQUDUDXAPA-MRXNPFEDSA-N | No-Effect | inactive |
| PMPA | BRD-A74387232 | O=C(O)CC[C@@H](CP(=O)(O)O)C(=O)O | ISEYJGQFXSTPMQ-BYPYZUCNSA-N | No-Effect | inactive |
| PNU-120596 | BRD-K71731651 | COc1cc(OC)c(NC(=O)Nc2cc(C)on2)cc1Cl | CEIIEALEIHQDBX-UHFFFAOYSA-N | No-Effect | inactive |
| PNU-142633 | BRD-K81616657 | CNC(=O)c1ccc2c(c1)CCO[C@H]2CCN1CCN(c2ccc(C(N)=O)cc2)CC1 | PNTVCCRNJOGKGA-QFIPXVFZSA-N | No-Effect | inactive |
| PNU-177864 | BRD-K04635846 | CCCNCCc1ccc(NS(=O)(=O)c2ccc(OC(F)(F)F)cc2)cc1 | JGGQWSXZZQPZTR-UHFFFAOYSA-N | No-Effect | inactive |
| PNU-22394 | BRD-K16551401 | Cn1c2c(c3ccccc31)CCNCC2 | ZBXDOQWPGBISAR-UHFFFAOYSA-N | No-Effect | inactive |
| PNU-282987 | BRD-K28863208 | O=C(N[C@H]1CN2CCC1CC2)c1ccc(Cl)cc1 | WECKJONDRAUFDD-ZDUSSCGKSA-N | No-Effect | inactive |
| PNU-37883 | BRD-K06341478 | C1CCC(N=C(NC23CC4CC(CC(C4)C2)C3)N2CCOCC2)CC1 | GKCGWXNLOYTUNC-UHFFFAOYSA-N | No-Effect | inactive |
| PNU-74654 | BRD-K30707190 | Cc1ccc(/C=N/NC(=O)c2ccccc2Oc2ccccc2)o1 | JJEDWBQZCRESJL-DEDYPNTBSA-N | No-Effect | inactive |

| | | | | | |
|---|---|---|---|---|---|
| PNU-89843 | BRD-K84507776 | Cc1cc2c(N3CCCC3)nc(N3CCCC3)nc2n1C | UADIKEUOAMACNA-UHFFFAOYSA-N | No-Effect | inactive |
| pocapavir | BRD-K00003464 | COc1ccc(OCc2ccc(COc3c(Cl)cccc3Cl)cc2)c(Cl)c1 | XXMDDBVNWRWNCW-UHFFFAOYSA-N | No-Effect | inactive |
| polaprezinc | BRD-K00003535 | O=C1CCN[Zn]OC(=O)[C@H](Cc2cnc[nH]2)N1 | VBHDYBCWTAXBNA-FJXQXJEOSA-M | No-Effect | inactive |
| polidocanol | BRD-K35798650 | CCCCCCCCCCCCOCCOCCOCCOCCOCCOCCOCCOCCOCCO | ONJQDTZCDSESIW-UHFFFAOYSA-N | No-Effect | inactive |
| polydatin | BRD-K43236057 | OC[C@H]1O[C@@H](Oc2cc(O)cc(/C=C/c3ccc(O)cc3)c2)[C@H](O)[C@@H](O)[C@@H]1O | HSTZMXCBWJGKHG-CUYWLFDKSA-N | No-Effect | inactive |
| polyinosine | BRD-A67531813 | O=c1nc[nH]c2c1ncn2[C@H]1O[C@H](COP(=O)(O)O)[C@H](O)[C@H]1O | GRSZFWQUAKGDAV-SQQSDITASA-N | No-Effect | inactive |
| polymyxin-B-sulfate | BRD-A08641215 | CC[C@@H](C)CCCCC(=O)N[C@@H](CCN)C(=O)N[C@H](C(=O)N[C@@H](CCN)C(=O)N[C@H]1CCNC(=O)[C@H]([C@@H](C)O)NC(=O)[C@@H](CCN)NC(=O)[C@@H](CCN)NC(=O)[C@@H](CC(C)C)NC(=O)[C@@H](Cc2ccccc2)NC(=O)[C@@H](CCN)NC1=O)[C@@H](C)O | WQVJHHACXVLGBL-NPZCUVBMSA-N | No-Effect | inactive |
| polythiazide | BRD-A15739803 | CN1[C@@H](CSCC(F)(F)F)Nc2cc(Cl)c(S(N)(=O)=O)cc2S1(=O)=O | CYLWJCABXYDINA-JTQLQIEISA-N | No-Effect | inactive |
| ponatinib | BRD-K44227013 | Cc1ccc(C(=O)Nc2ccc(CN3CCN(C)CC3)c(C(F)(F)F)c2)cc1C#Cc1cnc2cccnn12 | PHXJVRSECIGDHY-UHFFFAOYSA-N | No-Effect | inactive |
| ponesimod | BRD-K44030614 | CCC/N=C1\S/C(=C\c2ccc(OC[C@H](O)CO)c(Cl)c2)C(=O)N1c1ccccc1C | LPAUOXUZGSBGDU-STDDISTJSA-N | No-Effect | inactive |
| posaconazole | BRD-K06222852 | CC[C@@H]([C@H](C)O)n1ncn(-c2ccc(N3CCN(c4ccc(OC[C@@H]5CO[C@@](Cn6cncn6)(c6ccc(F)cc6F)C5)cc4)CC3)cc2)c1=O | RAGOYPUPXAKGKH-XAKZXMRKSA-N | No-Effect | inactive |
| potassium-canrenoate | BRD-K46556543 | C[C@]12CC[C@H]3[C@@H](C=CC4=CC(=O)CC[C@@]43C)[C@@H]1CC[C@@]2(O)CCC(=O)O | PBKZPPIHUVSDNM-WNHSNXHDSA-N | No-Effect | inactive |
| pozanicline | BRD-K79803033 | Cc1ncccc1OC[C@@H]1CCCN1 | YRVIKLBSVVNSHF-JTQLQIEISA-N | No-Effect | inactive |
| poziotinib | BRD-K50010139 | C=CC(=O)N1CCC(Oc2cc3c(Nc4ccc(Cl)c(Cl)c4F)ncnc3cc2OC)CC1 | LPFWVDIFUFFKJU-UHFFFAOYSA-N | No-Effect | inactive |
| PP-1 | BRD-K47598052 | Cc1ccc(-c2nn(C(C)(C)C)c3ncnc(N)c23)cc1 | ZVPDNRVYHLRXLX-UHFFFAOYSA-N | No-Effect | inactive |
| PP-121 | BRD-K81801188 | Nc1ncnc2c1c(-c1cnc3[nH]ccc3c1)nn2C1CCCC1 | NVRXTLZYXZNATH-UHFFFAOYSA-N | No-Effect | inactive |
| PP-2 | BRD-K95785537 | CC(C)(C)n1nc(-c2ccc(Cl)cc2)c2c(N)ncnc21 | PBBRWFOVCUAONR-UHFFFAOYSA-N | No-Effect | inactive |
| PP242 | BRD-K87112191 | CC(C)n1nc(-c2cc3cc(O)ccc3[nH]2)c2c(N)ncnc21 | MFAQYJIYDMLAIM-UHFFFAOYSA-N | No-Effect | inactive |
| PPY-A | BRD-K72514671 | COc1ccccc1-c1c[nH]c2ncc(-c3cncc(C(=O)N(C)C)c3)cc12 | GYQRHHQPEMOLKH-UHFFFAOYSA-N | No-Effect | inactive |
| PQ-401 | BRD-K67537649 | COc1ccc(Cl)cc1NC(=O)Nc1cc(C)nc2ccccc12 | YBLWOZUPHDKFOT-UHFFFAOYSA-N | No-Effect | inactive |
| PR-619 | BRD-K54395039 | N#CSc1cc(SC#N)c(N)nc1N | ZXOBLNBVNROVLC-UHFFFAOYSA-N | No-Effect | inactive |
| practolol | BRD-A41304429 | CC(=O)Nc1ccc(OC[C@H](O)CNC(C)C)cc1 | DURULFYMVIFBIR-CYBMUJFWSA-N | No-Effect | inactive |
| pradefovir | BRD-K00003119 | Nc1ncnc2c1ncn2CCOC[P@@]1(=O)OCC[C@@H](c2cccc(Cl)c2)O1 | GWNHAOBXDGOXRR-HJFSHJIFSA-N | No-Effect | inactive |
| pralidoxime | BRD-K84281997 | C[n+]1ccccc1/C=N/O | JBKPUQTUERUYQE-UHFFFAOYSA-O | No-Effect | inactive |
| pralidoxime-chloride | BRD-K24221957 | C[n+]1ccccc1CN=O | MHOAUIZFSQGCNM-UHFFFAOYSA-N | No-Effect | inactive |
| pralmorelin | BRD-K00004629 | C[C@H](NC(=O)[C@@H](Cc1ccc2ccccc2c1)NC(=O)[C@@H](C)N)C(=O)N[C@@H](Cc1c[nH]c2ccccc12)C(=O)N[C@H](Cc1ccccc1)C(=O)N[C@@H](CCCCN)C(N)=O | HRNLPPBUBKMZMT-RDRUQFPZSA-N | No-Effect | inactive |
| pramipexole | BRD-K06388322 | CCCN[C@H]1CCc2nc(N)sc2C1 | FASDKYOPVNHBLU-ZETCQYMHSA-N | No-Effect | inactive |
| pramiracetam | BRD-K21123171 | CC(C)N(CCNC(=O)CN1CCCC1=O)C(C)C | ZULJGOSFKWFVRX-UHFFFAOYSA-N | No-Effect | inactive |
| pramoxine | BRD-K46523383 | CCCCOc1ccc(OCCCN2CCOCC2)cc1 | DQKXQSGTHWVTAD-UHFFFAOYSA-N | No-Effect | inactive |
| pranidipine | BRD-A68083442 | COC(=O)C1=C(C)NC(C)=C(C(=O)OC/C=C/c2ccccc2)[C@@H]1c1cccc([N+](=O)[O-])c1 | XTFPDGZNWTZCMF-OXHGWOFKSA-N | No-Effect | inactive |
| pranlukast | BRD-K97045029 | O=C(Nc1cccc2c(=O)cc(-c3nn[nH]n3)oc12)c1ccc(OCCCCc2ccccc2)cc1 | NBQKINXMPLXUET-UHFFFAOYSA-N | No-Effect | inactive |
| pranoprofen | BRD-A64610707 | C[C@@H](C(=O)O)c1ccc2c(c1)Cc1cccnc1O2 | TVQZAMVBTVNYLA-SECBINFHSA-N | No-Effect | inactive |
| prasterone-acetate | BRD-K96527333 | CC(=O)O[C@H]1CC[C@@]2(C)C(=CC[C@H]3[C@@H]4CCC(=O)[C@@]4(C)CC[C@@H]32)C1 | NCMZQTLCXHGLOK-ZKHIMWLXSA-N | No-Effect | inactive |
| prasugrel | BRD-A83029242 | CC(=O)Oc1cc2c(s1)CCN([C@@H](C(=O)C1CC1)c1ccccc1F)C2 | DTGLZDAWLRGWQN-LJQANCHMSA-N | No-Effect | inactive |
| pravadoline | BRD-K46209126 | COc1ccc(C(=O)c2c(C)n(CCN3CCOCC3)c3ccccc23)cc1 | MEUQWHZOUDZXHH-UHFFFAOYSA-N | No-Effect | inactive |
| pravastatin | BRD-K60511616 | CC[C@H](C)C(=O)O[C@H]1C[C@H](O)C=C2C=C[C@H](C)[C@H](CC[C@@H](O)C[C@@H](O)CC(=O)O)[C@H]21 | TUZYXOIXSAXUGO-PZAWKZKUSA-N | No-Effect | inactive |
| praziquantel | BRD-A21858158 | O=C(C1CCCCC1)N1CC(=O)N2CCc3ccccc3[C@H]2C1 | FSVJFNAIGNNGKK-QGZVFWFLSA-N | No-Effect | inactive |
| prazosin | BRD-K49111258 | COc1cc2nc(N3CCN(C(=O)c4ccco4)CC3)nc(N)c2cc1OC | IENZQIKPVFGBNW-UHFFFAOYSA-N | No-Effect | inactive |
| preclamol | BRD-K02908048 | CCCN1CCC[C@H](c2cccc(O)c2)C1 | HTSNFXAICLXZMA-ZDUSSCGKSA-N | No-Effect | inactive |
| prednicarbate | BRD-A93963402 | CCOC(=O)O[C@]1(C(=O)COC(=O)CC)CC[C@H]2[C@H]3CCC4=CC(=O)C=C[C@]4(C)[C@@H]3[C@@H](O)C[C@@]21C | FNPXMHRZILFCKX-HGVFMWMISA-N | No-Effect | inactive |
| prednisolone | BRD-A27887842 | C[C@]12C[C@H](O)[C@H]3[C@@H](CCC4=CC(=O)C=C[C@@]43C)[C@@H]1CC[C@]2(O)C(=O)CO | OIGNJSKKLXVSLS-VWUMJDOOSA-N | No-Effect | inactive |
| prednisolone-acetate | BRD-K45664306 | CC(=O)OCC(=O)[C@@]1(O)CC[C@H]2[C@@H]3CCC4=CC(=O)C=C[C@]4(C)[C@H]3[C@@H](O)C[C@@]21C | LRJOMUJRLNCICJ-JZYPGELDSA-N | No-Effect | inactive |
| prednisolone-hemisuccinate | BRD-A78391468 | C[C@]12C[C@H](O)[C@H]3[C@@H](CCC4=CC(=O)C=C[C@@]43C)[C@@H]1CC[C@]2(O)C(=O)COC(=O)CCC(=O)O | APGDTXUMTIZLCJ-CGVGKPPMSA-N | No-Effect | inactive |
| prednisolone-sodium-phosphate | BRD-K00003654 | C[C@]12C[C@H](O)[C@H]3[C@@H](CCC4=CC(=O)C=C[C@@]43C)[C@@H]1CC[C@]2(O)C(=O)COP(=O)(O)O | JDOZJEUDSLGTLU-VWUMJDOOSA-N | No-Effect | inactive |
| prednisolone-tebutate | BRD-A53151380 | CC(C)(C)CC(=O)OCC(=O)[C@@]1(O)CC[C@H]2[C@H]3CCC4=CC(=O)C=C[C@]4(C)[C@@H]3[C@@H](O)C[C@@]21C | HUMXXHTVHHLNRO-HGVFMWMISA-N | No-Effect | inactive |
| prednisone | BRD-K85883481 | C[C@]12CC(=O)[C@H]3[C@@H](CCC4=CC(=O)C=C[C@@]43C)[C@@H]1CC[C@]2(O)C(=O)CO | XOFYZVNMUHMLCC-ZPOLXVRWSA-N | No-Effect | inactive |
| pregnenolone | BRD-K43880410 | CC(=O)[C@H]1CC[C@H]2[C@@H]3CC=C4C[C@@H](O)CC[C@]4(C)[C@H]3CC[C@]12C | ORNBQBCIOKFOEO-QGVNFLHTSA-N | No-Effect | inactive |
| pregnenolone-succinate | BRD-A35912562 | CC(=O)[C@H]1CC[C@H]2[C@H]3CC=C4C[C@@H](OC(=O)CCC(=O)O)CC[C@]4(C)[C@@H]3CC[C@]12C | OZZAYJQNMKMUSD-VQLNUFJUSA-N | No-Effect | inactive |
| preladenant | BRD-K46290096 | COCCOc1ccc(N2CCN(CCn3ncc4c3nc(N)n3nc(-c5ccco5)nc43)CC2)cc1 | DTYWJKSSUANMHD-UHFFFAOYSA-N | No-Effect | inactive |
| presatovir | BRD-K45906612 | Cc1cn2nc([C@@H]3CCCCN3C(=O)c3cc(Cl)ccc3NS(C)(=O)=O)cc2nc1N1CC[C@H](N)C1 | GOFXWTVKPWJNGD-UWJYYQICSA-N | No-Effect | inactive |
| pretomanid | BRD-K96671969 | O=[N+]([O-])c1cn2c(n1)OC[C@@H](OCc1ccc(OC(F)(F)F)cc1)C2 | ZLHZLMOSPGACSZ-NSHDSACASA-N | No-Effect | inactive |
| pridinol | BRD-K17565903 | OC(CCN1CCCCC1)(c1ccccc1)c1ccccc1 | RQXCLMGKHJWMOA-UHFFFAOYSA-N | No-Effect | inactive |
| pridopidine | BRD-K00003473 | CCCN1CCC(c2cccc(S(C)(=O)=O)c2)CC1 | YGKUEOZJFIXDGI-UHFFFAOYSA-N | No-Effect | inactive |
| PRIMA1 | BRD-K15318909 | O=C1C2CCN(CC2)C1(CO)CO | RFBVBRVVOPAAFS-UHFFFAOYSA-N | No-Effect | inactive |
| primaquine | BRD-K00003556 | COc1cc(N[C@@H](C)CCCN)c2ncccc2c1 | INDBQLZJXZLFIT-NSHDSACASA-N | No-Effect | inactive |
| primidone | BRD-K32247306 | CCC1(c2ccccc2)C(=O)NCNC1=O | DQMZLTXERSFNPB-UHFFFAOYSA-N | No-Effect | inactive |
| prinaberel | BRD-K87316765 | C=Cc1cc(O)cc2nc(-c3ccc(O)c(F)c3)oc12 | MQIMZDXIAHJKQP-UHFFFAOYSA-N | No-Effect | inactive |
| prinomastat | BRD-K00003122 | CC1(C)SCCN(S(=O)(=O)c2ccc(Oc3ccncc3)cc2)[C@H]1C(=O)NO | YKPYIPVDTNNYCN-INIZCTEOSA-N | No-Effect | inactive |
| priralfinamide | BRD-K01662324 | C[C@H](NCc1ccc(OCc2ccccc2F)cc1)C(N)=O | BHJIBOFHEFDSAU-LBPRGKRZSA-N | No-Effect | inactive |
| prisotinol | BRD-K00004547 | CC(C)N[C@H](C)Cc1ccc(O)cn1 | WQRPBKUCJBWQRT-SECBINFHSA-N | No-Effect | inactive |
| pritelivir | BRD-K25204779 | Cc1nc(N(C)C(=O)Cc2ccc(-c3ccccn3)cc2)sc1S(N)(=O)=O | IVZKZONQVYTCKC-UHFFFAOYSA-N | No-Effect | inactive |

| | | | | | |
|---|---|---|---|---|---|
| PRLX-93936 | BRD-K00004604 | CCOc1cccc1-n1c(CN2CCNCC2)nc2ccccc2c1=O | OBWOSMGNXYUDFB-UHFFFAOYSA-N | No-Effect | inactive |
| PRN1008 | BRD-K00004599 | CC(C)(/C=C(\C#N)C(=O)N1CCC[C@@H](n2nc(-c3ccc(Oc4ccccc4)cc3F)c3c(N)ncnc32)C1)N1CCN(C2COC2)CC1 | LCFFREMLXLZNHE-GBOLQPHISA-N | No-Effect | inactive |
| PRN1371 | BRD-K00003430 | C=CC(=O)N1CCN(CCCn2c(=O)c(-c3c(Cl)c(OC)cc(OC)c3Cl)cc3cnc(NC)nc32)CC1 | PUIXMSRTTHLNKI-UHFFFAOYSA-N | No-Effect | inactive |
| proacipimox | BRD-K53605365 | COCc1c[n+]([O-])c(C)cn1 | KKKHJYGCTVKQEM-UHFFFAOYSA-N | No-Effect | inactive |
| probenecid | BRD-K95237249 | CCCN(CCC)S(=O)(=O)c1ccc(C(=O)O)cc1 | DBABZHXKTCFAPX-UHFFFAOYSA-N | No-Effect | inactive |
| probucol | BRD-K72029282 | CC(C)(Sc1cc(C(C)(C)C)c(O)c(C(C)(C)C)c1)Sc1cc(C(C)(C)C)c(O)c(C(C)(C)C)c1 | FYPMFJGVHOHGLL-UHFFFAOYSA-N | No-Effect | inactive |
| procainamide | BRD-K75089421 | CCN(CC)CCNC(=O)c1ccc(N)cc1 | REQCZEXYDRLIBE-UHFFFAOYSA-N | No-Effect | inactive |
| procaine | BRD-K24616672 | CCN(CC)CCOC(=O)c1ccc(N)cc1 | MFDFERRIHVXMIY-UHFFFAOYSA-N | No-Effect | inactive |
| procaterol | BRD-K78666826 | CC[C@H](NC(C)C)[C@H](O)c1ccc(O)c2[nH]c(=O)ccc12 | FKNXQNWAXFXVNW-BLLLJJGKSA-N | No-Effect | inactive |
| procodazole | BRD-K54824716 | O=C(O)CCc1nc2ccccc2[nH]1 | XYWJNTOURDMTPI-UHFFFAOYSA-N | No-Effect | inactive |
| procyanidin-B-2 | BRD-K72093555 | Oc1cc(O)c2c(c1)O[C@H](c1ccc(O)c(O)c1)[C@H](O)[C@H]2c1c(O)cc(O)c2c1O[C@H](c1ccc(O)c(O)c1)[C@@H](O)C2 | XFZJEEAOWLFHDH-UKWJTHFESA-N | No-Effect | inactive |
| procyclidine | BRD-K01826750 | COc1cc(N[C@@H](C)CCCN)c2ncccc2c1 | INDBQLZJXZLFIT-NSHDSACASA-N | No-Effect | inactive |
| procysteine | BRD-K92413528 | O=C1N[C@H](C(=O)O)CS1 | BMLMGCPTLHPWPY-REOHCLBHSA-N | No-Effect | inactive |
| proflavine-hemisulfate | BRD-K68408467 | Nc1ccc2cc3ccc(N)cc3nc2c1 | WDVSHHCDHLJJJR-UHFFFAOYSA-N | No-Effect | inactive |
| proglumetacin | BRD-A19458515 | CCCN(CCC)C(=O)[C@@H](CCC(=O)OCCCN1CCN(CCOC(=O)Cc2c(C)n(C(=O)c3ccc(Cl)cc3)c3ccc(OC)cc23)CC1)NC(=O)c1ccccc1 | PTXGHCGBYMQQIG-RRHRGVEJSA-N | No-Effect | inactive |
| proglumide | BRD-K01826553 | CCCN(CCC)C(=O)[C@@H](CCC(=O)O)NC(=O)c1ccccc1 | DGMKFQYCZXERLX-OAHLLOKOSA-N | No-Effect | inactive |
| proguanil | BRD-K28183345 | CC(C)NC(=N)N=C(N)Nc1ccc(Cl)cc1 | SSOLNOMRVKKSON-UHFFFAOYSA-N | No-Effect | inactive |
| prolylleucylglycinamide | BRD-A09911125 | CC(C)C[C@@H](NC(=O)[C@H]1CCCN1)C(=O)NCC(N)=O | NOOJLZTTWSNHOX-NXEZZACHSA-N | No-Effect | inactive |
| promestriene | BRD-K75097983 | CCCOc1ccc2c(c1)CC[C@@H]1[C@@H]2CC[C@]2(C)[C@@H](OC)CC[C@@H]12 | IUWKNLFTJBHTSD-AANPDWTMSA-N | No-Effect | inactive |
| pronethalol | BRD-K01825804 | CC(C)NC[C@@H](O)c1ccc2ccccc2c1 | HRSANNODOVBCST-OAHLLOKOSA-N | No-Effect | inactive |
| propacetamol | BRD-K54496168 | CCN(CC)CC(=O)Oc1ccc(NC(C)=O)cc1 | QTGAJCQTLIRCFL-UHFFFAOYSA-N | No-Effect | inactive |
| propafenone | BRD-K01826541 | CCCNC[C@@H](O)COc1ccccc1C(=O)CCc1ccccc1 | JWHAUXFOSRPERK-GOSISDBHSA-N | No-Effect | inactive |
| propagermanium | BRD-K53382462 | O=C(O)CC[Ge](=O)O[Ge](=O)CCC(=O)O | XEABSBMNTNXEJM-UHFFFAOYSA-N | No-Effect | inactive |
| propantheline | BRD-K90885812 | CC(C)[N+](C)(CCOC(=O)C1c2ccccc2Oc2ccccc21)C(C)C | VVWYOYDLCMFIEM-UHFFFAOYSA-N | No-Effect | inactive |
| propatylnitrate | BRD-K56276053 | CCC(CO[N+](=O)[O-])(CO[N+](=O)[O-])CO[N+](=O)[O-] | YZZCJYJBCUJISI-UHFFFAOYSA-N | No-Effect | inactive |
| propentofylline | BRD-K59273480 | CCCn1cnc2c1c(=O)n(CCCCC(C)=O)c(=O)n2C | RBQOQRRFDPXAGN-UHFFFAOYSA-N | No-Effect | inactive |
| propidium-iodide | BRD-K14821965 | CC[N+](C)(CC)CCC[n+]1c(-c2ccccc2)c2cc(N)ccc2c2ccc(N)cc21 | ZDWVWKDAWBGPDN-UHFFFAOYSA-O | No-Effect | inactive |
| propiolactone | BRD-K76867903 | O=C1CCO1 | VEZXCJBBBCKRPI-UHFFFAOYSA-N | No-Effect | inactive |
| propofol | BRD-K82255054 | CC(C)c1cccc(C(C)C)c1O | OLBCVFGFOZPWHH-UHFFFAOYSA-N | No-Effect | inactive |
| propoxur | BRD-K08231299 | CNC(=O)Oc1ccccc1OC(C)C | ISRUGXGCCGIOQO-UHFFFAOYSA-N | No-Effect | inactive |
| propoxycaine | BRD-K18250272 | CCCOc1cc(N)ccc1C(=O)OCCN(CC)CC | CAJIGINSTLKQMM-UHFFFAOYSA-N | No-Effect | inactive |
| propranolol | BRD-A10070317 | CC(C)NCC(O)COc1cccc2ccccc12 | AQHHHDLHHXJYJD-UHFFFAOYSA-N | No-Effect | inactive |
| propranolol-(S) | BRD-K13994703 | CC(C)NC[C@H](O)COc1cccc2ccccc12 | AQHHHDLHHXJYJD-AWEZNQCLSA-N | No-Effect | inactive |
| propylene-glycol | BRD-A19232309 | C[C@@H](O)CO | DNIAPMSPPWPWGF-GSVOUGTGSA-N | No-Effect | inactive |
| propylhexedrine | BRD-K01825844 | CN[C@@H](C)CC1CCCCC1 | JCRIVQIOJSSCQD-VIFPVBQESA-N | No-Effect | inactive |
| propylparaben | BRD-K60783397 | CCCOC(=O)c1ccc(O)cc1 | QELSKZZBTMNZEB-UHFFFAOYSA-N | No-Effect | inactive |
| propylthiouracil | BRD-K48168960 | CCCc1cc(=O)[nH]c(=S)[nH]1 | KNAHARQHSZJURB-UHFFFAOYSA-N | No-Effect | inactive |
| propyphenazone | BRD-K06520168 | Cc1c(C(C)C)c(=O)n(-c2ccccc2)n1C | PXWLVJLKJGVOKE-UHFFFAOYSA-N | No-Effect | inactive |
| protionamide | BRD-K97863768 | CCCc1cc(C(N)=S)ccn1 | VRDIULHPQTYCLN-UHFFFAOYSA-N | No-Effect | inactive |
| protirelin | BRD-K00003716 | NC(=O)[C@@H]1CCCN1C(=O)[C@H](Cc1c[nH]cn1)NC(=O)[C@@H]1CCC(=O)N1 | XNSAINXGIQZQOO-SRVKXCTJSA-N | No-Effect | inactive |
| protriptyline | BRD-K42098891 | CNCCCC1c2ccccc2C=Cc2ccccc21 | BWPIARFWQZKAIA-UHFFFAOYSA-N | No-Effect | inactive |
| proxodolol | BRD-A29944538 | Cc1noc(COc2ccccc2OC[C@@H](O)CNC(C)(C)C)n1 | CEDZURSVVKNCSL-ZDUSSCGKSA-N | No-Effect | inactive |
| proxyfan | BRD-K77171813 | c1ccc(COCCCc2cnc[nH]2)cc1 | WNWALBVQAAIULR-UHFFFAOYSA-N | No-Effect | inactive |
| proxymetacaine | BRD-K79116891 | CCCOc1ccc(C(=O)OCCN(CC)CC)cc1N | KCLANYCVBBTKTO-UHFFFAOYSA-N | No-Effect | inactive |
| Proxyphylline | BRD-A28887267 | C[C@@H](O)Cn1cnc2c1c(=O)n(C)c(=O)n2C | KYHQZNGJUGFTGR-ZCFIWIBFSA-N | No-Effect | inactive |
| PRT062070 | BRD-K25835157 | CCS(=O)(=O)N1CCN(c2ccc(Nc3ncc(C(N)=O)c(NC4CC4)n3)cc2)CC1 | BGLPECHZZQDNCD-UHFFFAOYSA-N | No-Effect | inactive |
| PRT4165 | BRD-K32456483 | O=C1C(=Cc2cccnc2)C(=O)c2ccccc21 | OMHZFEWYVFWVLI-UHFFFAOYSA-N | No-Effect | inactive |
| prucalopride | BRD-K33681278 | COCCCN1CCC(NC(=O)c2cc(Cl)c(N)c3c2OCC3)CC1 | ZPMNHBXQOOVQJL-UHFFFAOYSA-N | No-Effect | inactive |
| prulifloxacin | BRD-A92341659 | Cc1oc(=O)oc1CN1CCN(c2cc3c(cc2F)c(=O)c(C(=O)O)c2n3[C@@H](C)S2)CC1 | PWNMXPDKBYZCOO-LLVKDONJSA-N | No-Effect | inactive |
| pruvanserin | BRD-K43837174 | N#Cc1c[nH]c2c(C(=O)N3CCN(CCc4ccc(F)cc4)CC3)cccc12 | AQRLDDAFYYAIJP-UHFFFAOYSA-N | No-Effect | inactive |
| PRX-07034 | BRD-K00004616 | COc1cc(Cl)cc([C@H](C)Nc2cc(N3CCNCC3)ccc2S(C)(=O)=O)c1OC | BSLXKMCHXRCBIH-AWEZNQCLSA-N | No-Effect | inactive |
| PRX-08066 | BRD-K16757695 | N#Cc1cc(CN2CCC(Nc3ncnc4sc(Cl)cc34)CC2)ccc1F | IENZFHBNCRQMNP-UHFFFAOYSA-N | No-Effect | inactive |
| PS178990 | BRD-K99092662 | Cc1c(N2C(=O)[C@@H]3[C@H](O)CCN3C2=O)ccc(C#N)c1Cl | KEJORAMIZFOODM-PWSUYJOCSA-N | No-Effect | inactive |
| PSB-06126 | BRD-K71266197 | Nc1c(S(=O)(=O)O)cc(Nc2cccc3ccccc23)c2c1C(=O)c1ccccc1C2=O | ZHQRPIRGSWEALJ-UHFFFAOYSA-N | No-Effect | inactive |
| PSB-1115 | BRD-K49027941 | CCCn1c(=O)[nH]c2nc(-c3ccc(S(=O)(=O)O)cc3)[nH]c2c1=O | UYDRRQPGDSIMNU-UHFFFAOYSA-N | No-Effect | inactive |
| PSB-36 | BRD-K00004243 | CCCCn1c(=O)c2[nH]c([C@]34C[C@@H]5C[C@H]3C[C@@H](C5)C4)nc2n(CCCO)c1=O | CIBIXJYFYPFMTN-FZUGUKJMSA-N | No-Effect | inactive |
| PSB-603 | BRD-K05419687 | CCCn1c(=O)[nH]c2nc(-c3ccc(S(=O)(=O)N4CCN(c5ccc(Cl)cc5)CC4)cc3)[nH]c2c1=O | OVHCTHHFOHMNFV-UHFFFAOYSA-N | No-Effect | inactive |
| PSI-6130 | BRD-K07798980 | C[C@@]1(F)[C@H](O)[C@@H](CO)O[C@H]1n1ccc(N)nc1=O | NYPIRLYMDJMKGW-VPCXQMTMSA-N | No-Effect | inactive |
| PSI-697 | BRD-K00003192 | O=C(O)c1c(O)c(Cc2ccc(Cl)cc2)nc2c3c(ccc12)CCCC3 | DIEPFYNZGUUVHD-UHFFFAOYSA-N | No-Effect | inactive |
| PSI-7976 | BRD-K05674516 | CC(C)OC(=O)[C@H](C)N[P@@](=O)(OC[C@H]1O[C@@H](n2ccc(=O)[nH]c2=O)[C@](C)(F)[C@@H]1O)Oc1ccccc1 | TTZHDVOVKQGIBA-YBSJRAAASA-N | No-Effect | inactive |
| PSN-375963 | BRD-K25524093 | CCCCC1CCC(c2nc(-c3ccncc3)no2)CC1 | OAVLEYPTWABFLF-UHFFFAOYSA-N | No-Effect | inactive |
| psoralen | BRD-K47264279 | O=c1ccc2cc3ccoc3cc2o1 | ZCCUUQDIBDJBTK-UHFFFAOYSA-N | No-Effect | inactive |
| PT-2385 | BRD-K20850787 | CS(=O)(=O)c1ccc(Oc2cc(F)cc(C#N)c2)c2c1[C@H](O)C(F)(F)C2 | ONBSHRSJOPSEGS-INIZCTEOSA-N | No-Effect | inactive |
| PTC-028 | BRD-K00010388 | Cc1nc2cc(F)c(F)cc2n1-c1ncc(Nc2ccc(C(F)(F)F)cc2)n1 | JEZGPBWIZWPDHP-UHFFFAOYSA-N | No-Effect | inactive |
| pterostilbene | BRD-K92870997 | COc1cc(/C=C/c2ccc(O)cc2)cc(OC)c1 | VLEUZFDZJKSGMX-ONEGZZNKSA-N | No-Effect | inactive |
| pumosetrag | BRD-K44164034 | O=C(N[C@H]1CN2CCC1CC2)c1c[nH]c2ccsc2c1=O | AFUWQWYPPZFWCO-LBPRGKRZSA-N | No-Effect | inactive |
| purmorphamine | BRD-K73397362 | c1ccc2c(Oc3nc(Nc4ccc(N5CCOCC5)cc4)c4ncn(C5CCCCC5)c4n3)cccc2c1 | FYBHCRQFSFYWPY-UHFFFAOYSA-N | No-Effect | inactive |
| puromycin | BRD-K36007650 | COc1ccc(C[C@H](N)C(=O)N[C@@H]2[C@@H](CO)O[C@@H](n3cnc4c(N(C)C)ncnc43)[C@@H]2O)cc1 | RXWNCPJZOCPEPQ-NVWDDTSBSA-N | No-Effect | inactive |
| purvalanol-A | BRD-K50836978 | CC(C)[C@H](CO)Nc1nc(Nc2cccc(Cl)c2)c2ncn(C(C)C)c2n1 | PMXCMJLOPOFPBT-HNNXBMFYSA-N | No-Effect | inactive |
| purvalanol-B | BRD-K41564320 | CC(C)[C@H](CO)Nc1nc(Nc2ccc(C(=O)O)c(Cl)c2)c2ncn(C(C)C)c2n1 | ZKDXRFMOHZVXSG-HNNXBMFYSA-N | No-Effect | inactive |
| putrescine | BRD-K75282878 | NCCCCN | KIDHWZJUCRJVML-UHFFFAOYSA-N | No-Effect | inactive |
| PX-12 | BRD-A56592690 | CC[C@@H](C)SSc1ncc[nH]1 | BPBPYQWMFCTCNG-ZCFIWIBFSA-N | No-Effect | inactive |
| PX-478 | BRD-A26835809 | N[C@@H](Cc1ccc([N+]([O-])(CCCl)CCCl)cc1)C(=O)O | GSKQMLGAUOTSKT-LBPRGKRZSA-N | No-Effect | inactive |

| | | | | | |
|---|---|---|---|---|---|
| PYM50028 | BRD-K62277907 | C[C@H]1CC[C@@]2(OC1)O[C@H]1C[C@H]3[C@@H]4CC[C@@H]5C[C@@H](O)CC[C@]5(C)[C@H]4CC[C@]3(C)[C@H]1[C@@H]2C | GMBQZIIUCVWOCD-WWASVFFGSA-N | No-Effect | inactive |
| PYR-41 | BRD-K66101666 | CCOC(=O)c1ccc(N2NC(=O)/C(=C/c3ccc([N+](=O)[O-])o3)C2=O)cc1 | ARGIPZKQJGFSGQ-LCYFTJDESA-N | No-Effect | inactive |
| pyrantel | BRD-K82118441 | O=C(O)c1cc2ccccc2c(Cc2c(O)c(C(=O)O)cc3ccccc23)c1O | WLJNZVDCPSBLRP-UHFFFAOYSA-N | No-Effect | inactive |
| pyrantel-pamoate | BRD-K20672254 | CN1CCCN=C1/C=C/c1cccs1 | YSAUAVHXTIETRK-AATRIKPKSA-N | No-Effect | inactive |
| pyrazinamide | BRD-K28667793 | NC(=O)c1cnccn1 | IPEHBUMCGVEMRF-UHFFFAOYSA-N | No-Effect | inactive |
| pyrazinoylguanidine | BRD-K61693562 | N=C(N)NC(=O)c1cnccn1 | WNBSDCKJFDZMHT-UHFFFAOYSA-N | No-Effect | inactive |
| pyrazolanthrone | BRD-K01567962 | O=C1c2ccccc2-c2[nH]nc3cccc1c23 | ACPOUJIDANTYHO-UHFFFAOYSA-N | No-Effect | inactive |
| pyrazoloacridine | BRD-K42607789 | COc1ccc2nc3c([N+](=O)[O-])ccc4c3c(c2c1)NN4CCCN(C)C | HVRZAFDJOWIDMM-UHFFFAOYSA-N | No-Effect | inactive |
| pyrethrins | BRD-K00010129 | C=C/C=C\CC1=C(C)[C@@H](OC(=O)[C@@H]2[C@@H](C=C(C)C)C2(C)C)CC1=O | ROVGZAWFACYCSP-VUMXUWRFSA-N | No-Effect | inactive |
| pyridoxal | BRD-K76204040 | Cc1ncc(CO)c(C=O)c1O | RADKZDMFGJYCBB-UHFFFAOYSA-N | No-Effect | inactive |
| pyridoxal-isonicotinoyl-hydrazone | BRD-K42019693 | Cc1ncc(CO)c(/C=N/NC(=O)c2ccncc2)c1O | BQYIXOPJPLGCRZ-REZTVBANSA-N | No-Effect | inactive |
| pyridoxamine | BRD-K96669468 | Cc1ncc(CO)c(CN)c1O | NHZMQXZHNVQTQA-UHFFFAOYSA-N | No-Effect | inactive |
| pyrimethamine | BRD-K88429204 | CCc1nc(N)nc(N)c1-c1ccc(Cl)cc1 | WKSAUQYGYAYLPV-UHFFFAOYSA-N | No-Effect | inactive |
| pyrithyldione | BRD-K36116267 | CCC1(CC)C(=O)C=CNC1=O | NZASCBIBXNPDMH-UHFFFAOYSA-N | No-Effect | inactive |
| pyritinol | BRD-K82181219 | Cc1ncc(CSSCc2cnc(C)c(O)c2CO)c(CO)c1O | SIXLXDIJGIWWFU-UHFFFAOYSA-N | No-Effect | inactive |
| pyrotinib | BRD-K00003463 | CCOc1cc2ncc(C#N)c(Nc3ccc(OCc4ccccn4)c(Cl)c3)c2cc1NC(=O)/C=C/[C@H]1CCCN1C | SADXACCFNXBCFY-IYNHSRRRSA-N | No-Effect | inactive |
| pyroxamide | BRD-K11663430 | O=C(CCCCCCC(=O)Nc1cccnc1)NO | PTJGLFIIZFVFJV-UHFFFAOYSA-N | No-Effect | inactive |
| pyrrolidine-dithiocarbamate | BRD-K80970344 | S=C(S)N1CCCC1 | VSWDORGPIHIGNW-UHFFFAOYSA-N | No-Effect | inactive |
| pyrvinium-pamoate | BRD-K39479472 | Cc1cc(/C=C/c2ccc3cc(N(C)C)ccc3[n+]2C)c(C)n1-c1ccccc1 | QMHSXPLYMTVAMK-UHFFFAOYSA-N | No-Effect | inactive |
| PZ-2891 | BRD-K00005336 | CC(C)c1ccc(CC(=O)N2CCN(c3ccc(C#N)nn3)CC2)cc1 | LGWDVWIZDPGCFG-UHFFFAOYSA-N | No-Effect | inactive |
| Q-203 | BRD-K59853741 | CCc1nc2ccc(Cl)cn2c1C(=O)NCc1ccc(N2CCC(c3ccc(OC(F)(F)F)cc3)CC2)cc1 | OJICYBSWSZGRFB-UHFFFAOYSA-N | No-Effect | inactive |
| quazinone | BRD-K63631219 | CC1C(=O)N=C2Nc3cccc(Cl)c3CN21 | BHZFZYLBVSWUMT-UHFFFAOYSA-N | No-Effect | inactive |
| quetiapine | BRD-K68867920 | OCCOCCN1CCN(C2=Nc3ccccc3Sc3ccccc32)CC1 | URKOMYMAXPYINW-UHFFFAOYSA-N | No-Effect | inactive |
| quiflapon | BRD-K39503511 | CC(C)(C)Sc1c(CC(C)(C)C(=O)O)n(Cc2ccc(Cl)cc2)c2ccc(OCc3ccc4ccccc4n3)cc12 | NZOONKHCNQFYCI-UHFFFAOYSA-N | No-Effect | inactive |
| quinagolide | BRD-K02995728 | CCCN1C[C@@H](NS(=O)(=O)N(CC)CC)C[C@@H]2Cc3c(O)cccc3C[C@H]21 | GDFGTRDCCWFXTG-ZIFCJYIRSA-N | No-Effect | inactive |
| quinapril | BRD-K72222507 | CCOC(=O)[C@H](CCc1ccccc1)N[C@@H](C)C(=O)N1Cc2ccccc2C[C@H]1C(=O)O | JSDRRTOADPPCHY-HSQYWUDLSA-N | No-Effect | inactive |
| quinaprilat | BRD-K42317111 | C[C@H](N[C@@H](CCc1ccccc1)C(=O)O)C(=O)N1Cc2ccccc2C[C@H]1C(=O)O | FLSLEGPOVLMJMN-YSSFQJQWSA-N | No-Effect | inactive |
| quinestrol | BRD-K63794707 | C#C[C@]1(O)CC[C@H]2[C@@H]3CCc4cc(OC5CCCC5)ccc4[C@H]3CC[C@@]21C | PWZUUYSISTUNDW-VAFBSOEGSA-N | No-Effect | inactive |
| quinethazone | BRD-A59303141 | CC[C@H]1NC(=O)c2cc(S(N)(=O)=O)c(Cl)cc2N1 | AGMMTXLNIQSRCG-SECBINFHSA-N | No-Effect | inactive |
| quinine | BRD-K07940445 | C=C[C@H]1CN2CC[C@H]1C[C@H]2[C@H](O)c1ccnc2ccc(OC)cc12 | LOUPRKONTZGTKE-WZBLMQSHSA-N | No-Effect | inactive |
| quinine-ethyl-carbonate | BRD-K97028990 | C=C[C@H]1CN2CC[C@H]1C[C@@H]2[C@@H](OC(=O)OCC)c1ccnc2ccc(OC)cc12 | NSBRKSWSLRQPJW-FRMGNDQPSA-N | No-Effect | inactive |
| quinolinic-acid | BRD-K51885093 | O=C(O)c1cccnc1C(=O)O | GJAWHXHKYYXBSV-UHFFFAOYSA-N | No-Effect | inactive |
| quinpirol-(-) | BRD-K26548821 | CCCN1CCC[C@@H]2Cc3[nH]ncc3C[C@H]21 | FTSUPYGMFAPCFZ-ZWNOBZJWSA-N | No-Effect | inactive |
| quipazine | BRD-K77925998 | c1ccc2nc(N3CCNCC3)ccc2c1 | XRXDAJYKGWNHTQ-UHFFFAOYSA-N | No-Effect | inactive |
| quizartinib | BRD-K93918653 | CC(C)(C)c1cc(NC(=O)Nc2ccc(-c3cn4c(n3)sc3cc(OCCN5CCOCC5)ccc34)cc2)no1 | CVWXJKQAOSCOAB-UHFFFAOYSA-N | No-Effect | inactive |
| QX-222 | BRD-K40782193 | Cc1cccc(C)c1NC(=O)C[N+](C)(C)C | QYUXPWDVDMSTKR-UHFFFAOYSA-O | No-Effect | inactive |
| QX-314 | BRD-K56596464 | CC[N+](CC)(CC)CC(=O)Nc1c(C)cccc1C | PYEBKOFMWAMBFV-UHFFFAOYSA-O | No-Effect | inactive |
| R-121919 | BRD-K00004717 | CCCN(CCC)c1cc(C)nc2c(-c3cnc(N(C)C)cc3C)c(C)nn12 | ANNRUWYFVIGKHA-UHFFFAOYSA-N | No-Effect | inactive |
| R-1479 | BRD-K28442028 | [N-]=[N+]=N[C@]1(CO)O[C@@H](n2ccc(N)nc2=O)[C@H](O)[C@@H]1O | ODLGMSQBFONGNG-JVZYCSMKSA-N | No-Effect | inactive |
| R-1485 | BRD-K96418224 | O=S(=O)(c1ccccc1F)N1CCOc2c(N3CCNCC3)cccc21 | UPROBLPRPWYJGN-UHFFFAOYSA-N | No-Effect | inactive |
| R-1487 | BRD-K00004710 | Cn1c(=O)c(Oc2ccc(F)cc2F)cc2cnc(NC3CCOCC3)nc21 | KKKRKRMVJRHDMG-UHFFFAOYSA-N | No-Effect | inactive |
| R-268712 | BRD-K83904070 | Cc1cccc(-c2n[nH]cc2-c2ccc(F)c(-c3cnn(CCO)c3)c2)n1 | JQGOCCALXFSRHZ-UHFFFAOYSA-N | No-Effect | inactive |
| R-59022 | BRD-K54665485 | Cc1nc2sccn2c(=O)c1CCN1CCC(=C(c2ccccc2)c2ccc(F)cc2)CC1 | MFVJXLPANKSLLD-UHFFFAOYSA-N | No-Effect | inactive |
| R-96544 | BRD-K15588452 | COc1cccc(CCc2ccccc2OCC[C@@H]2C[C@@H](O)CN2C)c1 | AFZLABYDOCWQBQ-WOJBJXKFSA-N | No-Effect | inactive |
| R112 | BRD-K23902832 | Oc1cccc(Nc2ncc(F)c(Nc3cccc(O)c3)n2)c1 | TVKGTSHBQZEFEE-UHFFFAOYSA-N | No-Effect | inactive |
| R406 | BRD-K20285085 | COc1cc(Nc2ncc(F)c(Nc3ccc4c(n3)NC(=O)C(C)(C)O4)n2)cc(OC)c1OC | NHHQJBCNYHBUSI-UHFFFAOYSA-N | No-Effect | inactive |
| rabeprazole | BRD-K01826765 | COCCCOc1ccnc(C[S@@](=O)c2nc3ccccc3[nH]2)c1C | YREYEVIYCVEVJK-RUZDIDTESA-N | No-Effect | inactive |
| rac-BHFF | BRD-A11069678 | CC(C)(C)c1cc(C(C)(C)C)c2c(c1)[C@](O)(C(F)(F)F)C(=O)O2 | RVNOANDLZIIFHB-MRXNPFEDSA-N | No-Effect | inactive |
| racecadotril | BRD-A08187463 | CC(=O)SC[C@@H](Cc1ccccc1)C(=O)NCC(=O)OCc1ccccc1 | ODUOJXZPIYUATO-LJQANCHMSA-N | No-Effect | inactive |
| raclopride | BRD-K04111260 | CCN1CCC[C@H]1CNC(=O)c1c(O)c(Cl)cc(Cl)c1OC | WAOQONBSWFLFPE-VIFPVBQESA-N | No-Effect | inactive |
| ractopamine | BRD-K01825837 | C[C@@H](CCc1ccc(O)cc1)NC[C@@H](O)c1ccc(O)cc1 | YJQZYXCXBBCEAQ-SCLBCKFNSA-N | No-Effect | inactive |
| RAD140 | BRD-K00003169 | Cc1c(N[C@@H](c2nnc(-c3ccc(C#N)cc3)o2)[C@H](C)O)ccc(C#N)c1Cl | XMBUPPIEVAFYHO-KPZWWZAWSA-N | No-Effect | inactive |
| radafaxine | BRD-K13394247 | C[C@@H]1NC(C)(C)CO[C@@]1(O)c1cccc(Cl)c1 | RCOBKSKAZMVBHT-TVQRCGJNSA-N | No-Effect | inactive |
| radezolid | BRD-K84683831 | CC(=O)NC[C@H]1CN(c2ccc(-c3ccc(CNCc4cn[nH]n4)cc3)c(F)c2)C(=O)O1 | BTTNOGHPGJANSW-IBGZPJMESA-N | No-Effect | inactive |
| radiprodil | BRD-K00005328 | O=C(Nc1ccc2[nH]c(=O)oc2c1)C(=O)N1CCC(Cc2ccc(F)cc2)CC1 | GKGRZLGAQZPEHO-UHFFFAOYSA-N | No-Effect | inactive |
| radotinib | BRD-K00004716 | Cc1cn(-c2cc(NC(=O)c3ccc(C)c(Nc4nccc(-c5cnccn5)n4)c3)cc(C(F)(F)F)c2)cn1 | DUPWHXBITIZIKZ-UHFFFAOYSA-N | No-Effect | inactive |
| RAF265 | BRD-K93123848 | Cn1c(Nc2ccc(C(F)(F)F)cc2)nc2cc(Oc3ccnc(-c4ncc(C(F)(F)F)[nH]4)c3)ccc21 | YABJJWZLRMPFSI-UHFFFAOYSA-N | No-Effect | inactive |
| RAF709 | BRD-K00010389 | Cc1ncc(NC(=O)c2cccc(C(F)(F)F)c2)cc1-c1cnc(OC2CCOCC2)c(N2CCOCC2)c1 | FYNMINFUAIDIFL-UHFFFAOYSA-N | No-Effect | inactive |
| ralinepag | BRD-K00003227 | O=C(O)COC[C@H]1CC[C@H](COC(=O)N(c2ccccc2)c2ccc(Cl)cc2)CC1 | NPDKXVKJRHPDQT-IYARVYRRSA-N | No-Effect | inactive |
| raltitrexed | BRD-K08151102 | Cc1nc(=O)c2cc(CN(C)c3ccc(C(=O)N[C@@H](CCC(=O)O)C(=O)O)s3)ccc2[nH]1 | IVTVGDXNLFLDRM-HNNXBMFYSA-N | No-Effect | inactive |
| ramifenazone | BRD-K85333151 | Cc1c(NC(C)C)c(=O)n(-c2ccccc2)n1C | XOZLRRYPUKAKMU-UHFFFAOYSA-N | No-Effect | inactive |
| ramipril | BRD-K89348303 | CCOC(=O)[C@H](CCc1ccccc1)N[C@@H](C)C(=O)N1[C@H](C(=O)O)C[C@@H]2CCC[C@@H]21 | HDACQVRGBOVJII-JBDAPHQKSA-N | No-Effect | inactive |
| ramosetron | BRD-K85046107 | Cn1cc(C(=O)[C@@H]2CCc3nc[nH]c3C2)c2ccccc21 | NTHPAPBPFQJABD-LLVKDONJSA-N | No-Effect | inactive |

| | | | | | |
|---|---|---|---|---|---|
| ranirestat | BRD-K50489389 | O=C1C[C@@]2(C(=O)N1)C(=O)N(Cc1ccc(Br)cc1F)C(=O)c1cccn12 | QCVNMNYRNIMDKV-QGZVFWFLSA-N | No-Effect | inactive |
| ranitidine | BRD-K30816563 | CN=C(C[N+](=O)[O-])NCCSCc1ccc(CN(C)C)o1 | SYNOVMDJYWJCON-UHFFFAOYSA-N | No-Effect | inactive |
| ranolazine | BRD-K01825102 | COc1ccccc1OC[C@@H](O)CN1CCN(CC(=O)Nc2c(C)cccc2C)CC1 | XKLMZUWKNUAPSZ-FQEVSTJZSA-N | No-Effect | inactive |
| rapastinel | BRD-K02206208 | C[C@@H](O)[C@H](N)C(=O)N1CCC[C@H]1C(=O)N1CCC[C@H]1C(=O)N[C@H](C(N)=O)[C@@H](C)O | GIBQQARAXHVEGD-BSOLPCOYSA-N | No-Effect | inactive |
| rasagiline | BRD-K58114536 | C#CCN[C@@H]1CCc2ccccc21 | RUOKEQAAGRXIBM-GFCCVEGCSA-N | No-Effect | inactive |
| rauwolscine | BRD-K77474816 | COC(=O)[C@@H]1[C@@H](O)CC[C@@H]2CN3CCc4c([nH]c5ccccc45)[C@@H]3C[C@@H]21 | BLGXFZZNTVWLAY-DIRVCLHFSA-N | No-Effect | inactive |
| ravoxertinib | BRD-K86309810 | Cn1nccc1Nc1nccc(-c2ccn([C@H](CO)c3ccc(Cl)c(F)c3)c(=O)c2)n1 | RZUOCXOYPYGSKL-GOSISDBHSA-N | No-Effect | inactive |
| ravuconazole | BRD-K03503561 | C[C@@H](c1nc(-c2ccc(C#N)cc2)cs1)[C@](O)(Cn1cncn1)c1ccc(F)cc1F | OPAHEYNNJWPQPX-RCDICMHDSA-N | No-Effect | inactive |
| RBC8 | BRD-A67036568 | COc1ccc(OC)c([C@H]2c3c(-c4ccc5ccccc5c4)n[nH]c3OC(=N)C2C#N)c1 | RFEAEBNMNYRJFD-VGAJERRHSA-N | No-Effect | inactive |
| rebamipide | BRD-A15909516 | O=C(N[C@H](Cc1cc(=O)[nH]c2ccccc12)C(=O)O)c1ccc(Cl)cc1 | ALLWOAVDORUJLA-MRXNPFEDSA-N | No-Effect | inactive |
| rebastinib | BRD-K76694128 | CNC(=O)c1cc(Oc2ccc(NC(=O)Nc3cc(C(C)(C)C)nn3-c3ccc4ncccc4c3)c(F)c2)ccn1 | WVXNSAVVKYZVOE-UHFFFAOYSA-N | No-Effect | inactive |
| reboxetine | BRD-K77224738 | CCOc1ccccc1O[C@H](c1ccccc1)[C@H]1CNCCO1 | CBQGYUDMJHNJBX-RTBURBONSA-N | No-Effect | inactive |
| Rec-15/2615 | BRD-K84924479 | COc1cc2nc(N3CCN(C(=O)COc4c(OC)cccc4C(C)C)CC3)nc(N)c2cc1OC | ZZMFJJDBKSYGRM-UHFFFAOYSA-N | No-Effect | inactive |
| refametinib | BRD-K85751432 | COc1cc(F)c(F)c(Nc2ccc(I)cc2F)c1NS(=O)(=O)C1(C[C@H](O)CO)CC1 | RDSACQWTXKSHJT-NSHDSACASA-N | No-Effect | inactive |
| regorafenib | BRD-K16730910 | CNC(=O)c1cc(Oc2ccc(NC(=O)Nc3ccc(Cl)c(C(F)(F)F)c3)c(F)c2)ccn1 | FNHKPVJBJVTLMP-UHFFFAOYSA-N | No-Effect | inactive |
| relcovaptan | BRD-K96720755 | COc1ccc(S(=O)(=O)N2c3ccc(Cl)cc3[C@](O)(c3ccccc3Cl)[C@@H]2C(=O)N2CCC[C@H]2C(N)=O)cc1OC | CEBYCSRFKCEUSW-NAYZPBBASA-N | No-Effect | inactive |
| relebactam | BRD-K63018010 | O=C(NC1CCNCC1)[C@@H]1CC[C@@H]2CN1C(=O)N2OS(=O)(=O)O | SMOBCLHAZXOKDQ-ZJUUUORDSA-N | No-Effect | inactive |
| relugolix | BRD-K00003216 | CONC(=O)Nc1ccc(-c2sc3c(c2CN(C)C)c(=O)n(-c2ccc(OC)nn2)c(=O)n3Cc2c(F)cccc2F)cc1 | AOMXMOCNKJTRQP-UHFFFAOYSA-N | No-Effect | inactive |
| remacemide | BRD-A36074203 | C[C@@](Cc1ccccc1)(NC(=O)CN)c1ccccc1 | YSGASDXSLKIKOD-KRWDZBQOSA-N | No-Effect | inactive |
| remetinostat | BRD-K00004584 | COC(=O)c1ccc(OC(=O)CCCCCCC(=O)NO)cc1 | XDZAHHULFQIBFE-UHFFFAOYSA-N | No-Effect | inactive |
| remimazolam | BRD-K89413285 | COC(=O)CC[C@@H]1N=C(c2ccccn2)c2cc(Br)ccc2-n2c(C)cnc21 | CYHWMBVXXDIZNZ-KRWDZBQOSA-N | No-Effect | inactive |
| remodelin | BRD-K75349868 | N#Cc1ccc(-c2csc(NN=C3CCCC3)n2)cc1 | XAEJIFARBQJLML-UHFFFAOYSA-N | No-Effect | inactive |
| remoxipride | BRD-K54094468 | CCN1CCC[C@H]1CNC(=O)c1c(OC)ccc(Br)c1OC | GUJRSXAPGDDABA-NSHDSACASA-N | No-Effect | inactive |
| reparixin | BRD-K58486055 | CC(C)Cc1ccc([C@@H](C)C(=O)NS(C)(=O)=O)cc1 | KQDRVXQXKZXMHP-LLVKDONJSA-N | No-Effect | inactive |
| repotrectinib | BRD-K00003447 | C[C@H]1CNC(=O)c2cnn3ccc(nc23)N[C@H](C)c2cc(F)ccc2O1 | FIKPXCOQUIZNHB-WDEREUQCSA-N | No-Effect | inactive |
| resatorvid | BRD-K61567297 | CCOC(=O)C1=CCCC[C@H]1S(=O)(=O)Nc1ccc(F)cc1Cl | LEEIJTHMHDMWLJ-CQSZACIVSA-N | No-Effect | inactive |
| reserpine | BRD-K95921201 | COC(=O)[C@H]1[C@H]2C[C@@H]3c4[nH]c5cc(OC)ccc5c4CCN3C[C@H]2C[C@@H](OC(=O)c2cc(OC)c(OC)c(OC)c2)[C@@H]1OC | QEVHRUUCFGRFIF-MDEJGZGSSA-N | No-Effect | inactive |
| resiquimod | BRD-K76210423 | CCOCc1nc2c(N)nc3ccccc3c2n1CC(C)(C)O | BXNMTOQRYBFHNZ-UHFFFAOYSA-N | No-Effect | inactive |
| resmetirom | BRD-K00003123 | CC(C)c1cc(Oc2c(Cl)cc(-n3nc(C#N)c(=O)[nH]c3=O)cc2Cl)nnc1O | FDBYIYFVSAHJLY-UHFFFAOYSA-N | No-Effect | inactive |
| resminostat | BRD-K28822270 | CN(C)Cc1ccc(S(=O)(=O)n2ccc(/C=C/C(=O)NO)c2)cc1 | FECGNJPYVFEKOD-VMPITWQZSA-N | No-Effect | inactive |
| resorcinol | BRD-K74190368 | Oc1cccc(O)c1 | GHMLBKRAJCXXBS-UHFFFAOYSA-N | No-Effect | inactive |
| resorcinol-monoacetate | BRD-K85603128 | CC(=O)Oc1cccc(O)c1 | ZZPKZRHERLGEKA-UHFFFAOYSA-N | No-Effect | inactive |
| resveratrol | BRD-K80738081 | Oc1ccc(/C=C/c2cc(O)cc(O)c2)cc1 | LUKBXSAWLPMMSZ-OWOJBTEDSA-N | No-Effect | inactive |
| retagliptin | BRD-K00003118 | COC(=O)c1nc(C(F)(F)F)n2c1CN(C(=O)C[C@H](N)Cc1cc(F)c(F)cc1F)CC2 | WIIAMRXFUJLYEF-SNVBAGLBSA-N | No-Effect | inactive |
| retapamulin | BRD-K33082088 | C=C[C@]1(C)C[C@@H](OC(=O)CS[C@H]2C[C@@H]3CC[C@H](C2)N3C)[C@]2(C)[C@H](C)CC[C@]3(CCC(=O)[C@H]32)[C@@H](C)[C@@H]1O | STZYTFJPGGDRJD-NHUWBDDWSA-N | No-Effect | inactive |
| retinaldehyde | BRD-K79884267 | CC1=C(/C=C/C(C)=C/C=C/C(C)=C/C=O)C(C)(C)CCC1 | NCYCYZXNIZJOKI-OVSJKPMPSA-N | No-Effect | inactive |
| retinol | BRD-K22429181 | CC1=C(/C=C/C(C)=C/C=C/C(C)=C/CO)C(C)(C)CCC1 | FPIPGXGPPPQFEQ-OVSJKPMPSA-N | No-Effect | inactive |
| retinyl-acetate | BRD-K65331431 | CC(=O)OC/C=C(C)/C=C/C=C(C)/C=C/C1=C(C)CCCC1(C)C | QGNJRVVDBSJHIZ-QHLGVNSISA-N | No-Effect | inactive |
| retinyl-palmitate | BRD-K80595380 | CCCCCCCCCCCCCCCC(=O)OC/C=C(C)/C=C/C=C(C)/C=C/C1=C(C)CCCC1(C)C | VYGQUTWHTHXGQB-FFHKNEKCSA-N | No-Effect | inactive |
| REV-5901 | BRD-A68281735 | CCCCC[C@@H](O)c1cccc(OCc2ccc3ccccc3n2)c1 | JRLOEMCOOZSCQP-JOCHJYFZSA-N | No-Effect | inactive |
| revaprazan | BRD-A51497544 | Cc1nc(Nc2ccc(F)cc2)nc(N2CCc3ccccc3[C@H]2C)c1C | LECZXZOBEZITCL-MRXNPFEDSA-N | No-Effect | inactive |
| revefenacin | BRD-K00003198 | CN(CCN1CCC(OC(=O)Nc2ccccc2-c2ccccc2)CC1)C(=O)c1ccc(CN2CCC(C(N)=O)CC2)cc1 | FYDWDCIFZSGNBU-UHFFFAOYSA-N | No-Effect | inactive |
| reversan | BRD-K79700588 | O=C(NCCCN1CCOCC1)c1cnn2c(-c3ccccc3)cc(-c3ccccc3)nc12 | JTRXWCLQFAZHGP-UHFFFAOYSA-N | No-Effect | inactive |
| reversine | BRD-K77286328 | c1nc2nc(Nc3ccc(N4CCOCC4)cc3)nc(NC3CCCCC3)c2[nH]1 | ZFLJHSQHILSNCM-UHFFFAOYSA-N | No-Effect | inactive |
| reynoutrin | BRD-K00005343 | O=c1c(O[C@@H]2OC[C@@H](O)[C@H](O)[C@H]2O)c(-c2ccc(O)c(O)c2)oc2cc(O)cc(O)c12 | PZZRDJXEMZMZFD-BWYUNELBSA-N | No-Effect | inactive |
| RG108 | BRD-K89391146 | O=C(O)[C@H](Cc1c[nH]c2ccccc12)N1C(=O)c2ccccc2C1=O | HPTXLHAHLXOAKV-INIZCTEOSA-N | No-Effect | inactive |
| RG1530 | BRD-K32942258 | COc1cc2c(cc1F)C(c1ccccc1Cl)=NC1C(C)=NNC1=N2 | XXAKBYBCWYYQTL-UHFFFAOYSA-N | No-Effect | inactive |
| RG2833 | BRD-K02389548 | Cc1ccc(C(=O)NCCCCCC(=O)Nc2ccccc2N)cc1 | VOPDXHFYDJAYNS-UHFFFAOYSA-N | No-Effect | inactive |
| RG4733 | BRD-K22024824 | CC(C)(C(=O)NCC(F)(F)C(F)(F)F)C(=O)N[C@@H]1C(=O)Nc2ccccc2-c2ccccc21 | OJPLJFIFUQPSJR-INIZCTEOSA-N | No-Effect | inactive |
| RG7112 | BRD-K13178532 | CCOc1cc(C(C)(C)C)ccc1C1=N[C@@](C)(c2ccc(Cl)cc2)[C@@](C)(c2ccc(Cl)cc2)N1C(=O)N1CCN(CCCS(C)(=O)=O)CC1 | QBGKPEROWUKSBK-QPPIDDCLSA-N | No-Effect | inactive |
| RG7800 | BRD-K00004590 | CCc1nc(C)cn2nc(-c3cc(=O)n4cc(C5CCN(C)CC5)cc(C)c4n3)cc12 | GYFRQCMDLBNZSF-UHFFFAOYSA-N | No-Effect | inactive |
| RG7834 | BRD-K00004573 | COCCCOc1cc2c(cc1OC)-c1cc(=O)c(C(=O)O)cn1[C@H](C(C)C)C2 | KBXLMOYQNDMHQT-KRWDZBQOSA-N | No-Effect | inactive |
| RGB-286638 | BRD-K20986415 | COCCN1CCN(Cc2ccc(-c3n[nH]c4c3C(=O)c3c(NC(=O)NN5CCOCC5)cccc3-4)cc2)CC1 | XLSYZSRXVVCHLS-UHFFFAOYSA-N | No-Effect | inactive |
| RGFP966 | BRD-K61688984 | Nc1cc(F)ccc1NC(=O)/C=C/c1cnn(/C=C/c2ccccc2)c1 | BLVQHYHDYFTPDV-VCABWLAWSA-N | No-Effect | inactive |
| RGX-104 | BRD-K00004597 | C[C@H](CCOc1cccc(CC(=O)O)c1)N(Cc1cccc(C(F)(F)F)c1Cl)CC(c1ccccc1)c1ccccc1 | ZLJZDYOBXVOTSA-XMMPIXPASA-N | No-Effect | inactive |
| RHC-80267 | BRD-K56047318 | O=C(NCCCCCCNC(=O)ON=C1CCCCC1)ON=C1CCCCC1 | RXSVYGIGWRDVQC-UHFFFAOYSA-N | No-Effect | inactive |
| rhein | BRD-K27335680 | O=C(O)c1cc(O)c2c(c1)C(=O)c1cccc(O)c1C2=O | FCDLCPWAQCPTKC-UHFFFAOYSA-N | No-Effect | inactive |
| rheochrysidin | BRD-K67772619 | COc1cc(O)c2c(c1)C(=O)c1cc(C)cc(O)c1C2=O | FFWOKTFYGVYKIR-UHFFFAOYSA-N | No-Effect | inactive |
| rhodamine-123 | BRD-K47169076 | COC(=O)c1ccccc1-c1c2ccc(N)cc2[o+]c2cc(N)ccc12 | ZWODABSNRKVFCK-UHFFFAOYSA-N | No-Effect | inactive |
| RI-1 | BRD-K79555887 | O=C1C(Cl)=C(N2CCOCC2)C(=O)N1c1ccc(Cl)c(Cl)c1 | MWSUIZKGNWELRF-UHFFFAOYSA-N | No-Effect | inactive |
| ribavirin | BRD-K60369935 | NC(=O)c1ncn([C@@H]2O[C@H](CO)[C@@H](O)[C@H]2O)n1 | IWUCXVSUMQZMFG-AFCXAGJDSA-N | No-Effect | inactive |
| ribitol | BRD-K49313711 | OC[C@H](O)[C@H](O)[C@H](O)CO | HEBKCHPVOIAQTA-ZXFHETKHSA-N | No-Effect | inactive |
| ribociclib | BRD-K36788280 | CN(C)C(=O)c1cc2cnc(Nc3ccc(N4CCNCC4)cn3)nc2n1C1CCCC1 | RHXHGRAEPCAFML-UHFFFAOYSA-N | No-Effect | inactive |
| riboflavin | BRD-K92760278 | Cc1cc2nc3c(=O)[nH]c(=O)nc-3n(C[C@H](O)[C@H](O)[C@H](O)CO)c2cc1C | AUNGANRZJHBGPY-SCRDCRAPSA-N | No-Effect | inactive |
| riboflavin-tetrabutyrate | BRD-K74227494 | CCCC(=O)OC[C@@H](OC(=O)CCC)[C@@H](OC(=O)CCC)[C@H](Cn1c2nc(=O)[nH]c(=O)c-2nc2cc(C)c(C)cc21)OC(=O)CCC | MJNIWUJSIGSWKK-BBANNHEPSA-N | No-Effect | inactive |

| | | | | | |
|---|---|---|---|---|---|
| ribostamycin | BRD-K55188888 | NC[C@H]1O[C@H](O[C@@H]2[C@@H](N)C[C@@H](N)[C@H](O)[C@H]2O[C@@H]2O[C@H](CO)[C@@H](O)[C@H]2O)[C@H](N)[C@@H](O)[C@@H]1O | NSKGQURZWSPSBC-VVPCINPTSA-N | No-Effect | inactive |
| ribostamycin-sulfate | BRD-K01826536 | NC[C@H]1O[C@H](O[C@@H]2[C@@H](N)C[C@@H](N)[C@H](O)[C@@H]2O[C@@H]2O[C@H](CO)[C@@H](O)[C@H]2O)[C@H](N)[C@@H](O)[C@@H]1O | NSKGQURZWSPSBC-NZJHKTKDSA-N | No-Effect | inactive |
| ricinoleic-acid | BRD-K33682646 | CCCCCC[C@@H](O)C/C=C\CCCCCCCC(=O)O | WBHHMMIMDMUBKC-QJWNTBNXSA-N | No-Effect | inactive |
| ridinilazole | BRD-K77897005 | c1cc(-c2nc3ccc(-c4ccc5nc(-c6ccncc6)[nH]c5c4)cc3[nH]2)ccn1 | UHQFBTAJFNVZIV-UHFFFAOYSA-N | No-Effect | inactive |
| rifabutin | BRD-K31484631 | CO[C@H]1/C=C/O[C@@]2(C)Oc3c(C)c(O)c4c(c3C2=O)C2=NC3(CCN(CC(C)C)CC3)NC2=C(NC(=O)/C(C)=C\C=C\[C@H](C)[C@H](O)[C@@H](C)[C@@H](O)[C@@H](C)[C@H](OC(C)=O)[C@@H]1C)C4=O | ATEBXHFBFRCZMA-VXTBVIBXSA-N | No-Effect | inactive |
| rifampin | BRD-K01507359 | CO[C@H]1/C=C/O[C@@]2(C)Oc3c(C)c(O)c4c(O)c(c(/C=N/N5CCN(C)CC5)c(O)c4c3C2=O)NC(=O)/C(C)=C\C=C\[C@H](C)[C@H](O)[C@@H](C)[C@@H](O)[C@@H](C)[C@H](OC(C)=O)[C@@H]1C | JQXXHWHPUNPDRT-WLSIYKJHSA-N | No-Effect | inactive |
| rifamycin | BRD-K38512030 | CO[C@H]1/C=C/O[C@@]2(C)Oc3c(C)c(O)c4c(O)c(cc(O)c4c3C2=O)NC(=O)/C(C)=C\C=C\[C@H](C)[C@H](O)[C@@H](C)[C@@H](O)[C@@H](C)[C@H](OC(C)=O)[C@@H]1C | HJYYPODYNSCCOU-ODRIEIDWSA-N | No-Effect | inactive |
| rifamycin-sv | BRD-K66048476 | CO[C@H]1/C=C/O[C@@]2(C)Oc3c(C)c(O)c4c(O)c(cc(O)c4c3C2=O)NC(=O)/C(C)=C\C=C\[C@H](C)[C@H](O)[C@@H](C)[C@@H](O)[C@@H](C)[C@H](OC(C)=O)[C@@H]1C | HJYYPODYNSCCOU-ODRIEIDWSA-N | No-Effect | inactive |
| rifapentine | BRD-K28346421 | CO[C@H]1/C=C/O[C@@]2(C)Oc3c(C)c(O)c4c(O)c(c(/C=N/N5CCN(C6CCCC6)CC5)c(O)c4c3C2=O)NC(=O)/C(C)=C\C=C\[C@H](C)[C@H](O)[C@@H](C)[C@@H](O)[C@@H](C)[C@H](OC(C)=O)[C@@H]1C | WDZCUPBHRAEYDL-GZAUEHORSA-N | No-Effect | inactive |
| rifaximin | BRD-K53484294 | CO[C@H]1/C=C/O[C@@]2(C)Oc3c(C)c(O)c4c(O)c(c5c(nc6cc(C)ccn65)c4c3C2=O)NC(=O)/C(C)=C\C=C\[C@H](C)[C@H](O)[C@@H](C)[C@@H](O)[C@@H](C)[C@H](OC(C)=O)[C@@H]1C | NZCRJKRKKOLAOJ-XRCRFVBUSA-N | No-Effect | inactive |
| rilapladib | BRD-K00004676 | COCCN1CCC(N(Cc2ccc(-c3ccc(C(F)(F)F)cc3)cc2)C(=O)Cn2c(SCc3cccc(F)c3F)cc(=O)c3ccccc32)CC1 | NNBGCSGCRSCFEA-UHFFFAOYSA-N | No-Effect | inactive |
| rimantadine | BRD-K01825586 | C[C@@H](N)C12CC3CC(CC(C3)C1)C2 | UBCHPRBFMUDMNC-AAWJQDODSA-N | No-Effect | inactive |
| rimegepant | BRD-K55781385 | N[C@@H]1c2cccnc2[C@H](OC(=O)N2CCC(n3c(=O)[nH]c4ncccc43)CC2)CC[C@H]1c1cccc(F)c1F | KRNAOFGYEFKHPB-ANJVHQHFSA-N | No-Effect | inactive |
| rimeporide | BRD-K00004645 | Cc1cc(S(C)(=O)=O)c(S(C)(=O)=O)cc1C(=O)NC(=N)N | GROMEQPXDKRRIE-UHFFFAOYSA-N | No-Effect | inactive |
| rimexolone | BRD-K31627533 | CCC(=O)[C@@]1(C)[C@H](C)C[C@H]2[C@@H]3CCC4=CC(=O)C=C[C@]4(C)[C@H]3[C@@H](O)C[C@@]21C | QTTRZHGPGKRAFB-OOKHYKNYSA-N | No-Effect | inactive |
| rimonabant | BRD-K13296708 | Cc1c(C(=O)NN2CCCCC2)nn(-c2ccc(Cl)cc2Cl)c1-c1ccc(Cl)cc1 | JZCPYUJPEARBJL-UHFFFAOYSA-N | No-Effect | inactive |
| riociguat | BRD-K84783599 | COC(=O)N(C)c1c(N)nc(-c2nn(Cc3ccccc3F)c3ncccc23)nc1N | WXXSNCNJFUAIDG-UHFFFAOYSA-N | No-Effect | inactive |
| riodipine | BRD-K36379990 | COC(=O)C1=C(C)NC(C)=C(C(=O)OC)C1c1ccccc1OC(F)F | AZVFIZVKGSPGPK-UHFFFAOYSA-N | No-Effect | inactive |
| riodoxol | BRD-K48963429 | Oc1c(I)cc(I)c(O)c1I | XKFZYVWWXHCHIX-UHFFFAOYSA-N | No-Effect | inactive |
| ripasudil | BRD-K53814070 | C[C@H]1CNCCCN1S(=O)(=O)c1cccc2cncc(F)c12 | QSKQVZWVLOIIEV-NSHDSACASA-N | No-Effect | inactive |
| ripretinib | BRD-K00005241 | CCn1c(=O)c(-c2cc(NC(=O)Nc3ccccc3)c(F)cc2Br)cc2cnc(NC)cc21 | CEFJVGZHQAGLHS-UHFFFAOYSA-N | No-Effect | inactive |
| risdiplam | BRD-K00003506 | Cc1cn2nc(-c3cc(=O)n4cc(N5CCNC6(CC6)C5)ccc4n3)cc(C)c2n1 | ASKZRYGFUPSJPN-UHFFFAOYSA-N | No-Effect | inactive |
| risedronate | BRD-K05185389 | O=P(O)(O)C(O)(Cc1cccnc1)P(=O)(O)O | IIDJRNMFWXDHID-UHFFFAOYSA-N | No-Effect | inactive |
| rislenemdaz | BRD-K00003471 | Cc1ccc(COC(=O)N2CC[C@H](CNc3ncccn3)[C@H](F)C2)cc1 | RECBFDWSXWAXHY-IAGOWNOFSA-N | No-Effect | inactive |
| risperidone | BRD-K53857191 | Cc1nc2n(c(=O)c1CCN1CCC(c3noc4cc(F)ccc34)CC1)CCCC2 | RAPZEAPATHNIPO-UHFFFAOYSA-N | No-Effect | inactive |
| RITA | BRD-K00317371 | OCc1ccc(-c2ccc(-c3ccc(CO)s3)o2)s1 | KZENBFUSKMWCJF-UHFFFAOYSA-N | No-Effect | inactive |
| ritanserin | BRD-K40887525 | Cc1nc2sccn2c(=O)c1CCN1CCC(=C(c2ccc(F)cc2)c2ccc(F)cc2)CC1 | JUQLTPCYUFPYKE-UHFFFAOYSA-N | No-Effect | inactive |
| ritodrine | BRD-K14003026 | C[C@@H](NCCc1ccc(O)cc1)[C@@H](O)c1ccc(O)cc1 | IOVGROKTTNBUGK-SJKOYZFVSA-N | No-Effect | inactive |
| ritonavir | BRD-K51485625 | CC(C)c1nc(CN(C)C(=O)N[C@H](C(=O)N[C@@H](Cc2ccccc2)C[C@H](O)[C@H](Cc2ccccc2)NC(=O)OCc2cncs2)C(C)C)cs1 | NCDNCNXCDXHOMX-XGKFQTDJSA-N | No-Effect | inactive |
| rivanicline | BRD-K79966802 | CNCC/C=C/c1cccnc1 | JUOSGGQXEBBCJB-GORDUTHDSA-N | No-Effect | inactive |
| rivaroxaban | BRD-K37130656 | O=C(NC[C@H]1CN(c2ccc(N3CCOCC3=O)cc2)C(=O)O1)c1ccc(Cl)s1 | KGFYHTZWPPHNLQ-AWEZNQCLSA-N | No-Effect | inactive |
| rivastigmine | BRD-K10706131 | CCN(C)C(=O)Oc1cccc([C@H](C)N(C)C)c1 | XSVMFMHYUFZWBK-NSHDSACASA-N | No-Effect | inactive |
| rizatriptan | BRD-K75699339 | CN(C)CCc1c[nH]c2ccc(Cn3cncn3)cc12 | ULFRLSNUDGIQQP-UHFFFAOYSA-N | No-Effect | inactive |
| RKI-1447 | BRD-K19333160 | O=C(NCc1cccc(O)c1)Nc1nc(-c2ccncc2)cs1 | GDVRVPIXWXOKQO-UHFFFAOYSA-N | No-Effect | inactive |
| RLX | BRD-K28773847 | O=c1c2ccccc2nc2n1CCCCC2 | HTLIIBRHXAULMB-UHFFFAOYSA-N | No-Effect | inactive |
| RMC-4550 | BRD-K00010386 | Cc1nc(N2CCC3(CC2)CO[C@@H](C)[C@H]3N)c(CO)nc1-c1cccc(Cl)c1Cl | IKUYEYLZXGGCRD-ORAYPTAESA-N | No-Effect | inactive |
| RN-1 | BRD-K41415459 | CN1CCN(C(=O)CN[C@@H]2C[C@H]2c2ccc(OCc3ccccc3)cc2)CC1 | YAMSXCOVJUUMCT-FCHUYYIVSA-N | No-Effect | inactive |
| RN-1734 | BRD-K95581532 | CC(C)NCCN(C(C)C)S(=O)(=O)c1ccc(Cl)cc1Cl | IHYZMEAZAIFMTN-UHFFFAOYSA-N | No-Effect | inactive |
| Ro-04-5595 | BRD-A63546914 | COc1cc2c(cc1O)[C@H](CCc1ccc(Cl)cc1)N(C)CC2 | DSTWURHEHMZWEI-KRWDZBQOSA-N | No-Effect | inactive |
| Ro-08-2750 | BRD-K00486786 | Cc1cc2nc3c(=O)[nH]c(=O)nc-3n(C)c2cc1C=O | JDEMVNYMYPJJIM-UHFFFAOYSA-N | No-Effect | inactive |
| Ro-15-4513 | BRD-K82823076 | CCOC(=O)c1ncn2c1CN(C)C(=O)c1cc(N=[N+]=[N-])ccc1-2 | CFSOJZTUTOQNIA-UHFFFAOYSA-N | No-Effect | inactive |
| Ro-19-4605 | BRD-K80778372 | CN1Cc2c(C(=O)OC(C)(C)C)ncn2-c2ccsc2C1=O | ZIGMMUKDYCABPW-UHFFFAOYSA-N | No-Effect | inactive |
| Ro-20-1724 | BRD-A07207424 | CCCCOc1cc(C[C@H]2CNC(=O)N2)ccc1OC | PDMUULPVBYQBBK-LBPRGKRZSA-N | No-Effect | inactive |
| Ro-25-6981 | BRD-K51541829 | C[C@@H](CN1CCC(Cc2ccccc2)CC1)[C@@H](O)c1ccc(O)cc1 | WVZSEUPGUDIELE-HTAPYJJXSA-N | No-Effect | inactive |
| Ro-28-1675 | BRD-K21672174 | CS(=O)(=O)c1ccc([C@@H](CC2CCCC2)C(=O)Nc2nccs2)cc1 | NEQSWPCDHDQINX-MRXNPFEDSA-N | No-Effect | inactive |
| Ro-3 | BRD-K00354853 | COc1cc(Cc2cnc(N)nc2N)c(C(C)C)cc1OC | PYNPWUIBJMVRIG-UHFFFAOYSA-N | No-Effect | inactive |
| Ro-3306 | BRD-K78373679 | O=C1N=C(NCc2cccs2)S/C1=C\c1ccc2ncccc2c1 | XOLMRFUGOINFDQ-YBEGLDIGSA-N | No-Effect | inactive |
| Ro-48-6791 | BRD-K00004572 | CCCN(CCC)Cc1nc(-c2ncn3c2CN(C)C(=O)c2cc(F)ccc2-3)no1 | NOQIYRGMEFBZTI-UHFFFAOYSA-N | No-Effect | inactive |
| Ro-48-8071 | BRD-K55192287 | C=CCN(C)CCCCCCOc1ccc(C(=O)c2ccc(Br)cc2)c(F)c1 | CMYCCJYVZIMDFU-UHFFFAOYSA-N | No-Effect | inactive |
| Ro-4987655 | BRD-K64538373 | O=C(NOCCO)c1cc(CN2OCCCC2=O)c(F)c(F)c1Nc1ccc(I)cc1F | FIMYFEGKMOCQKT-UHFFFAOYSA-N | No-Effect | inactive |
| Ro-5028442 | BRD-K00003149 | CN(C)CCn1cc(C(=O)N2CCC3(CC2)OCc2ccccc23)c2ccc(Cl)cc21 | QZXVLRCMAHJVIP-UHFFFAOYSA-N | No-Effect | inactive |
| Ro-5126766 | BRD-K82804538 | CNS(=O)(=O)Nc1nccc(Cc2c(C)c3ccc(Oc4ncccn4)cc3oc2=O)c1F | LMMJFBMMJUMSJS-UHFFFAOYSA-N | No-Effect | inactive |
| Ro-5263397 | BRD-K00004623 | Cc1c(F)cccc1[C@H]1COC(N)=N1 | IOHOUWIYOVWGHV-SECBINFHSA-N | No-Effect | inactive |
| Ro-60-0175 | BRD-K28007504 | C[C@H](N)Cn1ccc2cc(F)c(Cl)cc21 | XJJZQXUGLLXTHO-ZETCQYMHSA-N | No-Effect | inactive |
| Ro-61-8048 | BRD-K12639498 | COc1ccc(S(=O)(=O)Nc2nc(-c3cccc([N+](=O)[O-])c3)cs2)cc1OC | NDPBMCKQJOZAQX-UHFFFAOYSA-N | No-Effect | inactive |
| Ro-67-7476 | BRD-K91333421 | Cc1ccc(S(=O)(=O)N2CCC[C@H]2c2ccc(F)cc2)cc1 | DAEHFYNGSSBGSS-KRWDZBQOSA-N | No-Effect | inactive |
| Ro-90-7501 | BRD-K58299615 | Nc1ccc(-c2nc3ccc(-c4nc5ccc(N)cc5[nH]4)cc3[nH]2)cc1 | PAGZCEHLFCJSPV-UHFFFAOYSA-N | No-Effect | inactive |
| Ro-9187 | BRD-K57284066 | [N-]=[N+]=N[C@]1(CO)O[C@@H](n2ccc(N)nc2=O)[C@@H](O)[C@@H]1O | ODLGMSQBFONGNG-XZMZPDFPSA-N | No-Effect | inactive |
| robalzotan | BRD-K26169680 | NC(=O)c1ccc(F)c2c1C[C@@H](N(C1CCC1)C1CCC1)CO2 | MQTUXRKNJYPMCG-CYBMUJFWSA-N | No-Effect | inactive |
| roblitinib | BRD-K00003424 | COCCNc1cc(NC(=O)N2CCCc3cc(CN4CCN(C)CC4=O)c(C=O)nc32)ncc1C#N | BHKDKKZMPODMIQ-UHFFFAOYSA-N | No-Effect | inactive |
| rociletinib | BRD-K33610132 | C=CC(=O)Nc1cccc(Nc2nc(Nc3ccc(N4CCN(C(C)=O)CC4)cc3OC)ncc2C(F)(F)F)c1 | HUFOZJXAKZVRNJ-UHFFFAOYSA-N | No-Effect | inactive |

| | | | | | |
|---|---|---|---|---|---|
| rocuronium | BRD-K43305603 | C=CC[N+]1([C@H]2C[C@H]3[C@@H]4CC[C@H]5C[C@H](O)[C@@H](N6CCOCC6)C[C@]5(C)[C@H]4CC[C@]3(C)[C@H]2OC(C)=O)CCCC1 | YXRDKMPIGHSVRX-OOJCLDBCSA-N | No-Effect | inactive |
| rofecoxib | BRD-K21733600 | CS(=O)(=O)c1ccc(C2=C(c3ccccc3)C(=O)OC2)cc1 | RZJQGNCSTQAWON-UHFFFAOYSA-N | No-Effect | inactive |
| roflumilast | BRD-K03194791 | O=C(Nc1c(Cl)cncc1Cl)c1ccc(OC(F)F)c(OCC2CC2)c1 | MNDBXUUTURYVHR-UHFFFAOYSA-N | No-Effect | inactive |
| rogaratinib | BRD-K00003358 | COCc1c(-c2cc3cc(C)cc(OC)c3s2)c2c(N)ncnn2c1CN1CCNC(=O)C1 | HNLRRJSKGXOYNO-UHFFFAOYSA-N | No-Effect | inactive |
| rolapitant | BRD-K00003175 | C[C@@H](OC[C@@]1(c2ccccc2)CC[C@]2(CCC(=O)N2)CN1)c1cc(C(F)(F)F)cc(C(F)(F)F)c1 | FIVSJYGQAIEMOC-ZGNKEGEESA-N | No-Effect | inactive |
| rolipram | BRD-A34255068 | COc1ccc([C@H]2CNC(=O)C2)cc1OC1CCCC1 | HJORMJIFDVBMOB-GFCCVEGCSA-N | No-Effect | inactive |
| roluperidone | BRD-K00003283 | O=C(CN1CCC(CN2Cc3ccccc3C2=O)CC1)c1ccc(F)cc1 | RNRYULFRLCBRQS-UHFFFAOYSA-N | No-Effect | inactive |
| rolziracetam | BRD-K38564664 | O=C1CCC2CCC(=O)N12 | IEZDOKQWPWZVQF-UHFFFAOYSA-N | No-Effect | inactive |
| ropinirole | BRD-K15933101 | CCCN(CCC)CCc1cccc2c1CC(=O)N2 | UHSKFQJFRQCDBE-UHFFFAOYSA-N | No-Effect | inactive |
| ropivacaine | BRD-K50938786 | CCCN1CCCC[C@H]1C(=O)Nc1c(C)cccc1C | ZKMNUMMKYBVTFN-HNNXBMFYSA-N | No-Effect | inactive |
| roquinimex | BRD-K03384561 | CN(C(=O)c1c(O)c2ccccc2n(C)c1=O)c1ccccc1 | SGOOQMRIPALTEL-UHFFFAOYSA-N | No-Effect | inactive |
| rosamicin | BRD-K00004538 | CC[C@H]1OC(=O)C[C@@H](O)[C@H](C)[C@@H](O[C@@H]2O[C@H](C)C[C@H](N(C)C)[C@H]2O)[C@@H](CC=O)C[C@@H](C)C(=O)/C=C/[C@]2(C)O[C@H]2[C@@H]1C | IUPCWCLVECYZRV-JZMZINANSA-N | No-Effect | inactive |
| roscovitine | BRD-K07691486 | CC[C@H](CO)Nc1nc(NCc2ccccc2)c2ncn(C(C)C)c2n1 | BTIHMVBBUGXLCJ-OAHLLOKOSA-N | No-Effect | inactive |
| rose-bengal-lactone | BRD-K36163066 | O=C1OC2(c3cc(I)c(O)c(I)c3Oc3c2cc(I)c(O)c3I)c2c(Cl)c(Cl)c(Cl)c(Cl)c21 | IICCLYANAQEHCI-UHFFFAOYSA-N | No-Effect | inactive |
| rosiglitazone | BRD-K01826734 | CN(CCOc1ccc(C[C@H]2SC(=O)NC2=O)cc1)c1ccccn1 | YASAKCUCGLMORW-OAHLLOKOSA-N | No-Effect | inactive |
| rosiptor | BRD-K00003494 | C=C1CC[C@H]2[C@H](CN)[C@@H]([C@@]3(C)CC[C@H](O)C[C@@H]3CO)CC[C@]12C | MDEJTPWQNNMAQF-BVMLLJBZSA-N | No-Effect | inactive |
| rosmarinic-acid | BRD-K20313525 | O=C(/C=C/c1ccc(O)c(O)c1)O[C@H](Cc1ccc(O)c(O)c1)C(=O)O | DOUMFZQKYFQNTF-WUTVXBCWSA-N | No-Effect | inactive |
| rosoxacin | BRD-K49841478 | CCn1cc(C(=O)O)c(=O)c2ccc(-c3ccncc3)cc21 | XBPZXDSZHPDXQU-UHFFFAOYSA-N | No-Effect | inactive |
| rostafuroxine | BRD-K11672787 | C[C@]12CC[C@H](O)C[C@H]1CC[C@@H]1[C@@H]2CC[C@@]2(C)[C@]1(O)CC[C@]2(O)c1ccoc1 | AEAPORIZZWBIEX-DTBDINHYSA-N | No-Effect | inactive |
| rotigotine | BRD-K91111634 | CCCN(CCc1cccs1)[C@H]1CCc2c(O)cccc2C1 | KFQYTPMOWPVWEJ-INIZCTEOSA-N | No-Effect | inactive |
| rotundine | BRD-K35719256 | COc1cc2c(cc1OC)[C@@H]1Cc3ccc(OC)c(OC)c3CN1CC2 | AEQDJSLRWYMAQI-KRWDZBQOSA-N | No-Effect | inactive |
| rovazolac | BRD-K00003502 | CCOC(=O)Cn1nc(C(F)(F)F)cc1-c1ccc(-c2cccc(S(C)(=O)=O)c2)cc1 | ZUMNJDGBYXHASJ-UHFFFAOYSA-N | No-Effect | inactive |
| roxarsone | BRD-K22685337 | O=[N+]([O-])c1cc([As](=O)(O)O)ccc1O | XMVJITFPVVRMHC-UHFFFAOYSA-N | No-Effect | inactive |
| roxatidine-acetate | BRD-K81855038 | CC(=O)OCC(=O)NCCCOc1cccc(CN2CCCCC2)c1 | SMTZFNFIKUPEJC-UHFFFAOYSA-N | No-Effect | inactive |
| roxithromycin | BRD-K38684403 | CC[C@H]1OC(=O)[C@H](C)[C@@H](O[C@H]2C[C@@](C)(OC)[C@@H](O)[C@H](C)O2)[C@H](C)[C@@H](O[C@@H]2O[C@H](C)C[C@H](N(C)C)[C@H]2O)[C@](C)(O)C[C@@H](C)/C(=N\OCOCCOC)[C@H](C)[C@@H](O)[C@]1(C)O | RXZBMPWDPOLZGW-XMRMVWPWSA-N | No-Effect | inactive |
| RP-001 | BRD-A29148886 | CC(C)Oc1ccc(-c2nc(-c3cccc4c3CC[C@H]4NCCC(=O)O)no2)cc1C#N | WNDDDFYFTSGSEC-HXUWFJFHSA-N | No-Effect | inactive |
| RP67580 | BRD-K40767677 | COc1ccccc1CC(=N)N1C[C@@H]2C(=O)CCC(c3ccccc3)(c3ccccc3)[C@@H]2C1 | VWBOQFANCXZMAU-LOSJGSFVSA-N | No-Effect | inactive |
| RQ-00203078 | BRD-K87973152 | O=C(O)c1ccc(S(=O)(=O)N(Cc2ccc(OC(F)(F)F)cc2)c2ncc(C(F)(F)F)cc2Cl)cc1 | IJGQFZYYEHCCIZ-UHFFFAOYSA-N | No-Effect | inactive |
| RRx-001 | BRD-K06894065 | O=C(CBr)N1CC([N+](=O)[O-])([N+](=O)[O-])C1 | JODKFOVZURLVTG-UHFFFAOYSA-N | No-Effect | inactive |
| RS-0481 | BRD-K00004000 | C[C@@H](NC(=O)[C@@H]1CSCN1C(=O)c1ccccc1)c1ccccc1 | MEXWLVVJOPSWLN-PBHICJAKSA-N | No-Effect | inactive |
| RS-100329 | BRD-K08640512 | Cc1c[nH]c(=O)n(CCCN2CCN(c3ccccc3OCC(F)(F)F)CC2)c1=O | RCOBWVAGWYRNHZ-UHFFFAOYSA-N | No-Effect | inactive |
| RS-102221 | BRD-K87048468 | COc1cc(OC)c(C(=O)CCCCN2CCC3(CC2)NC(=O)NC3=O)cc1NS(=O)(=O)c1ccc(C(F)(F)F)cc1 | HZZZZODVDSHQRG-UHFFFAOYSA-N | No-Effect | inactive |
| RS-102895 | BRD-K83063356 | O=C1Nc2ccccc2C2(CCN(CCc3ccc(C(F)(F)F)cc3)CC2)O1 | HIDWEYPGMLIQSN-UHFFFAOYSA-N | No-Effect | inactive |
| RS-127445 | BRD-K90825648 | CC(C)c1cc(-c2ccc(F)c3ccccc23)nc(N)n1 | ZZZQXCUPAJFVBN-UHFFFAOYSA-N | No-Effect | inactive |
| RS-16566 | BRD-K00004221 | CC(C)n1cnc2c(C(=O)N[C@H]3C[N@]4CC[C@@H]3CC4)cc(Cl)cc21 | OGXMGJGFLBSEJW-HNNXBMFYSA-N | No-Effect | inactive |
| RS-23597-190 | BRD-K01868942 | COc1cc(N)c(Cl)cc1C(=O)OCCCN1CCCCC1 | RLSGBCUXLRMTPF-UHFFFAOYSA-N | No-Effect | inactive |
| RS-39604 | BRD-K20742498 | COc1cc(COc2cc(N)c(Cl)cc2C(=O)CCC2CCN(CCNS(C)(=O)=O)CC2)cc(OC)c1 | BKVIWGRFRKLFIO-UHFFFAOYSA-N | No-Effect | inactive |
| RS-45041-190 | BRD-K26160755 | Clc1cccc2c1CN(C1=NCCN1)C2 | IVCMOOBKRRDNHX-UHFFFAOYSA-N | No-Effect | inactive |
| RS-504393 | BRD-K87510569 | Cc1ccc2c(c1)C1(CCN(CCc3nc(-c4ccccc4)oc3C)CC1)OC(=O)N2 | ODNICNWASXKNNQ-UHFFFAOYSA-N | No-Effect | inactive |
| RS-56812 | BRD-K00004219 | Cn1cc(C(=O)C(=O)N[C@H]2C[N@]3CC[C@@H]2CC3)c2ccccc21 | ZNAPADWWBWFMCQ-HNNXBMFYSA-N | No-Effect | inactive |
| RS-67333 | BRD-K46142322 | CCCCN1CCC(CCC(=O)c2cc(Cl)c(N)cc2OC)CC1 | JBHLYIVFFLNISJ-UHFFFAOYSA-N | No-Effect | inactive |
| RS-67506 | BRD-K50018155 | COc1cc(N)c(Cl)cc1C(=O)CCC1CCN(CCNS(C)(=O)=O)CC1 | UATDVRSFFGFTAI-UHFFFAOYSA-N | No-Effect | inactive |
| RS-79948 | BRD-K86600316 | CCS(=O)(=O)N1CCC[C@@H]2CN3CCc4cc(OC)ccc4[C@@H]3C[C@@H]21 | UMGBFFAJXFXOIL-MNEFBYGVSA-N | No-Effect | inactive |
| RSV604 | BRD-K30159788 | O=C(Nc1ccccc1F)N[C@H]1N=C(c2ccccc2)c2ccccc2NC1=O | MTPVBMVUENFFLL-HXUWFJFHSA-N | No-Effect | inactive |
| RU-24969 | BRD-K00234327 | COc1ccc2[nH]cc(C3=CCNCC3)c2c1 | KRVMLPUDAOWOGN-UHFFFAOYSA-N | No-Effect | inactive |
| RU-28318 | BRD-K37687387 | CCC[C@@H]1CC2=CC(=O)CC[C@]2(C)[C@H]2CC[C@@]3(C)[C@@H](CC[C@@]3(O)CCC(=O)O)[C@@H]21 | DNHCHRGCTVRAFT-JEHIOXJOSA-N | No-Effect | inactive |
| RU-42173 | BRD-K85285268 | CC(C)N[C@@H]1CCn2c(=O)[nH]c3cccc(c32)[C@H]1O | ZSTCZWJCLIRCOJ-DGCLKSJQSA-N | No-Effect | inactive |
| RU-58841 | BRD-K62762455 | CC1(C)C(=O)N(c2ccc(C#N)c(C(F)(F)F)c2)C(=O)N1CCCCO | ARBYGDBJECGMGA-UHFFFAOYSA-N | No-Effect | inactive |
| RU-SKI-43 | BRD-A33603104 | CC[C@@H](C)CNCC(=O)N1CCc2sccc2[C@@H]1COc1cccc(C)c1 | AEENEMOEBJOKGN-UZLBHIALSA-N | No-Effect | inactive |
| rubitecan | BRD-K79821389 | CC[C@@]1(O)C(=O)OCc2c1cc1n(c2=O)Cc2cc3c([N+](=O)[O-])cccc3nc2-1 | VHXNKPBCCMUMSW-FQEVSTJZSA-N | No-Effect | inactive |
| rucaparib | BRD-K88560311 | CNCc1ccc(-c2[nH]c3cc(F)cc4c3c2CCNC4=O)cc1 | HMABYWSNWIZPAG-UHFFFAOYSA-N | No-Effect | inactive |
| rucinol | BRD-K61036791 | CCCCc1ccc(O)cc1O | CSHZYWUPJWVTMQ-UHFFFAOYSA-N | No-Effect | inactive |
| rufinamide | BRD-K20079257 | NC(=O)c1cn(Cc2c(F)cccc2F)nn1 | POGQSBRIGCQNEG-UHFFFAOYSA-N | No-Effect | inactive |
| rufloxacin | BRD-K62374253 | CN1CCN(c2c(F)cc3c(=O)c(C(=O)O)cn4c3c2SCC4)CC1 | NJCJBUHJQLFDSW-UHFFFAOYSA-N | No-Effect | inactive |
| rupatadine | BRD-K98004573 | Cc1cncc(CN2CCC(=C3c4ccc(Cl)cc4CCc4cccnc43)CC2)c1 | WUZYKBABMWJHDL-UHFFFAOYSA-N | No-Effect | inactive |
| rutaecarpine | BRD-K94723713 | O=c1c2ccccc2nc2n1CCc1c-2[nH]c2ccccc12 | ACVGWSKVRYFWRP-UHFFFAOYSA-N | No-Effect | inactive |
| rutin | BRD-K20482099 | C[C@@H]1O[C@@H](OC[C@H]2O[C@@H](Oc3c(-c4ccc(O)c(O)c4)oc4cc(O)cc(O)c4c3=O)[C@H](O)[C@@H](O)[C@@H]2O)[C@H](O)[C@H](O)[C@H]1O | IKGXIBQEEMLURG-NVPNHPEKSA-N | No-Effect | inactive |
| ruxolitinib | BRD-K53972329 | N#CC[C@H](C1CCCC1)n1cc(-c2ncnc3[nH]ccc23)cn1 | HFNKQEVNSGCOJV-OAHLLOKOSA-N | No-Effect | inactive |
| ruxolitinib-(S) | BRD-K19601669 | N#CC[C@@H](C1CCCC1)n1cc(-c2ncnc3[nH]ccc23)cn1 | HFNKQEVNSGCOJV-HNNXBMFYSA-N | No-Effect | inactive |
| RWJ-21757 | BRD-K00091711 | C=CCn1c(=O)n([C@@H]2O[C@H](CO)[C@@H](O)[C@H]2O)c2[nH]c(N)nc(=O)c21 | VDCRFBBZFHHYGT-IOSLPCCCSA-N | No-Effect | inactive |
| RWJ-50271 | BRD-K94623186 | Cc1c(C(=O)NCCCO)cnn1-c1nc(-c2cccc(C(F)(F)F)c2)cs1 | HOFGTYCLOKDAES-UHFFFAOYSA-N | No-Effect | inactive |
| RWJ-67657 | BRD-K07074923 | OCCC#Cc1nc(-c2ccc(F)cc2)c(-c2ccncc2)n1CCCc1ccccc1 | QSUSKMBNZQHHPA-UHFFFAOYSA-N | No-Effect | inactive |
| RX-3117 | BRD-K66213167 | Nc1ccn([C@@H]2C(F)=C(CO)[C@@H](O)[C@H]2O)c(=O)n1 | QLLGKCJUPWYJON-HLTSFMKQSA-N | No-Effect | inactive |
| RX-821002 | BRD-A65597028 | CO[C@@]1(C2=NCCN2)COc2ccccc2O1 | HQGWKNGAKBPTBX-LBPRGKRZSA-N | No-Effect | inactive |
| ryuvidine | BRD-K06426971 | Cc1ccc(NC2=CC(=O)c3sc(C)nc3C2=O)cc1 | HFPLHASLIOXVGS-UHFFFAOYSA-N | No-Effect | inactive |
| S-07662 | BRD-K37635256 | Cc1oc2ccccc2c1CNC(=O)NCc1cccs1 | UITFWLJMEZPXOF-UHFFFAOYSA-N | No-Effect | inactive |
| S-111 | BRD-K60005752 | CC(C)=CCC[C@](C)(O)[C@H]1CC[C@]2(C)[C@@H]1[C@H](O)C[C@@H]1[C@@]3(C)CC[C@H](O)C(C)(C)[C@@H]3CC[C@]12C | PYXFVCFISTUSOO-HKUCOEKDSA-N | No-Effect | inactive |

| | | | | | |
|---|---|---|---|---|---|
| S-14506 | BRD-K46766488 | COc1ccc2cccc(N3CCN(CCNC(=O)c4ccc(F)cc4)CC3)c2c1 | IFMQODYDAUKKEN-UHFFFAOYSA-N | No-Effect | inactive |
| S-17092 | BRD-K00004618 | O=C([C@@H]1C[C@@H]2CCCC[C@@H]2N1C(=O)[C@@H]1C[C@H]1c1ccccc1)N1CCSC1 | NXSXRIHXEQSYEZ-KNJMJIDISA-N | No-Effect | inactive |
| S-3304 | BRD-K00003475 | Cc1ccc(C#Cc2ccc(S(=O)(=O)N[C@H](Cc3c[nH]c4ccccc34)C(=O)O)s2)cc1 | YWCLDDLVLSQGSZ-JOCHJYFZSA-N | No-Effect | inactive |
| S-38093 | BRD-K00003455 | NC(=O)c1ccc(OCCCN2C[C@H]3CCC[C@H]3C2)cc1 | MRNMYWNBLVJWKG-GASCZTMLSA-N | No-Effect | inactive |
| S-methylcysteine | BRD-K63123526 | CSC[C@H](N)C(=O)O | IDIDJDIHTAOVLG-VKHMYHEASA-N | No-Effect | inactive |
| S-nitrosoglutathione | BRD-K00024724 | N[C@@H](CCC(=O)N[C@@H](CS[N+][O-])C(=O)NCC(=O)O)C(=O)O | HYHSBSXUHZOYLX-WDSKDSINSA-N | No-Effect | inactive |
| S-sulfo-L-cysteine | BRD-K59527975 | N[C@@H](CSS(=O)(=O)O)C(=O)O | NOKPBJYHPHHWAN-REOHCLBHSA-N | No-Effect | inactive |
| S-trityl-L-cysteine | BRD-K09774593 | N[C@H](CSC(c1ccccc1)(c1ccccc1)c1ccccc1)C(=O)O | DLMYFMLKORXJPO-HXUWFJFHSA-N | No-Effect | inactive |
| S15535 | BRD-K00004653 | c1ccc2c(c1)CC(N1CCN(c3cccc4c3OCCO4)CC1)C2 | QJPPEMXOOWNICQ-UHFFFAOYSA-N | No-Effect | inactive |
| S18986 | BRD-K02501528 | O=S1(=O)N[C@H]2CCCN2c2ccccc21 | MNTIJYGEITVWHU-SNVBAGLBSA-N | No-Effect | inactive |
| S1P1-agonist-III | BRD-K12745498 | COc1ccncc1C(=O)NC(=O)Nc1ccc(-c2ccccc2)c(C(F)(F)F)c1 | MLDQTQOMWDNTNN-UHFFFAOYSA-N | No-Effect | inactive |
| S26948 | BRD-K48826825 | COC(=O)C(Cc1ccc(OCCn2c(=O)sc3cc(C(=O)c4ccccc4)ccc32)cc1)C(=O)OC | NSMJEHGOMXSLCW-UHFFFAOYSA-N | No-Effect | inactive |
| S4 | BRD-K96798615 | Cc1cc(C)cc(NC(=O)Nc2ccc(OS(N)(=O)=O)cc2)c1 | HGVHSNXRZYOTPD-UHFFFAOYSA-N | No-Effect | inactive |
| S49076 | BRD-K00003147 | O=C1Nc2ccc(CN3C(=O)CSC3=O)cc2/C1=C/c1cc(CN2CCOCC2)c[nH]1 | AREYWCZYVPSHGS-NVMNQCDNSA-N | No-Effect | inactive |
| S63845 | BRD-K91876515 | Cc1c(-c2c(-c3ccc(F)o3)sc3ncnc(O[C@H](Cc4ccccc4OCc4ccnn4CC(F)(F)F)C(=O)O)c23)ccc(OCCN2CCN(C)CC2)c1Cl | ZFBHXVOCZBPADE-SSEXGKCCSA-N | No-Effect | inactive |
| SA-47 | BRD-K99819791 | CNC(=O)COC(=O)NCCC1CCN(c2cccc(C)n2)CC1 | HSYCMGWPPRTNKH-UHFFFAOYSA-N | No-Effect | inactive |
| saclofen | BRD-A24122750 | NC[C@@H](CS(=O)(=O)O)c1ccc(Cl)cc1 | JYLNVJYYQQXNEK-QMMMGPOBSA-N | No-Effect | inactive |
| sacubitril | BRD-K91456750 | CCOC(=O)[C@H](C)C[C@@H](Cc1ccc(-c2ccccc2)cc1)NC(=O)CCC(=O)O | PYNXFZCZUAOOQC-UTKZUKDTSA-N | No-Effect | inactive |
| sacubitrilat | BRD-K00003252 | C[C@H](C[C@@H](Cc1ccc(-c2ccccc2)cc1)NC(=O)CCC(=O)O)C(=O)O | DOBNVUFHFMVMDB-BEFAXECRSA-N | No-Effect | inactive |
| safflower-yellow | BRD-A96503329 | O=C1C(=C(O)/C=C/c2ccc(O)cc2)C(=O)[C@@](O)([C@H]2O[C@H](CO)[C@@H](O)[C@H](O)[C@H]2O)C(=O)/C1=C/[C@H]1C(=O)C(=C(O)/C=C/c2ccc(O)cc2)C(=O)[C@](O)([C@H]2O[C@H](CO)[C@@H](O)[C@H](O)[C@H]2O)C1=O | HLUBVUICXYRPPM-DVSWWXTKSA-N | No-Effect | inactive |
| safinamide | BRD-K92613113 | C[C@H](NCc1ccc(OCc2cccc(F)c2)cc1)C(N)=O | NEMGRZFTLSKBAP-LBPRGKRZSA-N | No-Effect | inactive |
| safingol | BRD-K93944989 | CCCCCCCCCCCCCCC[C@@H](O)[C@@H](N)CO | OTKJDMGTUTTYMP-ZWKOTPCHSA-N | No-Effect | inactive |
| safranal | BRD-K90582192 | CC1=C(C=O)C(C)(C)CC=C1 | SGAWOGXMMPSZPB-UHFFFAOYSA-N | No-Effect | inactive |
| SAG | BRD-K99433989 | CN[C@H]1CC[C@H](N(Cc2cccc(-c3ccncc3)c2)C(=O)c2sc3ccccc3c2Cl)CC1 | VFSUUTYAEQOIMW-YHBQERECSA-N | No-Effect | inactive |
| sal003 | BRD-K01828772 | O=C(/C=C/c1ccccc1)N[C@@H](NC(=S)Nc1ccc(Cl)cc1)C(Cl)(Cl)Cl | TVNBASWNLOIQML-RFKZRZAASA-N | No-Effect | inactive |
| salazodine | BRD-K15045219 | COc1ccc(NS(=O)(=O)c2ccc(N=Nc3ccc(O)c(C(=O)O)c3)cc2)nn1 | YFUATOCFMCWUEH-UHFFFAOYSA-N | No-Effect | inactive |
| salicin | BRD-K64614248 | OCc1ccccc1O[C@@H]1O[C@H](CO)[C@@H](O)[C@H](O)[C@H]1O | NGFMICBWJRZIBI-UJPOAAIJSA-N | No-Effect | inactive |
| salicyl-alcohol | BRD-K08493205 | OCc1ccccc1O | CQRYARSYNCAZFO-UHFFFAOYSA-N | No-Effect | inactive |
| salicylamide | BRD-K81130846 | NC(=O)c1ccccc1O | SKZKKFZAGNVIMN-UHFFFAOYSA-N | No-Effect | inactive |
| salicylanilide | BRD-K83036479 | O=C(Nc1ccccc1)c1ccccc1O | WKEDVNSFRWHDNR-UHFFFAOYSA-N | No-Effect | inactive |
| salidroside | BRD-K66030860 | OC[C@H]1O[C@@H](OCCc2ccc(O)cc2)[C@H](O)[C@@H](O)[C@@H]1O | ILRCGYURZSFMEG-RKQHYHRCSA-N | No-Effect | inactive |
| salirasib | BRD-K98453471 | CC(C)=CCC/C(C)=C/CC/C(C)=C/CSc1ccccc1C(=O)O | WUILNKCFCLNXOK-CFBAGHHKSA-N | No-Effect | inactive |
| salmeterol | BRD-K01826763 | OCc1cc([C@@H](O)CNCCCCCCOCCCCc2ccccc2)ccc1O | GIIZNNXWQWCKIB-VWLOTQADSA-N | No-Effect | inactive |
| salsolinol-1-carboxylic-acid | BRD-A49952467 | C[C@@]1(C(=O)O)NCCc2cc(O)c(O)cc21 | XHGLVMDBZZZXDP-LLVKDONJSA-N | No-Effect | inactive |
| salubrinal | BRD-A45107007 | O=C(/C=C/c1ccccc1)N[C@@H](NC(=S)Nc1cccc2cccnc12)C(Cl)(Cl)Cl | LCOIAYJMPKXARU-CVTTXWKISA-N | No-Effect | inactive |
| salvianolic-acid-A | BRD-K00005247 | O=C(/C=C/c1ccc(O)c(O)c1/C=C/c1ccc(O)c(O)c1)O[C@H](Cc1ccc(O)c(O)c1)C(=O)O | YMGFTDKNIWPMGF-UCPJVGPRSA-N | No-Effect | inactive |
| salvianolic-acid-B | BRD-K18898553 | O=C(/C=C/c1ccc(O)c2c1[C@@H](C(=O)O[C@H](Cc1ccc(O)c(O)c1)C(=O)O)[C@H](c1ccc(O)c(O)c1)O2)O[C@H](Cc1ccc(O)c(O)c1)C(=O)O | SNKFFCBZYFGCQN-PDVBOLEISA-N | No-Effect | inactive |
| salvinorin-A | BRD-K19284129 | COC(=O)[C@@H]1C[C@H](OC(C)=O)C(=O)[C@H]2[C@@]1(C)CC[C@H]1C(=O)O[C@H](c3ccoc3)C[C@]21C | OBSYBRPAKCASQB-AGQYDFLVSA-N | No-Effect | inactive |
| SAM-315 | BRD-K00004553 | O=S(=O)(c1cccc2ccccc12)c1n[nH]c2ccc(N3CCNCC3)cc12 | FYOVZCDHYOEKDE-UHFFFAOYSA-N | No-Effect | inactive |
| sameridine | BRD-K00004552 | CCCCCCN1CCC(C(=O)N(C)CC)(c2ccccc2)CC1 | TYWUGCGYWNSRPS-UHFFFAOYSA-N | No-Effect | inactive |
| sanazole | BRD-K37747025 | COCCNC(=O)Cn1cnc([N+](=O)[O-])n1 | YKDRHKCETNMLHL-UHFFFAOYSA-N | No-Effect | inactive |
| sancycline | BRD-K00003242 | CN(C)[C@@H]1C(=O)[C@@H](C(N)=O)C(=O)[C@@]2(O)C(=O)[C@H]3C(=O)c4c(O)cccc4C[C@H]3C[C@@H]12 | CRNIKQQPTHSPBA-IEPLVVIMSA-N | No-Effect | inactive |
| SANT-1 | BRD-K99741577 | Cc1nn(-c2ccccc2)c(C)c1/C=N/N1CCN(Cc2ccccc2)CC1 | FOORCIAZMIWALX-JJIBRWJFSA-N | No-Effect | inactive |
| SANT-2 | BRD-K35781423 | CCOc1cc(C(=O)Nc2ccc(Cl)c(-c3nc4ccccc4[nH]3)c2)cc(OCC)c1OCC | VQOJFGFKIVFMDH-UHFFFAOYSA-N | No-Effect | inactive |
| sapacitabine | BRD-K00003214 | CCCCCCCCCCCCCCCC(=O)Nc1ccn([C@@H]2O[C@H](CO)[C@@H](O)[C@@H]2C#N)c(=O)n1 | LBGFKUUHOPIEMA-PEARBKPGSA-N | No-Effect | inactive |
| sapropterin | BRD-K36859431 | C[C@H](O)[C@H](O)[C@H]1CNc2nc(N)nc(=O)c2N1 | FNKQXYHWGSIFBK-RPDRRWSUSA-N | No-Effect | inactive |
| saquinavir | BRD-K09963420 | CC(C)(C)NC(=O)[C@@H]1C[C@@H]2CCCC[C@@H]2CN1C[C@@H](O)[C@H](Cc1ccccc1)NC(=O)[C@H](CC(N)=O)NC(=O)c1ccc2ccccc2n1 | QWAXKHKRTORLEM-UGJKXSETSA-N | No-Effect | inactive |
| SAR-245409 | BRD-K75308783 | COc1cc(Nc2nc3ccccc3nc2NS(=O)(=O)c2ccc(NC(=O)c3ccc(C)c(OC)c3)cc2)cc(OC)c1 | HJSSPYJVWLTYHG-UHFFFAOYSA-N | No-Effect | inactive |
| SAR131675 | BRD-K38904395 | CCn1c(N)c(C(=O)NC)c(=O)c2ccc(C#C[C@@](C)(O)COC)nc21 | PFMPOBVAYMTUOX-GOSISDBHSA-N | No-Effect | inactive |
| SAR405838 | BRD-K45349653 | CC(C)(C)C[C@@H]1N[C@@H](C(=O)N[C@H]2CC[C@H](O)CC2)[C@H](c2cccc(Cl)c2F)[C@]12C(=O)Nc1cc(Cl)ccc12 | IDKAKZRYYDCJDU-HBMMIIHUSA-N | No-Effect | inactive |
| SAR407899 | BRD-K56912469 | O=c1[nH]ccc2cc(OC3CCNCC3)ccc12 | IPEXHQGMTHOKQV-UHFFFAOYSA-N | No-Effect | inactive |
| saracatinib | BRD-K19540840 | CN1CCN(CCOc2cc(OC3CCOCC3)c3c(Nc4c(Cl)ccc5c4OCO5)ncnc3c2)CC1 | OUKYUETWWIPKQR-UHFFFAOYSA-N | No-Effect | inactive |
| sarafloxacin | BRD-K08525451 | O=C(O)c1cn(-c2ccc(F)cc2)c2cc(N3CCNCC3)c(F)cc2c1=O | XBHBWNFJWIASRO-UHFFFAOYSA-N | No-Effect | inactive |
| sarcosine | BRD-K51263939 | CNCC(=O)O | FSYKKLYZXJSNPZ-UHFFFAOYSA-N | No-Effect | inactive |
| sardomozide | BRD-K00003165 | N=C(N)N/N=C1\CCc2c(C(=N)N)cccc21 | CYPGNVSXMAUSJY-CXUHLZMHSA-N | No-Effect | inactive |
| sari-59-801 | BRD-A92448440 | CCc1noc(C)c1-c1[nH]c2ccccc2c1[C@@H](O)CN(C)C | GMGOZCSXMVXXRS-HNNXBMFYSA-N | No-Effect | inactive |
| sarpogrelate | BRD-A34358106 | COc1cccc(CCc2ccccc2OC[C@@H](CN(C)C)OC(=O)CCC(=O)O)c1 | FFYNAVGJSYHHFO-OAQYLSRUSA-N | No-Effect | inactive |
| sasapyrine | BRD-K48892307 | O=C(Oc1ccccc1C(=O)O)c1ccccc1O | WVYADZUPLLSGPU-UHFFFAOYSA-N | No-Effect | inactive |
| satraplatin | BRD-K42497885 | CC(=O)O[Pt](N)(Cl)(Cl)(NC1CCCCC1)OC(C)=O | JWTNVKZACPRKPF-UHFFFAOYSA-J | No-Effect | inactive |
| savolitinib | BRD-K98028353 | C[C@@H](c1ccc2nccn2c1)n1nnc2ncc(-c3cnn(C)c3)nc21 | XYDNMOZJKOGZLS-NSHDSACASA-N | No-Effect | inactive |
| saxagliptin | BRD-K00003546 | N#C[C@@H]1C[C@@H]2C[C@@H]2N1C(=O)[C@@H](N)[C@@]12C[C@@H]3C[C@@H](C[C@@](O)(C3)C1)C2 | QGJUIPDUBHWZPV-YQBUGCKMSA-N | No-Effect | inactive |

| | | | | | |
|---|---|---|---|---|---|
| SB-200646 | BRD-K77677632 | Cn1ccc2cc(NC(=O)Nc3cccnc3)ccc21 | OJZZJTLBYXHUSJ-UHFFFAOYSA-N | No-Effect | inactive |
| SB-203186 | BRD-K27141178 | O=C(OCCN1CCCCC1)c1c[nH]c2ccccc12 | YGKPIROTKVQCCU-UHFFFAOYSA-N | No-Effect | inactive |
| SB-203580 | BRD-K01826796 | C[S@@](=O)c1ccc(-c2nc(-c3ccncc3)c(-c3ccc(F)cc3)[nH]2)cc1 | CDMGBJANTYXAIV-HHHXNRCGSA-N | No-Effect | inactive |
| SB-205384 | BRD-A22707317 | CC#CCOC(=O)c1c(C)nc2sc3c(c2c1N)CC[C@H](O)C3 | JDTZAGLGBRRCJT-JTQLQIEISA-N | No-Effect | inactive |
| SB-206553 | BRD-K36395411 | Cn1ccc2cc3c(cc21)CCN3C(=O)Nc1cccnc1 | QJQORSLQNXDVGE-UHFFFAOYSA-N | No-Effect | inactive |
| SB-216641 | BRD-K30867024 | COc1ccc(NC(=O)c2ccc(-c3ccc(-c4noc(C)n4)cc3C)cc2)cc1OCCN(C)C | JRNUKVFYILMMLX-UHFFFAOYSA-N | No-Effect | inactive |
| SB-216763 | BRD-K59184148 | Cn1cc(C2=C(c3ccc(Cl)cc3Cl)C(=O)NC2=O)c2ccccc21 | JCSGFHVFHSKIJH-UHFFFAOYSA-N | No-Effect | inactive |
| SB-218795 | BRD-K61097567 | COC(=O)[C@H](NC(=O)c1cc(-c2ccccc2)nc2ccccc12)c1ccccc1 | IUMQXQJZIHWLIN-HSZRJFAPSA-N | No-Effect | inactive |
| SB-221284 | BRD-K14807180 | CSc1cc2c(cc1C(F)(F)F)N(C(=O)Nc1cccnc1)CC2 | OQZOXHCRSXYSPM-UHFFFAOYSA-N | No-Effect | inactive |
| SB-225002 | BRD-K61323504 | O=C(Nc1ccc([N+](=O)[O-])cc1O)Nc1ccccc1Br | MQBZVUNNWUIPMK-UHFFFAOYSA-N | No-Effect | inactive |
| SB-228357 | BRD-K15424032 | COc1cc2c(cc1C(F)(F)F)N(C(=O)Nc1cc(F)cc(-c3cccnc3)c1)CC2 | RRJLJKRFFRZRAF-UHFFFAOYSA-N | No-Effect | inactive |
| SB-239063 | BRD-K82091397 | COc1nccc(-c2c(-c3ccc(F)cc3)ncn2[C@H]2CC[C@H](O)CC2)n1 | ZQUSFAUAYSEREK-WKILWMFISA-N | No-Effect | inactive |
| SB-242235 | BRD-K19412355 | COc1nccc(-c2c(-c3ccc(F)cc3)ncn2C2CCNCC2)n1 | PDTYLGXVBIWRIM-UHFFFAOYSA-N | No-Effect | inactive |
| SB-243213 | BRD-K99968849 | Cc1cc2c(cc1C(F)(F)F)N(C(=O)Nc1ccc(Oc3cccnc3C)nc1)CC2 | ZETBBVYSBABLHL-UHFFFAOYSA-N | No-Effect | inactive |
| SB-258585 | BRD-K80639402 | COc1ccc(NS(=O)(=O)c2ccc(I)cc2)cc1N1CCN(C)CC1 | BDHMSYNBSBZCAF-UHFFFAOYSA-N | No-Effect | inactive |
| SB-268262 | BRD-A91031770 | Cc1ccccc1N(C)C(=O)c1ccc([S@@](=O)c2nccs2)c([N+](=O)[O-])c1 | YXHLPJQQPOMPGE-HHHXNRCGSA-N | No-Effect | inactive |
| SB-269970 | BRD-K24201553 | CC1CCN(CC[C@H]2CCCN2S(=O)(=O)c2cccc(O)c2)CC1 | HWKROQUZSKPIKQ-MRXNPFEDSA-N | No-Effect | inactive |
| SB-297006 | BRD-K16107558 | CCOC(=O)[C@H](Cc1ccc([N+](=O)[O-])cc1)NC(=O)c1ccccc1 | BEZXGSZPWXRHIN-INIZCTEOSA-N | No-Effect | inactive |
| SB-328437 | BRD-K78720403 | COC(=O)[C@H](Cc1ccc([N+](=O)[O-])cc1)NC(=O)c1cccc2ccccc12 | VMFGCGRAIBLAFY-IBGZPJMESA-N | No-Effect | inactive |
| SB-334867 | BRD-K41567364 | Cc1nc2ccc(NC(=O)Nc3ccnc4cccnc34)cc2o1 | AKMNUCBQGHFICM-UHFFFAOYSA-N | No-Effect | inactive |
| SB-366791 | BRD-K61536336 | COc1cccc(NC(=O)/C=C/c2ccc(Cl)cc2)c1 | RYAMDQKWNKKFHD-JXMROGBWSA-N | No-Effect | inactive |
| SB-399885 | BRD-K03988082 | COc1ccc(S(=O)(=O)Nc2cc(Cl)cc(Cl)c2OC)cc1N1CCNCC1 | ATKZKAYWARYLBW-UHFFFAOYSA-N | No-Effect | inactive |
| SB-408124 | BRD-K53913732 | Cc1cc(NC(=O)Nc2ccc(N(C)C)cc2)c2cc(F)cc(F)c2n1 | JTARFZSNUAGHRB-UHFFFAOYSA-N | No-Effect | inactive |
| SB-415286 | BRD-K76805682 | O=C1NC(=O)C(c2ccccc2[N+](=O)[O-])=C1Nc1ccc(O)c(Cl)c1 | PQCXVIPXISBFPN-UHFFFAOYSA-N | No-Effect | inactive |
| SB-431542 | BRD-K67298865 | NC(=O)c1ccc(-c2nc(-c3ccccn3)c(-c3ccc4c(c3)OCO4)[nH]2)cc1 | FHYUGAJXYORMHI-UHFFFAOYSA-N | No-Effect | inactive |
| SB-452533 | BRD-K03665769 | CCN(CCNC(=O)Nc1ccccc1Br)c1cccc(C)c1 | IFJYEGJUQIBBQV-UHFFFAOYSA-N | No-Effect | inactive |
| SB-505124 | BRD-K40109029 | Cc1cccc(-c2nc(C(C)(C)C)[nH]c2-c2ccc3c(c2)OCO3)n1 | WGZOTBUYUFBEPZ-UHFFFAOYSA-N | No-Effect | inactive |
| SB-525334 | BRD-K37720887 | Cc1cccc(-c2nc(C(C)(C)C)[nH]c2-c2ccc3nccnc3c2)n1 | DKPQHFZUICCZHF-UHFFFAOYSA-N | No-Effect | inactive |
| SB-612111 | BRD-K08333979 | Cc1cccc2c1CC[C@H](CN1CCC(c3c(Cl)cccc3Cl)CC1)C[C@@H]2O | OHRDCQFCAWLDBP-SBUREZEXSA-N | No-Effect | inactive |
| SB-649868 | BRD-K00003485 | Cc1nc(C(=O)N2CCCC[C@H]2CNC(=O)c2cccc3occc23)c(-c2ccc(F)cc2)s1 | ZJXIUGNEAIHSBI-IBGZPJMESA-N | No-Effect | inactive |
| SB-657510 | BRD-K95292280 | COc1cc(Br)c(S(=O)(=O)Nc2ccc(Cl)c(O[C@@H]3CCN(C)C3)c2)cc1OC | KQCZCINJGIRLCD-CYBMUJFWSA-N | No-Effect | inactive |
| SB-683698 | BRD-K24968862 | COc1cccc(OC)c1-c1ccc(C[C@H](NC(=O)c2c(Cl)cccc2Cl)C(=O)O)cc1 | DRSJLVGDSNWQBI-SFHVURJKSA-N | No-Effect | inactive |
| SB-705498 | BRD-K65991129 | O=C(Nc1ccccc1Br)N[C@@H]1CCN(c2ccc(C(F)(F)F)cn2)C1 | JYILLRHXRVTRSH-GFCCVEGCSA-N | No-Effect | inactive |
| SB-706375 | BRD-K68633027 | COc1cc(Br)c(S(=O)(=O)Nc2ccc(C(F)(F)F)c(O[C@@H]3CCN(C)C3)c2)cc1OC | BPOWQJYAMDEAFF-CYBMUJFWSA-N | No-Effect | inactive |
| SB-742457 | BRD-K78941306 | O=S(=O)(c1ccccc1)c1cnc2c(N3CCNCC3)cccc2c1 | JJZFWROHYSMCMU-UHFFFAOYSA-N | No-Effect | inactive |
| SB-747651A | BRD-K49597925 | CCn1c(-c2nonc2N)nc2cncc(CNC3CCNCC3)c21 | MBCJUIJWPYUEBX-UHFFFAOYSA-N | No-Effect | inactive |
| SB-756050 | BRD-K45168550 | COc1ccc(S(=O)(=O)N2CCCN(S(=O)(=O)c3ccc(OC)c(OC)c3)CC2)cc1OC | GJUFPAZNBPFNRI-UHFFFAOYSA-N | No-Effect | inactive |
| SB-772077B | BRD-K49110935 | CCn1c(-c2nonc2N)nc2cncc(C(=O)N3CC[C@H](N)C3)c21 | PHWHNHYZZNJPLB-QMMMGPOBSA-N | No-Effect | inactive |
| SBE-13 | BRD-K71667152 | COc1ccc(CCNCc2ccc(OCc3ccc(Cl)nc3)c(OC)c2)cc1OC | XNPRQBVQOJZMMA-UHFFFAOYSA-N | No-Effect | inactive |
| SBHA | BRD-K54729176 | O=C(CCCCCCC(=O)NO)NO | IDQPVOFTURLJPT-UHFFFAOYSA-N | No-Effect | inactive |
| SBI-115 | BRD-K00005340 | CCS(=O)(=O)c1ncc(Cl)c(C(=O)Oc2cccc(C)c2)n1 | IJPXOPBVXVPPEW-UHFFFAOYSA-N | No-Effect | inactive |
| SC-10 | BRD-K05361803 | CCCCCCCNS(=O)(=O)c1cccc2c(Cl)cccc12 | MJMJERJFCHYXEM-UHFFFAOYSA-N | No-Effect | inactive |
| SC-12267 | BRD-K40797222 | COc1cccc(-c2ccc(NC(=O)C3=C(C(=O)O)CCC3)c(F)c2)c1 | XPRDUGXOWVXZLL-UHFFFAOYSA-N | No-Effect | inactive |
| SC-144 | BRD-K55305701 | O=C(NNc1nc2cc(F)ccc2n2cccc12)c1cnccn1 | UEADAWQSJOWXBK-UHFFFAOYSA-N | No-Effect | inactive |
| SC-19220 | BRD-K52512893 | CC(=O)NNC(=O)N1Cc2ccccc2Oc2ccc(Cl)cc21 | KNURFLJTOUGOOQ-UHFFFAOYSA-N | No-Effect | inactive |
| SC-236 | BRD-K76365178 | NS(=O)(=O)c1ccc(-n2nc(C(F)(F)F)cc2-c2ccc(Cl)cc2)cc1 | NSQNZEUFHPTJME-UHFFFAOYSA-N | No-Effect | inactive |
| SC-51089 | BRD-K62715847 | O=C(CCc1ccncc1)NNC(=O)N1Cc2ccccc2Oc2ccc(Cl)cc21 | XDJPQOUDGROSEU-UHFFFAOYSA-N | No-Effect | inactive |
| SC-560 | BRD-K14767410 | COc1ccc(-n2nc(C(F)(F)F)cc2-c2ccc(Cl)cc2)cc1 | PQUGCKBLVKJMNT-UHFFFAOYSA-N | No-Effect | inactive |
| SC-9 | BRD-K25875056 | O=S(=O)(NCCCCCCc1ccccc1)c1cccc2c(Cl)cccc12 | YUXMTDYHFPRPLG-UHFFFAOYSA-N | No-Effect | inactive |
| SCH-202676 | BRD-K29562203 | CN=c1nc(-c2ccccc2)n(-c2ccccc2)s1 | FFUBTEITUNMMOK-UHFFFAOYSA-N | No-Effect | inactive |
| SCH-221510 | BRD-K00004286 | Cc1ccccc1C(c1ccccc1C)N1[C@H]2CC[C@@H]1C[C@](O)(c1ccccc1)C2 | LOSJNRBXNQTUNT-FOURXJPJSA-N | No-Effect | inactive |
| SCH-51344 | BRD-K02460719 | COc1ccc2nc3n[nH]c(C)c3c(NCCOCCO)c2c1 | YWEGXZZAORIRQR-UHFFFAOYSA-N | No-Effect | inactive |
| SCH-58261 | BRD-K81225797 | Nc1nc2c(cnn2CCc2ccccc2)c2nc(-c3ccco3)nn12 | UTLPKQYUXOEJIL-UHFFFAOYSA-N | No-Effect | inactive |
| SCH-900776 | BRD-K75009076 | Cn1cc(-c2cnn3c(N)c(Br)c([C@@H]4CCCNC4)nc23)cn1 | GMIZZEXBPRLVIV-SECBINFHSA-N | No-Effect | inactive |
| schisandrol-B | BRD-K18878288 | COc1cc2c(c(OC)c1OC)-c1cc(cc3c(c1OC)OCO3)C[C@H](C)[C@@](C)(O)C2 | ZWRRJEICIPUPHZ-MYODQAERSA-N | No-Effect | inactive |
| sclareol | BRD-A23127772 | C=CC(C)(O)CC[C@@H]1[C@@]2(C)CCCC(C)(C)[C@@H]2CC[C@@]1(C)O | XVULBTBTFGYVRC-SVPXJYONSA-N | No-Effect | inactive |
| sclareolide | BRD-K72925150 | CC1(C)CCC[C@@]2(C)[C@H]1CC[C@@]1(C)OC(=O)C[C@H]21 | IMKJGXCIJJXALX-SHUKQUCYSA-N | No-Effect | inactive |
| SCMC-Lys | BRD-K33809398 | N[C@@H](CSCC(=O)O)C(=O)O | GBFLZEXEOZUWRN-VKHMYHEASA-N | No-Effect | inactive |
| scopine | BRD-K75923785 | CN1[C@H]2C[C@H](O)C[C@@H]1[C@H]1O[C@@H]21 | FIMXSEMBHGTNKT-RZVDLVGDSA-N | No-Effect | inactive |
| scopolamine-N-oxide | BRD-A49906757 | C[N@@+]1([O-])[C@H]2C[C@H](OC(=O)[C@H](CO)c3ccccc3)C[C@@H]1[C@H]1O[C@@H]21 | GSYQNAMOFFWAPF-SXWCTKSRSA-N | No-Effect | inactive |
| SCP-1 | BRD-K20548681 | O=C(CN1C(=O)c2ccccc2S1(=O)=O)Nc1ccc(O)cc1 | PUPNJSIFIXXJCH-UHFFFAOYSA-N | No-Effect | inactive |
| scriptaid | BRD-K22503835 | O=C(CCCCCN1C(=O)c2cccc3cccc(c23)C1=O)NO | JTDYUFSDZATMKU-UHFFFAOYSA-N | No-Effect | inactive |
| SCS | BRD-K78496197 | O=C(N/N=C/c1ccccc1O)c1ccccc1O | OMCYEZUIYGPHDJ-OQLLNIDSSA-N | No-Effect | inactive |
| SD-0006 | BRD-K00003093 | O=C(CO)N1CCC(c2[nH]nc(-c3ccc(Cl)cc3)c2-c2ccncn2)CC1 | CATQHDWESBRRQA-UHFFFAOYSA-N | No-Effect | inactive |
| SD-169 | BRD-K91904471 | NC(=O)c1ccc2[nH]ccc2c1 | GQMYQEAXTITUAE-UHFFFAOYSA-N | No-Effect | inactive |
| SD-208 | BRD-K24566386 | Fc1ccc(Cl)cc1-c1nc(Nc2ccncc2)c2nccnc2n1 | BERLXWPRSBJFHO-UHFFFAOYSA-N | No-Effect | inactive |
| SDM25N | BRD-K01826797 | C=C(C)CN1CC[C@]23c4c5ccc(O)c4O[C@H]2c2[nH]c4ccccc4c2C[C@@]3(O)[C@H]1C5 | ZFMVWAWSKLAVOE-IFKAHUTRSA-N | No-Effect | inactive |
| SDZ-205-557 | BRD-K15868788 | CCN(CC)CCOC(=O)c1cc(Cl)c(N)cc1OC | FFNWMBDISAYHDC-UHFFFAOYSA-N | No-Effect | inactive |
| SDZ-21009 | BRD-A15530910 | CC(C)OC(=O)c1cc2c(OC[C@@H](O)CNC(C)(C)C)cccc2[nH]1 | SJYFDORQYYEJLB-ZDUSSCGKSA-N | No-Effect | inactive |
| SDZ-220-040 | BRD-K82266792 | N[C@@H](Cc1cc(-c2ccc(Cl)cc2Cl)cc(CP(=O)(O)O)c1O)C(=O)O | NYZFUZCCDOSQBG-AWEZNQCLSA-N | No-Effect | inactive |
| SDZ-220-581 | BRD-K27168049 | N[C@@H](Cc1cc(CP(=O)(O)O)cc(-c2ccccc2Cl)c1)C(=O)O | VBRJFXSFCYEZMQ-HNNXBMFYSA-N | No-Effect | inactive |
| SDZ-NKT-343 | BRD-K65820248 | CN(Cc1ccccc1)C(=O)[C@H](Cc1ccc2ccccc2c1)NC(=O)[C@@H]1CCCN1C(=O)Nc1ccccc1[N+](=O)[O-] | PCVSIMQAFWRUEC-JDXGNMNLSA-N | No-Effect | inactive |
| SDZ-WAG-994 | BRD-K39558284 | CO[C@@H]1[C@H](O)[C@@H](CO)O[C@H]1n1cnc2c(NC3CCCCC3)ncnc21 | JAKAFSGZUXCHLF-LSCFUAHRSA-N | No-Effect | inactive |
| secalciferol | BRD-K02965577 | C=C1CC[C@H](O)C/C1=C/C=C1\CCC[C@@]2(C)[C@H]1CC[C@@H]2[C@H](C)CC[C@@H](O)C(C)(C)O | FCKJYANJHNLEEP-XRWYNYHCSA-N | No-Effect | inactive |
| seclazone | BRD-A53472930 | O=C1c2cc(Cl)ccc2O[C@H]2CCON12 | XWXVKXXKKLBDDJ-VIFPVBQESA-N | No-Effect | inactive |

| | | | | | |
|---|---|---|---|---|---|
| seclidemstat | BRD-K00003454 | C/C(=N\NC(=O)c1cccc(S(=O)(=O)N2CCN(C)CC2)c1)c1cc(Cl)ccc1O | MVSQDUZRRVBYLA-HYARGMPZSA-N | No-Effect | inactive |
| secnidazole | BRD-A70083328 | Cc1ncc([N+](=O)[O-])n1C[C@@H](C)O | KPQZUUQMTUIKBP-RXMQYKEDSA-N | No-Effect | inactive |
| secoisolariciresinol-(-) | BRD-K91733562 | COc1cc(C[C@@H](CO)[C@H](CO)Cc2ccc(O)c(OC)c2)ccc1O | PUETUDUXMCLALY-HOTGVXAUSA-N | No-Effect | inactive |
| secoisolariciresinol-diglucoside | BRD-K77320845 | COc1cc(C[C@@H](CO[C@@H]2O[C@H](CO)[C@@H](O)[C@H](O)[C@H]2O)[C@H](CO[C@@H]2O[C@H](CO)[C@@H](O)[C@H](O)[C@H]2O)Cc2ccc(O)c(OC)c2)ccc1O | SBVBJPHMDABKJV-PGCJWIIOSA-N | No-Effect | inactive |
| securinine | BRD-K49071277 | O=C1C=C2C=C[C@@H]3C[C@@]2(O1)[C@H]1CCCCN31 | SWZMSZQQJRKFBP-WZRBSPASSA-N | No-Effect | inactive |
| selamectin | BRD-K00003549 | CO[C@H]1C[C@H](O[C@@H]2/C(C)=C/C[C@@H]3C[C@@H](C[C@]4(CC[C@H](C)[C@@H](C5CCCCC5)O4)O3)OC(=O)[C@@H]3C=C(C)/C(=N/O)[C@H]4OC/C(=C\C=C\[C@@H]2C)[C@@]34O)O[C@@H](C)[C@@H]1O | AFJYYKSVHJGXSN-XHKIUTQPSA-N | No-Effect | inactive |
| selenomethionine | BRD-K74664543 | C[Se]CC[C@H](N)C(=O)O | RJFAYQIBOAGBLC-BYPYZUCNSA-N | No-Effect | inactive |
| seletalisib | BRD-K00003228 | [O-][n+]1cccc(-c2nc3c(Cl)cccc3cc2[C@@H](Nc2ncnc3cccnc23)C(F)(F)F)c1 | LNLJHGXOFYUARS-OAQYLSRUSA-N | No-Effect | inactive |
| selexipag | BRD-K21191422 | CC(C)N(CCCCOCC(=O)NS(C)(=O)=O)c1cnc(-c2ccccc2)c(-c2ccccc2)n1 | QXWZQTURMXZVHJ-UHFFFAOYSA-N | No-Effect | inactive |
| selfotel | BRD-K48738253 | O=C(O)[C@H]1C[C@@H](CP(=O)(O)O)CCN1 | LPMRCCNDNGONCD-NTSWFWBYSA-N | No-Effect | inactive |
| selinexor | BRD-K21361524 | O=C(/C=C\n1cnc(-c2cc(C(F)(F)F)cc(C(F)(F)F)c2)n1)NNc1cnccn1 | DEVSOMFAQLZNKR-RJRFIUFISA-N | No-Effect | inactive |
| selitrectinib | BRD-K00003441 | C[C@@H]1CCc2ncc(F)cc2[C@H]2CCCN2c2ccn3ncc(c3n2)C(=O)N1 | OEBIHOVSAMBXIB-SJKOYZFVSA-N | No-Effect | inactive |
| selonsertib | BRD-K00003274 | Cc1cc(F)c(C(=O)Nc2cccc(-c3nncn3C(C)C)n2)cc1-n1cnc(C2CC2)c1 | YIDDLAAKOYYGJG-UHFFFAOYSA-N | No-Effect | inactive |
| selumetinib | BRD-K57080016 | Cn1cnc2c(F)c(Nc3ccc(Br)cc3Cl)c(C(=O)NOCCO)cc21 | CYOHGALHFOKKQC-UHFFFAOYSA-N | No-Effect | inactive |
| semagacestat | BRD-K65592642 | CC(C)[C@H](O)C(=O)N[C@@H](C)C(=O)N[C@@H]1C(=O)N(C)CCc2ccccc21 | PKXWXXPNHIWQHW-RCBQFDQVSA-N | No-Effect | inactive |
| semaxanib | BRD-K63504947 | Cc1cc(C)c(/C=C2\C(=O)Nc3ccccc32)[nH]1 | WUWDLXZGHZSWQZ-WQLSENKSSA-N | No-Effect | inactive |
| semustine | BRD-K13281575 | CC1CCC(NC(=O)N(CCCl)[NH2+][O-])CC1 | JUUWEJHZVBABHQ-UHFFFAOYSA-N | No-Effect | inactive |
| SEN-1269 | BRD-K70614402 | CN(C)c1cccc(Oc2cnc(Nc3cccc(O)c3)nc2)c1 | MFPAAKZZUIRMKR-UHFFFAOYSA-N | No-Effect | inactive |
| senicapoc | BRD-K21295289 | NC(=O)C(c1ccccc1)(c1ccc(F)cc1)c1ccc(F)cc1 | SCTZUZTYRMOMKT-UHFFFAOYSA-N | No-Effect | inactive |
| sennoside-A | BRD-A30370660 | O=C(O)c1cc(O)c2c(O)c3c(O[C@H]4O[C@H](CO)[C@@H](O)[C@H](O)[C@H]4O)cccc3c(-c3c4cc(C(=O)O)cc(O)c4c(O)c4c(O[C@H]5O[C@H](CO)[C@@H](O)[C@@H](O)[C@H]5O)cccc34)c2c1 | QFJIQUOMKJNYLI-QSXPFVSBSA-N | No-Effect | inactive |
| sennoside-protonated | BRD-A53504839 | O=C(O)c1cc(O)c2c(O)c3c(O[C@@H]4O[C@H](CO)[C@@H](O)[C@H](O)[C@@H]4O)cccc3c(-c3c4cc(C(=O)O)cc(O)c4c(O)c4c(O[C@H]5O[C@@H](CO)[C@@H](O)[C@@H](O)[C@@H]5O)cccc34)c2c1 | QFJIQUOMKJNYLI-NGMZATEOSA-N | No-Effect | inactive |
| seocalcitol | BRD-K28456624 | C=C1/C(=C\C=C2/CCC[C@@]3(C)[C@H]2CC[C@@H]3[C@H](C)/C=C/C=C/C(O)(CC)CC)C[C@@H](O)C[C@@H]1O | LVLLALCJVJNGQQ-SEODYNFXSA-N | No-Effect | inactive |
| SEP-227900 | BRD-K49504902 | O=C(O)c1cc2occc2[nH]1 | MMAIBGHDBYQYDI-UHFFFAOYSA-N | No-Effect | inactive |
| seratrodast | BRD-A60765580 | CC1=C(C)C(=O)C([C@@H](CCCCCC(=O)O)c2ccccc2)=C(C)C1=O | ZBVKEHDGYSLCCC-SFHVURJKSA-N | No-Effect | inactive |
| serotonin | BRD-K38580127 | NCCc1c[nH]c2ccc(O)cc12 | QZAYGJVTTNCVMB-UHFFFAOYSA-N | No-Effect | inactive |
| sertaconazole | BRD-K01825659 | Clc1ccc([C@@H](Cn2ccnc2)OCc2csc3c(Cl)cccc23)c(Cl)c1 | JLGKQTAYUIMGRK-LJQANCHMSA-N | No-Effect | inactive |
| sesamin | BRD-K11631773 | c1cc2c(cc1[C@H]1OC[C@@H]3[C@@H](c4ccc5c(c4)OCO5)OC[C@H]13)OCO2 | PEYUIKBAABKQKQ-AFHBHXEDSA-N | No-Effect | inactive |
| setipiprant | BRD-K24497734 | O=C(O)Cn1c2c(c3cc(F)ccc31)CN(C(=O)c1cccc3ccccc13)CC2 | IHAXLPDVOWLUOS-UHFFFAOYSA-N | No-Effect | inactive |
| setiptiline | BRD-K36732695 | CN1CCC2=C(C1)c1ccccc1Cc1ccccc12 | GVPIXRLYKVFFMK-UHFFFAOYSA-N | No-Effect | inactive |
| setmelanotide | BRD-K00003298 | CC(=O)N[C@@H](CCCNC(=N)N)C(=O)N[C@H]1CSSC[C@@H](C(N)=O)NC(=O)[C@H](Cc2c[nH]c3ccccc23)NC(=O)[C@H](CCCNC(=N)N)NC(=O)[C@@H](Cc2ccccc2)NC(=O)[C@H](Cc2cnc[nH]2)NC(=O)[C@@H](C)NC1=O | HDHDTKMUACZDAA-PHNIDTBTSA-N | No-Effect | inactive |
| sevoflurane | BRD-K05638300 | FCOC(C(F)(F)F)C(F)(F)F | DFEYYRMXOJXZRJ-UHFFFAOYSA-N | No-Effect | inactive |
| SEW-2871 | BRD-K15601958 | FC(F)(F)c1cccc(-c2noc(-c3cc(-c4ccccc4)c(C(F)(F)F)s3)n2)c1 | OYMNPJXKQVTQTR-UHFFFAOYSA-N | No-Effect | inactive |
| SF-11 | BRD-K42629487 | CCOc1ccc(NC(=S)N2CCC(C(O)(c3ccccc3)c3ccccc3)CC2)cc1 | PMEQBGAGFZDWQX-UHFFFAOYSA-N | No-Effect | inactive |
| SGC-0946 | BRD-K69577666 | CC(C)N(CCCNC(=O)Nc1ccc(C(C)(C)C)cc1)C[C@H]1O[C@@H](n2cc(Br)c3c(N)ncnc32)[C@H](O)[C@@H]1O | IQCKJUKAQJINMK-HUBRGWSESA-N | No-Effect | inactive |
| SGC-707 | BRD-K86718786 | O=C(NCC(=O)N1CCCC1)Nc1ccc2cnccc2c1 | DMIDPTCQPIJYFE-UHFFFAOYSA-N | No-Effect | inactive |
| SGC-CBP30 | BRD-K24821794 | COc1ccc(CCc2nc3cc(-c4c(C)noc4C)ccc3n2C[C@H](C)N2CCOCC2)cc1Cl | GEPYBHCJBORHCE-SFHVURJKSA-N | No-Effect | inactive |
| SGI-1776 | BRD-K80700417 | CN1CCC(CNc2ccc3ncc(-c4cccc(OC(F)(F)F)c4)n3n2)CC1 | MHXGEROHKGDZGO-UHFFFAOYSA-N | No-Effect | inactive |
| SGS-742 | BRD-K00004650 | CCCCP(=O)(O)CCCN | ONNMDRQRSGKZCN-UHFFFAOYSA-N | No-Effect | inactive |
| SGX523 | BRD-K19477839 | Cn1cc(-c2ccc3nnc(Sc4ccc5ncccc5c4)n3n2)cn1 | BCZUAADEACICHN-UHFFFAOYSA-N | No-Effect | inactive |
| SHA-68 | BRD-A63344375 | O=C(NCc1ccc(F)cc1)N1CCN2C(=O)OC(c3ccccc3)(c3ccccc3)[C@@H]2C1 | SFRQIPRTNYHJHP-QHCPKHFHSA-N | No-Effect | inactive |
| SIB-1553A | BRD-K01826006 | CN1CCC[C@@H]1CCSc1ccc(O)cc1 | NVZGJSVPOOILDI-LLVKDONJSA-N | No-Effect | inactive |
| SIB-1757 | BRD-K98157055 | C=C1C=CC(=O)C(=NNc2ccccc2)N1 | UOJMXHLMOUOGPY-UHFFFAOYSA-N | No-Effect | inactive |
| SIB-1893 | BRD-K14561059 | Cc1cccc(/C=C/c2ccccc2)n1 | SISOFUCTXZKSOQ-ZHACJKMWSA-N | No-Effect | inactive |
| SID-7969543 | BRD-A43885598 | CCOC(=O)[C@@H](C)Oc1cccc2c(=O)n(CC(=O)Nc3ccc4c(c3)OCCO4)ccc12 | KWMBIIQCLUIHDI-OAHLLOKOSA-N | No-Effect | inactive |
| siguazodan | BRD-A62071884 | CN/C(=N\c1ccc(C2=NNC(=O)C[C@@H]2C)cc1)NC#N | NUHPODZZKHQQET-VIFPVBQESA-N | No-Effect | inactive |
| sildenafil | BRD-K79759585 | CCCc1nn(C)c2c(=O)[nH]c(-c3cc(S(=O)(=O)N4CCN(C)CC4)ccc3OCC)nc12 | BNRNXUUZRGQAQC-UHFFFAOYSA-N | No-Effect | inactive |
| silibinin | BRD-K80353138 | COc1cc([C@H]2Oc3cc([C@H]4Oc5cc(O)cc(O)c5C(=O)[C@@H]4O)ccc3O[C@@H]2CO)ccc1O | SEBFKMXJBCUCAI-HKTJVKLFSA-N | No-Effect | inactive |
| silodosin | BRD-K10906636 | C[C@H](Cc1cc2c(c(C(N)=O)c1)N(CCCO)CC2)NCCOc1ccccc1OCC(F)(F)F | PNCPYILNMDWPEY-QGZVFWFLSA-N | No-Effect | inactive |
| silver-sulfadiazine | BRD-K68379918 | Nc1ccc(S(=O)(=O)N([Ag])c2ncccn2)cc1 | UEJSSZHHYBHCEL-UHFFFAOYSA-N | No-Effect | inactive |
| simurosertib | BRD-K00004727 | Cc1n[nH]cc1-c1cc2[nH]c([C@H]3CC4CCN3CC4)nc(=O)c2s1 | XGVXKJKTISMIOW-CYBMUJFWSA-N | No-Effect | inactive |
| simvastatin | BRD-K22134346 | CCC(C)(C)C(=O)O[C@H]1C[C@@H](C)C=C2C=C[C@H](C)[C@H](CC[C@@H]3C[C@@H](O)CC(=O)O3)[C@H]21 | RYMZZMVNJRMUDD-HGQWONQESA-N | No-Effect | inactive |
| sincalide | BRD-K00003334 | CSCC[C@H](NC(=O)[C@H](Cc1ccc(OS(=O)(=O)O)cc1)NC(=O)[C@@H](N)CC(=O)O)C(=O)NCC(=O)N[C@@H](Cc1c[nH]c2ccccc12)C(=O)N[C@@H](CCSC)C(=O)N[C@@H](CC(=O)O)C(=O)N[C@@H](Cc1ccccc1)C(N)=O | IZTQOLKUZKXIRV-YRVFCXMDSA-N | No-Effect | inactive |
| sinefungin | BRD-K91100918 | Nc1ncnc2c1ncn2[C@@H]1O[C@H](C[C@@H](N)CC[C@H](N)C(=O)O)[C@@H](O)[C@H]1O | LMXOHSDXUQEUSF-YECHIGJVSA-N | No-Effect | inactive |
| sinomenine | BRD-K83459933 | COC1=C[C@@H]2[C@@H]3Cc4ccc(OC)c(O)c4[C@]2(CCN3C)CC1=O | INYYVPJSBIVGPH-QHRIQVFBSA-N | No-Effect | inactive |
| sipatrigine | BRD-K53195974 | CN1CCN(c2ncc(-c3cc(Cl)cc(Cl)c3Cl)c(N)n2)CC1 | PDOCBJADCWMDGL-UHFFFAOYSA-N | No-Effect | inactive |
| siponimod | BRD-K62384160 | CCc1cc(/C(C)=N/OCc2ccc(C3CCCCC3)c(C(F)(F)F)c2)ccc1CN1CC(C(=O)O)C1 | KIHYPELVXPAIDH-HNSNBQBZSA-N | No-Effect | inactive |

| | | | | | |
|---|---|---|---|---|---|
| siramesine | BRD-K51333959 | Fc1ccc(-n2cc(CCCCN3CCC4(CC3)OCc3ccccc34)c3ccccc32)cc1 | XWAONOGAGZNUSF-UHFFFAOYSA-N | No-Effect | inactive |
| siremadlin | BRD-K65480835 | COc1ncc(-c2nc3c(n2C(C)C)[C@H](c2ccc(Cl)cc2)N(c2cc(Cl)cn(C)c2=O)C3=O)c(OC)n1 | AGBSXNCBIWWLHD-FQEVSTJZSA-N | No-Effect | inactive |
| sirtinol | BRD-A23747351 | C[C@@H](NC(=O)c1ccccc1/N=C/c1c(O)ccc2ccccc12)c1ccccc1 | UXJFDYIHRJGPFS-OVKKOYPGSA-N | No-Effect | inactive |
| SIS3 | BRD-K38905597 | COc1cc2c(cc1OC)CN(C(=O)/C=C/c1c(-c3ccccc3)n(C)c3ncccc13)CC2 | IJYPHMXWKKKHGT-VAWYXSNFSA-N | No-Effect | inactive |
| sisomicin | BRD-K71772032 | CN[C@@H]1[C@@H](O)[C@@H](O[C@@H]2[C@@H](O)[C@H](O[C@H]3OC(CN)=CC[C@H]3N)[C@@H](N)C[C@H]2N)OC[C@]1(C)O | URWAJWIAIPFPJE-YFMIWBNJSA-N | No-Effect | inactive |
| sisomicin-sulfate | BRD-A68997559 | CN[C@@H]1[C@@H](O)[C@@H](O[C@@H]2[C@@H](O)[C@H](O[C@H]3OC(CN)=CC[C@H]3N)[C@@H](N)C[C@H]2N)OC[C@]1(C)O | URWAJWIAIPFPJE-YFMIWBNJSA-N | No-Effect | inactive |
| sitagliptin | BRD-K19416115 | N[C@@H](CC(=O)N1CCn2c(nnc2C(F)(F)F)C1)Cc1cc(F)c(F)cc1F | MFFMDFFZMYYVKS-SECBINFHSA-N | No-Effect | inactive |
| sitaxentan | BRD-K63806092 | Cc1cc2c(cc1CC(=O)c1sccc1S(=O)(=O)Nc1onc(C)c1Cl)OCO2 | PHWXUGHIIBDVKD-UHFFFAOYSA-N | No-Effect | inactive |
| sitosterol | BRD-K79402892 | CC[C@H](CC[C@@H](C)[C@H]1CC[C@H]2[C@@H]3CC=C4C[C@@H](O)CC[C@]4(C)[C@H]3CC[C@@]21C)C(C)C | KZJWDPNRJALLNS-VJSFXXLFSA-N | No-Effect | inactive |
| SJ-172550 | BRD-K35945299 | CCOc1cc(/C=C2/C(=O)N(c3ccccc3)N=C2C)cc(Cl)c1OCC(=O)OC | RKKFQJXGAQWHBZ-LICLKQGHSA-N | No-Effect | inactive |
| SJ-733 | BRD-K34798115 | N#Cc1cc(NC(=O)[C@H]2c3ccccc3C(=O)N(CC(F)(F)F)[C@@H]2c2cccnc2)ccc1F | VKCPFWKTFZAOTO-LEWJYISDSA-N | No-Effect | inactive |
| SJG-136 | BRD-K00003173 | C=C1C[C@H]2C=Nc3cc(OCCCOc4cc5c(cc4OC)C(=O)N4CC(=C)C[C@H]4C=N5)c(OC)cc3C(=O)N2C1 | RWZVMMQNDHPRQD-SFTDATJTSA-N | No-Effect | inactive |
| SK&F-10047-(+) | BRD-K00004226 | C=CCN1CC[C@]2(C)c3cc(O)ccc3C[C@H]1[C@H]2C | LGQCVMYAEFTEFN-DQYPLSBCSA-N | No-Effect | inactive |
| SKA-31 | BRD-K67304264 | Nc1nc2c(ccc3ccccc32)s1 | FECQXVPRUCCUIL-UHFFFAOYSA-N | No-Effect | inactive |
| skepinone-L | BRD-K83607951 | O=C1c2ccc(Nc3ccc(F)cc3F)cc2CCc2ccc(OC[C@H](O)CO)cc21 | HXMGCTFLLWPVFM-GOSISDBHSA-N | No-Effect | inactive |
| SKF-38393 | BRD-K01826815 | Oc1cc2c(cc1O)[C@H](c1ccccc1)CNCC2 | JUDKOGFHZYMDMF-AWEZNQCLSA-N | No-Effect | inactive |
| SKF-77434 | BRD-A64227845 | C=CCN1CCc2cc(O)c(O)cc2[C@H](c2ccccc2)C1 | QBUVZVXIRYFENV-KRWDZBQOSA-N | No-Effect | inactive |
| SKF-81297 | BRD-A09828896 | Oc1cc2c(c(Cl)c1O)CCNCC2c1ccccc1 | GHWJEDJMOVUXEC-UHFFFAOYSA-N | No-Effect | inactive |
| SKF-83566 | BRD-A52588987 | CN1CCc2cc(Br)c(O)cc2[C@H](c2ccccc2)C1 | XFTVOHWWEQGXLS-HNNXBMFYSA-N | No-Effect | inactive |
| SKF-86002 | BRD-K96809896 | Fc1ccc(-c2nc3n(c2-c2ccncc2)CCS3)cc1 | YOELZIQOLWZLQC-UHFFFAOYSA-N | No-Effect | inactive |
| SKF-89976A | BRD-K01826728 | O=C(O)[C@H]1CCCN(CCC=C(c2ccccc2)c2ccccc2)C1 | TXQKSMSLZVKQBI-FQEVSTJZSA-N | No-Effect | inactive |
| SKF-91488 | BRD-K49776472 | CN(C)CCCCSC(=N)N | UFYJLJINUGVUHO-UHFFFAOYSA-N | No-Effect | inactive |
| SKF-96365 | BRD-K01826819 | COc1ccc(CCCO[C@@H](Cn2ccnc2)c2ccc(OC)cc2)cc1 | HLMBXBGDBBCYII-QFIPXVFZSA-N | No-Effect | inactive |
| SKLB1002 | BRD-K29895144 | COc1cc2ncnc(Sc3nnc(C)s3)c2cc1OC | RQVGFDBMONQTBC-UHFFFAOYSA-N | No-Effect | inactive |
| SLV-319-(+/-) | BRD-K00004304 | C/N=C(/NS(=O)(=O)c1ccc(Cl)cc1)N1C[C@H](c2ccccc2)C(c2ccc(Cl)cc2)=N1 | AXJQVVLKUYCICH-OAQYLSRUSA-N | No-Effect | inactive |
| SM-164 | BRD-K06191411 | CN[C@@H](C)C(=O)N[C@H]1CCCC[C@H]2CC[C@@H](C(=O)N[C@@H](c3ccccc3)c3cn(CCCCc4ccc(CCCCn5cc([C@@H](NC(=O)[C@@H]6CC[C@@H]7CCCC[C@H](NC(=O)[C@H](C)NC)C(=O)N76)c6ccccc6)nn5)cc4)nn3)N2C1=O | LGYDZXNSSLRFJS-IOQQVAQYSA-N | No-Effect | inactive |
| SM-21 | BRD-K00004213 | CC[C@H](Oc1ccc(Cl)cc1)C(=O)O[C@@H]1C[C@@H]2CC[C@H](C1)N2C | WTPAXDRULIZRDJ-ABFRBSLYSA-N | No-Effect | inactive |
| SMER-28 | BRD-K08478539 | C=CCNc1ncnc2ccc(Br)cc12 | BCPOLXUSCUFDGE-UHFFFAOYSA-N | No-Effect | inactive |
| SMI-4a | BRD-K98396737 | O=C1NC(=O)/C(=C/c2cccc(C(F)(F)F)c2)S1 | NGJLOFCOEOHFKQ-YVMONPNESA-N | No-Effect | inactive |
| SN-2 | BRD-K00004310 | Cc1cc(C)c(C2=NO[C@@H]3[C@H]4CC[C@H](C4)[C@H]23)c(C)c1 | WKLZNTYMDOPBSE-IARIHHJXSA-N | No-Effect | inactive |
| SN-38 | BRD-K89561498 | CCc1c2c(nc3ccc(O)cc13)-c1cc3c(c(=O)n1C2)COC(=O)[C@]3(O)CC | FJHBVJOVLFPMQE-QFIPXVFZSA-N | No-Effect | inactive |
| SN-6 | BRD-K01825672 | CCOC(=O)[C@H]1CS[C@H](Cc2ccc(OCc3ccc([N+](=O)[O-])cc3)cc2)N1 | ZVYIJXLMBWCGHP-RTBURBONSA-N | No-Effect | inactive |
| SNAP-94847 | BRD-K49702221 | Cc1ccc(NC(=O)C(C)C)cc1C1CCN(Cc2ccc(Oc3ccc(F)c(F)c3)cc2)CC1 | VMLZFUVIKCGATC-UHFFFAOYSA-N | No-Effect | inactive |
| SNC-80 | BRD-K29546239 | C=CCN1C[C@H](C)N([C@H](c2ccc(C(=O)N(CC)CC)cc2)c2cccc(OC)c2)C[C@H]1C | KQWVAUSXZDRQPZ-UMTXDNHDSA-N | No-Effect | inactive |
| SNG-1153 | BRD-K75016267 | CC(C)=CCc1c(O)cc(O)c2c(=O)c(O)c(-c3ccc(C(F)(F)F)cc3)oc12 | OXHMDMVBLQNMGP-UHFFFAOYSA-N | No-Effect | inactive |
| SNS-314 | BRD-K71512533 | O=C(Nc1cccc(Cl)c1)Nc1ncc(CCNc2ncnc3ccsc23)s1 | FAYAUAZLLLJJGH-UHFFFAOYSA-N | No-Effect | inactive |
| SNX-5422 | BRD-K73197500 | CC1(C)CC(=O)c2c(C(F)(F)F)nn(-c3ccc(C(N)=O)c(N[C@H]4CC[C@H](OC(=O)CN)CC4)c3)c2C1 | AVDSOVJPJZVBTC-CTYIDZIISA-N | No-Effect | inactive |
| sobetirome | BRD-K85406984 | Cc1cc(OCC(=O)O)cc(C)c1Cc1ccc(O)c(C(C)C)c1 | QNAZTOHXCZPOSA-UHFFFAOYSA-N | No-Effect | inactive |
| sobrepin | BRD-K58262380 | CC1=CC[C@@H](C(C)(C)O)C[C@@H]1O | OMDMTHRBGUBUCO-BDAKNGLRSA-N | No-Effect | inactive |
| sobuzoxane | BRD-K87595680 | CC(C)COC(=O)OCN1C(=O)CN(CCN2CC(=O)N(COC(=O)OCC(C)C)C(=O)C2)CC1=O | OCOKWVBYZHBHLU-UHFFFAOYSA-N | No-Effect | inactive |
| sodium-butyrate | BRD-K05878375 | CCCC(=O)O | FERIUCNNQQJTOY-UHFFFAOYSA-N | No-Effect | inactive |
| sodium-dodecyl-sulfate | BRD-M58433298 | CCCCCCCCCCCCOS(=O)(=O)O.[Na] | FMWAXKQEIXRUTI-UHFFFAOYSA-N | No-Effect | inactive |
| sodium-gluconate | BRD-K93201765 | O=C(O)[C@H](O)[C@@H](O)[C@H](O)[C@H](O)CO | RGHNJXZEOKUKBD-SQOUGZDYSA-N | No-Effect | inactive |
| sodium-gualenate | BRD-K80043866 | Cc1cc(S(=O)(=O)O)c2c(C)ccc(C(C)C)cc1-2 | VIZXMHCBZLGUET-UHFFFAOYSA-N | No-Effect | inactive |
| sodium-monofluorophosphate | BRD-K99308954 | O=P(O)(O)F | DWYMPOCYEZONEA-UHFFFAOYSA-N | No-Effect | inactive |
| sodium-nitrite | BRD-K61230716 | [O-][NH2+][O-] | ROODZSDSPAHHIS-UHFFFAOYSA-N | No-Effect | inactive |
| sodium-nitroprusside | BRD-K00003606 | N#C[Fe-2](C#N)(C#N)(C#N)(C#N)[N+][O-] | WTISQLWAPBSCSF-UHFFFAOYSA-N | No-Effect | inactive |
| sodium-orthovanadate | BRD-K60600693 | O=[V](O)(O)O | WQEVDHBJGNOKKO-UHFFFAOYSA-K | No-Effect | inactive |
| sodium-picosulfate | BRD-K77251551 | O=S(=O)(O)Oc1ccc(C(c2ccc(OS(=O)(=O)O)cc2)c2ccccn2)cc1 | UJIDKYTZIQTXPM-UHFFFAOYSA-N | No-Effect | inactive |
| sodium-stibogluconate | BRD-K00003543 | O=C(O)[C@@H]1O[Sb]2(O)(O[Sb]34(O)O[C@H]([C@H](C(=O)O)O3)[C@@H]([C@H](O)CO)O4)O[C@H]1[C@@H]([C@H](O)CO)O2 | RPOBPHCDIPHDQL-XCCFGPONSA-L | No-Effect | inactive |
| sodium-tanshinone-II-A-sulfonate | BRD-K84798689 | Cc1c(S(=O)(=O)O)oc2c1C(=O)C(=O)c1c-2ccc2c1CCCC2(C)C | UJCACMLMPLLRGW-UHFFFAOYSA-N | No-Effect | inactive |
| sodium-tetradecyl-sulfate | BRD-A55664972 | CCCC[C@@H](CC)CC[C@@H](CC(C)C)OS(=O)(=O)O | GROJOWHVXQYQGN-KGLIPLIRSA-N | No-Effect | inactive |
| sofalcone | BRD-K01039503 | CC(C)=CCOc1ccc(/C=C/C(=O)c2ccc(OCC=C(C)C)cc2OCC(=O)O)cc1 | GFWRVVCDTLRWPK-KPKJPENVSA-N | No-Effect | inactive |
| sofosbuvir | BRD-K00026984 | CC(C)OC(=O)[C@H](C)N[P@](=O)(OC[C@H]1O[C@@H](n2ccc(=O)[nH]c2=O)[C@](C)(F)[C@@H]1O)Oc1ccccc1 | TTZHDVOVKQGIBA-IQWMDFIBSA-N | No-Effect | inactive |
| sofosbuvir-impurity-m | BRD-K00003527 | CC(C)OC(=O)[C@H](C)N[P@](=O)(OC[C@H]1O[C@@H](n2ccc(=O)[nH]c2=O)[C@@](C)(O)[C@@H]1O)Oc1ccccc1 | WAKQLZVLNOKKHE-BFQGEBBDSA-N | No-Effect | inactive |
| sograzepide | BRD-K00004608 | CNc1cccc(NC(=O)N[C@@H]2N=C(c3ccccn3)c3ccccc3N(CC(=O)C(C)(C)C)C2=O)c1 | YDZYKNJZCVIKPP-VWLOTQADSA-N | No-Effect | inactive |

| | | | | | |
|---|---|---|---|---|---|
| solamargine | BRD-A65562236 | C[C@@H]1O[C@H](O[C@@H]2[C@H](CO)O[C@H](O[C@@H]3CC[C@]4(C)C(=CC[C@@H]5[C@@H]6C[C@H]7O[C@]8(CC[C@@H](C)CN8)[C@@H](C)[C@@H]7[C@]6(C)CC[C@H]54)C3)[C@@H](O[C@H]3O[C@@H](C)[C@@H](O)[C@@H](O)[C@@H]3O)[C@@H]2O)[C@@H](O)[C@H](O)[C@@H]1O | MBWUSSKCCUMJHO-OXBNUUDKSA-N | No-Effect | inactive |
| solcitinib | BRD-K69215269 | CC1(C)CN(C(=O)c2ccc(-c3cccc4nc(NC(=O)C5CC5)nn34)cc2)C1 | MPYACSQFXVMWNO-UHFFFAOYSA-N | No-Effect | inactive |
| solifenacin-succinate | BRD-K00003600 | O=C(O[C@H]1C[N@]2CC[C@@H]1CC2)N1CCc2ccccc2[C@@H]1c1ccccc1 | FBOUYBDGKBSUES-VXKWHMMOSA-N | No-Effect | inactive |
| solithromycin | BRD-K32256160 | CC[C@H]1OC(=O)[C@@](C)(F)C(=O)[C@H](C)[C@@H](O[C@@H]2O[C@H](C)C[C@H](N(C)C)[C@H]2O)[C@](C)(OC)C[C@@H](C)C(=O)[C@H](C)[C@H]2N(CCCCn3cc(-c4cccc(N)c4)nn3)C(=O)O[C@]12C | IXXFZUPTQVDPPK-ZAWHAJPISA-N | No-Effect | inactive |
| solriamfetol | BRD-K00004707 | NC(=O)OC[C@H](N)Cc1ccccc1 | UCTRAOBQFUDCSR-SECBINFHSA-N | No-Effect | inactive |
| somantadine | BRD-K85764479 | CC(C)(N)CC12CC3CC(CC(C3)C1)C2 | OWKXRDQRMTVCEX-UHFFFAOYSA-N | No-Effect | inactive |
| sonepiprazole | BRD-K69857269 | NS(=O)(=O)c1ccc(N2CCN(CC[C@@H]3OCCc4ccccc43)CC2)cc1 | WNUQCGWXPNGORO-NRFANRHFSA-N | No-Effect | inactive |
| sonidegib | BRD-K19796430 | Cc1c(C(=O)Nc2ccc(N3C[C@H](C)O[C@H](C)C3)nc2)cccc1-c1ccc(OC(F)(F)F)cc1 | VZZJRYRQSPEMTK-CALCHBBNSA-N | No-Effect | inactive |
| sophocarpine | BRD-K66845263 | O=C1C=CC[C@@H]2[C@H]3CCCN4CCC[C@@H](CN12)[C@@H]34 | AAGFPTSOPGCENQ-JLNYLFASSA-N | No-Effect | inactive |
| sorafenib | BRD-K23984367 | CNC(=O)c1cc(Oc2ccc(NC(=O)Nc3ccc(Cl)c(C(F)(F)F)c3)cc2)ccn1 | MLDQJTXFUGDVEO-UHFFFAOYSA-N | No-Effect | inactive |
| soraprazan | BRD-K00004585 | COCCO[C@@H]1c2ccn3c(C)c(C)nc3c2N[C@H](c2ccccc2)[C@H]1O | PWILYDZRJORZDR-MISYRCLQSA-N | No-Effect | inactive |
| sorbic-acid | BRD-K88734090 | C/C=C/C=C/C(=O)O | WSWCOQWTEOXDQX-MQQKCMAXSA-N | No-Effect | inactive |
| sorbinil | BRD-A89164055 | O=C1NC(=O)[C@@]2(CCOc3ccc(F)cc32)N1 | LXANPKRCLVQAOG-NSHDSACASA-N | No-Effect | inactive |
| sorbitol | BRD-K66035042 | OC[C@H](O)[C@@H](O)[C@H](O)[C@H](O)CO | FBPFZTCFMRRESA-JGWLITMVSA-N | No-Effect | inactive |
| sotagliflozin | BRD-K05850076 | CCOc1ccc(Cc2cc([C@@H]3O[C@H](SC)[C@@H](O)[C@H](O)[C@H]3O)ccc2Cl)cc1 | QKDRXGFQVGOQKS-CRSSMBPESA-N | No-Effect | inactive |
| sotrastaurin | BRD-K16761703 | CN1CCN(c2nc(C3=C(c4c[nH]c5ccccc45)C(=O)NC3=O)c3ccccc3n2)CC1 | OAVGBZOFDPFGPJ-UHFFFAOYSA-N | No-Effect | inactive |
| SP-100030 | BRD-K71900389 | O=C(Nc1cc(C(F)(F)F)cc(C(F)(F)F)c1)c1cnc(Cl)nc1C(F)(F)F | GHMKQBWHPQMXSH-UHFFFAOYSA-N | No-Effect | inactive |
| SP-420 | BRD-K23005288 | COCCOCCOc1ccc(C2=N[C@@](C)(C(=O)O)CS2)c(O)c1 | YASYAEVZKXPYIZ-MRXNPFEDSA-N | No-Effect | inactive |
| sparfloxacin | BRD-K07612980 | C[C@H]1CN(c2c(F)c(N)c3c(=O)c(C(=O)O)cn(C4CC4)c3c2F)C[C@@H](C)N1 | DZZWHBIBMUVIIW-DTORHVGOSA-N | No-Effect | inactive |
| sparfosate | BRD-A81539525 | O=C(O)C[C@@H](NC(=O)CP(=O)(O)O)C(=O)O | ZZKNRXZVGOYGJT-GSVOUGTGSA-N | No-Effect | inactive |
| sparsentan | BRD-K00003253 | CCCCC1=NC2(CCCC2)C(=O)N1Cc1ccc(-c2ccccc2S(=O)(=O)Nc2noc(C)c2C)c(COCC)c1 | WRFHGDPIDHPWIQ-UHFFFAOYSA-N | No-Effect | inactive |
| sparteine-(-) | BRD-K86872788 | C1CCN2C[C@@H]3C[C@@H](CN4CCCC[C@H]34)[C@@H]2C1 | SLRCCWJSBJZJBV-ZQDZILKHSA-N | No-Effect | inactive |
| spautin-1 | BRD-K84964099 | Fc1ccc(CNc2ncnc3ccc(F)cc23)cc1 | AWIVHRPYFSSVOG-UHFFFAOYSA-N | No-Effect | inactive |
| spectinomycin | BRD-K11905747 | CN[C@H]1[C@@H](O)[C@@H](NC)[C@H](O)[C@H]2O[C@@H]3O[C@H](C)CC(=O)[C@]3(O)O[C@H]12 | UNFWWIHTNXNPBV-WXKVUWSESA-N | No-Effect | inactive |
| spermidine | BRD-K40438880 | NCCCCNCCCN | ATHGHQPFGPMSJY-UHFFFAOYSA-N | No-Effect | inactive |
| spermine | BRD-K18375018 | NCCCNCCCCNCCCN | PFNFFQXMRSDOHW-UHFFFAOYSA-N | No-Effect | inactive |
| sphingosylphosphorylcholine | BRD-K01825110 | CCCCCCCCCCCCC/C=C/[C@@H](O)[C@@H](N)COP(=O)(O)OCC[N+](C)(C)C | JLVSPVFPBBFMBE-HXSWCURESA-O | No-Effect | inactive |
| spiradoline | BRD-K28192623 | CN(C(=O)Cc1ccc(Cl)c(Cl)c1)[C@H]1CC[C@@]2(CCCO2)C[C@@H]1N1CCCC1 | NYKCGQQJNVPOLU-ONTIZHBOSA-N | No-Effect | inactive |
| spiramycin | BRD-K00003542 | CO[C@@H]1[C@@H](O[C@@H]2O[C@H](C)[C@@H](O[C@H]3C[C@@](C)(O)[C@@H](O)[C@H](C)O3)[C@H](N(C)C)[C@H]2O)[C@@H](CC=O)C[C@@H](C)[C@@H](O[C@H]2CC[C@H](N(C)C)[C@@H](C)O2)/C=C/C=C/C[C@@H](C)OC(=O)C[C@H]1O | ACTOXUHEUCPTEW-CEUOBAOPSA-N | No-Effect | inactive |
| spirobromin | BRD-K96684644 | O=C(CCBr)N1CC[N+]2(CC1)CC[N+]1(CCN(C(=O)CCBr)CC1)CC2 | JXEICPOBKSQAIU-UHFFFAOYSA-N | No-Effect | inactive |
| spironolactone | BRD-K90027355 | CC(=O)S[C@@H]1CC2=CC(=O)CC[C@]2(C)[C@H]2CC[C@@]3(C)[C@@H](CC[C@@]34CCC(=O)O4)[C@@H]21 | LXMSZDCAJNLERA-ZHYRCANASA-N | No-Effect | inactive |
| spiroxatrine | BRD-A72483914 | O=C1NCN(c2ccccc2)C12CCN(C[C@H]1COc3ccccc3O1)CC2 | JVGBTTIJPBFLTE-SFHVURJKSA-N | No-Effect | inactive |
| spizofurone | BRD-K94934625 | CC(=O)c1ccc2c(c1)C(=O)C1(CC1)O2 | SETMGIIITGNLAS-UHFFFAOYSA-N | No-Effect | inactive |
| splitomycin | BRD-K27710560 | O=C1CCc2c(ccc3ccccc23)O1 | ISFPDBUKMJDAJH-UHFFFAOYSA-N | No-Effect | inactive |
| SPP301 | BRD-K19203487 | COc1ccccc1Oc1c(NS(=O)(=O)c2ccc(C)cn2)nc(-c2ccncc2)nc1OC | YBWLTKFZAOSWSM-UHFFFAOYSA-N | No-Effect | inactive |
| SQ-22536 | BRD-A56987319 | Nc1ncnc2c1ncn2[C@H]1CCCO1 | UKHMZCMKHPHFOT-ZCFIWIBFSA-N | No-Effect | inactive |
| squalamine | BRD-K00003215 | CC(C)[C@@H](CC[C@@H](C)[C@H]1CC[C@H]2[C@H]3[C@H](CC[C@@]21C)[C@@]1(C)CC[C@H](NCCCNCCCCN)C[C@@H]1C[C@H]3O)OS(=O)(=O)O | UIRKNQLZZXALBI-MSVGPLKSSA-N | No-Effect | inactive |
| squaric-acid-dibutyl-ester | BRD-K00004684 | CCCCOc1c(OCCCC)c(=O)c1=O | XBRWELTXMQSEIN-UHFFFAOYSA-N | No-Effect | inactive |
| SR-11302 | BRD-K16229414 | CC1=C(/C=C/C(=C/C=C/C(C)=C/C(=O)O)c2ccc(C)cc2)C(C)(C)CCC1 | RQANARBNMTXCDM-DKOHIBGUSA-N | No-Effect | inactive |
| SR-140333 | BRD-K60931454 | CC(C)Oc1cccc(CC(=O)N2CCC[C@](CC[N+]34CCC(c5ccccc5)(CC3)CC4)(c3ccc(Cl)c(Cl)c3)C2)c1 | RPDFDSQFBCJTDY-GAQXSTBRSA-N | No-Effect | inactive |
| SR-1664 | BRD-K62814487 | Cc1c(C)n(Cc2ccc(-c3ccccc3C(=O)O)cc2)c2ccc(C(=O)N[C@@H](C)c3ccc([N+](=O)[O-])cc3)cc12 | IIJDFXNUWZTHIM-NRFANRHFSA-N | No-Effect | inactive |
| SR-2211 | BRD-K11641193 | OC(c1ccc(-c2ccc(CN3CCN(Cc4ccncc4)CC3)cc2)c(F)c1)(C(F)(F)F)C(F)(F)F | KVHKWAZUPPBMLL-UHFFFAOYSA-N | No-Effect | inactive |
| SR-2640 | BRD-K62562723 | O=C(O)c1ccccc1Nc1cccc(OCc2ccc3ccccc3n2)c1 | LMPZHLXYBWGGNT-UHFFFAOYSA-N | No-Effect | inactive |
| SR-27897 | BRD-K35629949 | O=C(O)Cn1c(C(=O)Nc2nc(-c3ccccc3Cl)cs2)cc2ccccc21 | ILNRQFBVVQUOLP-UHFFFAOYSA-N | No-Effect | inactive |
| SR-3306 | BRD-K60952143 | Cc1ccc(-c2ncn(-c3ccc(Nc4nccc(-c5cccc(N6CCOCC6)c5)n4)cc3)n2)cn1 | QDMXVKKPYBGIAS-UHFFFAOYSA-N | No-Effect | inactive |
| SR-33805 | BRD-K43002771 | COc1ccc(CCN(C)CCCOc2ccc(S(=O)(=O)c3c(C(C)C)c4ccccc4n3C)cc2)cc1OC | GCVFQRRBTCHIPK-UHFFFAOYSA-N | No-Effect | inactive |
| SR-3677 | BRD-A86468475 | CN(C)CCOc1cc(-c2cn[nH]c2)ccc1NC(=O)[C@H]1COc2ccccc2O1 | OQWZIAVXCYIZNN-OAQYLSRUSA-N | No-Effect | inactive |
| SR-57227A | BRD-K09397065 | NC1CCN(c2cccc(Cl)n2)CC1 | WPVVMKYQOMJPIN-UHFFFAOYSA-N | No-Effect | inactive |
| SR-59230A | BRD-K35430135 | CCc1ccccc1OC[C@@H](O)CN[C@H]1CCCc2ccccc21 | VFDHMSXXELYMRW-ICSRJNTNSA-N | No-Effect | inactive |
| SR-95639A | BRD-K19309090 | c1ccc2c(c1)CCCc1cc(NCCN3CCOCC3)nnc1-2 | RUTYAHFAKWVMNH-UHFFFAOYSA-N | No-Effect | inactive |
| SRC-kinase-inhibitor-I | BRD-K50495309 | COc1cc2ncnc(Nc3ccc(Oc4ccccc4)cc3)c2cc1OC | DMWVGXGXHPOEPT-UHFFFAOYSA-N | No-Effect | inactive |
| SRT2104 | BRD-K48452630 | Cc1nc(-c2cccnc2)sc1C(=O)Nc1ccccc1-c1cn2c(CN3CCOCC3)csc2n1 | LAMQVIQMVKWXOC-UHFFFAOYSA-N | No-Effect | inactive |
| SRT3190 | BRD-K34801930 | C[C@H](Nc1cc(NS(=O)(=O)N2CCC2)nc(SCc2cccc(F)c2F)n1)[C@@H](O)CO | QVKPEMXUBULFBM-FZMZJTMJSA-N | No-Effect | inactive |
| SSR128129E | BRD-K65890018 | COc1c(C)c(C(=O)c2ccc(N)c(C(=O)O)c2)n2ccccc12 | SNMCWIDCSPHZRD-UHFFFAOYSA-N | No-Effect | inactive |
| SSR180711 | BRD-K00004316 | O=C(Oc1ccc(Br)cc1)N1CC[N@]2CC[C@@H]1CC2 | RXLOZRCLQMJJLC-UHFFFAOYSA-N | No-Effect | inactive |

| | | | | | |
|---|---|---|---|---|---|
| SSR240612 | BRD-K00003186 | COc1ccc2cc(S(=O)(=O)N[C@H](CC(=O)N[C@H](Cc3ccc(CN4[C@@H](C)CCC[C@H]4C)cc3)C(=O)N(C)C(C)C)c3ccc4c(c3)OCO4)ccc2c1 | QGWIQIAWOCJRPI-WSCVWKGISA-N | No-Effect | inactive |
| ST-1535 | BRD-K00004706 | CCCCc1nc(N)c2nc(-n3nccn3)n(C)c2n1 | CYYQMAWUIRPCNW-UHFFFAOYSA-N | No-Effect | inactive |
| ST-1859 | BRD-K73524797 | Oc1ccc2ccccc2c1Cc1c(O)ccc2ccccc12 | ZPANWZBSGMDWON-UHFFFAOYSA-N | No-Effect | inactive |
| ST-2825 | BRD-K68470841 | NC(=O)[C@H]1CCS[C@@H]2C[C@]3(CCCN3C(=O)Cc3ccc(NC(=O)COc4ccc(Cl)cc4Cl)cc3)C(=O)N21 | HBLHLJXFIPCEMW-ZJSFPPFMSA-N | No-Effect | inactive |
| ST-91 | BRD-K18905250 | CCc1cccc(CC)c1NC1=NCCN1 | IALHTUPVRAQZFI-UHFFFAOYSA-N | No-Effect | inactive |
| STAT3-inhibitor-VI | BRD-K40853697 | Cc1ccc(S(=O)(=O)OCC(=O)Nc2ccc(C(=O)O)c(O)c2)cc1 | HWNUSGNZBAISFM-UHFFFAOYSA-N | No-Effect | inactive |
| statil | BRD-K68332390 | O=C(O)Cc1nn(Cc2ccc(Br)cc2F)c(=O)c2ccccc12 | LKBFFDOJUKLQNY-UHFFFAOYSA-N | No-Effect | inactive |
| stattic | BRD-K17061841 | O=[N+]([O-])c1ccc2c(c1)S(=O)(=O)C=C2 | ZRRGOUHITGRLBA-UHFFFAOYSA-N | No-Effect | inactive |
| stavudine | BRD-K93880783 | Cc1cn([C@H]2C=C[C@@H](CO)O2)c(=O)[nH]c1=O | XNKLLVCARDGLGL-JGVFFNPUSA-N | No-Effect | inactive |
| stemregenin-1 | BRD-K36739687 | CC(C)n1cnc2c(NCCc3ccc(O)cc3)nc(-c3csc4ccccc34)nc21 | BGFHMYJZJZLMHW-UHFFFAOYSA-N | No-Effect | inactive |
| stepronin | BRD-A04506385 | C[C@@H](SC(=O)c1cccs1)C(=O)NCC(=O)O | JNYSEDHQJCOWQU-ZCFIWIBFSA-N | No-Effect | inactive |
| stetaderm | BRD-K93900986 | CC(=O)[C@H]1CC[C@H]2[C@@H]3CCC4=CC(=O)CC[C@]4(C)[C@H]3[C@H](O)C[C@]12C | BFZHCUBIASXHPK-QJSKAATBSA-N | No-Effect | inactive |
| STF-083010 | BRD-K13049524 | O=S(=O)(/N=C/c1c(O)ccc2ccccc12)c1cccs1 | TVIVJHZHPKNDAQ-MHWRWJLKSA-N | No-Effect | inactive |
| STF-118804 | BRD-K39171998 | Cc1ccc(S(=O)(=O)Cc2nc(-c3ccc(C(=O)NCc4cccnc4)cc3)oc2C)cc1 | DLFCEZOMHBPDGI-UHFFFAOYSA-N | No-Effect | inactive |
| stigmasterol | BRD-K41422683 | CC[C@H](/C=C/[C@@H](C)[C@H]1CC[C@H]2[C@@H]3CC=C4C[C@@H](O)CC[C@]4(C)[C@H]3CC[C@@]21C)C(C)C | HCXVJBMSMIARIN-PHZDYDNGSA-N | No-Effect | inactive |
| stiripentol | BRD-A72441487 | CC(C)(C)[C@@H](O)/C=C/c1ccc2c(c1)OCO2 | IBLNKMRFIPWSOY-IYNGYCSASA-N | No-Effect | inactive |
| stobadine | BRD-K33839500 | Cc1ccc2c(c1)[C@H]1CN(C)CC[C@H]1N2 | CYJQCYXRNNCURD-DGCLKSJQSA-N | No-Effect | inactive |
| streptomycin | BRD-K07166362 | CN[C@@H]1[C@H](O[C@H]2[C@H](O[C@@H]3[C@@H](NC(=N)N)[C@H](O)[C@@H](NC(=N)N)[C@H](O)[C@H]3O)O[C@@H](C)[C@]2(O)C=O)O[C@@H](CO)[C@H](O)[C@H]1O | UCSJYZPVAKXKNQ-HZYVHMACSA-N | No-Effect | inactive |
| streptozotocin | BRD-K34411947 | CN([NH2+][O-])C(=O)N[C@H]1[C@@H](O)O[C@H](CO)[C@@H](O)[C@@H]1O | BKWHZSFINQBEQH-GKHCUFPYSA-N | No-Effect | inactive |
| strontium-ranelate | BRD-K83146339 | N#Cc1c(N2C=C(O)O[Sr]OC(O)=C2)sc2c1CC(=O)O[Sr]OC2=O | TWHPPRXFYWSOQF-UHFFFAOYSA-J | No-Effect | inactive |
| SU014813 | BRD-K00003547 | Cc1[nH]c(/C=C2\C(=O)Nc3ccc(F)cc32)c(C)c1C(=O)NC[C@H](O)CN1CCOCC1 | CTNPALGJUAXMMC-PMFHANACSA-N | No-Effect | inactive |
| SU11274 | BRD-K02965346 | Cc1[nH]c(/C=C2\C(=O)Nc3ccc(S(=O)(=O)N(C)c4cccc(Cl)c4)cc32)c(C)c1C(=O)N1CCN(C)CC1 | FPYJSJDOHRDAMT-KQWNVCNZSA-N | No-Effect | inactive |
| SU16f | BRD-K91106808 | Cc1[nH]c(/C=C2\C(=O)Nc3cc(-c4ccccc4)ccc32)c(C)c1CCC(=O)O | APYYTEJNOZQZNA-MOSHPQCFSA-N | No-Effect | inactive |
| SU3327 | BRD-K32730239 | Nc1nnc(Sc2ncc([N+](=O)[O-])s2)s1 | NQQBNZBOOHHVQP-UHFFFAOYSA-N | No-Effect | inactive |
| SU4312 | BRD-K20287671 | CN(C)c1ccc(/C=C2\C(=O)Nc3ccccc32)cc1 | UAKWLVYMKBWHMX-PTNGSMBKSA-N | No-Effect | inactive |
| SU9516 | BRD-K94883909 | COc1ccc2c(c1)/C(=C/c1cnc[nH]1)C(=O)N2 | QNUKRWAIZMBVCU-WCIBSUBMSA-N | No-Effect | inactive |
| substance-P | BRD-K23106423 | CSCC[C@H](NC(=O)[C@H](CC(C)C)NC(=O)CNC(=O)[C@H](Cc1ccccc1)NC(=O)[C@H](Cc1ccccc1)NC(=O)[C@H](CCC(N)=O)NC(=O)[C@H](CCC(N)=O)NC(=O)[C@@H]1CCCN1C(=O)[C@H](CCCCN)NC(=O)[C@@H]1CCCN1C(=O)[C@@H](N)CCCNC(=N)N)C(N)=O | ADNPLDHMAVUMIW-CUZNLEPHSA-N | No-Effect | inactive |
| succinic-acid | BRD-K45238818 | O=C(O)CCC(=O)O | KDYFGRWQOYBRFD-UHFFFAOYSA-N | No-Effect | inactive |
| succinylcholine-chloride | BRD-K45497762 | C[N+](C)(C)CCOC(=O)CCC(=O)OCC[N+](C)(C)C | AXOIZCJOOAYSMI-UHFFFAOYSA-N | No-Effect | inactive |
| succinylsulfathiazole | BRD-K01950558 | O=C(O)CCC(=O)Nc1ccc(S(=O)(=O)Nc2nccs2)cc1 | SKVLYVHULOWXTD-UHFFFAOYSA-N | No-Effect | inactive |
| sucralfate | BRD-M71167108 | O=S(=O)(OC[C@H]1O[C@H](O[C@]2(COS(=O)(=O)O[Al](O)O)O[C@H](COS(=O)(=O)O[Al](O)O)[C@@H](OS(=O)(=O)O[Al](O)O)[C@@H]2OS(=O)(=O)O[Al](O)O)[C@H](OS(=O)(=O)O[Al](O)O)[C@@H](OS(=O)(=O)O[Al](O)O)[C@@H]1OS(=O)(=O)O[Al](O)O)O[Al](O)O.O[Al](O)O.O[Al](O)O.O[Al](O)O.O[Al](O)O.O[Al](O)O.O[Al](O)O.O[Al](O)O | DTULOKYBGCTCFP-QRLOXDDHSA-A | No-Effect | inactive |
| sucralose | BRD-K58968598 | OC[C@H]1O[C@H](O[C@]2(CCl)O[C@H](CCl)[C@@H](O)[C@@H]2O)[C@H](O)[C@@H](O)[C@H]1Cl | BAQAVOSOZGMPRM-QBMZZYIRSA-N | No-Effect | inactive |
| sucrose | BRD-K74724920 | OC[C@H]1O[C@H](O[C@]2(CO)O[C@H](CO)[C@@H](O)[C@@H]2O)[C@H](O)[C@@H](O)[C@@H]1O | CZMRCDWAGMRECN-UGDNZRGBSA-N | No-Effect | inactive |
| sufugolix | BRD-K00004582 | CONC(=O)Nc1ccc(-c2sc3c(c2CN(C)Cc2ccccc2)c(=O)n(-c2ccccc2)c(=O)n3Cc2c(F)cccc2F)cc1 | UCQSBGOFELXYIN-UHFFFAOYSA-N | No-Effect | inactive |
| sugammadex | BRD-K00003322 | O=C(O)CCSC[C@@H]1O[C@H]2O[C@@H]3[C@@H](O)[C@H](O)[C@@H](O[C@H]3CSCCC(=O)O)O[C@@H]3[C@@H](O)[C@H](O)[C@@H](O[C@H]3CSCCC(=O)O)O[C@@H]3[C@@H](O)[C@H](O)[C@@H](O[C@H]3CSCCC(=O)O)O[C@@H]3[C@@H](O)[C@H](O)[C@@H](O[C@H]3CSCCC(=O)O)O[C@@H]3[C@@H](O)[C@H](O)[C@@H](O[C@H]3CSCCC(=O)O)O[C@@H]3[C@@H](O)[C@H](O)[C@@H](O[C@H]3CSCCC(=O)O)O[C@@H]3[C@@H](O)[C@H](O)[C@@H](O[C@@H]3CSCCC(=O)O)O[C@@H]1[C@@H](O)[C@@H]2O | WHRODDIHRRDWEW-WXUQZJQRSA-N | No-Effect | inactive |
| sulbactam | BRD-K44133266 | CC1(C)[C@H](C(=O)O)N2C(=O)C[C@H]2S1(=O)=O | FKENQMMABCRJMK-RITPCOANSA-N | No-Effect | inactive |
| sulbactam-pivoxil | BRD-A63431791 | CC(C)(C)C(=O)OCOC(=O)[C@H]1N2C(=O)C[C@H]2S(=O)(=O)C1(C)C | OHPVYKXTRACOSQ-NXEZZACHSA-N | No-Effect | inactive |
| sulbentine | BRD-K72384124 | S=C1SCN(Cc2ccccc2)CN1Cc1ccccc1 | QFVAWNPSRQWSDU-UHFFFAOYSA-N | No-Effect | inactive |
| sulbutiamine | BRD-K12539416 | C/C(=C(/CCOC(=O)C(C)C)SS/C(CCOC(=O)C(C)C)=C(\C)N(C=O)Cc1cnc(C)nc1N)N(C=O)Cc1cnc(C)nc1N | CKHJPWQVLKHBIH-GPAWKIAZSA-N | No-Effect | inactive |
| sulconazole | BRD-A70649075 | Clc1ccc(CS[C@@H](Cn2ccnc2)c2ccc(Cl)cc2Cl)cc1 | AFNXATANNDIXLG-SFHVURJKSA-N | No-Effect | inactive |
| sulfachlorpyridazine | BRD-K32021043 | Nc1ccc(S(=O)(=O)Nc2ccc(Cl)nn2)cc1 | XOXHILFPRYWFOD-UHFFFAOYSA-N | No-Effect | inactive |
| sulfadiazine | BRD-K32273377 | Nc1ccc(S(=O)(=O)Nc2ncccn2)cc1 | SEEPANYCNGTZFQ-UHFFFAOYSA-N | No-Effect | inactive |
| sulfadoxine | BRD-K55250441 | COc1ncnc(NS(=O)(=O)c2ccc(N)cc2)c1OC | PJSFRIWCGOHTNF-UHFFFAOYSA-N | No-Effect | inactive |
| sulfafurazole | BRD-K50859149 | Cc1noc(NS(=O)(=O)c2ccc(N)cc2)c1C | NHUHCSRWZMLRLA-UHFFFAOYSA-N | No-Effect | inactive |
| sulfaguanidine | BRD-K45086103 | N=C(N)NS(=O)(=O)c1ccc(N)cc1 | BRBKOPJOKNSWSG-UHFFFAOYSA-N | No-Effect | inactive |
| sulfamerazine | BRD-K93524252 | Cc1ccnc(NS(=O)(=O)c2ccc(N)cc2)n1 | QPPBRPIAZZHUNT-UHFFFAOYSA-N | No-Effect | inactive |
| sulfameter | BRD-K87492696 | COc1cnc(NS(=O)(=O)c2ccc(N)cc2)nc1 | GPTONYMQFTZPKC-UHFFFAOYSA-N | No-Effect | inactive |
| sulfamethazine | BRD-K11640013 | Cc1cc(C)nc(NS(=O)(=O)c2ccc(N)cc2)n1 | ASWVTGNCAZCNNR-UHFFFAOYSA-N | No-Effect | inactive |
| sulfamethizole | BRD-K31682896 | Cc1nnc(NS(=O)(=O)c2ccc(N)cc2)s1 | VACCAVUAMIDAGB-UHFFFAOYSA-N | No-Effect | inactive |
| sulfamethoxypyridazine | BRD-K00938507 | COc1ccc(NS(=O)(=O)c2ccc(N)cc2)nn1 | VLYWMPOKSSWJAL-UHFFFAOYSA-N | No-Effect | inactive |
| sulfametopyrazine | BRD-K17205817 | COc1nccnc1NS(=O)(=O)c1ccc(N)cc1 | KXRZBTAEDBELFD-UHFFFAOYSA-N | No-Effect | inactive |
| sulfamoxole | BRD-K19742012 | Cc1nc(NS(=O)(=O)c2ccc(N)cc2)oc1C | CYFLXLSBHQBMFT-UHFFFAOYSA-N | No-Effect | inactive |
| sulfanilate-zinc | BRD-K04254814 | Nc1ccc(S(=O)(=O)O[Zn]OS(=O)(=O)c2ccc(N)cc2)cc1 | RIUORJCWAHCMSA-UHFFFAOYSA-L | No-Effect | inactive |
| sulfanitran | BRD-K76845197 | CC(=O)Nc1ccc(S(=O)(=O)Nc2ccc([N+](=O)[O-])cc2)cc1 | GWBPFRGXNGPPMF-UHFFFAOYSA-N | No-Effect | inactive |
| sulfaphenazole | BRD-K10671814 | Nc1ccc(S(=O)(=O)Nc2ccnn2-c2ccccc2)cc1 | QWCJHSGMANYXCW-UHFFFAOYSA-N | No-Effect | inactive |
| sulfapyridine | BRD-K41406082 | Nc1ccc(S(=O)(=O)Nc2ccccn2)cc1 | GECHUMIMRBOMGK-UHFFFAOYSA-N | No-Effect | inactive |
| sulfaquinoxaline | BRD-K71133585 | Nc1ccc(S(=O)(=O)Nc2cnc3ccccc3n2)cc1 | NHZLNPMOSADWGC-UHFFFAOYSA-N | No-Effect | inactive |

| | | | | | |
|---|---|---|---|---|---|
| sulfasalazine | BRD-K10670311 | O=C(O)c1cc(/N=N/c2ccc(S(=O)(=O)Nc3ccccn3)cc2)ccc1O | NCEXYHBECQHGNR-QZQOTICOSA-N | No-Effect | inactive |
| sulfathiazole | BRD-K14705039 | Nc1ccc(S(=O)(=O)Nc2nccs2)cc1 | JNMRHUJNCSQMMB-UHFFFAOYSA-N | No-Effect | inactive |
| sulfinpyrazone | BRD-K00004525 | O=C1C(CC[S@](=O)c2ccccc2)C(=O)N(c2ccccc2)N1c1ccccc1 | MBGGBVCUIVRRBF-LJAQVGFWSA-N | No-Effect | inactive |
| sulfisomidin | BRD-K52416806 | Cc1cc(NS(=O)(=O)c2ccc(N)cc2)nc(C)n1 | YZMCKZRAOLZXAZ-UHFFFAOYSA-N | No-Effect | inactive |
| sulfisoxazole-acetyl | BRD-K84810336 | CC(=O)N(c1onc(C)c1C)S(=O)(=O)c1ccc(N)cc1 | JFNWFXVFBDDWCX-UHFFFAOYSA-N | No-Effect | inactive |
| sulindac | BRD-K01827279 | CC1=C(CC(=O)O)c2cc(F)ccc2/C1=C\c1ccc([S@@](C)=O)cc1 | MLKXDPUZXIRXEP-LQVWSKNFSA-N | No-Effect | inactive |
| sulisobenzone | BRD-K38602774 | COc1cc(O)c(C(=O)c2ccccc2)cc1S(=O)(=O)O | CXVGEDCSTKKODG-UHFFFAOYSA-N | No-Effect | inactive |
| sulmazole | BRD-K01824719 | COc1cc([S@@](C)=O)ccc1-c1nc2ncccc2[nH]1 | XMFCOYRWYYXZMY-HXUWFJFHSA-N | No-Effect | inactive |
| sulmetozine | BRD-K29570907 | COc1cc(C(=S)N2CCOCC2)cc(OC)c1OC | MVOUIYUWRXPNKD-UHFFFAOYSA-N | No-Effect | inactive |
| suloctidil | BRD-K42635745 | CCCCCCCCN[C@H](C)[C@@H](O)c1ccc(SC(C)C)cc1 | BFCDFTHTSVTWOG-YLJYHZDGSA-N | No-Effect | inactive |
| sulopenem | BRD-K00004713 | C[C@@H](O)[C@H]1C(=O)N2C(C(=O)O)=C(S[C@H]3CC[S@@](=O)C3)S[C@H]12 | FLSUCZWOEMTFAQ-PRBGKLEPSA-N | No-Effect | inactive |
| sulphadimethoxine | BRD-K71125014 | COc1cc(NS(=O)(=O)c2ccc(N)cc2)nc(OC)n1 | ZZORFUFYDOWNEF-UHFFFAOYSA-N | No-Effect | inactive |
| sultamicillin | BRD-K00004694 | CC1(C)S[C@@H]2[C@H](NC(=O)[C@H](N)c3ccccc3)C(=O)N2[C@H]1C(=O)OCOC(=O)[C@@H]1N2C(=O)C[C@H]2S(=O)(=O)C1(C)C | OPYGFNJSCUDTBT-PMLPCWDUSA-N | No-Effect | inactive |
| sultiame | BRD-K00003488 | NS(=O)(=O)c1ccc(N2CCCCS2(=O)=O)cc1 | HMHVCUVYZFYAJI-UHFFFAOYSA-N | No-Effect | inactive |
| sultopride | BRD-K00003312 | CCN1CCC[C@H]1CNC(=O)c1cc(S(=O)(=O)CC)ccc1OC | UNRHXEPDKXPRTM-ZDUSSCGKSA-N | No-Effect | inactive |
| sumanirole | BRD-K30485548 | CN[C@@H]1Cc2cccc3[nH]c(=O)n(c23)C1 | RKZSNTNMEFVBDT-MRVPVSSYSA-N | No-Effect | inactive |
| sumatriptan | BRD-K50938287 | CNS(=O)(=O)Cc1ccc2[nH]cc(CCN(C)C)c2c1 | KQKPFRSPSRPDEB-UHFFFAOYSA-N | No-Effect | inactive |
| SUN-11602 | BRD-K71250460 | Cc1c(C)c(NCC(=O)N(C)C2CCN(Cc3ccc(C(N)=O)cc3)CC2)c(C)c(C)c1N | KCODNOOPOPTZMO-UHFFFAOYSA-N | No-Effect | inactive |
| SUN-B-8155 | BRD-K36274133 | C/C(=N\c1ccccc1N)c1c(C)cc(=O)n(O)c1O | SGJLINMLDNLWOS-CXUHLZMHSA-N | No-Effect | inactive |
| sunitinib | BRD-K42828737 | CCN(CC)CCNC(=O)c1c(C)[nH]c(/C=C2\C(=O)Nc3ccc(F)cc32)c1C | WINHZLLDWRZWRT-ATVHPVEESA-N | No-Effect | inactive |
| suplatast | BRD-A98427211 | CCOC[C@@H](O)COc1ccc(NC(=O)CC[S+](C)C)cc1 | DYZJXZOQQRXDLE-CQSZACIVSA-O | No-Effect | inactive |
| suprofen | BRD-A34006693 | C[C@@H](C(=O)O)c1ccc(C(=O)c2cccs2)cc1 | MDKGKXOCJGEUJW-SECBINFHSA-N | No-Effect | inactive |
| suritozole | BRD-K98557884 | Cn1nc(-c2cccc(F)c2)n(C)c1=S | IWDUZEHNLHFBRZ-UHFFFAOYSA-N | No-Effect | inactive |
| sutezolid | BRD-K86670086 | CC(=O)NC[C@H]1CN(c2ccc(N3CCSCC3)c(F)c2)C(=O)O1 | FNDDDNOJWPQCBZ-ZDUSSCGKSA-N | No-Effect | inactive |
| suxibuzone | BRD-K78815826 | CCCCC1(COC(=O)CCC(=O)O)C(=O)N(c2ccccc2)N(c2ccccc2)C1=O | ONWXNHPOAGOMTG-UHFFFAOYSA-N | No-Effect | inactive |
| swainsonine | BRD-K15170068 | O[C@@H]1[C@H](O)CN2CCC[C@@H](O)[C@H]12 | FXUAIOOAOAVCGD-WCTZXXKLSA-N | No-Effect | inactive |
| SX-011 | BRD-K45385835 | CN(C)C(=O)C(=O)c1cn(C)c2cc(Cl)c(C(=O)N3CCC(Cc4ccc(F)cc4)CC3)cc12 | GUTYHDCSDBBMGW-UHFFFAOYSA-N | No-Effect | inactive |
| SY-640 | BRD-K00004665 | CC(=O)N(C)Cc1ccc2c(c1)OCO2 | JIWDGIYBLGMPRV-UHFFFAOYSA-N | No-Effect | inactive |
| SYM-2081 | BRD-K72106461 | C[C@H](C[C@H](N)C(=O)O)C(=O)O | KRKRAOXTGDJWNI-DMTCNVIQSA-N | No-Effect | inactive |
| SYM-2206 | BRD-A77218119 | CCCNC(=O)N1N=C(c2ccc(N)cc2)c2cc3c(cc2[C@@H]1C)OCO3 | OFUDZKKOKPGXOH-LBPRGKRZSA-N | No-Effect | inactive |
| symclosene | BRD-K33642804 | O=c1n(Cl)c(=O)n(Cl)c(=O)n1Cl | YRIZYWQGELRKNT-UHFFFAOYSA-N | No-Effect | inactive |
| synephrine | BRD-K01825585 | CNC[C@@H](O)c1ccc(O)cc1 | YRCWQPVGYLYSOX-SECBINFHSA-N | No-Effect | inactive |
| T-0156 | BRD-K63516691 | COC(=O)c1c(-c2cc(OC)c(OC)c(OC)c2)c2ccnc(OCc3ncccn3)c2c(=O)n1Cc1ccnc(C)c1 | JEMJAABFSYOLAP-UHFFFAOYSA-N | No-Effect | inactive |
| T-025 | BRD-K16605796 | CNc1nc(NCc2ncccn2)c2c(-c3ccc4ncccc4c3)c[nH]c2n1 | PMWVYONICYRLNY-UHFFFAOYSA-N | No-Effect | inactive |
| T-0901317 | BRD-K23383398 | O=S(=O)(c1ccccc1)N(CC(F)(F)F)c1ccc(C(O)(C(F)(F)F)C(F)(F)F)cc1 | SGIWFELWJPNFDH-UHFFFAOYSA-N | No-Effect | inactive |
| T-1095 | BRD-K00004631 | COC(=O)OC[C@H]1O[C@@H](Oc2cc(C)cc(O)c2C(=O)CCc2ccc3occc3c2)[C@H](O)[C@@H](O)[C@@H]1O | BXNCIERBDJYIQT-PRDVQWLOSA-N | No-Effect | inactive |
| T-5224 | BRD-K84925803 | O=C(O)CCc1cc(C(=O)c2ccc(OC3CCCC3)cc2O)ccc1OCc1ccc2c(=O)[nH]oc2c1 | DALCQQSLNPLQFZ-UHFFFAOYSA-N | No-Effect | inactive |
| T-5601640 | BRD-K83492904 | Cc1cc(C(=O)Nc2cccc(C(=O)Nc3cccc(C(F)(F)F)c3)c2)on1 | XVOKFRPKSAWELK-UHFFFAOYSA-N | No-Effect | inactive |
| T-62 | BRD-K09001147 | Nc1sc2c(c1C(=O)c1ccc(Cl)cc1)CCCC2 | OTZVBZFYMFTYKH-UHFFFAOYSA-N | No-Effect | inactive |
| TA-01 | BRD-K79856806 | Fc1ccc(-c2[nH]c(-c3c(F)cccc3F)nc2-c2ccncc2)cc1 | TWPJJZCYVFUOA-UHFFFAOYSA-N | No-Effect | inactive |
| tabimorelin | BRD-K53600683 | CNC(=O)[C@@H](Cc1ccccc1)N(C)C(=O)[C@@H](Cc1ccc2ccccc2c1)N(C)C(=O)/C=C/CC(C)(C)N | WURGZWOTGMLDJP-ZCYANPAGSA-N | No-Effect | inactive |
| tacalcitol | BRD-K73237276 | C=C1/C(=C\C=C2/CCC[C@@]3(C)[C@H]2CC[C@@H]3[C@H](C)CC[C@@H](O)C(C)C)C[C@@H](O)C[C@@H]1O | BJYLYJCXYAMOFT-RSFVBTMBSA-N | No-Effect | inactive |
| tacedinaline | BRD-K52313696 | CC(=O)Nc1ccc(C(=O)Nc2ccccc2N)cc1 | VAZAPHZUAVEOMC-UHFFFAOYSA-N | No-Effect | inactive |
| tacrolimus | BRD-K69608737 | C=CC[C@@H]1/C=C(\C)C[C@H](C)C[C@H](OC)[C@H]2O[C@@](O)(C(=O)C(=O)N3CCCC[C@H]3C(=O)O[C@H](/C(C)=C/[C@@H]3CC[C@@H](O)[C@H](OC)C3)[C@H](C)[C@@H](O)CC1=O)[C@H](C)C[C@@H]2OC | QJJXYPPXXYFBGM-LFZNUXCKSA-N | No-Effect | inactive |
| tadalafil | BRD-K93645900 | CN1CC(=O)N2[C@H](Cc3c([nH]c4ccccc34)[C@H]2c2ccc3c(c2)OCO3)C1=O | WOXKDUGGOYFFRN-IIBYNOLFSA-N | No-Effect | inactive |
| tafamidis-meglumine | BRD-K23728141 | O=C(O)c1ccc2nc(-c3cc(Cl)cc(Cl)c3)oc2c1 | TXEIIPDJKFWEEC-UHFFFAOYSA-N | No-Effect | inactive |
| tagatose | BRD-K18320600 | O=C(CO)[C@@H](O)[C@@H](O)[C@H](O)CO | BJHIKXHVCXFQLS-PQLUHFTBSA-N | No-Effect | inactive |
| TAK-063 | BRD-K57155378 | COc1cn(-c2ccc(-n3cccn3)cc2F)nc(-c2ccnn2-c2ccccc2)c1=O | KVHRYLNQDWXAGI-UHFFFAOYSA-N | No-Effect | inactive |
| TAK-220 | BRD-K38793468 | CC(=O)N1CCC(C(=O)N(CCCN2CCC(Cc3ccc(C(N)=O)cc3)CC2)c2ccc(C)c(Cl)c2)CC1 | ASSJTMUEFHUKMJ-UHFFFAOYSA-N | No-Effect | inactive |
| TAK-285 | BRD-K80343549 | CC(C)(O)CC(=O)NCCn1ccc2ncnc(Nc3ccc(Oc4cccc(C(F)(F)F)c4)c(Cl)c3)c21 | ZYQXEVJIFYIBHZ-UHFFFAOYSA-N | No-Effect | inactive |
| TAK-593 | BRD-K67262033 | Cc1cc(C(=O)Nc2cc(Oc3ccc4nc(NC(=O)C5CC5)cn4n3)ccc2C)n(C)n1 | DZFZXPPHBWCXPQ-UHFFFAOYSA-N | No-Effect | inactive |
| TAK-632 | BRD-K31866293 | N#Cc1c(Oc2ccc(F)c(NC(=O)Cc3cccc(C(F)(F)F)c3)c2)ccc2nc(NC(=O)C3CC3)sc12 | OJFKUJDRGJSAQB-UHFFFAOYSA-N | No-Effect | inactive |
| TAK-659 | BRD-K71500795 | Cn1cc(-c2nc(N[C@@H]3CCCC[C@@H]3N)c(F)c3c2C(=O)NC3)cn1 | MJHOMTRKVMKCNE-NWDGAFQWSA-N | No-Effect | inactive |
| TAK-715 | BRD-K52751261 | CCc1nc(-c2cccc(C)c2)c(-c2ccnc(NC(=O)c3ccccc3)c2)s1 | HEKAIDKUDLCBRU-UHFFFAOYSA-N | No-Effect | inactive |
| TAK-733 | BRD-K26667523 | Cn1c(=O)c(F)c(Nc2ccc(I)cc2F)c2c(=O)n(C[C@@H](O)CO)cnc21 | RCLQNICOARASSR-SECBINFHSA-N | No-Effect | inactive |
| TAK-779 | BRD-K00003155 | Cc1ccc(-c2ccc3c(c2)C=C(C(=O)Nc2ccc(C[N+](C)(C)C4CCOCC4)cc2)CCC3)cc1 | XNHZXMPLVSJQFK-UHFFFAOYSA-O | No-Effect | inactive |
| TAK-875 | BRD-K58501140 | Cc1cc(OCCCS(C)(=O)=O)cc(C)c1-c1cccc(COc2ccc3c(c2)OC[C@H]3CC(=O)O)c1 | BZCALJIHZVNMGJ-HSZRJFAPSA-N | No-Effect | inactive |
| TAK-901 | BRD-K01683783 | CCS(=O)(=O)c1cccc(-c2cc(C(=O)NC3CCN(C)CC3)c(C)c3[nH]c4ncc(C)cc4c23)c1 | WKDACQVEJIVHMZ-UHFFFAOYSA-N | No-Effect | inactive |
| talabostat | BRD-K37419961 | CC(C)[C@H](N)C(=O)N1CCC[C@H]1B(O)O | FKCMADOPPWWGNZ-YUMQZZPRSA-N | No-Effect | inactive |
| taladegib | BRD-K50498985 | CN(C(=O)c1ccc(F)cc1C(F)(F)F)C1CCN(c2nnc(-c3ccnn3C)c3ccccc23)CC1 | SZBGQDXLNMELTB-UHFFFAOYSA-N | No-Effect | inactive |
| talampanel | BRD-K80359953 | CC(=O)N1N=C(c2ccc(N)cc2)c2cc3c(cc2C[C@H]1C)OCO3 | JACAAXNEHGBPOQ-LLVKDONJSA-N | No-Effect | inactive |
| talarozole | BRD-A95755460 | CCC(CC)[C@@H](c1ccc(Nc2nc3ccccc3s2)cc1)n1cncn1 | SNFYYXUGUBUECJ-FQEVSTJZSA-N | No-Effect | inactive |
| talazoparib | BRD-K95142244 | Cn1ncnc1[C@H]1c2n[nH]c(=O)c3cc(F)cc(c23)N[C@@H]1c1ccc(F)cc1 | HWGQMRYQVZSGDQ-HZPDHXFCSA-N | No-Effect | inactive |
| talinolol | BRD-A94276798 | CC(C)(C)NC[C@@H](O)COc1ccc(NC(=O)NC2CCCCC2)cc1 | MXFWWQICDIZSOA-QGZVFWFLSA-N | No-Effect | inactive |
| talmapimod | BRD-K17555800 | C[C@H]1CN(C(=O)c2cc3c(C(=O)C(=O)N(C)C)cn(C)c3cc2Cl)[C@H](C)CN1Cc1ccc(F)cc1 | ZMELOYOKMZBMRB-DLBZAZTESA-N | No-Effect | inactive |

| | | | | | |
|---|---|---|---|---|---|
| talnetant | BRD-K47095176 | CC[C@H](NC(=O)c1c(O)c(-c2ccccc2)nc2ccccc12)c1ccccc1 | BIAVGWDGIJKWRM-FQEVSTJZSA-N | No-Effect | inactive |
| talniflumate | BRD-A98378129 | O=C1O[C@H](OC(=O)c2cccnc2Nc2cccc(C(F)(F)F)c2)c2ccccc21 | ANMLJLFWUCQGKZ-HXUWFJFHSA-N | No-Effect | inactive |
| talopram | BRD-A02532772 | CNCCC[C@@]1(c2ccccc2)OC(C)(C)c2ccccc21 | LJBBMCNHIUJBDU-FQEVSTJZSA-N | No-Effect | inactive |
| talsaclidine | BRD-K00004651 | C#CCO[C@H]1CN2CCC1CC2 | XVFJONKUSLSKSW-JTQLQIEISA-N | No-Effect | inactive |
| taltirelin | BRD-K93869735 | CN1C(=O)C[C@@H](C(=O)N[C@@H](Cc2cnc[nH]2)C(=O)N2CCC[C@H]2C(N)=O)NC1=O | LQZAIAZUDWIVPM-SRVKXCTJSA-N | No-Effect | inactive |
| TAME | BRD-K72231366 | COC(=O)[C@H](CCCNC(=N)N)NS(=O)(=O)c1ccc(C)cc1 | FKMJXALNHKIDOD-LBPRGKRZSA-N | No-Effect | inactive |
| tamibarotene | BRD-K36627727 | CC1(C)CCC(C)(C)c2cc(NC(=O)c3ccc(C(=O)O)cc3)ccc21 | MUTNCGKQJGXKEM-UHFFFAOYSA-N | No-Effect | inactive |
| taminadenant | BRD-K00004682 | Nc1nc(-n2cccn2)nc(-n2cccn2)c1Br | ATFXVNUWQOXRRU-UHFFFAOYSA-N | No-Effect | inactive |
| tamoxifen | BRD-K93754473 | CC/C(=C(\c1ccccc1)c1ccc(OCCN(C)C)cc1)c1ccccc1 | NKANXQFJJICGDU-QPLCGJKRSA-N | No-Effect | inactive |
| tanaproget | BRD-K99604664 | Cn1c(C#N)ccc1-c1ccc2c(c1)C(C)(C)OC(=S)N2 | PYVFWTPEBMRKSR-UHFFFAOYSA-N | No-Effect | inactive |
| tandospirone | BRD-K65904673 | O=C1[C@H]2[C@@H]3CC[C@@H](C3)[C@H]2C(=O)N1CCCCN1CCN(c2ncccn2)CC1 | CEIJFEGBUDEYSX-FZDBZEDMSA-N | No-Effect | inactive |
| tangeritin | BRD-K25186396 | COc1ccc(-c2cc(=O)c3c(OC)c(OC)c(OC)c(OC)c3o2)cc1 | ULSUXBXHSYSGDT-UHFFFAOYSA-N | No-Effect | inactive |
| tanshinone-I | BRD-K34073885 | Cc1coc2c1C(=O)C(=O)c1c-2ccc2c(C)cccc12 | AIGAZQPHXLWMOJ-UHFFFAOYSA-N | No-Effect | inactive |
| tanshinone-IIA | BRD-K00141480 | Cc1coc2c1C(=O)C(=O)c1c-2ccc2c1CCCC2(C)C | HYXITZLLTYIPOF-UHFFFAOYSA-N | No-Effect | inactive |
| tapinarof | BRD-K00003498 | CC(C)c1c(O)cc(/C=C/c2ccccc2)cc1O | ZISJNXNHJRQYJO-CMDGGOBGSA-N | No-Effect | inactive |
| taprenepag | BRD-K88854110 | O=C(O)COc1cccc(CN(Cc2ccc(-n3cccn3)cc2)S(=O)(=O)c2cccnc2)c1 | MFFBXYNKZHTCEY-UHFFFAOYSA-N | No-Effect | inactive |
| tarafenacin | BRD-K00003568 | O=C(O[C@H]1C[N@]2CC[C@@H]1CC2)N(Cc1cc(F)c(F)c(F)c1)c1cccc(F)c1 | UXZDMXYRRQJIBJ-IBGZPJMESA-N | No-Effect | inactive |
| taranabant | BRD-K35169477 | C[C@H](NC(=O)C(C)(C)Oc1ccc(C(F)(F)F)cn1)[C@@H](Cc1ccc(Cl)cc1)c1cccc(C#N)c1 | QLYKJCMUNUWAGO-GAJHUEQPSA-N | No-Effect | inactive |
| targinine | BRD-K22227508 | CNC(=N)NCCC[C@H](N)C(=O)O | NTNWOCRCBQPEKQ-YFKPBYRVSA-N | No-Effect | inactive |
| tariquidar | BRD-K09353376 | COc1cc2c(cc1OC)CN(CCc1ccc(NC(=O)c3cc(OC)c(OC)cc3NC(=O)c3cnc4ccccc4c3)cc1)CC2 | LGGHDPFKSSRQNS-UHFFFAOYSA-N | No-Effect | inactive |
| tartaric-acid | BRD-K36474365 | O=C(O)[C@H](O)[C@@H](O)C(=O)O | FEWJPZIEWOKRBE-JCYAYHJZSA-N | No-Effect | inactive |
| TAS-103 | BRD-K09426783 | CN(C)CCNc1nc2cc(O)ccc2c2c1C(=O)c1ccccc1-2 | ROWSTIYZUWEOMM-UHFFFAOYSA-N | No-Effect | inactive |
| tasisulam | BRD-K42495768 | O=C(NS(=O)(=O)c1ccc(Br)s1)c1ccc(Cl)cc1Cl | WWONFUQGBVOKOF-UHFFFAOYSA-N | No-Effect | inactive |
| tasquinimod | BRD-K14653796 | COc1cccc2c1c(O)c(C(=O)N(C)c1ccc(C(F)(F)F)cc1)c(=O)n2C | ONDYALNGTUAJDX-UHFFFAOYSA-N | No-Effect | inactive |
| tasuldine | BRD-K10415035 | c1cnc(SCc2cccnc2)nc1 | HMCTXMOKMWELFJ-UHFFFAOYSA-N | No-Effect | inactive |
| taurine | BRD-K51474575 | NCCS(=O)(=O)O | XOAAWQZATWQOTB-UHFFFAOYSA-N | No-Effect | inactive |
| taurocholate | BRD-K81062487 | C[C@H](CCC(=O)NCCS(=O)(=O)O)[C@H]1CC[C@H]2[C@H]3[C@H](C[C@H](O)[C@@]21C)[C@@]1(C)CC[C@@H](O)C[C@H]1C[C@H]3O | WBWWGRHZICKQGZ-HZAMXZRMSA-N | No-Effect | inactive |
| tauroglycocholate | BRD-K00003523 | C[C@H](CCC(=O)NCC(=O)NCCS(=O)(=O)O)[C@H]1CC[C@H]2[C@H]3[C@H](C[C@H](O)[C@@]21C)[C@@]1(C)CC[C@@H](O)C[C@H]1C[C@H]3O | URJQSMIFSMHWSP-VVHBOOHCSA-N | No-Effect | inactive |
| taurolidine | BRD-K98765504 | O=S1(=O)CCN(CN2CCS(=O)(=O)NC2)CN1 | AJKIRUJIDFJUKJ-UHFFFAOYSA-N | No-Effect | inactive |
| tavaborole | BRD-K76713213 | OB1OCc2cc(F)ccc21 | LFQDNHWZDQTITF-UHFFFAOYSA-N | No-Effect | inactive |
| tavilermide | BRD-K00003254 | NCCCC[C@@H]1NC(=O)[C@H](CCC(=O)O)NC(=O)c2cc([N+](=O)[O-])ccc2OCC[C@@H](C(=O)NCC(=O)O)NC1=O | DVJXNXPFYJIACK-ULQDDVLXSA-N | No-Effect | inactive |
| taxifolin | BRD-K94649603 | O=C1c2c(O)cc(O)cc2O[C@H](c2ccc(O)c(O)c2)[C@H]1O | CXQWRCVTCMQVQX-LSDHHAIUSA-N | No-Effect | inactive |
| tazemetostat | BRD-K11215326 | CCN(c1cc(-c2ccc(CN3CCOCC3)cc2)cc(C(=O)NCc2c(C)cc(C)[nH]c2=O)c1C)C1CCOCC1 | NSQSAUGJQHDYNO-UHFFFAOYSA-N | No-Effect | inactive |
| tazobactam | BRD-K14312455 | C[C@]1(Cn2ccnn2)[C@H](C(=O)O)N2C(=O)C[C@H]2S1(=O)=O | LPQZKKCYTLCDGQ-WEDXCCLWSA-N | No-Effect | inactive |
| TBA-354 | BRD-K56405978 | O=[N+]([O-])c1cn2c(n1)OC[C@@H](OCc1ccc(-c3ccc(OC(F)(F)F)cc3)nc1)C2 | ZXSGSFMORAILEY-HNNXBMFYSA-N | No-Effect | inactive |
| TBOA-(DL) | BRD-M77041484 | N[C@@H](C(=O)O)[C@@H](OCc1ccccc1)C(=O)O.N[C@H](C(=O)O)[C@H](OCc1ccccc1)C(=O)O | CSEZCYBVEGIYNG-MFBWXLJUSA-N | No-Effect | inactive |
| TC-A-2317 | BRD-A12147465 | Cc1cc(Nc2cc(C)c(C#N)c(N[C@@H](C)CCCC(C)(C)O)n2)[nH]n1 | XHHRGYCOEVNAKV-ZDUSSCGKSA-N | No-Effect | inactive |
| TC-ASK-10 | BRD-K29660062 | CC(C)(C)c1ccc(C(=O)Nc2cn3cc(-n4ccnc4)ccc3n2)cc1 | SMJAMOAYGIVKRI-UHFFFAOYSA-N | No-Effect | inactive |
| TC-E-5006 | BRD-A66521565 | CC1CCN([C@@H](c2ccc(F)cc2)c2ccc([C@@H](C)C(=O)O)cc2-c2ccc(C(F)(F)F)cc2)CC1 | ONYWALXTZFPKAS-WINIVTDRSA-N | No-Effect | inactive |
| TC-F-2 | BRD-K26041695 | CCn1c(=O)n([C@H]2CCCN(c3nccc(-c4cc5ccccc5o4)n3)C2)c2ccccc21 | NXTBLPPTZRPJCA-IBGZPJMESA-N | No-Effect | inactive |
| TC-FPR-43 | BRD-K16497916 | CC(C)c1c(NC(=O)Nc2ccc(Cl)cc2)c(=O)n(-c2ccccc2)n1C | PAEBEUZTAPIOIO-UHFFFAOYSA-N | No-Effect | inactive |
| TC-G-1000 | BRD-K27010807 | Cc1sc(C)c2c1CCC=C2c1cnc[nH]1 | RLKWQKZPCMICPS-UHFFFAOYSA-N | No-Effect | inactive |
| TC-G-1004 | BRD-K76989898 | COC1CCN(c2cccc(-c3cc(NC(C)=O)nc(-n4nc(C)cc4C)n3)n2)CC1 | JENSDTKXNVHSSN-UHFFFAOYSA-N | No-Effect | inactive |
| TC-G-1005 | BRD-K16939217 | Cc1ccc(C)c(Oc2ccncc2C(=O)N2CCN(C3CC3)c3ccccc32)c1 | JQULIQJSYPZQMA-UHFFFAOYSA-N | No-Effect | inactive |
| TC-G-1008 | BRD-K87199310 | CNc1nc(NCc2ccc(NS(C)(=O)=O)cc2Cl)cc(-c2ccccn2)n1 | DRSZMILOMUPIBJ-UHFFFAOYSA-N | No-Effect | inactive |
| TC-H-106 | BRD-K45044657 | Cc1ccc(NC(=O)CCCCCC(=O)Nc2ccccc2N)cc1 | WTKBRPXPNAKVEQ-UHFFFAOYSA-N | No-Effect | inactive |
| TC-I-15 | BRD-K17621556 | CC1(C)SCN(S(=O)(=O)c2ccccc2)[C@@H]1C(=O)N[C@@H](CNC(=O)NCc1ccccc1)C(=O)O | XKLHCUGVLCGKKX-RBUKOAKNSA-N | No-Effect | inactive |
| TC-I-2000 | BRD-A83792282 | O=C(Nc1ccc(F)cc1)N1CCc2ccccc2[C@@H]1c1ccc(C(F)(F)F)cc1 | ADCDUDFEGFKKQH-NRFANRHFSA-N | No-Effect | inactive |
| TC-I-2014 | BRD-K05262258 | FC(F)(F)c1ccccc1-c1cc(C(F)(F)F)c2[nH]c(C3=NOC4(CCCCC4)C3)nc2c1 | VSMFCZZEKTVDHM-UHFFFAOYSA-N | No-Effect | inactive |
| TC-LPA5-4 | BRD-K32031961 | COc1cccc(-n2nc(C(=O)O)cc2-c2cc(C3CCCCC3)c(Cl)c2)c1 | BNALUYKEGYUHQC-UHFFFAOYSA-N | No-Effect | inactive |
| TC-Mps1-12 | BRD-K76792630 | CC(C)(C)Nc1nc(Nc2ccc(C(N)=O)cc2)cc(N)c1C#N | XDEFNAWAKYQBQY-UHFFFAOYSA-N | No-Effect | inactive |
| TC-N-1752 | BRD-K78161388 | CC(=O)Nc1cccc(Nc2ncnc(N3CCC(OCc4ccc(OC(F)(F)F)cc4)CC3)n2)c1C | QLKAFHZJICDACE-UHFFFAOYSA-N | No-Effect | inactive |
| TC-OT-39 | BRD-K55013981 | Cc1cc(C(=O)N2Cc3cnn(C)c3Nc3ccccc32)ccc1CNC(=O)N1CCC[C@H]1C(=S)N1CCCN(C)CC1 | KSNHHKZYKYNBEI-NDEPHWFRSA-N | No-Effect | inactive |
| TC-S-7003 | BRD-K34362291 | Cc1cccc(C)c1-n1c(=O)c2cnc(Nc3ccc(N4CCN(C)CC4)cc3)nc2n2c3ccccc3nc12 | BHJJWVDKNXABFS-UHFFFAOYSA-N | No-Effect | inactive |
| TC-S-7006 | BRD-K51776215 | N#Cc1cnc2cnc(NCc3cccnc3)cc2c1Nc1ccc(F)c(Cl)c1 | NMEUKWOOQOHUNA-UHFFFAOYSA-N | No-Effect | inactive |
| TC-S-7009 | BRD-K21894922 | O=[N+]([O-])c1c(Nc2cc(F)cc(Cl)c2)ccc2nonc12 | CDQUJZKBRAFWNG-UHFFFAOYSA-N | No-Effect | inactive |
| TC-SP-14 | BRD-K87461578 | O=C(O)C1CN(Cc2ccc(-c3nc4cc(Cc5ccccc5F)ccc4s3)c(F)c2)C1 | GVXGVDIXINMAAL-UHFFFAOYSA-N | No-Effect | inactive |
| TC1 | BRD-K01825674 | O[C@@]12[C@@H]3[C@@H]4[C@H]5[C@H]6C[C@@H]([C@H]53)[C@@H]1[C@H]6[C@H]4N2CCc1cccc(F)c1 | FYGREZKTJIXWIH-APDWONGJSA-N | No-Effect | inactive |
| TC2559 | BRD-K24227647 | CCOc1cncc(/C=C/CCNC)c1 | HFZDMKMXPGRKCK-GQCTYLIASA-N | No-Effect | inactive |
| TCID | BRD-K27536248 | O=C1CC(=O)c2c(Cl)c(Cl)c(Cl)c(Cl)c21 | IDLAOWFFKWRNHB-UHFFFAOYSA-N | No-Effect | inactive |
| TCN201 | BRD-K09787262 | O=C(NNC(=O)c1ccc(CNS(=O)(=O)c2ccc(F)c(Cl)c2)cc1)c1ccccc1 | FYIBXBFDXNPBSF-UHFFFAOYSA-N | No-Effect | inactive |
| TCN238 | BRD-K77915400 | Nc1nccc(/C=C/c2ccccc2)n1 | LNUXNUNUGIHCPA-VOTSOKGWSA-N | No-Effect | inactive |
| TCS-2002 | BRD-K00004294 | Cc1nnc(-c2ccc3occ(-c4ccc([S@](C)(=O)cc4)c3c2)o1 | RCKYSTKYIVULEK-DEOSSOPVSA-N | No-Effect | inactive |
| TCS-21311 | BRD-K49461364 | CC(C)(O)C(=O)N1CCN(c2ccc(C(F)(F)F)c(C3=C(c4c[nH]c5ccccc45)C(=O)NC3=O)c2)CC1 | CLGRAWDGLMENOD-UHFFFAOYSA-N | No-Effect | inactive |
| TCS-2210 | BRD-K19958299 | O=C(NO)c1ccc2[nH]c(=O)c(CCCc3ccccc3)nc2c1 | ZOFZKGFBKISZRI-UHFFFAOYSA-N | No-Effect | inactive |
| TCS-2314 | BRD-K55216666 | Cc1ccccc1NC(=O)Nc1ccc(CC(=O)N2CCOC[C@H]2C(=O)N2CCC(CC(=O)O)CC2)cc1 | ITXAAOWFOURIHK-DEOSSOPVSA-N | No-Effect | inactive |

| | | | | | |
|---|---|---|---|---|---|
| TCS-3035 | BRD-K81935556 | O=C(O)COc1ccc(/C=C2\SC(=O)NC2=O)cc1 | WXMCOLGPDOYHNK-UITAMQMPSA-N | No-Effect | inactive |
| TCS-359 | BRD-K81376179 | COc1ccc(C(=O)Nc2sc3c(c2C(N)=O)CCCC3)cc1OC | FSPQCTGGIANIJZ-UHFFFAOYSA-N | No-Effect | inactive |
| TCS-5861528 | BRD-A16560607 | CC[C@@H](C)c1ccc(NC(=O)Cn2cnc3c2c(=O)n(C)c(=O)n3C)cc1 | ZUTUWJYMCADJHD-GFCCVEGCSA-N | No-Effect | inactive |
| TCS-OX2-29 | BRD-K29424831 | COc1cc2c(cc1OC)CN(C(=O)[C@@H](NCc1ccncc1)C(C)(C)C)CC2 | COFVZFLCAOUMJT-OAQYLSRUSA-N | No-Effect | inactive |
| TCS-PIM-1-1 | BRD-K08902330 | N#Cc1c(-c2ccccc2)cc(-c2cc(Br)ccc2O)[nH]c1=O | SVSYJTYGPLVUOZ-UHFFFAOYSA-N | No-Effect | inactive |
| TCV-309 | BRD-K00003278 | CCC[n+]1cc(Br)cc(C(=O)N(CCC(=O)NCCOC(=O)N2CCc3ccccc3C2)c2ccccc2)c1 | NWWZMECSQFUIGC-UHFFFAOYSA-O | No-Effect | inactive |
| TD139 | BRD-K00003308 | OC[C@H]1O[C@@H](S[C@@H]2O[C@H](CO)[C@H](O)[C@H](n3cc(-c4cccc(F)c4)nn3)[C@H]2O)[C@H](O)[C@@H](n2cc(-c3cccc(F)c3)nn2)[C@H]1O | YGIDGBAHDZEYMT-MQFIMZJJSA-N | No-Effect | inactive |
| tebipenem | BRD-K28224173 | C[C@@H](O)[C@H]1C(=O)N2C(C(=O)OCOC(=O)C(C)(C)C)=C(SC3CN(C4=NCCS4)C3)[C@H](C)[C@H]12 | SNUDIPVBUUXCDG-QHSBEEBCSA-N | No-Effect | inactive |
| tecadenoson | BRD-K00003289 | OC[C@H]1O[C@@H](n2cnc3c(N[C@@H]4CCOC4)ncnc32)[C@H](O)[C@@H]1O | OESBDSFYJMDRJY-BAYCTPFLSA-N | No-Effect | inactive |
| tecalcet | BRD-K41662846 | COc1cccc([C@@H](C)NCCCc2ccccc2Cl)c1 | ZVQUCWXZCKWZBP-CQSZACIVSA-N | No-Effect | inactive |
| tecastemizole | BRD-K48195801 | Fc1ccc(Cn2c(NC3CCNCC3)nc3ccccc32)cc1 | SFOVDSLXFUGAIV-UHFFFAOYSA-N | No-Effect | inactive |
| tecovirimat | BRD-K24312986 | O=C(NN1C(=O)[C@H]2[C@H]3C=C[C@H]([C@@H]4C[C@H]34)[C@H]2C1=O)c1ccc(C(F)(F)F)cc1 | CSKDFZIMJXRJGH-VWLPUNTISA-N | No-Effect | inactive |
| tedizolid | BRD-K59436580 | Cn1nnc(-c2ccc(-c3ccc(N4C[C@H](CO)OC4=O)cc3F)cn2)n1 | XFALPSLJIHVRKE-GFCCVEGCSA-N | No-Effect | inactive |
| tedizolid-phosphate | BRD-K89121095 | Cn1nnc(-c2ccc(-c3ccc(N4C[C@H](COP(=O)(O)O)OC4=O)cc3F)cn2)n1 | QCGUSIANLFXSGE-GFCCVEGCSA-N | No-Effect | inactive |
| tegaserod | BRD-K88743730 | CCCCCNC(=N)N/N=C/c1c[nH]c2ccc(OC)cc12 | IKBKZGMPCYNSLU-RGVLZGJSSA-N | No-Effect | inactive |
| tegobuvir | BRD-K26701552 | Fc1ccccc1-c1nc2ccn(Cc3ccc(-c4ccc(C(F)(F)F)cc4C(F)(F)F)nn3)cc-2n1 | XBEQSQDCBSKCHJ-UHFFFAOYSA-N | No-Effect | inactive |
| teicoplanin | BRD-K07712545 | CCCCCCCCCC(=O)N[C@H]1[C@H](Oc2c3cc4cc2Oc2ccc(cc2Cl)[C@@H](O[C@@H]2O[C@H](CO)[C@@H](O)[C@H](O)[C@H]2NC(C)=O)[C@@H]2NC(=O)[C@H](NC(=O)[C@@H]4NC(=O)[C@H]4NC(=O)[C@@H](Cc5ccc(c(Cl)c5)O3)NC(=O)[C@@H](N)c3ccc(O)c(c3)Oc3cc(O)cc4c3)c3ccc(O)c(c3)-c3c(O[C@H]4O[C@H](CO)[C@@H](O)[C@H](O)[C@@H]4O)cc(O)cc3[C@H](C(=O)O)NC2=O)O[C@H](CO)[C@@H](O)[C@@H]1O | BJNLLBUOHPVGFT-VPMXZCLFSA-N | No-Effect | inactive |
| teicoplanin-A2-1 | BRD-A09761158 | CCCCC/C=C/CCC(=O)N[C@H]1[C@H](Oc2c3cc4cc2Oc2ccc(cc2Cl)[C@@H](O[C@@H]2O[C@H](CO)[C@@H](O)[C@H](O)[C@H]2NC(C)=O)[C@@H]2NC(=O)[C@H](NC(=O)[C@@H]4NC(=O)[C@H]4NC(=O)[C@@H](Cc5ccc(c(Cl)c5)O3)NC(=O)[C@H](N)c3ccc(O)c(c3)Oc3cc(O)cc4c3)c3ccc(O)c(c3)-c3c(O[C@H]4O[C@H](CO)[C@@H](O)[C@H](O)[C@@H]4O)cc(O)cc3[C@@H](C(=O)O)NC2=O)O[C@H](CO)[C@@H](O)[C@@H]1O | YMWQIYMUNZBOOV-SFRGEMKNSA-N | No-Effect | inactive |
| teicoplanin-A2-3 | BRD-K96571715 | CC(=O)N[C@@H]1[C@H](O[C@@H]2c3ccc(c(Cl)c3)Oc3cc4cc(c3O[C@@H]3O[C@H](CO)[C@@H](O)[C@H](O)[C@@H]3NC(=O)CCCCCCCC(C)C)Oc3ccc(cc3Cl)C[C@H]3NC(=O)[C@H](N)c5ccc(O)c(c5)Oc5cc(O)cc(c5)[C@H](NC3=O)C(=O)N[C@@H]4C(=O)N[C@@H]3C(=O)N[C@@H]2C(=O)N[C@@H](C(=O)O)c2cc(O)cc(O[C@H]4O[C@H](CO)[C@@H](O)[C@H](O)[C@@H]4O)c2-c2cc3ccc2O)O[C@H](CO)[C@@H](O)[C@@H]1O | FHBQKTSCJKPYIO-UBGUEUOASA-N | No-Effect | inactive |
| teijin-compound-1 | BRD-K51528698 | O=C(CNC(=O)c1cccc(C(F)(F)F)c1)N[C@@H]1CCN(Cc2ccc(Cl)cc2)C1 | BAOQJSULMWXFRK-GOSISDBHSA-N | No-Effect | inactive |
| telatinib | BRD-K84564571 | CNC(=O)c1cc(COc2nnc(Nc3ccc(Cl)cc3)c3ccoc23)ccn1 | QFCXANHHBCGMAS-UHFFFAOYSA-N | No-Effect | inactive |
| telbivudine | BRD-K15976406 | Cc1cn([C@@H]2C[C@@H](O)[C@H](CO)O2)c(=O)[nH]c1=O | IQFYYKKMVGJFEH-CSMHCCOUSA-N | No-Effect | inactive |
| telcagepant | BRD-K91159026 | O=C(N[C@@H]1CC[C@@H](c2cccc(F)c2F)CN(CC(F)(F)F)C1=O)N1CCC(n2c(=O)[nH]c3ncccc32)CC1 | CGDZXLJGHVKVIE-DNVCBOLYSA-N | No-Effect | inactive |
| telenzepine | BRD-K06147391 | Cc1scc2c1N(C(=O)CN1CCN(C)CC1)c1ccccc1NC2=O | VSWPGAIWKHPTKX-UHFFFAOYSA-N | No-Effect | inactive |
| telithromycin | BRD-K82240271 | CC[C@H]1OC(=O)[C@H](C)C(=O)[C@H](C)[C@@H](O[C@@H]2O[C@H](C)C[C@H](N(C)C)[C@H]2O)[C@](C)(OC)C[C@@H](C)C(=O)[C@H](C)[C@H]2N(CCCCn3cnc(-c4cccnc4)c3)C(=O)O[C@]12C | LJVAJPDWBABPEJ-PNUFFHFMSA-N | No-Effect | inactive |
| telmisartan | BRD-K73999723 | CCCc1nc2c(C)cc(-c3nc4ccccc4n3C)cc2n1Cc1ccc(-c2ccccc2C(=O)O)cc1 | RMMXLENWKUUMAY-UHFFFAOYSA-N | No-Effect | inactive |
| telotristat | BRD-K63178091 | Cc1ccn(-c2cc(Cl)ccc2[C@@H](Oc2cc(-c3ccc(C[C@H](N)C(=O)O)cc3)nc(N)n2)C(F)(F)F)n1 | NCLGDOBQAWBXRA-PGRDOPGGSA-N | No-Effect | inactive |
| telratolimod | BRD-K00004596 | CCCCCCCCCCCCCCCCCC(=O)NCCCCOn1c(CCCC)nc2c(N)nc3ccccc3c21 | RRTPWQXEERTRRK-UHFFFAOYSA-N | No-Effect | inactive |
| temanogrel | BRD-K00003467 | COc1cccc(C(=O)Nc2ccc(OCCN3CCOCC3)c(-c3ccnn3C)c2)c1 | ZEOQUKRCASTCFR-UHFFFAOYSA-N | No-Effect | inactive |
| temefos | BRD-K51805276 | COP(=S)(OC)Oc1ccc(Sc2ccc(OP(=S)(OC)OC)cc2)cc1 | WWJZWCUNLNYYAU-UHFFFAOYSA-N | No-Effect | inactive |
| temocapril | BRD-K10784860 | CCOC(=O)[C@H](CCc1ccccc1)N[C@H]1CS[C@H](c2cccs2)CN(CC(=O)O)C1=O | FIQOFIRCTOWDOW-BJLQDIEVSA-N | No-Effect | inactive |
| temozolomide | BRD-K32107296 | Cn1nnc2c(C(N)=O)ncn2c1=O | BPEGJWRSRHCHSN-UHFFFAOYSA-N | No-Effect | inactive |
| tempol | BRD-K95053546 | CC1(C)CC(O)CC(C)(C)N1O | CSGAUKGQUCHWDP-UHFFFAOYSA-N | No-Effect | inactive |
| temsirolimus | BRD-K08177763 | CO[C@H]1C[C@@H]2CC[C@@H](C)[C@@](O)(O2)C(=O)C(=O)N2CCCC[C@H]2C(=O)O[C@H]([C@H](C)C[C@@H]2CC[C@@H](OC(=O)C(C)(CO)CO)[C@H](OC)C2)CC(=O)[C@H](C)/C=C(\C)[C@@H](O)[C@@H](OC)C(=O)[C@H](C)C[C@H](C)/C=C/C=C/C=C/1C | CBPNZQVSJQDFBE-FUXHJELOSA-N | No-Effect | inactive |
| tenalisib | BRD-K00003263 | CC[C@H](Nc1ncnc2nc[nH]c12)c1oc2ccccc2c(=O)c1-c1cccc(F)c1 | HDXDQPRPFRKGKZ-INIZCTEOSA-N | No-Effect | inactive |
| tenapanor | BRD-K00003201 | CN1Cc2c(Cl)cc(Cl)cc2[C@H](c2cccc(S(=O)(=O)NCCOCCOCCNC(=O)NCCCCNC(=O)NCCOCCOCCNS(=O)(=O)c3cccc([C@@H]4CN(C)Cc5c(Cl)cc(Cl)cc54)c3)c2)C1 | DNHPDWGIXIMXSA-CXNSMIOJSA-N | No-Effect | inactive |
| tenatoprazole | BRD-A42270467 | COc1ccc2[nH]c([S@@](=O)Cc3ncc(C)c(OC)c3C)nc2n1 | ZBFDAUIVDSSISP-DEOSSOPVSA-N | No-Effect | inactive |
| teneligliptin | BRD-K48068743 | Cc1cc(N2CCN([C@@H]3CN[C@H](C(=O)N4CCSC4)C3)CC2)n(-c2ccccc2)n1 | WGRQANOPCQRCME-PMACEKPBSA-N | No-Effect | inactive |
| tenidap | BRD-K00004279 | NC(=O)N1C(=O)[C@@H](C(=O)c2cccs2)c2cc(Cl)ccc21 | IGPDWKCUDHIIRL-LLVKDONJSA-N | No-Effect | inactive |
| tenilsetam | BRD-A41835910 | O=C1NCCN[C@@H]1c1cccs1 | BBACKDMGYMXYLE-SSDOTTSWSA-N | No-Effect | inactive |
| tenofovir | BRD-K15891719 | C[C@H](Cn1cnc2c(N)ncnc21)OCP(=O)(O)O | SGOIRFVFHAKUTI-ZCFIWIBFSA-N | No-Effect | inactive |
| tenofovir-alafenamide | BRD-A25988899 | CC(C)OC(=O)[C@@H](C)NP(=O)(CO[C@@H](C)Cn1cnc2c(N)ncnc21)Oc1ccccc1 | LDEKQSIMHVQZJK-FOCZCJQRSA-N | No-Effect | inactive |
| tenofovir-disoproxil | BRD-K61868322 | CC(C)OC(=O)OCOP(=O)(CO[C@H](C)Cn1cnc2c(N)ncnc21)OCOC(=O)OC(C)C | JFVZFKDSXNQEJW-CQSZACIVSA-N | No-Effect | inactive |
| tenovin-1 | BRD-K05977823 | CC(=O)Nc1ccc(NC(=S)NC(=O)c2ccc(C(C)(C)C)cc2)cc1 | WOWJIWFCOPZFGV-UHFFFAOYSA-N | No-Effect | inactive |
| tenylidone | BRD-K91298281 | O=C1/C(=C/c2cccs2)CCC/C1=C\c1cccs1 | KHXBULXCCPIELT-DCIPZJNNSA-N | No-Effect | inactive |
| tepoxalin | BRD-K89714990 | COc1ccc(-n2nc(CCC(=O)N(C)O)cc2-c2ccc(Cl)cc2)cc1 | XYKWNRUXCOIMFZ-UHFFFAOYSA-N | No-Effect | inactive |
| TEPP-46 | BRD-K01826017 | Cn1c2cc([S@@](C)=O)sc2c2cnn(Cc3cccc(N)c3)c(=O)c21 | ZWKJWVSEDISQIS-RUZDIDTESA-N | No-Effect | inactive |
| teprenone | BRD-K11510141 | CC(=O)CC/C=C(\C)CC/C=C(\C)CC/C=C(\C)CCC=C(C)C | HUCXKZBETONXFO-NJFMWZAGSA-N | No-Effect | inactive |

| | | | | | |
|---|---|---|---|---|---|
| terameprocol | BRD-K76788410 | COc1ccc(C[C@H](C)[C@H](C)Cc2ccc(OC)c(OC)c2)cc1OC | ORQFDHFZSMXRLM-IYBDPMFKSA-N | No-Effect | inactive |
| terazosin | BRD-K01826569 | COc1cc2nc(N3CCN(C(=O)[C@H]4CCCO4)CC3)nc(N)c2cc1OC | VCKUSRYTPJJLNI-CQSZACIVSA-N | No-Effect | inactive |
| terbinafine | BRD-K68132782 | CN(C/C=C/C#CC(C)(C)C)Cc1cccc2ccccc12 | DOMXUEMWDBAQBQ-WEVVVXLNSA-N | No-Effect | inactive |
| terbutaline | BRD-A50157456 | CC(C)(C)NC[C@@H](O)c1cc(O)cc(O)c1 | XWTYSIMOBUGWOL-LLVKDONJSA-N | No-Effect | inactive |
| terciprazine | BRD-K01826018 | C#CC1(OC[C@H](O)CN2CCN(c3cccc(C(F)(F)F)c3)CC2)CCCCC1 | SDKXUWMRKYQIFU-HXUWFJFHSA-N | No-Effect | inactive |
| terguride | BRD-A89207210 | CCN(CC)C(=O)N[C@H]1C[C@@H]2c3cccc4[nH]cc(c34)C[C@H]2N(C)C1 | JOAHPSVPXZTVEP-YXJHDRRASA-N | No-Effect | inactive |
| teriflunomide | BRD-A42699921 | CC(=O)[C@@H](C#N)C(=O)Nc1ccc(C(F)(F)F)cc1 | NABQBYDEMLGCEO-SNVBAGLBSA-N | No-Effect | inactive |
| terlipressin | BRD-K73451719 | NCCCC[C@H](NC(=O)[C@@H]1CCCN1C(=O)[C@@H]1CSSC[C@H](NC(=O)CNC(=O)CNC(=O)CN)C(=O)N[C@@H](Cc2ccc(O)cc2)C(=O)N[C@@H](Cc2ccccc2)C(=O)N[C@@H](CCC(N)=O)C(=O)N[C@@H](CC(N)=O)C(=O)N1)C(=O)NCC(N)=O | BENFXAYNYRLAIU-QSVFAHTRSA-N | No-Effect | inactive |
| terodiline | BRD-A33479817 | C[C@@H](CC(c1ccccc1)c1ccccc1)NC(C)(C)C | UISARWKNNNHPGI-INIZCTEOSA-N | No-Effect | inactive |
| teroxirone | BRD-A39935389 | O=c1n(C[C@H]2CO2)c(=O)n(C[C@H]2CO2)c(=O)n1C[C@H]1CO1 | OUPZKGBUJRBPGC-CIUDSAMLSA-N | No-Effect | inactive |
| terpene | BRD-K00003674 | CC(C)(O)[C@H]1CC[C@](C)(O)CC1 | RBNWAMSGVWEHFP-CZMCAQCFSA-N | No-Effect | inactive |
| terreic-acid-(-) | BRD-K00024725 | C[C@H]1C(=O)C(=O)[C@@H]2O[C@@H]2C1=O | NYFMSQHOGRWGFD-OOVQLBIQSA-N | No-Effect | inactive |
| terutroban | BRD-K43975632 | Cc1ccc2c(c1CCC(=O)O)CC[C@@H](NS(=O)(=O)c1ccc(Cl)cc1)C2 | HWEOXFSBSQIWSY-MRXNPFEDSA-N | No-Effect | inactive |
| tetomilast | BRD-K00004564 | CCOc1ccc(-c2nc(-c3cccc(C(=O)O)n3)cs2)cc1OCC | XDBHURGONHZNJF-UHFFFAOYSA-N | No-Effect | inactive |
| tetrabenazine | BRD-K95739795 | COc1cc2c(cc1OC)[C@@H]1CC(=O)[C@@H](CC(C)C)CN1CC2 | MKJIEFSOBYUXJB-HOCLYGCPSA-N | No-Effect | inactive |
| tetracaine | BRD-K45071273 | CCCCNc1ccc(C(=O)OCCN(C)C)cc1 | GKCBAIGFKIBETG-UHFFFAOYSA-N | No-Effect | inactive |
| tetracycline | BRD-K00003352 | CN(C)[C@@H]1C(=O)[C@H](C(N)=O)C(=O)[C@@]2(O)C(=O)[C@@H]3C(=O)c4c(O)cccc4[C@@](C)(O)[C@H]3C[C@@H]12 | MUHBBHLZPHKTTR-VHOYBAKPSA-N | No-Effect | inactive |
| tetradecylthioacetic-acid | BRD-K47539947 | CCCCCCCCCCCCCCSCC(=O)O | IPBCWPPBAWQYOO-UHFFFAOYSA-N | No-Effect | inactive |
| tetraethylenepentamine | BRD-K41298358 | NCCNCCNCCNCCN | FAGUFWYHJQFNRV-UHFFFAOYSA-N | No-Effect | inactive |
| tetrahydrofolic-acid | BRD-A36978621 | Nc1nc(=O)c2c([nH]1)NC[C@H](CNc1ccc(C(=O)N[C@@H](CCC(=O)O)C(=O)O)cc1)N2 | MSTNYGQPCMXVAQ-RYUDHWBXSA-N | No-Effect | inactive |
| tetrahydrouridine | BRD-A86248581 | O=C1N[C@H](O)CCN1[C@@H]1O[C@H](CO)[C@@H](O)[C@H]1O | UCKYOOZPSJFJIZ-FMDGEEDCSA-N | No-Effect | inactive |
| tetrahydrozoline | BRD-A28856712 | c1ccc2c(c1)CCC[C@@H]2C1=NCCN1 | BYJAVTDNIXVSPW-LBPRGKRZSA-N | No-Effect | inactive |
| tetramethylthiuram-monosulfide | BRD-K36889451 | CN(C)C(=S)SC(=S)N(C)C | REQPQFUJGGOFQL-UHFFFAOYSA-N | No-Effect | inactive |
| tetramisole | BRD-A46393198 | c1ccc([C@H]2CN3CCSC3=N2)cc1 | HLFSDGLLUJUHTE-SNVBAGLBSA-N | No-Effect | inactive |
| tetrazol-5-yl-glycine-(RS) | BRD-A14757586 | N[C@@H](C(=O)O)c1nn[nH]n1 | UKBRUIZWQZHXFL-PVQJCKRUSA-N | No-Effect | inactive |
| tetroquinone | BRD-K90177246 | O=C1C(=O)C(O)C(=O)C(=O)C1O | GQPHQLHUIDOQTA-UHFFFAOYSA-N | No-Effect | inactive |
| tezacaftor | BRD-K97101532 | CC(C)(CO)c1cc2cc(NC(=O)C3(c4ccc5c(c4)OC(F)(F)O5)CC3)c(F)cc2n1C[C@@H](O)CO | MJUVRTYWUMPBTR-MRXNPFEDSA-N | No-Effect | inactive |
| TFC-007 | BRD-K57708067 | O=C(Nc1ccc(N2CCC(C(=O)N3CCOCC3)CC2)cc1)c1cnc(Oc2ccccc2)nc1 | NLSSUSRERAMBTA-UHFFFAOYSA-N | No-Effect | inactive |
| TG-003 | BRD-K97072811 | CCN1/C(=C/C(C)=O)Sc2ccc(OC)cc21 | BGVLELSCIHASRV-QPEQYQDCSA-N | No-Effect | inactive |
| TG-100713 | BRD-K46392623 | Nc1nc(N)c2nc(-c3cccc(O)c3)cnc2n1 | UOORQSPLBHUQDQ-UHFFFAOYSA-N | No-Effect | inactive |
| TG-100801 | BRD-K12735121 | Cc1cc(-c2cc(OC(=O)c3ccccc3)ccc2Cl)cc2nnc(Nc3ccc(OCCN4CCCC4)cc3)nc12 | JMGXJHWTVBGOKG-UHFFFAOYSA-N | No-Effect | inactive |
| TG100-115 | BRD-K64785675 | Nc1nc(N)c2nc(-c3cccc(O)c3)c(-c3cccc(O)c3)nc2n1 | UJIAQDJKSXQLIT-UHFFFAOYSA-N | No-Effect | inactive |
| TGR-1202 | BRD-K52579889 | CC(C)Oc1ccc(-c2nn([C@@H](C)c3oc4ccc(F)cc4c(=O)c3-c3cccc(F)c3)c3ncnc(N)c23)cc1F | IUVCFHHAEHNCFT-INIZCTEOSA-N | No-Effect | inactive |
| TGX-221 | BRD-A41692738 | Cc1cc([C@@H](C)Nc2ccccc2)c2nc(N3CCOCC3)cc(=O)n2c1 | CPRAGQJXBLMUEL-MRXNPFEDSA-N | No-Effect | inactive |
| TH-302 | BRD-K20958582 | Cn1c(COP(=O)(NCCBr)NCCBr)cnc1[N+](=O)[O-] | UGJWRPJDTDGERK-UHFFFAOYSA-N | No-Effect | inactive |
| TH-588 | BRD-K01054709 | Nc1nc(NC2CC2)cc(-c2cccc(Cl)c2Cl)n1 | PNMYJIOQIAEYQL-UHFFFAOYSA-N | No-Effect | inactive |
| thalidomide | BRD-A93255169 | O=C1CC[C@H](N2C(=O)c3ccccc3C2=O)C(=O)N1 | UEJJHQNACJXSKW-VIFPVBQESA-N | No-Effect | inactive |
| theaflavin | BRD-K60842365 | O=c1c(O)cc([C@H]2Oc3cc(O)cc(O)c3C[C@H]2O)c2cc([C@H]3Oc4cc(O)cc(O)c4C[C@H]3O)cc(O)c(O)c1-2 | SPQUOZYCYRJYJU-ZKSIBHASSA-N | No-Effect | inactive |
| theobromine | BRD-K34888156 | Cn1cnc2c1c(=O)[nH]c(=O)n2C | YAPQBXQYLJRXSA-UHFFFAOYSA-N | No-Effect | inactive |
| thiamine | BRD-K95785345 | Cc1ncc(C[n+]2csc(CCO)c2C)c(N)n1 | JZRWCGZRTZMZEH-UHFFFAOYSA-N | No-Effect | inactive |
| thiamine-pyrophosphate | BRD-A09994331 | Cc1ncc(C[n+]2csc(CCOP(=O)(O)OP(=O)(O)O)c2C)c(N)n1 | AYEKOFBPNLCAJY-UHFFFAOYSA-O | No-Effect | inactive |
| thiamphenicol | BRD-K79711234 | CS(=O)(=O)c1ccc([C@@H](O)[C@@H](CO)NC(=O)C(Cl)Cl)cc1 | OTVAEFIXJLOWRX-NXEZZACHSA-N | No-Effect | inactive |
| thiamylal | BRD-A23970436 | C=CCC1([C@H](C)CCC)C(=O)NC(=S)NC1=O | XLOMZPUITCYLMJ-MRVPVSSYSA-N | No-Effect | inactive |
| thiazovivin | BRD-K03164761 | O=C(NCc1ccccc1)c1csc(Nc2ccncn2)n1 | DOBKQCZBPPCLEG-UHFFFAOYSA-N | No-Effect | inactive |
| thioctic-acid | BRD-A16347691 | O=C(O)CCCC[C@H]1CCSS1 | AGBQKNBQESQNJD-ZETCQYMHSA-N | No-Effect | inactive |
| thiodiglycol | BRD-K54521836 | OCCSCCO | YODZTKMDCQEPHD-UHFFFAOYSA-N | No-Effect | inactive |
| thiopental | BRD-A72603112 | CCC[C@@H](C)C1(CC)C(=O)NC(=S)NC1=O | IUJDSEJGGMCXSG-SSDOTTSWSA-N | No-Effect | inactive |
| thioperamide | BRD-A85424553 | S=C(NC1CCCCC1)N1CCC([C@H]2C=NC=N2)CC1 | JFAYRTHYLOZRNL-CQSZACIVSA-N | No-Effect | inactive |
| thiophanate | BRD-K90168339 | COC(=O)NC(=S)Nc1ccccc1NC(=S)NC(=O)OC | QGHREAKMXXNCOA-UHFFFAOYSA-N | No-Effect | inactive |
| thiorphan | BRD-A56012032 | O=C(O)CNC(=O)[C@@H](CS)Cc1ccccc1 | LJJKNPQAGWVLDQ-SNVBAGLBSA-N | No-Effect | inactive |
| thiotepa | BRD-K09631521 | S=P(N1CC1)(N1CC1)N1CC1 | FOCVUCIESVLUNU-UHFFFAOYSA-N | No-Effect | inactive |
| thiothixene | BRD-K04175245 | CN1CCN(CC/C=C2/c3ccccc3Sc3ccc(S(=O)(=O)N(C)C)cc32)CC1 | GFBKORZTTCHDGY-UWVJOHFNSA-N | No-Effect | inactive |
| thiram | BRD-K29254801 | CN(C)C(=S)SSC(=S)N(C)C | KUAZQDVKQLNFPE-UHFFFAOYSA-N | No-Effect | inactive |
| thonzonium | BRD-K13387373 | CCCCCCCCCCCCCCCC[N+](C)(C)CCN(Cc1ccc(OC)cc1)c1ncccn1 | IOYZYMQFUSNATM-UHFFFAOYSA-N | No-Effect | inactive |
| thonzylamine | BRD-K88405679 | COc1ccc(CN(CCN(C)C)c2ncccn2)cc1 | GULNIHOSWFYMRN-UHFFFAOYSA-N | No-Effect | inactive |
| threo-2-methylisocitrate-(DL) | BRD-M01344114 | CC(O)(C(=O)O)[C@@H](CC(=O)O)C(=O)O.CC(O)(C(=O)O)[C@H](CC(=O)O)C(=O)O.[NaH].[NaH].[NaH].[NaH].[NaH].[NaH] | ZSOGGZWFKNMGJN-WREOTHMNSA-N | No-Effect | inactive |
| thymalfasin | BRD-K00004602 | CC[C@H](C)[C@H](NC(=O)[C@H](CCC(=O)O)NC(=O)[C@H](CO)NC(=O)[C@H](CO)NC(=O)[C@@H](NC(=O)[C@H](CC(=O)O)NC(=O)[C@@H](NC(=O)[C@H](C)NC(=O)[C@H](C)NC(=O)[C@H](CC(=O)O)NC(=O)[C@H](CO)NC(C)=O)C(C)C)[C@@H](C)O)C(=O)N[C@H](C(=O)N[C@H](C(=O)N[C@@H](CCCCN)C(=O)N[C@@H](CC(=O)O)C(=O)N[C@@H](CC(C)C)C(=O)N[C@@H](CCCCN)C(=O)N[C@@H](CCC(=O)O)C(=O)N[C@@H](CCCCN)C(=O)N[C@@H](CCCCN)C(=O)N[C@@H](CCC(=O)O)C(=O)N[C@H](C(=O)N[C@H](C(=O)N[C@@H](CCC(=O)O)C(=O)N[C@@H](CCC(=O)O)C(=O)N[C@@H](C)C(=O)N[C@@H](CCC(=O)O)C(=O)N[C@@H](CC(N)=O)C(=O)O)C(C)C)C(C)C)[C@@H](C)O)[C@@H](C)O | NZVYCXVTEHPMHE-ZSUJOUNUSA-N | No-Effect | inactive |
| thymol-iodide | BRD-K36617478 | Cc1cc(OI)c(C(C)C)cc1-c1cc(C(C)C)c(OI)cc1C | SHOKWSLXDAIZPP-UHFFFAOYSA-N | No-Effect | inactive |
| thymopentin | BRD-K43977121 | CC(C)[C@H](NC(=O)[C@H](CC(=O)O)NC(=O)[C@H](CCCCN)NC(=O)[C@@H](N)CCCNC(=N)N)C(=O)N[C@@H](Cc1ccc(O)cc1)C(=O)O | PSWFFKRAVBDQEG-YGQNSOCVSA-N | No-Effect | inactive |

| | | | | | |
|---|---|---|---|---|---|
| THZ1 | BRD-K66401437 | CN(C)C/C=C/C(=O)Nc1ccc(C(=O)Nc2cccc(Nc3ncc(Cl)c(-c4c[nH]c5ccccc45)n3)c2)cc1 | OBJNFLYHUXWUPF-IZZDOVSWSA-N | No-Effect | inactive |
| THZ1-R | BRD-K02135007 | CN(C)CCCC(=O)Nc1ccc(C(=O)Nc2cccc(Nc3ncc(Cl)c(-c4c[nH]c5ccccc45)n3)c2)cc1 | TUERFPPIPKZNKE-UHFFFAOYSA-N | No-Effect | inactive |
| THZ2 | BRD-K95902121 | CN(C)C/C=C/C(=O)Nc1cccc(C(=O)Nc2cccc(Nc3ncc(Cl)c(-c4c[nH]c5ccccc45)n3)c2)c1 | FONRCZUZCHXWBD-VGOFMYFVSA-N | No-Effect | inactive |
| tiabendazole | BRD-K77695569 | c1ccc2[nH]c(-c3cscn3)nc2c1 | WJCNZQLZVWNLKY-UHFFFAOYSA-N | No-Effect | inactive |
| tiagabine | BRD-K60160658 | Cc1ccsc1C(=CCCN1CCC[C@@H](C(=O)O)C1)c1sccc1C | PBJUNZJWGZTSKL-MRXNPFEDSA-N | No-Effect | inactive |
| tianeptine | BRD-K01826304 | CN1c2ccccc2[C@@H](NCCCCCCC(=O)O)c2ccc(Cl)cc2S1(=O)=O | JICJBGPOMZQUBB-OAQYLSRUSA-N | No-Effect | inactive |
| tiapride | BRD-K57432881 | CCN(CC)CCNC(=O)c1cc(S(C)(=O)=O)ccc1OC | JTVPZMFULRWINT-UHFFFAOYSA-N | No-Effect | inactive |
| tiaprofenic-acid | BRD-A72988804 | C[C@@H](C(=O)O)c1ccc(C(=O)c2ccccc2)s1 | GUHPRPJDBZHYCJ-SECBINFHSA-N | No-Effect | inactive |
| tibolone | BRD-A82522119 | C#C[C@]1(O)CC[C@@H]2[C@@H]3[C@H](CC[C@@]21C)C1=C(CC(=O)CC1)C[C@H]3C | WZDGZWOAQTVYBX-TZTUDTKPSA-N | No-Effect | inactive |
| TIC10 | BRD-K19939891 | Cc1ccccc1CN1C2=NCCN2C(=O)C2=C1CCN(Cc1ccccc1)C2 | RSAQARAFWMUYLL-UHFFFAOYSA-N | No-Effect | inactive |
| ticlopidine | BRD-K00603606 | Clc1ccccc1CN1CCc2sccc2C1 | PHWBOXQYWZNQIN-UHFFFAOYSA-N | No-Effect | inactive |
| tideglusib | BRD-K81330143 | O=c1sn(-c2cccc3ccccc23)c(=O)n1Cc1ccccc1 | PMJIHLSCWIDGMD-UHFFFAOYSA-N | No-Effect | inactive |
| tie2-kinase-inhibitor | BRD-A92800748 | COc1ccc2cc(-c3[nH]c(-c4ccc([S@@](C)=O)cc4)nc3-c3ccncc3)ccc2c1 | SINQIEAULQKUPD-JGCGQSQUSA-N | No-Effect | inactive |
| tienilic-acid | BRD-K34098590 | O=C(O)COc1ccc(C(=O)c2cccs2)c(Cl)c1Cl | AGHANLSBXUWXTB-UHFFFAOYSA-N | No-Effect | inactive |
| tifenazoxide | BRD-K20568871 | CC1(NC2=NS(=O)(=O)c3sc(Cl)cc3N2)CC1 | KYSFUHHFTIGRJN-UHFFFAOYSA-N | No-Effect | inactive |
| tigecycline | BRD-K00003349 | CN(C)c1cc(NC(=O)CNC(C)(C)C)c(O)c2c1C[C@H]1C[C@H]3[C@H](N(C)C)C(=O)[C@@H](C(N)=O)C(=O)[C@@]3(O)C(=O)[C@H]1C2=O | CGZOGNUFXMNYEI-PBZRBWFISA-N | No-Effect | inactive |
| tilmicosin | BRD-K36796419 | CC[C@H]1OC(=O)C[C@@H](O)[C@H](C)[C@@H](O[C@@H]2O[C@H](C)[C@@H](O)[C@H](N(C)C)[C@H]2O)[C@@H](CCN2C[C@@H](C)C[C@@H](C)C2)C[C@@H](C)C(=O)/C=C/C(C)=C/[C@@H]1CO[C@@H]1O[C@H](C)[C@@H](O)[C@@H](OC)[C@H]1OC | JTSDBFGMPLKDCD-XVFHVFLVSA-N | No-Effect | inactive |
| tiludronate | BRD-K10845183 | O=P(O)(O)C(Sc1ccc(Cl)cc1)P(=O)(O)O | DKJJVAGXPKPDRL-UHFFFAOYSA-N | No-Effect | inactive |
| timofibrate | BRD-A64406146 | CC(C)(Oc1ccc(Cl)cc1)C(=O)N1CSC[C@@H]1C(=O)O | QSNOWCXXECLALL-LLVKDONJSA-N | No-Effect | inactive |
| timolol | BRD-K08806317 | CC(C)(C)NC[C@H](O)COc1nsnc1N1CCOCC1 | BLJRIMJGRPQVNF-JTQLQIEISA-N | No-Effect | inactive |
| timonacic | BRD-K00003683 | O=C(O)[C@H]1CSCN1 | DZLNHFMRPBPULJ-GSVOUGTGSA-N | No-Effect | inactive |
| tinazoline | BRD-K00004663 | c1ccc2c(SC3=NCCN3)c[nH]c2c1 | JENBDJGHNIOHAJ-UHFFFAOYSA-N | No-Effect | inactive |
| tinidazole | BRD-K89125793 | CCS(=O)(=O)CCn1c([N+](=O)[O-])cnc1C | HJLSLZFTEKNLFI-UHFFFAOYSA-N | No-Effect | inactive |
| tioconazole | BRD-K00003630 | Clc1ccc([C@H](Cn2ccnc2)OCc2ccsc2Cl)c(Cl)c1 | QXHHHPZILQDDPS-HNNXBMFYSA-N | No-Effect | inactive |
| tiopronin | BRD-A85295731 | C[C@@H](S)C(=O)NCC(=O)O | YTGJWQPHMWSCST-GSVOUGTGSA-N | No-Effect | inactive |
| tiotidine | BRD-K11107424 | CN=C(NC#N)NCCSCc1csc(NC(=N)N)n1 | YDDXVAXDYKBWDX-UHFFFAOYSA-N | No-Effect | inactive |
| tiotropium | BRD-K62782754 | C[N+]1(C)[C@H]2C[C@H](OC(=O)C(O)(c3cccs3)c3cccs3)C[C@@H]1[C@H]1O[C@@H]21 | LERNTVKEWCAPOY-DZZGSBJMSA-N | No-Effect | inactive |
| tipelukast | BRD-K00004627 | CCCc1c(SCCCOc2ccc(C(C)=O)c(OCCCC(=O)O)c2CCC)ccc(C(C)=O)c1O | KPWYNAGOBXLMSE-UHFFFAOYSA-N | No-Effect | inactive |
| tipifarnib | BRD-K62965247 | Cn1cncc1[C@@](N)(c1ccc(Cl)cc1)c1ccc2c(c1)c(-c1cccc(Cl)c1)cc(=O)n2C | PLHJCIYEEKOWNM-HHHXNRCGSA-N | No-Effect | inactive |
| tiplaxtinin | BRD-K30882994 | O=C(O)C(=O)c1cn(Cc2ccccc2)c2ccc(-c3ccc(OC(F)(F)F)cc3)cc12 | ODXQFEWQSHNQNI-UHFFFAOYSA-N | No-Effect | inactive |
| tipranavir | BRD-A10039652 | CCC[C@@]1(CCc2ccccc2)CC(=O)[C@@H]([C@H](CC)c2cccc(NS(=O)(=O)c3ccc(C(F)(F)F)cn3)c2)C(=O)O1 | MYXACKZCYFKMTE-WWKQIGMOSA-N | No-Effect | inactive |
| tiprenolol | BRD-A73566563 | CSc1ccccc1OC[C@@H](O)CNC(C)C | CSUNLSYSEQIDMO-NSHDSACASA-N | No-Effect | inactive |
| tiracizine | BRD-K75250001 | CCOC(=O)Nc1ccc2c(c1)N(C(=O)CN(C)C)c1ccccc1CC2 | KJAMZCVTJDTESW-UHFFFAOYSA-N | No-Effect | inactive |
| tirapazamine | BRD-K92968657 | Nc1n[n+]([O-])c2ccccc2[n+]1[O-] | ORYDPOVDJJZGHQ-UHFFFAOYSA-N | No-Effect | inactive |
| tirasemtiv | BRD-K51087630 | C#Cc1cnc2[nH]c(=O)n(C(CC)CC)c2n1 | RSQGZEAXODVTOL-UHFFFAOYSA-N | No-Effect | inactive |
| tiratricol | BRD-K61236135 | O=C(O)Cc1cc(I)c(Oc2ccc(O)c(I)c2)c(I)c1 | UOWZUVNAGUAEQC-UHFFFAOYSA-N | No-Effect | inactive |
| tirofiban | BRD-K37872686 | CCCCS(=O)(=O)N[C@@H](Cc1ccc(OCCCCC2CCNCC2)cc1)C(=O)O | COKMIXFXJJXBQG-NRFANRHFSA-N | No-Effect | inactive |
| titanocene-dichloride | BRD-K60284353 | Cl[Ti](Cl)(C1C=CC=C1)C1C=CC=C1 | XKLWATAZDMHTSH-UHFFFAOYSA-L | No-Effect | inactive |
| tivantinib | BRD-K33379087 | O=C1NC(=O)[C@@H](c2cn3c4c(cccc24)CCC3)[C@@H]1c1c[nH]c2ccccc12 | UCEQXRCJXIVODC-PMACEKPBSA-N | No-Effect | inactive |
| tivozanib | BRD-K53414658 | COc1cc2nccc(Oc3ccc(NC(=O)Nc4cc(C)on4)c(Cl)c3)c2cc1OC | SPMVMDHWKHCIDT-UHFFFAOYSA-N | No-Effect | inactive |
| tizanidine | BRD-K06335600 | Clc1ccc2nsnc2c1NC1=NCCN1 | XFYDIVBRZNQMJC-UHFFFAOYSA-N | No-Effect | inactive |
| TMC-353121 | BRD-K66097663 | Cc1ccc(CCCO)c(NCc2ccc3nc(NCCCN4CCOCC4)n(Cc4nc(C)ccc4O)c3c2)c1 | DKORMNNYNRPTBJ-UHFFFAOYSA-N | No-Effect | inactive |
| TMPH | BRD-K49119234 | CCCCCCC(=O)OC1CC(C)(C)NC(C)(C)C1 | PWWKJRFUJIFGGD-UHFFFAOYSA-N | No-Effect | inactive |
| TMS | BRD-K96233303 | COc1cc(/C=C/c2ccc(OC)cc2OC)cc(OC)c1 | JDBCWSHYEQUBLW-AATRIKPKSA-N | No-Effect | inactive |
| TNP-470 | BRD-K53597484 | CO[C@H]1[C@H]([C@@]2(C)O[C@@H]2CC=C(C)C)[C@]2(CC[C@H]1OC(=O)NC(=O)CCl)CO2 | MSHZHSPISPJWHW-PVDLLORBSA-N | No-Effect | inactive |
| tobramycin | BRD-K05619559 | NC[C@H]1O[C@H](O[C@@H]2[C@@H](N)C[C@@H](N)[C@H](O[C@H]3O[C@H](CO)[C@@H](O)[C@H](N)[C@H]3O)[C@H]2O)[C@H](N)C[C@@H]1O | NLVFBUXFDBBNBW-PBSUHMDJSA-N | No-Effect | inactive |
| tocainide | BRD-K01826767 | Cc1cccc(C)c1NC(=O)[C@@H](C)N | BUJAGSGYPOAWEI-SECBINFHSA-N | No-Effect | inactive |
| toceranib | BRD-K62040061 | Cc1[nH]c(/C=C2\C(=O)Nc3ccc(F)cc32)c(C)c1C(=O)NCCN1CCCC1 | SRSGVKWWVXWSJT-ATVHPVEESA-N | No-Effect | inactive |
| tocofersolan | BRD-K57305459 | Cc1c(C)c2c(c(C)c1OC(=O)CCC(=O)OCCO)CC[C@@](C)(CCC[C@H](C)CCC[C@H](C)CCCC(C)C)O2 | AOBORMOPSGHCAX-AZAGJHQNSA-N | No-Effect | inactive |
| tofacitinib | BRD-K31283835 | C[C@@H]1CCN(C(=O)CC#N)C[C@@H]1N(C)c1ncnc2[nH]ccc12 | UJLAWZDWDVHWOW-YPMHNXCESA-N | No-Effect | inactive |
| tofogliflozin | BRD-K96344439 | CCc1ccc(Cc2ccc3c(c2)[C@]2(OC3)O[C@H](CO)[C@@H](O)[C@H](O)[C@H]2O)cc1 | VWVKUNOPTJGDOB-BDHVOXNPSA-N | No-Effect | inactive |
| tolamolol | BRD-A60498477 | Cc1ccccc1OC[C@@H](O)CNCCOc1ccc(C(N)=O)cc1 | SKQDKFOTIPJUSV-INIZCTEOSA-N | No-Effect | inactive |
| tolazamide | BRD-K32164935 | Cc1ccc(S(=O)(=O)NC(=O)NN2CCCCCC2)cc1 | OUDSBRTVNLOZBN-UHFFFAOYSA-N | No-Effect | inactive |
| tolbutamide | BRD-K85119730 | CCCCNC(=O)NS(=O)(=O)c1ccc(C)cc1 | JLRGJRBPOGGCBT-UHFFFAOYSA-N | No-Effect | inactive |
| tolcapone | BRD-K10852020 | Cc1ccc(C(=O)c2cc(O)c(O)c([N+](=O)[O-])c2)cc1 | MIQPIUSUKVNLNT-UHFFFAOYSA-N | No-Effect | inactive |
| tolfenamic-acid | BRD-K50133271 | Cc1c(Cl)cccc1Nc1ccccc1C(=O)O | YEZNLOUZAIOMLT-UHFFFAOYSA-N | No-Effect | inactive |
| tolimidone | BRD-K41772907 | Cc1cccc(Oc2cnc(=O)[nH]c2)c1 | HJQILFPVRNHTIG-UHFFFAOYSA-N | No-Effect | inactive |
| tolmetin | BRD-K82562631 | Cc1ccc(C(=O)c2ccc(CC(=O)O)n2C)cc1 | UPSPUYADGBWSHF-UHFFFAOYSA-N | No-Effect | inactive |
| tolnaftate | BRD-K44273375 | Cc1cccc(N(C)C(=S)Oc2ccc3ccccc3c2)c1 | FUSNMLFNXJSCDI-UHFFFAOYSA-N | No-Effect | inactive |
| tolonidine | BRD-K82687598 | Cc1ccc(NC2=NCCN2)c(Cl)c1 | KWBTZIFLQYYPTH-UHFFFAOYSA-N | No-Effect | inactive |
| toloxatone | BRD-A67182178 | Cc1cccc(N2C[C@@H](CO)OC2=O)c1 | MXUNKHLAEDCYJL-JTQLQIEISA-N | No-Effect | inactive |
| tolperisone | BRD-A27732521 | Cc1ccc(C(=O)[C@@H](C)CN2CCCCC2)cc1 | FSKFPVLPFLJRQB-AWEZNQCLSA-N | No-Effect | inactive |
| tolterodine | BRD-K54316499 | Cc1ccc(O)c([C@H](CCN(C(C)C)C(C)C)c2ccccc2)c1 | OOGJQPCLVADCPB-HXUWFJFHSA-N | No-Effect | inactive |
| tolvaptan | BRD-A82035391 | Cc1ccccc1C(=O)Nc1ccc(C(=O)N2CCC[C@@H](O)c3cc(Cl)ccc32)c(C)c1 | GYHCTFXIZSNGJT-XMMPIXPASA-N | No-Effect | inactive |
| tomeglovir | BRD-K00003487 | CN(C)c1cccc2c(S(=O)(=O)Nc3ccc(NC(=O)C(C)(C)CO)cc3)cccc12 | OSQAKHSYTKBSPB-UHFFFAOYSA-N | No-Effect | inactive |
| tomelukast | BRD-K74765201 | CCCc1c(OCCCCc2nn[nH]n2)ccc(C(C)=O)c1O | MWYHLEQJTQJHSS-UHFFFAOYSA-N | No-Effect | inactive |
| tomivosertib | BRD-K00003361 | Cc1cc(Nc2cc(N)ncn2)c(=O)n2c1C(=O)NC21CCCCC1 | HKTBYUWLRDZAJK-UHFFFAOYSA-N | No-Effect | inactive |

| | | | | | |
|---|---|---|---|---|---|
| tonabersat | BRD-K36970462 | CC(=O)c1ccc2c(c1)[C@H](NC(=O)c1ccc(F)c(Cl)c1)[C@H](O)C(C)(C)O2 | XLIIRNOPGJTBJD-ROUUACIJSA-N | No-Effect | inactive |
| topilutamide | BRD-K00003325 | C[C@](O)(CNC(=O)C(F)(F)F)C(=O)Nc1ccc([N+](=O)[O-])c(C(F)(F)F)c1 | YCNCRLKXSLARFT-NSHDSACASA-N | No-Effect | inactive |
| topiroxostat | BRD-K77561571 | N#Cc1cc(-c2nnc(-c3ccncc3)[nH]2)ccn1 | UBVZQGOVTLIHLH-UHFFFAOYSA-N | No-Effect | inactive |
| topotecan | BRD-K55696337 | CC[C@@]1(O)C(=O)OCc2c1cc1n(c2=O)Cc2cc3c(CN(C)C)c(O)ccc3nc2-1 | UCFGDBYHRUNTLO-QHCPKHFHSA-N | No-Effect | inactive |
| torasemide | BRD-K30480208 | Cc1cccc(Nc2ccncc2S(=O)(=O)NC(=O)NC(C)C)c1 | NGBFQHCMQULJNZ-UHFFFAOYSA-N | No-Effect | inactive |
| torcetrapib | BRD-K55675242 | CCOC(=O)N1c2ccc(C(F)(F)F)cc2[C@@H](N(Cc2cc(C(F)(F)F)cc(C(F)(F)F)c2)C(=O)OC)C[C@H]1CC | CMSGWTNRGKRWGS-NQIIRXRSSA-N | No-Effect | inactive |
| torcitabine | BRD-K91822704 | Nc1ccn([C@H]2C[C@H](O)[C@@H](CO)O2)c(=O)n1 | CKTSBUTUHBMZGZ-SHYZEUOFSA-N | No-Effect | inactive |
| torin-1 | BRD-K40175214 | CCC(=O)N1CCN(c2ccc(-n3c(=O)ccc4cnc5ccc(-c6cnc7ccccc7c6)cc5c43)cc2C(F)(F)F)CC1 | AKCRNFFTGXBONI-UHFFFAOYSA-N | No-Effect | inactive |
| tosedostat | BRD-K92241597 | CC(C)C[C@@H](C(=O)N[C@H](C(=O)OC1CCCC1)c1ccccc1)[C@H](O)C(=O)NO | FWFGIHPGRQZWIW-SQNIBIBYSA-N | No-Effect | inactive |
| tosufloxacin | BRD-K01825158 | N[C@H]1CCN(c2nc3c(cc2F)c(=O)c(C(=O)O)cn3-c2ccc(F)cc2F)C1 | WUWFMDMBOJLQIV-JTQLQIEISA-N | No-Effect | inactive |
| tozadenant | BRD-K93045741 | COc1ccc(N2CCOCC2)c2sc(NC(=O)N3CCC(C)(O)CC3)nc12 | XNBRWUQWSKXMPW-UHFFFAOYSA-N | No-Effect | inactive |
| tozasertib | BRD-K59369769 | Cc1cc(Nc2cc(N3CCN(C)CC3)nc(Sc3ccc(NC(=O)C4CC4)cc3)n2)[nH]n1 | GCIKSSRWRFVXBI-UHFFFAOYSA-N | No-Effect | inactive |
| TP-003 | BRD-K87941707 | CC(C)(O)c1ccn2c(-c3ccc(F)c(-c4ccc(F)cc4C#N)c3)cnc2c1F | SJMMDWXJSSJHOQ-UHFFFAOYSA-N | No-Effect | inactive |
| TP-3654 | BRD-K00005239 | CC(C)(O)[C@H]1CC[C@H](Nc2ccc3ncc(-c4cccc(C(F)(F)F)c4)n3n2)CC1 | XRNVABDYQLHODA-JCNLHEQBSA-N | No-Effect | inactive |
| TPCA-1 | BRD-K51575138 | NC(=O)Nc1sc(-c2ccc(F)cc2)cc1C(N)=O | SAYGKHKXGCPTLX-UHFFFAOYSA-N | No-Effect | inactive |
| TPPS4 | BRD-K00004542 | O=S(=O)(O)c1ccc(-c2c3nc(c(-c4ccc(S(=O)(=O)O)cc4)c4ccc([nH]4)c(-c4ccc(S(=O)(=O)O)cc4)c4nc(c(-c5ccc(S(=O)(=O)O)cc5)c5ccc2[nH]5)C=C4)C=C3)cc1 | PBHVCRIXMXQXPD-UHFFFAOYSA-N | No-Effect | inactive |
| trabodenoson | BRD-K00004698 | O=[N+]([O-])OC[C@H]1O[C@@H](n2cnc3c(NC4CCCC4)ncnc32)[C@H](O)[C@@H]1O | AQLVRTWKJDTWQQ-SDBHATRESA-N | No-Effect | inactive |
| tracazolate | BRD-K34154330 | CCCCNc1c(C(=O)OCC)c(C)nc2c1cnn2CC | PCTRYMLLRKWXGF-UHFFFAOYSA-N | No-Effect | inactive |
| TRAM-34 | BRD-K30071764 | Clc1ccccc1C(c1ccccc1)(c1ccccc1)n1cccn1 | KBFUQFVFYYBHBT-UHFFFAOYSA-N | No-Effect | inactive |
| trametinib | BRD-K12343256 | CC(=O)Nc1cccc(-n2c(=O)n(C3CC3)c(=O)c3c(Nc4ccc(I)cc4F)n(C)c(=O)c(C)c32)c1 | LIRYPHYGHXZJBZ-UHFFFAOYSA-N | No-Effect | inactive |
| tramiprosate | BRD-K82234479 | NCCCS(=O)(=O)O | SNKZJIOFVMKAOJ-UHFFFAOYSA-N | No-Effect | inactive |
| trandolapril | BRD-K28550399 | CCOC(=O)[C@H](CCc1ccccc1)N[C@@H](C)C(=O)N1[C@H](C(=O)O)C[C@H]2CCCC[C@@H]21 | VXFJYXUZANRPDJ-WTNASJBWSA-N | No-Effect | inactive |
| tranexamic-acid | BRD-K15014948 | NC[C@H]1CC[C@H](C(=O)O)CC1 | GYDJEQRTZSCIOI-LJGSYFOKSA-N | No-Effect | inactive |
| tranilast | BRD-K17849083 | COc1ccc(/C=C/C(=O)Nc2ccccc2C(=O)O)cc1OC | NZHGWWWHIYHZNX-CSKARUKUSA-N | No-Effect | inactive |
| trans-10,cis-12-conjugated-linoleic-acid | BRD-K26222048 | CCCCC/C=C\C=C\CCCCCCCCC(=O)O | GKJZMAHZJGSBKD-NMMTYZSQSA-N | No-Effect | inactive |
| trans-2-undecenoic-acid | BRD-K42959922 | CCCCCCCC/C=C/C(=O)O | IGBBVTAVILYDIO-MDZDMXLPSA-N | No-Effect | inactive |
| trans-4-[8-(3-fluorophenyl)-1,7-naphthyridin-6-yl]cyclohexanecarboxylic-acid | BRD-K75080769 | O=C(O)C1CCC(c2cc3cccnc3c(-c3cccc(F)c3)n2)CC1 | XWVBOPKUSVZZPJ-UHFFFAOYSA-N | No-Effect | inactive |
| trans-4-hydroxycrotonic-acid | BRD-K91614056 | O=C(O)/C=C/CO | RMQJECWPWQIIPW-OWOJBTEDSA-N | No-Effect | inactive |
| trans-4-methoxycinnamic-acid | BRD-K26273521 | COc1ccc(/C=C/C(=O)O)cc1 | AFDXODALSZRGIH-QPJJXVBHSA-N | No-Effect | inactive |
| transcrocetinate | BRD-K00003217 | CC(/C=C/C=C(\C)C(=O)O)=C\C=C\C=C(C)\C=C\C=C(/C)C(=O)O | PANKHBYNKQNAHN-MQQNZMFNSA-N | No-Effect | inactive |
| tranylcypromine | BRD-K00003699 | N[C@H]1C[C@@H]1c1ccccc1 | AELCINSCMGFISI-BDAKNGLRSA-N | No-Effect | inactive |
| trap-101 | BRD-K69837166 | CCn1c(=O)n(C2=C(CO)CN(CC3CCCCCCC3)CC2)c2ccccc21 | DCRGHMJXEBSRQG-UHFFFAOYSA-N | No-Effect | inactive |
| trapidil | BRD-K95763993 | CCN(CC)c1cc(C)nc2ncnn12 | GSNOZLZNQMLSKJ-UHFFFAOYSA-N | No-Effect | inactive |
| travoprost | BRD-K23976833 | CC(C)OC(=O)CCC/C=C\C[C@H]1[C@@H](O)C[C@@H](O)[C@@H]1/C=C/[C@@H](O)COc1cccc(C(F)(F)F)c1 | MKPLKVHSHYCHOC-AHTXBMBWSA-N | No-Effect | inactive |
| traxoprodil | BRD-K38158531 | C[C@@H]([C@@H](O)c1ccc(O)cc1)N1CCC(O)(c2ccccc2)CC1 | QEMSVZNTSXPFJA-HNAYVOBHSA-N | No-Effect | inactive |
| trazodone | BRD-K70778732 | O=c1n(CCCN2CCN(c3cccc(Cl)c3)CC2)nc2ccccn12 | PHLBKPHSAVXXEF-UHFFFAOYSA-N | No-Effect | inactive |
| trebenzomine | BRD-A20686268 | C[C@H]1Oc2ccccc2C[C@@H]1N(C)C | FDJAKSHVOONYHX-KOLCDFICSA-N | No-Effect | inactive |
| trelagliptin | BRD-K36407194 | Cn1c(=O)cc(N2CCC[C@@H](N)C2)n(Cc2cc(F)ccc2C#N)c1=O | IWYJYHUNXVAVAA-OAHLLOKOSA-N | No-Effect | inactive |
| tremorine | BRD-K08619838 | C(#CCN1CCCC1)CN1CCCC1 | JSUAJTLKVREZHV-UHFFFAOYSA-N | No-Effect | inactive |
| treosulfan | BRD-K20902145 | CS(=O)(=O)OC[C@H](O)[C@@H](O)COS(C)(=O)=O | YCPOZVAOBBQLRI-WDSKDSINSA-N | No-Effect | inactive |
| trepibutone | BRD-K37543758 | CCOc1cc(OCC)c(C(=O)CCC(=O)O)cc1OCC | YPTFHLJNWSJXKG-UHFFFAOYSA-N | No-Effect | inactive |
| treprostinil | BRD-K19706299 | CCCCC[C@H](O)CC[C@@H]1[C@H]2Cc3cccc(OCC(=O)O)c3C[C@H]2C[C@H]1O | PAJMKGZZBBTTOY-ZFORQUDYSA-N | No-Effect | inactive |
| trequinsin | BRD-K84663978 | COc1cc2c(cc1OC)-c1c/c(=N\c3c(C)cc(C)cc3C)n(C)c(=O)n1CC2 | MCMSJVMUSBZUCN-YYDJUVGSSA-N | No-Effect | inactive |
| tretazicar | BRD-K08748705 | NC(=O)c1cc(N2CC2)c([N+](=O)[O-])cc1[N+](=O)[O-] | WOCXQMCIOTUMJV-UHFFFAOYSA-N | No-Effect | inactive |
| triacetin | BRD-K92844263 | CC(=O)OCC(COC(C)=O)OC(C)=O | URAYPUMNDPQOKB-UHFFFAOYSA-N | No-Effect | inactive |
| triamcinolone | BRD-K77554836 | C[C@]12C[C@H](O)[C@@]3(F)[C@@H](CCC4=CC(=O)C=C[C@@]43C)[C@@H]1C[C@@H](O)[C@]2(O)C(=O)CO | GFNANZIMVAIWHM-OBYCQNJPSA-N | No-Effect | inactive |
| triamcinolone-acetonide | BRD-K53790871 | CC1(C)O[C@@H]2C[C@H]3[C@@H]4CCC5=CC(=O)C=C[C@]5(C)[C@@]4(F)[C@@H](O)C[C@]3(C)[C@]2(C(=O)CO)O1 | YNDXUCZADRHECN-JNQJZLCISA-N | No-Effect | inactive |
| triamcinolone-hexacetonide | BRD-K00003338 | CC(C)(C)CC(=O)OCC(=O)[C@@]12OC(C)(C)O[C@@H]1C[C@H]1[C@@H]3CCC4=CC(=O)C=C[C@]4(C)[C@@]3(F)[C@@H](O)C[C@@]12C | TZIZWYVVGLXXFV-FLRHRWPCSA-N | No-Effect | inactive |
| triamterene | BRD-K92049597 | Nc1nc(N)c2nc(-c3ccccc3)c(N)nc2n1 | FNYLWPVRPXGIIP-UHFFFAOYSA-N | No-Effect | inactive |
| triapine | BRD-K01669786 | NC(=S)N/N=C/c1ncccc1N | XMYKNCNAZKMVQN-NYYWCZLTSA-N | No-Effect | inactive |
| tribenoside | BRD-A60294240 | CCO[C@H]1O[C@H]([C@@H](COCc2ccccc2)OCc2ccccc2)[C@H](OCc2ccccc2)[C@H]1O | ULLNJSBQMBKOJH-JPHCZMGXSA-N | No-Effect | inactive |
| tribomsalan | BRD-K09118247 | O=C(Nc1ccc(Br)cc1)c1cc(Br)cc(Br)c1O | KVSKGMLNBAPGKH-UHFFFAOYSA-N | No-Effect | inactive |
| tribromoethanol | BRD-K69632891 | OCC(Br)(Br)Br | YFDSDPIBEUFTMI-UHFFFAOYSA-N | No-Effect | inactive |
| tributyrin | BRD-K38969488 | CCCC(=O)OCC(COC(=O)CCC)OC(=O)CCC | UYXTWWCETRIEDR-UHFFFAOYSA-N | No-Effect | inactive |
| trichlormethiazide | BRD-A28940325 | NS(=O)(=O)c1cc2c(cc1Cl)N[C@H](C(Cl)Cl)NS2(=O)=O | LMJSLTNSBFUCMU-QMMMGPOBSA-N | No-Effect | inactive |
| trichloroacetic-acid | BRD-K42090719 | O=C(O)C(Cl)(Cl)Cl | YNJBWRMUSHSURL-UHFFFAOYSA-N | No-Effect | inactive |
| trichloroethylene | BRD-K46435528 | ClC=C(Cl)Cl | XSTXAVWGXDQKEL-UHFFFAOYSA-N | No-Effect | inactive |
| triciribine | BRD-K80431395 | CN1N=C(N)c2cn([C@@H]3O[C@H](CO)[C@@H](O)[C@H]3O)c3ncnc1c23 | HOGVTUZUJGHKPL-HTVVRFAVSA-N | No-Effect | inactive |

| | | | | | |
|---|---|---|---|---|---|
| triciribine-phosphate | BRD-K00004666 | CN1N=C(N)c2cn([C@@H]3O[C@H](COP(=O)(O)O)[C@@H](O)[C@H]3O)c3ncnc1c23 | URLYINUFLXOMHP-HTVVRFAVSA-N | No-Effect | inactive |
| triclabendazole | BRD-K81916719 | CSc1nc2cc(Cl)c(Oc3cccc(Cl)c3Cl)cc2[nH]1 | NQPDXQQQCQDHHW-UHFFFAOYSA-N | No-Effect | inactive |
| triclosan | BRD-K41731458 | Oc1cc(Cl)ccc1Oc1ccc(Cl)cc1Cl | XEFQLINVKFYRCS-UHFFFAOYSA-N | No-Effect | inactive |
| tridihexethyl | BRD-A22152765 | CC[N+](CC)(CC)CC[C@](O)(c1ccccc1)C1CCCCC1 | NPRHVSBSZMAEIN-NRFANRHFSA-N | No-Effect | inactive |
| trientine | BRD-K37111771 | NCCNCCNCCN | VILCJCGEZXAXTO-UHFFFAOYSA-N | No-Effect | inactive |
| triflupromazine | BRD-K63675182 | CN(C)CCCN1c2ccccc2Sc2ccc(C(F)(F)F)cc21 | XSCGXQMFQXDFCW-UHFFFAOYSA-N | No-Effect | inactive |
| triflusal | BRD-K71696703 | CC(=O)Oc1cc(C(F)(F)F)ccc1C(=O)O | RMWVZGDJPAKBDE-UHFFFAOYSA-N | No-Effect | inactive |
| trigonelline | BRD-K69231085 | C[n+]1cccc(C(=O)O)c1 | WWNNZCOKKKDOPX-UHFFFAOYSA-O | No-Effect | inactive |
| trihexyphenidyl | BRD-K00003691 | O[C@](CCN1CCCCC1)(c1ccccc1)C1CCCCC1 | HWHLPVGTWGOCJO-HXUWFJFHSA-N | No-Effect | inactive |
| trilaciclib | BRD-K00003412 | CN1CCN(c2ccc(Nc3ncc4cc5n(c4n3)C3(CCCCC3)CNC5=O)nc2)CC1 | PDGKHKMBHVFCMG-UHFFFAOYSA-N | No-Effect | inactive |
| trilostane | BRD-A04497688 | C[C@]12CC[C@H]3[C@@H](CC[C@@]45O[C@@H]4C(=O)[C@H](C#N)C[C@]35C)[C@@H]1CC[C@@H]2O | RLQVKDVIBJCQGE-PORJLZOZSA-N | No-Effect | inactive |
| trimebutine | BRD-K00003635 | CC[C@@](COC(=O)c1cc(OC)c(OC)c(OC)c1)(c1ccccc1)N(C)C | LORDFXWUHHSAQU-JOCHJYFZSA-N | No-Effect | inactive |
| trimetazidine | BRD-K88366685 | COc1ccc(CN2CCNCC2)c(OC)c1OC | UHWVSEOVJBQKBE-UHFFFAOYSA-N | No-Effect | inactive |
| trimethobenzamide | BRD-K34415467 | COc1cc(C(=O)NCc2ccc(OCCN(C)C)cc2)cc(OC)c1OC | FEZBIKUBAYAZIU-UHFFFAOYSA-N | No-Effect | inactive |
| trimethoprim | BRD-K07208025 | COc1cc(Cc2cnc(N)nc2N)cc(OC)c1OC | IEDVJHCEMCRBQM-UHFFFAOYSA-N | No-Effect | inactive |
| trimethoquinol | BRD-A56241705 | COc1cc(C[C@H]2NCCc3cc(O)c(O)cc32)cc(OC)c1OC | RGVPOXRFEPSFGH-CQSZACIVSA-N | No-Effect | inactive |
| trimethylolpropane-triacrylate | BRD-K24483849 | C=CC(=O)OCC(CC)(COC(=O)C=C)COC(=O)C=C | DAKWPKUUDNSNPN-UHFFFAOYSA-N | No-Effect | inactive |
| trimetozine | BRD-K53220666 | COc1cc(C(=O)N2CCOCC2)cc(OC)c1OC | XWVOEFLBOSSYGM-UHFFFAOYSA-N | No-Effect | inactive |
| trioxsalen | BRD-K54790157 | Cc1cc2cc3c(C)cc(=O)oc3c(C)c2o1 | FMHHVULEAZTJMA-UHFFFAOYSA-N | No-Effect | inactive |
| triprolidine | BRD-K11742128 | Cc1ccc(/C(=C\CN2CCCC2)c2ccccn2)cc1 | CBEQULMOCCWAQT-WOJGMQOQSA-N | No-Effect | inactive |
| triptan | BRD-K19427720 | CC(C)C(C)(C)C | ZISSAWUMDACLOM-UHFFFAOYSA-N | No-Effect | inactive |
| triptorelin | BRD-K62685538 | CC(C)C[C@H](NC(=O)[C@@H](Cc1c[nH]c2ccccc12)NC(=O)[C@H](Cc1ccc(O)cc1)NC(=O)[C@H](CO)NC(=O)[C@H](Cc1c[nH]c2ccccc12)NC(=O)[C@H](Cc1cnc[nH]1)NC(=O)[C@@H]1CCC(=O)N1)C(=O)N[C@@H](CCCNC(=N)N)C(=O)N1CCC[C@H]1C(=O)NCC(N)=O | VXKHXGOKWPXYNA-PGBVPBMZSA-N | No-Effect | inactive |
| troclosene | BRD-K76375510 | O=c1[nH]c(=O)n(Cl)c(=O)n1Cl | CEJLBZWIKQJOAT-UHFFFAOYSA-N | No-Effect | inactive |
| trodusquemine | BRD-K00003124 | CC(C)[C@@H](CC[C@@H](C)[C@H]1CC[C@H]2[C@H]3[C@H](CC[C@@]21C)[C@@]1(C)CC[C@H](NCCCNCCCCNCCCN)C[C@@H]1C[C@H]3O)OS(=O)(=O)O | WUJVPODXELZABP-FWJXURDUSA-N | No-Effect | inactive |
| trofinetide | BRD-K83502127 | NCC(=O)N1CCC[C@H]1C(=O)N[C@@H](CCC(=O)O)C(=O)O | JJGBXTYGTKWGAT-YUMQZZPRSA-N | No-Effect | inactive |
| trofosfamide | BRD-A91386192 | O=[P@]1(N(CCCl)CCCl)OCCCN1CCCl | UMKFEPPTGMDVMI-KRWDZBQOSA-N | No-Effect | inactive |
| troglitazone | BRD-A13084692 | Cc1c(C)c2c(c(C)c1O)CC[C@](C)(COc1ccc(C[C@H]3SC(=O)NC3=O)cc1)O2 | GXPHKUHSUJUWKP-NTKDMRAZSA-N | No-Effect | inactive |
| troleandomycin | BRD-K38310698 | CO[C@H]1C[C@H](O[C@@H]2[C@@H](C)C(=O)O[C@H](C)[C@H](C)[C@H](OC(C)=O)[C@@H](C)C(=O)[C@]3(CO3)C[C@H](C)[C@H](O[C@@H]3O[C@H](C)C[C@H](N(C)C)[C@H]3OC(C)=O)[C@H]2C)O[C@@H](C)[C@@H]1OC(C)=O | LQCLVBQBTUVCEQ-QTFUVMRISA-N | No-Effect | inactive |
| trolox | BRD-A17846016 | Cc1c(C)c2c(c(C)c1O)CC[C@@](C)(C(=O)O)O2 | GLEVLJDDWXEYCO-AWEZNQCLSA-N | No-Effect | inactive |
| tromaril | BRD-K91422571 | O=C(O)c1ccccc1NCCc1ccccc1 | HLNLBEFKHHCAMV-UHFFFAOYSA-N | No-Effect | inactive |
| trometamol | BRD-K47978074 | NC(CO)(CO)CO | LENZDBCJOHFCAS-UHFFFAOYSA-N | No-Effect | inactive |
| tropanyl-3,5-dimethylbenzoate | BRD-A84389633 | Cc1cc(C)cc(C(=O)O[C@@H]2C[C@@H]3CC[C@H](C2)N3C)c1 | HDDNYFLPWFSBLN-ZSHCYNCHSA-N | No-Effect | inactive |
| tropesin | BRD-A34603619 | COc1ccc2c(c1)c(CC(=O)OC[C@@H](C(=O)O)c1ccccc1)c(C)n2C(=O)c1ccc(Cl)cc1 | UCCJWNPWWPJKGL-XMMPIXPASA-N | No-Effect | inactive |
| tropicamide | BRD-A79672927 | CCN(Cc1ccncc1)C(=O)[C@@H](CO)c1ccccc1 | BGDKAVGWHJFAGW-INIZCTEOSA-N | No-Effect | inactive |
| tropifexor | BRD-K00003480 | O=C(O)c1cc(F)c2nc(N3[C@H]4CC[C@@H]3C[C@H](OCc3c(-c5ccccc5OC(F)(F)F)noc3C3CC3)C4)sc2c1 | VYLOOGHLKSNNEK-PIIMJCKOSA-N | No-Effect | inactive |
| tropisetron | BRD-K50866992 | CN1[C@H]2CC[C@@H]1C[C@H](OC(=O)c1c[nH]c3ccccc13)C2 | ZNRGQMMCGHDTEI-ITGUQSILSA-N | No-Effect | inactive |
| trospium | BRD-K00003641 | O=C(O[C@@H]1C[C@@H]2CC[C@H](C1)[N+]21CCCC1)C(O)(c1ccccc1)c1ccccc1 | OYYDSUSKLWTMMQ-JKHIJQBDSA-N | No-Effect | inactive |
| trovafloxacin | BRD-K00494077 | N[C@H]1[C@@H]2CN(c3nc4c(cc3F)c(=O)c(C(=O)O)cn4-c3ccc(F)cc3F)C[C@H]12 | WVPSKSLAZQPAKQ-CDMJZVDBSA-N | No-Effect | inactive |
| troventol | BRD-A35868371 | CCC(CO)(C(=O)OC1C[C@@H]2CC[C@H](C1)[N+]2(C)C)c1ccccc1 | QFWSRMACUIYPGS-ZVBHREPYSA-N | No-Effect | inactive |
| trovirdine | BRD-K00989047 | S=C(NCCc1ccccn1)Nc1ccc(Br)cn1 | HOCFDYZWQYGULA-UHFFFAOYSA-N | No-Effect | inactive |
| troxipide | BRD-A68589262 | COc1cc(C(=O)N[C@H]2CCCNC2)cc(OC)c1OC | YSIITVVESCNIPR-NSHDSACASA-N | No-Effect | inactive |
| TRV130 | BRD-K35626314 | COc1ccsc1CNCC[C@@]1(c2ccccn2)CCOC2(CCCC2)C1 | DMNOVGJWPASQDL-OAQYLSRUSA-N | No-Effect | inactive |
| tryptanthrin | BRD-K76973696 | O=C1c2ccccc2-n2c1nc1ccccc1c2=O | VQQVWGVXDIPORV-UHFFFAOYSA-N | No-Effect | inactive |
| TS-011 | BRD-K00004685 | [O-][N+]CNc1ccc(N2CCOCC2)c(Cl)c1 | CQMCOBQPZDLWIX-UHFFFAOYSA-N | No-Effect | inactive |
| TTNPB | BRD-K49685476 | C/C(=C\c1ccc(C(=O)O)cc1)c1ccc2c(c1)C(C)(C)CCC2(C)C | FOIVPCKZDPCJJY-JQIJEIRASA-N | No-Effect | inactive |
| TTP-22 | BRD-K64852730 | Cc1ccc(-c2csc3ncnc(SCCC(=O)O)c23)cc1 | RAOULLCLLOGTDA-UHFFFAOYSA-N | No-Effect | inactive |
| TU-2100 | BRD-K44590731 | CCOC(=O)c1ccccc1OC(=O)CCCCCCCC(=O)Oc1ccccc1C(=O)OCC | LIBAUACOIKWWKP-UHFFFAOYSA-N | No-Effect | inactive |
| tuaminoheptane | BRD-A19958636 | CCCCC[C@@H](C)N | VSRBKQFNFZQRBM-SSDOTTSWSA-N | No-Effect | inactive |
| tubastatin-A | BRD-K00627859 | CN1CCc2c(c3ccccc3n2Cc2ccc(C(=O)NO)cc2)C1 | GOVYBPLHWIEHEJ-UHFFFAOYSA-N | No-Effect | inactive |
| tucatinib | BRD-K26026438 | Cc1cc(Nc2ncnc3ccc(NC4=NC(C)(C)CO4)cc23)ccc1Oc1ccn2ncnc2c1 | SDEAXTCZPQIFQM-UHFFFAOYSA-N | No-Effect | inactive |
| tucidinostat | BRD-K95216408 | Nc1cc(F)ccc1NC(=O)c1ccc(CNC(=O)/C=C/c2cccnc2)cc1 | SZMJVTADHFNAIS-BJMVGYQFSA-N | No-Effect | inactive |
| tuftsin | BRD-K00003336 | C[C@@H](O)[C@H](N)C(=O)N[C@@H](CCCCN)C(=O)N1CCC[C@H]1C(=O)N[C@@H](CCCNC(=N)N)C(=O)O | IESDGNYHXIOKRW-YXMSTPNBSA-N | No-Effect | inactive |
| TUG-770 | BRD-K56086081 | N#CCc1ccccc1C#Cc1ccc(CCC(=O)O)c(F)c1 | KIZUBVPJNPVIIN-UHFFFAOYSA-N | No-Effect | inactive |
| TUG-891 | BRD-K24282351 | Cc1ccc(-c2ccc(F)cc2COc2ccc(CCC(=O)O)cc2)cc1 | LPGBXHWIQNZEJB-UHFFFAOYSA-N | No-Effect | inactive |
| tulobuterol | BRD-K00003719 | CC(C)(C)NC[C@H](O)c1ccccc1Cl | YREYLAVBNPACJM-NSHDSACASA-N | No-Effect | inactive |
| tulrampator | BRD-K00005240 | O=C1c2cc3nnn(CCc4cccc(F)c4)c(=O)c3cc2OCN1C1CC1 | JHCFQXNWYDLBOG-UHFFFAOYSA-N | No-Effect | inactive |
| TWS-119 | BRD-K94176593 | Nc1cccc(-c2cc3c(Oc4cccc(O)c4)ncnc3[nH]2)c1 | VPVLEBIVXZSOMQ-UHFFFAOYSA-N | No-Effect | inactive |
| TXA127 | BRD-K36040345 | CC[C@H](C)[C@H](NC(=O)[C@H](Cc1ccc(O)cc1)NC(=O)[C@@H](NC(=O)[C@H](CCCNC(=N)N)NC(=O)[C@@H](N)CC(=O)O)C(C)C)C(=O)N[C@@H](Cc1cnc[nH]1)C(=O)N1CCC[C@H]1C(=O)O | PVHLMTREZMEJCG-GDTLVBQBSA-N | No-Effect | inactive |
| TY-52156 | BRD-K66584608 | CC(C)(C)C(=O)C(=NNc1ccc(Cl)cc1)Nc1ccc(Cl)cc1 | XONRRGIRSGNWFP-UHFFFAOYSA-N | No-Effect | inactive |
| tylosin | BRD-K37753391 | CC[C@H]1OC(=O)C[C@@H](O)[C@H](C)[C@@H](O[C@@H]2O[C@H](C)[C@@H](O[C@H]3C[C@@](C)(O)[C@@H](O)[C@H](C)O3)[C@H](N(C)C)[C@H]2O)[C@@H](CC=O)C[C@@H](C)C(=O)/C=C/C(C)=C/[C@@H]1CO[C@@H]1O[C@H](C)[C@@H](O)[C@@H](OC)[C@H]1OC | WBPYTXDJUQJLPQ-VMXQISHHSA-N | No-Effect | inactive |
| tymazoline | BRD-K64414150 | Cc1ccc(C(C)C)c(OCC2=NCCN2)c1 | QRORCRWSRPKEHR-UHFFFAOYSA-N | No-Effect | inactive |
| tyrphostin-A9 | BRD-K40255344 | CC(C)(C)c1cc(C=C(C#N)C#N)cc(C(C)(C)C)c1O | MZOPWQKISXCCTP-UHFFFAOYSA-N | No-Effect | inactive |
| tyrphostin-AG-1296 | BRD-K76064317 | COc1cc2ncc(-c3ccccc3)nc2cc1OC | QNOXYUNHIGOWNY-UHFFFAOYSA-N | No-Effect | inactive |
| tyrphostin-AG-1478 | BRD-K68336408 | COc1cc2ncnc(Nc3cccc(Cl)c3)c2cc1OC | GFNNBHLJANVSQV-UHFFFAOYSA-N | No-Effect | inactive |

| | | | | | |
|---|---|---|---|---|---|
| tyrphostin-AG-18 | BRD-K49657628 | N#CC(C#N)=Cc1ccc(O)c(O)c1 | VTJXFTPMFYAJJU-UHFFFAOYSA-N | No-Effect | inactive |
| tyrphostin-AG-494 | BRD-K34533029 | N#C/C(=C\c1ccc(O)c(O)c1)C(=O)Nc1ccccc1 | HKHOVJYOELRGMV-XYOKQWHBSA-N | No-Effect | inactive |
| tyrphostin-AG-825 | BRD-K87919739 | COc1cc(/C=C(\C#N)C(N)=O)cc(CSc2nc3ccccc3s2)c1O | KXDONFLNGBQLTN-WUXMJOGZSA-N | No-Effect | inactive |
| tyrphostin-AG-879 | BRD-K59469039 | CC(C)(C)c1cc(/C=C(\C#N)C(N)=S)cc(C(C)(C)C)c1O | XRZYELWZLNAXGE-KPKJPENVSA-N | No-Effect | inactive |
| tyrphostin-AG-99 | BRD-K60184833 | N#C/C(=C\c1ccc(O)c(O)c1)C(N)=O | USOXQZNJFMKTKJ-XVNBXDOJSA-N | No-Effect | inactive |
| U-0124 | BRD-K03635852 | CSC(=N)C(C#N)C(C#N)C(=N)SC | GXGXSVFKGDVBMD-UHFFFAOYSA-N | No-Effect | inactive |
| U-0521 | BRD-K08953028 | CC(C)C(=O)c1ccc(O)c(O)c1 | VDQLKIBLTMPAHI-UHFFFAOYSA-N | No-Effect | inactive |
| U-104 | BRD-K77867004 | NS(=O)(=O)c1ccc(NC(=O)Nc2ccc(F)cc2)cc1 | YJQZNWPYLCNRLP-UHFFFAOYSA-N | No-Effect | inactive |
| U-50488-(-) | BRD-K53532120 | CN(C(=O)Cc1ccc(Cl)c(Cl)c1)[C@H]1CCCC[C@@H]1N1CCCC1 | VQLPLYSROCPWFF-ROUUACIJSA-N | No-Effect | inactive |
| U-54494A | BRD-K20995441 | CN(C(=O)c1ccc(Cl)c(Cl)c1)[C@@H]1CCCC[C@@H]1N1CCCC1 | JUBNVWGVNWIXMB-SJORKVTESA-N | No-Effect | inactive |
| U-75799E | BRD-K77070363 | C[C@H](NC(=O)[C@@H](Cc1c[nH]c2ccccc12)NC(=O)[C@@H](N)Cc1cnc[nH]1)C(=O)N[C@@H](Cc1c[nH]c2ccccc12)C(=O)N[C@H](Cc1ccccc1)C(=O)N[C@@H](CCCCN)C(N)=O | WZHKXNSOCOQYQX-FUAFALNISA-N | No-Effect | inactive |
| U-99194 | BRD-K70281171 | CCCN(CCC)C1Cc2cc(OC)c(OC)cc2C1 | UOLJKAPABHXFRE-UHFFFAOYSA-N | No-Effect | inactive |
| UAMC-00039 | BRD-K38966689 | N[C@@H](CCNCc1ccc(Cl)cc1)C(=O)N1CCCCC1 | DNVZYSMOLXCOLY-HNNXBMFYSA-N | No-Effect | inactive |
| UB-165 | BRD-A14574269 | Clc1ccc(C2=CCC[C@H]3CC[C@@H]2N3)cn1 | TVSNVPGPFGAKPT-JQWIXIFHSA-N | No-Effect | inactive |
| ubenimex | BRD-K59574735 | CC(C)C[C@H](NC(=O)[C@@H](O)[C@H](N)Cc1ccccc1)C(=O)O | VGGGPCQERPFHOB-RDBSUJKOSA-N | No-Effect | inactive |
| UBP-296 | BRD-A45499626 | N[C@@H](Cn1ccc(=O)n(Cc2ccccc2C(=O)O)c1=O)C(=O)O | UUIYULWYHDSXHL-NSHDSACASA-N | No-Effect | inactive |
| UBP-302 | BRD-K79753741 | N[C@@H](Cn1ccc(=O)n(Cc2ccccc2C(=O)O)c1=O)C(=O)O | UUIYULWYHDSXHL-NSHDSACASA-N | No-Effect | inactive |
| UBP-310 | BRD-K09597601 | Cc1cn(C[C@H](N)C(=O)O)c(=O)n(Cc2ccsc2C(=O)O)c1=O | ZTAZUCRXCRXNSU-VIFPVBQESA-N | No-Effect | inactive |
| UCL-2077 | BRD-K50325075 | c1ccc(C(NCc2cccnc2)(c2ccccc2)c2ccccc2)cc1 | PQFNWDHABGBCHB-UHFFFAOYSA-N | No-Effect | inactive |
| udenafil | BRD-K00003266 | CCCOc1ccc(S(=O)(=O)NCC[C@@H]2CCCN2C)cc1-c1nc2c(CCC)nn(C)c2c(=O)[nH]1 | IYFNEFQTYQPVOC-KRWDZBQOSA-N | No-Effect | inactive |
| UF-010 | BRD-K20480395 | CCCCNNC(=O)c1ccc(Br)cc1 | BVQCFCYPFJOOAV-UHFFFAOYSA-N | No-Effect | inactive |
| ufenamate | BRD-K84173274 | CCCCOC(=O)c1ccccc1Nc1cccc(C(F)(F)F)c1 | JDLSRXWHEBFHNC-UHFFFAOYSA-N | No-Effect | inactive |
| UH-232-(+) | BRD-K66715657 | CCCN(CCC)[C@@H]1CCc2c(OC)cccc2[C@@H]1C | BTOJYCTUJJHANF-WMLDXEAASA-N | No-Effect | inactive |
| UK-356618 | BRD-K57011718 | Cc1cc(CCC[C@H](CC(=O)NO)C(=O)N[C@H](C(=O)N[C@H](C)c2ccccc2)C(C)(C)C)ccc1-c1ccccc1 | JJHRUUKMPWUYIB-HVOSOHGQSA-N | No-Effect | inactive |
| UK-383367 | BRD-K06294530 | NC(=O)c1noc([C@H](CCCC2CCCCC2)CC(=O)NO)n1 | ARJCBSRIPGJMAD-LLVKDONJSA-N | No-Effect | inactive |
| UK-5099 | BRD-K19558579 | N#C/C(=C\c1cn(-c2ccccc2)c2ccccc12)C(=O)O | BIZNHCWFGNKBBZ-JLHYYAGUSA-N | No-Effect | inactive |
| uki-1 | BRD-K00003367 | CCOC(=O)N1CCN(C(=O)[C@H](Cc2cccc(C(=N)N)c2)NS(=O)(=O)c2c(C(C)C)cc(C(C)C)cc2C(C)C)CC1 | ISJSHQTWOHGCMM-NDEPHWFRSA-N | No-Effect | inactive |
| ulifloxacin | BRD-A61633223 | C[C@H]1Sc2c(C(=O)O)c(=O)c3cc(F)c(N4CCNCC4)cc3n21 | SUXQDLLXIBLQHW-MRVPVSSYSA-N | No-Effect | inactive |
| unbs-5162 | BRD-K30933884 | CN(C)CCN1C(=O)c2cccc3cc(NC(N)=O)cc(c23)C1=O | WCKZRLOUKYFJDY-UHFFFAOYSA-N | No-Effect | inactive |
| UNC0224 | BRD-K77428767 | COc1cc2c(NC3CCN(C)CC3)nc(N3CCCN(C)CC3)nc2cc1OCCCN(C)C | XIVUGRBSBIXXJE-UHFFFAOYSA-N | No-Effect | inactive |
| UNC0321 | BRD-K74236984 | COc1cc2c(NC3CCN(C)CC3)nc(N3CCCN(C)CC3)nc2cc1OCCOCCN(C)C | AULLUGALUBVBDD-UHFFFAOYSA-N | No-Effect | inactive |
| UNC0638 | BRD-K86856088 | COc1cc2c(NC3CCN(C(C)C)CC3)nc(C3CCCCC3)nc2cc1OCCCN1CCCC1 | QOECJCJVIMVJGX-UHFFFAOYSA-N | No-Effect | inactive |
| UNC0646 | BRD-K87465484 | COc1cc2c(NC3CCN(C4CCCCC4)CC3)nc(N3CCCN(C(C)C)CC3)nc2cc1OCCCN1CCCCC1 | OUKWLRHRXOPODD-UHFFFAOYSA-N | No-Effect | inactive |
| UNC1215 | BRD-K83189926 | O=C(c1ccc(C(=O)N2CCC(N3CCCC3)CC2)c(Nc2ccccc2)c1)N1CCC(N2CCCC2)CC1 | PQOOIERVZAXHBP-UHFFFAOYSA-N | No-Effect | inactive |
| UNC2250 | BRD-K01773560 | CCCCNc1ncc(-c2ccc(CN3CCOCC3)cn2)c(N[C@H]2CC[C@H](O)CC2)n1 | HSYSSKFCQHXOBP-MXVIHJGJSA-N | No-Effect | inactive |
| UNC2327 | BRD-K60635008 | O=C(NCC(=O)N1CCCCC1)Nc1ccc2nnsc2c1 | MYTRGTBDVGKKRO-UHFFFAOYSA-N | No-Effect | inactive |
| UNC669 | BRD-K21906513 | O=C(c1cncc(Br)c1)N1CCC(N2CCCC2)CC1 | CQERVFFAOOUFEQ-UHFFFAOYSA-N | No-Effect | inactive |
| UNC926 | BRD-K22229656 | O=C(c1cccc(Br)c1)N1CCC(N2CCCC2)CC1 | OWGLFIKZKQOYHZ-UHFFFAOYSA-N | No-Effect | inactive |
| undecylenic-acid | BRD-K19189338 | C=CCCCCCCCCC(=O)O | FRPZMMHWLSIFAZ-UHFFFAOYSA-N | No-Effect | inactive |
| upadacitinib | BRD-K00003286 | CC[C@@H]1CN(C(=O)NCC(F)(F)F)C[C@@H]1c1cnc2cnc3[nH]ccc3n12 | WYQFJHHDOKWSHR-MNOVXSKESA-N | No-Effect | inactive |
| upamostat | BRD-K00003218 | CCOC(=O)N1CCN(C(=O)[C@H](Cc2cccc(C(=N)NO)c2)NS(=O)(=O)c2c(C(C)C)cc(C(C)C)cc2C(C)C)CC1 | HUASEDVYRABWCV-NDEPHWFRSA-N | No-Effect | inactive |
| UPF-1069 | BRD-K77950083 | O=C(COc1cccc2c(=O)[nH]ccc12)c1ccccc1 | JJWMRRNGWSITSQ-UHFFFAOYSA-N | No-Effect | inactive |
| uprifosbuvir | BRD-K00003526 | CC(C)OC(=O)[C@H](C)N[P@](=O)(OC[C@H]1O[C@@H](n2ccc(=O)[nH]c2=O)[C@](C)(Cl)[C@@H]1O)Oc1ccccc1 | SFPFZQKYPOWCSI-IQWMDFIBSA-N | No-Effect | inactive |
| uprosertib | BRD-K38332599 | Cn1ncc(Cl)c1-c1cc(C(=O)N[C@H](CN)Cc2ccc(F)c(F)c2)oc1Cl | AXTAPYRUEKNRBA-JTQLQIEISA-N | No-Effect | inactive |
| uracil | BRD-K80129304 | O=c1cc[nH]c(=O)[nH]1 | ISAKRJDGNUQOIC-UHFFFAOYSA-N | No-Effect | inactive |
| urapidil | BRD-K73196317 | COc1ccccc1N1CCN(CCCNc2cc(=O)n(C)c(=O)n2C)CC1 | ICMGLRUYEQNHPF-UHFFFAOYSA-N | No-Effect | inactive |
| URB597 | BRD-K92169341 | NC(=O)c1cccc(-c2cccc(OC(=O)NC3CCCCC3)c2)c1 | ROFVXGGUISEHAM-UHFFFAOYSA-N | No-Effect | inactive |
| urea | BRD-K88052444 | NC(N)=O | XSQUKJJJFZCRTK-UHFFFAOYSA-N | No-Effect | inactive |
| urethane | BRD-K24297741 | CCOC(N)=O | JOYRKODLDBILNP-UHFFFAOYSA-N | No-Effect | inactive |
| uric-acid | BRD-K01295354 | O=c1[nH]c(=O)c2[nH]c(=O)[nH]c2[nH]1 | LEHOTFFKMJEONL-UHFFFAOYSA-N | No-Effect | inactive |
| uridine | BRD-K13050303 | O=c1ccn([C@@H]2O[C@H](CO)[C@@H](O)[C@H]2O)c(=O)[nH]1 | DRTQHJPVMGBUCF-XVFCMESISA-N | No-Effect | inactive |
| uridine-triacetate | BRD-K18323388 | CC(=O)OC[C@H]1O[C@@H](n2ccc(=O)[nH]c2=O)[C@H](OC(C)=O)[C@@H]1OC(C)=O | AUFUWRKPQLGTGF-FMKGYKFTSA-N | No-Effect | inactive |
| urolithin-A | BRD-K00003383 | O=c1oc2cc(O)ccc2c2ccc(O)cc12 | RIUPLDUFZCXCHM-UHFFFAOYSA-N | No-Effect | inactive |
| ursodeoxycholyltaurine | BRD-K45245728 | C[C@H](CCC(=O)NCCS(=O)(=O)O)[C@H]1CC[C@H]2[C@@H]3[C@@H](O)C[C@@H]4C[C@H](O)CC[C@]4(C)[C@H]3CC[C@@]21C | BHTRKEVKTKCXOH-LBSADWJPSA-N | No-Effect | inactive |
| ursodiol | BRD-K15697815 | C[C@H](CCC(=O)O)[C@H]1CC[C@H]2[C@@H]3[C@@H](O)C[C@@H]4C[C@H](O)CC[C@]4(C)[C@H]3CC[C@@]21C | RUDATBOHQWOJDD-UZVSRGJWSA-N | No-Effect | inactive |
| ursolic-acid | BRD-K68185022 | C[C@@H]1[C@H]2C3=CC[C@@H]4[C@@]5(C)CC[C@H](O)C(C)(C)[C@@H]5CC[C@@]4(C)[C@]3(C)CC[C@@]2(C(=O)O)CC[C@H]1C | WCGUUGGRBIKTOS-GPOJBZKASA-N | No-Effect | inactive |
| V-51 | BRD-K75052463 | COc1cc(CN2CCN(c3ccc(F)cc3)CC2)cc(OC)c1OC | KKCOGNFFBYOKMA-UHFFFAOYSA-N | No-Effect | inactive |
| vaborbactam | BRD-K00003306 | O=C(O)C[C@@H]1CC[C@H](NC(=O)Cc2cccs2)B(O)O1 | IOOWNWLVCOUUEX-WPRPVWTQSA-N | No-Effect | inactive |
| vadadustat | BRD-K00003407 | O=C(O)CNC(=O)c1ncc(-c2cccc(Cl)c2)cc1O | JGRXMPYUTJLTKT-UHFFFAOYSA-N | No-Effect | inactive |
| val-tyr | BRD-K00004630 | CC(C)[C@H](N)C(=O)N[C@@H](Cc1ccc(O)cc1)C(=O)O | VEYJKJORLPYVLO-RYUDHWBXSA-N | No-Effect | inactive |
| valaciclovir | BRD-K46435977 | CC(C)[C@H](N)C(=O)OCCOCn1cnc2c(=O)nc(N)[nH]c21 | HDOVUKNUBWVHOX-QMMMGPOBSA-N | No-Effect | inactive |
| valdecoxib | BRD-K12994359 | Cc1onc(-c2ccccc2)c1-c1ccc(S(N)(=O)=O)cc1 | LNPDTQAFDNKSHK-UHFFFAOYSA-N | No-Effect | inactive |
| valethamate | BRD-A56613951 | CC[C@@H](C)[C@@H](C(=O)OCC[N+](C)(CC)CC)c1ccccc1 | UPPMZCXMQRVMME-SJLPKXTDSA-N | No-Effect | inactive |
| valganciclovir | BRD-K01825663 | CC(C)[C@H](N)C(=O)OC[C@@H](CO)OCn1cnc2c(=O)nc(N)[nH]c21 | WPVFJKSGQUFQAP-BDAKNGLRSA-N | No-Effect | inactive |
| valnemulin | BRD-K22938173 | C=C[C@]1(C)C[C@@H](OC(=O)CSC(C)(C)CNC(=O)[C@H](N)C(C)C)[C@]2(C)[C@H](C)CC[C@]3(CCC(=O)[C@H]32)[C@@H](C)[C@@H]1O | LLYYNOVSVPBRGV-MVNKZKPCSA-N | No-Effect | inactive |
| valnoctamide | BRD-A96644976 | CC[C@@H](C)[C@@H](CC)C(N)=O | QRCJOCOSPZMDJY-RNFRBKRXSA-N | No-Effect | inactive |
| valproic-acid | BRD-K41260949 | CCCC(CCC)C(=O)O | NIJJYAXOARWZEE-UHFFFAOYSA-N | No-Effect | inactive |

| | | | | | |
|---|---|---|---|---|---|
| valpromide | BRD-K54698528 | CCCC(CCC)C(N)=O | OMOMUFTZPTXCHP-UHFFFAOYSA-N | No-Effect | inactive |
| valrocemide | BRD-K32804101 | CCCC(CCC)C(=O)NCC(N)=O | RALGCAOVRLYSMA-UHFFFAOYSA-N | No-Effect | inactive |
| valrubicin | BRD-K72951360 | CCCCC(=O)OCC(=O)[C@]1(O)Cc2c(O)c3c(c(O)c2[C@@H](O[C@H]2C[C@H](NC(=O)C(F)(F)F)[C@H](O)[C@H](C)O2)C1)C(=O)c1c(OC)cccc1C3=O | ZOCKGBMQLCSHFP-KQRAQHLDSA-N | No-Effect | inactive |
| valspodar | BRD-K19412342 | C/C=C/C[C@@H](C)C(=O)[C@H]1C(=O)N[C@@H](C(C)C)C(=O)N(C)CC(=O)N(C)[C@@H](CC(C)C)C(=O)N[C@@H](C(C)C)C(=O)N(C)[C@@H](CC(C)C)C(=O)N[C@@H](C)C(=O)N[C@H](C)C(=O)N(C)[C@@H](CC(C)C)C(=O)N(C)[C@@H](CC(C)C)C(=O)N(C)[C@@H](C(C)C)C(=O)N1C | YJDYDFNKCBANTM-QCWCSKBGSA-N | No-Effect | inactive |
| vancomycin | BRD-K91065602 | CN[C@H](CC(C)C)C(=O)N[C@H]1C(=O)N[C@@H](CC(N)=O)C(=O)N[C@H]2C(=O)N[C@H]3C(=O)N[C@H](C(=O)N[C@H](C(=O)O)c4cc(O)cc(O)c4-c4cc3ccc4O)[C@H](O)c3ccc(c(Cl)c3)Oc3cc2cc(c3O[C@@H]2O[C@H](CO)[C@@H](O)[C@H](O)[C@H]2O[C@H]2C[C@](C)(N)[C@H](O)[C@H](C)O2)Oc2ccc(cc2Cl)[C@H]1O | MYPYJXKWCTUITO-LYRMYLQWSA-N | No-Effect | inactive |
| vanillin | BRD-K24258499 | COc1cc(C=O)ccc1O | MWOOGOJBHIARFG-UHFFFAOYSA-N | No-Effect | inactive |
| vanillylacetone | BRD-K33510429 | COc1cc(CCC(C)=O)ccc1O | OJYLAHXKWMRDGS-UHFFFAOYSA-N | No-Effect | inactive |
| vaniprevir | BRD-K00003445 | CC[C@@H]1C[C@]1(NC(=O)[C@@H]1C[C@@H]2CN1C(=O)[C@H](C(C)(C)C)NC(=O)OCC(C)(C)CCCCc1cccc3c1CN(C3)C(=O)O2)C(=O)NS(=O)(=O)C1CC1 | KUQWGLQLLVFLSM-ONAXAZCASA-N | No-Effect | inactive |
| vanoxerine | BRD-K32501161 | Fc1ccc(C(OCCN2CCN(CCCc3ccccc3)CC2)c2ccc(F)cc2)cc1 | NAUWTFJOPJWYOT-UHFFFAOYSA-N | No-Effect | inactive |
| vapendavir | BRD-K00004577 | CCOc1noc2cc(OCCC3CCN(c4ccc(C)nn4)CC3)ccc12 | DKSVBVKHUICELN-UHFFFAOYSA-N | No-Effect | inactive |
| vardenafil | BRD-K13926615 | CCCc1nc(C)c2c(=O)nc(-c3cc(S(=O)(=O)N4CCN(CC)CC4)ccc3OCC)[nH]n12 | SECKRCOLJRRGGV-UHFFFAOYSA-N | No-Effect | inactive |
| varenicline | BRD-K18855837 | c1cnc2cc3c(cc2n1)[C@H]1CNC[C@@H]3C1 | JQSHBVHOMNKWFT-DTORHVGOSA-N | No-Effect | inactive |
| varespladib | BRD-K09711437 | CCc1c(C(=O)C(N)=O)c2c(OCC(=O)O)cccc2n1Cc1ccccc1 | BHLXTPHDSZUFHR-UHFFFAOYSA-N | No-Effect | inactive |
| vasopressin | BRD-K73028949 | N=C(N)NCCC[C@H](NC(=O)[C@@H]1CCCN1C(=O)[C@@H]1CSSC[C@H](N)C(=O)N[C@@H](Cc2ccc(O)cc2)C(=O)N[C@@H](Cc2ccccc2)C(=O)N[C@@H](CCC(N)=O)C(=O)N[C@@H](CC(N)=O)C(=O)N1)C(=O)NCC(N)=O | KBZOIRJILGZLEJ-LGYYRGKSSA-N | No-Effect | inactive |
| vatinoxan | BRD-K00003277 | CS(=O)(=O)NCCN1C(=O)NC[C@]12CCN1CCc3c(oc4ccccc34)[C@@H]1C2 | GTBKISRCRQUFNL-OXJNMPFZSA-N | No-Effect | inactive |
| VCH-916 | BRD-K75956464 | CO[C@H]1CC[C@H](N(c2cc(C3=CCCCC3)sc2C(=O)O)C(=O)[C@H]2CC[C@H](C)CC2)CC1 | LNYKGELUODVSNS-YSXIEMCBSA-N | No-Effect | inactive |
| vecabrutinib | BRD-K00003503 | NC(=O)[C@H]1CCN(c2ncnc(N)c2F)C[C@@H]1N1CCC[C@@H](Nc2cc(Cl)cc(C(F)(F)F)c2)C1=O | QLRRJMOBVVGXEJ-XHSDSOJGSA-N | No-Effect | inactive |
| vecuronium | BRD-K20168442 | CC(=O)O[C@H]1C[C@@H]2CC[C@@H]3[C@H](CC[C@@]4(C)[C@H]3C[C@H]([N+]3(C)CCCCC3)[C@@H]4OC(C)=O)[C@@]2(C)C[C@@H]1N1CCCCC1 | BGSZAXLLHYERSY-XQIGCQGXSA-N | No-Effect | inactive |
| vedaprofen | BRD-K00004714 | C[C@H](C(=O)O)c1ccc(C2CCCCC2)c2ccccc12 | VZUGVMQFWFVFBX-ZDUSSCGKSA-N | No-Effect | inactive |
| velaresol | BRD-K00004580 | O=Cc1c(O)cccc1OCCCCC(=O)O | NSUDGNLOXMLAEB-UHFFFAOYSA-N | No-Effect | inactive |
| veledimex | BRD-K00003235 | CCC[C@@H](N(NC(=O)c1cccc(OC)c1CC)C(=O)c1cc(C)cc(C)c1)C(C)(C)C | LZWZPGLVHLSWQX-XMMPIXPASA-N | No-Effect | inactive |
| veliflapon | BRD-K12852738 | O=C(O)[C@@H](c1ccc(OCc2ccc3ccccc3n2)cc1)C1CCCC1 | ZEYYDOLCHFETHQ-JOCHJYFZSA-N | No-Effect | inactive |
| veliparib | BRD-K87142802 | C[C@]1(c2nc3cccc(C(N)=O)c3[nH]2)CCCN1 | JNAHVYVRKWKWKQ-CYBMUJFWSA-N | No-Effect | inactive |
| velneperit | BRD-K06357403 | CC(C)(C)S(=O)(=O)N[C@H]1CC[C@H](C(=O)Nc2ccc(C(F)(F)F)cn2)CC1 | WGEWUYACXPEFPO-AULYBMBSSA-N | No-Effect | inactive |
| velpatasvir | BRD-K00003133 | COC[C@H]1C[C@@H](c2nc(-c3ccc4c(c3)COc3cc5c(ccc6[nH]c([C@@H]7CC[C@H](C)N7C(=O)[C@@H](NC(=O)OC)C(C)C)nc65)cc3-4)c[nH]2)N(C(=O)[C@H](NC(=O)OC)c2ccccc2)C1 | FHCUMDQMBHQXKK-CDIODLITSA-N | No-Effect | inactive |
| vemurafenib | BRD-K56343971 | CCCS(=O)(=O)Nc1ccc(F)c(C(=O)c2c[nH]c3ncc(-c4ccc(Cl)cc4)cc23)c1F | GPXBXXGIAQBQNI-UHFFFAOYSA-N | No-Effect | inactive |
| venetoclax | BRD-K62391742 | CC1(C)CCC(CN2CCN(c3ccc(C(=O)NS(=O)(=O)c4ccc(NCC5CCOCC5)c([N+](=O)[O-])c4)c(Oc4cnc5[nH]ccc5c4)c3)CC2)=C(c2ccc(Cl)cc2)C1 | LQBVNQSMGBZMKD-UHFFFAOYSA-N | No-Effect | inactive |
| VER-155008 | BRD-K32330832 | N#Cc1ccc(COC[C@H]2O[C@@H](n3c(NCc4ccc(Cl)c(Cl)c4)nc4c(N)ncnc43)[C@H](O)[C@@H]2O)cc1 | ZXGGCBQORXDVTE-UMCMBGNQSA-N | No-Effect | inactive |
| veralipride | BRD-K00003433 | C=CCN1CCC[C@H]1CNC(=O)c1cc(S(N)(=O)=O)cc(OC)c1OC | RYJXBGGBZJGVQF-LBPRGKRZSA-N | No-Effect | inactive |
| verapamil | BRD-A09533288 | COc1ccc(CCN(C)CCCC(C#N)(c2ccc(OC)c(OC)c2)C(C)C)cc1OC | SGTNSNPWRIOYBX-UHFFFAOYSA-N | No-Effect | inactive |
| verbascoside | BRD-K36201510 | C[C@@H]1O[C@@H](O[C@@H]2[C@@H](O)[C@H](OCCc3ccc(O)c(O)c3)O[C@H](CO)[C@H]2OC(=O)/C=C/c2ccc(O)c(O)c2)[C@H](O)[C@@H](O)[C@H]1O | FBSKJMQYURKNSU-ZLSOWSIRSA-N | No-Effect | inactive |
| vercirnon | BRD-K47929404 | CC(C)(C)c1ccc(S(=O)(=O)Nc2ccc(Cl)cc2C(=O)c2cc[n+]([O-])cc2)cc1 | JRWROCIMSDXGOZ-UHFFFAOYSA-N | No-Effect | inactive |
| verdinexor | BRD-K00004674 | O=C(/C=C\n1cnc(-c2cc(C(F)(F)F)cc(C(F)(F)F)c2)n1)NNc1ccccn1 | OPAKEJZFFCECPN-XQRVVYSFSA-N | No-Effect | inactive |
| verdiperstat | BRD-K00004612 | CC(C)OCCn1c(=S)[nH]c(=O)c2[nH]ccc21 | FVJCUZCRPIMVLB-UHFFFAOYSA-N | No-Effect | inactive |
| vericiguat | BRD-K00003234 | COC(=O)Nc1c(N)nc(-c2nn(Cc3ccccc3F)c3ncc(F)cc23)nc1N | QZFHIXARHDBPBY-UHFFFAOYSA-N | No-Effect | inactive |
| verinurad | BRD-K36741660 | CC(C)(Sc1ccncc1-c1ccc(C#N)c2ccccc12)C(=O)O | YYBOLPLTQDKXPM-UHFFFAOYSA-N | No-Effect | inactive |
| vernakalant | BRD-K19741547 | COc1ccc(CCO[C@@H]2CCCC[C@H]2N2CC[C@@H](O)C2)cc1OC | VBHQKCBVWWUUKN-KZNAEPCWSA-N | No-Effect | inactive |
| verubecestat | BRD-K00003229 | CN1C(N)=N[C@](C)(c2cc(NC(=O)c3ccc(F)cn3)ccc2F)CS1(=O)=O | YHYKUSGACIYRML-KRWDZBQOSA-N | No-Effect | inactive |
| verucerfont | BRD-K00003180 | CC[C@H](Nc1cc(C)nc2c(-c3ccc(OC)cc3C)c(C)nn12)c1nc(C)no1 | VKHVAUKFLBBZFJ-SFHVURJKSA-N | No-Effect | inactive |
| vesamicol | BRD-K14765469 | O[C@@H]1CCCC[C@H]1N1CCC(c2ccccc2)CC1 | YSSBJODGIYRAMI-IAGOWNOFSA-N | No-Effect | inactive |
| vesnarinone | BRD-K13414800 | COc1ccc(C(=O)N2CCN(c3ccc4c(c3)CCC(=O)N4)CC2)cc1OC | ZVNYJIZDIRKMBF-UHFFFAOYSA-N | No-Effect | inactive |
| VGX-1027 | BRD-A32269193 | O=C(O)C[C@H]1CC(c2ccccc2)=NO1 | MUFJHYRCIHHATF-SECBINFHSA-N | No-Effect | inactive |
| vicriviroc | BRD-K84748119 | COC[C@@H](c1ccc(C(F)(F)F)cc1)N1CCN(C2(C)CCN(C(=O)c3c(C)ncnc3C)CC2)C[C@@H]1C | CNPVJJQCETWNEU-CYFREDJKSA-N | No-Effect | inactive |
| vidarabine | BRD-K72093121 | Nc1ncnc2c1ncn2[C@@H]1O[C@H](CO)[C@@H](O)[C@@H]1O | OIRDTQYFTABQOQ-UHTZMRCNSA-N | No-Effect | inactive |
| vigabatrin | BRD-K01825700 | C=C[C@@H](N)CCC(=O)O | PJDFLNIOAUIZSL-RXMQYKEDSA-N | No-Effect | inactive |
| vilanterol | BRD-M36487418 | O=C(O)C(c1ccccc1)(c1ccccc1)c1ccccc1.OCc1cc([C@@H](O)CNCCCCCCOCCOCc2c(Cl)cccc2Cl)ccc1O | KLOLZALDXGTNQE-JIDHJSLPSA-N | No-Effect | inactive |
| vilazodone | BRD-K58010567 | N#Cc1ccc2[nH]cc(CCCCN3CCN(c4ccc5oc(C(N)=O)cc5c4)CC3)c2c1 | SGEGOXDYSFKCPT-UHFFFAOYSA-N | No-Effect | inactive |
| vildagliptin | BRD-K00003565 | N#C[C@@H]1CCCN1C(=O)CN[C@@]12C[C@@H]3C[C@H](C[C@@](O)(C3)C1)C2 | SYOKIDBDQMKNDQ-NHMCJKAESA-N | No-Effect | inactive |
| viloxazine | BRD-A59198242 | CCOc1ccccc1OC[C@H]1CNCCO1 | YWPHCCPCQOJSGZ-LLVKDONJSA-N | No-Effect | inactive |
| vinflunine | BRD-K64120610 | CC[C@]12C=CCN3CC[C@@]4(c5cc([C@@]6(C(=O)OC)C[C@@H]7C[C@@H](C(C)(F)F)CN(CCc8c6[nH]c6ccccc86)C7)c(OC)cc5N(C)[C@H]4[C@@](O)(C(=O)OC)[C@@H]1OC(C)=O)[C@@H]32 | NOSAJPUYIASWEH-USRBKURZSA-N | No-Effect | inactive |

| | | | | | |
|---|---|---|---|---|---|
| vinorelbine | BRD-K04269837 | CCC1=C[C@@H]2CN(C1)Cc1c([nH]c3ccccc13)[C@@](C(=O)OC)(c1cc3c(cc1OC)N(C)[C@H]1[C@@](O)(C(=O)OC)[C@H](OC(C)=O)[C@]4(CC)C=CCN5CC[C@]31[C@@H]54)C2 | GBABOYUKABKIAF-IELIFDKJSA-N | No-Effect | inactive |
| vinpocetine | BRD-K53318339 | CCOC(=O)C1=C[C@]2(CC)CCCN3CCc4c(n1c1ccccc41)[C@@H]32 | DDNCQMVWWZOMLN-IRLDBZIGSA-N | No-Effect | inactive |
| viomycin | BRD-A42379279 | NCCC[C@H](N)CC(=O)N[C@H]1CNC(=O)[C@H]([C@H]2C[C@H](O)N=C(N)N2)NC(=O)C(C=NC(N)=O)NC(=O)[C@H](CO)NC(=O)[C@H](CO)NC1=O | NSWGELWYFNCPHU-XJSZFKHMSA-N | No-Effect | inactive |
| vipadenant | BRD-K56565215 | Cc1cc(Cn2nnc3c(-c4ccco4)nc(N)nc32)ccc1N | HQSBCDPYXDGTCL-UHFFFAOYSA-N | No-Effect | inactive |
| visomitin | BRD-K00003375 | CC1=C(C)C(=O)C(CCCCCCCCCC[P+](c2ccccc2)(c2ccccc2)c2ccccc2)=CC1=O | ZFUDULLGKXSPPH-UHFFFAOYSA-N | No-Effect | inactive |
| vitamin-B12 | BRD-K00003629 | C/C1=C2/N=C(/C=C3\N=C(/C(C)=C4/[C@@H](CCC(N)=O)[C@](C)(CC(N)=O)[C@](C)([C@@H]5N=C1[C@](C)(CCC(=O)NC[C@@H](C)OP(=O)(O)O[C@@H]1[C@@H](CO)O[C@H](n6cnc7cc(C)c(C)cc76)[C@@H]1O)[C@H]5CC(N)=O)N4[Co+]C#N)[C@@](C)(CC(N)=O)[C@@H]3CCC(N)=O)C(C)(C)[C@@H]2CCC(N)=O | RMRCNWBMXRMIRW-WZHZPDAFSA-M | No-Effect | inactive |
| vitamin-E | BRD-K97091514 | Cc1c(C)c2c(c(C)c1O)CC[C@@](C)(CCC[C@H](C)CCC[C@H](C)CCCC(C)C)O2 | GVJHHUAWPYXKBD-IEOSBIPESA-N | No-Effect | inactive |
| vixotrigine | BRD-K00003141 | NC(=O)[C@@H]1CC[C@H](c2ccc(OCc3ccccc3F)cc2)N1 | JESCETIFNOFKEU-SJORKVTESA-N | No-Effect | inactive |
| VL285 | BRD-K00005338 | Cc1ncsc1-c1ccc(CNC(=O)[C@@H]2C[C@@H](O)CN2C(=O)[C@H](C(C)C)N2Cc3ccccc3C2=O)cc1 | HEDFFPYRFJKXQP-VJTSUQJLSA-N | No-Effect | inactive |
| VLX1570 | BRD-K00003343 | C=CC(=O)N1CC/C(=C\c2ccc(F)c([N+](=O)[O-])c2)C(=O)/C(=C/c2ccc(F)c([N+](=O)[O-])c2)C1 | SCKXBVLYWLLALY-CZCYGEDCSA-N | No-Effect | inactive |
| VLX600 | BRD-K26122255 | CC(=NNc1nnc2c(n1)[nH]c1c(C)cccc12)c1ccccn1.C/C(=N/Nc1nnc2c(n1)[nH]c1c(C)cccc12)c1ccccn1 | AEFWYHATFJIATC-KELDOBJCSA-N | No-Effect | inactive |
| voclosporin | BRD-K00003472 | C=C/C=C/C[C@@H](C)[C@@H](O)[C@H]1C(=O)N[C@@H](CC)C(=O)N(C)CC(=O)N(C)[C@@H](CC(C)C)C(=O)N[C@@H](C(C)C)C(=O)N(C)[C@@H](CC(C)C)C(=O)N[C@@H](C)C(=O)N[C@H](C)C(=O)N(C)[C@@H](CC(C)C)C(=O)N(C)[C@@H](CC(C)C)C(=O)N(C)[C@@H](C(C)C)C(=O)N1C | BICRTLVBTLFLRD-PTWUADNWSA-N | No-Effect | inactive |
| voglibose | BRD-K66850609 | OCC(CO)N[C@H]1C[C@](O)(CO)[C@@H](O)[C@H](O)[C@H]1O | FZNCGRZWXLXZSZ-CIQUZCHMSA-N | No-Effect | inactive |
| volinanserin | BRD-K23647661 | COc1cccc([C@H](O)C2CCN(CCc3ccc(F)cc3)CC2)c1OC | HXTGXYRHXAGCFP-OAQYLSRUSA-N | No-Effect | inactive |
| vonoprazan | BRD-K32768103 | CNCc1cc(-c2ccccc2F)n(S(=O)(=O)c2cccnc2)c1 | BFDBKMOZYNOTPK-UHFFFAOYSA-N | No-Effect | inactive |
| vorasidenib | BRD-K00003460 | C[C@@H](Nc1nc(N[C@H](C)C(F)(F)F)nc(-c2cccc(Cl)n2)n1)C(F)(F)F | QCZAWDGAVJMPTA-RNFRBKRXSA-N | No-Effect | inactive |
| voreloxin | BRD-K23677682 | CN[C@H]1CN(c2ccc3c(=O)c(C(=O)O)cn(-c4nccs4)c3n2)C[C@@H]1OC | XZAFZXJXZHRNAQ-STQMWFEESA-N | No-Effect | inactive |
| voriconazole | BRD-K06557128 | C[C@@H](c1ncncc1F)[C@](O)(Cn1cncn1)c1ccc(F)cc1F | BCEHBSKCWLPMDN-MGPLVRAMSA-N | No-Effect | inactive |
| vorinostat | BRD-K81418486 | O=C(CCCCCCC(=O)Nc1ccccc1)NO | WAEXFXRVDQXREF-UHFFFAOYSA-N | No-Effect | inactive |
| vortioxetine | BRD-K53963539 | Cc1ccc(Sc2ccccc2N2CCNCC2)c(C)c1 | YQNWZWMKLDQSAC-UHFFFAOYSA-N | No-Effect | inactive |
| voruciclib | BRD-K00004725 | CN1CC[C@@H](c2c(O)cc(O)c3c(=O)cc(-c4ccc(C(F)(F)F)cc4Cl)oc23)[C@@H]1CO | MRPGRAKIAJJGMM-OCCSQVGLSA-N | No-Effect | inactive |
| VS-4718 | BRD-K09877029 | CNC(=O)c1ccccc1Nc1cc(Nc2ccc(N3CCOCC3)cc2OC)ncc1C(F)(F)F | IGUBBWJDMLCRIK-UHFFFAOYSA-N | No-Effect | inactive |
| VT-464 | BRD-K72005722 | CC(C)[C@](O)(c1ccc2cc(OC(F)F)c(OC(F)F)cc2c1)c1cn[nH]n1 | ZBRAJOQFSNYJMF-SFHVURJKSA-N | No-Effect | inactive |
| VTP-27999 | BRD-K81694556 | CN[C@H](CNC(=O)N1CCC[C@@H]([C@@H](OCCNC(=O)OC)c2cccc(Cl)c2)C1)C[C@H]1CCCOC1 | NXWASIVXQMMPLM-ZXMXYHOLSA-N | No-Effect | inactive |
| VU0155069 | BRD-K48535733 | C[C@@H](CN1CCC(n2c(=O)[nH]c3cc(Cl)ccc32)CC1)NC(=O)c1ccc2ccccc2c1 | DRIMIUYGTDAQOX-KRWDZBQOSA-N | No-Effect | inactive |
| VU0238429 | BRD-K24187789 | COc1ccc(CN2C(=O)C(=O)c3cc(OC(F)(F)F)ccc32)cc1 | CKLGZXFOLMHCMC-UHFFFAOYSA-N | No-Effect | inactive |
| VU0240551 | BRD-K27074404 | Cc1csc(NC(=O)CSc2ccc(-c3ccccc3)nn2)n1 | WJRWSLORVIHRNX-UHFFFAOYSA-N | No-Effect | inactive |
| VU0357121 | BRD-K86109159 | CCCCOc1ccc(C(=O)Nc2ccc(F)cc2F)cc1 | AHCYOTLTLQTPSU-UHFFFAOYSA-N | No-Effect | inactive |
| VU0360172 | BRD-K83649660 | O=C(NC1CCC1)c1ccc(C#Cc2cccc(F)c2)nc1 | NEMHWVUKNWAKTM-UHFFFAOYSA-N | No-Effect | inactive |
| VU0364439 | BRD-K72687424 | O=C(Nc1ccc(S(=O)(=O)Nc2ccccc2Cl)c(Cl)c1)c1ccccn1 | IXHCGJXBIHHIEF-UHFFFAOYSA-N | No-Effect | inactive |
| VU0364770 | BRD-K66527818 | O=C(Nc1cccc(Cl)c1)c1ccccn1 | SUYUTNCKIOLMAJ-UHFFFAOYSA-N | No-Effect | inactive |
| VU0422288 | BRD-K68674163 | O=C(Nc1ccc(Oc2ccc(Cl)cn2)c(Cl)c1)c1ccccn1 | MZRLPXFGQKQHST-UHFFFAOYSA-N | No-Effect | inactive |
| VU10010 | BRD-K12483104 | Cc1cc(C)c2c(N)c(C(=O)NCc3ccc(Cl)cc3)sc2n1 | FPRULFHDSFKYBV-UHFFFAOYSA-N | No-Effect | inactive |
| VU152100 | BRD-K87727457 | COc1ccc(CNC(=O)c2sc3nc(C)cc(C)c3c2N)cc1 | MDNWGCQSCGNTKH-UHFFFAOYSA-N | No-Effect | inactive |
| VU1545 | BRD-K48371964 | O=C(Nc1cc(-c2ccccc2)nn1-c1ccccc1F)c1ccc([N+](=O)[O-])cc1 | BRAZLURTFMCAHU-UHFFFAOYSA-N | No-Effect | inactive |
| VU29 | BRD-K54783522 | O=C(Nc1cc(-c2ccccc2)nn1-c1ccccc1)c1ccc([N+](=O)[O-])cc1 | KIALCSMRIHRFPL-UHFFFAOYSA-N | No-Effect | inactive |
| VU591 | BRD-K72063794 | O=[N+]([O-])c1ccc2nc(COCc3nc4ccc([N+](=O)[O-])cc4[nH]3)[nH]c2c1 | ZQPXNYLXYNRFNP-UHFFFAOYSA-N | No-Effect | inactive |
| VUF10460 | BRD-K90082312 | CN1CCN(c2cc(-c3ccccc3)nc(N)n2)CC1 | NIJGWJIOMPHDBP-UHFFFAOYSA-N | No-Effect | inactive |
| VUF11207 | BRD-A93717708 | COc1cc(C(=O)N(CC[C@H]2CCCN2C)C/C(C)=C/c2ccccc2F)cc(OC)c1OC | JVRRRKAXHMGUHZ-JIKIWWBOSA-N | No-Effect | inactive |
| VX-11e | BRD-K93725829 | Cc1cnc(Nc2ccc(F)cc2Cl)nc1-c1c[nH]c(C(=O)N[C@H](CO)c2cccc(Cl)c2)c1 | WUTVMXLIGHTZJC-OAQYLSRUSA-N | No-Effect | inactive |
| VX-222 | BRD-K70610771 | CC(C)(C)C#Cc1cc(N(C(=O)[C@H]2CC[C@H](C)CC2)[C@H]2CC[C@@H](O)CC2)c(C(=O)O)s1 | WPMJNLCLKAKMLA-UGLKCIBTSA-N | No-Effect | inactive |
| VX-702 | BRD-K07736136 | NC(=O)c1ccc(N(C(N)=O)c2c(F)cccc2F)nc1-c1ccc(F)cc1F | FYSRKRZDBHOFAY-UHFFFAOYSA-N | No-Effect | inactive |
| VX-745 | BRD-K91900765 | O=c1ncn2nc(Sc3ccc(F)cc3F)ccc2c1-c1c(Cl)cccc1Cl | VEPKQEUBKLEPRA-UHFFFAOYSA-N | No-Effect | inactive |
| VX-765 | BRD-K78430341 | CCO[C@@H]1OC(=O)C[C@@H]1NC(=O)[C@@H]1CCCN1C(=O)[C@@H](NC(=O)c1ccc(N)c(Cl)c1)C(C)(C)C | SJDDOCKBXFJEJB-MOKWFATOSA-N | No-Effect | inactive |
| W-54011 | BRD-A00559310 | COc1ccc2c(c1)[C@H](C(=O)N(Cc1ccc(N(C)C)cc1)c1ccc(C(C)C)cc1)CCC2 | DVYASSBBADJRAS-MUUNZHRXSA-N | No-Effect | inactive |
| walrycin-B | BRD-K01129913 | Cn1nc(-c2ccc(C(F)(F)F)cc2)nc2c(=O)n(C)c(=O)nc1-2 | XRVMPTWWKLKLPB-UHFFFAOYSA-N | No-Effect | inactive |
| warfarin | BRD-K25991148 | CC(=O)C[C@@H](c1ccccc1)c1c(O)c2ccccc2oc1=O | PJVWKTKQMONHTI-HNNXBMFYSA-N | No-Effect | inactive |
| WAY-100635 | BRD-K69812556 | COc1ccccc1N1CCN(CCN(C(=O)C2CCCCC2)c2ccccn2)CC1 | SBPRIAGPYFYCRT-UHFFFAOYSA-N | No-Effect | inactive |
| WAY-161503 | BRD-A62021152 | O=C1Nc2cc(Cl)c(Cl)cc2N2CCNC[C@H]12 | PHGWDAICBXUJDU-SNVBAGLBSA-N | No-Effect | inactive |
| WAY-170523 | BRD-K36198571 | Cc1cc(C)c(N(Cc2ccccc2)S(=O)(=O)c2ccc(OCCNC(=O)c3cc4ccccc4o3)cc2)c(C(=O)NO)c1 | FARMEEAGJWMFSZ-UHFFFAOYSA-N | No-Effect | inactive |
| WAY-200070 | BRD-K08223200 | Oc1ccc(-c2nc3cc(O)cc(Br)c3o2)cc1 | BAAILVWEAXFTSF-UHFFFAOYSA-N | No-Effect | inactive |
| WAY-207024 | BRD-K08657056 | CC(C)(C)c1ccc(-c2nc3c(N4CCN(Cc5ccc6nccnc6c5)CC4)cccc3[nH]2)cc1 | AVEVJMZJJJDOCI-UHFFFAOYSA-N | No-Effect | inactive |
| WAY-208466 | BRD-K63483542 | CN(C)CCn1cc(S(=O)(=O)c2cccc(F)c2)c2cccnc21 | SFSFIDVAEMDPIP-UHFFFAOYSA-N | No-Effect | inactive |
| WAY-213613 | BRD-K96758128 | N[C@@H](CC(=O)Nc1ccc(Oc2cc(F)c(F)cc2Br)cc1)C(=O)O | BNYDDAAZMBUFRG-ZDUSSCGKSA-N | No-Effect | inactive |
| WAY-255348 | BRD-K00004556 | Cn1c(C#N)ccc1-c1cc(F)c2c(c1)C(C)(C)C(=O)N2 | KIOOLNRTWPFVHX-UHFFFAOYSA-N | No-Effect | inactive |
| WAY-362450 | BRD-K54640016 | CC(C)OC(=O)C1=CN(C(=O)c2ccc(F)c(F)c2)CC(C)(C)c2c1[nH]c1ccccc21 | INASOKQDNHHMRE-UHFFFAOYSA-N | No-Effect | inactive |
| WB-4101 | BRD-A01493904 | COc1cccc(OC)c1OCCNC[C@H]1COc2ccccc2O1 | GYSZUJHYXCZAKI-AWEZNQCLSA-N | No-Effect | inactive |
| WDR5-0103 | BRD-K47726410 | COC(=O)c1ccc(N2CCN(C)CC2)c(NC(=O)c2cccc(OC)c2)c1 | ZPLBXOVTSNRBFB-UHFFFAOYSA-N | No-Effect | inactive |

| | | | | | |
|---|---|---|---|---|---|
| WEHI-345-analog | BRD-K55487946 | Cc1ccc(-c2nn(C(C)(C)CNC(=O)c3ncccc3C)c3ncnc(N)c23)cc1 | ZCZGSEXBXXYQQC-UHFFFAOYSA-N | No-Effect | inactive |
| WHI-P154 | BRD-K07487750 | COc1cc2ncnc(Nc3ccc(O)c(Br)c3)c2cc1OC | CBIAKDAYHRWZCU-UHFFFAOYSA-N | No-Effect | inactive |
| WIKI4 | BRD-K19316653 | COc1ccc(-n2c(SCCCN3C(=O)c4cccc5cccc(c45)C3=O)nnc2-c2ccncc2)cc1 | RNUXIZKXJOGYQP-UHFFFAOYSA-N | No-Effect | inactive |
| WIN-18446 | BRD-K30930903 | O=C(NCCCCCCCCNC(=O)C(Cl)Cl)C(Cl)Cl | FAOMZVDZARKPFJ-UHFFFAOYSA-N | No-Effect | inactive |
| WIN-64338 | BRD-K22662557 | CCCC[P+](CCCC)(CCCC)Cc1ccc(NC(=O)[C@H](Cc2ccc3ccccc3c2)N/C(=N/C2CCCCC2)NC2CCCCC2)cc1 | UKMJWGFHXMGRNG-QLKFWGTOSA-O | No-Effect | inactive |
| wnt-c59 | BRD-K78455053 | Cc1cc(-c2ccc(CC(=O)Nc3ccc(-c4cccnc4)cc3)cc2)ccn1 | KHZOJCQBHJUJFY-UHFFFAOYSA-N | No-Effect | inactive |
| WP1130 | BRD-K43644456 | CCC[C@H](NC(=O)/C(C#N)=C/c1cccc(Br)n1)c1ccccc1 | LIDOPKHSVQTSJY-VMEIHUARSA-N | No-Effect | inactive |
| WP921 | BRD-K00005248 | O=C(O)Cn1nnnc1-c1ccsc1 | DDVAZOPRCKAYOV-UHFFFAOYSA-N | No-Effect | inactive |
| WR99210 | BRD-K45973863 | CC1(C)N=C(N)N=C(N)N1OCCCOc1cc(Cl)c(Cl)cc1Cl | MJZJYWCQPMNPRM-UHFFFAOYSA-N | No-Effect | inactive |
| WWL-113 | BRD-K49389158 | CCOC(=O)c1ccc(-c2ccc(OC(=O)N(C)Cc3cccc(-c4ccncc4)c3)cc2)cc1 | AKIIPHDGVCFVCC-UHFFFAOYSA-N | No-Effect | inactive |
| WWL-123 | BRD-K83176916 | CN(Cc1cccc(-c2ccccc2)c1)C(=O)Oc1ccc(-c2ccc(C(N)=O)cc2)cc1 | UUJGYRHAOHORFC-UHFFFAOYSA-N | No-Effect | inactive |
| WY-16922 | BRD-K43419711 | CCOC(=O)C(=O)Nc1cccc(OC)c1C(N)=O | GGLRYZACPLNCKB-UHFFFAOYSA-N | No-Effect | inactive |
| WZ-3146 | BRD-K73293050 | C=CC(=O)Nc1cccc(Oc2nc(Nc3ccc(N4CCN(C)CC4)cc3)ncc2Cl)c1 | APHGZZPEOCCYNO-UHFFFAOYSA-N | No-Effect | inactive |
| WZ4003 | BRD-K45909031 | CCC(=O)Nc1cccc(Oc2nc(Nc3ccc(N4CCN(C)CC4)cc3OC)ncc2Cl)c1 | SDGJBAUIGHSMRI-UHFFFAOYSA-N | No-Effect | inactive |
| WZ811 | BRD-K25359048 | c1ccc(NCc2ccc(CNc3ccccn3)cc2)nc1 | KBVFRXIGQQRMEF-UHFFFAOYSA-N | No-Effect | inactive |
| xaliproden | BRD-K88358234 | FC(F)(F)c1cccc(C2=CCN(CCc3ccc4ccccc4c3)CC2)c1 | WJJYZXPHLSLMGE-UHFFFAOYSA-N | No-Effect | inactive |
| xamoterol | BRD-A59337442 | O=C(NCCNCC(O)COc1ccc(O)cc1)N1CCOCC1 | DXPOSRCHIDYWHW-UHFFFAOYSA-N | No-Effect | inactive |
| xanomeline | BRD-K72815923 | CCCCCCOc1nsnc1C1=CCCN(C)C1 | JOLJIIDDOBNFHW-UHFFFAOYSA-N | No-Effect | inactive |
| xanthone | BRD-K27135764 | O=c1c2ccccc2oc2ccccc12 | JNELGWHKGNBSMD-UHFFFAOYSA-N | No-Effect | inactive |
| XAV-939 | BRD-K12762134 | O=c1nc(-c2ccc(C(F)(F)F)cc2)[nH]c2c1CSCC2 | KLGQSVMIPOVQAX-UHFFFAOYSA-N | No-Effect | inactive |
| XBD173 | BRD-K45875383 | CCN(Cc1ccccc1)C(=O)Cn1c(=O)n(C)c2cnc(-c3ccccc3)nc21 | NBMBIEOUVBHEBM-UHFFFAOYSA-N | No-Effect | inactive |
| XD-14 | BRD-K69211575 | CCc1c(C(=O)Nc2cc(S(=O)(=O)N(CC)CC)ccc2O)[nH]c(C)c1C(C)=O | DPBKLIVPNYGQQG-UHFFFAOYSA-N | No-Effect | inactive |
| XE-991 | BRD-K42748308 | O=C1c2ccccc2C(Cc2ccncc2)(Cc2ccncc2)c2ccccc21 | KHJFBUUFMUBONL-UHFFFAOYSA-N | No-Effect | inactive |
| xenalipin | BRD-K99506538 | O=C(O)c1ccccc1-c1ccc(C(F)(F)F)cc1 | IQOMYCGTGFGDFN-UHFFFAOYSA-N | No-Effect | inactive |
| xibenolol | BRD-A21446384 | Cc1cccc(OC[C@@H](O)CNC(C)(C)C)c1C | RKUQLAPSGZJLGP-ZDUSSCGKSA-N | No-Effect | inactive |
| xilobam | BRD-K79633074 | Cc1cccc(C)c1NC(=O)/N=C1/CCCN1C | PAXRPWSXCPTPCA-QINSGFPZSA-N | No-Effect | inactive |
| xipamide | BRD-K27393415 | Cc1cccc(C)c1NC(=O)c1cc(S(N)(=O)=O)c(Cl)cc1O | MTZBBNMLMNBNJL-UHFFFAOYSA-N | No-Effect | inactive |
| XL041 | BRD-K00003439 | CC(C)(O)c1cn(-c2ccc(-c3cc(F)c(CO)c(S(C)(=O)=O)c3)cc2F)c(C(C)(C)c2c(Cl)cccc2Cl)n1 | HNAJDMYOTDNOBK-UHFFFAOYSA-N | No-Effect | inactive |
| XL147 | BRD-K95901403 | Cc1ccc(S(=O)(=O)Nc2nc3ccccc3nc2Nc2ccc3nsnc3c2)cc1 | MQMKRQLTIWPEDM-UHFFFAOYSA-N | No-Effect | inactive |
| XL228 | BRD-K14274271 | CC(C)c1cc(CNc2nc(Nc3cc(C4CC4)n[nH]3)cc(N3CCN(C)CC3)n2)on1 | ALKJNCZNEOTEMP-UHFFFAOYSA-N | No-Effect | inactive |
| XL388 | BRD-K03549949 | Cc1c(C(=O)N2CCOc3ccc(-c4ccc(N)nc4)cc3C2)ccc(S(C)(=O)=O)c1F | LNFBAYSBVQBKFR-UHFFFAOYSA-N | No-Effect | inactive |
| xylazine | BRD-K21565985 | Cc1cccc(C)c1NC1=NCCCS1 | BPICBUSOMSTKRF-UHFFFAOYSA-N | No-Effect | inactive |
| xylometazoline | BRD-K08356259 | Cc1cc(C(C)(C)C)cc(C)c1CC1=NCCN1 | HUCJFAOMUPXHDK-UHFFFAOYSA-N | No-Effect | inactive |
| xylose | BRD-K10759144 | O=C[C@H](O)[C@@H](O)[C@H](O)CO | PYMYPHUHKUWMLA-VPENINKCSA-N | No-Effect | inactive |
| Y-11 | BRD-K09317629 | OCC[N+]12CN3CN(CN(C3)C1)C2 | SQWGHMPVSMSPNY-UHFFFAOYSA-N | No-Effect | inactive |
| Y-27152 | BRD-K42679050 | CC(=O)N(OCc1ccccc1)[C@@H]1c2cc(C#N)ccc2OC(C)(C)[C@H]1O | RHFUXPCCELGMFC-UXHICEINSA-N | No-Effect | inactive |
| Y-27632 | BRD-K44084986 | C[C@@H](N)[C@H]1CC[C@H](C(=O)Nc2ccncc2)CC1 | IYOZTVGMEWJPKR-IJLUTSLNSA-N | No-Effect | inactive |
| Y-39983 | BRD-K56751279 | C[C@@H](N)c1ccc(C(=O)Nc2ccnc3[nH]ccc23)cc1 | JTVBXQAYBIJXRP-SNVBAGLBSA-N | No-Effect | inactive |
| Y16 | BRD-K81865756 | Cc1cccc(COc2cccc(/C=C3\C(=O)NN(c4ccccc4)C3=O)c2)c1 | ITMLWGWTDWJSRZ-PXLXIMEGSA-N | No-Effect | inactive |
| YC-1 | BRD-K60476892 | OCc1ccc(-c2nn(Cc3ccccc3)c3ccccc23)o1 | OQQVFCKUDYMWGV-UHFFFAOYSA-N | No-Effect | inactive |
| YIL-781 | BRD-K91403886 | Cc1nc2ccc(Oc3ccc(F)cc3)cc2c(=O)n1C[C@H]1CCCN(C(C)C)C1 | FRKXOBMDEXCHHD-SFHVURJKSA-N | No-Effect | inactive |
| YK-4-279 | BRD-A62182663 | COc1ccc(C(=O)C[C@@]2(O)C(=O)Nc3c(Cl)ccc(Cl)c32)cc1 | HLXSCTYHLQHQDJ-KRWDZBQOSA-N | No-Effect | inactive |
| YL0919 | BRD-K00003390 | O=c1ccccn1CC1(O)CCN(Cc2ccccc2)CC1 | BGMVOROUOUVVRV-UHFFFAOYSA-N | No-Effect | inactive |
| YM-022 | BRD-K09436144 | Cc1cccc(NC(=O)N[C@@H]2N=C(c3ccccc3)c3ccccc3N(CC(=O)c3ccccc3C)C2=O)c1 | YCXFHPUBGMMWJQ-PMERELPUSA-N | No-Effect | inactive |
| YM-155 | BRD-K76703230 | COCCn1c2c([n+](Cc3cnccn3)c1C)C(=O)c1ccccc1C2=O | OTSOOHRUMBRSHZ-UHFFFAOYSA-N | No-Effect | inactive |
| YM-244769 | BRD-K52087522 | Nc1cccc(CNC(=O)c2ccc(Oc3ccc(OCc4cccc(F)c4)cc3)nc2)c1 | JZMLHJRKSJXARY-UHFFFAOYSA-N | No-Effect | inactive |
| YM-298198 | BRD-K59650319 | Cc1c(C(=O)N(C)C2CCCCC2)sc2nc3ccc(N)cc3n12 | KCBXOMYXOBVLED-UHFFFAOYSA-N | No-Effect | inactive |
| YM-298198-desmethyl | BRD-K48059230 | Cc1c(C(=O)NC2CCCCC2)sc2nc3ccc(N)cc3n12 | VWOMTTIMBHBPBI-UHFFFAOYSA-N | No-Effect | inactive |
| YM-511 | BRD-K38398115 | N#Cc1ccc(N(Cc2ccc(Br)cc2)n2cnnc2)cc1 | GGPPBTSXFROGAE-UHFFFAOYSA-N | No-Effect | inactive |
| YM-750 | BRD-K91716465 | Cc1cc(C)c(NC(=O)N(Cc2ccc3c(c2)Cc2ccccc2-3)C2CCCCCC2)c(C)c1 | FMLJREWZCZHGGW-UHFFFAOYSA-N | No-Effect | inactive |
| YM-90709 | BRD-K06712146 | COc1cc2c(cc1OC)-c1cc3nc4ccccc4nc3n1C(C)(C)C2 | HIXSPVQXXDULHS-UHFFFAOYSA-N | No-Effect | inactive |
| YO-01027 | BRD-K04622161 | C[C@H](NC(=O)Cc1cc(F)cc(F)c1)C(=O)N[C@@H]1C(=O)N(C)c2ccccc2-c2ccccc21 | QSHGISMANBKLQL-OWJWWREXSA-N | No-Effect | inactive |
| yoda-1 | BRD-K88570498 | Clc1cccc(Cl)c1CSc1nnc(-c2cnccn2)s1 | BQNXBSYSQXSXPT-UHFFFAOYSA-N | No-Effect | inactive |
| yohimbine | BRD-K35586044 | COC(=O)[C@H]1[C@@H](O)CC[C@H]2CN3CCc4c([nH]c5ccccc45)[C@@H]3C[C@@H]21 | BLGXFZZNTVWLAY-SCYLSFHTSA-N | No-Effect | inactive |
| YS-035 | BRD-K06208435 | COc1ccc(CCN(C)CCc2ccc(OC)c(OC)c2)cc1OC | OXZMQFKVNVQMLI-UHFFFAOYSA-N | No-Effect | inactive |
| YS-201 | BRD-K00004746 | CCOC(=O)C1=C(C)NC(C)=C(C(=O)OCCN2CCCCC2)[C@@H]1c1cccc([N+](=O)[O-])c1 | LBSRZVRITCRLIU-JOCHJYFZSA-N | No-Effect | inactive |
| YZ9 | BRD-K53438416 | CCOC(=O)c1cc2ccc(O)cc2oc1=O | IETDBZQIWIJQJG-UHFFFAOYSA-N | No-Effect | inactive |
| Z160 | BRD-K16037980 | O=C(CC(c1ccccc1)c1ccccc1)N1CCN(C(c2ccccc2)c2ccccc2)CC1 | VCPMZDWBEWTGNW-UHFFFAOYSA-N | No-Effect | inactive |
| zacopride | BRD-K01826799 | COc1cc(N)c(Cl)cc1C(=O)N[C@H]1CN2CCC1CC2 | FEROPKNOYKURCJ-ZDUSSCGKSA-N | No-Effect | inactive |
| zafirlukast | BRD-K71289571 | COc1cc(C(=O)NS(=O)(=O)c2ccccc2C)ccc1Cc1cn(C)c2ccc(NC(=O)OC3CCCC3)cc12 | YEEZWCHGZNKEEK-UHFFFAOYSA-N | No-Effect | inactive |
| zaldaride | BRD-A26032986 | C[C@@]1(CN2CCC(n3c(=O)[nH]c4ccccc43)CC2)OCc2ccccc2-n2cccc21 | HTGCJAAPDHZCHL-SANMLTNESA-N | No-Effect | inactive |
| zaltidine | BRD-K12261274 | Cc1nc(-c2csc(NC(=N)N)n2)c[nH]1 | GIMNAEMRNXUAQP-UHFFFAOYSA-N | No-Effect | inactive |
| zaltoprofen | BRD-A00578795 | C[C@@H](C(=O)O)c1ccc2c(c1)CC(=O)c1ccccc1S2 | MUXFZBHBYYYLTH-SNVBAGLBSA-N | No-Effect | inactive |
| ZAMI-633 | BRD-K11984358 | CCC(=O)NO | RSIPQHOWTCNEBI-UHFFFAOYSA-N | No-Effect | inactive |
| zamifenacin | BRD-K80451230 | c1ccc(C(O[C@@H]2CCCN(CCc3ccc4c(c3)OCO4)C2)c2ccccc2)cc1 | BDNFQGRSKSQXRI-XMMPIXPASA-N | No-Effect | inactive |
| zanamivir | BRD-K23067689 | CC(=O)N[C@@H]1[C@@H](NC(=N)N)C=C(C(=O)O)O[C@H]1[C@H](O)[C@H](O)CO | ARAIBEBZBOPLMB-UFGQHTETSA-N | No-Effect | inactive |
| zanubrutinib | BRD-K00003413 | C=CC(=O)N1CCC([C@@H]2CCNc3c(C(N)=O)c(-c4ccc(Oc5ccccc5)cc4)nn32)CC1 | RNOAOAWBMHREKO-QFIPXVFZSA-N | No-Effect | inactive |
| ZAPA | BRD-K44825564 | N=C(N)S/C=C\C(=O)O | QEYNZJBVNYDZKZ-UPHRSURJSA-N | No-Effect | inactive |
| zaprinast | BRD-K16542329 | CCCOc1ccccc1-c1nc(=O)c2n[nH]nc2[nH]1 | REZGGXNDEMKIQB-UHFFFAOYSA-N | No-Effect | inactive |
| zardaverine | BRD-K37561857 | COc1cc(-c2ccc(=O)[nH]n2)ccc1OC(F)F | HJMQDJPMQIHLPB-UHFFFAOYSA-N | No-Effect | inactive |
| zatebradine | BRD-K92446736 | COc1ccc(CCN(C)CCCN2CCc3cc(OC)c(OC)cc3CC2=O)cc1OC | KEDQCFRVSHYKLR-UHFFFAOYSA-N | No-Effect | inactive |
| ZD-0947 | BRD-K00004620 | N#Cc1cccc([C@@H]2C=C(C(F)(F)F)NC3=C2C(=O)CCC3)c1 | RUVMMEREJMHLOS-LBPRGKRZSA-N | No-Effect | inactive |
| ZD-2079 | BRD-K64157027 | O=C(O)Cc1ccc(OCCNC[C@H](O)c2ccccc2)cc1 | SRBPKVWITYPHQR-KRWDZBQOSA-N | No-Effect | inactive |
| ZD-7114 | BRD-K28971625 | COCCNC(=O)COc1ccc(OCCNC[C@H](O)COc2ccccc2)cc1 | RVMBDLSFFNKKLG-SFHVURJKSA-N | No-Effect | inactive |
| ZD-7155 | BRD-K11373525 | CCc1cc2c(c(CC)n1)CCC(=O)N2Cc1ccc(-c2ccccc2-c2nn[nH]n2)cc1 | BFVNEYDCFJNLGN-UHFFFAOYSA-N | No-Effect | inactive |
| ZD-7288 | BRD-K18678457 | CCN(c1ccccc1)c1cc(=NC)n(C)c(C)n1 | JABSKGQQWUDVRU-UHFFFAOYSA-N | No-Effect | inactive |

| | | | | | |
|---|---|---|---|---|---|
| zeatin | BRD-K90033078 | C/C(=C\CNc1ncnc2nc[nH]c12)CO | UZKQTCBAMSWPJD-FARCUNLSSA-N | No-Effect | inactive |
| zebularine | BRD-K87714311 | O=c1ncccn1[C@@H]1O[C@H](CO)[C@@H](O)[C@H]1O | RPQZTTQVRYEKCR-WCTZXXKLSA-N | No-Effect | inactive |
| zibotentan | BRD-K31553034 | COc1nc(C)cnc1NS(=O)(=O)c1cccnc1-c1ccc(-c2nnco2)cc1 | FJHHZXWJVIEFGJ-UHFFFAOYSA-N | No-Effect | inactive |
| zidovudine | BRD-K72903603 | Cc1cn([C@H]2C[C@H](N=[N+]=[N-])[C@@H](CO)O2)c(=O)[nH]c1=O | HBOMLICNUCNMMY-XLPZGREQSA-N | No-Effect | inactive |
| zileuton | BRD-A56359832 | C[C@H](c1cc2ccccc2s1)N(O)C(N)=O | MWLSOWXNZPKENC-SSDOTTSWSA-N | No-Effect | inactive |
| zilpaterol | BRD-K42816473 | CC(C)N[C@H]1CCn2c(=O)[nH]c3cccc(c32)[C@@H]1O | ZSTCZWJCLIRCOJ-AAEUAGOBSA-N | No-Effect | inactive |
| zimelidine | BRD-K46914825 | CN(C)C/C=C(/c1ccc(Br)cc1)c1cccnc1 | OYPPVKRFBIWMSX-SXGWCWSVSA-N | No-Effect | inactive |
| zinc-undecylenate | BRD-K13330768 | C=CCCCCCCCCC(=O)O[Zn]OC(=O)CCCCCCCCC=C | YMCOHQVWOBMDCZ-UHFFFAOYSA-L | No-Effect | inactive |
| zindotrine | BRD-K26598477 | Cc1cc(N2CCCCC2)nn2cnnc12 | YYHWOTFWBVRTQD-UHFFFAOYSA-N | No-Effect | inactive |
| ziprasidone | BRD-K29582115 | O=C1Cc2cc(CCN3CCN(c4nsc5ccccc45)CC3)c(Cl)cc2N1 | MVWVFYHBGMAFLY-UHFFFAOYSA-N | No-Effect | inactive |
| ZK-164015 | BRD-K05151076 | CCCCCS(=O)(=O)CCCCCCCCCCn1c(-c2ccc(O)cc2)c(C)c2cc(O)ccc21 | LYJSJVYJLZOMCD-UHFFFAOYSA-N | No-Effect | inactive |
| ZK-200775 | BRD-K67525671 | O=c1[nH]c2cc(C(F)(F)F)c(N3CCOCC3)cc2n(CP(=O)(O)O)c1=O | WZMQMKNCWDCCMT-UHFFFAOYSA-N | No-Effect | inactive |
| ZK-811752 | BRD-K91825936 | C[C@@H]1CN(Cc2ccc(F)cc2)CCN1C(=O)COc1ccc(Cl)cc1NC(N)=O | XQYASZNUFDVMFH-CQSZACIVSA-N | No-Effect | inactive |
| ZK-93423 | BRD-K33882852 | CCOC(=O)c1ncc2[nH]c3ccc(OCc4ccccc4)cc3c2c1COC | ALBKMJDFBZVHAK-UHFFFAOYSA-N | No-Effect | inactive |
| ZLN005 | BRD-K73073971 | CC(C)(C)c1ccc(-c2nc3ccccc3[nH]2)cc1 | LQUNNCQSFFKSSK-UHFFFAOYSA-N | No-Effect | inactive |
| ZLN024 | BRD-K64103579 | Cc1ccc(OCCSc2nccn2)c(Br)c1 | KWJRSHZSULRJHE-UHFFFAOYSA-N | No-Effect | inactive |
| ZM-226600 | BRD-A62209527 | C[C@@](O)(C(=O)Nc1ccc(S(=O)(=O)c2ccccc2)cc1)C(F)(F)F | LJLXQHHFAKVTNP-OAHLLOKOSA-N | No-Effect | inactive |
| ZM-241385 | BRD-K19605405 | Nc1nc(NCCc2ccc(O)cc2)nc2nc(-c3ccco3)nn12 | PWTBZOIUWZOPFT-UHFFFAOYSA-N | No-Effect | inactive |
| ZM-306416 | BRD-K41337261 | COc1cc2ncnc(Nc3ccc(Cl)cc3F)c2cc1OC | YHUIUSRCUKUUQA-UHFFFAOYSA-N | No-Effect | inactive |
| ZM-323881 | BRD-K67831364 | Cc1cc(F)c(Nc2ncnc3cc(OCc4ccccc4)ccc23)cc1O | NVBNDZZLJRYRPD-UHFFFAOYSA-N | No-Effect | inactive |
| ZM-336372 | BRD-K73789395 | Cc1ccc(NC(=O)c2cccc(N(C)C)c2)cc1NC(=O)c1ccc(O)cc1 | PYEFPDQFAZNXLI-UHFFFAOYSA-N | No-Effect | inactive |
| ZM-39923 | BRD-K40624912 | CC(C)N(CCC(=O)c1ccc2ccccc2c1)Cc1ccccc1 | JSASWRWALCMOQP-UHFFFAOYSA-N | No-Effect | inactive |
| ZM-447439 | BRD-K72703948 | COc1cc2c(Nc3ccc(NC(=O)c4ccccc4)cc3)ncnc2cc1OCCCN1CCOCC1 | OGNYUTNQZVRGMN-UHFFFAOYSA-N | No-Effect | inactive |
| zofenopril-calcium | BRD-K46654563 | C[C@H](CSC(=O)c1ccccc1)C(=O)N1C[C@@H](Sc2ccccc2)C[C@H]1C(=O)O | IAIDUHCBNLFXEF-MNEFBYGVSA-N | No-Effect | inactive |
| zolantidine | BRD-K08996725 | c1cc(CN2CCCCC2)cc(OCCCNc2nc3ccccc3s2)c1 | KUBONGDXTUOOLM-UHFFFAOYSA-N | No-Effect | inactive |
| zoledronic-acid | BRD-K84253429 | O=P(O)(O)C(O)(Cn1ccnc1)P(=O)(O)O | XRASPMIURGNCCH-UHFFFAOYSA-N | No-Effect | inactive |
| zoliflodacin | BRD-K00003264 | C[C@@H]1O[C@H](C)CN2c3c(cc4c(N5C(=O)OC[C@@H]5C)noc4c3F)CC3(C(=O)NC(=O)NC3=O)[C@@H]12 | ZSWMIFNWDQEXDT-ZESJGQACSA-N | No-Effect | inactive |
| zolimidine | BRD-K45125084 | CS(=O)(=O)c1ccc(-c2cn3ccccc3n2)cc1 | VSLIUWLPFRVCDL-UHFFFAOYSA-N | No-Effect | inactive |
| zometapine | BRD-K14571191 | Cc1nn(C)c2c1C(c1cccc(Cl)c1)=NCCN2 | YCWROMXNZZIJDQ-UHFFFAOYSA-N | No-Effect | inactive |
| zoniporide | BRD-K43992824 | NC(N)=NC(=O)c1cnn(-c2cccc3ncccc23)c1C1CC1 | GDXBRVCQGGKXJY-UHFFFAOYSA-N | No-Effect | inactive |
| zonisamide | BRD-K48300629 | NS(=O)(=O)Cc1noc2ccccc12 | UBQNRHZMVUUOMG-UHFFFAOYSA-N | No-Effect | inactive |
| zopolrestat | BRD-K47710001 | O=C(O)Cc1nn(Cc2nc3cc(C(F)(F)F)ccc3s2)c(=O)c2ccccc12 | BCSVCWVQNOXFGL-UHFFFAOYSA-N | No-Effect | inactive |
| zosuquidar | BRD-K70557564 | O[C@@H](COc1cccc2ncccc12)CN1CCN([C@@H]2c3ccccc3[C@@H]3[C@H](c4ccccc42)C3(F)F)CC1 | IHOVFYSQUDPMCN-DBEBIPAYSA-N | No-Effect | inactive |
| zotarolimus | BRD-K46843573 | CO[C@H]1C[C@@H]2CC[C@@H](C)[C@@](O)(O2)C(=O)C(=O)N2CCC[C@H]2C(=O)O[C@@H]([C@H](C)C[C@@H]2CC[C@H](n3cnnn3)[C@H](OC)C2)CC(=O)[C@H](C)/C=C(\C)[C@@H](O)[C@@H](OC)C(=O)[C@H](C)C[C@H](C)/C=C/C=C/C=C/1C | CGTADGCBEXYWNE-JUKNQOCSSA-N | No-Effect | inactive |
| ZSET1446 | BRD-K35984734 | O=C1N=C2C=CC=CN2C12Cc1ccccc1C2 | QZWYXEBIQWJXAR-UHFFFAOYSA-N | No-Effect | inactive |
| ZSTK-474 | BRD-K63068307 | FC(F)c1nc2ccccc2n1-c1nc(N2CCOCC2)nc(N2CCOCC2)n1 | HGVNLRPZOWWDKD-UHFFFAOYSA-N | No-Effect | inactive |
| zuclopenthixol | BRD-K28761384 | OCCN1CCN(CC/C=C2/c3ccccc3Sc3ccc(Cl)cc32)CC1 | WFPIAZLQTJBIFN-DVZOWYKESA-N | No-Effect | inactive |
| zuranolone | BRD-K00003448 | C[C@]12CC[C@@H]3[C@H]4CC[C@@](C)(O)C[C@H]4CC[C@H]3[C@@H]1CC[C@@H]2C(=O)Cn1cc(C#N)cn1 | HARRKNSQXBRBGZ-GVKWWOCJSA-N | No-Effect | inactive |
| 10-DEBC | BRD-K70792160 | CCN(CC)CCCCN1c2ccccc2Oc2ccc(Cl)cc21 | GYBXAGDWMCJZJK-UHFFFAOYSA-N | Strong | inactive |
| A-485 | BRD-K30233223 | CNC(=O)Nc1ccc2c(c1)CC[C@@]21OC(=O)N(CC(=O)N(Cc2ccc(F)cc2)[C@@H](C)C(F)(F)F)C1=O | VRVJKILQRBSEAG-LFPIHBKWSA-N | Strong | inactive |
| acepromazine | BRD-K37814297 | CC(=O)c1ccc2c(c1)N(CCCN(C)C)c1ccccc1S2 | NOSIYYJFMPDDSA-UHFFFAOYSA-N | Strong | inactive |
| actinomycin-d | BRD-K16547679 | Cc1c2oc3c(C)ccc(C(=O)N[C@@H]4C(=O)N[C@@H](C(C)C)C(=O)N5CCC[C@H]5C(=O)N(C)CC(=O)N(C)[C@@H](C(C)C)C(=O)O[C@@H]4C)c3nc-2c(C(=O)N[C@@H]2C(=O)N[C@@H](C(C)C)C(=O)N3CCC[C@H]3C(=O)N(C)CC(=O)N(C)[C@@H](C(C)C)C(=O)O[C@@H]2C)c(N)c1=O | RJURFGZVJUQBHK-DCUCITLNSA-N | Strong | inactive |
| afatinib | BRD-K66175015 | CN(C)C/C=C/C(=O)Nc1cc2c(Nc3ccc(F)c(Cl)c3)ncnc2cc1O[C@H]1CCOC1 | ULXXDDBFHOBEHA-CWDCEQMOSA-N | Strong | inactive |
| AGN-195183 | BRD-K32533226 | CC1(C)CCC(C)(C)c2c1cc(C(=O)Nc1cc(F)c(C(=O)O)c(F)c1)c(O)c2Cl | PNAWUIKCVQSLFG-UHFFFAOYSA-N | Strong | inactive |
| aloxistatin | BRD-K91188791 | CCOC(=O)[C@H]1O[C@@H]1C(=O)N[C@@H](CC(C)C)C(=O)NCCC(C)C | SRVFFFJZQVENJC-IHRRRGAJSA-N | Strong | inactive |
| amiodarone | BRD-K17561142 | CCCCc1oc2ccccc2c1C(=O)c1cc(I)c(OCCN(CC)CC)c(I)c1 | IYIKLHRQXLHMJQ-UHFFFAOYSA-N | Strong | inactive |
| amitriptyline | BRD-K53737926 | CN(C)CCC=C1c2ccccc2CCc2ccccc21 | KRMDCWKBEZIMAB-UHFFFAOYSA-N | Strong | inactive |
| amodiaquine | BRD-K91290917 | CCN(CC)Cc1cc(Nc2ccnc3cc(Cl)ccc23)ccc1O | OVCDSSHSILBFBN-UHFFFAOYSA-N | Strong | inactive |
| AQ-13 | BRD-K00003365 | CCN(CC)CCCNc1ccnc2cc(Cl)ccc12 | NCPLTAGJJVCHOW-UHFFFAOYSA-N | Strong | inactive |
| asenapine | BRD-K00003551 | CN1C[C@@H]2c3ccccc3Oc3ccc(Cl)cc3[C@H]2C1 | VSWBSWWIRNCQIJ-HUUCEWRRSA-N | Strong | inactive |
| atiprimod | BRD-K38527262 | CCCC1(CCC)CCC2(CCN(CCCN(CC)CC)C2)CC1 | SERHTTSLBVGRBY-UHFFFAOYSA-N | Strong | inactive |
| AZ5104 | BRD-K81958169 | C=CC(=O)Nc1cc(Nc2nccc(-c3c[nH]c4ccccc34)n2)c(OC)cc1N(C)CCN(C)C | IQNVEOMHJHBNHC-UHFFFAOYSA-N | Strong | inactive |
| AZ7550 | BRD-K00003515 | C=CC(=O)Nc1cc(Nc2nccc(-c3cn(C)c4ccccc34)n2)c(OC)cc1N(C)CCNC | ZROCWKZRGJYPTG-UHFFFAOYSA-N | Strong | inactive |
| AZD1390 | BRD-K00003509 | CC(C)n1c(=O)n(C)c2cnc3ccc(F)c(-c4ccc(OCCCN5CCCCC5)nc4)cc3c21 | VQSZIPCGAGVRRP-UHFFFAOYSA-N | Strong | inactive |
| bafilomycin-a1 | BRD-K00010387 | CO/C1=C\C(C)=C/[C@@H](C)[C@@H](O)[C@@H](C)C/C(C)=C/C=C/[C@H](OC)[C@@H]([C@@H](C)[C@@H](O)[C@H](C)[C@@]2(O)C[C@@H](O)[C@H](C)[C@@H](C(C)C)O2)OC1=O | XDHNQDDQEHDUTM-UTRUWJARSA-N | Strong | inactive |
| balicatib | BRD-K27204852 | CCCN1CCN(c2ccc(C(=O)NC3(C(=O)NCC#N)CCCCC3)cc2)CC1 | LLCRBOWRJOUJAE-UHFFFAOYSA-N | Strong | inactive |
| bardoxolone | BRD-K20545304 | CC1(C)CC[C@]2(C(=O)O)CC[C@]3(C)[C@H](C(=O)C=C4[C@@]3(C)CC[C@H]3C(C)(C)C(=O)C(C#N)=C[C@]43C)[C@@H]2C1 | TXGZJQLMVSIZEI-UQMAOPSPSA-N | Strong | inactive |
| BAX-channel-blocker | BRD-K01826533 | O[C@@H](CN1CCNCC1)Cn1c2ccc(Br)cc2c2cc(Br)ccc21 | KUUJEXLRLIPQQJ-HNNXBMFYSA-N | Strong | inactive |
| BAY-2402234 | BRD-K22379844 | CCn1c(CO)nn(-c2cc(O[C@@H](C)C(F)(F)F)c(C(=O)Nc3c(F)cccc3Cl)cc2F)c1=O | KNVJMHHAXCPZHF-JTQLQIEISA-N | Strong | inactive |
| BMS-566419 | BRD-K56932272 | CCN1CCN(c2ccc(C(C)(C)NC(=O)c3cc4[nH]c5ccccc5c(=O)c4cc3F)cn2)CC1 | XEVJUIZOZCFECP-UHFFFAOYSA-N | Strong | inactive |
| brequinar | BRD-K96123349 | Cc1c(-c2ccc(-c3ccccc3F)cc2)nc2ccc(F)cc2c1C(=O)O | PHEZJEYUWHETKO-UHFFFAOYSA-N | Strong | inactive |
| bruceantin | BRD-A36057565 | COC(=O)[C@]12OC[C@@]34[C@H]1[C@@H](OC(=O)/C=C(\C)C(C)C)C(=O)O[C@@H]3C[C@H]1[C@@H](C)C(=O)C(=O)C[C@]1(C)[C@H]4[C@@H](O)[C@@H]2O | KMCZIPGKKBSLEX-AOHBHZTBSA-N | Strong | inactive |
| buparvaquone | BRD-A07947329 | CC(C)(C)C1CCC(C[C@H]2C(=O)C(=O)c3ccccc3C2=O)CC1 | OMMKCJLAGTXGLC-MQBCKMQZSA-N | Strong | inactive |

| | | | | | |
|---|---|---|---|---|---|
| C-021 | BRD-K98179423 | COc1cc2nc(N3CCC(N4CCCCC4)CC3)nc(NC3CCCCCC3)c2cc1OC | WYVBISCFCHREDA-UHFFFAOYSA-N | Strong | inactive |
| calpeptin | BRD-K26134695 | CCCC[C@@H](C=O)NC(=O)[C@H](CC(C)C)NC(=O)OCc1ccccc1 | PGGUOGKHUUUWAF-ROUUACIJSA-N | Strong | inactive |
| CC-115 | BRD-K16755045 | CCN1C(=O)CN=C2NC=C(c3ccc(-c4nnc[nH]4)nc3C)N=C21 | GMYLVKUGJMYTFB-UHFFFAOYSA-N | Strong | inactive |
| cetaben | BRD-K86954340 | CCCCCCCCCCCCCCCCNc1ccc(C(=O)O)cc1 | QXWKHSSBFQDQPR-UHFFFAOYSA-N | Strong | inactive |
| chlorcyclizine | BRD-A57133233 | CN1CCN([C@@H](c2ccccc2)c2ccc(Cl)cc2)CC1 | WFNAKBGANONZEQ-SFHVURJKSA-N | Strong | inactive |
| chlorpromazine | BRD-K89997465 | CN(C)CCCN1c2ccccc2Sc2ccc(Cl)cc21 | ZPEIMTDSQAKGNT-UHFFFAOYSA-N | Strong | inactive |
| chlorquinaldol | BRD-K90374350 | Cc1ccc2c(Cl)cc(Cl)c(O)c2n1 | GPTXWRGISTZRIO-UHFFFAOYSA-N | Strong | inactive |
| cinobufotalin | BRD-K00003354 | CC(=O)O[C@@H]1[C@H](c2ccc(=O)oc2)[C@@]2(C)CC[C@H]3[C@@H](CC[C@]4(O)C[C@@H](O)CC[C@]34C)[C@]23O[C@H]13 | KBKUJJFDSHBPPA-ZNCGZLKOSA-N | Strong | inactive |
| clemastine | BRD-K30240666 | CN1CCC[C@@H]1CCO[C@](C)(c1ccccc1)c1ccc(Cl)cc1 | YNNUSGIPVFPVBX-NHCUHLMSSA-N | Strong | inactive |
| clomifene | BRD-K29950728 | CCN(CC)CCOc1ccc(/C(=C(\Cl)c2ccccc2)c2ccccc2)cc1 | GKIRPKYJQBWNGO-QPLCGJKRSA-N | Strong | inactive |
| clorotepine | BRD-A19053834 | CN1CCN([C@H]2Cc3ccccc3Sc3ccc(Cl)cc32)CC1 | XRYLGRGAWQSVQW-KRWDZBQOSA-N | Strong | inactive |
| cobimetinib | BRD-K03390685 | O=C(c1ccc(F)c(F)c1Nc1ccc(I)cc1F)N1CC(O)([C@@H]2CCCCN2)C1 | BSMCAPRUBJMWDF-KRWDZBQOSA-N | Strong | inactive |
| COH29 | BRD-K00003307 | O=C(Nc1nc(-c2ccc(O)c(O)c2)c(-c2ccccc2)s1)c1ccc(O)c(O)c1 | LGGDLPSXAGQFSG-UHFFFAOYSA-N | Strong | inactive |
| combretastatin-A-4 | BRD-K61195623 | COc1ccc(/C=C\c2cc(OC)c(OC)c(OC)c2)cc1O | HVXBOLULGPECHP-WAYWQWQTSA-N | Strong | inactive |
| convallatoxin | BRD-K00004559 | C[C@@H]1O[C@@H](O[C@H]2CC[C@]3(C=O)[C@H]4CC[C@]5(C)[C@@H](C6=CC(=O)OC6)CC[C@]5(O)[C@@H]4CC[C@]3(O)C2)[C@H](O)[C@H](O)[C@H]1O | HULMNSIAKWANQO-JQKSAQOKSA-N | Strong | inactive |
| CYM-5442 | BRD-A43005764 | CCOc1ccc(-c2nc(-c3cccc4c3CC[C@H]4NCCO)no2)cc1OCC | NUIKTBLZSPQGCP-LJQANCHMSA-N | Strong | inactive |
| D-3263 | BRD-K00003205 | COc1ccc2c(c1)n(CCN)c(=O)n2C(=O)[C@@H]1C[C@H](C)CC[C@H]1C(C)C | ZGTYTFYRCIRWBL-HYVNUMGLSA-N | Strong | inactive |
| dasotraline | BRD-K00003142 | N[C@@H]1CC[C@@H](c2ccc(Cl)c(Cl)c2)c2ccccc21 | SRPXSILJHWNFMK-MEDUHNTESA-N | Strong | inactive |
| Debio-0932 | BRD-K00003157 | CN(C)c1cc2c(cc1Sc1nc3c(N)nccc3n1CCNCC(C)(C)C)OCO2 | RVJIQAYFTOPTKK-UHFFFAOYSA-N | Strong | inactive |
| deltarasin | BRD-K90830492 | c1ccc(Cn2c(-c3ccc(OC[C@H](C4CCNCC4)n4c(-c5ccccc5)nc5ccccc54)cc3)nc3ccccc32)cc1 | LTZKEDSUXKTTTC-KXQOOQHDSA-N | Strong | inactive |
| deslanoside | BRD-K99919177 | C[C@H]1O[C@@H](O[C@H]2[C@@H](O)C[C@H](O[C@H]3CC[C@]4(C)[C@H]5C[C@@H](O)[C@]6(C)[C@@H](C7=CC(=O)OC7)CC[C@]6(O)[C@@H]5CC[C@@H]4C3)O[C@@H]2C)C[C@H](O)[C@@H]1O[C@H]1C[C@H](O)[C@H](O[C@@H]2O[C@H](CO)[C@@H](O)[C@H](O)[C@H]2O)[C@@H](C)O1 | OBATZBGFDSVCJD-LALPQLPRSA-N | Strong | inactive |
| digitoxin | BRD-K63668566 | C[C@H]1O[C@@H](O[C@H]2[C@@H](O)C[C@H](O[C@H]3[C@@H](O)C[C@H](O[C@H]4CC[C@]5(C)[C@H]6CC[C@]7(C)[C@@H](C8=CC(=O)OC8)CC[C@]7(O)[C@@H]6CC[C@@H]5C4)O[C@@H]3C)O[C@@H]2C)C[C@H](O)[C@@H]1O | WDJUZGPOPHTGOT-XUDUSOBPSA-N | Strong | inactive |
| digoxigenin | BRD-K18619710 | C[C@]12CC[C@H](O)C[C@H]1CC[C@@H]1[C@@H]2C[C@@H](O)[C@]2(C)[C@@H](C3=CC(=O)OC3)CC[C@]12O | SHIBSTMRCDJXLN-KCZCNTNESA-N | Strong | inactive |
| digoxin | BRD-K00004302 | C[C@H]1O[C@@H](O[C@H]2[C@@H](O)C[C@H](O[C@H]3[C@@H](O)C[C@H](O[C@H]4CC[C@]5(C)[C@H]6C[C@@H](O)[C@@]7(C)[C@@H](C8=CC(=O)OC8)CC[C@]7(O)[C@@H]6CC[C@@H]5C4)O[C@@H]3C)O[C@@H]2C)C[C@H](O)[C@@H]1O | LTMHDMANZUZIPE-SYUUTQQWSA-N | Strong | inactive |
| dolastatin-10 | BRD-K78567475 | CC[C@H](C)[C@@H]([C@@H](CC(=O)N1CCC[C@H]1[C@H](OC)[C@@H](C)C(=O)N[C@@H](Cc1ccccc1)c1nccs1)OC)N(C)C(=O)[C@@H](NC(=O)[C@H](C(C)C)N(C)C)C(C)C | OFDNQWIFNXBECV-VFSYNPLYSA-N | Strong | inactive |
| drofenine | BRD-A22267103 | CCN(CC)CCOC(=O)[C@H](c1ccccc1)C1CCCCC1 | AGJBLWCLQCKRJP-LJQANCHMSA-N | Strong | inactive |
| duloxetine | BRD-K71103788 | CNCC[C@H](Oc1cccc2ccccc12)c1cccs1 | ZEUITGRIYCTCEM-KRWDZBQOSA-N | Strong | inactive |
| elacestrant | BRD-K00003295 | CCNCCc1ccc(CN(CC)c2cc(OC)ccc2[C@@H]2CCc3cc(O)ccc3C2)cc1 | SIFNOOUKXBRGGB-AREMUKBSSA-N | Strong | inactive |
| eliglustat | BRD-K86410485 | CCCCCCCC(=O)N[C@H](CN1CCCC1)[C@H](O)c1ccc2c(c1)OCCO2 | FJZZPCZKBUKGGU-AUSIDOKSSA-N | Strong | inactive |
| ELN-441958 | BRD-K50699835 | O=C(c1cccc(N2Cc3cccc(Cl)c3C2=O)c1)N1CCC2(CC1)CCN(c1ccncc1)CC2 | ARYQHSWJGHCGJS-UHFFFAOYSA-N | Strong | inactive |
| emetine | BRD-K03067624 | CC[C@H]1CN2CCc3cc(OC)c(OC)cc3[C@@H]2C[C@@H]1C[C@H]1NCCc2cc(OC)c(OC)cc21 | AUVVAXYIELKVAI-CKBKHPSWSA-N | Strong | inactive |
| endoxifen | BRD-K62752296 | CC/C(=C(\c1ccc(O)cc1)c1ccc(OCCNC)cc1)c1ccccc1 | MHJBZVSGOZTKRH-IZHYLOQSSA-N | Strong | inactive |
| ferroquine | BRD-K00003281 | CN(C)CC1=CC=C[C@@]1(CNc1ccnc2cc(Cl)ccc12)[Fe]C1C=CC=C1 | UZTPLPOYSVSMBU-UHFFFAOYSA-N | Strong | inactive |
| firsocostat | BRD-K00003236 | COc1ccccc1[C@H](Cn1c(=O)n(C(C)(C)C(=O)O)c(=O)c2c(C)c(-c3ncco3)sc21)OC1CCOCC1 | ZZWWXIBKLBMSCS-FQEVSTJZSA-N | Strong | inactive |
| FLT3-IN-1 | BRD-K00003510 | CCCNc1nc(Nc2ccc(C#N)cc2)ncc1C#CCCCNC(=O)[C@H](C)N(C)C(=O)/C=C/CN(C)C | HJFSVYUFOXAVAA-YUAYGMJFSA-N | Strong | inactive |
| flumatinib | BRD-K68870568 | Cc1ncc(NC(=O)c2ccc(CN3CCN(C)CC3)c(C(F)(F)F)c2)cc1Nc1nccc(-c2cccnc2)n1 | BJCJYEYYYGBROF-UHFFFAOYSA-N | Strong | inactive |
| fosbretabulin | BRD-K06814349 | COc1ccc(/C=C\c2cc(OC)c(OC)c(OC)c2)cc1OP(=O)(O)O | WDOGQTQEKVLZIJ-WAYWQWQTSA-N | Strong | inactive |
| gamithromycin | BRD-K00003489 | CCCN1C[C@H](C)[C@@H](O)[C@](C)(O)[C@@H](CC)OC(=O)[C@H](C)[C@@H](O[C@H]2C[C@@](C)(OC)[C@@H](O)[C@H](C)O2)[C@H](C)[C@@H](O[C@@H]2O[C@H](C)C[C@H](N(C)C)[C@H]2O)[C@](C)(O)C[C@H]1C | VWAMTBXLZPEDQO-UZSBJOJWSA-N | Strong | inactive |
| GR-127935 | BRD-K11911061 | COc1ccc(NC(=O)c2ccc(-c3ccc(-c4noc(C)n4)cc3C)cc2)cc1N1CCN(C)CC1 | YDBCEBYHYKAFRX-UHFFFAOYSA-N | Strong | inactive |
| GSK2126458 | BRD-K07972848 | COc1ncc(-c2ccc3nccc(-c4ccnnc4)c3c2)cc1NS(=O)(=O)c1ccc(F)cc1F | CGBJSGAELGCMKE-UHFFFAOYSA-N | Strong | inactive |
| GSK2606414 | BRD-K81689258 | Cn1cc(-c2ccc3c(c2)CCN3C(=O)Cc2cccc(C(F)(F)F)c2)c2c(N)ncnc21 | SIXVRXARNAVBTC-UHFFFAOYSA-N | Strong | inactive |
| GSK369796 | BRD-K00003121 | CC(C)(C)NCc1ccc(Nc2ccnc3cc(Cl)ccc23)cc1O | ZVMMVSSEAMUNGI-UHFFFAOYSA-N | Strong | inactive |
| GSK461364 | BRD-K92428232 | C[C@@H](Oc1cc(-n2cnc3ccc(CN4CCN(C)CC4)cc32)sc1C(N)=O)c1ccccc1C(F)(F)F | ZHJGWYRLJUCMRT-QGZVFWFLSA-N | Strong | inactive |
| guanosine | BRD-K67644125 | Nc1nc(=O)c2ncn([C@@H]3O[C@H](CO)[C@@H](O)[C@H]3O)c2[nH]1 | NYHBQMYGNKIUIF-UUOKFMHZSA-N | Strong | inactive |
| GZD824 | BRD-K35329391 | Cc1ccc(C(=O)Nc2ccc(CN3CCN(C)CC3)c(C(F)(F)F)c2)cc1C#Cc1cnc2n[nH]cc2c1 | TZKBVRDEOITLRB-UHFFFAOYSA-N | Strong | inactive |
| halofuginone | BRD-K54247840 | O=C(C[C@H]1NCCC[C@@H]1O)Cn1cnc2cc(Br)c(Cl)cc2c1=O | LVASCWIMLIKXLA-CABCVRRESA-N | Strong | inactive |
| harringtonine | BRD-K00003715 | COC(=O)C[C@](O)(CCC(C)(C)O)C(=O)O[C@@H]1C(OC)=C[C@]23CCCN2CCc2cc4c(cc2[C@H]13)OCO4 | HAVJATCHLFRDHY-KSZYUSJVSA-N | Strong | inactive |
| homoharringtonine | BRD-K76674262 | COC(=O)C[C@](O)(CCCC(C)(C)O)C(=O)O[C@@H]1C(OC)=C[C@]23CCCN2CCc2cc4c(cc2[C@H]13)OCO4 | HYFHYPWGAURHIV-JFIAXGOJSA-N | Strong | inactive |
| hypoestoxide | BRD-K13881745 | C=C1C(=O)C[C@H]2C[C@@H]3O[C@@]3(C)CC[C@@H]3O[C@@]3(C)C[C@@H](OC(C)=O)[C@H]1C2(C)C | HNPAHGHFONBTLV-KSJQNFQUSA-N | Strong | inactive |
| idarubicin | BRD-K69650333 | CC(=O)[C@]1(O)Cc2c(O)c3c(c(O)c2[C@@H](O[C@H]2C[C@H](N)[C@@H](O)[C@H](C)O2)C1)C(=O)c1ccccc1C3=O | XDXDZDZNSLXDNA-TZNDIEGXSA-N | Strong | inactive |
| IPAG | BRD-K82421491 | N=C(Nc1ccc(I)cc1)NC1C2CC3CC(C2)CC1C3 | UUKPIWYXWLJPJF-UHFFFAOYSA-N | Strong | inactive |
| IT1t | BRD-K73499906 | CC1(C)CN2C(CS/C(=N/C3CCCCC3)NC3CCCCC3)=CSC2=N1 | OOSUDWRRWZVFEB-UHFFFAOYSA-N | Strong | inactive |

| | | | | | |
|---|---|---|---|---|---|
| ixazomib | BRD-K78659596 | CC(C)C[C@H](NC(=O)CNC(=O)c1cc(Cl)ccc1Cl)B(O)O | MXAYKZJJDUDWDS-LBPRGKRZSA-N | Strong | inactive |
| JHW-007 | BRD-K32892871 | CCCCN1[C@H]2CC[C@@H]1C[C@H](OC(c1ccc(F)cc1)c1ccc(F)cc1)C2 | HZEVDVGPBUFIAW-JKHIJQBDSA-N | Strong | inactive |
| JTC-801 | BRD-K17705806 | CCc1ccc(OCc2ccccc2C(=O)Nc2ccc3nc(C)cc(N)c3c2)cc1 | VTGBZWHPJFMTKS-UHFFFAOYSA-N | Strong | inactive |
| K-strophanthidin | BRD-K84595254 | C[C@]12CC[C@H]3[C@@H](CC[C@]4(O)C[C@@H](O)CC[C@]34C=O)[C@@]1(O)CC[C@@H]2C1=CC(=O)OC1 | ODJLBQGVINUMMR-HZXDTFASSA-N | Strong | inactive |
| KU-60019 | BRD-K36016295 | C[C@H]1CN(CC(=O)Nc2ccc3c(c2)Cc2cccc(-c4cc(=O)cc(N5CCOCC5)o4)c2S3)C[C@@H](C)O1 | SCELLOWTHJGVIC-BGYRXZFFSA-N | Strong | inactive |
| lasalocid | BRD-K10207760 | CC[C@@H](C(=O)[C@@H](C)[C@@H](O)[C@H](C)CCc1ccc(C)c(O)c1C(=O)O)[C@H]1O[C@](CC)([C@H]2CC[C@](O)(CC)[C@H](C)O2)C[C@@H]1C | BBMULGJBVDDDNI-OWKLGTHSSA-N | Strong | inactive |
| lasofoxifene | BRD-M93148412 | O=C(O)[C@@H](O)[C@H](O)C(=O)O.Oc1ccc2c(c1)CC[C@H](c1ccccc1)[C@@H]2c1ccc(OCCN2CCCC2)cc1 | INEHJXCWEVNEDZ-LUDNRVPPSA-N | Strong | inactive |
| latrunculin-B | BRD-K61292460 | C/C1=C/C(=O)O[C@@H]2C[C@@H](CC[C@H](C)/C=C\CC1)O[C@@](O)([C@@H]1CSC(=O)N1)C2 | NSHPHXHGRHSMIK-JRIKCGFMSA-N | Strong | inactive |
| leucomethylene-blue | BRD-K27460187 | CN(C)c1ccc2c(c1)Sc1cc(N(C)C)ccc1N2 | QTWZICCBKBYHDM-UHFFFAOYSA-N | Strong | inactive |
| levamlodipine | BRD-K00005329 | CCOC(=O)C1=C(COCCN)NC(C)=C(C(=O)OC)[C@@H]1c1ccccc1Cl | HTIQEAQVCYTUBX-KRWDZBQOSA-N | Strong | inactive |
| levomepromazine | BRD-K35559145 | COc1ccc2c(c1)N(C[C@H](C)CN(C)C)c1ccccc1S2 | VRQVVMDWGGWHTJ-CQSZACIVSA-N | Strong | inactive |
| LY2228820 | BRD-K59227464 | CC(C)(C)Cn1c(N)nc2ccc(-c3nc(C(C)(C)C)[nH]c3-c3ccc(F)cc3)nc21 | XPPBBJCBDOEXDN-UHFFFAOYSA-N | Strong | inactive |
| lylamine | BRD-K62289640 | CC(C)c1ccc2c(c1)CC[C@@H]1[C@]2(C)CCC[C@@]1(C)CN | JVVXZOOGOGPDRZ-SLFFLAALSA-N | Strong | inactive |
| MCOPPB | BRD-K73276278 | CC1(N2CCC(n3c([C@@H]4CCCNC4)nc4ccccc43)CC2)CCCCCCC1 | CYYNMPPFEJPBJD-OAQYLSRUSA-N | Strong | inactive |
| meglutol | BRD-K00495673 | CC(O)(CC(=O)O)CC(=O)O | NPOAOTPXWNWTSH-UHFFFAOYSA-N | Strong | inactive |
| metaphit | BRD-K95701999 | S=C=Nc1cccc(C2(N3CCCCC3)CCCCC2)c1 | FGSGBQAQSPSRJK-UHFFFAOYSA-N | Strong | inactive |
| methotrexate | BRD-K59456551 | CN(Cc1cnc2nc(N)nc(N)c2n1)c1ccc(C(=O)N[C@@H](CCC(=O)O)C(=O)O)cc1 | FBOZXECLQNJBKD-ZDUSSCGKSA-N | Strong | inactive |
| mianserin | BRD-A19661776 | CN1CCN2c3ccccc3Cc3ccccc3C2C1 | UEQUQVLFIPOEMF-UHFFFAOYSA-N | Strong | inactive |
| mibefradil | BRD-K09549677 | COCC(=O)O[C@]1(CCN(C)CCCc2nc3ccccc3[nH]2)CCc2cc(F)ccc2[C@@H]1C(C)C | HBNPJJILLOYFJU-VMPREFPWSA-N | Strong | inactive |
| MK-2206 | BRD-K68065987 | NC1(c2ccc(-c3nc4ccn5c(=O)[nH]nc5c4cc3-c3ccccc3)cc2)CCC1 | ULDXWLCXEDXJGE-UHFFFAOYSA-N | Strong | inactive |
| monensin | BRD-K55375480 | CC[C@@]1([C@@H]2O[C@@H]([C@H]3O[C@@](O)(CO)[C@H](C)C[C@@H]3C)C[C@@H]2C)CC[C@H]([C@]2(C)CC[C@]3(C[C@H](O)[C@@H](C)[C@@H]([C@@H](C)[C@@H](OC)[C@H](C)C(=O)O)O3)O2)O1 | GAOZTHIDHYLHMS-KEOBGNEYSA-N | Strong | inactive |
| MRX-2843 | BRD-K00003417 | CN1CCN(Cc2ccc(-c3cn([C@H]4CC[C@H](O)CC4)c4nc(NCCC5CC5)ncc34)cc2)CC1 | LBEJYFVJIPQSNX-SOAUALDESA-N | Strong | inactive |
| mycophenolate-mofetil | BRD-K92428153 | COc1c(C)c2c(c(O)c1C/C=C(\C)CCC(=O)OCCN1CCOCC1)C(=O)OC2 | RTGDFNSFWBGLEC-SYZQJQIISA-N | Strong | inactive |
| mycophenolic-acid | BRD-K63750851 | COc1c(C)c2c(c(O)c1C/C=C(\C)CCC(=O)O)C(=O)OC2 | HPNSFSBZBAHARI-RUDMXATFSA-N | Strong | inactive |
| nanchangmycin | BRD-A73871920 | CO[C@H]1CC[C@@H](O[C@H]2C[C@H]([C@]3(C)CC[C@]4(C[C@H](O)[C@@H](C)[C@@H]([C@@H](C)/C=C(\C)C(=O)[C@H](C)C[C@H](C)C(=O)O)O4)O3)O[C@]3(O[C@@H]([C@H]4O[C@@](O)(CO)[C@H](C)C[C@@H]4C)C[C@@H]3C)[C@@H]2C)O[C@@H]1C | FELYAZAWTURXNF-KWJIWYEDSA-N | Strong | inactive |
| naquotinib | BRD-K64114733 | C=CC(=O)N1CC[C@@H](Oc2nc(Nc3ccc(N4CCC(N5CCN(C)CC5)CC4)cc3)c(C(N)=O)nc2CC)C1 | QKDCLUARMDUUKN-XMMPIXPASA-N | Strong | inactive |
| narasin | BRD-A25608658 | CC[C@@H](C(=O)[C@@H](C)[C@@H](O)[C@H](C)[C@@H]1O[C@@H]([C@@H](CC)C(=O)O)[C@H](C)C[C@@H]1C)[C@H]1O[C@]2(C=C[C@@H](O)[C@]3(CC[C@@](C)([C@H]4CC[C@](O)(CC)[C@H](C)O4)O3)O2)[C@H](C)C[C@@H]1C | VHKXXVVRRDYCIK-OQMITHEGSA-N | Strong | inactive |
| nelfinavir | BRD-K74117820 | Cc1c(O)cccc1C(=O)N[C@@H](CSc1ccccc1)[C@H](O)CN1C[C@H]2CCCC[C@H]2C[C@H]1C(=O)NC(C)(C)C | QAGYKUNXZHXKMR-HKWSIXNMSA-N | Strong | inactive |
| NNC-55-0396 | BRD-K78122587 | CC(C)[C@H]1c2ccc(F)cc2CC[C@@]1(CCN(C)CCCc1nc2ccccc2[nH]1)OC(=O)C1CC1 | VHHVPDKNKPNKHY-JDXGNMNLSA-N | Strong | inactive |
| NSC-319726 | BRD-K99170882 | C/C(=N\NC(=S)N1CCC1)c1ccccn1 | XDHBUMNIQRLHGO-UKTHLTGXSA-N | Strong | inactive |
| NVP-ADW742 | BRD-K15179513 | Nc1ncnc2c1c(-c1cccc(OCc3ccccc3)c1)cn2[C@H]1C[C@H](CN2CCCC2)C1 | LSFLAQVDISHMNB-AFARHQOCSA-N | Strong | inactive |
| NVP-AEW541 | BRD-K24696047 | Nc1ncnc2c1c(-c1cccc(OCc3ccccc3)c1)cn2[C@H]1C[C@@H](CN2CCC2)C1 | AECDBHGVIIRMOI-GRGXKFILSA-N | Strong | inactive |
| NVP-BHG712 | BRD-K79259477 | Cc1ccc(C(=O)Nc2cccc(C(F)(F)F)c2)cc1Nc1nc(-c2cccnc2)nc2c1cnn2C | ZCCPLJOKGAACRT-UHFFFAOYSA-N | Strong | inactive |
| oleandrin | BRD-K98006098 | CO[C@H]1C[C@H](O[C@H]2CC[C@]3(C)[C@H]4CC[C@]5(C)[C@@H](C6=CC(=O)OC6)[C@@H](OC(C)=O)C[C@]5(O)[C@@H]4CC[C@@H]3C2)O[C@@H](C)[C@@H]1O | JLPDBLFIVFSOCC-XYXFTTADSA-N | Strong | inactive |
| osimertinib | BRD-K42805893 | C=CC(=O)Nc1cc(Nc2nccc(-c3cn(C)c4ccccc34)n2)c(OC)cc1N(C)CCN(C)C | DUYJMQONPNNFPI-UHFFFAOYSA-N | Strong | inactive |
| ouabain | BRD-K33277808 | C[C@@H]1O[C@@H](O[C@@H]2C[C@@H](O)[C@]3(CO)[C@H]4[C@H](O)C[C@]5(C)[C@@H](C6=CC(=O)OC6)CC[C@]5(O)[C@@H]4CC[C@]3(O)C2)[C@H](O)[C@H](O)[C@H]1O | LPMXVESGRSUGHW-VGFRVVOQSA-N | Strong | inactive |
| pacritinib | BRD-K19034817 | C1=C/COCc2cc(ccc2OCCN2CCCC2)Nc2nccc(n2)-c2cccc(c2)COC/1 | HWXVIOGONBBTBY-ONEGZZNKSA-N | Strong | inactive |
| perhexiline | BRD-A19633847 | C1CCC(C(C[C@H]2CCCCN2)C2CCCCC2)CC1 | CYXKNKQEMFBLER-GOSISDBHSA-N | Strong | inactive |
| pimavanserin | BRD-K83405785 | CC(C)COc1ccc(CNC(=O)N(Cc2ccc(F)cc2)C2CCN(C)CC2)cc1 | RKEWSXXUOLRFBX-UHFFFAOYSA-N | Strong | inactive |
| pimethixene | BRD-K88090157 | CN1CCC(=C2c3ccccc3Sc3ccccc32)CC1 | NZLVRVYNQYGMAB-UHFFFAOYSA-N | Strong | inactive |
| pitavastatin | BRD-K75958547 | O=C(O)C[C@H](O)C[C@H](O)/C=C/c1c(C2CC2)nc2ccccc2c1-c1ccc(F)cc1 | VGYFMXBACGZSIL-MCBHFWOFSA-N | Strong | inactive |
| plicamycin | BRD-K36368047 | CO[C@H](C(=O)[C@@H](O)[C@@H](C)O)[C@@H]1Cc2cc3cc(O[C@H]4C[C@@H](O[C@H]5C[C@@H](O)[C@H](O)[C@@H](C)O5)[C@@H](O)[C@@H](C)O4)c(C)c(O)c3c(O)c2C(=O)[C@H]1O[C@H]1C[C@@H](O[C@H]2C[C@@H](O[C@H]3C[C@](C)(O)[C@H](O)[C@@H](C)O3)[C@H](O)[C@@H](C)O2)[C@H](O)[C@@H](C)O1 | CFCUWKMKBJTWLW-BGLFSJPPSA-N | Strong | inactive |
| PPT | BRD-K00312224 | CCCc1c(-c2ccc(O)cc2)nn(-c2ccc(O)cc2)c1-c1ccc(O)cc1 | IOTXSIGGFRQYKW-UHFFFAOYSA-N | Strong | inactive |
| pralatrexate | BRD-A74914197 | C#CC[C@H](Cc1cnc2nc(N)nc(N)c2n1)c1ccc(C(=O)N[C@@H](CCC(=O)O)C(=O)O)cc1 | OGSBUKJUDHAQEA-ZBFHGGJFSA-N | Strong | inactive |
| prenylamine | BRD-A10662413 | C[C@@H](Cc1ccccc1)NCCC(c1ccccc1)c1ccccc1 | IFFPICMESYHZPQ-FQEVSTJZSA-N | Strong | inactive |
| proadifen | BRD-K46317332 | CCCC(C(=O)OCCN(CC)CC)(c1ccccc1)c1ccccc1 | SNTQPLDRUZOSDP-UHFFFAOYSA-N | Strong | inactive |
| prochlorperazine | BRD-K19352500 | CN1CCN(CCCN2c3ccccc3Sc3ccc(Cl)cc32)CC1 | WIKYUJGCLQQFNW-UHFFFAOYSA-N | Strong | inactive |
| proscillaridin-A | BRD-K00005354 | C[C@@H]1O[C@@H](O[C@@H]2C=C3CC[C@@H]4[C@H](CC[C@]5(C)[C@@H](c6ccc(=O)oc6)CC[C@]45O)[C@@]3(C)CC2)[C@H](O)[C@H](O)[C@H]1O | MYEJFUXQJGHEQK-ALRJYLEOSA-N | Strong | inactive |
| PRT062607 | BRD-K53734668 | NC(=O)c1cnc(N[C@@H]2CCCC[C@@H]2N)nc1Nc1cccc(-n2nccn2)c1 | TXGKRVFSSHPBAJ-JKSUJKDBSA-N | Strong | inactive |

| | | | | | |
|---|---|---|---|---|---|
| PTC-209 | BRD-K83834509 | COc1cc(Br)c(Nc2nc(-c3c(C)nc4ncccn34)cs2)c(Br)c1 | XVOOCQSWCCRVDY-UHFFFAOYSA-N | Strong | inactive |
| PU-H71 | BRD-K36529613 | CC(C)NCCCn1c(Sc2cc3c(cc2I)OCO3)nc2c(N)ncnc21 | SUPVGFZUWFMATN-UHFFFAOYSA-N | Strong | inactive |
| pyrithione-zinc | BRD-K16136380 | c1cc[n+]2c(c1)S[Zn]1(O2)O[n+]2ccccc2S1 | KTIHEFSORPLWRR-UHFFFAOYSA-L | Strong | inactive |
| quinidine | BRD-K59632282 | C=C[C@H]1CN2CC[C@H]1C[C@@H]2[C@@H](O)c1ccnc2ccc(OC)cc12 | LOUPRKONTZGTKE-LHHVKLHASA-N | Strong | inactive |
| R-428 | BRD-K14870255 | Nc1nc(Nc2ccc3c(c2)CC[C@@H](N2CCCC2)CC3)nn1-c1cc2c(nn1)-c1ccccc1CCC2 | KXMZDGSRSGHMMK-VWLOTQADSA-N | Strong | inactive |
| salinomycin | BRD-K23414042 | CC[C@@H](C(=O)[C@@H](C)[C@@H](O)[C@H](C)[C@@H]1O[C@@H]([C@@H](CC)C(=O)O)CC[C@@H]1C)[C@H]1O[C@]2(C=C[C@@H](O)[C@]3(CC[C@@](C)([C@H]4CC[C@](O)(CC)[C@H](C)O4)O3)O2)[C@H](C)C[C@@H]1C | KQXDHUJYNAXLNZ-XQSDOZFQSA-N | Strong | inactive |
| SAR405 | BRD-K72868067 | C[C@@H]1COCCN1c1cc(=O)n2c(n1)N(Cc1cncc(Cl)c1)[C@H](C(F)(F)F)CC2 | SPDQRCUBFSRAFI-DOMZBBRYSA-N | Strong | inactive |
| SB-743921 | BRD-K68532323 | Cc1ccc(C(=O)N(CCCN)[C@@H](c2oc3cc(Cl)ccc3c(=O)c2Cc2ccccc2)C(C)C)cc1 | PGXYIBJJCLWJST-MUUNZHRXSA-N | Strong | inactive |
| scabertopin | BRD-K00005327 | C=C1C(=O)O[C@@H]2/C=C(/C)C[C@H]3C=C(C[C@H](OC(=O)/C(C)=C\C)[C@@H]12)C(=O)O3 | FTPHYXGWMIZVMP-WQYRJXRXSA-N | Strong | inactive |
| serdemetan | BRD-K65924316 | c1ccc2c(CCNc3ccc(Nc4ccncc4)cc3)c[nH]c2c1 | CEGSUKYESLWKJP-UHFFFAOYSA-N | Strong | inactive |
| sertraline | BRD-K82036761 | CN[C@H]1CC[C@@H](c2ccc(Cl)c(Cl)c2)c2ccccc21 | VGKDLMBJGBXTGI-SJCJKPOMSA-N | Strong | inactive |
| SKI-II | BRD-K20755323 | Oc1ccc(Nc2nc(-c3ccc(Cl)cc3)cs2)cc1 | ZFGXZJKLOFCECI-UHFFFAOYSA-N | Strong | inactive |
| SNAP-5089 | BRD-A21428877 | COC(=O)C1=C(C)NC(C)=C(C(=O)NCCCN2CCC(c3ccccc3)(c3ccccc3)CC2)[C@@H]1c1ccc([N+](=O)[O-])cc1 | FIIXCJGBCCCOQQ-XIFFEERXSA-N | Strong | inactive |
| spiperone | BRD-K55468218 | O=C(CCCN1CCC2(CC1)C(=O)NCN2c1ccccc1)c1ccc(F)cc1 | DKGZKTPJOSAWFA-UHFFFAOYSA-N | Strong | inactive |
| SPP86 | BRD-K63253858 | CC(C)n1nc(C#Cc2ccccc2)c2c(N)ncnc21 | JQOIRTDBHMDWMT-UHFFFAOYSA-N | Strong | inactive |
| SRT1720 | BRD-K95926480 | O=C(Nc1ccccc1-c1cn2c(CN3CCNCC3)csc2n1)c1cnc2ccccc2n1 | IASPBORHOMBZMY-UHFFFAOYSA-N | Strong | inactive |
| STA-5326 | BRD-K41783120 | Cc1cccc(/C=N/Nc2cc(N3CCOCC3)nc(OCCc3ccccn3)n2)c1 | HSKAZIJJKRAJAV-KOEQRZSOSA-N | Strong | inactive |
| sulfatinib | BRD-K00003125 | Cc1cc2cc(Oc3ccnc(Nc4cccc(CS(=O)(=O)NCCN(C)C)c4)n3)ccc2[nH]1 | TTZSNFLLYPYKIL-UHFFFAOYSA-N | Strong | inactive |
| sulforaphane | BRD-A58955223 | C[S@@](=O)CCCCN=C=S | SUVMJBTUFCVSAD-SNVBAGLBSA-N | Strong | inactive |
| tafenoquine | BRD-A78965079 | COc1cc(C)c2c(Oc3cccc(C(F)(F)F)c3)c(OC)cc(N[C@@H](C)CCCN)c2n1 | LBHLFPGPEGDCJG-HNNXBMFYSA-N | Strong | inactive |
| tetrandrine | BRD-K08078237 | COc1ccc2cc1Oc1ccc(cc1)C[C@H]1c3cc(c(OC)cc3CCN1C)Oc1c(OC)c(OC)cc3c1[C@H](C2)N(C)CC3 | WVTKBKWTSCPRNU-KYJUHHDHSA-N | Strong | inactive |
| thaliblastine | BRD-K87548127 | COc1cc(C[C@H]2c3cc(OC)c(OC)cc3CCN2C)c(Oc2cc3c(cc2OC)-c2c(OC)c(OC)cc4c2[C@H](C3)N(C)CC4)cc1OC | ZCTJIMXXSXQXRI-KYJUHHDHSA-N | Strong | inactive |
| thioguanine | BRD-K49350383 | Nc1nc2nc[nH]c2c(=S)[nH]1 | WYWHKKSPHMUBEB-UHFFFAOYSA-N | Strong | inactive |
| thiomersal | BRD-K61443650 | CC[Hg]Sc1ccccc1C(=O)O | HXQVQGWHFRNKMS-UHFFFAOYSA-M | Strong | inactive |
| thioproperazine | BRD-K08619574 | CN1CCN(CCCN2c3ccccc3Sc3ccc(S(=O)(=O)N(C)C)cc32)CC1 | VZYCZNZBPPHOFY-UHFFFAOYSA-N | Strong | inactive |
| thioridazine | BRD-A84481105 | CSc1ccc2c(c1)N(CCC1CCCCN1C)c1ccccc1S2 | KLBQZWRITKRQQV-UHFFFAOYSA-N | Strong | inactive |
| tioguanine | BRD-K75649340 | Nc1nc(=S)c2ncn([C@@H]3O[C@H](CO)[C@@H](O)[C@H]3O)c2[nH]1 | OTDJAMXESTUWLO-UUOKFMHZSA-N | Strong | inactive |
| tolonium | BRD-K94882685 | Cc1cc2nc3ccc(=[N+](C)C)cc-3sc2cc1N | KZEUBCUXBNEMSQ-UHFFFAOYSA-O | Strong | inactive |
| trabectedin | BRD-K00003313 | COc1cc2c(cc1O)CCN[C@]21CS[C@@H]2c3c(OC(C)=O)c(C)c4c(c3[C@H](COC1=O)N1[C@@H](O)[C@@H]3Cc5cc(C)c(OC)c(O)c5[C@H]([C@H]21)N3C)OCO4 | PKVRCIRHQMSYJX-AIFWHQITSA-N | Strong | inactive |
| tradipitant | BRD-K00003222 | O=C(c1ccccc1Cl)c1cccnc1-c1nnn(Cc2cc(C(F)(F)F)cc(C(F)(F)F)c2)c1-c1ccncc1 | CAVRKWRKTNINFF-UHFFFAOYSA-N | Strong | inactive |
| trifloxystrobin | BRD-K00005342 | CO/N=C(/C(=O)OC)c1ccccc1CO/N=C(\C)c1cccc(C(F)(F)F)c1 | ONCZDRURRATYFI-TVJDWZFNSA-N | Strong | inactive |
| trimetrexate | BRD-K13642819 | COc1cc(NCc2ccc3nc(N)nc(N)c3c2C)cc(OC)c1OC | NOYPYLRCIDNJJB-UHFFFAOYSA-N | Strong | inactive |
| U-18666A | BRD-K22025381 | CCN(CC)CCO[C@H]1CC[C@@]2(C)C(=CC[C@H]3[C@@H]4CCC(=O)[C@@]4(C)CC[C@@H]32)C1 | DMZCCFMMPHJWQY-BKWLFHPQSA-N | Strong | inactive |
| UNC0631 | BRD-K13060017 | COc1cc2c(NC3CCN(CC4CCCCC4)CC3)nc(N3CCCN(C(C)C)CC3)nc2cc1OCCCN1CCCCC1 | XFAXSWXKPQWHDW-UHFFFAOYSA-N | Strong | inactive |
| UNC0737 | BRD-K02988401 | COc1cc2c(N(C)C3CCN(C(C)C)CC3)nc(C3CCCCC3)nc2cc1OCCCN1CCCC1 | ATORKVYCKKBNTF-UHFFFAOYSA-N | Strong | inactive |
| UNC2025 | BRD-K91283740 | CCCCNc1ncc2c(-c3ccc(CN4CCN(C)CC4)cc3)cn([C@H]3CC[C@H](O)CC3)c2n1 | MJSHVHLADKXCML-RQNOJGIXSA-N | Strong | inactive |
| URMC-099 | BRD-K66908976 | CN1CCN(Cc2ccc(-c3cnc4[nH]cc(-c5ccc6[nH]ccc6c5)c4c3)cc2)CC1 | QKKIWEILHCXECO-UHFFFAOYSA-N | Strong | inactive |
| vandetanib | BRD-K77625799 | COc1cc2c(Nc3ccc(Br)cc3F)ncnc2cc1OCC1CCN(C)CC1 | UHTHHESEBZOYNR-UHFFFAOYSA-N | Strong | inactive |
| VBY-825 | BRD-K36338614 | CC[C@H](NC(=O)[C@H](CS(=O)(=O)CC1CC1)N[C@@H](c1ccc(F)cc1)C(F)(F)F)C(=O)C(=O)NC1CC1 | PPUXXDKQNAHHON-BJLQDIEVSA-N | Strong | inactive |
| vinblastine | BRD-K06519765 | CC[C@]1(O)C[C@@H]2CN(CCc3c([nH]c4ccccc34)[C@@](C(=O)OC)(c3cc4c(cc3OC)N(C)[C@H]3[C@@](O)(C(=O)OC)[C@H](OC(C)=O)[C@]5(CC)C=CCN6CC[C@]43[C@@H]65)C2)C1 | JXLYSJRDGCGARV-CFWMRBGOSA-N | Strong | inactive |
| VU0364739 | BRD-K93317811 | O=C(NCCN1CCC2(CC1)C(=O)NCN2c1cccc(F)c1)c1ccc2ccccc2c1 | MSTXJJGAXXJCBY-UHFFFAOYSA-N | Strong | inactive |
| WAY-600 | BRD-K85920262 | c1cncc(CN2CCC(n3ncc4c(N5CCOCC5)nc(-c5ccc6[nH]ccc6c5)nc43)CC2)c1 | FPEIJQLXFHKLJV-UHFFFAOYSA-N | Strong | inactive |
| WP1066 | BRD-K05445342 | C[C@H](NC(=O)/C(C#N)=C/c1cccc(Br)n1)c1ccccc1 | VFUAJMPDXIRPKO-LQELWAHVSA-N | Strong | inactive |
| Y-134 | BRD-K94832621 | CC(C)N1CCN(c2ccc(C(=O)c3c(-c4ccc(O)cc4)sc4cc(O)ccc34)cc2)CC1 | LQEOPHGPHCWOAC-UHFFFAOYSA-N | Strong | inactive |
| Y-29794 | BRD-K48449073 | CC(C)c1ccc(C(=O)c2cccs2)c(SCCCCCCCCN(C)C)n1 | HAGHKJDWBDSFKC-UHFFFAOYSA-N | Strong | inactive |
| Y-320 | BRD-K96259238 | Cc1c(C(=O)Nc2ccc(N3CCC(N4CCOCC4)CC3)c(C#N)c2)cnn1-c1ccc(Cl)cc1 | BWZNJVZTAWBIFG-UHFFFAOYSA-N | Strong | inactive |
| 1,12-Besm | BRD-K04603573 | CCNCCCNCCCCNCCCNCC | PGMIWVSHZMWSSI-UHFFFAOYSA-N | Very Weak | inactive |
| AC-264613 | BRD-K81548480 | C/C(=N\NC(=O)[C@@H]1C(=O)NC[C@H]1c1ccccc1)c1cccc(Br)c1 | RQKXQCSEZPQBNZ-QSBCOWLBSA-N | Very Weak | inactive |
| AC-710 | BRD-K15819326 | CCN1C(C)(C)CC(Oc2ccc(C(=O)Nc3ccc(NC(=O)Nc4cc(C(C)(C)C)on4)cc3)nc2)CC1(C)C | JVCWPUFNLFSKFS-UHFFFAOYSA-N | Very Weak | inactive |
| acemetacin | BRD-K67563174 | COc1ccc2c(c1)c(CC(=O)OCC(=O)O)c(C)n2C(=O)c1ccc(Cl)cc1 | FSQKKOOTNAMONP-UHFFFAOYSA-N | Very Weak | inactive |
| acyclovir | BRD-K32318651 | Nc1nc(=O)c2ncn(COCCO)c2[nH]1 | MKUXAQIIEYXACX-UHFFFAOYSA-N | Very Weak | inactive |
| AEG3482 | BRD-K77634909 | NS(=O)(=O)c1nn2cc(-c3ccccc3)nc2s1 | MQUYTXDAVCOCMX-UHFFFAOYSA-N | Very Weak | inactive |
| alfuzosin | BRD-K01826519 | COc1cc2nc(N(C)CCCNC(=O)[C@H]3CCCO3)nc(N)c2cc1OC | WNMJYKCGWZFFKR-CQSZACIVSA-N | Very Weak | inactive |
| AM-1241 | BRD-A66116161 | CN1CCCC[C@@H]1Cn1cc(C(=O)c2cc([N+](=O)[O-])ccc2I)c2ccccc21 | ZUHIXXCLLBMBDW-MRXNPFEDSA-N | Very Weak | inactive |
| AM-281 | BRD-K59419204 | Cc1c(C(=O)NN2CCOCC2)nn(-c2ccc(Cl)cc2Cl)c1-c1ccc(I)cc1 | AJFFBPZYXRNAIC-UHFFFAOYSA-N | Very Weak | inactive |
| AMG-PERK-44 | BRD-K04374714 | Cc1ccc2cc(-c3c(C)ccc(C(=O)c4c(-c5ccccc5)n(C)n(-c5ccccc5)c4=O)c3N)ccc2n1 | YPPSMYLMSKHYAY-UHFFFAOYSA-N | Very Weak | inactive |
| aminopentamide | BRD-A69917777 | CC(CC(C(N)=O)(c1ccccc1)c1ccccc1)N(C)C | NARHAGIVSFTMIG-UHFFFAOYSA-N | Very Weak | inactive |
| ampicillin | BRD-K68432770 | CC1(C)S[C@@H]2[C@H](NC(=O)[C@H](N)c3ccccc3)C(=O)N2[C@H]1C(=O)O | AVKUERGKIZMTKX-NJBDSQKTSA-N | Very Weak | inactive |
| amrinone | BRD-K45924332 | Nc1cc(-c2ccncc2)c[nH]c1=O | RNLQIBCLLYYYFJ-UHFFFAOYSA-N | Very Weak | inactive |

| | | | | | |
|---|---|---|---|---|---|
| apigenin | BRD-K01493881 | O=c1cc(-c2ccc(O)cc2)oc2cc(O)cc(O)c12 | KZNIFHPLKGYRTM-UHFFFAOYSA-N | Very Weak | inactive |
| apraclonidine | BRD-K38911213 | Nc1cc(Cl)c(NC2=NCCN2)c(Cl)c1 | IEJXVRYNEISIKR-UHFFFAOYSA-N | Very Weak | inactive |
| arsenic-trioxide | BRD-K32786395 | O=[As]O[As]=O | IKWTVSLWAPBBKU-UHFFFAOYSA-N | Very Weak | inactive |
| AT-406 | BRD-K83030136 | CN[C@@H](C)C(=O)N[C@H]1CN(C(=O)CC(C)C)CC[C@H]2CC[C@@H](C(=O)NC(c3ccccc3)c3ccccc3)N2C1=O | LSXUTRRVVSPWDZ-MKKUMYSQSA-N | Very Weak | inactive |
| AT13387 | BRD-K58435339 | CC(C)c1cc(C(=O)N2Cc3ccc(CN4CCN(C)CC4)cc3C2)c(O)cc1O | IFRGXKKQHBVPCQ-UHFFFAOYSA-N | Very Weak | inactive |
| atracurium | BRD-K01826774 | COc1ccc(C[C@H]2c3cc(OC)c(OC)cc3CC[N@@+]2(C)CCC(=O)OCCCCCOC(=O)CC[N@@+]2(C)CCc3cc(OC)c(OC)cc3[C@@H]2Cc2ccc(OC)c(OC)c2)cc1OC | YXSLJKQTIDHPOT-MWAPPURBSA-N | Very Weak | inactive |
| AZD0156 | BRD-K69555850 | CN(C)CCCOc1ccc(-c2ccc3ncc4c(c3c2)n(C2CCOCC2)c(=O)n4C)cn1 | AOTRIQLYUAFVSC-UHFFFAOYSA-N | Very Weak | inactive |
| AZD2014 | BRD-K99113996 | CNC(=O)c1cccc(-c2ccc3c(N4CCOC[C@@H]4C)nc(N4CCOC[C@@H]4C)nc3n2)c1 | JUSFANSTBFGBAF-IRXDYDNUSA-N | Very Weak | inactive |
| azilsartan | BRD-K81074237 | CCOc1nc2cccc(C(=O)O)c2n1Cc1ccc(-c2ccccc2-c2nc(=O)o[nH]2)cc1 | KGSXMPPBFPAXLY-UHFFFAOYSA-N | Very Weak | inactive |
| azimilide | BRD-K38913877 | CN1CCN(CCCCN2C(=O)CN(/N=C/c3ccc(-c4ccc(Cl)cc4)o3)C2=O)CC1 | MREBEPTUUMTTIA-PCLIKHOPSA-N | Very Weak | inactive |
| BAG-956 | BRD-K00652956 | Cc1nc2cnc3ccc(C#Cc4cccnc4)cc3c2n1-c1ccc(C(C)(C)C#N)cc1 | GVPAGJWVBUZHNQ-UHFFFAOYSA-N | Very Weak | inactive |
| baicalin | BRD-K49962337 | O=C(O)[C@H]1O[C@@H](Oc2cc3oc(-c4ccccc4)cc(O)c-3c(=O)c2O)[C@H](O)[C@@H](O)[C@@H]1O | DAIGSUHCWOZEQG-ZFORQUDYSA-N | Very Weak | inactive |
| BAY-1895344 | BRD-K00003422 | C[C@@H]1COCCN1c1cc(-c2ccnn2C)c2ccnc(-c3ccn[nH]3)c2n1 | YBXRSCXGRPSTMW-CYBMUJFWSA-N | Very Weak | inactive |
| bempedoic-acid | BRD-K63177377 | CC(C)(CCCCCC(O)CCCCCC(C)(C)C(=O)O)C(=O)O | HYHMLYSLQUKXKP-UHFFFAOYSA-N | Very Weak | inactive |
| bentiromide | BRD-K58794715 | O=C(O)c1ccc(NC(=O)[C@H](Cc2ccc(O)cc2)NC(=O)c2ccccc2)cc1 | SPPTWHFVYKCNNK-FQEVSTJZSA-N | Very Weak | inactive |
| benzocaine | BRD-K75466013 | CCOC(=O)c1ccc(N)cc1 | BLFLLBZGZJTVJG-UHFFFAOYSA-N | Very Weak | inactive |
| BET-BAY-002 | BRD-K72735080 | Cc1nnc(C[C@@H]2C=C(c3ccc(Cl)cc3)c3ccccc3-n3c(C)nnc32)o1 | AGYIAWHWIUZNSD-INIZCTEOSA-N | Very Weak | inactive |
| betahistine | BRD-K91315211 | CNCCc1ccccn1 | UUQMNUMQCIQDMZ-UHFFFAOYSA-N | Very Weak | inactive |
| betamethasone-phosphate | BRD-K00005349 | C[C@H]1C[C@H]2[C@@H]3CCC4=CC(=O)C=C[C@]4(C)[C@@]3(F)[C@@H](O)C[C@]2(C)[C@@]1(O)C(=O)COP(=O)(O)O | VQODGRNSFPNSQE-DVTGEIKXSA-N | Very Weak | inactive |
| biperiden | BRD-K01824983 | O[C@](CCN1CCCCC1)(c1ccccc1)[C@@H]1C[C@H]2C=C[C@@H]1C2 | YSXKPIUOCJLQIE-UYWIDEMCSA-N | Very Weak | inactive |
| bismuth-subsalicylate | BRD-K80528473 | O=C1O[Bi](O)Oc2ccccc21 | ZREIPSZUJIFJNP-UHFFFAOYSA-K | Very Weak | inactive |
| BMS-833923 | BRD-K33245965 | CNCc1ccc(C)c(NC(=O)c2ccc(Nc3nc(-c4ccccc4)c4ccccc4n3)cc2)c1 | KLRRGBHZCJLIEL-UHFFFAOYSA-N | Very Weak | inactive |
| BNC105 | BRD-K20468903 | COc1cc(C(=O)c2c(C)oc3c(O)c(OC)ccc23)cc(OC)c1OC | RADMJHVVIZTENA-UHFFFAOYSA-N | Very Weak | inactive |
| bretazenil | BRD-K96936751 | CC(C)(C)OC(=O)c1ncn2c1[C@@H]1CCCN1C(=O)c1c(Br)cccc1-2 | LWUDDYHYYNNIQI-ZDUSSCGKSA-N | Very Weak | inactive |
| brigatinib | BRD-K56981171 | COc1cc(N2CCC(N3CCN(C)CC3)CC2)ccc1Nc1ncc(Cl)c(Nc2ccccc2P(C)(C)=O)n1 | AILRADAXUVEEIR-UHFFFAOYSA-N | Very Weak | inactive |
| bromosporine | BRD-K44665581 | CCOC(=O)Nc1cc(-c2ccc(C)c(NS(C)(=O)=O)c2)nn2c(C)nnc12 | UYBRROMMFMPJAN-UHFFFAOYSA-N | Very Weak | inactive |
| BTK-IN-1 | BRD-K00003437 | O=C(CNc1cccc(Cl)c1)N[C@@H]1CCCN(c2ncnc3[nH]ccc23)C1 | SPSNKQVUYDXOGY-OAHLLOKOSA-N | Very Weak | inactive |
| butalbital | BRD-K71350836 | C=CCC1(CC(C)C)C(=O)NC(=O)NC1=O | UZVHFVZFNXBMQJ-UHFFFAOYSA-N | Very Weak | inactive |
| butamben | BRD-K27217864 | CCCCOC(=O)c1ccc(N)cc1 | IUWVALYLNVXWKX-UHFFFAOYSA-N | Very Weak | inactive |
| calcitriol | BRD-K27316855 | C=C1/C(=C\C=C2/CCC[C@@]3(C)[C@H]2CC[C@@H]3[C@H](C)CCCC(C)(C)O)C[C@@H](O)C[C@@H]1O | GMRQFYUYWCNGIN-NKMMMXOESA-N | Very Weak | inactive |
| carbidopa | BRD-K78712176 | C[C@@](Cc1ccc(O)c(O)c1)(NN)C(=O)O | TZFNLOMSOLWIDK-JTQLQIEISA-N | Very Weak | inactive |
| carbimazole | BRD-K87156652 | CCOC(=O)n1ccn(C)c1=S | CFOYWRHIYXMDOT-UHFFFAOYSA-N | Very Weak | inactive |
| CBS-1114 | BRD-K91874254 | NC(=NNc1ccccc1)c1ccccc1 | ZOBKPJRHDMKXQG-UHFFFAOYSA-N | Very Weak | inactive |
| cefodizime | BRD-K22641081 | CO/N=C(\C(=O)N[C@@H]1C(=O)N2C(C(=O)O)=C(CSc3nc(C)c(CC(=O)O)s3)CS[C@H]12)c1csc(N)n1 | XDZKBRJLTGRPSS-BGZQYGJUSA-N | Very Weak | inactive |
| cefozopran | BRD-K46970505 | CO/N=C(\C(=O)N[C@@H]1C(=O)N2C(C(=O)O)=C(C[n+]3ccn4ncccc43)CS[C@H]12)c1nsc(N)n1 | QDUIJCOKQCCXQY-WHJQOFBOSA-O | Very Weak | inactive |
| cefprozil | BRD-K24871708 | C/C=C/C1=C(C(=O)O)N2C(=O)[C@@H](NC(=O)[C@H](N)c3ccc(O)cc3)[C@H]2SC1 | WDLWHQDACQUCJR-ZAMMOSSLSA-N | Very Weak | inactive |
| cetylpyridinium | BRD-K87700323 | CCCCCCCCCCCCCCCC[n+]1ccccc1 | NEUSVAOJNUQRTM-UHFFFAOYSA-N | Very Weak | inactive |
| chlorobutanol | BRD-K39495750 | CC(C)(O)C(Cl)(Cl)Cl | OSASVXMJTNOKOY-UHFFFAOYSA-N | Very Weak | inactive |
| chloroquine | BRD-A91699651 | CCN(CC)CCC[C@@H](C)Nc1ccnc2cc(Cl)ccc12 | WHTVZRBIWZFKQO-CQSZACIVSA-N | Very Weak | inactive |
| cholic-acid | BRD-K43164539 | C[C@H](CCC(=O)O)[C@H]1CC[C@H]2[C@H]3[C@H](C[C@H](O)[C@@]21C)[C@@]1(C)CC[C@@H](O)C[C@H]1C[C@H]3O | BHQCQFFYRZLCQQ-OELDTZBJSA-N | Very Weak | inactive |
| chromanol-293B-(-)-[3R,4S] | BRD-K27911943 | CCS(=O)(=O)N(C)[C@H]1c2cc(C#N)ccc2OC(C)(C)[C@@H]1O | HVSJHHXUORMCGK-UONOGXRCSA-N | Very Weak | inactive |
| CIM-0216 | BRD-A75514485 | Cc1cc(NC(=O)[C@H](c2ccccc2)N2CCCc3ccccc32)no1 | KSEXDSJYVSEVGF-FQEVSTJZSA-N | Very Weak | inactive |
| clenbuterol | BRD-K00003701 | CC(C)(C)NC[C@@H](O)c1cc(Cl)c(N)c(Cl)c1 | STJMRWALKKWQGH-SNVBAGLBSA-N | Very Weak | inactive |
| clomipramine | BRD-K52989797 | CN(C)CCCN1c2ccccc2CCc2ccc(Cl)cc21 | GDLIGKIOYRNHDA-UHFFFAOYSA-N | Very Weak | inactive |
| clozapine | BRD-K37289225 | CN1CCN(C2=Nc3cc(Cl)ccc3Nc3ccccc32)CC1 | QZUDBNBUXVUHMW-UHFFFAOYSA-N | Very Weak | inactive |
| CNX-774 | BRD-K48443249 | C=CC(=O)Nc1cccc(Nc2nc(Nc3ccc(Oc4ccnc(C(=O)NC)c4)cc3)ncc2F)c1 | VVLHQJDAUIPZFH-UHFFFAOYSA-N | Very Weak | inactive |
| crizotinib-(S) | BRD-K40308497 | C[C@H](Oc1cc(-c2cnn(C3CCNCC3)c2)cnc1N)c1c(Cl)ccc(F)c1Cl | KTEIFNKAUNYNJU-LBPRGKRZSA-N | Very Weak | inactive |
| cromoglicic-acid | BRD-K20920669 | O=C(O)c1cc(=O)c2c(OCC(O)COc3cccc4oc(C(=O)O)cc(=O)c34)cccc2o1 | IMZMKUWMOSJXDT-UHFFFAOYSA-N | Very Weak | inactive |
| CX-4945 | BRD-K81458380 | O=C(O)c1ccc2c(c1)nc(Nc1cccc(Cl)c1)c1ccncc12 | MUOKSQABCJCOPU-UHFFFAOYSA-N | Very Weak | inactive |
| cyclizine | BRD-K79501723 | CN1CCN(C(c2ccccc2)c2ccccc2)CC1 | UVKZSORBKUEBAZ-UHFFFAOYSA-N | Very Weak | inactive |
| cyclosporin-A | BRD-K13533483 | C/C=C/C[C@H](C)[C@H](O)[C@@H]1C(=O)N[C@H](CC)C(=O)N(C)CC(=O)N(C)[C@H](CC(C)C)C(=O)N[C@@H](C(C)C)C(=O)N(C)[C@H](CC(C)C)C(=O)N[C@H](C)C(=O)N[C@H](C)C(=O)N(C)[C@H](CC(C)C)C(=O)N(C)[C@H](CC(C)C)C(=O)N(C)[C@H](C(C)C)C(=O)N1C.C/C=C/C[C@@H](C)[C@@H](O)[C@H]1C(=O)N[C@@H](CC)C(=O)N(C)CC(=O)N(C)[C@@H](CC(C)C)C(=O)N[C@@H](C(C)C)C(=O)N(C)[C@@H](CC(C)C)C(=O)N[C@@H](C)C(=O)N[C@H](C)C(=O)N(C)[C@@H](CC(C)C)C(=O)N(C)[C@@H](CC(C)C)C(=O)N(C)[C@@H](C(C)C)C(=O)N1C | FXSYVLZSAPZXIH-IHNLIXADSA-N | Very Weak | inactive |
| cyclovalone | BRD-K59866480 | COc1cc(/C=C2\CCC/C(=C\c3ccc(O)c(OC)c3)C2=O)ccc1O | DHKKONBXGAAFTB-OTYYAQKOSA-N | Very Weak | inactive |
| dabrafenib | BRD-K09951645 | CC(C)(C)c1nc(-c2cccc(NS(=O)(=O)c3c(F)cccc3F)c2F)c(-c2ccnc(N)n2)s1 | BFSMGDJOXZAERB-UHFFFAOYSA-N | Very Weak | inactive |
| dapagliflozin | BRD-K58160573 | CCOc1ccc(Cc2cc([C@@H]3O[C@H](CO)[C@@H](O)[C@H](O)[C@H]3O)ccc2Cl)cc1 | JVHXJTBJCFBINQ-ADAARDCZSA-N | Very Weak | inactive |
| dehydrocholate-acid | BRD-K90976994 | C[C@H](CCC(=O)O)[C@H]1CC[C@H]2[C@@H]3C(=O)C[C@@H]4CC(=O)CC[C@]4(C)[C@H]3CC(=O)[C@@]21C | OHXPGWPVLFPUSM-KLRNGDHRSA-N | Very Weak | inactive |
| denopamine | BRD-K00004656 | COc1ccc(CCNC[C@H](O)c2ccc(O)cc2)cc1OC | VHSBBVZJABQOSG-INIZCTEOSA-N | Very Weak | inactive |
| deptropine | BRD-K00005351 | CN1[C@H]2CC[C@@H]1C[C@H](OC1c3ccccc3CCc3ccccc31)C2 | ZWPODSUQWXAZNC-PMOLBWCYSA-N | Very Weak | inactive |
| deserpidine | BRD-K29462201 | COC(=O)[C@H]1[C@H]2C[C@@H]3c4[nH]c5ccccc5c4CCN3C[C@H]2C[C@@H](OC(=O)c2cc(OC)c(OC)c(OC)c2)[C@@H]1OC | CVBMAZKKCSYWQR-WCGOZPBSSA-N | Very Weak | inactive |

| | | | | | |
|---|---|---|---|---|---|
| dichlorvos | BRD-K23779958 | COP(=O)(OC)OC=C(Cl)Cl | OEBRKCOSUFCWJD-UHFFFAOYSA-N | Very Weak | inactive |
| dicyclohexylamine | BRD-K57091285 | C1CCC(NC2CCCCC2)CC1 | XBPCUCUWBYBCDP-UHFFFAOYSA-N | Very Weak | inactive |
| dicycloverine | BRD-K68507560 | CCN(CC)CCOC(=O)C1(C2CCCCC2)CCCCC1 | CURUTKGFNZGFSE-UHFFFAOYSA-N | Very Weak | inactive |
| difenpiramide | BRD-K96206066 | O=C(Cc1ccc(-c2ccccc2)cc1)Nc1ccccn1 | PWHROYKAGRUWDQ-UHFFFAOYSA-N | Very Weak | inactive |
| dinitolmide | BRD-K93240442 | Cc1c(C(N)=O)cc([N+](=O)[O-])cc1[N+](=O)[O-] | ZEFNOZRLAWVAQF-UHFFFAOYSA-N | Very Weak | inactive |
| doconexent-ethyl-ester | BRD-K61937613 | CC/C=C\C/C=C\C/C=C\C/C=C\C/C=C\C/C=C\CCC(=O)OCC | ITNKVODZACVXDS-YNUSHXQLSA-N | Very Weak | inactive |
| dydrogesterone | BRD-K68620903 | CC(=O)[C@H]1CC[C@H]2[C@@H]3C=CC4=CC(=O)CC[C@@]4(C)[C@@H]3CC[C@]12C | JGMOKGBVKVMRFX-HQZYFCCVSA-N | Very Weak | inactive |
| elbasvir | BRD-K09735259 | COC(=O)N[C@H](C(=O)N1CCC[C@H]1c1nc(-c2ccc3c(c2)O[C@@H](c2ccccc2)n2c-3cc3cc(-c4c[nH]c([C@@H]5CCCN5C(=O)[C@@H](NC(=O)OC)C(C)C)n4)ccc32)c[nH]1)C(C)C | BVAZQCUMNICBAQ-PZHYSIFUSA-N | Very Weak | inactive |
| enasidenib | BRD-K70601687 | CC(C)(O)CNc1nc(Nc2ccnc(C(F)(F)F)c2)nc(-c2cccc(C(F)(F)F)n2)n1 | DYLUUSLLRIQKOE-UHFFFAOYSA-N | Very Weak | inactive |
| erbstatin-analog | BRD-K35573744 | COC(=O)/C=C/c1cc(O)ccc1O | BQCNSTFWSKOWMA-GORDUTHDSA-N | Very Weak | inactive |
| ERK5-IN-1 | BRD-K64857848 | COc1cc(N2CCN(C)CC2)ccc1Nc1ncc2c(n1)N(C)c1ccccc1C(=O)N2C | DDTPGANIPBKTNU-UHFFFAOYSA-N | Very Weak | inactive |
| estrone | BRD-K81839095 | C[C@]12CC[C@@H]3c4ccc(O)cc4CC[C@H]3[C@@H]1CCC2=O | DNXHEGUUPJUMQT-CBZIJGRNSA-N | Very Weak | inactive |
| etafenone | BRD-K84971356 | CCN(CC)CCOc1ccccc1C(=O)CCc1ccccc1 | OEGDFSLNGABBKJ-UHFFFAOYSA-N | Very Weak | inactive |
| ethionamide | BRD-K33710385 | CCc1cc(C(N)=S)ccn1 | AEOCXXJPGCBFJA-UHFFFAOYSA-N | Very Weak | inactive |
| ethisterone | BRD-K03981224 | C#C[C@]1(O)CC[C@H]2[C@@H]3CCC4=CC(=O)CC[C@]4(C)[C@H]3CC[C@@]21C | CHNXZKVNWQUJIB-CEGNMAFCSA-N | Very Weak | inactive |
| fludarabine | BRD-K66788707 | Nc1nc(F)nc2c1ncn2[C@@H]1O[C@H](CO)[C@@H](O)[C@@H]1O | HBUBKKRHXORPQB-FJFJXFQQSA-N | Very Weak | inactive |
| fludrocortisone-acetate | BRD-K06408664 | CC(=O)OCC(=O)[C@@]1(O)CC[C@H]2[C@@H]3CCC4=CC(=O)CC[C@]4(C)[C@@]3(F)[C@@H](O)C[C@@]21C | SYWHXTATXSMDSB-GSLJADNHSA-N | Very Weak | inactive |
| flumecinol | BRD-A44181516 | CC[C@@](O)(c1ccccc1)c1cccc(C(F)(F)F)c1 | DVASNQYQOZHAJN-OAHLLOKOSA-N | Very Weak | inactive |
| foropafant | BRD-K57774929 | CC(C)c1cc(C(C)C)c(-c2csc(N(CCN(C)C)Cc3cccnc3)n2)c(C(C)C)c1 | VVBFISAUNSXQGZ-UHFFFAOYSA-N | Very Weak | inactive |
| fusidic-acid | BRD-K78355918 | CC(=O)O[C@H]1C[C@@]2(C)[C@@H](C[C@@H](O)[C@H]3[C@@]4(C)CC[C@@H](O)[C@@H](C)[C@@H]4CC[C@@]32C)/C1=C(\CCC=C(C)C)C(=O)O | IECPWNUMDGFDKC-MZJAQBGESA-N | Very Weak | inactive |
| gabexate | BRD-K59256312 | CCOC(=O)c1ccc(OC(=O)CCCCCNC(=N)N)cc1 | YKGYIDJEEQRWQH-UHFFFAOYSA-N | Very Weak | inactive |
| galidesivir | BRD-K00003270 | Nc1ncnc2c([C@@H]3N[C@H](CO)[C@@H](O)[C@H]3O)c[nH]c12 | AMFDITJFBUXZQN-KUBHLMPHSA-N | Very Weak | inactive |
| gatifloxacin | BRD-K01825698 | COc1c(N2CCN[C@@H](C)C2)c(F)cc2c(=O)c(C(=O)O)cn(C3CC3)c12 | XUBOMFCQGDBHNK-JTQLQIEISA-N | Very Weak | inactive |
| genistein | BRD-K43797669 | O=c1c(-c2ccc(O)cc2)coc2cc(O)cc(O)c12 | TZBJGXHYKVUXJN-UHFFFAOYSA-N | Very Weak | inactive |
| GKT137831 | BRD-K79710622 | CN(C)c1cccc(-c2c3c(=O)n(-c4ccccc4Cl)[nH]c3cc(=O)n2C)c1 | RGYQPQARIQKJKH-UHFFFAOYSA-N | Very Weak | inactive |
| GSK2194069 | BRD-K95123482 | O=C(C1CC1)N1CC[C@@H](Cc2n[nH]c(=O)n2-c2ccc(-c3ccc4occc4c3)cc2)C1 | AQTPWCUIYUOEMG-INIZCTEOSA-N | Very Weak | inactive |
| GSK2636771 | BRD-K49215523 | Cc1c(Cn2c(C)nc3c(C(=O)O)cc(N4CCOCC4)cc32)cccc1C(F)(F)F | XTKLTGBKIDQGQL-UHFFFAOYSA-N | Very Weak | inactive |
| GSK2656157 | BRD-K41104902 | Cc1cccc(CC(=O)N2CCc3c2ccc(-c2cn(C)c4ncnc(N)c24)c3F)n1 | PRWSIEBRGXYXAJ-UHFFFAOYSA-N | Very Weak | inactive |
| guanadrel | BRD-A14262390 | N=C(N)NC[C@H]1COC2(CCCCC2)O1 | HPBNRIOWIXYZFK-QMMMGPOBSA-N | Very Weak | inactive |
| halobetasol-propionate | BRD-K71487404 | CCC(=O)O[C@]1(C(=O)CCl)[C@@H](C)C[C@H]2[C@@H]3C[C@H](F)C4=CC(=O)C=C[C@]4(C)[C@@]3(F)[C@@H](O)C[C@@]21C | BDSYKGHYMJNPAB-LICBFIPMSA-N | Very Weak | inactive |
| hematoxylin | BRD-K78342372 | Oc1cc2c(cc1O)[C@@H]1c3ccc(O)c(O)c3OC[C@]1(O)C2 | WZUVPPKBWHMQCE-XJKSGUPXSA-N | Very Weak | inactive |
| hydrocortisone | BRD-K93568044 | C[C@]12C[C@H](O)[C@H]3[C@@H](CCC4=CC(=O)CC[C@@]43C)[C@@H]1CC[C@]2(O)C(=O)CO | JYGXADMDTFJGBT-VWUMJDOOSA-N | Very Weak | inactive |
| I-BZA2 | BRD-K71698503 | CCN(CC)CCNC(=O)c1ccccc1I | WNSWOZIBBVUJBI-UHFFFAOYSA-N | Very Weak | inactive |
| icotinib | BRD-K31698212 | C#Cc1cccc(Nc2ncnc3cc4c(cc23)OCCOCCOCCO4)c1 | QQLKULDARVNMAL-UHFFFAOYSA-N | Very Weak | inactive |
| ilaprazole | BRD-K00003427 | COc1ccnc(C[S@@](=O)c2nc3ccc(-n4cccc4)cc3[nH]2)c1C | HRRXCXABAPSOCP-AREMUKBSSA-N | Very Weak | inactive |
| imatinib | BRD-K92723993 | Cc1ccc(NC(=O)c2ccc(CN3CCN(C)CC3)cc2)cc1Nc1nccc(-c2cccnc2)n1 | KTUFNOKKBVMGRW-UHFFFAOYSA-N | Very Weak | inactive |
| imipenem | BRD-K23200955 | C[C@@H](O)[C@H]1C(=O)N2C(C(=O)O)=C(SCCNC=N)C[C@H]12 | ZSKVGTPCRGIANV-ZXFLCMHBSA-N | Very Weak | inactive |
| INCB-057643 | BRD-K00003102 | CN1C(=O)C(C)(C)Oc2c(-c3cn(C)c(=O)c4[nH]ccc3-4)cc(S(C)(=O)=O)cc21 | ZNFWPFHSBWLNSO-UHFFFAOYSA-N | Very Weak | inactive |
| indinavir | BRD-K33226500 | CC(C)(C)NC(=O)[C@@H]1CN(Cc2cccnc2)CCN1C[C@@H](O)C[C@@H](Cc1ccccc1)C(=O)N[C@H]1c2ccccc2C[C@H]1O | CBVCZFGXHXORBI-PXQQMZJSSA-N | Very Weak | inactive |
| indocyanine-green | BRD-K80762071 | CC1(C)C(/C=C/C=C/C=C/C=C2/N(CCCCS(=O)(=O)O)c3ccc4ccccc4c3C2(C)C)=[N+](CCCCS(=O)(=O)O)c2ccc3ccccc3c21 | BDBMLMBYCXNVMC-UHFFFAOYSA-O | Very Weak | inactive |
| inositol | BRD-K52618540 | O[C@H]1[C@H](O)[C@@H](O)[C@H](O)[C@@H](O)[C@H]1O | CDAISMWEOUEBRE-GPIVLXJGSA-N | Very Weak | inactive |
| iopromide | BRD-A72075775 | COCC(=O)Nc1c(I)c(C(=O)NC[C@@H](O)CO)c(I)c(C(=O)N(C)C[C@@H](O)CO)c1I | DGAIEPBNLOQYER-RKDXNWHRSA-N | Very Weak | inactive |
| isometheptene-mucate | BRD-A64743628 | CN[C@@H](C)CCC=C(C)C | XVQUOJBERHHONY-VIFPVBQESA-N | Very Weak | inactive |
| JNJ-47965567 | BRD-K83547600 | O=C(NCC1(N2CCN(c3ccccc3)CC2)CCOCC1)c1cccnc1Sc1ccccc1 | XREFXUCWSYMIOG-UHFFFAOYSA-N | Very Weak | inactive |
| KB-R7943 | BRD-K17743697 | N=C(N)SCCc1ccc(OCc2ccc([N+](=O)[O-])cc2)cc1 | UMJUNUMSJDTYTM-UHFFFAOYSA-N | Very Weak | inactive |
| kifunensine | BRD-K02898090 | O=C1N[C@@H]2[C@@H](O)[C@@H](O)[C@H](O)[C@@H](CO)N2C1=O | OIURYJWYVIAOCW-PQMKYFCFSA-N | Very Weak | inactive |
| KPT-9274 | BRD-K00004701 | Nc1ccc(/C=C/C(=O)NCc2cc3cc(-c4ccc(C(=O)N5CCC(F)(F)CC5)cc4)cc(-c4ccc(F)cc4)c3o2)cn1 | MRFOPLWJZULAQD-SWGQDTFXSA-N | Very Weak | inactive |
| KW-3902 | BRD-A95683721 | CCCn1c(=O)c2[nH]c(C34C[C@@H]5C[C@H]3C[C@@H](C5)C4)nc2n(CCC)c1=O | PJBFVWGQFLYWCB-XEDLVCFGSA-N | Very Weak | inactive |
| KX2-391 | BRD-K29968218 | O=C(Cc1ccc(-c2ccc(OCCN3CCOCC3)cc2)cn1)NCc1ccccc1 | HUNGUWOZPQBXGX-UHFFFAOYSA-N | Very Weak | inactive |
| L-aspartic-acid | BRD-K97163964 | N[C@@H](CC(=O)O)C(=O)O | CKLJMWTZIZZHCS-REOHCLBHSA-N | Very Weak | inactive |
| L-cysteinesulfinic-acid | BRD-A25800195 | N[C@@H](C[S@](=O)O)C(=O)O | ADVPTQAUNPRNPO-REOHCLBHSA-N | Very Weak | inactive |
| L-cystine | BRD-K99679910 | N[C@@H](CSSC[C@H](N)C(=O)O)C(=O)O | LEVWYRKDKASIDU-IMJSIDKUSA-N | Very Weak | inactive |
| L-glutamic-acid | BRD-K62309135 | N[C@@H](CCC(=O)O)C(=O)O | WHUUTDBJXJRKMK-VKHMYHEASA-N | Very Weak | inactive |
| L-theanine | BRD-K04843140 | CCNC(=O)CC[C@H](N)C(=O)O | DATAGRPVKZEWHA-YFKPBYRVSA-N | Very Weak | inactive |
| latanoprost | BRD-K10961822 | CC(C)OC(=O)CCC/C=C\C[C@H]1[C@@H](O)C[C@@H](O)[C@@H]1CC[C@@H](O)CCc1ccccc1 | GGXICVAJURFBLW-CEYXHVGTSA-N | Very Weak | inactive |
| lenalidomide | BRD-A17883755 | Nc1cccc2c1CN([C@H]1CCC(=O)NC1=O)C2=O | GOTYRUGSSMKFNF-JTQLQIEISA-N | Very Weak | inactive |
| levamisole | BRD-K73107279 | c1ccc([C@H]2CN3CCSC3=N2)cc1 | HLFSDGLLUJUHTE-SNVBAGLBSA-N | Very Weak | inactive |
| levosulpiride | BRD-K51671335 | CCN1CCC[C@H]1CNC(=O)c1cc(S(N)(=O)=O)ccc1OC | BGRJTUBHPOOWDU-NSHDSACASA-N | Very Weak | inactive |
| levothyroxine | BRD-K30685142 | N[C@@H](Cc1cc(I)c(Oc2cc(I)c(O)c(I)c2)c(I)c1)C(=O)O | XUIIKFGFIJCVMT-LBPRGKRZSA-N | Very Weak | inactive |
| linaclotide | BRD-K00003592 | C[C@@H]1NC(=O)[C@@H]2CCCN2C(=O)[C@H](CC(N)=O)NC(=O)[C@@H]2CSSC[C@H](N)C(=O)N[C@H]3CSSC[C@H](NC1=O)C(=O)N[C@@H]([C@@H](C)O)C(=O)NCC(=O)N[C@H](C(=O)N[C@@H](Cc1ccc(O)cc1)C(=O)O)CSSC[C@H](NC(=O)[C@H](Cc1ccc(O)cc1)NC(=O)[C@H](CCC(=O)O)NC3=O)C(=O)N2 | KXGCNMMJRFDFNR-WDRJZQOASA-N | Very Weak | inactive |

| | | | | | |
|---|---|---|---|---|---|
| liraglutide | BRD-K00003717 | CCCCCCCCCCCCCCCC(=O)N[C@@H](CCC(=O)NCCCC[C@H](NC(=O)[C@H](C)NC(=O)[C@H](C)NC(=O)[C@H](CCC(N)=O)NC(=O)CNC(=O)[C@H](CCC(=O)O)NC(=O)[C@H](CC(C)C)NC(=O)[C@H](Cc1ccc(O)cc1)NC(=O)[C@H](CO)NC(=O)[C@H](CO)NC(=O)[C@@H](NC(=O)[C@H](CC(=O)O)NC(=O)[C@H](CO)NC(=O)[C@@H](NC(=O)[C@H](Cc1ccccc1)NC(=O)[C@@H](NC(=O)CNC(=O)[C@H](CCC(=O)O)NC(=O)[C@H](C)NC(=O)[C@@H](N)Cc1cnc[nH]1)[C@@H](C)O)[C@@H](C)O)C(C)C)C(=O)N[C@@H](CCC(=O)O)C(=O)N[C@@H](Cc1ccccc1)C(=O)N[C@H](C(=O)N[C@@H](C)C(=O)N[C@@H](Cc1c[nH]c2ccccc12)C(=O)N[C@@H](CC(C)C)C(=O)N[C@H](C(=O)N[C@@H](CCCNC(=N)N)C(=O)NCC(=O)N[C@@H](CCCNC(=N)N)C(=O)NCC(=O)O)C(C)C)[C@@H](C)CC)C(=O)O | YSDQQAXHVYUZIW-QCIJIYAXSA-N | Very Weak | inactive |
| lurbinectedin | BRD-K00003211 | COc1ccc2[nH]c3c(c2c1)CCN[C@]31CS[C@@H]2c3c(OC(C)=O)c(C)c4c(c3[C@H](COC1=O)N1[C@@H](O)[C@@H]3Cc5cc(C)c(OC)c(O)c5[C@H]([C@H]21)N3C)OCO4 | YDDMIZRDDREKEP-HWTBNCOESA-N | Very Weak | inactive |
| LY3023414 | BRD-K82967180 | CO[C@@H](C)Cn1c(=O)n(C)c2cnc3ccc(-c4cncc(C(C)(C)O)c4)cc3c21 | ACCFLVVUVBJNGT-AWEZNQCLSA-N | Very Weak | inactive |
| mefloquine | BRD-K40645748 | O[C@@H](c1cc(C(F)(F)F)nc2c(C(F)(F)F)cccc12)[C@H]1CCCCN1 | XEEQGYMUWCZPDN-DOMZBBRYSA-N | Very Weak | inactive |
| metacresol | BRD-K22518984 | Cc1cccc(O)c1 | RLSSMJSEOOYNOY-UHFFFAOYSA-N | Very Weak | inactive |
| metrizoic-acid | BRD-K99241550 | CC(=O)Nc1c(I)c(C(=O)O)c(I)c(N(C)C(C)=O)c1I | GGGDNPWHMNJRFN-UHFFFAOYSA-N | Very Weak | inactive |
| MI-14 | BRD-K26726854 | CC(=O)NCCNc1cc(Cl)nn2c(-c3cccc(S(=O)(=O)N(C)C)c3)c(C)nc12 | PSTAJIONBVMKLA-UHFFFAOYSA-N | Very Weak | inactive |
| mirodenafil | BRD-K82967685 | CCCOc1ccc(S(=O)(=O)N2CCN(CCO)CC2)cc1-c1nc2c(CCC)cn(CC)c2c(=O)[nH]1 | MIJFNYMSCFYZNY-UHFFFAOYSA-N | Very Weak | inactive |
| mitotane | BRD-A31204924 | Clc1ccc([C@@H](c2ccccc2Cl)C(Cl)Cl)cc1 | JWBOIMRXGHLCPP-ZDUSSCGKSA-N | Very Weak | inactive |
| MK-0893 | BRD-K16732600 | COc1ccc2cc(-c3cc(-c4cc(Cl)cc(Cl)c4)nn3[C@@H](C)c3ccc(C(=O)NCCC(=O)O)cc3)ccc2c1 | DNTVJEMGHBIUMW-IBGZPJMESA-N | Very Weak | inactive |
| morantel | BRD-K71454870 | Cc1ccsc1/C=C/C1=NCCCN1C | NVEPPWDVLBMNMB-SNAWJCMRSA-N | Very Weak | inactive |
| moxidectin | BRD-K00003660 | CO/N=C1\C[C@]2(C[C@@H]3C[C@@H](C/C=C(\C)C[C@@H](C)/C=C/C=C4\CO[C@@H]5[C@H](O)C(C)=C[C@@H](C(=O)O3)[C@]45O)O2)O[C@H](/C(C)=C/C(C)C)[C@H]1C | YZBLFMPOMVTDJY-LSGXYNIPSA-N | Very Weak | inactive |
| MSX-122 | BRD-K25143390 | c1cnc(NCc2ccc(CNc3ncccn3)cc2)nc1 | PXZXYRKDDXKDTK-UHFFFAOYSA-N | Very Weak | inactive |
| NE-100 | BRD-K11443721 | CCCN(CCC)CCc1ccc(OC)c(OCCc2ccccc2)c1 | YBLIQJGXRLZBCZ-UHFFFAOYSA-N | Very Weak | inactive |
| nebracetam | BRD-A51355958 | NC[C@@H]1CC(=O)N(Cc2ccccc2)C1 | LCAFGJGYCUMTGS-NSHDSACASA-N | Very Weak | inactive |
| nedaplatin | BRD-K73469236 | N[Pt]1(N)OC=C(O)O1 | AUZDMWWRYCUANI-UHFFFAOYSA-L | Very Weak | inactive |
| nedocromil | BRD-K64716392 | CCCc1c2oc(C(=O)O)cc(=O)c2cc2c(=O)cc(C(=O)O)n(CC)c12 | RQTOOFIXOKYGAN-UHFFFAOYSA-N | Very Weak | inactive |
| neostigmine | BRD-K18922609 | CN(C)C(=O)Oc1cccc([N+](C)(C)C)c1 | ALWKGYPQUAPLQC-UHFFFAOYSA-N | Very Weak | inactive |
| nepafenac | BRD-K04112579 | NC(=O)Cc1cccc(C(=O)c2ccccc2)c1N | QEFAQIPZVLVERP-UHFFFAOYSA-N | Very Weak | inactive |
| norfluoxetine | BRD-A79769222 | NCC[C@@H](Oc1ccc(C(F)(F)F)cc1)c1ccccc1 | WIQRCHMSJFFONW-OAHLLOKOSA-N | Very Weak | inactive |
| norgestrel | BRD-A14886633 | C#C[C@]1(O)CC[C@H]2[C@@H]3CCC4=CC(=O)CC[C@@H]4[C@H]3CC[C@@]21CC | WWYNJERNGUHSAO-XUDSTZEESA-N | Very Weak | inactive |
| NPS-2143 | BRD-K75506300 | CC(C)(Cc1ccc2ccccc2c1)NC[C@@H](O)COc1cccc(Cl)c1C#N | PZUJQWHTIRWCID-HXUWFJFHSA-N | Very Weak | inactive |
| NS-11021 | BRD-K94342292 | FC(F)(F)c1cc(NC(=S)Nc2ccc(Br)cc2-c2nn[nH]n2)cc(C(F)(F)F)c1 | MDKAFDIKYQMOMF-UHFFFAOYSA-N | Very Weak | inactive |
| NVP-AUY922 | BRD-K41859756 | CCNC(=O)c1noc(-c2cc(C(C)C)c(O)cc2O)c1-c1ccc(CN2CCOCC2)cc1 | NDAZATDQFDPQBD-UHFFFAOYSA-N | Very Weak | inactive |
| nystatin | BRD-K21656326 | C[C@@H]1OC(=O)C[C@H](O)C[C@H](O)C[C@H](O)CC[C@@H](O)[C@H](O)C[C@]2(O)C[C@H](O)[C@@H](C(=O)O)[C@H](C[C@@H](O[C@@H]3O[C@H](C)[C@@H](O)[C@H](N)[C@@H]3O)/C=C/C=C/C=C/C=C/CC/C=C/C=C/[C@H](C)[C@@H](O)[C@H]1C)O2 | VQOXZBDYSJBXMA-NQTDYLQESA-N | Very Weak | inactive |
| olaparib | BRD-K02113016 | O=C(c1cc(Cc2n[nH]c(=O)c3ccccc23)ccc1F)N1CCN(C(=O)C2CC2)CC1 | FDLYAMZZIXQODN-UHFFFAOYSA-N | Very Weak | inactive |
| olmutinib | BRD-K60421517 | C=CC(=O)Nc1cccc(Oc2nc(Nc3ccc(N4CCN(C)CC4)cc3)nc3ccsc23)c1 | FDMQDKQUTRLUBU-UHFFFAOYSA-N | Very Weak | inactive |
| omarigliptin | BRD-K14171291 | CS(=O)(=O)n1cc2c(n1)CN([C@H]1CO[C@H](c3cc(F)ccc3F)[C@@H](N)C1)C2 | MKMPWKUAHLTIBJ-ISTRZQFTSA-N | Very Weak | inactive |
| oritavancin | BRD-K00003704 | CN[C@H](CC(C)C)C(=O)N[C@H]1C(=O)N[C@@H](CC(N)=O)C(=O)N[C@H]2C(=O)N[C@H]3C(=O)N[C@H](C(=O)N[C@H](C(=O)O)c4cc(O)cc(O)c4-c4cc3ccc4O)[C@H](O[C@H]3C[C@](C)(N)[C@@H](O)[C@H](C)O3)c3ccc(c(Cl)c3)Oc3cc2cc(c3O[C@@H]2O[C@H](CO)[C@@H](O)[C@H](O)[C@H]2O[C@H]2C[C@](C)(NCc3ccc(-c4ccc(Cl)cc4)cc3)[C@@H](O)[C@H](C)O2)Oc2ccc(cc2Cl)[C@H]1O | VHFGEBVPHAGQPI-LXKZPTCJSA-N | Very Weak | inactive |
| ormeloxifene | BRD-K00004639 | COc1ccc2c(c1)OC(C)(C)[C@@H](c1ccccc1)[C@@H]2c1ccc(OCCN2CCCC2)cc1 | XZEUAXYWNKYKPL-WDYNHAJCSA-N | Very Weak | inactive |
| oxybuprocaine | BRD-K04185004 | CCCCOc1cc(C(=O)OCCN(CC)CC)ccc1N | CMHHMUWAYWTMGS-UHFFFAOYSA-N | Very Weak | inactive |
| ozolinone | BRD-A72297358 | CN1C(=O)[C@H](N2CCCCC2)S/C1=C\C(=O)O | NQFBZYYUAFJYNS-SKVAFPRGSA-N | Very Weak | inactive |
| pantoprazole | BRD-K00003243 | COc1ccnc(C[S@@](=O)c2nc3ccc(OC(F)F)cc3[nH]2)c1OC | IQPSEEYGBUAQFF-AREMUKBSSA-N | Very Weak | inactive |
| papaverine | BRD-K15567136 | COc1ccc(Cc2nccc3cc(OC)c(OC)cc23)cc1OC | XQYZDYMELSJDRZ-UHFFFAOYSA-N | Very Weak | inactive |
| PD-161570 | BRD-K97152684 | CCN(CC)CCCCNc1ncc2cc(-c3c(Cl)cccc3Cl)c(NC(=O)NC(C)(C)C)nc2n1 | MKVMEJKNLUWFSQ-UHFFFAOYSA-N | Very Weak | inactive |
| pemetrexed | BRD-K32842773 | Nc1nc(=O)c2c(CCc3ccc(C(=O)N[C@@H](CCC(=O)O)C(=O)O)cc3)c[nH]c2[nH]1 | WBXPDJSOTKVWSJ-ZDUSSCGKSA-N | Very Weak | inactive |
| PF-429242 | BRD-K11821338 | CCN(CC)Cc1ccc(C(=O)N(CCc2ccccc2OC)[C@@H]2CCNC2)cc1 | XKPJTOHUPQWSOJ-HSZRJFAPSA-N | Very Weak | inactive |
| phenothrin | BRD-A22106989 | CC(C)=C[C@H]1[C@@H](C(=O)OCc2cccc(Oc3ccccc3)c2)C1(C)C | SBNFWQZLDJGRLK-SFTDATJTSA-N | Very Weak | inactive |
| phenylephrine | BRD-K73381542 | CNC[C@H](O)c1cccc(O)c1 | SONNWYBIRXJNDC-VIFPVBQESA-N | Very Weak | inactive |
| pidotimod | BRD-K44442813 | O=C1CC[C@@H](C(=O)N2CSC[C@H]2C(=O)O)N1 | UUTKICFRNVKFRG-WDSKDSINSA-N | Very Weak | inactive |
| PIK-93 | BRD-K29395450 | CC(=O)Nc1nc(C)c(-c2ccc(Cl)c(S(=O)(=O)NCCO)c2)s1 | JFVNFXCESCXMBC-UHFFFAOYSA-N | Very Weak | inactive |
| pinitol | BRD-K87873585 | CO[C@@H]1[C@H](O)[C@@H](O)[C@H](O)[C@H](O)[C@H]1O | DSCFFEYYQKSRSV-KLJZZCKASA-N | Very Weak | inactive |
| PLX51107 | BRD-K00003099 | Cc1noc(C)c1-c1cnc2c(-c3ccc(C(=O)O)cc3)cn([C@@H](C)c3ccccn3)c2c1 | AMSUHYUVOVCWTP-INIZCTEOSA-N | Very Weak | inactive |
| potassium-p-aminobenzoate | BRD-K58497451 | Nc1ccc(C(=O)O)cc1 | ALYNCZNDIQEVRV-UHFFFAOYSA-N | Very Weak | inactive |
| promethazine | BRD-A46335897 | CC(CN1c2ccccc2Sc2ccccc21)N(C)C | PWWVAXIEGOYWEE-UHFFFAOYSA-N | Very Weak | inactive |
| quercetin | BRD-K97399794 | O=c1c(O)c(-c2ccc(O)c(O)c2)oc2cc(O)cc(O)c12 | REFJWTPEDVJJIY-UHFFFAOYSA-N | Very Weak | inactive |
| quinelorane | BRD-K13261168 | CCCN1CCC[C@@H]2Cc3nc(N)ncc3C[C@H]21 | TUFADSGTJUOBEH-ZWNOBZJWSA-N | Very Weak | inactive |
| raloxifene | BRD-K63828191 | O=C(c1ccc(OCCN2CCCCC2)cc1)c1c(-c2ccc(O)cc2)sc2cc(O)ccc12 | GZUITABIAKMVPG-UHFFFAOYSA-N | Very Weak | inactive |
| raltegravir | BRD-K05658747 | Cc1nnc(C(=O)NC(C)(C)c2nc(C(=O)NCc3ccc(F)cc3)c(O)c(=O)n2C)o1 | CZFFBEXEKNGXKS-UHFFFAOYSA-N | Very Weak | inactive |
| riboflavin-5-phosphate-sodium | BRD-K09229706 | Cc1cc2nc3c(=O)[nH]c(=O)nc-3n(C[C@H](O)[C@H](O)[C@H](O)COP(=O)(O)O)c2cc1C | FVTCRASFADXXNN-SCRDCRAPSA-N | Very Weak | inactive |

| | | | | | |
|---|---|---|---|---|---|
| riluzole | BRD-K21283037 | Nc1nc2ccc(OC(F)(F)F)cc2s1 | FTALBRSUTCGOEG-UHFFFAOYSA-N | Very Weak | inactive |
| Ro-106-9920 | BRD-A53134341 | O=[S@@](c1ccccc1)c1ccc2nnnnn2n1 | JFSXSNSCPNFCDM-KRWDZBQOSA-N | Very Weak | inactive |
| rolitetracycline | BRD-A62336573 | CN(C)[C@@H]1C(=O)C(C(=O)NCN2CCCC2)=C(O)[C@@]2(O)C(O)=C3C(=O)c4c(O)cccc4[C@@](C)(O)[C@H]3C[C@@H]12 | YEHCEZXUSLIKDU-IAHYZSEUSA-N | Very Weak | inactive |
| rose-bengal | BRD-K33623053 | O=C(O)c1c(Cl)c(Cl)c(Cl)c(Cl)c1-c1c2cc(I)c(=O)c(I)c-2oc2c(I)c(O)c(I)cc12 | VDNLFJGJEQUWRB-UHFFFAOYSA-N | Very Weak | inactive |
| RS-17053 | BRD-K76840893 | CC(C)(Cc1c[nH]c2ccc(Cl)cc12)NCCOc1ccccc1OCC1CC1 | XLZHXAXXJVKTFM-UHFFFAOYSA-N | Very Weak | inactive |
| RV1729 | BRD-K00004579 | COCCN(CCOC)C(=O)CCCC#Cc1cccc2nc(Cn3nc(-c4cccc(O)c4)c4c(N)ncnc43)n(Cc3ccccc3Cl)c(=O)c12 | UBLOHCIYTDRGJH-UHFFFAOYSA-N | Very Weak | inactive |
| saroglitazar | BRD-K79878800 | CCO[C@@H](Cc1ccc(OCCn2c(C)ccc2-c2ccc(SC)cc2)cc1)C(=O)O | MRWFZSLZNUJVQW-DEOSSOPVSA-N | Very Weak | inactive |
| SB-202190 | BRD-K54330070 | Oc1ccc(-c2nc(-c3ccncc3)c(-c3ccc(F)cc3)[nH]2)cc1 | QHKYPYXTTXKZST-UHFFFAOYSA-N | Very Weak | inactive |
| SB-222200 | BRD-K04414442 | CC[C@H](NC(=O)c1c(C)c(-c2ccccc2)nc2ccccc12)c1ccccc1 | MQNYRKWJSMQECI-QFIPXVFZSA-N | Very Weak | inactive |
| SB-2343 | BRD-K98795921 | Cc1nc2c(-c3cnc(N)nc3)nc(N3CCOCC3)nc2n1C(C)C | QYBGBLQCOOISAR-UHFFFAOYSA-N | Very Weak | inactive |
| SB-271046 | BRD-K82467063 | COc1ccc(NS(=O)(=O)c2sc3ccc(Cl)cc3c2C)cc1N1CCNCC1 | LOCQRDBFWSXQQI-UHFFFAOYSA-N | Very Weak | inactive |
| scopolamine | BRD-K89923877 | CN1[C@H]2C[C@H](OC(=O)[C@H](CO)c3ccccc3)C[C@@H]1[C@H]1O[C@@H]21 | STECJAGHUSJQJN-FWXGHANASA-N | Very Weak | inactive |
| seladelpar | BRD-K00003284 | CCO[C@H](COc1ccc(C(F)(F)F)cc1)CSc1ccc(OCC(=O)O)c(C)c1 | JWHYSEDOYMYMNM-QGZVFWFLSA-N | Very Weak | inactive |
| selegiline | BRD-K86434416 | C#CCN(C)[C@H](C)Cc1ccccc1 | MEZLKOACVSPNER-GFCCVEGCSA-N | Very Weak | inactive |
| SGI-1027 | BRD-K61228301 | Cc1cc(Nc2ccc(NC(=O)c3ccc(Nc4ccnc5ccccc45)cc3)cc2)nc(N)n1 | QSYLKMKIVWJAAK-UHFFFAOYSA-N | Very Weak | inactive |
| silymarin | BRD-A06629414 | COc1cc([C@H]2Oc3ccc([C@H]4Oc5cc(O)cc(O)c5C(=O)[C@@H]4O)cc3O[C@@H]2CO)ccc1O | FDQAOULAVFHKBX-HKTJVKLFSA-N | Very Weak | inactive |
| sirolimus | BRD-K84937637 | CO[C@H]1C[C@@H]2CC[C@@H](C)[C@@](O)(O2)C(=O)C(=O)N2CCCC[C@H]2C(=O)O[C@H]([C@H](C)C[C@@H]2CC[C@@H](O)[C@@H](OC)C2)CC(=O)[C@H](C)/C=C(\C)[C@@H](O)[C@@H](OC)C(=O)[C@H](C)C[C@H](C)/C=C/C=C/C=C/1C | QFJCIRLUMZQUOT-HPLJOQBZSA-N | Very Weak | inactive |
| sodium-ascorbate | BRD-A67265925 | O=C1O[C@H]([C@@H](O)CO)[C@@H](O)[C@@H]1O | SXZYCXMUPBBULW-NEEWWZBLSA-N | Very Weak | inactive |
| spaglumic-acid | BRD-K68405354 | CC(=O)N[C@@H](CC(=O)O)C(=O)N[C@@H](CCC(=O)O)C(=O)O | OPVPGKGADVGKTG-BQBZGAKWSA-N | Very Weak | inactive |
| sparteine-sulfate | BRD-K32847234 | C1CCN2C[C@H]3C[C@@H](CN4CCCC[C@H]34)[C@@H]2C1 | SLRCCWJSBJZJBV-NMWPEEMBSA-N | Very Weak | inactive |
| succinobucol | BRD-K00003183 | CC(C)(Sc1cc(C(C)(C)C)c(O)c(C(C)(C)C)c1)Sc1cc(C(C)(C)C)c(OC(=O)CCC(=O)O)c(C(C)(C)C)c1 | RKSMVPNZHBRNNS-UHFFFAOYSA-N | Very Weak | inactive |
| sulfacarbamide | BRD-K13486660 | NC(=O)NS(=O)(=O)c1ccc(N)cc1 | WVAKABMNNSMCDK-UHFFFAOYSA-N | Very Weak | inactive |
| sulfamethoxazole | BRD-K28494619 | Cc1cc(NS(=O)(=O)c2ccc(N)cc2)no1 | JLKIGFTWXXRPMT-UHFFFAOYSA-N | Very Weak | inactive |
| sulfanilamide | BRD-K02594908 | Nc1ccc(S(N)(=O)=O)cc1 | FDDDEECHVMSUSB-UHFFFAOYSA-N | Very Weak | inactive |
| sulpiride | BRD-A55272860 | CCN1CCC[C@@H]1CNC(=O)c1cc(S(N)(=O)=O)ccc1OC | BGRJTUBHPOOWDU-LLVKDONJSA-N | Very Weak | inactive |
| tacrine | BRD-K81473089 | Nc1c2c(nc3ccccc13)CCCC2 | YLJREFDVOIBQDA-UHFFFAOYSA-N | Very Weak | inactive |
| talipexole | BRD-K75615183 | C=CCN1CCc2nc(N)sc2CC1 | DHSSDEDRBUKTQY-UHFFFAOYSA-N | Very Weak | inactive |
| tandutinib | BRD-K89162000 | COc1cc2c(N3CCN(C(=O)Nc4ccc(OC(C)C)cc4)CC3)ncnc2cc1OCCCN1CCCCC1 | UXXQOJXBIDBUAC-UHFFFAOYSA-N | Very Weak | inactive |
| tanespimycin | BRD-K81473043 | C=CCNC1=C2C[C@@H](C)C[C@H](OC)[C@H](O)[C@@H](C)/C=C(\C)[C@H](OC(N)=O)[C@@H](OC)/C=C\C=C(/C)C(=O)NC(=CC1=O)C2=O | AYUNIORJHRXIBJ-TXHRRWQRSA-N | Very Weak | inactive |
| telaprevir | BRD-K72912098 | CCC[C@H](NC(=O)[C@@H]1[C@H]2CCC[C@H]2CN1C(=O)[C@@H](NC(=O)[C@@H](NC(=O)c1cnccn1)C1CCCCC1)C(C)(C)C)C(=O)C(=O)NC1CC1 | BBAWEDCPNXPBQM-GDEBMMAJSA-N | Very Weak | inactive |
| telotristat-ethyl | BRD-K94559059 | CCOC(=O)[C@@H](N)Cc1ccc(-c2cc(O[C@H](c3ccc(Cl)cc3-n3ccc(C)n3)C(F)(F)F)nc(N)n2)cc1 | MDSQOJYHHZBZKA-GBXCKJPGSA-N | Very Weak | inactive |
| temoporfin | BRD-K63712959 | Oc1cccc(-c2c3nc(c(-c4cccc(O)c4)c4[nH]c(c(-c5cccc(O)c5)c5ccc([nH]5)c(-c5cccc(O)c5)c5nc2C=C5)CC4)C=C3)c1 | JGLSWAGTNWZKQU-UHFFFAOYSA-N | Very Weak | inactive |
| tenovin-6 | BRD-K13949769 | CN(C)CCCCC(=O)Nc1ccc(NC(=S)NC(=O)c2ccc(C(C)(C)C)cc2)cc1 | BVJSXSQRIUSRCO-UHFFFAOYSA-N | Very Weak | inactive |
| terfenadine | BRD-A06352418 | CC(C)(C)c1ccc([C@@H](O)CCCN2CCC(C(O)(c3ccccc3)c3ccccc3)CC2)cc1 | GUGOEEXESWIERI-PMERELPUSA-N | Very Weak | inactive |
| thiostrepton | BRD-K58049875 | C=C(NC(=O)C(=C)NC(=O)c1csc(C2=N[C@@H]3c4csc(n4)[C@H]4NC(=O)c5csc(n5)[C@H]([C@](C)(O)[C@@H](C)O)NC(=O)[C@H]5CSC(=N5)/C(=C/C)NC(=O)[C@H]([C@@H](C)O)NC(=O)c5csc(n5)[C@]3(CC2)NC(=O)[C@H](C)NC(=O)C(=C)NC(=O)[C@H](C)NC(=O)[C@H]([C@@H](C)CC)N[C@@H]2C=Cc3c([C@H](C)O)cc(nc3[C@H]2O)C(=O)O[C@@H]4C)n1)C(N)=O | NSFFHOGKXHRQEW-AIHSUZKVSA-N | Very Weak | inactive |
| ticarcillin | BRD-K29229654 | CC1(C)S[C@@H]2[C@H](NC(=O)[C@H](C(=O)O)c3ccsc3)C(=O)N2[C@H]1C(=O)O | OHKOGUYZJXTSFX-KZFFXBSXSA-N | Very Weak | inactive |
| tioxolone | BRD-K41876534 | O=c1oc2cc(O)ccc2s1 | SLYPOVJCSQHITR-UHFFFAOYSA-N | Very Weak | inactive |
| tipiracil | BRD-K11973162 | N=C1CCCN1Cc1[nH]c(=O)[nH]c(=O)c1Cl | QQHMKNYGKVVGCZ-UHFFFAOYSA-N | Very Weak | inactive |
| tiquizium | BRD-K77552740 | C[N@+]12CCCC[C@@H]1CCC(=C(c1cccs1)c1cccs1)C2 | ZGSDGGRVFIYKKE-OXQOHEQNSA-N | Very Weak | inactive |
| todralazine | BRD-K68553471 | CCOC(=O)NNc1nncc2ccccc12 | WGZDBVOTUVNQFP-UHFFFAOYSA-N | Very Weak | inactive |
| tolrestat | BRD-K85693895 | COc1ccc2c(C(=S)N(C)CC(=O)O)cccc2c1C(F)(F)F | LUBHDINQXIHVLS-UHFFFAOYSA-N | Very Weak | inactive |
| TRAM-39 | BRD-K81341251 | N#CC(c1ccccc1)(c1ccccc1)c1ccccc1Cl | JHNRTPKGSCVKKC-UHFFFAOYSA-N | Very Weak | inactive |
| tretinoin | BRD-K71879491 | CC1=C(/C=C/C(C)=C/C=C/C(C)=C/C(=O)O)C(C)(C)CCC1 | SHGAZHPCJJPHSC-YCNIQYBTSA-N | Very Weak | inactive |
| triclocarban | BRD-K25029121 | O=C(Nc1ccc(Cl)cc1)Nc1ccc(Cl)c(Cl)c1 | ICUTUKXCWQYESQ-UHFFFAOYSA-N | Very Weak | inactive |
| tubocurarine | BRD-K99621550 | COc1cc2c3cc1Oc1cc(ccc1O)C[C@@H]1c4c(cc(OC)c(O)c4Oc4ccc(cc4)C[C@@H]3N(C)CC2)CC[N+]1(C)C | JFJZZMVDLULRGK-URLMMPGGSA-O | Very Weak | inactive |
| tyrphostin-AG-835 | BRD-K17415526 | C[C@H](NC(=O)/C(C#N)=C/c1ccc(O)c(O)c1)c1ccccc1 | UMGQVUWXNOJOSJ-DGGAMASNSA-N | Very Weak | inactive |
| U-0126 | BRD-K18787491 | N#CC(C(=N)Sc1ccccc1N)C(C#N)C(=N)Sc1ccccc1N | ZCHZSBUNJOCWCW-UHFFFAOYSA-N | Very Weak | inactive |
| umeclidinium | BRD-K46692793 | OC(c1ccccc1)(c1ccccc1)C12CC[N+](CCOCc3ccccc3)(CC1)CC2 | FVTWTVQXNAJTQP-UHFFFAOYSA-N | Very Weak | inactive |
| uracil-mustard | BRD-K10180294 | O=c1[nH]cc(N(CCCl)CCCl)c(=O)[nH]1 | IDPUKCWIGUEADI-UHFFFAOYSA-N | Very Weak | inactive |
| VER-49009 | BRD-K38548312 | CCNC(=O)c1[nH]nc(-c2cc(Cl)c(O)cc2O)c1-c1ccc(OC)cc1 | HUNAOTXNHVALTN-UHFFFAOYSA-N | Very Weak | inactive |
| verubulin | BRD-K42673188 | COc1ccc(N(C)c2nc(C)nc3ccccc23)cc1 | SNHCRNMVYDHVDT-UHFFFAOYSA-N | Very Weak | inactive |
| vismodegib | BRD-K44827188 | CS(=O)(=O)c1ccc(C(=O)Nc2ccc(Cl)c(-c3ccccn3)c2)c(Cl)c1 | BPQMGSKTAYIVFO-UHFFFAOYSA-N | Very Weak | inactive |
| vorapaxar | BRD-K93442924 | CCOC(=O)N[C@@H]1CC[C@@H]2[C@@H](C1)C[C@H]1C(=O)O[C@@H](C)[C@H]1[C@H]2/C=C/c1ccc(-c2cccc(F)c2)cn1 | ZBGXUVOIWDMMJE-QHNZEKIYSA-N | Very Weak | inactive |
| voxtalisib | BRD-K56291712 | CCn1c(=O)c(-c2cc[nH]n2)cc2c(C)nc(N)nc21 | RGHYDLZMTYDBDT-UHFFFAOYSA-N | Very Weak | inactive |
| VU0361737 | BRD-K45462423 | COc1cc(NC(=O)c2ccccn2)ccc1Cl | ARYUXFNGXHNNDM-UHFFFAOYSA-N | Very Weak | inactive |
| WYE-687 | BRD-K49900195 | COC(=O)Nc1ccc(-c2nc(N3CCOCC3)c3cnn(C4CCN(Cc5cccnc5)CC4)c3n2)cc1 | VDOCQQKGPJENHJ-UHFFFAOYSA-N | Very Weak | inactive |
| XL647 | BRD-K03765900 | COc1cc2c(Nc3ccc(Cl)c(Cl)c3F)ncnc2cc1OC[C@@H]1C[C@@H]2CN(C)C[C@@H]2C1 | HVXKQKFEHMGHSL-QKDCVEJESA-N | Very Weak | inactive |
| XL888 | BRD-K00003558 | CC[C@@H](C)Nc1cc(C(=O)N[C@@H]2C[C@@H]3CC[C@H](C2)N3c2ccc(C(=O)C3CC3)cn2)c(C)cc1C(N)=O | LHGWWAFKVCIILM-HLRQEUIKSA-N | Very Weak | inactive |
| XMD17-109 | BRD-K48449981 | CCOc1cc(C(=O)N2CCC(N3CCN(C)CC3)CC2)ccc1Nc1ncc2c(n1)N(C1CCCC1)c1ccccc1C(=O)N2C | XVBGRTMNFNMINE-UHFFFAOYSA-N | Very Weak | inactive |
| zoxazolamine | BRD-K66353228 | Nc1nc2cc(Cl)ccc2o1 | YGCODSQDUUUKIV-UHFFFAOYSA-N | Very Weak | inactive |

| | | | | | |
|---|---|---|---|---|---|
| 3,4-methylenedioxy-beta-nitrostyrene | BRD-K42205652 | O=[N+]([O-])/C=C/c1ccc2c(c1)OCO2 | KFLWBZPSJQPRDD-ONEGZZNKSA-N | Weak | inactive |
| 3'-fluorobenzylspiperone | BRD-K45662124 | O=C(CCCN1CCC2(CC1)C(=O)N(Cc1cccc(F)c1)CN2c1ccccc1)c1ccc(F)cc1 | DXZKTRNEOISOCA-UHFFFAOYSA-N | Weak | inactive |
| acolbifene | BRD-K00004720 | CC1=C(c2ccc(O)cc2)[C@H](c2ccc(OCCN3CCCCC3)cc2)Oc2cc(O)ccc21 | DUYNJNWVGIWJRI-LJAQVGFWSA-N | Weak | inactive |
| amineptine | BRD-K24219278 | O=C(O)CCCCCCNC1c2ccccc2CCc2ccccc21 | ONNOFKFOZAJDHT-UHFFFAOYSA-N | Weak | inactive |
| ammonium-glycyrrhizinate | BRD-K83486494 | CC1(C)[C@@H](O[C@H]2O[C@H](C(=O)O)[C@@H](O)[C@H](O)[C@H]2O[C@@H]2O[C@H](C(=O)O)[C@@H](O)[C@H](O)[C@H]2O)CC[C@@]2(C)[C@H]1CC[C@]1(C)[C@@H]2C(=O)C=C2[C@@H]3C[C@@](C)(C(=O)O)CC[C@]3(C)CC[C@]21C | LPLVUJXQOOQHMX-QWBHMCJMSA-N | Weak | inactive |
| APY0201 | BRD-K38627885 | Cc1cccc(/C=N/Nc2cc(N3CCOCC3)n3nc(-c4ccncc4)cc3n2)c1 | RFZQYGBLRIKROZ-PCLIKHOPSA-N | Weak | inactive |
| arbidol | BRD-K73881242 | CCOC(=O)c1c(CSc2ccccc2)n(C)c2cc(Br)c(O)c(CN(C)C)c12 | KCFYEAOKVJSACF-UHFFFAOYSA-N | Weak | inactive |
| ASP-9521 | BRD-K00003301 | COc1ccc2[nH]c(C(=O)N3CCC(CC(C)(C)O)CC3)cc2c1 | OXSCPDKUZWPWFR-UHFFFAOYSA-N | Weak | inactive |
| astemizole | BRD-K37249724 | COc1ccc(CCN2CCC(Nc3nc4ccccc4n3Cc3ccc(F)cc3)CC2)cc1 | GXDALQBWZGODGZ-UHFFFAOYSA-N | Weak | inactive |
| AY-9944 | BRD-K77012840 | Clc1ccccc1CNC[C@H]1CC[C@H](CNCc2ccccc2Cl)CC1 | RZRPZWGIOOZIBE-IYARVYRRSA-N | Weak | inactive |
| AZ3146 | BRD-K59146805 | COc1cc(OC2CCN(C)CC2)ccc1Nc1ncc2c(n1)n(C1CCCC1)c(=O)n2C | YUKWVHPTFRQHMF-UHFFFAOYSA-N | Weak | inactive |
| AZD3463 | BRD-K24593301 | COc1cc(N2CCC(N)CC2)ccc1Nc1ncc(Cl)c(-c2c[nH]c3ccccc23)n1 | GCYIGMXOIWJGBU-UHFFFAOYSA-N | Weak | inactive |
| AZD4573 | BRD-K00003106 | CC(=O)N[C@@H]1CCC[C@H](C(=O)Nc2cc(-c3cnn4c3CC(C)(C)C4)c(Cl)cn2)C1 | AVIWDYSJSPOOAR-LSDHHAIUSA-N | Weak | inactive |
| AZD5153 | BRD-K00003528 | COc1nnc2ccc(N3CCC(c4ccc(OCCN5CCN(C)C(=O)[C@H]5C)cc4)CC3)nn12 | RSMYFSPOTCDHHJ-GOSISDBHSA-N | Weak | inactive |
| AZD5438 | BRD-K72414522 | Cc1ncc(-c2ccnc(Nc3ccc(S(C)(=O)=O)cc3)n2)n1C(C)C | WJRRGYBTGDJBFX-UHFFFAOYSA-N | Weak | inactive |
| AZD9056 | BRD-K13699028 | O=C(NCC12CC3CC(CC(C3)C1)C2)c1cc(CCCNCCCO)ccc1Cl | HSQAARMBHJCUOK-UHFFFAOYSA-N | Weak | inactive |
| azilsartan-medoxomil | BRD-K25723200 | CCOc1nc2cccc(C(=O)OCc3oc(=O)oc3C)c2n1Cc1ccc(-c2ccccc2-c2nc(=O)o[nH]2)cc1 | QJFSABGVXDWMIW-UHFFFAOYSA-N | Weak | inactive |
| bafetinib | BRD-K82196541 | Cc1ccc(NC(=O)c2ccc(CN3CC[C@H](N(C)C)C3)c(C(F)(F)F)c2)cc1Nc1nccc(-c2cncnc2)n1 | ZGBAJMQHJDFTQJ-DEOSSOPVSA-N | Weak | inactive |
| BAY1125976 | BRD-K00003357 | NC(=O)c1ccc2nc(-c3ccc(C4(N)CCC4)cc3)c(-c3ccccc3)n2n1 | JBGYKRAZYDNCNV-UHFFFAOYSA-N | Weak | inactive |
| belizatinib | BRD-K00003245 | CC(C)NC(=O)[C@H]1CC[C@@H](n2c(NC(=O)c3ccc(F)cc3)nc3ccc(CN4CCC(C(C)(C)O)CC4)cc32)CC1 | WSTUJEXAPHIEIM-FEGDYQJNSA-N | Weak | inactive |
| BIBU-1361 | BRD-K49294207 | CCN(CC)CC1CCN(c2ncc3ncnc(Nc4ccc(F)c(Cl)c4)c3n2)CC1 | JUGSLZCFNSTKBN-UHFFFAOYSA-N | Weak | inactive |
| BIBX-1382 | BRD-K70914287 | CN1CCC(Nc2ncc3ncnc(Nc4ccc(F)c(Cl)c4)c3n2)CC1 | FTFRZXFNZVCRSK-UHFFFAOYSA-N | Weak | inactive |
| BIIB021 | BRD-K51967704 | COc1c(C)cnc(Cn2cnc3c(Cl)nc(N)nc32)c1C | QULDDKSCVCJTPV-UHFFFAOYSA-N | Weak | inactive |
| bithionol | BRD-K39120595 | Oc1c(Cl)cc(Cl)cc1Sc1cc(Cl)cc(Cl)c1O | JFIOVJDNOJYLKP-UHFFFAOYSA-N | Weak | inactive |
| BMS-986158 | BRD-K00003423 | Cc1nnn(C)c1-c1cnc2c3ccc(C(C)(C)O)cc3n([C@H](c3ccccc3)C3CCOCC3)c2c1 | KGERZPVQIRYWRK-GDLZYMKVSA-N | Weak | inactive |
| briciclib | BRD-K05531427 | COc1cc(OC)c(/C=C/S(=O)(=O)Cc2ccc(OC)c(OP(=O)(O)O)c2)c(OC)c1 | LXENKEWVEVKKGV-BQYQJAHWSA-N | Weak | inactive |
| brivudine | BRD-A63675168 | O=c1[nH]c(=O)n([C@H]2C[C@H](O)[C@@H](CO)O2)cc1/C=C/Br | ODZBBRURCPAEIQ-PIXDULNESA-N | Weak | inactive |
| buserelin | BRD-K00003161 | CCNC(=O)[C@@H]1CCCN1C(=O)[C@H](CCCN=C(N)N)NC(=O)[C@H](CC(C)C)NC(=O)[C@@H](COC(C)(C)C)NC(=O)[C@H](Cc1ccc(O)cc1)NC(=O)[C@H](CO)NC(=O)[C@H](Cc1c[nH]c2ccccc12)NC(=O)[C@H](Cc1c[nH]cn1)NC(=O)[C@@H]1CCC(=O)N1 | CUWODFFVMXJOKD-UVLQAERKSA-N | Weak | inactive |
| butein | BRD-K17497770 | O=C(/C=C/c1ccc(O)c(O)c1)c1ccc(O)cc1O | AYMYWHCQALZEGT-ORCRQEGFSA-N | Weak | inactive |
| butylphthalide | BRD-A59378440 | CCCC[C@H]1OC(=O)c2ccccc21 | HJXMNVQARNZTEE-LLVKDONJSA-N | Weak | inactive |
| camylofine-chlorhydrate | BRD-A91733352 | CCN(CC)CCN[C@@H](C(=O)OCCC(C)C)c1ccccc1 | RYOOHIUJEJZCFT-GOSISDBHSA-N | Weak | inactive |
| candesartan | BRD-K84091759 | CCOc1nc2cccc(C(=O)O)c2n1Cc1ccc(-c2ccccc2-c2nn[nH]n2)cc1 | HTQMVQVXFRQIKW-UHFFFAOYSA-N | Weak | inactive |
| carfilzomib | BRD-K15179879 | CC(C)C[C@H](NC(=O)[C@H](CCc1ccccc1)NC(=O)CN1CCOCC1)C(=O)N[C@@H](Cc1ccccc1)C(=O)N[C@@H](CC(C)C)C(=O)[C@@]1(C)CO1 | BLMPQMFVWMYDKT-NZTKNTHTSA-N | Weak | inactive |
| CCT-031374 | BRD-K90999434 | O=C(CN1C2=NCCN2c2ccccc21)c1ccc(-c2ccccc2)cc1 | JPCYSKNELCOPKJ-UHFFFAOYSA-N | Weak | inactive |
| CD-1530 | BRD-K09619322 | O=C(O)c1ccc(-c2ccc3cc(O)c(C45CC6CC(CC(C6)C4)C5)cc3c2)cc1 | VCQGNUWOMLYNNG-UHFFFAOYSA-N | Weak | inactive |
| ceftibuten | BRD-K11124378 | Nc1nc(/C(=C/CC(=O)O)C(=O)N[C@@H]2C(=O)N3C(C(=O)O)=CCS[C@H]23)cs1 | UNJFKXSSGBWRBZ-BJCIPQKHSA-N | Weak | inactive |
| celgosivir | BRD-K00003204 | CCCC(=O)O[C@H]1CN2CC[C@H](O)[C@@H]2[C@@H](O)[C@@H]1O | HTJGLYIJVSDQAE-VWNXEWBOSA-N | Weak | inactive |
| CEP-37440 | BRD-K72827473 | CNC(=O)c1ccccc1Nc1nc(Nc2ccc3c(c2OC)CCC[C@H](N2CCN(CCO)CC2)C3)ncc1Cl | BCSHRERPHLTPEE-NRFANRHFSA-N | Weak | inactive |
| chloramphenicol-palmitate | BRD-K33742430 | CCCCCCCCCCCCCCCC(=O)OC[C@@H](NC(=O)C(Cl)Cl)[C@H](O)c1ccc([N+](=O)[O-])cc1 | PXKHGMGELZGJQE-ILBGXUMGSA-N | Weak | inactive |
| chlorophyllin | BRD-K23728266 | C=CC1=C(C)c2cc3[nH]c(c(CC(=O)O)c4[nH]c(cc5nc(cc1n2)C(C)=C5CC)c(C)c4C(=O)O)[C@@H](CCC(=O)O)[C@@H]3C | HZDQUOWQHHSPIE-UWJYYQICSA-N | Weak | inactive |
| chlorothymol | BRD-K04358951 | Cc1cc(O)c(C(C)C)cc1Cl | KFZXVMNBUMVKLN-UHFFFAOYSA-N | Weak | inactive |
| chlorproguanil | BRD-K25494650 | CC(C)NC(=N)N=C(N)Nc1ccc(Cl)c(Cl)c1 | ISZNZKHCRKXXAU-UHFFFAOYSA-N | Weak | inactive |
| CHR-6494 | BRD-K76259716 | CCCNc1ccc2ncc(-c3ccc4n[nH]cc4c3)n2n1 | CZZCAOGIEGXMBZ-UHFFFAOYSA-N | Weak | inactive |
| cinchocaine | BRD-K99121711 | CCCCOc1cc(C(=O)NCCN(CC)CC)c2ccccc2n1 | PUFQVTATUTYEAL-UHFFFAOYSA-N | Weak | inactive |
| cinchophen | BRD-K72915123 | O=C(O)c1cc(-c2ccccc2)nc2ccccc12 | YTRMTPPVNRALON-UHFFFAOYSA-N | Weak | inactive |
| clobetasol-propionate | BRD-K10799896 | CCC(=O)O[C@]1(C(=O)CCl)[C@@H](C)C[C@H]2[C@@H]3CCC4=CC(=O)C=C[C@]4(C)[C@@]3(F)[C@@H](O)C[C@@]21C | CBGUOGMQLZIXBE-XGQKBEPLSA-N | Weak | inactive |
| clofazimine | BRD-K56614220 | CC(C)/N=c1\cc2n(-c3ccc(Cl)cc3)c3ccccc3nc-2cc1Nc1ccc(Cl)cc1 | WDQPAMHFFCXSNU-BGABXYSRSA-N | Weak | inactive |
| coenzyme-A | BRD-A67956599 | CC(C)(COP(=O)(O)OP(=O)(O)OC[C@H]1O[C@@H](n2cnc3c(N)ncnc32)[C@H](O)[C@@H]1OP(=O)(O)O)[C@@H](O)C(=O)NCCC(=O)NCCS | RGJOEKWQDUBAIZ-IBOSZNHHSA-N | Weak | inactive |
| colistimethate | BRD-K01666924 | CC(C)CCCC(=O)N[C@@H](CCNCS(=O)(=O)O)C(=O)N[C@H](C(=O)N[C@@H](CCNCS(=O)(=O)O)C(=O)N[C@H]1CCNC(=O)[C@H]([C@@H](C)O)NC(=O)[C@H](CCNCS(=O)(=O)O)NC(=O)[C@H](CCNCS(=O)(=O)O)NC(=O)[C@H](CC(C)C)NC(=O)[C@H](CC(C)C)NC(=O)[C@H](CCNCS(=O)(=O)O)NC1=O)[C@@H](C)O | PTQGBRPUALXNCZ-INRRNNIXSA-N | Weak | inactive |
| coumarin | BRD-K23913458 | O=c1ccc2ccccc2o1 | ZYGHJZDHTFUPRJ-UHFFFAOYSA-N | Weak | inactive |
| CP-640186 | BRD-K26921335 | O=C(c1c2ccccc2cc2ccccc12)N1CCC(N2CCC[C@@H](C(=O)N3CCOCC3)C2)CC1 | LDQKDRLEMKIYMC-XMMPIXPASA-N | Weak | inactive |
| CPI-0610 | BRD-K19871656 | Cc1noc2c1-c1ccccc1C(c1ccc(Cl)cc1)=N[C@H]2CC(N)=O | GCWIQUVXWZWCLE-INIZCTEOSA-N | Weak | inactive |
| cycloheximide | BRD-K36055864 | C[C@H]1C[C@H](C)C(=O)[C@H]([C@H](O)CC2CC(=O)NC(=O)C2)C1 | YPHMISFOHDHNIV-FSZOTQKASA-N | Weak | inactive |
| CYM-50358 | BRD-K80666802 | Cc1cc(CN)cc(C)c1NC(=O)c1ccc(-c2cc(Cl)ccc2Cl)o1 | QWJOPXDAQCDRRM-UHFFFAOYSA-N | Weak | inactive |
| D-157495 | BRD-K29066607 | Nc1nc(N)c2c(OCC3CCN(Cc4c(Cl)cccc4Cl)CC3)cccc2n1 | MNLHFGXIUJNDAF-UHFFFAOYSA-N | Weak | inactive |
| dalfampridine | BRD-K22482860 | Nc1ccncc1 | NUKYPUAOHBNCPY-UHFFFAOYSA-N | Weak | inactive |

| | | | | | |
|---|---|---|---|---|---|
| dantron | BRD-K10065684 | O=C1c2cccc(O)c2C(=O)c2c(O)cccc21 | QBPFLULOKWLNNW-UHFFFAOYSA-N | Weak | inactive |
| deazaguanine | BRD-K00004562 | Nc1cc2[nH]cnc2c(O)n1 | KXBCLNRMQPRVTP-UHFFFAOYSA-N | Weak | inactive |
| dihydroartemisinin | BRD-K62213621 | C[C@@H]1CC[C@H]2[C@@H](C)[C@@H](O)O[C@@H]3O[C@@]4(C)CC[C@@H]1[C@@]23OO4 | BJDCWCLMFKKGEE-ISOSDAIHSA-N | Weak | inactive |
| dimethisoquin | BRD-K73391359 | CCCCc1cc2ccccc2c(OCCN(C)C)n1 | XNMYNYSCEJBRPZ-UHFFFAOYSA-N | Weak | inactive |
| dorsomorphin | BRD-K54233340 | c1cc(-c2cnn3cc(-c4ccc(OCCN5CCCCC5)cc4)cnc23)ccn1 | XHBVYDAKJHETMP-UHFFFAOYSA-N | Weak | inactive |
| dyclonine | BRD-K72259270 | CCCCOc1ccc(C(=O)CCN2CCCCC2)cc1 | BZEWSEKUUPWQDQ-UHFFFAOYSA-N | Weak | inactive |
| EGF816 | BRD-K22514639 | Cc1cc(C(=O)Nc2nc3cccc(Cl)c3n2[C@@H]2CCCCN(C(=O)/C=C/CN(C)C)C2)ccn1 | IOMMMLWIABWRKL-WUTDNEBXSA-N | Weak | inactive |
| eletriptan | BRD-K30652337 | CN1CCC[C@@H]1Cc1c[nH]c2ccc(CCS(=O)(=O)c3ccccc3)cc12 | PWVXXGRKLHYWKM-LJQANCHMSA-N | Weak | inactive |
| eperisone | BRD-K01825729 | CCc1ccc(C(=O)[C@@H](C)CN2CCCCC2)cc1 | SQUNAWUMZGQQJD-AWEZNQCLSA-N | Weak | inactive |
| etifoxine | BRD-K01826330 | CCNC1=Nc2ccc(Cl)cc2[C@](C)(c2ccccc2)O1 | IBYCYJFUEJQSMK-KRWDZBQOSA-N | Weak | inactive |
| etoposide-phosphate | BRD-K72533376 | COc1cc([C@@H]2c3cc4c(cc3[C@@H](O[C@@H]3O[C@@H]5CO[C@@H](C)O[C@H]5[C@H](O)[C@H]3O)[C@H]3COC(=O)[C@H]23)OCO4)cc(OC)c1OP(=O)(O)O | LIQODXNTTZAGID-OCBXBXKTSA-N | Weak | inactive |
| flubendazole | BRD-K86003836 | COC(=O)Nc1nc2ccc(C(=O)c3ccc(F)cc3)cc2[nH]1 | CPEUVMUXAHMANV-UHFFFAOYSA-N | Weak | inactive |
| frovatriptan | BRD-K08255599 | CN[C@@H]1CCc2[nH]c3ccc(C(N)=O)cc3c2C1 | XPSQPHWEGNHMSK-SECBINFHSA-N | Weak | inactive |
| futibatinib | BRD-K00003392 | C=CC(=O)N1CC[C@H](n2nc(C#Cc3cc(OC)cc(OC)c3)c3c(N)ncnc32)C1 | KEIPNCCJPRMIAX-HNNXBMFYSA-N | Weak | inactive |
| G-749 | BRD-K00003126 | CN1CCC(Nc2nc(Nc3ccc(Oc4ccccc4)cc3)c3c(O)ncc(Br)c3n2)CC1 | SXWMIXPJPNCXQQ-UHFFFAOYSA-N | Weak | inactive |
| ganciclovir | BRD-K22662435 | Nc1nc(=O)c2ncn(COC(CO)CO)c2[nH]1 | IRSCQMHQWWYFCW-UHFFFAOYSA-N | Weak | inactive |
| ganetespib | BRD-K38852836 | CC(C)c1cc(-c2n[nH]c(=O)n2-c2ccc3c(ccn3C)c2)c(O)cc1O | RVAQIUULWULRNW-UHFFFAOYSA-N | Weak | inactive |
| GDC-0084 | BRD-K00003309 | CC1(C)OCCn2c1nc1c(N3CCOCC3)nc(-c3cnc(N)nc3)nc12 | LGWACEZVCMBSKW-UHFFFAOYSA-N | Weak | inactive |
| GDC-0941 | BRD-K52911425 | CS(=O)(=O)N1CCN(Cc2cc3nc(-c4cccc5n[nH]cc45)nc(N4CCOCC4)c3s2)CC1 | LHNIIDJUOCFXAP-UHFFFAOYSA-N | Weak | inactive |
| GDC-0980 | BRD-K23228615 | Cc1c(CN2CCN(C(=O)[C@H](C)O)CC2)sc2c(N3CCOCC3)nc(-c3cnc(N)nc3)nc12 | YOVVNQKCSKSHKT-HNNXBMFYSA-N | Weak | inactive |
| geldanamycin | BRD-K43570081 | COC1=C2C[C@@H](C)C[C@H](OC)[C@H](O)[C@@H](C)/C=C(\C)[C@H](OC(N)=O)[C@@H](OC)/C=C/C=C(\C)C(=O)NC(=CC1=O)C2=O | QTQAWLPCGQOSGP-PHLMVCJGSA-N | Weak | inactive |
| ginsenoside-RE3 | BRD-A44511856 | CC(C)=CCC[C@@](C)(O)[C@H]1CC[C@]2(C)[C@@H]1[C@H](O)C[C@@H]1[C@@]3(C)CC[C@H](O[C@@H]4O[C@H](CO)[C@@H](O)[C@H](O)[C@H]4O[C@@H]4O[C@H](CO)[C@@H](O)[C@H](O)[C@H]4O)C(C)(C)[C@@H]3CC[C@]12C | RWXIFXNRCLMQCD-CZIWJLDFSA-N | Weak | inactive |
| golvatinib | BRD-K69001009 | CN1CCN(C2CCN(C(=O)Nc3cc(Oc4ccc(NC(=O)C5(C(=O)Nc6ccc(F)cc6)CC5)c(F)c4)ccn3)CC2)CC1 | UQRCJCNVNUFYDX-UHFFFAOYSA-N | Weak | inactive |
| GPP-78 | BRD-K83289131 | O=C(CCCCCCn1cc(-c2cccnc2)nn1)Nc1ccccc1-c1ccccc1 | SJOLTIOPWDLDEB-UHFFFAOYSA-N | Weak | inactive |
| GSK1562590 | BRD-K36697397 | CN(C(=O)CN1C(=O)COc2cc(Cl)c(Cl)cc21)[C@@H](CN1CCCC1)c1ccc(-c2cccc(C(N)=O)c2)cc1 | QDUMEBKFKOODFN-SANMLTNESA-N | Weak | inactive |
| hexachlorophene | BRD-K99792991 | Oc1c(Cl)cc(Cl)c(Cl)c1Cc1c(O)c(Cl)cc(Cl)c1Cl | ACGUYXCXAPNIKK-UHFFFAOYSA-N | Weak | inactive |
| hexylresorcinol | BRD-K99946902 | CCCCCCc1ccc(O)cc1O | WFJIVOKAWHGMBH-UHFFFAOYSA-N | Weak | inactive |
| I-BET-762 | BRD-K08109215 | CCNC(=O)C[C@@H]1N=C(c2ccc(Cl)cc2)c2cc(OC)ccc2-n2c(C)nnc21 | AAAQFGUYHFJNHI-SFHVURJKSA-N | Weak | inactive |
| ibuprofen | BRD-A17655518 | CC(C)Cc1ccc([C@@H](C)C(=O)O)cc1 | HEFNNWSXXWATRW-SNVBAGLBSA-N | Weak | inactive |
| IKK-2-inhibitor-V | BRD-K74305673 | O=C(Nc1cc(C(F)(F)F)cc(C(F)(F)F)c1)c1cc(Cl)ccc1O | CHILCFMQWMQVAL-UHFFFAOYSA-N | Weak | inactive |
| imipramine | BRD-K38436528 | CN(C)CCCN1c2ccccc2CCc2ccccc21 | BCGWQEUPMDMJNV-UHFFFAOYSA-N | Weak | inactive |
| indatraline | BRD-K01649396 | CN[C@@H]1C[C@@H](c2ccc(Cl)c(Cl)c2)c2ccccc21 | SVFXPTLYMIXFRX-XJKSGUPXSA-N | Weak | inactive |
| ingenol-mebutate | BRD-K93779381 | C/C=C(/C)C(=O)O[C@H]1C(C)=C[C@]23C(=O)[C@@H](C=C(CO)[C@@H](O)[C@]12O)[C@H]1[C@@H](C[C@H]3C)C1(C)C | VDJHFHXMUKFKET-WDUFCVPESA-N | Weak | inactive |
| iodipamide | BRD-K67261995 | O=C(CCCCC(=O)Nc1c(I)cc(I)c(C(=O)O)c1I)Nc1c(I)cc(I)c(C(=O)O)c1I | FFINMCNLQNTKLU-UHFFFAOYSA-N | Weak | inactive |
| istaroxime | BRD-K95412502 | C[C@]12CC[C@H]3[C@@H](CC(=O)[C@H]4C/C(=N/OCCN)CC[C@@]43C)[C@@H]1CCC2=O | MPYLDWFDPHRTEG-PAAYLBSLSA-N | Weak | inactive |
| K02288 | BRD-K58568447 | COc1cc(-c2cc(-c3cccc(O)c3)cnc2N)cc(OC)c1OC | CJLMANFTWLNAKC-UHFFFAOYSA-N | Weak | inactive |
| L-alanine | BRD-K25125382 | C[C@H](N)C(=O)O | QNAYBMKLOCPYGJ-REOHCLBHSA-N | Weak | inactive |
| lanatoside-C | BRD-A64242993 | CC(=O)O[C@H]1C[C@H](O[C@H]2[C@@H](O)C[C@H](O[C@H]3[C@@H](O)C[C@H](O[C@H]4CC[C@@]5(C)[C@@H](CC[C@H]6[C@@H]5C[C@@H](O)[C@]5(C)[C@@H](C7=CC(=O)OC7)CC[C@]65O)C4)O[C@@H]3C)O[C@@H]2C)O[C@H](C)[C@H]1O[C@@H]1O[C@H](CO)[C@@H](O)[C@H](O)[C@H]1O | JAYAGJDXJIDEKI-FUVOQQLASA-N | Weak | inactive |
| levallorphan | BRD-K30634073 | C=CCN1CC[C@]23CCCC[C@H]2[C@H]1Cc1ccc(O)cc13 | OZYUPQUCAUTOBP-QXAKKESOSA-N | Weak | inactive |
| levetiracetam | BRD-K49404994 | CC[C@@H](C(N)=O)N1CCCC1=O | HPHUVLMMVZITSG-LURJTMIESA-N | Weak | inactive |
| levomequitazine | BRD-A67616246 | c1ccc2c(c1)Sc1ccccc1N2C[C@H]1CN2CCC1CC2 | HOKDBMAJZXIPGC-MRXNPFEDSA-N | Weak | inactive |
| lometrexol | BRD-K66036108 | Nc1nc(=O)c2c([nH]1)NC[C@H](CCc1ccc(C(=O)N[C@@H](CCC(=O)O)C(=O)O)cc1)C2 | ZUQBAQVRAURMCL-DOMZBBRYSA-N | Weak | inactive |
| LUF-5834 | BRD-K67079690 | N#Cc1c(N)nc(SCc2ncc[nH]2)c(C#N)c1-c1ccc(O)cc1 | OFHKDLYFKPBXER-UHFFFAOYSA-N | Weak | inactive |
| lumateperone | BRD-K10031916 | CN1CCN2c3c(cccc31)[C@@H]1CN(CCCC(=O)c3ccc(F)cc3)CC[C@@H]12 | HOIIHACBCFLJET-SFTDATJTSA-N | Weak | inactive |
| LXH254 | BRD-K00003107 | Cc1ccc(NC(=O)c2ccnc(C(F)(F)F)c2)cc1-c1cc(OCCO)nc(N2CCOCC2)c1 | UEPXBTCUIIGYCY-UHFFFAOYSA-N | Weak | inactive |
| LXS196 | BRD-K00003425 | CC1(N)CCN(c2cccnc2NC(=O)c2nc(-c3ncccc3C(F)(F)F)cnc2N)CC1 | XXJXHXJWQSCNPX-UHFFFAOYSA-N | Weak | inactive |
| LY2857785 | BRD-K15382985 | CC(C)c1c2cc(-c3ccnc(N[C@H]4CC[C@H](NC5CCOCC5)CC4)n3)ccc2nn1C | LHIUZPIDLZYPRL-MXVIHJGJSA-N | Weak | inactive |
| LY334370 | BRD-K00003451 | CN1CCC(c2c[nH]c3ccc(NC(=O)c4ccc(F)cc4)cc23)CC1 | MDMJLMDBRQXOOI-UHFFFAOYSA-N | Weak | inactive |
| marbofloxacin | BRD-K71926323 | CN1CCN(c2c(F)cc3c(=O)c(C(=O)O)cn4c3c2OCN4C)CC1 | BPFYOAJNDMUVBL-UHFFFAOYSA-N | Weak | inactive |
| maropitant | BRD-K76594255 | COc1ccc(C(C)(C)C)cc1CN[C@H]1C2CCN(CC2)[C@H]1C(c1ccccc1)c1ccccc1 | OMPCVMLFFSQFIX-CONSDPRKSA-N | Weak | inactive |
| masitinib | BRD-K71035033 | Cc1ccc(NC(=O)c2ccc(CN3CCN(C)CC3)cc2)cc1Nc1nc(-c2cccnc2)cs1 | WJEOLQLKVOPQFV-UHFFFAOYSA-N | Weak | inactive |
| meclizine | BRD-K01826560 | Cc1cccc(CN2CCN([C@@H](c3ccccc3)c3ccc(Cl)cc3)CC2)c1 | OCJYIGYOJCODJL-VWLOTQADSA-N | Weak | inactive |
| medronic-acid | BRD-K94072573 | O=P(O)(O)CP(=O)(O)O | MBKDYNNUVRNNRF-UHFFFAOYSA-N | Weak | inactive |
| mertansine | BRD-K00003293 | COc1cc2cc(c1Cl)N(C)C(=O)C[C@H](OC(=O)[C@H](C)N(C)C(=O)CCS)[C@]1(C)O[C@H]1[C@H](C)[C@@H]1C[C@@](O)(NC(=O)O1)[C@H](OC)/C=C/C=C(\C)C2 | ANZJBCHSOXCCRQ-FKUXLPTCSA-N | Weak | inactive |
| methylene-blue | BRD-K54770538 | CN(C)c1ccc2nc3ccc(=[N+](C)C)cc-3sc2c1 | RBTBFTRPCNLSDE-UHFFFAOYSA-N | Weak | inactive |
| methylparaben | BRD-K78321730 | COC(=O)c1ccc(O)cc1 | LXCFILQKKLGQFO-UHFFFAOYSA-N | Weak | inactive |
| metitepine | BRD-K01826545 | CSc1ccc2c(c1)[C@H](N1CCN(C)CC1)Cc1ccccc1S2 | RLJFTICUTYVZDG-GOSISDBHSA-N | Weak | inactive |
| metixene | BRD-K01826531 | CN1CCC[C@@H](CC2c3ccccc3Sc3ccccc32)C1 | MJFJKKXQDNNUJF-HNNXBMFYSA-N | Weak | inactive |
| miglustat | BRD-K33251802 | CCCCN1C[C@H](O)[C@@H](O)[C@H](O)[C@H]1CO | UQRORFVVSGFNRO-UTINFBMNSA-N | Weak | inactive |
| mivebresib | BRD-K00003356 | CCS(=O)(=O)Nc1ccc(Oc2ccc(F)cc2F)c(-c2cn(C)c(=O)c3[nH]ccc23)c1 | RDONXGFGWSSFMY-UHFFFAOYSA-N | Weak | inactive |

| | | | | | |
|---|---|---|---|---|---|
| MK-4074 | BRD-K00003482 | COc1cc(C(=O)N2CCC3(CC2)CC(=O)c2cc(-c4cncc(C(=O)O)c4)ccc2O3)cc2c1c(C)cn2C1CC1 | WDBNGXLHMZSUEI-UHFFFAOYSA-N | Weak | inactive |
| moxifloxacin | BRD-K66615216 | COc1c(N2C[C@@H]3CCCN[C@@H]3C2)c(F)cc2c(=O)c(C(=O)O)cn(C3CC3)c12 | FABPRXSRWADJSP-MEDUHNTESA-N | Weak | inactive |
| napabucasin | BRD-K22064724 | CC(=O)c1cc2c(o1)C(=O)c1ccccc1C2=O | DPHUWDIXHNQOSY-UHFFFAOYSA-N | Weak | inactive |
| navitoclax | BRD-K82746043 | CC1(C)CCC(c2ccc(Cl)cc2)=C(CN2CCN(c3ccc(C(=O)NS(=O)(=O)c4ccc(N[C@H](CCN5CCOCC5)CSc5ccccc5)c(S(=O)(=O)C(F)(F)F)c4)cc3)CC2)C1 | JLYAXFNOILIKPP-KXQOOQHDSA-N | Weak | inactive |
| NECA | BRD-K14395568 | CCNC(=O)[C@H]1O[C@@H](n2cnc3c(N)ncnc32)[C@H](O)[C@@H]1O | JADDQZYHOWSFJD-FLNNQWSLSA-N | Weak | inactive |
| nemiralisib | BRD-K00003285 | CC(C)N1CCN(Cc2cnc(-c3cc(-c4cccc5[nH]ccc45)cc4[nH]ncc34)o2)CC1 | MCIDWGZGWVSZMK-UHFFFAOYSA-N | Weak | inactive |
| nemorubicin | BRD-K51911221 | COc1cccc2c1C(=O)c1c(O)c3c(c(O)c1C2=O)C[C@@](O)(C(=O)CO)C[C@@H]3O[C@H]1C[C@H](N2CCO[C@H](OC)C2)[C@H](O)[C@H](C)O1 | CTMCWCONSULRHO-UHQPFXKFSA-N | Weak | inactive |
| NG-nitro-arginine | BRD-K89223384 | NC(=N[N+](=O)[O-])NCCC[C@H](N)C(=O)O | MRAUNPAHJZDYCK-BYPYZUCNSA-N | Weak | inactive |
| niclosamide | BRD-K35960502 | O=C(Nc1ccc([N+](=O)[O-])cc1Cl)c1cc(Cl)ccc1O | RJMUSRYZPJIFPJ-UHFFFAOYSA-N | Weak | inactive |
| nitazoxanide | BRD-K96471533 | CC(=O)Oc1ccccc1C(=O)Nc1ncc([N+](=O)[O-])s1 | YQNQNVDNTFHQSW-UHFFFAOYSA-N | Weak | inactive |
| NNC-05-2090 | BRD-K85015012 | COc1ccccc1C1(O)CCN(CCCn2c3ccccc3c3ccccc32)CC1 | XZZYCJOGZYEPPW-UHFFFAOYSA-N | Weak | inactive |
| nocodazole | BRD-K12539581 | COC(=O)Nc1nc2ccc(C(=O)c3cccs3)cc2[nH]1 | KYRVNWMVYQXFEU-UHFFFAOYSA-N | Weak | inactive |
| norelgestromin | BRD-K42504537 | C#C[C@]1(O)CC[C@H]2[C@@H]3CCC4=C/C(=N/O)CC[C@@H]4[C@H]3CC[C@@]21CC | ISHXLNHNDMZNMC-VTKCIJPMSA-N | Weak | inactive |
| norethisterone-enanthate | BRD-K07625016 | C#C[C@]1(OC(=O)CCCCCC)CC[C@H]2[C@@H]3CCC4=CC(=O)CC[C@@H]4[C@H]3CC[C@@]21C | APTGJECXMIKIET-WOSSHHRXSA-N | Weak | inactive |
| NTNCB | BRD-A40413915 | O=[N+]([O-])c1ccccc1S(=O)(=O)NC[C@H]1CC[C@H](CNC[C@H]2CCc3ccccc32)CC1 | ISSSVYKMIIAPTH-ACRUOGEOSA-N | Weak | inactive |
| NVP-HSP990 | BRD-K67915505 | COc1cccc(-c2cc(F)ccc2[C@H]2Cc3nc(N)nc(C)c3C(=O)N2)n1 | WSMQUUGTQYPVPD-OAHLLOKOSA-N | Weak | inactive |
| octocrylene | BRD-K00003605 | CCCC[C@@H](CC)COC(=O)C(C#N)=C(c1ccccc1)c1ccccc1 | FMJSMJQBSVNSBF-LJQANCHMSA-N | Weak | inactive |
| oglemilast | BRD-K92303087 | CS(=O)(=O)Nc1ccc2oc3c(OC(F)F)ccc(C(=O)Nc4c(Cl)cncc4Cl)c3c2c1 | OKFDRAHPFKMAJH-UHFFFAOYSA-N | Weak | inactive |
| olumacostat-glasaretil | BRD-K00004611 | CCCCCCCCCCCCCCOc1ccc(C(=O)OCC(=O)N(C)CC(=O)OCC)o1 | FYJLDICZGDFWKP-UHFFFAOYSA-N | Weak | inactive |
| ombrabulin | BRD-K00003267 | COc1ccc(/C=C\c2cc(OC)c(OC)c(OC)c2)cc1NC(=O)[C@@H](N)CO | IXWNTLSTOZFSCM-YVACAVLKSA-N | Weak | inactive |
| oridonin | BRD-K00003710 | C=C1C(=O)[C@@]23[C@@H](CC[C@@H]1[C@H]2O)[C@@]12CO[C@]3(O)[C@@H](O)[C@@H]1C(C)(C)CC[C@@H]2O | SDHTXBWLVGWJFT-XKCURVIJSA-N | Weak | inactive |
| OSI-027 | BRD-K87124298 | COc1cccc2cc(-c3nc([C@H]4CC[C@H](C(=O)O)CC4)n4ncnc(N)c34)[nH]c12 | JROFGZPOBKIAEW-HAQNSBGRSA-N | Weak | inactive |
| OTS167 | BRD-K53417444 | CC(=O)c1cnc2ccc(-c3cc(Cl)c(O)c(Cl)c3)nc2c1N[C@H]1CC[C@H](CN(C)C)CC1 | DKZYXHCYPUVGAF-JCNLHEQBSA-N | Weak | inactive |
| oxelaidin | BRD-K17977438 | CCN(CC)CCOCCOC(=O)C(CC)(CC)c1ccccc1 | IQADUMSPOQKAAO-UHFFFAOYSA-N | Weak | inactive |
| OXF-BD-02 | BRD-A73891132 | Cc1noc(C)c1-c1cc(O)cc([C@@H](O)c2ccccc2)c1 | FEQUIPXIENTMJN-SFHVURJKSA-N | Weak | inactive |
| oxybenzone | BRD-K59037100 | COc1ccc(C(=O)c2ccccc2)c(O)c1 | DXGLGDHPHMLXJC-UHFFFAOYSA-N | Weak | inactive |
| oxyphenbutazone | BRD-A33749298 | CCCC[C@H]1C(=O)N(c2ccccc2)N(c2ccc(O)cc2)C1=O | HFHZKZSRXITVMK-KRWDZBQOSA-N | Weak | inactive |
| PD-153035 | BRD-K26603252 | COc1cc2ncnc(Nc3cccc(Br)c3)c2cc1OC | LSPANGZZENHZNJ-UHFFFAOYSA-N | Weak | inactive |
| PD-168393 | BRD-K17702546 | C=CC(=O)Nc1ccc2ncnc(Nc3cccc(Br)c3)c2c1 | HTUBKQUPEREOGA-UHFFFAOYSA-N | Weak | inactive |
| perphenazine | BRD-K10995081 | OCCN1CCN(CCCN2c3ccccc3Sc3ccc(Cl)cc32)CC1 | RGCVKNLCSQQDEP-UHFFFAOYSA-N | Weak | inactive |
| PF-04691502 | BRD-K07310275 | COc1ccc(-c2cc3c(C)nc(N)nc3n([C@H]3CC[C@H](OCCO)CC3)c2=O)cn1 | XDLYKKIQACFMJG-WKILWMFISA-N | Weak | inactive |
| PF-05175157 | BRD-K00002423 | Cc1nc2ccc(C(=O)N3CCC4(CC3)CC(=O)c3c(cnn3C(C)(C)C)C4)cc2[nH]1 | BDXXSFOJPYSYOC-UHFFFAOYSA-N | Weak | inactive |
| PFI-1 | BRD-K13094524 | COc1ccccc1S(=O)(=O)Nc1ccc2c(c1)CN(C)C(=O)N2 | TXZPMHLMPKIUGK-UHFFFAOYSA-N | Weak | inactive |
| phthalylsulfathiazole | BRD-K64659768 | O=C(O)c1ccccc1C(=O)Nc1ccc(S(=O)(=O)Nc2nccs2)cc1 | PBMSWVPMRUJMPE-UHFFFAOYSA-N | Weak | inactive |
| PI-103 | BRD-K67868012 | Oc1cccc(-c2nc(N3CCOCC3)c3oc4ncccc4c3n2)c1 | TUVCWJQQGGETHL-UHFFFAOYSA-N | Weak | inactive |
| PI-828 | BRD-K97365803 | Nc1ccc(-c2cccc3c(=O)cc(N4CCOCC4)oc23)cc1 | WUKMIBOGGXMBAC-UHFFFAOYSA-N | Weak | inactive |
| pifithrin-mu | BRD-K96799727 | NS(=O)(=O)C#Cc1ccccc1 | ZZUZYEMRHCMVTB-UHFFFAOYSA-N | Weak | inactive |
| pitolisant | BRD-K21673112 | Clc1ccc(CCCOCCCN2CCCCC2)cc1 | NNACHAUCXXVJSP-UHFFFAOYSA-N | Weak | inactive |
| plinabulin | BRD-K99498722 | CC(C)(C)c1nc[nH]c1/C=c1\[nH]c(=O)/c(=C/c2ccccc2)[nH]c1=O | UNRCMCRRFYFGFX-TYPNBTCFSA-N | Weak | inactive |
| procarbazine | BRD-K13032584 | CNNCc1ccc(C(=O)NC(C)C)cc1 | CPTBDICYNRMXFX-UHFFFAOYSA-N | Weak | inactive |
| profenamine | BRD-A16311756 | CCN(CC)[C@@H](C)CN1c2ccccc2Sc2ccccc21 | CDOZDBSBBXSXLB-HNNXBMFYSA-N | Weak | inactive |
| progesterone | BRD-K64994968 | CC(=O)[C@H]1CC[C@H]2[C@@H]3CCC4=CC(=O)CC[C@]4(C)[C@H]3CC[C@]12C | RJKFOVLPORLFTN-LEKSSAKUSA-N | Weak | inactive |
| promazine | BRD-K06980535 | CN(C)CCCN1c2ccccc2Sc2ccccc21 | ZGUGWUXLJSTTMA-UHFFFAOYSA-N | Weak | inactive |
| propiverine | BRD-K91244729 | CCCOC(C(=O)OC1CCN(C)CC1)(c1ccccc1)c1ccccc1 | QPCVHQBVMYCJOM-UHFFFAOYSA-N | Weak | inactive |
| protoporphyrin-IX | BRD-K26813314 | C=CC1=C(C)c2cc3[nH]c(cc4[nH]c(cc5nc(cc1n2)C(C)=C5C=C)c(C)c4CCC(=O)O)c(CCC(=O)O)c3C | HGXFBVDWWKPUKZ-UHFFFAOYSA-N | Weak | inactive |
| puerarin | BRD-K89048337 | O=c1c(-c2ccc(O)cc2)coc2c([C@@H]3O[C@H](CO)[C@@H](O)[C@H](O)[C@H]3O)c(O)ccc12 | HKEAFJYKMMKDOR-VPRICQMDSA-N | Weak | inactive |
| pyronaridine | BRD-K73132780 | COc1ccc2nc3cc(Cl)ccc3c(Nc3cc(CN4CCCC4)c(O)c(CN4CCCC4)c3)c2n1 | DJUFPMUQJKWIJB-UHFFFAOYSA-N | Weak | inactive |
| regadenoson | BRD-K00003609 | CNC(=O)c1cnn(-c2nc(N)c3ncn([C@@H]4O[C@H](CO)[C@@H](O)[C@H]4O)c3n2)c1 | LZPZPHGJDAGEJZ-AKAIJSEGSA-N | Weak | inactive |
| retaspimycin | BRD-K67935717 | C=CCNc1c(O)cc2c(O)c1C[C@@H](C)C[C@H](OC)[C@H](O)[C@@H](C)/C=C(\C)[C@H](OC(N)=O)[C@@H](OC)/C=C\C=C(/C)C(=O)N2 | OAKGNIRUXAZDQF-TXHRRWQRSA-N | Weak | inactive |
| rimcazole | BRD-K70490179 | C[C@H]1CN(CCCn2c3ccccc3c3ccccc32)C[C@@H](C)N1 | GUDVQJXODNJRIJ-CALCHBBNSA-N | Weak | inactive |
| sangivamycin | BRD-K36169695 | NC(=O)c1cn([C@@H]2O[C@H](CO)[C@@H](O)[C@H]2O)c2ncnc(N)c12 | OBZJZDHRXBKKTJ-JTFADIMSSA-N | Weak | inactive |
| SB-590885 | BRD-K78809024 | CN(C)CCOc1ccc(-c2nc(-c3ccncc3)c(-c3ccc4c(c3)CC/C4=N\O)[nH]2)cc1 | MLSAQOINCGAULQ-QFMPWRQOSA-N | Weak | inactive |
| semapimod | BRD-K21237892 | CC(=NNC(=N)N)c1cc(NC(=O)CCCCCCCCC(=O)Nc2cc(C(C)=NNC(=N)N)cc(C(C)=NNC(=N)N)c2)cc(C(C)=NNC(=N)N)c1 | PWDYHMBTPGXCSN-UHFFFAOYSA-N | Weak | inactive |
| SKLB-1028 | BRD-K00004690 | CC(C)n1c(Nc2cccnc2)nc2cnc(Nc3ccc(N4CCN(C)CC4)cc3)nc21 | WSOHOUHPUOAXIN-UHFFFAOYSA-N | Weak | inactive |
| SNX-2112 | BRD-K71281111 | CC1(C)CC(=O)c2c(C(F)(F)F)nn(-c3ccc(C(N)=O)c(N[C@H]4CC[C@H](O)CC4)c3)c2C1 | ZFVRYNYOPQZKDG-MQMHXKEQSA-N | Weak | inactive |
| soblidotin | BRD-K00003174 | CC[C@H](C)[C@@H]([C@@H](CC(=O)N1CCC[C@H]1[C@H](OC)[C@@H](C)C(=O)NCCc1ccccc1)OC)N(C)C(=O)[C@@H](NC(=O)[C@H](C(C)C)N(C)C)C(C)C | DZMVCVHATYROOS-ZBFGKEHZSA-N | Weak | inactive |

| | | | | | |
|---|---|---|---|---|---|
| somatostatin | BRD-K14681867 | C[C@H](N)C(=O)NCC(=O)N[C@H]1CSSC[C@@H](C(=O)O)NC(=O)[C@H](CO)NC(=O)[C@H]([C@@H](C)O)NC(=O)[C@H](Cc2ccccc2)NC(=O)[C@H]([C@@H](C)O)NC(=O)[C@H](CCCCN)NC(=O)[C@H](Cc2c[nH]c3ccccc23)NC(=O)[C@H](Cc2ccccc2)NC(=O)[C@H](Cc2ccccc2)NC(=O)[C@H](CC(N)=O)NC(=O)[C@H](CCCCN)NC1=O | NHXLMOGPVYXJNR-ATOGVRKGSA-N | Weak | inactive |
| sotalol | BRD-A33168282 | CC(C)NCC(O)c1ccc(NS(C)(=O)=O)cc1 | ZBMZVLHSJCTVON-UHFFFAOYSA-N | Weak | inactive |
| STF-31 | BRD-K22477529 | CC(C)(C)c1ccc(S(=O)(=O)NCc2ccc(C(=O)Nc3cccnc3)cc2)cc1 | NGQPRVWTFNBUHA-UHFFFAOYSA-N | Weak | inactive |
| sulfamonomethoxine | BRD-K14116214 | COc1cc(NS(=O)(=O)c2ccc(N)cc2)ncn1 | WMPXPUYPYQKQCX-UHFFFAOYSA-N | Weak | inactive |
| T-0070907 | BRD-K31843556 | O=C(Nc1ccncc1)c1cc([N+](=O)[O-])ccc1Cl | FRPJSHKMZHWJBE-UHFFFAOYSA-N | Weak | inactive |
| T-807 | BRD-K00003402 | Fc1ccc(-c2ccc3c(c2)[nH]c2ccncc23)cn1 | GETAAWDSFUCLBS-UHFFFAOYSA-N | Weak | inactive |
| TAK-243 | BRD-K00003376 | NS(=O)(=O)OC[C@H]1C[C@@H](Nc2ccnc3cc(-c4cccc(SC(F)(F)F)c4)nn23)[C@H](O)[C@@H]1O | KJDAGXLMHXUAGV-DGWLBADLSA-N | Weak | inactive |
| taltobulin | BRD-K87512222 | CN[C@H](C(=O)N[C@H](C(=O)N(C)[C@H](/C=C(\C)C(=O)O)C(C)C)C(C)(C)C)C(C)(C)c1ccccc1 | CNTMOLDWXSVYKD-PSRNMDMQSA-N | Weak | inactive |
| taselisib | BRD-K44844162 | Cc1nc(-c2cn3c(n2)-c2ccc(-c4cnn(C(C)(C)C(N)=O)c4)cc2OCC3)n(C(C)C)n1 | BEUQXVWXFDOSAQ-UHFFFAOYSA-N | Weak | inactive |
| tasimelteon | BRD-K62971431 | CCC(=O)NC[C@@H]1C[C@H]1c1cccc2c1CCO2 | PTOIAAWZLUQTIO-GXFFZTMASA-N | Weak | inactive |
| tenoxicam | BRD-A22844106 | CN1C(C(=O)Nc2ccccn2)=C(O)c2sccc2S1(=O)=O | LZNWYQJJBLGYLT-UHFFFAOYSA-N | Weak | inactive |
| terconazole | BRD-K86204871 | CC(C)N1CCN(c2ccc(OC[C@H]3CO[C@](Cn4cncn4)(c4ccc(Cl)cc4Cl)O3)cc2)CC1 | BLSQLHNBWJLIBQ-OZXSUGGESA-N | Weak | inactive |
| tetrindole | BRD-A52660433 | c1cc2c(cc1C1CCCCC1)c1c3n2CCNC3CCC1 | AUXCHYJDVJZEPG-UHFFFAOYSA-N | Weak | inactive |
| TG-100572 | BRD-K18424115 | Cc1cc(-c2cc(O)ccc2Cl)cc2nnc(Nc3ccc(OCCN4CCCC4)cc3)nc12 | AQSSMEORRLJZLU-UHFFFAOYSA-N | Weak | inactive |
| thiethylperazine | BRD-K69600043 | CCSc1ccc2c(c1)N(CCCN1CCN(C)CC1)c1ccccc1S2 | XCTYLCDETUVOIP-UHFFFAOYSA-N | Weak | inactive |
| tilorone | BRD-K34568562 | CCN(CC)CCOc1ccc2c(c1)C(=O)c1cc(OCCN(CC)CC)ccc1-2 | MPMFCABZENCRHV-UHFFFAOYSA-N | Weak | inactive |
| toltrazuril | BRD-K64514229 | Cc1cc(-n2c(=O)[nH]c(=O)n(C)c2=O)ccc1Oc1ccc(SC(F)(F)F)cc1 | OCINXEZVIIVXFU-UHFFFAOYSA-N | Weak | inactive |
| toremifene | BRD-K51350053 | CN(C)CCOc1ccc(/C(=C(/CCCl)c2ccccc2)c2ccccc2)cc1 | XFCLJVABOIYOMF-QPLCGJKRSA-N | Weak | inactive |
| toyocamycin | BRD-K26420709 | N#Cc1cn([C@@H]2O[C@H](CO)[C@@H](O)[C@H]2O)c2ncnc(N)c12 | XOKJUSAYZUAMGJ-WOUKDFQISA-N | Weak | inactive |
| trifluoperazine | BRD-K89732114 | CN1CCN(CCCN2c3ccccc3Sc3ccc(C(F)(F)F)cc32)CC1 | ZEWQUBUPAILYHI-UHFFFAOYSA-N | Weak | inactive |
| triptolide | BRD-K39484304 | CC(C)[C@]12O[C@H]1[C@@H]1O[C@@]13[C@@]1(C)CCC4=C(COC4=O)[C@@H]1C[C@@H]1O[C@@]13[C@@H]2O | DFBIRQPKNDILPW-CIVMWXNOSA-N | Weak | inactive |
| troxerutin | BRD-K73529671 | C[C@@H]1O[C@@H](OC[C@H]2O[C@@H](Oc3c(-c4ccc(OCCO)c(OCCO)c4)oc4cc(OCCO)cc(O)c4c3=O)[C@H](O)[C@@H](O)[C@@H]2O)[C@H](O)[C@H](O)[C@H]1O | IYVFNTXFRYQLRP-VVSTWUKXSA-N | Weak | inactive |
| UC-112 | BRD-K73610817 | Oc1c(CN2CCCC2)cc(COCc2ccccc2)c2cccnc12 | LTGLGIQQZXSLLF-UHFFFAOYSA-N | Weak | inactive |
| ulipristal | BRD-K64381438 | CC(=O)O[C@]1(C(C)=O)CC[C@H]2[C@@H]3CCC4=CC(=O)CCC4=C3[C@@H](c3ccc(N(C)C)cc3)C[C@@]21C | OOLLAFOLCSJHRE-ZHAKMVSLSA-N | Weak | inactive |
| UNC0642 | BRD-K60130390 | COc1cc2c(NC3CCN(C(C)C)CC3)nc(N3CCC(F)(F)CC3)nc2cc1OCCCN1CCCC1 | RNAMYOYQYRYFQY-UHFFFAOYSA-N | Weak | inactive |
| UNC1999 | BRD-K26989966 | CCCc1cc(C)[nH]c(=O)c1CNC(=O)c1cc(-c2ccc(N3CCN(C(C)C)CC3)nc2)cc2c1cnn2C(C)C | DPJNKUOXBZSZAI-UHFFFAOYSA-N | Weak | inactive |
| UNC3230 | BRD-K46344388 | NC(=O)c1nc(Nc2ccccc2)sc1NC(=O)C1CCCCC1 | RZCNASHHHSKTGP-UHFFFAOYSA-N | Weak | inactive |
| uridine-5'-triphosphate | BRD-K01825013 | O=c1ccn([C@@H]2O[C@H](COP(=O)(O)OP(=O)(O)OP(=O)(O)O)[C@@H](O)[C@H]2O)c(=O)[nH]1 | PGAVKCOVUIYSFO-XVFCMESISA-N | Weak | inactive |
| valsartan | BRD-K45158365 | CCCCC(=O)N(Cc1ccc(-c2ccccc2-c2nn[nH]n2)cc1)[C@H](C(=O)O)C(C)C | ACWBQPMHZXGDFX-QFIPXVFZSA-N | Weak | inactive |
| vatalanib | BRD-K70163715 | Clc1ccc(Nc2nnc(Cc3ccncc3)c3ccccc23)cc1 | YCOYDOIWSSHVCK-UHFFFAOYSA-N | Weak | inactive |
| VE-822 | BRD-K04701033 | CNCc1ccc(-c2cc(-c3nc(-c4ccc(S(=O)(=O)C(C)C)cc4)cnc3N)on2)cc1 | JZCWLJDSIRUGIN-UHFFFAOYSA-N | Weak | inactive |
| venlafaxine | BRD-K01826548 | COc1ccc([C@@H](CN(C)C)C2(O)CCCCC2)cc1 | PNVNVHUZROJLTJ-MRXNPFEDSA-N | Weak | inactive |
| verteporfin | BRD-K81849500 | C=CC1=C(C)c2cc3[nH]c(cc4[nH]c(cc5nc(cc1n2)C(C)=C5CCC(=O)O)c(CCC(=O)OC)c4C)[C@@]1(C)C3=CC=C(C(=O)OC)[C@H]1C(=O)OC | VLGTZPSSBIXGAI-ZSFNYQMMSA-N | Weak | inactive |
| vincristine | BRD-K12251893 | CC[C@]1(O)C[C@@H]2CN(CCc3c([nH]c4ccccc34)[C@@](C(=O)OC)(c3cc4c(cc3OC)N(C=O)[C@H]3[C@@](O)(C(=O)OC)[C@H](OC(C)=O)[C@]5(CC)C=CCN6CC[C@]43[C@@H]65)C2)C1 | OGWKCGZFUXNPDA-CFWMRBGOSA-N | Weak | inactive |
| vindesine | BRD-K59753975 | CC[C@]1(O)C[C@H]2CN(CCc3c([nH]c4ccccc34)[C@@](C(=O)OC)(c3cc4c(cc3OC)N(C)[C@H]3[C@@](O)(C(N)=O)[C@H](O)[C@]5(CC)C=CCN6CC[C@]43[C@@H]65)C2)C1 | HHJUWIANJFBDHT-KOTLKJBCSA-N | Weak | inactive |
| voxelotor | BRD-K43639297 | CC(C)n1nccc1-c1ncccc1COc1cccc(O)c1C=O | FWCVZAQENIZVMY-UHFFFAOYSA-N | Weak | inactive |
| WH-4-023 | BRD-K50199947 | COc1ccc(N(C(=O)Oc2c(C)cccc2C)c2ccnc(Nc3ccc(N4CCN(C)CC4)cc3)n2)c(OC)c1 | NBTNHSGBRGTFJS-UHFFFAOYSA-N | Weak | inactive |
| wiskostatin | BRD-A18579359 | CN(C)C[C@@H](O)Cn1c2ccc(Br)cc2c2cc(Br)ccc21 | XUBJEDZHBUPBKL-CYBMUJFWSA-N | Weak | inactive |
| WYE-125132 | BRD-K00003540 | CNC(=O)Nc1ccc(-c2nc(N3C[C@@H]4CC[C@H](C3)O4)c3cnn(C4CCC5(CC4)OCCO5)c3n2)cc1 | QLHHRYZMBGPBJG-OYRHEFFESA-N | Weak | inactive |
| WYE-354 | BRD-K77008974 | COC(=O)Nc1ccc(-c2nc(N3CCOCC3)c3cnn(C4CCN(C(=O)OC)CC4)c3n2)cc1 | IMXHGCRIEAKIBU-UHFFFAOYSA-N | Weak | inactive |
| XL019 | BRD-K04266228 | O=C(Nc1ccc(-c2ccnc(Nc3ccc(N4CCOCC4)cc3)n2)cc1)[C@@H]1CCCN1 | ISOCDPQFIXDIMS-QHCPKHFHSA-N | Weak | inactive |
| XMD8-92 | BRD-K50387473 | CCOc1cc(N2CCC(O)CC2)ccc1Nc1ncc2c(n1)N(C)c1ccccc1C(=O)N2C | QAPAJIZPZGWAND-UHFFFAOYSA-N | Weak | inactive |
| YM-201636 | BRD-K48488978 | Nc1ccc(C(=O)Nc2cccc(-c3nc(N4CCOCC4)c4oc5ncccc5c4n3)c2)cn1 | YBPIBGNBHHGLEB-UHFFFAOYSA-N | Weak | inactive |
| zalcitabine | BRD-K85925969 | Nc1ccn([C@H]2CC[C@@H](CO)O2)c(=O)n1 | WREGKURFCTUGRC-POYBYMJQSA-N | Weak | inactive |
| zolmitriptan | BRD-K54314721 | CN(C)CCc1c[nH]c2ccc(C[C@H]3COC(=O)N3)cc12 | ULSDMUVEXKOYBU-ZDUSSCGKSA-N | Weak | inactive |
