## Supplemental Table 3 for "Identification of druggable host targets needed for SARS-CoV-2 infection by combined pharmacological evaluation and cellular network directed prioritization both in vitro and in vivo"

| | FragmentsBi | AvgNumNon | StdNumNonI | countStrong | p-valueStron | countWeak | p-valueWeak | countVeryWe | p-valueVery | countLow | p-valueLow | countCyto | p-valueCyto | countNE | p-valueNE | hStrongBH | p-valueStron | hWeakBH | p-valueWeak | hVeryWeakB | p-valueVery | hLowBH | p-valueLowB | hCytoBH | p-valueCytoB | hNEBH | p-valueNEBH |
|---|---|---|---|---|---|---|---|---|---|---|---|---|---|---|---|---|---|---|---|---|---|---|---|---|---|---|---|
| 0 | 0 | 8.77272727 | 2.1091105 | 0 | 1 | 0 | 1 | 0 | 1 | 2 | 0.26791271 | 1 | 0.30529951 | 19 | 0.54233419 | FALSE | 1 | FALSE | 1 | FALSE | 1 | FALSE | 1 | FALSE | 1 | FALSE | 1 |
| 1 | 1 | 7.84615385 | 2.08460119 | 4 | 0.08106144 | 1 | 0.87929209 | 4 | 0.19706565 | 4 | 0.33842086 | 3 | 0.08717933 | 48 | 0.98341919 | FALSE | 0.79236883 | FALSE | 1 | FALSE | 1 | FALSE | 1 | FALSE | 1 | FALSE | 1 |
| 2 | 2 | 10.6206897 | 2.2501156 | 1 | 0.52988231 | 0 | 1 | 2 | 0.27983822 | 2 | 0.3871811 | 0 | 1 | 24 | 0.70563334 | FALSE | 1 | FALSE | 1 | FALSE | 1 | FALSE | 1 | FALSE | 1 | FALSE | 1 |
| 3 | 3 | 10.2857143 | 1.48461498 | 0 | 1 | 1 | 0.20567256 | 0 | 1 | 0 | 1 | 0 | 1 | 6 | 0.69920685 | FALSE | 1 | FALSE | 1 | FALSE | 1 | FALSE | 1 | FALSE | 1 | FALSE | 1 |
| 4 | 4 | 9.09090909 | 2.23421922 | 0 | 1 | 0 | 1 | 1 | 0.33149969 | 0 | 1 | 0 | 1 | 10 | 0.46859937 | FALSE | 1 | FALSE | 1 | FALSE | 1 | FALSE | 1 | FALSE | 1 | FALSE | 1 |
| 5 | 5 | 4.57142857 | 1.79694879 | 2 | 0.9579149 | 4 | 0.86657341 | 11 | 0.0775154 | 9 | 0.50446879 | 5 | 0.19799787 | 158 | 0.66275545 | FALSE | 1 | FALSE | 1 | FALSE | 1 | FALSE | 1 | FALSE | 1 | FALSE | 1 |
| 6 | 6 | 9.5 | 3.13391585 | 0 | 1 | 0 | 1 | 1 | 0.40110418 | 3 | 0.0239258 | 0 | 1 | 10 | 0.94579623 | FALSE | 1 | FALSE | 1 | FALSE | 1 | FALSE | 1 | FALSE | 1 | FALSE | 1 |
| 7 | 7 | 10.3461539 | 1.62770031 | 1 | 0.71380452 | 2 | 0.46321244 | 2 | 0.51909752 | 0 | 1 | 1 | 0.54902428 | 42 | 0.36011057 | FALSE | 1 | FALSE | 1 | FALSE | 1 | FALSE | 1 | FALSE | 1 | FALSE | 1 |
| 8 | 8 | 11.5517241 | 1.65229635 | 2 | 0.16962005 | 1 | 0.61539065 | 1 | 0.65463982 | 2 | 0.3871811 | 1 | 0.38148475 | 22 | 0.92902102 | FALSE | 1 | FALSE | 1 | FALSE | 1 | FALSE | 1 | FALSE | 1 | FALSE | 1 |
| 9 | 9 | 10.3529412 | 1.53167254 | 3 | 0.05546921 | 2 | 0.30150529 | 3 | 0.12152681 | 1 | 0.7984848 | 0 | 1 | 25 | 0.9693929 | FALSE | 0.68849059 | FALSE | 1 | FALSE | 1 | FALSE | 1 | FALSE | 1 | FALSE | 1 |
| 10 | 10 | 8 | 2.3204774 | 0 | 1 | 0 | 1 | 0 | 1 | 0 | 1 | 0 | 1 | 13 | 0.10967906 | FALSE | 1 | FALSE | 1 | FALSE | 1 | FALSE | 1 | FALSE | 1 | FALSE | 1 |
| 11 | 11 | 7.45238095 | 2.2168444 | 1 | 0.66519151 | 1 | 0.74973543 | 0 | 1 | 2 | 0.5811115 | 3 | 0.03099591 | 35 | 0.67024784 | FALSE | 1 | FALSE | 1 | FALSE | 1 | FALSE | 1 | FALSE | 0.82559785 | FALSE | 1 |
| 12 | 12 | 10.7 | 1.61554944 | 1 | 0.40558521 | 0 | 1 | 0 | 1 | 1 | 0.60987146 | 0 | 1 | 18 | 0.37504645 | FALSE | 1 | FALSE | 1 | FALSE | 1 | FALSE | 1 | FALSE | 1 | FALSE | 1 |
| 13 | 13 | 3.32321041 | 1.35374603 | 16 | 0.12675623 | 20 | 0.10317716 | 20 | 0.21004268 | 28 | 0.07132865 | 5 | 0.87952001 | 369 | 0.99032617 | FALSE | 0.9659414 | FALSE | 1 | FALSE | 1 | FALSE | 1 | FALSE | 1 | FALSE | 1 |
| 14 | 14 | 6.65432099 | 1.7579484 | 14 | 6.34E-05 | 8 | 0.15374722 | 11 | 0.03108868 | 3 | 0.98174209 | 10 | 0.00030001 | 116 | 0.99999086 | TRUE | 0.01330875 | FALSE | 1 | FALSE | 1 | FALSE | 1 | FALSE | 0.0910253 | FALSE | 1 |
| 15 | 15 | 10 | 2.21880079 | 1 | 0.28675951 | 0 | 1 | 0 | 1 | 0 | 1 | 0 | 1 | 12 | 0.374198 | FALSE | 1 | FALSE | 1 | FALSE | 1 | FALSE | 1 | FALSE | 1 | FALSE | 1 |
| 16 | 16 | 8.88 | 3.43689783 | 2 | 0.57749853 | 4 | 0.22394338 | 4 | 0.2829119 | 0 | 1 | 0 | 1 | 65 | 0.36047181 | FALSE | 1 | FALSE | 1 | FALSE | 1 | FALSE | 1 | FALSE | 1 | FALSE | 1 |
| 17 | 17 | 9.45833333 | 2.23567974 | 1 | 0.46441868 | 1 | 0.54637326 | 2 | 0.21260257 | 0 | 1 | 1 | 0.32796862 | 19 | 0.83975208 | FALSE | 1 | FALSE | 1 | FALSE | 1 | FALSE | 1 | FALSE | 1 | FALSE | 1 |
| 18 | 18 | 9.8125 | 2.24217389 | 0 | 1 | 0 | 1 | 1 | 0.44344817 | 0 | 1 | 0 | 1 | 15 | 0.26127859 | FALSE | 1 | FALSE | 1 | FALSE | 1 | FALSE | 1 | FALSE | 1 | FALSE | 1 |
| 19 | 19 | 9.33333333 | 1.56347192 | 0 | 1 | 0 | 1 | 0 | 1 | 0 | 1 | 0 | 1 | 9 | 0.21661397 | FALSE | 1 | FALSE | 1 | FALSE | 1 | FALSE | 1 | FALSE | 1 | FALSE | 1 |
| 20 | 20 | 9.09302326 | 3.21922967 | 0 | 1 | 2 | 0.40761584 | 1 | 0.79363407 | 5 | 0.0456107 | 2 | 0.1563365 | 33 | 0.93815092 | FALSE | 1 | FALSE | 1 | FALSE | 1 | FALSE | 1 | FALSE | 1 | FALSE | 1 |
| 21 | 21 | 8.95454546 | 2.70483192 | 2 | 0.10809809 | 2 | 0.15794013 | 1 | 0.55340851 | 0 | 1 | 1 | 0.30529951 | 16 | 0.95553951 | FALSE | 0.88820416 | FALSE | 1 | FALSE | 1 | FALSE | 1 | FALSE | 1 | FALSE | 1 |
| 22 | 22 | 9.27272727 | 2.63165724 | 0 | 1 | 2 | 0.15794013 | 1 | 0.55340851 | 1 | 0.64497439 | 0 | 1 | 18 | 0.74744813 | FALSE | 1 | FALSE | 1 | FALSE | 1 | FALSE | 1 | FALSE | 1 | FALSE | 1 |
| 23 | 23 | 10.0588235 | 2.79643866 | 0 | 1 | 3 | 0.01622926 | 1 | 0.46348836 | 0 | 1 | 0 | 1 | 13 | 0.88719222 | FALSE | 1 | FALSE | 0.7256023 | FALSE | 1 | FALSE | 1 | FALSE | 1 | FALSE | 1 |
| 24 | 24 | 9.90909091 | 2.31416765 | 1 | 0.24866461 | 0 | 1 | 0 | 1 | 0 | 1 | 0 | 1 | 10 | 0.46859937 | FALSE | 1 | FALSE | 1 | FALSE | 1 | FALSE | 1 | FALSE | 1 | FALSE | 1 |
| 25 | 25 | 6.26851852 | 1.19881486 | 2 | 0.77000471 | 3 | 0.6846759 | 3 | 0.75139368 | 3 | 0.87972658 | 6 | 0.00838282 | 91 | 0.57807597 | FALSE | 1 | FALSE | 1 | FALSE | 1 | FALSE | 1 | FALSE | 0.50674765 | FALSE | 1 |
| 26 | 26 | 6.57142857 | 2.82120252 | 0 | 1 | 0 | 1 | 0 | 1 | 0 | 1 | 0 | 1 | 7 | 0.3043657 | FALSE | 1 | FALSE | 1 | FALSE | 1 | FALSE | 1 | FALSE | 1 | FALSE | 1 |
| 27 | 27 | 7.5 | 3.43823044 | 0 | 1 | 1 | 0.36919824 | 0 | 1 | 3 | 0.0239258 | 0 | 1 | 10 | 0.94579623 | FALSE | 1 | FALSE | 1 | FALSE | 1 | FALSE | 1 | FALSE | 1 | FALSE | 1 |
| 28 | 28 | 6.63888889 | 3.59901821 | 1 | 0.60837712 | 1 | 0.69480472 | 2 | 0.37289966 | 3 | 0.22784656 | 1 | 0.44938266 | 28 | 0.90259714 | FALSE | 1 | FALSE | 1 | FALSE | 1 | FALSE | 1 | FALSE | 1 | FALSE | 1 |
| 29 | 29 | 9.33333333 | 2.80871659 | 2 | 0.16962005 | 0 | 1 | 1 | 0.65463982 | 1 | 0.74482229 | 1 | 0.38148475 | 24 | 0.70563334 | FALSE | 1 | FALSE | 1 | FALSE | 1 | FALSE | 1 | FALSE | 1 | FALSE | 1 |
| 30 | 30 | 9.63333333 | 2.05723655 | 1 | 0.54198714 | 2 | 0.25306331 | 2 | 0.29330414 | 0 | 1 | 2 | 0.08611595 | 23 | 0.9158696 | FALSE | 1 | FALSE | 1 | FALSE | 1 | FALSE | 1 | FALSE | 1 | FALSE | 1 |
| 31 | 31 | 9.75 | 3.19178633 | 1 | 0.34033054 | 1 | 0.40942991 | 1 | 0.44344817 | 0 | 1 | 0 | 1 | 13 | 0.76799161 | FALSE | 1 | FALSE | 1 | FALSE | 1 | FALSE | 1 | FALSE | 1 | FALSE | 1 |
| 32 | 32 | 7.17283951 | 0.78236936 | 8 | 0.57398961 | 4 | 0.99331019 | 10 | 0.71390483 | 13 | 0.70896383 | 8 | 0.14996266 | 275 | 0.16436059 | FALSE | 1 | FALSE | 1 | FALSE | 1 | FALSE | 1 | FALSE | 1 | FALSE | 1 |
| 33 | 33 | 11.0909091 | 4.52175641 | 0 | 1 | 0 | 1 | 1 | 0.48280983 | 0 | 1 | 1 | 0.25766374 | 16 | 0.44998001 | FALSE | 1 | FALSE | 1 | FALSE | 1 | FALSE | 1 | FALSE | 1 | FALSE | 1 |
| 34 | 34 | 7.79069767 | 1.89930391 | 2 | 0.30241117 | 1 | 0.75788114 | 0 | 1 | 2 | 0.59404271 | 2 | 0.1563365 | 36 | 0.6451239 | FALSE | 1 | FALSE | 1 | FALSE | 1 | FALSE | 1 | FALSE | 1 | FALSE | 1 |
| 35 | 35 | 9.69230769 | 1.13575562 | 0 | 1 | 0 | 1 | 1 | 0.37874417 | 2 | 0.1174583 | 0 | 1 | 10 | 0.86791642 | FALSE | 1 | FALSE | 1 | FALSE | 1 | FALSE | 1 | FALSE | 1 | FALSE | 1 |
| 36 | 36 | 10.5 | 2.29128785 | 1 | 0.4357638 | 2 | 0.15794013 | 0 | 1 | 3 | 0.07743186 | 0 | 1 | 16 | 0.95553951 | FALSE | 1 | FALSE | 1 | FALSE | 1 | FALSE | 1 | FALSE | 1 | FALSE | 1 |
| 37 | 37 | 8.625 | 2.9973947 | 1 | 0.34033054 | 3 | 0.01368613 | 1 | 0.44344817 | 0 | 1 | 0 | 1 | 11 | 0.9718531 | FALSE | 1 | FALSE | 0.69207862 | FALSE | 1 | FALSE | 1 | FALSE | 1 | FALSE | 1 |
| 38 | 38 | 10.7826087 | 2.76557285 | 1 | 0.45027683 | 0 | 1 | 3 | 0.04772439 | 1 | 0.6613267 | 0 | 1 | 18 | 0.86273155 | FALSE | 1 | FALSE | 1 | FALSE | 1 | FALSE | 1 | FALSE | 1 | FALSE | 1 |
| 39 | 39 | 6.87804878 | 1.97786682 | 1 | 0.65632651 | 1 | 0.74131698 | 0 | 1 | 1 | 0.85524728 | 1 | 0.49330646 | 37 | 0.21027077 | FALSE | 1 | FALSE | 1 | FALSE | 1 | FALSE | 1 | FALSE | 1 | FALSE | 1 |
| 40 | 40 | 9.25 | 1.39194109 | 1 | 0.18769745 | 0 | 1 | 1 | 0.25384309 | 0 | 1 | 1 | 0.12394282 | 5 | 0.97551477 | FALSE | 1 | FALSE | 1 | FALSE | 1 | FALSE | 1 | FALSE | 1 | FALSE | 1 |
| 41 | 41 | 4.86769231 | 2.22939162 | 4 | 0.96530641 | 10 | 0.57541473 | 9 | 0.80908096 | 15 | 0.48558714 | 7 | 0.26058953 | 271 | 0.27341267 | FALSE | 1 | FALSE | 1 | FALSE | 1 | FALSE | 1 | FALSE | 1 | FALSE | 1 |
| 42 | 42 | 8 | 3.1896572 | 0 | 1 | 1 | 0.53115654 | 2 | 0.19931429 | 0 | 1 | 1 | 0.31672722 | 19 | 0.71542496 | FALSE | 1 | FALSE | 1 | FALSE | 1 | FALSE | 1 | FALSE | 1 | FALSE | 1 |
| 43 | 43 | 10.2580645 | 3.20249643 | 0 | 1 | 1 | 0.63997468 | 1 | 0.67911179 | 0 | 1 | 0 | 1 | 29 | 0.11661149 | FALSE | 1 | FALSE | 1 | FALSE | 1 | FALSE | 1 | FALSE | 1 | FALSE | 1 |
| 44 | 44 | 7.875 | 2.20439901 | 1 | 0.18769745 | 1 | 0.23139161 | 0 | 1 | 0 | 1 | 0 | 1 | 6 | 0.88425551 | FALSE | 1 | FALSE | 1 | FALSE | 1 | FALSE | 1 | FALSE | 1 | FALSE | 1 |
| 45 | 45 | 8.39393939 | 1.9453035 | 5 | 0.02655059 | 2 | 0.63534334 | 3 | 0.4248691 | 3 | 0.58996791 | 5 | 0.00436843 | 48 | 0.99535371 | FALSE | 0.50937328 | FALSE | 1 | FALSE | 1 | FALSE | 1 | FALSE | 0.39762413 | FALSE | 1 |
| 46 | 46 | 8.48 | 3.0215228 | 2 | 0.13369621 | 4 | 0.00789279 | 1 | 0.59998149 | 2 | 0.31968852 | 0 | 1 | 16 | 0.99713666 | FALSE | 0.97964162 | FALSE | 0.6167796 | FALSE | 1 | FALSE | 1 | FALSE | 1 | FALSE | 1 |
| 47 | 47 | 10.2647059 | 3.45822824 | 0 | 1 | 0 | 1 | 5 | 0.00597743 | 1 | 0.78873814 | 0 | 1 | 27 | 0.75124583 | FALSE | 1 | FALSE | 1 | FALSE | 1 | FALSE | 1 | FALSE | 1 | FALSE | 1 |
| 48 | 48 | 9.23809524 | 1.94947519 | 1 | 0.42086991 | 2 | 0.14655134 | 1 | 0.53671415 | 0 | 1 | 0 | 1 | 17 | 0.77849076 | FALSE | 1 | FALSE | 1 | FALSE | 1 | FALSE | 1 | FALSE | 1 | FALSE | 1 |
| 49 | 49 | 9 | 2.43086217 | 0 | 1 | 1 | 0.49918218 | 0 | 1 | 1 | 0.62783522 | 2 | 0.04568221 | 17 | 0.77849076 | FALSE | 1 | FALSE | 1 | FALSE | 1 | FALSE | 1 | FALSE | 0.95710659 | FALSE | 1 |
| 50 | 50 | 9.26086957 | 1.72494212 | 0 | 1 | 0 | 1 | 1 | 0.5695037 | 4 | 0.01932195 | 0 | 1 | 18 | 0.86273155 | FALSE | 1 | FALSE | 1 | FALSE | 1 | FALSE | 1 | FALSE | 1 | FALSE | 1 |
| 51 | 51 | 8.64285714 | 1.34202102 | 1 | 0.3050805 | 1 | 0.36919824 | 0 | 1 | 0 | 1 | 0 | 1 | 12 | 0.62236227 | FALSE | 1 | FALSE | 1 | FALSE | 1 | FALSE | 1 | FALSE | 1 | FALSE | 1 |
| 52 | 52 | 8.43478261 | 1.95216907 | 0 | 1 | 4 | 0.06038631 | 1 | 0.81522035 | 1 | 0.88573774 | 1 | 0.53375569 | 39 | 0.56897353 | FALSE | 1 | FALSE | 1 | FALSE | 1 | FALSE | 1 | FALSE | 1 | FALSE | 1 |
| 53 | 53 | 9.13043478 | 2.87877311 | 0 | 1 | 0 | 1 | 4 | 0.00713063 | 1 | 0.64497439 | 0 | 1 | 17 | 0.8840357 | FALSE | 1 | FALSE | 1 | FALSE | 1 | FALSE | 1 | FALSE | 1 | FALSE | 1 |
| 54 | 54 | 8.06666667 | 2.59401019 | 2 | 0.17884234 | 1 | 0.6278847 | 0 | 1 | 3 | 0.15707004 | 0 | 1 | 24 | 0.82294782 | FALSE | 1 | FALSE | 1 | FALSE | 1 | FALSE | 1 | FALSE | 1 | FALSE | 1 |
| 55 | 55 | 9.42857143 | 1.17803018 | 1 | 0.16630142 | 0 | 1 | 1 | 0.22601089 | 0 | 1 | 0 | 1 | 5 | 0.91847773 | FALSE | 1 | FALSE | 1 | FALSE | 1 | FALSE | 1 | FALSE | 1 | FALSE | 1 |
| 56 | 56 | 9.57142857 | 2.77010278 | 0 | 1 | 0 | 1 | 0 | 1 | 0 | 1 | 0 | 1 | 7 | 0.3043657 | FALSE | 1 | FALSE | 1 | FALSE | 1 | FALSE | 1 | FALSE | 1 | FALSE | 1 |
| 57 | 57 | 9.04761905 | 2.49716392 | 0 | 1 | 2 | 0.14655134 | 0 | 1 | 2 | 0.25059627 | 0 | 1 | 17 | 0.77849076 | FALSE | 1 | FALSE | 1 | FALSE | 1 | FALSE | 1 | FALSE | 1 | FALSE | 1 |
| 58 | 58 | 8.79411765 | 1.36725168 | 0 | 1 | 3 | 0.37822937 | 2 | 0.70787008 | 6 | 0.09079991 | 2 | 0.30695409 | 55 | 0.83409155 | FALSE | 1 | FALSE | 1 | FALSE | 1 | FALSE | 1 | FALSE | 1 | FALSE | 1 |
| 59 | 59 | 11.0526316 | 2.01242677 | 1 | 0.38989951 | 0 | 1 | 0 | 1 | 0 | 1 | 1 | 0.26986775 | 17 | 0.41141543 | FALSE | 1 | FALSE | 1 | FALSE | 1 | FALSE | 1 | FALSE | 1 | FALSE | 1 |
| 60 | 60 | 10.2727273 | 2.48964799 | 0 | 1 | 0 | 1 | 0 | 1 | 1 | 0.40390032 | 0 | 1 | 10 | 0.46859937 | FALSE | 1 | FALSE | 1 | FALSE | 1 | FALSE | 1 | FALSE | 1 | FALSE | 1 |
| 61 | 61 | 10 | 2.64575131 | 0 | 1 | 0 | 1 | 1 | 0.30655256 | 0 | 1 | 2 | 0.01100138 | 7 | 0.94326215 | FALSE | 1 | FALSE | 1 | FALSE | 1 | FALSE | 1 | FALSE | 0.57403367 | FALSE | 1 |
| 62 | 62 | 7.0483871 | 3.87892328 | 0 | 1 | 5 | 0.04934001 | 2 | 0.65873577 | 2 | 0.78520976 | 2 | 0.27036738 | 51 | 0.74578096 | FALSE | 1 | FALSE | 1 | FALSE | 1 | FALSE | 1 | FALSE | 1 | FALSE | 1 |
| 63 | 63 | 9.31578947 | 2.75399483 | 0 | 1 | 1 | 0.46503795 | 1 | 0.50143826 | 0 | 1 | 0 | 1 | 17 | 0.41141543 | FALSE | 1 | FALSE | 1 | FALSE | 1 | FALSE | 1 | FALSE | 1 | FALSE | 1 |
| 64 | 64 | 11.75 | 2.2220486 | 0 | 1 | 2 | 0.09285443 | 0 | 1 | 0 | 1 | 0 | 1 | 14 | 0.53308261 | FALSE | 1 | FALSE | 1 | FALSE | 1 | FALSE | 1 | FALSE | 1 | FALSE | 1 |
| 65 | 65 | 8.8 | 1.2489996 | 0 | 1 | 0 | 1 | 0 | 1 | 0 | 1 | 0 | 1 | 10 | 0.18273166 | FALSE | 1 | FALSE | 1 | FALSE | 1 | FALSE | 1 | FALSE | 1 | FALSE | 1 |
| 66 | 66 | 8.96774194 | 2.11726956 | 0 | 1 | 1 | 0.63997468 | 3 | 0.09854319 | 2 | 0.41984187 | 1 | 0.4016888 | 24 | 0.90138666 | FALSE | 1 | FALSE | 1 | FALSE | 1 | FALSE | 1 | FALSE | 1 | FALSE | 1 |
| 67 | 67 | 7.97142857 | 3.67606711 | 2 | 0.21638713 | 2 | 0.30150529 | 1 | 0.71261921 | 4 | 0.06848449 | 1 | 0.43077488 | 24 | 0.98848472 | FALSE | 1 | FALSE | 1 | FALSE | 1 | FALSE | 1 | FALSE | 1 | FALSE | 1 |
| 68 | 68 | 9.64285714 | 3.68601881 | 3 | 0.03391597 | 2 | 0.2288771 | 0 | 1 | 2 | 0.3705495 | 1 | 0.37113067 | 20 | 0.97757517 | FALSE | 0.56934354 | FALSE | 1 | FALSE | 1 | FALSE | 1 | FALSE | 1 | FALSE | 1 |
| 69 | 69 | 5.61386139 | 2.11533906 | 4 | 0.76377687 | 5 | 0.78527295 | 4 | 0.93534132 | 13 | 0.13293166 | 7 | 0.04653437 | 168 | 0.66716773 | FALSE | 1 | FALSE | 1 | FALSE | 1 | FALSE | 1 | FALSE | 0.96999891 | FALSE | 1 |
| 70 | 70 | 9.83333333 | 2.19215774 | 0 | 1 | 0 | 1 | 0 | 1 | 0 | 1 | 0 | 1 | 6 | 0.36077158 | FALSE | 1 | FALSE | 1 | FALSE | 1 | FALSE | 1 | FALSE | 1 | FALSE | 1 |
| 71 | 71 | 8.29032258 | 2.55511566 | 0 | 1 | 0 | 1 | 1 | 0.67911179 | 1 | 0.76781284 | 0 | 1 | 29 | 0.11661149 | FALSE | 1 | FALSE | 1 | FALSE | 1 | FALSE | 1 | FALSE | 1 | FALSE | 1 |
| 72 | 72 | 9.30769231 | 2.83886801 | 0 | 1 | 2 | 0.06426041 | 1 | 0.37874417 | 2 | 0.1174583 | 0 | 1 | 8 | 0.99085298 | FALSE | 1 | FALSE | 1 | FALSE | 1 | FALSE | 1 | FALSE | 1 | FALSE | 1 |
| 73 | 73 | 9.31578947 | 1.21623369 | 0 | 1 | 4 | 0.00281443 | 0 | 1 | 1 | 0.59104357 | 0 | 1 | 14 | 0.93677253 | FALSE | 1 | FALSE | 0.46409366 | FALSE | 1 | FALSE | 1 | FALSE | 1 | FALSE | 1 |
| 74 | 74 | 9.71428571 | 2.27078458 | 1 | 0.40558521 | 1 | 0.48239024 | 0 | 1 | 0 | 1 | 0 | 1 | 18 | 0.37504645 | FALSE | 1 | FALSE | 1 | FALSE | 1 | FALSE | 1 | FALSE | 1 | FALSE | 1 |
| 75 | 75 | 7.55454546 | 1.1953422 | 3 | 0.52695321 | 6 | 0.13654729 | 1 | 0.98137785 | 1 | 0.99400862 | 2 | 0.5321878 | 95 | 0.18526068 | FALSE | 1 | FALSE | 1 | FALSE | 1 | FALSE | 1 | FALSE | 1 | FALSE | 1 |
| 76 | 76 | 6.61538462 | 2.10756129 | 2 | 0.26413583 | 2 | 0.36122049 | 1 | 0.76089851 | 1 | 0.84089119 | 0 | 1 | 33 | 0.59061307 | FALSE | 1 | FALSE | 1 | FALSE | 1 | FALSE | 1 | FALSE | 1 | FALSE | 1 |
| 77 | 77 | 9.38235294 | 2.98047278 | 1 | 0.58738519 | 1 | 0.6739483 | 0 | 1 | 2 | 0.46709861 | 1 | 0.43077488 | 29 | 0.55707911 | FALSE | 1 | FALSE | 1 | FALSE | 1 | FALSE | 1 | FALSE | 1 | FALSE | 1 |
| 78 | 78 | 9.5 | 1.63663418 | 1 | 0.3050805 | 0 | 1 | 1 | 0.40110418 | 0 | 1 | 1 | 0.20679328 | 11 | 0.83676449 | FALSE | 1 | FALSE | 1 | FALSE | 1 | FALSE | 1 | FALSE | 1 | FALSE | 1 |
| 79 | 79 | 10 | 2.50713268 | 0 | 1 | 0 | 1 | 0 | 1 | 0 | 1 | 0 | 1 | 14 | 0.09251304 | FALSE | 1 | FALSE | 1 | FALSE | 1 | FALSE | 1 | FALSE | 1 | FALSE | 1 |
| 80 | 80 | 9.17647059 | 3.03383458 | 0 | 1 | 1 | 0.42857733 | 0 | 1 | 1 | 0.55062772 | 0 | 1 | 15 | 0.49060435 | FALSE | 1 | FALSE | 1 | FALSE | 1 | FALSE | 1 | FALSE | 1 | FALSE | 1 |
| 81 | 81 | 8.11111111 | 3.49426338 | 2 | 0.07645454 | 1 | 0.44710673 | 1 | 0.48280983 | 0 | 1 | 1 | 0.25766374 | 13 | 0.95041913 | FALSE | 0.77424195 | FALSE | 1 | FALSE | 1 | FALSE | 1 | FALSE | 1 | FALSE | 1 |
| 82 | 82 | 10.6818182 | 2.07553637 | 4 | 0.00212601 | 2 | 0.15794013 | 0 | 1 | 1 | 0.64497439 | 0 | 1 | 15 | 0.98570275 | FALSE | 0.12179217 | FALSE | 1 | FALSE | 1 | FALSE | 1 | FALSE | 1 | FALSE | 1 |
| 83 | 83 | 8 | 2.76887462 | 0 | 1 | 0 | 1 | 0 | 1 | 0 | 1 | 0 | 1 | 24 | 0.01683964 | FALSE | 1 | FALSE | 1 | FALSE | 1 | FALSE | 1 | FALSE | 1 | FALSE | 1 |
| 84 | 84 | 7.41818182 | 2.26186495 | 4 | 0.05176316 | 3 | 0.26245929 | 2 | 0.59334725 | 0 | 1 | 0 | 1 | 46 | 0.64645216 | FALSE | 0.67847005 | FALSE | 1 | FALSE | 1 | FALSE | 1 | FALSE | 1 | FALSE | 1 |
| 85 | 85 | 5.15384615 | 2.38089039 | 0 | 1 | 1 | 0.56109832 | 0 | 1 | 2 | 0.31968852 | 1 | 0.33902672 | 21 | 0.64957167 | FALSE | 1 | FALSE | 1 | FALSE | 1 | FALSE | 1 | FALSE | 1 | FALSE | 1 |
| 86 | 86 | 9.43478261 | 1.9962157 | 0 | 1 | 1 | 0.53115654 | 2 | 0.19931429 | 1 | 0.6613267 | 0 | 1 | 19 | 0.71542496 | FALSE | 1 | FALSE | 1 | FALSE | 1 | FALSE | 1 | FALSE | 1 | FALSE | 1 |
| 87 | 87 | 7.765625 | 1.28381983 | 3 | 0.22550017 | 1 | 0.87929209 | 1 | 0.90487646 | 4 | 0.33842086 | 1 | 0.65460803 | 54 | 0.58381086 | FALSE | 1 | FALSE | 1 | FALSE | 1 | FALSE | 1 | FALSE | 1 | FALSE | 1 |
| 88 | 88 | 8.95454546 | 3.78366432 | 0 | 1 | 0 | 1 | 2 | 0.18613256 | 0 | 1 | 0 | 1 | 20 | 0.30915202 | FALSE | 1 | FALSE | 1 | FALSE | 1 | FALSE | 1 | FALSE | 1 | FALSE | 1 |
| 89 | 89 | 6.95238095 | 3.4153183 | 1 | 0.40558521 | 1 | 0.48239024 | 1 | 0.51939841 | 1 | 0.60987146 | 0 | 1 | 16 | 0.80826361 | FALSE | 1 | FALSE | 1 | FALSE | 1 | FALSE | 1 | FALSE | 1 | FALSE | 1 |
| 90 | 90 | 8.53125 | 3.04122072 | 1 | 0.56527493 | 2 | 0.27730767 | 3 | 0.10599423 | 4 | 0.05697506 | 0 | 1 | 22 | 0.99301316 | FALSE | 1 | FALSE | 1 | FALSE | 1 | FALSE | 1 | FALSE | 1 | FALSE | 1 |
| 91 | 91 | 9.07142857 | 5.41775238 | 0 | 1 | 6 | 0.0021153 | 0 | 1 | 2 | 0.5811115 | 2 | 0.15059682 | 32 | 0.94677277 | FALSE | 1 | FALSE | 0.42264619 | FALSE | 1 | FALSE | 1 | FALSE | 1 | FALSE | 1 |
| 92 | 92 | 4.99206349 | 2.58351315 | 3 | 0.62666532 | 5 | 0.3797149 | 7 | 0.16226546 | 7 | 0.35028332 | 1 | 0.87581372 | 102 | 0.83891146 | FALSE | 1 | FALSE | 1 | FALSE | 1 | FALSE | 1 | FALSE | 1 | FALSE | 1 |

| | | | | | | | | | | | | | | | | | | | | | | | | | | | |
|---|---|---|---|---|---|---|---|---|---|---|---|---|---|---|---|---|---|---|---|---|---|---|---|---|---|---|---|
| 93 | 93 | 8 | 2.82842713 | 0 | 1 | 2 | 0.1135776 | 0 | 1 | 1 | 0.57131013 | 0 | 1 | 15 | 0.69372602 | FALSE | 1 | FALSE | 1 | FALSE | 1 | FALSE | 1 | FALSE | 1 | FALSE | 1 |
| 94 | 94 | 6.21235521 | 1.00446967 | 6 | 0.65850749 | 8 | 0.60491261 | 12 | 0.22059163 | 11 | 0.64989339 | 4 | 0.61950435 | 218 | 0.57914159 | FALSE | 1 | FALSE | 1 | FALSE | 1 | FALSE | 1 | FALSE | 1 | FALSE | 1 |
| 95 | 95 | 9.39285714 | 2.36551543 | 4 | 0.00528025 | 1 | 0.60247911 | 1 | 0.64171563 | 2 | 0.3705495 | 0 | 1 | 20 | 0.97757517 | FALSE | 0.2089654 | FALSE | 1 | FALSE | 1 | FALSE | 1 | FALSE | 1 | FALSE | 1 |
| 96 | 96 | 9.2 | 2.63818119 | 0 | 1 | 0 | 1 | 1 | 0.30655256 | 2 | 0.0740981 | 0 | 1 | 7 | 0.94326215 | FALSE | 1 | FALSE | 1 | FALSE | 1 | FALSE | 1 | FALSE | 1 | FALSE | 1 |
| 97 | 97 | 3.192 | 4.08058035 | 7 | 0.03820623 | 0 | 1 | 7 | 0.15312344 | 4 | 0.82348102 | 0 | 1 | 105 | 0.43974013 | FALSE | 0.59844257 | FALSE | 1 | FALSE | 1 | FALSE | 1 | FALSE | 1 | FALSE | 1 |
| 98 | 98 | 9 | 2.1821789 | 0 | 1 | 2 | 0.14655134 | 0 | 1 | 3 | 0.06912908 | 1 | 0.29368245 | 15 | 0.96513691 | FALSE | 1 | FALSE | 1 | FALSE | 1 | FALSE | 1 | FALSE | 1 | FALSE | 1 |
| 99 | 99 | 7.75675676 | 1.61756249 | 1 | 0.59801736 | 0 | 1 | 0 | 1 | 2 | 0.48233989 | 1 | 0.44015537 | 31 | 0.34231756 | FALSE | 1 | FALSE | 1 | FALSE | 1 | FALSE | 1 | FALSE | 1 | FALSE | 1 |
| 100 | 100 | 9.51923077 | 0.77187047 | 3 | 0.14187072 | 4 | 0.08182743 | 1 | 0.84631478 | 2 | 0.68678423 | 1 | 0.57100288 | 40 | 0.90979546 | FALSE | 1 | FALSE | 1 | FALSE | 1 | FALSE | 1 | FALSE | 1 | FALSE | 1 |
| 101 | 101 | 8.45454546 | 1.98617813 | 0 | 1 | 1 | 0.65167362 | 3 | 0.10599423 | 4 | 0.05697506 | 1 | 0.41154418 | 23 | 0.97984382 | FALSE | 1 | FALSE | 1 | FALSE | 1 | FALSE | 1 | FALSE | 1 | FALSE | 1 |
| 102 | 102 | 6.06049822 | 1.63150863 | 10 | 0.17851255 | 11 | 0.2923223 | 10 | 0.54985724 | 15 | 0.29987547 | 3 | 0.84273268 | 230 | 0.84071751 | FALSE | 1 | FALSE | 1 | FALSE | 1 | FALSE | 1 | FALSE | 1 | FALSE | 1 |
| 103 | 103 | 11.5625 | 2.57314861 | 0 | 1 | 0 | 1 | 1 | 0.44344817 | 0 | 1 | 1 | 0.23264612 | 14 | 0.53308261 | FALSE | 1 | FALSE | 1 | FALSE | 1 | FALSE | 1 | FALSE | 1 | FALSE | 1 |
| 104 | 104 | 10.3125 | 1.3564084 | 1 | 0.34033054 | 1 | 0.40942991 | 0 | 1 | 0 | 1 | 0 | 1 | 14 | 0.53308261 | FALSE | 1 | FALSE | 1 | FALSE | 1 | FALSE | 1 | FALSE | 1 | FALSE | 1 |
| 105 | 105 | 11.0217391 | 1.78750653 | 1 | 0.66519151 | 2 | 0.39616059 | 3 | 0.19060367 | 5 | 0.04181362 | 0 | 1 | 31 | 0.97647435 | FALSE | 1 | FALSE | 1 | FALSE | 1 | FALSE | 1 | FALSE | 1 | FALSE | 1 |
| 106 | 106 | 8 | 1.78885438 | 1 | 0.85897707 | 3 | 0.43942904 | 3 | 0.50898113 | 2 | 0.86569532 | 1 | 0.71258573 | 65 | 0.36047181 | FALSE | 1 | FALSE | 1 | FALSE | 1 | FALSE | 1 | FALSE | 1 | FALSE | 1 |
| 107 | 107 | 9.39473684 | 1.8287107 | 2 | 0.25455533 | 0 | 1 | 0 | 1 | 1 | 0.83318955 | 0 | 1 | 35 | 0.13407055 | FALSE | 1 | FALSE | 1 | FALSE | 1 | FALSE | 1 | FALSE | 1 | FALSE | 1 |
| 108 | 108 | 4.90384615 | 1.9538878 | 0 | 1 | 2 | 0.85560113 | 2 | 0.89330417 | 7 | 0.20017129 | 0 | 1 | 93 | 0.09390327 | FALSE | 1 | FALSE | 1 | FALSE | 1 | FALSE | 1 | FALSE | 1 | FALSE | 1 |
| 109 | 109 | 11.2978723 | 1.69999867 | 3 | 0.11868304 | 1 | 0.78790827 | 4 | 0.08716543 | 2 | 0.64277396 | 0 | 1 | 37 | 0.89489966 | FALSE | 0.94393345 | FALSE | 1 | FALSE | 1 | FALSE | 1 | FALSE | 1 | FALSE | 1 |
| 110 | 110 | 9.27272727 | 1.54277843 | 0 | 1 | 0 | 1 | 0 | 1 | 1 | 0.40390032 | 0 | 1 | 10 | 0.46859937 | FALSE | 1 | FALSE | 1 | FALSE | 1 | FALSE | 1 | FALSE | 1 | FALSE | 1 |
| 111 | 111 | 9.65217391 | 3.29173795 | 0 | 1 | 1 | 0.53115654 | 1 | 0.5695037 | 0 | 1 | 0 | 1 | 21 | 0.27964072 | FALSE | 1 | FALSE | 1 | FALSE | 1 | FALSE | 1 | FALSE | 1 | FALSE | 1 |
| 112 | 112 | 9.625 | 2.59506744 | 1 | 0.46441868 | 1 | 0.54637326 | 2 | 0.21260257 | 0 | 1 | 0 | 1 | 20 | 0.68270729 | FALSE | 1 | FALSE | 1 | FALSE | 1 | FALSE | 1 | FALSE | 1 | FALSE | 1 |
| 113 | 113 | 9.36363636 | 1.55345523 | 0 | 1 | 0 | 1 | 1 | 0.33149969 | 0 | 1 | 0 | 1 | 10 | 0.46859937 | FALSE | 1 | FALSE | 1 | FALSE | 1 | FALSE | 1 | FALSE | 1 | FALSE | 1 |
| 114 | 114 | 8.54347826 | 2.74025167 | 1 | 0.69843502 | 3 | 0.18548475 | 3 | 0.2282043 | 3 | 0.35428318 | 3 | 0.03911573 | 33 | 0.99158534 | FALSE | 1 | FALSE | 1 | FALSE | 1 | FALSE | 1 | FALSE | 0.88763447 | FALSE | 1 |
| 115 | 115 | 5.22368421 | 2.78434857 | 1 | 0.80677819 | 3 | 0.33367983 | 3 | 0.39570451 | 4 | 0.3278565 | 0 | 1 | 52 | 0.72716859 | FALSE | 1 | FALSE | 1 | FALSE | 1 | FALSE | 1 | FALSE | 1 | FALSE | 1 |
| 116 | 116 | 8.65384615 | 2.54108256 | 2 | 0.14250978 | 1 | 0.57534752 | 1 | 0.61440466 | 1 | 0.70601446 | 0 | 1 | 21 | 0.78939668 | FALSE | 1 | FALSE | 1 | FALSE | 1 | FALSE | 1 | FALSE | 1 | FALSE | 1 |
| 117 | 117 | 3.34242424 | 1.51187828 | 10 | 0.33559573 | 12 | 0.37606547 | 10 | 0.75254391 | 18 | 0.25180855 | 6 | 0.45507896 | 273 | 0.78853336 | FALSE | 1 | FALSE | 1 | FALSE | 1 | FALSE | 1 | FALSE | 1 | FALSE | 1 |
| 118 | 118 | 8.4 | 3.41174442 | 0 | 1 | 0 | 1 | 0 | 1 | 0 | 1 | 1 | 0.15247062 | 9 | 0.5215547 | FALSE | 1 | FALSE | 1 | FALSE | 1 | FALSE | 1 | FALSE | 1 | FALSE | 1 |
| 119 | 119 | 7.64 | 2.09532814 | 1 | 0.47819878 | 1 | 0.56109832 | 1 | 0.59998149 | 0 | 1 | 1 | 0.33902672 | 21 | 0.64957167 | FALSE | 1 | FALSE | 1 | FALSE | 1 | FALSE | 1 | FALSE | 1 | FALSE | 1 |
| 120 | 120 | 7.94117647 | 2.97974709 | 1 | 0.35728305 | 0 | 1 | 0 | 1 | 1 | 0.55062772 | 1 | 0.24525764 | 14 | 0.7313145 | FALSE | 1 | FALSE | 1 | FALSE | 1 | FALSE | 1 | FALSE | 1 | FALSE | 1 |
| 121 | 121 | 10.92 | 2.36507928 | 1 | 0.47819878 | 1 | 0.56109832 | 0 | 1 | 0 | 1 | 0 | 1 | 23 | 0.22727183 | FALSE | 1 | FALSE | 1 | FALSE | 1 | FALSE | 1 | FALSE | 1 | FALSE | 1 |
| 122 | 122 | 7.02631579 | 3.86949502 | 1 | 0.62830697 | 2 | 0.3494045 | 1 | 0.75193685 | 1 | 0.83318955 | 1 | 0.4673875 | 32 | 0.61800916 | FALSE | 1 | FALSE | 1 | FALSE | 1 | FALSE | 1 | FALSE | 1 | FALSE | 1 |
| 123 | 123 | 9.18181818 | 2.18749262 | 0 | 1 | 1 | 0.5154318 | 0 | 1 | 2 | 0.26791271 | 1 | 0.30529951 | 18 | 0.74744813 | FALSE | 1 | FALSE | 1 | FALSE | 1 | FALSE | 1 | FALSE | 1 | FALSE | 1 |
| 124 | 124 | 9.7 | 2.41039416 | 0 | 1 | 0 | 1 | 1 | 0.30655256 | 2 | 0.0740981 | 0 | 1 | 7 | 0.94326215 | FALSE | 1 | FALSE | 1 | FALSE | 1 | FALSE | 1 | FALSE | 1 | FALSE | 1 |
| 125 | 125 | 10.4545455 | 2.53569558 | 1 | 0.24866461 | 0 | 1 | 2 | 0.05708125 | 2 | 0.08789566 | 0 | 1 | 6 | 0.99674012 | FALSE | 1 | FALSE | 1 | FALSE | 1 | FALSE | 1 | FALSE | 1 | FALSE | 1 |
| 126 | 126 | 9.09090909 | 2.06505758 | 0 | 1 | 2 | 0.04725998 | 0 | 1 | 0 | 1 | 0 | 1 | 9 | 0.75985366 | FALSE | 1 | FALSE | 1 | FALSE | 1 | FALSE | 1 | FALSE | 1 | FALSE | 1 |
| 127 | 127 | 9.23529412 | 2.55594835 | 0 | 1 | 1 | 0.42857733 | 0 | 1 | 3 | 0.040397 | 1 | 0.24525764 | 12 | 0.962093 | FALSE | 1 | FALSE | 1 | FALSE | 1 | FALSE | 1 | FALSE | 1 | FALSE | 1 |
| 128 | 128 | 10.4285714 | 2.32115383 | 0 | 1 | 0 | 1 | 1 | 0.40110418 | 1 | 0.48242099 | 0 | 1 | 12 | 0.62236227 | FALSE | 1 | FALSE | 1 | FALSE | 1 | FALSE | 1 | FALSE | 1 | FALSE | 1 |
| 129 | 129 | 8.5 | 3.09155176 | 0 | 1 | 0 | 1 | 1 | 0.59998149 | 2 | 0.31968852 | 0 | 1 | 22 | 0.43642846 | FALSE | 1 | FALSE | 1 | FALSE | 1 | FALSE | 1 | FALSE | 1 | FALSE | 1 |
| 130 | 130 | 10.0833333 | 2.46503324 | 0 | 1 | 0 | 1 | 0 | 1 | 1 | 0.67692816 | 0 | 1 | 23 | 0.09196389 | FALSE | 1 | FALSE | 1 | FALSE | 1 | FALSE | 1 | FALSE | 1 | FALSE | 1 |
| 131 | 131 | 6.48148148 | 3.41966395 | 0 | 1 | 4 | 0.09626401 | 4 | 0.12796328 | 2 | 0.71679995 | 3 | 0.05824229 | 41 | 0.96572325 | FALSE | 1 | FALSE | 1 | FALSE | 1 | FALSE | 1 | FALSE | 1 | FALSE | 1 |
| 132 | 132 | 9.32432432 | 3.23873561 | 2 | 0.22589135 | 1 | 0.68454804 | 2 | 0.35984046 | 1 | 0.80778317 | 1 | 0.44015537 | 28 | 0.83079716 | FALSE | 1 | FALSE | 1 | FALSE | 1 | FALSE | 1 | FALSE | 1 | FALSE | 1 |
| 133 | 133 | 4.38888889 | 1.35286619 | 1 | 0.75539413 | 5 | 0.02959485 | 4 | 0.12796328 | 0 | 1 | 3 | 0.05824229 | 41 | 0.96572325 | FALSE | 1 | FALSE | 0.89461624 | FALSE | 1 | FALSE | 1 | FALSE | 1 | FALSE | 1 |
| 134 | 134 | 9.57142857 | 2.49897938 | 1 | 0.42086991 | 1 | 0.49918218 | 0 | 1 | 1 | 0.62783522 | 0 | 1 | 18 | 0.57962763 | FALSE | 1 | FALSE | 1 | FALSE | 1 | FALSE | 1 | FALSE | 1 | FALSE | 1 |
| 135 | 135 | 7.19047619 | 3.59389757 | 1 | 0.42086991 | 0 | 1 | 0 | 1 | 1 | 0.62783522 | 1 | 0.29368245 | 18 | 0.57962763 | FALSE | 1 | FALSE | 1 | FALSE | 1 | FALSE | 1 | FALSE | 1 | FALSE | 1 |
| 136 | 136 | 9.25 | 3.17214439 | 0 | 1 | 0 | 1 | 0 | 1 | 1 | 0.52895088 | 1 | 0.23264612 | 14 | 0.53308261 | FALSE | 1 | FALSE | 1 | FALSE | 1 | FALSE | 1 | FALSE | 1 | FALSE | 1 |
| 137 | 137 | 10.7058824 | 2.65163004 | 0 | 1 | 0 | 1 | 1 | 0.46348836 | 1 | 0.55062772 | 0 | 1 | 15 | 0.49060435 | FALSE | 1 | FALSE | 1 | FALSE | 1 | FALSE | 1 | FALSE | 1 | FALSE | 1 |
| 138 | 138 | 11.875 | 4.04467242 | 1 | 0.34033054 | 0 | 1 | 1 | 0.44344817 | 0 | 1 | 0 | 1 | 14 | 0.53308261 | FALSE | 1 | FALSE | 1 | FALSE | 1 | FALSE | 1 | FALSE | 1 | FALSE | 1 |
| 139 | 139 | 9.75 | 3.91843932 | 0 | 1 | 1 | 0.32623933 | 2 | 0.06691409 | 2 | 0.1023796 | 0 | 1 | 7 | 0.99433481 | FALSE | 1 | FALSE | 1 | FALSE | 1 | FALSE | 1 | FALSE | 1 | FALSE | 1 |
| 140 | 140 | 2.25721154 | 1.55776464 | 4 | 0.99494617 | 17 | 0.18376435 | 16 | 0.41792556 | 19 | 0.53610123 | 4 | 0.91518357 | 354 | 0.28319308 | FALSE | 1 | FALSE | 1 | FALSE | 1 | FALSE | 1 | FALSE | 1 | FALSE | 1 |
| 141 | 141 | 10.6666667 | 1.92064804 | 0 | 1 | 1 | 0.38964394 | 1 | 0.42266264 | 1 | 0.50623194 | 1 | 0.2198258 | 11 | 0.92879412 | FALSE | 1 | FALSE | 1 | FALSE | 1 | FALSE | 1 | FALSE | 1 | FALSE | 1 |
| 142 | 142 | 7.0754717 | 2.86717965 | 1 | 0.74890551 | 3 | 0.24492539 | 2 | 0.57303317 | 1 | 0.91797304 | 1 | 0.58506217 | 45 | 0.5504765 | FALSE | 1 | FALSE | 1 | FALSE | 1 | FALSE | 1 | FALSE | 1 | FALSE | 1 |
| 143 | 143 | 9.46153846 | 3.00295712 | 0 | 1 | 1 | 0.3480709 | 0 | 1 | 0 | 1 | 0 | 1 | 12 | 0.374198 | FALSE | 1 | FALSE | 1 | FALSE | 1 | FALSE | 1 | FALSE | 1 | FALSE | 1 |
| 144 | 144 | 10 | 1.37840488 | 5 | 0.00332431 | 1 | 0.7326167 | 3 | 0.17245215 | 1 | 0.84823834 | 1 | 0.48480878 | 29 | 0.98400908 | FALSE | 0.15832977 | FALSE | 1 | FALSE | 1 | FALSE | 1 | FALSE | 1 | FALSE | 1 |
| 145 | 145 | 9.125 | 2.47171499 | 1 | 0.34033054 | 1 | 0.40942991 | 2 | 0.11087977 | 0 | 1 | 0 | 1 | 12 | 0.90922186 | FALSE | 1 | FALSE | 1 | FALSE | 1 | FALSE | 1 | FALSE | 1 | FALSE | 1 |
| 146 | 146 | 4.77083333 | 2.34733522 | 0 | 1 | 3 | 0.20208567 | 1 | 0.8283465 | 5 | 0.06758937 | 0 | 1 | 39 | 0.79274273 | FALSE | 1 | FALSE | 1 | FALSE | 1 | FALSE | 1 | FALSE | 1 | FALSE | 1 |
| 147 | 147 | 10.1489362 | 2.47522811 | 1 | 0.69843502 | 1 | 0.78076835 | 1 | 0.81522035 | 4 | 0.1589264 | 0 | 1 | 39 | 0.56897353 | FALSE | 1 | FALSE | 1 | FALSE | 1 | FALSE | 1 | FALSE | 1 | FALSE | 1 |
| 148 | 148 | 9.54166667 | 1.82526634 | 1 | 0.46441868 | 1 | 0.54637326 | 3 | 0.05314317 | 0 | 1 | 1 | 0.32796862 | 18 | 0.93166998 | FALSE | 1 | FALSE | 1 | FALSE | 1 | FALSE | 1 | FALSE | 1 | FALSE | 1 |
| 149 | 149 | 10.7647059 | 2.57886003 | 0 | 1 | 0 | 1 | 0 | 1 | 1 | 0.55062772 | 0 | 1 | 16 | 0.23070182 | FALSE | 1 | FALSE | 1 | FALSE | 1 | FALSE | 1 | FALSE | 1 | FALSE | 1 |
| 150 | 150 | 8.61538462 | 2.45069728 | 1 | 0.74224579 | 1 | 0.82028131 | 1 | 0.85187773 | 4 | 0.21499032 | 0 | 1 | 45 | 0.4221244 | FALSE | 1 | FALSE | 1 | FALSE | 1 | FALSE | 1 | FALSE | 1 | FALSE | 1 |
| 151 | 151 | 5.94736842 | 3.27361088 | 3 | 0.17914398 | 3 | 0.28014665 | 2 | 0.61293419 | 2 | 0.74436726 | 0 | 1 | 47 | 0.72965359 | FALSE | 1 | FALSE | 1 | FALSE | 1 | FALSE | 1 | FALSE | 1 | FALSE | 1 |
| 152 | 152 | 8.85714286 | 3.44045559 | 0 | 1 | 0 | 1 | 1 | 0.37874417 | 0 | 1 | 2 | 0.01846832 | 10 | 0.86791642 | FALSE | 1 | FALSE | 1 | FALSE | 1 | FALSE | 1 | FALSE | 0.69720022 | FALSE | 1 |
| 153 | 153 | 8.4 | 0.8 | 0 | 1 | 0 | 1 | 0 | 1 | 0 | 1 | 0 | 1 | 5 | 0.42761893 | FALSE | 1 | FALSE | 1 | FALSE | 1 | FALSE | 1 | FALSE | 1 | FALSE | 1 |
| 154 | 154 | 8.27272727 | 1.28564869 | 0 | 1 | 1 | 0.30368014 | 0 | 1 | 1 | 0.40390032 | 1 | 0.16638736 | 8 | 0.92152993 | FALSE | 1 | FALSE | 1 | FALSE | 1 | FALSE | 1 | FALSE | 1 | FALSE | 1 |
| 155 | 155 | 8.58333333 | 1.381927 | 0 | 1 | 2 | 0.32556427 | 4 | 0.03888854 | 0 | 1 | 1 | 0.44938266 | 29 | 0.81028289 | FALSE | 1 | FALSE | 1 | FALSE | 1 | FALSE | 1 | FALSE | 1 | FALSE | 1 |
| 156 | 156 | 11.6470588 | 1.02562328 | 1 | 0.32293355 | 0 | 1 | 1 | 0.42266264 | 2 | 0.14906763 | 0 | 1 | 11 | 0.92879412 | FALSE | 1 | FALSE | 1 | FALSE | 1 | FALSE | 1 | FALSE | 1 | FALSE | 1 |
| 157 | 157 | 5.59649123 | 3.4328631 | 3 | 0.16642539 | 2 | 0.5356583 | 2 | 0.59334725 | 5 | 0.10658542 | 0 | 1 | 43 | 0.92240996 | FALSE | 1 | FALSE | 1 | FALSE | 1 | FALSE | 1 | FALSE | 1 | FALSE | 1 |
| 158 | 158 | 5.95789474 | 2.58707634 | 2 | 0.70534724 | 1 | 0.95697569 | 4 | 0.44662765 | 4 | 0.64110971 | 1 | 0.79437879 | 83 | 0.25859323 | FALSE | 1 | FALSE | 1 | FALSE | 1 | FALSE | 1 | FALSE | 1 | FALSE | 1 |
| 159 | 159 | 9.62962963 | 3.10493214 | 2 | 0.15144274 | 2 | 0.21684246 | 1 | 0.62830986 | 1 | 0.71956339 | 0 | 1 | 21 | 0.88395936 | FALSE | 1 | FALSE | 1 | FALSE | 1 | FALSE | 1 | FALSE | 1 | FALSE | 1 |
| 160 | 160 | 7.90909091 | 2.10862064 | 1 | 0.4357638 | 1 | 0.5154318 | 0 | 1 | 1 | 0.64497439 | 0 | 1 | 19 | 0.54233419 | FALSE | 1 | FALSE | 1 | FALSE | 1 | FALSE | 1 | FALSE | 1 | FALSE | 1 |
| 161 | 161 | 6.80645161 | 1.5947364 | 3 | 0.04402911 | 1 | 0.63997468 | 3 | 0.09854319 | 2 | 0.41984187 | 2 | 0.09110253 | 20 | 0.9984454 | FALSE | 0.64293489 | FALSE | 1 | FALSE | 1 | FALSE | 1 | FALSE | 1 | FALSE | 1 |
| 162 | 162 | 3.69506727 | 2.14210886 | 6 | 0.51142422 | 5 | 0.85436185 | 9 | 0.40886765 | 9 | 0.70297841 | 7 | 0.07296038 | 187 | 0.6294654 | FALSE | 1 | FALSE | 1 | FALSE | 1 | FALSE | 1 | FALSE | 1 | FALSE | 1 |
| 163 | 163 | 8.5 | 2.20227156 | 0 | 1 | 1 | 0.48239024 | 3 | 0.03322143 | 1 | 0.60987146 | 1 | 0.28187291 | 14 | 0.97325593 | FALSE | 1 | FALSE | 1 | FALSE | 1 | FALSE | 1 | FALSE | 1 | FALSE | 1 |
| 164 | 164 | 9.23529412 | 1.86387994 | 3 | 0.00725145 | 1 | 0.40942991 | 0 | 1 | 2 | 0.16544789 | 0 | 1 | 10 | 0.99304605 | FALSE | 0.25171133 | FALSE | 1 | FALSE | 1 | FALSE | 1 | FALSE | 1 | FALSE | 1 |
| 165 | 165 | 6.125 | 1.1110243 | 0 | 1 | 0 | 1 | 0 | 1 | 2 | 0.16544789 | 0 | 1 | 14 | 0.53308261 | FALSE | 1 | FALSE | 1 | FALSE | 1 | FALSE | 1 | FALSE | 1 | FALSE | 1 |
| 166 | 166 | 6.80769231 | 2.30416397 | 1 | 0.49162634 | 0 | 1 | 2 | 0.23940226 | 1 | 0.70601446 | 0 | 1 | 22 | 0.61628 | FALSE | 1 | FALSE | 1 | FALSE | 1 | FALSE | 1 | FALSE | 1 | FALSE | 1 |
| 167 | 167 | 7.21428571 | 2.17710724 | 1 | 0.3050805 | 0 | 1 | 0 | 1 | 0 | 1 | 0 | 1 | 13 | 0.33283729 | FALSE | 1 | FALSE | 1 | FALSE | 1 | FALSE | 1 | FALSE | 1 | FALSE | 1 |
| 168 | 168 | 10.4285714 | 2.49897938 | 2 | 0.04858661 | 1 | 0.36919824 | 1 | 0.40110418 | 2 | 0.13304705 | 0 | 1 | 8 | 0.99722299 | FALSE | 0.649301 | FALSE | 1 | FALSE | 1 | FALSE | 1 | FALSE | 1 | FALSE | 1 |
| 169 | 169 | 8.28125 | 2.60089378 | 1 | 0.56527493 | 0 | 1 | 1 | 0.69069222 | 1 | 0.77852168 | 0 | 1 | 29 | 0.24088541 | FALSE | 1 | FALSE | 1 | FALSE | 1 | FALSE | 1 | FALSE | 1 | FALSE | 1 |
| 170 | 170 | 10.4117647 | 2.42678251 | 0 | 1 | 0 | 1 | 0 | 1 | 0 | 1 | 1 | 0.24525764 | 16 | 0.23070182 | FALSE | 1 | FALSE | 1 | FALSE | 1 | FALSE | 1 | FALSE | 1 | FALSE | 1 |
| 171 | 171 | 9.92857143 | 1.66751679 | 2 | 0.16048326 | 0 | 1 | 0 | 1 | 0 | 1 | 1 | 0.37113067 | 25 | 0.34293 | FALSE | 1 | FALSE | 1 | FALSE | 1 | FALSE | 1 | FALSE | 1 | FALSE | 1 |
| 172 | 172 | 7.77777778 | 2.92287699 | 1 | 0.50471032 | 0 | 1 | 1 | 0.62830986 | 3 | 0.12475952 | 1 | 0.36060486 | 21 | 0.88395936 | FALSE | 1 | FALSE | 1 | FALSE | 1 | FALSE | 1 | FALSE | 1 | FALSE | 1 |
| 173 | 173 | 7.92307692 | 2.7021797 | 1 | 0.63789042 | 0 | 1 | 2 | 0.41139386 | 2 | 0.54052736 | 0 | 1 | 34 | 0.41570781 | FALSE | 1 | FALSE | 1 | FALSE | 1 | FALSE | 1 | FALSE | 1 | FALSE | 1 |
| 174 | 174 | 10.0571429 | 1.94117574 | 2 | 0.22589135 | 0 | 1 | 1 | 0.72299506 | 5 | 0.02066043 | 1 | 0.44015537 | 26 | 0.96294373 | FALSE | 1 | FALSE | 1 | FALSE | 1 | FALSE | 1 | FALSE | 1 | FALSE | 1 |
| 175 | 175 | 8.06666667 | 2.5157283 | 1 | 0.32293355 | 0 | 1 | 1 | 0.42266264 | 2 | 0.14906763 | 2 | 0.02433796 | 9 | 0.995488 | FALSE | 1 | FALSE | 1 | FALSE | 1 | FALSE | 1 | FALSE | 0.77578444 | FALSE | 1 |
| 176 | 176 | 9.42857143 | 1.80136003 | 0 | 1 | 0 | 1 | 0 | 1 | 4 | 0.03737468 | 0 | 1 | 24 | 0.55018107 | FALSE | 1 | FALSE | 1 | FALSE | 1 | FALSE | 1 | FALSE | 1 | FALSE | 1 |
| 177 | 177 | 9.92857143 | 2.46299138 | 0 | 1 | 0 | 1 | 0 | 1 | 0 | 1 | 0 | 1 | 14 | 0.09251304 | FALSE | 1 | FALSE | 1 | FALSE | 1 | FALSE | 1 | FALSE | 1 | FALSE | 1 |
| 178 | 178 | 9.7 | 2.49198716 | 1 | 0.22886536 | 1 | 0.28036922 | 0 | 1 | 3 | 0.00903718 | 0 | 1 | 5 | 0.99829382 | FALSE | 1 | FALSE | 1 | FALSE | 1 | FALSE | 1 | FALSE | 1 | FALSE | 1 |
| 179 | 179 | 7.32352941 | 3.3669727 | 1 | 0.57647348 | 0 | 1 | 1 | 0.70185639 | 3 | 0.19161876 | 1 | 0.42123867 | 27 | 0.75124583 | FALSE | 1 | FALSE | 1 | FALSE | 1 | FALSE | 1 | FALSE | 1 | FALSE | 1 |
| 180 | 180 | 7.21428571 | 1.52027119 | 1 | 0.3050805 | 0 | 1 | 0 | 1 | 1 | 0.48242099 | 0 | 1 | 12 | 0.62236227 | FALSE | 1 | FALSE | 1 | FALSE | 1 | FALSE | 1 | FALSE | 1 | FALSE | 1 |
| 181 | 181 | 9.92307692 | 2.89459811 | 0 | 1 | 0 | 1 | 1 | 0.37874417 | 0 | 1 | 0 | 1 | 12 | 0.374198 | FALSE | 1 | FALSE | 1 | FALSE | 1 | FALSE | 1 | FALSE | 1 | FALSE | 1 |
| 182 | 182 | 10.7272727 | 1.76279268 | 0 | 1 | 0 | 1 | 1 | 0.33149969 | 0 | 1 | 1 | 0.16638736 | 9 | 0.75985366 | FALSE | 1 | FALSE | 1 | FALSE | 1 | FALSE | 1 | FALSE | 1 | FALSE | 1 |
| 183 | 183 | 11 | 2.94392029 | 0 | 1 | 0 | 1 | 0 | 1 | 0 | 1 | 0 | 1 | 6 | 0.36077158 | FALSE | 1 | FALSE | 1 | FALSE | 1 | FALSE | 1 | FALSE | 1 | FALSE | 1 |
| 184 | 184 | 8.91176471 | 1.52175686 | 1 | 0.58738519 | 0 | 1 | 1 | 0.71261921 | 4 | 0.06848449 | 1 | 0.43077488 | 27 | 0.85023803 | FALSE | 1 | FALSE | 1 | FALSE | 1 | FALSE | 1 | FALSE | 1 | FALSE | 1 |
| 185 | 185 | 8.91111111 | 2.24916994 | 5 | 0.00556946 | 4 | 0.05650644 | 4 | 0.07687709 | 4 | 0.15016749 | 2 | 0.16793736 | 26 | 0.99999605 | FALSE | 0.21726186 | FALSE | 1 | FALSE | 1 | FALSE | 1 | FALSE | 1 | FALSE | 1 |
| 186 | 186 | 7.78947368 | 2.54597613 | 0 | 1 | 0 | 1 | 1 | 0.50143826 | 0 | 1 | 1 | 0.26986775 | 17 | 0.41141543 | FALSE | 1 | FALSE | 1 | FALSE | 1 | FALSE | 1 | FALSE | 1 | FALSE | 1 |

| | | | | | | | | | | | | | | | | | | | | | | | | | | | |
|---|---|---|---|---|---|---|---|---|---|---|---|---|---|---|---|---|---|---|---|---|---|---|---|---|---|---|---|
| 187 | 187 | 9.03846154 | 1.95095182 | 0 | 1 | 0 | 1 | 2 | 0.23940226 | 3 | 0.11458571 | 1 | 0.3499045 | 20 | 0.90142689 | FALSE | 1 | FALSE | 1 | FALSE | 1 | FALSE | 1 | FALSE | 1 | FALSE | 1 |
| 188 | 188 | 10.1052632 | 1.58594939 | 0 | 1 | 1 | 0.71433277 | 0 | 1 | 3 | 0.25266879 | 1 | 0.4673875 | 33 | 0.44261631 | FALSE | 1 | FALSE | 1 | FALSE | 1 | FALSE | 1 | FALSE | 1 | FALSE | 1 |
| 189 | 189 | 9.32 | 1.40627167 | 0 | 1 | 0 | 1 | 2 | 0.22597287 | 0 | 1 | 0 | 1 | 23 | 0.22727183 | FALSE | 1 | FALSE | 1 | FALSE | 1 | FALSE | 1 | FALSE | 1 | FALSE | 1 |
| 190 | 190 | 9.63888889 | 2.07032979 | 0 | 1 | 1 | 0.69480472 | 4 | 0.03888854 | 1 | 0.81665383 | 0 | 1 | 30 | 0.67248271 | FALSE | 1 | FALSE | 1 | FALSE | 1 | FALSE | 1 | FALSE | 1 | FALSE | 1 |
| 191 | 191 | 8.23076923 | 2.93922664 | 0 | 1 | 1 | 0.57534752 | 1 | 0.61440466 | 0 | 1 | 0 | 1 | 24 | 0.20425579 | FALSE | 1 | FALSE | 1 | FALSE | 1 | FALSE | 1 | FALSE | 1 | FALSE | 1 |
| 192 | 192 | 9.04 | 2.23571018 | 0 | 1 | 4 | 0.00789279 | 1 | 0.59998149 | 0 | 1 | 0 | 1 | 20 | 0.8152498 | FALSE | 1 | FALSE | 0.6167796 | FALSE | 1 | FALSE | 1 | FALSE | 1 | FALSE | 1 |
| 193 | 193 | 8 | 1.41421356 | 0 | 1 | 0 | 1 | 1 | 0.33149969 | 0 | 1 | 0 | 1 | 10 | 0.46859937 | FALSE | 1 | FALSE | 1 | FALSE | 1 | FALSE | 1 | FALSE | 1 | FALSE | 1 |
| 194 | 194 | 8.04444444 | 2.47615792 | 0 | 1 | 4 | 0.05276567 | 2 | 0.47281647 | 2 | 0.60667406 | 1 | 0.51797501 | 35 | 0.8620983 | FALSE | 1 | FALSE | 1 | FALSE | 1 | FALSE | 1 | FALSE | 1 | FALSE | 1 |
| 195 | 195 | 7.81481482 | 2.84197424 | 4 | 0.00461682 | 2 | 0.21684246 | 0 | 1 | 0 | 1 | 1 | 0.36060486 | 20 | 0.95133411 | FALSE | 0.19698443 | FALSE | 1 | FALSE | 1 | FALSE | 1 | FALSE | 1 | FALSE | 1 |
| 196 | 196 | 9.72727273 | 2.37758124 | 0 | 1 | 0 | 1 | 0 | 1 | 0 | 1 | 0 | 1 | 11 | 0.15414489 | FALSE | 1 | FALSE | 1 | FALSE | 1 | FALSE | 1 | FALSE | 1 | FALSE | 1 |
| 197 | 197 | 3.22929936 | 1.25449144 | 7 | 0.96186579 | 11 | 0.90175005 | 19 | 0.32525552 | 21 | 0.58210932 | 7 | 0.65717054 | 404 | 0.15279768 | FALSE | 1 | FALSE | 1 | FALSE | 1 | FALSE | 1 | FALSE | 1 | FALSE | 1 |
| 198 | 198 | 9.15 | 1.60545944 | 0 | 1 | 1 | 0.7326167 | 1 | 0.76953771 | 1 | 0.84823834 | 3 | 0.02730775 | 34 | 0.56328904 | FALSE | 1 | FALSE | 1 | FALSE | 1 | FALSE | 1 | FALSE | 0.79106376 | FALSE | 1 |
| 199 | 199 | 9.25 | 3.25384388 | 0 | 1 | 1 | 0.48239024 | 1 | 0.51939841 | 2 | 0.23331368 | 0 | 1 | 16 | 0.80826361 | FALSE | 1 | FALSE | 1 | FALSE | 1 | FALSE | 1 | FALSE | 1 | FALSE | 1 |
| 200 | 200 | 8.875 | 3.03881145 | 0 | 1 | 1 | 0.40942991 | 2 | 0.11087977 | 0 | 1 | 0 | 1 | 13 | 0.76799161 | FALSE | 1 | FALSE | 1 | FALSE | 1 | FALSE | 1 | FALSE | 1 | FALSE | 1 |
| 201 | 201 | 6.52777778 | 2.56565035 | 4 | 0.01296492 | 3 | 0.10944025 | 1 | 0.73299778 | 2 | 0.49731009 | 0 | 1 | 26 | 0.98200522 | FALSE | 0.35760478 | FALSE | 1 | FALSE | 1 | FALSE | 1 | FALSE | 1 | FALSE | 1 |
| 202 | 202 | 10.525 | 2.40818915 | 0 | 1 | 2 | 0.37295654 | 0 | 1 | 0 | 1 | 0 | 1 | 38 | 0.03896014 | FALSE | 1 | FALSE | 1 | FALSE | 1 | FALSE | 1 | FALSE | 1 | FALSE | 1 |
| 203 | 203 | 8.95454546 | 2.77123656 | 0 | 1 | 0 | 1 | 0 | 1 | 1 | 0.62783522 | 0 | 1 | 20 | 0.13763971 | FALSE | 1 | FALSE | 1 | FALSE | 1 | FALSE | 1 | FALSE | 1 | FALSE | 1 |
| 204 | 204 | 8.69387755 | 2.27859679 | 3 | 0.12433927 | 2 | 0.46321244 | 6 | 0.00699668 | 3 | 0.37966044 | 2 | 0.18560062 | 32 | 0.99940757 | FALSE | 0.95622564 | FALSE | 1 | FALSE | 1 | FALSE | 1 | FALSE | 1 | FALSE | 1 |
| 205 | 205 | 8 | 2.57460122 | 2 | 0.22589135 | 2 | 0.31355738 | 1 | 0.72299506 | 2 | 0.48233989 | 0 | 1 | 28 | 0.83079716 | FALSE | 1 | FALSE | 1 | FALSE | 1 | FALSE | 1 | FALSE | 1 | FALSE | 1 |
| 206 | 206 | 10.6470588 | 3.28886135 | 0 | 1 | 1 | 0.40942991 | 1 | 0.44344817 | 3 | 0.03440583 | 2 | 0.02752114 | 9 | 0.99862766 | FALSE | 1 | FALSE | 1 | FALSE | 1 | FALSE | 1 | FALSE | 0.79106376 | FALSE | 1 |
| 207 | 207 | 8.75 | 2.1318168 | 0 | 1 | 0 | 1 | 2 | 0.23940226 | 2 | 0.33678042 | 0 | 1 | 22 | 0.61628 | FALSE | 1 | FALSE | 1 | FALSE | 1 | FALSE | 1 | FALSE | 1 | FALSE | 1 |
| 208 | 208 | 9.53424658 | 2.9751583 | 3 | 0.28086759 | 5 | 0.08277437 | 5 | 0.11589609 | 2 | 0.85003314 | 3 | 0.11388306 | 54 | 0.98741896 | FALSE | 1 | FALSE | 1 | FALSE | 1 | FALSE | 1 | FALSE | 1 | FALSE | 1 |
| 209 | 209 | 9.95238095 | 2.27776395 | 0 | 1 | 1 | 0.48239024 | 1 | 0.51939841 | 1 | 0.60987146 | 0 | 1 | 17 | 0.61750632 | FALSE | 1 | FALSE | 1 | FALSE | 1 | FALSE | 1 | FALSE | 1 | FALSE | 1 |
| 210 | 210 | 7.7 | 3.01827766 | 0 | 1 | 0 | 1 | 0 | 1 | 2 | 0.23331368 | 0 | 1 | 18 | 0.37504645 | FALSE | 1 | FALSE | 1 | FALSE | 1 | FALSE | 1 | FALSE | 1 | FALSE | 1 |
| 211 | 211 | 8.57894737 | 2.2317279 | 1 | 0.38989951 | 0 | 1 | 1 | 0.50143826 | 1 | 0.59104357 | 0 | 1 | 16 | 0.65565778 | FALSE | 1 | FALSE | 1 | FALSE | 1 | FALSE | 1 | FALSE | 1 | FALSE | 1 |
| 212 | 212 | 8.1 | 1.5132746 | 0 | 1 | 0 | 1 | 1 | 0.51939841 | 2 | 0.23331368 | 0 | 1 | 17 | 0.61750632 | FALSE | 1 | FALSE | 1 | FALSE | 1 | FALSE | 1 | FALSE | 1 | FALSE | 1 |
| 213 | 213 | 9.88235294 | 3.73980737 | 1 | 0.35728305 | 1 | 0.42857733 | 0 | 1 | 0 | 1 | 0 | 1 | 15 | 0.49060435 | FALSE | 1 | FALSE | 1 | FALSE | 1 | FALSE | 1 | FALSE | 1 | FALSE | 1 |
| 214 | 214 | 9 | 1.63299316 | 0 | 1 | 3 | 0.02888061 | 1 | 0.53671415 | 1 | 0.62783522 | 1 | 0.29368245 | 15 | 0.96513691 | FALSE | 1 | FALSE | 0.88012077 | FALSE | 1 | FALSE | 1 | FALSE | 1 | FALSE | 1 |
| 215 | 215 | 8.625 | 2.49687304 | 0 | 1 | 0 | 1 | 1 | 0.58502114 | 1 | 0.67692816 | 2 | 0.058175 | 20 | 0.68270729 | FALSE | 1 | FALSE | 1 | FALSE | 1 | FALSE | 1 | FALSE | 1 | FALSE | 1 |
| 216 | 216 | 8 | 2.61861468 | 2 | 0.09187136 | 1 | 0.48239024 | 0 | 1 | 1 | 0.60987146 | 0 | 1 | 16 | 0.80826361 | FALSE | 0.83248965 | FALSE | 1 | FALSE | 1 | FALSE | 1 | FALSE | 1 | FALSE | 1 |
| 217 | 217 | 8.57894737 | 2.15981942 | 0 | 1 | 0 | 1 | 1 | 0.50143826 | 0 | 1 | 0 | 1 | 18 | 0.17880694 | FALSE | 1 | FALSE | 1 | FALSE | 1 | FALSE | 1 | FALSE | 1 | FALSE | 1 |
| 218 | 218 | 5.79166667 | 2.13656981 | 1 | 0.95577586 | 2 | 0.90254753 | 2 | 0.93178691 | 5 | 0.64416638 | 1 | 0.86261071 | 108 | 0.02962185 | FALSE | 1 | FALSE | 1 | FALSE | 1 | FALSE | 1 | FALSE | 1 | FALSE | 1 |
| 219 | 219 | 10.05 | 2.15580611 | 3 | 0.01186104 | 2 | 0.12434464 | 0 | 1 | 1 | 0.59104357 | 0 | 1 | 13 | 0.97998676 | FALSE | 0.3373808 | FALSE | 1 | FALSE | 1 | FALSE | 1 | FALSE | 1 | FALSE | 1 |
| 220 | 220 | 9.11111111 | 1.96889391 | 0 | 1 | 1 | 0.44710673 | 1 | 0.48280983 | 1 | 0.57131013 | 0 | 1 | 15 | 0.69372602 | FALSE | 1 | FALSE | 1 | FALSE | 1 | FALSE | 1 | FALSE | 1 | FALSE | 1 |
| 221 | 221 | 5.28395062 | 2.42535255 | 2 | 0.539854 | 2 | 0.66721401 | 1 | 0.92379401 | 2 | 0.83868849 | 1 | 0.68753645 | 62 | 0.21402731 | FALSE | 1 | FALSE | 1 | FALSE | 1 | FALSE | 1 | FALSE | 1 | FALSE | 1 |
| 222 | 222 | 8.21875 | 2.0575224 | 0 | 1 | 1 | 0.63997468 | 1 | 0.67911179 | 1 | 0.76781284 | 0 | 1 | 28 | 0.26393996 | FALSE | 1 | FALSE | 1 | FALSE | 1 | FALSE | 1 | FALSE | 1 | FALSE | 1 |
| 223 | 223 | 8.95454546 | 2.28587108 | 1 | 0.68224519 | 2 | 0.41896557 | 2 | 0.47281647 | 1 | 0.87439647 | 0 | 1 | 38 | 0.45705039 | FALSE | 1 | FALSE | 1 | FALSE | 1 | FALSE | 1 | FALSE | 1 | FALSE | 1 |
| 224 | 224 | 7.93548387 | 2.15429779 | 0 | 1 | 0 | 1 | 3 | 0.09854319 | 0 | 1 | 0 | 1 | 28 | 0.26393996 | FALSE | 1 | FALSE | 1 | FALSE | 1 | FALSE | 1 | FALSE | 1 | FALSE | 1 |
| 225 | 225 | 7.55555556 | 1.58733766 | 6 | 0.39768943 | 6 | 0.62342573 | 9 | 0.28163146 | 18 | 0.00416558 | 1 | 0.96394579 | 158 | 0.96810024 | FALSE | 1 | FALSE | 1 | FALSE | 1 | FALSE | 1 | FALSE | 1 | FALSE | 1 |
| 226 | 226 | 5.90196078 | 3.11388353 | 2 | 0.37762121 | 4 | 0.08182743 | 0 | 1 | 3 | 0.41729179 | 1 | 0.57100288 | 41 | 0.8377026 | FALSE | 1 | FALSE | 1 | FALSE | 1 | FALSE | 1 | FALSE | 1 | FALSE | 1 |
| 227 | 227 | 9 | 2.3741027 | 0 | 1 | 0 | 1 | 1 | 0.55340851 | 1 | 0.64497439 | 0 | 1 | 20 | 0.30915202 | FALSE | 1 | FALSE | 1 | FALSE | 1 | FALSE | 1 | FALSE | 1 | FALSE | 1 |
| 228 | 228 | 9.88235294 | 1.74498788 | 0 | 1 | 0 | 1 | 0 | 1 | 1 | 0.55062772 | 1 | 0.24525764 | 15 | 0.49060435 | FALSE | 1 | FALSE | 1 | FALSE | 1 | FALSE | 1 | FALSE | 1 | FALSE | 1 |
| 229 | 229 | 2.73684211 | 3.74646648 | 0 | 1 | 2 | 0.3494045 | 0 | 1 | 1 | 0.83318955 | 0 | 1 | 35 | 0.13407055 | FALSE | 1 | FALSE | 1 | FALSE | 1 | FALSE | 1 | FALSE | 1 | FALSE | 1 |
| 230 | 230 | 11 | 2.07549809 | 0 | 1 | 2 | 0.20486827 | 0 | 1 | 1 | 0.70601446 | 0 | 1 | 23 | 0.40372818 | FALSE | 1 | FALSE | 1 | FALSE | 1 | FALSE | 1 | FALSE | 1 | FALSE | 1 |
| 231 | 231 | 4.51115619 | 2.23433862 | 14 | 0.35873469 | 18 | 0.30189042 | 24 | 0.06533774 | 17 | 0.90545551 | 7 | 0.68968989 | 404 | 0.74015148 | FALSE | 1 | FALSE | 1 | FALSE | 1 | FALSE | 1 | FALSE | 1 | FALSE | 1 |
| 232 | 232 | 9.88888889 | 1.79161283 | 1 | 0.69044531 | 0 | 1 | 2 | 0.48463745 | 3 | 0.34154252 | 0 | 1 | 39 | 0.4317533 | FALSE | 1 | FALSE | 1 | FALSE | 1 | FALSE | 1 | FALSE | 1 | FALSE | 1 |
| 233 | 233 | 9.39285714 | 2.17681428 | 0 | 1 | 2 | 0.2288771 | 1 | 0.64171563 | 0 | 1 | 2 | 0.07640798 | 23 | 0.73439118 | FALSE | 1 | FALSE | 1 | FALSE | 1 | FALSE | 1 | FALSE | 1 | FALSE | 1 |
| 234 | 234 | 10.8333333 | 3.07769755 | 0 | 1 | 0 | 1 | 0 | 1 | 0 | 1 | 0 | 1 | 18 | 0.04681656 | FALSE | 1 | FALSE | 1 | FALSE | 1 | FALSE | 1 | FALSE | 1 | FALSE | 1 |
| 235 | 235 | 6.53719008 | 1.61624666 | 3 | 0.59315467 | 2 | 0.90254753 | 0 | 1 | 6 | 0.46726758 | 1 | 0.86261071 | 107 | 0.05478417 | FALSE | 1 | FALSE | 1 | FALSE | 1 | FALSE | 1 | FALSE | 1 | FALSE | 1 |
| 236 | 236 | 9.57575758 | 1.34908056 | 0 | 1 | 0 | 1 | 2 | 0.33343871 | 4 | 0.06258966 | 0 | 1 | 27 | 0.75124583 | FALSE | 1 | FALSE | 1 | FALSE | 1 | FALSE | 1 | FALSE | 1 | FALSE | 1 |
| 237 | 237 | 5.34482759 | 2.45805075 | 3 | 0.38658756 | 3 | 0.53828446 | 4 | 0.38141809 | 7 | 0.10374527 | 0 | 1 | 70 | 0.87568001 | FALSE | 1 | FALSE | 1 | FALSE | 1 | FALSE | 1 | FALSE | 1 | FALSE | 1 |
| 238 | 238 | 8.66666667 | 2.47206616 | 4 | 0.00096345 | 0 | 1 | 0 | 1 | 1 | 0.57131013 | 0 | 1 | 13 | 0.95041913 | FALSE | 0.07376244 | FALSE | 1 | FALSE | 1 | FALSE | 1 | FALSE | 1 | FALSE | 1 |
| 239 | 239 | 8.05 | 2.2130296 | 2 | 0.27371785 | 1 | 0.7326167 | 1 | 0.76953771 | 1 | 0.84823834 | 2 | 0.13925369 | 33 | 0.71942635 | FALSE | 1 | FALSE | 1 | FALSE | 1 | FALSE | 1 | FALSE | 1 | FALSE | 1 |
| 240 | 240 | 6.94366197 | 2.26353409 | 1 | 0.84335867 | 1 | 0.90433446 | 1 | 0.92656034 | 3 | 0.63976353 | 0 | 1 | 65 | 0.05811248 | FALSE | 1 | FALSE | 1 | FALSE | 1 | FALSE | 1 | FALSE | 1 | FALSE | 1 |
| 241 | 241 | 9.64285714 | 3.22062866 | 0 | 1 | 0 | 1 | 1 | 0.40110418 | 0 | 1 | 0 | 1 | 13 | 0.33283729 | FALSE | 1 | FALSE | 1 | FALSE | 1 | FALSE | 1 | FALSE | 1 | FALSE | 1 |
| 242 | 242 | 5.24666667 | 2.51379306 | 2 | 0.90202666 | 3 | 0.86947639 | 5 | 0.63139269 | 8 | 0.38363278 | 1 | 0.91856551 | 131 | 0.18686143 | FALSE | 1 | FALSE | 1 | FALSE | 1 | FALSE | 1 | FALSE | 1 | FALSE | 1 |
| 243 | 243 | 9.08333333 | 2.53174292 | 1 | 0.26795838 | 0 | 1 | 0 | 1 | 1 | 0.43131129 | 0 | 1 | 10 | 0.71436964 | FALSE | 1 | FALSE | 1 | FALSE | 1 | FALSE | 1 | FALSE | 1 | FALSE | 1 |
| 244 | 244 | 10.0555556 | 1.68233377 | 0 | 1 | 1 | 0.44710673 | 0 | 1 | 0 | 1 | 2 | 0.03435384 | 15 | 0.69372602 | FALSE | 1 | FALSE | 1 | FALSE | 1 | FALSE | 1 | FALSE | 0.86063198 | FALSE | 1 |
| 245 | 245 | 4.58730159 | 2.20486418 | 0 | 1 | 1 | 0.98474009 | 5 | 0.47556774 | 10 | 0.06342794 | 1 | 0.87788852 | 109 | 0.30130908 | FALSE | 1 | FALSE | 1 | FALSE | 1 | FALSE | 1 | FALSE | 1 | FALSE | 1 |
| 246 | 246 | 8.73684211 | 2.22052875 | 0 | 1 | 2 | 0.12434464 | 1 | 0.50143826 | 2 | 0.21610812 | 0 | 1 | 14 | 0.93677253 | FALSE | 1 | FALSE | 1 | FALSE | 1 | FALSE | 1 | FALSE | 1 | FALSE | 1 |
| 247 | 247 | 8.14285714 | 2.97266458 | 0 | 1 | 0 | 1 | 1 | 0.40110418 | 1 | 0.48242099 | 0 | 1 | 12 | 0.62236227 | FALSE | 1 | FALSE | 1 | FALSE | 1 | FALSE | 1 | FALSE | 1 | FALSE | 1 |
| 248 | 248 | 4.07142857 | 2.60738746 | 2 | 0.7830614 | 5 | 0.28966038 | 5 | 0.36853505 | 2 | 0.96659807 | 0 | 1 | 97 | 0.2310598 | FALSE | 1 | FALSE | 1 | FALSE | 1 | FALSE | 1 | FALSE | 1 | FALSE | 1 |
| 249 | 249 | 9.92857143 | 2.60396088 | 0 | 1 | 1 | 0.60247911 | 1 | 0.64171563 | 1 | 0.73248983 | 0 | 1 | 25 | 0.34293 | FALSE | 1 | FALSE | 1 | FALSE | 1 | FALSE | 1 | FALSE | 1 | FALSE | 1 |
| 250 | 250 | 8.31578947 | 3.24528211 | 2 | 0.08405377 | 0 | 1 | 2 | 0.14749808 | 0 | 1 | 0 | 1 | 15 | 0.83648191 | FALSE | 0.79236883 | FALSE | 1 | FALSE | 1 | FALSE | 1 | FALSE | 1 | FALSE | 1 |
| 251 | 251 | 7.88888889 | 2.76664436 | 1 | 0.50471032 | 0 | 1 | 4 | 0.01487236 | 0 | 1 | 0 | 1 | 22 | 0.76237877 | FALSE | 1 | FALSE | 1 | FALSE | 1 | FALSE | 1 | FALSE | 1 | FALSE | 1 |
| 252 | 252 | 8.28571429 | 1.9552824 | 0 | 1 | 0 | 1 | 1 | 0.50143826 | 0 | 1 | 0 | 1 | 18 | 0.17880694 | FALSE | 1 | FALSE | 1 | FALSE | 1 | FALSE | 1 | FALSE | 1 | FALSE | 1 |
| 253 | 253 | 6.78947368 | 3.0934954 | 5 | 0.01390429 | 0 | 1 | 4 | 0.14086741 | 4 | 0.25497934 | 3 | 0.06360074 | 40 | 0.9960055 | FALSE | 0.36625055 | FALSE | 1 | FALSE | 1 | FALSE | 1 | FALSE | 1 | FALSE | 1 |
| 254 | 254 | 6.82352941 | 2.33226207 | 1 | 0.35728305 | 0 | 1 | 0 | 1 | 0 | 1 | 0 | 1 | 16 | 0.23070182 | FALSE | 1 | FALSE | 1 | FALSE | 1 | FALSE | 1 | FALSE | 1 | FALSE | 1 |
| 255 | 255 | 7.73333333 | 3.49221356 | 0 | 1 | 0 | 1 | 1 | 0.42266264 | 1 | 0.50623194 | 1 | 0.2198258 | 12 | 0.80329917 | FALSE | 1 | FALSE | 1 | FALSE | 1 | FALSE | 1 | FALSE | 1 | FALSE | 1 |
| 256 | 256 | 8.57142857 | 2.90437156 | 0 | 1 | 1 | 0.49918218 | 0 | 1 | 0 | 1 | 0 | 1 | 20 | 0.13763971 | FALSE | 1 | FALSE | 1 | FALSE | 1 | FALSE | 1 | FALSE | 1 | FALSE | 1 |
| 257 | 257 | 7.41509434 | 2.26901992 | 2 | 0.39594061 | 6 | 0.00689664 | 4 | 0.12170726 | 1 | 0.91797304 | 1 | 0.58506217 | 39 | 0.98676224 | FALSE | 1 | FALSE | 1 | FALSE | 1 | FALSE | 1 | FALSE | 0.60103504 | FALSE | 1 |
| 258 | 258 | 12.1363636 | 2.02922857 | 0 | 1 | 0 | 1 | 1 | 0.51939841 | 1 | 0.60987146 | 1 | 0.28187291 | 17 | 0.61750632 | FALSE | 1 | FALSE | 1 | FALSE | 1 | FALSE | 1 | FALSE | 1 | FALSE | 1 |
| 259 | 259 | 8.25 | 2.384848 | 0 | 1 | 0 | 1 | 0 | 1 | 0 | 1 | 4 | 4.55E-06 | 4 | 0.99658732 | FALSE | 1 | FALSE | 1 | FALSE | 1 | FALSE | 1 | FALSE | 1 | FALSE | 1 |
| 260 | 260 | 7.19047619 | 2.78805356 | 2 | 0.26413583 | 0 | 1 | 0 | 1 | 1 | 0.84089119 | 1 | 0.4761699 | 35 | 0.24915769 | FALSE | 1 | FALSE | 1 | FALSE | 1 | FALSE | 1 | TRUE | 0.00373057 | FALSE | 1 |
| 261 | 261 | 8.84615385 | 2.38089039 | 0 | 1 | 0 | 1 | 1 | 0.37874417 | 0 | 1 | 2 | 0.01846832 | 10 | 0.86791642 | FALSE | 1 | FALSE | 1 | FALSE | 1 | FALSE | 1 | FALSE | 0.69720022 | FALSE | 1 |
| 262 | 262 | 8.65116279 | 1.58310441 | 4 | 0.02366788 | 5 | 0.01196839 | 2 | 0.46083448 | 1 | 0.86831187 | 0 | 1 | 31 | 0.98840896 | FALSE | 0.48961422 | FALSE | 0.67617265 | FALSE | 1 | FALSE | 1 | FALSE | 1 | FALSE | 1 |
| 263 | 263 | 8.88888889 | 2.33068633 | 0 | 1 | 2 | 0.1135776 | 0 | 1 | 0 | 1 | 0 | 1 | 16 | 0.44998001 | FALSE | 1 | FALSE | 1 | FALSE | 1 | FALSE | 1 | FALSE | 1 | FALSE | 1 |
| 264 | 264 | 6.90625 | 2.49197932 | 3 | 0.0476977 | 1 | 0.65167362 | 1 | 0.69069222 | 3 | 0.17988885 | 0 | 1 | 24 | 0.94876798 | FALSE | 0.649301 | FALSE | 1 | FALSE | 1 | FALSE | 1 | FALSE | 1 | FALSE | 1 |
| 265 | 265 | 7.14285714 | 2.55017673 | 2 | 0.09989125 | 1 | 0.49918218 | 0 | 1 | 0 | 1 | 0 | 1 | 18 | 0.57962763 | FALSE | 0.86047225 | FALSE | 1 | FALSE | 1 | FALSE | 1 | FALSE | 1 | FALSE | 1 |
| 266 | 266 | 11.46875 | 1.76749072 | 2 | 0.1881398 | 1 | 0.63997468 | 1 | 0.67911179 | 4 | 0.05164411 | 1 | 0.4016888 | 22 | 0.98395924 | FALSE | 1 | FALSE | 1 | FALSE | 1 | FALSE | 1 | FALSE | 1 | FALSE | 1 |
| 267 | 267 | 11.5555556 | 2.00616334 | 0 | 1 | 1 | 0.25628163 | 1 | 0.28067846 | 0 | 1 | 0 | 1 | 7 | 0.84574168 | FALSE | 1 | FALSE | 1 | FALSE | 1 | FALSE | 1 | FALSE | 1 | FALSE | 1 |
| 268 | 268 | 8.625 | 2.23257139 | 0 | 1 | 1 | 0.40942991 | 0 | 1 | 2 | 0.16544789 | 1 | 0.23264612 | 12 | 0.90922186 | FALSE | 1 | FALSE | 1 | FALSE | 1 | FALSE | 1 | FALSE | 1 | FALSE | 1 |
| 269 | 269 | 8.42857143 | 3.02126047 | 0 | 1 | 1 | 0.9134145 | 4 | 0.27483745 | 2 | 0.86064871 | 1 | 0.70774043 | 66 | 0.16217093 | FALSE | 1 | FALSE | 1 | FALSE | 1 | FALSE | 1 | FALSE | 1 | FALSE | 1 |
| 270 | 270 | 6.5 | 3.27363203 | 0 | 1 | 0 | 1 | 0 | 1 | 1 | 0.75658797 | 0 | 1 | 29 | 0.03983039 | FALSE | 1 | FALSE | 1 | FALSE | 1 | FALSE | 1 | FALSE | 1 | FALSE | 1 |
| 271 | 271 | 8.40909091 | 2.67410292 | 2 | 0.10809809 | 1 | 0.5154318 | 1 | 0.55340851 | 1 | 0.64497439 | 0 | 1 | 17 | 0.8840357 | FALSE | 0.88820416 | FALSE | 1 | FALSE | 1 | FALSE | 1 | FALSE | 1 | FALSE | 1 |
| 272 | 272 | 9 | 2.56904652 | 1 | 0.54198714 | 2 | 0.25306331 | 1 | 0.66709967 | 4 | 0.04659941 | 2 | 0.08611595 | 20 | 0.99601577 | FALSE | 1 | FALSE | 1 | FALSE | 1 | FALSE | 1 | FALSE | 1 | FALSE | 1 |
| 273 | 273 | 9.54545455 | 2.33018284 | 0 | 1 | 1 | 0.49918218 | 2 | 0.17308361 | 0 | 1 | 0 | 1 | 18 | 0.57962763 | FALSE | 1 | FALSE | 1 | FALSE | 1 | FALSE | 1 | FALSE | 1 | FALSE | 1 |
| 274 | 274 | 9.26666667 | 1.69181034 | 0 | 1 | 1 | 0.38964394 | 0 | 1 | 4 | 0.00397592 | 0 | 1 | 10 | 0.97980046 | FALSE | 1 | FALSE | 1 | FALSE | 1 | FALSE | 1 | FALSE | 1 | FALSE | 1 |
| 275 | 275 | 11.0178571 | 1.72679587 | 0 | 1 | 1 | 0.82613965 | 1 | 0.85724013 | 3 | 0.44196866 | 0 | 1 | 48 | 0.14364394 | FALSE | 1 | FALSE | 1 | FALSE | 1 | FALSE | 1 | FALSE | 1 | FALSE | 1 |
| 276 | 276 | 4.67307692 | 1.84725528 | 4 | 0.03851986 | 0 | 1 | 2 | 0.54120634 | 4 | 0.19569978 | 0 | 1 | 40 | 0.85325204 | FALSE | 0.60156825 | FALSE | 1 | FALSE | 1 | FALSE | 1 | FALSE | 1 | FALSE | 1 |
| 277 | 277 | 7.87356322 | 2.32331807 | 1 | 0.89439119 | 1 | 0.94192298 | 5 | 0.1959611 | 3 | 0.76292097 | 2 | 0.41367714 | 74 | 0.40225767 | FALSE | 1 | FALSE | 1 | FALSE | 1 | FALSE | 1 | FALSE | 1 | FALSE | 1 |
| 278 | 278 | 8.45 | 2.31246622 | 1 | 0.38989951 | 1 | 0.46503795 | 0 | 1 | 1 | 0.59104357 | 0 | 1 | 16 | 0.65565778 | FALSE | 1 | FALSE | 1 | FALSE | 1 | FALSE | 1 | FALSE | 1 | FALSE | 1 |
| 279 | 279 | 9.08333333 | 2.30789707 | 2 | 0.12501436 | 0 | 1 | 1 | 0.58502114 | 1 | 0.67692816 | 0 | 1 | 20 | 0.68270729 | FALSE | 0.95622564 | FALSE | 1 | FALSE | 1 | FALSE | 1 | FALSE | 1 | FALSE | 1 |
| 280 | 280 | 9.59375 | 2.88161603 | 3 | 0.0476977 | 0 | 1 | 0 | 1 | 0 | 1 | 0 | 1 | 29 | 0.24088541 | FALSE | 0.649301 | FALSE | 1 | FALSE | 1 | FALSE | 1 | FALSE | 1 | FALSE | 1 |

| | | | | | | | | | | | | | | | | | | | | | | | | | | | |
|---|---|---|---|---|---|---|---|---|---|---|---|---|---|---|---|---|---|---|---|---|---|---|---|---|---|---|---|
| 281 | 281 | 9.26086957 | 1.69954959 | 1 | 0.45027683 | 1 | 0.53115654 | 1 | 0.5695037 | 1 | 0.6613267 | 0 | 1 | 19 | 0.71542496 | FALSE | 1 | FALSE | 1 | FALSE | 1 | FALSE | 1 | FALSE | 1 | FALSE | 1 |
| 282 | 282 | 9.33333333 | 2.88675135 | 0 | 1 | 0 | 1 | 1 | 0.46348836 | 2 | 0.18212134 | 0 | 1 | 14 | 0.7313145 | FALSE | 1 | FALSE | 1 | FALSE | 1 | FALSE | 1 | FALSE | 1 | FALSE | 1 |
| 283 | 283 | 9.33333333 | 1.79505494 | 0 | 1 | 1 | 0.32623933 | 0 | 1 | 0 | 1 | 2 | 0.01579445 | 9 | 0.89630573 | FALSE | 1 | FALSE | 1 | FALSE | 1 | FALSE | 1 | FALSE | 0.66313193 | FALSE | 1 |
| 284 | 284 | 10.95 | 1.90984293 | 0 | 1 | 2 | 0.1353446 | 1 | 0.51939841 | 1 | 0.60987146 | 1 | 0.28187291 | 15 | 0.92113679 | FALSE | 1 | FALSE | 1 | FALSE | 1 | FALSE | 1 | FALSE | 1 | FALSE | 1 |
| 285 | 285 | 8.86363636 | 2.56365248 | 1 | 0.4357638 | 0 | 1 | 1 | 0.55340851 | 0 | 1 | 0 | 1 | 20 | 0.30915202 | FALSE | 1 | FALSE | 1 | FALSE | 1 | FALSE | 1 | FALSE | 1 | FALSE | 1 |
| 286 | 286 | 8.95833333 | 2.96478171 | 0 | 1 | 2 | 0.18117183 | 0 | 1 | 1 | 0.67692816 | 0 | 1 | 21 | 0.47053137 | FALSE | 1 | FALSE | 1 | FALSE | 1 | FALSE | 1 | FALSE | 1 | FALSE | 1 |
| 287 | 287 | 9.25 | 3.29456623 | 0 | 1 | 1 | 0.32623933 | 0 | 1 | 0 | 1 | 0 | 1 | 11 | 0.41944532 | FALSE | 1 | FALSE | 1 | FALSE | 1 | FALSE | 1 | FALSE | 1 | FALSE | 1 |
| 288 | 288 | 10 | 2.80975744 | 1 | 0.38989951 | 1 | 0.46503795 | 2 | 0.14749808 | 1 | 0.59104357 | 1 | 0.26986775 | 13 | 0.97998676 | FALSE | 1 | FALSE | 1 | FALSE | 1 | FALSE | 1 | FALSE | 1 | FALSE | 1 |
| 289 | 289 | 6.51785714 | 1.18760069 | 7 | 0.35110673 | 14 | 0.01353108 | 12 | 0.10762857 | 8 | 0.81513269 | 4 | 0.50365467 | 179 | 0.97241789 | FALSE | 1 | FALSE | 0.69207862 | FALSE | 1 | FALSE | 1 | FALSE | 1 | FALSE | 1 |
| 290 | 290 | 8 | 3.24893145 | 2 | 0.06909038 | 3 | 0.01622926 | 0 | 1 | 0 | 1 | 1 | 0.24525764 | 11 | 0.98974663 | FALSE | 0.74276693 | FALSE | 0.7256023 | FALSE | 1 | FALSE | 1 | FALSE | 1 | FALSE | 1 |
| 291 | 291 | 5.51020408 | 1.96003383 | 1 | 0.71380452 | 0 | 1 | 1 | 0.8283465 | 2 | 0.65421368 | 1 | 0.54902428 | 43 | 0.21763056 | FALSE | 1 | FALSE | 1 | FALSE | 1 | FALSE | 1 | FALSE | 1 | FALSE | 1 |
| 292 | 292 | 10.2307692 | 2.18926915 | 0 | 1 | 0 | 1 | 0 | 1 | 0 | 1 | 1 | 0.19354507 | 12 | 0.374198 | FALSE | 1 | FALSE | 1 | FALSE | 1 | FALSE | 1 | FALSE | 1 | FALSE | 1 |
| 293 | 293 | 9.16 | 2.72293959 | 1 | 0.72119464 | 1 | 0.80150109 | 5 | 0.03025969 | 3 | 0.39227316 | 1 | 0.55647171 | 38 | 0.92985909 | FALSE | 1 | FALSE | 1 | FALSE | 1 | FALSE | 1 | FALSE | 1 | FALSE | 1 |
| 294 | 294 | 9.26666667 | 1.12348664 | 0 | 1 | 0 | 1 | 1 | 0.42266264 | 0 | 1 | 1 | 0.2198258 | 13 | 0.57712805 | FALSE | 1 | FALSE | 1 | FALSE | 1 | FALSE | 1 | FALSE | 1 | FALSE | 1 |
| 295 | 295 | 8.9 | 1.92093727 | 0 | 1 | 1 | 0.7326167 | 0 | 1 | 2 | 0.5543522 | 0 | 1 | 37 | 0.10889911 | FALSE | 1 | FALSE | 1 | FALSE | 1 | FALSE | 1 | FALSE | 1 | FALSE | 1 |
| 296 | 296 | 6.125 | 2.89125838 | 1 | 0.76171602 | 2 | 0.5356583 | 1 | 0.8673918 | 5 | 0.10658542 | 0 | 1 | 46 | 0.64645216 | FALSE | 1 | FALSE | 1 | FALSE | 1 | FALSE | 1 | FALSE | 1 | FALSE | 1 |
| 297 | 297 | 10.375 | 2.9553976 | 1 | 0.18769745 | 0 | 1 | 0 | 1 | 0 | 1 | 0 | 1 | 7 | 0.63752397 | FALSE | 1 | FALSE | 1 | FALSE | 1 | FALSE | 1 | FALSE | 1 | FALSE | 1 |
| 298 | 298 | 8.42307692 | 1.86409551 | 0 | 1 | 0 | 1 | 1 | 0.61440466 | 3 | 0.11458571 | 0 | 1 | 22 | 0.61628 | FALSE | 1 | FALSE | 1 | FALSE | 1 | FALSE | 1 | FALSE | 1 | FALSE | 1 |
| 299 | 299 | 8.73913044 | 2.39959039 | 0 | 1 | 1 | 0.53115654 | 0 | 1 | 2 | 0.2852236 | 0 | 1 | 20 | 0.50589402 | FALSE | 1 | FALSE | 1 | FALSE | 1 | FALSE | 1 | FALSE | 1 | FALSE | 1 |
| 300 | 300 | 7.94117647 | 1.21125061 | 1 | 0.35728305 | 0 | 1 | 0 | 1 | 2 | 0.18212134 | 0 | 1 | 14 | 0.7313145 | FALSE | 1 | FALSE | 1 | FALSE | 1 | FALSE | 1 | FALSE | 1 | FALSE | 1 |
| 301 | 301 | 6.38095238 | 3.4568936 | 1 | 0.42086991 | 2 | 0.14655134 | 1 | 0.53671415 | 1 | 0.62783522 | 1 | 0.29368245 | 15 | 0.96513691 | FALSE | 1 | FALSE | 1 | FALSE | 1 | FALSE | 1 | FALSE | 1 | FALSE | 1 |
| 302 | 302 | 8.42857143 | 1.59078982 | 0 | 1 | 0 | 1 | 0 | 1 | 0 | 1 | 0 | 1 | 7 | 0.3043657 | FALSE | 1 | FALSE | 1 | FALSE | 1 | FALSE | 1 | FALSE | 1 | FALSE | 1 |
| 303 | 303 | 11.3125 | 2.77756076 | 1 | 0.34033054 | 1 | 0.40942991 | 0 | 1 | 0 | 1 | 0 | 1 | 14 | 0.53308261 | FALSE | 1 | FALSE | 1 | FALSE | 1 | FALSE | 1 | FALSE | 1 | FALSE | 1 |
| 304 | 304 | 8.375 | 1.86664807 | 0 | 1 | 0 | 1 | 0 | 1 | 0 | 1 | 0 | 1 | 8 | 0.25677166 | FALSE | 1 | FALSE | 1 | FALSE | 1 | FALSE | 1 | FALSE | 1 | FALSE | 1 |
| 305 | 305 | 9.45454546 | 2.06105165 | 0 | 1 | 0 | 1 | 0 | 1 | 1 | 0.40390032 | 0 | 1 | 10 | 0.46859937 | FALSE | 1 | FALSE | 1 | FALSE | 1 | FALSE | 1 | FALSE | 1 | FALSE | 1 |
| 306 | 306 | 10.1363636 | 3.2374998 | 0 | 1 | 2 | 0.15794013 | 0 | 1 | 0 | 1 | 0 | 1 | 20 | 0.30915202 | FALSE | 1 | FALSE | 1 | FALSE | 1 | FALSE | 1 | FALSE | 1 | FALSE | 1 |
| 307 | 307 | 9.70588235 | 2.69563276 | 3 | 0.05546921 | 2 | 0.30150529 | 1 | 0.71261921 | 0 | 1 | 2 | 0.1065437 | 26 | 0.92796052 | FALSE | 0.68849059 | FALSE | 1 | FALSE | 1 | FALSE | 1 | FALSE | 1 | FALSE | 1 |
| 308 | 308 | 6.21212121 | 3.24511358 | 2 | 0.1881398 | 1 | 0.63997468 | 2 | 0.30673593 | 1 | 0.76781284 | 1 | 0.4016888 | 24 | 0.90138666 | FALSE | 1 | FALSE | 1 | FALSE | 1 | FALSE | 1 | FALSE | 1 | FALSE | 1 |
| 309 | 309 | 6.92307692 | 2.23474447 | 2 | 0.04234484 | 1 | 0.3480709 | 1 | 0.37874417 | 0 | 1 | 0 | 1 | 9 | 0.96018168 | FALSE | 0.62842194 | FALSE | 1 | FALSE | 1 | FALSE | 1 | FALSE | 1 | FALSE | 1 |
| 310 | 310 | 6.56 | 2.6545056 | 0 | 1 | 1 | 0.80150109 | 2 | 0.53023996 | 0 | 1 | 0 | 1 | 46 | 0.04012052 | FALSE | 1 | FALSE | 1 | FALSE | 1 | FALSE | 1 | FALSE | 1 | FALSE | 1 |
| 311 | 311 | 10.8 | 1.98997487 | 0 | 1 | 0 | 1 | 0 | 1 | 1 | 0.37517262 | 0 | 1 | 9 | 0.5215547 | FALSE | 1 | FALSE | 1 | FALSE | 1 | FALSE | 1 | FALSE | 1 | FALSE | 1 |
| 312 | 312 | 9.66666667 | 3.05505046 | 0 | 1 | 0 | 1 | 0 | 1 | 0 | 1 | 0 | 1 | 9 | 0.21661397 | FALSE | 1 | FALSE | 1 | FALSE | 1 | FALSE | 1 | FALSE | 1 | FALSE | 1 |
| 313 | 313 | 9.44444444 | 2.24158233 | 3 | 0.01017712 | 1 | 0.44710673 | 0 | 1 | 1 | 0.57131013 | 0 | 1 | 13 | 0.95041913 | FALSE | 0.30467829 | FALSE | 1 | FALSE | 1 | FALSE | 1 | FALSE | 1 | FALSE | 1 |
| 314 | 314 | 10.4857143 | 0.99631976 | 0 | 1 | 1 | 0.68454804 | 3 | 0.12958859 | 2 | 0.48233989 | 0 | 1 | 29 | 0.69927719 | FALSE | 1 | FALSE | 1 | FALSE | 1 | FALSE | 1 | FALSE | 1 | FALSE | 1 |
| 315 | 315 | 8.65217391 | 2.31457942 | 0 | 1 | 2 | 0.16948764 | 0 | 1 | 0 | 1 | 2 | 0.05389003 | 19 | 0.71542496 | FALSE | 1 | FALSE | 1 | FALSE | 1 | FALSE | 1 | FALSE | 1 | FALSE | 1 |
| 316 | 316 | 6.37096774 | 4.94561999 | 0 | 1 | 2 | 0.60114557 | 4 | 0.18240181 | 5 | 0.15434169 | 0 | 1 | 51 | 0.74578096 | FALSE | 1 | FALSE | 1 | FALSE | 1 | FALSE | 1 | FALSE | 1 | FALSE | 1 |
| 317 | 317 | 9.28571429 | 2.43276948 | 1 | 0.51745948 | 3 | 0.06039219 | 1 | 0.64171563 | 1 | 0.73248983 | 3 | 0.0104283 | 19 | 0.99262381 | FALSE | 1 | FALSE | 1 | FALSE | 1 | FALSE | 1 | FALSE | 0.5620304 | FALSE | 1 |
| 318 | 318 | 6.04255319 | 4.46240774 | 1 | 0.70621968 | 2 | 0.45233216 | 1 | 0.82190381 | 1 | 0.89101956 | 0 | 1 | 42 | 0.23544056 | FALSE | 1 | FALSE | 1 | FALSE | 1 | FALSE | 1 | FALSE | 1 | FALSE | 1 |
| 319 | 319 | 5.67708333 | 2.5799499 | 4 | 0.23177249 | 2 | 0.82287436 | 3 | 0.67648354 | 4 | 0.64930879 | 4 | 0.07228874 | 79 | 0.76570638 | FALSE | 1 | FALSE | 1 | FALSE | 1 | FALSE | 1 | FALSE | 1 | FALSE | 1 |
| 320 | 320 | 7.21428571 | 2.78204887 | 0 | 1 | 2 | 0.07341559 | 1 | 0.40110418 | 2 | 0.13304705 | 0 | 1 | 9 | 0.98607548 | FALSE | 1 | FALSE | 1 | FALSE | 1 | FALSE | 1 | FALSE | 1 | FALSE | 1 |
| 321 | 321 | 9.64516129 | 2.81182282 | 3 | 0.04402911 | 1 | 0.63997468 | 0 | 1 | 1 | 0.76781284 | 1 | 0.4016888 | 25 | 0.80006983 | FALSE | 0.64293489 | FALSE | 1 | FALSE | 1 | FALSE | 1 | FALSE | 1 | FALSE | 1 |
| 322 | 322 | 10.3333333 | 2.98142397 | 0 | 1 | 0 | 1 | 1 | 0.35555293 | 1 | 0.43131129 | 0 | 1 | 10 | 0.71436964 | FALSE | 1 | FALSE | 1 | FALSE | 1 | FALSE | 1 | FALSE | 1 | FALSE | 1 |
| 323 | 323 | 9.28571429 | 2.0734801 | 0 | 1 | 2 | 0.14655134 | 3 | 0.03775825 | 0 | 1 | 1 | 0.29368245 | 15 | 0.96513691 | FALSE | 1 | FALSE | 1 | FALSE | 1 | FALSE | 1 | FALSE | 1 | FALSE | 1 |
| 324 | 324 | 8.22222222 | 1.71773609 | 0 | 1 | 0 | 1 | 2 | 0.13502329 | 0 | 1 | 1 | 0.25766374 | 15 | 0.69372602 | FALSE | 1 | FALSE | 1 | FALSE | 1 | FALSE | 1 | FALSE | 1 | FALSE | 1 |
| 325 | 325 | 4.31372549 | 2.64073318 | 1 | 0.98106619 | 2 | 0.95935609 | 8 | 0.17549538 | 6 | 0.69931079 | 1 | 0.91993124 | 133 | 0.1222964 | FALSE | 1 | FALSE | 1 | FALSE | 1 | FALSE | 1 | FALSE | 1 | FALSE | 1 |
| 326 | 326 | 7.44444444 | 3.11903752 | 2 | 0.15144274 | 1 | 0.58913618 | 1 | 0.62830986 | 1 | 0.71956339 | 1 | 0.36060486 | 21 | 0.88395936 | FALSE | 1 | FALSE | 1 | FALSE | 1 | FALSE | 1 | FALSE | 1 | FALSE | 1 |
| 327 | 327 | 6.54285714 | 1.53702602 | 1 | 0.59801736 | 1 | 0.68454804 | 1 | 0.72299506 | 2 | 0.48233989 | 6 | 1.87E-05 | 24 | 0.99496229 | FALSE | 1 | FALSE | 1 | FALSE | 1 | FALSE | 1 | TRUE | 0.01092769 | FALSE | 1 |
| 328 | 328 | 7.9765625 | 2.06709656 | 2 | 0.84227743 | 7 | 0.11675455 | 5 | 0.48253777 | 7 | 0.36545349 | 1 | 0.87992897 | 105 | 0.74926772 | FALSE | 1 | FALSE | 1 | FALSE | 1 | FALSE | 1 | FALSE | 1 | FALSE | 1 |
| 329 | 329 | 8.27777778 | 3.10564997 | 0 | 1 | 0 | 1 | 1 | 0.48280983 | 0 | 1 | 1 | 0.25766374 | 16 | 0.44998001 | FALSE | 1 | FALSE | 1 | FALSE | 1 | FALSE | 1 | FALSE | 1 | FALSE | 1 |
| 330 | 330 | 10.3333333 | 2.4267033 | 0 | 1 | 1 | 0.32623933 | 0 | 1 | 0 | 1 | 0 | 1 | 11 | 0.41944532 | FALSE | 1 | FALSE | 1 | FALSE | 1 | FALSE | 1 | FALSE | 1 | FALSE | 1 |
| 331 | 331 | 10.2790698 | 3.08277722 | 1 | 0.67382914 | 1 | 0.75788114 | 1 | 0.79363407 | 2 | 0.59404271 | 1 | 0.50988734 | 37 | 0.48294184 | FALSE | 1 | FALSE | 1 | FALSE | 1 | FALSE | 1 | FALSE | 1 | FALSE | 1 |
| 332 | 332 | 8.15789474 | 3.18323169 | 0 | 1 | 1 | 0.46503795 | 0 | 1 | 2 | 0.21610812 | 0 | 1 | 16 | 0.65565778 | FALSE | 1 | FALSE | 1 | FALSE | 1 | FALSE | 1 | FALSE | 1 | FALSE | 1 |
| 333 | 333 | 10.8461539 | 2.31537184 | 0 | 1 | 0 | 1 | 0 | 1 | 0 | 1 | 0 | 1 | 13 | 0.10967906 | FALSE | 1 | FALSE | 1 | FALSE | 1 | FALSE | 1 | FALSE | 1 | FALSE | 1 |
| 334 | 334 | 6.55263158 | 2.52041526 | 0 | 1 | 1 | 0.71433277 | 0 | 1 | 4 | 0.09477133 | 0 | 1 | 33 | 0.44261631 | FALSE | 1 | FALSE | 1 | FALSE | 1 | FALSE | 1 | FALSE | 1 | FALSE | 1 |
| 335 | 335 | 7.2247191 | 1.97030089 | 3 | 0.40052974 | 4 | 0.32490044 | 5 | 0.21518975 | 0 | 1 | 0 | 1 | 77 | 0.35047347 | FALSE | 1 | FALSE | 1 | FALSE | 1 | FALSE | 1 | FALSE | 1 | FALSE | 1 |
| 336 | 336 | 7.5 | 1.58113883 | 0 | 1 | 0 | 1 | 0 | 1 | 0 | 1 | 0 | 1 | 8 | 0.25677166 | FALSE | 1 | FALSE | 1 | FALSE | 1 | FALSE | 1 | FALSE | 1 | FALSE | 1 |
| 337 | 337 | 8.82352941 | 3.0723682 | 4 | 0.00076452 | 2 | 0.10307112 | 0 | 1 | 0 | 1 | 0 | 1 | 11 | 0.98974663 | FALSE | 0.06390799 | FALSE | 1 | FALSE | 1 | FALSE | 1 | FALSE | 1 | FALSE | 1 |
| 338 | 338 | 6.71428571 | 2.76272566 | 2 | 0.15144274 | 0 | 1 | 0 | 1 | 2 | 0.35374209 | 0 | 1 | 23 | 0.58307573 | FALSE | 1 | FALSE | 1 | FALSE | 1 | FALSE | 1 | FALSE | 1 | FALSE | 1 |
| 339 | 339 | 6.6875 | 3.08663956 | 1 | 0.56527493 | 1 | 0.65167362 | 0 | 1 | 1 | 0.77852168 | 1 | 0.41154418 | 28 | 0.42532177 | FALSE | 1 | FALSE | 1 | FALSE | 1 | FALSE | 1 | FALSE | 1 | FALSE | 1 |
| 340 | 340 | 9.28571429 | 2.02232439 | 1 | 0.59801736 | 1 | 0.68454804 | 3 | 0.12958859 | 1 | 0.80778317 | 1 | 0.44015537 | 28 | 0.83079716 | FALSE | 1 | FALSE | 1 | FALSE | 1 | FALSE | 1 | FALSE | 1 | FALSE | 1 |
| 341 | 341 | 11.5 | 2.08348484 | 0 | 1 | 0 | 1 | 0 | 1 | 2 | 0.23331368 | 0 | 1 | 18 | 0.37504645 | FALSE | 1 | FALSE | 1 | FALSE | 1 | FALSE | 1 | FALSE | 1 | FALSE | 1 |
| 342 | 342 | 10.5384615 | 2.95128094 | 0 | 1 | 0 | 1 | 0 | 1 | 1 | 0.43131129 | 0 | 1 | 11 | 0.41944532 | FALSE | 1 | FALSE | 1 | FALSE | 1 | FALSE | 1 | FALSE | 1 | FALSE | 1 |
| 343 | 343 | 8.02631579 | 1.58645874 | 5 | 0.16803892 | 4 | 0.50674923 | 6 | 0.22590386 | 4 | 0.7753231 | 0 | 1 | 95 | 0.67842369 | FALSE | 1 | FALSE | 1 | FALSE | 1 | FALSE | 1 | FALSE | 1 | FALSE | 1 |
| 344 | 344 | 5.48888889 | 3.05957312 | 4 | 0.02747663 | 0 | 1 | 1 | 0.80828716 | 4 | 0.15016749 | 0 | 1 | 36 | 0.84604146 | FALSE | 0.50964275 | FALSE | 1 | FALSE | 1 | FALSE | 1 | FALSE | 1 | FALSE | 1 |
| 345 | 345 | 4.19178082 | 2.75808915 | 1 | 0.85137169 | 3 | 0.42214364 | 3 | 0.49081265 | 2 | 0.8554296 | 1 | 0.7028142 | 63 | 0.39803869 | FALSE | 1 | FALSE | 1 | FALSE | 1 | FALSE | 1 | FALSE | 1 | FALSE | 1 |
| 346 | 346 | 10.3846154 | 3.17534884 | 0 | 1 | 1 | 0.3480709 | 0 | 1 | 0 | 1 | 1 | 0.19354507 | 11 | 0.6683056 | FALSE | 1 | FALSE | 1 | FALSE | 1 | FALSE | 1 | FALSE | 1 | FALSE | 1 |
| 347 | 347 | 7.81481482 | 2.82891207 | 0 | 1 | 1 | 0.57534752 | 0 | 1 | 1 | 0.70601446 | 1 | 0.3499045 | 23 | 0.40372818 | FALSE | 1 | FALSE | 1 | FALSE | 1 | FALSE | 1 | FALSE | 1 | FALSE | 1 |
| 348 | 348 | 6.73333333 | 3.27549827 | 0 | 1 | 1 | 0.6278847 | 0 | 1 | 0 | 1 | 0 | 1 | 29 | 0.03983039 | FALSE | 1 | FALSE | 1 | FALSE | 1 | FALSE | 1 | FALSE | 1 | FALSE | 1 |
| 349 | 349 | 8.92307692 | 1.73034183 | 0 | 1 | 0 | 1 | 2 | 0.07725803 | 0 | 1 | 0 | 1 | 11 | 0.6683056 | FALSE | 1 | FALSE | 1 | FALSE | 1 | FALSE | 1 | FALSE | 1 | FALSE | 1 |
| 350 | 350 | 1.37162162 | 1.91995357 | 6 | 0.99833833 | 17 | 0.72911549 | 18 | 0.80182356 | 27 | 0.5378966 | 14 | 0.10101834 | 508 | 0.12480581 | FALSE | 1 | FALSE | 1 | FALSE | 1 | FALSE | 1 | FALSE | 1 | FALSE | 1 |
| 351 | 351 | 6.81818182 | 2.97315816 | 2 | 0.41401181 | 3 | 0.26245929 | 3 | 0.31652594 | 3 | 0.46623289 | 0 | 1 | 44 | 0.86014424 | FALSE | 1 | FALSE | 1 | FALSE | 1 | FALSE | 1 | FALSE | 1 | FALSE | 1 |
| 352 | 352 | 9.8125 | 1.97543508 | 0 | 1 | 1 | 0.40942991 | 2 | 0.11087977 | 0 | 1 | 0 | 1 | 13 | 0.76799161 | FALSE | 1 | FALSE | 1 | FALSE | 1 | FALSE | 1 | FALSE | 1 | FALSE | 1 |
| 353 | 353 | 9.5 | 2.32737334 | 0 | 1 | 0 | 1 | 0 | 1 | 1 | 0.43131129 | 0 | 1 | 11 | 0.41944532 | FALSE | 1 | FALSE | 1 | FALSE | 1 | FALSE | 1 | FALSE | 1 | FALSE | 1 |
| 354 | 354 | 9.61538462 | 3.15103064 | 0 | 1 | 0 | 1 | 1 | 0.37874417 | 0 | 1 | 0 | 1 | 12 | 0.374198 | FALSE | 1 | FALSE | 1 | FALSE | 1 | FALSE | 1 | FALSE | 1 | FALSE | 1 |
| 355 | 355 | 10.0909091 | 2.74539434 | 0 | 1 | 1 | 0.30368014 | 0 | 1 | 0 | 1 | 0 | 1 | 10 | 0.46859937 | FALSE | 1 | FALSE | 1 | FALSE | 1 | FALSE | 1 | FALSE | 1 | FALSE | 1 |
| 356 | 356 | 10.375 | 1.2520816 | 0 | 1 | 0 | 1 | 3 | 0.05314317 | 2 | 0.30249307 | 0 | 1 | 19 | 0.83975208 | FALSE | 1 | FALSE | 1 | FALSE | 1 | FALSE | 1 | FALSE | 1 | FALSE | 1 |
| 357 | 357 | 9.17073171 | 1.89868297 | 2 | 0.28329504 | 3 | 0.14580275 | 0 | 1 | 0 | 1 | 0 | 1 | 36 | 0.36455032 | FALSE | 1 | FALSE | 1 | FALSE | 1 | FALSE | 1 | FALSE | 1 | FALSE | 1 |
| 358 | 358 | 8.79166667 | 2.08124896 | 1 | 0.46441868 | 1 | 0.54637326 | 2 | 0.21260257 | 2 | 0.30249307 | 0 | 1 | 18 | 0.93166998 | FALSE | 1 | FALSE | 1 | FALSE | 1 | FALSE | 1 | FALSE | 1 | FALSE | 1 |
| 359 | 359 | 8.05128205 | 2.06246949 | 0 | 1 | 0 | 1 | 1 | 0.75193685 | 4 | 0.09477133 | 2 | 0.12811285 | 31 | 0.76640548 | FALSE | 1 | FALSE | 1 | FALSE | 1 | FALSE | 1 | FALSE | 1 | FALSE | 1 |
| 360 | 360 | 8.41176471 | 3.53235172 | 3 | 0.00864151 | 3 | 0.01622926 | 1 | 0.46348836 | 0 | 1 | 0 | 1 | 10 | 0.99775952 | FALSE | 0.27019545 | FALSE | 0.7256023 | FALSE | 1 | FALSE | 1 | FALSE | 1 | FALSE | 1 |
| 361 | 361 | 6.29113924 | 2.15331301 | 1 | 0.87304634 | 3 | 0.47338398 | 0 | 1 | 3 | 0.71030908 | 0 | 1 | 72 | 0.05857603 | FALSE | 1 | FALSE | 1 | FALSE | 1 | FALSE | 1 | FALSE | 1 | FALSE | 1 |
| 362 | 362 | 12.4166667 | 2.25308431 | 0 | 1 | 0 | 1 | 0 | 1 | 2 | 0.1023796 | 0 | 1 | 10 | 0.71436964 | FALSE | 1 | FALSE | 1 | FALSE | 1 | FALSE | 1 | FALSE | 1 | FALSE | 1 |
| 363 | 363 | 7.4516129 | 2.76893726 | 1 | 0.55378205 | 1 | 0.63997468 | 2 | 0.30673593 | 2 | 0.41984187 | 2 | 0.09110253 | 23 | 0.95750806 | FALSE | 1 | FALSE | 1 | FALSE | 1 | FALSE | 1 | FALSE | 1 | FALSE | 1 |
| 364 | 364 | 8.625 | 1.97510548 | 1 | 0.46441868 | 0 | 1 | 2 | 0.21260257 | 1 | 0.67692816 | 0 | 1 | 20 | 0.68270729 | FALSE | 1 | FALSE | 1 | FALSE | 1 | FALSE | 1 | FALSE | 1 | FALSE | 1 |
| 365 | 365 | 8.25 | 2.04633819 | 0 | 1 | 2 | 0.09285443 | 0 | 1 | 1 | 0.52895088 | 0 | 1 | 13 | 0.76799161 | FALSE | 1 | FALSE | 1 | FALSE | 1 | FALSE | 1 | FALSE | 1 | FALSE | 1 |
| 366 | 366 | 7.74358974 | 3.34141033 | 1 | 0.63789042 | 2 | 0.36122049 | 3 | 0.16357218 | 2 | 0.54052736 | 0 | 1 | 31 | 0.85613899 | FALSE | 1 | FALSE | 1 | FALSE | 1 | FALSE | 1 | FALSE | 1 | FALSE | 1 |
| 367 | 367 | 8.83333333 | 1.58989867 | 0 | 1 | 0 | 1 | 0 | 1 | 2 | 0.49731009 | 0 | 1 | 34 | 0.06417017 | FALSE | 1 | FALSE | 1 | FALSE | 1 | FALSE | 1 | FALSE | 1 | FALSE | 1 |
| 368 | 368 | 10 | 1.60727513 | 2 | 0.11647716 | 0 | 1 | 0 | 1 | 2 | 0.2852236 | 1 | 0.31672722 | 18 | 0.86273155 | FALSE | 0.93181727 | FALSE | 1 | FALSE | 1 | FALSE | 1 | FALSE | 1 | FALSE | 1 |
| 369 | 369 | 9.84615385 | 2.10662522 | 1 | 0.26795838 | 1 | 0.32623933 | 0 | 1 | 0 | 1 | 1 | 0.18007762 | 9 | 0.89630573 | FALSE | 1 | FALSE | 1 | FALSE | 1 | FALSE | 1 | FALSE | 1 | FALSE | 1 |
| 370 | 370 | 8 | 3.30943816 | 2 | 0.09187136 | 0 | 1 | 1 | 0.51939841 | 0 | 1 | 0 | 1 | 17 | 0.61750632 | FALSE | 0.83248965 | FALSE | 1 | FALSE | 1 | FALSE | 1 | FALSE | 1 | FALSE | 1 |
| 371 | 371 | 10.2352941 | 3.47307986 | 0 | 1 | 0 | 1 | 1 | 0.46348836 | 1 | 0.55062772 | 0 | 1 | 15 | 0.49060435 | FALSE | 1 | FALSE | 1 | FALSE | 1 | FALSE | 1 | FALSE | 1 | FALSE | 1 |
| 372 | 372 | 9.15789474 | 3.60209234 | 0 | 1 | 1 | 0.46503795 | 1 | 0.50143826 | 1 | 0.59104357 | 0 | 1 | 16 | 0.65565778 | FALSE | 1 | FALSE | 1 | FALSE | 1 | FALSE | 1 | FALSE | 1 | FALSE | 1 |
| 373 | 373 | 9.09375 | 2.71982737 | 3 | 0.0476977 | 0 | 1 | 2 | 0.32011871 | 2 | 0.43583874 | 1 | 0.41154418 | 24 | 0.94876798 | FALSE | 0.649301 | FALSE | 1 | FALSE | 1 | FALSE | 1 | FALSE | 1 | FALSE | 1 |
| 374 | 374 | 8.75 | 1.94615005 | 0 | 1 | 1 | 0.46503795 | 0 | 1 | 0 | 1 | 0 | 1 | 18 | 0.17880694 | FALSE | 1 | FALSE | 1 | FALSE | 1 | FALSE | 1 | FALSE | 1 | FALSE | 1 |

| | | | | | | | | | | | | | | | | | | | | | | | | | | | |
|---|---|---|---|---|---|---|---|---|---|---|---|---|---|---|---|---|---|---|---|---|---|---|---|---|---|---|---|
| 375 | 375 | 9.05128205 | 1.73868087 | 0 | 1 | 2 | 0.36122049 | 4 | 0.05000009 | 0 | 1 | 2 | 0.13365639 | 31 | 0.85613899 | FALSE | 1 | FALSE | 1 | FALSE | 1 | FALSE | 1 | FALSE | 1 | FALSE | 1 |
| 376 | 376 | 9.28571429 | 1.77664363 | 0 | 1 | 2 | 0.14655134 | 2 | 0.17308361 | 1 | 0.62783522 | 0 | 1 | 16 | 0.90353674 | FALSE | 1 | FALSE | 1 | FALSE | 1 | FALSE | 1 | FALSE | 1 | FALSE | 1 |
| 377 | 377 | 9.44444444 | 2.54344959 | 0 | 1 | 0 | 1 | 0 | 1 | 1 | 0.34506517 | 0 | 1 | 8 | 0.5780332 | FALSE | 1 | FALSE | 1 | FALSE | 1 | FALSE | 1 | FALSE | 1 | FALSE | 1 |
| 378 | 378 | 9.66666667 | 1.95505044 | 0 | 1 | 0 | 1 | 1 | 0.42266264 | 0 | 1 | 0 | 1 | 14 | 0.29525409 | FALSE | 1 | FALSE | 1 | FALSE | 1 | FALSE | 1 | FALSE | 1 | FALSE | 1 |
| 379 | 379 | 9.47727273 | 2.10506571 | 3 | 0.09709068 | 1 | 0.75788114 | 1 | 0.79363407 | 3 | 0.31602175 | 1 | 0.50988734 | 34 | 0.87722689 | FALSE | 0.86047225 | FALSE | 1 | FALSE | 1 | FALSE | 1 | FALSE | 1 | FALSE | 1 |
| 380 | 380 | 7.16666667 | 3.37062474 | 0 | 1 | 0 | 1 | 0 | 1 | 0 | 1 | 0 | 1 | 18 | 0.04681656 | FALSE | 1 | FALSE | 1 | FALSE | 1 | FALSE | 1 | FALSE | 1 | FALSE | 1 |
| 381 | 381 | 10 | 2.05480467 | 1 | 0.20854748 | 2 | 0.03227145 | 0 | 1 | 1 | 0.34506517 | 0 | 1 | 5 | 0.99330426 | FALSE | 1 | FALSE | 0.92268438 | FALSE | 1 | FALSE | 1 | FALSE | 1 | FALSE | 1 |
| 382 | 382 | 7.71428571 | 3.10365157 | 0 | 1 | 1 | 0.36919824 | 1 | 0.40110418 | 1 | 0.48242099 | 0 | 1 | 11 | 0.83676449 | FALSE | 1 | FALSE | 1 | FALSE | 1 | FALSE | 1 | FALSE | 1 | FALSE | 1 |
| 383 | 383 | 8.93939394 | 3.19032053 | 0 | 1 | 2 | 0.28941836 | 2 | 0.33343871 | 2 | 0.45159487 | 4 | 0.00194079 | 23 | 0.99096587 | FALSE | 1 | FALSE | 1 | FALSE | 1 | FALSE | 1 | FALSE | 0.2523647 | FALSE | 1 |
| 384 | 384 | 6.65306122 | 4.42419609 | 3 | 0.13009168 | 2 | 0.47396556 | 3 | 0.25725031 | 1 | 0.90086419 | 0 | 1 | 40 | 0.77344956 | FALSE | 0.97964162 | FALSE | 1 | FALSE | 1 | FALSE | 1 | FALSE | 1 | FALSE | 1 |
| 385 | 385 | 4.88607595 | 2.2557232 | 3 | 0.33025289 | 4 | 0.25205408 | 3 | 0.54427311 | 3 | 0.71030908 | 1 | 0.73118367 | 65 | 0.75562402 | FALSE | 1 | FALSE | 1 | FALSE | 1 | FALSE | 1 | FALSE | 1 | FALSE | 1 |
| 386 | 386 | 7.83333333 | 2.19215774 | 0 | 1 | 0 | 1 | 0 | 1 | 0 | 1 | 0 | 1 | 6 | 0.36077158 | FALSE | 1 | FALSE | 1 | FALSE | 1 | FALSE | 1 | FALSE | 1 | FALSE | 1 |
| 387 | 387 | 11.1481482 | 3.26262453 | 1 | 0.50471032 | 2 | 0.21684246 | 0 | 1 | 0 | 1 | 1 | 0.36060486 | 23 | 0.58307573 | FALSE | 1 | FALSE | 1 | FALSE | 1 | FALSE | 1 | FALSE | 1 | FALSE | 1 |
| 388 | 388 | 10.3333333 | 2.02758751 | 2 | 0.06909038 | 0 | 1 | 0 | 1 | 0 | 1 | 0 | 1 | 15 | 0.49060435 | FALSE | 0.74276693 | FALSE | 1 | FALSE | 1 | FALSE | 1 | FALSE | 1 | FALSE | 1 |
| 389 | 389 | 11.25 | 2.34964536 | 0 | 1 | 0 | 1 | 0 | 1 | 0 | 1 | 0 | 1 | 12 | 0.13002667 | FALSE | 1 | FALSE | 1 | FALSE | 1 | FALSE | 1 | FALSE | 1 | FALSE | 1 |
| 390 | 390 | 10.2222222 | 2.1227748 | 1 | 0.37380237 | 0 | 1 | 0 | 1 | 0 | 1 | 1 | 0.25766374 | 16 | 0.44998001 | FALSE | 1 | FALSE | 1 | FALSE | 1 | FALSE | 1 | FALSE | 1 | FALSE | 1 |
| 391 | 391 | 8 | 2.0519567 | 0 | 1 | 0 | 1 | 1 | 0.50143826 | 1 | 0.59104357 | 0 | 1 | 17 | 0.41141543 | FALSE | 1 | FALSE | 1 | FALSE | 1 | FALSE | 1 | FALSE | 1 | FALSE | 1 |
| 392 | 392 | 11.2333333 | 3.05159339 | 1 | 0.51745948 | 3 | 0.06039219 | 0 | 1 | 2 | 0.3705495 | 0 | 1 | 22 | 0.86499992 | FALSE | 1 | FALSE | 1 | FALSE | 1 | FALSE | 1 | FALSE | 1 | FALSE | 1 |
| 393 | 393 | 10.7857143 | 3.34191073 | 0 | 1 | 0 | 1 | 2 | 0.08806398 | 0 | 1 | 0 | 1 | 12 | 0.62236227 | FALSE | 1 | FALSE | 1 | FALSE | 1 | FALSE | 1 | FALSE | 1 | FALSE | 1 |
| 394 | 394 | 5.25 | 3.93092939 | 1 | 0.83052698 | 3 | 0.37822937 | 4 | 0.22739663 | 4 | 0.38072365 | 0 | 1 | 56 | 0.74314178 | FALSE | 1 | FALSE | 1 | FALSE | 1 | FALSE | 1 | FALSE | 1 | FALSE | 1 |
| 395 | 395 | 10.4 | 1.9078784 | 0 | 1 | 0 | 1 | 1 | 0.30655256 | 1 | 0.37517262 | 0 | 1 | 8 | 0.80394719 | FALSE | 1 | FALSE | 1 | FALSE | 1 | FALSE | 1 | FALSE | 1 | FALSE | 1 |
| 396 | 396 | 9.28571429 | 2.84760312 | 0 | 1 | 0 | 1 | 1 | 0.53671415 | 2 | 0.25059627 | 0 | 1 | 18 | 0.57962763 | FALSE | 1 | FALSE | 1 | FALSE | 1 | FALSE | 1 | FALSE | 1 | FALSE | 1 |
| 397 | 397 | 10.8333333 | 1.99304346 | 0 | 1 | 0 | 1 | 0 | 1 | 1 | 0.43131129 | 1 | 0.18007762 | 10 | 0.71436964 | FALSE | 1 | FALSE | 1 | FALSE | 1 | FALSE | 1 | FALSE | 1 | FALSE | 1 |
| 398 | 398 | 9.68421053 | 1.80795411 | 0 | 1 | 1 | 0.44710673 | 0 | 1 | 2 | 0.19902682 | 0 | 1 | 15 | 0.69372602 | FALSE | 1 | FALSE | 1 | FALSE | 1 | FALSE | 1 | FALSE | 1 | FALSE | 1 |
| 399 | 399 | 8.43478261 | 2.9756415 | 0 | 1 | 1 | 0.53115654 | 0 | 1 | 1 | 0.6613267 | 1 | 0.31672722 | 20 | 0.50589402 | FALSE | 1 | FALSE | 1 | FALSE | 1 | FALSE | 1 | FALSE | 1 | FALSE | 1 |
| 400 | 400 | 9.51351351 | 2.56400891 | 3 | 0.0681763 | 1 | 0.70472939 | 1 | 0.74264075 | 0 | 1 | 2 | 0.12262638 | 30 | 0.78878581 | FALSE | 0.74276693 | FALSE | 1 | FALSE | 1 | FALSE | 1 | FALSE | 1 | FALSE | 1 |
| 401 | 401 | 10.5416667 | 2.51626652 | 0 | 1 | 1 | 0.54637326 | 0 | 1 | 2 | 0.30249307 | 1 | 0.32796862 | 20 | 0.68270729 | FALSE | 1 | FALSE | 1 | FALSE | 1 | FALSE | 1 | FALSE | 1 | FALSE | 1 |
| 402 | 402 | 11.1666667 | 2.79384244 | 0 | 1 | 0 | 1 | 1 | 0.48280983 | 0 | 1 | 0 | 1 | 17 | 0.20329205 | FALSE | 1 | FALSE | 1 | FALSE | 1 | FALSE | 1 | FALSE | 1 | FALSE | 1 |
| 403 | 403 | 10 | 2.79610118 | 1 | 0.24866461 | 1 | 0.30368014 | 0 | 1 | 0 | 1 | 0 | 1 | 9 | 0.75985366 | FALSE | 1 | FALSE | 1 | FALSE | 1 | FALSE | 1 | FALSE | 1 | FALSE | 1 |
| 404 | 404 | 11.2083333 | 3.65124598 | 0 | 1 | 3 | 0.04096158 | 0 | 1 | 1 | 0.67692816 | 0 | 1 | 20 | 0.68270729 | FALSE | 1 | FALSE | 0.9698186 | FALSE | 1 | FALSE | 1 | FALSE | 1 | FALSE | 1 |
| 405 | 405 | 8.47058824 | 1.57620718 | 0 | 1 | 1 | 0.42857733 | 1 | 0.46348836 | 0 | 1 | 0 | 1 | 15 | 0.49060435 | FALSE | 1 | FALSE | 1 | FALSE | 1 | FALSE | 1 | FALSE | 1 | FALSE | 1 |
| 406 | 406 | 7.70588235 | 3.90722868 | 0 | 1 | 0 | 1 | 1 | 0.44344817 | 2 | 0.16544789 | 1 | 0.23264612 | 12 | 0.90922186 | FALSE | 1 | FALSE | 1 | FALSE | 1 | FALSE | 1 | FALSE | 1 | FALSE | 1 |
| 407 | 407 | 10.7619048 | 2.22335572 | 0 | 1 | 2 | 0.14655134 | 0 | 1 | 1 | 0.62783522 | 0 | 1 | 18 | 0.57962763 | FALSE | 1 | FALSE | 1 | FALSE | 1 | FALSE | 1 | FALSE | 1 | FALSE | 1 |
| 408 | 408 | 10.5833333 | 3.90423559 | 3 | 0.00307177 | 0 | 1 | 1 | 0.35555293 | 1 | 0.43131129 | 0 | 1 | 7 | 0.99433481 | FALSE | 0.14802295 | FALSE | 1 | FALSE | 1 | FALSE | 1 | FALSE | 1 | FALSE | 1 |
| 409 | 409 | 9.42857143 | 2.82120252 | 0 | 1 | 0 | 1 | 2 | 0.08806398 | 0 | 1 | 1 | 0.20679328 | 11 | 0.83676449 | FALSE | 1 | FALSE | 1 | FALSE | 1 | FALSE | 1 | FALSE | 1 | FALSE | 1 |
| 410 | 410 | 8.90384615 | 2.36366417 | 1 | 0.68224519 | 2 | 0.41896557 | 2 | 0.47281647 | 0 | 1 | 1 | 0.51797501 | 38 | 0.45705039 | FALSE | 1 | FALSE | 1 | FALSE | 1 | FALSE | 1 | FALSE | 1 | FALSE | 1 |
| 411 | 411 | 9.90909091 | 2.10862064 | 2 | 0.03087289 | 0 | 1 | 2 | 0.05708125 | 0 | 1 | 0 | 1 | 7 | 0.98129353 | FALSE | 0.53356688 | FALSE | 1 | FALSE | 1 | FALSE | 1 | FALSE | 1 | FALSE | 1 |
| 412 | 412 | 9.53846154 | 2.08592541 | 2 | 0.26413583 | 0 | 1 | 1 | 0.76089851 | 2 | 0.54052736 | 0 | 1 | 34 | 0.41570781 | FALSE | 1 | FALSE | 1 | FALSE | 1 | FALSE | 1 | FALSE | 1 | FALSE | 1 |
| 413 | 413 | 8.70588235 | 2.74964609 | 0 | 1 | 1 | 0.6739483 | 0 | 1 | 1 | 0.7984848 | 0 | 1 | 32 | 0.08180309 | FALSE | 1 | FALSE | 1 | FALSE | 1 | FALSE | 1 | FALSE | 1 | FALSE | 1 |
| 414 | 414 | 9.5 | 1.64455023 | 1 | 0.4357638 | 0 | 1 | 2 | 0.18613256 | 0 | 1 | 1 | 0.30529951 | 18 | 0.74744813 | FALSE | 1 | FALSE | 1 | FALSE | 1 | FALSE | 1 | FALSE | 1 | FALSE | 1 |
| 415 | 415 | 8.42857143 | 2.38190457 | 0 | 1 | 1 | 0.20567256 | 0 | 1 | 0 | 1 | 0 | 1 | 6 | 0.69920685 | FALSE | 1 | FALSE | 1 | FALSE | 1 | FALSE | 1 | FALSE | 1 | FALSE | 1 |
| 416 | 416 | 11.25 | 1.83143842 | 0 | 1 | 0 | 1 | 1 | 0.58502114 | 1 | 0.67692816 | 0 | 1 | 22 | 0.25237063 | FALSE | 1 | FALSE | 1 | FALSE | 1 | FALSE | 1 | FALSE | 1 | FALSE | 1 |
| 417 | 417 | 9.39285714 | 1.29115911 | 0 | 1 | 0 | 1 | 1 | 0.64171563 | 0 | 1 | 0 | 1 | 27 | 0.05284755 | FALSE | 1 | FALSE | 1 | FALSE | 1 | FALSE | 1 | FALSE | 1 | FALSE | 1 |
| 418 | 418 | 9.25 | 2.65753645 | 0 | 1 | 0 | 1 | 1 | 0.44344817 | 1 | 0.52895088 | 0 | 1 | 14 | 0.53308261 | FALSE | 1 | FALSE | 1 | FALSE | 1 | FALSE | 1 | FALSE | 1 | FALSE | 1 |
| 419 | 419 | 10.0645161 | 2.29916512 | 1 | 0.55378205 | 0 | 1 | 0 | 1 | 2 | 0.41984187 | 0 | 1 | 28 | 0.26393996 | FALSE | 1 | FALSE | 1 | FALSE | 1 | FALSE | 1 | FALSE | 1 | FALSE | 1 |
| 420 | 420 | 8.13333333 | 1.01105006 | 2 | 0.55518392 | 1 | 0.90746182 | 5 | 0.11589609 | 3 | 0.64919942 | 3 | 0.11388306 | 58 | 0.85533855 | FALSE | 1 | FALSE | 1 | FALSE | 1 | FALSE | 1 | FALSE | 1 | FALSE | 1 |
| 421 | 421 | 11.1538462 | 3.25449166 | 0 | 1 | 1 | 0.3480709 | 1 | 0.37874417 | 2 | 0.1174583 | 0 | 1 | 9 | 0.96018168 | FALSE | 1 | FALSE | 1 | FALSE | 1 | FALSE | 1 | FALSE | 1 | FALSE | 1 |
| 422 | 422 | 8.64 | 1.6218508 | 0 | 1 | 0 | 1 | 1 | 0.59998149 | 1 | 0.69181315 | 0 | 1 | 23 | 0.22727183 | FALSE | 1 | FALSE | 1 | FALSE | 1 | FALSE | 1 | FALSE | 1 | FALSE | 1 |
| 423 | 423 | 5 | 1.78376517 | 0 | 1 | 0 | 1 | 1 | 0.46348836 | 0 | 1 | 0 | 1 | 16 | 0.23070182 | FALSE | 1 | FALSE | 1 | FALSE | 1 | FALSE | 1 | FALSE | 1 | FALSE | 1 |
| 424 | 424 | 5.91891892 | 2.4973324 | 1 | 0.6184714 | 0 | 1 | 0 | 1 | 0 | 1 | 0 | 1 | 36 | 0.01445326 | FALSE | 1 | FALSE | 1 | FALSE | 1 | FALSE | 1 | FALSE | 1 | FALSE | 1 |
| 425 | 425 | 10.3125 | 2.1713691 | 0 | 1 | 1 | 0.40942991 | 2 | 0.11087977 | 0 | 1 | 0 | 1 | 13 | 0.76799161 | FALSE | 1 | FALSE | 1 | FALSE | 1 | FALSE | 1 | FALSE | 1 | FALSE | 1 |
| 426 | 426 | 7.48648649 | 1.71828987 | 0 | 1 | 1 | 0.69480472 | 0 | 1 | 5 | 0.02311989 | 0 | 1 | 30 | 0.67248271 | FALSE | 1 | FALSE | 1 | FALSE | 1 | FALSE | 1 | FALSE | 1 | FALSE | 1 |
| 427 | 427 | 9.4 | 2.62805378 | 0 | 1 | 1 | 0.38964394 | 1 | 0.42266264 | 0 | 1 | 0 | 1 | 13 | 0.57712805 | FALSE | 1 | FALSE | 1 | FALSE | 1 | FALSE | 1 | FALSE | 1 | FALSE | 1 |
| 428 | 428 | 8.38888889 | 3.66877044 | 0 | 1 | 1 | 0.44710673 | 1 | 0.48280983 | 0 | 1 | 0 | 1 | 16 | 0.44998001 | FALSE | 1 | FALSE | 1 | FALSE | 1 | FALSE | 1 | FALSE | 1 | FALSE | 1 |
| 429 | 429 | 9.26666667 | 1.69181034 | 0 | 1 | 1 | 0.38964394 | 0 | 1 | 1 | 0.50623194 | 0 | 1 | 13 | 0.57712805 | FALSE | 1 | FALSE | 1 | FALSE | 1 | FALSE | 1 | FALSE | 1 | FALSE | 1 |
| 430 | 430 | 4.89361702 | 3.41606992 | 3 | 0.11868304 | 5 | 0.01719421 | 4 | 0.08716543 | 4 | 0.1678689 | 0 | 1 | 31 | 0.99955348 | FALSE | 0.94393345 | FALSE | 0.74134176 | FALSE | 1 | FALSE | 1 | FALSE | 1 | FALSE | 1 |
| 431 | 431 | 8.66666667 | 1.84992492 | 0 | 1 | 0 | 1 | 1 | 0.42266264 | 0 | 1 | 0 | 1 | 14 | 0.29525409 | FALSE | 1 | FALSE | 1 | FALSE | 1 | FALSE | 1 | FALSE | 1 | FALSE | 1 |
| 432 | 432 | 9.35294118 | 2.61086365 | 0 | 1 | 0 | 1 | 1 | 0.46348836 | 0 | 1 | 1 | 0.24525764 | 15 | 0.49060435 | FALSE | 1 | FALSE | 1 | FALSE | 1 | FALSE | 1 | FALSE | 1 | FALSE | 1 |
| 433 | 433 | 7.48148148 | 2.02521824 | 0 | 1 | 0 | 1 | 1 | 0.62830986 | 2 | 0.35374209 | 0 | 1 | 24 | 0.3725373 | FALSE | 1 | FALSE | 1 | FALSE | 1 | FALSE | 1 | FALSE | 1 | FALSE | 1 |
| 434 | 434 | 8.31578947 | 2.34077431 | 0 | 1 | 2 | 0.12434464 | 3 | 0.02898764 | 1 | 0.59104357 | 4 | 0.00021933 | 9 | 0.99997115 | FALSE | 1 | FALSE | 1 | FALSE | 1 | FALSE | 1 | FALSE | 0.07486439 | FALSE | 1 |
| 435 | 435 | 7.5 | 1.6583124 | 0 | 1 | 2 | 0.38460545 | 2 | 0.43640583 | 3 | 0.29054213 | 1 | 0.49330646 | 33 | 0.81996213 | FALSE | 1 | FALSE | 1 | FALSE | 1 | FALSE | 1 | FALSE | 1 | FALSE | 1 |
| 436 | 436 | 9.65 | 2.15116248 | 1 | 0.40558521 | 0 | 1 | 0 | 1 | 0 | 1 | 0 | 1 | 19 | 0.15700255 | FALSE | 1 | FALSE | 1 | FALSE | 1 | FALSE | 1 | FALSE | 1 | FALSE | 1 |
| 437 | 437 | 9.68421053 | 2.24903796 | 3 | 0.01186104 | 0 | 1 | 0 | 1 | 0 | 1 | 1 | 0.26986775 | 15 | 0.83648191 | FALSE | 0.3373808 | FALSE | 1 | FALSE | 1 | FALSE | 1 | FALSE | 1 | FALSE | 1 |
| 438 | 438 | 6.73015873 | 3.06131183 | 2 | 0.47501604 | 2 | 0.60114557 | 1 | 0.89758397 | 1 | 0.94646288 | 0 | 1 | 56 | 0.12847218 | FALSE | 1 | FALSE | 1 | FALSE | 1 | FALSE | 1 | FALSE | 1 | FALSE | 1 |
| 439 | 439 | 8.77777778 | 2.5724082 | 2 | 0.23542589 | 0 | 1 | 1 | 0.73299778 | 2 | 0.49731009 | 0 | 1 | 31 | 0.49869404 | FALSE | 1 | FALSE | 1 | FALSE | 1 | FALSE | 1 | FALSE | 1 | FALSE | 1 |
| 440 | 440 | 7.70512821 | 3.6448382 | 4 | 0.1347453 | 5 | 0.10306379 | 5 | 0.14243046 | 2 | 0.87529005 | 5 | 0.00838171 | 56 | 0.99731945 | FALSE | 0.98380879 | FALSE | 1 | FALSE | 1 | FALSE | 1 | FALSE | 0.50674765 | FALSE | 1 |
| 441 | 441 | 9.675 | 3.18109651 | 1 | 0.63789042 | 2 | 0.36122049 | 0 | 1 | 2 | 0.54052736 | 1 | 0.4761699 | 33 | 0.59061307 | FALSE | 1 | FALSE | 1 | FALSE | 1 | FALSE | 1 | FALSE | 1 | FALSE | 1 |
| 442 | 442 | 10.65 | 3.45362129 | 1 | 0.40558521 | 0 | 1 | 0 | 1 | 1 | 0.60987146 | 0 | 1 | 18 | 0.37504645 | FALSE | 1 | FALSE | 1 | FALSE | 1 | FALSE | 1 | FALSE | 1 | FALSE | 1 |
| 443 | 443 | 10.8636364 | 2.34124005 | 2 | 0.10809809 | 0 | 1 | 3 | 0.04259419 | 1 | 0.64497439 | 1 | 0.30529951 | 15 | 0.98570275 | FALSE | 0.88820416 | FALSE | 1 | FALSE | 1 | FALSE | 1 | FALSE | 1 | FALSE | 1 |
| 444 | 444 | 5.35 | 2.16275288 | 0 | 1 | 1 | 0.7326167 | 3 | 0.17245215 | 3 | 0.27785536 | 0 | 1 | 33 | 0.71942635 | FALSE | 1 | FALSE | 1 | FALSE | 1 | FALSE | 1 | FALSE | 1 | FALSE | 1 |
| 445 | 445 | 8.71428571 | 2.31234487 | 0 | 1 | 0 | 1 | 2 | 0.08806398 | 2 | 0.13304705 | 0 | 1 | 10 | 0.94579623 | FALSE | 1 | FALSE | 1 | FALSE | 1 | FALSE | 1 | FALSE | 1 | FALSE | 1 |
| 446 | 446 | 9.7826087 | 2.10499451 | 1 | 0.45027683 | 1 | 0.53115654 | 0 | 1 | 1 | 0.6613267 | 0 | 1 | 20 | 0.50589402 | FALSE | 1 | FALSE | 1 | FALSE | 1 | FALSE | 1 | FALSE | 1 | FALSE | 1 |
| 447 | 447 | 11.1428571 | 1.64130361 | 0 | 1 | 0 | 1 | 1 | 0.22601089 | 0 | 1 | 0 | 1 | 6 | 0.69920685 | FALSE | 1 | FALSE | 1 | FALSE | 1 | FALSE | 1 | FALSE | 1 | FALSE | 1 |
| 448 | 448 | 6.88888889 | 3.33148097 | 1 | 0.37380237 | 2 | 0.1135776 | 0 | 1 | 0 | 1 | 0 | 1 | 15 | 0.69372602 | FALSE | 1 | FALSE | 1 | FALSE | 1 | FALSE | 1 | FALSE | 1 | FALSE | 1 |
| 449 | 449 | 9.85294118 | 2.3405926 | 0 | 1 | 2 | 0.30150529 | 0 | 1 | 3 | 0.20353499 | 0 | 1 | 29 | 0.55707911 | FALSE | 1 | FALSE | 1 | FALSE | 1 | FALSE | 1 | FALSE | 1 | FALSE | 1 |
| 450 | 450 | 10.5333333 | 1.78387842 | 7 | 3.49E-08 | 2 | 0.08295839 | 0 | 1 | 1 | 0.50623194 | 0 | 1 | 5 | 0.99999904 | TRUE | 3.57E-05 | FALSE | 1 | FALSE | 1 | FALSE | 1 | FALSE | 1 | FALSE | 1 |
| 451 | 451 | 10.4444444 | 2.1659543 | 1 | 0.20854748 | 0 | 1 | 0 | 1 | 0 | 1 | 0 | 1 | 8 | 0.5780332 | FALSE | 1 | FALSE | 1 | FALSE | 1 | FALSE | 1 | FALSE | 1 | FALSE | 1 |
| 452 | 452 | 10.44 | 1.23547562 | 2 | 0.13369621 | 0 | 1 | 1 | 0.59998149 | 1 | 0.69181315 | 0 | 1 | 21 | 0.64957167 | FALSE | 0.97964162 | FALSE | 1 | FALSE | 1 | FALSE | 1 | FALSE | 1 | FALSE | 1 |
| 453 | 453 | 9.43636364 | 1.14039286 | 0 | 1 | 4 | 0.10132561 | 2 | 0.59334725 | 5 | 0.10658542 | 0 | 1 | 44 | 0.86014424 | FALSE | 1 | FALSE | 1 | FALSE | 1 | FALSE | 1 | FALSE | 1 | FALSE | 1 |
| 454 | 454 | 8.05494506 | 1.09306141 | 2 | 0.6707556 | 6 | 0.06764647 | 3 | 0.62570603 | 1 | 0.98516783 | 1 | 0.77262299 | 76 | 0.46596844 | FALSE | 1 | FALSE | 1 | FALSE | 1 | FALSE | 1 | FALSE | 1 | FALSE | 1 |
| 455 | 455 | 12.4 | 1.0198039 | 0 | 1 | 0 | 1 | 0 | 1 | 0 | 1 | 0 | 1 | 5 | 0.42761893 | FALSE | 1 | FALSE | 1 | FALSE | 1 | FALSE | 1 | FALSE | 1 | FALSE | 1 |
| 456 | 456 | 10.0344828 | 1.65013602 | 3 | 0.03713651 | 2 | 0.24095584 | 2 | 0.27983822 | 0 | 1 | 3 | 0.01149488 | 19 | 0.99707037 | FALSE | 0.58392 | FALSE | 1 | FALSE | 1 | FALSE | 1 | FALSE | 0.57972935 | FALSE | 1 |
| 457 | 457 | 10.3076923 | 2.49259258 | 0 | 1 | 0 | 1 | 0 | 1 | 1 | 0.45746571 | 0 | 1 | 12 | 0.374198 | FALSE | 1 | FALSE | 1 | FALSE | 1 | FALSE | 1 | FALSE | 1 | FALSE | 1 |
| 458 | 458 | 3.60227273 | 1.69584903 | 1 | 0.89980571 | 3 | 0.5460764 | 4 | 0.38964212 | 3 | 0.77645479 | 1 | 0.76878158 | 76 | 0.36734352 | FALSE | 1 | FALSE | 1 | FALSE | 1 | FALSE | 1 | FALSE | 1 | FALSE | 1 |
| 459 | 459 | 8.6875 | 2.39057394 | 2 | 0.19750261 | 0 | 1 | 1 | 0.69069222 | 0 | 1 | 0 | 1 | 29 | 0.24088541 | FALSE | 1 | FALSE | 1 | FALSE | 1 | FALSE | 1 | FALSE | 1 | FALSE | 1 |
| 460 | 460 | 7.21052632 | 3.90113273 | 1 | 0.62830697 | 2 | 0.3494045 | 1 | 0.75193685 | 1 | 0.83318955 | 1 | 0.4673875 | 32 | 0.61800916 | FALSE | 1 | FALSE | 1 | FALSE | 1 | FALSE | 1 | FALSE | 1 | FALSE | 1 |
| 461 | 461 | 9.9 | 1.81383572 | 0 | 1 | 0 | 1 | 2 | 0.16019548 | 1 | 0.60987146 | 0 | 1 | 17 | 0.61750632 | FALSE | 1 | FALSE | 1 | FALSE | 1 | FALSE | 1 | FALSE | 1 | FALSE | 1 |
| 462 | 462 | 10.0333333 | 2.25807194 | 0 | 1 | 0 | 1 | 1 | 0.65463982 | 0 | 1 | 0 | 1 | 28 | 0.04589964 | FALSE | 1 | FALSE | 1 | FALSE | 1 | FALSE | 1 | FALSE | 1 | FALSE | 1 |
| 463 | 463 | 9.06666667 | 2.76807193 | 1 | 0.32293355 | 0 | 1 | 0 | 1 | 0 | 1 | 0 | 1 | 14 | 0.29525409 | FALSE | 1 | FALSE | 1 | FALSE | 1 | FALSE | 1 | FALSE | 1 | FALSE | 1 |
| 464 | 464 | 3.6192053 | 1.97434591 | 8 | 0.51433781 | 12 | 0.27158566 | 9 | 0.76513756 | 11 | 0.82744434 | 6 | 0.37516769 | 256 | 0.46309943 | FALSE | 1 | FALSE | 1 | FALSE | 1 | FALSE | 1 | FALSE | 1 | FALSE | 1 |
| 465 | 465 | 8.88461539 | 2.8328692 | 0 | 1 | 0 | 1 | 1 | 0.61440466 | 2 | 0.33678042 | 0 | 1 | 23 | 0.40372818 | FALSE | 1 | FALSE | 1 | FALSE | 1 | FALSE | 1 | FALSE | 1 | FALSE | 1 |
| 466 | 466 | 7 | 0.70710678 | 0 | 1 | 0 | 1 | 0 | 1 | 0 | 1 | 0 | 1 | 20 | 0.03329856 | FALSE | 1 | FALSE | 1 | FALSE | 1 | FALSE | 1 | FALSE | 1 | FALSE | 1 |
| 467 | 467 | 10.4615385 | 1.88579241 | 0 | 1 | 1 | 0.56109832 | 1 | 0.59998149 | 1 | 0.69181315 | 0 | 1 | 22 | 0.43642846 | FALSE | 1 | FALSE | 1 | FALSE | 1 | FALSE | 1 | FALSE | 1 | FALSE | 1 |
| 468 | 468 | 8.08333333 | 2.6127359 | 1 | 0.46441868 | 0 | 1 | 0 | 1 | 0 | 1 | 2 | 0.058175 | 21 | 0.47053137 | FALSE | 1 | FALSE | 1 | FALSE | 1 | FALSE | 1 | FALSE | 1 | FALSE | 1 |

| | | | | | | | | | | | | | | | | | | | | | | | | | | | |
|---|---|---|---|---|---|---|---|---|---|---|---|---|---|---|---|---|---|---|---|---|---|---|---|---|---|---|---|
| 469 | 469 | 7.40909091 | 2.9643195 | 0 | 1 | 0 | 0.5154318 | 0 | 1 | 0 | 1 | 2 | 0.04972408 | 20 | 0.30915202 | FALSE | 1 | FALSE | 1 | FALSE | 1 | FALSE | 1 | FALSE | 0.98629461 | FALSE | 1 |
| 470 | 470 | 8.27272727 | 1.57196513 | 1 | 0.4357638 | 1 | 0.5154318 | 1 | 0.55340851 | 0 | 1 | 0 | 1 | 19 | 0.54233419 | FALSE | 1 | FALSE | 1 | FALSE | 1 | FALSE | 1 | FALSE | 1 | FALSE | 1 |
| 471 | 471 | 11.1724138 | 1.98388274 | 0 | 1 | 1 | 0.61539065 | 3 | 0.08431992 | 0 | 1 | 2 | 0.08121636 | 23 | 0.84462957 | FALSE | 1 | FALSE | 1 | FALSE | 1 | FALSE | 1 | FALSE | 1 | FALSE | 1 |
| 472 | 472 | 6.60416667 | 2.47268059 | 0 | 1 | 0 | 1 | 6 | 0.00699668 | 1 | 0.89605799 | 0 | 1 | 41 | 0.51861324 | FALSE | 1 | FALSE | 1 | FALSE | 1 | FALSE | 1 | FALSE | 1 | FALSE | 1 |
| 473 | 473 | 8.47368421 | 2.7408877 | 2 | 0.08405377 | 0 | 1 | 1 | 0.50143826 | 0 | 1 | 0 | 1 | 16 | 0.65565778 | FALSE | 0.79236883 | FALSE | 1 | FALSE | 1 | FALSE | 1 | FALSE | 1 | FALSE | 1 |
| 474 | 474 | 10.28 | 2.55374235 | 1 | 0.47819878 | 1 | 0.56109832 | 1 | 0.59998149 | 3 | 0.10474541 | 0 | 1 | 19 | 0.91734261 | FALSE | 1 | FALSE | 1 | FALSE | 1 | FALSE | 1 | FALSE | 1 | FALSE | 1 |
| 475 | 475 | 10.3255814 | 2.54924458 | 1 | 0.67382914 | 2 | 0.40761584 | 1 | 0.79363407 | 0 | 1 | 1 | 0.50988734 | 38 | 0.31730615 | FALSE | 1 | FALSE | 1 | FALSE | 1 | FALSE | 1 | FALSE | 1 | FALSE | 1 |
| 476 | 476 | 8.41666667 | 2.01900691 | 0 | 1 | 0 | 1 | 2 | 0.19931429 | 1 | 0.6613267 | 0 | 1 | 20 | 0.50589402 | FALSE | 1 | FALSE | 1 | FALSE | 1 | FALSE | 1 | FALSE | 1 | FALSE | 1 |
| 477 | 477 | 9 | 2.23606798 | 0 | 1 | 0 | 1 | 0 | 1 | 1 | 0.31351192 | 0 | 1 | 7 | 0.63752397 | FALSE | 1 | FALSE | 1 | FALSE | 1 | FALSE | 1 | FALSE | 1 | FALSE | 1 |
| 478 | 478 | 5.4125 | 2.76899689 | 2 | 0.51618914 | 3 | 0.36935445 | 0 | 1 | 2 | 0.82021683 | 1 | 0.6714808 | 59 | 0.26021394 | FALSE | 1 | FALSE | 1 | FALSE | 1 | FALSE | 1 | FALSE | 1 | FALSE | 1 |
| 479 | 479 | 9.13333333 | 3.24276974 | 0 | 1 | 1 | 0.38964394 | 1 | 0.42266264 | 0 | 1 | 0 | 1 | 13 | 0.57712805 | FALSE | 1 | FALSE | 1 | FALSE | 1 | FALSE | 1 | FALSE | 1 | FALSE | 1 |
| 480 | 480 | 8.93939394 | 1.17909554 | 0 | 1 | 1 | 0.6629941 | 1 | 0.70185639 | 2 | 0.45159487 | 0 | 1 | 29 | 0.39648374 | FALSE | 1 | FALSE | 1 | FALSE | 1 | FALSE | 1 | FALSE | 1 | FALSE | 1 |
| 481 | 481 | 7.61904762 | 2.57231025 | 0 | 1 | 2 | 0.14655134 | 0 | 1 | 0 | 1 | 4 | 0.00033021 | 15 | 0.96513691 | FALSE | 1 | FALSE | 1 | FALSE | 1 | FALSE | 1 | FALSE | 0.09327956 | FALSE | 1 |
| 482 | 482 | 10.3076923 | 3.12273587 | 1 | 0.28675951 | 0 | 1 | 1 | 0.37874417 | 0 | 1 | 1 | 0.19354507 | 10 | 0.86791642 | FALSE | 1 | FALSE | 1 | FALSE | 1 | FALSE | 1 | FALSE | 1 | FALSE | 1 |
| 483 | 483 | 9.13888889 | 3.21587018 | 0 | 1 | 2 | 0.32556427 | 0 | 1 | 2 | 0.49731009 | 0 | 1 | 32 | 0.31709555 | FALSE | 1 | FALSE | 1 | FALSE | 1 | FALSE | 1 | FALSE | 1 | FALSE | 1 |
| 484 | 484 | 6.44827586 | 1.22012455 | 1 | 0.52988231 | 1 | 0.61539065 | 1 | 0.65463982 | 2 | 0.3871811 | 0 | 1 | 24 | 0.70563334 | FALSE | 1 | FALSE | 1 | FALSE | 1 | FALSE | 1 | FALSE | 1 | FALSE | 1 |
| 485 | 485 | 9.78125 | 2.54625184 | 0 | 1 | 2 | 0.27730767 | 4 | 0.02651644 | 1 | 0.77852168 | 1 | 0.41154418 | 24 | 0.94876798 | FALSE | 1 | FALSE | 1 | FALSE | 1 | FALSE | 1 | FALSE | 1 | FALSE | 1 |
| 486 | 486 | 5.18867925 | 3.10845836 | 3 | 0.15399149 | 0 | 1 | 1 | 0.85724013 | 4 | 0.22482652 | 0 | 1 | 45 | 0.5504765 | FALSE | 1 | FALSE | 1 | FALSE | 1 | FALSE | 1 | FALSE | 1 | FALSE | 1 |
| 487 | 487 | 4 | 2.49443826 | 0 | 1 | 1 | 0.57534752 | 0 | 1 | 4 | 0.02930308 | 1 | 0.3499045 | 20 | 0.90142689 | FALSE | 1 | FALSE | 1 | FALSE | 1 | FALSE | 1 | FALSE | 1 | FALSE | 1 |
| 488 | 488 | 9.21568628 | 1.96352749 | 0 | 1 | 2 | 0.49507792 | 1 | 0.84631478 | 4 | 0.20527806 | 0 | 1 | 44 | 0.44552698 | FALSE | 1 | FALSE | 1 | FALSE | 1 | FALSE | 1 | FALSE | 1 | FALSE | 1 |
| 489 | 489 | 10.5333333 | 1.54344492 | 0 | 1 | 0 | 1 | 0 | 1 | 2 | 0.14906763 | 0 | 1 | 13 | 0.57712805 | FALSE | 1 | FALSE | 1 | FALSE | 1 | FALSE | 1 | FALSE | 1 | FALSE | 1 |
| 490 | 490 | 8.54166667 | 2.36254042 | 0 | 1 | 3 | 0.04096158 | 1 | 0.58502114 | 1 | 0.67692816 | 0 | 1 | 19 | 0.83975208 | FALSE | 1 | FALSE | 0.9698186 | FALSE | 1 | FALSE | 1 | FALSE | 1 | FALSE | 1 |
| 491 | 491 | 3.32857143 | 1.461129 | 18 | 0.00326263 | 20 | 0.00834152 | 15 | 0.26465537 | 11 | 0.93024295 | 19 | 2.84E-06 | 264 | 0.9999874 | FALSE | 0.15630083 | FALSE | 0.62691516 | FALSE | 1 | FALSE | 1 | TRUE | 0.00271857 | FALSE | 1 |
| 492 | 492 | 9 | 1.34164079 | 2 | 0.0256865 | 1 | 0.28036922 | 0 | 1 | 0 | 1 | 0 | 1 | 7 | 0.94326215 | FALSE | 0.4974559 | FALSE | 1 | FALSE | 1 | FALSE | 1 | FALSE | 1 | FALSE | 1 |
| 493 | 493 | 7.34482759 | 2.59011199 | 0 | 1 | 2 | 0.56459951 | 4 | 0.15426575 | 5 | 0.12605748 | 1 | 0.61824432 | 46 | 0.89115248 | FALSE | 1 | FALSE | 1 | FALSE | 1 | FALSE | 1 | FALSE | 1 | FALSE | 1 |
| 494 | 494 | 8.14634146 | 2.25843649 | 0 | 1 | 0 | 1 | 1 | 0.77786601 | 0 | 1 | 0 | 1 | 40 | 0.00798922 | FALSE | 1 | FALSE | 1 | FALSE | 1 | FALSE | 1 | FALSE | 1 | FALSE | 1 |
| 495 | 495 | 9 | 1.91942974 | 1 | 0.38989951 | 1 | 0.46503795 | 0 | 1 | 1 | 0.59104357 | 1 | 0.26986775 | 15 | 0.83648191 | FALSE | 1 | FALSE | 1 | FALSE | 1 | FALSE | 1 | FALSE | 1 | FALSE | 1 |
| 496 | 496 | 8.2173913 | 1.23792212 | 0 | 1 | 4 | 0.18354682 | 2 | 0.71546205 | 0 | 1 | 1 | 0.6822729 | 62 | 0.13493316 | FALSE | 1 | FALSE | 1 | FALSE | 1 | FALSE | 1 | FALSE | 1 | FALSE | 1 |
| 497 | 497 | 8.49425287 | 1.71453263 | 1 | 0.87304634 | 5 | 0.11181294 | 4 | 0.31552613 | 3 | 0.71030908 | 0 | 1 | 66 | 0.65229617 | FALSE | 1 | FALSE | 1 | FALSE | 1 | FALSE | 1 | FALSE | 1 | FALSE | 1 |
| 498 | 498 | 9.1 | 2.02237484 | 0 | 1 | 0 | 1 | 0 | 1 | 0 | 1 | 0 | 1 | 10 | 0.18273166 | FALSE | 1 | FALSE | 1 | FALSE | 1 | FALSE | 1 | FALSE | 1 | FALSE | 1 |
| 499 | 499 | 8.88 | 1.76227126 | 0 | 1 | 1 | 0.80796869 | 0 | 1 | 1 | 0.90544885 | 1 | 0.56379725 | 47 | 0.03571077 | FALSE | 1 | FALSE | 1 | FALSE | 1 | FALSE | 1 | FALSE | 1 | FALSE | 1 |
| 500 | 500 | 10.3968254 | 2.27156112 | 2 | 0.47501604 | 6 | 0.01453703 | 3 | 0.3858942 | 5 | 0.15434169 | 0 | 1 | 46 | 0.98794737 | FALSE | 1 | FALSE | 0.71309764 | FALSE | 1 | FALSE | 1 | FALSE | 1 | FALSE | 1 |
| 501 | 501 | 9.81818182 | 2.85460336 | 0 | 1 | 2 | 0.28941836 | 1 | 0.70185639 | 2 | 0.45159487 | 0 | 1 | 28 | 0.58681477 | FALSE | 1 | FALSE | 1 | FALSE | 1 | FALSE | 1 | FALSE | 1 | FALSE | 1 |
| 502 | 502 | 8.36363636 | 3.45813211 | 0 | 1 | 0 | 1 | 0 | 1 | 1 | 0.59104357 | 2 | 0.03799168 | 16 | 0.65565778 | FALSE | 1 | FALSE | 1 | FALSE | 1 | FALSE | 1 | FALSE | 0.87669812 | FALSE | 1 |
| 503 | 503 | 10.0909091 | 2.02055553 | 0 | 1 | 0 | 1 | 0 | 1 | 0 | 1 | 1 | 0.16638736 | 10 | 0.46859937 | FALSE | 1 | FALSE | 1 | FALSE | 1 | FALSE | 1 | FALSE | 1 | FALSE | 1 |
| 504 | 504 | 9.07692308 | 2.44828161 | 0 | 1 | 1 | 0.57534752 | 1 | 0.61440466 | 1 | 0.70601446 | 1 | 0.3499045 | 22 | 0.61628 | FALSE | 1 | FALSE | 1 | FALSE | 1 | FALSE | 1 | FALSE | 1 | FALSE | 1 |
| 505 | 505 | 9.53333333 | 1.49962958 | 1 | 0.32293355 | 1 | 0.38964394 | 0 | 1 | 0 | 1 | 0 | 1 | 13 | 0.57712805 | FALSE | 1 | FALSE | 1 | FALSE | 1 | FALSE | 1 | FALSE | 1 | FALSE | 1 |
| 506 | 506 | 8.5 | 1.83030052 | 0 | 1 | 2 | 0.1353446 | 1 | 0.51939841 | 2 | 0.23331368 | 0 | 1 | 15 | 0.92113679 | FALSE | 1 | FALSE | 1 | FALSE | 1 | FALSE | 1 | FALSE | 1 | FALSE | 1 |
| 507 | 507 | 7.41666667 | 2.16506351 | 1 | 0.59801736 | 1 | 0.68454804 | 2 | 0.35984046 | 4 | 0.07465533 | 0 | 1 | 27 | 0.91585095 | FALSE | 1 | FALSE | 1 | FALSE | 1 | FALSE | 1 | FALSE | 1 | FALSE | 1 |
| 508 | 508 | 8.3 | 3.22645316 | 1 | 0.22886536 | 1 | 0.28036922 | 1 | 0.30655256 | 0 | 1 | 0 | 1 | 7 | 0.94326215 | FALSE | 1 | FALSE | 1 | FALSE | 1 | FALSE | 1 | FALSE | 1 | FALSE | 1 |
| 509 | 509 | 9.81818182 | 2.51606409 | 0 | 1 | 1 | 0.5154318 | 0 | 1 | 1 | 0.64497439 | 0 | 1 | 20 | 0.30915202 | FALSE | 1 | FALSE | 1 | FALSE | 1 | FALSE | 1 | FALSE | 1 | FALSE | 1 |
| 510 | 510 | 6.42857143 | 2.87139304 | 1 | 0.66519151 | 0 | 1 | 0 | 1 | 2 | 0.5811115 | 1 | 0.50166523 | 38 | 0.19268329 | FALSE | 1 | FALSE | 1 | FALSE | 1 | FALSE | 1 | FALSE | 1 | FALSE | 1 |
| 511 | 511 | 7.40909091 | 3.05493775 | 1 | 0.4357638 | 1 | 0.5154318 | 0 | 1 | 1 | 0.64497439 | 0 | 1 | 19 | 0.54233419 | FALSE | 1 | FALSE | 1 | FALSE | 1 | FALSE | 1 | FALSE | 1 | FALSE | 1 |
| 512 | 512 | 7.21428571 | 4.16067597 | 2 | 0.16048326 | 4 | 0.01184672 | 0 | 1 | 2 | 0.3705495 | 0 | 1 | 20 | 0.97757517 | FALSE | 1 | FALSE | 0.67617265 | FALSE | 1 | FALSE | 1 | FALSE | 1 | FALSE | 1 |
| 513 | 513 | 10.4375 | 2.14967294 | 3 | 0.00725145 | 0 | 1 | 0 | 1 | 0 | 1 | 1 | 0.23264612 | 12 | 0.90922186 | FALSE | 0.25171133 | FALSE | 1 | FALSE | 1 | FALSE | 1 | FALSE | 1 | FALSE | 1 |
| 514 | 514 | 10 | 3.43511281 | 0 | 1 | 0 | 1 | 1 | 0.51939841 | 2 | 0.23331368 | 0 | 1 | 17 | 0.61750632 | FALSE | 1 | FALSE | 1 | FALSE | 1 | FALSE | 1 | FALSE | 1 | FALSE | 1 |
| 515 | 515 | 8.11538462 | 1.90801796 | 1 | 0.49162634 | 0 | 1 | 1 | 0.61440466 | 0 | 1 | 1 | 0.3499045 | 23 | 0.40372818 | FALSE | 1 | FALSE | 1 | FALSE | 1 | FALSE | 1 | FALSE | 1 | FALSE | 1 |
| 516 | 516 | 8.7027027 | 2.24047371 | 3 | 0.0681763 | 4 | 0.0305219 | 1 | 0.74264075 | 0 | 1 | 0 | 1 | 29 | 0.88820978 | FALSE | 0.74276693 | FALSE | 0.90921954 | FALSE | 1 | FALSE | 1 | FALSE | 1 | FALSE | 1 |
| 517 | 517 | 8.55555556 | 1.94999209 | 1 | 0.35728305 | 0 | 1 | 2 | 0.12280513 | 1 | 0.55062772 | 1 | 0.24525764 | 12 | 0.962093 | FALSE | 1 | FALSE | 1 | FALSE | 1 | FALSE | 1 | FALSE | 1 | FALSE | 1 |
| 518 | 518 | 8.24 | 2.8464715 | 1 | 0.47819878 | 0 | 1 | 1 | 0.59998149 | 2 | 0.31968852 | 0 | 1 | 21 | 0.64957167 | FALSE | 1 | FALSE | 1 | FALSE | 1 | FALSE | 1 | FALSE | 1 | FALSE | 1 |
| 519 | 519 | 10.9166667 | 2.51523138 | 1 | 0.46441868 | 1 | 0.54637326 | 1 | 0.58502114 | 3 | 0.09525947 | 0 | 1 | 18 | 0.93166998 | FALSE | 1 | FALSE | 1 | FALSE | 1 | FALSE | 1 | FALSE | 1 | FALSE | 1 |
| 520 | 520 | 8.85714286 | 2.09956264 | 1 | 0.42086991 | 1 | 0.49918218 | 0 | 1 | 1 | 0.62783522 | 2 | 0.04568221 | 16 | 0.90353674 | FALSE | 1 | FALSE | 1 | FALSE | 1 | FALSE | 1 | FALSE | 0.95710659 | FALSE | 1 |
| 521 | 521 | 8.37142857 | 2.59764415 | 1 | 0.59801736 | 0 | 1 | 2 | 0.35984046 | 3 | 0.21561745 | 1 | 0.44015537 | 28 | 0.83079716 | FALSE | 1 | FALSE | 1 | FALSE | 1 | FALSE | 1 | FALSE | 1 | FALSE | 1 |
| 522 | 522 | 10 | 1.8547237 | 2 | 0.13369621 | 2 | 0.19297191 | 0 | 1 | 2 | 0.31968852 | 0 | 1 | 19 | 0.91734261 | FALSE | 0.97964162 | FALSE | 1 | FALSE | 1 | FALSE | 1 | FALSE | 1 | FALSE | 1 |
| 523 | 523 | 8.23529412 | 2.10124366 | 1 | 0.35728305 | 0 | 1 | 1 | 0.46348836 | 0 | 1 | 0 | 1 | 15 | 0.49060435 | FALSE | 1 | FALSE | 1 | FALSE | 1 | FALSE | 1 | FALSE | 1 | FALSE | 1 |
| 524 | 524 | 9.71428571 | 3.41066754 | 0 | 1 | 2 | 0.14655134 | 2 | 0.17308361 | 0 | 1 | 0 | 1 | 17 | 0.77849076 | FALSE | 1 | FALSE | 1 | FALSE | 1 | FALSE | 1 | FALSE | 1 | FALSE | 1 |
| 525 | 525 | 8.17647059 | 2.74901681 | 0 | 1 | 1 | 0.42857733 | 0 | 1 | 0 | 1 | 0 | 1 | 16 | 0.23070182 | FALSE | 1 | FALSE | 1 | FALSE | 1 | FALSE | 1 | FALSE | 1 | FALSE | 1 |
| 526 | 526 | 8.41414141 | 1.30275622 | 5 | 0.09306405 | 4 | 0.36195667 | 3 | 0.66251153 | 5 | 0.43432519 | 2 | 0.45858138 | 75 | 0.91227221 | FALSE | 0.84055203 | FALSE | 1 | FALSE | 1 | FALSE | 1 | FALSE | 1 | FALSE | 1 |
| 527 | 527 | 4.95918367 | 2.22654838 | 5 | 0.10296654 | 5 | 0.20465359 | 2 | 0.86915992 | 2 | 0.9418724 | 4 | 0.07448782 | 79 | 0.82835389 | FALSE | 0.87864781 | FALSE | 1 | FALSE | 1 | FALSE | 1 | FALSE | 1 | FALSE | 1 |
| 528 | 528 | 8.8 | 2.31516738 | 0 | 1 | 0 | 1 | 0 | 1 | 0 | 1 | 1 | 0.2198258 | 14 | 0.29525409 | FALSE | 1 | FALSE | 1 | FALSE | 1 | FALSE | 1 | FALSE | 1 | FALSE | 1 |
| 529 | 529 | 9.42857143 | 1.63507474 | 0 | 1 | 1 | 0.36919824 | 0 | 1 | 0 | 1 | 0 | 1 | 13 | 0.33283729 | FALSE | 1 | FALSE | 1 | FALSE | 1 | FALSE | 1 | FALSE | 1 | FALSE | 1 |
| 530 | 530 | 11.0434783 | 2.17652018 | 0 | 1 | 1 | 0.53115654 | 1 | 0.5695037 | 1 | 0.6613267 | 0 | 1 | 20 | 0.50589402 | FALSE | 1 | FALSE | 1 | FALSE | 1 | FALSE | 1 | FALSE | 1 | FALSE | 1 |
| 531 | 531 | 11.5 | 1.5 | 0 | 1 | 1 | 0.56109832 | 3 | 0.05884411 | 2 | 0.31968852 | 0 | 1 | 19 | 0.91734261 | FALSE | 1 | FALSE | 1 | FALSE | 1 | FALSE | 1 | FALSE | 1 | FALSE | 1 |
| 532 | 532 | 8.2631579 | 0.84865871 | 0 | 1 | 0 | 1 | 0 | 1 | 1 | 0.59104357 | 1 | 0.26986775 | 17 | 0.41141543 | FALSE | 1 | FALSE | 1 | FALSE | 1 | FALSE | 1 | FALSE | 1 | FALSE | 1 |
| 533 | 533 | 11.2 | 2.18174242 | 0 | 1 | 1 | 0.28036922 | 0 | 1 | 1 | 0.37517262 | 0 | 1 | 8 | 0.80394719 | FALSE | 1 | FALSE | 1 | FALSE | 1 | FALSE | 1 | FALSE | 1 | FALSE | 1 |
| 534 | 534 | 5.72093023 | 3.14254808 | 2 | 0.30241117 | 1 | 0.75788114 | 3 | 0.19985581 | 1 | 0.86831187 | 1 | 0.50988734 | 35 | 0.7801651 | FALSE | 1 | FALSE | 1 | FALSE | 1 | FALSE | 1 | FALSE | 1 | FALSE | 1 |
| 535 | 535 | 11.1 | 3.01496269 | 1 | 0.22886536 | 0 | 1 | 1 | 0.30655256 | 1 | 0.37517262 | 0 | 1 | 7 | 0.94326215 | FALSE | 1 | FALSE | 1 | FALSE | 1 | FALSE | 1 | FALSE | 1 | FALSE | 1 |
| 536 | 536 | 9.33333333 | 1.97202659 | 0 | 1 | 1 | 0.179097 | 0 | 1 | 1 | 0.24578796 | 0 | 1 | 4 | 0.94743772 | FALSE | 1 | FALSE | 1 | FALSE | 1 | FALSE | 1 | FALSE | 1 | FALSE | 1 |
| 537 | 537 | 10.5625 | 2.69185693 | 2 | 0.06197875 | 1 | 0.40942991 | 1 | 0.44344817 | 0 | 1 | 1 | 0.23264612 | 11 | 0.9718531 | FALSE | 0.72532843 | FALSE | 1 | FALSE | 1 | FALSE | 1 | FALSE | 1 | FALSE | 1 |
| 538 | 538 | 10.1538462 | 1.16659622 | 0 | 1 | 0 | 1 | 1 | 0.37874417 | 1 | 0.45746571 | 1 | 0.19354507 | 10 | 0.86791642 | FALSE | 1 | FALSE | 1 | FALSE | 1 | FALSE | 1 | FALSE | 1 | FALSE | 1 |
| 539 | 539 | 9.625 | 2.42061459 | 0 | 1 | 0 | 1 | 1 | 0.44344817 | 1 | 0.52895088 | 0 | 1 | 14 | 0.53308261 | FALSE | 1 | FALSE | 1 | FALSE | 1 | FALSE | 1 | FALSE | 1 | FALSE | 1 |
| 540 | 540 | 9.65 | 3.33579076 | 0 | 1 | 1 | 0.46503795 | 0 | 1 | 1 | 0.59104357 | 0 | 1 | 17 | 0.41141543 | FALSE | 1 | FALSE | 1 | FALSE | 1 | FALSE | 1 | FALSE | 1 | FALSE | 1 |
| 541 | 541 | 8.65789474 | 2.06826034 | 3 | 0.07268126 | 3 | 0.12352882 | 3 | 0.15483587 | 1 | 0.83318955 | 0 | 1 | 28 | 0.97314817 | FALSE | 0.76727438 | FALSE | 1 | FALSE | 1 | FALSE | 1 | FALSE | 1 | FALSE | 1 |
| 542 | 542 | 7.90322581 | 4.54621316 | 0 | 1 | 2 | 0.2651851 | 0 | 1 | 3 | 0.16836568 | 0 | 1 | 26 | 0.64660109 | FALSE | 1 | FALSE | 1 | FALSE | 1 | FALSE | 1 | FALSE | 1 | FALSE | 1 |
| 543 | 543 | 9.94444444 | 1.44551243 | 1 | 0.74890551 | 2 | 0.51564773 | 5 | 0.04061975 | 1 | 0.91797304 | 1 | 0.58506217 | 43 | 0.80430518 | FALSE | 1 | FALSE | 1 | FALSE | 1 | FALSE | 1 | FALSE | 1 | FALSE | 1 |
| 544 | 544 | 9.07142857 | 1.86946462 | 0 | 1 | 0 | 1 | 0 | 1 | 3 | 0.13524571 | 1 | 0.37113067 | 24 | 0.55018107 | FALSE | 1 | FALSE | 1 | FALSE | 1 | FALSE | 1 | FALSE | 1 | FALSE | 1 |
| 545 | 545 | 9.375 | 1.37878268 | 3 | 0.02256792 | 2 | 0.18117183 | 0 | 1 | 1 | 0.67692816 | 0 | 1 | 18 | 0.93166998 | FALSE | 0.4692294 | FALSE | 1 | FALSE | 1 | FALSE | 1 | FALSE | 1 | FALSE | 1 |
| 546 | 546 | 10.8 | 2.03960781 | 0 | 1 | 1 | 0.12326188 | 1 | 0.13615783 | 0 | 1 | 0 | 1 | 2 | 0.98656791 | FALSE | 1 | FALSE | 1 | FALSE | 1 | FALSE | 1 | FALSE | 1 | FALSE | 1 |
| 547 | 547 | 3.87022901 | 2.29635272 | 2 | 0.83905117 | 4 | 0.58644376 | 6 | 0.29917534 | 8 | 0.22146501 | 2 | 0.61646357 | 104 | 0.76178277 | FALSE | 1 | FALSE | 1 | FALSE | 1 | FALSE | 1 | FALSE | 1 | FALSE | 1 |
| 548 | 548 | 5.74736842 | 3.63598512 | 3 | 0.44178082 | 0 | 1 | 3 | 0.66955123 | 1 | 0.98885678 | 0 | 1 | 88 | 0.01275701 | FALSE | 1 | FALSE | 1 | FALSE | 1 | FALSE | 1 | FALSE | 1 | FALSE | 1 |
| 549 | 549 | 6.52307692 | 2.43757567 | 2 | 0.49997462 | 5 | 0.05834767 | 3 | 0.41519671 | 3 | 0.57949569 | 0 | 1 | 52 | 0.87296036 | FALSE | 1 | FALSE | 1 | FALSE | 1 | FALSE | 1 | FALSE | 1 | FALSE | 1 |
| 550 | 550 | 9.14285714 | 2.06526176 | 3 | 0.0048942 | 0 | 1 | 1 | 0.40110418 | 2 | 0.13304705 | 0 | 1 | 8 | 0.99722299 | FALSE | 0.19848171 | FALSE | 1 | FALSE | 1 | FALSE | 1 | FALSE | 1 | FALSE | 1 |
| 551 | 551 | 8.53333333 | 1.08730043 | 0 | 1 | 1 | 0.38964394 | 0 | 1 | 1 | 0.50623194 | 1 | 0.2198258 | 12 | 0.80329917 | FALSE | 1 | FALSE | 1 | FALSE | 1 | FALSE | 1 | FALSE | 1 | FALSE | 1 |
| 552 | 552 | 3.62295082 | 1.60085975 | 4 | 0.3819299 | 2 | 0.91004611 | 4 | 0.64412798 | 5 | 0.66589028 | 1 | 0.86937741 | 106 | 0.26537725 | FALSE | 1 | FALSE | 1 | FALSE | 1 | FALSE | 1 | FALSE | 1 | FALSE | 1 |
| 553 | 553 | 8.57142857 | 2.55550626 | 1 | 0.42086991 | 1 | 0.49918218 | 1 | 0.53671415 | 1 | 0.62783522 | 0 | 1 | 17 | 0.77849076 | FALSE | 1 | FALSE | 1 | FALSE | 1 | FALSE | 1 | FALSE | 1 | FALSE | 1 |
| 554 | 554 | 6.66666667 | 2.59629366 | 2 | 0.15144274 | 2 | 0.21684246 | 1 | 0.62830986 | 3 | 0.12475952 | 0 | 1 | 19 | 0.98241223 | FALSE | 1 | FALSE | 1 | FALSE | 1 | FALSE | 1 | FALSE | 1 | FALSE | 1 |
| 555 | 555 | 7.47368421 | 1.25803053 | 5 | 0.01493383 | 7 | 0.00224617 | 4 | 0.147507 | 0 | 1 | 4 | 0.01388046 | 37 | 0.99992793 | FALSE | 0.38230603 | FALSE | 0.42431176 | FALSE | 1 | FALSE | 1 | FALSE | 0.64607242 | FALSE | 1 |
| 556 | 556 | 9.66666667 | 2.76887462 | 0 | 1 | 0 | 1 | 0 | 1 | 1 | 0.57131013 | 0 | 1 | 17 | 0.20329205 | FALSE | 1 | FALSE | 1 | FALSE | 1 | FALSE | 1 | FALSE | 1 | FALSE | 1 |
| 557 | 557 | 7.90909091 | 1.67628081 | 0 | 1 | 0 | 1 | 2 | 0.05708125 | 0 | 1 | 0 | 1 | 9 | 0.75985366 | FALSE | 1 | FALSE | 1 | FALSE | 1 | FALSE | 1 | FALSE | 1 | FALSE | 1 |
| 558 | 558 | 9 | 3.19089614 | 1 | 0.4357638 | 1 | 0.5154318 | 1 | 0.55340851 | 0 | 1 | 1 | 0.30529951 | 18 | 0.74744813 | FALSE | 1 | FALSE | 1 | FALSE | 1 | FALSE | 1 | FALSE | 1 | FALSE | 1 |
| 559 | 559 | 9.0625 | 2.78318051 | 1 | 0.81177881 | 4 | 0.15197537 | 3 | 0.40547364 | 4 | 0.33842086 | 1 | 0.65460803 | 51 | 0.88464035 | FALSE | 1 | FALSE | 1 | FALSE | 1 | FALSE | 1 | FALSE | 1 | FALSE | 1 |
| 560 | 560 | 7.09302326 | 3.75910648 | 1 | 0.64722818 | 0 | 1 | 0 | 1 | 1 | 0.84823834 | 1 | 0.48480878 | 37 | 0.10889911 | FALSE | 1 | FALSE | 1 | FALSE | 1 | FALSE | 1 | FALSE | 1 | FALSE | 1 |
| 561 | 561 | 3.21302999 | 1.23131991 | 29 | 0.19839898 | 36 | 0.19414769 | 36 | 0.42475579 | 43 | 0.6055609 | 19 | 0.22435228 | 800 | 0.89426102 | FALSE | 1 | FALSE | 1 | FALSE | 1 | FALSE | 1 | FALSE | 1 | FALSE | 1 |
| 562 | 562 | 6.12048193 | 0.99876539 | 5 | 0.05627772 | 4 | 0.26638189 | 4 | 0.33196391 | 5 | 0.31464362 | 0 | 1 | 63 | 0.95896498 | FALSE | 0.69536509 | FALSE | 1 | FALSE | 1 | FALSE | 1 | FALSE | 1 | FALSE | 1 |

| | | | | | | | | | | | | | | | | | | | | | | | | | | | |
|---|---|---|---|---|---|---|---|---|---|---|---|---|---|---|---|---|---|---|---|---|---|---|---|---|---|---|---|
| 563 | 563 | 7.54166667 | 3.95789471 | 0 | 1 | 0 | 1 | 0 | 1 | 0 | 1 | 0 | 1 | 24 | 0.01683964 | FALSE | 1 | FALSE | 1 | FALSE | 1 | FALSE | 1 | FALSE | 1 | FALSE | 1 |
| 564 | 564 | 6.01219512 | 2.35037051 | 2 | 0.59898979 | 4 | 0.24495326 | 2 | 0.77659406 | 3 | 0.7021069 | 1 | 0.72664912 | 66 | 0.55329149 | FALSE | 1 | FALSE | 1 | FALSE | 1 | FALSE | 1 | FALSE | 1 | FALSE | 1 |
| 565 | 565 | 6.89285714 | 3.44693742 | 1 | 0.51745948 | 1 | 0.60247911 | 1 | 0.64171563 | 1 | 0.73248983 | 0 | 1 | 24 | 0.55018107 | FALSE | 1 | FALSE | 1 | FALSE | 1 | FALSE | 1 | FALSE | 1 | FALSE | 1 |
| 566 | 566 | 10.125 | 2.75850956 | 0 | 1 | 1 | 0.23139161 | 0 | 1 | 0 | 1 | 0 | 1 | 7 | 0.63752397 | FALSE | 1 | FALSE | 1 | FALSE | 1 | FALSE | 1 | FALSE | 1 | FALSE | 1 |
| 567 | 567 | 9.42857143 | 2.4215585 | 0 | 1 | 0 | 1 | 0 | 1 | 0 | 1 | 1 | 0.29368245 | 20 | 0.13763971 | FALSE | 1 | FALSE | 1 | FALSE | 1 | FALSE | 1 | FALSE | 1 | FALSE | 1 |
| 568 | 568 | 12 | 2.74873708 | 1 | 0.20854748 | 0 | 1 | 0 | 1 | 1 | 0.34506517 | 0 | 1 | 7 | 0.84574168 | FALSE | 1 | FALSE | 1 | FALSE | 1 | FALSE | 1 | FALSE | 1 | FALSE | 1 |
| 569 | 569 | 9.14285714 | 2.69542318 | 0 | 1 | 0 | 1 | 1 | 0.53671415 | 0 | 1 | 0 | 1 | 20 | 0.13763971 | FALSE | 1 | FALSE | 1 | FALSE | 1 | FALSE | 1 | FALSE | 1 | FALSE | 1 |
| 570 | 570 | 8 | 4.01467895 | 0 | 1 | 0 | 1 | 0 | 1 | 1 | 0.55062772 | 0 | 1 | 16 | 0.23070182 | FALSE | 1 | FALSE | 1 | FALSE | 1 | FALSE | 1 | FALSE | 1 | FALSE | 1 |
| 571 | 571 | 10.45 | 2.06094639 | 0 | 1 | 1 | 0.48239024 | 0 | 1 | 0 | 1 | 1 | 0.28187291 | 18 | 0.37504645 | FALSE | 1 | FALSE | 1 | FALSE | 1 | FALSE | 1 | FALSE | 1 | FALSE | 1 |
| 572 | 572 | 10.375 | 2.0577597 | 1 | 0.3050805 | 0 | 1 | 1 | 0.40110418 | 0 | 1 | 0 | 1 | 12 | 0.62236227 | FALSE | 1 | FALSE | 1 | FALSE | 1 | FALSE | 1 | FALSE | 1 | FALSE | 1 |
| 573 | 573 | 10.2692308 | 2.8360006 | 0 | 1 | 0 | 1 | 2 | 0.23940226 | 0 | 1 | 0 | 1 | 24 | 0.20425579 | FALSE | 1 | FALSE | 1 | FALSE | 1 | FALSE | 1 | FALSE | 1 | FALSE | 1 |
| 574 | 574 | 7.79411765 | 3.55444553 | 0 | 1 | 2 | 0.30150529 | 1 | 0.71261921 | 1 | 0.7984848 | 1 | 0.43077488 | 29 | 0.55707911 | FALSE | 1 | FALSE | 1 | FALSE | 1 | FALSE | 1 | FALSE | 1 | FALSE | 1 |
| 575 | 575 | 7.04347826 | 2.32151114 | 1 | 0.69843502 | 2 | 0.44132825 | 1 | 0.81522035 | 2 | 0.63103861 | 3 | 0.03911573 | 37 | 0.82909673 | FALSE | 1 | FALSE | 1 | FALSE | 1 | FALSE | 1 | FALSE | 0.88763447 | FALSE | 1 |
| 576 | 576 | 7.1 | 3.9694668 | 0 | 1 | 1 | 0.6278847 | 1 | 0.66709967 | 1 | 0.75658797 | 0 | 1 | 27 | 0.28862208 | FALSE | 1 | FALSE | 1 | FALSE | 1 | FALSE | 1 | FALSE | 1 | FALSE | 1 |
| 577 | 577 | 8.36 | 3.50861796 | 1 | 0.47819878 | 2 | 0.19297191 | 3 | 0.05884411 | 1 | 0.69181315 | 0 | 1 | 18 | 0.96851178 | FALSE | 1 | FALSE | 1 | FALSE | 1 | FALSE | 1 | FALSE | 1 | FALSE | 1 |
| 578 | 578 | 9.92 | 2.93829883 | 1 | 0.47819878 | 4 | 0.00789279 | 2 | 0.22597287 | 3 | 0.10474541 | 1 | 0.33902672 | 14 | 0.99985788 | FALSE | 1 | FALSE | 0.6167796 | FALSE | 1 | FALSE | 1 | FALSE | 1 | FALSE | 1 |
| 579 | 579 | 7.15625 | 3.31765519 | 3 | 0.0476977 | 2 | 0.27730767 | 3 | 0.10599423 | 3 | 0.17988885 | 2 | 0.096172 | 19 | 0.9998606 | FALSE | 0.649301 | FALSE | 1 | FALSE | 1 | FALSE | 1 | FALSE | 1 | FALSE | 1 |
| 580 | 580 | 10.4137931 | 2.2971833 | 2 | 0.16962005 | 2 | 0.24095584 | 1 | 0.65463982 | 2 | 0.3871811 | 0 | 1 | 22 | 0.92902102 | FALSE | 1 | FALSE | 1 | FALSE | 1 | FALSE | 1 | FALSE | 1 | FALSE | 1 |
| 581 | 581 | 10.7368421 | 2.61256287 | 1 | 0.38989951 | 1 | 0.46503795 | 0 | 1 | 2 | 0.21610812 | 0 | 1 | 15 | 0.83648191 | FALSE | 1 | FALSE | 1 | FALSE | 1 | FALSE | 1 | FALSE | 1 | FALSE | 1 |
| 582 | 582 | 10.0645161 | 2.15429779 | 1 | 0.55378205 | 4 | 0.01688692 | 3 | 0.09854319 | 1 | 0.76781284 | 2 | 0.09110253 | 20 | 0.9984454 | FALSE | 1 | FALSE | 0.73194511 | FALSE | 1 | FALSE | 1 | FALSE | 1 | FALSE | 1 |
| 583 | 583 | 11.5 | 3.70809924 | 1 | 0.34033054 | 1 | 0.40942991 | 2 | 0.11087977 | 1 | 0.52895088 | 0 | 1 | 11 | 0.9718531 | FALSE | 1 | FALSE | 1 | FALSE | 1 | FALSE | 1 | FALSE | 1 | FALSE | 1 |
| 584 | 584 | 9.90909091 | 2.46630181 | 1 | 0.22886536 | 0 | 1 | 0 | 1 | 0 | 1 | 0 | 1 | 9 | 0.5215547 | FALSE | 1 | FALSE | 1 | FALSE | 1 | FALSE | 1 | FALSE | 1 | FALSE | 1 |
| 585 | 585 | 12.0833333 | 3.4268142 | 0 | 1 | 1 | 0.32623933 | 0 | 1 | 0 | 1 | 1 | 0.18007762 | 10 | 0.71436964 | FALSE | 1 | FALSE | 1 | FALSE | 1 | FALSE | 1 | FALSE | 1 | FALSE | 1 |
| 586 | 586 | 10.6923077 | 3.36005072 | 0 | 1 | 0 | 1 | 0 | 1 | 0 | 1 | 0 | 1 | 13 | 0.10967906 | FALSE | 1 | FALSE | 1 | FALSE | 1 | FALSE | 1 | FALSE | 1 | FALSE | 1 |
| 587 | 587 | 8.93333333 | 2.64491126 | 0 | 1 | 1 | 0.38964394 | 0 | 1 | 0 | 1 | 1 | 0.2198258 | 13 | 0.57712805 | FALSE | 1 | FALSE | 1 | FALSE | 1 | FALSE | 1 | FALSE | 1 | FALSE | 1 |
| 588 | 588 | 9.6 | 1.74355958 | 0 | 1 | 0 | 1 | 0 | 1 | 0 | 1 | 0 | 1 | 5 | 0.42761893 | FALSE | 1 | FALSE | 1 | FALSE | 1 | FALSE | 1 | FALSE | 1 | FALSE | 1 |
| 589 | 589 | 7.17679558 | 1.15711238 | 6 | 0.3194466 | 5 | 0.70279906 | 6 | 0.63819571 | 9 | 0.45206534 | 1 | 0.95185195 | 154 | 0.44496718 | FALSE | 1 | FALSE | 1 | FALSE | 1 | FALSE | 1 | FALSE | 1 | FALSE | 1 |
| 590 | 590 | 9.28571429 | 2.38532931 | 0 | 1 | 1 | 0.68454804 | 1 | 0.72299506 | 2 | 0.48233989 | 1 | 0.44015537 | 30 | 0.52765882 | FALSE | 1 | FALSE | 1 | FALSE | 1 | FALSE | 1 | FALSE | 1 | FALSE | 1 |
| 591 | 591 | 9.43396226 | 1.9475135 | 4 | 0.04620161 | 2 | 0.51564773 | 1 | 0.85724013 | 1 | 0.91797304 | 0 | 1 | 45 | 0.5504765 | FALSE | 0.649301 | FALSE | 1 | FALSE | 1 | FALSE | 1 | FALSE | 1 | FALSE | 1 |
| 592 | 592 | 3.49350649 | 3.28958757 | 2 | 0.59191779 | 1 | 0.92163631 | 2 | 0.77041438 | 3 | 0.69372972 | 1 | 0.72203879 | 68 | 0.21618945 | FALSE | 1 | FALSE | 1 | FALSE | 1 | FALSE | 1 | FALSE | 1 | FALSE | 1 |
| 593 | 593 | 10.9047619 | 2.20183391 | 0 | 1 | 0 | 1 | 0 | 1 | 4 | 0.01177595 | 0 | 1 | 16 | 0.80826361 | FALSE | 1 | FALSE | 1 | FALSE | 1 | FALSE | 1 | FALSE | 1 | FALSE | 1 |
| 594 | 594 | 9.28125 | 2.00365876 | 0 | 1 | 3 | 0.08334648 | 0 | 1 | 1 | 0.77852168 | 0 | 1 | 28 | 0.42532177 | FALSE | 1 | FALSE | 1 | FALSE | 1 | FALSE | 1 | FALSE | 1 | FALSE | 1 |
| 595 | 595 | 7.6 | 1.42828569 | 0 | 1 | 1 | 0.48239024 | 0 | 1 | 0 | 1 | 0 | 1 | 19 | 0.15700255 | FALSE | 1 | FALSE | 1 | FALSE | 1 | FALSE | 1 | FALSE | 1 | FALSE | 1 |
| 596 | 596 | 11.8823529 | 3.47905249 | 1 | 0.35728305 | 2 | 0.10307112 | 1 | 0.46348836 | 1 | 0.55062772 | 0 | 1 | 12 | 0.962093 | FALSE | 1 | FALSE | 1 | FALSE | 1 | FALSE | 1 | FALSE | 1 | FALSE | 1 |
| 597 | 597 | 9.5 | 2.40947205 | 0 | 1 | 0 | 1 | 0 | 1 | 2 | 0.19902682 | 0 | 1 | 16 | 0.44998001 | FALSE | 1 | FALSE | 1 | FALSE | 1 | FALSE | 1 | FALSE | 1 | FALSE | 1 |
| 598 | 598 | 9.64285714 | 3.22062866 | 0 | 1 | 1 | 0.36919824 | 1 | 0.40110418 | 1 | 0.48242099 | 1 | 0.20679328 | 10 | 0.94579623 | FALSE | 1 | FALSE | 1 | FALSE | 1 | FALSE | 1 | FALSE | 1 | FALSE | 1 |
| 599 | 599 | 8.6 | 2.67207784 | 2 | 0.09187136 | 0 | 1 | 1 | 0.51939841 | 1 | 0.60987146 | 1 | 0.28187291 | 15 | 0.92113679 | FALSE | 0.83248965 | FALSE | 1 | FALSE | 1 | FALSE | 1 | FALSE | 1 | FALSE | 1 |
| 600 | 600 | 10.4347826 | 3.26810791 | 1 | 0.45027683 | 2 | 0.16948764 | 1 | 0.5695037 | 0 | 1 | 0 | 1 | 19 | 0.71542496 | FALSE | 1 | FALSE | 1 | FALSE | 1 | FALSE | 1 | FALSE | 1 | FALSE | 1 |
| 601 | 601 | 11.0666667 | 2.29395341 | 0 | 1 | 0 | 1 | 0 | 1 | 0 | 1 | 0 | 1 | 15 | 0.07803154 | FALSE | 1 | FALSE | 1 | FALSE | 1 | FALSE | 1 | FALSE | 1 | FALSE | 1 |
| 602 | 602 | 8.63157895 | 5.07014509 | 1 | 0.38989951 | 1 | 0.46503795 | 0 | 1 | 0 | 1 | 0 | 1 | 17 | 0.41141543 | FALSE | 1 | FALSE | 1 | FALSE | 1 | FALSE | 1 | FALSE | 1 | FALSE | 1 |
| 603 | 603 | 10.3103448 | 3.24943972 | 1 | 0.52988231 | 3 | 0.06581417 | 1 | 0.65463982 | 1 | 0.74482229 | 0 | 1 | 23 | 0.84462957 | FALSE | 1 | FALSE | 1 | FALSE | 1 | FALSE | 1 | FALSE | 1 | FALSE | 1 |
| 604 | 604 | 10.3333333 | 2.30940108 | 0 | 1 | 1 | 0.44710673 | 1 | 0.48280983 | 0 | 1 | 0 | 1 | 16 | 0.44998001 | FALSE | 1 | FALSE | 1 | FALSE | 1 | FALSE | 1 | FALSE | 1 | FALSE | 1 |
| 605 | 605 | 10.3478261 | 2.13884763 | 1 | 0.45027683 | 0 | 1 | 1 | 0.5695037 | 0 | 1 | 0 | 1 | 21 | 0.27964072 | FALSE | 1 | FALSE | 1 | FALSE | 1 | FALSE | 1 | FALSE | 1 | FALSE | 1 |
| 606 | 606 | 9.56756757 | 3.12393152 | 1 | 0.6184714 | 0 | 1 | 1 | 0.74264075 | 2 | 0.51200191 | 1 | 0.45845922 | 32 | 0.47030992 | FALSE | 1 | FALSE | 1 | FALSE | 1 | FALSE | 1 | FALSE | 1 | FALSE | 1 |
| 607 | 607 | 10.125 | 2.89125838 | 1 | 0.34033054 | 0 | 1 | 0 | 1 | 1 | 0.52895088 | 0 | 1 | 14 | 0.53308261 | FALSE | 1 | FALSE | 1 | FALSE | 1 | FALSE | 1 | FALSE | 1 | FALSE | 1 |
| 608 | 608 | 5.46153846 | 2.58046064 | 1 | 0.63789042 | 0 | 1 | 2 | 0.41139386 | 5 | 0.03160407 | 1 | 0.4761699 | 30 | 0.92795554 | FALSE | 1 | FALSE | 1 | FALSE | 1 | FALSE | 1 | FALSE | 1 | FALSE | 1 |
| 609 | 609 | 4.16326531 | 2.50181109 | 12 | 6.57E-06 | 6 | 0.09708054 | 2 | 0.87289187 | 1 | 0.99034238 | 2 | 0.48029053 | 75 | 0.98575989 | TRUE | 0.00299395 | FALSE | 1 | FALSE | 1 | FALSE | 1 | FALSE | 1 | FALSE | 1 |
| 610 | 610 | 8.81818182 | 3.0989201 | 1 | 0.4357638 | 0 | 1 | 1 | 0.55340851 | 1 | 0.64497439 | 0 | 1 | 19 | 0.54233419 | FALSE | 1 | FALSE | 1 | FALSE | 1 | FALSE | 1 | FALSE | 1 | FALSE | 1 |
| 611 | 611 | 7.81818182 | 2.93313924 | 2 | 0.10809809 | 1 | 0.5154318 | 0 | 1 | 1 | 0.64497439 | 0 | 1 | 18 | 0.74744813 | FALSE | 0.88820416 | FALSE | 1 | FALSE | 1 | FALSE | 1 | FALSE | 1 | FALSE | 1 |
| 612 | 612 | 8.4375 | 1.36787929 | 0 | 1 | 1 | 0.40942991 | 1 | 0.44344817 | 0 | 1 | 1 | 0.23264612 | 13 | 0.76799161 | FALSE | 1 | FALSE | 1 | FALSE | 1 | FALSE | 1 | FALSE | 1 | FALSE | 1 |
| 613 | 613 | 9.05 | 3.20117166 | 1 | 0.40558521 | 1 | 0.48239024 | 0 | 1 | 0 | 1 | 0 | 1 | 18 | 0.37504645 | FALSE | 1 | FALSE | 1 | FALSE | 1 | FALSE | 1 | FALSE | 1 | FALSE | 1 |
| 614 | 614 | 11.7777778 | 1.87247773 | 2 | 0.02089722 | 0 | 1 | 0 | 1 | 3 | 0.00654627 | 0 | 1 | 4 | 0.99921376 | FALSE | 0.45408499 | FALSE | 1 | FALSE | 1 | FALSE | 1 | FALSE | 1 | FALSE | 1 |
| 615 | 615 | 9.73684211 | 2.69193934 | 0 | 1 | 0 | 1 | 1 | 0.50143826 | 0 | 1 | 0 | 1 | 18 | 0.17880694 | FALSE | 1 | FALSE | 1 | FALSE | 1 | FALSE | 1 | FALSE | 1 | FALSE | 1 |
| 616 | 616 | 9.76923077 | 1.42255708 | 0 | 1 | 1 | 0.3480709 | 1 | 0.37874417 | 1 | 0.45746571 | 0 | 1 | 10 | 0.86791642 | FALSE | 1 | FALSE | 1 | FALSE | 1 | FALSE | 1 | FALSE | 1 | FALSE | 1 |
| 617 | 617 | 9.09090909 | 2.46630181 | 2 | 0.03087289 | 0 | 1 | 0 | 1 | 0 | 1 | 0 | 1 | 9 | 0.75985366 | FALSE | 0.53356688 | FALSE | 1 | FALSE | 1 | FALSE | 1 | FALSE | 1 | FALSE | 1 |
| 618 | 618 | 9.88888889 | 1.91162784 | 0 | 1 | 0 | 1 | 0 | 1 | 0 | 1 | 0 | 1 | 9 | 0.21661397 | FALSE | 1 | FALSE | 1 | FALSE | 1 | FALSE | 1 | FALSE | 1 | FALSE | 1 |
| 619 | 619 | 7.73684211 | 2.12360432 | 0 | 1 | 0 | 1 | 1 | 0.48280983 | 0 | 1 | 0 | 1 | 17 | 0.20329205 | FALSE | 1 | FALSE | 1 | FALSE | 1 | FALSE | 1 | FALSE | 1 | FALSE | 1 |
| 620 | 620 | 8.82352941 | 2.35734881 | 0 | 1 | 0 | 1 | 1 | 0.46348836 | 0 | 1 | 0 | 1 | 16 | 0.23070182 | FALSE | 1 | FALSE | 1 | FALSE | 1 | FALSE | 1 | FALSE | 1 | FALSE | 1 |
| 621 | 621 | 9.04347826 | 3.96153529 | 0 | 1 | 1 | 0.53115654 | 1 | 0.5695037 | 0 | 1 | 0 | 1 | 21 | 0.27964072 | FALSE | 1 | FALSE | 1 | FALSE | 1 | FALSE | 1 | FALSE | 1 | FALSE | 1 |
| 622 | 622 | 4.60309278 | 1.84228419 | 2 | 0.9579149 | 6 | 0.57768024 | 8 | 0.36877444 | 6 | 0.8727896 | 4 | 0.37532838 | 163 | 0.27410001 | FALSE | 1 | FALSE | 1 | FALSE | 1 | FALSE | 1 | FALSE | 1 | FALSE | 1 |
| 623 | 623 | 8.175 | 1.84238297 | 3 | 0.08207756 | 1 | 0.7326167 | 1 | 0.76953771 | 4 | 0.10945471 | 0 | 1 | 31 | 0.91676852 | FALSE | 0.79236883 | FALSE | 1 | FALSE | 1 | FALSE | 1 | FALSE | 1 | FALSE | 1 |
| 624 | 624 | 8.17647059 | 2.35734881 | 0 | 1 | 1 | 0.42857733 | 2 | 0.12280513 | 1 | 0.55062772 | 0 | 1 | 13 | 0.88719222 | FALSE | 1 | FALSE | 1 | FALSE | 1 | FALSE | 1 | FALSE | 1 | FALSE | 1 |
| 625 | 625 | 9.43037975 | 2.33673766 | 0 | 1 | 3 | 0.47338398 | 2 | 0.78262837 | 2 | 0.8842511 | 0 | 1 | 72 | 0.05857603 | FALSE | 1 | FALSE | 1 | FALSE | 1 | FALSE | 1 | FALSE | 1 | FALSE | 1 |
| 626 | 626 | 7.24242424 | 2.65181817 | 3 | 0.0476977 | 0 | 1 | 1 | 0.69069222 | 1 | 0.77852168 | 0 | 1 | 27 | 0.61671133 | FALSE | 0.649301 | FALSE | 1 | FALSE | 1 | FALSE | 1 | FALSE | 1 | FALSE | 1 |
| 627 | 627 | 10.375 | 3.03881145 | 0 | 1 | 0 | 1 | 0 | 1 | 0 | 1 | 0 | 1 | 8 | 0.25677166 | FALSE | 1 | FALSE | 1 | FALSE | 1 | FALSE | 1 | FALSE | 1 | FALSE | 1 |
| 628 | 628 | 9.91666667 | 1.70579079 | 0 | 1 | 0 | 1 | 0 | 1 | 1 | 0.43131129 | 0 | 1 | 11 | 0.41944532 | FALSE | 1 | FALSE | 1 | FALSE | 1 | FALSE | 1 | FALSE | 1 | FALSE | 1 |
| 629 | 629 | 10.9090909 | 3.3336088 | 0 | 1 | 0 | 1 | 1 | 0.70185639 | 1 | 0.78873814 | 2 | 0.10132035 | 29 | 0.39648374 | FALSE | 1 | FALSE | 1 | FALSE | 1 | FALSE | 1 | FALSE | 1 | FALSE | 1 |
| 630 | 630 | 8.38888889 | 3.33564735 | 1 | 0.37380237 | 0 | 1 | 1 | 0.48280983 | 2 | 0.19902682 | 0 | 1 | 14 | 0.86287418 | FALSE | 1 | FALSE | 1 | FALSE | 1 | FALSE | 1 | FALSE | 1 | FALSE | 1 |
| 631 | 631 | 9.83333333 | 2.19215774 | 0 | 1 | 0 | 1 | 0 | 1 | 1 | 0.40390032 | 0 | 1 | 10 | 0.46859937 | FALSE | 1 | FALSE | 1 | FALSE | 1 | FALSE | 1 | FALSE | 1 | FALSE | 1 |
| 632 | 632 | 7.95454546 | 3.18344115 | 1 | 0.4357638 | 0 | 1 | 2 | 0.18613256 | 0 | 1 | 0 | 1 | 19 | 0.54233419 | FALSE | 1 | FALSE | 1 | FALSE | 1 | FALSE | 1 | FALSE | 1 | FALSE | 1 |
| 633 | 633 | 9.38461539 | 1.98216904 | 0 | 1 | 0 | 1 | 1 | 0.37874417 | 0 | 1 | 0 | 1 | 12 | 0.374198 | FALSE | 1 | FALSE | 1 | FALSE | 1 | FALSE | 1 | FALSE | 1 | FALSE | 1 |
| 634 | 634 | 8.86363636 | 2.4363806 | 0 | 1 | 1 | 0.5154318 | 1 | 0.55340851 | 0 | 1 | 1 | 0.30529951 | 19 | 0.54233419 | FALSE | 1 | FALSE | 1 | FALSE | 1 | FALSE | 1 | FALSE | 1 | FALSE | 1 |
| 635 | 635 | 8.11627907 | 2.09328164 | 0 | 1 | 5 | 0.01196839 | 0 | 1 | 1 | 0.86831187 | 3 | 0.03293335 | 34 | 0.87722689 | FALSE | 1 | FALSE | 0.67617265 | FALSE | 1 | FALSE | 1 | FALSE | 0.86063198 | FALSE | 1 |
| 636 | 636 | 9.44444444 | 2.58676594 | 0 | 1 | 0 | 1 | 0 | 1 | 1 | 0.34506517 | 1 | 0.1383237 | 7 | 0.84574168 | FALSE | 1 | FALSE | 1 | FALSE | 1 | FALSE | 1 | FALSE | 1 | FALSE | 1 |
| 637 | 637 | 9.60526316 | 4.01667232 | 3 | 0.07268126 | 3 | 0.12352882 | 1 | 0.75193685 | 0 | 1 | 2 | 0.12811285 | 29 | 0.93814695 | FALSE | 0.76727438 | FALSE | 1 | FALSE | 1 | FALSE | 1 | FALSE | 1 | FALSE | 1 |
| 638 | 638 | 7.3974359 | 1.58793321 | 1 | 0.86966563 | 3 | 0.46497905 | 5 | 0.14803148 | 0 | 1 | 3 | 0.13569451 | 66 | 0.55329149 | FALSE | 1 | FALSE | 1 | FALSE | 1 | FALSE | 1 | FALSE | 1 | FALSE | 1 |
| 639 | 639 | 9.41666667 | 2.56444709 | 0 | 1 | 1 | 0.32623933 | 0 | 1 | 1 | 0.43131129 | 0 | 1 | 10 | 0.71436964 | FALSE | 1 | FALSE | 1 | FALSE | 1 | FALSE | 1 | FALSE | 1 | FALSE | 1 |
| 640 | 640 | 6.89285714 | 3.41571249 | 2 | 0.15144274 | 0 | 1 | 0 | 1 | 2 | 0.35374209 | 2 | 0.07169515 | 21 | 0.88395936 | FALSE | 1 | FALSE | 1 | FALSE | 1 | FALSE | 1 | FALSE | 1 | FALSE | 1 |
| 641 | 641 | 8.94594595 | 0.98491174 | 1 | 0.60837712 | 3 | 0.10944025 | 2 | 0.37289966 | 2 | 0.49731009 | 1 | 0.44938266 | 27 | 0.95561238 | FALSE | 1 | FALSE | 1 | FALSE | 1 | FALSE | 1 | FALSE | 1 | FALSE | 1 |
| 642 | 642 | 8.83870968 | 3.10147493 | 0 | 1 | 1 | 0.63997468 | 2 | 0.30673593 | 2 | 0.41984187 | 0 | 1 | 26 | 0.64660109 | FALSE | 1 | FALSE | 1 | FALSE | 1 | FALSE | 1 | FALSE | 1 | FALSE | 1 |
| 643 | 643 | 7.26666667 | 1.28927197 | 0 | 1 | 0 | 1 | 1 | 0.42266264 | 2 | 0.14906763 | 0 | 1 | 12 | 0.80329917 | FALSE | 1 | FALSE | 1 | FALSE | 1 | FALSE | 1 | FALSE | 1 | FALSE | 1 |
| 644 | 644 | 10.2222222 | 2.52884593 | 1 | 0.20854748 | 0 | 1 | 0 | 1 | 1 | 0.34506517 | 0 | 1 | 7 | 0.84574168 | FALSE | 1 | FALSE | 1 | FALSE | 1 | FALSE | 1 | FALSE | 1 | FALSE | 1 |
| 645 | 645 | 8.4 | 1.40475383 | 1 | 0.54198714 | 3 | 0.07145121 | 0 | 1 | 1 | 0.75658797 | 0 | 1 | 25 | 0.67630471 | FALSE | 1 | FALSE | 1 | FALSE | 1 | FALSE | 1 | FALSE | 1 | FALSE | 1 |
| 646 | 646 | 10.1538462 | 1.83328851 | 0 | 1 | 0 | 1 | 0 | 1 | 2 | 0.1174583 | 1 | 0.19354507 | 10 | 0.86791642 | FALSE | 1 | FALSE | 1 | FALSE | 1 | FALSE | 1 | FALSE | 1 | FALSE | 1 |
| 647 | 647 | 7.84444444 | 2.65794871 | 0 | 1 | 0 | 1 | 2 | 0.48463745 | 4 | 0.15016749 | 0 | 1 | 39 | 0.4317533 | FALSE | 1 | FALSE | 1 | FALSE | 1 | FALSE | 1 | FALSE | 1 | FALSE | 1 |
| 648 | 648 | 12.2692308 | 2.52003804 | 1 | 0.49162634 | 1 | 0.57534752 | 1 | 0.61440466 | 1 | 0.70601446 | 1 | 0.3499045 | 21 | 0.78939668 | FALSE | 1 | FALSE | 1 | FALSE | 1 | FALSE | 1 | FALSE | 1 | FALSE | 1 |
| 649 | 649 | 7.06779661 | 2.99640693 | 3 | 0.19211919 | 1 | 0.85750546 | 3 | 0.35626822 | 2 | 0.76144533 | 3 | 0.07204991 | 47 | 0.87928796 | FALSE | 1 | FALSE | 1 | FALSE | 1 | FALSE | 1 | FALSE | 1 | FALSE | 1 |
| 650 | 650 | 9.55555556 | 3.35364184 | 3 | 0.01017712 | 1 | 0.44710673 | 0 | 1 | 0 | 1 | 1 | 0.25766374 | 13 | 0.95041913 | FALSE | 0.30467829 | FALSE | 1 | FALSE | 1 | FALSE | 1 | FALSE | 1 | FALSE | 1 |
| 651 | 651 | 9.14285714 | 1.55182578 | 0 | 1 | 1 | 0.20567256 | 0 | 1 | 1 | 0.28044368 | 0 | 1 | 5 | 0.91847773 | FALSE | 1 | FALSE | 1 | FALSE | 1 | FALSE | 1 | FALSE | 1 | FALSE | 1 |
| 652 | 652 | 6.93617021 | 2.38259875 | 1 | 0.91216464 | 8 | 0.01013782 | 3 | 0.65536448 | 5 | 0.42517356 | 2 | 0.45307384 | 74 | 0.91970269 | FALSE | 1 | FALSE | 0.67617265 | FALSE | 1 | FALSE | 1 | FALSE | 1 | FALSE | 1 |
| 653 | 653 | 8 | 3.7583241 | 0 | 1 | 0 | 1 | 1 | 0.44344817 | 2 | 0.16544789 | 1 | 0.23264612 | 12 | 0.90922186 | FALSE | 1 | FALSE | 1 | FALSE | 1 | FALSE | 1 | FALSE | 1 | FALSE | 1 |
| 654 | 654 | 11.0588235 | 2.04279475 | 0 | 1 | 0 | 1 | 3 | 0.02143959 | 0 | 1 | 0 | 1 | 14 | 0.7313145 | FALSE | 1 | FALSE | 1 | FALSE | 1 | FALSE | 1 | FALSE | 1 | FALSE | 1 |
| 655 | 655 | 8 | 2.51661148 | 1 | 0.26795838 | 0 | 1 | 2 | 0.06691409 | 0 | 1 | 0 | 1 | 9 | 0.89630573 | FALSE | 1 | FALSE | 1 | FALSE | 1 | FALSE | 1 | FALSE | 1 | FALSE | 1 |
| 656 | 656 | 1.08980419 | 0.86536478 | 52 | 0.00712042 | 56 | 0.10347394 | 55 | 0.41125687 | 72 | 0.30536962 | 27 | 0.29701103 | 1218 | 0.99401611 | FALSE | 0.25171133 | FALSE | 1 | FALSE | 1 | FALSE | 1 | FALSE | 1 | FALSE | 1 |

| | | | | | | | | | | | | | | | | | | | | | | | | | | | |
|---|---|---|---|---|---|---|---|---|---|---|---|---|---|---|---|---|---|---|---|---|---|---|---|---|---|---|---|
| 657 | 657 | 12.25 | 2.98956519 | 0 | 1 | 0 | 1 | 0 | 1 | 0 | 1 | 0 | 1 | 8 | 0.25677166 | FALSE | 1 | FALSE | 1 | FALSE | 1 | FALSE | 1 | FALSE | 1 | FALSE | 1 |
| 658 | 658 | 6.20238095 | 1.07769333 | 1 | 0.88571933 | 0 | 1 | 2 | 0.80536046 | 3 | 0.74138858 | 1 | 0.74858842 | 76 | 0.04081905 | FALSE | 1 | FALSE | 1 | FALSE | 1 | FALSE | 1 | FALSE | 1 | FALSE | 1 |
| 659 | 659 | 9.61538462 | 2.78839523 | 0 | 1 | 1 | 0.3480709 | 0 | 1 | 0 | 1 | 0 | 1 | 12 | 0.374198 | FALSE | 1 | FALSE | 1 | FALSE | 1 | FALSE | 1 | FALSE | 1 | FALSE | 1 |
| 660 | 660 | 6.51851852 | 2.25043815 | 0 | 1 | 2 | 0.21684246 | 0 | 1 | 1 | 0.71956339 | 0 | 1 | 24 | 0.3725373 | FALSE | 1 | FALSE | 1 | FALSE | 1 | FALSE | 1 | FALSE | 1 | FALSE | 1 |
| 661 | 661 | 9.3 | 2.23830293 | 2 | 0.17884234 | 2 | 0.25306331 | 1 | 0.66709967 | 1 | 0.75658797 | 0 | 1 | 24 | 0.82294782 | FALSE | 1 | FALSE | 1 | FALSE | 1 | FALSE | 1 | FALSE | 1 | FALSE | 1 |
| 662 | 662 | 10.3125 | 1.6851836 | 0 | 1 | 0 | 1 | 1 | 0.44344817 | 1 | 0.52895088 | 0 | 1 | 14 | 0.53308261 | FALSE | 1 | FALSE | 1 | FALSE | 1 | FALSE | 1 | FALSE | 1 | FALSE | 1 |
| 663 | 663 | 7.25 | 3.526684 | 0 | 1 | 0 | 1 | 2 | 0.03118552 | 0 | 1 | 0 | 1 | 6 | 0.88425551 | FALSE | 1 | FALSE | 1 | FALSE | 1 | FALSE | 1 | FALSE | 1 | FALSE | 1 |
| 664 | 664 | 4.2972973 | 2.91190348 | 3 | 0.2949342 | 4 | 0.21704891 | 6 | 0.04913907 | 3 | 0.66754163 | 1 | 0.70774043 | 57 | 0.96691729 | FALSE | 1 | FALSE | 1 | FALSE | 1 | FALSE | 1 | FALSE | 1 | FALSE | 1 |
| 665 | 665 | 7.84615385 | 4.05436429 | 1 | 0.49162634 | 0 | 1 | 1 | 0.61440466 | 1 | 0.70601446 | 1 | 0.3499045 | 22 | 0.61628 | FALSE | 1 | FALSE | 1 | FALSE | 1 | FALSE | 1 | FALSE | 1 | FALSE | 1 |
| 666 | 666 | 10.0476191 | 3.16944025 | 0 | 1 | 1 | 0.49918218 | 0 | 1 | 3 | 0.06912908 | 0 | 1 | 17 | 0.77849076 | FALSE | 1 | FALSE | 1 | FALSE | 1 | FALSE | 1 | FALSE | 1 | FALSE | 1 |
| 667 | 667 | 9.32142857 | 2.1389846 | 0 | 1 | 0 | 1 | 3 | 0.07106362 | 1 | 0.71956339 | 1 | 0.36060486 | 22 | 0.76237877 | FALSE | 1 | FALSE | 1 | FALSE | 1 | FALSE | 1 | FALSE | 1 | FALSE | 1 |
| 668 | 668 | 7.54 | 1.71125685 | 1 | 0.92696133 | 4 | 0.40632801 | 1 | 0.97492961 | 7 | 0.17487335 | 2 | 0.49094677 | 85 | 0.49905328 | FALSE | 1 | FALSE | 1 | FALSE | 1 | FALSE | 1 | FALSE | 1 | FALSE | 1 |
| 669 | 669 | 11.6666667 | 2.05480467 | 0 | 1 | 1 | 0.38964394 | 0 | 1 | 2 | 0.14906763 | 0 | 1 | 12 | 0.80329917 | FALSE | 1 | FALSE | 1 | FALSE | 1 | FALSE | 1 | FALSE | 1 | FALSE | 1 |
| 670 | 670 | 10.1538462 | 4.27598219 | 1 | 0.28675951 | 0 | 1 | 1 | 0.37874417 | 2 | 0.1174583 | 0 | 1 | 9 | 0.96018168 | FALSE | 1 | FALSE | 1 | FALSE | 1 | FALSE | 1 | FALSE | 1 | FALSE | 1 |
| 671 | 671 | 7.86956522 | 3.36612728 | 0 | 1 | 2 | 0.16948764 | 1 | 0.5695037 | 0 | 1 | 1 | 0.31672722 | 19 | 0.71542496 | FALSE | 1 | FALSE | 1 | FALSE | 1 | FALSE | 1 | FALSE | 1 | FALSE | 1 |
| 672 | 672 | 10.3 | 1.48660688 | 0 | 1 | 2 | 0.03949327 | 0 | 1 | 2 | 0.0740981 | 0 | 1 | 6 | 0.98831471 | FALSE | 1 | FALSE | 0.96864929 | FALSE | 1 | FALSE | 1 | FALSE | 1 | FALSE | 1 |
| 673 | 673 | 10.5 | 2.39791576 | 4 | 0.00059649 | 0 | 1 | 1 | 0.44344817 | 1 | 0.52895088 | 0 | 1 | 10 | 0.99304605 | FALSE | 0.05254197 | FALSE | 1 | FALSE | 1 | FALSE | 1 | FALSE | 1 | FALSE | 1 |
| 674 | 674 | 11.4285714 | 2.75285875 | 0 | 1 | 2 | 0.14655134 | 0 | 1 | 0 | 1 | 0 | 1 | 19 | 0.34094959 | FALSE | 1 | FALSE | 1 | FALSE | 1 | FALSE | 1 | FALSE | 1 | FALSE | 1 |
| 675 | 675 | 7.29850746 | 3.95879872 | 0 | 1 | 2 | 0.62701348 | 2 | 0.68408105 | 2 | 0.80687302 | 2 | 0.28868487 | 57 | 0.2945347 | FALSE | 1 | FALSE | 1 | FALSE | 1 | FALSE | 1 | FALSE | 1 | FALSE | 1 |
| 676 | 676 | 7.71794872 | 3.59450234 | 1 | 0.63789042 | 2 | 0.36122049 | 2 | 0.41139386 | 0 | 1 | 0 | 1 | 34 | 0.41570781 | FALSE | 1 | FALSE | 1 | FALSE | 1 | FALSE | 1 | FALSE | 1 | FALSE | 1 |
| 677 | 677 | 11.6428571 | 1.79710652 | 0 | 1 | 0 | 1 | 2 | 0.08806398 | 0 | 1 | 0 | 1 | 12 | 0.62236227 | FALSE | 1 | FALSE | 1 | FALSE | 1 | FALSE | 1 | FALSE | 1 | FALSE | 1 |
| 678 | 678 | 9.07692308 | 2.96461711 | 2 | 0.26413583 | 0 | 1 | 2 | 0.41139386 | 1 | 0.84089119 | 1 | 0.4761699 | 33 | 0.59061307 | FALSE | 1 | FALSE | 1 | FALSE | 1 | FALSE | 1 | FALSE | 1 | FALSE | 1 |
| 679 | 679 | 8.57894737 | 3.18410178 | 0 | 1 | 1 | 0.46503795 | 1 | 0.50143826 | 2 | 0.21610812 | 0 | 1 | 15 | 0.83648191 | FALSE | 1 | FALSE | 1 | FALSE | 1 | FALSE | 1 | FALSE | 1 | FALSE | 1 |
| 680 | 680 | 11 | 2.70185122 | 0 | 1 | 0 | 1 | 0 | 1 | 1 | 0.60987146 | 2 | 0.04176963 | 17 | 0.61750632 | FALSE | 1 | FALSE | 1 | FALSE | 1 | FALSE | 1 | FALSE | 0.90763079 | FALSE | 1 |
| 681 | 681 | 8.76470588 | 2.12905424 | 2 | 0.06909038 | 0 | 1 | 0 | 1 | 0 | 1 | 0 | 1 | 15 | 0.49060435 | FALSE | 0.74276693 | FALSE | 1 | FALSE | 1 | FALSE | 1 | FALSE | 1 | FALSE | 1 |
| 682 | 682 | 11 | 3.10424929 | 0 | 1 | 0 | 1 | 1 | 0.33149969 | 2 | 0.08789566 | 0 | 1 | 8 | 0.92152993 | FALSE | 1 | FALSE | 1 | FALSE | 1 | FALSE | 1 | FALSE | 1 | FALSE | 1 |
| 683 | 683 | 9.61290323 | 1.20956989 | 0 | 1 | 1 | 0.63997468 | 5 | 0.00453596 | 2 | 0.41984187 | 0 | 1 | 23 | 0.95750806 | FALSE | 1 | FALSE | 1 | FALSE | 1 | FALSE | 1 | FALSE | 1 | FALSE | 1 |
| 684 | 684 | 9.08108108 | 2.82248097 | 0 | 1 | 0 | 1 | 3 | 0.13783405 | 2 | 0.49731009 | 0 | 1 | 31 | 0.49869404 | FALSE | 1 | FALSE | 1 | FALSE | 1 | FALSE | 1 | FALSE | 1 | FALSE | 1 |
| 685 | 685 | 10.1111111 | 2.99794168 | 0 | 1 | 1 | 0.25628163 | 1 | 0.28067846 | 0 | 1 | 0 | 1 | 7 | 0.84574168 | FALSE | 1 | FALSE | 1 | FALSE | 1 | FALSE | 1 | FALSE | 1 | FALSE | 1 |
| 686 | 686 | 9.625 | 2.39465551 | 0 | 1 | 0 | 1 | 2 | 0.11087977 | 0 | 1 | 0 | 1 | 14 | 0.53308261 | FALSE | 1 | FALSE | 1 | FALSE | 1 | FALSE | 1 | FALSE | 1 | FALSE | 1 |
| 687 | 687 | 8.28571429 | 2.44740596 | 2 | 0.16048326 | 0 | 1 | 4 | 0.01687379 | 1 | 0.73248983 | 1 | 0.37113067 | 20 | 0.97757517 | FALSE | 1 | FALSE | 1 | FALSE | 1 | FALSE | 1 | FALSE | 1 | FALSE | 1 |
| 688 | 688 | 8 | 1.78376517 | 1 | 0.4357638 | 0 | 1 | 0 | 1 | 2 | 0.26791271 | 0 | 1 | 19 | 0.54233419 | FALSE | 1 | FALSE | 1 | FALSE | 1 | FALSE | 1 | FALSE | 1 | FALSE | 1 |
| 689 | 689 | 9.52941177 | 1.97387785 | 0 | 1 | 0 | 1 | 0 | 1 | 0 | 1 | 0 | 1 | 17 | 0.05550965 | FALSE | 1 | FALSE | 1 | FALSE | 1 | FALSE | 1 | FALSE | 1 | FALSE | 1 |
| 690 | 690 | 4.92045455 | 3.02347479 | 3 | 0.39356864 | 2 | 0.78367607 | 3 | 0.61802554 | 4 | 0.58033022 | 3 | 0.17486078 | 73 | 0.7061576 | FALSE | 1 | FALSE | 1 | FALSE | 1 | FALSE | 1 | FALSE | 1 | FALSE | 1 |
| 691 | 691 | 7.05660377 | 3.08627573 | 1 | 0.74224579 | 2 | 0.50543179 | 3 | 0.28676219 | 1 | 0.91399378 | 0 | 1 | 45 | 0.4221244 | FALSE | 1 | FALSE | 1 | FALSE | 1 | FALSE | 1 | FALSE | 1 | FALSE | 1 |
| 692 | 692 | 3.11864407 | 2.23038655 | 0 | 1 | 6 | 0.51277283 | 5 | 0.76880524 | 5 | 0.91533579 | 6 | 0.07009621 | 155 | 0.13888892 | FALSE | 1 | FALSE | 1 | FALSE | 1 | FALSE | 1 | FALSE | 1 | FALSE | 1 |
| 693 | 693 | 6.72413793 | 2.44803301 | 0 | 1 | 1 | 0.61539065 | 1 | 0.65463982 | 4 | 0.04184267 | 0 | 1 | 23 | 0.84462957 | FALSE | 1 | FALSE | 1 | FALSE | 1 | FALSE | 1 | FALSE | 1 | FALSE | 1 |
| 694 | 694 | 11.5333333 | 2.15612822 | 0 | 1 | 0 | 1 | 0 | 1 | 1 | 0.50623194 | 0 | 1 | 14 | 0.29525409 | FALSE | 1 | FALSE | 1 | FALSE | 1 | FALSE | 1 | FALSE | 1 | FALSE | 1 |
| 695 | 695 | 1.04964539 | 0.70675695 | 77 | 0.03521477 | 86 | 0.31236771 | 93 | 0.42668174 | 120 | 0.35839335 | 48 | 0.12206733 | 2114 | 0.97378608 | FALSE | 0.58392 | FALSE | 1 | FALSE | 1 | FALSE | 1 | FALSE | 1 | FALSE | 1 |
| 696 | 696 | 9.66666667 | 1.65998661 | 2 | 0.05513784 | 0 | 1 | 1 | 0.42266264 | 0 | 1 | 0 | 1 | 12 | 0.80329917 | FALSE | 0.68849059 | FALSE | 1 | FALSE | 1 | FALSE | 1 | FALSE | 1 | FALSE | 1 |
| 697 | 697 | 10.3243243 | 2.52612356 | 2 | 0.24498301 | 0 | 1 | 1 | 0.74264075 | 2 | 0.51200191 | 1 | 0.45845922 | 31 | 0.64534533 | FALSE | 1 | FALSE | 1 | FALSE | 1 | FALSE | 1 | FALSE | 1 | FALSE | 1 |
| 698 | 698 | 11.1 | 2.07123152 | 0 | 1 | 1 | 0.28036922 | 2 | 0.04781153 | 1 | 0.37517262 | 0 | 1 | 6 | 0.98831471 | FALSE | 1 | FALSE | 1 | FALSE | 1 | FALSE | 1 | FALSE | 1 | FALSE | 1 |
| 699 | 699 | 10.5454546 | 2.38798646 | 0 | 1 | 1 | 0.30368014 | 0 | 1 | 0 | 1 | 0 | 1 | 10 | 0.46859937 | FALSE | 1 | FALSE | 1 | FALSE | 1 | FALSE | 1 | FALSE | 1 | FALSE | 1 |
| 700 | 700 | 9.22413793 | 2.1738501 | 3 | 0.18560124 | 3 | 0.28903336 | 2 | 0.62245518 | 4 | 0.27551999 | 2 | 0.24594209 | 44 | 0.97031903 | FALSE | 1 | FALSE | 1 | FALSE | 1 | FALSE | 1 | FALSE | 1 | FALSE | 1 |
| 701 | 701 | 8.63636364 | 1.39952767 | 0 | 1 | 1 | 0.5154318 | 3 | 0.04259419 | 1 | 0.64497439 | 0 | 1 | 17 | 0.8840357 | FALSE | 1 | FALSE | 1 | FALSE | 1 | FALSE | 1 | FALSE | 1 | FALSE | 1 |
| 702 | 702 | 7.58823529 | 3.4137925 | 1 | 0.35728305 | 1 | 0.42857733 | 1 | 0.46348836 | 2 | 0.18212134 | 0 | 1 | 12 | 0.962093 | FALSE | 1 | FALSE | 1 | FALSE | 1 | FALSE | 1 | FALSE | 1 | FALSE | 1 |
| 703 | 703 | 10.0714286 | 2.08615455 | 0 | 1 | 1 | 0.60247911 | 2 | 0.2663542 | 1 | 0.73248983 | 1 | 0.37113067 | 23 | 0.73439118 | FALSE | 1 | FALSE | 1 | FALSE | 1 | FALSE | 1 | FALSE | 1 | FALSE | 1 |
| 704 | 704 | 7.53571429 | 2.79645366 | 4 | 0.00528025 | 0 | 1 | 1 | 0.64171563 | 0 | 1 | 0 | 1 | 23 | 0.73439118 | FALSE | 0.2089654 | FALSE | 1 | FALSE | 1 | FALSE | 1 | FALSE | 1 | FALSE | 1 |
| 705 | 705 | 9.05263158 | 3.34779871 | 0 | 1 | 1 | 0.46503795 | 1 | 0.50143826 | 2 | 0.21610812 | 0 | 1 | 15 | 0.83648191 | FALSE | 1 | FALSE | 1 | FALSE | 1 | FALSE | 1 | FALSE | 1 | FALSE | 1 |
| 706 | 706 | 9.45454546 | 3.22951267 | 2 | 0.03087289 | 1 | 0.30368014 | 0 | 1 | 0 | 1 | 0 | 1 | 8 | 0.92152993 | FALSE | 0.53356688 | FALSE | 1 | FALSE | 1 | FALSE | 1 | FALSE | 1 | FALSE | 1 |
| 707 | 707 | 9.23529412 | 2.76595716 | 1 | 0.56527493 | 3 | 0.08334648 | 4 | 0.02651644 | 2 | 0.43583874 | 0 | 1 | 22 | 0.99301316 | FALSE | 1 | FALSE | 1 | FALSE | 1 | FALSE | 1 | FALSE | 1 | FALSE | 1 |
| 708 | 708 | 4.31623932 | 1.49156931 | 10 | 0.07583286 | 9 | 0.34070193 | 8 | 0.60386997 | 6 | 0.9610208 | 7 | 0.08742973 | 193 | 0.77702661 | FALSE | 0.77424195 | FALSE | 1 | FALSE | 1 | FALSE | 1 | FALSE | 1 | FALSE | 1 |
| 709 | 709 | 9.41666667 | 2.56444709 | 2 | 0.12501436 | 0 | 1 | 2 | 0.21260257 | 1 | 0.67692816 | 0 | 1 | 19 | 0.83975208 | FALSE | 0.95622564 | FALSE | 1 | FALSE | 1 | FALSE | 1 | FALSE | 1 | FALSE | 1 |
| 710 | 710 | 9.60606061 | 2.52189676 | 1 | 0.57647348 | 0 | 1 | 1 | 0.70185639 | 1 | 0.78873814 | 0 | 1 | 30 | 0.21942357 | FALSE | 1 | FALSE | 1 | FALSE | 1 | FALSE | 1 | FALSE | 1 | FALSE | 1 |
| 711 | 711 | 8.10526316 | 2.14953451 | 4 | 0.00119617 | 2 | 0.12434464 | 0 | 1 | 0 | 1 | 1 | 0.26986775 | 12 | 0.99479039 | FALSE | 0.08447413 | FALSE | 1 | FALSE | 1 | FALSE | 1 | FALSE | 1 | FALSE | 1 |
| 712 | 712 | 9.375 | 2.46327188 | 1 | 0.46441868 | 1 | 0.54637326 | 0 | 1 | 1 | 0.67692816 | 0 | 1 | 21 | 0.47053137 | FALSE | 1 | FALSE | 1 | FALSE | 1 | FALSE | 1 | FALSE | 1 | FALSE | 1 |
| 713 | 713 | 3.45794393 | 1.8458358 | 2 | 0.76550375 | 3 | 0.67855912 | 2 | 0.90233169 | 6 | 0.36804259 | 0 | 1 | 94 | 0.1961492 | FALSE | 1 | FALSE | 1 | FALSE | 1 | FALSE | 1 | FALSE | 1 | FALSE | 1 |
| 714 | 714 | 8.5 | 1.59239263 | 0 | 1 | 1 | 0.36919824 | 2 | 0.08806398 | 0 | 1 | 2 | 0.02131828 | 9 | 0.98607548 | FALSE | 1 | FALSE | 1 | FALSE | 1 | FALSE | 1 | FALSE | 0.73999729 | FALSE | 1 |
| 715 | 715 | 10.4 | 0.66332496 | 0 | 1 | 0 | 1 | 0 | 1 | 1 | 0.37517262 | 0 | 1 | 9 | 0.5215547 | FALSE | 1 | FALSE | 1 | FALSE | 1 | FALSE | 1 | FALSE | 1 | FALSE | 1 |
| 716 | 716 | 5.68421053 | 2.56602563 | 3 | 0.53942587 | 4 | 0.47871467 | 3 | 0.76242935 | 3 | 0.88727759 | 2 | 0.54214185 | 95 | 0.3367471 | FALSE | 1 | FALSE | 1 | FALSE | 1 | FALSE | 1 | FALSE | 1 | FALSE | 1 |
| 717 | 717 | 10.6 | 2.0976177 | 0 | 1 | 2 | 0.19297191 | 0 | 1 | 2 | 0.31968852 | 1 | 0.33902672 | 20 | 0.8152498 | FALSE | 1 | FALSE | 1 | FALSE | 1 | FALSE | 1 | FALSE | 1 | FALSE | 1 |
| 718 | 718 | 7.07142857 | 2.24290264 | 2 | 0.41401181 | 3 | 0.26245929 | 2 | 0.59334725 | 1 | 0.92538938 | 0 | 1 | 47 | 0.50345693 | FALSE | 1 | FALSE | 1 | FALSE | 1 | FALSE | 1 | FALSE | 1 | FALSE | 1 |
| 719 | 719 | 12.1 | 1.57797338 | 0 | 1 | 2 | 0.03949327 | 0 | 1 | 0 | 1 | 0 | 1 | 8 | 0.80394719 | FALSE | 1 | FALSE | 0.96864929 | FALSE | 1 | FALSE | 1 | FALSE | 1 | FALSE | 1 |
| 720 | 720 | 7.4 | 1.38564065 | 1 | 0.47819878 | 2 | 0.19297191 | 0 | 1 | 2 | 0.31968852 | 0 | 1 | 20 | 0.8152498 | FALSE | 1 | FALSE | 1 | FALSE | 1 | FALSE | 1 | FALSE | 1 | FALSE | 1 |
| 721 | 721 | 8.73214286 | 2.4675832 | 1 | 0.74890551 | 6 | 0.00689664 | 2 | 0.57303317 | 3 | 0.44196866 | 4 | 0.01081442 | 37 | 0.99791276 | FALSE | 1 | FALSE | 0.60103504 | FALSE | 1 | FALSE | 1 | FALSE | 0.57142086 | FALSE | 1 |
| 722 | 722 | 8.57142857 | 2.49897938 | 0 | 1 | 1 | 0.36919824 | 1 | 0.40110418 | 1 | 0.48242099 | 0 | 1 | 11 | 0.83676449 | FALSE | 1 | FALSE | 1 | FALSE | 1 | FALSE | 1 | FALSE | 1 | FALSE | 1 |
| 723 | 723 | 12.15 | 2.21979729 | 1 | 0.40558521 | 1 | 0.48239024 | 2 | 0.16019548 | 0 | 1 | 1 | 0.28187291 | 15 | 0.92113679 | FALSE | 1 | FALSE | 1 | FALSE | 1 | FALSE | 1 | FALSE | 1 | FALSE | 1 |
| 724 | 724 | 4.63551402 | 1.81577986 | 3 | 0.52064823 | 2 | 0.86638697 | 4 | 0.53987568 | 6 | 0.36804259 | 1 | 0.83189113 | 91 | 0.48903513 | FALSE | 1 | FALSE | 1 | FALSE | 1 | FALSE | 1 | FALSE | 1 | FALSE | 1 |
| 725 | 725 | 7.51515152 | 3.01633143 | 0 | 1 | 1 | 0.58913618 | 1 | 0.62830986 | 2 | 0.35374209 | 0 | 1 | 23 | 0.58307573 | FALSE | 1 | FALSE | 1 | FALSE | 1 | FALSE | 1 | FALSE | 1 | FALSE | 1 |
| 726 | 726 | 10 | 3.34034351 | 1 | 0.38989951 | 0 | 1 | 1 | 0.50143826 | 3 | 0.05383577 | 0 | 1 | 14 | 0.93677253 | FALSE | 1 | FALSE | 1 | FALSE | 1 | FALSE | 1 | FALSE | 1 | FALSE | 1 |
| 727 | 727 | 5.93548387 | 2.38185999 | 0 | 1 | 0 | 1 | 0 | 1 | 3 | 0.15707004 | 1 | 0.39166989 | 26 | 0.48609328 | FALSE | 1 | FALSE | 1 | FALSE | 1 | FALSE | 1 | FALSE | 1 | FALSE | 1 |
| 728 | 728 | 3.07253886 | 2.53982217 | 6 | 0.37456823 | 6 | 0.59835315 | 5 | 0.82977877 | 4 | 0.98005486 | 2 | 0.83065424 | 170 | 0.08729665 | FALSE | 1 | FALSE | 1 | FALSE | 1 | FALSE | 1 | FALSE | 1 | FALSE | 1 |
| 729 | 729 | 8.55263158 | 2.1299539 | 0 | 1 | 3 | 0.44799895 | 2 | 0.76408693 | 3 | 0.68517679 | 4 | 0.03564889 | 64 | 0.59240184 | FALSE | 1 | FALSE | 1 | FALSE | 1 | FALSE | 1 | FALSE | 0.87669812 | FALSE | 1 |
| 730 | 730 | 8.41666667 | 1.65621724 | 0 | 1 | 0 | 1 | 2 | 0.06691409 | 0 | 1 | 0 | 1 | 10 | 0.71436964 | FALSE | 1 | FALSE | 1 | FALSE | 1 | FALSE | 1 | FALSE | 1 | FALSE | 1 |
| 731 | 731 | 8.22727273 | 2.62733564 | 1 | 0.89713386 | 1 | 0.94382585 | 2 | 0.82596174 | 9 | 0.01777745 | 1 | 0.76487586 | 73 | 0.61830614 | FALSE | 1 | FALSE | 1 | FALSE | 1 | FALSE | 1 | FALSE | 1 | FALSE | 1 |
| 732 | 732 | 9.575 | 2.9401318 | 2 | 0.27371785 | 0 | 1 | 2 | 0.42396974 | 2 | 0.5543522 | 1 | 0.48480878 | 33 | 0.71942635 | FALSE | 1 | FALSE | 1 | FALSE | 1 | FALSE | 1 | FALSE | 1 | FALSE | 1 |
| 733 | 733 | 10.9 | 2.66270539 | 0 | 1 | 0 | 1 | 0 | 1 | 0 | 1 | 0 | 1 | 10 | 0.18273166 | FALSE | 1 | FALSE | 1 | FALSE | 1 | FALSE | 1 | FALSE | 1 | FALSE | 1 |
| 734 | 734 | 10.4444444 | 2.06080411 | 0 | 1 | 0 | 1 | 0 | 1 | 0 | 1 | 0 | 1 | 9 | 0.21661397 | FALSE | 1 | FALSE | 1 | FALSE | 1 | FALSE | 1 | FALSE | 1 | FALSE | 1 |
| 735 | 735 | 8.21428571 | 1.95223577 | 1 | 0.51745948 | 0 | 1 | 3 | 0.07756639 | 0 | 1 | 0 | 1 | 24 | 0.55018107 | FALSE | 1 | FALSE | 1 | FALSE | 1 | FALSE | 1 | FALSE | 1 | FALSE | 1 |
| 736 | 736 | 4.49173554 | 1.84131137 | 5 | 0.66724277 | 6 | 0.71838968 | 9 | 0.38810777 | 14 | 0.13066817 | 4 | 0.48588925 | 181 | 0.7936038 | FALSE | 1 | FALSE | 1 | FALSE | 1 | FALSE | 1 | FALSE | 1 | FALSE | 1 |
| 737 | 737 | 8.81818182 | 2.70750411 | 0 | 1 | 2 | 0.15794013 | 0 | 1 | 0 | 1 | 0 | 1 | 20 | 0.30915202 | FALSE | 1 | FALSE | 1 | FALSE | 1 | FALSE | 1 | FALSE | 1 | FALSE | 1 |
| 738 | 738 | 8.42857143 | 2.34157921 | 2 | 0.09989125 | 1 | 0.49918218 | 1 | 0.53671415 | 0 | 1 | 0 | 1 | 17 | 0.77849076 | FALSE | 0.86047225 | FALSE | 1 | FALSE | 1 | FALSE | 1 | FALSE | 1 | FALSE | 1 |
| 739 | 739 | 8.07142857 | 3.4530377 | 0 | 1 | 0 | 1 | 2 | 0.08806398 | 2 | 0.13304705 | 0 | 1 | 10 | 0.94579623 | FALSE | 1 | FALSE | 1 | FALSE | 1 | FALSE | 1 | FALSE | 1 | FALSE | 1 |
| 740 | 740 | 7.3627451 | 1.28176667 | 2 | 0.74184548 | 3 | 0.6466734 | 3 | 0.71583185 | 4 | 0.69586384 | 1 | 0.81716516 | 89 | 0.25743428 | FALSE | 1 | FALSE | 1 | FALSE | 1 | FALSE | 1 | FALSE | 1 | FALSE | 1 |
| 741 | 741 | 11.5185185 | 2.85929764 | 1 | 0.50471032 | 1 | 0.58913618 | 0 | 1 | 1 | 0.71956339 | 0 | 1 | 24 | 0.3725373 | FALSE | 1 | FALSE | 1 | FALSE | 1 | FALSE | 1 | FALSE | 1 | FALSE | 1 |
| 742 | 742 | 10.1 | 2.3430749 | 1 | 0.22886536 | 0 | 1 | 0 | 1 | 0 | 1 | 0 | 1 | 9 | 0.5215547 | FALSE | 1 | FALSE | 1 | FALSE | 1 | FALSE | 1 | FALSE | 1 | FALSE | 1 |
| 743 | 743 | 7.96875 | 2.56763966 | 1 | 0.56527493 | 0 | 1 | 2 | 0.32011871 | 2 | 0.43583874 | 1 | 0.41154418 | 26 | 0.77612337 | FALSE | 1 | FALSE | 1 | FALSE | 1 | FALSE | 1 | FALSE | 1 | FALSE | 1 |
| 744 | 744 | 9.10204082 | 2.78659252 | 1 | 0.72119464 | 4 | 0.07284863 | 2 | 0.53023996 | 3 | 0.39227316 | 0 | 1 | 39 | 0.86798612 | FALSE | 1 | FALSE | 1 | FALSE | 1 | FALSE | 1 | FALSE | 1 | FALSE | 1 |
| 745 | 745 | 4.31100479 | 2.81552828 | 9 | 0.08795309 | 4 | 0.91193307 | 6 | 0.76870091 | 9 | 0.62737993 | 6 | 0.12794878 | 175 | 0.64674954 | FALSE | 0.82250193 | FALSE | 1 | FALSE | 1 | FALSE | 1 | FALSE | 1 | FALSE | 1 |
| 746 | 746 | 9.53333333 | 3.53773311 | 0 | 1 | 2 | 0.07341559 | 0 | 1 | 0 | 1 | 1 | 0.20679328 | 11 | 0.83676449 | FALSE | 1 | FALSE | 1 | FALSE | 1 | FALSE | 1 | FALSE | 1 | FALSE | 1 |
| 747 | 747 | 8.41071429 | 1.16153817 | 4 | 0.05467538 | 2 | 0.54544984 | 4 | 0.14086741 | 2 | 0.73544289 | 0 | 1 | 44 | 0.91271545 | FALSE | 0.68849059 | FALSE | 1 | FALSE | 1 | FALSE | 1 | FALSE | 1 | FALSE | 1 |
| 748 | 748 | 10.1891892 | 1.46744562 | 1 | 0.6184714 | 0 | 1 | 3 | 0.14625315 | 1 | 0.82511639 | 1 | 0.45845922 | 31 | 0.64534533 | FALSE | 1 | FALSE | 1 | FALSE | 1 | FALSE | 1 | FALSE | 1 | FALSE | 1 |
| 749 | 749 | 8.25925926 | 1.69077218 | 2 | 0.40500867 | 0 | 1 | 4 | 0.12796328 | 1 | 0.92176876 | 2 | 0.22161752 | 45 | 0.66857827 | FALSE | 1 | FALSE | 1 | FALSE | 1 | FALSE | 1 | FALSE | 1 | FALSE | 1 |
| 750 | 750 | 3.88292683 | 2.69886177 | 9 | 0.07646017 | 4 | 0.90000978 | 9 | 0.30634569 | 9 | 0.59210468 | 7 | 0.04863345 | 165 | 0.9063806 | FALSE | 0.77424195 | FALSE | 1 | FALSE | 1 | FALSE | 1 | FALSE | 0.98629461 | FALSE | 1 |

| | | | | | | | | | | | | | | | | | | | | | | | | | | | |
|---|---|---|---|---|---|---|---|---|---|---|---|---|---|---|---|---|---|---|---|---|---|---|---|---|---|---|---|
| 751 | 751 | 9.17647059 | 2.61748182 | 1 | 0.35728305 | 0 | 1 | 1 | 0.46348836 | 0 | 1 | 0 | 1 | 15 | 0.49060435 | FALSE | 1 | FALSE | 1 | FALSE | 1 | FALSE | 1 | FALSE | 1 | FALSE | 1 |
| 752 | 752 | 8.51851852 | 2.92334626 | 0 | 1 | 2 | 0.21684246 | 1 | 0.62830986 | 3 | 0.12475952 | 0 | 1 | 21 | 0.88395936 | FALSE | 1 | FALSE | 1 | FALSE | 1 | FALSE | 1 | FALSE | 1 | FALSE | 1 |
| 753 | 753 | 8.15384615 | 0.68081631 | 1 | 0.86966563 | 1 | 0.92420038 | 5 | 0.14803148 | 4 | 0.48430022 | 0 | 1 | 67 | 0.4284743 | FALSE | 1 | FALSE | 1 | FALSE | 1 | FALSE | 1 | FALSE | 1 | FALSE | 1 |
| 754 | 754 | 7.93617021 | 1.56292491 | 2 | 0.34034075 | 1 | 0.78790827 | 1 | 0.82190381 | 1 | 0.89101956 | 0 | 1 | 42 | 0.23544056 | FALSE | 1 | FALSE | 1 | FALSE | 1 | FALSE | 1 | FALSE | 1 | FALSE | 1 |
| 755 | 755 | 5.90476191 | 2.67981453 | 0 | 1 | 1 | 0.74973543 | 0 | 1 | 2 | 0.5811115 | 2 | 0.15059682 | 37 | 0.34041811 | FALSE | 1 | FALSE | 1 | FALSE | 1 | FALSE | 1 | FALSE | 1 | FALSE | 1 |
| 756 | 756 | 10.8235294 | 2.47897735 | 0 | 1 | 0 | 1 | 2 | 0.12280513 | 1 | 0.55062772 | 0 | 1 | 14 | 0.7313145 | FALSE | 1 | FALSE | 1 | FALSE | 1 | FALSE | 1 | FALSE | 1 | FALSE | 1 |
| 757 | 757 | 9.66666667 | 2.82842713 | 0 | 1 | 1 | 0.25628163 | 0 | 1 | 0 | 1 | 1 | 0.1383237 | 7 | 0.84574168 | FALSE | 1 | FALSE | 1 | FALSE | 1 | FALSE | 1 | FALSE | 1 | FALSE | 1 |
| 758 | 758 | 8.42857143 | 1.84058553 | 1 | 0.51745948 | 2 | 0.2288771 | 1 | 0.64171563 | 2 | 0.3705495 | 1 | 0.37113067 | 21 | 0.94083768 | FALSE | 1 | FALSE | 1 | FALSE | 1 | FALSE | 1 | FALSE | 1 | FALSE | 1 |
| 759 | 759 | 9.55555556 | 1.80191942 | 0 | 1 | 0 | 1 | 2 | 0.13502329 | 2 | 0.19902682 | 3 | 0.00292491 | 11 | 0.99651304 | FALSE | 1 | FALSE | 1 | FALSE | 1 | FALSE | 1 | FALSE | 0.29951111 | FALSE | 1 |
| 760 | 760 | 9.29411765 | 2.3203627 | 1 | 0.35728305 | 1 | 0.42857733 | 0 | 1 | 3 | 0.040397 | 0 | 1 | 12 | 0.962093 | FALSE | 1 | FALSE | 1 | FALSE | 1 | FALSE | 1 | FALSE | 1 | FALSE | 1 |
| 761 | 761 | 8.63636364 | 0.88139634 | 1 | 0.24866461 | 2 | 0.04725998 | 0 | 1 | 0 | 1 | 0 | 1 | 8 | 0.92152993 | FALSE | 1 | FALSE | 1 | FALSE | 1 | FALSE | 1 | FALSE | 1 | FALSE | 1 |
| 762 | 762 | 8.25 | 2.2107691 | 0 | 1 | 1 | 0.48239024 | 0 | 1 | 2 | 0.23331368 | 0 | 1 | 17 | 0.61750632 | FALSE | 1 | FALSE | 1 | FALSE | 1 | FALSE | 1 | FALSE | 1 | FALSE | 1 |
| 763 | 763 | 4.85 | 1.31434394 | 1 | 0.40558521 | 1 | 0.48239024 | 1 | 0.51939841 | 2 | 0.23331368 | 1 | 0.28187291 | 14 | 0.97325593 | FALSE | 1 | FALSE | 1 | FALSE | 1 | FALSE | 1 | FALSE | 1 | FALSE | 1 |
| 764 | 764 | 8.96666667 | 2.21334337 | 1 | 0.54198714 | 0 | 1 | 3 | 0.09131526 | 0 | 1 | 0 | 1 | 26 | 0.48609328 | FALSE | 1 | FALSE | 1 | FALSE | 1 | FALSE | 1 | FALSE | 1 | FALSE | 1 |
| 765 | 765 | 8.90625 | 3.49427545 | 3 | 0.03713651 | 1 | 0.61539065 | 3 | 0.08431992 | 3 | 0.14602298 | 2 | 0.08121636 | 17 | 0.99982101 | FALSE | 0.58392 | FALSE | 1 | FALSE | 1 | FALSE | 1 | FALSE | 1 | FALSE | 1 |
| 766 | 766 | 9.83333333 | 2.54405625 | 0 | 1 | 0 | 1 | 0 | 1 | 0 | 1 | 0 | 1 | 6 | 0.36077158 | FALSE | 1 | FALSE | 1 | FALSE | 1 | FALSE | 1 | FALSE | 1 | FALSE | 1 |
| 767 | 767 | 12.0909091 | 3.77683238 | 0 | 1 | 0 | 1 | 0 | 1 | 0 | 1 | 0 | 1 | 11 | 0.15414489 | FALSE | 1 | FALSE | 1 | FALSE | 1 | FALSE | 1 | FALSE | 1 | FALSE | 1 |
| 768 | 768 | 8.2962963 | 2.27408614 | 1 | 0.75539413 | 0 | 1 | 1 | 0.86240917 | 4 | 0.2347767 | 3 | 0.05824229 | 45 | 0.66857827 | FALSE | 1 | FALSE | 1 | FALSE | 1 | FALSE | 1 | FALSE | 1 | FALSE | 1 |
| 769 | 769 | 10.4285714 | 2.44114393 | 0 | 1 | 0 | 1 | 0 | 1 | 0 | 1 | 0 | 1 | 7 | 0.3043657 | FALSE | 1 | FALSE | 1 | FALSE | 1 | FALSE | 1 | FALSE | 1 | FALSE | 1 |
| 770 | 770 | 6.37931035 | 4.83103325 | 1 | 0.52988231 | 1 | 0.61539065 | 0 | 1 | 3 | 0.14602298 | 0 | 1 | 24 | 0.70563334 | FALSE | 1 | FALSE | 1 | FALSE | 1 | FALSE | 1 | FALSE | 1 | FALSE | 1 |
| 771 | 771 | 4.66666667 | 1.62731317 | 4 | 0.29417686 | 3 | 0.67855912 | 4 | 0.53987568 | 4 | 0.73116831 | 1 | 0.83189113 | 91 | 0.48903513 | FALSE | 1 | FALSE | 1 | FALSE | 1 | FALSE | 1 | FALSE | 1 | FALSE | 1 |
| 772 | 772 | 8.86666667 | 2.07203604 | 1 | 0.69044531 | 1 | 0.77338922 | 1 | 0.80828716 | 4 | 0.15016749 | 1 | 0.52593041 | 37 | 0.73736034 | FALSE | 1 | FALSE | 1 | FALSE | 1 | FALSE | 1 | FALSE | 1 | FALSE | 1 |
| 773 | 773 | 7.45833333 | 1.99956593 | 0 | 1 | 2 | 0.18117183 | 0 | 1 | 2 | 0.30249307 | 0 | 1 | 20 | 0.68270729 | FALSE | 1 | FALSE | 1 | FALSE | 1 | FALSE | 1 | FALSE | 1 | FALSE | 1 |
| 774 | 774 | 9.09090909 | 1.975051 | 0 | 1 | 0 | 1 | 1 | 0.50143826 | 4 | 0.00976682 | 0 | 1 | 14 | 0.93677253 | FALSE | 1 | FALSE | 1 | FALSE | 1 | FALSE | 1 | FALSE | 1 | FALSE | 1 |
| 775 | 775 | 6.8372093 | 2.29184837 | 3 | 0.09709068 | 0 | 1 | 1 | 0.79363407 | 3 | 0.31602175 | 1 | 0.50988734 | 35 | 0.7801651 | FALSE | 0.86047225 | FALSE | 1 | FALSE | 1 | FALSE | 1 | FALSE | 1 | FALSE | 1 |
| 776 | 776 | 8.75 | 1.95256242 | 1 | 0.34033054 | 0 | 1 | 1 | 0.44344817 | 1 | 0.52895088 | 0 | 1 | 13 | 0.76799161 | FALSE | 1 | FALSE | 1 | FALSE | 1 | FALSE | 1 | FALSE | 1 | FALSE | 1 |
| 777 | 777 | 8.9 | 2.84429253 | 2 | 0.17884234 | 1 | 0.6278847 | 1 | 0.66709967 | 0 | 1 | 1 | 0.39166989 | 25 | 0.67630471 | FALSE | 1 | FALSE | 1 | FALSE | 1 | FALSE | 1 | FALSE | 1 | FALSE | 1 |
| 778 | 778 | 8.44444444 | 2.40883149 | 1 | 0.20854748 | 0 | 1 | 0 | 1 | 0 | 1 | 0 | 1 | 8 | 0.5780332 | FALSE | 1 | FALSE | 1 | FALSE | 1 | FALSE | 1 | FALSE | 1 | FALSE | 1 |
| 779 | 779 | 9.96774194 | 2.68188512 | 1 | 0.55378205 | 2 | 0.2651851 | 0 | 1 | 2 | 0.41984187 | 0 | 1 | 26 | 0.64660109 | FALSE | 1 | FALSE | 1 | FALSE | 1 | FALSE | 1 | FALSE | 1 | FALSE | 1 |
| 780 | 780 | 10.5238095 | 1.13888198 | 0 | 1 | 2 | 0.14655134 | 1 | 0.53671415 | 0 | 1 | 0 | 1 | 18 | 0.57962763 | FALSE | 1 | FALSE | 1 | FALSE | 1 | FALSE | 1 | FALSE | 1 | FALSE | 1 |
| 781 | 781 | 9.57142857 | 2.39386972 | 1 | 0.59801736 | 1 | 0.68454804 | 2 | 0.35984046 | 1 | 0.80778317 | 1 | 0.44015537 | 29 | 0.69927719 | FALSE | 1 | FALSE | 1 | FALSE | 1 | FALSE | 1 | FALSE | 1 | FALSE | 1 |
| 782 | 782 | 4.11458333 | 2.54079135 | 3 | 0.44855528 | 2 | 0.82287436 | 0 | 1 | 4 | 0.64930879 | 4 | 0.07228874 | 83 | 0.3457435 | FALSE | 1 | FALSE | 1 | FALSE | 1 | FALSE | 1 | FALSE | 1 | FALSE | 1 |
| 783 | 783 | 8.11627907 | 4.50430958 | 0 | 1 | 1 | 0.75788114 | 1 | 0.79363407 | 3 | 0.31602175 | 1 | 0.50988734 | 37 | 0.48294184 | FALSE | 1 | FALSE | 1 | FALSE | 1 | FALSE | 1 | FALSE | 1 | FALSE | 1 |
| 784 | 784 | 9.60869565 | 3.00346366 | 0 | 1 | 0 | 1 | 0 | 1 | 0 | 1 | 0 | 1 | 23 | 0.01996981 | FALSE | 1 | FALSE | 1 | FALSE | 1 | FALSE | 1 | FALSE | 1 | FALSE | 1 |
| 785 | 785 | 12.3125 | 3.13685252 | 2 | 0.1881398 | 0 | 1 | 1 | 0.67911179 | 2 | 0.41984187 | 0 | 1 | 26 | 0.64660109 | FALSE | 1 | FALSE | 1 | FALSE | 1 | FALSE | 1 | FALSE | 1 | FALSE | 1 |
| 786 | 786 | 7.36363636 | 2.05703791 | 0 | 1 | 1 | 0.5154318 | 1 | 0.55340851 | 1 | 0.64497439 | 0 | 1 | 19 | 0.54233419 | FALSE | 1 | FALSE | 1 | FALSE | 1 | FALSE | 1 | FALSE | 1 | FALSE | 1 |
| 787 | 787 | 10.2325581 | 2.76890718 | 3 | 0.09709068 | 4 | 0.04916488 | 2 | 0.46083448 | 3 | 0.31602175 | 0 | 1 | 31 | 0.98840896 | FALSE | 0.86047225 | FALSE | 1 | FALSE | 1 | FALSE | 1 | FALSE | 1 | FALSE | 1 |
| 788 | 788 | 7.1875 | 4.20146924 | 3 | 0.00725145 | 1 | 0.40942991 | 1 | 0.44344817 | 0 | 1 | 0 | 1 | 11 | 0.9718531 | FALSE | 0.25171133 | FALSE | 1 | FALSE | 1 | FALSE | 1 | FALSE | 1 | FALSE | 1 |
| 789 | 789 | 10 | 1.47196014 | 0 | 1 | 1 | 0.32623933 | 0 | 1 | 0 | 1 | 0 | 1 | 11 | 0.41944532 | FALSE | 1 | FALSE | 1 | FALSE | 1 | FALSE | 1 | FALSE | 1 | FALSE | 1 |
| 790 | 790 | 8.51851852 | 2.21727845 | 0 | 1 | 2 | 0.21684246 | 0 | 1 | 1 | 0.71956339 | 0 | 1 | 24 | 0.3725373 | FALSE | 1 | FALSE | 1 | FALSE | 1 | FALSE | 1 | FALSE | 1 | FALSE | 1 |
| 791 | 791 | 7.90476191 | 3.9146217 | 1 | 0.42086991 | 2 | 0.14655134 | 0 | 1 | 1 | 0.62783522 | 1 | 0.29368245 | 16 | 0.90353674 | FALSE | 1 | FALSE | 1 | FALSE | 1 | FALSE | 1 | FALSE | 1 | FALSE | 1 |
| 792 | 792 | 9 | 2.52677965 | 0 | 1 | 0 | 1 | 1 | 0.61440466 | 3 | 0.11458571 | 0 | 1 | 22 | 0.61628 | FALSE | 1 | FALSE | 1 | FALSE | 1 | FALSE | 1 | FALSE | 1 | FALSE | 1 |
| 793 | 793 | 8.22222222 | 2.71938628 | 3 | 0.01017712 | 1 | 0.44710673 | 0 | 1 | 1 | 0.57131013 | 2 | 0.03435384 | 11 | 0.99651304 | FALSE | 0.30467829 | FALSE | 1 | FALSE | 1 | FALSE | 1 | FALSE | 0.86063198 | FALSE | 1 |
| 794 | 794 | 8.66666667 | 2.153808 | 0 | 1 | 0 | 1 | 2 | 0.21260257 | 0 | 1 | 0 | 1 | 22 | 0.25237063 | FALSE | 1 | FALSE | 1 | FALSE | 1 | FALSE | 1 | FALSE | 1 | FALSE | 1 |
| 795 | 795 | 8.05882353 | 1.60772357 | 4 | 0.01061864 | 0 | 1 | 1 | 0.71261921 | 4 | 0.06848449 | 1 | 0.43077488 | 24 | 0.98848472 | FALSE | 0.31290615 | FALSE | 1 | FALSE | 1 | FALSE | 1 | FALSE | 1 | FALSE | 1 |
| 796 | 796 | 8.875 | 1.45236876 | 0 | 1 | 0 | 1 | 0 | 1 | 1 | 0.67692816 | 0 | 1 | 23 | 0.09196389 | FALSE | 1 | FALSE | 1 | FALSE | 1 | FALSE | 1 | FALSE | 1 | FALSE | 1 |
| 797 | 797 | 8.86111111 | 2.16221052 | 0 | 1 | 2 | 0.32556427 | 0 | 1 | 0 | 1 | 0 | 1 | 34 | 0.06417017 | FALSE | 1 | FALSE | 1 | FALSE | 1 | FALSE | 1 | FALSE | 1 | FALSE | 1 |
| 798 | 798 | 5.51239669 | 2.60994824 | 6 | 0.07447161 | 6 | 0.16801732 | 6 | 0.23178146 | 4 | 0.78114102 | 3 | 0.29179524 | 90 | 0.97037573 | FALSE | 0.77424195 | FALSE | 1 | FALSE | 1 | FALSE | 1 | FALSE | 1 | FALSE | 1 |
| 799 | 799 | 8.46428571 | 1.8609165 | 3 | 0.03084786 | 1 | 0.58913618 | 0 | 1 | 2 | 0.35374209 | 1 | 0.36060486 | 20 | 0.95133411 | FALSE | 0.53356688 | FALSE | 1 | FALSE | 1 | FALSE | 1 | FALSE | 1 | FALSE | 1 |
| 800 | 800 | 7.22222222 | 2.18540937 | 2 | 0.48341887 | 3 | 0.33367983 | 2 | 0.66735975 | 3 | 0.55805561 | 2 | 0.27647607 | 51 | 0.82488355 | FALSE | 1 | FALSE | 1 | FALSE | 1 | FALSE | 1 | FALSE | 1 | FALSE | 1 |
| 801 | 801 | 9.76190476 | 1.84918932 | 0 | 1 | 0 | 1 | 2 | 0.17308361 | 1 | 0.62783522 | 1 | 0.29368245 | 17 | 0.77849076 | FALSE | 1 | FALSE | 1 | FALSE | 1 | FALSE | 1 | FALSE | 1 | FALSE | 1 |
| 802 | 802 | 4.94392523 | 2.27678974 | 9 | 0.09406713 | 6 | 0.6886738 | 10 | 0.2312291 | 7 | 0.86228256 | 6 | 0.13430753 | 174 | 0.84949433 | FALSE | 0.84867611 | FALSE | 1 | FALSE | 1 | FALSE | 1 | FALSE | 1 | FALSE | 1 |
| 803 | 803 | 5.34615385 | 3.12486686 | 1 | 0.49162634 | 0 | 1 | 1 | 0.61440466 | 1 | 0.70601446 | 0 | 1 | 23 | 0.40372818 | FALSE | 1 | FALSE | 1 | FALSE | 1 | FALSE | 1 | FALSE | 1 | FALSE | 1 |
| 804 | 804 | 11 | 3.10912635 | 0 | 1 | 0 | 1 | 0 | 1 | 0 | 1 | 0 | 1 | 6 | 0.36077158 | FALSE | 1 | FALSE | 1 | FALSE | 1 | FALSE | 1 | FALSE | 1 | FALSE | 1 |
| 805 | 805 | 8.14583333 | 2.80616571 | 1 | 0.71380452 | 2 | 0.46321244 | 1 | 0.8283465 | 4 | 0.17698522 | 0 | 1 | 40 | 0.66905473 | FALSE | 1 | FALSE | 1 | FALSE | 1 | FALSE | 1 | FALSE | 1 | FALSE | 1 |
| 806 | 806 | 9.14814815 | 2.47511621 | 0 | 1 | 2 | 0.21684246 | 1 | 0.62830986 | 0 | 1 | 1 | 0.36060486 | 23 | 0.58307573 | FALSE | 1 | FALSE | 1 | FALSE | 1 | FALSE | 1 | FALSE | 1 | FALSE | 1 |
| 807 | 807 | 9.35 | 4.31595876 | 0 | 1 | 0 | 1 | 2 | 0.14749808 | 3 | 0.05383577 | 0 | 1 | 14 | 0.93677253 | FALSE | 1 | FALSE | 1 | FALSE | 1 | FALSE | 1 | FALSE | 1 | FALSE | 1 |
| 808 | 808 | 11.0714286 | 1.1627729 | 0 | 1 | 0 | 1 | 1 | 0.40110418 | 1 | 0.48242099 | 0 | 1 | 12 | 0.62236227 | FALSE | 1 | FALSE | 1 | FALSE | 1 | FALSE | 1 | FALSE | 1 | FALSE | 1 |
| 809 | 809 | 8.27777778 | 1.81981548 | 2 | 0.20692137 | 1 | 0.6629941 | 1 | 0.70185639 | 2 | 0.45159487 | 1 | 0.42123867 | 26 | 0.86852567 | FALSE | 1 | FALSE | 1 | FALSE | 1 | FALSE | 1 | FALSE | 1 | FALSE | 1 |
| 810 | 810 | 5.70833333 | 4.37301542 | 0 | 1 | 1 | 0.54637326 | 1 | 0.58502114 | 1 | 0.67692816 | 0 | 1 | 21 | 0.47053137 | FALSE | 1 | FALSE | 1 | FALSE | 1 | FALSE | 1 | FALSE | 1 | FALSE | 1 |
| 811 | 811 | 8.10526316 | 2.61362295 | 1 | 0.38989951 | 0 | 1 | 1 | 0.50143826 | 1 | 0.59104357 | 0 | 1 | 16 | 0.65565778 | FALSE | 1 | FALSE | 1 | FALSE | 1 | FALSE | 1 | FALSE | 1 | FALSE | 1 |
| 812 | 812 | 9.77777778 | 2.73973956 | 0 | 1 | 1 | 0.25628163 | 0 | 1 | 0 | 1 | 1 | 0.1383237 | 7 | 0.84574168 | FALSE | 1 | FALSE | 1 | FALSE | 1 | FALSE | 1 | FALSE | 1 | FALSE | 1 |
| 813 | 813 | 9.11111111 | 1.2862041 | 1 | 0.20854748 | 0 | 1 | 1 | 0.28067846 | 0 | 1 | 0 | 1 | 7 | 0.84574168 | FALSE | 1 | FALSE | 1 | FALSE | 1 | FALSE | 1 | FALSE | 1 | FALSE | 1 |
| 814 | 814 | 9.21052632 | 2.1662227 | 0 | 1 | 1 | 0.46503795 | 1 | 0.50143826 | 0 | 1 | 1 | 0.26986775 | 16 | 0.65565778 | FALSE | 1 | FALSE | 1 | FALSE | 1 | FALSE | 1 | FALSE | 1 | FALSE | 1 |
| 815 | 815 | 11.7619048 | 1.97375065 | 3 | 0.01568237 | 1 | 0.49918218 | 1 | 0.53671415 | 1 | 0.62783522 | 0 | 1 | 15 | 0.96513691 | FALSE | 0.38695764 | FALSE | 1 | FALSE | 1 | FALSE | 1 | FALSE | 1 | FALSE | 1 |
| 816 | 816 | 9 | 3.31662479 | 0 | 1 | 0 | 1 | 0 | 1 | 0 | 1 | 1 | 0.09446378 | 5 | 0.76185568 | FALSE | 1 | FALSE | 1 | FALSE | 1 | FALSE | 1 | FALSE | 1 | FALSE | 1 |
| 817 | 817 | 10.0666667 | 0.77172246 | 1 | 0.32293355 | 0 | 1 | 1 | 0.42266264 | 1 | 0.50623194 | 0 | 1 | 12 | 0.80329917 | FALSE | 1 | FALSE | 1 | FALSE | 1 | FALSE | 1 | FALSE | 1 | FALSE | 1 |
| 818 | 818 | 8 | 2.76026224 | 1 | 0.42086991 | 0 | 1 | 1 | 0.53671415 | 1 | 0.62783522 | 0 | 1 | 18 | 0.57962763 | FALSE | 1 | FALSE | 1 | FALSE | 1 | FALSE | 1 | FALSE | 1 | FALSE | 1 |
| 819 | 819 | 8.04 | 2.64544892 | 0 | 1 | 0 | 1 | 0 | 1 | 2 | 0.31968852 | 0 | 1 | 23 | 0.22727183 | FALSE | 1 | FALSE | 1 | FALSE | 1 | FALSE | 1 | FALSE | 1 | FALSE | 1 |
| 820 | 820 | 10.9375 | 2.35766701 | 1 | 0.34033054 | 2 | 0.09285443 | 0 | 1 | 0 | 1 | 0 | 1 | 13 | 0.76799161 | FALSE | 1 | FALSE | 1 | FALSE | 1 | FALSE | 1 | FALSE | 1 | FALSE | 1 |
| 821 | 821 | 8.76190476 | 2.34834819 | 0 | 1 | 0 | 1 | 1 | 0.53671415 | 0 | 1 | 0 | 1 | 20 | 0.13763971 | FALSE | 1 | FALSE | 1 | FALSE | 1 | FALSE | 1 | FALSE | 1 | FALSE | 1 |
| 822 | 822 | 10.5454546 | 3.47363149 | 0 | 1 | 1 | 0.30368014 | 1 | 0.33149969 | 0 | 1 | 0 | 1 | 9 | 0.75985366 | FALSE | 1 | FALSE | 1 | FALSE | 1 | FALSE | 1 | FALSE | 1 | FALSE | 1 |
| 823 | 823 | 10.75 | 1.98431348 | 1 | 0.18769745 | 0 | 1 | 0 | 1 | 0 | 1 | 0 | 1 | 7 | 0.63752397 | FALSE | 1 | FALSE | 1 | FALSE | 1 | FALSE | 1 | FALSE | 1 | FALSE | 1 |
| 824 | 824 | 10.3125 | 2.51790464 | 0 | 1 | 0 | 1 | 1 | 0.42266264 | 0 | 1 | 0 | 1 | 14 | 0.29525409 | FALSE | 1 | FALSE | 1 | FALSE | 1 | FALSE | 1 | FALSE | 1 | FALSE | 1 |
| 825 | 825 | 9.70588235 | 2.53828812 | 0 | 1 | 0 | 1 | 0 | 1 | 2 | 0.18212134 | 0 | 1 | 15 | 0.49060435 | FALSE | 1 | FALSE | 1 | FALSE | 1 | FALSE | 1 | FALSE | 1 | FALSE | 1 |
| 826 | 826 | 8.4 | 2.69602883 | 2 | 0.22589135 | 0 | 1 | 1 | 0.72299506 | 2 | 0.48233989 | 1 | 0.44015537 | 29 | 0.69927719 | FALSE | 1 | FALSE | 1 | FALSE | 1 | FALSE | 1 | FALSE | 1 | FALSE | 1 |
| 827 | 827 | 10.64 | 1.49345238 | 1 | 0.47819878 | 2 | 0.19297191 | 1 | 0.59998149 | 0 | 1 | 1 | 0.33902672 | 20 | 0.8152498 | FALSE | 1 | FALSE | 1 | FALSE | 1 | FALSE | 1 | FALSE | 1 | FALSE | 1 |
| 828 | 828 | 6.7027027 | 1.75321889 | 1 | 0.85522402 | 3 | 0.43080961 | 2 | 0.75097925 | 2 | 0.86064871 | 1 | 0.70774043 | 65 | 0.2604438 | FALSE | 1 | FALSE | 1 | FALSE | 1 | FALSE | 1 | FALSE | 1 | FALSE | 1 |
| 829 | 829 | 7.84848485 | 2.46332139 | 0 | 1 | 2 | 0.28941836 | 1 | 0.70185639 | 1 | 0.78873814 | 0 | 1 | 29 | 0.39648374 | FALSE | 1 | FALSE | 1 | FALSE | 1 | FALSE | 1 | FALSE | 1 | FALSE | 1 |
| 830 | 830 | 10.0869565 | 1.86298133 | 2 | 0.11647716 | 1 | 0.53115654 | 3 | 0.04772439 | 1 | 0.6613267 | 2 | 0.05389003 | 14 | 0.99865094 | FALSE | 0.93181727 | FALSE | 1 | FALSE | 1 | FALSE | 1 | FALSE | 1 | FALSE | 1 |
| 831 | 831 | 10.5238095 | 2.06141532 | 3 | 0.01568237 | 0 | 1 | 0 | 1 | 1 | 0.62783522 | 0 | 1 | 17 | 0.77849076 | FALSE | 0.38695764 | FALSE | 1 | FALSE | 1 | FALSE | 1 | FALSE | 1 | FALSE | 1 |
| 832 | 832 | 8.6 | 2.10713075 | 0 | 1 | 0 | 1 | 0 | 1 | 0 | 1 | 0 | 1 | 10 | 0.18273166 | FALSE | 1 | FALSE | 1 | FALSE | 1 | FALSE | 1 | FALSE | 1 | FALSE | 1 |
| 833 | 833 | 10.5 | 1.6583124 | 0 | 1 | 0 | 1 | 0 | 1 | 2 | 0.048966 | 0 | 1 | 6 | 0.88425551 | FALSE | 1 | FALSE | 1 | FALSE | 1 | FALSE | 1 | FALSE | 1 | FALSE | 1 |
| 834 | 834 | 6.23684211 | 2.10180637 | 2 | 0.58475401 | 4 | 0.23089453 | 0 | 1 | 4 | 0.46403589 | 1 | 0.71735141 | 65 | 0.46718352 | FALSE | 1 | FALSE | 1 | FALSE | 1 | FALSE | 1 | FALSE | 1 | FALSE | 1 |
| 835 | 835 | 6.22641509 | 3.39056131 | 0 | 1 | 2 | 0.50543179 | 3 | 0.28676219 | 4 | 0.21499032 | 0 | 1 | 43 | 0.71191105 | FALSE | 1 | FALSE | 1 | FALSE | 1 | FALSE | 1 | FALSE | 1 | FALSE | 1 |
| 836 | 836 | 9.28571429 | 1.57790872 | 0 | 1 | 0 | 1 | 0 | 1 | 0 | 1 | 1 | 0.20679328 | 13 | 0.33283729 | FALSE | 1 | FALSE | 1 | FALSE | 1 | FALSE | 1 | FALSE | 1 | FALSE | 1 |
| 837 | 837 | 7.42424242 | 1.27992194 | 1 | 0.8213973 | 1 | 0.88704765 | 2 | 0.69218142 | 3 | 0.58996791 | 1 | 0.66594933 | 58 | 0.27702369 | FALSE | 1 | FALSE | 1 | FALSE | 1 | FALSE | 1 | FALSE | 1 | FALSE | 1 |
| 838 | 838 | 9.88888889 | 2.2825154 | 0 | 1 | 0 | 1 | 0 | 1 | 0 | 1 | 0 | 1 | 9 | 0.21661397 | FALSE | 1 | FALSE | 1 | FALSE | 1 | FALSE | 1 | FALSE | 1 | FALSE | 1 |
| 839 | 839 | 8 | 2.1821789 | 0 | 1 | 0 | 1 | 0 | 1 | 0 | 1 | 1 | 0.29368245 | 20 | 0.13763971 | FALSE | 1 | FALSE | 1 | FALSE | 1 | FALSE | 1 | FALSE | 1 | FALSE | 1 |
| 840 | 840 | 7.83333333 | 2.05480467 | 0 | 1 | 0 | 1 | 0 | 1 | 1 | 0.67692816 | 0 | 1 | 23 | 0.09196389 | FALSE | 1 | FALSE | 1 | FALSE | 1 | FALSE | 1 | FALSE | 1 | FALSE | 1 |
| 841 | 841 | 2.13862333 | 1.05982593 | 40 | 0.00444164 | 40 | 0.13272086 | 43 | 0.17492671 | 40 | 0.91044897 | 29 | 0.00215178 | 848 | 0.99698283 | FALSE | 0.19457727 | FALSE | 1 | FALSE | 1 | FALSE | 1 | FALSE | 0.26309483 | FALSE | 1 |
| 842 | 842 | 9 | 2.61116484 | 1 | 0.4357638 | 0 | 1 | 0 | 1 | 1 | 0.64497439 | 1 | 0.30529951 | 19 | 0.54233419 | FALSE | 1 | FALSE | 1 | FALSE | 1 | FALSE | 1 | FALSE | 1 | FALSE | 1 |
| 843 | 843 | 9.8 | 1.6 | 2 | 0.04858661 | 0 | 1 | 0 | 1 | 0 | 1 | 0 | 1 | 12 | 0.62236227 | FALSE | 0.649301 | FALSE | 1 | FALSE | 1 | FALSE | 1 | FALSE | 1 | FALSE | 1 |
| 844 | 844 | 9.92307692 | 2.15293021 | 1 | 0.63789042 | 0 | 1 | 2 | 0.41139386 | 2 | 0.54052736 | 1 | 0.4761699 | 33 | 0.59061307 | FALSE | 1 | FALSE | 1 | FALSE | 1 | FALSE | 1 | FALSE | 1 | FALSE | 1 |

| | | | | | | | | | | | | | | | | | | | | | | | | | | | |
|---|---|---|---|---|---|---|---|---|---|---|---|---|---|---|---|---|---|---|---|---|---|---|---|---|---|---|---|
| 845 | 845 | 8.53846154 | 2.5605103 | 1 | 0.47819878 | 1 | 0.56109832 | 0 | 1 | 0 | 1 | 0 | 1 | 23 | 0.22727183 | FALSE | 1 | FALSE | 1 | FALSE | 1 | FALSE | 1 | FALSE | 1 | FALSE | 1 |
| 846 | 846 | 9.13157895 | 2.62749683 | 1 | 0.62830697 | 0 | 1 | 1 | 0.75193685 | 1 | 0.83318955 | 0 | 1 | 35 | 0.13407055 | FALSE | 1 | FALSE | 1 | FALSE | 1 | FALSE | 1 | FALSE | 1 | FALSE | 1 |
| 847 | 847 | 7.51428571 | 1.27327314 | 5 | 0.03316372 | 6 | 0.02509509 | 1 | 0.92379401 | 1 | 0.9633793 | 1 | 0.68753645 | 56 | 0.87889835 | FALSE | 0.56836235 | FALSE | 0.82367276 | FALSE | 1 | FALSE | 1 | FALSE | 1 | FALSE | 1 |
| 848 | 848 | 10.8571429 | 2.09956264 | 0 | 1 | 0 | 1 | 0 | 1 | 1 | 0.28044368 | 0 | 1 | 6 | 0.69920655 | FALSE | 1 | FALSE | 1 | FALSE | 1 | FALSE | 1 | FALSE | 1 | FALSE | 1 |
| 849 | 849 | 8.61111111 | 1.76820339 | 2 | 0.07645454 | 4 | 0.00227902 | 0 | 1 | 1 | 0.57131013 | 1 | 0.25766374 | 10 | 0.99931761 | FALSE | 0.77424195 | FALSE | 0.42431176 | FALSE | 1 | FALSE | 1 | FALSE | 1 | FALSE | 1 |
| 850 | 850 | 5.51351351 | 2.43423357 | 1 | 0.6184714 | 1 | 0.70472939 | 3 | 0.14625315 | 4 | 0.08780467 | 1 | 0.45845922 | 27 | 0.97789976 | FALSE | 1 | FALSE | 1 | FALSE | 1 | FALSE | 1 | FALSE | 1 | FALSE | 1 |
| 851 | 851 | 11.375 | 1.9324531 | 2 | 0.12501436 | 1 | 0.54637326 | 0 | 1 | 0 | 1 | 0 | 1 | 21 | 0.47053137 | FALSE | 0.95622564 | FALSE | 1 | FALSE | 1 | FALSE | 1 | FALSE | 1 | FALSE | 1 |
| 852 | 852 | 9.875 | 2.52178013 | 1 | 0.32293355 | 0 | 1 | 0 | 1 | 0 | 1 | 0 | 1 | 14 | 0.29525409 | FALSE | 1 | FALSE | 1 | FALSE | 1 | FALSE | 1 | FALSE | 1 | FALSE | 1 |
| 853 | 853 | 11.16 | 2.34401365 | 1 | 0.47819878 | 1 | 0.56109832 | 3 | 0.05884411 | 1 | 0.69181315 | 0 | 1 | 19 | 0.91734261 | FALSE | 1 | FALSE | 1 | FALSE | 1 | FALSE | 1 | FALSE | 1 | FALSE | 1 |
| 854 | 854 | 9.8 | 2.87981481 | 0 | 1 | 0 | 1 | 1 | 0.42266264 | 0 | 1 | 0 | 1 | 14 | 0.29525409 | FALSE | 1 | FALSE | 1 | FALSE | 1 | FALSE | 1 | FALSE | 1 | FALSE | 1 |
| 855 | 855 | 5.84615385 | 3.02161836 | 3 | 0.6427009 | 4 | 0.59899011 | 7 | 0.17642727 | 5 | 0.70669001 | 2 | 0.62508358 | 107 | 0.65291815 | FALSE | 1 | FALSE | 1 | FALSE | 1 | FALSE | 1 | FALSE | 1 | FALSE | 1 |
| 856 | 856 | 8.38888889 | 1.49587911 | 0 | 1 | 1 | 0.68454804 | 0 | 1 | 3 | 0.21561745 | 0 | 1 | 31 | 0.34231756 | FALSE | 1 | FALSE | 1 | FALSE | 1 | FALSE | 1 | FALSE | 1 | FALSE | 1 |
| 857 | 857 | 8.66666667 | 2.13437475 | 0 | 1 | 0 | 1 | 1 | 0.33149969 | 1 | 0.40390032 | 0 | 1 | 9 | 0.75985366 | FALSE | 1 | FALSE | 1 | FALSE | 1 | FALSE | 1 | FALSE | 1 | FALSE | 1 |
| 858 | 858 | 9.05882353 | 1.78129458 | 1 | 0.58738519 | 0 | 1 | 2 | 0.34668321 | 2 | 0.46709861 | 1 | 0.43077488 | 28 | 0.72558119 | FALSE | 1 | FALSE | 1 | FALSE | 1 | FALSE | 1 | FALSE | 1 | FALSE | 1 |
| 859 | 859 | 9.77777778 | 3.29234053 | 0 | 1 | 0 | 1 | 1 | 0.28067846 | 0 | 1 | 0 | 1 | 8 | 0.5780332 | FALSE | 1 | FALSE | 1 | FALSE | 1 | FALSE | 1 | FALSE | 1 | FALSE | 1 |
| 860 | 860 | 8.05 | 2.78343313 | 0 | 1 | 2 | 0.12434464 | 2 | 0.14749808 | 1 | 0.59104357 | 0 | 1 | 14 | 0.93677253 | FALSE | 1 | FALSE | 1 | FALSE | 1 | FALSE | 1 | FALSE | 1 | FALSE | 1 |
| 861 | 861 | 6.45714286 | 3.77467393 | 0 | 1 | 0 | 1 | 0 | 1 | 4 | 0.07465533 | 0 | 1 | 31 | 0.34231756 | FALSE | 1 | FALSE | 1 | FALSE | 1 | FALSE | 1 | FALSE | 1 | FALSE | 1 |
| 862 | 862 | 8.82352941 | 1.72303747 | 0 | 1 | 2 | 0.10307112 | 0 | 1 | 1 | 0.55062772 | 0 | 1 | 14 | 0.7313145 | FALSE | 1 | FALSE | 1 | FALSE | 1 | FALSE | 1 | FALSE | 1 | FALSE | 1 |
| 863 | 863 | 8.05882353 | 1.69673001 | 1 | 0.35728305 | 0 | 1 | 0 | 1 | 2 | 0.18212134 | 0 | 1 | 14 | 0.7313145 | FALSE | 1 | FALSE | 1 | FALSE | 1 | FALSE | 1 | FALSE | 1 | FALSE | 1 |
| 864 | 864 | 4.90740741 | 3.8213267 | 1 | 0.74890551 | 1 | 0.82613965 | 2 | 0.57303317 | 1 | 0.91797304 | 0 | 1 | 48 | 0.14364394 | FALSE | 1 | FALSE | 1 | FALSE | 1 | FALSE | 1 | FALSE | 1 | FALSE | 1 |
| 865 | 865 | 9.36363636 | 2.53243423 | 0 | 1 | 0 | 1 | 0 | 1 | 2 | 0.08789566 | 0 | 1 | 9 | 0.75985366 | FALSE | 1 | FALSE | 1 | FALSE | 1 | FALSE | 1 | FALSE | 1 | FALSE | 1 |
| 866 | 866 | 10.1666667 | 1.72401341 | 2 | 0.12501436 | 0 | 1 | 0 | 1 | 2 | 0.30249307 | 0 | 1 | 20 | 0.68270729 | FALSE | 0.95622564 | FALSE | 1 | FALSE | 1 | FALSE | 1 | FALSE | 1 | FALSE | 1 |
| 867 | 867 | 11.2857143 | 2.18529408 | 2 | 0.04858661 | 0 | 1 | 0 | 1 | 0 | 1 | 0 | 1 | 12 | 0.62236227 | FALSE | 0.649301 | FALSE | 1 | FALSE | 1 | FALSE | 1 | FALSE | 1 | FALSE | 1 |
| 868 | 868 | 10.8947368 | 2.57303296 | 0 | 1 | 0 | 1 | 2 | 0.14749808 | 0 | 1 | 0 | 1 | 17 | 0.41141543 | FALSE | 1 | FALSE | 1 | FALSE | 1 | FALSE | 1 | FALSE | 1 | FALSE | 1 |
| 869 | 869 | 8.24719101 | 0.6915363 | 8 | 0.00142178 | 4 | 0.29545046 | 2 | 0.81591765 | 7 | 0.09449282 | 1 | 0.75686725 | 63 | 0.99538493 | FALSE | 0.09317759 | FALSE | 1 | FALSE | 1 | FALSE | 1 | FALSE | 1 | FALSE | 1 |
| 870 | 870 | 5.80952381 | 3.06468436 | 0 | 1 | 1 | 0.49918218 | 1 | 0.53671415 | 1 | 0.62783522 | 0 | 1 | 18 | 0.57962763 | FALSE | 1 | FALSE | 1 | FALSE | 1 | FALSE | 1 | FALSE | 1 | FALSE | 1 |
| 871 | 871 | 8.85714286 | 2.74791201 | 1 | 0.16630142 | 0 | 1 | 1 | 0.22601089 | 1 | 0.28044368 | 0 | 1 | 4 | 0.98605104 | FALSE | 1 | FALSE | 1 | FALSE | 1 | FALSE | 1 | FALSE | 1 | FALSE | 1 |
| 872 | 872 | 9.70588235 | 2.58422149 | 0 | 1 | 0 | 1 | 0 | 1 | 0 | 1 | 1 | 0.24525764 | 16 | 0.23070182 | FALSE | 1 | FALSE | 1 | FALSE | 1 | FALSE | 1 | FALSE | 1 | FALSE | 1 |
| 873 | 873 | 6.25 | 2.94745653 | 1 | 0.40558521 | 1 | 0.48239024 | 1 | 0.51939841 | 1 | 0.60987146 | 1 | 0.28187291 | 15 | 0.92113679 | FALSE | 1 | FALSE | 1 | FALSE | 1 | FALSE | 1 | FALSE | 1 | FALSE | 1 |
| 874 | 874 | 10.25 | 1.29903811 | 0 | 1 | 0 | 1 | 0 | 1 | 0 | 1 | 1 | 0.06400211 | 3 | 0.8823539 | FALSE | 1 | FALSE | 1 | FALSE | 1 | FALSE | 1 | FALSE | 1 | FALSE | 1 |
| 875 | 875 | 3.09667467 | 0.71295802 | 90 | 0.00942607 | 115 | 0.00208258 | 117 | 0.05219592 | 118 | 0.96808904 | 61 | 0.00624965 | 2400 | 0.99946783 | FALSE | 0.29138989 | FALSE | 0.42264619 | FALSE | 1 | FALSE | 1 | FALSE | 0.46542874 | FALSE | 1 |
| 876 | 876 | 9.38888889 | 2.28859221 | 1 | 0.37380237 | 1 | 0.44710673 | 2 | 0.13502329 | 1 | 0.57131013 | 0 | 1 | 13 | 0.95041913 | FALSE | 1 | FALSE | 1 | FALSE | 1 | FALSE | 1 | FALSE | 1 | FALSE | 1 |
| 877 | 877 | 12 | 3.33809184 | 0 | 1 | 0 | 1 | 2 | 0.02394967 | 0 | 1 | 0 | 1 | 5 | 0.91847773 | FALSE | 1 | FALSE | 1 | FALSE | 1 | FALSE | 1 | FALSE | 1 | FALSE | 1 |
| 878 | 878 | 9.86666667 | 3.11697859 | 1 | 0.32293355 | 0 | 1 | 0 | 1 | 1 | 0.50623194 | 0 | 1 | 13 | 0.57712805 | FALSE | 1 | FALSE | 1 | FALSE | 1 | FALSE | 1 | FALSE | 1 | FALSE | 1 |
| 879 | 879 | 10.6470588 | 3.23475931 | 0 | 1 | 1 | 0.40942991 | 0 | 1 | 1 | 0.52895088 | 2 | 0.02752114 | 12 | 0.90922186 | FALSE | 1 | FALSE | 1 | FALSE | 1 | FALSE | 1 | FALSE | 0.79106376 | FALSE | 1 |
| 880 | 880 | 9.2 | 0.9797959 | 0 | 1 | 0 | 1 | 0 | 1 | 1 | 0.69181315 | 0 | 1 | 24 | 0.08019798 | FALSE | 1 | FALSE | 1 | FALSE | 1 | FALSE | 1 | FALSE | 1 | FALSE | 1 |
| 881 | 881 | 9.37037037 | 3.88871252 | 0 | 1 | 1 | 0.57534752 | 1 | 0.61440466 | 0 | 1 | 1 | 0.3499045 | 23 | 0.40372818 | FALSE | 1 | FALSE | 1 | FALSE | 1 | FALSE | 1 | FALSE | 1 | FALSE | 1 |
| 882 | 882 | 10.9 | 2.38537209 | 1 | 0.38989951 | 2 | 0.12434464 | 0 | 1 | 0 | 1 | 2 | 0.03799168 | 14 | 0.93677253 | FALSE | 1 | FALSE | 1 | FALSE | 1 | FALSE | 1 | FALSE | 0.87669812 | FALSE | 1 |
| 883 | 883 | 6.34090909 | 3.26790997 | 0 | 1 | 0 | 1 | 3 | 0.20921284 | 2 | 0.60667406 | 1 | 0.51797501 | 38 | 0.45705039 | FALSE | 1 | FALSE | 1 | FALSE | 1 | FALSE | 1 | FALSE | 1 | FALSE | 1 |
| 884 | 884 | 8.37209302 | 2.53370999 | 1 | 0.67382914 | 5 | 0.01196839 | 1 | 0.79363407 | 0 | 1 | 1 | 0.50988734 | 35 | 0.7801651 | FALSE | 1 | FALSE | 0.67617265 | FALSE | 1 | FALSE | 1 | FALSE | 1 | FALSE | 1 |
| 885 | 885 | 5.69411765 | 2.71823269 | 2 | 0.62636022 | 1 | 0.93364809 | 5 | 0.17133706 | 9 | 0.01238625 | 1 | 0.74434486 | 64 | 0.95410071 | FALSE | 1 | FALSE | 1 | FALSE | 1 | FALSE | 1 | FALSE | 1 | FALSE | 1 |
| 886 | 886 | 9.83333333 | 2.29613201 | 1 | 0.52988231 | 2 | 0.24095584 | 2 | 0.27983822 | 2 | 0.3871811 | 1 | 0.38148475 | 21 | 0.97185642 | FALSE | 1 | FALSE | 1 | FALSE | 1 | FALSE | 1 | FALSE | 1 | FALSE | 1 |
| 887 | 887 | 8.83333333 | 1.21335165 | 1 | 0.14434519 | 1 | 0.179097 | 0 | 1 | 1 | 0.24578796 | 0 | 1 | 3 | 0.9931827 | FALSE | 1 | FALSE | 1 | FALSE | 1 | FALSE | 1 | FALSE | 1 | FALSE | 1 |
| 888 | 888 | 6.56756757 | 2.76112531 | 0 | 1 | 3 | 0.43080961 | 5 | 0.12620681 | 4 | 0.4435008 | 3 | 0.1209957 | 59 | 0.89469615 | FALSE | 1 | FALSE | 1 | FALSE | 1 | FALSE | 1 | FALSE | 1 | FALSE | 1 |
| 889 | 889 | 8.94117647 | 3.09592879 | 1 | 0.34033054 | 0 | 1 | 1 | 0.44344817 | 0 | 1 | 0 | 1 | 14 | 0.53308261 | FALSE | 1 | FALSE | 1 | FALSE | 1 | FALSE | 1 | FALSE | 1 | FALSE | 1 |
| 890 | 890 | 6.875 | 3.71609674 | 2 | 0.26413583 | 0 | 1 | 3 | 0.16357218 | 3 | 0.26522531 | 0 | 1 | 31 | 0.85613899 | FALSE | 1 | FALSE | 1 | FALSE | 1 | FALSE | 1 | FALSE | 1 | FALSE | 1 |
| 891 | 891 | 9.63157895 | 1.86971786 | 0 | 1 | 0 | 1 | 0 | 1 | 1 | 0.59104357 | 1 | 0.26986775 | 17 | 0.41141543 | FALSE | 1 | FALSE | 1 | FALSE | 1 | FALSE | 1 | FALSE | 1 | FALSE | 1 |
| 892 | 892 | 5.4057971 | 2.81465544 | 1 | 0.83491628 | 0 | 1 | 2 | 0.71546205 | 4 | 0.39127216 | 0 | 1 | 62 | 0.13493316 | FALSE | 1 | FALSE | 1 | FALSE | 1 | FALSE | 1 | FALSE | 1 | FALSE | 1 |
| 893 | 893 | 8.375 | 2.19492787 | 0 | 1 | 0 | 1 | 0 | 1 | 2 | 0.30249307 | 1 | 0.32796862 | 21 | 0.47053137 | FALSE | 1 | FALSE | 1 | FALSE | 1 | FALSE | 1 | FALSE | 1 | FALSE | 1 |
| 894 | 894 | 9.8 | 3.81575681 | 0 | 1 | 1 | 0.38964394 | 2 | 0.09928574 | 1 | 0.50623194 | 0 | 1 | 11 | 0.92879412 | FALSE | 1 | FALSE | 1 | FALSE | 1 | FALSE | 1 | FALSE | 1 | FALSE | 1 |
| 895 | 895 | 8.06976744 | 2.81520249 | 2 | 0.30241117 | 1 | 0.75788114 | 2 | 0.46083448 | 0 | 1 | 2 | 0.1563365 | 36 | 0.6451239 | FALSE | 1 | FALSE | 1 | FALSE | 1 | FALSE | 1 | FALSE | 1 | FALSE | 1 |
| 896 | 896 | 9.11538462 | 3.14186279 | 1 | 0.49162634 | 0 | 1 | 0 | 1 | 0 | 1 | 0 | 1 | 25 | 0.06985983 | FALSE | 1 | FALSE | 1 | FALSE | 1 | FALSE | 1 | FALSE | 1 | FALSE | 1 |
| 897 | 897 | 10.2 | 1.76635217 | 0 | 1 | 0 | 1 | 2 | 0.22597287 | 2 | 0.31968852 | 0 | 1 | 21 | 0.64957167 | FALSE | 1 | FALSE | 1 | FALSE | 1 | FALSE | 1 | FALSE | 1 | FALSE | 1 |
| 898 | 898 | 8.74074074 | 2.8099373 | 0 | 1 | 2 | 0.21684246 | 2 | 0.25286935 | 0 | 1 | 1 | 0.36060486 | 22 | 0.76237877 | FALSE | 1 | FALSE | 1 | FALSE | 1 | FALSE | 1 | FALSE | 1 | FALSE | 1 |
| 899 | 899 | 7.5625 | 3.25900196 | 0 | 1 | 2 | 0.2651851 | 0 | 1 | 0 | 1 | 0 | 1 | 29 | 0.11661149 | FALSE | 1 | FALSE | 1 | FALSE | 1 | FALSE | 1 | FALSE | 1 | FALSE | 1 |
| 900 | 900 | 9.8 | 2.85657137 | 1 | 0.22886536 | 0 | 1 | 0 | 1 | 0 | 1 | 0 | 1 | 9 | 0.5215547 | FALSE | 1 | FALSE | 1 | FALSE | 1 | FALSE | 1 | FALSE | 1 | FALSE | 1 |
| 901 | 901 | 8.51807229 | 1.6233347 | 1 | 0.88571933 | 2 | 0.75548815 | 3 | 0.57805941 | 2 | 0.90041492 | 3 | 0.15488013 | 72 | 0.33817167 | FALSE | 1 | FALSE | 1 | FALSE | 1 | FALSE | 1 | FALSE | 1 | FALSE | 1 |
| 902 | 902 | 8.66666667 | 3.04776785 | 0 | 1 | 0 | 1 | 1 | 0.42266264 | 1 | 0.50623194 | 1 | 0.2198258 | 12 | 0.80329917 | FALSE | 1 | FALSE | 1 | FALSE | 1 | FALSE | 1 | FALSE | 1 | FALSE | 1 |
| 903 | 903 | 8.49090909 | 1.89614659 | 6 | 0.00266387 | 3 | 0.26245929 | 3 | 0.31652594 | 1 | 0.92538938 | 0 | 1 | 42 | 0.96038295 | FALSE | 0.13988708 | FALSE | 1 | FALSE | 1 | FALSE | 1 | FALSE | 1 | FALSE | 1 |
| 904 | 904 | 9.325 | 1.87599973 | 2 | 0.25455533 | 3 | 0.12352882 | 0 | 1 | 1 | 0.83318955 | 0 | 1 | 32 | 0.61800916 | FALSE | 1 | FALSE | 1 | FALSE | 1 | FALSE | 1 | FALSE | 1 | FALSE | 1 |
| 905 | 905 | 10.4615385 | 1.49950682 | 2 | 0.04234484 | 2 | 0.06426041 | 0 | 1 | 0 | 1 | 0 | 1 | 9 | 0.96018168 | FALSE | 0.62842194 | FALSE | 1 | FALSE | 1 | FALSE | 1 | FALSE | 1 | FALSE | 1 |
| 906 | 906 | 7.94117647 | 1.92416791 | 0 | 1 | 0 | 1 | 0 | 1 | 0 | 1 | 1 | 0.23264612 | 15 | 0.26127859 | FALSE | 1 | FALSE | 1 | FALSE | 1 | FALSE | 1 | FALSE | 1 | FALSE | 1 |
| 907 | 907 | 9.25 | 2.89035753 | 0 | 1 | 0 | 1 | 1 | 0.35555293 | 2 | 0.1023796 | 0 | 1 | 9 | 0.89630573 | FALSE | 1 | FALSE | 1 | FALSE | 1 | FALSE | 1 | FALSE | 1 | FALSE | 1 |
| 908 | 908 | 5.4375 | 3.22041826 | 6 | 0.005277 | 4 | 0.14592986 | 2 | 0.66735975 | 0 | 1 | 1 | 0.64879513 | 50 | 0.89566304 | FALSE | 0.2089654 | FALSE | 1 | FALSE | 1 | FALSE | 1 | FALSE | 1 | FALSE | 1 |
| 909 | 909 | 5.375 | 3.63790805 | 0 | 1 | 2 | 0.27730767 | 2 | 0.32011871 | 0 | 1 | 0 | 1 | 28 | 0.42532177 | FALSE | 1 | FALSE | 1 | FALSE | 1 | FALSE | 1 | FALSE | 1 | FALSE | 1 |
| 910 | 910 | 10.1 | 3.6180105 | 1 | 0.22886536 | 1 | 0.28036922 | 0 | 1 | 1 | 0.37517262 | 0 | 1 | 7 | 0.94326215 | FALSE | 1 | FALSE | 1 | FALSE | 1 | FALSE | 1 | FALSE | 1 | FALSE | 1 |
| 911 | 911 | 7.4 | 3.87126164 | 3 | 0.28789392 | 4 | 0.21021455 | 2 | 0.74419436 | 3 | 0.65845889 | 1 | 0.7028142 | 60 | 0.75780737 | FALSE | 1 | FALSE | 1 | FALSE | 1 | FALSE | 1 | FALSE | 1 | FALSE | 1 |
| 912 | 912 | 9.16 | 2.46056904 | 2 | 0.13369621 | 1 | 0.56109832 | 0 | 1 | 1 | 0.69181315 | 2 | 0.06257402 | 19 | 0.91734261 | FALSE | 0.97964162 | FALSE | 1 | FALSE | 1 | FALSE | 1 | FALSE | 1 | FALSE | 1 |
| 913 | 913 | 5.13114754 | 3.55968045 | 0 | 1 | 0 | 1 | 0 | 1 | 0 | 1 | 2 | 0.26425795 | 59 | 0.00229841 | FALSE | 1 | FALSE | 1 | FALSE | 1 | FALSE | 1 | FALSE | 1 | FALSE | 1 |
| 914 | 914 | 9.42105263 | 2.58163126 | 2 | 0.08405377 | 0 | 1 | 0 | 1 | 1 | 0.59104357 | 0 | 1 | 16 | 0.65565778 | FALSE | 0.79236883 | FALSE | 1 | FALSE | 1 | FALSE | 1 | FALSE | 1 | FALSE | 1 |
| 915 | 915 | 10.4285714 | 2.61080956 | 2 | 0.15144274 | 2 | 0.21684246 | 4 | 0.01487236 | 0 | 1 | 1 | 0.36060486 | 18 | 0.99450398 | FALSE | 1 | FALSE | 1 | FALSE | 1 | FALSE | 1 | FALSE | 1 | FALSE | 1 |
| 916 | 916 | 7.90909091 | 3.77683238 | 0 | 1 | 1 | 0.5154318 | 0 | 1 | 2 | 0.26791271 | 2 | 0.04972408 | 17 | 0.8840357 | FALSE | 1 | FALSE | 1 | FALSE | 1 | FALSE | 1 | FALSE | 0.98629461 | FALSE | 1 |
| 917 | 917 | 7.97368421 | 2.91535718 | 0 | 1 | 1 | 0.71433277 | 3 | 0.15483587 | 4 | 0.09477133 | 0 | 1 | 30 | 0.87271204 | FALSE | 1 | FALSE | 1 | FALSE | 1 | FALSE | 1 | FALSE | 1 | FALSE | 1 |
| 918 | 918 | 9.26086957 | 1.072072 | 1 | 0.69843502 | 2 | 0.44132825 | 0 | 1 | 3 | 0.35428318 | 0 | 1 | 40 | 0.40712403 | FALSE | 1 | FALSE | 1 | FALSE | 1 | FALSE | 1 | FALSE | 1 | FALSE | 1 |
| 919 | 919 | 8.0952381 | 3.05356563 | 0 | 1 | 0 | 1 | 2 | 0.16019548 | 2 | 0.23331368 | 0 | 1 | 16 | 0.80826361 | FALSE | 1 | FALSE | 1 | FALSE | 1 | FALSE | 1 | FALSE | 1 | FALSE | 1 |
| 920 | 920 | 11.1428571 | 2.79941685 | 1 | 0.3050805 | 0 | 1 | 0 | 1 | 0 | 1 | 0 | 1 | 13 | 0.33283729 | FALSE | 1 | FALSE | 1 | FALSE | 1 | FALSE | 1 | FALSE | 1 | FALSE | 1 |
| 921 | 921 | 10.2285714 | 2.76803917 | 2 | 0.21638713 | 3 | 0.09602744 | 0 | 1 | 4 | 0.06848449 | 0 | 1 | 25 | 0.9693929 | FALSE | 1 | FALSE | 1 | FALSE | 1 | FALSE | 1 | FALSE | 1 | FALSE | 1 |
| 922 | 922 | 9.7037037 | 2.732721 | 0 | 1 | 1 | 0.58913618 | 2 | 0.25286935 | 0 | 1 | 1 | 0.36060486 | 23 | 0.58307573 | FALSE | 1 | FALSE | 1 | FALSE | 1 | FALSE | 1 | FALSE | 1 | FALSE | 1 |
| 923 | 923 | 9.38461539 | 1.07692308 | 0 | 1 | 1 | 0.3480709 | 0 | 1 | 0 | 1 | 0 | 1 | 12 | 0.374198 | FALSE | 1 | FALSE | 1 | FALSE | 1 | FALSE | 1 | FALSE | 1 | FALSE | 1 |
| 924 | 924 | 11.2857143 | 1.74963553 | 0 | 1 | 0 | 1 | 0 | 1 | 0 | 1 | 0 | 1 | 14 | 0.09251304 | FALSE | 1 | FALSE | 1 | FALSE | 1 | FALSE | 1 | FALSE | 1 | FALSE | 1 |
| 925 | 925 | 7 | 1.45773797 | 0 | 1 | 0 | 1 | 0 | 1 | 2 | 0.16544789 | 1 | 0.23264612 | 13 | 0.76799161 | FALSE | 1 | FALSE | 1 | FALSE | 1 | FALSE | 1 | FALSE | 1 | FALSE | 1 |
| 926 | 926 | 10.6 | 2.30362034 | 0 | 1 | 1 | 0.38964394 | 0 | 1 | 0 | 1 | 0 | 1 | 14 | 0.29525409 | FALSE | 1 | FALSE | 1 | FALSE | 1 | FALSE | 1 | FALSE | 1 | FALSE | 1 |
| 927 | 927 | 9.78571429 | 2.11046953 | 0 | 1 | 0 | 1 | 0 | 1 | 0 | 1 | 0 | 1 | 14 | 0.09251304 | FALSE | 1 | FALSE | 1 | FALSE | 1 | FALSE | 1 | FALSE | 1 | FALSE | 1 |
| 928 | 928 | 9.23529412 | 2.55594835 | 1 | 0.35728305 | 3 | 0.01622926 | 0 | 1 | 1 | 0.55062772 | 0 | 1 | 12 | 0.962093 | FALSE | 1 | FALSE | 0.7256023 | FALSE | 1 | FALSE | 1 | FALSE | 1 | FALSE | 1 |
| 929 | 929 | 9.69047619 | 2.24125916 | 2 | 0.29286138 | 1 | 0.74973543 | 1 | 0.78589454 | 0 | 1 | 2 | 0.15059682 | 36 | 0.50934357 | FALSE | 1 | FALSE | 1 | FALSE | 1 | FALSE | 1 | FALSE | 1 | FALSE | 1 |
| 930 | 930 | 10.2083333 | 2.7984991 | 1 | 0.46441868 | 0 | 1 | 1 | 0.58502114 | 0 | 1 | 0 | 1 | 22 | 0.25237063 | FALSE | 1 | FALSE | 1 | FALSE | 1 | FALSE | 1 | FALSE | 1 | FALSE | 1 |
| 931 | 931 | 6.84210526 | 3.77335754 | 2 | 0.25455533 | 1 | 0.71433277 | 1 | 0.75193685 | 0 | 1 | 0 | 1 | 34 | 0.27051027 | FALSE | 1 | FALSE | 1 | FALSE | 1 | FALSE | 1 | FALSE | 1 | FALSE | 1 |
| 932 | 932 | 3.90225564 | 2.533578 | 10 | 0.00210194 | 7 | 0.13900139 | 8 | 0.1051942 | 3 | 0.94813262 | 6 | 0.02166317 | 99 | 0.99905759 | FALSE | 0.12179217 | FALSE | 1 | FALSE | 1 | FALSE | 1 | FALSE | 0.74564993 | FALSE | 1 |
| 933 | 933 | 9.625 | 3.13996417 | 0 | 1 | 1 | 0.40942991 | 1 | 0.44344817 | 1 | 0.52895088 | 0 | 1 | 13 | 0.76799161 | FALSE | 1 | FALSE | 1 | FALSE | 1 | FALSE | 1 | FALSE | 1 | FALSE | 1 |
| 934 | 934 | 9.44 | 1.29861465 | 1 | 0.47819878 | 4 | 0.00789279 | 3 | 0.05884411 | 2 | 0.31968852 | 1 | 0.33902672 | 14 | 0.99985788 | FALSE | 1 | FALSE | 0.6167796 | FALSE | 1 | FALSE | 1 | FALSE | 1 | FALSE | 1 |
| 935 | 935 | 6.24390244 | 2.9031563 | 2 | 0.28329504 | 2 | 0.38460545 | 0 | 1 | 4 | 0.11715635 | 0 | 1 | 33 | 0.81996213 | FALSE | 1 | FALSE | 1 | FALSE | 1 | FALSE | 1 | FALSE | 1 | FALSE | 1 |
| 936 | 936 | 6.82278481 | 1.72663054 | 1 | 0.87304634 | 1 | 0.92668093 | 5 | 0.15372513 | 5 | 0.29645796 | 3 | 0.13946252 | 64 | 0.83834292 | FALSE | 1 | FALSE | 1 | FALSE | 1 | FALSE | 1 | FALSE | 1 | FALSE | 1 |
| 937 | 937 | 5.59589041 | 2.92504248 | 4 | 0.51844612 | 4 | 0.70117193 | 9 | 0.07920225 | 2 | 0.99205936 | 8 | 0.00261863 | 119 | 0.85967389 | FALSE | 1 | FALSE | 1 | FALSE | 1 | FALSE | 1 | FALSE | 0.29784427 | FALSE | 1 |
| 938 | 938 | 8.75 | 2.04633819 | 0 | 1 | 0 | 1 | 0 | 1 | 0 | 1 | 0 | 1 | 8 | 0.25677166 | FALSE | 1 | FALSE | 1 | FALSE | 1 | FALSE | 1 | FALSE | 1 | FALSE | 1 |

| | | | | | | | | | | | | | | | | | | | | | | | | | | | |
|---|---|---|---|---|---|---|---|---|---|---|---|---|---|---|---|---|---|---|---|---|---|---|---|---|---|---|---|
| 939 | 939 | 9.52941177 | 2.59224289 | 1 | 0.35728305 | 1 | 0.42857733 | 1 | 0.46348836 | 1 | 0.55062772 | 1 | 0.24525764 | 12 | 0.962093 | FALSE | 1 | FALSE | 1 | FALSE | 1 | FALSE | 1 | FALSE | 1 | FALSE | 1 |
| 940 | 940 | 3.90909091 | 2.45151328 | 0 | 1 | 3 | 0.26245929 | 4 | 0.1343514 | 0 | 1 | 1 | 0.59866474 | 47 | 0.50345693 | FALSE | 1 | FALSE | 1 | FALSE | 1 | FALSE | 1 | FALSE | 1 | FALSE | 1 |
| 941 | 941 | 9.56666667 | 2.94033256 | 2 | 0.17884234 | 1 | 0.6278847 | 1 | 0.66709967 | 1 | 0.75658797 | 0 | 1 | 25 | 0.67630471 | FALSE | 1 | FALSE | 1 | FALSE | 1 | FALSE | 1 | FALSE | 1 | FALSE | 1 |
| 942 | 942 | 9.54545455 | 1.92418277 | 0 | 1 | 1 | 0.5154318 | 0 | 1 | 1 | 0.64497439 | 1 | 0.30529951 | 19 | 0.54233419 | FALSE | 1 | FALSE | 1 | FALSE | 1 | FALSE | 1 | FALSE | 1 | FALSE | 1 |
| 943 | 943 | 10.6666667 | 2.6148818 | 2 | 0.26413583 | 3 | 0.13080858 | 2 | 0.41139386 | 2 | 0.54052736 | 0 | 1 | 30 | 0.92795554 | FALSE | 1 | FALSE | 1 | FALSE | 1 | FALSE | 1 | FALSE | 1 | FALSE | 1 |
| 944 | 944 | 10.0714286 | 1.94437966 | 2 | 0.53205439 | 5 | 0.07172628 | 3 | 0.45353909 | 4 | 0.39127216 | 2 | 0.31302643 | 53 | 0.96638194 | FALSE | 1 | FALSE | 1 | FALSE | 1 | FALSE | 1 | FALSE | 1 | FALSE | 1 |
| 945 | 945 | 8.3 | 1.97737199 | 1 | 0.40558521 | 0 | 1 | 2 | 0.16019548 | 0 | 1 | 0 | 1 | 17 | 0.61750632 | FALSE | 1 | FALSE | 1 | FALSE | 1 | FALSE | 1 | FALSE | 1 | FALSE | 1 |
| 946 | 946 | 7.42857143 | 1.74086537 | 5 | 0.01390429 | 3 | 0.27128673 | 5 | 0.04964325 | 1 | 0.92884295 | 2 | 0.23375968 | 40 | 0.9960055 | FALSE | 0.36625055 | FALSE | 1 | FALSE | 1 | FALSE | 1 | FALSE | 1 | FALSE | 1 |
| 947 | 947 | 8.69230769 | 1.87635553 | 1 | 0.49162634 | 2 | 0.20486827 | 1 | 0.61440466 | 3 | 0.11458571 | 1 | 0.3499045 | 18 | 0.98644281 | FALSE | 1 | FALSE | 1 | FALSE | 1 | FALSE | 1 | FALSE | 1 | FALSE | 1 |
| 948 | 948 | 9.9 | 1.22065556 | 0 | 1 | 1 | 0.28036922 | 1 | 0.30655256 | 0 | 1 | 0 | 1 | 8 | 0.80394719 | FALSE | 1 | FALSE | 1 | FALSE | 1 | FALSE | 1 | FALSE | 1 | FALSE | 1 |
| 949 | 949 | 8.18181818 | 1.33608531 | 0 | 1 | 0 | 1 | 0 | 1 | 0 | 1 | 0 | 1 | 11 | 0.15414489 | FALSE | 1 | FALSE | 1 | FALSE | 1 | FALSE | 1 | FALSE | 1 | FALSE | 1 |
| 950 | 950 | 3.54153846 | 2.12290746 | 9 | 0.44425977 | 19 | 0.0079055 | 12 | 0.49086461 | 20 | 0.10237031 | 10 | 0.03779006 | 252 | 0.99886779 | FALSE | 1 | FALSE | 0.6167796 | FALSE | 1 | FALSE | 1 | FALSE | 0.87669812 | FALSE | 1 |
| 951 | 951 | 8.90909091 | 2.50289089 | 1 | 0.4357638 | 1 | 0.5154318 | 2 | 0.18613256 | 1 | 0.64497439 | 1 | 0.30529951 | 16 | 0.95553951 | FALSE | 1 | FALSE | 1 | FALSE | 1 | FALSE | 1 | FALSE | 1 | FALSE | 1 |
| 952 | 952 | 8.77777778 | 2.01143118 | 4 | 0.07746936 | 2 | 0.60991494 | 2 | 0.66735975 | 3 | 0.55805561 | 1 | 0.64879513 | 51 | 0.82488355 | FALSE | 0.77583006 | FALSE | 1 | FALSE | 1 | FALSE | 1 | FALSE | 1 | FALSE | 1 |
| 953 | 953 | 6.75862069 | 2.52784728 | 2 | 0.44060625 | 0 | 1 | 1 | 0.88128698 | 2 | 0.75303313 | 1 | 0.61824432 | 52 | 0.17733928 | FALSE | 1 | FALSE | 1 | FALSE | 1 | FALSE | 1 | FALSE | 1 | FALSE | 1 |
| 954 | 954 | 8.3 | 2.32594067 | 0 | 1 | 0 | 1 | 2 | 0.16019548 | 1 | 0.60987146 | 1 | 0.28187291 | 16 | 0.80826361 | FALSE | 1 | FALSE | 1 | FALSE | 1 | FALSE | 1 | FALSE | 1 | FALSE | 1 |
| 955 | 955 | 7.02631579 | 2.41145121 | 1 | 0.62830697 | 0 | 1 | 0 | 1 | 0 | 1 | 3 | 0.02386978 | 34 | 0.27051027 | FALSE | 1 | FALSE | 1 | FALSE | 1 | FALSE | 1 | FALSE | 0.77578444 | FALSE | 1 |
| 956 | 956 | 8.73333333 | 1.73076733 | 1 | 0.3050805 | 1 | 0.36919824 | 0 | 1 | 1 | 0.48242099 | 0 | 1 | 11 | 0.83676449 | FALSE | 1 | FALSE | 1 | FALSE | 1 | FALSE | 1 | FALSE | 1 | FALSE | 1 |
| 957 | 957 | 9.66666667 | 3.06956385 | 3 | 0.0405081 | 0 | 1 | 0 | 1 | 1 | 0.75658797 | 0 | 1 | 26 | 0.48609328 | FALSE | 0.62493854 | FALSE | 1 | FALSE | 1 | FALSE | 1 | FALSE | 1 | FALSE | 1 |
| 958 | 958 | 11.0769231 | 3.14915223 | 0 | 1 | 0 | 1 | 1 | 0.35555293 | 0 | 1 | 1 | 0.18007762 | 10 | 0.71436964 | FALSE | 1 | FALSE | 1 | FALSE | 1 | FALSE | 1 | FALSE | 1 | FALSE | 1 |
| 959 | 959 | 5.64814815 | 4.34680559 | 4 | 0.04893847 | 2 | 0.52572379 | 2 | 0.58328102 | 2 | 0.71679995 | 2 | 0.22161752 | 42 | 0.93138539 | FALSE | 0.65293798 | FALSE | 1 | FALSE | 1 | FALSE | 1 | FALSE | 1 | FALSE | 1 |
| 960 | 960 | 5.97402597 | 2.10116867 | 3 | 0.2949342 | 4 | 0.21704891 | 3 | 0.49993836 | 1 | 0.96971789 | 3 | 0.1209957 | 60 | 0.82945515 | FALSE | 1 | FALSE | 1 | FALSE | 1 | FALSE | 1 | FALSE | 1 | FALSE | 1 |
| 961 | 961 | 8.28571429 | 1.48461498 | 1 | 0.3050805 | 1 | 0.36919824 | 0 | 1 | 0 | 1 | 1 | 0.20679328 | 11 | 0.83676449 | FALSE | 1 | FALSE | 1 | FALSE | 1 | FALSE | 1 | FALSE | 1 | FALSE | 1 |
| 962 | 962 | 11.45 | 1.6874537 | 0 | 1 | 0 | 1 | 0 | 1 | 1 | 0.60987146 | 0 | 1 | 19 | 0.15700255 | FALSE | 1 | FALSE | 1 | FALSE | 1 | FALSE | 1 | FALSE | 1 | FALSE | 1 |
| 963 | 963 | 9.42857143 | 2.32115383 | 1 | 0.16630142 | 0 | 1 | 2 | 0.02394967 | 0 | 1 | 0 | 1 | 4 | 0.98605104 | FALSE | 1 | FALSE | 1 | FALSE | 1 | FALSE | 1 | FALSE | 1 | FALSE | 1 |
| 964 | 964 | 7.81818182 | 1.24205464 | 2 | 0.19750261 | 1 | 0.65167362 | 1 | 0.69069222 | 2 | 0.43583874 | 0 | 1 | 26 | 0.77612337 | FALSE | 1 | FALSE | 1 | FALSE | 1 | FALSE | 1 | FALSE | 1 | FALSE | 1 |
| 965 | 965 | 9.03448276 | 2.70991486 | 2 | 0.16962005 | 2 | 0.24095584 | 2 | 0.27983822 | 2 | 0.3871811 | 0 | 1 | 21 | 0.97185642 | FALSE | 1 | FALSE | 1 | FALSE | 1 | FALSE | 1 | FALSE | 1 | FALSE | 1 |
| 966 | 966 | 9.08333333 | 2.99188718 | 1 | 0.70621968 | 1 | 0.78790827 | 0 | 1 | 1 | 0.89101956 | 3 | 0.0412987 | 41 | 0.38322554 | FALSE | 1 | FALSE | 1 | FALSE | 1 | FALSE | 1 | FALSE | 0.90763079 | FALSE | 1 |
| 967 | 967 | 6.90322581 | 3.79604921 | 2 | 0.17884234 | 0 | 1 | 2 | 0.29330414 | 2 | 0.4036176 | 0 | 1 | 24 | 0.82294782 | FALSE | 1 | FALSE | 1 | FALSE | 1 | FALSE | 1 | FALSE | 1 | FALSE | 1 |
| 968 | 968 | 12.0769231 | 2.94726832 | 1 | 0.26795838 | 2 | 0.05552913 | 0 | 1 | 0 | 1 | 0 | 1 | 9 | 0.89630573 | FALSE | 1 | FALSE | 1 | FALSE | 1 | FALSE | 1 | FALSE | 1 | FALSE | 1 |
| 969 | 969 | 9.51851852 | 2.06146964 | 0 | 1 | 1 | 0.58913618 | 1 | 0.62830986 | 2 | 0.35374209 | 0 | 1 | 23 | 0.58307573 | FALSE | 1 | FALSE | 1 | FALSE | 1 | FALSE | 1 | FALSE | 1 | FALSE | 1 |
| 970 | 970 | 8.20689655 | 2.39797775 | 0 | 1 | 3 | 0.06581417 | 2 | 0.27983822 | 2 | 0.3871811 | 0 | 1 | 22 | 0.92902102 | FALSE | 1 | FALSE | 1 | FALSE | 1 | FALSE | 1 | FALSE | 1 | FALSE | 1 |
| 971 | 971 | 10.8571429 | 2.72179413 | 2 | 0.04858661 | 1 | 0.36919824 | 1 | 0.40110418 | 1 | 0.48242099 | 0 | 1 | 9 | 0.98607548 | FALSE | 0.649301 | FALSE | 1 | FALSE | 1 | FALSE | 1 | FALSE | 1 | FALSE | 1 |
| 972 | 972 | 10.6538462 | 2.55617364 | 0 | 1 | 1 | 0.57534752 | 1 | 0.61440466 | 2 | 0.33678042 | 1 | 0.3499045 | 21 | 0.78939668 | FALSE | 1 | FALSE | 1 | FALSE | 1 | FALSE | 1 | FALSE | 1 | FALSE | 1 |
| 973 | 973 | 6.5 | 2.22361068 | 1 | 0.84741747 | 2 | 0.68229183 | 1 | 0.92922664 | 5 | 0.2346015 | 1 | 0.69780571 | 62 | 0.41748931 | FALSE | 1 | FALSE | 1 | FALSE | 1 | FALSE | 1 | FALSE | 1 | FALSE | 1 |
| 974 | 974 | 7.19565217 | 1.09583332 | 4 | 0.47418487 | 3 | 0.82980188 | 6 | 0.37622245 | 4 | 0.88466269 | 4 | 0.190269 | 117 | 0.50509684 | FALSE | 1 | FALSE | 1 | FALSE | 1 | FALSE | 1 | FALSE | 1 | FALSE | 1 |
| 975 | 975 | 9.52173913 | 1.28094955 | 4 | 0.00252272 | 1 | 0.53115654 | 2 | 0.19931429 | 2 | 0.2852236 | 0 | 1 | 14 | 0.99865094 | FALSE | 0.13507246 | FALSE | 1 | FALSE | 1 | FALSE | 1 | FALSE | 1 | FALSE | 1 |
| 976 | 976 | 7.75 | 2.58602011 | 0 | 1 | 0 | 1 | 1 | 0.44344817 | 0 | 1 | 0 | 1 | 15 | 0.26127859 | FALSE | 1 | FALSE | 1 | FALSE | 1 | FALSE | 1 | FALSE | 1 | FALSE | 1 |
| 977 | 977 | 8.17857143 | 2.15561677 | 0 | 1 | 0 | 1 | 3 | 0.07756639 | 1 | 0.73248983 | 1 | 0.37113067 | 23 | 0.73439118 | FALSE | 1 | FALSE | 1 | FALSE | 1 | FALSE | 1 | FALSE | 1 | FALSE | 1 |
| 978 | 978 | 9.63265306 | 1.58685773 | 1 | 0.72119464 | 4 | 0.07284863 | 1 | 0.83455705 | 4 | 0.1862655 | 0 | 1 | 39 | 0.86798612 | FALSE | 1 | FALSE | 1 | FALSE | 1 | FALSE | 1 | FALSE | 1 | FALSE | 1 |
| 979 | 979 | 8.46666667 | 2.15612822 | 2 | 0.05513784 | 0 | 1 | 0 | 1 | 0 | 1 | 1 | 0.2198258 | 12 | 0.80329917 | FALSE | 0.68849059 | FALSE | 1 | FALSE | 1 | FALSE | 1 | FALSE | 1 | FALSE | 1 |
| 980 | 980 | 4.48387097 | 1.48550635 | 13 | 0.02523827 | 21 | 0.00022373 | 13 | 0.19601685 | 10 | 0.82647313 | 7 | 0.16865474 | 213 | 0.99968249 | FALSE | 0.4974559 | FALSE | 0.15723145 | FALSE | 1 | FALSE | 1 | FALSE | 1 | FALSE | 1 |
| 981 | 981 | 2.56451613 | 2.08389344 | 5 | 0.52176149 | 4 | 0.85827754 | 3 | 0.96646237 | 8 | 0.62685614 | 2 | 0.81470297 | 164 | 0.08636593 | FALSE | 1 | FALSE | 1 | FALSE | 1 | FALSE | 1 | FALSE | 1 | FALSE | 1 |
| 982 | 982 | 10.7272727 | 2.17802701 | 0 | 1 | 0 | 1 | 1 | 0.55340851 | 3 | 0.07743186 | 0 | 1 | 18 | 0.74744813 | FALSE | 1 | FALSE | 1 | FALSE | 1 | FALSE | 1 | FALSE | 1 | FALSE | 1 |
| 983 | 983 | 6.38235294 | 1.69800412 | 1 | 0.83052698 | 0 | 1 | 2 | 0.70787008 | 2 | 0.82657478 | 0 | 1 | 63 | 0.03435075 | FALSE | 1 | FALSE | 1 | FALSE | 1 | FALSE | 1 | FALSE | 1 | FALSE | 1 |
| 984 | 984 | 6.18575851 | 0.84541101 | 9 | 0.4182978 | 15 | 0.08504634 | 8 | 0.88634307 | 15 | 0.48052577 | 5 | 0.5942769 | 263 | 0.70388064 | FALSE | 1 | FALSE | 1 | FALSE | 1 | FALSE | 1 | FALSE | 1 | FALSE | 1 |
| 985 | 985 | 9.77777778 | 2.52884593 | 0 | 1 | 0 | 1 | 1 | 0.28067846 | 0 | 1 | 0 | 1 | 8 | 0.5780332 | FALSE | 1 | FALSE | 1 | FALSE | 1 | FALSE | 1 | FALSE | 1 | FALSE | 1 |
| 986 | 986 | 10 | 2.48069469 | 0 | 1 | 0 | 1 | 1 | 0.37874417 | 0 | 1 | 1 | 0.19354507 | 11 | 0.6683056 | FALSE | 1 | FALSE | 1 | FALSE | 1 | FALSE | 1 | FALSE | 1 | FALSE | 1 |
| 987 | 987 | 11.9230769 | 1.97918159 | 0 | 1 | 0 | 1 | 2 | 0.07725803 | 2 | 0.1174583 | 0 | 1 | 9 | 0.96018168 | FALSE | 1 | FALSE | 1 | FALSE | 1 | FALSE | 1 | FALSE | 1 | FALSE | 1 |
| 988 | 988 | 8.53846154 | 2.18791733 | 0 | 1 | 0 | 1 | 0 | 1 | 2 | 0.33678042 | 0 | 1 | 24 | 0.20425579 | FALSE | 1 | FALSE | 1 | FALSE | 1 | FALSE | 1 | FALSE | 1 | FALSE | 1 |
| 989 | 989 | 5 | 1.82093094 | 1 | 0.86263338 | 0 | 1 | 3 | 0.51793834 | 5 | 0.26954615 | 0 | 1 | 67 | 0.23030366 | FALSE | 1 | FALSE | 1 | FALSE | 1 | FALSE | 1 | FALSE | 1 | FALSE | 1 |
| 990 | 990 | 7.93181818 | 1.55793755 | 1 | 0.67382914 | 4 | 0.04916488 | 3 | 0.19985581 | 1 | 0.86831187 | 2 | 0.1563365 | 32 | 0.9718545 | FALSE | 1 | FALSE | 1 | FALSE | 1 | FALSE | 1 | FALSE | 1 | FALSE | 1 |
| 991 | 991 | 8.26666667 | 3.72320054 | 1 | 0.54198714 | 1 | 0.6278847 | 3 | 0.09131526 | 3 | 0.15707004 | 1 | 0.39166989 | 21 | 0.98741738 | FALSE | 1 | FALSE | 1 | FALSE | 1 | FALSE | 1 | FALSE | 1 | FALSE | 1 |
| 992 | 992 | 9.125 | 1.89983552 | 0 | 1 | 0 | 1 | 0 | 1 | 3 | 0.03440583 | 0 | 1 | 13 | 0.76799161 | FALSE | 1 | FALSE | 1 | FALSE | 1 | FALSE | 1 | FALSE | 1 | FALSE | 1 |
| 993 | 993 | 9.05882353 | 1.69673001 | 0 | 1 | 0 | 1 | 0 | 1 | 0 | 1 | 0 | 1 | 17 | 0.05550965 | FALSE | 1 | FALSE | 1 | FALSE | 1 | FALSE | 1 | FALSE | 1 | FALSE | 1 |
| 994 | 994 | 8.5 | 2.29128785 | 0 | 1 | 0 | 1 | 0 | 1 | 1 | 0.37517262 | 0 | 1 | 9 | 0.5215547 | FALSE | 1 | FALSE | 1 | FALSE | 1 | FALSE | 1 | FALSE | 1 | FALSE | 1 |
| 995 | 995 | 9.4375 | 3.06122422 | 1 | 0.34033054 | 0 | 1 | 0 | 1 | 2 | 0.16544789 | 0 | 1 | 13 | 0.76799161 | FALSE | 1 | FALSE | 1 | FALSE | 1 | FALSE | 1 | FALSE | 1 | FALSE | 1 |
| 996 | 996 | 8.22 | 1.87925517 | 1 | 0.72839502 | 1 | 0.80796869 | 3 | 0.26704679 | 1 | 0.90544885 | 0 | 1 | 44 | 0.31639336 | FALSE | 1 | FALSE | 1 | FALSE | 1 | FALSE | 1 | FALSE | 1 | FALSE | 1 |
| 997 | 997 | 9.42105263 | 2.56116317 | 1 | 0.38989951 | 0 | 1 | 1 | 0.50143826 | 0 | 1 | 0 | 1 | 17 | 0.41141543 | FALSE | 1 | FALSE | 1 | FALSE | 1 | FALSE | 1 | FALSE | 1 | FALSE | 1 |
| 998 | 998 | 10.6428571 | 3.37230618 | 2 | 0.04858661 | 0 | 1 | 0 | 1 | 3 | 0.0239258 | 0 | 1 | 9 | 0.98607548 | FALSE | 0.649301 | FALSE | 1 | FALSE | 1 | FALSE | 1 | FALSE | 1 | FALSE | 1 |
| 999 | 999 | 11.3888889 | 2.42988086 | 2 | 0.07645454 | 1 | 0.44710673 | 2 | 0.13502329 | 2 | 0.19902682 | 0 | 1 | 11 | 0.99651304 | FALSE | 0.77424195 | FALSE | 1 | FALSE | 1 | FALSE | 1 | FALSE | 1 | FALSE | 1 |
| 1000 | 1000 | 11.3888889 | 2.94654014 | 0 | 1 | 1 | 0.44710673 | 1 | 0.48280983 | 0 | 1 | 2 | 0.03435384 | 14 | 0.86287418 | FALSE | 1 | FALSE | 1 | FALSE | 1 | FALSE | 1 | FALSE | 0.86063198 | FALSE | 1 |
| 1001 | 1001 | 9.1 | 2.46779254 | 1 | 0.22886536 | 1 | 0.28036922 | 2 | 0.04781153 | 0 | 1 | 0 | 1 | 6 | 0.98831471 | FALSE | 1 | FALSE | 1 | FALSE | 1 | FALSE | 1 | FALSE | 1 | FALSE | 1 |
| 1002 | 1002 | 9.76470588 | 2.10124366 | 0 | 1 | 0 | 1 | 1 | 0.44344817 | 1 | 0.52895088 | 1 | 0.23264612 | 13 | 0.76799161 | FALSE | 1 | FALSE | 1 | FALSE | 1 | FALSE | 1 | FALSE | 1 | FALSE | 1 |
| 1003 | 1003 | 9.77272727 | 0.99689601 | 1 | 0.4357638 | 1 | 0.5154318 | 1 | 0.55340851 | 0 | 1 | 1 | 0.30529951 | 19 | 0.54233419 | FALSE | 1 | FALSE | 1 | FALSE | 1 | FALSE | 1 | FALSE | 1 | FALSE | 1 |
| 1004 | 1004 | 4.94067797 | 3.20035055 | 5 | 0.18092215 | 5 | 0.32796018 | 3 | 0.79796678 | 4 | 0.79241928 | 3 | 0.3007947 | 97 | 0.72247551 | FALSE | 1 | FALSE | 1 | FALSE | 1 | FALSE | 1 | FALSE | 1 | FALSE | 1 |
| 1005 | 1005 | 6.8515625 | 3.57309793 | 4 | 0.3057853 | 1 | 0.9730424 | 3 | 0.75696175 | 8 | 0.12709589 | 1 | 0.83744681 | 92 | 0.56157078 | FALSE | 1 | FALSE | 1 | FALSE | 1 | FALSE | 1 | FALSE | 1 | FALSE | 1 |
| 1006 | 1006 | 7.25925926 | 1.87759879 | 0 | 1 | 4 | 0.00909592 | 0 | 1 | 1 | 0.70601446 | 0 | 1 | 21 | 0.78939668 | FALSE | 1 | FALSE | 0.63686998 | FALSE | 1 | FALSE | 1 | FALSE | 1 | FALSE | 1 |
| 1007 | 1007 | 9.81818182 | 2.32840881 | 0 | 1 | 2 | 0.15794013 | 1 | 0.55340851 | 0 | 1 | 1 | 0.30529951 | 18 | 0.74744813 | FALSE | 1 | FALSE | 1 | FALSE | 1 | FALSE | 1 | FALSE | 1 | FALSE | 1 |
| 1008 | 1008 | 9.46153846 | 1.64622574 | 1 | 0.28675951 | 0 | 1 | 0 | 1 | 1 | 0.45746571 | 1 | 0.19354507 | 10 | 0.86791642 | FALSE | 1 | FALSE | 1 | FALSE | 1 | FALSE | 1 | FALSE | 1 | FALSE | 1 |
| 1009 | 1009 | 10.9090909 | 1.37870463 | 0 | 1 | 0 | 1 | 1 | 0.33149969 | 1 | 0.40390032 | 0 | 1 | 9 | 0.75985366 | FALSE | 1 | FALSE | 1 | FALSE | 1 | FALSE | 1 | FALSE | 1 | FALSE | 1 |
| 1010 | 1010 | 6.48672566 | 1.63000106 | 3 | 0.55171054 | 5 | 0.29599168 | 6 | 0.21430103 | 6 | 0.40959577 | 3 | 0.27833774 | 89 | 0.93825831 | FALSE | 1 | FALSE | 1 | FALSE | 1 | FALSE | 1 | FALSE | 1 | FALSE | 1 |
| 1011 | 1011 | 5.75531915 | 1.81416513 | 1 | 0.91444814 | 2 | 0.81371683 | 4 | 0.43857051 | 7 | 0.13978433 | 3 | 0.19972697 | 77 | 0.79315074 | FALSE | 1 | FALSE | 1 | FALSE | 1 | FALSE | 1 | FALSE | 1 | FALSE | 1 |
| 1012 | 1012 | 8.66666667 | 2.78316937 | 0 | 1 | 2 | 0.14655134 | 0 | 1 | 3 | 0.06912908 | 0 | 1 | 16 | 0.90353674 | FALSE | 1 | FALSE | 1 | FALSE | 1 | FALSE | 1 | FALSE | 1 | FALSE | 1 |
| 1013 | 1013 | 9.57142857 | 1.84058553 | 0 | 1 | 1 | 0.49918218 | 0 | 1 | 0 | 1 | 0 | 1 | 20 | 0.13763971 | FALSE | 1 | FALSE | 1 | FALSE | 1 | FALSE | 1 | FALSE | 1 | FALSE | 1 |
| 1014 | 1014 | 8.94871795 | 2.18325466 | 0 | 1 | 3 | 0.13080858 | 3 | 0.16357218 | 0 | 1 | 0 | 1 | 33 | 0.59061307 | FALSE | 1 | FALSE | 1 | FALSE | 1 | FALSE | 1 | FALSE | 1 | FALSE | 1 |
| 1015 | 1015 | 8.76190476 | 3.03793103 | 2 | 0.09187136 | 1 | 0.48239024 | 0 | 1 | 2 | 0.23331368 | 0 | 1 | 15 | 0.92113679 | FALSE | 0.83248965 | FALSE | 1 | FALSE | 1 | FALSE | 1 | FALSE | 1 | FALSE | 1 |
| 1016 | 1016 | 9.66666667 | 2.2852182 | 0 | 1 | 0 | 1 | 0 | 1 | 2 | 0.1023796 | 0 | 1 | 10 | 0.71436964 | FALSE | 1 | FALSE | 1 | FALSE | 1 | FALSE | 1 | FALSE | 1 | FALSE | 1 |
| 1017 | 1017 | 9 | 2.05480467 | 1 | 0.35728305 | 1 | 0.42857733 | 0 | 1 | 0 | 1 | 0 | 1 | 15 | 0.49060435 | FALSE | 1 | FALSE | 1 | FALSE | 1 | FALSE | 1 | FALSE | 1 | FALSE | 1 |
| 1018 | 1018 | 8.90909091 | 1.83176743 | 1 | 0.57647348 | 2 | 0.28941836 | 3 | 0.11365871 | 1 | 0.78873814 | 0 | 1 | 26 | 0.86852567 | FALSE | 1 | FALSE | 1 | FALSE | 1 | FALSE | 1 | FALSE | 1 | FALSE | 1 |
| 1019 | 1019 | 10.1304348 | 5.19451507 | 0 | 1 | 1 | 0.53115654 | 0 | 1 | 1 | 0.6613267 | 0 | 1 | 21 | 0.27964072 | FALSE | 1 | FALSE | 1 | FALSE | 1 | FALSE | 1 | FALSE | 1 | FALSE | 1 |
| 1020 | 1020 | 8.525 | 2.55917467 | 1 | 0.63789042 | 3 | 0.13080858 | 3 | 0.16357218 | 2 | 0.54052736 | 0 | 1 | 30 | 0.92795554 | FALSE | 1 | FALSE | 1 | FALSE | 1 | FALSE | 1 | FALSE | 1 | FALSE | 1 |
| 1021 | 1021 | 9.5 | 2.87228132 | 0 | 1 | 1 | 0.5154318 | 1 | 0.55340851 | 2 | 0.26791271 | 0 | 1 | 18 | 0.74744813 | FALSE | 1 | FALSE | 1 | FALSE | 1 | FALSE | 1 | FALSE | 1 | FALSE | 1 |
| 1022 | 1022 | 8.08333333 | 1.60511336 | 0 | 1 | 0 | 1 | 0 | 1 | 0 | 1 | 0 | 1 | 12 | 0.13002667 | FALSE | 1 | FALSE | 1 | FALSE | 1 | FALSE | 1 | FALSE | 1 | FALSE | 1 |
| 1023 | 1023 | 10.2857143 | 2.78846019 | 0 | 1 | 2 | 0.07341559 | 0 | 1 | 0 | 1 | 0 | 1 | 12 | 0.62236227 | FALSE | 1 | FALSE | 1 | FALSE | 1 | FALSE | 1 | FALSE | 1 | FALSE | 1 |
| 1024 | 1024 | 7.69387755 | 2.58096708 | 0 | 1 | 2 | 0.47396556 | 1 | 0.83455705 | 1 | 0.90086419 | 0 | 1 | 45 | 0.10025985 | FALSE | 1 | FALSE | 1 | FALSE | 1 | FALSE | 1 | FALSE | 1 | FALSE | 1 |
| 1025 | 1025 | 9.57142857 | 2.49897938 | 0 | 1 | 0 | 1 | 2 | 0.17308361 | 1 | 0.62783522 | 0 | 1 | 18 | 0.57962763 | FALSE | 1 | FALSE | 1 | FALSE | 1 | FALSE | 1 | FALSE | 1 | FALSE | 1 |
| 1026 | 1026 | 7.45454546 | 2.31059364 | 0 | 1 | 2 | 0.41896557 | 1 | 0.80109494 | 1 | 0.87439647 | 2 | 0.16211764 | 38 | 0.45705039 | FALSE | 1 | FALSE | 1 | FALSE | 1 | FALSE | 1 | FALSE | 1 | FALSE | 1 |
| 1027 | 1027 | 8.33333333 | 4.18993503 | 0 | 1 | 0 | 1 | 1 | 0.5695037 | 2 | 0.2852236 | 2 | 0.05389003 | 18 | 0.86273155 | FALSE | 1 | FALSE | 1 | FALSE | 1 | FALSE | 1 | FALSE | 1 | FALSE | 1 |
| 1028 | 1028 | 11.5833333 | 2.81242284 | 0 | 1 | 2 | 0.05552913 | 0 | 1 | 0 | 1 | 1 | 0.18007762 | 9 | 0.89630573 | FALSE | 1 | FALSE | 1 | FALSE | 1 | FALSE | 1 | FALSE | 1 | FALSE | 1 |
| 1029 | 1029 | 8.30434783 | 2.07332505 | 0 | 1 | 1 | 0.53115654 | 0 | 1 | 1 | 0.6613267 | 0 | 1 | 21 | 0.27964072 | FALSE | 1 | FALSE | 1 | FALSE | 1 | FALSE | 1 | FALSE | 1 | FALSE | 1 |
| 1030 | 1030 | 8.57142857 | 2.19461307 | 0 | 1 | 0 | 1 | 0 | 1 | 0 | 1 | 0 | 1 | 14 | 0.09251304 | FALSE | 1 | FALSE | 1 | FALSE | 1 | FALSE | 1 | FALSE | 1 | FALSE | 1 |
| 1031 | 1031 | 10.0212766 | 1.56408306 | 4 | 0.02951043 | 2 | 0.44132825 | 2 | 0.4962935 | 3 | 0.35428318 | 1 | 0.53375569 | 34 | 0.97940583 | FALSE | 0.53131751 | FALSE | 1 | FALSE | 1 | FALSE | 1 | FALSE | 1 | FALSE | 1 |
| 1032 | 1032 | 6.97894737 | 2.09751205 | 2 | 0.68844298 | 4 | 0.34712171 | 3 | 0.64811016 | 4 | 0.61577546 | 2 | 0.44753511 | 77 | 0.63869823 | FALSE | 1 | FALSE | 1 | FALSE | 1 | FALSE | 1 | FALSE | 1 | FALSE | 1 |

| | | | | | | | | | | | | | | | | | | | | | | | | | | | |
|---|---|---|---|---|---|---|---|---|---|---|---|---|---|---|---|---|---|---|---|---|---|---|---|---|---|---|---|
| 1033 | 1033 | 9 | 2.51661148 | 2 | 0.07645454 | 0 | 1 | 0 | 1 | 3 | 0.04687752 | 0 | 1 | 13 | 0.95041913 | FALSE | 0.77424195 | FALSE | 1 | FALSE | 1 | FALSE | 1 | FALSE | 1 | FALSE | 1 |
| 1034 | 1034 | 7.53125 | 1.93623951 | 2 | 0.19750261 | 2 | 0.27730767 | 3 | 0.10599423 | 2 | 0.43583874 | 0 | 1 | 23 | 0.97984382 | FALSE | 1 | FALSE | 1 | FALSE | 1 | FALSE | 1 | FALSE | 1 | FALSE | 1 |
| 1035 | 1035 | 9.90322581 | 2.54450501 | 0 | 1 | 0 | 1 | 1 | 0.67911179 | 0 | 1 | 1 | 0.4016888 | 29 | 0.11661149 | FALSE | 1 | FALSE | 1 | FALSE | 1 | FALSE | 1 | FALSE | 1 | FALSE | 1 |
| 1036 | 1036 | 9.89473684 | 3.58204201 | 0 | 1 | 1 | 0.44710673 | 0 | 1 | 0 | 1 | 1 | 0.25766374 | 16 | 0.44998001 | FALSE | 1 | FALSE | 1 | FALSE | 1 | FALSE | 1 | FALSE | 1 | FALSE | 1 |
| 1037 | 1037 | 8.35833333 | 1.25030552 | 5 | 0.13951176 | 5 | 0.2646113 | 6 | 0.1862724 | 8 | 0.11766557 | 1 | 0.83189113 | 82 | 0.98778349 | FALSE | 1 | FALSE | 1 | FALSE | 1 | FALSE | 1 | FALSE | 1 | FALSE | 1 |
| 1038 | 1038 | 8 | 2.84800125 | 0 | 1 | 1 | 0.44710673 | 3 | 0.02505979 | 1 | 0.57131013 | 0 | 1 | 13 | 0.95041913 | FALSE | 1 | FALSE | 1 | FALSE | 1 | FALSE | 1 | FALSE | 1 | FALSE | 1 |
| 1039 | 1039 | 10 | 2.43373723 | 0 | 1 | 1 | 0.57534752 | 1 | 0.61440466 | 3 | 0.11458571 | 0 | 1 | 21 | 0.78939668 | FALSE | 1 | FALSE | 1 | FALSE | 1 | FALSE | 1 | FALSE | 1 | FALSE | 1 |
| 1040 | 1040 | 10.3076923 | 3.12273587 | 1 | 0.28675951 | 0 | 1 | 0 | 1 | 0 | 1 | 0 | 1 | 12 | 0.374198 | FALSE | 1 | FALSE | 1 | FALSE | 1 | FALSE | 1 | FALSE | 1 | FALSE | 1 |
| 1041 | 1041 | 8.08333333 | 2.45373864 | 1 | 0.60837712 | 3 | 0.10944025 | 0 | 1 | 0 | 1 | 3 | 0.0206828 | 29 | 0.81028289 | FALSE | 1 | FALSE | 1 | FALSE | 1 | FALSE | 1 | FALSE | 0.7366672 | FALSE | 1 |
| 1042 | 1042 | 9.63157895 | 1.97911533 | 2 | 0.08405377 | 2 | 0.12434464 | 2 | 0.14749808 | 2 | 0.21610812 | 0 | 1 | 11 | 0.99888169 | FALSE | 0.79236883 | FALSE | 1 | FALSE | 1 | FALSE | 1 | FALSE | 1 | FALSE | 1 |
| 1043 | 1043 | 9.81818182 | 2.22866376 | 1 | 0.4357638 | 0 | 1 | 1 | 0.55340851 | 1 | 0.64497439 | 2 | 0.04972408 | 17 | 0.8840357 | FALSE | 1 | FALSE | 1 | FALSE | 1 | FALSE | 1 | FALSE | 0.98629461 | FALSE | 1 |
| 1044 | 1044 | 9.03703704 | 2.51633893 | 3 | 0.03084786 | 1 | 0.58913618 | 1 | 0.62830986 | 1 | 0.71956339 | 1 | 0.36060486 | 20 | 0.95133411 | FALSE | 0.53356688 | FALSE | 1 | FALSE | 1 | FALSE | 1 | FALSE | 1 | FALSE | 1 |
| 1045 | 1045 | 7.33950617 | 1.31545223 | 1 | 0.98586902 | 6 | 0.42716587 | 5 | 0.69789705 | 9 | 0.32651646 | 2 | 0.74956518 | 139 | 0.35384787 | FALSE | 1 | FALSE | 1 | FALSE | 1 | FALSE | 1 | FALSE | 1 | FALSE | 1 |
| 1046 | 1046 | 10 | 3.48807492 | 1 | 0.26795838 | 1 | 0.32623933 | 0 | 1 | 1 | 0.43131129 | 1 | 0.18007762 | 8 | 0.97197833 | FALSE | 1 | FALSE | 1 | FALSE | 1 | FALSE | 1 | FALSE | 1 | FALSE | 1 |
| 1047 | 1047 | 5.04444444 | 2.31857768 | 2 | 0.32143972 | 1 | 0.77338922 | 0 | 1 | 0 | 1 | 0 | 1 | 42 | 0.06326314 | FALSE | 1 | FALSE | 1 | FALSE | 1 | FALSE | 1 | FALSE | 1 | FALSE | 1 |
| 1048 | 1048 | 8.625 | 2.42061459 | 0 | 1 | 2 | 0.27730767 | 2 | 0.32011871 | 1 | 0.77852168 | 1 | 0.41154418 | 26 | 0.77612337 | FALSE | 1 | FALSE | 1 | FALSE | 1 | FALSE | 1 | FALSE | 1 | FALSE | 1 |
| 1049 | 1049 | 8.66666667 | 2.3570226 | 1 | 0.37380237 | 0 | 1 | 0 | 1 | 2 | 0.19902682 | 1 | 0.25766374 | 14 | 0.86287418 | FALSE | 1 | FALSE | 1 | FALSE | 1 | FALSE | 1 | FALSE | 1 | FALSE | 1 |
| 1050 | 1050 | 5.328125 | 2.96887335 | 3 | 0.22550017 | 1 | 0.87929209 | 3 | 0.40547364 | 3 | 0.56885732 | 0 | 1 | 54 | 0.58381086 | FALSE | 1 | FALSE | 1 | FALSE | 1 | FALSE | 1 | FALSE | 1 | FALSE | 1 |
| 1051 | 1051 | 7.84210526 | 3.52827911 | 0 | 1 | 0 | 1 | 0 | 1 | 2 | 0.21610812 | 0 | 1 | 17 | 0.41141543 | FALSE | 1 | FALSE | 1 | FALSE | 1 | FALSE | 1 | FALSE | 1 | FALSE | 1 |
| 1052 | 1052 | 8.8 | 3.50618501 | 0 | 1 | 1 | 0.38964394 | 0 | 1 | 1 | 0.50623194 | 0 | 1 | 13 | 0.57712805 | FALSE | 1 | FALSE | 1 | FALSE | 1 | FALSE | 1 | FALSE | 1 | FALSE | 1 |
| 1053 | 1053 | 10.1578947 | 2.62103149 | 1 | 0.38989951 | 0 | 1 | 1 | 0.50143826 | 3 | 0.05383577 | 1 | 0.26986775 | 13 | 0.97998676 | FALSE | 1 | FALSE | 1 | FALSE | 1 | FALSE | 1 | FALSE | 1 | FALSE | 1 |
| 1054 | 1054 | 10.3333333 | 4.45969605 | 1 | 0.37380237 | 1 | 0.44710673 | 0 | 1 | 0 | 1 | 0 | 1 | 16 | 0.44998001 | FALSE | 1 | FALSE | 1 | FALSE | 1 | FALSE | 1 | FALSE | 1 | FALSE | 1 |
| 1055 | 1055 | 8.14492754 | 1.95807775 | 7 | 0.00182332 | 7 | 0.00661418 | 4 | 0.23516085 | 3 | 0.6203651 | 2 | 0.31302643 | 46 | 0.99993388 | FALSE | 0.11146758 | FALSE | 0.60103504 | FALSE | 1 | FALSE | 1 | FALSE | 1 | FALSE | 1 |
| 1056 | 1056 | 7.57142857 | 3.08386188 | 1 | 0.71380452 | 1 | 0.79481669 | 2 | 0.51909752 | 4 | 0.17698522 | 0 | 1 | 40 | 0.66905473 | FALSE | 1 | FALSE | 1 | FALSE | 1 | FALSE | 1 | FALSE | 1 | FALSE | 1 |
| 1057 | 1057 | 10.4 | 2.83548938 | 0 | 1 | 2 | 0.12434464 | 1 | 0.50143826 | 0 | 1 | 1 | 0.26986775 | 15 | 0.83648191 | FALSE | 1 | FALSE | 1 | FALSE | 1 | FALSE | 1 | FALSE | 1 | FALSE | 1 |
| 1058 | 1058 | 7 | 1.93649167 | 1 | 0.55378205 | 2 | 0.2651851 | 1 | 0.67911179 | 0 | 1 | 0 | 1 | 27 | 0.45522639 | FALSE | 1 | FALSE | 1 | FALSE | 1 | FALSE | 1 | FALSE | 1 | FALSE | 1 |
| 1059 | 1059 | 9.29166667 | 3.02047412 | 0 | 1 | 0 | 1 | 0 | 1 | 1 | 0.67692816 | 3 | 0.00675546 | 20 | 0.68270729 | FALSE | 1 | FALSE | 1 | FALSE | 1 | FALSE | 1 | FALSE | 0.47461742 | FALSE | 1 |
| 1060 | 1060 | 8.47619048 | 2.87218264 | 2 | 0.09989125 | 3 | 0.02888061 | 0 | 1 | 2 | 0.25059627 | 2 | 0.04568221 | 12 | 0.99943951 | FALSE | 0.86047225 | FALSE | 0.88012077 | FALSE | 1 | FALSE | 1 | FALSE | 0.95710659 | FALSE | 1 |
| 1061 | 1061 | 8.875 | 3.15980617 | 0 | 1 | 1 | 0.40942991 | 0 | 1 | 0 | 1 | 0 | 1 | 15 | 0.26127859 | FALSE | 1 | FALSE | 1 | FALSE | 1 | FALSE | 1 | FALSE | 1 | FALSE | 1 |
| 1062 | 1062 | 11.2307692 | 2.96528235 | 0 | 1 | 0 | 1 | 0 | 1 | 1 | 0.45746571 | 0 | 1 | 12 | 0.374198 | FALSE | 1 | FALSE | 1 | FALSE | 1 | FALSE | 1 | FALSE | 1 | FALSE | 1 |
| 1063 | 1063 | 5.775 | 3.08615862 | 1 | 0.63789042 | 1 | 0.7236252 | 1 | 0.76089851 | 0 | 1 | 0 | 1 | 36 | 0.12091774 | FALSE | 1 | FALSE | 1 | FALSE | 1 | FALSE | 1 | FALSE | 1 | FALSE | 1 |
| 1064 | 1064 | 10.4285714 | 2.96922996 | 1 | 0.16630142 | 0 | 1 | 0 | 1 | 0 | 1 | 0 | 1 | 6 | 0.69920685 | FALSE | 1 | FALSE | 1 | FALSE | 1 | FALSE | 1 | FALSE | 1 | FALSE | 1 |
| 1065 | 1065 | 7.25 | 2.13600094 | 1 | 0.34033054 | 0 | 1 | 1 | 0.44344817 | 3 | 0.03440583 | 0 | 1 | 11 | 0.9718531 | FALSE | 1 | FALSE | 1 | FALSE | 1 | FALSE | 1 | FALSE | 1 | FALSE | 1 |
| 1066 | 1066 | 6.1509434 | 0.96438907 | 12 | 0.40436524 | 7 | 0.9866369 | 13 | 0.76407656 | 16 | 0.82860531 | 12 | 0.04410272 | 364 | 0.2156222 | FALSE | 1 | FALSE | 1 | FALSE | 1 | FALSE | 1 | FALSE | 0.95327047 | FALSE | 1 |
| 1067 | 1067 | 9.23529412 | 2.27594613 | 2 | 0.19750261 | 1 | 0.65167362 | 1 | 0.69069222 | 2 | 0.43583874 | 2 | 0.096172 | 24 | 0.94876798 | FALSE | 1 | FALSE | 1 | FALSE | 1 | FALSE | 1 | FALSE | 1 | FALSE | 1 |
| 1068 | 1068 | 9.80769231 | 1.77632358 | 1 | 0.46441868 | 1 | 0.54637326 | 1 | 0.58502114 | 0 | 1 | 0 | 1 | 21 | 0.47053137 | FALSE | 1 | FALSE | 1 | FALSE | 1 | FALSE | 1 | FALSE | 1 | FALSE | 1 |
| 1069 | 1069 | 7.29032258 | 3.07113038 | 5 | 0.00103368 | 1 | 0.63997468 | 2 | 0.30673593 | 1 | 0.76781284 | 1 | 0.4016888 | 21 | 0.99467947 | FALSE | 0.07493697 | FALSE | 1 | FALSE | 1 | FALSE | 1 | FALSE | 1 | FALSE | 1 |
| 1070 | 1070 | 8.16129032 | 2.31540109 | 0 | 1 | 2 | 0.25306331 | 0 | 1 | 1 | 0.75658797 | 0 | 1 | 27 | 0.28862208 | FALSE | 1 | FALSE | 1 | FALSE | 1 | FALSE | 1 | FALSE | 1 | FALSE | 1 |
| 1071 | 1071 | 7.74468085 | 3.56381391 | 1 | 0.70621968 | 2 | 0.45233216 | 1 | 0.82190381 | 2 | 0.64277396 | 1 | 0.54145295 | 40 | 0.54368682 | FALSE | 1 | FALSE | 1 | FALSE | 1 | FALSE | 1 | FALSE | 1 | FALSE | 1 |
| 1072 | 1072 | 9.84615385 | 1.85414935 | 2 | 0.12501436 | 1 | 0.54637326 | 3 | 0.05314317 | 1 | 0.67692816 | 2 | 0.058175 | 15 | 0.99800699 | FALSE | 0.95622564 | FALSE | 1 | FALSE | 1 | FALSE | 1 | FALSE | 1 | FALSE | 1 |
| 1073 | 1073 | 10 | 2.44948974 | 1 | 0.32293355 | 0 | 1 | 2 | 0.09928574 | 0 | 1 | 0 | 1 | 12 | 0.80329917 | FALSE | 1 | FALSE | 1 | FALSE | 1 | FALSE | 1 | FALSE | 1 | FALSE | 1 |
| 1074 | 1074 | 8.18181818 | 1.11340443 | 0 | 1 | 0 | 1 | 0 | 1 | 3 | 0.01200907 | 1 | 0.16638736 | 7 | 0.98129353 | FALSE | 1 | FALSE | 1 | FALSE | 1 | FALSE | 1 | FALSE | 1 | FALSE | 1 |
| 1075 | 1075 | 6.84 | 2.82389802 | 1 | 0.72119464 | 1 | 0.80150109 | 1 | 0.83455705 | 4 | 0.1862655 | 0 | 1 | 42 | 0.493842 | FALSE | 1 | FALSE | 1 | FALSE | 1 | FALSE | 1 | FALSE | 1 | FALSE | 1 |
| 1076 | 1076 | 9.4516129 | 1.1024521 | 1 | 0.55378205 | 2 | 0.2651851 | 0 | 1 | 0 | 1 | 1 | 0.4016888 | 27 | 0.45522639 | FALSE | 1 | FALSE | 1 | FALSE | 1 | FALSE | 1 | FALSE | 1 | FALSE | 1 |
| 1077 | 1077 | 7.10152284 | 0.96636202 | 1 | 0.99444827 | 6 | 0.61848207 | 7 | 0.56688065 | 9 | 0.55535368 | 5 | 0.22145483 | 169 | 0.33167181 | FALSE | 1 | FALSE | 1 | FALSE | 1 | FALSE | 1 | FALSE | 1 | FALSE | 1 |
| 1078 | 1078 | 12.2222222 | 1.39664501 | 1 | 0.20854748 | 0 | 1 | 0 | 1 | 0 | 1 | 0 | 1 | 8 | 0.5780332 | FALSE | 1 | FALSE | 1 | FALSE | 1 | FALSE | 1 | FALSE | 1 | FALSE | 1 |
| 1079 | 1079 | 6.96969697 | 2.51642903 | 1 | 0.57647348 | 3 | 0.08959205 | 1 | 0.70185639 | 2 | 0.45159487 | 0 | 1 | 26 | 0.86852567 | FALSE | 1 | FALSE | 1 | FALSE | 1 | FALSE | 1 | FALSE | 1 | FALSE | 1 |
| 1080 | 1080 | 7.67647059 | 2.68679328 | 1 | 0.58738519 | 1 | 0.6739483 | 0 | 1 | 1 | 0.7984848 | 3 | 0.01774672 | 28 | 0.72558119 | FALSE | 1 | FALSE | 1 | FALSE | 1 | FALSE | 1 | FALSE | 0.69720022 | FALSE | 1 |
| 1081 | 1081 | 10.15 | 1.66658333 | 2 | 0.27371785 | 1 | 0.7326167 | 0 | 1 | 1 | 0.84823834 | 0 | 1 | 36 | 0.22908535 | FALSE | 1 | FALSE | 1 | FALSE | 1 | FALSE | 1 | FALSE | 1 | FALSE | 1 |
| 1082 | 1082 | 6.57627119 | 2.75067566 | 1 | 0.78541994 | 1 | 0.85750546 | 0 | 1 | 4 | 0.28589315 | 0 | 1 | 53 | 0.16392156 | FALSE | 1 | FALSE | 1 | FALSE | 1 | FALSE | 1 | FALSE | 1 | FALSE | 1 |
| 1083 | 1083 | 10.5128205 | 2.2288527 | 1 | 0.63789042 | 1 | 0.7236252 | 2 | 0.41139386 | 2 | 0.54052736 | 1 | 0.4761699 | 32 | 0.74324845 | FALSE | 1 | FALSE | 1 | FALSE | 1 | FALSE | 1 | FALSE | 1 | FALSE | 1 |
| 1084 | 1084 | 7.9 | 2.25610284 | 0 | 1 | 0 | 1 | 0 | 1 | 0 | 1 | 0 | 1 | 10 | 0.18273166 | FALSE | 1 | FALSE | 1 | FALSE | 1 | FALSE | 1 | FALSE | 1 | FALSE | 1 |
| 1085 | 1085 | 6.91111111 | 2.02014545 | 5 | 0.00556946 | 4 | 0.05650644 | 2 | 0.48463745 | 0 | 1 | 4 | 0.00607992 | 30 | 0.99918792 | FALSE | 0.21726186 | FALSE | 1 | FALSE | 1 | FALSE | 1 | FALSE | 0.45694206 | FALSE | 1 |
| 1086 | 1086 | 10 | 1.69030851 | 0 | 1 | 0 | 1 | 0 | 1 | 1 | 0.28044368 | 0 | 1 | 6 | 0.69920685 | FALSE | 1 | FALSE | 1 | FALSE | 1 | FALSE | 1 | FALSE | 1 | FALSE | 1 |
| 1087 | 1087 | 10.5333333 | 2.60427 | 0 | 1 | 2 | 0.08295839 | 1 | 0.42266264 | 0 | 1 | 0 | 1 | 12 | 0.80329917 | FALSE | 1 | FALSE | 1 | FALSE | 1 | FALSE | 1 | FALSE | 1 | FALSE | 1 |
| 1088 | 1088 | 6.0483871 | 3.10788497 | 4 | 0.07395961 | 2 | 0.60114557 | 2 | 0.65873577 | 1 | 0.94646288 | 3 | 0.08097514 | 50 | 0.83948885 | FALSE | 0.77424195 | FALSE | 1 | FALSE | 1 | FALSE | 1 | FALSE | 1 | FALSE | 1 |
| 1089 | 1089 | 5.75 | 2.40134283 | 1 | 0.86263338 | 2 | 0.71077581 | 3 | 0.51793834 | 2 | 0.8705742 | 1 | 0.71735141 | 67 | 0.23030366 | FALSE | 1 | FALSE | 1 | FALSE | 1 | FALSE | 1 | FALSE | 1 | FALSE | 1 |
| 1090 | 1090 | 8.96774194 | 2.30684635 | 2 | 0.47501604 | 3 | 0.32473935 | 3 | 0.3858942 | 0 | 1 | 1 | 0.64288531 | 53 | 0.4900821 | FALSE | 1 | FALSE | 1 | FALSE | 1 | FALSE | 1 | FALSE | 1 | FALSE | 1 |
| 1091 | 1091 | 9.28571429 | 3.04166084 | 0 | 1 | 1 | 0.49918218 | 2 | 0.17308361 | 1 | 0.62783522 | 2 | 0.04568221 | 15 | 0.96513691 | FALSE | 1 | FALSE | 1 | FALSE | 1 | FALSE | 1 | FALSE | 0.95710659 | FALSE | 1 |
| 1092 | 1092 | 10.625 | 2.66634113 | 2 | 0.06197875 | 0 | 1 | 2 | 0.11087977 | 1 | 0.52895088 | 1 | 0.23264612 | 10 | 0.99304605 | FALSE | 0.72532843 | FALSE | 1 | FALSE | 1 | FALSE | 1 | FALSE | 1 | FALSE | 1 |
| 1093 | 1093 | 9.3 | 3.19530906 | 0 | 1 | 0 | 1 | 2 | 0.14749808 | 1 | 0.59104357 | 0 | 1 | 16 | 0.65565778 | FALSE | 1 | FALSE | 1 | FALSE | 1 | FALSE | 1 | FALSE | 1 | FALSE | 1 |
| 1094 | 1094 | 10.4761905 | 3.34691112 | 0 | 1 | 2 | 0.14655134 | 2 | 0.17308361 | 0 | 1 | 2 | 0.04568221 | 15 | 0.96513691 | FALSE | 1 | FALSE | 1 | FALSE | 1 | FALSE | 1 | FALSE | 0.95710659 | FALSE | 1 |
| 1095 | 1095 | 8.85714286 | 3.4611547 | 0 | 1 | 0 | 1 | 1 | 0.35555293 | 1 | 0.43131129 | 0 | 1 | 10 | 0.71436964 | FALSE | 1 | FALSE | 1 | FALSE | 1 | FALSE | 1 | FALSE | 1 | FALSE | 1 |
| 1096 | 1096 | 6.34146342 | 3.06522395 | 1 | 0.65632651 | 1 | 0.74131698 | 2 | 0.43640583 | 3 | 0.29054213 | 0 | 1 | 34 | 0.69505404 | FALSE | 1 | FALSE | 1 | FALSE | 1 | FALSE | 1 | FALSE | 1 | FALSE | 1 |
| 1097 | 1097 | 9.11111111 | 3.39752997 | 0 | 1 | 0 | 1 | 1 | 0.48280983 | 0 | 1 | 0 | 1 | 17 | 0.20329205 | FALSE | 1 | FALSE | 1 | FALSE | 1 | FALSE | 1 | FALSE | 1 | FALSE | 1 |
| 1098 | 1098 | 10.2173913 | 3.30949256 | 2 | 0.11647716 | 0 | 1 | 0 | 1 | 0 | 1 | 0 | 1 | 21 | 0.27964072 | FALSE | 0.93181727 | FALSE | 1 | FALSE | 1 | FALSE | 1 | FALSE | 1 | FALSE | 1 |
| 1099 | 1099 | 10.6956522 | 2.61159259 | 0 | 1 | 0 | 1 | 3 | 0.2282043 | 2 | 0.63103861 | 2 | 0.17379289 | 39 | 0.56897353 | FALSE | 1 | FALSE | 1 | FALSE | 1 | FALSE | 1 | FALSE | 1 | FALSE | 1 |
| 1100 | 1100 | 10.125 | 1.79843682 | 1 | 0.34033054 | 1 | 0.40942991 | 1 | 0.44344817 | 2 | 0.16544789 | 0 | 1 | 11 | 0.9718531 | FALSE | 1 | FALSE | 1 | FALSE | 1 | FALSE | 1 | FALSE | 1 | FALSE | 1 |
| 1101 | 1101 | 9.69047619 | 3.61536467 | 0 | 1 | 0 | 1 | 3 | 0.19060367 | 1 | 0.86193352 | 0 | 1 | 38 | 0.19268329 | FALSE | 1 | FALSE | 1 | FALSE | 1 | FALSE | 1 | FALSE | 1 | FALSE | 1 |
| 1102 | 1102 | 8.70175439 | 2.37652933 | 3 | 0.17914398 | 4 | 0.11180811 | 4 | 0.147507 | 1 | 0.93213716 | 0 | 1 | 45 | 0.90229646 | FALSE | 1 | FALSE | 1 | FALSE | 1 | FALSE | 1 | FALSE | 1 | FALSE | 1 |
| 1103 | 1103 | 9.38888889 | 1.56839936 | 0 | 1 | 1 | 0.40942991 | 0 | 1 | 1 | 0.52895088 | 0 | 1 | 14 | 0.53308261 | FALSE | 1 | FALSE | 1 | FALSE | 1 | FALSE | 1 | FALSE | 1 | FALSE | 1 |
| 1104 | 1104 | 9.7 | 2.43789527 | 1 | 0.52988231 | 0 | 1 | 1 | 0.65463982 | 2 | 0.3871811 | 2 | 0.08121636 | 23 | 0.84462957 | FALSE | 1 | FALSE | 1 | FALSE | 1 | FALSE | 1 | FALSE | 1 | FALSE | 1 |
| 1105 | 1105 | 9.85714286 | 1.68426088 | 1 | 0.3050805 | 0 | 1 | 1 | 0.40110418 | 1 | 0.48242099 | 0 | 1 | 11 | 0.83676449 | FALSE | 1 | FALSE | 1 | FALSE | 1 | FALSE | 1 | FALSE | 1 | FALSE | 1 |
| 1106 | 1106 | 9.88709677 | 2.31478306 | 5 | 0.02084841 | 2 | 0.60114557 | 3 | 0.3858942 | 2 | 0.78520976 | 1 | 0.64288531 | 49 | 0.90602303 | FALSE | 0.45408499 | FALSE | 1 | FALSE | 1 | FALSE | 1 | FALSE | 1 | FALSE | 1 |
| 1107 | 1107 | 10.0434783 | 3.91738889 | 1 | 0.45027683 | 1 | 0.53115654 | 0 | 1 | 0 | 1 | 0 | 1 | 21 | 0.27964072 | FALSE | 1 | FALSE | 1 | FALSE | 1 | FALSE | 1 | FALSE | 1 | FALSE | 1 |
| 1108 | 1108 | 7.53333333 | 1.35973854 | 0 | 1 | 2 | 0.24095584 | 4 | 0.01903587 | 0 | 1 | 1 | 0.38148475 | 22 | 0.92902102 | FALSE | 1 | FALSE | 1 | FALSE | 1 | FALSE | 1 | FALSE | 1 | FALSE | 1 |
| 1109 | 1109 | 5.92307692 | 4.01653388 | 0 | 1 | 2 | 0.60991494 | 4 | 0.18968799 | 0 | 1 | 3 | 0.08405237 | 54 | 0.46836572 | FALSE | 1 | FALSE | 1 | FALSE | 1 | FALSE | 1 | FALSE | 1 | FALSE | 1 |
| 1110 | 1110 | 6.86585366 | 2.11130635 | 2 | 0.62636022 | 2 | 0.74948557 | 1 | 0.95112069 | 4 | 0.52386423 | 1 | 0.74434486 | 72 | 0.24542477 | FALSE | 1 | FALSE | 1 | FALSE | 1 | FALSE | 1 | FALSE | 1 | FALSE | 1 |
| 1111 | 1111 | 11.8076923 | 2.00036979 | 1 | 0.49162634 | 2 | 0.20486827 | 0 | 1 | 1 | 0.70601446 | 1 | 0.3499045 | 21 | 0.78939668 | FALSE | 1 | FALSE | 1 | FALSE | 1 | FALSE | 1 | FALSE | 1 | FALSE | 1 |
| 1112 | 1112 | 9.52941177 | 1.68444954 | 2 | 0.06909038 | 0 | 1 | 0 | 1 | 0 | 1 | 0 | 1 | 15 | 0.49060435 | FALSE | 0.74276693 | FALSE | 1 | FALSE | 1 | FALSE | 1 | FALSE | 1 | FALSE | 1 |
| 1113 | 1113 | 9.16129032 | 1.96906354 | 0 | 1 | 1 | 0.6278847 | 1 | 0.66709967 | 4 | 0.04659941 | 0 | 1 | 24 | 0.82294782 | FALSE | 1 | FALSE | 1 | FALSE | 1 | FALSE | 1 | FALSE | 1 | FALSE | 1 |
| 1114 | 1114 | 9.68421053 | 2.00138456 | 1 | 0.38989951 | 1 | 0.46503795 | 3 | 0.02898764 | 0 | 1 | 0 | 1 | 14 | 0.93677253 | FALSE | 1 | FALSE | 1 | FALSE | 1 | FALSE | 1 | FALSE | 1 | FALSE | 1 |
| 1115 | 1115 | 8.64705882 | 2.14040064 | 0 | 1 | 3 | 0.01622926 | 1 | 0.46348836 | 3 | 0.040397 | 0 | 1 | 10 | 0.99775952 | FALSE | 1 | FALSE | 0.7256023 | FALSE | 1 | FALSE | 1 | FALSE | 1 | FALSE | 1 |
| 1116 | 1116 | 7.70588235 | 3.57712842 | 0 | 1 | 1 | 0.42857733 | 0 | 1 | 0 | 1 | 1 | 0.24525764 | 15 | 0.49060435 | FALSE | 1 | FALSE | 1 | FALSE | 1 | FALSE | 1 | FALSE | 1 | FALSE | 1 |
| 1117 | 1117 | 9.46666667 | 1.40791414 | 1 | 0.32293355 | 3 | 0.01138739 | 0 | 1 | 1 | 0.50623194 | 0 | 1 | 10 | 0.97980046 | FALSE | 1 | FALSE | 0.67617265 | FALSE | 1 | FALSE | 1 | FALSE | 1 | FALSE | 1 |
| 1118 | 1118 | 5.4 | 2.34459866 | 1 | 0.57647348 | 0 | 1 | 1 | 0.70185639 | 0 | 1 | 2 | 0.10132035 | 29 | 0.39648374 | FALSE | 1 | FALSE | 1 | FALSE | 1 | FALSE | 1 | FALSE | 1 | FALSE | 1 |
| 1119 | 1119 | 10 | 2.19848433 | 3 | 0.02256792 | 1 | 0.54637326 | 0 | 1 | 2 | 0.30249307 | 2 | 0.058175 | 16 | 0.99240722 | FALSE | 0.4692294 | FALSE | 1 | FALSE | 1 | FALSE | 1 | FALSE | 1 | FALSE | 1 |
| 1120 | 1120 | 10.0952381 | 2.34834819 | 0 | 1 | 2 | 0.14655134 | 0 | 1 | 3 | 0.06912908 | 0 | 1 | 16 | 0.90353674 | FALSE | 1 | FALSE | 1 | FALSE | 1 | FALSE | 1 | FALSE | 1 | FALSE | 1 |
| 1121 | 1121 | 9.52631579 | 3.16840357 | 0 | 1 | 1 | 0.46503795 | 0 | 1 | 2 | 0.21610812 | 0 | 1 | 16 | 0.65565778 | FALSE | 1 | FALSE | 1 | FALSE | 1 | FALSE | 1 | FALSE | 1 | FALSE | 1 |
| 1122 | 1122 | 9.5 | 3.14245127 | 2 | 0.06197875 | 0 | 1 | 0 | 1 | 0 | 1 | 1 | 0.23264612 | 13 | 0.76799161 | FALSE | 0.72532843 | FALSE | 1 | FALSE | 1 | FALSE | 1 | FALSE | 1 | FALSE | 1 |
| 1123 | 1123 | 9.28571429 | 1.97948664 | 0 | 1 | 1 | 0.20567256 | 0 | 1 | 0 | 1 | 0 | 1 | 6 | 0.69920685 | FALSE | 1 | FALSE | 1 | FALSE | 1 | FALSE | 1 | FALSE | 1 | FALSE | 1 |
| 1124 | 1124 | 10.4 | 1.56204994 | 0 | 1 | 1 | 0.28036922 | 0 | 1 | 0 | 1 | 0 | 1 | 9 | 0.5215547 | FALSE | 1 | FALSE | 1 | FALSE | 1 | FALSE | 1 | FALSE | 1 | FALSE | 1 |
| 1125 | 1125 | 9.53846154 | 2.061194 | 0 | 1 | 0 | 1 | 1 | 0.37874417 | 1 | 0.45746571 | 1 | 0.19354507 | 10 | 0.86791642 | FALSE | 1 | FALSE | 1 | FALSE | 1 | FALSE | 1 | FALSE | 1 | FALSE | 1 |
| 1126 | 1126 | 3.72850679 | 2.01769369 | 9 | 0.1138689 | 12 | 0.05415619 | 5 | 0.90479638 | 8 | 0.80347378 | 8 | 0.02822118 | 179 | 0.9306926 | FALSE | 0.92909767 | FALSE | 1 | FALSE | 1 | FALSE | 1 | FALSE | 0.80834928 | FALSE | 1 |

| | | | | | | | | | | | | | | | | | | | | | | | | | | | |
|---|---|---|---|---|---|---|---|---|---|---|---|---|---|---|---|---|---|---|---|---|---|---|---|---|---|---|---|
| 1127 | 1127 | 8.90909091 | 4.37876167 | 0 | 1 | 0 | 1 | 2 | 0.05708125 | 0 | 1 | 0 | 1 | 9 | 0.75985366 | FALSE | 1 | FALSE | 1 | FALSE | 1 | FALSE | 1 | FALSE | 1 | FALSE | 1 |
| 1128 | 1128 | 8.2631579 | 2.82548748 | 4 | 0.22626568 | 3 | 0.59841294 | 3 | 0.66955123 | 5 | 0.44344573 | 0 | 1 | 80 | 0.58645446 | FALSE | 1 | FALSE | 1 | FALSE | 1 | FALSE | 1 | FALSE | 1 | FALSE | 1 |
| 1129 | 1129 | 7.47368421 | 2.72161757 | 1 | 0.62830697 | 1 | 0.71433277 | 1 | 0.75193685 | 0 | 1 | 1 | 0.4673875 | 34 | 0.27051027 | FALSE | 1 | FALSE | 1 | FALSE | 1 | FALSE | 1 | FALSE | 1 | FALSE | 1 |
| 1130 | 1130 | 6.92156863 | 3.1238685 | 1 | 0.72119464 | 3 | 0.21051417 | 2 | 0.53023996 | 1 | 0.90086419 | 2 | 0.19154766 | 40 | 0.77344956 | FALSE | 1 | FALSE | 1 | FALSE | 1 | FALSE | 1 | FALSE | 1 | FALSE | 1 |
| 1131 | 1131 | 9.66666667 | 2.38513918 | 0 | 1 | 2 | 0.08295839 | 1 | 0.42266264 | 1 | 0.50623194 | 0 | 1 | 11 | 0.92879412 | FALSE | 1 | FALSE | 1 | FALSE | 1 | FALSE | 1 | FALSE | 1 | FALSE | 1 |
| 1132 | 1132 | 7.3 | 3.14271645 | 1 | 0.54198714 | 0 | 1 | 2 | 0.29330414 | 1 | 0.75658797 | 0 | 1 | 26 | 0.48609328 | FALSE | 1 | FALSE | 1 | FALSE | 1 | FALSE | 1 | FALSE | 1 | FALSE | 1 |
| 1133 | 1133 | 9.41176471 | 2.97160728 | 0 | 1 | 1 | 0.42857733 | 0 | 1 | 1 | 0.55062772 | 0 | 1 | 15 | 0.49060435 | FALSE | 1 | FALSE | 1 | FALSE | 1 | FALSE | 1 | FALSE | 1 | FALSE | 1 |
| 1134 | 1134 | 6.9 | 2.50798724 | 0 | 1 | 0 | 1 | 1 | 0.51939841 | 3 | 0.0612581 | 0 | 1 | 16 | 0.80826361 | FALSE | 1 | FALSE | 1 | FALSE | 1 | FALSE | 1 | FALSE | 1 | FALSE | 1 |
| 1135 | 1135 | 7.41269841 | 3.24470225 | 2 | 0.48341887 | 3 | 0.33367983 | 2 | 0.66735975 | 0 | 1 | 1 | 0.64879513 | 55 | 0.33161592 | FALSE | 1 | FALSE | 1 | FALSE | 1 | FALSE | 1 | FALSE | 1 | FALSE | 1 |
| 1136 | 1136 | 3.53719008 | 1.79550091 | 2 | 0.82201398 | 6 | 0.19719106 | 2 | 0.93579741 | 5 | 0.65874557 | 1 | 0.86715937 | 105 | 0.27807773 | FALSE | 1 | FALSE | 1 | FALSE | 1 | FALSE | 1 | FALSE | 1 | FALSE | 1 |
| 1137 | 1137 | 10.52 | 2.09990476 | 0 | 1 | 0 | 1 | 1 | 0.59998149 | 1 | 0.69181315 | 2 | 0.06257402 | 21 | 0.64957167 | FALSE | 1 | FALSE | 1 | FALSE | 1 | FALSE | 1 | FALSE | 1 | FALSE | 1 |
| 1138 | 1138 | 7.33333333 | 1.98885785 | 0 | 1 | 0 | 1 | 1 | 0.42266264 | 1 | 0.50623194 | 1 | 0.2198258 | 12 | 0.80329917 | FALSE | 1 | FALSE | 1 | FALSE | 1 | FALSE | 1 | FALSE | 1 | FALSE | 1 |
| 1139 | 1139 | 6.96428571 | 3.95042622 | 0 | 1 | 0 | 1 | 2 | 0.25286935 | 2 | 0.35374209 | 0 | 1 | 23 | 0.58307573 | FALSE | 1 | FALSE | 1 | FALSE | 1 | FALSE | 1 | FALSE | 1 | FALSE | 1 |
| 1140 | 1140 | 9.72222222 | 1.57429192 | 1 | 0.60837712 | 1 | 0.69480472 | 1 | 0.73299778 | 2 | 0.49731009 | 0 | 1 | 31 | 0.49869404 | FALSE | 1 | FALSE | 1 | FALSE | 1 | FALSE | 1 | FALSE | 1 | FALSE | 1 |
| 1141 | 1141 | 5.84482759 | 2.04940466 | 1 | 0.77972342 | 1 | 0.85269953 | 4 | 0.15426575 | 3 | 0.50172293 | 2 | 0.24594209 | 47 | 0.8149836 | FALSE | 1 | FALSE | 1 | FALSE | 1 | FALSE | 1 | FALSE | 1 | FALSE | 1 |
| 1142 | 1142 | 8.21276596 | 2.3057391 | 1 | 0.70621968 | 1 | 0.78790827 | 2 | 0.50778118 | 3 | 0.36699324 | 0 | 1 | 40 | 0.54368682 | FALSE | 1 | FALSE | 1 | FALSE | 1 | FALSE | 1 | FALSE | 1 | FALSE | 1 |
| 1143 | 1143 | 7.39393939 | 1.33574163 | 2 | 0.50812508 | 7 | 0.00517922 | 1 | 0.9116517 | 3 | 0.58996791 | 2 | 0.29478192 | 51 | 0.95588332 | FALSE | 1 | FALSE | 0.58120789 | FALSE | 1 | FALSE | 1 | FALSE | 1 | FALSE | 1 |
| 1144 | 1144 | 10.35 | 2.43464577 | 0 | 1 | 0 | 1 | 1 | 0.51939841 | 1 | 0.60987146 | 0 | 1 | 18 | 0.37504645 | FALSE | 1 | FALSE | 1 | FALSE | 1 | FALSE | 1 | FALSE | 1 | FALSE | 1 |
| 1145 | 1145 | 3.11206897 | 0.87613524 | 22 | 0.00183827 | 21 | 0.04601385 | 13 | 0.80742968 | 23 | 0.28643736 | 6 | 0.73698213 | 355 | 0.98703898 | FALSE | 0.11154888 | FALSE | 1 | FALSE | 1 | FALSE | 1 | FALSE | 1 | FALSE | 1 |
| 1146 | 1146 | 8.48275862 | 2.26853575 | 0 | 1 | 2 | 0.24095584 | 0 | 1 | 1 | 0.74482229 | 0 | 1 | 26 | 0.31495157 | FALSE | 1 | FALSE | 1 | FALSE | 1 | FALSE | 1 | FALSE | 1 | FALSE | 1 |
| 1147 | 1147 | 9.2 | 1.79629248 | 1 | 0.32293355 | 0 | 1 | 0 | 1 | 2 | 0.14906763 | 0 | 1 | 12 | 0.80329917 | FALSE | 1 | FALSE | 1 | FALSE | 1 | FALSE | 1 | FALSE | 1 | FALSE | 1 |
| 1148 | 1148 | 10.5789474 | 2.92567202 | 2 | 0.08405377 | 0 | 1 | 0 | 1 | 0 | 1 | 0 | 1 | 17 | 0.41141543 | FALSE | 0.79236883 | FALSE | 1 | FALSE | 1 | FALSE | 1 | FALSE | 1 | FALSE | 1 |
| 1149 | 1149 | 10.1176471 | 1.99653679 | 1 | 0.35728305 | 2 | 0.10307112 | 2 | 0.12280513 | 0 | 1 | 1 | 0.24525764 | 11 | 0.98974663 | FALSE | 1 | FALSE | 1 | FALSE | 1 | FALSE | 1 | FALSE | 1 | FALSE | 1 |
| 1150 | 1150 | 7.25531915 | 3.65518044 | 2 | 0.69979848 | 3 | 0.59118008 | 5 | 0.2484594 | 1 | 0.98831261 | 1 | 0.79090173 | 82 | 0.2727315 | FALSE | 1 | FALSE | 1 | FALSE | 1 | FALSE | 1 | FALSE | 1 | FALSE | 1 |
| 1151 | 1151 | 10.4615385 | 2.17026707 | 1 | 0.28675951 | 0 | 1 | 1 | 0.37874417 | 2 | 0.1174583 | 0 | 1 | 9 | 0.96018168 | FALSE | 1 | FALSE | 1 | FALSE | 1 | FALSE | 1 | FALSE | 1 | FALSE | 1 |
| 1152 | 1152 | 4.90410959 | 2.66465646 | 1 | 0.85137169 | 1 | 0.9104874 | 1 | 0.93179654 | 3 | 0.65845889 | 1 | 0.7028142 | 66 | 0.09785941 | FALSE | 1 | FALSE | 1 | FALSE | 1 | FALSE | 1 | FALSE | 1 | FALSE | 1 |
| 1153 | 1153 | 9.95454546 | 3.0965857 | 1 | 0.4357638 | 0 | 1 | 3 | 0.04259419 | 1 | 0.64497439 | 2 | 0.04972408 | 15 | 0.98570275 | FALSE | 1 | FALSE | 1 | FALSE | 1 | FALSE | 1 | FALSE | 0.98629461 | FALSE | 1 |
| 1154 | 1154 | 10.7222222 | 2.70402078 | 0 | 1 | 1 | 0.69480472 | 0 | 1 | 1 | 0.81665383 | 0 | 1 | 34 | 0.06417017 | FALSE | 1 | FALSE | 1 | FALSE | 1 | FALSE | 1 | FALSE | 1 | FALSE | 1 |
| 1155 | 1155 | 11.0769231 | 1.63902121 | 0 | 1 | 0 | 1 | 1 | 0.33149969 | 0 | 1 | 0 | 1 | 10 | 0.46859937 | FALSE | 1 | FALSE | 1 | FALSE | 1 | FALSE | 1 | FALSE | 1 | FALSE | 1 |
| 1156 | 1156 | 10.5789474 | 2.68162931 | 0 | 1 | 1 | 0.46503795 | 0 | 1 | 0 | 1 | 1 | 0.26986775 | 17 | 0.41141543 | FALSE | 1 | FALSE | 1 | FALSE | 1 | FALSE | 1 | FALSE | 1 | FALSE | 1 |
| 1157 | 1157 | 7.52702703 | 1.2544297 | 2 | 0.57015153 | 2 | 0.69680912 | 1 | 0.93427349 | 1 | 0.96971789 | 0 | 1 | 68 | 0.04378964 | FALSE | 1 | FALSE | 1 | FALSE | 1 | FALSE | 1 | FALSE | 1 | FALSE | 1 |
| 1158 | 1158 | 9.35294118 | 2.0505717 | 1 | 0.8826742 | 5 | 0.12558433 | 4 | 0.34020118 | 4 | 0.52386423 | 2 | 0.39055391 | 66 | 0.86930658 | FALSE | 1 | FALSE | 1 | FALSE | 1 | FALSE | 1 | FALSE | 1 | FALSE | 1 |
| 1159 | 1159 | 10 | 1.93649167 | 0 | 1 | 0 | 1 | 0 | 1 | 1 | 0.31351192 | 0 | 1 | 7 | 0.63752397 | FALSE | 1 | FALSE | 1 | FALSE | 1 | FALSE | 1 | FALSE | 1 | FALSE | 1 |
| 1160 | 1160 | 4.24534884 | 1.14892842 | 24 | 0.34845578 | 48 | 6.78E-05 | 44 | 0.00787572 | 26 | 0.99413807 | 21 | 0.03599426 | 692 | 0.99844438 | FALSE | 1 | FALSE | 0.10861097 | FALSE | 1 | FALSE | 1 | FALSE | 0.87669812 | FALSE | 1 |
| 1161 | 1161 | 8.86956522 | 0.99145689 | 0 | 1 | 1 | 0.53115654 | 0 | 1 | 3 | 0.08614834 | 0 | 1 | 19 | 0.71542496 | FALSE | 1 | FALSE | 1 | FALSE | 1 | FALSE | 1 | FALSE | 1 | FALSE | 1 |
| 1162 | 1162 | 10.4464286 | 1.58023103 | 3 | 0.17275086 | 1 | 0.84259834 | 2 | 0.60323163 | 0 | 1 | 2 | 0.23375968 | 48 | 0.48035049 | FALSE | 1 | FALSE | 1 | FALSE | 1 | FALSE | 1 | FALSE | 1 | FALSE | 1 |
| 1163 | 1163 | 2.28915663 | 1.53928571 | 22 | 0.00080595 | 21 | 0.02554157 | 18 | 0.22812819 | 16 | 0.7969273 | 10 | 0.13854878 | 326 | 0.99904375 | FALSE | 0.06668996 | FALSE | 0.82367276 | FALSE | 1 | FALSE | 1 | FALSE | 1 | FALSE | 1 |
| 1164 | 1164 | 10.2758621 | 1.89184369 | 1 | 0.52988231 | 1 | 0.61539065 | 1 | 0.65463982 | 1 | 0.74482229 | 1 | 0.38148475 | 24 | 0.70563334 | FALSE | 1 | FALSE | 1 | FALSE | 1 | FALSE | 1 | FALSE | 1 | FALSE | 1 |
| 1165 | 1165 | 9 | 3.34165628 | 1 | 0.26795838 | 0 | 1 | 0 | 1 | 0 | 1 | 0 | 1 | 11 | 0.41944532 | FALSE | 1 | FALSE | 1 | FALSE | 1 | FALSE | 1 | FALSE | 1 | FALSE | 1 |
| 1166 | 1166 | 8.92307692 | 2.99901365 | 1 | 0.28675951 | 0 | 1 | 0 | 1 | 0 | 1 | 0 | 1 | 12 | 0.374198 | FALSE | 1 | FALSE | 1 | FALSE | 1 | FALSE | 1 | FALSE | 1 | FALSE | 1 |
| 1167 | 1167 | 8.90277778 | 2.12236583 | 7 | 0.0023347 | 7 | 0.00832601 | 4 | 0.25881411 | 3 | 0.64919942 | 2 | 0.33117292 | 49 | 0.99985771 | FALSE | 0.12737083 | FALSE | 0.62691516 | FALSE | 1 | FALSE | 1 | FALSE | 1 | FALSE | 1 |
| 1168 | 1168 | 9.15384615 | 3.05956643 | 1 | 0.28675951 | 0 | 1 | 0 | 1 | 1 | 0.45746571 | 0 | 1 | 11 | 0.6683056 | FALSE | 1 | FALSE | 1 | FALSE | 1 | FALSE | 1 | FALSE | 1 | FALSE | 1 |
| 1169 | 1169 | 10.9285714 | 1.48633229 | 0 | 1 | 1 | 0.36919824 | 0 | 1 | 0 | 1 | 2 | 0.02131828 | 11 | 0.83676449 | FALSE | 1 | FALSE | 1 | FALSE | 1 | FALSE | 1 | FALSE | 0.73999729 | FALSE | 1 |
| 1170 | 1170 | 9.88888889 | 2.42415825 | 1 | 0.20854748 | 0 | 1 | 0 | 1 | 0 | 1 | 1 | 0.1383237 | 7 | 0.84574168 | FALSE | 1 | FALSE | 1 | FALSE | 1 | FALSE | 1 | FALSE | 1 | FALSE | 1 |
| 1171 | 1171 | 9.13333333 | 2.12498366 | 1 | 0.32293355 | 0 | 1 | 3 | 0.01512258 | 2 | 0.14906763 | 0 | 1 | 9 | 0.995488 | FALSE | 1 | FALSE | 1 | FALSE | 1 | FALSE | 1 | FALSE | 1 | FALSE | 1 |
| 1172 | 1172 | 10.0909091 | 1.62114132 | 0 | 1 | 0 | 1 | 0 | 1 | 0 | 1 | 0 | 1 | 11 | 0.15414489 | FALSE | 1 | FALSE | 1 | FALSE | 1 | FALSE | 1 | FALSE | 1 | FALSE | 1 |
| 1173 | 1173 | 8.24 | 2.6272419 | 0 | 1 | 0 | 1 | 1 | 0.59998149 | 0 | 1 | 1 | 0.33902672 | 23 | 0.22727183 | FALSE | 1 | FALSE | 1 | FALSE | 1 | FALSE | 1 | FALSE | 1 | FALSE | 1 |
| 1174 | 1174 | 8.04545455 | 2.34476734 | 3 | 0.10232687 | 0 | 1 | 2 | 0.47281647 | 1 | 0.87439647 | 2 | 0.16211764 | 36 | 0.75909657 | FALSE | 0.87592659 | FALSE | 1 | FALSE | 1 | FALSE | 1 | FALSE | 1 | FALSE | 1 |
| 1175 | 1175 | 8.89705882 | 1.84820824 | 2 | 0.49997462 | 3 | 0.35154359 | 3 | 0.41519671 | 1 | 0.95356658 | 2 | 0.28868487 | 54 | 0.68873256 | FALSE | 1 | FALSE | 1 | FALSE | 1 | FALSE | 1 | FALSE | 1 | FALSE | 1 |
| 1176 | 1176 | 9.95121951 | 1.44747416 | 0 | 1 | 1 | 0.74131698 | 1 | 0.77786601 | 3 | 0.29054213 | 0 | 1 | 36 | 0.36455032 | FALSE | 1 | FALSE | 1 | FALSE | 1 | FALSE | 1 | FALSE | 1 | FALSE | 1 |
| 1177 | 1177 | 8.95238095 | 2.10387829 | 0 | 1 | 0 | 1 | 1 | 0.53671415 | 1 | 0.62783522 | 2 | 0.04568221 | 17 | 0.77849076 | FALSE | 1 | FALSE | 1 | FALSE | 1 | FALSE | 1 | FALSE | 0.95710659 | FALSE | 1 |
| 1178 | 1178 | 10.2631579 | 2.74996852 | 0 | 1 | 1 | 0.46503795 | 2 | 0.14749808 | 1 | 0.59104357 | 0 | 1 | 15 | 0.83648191 | FALSE | 1 | FALSE | 1 | FALSE | 1 | FALSE | 1 | FALSE | 1 | FALSE | 1 |
| 1179 | 1179 | 9.55555556 | 2.49938264 | 0 | 1 | 0 | 1 | 1 | 0.48280983 | 1 | 0.57131013 | 0 | 1 | 16 | 0.44998001 | FALSE | 1 | FALSE | 1 | FALSE | 1 | FALSE | 1 | FALSE | 1 | FALSE | 1 |
| 1180 | 1180 | 9.3030303 | 1.9615215 | 1 | 0.56527493 | 0 | 1 | 1 | 0.69069222 | 1 | 0.77852168 | 0 | 1 | 29 | 0.24088541 | FALSE | 1 | FALSE | 1 | FALSE | 1 | FALSE | 1 | FALSE | 1 | FALSE | 1 |
| 1181 | 1181 | 6.15555556 | 3.44451613 | 6 | 0.00081822 | 0 | 1 | 4 | 0.07197653 | 3 | 0.32878418 | 0 | 1 | 31 | 0.99454415 | FALSE | 0.06702889 | FALSE | 1 | FALSE | 1 | FALSE | 1 | FALSE | 1 | FALSE | 1 |
| 1182 | 1182 | 4.65116279 | 2.95220628 | 5 | 0.06651199 | 1 | 0.93995595 | 2 | 0.81591765 | 5 | 0.35139151 | 0 | 1 | 72 | 0.54001499 | FALSE | 0.74276693 | FALSE | 1 | FALSE | 1 | FALSE | 1 | FALSE | 1 | FALSE | 1 |
| 1183 | 1183 | 8.9 | 1.97230829 | 0 | 1 | 0 | 1 | 1 | 0.30655256 | 1 | 0.37517262 | 0 | 1 | 8 | 0.80394719 | FALSE | 1 | FALSE | 1 | FALSE | 1 | FALSE | 1 | FALSE | 1 | FALSE | 1 |
| 1184 | 1184 | 6.01666667 | 1.52196875 | 2 | 0.45796754 | 2 | 0.58316597 | 0 | 1 | 3 | 0.52470434 | 0 | 1 | 53 | 0.25931015 | FALSE | 1 | FALSE | 1 | FALSE | 1 | FALSE | 1 | FALSE | 1 | FALSE | 1 |
| 1185 | 1185 | 9.25 | 2.91904665 | 0 | 1 | 1 | 0.32623933 | 0 | 1 | 2 | 0.1023796 | 1 | 0.18007762 | 8 | 0.97197833 | FALSE | 1 | FALSE | 1 | FALSE | 1 | FALSE | 1 | FALSE | 1 | FALSE | 1 |
| 1186 | 1186 | 8.29166667 | 2.79104471 | 6 | 2.39E-05 | 0 | 1 | 0 | 1 | 0 | 1 | 1 | 0.32796862 | 17 | 0.97530702 | TRUE | 0.00651577 | FALSE | 1 | FALSE | 1 | FALSE | 1 | FALSE | 1 | FALSE | 1 |
| 1187 | 1187 | 8.52941177 | 2.26527877 | 3 | 0.05151181 | 1 | 0.6629941 | 1 | 0.70185639 | 0 | 1 | 0 | 1 | 28 | 0.58681477 | FALSE | 0.67646333 | FALSE | 1 | FALSE | 1 | FALSE | 1 | FALSE | 1 | FALSE | 1 |
| 1188 | 1188 | 11.175 | 1.9220757 | 0 | 1 | 0 | 1 | 2 | 0.42396974 | 4 | 0.10945471 | 0 | 1 | 34 | 0.56328904 | FALSE | 1 | FALSE | 1 | FALSE | 1 | FALSE | 1 | FALSE | 1 | FALSE | 1 |
| 1189 | 1189 | 9.83333333 | 3.16315595 | 1 | 0.54198714 | 0 | 1 | 0 | 1 | 2 | 0.4036176 | 2 | 0.08611595 | 25 | 0.67630471 | FALSE | 1 | FALSE | 1 | FALSE | 1 | FALSE | 1 | FALSE | 1 | FALSE | 1 |
| 1190 | 1190 | 9.40909091 | 2.28948368 | 0 | 1 | 0 | 1 | 1 | 0.55340851 | 2 | 0.26791271 | 0 | 1 | 19 | 0.54233419 | FALSE | 1 | FALSE | 1 | FALSE | 1 | FALSE | 1 | FALSE | 1 | FALSE | 1 |
| 1191 | 1191 | 8.5 | 2.01246118 | 0 | 1 | 0 | 1 | 0 | 1 | 0 | 1 | 0 | 1 | 10 | 0.18273166 | FALSE | 1 | FALSE | 1 | FALSE | 1 | FALSE | 1 | FALSE | 1 | FALSE | 1 |
| 1192 | 1192 | 7.07142857 | 2.76364888 | 0 | 1 | 0 | 1 | 2 | 0.08806398 | 3 | 0.0239258 | 0 | 1 | 9 | 0.98607548 | FALSE | 1 | FALSE | 1 | FALSE | 1 | FALSE | 1 | FALSE | 1 | FALSE | 1 |
| 1193 | 1193 | 9.76923077 | 1.55186469 | 0 | 1 | 0 | 1 | 0 | 1 | 3 | 0.11458571 | 0 | 1 | 23 | 0.40372818 | FALSE | 1 | FALSE | 1 | FALSE | 1 | FALSE | 1 | FALSE | 1 | FALSE | 1 |
| 1194 | 1194 | 8.30434783 | 2.07332505 | 0 | 1 | 2 | 0.15794013 | 0 | 1 | 0 | 1 | 0 | 1 | 20 | 0.30915202 | FALSE | 1 | FALSE | 1 | FALSE | 1 | FALSE | 1 | FALSE | 1 | FALSE | 1 |
| 1195 | 1195 | 9.17857143 | 4.24309162 | 0 | 1 | 1 | 0.60247911 | 2 | 0.2663542 | 2 | 0.3705495 | 0 | 1 | 23 | 0.73439118 | FALSE | 1 | FALSE | 1 | FALSE | 1 | FALSE | 1 | FALSE | 1 | FALSE | 1 |
| 1196 | 1196 | 6.875 | 2.26039267 | 1 | 0.46441868 | 1 | 0.54637326 | 1 | 0.58502114 | 1 | 0.67692816 | 2 | 0.058175 | 18 | 0.93166998 | FALSE | 1 | FALSE | 1 | FALSE | 1 | FALSE | 1 | FALSE | 1 | FALSE | 1 |
| 1197 | 1197 | 9.91666667 | 1.60511336 | 0 | 1 | 0 | 1 | 0 | 1 | 0 | 1 | 0 | 1 | 11 | 0.15414489 | FALSE | 1 | FALSE | 1 | FALSE | 1 | FALSE | 1 | FALSE | 1 | FALSE | 1 |
| 1198 | 1198 | 10.3333333 | 3.24893145 | 1 | 0.26795838 | 0 | 1 | 1 | 0.35555293 | 0 | 1 | 0 | 1 | 10 | 0.71436964 | FALSE | 1 | FALSE | 1 | FALSE | 1 | FALSE | 1 | FALSE | 1 | FALSE | 1 |
| 1199 | 1199 | 11.4615385 | 4.43092948 | 0 | 1 | 0 | 1 | 1 | 0.35555293 | 0 | 1 | 0 | 1 | 11 | 0.41944532 | FALSE | 1 | FALSE | 1 | FALSE | 1 | FALSE | 1 | FALSE | 1 | FALSE | 1 |
| 1200 | 1200 | 7.63636364 | 3.74827784 | 0 | 1 | 0 | 1 | 1 | 0.70185639 | 0 | 1 | 0 | 1 | 32 | 0.02590314 | FALSE | 1 | FALSE | 1 | FALSE | 1 | FALSE | 1 | FALSE | 1 | FALSE | 1 |
| 1201 | 1201 | 8.6 | 2.82370915 | 0 | 1 | 0 | 1 | 1 | 0.42266264 | 0 | 1 | 0 | 1 | 14 | 0.29525409 | FALSE | 1 | FALSE | 1 | FALSE | 1 | FALSE | 1 | FALSE | 1 | FALSE | 1 |
| 1202 | 1202 | 10.1111111 | 2.0245408 | 0 | 1 | 0 | 1 | 0 | 1 | 0 | 1 | 1 | 0.25766374 | 17 | 0.20329205 | FALSE | 1 | FALSE | 1 | FALSE | 1 | FALSE | 1 | FALSE | 1 | FALSE | 1 |
| 1203 | 1203 | 7.54166667 | 2.23567974 | 0 | 1 | 1 | 0.54637326 | 0 | 1 | 1 | 0.67692816 | 0 | 1 | 22 | 0.25237063 | FALSE | 1 | FALSE | 1 | FALSE | 1 | FALSE | 1 | FALSE | 1 | FALSE | 1 |
| 1204 | 1204 | 10.6470588 | 2.42392915 | 1 | 0.35728305 | 0 | 1 | 1 | 0.46348836 | 0 | 1 | 1 | 0.24525764 | 14 | 0.7313145 | FALSE | 1 | FALSE | 1 | FALSE | 1 | FALSE | 1 | FALSE | 1 | FALSE | 1 |
| 1205 | 1205 | 7.83333333 | 2.33927814 | 0 | 1 | 1 | 0.44710673 | 2 | 0.13502329 | 1 | 0.57131013 | 0 | 1 | 14 | 0.86287418 | FALSE | 1 | FALSE | 1 | FALSE | 1 | FALSE | 1 | FALSE | 1 | FALSE | 1 |
| 1206 | 1206 | 9.83333333 | 2.67186992 | 0 | 1 | 0 | 1 | 0 | 1 | 1 | 0.43131129 | 0 | 1 | 11 | 0.41944532 | FALSE | 1 | FALSE | 1 | FALSE | 1 | FALSE | 1 | FALSE | 1 | FALSE | 1 |
| 1207 | 1207 | 10.8636364 | 2.0515062 | 0 | 1 | 0 | 1 | 1 | 0.55340851 | 1 | 0.64497439 | 0 | 1 | 20 | 0.30915202 | FALSE | 1 | FALSE | 1 | FALSE | 1 | FALSE | 1 | FALSE | 1 | FALSE | 1 |
| 1208 | 1208 | 5.17924528 | 0.89860657 | 5 | 0.13178753 | 7 | 0.05309476 | 4 | 0.52485421 | 3 | 0.86756002 | 1 | 0.82614736 | 85 | 0.86552172 | FALSE | 0.97964162 | FALSE | 1 | FALSE | 1 | FALSE | 1 | FALSE | 1 | FALSE | 1 |
| 1209 | 1209 | 11.25 | 2.2220486 | 1 | 0.18769745 | 0 | 1 | 0 | 1 | 0 | 1 | 1 | 0.12394282 | 6 | 0.88425551 | FALSE | 1 | FALSE | 1 | FALSE | 1 | FALSE | 1 | FALSE | 1 | FALSE | 1 |
| 1210 | 1210 | 6.96969697 | 2.36751818 | 1 | 0.57647348 | 0 | 1 | 0 | 1 | 1 | 0.78873814 | 1 | 0.42123867 | 30 | 0.21942357 | FALSE | 1 | FALSE | 1 | FALSE | 1 | FALSE | 1 | FALSE | 1 | FALSE | 1 |
| 1211 | 1211 | 9.34210526 | 1.93684926 | 0 | 1 | 1 | 0.71433277 | 3 | 0.15483587 | 3 | 0.25266879 | 0 | 1 | 31 | 0.76640548 | FALSE | 1 | FALSE | 1 | FALSE | 1 | FALSE | 1 | FALSE | 1 | FALSE | 1 |
| 1212 | 1212 | 9.41666667 | 2.46503324 | 0 | 1 | 0 | 1 | 0 | 1 | 0 | 1 | 1 | 0.18007762 | 11 | 0.41944532 | FALSE | 1 | FALSE | 1 | FALSE | 1 | FALSE | 1 | FALSE | 1 | FALSE | 1 |
| 1213 | 1213 | 8.2 | 1.93907194 | 0 | 1 | 0 | 1 | 0 | 1 | 0 | 1 | 0 | 1 | 10 | 0.18273166 | FALSE | 1 | FALSE | 1 | FALSE | 1 | FALSE | 1 | FALSE | 1 | FALSE | 1 |
| 1214 | 1214 | 7.93478261 | 1.93819923 | 1 | 0.68224519 | 0 | 1 | 2 | 0.47281647 | 2 | 0.60667406 | 0 | 1 | 39 | 0.29524127 | FALSE | 1 | FALSE | 1 | FALSE | 1 | FALSE | 1 | FALSE | 1 | FALSE | 1 |
| 1215 | 1215 | 10.4545455 | 1.8763425 | 0 | 1 | 0 | 1 | 0 | 1 | 1 | 0.40390032 | 0 | 1 | 10 | 0.46859937 | FALSE | 1 | FALSE | 1 | FALSE | 1 | FALSE | 1 | FALSE | 1 | FALSE | 1 |
| 1216 | 1216 | 8.82352941 | 3.25820763 | 0 | 1 | 0 | 1 | 0 | 1 | 0 | 1 | 0 | 1 | 17 | 0.05550965 | FALSE | 1 | FALSE | 1 | FALSE | 1 | FALSE | 1 | FALSE | 1 | FALSE | 1 |
| 1217 | 1217 | 9.75 | 2.71185668 | 1 | 0.26795838 | 0 | 1 | 1 | 0.35555293 | 0 | 1 | 0 | 1 | 10 | 0.71436964 | FALSE | 1 | FALSE | 1 | FALSE | 1 | FALSE | 1 | FALSE | 1 | FALSE | 1 |
| 1218 | 1218 | 10.7142857 | 3.10365157 | 1 | 0.3050805 | 1 | 0.36919824 | 0 | 1 | 1 | 0.48242099 | 0 | 1 | 11 | 0.83676449 | FALSE | 1 | FALSE | 1 | FALSE | 1 | FALSE | 1 | FALSE | 1 | FALSE | 1 |
| 1219 | 1219 | 9.6097561 | 1.60605635 | 0 | 1 | 1 | 0.74131698 | 2 | 0.43640583 | 1 | 0.85524728 | 0 | 1 | 37 | 0.21027077 | FALSE | 1 | FALSE | 1 | FALSE | 1 | FALSE | 1 | FALSE | 1 | FALSE | 1 |
| 1220 | 1220 | 8.19736842 | 0.93216975 | 5 | 0.04276052 | 5 | 0.09467212 | 2 | 0.75760933 | 0 | 1 | 2 | 0.34918837 | 61 | 0.81568423 | FALSE | 0.63230002 | FALSE | 1 | FALSE | 1 | FALSE | 1 | FALSE | 1 | FALSE | 1 |

| | | | | | | | | | | | | | | | | | | | | | | | | | | | |
|---|---|---|---|---|---|---|---|---|---|---|---|---|---|---|---|---|---|---|---|---|---|---|---|---|---|---|---|
| 1221 | 1221 | 8.5 | 2.64145277 | 1 | 0.8213973 | 3 | 0.3604578 | 3 | 0.4248691 | 4 | 0.35957896 | 2 | 0.29478192 | 53 | 0.86063256 | FALSE | 1 | FALSE | 1 | FALSE | 1 | FALSE | 1 | FALSE | 1 | FALSE | 1 |
| 1222 | 1222 | 9.36363636 | 1.74630661 | 1 | 0.68224519 | 1 | 0.76576289 | 1 | 0.80109494 | 4 | 0.14160176 | 2 | 0.16211764 | 35 | 0.8620983 | FALSE | 1 | FALSE | 1 | FALSE | 1 | FALSE | 1 | FALSE | 1 | FALSE | 1 |
| 1223 | 1223 | 11.5 | 2.51811618 | 2 | 0.10809809 | 2 | 0.15794013 | 1 | 0.55340851 | 1 | 0.64497439 | 0 | 1 | 16 | 0.95553951 | FALSE | 0.88820416 | FALSE | 1 | FALSE | 1 | FALSE | 1 | FALSE | 1 | FALSE | 1 |
| 1224 | 1224 | 4.24739583 | 1.12904066 | 18 | 0.00899941 | 13 | 0.47023332 | 13 | 0.62587902 | 14 | 0.84857647 | 8 | 0.29126138 | 317 | 0.83355909 | FALSE | 0.2803162 | FALSE | 1 | FALSE | 1 | FALSE | 1 | FALSE | 1 | FALSE | 1 |
| 1225 | 1225 | 10.35 | 2.30813778 | 0 | 1 | 2 | 0.1353446 | 0 | 1 | 1 | 0.60987146 | 0 | 1 | 17 | 0.61750632 | FALSE | 1 | FALSE | 1 | FALSE | 1 | FALSE | 1 | FALSE | 1 | FALSE | 1 |
| 1226 | 1226 | 6.84615385 | 1.91536917 | 0 | 1 | 1 | 0.7236252 | 0 | 1 | 2 | 0.54052736 | 1 | 0.4761699 | 35 | 0.24915769 | FALSE | 1 | FALSE | 1 | FALSE | 1 | FALSE | 1 | FALSE | 1 | FALSE | 1 |
| 1227 | 1227 | 9.52 | 1.83564703 | 2 | 0.13369621 | 0 | 1 | 2 | 0.22597287 | 1 | 0.69181315 | 0 | 1 | 20 | 0.8152498 | FALSE | 0.97964162 | FALSE | 1 | FALSE | 1 | FALSE | 1 | FALSE | 1 | FALSE | 1 |
| 1228 | 1228 | 8.6969697 | 2.57593582 | 0 | 1 | 2 | 0.28941836 | 0 | 1 | 1 | 0.78873814 | 1 | 0.42123867 | 29 | 0.39648374 | FALSE | 1 | FALSE | 1 | FALSE | 1 | FALSE | 1 | FALSE | 1 | FALSE | 1 |
| 1229 | 1229 | 8.75 | 2.48746859 | 0 | 1 | 0 | 1 | 1 | 0.44344817 | 1 | 0.52895088 | 0 | 1 | 14 | 0.53308261 | FALSE | 1 | FALSE | 1 | FALSE | 1 | FALSE | 1 | FALSE | 1 | FALSE | 1 |
| 1230 | 1230 | 8.94736842 | 1.73125097 | 0 | 1 | 0 | 1 | 0 | 1 | 2 | 0.21610812 | 0 | 1 | 17 | 0.41141543 | FALSE | 1 | FALSE | 1 | FALSE | 1 | FALSE | 1 | FALSE | 1 | FALSE | 1 |
| 1231 | 1231 | 8.67857143 | 1.3377372 | 1 | 0.50471032 | 2 | 0.21684246 | 1 | 0.62830986 | 3 | 0.12475952 | 0 | 1 | 20 | 0.95133411 | FALSE | 1 | FALSE | 1 | FALSE | 1 | FALSE | 1 | FALSE | 1 | FALSE | 1 |
| 1232 | 1232 | 7.95121951 | 2.43902439 | 0 | 1 | 1 | 0.74131698 | 2 | 0.43640583 | 0 | 1 | 0 | 1 | 38 | 0.09794002 | FALSE | 1 | FALSE | 1 | FALSE | 1 | FALSE | 1 | FALSE | 1 | FALSE | 1 |
| 1233 | 1233 | 9.7826087 | 2.14590655 | 0 | 1 | 1 | 0.53115654 | 0 | 1 | 0 | 1 | 1 | 0.31672722 | 21 | 0.27964072 | FALSE | 1 | FALSE | 1 | FALSE | 1 | FALSE | 1 | FALSE | 1 | FALSE | 1 |
| 1234 | 1234 | 9.47368421 | 2.0867608 | 1 | 0.37380237 | 0 | 1 | 1 | 0.48280983 | 0 | 1 | 1 | 0.25766374 | 15 | 0.69372602 | FALSE | 1 | FALSE | 1 | FALSE | 1 | FALSE | 1 | FALSE | 1 | FALSE | 1 |
| 1235 | 1235 | 9.5 | 2.74912574 | 2 | 0.13369621 | 0 | 1 | 0 | 1 | 1 | 0.69181315 | 0 | 1 | 22 | 0.43642846 | FALSE | 0.97964162 | FALSE | 1 | FALSE | 1 | FALSE | 1 | FALSE | 1 | FALSE | 1 |
| 1236 | 1236 | 4.28676471 | 1.19172857 | 23 | 0.00023523 | 10 | 0.85093623 | 13 | 0.70133623 | 18 | 0.58927342 | 4 | 0.90535574 | 336 | 0.77881481 | TRUE | 0.03045702 | FALSE | 1 | FALSE | 1 | FALSE | 1 | FALSE | 1 | FALSE | 1 |
| 1237 | 1237 | 12 | 3.082207 | 1 | 0.18769745 | 1 | 0.23139161 | 0 | 1 | 1 | 0.31351192 | 1 | 0.12394282 | 4 | 0.99658732 | FALSE | 1 | FALSE | 1 | FALSE | 1 | FALSE | 1 | FALSE | 1 | FALSE | 1 |
| 1238 | 1238 | 8.70967742 | 3.27447162 | 7 | 1.01E-05 | 5 | 0.00288926 | 2 | 0.30673593 | 0 | 1 | 1 | 0.4016888 | 16 | 0.99999671 | TRUE | 0.00393026 | FALSE | 0.46409366 | FALSE | 1 | FALSE | 1 | FALSE | 1 | FALSE | 1 |
| 1239 | 1239 | 9.71428571 | 2.68404203 | 0 | 1 | 2 | 0.07341559 | 0 | 1 | 0 | 1 | 0 | 1 | 12 | 0.62236227 | FALSE | 1 | FALSE | 1 | FALSE | 1 | FALSE | 1 | FALSE | 1 | FALSE | 1 |
| 1240 | 1240 | 9.97142857 | 3.15762763 | 0 | 1 | 1 | 0.68454804 | 1 | 0.72299506 | 1 | 0.80778317 | 1 | 0.44015537 | 31 | 0.34231756 | FALSE | 1 | FALSE | 1 | FALSE | 1 | FALSE | 1 | FALSE | 1 | FALSE | 1 |
| 1241 | 1241 | 9.34482759 | 2.70728089 | 1 | 0.52988231 | 0 | 1 | 1 | 0.65463982 | 1 | 0.74482229 | 0 | 1 | 26 | 0.31495157 | FALSE | 1 | FALSE | 1 | FALSE | 1 | FALSE | 1 | FALSE | 1 | FALSE | 1 |
| 1242 | 1242 | 9.6 | 2.33238076 | 1 | 0.22886536 | 0 | 1 | 0 | 1 | 0 | 1 | 0 | 1 | 9 | 0.5215547 | FALSE | 1 | FALSE | 1 | FALSE | 1 | FALSE | 1 | FALSE | 1 | FALSE | 1 |
| 1243 | 1243 | 7.42105263 | 2.49875315 | 0 | 1 | 1 | 0.46503795 | 0 | 1 | 0 | 1 | 1 | 0.26986775 | 17 | 0.41141543 | FALSE | 1 | FALSE | 1 | FALSE | 1 | FALSE | 1 | FALSE | 1 | FALSE | 1 |
| 1244 | 1244 | 10.8571429 | 2.91197387 | 1 | 0.51745948 | 1 | 0.60247911 | 0 | 1 | 2 | 0.3705495 | 0 | 1 | 24 | 0.55018107 | FALSE | 1 | FALSE | 1 | FALSE | 1 | FALSE | 1 | FALSE | 1 | FALSE | 1 |
| 1245 | 1245 | 9.02777778 | 1.73982935 | 5 | 0.0020683 | 3 | 0.10944025 | 0 | 1 | 2 | 0.49731009 | 1 | 0.44938266 | 25 | 0.99349451 | FALSE | 0.12179217 | FALSE | 1 | FALSE | 1 | FALSE | 1 | FALSE | 1 | FALSE | 1 |
| 1246 | 1246 | 8.73684211 | 2.63263137 | 0 | 1 | 1 | 0.46503795 | 1 | 0.50143826 | 0 | 1 | 1 | 0.26986775 | 16 | 0.65565778 | FALSE | 1 | FALSE | 1 | FALSE | 1 | FALSE | 1 | FALSE | 1 | FALSE | 1 |
| 1247 | 1247 | 10.7 | 3.19530906 | 0 | 1 | 0 | 1 | 0 | 1 | 0 | 1 | 0 | 1 | 10 | 0.18273166 | FALSE | 1 | FALSE | 1 | FALSE | 1 | FALSE | 1 | FALSE | 1 | FALSE | 1 |
| 1248 | 1248 | 9.29166667 | 2.69998714 | 0 | 1 | 2 | 0.16948764 | 1 | 0.5695037 | 0 | 1 | 1 | 0.31672722 | 19 | 0.71542496 | FALSE | 1 | FALSE | 1 | FALSE | 1 | FALSE | 1 | FALSE | 1 | FALSE | 1 |
| 1249 | 1249 | 4.89130435 | 3.402226 | 2 | 0.3309085 | 4 | 0.06038631 | 3 | 0.2282043 | 0 | 1 | 5 | 0.00086811 | 32 | 0.99686402 | FALSE | 1 | FALSE | 1 | FALSE | 1 | FALSE | 1 | FALSE | 0.16162556 | FALSE | 1 |
| 1250 | 1250 | 5.27272727 | 2.50783895 | 2 | 0.38681045 | 1 | 0.82028131 | 1 | 0.85187773 | 5 | 0.08873102 | 1 | 0.57809055 | 42 | 0.82137318 | FALSE | 1 | FALSE | 1 | FALSE | 1 | FALSE | 1 | FALSE | 1 | FALSE | 1 |
| 1251 | 1251 | 9.66666667 | 3.6270588 | 1 | 0.32293355 | 1 | 0.38964394 | 1 | 0.42266264 | 0 | 1 | 1 | 0.2198258 | 11 | 0.92879412 | FALSE | 1 | FALSE | 1 | FALSE | 1 | FALSE | 1 | FALSE | 1 | FALSE | 1 |
| 1252 | 1252 | 7.80952381 | 2.16653584 | 0 | 1 | 2 | 0.60991494 | 2 | 0.66735975 | 2 | 0.79265743 | 0 | 1 | 57 | 0.11815964 | FALSE | 1 | FALSE | 1 | FALSE | 1 | FALSE | 1 | FALSE | 1 | FALSE | 1 |
| 1253 | 1253 | 9.1875 | 1.52282974 | 0 | 1 | 3 | 0.20208567 | 3 | 0.24750546 | 0 | 1 | 0 | 1 | 42 | 0.36011057 | FALSE | 1 | FALSE | 1 | FALSE | 1 | FALSE | 1 | FALSE | 1 | FALSE | 1 |
| 1254 | 1254 | 10.5 | 2.6925824 | 1 | 0.37380237 | 0 | 1 | 0 | 1 | 1 | 0.57131013 | 0 | 1 | 16 | 0.44998001 | FALSE | 1 | FALSE | 1 | FALSE | 1 | FALSE | 1 | FALSE | 1 | FALSE | 1 |
| 1255 | 1255 | 7.51351351 | 3.30128248 | 1 | 0.6184714 | 2 | 0.3375163 | 3 | 0.14625315 | 1 | 0.82511639 | 0 | 1 | 30 | 0.78878581 | FALSE | 1 | FALSE | 1 | FALSE | 1 | FALSE | 1 | FALSE | 1 | FALSE | 1 |
| 1256 | 1256 | 9.38095238 | 1.49526843 | 2 | 0.09989125 | 2 | 0.14655134 | 0 | 1 | 1 | 0.62783522 | 0 | 1 | 16 | 0.90353674 | FALSE | 0.86047225 | FALSE | 1 | FALSE | 1 | FALSE | 1 | FALSE | 1 | FALSE | 1 |
| 1257 | 1257 | 2.62019915 | 2.47134162 | 26 | 0.03390081 | 32 | 0.02777344 | 28 | 0.30534869 | 43 | 0.02857621 | 15 | 0.1721634 | 558 | 0.99989628 | FALSE | 0.56934354 | FALSE | 0.87929047 | FALSE | 1 | FALSE | 1 | FALSE | 1 | FALSE | 1 |
| 1258 | 1258 | 9 | 4.28403328 | 0 | 1 | 0 | 1 | 0 | 1 | 0 | 1 | 0 | 1 | 17 | 0.05550965 | FALSE | 1 | FALSE | 1 | FALSE | 1 | FALSE | 1 | FALSE | 1 | FALSE | 1 |
| 1259 | 1259 | 9.15 | 4.16263138 | 0 | 1 | 0 | 1 | 2 | 0.16019548 | 1 | 0.60987146 | 0 | 1 | 17 | 0.61750632 | FALSE | 1 | FALSE | 1 | FALSE | 1 | FALSE | 1 | FALSE | 1 | FALSE | 1 |
| 1260 | 1260 | 8.875 | 1.98037244 | 2 | 0.19750261 | 0 | 1 | 0 | 1 | 3 | 0.17988885 | 1 | 0.41154418 | 26 | 0.77612337 | FALSE | 1 | FALSE | 1 | FALSE | 1 | FALSE | 1 | FALSE | 1 | FALSE | 1 |
| 1261 | 1261 | 6.39473684 | 3.14173497 | 0 | 1 | 3 | 0.12352882 | 2 | 0.39868506 | 1 | 0.83318955 | 0 | 1 | 32 | 0.61800916 | FALSE | 1 | FALSE | 1 | FALSE | 1 | FALSE | 1 | FALSE | 1 | FALSE | 1 |
| 1262 | 1262 | 7.4516129 | 2.42085975 | 1 | 0.55378205 | 0 | 1 | 0 | 1 | 1 | 0.76781284 | 3 | 0.01381051 | 26 | 0.64660109 | FALSE | 1 | FALSE | 1 | FALSE | 1 | FALSE | 1 | FALSE | 0.64607242 | FALSE | 1 |
| 1263 | 1263 | 5.81481482 | 2.41962711 | 0 | 1 | 1 | 0.58913618 | 1 | 0.62830986 | 1 | 0.71956339 | 0 | 1 | 24 | 0.3725373 | FALSE | 1 | FALSE | 1 | FALSE | 1 | FALSE | 1 | FALSE | 1 | FALSE | 1 |
| 1264 | 1264 | 10.1428571 | 1.55182578 | 1 | 0.3050805 | 1 | 0.36919824 | 0 | 1 | 1 | 0.48242099 | 0 | 1 | 11 | 0.83676449 | FALSE | 1 | FALSE | 1 | FALSE | 1 | FALSE | 1 | FALSE | 1 | FALSE | 1 |
| 1265 | 1265 | 10.5833333 | 2.92854723 | 1 | 0.26795838 | 0 | 1 | 1 | 0.35555293 | 1 | 0.43131129 | 1 | 0.18007762 | 8 | 0.97197833 | FALSE | 1 | FALSE | 1 | FALSE | 1 | FALSE | 1 | FALSE | 1 | FALSE | 1 |
| 1266 | 1266 | 8.38461539 | 1.54613471 | 0 | 1 | 1 | 0.3480709 | 1 | 0.37874417 | 0 | 1 | 0 | 1 | 11 | 0.6683056 | FALSE | 1 | FALSE | 1 | FALSE | 1 | FALSE | 1 | FALSE | 1 | FALSE | 1 |
| 1267 | 1267 | 9.07407407 | 2.41792574 | 0 | 1 | 1 | 0.58913618 | 1 | 0.62830986 | 2 | 0.35374209 | 0 | 1 | 23 | 0.58307573 | FALSE | 1 | FALSE | 1 | FALSE | 1 | FALSE | 1 | FALSE | 1 | FALSE | 1 |
| 1268 | 1268 | 11.8461539 | 1.45951277 | 0 | 1 | 1 | 0.3480709 | 1 | 0.37874417 | 0 | 1 | 0 | 1 | 11 | 0.6683056 | FALSE | 1 | FALSE | 1 | FALSE | 1 | FALSE | 1 | FALSE | 1 | FALSE | 1 |
| 1269 | 1269 | 10.8 | 1.77763888 | 0 | 1 | 0 | 1 | 0 | 1 | 2 | 0.0740981 | 0 | 1 | 8 | 0.80394719 | FALSE | 1 | FALSE | 1 | FALSE | 1 | FALSE | 1 | FALSE | 1 | FALSE | 1 |
| 1270 | 1270 | 3.64233577 | 2.0388104 | 1 | 0.97254344 | 7 | 0.15490136 | 4 | 0.73168489 | 4 | 0.88124375 | 2 | 0.66204068 | 119 | 0.24964601 | FALSE | 1 | FALSE | 1 | FALSE | 1 | FALSE | 1 | FALSE | 1 | FALSE | 1 |
| 1271 | 1271 | 9.63636364 | 2.18560278 | 0 | 1 | 0 | 1 | 2 | 0.33343871 | 3 | 0.19161876 | 0 | 1 | 28 | 0.58681477 | FALSE | 1 | FALSE | 1 | FALSE | 1 | FALSE | 1 | FALSE | 1 | FALSE | 1 |
| 1272 | 1272 | 10.9310345 | 2.31830846 | 0 | 1 | 0 | 1 | 1 | 0.64171563 | 0 | 1 | 0 | 1 | 27 | 0.05284755 | FALSE | 1 | FALSE | 1 | FALSE | 1 | FALSE | 1 | FALSE | 1 | FALSE | 1 |
| 1273 | 1273 | 8.2962963 | 2.4767783 | 0 | 1 | 3 | 0.04547096 | 2 | 0.22597287 | 1 | 0.69181315 | 0 | 1 | 19 | 0.91734261 | FALSE | 1 | FALSE | 1 | FALSE | 1 | FALSE | 1 | FALSE | 1 | FALSE | 1 |
| 1274 | 1274 | 10.7333333 | 1.94821856 | 0 | 1 | 0 | 1 | 2 | 0.09928574 | 1 | 0.50623194 | 0 | 1 | 12 | 0.80329917 | FALSE | 1 | FALSE | 1 | FALSE | 1 | FALSE | 1 | FALSE | 1 | FALSE | 1 |
| 1275 | 1275 | 8.45454546 | 3.47363149 | 0 | 1 | 2 | 0.15794013 | 0 | 1 | 2 | 0.26791271 | 0 | 1 | 18 | 0.74744813 | FALSE | 1 | FALSE | 1 | FALSE | 1 | FALSE | 1 | FALSE | 1 | FALSE | 1 |
| 1276 | 1276 | 8.41176471 | 2.14363143 | 1 | 0.35728305 | 1 | 0.42857733 | 0 | 1 | 0 | 1 | 0 | 1 | 15 | 0.49060435 | FALSE | 1 | FALSE | 1 | FALSE | 1 | FALSE | 1 | FALSE | 1 | FALSE | 1 |
| 1277 | 1277 | 9.69230769 | 2.91902553 | 0 | 1 | 2 | 0.06426041 | 0 | 1 | 2 | 0.1174583 | 0 | 1 | 9 | 0.96018168 | FALSE | 1 | FALSE | 1 | FALSE | 1 | FALSE | 1 | FALSE | 1 | FALSE | 1 |
| 1278 | 1278 | 9.22727273 | 2.55719694 | 0 | 1 | 1 | 0.5154318 | 3 | 0.04259419 | 1 | 0.64497439 | 0 | 1 | 17 | 0.8840357 | FALSE | 1 | FALSE | 1 | FALSE | 1 | FALSE | 1 | FALSE | 1 | FALSE | 1 |
| 1279 | 1279 | 4.43598616 | 1.7871763 | 6 | 0.75500615 | 7 | 0.83075496 | 13 | 0.23535258 | 16 | 0.24869834 | 4 | 0.70186779 | 242 | 0.60529048 | FALSE | 1 | FALSE | 1 | FALSE | 1 | FALSE | 1 | FALSE | 1 | FALSE | 1 |
| 1280 | 1280 | 10.6363636 | 2.14360475 | 0 | 1 | 0 | 1 | 1 | 0.33149969 | 0 | 1 | 1 | 0.16638736 | 9 | 0.75985366 | FALSE | 1 | FALSE | 1 | FALSE | 1 | FALSE | 1 | FALSE | 1 | FALSE | 1 |
| 1281 | 1281 | 8.30434783 | 2.11484983 | 0 | 1 | 2 | 0.16948764 | 0 | 1 | 1 | 0.6613267 | 0 | 1 | 20 | 0.50589402 | FALSE | 1 | FALSE | 1 | FALSE | 1 | FALSE | 1 | FALSE | 1 | FALSE | 1 |
| 1282 | 1282 | 9.13636364 | 2.66791465 | 0 | 1 | 1 | 0.5154318 | 0 | 1 | 2 | 0.26791271 | 3 | 0.00526351 | 16 | 0.95553951 | FALSE | 1 | FALSE | 1 | FALSE | 1 | FALSE | 1 | FALSE | 0.43998669 | FALSE | 1 |
| 1283 | 1283 | 9.6 | 2.4979992 | 0 | 1 | 0 | 1 | 0 | 1 | 0 | 1 | 0 | 1 | 5 | 0.42761893 | FALSE | 1 | FALSE | 1 | FALSE | 1 | FALSE | 1 | FALSE | 1 | FALSE | 1 |
| 1284 | 1284 | 9.85 | 3.92778564 | 0 | 1 | 1 | 0.48239024 | 0 | 1 | 0 | 1 | 0 | 1 | 19 | 0.15700255 | FALSE | 1 | FALSE | 1 | FALSE | 1 | FALSE | 1 | FALSE | 1 | FALSE | 1 |
| 1285 | 1285 | 9.05882353 | 3.3161031 | 2 | 0.06909038 | 0 | 1 | 3 | 0.02143959 | 1 | 0.55062772 | 1 | 0.24525764 | 10 | 0.99775952 | FALSE | 0.74276693 | FALSE | 1 | FALSE | 1 | FALSE | 1 | FALSE | 1 | FALSE | 1 |
| 1286 | 1286 | 8.08163265 | 2.2023803 | 0 | 1 | 1 | 0.80150109 | 4 | 0.09808429 | 2 | 0.66536008 | 0 | 1 | 42 | 0.493842 | FALSE | 1 | FALSE | 1 | FALSE | 1 | FALSE | 1 | FALSE | 1 | FALSE | 1 |
| 1287 | 1287 | 10.5 | 2.11476292 | 2 | 0.07645454 | 2 | 0.1135776 | 1 | 0.48280983 | 1 | 0.57131013 | 0 | 1 | 12 | 0.98544073 | FALSE | 0.77424195 | FALSE | 1 | FALSE | 1 | FALSE | 1 | FALSE | 1 | FALSE | 1 |
| 1288 | 1288 | 10.6521739 | 2.42466843 | 0 | 1 | 0 | 1 | 2 | 0.19931429 | 1 | 0.6613267 | 0 | 1 | 20 | 0.50589402 | FALSE | 1 | FALSE | 1 | FALSE | 1 | FALSE | 1 | FALSE | 1 | FALSE | 1 |
| 1289 | 1289 | 9.65217391 | 2.79142625 | 7 | 0.0001483 | 4 | 0.06038631 | 1 | 0.81522035 | 4 | 0.1589264 | 0 | 1 | 30 | 0.99966734 | TRUE | 0.02249687 | FALSE | 1 | FALSE | 1 | FALSE | 1 | FALSE | 1 | FALSE | 1 |
| 1290 | 1290 | 5.34883721 | 3.61017332 | 5 | 0.2262986 | 6 | 0.22810486 | 5 | 0.48253777 | 6 | 0.53119536 | 3 | 0.34592468 | 102 | 0.91550935 | FALSE | 1 | FALSE | 1 | FALSE | 1 | FALSE | 1 | FALSE | 1 | FALSE | 1 |
| 1291 | 1291 | 9.32 | 2.46121921 | 1 | 0.47819878 | 1 | 0.56109832 | 0 | 1 | 1 | 0.69181315 | 0 | 1 | 22 | 0.43642846 | FALSE | 1 | FALSE | 1 | FALSE | 1 | FALSE | 1 | FALSE | 1 | FALSE | 1 |
| 1292 | 1292 | 3.20662768 | 1.22839811 | 25 | 0.00117483 | 17 | 0.48572436 | 23 | 0.1505265 | 33 | 0.02686208 | 9 | 0.46012505 | 403 | 0.99967538 | FALSE | 0.0844226 | FALSE | 1 | FALSE | 1 | FALSE | 1 | FALSE | 1 | FALSE | 1 |
| 1293 | 1293 | 9.33333333 | 1.63299316 | 0 | 1 | 0 | 1 | 1 | 0.28067846 | 1 | 0.34506517 | 0 | 1 | 7 | 0.84574168 | FALSE | 1 | FALSE | 1 | FALSE | 1 | FALSE | 1 | FALSE | 1 | FALSE | 1 |
| 1294 | 1294 | 8.5 | 2.64575131 | 1 | 0.46441868 | 0 | 1 | 2 | 0.21260257 | 1 | 0.67692816 | 0 | 1 | 20 | 0.68270729 | FALSE | 1 | FALSE | 1 | FALSE | 1 | FALSE | 1 | FALSE | 1 | FALSE | 1 |
| 1295 | 1295 | 9.71428571 | 3.76883027 | 0 | 1 | 0 | 1 | 1 | 0.40110418 | 1 | 0.48242099 | 0 | 1 | 12 | 0.62236227 | FALSE | 1 | FALSE | 1 | FALSE | 1 | FALSE | 1 | FALSE | 1 | FALSE | 1 |
| 1296 | 1296 | 10.9090909 | 1.92847304 | 0 | 1 | 0 | 1 | 0 | 1 | 0 | 1 | 0 | 1 | 11 | 0.15414489 | FALSE | 1 | FALSE | 1 | FALSE | 1 | FALSE | 1 | FALSE | 1 | FALSE | 1 |
| 1297 | 1297 | 9.625 | 1.89983552 | 0 | 1 | 0 | 1 | 0 | 1 | 2 | 0.16544789 | 0 | 1 | 14 | 0.53308261 | FALSE | 1 | FALSE | 1 | FALSE | 1 | FALSE | 1 | FALSE | 1 | FALSE | 1 |
| 1298 | 1298 | 7.62790698 | 2.2415034 | 2 | 0.28329504 | 4 | 0.04238571 | 2 | 0.43640583 | 4 | 0.11715635 | 0 | 1 | 29 | 0.99246133 | FALSE | 1 | FALSE | 0.99776943 | FALSE | 1 | FALSE | 1 | FALSE | 1 | FALSE | 1 |
| 1299 | 1299 | 8.4516129 | 2.18998762 | 3 | 0.6157068 | 7 | 0.10304651 | 3 | 0.82480276 | 3 | 0.92683786 | 1 | 0.87155875 | 106 | 0.34225586 | FALSE | 1 | FALSE | 1 | FALSE | 1 | FALSE | 1 | FALSE | 1 | FALSE | 1 |
| 1300 | 1300 | 9.17647059 | 1.88602583 | 0 | 1 | 1 | 0.42857733 | 1 | 0.46348836 | 0 | 1 | 1 | 0.24525764 | 14 | 0.7313145 | FALSE | 1 | FALSE | 1 | FALSE | 1 | FALSE | 1 | FALSE | 1 | FALSE | 1 |
| 1301 | 1301 | 12.1538462 | 3.32464351 | 0 | 1 | 1 | 0.3480709 | 0 | 1 | 1 | 0.45746571 | 1 | 0.19354507 | 10 | 0.86791642 | FALSE | 1 | FALSE | 1 | FALSE | 1 | FALSE | 1 | FALSE | 1 | FALSE | 1 |
| 1302 | 1302 | 8.52631579 | 1.40930819 | 2 | 0.24498301 | 7 | 0.00015128 | 5 | 0.00977427 | 3 | 0.2402032 | 2 | 0.12262638 | 18 | 0.99999993 | FALSE | 1 | FALSE | 0.15490599 | FALSE | 1 | FALSE | 1 | FALSE | 1 | FALSE | 1 |
| 1303 | 1303 | 8.40625 | 2.3830046 | 0 | 1 | 2 | 0.27730767 | 0 | 1 | 1 | 0.77852168 | 0 | 1 | 29 | 0.24088541 | FALSE | 1 | FALSE | 1 | FALSE | 1 | FALSE | 1 | FALSE | 1 | FALSE | 1 |
| 1304 | 1304 | 9.23529412 | 2.21038756 | 2 | 0.21638713 | 1 | 0.6739483 | 1 | 0.71261921 | 1 | 0.7984848 | 0 | 1 | 29 | 0.55707911 | FALSE | 1 | FALSE | 1 | FALSE | 1 | FALSE | 1 | FALSE | 1 | FALSE | 1 |
| 1305 | 1305 | 7.68181818 | 2.60046086 | 0 | 1 | 0 | 1 | 0 | 1 | 0 | 1 | 0 | 1 | 22 | 0.02368118 | FALSE | 1 | FALSE | 1 | FALSE | 1 | FALSE | 1 | FALSE | 1 | FALSE | 1 |
| 1306 | 1306 | 8.93023256 | 0.87325504 | 0 | 1 | 2 | 0.40761584 | 0 | 1 | 0 | 1 | 1 | 0.50988734 | 40 | 0.07890873 | FALSE | 1 | FALSE | 1 | FALSE | 1 | FALSE | 1 | FALSE | 1 | FALSE | 1 |
| 1307 | 1307 | 8.72 | 2.14513403 | 1 | 0.47819878 | 0 | 1 | 1 | 0.59998149 | 0 | 1 | 0 | 1 | 23 | 0.22727183 | FALSE | 1 | FALSE | 1 | FALSE | 1 | FALSE | 1 | FALSE | 1 | FALSE | 1 |
| 1308 | 1308 | 3.96212121 | 2.4414517 | 5 | 0.25016225 | 1 | 0.98752114 | 3 | 0.85933994 | 6 | 0.56950866 | 4 | 0.17065314 | 113 | 0.40290099 | FALSE | 1 | FALSE | 1 | FALSE | 1 | FALSE | 1 | FALSE | 1 | FALSE | 1 |
| 1309 | 1309 | 2.35261708 | 1.60136393 | 22 | 0.00012266 | 15 | 0.19057917 | 17 | 0.15211667 | 20 | 0.22003278 | 11 | 0.03379963 | 276 | 0.99998091 | TRUE | 0.01970244 | FALSE | 1 | FALSE | 1 | FALSE | 1 | FALSE | 0.86063198 | FALSE | 1 |
| 1310 | 1310 | 9.33333333 | 2.72437557 | 1 | 0.32293355 | 0 | 1 | 0 | 1 | 1 | 0.50623194 | 0 | 1 | 13 | 0.57712805 | FALSE | 1 | FALSE | 1 | FALSE | 1 | FALSE | 1 | FALSE | 1 | FALSE | 1 |
| 1311 | 1311 | 9.22222222 | 2.24982853 | 1 | 0.37380237 | 0 | 1 | 0 | 1 | 2 | 0.19902682 | 1 | 0.25766374 | 14 | 0.86287418 | FALSE | 1 | FALSE | 1 | FALSE | 1 | FALSE | 1 | FALSE | 1 | FALSE | 1 |
| 1312 | 1312 | 5.2238806 | 3.75090395 | 4 | 0.45149984 | 6 | 0.26597002 | 1 | 0.99293277 | 7 | 0.41877501 | 2 | 0.6500531 | 114 | 0.46918017 | FALSE | 1 | FALSE | 1 | FALSE | 1 | FALSE | 1 | FALSE | 1 | FALSE | 1 |
| 1313 | 1313 | 6.49166667 | 1.69556202 | 7 | 0.41467797 | 11 | 0.15093763 | 13 | 0.08840188 | 8 | 0.86564468 | 3 | 0.75741735 | 197 | 0.8264056 | FALSE | 1 | FALSE | 1 | FALSE | 1 | FALSE | 1 | FALSE | 1 | FALSE | 1 |
| 1314 | 1314 | 6.86666667 | 3.18573208 | 0 | 1 | 2 | 0.50543179 | 0 | 1 | 3 | 0.4296776 | 1 | 0.57809055 | 46 | 0.27620659 | FALSE | 1 | FALSE | 1 | FALSE | 1 | FALSE | 1 | FALSE | 1 | FALSE | 1 |

| | | | | | | | | | | | | | | | | | | | | | | | | | | | |
|---|---|---|---|---|---|---|---|---|---|---|---|---|---|---|---|---|---|---|---|---|---|---|---|---|---|---|---|
| 1315 | 1315 | 10.8235294 | 2.63985949 | 2 | 0.06909038 | 2 | 0.10307112 | 2 | 0.12280513 | 0 | 1 | 0 | 1 | 11 | 0.98974663 | FALSE | 0.74276693 | FALSE | 1 | FALSE | 1 | FALSE | 1 | FALSE | 1 | FALSE | 1 |
| 1316 | 1316 | 8.86666667 | 3.18747549 | 2 | 0.32143972 | 4 | 0.05650644 | 1 | 0.80828716 | 1 | 0.8802008 | 0 | 1 | 37 | 0.73736034 | FALSE | 1 | FALSE | 1 | FALSE | 1 | FALSE | 1 | FALSE | 1 | FALSE | 1 |
| 1317 | 1317 | 8.44444444 | 3.31476309 | 1 | 0.50471032 | 0 | 1 | 1 | 0.62830986 | 1 | 0.71956339 | 1 | 0.36060486 | 23 | 0.58307573 | FALSE | 1 | FALSE | 1 | FALSE | 1 | FALSE | 1 | FALSE | 1 | FALSE | 1 |
| 1318 | 1318 | 9.5862069 | 2.41428566 | 0 | 1 | 0 | 1 | 2 | 0.2663542 | 1 | 0.73248983 | 0 | 1 | 25 | 0.34293 | FALSE | 1 | FALSE | 1 | FALSE | 1 | FALSE | 1 | FALSE | 1 | FALSE | 1 |
| 1319 | 1319 | 13.1428571 | 3.68117107 | 0 | 1 | 1 | 0.20567256 | 0 | 1 | 0 | 1 | 0 | 1 | 6 | 0.69920685 | FALSE | 1 | FALSE | 1 | FALSE | 1 | FALSE | 1 | FALSE | 1 | FALSE | 1 |
| 1320 | 1320 | 10.3333333 | 1.24721913 | 0 | 1 | 1 | 0.32623933 | 0 | 1 | 0 | 1 | 0 | 1 | 11 | 0.41944532 | FALSE | 1 | FALSE | 1 | FALSE | 1 | FALSE | 1 | FALSE | 1 | FALSE | 1 |
| 1321 | 1321 | 7.96363636 | 1.73691091 | 1 | 0.76171602 | 2 | 0.5356583 | 0 | 1 | 3 | 0.46623289 | 0 | 1 | 49 | 0.22274213 | FALSE | 1 | FALSE | 1 | FALSE | 1 | FALSE | 1 | FALSE | 1 | FALSE | 1 |
| 1322 | 1322 | 6.79545455 | 2.60750268 | 0 | 1 | 1 | 0.76576289 | 2 | 0.47281647 | 1 | 0.87439647 | 0 | 1 | 40 | 0.16103481 | FALSE | 1 | FALSE | 1 | FALSE | 1 | FALSE | 1 | FALSE | 1 | FALSE | 1 |
| 1323 | 1323 | 10.3809524 | 3.98181809 | 0 | 1 | 1 | 0.49918218 | 1 | 0.53671415 | 1 | 0.62783522 | 1 | 0.29368245 | 17 | 0.77849076 | FALSE | 1 | FALSE | 1 | FALSE | 1 | FALSE | 1 | FALSE | 1 | FALSE | 1 |
| 1324 | 1324 | 9.43243243 | 2.21225751 | 0 | 1 | 1 | 0.70472939 | 1 | 0.74264075 | 2 | 0.51200191 | 0 | 1 | 33 | 0.2931558 | FALSE | 1 | FALSE | 1 | FALSE | 1 | FALSE | 1 | FALSE | 1 | FALSE | 1 |
| 1325 | 1325 | 3.13342599 | 1.01359152 | 59 | 4.08E-05 | 59 | 0.01995404 | 45 | 0.85764998 | 68 | 0.37655247 | 23 | 0.5706351 | 1170 | 0.99542863 | TRUE | 0.01013098 | FALSE | 0.79351225 | FALSE | 1 | FALSE | 1 | FALSE | 1 | FALSE | 1 |
| 1326 | 1326 | 5.27027027 | 2.65745915 | 4 | 0.01425247 | 1 | 0.70472939 | 4 | 0.04241754 | 1 | 0.82511639 | 0 | 1 | 27 | 0.97789976 | FALSE | 0.37183513 | FALSE | 1 | FALSE | 1 | FALSE | 1 | FALSE | 1 | FALSE | 1 |
| 1327 | 1327 | 9.92857143 | 2.18646112 | 0 | 1 | 0 | 1 | 0 | 1 | 0 | 1 | 1 | 0.19354507 | 12 | 0.374198 | FALSE | 1 | FALSE | 1 | FALSE | 1 | FALSE | 1 | FALSE | 1 | FALSE | 1 |
| 1328 | 1328 | 7.0952381 | 3.74044512 | 1 | 0.42086991 | 1 | 0.49918218 | 0 | 1 | 0 | 1 | 1 | 0.29368245 | 18 | 0.57962763 | FALSE | 1 | FALSE | 1 | FALSE | 1 | FALSE | 1 | FALSE | 1 | FALSE | 1 |
| 1329 | 1329 | 6.78378378 | 3.66970899 | 2 | 0.24498301 | 1 | 0.70472939 | 1 | 0.74264075 | 0 | 1 | 0 | 1 | 33 | 0.2931558 | FALSE | 1 | FALSE | 1 | FALSE | 1 | FALSE | 1 | FALSE | 1 | FALSE | 1 |
| 1330 | 1330 | 10.6 | 3.77359245 | 1 | 0.22886536 | 0 | 1 | 0 | 1 | 0 | 1 | 1 | 0.15247062 | 8 | 0.80394719 | FALSE | 1 | FALSE | 1 | FALSE | 1 | FALSE | 1 | FALSE | 1 | FALSE | 1 |
| 1331 | 1331 | 9.0625 | 3.5963306 | 0 | 1 | 1 | 0.40942991 | 0 | 1 | 0 | 1 | 0 | 1 | 15 | 0.26127859 | FALSE | 1 | FALSE | 1 | FALSE | 1 | FALSE | 1 | FALSE | 1 | FALSE | 1 |
| 1332 | 1332 | 6.81967213 | 2.19916673 | 2 | 0.46653175 | 2 | 0.59222924 | 2 | 0.64993443 | 5 | 0.14703607 | 0 | 1 | 50 | 0.76391797 | FALSE | 1 | FALSE | 1 | FALSE | 1 | FALSE | 1 | FALSE | 1 | FALSE | 1 |
| 1333 | 1333 | 9.31578947 | 2.24903796 | 3 | 0.01186104 | 2 | 0.12434464 | 0 | 1 | 0 | 1 | 1 | 0.26986775 | 13 | 0.97998676 | FALSE | 0.3373808 | FALSE | 1 | FALSE | 1 | FALSE | 1 | FALSE | 1 | FALSE | 1 |
| 1334 | 1334 | 9.53333333 | 1.8208667 | 1 | 0.32293355 | 1 | 0.38964394 | 1 | 0.42266264 | 0 | 1 | 0 | 1 | 12 | 0.80329917 | FALSE | 1 | FALSE | 1 | FALSE | 1 | FALSE | 1 | FALSE | 1 | FALSE | 1 |
| 1335 | 1335 | 7.13333333 | 4.2089851 | 0 | 1 | 2 | 0.25306331 | 1 | 0.66709967 | 1 | 0.75658797 | 0 | 1 | 26 | 0.48609328 | FALSE | 1 | FALSE | 1 | FALSE | 1 | FALSE | 1 | FALSE | 1 | FALSE | 1 |
| 1336 | 1336 | 8.78787879 | 1.38733614 | 0 | 1 | 1 | 0.6629941 | 0 | 1 | 3 | 0.19161876 | 0 | 1 | 29 | 0.39648374 | FALSE | 1 | FALSE | 1 | FALSE | 1 | FALSE | 1 | FALSE | 1 | FALSE | 1 |
| 1337 | 1337 | 10.2941177 | 2.02236629 | 0 | 1 | 1 | 0.42857733 | 0 | 1 | 0 | 1 | 0 | 1 | 16 | 0.23070182 | FALSE | 1 | FALSE | 1 | FALSE | 1 | FALSE | 1 | FALSE | 1 | FALSE | 1 |
| 1338 | 1338 | 9.4 | 1.41421356 | 0 | 1 | 0 | 1 | 2 | 0.22597287 | 2 | 0.31968852 | 0 | 1 | 21 | 0.64957167 | FALSE | 1 | FALSE | 1 | FALSE | 1 | FALSE | 1 | FALSE | 1 | FALSE | 1 |
| 1339 | 1339 | 8.91666667 | 1.84654332 | 0 | 1 | 0 | 1 | 1 | 0.35555293 | 1 | 0.43131129 | 0 | 1 | 10 | 0.71436964 | FALSE | 1 | FALSE | 1 | FALSE | 1 | FALSE | 1 | FALSE | 1 | FALSE | 1 |
| 1340 | 1340 | 11.0769231 | 3.26900452 | 0 | 1 | 1 | 0.3480709 | 0 | 1 | 1 | 0.45746571 | 1 | 0.19354507 | 10 | 0.86791642 | FALSE | 1 | FALSE | 1 | FALSE | 1 | FALSE | 1 | FALSE | 1 | FALSE | 1 |
| 1341 | 1341 | 9.85714286 | 3.97953951 | 0 | 1 | 2 | 0.07341559 | 1 | 0.40110418 | 0 | 1 | 0 | 1 | 11 | 0.83676449 | FALSE | 1 | FALSE | 1 | FALSE | 1 | FALSE | 1 | FALSE | 1 | FALSE | 1 |
| 1342 | 1342 | 8.5 | 1.5 | 0 | 1 | 0 | 1 | 0 | 1 | 1 | 0.67692816 | 0 | 1 | 23 | 0.09196389 | FALSE | 1 | FALSE | 1 | FALSE | 1 | FALSE | 1 | FALSE | 1 | FALSE | 1 |
| 1343 | 1343 | 8.81818182 | 1.56581948 | 1 | 0.57647348 | 3 | 0.08959205 | 1 | 0.70185639 | 3 | 0.19161876 | 2 | 0.10132035 | 23 | 0.99096587 | FALSE | 1 | FALSE | 1 | FALSE | 1 | FALSE | 1 | FALSE | 1 | FALSE | 1 |
| 1344 | 1344 | 6.17073171 | 3.34884355 | 1 | 0.65632651 | 2 | 0.38460545 | 0 | 1 | 6 | 0.01031399 | 0 | 1 | 32 | 0.90458039 | FALSE | 1 | FALSE | 1 | FALSE | 1 | FALSE | 1 | FALSE | 1 | FALSE | 1 |
| 1345 | 1345 | 8.15789474 | 2.34550315 | 0 | 1 | 0 | 1 | 1 | 0.50143826 | 0 | 1 | 0 | 1 | 18 | 0.17880694 | FALSE | 1 | FALSE | 1 | FALSE | 1 | FALSE | 1 | FALSE | 1 | FALSE | 1 |
| 1346 | 1346 | 9.05405405 | 3.04866782 | 1 | 0.6184714 | 0 | 1 | 1 | 0.74264075 | 2 | 0.51200191 | 0 | 1 | 33 | 0.2931558 | FALSE | 1 | FALSE | 1 | FALSE | 1 | FALSE | 1 | FALSE | 1 | FALSE | 1 |
| 1347 | 1347 | 8.96551724 | 2.85853542 | 1 | 0.52988231 | 0 | 1 | 1 | 0.65463982 | 0 | 1 | 0 | 1 | 27 | 0.14664438 | FALSE | 1 | FALSE | 1 | FALSE | 1 | FALSE | 1 | FALSE | 1 | FALSE | 1 |
| 1348 | 1348 | 9 | 1.27342908 | 6 | 0.00031256 | 2 | 0.3375163 | 0 | 1 | 3 | 0.2402032 | 3 | 0.0222449 | 23 | 0.99977226 | TRUE | 0.03507515 | FALSE | 1 | FALSE | 1 | FALSE | 1 | FALSE | 0.74991845 | FALSE | 1 |
| 1349 | 1349 | 9.57142857 | 1.9166297 | 0 | 1 | 0 | 1 | 0 | 1 | 0 | 1 | 0 | 1 | 7 | 0.3043657 | FALSE | 1 | FALSE | 1 | FALSE | 1 | FALSE | 1 | FALSE | 1 | FALSE | 1 |
| 1350 | 1350 | 10.5 | 1.93649167 | 1 | 0.18769745 | 1 | 0.23139161 | 1 | 0.25384309 | 0 | 1 | 0 | 1 | 5 | 0.97551477 | FALSE | 1 | FALSE | 1 | FALSE | 1 | FALSE | 1 | FALSE | 1 | FALSE | 1 |
| 1351 | 1351 | 9.9375 | 2.43589691 | 0 | 1 | 0 | 1 | 2 | 0.11087977 | 0 | 1 | 1 | 0.23264612 | 13 | 0.76799161 | FALSE | 1 | FALSE | 1 | FALSE | 1 | FALSE | 1 | FALSE | 1 | FALSE | 1 |
| 1352 | 1352 | 9.30769231 | 1.43498139 | 1 | 0.28675951 | 3 | 0.00750707 | 1 | 0.37874417 | 2 | 0.1174583 | 0 | 1 | 6 | 0.99978701 | FALSE | 1 | FALSE | 0.6167796 | FALSE | 1 | FALSE | 1 | FALSE | 1 | FALSE | 1 |
| 1353 | 1353 | 6.72727273 | 1.88732177 | 1 | 0.8213973 | 0 | 1 | 2 | 0.69218142 | 2 | 0.8136513 | 1 | 0.66594933 | 60 | 0.09129047 | FALSE | 1 | FALSE | 1 | FALSE | 1 | FALSE | 1 | FALSE | 1 | FALSE | 1 |
| 1354 | 1354 | 9.58333333 | 2.32588096 | 1 | 0.24866461 | 0 | 1 | 0 | 1 | 1 | 0.40390032 | 2 | 0.01330324 | 7 | 0.98129353 | FALSE | 1 | FALSE | 1 | FALSE | 1 | FALSE | 1 | FALSE | 0.63731057 | FALSE | 1 |
| 1355 | 1355 | 10 | 2.5819889 | 1 | 0.3050805 | 2 | 0.07341559 | 2 | 0.08806398 | 0 | 1 | 0 | 1 | 9 | 0.98607548 | FALSE | 1 | FALSE | 1 | FALSE | 1 | FALSE | 1 | FALSE | 1 | FALSE | 1 |
| 1356 | 1356 | 10.1111111 | 2.96064724 | 0 | 1 | 0 | 1 | 0 | 1 | 1 | 0.34506517 | 0 | 1 | 8 | 0.5780332 | FALSE | 1 | FALSE | 1 | FALSE | 1 | FALSE | 1 | FALSE | 1 | FALSE | 1 |
| 1357 | 1357 | 6.81081081 | 2.86486487 | 1 | 0.60837712 | 1 | 0.69480472 | 1 | 0.73299778 | 4 | 0.08109727 | 0 | 1 | 29 | 0.81028289 | FALSE | 1 | FALSE | 1 | FALSE | 1 | FALSE | 1 | FALSE | 1 | FALSE | 1 |
| 1358 | 1358 | 10.125 | 1.99608993 | 1 | 0.34033054 | 2 | 0.09285443 | 0 | 1 | 0 | 1 | 0 | 1 | 13 | 0.76799161 | FALSE | 1 | FALSE | 1 | FALSE | 1 | FALSE | 1 | FALSE | 1 | FALSE | 1 |
| 1359 | 1359 | 11.4166667 | 1.93469779 | 0 | 1 | 0 | 1 | 1 | 0.35555293 | 2 | 0.1023796 | 0 | 1 | 9 | 0.89630573 | FALSE | 1 | FALSE | 1 | FALSE | 1 | FALSE | 1 | FALSE | 1 | FALSE | 1 |
| 1360 | 1360 | 7.77272727 | 1.76922731 | 8 | 1.28E-05 | 1 | 0.76576289 | 1 | 0.80109494 | 2 | 0.60667406 | 0 | 1 | 32 | 0.98583614 | TRUE | 0.00418892 | FALSE | 1 | FALSE | 1 | FALSE | 1 | FALSE | 1 | FALSE | 1 |
| 1361 | 1361 | 9.96666667 | 2.34497098 | 2 | 0.17884234 | 1 | 0.6278847 | 1 | 0.66709967 | 1 | 0.75658797 | 1 | 0.39166989 | 24 | 0.82294782 | FALSE | 1 | FALSE | 1 | FALSE | 1 | FALSE | 1 | FALSE | 1 | FALSE | 1 |
| 1362 | 1362 | 9.57894737 | 3.61744006 | 0 | 1 | 0 | 1 | 1 | 0.48280983 | 0 | 1 | 0 | 1 | 17 | 0.20329205 | FALSE | 1 | FALSE | 1 | FALSE | 1 | FALSE | 1 | FALSE | 1 | FALSE | 1 |
| 1363 | 1363 | 8.14705882 | 2.18460325 | 0 | 1 | 0 | 1 | 2 | 0.34668321 | 3 | 0.20353499 | 0 | 1 | 29 | 0.55707911 | FALSE | 1 | FALSE | 1 | FALSE | 1 | FALSE | 1 | FALSE | 1 | FALSE | 1 |
| 1364 | 1364 | 9.17241379 | 3.1301533 | 2 | 0.16962005 | 1 | 0.61539065 | 1 | 0.65463982 | 2 | 0.3871811 | 0 | 1 | 23 | 0.84462957 | FALSE | 1 | FALSE | 1 | FALSE | 1 | FALSE | 1 | FALSE | 1 | FALSE | 1 |
| 1365 | 1365 | 4.2014742 | 1.14273597 | 9 | 0.71953412 | 10 | 0.85300744 | 17 | 0.28633472 | 23 | 0.16827342 | 4 | 0.90638142 | 342 | 0.51894786 | FALSE | 1 | FALSE | 1 | FALSE | 1 | FALSE | 1 | FALSE | 1 | FALSE | 1 |
| 1366 | 1366 | 7.10714286 | 3.29869331 | 4 | 0.00528025 | 3 | 0.06039219 | 3 | 0.07756639 | 2 | 0.3705495 | 1 | 0.37113067 | 15 | 0.99997806 | FALSE | 0.2089654 | FALSE | 1 | FALSE | 1 | FALSE | 1 | FALSE | 1 | FALSE | 1 |
| 1367 | 1367 | 8.13793103 | 2.72566522 | 2 | 0.16962005 | 0 | 1 | 1 | 0.65463982 | 2 | 0.3871811 | 1 | 0.38148475 | 23 | 0.84462957 | FALSE | 1 | FALSE | 1 | FALSE | 1 | FALSE | 1 | FALSE | 1 | FALSE | 1 |
| 1368 | 1368 | 8.72 | 2.67611659 | 2 | 0.13369621 | 0 | 1 | 0 | 1 | 1 | 0.69181315 | 1 | 0.33902672 | 21 | 0.64957167 | FALSE | 0.97964162 | FALSE | 1 | FALSE | 1 | FALSE | 1 | FALSE | 1 | FALSE | 1 |
| 1369 | 1369 | 11.3157895 | 2.15339711 | 1 | 0.37380237 | 0 | 1 | 0 | 1 | 1 | 0.57131013 | 0 | 1 | 16 | 0.44998001 | FALSE | 1 | FALSE | 1 | FALSE | 1 | FALSE | 1 | FALSE | 1 | FALSE | 1 |
| 1370 | 1370 | 9.34375 | 2.11555807 | 0 | 1 | 0 | 1 | 0 | 1 | 3 | 0.17988885 | 0 | 1 | 29 | 0.24088541 | FALSE | 1 | FALSE | 1 | FALSE | 1 | FALSE | 1 | FALSE | 1 | FALSE | 1 |
| 1371 | 1371 | 9.18181818 | 3.36609247 | 1 | 0.4357638 | 0 | 1 | 0 | 1 | 0 | 1 | 3 | 0.00526351 | 18 | 0.74744813 | FALSE | 1 | FALSE | 1 | FALSE | 1 | FALSE | 1 | FALSE | 0.43998669 | FALSE | 1 |
| 1372 | 1372 | 12.2666667 | 4.21847787 | 0 | 1 | 0 | 1 | 1 | 0.42266264 | 0 | 1 | 1 | 0.2198258 | 13 | 0.57712805 | FALSE | 1 | FALSE | 1 | FALSE | 1 | FALSE | 1 | FALSE | 1 | FALSE | 1 |
| 1373 | 1373 | 6.73846154 | 3.31094652 | 1 | 0.81177881 | 4 | 0.15197537 | 2 | 0.67580769 | 1 | 0.95130976 | 3 | 0.08717933 | 53 | 0.70813409 | FALSE | 1 | FALSE | 1 | FALSE | 1 | FALSE | 1 | FALSE | 1 | FALSE | 1 |
| 1374 | 1374 | 8.19047619 | 2.32213054 | 2 | 0.09989125 | 0 | 1 | 1 | 0.53671415 | 1 | 0.62783522 | 0 | 1 | 17 | 0.77849076 | FALSE | 0.86047225 | FALSE | 1 | FALSE | 1 | FALSE | 1 | FALSE | 1 | FALSE | 1 |
| 1375 | 1375 | 6.86363636 | 2.26224857 | 3 | 0.01782284 | 2 | 0.15794013 | 0 | 1 | 1 | 0.64497439 | 0 | 1 | 16 | 0.95553951 | FALSE | 0.41596787 | FALSE | 1 | FALSE | 1 | FALSE | 1 | FALSE | 1 | FALSE | 1 |
| 1376 | 1376 | 10.3157895 | 1.92087238 | 0 | 1 | 2 | 0.12434464 | 1 | 0.50143826 | 0 | 1 | 1 | 0.26986775 | 15 | 0.83648191 | FALSE | 1 | FALSE | 1 | FALSE | 1 | FALSE | 1 | FALSE | 1 | FALSE | 1 |
| 1377 | 1377 | 8.97222222 | 2.49984567 | 1 | 0.59801736 | 1 | 0.68454804 | 2 | 0.35984046 | 1 | 0.80778317 | 0 | 1 | 30 | 0.52765882 | FALSE | 1 | FALSE | 1 | FALSE | 1 | FALSE | 1 | FALSE | 1 | FALSE | 1 |
| 1378 | 1378 | 8.42857143 | 2.64703659 | 0 | 1 | 0 | 1 | 0 | 1 | 0 | 1 | 0 | 1 | 21 | 0.02808151 | FALSE | 1 | FALSE | 1 | FALSE | 1 | FALSE | 1 | FALSE | 1 | FALSE | 1 |
| 1379 | 1379 | 9.64285714 | 1.71577316 | 2 | 0.04858661 | 0 | 1 | 0 | 1 | 2 | 0.13304705 | 0 | 1 | 10 | 0.94579623 | FALSE | 0.649301 | FALSE | 1 | FALSE | 1 | FALSE | 1 | FALSE | 1 | FALSE | 1 |
| 1380 | 1380 | 6.36363636 | 1.22643069 | 0 | 1 | 0 | 1 | 1 | 0.65463982 | 1 | 0.74482229 | 1 | 0.38148475 | 26 | 0.31495157 | FALSE | 1 | FALSE | 1 | FALSE | 1 | FALSE | 1 | FALSE | 1 | FALSE | 1 |
| 1381 | 1381 | 7.5 | 4.5 | 1 | 0.0986934 | 0 | 1 | 0 | 1 | 0 | 1 | 0 | 1 | 3 | 0.8823539 | FALSE | 0.86047225 | FALSE | 1 | FALSE | 1 | FALSE | 1 | FALSE | 1 | FALSE | 1 |
| 1382 | 1382 | 8.90476191 | 2.24467119 | 0 | 1 | 0 | 1 | 1 | 0.53671415 | 2 | 0.25059627 | 0 | 1 | 18 | 0.57962763 | FALSE | 1 | FALSE | 1 | FALSE | 1 | FALSE | 1 | FALSE | 1 | FALSE | 1 |
| 1383 | 1383 | 9.56666667 | 2.17076535 | 0 | 1 | 1 | 0.6278847 | 3 | 0.09131526 | 0 | 1 | 1 | 0.39166989 | 25 | 0.67630471 | FALSE | 1 | FALSE | 1 | FALSE | 1 | FALSE | 1 | FALSE | 1 | FALSE | 1 |
| 1384 | 1384 | 11.98 | 2.54943131 | 3 | 0.13593669 | 0 | 1 | 4 | 0.1037722 | 6 | 0.02587968 | 0 | 1 | 37 | 0.98198136 | FALSE | 0.98986074 | FALSE | 1 | FALSE | 1 | FALSE | 1 | FALSE | 1 | FALSE | 1 |
| 1385 | 1385 | 10.75 | 1.69926455 | 0 | 1 | 0 | 1 | 0 | 1 | 0 | 1 | 0 | 1 | 20 | 0.03329856 | FALSE | 1 | FALSE | 1 | FALSE | 1 | FALSE | 1 | FALSE | 1 | FALSE | 1 |
| 1386 | 1386 | 11.4705882 | 2.30390071 | 0 | 1 | 1 | 0.42857733 | 0 | 1 | 0 | 1 | 0 | 1 | 16 | 0.23070182 | FALSE | 1 | FALSE | 1 | FALSE | 1 | FALSE | 1 | FALSE | 1 | FALSE | 1 |
| 1387 | 1387 | 7.15517241 | 3.13930436 | 1 | 0.77972342 | 1 | 0.85269953 | 1 | 0.88128698 | 2 | 0.75303313 | 1 | 0.61824432 | 52 | 0.17733928 | FALSE | 1 | FALSE | 1 | FALSE | 1 | FALSE | 1 | FALSE | 1 | FALSE | 1 |
| 1388 | 1388 | 10.3888889 | 2.66955862 | 0 | 1 | 0 | 1 | 1 | 0.46348836 | 1 | 0.55062772 | 0 | 1 | 15 | 0.49060435 | FALSE | 1 | FALSE | 1 | FALSE | 1 | FALSE | 1 | FALSE | 1 | FALSE | 1 |
| 1389 | 1389 | 10.1578947 | 2.36783619 | 1 | 0.37380237 | 2 | 0.1135776 | 0 | 1 | 1 | 0.57131013 | 0 | 1 | 14 | 0.86287418 | FALSE | 1 | FALSE | 1 | FALSE | 1 | FALSE | 1 | FALSE | 1 | FALSE | 1 |
| 1390 | 1390 | 10.1111111 | 3.76468319 | 2 | 0.14250978 | 3 | 0.05021553 | 3 | 0.06482014 | 1 | 0.70601446 | 0 | 1 | 17 | 0.99598795 | FALSE | 1 | FALSE | 1 | FALSE | 1 | FALSE | 1 | FALSE | 1 | FALSE | 1 |
| 1391 | 1391 | 8.08695652 | 2.73325803 | 0 | 1 | 0 | 1 | 1 | 0.5695037 | 0 | 1 | 0 | 1 | 22 | 0.10533112 | FALSE | 1 | FALSE | 1 | FALSE | 1 | FALSE | 1 | FALSE | 1 | FALSE | 1 |
| 1392 | 1392 | 8.25925926 | 1.97376483 | 3 | 0.0279333 | 0 | 1 | 1 | 0.61440466 | 0 | 1 | 1 | 0.3499045 | 21 | 0.78939668 | FALSE | 0.50964275 | FALSE | 1 | FALSE | 1 | FALSE | 1 | FALSE | 1 | FALSE | 1 |
| 1393 | 1393 | 6.58571429 | 1.50705822 | 0 | 1 | 1 | 0.89430705 | 1 | 0.91794622 | 6 | 0.09079991 | 0 | 1 | 60 | 0.24411098 | FALSE | 1 | FALSE | 1 | FALSE | 1 | FALSE | 1 | FALSE | 1 | FALSE | 1 |
| 1394 | 1394 | 9.64285714 | 2.68879003 | 0 | 1 | 0 | 1 | 0 | 1 | 1 | 0.48242099 | 0 | 1 | 13 | 0.33283729 | FALSE | 1 | FALSE | 1 | FALSE | 1 | FALSE | 1 | FALSE | 1 | FALSE | 1 |
| 1395 | 1395 | 9.58333333 | 3.25213605 | 0 | 1 | 0 | 1 | 1 | 0.35555293 | 1 | 0.43131129 | 0 | 1 | 10 | 0.71436964 | FALSE | 1 | FALSE | 1 | FALSE | 1 | FALSE | 1 | FALSE | 1 | FALSE | 1 |
| 1396 | 1396 | 10.4444444 | 3.67003213 | 0 | 1 | 1 | 0.44710673 | 1 | 0.48280983 | 0 | 1 | 0 | 1 | 16 | 0.44998001 | FALSE | 1 | FALSE | 1 | FALSE | 1 | FALSE | 1 | FALSE | 1 | FALSE | 1 |
| 1397 | 1397 | 9.53333333 | 2.06128331 | 2 | 0.05513784 | 0 | 1 | 0 | 1 | 1 | 0.50623194 | 0 | 1 | 12 | 0.80329917 | FALSE | 0.68849059 | FALSE | 1 | FALSE | 1 | FALSE | 1 | FALSE | 1 | FALSE | 1 |
| 1398 | 1398 | 8.75 | 2.91904665 | 0 | 1 | 2 | 0.05552913 | 0 | 1 | 0 | 1 | 0 | 1 | 10 | 0.71436964 | FALSE | 1 | FALSE | 1 | FALSE | 1 | FALSE | 1 | FALSE | 1 | FALSE | 1 |
| 1399 | 1399 | 6.58490566 | 1.63030223 | 0 | 1 | 3 | 0.24492539 | 2 | 0.57303317 | 5 | 0.0944968 | 1 | 0.58506217 | 42 | 0.88654191 | FALSE | 1 | FALSE | 1 | FALSE | 1 | FALSE | 1 | FALSE | 1 | FALSE | 1 |
| 1400 | 1400 | 11.8823529 | 3.23368944 | 1 | 0.35728305 | 0 | 1 | 0 | 1 | 0 | 1 | 0 | 1 | 16 | 0.23070182 | FALSE | 1 | FALSE | 1 | FALSE | 1 | FALSE | 1 | FALSE | 1 | FALSE | 1 |
| 1401 | 1401 | 7.68965517 | 1.96702906 | 0 | 1 | 2 | 0.24095584 | 0 | 1 | 1 | 0.74482229 | 0 | 1 | 26 | 0.31495157 | FALSE | 1 | FALSE | 1 | FALSE | 1 | FALSE | 1 | FALSE | 1 | FALSE | 1 |
| 1402 | 1402 | 8.76190476 | 4.44977005 | 1 | 0.40558521 | 0 | 1 | 1 | 0.51939841 | 1 | 0.60987146 | 0 | 1 | 17 | 0.61750632 | FALSE | 1 | FALSE | 1 | FALSE | 1 | FALSE | 1 | FALSE | 1 | FALSE | 1 |
| 1403 | 1403 | 5.29850746 | 2.94737387 | 1 | 0.81177881 | 1 | 0.87929209 | 0 | 1 | 4 | 0.33842086 | 1 | 0.65460803 | 57 | 0.19643469 | FALSE | 1 | FALSE | 1 | FALSE | 1 | FALSE | 1 | FALSE | 1 | FALSE | 1 |
| 1404 | 1404 | 12 | 1.52752523 | 0 | 1 | 1 | 0.30368014 | 2 | 0.05708125 | 1 | 0.40390032 | 0 | 1 | 7 | 0.98129353 | FALSE | 1 | FALSE | 1 | FALSE | 1 | FALSE | 1 | FALSE | 1 | FALSE | 1 |
| 1405 | 1405 | 7.2 | 2.92574777 | 1 | 0.40558521 | 0 | 1 | 1 | 0.51939841 | 0 | 1 | 0 | 1 | 18 | 0.37504645 | FALSE | 1 | FALSE | 1 | FALSE | 1 | FALSE | 1 | FALSE | 1 | FALSE | 1 |
| 1406 | 1406 | 10.7857143 | 2.11046953 | 2 | 0.04858661 | 1 | 0.36919824 | 1 | 0.40110418 | 1 | 0.48242099 | 0 | 1 | 9 | 0.98607548 | FALSE | 0.649301 | FALSE | 1 | FALSE | 1 | FALSE | 1 | FALSE | 1 | FALSE | 1 |
| 1407 | 1407 | 8.35 | 1.10792599 | 1 | 0.40558521 | 0 | 1 | 0 | 1 | 2 | 0.23331368 | 0 | 1 | 17 | 0.61750632 | FALSE | 1 | FALSE | 1 | FALSE | 1 | FALSE | 1 | FALSE | 1 | FALSE | 1 |
| 1408 | 1408 | 9.33333333 | 2.21108319 | 0 | 1 | 1 | 0.179097 | 0 | 1 | 0 | 1 | 0 | 1 | 5 | 0.76185568 | FALSE | 1 | FALSE | 1 | FALSE | 1 | FALSE | 1 | FALSE | 1 | FALSE | 1 |

| | | | | | | | | | | | | | | | | | | | | | | | | | | | |
|---|---|---|---|---|---|---|---|---|---|---|---|---|---|---|---|---|---|---|---|---|---|---|---|---|---|---|---|
| 1409 | 1409 | 11.1666667 | 1.95078332 | 0 | 1 | 0 | 1 | 0 | 1 | 0 | 1 | 1 | 0.09446378 | 5 | 0.76185568 | FALSE | 1 | FALSE | 1 | FALSE | 1 | FALSE | 1 | FALSE | 1 | FALSE | 1 |
| 1410 | 1410 | 11.0454546 | 2.56687416 | 4 | 0.00212601 | 0 | 1 | 1 | 0.55340851 | 1 | 0.64497439 | 0 | 1 | 16 | 0.95553951 | FALSE | 0.12179217 | FALSE | 1 | FALSE | 1 | FALSE | 1 | FALSE | 1 | FALSE | 1 |
| 1411 | 1411 | 6.93181818 | 3.63941633 | 3 | 0.09709068 | 2 | 0.40761584 | 0 | 1 | 2 | 0.59404271 | 0 | 1 | 36 | 0.6451239 | FALSE | 0.86047225 | FALSE | 1 | FALSE | 1 | FALSE | 1 | FALSE | 1 | FALSE | 1 |
| 1412 | 1412 | 8.35135135 | 1.89189189 | 0 | 1 | 0 | 1 | 0 | 1 | 0 | 1 | 1 | 0.45845922 | 36 | 0.01445326 | FALSE | 1 | FALSE | 1 | FALSE | 1 | FALSE | 1 | FALSE | 1 | FALSE | 1 |
| 1413 | 1413 | 8.8 | 2.22710575 | 0 | 1 | 0 | 1 | 0 | 1 | 0 | 1 | 0 | 1 | 5 | 0.42761893 | FALSE | 1 | FALSE | 1 | FALSE | 1 | FALSE | 1 | FALSE | 1 | FALSE | 1 |
| 1414 | 1414 | 8.13793103 | 2.23952178 | 1 | 0.50471032 | 0 | 1 | 2 | 0.25286935 | 2 | 0.35374209 | 0 | 1 | 22 | 0.76237877 | FALSE | 1 | FALSE | 1 | FALSE | 1 | FALSE | 1 | FALSE | 1 | FALSE | 1 |
| 1415 | 1415 | 8.5 | 3.07112653 | 0 | 1 | 0 | 1 | 0 | 1 | 2 | 0.26791271 | 0 | 1 | 20 | 0.30915202 | FALSE | 1 | FALSE | 1 | FALSE | 1 | FALSE | 1 | FALSE | 1 | FALSE | 1 |
| 1416 | 1416 | 11.2307692 | 2.48546068 | 0 | 1 | 2 | 0.06426041 | 2 | 0.07725803 | 0 | 1 | 0 | 1 | 9 | 0.96018168 | FALSE | 1 | FALSE | 1 | FALSE | 1 | FALSE | 1 | FALSE | 1 | FALSE | 1 |
| 1417 | 1417 | 4.22666667 | 1.12928689 | 9 | 0.35539886 | 10 | 0.49929249 | 12 | 0.38392862 | 18 | 0.14055343 | 2 | 0.95984547 | 247 | 0.79217214 | FALSE | 1 | FALSE | 1 | FALSE | 1 | FALSE | 1 | FALSE | 1 | FALSE | 1 |
| 1418 | 1418 | 7.25 | 3.96074488 | 2 | 0.27371785 | 0 | 1 | 2 | 0.42396974 | 1 | 0.84823834 | 0 | 1 | 35 | 0.38966407 | FALSE | 1 | FALSE | 1 | FALSE | 1 | FALSE | 1 | FALSE | 1 | FALSE | 1 |
| 1419 | 1419 | 9.13043478 | 2.32761023 | 0 | 1 | 1 | 0.53115654 | 0 | 1 | 1 | 0.6613267 | 0 | 1 | 21 | 0.27964072 | FALSE | 1 | FALSE | 1 | FALSE | 1 | FALSE | 1 | FALSE | 1 | FALSE | 1 |
| 1420 | 1420 | 9.53571429 | 2.42725805 | 2 | 0.15144274 | 0 | 1 | 2 | 0.25286935 | 0 | 1 | 1 | 0.36060486 | 22 | 0.76237877 | FALSE | 1 | FALSE | 1 | FALSE | 1 | FALSE | 1 | FALSE | 1 | FALSE | 1 |
| 1421 | 1421 | 8 | 2.33549683 | 1 | 0.4357638 | 0 | 1 | 0 | 1 | 2 | 0.26791271 | 0 | 1 | 19 | 0.54233419 | FALSE | 1 | FALSE | 1 | FALSE | 1 | FALSE | 1 | FALSE | 1 | FALSE | 1 |
| 1422 | 1422 | 5.10810811 | 2.99354065 | 1 | 0.6184714 | 2 | 0.3375163 | 5 | 0.00977427 | 2 | 0.51200191 | 0 | 1 | 27 | 0.97789976 | FALSE | 1 | FALSE | 1 | FALSE | 1 | FALSE | 1 | FALSE | 1 | FALSE | 1 |
| 1423 | 1423 | 7.86666667 | 2.05318181 | 4 | 0.0671906 | 7 | 0.00302356 | 2 | 0.64095454 | 2 | 0.76960882 | 2 | 0.25814944 | 43 | 0.99642726 | FALSE | 0.74276693 | FALSE | 0.46839856 | FALSE | 1 | FALSE | 1 | FALSE | 1 | FALSE | 1 |
| 1424 | 1424 | 7.72 | 4.05482429 | 0 | 1 | 0 | 1 | 2 | 0.22597287 | 1 | 0.69181315 | 1 | 0.33902672 | 21 | 0.64957167 | FALSE | 1 | FALSE | 1 | FALSE | 1 | FALSE | 1 | FALSE | 1 | FALSE | 1 |
| 1425 | 1425 | 9.2 | 2.90615111 | 1 | 0.59801736 | 0 | 1 | 0 | 1 | 0 | 1 | 0 | 1 | 34 | 0.01937385 | FALSE | 1 | FALSE | 1 | FALSE | 1 | FALSE | 1 | FALSE | 1 | FALSE | 1 |
| 1426 | 1426 | 9.13333333 | 2.70472837 | 0 | 1 | 0 | 1 | 1 | 0.40110418 | 1 | 0.48242099 | 0 | 1 | 12 | 0.62236227 | FALSE | 1 | FALSE | 1 | FALSE | 1 | FALSE | 1 | FALSE | 1 | FALSE | 1 |
| 1427 | 1427 | 7.95833333 | 1.61964931 | 2 | 0.12501436 | 0 | 1 | 2 | 0.21260257 | 1 | 0.67692816 | 0 | 1 | 19 | 0.83975208 | FALSE | 0.95622564 | FALSE | 1 | FALSE | 1 | FALSE | 1 | FALSE | 1 | FALSE | 1 |
| 1428 | 1428 | 8.87878788 | 2.78260371 | 2 | 0.20692137 | 0 | 1 | 3 | 0.11365871 | 2 | 0.45159487 | 2 | 0.10132035 | 24 | 0.97500814 | FALSE | 1 | FALSE | 1 | FALSE | 1 | FALSE | 1 | FALSE | 1 | FALSE | 1 |
| 1429 | 1429 | 8.72727273 | 1.21287855 | 0 | 1 | 0 | 1 | 0 | 1 | 0 | 1 | 1 | 0.16638736 | 10 | 0.46859937 | FALSE | 1 | FALSE | 1 | FALSE | 1 | FALSE | 1 | FALSE | 1 | FALSE | 1 |
| 1430 | 1430 | 5.33043478 | 3.85547998 | 1 | 0.95084309 | 2 | 0.89164461 | 4 | 0.59743048 | 7 | 0.27601945 | 2 | 0.56638749 | 99 | 0.36165879 | FALSE | 1 | FALSE | 1 | FALSE | 1 | FALSE | 1 | FALSE | 1 | FALSE | 1 |
| 1431 | 1431 | 5.39597315 | 2.7017737 | 5 | 0.2945697 | 4 | 0.67478629 | 6 | 0.39568129 | 10 | 0.11294592 | 0 | 1 | 116 | 0.79490007 | FALSE | 1 | FALSE | 1 | FALSE | 1 | FALSE | 1 | FALSE | 1 | FALSE | 1 |
| 1432 | 1432 | 9.14814815 | 2.90215378 | 0 | 1 | 1 | 0.58913618 | 0 | 1 | 1 | 0.71956339 | 0 | 1 | 25 | 0.1832206 | FALSE | 1 | FALSE | 1 | FALSE | 1 | FALSE | 1 | FALSE | 1 | FALSE | 1 |
| 1433 | 1433 | 8.91304348 | 1.99810875 | 0 | 1 | 2 | 0.16948764 | 2 | 0.19931429 | 1 | 0.6613267 | 0 | 1 | 18 | 0.86273155 | FALSE | 1 | FALSE | 1 | FALSE | 1 | FALSE | 1 | FALSE | 1 | FALSE | 1 |
| 1434 | 1434 | 9.8 | 2.22710575 | 2 | 0.0256865 | 1 | 0.28036922 | 0 | 1 | 0 | 1 | 0 | 1 | 7 | 0.94326215 | FALSE | 0.4974559 | FALSE | 1 | FALSE | 1 | FALSE | 1 | FALSE | 1 | FALSE | 1 |
| 1435 | 1435 | 7.4375 | 1.53983562 | 2 | 0.34973212 | 2 | 0.46321244 | 3 | 0.24750546 | 3 | 0.37966044 | 2 | 0.18560062 | 36 | 0.97101084 | FALSE | 1 | FALSE | 1 | FALSE | 1 | FALSE | 1 | FALSE | 1 | FALSE | 1 |
| 1436 | 1436 | 5.14035088 | 1.33032903 | 8 | 0.0628901 | 6 | 0.45032499 | 8 | 0.24643487 | 14 | 0.01998979 | 1 | 0.93789185 | 129 | 0.99185286 | FALSE | 0.73494389 | FALSE | 1 | FALSE | 1 | FALSE | 1 | FALSE | 1 | FALSE | 1 |
| 1437 | 1437 | 7.68421053 | 1.85110583 | 3 | 0.07268126 | 0 | 1 | 1 | 0.75193685 | 2 | 0.52640931 | 0 | 1 | 32 | 0.61800916 | FALSE | 0.76727438 | FALSE | 1 | FALSE | 1 | FALSE | 1 | FALSE | 1 | FALSE | 1 |
| 1438 | 1438 | 8.52631579 | 3.83019061 | 1 | 0.38989951 | 1 | 0.46503795 | 0 | 1 | 0 | 1 | 0 | 1 | 17 | 0.41141543 | FALSE | 1 | FALSE | 1 | FALSE | 1 | FALSE | 1 | FALSE | 1 | FALSE | 1 |
| 1439 | 1439 | 9.8125 | 2.98367957 | 0 | 1 | 1 | 0.40942991 | 0 | 1 | 0 | 1 | 0 | 1 | 15 | 0.26127859 | FALSE | 1 | FALSE | 1 | FALSE | 1 | FALSE | 1 | FALSE | 1 | FALSE | 1 |
| 1440 | 1440 | 9.05 | 1.35922772 | 0 | 1 | 1 | 0.48239024 | 3 | 0.03322143 | 2 | 0.23331368 | 0 | 1 | 14 | 0.97325593 | FALSE | 1 | FALSE | 1 | FALSE | 1 | FALSE | 1 | FALSE | 1 | FALSE | 1 |
| 1441 | 1441 | 7.49275362 | 0.91085544 | 0 | 1 | 2 | 0.65946213 | 3 | 0.45353909 | 4 | 0.39127216 | 0 | 1 | 60 | 0.34674272 | FALSE | 1 | FALSE | 1 | FALSE | 1 | FALSE | 1 | FALSE | 1 | FALSE | 1 |
| 1442 | 1442 | 11 | 2.93056911 | 2 | 0.06909038 | 0 | 1 | 0 | 1 | 0 | 1 | 0 | 1 | 15 | 0.49060435 | FALSE | 0.74276693 | FALSE | 1 | FALSE | 1 | FALSE | 1 | FALSE | 1 | FALSE | 1 |
| 1443 | 1443 | 5.89473684 | 3.82367606 | 0 | 1 | 0 | 1 | 2 | 0.37289966 | 3 | 0.22784656 | 1 | 0.44938266 | 30 | 0.67248271 | FALSE | 1 | FALSE | 1 | FALSE | 1 | FALSE | 1 | FALSE | 1 | FALSE | 1 |
| 1444 | 1444 | 9.67647059 | 1.52856311 | 1 | 0.58738519 | 1 | 0.6739483 | 1 | 0.71261921 | 2 | 0.46709861 | 1 | 0.43077488 | 28 | 0.72558119 | FALSE | 1 | FALSE | 1 | FALSE | 1 | FALSE | 1 | FALSE | 1 | FALSE | 1 |
| 1445 | 1445 | 9.23076923 | 3.19022175 | 1 | 0.28675951 | 1 | 0.3480709 | 0 | 1 | 0 | 1 | 0 | 1 | 11 | 0.6683056 | FALSE | 1 | FALSE | 1 | FALSE | 1 | FALSE | 1 | FALSE | 1 | FALSE | 1 |
| 1446 | 1446 | 7.88888889 | 2.07869855 | 0 | 1 | 0 | 1 | 1 | 0.62830986 | 2 | 0.35374209 | 0 | 1 | 24 | 0.3725373 | FALSE | 1 | FALSE | 1 | FALSE | 1 | FALSE | 1 | FALSE | 1 | FALSE | 1 |
| 1447 | 1447 | 9.5 | 3.13847097 | 1 | 0.40558521 | 1 | 0.48239024 | 2 | 0.16019548 | 0 | 1 | 0 | 1 | 16 | 0.80826361 | FALSE | 1 | FALSE | 1 | FALSE | 1 | FALSE | 1 | FALSE | 1 | FALSE | 1 |
| 1448 | 1448 | 7.4 | 3.28071287 | 0 | 1 | 2 | 0.62701348 | 1 | 0.90832639 | 3 | 0.57949569 | 1 | 0.66032558 | 58 | 0.18266626 | FALSE | 1 | FALSE | 1 | FALSE | 1 | FALSE | 1 | FALSE | 1 | FALSE | 1 |
| 1449 | 1449 | 8.8 | 3.26190129 | 0 | 1 | 0 | 1 | 2 | 0.22597287 | 0 | 1 | 1 | 0.33902672 | 22 | 0.43642846 | FALSE | 1 | FALSE | 1 | FALSE | 1 | FALSE | 1 | FALSE | 1 | FALSE | 1 |
| 1450 | 1450 | 5.65957447 | 2.88264503 | 1 | 0.70621968 | 1 | 0.78790827 | 1 | 0.82190381 | 1 | 0.89101956 | 0 | 1 | 43 | 0.12168195 | FALSE | 1 | FALSE | 1 | FALSE | 1 | FALSE | 1 | FALSE | 1 | FALSE | 1 |
| 1451 | 1451 | 7.88461539 | 1.73929524 | 1 | 0.49162634 | 3 | 0.05021553 | 0 | 1 | 1 | 0.70601446 | 0 | 1 | 21 | 0.78939668 | FALSE | 1 | FALSE | 1 | FALSE | 1 | FALSE | 1 | FALSE | 1 | FALSE | 1 |
| 1452 | 1452 | 8.61904762 | 2.99962205 | 0 | 1 | 2 | 0.14655134 | 1 | 0.53671415 | 1 | 0.62783522 | 2 | 0.04568221 | 15 | 0.96513691 | FALSE | 1 | FALSE | 1 | FALSE | 1 | FALSE | 1 | FALSE | 0.95710659 | FALSE | 1 |
| 1453 | 1453 | 9.52941177 | 4.28564838 | 0 | 1 | 1 | 0.42857733 | 1 | 0.46348836 | 0 | 1 | 0 | 1 | 15 | 0.49060435 | FALSE | 1 | FALSE | 1 | FALSE | 1 | FALSE | 1 | FALSE | 1 | FALSE | 1 |
| 1454 | 1454 | 5.75714286 | 3.31419335 | 2 | 0.539854 | 0 | 1 | 3 | 0.46296673 | 2 | 0.83868849 | 3 | 0.10693796 | 60 | 0.45755253 | FALSE | 1 | FALSE | 1 | FALSE | 1 | FALSE | 1 | FALSE | 1 | FALSE | 1 |
| 1455 | 1455 | 8.80952381 | 1.78936136 | 0 | 1 | 1 | 0.49918218 | 0 | 1 | 0 | 1 | 0 | 1 | 20 | 0.13763971 | FALSE | 1 | FALSE | 1 | FALSE | 1 | FALSE | 1 | FALSE | 1 | FALSE | 1 |
| 1456 | 1456 | 9.46875 | 2.78370678 | 1 | 0.55378205 | 0 | 1 | 0 | 1 | 3 | 0.16836568 | 2 | 0.09110253 | 25 | 0.80006983 | FALSE | 1 | FALSE | 1 | FALSE | 1 | FALSE | 1 | FALSE | 1 | FALSE | 1 |
| 1457 | 1457 | 2.25938567 | 1.24791524 | 17 | 0.32967802 | 27 | 0.03670513 | 36 | 0.00086957 | 24 | 0.75189225 | 9 | 0.62982655 | 472 | 0.99500864 | FALSE | 1 | FALSE | 0.94259695 | FALSE | 1 | FALSE | 1 | FALSE | 1 | FALSE | 1 |
| 1458 | 1458 | 7.94642857 | 3.00249684 | 3 | 0.16642539 | 3 | 0.26245929 | 3 | 0.31652594 | 5 | 0.10658542 | 1 | 0.59866474 | 40 | 0.99191141 | FALSE | 1 | FALSE | 1 | FALSE | 1 | FALSE | 1 | FALSE | 1 | FALSE | 1 |
| 1459 | 1459 | 9.29411765 | 1.90246607 | 0 | 1 | 0 | 1 | 1 | 0.46348836 | 1 | 0.55062772 | 1 | 0.24525764 | 14 | 0.7313145 | FALSE | 1 | FALSE | 1 | FALSE | 1 | FALSE | 1 | FALSE | 1 | FALSE | 1 |
| 1460 | 1460 | 8.26829268 | 2.29553533 | 0 | 1 | 2 | 0.36122049 | 1 | 0.76089851 | 2 | 0.54052736 | 1 | 0.4761699 | 33 | 0.59061307 | FALSE | 1 | FALSE | 1 | FALSE | 1 | FALSE | 1 | FALSE | 1 | FALSE | 1 |
| 1461 | 1461 | 8.73684211 | 2.38067008 | 1 | 0.38989951 | 0 | 1 | 1 | 0.50143826 | 0 | 1 | 0 | 1 | 17 | 0.41141543 | FALSE | 1 | FALSE | 1 | FALSE | 1 | FALSE | 1 | FALSE | 1 | FALSE | 1 |
| 1462 | 1462 | 8.47619048 | 2.34254741 | 1 | 0.42086991 | 0 | 1 | 1 | 0.53671415 | 0 | 1 | 0 | 1 | 19 | 0.34094959 | FALSE | 1 | FALSE | 1 | FALSE | 1 | FALSE | 1 | FALSE | 1 | FALSE | 1 |
| 1463 | 1463 | 7.81578947 | 2.46934109 | 0 | 1 | 1 | 0.71433277 | 0 | 1 | 4 | 0.09477133 | 0 | 1 | 33 | 0.44261631 | FALSE | 1 | FALSE | 1 | FALSE | 1 | FALSE | 1 | FALSE | 1 | FALSE | 1 |
| 1464 | 1464 | 13.2857143 | 1.97948664 | 0 | 1 | 0 | 1 | 0 | 1 | 0 | 1 | 0 | 1 | 7 | 0.3043657 | FALSE | 1 | FALSE | 1 | FALSE | 1 | FALSE | 1 | FALSE | 1 | FALSE | 1 |
| 1465 | 1465 | 9.69090909 | 3.15002296 | 2 | 0.41401181 | 2 | 0.5356583 | 0 | 1 | 4 | 0.24483091 | 1 | 0.59866474 | 46 | 0.64645216 | FALSE | 1 | FALSE | 1 | FALSE | 1 | FALSE | 1 | FALSE | 1 | FALSE | 1 |
| 1466 | 1466 | 8 | 1.67332005 | 0 | 1 | 0 | 1 | 1 | 0.16720719 | 0 | 1 | 1 | 0.07935778 | 3 | 0.97031021 | FALSE | 1 | FALSE | 1 | FALSE | 1 | FALSE | 1 | FALSE | 1 | FALSE | 1 |
| 1467 | 1467 | 9.82222222 | 1.01737984 | 1 | 0.69044531 | 1 | 0.77338922 | 0 | 1 | 2 | 0.61900576 | 3 | 0.03699359 | 38 | 0.59437745 | FALSE | 1 | FALSE | 1 | FALSE | 1 | FALSE | 1 | FALSE | 0.87669812 | FALSE | 1 |
| 1468 | 1468 | 7.16666667 | 2.88193608 | 1 | 0.71380452 | 1 | 0.79481669 | 4 | 0.09254759 | 2 | 0.65421368 | 1 | 0.54902428 | 39 | 0.79274273 | FALSE | 1 | FALSE | 1 | FALSE | 1 | FALSE | 1 | FALSE | 1 | FALSE | 1 |
| 1469 | 1469 | 8.52631579 | 2.26804857 | 0 | 1 | 3 | 0.12352882 | 3 | 0.15483587 | 1 | 0.83318955 | 2 | 0.12811285 | 29 | 0.93814695 | FALSE | 1 | FALSE | 1 | FALSE | 1 | FALSE | 1 | FALSE | 1 | FALSE | 1 |
| 1470 | 1470 | 10.0769231 | 1.73034183 | 0 | 1 | 1 | 0.3480709 | 0 | 1 | 0 | 1 | 0 | 1 | 12 | 0.374198 | FALSE | 1 | FALSE | 1 | FALSE | 1 | FALSE | 1 | FALSE | 1 | FALSE | 1 |
| 1471 | 1471 | 8.35483871 | 1.99270261 | 3 | 0.21200292 | 4 | 0.13998159 | 1 | 0.89758397 | 3 | 0.54709375 | 3 | 0.08097514 | 48 | 0.94896265 | FALSE | 1 | FALSE | 1 | FALSE | 1 | FALSE | 1 | FALSE | 1 | FALSE | 1 |
| 1472 | 1472 | 8.41538462 | 1.09393162 | 2 | 0.49997462 | 7 | 0.00475737 | 2 | 0.68408105 | 3 | 0.57949569 | 3 | 0.09035518 | 48 | 0.99106774 | FALSE | 1 | FALSE | 0.58120789 | FALSE | 1 | FALSE | 1 | FALSE | 1 | FALSE | 1 |
| 1473 | 1473 | 10.6216216 | 2.96238902 | 1 | 0.56527493 | 1 | 0.65167362 | 3 | 0.10599423 | 3 | 0.17988885 | 0 | 1 | 24 | 0.94876798 | FALSE | 1 | FALSE | 1 | FALSE | 1 | FALSE | 1 | FALSE | 1 | FALSE | 1 |
| 1474 | 1474 | 8.82352941 | 1.2478355 | 1 | 0.35728305 | 0 | 1 | 1 | 0.46348836 | 2 | 0.18212134 | 0 | 1 | 13 | 0.88719222 | FALSE | 1 | FALSE | 1 | FALSE | 1 | FALSE | 1 | FALSE | 1 | FALSE | 1 |
| 1475 | 1475 | 6.4 | 1.70680132 | 2 | 0.96989994 | 5 | 0.79597743 | 9 | 0.31135075 | 6 | 0.91330276 | 1 | 0.96745086 | 181 | 0.04645154 | FALSE | 1 | FALSE | 1 | FALSE | 1 | FALSE | 1 | FALSE | 1 | FALSE | 1 |
| 1476 | 1476 | 8.86363636 | 3.2374998 | 1 | 0.4357638 | 1 | 0.5154318 | 1 | 0.55340851 | 0 | 1 | 0 | 1 | 19 | 0.54233419 | FALSE | 1 | FALSE | 1 | FALSE | 1 | FALSE | 1 | FALSE | 1 | FALSE | 1 |
| 1477 | 1477 | 9.93939394 | 1.70452862 | 0 | 1 | 2 | 0.28941836 | 2 | 0.33343871 | 1 | 0.78873814 | 4 | 0.00194079 | 24 | 0.97500814 | FALSE | 1 | FALSE | 1 | FALSE | 1 | FALSE | 1 | FALSE | 0.2523647 | FALSE | 1 |
| 1478 | 1478 | 9.5 | 1.41421356 | 0 | 1 | 0 | 1 | 0 | 1 | 0 | 1 | 0 | 1 | 7 | 0.3043657 | FALSE | 1 | FALSE | 1 | FALSE | 1 | FALSE | 1 | FALSE | 1 | FALSE | 1 |
| 1479 | 1479 | 9.07142857 | 0.79859571 | 0 | 1 | 1 | 0.36919824 | 1 | 0.40110418 | 1 | 0.48242099 | 0 | 1 | 11 | 0.83676449 | FALSE | 1 | FALSE | 1 | FALSE | 1 | FALSE | 1 | FALSE | 1 | FALSE | 1 |
| 1480 | 1480 | 7.92 | 2.05757138 | 1 | 0.47819878 | 0 | 1 | 1 | 0.59998149 | 2 | 0.31968852 | 2 | 0.06257402 | 19 | 0.91734261 | FALSE | 1 | FALSE | 1 | FALSE | 1 | FALSE | 1 | FALSE | 1 | FALSE | 1 |
| 1481 | 1481 | 7.90909091 | 1.83176743 | 0 | 1 | 0 | 1 | 0 | 1 | 1 | 0.40390032 | 0 | 1 | 10 | 0.46859937 | FALSE | 1 | FALSE | 1 | FALSE | 1 | FALSE | 1 | FALSE | 1 | FALSE | 1 |
| 1482 | 1482 | 7.40476191 | 1.94641947 | 3 | 0.34438864 | 3 | 0.49001146 | 4 | 0.33196391 | 3 | 0.72619227 | 1 | 0.74003033 | 67 | 0.72302002 | FALSE | 1 | FALSE | 1 | FALSE | 1 | FALSE | 1 | FALSE | 1 | FALSE | 1 |
| 1483 | 1483 | 10.625 | 2.86956007 | 1 | 0.18769745 | 0 | 1 | 0 | 1 | 0 | 1 | 1 | 0.12394282 | 6 | 0.88425551 | FALSE | 1 | FALSE | 1 | FALSE | 1 | FALSE | 1 | FALSE | 1 | FALSE | 1 |
| 1484 | 1484 | 8.30882353 | 2.8761563 | 1 | 0.82602167 | 5 | 0.06484224 | 3 | 0.43448639 | 4 | 0.3701568 | 1 | 0.6714808 | 53 | 0.90970332 | FALSE | 1 | FALSE | 1 | FALSE | 1 | FALSE | 1 | FALSE | 1 | FALSE | 1 |
| 1485 | 1485 | 9.63636364 | 1.77214443 | 0 | 1 | 0 | 1 | 1 | 0.33149969 | 0 | 1 | 0 | 1 | 10 | 0.46859937 | FALSE | 1 | FALSE | 1 | FALSE | 1 | FALSE | 1 | FALSE | 1 | FALSE | 1 |
| 1486 | 1486 | 9.6 | 2.15406592 | 0 | 1 | 0 | 1 | 0 | 1 | 0 | 1 | 0 | 1 | 9 | 0.21661397 | FALSE | 1 | FALSE | 1 | FALSE | 1 | FALSE | 1 | FALSE | 1 | FALSE | 1 |
| 1487 | 1487 | 6.13690476 | 1.23406797 | 5 | 0.42729556 | 3 | 0.91172387 | 3 | 0.943354 | 8 | 0.50305898 | 3 | 0.51937148 | 145 | 0.22222643 | FALSE | 1 | FALSE | 1 | FALSE | 1 | FALSE | 1 | FALSE | 1 | FALSE | 1 |
| 1488 | 1488 | 7.91666667 | 1.46011796 | 1 | 0.60837712 | 2 | 0.32556427 | 0 | 1 | 3 | 0.22784656 | 0 | 1 | 30 | 0.67248271 | FALSE | 1 | FALSE | 1 | FALSE | 1 | FALSE | 1 | FALSE | 1 | FALSE | 1 |
| 1489 | 1489 | 11 | 2.33549683 | 1 | 0.24866461 | 1 | 0.30368014 | 0 | 1 | 2 | 0.08789566 | 0 | 1 | 7 | 0.98129353 | FALSE | 1 | FALSE | 1 | FALSE | 1 | FALSE | 1 | FALSE | 1 | FALSE | 1 |
| 1490 | 1490 | 9.5106383 | 2.37784401 | 5 | 0.00612174 | 4 | 0.06038631 | 1 | 0.81522035 | 2 | 0.63103861 | 2 | 0.17379289 | 32 | 0.99686402 | FALSE | 0.23004259 | FALSE | 1 | FALSE | 1 | FALSE | 1 | FALSE | 1 | FALSE | 1 |
| 1491 | 1491 | 9.29411765 | 2.67372192 | 2 | 0.06909038 | 1 | 0.42857733 | 0 | 1 | 1 | 0.55062772 | 0 | 1 | 13 | 0.88719222 | FALSE | 0.74276693 | FALSE | 1 | FALSE | 1 | FALSE | 1 | FALSE | 1 | FALSE | 1 |
| 1492 | 1492 | 8.75 | 3.05845822 | 0 | 1 | 1 | 0.32623933 | 0 | 1 | 0 | 1 | 0 | 1 | 11 | 0.41944532 | FALSE | 1 | FALSE | 1 | FALSE | 1 | FALSE | 1 | FALSE | 1 | FALSE | 1 |
| 1493 | 1493 | 9.11111111 | 2.73973956 | 1 | 0.50471032 | 0 | 1 | 3 | 0.07106362 | 0 | 1 | 0 | 1 | 23 | 0.58307573 | FALSE | 1 | FALSE | 1 | FALSE | 1 | FALSE | 1 | FALSE | 1 | FALSE | 1 |
| 1494 | 1494 | 9.72727273 | 2.57932017 | 1 | 0.4357638 | 3 | 0.03266339 | 1 | 0.55340851 | 1 | 0.64497439 | 0 | 1 | 16 | 0.95553951 | FALSE | 1 | FALSE | 0.92268438 | FALSE | 1 | FALSE | 1 | FALSE | 1 | FALSE | 1 |
| 1495 | 1495 | 9.32 | 2.41197015 | 0 | 1 | 1 | 0.54637326 | 0 | 1 | 0 | 1 | 1 | 0.32796862 | 22 | 0.25237063 | FALSE | 1 | FALSE | 1 | FALSE | 1 | FALSE | 1 | FALSE | 1 | FALSE | 1 |
| 1496 | 1496 | 5.33870968 | 2.19929276 | 1 | 0.79637728 | 4 | 0.13413374 | 0 | 1 | 1 | 0.94386203 | 0 | 1 | 55 | 0.13951638 | FALSE | 1 | FALSE | 1 | FALSE | 1 | FALSE | 1 | FALSE | 1 | FALSE | 1 |
| 1497 | 1497 | 9.625 | 1.49478259 | 0 | 1 | 1 | 0.40942991 | 0 | 1 | 0 | 1 | 0 | 1 | 15 | 0.26127859 | FALSE | 1 | FALSE | 1 | FALSE | 1 | FALSE | 1 | FALSE | 1 | FALSE | 1 |
| 1498 | 1498 | 10.5428571 | 2.85199536 | 0 | 1 | 1 | 0.68454804 | 1 | 0.72299506 | 2 | 0.48233989 | 0 | 1 | 31 | 0.34231756 | FALSE | 1 | FALSE | 1 | FALSE | 1 | FALSE | 1 | FALSE | 1 | FALSE | 1 |
| 1499 | 1499 | 7.97142857 | 1.66451562 | 1 | 0.58738519 | 2 | 0.30150529 | 1 | 0.71261921 | 2 | 0.46709861 | 2 | 0.1065437 | 26 | 0.92796052 | FALSE | 1 | FALSE | 1 | FALSE | 1 | FALSE | 1 | FALSE | 1 | FALSE | 1 |
| 1500 | 1500 | 8.25 | 3.56195171 | 1 | 0.26795838 | 0 | 1 | 0 | 1 | 0 | 1 | 0 | 1 | 11 | 0.41944532 | FALSE | 1 | FALSE | 1 | FALSE | 1 | FALSE | 1 | FALSE | 1 | FALSE | 1 |
| 1501 | 1501 | 9.89473684 | 2.29174116 | 0 | 1 | 0 | 1 | 1 | 0.50143826 | 3 | 0.05383577 | 0 | 1 | 15 | 0.83648191 | FALSE | 1 | FALSE | 1 | FALSE | 1 | FALSE | 1 | FALSE | 1 | FALSE | 1 |
| 1502 | 1502 | 8.27777778 | 2.29263448 | 1 | 0.60837712 | 0 | 1 | 0 | 1 | 2 | 0.49731009 | 0 | 1 | 33 | 0.16407838 | FALSE | 1 | FALSE | 1 | FALSE | 1 | FALSE | 1 | FALSE | 1 | FALSE | 1 |

| | | | | | | | | | | | | | | | | | | | | | | | | | | | |
|---|---|---|---|---|---|---|---|---|---|---|---|---|---|---|---|---|---|---|---|---|---|---|---|---|---|---|---|
| 1503 | 1503 | 10.2592593 | 2.04744814 | 0 | 1 | 0 | 1 | 3 | 0.07106362 | 1 | 0.71956339 | 1 | 0.36060486 | 22 | 0.76237877 | FALSE | 1 | FALSE | 1 | FALSE | 1 | FALSE | 1 | FALSE | 1 | FALSE | 1 |
| 1504 | 1504 | 9.73684211 | 1.7424153 | 0 | 1 | 1 | 0.46503795 | 0 | 1 | 0 | 1 | 0 | 1 | 18 | 0.17880694 | FALSE | 1 | FALSE | 1 | FALSE | 1 | FALSE | 1 | FALSE | 1 | FALSE | 1 |
| 1505 | 1505 | 11.44 | 1.16893114 | 0 | 1 | 0 | 1 | 0 | 1 | 1 | 0.69181315 | 0 | 1 | 24 | 0.08019798 | FALSE | 1 | FALSE | 1 | FALSE | 1 | FALSE | 1 | FALSE | 1 | FALSE | 1 |
| 1506 | 1506 | 10.8571429 | 3.1590492 | 0 | 1 | 0 | 1 | 1 | 0.40110418 | 1 | 0.48242099 | 0 | 1 | 12 | 0.62236227 | FALSE | 1 | FALSE | 1 | FALSE | 1 | FALSE | 1 | FALSE | 1 | FALSE | 1 |
| 1507 | 1507 | 7.65811966 | 1.80295291 | 2 | 0.79946155 | 4 | 0.51365485 | 5 | 0.39735161 | 3 | 0.90433751 | 5 | 0.04008201 | 96 | 0.66343565 | FALSE | 1 | FALSE | 1 | FALSE | 1 | FALSE | 1 | FALSE | 0.90455059 | FALSE | 1 |
| 1508 | 1508 | 8.5 | 2.43926218 | 1 | 0.64722818 | 0 | 1 | 0 | 1 | 1 | 0.84823834 | 0 | 1 | 38 | 0.03896014 | FALSE | 1 | FALSE | 1 | FALSE | 1 | FALSE | 1 | FALSE | 1 | FALSE | 1 |
| 1509 | 1509 | 5.59701493 | 3.02753385 | 9 | 1.86E-05 | 2 | 0.58316597 | 2 | 0.64095454 | 0 | 1 | 4 | 0.016509 | 43 | 0.99642726 | TRUE | 0.00545372 | FALSE | 1 | FALSE | 1 | FALSE | 1 | FALSE | 0.66951368 | FALSE | 1 |
| 1510 | 1510 | 8.66666667 | 2.5819889 | 0 | 1 | 0 | 1 | 0 | 1 | 0 | 1 | 0 | 1 | 9 | 0.21661397 | FALSE | 1 | FALSE | 1 | FALSE | 1 | FALSE | 1 | FALSE | 1 | FALSE | 1 |
| 1511 | 1511 | 4.67857143 | 1.73315508 | 0 | 1 | 2 | 0.2288771 | 1 | 0.64171563 | 4 | 0.03737468 | 1 | 0.37113067 | 20 | 0.97757517 | FALSE | 1 | FALSE | 1 | FALSE | 1 | FALSE | 1 | FALSE | 1 | FALSE | 1 |
| 1512 | 1512 | 10.1851852 | 2.09119914 | 0 | 1 | 0 | 1 | 1 | 0.62830986 | 1 | 0.71956339 | 0 | 1 | 25 | 0.1832206 | FALSE | 1 | FALSE | 1 | FALSE | 1 | FALSE | 1 | FALSE | 1 | FALSE | 1 |
| 1513 | 1513 | 9.44444444 | 3.14858385 | 0 | 1 | 2 | 0.1135776 | 1 | 0.48280983 | 0 | 1 | 0 | 1 | 15 | 0.69372602 | FALSE | 1 | FALSE | 1 | FALSE | 1 | FALSE | 1 | FALSE | 1 | FALSE | 1 |
| 1514 | 1514 | 10.5365854 | 2.98040398 | 1 | 0.65632651 | 2 | 0.38460545 | 3 | 0.1814659 | 1 | 0.85524728 | 5 | 0.00050647 | 29 | 0.99246133 | FALSE | 1 | FALSE | 1 | FALSE | 1 | FALSE | 1 | FALSE | 0.11213569 | FALSE | 1 |
| 1515 | 1515 | 9.5 | 1.60727513 | 0 | 1 | 0 | 1 | 0 | 1 | 0 | 1 | 0 | 1 | 12 | 0.13002667 | FALSE | 1 | FALSE | 1 | FALSE | 1 | FALSE | 1 | FALSE | 1 | FALSE | 1 |
| 1516 | 1516 | 7.87234043 | 1.82904488 | 1 | 0.69843502 | 4 | 0.06038631 | 3 | 0.2282043 | 1 | 0.88573774 | 2 | 0.17379289 | 35 | 0.95410769 | FALSE | 1 | FALSE | 1 | FALSE | 1 | FALSE | 1 | FALSE | 1 | FALSE | 1 |
| 1517 | 1517 | 7.69230769 | 2.44586355 | 0 | 1 | 0 | 1 | 0 | 1 | 1 | 0.91399378 | 0 | 1 | 51 | 0.00150735 | FALSE | 1 | FALSE | 1 | FALSE | 1 | FALSE | 1 | FALSE | 1 | FALSE | 1 |
| 1518 | 1518 | 9.64285714 | 2.68879003 | 0 | 1 | 0 | 1 | 2 | 0.2663542 | 2 | 0.3705495 | 0 | 1 | 24 | 0.55018107 | FALSE | 1 | FALSE | 1 | FALSE | 1 | FALSE | 1 | FALSE | 1 | FALSE | 1 |
| 1519 | 1519 | 8.75 | 1.60078106 | 2 | 0.06197875 | 1 | 0.40942991 | 0 | 1 | 0 | 1 | 0 | 1 | 13 | 0.76799161 | FALSE | 0.72532843 | FALSE | 1 | FALSE | 1 | FALSE | 1 | FALSE | 1 | FALSE | 1 |
| 1520 | 1520 | 6.625 | 1.38631706 | 1 | 0.56527493 | 0 | 1 | 1 | 0.69069222 | 1 | 0.77852168 | 0 | 1 | 29 | 0.24088541 | FALSE | 1 | FALSE | 1 | FALSE | 1 | FALSE | 1 | FALSE | 1 | FALSE | 1 |
| 1521 | 1521 | 10.2173913 | 2.66957938 | 3 | 0.02011786 | 1 | 0.53115654 | 1 | 0.5695037 | 0 | 1 | 4 | 0.00047637 | 14 | 0.99865094 | FALSE | 0.44482142 | FALSE | 1 | FALSE | 1 | FALSE | 1 | FALSE | 0.10840111 | FALSE | 1 |
| 1522 | 1522 | 7.42 | 1.42954538 | 0 | 1 | 0 | 1 | 5 | 0.03267327 | 5 | 0.07777122 | 0 | 1 | 40 | 0.85325204 | FALSE | 1 | FALSE | 1 | FALSE | 1 | FALSE | 1 | FALSE | 1 | FALSE | 1 |
| 1523 | 1523 | 8.0625 | 2.22116946 | 0 | 1 | 1 | 0.40942991 | 0 | 1 | 0 | 1 | 0 | 1 | 15 | 0.26127859 | FALSE | 1 | FALSE | 1 | FALSE | 1 | FALSE | 1 | FALSE | 1 | FALSE | 1 |
| 1524 | 1524 | 6.91735537 | 1.32731492 | 8 | 0.01112945 | 11 | 0.00154348 | 6 | 0.25575376 | 8 | 0.17975796 | 8 | 0.00070197 | 78 | 0.99999992 | FALSE | 0.32445698 | FALSE | 0.40311621 | FALSE | 1 | FALSE | 1 | FALSE | 0.14103898 | FALSE | 1 |
| 1525 | 1525 | 9.5 | 3.04138127 | 1 | 0.22886536 | 0 | 1 | 1 | 0.30655256 | 1 | 0.37517262 | 0 | 1 | 7 | 0.94326215 | FALSE | 1 | FALSE | 1 | FALSE | 1 | FALSE | 1 | FALSE | 1 | FALSE | 1 |
| 1526 | 1526 | 7.75 | 2.45990576 | 2 | 0.31193899 | 2 | 0.41896557 | 1 | 0.80109494 | 6 | 0.0144401 | 1 | 0.51797501 | 32 | 0.98583614 | FALSE | 1 | FALSE | 1 | FALSE | 1 | FALSE | 1 | FALSE | 1 | FALSE | 1 |
| 1527 | 1527 | 10.5833333 | 2.77263413 | 2 | 0.23542589 | 0 | 1 | 0 | 1 | 3 | 0.22784656 | 0 | 1 | 31 | 0.49869404 | FALSE | 1 | FALSE | 1 | FALSE | 1 | FALSE | 1 | FALSE | 1 | FALSE | 1 |
| 1528 | 1528 | 9.7037037 | 2.82600116 | 1 | 0.50471032 | 1 | 0.58913618 | 4 | 0.01487236 | 2 | 0.35374209 | 2 | 0.07169515 | 17 | 0.99851071 | FALSE | 1 | FALSE | 1 | FALSE | 1 | FALSE | 1 | FALSE | 1 | FALSE | 1 |
| 1529 | 1529 | 9.69230769 | 2.03373909 | 1 | 0.49162634 | 0 | 1 | 1 | 0.61440466 | 0 | 1 | 1 | 0.3499045 | 23 | 0.40372818 | FALSE | 1 | FALSE | 1 | FALSE | 1 | FALSE | 1 | FALSE | 1 | FALSE | 1 |
| 1530 | 1530 | 8.22727273 | 2.19362337 | 3 | 0.09196816 | 3 | 0.1535026 | 1 | 0.78589454 | 2 | 0.5811115 | 0 | 1 | 33 | 0.89139527 | FALSE | 0.83248965 | FALSE | 1 | FALSE | 1 | FALSE | 1 | FALSE | 1 | FALSE | 1 |
| 1531 | 1531 | 8.77777778 | 2.04275292 | 0 | 1 | 0 | 1 | 1 | 0.48280983 | 1 | 0.57131013 | 0 | 1 | 16 | 0.44998001 | FALSE | 1 | FALSE | 1 | FALSE | 1 | FALSE | 1 | FALSE | 1 | FALSE | 1 |
| 1532 | 1532 | 9.11764706 | 2.16611208 | 0 | 1 | 0 | 1 | 0 | 1 | 1 | 0.55062772 | 1 | 0.24525764 | 15 | 0.49060435 | FALSE | 1 | FALSE | 1 | FALSE | 1 | FALSE | 1 | FALSE | 1 | FALSE | 1 |
| 1533 | 1533 | 9.88461539 | 2.91319179 | 0 | 1 | 0 | 1 | 0 | 1 | 1 | 0.70601446 | 0 | 1 | 25 | 0.06985983 | FALSE | 1 | FALSE | 1 | FALSE | 1 | FALSE | 1 | FALSE | 1 | FALSE | 1 |
| 1534 | 1534 | 7.71428571 | 1.88080168 | 1 | 0.42086991 | 2 | 0.14655134 | 2 | 0.17308361 | 1 | 0.62783522 | 0 | 1 | 15 | 0.96513691 | FALSE | 1 | FALSE | 1 | FALSE | 1 | FALSE | 1 | FALSE | 1 | FALSE | 1 |
| 1535 | 1535 | 3.09517767 | 0.78035986 | 19 | 0.64709655 | 38 | 0.00686774 | 25 | 0.77602326 | 40 | 0.26630219 | 31 | 1.86E-06 | 634 | 0.99909222 | FALSE | 1 | FALSE | 0.60103504 | FALSE | 1 | FALSE | 1 | TRUE | 0.00217217 | FALSE | 1 |
| 1536 | 1536 | 3.20603675 | 1.26358873 | 29 | 0.01627684 | 33 | 0.0429806 | 30 | 0.30414198 | 32 | 0.71529806 | 23 | 0.00224332 | 608 | 0.99893266 | FALSE | 0.39708985 | FALSE | 1 | FALSE | 1 | FALSE | 1 | FALSE | 0.27025449 | FALSE | 1 |
| 1537 | 1537 | 8.65 | 2.04389334 | 1 | 0.64722818 | 2 | 0.37295654 | 2 | 0.42396974 | 2 | 0.5543522 | 1 | 0.48480878 | 32 | 0.83853687 | FALSE | 1 | FALSE | 1 | FALSE | 1 | FALSE | 1 | FALSE | 1 | FALSE | 1 |
| 1538 | 1538 | 9.9375 | 3.00974978 | 1 | 0.34033054 | 0 | 1 | 0 | 1 | 1 | 0.52895088 | 0 | 1 | 14 | 0.53308261 | FALSE | 1 | FALSE | 1 | FALSE | 1 | FALSE | 1 | FALSE | 1 | FALSE | 1 |
| 1539 | 1539 | 9.86666667 | 2.30554886 | 2 | 0.05513784 | 0 | 1 | 1 | 0.42266264 | 0 | 1 | 0 | 1 | 12 | 0.80329917 | FALSE | 0.68849059 | FALSE | 1 | FALSE | 1 | FALSE | 1 | FALSE | 1 | FALSE | 1 |
| 1540 | 1540 | 8.88 | 2.65811964 | 0 | 1 | 1 | 0.56109832 | 1 | 0.59998149 | 0 | 1 | 2 | 0.06257402 | 21 | 0.64957167 | FALSE | 1 | FALSE | 1 | FALSE | 1 | FALSE | 1 | FALSE | 1 | FALSE | 1 |
| 1541 | 1541 | 10.8571429 | 3.13635691 | 1 | 0.16630142 | 0 | 1 | 0 | 1 | 0 | 1 | 0 | 1 | 6 | 0.69920685 | FALSE | 1 | FALSE | 1 | FALSE | 1 | FALSE | 1 | FALSE | 1 | FALSE | 1 |
| 1542 | 1542 | 7.07142857 | 1.98077495 | 0 | 1 | 0 | 1 | 0 | 1 | 0 | 1 | 0 | 1 | 14 | 0.09251304 | FALSE | 1 | FALSE | 1 | FALSE | 1 | FALSE | 1 | FALSE | 1 | FALSE | 1 |
| 1543 | 1543 | 9.49275362 | 1.05807028 | 0 | 1 | 4 | 0.18354682 | 2 | 0.71546205 | 4 | 0.39127216 | 0 | 1 | 59 | 0.47808459 | FALSE | 1 | FALSE | 1 | FALSE | 1 | FALSE | 1 | FALSE | 1 | FALSE | 1 |
| 1544 | 1544 | 8.92857143 | 2.34411987 | 1 | 0.51745948 | 0 | 1 | 1 | 0.64171563 | 2 | 0.3705495 | 0 | 1 | 24 | 0.55018107 | FALSE | 1 | FALSE | 1 | FALSE | 1 | FALSE | 1 | FALSE | 1 | FALSE | 1 |
| 1545 | 1545 | 7.92307692 | 2.94726832 | 0 | 1 | 0 | 1 | 1 | 0.37874417 | 1 | 0.45746571 | 1 | 0.19354507 | 10 | 0.86791642 | FALSE | 1 | FALSE | 1 | FALSE | 1 | FALSE | 1 | FALSE | 1 | FALSE | 1 |
| 1546 | 1546 | 10.3125 | 3.80326751 | 0 | 1 | 0 | 1 | 1 | 0.44344817 | 0 | 1 | 1 | 0.23264612 | 14 | 0.53308261 | FALSE | 1 | FALSE | 1 | FALSE | 1 | FALSE | 1 | FALSE | 1 | FALSE | 1 |
| 1547 | 1547 | 9.45454546 | 2.89556515 | 2 | 0.31193899 | 3 | 0.16927151 | 1 | 0.80109494 | 5 | 0.04960693 | 1 | 0.51797501 | 32 | 0.98583614 | FALSE | 1 | FALSE | 1 | FALSE | 1 | FALSE | 1 | FALSE | 1 | FALSE | 1 |
| 1548 | 1548 | 7.90909091 | 2.24436853 | 1 | 0.68224519 | 0 | 1 | 0 | 1 | 4 | 0.14160176 | 1 | 0.51797501 | 38 | 0.45705039 | FALSE | 1 | FALSE | 1 | FALSE | 1 | FALSE | 1 | FALSE | 1 | FALSE | 1 |
| 1549 | 1549 | 8.47368421 | 3.60363006 | 3 | 0.01017712 | 0 | 1 | 0 | 1 | 2 | 0.19902682 | 0 | 1 | 13 | 0.95041913 | FALSE | 0.30467829 | FALSE | 1 | FALSE | 1 | FALSE | 1 | FALSE | 1 | FALSE | 1 |
| 1550 | 1550 | 6.66666667 | 2.82842713 | 0 | 1 | 0 | 1 | 1 | 0.62830986 | 1 | 0.71956339 | 0 | 1 | 25 | 0.1832206 | FALSE | 1 | FALSE | 1 | FALSE | 1 | FALSE | 1 | FALSE | 1 | FALSE | 1 |
| 1551 | 1551 | 8.2 | 2.4 | 0 | 1 | 0 | 1 | 1 | 0.42266264 | 0 | 1 | 1 | 0.2198258 | 13 | 0.57712805 | FALSE | 1 | FALSE | 1 | FALSE | 1 | FALSE | 1 | FALSE | 1 | FALSE | 1 |
| 1552 | 1552 | 10.75 | 2.75 | 0 | 1 | 0 | 1 | 1 | 0.44344817 | 0 | 1 | 0 | 1 | 15 | 0.26127859 | FALSE | 1 | FALSE | 1 | FALSE | 1 | FALSE | 1 | FALSE | 1 | FALSE | 1 |
| 1553 | 1553 | 9.11111111 | 1.9969112 | 0 | 1 | 0 | 1 | 0 | 1 | 2 | 0.19902682 | 0 | 1 | 16 | 0.44998001 | FALSE | 1 | FALSE | 1 | FALSE | 1 | FALSE | 1 | FALSE | 1 | FALSE | 1 |
| 1554 | 1554 | 10.1538462 | 2.90480117 | 1 | 0.28675951 | 1 | 0.3480709 | 0 | 1 | 1 | 0.45746571 | 1 | 0.19354507 | 9 | 0.96018168 | FALSE | 1 | FALSE | 1 | FALSE | 1 | FALSE | 1 | FALSE | 1 | FALSE | 1 |
| 1555 | 1555 | 9.78571429 | 2.67738069 | 2 | 0.04858661 | 0 | 1 | 1 | 0.40110418 | 0 | 1 | 0 | 1 | 11 | 0.83676449 | FALSE | 0.649301 | FALSE | 1 | FALSE | 1 | FALSE | 1 | FALSE | 1 | FALSE | 1 |
| 1556 | 1556 | 9.45238095 | 1.19925336 | 0 | 1 | 1 | 0.74973543 | 3 | 0.19060367 | 0 | 1 | 1 | 0.50166523 | 37 | 0.34041811 | FALSE | 1 | FALSE | 1 | FALSE | 1 | FALSE | 1 | FALSE | 1 | FALSE | 1 |
| 1557 | 1557 | 10.4615385 | 2.6199056 | 1 | 0.28675951 | 0 | 1 | 0 | 1 | 0 | 1 | 0 | 1 | 12 | 0.374198 | FALSE | 1 | FALSE | 1 | FALSE | 1 | FALSE | 1 | FALSE | 1 | FALSE | 1 |
| 1558 | 1558 | 9.82352941 | 3.39957254 | 2 | 0.06909038 | 1 | 0.42857733 | 0 | 1 | 0 | 1 | 0 | 1 | 14 | 0.7313145 | FALSE | 0.74276693 | FALSE | 1 | FALSE | 1 | FALSE | 1 | FALSE | 1 | FALSE | 1 |
| 1559 | 1559 | 9.52941177 | 1.68444954 | 0 | 1 | 1 | 0.6739483 | 0 | 1 | 1 | 0.7984848 | 0 | 1 | 32 | 0.08180309 | FALSE | 1 | FALSE | 1 | FALSE | 1 | FALSE | 1 | FALSE | 1 | FALSE | 1 |
| 1560 | 1560 | 10.6 | 2.15406592 | 0 | 1 | 0 | 1 | 0 | 1 | 0 | 1 | 0 | 1 | 5 | 0.42761893 | FALSE | 1 | FALSE | 1 | FALSE | 1 | FALSE | 1 | FALSE | 1 | FALSE | 1 |
| 1561 | 1561 | 9.57142857 | 2.77010278 | 0 | 1 | 1 | 0.36919824 | 1 | 0.40110418 | 0 | 1 | 0 | 1 | 12 | 0.62236227 | FALSE | 1 | FALSE | 1 | FALSE | 1 | FALSE | 1 | FALSE | 1 | FALSE | 1 |
| 1562 | 1562 | 10.4722222 | 3.32904817 | 0 | 1 | 1 | 0.69480472 | 2 | 0.37289966 | 3 | 0.22784656 | 0 | 1 | 30 | 0.67248271 | FALSE | 1 | FALSE | 1 | FALSE | 1 | FALSE | 1 | FALSE | 1 | FALSE | 1 |
| 1563 | 1563 | 9.14285714 | 2.48715521 | 4 | 0.07746936 | 4 | 0.14592986 | 0 | 1 | 2 | 0.79265743 | 1 | 0.64879513 | 52 | 0.72716859 | FALSE | 0.77583006 | FALSE | 1 | FALSE | 1 | FALSE | 1 | FALSE | 1 | FALSE | 1 |
| 1564 | 1564 | 10.125 | 2.57087048 | 0 | 1 | 0 | 1 | 1 | 0.25384309 | 0 | 1 | 0 | 1 | 7 | 0.63752397 | FALSE | 1 | FALSE | 1 | FALSE | 1 | FALSE | 1 | FALSE | 1 | FALSE | 1 |
| 1565 | 1565 | 9.48 | 2.15629312 | 0 | 1 | 4 | 0.00789279 | 2 | 0.22597287 | 0 | 1 | 3 | 0.00758617 | 16 | 0.99713666 | FALSE | 1 | FALSE | 0.6167796 | FALSE | 1 | FALSE | 1 | FALSE | 0.49339011 | FALSE | 1 |
| 1566 | 1566 | 8.64705882 | 1.28067889 | 1 | 0.35728305 | 0 | 1 | 0 | 1 | 0 | 1 | 0 | 1 | 16 | 0.23070182 | FALSE | 1 | FALSE | 1 | FALSE | 1 | FALSE | 1 | FALSE | 1 | FALSE | 1 |
| 1567 | 1567 | 8.57692308 | 2.00332564 | 0 | 1 | 1 | 0.57534752 | 0 | 1 | 3 | 0.11458571 | 1 | 0.3499045 | 21 | 0.78939668 | FALSE | 1 | FALSE | 1 | FALSE | 1 | FALSE | 1 | FALSE | 1 | FALSE | 1 |
| 1568 | 1568 | 7.80952381 | 3.41730955 | 2 | 0.09989125 | 1 | 0.49918218 | 0 | 1 | 1 | 0.62783522 | 0 | 1 | 17 | 0.77849076 | FALSE | 0.86047225 | FALSE | 1 | FALSE | 1 | FALSE | 1 | FALSE | 1 | FALSE | 1 |
| 1569 | 1569 | 6.5 | 3.5152728 | 2 | 0.41401181 | 2 | 0.5356583 | 1 | 0.8673918 | 5 | 0.10658542 | 0 | 1 | 45 | 0.76814578 | FALSE | 1 | FALSE | 1 | FALSE | 1 | FALSE | 1 | FALSE | 1 | FALSE | 1 |
| 1570 | 1570 | 6.14705882 | 3.20048115 | 3 | 0.48847083 | 7 | 0.04665336 | 2 | 0.88686294 | 8 | 0.09581403 | 1 | 0.81716516 | 81 | 0.93271075 | FALSE | 1 | FALSE | 1 | FALSE | 1 | FALSE | 1 | FALSE | 1 | FALSE | 1 |
| 1571 | 1571 | 7.95121951 | 3.19260746 | 3 | 0.08696268 | 0 | 1 | 0 | 1 | 2 | 0.56788092 | 1 | 0.49330646 | 35 | 0.53616095 | FALSE | 0.8160347 | FALSE | 1 | FALSE | 1 | FALSE | 1 | FALSE | 1 | FALSE | 1 |
| 1572 | 1572 | 7.06862745 | 2.28490273 | 7 | 0.0147058 | 8 | 0.01616877 | 3 | 0.70953966 | 2 | 0.95031014 | 1 | 0.81407008 | 80 | 0.93855511 | FALSE | 0.38123399 | FALSE | 0.7256023 | FALSE | 1 | FALSE | 1 | FALSE | 1 | FALSE | 1 |
| 1573 | 1573 | 7.4375 | 2.59732434 | 0 | 1 | 0 | 1 | 2 | 0.51909752 | 1 | 0.89605799 | 0 | 1 | 45 | 0.04502985 | FALSE | 1 | FALSE | 1 | FALSE | 1 | FALSE | 1 | FALSE | 1 | FALSE | 1 |
| 1574 | 1574 | 7.87878788 | 4.20503 | 3 | 0.05151181 | 0 | 1 | 1 | 0.70185639 | 1 | 0.78873814 | 2 | 0.10132035 | 26 | 0.86852567 | FALSE | 0.67646333 | FALSE | 1 | FALSE | 1 | FALSE | 1 | FALSE | 1 | FALSE | 1 |
| 1575 | 1575 | 5.21126761 | 1.71090531 | 0 | 1 | 3 | 0.82607155 | 5 | 0.5502572 | 6 | 0.60625053 | 4 | 0.18694959 | 119 | 0.24964601 | FALSE | 1 | FALSE | 1 | FALSE | 1 | FALSE | 1 | FALSE | 1 | FALSE | 1 |
| 1576 | 1576 | 9.94736842 | 2.81861639 | 0 | 1 | 0 | 1 | 0 | 1 | 0 | 1 | 0 | 1 | 18 | 0.04681656 | FALSE | 1 | FALSE | 1 | FALSE | 1 | FALSE | 1 | FALSE | 1 | FALSE | 1 |
| 1577 | 1577 | 9.66666667 | 2.45596133 | 1 | 0.42086991 | 2 | 0.14655134 | 0 | 1 | 0 | 1 | 0 | 1 | 18 | 0.57962763 | FALSE | 1 | FALSE | 1 | FALSE | 1 | FALSE | 1 | FALSE | 1 | FALSE | 1 |
| 1578 | 1578 | 10.4 | 1.356466 | 0 | 1 | 0 | 1 | 0 | 1 | 1 | 0.2094688 | 0 | 1 | 4 | 0.82371636 | FALSE | 1 | FALSE | 1 | FALSE | 1 | FALSE | 1 | FALSE | 1 | FALSE | 1 |
| 1579 | 1579 | 9.95652174 | 1.9886255 | 0 | 1 | 1 | 0.53115654 | 0 | 1 | 0 | 1 | 0 | 1 | 22 | 0.10533112 | FALSE | 1 | FALSE | 1 | FALSE | 1 | FALSE | 1 | FALSE | 1 | FALSE | 1 |
| 1580 | 1580 | 9.26666667 | 2.23507395 | 0 | 1 | 0 | 1 | 0 | 1 | 0 | 1 | 0 | 1 | 15 | 0.07803154 | FALSE | 1 | FALSE | 1 | FALSE | 1 | FALSE | 1 | FALSE | 1 | FALSE | 1 |
| 1581 | 1581 | 8.19354839 | 2.19117519 | 4 | 0.21535847 | 7 | 0.03042764 | 6 | 0.11686853 | 4 | 0.62434183 | 2 | 0.45307384 | 70 | 0.9926682 | FALSE | 1 | FALSE | 0.90921954 | FALSE | 1 | FALSE | 1 | FALSE | 1 | FALSE | 1 |
| 1582 | 1582 | 4.8277512 | 2.67524452 | 10 | 0.03855265 | 14 | 0.00672433 | 11 | 0.12194936 | 7 | 0.84213832 | 18 | 4.64E-09 | 146 | 0.99999981 | FALSE | 0.60156825 | FALSE | 0.60103504 | FALSE | 1 | FALSE | 1 | TRUE | 1.27E-05 | FALSE | 1 |
| 1583 | 1583 | 9.47619048 | 2.42249473 | 0 | 1 | 1 | 0.49918218 | 0 | 1 | 2 | 0.25059627 | 0 | 1 | 18 | 0.57962763 | FALSE | 1 | FALSE | 1 | FALSE | 1 | FALSE | 1 | FALSE | 1 | FALSE | 1 |
| 1584 | 1584 | 9.08823529 | 2.2277349 | 0 | 1 | 2 | 0.30150529 | 1 | 0.71261921 | 2 | 0.46709861 | 0 | 1 | 29 | 0.55707911 | FALSE | 1 | FALSE | 1 | FALSE | 1 | FALSE | 1 | FALSE | 1 | FALSE | 1 |
| 1585 | 1585 | 9.78787879 | 2.33234926 | 2 | 0.19750261 | 3 | 0.08334648 | 2 | 0.32011871 | 2 | 0.43583874 | 0 | 1 | 23 | 0.97984382 | FALSE | 1 | FALSE | 1 | FALSE | 1 | FALSE | 1 | FALSE | 1 | FALSE | 1 |
| 1586 | 1586 | 5.97767857 | 1.57956876 | 7 | 0.34689586 | 9 | 0.29538344 | 11 | 0.17806277 | 11 | 0.4477466 | 14 | 1.45E-05 | 171 | 0.9991882 | FALSE | 1 | FALSE | 1 | FALSE | 1 | FALSE | 1 | TRUE | 0.00911067 | FALSE | 1 |
| 1587 | 1587 | 10.3809524 | 1.67548687 | 2 | 0.48341887 | 4 | 0.14592986 | 2 | 0.66735975 | 1 | 0.94894362 | 2 | 0.27647607 | 52 | 0.72716859 | FALSE | 1 | FALSE | 1 | FALSE | 1 | FALSE | 1 | FALSE | 1 | FALSE | 1 |
| 1588 | 1588 | 8.60869565 | 2.31702828 | 1 | 0.45027683 | 0 | 1 | 1 | 0.5695037 | 0 | 1 | 2 | 0.05389003 | 19 | 0.71542496 | FALSE | 1 | FALSE | 1 | FALSE | 1 | FALSE | 1 | FALSE | 1 | FALSE | 1 |
| 1589 | 1589 | 9.85714286 | 2.5872529 | 0 | 1 | 0 | 1 | 0 | 1 | 0 | 1 | 0 | 1 | 14 | 0.09251304 | FALSE | 1 | FALSE | 1 | FALSE | 1 | FALSE | 1 | FALSE | 1 | FALSE | 1 |
| 1590 | 1590 | 7.63636364 | 2.86904252 | 1 | 0.24866461 | 0 | 1 | 0 | 1 | 1 | 0.40390032 | 0 | 1 | 9 | 0.75985366 | FALSE | 1 | FALSE | 1 | FALSE | 1 | FALSE | 1 | FALSE | 1 | FALSE | 1 |
| 1591 | 1591 | 8.56521739 | 2.12287924 | 0 | 1 | 1 | 0.53115654 | 1 | 0.5695037 | 1 | 0.6613267 | 1 | 0.31672722 | 19 | 0.71542496 | FALSE | 1 | FALSE | 1 | FALSE | 1 | FALSE | 1 | FALSE | 1 | FALSE | 1 |
| 1592 | 1592 | 9.46153846 | 3.56511699 | 0 | 1 | 1 | 0.3480709 | 2 | 0.07725803 | 2 | 0.1174583 | 0 | 1 | 8 | 0.99085298 | FALSE | 1 | FALSE | 1 | FALSE | 1 | FALSE | 1 | FALSE | 1 | FALSE | 1 |
| 1593 | 1593 | 8.66666667 | 2.4267033 | 0 | 1 | 2 | 0.1135776 | 0 | 1 | 1 | 0.57131013 | 1 | 0.25766374 | 14 | 0.86287418 | FALSE | 1 | FALSE | 1 | FALSE | 1 | FALSE | 1 | FALSE | 1 | FALSE | 1 |
| 1594 | 1594 | 8.13333333 | 3.64904858 | 0 | 1 | 1 | 0.38964394 | 0 | 1 | 2 | 0.14906763 | 0 | 1 | 12 | 0.80329917 | FALSE | 1 | FALSE | 1 | FALSE | 1 | FALSE | 1 | FALSE | 1 | FALSE | 1 |
| 1595 | 1595 | 11.4736842 | 2.95347113 | 2 | 0.25455533 | 3 | 0.12352882 | 1 | 0.75193685 | 0 | 1 | 0 | 1 | 32 | 0.61800916 | FALSE | 1 | FALSE | 1 | FALSE | 1 | FALSE | 1 | FALSE | 1 | FALSE | 1 |
| 1596 | 1596 | 10.6 | 2.53771551 | 0 | 1 | 0 | 1 | 1 | 0.30655256 | 0 | 1 | 1 | 0.15247062 | 8 | 0.80394719 | FALSE | 1 | FALSE | 1 | FALSE | 1 | FALSE | 1 | FALSE | 1 | FALSE | 1 |

| | | | | | | | | | | | | | | | | | | | | | | | | | | | |
|---|---|---|---|---|---|---|---|---|---|---|---|---|---|---|---|---|---|---|---|---|---|---|---|---|---|---|---|
| 1597 | 1597 | 6.3030303 | 0.98426556 | 3 | 0.66332479 | 5 | 0.42496778 | 5 | 0.51688507 | 7 | 0.40353836 | 4 | 0.17065314 | 108 | 0.82728985 | FALSE | 1 | FALSE | 1 | FALSE | 1 | FALSE | 1 | FALSE | 1 | FALSE | 1 |
| 1598 | 1598 | 7.23076923 | 3.51160376 | 7 | 4.96E-05 | 2 | 0.36122049 | 2 | 0.41139386 | 2 | 0.54052736 | 2 | 0.13365639 | 24 | 0.99988054 | TRUE | 0.01105799 | FALSE | 1 | FALSE | 1 | FALSE | 1 | FALSE | 1 | FALSE | 1 |
| 1599 | 1599 | 8.13333333 | 1.7074998 | 0 | 1 | 1 | 0.38964394 | 1 | 0.42266264 | 1 | 0.50623194 | 0 | 1 | 12 | 0.80329917 | FALSE | 1 | FALSE | 1 | FALSE | 1 | FALSE | 1 | FALSE | 1 | FALSE | 1 |
| 1600 | 1600 | 8.88461539 | 2.05364447 | 3 | 0.14789018 | 4 | 0.08651261 | 0 | 1 | 1 | 0.91399378 | 0 | 1 | 44 | 0.57424118 | FALSE | 1 | FALSE | 1 | FALSE | 1 | FALSE | 1 | FALSE | 1 | FALSE | 1 |
| 1601 | 1601 | 8.79310345 | 2.65718688 | 1 | 0.52988231 | 1 | 0.61539065 | 0 | 1 | 2 | 0.3871811 | 1 | 0.38148475 | 24 | 0.70563334 | FALSE | 1 | FALSE | 1 | FALSE | 1 | FALSE | 1 | FALSE | 1 | FALSE | 1 |
| 1602 | 1602 | 9.4375 | 2.03004772 | 0 | 1 | 0 | 1 | 0 | 1 | 1 | 0.52895088 | 0 | 1 | 15 | 0.26127859 | FALSE | 1 | FALSE | 1 | FALSE | 1 | FALSE | 1 | FALSE | 1 | FALSE | 1 |
| 1603 | 1603 | 7.20869565 | 0.86973911 | 2 | 0.79546698 | 2 | 0.88874783 | 2 | 0.92070419 | 10 | 0.03600902 | 3 | 0.28730314 | 95 | 0.67842369 | FALSE | 1 | FALSE | 1 | FALSE | 1 | FALSE | 1 | FALSE | 1 | FALSE | 1 |
| 1604 | 1604 | 9.92857143 | 2.43381786 | 0 | 1 | 0 | 1 | 2 | 0.08806398 | 0 | 1 | 0 | 1 | 12 | 0.62236227 | FALSE | 1 | FALSE | 1 | FALSE | 1 | FALSE | 1 | FALSE | 1 | FALSE | 1 |
| 1605 | 1605 | 8.90909091 | 1.40051152 | 0 | 1 | 3 | 0.08959205 | 2 | 0.33343871 | 0 | 1 | 0 | 1 | 28 | 0.58681477 | FALSE | 1 | FALSE | 1 | FALSE | 1 | FALSE | 1 | FALSE | 1 | FALSE | 1 |
| 1606 | 1606 | 9.8 | 1.8547237 | 0 | 1 | 1 | 0.56109832 | 2 | 0.22597287 | 1 | 0.69181315 | 1 | 0.33902672 | 20 | 0.8152498 | FALSE | 1 | FALSE | 1 | FALSE | 1 | FALSE | 1 | FALSE | 1 | FALSE | 1 |
| 1607 | 1607 | 8.92771084 | 1.09525958 | 3 | 0.35850108 | 2 | 0.75548815 | 3 | 0.57805941 | 6 | 0.1806169 | 1 | 0.74858842 | 68 | 0.78384481 | FALSE | 1 | FALSE | 1 | FALSE | 1 | FALSE | 1 | FALSE | 1 | FALSE | 1 |
| 1608 | 1608 | 5.4137931 | 2.23633384 | 3 | 0.03713651 | 3 | 0.06581417 | 1 | 0.65463982 | 1 | 0.74482229 | 1 | 0.38148475 | 20 | 0.99028351 | FALSE | 0.58392 | FALSE | 1 | FALSE | 1 | FALSE | 1 | FALSE | 1 | FALSE | 1 |
| 1609 | 1609 | 9.19047619 | 2.73633484 | 0 | 1 | 0 | 1 | 2 | 0.17308361 | 3 | 0.06912908 | 0 | 1 | 16 | 0.90353674 | FALSE | 1 | FALSE | 1 | FALSE | 1 | FALSE | 1 | FALSE | 1 | FALSE | 1 |
| 1610 | 1610 | 8.625 | 2.42920598 | 1 | 0.46441868 | 1 | 0.54637326 | 1 | 0.58502114 | 3 | 0.09525947 | 0 | 1 | 18 | 0.93166998 | FALSE | 1 | FALSE | 1 | FALSE | 1 | FALSE | 1 | FALSE | 1 | FALSE | 1 |
| 1611 | 1611 | 8.05714286 | 3.64058753 | 0 | 1 | 1 | 0.68454804 | 0 | 1 | 4 | 0.07465533 | 0 | 1 | 30 | 0.52765882 | FALSE | 1 | FALSE | 1 | FALSE | 1 | FALSE | 1 | FALSE | 1 | FALSE | 1 |
| 1612 | 1612 | 9.08695652 | 2.53519648 | 2 | 0.11647716 | 1 | 0.53115654 | 0 | 1 | 0 | 1 | 3 | 0.00598158 | 17 | 0.94439745 | FALSE | 0.93181727 | FALSE | 1 | FALSE | 1 | FALSE | 1 | FALSE | 0.45694206 | FALSE | 1 |
| 1613 | 1613 | 8.43478261 | 3.06205511 | 4 | 0.00212601 | 1 | 0.5154318 | 0 | 1 | 1 | 0.64497439 | 0 | 1 | 16 | 0.95553951 | FALSE | 0.12179217 | FALSE | 1 | FALSE | 1 | FALSE | 1 | FALSE | 1 | FALSE | 1 |
| 1614 | 1614 | 8.92307692 | 2.33572715 | 0 | 1 | 0 | 1 | 1 | 0.37874417 | 0 | 1 | 0 | 1 | 12 | 0.374198 | FALSE | 1 | FALSE | 1 | FALSE | 1 | FALSE | 1 | FALSE | 1 | FALSE | 1 |
| 1615 | 1615 | 10.7692308 | 2.438595 | 2 | 0.14250978 | 0 | 1 | 2 | 0.23940226 | 0 | 1 | 0 | 1 | 22 | 0.61628 | FALSE | 1 | FALSE | 1 | FALSE | 1 | FALSE | 1 | FALSE | 1 | FALSE | 1 |
| 1616 | 1616 | 8.73333333 | 2.80396545 | 0 | 1 | 0 | 1 | 1 | 0.66709967 | 2 | 0.4036176 | 0 | 1 | 27 | 0.28862208 | FALSE | 1 | FALSE | 1 | FALSE | 1 | FALSE | 1 | FALSE | 1 | FALSE | 1 |
| 1617 | 1617 | 10.0909091 | 2.2611828 | 0 | 1 | 1 | 0.6629941 | 1 | 0.70185639 | 1 | 0.78873814 | 1 | 0.42123867 | 29 | 0.39648374 | FALSE | 1 | FALSE | 1 | FALSE | 1 | FALSE | 1 | FALSE | 1 | FALSE | 1 |
| 1618 | 1618 | 7.5 | 2.26295553 | 3 | 0.20532341 | 4 | 0.13413374 | 1 | 0.89373158 | 4 | 0.30679865 | 1 | 0.63687695 | 48 | 0.91571908 | FALSE | 1 | FALSE | 1 | FALSE | 1 | FALSE | 1 | FALSE | 1 | FALSE | 1 |
| 1619 | 1619 | 9.88888889 | 2.90380763 | 2 | 0.07645454 | 0 | 1 | 0 | 1 | 0 | 1 | 0 | 1 | 16 | 0.44998001 | FALSE | 0.77424195 | FALSE | 1 | FALSE | 1 | FALSE | 1 | FALSE | 1 | FALSE | 1 |
| 1620 | 1620 | 8.77272727 | 1.18443766 | 2 | 0.31193899 | 1 | 0.76576289 | 2 | 0.47281647 | 7 | 0.00355513 | 0 | 1 | 32 | 0.98583614 | FALSE | 1 | FALSE | 1 | FALSE | 1 | FALSE | 1 | FALSE | 1 | FALSE | 1 |
| 1621 | 1621 | 9.29411765 | 2.29487167 | 0 | 1 | 2 | 0.10307112 | 0 | 1 | 1 | 0.55062772 | 0 | 1 | 14 | 0.7313145 | FALSE | 1 | FALSE | 1 | FALSE | 1 | FALSE | 1 | FALSE | 1 | FALSE | 1 |
| 1622 | 1622 | 10.6944444 | 2.01134897 | 0 | 1 | 1 | 0.69480472 | 4 | 0.03888854 | 1 | 0.81665383 | 0 | 1 | 30 | 0.67248271 | FALSE | 1 | FALSE | 1 | FALSE | 1 | FALSE | 1 | FALSE | 1 | FALSE | 1 |
| 1623 | 1623 | 8.625 | 1.72753437 | 0 | 1 | 2 | 0.09285443 | 2 | 0.11087977 | 1 | 0.52895088 | 2 | 0.02752114 | 9 | 0.99862766 | FALSE | 1 | FALSE | 1 | FALSE | 1 | FALSE | 1 | FALSE | 0.79106376 | FALSE | 1 |
| 1624 | 1624 | 8.72727273 | 1.86308196 | 0 | 1 | 0 | 1 | 1 | 0.33149969 | 1 | 0.40390032 | 0 | 1 | 9 | 0.75985366 | FALSE | 1 | FALSE | 1 | FALSE | 1 | FALSE | 1 | FALSE | 1 | FALSE | 1 |
| 1625 | 1625 | 8.5 | 1.31021627 | 0 | 1 | 0 | 1 | 1 | 0.66709967 | 1 | 0.75658797 | 0 | 1 | 28 | 0.13086908 | FALSE | 1 | FALSE | 1 | FALSE | 1 | FALSE | 1 | FALSE | 1 | FALSE | 1 |
| 1626 | 1626 | 5.02272727 | 3.80483327 | 1 | 0.68224519 | 3 | 0.16927151 | 2 | 0.47281647 | 2 | 0.60667406 | 0 | 1 | 36 | 0.75909657 | FALSE | 1 | FALSE | 1 | FALSE | 1 | FALSE | 1 | FALSE | 1 | FALSE | 1 |
| 1627 | 1627 | 5.01428571 | 2.49280598 | 1 | 0.83919255 | 1 | 0.90110191 | 3 | 0.46296673 | 4 | 0.40179526 | 0 | 1 | 61 | 0.32827441 | FALSE | 1 | FALSE | 1 | FALSE | 1 | FALSE | 1 | FALSE | 1 | FALSE | 1 |
| 1628 | 1628 | 12.1818182 | 2.83863545 | 0 | 1 | 1 | 0.5154318 | 0 | 1 | 2 | 0.26791271 | 0 | 1 | 19 | 0.54233419 | FALSE | 1 | FALSE | 1 | FALSE | 1 | FALSE | 1 | FALSE | 1 | FALSE | 1 |
| 1629 | 1629 | 8.95454546 | 1.90638336 | 2 | 0.31193899 | 1 | 0.76576289 | 0 | 1 | 1 | 0.87439647 | 0 | 1 | 40 | 0.16103481 | FALSE | 1 | FALSE | 1 | FALSE | 1 | FALSE | 1 | FALSE | 1 | FALSE | 1 |
| 1630 | 1630 | 9.4 | 2.91204396 | 2 | 0.13369621 | 1 | 0.56109832 | 1 | 0.59998149 | 1 | 0.69181315 | 1 | 0.33902672 | 19 | 0.91734261 | FALSE | 0.97964162 | FALSE | 1 | FALSE | 1 | FALSE | 1 | FALSE | 1 | FALSE | 1 |
| 1631 | 1631 | 4.17058824 | 3.18457028 | 5 | 0.44253041 | 6 | 0.47328187 | 4 | 0.86614383 | 2 | 0.99712931 | 2 | 0.77318603 | 151 | 0.06116219 | FALSE | 1 | FALSE | 1 | FALSE | 1 | FALSE | 1 | FALSE | 1 | FALSE | 1 |
| 1632 | 1632 | 10.2352941 | 2.14282419 | 3 | 0.05546921 | 2 | 0.30150529 | 0 | 1 | 2 | 0.46709861 | 0 | 1 | 27 | 0.85023803 | FALSE | 0.68849059 | FALSE | 1 | FALSE | 1 | FALSE | 1 | FALSE | 1 | FALSE | 1 |
| 1633 | 1633 | 9.86206897 | 2.27010767 | 1 | 0.52988231 | 0 | 1 | 2 | 0.27983822 | 1 | 0.74482229 | 0 | 1 | 25 | 0.51779518 | FALSE | 1 | FALSE | 1 | FALSE | 1 | FALSE | 1 | FALSE | 1 | FALSE | 1 |
| 1634 | 1634 | 9.36363636 | 3.21283583 | 0 | 1 | 0 | 1 | 0 | 1 | 1 | 0.64497439 | 0 | 1 | 21 | 0.12048848 | FALSE | 1 | FALSE | 1 | FALSE | 1 | FALSE | 1 | FALSE | 1 | FALSE | 1 |
| 1635 | 1635 | 10.85 | 1.55804365 | 0 | 1 | 0 | 1 | 0 | 1 | 1 | 0.60987146 | 0 | 1 | 19 | 0.15700255 | FALSE | 1 | FALSE | 1 | FALSE | 1 | FALSE | 1 | FALSE | 1 | FALSE | 1 |
| 1636 | 1636 | 9.12765957 | 2.51478201 | 0 | 1 | 1 | 0.78790827 | 4 | 0.08716543 | 2 | 0.64277396 | 3 | 0.0412987 | 37 | 0.89489966 | FALSE | 1 | FALSE | 1 | FALSE | 1 | FALSE | 1 | FALSE | 0.90763079 | FALSE | 1 |
| 1637 | 1637 | 8.78787879 | 3.08228148 | 0 | 1 | 4 | 0.02089081 | 2 | 0.33343871 | 1 | 0.78873814 | 1 | 0.42123867 | 25 | 0.93892791 | FALSE | 1 | FALSE | 0.8034624 | FALSE | 1 | FALSE | 1 | FALSE | 1 | FALSE | 1 |
| 1638 | 1638 | 6.66153846 | 2.5375756 | 4 | 0.0847349 | 2 | 0.62701348 | 5 | 0.08323317 | 3 | 0.57949569 | 0 | 1 | 51 | 0.92718889 | FALSE | 0.79787163 | FALSE | 1 | FALSE | 1 | FALSE | 1 | FALSE | 1 | FALSE | 1 |
| 1639 | 1639 | 9.15384615 | 2.8105886 | 1 | 0.49162634 | 0 | 1 | 2 | 0.23940226 | 2 | 0.33678042 | 0 | 1 | 21 | 0.78939668 | FALSE | 1 | FALSE | 1 | FALSE | 1 | FALSE | 1 | FALSE | 1 | FALSE | 1 |
| 1640 | 1640 | 7.05555556 | 3.09968139 | 0 | 1 | 1 | 0.44710673 | 0 | 1 | 0 | 1 | 0 | 1 | 17 | 0.20329205 | FALSE | 1 | FALSE | 1 | FALSE | 1 | FALSE | 1 | FALSE | 1 | FALSE | 1 |
| 1641 | 1641 | 12.4285714 | 3.75907066 | 1 | 0.59801736 | 1 | 0.68454804 | 1 | 0.72299506 | 4 | 0.07465533 | 0 | 1 | 28 | 0.83079716 | FALSE | 1 | FALSE | 1 | FALSE | 1 | FALSE | 1 | FALSE | 1 | FALSE | 1 |
| 1642 | 1642 | 10.5384615 | 2.97723115 | 0 | 1 | 1 | 0.3480709 | 0 | 1 | 1 | 0.45746571 | 0 | 1 | 11 | 0.6683056 | FALSE | 1 | FALSE | 1 | FALSE | 1 | FALSE | 1 | FALSE | 1 | FALSE | 1 |
| 1643 | 1643 | 10.04 | 2.5215868 | 1 | 0.46441868 | 0 | 1 | 1 | 0.58502114 | 0 | 1 | 2 | 0.058175 | 20 | 0.68270729 | FALSE | 1 | FALSE | 1 | FALSE | 1 | FALSE | 1 | FALSE | 1 | FALSE | 1 |
| 1644 | 1644 | 7.73076923 | 3.55784823 | 0 | 1 | 2 | 0.20486827 | 2 | 0.23940226 | 0 | 1 | 1 | 0.3499045 | 21 | 0.78939668 | FALSE | 1 | FALSE | 1 | FALSE | 1 | FALSE | 1 | FALSE | 1 | FALSE | 1 |
| 1645 | 1645 | 9.10526316 | 3.71192583 | 0 | 1 | 1 | 0.46503795 | 0 | 1 | 2 | 0.21610812 | 1 | 0.26986775 | 15 | 0.83648191 | FALSE | 1 | FALSE | 1 | FALSE | 1 | FALSE | 1 | FALSE | 1 | FALSE | 1 |
| 1646 | 1646 | 8.425 | 1.41222342 | 7 | 4.96E-05 | 1 | 0.7236252 | 2 | 0.41139386 | 0 | 1 | 1 | 0.4761699 | 28 | 0.98701111 | TRUE | 0.01105799 | FALSE | 1 | FALSE | 1 | FALSE | 1 | FALSE | 1 | FALSE | 1 |
| 1647 | 1647 | 8.11428571 | 1.43967116 | 1 | 0.83919255 | 1 | 0.90110191 | 1 | 0.92379401 | 4 | 0.40179526 | 0 | 1 | 63 | 0.12472747 | FALSE | 1 | FALSE | 1 | FALSE | 1 | FALSE | 1 | FALSE | 1 | FALSE | 1 |
| 1648 | 1648 | 9.07843137 | 2.11292138 | 0 | 1 | 5 | 0.02197853 | 2 | 0.54120634 | 4 | 0.19569978 | 0 | 1 | 39 | 0.92022511 | FALSE | 1 | FALSE | 0.81029974 | FALSE | 1 | FALSE | 1 | FALSE | 1 | FALSE | 1 |
| 1649 | 1649 | 7.6 | 2.27449628 | 1 | 0.69044531 | 2 | 0.43020462 | 1 | 0.80828716 | 1 | 0.8802008 | 2 | 0.16793736 | 38 | 0.59437745 | FALSE | 1 | FALSE | 1 | FALSE | 1 | FALSE | 1 | FALSE | 1 | FALSE | 1 |
| 1650 | 1650 | 8.52631579 | 2.41417788 | 0 | 1 | 0 | 1 | 2 | 0.14749808 | 1 | 0.59104357 | 0 | 1 | 16 | 0.65565778 | FALSE | 1 | FALSE | 1 | FALSE | 1 | FALSE | 1 | FALSE | 1 | FALSE | 1 |
| 1651 | 1651 | 10.25 | 2.58602011 | 0 | 1 | 0 | 1 | 2 | 0.06691409 | 1 | 0.43131129 | 0 | 1 | 9 | 0.89630573 | FALSE | 1 | FALSE | 1 | FALSE | 1 | FALSE | 1 | FALSE | 1 | FALSE | 1 |
| 1652 | 1652 | 7.54545455 | 0.9875255 | 1 | 0.24866461 | 1 | 0.30368014 | 1 | 0.33149969 | 0 | 1 | 0 | 1 | 8 | 0.92152993 | FALSE | 1 | FALSE | 1 | FALSE | 1 | FALSE | 1 | FALSE | 1 | FALSE | 1 |
| 1653 | 1653 | 10.4090909 | 2.55234457 | 0 | 1 | 0 | 1 | 2 | 0.18613256 | 2 | 0.26791271 | 0 | 1 | 18 | 0.74744813 | FALSE | 1 | FALSE | 1 | FALSE | 1 | FALSE | 1 | FALSE | 1 | FALSE | 1 |
| 1654 | 1654 | 8.125 | 2.75850956 | 0 | 1 | 0 | 1 | 1 | 0.5695037 | 1 | 0.6613267 | 0 | 1 | 21 | 0.27964072 | FALSE | 1 | FALSE | 1 | FALSE | 1 | FALSE | 1 | FALSE | 1 | FALSE | 1 |
| 1655 | 1655 | 7.56730769 | 2.00846489 | 3 | 0.50146864 | 1 | 0.9681402 | 3 | 0.72810072 | 6 | 0.34318392 | 4 | 0.09081468 | 87 | 0.64353601 | FALSE | 1 | FALSE | 1 | FALSE | 1 | FALSE | 1 | FALSE | 1 | FALSE | 1 |
| 1656 | 1656 | 7 | 2.32590543 | 0 | 1 | 1 | 0.85750546 | 4 | 0.16113918 | 3 | 0.51328524 | 0 | 1 | 51 | 0.41341988 | FALSE | 1 | FALSE | 1 | FALSE | 1 | FALSE | 1 | FALSE | 1 | FALSE | 1 |
| 1657 | 1657 | 10.5625 | 2.49921863 | 0 | 1 | 0 | 1 | 1 | 0.44344817 | 1 | 0.52895088 | 1 | 0.23264612 | 13 | 0.76799161 | FALSE | 1 | FALSE | 1 | FALSE | 1 | FALSE | 1 | FALSE | 1 | FALSE | 1 |
| 1658 | 1658 | 7.73684211 | 2.4353127 | 1 | 0.62830697 | 2 | 0.3494045 | 1 | 0.75193685 | 2 | 0.52640931 | 0 | 1 | 32 | 0.61800916 | FALSE | 1 | FALSE | 1 | FALSE | 1 | FALSE | 1 | FALSE | 1 | FALSE | 1 |
| 1659 | 1659 | 9.85714286 | 1.72614943 | 1 | 0.3050805 | 0 | 1 | 1 | 0.40110418 | 0 | 1 | 0 | 1 | 12 | 0.62236227 | FALSE | 1 | FALSE | 1 | FALSE | 1 | FALSE | 1 | FALSE | 1 | FALSE | 1 |
| 1660 | 1660 | 8.41573034 | 1.45963662 | 9 | 0.00032756 | 2 | 0.77275903 | 7 | 0.03432404 | 4 | 0.56192759 | 0 | 1 | 64 | 0.99460611 | TRUE | 0.03626121 | FALSE | 1 | FALSE | 1 | FALSE | 1 | FALSE | 1 | FALSE | 1 |
| 1661 | 1661 | 8.95454546 | 3.14031372 | 0 | 1 | 0 | 1 | 1 | 0.55340851 | 0 | 1 | 1 | 0.30529951 | 20 | 0.30915202 | FALSE | 1 | FALSE | 1 | FALSE | 1 | FALSE | 1 | FALSE | 1 | FALSE | 1 |
| 1662 | 1662 | 9.71428571 | 3.45229885 | 2 | 0.0126078 | 0 | 1 | 1 | 0.22601089 | 0 | 1 | 1 | 0.10932414 | 3 | 0.99853144 | FALSE | 0.3525019 | FALSE | 1 | FALSE | 1 | FALSE | 1 | FALSE | 1 | FALSE | 1 |
| 1663 | 1663 | 8.41176471 | 1.84896866 | 0 | 1 | 0 | 1 | 0 | 1 | 2 | 0.18212134 | 1 | 0.24525764 | 14 | 0.7313145 | FALSE | 1 | FALSE | 1 | FALSE | 1 | FALSE | 1 | FALSE | 1 | FALSE | 1 |
| 1664 | 1664 | 8.22222222 | 2.0245408 | 1 | 0.50471032 | 2 | 0.21684246 | 2 | 0.25286935 | 0 | 1 | 0 | 1 | 22 | 0.76237877 | FALSE | 1 | FALSE | 1 | FALSE | 1 | FALSE | 1 | FALSE | 1 | FALSE | 1 |
| 1665 | 1665 | 7.41071429 | 2.48406914 | 3 | 0.17275086 | 6 | 0.00900454 | 6 | 0.01460362 | 4 | 0.25497934 | 1 | 0.60529938 | 36 | 0.99994677 | FALSE | 1 | FALSE | 0.63686998 | FALSE | 1 | FALSE | 1 | FALSE | 1 | FALSE | 1 |
| 1666 | 1666 | 8.4375 | 1.86978441 | 1 | 0.34033054 | 2 | 0.09285443 | 0 | 1 | 2 | 0.16544789 | 1 | 0.23264612 | 10 | 0.99304605 | FALSE | 1 | FALSE | 1 | FALSE | 1 | FALSE | 1 | FALSE | 1 | FALSE | 1 |
| 1667 | 1667 | 7.85 | 3.55351938 | 1 | 0.40558521 | 0 | 1 | 0 | 1 | 1 | 0.60987146 | 0 | 1 | 18 | 0.37504645 | FALSE | 1 | FALSE | 1 | FALSE | 1 | FALSE | 1 | FALSE | 1 | FALSE | 1 |
| 1668 | 1668 | 7.44 | 1.87075742 | 7 | 0.00295027 | 3 | 0.43942904 | 4 | 0.2829119 | 4 | 0.45379959 | 3 | 0.12461258 | 54 | 0.99814335 | FALSE | 0.14653312 | FALSE | 1 | FALSE | 1 | FALSE | 1 | FALSE | 1 | FALSE | 1 |
| 1669 | 1669 | 9.88 | 2.51904744 | 1 | 0.47819878 | 1 | 0.56109832 | 2 | 0.22597287 | 3 | 0.10474541 | 0 | 1 | 18 | 0.96851178 | FALSE | 1 | FALSE | 1 | FALSE | 1 | FALSE | 1 | FALSE | 1 | FALSE | 1 |
| 1670 | 1670 | 4.1779661 | 0.95932553 | 12 | 0.18895005 | 10 | 0.70687295 | 6 | 0.98898647 | 18 | 0.34687122 | 3 | 0.93288771 | 302 | 0.21204025 | FALSE | 1 | FALSE | 1 | FALSE | 1 | FALSE | 1 | FALSE | 1 | FALSE | 1 |
| 1671 | 1671 | 8.72727273 | 2.45286138 | 0 | 1 | 0 | 1 | 0 | 1 | 1 | 0.37517262 | 0 | 1 | 9 | 0.5215547 | FALSE | 1 | FALSE | 1 | FALSE | 1 | FALSE | 1 | FALSE | 1 | FALSE | 1 |
| 1672 | 1672 | 7.7826087 | 2.68581656 | 0 | 1 | 0 | 1 | 1 | 0.5695037 | 1 | 0.6613267 | 0 | 1 | 21 | 0.27964072 | FALSE | 1 | FALSE | 1 | FALSE | 1 | FALSE | 1 | FALSE | 1 | FALSE | 1 |
| 1673 | 1673 | 8.18367347 | 2.70806407 | 1 | 0.72119464 | 1 | 0.80150109 | 0 | 1 | 1 | 0.90086419 | 1 | 0.55647171 | 45 | 0.10025985 | FALSE | 1 | FALSE | 1 | FALSE | 1 | FALSE | 1 | FALSE | 1 | FALSE | 1 |
| 1674 | 1674 | 10.0952381 | 1.79694879 | 2 | 0.09989125 | 1 | 0.49918218 | 0 | 1 | 0 | 1 | 1 | 0.29368245 | 17 | 0.77849076 | FALSE | 0.86047225 | FALSE | 1 | FALSE | 1 | FALSE | 1 | FALSE | 1 | FALSE | 1 |
| 1675 | 1675 | 8.90476191 | 3.08459709 | 1 | 0.42086991 | 1 | 0.49918218 | 0 | 1 | 0 | 1 | 0 | 1 | 19 | 0.34094959 | FALSE | 1 | FALSE | 1 | FALSE | 1 | FALSE | 1 | FALSE | 1 | FALSE | 1 |
| 1676 | 1676 | 9.6 | 2.10713075 | 1 | 0.22886536 | 0 | 1 | 0 | 1 | 0 | 1 | 0 | 1 | 9 | 0.5215547 | FALSE | 1 | FALSE | 1 | FALSE | 1 | FALSE | 1 | FALSE | 1 | FALSE | 1 |
| 1677 | 1677 | 9 | 2.8654016 | 0 | 1 | 0 | 1 | 2 | 0.14749808 | 0 | 1 | 1 | 0.26986775 | 16 | 0.65565778 | FALSE | 1 | FALSE | 1 | FALSE | 1 | FALSE | 1 | FALSE | 1 | FALSE | 1 |
| 1678 | 1678 | 8 | 2.30007013 | 2 | 0.1881398 | 1 | 0.63997468 | 0 | 1 | 2 | 0.41984187 | 3 | 0.01381051 | 23 | 0.95750806 | FALSE | 1 | FALSE | 1 | FALSE | 1 | FALSE | 1 | FALSE | 0.64607242 | FALSE | 1 |
| 1679 | 1679 | 8.41666667 | 2.37317181 | 2 | 0.23542589 | 2 | 0.32556427 | 3 | 0.13783405 | 0 | 1 | 0 | 1 | 29 | 0.81028289 | FALSE | 1 | FALSE | 1 | FALSE | 1 | FALSE | 1 | FALSE | 1 | FALSE | 1 |
| 1680 | 1680 | 8.6 | 1.70489491 | 1 | 0.32293355 | 0 | 1 | 1 | 0.42266264 | 0 | 1 | 1 | 0.2198258 | 12 | 0.80329917 | FALSE | 1 | FALSE | 1 | FALSE | 1 | FALSE | 1 | FALSE | 1 | FALSE | 1 |
| 1681 | 1681 | 8.77777778 | 2.92287699 | 0 | 1 | 0 | 1 | 0 | 1 | 2 | 0.35374209 | 0 | 1 | 25 | 0.1832206 | FALSE | 1 | FALSE | 1 | FALSE | 1 | FALSE | 1 | FALSE | 1 | FALSE | 1 |
| 1682 | 1682 | 9.6875 | 2.19996449 | 0 | 1 | 1 | 0.40942991 | 0 | 1 | 1 | 0.52895088 | 0 | 1 | 14 | 0.53308261 | FALSE | 1 | FALSE | 1 | FALSE | 1 | FALSE | 1 | FALSE | 1 | FALSE | 1 |
| 1683 | 1683 | 2.11376673 | 0.93858329 | 27 | 0.51324553 | 38 | 0.23555993 | 40 | 0.35265836 | 44 | 0.75938139 | 20 | 0.25800172 | 875 | 0.7258007 | FALSE | 1 | FALSE | 1 | FALSE | 1 | FALSE | 1 | FALSE | 1 | FALSE | 1 |
| 1684 | 1684 | 8.22222222 | 2.80981525 | 0 | 1 | 1 | 0.69480472 | 0 | 1 | 2 | 0.49731009 | 0 | 1 | 33 | 0.16407838 | FALSE | 1 | FALSE | 1 | FALSE | 1 | FALSE | 1 | FALSE | 1 | FALSE | 1 |
| 1685 | 1685 | 7.52941177 | 2.97277148 | 1 | 0.35728305 | 0 | 1 | 0 | 1 | 0 | 1 | 0 | 1 | 16 | 0.23070182 | FALSE | 1 | FALSE | 1 | FALSE | 1 | FALSE | 1 | FALSE | 1 | FALSE | 1 |
| 1686 | 1686 | 7.5862069 | 2.42852646 | 1 | 0.52988231 | 2 | 0.24095584 | 0 | 1 | 2 | 0.3871811 | 1 | 0.38148475 | 23 | 0.84462957 | FALSE | 1 | FALSE | 1 | FALSE | 1 | FALSE | 1 | FALSE | 1 | FALSE | 1 |
| 1687 | 1687 | 7.30434783 | 2.13530942 | 0 | 1 | 0 | 1 | 1 | 0.5695037 | 1 | 0.6613267 | 1 | 0.31672722 | 20 | 0.50589402 | FALSE | 1 | FALSE | 1 | FALSE | 1 | FALSE | 1 | FALSE | 1 | FALSE | 1 |
| 1688 | 1688 | 8.88888889 | 2.0245408 | 1 | 0.20854748 | 1 | 0.25628163 | 0 | 1 | 1 | 0.34506517 | 0 | 1 | 6 | 0.96128317 | FALSE | 1 | FALSE | 1 | FALSE | 1 | FALSE | 1 | FALSE | 1 | FALSE | 1 |
| 1689 | 1689 | 8.94117647 | 2.43674296 | 0 | 1 | 2 | 0.10307112 | 0 | 1 | 0 | 1 | 0 | 1 | 15 | 0.49060435 | FALSE | 1 | FALSE | 1 | FALSE | 1 | FALSE | 1 | FALSE | 1 | FALSE | 1 |
| 1690 | 1690 | 10.4117647 | 5.04169467 | 0 | 1 | 0 | 1 | 0 | 1 | 1 | 0.55062772 | 0 | 1 | 16 | 0.23070182 | FALSE | 1 | FALSE | 1 | FALSE | 1 | FALSE | 1 | FALSE | 1 | FALSE | 1 |

| | | | | | | | | | | | | | | | | | | | | | | | | | | | |
|---|---|---|---|---|---|---|---|---|---|---|---|---|---|---|---|---|---|---|---|---|---|---|---|---|---|---|---|
| 1691 | 1691 | 6.625 | 4.18889902 | 0 | 1 | 1 | 0.65167362 | 3 | 0.10599423 | 2 | 0.43583874 | 0 | 1 | 26 | 0.77612337 | FALSE | 1 | FALSE | 1 | FALSE | 1 | FALSE | 1 | FALSE | 1 | FALSE | 1 |
| 1692 | 1692 | 7.54545455 | 3.24355687 | 0 | 1 | 1 | 0.5154318 | 1 | 0.55340851 | 1 | 0.64497439 | 1 | 0.30529951 | 18 | 0.74744813 | FALSE | 1 | FALSE | 1 | FALSE | 1 | FALSE | 1 | FALSE | 1 | FALSE | 1 |
| 1693 | 1693 | 6.04054054 | 2.89192347 | 1 | 0.85522402 | 1 | 0.9134145 | 5 | 0.12620681 | 5 | 0.25191953 | 1 | 0.70774043 | 61 | 0.74108955 | FALSE | 1 | FALSE | 1 | FALSE | 1 | FALSE | 1 | FALSE | 1 | FALSE | 1 |
| 1694 | 1694 | 10.4347826 | 2.24239947 | 1 | 0.4357638 | 1 | 0.5154318 | 1 | 0.55340851 | 1 | 0.64497439 | 0 | 1 | 18 | 0.74744813 | FALSE | 1 | FALSE | 1 | FALSE | 1 | FALSE | 1 | FALSE | 1 | FALSE | 1 |
| 1695 | 1695 | 11.2 | 2.4 | 0 | 1 | 0 | 1 | 0 | 1 | 1 | 0.37517262 | 0 | 1 | 9 | 0.5215547 | FALSE | 1 | FALSE | 1 | FALSE | 1 | FALSE | 1 | FALSE | 1 | FALSE | 1 |
| 1696 | 1696 | 4.4295082 | 1.46066954 | 11 | 0.15414876 | 16 | 0.03592936 | 9 | 0.76849506 | 12 | 0.74398616 | 8 | 0.12339972 | 247 | 0.92876779 | FALSE | 1 | FALSE | 0.94259695 | FALSE | 1 | FALSE | 1 | FALSE | 1 | FALSE | 1 |
| 1697 | 1697 | 6.0334728 | 0.54015424 | 17 | 0.10319245 | 21 | 0.09057393 | 20 | 0.26588442 | 20 | 0.69957994 | 9 | 0.38115591 | 390 | 0.95327855 | FALSE | 0.87965924 | FALSE | 1 | FALSE | 1 | FALSE | 1 | FALSE | 1 | FALSE | 1 |
| 1698 | 1698 | 9.4 | 2.37486842 | 0 | 1 | 0 | 1 | 1 | 0.51939841 | 0 | 1 | 0 | 1 | 19 | 0.15700255 | FALSE | 1 | FALSE | 1 | FALSE | 1 | FALSE | 1 | FALSE | 1 | FALSE | 1 |
| 1699 | 1699 | 9.625 | 2.23257139 | 0 | 1 | 0 | 1 | 1 | 0.25384309 | 0 | 1 | 0 | 1 | 7 | 0.63752397 | FALSE | 1 | FALSE | 1 | FALSE | 1 | FALSE | 1 | FALSE | 1 | FALSE | 1 |
| 1700 | 1700 | 11.0625 | 2.77192961 | 0 | 1 | 0 | 1 | 1 | 0.44344817 | 0 | 1 | 0 | 1 | 15 | 0.26127859 | FALSE | 1 | FALSE | 1 | FALSE | 1 | FALSE | 1 | FALSE | 1 | FALSE | 1 |
| 1701 | 1701 | 9.29411765 | 2.92584238 | 0 | 1 | 0 | 1 | 1 | 0.46348836 | 1 | 0.55062772 | 0 | 1 | 15 | 0.49060435 | FALSE | 1 | FALSE | 1 | FALSE | 1 | FALSE | 1 | FALSE | 1 | FALSE | 1 |
| 1702 | 1702 | 7.66666667 | 3 | 0 | 1 | 0 | 1 | 2 | 0.13502329 | 2 | 0.19902682 | 0 | 1 | 14 | 0.86287418 | FALSE | 1 | FALSE | 1 | FALSE | 1 | FALSE | 1 | FALSE | 1 | FALSE | 1 |
| 1703 | 1703 | 11.0833333 | 2.59673941 | 1 | 0.24866461 | 0 | 1 | 0 | 1 | 0 | 1 | 0 | 1 | 10 | 0.46859937 | FALSE | 1 | FALSE | 1 | FALSE | 1 | FALSE | 1 | FALSE | 1 | FALSE | 1 |
| 1704 | 1704 | 8.90909091 | 2.23421922 | 0 | 1 | 0 | 1 | 1 | 0.33149969 | 1 | 0.40390032 | 0 | 1 | 9 | 0.75985366 | FALSE | 1 | FALSE | 1 | FALSE | 1 | FALSE | 1 | FALSE | 1 | FALSE | 1 |
| 1705 | 1705 | 10.4 | 2.45764115 | 0 | 1 | 3 | 0.02534497 | 0 | 1 | 1 | 0.60987146 | 1 | 0.28187291 | 15 | 0.92113679 | FALSE | 1 | FALSE | 0.82367276 | FALSE | 1 | FALSE | 1 | FALSE | 1 | FALSE | 1 |
| 1706 | 1706 | 9.9 | 2.18860686 | 0 | 1 | 2 | 0.1135776 | 0 | 1 | 1 | 0.57131013 | 0 | 1 | 15 | 0.69372602 | FALSE | 1 | FALSE | 1 | FALSE | 1 | FALSE | 1 | FALSE | 1 | FALSE | 1 |
| 1707 | 1707 | 7.66666667 | 2.62466929 | 0 | 1 | 0 | 1 | 0 | 1 | 0 | 1 | 0 | 1 | 3 | 0.60071728 | FALSE | 1 | FALSE | 1 | FALSE | 1 | FALSE | 1 | FALSE | 1 | FALSE | 1 |
| 1708 | 1708 | 10.7857143 | 2.75625448 | 0 | 1 | 0 | 1 | 1 | 0.40110418 | 1 | 0.48242099 | 1 | 0.20679328 | 11 | 0.83676449 | FALSE | 1 | FALSE | 1 | FALSE | 1 | FALSE | 1 | FALSE | 1 | FALSE | 1 |
| 1709 | 1709 | 9.13157895 | 2.93052046 | 1 | 0.6184714 | 1 | 0.70472939 | 1 | 0.74264075 | 1 | 0.82511639 | 1 | 0.45845922 | 32 | 0.47030992 | FALSE | 1 | FALSE | 1 | FALSE | 1 | FALSE | 1 | FALSE | 1 | FALSE | 1 |
| 1710 | 1710 | 7.43636364 | 1.41759895 | 3 | 0.53321268 | 5 | 0.27707647 | 3 | 0.75696175 | 4 | 0.74440481 | 1 | 0.83744681 | 93 | 0.45592412 | FALSE | 1 | FALSE | 1 | FALSE | 1 | FALSE | 1 | FALSE | 1 | FALSE | 1 |
| 1711 | 1711 | 7.48148148 | 2.14942491 | 1 | 0.50471032 | 2 | 0.21684246 | 1 | 0.62830986 | 2 | 0.35374209 | 0 | 1 | 21 | 0.88395936 | FALSE | 1 | FALSE | 1 | FALSE | 1 | FALSE | 1 | FALSE | 1 | FALSE | 1 |
| 1712 | 1712 | 9.34615385 | 2.46424131 | 0 | 1 | 1 | 0.57534752 | 2 | 0.23940226 | 0 | 1 | 1 | 0.3499045 | 22 | 0.61628 | FALSE | 1 | FALSE | 1 | FALSE | 1 | FALSE | 1 | FALSE | 1 | FALSE | 1 |
| 1713 | 1713 | 8.13333333 | 2.47296493 | 1 | 0.54198714 | 0 | 1 | 0 | 1 | 1 | 0.75658797 | 0 | 1 | 28 | 0.13086908 | FALSE | 1 | FALSE | 1 | FALSE | 1 | FALSE | 1 | FALSE | 1 | FALSE | 1 |
| 1714 | 1714 | 10.2857143 | 4.29997627 | 0 | 1 | 1 | 0.74973543 | 2 | 0.44869594 | 6 | 0.01158182 | 0 | 1 | 33 | 0.89139527 | FALSE | 1 | FALSE | 1 | FALSE | 1 | FALSE | 1 | FALSE | 1 | FALSE | 1 |
| 1715 | 1715 | 7.54166667 | 3.00664773 | 2 | 0.34973212 | 1 | 0.79481669 | 2 | 0.51909752 | 4 | 0.17698522 | 1 | 0.54902428 | 38 | 0.88187597 | FALSE | 1 | FALSE | 1 | FALSE | 1 | FALSE | 1 | FALSE | 1 | FALSE | 1 |
| 1716 | 1716 | 9.72222222 | 2.07647018 | 0 | 1 | 1 | 0.44710673 | 1 | 0.48280983 | 0 | 1 | 2 | 0.03435384 | 14 | 0.86287418 | FALSE | 1 | FALSE | 1 | FALSE | 1 | FALSE | 1 | FALSE | 0.86063198 | FALSE | 1 |
| 1717 | 1717 | 10.2 | 1.93907194 | 0 | 1 | 1 | 0.28036922 | 2 | 0.04781153 | 0 | 1 | 0 | 1 | 7 | 0.94326215 | FALSE | 1 | FALSE | 1 | FALSE | 1 | FALSE | 1 | FALSE | 1 | FALSE | 1 |
| 1718 | 1718 | 6.57954546 | 3.66408115 | 4 | 0.1887901 | 4 | 0.31751295 | 3 | 0.61802554 | 4 | 0.58033022 | 2 | 0.42507851 | 71 | 0.865291 | FALSE | 1 | FALSE | 1 | FALSE | 1 | FALSE | 1 | FALSE | 1 | FALSE | 1 |
| 1719 | 1719 | 8.28 | 3.05312954 | 0 | 1 | 1 | 0.54637326 | 1 | 0.58502114 | 0 | 1 | 1 | 0.32796862 | 21 | 0.47053137 | FALSE | 1 | FALSE | 1 | FALSE | 1 | FALSE | 1 | FALSE | 1 | FALSE | 1 |
| 1720 | 1720 | 9.0625 | 1.93548282 | 1 | 0.56527493 | 1 | 0.65167362 | 1 | 0.69069222 | 1 | 0.77852168 | 0 | 1 | 28 | 0.42532177 | FALSE | 1 | FALSE | 1 | FALSE | 1 | FALSE | 1 | FALSE | 1 | FALSE | 1 |
| 1721 | 1721 | 9.05 | 2.57827462 | 0 | 1 | 1 | 0.48239024 | 2 | 0.16019548 | 1 | 0.60987146 | 0 | 1 | 16 | 0.80826361 | FALSE | 1 | FALSE | 1 | FALSE | 1 | FALSE | 1 | FALSE | 1 | FALSE | 1 |
| 1722 | 1722 | 9.14285714 | 1.98292027 | 1 | 0.40558521 | 1 | 0.48239024 | 1 | 0.51939841 | 0 | 1 | 0 | 1 | 17 | 0.61750632 | FALSE | 1 | FALSE | 1 | FALSE | 1 | FALSE | 1 | FALSE | 1 | FALSE | 1 |
| 1723 | 1723 | 9.66666667 | 2.53859104 | 0 | 1 | 0 | 1 | 1 | 0.28067846 | 0 | 1 | 0 | 1 | 8 | 0.5780332 | FALSE | 1 | FALSE | 1 | FALSE | 1 | FALSE | 1 | FALSE | 1 | FALSE | 1 |
| 1724 | 1724 | 9.39130435 | 1.97239934 | 0 | 1 | 1 | 0.5154318 | 2 | 0.18613256 | 1 | 0.64497439 | 0 | 1 | 18 | 0.74744813 | FALSE | 1 | FALSE | 1 | FALSE | 1 | FALSE | 1 | FALSE | 1 | FALSE | 1 |
| 1725 | 1725 | 7.77777778 | 3.02866959 | 1 | 0.37380237 | 0 | 1 | 1 | 0.48280983 | 1 | 0.57131013 | 0 | 1 | 15 | 0.69372602 | FALSE | 1 | FALSE | 1 | FALSE | 1 | FALSE | 1 | FALSE | 1 | FALSE | 1 |
| 1726 | 1726 | 10.3695652 | 3.10930368 | 0 | 1 | 1 | 0.78076835 | 2 | 0.4962935 | 2 | 0.63103861 | 3 | 0.03911573 | 38 | 0.71504609 | FALSE | 1 | FALSE | 1 | FALSE | 1 | FALSE | 1 | FALSE | 0.88763447 | FALSE | 1 |
| 1727 | 1727 | 12.3333333 | 2.62466929 | 0 | 1 | 0 | 1 | 0 | 1 | 0 | 1 | 0 | 1 | 3 | 0.60071728 | FALSE | 1 | FALSE | 1 | FALSE | 1 | FALSE | 1 | FALSE | 1 | FALSE | 1 |
| 1728 | 1728 | 9.32142857 | 2.63294473 | 0 | 1 | 3 | 0.06039219 | 2 | 0.2663542 | 3 | 0.13524571 | 0 | 1 | 20 | 0.97757517 | FALSE | 1 | FALSE | 1 | FALSE | 1 | FALSE | 1 | FALSE | 1 | FALSE | 1 |
| 1729 | 1729 | 8.34328358 | 1.6073733 | 0 | 1 | 2 | 0.64352762 | 2 | 0.7001105 | 4 | 0.3701568 | 1 | 0.6714808 | 58 | 0.38535314 | FALSE | 1 | FALSE | 1 | FALSE | 1 | FALSE | 1 | FALSE | 1 | FALSE | 1 |
| 1730 | 1730 | 8.11111111 | 3.47832797 | 1 | 0.37380237 | 1 | 0.44710673 | 1 | 0.48280983 | 3 | 0.04687752 | 0 | 1 | 12 | 0.98544073 | FALSE | 1 | FALSE | 1 | FALSE | 1 | FALSE | 1 | FALSE | 1 | FALSE | 1 |
| 1731 | 1731 | 9.75 | 1.41421356 | 1 | 0.81177881 | 3 | 0.34261611 | 2 | 0.67580769 | 5 | 0.16939111 | 1 | 0.65460803 | 52 | 0.80965143 | FALSE | 1 | FALSE | 1 | FALSE | 1 | FALSE | 1 | FALSE | 1 | FALSE | 1 |
| 1732 | 1732 | 10.4375 | 2.34437492 | 0 | 1 | 1 | 0.40942991 | 0 | 1 | 0 | 1 | 0 | 1 | 15 | 0.26127859 | FALSE | 1 | FALSE | 1 | FALSE | 1 | FALSE | 1 | FALSE | 1 | FALSE | 1 |
| 1733 | 1733 | 6.66666667 | 3.60995097 | 1 | 0.66519151 | 0 | 1 | 2 | 0.44869594 | 1 | 0.86193352 | 0 | 1 | 38 | 0.19268329 | FALSE | 1 | FALSE | 1 | FALSE | 1 | FALSE | 1 | FALSE | 1 | FALSE | 1 |
| 1734 | 1734 | 6.41935484 | 2.25437487 | 1 | 0.8016455 | 1 | 0.87100665 | 5 | 0.07095719 | 2 | 0.78520976 | 2 | 0.27036738 | 51 | 0.74578096 | FALSE | 1 | FALSE | 1 | FALSE | 1 | FALSE | 1 | FALSE | 1 | FALSE | 1 |
| 1735 | 1735 | 7.8030303 | 1.79409551 | 1 | 0.8213973 | 1 | 0.88704765 | 3 | 0.4248691 | 5 | 0.18498818 | 1 | 0.66594933 | 55 | 0.6690204 | FALSE | 1 | FALSE | 1 | FALSE | 1 | FALSE | 1 | FALSE | 1 | FALSE | 1 |
| 1736 | 1736 | 7.175 | 2.07228738 | 3 | 0.08207756 | 3 | 0.13823565 | 2 | 0.42396974 | 1 | 0.84823834 | 2 | 0.13925369 | 29 | 0.98400908 | FALSE | 0.79236883 | FALSE | 1 | FALSE | 1 | FALSE | 1 | FALSE | 1 | FALSE | 1 |
| 1737 | 1737 | 7.84615385 | 2.01326959 | 0 | 1 | 0 | 1 | 1 | 0.61440466 | 2 | 0.33678042 | 0 | 1 | 23 | 0.40372818 | FALSE | 1 | FALSE | 1 | FALSE | 1 | FALSE | 1 | FALSE | 1 | FALSE | 1 |
| 1738 | 1738 | 8.25806452 | 2.40922611 | 0 | 1 | 1 | 0.63997468 | 1 | 0.67911179 | 2 | 0.41984187 | 0 | 1 | 27 | 0.45522639 | FALSE | 1 | FALSE | 1 | FALSE | 1 | FALSE | 1 | FALSE | 1 | FALSE | 1 |
| 1739 | 1739 | 10.1904762 | 3.78743597 | 0 | 1 | 0 | 1 | 1 | 0.53671415 | 0 | 1 | 0 | 1 | 20 | 0.13763971 | FALSE | 1 | FALSE | 1 | FALSE | 1 | FALSE | 1 | FALSE | 1 | FALSE | 1 |
| 1740 | 1740 | 8.91666667 | 2.98491578 | 0 | 1 | 0 | 1 | 0 | 1 | 0 | 1 | 0 | 1 | 12 | 0.13002667 | FALSE | 1 | FALSE | 1 | FALSE | 1 | FALSE | 1 | FALSE | 1 | FALSE | 1 |
| 1741 | 1741 | 8.84090909 | 3.04469122 | 0 | 1 | 1 | 0.76576289 | 1 | 0.80109494 | 1 | 0.87439647 | 2 | 0.16211764 | 39 | 0.29524127 | FALSE | 1 | FALSE | 1 | FALSE | 1 | FALSE | 1 | FALSE | 1 | FALSE | 1 |
| 1742 | 1742 | 9.22222222 | 2.93552107 | 0 | 1 | 0 | 1 | 0 | 1 | 0 | 1 | 0 | 1 | 9 | 0.21661397 | FALSE | 1 | FALSE | 1 | FALSE | 1 | FALSE | 1 | FALSE | 1 | FALSE | 1 |
| 1743 | 1743 | 11.0588235 | 2.68920693 | 1 | 0.35728305 | 0 | 1 | 1 | 0.46348836 | 1 | 0.55062772 | 0 | 1 | 14 | 0.7313145 | FALSE | 1 | FALSE | 1 | FALSE | 1 | FALSE | 1 | FALSE | 1 | FALSE | 1 |
| 1744 | 1744 | 8.6875 | 2.05300846 | 0 | 1 | 0 | 1 | 0 | 1 | 1 | 0.52895088 | 2 | 0.02752114 | 13 | 0.76799161 | FALSE | 1 | FALSE | 1 | FALSE | 1 | FALSE | 1 | FALSE | 0.79106376 | FALSE | 1 |
| 1745 | 1745 | 6.82608696 | 4.75472769 | 0 | 1 | 5 | 0.05834767 | 5 | 0.08323317 | 3 | 0.57949569 | 0 | 1 | 52 | 0.87296036 | FALSE | 1 | FALSE | 1 | FALSE | 1 | FALSE | 1 | FALSE | 1 | FALSE | 1 |
| 1746 | 1746 | 10.1428571 | 2.29462549 | 0 | 1 | 1 | 0.49918218 | 4 | 0.0060005 | 1 | 0.62783522 | 0 | 1 | 15 | 0.96513691 | FALSE | 1 | FALSE | 1 | FALSE | 1 | FALSE | 1 | FALSE | 1 | FALSE | 1 |
| 1747 | 1747 | 4.58167331 | 2.16185299 | 10 | 0.1039238 | 7 | 0.69789617 | 13 | 0.10955098 | 8 | 0.89018426 | 11 | 0.00236384 | 199 | 0.96951182 | FALSE | 0.88497269 | FALSE | 1 | FALSE | 1 | FALSE | 1 | FALSE | 0.2806458 | FALSE | 1 |
| 1748 | 1748 | 10 | 2.64575131 | 0 | 1 | 0 | 1 | 1 | 0.40110418 | 2 | 0.13304705 | 1 | 0.20679328 | 10 | 0.94579623 | FALSE | 1 | FALSE | 1 | FALSE | 1 | FALSE | 1 | FALSE | 1 | FALSE | 1 |
| 1749 | 1749 | 9 | 2.20794022 | 1 | 0.34033054 | 0 | 1 | 0 | 1 | 0 | 1 | 0 | 1 | 15 | 0.26127859 | FALSE | 1 | FALSE | 1 | FALSE | 1 | FALSE | 1 | FALSE | 1 | FALSE | 1 |
| 1750 | 1750 | 12.1111111 | 3.41384256 | 0 | 1 | 0 | 1 | 0 | 1 | 0 | 1 | 1 | 0.1383237 | 8 | 0.5780332 | FALSE | 1 | FALSE | 1 | FALSE | 1 | FALSE | 1 | FALSE | 1 | FALSE | 1 |
| 1751 | 1751 | 10.0666667 | 2.08059821 | 3 | 0.00600364 | 1 | 0.38964394 | 0 | 1 | 1 | 0.50623194 | 1 | 0.2198258 | 9 | 0.995488 | FALSE | 0.22773046 | FALSE | 1 | FALSE | 1 | FALSE | 1 | FALSE | 1 | FALSE | 1 |
| 1752 | 1752 | 8.61111111 | 2.62760747 | 0 | 1 | 1 | 0.68454804 | 0 | 1 | 2 | 0.48233989 | 0 | 1 | 32 | 0.18108028 | FALSE | 1 | FALSE | 1 | FALSE | 1 | FALSE | 1 | FALSE | 1 | FALSE | 1 |
| 1753 | 1753 | 9.04761905 | 1.67548687 | 0 | 1 | 0 | 1 | 2 | 0.17308361 | 0 | 1 | 1 | 0.29368245 | 18 | 0.57962763 | FALSE | 1 | FALSE | 1 | FALSE | 1 | FALSE | 1 | FALSE | 1 | FALSE | 1 |
| 1754 | 1754 | 5.92682927 | 3.11871021 | 0 | 1 | 4 | 0.03920804 | 2 | 0.42396974 | 1 | 0.84823834 | 0 | 1 | 33 | 0.71942635 | FALSE | 1 | FALSE | 0.96864929 | FALSE | 1 | FALSE | 1 | FALSE | 1 | FALSE | 1 |
| 1755 | 1755 | 6.87142857 | 2.66684098 | 0 | 1 | 0 | 1 | 3 | 0.45353909 | 4 | 0.39127216 | 0 | 1 | 62 | 0.13493316 | FALSE | 1 | FALSE | 1 | FALSE | 1 | FALSE | 1 | FALSE | 1 | FALSE | 1 |
| 1756 | 1756 | 9.64285714 | 2.3786894 | 0 | 1 | 0 | 1 | 2 | 0.06691409 | 1 | 0.43131129 | 1 | 0.18007762 | 8 | 0.97197833 | FALSE | 1 | FALSE | 1 | FALSE | 1 | FALSE | 1 | FALSE | 1 | FALSE | 1 |
| 1757 | 1757 | 8.5 | 2.98208939 | 0 | 1 | 2 | 0.2288771 | 0 | 1 | 1 | 0.73248983 | 0 | 1 | 25 | 0.34293 | FALSE | 1 | FALSE | 1 | FALSE | 1 | FALSE | 1 | FALSE | 1 | FALSE | 1 |
| 1758 | 1758 | 9.05 | 2.80133897 | 0 | 1 | 1 | 0.48239024 | 1 | 0.51939841 | 0 | 1 | 0 | 1 | 18 | 0.37504645 | FALSE | 1 | FALSE | 1 | FALSE | 1 | FALSE | 1 | FALSE | 1 | FALSE | 1 |
| 1759 | 1759 | 10.2142857 | 1.73939938 | 0 | 1 | 0 | 1 | 0 | 1 | 3 | 0.0239258 | 0 | 1 | 11 | 0.83676449 | FALSE | 1 | FALSE | 1 | FALSE | 1 | FALSE | 1 | FALSE | 1 | FALSE | 1 |
| 1760 | 1760 | 5.81188119 | 2.07155593 | 1 | 0.92501001 | 1 | 0.96235137 | 1 | 0.97398096 | 4 | 0.67315543 | 0 | 1 | 92 | 0.00845635 | FALSE | 1 | FALSE | 1 | FALSE | 1 | FALSE | 1 | FALSE | 1 | FALSE | 1 |
| 1761 | 1761 | 6.16666667 | 3.57848509 | 0 | 1 | 1 | 0.54637326 | 1 | 0.58502114 | 0 | 1 | 0 | 1 | 22 | 0.25237063 | FALSE | 1 | FALSE | 1 | FALSE | 1 | FALSE | 1 | FALSE | 1 | FALSE | 1 |
| 1762 | 1762 | 9.47368421 | 2.92661869 | 2 | 0.08405377 | 1 | 0.46503795 | 1 | 0.50143826 | 0 | 1 | 1 | 0.26986775 | 14 | 0.93677253 | FALSE | 0.79236883 | FALSE | 1 | FALSE | 1 | FALSE | 1 | FALSE | 1 | FALSE | 1 |
| 1763 | 1763 | 8.72413793 | 2.3177955 | 1 | 0.52988231 | 1 | 0.61539065 | 1 | 0.65463982 | 3 | 0.14602298 | 0 | 1 | 23 | 0.84462957 | FALSE | 1 | FALSE | 1 | FALSE | 1 | FALSE | 1 | FALSE | 1 | FALSE | 1 |
| 1764 | 1764 | 7.28985507 | 2.03663156 | 2 | 0.53205439 | 2 | 0.65946213 | 1 | 0.92092393 | 5 | 0.20931033 | 0 | 1 | 59 | 0.47808459 | FALSE | 1 | FALSE | 1 | FALSE | 1 | FALSE | 1 | FALSE | 1 | FALSE | 1 |
| 1765 | 1765 | 3.43973941 | 2.01107291 | 11 | 0.15899239 | 16 | 0.03781951 | 10 | 0.66361937 | 9 | 0.94621659 | 3 | 0.88324918 | 256 | 0.62358521 | FALSE | 1 | FALSE | 0.9681794 | FALSE | 1 | FALSE | 1 | FALSE | 1 | FALSE | 1 |
| 1766 | 1766 | 8.96428571 | 2.33730833 | 0 | 1 | 0 | 1 | 1 | 0.64171563 | 1 | 0.73248983 | 0 | 1 | 26 | 0.16405547 | FALSE | 1 | FALSE | 1 | FALSE | 1 | FALSE | 1 | FALSE | 1 | FALSE | 1 |
| 1767 | 1767 | 10.8125 | 2.81111077 | 2 | 0.06197875 | 0 | 1 | 2 | 0.11087977 | 0 | 1 | 0 | 1 | 12 | 0.90922186 | FALSE | 0.72532843 | FALSE | 1 | FALSE | 1 | FALSE | 1 | FALSE | 1 | FALSE | 1 |
| 1768 | 1768 | 5.18518519 | 2.55367617 | 0 | 1 | 2 | 0.21684246 | 1 | 0.62830986 | 4 | 0.03319522 | 1 | 0.36060486 | 19 | 0.98241223 | FALSE | 1 | FALSE | 1 | FALSE | 1 | FALSE | 1 | FALSE | 1 | FALSE | 1 |
| 1769 | 1769 | 8.87096774 | 2.05946964 | 1 | 0.55378205 | 2 | 0.2651851 | 2 | 0.30673593 | 1 | 0.76781284 | 2 | 0.09110253 | 23 | 0.95750806 | FALSE | 1 | FALSE | 1 | FALSE | 1 | FALSE | 1 | FALSE | 1 | FALSE | 1 |
| 1770 | 1770 | 8.97142857 | 1.53967927 | 0 | 1 | 2 | 0.31355738 | 0 | 1 | 4 | 0.07465533 | 0 | 1 | 29 | 0.69927719 | FALSE | 1 | FALSE | 1 | FALSE | 1 | FALSE | 1 | FALSE | 1 | FALSE | 1 |
| 1771 | 1771 | 10.5555556 | 3.022549 | 0 | 1 | 0 | 1 | 0 | 1 | 0 | 1 | 0 | 1 | 9 | 0.21661397 | FALSE | 1 | FALSE | 1 | FALSE | 1 | FALSE | 1 | FALSE | 1 | FALSE | 1 |
| 1772 | 1772 | 9.14285714 | 1.68426088 | 1 | 0.3050805 | 0 | 1 | 1 | 0.40110418 | 0 | 1 | 0 | 1 | 12 | 0.62236227 | FALSE | 1 | FALSE | 1 | FALSE | 1 | FALSE | 1 | FALSE | 1 | FALSE | 1 |
| 1773 | 1773 | 8.7704918 | 3.11207823 | 2 | 0.4493251 | 2 | 0.57395594 | 3 | 0.35626822 | 1 | 0.93827647 | 0 | 1 | 51 | 0.41341988 | FALSE | 1 | FALSE | 1 | FALSE | 1 | FALSE | 1 | FALSE | 1 | FALSE | 1 |
| 1774 | 1774 | 11 | 1.08012345 | 0 | 1 | 0 | 1 | 2 | 0.06691409 | 0 | 1 | 0 | 1 | 10 | 0.71436964 | FALSE | 1 | FALSE | 1 | FALSE | 1 | FALSE | 1 | FALSE | 1 | FALSE | 1 |
| 1775 | 1775 | 6.78947368 | 2.17833706 | 2 | 0.25455533 | 1 | 0.71433277 | 2 | 0.39868506 | 0 | 1 | 0 | 1 | 33 | 0.44261631 | FALSE | 1 | FALSE | 1 | FALSE | 1 | FALSE | 1 | FALSE | 1 | FALSE | 1 |
| 1776 | 1776 | 8.6 | 1.6653328 | 1 | 0.32293355 | 0 | 1 | 0 | 1 | 0 | 1 | 0 | 1 | 14 | 0.29525409 | FALSE | 1 | FALSE | 1 | FALSE | 1 | FALSE | 1 | FALSE | 1 | FALSE | 1 |
| 1777 | 1777 | 8.73809524 | 2.1937088 | 3 | 0.09196816 | 2 | 0.39616059 | 1 | 0.78589454 | 1 | 0.86193352 | 1 | 0.50166523 | 34 | 0.80048036 | FALSE | 0.83248965 | FALSE | 1 | FALSE | 1 | FALSE | 1 | FALSE | 1 | FALSE | 1 |
| 1778 | 1778 | 9.25 | 2.2776084 | 0 | 1 | 0 | 1 | 0 | 1 | 1 | 0.43131129 | 0 | 1 | 11 | 0.41944532 | FALSE | 1 | FALSE | 1 | FALSE | 1 | FALSE | 1 | FALSE | 1 | FALSE | 1 |
| 1779 | 1779 | 8.12 | 1.79599555 | 0 | 1 | 0 | 1 | 2 | 0.19931429 | 2 | 0.2852236 | 0 | 1 | 19 | 0.71542496 | FALSE | 1 | FALSE | 1 | FALSE | 1 | FALSE | 1 | FALSE | 1 | FALSE | 1 |
| 1780 | 1780 | 6.91176471 | 3.45872849 | 1 | 0.58738519 | 3 | 0.09602744 | 2 | 0.34668321 | 1 | 0.7984848 | 1 | 0.43077488 | 26 | 0.92796052 | FALSE | 1 | FALSE | 1 | FALSE | 1 | FALSE | 1 | FALSE | 1 | FALSE | 1 |
| 1781 | 1781 | 10.2608696 | 2.28829519 | 1 | 0.45027683 | 0 | 1 | 0 | 1 | 1 | 0.6613267 | 0 | 1 | 21 | 0.27964072 | FALSE | 1 | FALSE | 1 | FALSE | 1 | FALSE | 1 | FALSE | 1 | FALSE | 1 |
| 1782 | 1782 | 7.93333333 | 2.32283352 | 0 | 1 | 2 | 0.25306331 | 1 | 0.66709967 | 1 | 0.75658797 | 0 | 1 | 26 | 0.48609328 | FALSE | 1 | FALSE | 1 | FALSE | 1 | FALSE | 1 | FALSE | 1 | FALSE | 1 |
| 1783 | 1783 | 10.8095238 | 2.10710923 | 2 | 0.09989125 | 1 | 0.49918218 | 1 | 0.53671415 | 0 | 1 | 0 | 1 | 17 | 0.77849076 | FALSE | 0.86047225 | FALSE | 1 | FALSE | 1 | FALSE | 1 | FALSE | 1 | FALSE | 1 |
| 1784 | 1784 | 9.75 | 1.78535711 | 0 | 1 | 0 | 1 | 0 | 1 | 0 | 1 | 0 | 1 | 4 | 0.50683841 | FALSE | 1 | FALSE | 1 | FALSE | 1 | FALSE | 1 | FALSE | 1 | FALSE | 1 |

| | | | | | | | | | | | | | | | | | | | | | | | | | | | |
|---|---|---|---|---|---|---|---|---|---|---|---|---|---|---|---|---|---|---|---|---|---|---|---|---|---|---|---|
| 1785 | 1785 | 8.5625 | 1.32139841 | 0 | 1 | 0 | 1 | 0 | 1 | 0 | 1 | 0 | 1 | 16 | 0.06581507 | FALSE | 1 | FALSE | 1 | FALSE | 1 | FALSE | 1 | FALSE | 1 | FALSE | 1 |
| 1786 | 1786 | 7.8 | 2.78567766 | 0 | 1 | 1 | 0.53115654 | 0 | 1 | 4 | 0.01932195 | 0 | 1 | 18 | 0.86273155 | FALSE | 1 | FALSE | 1 | FALSE | 1 | FALSE | 1 | FALSE | 1 | FALSE | 1 |
| 1787 | 1787 | 9.9375 | 3.07141559 | 0 | 1 | 1 | 0.40942991 | 1 | 0.44344817 | 2 | 0.16544789 | 1 | 0.23264612 | 11 | 0.9718531 | FALSE | 1 | FALSE | 1 | FALSE | 1 | FALSE | 1 | FALSE | 1 | FALSE | 1 |
| 1788 | 1788 | 9.35714286 | 2.52437101 | 0 | 1 | 0 | 1 | 1 | 0.40110418 | 1 | 0.48242099 | 0 | 1 | 12 | 0.62236227 | FALSE | 1 | FALSE | 1 | FALSE | 1 | FALSE | 1 | FALSE | 1 | FALSE | 1 |
| 1789 | 1789 | 9.75 | 2.2776084 | 1 | 0.34033054 | 3 | 0.01368613 | 0 | 1 | 0 | 1 | 1 | 0.23264612 | 11 | 0.9718531 | FALSE | 1 | FALSE | 0.69207862 | FALSE | 1 | FALSE | 1 | FALSE | 1 | FALSE | 1 |
| 1790 | 1790 | 10.4117647 | 2.40242092 | 0 | 1 | 1 | 0.40942991 | 0 | 1 | 1 | 0.52895088 | 0 | 1 | 14 | 0.53308261 | FALSE | 1 | FALSE | 1 | FALSE | 1 | FALSE | 1 | FALSE | 1 | FALSE | 1 |
| 1791 | 1791 | 8.04166667 | 2.05099908 | 1 | 0.46441868 | 2 | 0.18117183 | 0 | 1 | 0 | 1 | 0 | 1 | 21 | 0.47053137 | FALSE | 1 | FALSE | 1 | FALSE | 1 | FALSE | 1 | FALSE | 1 | FALSE | 1 |
| 1792 | 1792 | 8.76190476 | 2.23403888 | 2 | 0.28329504 | 3 | 0.14580275 | 0 | 1 | 4 | 0.11715635 | 0 | 1 | 32 | 0.90458039 | FALSE | 1 | FALSE | 1 | FALSE | 1 | FALSE | 1 | FALSE | 1 | FALSE | 1 |
| 1793 | 1793 | 9.80952381 | 1.62184632 | 1 | 0.42086991 | 3 | 0.02888061 | 0 | 1 | 1 | 0.62783522 | 3 | 0.00460011 | 13 | 0.9973503 | FALSE | 1 | FALSE | 0.88012077 | FALSE | 1 | FALSE | 1 | FALSE | 0.40960939 | FALSE | 1 |
| 1794 | 1794 | 9.26415094 | 1.519073 | 0 | 1 | 1 | 0.82613965 | 3 | 0.29666549 | 4 | 0.22482652 | 0 | 1 | 45 | 0.5504765 | FALSE | 1 | FALSE | 1 | FALSE | 1 | FALSE | 1 | FALSE | 1 | FALSE | 1 |
| 1795 | 1795 | 9.60869565 | 2.58320883 | 1 | 0.45027683 | 1 | 0.53115654 | 0 | 1 | 1 | 0.6613267 | 1 | 0.31672722 | 19 | 0.71542496 | FALSE | 1 | FALSE | 1 | FALSE | 1 | FALSE | 1 | FALSE | 1 | FALSE | 1 |
| 1796 | 1796 | 8.21428571 | 2.89528117 | 2 | 0.16048326 | 3 | 0.06039219 | 0 | 1 | 0 | 1 | 2 | 0.07640798 | 21 | 0.94083768 | FALSE | 1 | FALSE | 1 | FALSE | 1 | FALSE | 1 | FALSE | 1 | FALSE | 1 |
| 1797 | 1797 | 10.7142857 | 2.83922956 | 0 | 1 | 0 | 1 | 0 | 1 | 2 | 0.13304705 | 0 | 1 | 12 | 0.62236227 | FALSE | 1 | FALSE | 1 | FALSE | 1 | FALSE | 1 | FALSE | 1 | FALSE | 1 |
| 1798 | 1798 | 9.2 | 2.16641024 | 0 | 1 | 0 | 1 | 1 | 0.42266264 | 1 | 0.50623194 | 0 | 1 | 13 | 0.57712805 | FALSE | 1 | FALSE | 1 | FALSE | 1 | FALSE | 1 | FALSE | 1 | FALSE | 1 |
| 1799 | 1799 | 10.0833333 | 2.13924961 | 0 | 1 | 0 | 1 | 0 | 1 | 0 | 1 | 0 | 1 | 12 | 0.13002667 | FALSE | 1 | FALSE | 1 | FALSE | 1 | FALSE | 1 | FALSE | 1 | FALSE | 1 |
| 1800 | 1800 | 7.625 | 2.76878912 | 5 | 0.0748259 | 3 | 0.5460764 | 1 | 0.96086551 | 5 | 0.37910615 | 5 | 0.01443138 | 69 | 0.95054836 | FALSE | 0.77424195 | FALSE | 1 | FALSE | 1 | FALSE | 1 | FALSE | 0.66313193 | FALSE | 1 |
| 1801 | 1801 | 12.0555556 | 3.5819334 | 0 | 1 | 2 | 0.32556427 | 1 | 0.73299778 | 2 | 0.49731009 | 1 | 0.44938266 | 30 | 0.67248271 | FALSE | 1 | FALSE | 1 | FALSE | 1 | FALSE | 1 | FALSE | 1 | FALSE | 1 |
| 1802 | 1802 | 9.03278689 | 2.37450401 | 1 | 0.79637728 | 5 | 0.04653332 | 0 | 1 | 1 | 0.94386203 | 0 | 1 | 54 | 0.24240316 | FALSE | 1 | FALSE | 1 | FALSE | 1 | FALSE | 1 | FALSE | 1 | FALSE | 1 |
| 1803 | 1803 | 6.97959184 | 2.62242913 | 2 | 0.35907852 | 2 | 0.47396556 | 1 | 0.83455705 | 0 | 1 | 0 | 1 | 44 | 0.20085966 | FALSE | 1 | FALSE | 1 | FALSE | 1 | FALSE | 1 | FALSE | 1 | FALSE | 1 |
| 1804 | 1804 | 7.12 | 2.02622802 | 2 | 0.13369621 | 0 | 1 | 1 | 0.59998149 | 1 | 0.69181315 | 1 | 0.33902672 | 20 | 0.8152498 | FALSE | 0.97964162 | FALSE | 1 | FALSE | 1 | FALSE | 1 | FALSE | 1 | FALSE | 1 |
| 1805 | 1805 | 8.44444444 | 1.46143849 | 1 | 0.37380237 | 1 | 0.44710673 | 1 | 0.48280983 | 1 | 0.57131013 | 0 | 1 | 14 | 0.86287418 | FALSE | 1 | FALSE | 1 | FALSE | 1 | FALSE | 1 | FALSE | 1 | FALSE | 1 |
| 1806 | 1806 | 10.0454546 | 2.54910453 | 0 | 1 | 0 | 1 | 0 | 1 | 3 | 0.07743186 | 1 | 0.30529951 | 18 | 0.74744813 | FALSE | 1 | FALSE | 1 | FALSE | 1 | FALSE | 1 | FALSE | 1 | FALSE | 1 |
| 1807 | 1807 | 9.44 | 2.13691366 | 2 | 0.13369621 | 0 | 1 | 1 | 0.59998149 | 1 | 0.69181315 | 0 | 1 | 21 | 0.64957167 | FALSE | 0.97964162 | FALSE | 1 | FALSE | 1 | FALSE | 1 | FALSE | 1 | FALSE | 1 |
| 1808 | 1808 | 7.65 | 1.65151446 | 0 | 1 | 0 | 1 | 3 | 0.17245215 | 2 | 0.5543522 | 0 | 1 | 35 | 0.38966407 | FALSE | 1 | FALSE | 1 | FALSE | 1 | FALSE | 1 | FALSE | 1 | FALSE | 1 |
| 1809 | 1809 | 10 | 1.62221421 | 0 | 1 | 0 | 1 | 3 | 0.02898764 | 0 | 1 | 0 | 1 | 16 | 0.65565778 | FALSE | 1 | FALSE | 1 | FALSE | 1 | FALSE | 1 | FALSE | 1 | FALSE | 1 |
| 1810 | 1810 | 9.33333333 | 2.33673222 | 0 | 1 | 0 | 1 | 0 | 1 | 2 | 0.25059627 | 1 | 0.29368245 | 18 | 0.57962763 | FALSE | 1 | FALSE | 1 | FALSE | 1 | FALSE | 1 | FALSE | 1 | FALSE | 1 |
| 1811 | 1811 | 7.09259259 | 3.2675085 | 2 | 0.40500867 | 0 | 1 | 1 | 0.86240917 | 0 | 1 | 2 | 0.22161752 | 49 | 0.13164269 | FALSE | 1 | FALSE | 1 | FALSE | 1 | FALSE | 1 | FALSE | 1 | FALSE | 1 |
| 1812 | 1812 | 11 | 1.26491106 | 0 | 1 | 0 | 1 | 0 | 1 | 1 | 0.34506517 | 0 | 1 | 8 | 0.5780332 | FALSE | 1 | FALSE | 1 | FALSE | 1 | FALSE | 1 | FALSE | 1 | FALSE | 1 |
| 1813 | 1813 | 9.42857143 | 3.47194784 | 0 | 1 | 0 | 1 | 1 | 0.53671415 | 1 | 0.62783522 | 0 | 1 | 19 | 0.34094959 | FALSE | 1 | FALSE | 1 | FALSE | 1 | FALSE | 1 | FALSE | 1 | FALSE | 1 |
| 1814 | 1814 | 8.71428571 | 3.36852175 | 1 | 0.51745948 | 3 | 0.06039219 | 2 | 0.2663542 | 4 | 0.03737468 | 0 | 1 | 18 | 0.9978883 | FALSE | 1 | FALSE | 1 | FALSE | 1 | FALSE | 1 | FALSE | 1 | FALSE | 1 |
| 1815 | 1815 | 12.6923077 | 3.91023539 | 0 | 1 | 0 | 1 | 0 | 1 | 0 | 1 | 0 | 1 | 13 | 0.10967906 | FALSE | 1 | FALSE | 1 | FALSE | 1 | FALSE | 1 | FALSE | 1 | FALSE | 1 |
| 1816 | 1816 | 9.05882353 | 2.66724331 | 0 | 1 | 0 | 1 | 1 | 0.46348836 | 0 | 1 | 0 | 1 | 16 | 0.23070182 | FALSE | 1 | FALSE | 1 | FALSE | 1 | FALSE | 1 | FALSE | 1 | FALSE | 1 |
| 1817 | 1817 | 8.64516129 | 2.753864 | 0 | 1 | 1 | 0.63997468 | 1 | 0.67911179 | 2 | 0.41984187 | 0 | 1 | 27 | 0.45522639 | FALSE | 1 | FALSE | 1 | FALSE | 1 | FALSE | 1 | FALSE | 1 | FALSE | 1 |
| 1818 | 1818 | 10.3571429 | 3.08468898 | 0 | 1 | 0 | 1 | 0 | 1 | 0 | 1 | 0 | 1 | 14 | 0.09251304 | FALSE | 1 | FALSE | 1 | FALSE | 1 | FALSE | 1 | FALSE | 1 | FALSE | 1 |
| 1819 | 1819 | 5.5 | 2.17178882 | 6 | 0.00292136 | 2 | 0.54544984 | 2 | 0.60323163 | 3 | 0.47819064 | 3 | 0.06360074 | 40 | 0.9960055 | FALSE | 0.14653312 | FALSE | 1 | FALSE | 1 | FALSE | 1 | FALSE | 1 | FALSE | 1 |
| 1820 | 1820 | 10.125 | 2.14694551 | 0 | 1 | 0 | 1 | 0 | 1 | 0 | 1 | 0 | 1 | 8 | 0.25677166 | FALSE | 1 | FALSE | 1 | FALSE | 1 | FALSE | 1 | FALSE | 1 | FALSE | 1 |
| 1821 | 1821 | 9.78947368 | 3.12172266 | 2 | 0.08405377 | 1 | 0.46503795 | 0 | 1 | 2 | 0.21610812 | 1 | 0.26986775 | 13 | 0.97998676 | FALSE | 0.79236883 | FALSE | 1 | FALSE | 1 | FALSE | 1 | FALSE | 1 | FALSE | 1 |
| 1822 | 1822 | 9.21052632 | 2.04112836 | 0 | 1 | 0 | 1 | 0 | 1 | 1 | 0.57131013 | 0 | 1 | 17 | 0.20329205 | FALSE | 1 | FALSE | 1 | FALSE | 1 | FALSE | 1 | FALSE | 1 | FALSE | 1 |
| 1823 | 1823 | 7.5 | 2.61725047 | 0 | 1 | 0 | 1 | 1 | 0.30655256 | 2 | 0.0740981 | 0 | 1 | 7 | 0.94326215 | FALSE | 1 | FALSE | 1 | FALSE | 1 | FALSE | 1 | FALSE | 1 | FALSE | 1 |
| 1824 | 1824 | 9.75 | 2.49861073 | 0 | 1 | 2 | 0.32556427 | 1 | 0.73299778 | 2 | 0.49731009 | 0 | 1 | 31 | 0.49869404 | FALSE | 1 | FALSE | 1 | FALSE | 1 | FALSE | 1 | FALSE | 1 | FALSE | 1 |
| 1825 | 1825 | 9.18518519 | 2.07341259 | 3 | 0.03084786 | 3 | 0.05519089 | 1 | 0.62830986 | 2 | 0.35374209 | 1 | 0.36060486 | 17 | 0.99851071 | FALSE | 0.53356688 | FALSE | 1 | FALSE | 1 | FALSE | 1 | FALSE | 1 | FALSE | 1 |
| 1826 | 1826 | 9.83333333 | 2.03442594 | 0 | 1 | 1 | 0.44710673 | 0 | 1 | 1 | 0.57131013 | 0 | 1 | 16 | 0.44998001 | FALSE | 1 | FALSE | 1 | FALSE | 1 | FALSE | 1 | FALSE | 1 | FALSE | 1 |
| 1827 | 1827 | 9.33333333 | 2.7080128 | 0 | 1 | 0 | 1 | 1 | 0.28067846 | 0 | 1 | 1 | 0.1383237 | 7 | 0.84574168 | FALSE | 1 | FALSE | 1 | FALSE | 1 | FALSE | 1 | FALSE | 1 | FALSE | 1 |
| 1828 | 1828 | 8.36363636 | 3.52324043 | 1 | 0.24866461 | 1 | 0.30368014 | 0 | 1 | 0 | 1 | 0 | 1 | 9 | 0.75985366 | FALSE | 1 | FALSE | 1 | FALSE | 1 | FALSE | 1 | FALSE | 1 | FALSE | 1 |
| 1829 | 1829 | 9.38888889 | 2.58497558 | 0 | 1 | 1 | 0.44710673 | 1 | 0.48280983 | 0 | 1 | 0 | 1 | 16 | 0.44998001 | FALSE | 1 | FALSE | 1 | FALSE | 1 | FALSE | 1 | FALSE | 1 | FALSE | 1 |
| 1830 | 1830 | 8.94736842 | 3.18671063 | 0 | 1 | 2 | 0.12434464 | 1 | 0.50143826 | 0 | 1 | 0 | 1 | 16 | 0.65565778 | FALSE | 1 | FALSE | 1 | FALSE | 1 | FALSE | 1 | FALSE | 1 | FALSE | 1 |
| 1831 | 1831 | 7.40425532 | 2.67103912 | 1 | 0.69044531 | 1 | 0.77338922 | 2 | 0.48463745 | 1 | 0.8802008 | 0 | 1 | 40 | 0.27423942 | FALSE | 1 | FALSE | 1 | FALSE | 1 | FALSE | 1 | FALSE | 1 | FALSE | 1 |
| 1832 | 1832 | 11.3157895 | 2.15339711 | 0 | 1 | 1 | 0.46503795 | 1 | 0.50143826 | 0 | 1 | 0 | 1 | 17 | 0.41141543 | FALSE | 1 | FALSE | 1 | FALSE | 1 | FALSE | 1 | FALSE | 1 | FALSE | 1 |
| 1833 | 1833 | 8.68181818 | 2.88233434 | 0 | 1 | 2 | 0.15794013 | 0 | 1 | 2 | 0.26791271 | 0 | 1 | 18 | 0.74744813 | FALSE | 1 | FALSE | 1 | FALSE | 1 | FALSE | 1 | FALSE | 1 | FALSE | 1 |
| 1834 | 1834 | 9.9 | 1.64012195 | 0 | 1 | 0 | 1 | 0 | 1 | 0 | 1 | 0 | 1 | 10 | 0.18273166 | FALSE | 1 | FALSE | 1 | FALSE | 1 | FALSE | 1 | FALSE | 1 | FALSE | 1 |
| 1835 | 1835 | 7.58333333 | 3.4268142 | 1 | 0.26795838 | 0 | 1 | 0 | 1 | 0 | 1 | 0 | 1 | 11 | 0.41944532 | FALSE | 1 | FALSE | 1 | FALSE | 1 | FALSE | 1 | FALSE | 1 | FALSE | 1 |
| 1836 | 1836 | 9.92857143 | 3.12739704 | 0 | 1 | 0 | 1 | 1 | 0.40110418 | 0 | 1 | 1 | 0.20679328 | 12 | 0.62236227 | FALSE | 1 | FALSE | 1 | FALSE | 1 | FALSE | 1 | FALSE | 1 | FALSE | 1 |
| 1837 | 1837 | 9.77777778 | 2.65739128 | 0 | 1 | 0 | 1 | 0 | 1 | 0 | 1 | 1 | 0.1383237 | 8 | 0.5780332 | FALSE | 1 | FALSE | 1 | FALSE | 1 | FALSE | 1 | FALSE | 1 | FALSE | 1 |
| 1838 | 1838 | 10.2380952 | 2.84361879 | 0 | 1 | 0 | 1 | 2 | 0.17308361 | 2 | 0.25059627 | 0 | 1 | 17 | 0.77849076 | FALSE | 1 | FALSE | 1 | FALSE | 1 | FALSE | 1 | FALSE | 1 | FALSE | 1 |
| 1839 | 1839 | 9.05882353 | 1.79773021 | 0 | 1 | 0 | 1 | 0 | 1 | 0 | 1 | 0 | 1 | 16 | 0.06581507 | FALSE | 1 | FALSE | 1 | FALSE | 1 | FALSE | 1 | FALSE | 1 | FALSE | 1 |
| 1840 | 1840 | 7.95238095 | 1.29011592 | 0 | 1 | 0 | 1 | 1 | 0.90129726 | 2 | 0.79265743 | 0 | 1 | 60 | 0.00727099 | FALSE | 1 | FALSE | 1 | FALSE | 1 | FALSE | 1 | FALSE | 1 | FALSE | 1 |
| 1841 | 1841 | 10 | 2.56038192 | 0 | 1 | 0 | 1 | 0 | 1 | 2 | 0.19902682 | 0 | 1 | 16 | 0.44998001 | FALSE | 1 | FALSE | 1 | FALSE | 1 | FALSE | 1 | FALSE | 1 | FALSE | 1 |
| 1842 | 1842 | 9.90566038 | 2.13916004 | 1 | 0.74890551 | 1 | 0.82613965 | 4 | 0.12170726 | 3 | 0.44196866 | 2 | 0.21556658 | 42 | 0.88654191 | FALSE | 1 | FALSE | 1 | FALSE | 1 | FALSE | 1 | FALSE | 1 | FALSE | 1 |
| 1843 | 1843 | 10.84 | 2.92136954 | 3 | 0.02517314 | 1 | 0.56109832 | 1 | 0.59998149 | 2 | 0.31968852 | 0 | 1 | 18 | 0.96851178 | FALSE | 0.4974559 | FALSE | 1 | FALSE | 1 | FALSE | 1 | FALSE | 1 | FALSE | 1 |
| 1844 | 1844 | 5.6553288 | 2.06669408 | 11 | 0.53895511 | 13 | 0.6317617 | 16 | 0.46820114 | 20 | 0.49712943 | 2 | 0.99415632 | 365 | 0.28455062 | FALSE | 1 | FALSE | 1 | FALSE | 1 | FALSE | 1 | FALSE | 1 | FALSE | 1 |
| 1845 | 1845 | 10.1428571 | 2.29462549 | 0 | 1 | 0 | 1 | 0 | 1 | 2 | 0.13304705 | 0 | 1 | 12 | 0.62236227 | FALSE | 1 | FALSE | 1 | FALSE | 1 | FALSE | 1 | FALSE | 1 | FALSE | 1 |
| 1846 | 1846 | 9.72222222 | 3.47699672 | 1 | 0.37380237 | 2 | 0.1135776 | 0 | 1 | 1 | 0.57131013 | 3 | 0.00292491 | 11 | 0.99651304 | FALSE | 1 | FALSE | 1 | FALSE | 1 | FALSE | 1 | FALSE | 0.29951111 | FALSE | 1 |
| 1847 | 1847 | 15.125 | 4.53975495 | 1 | 0.34033054 | 0 | 1 | 1 | 0.44344817 | 2 | 0.16544789 | 1 | 0.23264612 | 11 | 0.9718531 | FALSE | 1 | FALSE | 1 | FALSE | 1 | FALSE | 1 | FALSE | 1 | FALSE | 1 |
| 1848 | 1848 | 8.03225807 | 1.46977881 | 1 | 0.55378205 | 1 | 0.63997468 | 0 | 1 | 3 | 0.16836568 | 0 | 1 | 26 | 0.64660109 | FALSE | 1 | FALSE | 1 | FALSE | 1 | FALSE | 1 | FALSE | 1 | FALSE | 1 |
| 1849 | 1849 | 4.48648649 | 1.52229595 | 1 | 0.9789433 | 2 | 0.95455861 | 9 | 0.08186367 | 5 | 0.81236931 | 0 | 1 | 130 | 0.10163053 | FALSE | 1 | FALSE | 1 | FALSE | 1 | FALSE | 1 | FALSE | 1 | FALSE | 1 |
| 1850 | 1850 | 8.90476191 | 3.26459743 | 0 | 1 | 1 | 0.49918218 | 1 | 0.53671415 | 3 | 0.06912908 | 0 | 1 | 16 | 0.90353674 | FALSE | 1 | FALSE | 1 | FALSE | 1 | FALSE | 1 | FALSE | 1 | FALSE | 1 |
| 1851 | 1851 | 10.08 | 1.49452334 | 2 | 0.13369621 | 0 | 1 | 1 | 0.59998149 | 0 | 1 | 1 | 0.33902672 | 21 | 0.64957167 | FALSE | 0.97964162 | FALSE | 1 | FALSE | 1 | FALSE | 1 | FALSE | 1 | FALSE | 1 |
| 1852 | 1852 | 9.75 | 3.05845822 | 0 | 1 | 1 | 0.54637326 | 1 | 0.58502114 | 0 | 1 | 0 | 1 | 22 | 0.25237063 | FALSE | 1 | FALSE | 1 | FALSE | 1 | FALSE | 1 | FALSE | 1 | FALSE | 1 |
| 1853 | 1853 | 10.5625 | 2.62127712 | 0 | 1 | 0 | 1 | 2 | 0.11087977 | 1 | 0.52895088 | 0 | 1 | 13 | 0.76799161 | FALSE | 1 | FALSE | 1 | FALSE | 1 | FALSE | 1 | FALSE | 1 | FALSE | 1 |
| 1854 | 1854 | 7.1025641 | 3.45516983 | 1 | 0.63789042 | 0 | 1 | 3 | 0.16357218 | 0 | 1 | 0 | 1 | 35 | 0.24915769 | FALSE | 1 | FALSE | 1 | FALSE | 1 | FALSE | 1 | FALSE | 1 | FALSE | 1 |
| 1855 | 1855 | 3.07724601 | 0.66764833 | 51 | 6.91E-05 | 61 | 7.95E-05 | 52 | 0.06721371 | 49 | 0.82143696 | 35 | 0.0002108 | 940 | 0.99999996 | TRUE | 0.01342173 | FALSE | 0.10861097 | FALSE | 1 | FALSE | 1 | FALSE | 0.07486439 | FALSE | 1 |
| 1856 | 1856 | 12.1428571 | 2.29462549 | 0 | 1 | 0 | 1 | 0 | 1 | 1 | 0.28044368 | 0 | 1 | 6 | 0.69920685 | FALSE | 1 | FALSE | 1 | FALSE | 1 | FALSE | 1 | FALSE | 1 | FALSE | 1 |
| 1857 | 1857 | 9.28571429 | 3.04166084 | 4 | 0.00146558 | 1 | 0.48239024 | 0 | 1 | 2 | 0.23331368 | 0 | 1 | 13 | 0.99248686 | FALSE | 0.09453559 | FALSE | 1 | FALSE | 1 | FALSE | 1 | FALSE | 1 | FALSE | 1 |
| 1858 | 1858 | 11.2142857 | 2.6234811 | 3 | 0.0048942 | 1 | 0.36919824 | 0 | 1 | 1 | 0.48242099 | 0 | 1 | 9 | 0.98607548 | FALSE | 0.19848171 | FALSE | 1 | FALSE | 1 | FALSE | 1 | FALSE | 1 | FALSE | 1 |
| 1859 | 1859 | 8.70731707 | 4.15694484 | 1 | 0.59801736 | 1 | 0.68454804 | 3 | 0.12958859 | 2 | 0.48233989 | 0 | 1 | 28 | 0.83079716 | FALSE | 1 | FALSE | 1 | FALSE | 1 | FALSE | 1 | FALSE | 1 | FALSE | 1 |
| 1860 | 1860 | 5.02040816 | 2.34729367 | 2 | 0.72148567 | 3 | 0.61960793 | 3 | 0.69002641 | 4 | 0.66533238 | 1 | 0.80446991 | 85 | 0.31459839 | FALSE | 1 | FALSE | 1 | FALSE | 1 | FALSE | 1 | FALSE | 1 | FALSE | 1 |
| 1861 | 1861 | 9.51428571 | 2.62266417 | 1 | 0.59801736 | 0 | 1 | 2 | 0.35984046 | 1 | 0.80778317 | 1 | 0.44015537 | 30 | 0.52765882 | FALSE | 1 | FALSE | 1 | FALSE | 1 | FALSE | 1 | FALSE | 1 | FALSE | 1 |
| 1862 | 1862 | 5.16901409 | 2.70620526 | 3 | 0.26686779 | 1 | 0.90110191 | 1 | 0.92379401 | 2 | 0.83868849 | 0 | 1 | 63 | 0.12472747 | FALSE | 1 | FALSE | 1 | FALSE | 1 | FALSE | 1 | FALSE | 1 | FALSE | 1 |
| 1863 | 1863 | 6.58823529 | 3.67917548 | 1 | 0.35728305 | 2 | 0.10307112 | 1 | 0.46348836 | 0 | 1 | 0 | 1 | 13 | 0.88719222 | FALSE | 1 | FALSE | 1 | FALSE | 1 | FALSE | 1 | FALSE | 1 | FALSE | 1 |
| 1864 | 1864 | 8.34782609 | 3.35712993 | 2 | 0.11647716 | 2 | 0.16948764 | 1 | 0.5695037 | 0 | 1 | 0 | 1 | 18 | 0.86273155 | FALSE | 0.93181727 | FALSE | 1 | FALSE | 1 | FALSE | 1 | FALSE | 1 | FALSE | 1 |
| 1865 | 1865 | 7.31914894 | 2.26073216 | 1 | 0.69843502 | 2 | 0.44132825 | 2 | 0.4962935 | 1 | 0.88573774 | 0 | 1 | 40 | 0.40712403 | FALSE | 1 | FALSE | 1 | FALSE | 1 | FALSE | 1 | FALSE | 1 | FALSE | 1 |
| 1866 | 1866 | 10 | 3.53553391 | 0 | 1 | 1 | 0.12326188 | 0 | 1 | 0 | 1 | 0 | 1 | 3 | 0.8823539 | FALSE | 1 | FALSE | 1 | FALSE | 1 | FALSE | 1 | FALSE | 1 | FALSE | 1 |
| 1867 | 1867 | 9.22222222 | 2.27438772 | 0 | 1 | 2 | 0.1135776 | 1 | 0.48280983 | 0 | 1 | 0 | 1 | 15 | 0.69372602 | FALSE | 1 | FALSE | 1 | FALSE | 1 | FALSE | 1 | FALSE | 1 | FALSE | 1 |
| 1868 | 1868 | 8.875 | 1.45236876 | 0 | 1 | 0 | 1 | 0 | 1 | 0 | 1 | 0 | 1 | 8 | 0.25677166 | FALSE | 1 | FALSE | 1 | FALSE | 1 | FALSE | 1 | FALSE | 1 | FALSE | 1 |
| 1869 | 1869 | 9.72727273 | 2.17802701 | 1 | 0.24866461 | 0 | 1 | 1 | 0.33149969 | 0 | 1 | 1 | 0.16638736 | 8 | 0.92152993 | FALSE | 1 | FALSE | 1 | FALSE | 1 | FALSE | 1 | FALSE | 1 | FALSE | 1 |
| 1870 | 1870 | 9.31818182 | 1.60641791 | 0 | 1 | 3 | 0.03266339 | 1 | 0.55340851 | 0 | 1 | 0 | 1 | 18 | 0.74744813 | FALSE | 1 | FALSE | 0.92268438 | FALSE | 1 | FALSE | 1 | FALSE | 1 | FALSE | 1 |
| 1871 | 1871 | 10.5 | 1.5 | 0 | 1 | 0 | 1 | 0 | 1 | 0 | 1 | 0 | 1 | 4 | 0.50683841 | FALSE | 1 | FALSE | 1 | FALSE | 1 | FALSE | 1 | FALSE | 1 | FALSE | 1 |
| 1872 | 1872 | 8.72413793 | 1.98088367 | 0 | 1 | 0 | 1 | 0 | 1 | 3 | 0.14602298 | 0 | 1 | 26 | 0.31495157 | FALSE | 1 | FALSE | 1 | FALSE | 1 | FALSE | 1 | FALSE | 1 | FALSE | 1 |
| 1873 | 1873 | 7.07894737 | 2.37178124 | 0 | 1 | 3 | 0.44799895 | 4 | 0.29102231 | 8 | 0.02253952 | 2 | 0.35515917 | 59 | 0.957857 | FALSE | 1 | FALSE | 1 | FALSE | 1 | FALSE | 1 | FALSE | 1 | FALSE | 1 |
| 1874 | 1874 | 8.82608696 | 2.72910519 | 1 | 0.45027683 | 0 | 1 | 1 | 0.5695037 | 1 | 0.6613267 | 0 | 1 | 20 | 0.50589402 | FALSE | 1 | FALSE | 1 | FALSE | 1 | FALSE | 1 | FALSE | 1 | FALSE | 1 |
| 1875 | 1875 | 9.5 | 3.82970843 | 0 | 1 | 0 | 1 | 1 | 0.58502114 | 3 | 0.09525947 | 0 | 1 | 20 | 0.68270729 | FALSE | 1 | FALSE | 1 | FALSE | 1 | FALSE | 1 | FALSE | 1 | FALSE | 1 |
| 1876 | 1876 | 9.35483871 | 5.07713653 | 3 | 0.04402911 | 2 | 0.2651851 | 1 | 0.67911179 | 0 | 1 | 0 | 1 | 25 | 0.80006983 | FALSE | 0.64293489 | FALSE | 1 | FALSE | 1 | FALSE | 1 | FALSE | 1 | FALSE | 1 |
| 1877 | 1877 | 9.81818182 | 4.74472304 | 0 | 1 | 0 | 1 | 0 | 1 | 1 | 0.40390032 | 0 | 1 | 10 | 0.46859937 | FALSE | 1 | FALSE | 1 | FALSE | 1 | FALSE | 1 | FALSE | 1 | FALSE | 1 |
| 1878 | 1878 | 10.6666667 | 2.37713981 | 0 | 1 | 0 | 1 | 1 | 0.53671415 | 0 | 1 | 0 | 1 | 20 | 0.13763971 | FALSE | 1 | FALSE | 1 | FALSE | 1 | FALSE | 1 | FALSE | 1 | FALSE | 1 |

| | | | | | | | | | | | | | | | | | | | | | | | | | | | |
|---|---|---|---|---|---|---|---|---|---|---|---|---|---|---|---|---|---|---|---|---|---|---|---|---|---|---|---|
| 1879 | 1879 | 11 | 2.80109869 | 0 | 1 | 0 | 1 | 1 | 0.35555293 | 0 | 1 | 0 | 1 | 11 | 0.41944532 | FALSE | 1 | FALSE | 1 | FALSE | 1 | FALSE | 1 | FALSE | 1 | FALSE | 1 |
| 1880 | 1880 | 9.33333333 | 1.61932771 | 1 | 0.32293355 | 0 | 1 | 2 | 0.09928574 | 1 | 0.50623194 | 2 | 0.02433796 | 9 | 0.995488 | FALSE | 1 | FALSE | 1 | FALSE | 1 | FALSE | 1 | FALSE | 0.77578444 | FALSE | 1 |
| 1881 | 1881 | 10 | 2.9011492 | 3 | 0.02256792 | 1 | 0.54637326 | 0 | 1 | 3 | 0.09525947 | 2 | 0.058175 | 15 | 0.99800699 | FALSE | 0.4692294 | FALSE | 1 | FALSE | 1 | FALSE | 1 | FALSE | 1 | FALSE | 1 |
| 1882 | 1882 | 8.88888889 | 1.6960375 | 0 | 1 | 1 | 0.44710673 | 2 | 0.13502329 | 0 | 1 | 1 | 0.25766374 | 14 | 0.86287418 | FALSE | 1 | FALSE | 1 | FALSE | 1 | FALSE | 1 | FALSE | 1 | FALSE | 1 |
| 1883 | 1883 | 10 | 1.89181061 | 1 | 0.38989951 | 0 | 1 | 0 | 1 | 0 | 1 | 1 | 0.26986775 | 17 | 0.41141543 | FALSE | 1 | FALSE | 1 | FALSE | 1 | FALSE | 1 | FALSE | 1 | FALSE | 1 |
| 1884 | 1884 | 10.6666667 | 2.82842713 | 1 | 0.20854748 | 1 | 0.25628163 | 0 | 1 | 0 | 1 | 0 | 1 | 7 | 0.84574168 | FALSE | 1 | FALSE | 1 | FALSE | 1 | FALSE | 1 | FALSE | 1 | FALSE | 1 |
| 1885 | 1885 | 9.625 | 3.07967125 | 1 | 0.34033054 | 2 | 0.09285443 | 1 | 0.44344817 | 1 | 0.52895088 | 0 | 1 | 11 | 0.9718531 | FALSE | 1 | FALSE | 1 | FALSE | 1 | FALSE | 1 | FALSE | 1 | FALSE | 1 |
| 1886 | 1886 | 9.28125 | 2.18280747 | 1 | 0.56527493 | 4 | 0.0188228 | 1 | 0.69069222 | 0 | 1 | 1 | 0.41154418 | 25 | 0.88559288 | FALSE | 1 | FALSE | 0.77515497 | FALSE | 1 | FALSE | 1 | FALSE | 1 | FALSE | 1 |
| 1887 | 1887 | 9.55813954 | 2.87169285 | 4 | 0.02189152 | 0 | 1 | 0 | 1 | 2 | 0.5811115 | 0 | 1 | 36 | 0.50934357 | FALSE | 0.46823848 | FALSE | 1 | FALSE | 1 | FALSE | 1 | FALSE | 1 | FALSE | 1 |
| 1888 | 1888 | 8.82857143 | 2.82322728 | 0 | 1 | 2 | 0.31355738 | 3 | 0.12958859 | 2 | 0.48233989 | 2 | 0.11183826 | 26 | 0.96294373 | FALSE | 1 | FALSE | 1 | FALSE | 1 | FALSE | 1 | FALSE | 1 | FALSE | 1 |
| 1889 | 1889 | 7.5 | 2.53610295 | 0 | 1 | 1 | 0.5154318 | 1 | 0.55340851 | 0 | 1 | 1 | 0.30529951 | 19 | 0.54233419 | FALSE | 1 | FALSE | 1 | FALSE | 1 | FALSE | 1 | FALSE | 1 | FALSE | 1 |
| 1890 | 1890 | 8 | 2.90593263 | 0 | 1 | 0 | 1 | 1 | 0.28067846 | 0 | 1 | 0 | 1 | 8 | 0.5780332 | FALSE | 1 | FALSE | 1 | FALSE | 1 | FALSE | 1 | FALSE | 1 | FALSE | 1 |
| 1891 | 1891 | 8.35294118 | 2.74271605 | 0 | 1 | 0 | 1 | 0 | 1 | 0 | 1 | 2 | 0.03086174 | 15 | 0.49060435 | FALSE | 1 | FALSE | 1 | FALSE | 1 | FALSE | 1 | FALSE | 0.82559785 | FALSE | 1 |
| 1892 | 1892 | 7.53333333 | 1.96185853 | 2 | 0.05513784 | 0 | 1 | 1 | 0.42266264 | 0 | 1 | 0 | 1 | 12 | 0.80329917 | FALSE | 0.68849059 | FALSE | 1 | FALSE | 1 | FALSE | 1 | FALSE | 1 | FALSE | 1 |
| 1893 | 1893 | 8.95 | 2.31246622 | 3 | 0.01369554 | 2 | 0.1353446 | 1 | 0.51939841 | 0 | 1 | 0 | 1 | 14 | 0.97325593 | FALSE | 0.36625055 | FALSE | 1 | FALSE | 1 | FALSE | 1 | FALSE | 1 | FALSE | 1 |
| 1894 | 1894 | 10.5333333 | 2.15612822 | 0 | 1 | 1 | 0.38964394 | 1 | 0.42266264 | 3 | 0.02891327 | 1 | 0.2198258 | 9 | 0.995488 | FALSE | 1 | FALSE | 1 | FALSE | 1 | FALSE | 1 | FALSE | 1 | FALSE | 1 |
| 1895 | 1895 | 6.33333333 | 2.92498813 | 0 | 1 | 2 | 0.05552913 | 0 | 1 | 3 | 0.01547597 | 0 | 1 | 7 | 0.99433481 | FALSE | 1 | FALSE | 1 | FALSE | 1 | FALSE | 1 | FALSE | 1 | FALSE | 1 |
| 1896 | 1896 | 10.4615385 | 2.6345452 | 1 | 0.49162634 | 1 | 0.57534752 | 2 | 0.23940226 | 1 | 0.70601446 | 0 | 1 | 21 | 0.78939668 | FALSE | 1 | FALSE | 1 | FALSE | 1 | FALSE | 1 | FALSE | 1 | FALSE | 1 |
| 1897 | 1897 | 8.66666667 | 3.56348323 | 0 | 1 | 2 | 0.14655134 | 0 | 1 | 1 | 0.62783522 | 0 | 1 | 18 | 0.57962763 | FALSE | 1 | FALSE | 1 | FALSE | 1 | FALSE | 1 | FALSE | 1 | FALSE | 1 |
| 1898 | 1898 | 9.22222222 | 2.91017797 | 1 | 0.50471032 | 0 | 1 | 0 | 1 | 2 | 0.35374209 | 1 | 0.36060486 | 23 | 0.58307573 | FALSE | 1 | FALSE | 1 | FALSE | 1 | FALSE | 1 | FALSE | 1 | FALSE | 1 |
| 1899 | 1899 | 7.26086957 | 2.04763684 | 0 | 1 | 0 | 1 | 1 | 0.55340851 | 0 | 1 | 0 | 1 | 21 | 0.12048848 | FALSE | 1 | FALSE | 1 | FALSE | 1 | FALSE | 1 | FALSE | 1 | FALSE | 1 |
| 1900 | 1900 | 8.73913044 | 3.45426506 | 0 | 1 | 0 | 1 | 0 | 1 | 0 | 1 | 1 | 0.30529951 | 21 | 0.12048848 | FALSE | 1 | FALSE | 1 | FALSE | 1 | FALSE | 1 | FALSE | 1 | FALSE | 1 |
| 1901 | 1901 | 10.25 | 2.04633819 | 0 | 1 | 0 | 1 | 0 | 1 | 0 | 1 | 1 | 0.18007762 | 11 | 0.41944532 | FALSE | 1 | FALSE | 1 | FALSE | 1 | FALSE | 1 | FALSE | 1 | FALSE | 1 |
| 1902 | 1902 | 9.44 | 1.94072152 | 0 | 1 | 0 | 1 | 1 | 0.58502114 | 0 | 1 | 0 | 1 | 23 | 0.09196389 | FALSE | 1 | FALSE | 1 | FALSE | 1 | FALSE | 1 | FALSE | 1 | FALSE | 1 |
| 1903 | 1903 | 8.70588235 | 1.88694293 | 1 | 0.58738519 | 1 | 0.6739483 | 1 | 0.71261921 | 0 | 1 | 1 | 0.43077488 | 30 | 0.36879488 | FALSE | 1 | FALSE | 1 | FALSE | 1 | FALSE | 1 | FALSE | 1 | FALSE | 1 |
| 1904 | 1904 | 10.2727273 | 3.27777389 | 0 | 1 | 1 | 0.30368014 | 0 | 1 | 0 | 1 | 0 | 1 | 10 | 0.46859937 | FALSE | 1 | FALSE | 1 | FALSE | 1 | FALSE | 1 | FALSE | 1 | FALSE | 1 |
| 1905 | 1905 | 10.4285714 | 3.18585308 | 2 | 0.09989125 | 2 | 0.14655134 | 2 | 0.17308361 | 1 | 0.62783522 | 0 | 1 | 14 | 0.98949398 | FALSE | 0.86047225 | FALSE | 1 | FALSE | 1 | FALSE | 1 | FALSE | 1 | FALSE | 1 |
| 1906 | 1906 | 9.14285714 | 2.1221219 | 1 | 0.42086991 | 2 | 0.14655134 | 0 | 1 | 0 | 1 | 0 | 1 | 18 | 0.57962763 | FALSE | 1 | FALSE | 1 | FALSE | 1 | FALSE | 1 | FALSE | 1 | FALSE | 1 |
| 1907 | 1907 | 7.25 | 2.51246891 | 0 | 1 | 1 | 0.40942991 | 1 | 0.44344817 | 1 | 0.52895088 | 2 | 0.02752114 | 11 | 0.9718531 | FALSE | 1 | FALSE | 1 | FALSE | 1 | FALSE | 1 | FALSE | 0.79106376 | FALSE | 1 |
| 1908 | 1908 | 8.36363636 | 1.89605942 | 1 | 0.4357638 | 0 | 1 | 2 | 0.18613256 | 1 | 0.64497439 | 0 | 1 | 18 | 0.74744813 | FALSE | 1 | FALSE | 1 | FALSE | 1 | FALSE | 1 | FALSE | 1 | FALSE | 1 |
| 1909 | 1909 | 10.1764706 | 2.85400223 | 0 | 1 | 0 | 1 | 2 | 0.12280513 | 1 | 0.55062772 | 1 | 0.24525764 | 13 | 0.88719222 | FALSE | 1 | FALSE | 1 | FALSE | 1 | FALSE | 1 | FALSE | 1 | FALSE | 1 |
| 1910 | 1910 | 4.39705882 | 1.52886963 | 12 | 0.00260908 | 13 | 0.00647786 | 10 | 0.12527692 | 6 | 0.85617949 | 4 | 0.35655526 | 139 | 0.99943992 | FALSE | 0.13878934 | FALSE | 0.60103504 | FALSE | 1 | FALSE | 1 | FALSE | 1 | FALSE | 1 |
| 1911 | 1911 | 8.60714286 | 2.69044985 | 2 | 0.16048326 | 1 | 0.60247911 | 0 | 1 | 2 | 0.3705495 | 2 | 0.07640798 | 21 | 0.94083768 | FALSE | 1 | FALSE | 1 | FALSE | 1 | FALSE | 1 | FALSE | 1 | FALSE | 1 |
| 1912 | 1912 | 7.21568628 | 2.86502729 | 4 | 0.04099222 | 3 | 0.22759346 | 1 | 0.84631478 | 0 | 1 | 1 | 0.57100288 | 42 | 0.73296146 | FALSE | 0.62767905 | FALSE | 1 | FALSE | 1 | FALSE | 1 | FALSE | 1 | FALSE | 1 |
| 1913 | 1913 | 9.10714286 | 1.61110622 | 2 | 0.16048326 | 0 | 1 | 0 | 1 | 1 | 0.73248983 | 0 | 1 | 25 | 0.34293 | FALSE | 1 | FALSE | 1 | FALSE | 1 | FALSE | 1 | FALSE | 1 | FALSE | 1 |
| 1914 | 1914 | 13.1111111 | 3.12496914 | 0 | 1 | 2 | 0.1135776 | 0 | 1 | 0 | 1 | 0 | 1 | 16 | 0.44998001 | FALSE | 1 | FALSE | 1 | FALSE | 1 | FALSE | 1 | FALSE | 1 | FALSE | 1 |
| 1915 | 1915 | 7.98717949 | 1.60522855 | 4 | 0.1347453 | 4 | 0.23789896 | 1 | 0.94117955 | 1 | 0.97374287 | 1 | 0.72203879 | 66 | 0.44767694 | FALSE | 0.98380879 | FALSE | 1 | FALSE | 1 | FALSE | 1 | FALSE | 1 | FALSE | 1 |
| 1916 | 1916 | 8.43636364 | 2.86523765 | 1 | 0.63789042 | 1 | 0.7236252 | 1 | 0.76089851 | 3 | 0.26522531 | 0 | 1 | 33 | 0.59061307 | FALSE | 1 | FALSE | 1 | FALSE | 1 | FALSE | 1 | FALSE | 1 | FALSE | 1 |
| 1917 | 1917 | 2.04759285 | 0.53611431 | 80 | 0.97967429 | 112 | 0.77658796 | 130 | 0.52316564 | 173 | 0.22541154 | 60 | 0.48590053 | 3062 | 0.26058438 | FALSE | 1 | FALSE | 1 | FALSE | 1 | FALSE | 1 | FALSE | 1 | FALSE | 1 |
| 1918 | 1918 | 9 | 2.44948974 | 2 | 0.01652925 | 1 | 0.23139161 | 0 | 1 | 0 | 1 | 0 | 1 | 5 | 0.97551477 | FALSE | 0.39708985 | FALSE | 1 | FALSE | 1 | FALSE | 1 | FALSE | 1 | FALSE | 1 |
| 1919 | 1919 | 10.2962963 | 2.60789137 | 0 | 1 | 0 | 1 | 1 | 0.62830986 | 3 | 0.12475952 | 0 | 1 | 23 | 0.58307573 | FALSE | 1 | FALSE | 1 | FALSE | 1 | FALSE | 1 | FALSE | 1 | FALSE | 1 |
| 1920 | 1920 | 7.12162162 | 2.56247717 | 2 | 0.57015153 | 0 | 1 | 0 | 1 | 1 | 0.96971789 | 2 | 0.34319968 | 69 | 0.01821645 | FALSE | 1 | FALSE | 1 | FALSE | 1 | FALSE | 1 | FALSE | 1 | FALSE | 1 |
| 1921 | 1921 | 3.36434109 | 1.42660935 | 20 | 0.00202379 | 18 | 0.07539418 | 6 | 0.99560665 | 22 | 0.17370494 | 10 | 0.10156027 | 311 | 0.98803837 | FALSE | 0.12179217 | FALSE | 1 | FALSE | 1 | FALSE | 1 | FALSE | 1 | FALSE | 1 |
| 1922 | 1922 | 5.78787879 | 2.84186853 | 0 | 1 | 2 | 0.93132803 | 8 | 0.10184066 | 5 | 0.73189483 | 7 | 0.00577322 | 110 | 0.68305172 | FALSE | 1 | FALSE | 1 | FALSE | 1 | FALSE | 1 | FALSE | 0.45694206 | FALSE | 1 |
| 1923 | 1923 | 7.16216216 | 2.70648384 | 1 | 0.6184714 | 2 | 0.3375163 | 4 | 0.04241754 | 2 | 0.51200191 | 3 | 0.0222449 | 25 | 0.99721423 | FALSE | 1 | FALSE | 1 | FALSE | 1 | FALSE | 1 | FALSE | 0.74991845 | FALSE | 1 |
| 1924 | 1924 | 9.8 | 1.83303028 | 0 | 1 | 0 | 1 | 1 | 0.30655256 | 0 | 1 | 0 | 1 | 9 | 0.5215547 | FALSE | 1 | FALSE | 1 | FALSE | 1 | FALSE | 1 | FALSE | 1 | FALSE | 1 |
| 1925 | 1925 | 8.60869565 | 1.34290828 | 0 | 1 | 3 | 0.03669128 | 0 | 1 | 0 | 1 | 1 | 0.31672722 | 19 | 0.71542496 | FALSE | 1 | FALSE | 0.94259695 | FALSE | 1 | FALSE | 1 | FALSE | 1 | FALSE | 1 |
| 1926 | 1926 | 9.10526316 | 2.78897712 | 0 | 1 | 0 | 1 | 3 | 0.02898764 | 3 | 0.05383577 | 0 | 1 | 13 | 0.97998676 | FALSE | 1 | FALSE | 1 | FALSE | 1 | FALSE | 1 | FALSE | 1 | FALSE | 1 |
| 1927 | 1927 | 10.3043478 | 3.1268095 | 0 | 1 | 2 | 0.16948764 | 0 | 1 | 0 | 1 | 0 | 1 | 21 | 0.27964072 | FALSE | 1 | FALSE | 1 | FALSE | 1 | FALSE | 1 | FALSE | 1 | FALSE | 1 |
| 1928 | 1928 | 10.3125 | 3.0561158 | 1 | 0.34033054 | 0 | 1 | 0 | 1 | 1 | 0.52895088 | 2 | 0.02752114 | 12 | 0.90922186 | FALSE | 1 | FALSE | 1 | FALSE | 1 | FALSE | 1 | FALSE | 0.79106376 | FALSE | 1 |
| 1929 | 1929 | 7.90566038 | 1.79374333 | 1 | 0.74224579 | 4 | 0.08651261 | 1 | 0.85187773 | 3 | 0.4296776 | 0 | 1 | 43 | 0.71191105 | FALSE | 1 | FALSE | 1 | FALSE | 1 | FALSE | 1 | FALSE | 1 | FALSE | 1 |
| 1930 | 1930 | 7.0625 | 1.91926907 | 1 | 0.34033054 | 2 | 0.09285443 | 1 | 0.44344817 | 2 | 0.16544789 | 0 | 1 | 10 | 0.99304605 | FALSE | 1 | FALSE | 1 | FALSE | 1 | FALSE | 1 | FALSE | 1 | FALSE | 1 |
| 1931 | 1931 | 11.6666667 | 1.33333333 | 0 | 1 | 0 | 1 | 0 | 1 | 2 | 0.06108592 | 0 | 1 | 7 | 0.84574168 | FALSE | 1 | FALSE | 1 | FALSE | 1 | FALSE | 1 | FALSE | 1 | FALSE | 1 |
| 1932 | 1932 | 9.84615385 | 2.47591953 | 1 | 0.28675951 | 0 | 1 | 1 | 0.37874417 | 2 | 0.1174583 | 0 | 1 | 9 | 0.96018168 | FALSE | 1 | FALSE | 1 | FALSE | 1 | FALSE | 1 | FALSE | 1 | FALSE | 1 |
| 1933 | 1933 | 9.78571429 | 1.77999083 | 0 | 1 | 1 | 0.36919824 | 0 | 1 | 0 | 1 | 0 | 1 | 13 | 0.33283729 | FALSE | 1 | FALSE | 1 | FALSE | 1 | FALSE | 1 | FALSE | 1 | FALSE | 1 |
| 1934 | 1934 | 7.65 | 1.54730949 | 2 | 0.4493251 | 3 | 0.2979413 | 3 | 0.35626822 | 1 | 0.93827647 | 4 | 0.01560072 | 46 | 0.93313465 | FALSE | 1 | FALSE | 1 | FALSE | 1 | FALSE | 1 | FALSE | 0.66313193 | FALSE | 1 |
| 1935 | 1935 | 9.31818182 | 2.63518783 | 1 | 0.4357638 | 1 | 0.5154318 | 1 | 0.55340851 | 1 | 0.64497439 | 0 | 1 | 18 | 0.74744813 | FALSE | 1 | FALSE | 1 | FALSE | 1 | FALSE | 1 | FALSE | 1 | FALSE | 1 |
| 1936 | 1936 | 7.18 | 2.58990347 | 1 | 0.71380452 | 1 | 0.79481669 | 3 | 0.24750546 | 1 | 0.89605799 | 1 | 0.54902428 | 41 | 0.51861324 | FALSE | 1 | FALSE | 1 | FALSE | 1 | FALSE | 1 | FALSE | 1 | FALSE | 1 |
| 1937 | 1937 | 6.61151079 | 1.69430188 | 6 | 0.14626155 | 7 | 0.16315538 | 2 | 0.96310807 | 8 | 0.30677177 | 1 | 0.90193101 | 115 | 0.74941026 | FALSE | 1 | FALSE | 1 | FALSE | 1 | FALSE | 1 | FALSE | 1 | FALSE | 1 |
| 1938 | 1938 | 7.87096774 | 3.88331365 | 2 | 0.1881398 | 0 | 1 | 5 | 0.00453596 | 1 | 0.76781284 | 1 | 0.4016888 | 22 | 0.98395924 | FALSE | 1 | FALSE | 1 | FALSE | 1 | FALSE | 1 | FALSE | 1 | FALSE | 1 |
| 1939 | 1939 | 6.72413793 | 1.40556902 | 4 | 0.34086006 | 5 | 0.31511843 | 4 | 0.59743048 | 5 | 0.61387583 | 1 | 0.85304535 | 96 | 0.66343565 | FALSE | 1 | FALSE | 1 | FALSE | 1 | FALSE | 1 | FALSE | 1 | FALSE | 1 |
| 1940 | 1940 | 8.65384615 | 1.96002959 | 0 | 1 | 2 | 0.50543179 | 1 | 0.85187773 | 2 | 0.6970685 | 0 | 1 | 47 | 0.15653036 | FALSE | 1 | FALSE | 1 | FALSE | 1 | FALSE | 1 | FALSE | 1 | FALSE | 1 |
| 1941 | 1941 | 10.875 | 2.49687304 | 1 | 0.34033054 | 0 | 1 | 0 | 1 | 1 | 0.52895088 | 0 | 1 | 14 | 0.53308261 | FALSE | 1 | FALSE | 1 | FALSE | 1 | FALSE | 1 | FALSE | 1 | FALSE | 1 |
| 1942 | 1942 | 6.97222222 | 3.91213292 | 0 | 1 | 0 | 1 | 0 | 1 | 0 | 1 | 1 | 0.44938266 | 35 | 0.01673877 | FALSE | 1 | FALSE | 1 | FALSE | 1 | FALSE | 1 | FALSE | 1 | FALSE | 1 |
| 1943 | 1943 | 9.72093023 | 2.45257891 | 1 | 0.67382914 | 2 | 0.40761584 | 3 | 0.19985581 | 1 | 0.86831187 | 1 | 0.50988734 | 35 | 0.7801651 | FALSE | 1 | FALSE | 1 | FALSE | 1 | FALSE | 1 | FALSE | 1 | FALSE | 1 |
| 1944 | 1944 | 12.25 | 1.23322072 | 0 | 1 | 1 | 0.32623933 | 0 | 1 | 2 | 0.1023796 | 0 | 1 | 9 | 0.89630573 | FALSE | 1 | FALSE | 1 | FALSE | 1 | FALSE | 1 | FALSE | 1 | FALSE | 1 |
| 1945 | 1945 | 8.94444444 | 2.39147147 | 3 | 0.16017108 | 1 | 0.83180787 | 1 | 0.86240917 | 1 | 0.92176876 | 0 | 1 | 48 | 0.23965388 | FALSE | 1 | FALSE | 1 | FALSE | 1 | FALSE | 1 | FALSE | 1 | FALSE | 1 |
| 1946 | 1946 | 9.46341463 | 0.88614157 | 0 | 1 | 2 | 0.38460545 | 2 | 0.43640583 | 1 | 0.85524728 | 0 | 1 | 36 | 0.36455032 | FALSE | 1 | FALSE | 1 | FALSE | 1 | FALSE | 1 | FALSE | 1 | FALSE | 1 |
| 1947 | 1947 | 6.08121331 | 0.76253985 | 23 | 0.78434284 | 29 | 0.80793311 | 37 | 0.50758924 | 49 | 0.39257264 | 17 | 0.51533081 | 867 | 0.35142555 | FALSE | 1 | FALSE | 1 | FALSE | 1 | FALSE | 1 | FALSE | 1 | FALSE | 1 |
| 1948 | 1948 | 5.55319149 | 2.59957861 | 0 | 1 | 4 | 0.06440418 | 4 | 0.08716543 | 2 | 0.64277396 | 0 | 1 | 37 | 0.89489966 | FALSE | 1 | FALSE | 1 | FALSE | 1 | FALSE | 1 | FALSE | 1 | FALSE | 1 |
| 1949 | 1949 | 8.54166667 | 1.82526634 | 0 | 1 | 1 | 0.54637326 | 2 | 0.21260257 | 0 | 1 | 1 | 0.32796862 | 20 | 0.68270729 | FALSE | 1 | FALSE | 1 | FALSE | 1 | FALSE | 1 | FALSE | 1 | FALSE | 1 |
| 1950 | 1950 | 10.4285714 | 2.77010278 | 0 | 1 | 0 | 1 | 0 | 1 | 0 | 1 | 1 | 0.10932414 | 6 | 0.69920685 | FALSE | 1 | FALSE | 1 | FALSE | 1 | FALSE | 1 | FALSE | 1 | FALSE | 1 |
| 1951 | 1951 | 7.5625 | 2.7226385 | 0 | 1 | 1 | 0.78790827 | 3 | 0.23782059 | 3 | 0.36699324 | 0 | 1 | 40 | 0.54368682 | FALSE | 1 | FALSE | 1 | FALSE | 1 | FALSE | 1 | FALSE | 1 | FALSE | 1 |
| 1952 | 1952 | 9.375 | 2.68967935 | 1 | 0.34033054 | 1 | 0.40942991 | 0 | 1 | 1 | 0.52895088 | 0 | 1 | 13 | 0.76799161 | FALSE | 1 | FALSE | 1 | FALSE | 1 | FALSE | 1 | FALSE | 1 | FALSE | 1 |
| 1953 | 1953 | 6.97916667 | 2.22195094 | 2 | 0.34973212 | 1 | 0.79481669 | 0 | 1 | 0 | 1 | 2 | 0.18560062 | 43 | 0.21763056 | FALSE | 1 | FALSE | 1 | FALSE | 1 | FALSE | 1 | FALSE | 1 | FALSE | 1 |
| 1954 | 1954 | 10.7692308 | 1.80400606 | 2 | 0.04234484 | 1 | 0.3480709 | 0 | 1 | 1 | 0.45746571 | 0 | 1 | 9 | 0.96018168 | FALSE | 0.62842194 | FALSE | 1 | FALSE | 1 | FALSE | 1 | FALSE | 1 | FALSE | 1 |
| 1955 | 1955 | 8.6 | 3.90384426 | 0 | 1 | 1 | 0.48239024 | 1 | 0.51939841 | 0 | 1 | 0 | 1 | 18 | 0.37504645 | FALSE | 1 | FALSE | 1 | FALSE | 1 | FALSE | 1 | FALSE | 1 | FALSE | 1 |
| 1956 | 1956 | 3.78346457 | 2.00500413 | 5 | 0.78662995 | 11 | 0.19925706 | 11 | 0.30469648 | 9 | 0.83282503 | 3 | 0.79301509 | 215 | 0.49504207 | FALSE | 1 | FALSE | 1 | FALSE | 1 | FALSE | 1 | FALSE | 1 | FALSE | 1 |
| 1957 | 1957 | 9.85 | 2.47537876 | 0 | 1 | 2 | 0.37295654 | 1 | 0.76953771 | 0 | 1 | 0 | 1 | 37 | 0.10889911 | FALSE | 1 | FALSE | 1 | FALSE | 1 | FALSE | 1 | FALSE | 1 | FALSE | 1 |
| 1958 | 1958 | 9.25 | 2.16506351 | 0 | 1 | 0 | 1 | 0 | 1 | 0 | 1 | 0 | 1 | 12 | 0.13002667 | FALSE | 1 | FALSE | 1 | FALSE | 1 | FALSE | 1 | FALSE | 1 | FALSE | 1 |
| 1959 | 1959 | 7.29166667 | 3.372427 | 2 | 0.34034075 | 3 | 0.19374024 | 1 | 0.82190381 | 1 | 0.89101956 | 1 | 0.54145295 | 39 | 0.69224623 | FALSE | 1 | FALSE | 1 | FALSE | 1 | FALSE | 1 | FALSE | 1 | FALSE | 1 |
| 1960 | 1960 | 9.20588235 | 3.79457362 | 0 | 1 | 0 | 1 | 3 | 0.11365871 | 0 | 1 | 0 | 1 | 30 | 0.21942357 | FALSE | 1 | FALSE | 1 | FALSE | 1 | FALSE | 1 | FALSE | 1 | FALSE | 1 |
| 1961 | 1961 | 9.45454546 | 1.85195898 | 1 | 0.4357638 | 2 | 0.15794013 | 0 | 1 | 0 | 1 | 0 | 1 | 19 | 0.54233419 | FALSE | 1 | FALSE | 1 | FALSE | 1 | FALSE | 1 | FALSE | 1 | FALSE | 1 |
| 1962 | 1962 | 9.05555556 | 3.3245254 | 1 | 0.37380237 | 2 | 0.1135776 | 0 | 1 | 1 | 0.57131013 | 0 | 1 | 14 | 0.86287418 | FALSE | 1 | FALSE | 1 | FALSE | 1 | FALSE | 1 | FALSE | 1 | FALSE | 1 |
| 1963 | 1963 | 2.13333333 | 3.05993049 | 2 | 0.97371239 | 9 | 0.23919345 | 5 | 0.87961572 | 6 | 0.92607714 | 2 | 0.86440123 | 186 | 0.05032176 | FALSE | 1 | FALSE | 1 | FALSE | 1 | FALSE | 1 | FALSE | 1 | FALSE | 1 |
| 1964 | 1964 | 8.44 | 1.66084316 | 2 | 0.83576566 | 5 | 0.3797149 | 7 | 0.16226546 | 4 | 0.83294623 | 0 | 1 | 107 | 0.40944656 | FALSE | 1 | FALSE | 1 | FALSE | 1 | FALSE | 1 | FALSE | 1 | FALSE | 1 |
| 1965 | 1965 | 9.31818182 | 2.80237503 | 0 | 1 | 1 | 0.5154318 | 2 | 0.18613256 | 0 | 1 | 0 | 1 | 19 | 0.54233419 | FALSE | 1 | FALSE | 1 | FALSE | 1 | FALSE | 1 | FALSE | 1 | FALSE | 1 |
| 1966 | 1966 | 10.2666667 | 2.54209013 | 0 | 1 | 1 | 0.38964394 | 1 | 0.42266264 | 0 | 1 | 0 | 1 | 13 | 0.57712805 | FALSE | 1 | FALSE | 1 | FALSE | 1 | FALSE | 1 | FALSE | 1 | FALSE | 1 |
| 1967 | 1967 | 9 | 2.62851496 | 0 | 1 | 0 | 1 | 0 | 1 | 0 | 1 | 0 | 1 | 11 | 0.15414489 | FALSE | 1 | FALSE | 1 | FALSE | 1 | FALSE | 1 | FALSE | 1 | FALSE | 1 |
| 1968 | 1968 | 10.5625 | 2.34437492 | 0 | 1 | 1 | 0.40942991 | 0 | 1 | 1 | 0.52895088 | 1 | 0.23264612 | 13 | 0.76799161 | FALSE | 1 | FALSE | 1 | FALSE | 1 | FALSE | 1 | FALSE | 1 | FALSE | 1 |
| 1969 | 1969 | 8.5 | 2.63899332 | 2 | 0.04234484 | 1 | 0.3480709 | 0 | 1 | 0 | 1 | 0 | 1 | 10 | 0.86791642 | FALSE | 0.62842194 | FALSE | 1 | FALSE | 1 | FALSE | 1 | FALSE | 1 | FALSE | 1 |
| 1970 | 1970 | 12.2083333 | 1.95744661 | 1 | 0.46441868 | 1 | 0.54637326 | 1 | 0.58502114 | 0 | 1 | 2 | 0.058175 | 19 | 0.83975208 | FALSE | 1 | FALSE | 1 | FALSE | 1 | FALSE | 1 | FALSE | 1 | FALSE | 1 |
| 1971 | 1971 | 9.92592593 | 3.52689062 | 0 | 1 | 0 | 1 | 0 | 1 | 0 | 1 | 1 | 0.36060486 | 26 | 0.06079097 | FALSE | 1 | FALSE | 1 | FALSE | 1 | FALSE | 1 | FALSE | 1 | FALSE | 1 |
| 1972 | 1972 | 6.975 | 2.89385124 | 3 | 0.08207756 | 0 | 1 | 1 | 0.76953771 | 4 | 0.10945471 | 2 | 0.13925369 | 30 | 0.96151659 | FALSE | 0.79236883 | FALSE | 1 | FALSE | 1 | FALSE | 1 | FALSE | 1 | FALSE | 1 |

| | | | | | | | | | | | | | | | | | | | | | | | | | | | |
|---|---|---|---|---|---|---|---|---|---|---|---|---|---|---|---|---|---|---|---|---|---|---|---|---|---|---|---|
| 1973 | 1973 | 5.69047619 | 2.85763885 | 1 | 0.66519151 | 5 | 0.01085499 | 1 | 0.78589454 | 3 | 0.30326943 | 2 | 0.15059682 | 30 | 0.99060471 | FALSE | 1 | FALSE | 0.67617265 | FALSE | 1 | FALSE | 1 | FALSE | 1 | FALSE | 1 |
| 1974 | 1974 | 9.65625 | 1.13493874 | 0 | 1 | 2 | 0.27730767 | 2 | 0.32011871 | 2 | 0.43583874 | 1 | 0.41154418 | 25 | 0.88559288 | FALSE | 1 | FALSE | 1 | FALSE | 1 | FALSE | 1 | FALSE | 1 | FALSE | 1 |
| 1975 | 1975 | 9.3 | 2.37907545 | 1 | 0.64722818 | 1 | 0.7326167 | 1 | 0.76953771 | 1 | 0.84823834 | 2 | 0.13925369 | 34 | 0.56328904 | FALSE | 1 | FALSE | 1 | FALSE | 1 | FALSE | 1 | FALSE | 1 | FALSE | 1 |
| 1976 | 1976 | 10.1764706 | 3.18517326 | 0 | 1 | 0 | 1 | 1 | 0.46348836 | 3 | 0.040397 | 0 | 1 | 13 | 0.88719222 | FALSE | 1 | FALSE | 1 | FALSE | 1 | FALSE | 1 | FALSE | 1 | FALSE | 1 |
| 1977 | 1977 | 5.21176471 | 1.94085559 | 1 | 0.89157583 | 0 | 1 | 3 | 0.59435517 | 3 | 0.75590957 | 0 | 1 | 78 | 0.03389538 | FALSE | 1 | FALSE | 1 | FALSE | 1 | FALSE | 1 | FALSE | 1 | FALSE | 1 |
| 1978 | 1978 | 7.79166667 | 2.9857881 | 0 | 1 | 2 | 0.18117183 | 0 | 1 | 1 | 0.67692816 | 1 | 0.32796862 | 20 | 0.68270729 | FALSE | 1 | FALSE | 1 | FALSE | 1 | FALSE | 1 | FALSE | 1 | FALSE | 1 |
| 1979 | 1979 | 7.45652174 | 3.60980863 | 3 | 0.11312654 | 2 | 0.44132825 | 1 | 0.81522035 | 3 | 0.35428318 | 4 | 0.00657628 | 33 | 0.99158534 | FALSE | 0.92488288 | FALSE | 1 | FALSE | 1 | FALSE | 1 | FALSE | 0.47461742 | FALSE | 1 |
| 1980 | 1980 | 11.6470588 | 2.31214635 | 4 | 0.01061864 | 2 | 0.30150529 | 2 | 0.34668321 | 0 | 1 | 0 | 1 | 26 | 0.92796052 | FALSE | 0.31290615 | FALSE | 1 | FALSE | 1 | FALSE | 1 | FALSE | 1 | FALSE | 1 |
| 1981 | 1981 | 9.85714286 | 1.35526185 | 0 | 1 | 1 | 0.20567256 | 0 | 1 | 0 | 1 | 0 | 1 | 6 | 0.69920685 | FALSE | 1 | FALSE | 1 | FALSE | 1 | FALSE | 1 | FALSE | 1 | FALSE | 1 |
| 1982 | 1982 | 8.66666667 | 2.49443826 | 0 | 1 | 0 | 1 | 0 | 1 | 0 | 1 | 0 | 1 | 15 | 0.07803154 | FALSE | 1 | FALSE | 1 | FALSE | 1 | FALSE | 1 | FALSE | 1 | FALSE | 1 |
| 1983 | 1983 | 10.1666667 | 2.76385399 | 0 | 1 | 0 | 1 | 0 | 1 | 0 | 1 | 1 | 0.18007762 | 11 | 0.41944532 | FALSE | 1 | FALSE | 1 | FALSE | 1 | FALSE | 1 | FALSE | 1 | FALSE | 1 |
| 1984 | 1984 | 11.0625 | 1.61897923 | 3 | 0.22550017 | 3 | 0.34261611 | 2 | 0.67580769 | 1 | 0.95130976 | 3 | 0.08717933 | 52 | 0.80965143 | FALSE | 1 | FALSE | 1 | FALSE | 1 | FALSE | 1 | FALSE | 1 | FALSE | 1 |
| 1985 | 1985 | 9.72 | 1.82252572 | 1 | 0.47819878 | 1 | 0.56109832 | 0 | 1 | 1 | 0.69181315 | 1 | 0.33902672 | 21 | 0.64957167 | FALSE | 1 | FALSE | 1 | FALSE | 1 | FALSE | 1 | FALSE | 1 | FALSE | 1 |
| 1986 | 1986 | 10.952381 | 2.05811265 | 0 | 1 | 0 | 1 | 0 | 1 | 0 | 1 | 0 | 1 | 21 | 0.02808151 | FALSE | 1 | FALSE | 1 | FALSE | 1 | FALSE | 1 | FALSE | 1 | FALSE | 1 |
| 1987 | 1987 | 11 | 2.26778684 | 0 | 1 | 0 | 1 | 1 | 0.22601089 | 0 | 1 | 0 | 1 | 6 | 0.69920685 | FALSE | 1 | FALSE | 1 | FALSE | 1 | FALSE | 1 | FALSE | 1 | FALSE | 1 |
| 1988 | 1988 | 9.625 | 0.69597055 | 0 | 1 | 1 | 0.23139161 | 1 | 0.25384309 | 0 | 1 | 0 | 1 | 6 | 0.88425551 | FALSE | 1 | FALSE | 1 | FALSE | 1 | FALSE | 1 | FALSE | 1 | FALSE | 1 |
| 1989 | 1989 | 8.31578947 | 1.68749679 | 0 | 1 | 0 | 1 | 0 | 1 | 0 | 1 | 1 | 0.26986775 | 18 | 0.17880694 | FALSE | 1 | FALSE | 1 | FALSE | 1 | FALSE | 1 | FALSE | 1 | FALSE | 1 |
| 1990 | 1990 | 9.88888889 | 3.91420332 | 0 | 1 | 0 | 1 | 1 | 0.48280983 | 0 | 1 | 0 | 1 | 17 | 0.20329205 | FALSE | 1 | FALSE | 1 | FALSE | 1 | FALSE | 1 | FALSE | 1 | FALSE | 1 |
| 1991 | 1991 | 6.42857143 | 3.13517361 | 2 | 0.58475401 | 2 | 0.71077581 | 0 | 1 | 3 | 0.68517679 | 4 | 0.03564889 | 65 | 0.46718352 | FALSE | 1 | FALSE | 1 | FALSE | 1 | FALSE | 1 | FALSE | 0.87669812 | FALSE | 1 |
| 1992 | 1992 | 10.9285714 | 2.57638411 | 2 | 0.04858661 | 0 | 1 | 1 | 0.40110418 | 2 | 0.13304705 | 1 | 0.20679328 | 8 | 0.99722299 | FALSE | 0.649301 | FALSE | 1 | FALSE | 1 | FALSE | 1 | FALSE | 1 | FALSE | 1 |
| 1993 | 1993 | 10.1724138 | 2.27533954 | 3 | 0.03713651 | 0 | 1 | 1 | 0.65463982 | 2 | 0.3871811 | 1 | 0.38148475 | 22 | 0.92902102 | FALSE | 0.58392 | FALSE | 1 | FALSE | 1 | FALSE | 1 | FALSE | 1 | FALSE | 1 |
| 1994 | 1994 | 7.63157895 | 1.69241423 | 0 | 1 | 1 | 0.46503795 | 1 | 0.50143826 | 0 | 1 | 0 | 1 | 17 | 0.41141543 | FALSE | 1 | FALSE | 1 | FALSE | 1 | FALSE | 1 | FALSE | 1 | FALSE | 1 |
| 1995 | 1995 | 8.83333333 | 0.68718427 | 0 | 1 | 0 | 1 | 0 | 1 | 0 | 1 | 0 | 1 | 6 | 0.36077158 | FALSE | 1 | FALSE | 1 | FALSE | 1 | FALSE | 1 | FALSE | 1 | FALSE | 1 |
| 1996 | 1996 | 9.72222222 | 2.37593386 | 2 | 0.07645454 | 0 | 1 | 0 | 1 | 1 | 0.57131013 | 0 | 1 | 15 | 0.69372602 | FALSE | 0.77424195 | FALSE | 1 | FALSE | 1 | FALSE | 1 | FALSE | 1 | FALSE | 1 |
| 1997 | 1997 | 11.4117647 | 2.03090312 | 1 | 0.57647348 | 2 | 0.28941836 | 3 | 0.11365871 | 4 | 0.06258966 | 0 | 1 | 23 | 0.99096587 | FALSE | 1 | FALSE | 1 | FALSE | 1 | FALSE | 1 | FALSE | 1 | FALSE | 1 |
| 1998 | 1998 | 9.66666667 | 0.9860133 | 0 | 1 | 0 | 1 | 1 | 0.58502114 | 2 | 0.30249307 | 2 | 0.058175 | 19 | 0.83975208 | FALSE | 1 | FALSE | 1 | FALSE | 1 | FALSE | 1 | FALSE | 1 | FALSE | 1 |
| 1999 | 1999 | 9.92857143 | 2.63124864 | 2 | 0.04858661 | 0 | 1 | 0 | 1 | 1 | 0.48242099 | 1 | 0.20679328 | 10 | 0.94579623 | FALSE | 0.649301 | FALSE | 1 | FALSE | 1 | FALSE | 1 | FALSE | 1 | FALSE | 1 |
| 2000 | 2000 | 8.72 | 2.42520102 | 0 | 1 | 2 | 0.19297191 | 2 | 0.22597287 | 0 | 1 | 1 | 0.33902672 | 20 | 0.8152498 | FALSE | 1 | FALSE | 1 | FALSE | 1 | FALSE | 1 | FALSE | 1 | FALSE | 1 |
| 2001 | 2001 | 10.4761905 | 1.76254815 | 0 | 1 | 1 | 0.49918218 | 0 | 1 | 0 | 1 | 0 | 1 | 20 | 0.13763971 | FALSE | 1 | FALSE | 1 | FALSE | 1 | FALSE | 1 | FALSE | 1 | FALSE | 1 |
| 2002 | 2002 | 6.88888889 | 2.88461222 | 3 | 0.01017712 | 2 | 0.1135776 | 0 | 1 | 2 | 0.19902682 | 0 | 1 | 11 | 0.99651304 | FALSE | 0.30467829 | FALSE | 1 | FALSE | 1 | FALSE | 1 | FALSE | 1 | FALSE | 1 |
| 2003 | 2003 | 9.25 | 2.04633819 | 1 | 0.40558521 | 0 | 1 | 0 | 1 | 2 | 0.23331368 | 0 | 1 | 17 | 0.61750632 | FALSE | 1 | FALSE | 1 | FALSE | 1 | FALSE | 1 | FALSE | 1 | FALSE | 1 |
| 2004 | 2004 | 4.33606557 | 3.10897868 | 2 | 0.82554509 | 4 | 0.56068985 | 6 | 0.27416206 | 9 | 0.10739474 | 0 | 1 | 101 | 0.73627522 | FALSE | 1 | FALSE | 1 | FALSE | 1 | FALSE | 1 | FALSE | 1 | FALSE | 1 |
| 2005 | 2005 | 7.04878049 | 3.52656275 | 0 | 1 | 2 | 0.38460545 | 5 | 0.01496777 | 0 | 1 | 3 | 0.02912063 | 31 | 0.95455175 | FALSE | 1 | FALSE | 1 | FALSE | 1 | FALSE | 1 | FALSE | 0.81978088 | FALSE | 1 |
| 2006 | 2006 | 10.2777778 | 2.95908311 | 2 | 0.07645454 | 1 | 0.44710673 | 0 | 1 | 0 | 1 | 0 | 1 | 15 | 0.69372602 | FALSE | 0.77424195 | FALSE | 1 | FALSE | 1 | FALSE | 1 | FALSE | 1 | FALSE | 1 |
| 2007 | 2007 | 8.25 | 3.15803963 | 0 | 1 | 1 | 0.58913618 | 1 | 0.62830986 | 2 | 0.35374209 | 1 | 0.36060486 | 22 | 0.76237877 | FALSE | 1 | FALSE | 1 | FALSE | 1 | FALSE | 1 | FALSE | 1 | FALSE | 1 |
| 2008 | 2008 | 8.5625 | 2.09071609 | 1 | 0.67382914 | 0 | 1 | 4 | 0.06724173 | 3 | 0.31602175 | 0 | 1 | 35 | 0.7801651 | FALSE | 1 | FALSE | 1 | FALSE | 1 | FALSE | 1 | FALSE | 1 | FALSE | 1 |
| 2009 | 2009 | 6.92307692 | 4.1779925 | 0 | 1 | 0 | 1 | 0 | 1 | 0 | 1 | 0 | 1 | 26 | 0.0119733 | FALSE | 1 | FALSE | 1 | FALSE | 1 | FALSE | 1 | FALSE | 1 | FALSE | 1 |
| 2010 | 2010 | 8.27586207 | 2.81494229 | 0 | 1 | 0 | 1 | 1 | 0.65463982 | 1 | 0.74482229 | 0 | 1 | 27 | 0.14664438 | FALSE | 1 | FALSE | 1 | FALSE | 1 | FALSE | 1 | FALSE | 1 | FALSE | 1 |
| 2011 | 2011 | 11.3846154 | 1.21138583 | 2 | 0.14250978 | 0 | 1 | 2 | 0.23940226 | 3 | 0.11458571 | 1 | 0.3499045 | 18 | 0.98644281 | FALSE | 1 | FALSE | 1 | FALSE | 1 | FALSE | 1 | FALSE | 1 | FALSE | 1 |
| 2012 | 2012 | 9.5 | 3.58236421 | 0 | 1 | 0 | 1 | 0 | 1 | 1 | 0.67692816 | 0 | 1 | 23 | 0.09196389 | FALSE | 1 | FALSE | 1 | FALSE | 1 | FALSE | 1 | FALSE | 1 | FALSE | 1 |
| 2013 | 2013 | 11 | 2 | 0 | 1 | 1 | 0.20567256 | 1 | 0.22601089 | 0 | 1 | 0 | 1 | 5 | 0.91847773 | FALSE | 1 | FALSE | 1 | FALSE | 1 | FALSE | 1 | FALSE | 1 | FALSE | 1 |
| 2014 | 2014 | 9.8125 | 1.91111061 | 1 | 0.55378205 | 1 | 0.63997468 | 2 | 0.30673593 | 2 | 0.41984187 | 2 | 0.09110253 | 23 | 0.95750806 | FALSE | 1 | FALSE | 1 | FALSE | 1 | FALSE | 1 | FALSE | 1 | FALSE | 1 |
| 2015 | 2015 | 9.5483871 | 2.56324781 | 1 | 0.55378205 | 2 | 0.2651851 | 1 | 0.67911179 | 1 | 0.76781284 | 0 | 1 | 26 | 0.64660109 | FALSE | 1 | FALSE | 1 | FALSE | 1 | FALSE | 1 | FALSE | 1 | FALSE | 1 |
| 2016 | 2016 | 9.33333333 | 2.51661148 | 0 | 1 | 2 | 0.1135776 | 0 | 1 | 0 | 1 | 0 | 1 | 16 | 0.44998001 | FALSE | 1 | FALSE | 1 | FALSE | 1 | FALSE | 1 | FALSE | 1 | FALSE | 1 |
| 2017 | 2017 | 9 | 2.28035085 | 0 | 1 | 0 | 1 | 0 | 1 | 0 | 1 | 1 | 0.15247062 | 9 | 0.5215547 | FALSE | 1 | FALSE | 1 | FALSE | 1 | FALSE | 1 | FALSE | 1 | FALSE | 1 |
| 2018 | 2018 | 9.13333333 | 2.30554886 | 0 | 1 | 2 | 0.08295839 | 0 | 1 | 1 | 0.50623194 | 2 | 0.02433796 | 10 | 0.97980046 | FALSE | 1 | FALSE | 1 | FALSE | 1 | FALSE | 1 | FALSE | 0.77578444 | FALSE | 1 |
| 2019 | 2019 | 9.34782609 | 2.52997157 | 1 | 0.4357638 | 1 | 0.5154318 | 0 | 1 | 0 | 1 | 0 | 1 | 20 | 0.30915202 | FALSE | 1 | FALSE | 1 | FALSE | 1 | FALSE | 1 | FALSE | 1 | FALSE | 1 |
| 2020 | 2020 | 5.84745763 | 1.90283962 | 2 | 0.4493251 | 2 | 0.57395594 | 3 | 0.35626822 | 2 | 0.76144533 | 1 | 0.62455813 | 49 | 0.68931377 | FALSE | 1 | FALSE | 1 | FALSE | 1 | FALSE | 1 | FALSE | 1 | FALSE | 1 |
| 2021 | 2021 | 6.93495935 | 1.82903341 | 5 | 0.20317587 | 3 | 0.76134863 | 4 | 0.64412798 | 3 | 0.92431696 | 3 | 0.32335088 | 104 | 0.45516479 | FALSE | 1 | FALSE | 1 | FALSE | 1 | FALSE | 1 | FALSE | 1 | FALSE | 1 |
| 2022 | 2022 | 9.5 | 2.26936114 | 2 | 0.09187136 | 2 | 0.1353446 | 1 | 0.51939841 | 1 | 0.60987146 | 0 | 1 | 14 | 0.97325593 | FALSE | 0.83248965 | FALSE | 1 | FALSE | 1 | FALSE | 1 | FALSE | 1 | FALSE | 1 |
| 2023 | 2023 | 7.94736842 | 2.62525556 | 0 | 1 | 1 | 0.46503795 | 0 | 1 | 0 | 1 | 1 | 0.26986775 | 17 | 0.41141543 | FALSE | 1 | FALSE | 1 | FALSE | 1 | FALSE | 1 | FALSE | 1 | FALSE | 1 |
| 2024 | 2024 | 9.6875 | 1.60929915 | 3 | 0.00725145 | 1 | 0.40942991 | 1 | 0.44344817 | 1 | 0.52895088 | 0 | 1 | 10 | 0.99304605 | FALSE | 0.25171133 | FALSE | 1 | FALSE | 1 | FALSE | 1 | FALSE | 1 | FALSE | 1 |
| 2025 | 2025 | 11.375 | 2.0577597 | 1 | 0.18769745 | 0 | 1 | 0 | 1 | 0 | 1 | 0 | 1 | 7 | 0.63752397 | FALSE | 1 | FALSE | 1 | FALSE | 1 | FALSE | 1 | FALSE | 1 | FALSE | 1 |
| 2026 | 2026 | 7.6 | 3.23728281 | 1 | 0.47819878 | 0 | 1 | 2 | 0.22597287 | 1 | 0.69181315 | 1 | 0.33902672 | 20 | 0.8152498 | FALSE | 1 | FALSE | 1 | FALSE | 1 | FALSE | 1 | FALSE | 1 | FALSE | 1 |
| 2027 | 2027 | 10.6 | 2.07846097 | 0 | 1 | 2 | 0.19297191 | 0 | 1 | 1 | 0.69181315 | 2 | 0.06257402 | 20 | 0.8152498 | FALSE | 1 | FALSE | 1 | FALSE | 1 | FALSE | 1 | FALSE | 1 | FALSE | 1 |
| 2028 | 2028 | 7.73529412 | 2.06239187 | 3 | 0.14789018 | 3 | 0.23623106 | 1 | 0.85187773 | 5 | 0.08873102 | 0 | 1 | 40 | 0.94722426 | FALSE | 1 | FALSE | 1 | FALSE | 1 | FALSE | 1 | FALSE | 1 | FALSE | 1 |
| 2029 | 2029 | 9.9 | 3.11287648 | 0 | 1 | 0 | 1 | 1 | 0.30655256 | 1 | 0.37517262 | 1 | 0.15247062 | 7 | 0.94326215 | FALSE | 1 | FALSE | 1 | FALSE | 1 | FALSE | 1 | FALSE | 1 | FALSE | 1 |
| 2030 | 2030 | 11.0769231 | 2.33572715 | 0 | 1 | 0 | 1 | 1 | 0.37874417 | 1 | 0.45746571 | 1 | 0.19354507 | 10 | 0.86791642 | FALSE | 1 | FALSE | 1 | FALSE | 1 | FALSE | 1 | FALSE | 1 | FALSE | 1 |
| 2031 | 2031 | 11.8571429 | 2.35606036 | 2 | 0.04234484 | 0 | 1 | 1 | 0.37874417 | 0 | 1 | 0 | 1 | 10 | 0.86791642 | FALSE | 0.62842194 | FALSE | 1 | FALSE | 1 | FALSE | 1 | FALSE | 1 | FALSE | 1 |
| 2032 | 2032 | 10.2 | 2.46170673 | 0 | 1 | 2 | 0.1353446 | 0 | 1 | 2 | 0.23331368 | 1 | 0.28187291 | 15 | 0.92113679 | FALSE | 1 | FALSE | 1 | FALSE | 1 | FALSE | 1 | FALSE | 1 | FALSE | 1 |
| 2033 | 2033 | 8.85714286 | 2.35606036 | 0 | 1 | 0 | 1 | 0 | 1 | 1 | 0.28044368 | 1 | 0.10932414 | 5 | 0.91847773 | FALSE | 1 | FALSE | 1 | FALSE | 1 | FALSE | 1 | FALSE | 1 | FALSE | 1 |
| 2034 | 2034 | 9.33333333 | 2.24433443 | 3 | 0.0279333 | 0 | 1 | 2 | 0.23940226 | 2 | 0.33678042 | 0 | 1 | 19 | 0.96054244 | FALSE | 0.50964275 | FALSE | 1 | FALSE | 1 | FALSE | 1 | FALSE | 1 | FALSE | 1 |
| 2035 | 2035 | 8.46153846 | 2.24003381 | 0 | 1 | 3 | 0.13080858 | 0 | 1 | 0 | 1 | 0 | 1 | 36 | 0.12091774 | FALSE | 1 | FALSE | 1 | FALSE | 1 | FALSE | 1 | FALSE | 1 | FALSE | 1 |
| 2036 | 2036 | 8.77272727 | 2.95314652 | 0 | 1 | 1 | 0.5154318 | 2 | 0.18613256 | 2 | 0.26791271 | 0 | 1 | 17 | 0.8840357 | FALSE | 1 | FALSE | 1 | FALSE | 1 | FALSE | 1 | FALSE | 1 | FALSE | 1 |
| 2037 | 2037 | 10.25 | 2.58602011 | 1 | 0.0986934 | 0 | 1 | 0 | 1 | 0 | 1 | 0 | 1 | 3 | 0.8823539 | FALSE | 0.86047225 | FALSE | 1 | FALSE | 1 | FALSE | 1 | FALSE | 1 | FALSE | 1 |
| 2038 | 2038 | 9.76190476 | 3.40734168 | 0 | 1 | 1 | 0.49918218 | 0 | 1 | 1 | 0.62783522 | 0 | 1 | 19 | 0.34094959 | FALSE | 1 | FALSE | 1 | FALSE | 1 | FALSE | 1 | FALSE | 1 | FALSE | 1 |
| 2039 | 2039 | 9.38095238 | 2.4780213 | 0 | 1 | 4 | 0.00413111 | 2 | 0.17308361 | 0 | 1 | 1 | 0.29368245 | 14 | 0.98949398 | FALSE | 1 | FALSE | 0.58120789 | FALSE | 1 | FALSE | 1 | FALSE | 1 | FALSE | 1 |
| 2040 | 2040 | 9.95454546 | 3.57360618 | 1 | 0.40558521 | 2 | 0.1353446 | 1 | 0.51939841 | 2 | 0.23331368 | 0 | 1 | 14 | 0.97325593 | FALSE | 1 | FALSE | 1 | FALSE | 1 | FALSE | 1 | FALSE | 1 | FALSE | 1 |
| 2041 | 2041 | 9.31578947 | 2.47200391 | 2 | 0.08405377 | 1 | 0.46503795 | 1 | 0.50143826 | 1 | 0.59104357 | 0 | 1 | 14 | 0.93677253 | FALSE | 0.79236883 | FALSE | 1 | FALSE | 1 | FALSE | 1 | FALSE | 1 | FALSE | 1 |
| 2042 | 2042 | 7.64583333 | 1.62646835 | 3 | 0.44855528 | 3 | 0.6055623 | 1 | 0.97091514 | 5 | 0.45253098 | 1 | 0.79779856 | 83 | 0.3457435 | FALSE | 1 | FALSE | 1 | FALSE | 1 | FALSE | 1 | FALSE | 1 | FALSE | 1 |
| 2043 | 2043 | 8.15151515 | 2.21730402 | 2 | 0.19750261 | 1 | 0.65167362 | 2 | 0.32011871 | 1 | 0.77852168 | 1 | 0.41154418 | 25 | 0.88559288 | FALSE | 1 | FALSE | 1 | FALSE | 1 | FALSE | 1 | FALSE | 1 | FALSE | 1 |
| 2044 | 2044 | 10.75 | 2.53722289 | 0 | 1 | 0 | 1 | 0 | 1 | 1 | 0.31351192 | 0 | 1 | 7 | 0.63752397 | FALSE | 1 | FALSE | 1 | FALSE | 1 | FALSE | 1 | FALSE | 1 | FALSE | 1 |
| 2045 | 2045 | 8.8 | 2.11187121 | 1 | 0.40558521 | 0 | 1 | 2 | 0.16019548 | 0 | 1 | 0 | 1 | 17 | 0.61750632 | FALSE | 1 | FALSE | 1 | FALSE | 1 | FALSE | 1 | FALSE | 1 | FALSE | 1 |
| 2046 | 2046 | 8.30555556 | 1.56026074 | 1 | 0.60837712 | 4 | 0.02790717 | 2 | 0.37289966 | 0 | 1 | 0 | 1 | 29 | 0.81028289 | FALSE | 1 | FALSE | 0.87929047 | FALSE | 1 | FALSE | 1 | FALSE | 1 | FALSE | 1 |
| 2047 | 2047 | 7.6 | 2.00997512 | 1 | 0.6184714 | 0 | 1 | 1 | 0.74264075 | 0 | 1 | 1 | 0.45845922 | 34 | 0.14843263 | FALSE | 1 | FALSE | 1 | FALSE | 1 | FALSE | 1 | FALSE | 1 | FALSE | 1 |
| 2048 | 2048 | 11.8181818 | 1.79990817 | 0 | 1 | 1 | 0.30368014 | 0 | 1 | 2 | 0.08789566 | 0 | 1 | 8 | 0.92152993 | FALSE | 1 | FALSE | 1 | FALSE | 1 | FALSE | 1 | FALSE | 1 | FALSE | 1 |
| 2049 | 2049 | 1.20477816 | 1.32883482 | 52 | 0.12054998 | 52 | 0.78456043 | 67 | 0.28711067 | 89 | 0.13675598 | 30 | 0.42065269 | 1458 | 0.90883495 | FALSE | 0.95622564 | FALSE | 1 | FALSE | 1 | FALSE | 1 | FALSE | 1 | FALSE | 1 |
| 2050 | 2050 | 9.31578947 | 2.49320406 | 1 | 0.38989951 | 1 | 0.46503795 | 0 | 1 | 2 | 0.21610812 | 0 | 1 | 15 | 0.83648191 | FALSE | 1 | FALSE | 1 | FALSE | 1 | FALSE | 1 | FALSE | 1 | FALSE | 1 |
| 2051 | 2051 | 7.5 | 1.9009567 | 0 | 1 | 0 | 1 | 0 | 1 | 2 | 0.26791271 | 0 | 1 | 20 | 0.30915202 | FALSE | 1 | FALSE | 1 | FALSE | 1 | FALSE | 1 | FALSE | 1 | FALSE | 1 |
| 2052 | 2052 | 9.76923077 | 2.51621957 | 0 | 1 | 1 | 0.3480709 | 1 | 0.37874417 | 1 | 0.45746571 | 0 | 1 | 10 | 0.86791642 | FALSE | 1 | FALSE | 1 | FALSE | 1 | FALSE | 1 | FALSE | 1 | FALSE | 1 |
| 2053 | 2053 | 7.3442623 | 2.97467588 | 1 | 0.79637728 | 2 | 0.59222924 | 1 | 0.89373158 | 4 | 0.30679865 | 2 | 0.26425795 | 51 | 0.64764523 | FALSE | 1 | FALSE | 1 | FALSE | 1 | FALSE | 1 | FALSE | 1 | FALSE | 1 |
| 2054 | 2054 | 10.4666667 | 3.05213951 | 0 | 1 | 0 | 1 | 0 | 1 | 1 | 0.50623194 | 0 | 1 | 14 | 0.29525409 | FALSE | 1 | FALSE | 1 | FALSE | 1 | FALSE | 1 | FALSE | 1 | FALSE | 1 |
| 2055 | 2055 | 8.90476191 | 2.44763758 | 0 | 1 | 0 | 1 | 1 | 0.53671415 | 1 | 0.62783522 | 0 | 1 | 19 | 0.34094959 | FALSE | 1 | FALSE | 1 | FALSE | 1 | FALSE | 1 | FALSE | 1 | FALSE | 1 |
| 2056 | 2056 | 6.88764045 | 1.83252881 | 6 | 0.02624524 | 1 | 0.9474474 | 2 | 0.83551221 | 5 | 0.38834525 | 2 | 0.43073682 | 73 | 0.78154321 | FALSE | 0.50496189 | FALSE | 1 | FALSE | 1 | FALSE | 1 | FALSE | 1 | FALSE | 1 |
| 2057 | 2057 | 6.92647059 | 3.10273208 | 2 | 0.52416586 | 5 | 0.06823576 | 3 | 0.44404438 | 2 | 0.82657478 | 1 | 0.6769215 | 55 | 0.83409155 | FALSE | 1 | FALSE | 1 | FALSE | 1 | FALSE | 1 | FALSE | 1 | FALSE | 1 |
| 2058 | 2058 | 9.08333333 | 3.22641011 | 0 | 1 | 0 | 1 | 0 | 1 | 0 | 1 | 0 | 1 | 12 | 0.13002667 | FALSE | 1 | FALSE | 1 | FALSE | 1 | FALSE | 1 | FALSE | 1 | FALSE | 1 |
| 2059 | 2059 | 8.93333333 | 2.54209013 | 1 | 0.54198714 | 2 | 0.25306331 | 1 | 0.66709967 | 4 | 0.04659941 | 2 | 0.08611595 | 20 | 0.99601577 | FALSE | 1 | FALSE | 1 | FALSE | 1 | FALSE | 1 | FALSE | 1 | FALSE | 1 |
| 2060 | 2060 | 8.4 | 2.05912603 | 0 | 1 | 0 | 1 | 0 | 1 | 0 | 1 | 0 | 1 | 10 | 0.18273166 | FALSE | 1 | FALSE | 1 | FALSE | 1 | FALSE | 1 | FALSE | 1 | FALSE | 1 |
| 2061 | 2061 | 7.91304348 | 2.48321397 | 0 | 1 | 2 | 0.16948764 | 0 | 1 | 1 | 0.6613267 | 1 | 0.31672722 | 19 | 0.71542496 | FALSE | 1 | FALSE | 1 | FALSE | 1 | FALSE | 1 | FALSE | 1 | FALSE | 1 |
| 2062 | 2062 | 8.79166667 | 2.66112706 | 0 | 1 | 1 | 0.53115654 | 0 | 1 | 1 | 0.6613267 | 0 | 1 | 21 | 0.27964072 | FALSE | 1 | FALSE | 1 | FALSE | 1 | FALSE | 1 | FALSE | 1 | FALSE | 1 |
| 2063 | 2063 | 10.8571429 | 2.4453204 | 0 | 1 | 1 | 0.36919824 | 1 | 0.40110418 | 0 | 1 | 1 | 0.20679328 | 11 | 0.83676449 | FALSE | 1 | FALSE | 1 | FALSE | 1 | FALSE | 1 | FALSE | 1 | FALSE | 1 |
| 2064 | 2064 | 10.7333333 | 3.2550815 | 0 | 1 | 0 | 1 | 0 | 1 | 0 | 1 | 0 | 1 | 15 | 0.07803154 | FALSE | 1 | FALSE | 1 | FALSE | 1 | FALSE | 1 | FALSE | 1 | FALSE | 1 |
| 2065 | 2065 | 11.0833333 | 3.70716274 | 0 | 1 | 0 | 1 | 0 | 1 | 0 | 1 | 1 | 0.18007762 | 11 | 0.41944532 | FALSE | 1 | FALSE | 1 | FALSE | 1 | FALSE | 1 | FALSE | 1 | FALSE | 1 |
| 2066 | 2066 | 8.15555556 | 2.58160635 | 1 | 0.68224519 | 0 | 1 | 2 | 0.47281647 | 2 | 0.60667406 | 0 | 1 | 39 | 0.29524127 | FALSE | 1 | FALSE | 1 | FALSE | 1 | FALSE | 1 | FALSE | 1 | FALSE | 1 |

| | | | | | | | | | | | | | | | | | | | | | | | | | | | |
|---|---|---|---|---|---|---|---|---|---|---|---|---|---|---|---|---|---|---|---|---|---|---|---|---|---|---|---|
| 2067 | 2067 | 7.98305085 | 2.02100433 | 2 | 0.4493251 | 0 | 1 | 2 | 0.63179508 | 2 | 0.76144533 | 2 | 0.25204357 | 51 | 0.41341988 | FALSE | 1 | FALSE | 1 | FALSE | 1 | FALSE | 1 | FALSE | 1 | FALSE | 1 |
| 2068 | 2068 | 9.83333333 | 2.49109525 | 0 | 1 | 2 | 0.25306331 | 1 | 0.66709967 | 0 | 1 | 2 | 0.08611595 | 25 | 0.67630471 | FALSE | 1 | FALSE | 1 | FALSE | 1 | FALSE | 1 | FALSE | 1 | FALSE | 1 |
| 2069 | 2069 | 9.33333333 | 2.40370085 | 1 | 0.20854748 | 0 | 1 | 0 | 1 | 0 | 1 | 0 | 1 | 8 | 0.5780332 | FALSE | 1 | FALSE | 1 | FALSE | 1 | FALSE | 1 | FALSE | 1 | FALSE | 1 |
| 2070 | 2070 | 9.7037037 | 2.03332996 | 0 | 1 | 0 | 1 | 1 | 0.62830986 | 1 | 0.71956339 | 1 | 0.36060486 | 24 | 0.3725373 | FALSE | 1 | FALSE | 1 | FALSE | 1 | FALSE | 1 | FALSE | 1 | FALSE | 1 |
| 2071 | 2071 | 9.82608696 | 1.57844651 | 0 | 1 | 0 | 1 | 0 | 1 | 2 | 0.2852236 | 1 | 0.31672722 | 20 | 0.50589402 | FALSE | 1 | FALSE | 1 | FALSE | 1 | FALSE | 1 | FALSE | 1 | FALSE | 1 |
| 2072 | 2072 | 10.2857143 | 2.76272566 | 0 | 1 | 0 | 1 | 0 | 1 | 1 | 0.28044368 | 0 | 1 | 6 | 0.69920685 | FALSE | 1 | FALSE | 1 | FALSE | 1 | FALSE | 1 | FALSE | 1 | FALSE | 1 |
| 2073 | 2073 | 9.12121212 | 3.5484318 | 0 | 1 | 1 | 0.6629941 | 1 | 0.70185639 | 3 | 0.19161876 | 0 | 1 | 28 | 0.58681477 | FALSE | 1 | FALSE | 1 | FALSE | 1 | FALSE | 1 | FALSE | 1 | FALSE | 1 |
| 2074 | 2074 | 8.47826087 | 2.91725847 | 1 | 0.45027683 | 1 | 0.53115654 | 1 | 0.5695037 | 0 | 1 | 2 | 0.05389003 | 18 | 0.86273155 | FALSE | 1 | FALSE | 1 | FALSE | 1 | FALSE | 1 | FALSE | 1 | FALSE | 1 |
| 2075 | 2075 | 8 | 2.45967478 | 3 | 0.08207756 | 3 | 0.13823565 | 2 | 0.42396974 | 1 | 0.84823834 | 1 | 0.48480878 | 30 | 0.96151659 | FALSE | 0.79236883 | FALSE | 1 | FALSE | 1 | FALSE | 1 | FALSE | 1 | FALSE | 1 |
| 2076 | 2076 | 9.8125 | 1.81034355 | 1 | 0.34033054 | 1 | 0.40942991 | 1 | 0.44344817 | 0 | 1 | 0 | 1 | 13 | 0.76799161 | FALSE | 1 | FALSE | 1 | FALSE | 1 | FALSE | 1 | FALSE | 1 | FALSE | 1 |
| 2077 | 2077 | 8.14583333 | 2.96500129 | 1 | 0.71380452 | 2 | 0.46321244 | 3 | 0.24750546 | 4 | 0.17698522 | 1 | 0.54902428 | 37 | 0.93870662 | FALSE | 1 | FALSE | 1 | FALSE | 1 | FALSE | 1 | FALSE | 1 | FALSE | 1 |
| 2078 | 2078 | 8.61538462 | 3.07692308 | 0 | 1 | 0 | 1 | 0 | 1 | 0 | 1 | 0 | 1 | 13 | 0.10967906 | FALSE | 1 | FALSE | 1 | FALSE | 1 | FALSE | 1 | FALSE | 1 | FALSE | 1 |
| 2079 | 2079 | 9.57894737 | 3.85685704 | 0 | 1 | 0 | 1 | 1 | 0.50143826 | 1 | 0.59104357 | 1 | 0.26986775 | 16 | 0.65565778 | FALSE | 1 | FALSE | 1 | FALSE | 1 | FALSE | 1 | FALSE | 1 | FALSE | 1 |
| 2080 | 2080 | 9.10344828 | 3.26258653 | 1 | 0.52988231 | 2 | 0.24095584 | 0 | 1 | 1 | 0.74482229 | 0 | 1 | 25 | 0.51779518 | FALSE | 1 | FALSE | 1 | FALSE | 1 | FALSE | 1 | FALSE | 1 | FALSE | 1 |
| 2081 | 2081 | 7.44285714 | 1.70407674 | 2 | 0.539854 | 4 | 0.19010643 | 2 | 0.72288838 | 1 | 0.9633793 | 0 | 1 | 61 | 0.32827441 | FALSE | 1 | FALSE | 1 | FALSE | 1 | FALSE | 1 | FALSE | 1 | FALSE | 1 |
| 2082 | 2082 | 8.36363636 | 2.20630202 | 1 | 0.4357638 | 3 | 0.03266339 | 1 | 0.55340851 | 1 | 0.64497439 | 2 | 0.04972408 | 14 | 0.99612862 | FALSE | 1 | FALSE | 0.92268438 | FALSE | 1 | FALSE | 1 | FALSE | 0.98629461 | FALSE | 1 |
| 2083 | 2083 | 8.72413793 | 3.55155674 | 1 | 0.51745948 | 0 | 1 | 0 | 1 | 0 | 1 | 0 | 1 | 27 | 0.05284755 | FALSE | 1 | FALSE | 1 | FALSE | 1 | FALSE | 1 | FALSE | 1 | FALSE | 1 |
| 2084 | 2084 | 8.43478261 | 1.89567212 | 4 | 0.02951043 | 3 | 0.18548475 | 5 | 0.02370648 | 1 | 0.88573774 | 1 | 0.53375569 | 32 | 0.99686402 | FALSE | 0.53131751 | FALSE | 1 | FALSE | 1 | FALSE | 1 | FALSE | 1 | FALSE | 1 |
| 2085 | 2085 | 9.20833333 | 2.87197909 | 0 | 1 | 0 | 1 | 1 | 0.58502114 | 1 | 0.67692816 | 0 | 1 | 22 | 0.25237063 | FALSE | 1 | FALSE | 1 | FALSE | 1 | FALSE | 1 | FALSE | 1 | FALSE | 1 |
| 2086 | 2086 | 9.02777778 | 3.63230957 | 0 | 1 | 0 | 1 | 1 | 0.73299778 | 3 | 0.22784656 | 0 | 1 | 32 | 0.31709555 | FALSE | 1 | FALSE | 1 | FALSE | 1 | FALSE | 1 | FALSE | 1 | FALSE | 1 |
| 2087 | 2087 | 9.52941177 | 3.21974875 | 1 | 0.35728305 | 0 | 1 | 1 | 0.46348836 | 1 | 0.55062772 | 0 | 1 | 14 | 0.7313145 | FALSE | 1 | FALSE | 1 | FALSE | 1 | FALSE | 1 | FALSE | 1 | FALSE | 1 |
| 2088 | 2088 | 10.0769231 | 1.73034183 | 3 | 0.07731604 | 0 | 1 | 2 | 0.41139386 | 2 | 0.54052736 | 1 | 0.4761699 | 31 | 0.85613899 | FALSE | 0.77583006 | FALSE | 1 | FALSE | 1 | FALSE | 1 | FALSE | 1 | FALSE | 1 |
| 2089 | 2089 | 8.95238095 | 1.96338363 | 0 | 1 | 3 | 0.02888061 | 2 | 0.17308361 | 1 | 0.62783522 | 0 | 1 | 15 | 0.96513691 | FALSE | 1 | FALSE | 0.88012077 | FALSE | 1 | FALSE | 1 | FALSE | 1 | FALSE | 1 |
| 2090 | 2090 | 9.08333333 | 1.99826314 | 1 | 0.46441868 | 2 | 0.18117183 | 0 | 1 | 0 | 1 | 0 | 1 | 21 | 0.47053137 | FALSE | 1 | FALSE | 1 | FALSE | 1 | FALSE | 1 | FALSE | 1 | FALSE | 1 |
| 2091 | 2091 | 5.79166667 | 2.7984991 | 0 | 1 | 1 | 0.54637326 | 2 | 0.21260257 | 1 | 0.67692816 | 1 | 0.32796862 | 19 | 0.83975208 | FALSE | 1 | FALSE | 1 | FALSE | 1 | FALSE | 1 | FALSE | 1 | FALSE | 1 |
| 2092 | 2092 | 8.83333333 | 3.3374974 | 0 | 1 | 0 | 1 | 2 | 0.13502329 | 0 | 1 | 0 | 1 | 16 | 0.44998001 | FALSE | 1 | FALSE | 1 | FALSE | 1 | FALSE | 1 | FALSE | 1 | FALSE | 1 |
| 2093 | 2093 | 9.08333333 | 1.552328 | 0 | 1 | 0 | 1 | 1 | 0.35555293 | 0 | 1 | 0 | 1 | 11 | 0.41944532 | FALSE | 1 | FALSE | 1 | FALSE | 1 | FALSE | 1 | FALSE | 1 | FALSE | 1 |
| 2094 | 2094 | 8.69230769 | 1.89674277 | 0 | 1 | 1 | 0.3480709 | 2 | 0.07725803 | 0 | 1 | 2 | 0.01846832 | 8 | 0.99085298 | FALSE | 1 | FALSE | 1 | FALSE | 1 | FALSE | 1 | FALSE | 0.69720022 | FALSE | 1 |
| 2095 | 2095 | 11.3333333 | 3.23178657 | 0 | 1 | 0 | 1 | 1 | 0.62830986 | 1 | 0.71956339 | 1 | 0.36060486 | 24 | 0.3725373 | FALSE | 1 | FALSE | 1 | FALSE | 1 | FALSE | 1 | FALSE | 1 | FALSE | 1 |
| 2096 | 2096 | 8.33333333 | 2.86744176 | 0 | 1 | 1 | 0.38964394 | 0 | 1 | 0 | 1 | 0 | 1 | 14 | 0.29525409 | FALSE | 1 | FALSE | 1 | FALSE | 1 | FALSE | 1 | FALSE | 1 | FALSE | 1 |
| 2097 | 2097 | 8.94117647 | 3.1895157 | 2 | 0.06909038 | 0 | 1 | 0 | 1 | 0 | 1 | 0 | 1 | 15 | 0.49060435 | FALSE | 0.74276693 | FALSE | 1 | FALSE | 1 | FALSE | 1 | FALSE | 1 | FALSE | 1 |
| 2098 | 2098 | 8.69047619 | 2.10993224 | 1 | 0.66519151 | 0 | 1 | 1 | 0.78589454 | 3 | 0.30326943 | 0 | 1 | 37 | 0.34041811 | FALSE | 1 | FALSE | 1 | FALSE | 1 | FALSE | 1 | FALSE | 1 | FALSE | 1 |
| 2099 | 2099 | 10 | 2.9612887 | 1 | 0.28675951 | 0 | 1 | 1 | 0.37874417 | 0 | 1 | 0 | 1 | 11 | 0.6683056 | FALSE | 1 | FALSE | 1 | FALSE | 1 | FALSE | 1 | FALSE | 1 | FALSE | 1 |
| 2100 | 2100 | 9.55172414 | 2.82337786 | 0 | 1 | 2 | 0.24095584 | 0 | 1 | 2 | 0.3871811 | 4 | 0.00118463 | 21 | 0.97185642 | FALSE | 1 | FALSE | 1 | FALSE | 1 | FALSE | 1 | FALSE | 0.20625174 | FALSE | 1 |
| 2101 | 2101 | 10.5294118 | 1.45996161 | 0 | 1 | 2 | 0.10307112 | 1 | 0.46348836 | 1 | 0.55062772 | 0 | 1 | 13 | 0.88719222 | FALSE | 1 | FALSE | 1 | FALSE | 1 | FALSE | 1 | FALSE | 1 | FALSE | 1 |
| 2102 | 2102 | 9.25 | 4.1215895 | 1 | 0.40558521 | 0 | 1 | 1 | 0.51939841 | 1 | 0.60987146 | 1 | 0.28187291 | 16 | 0.80826361 | FALSE | 1 | FALSE | 1 | FALSE | 1 | FALSE | 1 | FALSE | 1 | FALSE | 1 |
| 2103 | 2103 | 10.3076923 | 2.5535681 | 0 | 1 | 0 | 1 | 0 | 1 | 1 | 0.45746571 | 1 | 0.19354507 | 11 | 0.6683056 | FALSE | 1 | FALSE | 1 | FALSE | 1 | FALSE | 1 | FALSE | 1 | FALSE | 1 |
| 2104 | 2104 | 7.23809524 | 2.97457099 | 1 | 0.42086991 | 0 | 1 | 0 | 1 | 3 | 0.06912908 | 1 | 0.29368245 | 16 | 0.90353674 | FALSE | 1 | FALSE | 1 | FALSE | 1 | FALSE | 1 | FALSE | 1 | FALSE | 1 |
| 2105 | 2105 | 7.79220779 | 3.02945566 | 9 | 0.00013999 | 5 | 0.10306379 | 7 | 0.0201002 | 3 | 0.69372972 | 0 | 1 | 53 | 0.99983245 | TRUE | 0.02163774 | FALSE | 1 | FALSE | 1 | FALSE | 1 | FALSE | 1 | FALSE | 1 |
| 2106 | 2106 | 10.7727273 | 2.13055303 | 0 | 1 | 0 | 1 | 0 | 1 | 1 | 0.64497439 | 0 | 1 | 21 | 0.12048848 | FALSE | 1 | FALSE | 1 | FALSE | 1 | FALSE | 1 | FALSE | 1 | FALSE | 1 |
| 2107 | 2107 | 7.09756098 | 2.75685291 | 1 | 0.65632651 | 1 | 0.74131698 | 1 | 0.77786601 | 6 | 0.01031399 | 1 | 0.49330646 | 31 | 0.95455175 | FALSE | 1 | FALSE | 1 | FALSE | 1 | FALSE | 1 | FALSE | 1 | FALSE | 1 |
| 2108 | 2108 | 9 | 2.7080128 | 0 | 1 | 1 | 0.49918218 | 1 | 0.53671415 | 0 | 1 | 1 | 0.29368245 | 18 | 0.57962763 | FALSE | 1 | FALSE | 1 | FALSE | 1 | FALSE | 1 | FALSE | 1 | FALSE | 1 |
| 2109 | 2109 | 6.64556962 | 1.56737283 | 5 | 0.05152748 | 3 | 0.47338398 | 5 | 0.15372513 | 6 | 0.15385579 | 2 | 0.37295427 | 58 | 0.99620676 | FALSE | 0.67646333 | FALSE | 1 | FALSE | 1 | FALSE | 1 | FALSE | 1 | FALSE | 1 |
| 2110 | 2110 | 9.1875 | 2.53028531 | 2 | 0.06197875 | 1 | 0.40942991 | 0 | 1 | 0 | 1 | 0 | 1 | 13 | 0.76799161 | FALSE | 0.72532843 | FALSE | 1 | FALSE | 1 | FALSE | 1 | FALSE | 1 | FALSE | 1 |
| 2111 | 2111 | 8.73333333 | 3.95755256 | 0 | 1 | 0 | 1 | 0 | 1 | 2 | 0.13304705 | 0 | 1 | 12 | 0.62236227 | FALSE | 1 | FALSE | 1 | FALSE | 1 | FALSE | 1 | FALSE | 1 | FALSE | 1 |
| 2112 | 2112 | 8.69230769 | 2.64239447 | 0 | 1 | 3 | 0.05021553 | 1 | 0.61440466 | 1 | 0.70601446 | 0 | 1 | 21 | 0.78939668 | FALSE | 1 | FALSE | 1 | FALSE | 1 | FALSE | 1 | FALSE | 1 | FALSE | 1 |
| 2113 | 2113 | 8.77777778 | 1.68508343 | 0 | 1 | 3 | 0.10944025 | 3 | 0.13783405 | 1 | 0.81665383 | 4 | 0.00269049 | 25 | 0.99349451 | FALSE | 1 | FALSE | 1 | FALSE | 1 | FALSE | 1 | FALSE | 0.29784427 | FALSE | 1 |
| 2114 | 2114 | 9.92857143 | 2.9390058 | 0 | 1 | 1 | 0.36919824 | 0 | 1 | 1 | 0.48242099 | 1 | 0.20679328 | 11 | 0.83676449 | FALSE | 1 | FALSE | 1 | FALSE | 1 | FALSE | 1 | FALSE | 1 | FALSE | 1 |
| 2115 | 2115 | 8.7826087 | 2.60361825 | 7 | 1.12E-06 | 2 | 0.16948764 | 1 | 0.5695037 | 1 | 0.6613267 | 0 | 1 | 12 | 0.9999499 | TRUE | 0.00077787 | FALSE | 1 | FALSE | 1 | FALSE | 1 | FALSE | 1 | FALSE | 1 |
| 2116 | 2116 | 9.18181818 | 2.08100421 | 4 | 0.02552933 | 5 | 0.01315694 | 0 | 1 | 2 | 0.60667406 | 0 | 1 | 33 | 0.96661138 | FALSE | 0.4974559 | FALSE | 0.69207862 | FALSE | 1 | FALSE | 1 | FALSE | 1 | FALSE | 1 |
| 2117 | 2117 | 9.5 | 2.34520788 | 0 | 1 | 1 | 0.23139161 | 0 | 1 | 0 | 1 | 1 | 0.12394282 | 6 | 0.88425551 | FALSE | 1 | FALSE | 1 | FALSE | 1 | FALSE | 1 | FALSE | 1 | FALSE | 1 |
| 2118 | 2118 | 10.1666667 | 3.46008992 | 0 | 1 | 2 | 0.05552913 | 0 | 1 | 1 | 0.43131129 | 1 | 0.18007762 | 8 | 0.97197833 | FALSE | 1 | FALSE | 1 | FALSE | 1 | FALSE | 1 | FALSE | 1 | FALSE | 1 |
| 2119 | 2119 | 8.75 | 2.53722289 | 0 | 1 | 0 | 1 | 0 | 1 | 1 | 0.31351192 | 1 | 0.12394282 | 6 | 0.88425551 | FALSE | 1 | FALSE | 1 | FALSE | 1 | FALSE | 1 | FALSE | 1 | FALSE | 1 |
| 2120 | 2120 | 9.71428571 | 2.7454353 | 0 | 1 | 0 | 1 | 1 | 0.53671415 | 0 | 1 | 0 | 1 | 20 | 0.13763971 | FALSE | 1 | FALSE | 1 | FALSE | 1 | FALSE | 1 | FALSE | 1 | FALSE | 1 |
| 2121 | 2121 | 9.12068966 | 2.09331931 | 1 | 0.77387657 | 5 | 0.03627929 | 2 | 0.61293419 | 1 | 0.93213716 | 3 | 0.06636297 | 45 | 0.90229646 | FALSE | 1 | FALSE | 0.94259695 | FALSE | 1 | FALSE | 1 | FALSE | 1 | FALSE | 1 |
| 2122 | 2122 | 5.48188406 | 1.98992883 | 11 | 0.09125829 | 11 | 0.26882215 | 10 | 0.5216244 | 10 | 0.81287132 | 7 | 0.16016982 | 224 | 0.87723855 | FALSE | 0.83248965 | FALSE | 1 | FALSE | 1 | FALSE | 1 | FALSE | 1 | FALSE | 1 |
| 2123 | 2123 | 8.66666667 | 3.34995854 | 1 | 0.46441868 | 0 | 1 | 0 | 1 | 2 | 0.30249307 | 0 | 1 | 21 | 0.47053137 | FALSE | 1 | FALSE | 1 | FALSE | 1 | FALSE | 1 | FALSE | 1 | FALSE | 1 |
| 2124 | 2124 | 10.4761905 | 1.53123192 | 0 | 1 | 0 | 1 | 1 | 0.53671415 | 0 | 1 | 0 | 1 | 20 | 0.13763971 | FALSE | 1 | FALSE | 1 | FALSE | 1 | FALSE | 1 | FALSE | 1 | FALSE | 1 |
| 2125 | 2125 | 8.15789474 | 4.4634558 | 2 | 0.08405377 | 1 | 0.46503795 | 0 | 1 | 1 | 0.59104357 | 1 | 0.26986775 | 14 | 0.93677253 | FALSE | 0.79236883 | FALSE | 1 | FALSE | 1 | FALSE | 1 | FALSE | 1 | FALSE | 1 |
| 2126 | 2126 | 9.92857143 | 1.98976975 | 0 | 1 | 1 | 0.83180787 | 3 | 0.30658915 | 4 | 0.2347767 | 0 | 1 | 46 | 0.52685615 | FALSE | 1 | FALSE | 1 | FALSE | 1 | FALSE | 1 | FALSE | 1 | FALSE | 1 |
| 2127 | 2127 | 5 | 0 | 1 | 0.28675951 | 0 | 1 | 0 | 1 | 1 | 0.45746571 | 0 | 1 | 11 | 0.6683056 | FALSE | 1 | FALSE | 1 | FALSE | 1 | FALSE | 1 | FALSE | 1 | FALSE | 1 |
| 2128 | 2128 | 7.48837209 | 1.90924426 | 1 | 0.66519151 | 4 | 0.04570474 | 2 | 0.44869594 | 5 | 0.04181362 | 0 | 1 | 30 | 0.99060471 | FALSE | 1 | FALSE | 1 | FALSE | 1 | FALSE | 1 | FALSE | 1 | FALSE | 1 |
| 2129 | 2129 | 8.07317073 | 2.88300524 | 0 | 1 | 1 | 0.7326167 | 2 | 0.42396974 | 1 | 0.84823834 | 1 | 0.48480878 | 35 | 0.38966407 | FALSE | 1 | FALSE | 1 | FALSE | 1 | FALSE | 1 | FALSE | 1 | FALSE | 1 |
| 2130 | 2130 | 8.63636364 | 3.01579862 | 3 | 0.10232687 | 2 | 0.41896557 | 0 | 1 | 2 | 0.60667406 | 1 | 0.51797501 | 36 | 0.75909657 | FALSE | 0.87592659 | FALSE | 1 | FALSE | 1 | FALSE | 1 | FALSE | 1 | FALSE | 1 |
| 2131 | 2131 | 8.33333333 | 3.6968455 | 1 | 0.60837712 | 2 | 0.32556427 | 3 | 0.13783405 | 3 | 0.22784656 | 1 | 0.44938266 | 26 | 0.98200522 | FALSE | 1 | FALSE | 1 | FALSE | 1 | FALSE | 1 | FALSE | 1 | FALSE | 1 |
| 2132 | 2132 | 4.16666667 | 2.41331185 | 12 | 0.00837622 | 8 | 0.38416093 | 10 | 0.23554084 | 19 | 0.00404287 | 2 | 0.86968199 | 162 | 0.99953586 | FALSE | 0.27019545 | FALSE | 1 | FALSE | 1 | FALSE | 1 | FALSE | 1 | FALSE | 1 |
| 2133 | 2133 | 9.77777778 | 2.48451998 | 0 | 1 | 0 | 1 | 0 | 1 | 2 | 0.06108592 | 0 | 1 | 7 | 0.84574168 | FALSE | 1 | FALSE | 1 | FALSE | 1 | FALSE | 1 | FALSE | 1 | FALSE | 1 |
| 2134 | 2134 | 9.72727273 | 3.7195041 | 0 | 1 | 0 | 1 | 0 | 1 | 2 | 0.08789566 | 0 | 1 | 9 | 0.75985366 | FALSE | 1 | FALSE | 1 | FALSE | 1 | FALSE | 1 | FALSE | 1 | FALSE | 1 |
| 2135 | 2135 | 6.84615385 | 2.34835962 | 1 | 0.28675951 | 0 | 1 | 0 | 1 | 0 | 1 | 0 | 1 | 12 | 0.374198 | FALSE | 1 | FALSE | 1 | FALSE | 1 | FALSE | 1 | FALSE | 1 | FALSE | 1 |
| 2136 | 2136 | 8.36 | 1.24515059 | 1 | 0.72839502 | 1 | 0.80796869 | 3 | 0.26704679 | 2 | 0.6762159 | 1 | 0.56379725 | 42 | 0.62187327 | FALSE | 1 | FALSE | 1 | FALSE | 1 | FALSE | 1 | FALSE | 1 | FALSE | 1 |
| 2137 | 2137 | 9.64705882 | 1.87683013 | 1 | 0.35728305 | 0 | 1 | 3 | 0.02143959 | 0 | 1 | 0 | 1 | 13 | 0.88719222 | FALSE | 1 | FALSE | 1 | FALSE | 1 | FALSE | 1 | FALSE | 1 | FALSE | 1 |
| 2138 | 2138 | 8.57575758 | 3.16256803 | 1 | 0.56527493 | 2 | 0.27730767 | 0 | 1 | 2 | 0.43583874 | 1 | 0.41154418 | 26 | 0.77612337 | FALSE | 1 | FALSE | 1 | FALSE | 1 | FALSE | 1 | FALSE | 1 | FALSE | 1 |
| 2139 | 2139 | 9.28571429 | 2.08493136 | 0 | 1 | 2 | 0.07341559 | 0 | 1 | 1 | 0.48242099 | 0 | 1 | 11 | 0.83676449 | FALSE | 1 | FALSE | 1 | FALSE | 1 | FALSE | 1 | FALSE | 1 | FALSE | 1 |
| 2140 | 2140 | 7.11111111 | 3.18658453 | 3 | 0.0638041 | 5 | 0.0056244 | 1 | 0.73299778 | 2 | 0.49731009 | 0 | 1 | 25 | 0.99349451 | FALSE | 0.73766 | FALSE | 0.59070661 | FALSE | 1 | FALSE | 1 | FALSE | 1 | FALSE | 1 |
| 2141 | 2141 | 9.75 | 2.20321734 | 1 | 0.26795838 | 0 | 1 | 1 | 0.35555293 | 0 | 1 | 0 | 1 | 10 | 0.71436964 | FALSE | 1 | FALSE | 1 | FALSE | 1 | FALSE | 1 | FALSE | 1 | FALSE | 1 |
| 2142 | 2142 | 8.75 | 1.82002747 | 0 | 1 | 1 | 0.40942991 | 2 | 0.11087977 | 0 | 1 | 0 | 1 | 13 | 0.76799161 | FALSE | 1 | FALSE | 1 | FALSE | 1 | FALSE | 1 | FALSE | 1 | FALSE | 1 |
| 2143 | 2143 | 7.56 | 3.66693332 | 2 | 0.13369621 | 1 | 0.56109832 | 1 | 0.59998149 | 1 | 0.69181315 | 0 | 1 | 20 | 0.8152498 | FALSE | 0.97964162 | FALSE | 1 | FALSE | 1 | FALSE | 1 | FALSE | 1 | FALSE | 1 |
| 2144 | 2144 | 9.55555556 | 1.4229165 | 0 | 1 | 1 | 0.42857733 | 1 | 0.46348836 | 1 | 0.55062772 | 0 | 1 | 14 | 0.7313145 | FALSE | 1 | FALSE | 1 | FALSE | 1 | FALSE | 1 | FALSE | 1 | FALSE | 1 |
| 2145 | 2145 | 8.8974359 | 2.79922033 | 0 | 1 | 0 | 1 | 3 | 0.16357218 | 0 | 1 | 0 | 1 | 36 | 0.12091774 | FALSE | 1 | FALSE | 1 | FALSE | 1 | FALSE | 1 | FALSE | 1 | FALSE | 1 |
| 2146 | 2146 | 9.0625 | 1.88642353 | 0 | 1 | 0 | 1 | 1 | 0.44344817 | 1 | 0.52895088 | 1 | 0.23264612 | 13 | 0.76799161 | FALSE | 1 | FALSE | 1 | FALSE | 1 | FALSE | 1 | FALSE | 1 | FALSE | 1 |
| 2147 | 2147 | 8.23809524 | 2.7929292 | 1 | 0.42086991 | 0 | 1 | 0 | 1 | 3 | 0.06912908 | 1 | 0.29368245 | 16 | 0.90353674 | FALSE | 1 | FALSE | 1 | FALSE | 1 | FALSE | 1 | FALSE | 1 | FALSE | 1 |
| 2148 | 2148 | 12.6666667 | 0.47140452 | 0 | 1 | 0 | 1 | 0 | 1 | 0 | 1 | 0 | 1 | 3 | 0.60071728 | FALSE | 1 | FALSE | 1 | FALSE | 1 | FALSE | 1 | FALSE | 1 | FALSE | 1 |
| 2149 | 2149 | 8.23809524 | 2.06800483 | 0 | 1 | 0 | 1 | 1 | 0.53671415 | 0 | 1 | 0 | 1 | 20 | 0.13763971 | FALSE | 1 | FALSE | 1 | FALSE | 1 | FALSE | 1 | FALSE | 1 | FALSE | 1 |
| 2150 | 2150 | 9.46875 | 1.9840674 | 1 | 0.56527493 | 1 | 0.65167362 | 0 | 1 | 2 | 0.43583874 | 0 | 1 | 28 | 0.42532177 | FALSE | 1 | FALSE | 1 | FALSE | 1 | FALSE | 1 | FALSE | 1 | FALSE | 1 |
| 2151 | 2151 | 10.0714286 | 2.54850897 | 1 | 0.3050805 | 0 | 1 | 0 | 1 | 0 | 1 | 0 | 1 | 13 | 0.33283729 | FALSE | 1 | FALSE | 1 | FALSE | 1 | FALSE | 1 | FALSE | 1 | FALSE | 1 |
| 2152 | 2152 | 7.68965517 | 2.05281049 | 1 | 0.51745948 | 0 | 1 | 1 | 0.64171563 | 2 | 0.3705495 | 3 | 0.0104283 | 21 | 0.94083768 | FALSE | 1 | FALSE | 1 | FALSE | 1 | FALSE | 1 | FALSE | 0.5620304 | FALSE | 1 |
| 2153 | 2153 | 9.53846154 | 1.90607872 | 1 | 0.28675951 | 1 | 0.3480709 | 1 | 0.37874417 | 1 | 0.45746571 | 0 | 1 | 9 | 0.96018168 | FALSE | 1 | FALSE | 1 | FALSE | 1 | FALSE | 1 | FALSE | 1 | FALSE | 1 |
| 2154 | 2154 | 9.97142857 | 2.67779989 | 0 | 1 | 1 | 0.6629941 | 4 | 0.0293494 | 1 | 0.78873814 | 0 | 1 | 27 | 0.75124583 | FALSE | 1 | FALSE | 1 | FALSE | 1 | FALSE | 1 | FALSE | 1 | FALSE | 1 |
| 2155 | 2155 | 7.14864865 | 1.59121263 | 3 | 0.2949342 | 1 | 0.9134145 | 2 | 0.75097925 | 2 | 0.86064871 | 1 | 0.70774043 | 65 | 0.2604438 | FALSE | 1 | FALSE | 1 | FALSE | 1 | FALSE | 1 | FALSE | 1 | FALSE | 1 |
| 2156 | 2156 | 10.4285714 | 2.71804251 | 0 | 1 | 0 | 1 | 1 | 0.40110418 | 1 | 0.48242099 | 0 | 1 | 12 | 0.62236227 | FALSE | 1 | FALSE | 1 | FALSE | 1 | FALSE | 1 | FALSE | 1 | FALSE | 1 |
| 2157 | 2157 | 10.5 | 2.72946881 | 0 | 1 | 0 | 1 | 0 | 1 | 1 | 0.37517262 | 0 | 1 | 9 | 0.5215547 | FALSE | 1 | FALSE | 1 | FALSE | 1 | FALSE | 1 | FALSE | 1 | FALSE | 1 |
| 2158 | 2158 | 9.59375 | 3.15098571 | 1 | 0.56527493 | 1 | 0.65167362 | 1 | 0.69069222 | 3 | 0.17988885 | 1 | 0.41154418 | 25 | 0.88559288 | FALSE | 1 | FALSE | 1 | FALSE | 1 | FALSE | 1 | FALSE | 1 | FALSE | 1 |
| 2159 | 2159 | 10.625 | 1.9324531 | 0 | 1 | 0 | 1 | 1 | 0.25384309 | 1 | 0.31351192 | 0 | 1 | 6 | 0.88425551 | FALSE | 1 | FALSE | 1 | FALSE | 1 | FALSE | 1 | FALSE | 1 | FALSE | 1 |
| 2160 | 2160 | 9.10526316 | 2.8264677 | 1 | 0.38989951 | 1 | 0.46503795 | 1 | 0.50143826 | 1 | 0.59104357 | 1 | 0.26986775 | 14 | 0.93677253 | FALSE | 1 | FALSE | 1 | FALSE | 1 | FALSE | 1 | FALSE | 1 | FALSE | 1 |

| | | | | | | | | | | | | | | | | | | | | | | | | | | | |
|---|---|---|---|---|---|---|---|---|---|---|---|---|---|---|---|---|---|---|---|---|---|---|---|---|---|---|---|
| 2161 | 2161 | 8.06666667 | 3.37573037 | 1 | 0.32293355 | 0 | 1 | 0 | 1 | 0 | 1 | 0 | 1 | 14 | 0.29525409 | FALSE | 1 | FALSE | 1 | FALSE | 1 | FALSE | 1 | FALSE | 1 | FALSE | 1 |
| 2162 | 2162 | 9.28571429 | 2.7454353 | 1 | 0.42086991 | 1 | 0.49918218 | 1 | 0.53671415 | 5 | 0.00218688 | 0 | 1 | 13 | 0.9973503 | FALSE | 1 | FALSE | 1 | FALSE | 1 | FALSE | 1 | FALSE | 1 | FALSE | 1 |
| 2163 | 2163 | 8.26666667 | 2.04830553 | 1 | 0.32293355 | 0 | 1 | 0 | 1 | 1 | 0.50623194 | 0 | 1 | 13 | 0.57712805 | FALSE | 1 | FALSE | 1 | FALSE | 1 | FALSE | 1 | FALSE | 1 | FALSE | 1 |
| 2164 | 2164 | 5.90625 | 3.29165019 | 1 | 0.56527493 | 2 | 0.27730767 | 0 | 1 | 1 | 0.77852168 | 0 | 1 | 28 | 0.42532177 | FALSE | 1 | FALSE | 1 | FALSE | 1 | FALSE | 1 | FALSE | 1 | FALSE | 1 |
| 2165 | 2165 | 7.31818182 | 3.88922011 | 3 | 0.01782284 | 1 | 0.5154318 | 1 | 0.55340851 | 3 | 0.07743186 | 0 | 1 | 14 | 0.99612862 | FALSE | 0.41596787 | FALSE | 1 | FALSE | 1 | FALSE | 1 | FALSE | 1 | FALSE | 1 |
| 2166 | 2166 | 6.82142857 | 2.29990018 | 1 | 0.50471032 | 0 | 1 | 0 | 1 | 3 | 0.12475952 | 1 | 0.36060486 | 22 | 0.76237877 | FALSE | 1 | FALSE | 1 | FALSE | 1 | FALSE | 1 | FALSE | 1 | FALSE | 1 |
| 2167 | 2167 | 8.26666667 | 2.26470503 | 1 | 0.68224519 | 0 | 1 | 1 | 0.80109494 | 3 | 0.32878418 | 2 | 0.16211764 | 37 | 0.61979664 | FALSE | 1 | FALSE | 1 | FALSE | 1 | FALSE | 1 | FALSE | 1 | FALSE | 1 |
| 2168 | 2168 | 9.36363636 | 1.82272161 | 0 | 1 | 1 | 0.5154318 | 0 | 1 | 0 | 1 | 0 | 1 | 21 | 0.12048848 | FALSE | 1 | FALSE | 1 | FALSE | 1 | FALSE | 1 | FALSE | 1 | FALSE | 1 |
| 2169 | 2169 | 5.59770115 | 2.84649319 | 2 | 0.65852142 | 4 | 0.31014012 | 4 | 0.38141809 | 4 | 0.57118367 | 4 | 0.05404971 | 69 | 0.92352338 | FALSE | 1 | FALSE | 1 | FALSE | 1 | FALSE | 1 | FALSE | 1 | FALSE | 1 |
| 2170 | 2170 | 11.8064516 | 2.19117519 | 0 | 1 | 0 | 1 | 2 | 0.30673593 | 1 | 0.76781284 | 0 | 1 | 28 | 0.26393996 | FALSE | 1 | FALSE | 1 | FALSE | 1 | FALSE | 1 | FALSE | 1 | FALSE | 1 |
| 2171 | 2171 | 10.5925926 | 2.00479535 | 0 | 1 | 1 | 0.58913618 | 0 | 1 | 0 | 1 | 0 | 1 | 26 | 0.06079097 | FALSE | 1 | FALSE | 1 | FALSE | 1 | FALSE | 1 | FALSE | 1 | FALSE | 1 |
| 2172 | 2172 | 10.6111111 | 1.2969575 | 1 | 0.37380237 | 0 | 1 | 1 | 0.48280983 | 0 | 1 | 0 | 1 | 16 | 0.44998001 | FALSE | 1 | FALSE | 1 | FALSE | 1 | FALSE | 1 | FALSE | 1 | FALSE | 1 |
| 2173 | 2173 | 6.11111111 | 1.74129227 | 2 | 0.32143972 | 1 | 0.77338922 | 1 | 0.80828716 | 3 | 0.34154252 | 1 | 0.52593041 | 37 | 0.73736034 | FALSE | 1 | FALSE | 1 | FALSE | 1 | FALSE | 1 | FALSE | 1 | FALSE | 1 |
| 2174 | 2174 | 8.18918919 | 2.50229989 | 0 | 1 | 2 | 0.3375163 | 0 | 1 | 1 | 0.82511639 | 0 | 1 | 34 | 0.14843263 | FALSE | 1 | FALSE | 1 | FALSE | 1 | FALSE | 1 | FALSE | 1 | FALSE | 1 |
| 2175 | 2175 | 8.68181818 | 2.30387744 | 2 | 0.10809809 | 0 | 1 | 0 | 1 | 1 | 0.64497439 | 0 | 1 | 19 | 0.54233419 | FALSE | 0.88820416 | FALSE | 1 | FALSE | 1 | FALSE | 1 | FALSE | 1 | FALSE | 1 |
| 2176 | 2176 | 10.3684211 | 2.53834854 | 1 | 0.38989951 | 0 | 1 | 0 | 1 | 0 | 1 | 1 | 0.26986775 | 17 | 0.41141543 | FALSE | 1 | FALSE | 1 | FALSE | 1 | FALSE | 1 | FALSE | 1 | FALSE | 1 |
| 2177 | 2177 | 8.41666667 | 1.89113429 | 0 | 1 | 0 | 1 | 0 | 1 | 0 | 1 | 1 | 0.18007762 | 11 | 0.41944532 | FALSE | 1 | FALSE | 1 | FALSE | 1 | FALSE | 1 | FALSE | 1 | FALSE | 1 |
| 2178 | 2178 | 9.31818182 | 2.5115436 | 2 | 0.10809809 | 1 | 0.5154318 | 1 | 0.55340851 | 0 | 1 | 0 | 1 | 18 | 0.74744813 | FALSE | 0.88820416 | FALSE | 1 | FALSE | 1 | FALSE | 1 | FALSE | 1 | FALSE | 1 |
| 2179 | 2179 | 8.18518519 | 2.59682195 | 0 | 1 | 1 | 0.58913618 | 0 | 1 | 3 | 0.12475952 | 1 | 0.36060486 | 22 | 0.76237877 | FALSE | 1 | FALSE | 1 | FALSE | 1 | FALSE | 1 | FALSE | 1 | FALSE | 1 |
| 2180 | 2180 | 6.75757576 | 2.90277091 | 0 | 1 | 3 | 0.08959205 | 2 | 0.33343871 | 1 | 0.78873814 | 1 | 0.42123867 | 26 | 0.86852567 | FALSE | 1 | FALSE | 1 | FALSE | 1 | FALSE | 1 | FALSE | 1 | FALSE | 1 |
| 2181 | 2181 | 10.0769231 | 2.40069024 | 0 | 1 | 1 | 0.3480709 | 0 | 1 | 1 | 0.45746571 | 0 | 1 | 11 | 0.6683056 | FALSE | 1 | FALSE | 1 | FALSE | 1 | FALSE | 1 | FALSE | 1 | FALSE | 1 |
| 2182 | 2182 | 5.68493151 | 2.82782996 | 1 | 0.85137169 | 2 | 0.68961994 | 3 | 0.49081265 | 1 | 0.96824399 | 0 | 1 | 66 | 0.09785941 | FALSE | 1 | FALSE | 1 | FALSE | 1 | FALSE | 1 | FALSE | 1 | FALSE | 1 |
| 2183 | 2183 | 11.0476191 | 2.68108348 | 1 | 0.42086991 | 1 | 0.49918218 | 0 | 1 | 2 | 0.25059627 | 0 | 1 | 17 | 0.77849076 | FALSE | 1 | FALSE | 1 | FALSE | 1 | FALSE | 1 | FALSE | 1 | FALSE | 1 |
| 2184 | 2184 | 11.2941177 | 2.65163004 | 0 | 1 | 0 | 1 | 0 | 1 | 1 | 0.55062772 | 0 | 1 | 16 | 0.23070182 | FALSE | 1 | FALSE | 1 | FALSE | 1 | FALSE | 1 | FALSE | 1 | FALSE | 1 |
| 2185 | 2185 | 8.85714286 | 1.45686272 | 0 | 1 | 0 | 1 | 1 | 0.22601089 | 0 | 1 | 1 | 0.10932414 | 5 | 0.91847773 | FALSE | 1 | FALSE | 1 | FALSE | 1 | FALSE | 1 | FALSE | 1 | FALSE | 1 |
| 2186 | 2186 | 7.11627907 | 1.75485111 | 2 | 0.30241117 | 1 | 0.75788114 | 0 | 1 | 2 | 0.59404271 | 2 | 0.1563365 | 36 | 0.6451239 | FALSE | 1 | FALSE | 1 | FALSE | 1 | FALSE | 1 | FALSE | 1 | FALSE | 1 |
| 2187 | 2187 | 9.875 | 3.65504788 | 0 | 1 | 0 | 1 | 1 | 0.25384309 | 1 | 0.31351192 | 0 | 1 | 6 | 0.88425551 | FALSE | 1 | FALSE | 1 | FALSE | 1 | FALSE | 1 | FALSE | 1 | FALSE | 1 |
| 2188 | 2188 | 9.1038961 | 0.84645043 | 1 | 0.86619543 | 5 | 0.10306379 | 2 | 0.77041438 | 4 | 0.47420444 | 1 | 0.72203879 | 64 | 0.68868587 | FALSE | 1 | FALSE | 1 | FALSE | 1 | FALSE | 1 | FALSE | 1 | FALSE | 1 |
| 2189 | 2189 | 8.12 | 1.81813091 | 2 | 0.13369621 | 1 | 0.56109832 | 0 | 1 | 1 | 0.69181315 | 1 | 0.33902672 | 20 | 0.8152498 | FALSE | 0.97964162 | FALSE | 1 | FALSE | 1 | FALSE | 1 | FALSE | 1 | FALSE | 1 |
| 2190 | 2190 | 6.36111111 | 2.77040689 | 1 | 0.60837712 | 1 | 0.69480472 | 1 | 0.73299778 | 2 | 0.49731009 | 0 | 1 | 31 | 0.49869404 | FALSE | 1 | FALSE | 1 | FALSE | 1 | FALSE | 1 | FALSE | 1 | FALSE | 1 |
| 2191 | 2191 | 14.6363636 | 2.3845231 | 0 | 1 | 0 | 1 | 0 | 1 | 0 | 1 | 0 | 1 | 11 | 0.15414489 | FALSE | 1 | FALSE | 1 | FALSE | 1 | FALSE | 1 | FALSE | 1 | FALSE | 1 |
| 2192 | 2192 | 8 | 3.61033001 | 1 | 0.52988231 | 1 | 0.61539065 | 0 | 1 | 2 | 0.3871811 | 0 | 1 | 25 | 0.51779518 | FALSE | 1 | FALSE | 1 | FALSE | 1 | FALSE | 1 | FALSE | 1 | FALSE | 1 |
| 2193 | 2193 | 8.22222222 | 1.88430817 | 0 | 1 | 0 | 1 | 1 | 0.80828716 | 4 | 0.15016749 | 0 | 1 | 40 | 0.27423942 | FALSE | 1 | FALSE | 1 | FALSE | 1 | FALSE | 1 | FALSE | 1 | FALSE | 1 |
| 2194 | 2194 | 7.57692308 | 2.71991342 | 1 | 0.49162634 | 2 | 0.20486827 | 0 | 1 | 1 | 0.70601446 | 0 | 1 | 22 | 0.61628 | FALSE | 1 | FALSE | 1 | FALSE | 1 | FALSE | 1 | FALSE | 1 | FALSE | 1 |
| 2195 | 2195 | 5.46616541 | 2.86684147 | 1 | 0.96605914 | 3 | 0.79366541 | 3 | 0.84854739 | 7 | 0.38067194 | 5 | 0.06022929 | 110 | 0.4472263 | FALSE | 1 | FALSE | 1 | FALSE | 1 | FALSE | 1 | FALSE | 1 | FALSE | 1 |
| 2196 | 2196 | 8.92307692 | 3.68910117 | 0 | 1 | 0 | 1 | 0 | 1 | 2 | 0.1174583 | 1 | 0.19354507 | 10 | 0.86791642 | FALSE | 1 | FALSE | 1 | FALSE | 1 | FALSE | 1 | FALSE | 1 | FALSE | 1 |
| 2197 | 2197 | 9.33333333 | 2.87799032 | 0 | 1 | 1 | 0.6629941 | 1 | 0.70185639 | 0 | 1 | 1 | 0.42123867 | 30 | 0.21942357 | FALSE | 1 | FALSE | 1 | FALSE | 1 | FALSE | 1 | FALSE | 1 | FALSE | 1 |
| 2198 | 2198 | 9.625 | 1.65359457 | 0 | 1 | 1 | 0.23139161 | 1 | 0.25384309 | 0 | 1 | 0 | 1 | 6 | 0.88425551 | FALSE | 1 | FALSE | 1 | FALSE | 1 | FALSE | 1 | FALSE | 1 | FALSE | 1 |
| 2199 | 2199 | 9 | 2.52487624 | 0 | 1 | 1 | 0.40942991 | 1 | 0.44344817 | 0 | 1 | 0 | 1 | 14 | 0.53308261 | FALSE | 1 | FALSE | 1 | FALSE | 1 | FALSE | 1 | FALSE | 1 | FALSE | 1 |
| 2200 | 2200 | 10 | 2.32992949 | 0 | 1 | 0 | 1 | 2 | 0.08806398 | 0 | 1 | 1 | 0.20679328 | 11 | 0.83676449 | FALSE | 1 | FALSE | 1 | FALSE | 1 | FALSE | 1 | FALSE | 1 | FALSE | 1 |
| 2201 | 2201 | 10.5 | 1.96214169 | 0 | 1 | 1 | 0.28036922 | 0 | 1 | 1 | 0.37517262 | 0 | 1 | 8 | 0.80394719 | FALSE | 1 | FALSE | 1 | FALSE | 1 | FALSE | 1 | FALSE | 1 | FALSE | 1 |
| 2202 | 2202 | 9.66666667 | 3.02581486 | 0 | 1 | 1 | 0.36919824 | 1 | 0.40110418 | 1 | 0.48242099 | 0 | 1 | 11 | 0.83676449 | FALSE | 1 | FALSE | 1 | FALSE | 1 | FALSE | 1 | FALSE | 1 | FALSE | 1 |
| 2203 | 2203 | 8.16666667 | 1.74005109 | 1 | 0.34033054 | 1 | 0.40942991 | 1 | 0.44344817 | 1 | 0.52895088 | 1 | 0.23264612 | 11 | 0.9718531 | FALSE | 1 | FALSE | 1 | FALSE | 1 | FALSE | 1 | FALSE | 1 | FALSE | 1 |
| 2204 | 2204 | 9.76923077 | 2.00590843 | 1 | 0.49162634 | 2 | 0.20486827 | 0 | 1 | 1 | 0.70601446 | 0 | 1 | 22 | 0.61628 | FALSE | 1 | FALSE | 1 | FALSE | 1 | FALSE | 1 | FALSE | 1 | FALSE | 1 |
| 2205 | 2205 | 6.48717949 | 2.92532527 | 1 | 0.62830697 | 5 | 0.0071078 | 1 | 0.75193685 | 1 | 0.83318955 | 1 | 0.4673875 | 29 | 0.93814695 | FALSE | 1 | FALSE | 0.61291637 | FALSE | 1 | FALSE | 1 | FALSE | 1 | FALSE | 1 |
| 2206 | 2206 | 7.9 | 2.54754784 | 0 | 1 | 2 | 0.1353446 | 0 | 1 | 1 | 0.60987146 | 1 | 0.28187291 | 16 | 0.80826361 | FALSE | 1 | FALSE | 1 | FALSE | 1 | FALSE | 1 | FALSE | 1 | FALSE | 1 |
| 2207 | 2207 | 10.8333333 | 0.89752747 | 0 | 1 | 0 | 1 | 1 | 0.33149969 | 0 | 1 | 0 | 1 | 10 | 0.46859937 | FALSE | 1 | FALSE | 1 | FALSE | 1 | FALSE | 1 | FALSE | 1 | FALSE | 1 |
| 2208 | 2208 | 8.23333333 | 2.0604746 | 0 | 1 | 0 | 1 | 0 | 1 | 1 | 0.75658797 | 0 | 1 | 29 | 0.03983039 | FALSE | 1 | FALSE | 1 | FALSE | 1 | FALSE | 1 | FALSE | 1 | FALSE | 1 |
| 2209 | 2209 | 9.625 | 1.79843682 | 1 | 0.34033054 | 1 | 0.40942991 | 0 | 1 | 0 | 1 | 1 | 0.23264612 | 13 | 0.76799161 | FALSE | 1 | FALSE | 1 | FALSE | 1 | FALSE | 1 | FALSE | 1 | FALSE | 1 |
| 2210 | 2210 | 10.5833333 | 2.7220805 | 0 | 1 | 0 | 1 | 1 | 0.35555293 | 0 | 1 | 0 | 1 | 11 | 0.41944532 | FALSE | 1 | FALSE | 1 | FALSE | 1 | FALSE | 1 | FALSE | 1 | FALSE | 1 |
| 2211 | 2211 | 9 | 2.67889354 | 0 | 1 | 0 | 1 | 1 | 0.44344817 | 3 | 0.03440583 | 1 | 0.23264612 | 11 | 0.9718531 | FALSE | 1 | FALSE | 1 | FALSE | 1 | FALSE | 1 | FALSE | 1 | FALSE | 1 |
| 2212 | 2212 | 9.5 | 2.20227156 | 0 | 1 | 0 | 1 | 0 | 1 | 1 | 0.37517262 | 0 | 1 | 9 | 0.5215547 | FALSE | 1 | FALSE | 1 | FALSE | 1 | FALSE | 1 | FALSE | 1 | FALSE | 1 |
| 2213 | 2213 | 7.01941748 | 0.35473448 | 11 | 0.01650824 | 7 | 0.50221329 | 10 | 0.20603034 | 10 | 0.47484245 | 5 | 0.24880938 | 163 | 0.98373338 | FALSE | 0.39708985 | FALSE | 1 | FALSE | 1 | FALSE | 1 | FALSE | 1 | FALSE | 1 |
| 2214 | 2214 | 11.7272727 | 2.83134755 | 1 | 0.24866461 | 0 | 1 | 0 | 1 | 1 | 0.40390032 | 0 | 1 | 9 | 0.75985366 | FALSE | 1 | FALSE | 1 | FALSE | 1 | FALSE | 1 | FALSE | 1 | FALSE | 1 |
| 2215 | 2215 | 9.83333333 | 3.50792753 | 0 | 1 | 1 | 0.32623933 | 2 | 0.06691409 | 0 | 1 | 0 | 1 | 9 | 0.89630573 | FALSE | 1 | FALSE | 1 | FALSE | 1 | FALSE | 1 | FALSE | 1 | FALSE | 1 |
| 2216 | 2216 | 7.81538462 | 2.3980268 | 2 | 0.49997462 | 3 | 0.35154359 | 3 | 0.41519671 | 2 | 0.80687302 | 3 | 0.09035518 | 52 | 0.87296036 | FALSE | 1 | FALSE | 1 | FALSE | 1 | FALSE | 1 | FALSE | 1 | FALSE | 1 |
| 2217 | 2217 | 7.875 | 3.58599707 | 1 | 0.46441868 | 1 | 0.54637326 | 0 | 1 | 0 | 1 | 0 | 1 | 22 | 0.25237063 | FALSE | 1 | FALSE | 1 | FALSE | 1 | FALSE | 1 | FALSE | 1 | FALSE | 1 |
| 2218 | 2218 | 8.45833333 | 2.02029632 | 0 | 1 | 0 | 1 | 1 | 0.58502114 | 2 | 0.30249307 | 0 | 1 | 21 | 0.47053137 | FALSE | 1 | FALSE | 1 | FALSE | 1 | FALSE | 1 | FALSE | 1 | FALSE | 1 |
| 2219 | 2219 | 8.75 | 2.24602824 | 0 | 1 | 0 | 1 | 3 | 0.07756639 | 0 | 1 | 1 | 0.37113067 | 24 | 0.55018107 | FALSE | 1 | FALSE | 1 | FALSE | 1 | FALSE | 1 | FALSE | 1 | FALSE | 1 |
| 2220 | 2220 | 10.1333333 | 1.92757764 | 2 | 0.05513784 | 0 | 1 | 0 | 1 | 1 | 0.50623194 | 1 | 0.2198258 | 11 | 0.92879412 | FALSE | 0.68849059 | FALSE | 1 | FALSE | 1 | FALSE | 1 | FALSE | 1 | FALSE | 1 |
| 2221 | 2221 | 9.27272727 | 2.13588002 | 0 | 1 | 1 | 0.30368014 | 0 | 1 | 0 | 1 | 1 | 0.16638736 | 9 | 0.75985366 | FALSE | 1 | FALSE | 1 | FALSE | 1 | FALSE | 1 | FALSE | 1 | FALSE | 1 |
| 2222 | 2222 | 11 | 2.44948974 | 0 | 1 | 0 | 1 | 1 | 0.62830986 | 0 | 1 | 0 | 1 | 26 | 0.06079097 | FALSE | 1 | FALSE | 1 | FALSE | 1 | FALSE | 1 | FALSE | 1 | FALSE | 1 |
| 2223 | 2223 | 9.38636364 | 1.4336208 | 0 | 1 | 2 | 0.41896557 | 1 | 0.80109494 | 2 | 0.60667406 | 0 | 1 | 39 | 0.29524127 | FALSE | 1 | FALSE | 1 | FALSE | 1 | FALSE | 1 | FALSE | 1 | FALSE | 1 |
| 2224 | 2224 | 10.3809524 | 1.78555555 | 2 | 0.09187136 | 1 | 0.48239024 | 2 | 0.16019548 | 2 | 0.23331368 | 4 | 0.00027072 | 9 | 0.99999262 | FALSE | 0.83248965 | FALSE | 1 | FALSE | 1 | FALSE | 1 | FALSE | 0.08567695 | FALSE | 1 |
| 2225 | 2225 | 8.60714286 | 2.59586602 | 0 | 1 | 0 | 1 | 1 | 0.64171563 | 2 | 0.3705495 | 1 | 0.37113067 | 24 | 0.55018107 | FALSE | 1 | FALSE | 1 | FALSE | 1 | FALSE | 1 | FALSE | 1 | FALSE | 1 |
| 2226 | 2226 | 10.875 | 2.75850956 | 0 | 1 | 0 | 1 | 0 | 1 | 1 | 0.31351192 | 0 | 1 | 7 | 0.63752397 | FALSE | 1 | FALSE | 1 | FALSE | 1 | FALSE | 1 | FALSE | 1 | FALSE | 1 |
| 2227 | 2227 | 6.90476191 | 4.06968774 | 0 | 1 | 0 | 1 | 1 | 0.53671415 | 1 | 0.62783522 | 0 | 1 | 19 | 0.34094959 | FALSE | 1 | FALSE | 1 | FALSE | 1 | FALSE | 1 | FALSE | 1 | FALSE | 1 |
| 2228 | 2228 | 10.5 | 2.36290781 | 0 | 1 | 0 | 1 | 1 | 0.48280983 | 0 | 1 | 0 | 1 | 17 | 0.20329205 | FALSE | 1 | FALSE | 1 | FALSE | 1 | FALSE | 1 | FALSE | 1 | FALSE | 1 |
| 2229 | 2229 | 9.44444444 | 2.34652357 | 2 | 0.15144274 | 2 | 0.21684246 | 1 | 0.62830986 | 1 | 0.71956339 | 1 | 0.36060486 | 20 | 0.95133411 | FALSE | 1 | FALSE | 1 | FALSE | 1 | FALSE | 1 | FALSE | 1 | FALSE | 1 |
| 2230 | 2230 | 7 | 1.87923696 | 1 | 0.94536367 | 7 | 0.06758379 | 8 | 0.04571544 | 5 | 0.5821433 | 1 | 0.8428205 | 89 | 0.90990553 | FALSE | 1 | FALSE | 1 | FALSE | 1 | FALSE | 1 | FALSE | 1 | FALSE | 1 |
| 2231 | 2231 | 10.1428571 | 2.72179413 | 0 | 1 | 1 | 0.60247911 | 0 | 1 | 1 | 0.73248983 | 1 | 0.37113067 | 25 | 0.34293 | FALSE | 1 | FALSE | 1 | FALSE | 1 | FALSE | 1 | FALSE | 1 | FALSE | 1 |
| 2232 | 2232 | 7.43604651 | 1.23490069 | 7 | 0.15233704 | 7 | 0.32178233 | 7 | 0.42316075 | 6 | 0.80906761 | 4 | 0.31160882 | 141 | 0.83817723 | FALSE | 1 | FALSE | 1 | FALSE | 1 | FALSE | 1 | FALSE | 1 | FALSE | 1 |
| 2233 | 2233 | 9.16666667 | 1.81811869 | 1 | 0.26795838 | 0 | 1 | 2 | 0.06691409 | 0 | 1 | 0 | 1 | 9 | 0.89630573 | FALSE | 1 | FALSE | 1 | FALSE | 1 | FALSE | 1 | FALSE | 1 | FALSE | 1 |
| 2234 | 2234 | 5.95555556 | 3.62678649 | 0 | 1 | 0 | 1 | 3 | 0.21866539 | 1 | 0.8802008 | 0 | 1 | 41 | 0.14688423 | FALSE | 1 | FALSE | 1 | FALSE | 1 | FALSE | 1 | FALSE | 1 | FALSE | 1 |
| 2235 | 2235 | 9.51851852 | 2.5729414 | 1 | 0.50471032 | 0 | 1 | 2 | 0.25286935 | 2 | 0.35374209 | 0 | 1 | 22 | 0.76237877 | FALSE | 1 | FALSE | 1 | FALSE | 1 | FALSE | 1 | FALSE | 1 | FALSE | 1 |
| 2236 | 2236 | 7.5 | 2.83725219 | 2 | 0.17884234 | 3 | 0.07145121 | 1 | 0.66709967 | 1 | 0.75658797 | 1 | 0.39166989 | 22 | 0.96518743 | FALSE | 1 | FALSE | 1 | FALSE | 1 | FALSE | 1 | FALSE | 1 | FALSE | 1 |
| 2237 | 2237 | 8.9375 | 2.12039943 | 3 | 0.0476977 | 1 | 0.65167362 | 0 | 1 | 2 | 0.43583874 | 1 | 0.41154418 | 25 | 0.88559288 | FALSE | 0.649301 | FALSE | 1 | FALSE | 1 | FALSE | 1 | FALSE | 1 | FALSE | 1 |
| 2238 | 2238 | 9.4 | 1.8547237 | 0 | 1 | 0 | 1 | 0 | 1 | 1 | 0.2094688 | 0 | 1 | 4 | 0.82371636 | FALSE | 1 | FALSE | 1 | FALSE | 1 | FALSE | 1 | FALSE | 1 | FALSE | 1 |
| 2239 | 2239 | 8.22222222 | 3.90946935 | 0 | 1 | 1 | 0.44710673 | 0 | 1 | 0 | 1 | 0 | 1 | 17 | 0.20329205 | FALSE | 1 | FALSE | 1 | FALSE | 1 | FALSE | 1 | FALSE | 1 | FALSE | 1 |
| 2240 | 2240 | 2.5359116 | 2.78414669 | 1 | 0.99077508 | 6 | 0.51832838 | 4 | 0.88837507 | 5 | 0.91768789 | 1 | 0.94933441 | 161 | 0.01194149 | FALSE | 1 | FALSE | 1 | FALSE | 1 | FALSE | 1 | FALSE | 1 | FALSE | 1 |
| 2241 | 2241 | 6.67857143 | 2.38627382 | 4 | 0.16836665 | 1 | 0.93792263 | 1 | 0.95461143 | 5 | 0.34217166 | 1 | 0.75276217 | 72 | 0.43856934 | FALSE | 1 | FALSE | 1 | FALSE | 1 | FALSE | 1 | FALSE | 1 | FALSE | 1 |
| 2242 | 2242 | 9.76190476 | 2.20183391 | 1 | 0.42086991 | 1 | 0.49918218 | 1 | 0.53671415 | 2 | 0.25059627 | 0 | 1 | 16 | 0.90353674 | FALSE | 1 | FALSE | 1 | FALSE | 1 | FALSE | 1 | FALSE | 1 | FALSE | 1 |
| 2243 | 2243 | 10 | 2.04301567 | 0 | 1 | 1 | 0.53115654 | 2 | 0.19931429 | 0 | 1 | 0 | 1 | 20 | 0.50589402 | FALSE | 1 | FALSE | 1 | FALSE | 1 | FALSE | 1 | FALSE | 1 | FALSE | 1 |
| 2244 | 2244 | 5.3255814 | 2.55835094 | 0 | 1 | 1 | 0.75788114 | 0 | 1 | 0 | 1 | 0 | 1 | 42 | 0.00592178 | FALSE | 1 | FALSE | 1 | FALSE | 1 | FALSE | 1 | FALSE | 1 | FALSE | 1 |
| 2245 | 2245 | 11.3571429 | 3.01442112 | 0 | 1 | 1 | 0.36919824 | 0 | 1 | 0 | 1 | 0 | 1 | 13 | 0.33283729 | FALSE | 1 | FALSE | 1 | FALSE | 1 | FALSE | 1 | FALSE | 1 | FALSE | 1 |
| 2246 | 2246 | 10.7222222 | 2.57780172 | 5 | 6.83E-05 | 2 | 0.1135776 | 1 | 0.48280983 | 1 | 0.57131013 | 0 | 1 | 9 | 0.99989101 | TRUE | 0.01342173 | FALSE | 1 | FALSE | 1 | FALSE | 1 | FALSE | 1 | FALSE | 1 |
| 2247 | 2247 | 9.75 | 2.29582955 | 0 | 1 | 2 | 0.18117183 | 0 | 1 | 2 | 0.30249307 | 0 | 1 | 20 | 0.68270729 | FALSE | 1 | FALSE | 1 | FALSE | 1 | FALSE | 1 | FALSE | 1 | FALSE | 1 |
| 2248 | 2248 | 9.41176471 | 1.7170023 | 1 | 0.35728305 | 3 | 0.01622926 | 1 | 0.46348836 | 1 | 0.55062772 | 0 | 1 | 11 | 0.98974663 | FALSE | 1 | FALSE | 0.7256023 | FALSE | 1 | FALSE | 1 | FALSE | 1 | FALSE | 1 |
| 2249 | 2249 | 10.4 | 2.57681975 | 0 | 1 | 0 | 1 | 0 | 1 | 0 | 1 | 0 | 1 | 20 | 0.03329856 | FALSE | 1 | FALSE | 1 | FALSE | 1 | FALSE | 1 | FALSE | 1 | FALSE | 1 |
| 2250 | 2250 | 8.4 | 2.56385127 | 1 | 0.78541994 | 4 | 0.12275177 | 1 | 0.88558856 | 2 | 0.76144533 | 0 | 1 | 51 | 0.41341988 | FALSE | 1 | FALSE | 1 | FALSE | 1 | FALSE | 1 | FALSE | 1 | FALSE | 1 |
| 2251 | 2251 | 10.4615385 | 2.24003381 | 0 | 1 | 0 | 1 | 2 | 0.07725803 | 0 | 1 | 0 | 1 | 11 | 0.6683056 | FALSE | 1 | FALSE | 1 | FALSE | 1 | FALSE | 1 | FALSE | 1 | FALSE | 1 |
| 2252 | 2252 | 10.7407407 | 1.40350336 | 0 | 1 | 0 | 1 | 0 | 1 | 3 | 0.12475952 | 0 | 1 | 24 | 0.3725373 | FALSE | 1 | FALSE | 1 | FALSE | 1 | FALSE | 1 | FALSE | 1 | FALSE | 1 |
| 2253 | 2253 | 8.8 | 0.74833148 | 0 | 1 | 0 | 1 | 0 | 1 | 1 | 0.2094688 | 0 | 1 | 4 | 0.82371636 | FALSE | 1 | FALSE | 1 | FALSE | 1 | FALSE | 1 | FALSE | 1 | FALSE | 1 |
| 2254 | 2254 | 11.7647059 | 2.53282943 | 0 | 1 | 1 | 0.42857733 | 1 | 0.46348836 | 2 | 0.18212134 | 0 | 1 | 13 | 0.88719222 | FALSE | 1 | FALSE | 1 | FALSE | 1 | FALSE | 1 | FALSE | 1 | FALSE | 1 |

| | | | | | | | | | | | | | | | | | | | | | | | | | | | |
|---|---|---|---|---|---|---|---|---|---|---|---|---|---|---|---|---|---|---|---|---|---|---|---|---|---|---|---|
| 2255 | 2255 | 4.57 | 2.00626519 | 18 | 2.85E-06 | 14 | 0.00497488 | 11 | 0.10219356 | 4 | 0.98376963 | 12 | 9.14E-05 | 140 | 0.99999988 | TRUE | 0.00179363 | FALSE | 0.58120789 | FALSE | 1 | FALSE | 1 | TRUE | 0.04046262 | FALSE | 1 |
| 2256 | 2256 | 8.70588235 | 1.56298003 | 1 | 0.58738519 | 3 | 0.09602744 | 0 | 1 | 3 | 0.20353499 | 0 | 1 | 27 | 0.85023803 | FALSE | 1 | FALSE | 1 | FALSE | 1 | FALSE | 1 | FALSE | 1 | FALSE | 1 |
| 2257 | 2257 | 7.05263158 | 3.08602426 | 1 | 0.6184714 | 0 | 1 | 0 | 1 | 0 | 1 | 0 | 1 | 36 | 0.01445326 | FALSE | 1 | FALSE | 1 | FALSE | 1 | FALSE | 1 | FALSE | 1 | FALSE | 1 |
| 2258 | 2258 | 8.71428571 | 0.79539491 | 0 | 1 | 0 | 1 | 1 | 0.40110418 | 3 | 0.0239258 | 0 | 1 | 10 | 0.94579623 | FALSE | 1 | FALSE | 1 | FALSE | 1 | FALSE | 1 | FALSE | 1 | FALSE | 1 |
| 2259 | 2259 | 8.96 | 3.01303833 | 1 | 0.47819878 | 0 | 1 | 0 | 1 | 0 | 1 | 2 | 0.06257402 | 22 | 0.43642846 | FALSE | 1 | FALSE | 1 | FALSE | 1 | FALSE | 1 | FALSE | 1 | FALSE | 1 |
| 2260 | 2260 | 9.41666667 | 1.89113429 | 0 | 1 | 0 | 1 | 0 | 1 | 1 | 0.43131129 | 0 | 1 | 11 | 0.41944532 | FALSE | 1 | FALSE | 1 | FALSE | 1 | FALSE | 1 | FALSE | 1 | FALSE | 1 |
| 2261 | 2261 | 6.21875 | 3.03865076 | 0 | 1 | 0 | 1 | 1 | 0.69069222 | 0 | 1 | 0 | 1 | 31 | 0.02992045 | FALSE | 1 | FALSE | 1 | FALSE | 1 | FALSE | 1 | FALSE | 1 | FALSE | 1 |
| 2262 | 2262 | 10.05 | 3.00790625 | 0 | 1 | 1 | 0.48239024 | 1 | 0.51939841 | 1 | 0.60987146 | 0 | 1 | 17 | 0.61750632 | FALSE | 1 | FALSE | 1 | FALSE | 1 | FALSE | 1 | FALSE | 1 | FALSE | 1 |
| 2263 | 2263 | 12.2666667 | 2.56818137 | 1 | 0.32293355 | 0 | 1 | 0 | 1 | 0 | 1 | 0 | 1 | 14 | 0.29525409 | FALSE | 1 | FALSE | 1 | FALSE | 1 | FALSE | 1 | FALSE | 1 | FALSE | 1 |
| 2264 | 2264 | 6.95294118 | 2.39069708 | 2 | 0.64592863 | 2 | 0.76712318 | 4 | 0.36494003 | 8 | 0.04050293 | 1 | 0.75686725 | 68 | 0.89498314 | FALSE | 1 | FALSE | 1 | FALSE | 1 | FALSE | 1 | FALSE | 1 | FALSE | 1 |
| 2265 | 2265 | 10.0625 | 3.09169755 | 0 | 1 | 1 | 0.40942991 | 2 | 0.11087977 | 1 | 0.52895088 | 1 | 0.23264612 | 11 | 0.9718531 | FALSE | 1 | FALSE | 1 | FALSE | 1 | FALSE | 1 | FALSE | 1 | FALSE | 1 |
| 2266 | 2266 | 9 | 2.08637034 | 0 | 1 | 0 | 1 | 1 | 0.46348836 | 0 | 1 | 0 | 1 | 16 | 0.23070182 | FALSE | 1 | FALSE | 1 | FALSE | 1 | FALSE | 1 | FALSE | 1 | FALSE | 1 |
| 2267 | 2267 | 8.65384615 | 2.25221784 | 0 | 1 | 1 | 0.56109832 | 2 | 0.22597287 | 1 | 0.69181315 | 0 | 1 | 21 | 0.64957167 | FALSE | 1 | FALSE | 1 | FALSE | 1 | FALSE | 1 | FALSE | 1 | FALSE | 1 |
| 2268 | 2268 | 9.45454546 | 3.05595206 | 3 | 0.00234778 | 1 | 0.30368014 | 0 | 1 | 2 | 0.08789566 | 0 | 1 | 5 | 0.99958857 | FALSE | 0.12737083 | FALSE | 1 | FALSE | 1 | FALSE | 1 | FALSE | 1 | FALSE | 1 |
| 2269 | 2269 | 7.875 | 1.26861145 | 0 | 1 | 2 | 0.02563923 | 0 | 1 | 0 | 1 | 0 | 1 | 6 | 0.88425551 | FALSE | 1 | FALSE | 0.82367276 | FALSE | 1 | FALSE | 1 | FALSE | 1 | FALSE | 1 |
| 2270 | 2270 | 2.87755102 | 2.17583947 | 13 | 0.90199921 | 27 | 0.15333282 | 18 | 0.93990718 | 36 | 0.20432487 | 7 | 0.93951143 | 580 | 0.29601805 | FALSE | 1 | FALSE | 1 | FALSE | 1 | FALSE | 1 | FALSE | 1 | FALSE | 1 |
| 2271 | 2271 | 9.77142857 | 1.91364595 | 0 | 1 | 0 | 1 | 4 | 0.0355345 | 1 | 0.80778317 | 2 | 0.11183826 | 28 | 0.83079716 | FALSE | 1 | FALSE | 1 | FALSE | 1 | FALSE | 1 | FALSE | 1 | FALSE | 1 |
| 2272 | 2272 | 9.625 | 3.4255474 | 2 | 0.01652925 | 0 | 1 | 0 | 1 | 1 | 0.31351192 | 0 | 1 | 5 | 0.97551477 | FALSE | 0.39708985 | FALSE | 1 | FALSE | 1 | FALSE | 1 | FALSE | 1 | FALSE | 1 |
| 2273 | 2273 | 8.91891892 | 4.0893955 | 3 | 0.0681763 | 0 | 1 | 2 | 0.38585089 | 3 | 0.2402032 | 0 | 1 | 29 | 0.88820978 | FALSE | 0.74276693 | FALSE | 1 | FALSE | 1 | FALSE | 1 | FALSE | 1 | FALSE | 1 |
| 2274 | 2274 | 8.05454546 | 2.51119806 | 2 | 0.41401181 | 7 | 0.00182072 | 3 | 0.31652594 | 3 | 0.46623289 | 0 | 1 | 40 | 0.99191141 | FALSE | 1 | FALSE | 0.40311621 | FALSE | 1 | FALSE | 1 | FALSE | 1 | FALSE | 1 |
| 2275 | 2275 | 9.53846154 | 2.60518373 | 2 | 0.13369621 | 2 | 0.19297191 | 0 | 1 | 1 | 0.69181315 | 2 | 0.06257402 | 18 | 0.96851178 | FALSE | 0.97964162 | FALSE | 1 | FALSE | 1 | FALSE | 1 | FALSE | 1 | FALSE | 1 |
| 2276 | 2276 | 8.75 | 2.00520157 | 0 | 1 | 1 | 0.32623933 | 1 | 0.35555293 | 0 | 1 | 0 | 1 | 10 | 0.71436964 | FALSE | 1 | FALSE | 1 | FALSE | 1 | FALSE | 1 | FALSE | 1 | FALSE | 1 |
| 2277 | 2277 | 11.2222222 | 2.93552107 | 2 | 0.07645454 | 0 | 1 | 0 | 1 | 1 | 0.57131013 | 0 | 1 | 15 | 0.69372602 | FALSE | 0.77424195 | FALSE | 1 | FALSE | 1 | FALSE | 1 | FALSE | 1 | FALSE | 1 |
| 2278 | 2278 | 9.56666667 | 2.30482586 | 0 | 1 | 0 | 1 | 1 | 0.66709967 | 3 | 0.15707004 | 1 | 0.39166989 | 25 | 0.67630471 | FALSE | 1 | FALSE | 1 | FALSE | 1 | FALSE | 1 | FALSE | 1 | FALSE | 1 |
| 2279 | 2279 | 7.62962963 | 2.17133085 | 2 | 0.38681045 | 1 | 0.82028131 | 5 | 0.03785225 | 2 | 0.6970685 | 1 | 0.57809055 | 41 | 0.89856682 | FALSE | 1 | FALSE | 1 | FALSE | 1 | FALSE | 1 | FALSE | 1 | FALSE | 1 |
| 2280 | 2280 | 10.6 | 1.92527054 | 0 | 1 | 1 | 0.38964394 | 1 | 0.42266264 | 0 | 1 | 0 | 1 | 13 | 0.57712805 | FALSE | 1 | FALSE | 1 | FALSE | 1 | FALSE | 1 | FALSE | 1 | FALSE | 1 |
| 2281 | 2281 | 5.14563107 | 2.30410258 | 2 | 0.74184548 | 2 | 0.84797708 | 7 | 0.07345885 | 4 | 0.69586384 | 2 | 0.50146696 | 85 | 0.67551735 | FALSE | 1 | FALSE | 1 | FALSE | 1 | FALSE | 1 | FALSE | 1 | FALSE | 1 |
| 2282 | 2282 | 10.4 | 2.97321375 | 0 | 1 | 1 | 0.28036922 | 1 | 0.30655256 | 0 | 1 | 0 | 1 | 8 | 0.80394719 | FALSE | 1 | FALSE | 1 | FALSE | 1 | FALSE | 1 | FALSE | 1 | FALSE | 1 |
| 2283 | 2283 | 9.8 | 2.28619043 | 0 | 1 | 0 | 1 | 0 | 1 | 1 | 0.50623194 | 0 | 1 | 14 | 0.29525409 | FALSE | 1 | FALSE | 1 | FALSE | 1 | FALSE | 1 | FALSE | 1 | FALSE | 1 |
| 2284 | 2284 | 10.9285714 | 2.12011744 | 0 | 1 | 0 | 1 | 0 | 1 | 0 | 1 | 0 | 1 | 14 | 0.09251304 | FALSE | 1 | FALSE | 1 | FALSE | 1 | FALSE | 1 | FALSE | 1 | FALSE | 1 |
| 2285 | 2285 | 12.6666667 | 2.3570226 | 1 | 0.14434519 | 0 | 1 | 0 | 1 | 0 | 1 | 0 | 1 | 5 | 0.76185568 | FALSE | 1 | FALSE | 1 | FALSE | 1 | FALSE | 1 | FALSE | 1 | FALSE | 1 |
| 2286 | 2286 | 9.94736842 | 2.32772046 | 0 | 1 | 1 | 0.44710673 | 1 | 0.48280983 | 3 | 0.04687752 | 0 | 1 | 13 | 0.95041913 | FALSE | 1 | FALSE | 1 | FALSE | 1 | FALSE | 1 | FALSE | 1 | FALSE | 1 |
| 2287 | 2287 | 7.55319149 | 1.73583643 | 2 | 0.3309085 | 4 | 0.06038631 | 2 | 0.4962935 | 3 | 0.35428318 | 0 | 1 | 35 | 0.95410769 | FALSE | 1 | FALSE | 1 | FALSE | 1 | FALSE | 1 | FALSE | 1 | FALSE | 1 |
| 2288 | 2288 | 9.8125 | 2.55486668 | 0 | 1 | 2 | 0.09285443 | 0 | 1 | 0 | 1 | 0 | 1 | 14 | 0.53308261 | FALSE | 1 | FALSE | 1 | FALSE | 1 | FALSE | 1 | FALSE | 1 | FALSE | 1 |
| 2289 | 2289 | 10.25 | 3.73329613 | 0 | 1 | 0 | 1 | 1 | 0.44344817 | 2 | 0.16544789 | 0 | 1 | 13 | 0.76799161 | FALSE | 1 | FALSE | 1 | FALSE | 1 | FALSE | 1 | FALSE | 1 | FALSE | 1 |
| 2290 | 2290 | 7.92857143 | 2.48224991 | 0 | 1 | 1 | 0.74131698 | 0 | 1 | 0 | 1 | 1 | 0.49330646 | 39 | 0.03430384 | FALSE | 1 | FALSE | 1 | FALSE | 1 | FALSE | 1 | FALSE | 1 | FALSE | 1 |
| 2291 | 2291 | 10.0416667 | 3.07515808 | 1 | 0.45027683 | 0 | 1 | 5 | 0.00113518 | 0 | 1 | 0 | 1 | 17 | 0.94439745 | FALSE | 1 | FALSE | 1 | FALSE | 1 | FALSE | 1 | FALSE | 1 | FALSE | 1 |
| 2292 | 2292 | 9.46666667 | 3.34398299 | 0 | 1 | 1 | 0.38964394 | 0 | 1 | 3 | 0.02891327 | 0 | 1 | 11 | 0.92879412 | FALSE | 1 | FALSE | 1 | FALSE | 1 | FALSE | 1 | FALSE | 1 | FALSE | 1 |
| 2293 | 2293 | 9.73333333 | 2.88598144 | 2 | 0.05513784 | 0 | 1 | 1 | 0.42266264 | 0 | 1 | 1 | 0.2198258 | 11 | 0.92879412 | FALSE | 0.68849059 | FALSE | 1 | FALSE | 1 | FALSE | 1 | FALSE | 1 | FALSE | 1 |
| 2294 | 2294 | 8.84 | 1.66565303 | 1 | 0.47819878 | 2 | 0.19297191 | 1 | 0.59998149 | 1 | 0.69181315 | 1 | 0.33902672 | 19 | 0.91734261 | FALSE | 1 | FALSE | 1 | FALSE | 1 | FALSE | 1 | FALSE | 1 | FALSE | 1 |
| 2295 | 2295 | 10.6944444 | 1.95532669 | 0 | 1 | 1 | 0.69480472 | 0 | 1 | 2 | 0.49731009 | 0 | 1 | 33 | 0.16407838 | FALSE | 1 | FALSE | 1 | FALSE | 1 | FALSE | 1 | FALSE | 1 | FALSE | 1 |
| 2296 | 2296 | 8.24137931 | 2.23845964 | 0 | 1 | 1 | 0.60247911 | 2 | 0.2663542 | 3 | 0.13524571 | 0 | 1 | 22 | 0.86499992 | FALSE | 1 | FALSE | 1 | FALSE | 1 | FALSE | 1 | FALSE | 1 | FALSE | 1 |
| 2297 | 2297 | 10.661017 | 0.93174103 | 0 | 1 | 0 | 1 | 3 | 0.35626822 | 3 | 0.51328524 | 1 | 0.62455813 | 52 | 0.27701375 | FALSE | 1 | FALSE | 1 | FALSE | 1 | FALSE | 1 | FALSE | 1 | FALSE | 1 |
| 2298 | 2298 | 6.34782609 | 2.82240563 | 0 | 1 | 1 | 0.74131698 | 3 | 0.1814659 | 3 | 0.29054213 | 0 | 1 | 34 | 0.69505404 | FALSE | 1 | FALSE | 1 | FALSE | 1 | FALSE | 1 | FALSE | 1 | FALSE | 1 |
| 2299 | 2299 | 4.29268293 | 3.60043292 | 1 | 0.8826742 | 3 | 0.49822953 | 4 | 0.34020118 | 7 | 0.08151801 | 1 | 0.74434486 | 66 | 0.86930658 | FALSE | 1 | FALSE | 1 | FALSE | 1 | FALSE | 1 | FALSE | 1 | FALSE | 1 |
| 2300 | 2300 | 9.94444444 | 2.27235126 | 0 | 1 | 0 | 1 | 0 | 1 | 1 | 0.57131013 | 0 | 1 | 17 | 0.20329205 | FALSE | 1 | FALSE | 1 | FALSE | 1 | FALSE | 1 | FALSE | 1 | FALSE | 1 |
| 2301 | 2301 | 7.98387097 | 2.23961356 | 6 | 0.0048757 | 6 | 0.01453703 | 1 | 0.89758397 | 4 | 0.31731292 | 0 | 1 | 45 | 0.99474796 | FALSE | 0.19848171 | FALSE | 0.71309764 | FALSE | 1 | FALSE | 1 | FALSE | 1 | FALSE | 1 |
| 2302 | 2302 | 8.30769231 | 2.24926023 | 2 | 0.14250978 | 1 | 0.57534752 | 0 | 1 | 6 | 0.00094105 | 0 | 1 | 17 | 0.99598795 | FALSE | 1 | FALSE | 1 | FALSE | 1 | FALSE | 1 | FALSE | 1 | FALSE | 1 |
| 2303 | 2303 | 9.32941177 | 1.46639402 | 1 | 0.89157583 | 6 | 0.05649557 | 1 | 0.95626254 | 4 | 0.55256464 | 1 | 0.75686725 | 72 | 0.54001499 | FALSE | 1 | FALSE | 1 | FALSE | 1 | FALSE | 1 | FALSE | 1 | FALSE | 1 |
| 2304 | 2304 | 11.4285714 | 2.52740086 | 1 | 0.3050805 | 1 | 0.36919824 | 0 | 1 | 0 | 1 | 0 | 1 | 12 | 0.62236227 | FALSE | 1 | FALSE | 1 | FALSE | 1 | FALSE | 1 | FALSE | 1 | FALSE | 1 |
| 2305 | 2305 | 10.5833333 | 3.40240928 | 2 | 0.03643311 | 0 | 1 | 0 | 1 | 0 | 1 | 0 | 1 | 10 | 0.71436964 | FALSE | 0.58392 | FALSE | 1 | FALSE | 1 | FALSE | 1 | FALSE | 1 | FALSE | 1 |
| 2306 | 2306 | 10.3125 | 1.3564084 | 1 | 0.34033054 | 1 | 0.40942991 | 1 | 0.44344817 | 1 | 0.52895088 | 1 | 0.23264612 | 11 | 0.9718531 | FALSE | 1 | FALSE | 1 | FALSE | 1 | FALSE | 1 | FALSE | 1 | FALSE | 1 |
| 2307 | 2307 | 8.72727273 | 1.68365993 | 1 | 0.4357638 | 1 | 0.5154318 | 0 | 1 | 3 | 0.07743186 | 0 | 1 | 17 | 0.8840357 | FALSE | 1 | FALSE | 1 | FALSE | 1 | FALSE | 1 | FALSE | 1 | FALSE | 1 |
| 2308 | 2308 | 8.6 | 2.33238076 | 1 | 0.32293355 | 0 | 1 | 1 | 0.42266264 | 0 | 1 | 0 | 1 | 13 | 0.57712805 | FALSE | 1 | FALSE | 1 | FALSE | 1 | FALSE | 1 | FALSE | 1 | FALSE | 1 |
| 2309 | 2309 | 8.9 | 2.84429253 | 0 | 1 | 0 | 1 | 0 | 1 | 0 | 1 | 0 | 1 | 10 | 0.18273166 | FALSE | 1 | FALSE | 1 | FALSE | 1 | FALSE | 1 | FALSE | 1 | FALSE | 1 |
| 2310 | 2310 | 10.1176471 | 2.34852527 | 0 | 1 | 1 | 0.42857733 | 1 | 0.46348836 | 0 | 1 | 0 | 1 | 15 | 0.49060435 | FALSE | 1 | FALSE | 1 | FALSE | 1 | FALSE | 1 | FALSE | 1 | FALSE | 1 |
| 2311 | 2311 | 9.9375 | 2.65680141 | 4 | 0.00059649 | 2 | 0.09285443 | 0 | 1 | 2 | 0.16544789 | 0 | 1 | 8 | 0.99978399 | FALSE | 0.05254197 | FALSE | 1 | FALSE | 1 | FALSE | 1 | FALSE | 1 | FALSE | 1 |
| 2312 | 2312 | 9.28571429 | 2.77990848 | 0 | 1 | 0 | 1 | 0 | 1 | 0 | 1 | 0 | 1 | 21 | 0.02808151 | FALSE | 1 | FALSE | 1 | FALSE | 1 | FALSE | 1 | FALSE | 1 | FALSE | 1 |
| 2313 | 2313 | 10.4 | 3.24139888 | 1 | 0.32293355 | 0 | 1 | 2 | 0.09928574 | 0 | 1 | 1 | 0.2198258 | 11 | 0.92879412 | FALSE | 1 | FALSE | 1 | FALSE | 1 | FALSE | 1 | FALSE | 1 | FALSE | 1 |
| 2314 | 2314 | 8.1 | 1.92093727 | 0 | 1 | 0 | 1 | 0 | 1 | 1 | 0.37517262 | 0 | 1 | 9 | 0.5215547 | FALSE | 1 | FALSE | 1 | FALSE | 1 | FALSE | 1 | FALSE | 1 | FALSE | 1 |
| 2315 | 2315 | 8.03846154 | 2.57921827 | 1 | 0.49162634 | 0 | 1 | 0 | 1 | 3 | 0.11458571 | 1 | 0.3499045 | 21 | 0.78939668 | FALSE | 1 | FALSE | 1 | FALSE | 1 | FALSE | 1 | FALSE | 1 | FALSE | 1 |
| 2316 | 2316 | 8.04166667 | 2.97880243 | 1 | 0.46441868 | 0 | 1 | 2 | 0.21260257 | 2 | 0.30249307 | 0 | 1 | 19 | 0.83975208 | FALSE | 1 | FALSE | 1 | FALSE | 1 | FALSE | 1 | FALSE | 1 | FALSE | 1 |
| 2317 | 2317 | 10.2222222 | 1.98761598 | 1 | 0.20854748 | 0 | 1 | 0 | 1 | 0 | 1 | 0 | 1 | 8 | 0.5780332 | FALSE | 1 | FALSE | 1 | FALSE | 1 | FALSE | 1 | FALSE | 1 | FALSE | 1 |
| 2318 | 2318 | 6.97297297 | 1.46096019 | 1 | 0.6184714 | 1 | 0.70472939 | 2 | 0.38585089 | 4 | 0.08780467 | 0 | 1 | 29 | 0.88820978 | FALSE | 1 | FALSE | 1 | FALSE | 1 | FALSE | 1 | FALSE | 1 | FALSE | 1 |
| 2319 | 2319 | 7.07692308 | 4.77914602 | 2 | 0.14250978 | 0 | 1 | 2 | 0.23940226 | 1 | 0.70601446 | 2 | 0.0670823 | 19 | 0.96054244 | FALSE | 1 | FALSE | 1 | FALSE | 1 | FALSE | 1 | FALSE | 1 | FALSE | 1 |
| 2320 | 2320 | 9.29411765 | 1.67207887 | 0 | 1 | 2 | 0.10307112 | 0 | 1 | 0 | 1 | 0 | 1 | 15 | 0.49060435 | FALSE | 1 | FALSE | 1 | FALSE | 1 | FALSE | 1 | FALSE | 1 | FALSE | 1 |
| 2321 | 2321 | 7.8125 | 2.39057394 | 1 | 0.56527493 | 2 | 0.27730767 | 4 | 0.02651644 | 0 | 1 | 4 | 0.00172712 | 21 | 0.9978606 | FALSE | 1 | FALSE | 1 | FALSE | 1 | FALSE | 1 | FALSE | 0.23980666 | FALSE | 1 |
| 2322 | 2322 | 8.54545455 | 3.60211141 | 1 | 0.24866461 | 0 | 1 | 0 | 1 | 1 | 0.40390032 | 0 | 1 | 9 | 0.75985366 | FALSE | 1 | FALSE | 1 | FALSE | 1 | FALSE | 1 | FALSE | 1 | FALSE | 1 |
| 2323 | 2323 | 10.2941177 | 2.3203627 | 2 | 0.06909038 | 0 | 1 | 0 | 1 | 0 | 1 | 0 | 1 | 15 | 0.49060435 | FALSE | 0.74276693 | FALSE | 1 | FALSE | 1 | FALSE | 1 | FALSE | 1 | FALSE | 1 |
| 2324 | 2324 | 7 | 2.13497999 | 1 | 0.62830697 | 2 | 0.3494045 | 2 | 0.39868506 | 1 | 0.83318955 | 0 | 1 | 32 | 0.61800916 | FALSE | 1 | FALSE | 1 | FALSE | 1 | FALSE | 1 | FALSE | 1 | FALSE | 1 |
| 2325 | 2325 | 8.4 | 2.1725561 | 0 | 1 | 0 | 1 | 1 | 0.59998149 | 1 | 0.69181315 | 0 | 1 | 23 | 0.22727183 | FALSE | 1 | FALSE | 1 | FALSE | 1 | FALSE | 1 | FALSE | 1 | FALSE | 1 |
| 2326 | 2326 | 8 | 2.88230677 | 2 | 0.14250978 | 4 | 0.00909592 | 2 | 0.23940226 | 0 | 1 | 1 | 0.3499045 | 17 | 0.99598795 | FALSE | 1 | FALSE | 0.63686998 | FALSE | 1 | FALSE | 1 | FALSE | 1 | FALSE | 1 |
| 2327 | 2327 | 9.41666667 | 1.96885472 | 4 | 0.0671906 | 5 | 0.04382428 | 4 | 0.16812274 | 0 | 1 | 1 | 0.63076845 | 46 | 0.96067108 | FALSE | 0.74276693 | FALSE | 1 | FALSE | 1 | FALSE | 1 | FALSE | 1 | FALSE | 1 |
| 2328 | 2328 | 8.6 | 3.44093011 | 0 | 1 | 0 | 1 | 1 | 0.51939841 | 1 | 0.60987146 | 0 | 1 | 18 | 0.37504645 | FALSE | 1 | FALSE | 1 | FALSE | 1 | FALSE | 1 | FALSE | 1 | FALSE | 1 |
| 2329 | 2329 | 11.0714286 | 2.52032554 | 0 | 1 | 0 | 1 | 0 | 1 | 0 | 1 | 1 | 0.20679328 | 13 | 0.33283729 | FALSE | 1 | FALSE | 1 | FALSE | 1 | FALSE | 1 | FALSE | 1 | FALSE | 1 |
| 2330 | 2330 | 7.94444444 | 1.87000363 | 1 | 0.60837712 | 2 | 0.32556427 | 1 | 0.73299778 | 2 | 0.49731009 | 0 | 1 | 30 | 0.67248271 | FALSE | 1 | FALSE | 1 | FALSE | 1 | FALSE | 1 | FALSE | 1 | FALSE | 1 |
| 2331 | 2331 | 2.12334802 | 0.99236348 | 34 | 0.01337507 | 35 | 0.15047725 | 31 | 0.65084209 | 48 | 0.16048752 | 24 | 0.01088155 | 736 | 0.99843286 | FALSE | 0.36625055 | FALSE | 1 | FALSE | 1 | FALSE | 1 | FALSE | 0.57142086 | FALSE | 1 |
| 2332 | 2332 | 9.36585366 | 2.58283355 | 4 | 0.01859058 | 2 | 0.37295654 | 1 | 0.76953771 | 1 | 0.84823834 | 1 | 0.48480878 | 31 | 0.91676852 | FALSE | 0.43265346 | FALSE | 1 | FALSE | 1 | FALSE | 1 | FALSE | 1 | FALSE | 1 |
| 2333 | 2333 | 8.61111111 | 2.54162872 | 0 | 1 | 0 | 1 | 1 | 0.48280983 | 1 | 0.57131013 | 0 | 1 | 16 | 0.44998001 | FALSE | 1 | FALSE | 1 | FALSE | 1 | FALSE | 1 | FALSE | 1 | FALSE | 1 |
| 2334 | 2334 | 8.65384615 | 2.94751927 | 0 | 1 | 0 | 1 | 0 | 1 | 1 | 0.70601446 | 1 | 0.3499045 | 24 | 0.20425579 | FALSE | 1 | FALSE | 1 | FALSE | 1 | FALSE | 1 | FALSE | 1 | FALSE | 1 |
| 2335 | 2335 | 8.25 | 2.2220486 | 0 | 1 | 2 | 0.09285443 | 0 | 1 | 0 | 1 | 0 | 1 | 14 | 0.53308261 | FALSE | 1 | FALSE | 1 | FALSE | 1 | FALSE | 1 | FALSE | 1 | FALSE | 1 |
| 2336 | 2336 | 9.94444444 | 2.34454976 | 1 | 0.37380237 | 0 | 1 | 0 | 1 | 2 | 0.19902682 | 1 | 0.25766374 | 14 | 0.86287418 | FALSE | 1 | FALSE | 1 | FALSE | 1 | FALSE | 1 | FALSE | 1 | FALSE | 1 |
| 2337 | 2337 | 5.07575758 | 2.36969891 | 3 | 0.2187296 | 3 | 0.33367983 | 4 | 0.18968799 | 6 | 0.06788138 | 2 | 0.27647607 | 45 | 0.99738502 | FALSE | 1 | FALSE | 1 | FALSE | 1 | FALSE | 1 | FALSE | 1 | FALSE | 1 |
| 2338 | 2338 | 8.27272727 | 1.65644247 | 0 | 1 | 0 | 1 | 0 | 1 | 0 | 1 | 0 | 1 | 11 | 0.15414489 | FALSE | 1 | FALSE | 1 | FALSE | 1 | FALSE | 1 | FALSE | 1 | FALSE | 1 |
| 2339 | 2339 | 9.47368421 | 2.66297066 | 1 | 0.38989951 | 1 | 0.46503795 | 0 | 1 | 0 | 1 | 0 | 1 | 17 | 0.41141543 | FALSE | 1 | FALSE | 1 | FALSE | 1 | FALSE | 1 | FALSE | 1 | FALSE | 1 |
| 2340 | 2340 | 10.3684211 | 2.34432184 | 2 | 0.08405377 | 0 | 1 | 0 | 1 | 1 | 0.59104357 | 0 | 1 | 16 | 0.65565778 | FALSE | 0.79236883 | FALSE | 1 | FALSE | 1 | FALSE | 1 | FALSE | 1 | FALSE | 1 |
| 2341 | 2341 | 9.2 | 2.73130006 | 0 | 1 | 1 | 0.48239024 | 1 | 0.51939841 | 1 | 0.60987146 | 0 | 1 | 17 | 0.61750632 | FALSE | 1 | FALSE | 1 | FALSE | 1 | FALSE | 1 | FALSE | 1 | FALSE | 1 |
| 2342 | 2342 | 7.92 | 1.97828208 | 0 | 1 | 2 | 0.19297191 | 0 | 1 | 1 | 0.69181315 | 0 | 1 | 22 | 0.43642846 | FALSE | 1 | FALSE | 1 | FALSE | 1 | FALSE | 1 | FALSE | 1 | FALSE | 1 |
| 2343 | 2343 | 8.77777778 | 3.22413736 | 0 | 1 | 0 | 1 | 0 | 1 | 1 | 0.57131013 | 0 | 1 | 17 | 0.20329205 | FALSE | 1 | FALSE | 1 | FALSE | 1 | FALSE | 1 | FALSE | 1 | FALSE | 1 |
| 2344 | 2344 | 9.38461539 | 2.45069728 | 1 | 0.49162634 | 0 | 1 | 2 | 0.23940226 | 1 | 0.70601446 | 1 | 0.3499045 | 21 | 0.78939668 | FALSE | 1 | FALSE | 1 | FALSE | 1 | FALSE | 1 | FALSE | 1 | FALSE | 1 |
| 2345 | 2345 | 9.52173913 | 2.28167683 | 1 | 0.4357638 | 0 | 1 | 1 | 0.55340851 | 1 | 0.64497439 | 0 | 1 | 19 | 0.54233419 | FALSE | 1 | FALSE | 1 | FALSE | 1 | FALSE | 1 | FALSE | 1 | FALSE | 1 |
| 2346 | 2346 | 8 | 2.39918685 | 0 | 1 | 3 | 0.13823565 | 1 | 0.76953771 | 1 | 0.84823834 | 2 | 0.13925369 | 33 | 0.71942635 | FALSE | 1 | FALSE | 1 | FALSE | 1 | FALSE | 1 | FALSE | 1 | FALSE | 1 |
| 2347 | 2347 | 7.57142857 | 3.53986048 | 1 | 0.16630142 | 0 | 1 | 2 | 0.02394967 | 0 | 1 | 1 | 0.10932414 | 3 | 0.99853144 | FALSE | 1 | FALSE | 1 | FALSE | 1 | FALSE | 1 | FALSE | 1 | FALSE | 1 |
| 2348 | 2348 | 10.2666667 | 2.32283352 | 3 | 0.00600364 | 0 | 1 | 0 | 1 | 1 | 0.50623194 | 0 | 1 | 11 | 0.92879412 | FALSE | 0.22773046 | FALSE | 1 | FALSE | 1 | FALSE | 1 | FALSE | 1 | FALSE | 1 |

| | | | | | | | | | | | | | | | | | | | | | | | | | | | |
|---|---|---|---|---|---|---|---|---|---|---|---|---|---|---|---|---|---|---|---|---|---|---|---|---|---|---|---|
| 2349 | 2349 | 11.0952381 | 3.99886605 | 0 | 1 | 1 | 0.49918218 | 1 | 0.53671415 | 3 | 0.06912908 | 0 | 1 | 16 | 0.90353674 | FALSE | 1 | FALSE | 1 | FALSE | 1 | FALSE | 1 | FALSE | 1 | FALSE | 1 |
| 2350 | 2350 | 9.26666667 | 1.12348664 | 0 | 1 | 1 | 0.38964394 | 0 | 1 | 2 | 0.14906763 | 0 | 1 | 12 | 0.80329917 | FALSE | 1 | FALSE | 1 | FALSE | 1 | FALSE | 1 | FALSE | 1 | FALSE | 1 |
| 2351 | 2351 | 7.89473684 | 1.02056418 | 0 | 1 | 1 | 0.46503795 | 1 | 0.50143826 | 1 | 0.59104357 | 1 | 0.26986775 | 15 | 0.83648191 | FALSE | 1 | FALSE | 1 | FALSE | 1 | FALSE | 1 | FALSE | 1 | FALSE | 1 |
| 2352 | 2352 | 7.23076923 | 1.47564047 | 0 | 1 | 1 | 0.57534752 | 2 | 0.23940226 | 2 | 0.33678042 | 0 | 1 | 21 | 0.78939668 | FALSE | 1 | FALSE | 1 | FALSE | 1 | FALSE | 1 | FALSE | 1 | FALSE | 1 |
| 2353 | 2353 | 8.97826087 | 2.50641898 | 0 | 1 | 3 | 0.18548475 | 1 | 0.81522035 | 1 | 0.88573774 | 0 | 1 | 41 | 0.25430673 | FALSE | 1 | FALSE | 1 | FALSE | 1 | FALSE | 1 | FALSE | 1 | FALSE | 1 |
| 2354 | 2354 | 8.7037037 | 1.67364793 | 3 | 0.03084786 | 0 | 1 | 0 | 1 | 1 | 0.71956339 | 1 | 0.36060486 | 22 | 0.76237877 | FALSE | 0.53356688 | FALSE | 1 | FALSE | 1 | FALSE | 1 | FALSE | 1 | FALSE | 1 |
| 2355 | 2355 | 9.85714286 | 2.29462549 | 0 | 1 | 0 | 1 | 0 | 1 | 2 | 0.03785362 | 0 | 1 | 5 | 0.91847773 | FALSE | 1 | FALSE | 1 | FALSE | 1 | FALSE | 1 | FALSE | 1 | FALSE | 1 |
| 2356 | 2356 | 9.90476191 | 2.65302634 | 1 | 0.42086991 | 0 | 1 | 0 | 1 | 2 | 0.25059627 | 0 | 1 | 18 | 0.57962763 | FALSE | 1 | FALSE | 1 | FALSE | 1 | FALSE | 1 | FALSE | 1 | FALSE | 1 |
| 2357 | 2357 | 11.25 | 2.94745653 | 5 | 0.00011852 | 2 | 0.1353446 | 0 | 1 | 1 | 0.60987146 | 0 | 1 | 12 | 0.99824567 | TRUE | 0.01941834 | FALSE | 1 | FALSE | 1 | FALSE | 1 | FALSE | 1 | FALSE | 1 |
| 2358 | 2358 | 6.13907285 | 0.86153825 | 8 | 0.98955116 | 18 | 0.67141638 | 25 | 0.24670676 | 22 | 0.89632448 | 12 | 0.2793051 | 516 | 0.16187761 | FALSE | 1 | FALSE | 1 | FALSE | 1 | FALSE | 1 | FALSE | 1 | FALSE | 1 |
| 2359 | 2359 | 11.8571429 | 1.84612114 | 1 | 0.3050805 | 1 | 0.36919824 | 0 | 1 | 1 | 0.48242099 | 0 | 1 | 11 | 0.83676449 | FALSE | 1 | FALSE | 1 | FALSE | 1 | FALSE | 1 | FALSE | 1 | FALSE | 1 |
| 2360 | 2360 | 6.90909091 | 1.85641617 | 0 | 1 | 0 | 1 | 0 | 1 | 0 | 1 | 1 | 0.30529951 | 21 | 0.12048848 | FALSE | 1 | FALSE | 1 | FALSE | 1 | FALSE | 1 | FALSE | 1 | FALSE | 1 |
| 2361 | 2361 | 8.72222222 | 2.62055031 | 1 | 0.37380237 | 1 | 0.44710673 | 1 | 0.48280983 | 2 | 0.19902682 | 0 | 1 | 13 | 0.95041913 | FALSE | 1 | FALSE | 1 | FALSE | 1 | FALSE | 1 | FALSE | 1 | FALSE | 1 |
| 2362 | 2362 | 10.0769231 | 0.99703703 | 1 | 0.28675951 | 1 | 0.3480709 | 0 | 1 | 0 | 1 | 0 | 1 | 11 | 0.6683056 | FALSE | 1 | FALSE | 1 | FALSE | 1 | FALSE | 1 | FALSE | 1 | FALSE | 1 |
| 2363 | 2363 | 7.09722222 | 2.94938658 | 3 | 0.28086759 | 2 | 0.68229183 | 3 | 0.48160679 | 2 | 0.85003314 | 0 | 1 | 62 | 0.41748931 | FALSE | 1 | FALSE | 1 | FALSE | 1 | FALSE | 1 | FALSE | 1 | FALSE | 1 |
| 2364 | 2364 | 9.57692308 | 2.00332564 | 1 | 0.49162634 | 1 | 0.57534752 | 3 | 0.06482014 | 2 | 0.33678042 | 0 | 1 | 19 | 0.96054244 | FALSE | 1 | FALSE | 1 | FALSE | 1 | FALSE | 1 | FALSE | 1 | FALSE | 1 |
| 2365 | 2365 | 10.0217391 | 3.15876373 | 0 | 1 | 1 | 0.78076835 | 0 | 1 | 1 | 0.88573774 | 0 | 1 | 44 | 0.01791373 | FALSE | 1 | FALSE | 1 | FALSE | 1 | FALSE | 1 | FALSE | 1 | FALSE | 1 |
| 2366 | 2366 | 5.47826087 | 2.42336869 | 1 | 0.83491628 | 0 | 1 | 3 | 0.45353909 | 3 | 0.6203651 | 3 | 0.1035305 | 59 | 0.47808459 | FALSE | 1 | FALSE | 1 | FALSE | 1 | FALSE | 1 | FALSE | 1 | FALSE | 1 |
| 2367 | 2367 | 4.3984375 | 1.74928487 | 2 | 0.99046366 | 8 | 0.58256756 | 9 | 0.56605271 | 14 | 0.27713433 | 7 | 0.12275609 | 214 | 0.56484799 | FALSE | 1 | FALSE | 1 | FALSE | 1 | FALSE | 1 | FALSE | 1 | FALSE | 1 |
| 2368 | 2368 | 9.90476191 | 2.15814651 | 1 | 0.42086991 | 4 | 0.00413111 | 0 | 1 | 2 | 0.25059627 | 1 | 0.29368245 | 13 | 0.9973503 | FALSE | 1 | FALSE | 0.58120789 | FALSE | 1 | FALSE | 1 | FALSE | 1 | FALSE | 1 |
| 2369 | 2369 | 10 | 3.05505046 | 0 | 1 | 1 | 0.38964394 | 0 | 1 | 2 | 0.14906763 | 0 | 1 | 12 | 0.80329917 | FALSE | 1 | FALSE | 1 | FALSE | 1 | FALSE | 1 | FALSE | 1 | FALSE | 1 |
| 2370 | 2370 | 10.0952381 | 3.29364123 | 0 | 1 | 1 | 0.49918218 | 1 | 0.53671415 | 0 | 1 | 1 | 0.29368245 | 18 | 0.57962763 | FALSE | 1 | FALSE | 1 | FALSE | 1 | FALSE | 1 | FALSE | 1 | FALSE | 1 |
| 2371 | 2371 | 4.28571429 | 1.49623316 | 8 | 0.24077671 | 7 | 0.62481412 | 10 | 0.31758275 | 13 | 0.26273355 | 3 | 0.73652732 | 190 | 0.84122099 | FALSE | 1 | FALSE | 1 | FALSE | 1 | FALSE | 1 | FALSE | 1 | FALSE | 1 |
| 2372 | 2372 | 8.13333333 | 2.72926527 | 4 | 0.00045644 | 2 | 0.08295839 | 2 | 0.09928574 | 0 | 1 | 1 | 0.2198258 | 6 | 0.99998825 | TRUE | 0.0445135 | FALSE | 1 | FALSE | 1 | FALSE | 1 | FALSE | 1 | FALSE | 1 |
| 2373 | 2373 | 8.10344828 | 2.6043045 | 2 | 0.16962005 | 0 | 1 | 1 | 0.65463982 | 2 | 0.3871811 | 0 | 1 | 24 | 0.70563334 | FALSE | 1 | FALSE | 1 | FALSE | 1 | FALSE | 1 | FALSE | 1 | FALSE | 1 |
| 2374 | 2374 | 9.61538462 | 2.90988928 | 0 | 1 | 1 | 0.56109832 | 0 | 1 | 1 | 0.69181315 | 1 | 0.33902672 | 22 | 0.43642846 | FALSE | 1 | FALSE | 1 | FALSE | 1 | FALSE | 1 | FALSE | 1 | FALSE | 1 |
| 2375 | 2375 | 9.83333333 | 2.9178569 | 1 | 0.71380452 | 2 | 0.46321244 | 1 | 0.8283465 | 2 | 0.65421368 | 2 | 0.18560062 | 40 | 0.66905473 | FALSE | 1 | FALSE | 1 | FALSE | 1 | FALSE | 1 | FALSE | 1 | FALSE | 1 |
| 2376 | 2376 | 10.1333333 | 3.20138859 | 0 | 1 | 1 | 0.38964394 | 1 | 0.42266264 | 0 | 1 | 0 | 1 | 13 | 0.57712805 | FALSE | 1 | FALSE | 1 | FALSE | 1 | FALSE | 1 | FALSE | 1 | FALSE | 1 |
| 2377 | 2377 | 7.17032967 | 1.14287035 | 8 | 0.09525445 | 7 | 0.37485369 | 4 | 0.8982367 | 8 | 0.60213399 | 2 | 0.80500336 | 153 | 0.59590307 | FALSE | 0.85844272 | FALSE | 1 | FALSE | 1 | FALSE | 1 | FALSE | 1 | FALSE | 1 |
| 2378 | 2378 | 9.17647059 | 3.0723682 | 0 | 1 | 2 | 0.30150529 | 2 | 0.34668321 | 2 | 0.46709861 | 0 | 1 | 28 | 0.72558119 | FALSE | 1 | FALSE | 1 | FALSE | 1 | FALSE | 1 | FALSE | 1 | FALSE | 1 |
| 2379 | 2379 | 8.91666667 | 2.56444709 | 1 | 0.26795838 | 1 | 0.32623933 | 0 | 1 | 1 | 0.43131129 | 0 | 1 | 9 | 0.89630573 | FALSE | 1 | FALSE | 1 | FALSE | 1 | FALSE | 1 | FALSE | 1 | FALSE | 1 |
| 2380 | 2380 | 10.5 | 2.25989886 | 0 | 1 | 0 | 1 | 0 | 1 | 0 | 1 | 1 | 0.20679328 | 13 | 0.33283729 | FALSE | 1 | FALSE | 1 | FALSE | 1 | FALSE | 1 | FALSE | 1 | FALSE | 1 |
| 2381 | 2381 | 8.25 | 2.50748878 | 0 | 1 | 0 | 1 | 1 | 0.51939841 | 1 | 0.60987146 | 1 | 0.28187291 | 17 | 0.61750632 | FALSE | 1 | FALSE | 1 | FALSE | 1 | FALSE | 1 | FALSE | 1 | FALSE | 1 |
| 2382 | 2382 | 10.2307692 | 2.54660698 | 0 | 1 | 0 | 1 | 1 | 0.35555293 | 0 | 1 | 0 | 1 | 11 | 0.41944532 | FALSE | 1 | FALSE | 1 | FALSE | 1 | FALSE | 1 | FALSE | 1 | FALSE | 1 |
| 2383 | 2383 | 6.33333333 | 2.51818781 | 3 | 0.3655431 | 4 | 0.28813976 | 4 | 0.35669301 | 5 | 0.34217166 | 2 | 0.40216708 | 66 | 0.94315756 | FALSE | 1 | FALSE | 1 | FALSE | 1 | FALSE | 1 | FALSE | 1 | FALSE | 1 |
| 2384 | 2384 | 9.4375 | 2.26298337 | 0 | 1 | 0 | 1 | 0 | 1 | 2 | 0.16544789 | 0 | 1 | 14 | 0.53308261 | FALSE | 1 | FALSE | 1 | FALSE | 1 | FALSE | 1 | FALSE | 1 | FALSE | 1 |
| 2385 | 2385 | 10.2 | 2.44131112 | 0 | 1 | 1 | 0.25628163 | 0 | 1 | 0 | 1 | 0 | 1 | 8 | 0.5780332 | FALSE | 1 | FALSE | 1 | FALSE | 1 | FALSE | 1 | FALSE | 1 | FALSE | 1 |
| 2386 | 2386 | 9.08333333 | 2.8710722 | 0 | 1 | 0 | 1 | 1 | 0.35555293 | 1 | 0.43131129 | 0 | 1 | 10 | 0.71436964 | FALSE | 1 | FALSE | 1 | FALSE | 1 | FALSE | 1 | FALSE | 1 | FALSE | 1 |
| 2387 | 2387 | 6.52941177 | 1.89791362 | 1 | 0.58738519 | 0 | 1 | 2 | 0.34668321 | 3 | 0.20353499 | 0 | 1 | 28 | 0.72558119 | FALSE | 1 | FALSE | 1 | FALSE | 1 | FALSE | 1 | FALSE | 1 | FALSE | 1 |
| 2388 | 2388 | 8.73684211 | 2.3360359 | 1 | 0.38989951 | 0 | 1 | 1 | 0.50143826 | 0 | 1 | 0 | 1 | 17 | 0.41141543 | FALSE | 1 | FALSE | 1 | FALSE | 1 | FALSE | 1 | FALSE | 1 | FALSE | 1 |
| 2389 | 2389 | 10.1428571 | 2.23149991 | 0 | 1 | 0 | 1 | 0 | 1 | 0 | 1 | 0 | 1 | 7 | 0.3043657 | FALSE | 1 | FALSE | 1 | FALSE | 1 | FALSE | 1 | FALSE | 1 | FALSE | 1 |
| 2390 | 2390 | 8.77777778 | 2.8781853 | 0 | 1 | 0 | 1 | 1 | 0.73299778 | 0 | 1 | 2 | 0.11720035 | 33 | 0.16407838 | FALSE | 1 | FALSE | 1 | FALSE | 1 | FALSE | 1 | FALSE | 1 | FALSE | 1 |
| 2391 | 2391 | 9.8 | 1.80554701 | 6 | 0.00048425 | 6 | 0.00163322 | 1 | 0.76953771 | 3 | 0.27785536 | 0 | 1 | 24 | 0.99995724 | TRUE | 0.0450788 | FALSE | 0.40311621 | FALSE | 1 | FALSE | 1 | FALSE | 1 | FALSE | 1 |
| 2392 | 2392 | 10.5333333 | 3.20138859 | 0 | 1 | 0 | 1 | 0 | 1 | 0 | 1 | 1 | 0.20679328 | 13 | 0.33283729 | FALSE | 1 | FALSE | 1 | FALSE | 1 | FALSE | 1 | FALSE | 1 | FALSE | 1 |
| 2393 | 2393 | 8.85714286 | 1.28769688 | 1 | 0.74890551 | 0 | 1 | 1 | 0.85724013 | 3 | 0.44196866 | 1 | 0.58506217 | 47 | 0.25747403 | FALSE | 1 | FALSE | 1 | FALSE | 1 | FALSE | 1 | FALSE | 1 | FALSE | 1 |
| 2394 | 2394 | 11.2 | 3.34065862 | 0 | 1 | 1 | 0.28036922 | 0 | 1 | 0 | 1 | 1 | 0.15247062 | 8 | 0.80394719 | FALSE | 1 | FALSE | 1 | FALSE | 1 | FALSE | 1 | FALSE | 1 | FALSE | 1 |
| 2395 | 2395 | 9.2 | 1.72046505 | 0 | 1 | 0 | 1 | 0 | 1 | 0 | 1 | 0 | 1 | 5 | 0.42761893 | FALSE | 1 | FALSE | 1 | FALSE | 1 | FALSE | 1 | FALSE | 1 | FALSE | 1 |
| 2396 | 2396 | 9.07692308 | 3.5833276 | 1 | 0.49162634 | 0 | 1 | 0 | 1 | 2 | 0.33678042 | 0 | 1 | 23 | 0.40372818 | FALSE | 1 | FALSE | 1 | FALSE | 1 | FALSE | 1 | FALSE | 1 | FALSE | 1 |
| 2397 | 2397 | 11.1190476 | 4.81170389 | 2 | 0.28329504 | 1 | 0.74131698 | 2 | 0.43640583 | 3 | 0.29054213 | 0 | 1 | 33 | 0.81996213 | FALSE | 1 | FALSE | 1 | FALSE | 1 | FALSE | 1 | FALSE | 1 | FALSE | 1 |
| 2398 | 2398 | 5.62790698 | 2.86943197 | 2 | 0.30241117 | 2 | 0.40761584 | 1 | 0.79363407 | 2 | 0.59404271 | 1 | 0.50988734 | 35 | 0.7801651 | FALSE | 1 | FALSE | 1 | FALSE | 1 | FALSE | 1 | FALSE | 1 | FALSE | 1 |
| 2399 | 2399 | 9.17647059 | 1.42396688 | 3 | 0.00864151 | 0 | 1 | 0 | 1 | 0 | 1 | 0 | 1 | 14 | 0.7313145 | FALSE | 0.27019545 | FALSE | 1 | FALSE | 1 | FALSE | 1 | FALSE | 1 | FALSE | 1 |
| 2400 | 2400 | 4.20285714 | 0.98212182 | 25 | 0.04589586 | 35 | 0.00432922 | 24 | 0.59682733 | 34 | 0.35228296 | 13 | 0.33888856 | 558 | 0.99520197 | FALSE | 0.649301 | FALSE | 0.58120789 | FALSE | 1 | FALSE | 1 | FALSE | 1 | FALSE | 1 |
| 2401 | 2401 | 8.5 | 3.32080981 | 0 | 1 | 0 | 1 | 0 | 1 | 1 | 0.57131013 | 0 | 1 | 17 | 0.20329205 | FALSE | 1 | FALSE | 1 | FALSE | 1 | FALSE | 1 | FALSE | 1 | FALSE | 1 |
| 2402 | 2402 | 7.46153846 | 3.44440203 | 2 | 0.14250978 | 2 | 0.20486827 | 0 | 1 | 2 | 0.33678042 | 0 | 1 | 20 | 0.90142689 | FALSE | 1 | FALSE | 1 | FALSE | 1 | FALSE | 1 | FALSE | 1 | FALSE | 1 |
| 2403 | 2403 | 10 | 2.39791576 | 1 | 0.18769745 | 0 | 1 | 0 | 1 | 0 | 1 | 1 | 0.12394282 | 6 | 0.88425551 | FALSE | 1 | FALSE | 1 | FALSE | 1 | FALSE | 1 | FALSE | 1 | FALSE | 1 |
| 2404 | 2404 | 9.4 | 1.2 | 1 | 0.54198714 | 0 | 1 | 1 | 0.66709967 | 0 | 1 | 0 | 1 | 28 | 0.13086908 | FALSE | 1 | FALSE | 1 | FALSE | 1 | FALSE | 1 | FALSE | 1 | FALSE | 1 |
| 2405 | 2405 | 8.11111111 | 3.26976422 | 1 | 0.50471032 | 0 | 1 | 3 | 0.07106362 | 0 | 1 | 0 | 1 | 23 | 0.58307573 | FALSE | 1 | FALSE | 1 | FALSE | 1 | FALSE | 1 | FALSE | 1 | FALSE | 1 |
| 2406 | 2406 | 11.1428571 | 2.82481713 | 0 | 1 | 1 | 0.36919824 | 0 | 1 | 3 | 0.0239258 | 0 | 1 | 10 | 0.94579623 | FALSE | 1 | FALSE | 1 | FALSE | 1 | FALSE | 1 | FALSE | 1 | FALSE | 1 |
| 2407 | 2407 | 11.4117647 | 2.35294118 | 0 | 1 | 0 | 1 | 1 | 0.44344817 | 2 | 0.16544789 | 0 | 1 | 13 | 0.76799161 | FALSE | 1 | FALSE | 1 | FALSE | 1 | FALSE | 1 | FALSE | 1 | FALSE | 1 |
| 2408 | 2408 | 8.94871795 | 2.1116128 | 3 | 0.07731604 | 0 | 1 | 2 | 0.41139386 | 2 | 0.54052736 | 1 | 0.4761699 | 31 | 0.85613899 | FALSE | 0.77583006 | FALSE | 1 | FALSE | 1 | FALSE | 1 | FALSE | 1 | FALSE | 1 |
| 2409 | 2409 | 10.0952381 | 2.28670613 | 0 | 1 | 2 | 0.1353446 | 0 | 1 | 0 | 1 | 1 | 0.28187291 | 17 | 0.61750632 | FALSE | 1 | FALSE | 1 | FALSE | 1 | FALSE | 1 | FALSE | 1 | FALSE | 1 |
| 2410 | 2410 | 9.22727273 | 1.97557398 | 1 | 0.81665075 | 0 | 1 | 6 | 0.02850192 | 2 | 0.80687302 | 1 | 0.66032558 | 55 | 0.56243792 | FALSE | 1 | FALSE | 1 | FALSE | 1 | FALSE | 1 | FALSE | 1 | FALSE | 1 |
| 2411 | 2411 | 9.48148148 | 2.33127481 | 0 | 1 | 0 | 1 | 2 | 0.25286935 | 0 | 1 | 1 | 0.36060486 | 24 | 0.3725373 | FALSE | 1 | FALSE | 1 | FALSE | 1 | FALSE | 1 | FALSE | 1 | FALSE | 1 |
| 2412 | 2412 | 6.56321839 | 1.7662295 | 2 | 0.65852142 | 5 | 0.15016498 | 3 | 0.61023975 | 4 | 0.57118367 | 1 | 0.76487586 | 72 | 0.72239496 | FALSE | 1 | FALSE | 1 | FALSE | 1 | FALSE | 1 | FALSE | 1 | FALSE | 1 |
| 2413 | 2413 | 10.0555556 | 3.08170628 | 1 | 0.37380237 | 1 | 0.44710673 | 1 | 0.48280983 | 0 | 1 | 0 | 1 | 15 | 0.69372602 | FALSE | 1 | FALSE | 1 | FALSE | 1 | FALSE | 1 | FALSE | 1 | FALSE | 1 |
| 2414 | 2414 | 5.37371663 | 1.68699797 | 10 | 0.81092704 | 22 | 0.0680905 | 23 | 0.10569742 | 18 | 0.86426986 | 8 | 0.55122544 | 406 | 0.76106467 | FALSE | 1 | FALSE | 1 | FALSE | 1 | FALSE | 1 | FALSE | 1 | FALSE | 1 |
| 2415 | 2415 | 7.88888889 | 3.69517537 | 1 | 0.20854748 | 0 | 1 | 0 | 1 | 1 | 0.34506517 | 0 | 1 | 7 | 0.84574168 | FALSE | 1 | FALSE | 1 | FALSE | 1 | FALSE | 1 | FALSE | 1 | FALSE | 1 |
| 2416 | 2416 | 8.01818182 | 2.10186851 | 1 | 0.76171602 | 1 | 0.83729212 | 0 | 1 | 4 | 0.24483091 | 0 | 1 | 49 | 0.22274213 | FALSE | 1 | FALSE | 1 | FALSE | 1 | FALSE | 1 | FALSE | 1 | FALSE | 1 |
| 2417 | 2417 | 10.4 | 2.05912603 | 1 | 0.40558521 | 0 | 1 | 0 | 1 | 0 | 1 | 0 | 1 | 19 | 0.15700255 | FALSE | 1 | FALSE | 1 | FALSE | 1 | FALSE | 1 | FALSE | 1 | FALSE | 1 |
| 2418 | 2418 | 8.69230769 | 2.46153846 | 1 | 0.49162634 | 1 | 0.57534752 | 0 | 1 | 1 | 0.70601446 | 1 | 0.3499045 | 22 | 0.61628 | FALSE | 1 | FALSE | 1 | FALSE | 1 | FALSE | 1 | FALSE | 1 | FALSE | 1 |
| 2419 | 2419 | 6.96753247 | 1.89469126 | 8 | 0.04248241 | 7 | 0.22598342 | 3 | 0.91771138 | 11 | 0.09275085 | 0 | 1 | 124 | 0.89424236 | FALSE | 0.62932357 | FALSE | 1 | FALSE | 1 | FALSE | 1 | FALSE | 1 | FALSE | 1 |
| 2420 | 2420 | 7.90909091 | 3.44975745 | 2 | 0.03087289 | 0 | 1 | 2 | 0.05708125 | 0 | 1 | 0 | 1 | 7 | 0.98129353 | FALSE | 0.53356688 | FALSE | 1 | FALSE | 1 | FALSE | 1 | FALSE | 1 | FALSE | 1 |
| 2421 | 2421 | 6.94444444 | 1.54460431 | 1 | 0.37380237 | 0 | 1 | 0 | 1 | 1 | 0.57131013 | 0 | 1 | 16 | 0.44998001 | FALSE | 1 | FALSE | 1 | FALSE | 1 | FALSE | 1 | FALSE | 1 | FALSE | 1 |
| 2422 | 2422 | 10.2068966 | 2.12342129 | 0 | 1 | 1 | 0.61539065 | 2 | 0.27983822 | 1 | 0.74482229 | 0 | 1 | 25 | 0.51779518 | FALSE | 1 | FALSE | 1 | FALSE | 1 | FALSE | 1 | FALSE | 1 | FALSE | 1 |
| 2423 | 2423 | 9.85714286 | 1.88441514 | 0 | 1 | 0 | 1 | 0 | 1 | 0 | 1 | 0 | 1 | 14 | 0.09251304 | FALSE | 1 | FALSE | 1 | FALSE | 1 | FALSE | 1 | FALSE | 1 | FALSE | 1 |
| 2424 | 2424 | 10.375 | 1.86664807 | 0 | 1 | 0 | 1 | 0 | 1 | 3 | 0.00451649 | 0 | 1 | 5 | 0.97551477 | FALSE | 1 | FALSE | 1 | FALSE | 1 | FALSE | 1 | FALSE | 1 | FALSE | 1 |
| 2425 | 2425 | 7.39393939 | 1.82473566 | 2 | 0.20692137 | 0 | 1 | 1 | 0.70185639 | 2 | 0.45159487 | 1 | 0.42123867 | 27 | 0.75124583 | FALSE | 1 | FALSE | 1 | FALSE | 1 | FALSE | 1 | FALSE | 1 | FALSE | 1 |
| 2426 | 2426 | 10 | 2.23606798 | 0 | 1 | 0 | 1 | 1 | 0.35555293 | 0 | 1 | 0 | 1 | 11 | 0.41944532 | FALSE | 1 | FALSE | 1 | FALSE | 1 | FALSE | 1 | FALSE | 1 | FALSE | 1 |
| 2427 | 2427 | 8.95454546 | 1.94181903 | 1 | 0.4357638 | 2 | 0.15794013 | 0 | 1 | 2 | 0.26791271 | 0 | 1 | 17 | 0.8840357 | FALSE | 1 | FALSE | 1 | FALSE | 1 | FALSE | 1 | FALSE | 1 | FALSE | 1 |
| 2428 | 2428 | 5.91891892 | 2.45366112 | 1 | 0.6184714 | 1 | 0.70472939 | 0 | 1 | 1 | 0.82511639 | 0 | 1 | 34 | 0.14843263 | FALSE | 1 | FALSE | 1 | FALSE | 1 | FALSE | 1 | FALSE | 1 | FALSE | 1 |
| 2429 | 2429 | 6.48739496 | 1.76742488 | 3 | 0.59315467 | 7 | 0.09027056 | 8 | 0.06387979 | 3 | 0.91627151 | 1 | 0.86261071 | 97 | 0.84081583 | FALSE | 1 | FALSE | 1 | FALSE | 1 | FALSE | 1 | FALSE | 1 | FALSE | 1 |
| 2430 | 2430 | 12.05 | 1.56444879 | 0 | 1 | 0 | 1 | 0 | 1 | 0 | 1 | 0 | 1 | 20 | 0.03329856 | FALSE | 1 | FALSE | 1 | FALSE | 1 | FALSE | 1 | FALSE | 1 | FALSE | 1 |
| 2431 | 2431 | 6.81690141 | 3.20786823 | 2 | 0.54756403 | 1 | 0.90433446 | 2 | 0.73015113 | 6 | 0.10633312 | 2 | 0.3251371 | 58 | 0.7898837 | FALSE | 1 | FALSE | 1 | FALSE | 1 | FALSE | 1 | FALSE | 1 | FALSE | 1 |
| 2432 | 2432 | 8.48717949 | 2.2403273 | 1 | 0.63789042 | 0 | 1 | 2 | 0.41139386 | 1 | 0.84089119 | 0 | 1 | 35 | 0.24915769 | FALSE | 1 | FALSE | 1 | FALSE | 1 | FALSE | 1 | FALSE | 1 | FALSE | 1 |
| 2433 | 2433 | 10.1666667 | 2.79384244 | 1 | 0.14434519 | 0 | 1 | 0 | 1 | 0 | 1 | 0 | 1 | 5 | 0.76185568 | FALSE | 1 | FALSE | 1 | FALSE | 1 | FALSE | 1 | FALSE | 1 | FALSE | 1 |
| 2434 | 2434 | 12.3913044 | 2.18172509 | 1 | 0.45027683 | 0 | 1 | 0 | 1 | 0 | 1 | 0 | 1 | 22 | 0.10533112 | FALSE | 1 | FALSE | 1 | FALSE | 1 | FALSE | 1 | FALSE | 1 | FALSE | 1 |
| 2435 | 2435 | 9.64285714 | 2.5803417 | 3 | 0.0048942 | 0 | 1 | 0 | 1 | 0 | 1 | 0 | 1 | 11 | 0.83676449 | FALSE | 0.19848171 | FALSE | 1 | FALSE | 1 | FALSE | 1 | FALSE | 1 | FALSE | 1 |
| 2436 | 2436 | 10.28 | 2.30685934 | 1 | 0.47819878 | 0 | 1 | 0 | 1 | 2 | 0.31968852 | 0 | 1 | 22 | 0.43642846 | FALSE | 1 | FALSE | 1 | FALSE | 1 | FALSE | 1 | FALSE | 1 | FALSE | 1 |
| 2437 | 2437 | 2.44525547 | 1.60861296 | 21 | 0.98862766 | 23 | 0.99943001 | 48 | 0.27487347 | 59 | 0.36417764 | 7 | 0.99992693 | 1068 | 0.00173338 | FALSE | 1 | FALSE | 1 | FALSE | 1 | FALSE | 1 | FALSE | 1 | FALSE | 1 |
| 2438 | 2438 | 9.88461539 | 2.80558404 | 0 | 1 | 0 | 1 | 2 | 0.23940226 | 1 | 0.70601446 | 0 | 1 | 23 | 0.40372818 | FALSE | 1 | FALSE | 1 | FALSE | 1 | FALSE | 1 | FALSE | 1 | FALSE | 1 |
| 2439 | 2439 | 11.6363636 | 1.66639116 | 0 | 1 | 1 | 0.30368014 | 0 | 1 | 2 | 0.08789566 | 0 | 1 | 8 | 0.92152993 | FALSE | 1 | FALSE | 1 | FALSE | 1 | FALSE | 1 | FALSE | 1 | FALSE | 1 |
| 2440 | 2440 | 8.74074074 | 2.54722205 | 0 | 1 | 0 | 1 | 1 | 0.62830986 | 3 | 0.12475952 | 0 | 1 | 23 | 0.58307573 | FALSE | 1 | FALSE | 1 | FALSE | 1 | FALSE | 1 | FALSE | 1 | FALSE | 1 |
| 2441 | 2441 | 9.32142857 | 2.03634076 | 0 | 1 | 2 | 0.2288771 | 1 | 0.64171563 | 1 | 0.73248983 | 1 | 0.37113067 | 23 | 0.73439118 | FALSE | 1 | FALSE | 1 | FALSE | 1 | FALSE | 1 | FALSE | 1 | FALSE | 1 |
| 2442 | 2442 | 6.73684211 | 2.26743781 | 2 | 0.24498301 | 0 | 1 | 2 | 0.38585089 | 1 | 0.82511639 | 2 | 0.12262638 | 30 | 0.78878581 | FALSE | 1 | FALSE | 1 | FALSE | 1 | FALSE | 1 | FALSE | 1 | FALSE | 1 |

| | | | | | | | | | | | | | | | | | | | | | | | | | | | |
|---|---|---|---|---|---|---|---|---|---|---|---|---|---|---|---|---|---|---|---|---|---|---|---|---|---|---|---|
| 2443 | 2443 | 7.91666667 | 3.4990078 | 0 | 1 | 0 | 1 | 0 | 1 | 0 | 1 | 0 | 1 | 12 | 0.13002667 | FALSE | 1 | FALSE | 1 | FALSE | 1 | FALSE | 1 | FALSE | 1 | FALSE | 1 |
| 2444 | 2444 | 8.8125 | 2.81111077 | 1 | 0.34033054 | 0 | 1 | 0 | 1 | 3 | 0.03440583 | 0 | 1 | 12 | 0.90922186 | FALSE | 1 | FALSE | 1 | FALSE | 1 | FALSE | 1 | FALSE | 1 | FALSE | 1 |
| 2445 | 2445 | 7.95238095 | 2.39992441 | 0 | 1 | 1 | 0.49918218 | 1 | 0.53671415 | 0 | 1 | 0 | 1 | 19 | 0.34094959 | FALSE | 1 | FALSE | 1 | FALSE | 1 | FALSE | 1 | FALSE | 1 | FALSE | 1 |
| 2446 | 2446 | 8.02439024 | 0.15425745 | 8 | 7.36E-06 | 3 | 0.14580275 | 5 | 0.01496777 | 1 | 0.85524728 | 2 | 0.14490154 | 22 | 0.99999937 | TRUE | 0.0030143 | FALSE | 1 | FALSE | 1 | FALSE | 1 | FALSE | 1 | FALSE | 1 |
| 2447 | 2447 | 10.5333333 | 2.70472837 | 0 | 1 | 1 | 0.38964394 | 0 | 1 | 2 | 0.14906763 | 0 | 1 | 12 | 0.80329917 | FALSE | 1 | FALSE | 1 | FALSE | 1 | FALSE | 1 | FALSE | 1 | FALSE | 1 |
| 2448 | 2448 | 7.64285714 | 2.31785439 | 0 | 1 | 1 | 0.36919824 | 0 | 1 | 1 | 0.48242099 | 0 | 1 | 12 | 0.62236227 | FALSE | 1 | FALSE | 1 | FALSE | 1 | FALSE | 1 | FALSE | 1 | FALSE | 1 |
| 2449 | 2449 | 8.28571429 | 2.93002287 | 0 | 1 | 0 | 1 | 0 | 1 | 1 | 0.62783522 | 0 | 1 | 20 | 0.13763971 | FALSE | 1 | FALSE | 1 | FALSE | 1 | FALSE | 1 | FALSE | 1 | FALSE | 1 |
| 2450 | 2450 | 11.8076923 | 3.36291118 | 0 | 1 | 2 | 0.20486827 | 2 | 0.23940226 | 1 | 0.70601446 | 0 | 1 | 21 | 0.78939668 | FALSE | 1 | FALSE | 1 | FALSE | 1 | FALSE | 1 | FALSE | 1 | FALSE | 1 |
| 2451 | 2451 | 6.66666667 | 3.56682243 | 0 | 1 | 0 | 1 | 1 | 0.58502114 | 2 | 0.30249307 | 0 | 1 | 21 | 0.47053137 | FALSE | 1 | FALSE | 1 | FALSE | 1 | FALSE | 1 | FALSE | 1 | FALSE | 1 |
| 2452 | 2452 | 7.10526316 | 4.78918449 | 1 | 0.38989951 | 0 | 1 | 1 | 0.50143826 | 1 | 0.59104357 | 0 | 1 | 16 | 0.65565778 | FALSE | 1 | FALSE | 1 | FALSE | 1 | FALSE | 1 | FALSE | 1 | FALSE | 1 |
| 2453 | 2453 | 9.33333333 | 1.69967317 | 0 | 1 | 0 | 1 | 0 | 1 | 0 | 1 | 0 | 1 | 6 | 0.36077158 | FALSE | 1 | FALSE | 1 | FALSE | 1 | FALSE | 1 | FALSE | 1 | FALSE | 1 |
| 2454 | 2454 | 9.66666667 | 3.74165739 | 0 | 1 | 0 | 1 | 0 | 1 | 3 | 0.00654627 | 1 | 0.1383237 | 5 | 0.99330426 | FALSE | 1 | FALSE | 1 | FALSE | 1 | FALSE | 1 | FALSE | 1 | FALSE | 1 |
| 2455 | 2455 | 4.19626168 | 1.02482506 | 13 | 0.30001996 | 24 | 0.00541911 | 15 | 0.57544198 | 15 | 0.89226965 | 14 | 0.00960705 | 346 | 0.97722882 | FALSE | 1 | FALSE | 0.59070661 | FALSE | 1 | FALSE | 1 | FALSE | 0.53904735 | FALSE | 1 |
| 2456 | 2456 | 11.25 | 2.04633819 | 0 | 1 | 0 | 1 | 2 | 0.06691409 | 0 | 1 | 0 | 1 | 10 | 0.71436964 | FALSE | 1 | FALSE | 1 | FALSE | 1 | FALSE | 1 | FALSE | 1 | FALSE | 1 |
| 2457 | 2457 | 5.96 | 1.78840711 | 0 | 1 | 2 | 0.48458833 | 1 | 0.8405438 | 2 | 0.6762159 | 0 | 1 | 45 | 0.18510506 | FALSE | 1 | FALSE | 1 | FALSE | 1 | FALSE | 1 | FALSE | 1 | FALSE | 1 |
| 2458 | 2458 | 9.91666667 | 2.19690439 | 0 | 1 | 0 | 1 | 2 | 0.21260257 | 1 | 0.67692816 | 0 | 1 | 21 | 0.47053137 | FALSE | 1 | FALSE | 1 | FALSE | 1 | FALSE | 1 | FALSE | 1 | FALSE | 1 |
| 2459 | 2459 | 8.6122449 | 1.58843173 | 2 | 0.34973212 | 1 | 0.79481669 | 1 | 0.8283465 | 7 | 0.00584147 | 0 | 1 | 37 | 0.93870662 | FALSE | 1 | FALSE | 1 | FALSE | 1 | FALSE | 1 | FALSE | 1 | FALSE | 1 |
| 2460 | 2460 | 7.4137931 | 2.32701142 | 0 | 1 | 1 | 0.61539065 | 0 | 1 | 1 | 0.74482229 | 0 | 1 | 27 | 0.14664438 | FALSE | 1 | FALSE | 1 | FALSE | 1 | FALSE | 1 | FALSE | 1 | FALSE | 1 |
| 2461 | 2461 | 8.39130435 | 3.22677832 | 1 | 0.45027683 | 3 | 0.03669128 | 0 | 1 | 1 | 0.6613267 | 0 | 1 | 18 | 0.86273155 | FALSE | 1 | FALSE | 0.94259695 | FALSE | 1 | FALSE | 1 | FALSE | 1 | FALSE | 1 |
| 2462 | 2462 | 9.5 | 1.5 | 0 | 1 | 0 | 1 | 1 | 0.197145 | 0 | 1 | 0 | 1 | 5 | 0.76185568 | FALSE | 1 | FALSE | 1 | FALSE | 1 | FALSE | 1 | FALSE | 1 | FALSE | 1 |
| 2463 | 2463 | 9.85 | 2.19715725 | 1 | 0.40558521 | 1 | 0.48239024 | 1 | 0.51939841 | 1 | 0.60987146 | 0 | 1 | 16 | 0.80826361 | FALSE | 1 | FALSE | 1 | FALSE | 1 | FALSE | 1 | FALSE | 1 | FALSE | 1 |
| 2464 | 2464 | 10.6363636 | 1.96666433 | 0 | 1 | 1 | 0.30368014 | 0 | 1 | 1 | 0.40390032 | 0 | 1 | 9 | 0.75985366 | FALSE | 1 | FALSE | 1 | FALSE | 1 | FALSE | 1 | FALSE | 1 | FALSE | 1 |
| 2465 | 2465 | 9.9375 | 2.24913178 | 0 | 1 | 0 | 1 | 1 | 0.44344817 | 0 | 1 | 2 | 0.02752114 | 13 | 0.76799161 | FALSE | 1 | FALSE | 1 | FALSE | 1 | FALSE | 1 | FALSE | 0.79106376 | FALSE | 1 |
| 2466 | 2466 | 6.91428571 | 2.84224688 | 1 | 0.59801736 | 3 | 0.10264583 | 2 | 0.35984046 | 3 | 0.21561745 | 0 | 1 | 26 | 0.96294373 | FALSE | 1 | FALSE | 1 | FALSE | 1 | FALSE | 1 | FALSE | 1 | FALSE | 1 |
| 2467 | 2467 | 9.4375 | 2.44869225 | 0 | 1 | 2 | 0.09285443 | 0 | 1 | 0 | 1 | 0 | 1 | 14 | 0.53308261 | FALSE | 1 | FALSE | 1 | FALSE | 1 | FALSE | 1 | FALSE | 1 | FALSE | 1 |
| 2468 | 2468 | 9.05405405 | 2.39307499 | 0 | 1 | 2 | 0.3375163 | 2 | 0.38585089 | 2 | 0.51200191 | 2 | 0.12262638 | 29 | 0.88820978 | FALSE | 1 | FALSE | 1 | FALSE | 1 | FALSE | 1 | FALSE | 1 | FALSE | 1 |
| 2469 | 2469 | 11.2352941 | 2.04448791 | 1 | 0.35728305 | 1 | 0.42857733 | 0 | 1 | 0 | 1 | 1 | 0.24525764 | 14 | 0.7313145 | FALSE | 1 | FALSE | 1 | FALSE | 1 | FALSE | 1 | FALSE | 1 | FALSE | 1 |
| 2470 | 2470 | 5.46666667 | 1.30979218 | 2 | 0.17884234 | 0 | 1 | 2 | 0.29330414 | 1 | 0.75658797 | 0 | 1 | 25 | 0.67630471 | FALSE | 1 | FALSE | 1 | FALSE | 1 | FALSE | 1 | FALSE | 1 | FALSE | 1 |
| 2471 | 2471 | 6.90476191 | 3.30806751 | 0 | 1 | 0 | 1 | 0 | 1 | 1 | 0.62783522 | 0 | 1 | 20 | 0.13763971 | FALSE | 1 | FALSE | 1 | FALSE | 1 | FALSE | 1 | FALSE | 1 | FALSE | 1 |
| 2472 | 2472 | 10.3157895 | 3.17974894 | 1 | 0.38989951 | 1 | 0.46503795 | 1 | 0.50143826 | 1 | 0.59104357 | 0 | 1 | 15 | 0.83648191 | FALSE | 1 | FALSE | 1 | FALSE | 1 | FALSE | 1 | FALSE | 1 | FALSE | 1 |
| 2473 | 2473 | 11.75 | 3.10516432 | 1 | 0.68224519 | 2 | 0.41896557 | 1 | 0.80109494 | 2 | 0.60667406 | 0 | 1 | 38 | 0.45705039 | FALSE | 1 | FALSE | 1 | FALSE | 1 | FALSE | 1 | FALSE | 1 | FALSE | 1 |
| 2474 | 2474 | 10.1666667 | 1.4043583 | 0 | 1 | 0 | 1 | 2 | 0.06691409 | 0 | 1 | 0 | 1 | 10 | 0.71436964 | FALSE | 1 | FALSE | 1 | FALSE | 1 | FALSE | 1 | FALSE | 1 | FALSE | 1 |
| 2475 | 2475 | 10.625 | 2.78107443 | 0 | 1 | 0 | 1 | 0 | 1 | 0 | 1 | 2 | 0.00699343 | 6 | 0.88425551 | FALSE | 1 | FALSE | 1 | FALSE | 1 | FALSE | 1 | FALSE | 0.4774184 | FALSE | 1 |
| 2476 | 2476 | 7.29824561 | 1.13886543 | 0 | 1 | 3 | 0.28014665 | 4 | 0.147507 | 3 | 0.49002278 | 2 | 0.23984683 | 45 | 0.90229646 | FALSE | 1 | FALSE | 1 | FALSE | 1 | FALSE | 1 | FALSE | 1 | FALSE | 1 |
| 2477 | 2477 | 7.75 | 2.2776084 | 0 | 1 | 0 | 1 | 0 | 1 | 1 | 0.60987146 | 0 | 1 | 19 | 0.15700255 | FALSE | 1 | FALSE | 1 | FALSE | 1 | FALSE | 1 | FALSE | 1 | FALSE | 1 |
| 2478 | 2478 | 8.95 | 1.59608897 | 0 | 1 | 3 | 0.02534497 | 2 | 0.16019548 | 1 | 0.60987146 | 0 | 1 | 14 | 0.97325593 | FALSE | 1 | FALSE | 0.82367276 | FALSE | 1 | FALSE | 1 | FALSE | 1 | FALSE | 1 |
| 2479 | 2479 | 8.91666667 | 2.7220805 | 2 | 0.03643311 | 0 | 1 | 0 | 1 | 0 | 1 | 0 | 1 | 10 | 0.71436964 | FALSE | 0.58392 | FALSE | 1 | FALSE | 1 | FALSE | 1 | FALSE | 1 | FALSE | 1 |
| 2480 | 2480 | 8.48148148 | 2.13212414 | 0 | 1 | 1 | 0.58913618 | 0 | 1 | 1 | 0.71956339 | 1 | 0.36060486 | 24 | 0.3725373 | FALSE | 1 | FALSE | 1 | FALSE | 1 | FALSE | 1 | FALSE | 1 | FALSE | 1 |
| 2481 | 2481 | 10.4375 | 1.2732218 | 1 | 0.34033054 | 0 | 1 | 0 | 1 | 0 | 1 | 0 | 1 | 15 | 0.26127859 | FALSE | 1 | FALSE | 1 | FALSE | 1 | FALSE | 1 | FALSE | 1 | FALSE | 1 |
| 2482 | 2482 | 11 | 1.52177182 | 0 | 1 | 1 | 0.46503795 | 1 | 0.50143826 | 2 | 0.21610812 | 0 | 1 | 15 | 0.83648191 | FALSE | 1 | FALSE | 1 | FALSE | 1 | FALSE | 1 | FALSE | 1 | FALSE | 1 |
| 2483 | 2483 | 9.36 | 3.87690598 | 2 | 0.13369621 | 0 | 1 | 1 | 0.59998149 | 0 | 1 | 1 | 0.33902672 | 21 | 0.64957167 | FALSE | 0.97964162 | FALSE | 1 | FALSE | 1 | FALSE | 1 | FALSE | 1 | FALSE | 1 |
| 2484 | 2484 | 10.8275862 | 1.3148742 | 0 | 1 | 1 | 0.61539065 | 1 | 0.65463982 | 2 | 0.3871811 | 1 | 0.38148475 | 24 | 0.70563334 | FALSE | 1 | FALSE | 1 | FALSE | 1 | FALSE | 1 | FALSE | 1 | FALSE | 1 |
| 2485 | 2485 | 5.90350877 | 2.1519852 | 3 | 0.56380187 | 4 | 0.50674923 | 6 | 0.22590386 | 7 | 0.2688136 | 2 | 0.56161197 | 92 | 0.8867997 | FALSE | 1 | FALSE | 1 | FALSE | 1 | FALSE | 1 | FALSE | 1 | FALSE | 1 |
| 2486 | 2486 | 10.6 | 1.2 | 1 | 0.22886536 | 0 | 1 | 2 | 0.04781153 | 3 | 0.00903718 | 0 | 1 | 4 | 0.99982705 | FALSE | 1 | FALSE | 1 | FALSE | 1 | FALSE | 1 | FALSE | 1 | FALSE | 1 |
| 2487 | 2487 | 10.952381 | 2.05811265 | 1 | 0.42086991 | 2 | 0.14655134 | 1 | 0.53671415 | 2 | 0.25059627 | 0 | 1 | 15 | 0.96513691 | FALSE | 1 | FALSE | 1 | FALSE | 1 | FALSE | 1 | FALSE | 1 | FALSE | 1 |
| 2488 | 2488 | 9.85185185 | 2.77159806 | 1 | 0.50471032 | 0 | 1 | 2 | 0.25286935 | 2 | 0.35374209 | 1 | 0.36060486 | 21 | 0.88395936 | FALSE | 1 | FALSE | 1 | FALSE | 1 | FALSE | 1 | FALSE | 1 | FALSE | 1 |
| 2489 | 2489 | 8.65217391 | 2.72841243 | 1 | 0.4357638 | 0 | 1 | 0 | 1 | 1 | 0.64497439 | 0 | 1 | 20 | 0.30915202 | FALSE | 1 | FALSE | 1 | FALSE | 1 | FALSE | 1 | FALSE | 1 | FALSE | 1 |
| 2490 | 2490 | 9.59090909 | 2.12472653 | 2 | 0.10809809 | 4 | 0.00492303 | 0 | 1 | 1 | 0.64497439 | 0 | 1 | 15 | 0.98570275 | FALSE | 0.88820416 | FALSE | 0.58120789 | FALSE | 1 | FALSE | 1 | FALSE | 1 | FALSE | 1 |
| 2491 | 2491 | 10.2222222 | 1.82743158 | 6 | 0.00026749 | 1 | 0.69480472 | 1 | 0.73299778 | 0 | 1 | 3 | 0.0206828 | 25 | 0.99349451 | TRUE | 0.03175788 | FALSE | 1 | FALSE | 1 | FALSE | 1 | FALSE | 0.7366672 | FALSE | 1 |
| 2492 | 2492 | 9.25 | 4.06970515 | 1 | 0.34033054 | 3 | 0.01368613 | 1 | 0.44344817 | 2 | 0.16544789 | 0 | 1 | 9 | 0.99862766 | FALSE | 1 | FALSE | 0.69207862 | FALSE | 1 | FALSE | 1 | FALSE | 1 | FALSE | 1 |
| 2493 | 2493 | 9.375 | 3.72281815 | 1 | 0.56527493 | 1 | 0.65167362 | 2 | 0.32011871 | 1 | 0.77852168 | 0 | 1 | 27 | 0.61671133 | FALSE | 1 | FALSE | 1 | FALSE | 1 | FALSE | 1 | FALSE | 1 | FALSE | 1 |
| 2494 | 2494 | 5.2010582 | 1.37313566 | 5 | 0.53134868 | 6 | 0.57243098 | 8 | 0.36327078 | 8 | 0.63890292 | 1 | 0.95725437 | 160 | 0.43838574 | FALSE | 1 | FALSE | 1 | FALSE | 1 | FALSE | 1 | FALSE | 1 | FALSE | 1 |
| 2495 | 2495 | 8.0625 | 1.91926907 | 2 | 0.06197875 | 0 | 1 | 1 | 0.44344817 | 0 | 1 | 1 | 0.23264612 | 12 | 0.90922186 | FALSE | 0.72532843 | FALSE | 1 | FALSE | 1 | FALSE | 1 | FALSE | 1 | FALSE | 1 |
| 2496 | 2496 | 8.7 | 2.28254244 | 3 | 0.01369554 | 2 | 0.1353446 | 0 | 1 | 0 | 1 | 0 | 1 | 15 | 0.92113679 | FALSE | 0.36625055 | FALSE | 1 | FALSE | 1 | FALSE | 1 | FALSE | 1 | FALSE | 1 |
| 2497 | 2497 | 9 | 2.25831796 | 1 | 0.63789042 | 3 | 0.13080858 | 1 | 0.76089851 | 0 | 1 | 1 | 0.4761699 | 33 | 0.59061307 | FALSE | 1 | FALSE | 1 | FALSE | 1 | FALSE | 1 | FALSE | 1 | FALSE | 1 |
| 2498 | 2498 | 11 | 1.5666989 | 1 | 0.4357638 | 1 | 0.5154318 | 0 | 1 | 0 | 1 | 0 | 1 | 20 | 0.30915202 | FALSE | 1 | FALSE | 1 | FALSE | 1 | FALSE | 1 | FALSE | 1 | FALSE | 1 |
| 2499 | 2499 | 9.61904762 | 2.51616092 | 0 | 1 | 1 | 0.49918218 | 1 | 0.53671415 | 1 | 0.62783522 | 1 | 0.29368245 | 17 | 0.77849076 | FALSE | 1 | FALSE | 1 | FALSE | 1 | FALSE | 1 | FALSE | 1 | FALSE | 1 |
| 2500 | 2500 | 9.46153846 | 1.94601702 | 0 | 1 | 1 | 0.3480709 | 0 | 1 | 0 | 1 | 1 | 0.19354507 | 11 | 0.6683056 | FALSE | 1 | FALSE | 1 | FALSE | 1 | FALSE | 1 | FALSE | 1 | FALSE | 1 |
| 2501 | 2501 | 10 | 3.33809184 | 0 | 1 | 0 | 1 | 0 | 1 | 1 | 0.28044368 | 0 | 1 | 6 | 0.69920685 | FALSE | 1 | FALSE | 1 | FALSE | 1 | FALSE | 1 | FALSE | 1 | FALSE | 1 |
| 2502 | 2502 | 8.9375 | 1.91926907 | 2 | 0.19750261 | 2 | 0.27730767 | 2 | 0.32011871 | 1 | 0.77852168 | 0 | 1 | 25 | 0.88559288 | FALSE | 1 | FALSE | 1 | FALSE | 1 | FALSE | 1 | FALSE | 1 | FALSE | 1 |
| 2503 | 2503 | 4.57657658 | 1.63050619 | 2 | 0.77878267 | 6 | 0.14523534 | 4 | 0.56195051 | 11 | 0.0118055 | 2 | 0.54214185 | 85 | 0.9826298 | FALSE | 1 | FALSE | 1 | FALSE | 1 | FALSE | 1 | FALSE | 1 | FALSE | 1 |
| 2504 | 2504 | 9.90909091 | 2.02055553 | 0 | 1 | 0 | 1 | 0 | 1 | 2 | 0.08789566 | 0 | 1 | 9 | 0.75985366 | FALSE | 1 | FALSE | 1 | FALSE | 1 | FALSE | 1 | FALSE | 1 | FALSE | 1 |
| 2505 | 2505 | 7.79069767 | 2.28806956 | 1 | 0.67382914 | 0 | 1 | 3 | 0.19985581 | 1 | 0.86831187 | 2 | 0.1563365 | 36 | 0.6451239 | FALSE | 1 | FALSE | 1 | FALSE | 1 | FALSE | 1 | FALSE | 1 | FALSE | 1 |
| 2506 | 2506 | 7.47107438 | 1.2535858 | 6 | 0.0873213 | 9 | 0.01538996 | 5 | 0.43318786 | 6 | 0.47540116 | 14 | 6.92E-09 | 80 | 0.99999967 | FALSE | 0.81752699 | FALSE | 0.7256023 | FALSE | 1 | FALSE | 1 | TRUE | 1.42E-05 | FALSE | 1 |
| 2507 | 2507 | 9.375 | 3.90312375 | 1 | 0.34033054 | 1 | 0.40942991 | 0 | 1 | 0 | 1 | 1 | 0.23264612 | 13 | 0.76799161 | FALSE | 1 | FALSE | 1 | FALSE | 1 | FALSE | 1 | FALSE | 1 | FALSE | 1 |
| 2508 | 2508 | 9.35 | 2.12779228 | 2 | 0.27371785 | 2 | 0.37295654 | 2 | 0.42396974 | 4 | 0.10945471 | 0 | 1 | 30 | 0.96151659 | FALSE | 1 | FALSE | 1 | FALSE | 1 | FALSE | 1 | FALSE | 1 | FALSE | 1 |
| 2509 | 2509 | 6.796875 | 2.83493655 | 4 | 0.07746936 | 3 | 0.33367983 | 3 | 0.39570451 | 3 | 0.55805561 | 0 | 1 | 50 | 0.89566304 | FALSE | 0.77583006 | FALSE | 1 | FALSE | 1 | FALSE | 1 | FALSE | 1 | FALSE | 1 |
| 2510 | 2510 | 9.05555556 | 2.01307455 | 0 | 1 | 1 | 0.44710673 | 2 | 0.13502329 | 2 | 0.19902682 | 0 | 1 | 13 | 0.95041913 | FALSE | 1 | FALSE | 1 | FALSE | 1 | FALSE | 1 | FALSE | 1 | FALSE | 1 |
| 2511 | 2511 | 9.03448276 | 2.26643817 | 0 | 1 | 1 | 0.61539065 | 2 | 0.27983822 | 1 | 0.74482229 | 0 | 1 | 25 | 0.51779518 | FALSE | 1 | FALSE | 1 | FALSE | 1 | FALSE | 1 | FALSE | 1 | FALSE | 1 |
| 2512 | 2512 | 7.73684211 | 2.0986739 | 1 | 0.38989951 | 3 | 0.02205772 | 0 | 1 | 1 | 0.59104357 | 5 | 1.05E-05 | 9 | 0.99997115 | FALSE | 1 | FALSE | 0.81029974 | FALSE | 1 | FALSE | 1 | TRUE | 0.00740799 | FALSE | 1 |
| 2513 | 2513 | 11.3 | 2.32594067 | 0 | 1 | 0 | 1 | 0 | 1 | 1 | 0.37517262 | 0 | 1 | 9 | 0.5215547 | FALSE | 1 | FALSE | 1 | FALSE | 1 | FALSE | 1 | FALSE | 1 | FALSE | 1 |
| 2514 | 2514 | 7.69230769 | 2.64239447 | 0 | 1 | 0 | 1 | 1 | 0.37874417 | 0 | 1 | 0 | 1 | 12 | 0.374198 | FALSE | 1 | FALSE | 1 | FALSE | 1 | FALSE | 1 | FALSE | 1 | FALSE | 1 |
| 2515 | 2515 | 11.8571429 | 1.64130361 | 1 | 0.16630142 | 0 | 1 | 0 | 1 | 0 | 1 | 0 | 1 | 6 | 0.69920685 | FALSE | 1 | FALSE | 1 | FALSE | 1 | FALSE | 1 | FALSE | 1 | FALSE | 1 |
| 2516 | 2516 | 9.52380952 | 2.49988662 | 0 | 1 | 0 | 1 | 1 | 0.53671415 | 0 | 1 | 0 | 1 | 20 | 0.13763971 | FALSE | 1 | FALSE | 1 | FALSE | 1 | FALSE | 1 | FALSE | 1 | FALSE | 1 |
| 2517 | 2517 | 8.66666667 | 1.64991582 | 1 | 0.46441868 | 1 | 0.54637326 | 1 | 0.58502114 | 2 | 0.30249307 | 0 | 1 | 19 | 0.83975208 | FALSE | 1 | FALSE | 1 | FALSE | 1 | FALSE | 1 | FALSE | 1 | FALSE | 1 |
| 2518 | 2518 | 4.29885058 | 2.6614098 | 2 | 0.65852142 | 2 | 0.77827608 | 2 | 0.82596174 | 3 | 0.76976873 | 3 | 0.17080654 | 75 | 0.38461264 | FALSE | 1 | FALSE | 1 | FALSE | 1 | FALSE | 1 | FALSE | 1 | FALSE | 1 |
| 2519 | 2519 | 9.45 | 2.39739442 | 0 | 1 | 0 | 1 | 1 | 0.51939841 | 1 | 0.60987146 | 0 | 1 | 18 | 0.37504645 | FALSE | 1 | FALSE | 1 | FALSE | 1 | FALSE | 1 | FALSE | 1 | FALSE | 1 |
| 2520 | 2520 | 8.21428571 | 3.66798059 | 1 | 0.3050805 | 1 | 0.36919824 | 0 | 1 | 0 | 1 | 0 | 1 | 12 | 0.62236227 | FALSE | 1 | FALSE | 1 | FALSE | 1 | FALSE | 1 | FALSE | 1 | FALSE | 1 |
| 2521 | 2521 | 8.90476191 | 3.35097373 | 2 | 0.09989125 | 1 | 0.49918218 | 3 | 0.03775825 | 0 | 1 | 0 | 1 | 15 | 0.96513691 | FALSE | 0.86047225 | FALSE | 1 | FALSE | 1 | FALSE | 1 | FALSE | 1 | FALSE | 1 |
| 2522 | 2522 | 7.375 | 3.00928622 | 1 | 0.76171602 | 0 | 1 | 0 | 1 | 3 | 0.46623289 | 0 | 1 | 51 | 0.05446441 | FALSE | 1 | FALSE | 1 | FALSE | 1 | FALSE | 1 | FALSE | 1 | FALSE | 1 |
| 2523 | 2523 | 8.85294118 | 2.62490731 | 0 | 1 | 1 | 0.6629941 | 1 | 0.70185639 | 1 | 0.78873814 | 0 | 1 | 30 | 0.21942357 | FALSE | 1 | FALSE | 1 | FALSE | 1 | FALSE | 1 | FALSE | 1 | FALSE | 1 |
| 2524 | 2524 | 9.32 | 1.92291446 | 2 | 0.36837609 | 3 | 0.21901899 | 1 | 0.8405438 | 2 | 0.6762159 | 1 | 0.56379725 | 41 | 0.75349894 | FALSE | 1 | FALSE | 1 | FALSE | 1 | FALSE | 1 | FALSE | 1 | FALSE | 1 |
| 2525 | 2525 | 9.71428571 | 2.65729646 | 1 | 0.3050805 | 0 | 1 | 0 | 1 | 1 | 0.48242099 | 1 | 0.20679328 | 11 | 0.83676449 | FALSE | 1 | FALSE | 1 | FALSE | 1 | FALSE | 1 | FALSE | 1 | FALSE | 1 |
| 2526 | 2526 | 10.65 | 2.03162497 | 1 | 0.40558521 | 0 | 1 | 0 | 1 | 2 | 0.23331368 | 0 | 1 | 17 | 0.61750632 | FALSE | 1 | FALSE | 1 | FALSE | 1 | FALSE | 1 | FALSE | 1 | FALSE | 1 |
| 2527 | 2527 | 7.29411765 | 2.59557782 | 0 | 1 | 0 | 1 | 2 | 0.34668321 | 1 | 0.7984848 | 1 | 0.43077488 | 30 | 0.36879488 | FALSE | 1 | FALSE | 1 | FALSE | 1 | FALSE | 1 | FALSE | 1 | FALSE | 1 |
| 2528 | 2528 | 7.91666667 | 2.25308431 | 1 | 0.46441868 | 2 | 0.18117183 | 0 | 1 | 0 | 1 | 0 | 1 | 21 | 0.47053137 | FALSE | 1 | FALSE | 1 | FALSE | 1 | FALSE | 1 | FALSE | 1 | FALSE | 1 |
| 2529 | 2529 | 4.2195122 | 2.68248333 | 1 | 0.64722818 | 2 | 0.37295654 | 2 | 0.42396974 | 0 | 1 | 0 | 1 | 35 | 0.38966407 | FALSE | 1 | FALSE | 1 | FALSE | 1 | FALSE | 1 | FALSE | 1 | FALSE | 1 |
| 2530 | 2530 | 3.78767123 | 2.57013646 | 3 | 0.72868611 | 3 | 0.85725387 | 5 | 0.60737487 | 6 | 0.66783939 | 0 | 1 | 129 | 0.10781104 | FALSE | 1 | FALSE | 1 | FALSE | 1 | FALSE | 1 | FALSE | 1 | FALSE | 1 |
| 2531 | 2531 | 11.25 | 1.08972474 | 0 | 1 | 1 | 0.12326188 | 0 | 1 | 2 | 0.01185256 | 0 | 1 | 1 | 0.99940747 | FALSE | 1 | FALSE | 1 | FALSE | 1 | FALSE | 1 | FALSE | 1 | FALSE | 1 |
| 2532 | 2532 | 10.25 | 3.09232922 | 0 | 1 | 1 | 0.40942991 | 1 | 0.44344817 | 1 | 0.52895088 | 0 | 1 | 13 | 0.76799161 | FALSE | 1 | FALSE | 1 | FALSE | 1 | FALSE | 1 | FALSE | 1 | FALSE | 1 |
| 2533 | 2533 | 8.24675325 | 2.94166595 | 6 | 0.01374005 | 3 | 0.45651652 | 4 | 0.29916433 | 3 | 0.69372972 | 3 | 0.13196291 | 58 | 0.9872953 | FALSE | 0.36625055 | FALSE | 1 | FALSE | 1 | FALSE | 1 | FALSE | 1 | FALSE | 1 |
| 2534 | 2534 | 3.47826087 | 3.0320994 | 2 | 0.92246894 | 6 | 0.42135278 | 5 | 0.69268343 | 7 | 0.61373612 | 0 | 1 | 141 | 0.15300405 | FALSE | 1 | FALSE | 1 | FALSE | 1 | FALSE | 1 | FALSE | 1 | FALSE | 1 |
| 2535 | 2535 | 11.4666667 | 3.38362068 | 0 | 1 | 1 | 0.38964394 | 0 | 1 | 1 | 0.50623194 | 1 | 0.2198258 | 12 | 0.80329917 | FALSE | 1 | FALSE | 1 | FALSE | 1 | FALSE | 1 | FALSE | 1 | FALSE | 1 |
| 2536 | 2536 | 9.57142857 | 1.87899235 | 1 | 0.3050805 | 0 | 1 | 1 | 0.40110418 | 1 | 0.48242099 | 1 | 0.20679328 | 10 | 0.94579623 | FALSE | 1 | FALSE | 1 | FALSE | 1 | FALSE | 1 | FALSE | 1 | FALSE | 1 |

| | | | | | | | | | | | | | | | | | | | | | | | | | | | |
|---|---|---|---|---|---|---|---|---|---|---|---|---|---|---|---|---|---|---|---|---|---|---|---|---|---|---|---|
| 2537 | 2537 | 4.55142857 | 2.50745989 | 11 | 0.24907395 | 18 | 0.02504184 | 8 | 0.92557716 | 7 | 0.99599305 | 3 | 0.92126998 | 291 | 0.20917786 | FALSE | 1 | FALSE | 0.82367276 | FALSE | 1 | FALSE | 1 | FALSE | 1 | FALSE | 1 |
| 2538 | 2538 | 8.33333333 | 2.28137642 | 5 | 0.01493383 | 1 | 0.84773229 | 2 | 0.61293419 | 1 | 0.93213716 | 0 | 1 | 48 | 0.60167081 | FALSE | 0.38230603 | FALSE | 1 | FALSE | 1 | FALSE | 1 | FALSE | 1 | FALSE | 1 |
| 2539 | 2539 | 8.63636364 | 1.55345523 | 0 | 1 | 0 | 1 | 0 | 1 | 0 | 1 | 0 | 1 | 22 | 0.02368118 | FALSE | 1 | FALSE | 1 | FALSE | 1 | FALSE | 1 | FALSE | 1 | FALSE | 1 |
| 2540 | 2540 | 10 | 3.16227766 | 0 | 1 | 0 | 1 | 0 | 1 | 0 | 1 | 0 | 1 | 5 | 0.42761893 | FALSE | 1 | FALSE | 1 | FALSE | 1 | FALSE | 1 | FALSE | 1 | FALSE | 1 |
| 2541 | 2541 | 8.70588235 | 2.3952086 | 0 | 1 | 0 | 1 | 0 | 1 | 1 | 0.55062772 | 0 | 1 | 16 | 0.23070182 | FALSE | 1 | FALSE | 1 | FALSE | 1 | FALSE | 1 | FALSE | 1 | FALSE | 1 |
| 2542 | 2542 | 8.79545455 | 2.97397458 | 1 | 0.68224519 | 3 | 0.16927151 | 4 | 0.07197653 | 1 | 0.87439647 | 0 | 1 | 35 | 0.8620983 | FALSE | 1 | FALSE | 1 | FALSE | 1 | FALSE | 1 | FALSE | 1 | FALSE | 1 |
| 2543 | 2543 | 7.57142857 | 2.96922996 | 0 | 1 | 1 | 0.49918218 | 1 | 0.53671415 | 1 | 0.62783522 | 0 | 1 | 18 | 0.57962763 | FALSE | 1 | FALSE | 1 | FALSE | 1 | FALSE | 1 | FALSE | 1 | FALSE | 1 |
| 2544 | 2544 | 9.18181818 | 2.46127025 | 0 | 1 | 0 | 1 | 0 | 1 | 2 | 0.26791271 | 0 | 1 | 20 | 0.30915202 | FALSE | 1 | FALSE | 1 | FALSE | 1 | FALSE | 1 | FALSE | 1 | FALSE | 1 |
| 2545 | 2545 | 9.36363636 | 1.72008072 | 0 | 1 | 0 | 1 | 2 | 0.04781153 | 1 | 0.37517262 | 0 | 1 | 7 | 0.94326215 | FALSE | 1 | FALSE | 1 | FALSE | 1 | FALSE | 1 | FALSE | 1 | FALSE | 1 |
| 2546 | 2546 | 7.08441558 | 0.58054272 | 3 | 0.76136663 | 4 | 0.74013817 | 6 | 0.47921491 | 6 | 0.71721273 | 3 | 0.46520079 | 132 | 0.37169225 | FALSE | 1 | FALSE | 1 | FALSE | 1 | FALSE | 1 | FALSE | 1 | FALSE | 1 |
| 2547 | 2547 | 10.3076923 | 3.0980047 | 0 | 1 | 0 | 1 | 2 | 0.23940226 | 1 | 0.70601446 | 1 | 0.3499045 | 22 | 0.61628 | FALSE | 1 | FALSE | 1 | FALSE | 1 | FALSE | 1 | FALSE | 1 | FALSE | 1 |
| 2548 | 2548 | 9.36363636 | 2.22680886 | 0 | 1 | 0 | 1 | 0 | 1 | 3 | 0.19161876 | 0 | 1 | 30 | 0.21942357 | FALSE | 1 | FALSE | 1 | FALSE | 1 | FALSE | 1 | FALSE | 1 | FALSE | 1 |
| 2549 | 2549 | 9.03571429 | 2.55625483 | 0 | 1 | 3 | 0.06039219 | 1 | 0.64171563 | 1 | 0.73248983 | 1 | 0.37113067 | 22 | 0.86499992 | FALSE | 1 | FALSE | 1 | FALSE | 1 | FALSE | 1 | FALSE | 1 | FALSE | 1 |
| 2550 | 2550 | 7.65625 | 2.10073462 | 0 | 1 | 1 | 0.65167362 | 2 | 0.32011871 | 0 | 1 | 1 | 0.41154418 | 28 | 0.42532177 | FALSE | 1 | FALSE | 1 | FALSE | 1 | FALSE | 1 | FALSE | 1 | FALSE | 1 |
| 2551 | 2551 | 8.38461539 | 2.07692308 | 1 | 0.74224579 | 2 | 0.50543179 | 0 | 1 | 4 | 0.21499032 | 1 | 0.57809055 | 44 | 0.57424118 | FALSE | 1 | FALSE | 1 | FALSE | 1 | FALSE | 1 | FALSE | 1 | FALSE | 1 |
| 2552 | 2552 | 4.25 | 1.29632471 | 6 | 0.74297508 | 4 | 0.98432982 | 11 | 0.44308751 | 8 | 0.95425368 | 3 | 0.85135188 | 252 | 0.02062311 | FALSE | 1 | FALSE | 1 | FALSE | 1 | FALSE | 1 | FALSE | 1 | FALSE | 1 |
| 2553 | 2553 | 9.94736842 | 3.051724 | 2 | 0.24498301 | 4 | 0.0305219 | 0 | 1 | 1 | 0.82511639 | 2 | 0.12262638 | 28 | 0.9473578 | FALSE | 1 | FALSE | 0.90921954 | FALSE | 1 | FALSE | 1 | FALSE | 1 | FALSE | 1 |
| 2554 | 2554 | 9.95238095 | 1.49526843 | 2 | 0.09989125 | 0 | 1 | 0 | 1 | 1 | 0.62783522 | 2 | 0.04568221 | 16 | 0.90353674 | FALSE | 0.86047225 | FALSE | 1 | FALSE | 1 | FALSE | 1 | FALSE | 0.95710659 | FALSE | 1 |
| 2555 | 2555 | 9.27027027 | 2.35554147 | 2 | 0.24498301 | 0 | 1 | 0 | 1 | 3 | 0.2402032 | 1 | 0.45845922 | 31 | 0.64534533 | FALSE | 1 | FALSE | 1 | FALSE | 1 | FALSE | 1 | FALSE | 1 | FALSE | 1 |
| 2556 | 2556 | 9.65 | 2.41402154 | 1 | 0.40558521 | 1 | 0.48239024 | 0 | 1 | 0 | 1 | 0 | 1 | 18 | 0.37504645 | FALSE | 1 | FALSE | 1 | FALSE | 1 | FALSE | 1 | FALSE | 1 | FALSE | 1 |
| 2557 | 2557 | 10 | 1.95789002 | 1 | 0.26795838 | 1 | 0.32623933 | 1 | 0.35555293 | 0 | 1 | 0 | 1 | 9 | 0.89630573 | FALSE | 1 | FALSE | 1 | FALSE | 1 | FALSE | 1 | FALSE | 1 | FALSE | 1 |
| 2558 | 2558 | 8.33333333 | 3.6270588 | 0 | 1 | 0 | 1 | 1 | 0.42266264 | 0 | 1 | 1 | 0.2198258 | 13 | 0.57712805 | FALSE | 1 | FALSE | 1 | FALSE | 1 | FALSE | 1 | FALSE | 1 | FALSE | 1 |
| 2559 | 2559 | 10.3181818 | 3.74634808 | 0 | 1 | 0 | 1 | 1 | 0.53671415 | 0 | 1 | 0 | 1 | 20 | 0.13763971 | FALSE | 1 | FALSE | 1 | FALSE | 1 | FALSE | 1 | FALSE | 1 | FALSE | 1 |
| 2560 | 2560 | 10.2352941 | 3.2814884 | 0 | 1 | 0 | 1 | 0 | 1 | 1 | 0.55062772 | 0 | 1 | 16 | 0.23070182 | FALSE | 1 | FALSE | 1 | FALSE | 1 | FALSE | 1 | FALSE | 1 | FALSE | 1 |
| 2561 | 2561 | 10 | 3.7032804 | 1 | 0.3050805 | 0 | 1 | 1 | 0.40110418 | 0 | 1 | 1 | 0.20679328 | 11 | 0.83676449 | FALSE | 1 | FALSE | 1 | FALSE | 1 | FALSE | 1 | FALSE | 1 | FALSE | 1 |
| 2562 | 2562 | 10.5 | 2.41867732 | 0 | 1 | 0 | 1 | 0 | 1 | 1 | 0.37517262 | 0 | 1 | 9 | 0.5215547 | FALSE | 1 | FALSE | 1 | FALSE | 1 | FALSE | 1 | FALSE | 1 | FALSE | 1 |
| 2563 | 2563 | 6.4137931 | 2.61978206 | 2 | 0.16962005 | 2 | 0.24095584 | 2 | 0.27983822 | 1 | 0.74482229 | 0 | 1 | 22 | 0.92902102 | FALSE | 1 | FALSE | 1 | FALSE | 1 | FALSE | 1 | FALSE | 1 | FALSE | 1 |
| 2564 | 2564 | 8.23076923 | 2.63566795 | 0 | 1 | 0 | 1 | 1 | 0.61440466 | 1 | 0.70601446 | 2 | 0.0670823 | 22 | 0.61628 | FALSE | 1 | FALSE | 1 | FALSE | 1 | FALSE | 1 | FALSE | 1 | FALSE | 1 |
| 2565 | 2565 | 8.45 | 1.85674446 | 0 | 1 | 0 | 1 | 1 | 0.51939841 | 1 | 0.60987146 | 0 | 1 | 18 | 0.37504645 | FALSE | 1 | FALSE | 1 | FALSE | 1 | FALSE | 1 | FALSE | 1 | FALSE | 1 |
| 2566 | 2566 | 11.625 | 2.0577597 | 0 | 1 | 0 | 1 | 1 | 0.25384309 | 0 | 1 | 0 | 1 | 7 | 0.63752397 | FALSE | 1 | FALSE | 1 | FALSE | 1 | FALSE | 1 | FALSE | 1 | FALSE | 1 |
| 2567 | 2567 | 10.375 | 1.86664807 | 0 | 1 | 0 | 1 | 0 | 1 | 0 | 1 | 1 | 0.23264612 | 15 | 0.26127859 | FALSE | 1 | FALSE | 1 | FALSE | 1 | FALSE | 1 | FALSE | 1 | FALSE | 1 |
| 2568 | 2568 | 10 | 2.68900897 | 0 | 1 | 2 | 0.06426041 | 0 | 1 | 0 | 1 | 1 | 0.19354507 | 10 | 0.86791642 | FALSE | 1 | FALSE | 1 | FALSE | 1 | FALSE | 1 | FALSE | 1 | FALSE | 1 |
| 2569 | 2569 | 7.93333333 | 1.48174072 | 1 | 0.32293355 | 0 | 1 | 1 | 0.42266264 | 0 | 1 | 0 | 1 | 13 | 0.57712805 | FALSE | 1 | FALSE | 1 | FALSE | 1 | FALSE | 1 | FALSE | 1 | FALSE | 1 |
| 2570 | 2570 | 9.07142857 | 1.86946462 | 0 | 1 | 0 | 1 | 0 | 1 | 1 | 0.48242099 | 0 | 1 | 13 | 0.33283729 | FALSE | 1 | FALSE | 1 | FALSE | 1 | FALSE | 1 | FALSE | 1 | FALSE | 1 |
| 2571 | 2571 | 8.5 | 1.72171011 | 0 | 1 | 0 | 1 | 1 | 0.40110418 | 0 | 1 | 0 | 1 | 13 | 0.33283729 | FALSE | 1 | FALSE | 1 | FALSE | 1 | FALSE | 1 | FALSE | 1 | FALSE | 1 |
| 2572 | 2572 | 8.23529412 | 3.07799422 | 0 | 1 | 1 | 0.42857733 | 0 | 1 | 0 | 1 | 0 | 1 | 16 | 0.23070182 | FALSE | 1 | FALSE | 1 | FALSE | 1 | FALSE | 1 | FALSE | 1 | FALSE | 1 |
| 2573 | 2573 | 4.7734375 | 2.22280385 | 3 | 0.63740957 | 4 | 0.59274513 | 5 | 0.48253777 | 3 | 0.93617141 | 3 | 0.34592468 | 109 | 0.38001585 | FALSE | 1 | FALSE | 1 | FALSE | 1 | FALSE | 1 | FALSE | 1 | FALSE | 1 |
| 2574 | 2574 | 8.42857143 | 2.04290709 | 1 | 0.51745948 | 0 | 1 | 1 | 0.64171563 | 1 | 0.73248983 | 1 | 0.37113067 | 24 | 0.55018107 | FALSE | 1 | FALSE | 1 | FALSE | 1 | FALSE | 1 | FALSE | 1 | FALSE | 1 |
| 2575 | 2575 | 10.1818182 | 1.79990817 | 1 | 0.24866461 | 0 | 1 | 0 | 1 | 1 | 0.40390032 | 0 | 1 | 9 | 0.75985366 | FALSE | 1 | FALSE | 1 | FALSE | 1 | FALSE | 1 | FALSE | 1 | FALSE | 1 |
| 2576 | 2576 | 8.38095238 | 2.90280965 | 3 | 0.01568237 | 0 | 1 | 2 | 0.17308361 | 1 | 0.62783522 | 0 | 1 | 15 | 0.96513691 | FALSE | 0.38695764 | FALSE | 1 | FALSE | 1 | FALSE | 1 | FALSE | 1 | FALSE | 1 |
| 2577 | 2577 | 10.7777778 | 2.59391501 | 1 | 0.37380237 | 3 | 0.01901931 | 0 | 1 | 1 | 0.57131013 | 1 | 0.25766374 | 12 | 0.98544073 | FALSE | 1 | FALSE | 0.77515497 | FALSE | 1 | FALSE | 1 | FALSE | 1 | FALSE | 1 |
| 2578 | 2578 | 8.45 | 2.57180222 | 1 | 0.79096998 | 3 | 0.30686508 | 1 | 0.8897349 | 0 | 1 | 0 | 1 | 55 | 0.07598467 | FALSE | 1 | FALSE | 1 | FALSE | 1 | FALSE | 1 | FALSE | 1 | FALSE | 1 |
| 2579 | 2579 | 8.46666667 | 1.76509364 | 0 | 1 | 1 | 0.6278847 | 0 | 1 | 3 | 0.15707004 | 1 | 0.39166989 | 25 | 0.67630471 | FALSE | 1 | FALSE | 1 | FALSE | 1 | FALSE | 1 | FALSE | 1 | FALSE | 1 |
| 2580 | 2580 | 8.82352941 | 2.93528822 | 0 | 1 | 0 | 1 | 0 | 1 | 0 | 1 | 0 | 1 | 17 | 0.05550965 | FALSE | 1 | FALSE | 1 | FALSE | 1 | FALSE | 1 | FALSE | 1 | FALSE | 1 |
| 2581 | 2581 | 6.64150943 | 2.21567605 | 6 | 0.05427402 | 3 | 0.67235559 | 5 | 0.33257943 | 10 | 0.02316714 | 2 | 0.52208951 | 80 | 0.99440721 | FALSE | 0.68849059 | FALSE | 1 | FALSE | 1 | FALSE | 1 | FALSE | 1 | FALSE | 1 |
| 2582 | 2582 | 9.22222222 | 2.29868454 | 0 | 1 | 1 | 0.44710673 | 2 | 0.13502329 | 0 | 1 | 0 | 1 | 15 | 0.69372602 | FALSE | 1 | FALSE | 1 | FALSE | 1 | FALSE | 1 | FALSE | 1 | FALSE | 1 |
| 2583 | 2583 | 10.9166667 | 3.40240928 | 1 | 0.26795838 | 0 | 1 | 1 | 0.35555293 | 0 | 1 | 0 | 1 | 10 | 0.71436964 | FALSE | 1 | FALSE | 1 | FALSE | 1 | FALSE | 1 | FALSE | 1 | FALSE | 1 |
| 2584 | 2584 | 11.6666667 | 1.77638835 | 0 | 1 | 2 | 0.08295839 | 0 | 1 | 0 | 1 | 0 | 1 | 13 | 0.57712805 | FALSE | 1 | FALSE | 1 | FALSE | 1 | FALSE | 1 | FALSE | 1 | FALSE | 1 |
| 2585 | 2585 | 9 | 2.73861279 | 0 | 1 | 0 | 1 | 1 | 0.51939841 | 0 | 1 | 0 | 1 | 19 | 0.15700255 | FALSE | 1 | FALSE | 1 | FALSE | 1 | FALSE | 1 | FALSE | 1 | FALSE | 1 |
| 2586 | 2586 | 8.77777778 | 2.8196839 | 1 | 0.37380237 | 0 | 1 | 1 | 0.48280983 | 1 | 0.57131013 | 0 | 1 | 15 | 0.69372602 | FALSE | 1 | FALSE | 1 | FALSE | 1 | FALSE | 1 | FALSE | 1 | FALSE | 1 |
| 2587 | 2587 | 8 | 2.05480467 | 1 | 0.50471032 | 1 | 0.58913618 | 0 | 1 | 3 | 0.12475952 | 0 | 1 | 22 | 0.76237877 | FALSE | 1 | FALSE | 1 | FALSE | 1 | FALSE | 1 | FALSE | 1 | FALSE | 1 |
| 2588 | 2588 | 9.08333333 | 2.69129254 | 0 | 1 | 0 | 1 | 1 | 0.35555293 | 2 | 0.1023796 | 0 | 1 | 9 | 0.89630573 | FALSE | 1 | FALSE | 1 | FALSE | 1 | FALSE | 1 | FALSE | 1 | FALSE | 1 |
| 2589 | 2589 | 8.1875 | 2.15692924 | 0 | 1 | 2 | 0.27730767 | 1 | 0.69069222 | 2 | 0.43583874 | 0 | 1 | 27 | 0.61671133 | FALSE | 1 | FALSE | 1 | FALSE | 1 | FALSE | 1 | FALSE | 1 | FALSE | 1 |
| 2590 | 2590 | 8.59090909 | 3.22855288 | 0 | 1 | 0 | 1 | 1 | 0.55340851 | 1 | 0.64497439 | 0 | 1 | 20 | 0.30915202 | FALSE | 1 | FALSE | 1 | FALSE | 1 | FALSE | 1 | FALSE | 1 | FALSE | 1 |
| 2591 | 2591 | 8.73170732 | 2.74993916 | 3 | 0.08696268 | 1 | 0.74131698 | 2 | 0.43640583 | 1 | 0.85524728 | 0 | 1 | 34 | 0.69505404 | FALSE | 0.8160347 | FALSE | 1 | FALSE | 1 | FALSE | 1 | FALSE | 1 | FALSE | 1 |
| 2592 | 2592 | 9.67647059 | 1.8266103 | 2 | 0.17884234 | 1 | 0.6278847 | 1 | 0.66709967 | 1 | 0.75658797 | 0 | 1 | 25 | 0.67630471 | FALSE | 1 | FALSE | 1 | FALSE | 1 | FALSE | 1 | FALSE | 1 | FALSE | 1 |
| 2593 | 2593 | 6.89361702 | 2.84518288 | 0 | 1 | 3 | 0.19374024 | 1 | 0.82190381 | 3 | 0.36699324 | 0 | 1 | 40 | 0.54368682 | FALSE | 1 | FALSE | 1 | FALSE | 1 | FALSE | 1 | FALSE | 1 | FALSE | 1 |
| 2594 | 2594 | 8.08571429 | 2.25985371 | 2 | 0.22589135 | 2 | 0.31355738 | 0 | 1 | 0 | 1 | 1 | 0.44015537 | 30 | 0.52765882 | FALSE | 1 | FALSE | 1 | FALSE | 1 | FALSE | 1 | FALSE | 1 | FALSE | 1 |
| 2595 | 2595 | 9.66666667 | 2.10818511 | 1 | 0.37380237 | 1 | 0.44710673 | 0 | 1 | 0 | 1 | 1 | 0.25766374 | 15 | 0.69372602 | FALSE | 1 | FALSE | 1 | FALSE | 1 | FALSE | 1 | FALSE | 1 | FALSE | 1 |
| 2596 | 2596 | 11.5357143 | 2.36767129 | 0 | 1 | 1 | 0.60247911 | 0 | 1 | 2 | 0.3705495 | 0 | 1 | 25 | 0.34293 | FALSE | 1 | FALSE | 1 | FALSE | 1 | FALSE | 1 | FALSE | 1 | FALSE | 1 |
| 2597 | 2597 | 9.64285714 | 2.81848852 | 0 | 1 | 1 | 0.36919824 | 1 | 0.40110418 | 1 | 0.48242099 | 0 | 1 | 11 | 0.83676449 | FALSE | 1 | FALSE | 1 | FALSE | 1 | FALSE | 1 | FALSE | 1 | FALSE | 1 |
| 2598 | 2598 | 9.125 | 1.16592238 | 0 | 1 | 0 | 1 | 0 | 1 | 1 | 0.31351192 | 0 | 1 | 7 | 0.63752397 | FALSE | 1 | FALSE | 1 | FALSE | 1 | FALSE | 1 | FALSE | 1 | FALSE | 1 |
| 2599 | 2599 | 9.46428571 | 1.97270921 | 3 | 0.0279333 | 3 | 0.05021553 | 1 | 0.61440466 | 1 | 0.70601446 | 0 | 1 | 18 | 0.98644281 | FALSE | 0.50964275 | FALSE | 1 | FALSE | 1 | FALSE | 1 | FALSE | 1 | FALSE | 1 |
| 2600 | 2600 | 6.30882353 | 2.4086691 | 4 | 0.09232186 | 1 | 0.89073735 | 4 | 0.21970029 | 0 | 1 | 3 | 0.09685018 | 55 | 0.76052924 | FALSE | 0.83476892 | FALSE | 1 | FALSE | 1 | FALSE | 1 | FALSE | 1 | FALSE | 1 |
| 2601 | 2601 | 8.90909091 | 3.2320707 | 0 | 1 | 0 | 1 | 0 | 1 | 1 | 0.40390032 | 0 | 1 | 10 | 0.46859937 | FALSE | 1 | FALSE | 1 | FALSE | 1 | FALSE | 1 | FALSE | 1 | FALSE | 1 |
| 2602 | 2602 | 6.98039216 | 2.76880519 | 1 | 0.72839502 | 1 | 0.80796869 | 2 | 0.54120634 | 1 | 0.90544885 | 2 | 0.19752014 | 43 | 0.46945509 | FALSE | 1 | FALSE | 1 | FALSE | 1 | FALSE | 1 | FALSE | 1 | FALSE | 1 |
| 2603 | 2603 | 7.57777778 | 2.67073763 | 1 | 0.69044531 | 1 | 0.77338922 | 4 | 0.07687709 | 5 | 0.05380295 | 1 | 0.52593041 | 33 | 0.98284787 | FALSE | 1 | FALSE | 1 | FALSE | 1 | FALSE | 1 | FALSE | 1 | FALSE | 1 |
| 2604 | 2604 | 9.12 | 2.48708665 | 0 | 1 | 0 | 1 | 1 | 0.59998149 | 0 | 1 | 1 | 0.33902672 | 23 | 0.22727183 | FALSE | 1 | FALSE | 1 | FALSE | 1 | FALSE | 1 | FALSE | 1 | FALSE | 1 |
| 2605 | 2605 | 11.7037037 | 3.00388409 | 0 | 1 | 1 | 0.58913618 | 1 | 0.62830986 | 1 | 0.71956339 | 1 | 0.36060486 | 23 | 0.58307573 | FALSE | 1 | FALSE | 1 | FALSE | 1 | FALSE | 1 | FALSE | 1 | FALSE | 1 |
| 2606 | 2606 | 10.1052632 | 2.0747793 | 0 | 1 | 1 | 0.46503795 | 0 | 1 | 0 | 1 | 0 | 1 | 18 | 0.17880694 | FALSE | 1 | FALSE | 1 | FALSE | 1 | FALSE | 1 | FALSE | 1 | FALSE | 1 |
| 2607 | 2607 | 7.65 | 3.22916398 | 1 | 0.40558521 | 2 | 0.1353446 | 0 | 1 | 0 | 1 | 1 | 0.28187291 | 16 | 0.80826361 | FALSE | 1 | FALSE | 1 | FALSE | 1 | FALSE | 1 | FALSE | 1 | FALSE | 1 |
| 2608 | 2608 | 7.88 | 2.79742739 | 1 | 0.72839502 | 2 | 0.48458833 | 1 | 0.8405438 | 4 | 0.19569978 | 1 | 0.56379725 | 41 | 0.75349894 | FALSE | 1 | FALSE | 1 | FALSE | 1 | FALSE | 1 | FALSE | 1 | FALSE | 1 |
| 2609 | 2609 | 9.8 | 1.97315315 | 2 | 0.05513784 | 1 | 0.38964394 | 0 | 1 | 0 | 1 | 0 | 1 | 12 | 0.80329917 | FALSE | 0.68849059 | FALSE | 1 | FALSE | 1 | FALSE | 1 | FALSE | 1 | FALSE | 1 |
| 2610 | 2610 | 6.22147651 | 0.91845828 | 3 | 0.73716798 | 4 | 0.71128806 | 3 | 0.90624821 | 3 | 0.96953633 | 3 | 0.43933641 | 132 | 0.06002961 | FALSE | 1 | FALSE | 1 | FALSE | 1 | FALSE | 1 | FALSE | 1 | FALSE | 1 |
| 2611 | 2611 | 8.66666667 | 2.29561348 | 1 | 0.42086991 | 0 | 1 | 1 | 0.53671415 | 2 | 0.25059627 | 0 | 1 | 17 | 0.77849076 | FALSE | 1 | FALSE | 1 | FALSE | 1 | FALSE | 1 | FALSE | 1 | FALSE | 1 |
| 2612 | 2612 | 9.43478261 | 1.78948832 | 0 | 1 | 0 | 1 | 2 | 0.18613256 | 0 | 1 | 1 | 0.30529951 | 19 | 0.54233419 | FALSE | 1 | FALSE | 1 | FALSE | 1 | FALSE | 1 | FALSE | 1 | FALSE | 1 |
| 2613 | 2613 | 9.80487805 | 2.9402133 | 0 | 1 | 2 | 0.37295654 | 3 | 0.17245215 | 1 | 0.84823834 | 2 | 0.13925369 | 32 | 0.83853687 | FALSE | 1 | FALSE | 1 | FALSE | 1 | FALSE | 1 | FALSE | 1 | FALSE | 1 |
| 2614 | 2614 | 7.46666667 | 2.57854395 | 0 | 1 | 0 | 1 | 1 | 0.42266264 | 1 | 0.50623194 | 0 | 1 | 13 | 0.57712805 | FALSE | 1 | FALSE | 1 | FALSE | 1 | FALSE | 1 | FALSE | 1 | FALSE | 1 |
| 2615 | 2615 | 7.78125 | 2.57068054 | 0 | 1 | 0 | 1 | 1 | 0.69069222 | 6 | 0.00293693 | 0 | 1 | 25 | 0.88559288 | FALSE | 1 | FALSE | 1 | FALSE | 1 | FALSE | 1 | FALSE | 1 | FALSE | 1 |
| 2616 | 2616 | 9.81481482 | 2.22838651 | 1 | 0.50471032 | 0 | 1 | 2 | 0.25286935 | 2 | 0.35374209 | 1 | 0.36060486 | 21 | 0.88395936 | FALSE | 1 | FALSE | 1 | FALSE | 1 | FALSE | 1 | FALSE | 1 | FALSE | 1 |
| 2617 | 2617 | 8.58333333 | 1.76973005 | 2 | 0.22589135 | 1 | 0.68454804 | 1 | 0.72299506 | 0 | 1 | 1 | 0.44015537 | 30 | 0.52765882 | FALSE | 1 | FALSE | 1 | FALSE | 1 | FALSE | 1 | FALSE | 1 | FALSE | 1 |
| 2618 | 2618 | 11.3333333 | 2.98142397 | 2 | 0.09989125 | 1 | 0.49918218 | 0 | 1 | 2 | 0.25059627 | 0 | 1 | 16 | 0.90353674 | FALSE | 0.86047225 | FALSE | 1 | FALSE | 1 | FALSE | 1 | FALSE | 1 | FALSE | 1 |
| 2619 | 2619 | 8.57142857 | 3.84919817 | 0 | 1 | 0 | 1 | 1 | 0.62830986 | 3 | 0.12475952 | 0 | 1 | 23 | 0.58307573 | FALSE | 1 | FALSE | 1 | FALSE | 1 | FALSE | 1 | FALSE | 1 | FALSE | 1 |
| 2620 | 2620 | 9.91666667 | 3.0265033 | 0 | 1 | 0 | 1 | 1 | 0.5695037 | 1 | 0.6613267 | 0 | 1 | 21 | 0.27964072 | FALSE | 1 | FALSE | 1 | FALSE | 1 | FALSE | 1 | FALSE | 1 | FALSE | 1 |
| 2621 | 2621 | 9.0625 | 2.24913178 | 0 | 1 | 0 | 1 | 0 | 1 | 2 | 0.16544789 | 1 | 0.23264612 | 13 | 0.76799161 | FALSE | 1 | FALSE | 1 | FALSE | 1 | FALSE | 1 | FALSE | 1 | FALSE | 1 |
| 2622 | 2622 | 9.20833333 | 1.73154956 | 2 | 0.12501436 | 0 | 1 | 0 | 1 | 0 | 1 | 0 | 1 | 22 | 0.25237063 | FALSE | 0.95622564 | FALSE | 1 | FALSE | 1 | FALSE | 1 | FALSE | 1 | FALSE | 1 |
| 2623 | 2623 | 8.75 | 2.384848 | 1 | 0.0986934 | 0 | 1 | 1 | 0.13615783 | 0 | 1 | 0 | 1 | 2 | 0.98656791 | FALSE | 0.86047225 | FALSE | 1 | FALSE | 1 | FALSE | 1 | FALSE | 1 | FALSE | 1 |
| 2624 | 2624 | 8 | 3.50509833 | 0 | 1 | 0 | 1 | 0 | 1 | 0 | 1 | 0 | 1 | 28 | 0.0085123 | FALSE | 1 | FALSE | 1 | FALSE | 1 | FALSE | 1 | FALSE | 1 | FALSE | 1 |
| 2625 | 2625 | 8.38235294 | 2.36412782 | 2 | 0.21638713 | 1 | 0.6739483 | 0 | 1 | 1 | 0.7984848 | 2 | 0.1065437 | 28 | 0.72558119 | FALSE | 1 | FALSE | 1 | FALSE | 1 | FALSE | 1 | FALSE | 1 | FALSE | 1 |
| 2626 | 2626 | 8.88888889 | 1.6292087 | 2 | 0.06909038 | 1 | 0.42857733 | 0 | 1 | 0 | 1 | 1 | 0.24525764 | 13 | 0.88719222 | FALSE | 0.74276693 | FALSE | 1 | FALSE | 1 | FALSE | 1 | FALSE | 1 | FALSE | 1 |
| 2627 | 2627 | 9.63157895 | 2.79493012 | 1 | 0.37380237 | 1 | 0.44710673 | 0 | 1 | 0 | 1 | 0 | 1 | 16 | 0.44998001 | FALSE | 1 | FALSE | 1 | FALSE | 1 | FALSE | 1 | FALSE | 1 | FALSE | 1 |
| 2628 | 2628 | 9.0625 | 3.02011486 | 1 | 0.56527493 | 2 | 0.27730767 | 1 | 0.69069222 | 0 | 1 | 0 | 1 | 28 | 0.42532177 | FALSE | 1 | FALSE | 1 | FALSE | 1 | FALSE | 1 | FALSE | 1 | FALSE | 1 |
| 2629 | 2629 | 7.61538462 | 1.68880757 | 1 | 0.24866461 | 0 | 1 | 0 | 1 | 0 | 1 | 0 | 1 | 10 | 0.46859937 | FALSE | 1 | FALSE | 1 | FALSE | 1 | FALSE | 1 | FALSE | 1 | FALSE | 1 |
| 2630 | 2630 | 8.87719298 | 2.24053702 | 0 | 1 | 1 | 0.83180787 | 5 | 0.04350703 | 2 | 0.71679995 | 0 | 1 | 46 | 0.52685615 | FALSE | 1 | FALSE | 1 | FALSE | 1 | FALSE | 1 | FALSE | 1 | FALSE | 1 |

| | | | | | | | | | | | | | | | | | | | | | | | | | | | |
|---|---|---|---|---|---|---|---|---|---|---|---|---|---|---|---|---|---|---|---|---|---|---|---|---|---|---|---|
| 2631 | 2631 | 10.137931 | 3.08191765 | 3 | 0.03713651 | 0 | 1 | 0 | 1 | 0 | 1 | 0 | 1 | 26 | 0.31495157 | FALSE | 0.58392 | FALSE | 1 | FALSE | 1 | FALSE | 1 | FALSE | 1 | FALSE | 1 |
| 2632 | 2632 | 10.6666667 | 2.41906012 | 4 | 0.00461682 | 4 | 0.01041264 | 2 | 0.25286935 | 1 | 0.71956339 | 1 | 0.36060486 | 15 | 0.99992818 | FALSE | 0.19698443 | FALSE | 0.67617265 | FALSE | 1 | FALSE | 1 | FALSE | 1 | FALSE | 1 |
| 2633 | 2633 | 4.91666667 | 2.5167369 | 3 | 0.62121274 | 3 | 0.77097552 | 6 | 0.28660842 | 4 | 0.82826751 | 0 | 1 | 108 | 0.24109314 | FALSE | 1 | FALSE | 1 | FALSE | 1 | FALSE | 1 | FALSE | 1 | FALSE | 1 |
| 2634 | 2634 | 8.7 | 1.84661853 | 2 | 0.09187136 | 1 | 0.48239024 | 2 | 0.16019548 | 0 | 1 | 0 | 1 | 15 | 0.92113679 | FALSE | 0.83248965 | FALSE | 1 | FALSE | 1 | FALSE | 1 | FALSE | 1 | FALSE | 1 |
| 2635 | 2635 | 8.39534884 | 1.16418521 | 1 | 0.66519151 | 3 | 0.1535026 | 0 | 1 | 0 | 1 | 3 | 0.03099591 | 35 | 0.67024784 | FALSE | 1 | FALSE | 1 | FALSE | 1 | FALSE | 1 | FALSE | 0.82559785 | FALSE | 1 |
| 2636 | 2636 | 4.67592593 | 1.95707976 | 9 | 0.00172182 | 9 | 0.00801754 | 5 | 0.34693545 | 6 | 0.37635117 | 0 | 1 | 79 | 0.99916965 | FALSE | 0.10850113 | FALSE | 0.61961954 | FALSE | 1 | FALSE | 1 | FALSE | 1 | FALSE | 1 |
| 2637 | 2637 | 7.78378378 | 4.52700686 | 1 | 0.6184714 | 1 | 0.70472939 | 1 | 0.74264075 | 2 | 0.51200191 | 0 | 1 | 32 | 0.47030992 | FALSE | 1 | FALSE | 1 | FALSE | 1 | FALSE | 1 | FALSE | 1 | FALSE | 1 |
| 2638 | 2638 | 4.62727273 | 2.05319747 | 6 | 0.05843558 | 2 | 0.86981548 | 3 | 0.75139368 | 8 | 0.1223324 | 3 | 0.26050233 | 86 | 0.9296732 | FALSE | 0.71985605 | FALSE | 1 | FALSE | 1 | FALSE | 1 | FALSE | 1 | FALSE | 1 |
| 2639 | 2639 | 7.78787879 | 2.91035326 | 1 | 0.57647348 | 0 | 1 | 2 | 0.33343871 | 1 | 0.78873814 | 1 | 0.42123867 | 28 | 0.58681477 | FALSE | 1 | FALSE | 1 | FALSE | 1 | FALSE | 1 | FALSE | 1 | FALSE | 1 |
| 2640 | 2640 | 7.77777778 | 1.8872542 | 0 | 1 | 1 | 0.90746182 | 2 | 0.73725239 | 4 | 0.42273814 | 2 | 0.33117292 | 63 | 0.2931215 | FALSE | 1 | FALSE | 1 | FALSE | 1 | FALSE | 1 | FALSE | 1 | FALSE | 1 |
| 2641 | 2641 | 8.65789474 | 2.78413093 | 1 | 0.6184714 | 3 | 0.11640364 | 1 | 0.74264075 | 2 | 0.51200191 | 0 | 1 | 30 | 0.78878581 | FALSE | 1 | FALSE | 1 | FALSE | 1 | FALSE | 1 | FALSE | 1 | FALSE | 1 |
| 2642 | 2642 | 9.4 | 2.87054002 | 1 | 0.32293355 | 0 | 1 | 0 | 1 | 0 | 1 | 1 | 0.2198258 | 13 | 0.57712805 | FALSE | 1 | FALSE | 1 | FALSE | 1 | FALSE | 1 | FALSE | 1 | FALSE | 1 |
| 2643 | 2643 | 8.52380952 | 2.43556416 | 2 | 0.45796754 | 3 | 0.30686508 | 3 | 0.36617062 | 3 | 0.52470434 | 2 | 0.25814944 | 47 | 0.924754 | FALSE | 1 | FALSE | 1 | FALSE | 1 | FALSE | 1 | FALSE | 1 | FALSE | 1 |
| 2644 | 2644 | 11.4375 | 2.17855313 | 0 | 1 | 2 | 0.09285443 | 0 | 1 | 1 | 0.52895088 | 0 | 1 | 13 | 0.76799161 | FALSE | 1 | FALSE | 1 | FALSE | 1 | FALSE | 1 | FALSE | 1 | FALSE | 1 |
| 2645 | 2645 | 7.94444444 | 2.39147147 | 0 | 1 | 0 | 1 | 1 | 0.48280983 | 2 | 0.19902682 | 0 | 1 | 15 | 0.69372602 | FALSE | 1 | FALSE | 1 | FALSE | 1 | FALSE | 1 | FALSE | 1 | FALSE | 1 |
| 2646 | 2646 | 4.38513514 | 2.22250057 | 3 | 0.73716798 | 3 | 0.86348461 | 4 | 0.78496075 | 6 | 0.68067078 | 4 | 0.22443756 | 128 | 0.27992276 | FALSE | 1 | FALSE | 1 | FALSE | 1 | FALSE | 1 | FALSE | 1 | FALSE | 1 |
| 2647 | 2647 | 6.58 | 3.10541463 | 3 | 0.13593669 | 2 | 0.48458833 | 0 | 1 | 3 | 0.4048204 | 0 | 1 | 42 | 0.62187327 | FALSE | 0.98986074 | FALSE | 1 | FALSE | 1 | FALSE | 1 | FALSE | 1 | FALSE | 1 |
| 2648 | 2648 | 7.62162162 | 2.50784015 | 2 | 0.24498301 | 1 | 0.70472939 | 0 | 1 | 6 | 0.00621583 | 0 | 1 | 28 | 0.9473578 | FALSE | 1 | FALSE | 1 | FALSE | 1 | FALSE | 1 | FALSE | 1 | FALSE | 1 |
| 2649 | 2649 | 8.93103448 | 2.53160757 | 1 | 0.52988231 | 1 | 0.61539065 | 1 | 0.65463982 | 1 | 0.74482229 | 0 | 1 | 25 | 0.51779518 | FALSE | 1 | FALSE | 1 | FALSE | 1 | FALSE | 1 | FALSE | 1 | FALSE | 1 |
| 2650 | 2650 | 5.24590164 | 3.48606471 | 7 | 0.00087526 | 5 | 0.04653332 | 2 | 0.64993443 | 2 | 0.7775286 | 2 | 0.26425795 | 43 | 0.99827461 | FALSE | 0.07029574 | FALSE | 1 | FALSE | 1 | FALSE | 1 | FALSE | 1 | FALSE | 1 |
| 2651 | 2651 | 9.61538462 | 2.16753889 | 1 | 0.49162634 | 2 | 0.20486827 | 0 | 1 | 1 | 0.70601446 | 0 | 1 | 22 | 0.61628 | FALSE | 1 | FALSE | 1 | FALSE | 1 | FALSE | 1 | FALSE | 1 | FALSE | 1 |
| 2652 | 2652 | 9.15 | 2.77983813 | 0 | 1 | 0 | 1 | 1 | 0.51939841 | 1 | 0.60987146 | 0 | 1 | 18 | 0.37504645 | FALSE | 1 | FALSE | 1 | FALSE | 1 | FALSE | 1 | FALSE | 1 | FALSE | 1 |
| 2653 | 2653 | 7.15 | 2.77983813 | 1 | 0.40558521 | 1 | 0.48239024 | 1 | 0.51939841 | 0 | 1 | 0 | 1 | 17 | 0.61750632 | FALSE | 1 | FALSE | 1 | FALSE | 1 | FALSE | 1 | FALSE | 1 | FALSE | 1 |
| 2654 | 2654 | 4.35483871 | 2.45232022 | 0 | 1 | 8 | 0.24487724 | 6 | 0.65402337 | 2 | 0.99842943 | 2 | 0.80990767 | 166 | 0.01378336 | FALSE | 1 | FALSE | 1 | FALSE | 1 | FALSE | 1 | FALSE | 1 | FALSE | 1 |
| 2655 | 2655 | 8.72727273 | 2.16406922 | 1 | 0.57647348 | 0 | 1 | 4 | 0.0293494 | 1 | 0.78873814 | 1 | 0.42123867 | 26 | 0.86852567 | FALSE | 1 | FALSE | 1 | FALSE | 1 | FALSE | 1 | FALSE | 1 | FALSE | 1 |
| 2656 | 2656 | 9.61111111 | 1.63771144 | 1 | 0.34033054 | 3 | 0.01368613 | 0 | 1 | 0 | 1 | 0 | 1 | 12 | 0.90922186 | FALSE | 1 | FALSE | 0.69207862 | FALSE | 1 | FALSE | 1 | FALSE | 1 | FALSE | 1 |
| 2657 | 2657 | 11.9 | 2.54754784 | 0 | 1 | 0 | 1 | 0 | 1 | 0 | 1 | 0 | 1 | 10 | 0.18273166 | FALSE | 1 | FALSE | 1 | FALSE | 1 | FALSE | 1 | FALSE | 1 | FALSE | 1 |
| 2658 | 2658 | 10.7647059 | 2.10124366 | 2 | 0.06909038 | 0 | 1 | 1 | 0.46348836 | 0 | 1 | 0 | 1 | 14 | 0.7313145 | FALSE | 0.74276693 | FALSE | 1 | FALSE | 1 | FALSE | 1 | FALSE | 1 | FALSE | 1 |
| 2659 | 2659 | 5.71698113 | 2.22657499 | 1 | 0.74890551 | 4 | 0.09132555 | 1 | 0.85724013 | 2 | 0.70707241 | 0 | 1 | 45 | 0.5504765 | FALSE | 1 | FALSE | 1 | FALSE | 1 | FALSE | 1 | FALSE | 1 | FALSE | 1 |
| 2660 | 2660 | 9 | 3.43511281 | 1 | 0.40558521 | 0 | 1 | 1 | 0.51939841 | 0 | 1 | 1 | 0.28187291 | 17 | 0.61750632 | FALSE | 1 | FALSE | 1 | FALSE | 1 | FALSE | 1 | FALSE | 1 | FALSE | 1 |
| 2661 | 2661 | 10.6842105 | 1.92087238 | 1 | 0.38989951 | 0 | 1 | 1 | 0.50143826 | 2 | 0.21610812 | 2 | 0.03799168 | 13 | 0.97998676 | FALSE | 1 | FALSE | 1 | FALSE | 1 | FALSE | 1 | FALSE | 0.87669812 | FALSE | 1 |
| 2662 | 2662 | 10.1428571 | 1.95875846 | 0 | 1 | 1 | 0.36919824 | 0 | 1 | 0 | 1 | 0 | 1 | 13 | 0.33283729 | FALSE | 1 | FALSE | 1 | FALSE | 1 | FALSE | 1 | FALSE | 1 | FALSE | 1 |
| 2663 | 2663 | 9.88888889 | 2.18298697 | 0 | 1 | 0 | 1 | 0 | 1 | 1 | 0.34506517 | 0 | 1 | 8 | 0.5780332 | FALSE | 1 | FALSE | 1 | FALSE | 1 | FALSE | 1 | FALSE | 1 | FALSE | 1 |
| 2664 | 2664 | 9.6 | 1.59373775 | 0 | 1 | 0 | 1 | 0 | 1 | 0 | 1 | 0 | 1 | 20 | 0.03329856 | FALSE | 1 | FALSE | 1 | FALSE | 1 | FALSE | 1 | FALSE | 1 | FALSE | 1 |
| 2665 | 2665 | 9.05882353 | 2.33818905 | 0 | 1 | 0 | 1 | 1 | 0.46348836 | 0 | 1 | 1 | 0.24525764 | 15 | 0.49060435 | FALSE | 1 | FALSE | 1 | FALSE | 1 | FALSE | 1 | FALSE | 1 | FALSE | 1 |
| 2666 | 2666 | 9.56097561 | 2.62305706 | 1 | 0.65632651 | 1 | 0.74131698 | 3 | 0.1814659 | 2 | 0.56788092 | 0 | 1 | 34 | 0.69505404 | FALSE | 1 | FALSE | 1 | FALSE | 1 | FALSE | 1 | FALSE | 1 | FALSE | 1 |
| 2667 | 2667 | 11.125 | 4.19635258 | 0 | 1 | 0 | 1 | 1 | 0.44344817 | 2 | 0.16544789 | 0 | 1 | 13 | 0.76799161 | FALSE | 1 | FALSE | 1 | FALSE | 1 | FALSE | 1 | FALSE | 1 | FALSE | 1 |
| 2668 | 2668 | 10.1666667 | 4.19324854 | 1 | 0.37380237 | 1 | 0.44710673 | 0 | 1 | 1 | 0.57131013 | 0 | 1 | 15 | 0.69372602 | FALSE | 1 | FALSE | 1 | FALSE | 1 | FALSE | 1 | FALSE | 1 | FALSE | 1 |
| 2669 | 2669 | 10.2142857 | 1.85851598 | 0 | 1 | 0 | 1 | 0 | 1 | 1 | 0.48242099 | 0 | 1 | 13 | 0.33283729 | FALSE | 1 | FALSE | 1 | FALSE | 1 | FALSE | 1 | FALSE | 1 | FALSE | 1 |
| 2670 | 2670 | 9.80952381 | 2.23860177 | 0 | 1 | 1 | 0.48239024 | 2 | 0.16019548 | 1 | 0.60987146 | 1 | 0.28187291 | 15 | 0.92113679 | FALSE | 1 | FALSE | 1 | FALSE | 1 | FALSE | 1 | FALSE | 1 | FALSE | 1 |
| 2671 | 2671 | 2.53092784 | 3.46433555 | 9 | 0.05639971 | 5 | 0.74648259 | 7 | 0.53363191 | 9 | 0.51736146 | 2 | 0.82622619 | 159 | 0.71023741 | FALSE | 0.69582293 | FALSE | 1 | FALSE | 1 | FALSE | 1 | FALSE | 1 | FALSE | 1 |
| 2672 | 2672 | 8.78947368 | 2.26132116 | 1 | 0.38989951 | 0 | 1 | 1 | 0.50143826 | 0 | 1 | 1 | 0.26986775 | 16 | 0.65565778 | FALSE | 1 | FALSE | 1 | FALSE | 1 | FALSE | 1 | FALSE | 1 | FALSE | 1 |
| 2673 | 2673 | 9.72727273 | 2.45286138 | 0 | 1 | 0 | 1 | 0 | 1 | 0 | 1 | 0 | 1 | 22 | 0.02368118 | FALSE | 1 | FALSE | 1 | FALSE | 1 | FALSE | 1 | FALSE | 1 | FALSE | 1 |
| 2674 | 2674 | 8.62068966 | 2.04119291 | 2 | 0.16962005 | 0 | 1 | 4 | 0.01903587 | 3 | 0.14602298 | 0 | 1 | 20 | 0.99028351 | FALSE | 1 | FALSE | 1 | FALSE | 1 | FALSE | 1 | FALSE | 1 | FALSE | 1 |
| 2675 | 2675 | 9.0625 | 2.94679381 | 0 | 1 | 0 | 1 | 0 | 1 | 0 | 1 | 0 | 1 | 16 | 0.06581507 | FALSE | 1 | FALSE | 1 | FALSE | 1 | FALSE | 1 | FALSE | 1 | FALSE | 1 |
| 2676 | 2676 | 4.64864865 | 4.30409118 | 3 | 0.2949342 | 0 | 1 | 2 | 0.75097925 | 1 | 0.96971789 | 3 | 0.1209957 | 65 | 0.2604438 | FALSE | 1 | FALSE | 1 | FALSE | 1 | FALSE | 1 | FALSE | 1 | FALSE | 1 |
| 2677 | 2677 | 10.0909091 | 2.77831033 | 0 | 1 | 0 | 1 | 0 | 1 | 1 | 0.40390032 | 0 | 1 | 10 | 0.46859937 | FALSE | 1 | FALSE | 1 | FALSE | 1 | FALSE | 1 | FALSE | 1 | FALSE | 1 |
| 2678 | 2678 | 9.60869565 | 2.88533218 | 3 | 0.02011786 | 0 | 1 | 1 | 0.5695037 | 3 | 0.08614834 | 0 | 1 | 16 | 0.98100709 | FALSE | 0.44482142 | FALSE | 1 | FALSE | 1 | FALSE | 1 | FALSE | 1 | FALSE | 1 |
| 2679 | 2679 | 10.7 | 2.57099203 | 1 | 0.22886536 | 1 | 0.28036922 | 0 | 1 | 0 | 1 | 1 | 0.15247062 | 7 | 0.94326215 | FALSE | 1 | FALSE | 1 | FALSE | 1 | FALSE | 1 | FALSE | 1 | FALSE | 1 |
| 2680 | 2680 | 8.42372881 | 2.3084472 | 3 | 0.18560124 | 1 | 0.85269953 | 3 | 0.34634641 | 1 | 0.93527934 | 1 | 0.61824432 | 49 | 0.57916832 | FALSE | 1 | FALSE | 1 | FALSE | 1 | FALSE | 1 | FALSE | 1 | FALSE | 1 |
| 2681 | 2681 | 9 | 2.79880927 | 0 | 1 | 0 | 1 | 0 | 1 | 0 | 1 | 0 | 1 | 12 | 0.13002667 | FALSE | 1 | FALSE | 1 | FALSE | 1 | FALSE | 1 | FALSE | 1 | FALSE | 1 |
| 2682 | 2682 | 8.8 | 2.71293199 | 0 | 1 | 1 | 0.15163655 | 0 | 1 | 0 | 1 | 0 | 1 | 4 | 0.82371636 | FALSE | 1 | FALSE | 1 | FALSE | 1 | FALSE | 1 | FALSE | 1 | FALSE | 1 |
| 2683 | 2683 | 8 | 3.21182027 | 4 | 0.01561835 | 2 | 0.3494045 | 1 | 0.75193685 | 1 | 0.83318955 | 0 | 1 | 30 | 0.87271204 | FALSE | 0.38695764 | FALSE | 1 | FALSE | 1 | FALSE | 1 | FALSE | 1 | FALSE | 1 |
| 2684 | 2684 | 10.0833333 | 3.4268142 | 0 | 1 | 1 | 0.54637326 | 1 | 0.58502114 | 3 | 0.09525947 | 2 | 0.058175 | 17 | 0.97530702 | FALSE | 1 | FALSE | 1 | FALSE | 1 | FALSE | 1 | FALSE | 1 | FALSE | 1 |
| 2685 | 2685 | 7.09848485 | 0.85155694 | 12 | 0.00013241 | 8 | 0.06368144 | 9 | 0.04751705 | 4 | 0.86280699 | 8 | 0.00138268 | 91 | 0.99999791 | TRUE | 0.02085992 | FALSE | 1 | FALSE | 1 | FALSE | 1 | FALSE | 0.20625174 | FALSE | 1 |
| 2686 | 2686 | 9.8 | 2.31516738 | 1 | 0.40558521 | 0 | 1 | 4 | 0.00499622 | 1 | 0.60987146 | 0 | 1 | 14 | 0.97325593 | FALSE | 1 | FALSE | 1 | FALSE | 1 | FALSE | 1 | FALSE | 1 | FALSE | 1 |
| 2687 | 2687 | 7.42307692 | 2.44435105 | 0 | 1 | 0 | 1 | 3 | 0.26704679 | 5 | 0.07777122 | 0 | 1 | 42 | 0.62187327 | FALSE | 1 | FALSE | 1 | FALSE | 1 | FALSE | 1 | FALSE | 1 | FALSE | 1 |
| 2688 | 2688 | 8.55 | 4.05555175 | 0 | 1 | 0 | 1 | 0 | 1 | 1 | 0.60987146 | 0 | 1 | 19 | 0.15700255 | FALSE | 1 | FALSE | 1 | FALSE | 1 | FALSE | 1 | FALSE | 1 | FALSE | 1 |
| 2689 | 2689 | 10.4285714 | 3.048363 | 1 | 0.42086991 | 0 | 1 | 1 | 0.53671415 | 1 | 0.62783522 | 0 | 1 | 18 | 0.57962763 | FALSE | 1 | FALSE | 1 | FALSE | 1 | FALSE | 1 | FALSE | 1 | FALSE | 1 |
| 2690 | 2690 | 8.24390244 | 1.74998938 | 1 | 0.63789042 | 1 | 0.7236252 | 1 | 0.76089851 | 3 | 0.26522531 | 0 | 1 | 33 | 0.59061307 | FALSE | 1 | FALSE | 1 | FALSE | 1 | FALSE | 1 | FALSE | 1 | FALSE | 1 |
| 2691 | 2691 | 5.60869565 | 3.64440167 | 0 | 1 | 1 | 0.53115654 | 1 | 0.5695037 | 0 | 1 | 0 | 1 | 21 | 0.27964072 | FALSE | 1 | FALSE | 1 | FALSE | 1 | FALSE | 1 | FALSE | 1 | FALSE | 1 |
| 2692 | 2692 | 9.25 | 2.56661255 | 2 | 0.09187136 | 1 | 0.48239024 | 1 | 0.51939841 | 0 | 1 | 2 | 0.04176963 | 14 | 0.97325593 | FALSE | 0.83248965 | FALSE | 1 | FALSE | 1 | FALSE | 1 | FALSE | 0.90763079 | FALSE | 1 |
| 2693 | 2693 | 7.64285714 | 2.37117038 | 5 | 0.01390429 | 2 | 0.54544984 | 5 | 0.04964325 | 2 | 0.73544289 | 0 | 1 | 42 | 0.97810254 | FALSE | 0.36625055 | FALSE | 1 | FALSE | 1 | FALSE | 1 | FALSE | 1 | FALSE | 1 |
| 2694 | 2694 | 8.7 | 2.16256021 | 1 | 0.52988231 | 2 | 0.24095584 | 1 | 0.65463982 | 1 | 0.74482229 | 2 | 0.08121636 | 22 | 0.92902102 | FALSE | 1 | FALSE | 1 | FALSE | 1 | FALSE | 1 | FALSE | 1 | FALSE | 1 |
| 2695 | 2695 | 9.33333333 | 1.69967317 | 0 | 1 | 0 | 1 | 1 | 0.35555293 | 0 | 1 | 2 | 0.01579445 | 9 | 0.89630573 | FALSE | 1 | FALSE | 1 | FALSE | 1 | FALSE | 1 | FALSE | 0.66313193 | FALSE | 1 |
| 2696 | 2696 | 7.97959184 | 1.74367423 | 0 | 1 | 2 | 0.44132825 | 1 | 0.81522035 | 2 | 0.63103861 | 0 | 1 | 41 | 0.25430673 | FALSE | 1 | FALSE | 1 | FALSE | 1 | FALSE | 1 | FALSE | 1 | FALSE | 1 |
| 2697 | 2697 | 8.28947368 | 1.89052895 | 1 | 0.62830697 | 1 | 0.71433277 | 1 | 0.75193685 | 0 | 1 | 0 | 1 | 35 | 0.13407055 | FALSE | 1 | FALSE | 1 | FALSE | 1 | FALSE | 1 | FALSE | 1 | FALSE | 1 |
| 2698 | 2698 | 7.83928571 | 2.86466643 | 1 | 0.76787545 | 2 | 0.54544984 | 1 | 0.87219471 | 5 | 0.11290142 | 1 | 0.60529938 | 46 | 0.74916201 | FALSE | 1 | FALSE | 1 | FALSE | 1 | FALSE | 1 | FALSE | 1 | FALSE | 1 |
| 2699 | 2699 | 7.625 | 3.63790805 | 2 | 0.12501436 | 2 | 0.18117183 | 0 | 1 | 0 | 1 | 0 | 1 | 20 | 0.68270729 | FALSE | 0.95622564 | FALSE | 1 | FALSE | 1 | FALSE | 1 | FALSE | 1 | FALSE | 1 |
| 2700 | 2700 | 4.47741936 | 1.86421473 | 12 | 0.00059051 | 12 | 0.00427511 | 5 | 0.66013738 | 8 | 0.4190394 | 1 | 0.92517085 | 117 | 0.99873904 | FALSE | 0.05254197 | FALSE | 0.58120789 | FALSE | 1 | FALSE | 1 | FALSE | 1 | FALSE | 1 |
| 2701 | 2701 | 6.40883978 | 2.21215359 | 11 | 0.0066239 | 9 | 0.13168642 | 6 | 0.63819571 | 10 | 0.31897316 | 2 | 0.80250974 | 143 | 0.98010203 | FALSE | 0.24562356 | FALSE | 1 | FALSE | 1 | FALSE | 1 | FALSE | 1 | FALSE | 1 |
| 2702 | 2702 | 11.7777778 | 2.52884593 | 0 | 1 | 0 | 1 | 0 | 1 | 1 | 0.34506517 | 0 | 1 | 8 | 0.5780332 | FALSE | 1 | FALSE | 1 | FALSE | 1 | FALSE | 1 | FALSE | 1 | FALSE | 1 |
| 2703 | 2703 | 10.3684211 | 2.20550809 | 0 | 1 | 1 | 0.44710673 | 1 | 0.48280983 | 1 | 0.57131013 | 3 | 0.00292491 | 12 | 0.98544073 | FALSE | 1 | FALSE | 1 | FALSE | 1 | FALSE | 1 | FALSE | 0.29951111 | FALSE | 1 |
| 2704 | 2704 | 10.08 | 1.97828208 | 0 | 1 | 1 | 0.54637326 | 3 | 0.05314317 | 1 | 0.67692816 | 0 | 1 | 19 | 0.83975208 | FALSE | 1 | FALSE | 1 | FALSE | 1 | FALSE | 1 | FALSE | 1 | FALSE | 1 |
| 2705 | 2705 | 8.91304348 | 0.92844159 | 0 | 1 | 0 | 1 | 3 | 0.04772439 | 0 | 1 | 0 | 1 | 20 | 0.50589402 | FALSE | 1 | FALSE | 1 | FALSE | 1 | FALSE | 1 | FALSE | 1 | FALSE | 1 |
| 2706 | 2706 | 10.1818182 | 1.58505416 | 1 | 0.24866461 | 1 | 0.30368014 | 0 | 1 | 0 | 1 | 0 | 1 | 9 | 0.75985366 | FALSE | 1 | FALSE | 1 | FALSE | 1 | FALSE | 1 | FALSE | 1 | FALSE | 1 |
| 2707 | 2707 | 9.6 | 3.29241553 | 0 | 1 | 0 | 1 | 0 | 1 | 2 | 0.0740981 | 0 | 1 | 8 | 0.80394719 | FALSE | 1 | FALSE | 1 | FALSE | 1 | FALSE | 1 | FALSE | 1 | FALSE | 1 |
| 2708 | 2708 | 11.2857143 | 1.57790872 | 0 | 1 | 0 | 1 | 1 | 0.40110418 | 0 | 1 | 0 | 1 | 13 | 0.33283729 | FALSE | 1 | FALSE | 1 | FALSE | 1 | FALSE | 1 | FALSE | 1 | FALSE | 1 |
| 2709 | 2709 | 9.61111111 | 2.47518549 | 2 | 0.07645454 | 0 | 1 | 1 | 0.48280983 | 1 | 0.57131013 | 1 | 0.25766374 | 13 | 0.95041913 | FALSE | 0.77424195 | FALSE | 1 | FALSE | 1 | FALSE | 1 | FALSE | 1 | FALSE | 1 |
| 2710 | 2710 | 9.09090909 | 1.92847304 | 1 | 0.24866461 | 0 | 1 | 0 | 1 | 0 | 1 | 1 | 0.16638736 | 9 | 0.75985366 | FALSE | 1 | FALSE | 1 | FALSE | 1 | FALSE | 1 | FALSE | 1 | FALSE | 1 |
| 2711 | 2711 | 10.0833333 | 2.39646731 | 0 | 1 | 0 | 1 | 0 | 1 | 2 | 0.1023796 | 0 | 1 | 10 | 0.71436964 | FALSE | 1 | FALSE | 1 | FALSE | 1 | FALSE | 1 | FALSE | 1 | FALSE | 1 |
| 2712 | 2712 | 8.95454546 | 2.9461419 | 1 | 0.4357638 | 1 | 0.5154318 | 0 | 1 | 2 | 0.26791271 | 0 | 1 | 18 | 0.74744813 | FALSE | 1 | FALSE | 1 | FALSE | 1 | FALSE | 1 | FALSE | 1 | FALSE | 1 |
| 2713 | 2713 | 9.90909091 | 1.72966342 | 0 | 1 | 0 | 1 | 0 | 1 | 0 | 1 | 0 | 1 | 11 | 0.15414489 | FALSE | 1 | FALSE | 1 | FALSE | 1 | FALSE | 1 | FALSE | 1 | FALSE | 1 |
| 2714 | 2714 | 10.5384615 | 2.49970413 | 0 | 1 | 0 | 1 | 1 | 0.35555293 | 1 | 0.43131129 | 0 | 1 | 10 | 0.71436964 | FALSE | 1 | FALSE | 1 | FALSE | 1 | FALSE | 1 | FALSE | 1 | FALSE | 1 |
| 2715 | 2715 | 8.19047619 | 1.49981102 | 0 | 1 | 0 | 1 | 1 | 0.53671415 | 0 | 1 | 0 | 1 | 20 | 0.13763971 | FALSE | 1 | FALSE | 1 | FALSE | 1 | FALSE | 1 | FALSE | 1 | FALSE | 1 |
| 2716 | 2716 | 10.75 | 2.94745653 | 0 | 1 | 1 | 0.48239024 | 2 | 0.16019548 | 2 | 0.23331368 | 0 | 1 | 15 | 0.92113679 | FALSE | 1 | FALSE | 1 | FALSE | 1 | FALSE | 1 | FALSE | 1 | FALSE | 1 |
| 2717 | 2717 | 10.6666667 | 2.74873708 | 0 | 1 | 0 | 1 | 0 | 1 | 0 | 1 | 0 | 1 | 18 | 0.04681656 | FALSE | 1 | FALSE | 1 | FALSE | 1 | FALSE | 1 | FALSE | 1 | FALSE | 1 |
| 2718 | 2718 | 9.39130435 | 3.07499212 | 0 | 1 | 1 | 0.53115654 | 1 | 0.5695037 | 1 | 0.6613267 | 1 | 0.31672722 | 19 | 0.71542496 | FALSE | 1 | FALSE | 1 | FALSE | 1 | FALSE | 1 | FALSE | 1 | FALSE | 1 |
| 2719 | 2719 | 9.94736842 | 2.06407069 | 0 | 1 | 0 | 1 | 0 | 1 | 1 | 0.59104357 | 0 | 1 | 18 | 0.17880694 | FALSE | 1 | FALSE | 1 | FALSE | 1 | FALSE | 1 | FALSE | 1 | FALSE | 1 |
| 2720 | 2720 | 9.19444444 | 1.89765125 | 1 | 0.60837712 | 3 | 0.10944025 | 0 | 1 | 1 | 0.81665383 | 0 | 1 | 31 | 0.49869404 | FALSE | 1 | FALSE | 1 | FALSE | 1 | FALSE | 1 | FALSE | 1 | FALSE | 1 |
| 2721 | 2721 | 7.14285714 | 4.099776 | 0 | 1 | 0 | 1 | 0 | 1 | 2 | 0.45159487 | 0 | 1 | 31 | 0.09218876 | FALSE | 1 | FALSE | 1 | FALSE | 1 | FALSE | 1 | FALSE | 1 | FALSE | 1 |
| 2722 | 2722 | 11.0714286 | 3.03466368 | 2 | 0.04858661 | 1 | 0.36919824 | 1 | 0.40110418 | 0 | 1 | 0 | 1 | 10 | 0.94579623 | FALSE | 0.649301 | FALSE | 1 | FALSE | 1 | FALSE | 1 | FALSE | 1 | FALSE | 1 |
| 2723 | 2723 | 9.41666667 | 3.14797113 | 2 | 0.03643311 | 1 | 0.32623933 | 1 | 0.35555293 | 0 | 1 | 0 | 1 | 8 | 0.97197833 | FALSE | 0.58392 | FALSE | 1 | FALSE | 1 | FALSE | 1 | FALSE | 1 | FALSE | 1 |
| 2724 | 2724 | 8.4 | 1.74355958 | 0 | 1 | 0 | 1 | 0 | 1 | 0 | 1 | 0 | 1 | 5 | 0.42761893 | FALSE | 1 | FALSE | 1 | FALSE | 1 | FALSE | 1 | FALSE | 1 | FALSE | 1 |

| | | | | | | | | | | | | | | | | | | | | | | | | | | | |
|---|---|---|---|---|---|---|---|---|---|---|---|---|---|---|---|---|---|---|---|---|---|---|---|---|---|---|---|
| 2725 | 2725 | 7.015625 | 1.94045893 | 2 | 0.49173883 | 1 | 0.87929209 | 3 | 0.40547364 | 5 | 0.16939111 | 0 | 1 | 53 | 0.70813409 | FALSE | 1 | FALSE | 1 | FALSE | 1 | FALSE | 1 | FALSE | 1 | FALSE | 1 |
| 2726 | 2726 | 6.66666667 | 1.90515869 | 0 | 1 | 0 | 1 | 2 | 0.25286935 | 3 | 0.12475952 | 1 | 0.36060486 | 21 | 0.88395936 | FALSE | 1 | FALSE | 1 | FALSE | 1 | FALSE | 1 | FALSE | 1 | FALSE | 1 |
| 2727 | 2727 | 3.4787234 | 1.75682291 | 16 | 0.00218994 | 9 | 0.56133888 | 11 | 0.42907101 | 18 | 0.09435963 | 6 | 0.31335749 | 221 | 0.99638882 | FALSE | 0.12457305 | FALSE | 1 | FALSE | 1 | FALSE | 1 | FALSE | 1 | FALSE | 1 |
| 2728 | 2728 | 4.40677966 | 2.41898315 | 6 | 0.08204291 | 3 | 0.74112112 | 3 | 0.80266463 | 4 | 0.79788193 | 4 | 0.12806481 | 98 | 0.70851935 | FALSE | 0.79236883 | FALSE | 1 | FALSE | 1 | FALSE | 1 | FALSE | 1 | FALSE | 1 |
| 2729 | 2729 | 10.5217391 | 2.90231639 | 4 | 0.00252272 | 2 | 0.16948764 | 3 | 0.04772439 | 0 | 1 | 0 | 1 | 14 | 0.99865094 | FALSE | 0.13507246 | FALSE | 1 | FALSE | 1 | FALSE | 1 | FALSE | 1 | FALSE | 1 |
| 2730 | 2730 | 8.70833333 | 3.51756505 | 0 | 1 | 1 | 0.54637326 | 1 | 0.58502114 | 2 | 0.30249307 | 0 | 1 | 20 | 0.68270729 | FALSE | 1 | FALSE | 1 | FALSE | 1 | FALSE | 1 | FALSE | 1 | FALSE | 1 |
| 2731 | 2731 | 10 | 2.13347701 | 2 | 0.16962005 | 2 | 0.24095584 | 0 | 1 | 2 | 0.3871811 | 0 | 1 | 23 | 0.84462957 | FALSE | 1 | FALSE | 1 | FALSE | 1 | FALSE | 1 | FALSE | 1 | FALSE | 1 |
| 2732 | 2732 | 4.56382979 | 2.3089028 | 3 | 0.4212756 | 1 | 0.95244851 | 6 | 0.1125 | 3 | 0.80162372 | 4 | 0.06383454 | 75 | 0.81898498 | FALSE | 1 | FALSE | 1 | FALSE | 1 | FALSE | 1 | FALSE | 1 | FALSE | 1 |
| 2733 | 2733 | 9.93333333 | 1.69181034 | 1 | 0.32293355 | 1 | 0.38964394 | 1 | 0.42266264 | 1 | 0.50623194 | 0 | 1 | 11 | 0.92879412 | FALSE | 1 | FALSE | 1 | FALSE | 1 | FALSE | 1 | FALSE | 1 | FALSE | 1 |
| 2734 | 2734 | 10.5909091 | 1.92257145 | 1 | 0.4357638 | 1 | 0.5154318 | 1 | 0.55340851 | 1 | 0.64497439 | 0 | 1 | 18 | 0.74744813 | FALSE | 1 | FALSE | 1 | FALSE | 1 | FALSE | 1 | FALSE | 1 | FALSE | 1 |
| 2735 | 2735 | 7.77011494 | 0.96133467 | 12 | 0.0010292 | 9 | 0.08627167 | 6 | 0.54714381 | 4 | 0.94961602 | 8 | 0.0055196 | 126 | 0.99767808 | FALSE | 0.07493697 | FALSE | 1 | FALSE | 1 | FALSE | 1 | FALSE | 0.45257452 | FALSE | 1 |
| 2736 | 2736 | 9.35294118 | 2.88894804 | 0 | 1 | 0 | 1 | 0 | 1 | 2 | 0.18212134 | 0 | 1 | 15 | 0.49060435 | FALSE | 1 | FALSE | 1 | FALSE | 1 | FALSE | 1 | FALSE | 1 | FALSE | 1 |
| 2737 | 2737 | 8.875 | 1.83285979 | 0 | 1 | 0 | 1 | 0 | 1 | 0 | 1 | 0 | 1 | 8 | 0.25677166 | FALSE | 1 | FALSE | 1 | FALSE | 1 | FALSE | 1 | FALSE | 1 | FALSE | 1 |
| 2738 | 2738 | 9.02325581 | 1.48619026 | 0 | 1 | 1 | 0.75788114 | 2 | 0.46083448 | 4 | 0.13323859 | 1 | 0.50988734 | 35 | 0.7801651 | FALSE | 1 | FALSE | 1 | FALSE | 1 | FALSE | 1 | FALSE | 1 | FALSE | 1 |
| 2739 | 2739 | 7.84313726 | 1.40548713 | 1 | 0.72119464 | 2 | 0.47396556 | 0 | 1 | 3 | 0.39227316 | 0 | 1 | 43 | 0.337822 | FALSE | 1 | FALSE | 1 | FALSE | 1 | FALSE | 1 | FALSE | 1 | FALSE | 1 |
| 2740 | 2740 | 9.34375 | 3.12859169 | 1 | 0.56527493 | 0 | 1 | 0 | 1 | 2 | 0.43583874 | 0 | 1 | 29 | 0.24088541 | FALSE | 1 | FALSE | 1 | FALSE | 1 | FALSE | 1 | FALSE | 1 | FALSE | 1 |
| 2741 | 2741 | 8.3125 | 1.8946883 | 0 | 1 | 0 | 1 | 1 | 0.44344817 | 4 | 0.00511215 | 0 | 1 | 11 | 0.9718531 | FALSE | 1 | FALSE | 1 | FALSE | 1 | FALSE | 1 | FALSE | 1 | FALSE | 1 |
| 2742 | 2742 | 11.9166667 | 2.8710722 | 0 | 1 | 1 | 0.32623933 | 0 | 1 | 1 | 0.43131129 | 0 | 1 | 10 | 0.71436964 | FALSE | 1 | FALSE | 1 | FALSE | 1 | FALSE | 1 | FALSE | 1 | FALSE | 1 |
| 2743 | 2743 | 8.71428571 | 2.05038573 | 0 | 1 | 1 | 0.49918218 | 0 | 1 | 0 | 1 | 0 | 1 | 20 | 0.13763971 | FALSE | 1 | FALSE | 1 | FALSE | 1 | FALSE | 1 | FALSE | 1 | FALSE | 1 |
| 2744 | 2744 | 8.26829268 | 1.71114553 | 0 | 1 | 1 | 0.7326167 | 2 | 0.42396974 | 2 | 0.5543522 | 0 | 1 | 35 | 0.38966407 | FALSE | 1 | FALSE | 1 | FALSE | 1 | FALSE | 1 | FALSE | 1 | FALSE | 1 |
| 2745 | 2745 | 5.81632653 | 2.47166365 | 3 | 0.13009168 | 3 | 0.21051417 | 2 | 0.53023996 | 2 | 0.66536008 | 1 | 0.55647171 | 38 | 0.92985909 | FALSE | 0.97964162 | FALSE | 1 | FALSE | 1 | FALSE | 1 | FALSE | 1 | FALSE | 1 |
| 2746 | 2746 | 9.43103448 | 1.52125254 | 1 | 0.77972342 | 1 | 0.85269953 | 3 | 0.34634641 | 5 | 0.12605748 | 0 | 1 | 48 | 0.70968273 | FALSE | 1 | FALSE | 1 | FALSE | 1 | FALSE | 1 | FALSE | 1 | FALSE | 1 |
| 2747 | 2747 | 9.53846154 | 1.54995705 | 0 | 1 | 0 | 1 | 1 | 0.37874417 | 1 | 0.45746571 | 0 | 1 | 11 | 0.6683056 | FALSE | 1 | FALSE | 1 | FALSE | 1 | FALSE | 1 | FALSE | 1 | FALSE | 1 |
| 2748 | 2748 | 9.25 | 1.86748494 | 0 | 1 | 1 | 0.48239024 | 3 | 0.03322143 | 0 | 1 | 0 | 1 | 16 | 0.80826361 | FALSE | 1 | FALSE | 1 | FALSE | 1 | FALSE | 1 | FALSE | 1 | FALSE | 1 |
| 2749 | 2749 | 7.08938548 | 0.54192824 | 8 | 0.08857042 | 8 | 0.22267763 | 3 | 0.95923034 | 9 | 0.43882028 | 1 | 0.95018775 | 150 | 0.63354399 | FALSE | 0.82638822 | FALSE | 1 | FALSE | 1 | FALSE | 1 | FALSE | 1 | FALSE | 1 |
| 2750 | 2750 | 7.89189189 | 1.73647334 | 3 | 0.0681763 | 2 | 0.3375163 | 1 | 0.74264075 | 2 | 0.51200191 | 2 | 0.12262638 | 27 | 0.97789976 | FALSE | 0.74276693 | FALSE | 1 | FALSE | 1 | FALSE | 1 | FALSE | 1 | FALSE | 1 |
| 2751 | 2751 | 6.43258427 | 1.34015556 | 3 | 0.83792675 | 8 | 0.21401708 | 10 | 0.10397021 | 10 | 0.29486636 | 8 | 0.00833541 | 138 | 0.99154957 | FALSE | 1 | FALSE | 1 | FALSE | 1 | FALSE | 1 | FALSE | 0.50674765 | FALSE | 1 |
| 2752 | 2752 | 8.14285714 | 1.95875846 | 0 | 1 | 0 | 1 | 1 | 0.40110418 | 1 | 0.48242099 | 0 | 1 | 12 | 0.62236227 | FALSE | 1 | FALSE | 1 | FALSE | 1 | FALSE | 1 | FALSE | 1 | FALSE | 1 |
| 2753 | 2753 | 9 | 2.20047842 | 1 | 0.38989951 | 3 | 0.02205772 | 0 | 1 | 0 | 1 | 1 | 0.26986775 | 14 | 0.93677253 | FALSE | 1 | FALSE | 0.81029974 | FALSE | 1 | FALSE | 1 | FALSE | 1 | FALSE | 1 |
| 2754 | 2754 | 6.05882353 | 3.35139283 | 9 | 4.62E-05 | 3 | 0.36935445 | 1 | 0.91485686 | 2 | 0.82021683 | 1 | 0.6714808 | 51 | 0.97428925 | TRUE | 0.01105799 | FALSE | 1 | FALSE | 1 | FALSE | 1 | FALSE | 1 | FALSE | 1 |
| 2755 | 2755 | 8.7 | 1.45258391 | 2 | 0.09187136 | 0 | 1 | 2 | 0.16019548 | 1 | 0.60987146 | 0 | 1 | 15 | 0.92113679 | FALSE | 0.83248965 | FALSE | 1 | FALSE | 1 | FALSE | 1 | FALSE | 1 | FALSE | 1 |
| 2756 | 2756 | 10.3333333 | 2.89059779 | 0 | 1 | 0 | 1 | 1 | 0.42266264 | 2 | 0.14906763 | 0 | 1 | 12 | 0.80329917 | FALSE | 1 | FALSE | 1 | FALSE | 1 | FALSE | 1 | FALSE | 1 | FALSE | 1 |
| 2757 | 2757 | 9.66666667 | 3.06024172 | 2 | 0.09989125 | 2 | 0.14655134 | 1 | 0.53671415 | 3 | 0.06912908 | 2 | 0.04568221 | 11 | 0.99990054 | FALSE | 0.86047225 | FALSE | 1 | FALSE | 1 | FALSE | 1 | FALSE | 0.95710659 | FALSE | 1 |
| 2758 | 2758 | 10 | 2.75680975 | 0 | 1 | 0 | 1 | 0 | 1 | 0 | 1 | 0 | 1 | 5 | 0.42761893 | FALSE | 1 | FALSE | 1 | FALSE | 1 | FALSE | 1 | FALSE | 1 | FALSE | 1 |
| 2759 | 2759 | 8.31914894 | 2.61017961 | 3 | 0.11868304 | 1 | 0.78790827 | 2 | 0.50778118 | 2 | 0.64277396 | 0 | 1 | 39 | 0.69224623 | FALSE | 0.94393345 | FALSE | 1 | FALSE | 1 | FALSE | 1 | FALSE | 1 | FALSE | 1 |
| 2760 | 2760 | 8.93548387 | 2.07808044 | 0 | 1 | 3 | 0.07145121 | 2 | 0.29330414 | 0 | 1 | 0 | 1 | 25 | 0.67630471 | FALSE | 1 | FALSE | 1 | FALSE | 1 | FALSE | 1 | FALSE | 1 | FALSE | 1 |
| 2761 | 2761 | 7.34146342 | 1.71236189 | 4 | 0.1584689 | 1 | 0.93364809 | 2 | 0.79988333 | 7 | 0.08151801 | 3 | 0.15097573 | 65 | 0.92022824 | FALSE | 1 | FALSE | 1 | FALSE | 1 | FALSE | 1 | FALSE | 1 | FALSE | 1 |
| 2762 | 2762 | 7.02777778 | 1.90738493 | 2 | 0.22589135 | 1 | 0.68454804 | 2 | 0.35984046 | 3 | 0.21561745 | 3 | 0.01918344 | 24 | 0.99496229 | FALSE | 1 | FALSE | 1 | FALSE | 1 | FALSE | 1 | FALSE | 0.70156566 | FALSE | 1 |
| 2763 | 2763 | 8.2173913 | 2.65324284 | 1 | 0.45027683 | 1 | 0.53115654 | 0 | 1 | 1 | 0.6613267 | 1 | 0.31672722 | 19 | 0.71542496 | FALSE | 1 | FALSE | 1 | FALSE | 1 | FALSE | 1 | FALSE | 1 | FALSE | 1 |
| 2764 | 2764 | 9.19230769 | 2.63145515 | 0 | 1 | 0 | 1 | 1 | 0.61440466 | 1 | 0.70601446 | 0 | 1 | 24 | 0.20425579 | FALSE | 1 | FALSE | 1 | FALSE | 1 | FALSE | 1 | FALSE | 1 | FALSE | 1 |
| 2765 | 2765 | 9.05555556 | 3.08170628 | 0 | 1 | 1 | 0.44710673 | 2 | 0.13502329 | 0 | 1 | 0 | 1 | 15 | 0.69372602 | FALSE | 1 | FALSE | 1 | FALSE | 1 | FALSE | 1 | FALSE | 1 | FALSE | 1 |
| 2766 | 2766 | 7.46153846 | 2.41625005 | 0 | 1 | 3 | 0.13080858 | 0 | 1 | 4 | 0.10199045 | 0 | 1 | 32 | 0.74324845 | FALSE | 1 | FALSE | 1 | FALSE | 1 | FALSE | 1 | FALSE | 1 | FALSE | 1 |
| 2767 | 2767 | 6.34482759 | 3.31429362 | 0 | 1 | 1 | 0.61539065 | 1 | 0.65463982 | 2 | 0.3871811 | 0 | 1 | 25 | 0.51779518 | FALSE | 1 | FALSE | 1 | FALSE | 1 | FALSE | 1 | FALSE | 1 | FALSE | 1 |
| 2768 | 2768 | 9.03448276 | 2.31163114 | 0 | 1 | 3 | 0.06581417 | 0 | 1 | 1 | 0.74482229 | 0 | 1 | 25 | 0.51779518 | FALSE | 1 | FALSE | 1 | FALSE | 1 | FALSE | 1 | FALSE | 1 | FALSE | 1 |
| 2769 | 2769 | 6.2 | 2.90860791 | 0 | 1 | 1 | 0.7326167 | 4 | 0.05405259 | 1 | 0.84823834 | 0 | 1 | 34 | 0.56328904 | FALSE | 1 | FALSE | 1 | FALSE | 1 | FALSE | 1 | FALSE | 1 | FALSE | 1 |
| 2770 | 2770 | 5.408 | 2.69843214 | 16 | 2.21E-08 | 4 | 0.49980093 | 5 | 0.382949 | 5 | 0.59818382 | 5 | 0.03759894 | 78 | 0.99998924 | TRUE | 2.93E-05 | FALSE | 1 | FALSE | 1 | FALSE | 1 | FALSE | 0.87669812 | FALSE | 1 |
| 2771 | 2771 | 6.47826087 | 3.13747196 | 3 | 0.25989998 | 3 | 0.38707857 | 6 | 0.03679232 | 1 | 0.96159801 | 3 | 0.1035305 | 53 | 0.96638194 | FALSE | 1 | FALSE | 1 | FALSE | 1 | FALSE | 1 | FALSE | 1 | FALSE | 1 |
| 2772 | 2772 | 8.58823529 | 2.5218766 | 2 | 0.06909038 | 0 | 1 | 0 | 1 | 1 | 0.55062772 | 0 | 1 | 14 | 0.7313145 | FALSE | 0.74276693 | FALSE | 1 | FALSE | 1 | FALSE | 1 | FALSE | 1 | FALSE | 1 |
| 2773 | 2773 | 8.25 | 3.07205143 | 1 | 0.34033054 | 0 | 1 | 0 | 1 | 1 | 0.52895088 | 1 | 0.23264612 | 13 | 0.76799161 | FALSE | 1 | FALSE | 1 | FALSE | 1 | FALSE | 1 | FALSE | 1 | FALSE | 1 |
| 2774 | 2774 | 8.06818182 | 2.00451453 | 4 | 0.02552933 | 3 | 0.16927151 | 2 | 0.47281647 | 0 | 1 | 1 | 0.51797501 | 34 | 0.9286641 | FALSE | 0.4974559 | FALSE | 1 | FALSE | 1 | FALSE | 1 | FALSE | 1 | FALSE | 1 |
| 2775 | 2775 | 8.43181818 | 3.42051117 | 0 | 1 | 1 | 0.70472939 | 3 | 0.14625315 | 0 | 1 | 1 | 0.45845922 | 32 | 0.47030992 | FALSE | 1 | FALSE | 1 | FALSE | 1 | FALSE | 1 | FALSE | 1 | FALSE | 1 |
| 2776 | 2776 | 7.63461539 | 2.87581992 | 3 | 0.14789018 | 5 | 0.02560527 | 0 | 1 | 3 | 0.4296776 | 0 | 1 | 41 | 0.89856682 | FALSE | 1 | FALSE | 0.82367276 | FALSE | 1 | FALSE | 1 | FALSE | 1 | FALSE | 1 |
| 2777 | 2777 | 10 | 2.86356421 | 1 | 0.40558521 | 2 | 0.1353446 | 1 | 0.51939841 | 1 | 0.60987146 | 0 | 1 | 15 | 0.92113679 | FALSE | 1 | FALSE | 1 | FALSE | 1 | FALSE | 1 | FALSE | 1 | FALSE | 1 |
| 2778 | 2778 | 7.67816092 | 1.71198784 | 2 | 0.65227007 | 3 | 0.53041741 | 2 | 0.82100261 | 5 | 0.36062409 | 0 | 1 | 74 | 0.40225767 | FALSE | 1 | FALSE | 1 | FALSE | 1 | FALSE | 1 | FALSE | 1 | FALSE | 1 |
| 2779 | 2779 | 10.0476191 | 2.41968496 | 0 | 1 | 0 | 1 | 1 | 0.53671415 | 0 | 1 | 1 | 0.29368245 | 19 | 0.34094959 | FALSE | 1 | FALSE | 1 | FALSE | 1 | FALSE | 1 | FALSE | 1 | FALSE | 1 |
| 2780 | 2780 | 8.95833333 | 3.06157866 | 0 | 1 | 2 | 0.18117183 | 1 | 0.58502114 | 2 | 0.30249307 | 1 | 0.32796862 | 18 | 0.93166998 | FALSE | 1 | FALSE | 1 | FALSE | 1 | FALSE | 1 | FALSE | 1 | FALSE | 1 |
| 2781 | 2781 | 9.2173913 | 2.57000291 | 0 | 1 | 1 | 0.53115654 | 0 | 1 | 1 | 0.6613267 | 0 | 1 | 21 | 0.27964072 | FALSE | 1 | FALSE | 1 | FALSE | 1 | FALSE | 1 | FALSE | 1 | FALSE | 1 |
| 2782 | 2782 | 9.64285714 | 2.02156739 | 0 | 1 | 1 | 0.60247911 | 3 | 0.07756639 | 2 | 0.3705495 | 1 | 0.37113067 | 21 | 0.94083768 | FALSE | 1 | FALSE | 1 | FALSE | 1 | FALSE | 1 | FALSE | 1 | FALSE | 1 |
| 2783 | 2783 | 10.0526316 | 2.01242677 | 2 | 0.08405377 | 2 | 0.12434464 | 1 | 0.50143826 | 2 | 0.21610812 | 2 | 0.03799168 | 10 | 0.99980197 | FALSE | 0.79236883 | FALSE | 1 | FALSE | 1 | FALSE | 1 | FALSE | 0.87669812 | FALSE | 1 |
| 2784 | 2784 | 10.3703704 | 2.626759 | 1 | 0.50471032 | 0 | 1 | 2 | 0.25286935 | 1 | 0.71956339 | 1 | 0.36060486 | 22 | 0.76237877 | FALSE | 1 | FALSE | 1 | FALSE | 1 | FALSE | 1 | FALSE | 1 | FALSE | 1 |
| 2785 | 2785 | 8 | 4.5607017 | 0 | 1 | 0 | 1 | 0 | 1 | 0 | 1 | 0 | 1 | 10 | 0.18273166 | FALSE | 1 | FALSE | 1 | FALSE | 1 | FALSE | 1 | FALSE | 1 | FALSE | 1 |
| 2786 | 2786 | 9.62068966 | 1.34968178 | 0 | 1 | 0 | 1 | 0 | 1 | 1 | 0.74482229 | 1 | 0.38148475 | 27 | 0.14664438 | FALSE | 1 | FALSE | 1 | FALSE | 1 | FALSE | 1 | FALSE | 1 | FALSE | 1 |
| 2787 | 2787 | 10.6666667 | 2.1602469 | 1 | 0.20854748 | 0 | 1 | 0 | 1 | 0 | 1 | 0 | 1 | 8 | 0.5780332 | FALSE | 1 | FALSE | 1 | FALSE | 1 | FALSE | 1 | FALSE | 1 | FALSE | 1 |
| 2788 | 2788 | 8.09090909 | 2.77831033 | 0 | 1 | 1 | 0.30368014 | 0 | 1 | 0 | 1 | 1 | 0.16638736 | 9 | 0.75985366 | FALSE | 1 | FALSE | 1 | FALSE | 1 | FALSE | 1 | FALSE | 1 | FALSE | 1 |
| 2789 | 2789 | 8.35135135 | 0.96467937 | 1 | 0.6184714 | 0 | 1 | 2 | 0.38585089 | 3 | 0.2402032 | 1 | 0.45845922 | 30 | 0.78878581 | FALSE | 1 | FALSE | 1 | FALSE | 1 | FALSE | 1 | FALSE | 1 | FALSE | 1 |
| 2790 | 2790 | 9.04347826 | 2.27420814 | 1 | 0.45027683 | 3 | 0.03669128 | 1 | 0.5695037 | 1 | 0.6613267 | 0 | 1 | 17 | 0.94439745 | FALSE | 1 | FALSE | 0.94259695 | FALSE | 1 | FALSE | 1 | FALSE | 1 | FALSE | 1 |
| 2791 | 2791 | 9.81481482 | 1.49163189 | 1 | 0.50471032 | 5 | 0.00152482 | 1 | 0.62830986 | 1 | 0.71956339 | 0 | 1 | 19 | 0.98241223 | FALSE | 1 | FALSE | 0.40311621 | FALSE | 1 | FALSE | 1 | FALSE | 1 | FALSE | 1 |
| 2792 | 2792 | 7.18181818 | 1.91017285 | 9 | 0.00038952 | 9 | 0.00204035 | 1 | 0.96086551 | 8 | 0.04815327 | 0 | 1 | 61 | 0.99989658 | TRUE | 0.04090948 | FALSE | 0.42264619 | FALSE | 1 | FALSE | 1 | FALSE | 1 | FALSE | 1 |
| 2793 | 2793 | 7.4516129 | 4.29432266 | 2 | 0.16962005 | 2 | 0.24095584 | 1 | 0.65463982 | 2 | 0.3871811 | 3 | 0.01149488 | 19 | 0.99707037 | FALSE | 1 | FALSE | 1 | FALSE | 1 | FALSE | 1 | FALSE | 0.57972935 | FALSE | 1 |
| 2794 | 2794 | 9.5625 | 1.7309228 | 0 | 1 | 0 | 1 | 0 | 1 | 0 | 1 | 1 | 0.19354507 | 12 | 0.374198 | FALSE | 1 | FALSE | 1 | FALSE | 1 | FALSE | 1 | FALSE | 1 | FALSE | 1 |
| 2795 | 2795 | 4.55555556 | 1.58542747 | 2 | 0.32143972 | 1 | 0.77338922 | 2 | 0.80828716 | 1 | 0.8802008 | 2 | 0.16793736 | 38 | 0.59437745 | FALSE | 1 | FALSE | 1 | FALSE | 1 | FALSE | 1 | FALSE | 1 | FALSE | 1 |
| 2796 | 2796 | 6.98461539 | 2.53888108 | 3 | 0.23231139 | 3 | 0.35154359 | 7 | 0.00834811 | 6 | 0.07659646 | 0 | 1 | 46 | 0.9984417 | FALSE | 1 | FALSE | 1 | FALSE | 1 | FALSE | 1 | FALSE | 1 | FALSE | 1 |
| 2797 | 2797 | 8.17647059 | 2.68405519 | 0 | 1 | 2 | 0.30150529 | 2 | 0.34668321 | 1 | 0.7984848 | 1 | 0.43077488 | 28 | 0.72558119 | FALSE | 1 | FALSE | 1 | FALSE | 1 | FALSE | 1 | FALSE | 1 | FALSE | 1 |
| 2798 | 2798 | 4.94174757 | 2.25045947 | 0 | 1 | 4 | 0.42831767 | 6 | 0.16497789 | 2 | 0.95407875 | 2 | 0.50667524 | 89 | 0.34117234 | FALSE | 1 | FALSE | 1 | FALSE | 1 | FALSE | 1 | FALSE | 1 | FALSE | 1 |
| 2799 | 2799 | 8.33333333 | 2.6119384 | 2 | 0.17884234 | 1 | 0.6278847 | 0 | 1 | 0 | 1 | 4 | 0.00134977 | 23 | 0.9158696 | FALSE | 1 | FALSE | 1 | FALSE | 1 | FALSE | 1 | FALSE | 0.20625174 | FALSE | 1 |
| 2800 | 2800 | 9.64 | 1.22898332 | 0 | 1 | 0 | 1 | 1 | 0.59998149 | 2 | 0.31968852 | 0 | 1 | 22 | 0.43642846 | FALSE | 1 | FALSE | 1 | FALSE | 1 | FALSE | 1 | FALSE | 1 | FALSE | 1 |
| 2801 | 2801 | 3.77824268 | 3.19469151 | 2 | 0.98583246 | 7 | 0.65168128 | 6 | 0.86083541 | 6 | 0.96527859 | 2 | 0.90563009 | 214 | 0.00488447 | FALSE | 1 | FALSE | 1 | FALSE | 1 | FALSE | 1 | FALSE | 1 | FALSE | 1 |
| 2802 | 2802 | 12.08 | 3.77274436 | 2 | 0.13369621 | 1 | 0.56109832 | 3 | 0.05884411 | 1 | 0.69181315 | 0 | 1 | 18 | 0.96851178 | FALSE | 0.97964162 | FALSE | 1 | FALSE | 1 | FALSE | 1 | FALSE | 1 | FALSE | 1 |
| 2803 | 2803 | 8.72727273 | 2.68297356 | 2 | 0.10809809 | 0 | 1 | 1 | 0.55340851 | 2 | 0.26791271 | 1 | 0.30529951 | 16 | 0.95553951 | FALSE | 0.88820416 | FALSE | 1 | FALSE | 1 | FALSE | 1 | FALSE | 1 | FALSE | 1 |
| 2804 | 2804 | 8.20689655 | 0.84605822 | 2 | 0.16962005 | 3 | 0.06581417 | 5 | 0.0033608 | 1 | 0.74482229 | 0 | 1 | 18 | 0.99922671 | FALSE | 1 | FALSE | 1 | FALSE | 1 | FALSE | 1 | FALSE | 1 | FALSE | 1 |
| 2805 | 2805 | 7.84848485 | 3.15384529 | 0 | 1 | 0 | 1 | 1 | 0.70185639 | 1 | 0.78873814 | 0 | 1 | 31 | 0.09218876 | FALSE | 1 | FALSE | 1 | FALSE | 1 | FALSE | 1 | FALSE | 1 | FALSE | 1 |
| 2806 | 2806 | 8.41666667 | 1.44096804 | 1 | 0.45027683 | 1 | 0.53115654 | 3 | 0.04772439 | 0 | 1 | 1 | 0.31672722 | 17 | 0.94439745 | FALSE | 1 | FALSE | 1 | FALSE | 1 | FALSE | 1 | FALSE | 1 | FALSE | 1 |
| 2807 | 2807 | 6.5 | 3.14661132 | 4 | 0.17847815 | 2 | 0.77275903 | 3 | 0.60234934 | 1 | 0.98288967 | 4 | 0.05219855 | 72 | 0.63625249 | FALSE | 1 | FALSE | 1 | FALSE | 1 | FALSE | 1 | FALSE | 1 | FALSE | 1 |
| 2808 | 2808 | 6.09565217 | 3.43631226 | 6 | 0.07204072 | 4 | 0.50674923 | 4 | 0.59047122 | 8 | 0.1523207 | 3 | 0.28730314 | 89 | 0.97324198 | FALSE | 0.76727438 | FALSE | 1 | FALSE | 1 | FALSE | 1 | FALSE | 1 | FALSE | 1 |
| 2809 | 2809 | 10.0909091 | 2.42916168 | 1 | 0.24866461 | 0 | 1 | 1 | 0.33149969 | 0 | 1 | 1 | 0.16638736 | 8 | 0.92152993 | FALSE | 1 | FALSE | 1 | FALSE | 1 | FALSE | 1 | FALSE | 1 | FALSE | 1 |
| 2810 | 2810 | 3.90909091 | 2.26653702 | 1 | 0.76171602 | 4 | 0.10132561 | 1 | 0.8673918 | 2 | 0.72625531 | 0 | 1 | 47 | 0.50345693 | FALSE | 1 | FALSE | 1 | FALSE | 1 | FALSE | 1 | FALSE | 1 | FALSE | 1 |
| 2811 | 2811 | 9 | 2.1725561 | 0 | 1 | 0 | 1 | 1 | 0.59998149 | 0 | 1 | 0 | 1 | 24 | 0.08019798 | FALSE | 1 | FALSE | 1 | FALSE | 1 | FALSE | 1 | FALSE | 1 | FALSE | 1 |
| 2812 | 2812 | 7.81818182 | 2.08100421 | 0 | 1 | 0 | 1 | 0 | 1 | 0 | 1 | 0 | 1 | 11 | 0.15414489 | FALSE | 1 | FALSE | 1 | FALSE | 1 | FALSE | 1 | FALSE | 1 | FALSE | 1 |
| 2813 | 2813 | 7.84 | 3.5290792 | 0 | 1 | 0 | 1 | 0 | 1 | 1 | 0.69181315 | 1 | 0.33902672 | 23 | 0.22727183 | FALSE | 1 | FALSE | 1 | FALSE | 1 | FALSE | 1 | FALSE | 1 | FALSE | 1 |
| 2814 | 2814 | 7.61538462 | 3.91326072 | 0 | 1 | 0 | 1 | 2 | 0.07725803 | 0 | 1 | 0 | 1 | 11 | 0.6683056 | FALSE | 1 | FALSE | 1 | FALSE | 1 | FALSE | 1 | FALSE | 1 | FALSE | 1 |
| 2815 | 2815 | 8.14814815 | 3.6281176 | 1 | 0.49162634 | 1 | 0.57534752 | 2 | 0.23940226 | 1 | 0.70601446 | 2 | 0.0670823 | 19 | 0.96054244 | FALSE | 1 | FALSE | 1 | FALSE | 1 | FALSE | 1 | FALSE | 1 | FALSE | 1 |
| 2816 | 2816 | 9.46428571 | 2.59783073 | 0 | 1 | 0 | 1 | 1 | 0.64171563 | 1 | 0.73248983 | 2 | 0.07640798 | 24 | 0.55018107 | FALSE | 1 | FALSE | 1 | FALSE | 1 | FALSE | 1 | FALSE | 1 | FALSE | 1 |
| 2817 | 2817 | 10.6111111 | 2.60637869 | 0 | 1 | 0 | 1 | 1 | 0.48280983 | 0 | 1 | 2 | 0.03435384 | 15 | 0.69372602 | FALSE | 1 | FALSE | 1 | FALSE | 1 | FALSE | 1 | FALSE | 0.86063198 | FALSE | 1 |
| 2818 | 2818 | 7.61764706 | 2.33911378 | 1 | 0.58738519 | 1 | 0.6739483 | 4 | 0.03235504 | 2 | 0.46709861 | 3 | 0.01774672 | 23 | 0.99615372 | FALSE | 1 | FALSE | 1 | FALSE | 1 | FALSE | 1 | FALSE | 0.69720022 | FALSE | 1 |

| | | | | | | | | | | | | | | | | | | | | | | | | | | | |
|---|---|---|---|---|---|---|---|---|---|---|---|---|---|---|---|---|---|---|---|---|---|---|---|---|---|---|---|
| 2819 | 2819 | 9 | 3.10912635 | 0 | 1 | 0 | 1 | 1 | 0.35555293 | 1 | 0.43131129 | 0 | 1 | 10 | 0.71436964 | FALSE | 1 | FALSE | 1 | FALSE | 1 | FALSE | 1 | FALSE | 1 | FALSE | 1 |
| 2820 | 2820 | 7.16455696 | 2.02159468 | 2 | 0.60596997 | 1 | 0.92668093 | 0 | 1 | 3 | 0.71030908 | 0 | 1 | 73 | 0.02683162 | FALSE | 1 | FALSE | 1 | FALSE | 1 | FALSE | 1 | FALSE | 1 | FALSE | 1 |
| 2821 | 2821 | 8.63636364 | 2.70597747 | 0 | 1 | 0 | 1 | 1 | 0.33149969 | 0 | 1 | 0 | 1 | 10 | 0.46859937 | FALSE | 1 | FALSE | 1 | FALSE | 1 | FALSE | 1 | FALSE | 1 | FALSE | 1 |
| 2822 | 2822 | 11.05 | 1.88348082 | 0 | 1 | 0 | 1 | 0 | 1 | 0 | 1 | 0 | 1 | 19 | 0.03948376 | FALSE | 1 | FALSE | 1 | FALSE | 1 | FALSE | 1 | FALSE | 1 | FALSE | 1 |
| 2823 | 2823 | 8.53333333 | 2.39072281 | 0 | 1 | 1 | 0.38964394 | 0 | 1 | 0 | 1 | 0 | 1 | 14 | 0.29525409 | FALSE | 1 | FALSE | 1 | FALSE | 1 | FALSE | 1 | FALSE | 1 | FALSE | 1 |
| 2824 | 2824 | 8.64705882 | 3.41176471 | 2 | 0.06909038 | 0 | 1 | 1 | 0.46348836 | 0 | 1 | 0 | 1 | 14 | 0.7313145 | FALSE | 0.74276693 | FALSE | 1 | FALSE | 1 | FALSE | 1 | FALSE | 1 | FALSE | 1 |
| 2825 | 2825 | 7.73684211 | 2.46751751 | 0 | 1 | 0 | 1 | 0 | 1 | 0 | 1 | 2 | 0.03799168 | 17 | 0.41141543 | FALSE | 1 | FALSE | 1 | FALSE | 1 | FALSE | 1 | FALSE | 0.87669812 | FALSE | 1 |
| 2826 | 2826 | 8.96296296 | 1.87540576 | 0 | 1 | 0 | 1 | 2 | 0.25286935 | 2 | 0.35374209 | 2 | 0.07169515 | 21 | 0.88395936 | FALSE | 1 | FALSE | 1 | FALSE | 1 | FALSE | 1 | FALSE | 1 | FALSE | 1 |
| 2827 | 2827 | 11.2142857 | 2.14404729 | 0 | 1 | 2 | 0.07341559 | 0 | 1 | 0 | 1 | 1 | 0.20679328 | 11 | 0.83676449 | FALSE | 1 | FALSE | 1 | FALSE | 1 | FALSE | 1 | FALSE | 1 | FALSE | 1 |
| 2828 | 2828 | 8.93548387 | 2.83907622 | 1 | 0.55378205 | 1 | 0.63997468 | 0 | 1 | 3 | 0.16836568 | 0 | 1 | 26 | 0.64660109 | FALSE | 1 | FALSE | 1 | FALSE | 1 | FALSE | 1 | FALSE | 1 | FALSE | 1 |
| 2829 | 2829 | 8.20689655 | 2.75905169 | 2 | 0.16962005 | 0 | 1 | 2 | 0.27983822 | 4 | 0.04184267 | 0 | 1 | 21 | 0.97185642 | FALSE | 1 | FALSE | 1 | FALSE | 1 | FALSE | 1 | FALSE | 1 | FALSE | 1 |
| 2830 | 2830 | 10.2 | 1.93907194 | 1 | 0.22886536 | 1 | 0.28036922 | 0 | 1 | 0 | 1 | 0 | 1 | 8 | 0.80394719 | FALSE | 1 | FALSE | 1 | FALSE | 1 | FALSE | 1 | FALSE | 1 | FALSE | 1 |
| 2831 | 2831 | 10.5714286 | 2.29017422 | 0 | 1 | 0 | 1 | 2 | 0.08806398 | 2 | 0.13304705 | 0 | 1 | 10 | 0.94579623 | FALSE | 1 | FALSE | 1 | FALSE | 1 | FALSE | 1 | FALSE | 1 | FALSE | 1 |
| 2832 | 2832 | 7.21008403 | 0.78722947 | 9 | 0.00334987 | 10 | 0.00499748 | 4 | 0.62455469 | 6 | 0.46726758 | 2 | 0.5851187 | 88 | 0.99896326 | FALSE | 0.15862482 | FALSE | 0.58120789 | FALSE | 1 | FALSE | 1 | FALSE | 1 | FALSE | 1 |
| 2833 | 2833 | 9.25 | 1.82002747 | 2 | 0.06197875 | 0 | 1 | 1 | 0.44344817 | 1 | 0.52895088 | 0 | 1 | 12 | 0.90922186 | FALSE | 0.72532843 | FALSE | 1 | FALSE | 1 | FALSE | 1 | FALSE | 1 | FALSE | 1 |
| 2834 | 2834 | 7.86666667 | 2.12498366 | 2 | 0.17884234 | 1 | 0.6278847 | 3 | 0.09131526 | 0 | 1 | 0 | 1 | 24 | 0.82294782 | FALSE | 1 | FALSE | 1 | FALSE | 1 | FALSE | 1 | FALSE | 1 | FALSE | 1 |
| 2835 | 2835 | 9.10344828 | 2.33924284 | 0 | 1 | 2 | 0.24095584 | 0 | 1 | 0 | 1 | 0 | 1 | 27 | 0.14664438 | FALSE | 1 | FALSE | 1 | FALSE | 1 | FALSE | 1 | FALSE | 1 | FALSE | 1 |
| 2836 | 2836 | 11.25 | 0.8291562 | 1 | 0.34033054 | 0 | 1 | 2 | 0.11087977 | 0 | 1 | 0 | 1 | 13 | 0.76799161 | FALSE | 1 | FALSE | 1 | FALSE | 1 | FALSE | 1 | FALSE | 1 | FALSE | 1 |
| 2837 | 2837 | 9.41176471 | 2.47478635 | 0 | 1 | 0 | 1 | 1 | 0.46348836 | 2 | 0.18212134 | 0 | 1 | 14 | 0.7313145 | FALSE | 1 | FALSE | 1 | FALSE | 1 | FALSE | 1 | FALSE | 1 | FALSE | 1 |
| 2838 | 2838 | 2.91544118 | 2.83171063 | 6 | 0.70103235 | 3 | 0.99387335 | 6 | 0.92992101 | 12 | 0.59478202 | 3 | 0.828056 | 241 | 0.01848523 | FALSE | 1 | FALSE | 1 | FALSE | 1 | FALSE | 1 | FALSE | 1 | FALSE | 1 |
| 2839 | 2839 | 10.1666667 | 1.77169097 | 0 | 1 | 1 | 0.179097 | 0 | 1 | 1 | 0.24578796 | 0 | 1 | 4 | 0.94743772 | FALSE | 1 | FALSE | 1 | FALSE | 1 | FALSE | 1 | FALSE | 1 | FALSE | 1 |
| 2840 | 2840 | 9.33333333 | 3.0446414 | 1 | 0.42086991 | 0 | 1 | 2 | 0.17308361 | 3 | 0.06912908 | 0 | 1 | 15 | 0.96513691 | FALSE | 1 | FALSE | 1 | FALSE | 1 | FALSE | 1 | FALSE | 1 | FALSE | 1 |
| 2841 | 2841 | 9.5 | 2 | 0 | 1 | 0 | 1 | 1 | 0.58502114 | 2 | 0.30249307 | 0 | 1 | 21 | 0.47053137 | FALSE | 1 | FALSE | 1 | FALSE | 1 | FALSE | 1 | FALSE | 1 | FALSE | 1 |
| 2842 | 2842 | 9.24 | 1.49523688 | 4 | 0.12569732 | 3 | 0.43942904 | 3 | 0.50898113 | 2 | 0.86569532 | 0 | 1 | 63 | 0.61191651 | FALSE | 0.95955826 | FALSE | 1 | FALSE | 1 | FALSE | 1 | FALSE | 1 | FALSE | 1 |
| 2843 | 2843 | 10.0476191 | 2.2141577 | 1 | 0.42086991 | 1 | 0.49918218 | 2 | 0.17308361 | 1 | 0.62783522 | 0 | 1 | 16 | 0.90353674 | FALSE | 1 | FALSE | 1 | FALSE | 1 | FALSE | 1 | FALSE | 1 | FALSE | 1 |
| 2844 | 2844 | 9.5 | 3.47953199 | 5 | 0.00412436 | 4 | 0.04570474 | 2 | 0.44869594 | 2 | 0.5811115 | 1 | 0.50166523 | 28 | 0.99888598 | FALSE | 0.18198109 | FALSE | 1 | FALSE | 1 | FALSE | 1 | FALSE | 1 | FALSE | 1 |
| 2845 | 2845 | 8.2173913 | 2.4664095 | 1 | 0.45027683 | 1 | 0.53115654 | 1 | 0.5695037 | 0 | 1 | 1 | 0.31672722 | 19 | 0.71542496 | FALSE | 1 | FALSE | 1 | FALSE | 1 | FALSE | 1 | FALSE | 1 | FALSE | 1 |
| 2846 | 2846 | 9.01754386 | 2.61903443 | 2 | 0.4229474 | 2 | 0.54544984 | 3 | 0.32646887 | 2 | 0.73544289 | 0 | 1 | 47 | 0.62412393 | FALSE | 1 | FALSE | 1 | FALSE | 1 | FALSE | 1 | FALSE | 1 | FALSE | 1 |
| 2847 | 2847 | 9.30769231 | 3.5601343 | 0 | 1 | 0 | 1 | 1 | 0.37874417 | 0 | 1 | 0 | 1 | 12 | 0.374198 | FALSE | 1 | FALSE | 1 | FALSE | 1 | FALSE | 1 | FALSE | 1 | FALSE | 1 |
| 2848 | 2848 | 8.51851852 | 2.69938773 | 1 | 0.50471032 | 0 | 1 | 1 | 0.62830986 | 0 | 1 | 0 | 1 | 25 | 0.1832206 | FALSE | 1 | FALSE | 1 | FALSE | 1 | FALSE | 1 | FALSE | 1 | FALSE | 1 |
| 2849 | 2849 | 10.7037037 | 1.90155521 | 0 | 1 | 2 | 0.21684246 | 2 | 0.25286935 | 1 | 0.71956339 | 0 | 1 | 22 | 0.76237877 | FALSE | 1 | FALSE | 1 | FALSE | 1 | FALSE | 1 | FALSE | 1 | FALSE | 1 |
| 2850 | 2850 | 4.25242718 | 2.93858974 | 2 | 0.7467337 | 3 | 0.65322408 | 3 | 0.72201868 | 5 | 0.51481541 | 3 | 0.23844883 | 87 | 0.55659837 | FALSE | 1 | FALSE | 1 | FALSE | 1 | FALSE | 1 | FALSE | 1 | FALSE | 1 |
| 2851 | 2851 | 9.58333333 | 2.39646731 | 3 | 0.0638041 | 2 | 0.32556427 | 0 | 1 | 0 | 1 | 3 | 0.0206828 | 28 | 0.90259714 | FALSE | 0.73766 | FALSE | 1 | FALSE | 1 | FALSE | 1 | FALSE | 0.7366672 | FALSE | 1 |
| 2852 | 2852 | 5.94230769 | 1.57398883 | 7 | 0.00025388 | 6 | 0.00517174 | 2 | 0.54120634 | 3 | 0.4048204 | 0 | 1 | 32 | 0.99990252 | TRUE | 0.03104128 | FALSE | 0.58120789 | FALSE | 1 | FALSE | 1 | FALSE | 1 | FALSE | 1 |
| 2853 | 2853 | 11.8235294 | 2.25532818 | 0 | 1 | 1 | 0.42857733 | 0 | 1 | 0 | 1 | 1 | 0.24525764 | 15 | 0.49060435 | FALSE | 1 | FALSE | 1 | FALSE | 1 | FALSE | 1 | FALSE | 1 | FALSE | 1 |
| 2854 | 2854 | 8.22222222 | 1.25707872 | 1 | 0.75539413 | 2 | 0.52572379 | 4 | 0.12796328 | 2 | 0.71679995 | 4 | 0.01153514 | 41 | 0.96572325 | FALSE | 1 | FALSE | 1 | FALSE | 1 | FALSE | 1 | FALSE | 0.57972935 | FALSE | 1 |
| 2855 | 2855 | 1.04503797 | 0.63876958 | 85 | 0.97887986 | 115 | 0.88440547 | 138 | 0.45990076 | 179 | 0.32919427 | 60 | 0.71425352 | 3229 | 0.12395847 | FALSE | 1 | FALSE | 1 | FALSE | 1 | FALSE | 1 | FALSE | 1 | FALSE | 1 |
| 2856 | 2856 | 8.36842105 | 1.56307815 | 0 | 1 | 0 | 1 | 0 | 1 | 2 | 0.21610812 | 0 | 1 | 17 | 0.41141543 | FALSE | 1 | FALSE | 1 | FALSE | 1 | FALSE | 1 | FALSE | 1 | FALSE | 1 |
| 2857 | 2857 | 5.10402685 | 0.83500426 | 8 | 0.49900285 | 7 | 0.85560128 | 8 | 0.84747844 | 15 | 0.3940202 | 3 | 0.87335417 | 257 | 0.20643046 | FALSE | 1 | FALSE | 1 | FALSE | 1 | FALSE | 1 | FALSE | 1 | FALSE | 1 |
| 2858 | 2858 | 11.1818182 | 2.7901835 | 0 | 1 | 0 | 1 | 0 | 1 | 3 | 0.07743186 | 0 | 1 | 19 | 0.54233419 | FALSE | 1 | FALSE | 1 | FALSE | 1 | FALSE | 1 | FALSE | 1 | FALSE | 1 |
| 2859 | 2859 | 10.2142857 | 1.9338552 | 0 | 1 | 0 | 1 | 0 | 1 | 1 | 0.48242099 | 0 | 1 | 13 | 0.33283729 | FALSE | 1 | FALSE | 1 | FALSE | 1 | FALSE | 1 | FALSE | 1 | FALSE | 1 |
| 2860 | 2860 | 7.11538462 | 2.56195423 | 2 | 0.36837609 | 0 | 1 | 3 | 0.26704679 | 5 | 0.07777122 | 4 | 0.00882912 | 36 | 0.99249228 | FALSE | 1 | FALSE | 1 | FALSE | 1 | FALSE | 1 | FALSE | 0.52052899 | FALSE | 1 |
| 2861 | 2861 | 11 | 4.163332 | 0 | 1 | 1 | 0.54637326 | 0 | 1 | 3 | 0.09525947 | 2 | 0.058175 | 18 | 0.93166998 | FALSE | 1 | FALSE | 1 | FALSE | 1 | FALSE | 1 | FALSE | 1 | FALSE | 1 |
| 2862 | 2862 | 6.21052632 | 3.22129342 | 4 | 0.01561835 | 2 | 0.3494045 | 0 | 1 | 2 | 0.52640931 | 1 | 0.4673875 | 29 | 0.93814695 | FALSE | 0.38695764 | FALSE | 1 | FALSE | 1 | FALSE | 1 | FALSE | 1 | FALSE | 1 |
| 2863 | 2863 | 9.88888889 | 2.96064724 | 1 | 0.20854748 | 2 | 0.03227145 | 0 | 1 | 0 | 1 | 0 | 1 | 6 | 0.96128317 | FALSE | 1 | FALSE | 0.92268438 | FALSE | 1 | FALSE | 1 | FALSE | 1 | FALSE | 1 |
| 2864 | 2864 | 9.9 | 1.3 | 0 | 1 | 0 | 1 | 0 | 1 | 0 | 1 | 0 | 1 | 10 | 0.18273166 | FALSE | 1 | FALSE | 1 | FALSE | 1 | FALSE | 1 | FALSE | 1 | FALSE | 1 |
| 2865 | 2865 | 9.42307692 | 3.02773179 | 0 | 1 | 1 | 0.57534752 | 0 | 1 | 0 | 1 | 0 | 1 | 25 | 0.06985983 | FALSE | 1 | FALSE | 1 | FALSE | 1 | FALSE | 1 | FALSE | 1 | FALSE | 1 |
| 2866 | 2866 | 11.4137931 | 3.0458252 | 0 | 1 | 1 | 0.58913618 | 1 | 0.62830986 | 0 | 1 | 0 | 1 | 25 | 0.1832206 | FALSE | 1 | FALSE | 1 | FALSE | 1 | FALSE | 1 | FALSE | 1 | FALSE | 1 |
| 2867 | 2867 | 10 | 2.49443826 | 0 | 1 | 0 | 1 | 1 | 0.28067846 | 0 | 1 | 1 | 0.1383237 | 7 | 0.84574168 | FALSE | 1 | FALSE | 1 | FALSE | 1 | FALSE | 1 | FALSE | 1 | FALSE | 1 |
| 2868 | 2868 | 9.55172414 | 2.2982183 | 2 | 0.16048326 | 0 | 1 | 2 | 0.2663542 | 1 | 0.73248983 | 2 | 0.07640798 | 21 | 0.94083768 | FALSE | 1 | FALSE | 1 | FALSE | 1 | FALSE | 1 | FALSE | 1 | FALSE | 1 |
| 2869 | 2869 | 11.15 | 3.03768004 | 0 | 1 | 3 | 0.02534497 | 1 | 0.51939841 | 0 | 1 | 2 | 0.04176963 | 14 | 0.97325593 | FALSE | 1 | FALSE | 0.82367276 | FALSE | 1 | FALSE | 1 | FALSE | 0.90763079 | FALSE | 1 |
| 2870 | 2870 | 9.86956522 | 2.11216657 | 1 | 0.45027683 | 0 | 1 | 2 | 0.19931429 | 0 | 1 | 1 | 0.31672722 | 19 | 0.71542496 | FALSE | 1 | FALSE | 1 | FALSE | 1 | FALSE | 1 | FALSE | 1 | FALSE | 1 |
| 2871 | 2871 | 9.8125 | 1.91111061 | 1 | 0.34033054 | 1 | 0.40942991 | 0 | 1 | 0 | 1 | 1 | 0.23264612 | 13 | 0.76799161 | FALSE | 1 | FALSE | 1 | FALSE | 1 | FALSE | 1 | FALSE | 1 | FALSE | 1 |
| 2872 | 2872 | 8.46153846 | 3.94738572 | 2 | 0.26413583 | 1 | 0.7236252 | 2 | 0.41139386 | 0 | 1 | 1 | 0.4761699 | 33 | 0.59061307 | FALSE | 1 | FALSE | 1 | FALSE | 1 | FALSE | 1 | FALSE | 1 | FALSE | 1 |
| 2873 | 2873 | 13.44 | 4.94837347 | 1 | 0.45027683 | 0 | 1 | 0 | 1 | 2 | 0.2852236 | 1 | 0.31672722 | 19 | 0.71542496 | FALSE | 1 | FALSE | 1 | FALSE | 1 | FALSE | 1 | FALSE | 1 | FALSE | 1 |
| 2874 | 2874 | 8.63461539 | 2.08500919 | 3 | 0.14789018 | 3 | 0.23623106 | 1 | 0.85187773 | 3 | 0.4296776 | 2 | 0.2095319 | 40 | 0.94722426 | FALSE | 1 | FALSE | 1 | FALSE | 1 | FALSE | 1 | FALSE | 1 | FALSE | 1 |
| 2875 | 2875 | 9.66666667 | 2.78886676 | 0 | 1 | 3 | 0.01901931 | 1 | 0.48280983 | 0 | 1 | 0 | 1 | 14 | 0.86287418 | FALSE | 1 | FALSE | 0.77515497 | FALSE | 1 | FALSE | 1 | FALSE | 1 | FALSE | 1 |
| 2876 | 2876 | 7.42307692 | 1.82236273 | 2 | 0.14250978 | 1 | 0.57534752 | 1 | 0.61440466 | 2 | 0.33678042 | 0 | 1 | 20 | 0.90142689 | FALSE | 1 | FALSE | 1 | FALSE | 1 | FALSE | 1 | FALSE | 1 | FALSE | 1 |
| 2877 | 2877 | 6.52941177 | 3.51245929 | 7 | 0.00057802 | 0 | 1 | 2 | 0.61293419 | 2 | 0.74436726 | 0 | 1 | 46 | 0.83074268 | FALSE | 0.05254197 | FALSE | 1 | FALSE | 1 | FALSE | 1 | FALSE | 1 | FALSE | 1 |
| 2878 | 2878 | 9.14285714 | 1.22224284 | 0 | 1 | 1 | 0.68454804 | 1 | 0.72299506 | 4 | 0.07465533 | 1 | 0.44015537 | 28 | 0.83079716 | FALSE | 1 | FALSE | 1 | FALSE | 1 | FALSE | 1 | FALSE | 1 | FALSE | 1 |
| 2879 | 2879 | 6.97278912 | 1.52505416 | 0 | 1 | 3 | 0.84058578 | 3 | 0.88778543 | 6 | 0.6343804 | 4 | 0.20033966 | 125 | 0.09400764 | FALSE | 1 | FALSE | 1 | FALSE | 1 | FALSE | 1 | FALSE | 1 | FALSE | 1 |
| 2880 | 2880 | 3.81060606 | 2.37133314 | 4 | 0.44004265 | 7 | 0.1351573 | 3 | 0.85933994 | 7 | 0.40353836 | 1 | 0.8896353 | 110 | 0.68305172 | FALSE | 1 | FALSE | 1 | FALSE | 1 | FALSE | 1 | FALSE | 1 | FALSE | 1 |
| 2881 | 2881 | 8.92307692 | 1.97918159 | 2 | 0.14250978 | 0 | 1 | 0 | 1 | 0 | 1 | 3 | 0.00847466 | 21 | 0.78939668 | FALSE | 1 | FALSE | 1 | FALSE | 1 | FALSE | 1 | FALSE | 0.50674765 | FALSE | 1 |
| 2882 | 2882 | 12.9 | 3.44818793 | 0 | 1 | 0 | 1 | 0 | 1 | 0 | 1 | 0 | 1 | 10 | 0.18273166 | FALSE | 1 | FALSE | 1 | FALSE | 1 | FALSE | 1 | FALSE | 1 | FALSE | 1 |
| 2883 | 2883 | 4.31132076 | 2.59848725 | 4 | 0.78673083 | 7 | 0.51254122 | 10 | 0.21429748 | 11 | 0.35836698 | 9 | 0.00689637 | 167 | 0.95643148 | FALSE | 1 | FALSE | 1 | FALSE | 1 | FALSE | 1 | FALSE | 0.4774184 | FALSE | 1 |
| 2884 | 2884 | 11.7777778 | 2.63640234 | 0 | 1 | 1 | 0.44710673 | 0 | 1 | 2 | 0.19902682 | 0 | 1 | 15 | 0.69372602 | FALSE | 1 | FALSE | 1 | FALSE | 1 | FALSE | 1 | FALSE | 1 | FALSE | 1 |
| 2885 | 2885 | 10.0714286 | 3.36927898 | 0 | 1 | 0 | 1 | 0 | 1 | 0 | 1 | 0 | 1 | 13 | 0.10967906 | FALSE | 1 | FALSE | 1 | FALSE | 1 | FALSE | 1 | FALSE | 1 | FALSE | 1 |
| 2886 | 2886 | 10.8275862 | 2.6918647 | 1 | 0.52988231 | 3 | 0.06581417 | 3 | 0.08431992 | 1 | 0.74482229 | 0 | 1 | 21 | 0.97185642 | FALSE | 1 | FALSE | 1 | FALSE | 1 | FALSE | 1 | FALSE | 1 | FALSE | 1 |
| 2887 | 2887 | 9.18181818 | 3.37956915 | 1 | 0.4357638 | 1 | 0.5154318 | 2 | 0.18613256 | 0 | 1 | 0 | 1 | 18 | 0.74744813 | FALSE | 1 | FALSE | 1 | FALSE | 1 | FALSE | 1 | FALSE | 1 | FALSE | 1 |
| 2888 | 2888 | 5 | 2.03809866 | 0 | 1 | 2 | 0.19297191 | 1 | 0.59998149 | 1 | 0.69181315 | 1 | 0.33902672 | 20 | 0.8152498 | FALSE | 1 | FALSE | 1 | FALSE | 1 | FALSE | 1 | FALSE | 1 | FALSE | 1 |
| 2889 | 2889 | 9.48387097 | 2.81552113 | 1 | 0.55378205 | 4 | 0.01688692 | 2 | 0.30673593 | 0 | 1 | 1 | 0.4016888 | 23 | 0.95750806 | FALSE | 1 | FALSE | 0.73194511 | FALSE | 1 | FALSE | 1 | FALSE | 1 | FALSE | 1 |
| 2890 | 2890 | 10.6041667 | 1.22030705 | 5 | 0.00733988 | 3 | 0.20208567 | 0 | 1 | 3 | 0.37966044 | 0 | 1 | 37 | 0.93870662 | FALSE | 0.2537058 | FALSE | 1 | FALSE | 1 | FALSE | 1 | FALSE | 1 | FALSE | 1 |
| 2891 | 2891 | 9.71875 | 3.02318515 | 0 | 1 | 1 | 0.65167362 | 2 | 0.32011871 | 2 | 0.43583874 | 0 | 1 | 27 | 0.61671133 | FALSE | 1 | FALSE | 1 | FALSE | 1 | FALSE | 1 | FALSE | 1 | FALSE | 1 |
| 2892 | 2892 | 5.12903226 | 4.47027412 | 1 | 0.55378205 | 1 | 0.63997468 | 3 | 0.09854319 | 0 | 1 | 1 | 0.4016888 | 25 | 0.80006983 | FALSE | 1 | FALSE | 1 | FALSE | 1 | FALSE | 1 | FALSE | 1 | FALSE | 1 |
| 2893 | 2893 | 8.125 | 2.58057649 | 3 | 0.08207756 | 4 | 0.03920804 | 2 | 0.42396974 | 2 | 0.5543522 | 0 | 1 | 29 | 0.98400908 | FALSE | 0.79236883 | FALSE | 0.96864929 | FALSE | 1 | FALSE | 1 | FALSE | 1 | FALSE | 1 |
| 2894 | 2894 | 10.5 | 2.29128785 | 0 | 1 | 0 | 1 | 1 | 0.50143826 | 1 | 0.59104357 | 0 | 1 | 17 | 0.41141543 | FALSE | 1 | FALSE | 1 | FALSE | 1 | FALSE | 1 | FALSE | 1 | FALSE | 1 |
| 2895 | 2895 | 6.74666667 | 1.6255734 | 6 | 0.01216612 | 5 | 0.09467212 | 6 | 0.05188029 | 0 | 1 | 2 | 0.34918837 | 56 | 0.99057646 | FALSE | 0.3424908 | FALSE | 1 | FALSE | 1 | FALSE | 1 | FALSE | 1 | FALSE | 1 |
| 2896 | 2896 | 8.25 | 1.95256242 | 0 | 1 | 1 | 0.40942991 | 1 | 0.44344817 | 1 | 0.52895088 | 0 | 1 | 13 | 0.76799161 | FALSE | 1 | FALSE | 1 | FALSE | 1 | FALSE | 1 | FALSE | 1 | FALSE | 1 |
| 2897 | 2897 | 10 | 2.47487373 | 1 | 0.34033054 | 1 | 0.40942991 | 0 | 1 | 0 | 1 | 1 | 0.23264612 | 13 | 0.76799161 | FALSE | 1 | FALSE | 1 | FALSE | 1 | FALSE | 1 | FALSE | 1 | FALSE | 1 |
| 2898 | 2898 | 8.41935484 | 1.99635464 | 0 | 1 | 1 | 0.63997468 | 1 | 0.67911179 | 3 | 0.16836568 | 0 | 1 | 26 | 0.64660109 | FALSE | 1 | FALSE | 1 | FALSE | 1 | FALSE | 1 | FALSE | 1 | FALSE | 1 |
| 2899 | 2899 | 10.8181818 | 2.16661369 | 0 | 1 | 1 | 0.30368014 | 1 | 0.33149969 | 0 | 1 | 0 | 1 | 9 | 0.75985366 | FALSE | 1 | FALSE | 1 | FALSE | 1 | FALSE | 1 | FALSE | 1 | FALSE | 1 |
| 2900 | 2900 | 7.875 | 1.64094739 | 0 | 1 | 1 | 0.54637326 | 0 | 1 | 2 | 0.30249307 | 0 | 1 | 21 | 0.47053137 | FALSE | 1 | FALSE | 1 | FALSE | 1 | FALSE | 1 | FALSE | 1 | FALSE | 1 |
| 2901 | 2901 | 8.7 | 4.3829214 | 0 | 1 | 0 | 1 | 0 | 1 | 0 | 1 | 0 | 1 | 10 | 0.18273166 | FALSE | 1 | FALSE | 1 | FALSE | 1 | FALSE | 1 | FALSE | 1 | FALSE | 1 |
| 2902 | 2902 | 8.18918919 | 2.50229989 | 1 | 0.6184714 | 1 | 0.70472939 | 0 | 1 | 2 | 0.51200191 | 0 | 1 | 33 | 0.2931558 | FALSE | 1 | FALSE | 1 | FALSE | 1 | FALSE | 1 | FALSE | 1 | FALSE | 1 |
| 2903 | 2903 | 9.83333333 | 2.19215774 | 0 | 1 | 1 | 0.44710673 | 3 | 0.02505979 | 0 | 1 | 2 | 0.03435384 | 12 | 0.98544073 | FALSE | 1 | FALSE | 1 | FALSE | 1 | FALSE | 1 | FALSE | 0.86063198 | FALSE | 1 |
| 2904 | 2904 | 7.57142857 | 3.25973164 | 0 | 1 | 0 | 1 | 1 | 0.53671415 | 0 | 1 | 0 | 1 | 20 | 0.13763971 | FALSE | 1 | FALSE | 1 | FALSE | 1 | FALSE | 1 | FALSE | 1 | FALSE | 1 |
| 2905 | 2905 | 6.89570552 | 0.62281271 | 2 | 0.92411467 | 5 | 0.60643345 | 7 | 0.36371237 | 7 | 0.62030147 | 3 | 0.49886688 | 138 | 0.43918548 | FALSE | 1 | FALSE | 1 | FALSE | 1 | FALSE | 1 | FALSE | 1 | FALSE | 1 |
| 2906 | 2906 | 8.52 | 1.89989473 | 0 | 1 | 0 | 1 | 1 | 0.59998149 | 1 | 0.69181315 | 0 | 1 | 23 | 0.22727183 | FALSE | 1 | FALSE | 1 | FALSE | 1 | FALSE | 1 | FALSE | 1 | FALSE | 1 |
| 2907 | 2907 | 9.42857143 | 1.22775313 | 3 | 0.13009168 | 2 | 0.47396556 | 1 | 0.83455705 | 3 | 0.39227316 | 0 | 1 | 40 | 0.77344956 | FALSE | 0.97964162 | FALSE | 1 | FALSE | 1 | FALSE | 1 | FALSE | 1 | FALSE | 1 |
| 2908 | 2908 | 9.14814815 | 1.73640121 | 0 | 1 | 2 | 0.21684246 | 0 | 1 | 1 | 0.71956339 | 0 | 1 | 24 | 0.3725373 | FALSE | 1 | FALSE | 1 | FALSE | 1 | FALSE | 1 | FALSE | 1 | FALSE | 1 |
| 2909 | 2909 | 8.04347826 | 3.47009912 | 2 | 0.11647716 | 2 | 0.16948764 | 0 | 1 | 0 | 1 | 1 | 0.31672722 | 18 | 0.86273155 | FALSE | 0.93181727 | FALSE | 1 | FALSE | 1 | FALSE | 1 | FALSE | 1 | FALSE | 1 |
| 2910 | 2910 | 7.29166667 | 3.98935215 | 3 | 0.12433927 | 5 | 0.01870362 | 1 | 0.8283465 | 2 | 0.65421368 | 1 | 0.54902428 | 36 | 0.97101084 | FALSE | 0.95622564 | FALSE | 0.77515497 | FALSE | 1 | FALSE | 1 | FALSE | 1 | FALSE | 1 |
| 2911 | 2911 | 9.36842105 | 2.59980822 | 0 | 1 | 3 | 0.02205772 | 0 | 1 | 2 | 0.21610812 | 0 | 1 | 14 | 0.93677253 | FALSE | 1 | FALSE | 0.81029974 | FALSE | 1 | FALSE | 1 | FALSE | 1 | FALSE | 1 |
| 2912 | 2912 | 9.55 | 3.24769149 | 4 | 0.00146558 | 1 | 0.48239024 | 1 | 0.51939841 | 1 | 0.60987146 | 0 | 1 | 13 | 0.99248686 | FALSE | 0.09453559 | FALSE | 1 | FALSE | 1 | FALSE | 1 | FALSE | 1 | FALSE | 1 |

| | | | | | | | | | | | | | | | | | | | | | | | | | | | |
|---|---|---|---|---|---|---|---|---|---|---|---|---|---|---|---|---|---|---|---|---|---|---|---|---|---|---|---|
| 2913 | 2913 | 8.3 | 2.05182845 | 0 | 1 | 0 | 1 | 3 | 0.09131526 | 2 | 0.4036176 | 0 | 1 | 25 | 0.67630471 | FALSE | 1 | FALSE | 1 | FALSE | 1 | FALSE | 1 | FALSE | 1 | FALSE | 1 |
| 2914 | 2914 | 7.36666667 | 2.54929184 | 1 | 0.54198714 | 0 | 1 | 0 | 1 | 1 | 0.75658797 | 1 | 0.39166989 | 27 | 0.28862208 | FALSE | 1 | FALSE | 1 | FALSE | 1 | FALSE | 1 | FALSE | 1 | FALSE | 1 |
| 2915 | 2915 | 9.51351351 | 1.73394756 | 1 | 0.6184714 | 2 | 0.3375163 | 1 | 0.74264075 | 1 | 0.82511639 | 1 | 0.45845922 | 31 | 0.64534533 | FALSE | 1 | FALSE | 1 | FALSE | 1 | FALSE | 1 | FALSE | 1 | FALSE | 1 |
| 2916 | 2916 | 9.1 | 2.3430749 | 0 | 1 | 0 | 1 | 2 | 0.29330414 | 1 | 0.75658797 | 0 | 1 | 27 | 0.28862208 | FALSE | 1 | FALSE | 1 | FALSE | 1 | FALSE | 1 | FALSE | 1 | FALSE | 1 |
| 2917 | 2917 | 10.0909091 | 2.10862064 | 1 | 0.24866461 | 0 | 1 | 0 | 1 | 0 | 1 | 0 | 1 | 10 | 0.46859937 | FALSE | 1 | FALSE | 1 | FALSE | 1 | FALSE | 1 | FALSE | 1 | FALSE | 1 |
| 2918 | 2918 | 9.82608696 | 2.40745534 | 0 | 1 | 1 | 0.5154318 | 0 | 1 | 0 | 1 | 1 | 0.30529951 | 20 | 0.30915202 | FALSE | 1 | FALSE | 1 | FALSE | 1 | FALSE | 1 | FALSE | 1 | FALSE | 1 |
| 2919 | 2919 | 8.90909091 | 3.31537864 | 0 | 1 | 0 | 1 | 0 | 1 | 1 | 0.78873814 | 0 | 1 | 32 | 0.02590314 | FALSE | 1 | FALSE | 1 | FALSE | 1 | FALSE | 1 | FALSE | 1 | FALSE | 1 |
| 2920 | 2920 | 8.95454546 | 3.36394348 | 0 | 1 | 0 | 1 | 0 | 1 | 0 | 1 | 0 | 1 | 22 | 0.02368118 | FALSE | 1 | FALSE | 1 | FALSE | 1 | FALSE | 1 | FALSE | 1 | FALSE | 1 |
| 2921 | 2921 | 10.6153846 | 3.28255205 | 1 | 0.49162634 | 3 | 0.05021553 | 2 | 0.23940226 | 0 | 1 | 0 | 1 | 20 | 0.90142689 | FALSE | 1 | FALSE | 1 | FALSE | 1 | FALSE | 1 | FALSE | 1 | FALSE | 1 |
| 2922 | 2922 | 8.66666667 | 1.49071199 | 1 | 0.63789042 | 1 | 0.7236252 | 1 | 0.76089851 | 1 | 0.84089119 | 0 | 1 | 35 | 0.24915769 | FALSE | 1 | FALSE | 1 | FALSE | 1 | FALSE | 1 | FALSE | 1 | FALSE | 1 |
| 2923 | 2923 | 9.14285714 | 1.50509339 | 1 | 0.3050805 | 1 | 0.36919824 | 0 | 1 | 0 | 1 | 0 | 1 | 12 | 0.62236227 | FALSE | 1 | FALSE | 1 | FALSE | 1 | FALSE | 1 | FALSE | 1 | FALSE | 1 |
| 2924 | 2924 | 8.25 | 1.86748494 | 0 | 1 | 1 | 0.48239024 | 0 | 1 | 0 | 1 | 1 | 0.28187291 | 18 | 0.37504645 | FALSE | 1 | FALSE | 1 | FALSE | 1 | FALSE | 1 | FALSE | 1 | FALSE | 1 |
| 2925 | 2925 | 10.5416667 | 1.87036464 | 0 | 1 | 1 | 0.54637326 | 2 | 0.21260257 | 1 | 0.67692816 | 0 | 1 | 20 | 0.68270729 | FALSE | 1 | FALSE | 1 | FALSE | 1 | FALSE | 1 | FALSE | 1 | FALSE | 1 |
| 2926 | 2926 | 7.008 | 2.01790386 | 6 | 0.0873213 | 2 | 0.90510937 | 1 | 0.98808666 | 4 | 0.80846044 | 4 | 0.13384634 | 103 | 0.38590301 | FALSE | 0.81752699 | FALSE | 1 | FALSE | 1 | FALSE | 1 | FALSE | 1 | FALSE | 1 |
| 2927 | 2927 | 10.5333333 | 3.13829395 | 1 | 0.32293355 | 0 | 1 | 2 | 0.09928574 | 0 | 1 | 0 | 1 | 12 | 0.80329917 | FALSE | 1 | FALSE | 1 | FALSE | 1 | FALSE | 1 | FALSE | 1 | FALSE | 1 |
| 2928 | 2928 | 9.5625 | 3.62230503 | 2 | 0.19750261 | 0 | 1 | 2 | 0.32011871 | 4 | 0.05697506 | 2 | 0.096172 | 22 | 0.99301316 | FALSE | 1 | FALSE | 1 | FALSE | 1 | FALSE | 1 | FALSE | 1 | FALSE | 1 |
| 2929 | 2929 | 1.39295775 | 1.9023206 | 35 | 9.40E-05 | 29 | 0.10943265 | 28 | 0.32754766 | 26 | 0.91417035 | 17 | 0.06978185 | 575 | 0.99576742 | TRUE | 0.01750963 | FALSE | 1 | FALSE | 1 | FALSE | 1 | FALSE | 1 | FALSE | 1 |
| 2930 | 2930 | 7.18421053 | 1.61172126 | 2 | 0.58475401 | 7 | 0.01108961 | 6 | 0.05471327 | 8 | 0.02253952 | 2 | 0.35515917 | 51 | 0.99995398 | FALSE | 1 | FALSE | 0.67617265 | FALSE | 1 | FALSE | 1 | FALSE | 1 | FALSE | 1 |
| 2931 | 2931 | 9.36363636 | 2.68912717 | 0 | 1 | 2 | 0.15794013 | 0 | 1 | 1 | 0.64497439 | 2 | 0.04972408 | 17 | 0.8840357 | FALSE | 1 | FALSE | 1 | FALSE | 1 | FALSE | 1 | FALSE | 0.98629461 | FALSE | 1 |
| 2932 | 2932 | 6.92156863 | 3.65600842 | 1 | 0.7354105 | 3 | 0.22759346 | 2 | 0.55199489 | 1 | 0.90982214 | 2 | 0.20351566 | 42 | 0.73296146 | FALSE | 1 | FALSE | 1 | FALSE | 1 | FALSE | 1 | FALSE | 1 | FALSE | 1 |
| 2933 | 2933 | 10.4848485 | 2.51204628 | 2 | 0.20692137 | 2 | 0.28941836 | 0 | 1 | 0 | 1 | 3 | 0.01637249 | 26 | 0.86852567 | FALSE | 1 | FALSE | 1 | FALSE | 1 | FALSE | 1 | FALSE | 0.66728076 | FALSE | 1 |
| 2934 | 2934 | 11 | 3.82099464 | 0 | 1 | 0 | 1 | 1 | 0.30655256 | 1 | 0.37517262 | 1 | 0.15247062 | 7 | 0.94326215 | FALSE | 1 | FALSE | 1 | FALSE | 1 | FALSE | 1 | FALSE | 1 | FALSE | 1 |
| 2935 | 2935 | 8.40909091 | 2.82294319 | 0 | 1 | 0 | 1 | 0 | 1 | 1 | 0.64497439 | 0 | 1 | 21 | 0.12048848 | FALSE | 1 | FALSE | 1 | FALSE | 1 | FALSE | 1 | FALSE | 1 | FALSE | 1 |
| 2936 | 2936 | 7.35714286 | 2.81123838 | 3 | 0.46200485 | 6 | 0.09708054 | 4 | 0.47057639 | 6 | 0.29412863 | 5 | 0.02201494 | 74 | 0.99265016 | FALSE | 1 | FALSE | 1 | FALSE | 1 | FALSE | 1 | FALSE | 0.74991845 | FALSE | 1 |
| 2937 | 2937 | 9.87179487 | 2.5437655 | 0 | 1 | 2 | 0.36122049 | 1 | 0.76089851 | 0 | 1 | 1 | 0.4761699 | 35 | 0.24915769 | FALSE | 1 | FALSE | 1 | FALSE | 1 | FALSE | 1 | FALSE | 1 | FALSE | 1 |
| 2938 | 2938 | 5.28813559 | 1.41781459 | 10 | 0.08102084 | 5 | 0.8857741 | 10 | 0.34150534 | 6 | 0.96425576 | 8 | 0.03940625 | 197 | 0.68997151 | FALSE | 0.79236883 | FALSE | 1 | FALSE | 1 | FALSE | 1 | FALSE | 0.89175691 | FALSE | 1 |
| 2939 | 2939 | 8.6 | 1.356466 | 0 | 1 | 0 | 1 | 0 | 1 | 0 | 1 | 0 | 1 | 5 | 0.42761893 | FALSE | 1 | FALSE | 1 | FALSE | 1 | FALSE | 1 | FALSE | 1 | FALSE | 1 |
| 2940 | 2940 | 7.72340426 | 1.59389964 | 1 | 0.70621968 | 0 | 1 | 5 | 0.02577785 | 2 | 0.64277396 | 0 | 1 | 39 | 0.69224623 | FALSE | 1 | FALSE | 1 | FALSE | 1 | FALSE | 1 | FALSE | 1 | FALSE | 1 |
| 2941 | 2941 | 10 | 2.12132034 | 0 | 1 | 1 | 0.23139161 | 1 | 0.25384309 | 0 | 1 | 0 | 1 | 6 | 0.88425551 | FALSE | 1 | FALSE | 1 | FALSE | 1 | FALSE | 1 | FALSE | 1 | FALSE | 1 |
| 2942 | 2942 | 8.5 | 2.5 | 2 | 0.17884234 | 1 | 0.6278847 | 0 | 1 | 1 | 0.75658797 | 1 | 0.39166989 | 25 | 0.67630471 | FALSE | 1 | FALSE | 1 | FALSE | 1 | FALSE | 1 | FALSE | 1 | FALSE | 1 |
| 2943 | 2943 | 9.18918919 | 2.2036559 | 1 | 0.60837712 | 0 | 1 | 2 | 0.37289966 | 4 | 0.08109727 | 1 | 0.44938266 | 28 | 0.90259714 | FALSE | 1 | FALSE | 1 | FALSE | 1 | FALSE | 1 | FALSE | 1 | FALSE | 1 |
| 2944 | 2944 | 6.09193246 | 0.68554536 | 18 | 0.13429715 | 28 | 0.00653714 | 28 | 0.02473592 | 16 | 0.97775075 | 13 | 0.0932102 | 428 | 0.99380256 | FALSE | 0.98316554 | FALSE | 0.60103504 | FALSE | 1 | FALSE | 1 | FALSE | 1 | FALSE | 1 |
| 2945 | 2945 | 8.03773585 | 4.00924528 | 1 | 0.74890551 | 5 | 0.02755409 | 1 | 0.85724013 | 2 | 0.70707241 | 2 | 0.21556658 | 42 | 0.88654191 | FALSE | 1 | FALSE | 0.87929047 | FALSE | 1 | FALSE | 1 | FALSE | 1 | FALSE | 1 |
| 2946 | 2946 | 4.7184466 | 1.9228694 | 2 | 0.7467337 | 4 | 0.42831767 | 6 | 0.16497789 | 7 | 0.19371537 | 1 | 0.82020918 | 83 | 0.88406659 | FALSE | 1 | FALSE | 1 | FALSE | 1 | FALSE | 1 | FALSE | 1 | FALSE | 1 |
| 2947 | 2947 | 7.65853659 | 2.47533941 | 2 | 0.28329504 | 2 | 0.38460545 | 1 | 0.77786601 | 1 | 0.85524728 | 0 | 1 | 35 | 0.53616095 | FALSE | 1 | FALSE | 1 | FALSE | 1 | FALSE | 1 | FALSE | 1 | FALSE | 1 |
| 2948 | 2948 | 6.55769231 | 3.50501543 | 2 | 0.38681045 | 4 | 0.08651261 | 1 | 0.85187773 | 3 | 0.4296776 | 0 | 1 | 42 | 0.82137318 | FALSE | 1 | FALSE | 1 | FALSE | 1 | FALSE | 1 | FALSE | 1 | FALSE | 1 |
| 2949 | 2949 | 6.86842105 | 3.96146605 | 0 | 1 | 0 | 1 | 1 | 0.75193685 | 2 | 0.52640931 | 1 | 0.4673875 | 34 | 0.27051027 | FALSE | 1 | FALSE | 1 | FALSE | 1 | FALSE | 1 | FALSE | 1 | FALSE | 1 |
| 2950 | 2950 | 9.61538462 | 2.84303363 | 0 | 1 | 0 | 1 | 0 | 1 | 2 | 0.1174583 | 0 | 1 | 11 | 0.6683056 | FALSE | 1 | FALSE | 1 | FALSE | 1 | FALSE | 1 | FALSE | 1 | FALSE | 1 |
| 2951 | 2951 | 7.47619048 | 4.66618073 | 1 | 0.42086991 | 0 | 1 | 0 | 1 | 2 | 0.25059627 | 0 | 1 | 18 | 0.57962763 | FALSE | 1 | FALSE | 1 | FALSE | 1 | FALSE | 1 | FALSE | 1 | FALSE | 1 |
| 2952 | 2952 | 8.66666667 | 1.91070501 | 3 | 0.01568237 | 2 | 0.14655134 | 0 | 1 | 2 | 0.25059627 | 1 | 0.29368245 | 13 | 0.9973503 | FALSE | 0.38695764 | FALSE | 1 | FALSE | 1 | FALSE | 1 | FALSE | 1 | FALSE | 1 |
| 2953 | 2953 | 8.29166667 | 3.27209371 | 1 | 0.45027683 | 1 | 0.53115654 | 2 | 0.19931429 | 2 | 0.2852236 | 1 | 0.31672722 | 16 | 0.98100709 | FALSE | 1 | FALSE | 1 | FALSE | 1 | FALSE | 1 | FALSE | 1 | FALSE | 1 |
| 2954 | 2954 | 6.44736842 | 2.33618412 | 4 | 0.13018824 | 2 | 0.71077581 | 0 | 1 | 3 | 0.68517679 | 0 | 1 | 67 | 0.23030366 | FALSE | 0.97964162 | FALSE | 1 | FALSE | 1 | FALSE | 1 | FALSE | 1 | FALSE | 1 |
| 2955 | 2955 | 9.46153846 | 3.38811008 | 0 | 1 | 2 | 0.20486827 | 0 | 1 | 1 | 0.70601446 | 1 | 0.3499045 | 22 | 0.61628 | FALSE | 1 | FALSE | 1 | FALSE | 1 | FALSE | 1 | FALSE | 1 | FALSE | 1 |
| 2956 | 2956 | 10.0625 | 2.9889118 | 0 | 1 | 0 | 1 | 0 | 1 | 1 | 0.52895088 | 0 | 1 | 15 | 0.26127859 | FALSE | 1 | FALSE | 1 | FALSE | 1 | FALSE | 1 | FALSE | 1 | FALSE | 1 |
| 2957 | 2957 | 10 | 2.71569512 | 0 | 1 | 1 | 0.40942991 | 0 | 1 | 0 | 1 | 0 | 1 | 15 | 0.26127859 | FALSE | 1 | FALSE | 1 | FALSE | 1 | FALSE | 1 | FALSE | 1 | FALSE | 1 |
| 2958 | 2958 | 9.83333333 | 3.14907394 | 0 | 1 | 1 | 0.44710673 | 0 | 1 | 1 | 0.57131013 | 0 | 1 | 16 | 0.44998001 | FALSE | 1 | FALSE | 1 | FALSE | 1 | FALSE | 1 | FALSE | 1 | FALSE | 1 |
| 2959 | 2959 | 9.12 | 2.0458739 | 1 | 0.47819878 | 2 | 0.19297191 | 0 | 1 | 2 | 0.31968852 | 1 | 0.33902672 | 19 | 0.91734261 | FALSE | 1 | FALSE | 1 | FALSE | 1 | FALSE | 1 | FALSE | 1 | FALSE | 1 |
| 2960 | 2960 | 8.16129032 | 1.77975667 | 0 | 1 | 2 | 0.2651851 | 2 | 0.30673593 | 2 | 0.41984187 | 1 | 0.4016888 | 24 | 0.90138666 | FALSE | 1 | FALSE | 1 | FALSE | 1 | FALSE | 1 | FALSE | 1 | FALSE | 1 |
| 2961 | 2961 | 11.5517241 | 4.11545621 | 3 | 0.03713651 | 2 | 0.24095584 | 1 | 0.65463982 | 1 | 0.74482229 | 0 | 1 | 22 | 0.92902102 | FALSE | 0.58392 | FALSE | 1 | FALSE | 1 | FALSE | 1 | FALSE | 1 | FALSE | 1 |
| 2962 | 2962 | 10.2608696 | 3.21974062 | 5 | 0.00024166 | 1 | 0.53115654 | 1 | 0.5695037 | 3 | 0.08614834 | 1 | 0.31672722 | 12 | 0.9999499 | TRUE | 0.03045702 | FALSE | 1 | FALSE | 1 | FALSE | 1 | FALSE | 1 | FALSE | 1 |
| 2963 | 2963 | 3.75714286 | 3.65839822 | 2 | 0.53205439 | 5 | 0.07172628 | 5 | 0.1012281 | 0 | 1 | 0 | 1 | 57 | 0.72531896 | FALSE | 1 | FALSE | 1 | FALSE | 1 | FALSE | 1 | FALSE | 1 | FALSE | 1 |
| 2964 | 2964 | 9.76923077 | 1.84615385 | 0 | 1 | 0 | 1 | 0 | 1 | 0 | 1 | 0 | 1 | 13 | 0.10967906 | FALSE | 1 | FALSE | 1 | FALSE | 1 | FALSE | 1 | FALSE | 1 | FALSE | 1 |
| 2965 | 2965 | 5.43636364 | 2.53530444 | 1 | 0.76171602 | 3 | 0.26245929 | 5 | 0.04651472 | 2 | 0.72625531 | 2 | 0.22768257 | 42 | 0.96038295 | FALSE | 1 | FALSE | 1 | FALSE | 1 | FALSE | 1 | FALSE | 1 | FALSE | 1 |
| 2966 | 2966 | 4.05063291 | 2.50012017 | 3 | 0.32318196 | 4 | 0.24495326 | 4 | 0.30733368 | 6 | 0.14745982 | 3 | 0.13569451 | 58 | 0.99295041 | FALSE | 1 | FALSE | 1 | FALSE | 1 | FALSE | 1 | FALSE | 1 | FALSE | 1 |
| 2967 | 2967 | 9.8 | 3.18747549 | 1 | 0.47819878 | 2 | 0.19297191 | 0 | 1 | 2 | 0.31968852 | 1 | 0.33902672 | 19 | 0.91734261 | FALSE | 1 | FALSE | 1 | FALSE | 1 | FALSE | 1 | FALSE | 1 | FALSE | 1 |
| 2968 | 2968 | 7.57142857 | 2.71804251 | 0 | 1 | 0 | 1 | 0 | 1 | 0 | 1 | 0 | 1 | 21 | 0.02808151 | FALSE | 1 | FALSE | 1 | FALSE | 1 | FALSE | 1 | FALSE | 1 | FALSE | 1 |
| 2969 | 2969 | 7.7 | 2.18174242 | 1 | 0.64722818 | 1 | 0.7326167 | 0 | 1 | 2 | 0.5543522 | 0 | 1 | 36 | 0.22908535 | FALSE | 1 | FALSE | 1 | FALSE | 1 | FALSE | 1 | FALSE | 1 | FALSE | 1 |
| 2970 | 2970 | 9.625 | 2.59506744 | 1 | 0.46441868 | 1 | 0.54637326 | 2 | 0.21260257 | 0 | 1 | 0 | 1 | 20 | 0.68270729 | FALSE | 1 | FALSE | 1 | FALSE | 1 | FALSE | 1 | FALSE | 1 | FALSE | 1 |
| 2971 | 2971 | 8.53846154 | 1.98515199 | 1 | 0.28675951 | 0 | 1 | 0 | 1 | 1 | 0.45746571 | 0 | 1 | 11 | 0.6683056 | FALSE | 1 | FALSE | 1 | FALSE | 1 | FALSE | 1 | FALSE | 1 | FALSE | 1 |
| 2972 | 2972 | 8.12121212 | 3.30066638 | 1 | 0.57647348 | 0 | 1 | 0 | 1 | 2 | 0.45159487 | 0 | 1 | 30 | 0.21942357 | FALSE | 1 | FALSE | 1 | FALSE | 1 | FALSE | 1 | FALSE | 1 | FALSE | 1 |
| 2973 | 2973 | 8.52 | 3.88967865 | 0 | 1 | 1 | 0.54637326 | 2 | 0.21260257 | 3 | 0.09525947 | 0 | 1 | 18 | 0.93166998 | FALSE | 1 | FALSE | 1 | FALSE | 1 | FALSE | 1 | FALSE | 1 | FALSE | 1 |
| 2974 | 2974 | 6.5625 | 2.97143519 | 0 | 1 | 1 | 0.74131698 | 1 | 0.77786601 | 1 | 0.85524728 | 1 | 0.49330646 | 37 | 0.21027077 | FALSE | 1 | FALSE | 1 | FALSE | 1 | FALSE | 1 | FALSE | 1 | FALSE | 1 |
| 2975 | 2975 | 4.9875 | 2.81732919 | 3 | 0.32318196 | 1 | 0.92420038 | 3 | 0.53558651 | 4 | 0.48430022 | 1 | 0.72664912 | 66 | 0.55329149 | FALSE | 1 | FALSE | 1 | FALSE | 1 | FALSE | 1 | FALSE | 1 | FALSE | 1 |
| 2976 | 2976 | 9.6 | 1.356466 | 1 | 0.40558521 | 2 | 0.1353446 | 1 | 0.51939841 | 1 | 0.60987146 | 0 | 1 | 15 | 0.92113679 | FALSE | 1 | FALSE | 1 | FALSE | 1 | FALSE | 1 | FALSE | 1 | FALSE | 1 |
| 2977 | 2977 | 9.52631579 | 2.39227751 | 0 | 1 | 0 | 1 | 0 | 1 | 1 | 0.57131013 | 0 | 1 | 17 | 0.20329205 | FALSE | 1 | FALSE | 1 | FALSE | 1 | FALSE | 1 | FALSE | 1 | FALSE | 1 |
| 2978 | 2978 | 9.5 | 2.2654697 | 0 | 1 | 0 | 1 | 0 | 1 | 4 | 0.06848449 | 0 | 1 | 30 | 0.36879488 | FALSE | 1 | FALSE | 1 | FALSE | 1 | FALSE | 1 | FALSE | 1 | FALSE | 1 |
| 2979 | 2979 | 8.23809524 | 1.82325616 | 0 | 1 | 0 | 1 | 1 | 0.53671415 | 1 | 0.62783522 | 0 | 1 | 19 | 0.34094959 | FALSE | 1 | FALSE | 1 | FALSE | 1 | FALSE | 1 | FALSE | 1 | FALSE | 1 |
| 2980 | 2980 | 10.5 | 3.39116499 | 0 | 1 | 0 | 1 | 1 | 0.25384309 | 0 | 1 | 0 | 1 | 7 | 0.63752397 | FALSE | 1 | FALSE | 1 | FALSE | 1 | FALSE | 1 | FALSE | 1 | FALSE | 1 |
| 2981 | 2981 | 10 | 2.15058132 | 0 | 1 | 0 | 1 | 0 | 1 | 2 | 0.16544789 | 0 | 1 | 14 | 0.53308261 | FALSE | 1 | FALSE | 1 | FALSE | 1 | FALSE | 1 | FALSE | 1 | FALSE | 1 |
| 2982 | 2982 | 9.97222222 | 3.73784037 | 1 | 0.59801736 | 2 | 0.31355738 | 2 | 0.35984046 | 4 | 0.07465533 | 0 | 1 | 26 | 0.96294373 | FALSE | 1 | FALSE | 1 | FALSE | 1 | FALSE | 1 | FALSE | 1 | FALSE | 1 |
| 2983 | 2983 | 9 | 2.47338777 | 0 | 1 | 1 | 0.42857733 | 1 | 0.46348836 | 0 | 1 | 1 | 0.24525764 | 14 | 0.7313145 | FALSE | 1 | FALSE | 1 | FALSE | 1 | FALSE | 1 | FALSE | 1 | FALSE | 1 |
| 2984 | 2984 | 8.36842105 | 2.4540091 | 0 | 1 | 0 | 1 | 0 | 1 | 0 | 1 | 1 | 0.26986775 | 18 | 0.17880694 | FALSE | 1 | FALSE | 1 | FALSE | 1 | FALSE | 1 | FALSE | 1 | FALSE | 1 |
| 2985 | 2985 | 8.375 | 3.39806931 | 0 | 1 | 1 | 0.65167362 | 0 | 1 | 4 | 0.05697506 | 2 | 0.096172 | 25 | 0.88559288 | FALSE | 1 | FALSE | 1 | FALSE | 1 | FALSE | 1 | FALSE | 1 | FALSE | 1 |
| 2986 | 2986 | 9.71428571 | 3.53409054 | 0 | 1 | 0 | 1 | 1 | 0.22601089 | 1 | 0.28044368 | 0 | 1 | 5 | 0.91847773 | FALSE | 1 | FALSE | 1 | FALSE | 1 | FALSE | 1 | FALSE | 1 | FALSE | 1 |
| 2987 | 2987 | 8.13333333 | 3.28362943 | 2 | 0.05513784 | 1 | 0.38964394 | 0 | 1 | 2 | 0.14906763 | 0 | 1 | 10 | 0.97980046 | FALSE | 0.68849059 | FALSE | 1 | FALSE | 1 | FALSE | 1 | FALSE | 1 | FALSE | 1 |
| 2988 | 2988 | 9 | 1.69967317 | 0 | 1 | 0 | 1 | 0 | 1 | 1 | 0.34506517 | 0 | 1 | 8 | 0.5780332 | FALSE | 1 | FALSE | 1 | FALSE | 1 | FALSE | 1 | FALSE | 1 | FALSE | 1 |
| 2989 | 2989 | 6.22093023 | 3.38396067 | 4 | 0.17847815 | 2 | 0.77275903 | 5 | 0.1959611 | 2 | 0.91113247 | 8 | 7.43E-05 | 65 | 0.98889609 | FALSE | 1 | FALSE | 1 | FALSE | 1 | FALSE | 1 | TRUE | 0.03578309 | FALSE | 1 |
| 2990 | 2990 | 9.31818182 | 1.76805912 | 0 | 1 | 0 | 1 | 1 | 0.55340851 | 0 | 1 | 0 | 1 | 21 | 0.12048848 | FALSE | 1 | FALSE | 1 | FALSE | 1 | FALSE | 1 | FALSE | 1 | FALSE | 1 |
| 2991 | 2991 | 8.27777778 | 3.15885977 | 0 | 1 | 1 | 0.44710673 | 0 | 1 | 0 | 1 | 0 | 1 | 17 | 0.20329205 | FALSE | 1 | FALSE | 1 | FALSE | 1 | FALSE | 1 | FALSE | 1 | FALSE | 1 |
| 2992 | 2992 | 9 | 2.25831796 | 1 | 0.40558521 | 0 | 1 | 0 | 1 | 1 | 0.60987146 | 1 | 0.28187291 | 17 | 0.61750632 | FALSE | 1 | FALSE | 1 | FALSE | 1 | FALSE | 1 | FALSE | 1 | FALSE | 1 |
| 2993 | 2993 | 9.29166667 | 1.24093938 | 0 | 1 | 3 | 0.04096158 | 1 | 0.58502114 | 1 | 0.67692816 | 2 | 0.058175 | 17 | 0.97530702 | FALSE | 1 | FALSE | 0.9698186 | FALSE | 1 | FALSE | 1 | FALSE | 1 | FALSE | 1 |
| 2994 | 2994 | 10.1538462 | 3.18279398 | 0 | 1 | 1 | 0.3480709 | 0 | 1 | 0 | 1 | 1 | 0.19354507 | 11 | 0.6683056 | FALSE | 1 | FALSE | 1 | FALSE | 1 | FALSE | 1 | FALSE | 1 | FALSE | 1 |
| 2995 | 2995 | 9.16666667 | 2.47767813 | 0 | 1 | 0 | 1 | 2 | 0.21260257 | 0 | 1 | 0 | 1 | 22 | 0.25237063 | FALSE | 1 | FALSE | 1 | FALSE | 1 | FALSE | 1 | FALSE | 1 | FALSE | 1 |
| 2996 | 2996 | 10.5263158 | 1.98330986 | 2 | 0.08405377 | 2 | 0.12434464 | 2 | 0.14749808 | 0 | 1 | 0 | 1 | 13 | 0.97998676 | FALSE | 0.79236883 | FALSE | 1 | FALSE | 1 | FALSE | 1 | FALSE | 1 | FALSE | 1 |
| 2997 | 2997 | 7.32432432 | 2.82894359 | 0 | 1 | 1 | 0.69480472 | 0 | 1 | 2 | 0.49731009 | 1 | 0.44938266 | 32 | 0.31709555 | FALSE | 1 | FALSE | 1 | FALSE | 1 | FALSE | 1 | FALSE | 1 | FALSE | 1 |
| 2998 | 2998 | 9.47826087 | 2.53445072 | 1 | 0.45027683 | 1 | 0.53115654 | 1 | 0.5695037 | 1 | 0.6613267 | 2 | 0.05389003 | 17 | 0.94439745 | FALSE | 1 | FALSE | 1 | FALSE | 1 | FALSE | 1 | FALSE | 1 | FALSE | 1 |
| 2999 | 2999 | 8 | 1.58943883 | 0 | 1 | 0 | 1 | 0 | 1 | 0 | 1 | 0 | 1 | 38 | 0.00154342 | FALSE | 1 | FALSE | 1 | FALSE | 1 | FALSE | 1 | FALSE | 1 | FALSE | 1 |
| 3000 | 3000 | 12.4285714 | 2.0946969 | 0 | 1 | 0 | 1 | 1 | 0.40110418 | 1 | 0.48242099 | 0 | 1 | 12 | 0.62236227 | FALSE | 1 | FALSE | 1 | FALSE | 1 | FALSE | 1 | FALSE | 1 | FALSE | 1 |
| 3001 | 3001 | 8.41176471 | 2.47478635 | 0 | 1 | 1 | 0.6739483 | 3 | 0.12152681 | 1 | 0.7984848 | 0 | 1 | 29 | 0.55707911 | FALSE | 1 | FALSE | 1 | FALSE | 1 | FALSE | 1 | FALSE | 1 | FALSE | 1 |
| 3002 | 3002 | 8.68421053 | 2.53507254 | 0 | 1 | 1 | 0.46503795 | 1 | 0.50143826 | 2 | 0.21610812 | 2 | 0.03799168 | 13 | 0.97998676 | FALSE | 1 | FALSE | 1 | FALSE | 1 | FALSE | 1 | FALSE | 0.87669812 | FALSE | 1 |
| 3003 | 3003 | 6.64285714 | 1.92008716 | 2 | 0.539854 | 3 | 0.39589822 | 6 | 0.0390825 | 5 | 0.21764111 | 0 | 1 | 54 | 0.9618177 | FALSE | 1 | FALSE | 1 | FALSE | 1 | FALSE | 1 | FALSE | 1 | FALSE | 1 |
| 3004 | 3004 | 8.2 | 3.6 | 1 | 0.40558521 | 0 | 1 | 0 | 1 | 1 | 0.60987146 | 0 | 1 | 18 | 0.37504645 | FALSE | 1 | FALSE | 1 | FALSE | 1 | FALSE | 1 | FALSE | 1 | FALSE | 1 |
| 3005 | 3005 | 7.90909091 | 1.79837699 | 0 | 1 | 1 | 0.65167362 | 1 | 0.69069222 | 2 | 0.43583874 | 0 | 1 | 28 | 0.42532177 | FALSE | 1 | FALSE | 1 | FALSE | 1 | FALSE | 1 | FALSE | 1 | FALSE | 1 |
| 3006 | 3006 | 8.33333333 | 2.4267033 | 1 | 0.26795838 | 0 | 1 | 0 | 1 | 0 | 1 | 0 | 1 | 11 | 0.41944532 | FALSE | 1 | FALSE | 1 | FALSE | 1 | FALSE | 1 | FALSE | 1 | FALSE | 1 |

| | | | | | | | | | | | | | | | | | | | | | | | | | | | |
|---|---|---|---|---|---|---|---|---|---|---|---|---|---|---|---|---|---|---|---|---|---|---|---|---|---|---|---|
| 3007 | 3007 | 9.16666667 | 2.88771344 | 2 | 0.17884234 | 2 | 0.25306331 | 3 | 0.09131526 | 1 | 0.75658797 | 0 | 1 | 22 | 0.96518743 | FALSE | 1 | FALSE | 1 | FALSE | 1 | FALSE | 1 | FALSE | 1 | FALSE | 1 |
| 3008 | 3008 | 9.34615385 | 2.35245057 | 0 | 1 | 1 | 0.56109832 | 0 | 1 | 0 | 1 | 0 | 1 | 24 | 0.08019798 | FALSE | 1 | FALSE | 1 | FALSE | 1 | FALSE | 1 | FALSE | 1 | FALSE | 1 |
| 3009 | 3009 | 7.29166667 | 2.82074409 | 2 | 0.12501436 | 2 | 0.18117183 | 0 | 1 | 0 | 1 | 1 | 0.32796862 | 19 | 0.83975208 | FALSE | 0.95622564 | FALSE | 1 | FALSE | 1 | FALSE | 1 | FALSE | 1 | FALSE | 1 |
| 3010 | 3010 | 9.26666667 | 2.97694848 | 1 | 0.32293355 | 1 | 0.38964394 | 1 | 0.42266264 | 1 | 0.50623194 | 1 | 0.2198258 | 10 | 0.97980046 | FALSE | 1 | FALSE | 1 | FALSE | 1 | FALSE | 1 | FALSE | 1 | FALSE | 1 |
| 3011 | 3011 | 8.95 | 1.9185498 | 0 | 1 | 2 | 0.58316597 | 3 | 0.36617062 | 4 | 0.29632241 | 0 | 1 | 51 | 0.5343051 | FALSE | 1 | FALSE | 1 | FALSE | 1 | FALSE | 1 | FALSE | 1 | FALSE | 1 |
| 3012 | 3012 | 8.66666667 | 1.81811869 | 0 | 1 | 1 | 0.54637326 | 2 | 0.21260257 | 2 | 0.30249307 | 1 | 0.32796862 | 18 | 0.93166998 | FALSE | 1 | FALSE | 1 | FALSE | 1 | FALSE | 1 | FALSE | 1 | FALSE | 1 |
| 3013 | 3013 | 11.0625 | 2.04538352 | 0 | 1 | 0 | 1 | 0 | 1 | 1 | 0.52895088 | 0 | 1 | 15 | 0.26127859 | FALSE | 1 | FALSE | 1 | FALSE | 1 | FALSE | 1 | FALSE | 1 | FALSE | 1 |
| 3014 | 3014 | 7.55555556 | 2.72618759 | 1 | 0.50471032 | 1 | 0.58913618 | 1 | 0.62830986 | 0 | 1 | 2 | 0.07169515 | 22 | 0.76237877 | FALSE | 1 | FALSE | 1 | FALSE | 1 | FALSE | 1 | FALSE | 1 | FALSE | 1 |
| 3015 | 3015 | 8.92857143 | 2.78937488 | 1 | 0.3050805 | 0 | 1 | 0 | 1 | 4 | 0.0030237 | 0 | 1 | 9 | 0.98607548 | FALSE | 1 | FALSE | 1 | FALSE | 1 | FALSE | 1 | FALSE | 1 | FALSE | 1 |
| 3016 | 3016 | 8.18918919 | 2.75914046 | 1 | 0.6184714 | 0 | 1 | 3 | 0.14625315 | 2 | 0.51200191 | 1 | 0.45845922 | 30 | 0.78878581 | FALSE | 1 | FALSE | 1 | FALSE | 1 | FALSE | 1 | FALSE | 1 | FALSE | 1 |
| 3017 | 3017 | 10.125 | 1.92164209 | 3 | 0.01782284 | 2 | 0.15794013 | 0 | 1 | 1 | 0.64497439 | 0 | 1 | 16 | 0.95553951 | FALSE | 0.41596787 | FALSE | 1 | FALSE | 1 | FALSE | 1 | FALSE | 1 | FALSE | 1 |
| 3018 | 3018 | 5.91044776 | 2.79515222 | 0 | 1 | 0 | 1 | 3 | 0.43448639 | 4 | 0.3701568 | 1 | 0.6714808 | 59 | 0.26021394 | FALSE | 1 | FALSE | 1 | FALSE | 1 | FALSE | 1 | FALSE | 1 | FALSE | 1 |
| 3019 | 3019 | 8.12903226 | 4.22543707 | 0 | 1 | 1 | 0.63997468 | 2 | 0.30673593 | 2 | 0.41984187 | 0 | 1 | 26 | 0.64660109 | FALSE | 1 | FALSE | 1 | FALSE | 1 | FALSE | 1 | FALSE | 1 | FALSE | 1 |
| 3020 | 3020 | 10.1818182 | 3.90380192 | 0 | 1 | 0 | 1 | 1 | 0.33149969 | 1 | 0.40390032 | 0 | 1 | 9 | 0.75985366 | FALSE | 1 | FALSE | 1 | FALSE | 1 | FALSE | 1 | FALSE | 1 | FALSE | 1 |
| 3021 | 3021 | 12.2222222 | 1.87247773 | 1 | 0.20854748 | 1 | 0.25628163 | 1 | 0.28067846 | 0 | 1 | 0 | 1 | 6 | 0.96128317 | FALSE | 1 | FALSE | 1 | FALSE | 1 | FALSE | 1 | FALSE | 1 | FALSE | 1 |
| 3022 | 3022 | 4.83333333 | 3.72677996 | 1 | 0.84741747 | 3 | 0.41343422 | 5 | 0.11589609 | 4 | 0.42273814 | 0 | 1 | 59 | 0.77408425 | FALSE | 1 | FALSE | 1 | FALSE | 1 | FALSE | 1 | FALSE | 1 | FALSE | 1 |
| 3023 | 3023 | 9.66666667 | 2.65622958 | 1 | 0.46441868 | 2 | 0.18117183 | 1 | 0.58502114 | 1 | 0.67692816 | 0 | 1 | 19 | 0.83975208 | FALSE | 1 | FALSE | 1 | FALSE | 1 | FALSE | 1 | FALSE | 1 | FALSE | 1 |
| 3024 | 3024 | 9.5 | 2.31104069 | 0 | 1 | 0 | 1 | 2 | 0.18613256 | 2 | 0.26791271 | 0 | 1 | 18 | 0.74744813 | FALSE | 1 | FALSE | 1 | FALSE | 1 | FALSE | 1 | FALSE | 1 | FALSE | 1 |
| 3025 | 3025 | 8.52173913 | 2.73256632 | 3 | 0.02011786 | 1 | 0.53115654 | 1 | 0.5695037 | 0 | 1 | 0 | 1 | 18 | 0.86273155 | FALSE | 0.44482142 | FALSE | 1 | FALSE | 1 | FALSE | 1 | FALSE | 1 | FALSE | 1 |
| 3026 | 3026 | 6.625 | 1.88568385 | 0 | 1 | 2 | 0.54544984 | 0 | 1 | 2 | 0.73544289 | 0 | 1 | 52 | 0.04900615 | FALSE | 1 | FALSE | 1 | FALSE | 1 | FALSE | 1 | FALSE | 1 | FALSE | 1 |
| 3027 | 3027 | 11.85 | 3.33579076 | 0 | 1 | 0 | 1 | 1 | 0.51939841 | 1 | 0.60987146 | 1 | 0.28187291 | 17 | 0.61750632 | FALSE | 1 | FALSE | 1 | FALSE | 1 | FALSE | 1 | FALSE | 1 | FALSE | 1 |
| 3028 | 3028 | 5.78571429 | 2.18529408 | 0 | 1 | 3 | 0.25367018 | 1 | 0.86240917 | 2 | 0.71679995 | 0 | 1 | 48 | 0.23965388 | FALSE | 1 | FALSE | 1 | FALSE | 1 | FALSE | 1 | FALSE | 1 | FALSE | 1 |
| 3029 | 3029 | 8.55555556 | 3.59354703 | 0 | 1 | 0 | 1 | 1 | 0.28067846 | 0 | 1 | 0 | 1 | 8 | 0.5780332 | FALSE | 1 | FALSE | 1 | FALSE | 1 | FALSE | 1 | FALSE | 1 | FALSE | 1 |
| 3030 | 3030 | 8.27272727 | 2.89485151 | 0 | 1 | 1 | 0.5154318 | 1 | 0.55340851 | 1 | 0.64497439 | 1 | 0.30529951 | 18 | 0.74744813 | FALSE | 1 | FALSE | 1 | FALSE | 1 | FALSE | 1 | FALSE | 1 | FALSE | 1 |
| 3031 | 3031 | 9.625 | 2.36841192 | 2 | 0.06197875 | 1 | 0.40942991 | 0 | 1 | 1 | 0.52895088 | 0 | 1 | 12 | 0.90922186 | FALSE | 0.72532843 | FALSE | 1 | FALSE | 1 | FALSE | 1 | FALSE | 1 | FALSE | 1 |
| 3032 | 3032 | 7.92592593 | 3.57901213 | 0 | 1 | 4 | 0.01041264 | 1 | 0.62830986 | 2 | 0.35374209 | 1 | 0.36060486 | 19 | 0.98241223 | FALSE | 1 | FALSE | 0.67617265 | FALSE | 1 | FALSE | 1 | FALSE | 1 | FALSE | 1 |
| 3033 | 3033 | 9.625 | 1.72753437 | 0 | 1 | 0 | 1 | 0 | 1 | 0 | 1 | 0 | 1 | 24 | 0.01683964 | FALSE | 1 | FALSE | 1 | FALSE | 1 | FALSE | 1 | FALSE | 1 | FALSE | 1 |
| 3034 | 3034 | 9.25 | 3.03108891 | 0 | 1 | 0 | 1 | 0 | 1 | 1 | 0.17140665 | 0 | 1 | 3 | 0.8823539 | FALSE | 1 | FALSE | 1 | FALSE | 1 | FALSE | 1 | FALSE | 1 | FALSE | 1 |
| 3035 | 3035 | 5.90291262 | 2.26203175 | 0 | 1 | 3 | 0.62650314 | 5 | 0.28294346 | 4 | 0.67315543 | 6 | 0.00552459 | 81 | 0.80397724 | FALSE | 1 | FALSE | 1 | FALSE | 1 | FALSE | 1 | FALSE | 0.45257452 | FALSE | 1 |
| 3036 | 3036 | 11.5 | 2.59807621 | 0 | 1 | 0 | 1 | 2 | 0.16019548 | 1 | 0.60987146 | 0 | 1 | 17 | 0.61750632 | FALSE | 1 | FALSE | 1 | FALSE | 1 | FALSE | 1 | FALSE | 1 | FALSE | 1 |
| 3037 | 3037 | 8.79310345 | 1.58395643 | 0 | 1 | 0 | 1 | 3 | 0.08431992 | 2 | 0.3871811 | 0 | 1 | 24 | 0.70563334 | FALSE | 1 | FALSE | 1 | FALSE | 1 | FALSE | 1 | FALSE | 1 | FALSE | 1 |
| 3038 | 3038 | 8.91666667 | 3.7518514 | 1 | 0.26795838 | 0 | 1 | 1 | 0.35555293 | 0 | 1 | 0 | 1 | 10 | 0.71436964 | FALSE | 1 | FALSE | 1 | FALSE | 1 | FALSE | 1 | FALSE | 1 | FALSE | 1 |
| 3039 | 3039 | 8.4 | 2.23223143 | 2 | 0.21638713 | 3 | 0.09602744 | 0 | 1 | 2 | 0.46709861 | 1 | 0.43077488 | 26 | 0.92796052 | FALSE | 1 | FALSE | 1 | FALSE | 1 | FALSE | 1 | FALSE | 1 | FALSE | 1 |
| 3040 | 3040 | 8.125 | 1.9324531 | 1 | 0.56527493 | 0 | 1 | 2 | 0.32011871 | 2 | 0.43583874 | 3 | 0.01506052 | 24 | 0.94876798 | FALSE | 1 | FALSE | 1 | FALSE | 1 | FALSE | 1 | FALSE | 0.66313193 | FALSE | 1 |
| 3041 | 3041 | 8.7826087 | 2.65324284 | 1 | 0.45027683 | 1 | 0.53115654 | 0 | 1 | 4 | 0.01932195 | 1 | 0.31672722 | 16 | 0.98100709 | FALSE | 1 | FALSE | 1 | FALSE | 1 | FALSE | 1 | FALSE | 1 | FALSE | 1 |
| 3042 | 3042 | 8.89473684 | 2.29174116 | 0 | 1 | 0 | 1 | 0 | 1 | 0 | 1 | 0 | 1 | 19 | 0.03948376 | FALSE | 1 | FALSE | 1 | FALSE | 1 | FALSE | 1 | FALSE | 1 | FALSE | 1 |
| 3043 | 3043 | 11 | 2.52982213 | 0 | 1 | 0 | 1 | 0 | 1 | 0 | 1 | 0 | 1 | 5 | 0.42761893 | FALSE | 1 | FALSE | 1 | FALSE | 1 | FALSE | 1 | FALSE | 1 | FALSE | 1 |
| 3044 | 3044 | 9.86666667 | 1.96185853 | 0 | 1 | 0 | 1 | 2 | 0.09928574 | 1 | 0.50623194 | 3 | 0.00169035 | 9 | 0.995488 | FALSE | 1 | FALSE | 1 | FALSE | 1 | FALSE | 1 | FALSE | 0.23874659 | FALSE | 1 |
| 3045 | 3045 | 7.88235294 | 2.86730767 | 1 | 0.72839502 | 4 | 0.07727212 | 2 | 0.54120634 | 2 | 0.6762159 | 2 | 0.19752014 | 39 | 0.92022511 | FALSE | 1 | FALSE | 1 | FALSE | 1 | FALSE | 1 | FALSE | 1 | FALSE | 1 |
| 3046 | 3046 | 9.64705882 | 2.37489756 | 0 | 1 | 1 | 0.42857733 | 1 | 0.46348836 | 0 | 1 | 0 | 1 | 15 | 0.49060435 | FALSE | 1 | FALSE | 1 | FALSE | 1 | FALSE | 1 | FALSE | 1 | FALSE | 1 |
| 3047 | 3047 | 9.61904762 | 2.35990699 | 0 | 1 | 1 | 0.49918218 | 1 | 0.53671415 | 2 | 0.25059627 | 0 | 1 | 17 | 0.77849076 | FALSE | 1 | FALSE | 1 | FALSE | 1 | FALSE | 1 | FALSE | 1 | FALSE | 1 |
| 3048 | 3048 | 9.22580645 | 1.40757576 | 0 | 1 | 3 | 0.07729739 | 1 | 0.67911179 | 1 | 0.76781284 | 1 | 0.4016888 | 25 | 0.80006983 | FALSE | 1 | FALSE | 1 | FALSE | 1 | FALSE | 1 | FALSE | 1 | FALSE | 1 |
| 3049 | 3049 | 10.1818182 | 3.40636255 | 0 | 1 | 0 | 1 | 0 | 1 | 2 | 0.08789566 | 0 | 1 | 9 | 0.75985366 | FALSE | 1 | FALSE | 1 | FALSE | 1 | FALSE | 1 | FALSE | 1 | FALSE | 1 |
| 3050 | 3050 | 10.6363636 | 1.36666331 | 0 | 1 | 0 | 1 | 1 | 0.33149969 | 0 | 1 | 0 | 1 | 10 | 0.46859937 | FALSE | 1 | FALSE | 1 | FALSE | 1 | FALSE | 1 | FALSE | 1 | FALSE | 1 |
| 3051 | 3051 | 9.20833333 | 3.3037752 | 0 | 1 | 2 | 0.18117183 | 0 | 1 | 2 | 0.30249307 | 1 | 0.32796862 | 19 | 0.83975208 | FALSE | 1 | FALSE | 1 | FALSE | 1 | FALSE | 1 | FALSE | 1 | FALSE | 1 |
| 3052 | 3052 | 3.2 | 2.88870479 | 3 | 0.23231139 | 2 | 0.62701348 | 0 | 1 | 3 | 0.57949569 | 2 | 0.28868487 | 55 | 0.56243792 | FALSE | 1 | FALSE | 1 | FALSE | 1 | FALSE | 1 | FALSE | 1 | FALSE | 1 |
| 3053 | 3053 | 11.25 | 2.384848 | 0 | 1 | 0 | 1 | 1 | 0.25384309 | 0 | 1 | 0 | 1 | 7 | 0.63752397 | FALSE | 1 | FALSE | 1 | FALSE | 1 | FALSE | 1 | FALSE | 1 | FALSE | 1 |
| 3054 | 3054 | 8.17647059 | 3.43400459 | 0 | 1 | 1 | 0.40942991 | 0 | 1 | 0 | 1 | 1 | 0.23264612 | 14 | 0.53308261 | FALSE | 1 | FALSE | 1 | FALSE | 1 | FALSE | 1 | FALSE | 1 | FALSE | 1 |
| 3055 | 3055 | 9.83333333 | 2.03442594 | 0 | 1 | 0 | 1 | 1 | 0.35555293 | 1 | 0.43131129 | 0 | 1 | 10 | 0.71436964 | FALSE | 1 | FALSE | 1 | FALSE | 1 | FALSE | 1 | FALSE | 1 | FALSE | 1 |
| 3056 | 3056 | 8.42424242 | 3.24765933 | 0 | 1 | 0 | 1 | 3 | 0.11365871 | 0 | 1 | 1 | 0.42123867 | 29 | 0.39648374 | FALSE | 1 | FALSE | 1 | FALSE | 1 | FALSE | 1 | FALSE | 1 | FALSE | 1 |
| 3057 | 3057 | 5.14457831 | 2.10067419 | 1 | 0.88571933 | 3 | 0.50638104 | 1 | 0.95289825 | 5 | 0.33297 | 0 | 1 | 73 | 0.2312551 | FALSE | 1 | FALSE | 1 | FALSE | 1 | FALSE | 1 | FALSE | 1 | FALSE | 1 |
| 3058 | 3058 | 10.9354839 | 2.42214892 | 0 | 1 | 0 | 1 | 0 | 1 | 1 | 0.75658797 | 0 | 1 | 29 | 0.03983039 | FALSE | 1 | FALSE | 1 | FALSE | 1 | FALSE | 1 | FALSE | 1 | FALSE | 1 |
| 3059 | 3059 | 8.6 | 2.39582971 | 2 | 0.09187136 | 1 | 0.48239024 | 1 | 0.51939841 | 0 | 1 | 1 | 0.28187291 | 15 | 0.92113679 | FALSE | 0.83248965 | FALSE | 1 | FALSE | 1 | FALSE | 1 | FALSE | 1 | FALSE | 1 |
| 3060 | 3060 | 14.0714286 | 5.18878304 | 0 | 1 | 0 | 1 | 0 | 1 | 1 | 0.48242099 | 0 | 1 | 13 | 0.33283729 | FALSE | 1 | FALSE | 1 | FALSE | 1 | FALSE | 1 | FALSE | 1 | FALSE | 1 |
| 3061 | 3061 | 9 | 2.07744783 | 0 | 1 | 1 | 0.46503795 | 1 | 0.50143826 | 1 | 0.59104357 | 0 | 1 | 16 | 0.65565778 | FALSE | 1 | FALSE | 1 | FALSE | 1 | FALSE | 1 | FALSE | 1 | FALSE | 1 |
| 3062 | 3062 | 6.65333333 | 1.73199949 | 3 | 0.30198574 | 11 | 2.49E-05 | 4 | 0.2829119 | 4 | 0.45379959 | 4 | 0.03419047 | 49 | 0.99998858 | FALSE | 1 | FALSE | 0.07208458 | FALSE | 1 | FALSE | 1 | FALSE | 0.86063198 | FALSE | 1 |
| 3063 | 3063 | 9.85714286 | 2.18840482 | 3 | 0.09196816 | 0 | 1 | 2 | 0.44869594 | 0 | 1 | 1 | 0.50166523 | 36 | 0.50934357 | FALSE | 0.83248965 | FALSE | 1 | FALSE | 1 | FALSE | 1 | FALSE | 1 | FALSE | 1 |
| 3064 | 3064 | 9.57142857 | 2.19461307 | 0 | 1 | 1 | 0.36919824 | 1 | 0.40110418 | 0 | 1 | 0 | 1 | 12 | 0.62236227 | FALSE | 1 | FALSE | 1 | FALSE | 1 | FALSE | 1 | FALSE | 1 | FALSE | 1 |
| 3065 | 3065 | 11.4117647 | 1.97212407 | 0 | 1 | 2 | 0.10307112 | 0 | 1 | 0 | 1 | 1 | 0.24525764 | 14 | 0.7313145 | FALSE | 1 | FALSE | 1 | FALSE | 1 | FALSE | 1 | FALSE | 1 | FALSE | 1 |
| 3066 | 3066 | 10.6190476 | 2.88635856 | 4 | 0.00177457 | 0 | 1 | 2 | 0.17308361 | 0 | 1 | 0 | 1 | 15 | 0.96513691 | FALSE | 0.10930302 | FALSE | 1 | FALSE | 1 | FALSE | 1 | FALSE | 1 | FALSE | 1 |
| 3067 | 3067 | 1.14802131 | 1.02246823 | 86 | 0.00120907 | 100 | 0.01207464 | 91 | 0.61566435 | 130 | 0.09386214 | 45 | 0.33107586 | 2129 | 0.99965357 | FALSE | 0.08465577 | FALSE | 0.67750333 | FALSE | 1 | FALSE | 1 | FALSE | 1 | FALSE | 1 |
| 3068 | 3068 | 7.69565217 | 2.59489101 | 0 | 1 | 2 | 0.14655134 | 1 | 0.53671415 | 0 | 1 | 0 | 1 | 18 | 0.57962763 | FALSE | 1 | FALSE | 1 | FALSE | 1 | FALSE | 1 | FALSE | 1 | FALSE | 1 |
| 3069 | 3069 | 8.79166667 | 2.6454232 | 2 | 0.12501436 | 0 | 1 | 0 | 1 | 2 | 0.30249307 | 0 | 1 | 20 | 0.68270729 | FALSE | 0.95622564 | FALSE | 1 | FALSE | 1 | FALSE | 1 | FALSE | 1 | FALSE | 1 |
| 3070 | 3070 | 4.21212121 | 2.91035326 | 0 | 1 | 1 | 0.6629941 | 1 | 0.70185639 | 2 | 0.45159487 | 2 | 0.10132035 | 27 | 0.75124583 | FALSE | 1 | FALSE | 1 | FALSE | 1 | FALSE | 1 | FALSE | 1 | FALSE | 1 |
| 3071 | 3071 | 9.17142857 | 2.02111305 | 1 | 0.50471032 | 1 | 0.58913618 | 1 | 0.62830986 | 3 | 0.12475952 | 0 | 1 | 21 | 0.88395936 | FALSE | 1 | FALSE | 1 | FALSE | 1 | FALSE | 1 | FALSE | 1 | FALSE | 1 |
| 3072 | 3072 | 10.5384615 | 3.8353695 | 2 | 0.04234484 | 2 | 0.06426041 | 0 | 1 | 0 | 1 | 0 | 1 | 9 | 0.96018168 | FALSE | 0.62842194 | FALSE | 1 | FALSE | 1 | FALSE | 1 | FALSE | 1 | FALSE | 1 |
| 3073 | 3073 | 8.72580645 | 2.08039508 | 4 | 0.07395961 | 2 | 0.60114557 | 2 | 0.65873577 | 3 | 0.54709375 | 0 | 1 | 51 | 0.74578096 | FALSE | 0.77424195 | FALSE | 1 | FALSE | 1 | FALSE | 1 | FALSE | 1 | FALSE | 1 |
| 3074 | 3074 | 9.5 | 2.54950976 | 0 | 1 | 0 | 1 | 1 | 0.25384309 | 0 | 1 | 0 | 1 | 7 | 0.63752397 | FALSE | 1 | FALSE | 1 | FALSE | 1 | FALSE | 1 | FALSE | 1 | FALSE | 1 |
| 3075 | 3075 | 10.6923077 | 2.43008754 | 0 | 1 | 0 | 1 | 0 | 1 | 0 | 1 | 1 | 0.19354507 | 12 | 0.374198 | FALSE | 1 | FALSE | 1 | FALSE | 1 | FALSE | 1 | FALSE | 1 | FALSE | 1 |
| 3076 | 3076 | 5.27272727 | 2.48558747 | 1 | 0.92501001 | 4 | 0.3989613 | 5 | 0.28294346 | 1 | 0.99079232 | 5 | 0.02289208 | 83 | 0.62438251 | FALSE | 1 | FALSE | 1 | FALSE | 1 | FALSE | 1 | FALSE | 0.7685735 | FALSE | 1 |
| 3077 | 3077 | 8.94736842 | 1.82017015 | 0 | 1 | 1 | 0.46503795 | 0 | 1 | 1 | 0.59104357 | 0 | 1 | 17 | 0.41141543 | FALSE | 1 | FALSE | 1 | FALSE | 1 | FALSE | 1 | FALSE | 1 | FALSE | 1 |
| 3078 | 3078 | 7.73333333 | 1.91369335 | 0 | 1 | 1 | 0.38964394 | 2 | 0.09928574 | 1 | 0.50623194 | 1 | 0.2198258 | 10 | 0.97980046 | FALSE | 1 | FALSE | 1 | FALSE | 1 | FALSE | 1 | FALSE | 1 | FALSE | 1 |
| 3079 | 3079 | 9.27777778 | 2.67302483 | 0 | 1 | 1 | 0.69480472 | 1 | 0.73299778 | 5 | 0.02311989 | 0 | 1 | 29 | 0.81028289 | FALSE | 1 | FALSE | 1 | FALSE | 1 | FALSE | 1 | FALSE | 1 | FALSE | 1 |
| 3080 | 3080 | 8.5 | 2.16899642 | 2 | 0.30241117 | 1 | 0.75788114 | 0 | 1 | 4 | 0.13323859 | 1 | 0.50988734 | 35 | 0.7801651 | FALSE | 1 | FALSE | 1 | FALSE | 1 | FALSE | 1 | FALSE | 1 | FALSE | 1 |
| 3081 | 3081 | 9.57142857 | 1.52975032 | 1 | 0.42086991 | 1 | 0.49918218 | 1 | 0.53671415 | 0 | 1 | 0 | 1 | 18 | 0.57962763 | FALSE | 1 | FALSE | 1 | FALSE | 1 | FALSE | 1 | FALSE | 1 | FALSE | 1 |
| 3082 | 3082 | 5.11403509 | 2.47002654 | 1 | 0.94952673 | 5 | 0.30872273 | 6 | 0.22590386 | 9 | 0.07781938 | 2 | 0.56161197 | 91 | 0.92683473 | FALSE | 1 | FALSE | 1 | FALSE | 1 | FALSE | 1 | FALSE | 1 | FALSE | 1 |
| 3083 | 3083 | 6.41085271 | 1.21113953 | 3 | 0.64793818 | 2 | 0.92549961 | 5 | 0.49638206 | 9 | 0.13780368 | 0 | 1 | 110 | 0.4472263 | FALSE | 1 | FALSE | 1 | FALSE | 1 | FALSE | 1 | FALSE | 1 | FALSE | 1 |
| 3084 | 3084 | 9.75862069 | 2.48658709 | 0 | 1 | 0 | 1 | 2 | 0.27983822 | 1 | 0.74482229 | 0 | 1 | 26 | 0.31495157 | FALSE | 1 | FALSE | 1 | FALSE | 1 | FALSE | 1 | FALSE | 1 | FALSE | 1 |
| 3085 | 3085 | 12 | 4.30945804 | 1 | 0.3050805 | 0 | 1 | 1 | 0.40110418 | 1 | 0.48242099 | 0 | 1 | 11 | 0.83676449 | FALSE | 1 | FALSE | 1 | FALSE | 1 | FALSE | 1 | FALSE | 1 | FALSE | 1 |
| 3086 | 3086 | 9 | 3.09232922 | 3 | 0.0476977 | 1 | 0.65167362 | 0 | 1 | 3 | 0.17988885 | 1 | 0.41154418 | 24 | 0.94876798 | FALSE | 0.649301 | FALSE | 1 | FALSE | 1 | FALSE | 1 | FALSE | 1 | FALSE | 1 |
| 3087 | 3087 | 5.06621773 | 0.63836663 | 19 | 0.83156867 | 25 | 0.7998229 | 31 | 0.59336284 | 35 | 0.85707462 | 19 | 0.13096259 | 757 | 0.18882095 | FALSE | 1 | FALSE | 1 | FALSE | 1 | FALSE | 1 | FALSE | 1 | FALSE | 1 |
| 3088 | 3088 | 9.64705882 | 2.34999816 | 0 | 1 | 3 | 0.01622926 | 0 | 1 | 0 | 1 | 1 | 0.24525764 | 13 | 0.88719222 | FALSE | 1 | FALSE | 0.7256023 | FALSE | 1 | FALSE | 1 | FALSE | 1 | FALSE | 1 |
| 3089 | 3089 | 9.85714286 | 2.79941685 | 1 | 0.3050805 | 0 | 1 | 0 | 1 | 2 | 0.13304705 | 0 | 1 | 11 | 0.83676449 | FALSE | 1 | FALSE | 1 | FALSE | 1 | FALSE | 1 | FALSE | 1 | FALSE | 1 |
| 3090 | 3090 | 10.1290323 | 1.60124821 | 4 | 0.00764535 | 1 | 0.63997468 | 2 | 0.30673593 | 6 | 0.00248013 | 0 | 1 | 18 | 0.99990857 | FALSE | 0.2566832 | FALSE | 1 | FALSE | 1 | FALSE | 1 | FALSE | 1 | FALSE | 1 |
| 3091 | 3091 | 9.78378378 | 2.09420223 | 0 | 1 | 1 | 0.70472939 | 2 | 0.38585089 | 2 | 0.51200191 | 1 | 0.45845922 | 31 | 0.64534533 | FALSE | 1 | FALSE | 1 | FALSE | 1 | FALSE | 1 | FALSE | 1 | FALSE | 1 |
| 3092 | 3092 | 10.0344828 | 2.35595736 | 5 | 0.00075302 | 2 | 0.24095584 | 1 | 0.65463982 | 0 | 1 | 1 | 0.38148475 | 20 | 0.99028351 | FALSE | 0.06390799 | FALSE | 1 | FALSE | 1 | FALSE | 1 | FALSE | 1 | FALSE | 1 |
| 3093 | 3093 | 9.5 | 3.04138127 | 0 | 1 | 0 | 1 | 0 | 1 | 1 | 0.31351192 | 0 | 1 | 7 | 0.63752397 | FALSE | 1 | FALSE | 1 | FALSE | 1 | FALSE | 1 | FALSE | 1 | FALSE | 1 |
| 3094 | 3094 | 9.43478261 | 2.33733566 | 1 | 0.45027683 | 1 | 0.53115654 | 2 | 0.19931429 | 1 | 0.6613267 | 1 | 0.31672722 | 17 | 0.94439745 | FALSE | 1 | FALSE | 1 | FALSE | 1 | FALSE | 1 | FALSE | 1 | FALSE | 1 |
| 3095 | 3095 | 9.21428571 | 2.6234811 | 0 | 1 | 1 | 0.36919824 | 0 | 1 | 1 | 0.48242099 | 1 | 0.20679328 | 11 | 0.83676449 | FALSE | 1 | FALSE | 1 | FALSE | 1 | FALSE | 1 | FALSE | 1 | FALSE | 1 |
| 3096 | 3096 | 7.56521739 | 2.6674542 | 0 | 1 | 0 | 1 | 0 | 1 | 0 | 1 | 0 | 1 | 23 | 0.01996981 | FALSE | 1 | FALSE | 1 | FALSE | 1 | FALSE | 1 | FALSE | 1 | FALSE | 1 |
| 3097 | 3097 | 10.5555556 | 1.77081972 | 0 | 1 | 0 | 1 | 0 | 1 | 0 | 1 | 0 | 1 | 9 | 0.21661397 | FALSE | 1 | FALSE | 1 | FALSE | 1 | FALSE | 1 | FALSE | 1 | FALSE | 1 |
| 3098 | 3098 | 10.4583333 | 1.25761569 | 0 | 1 | 1 | 0.54637326 | 2 | 0.21260257 | 1 | 0.67692816 | 1 | 0.32796862 | 19 | 0.83975208 | FALSE | 1 | FALSE | 1 | FALSE | 1 | FALSE | 1 | FALSE | 1 | FALSE | 1 |
| 3099 | 3099 | 11.2 | 3.01551543 | 1 | 0.32293355 | 1 | 0.38964394 | 0 | 1 | 0 | 1 | 1 | 0.2198258 | 12 | 0.80329917 | FALSE | 1 | FALSE | 1 | FALSE | 1 | FALSE | 1 | FALSE | 1 | FALSE | 1 |
| 3100 | 3100 | 9 | 2.49443826 | 1 | 0.20854748 | 0 | 1 | 0 | 1 | 2 | 0.06108592 | 0 | 1 | 6 | 0.96128317 | FALSE | 1 | FALSE | 1 | FALSE | 1 | FALSE | 1 | FALSE | 1 | FALSE | 1 |

| | | | | | | | | | | | | | | | | | | | | | | | | | | | |
|---|---|---|---|---|---|---|---|---|---|---|---|---|---|---|---|---|---|---|---|---|---|---|---|---|---|---|---|
| 3101 | 3101 | 8.33333333 | 1.84992492 | 0 | 1 | 0 | 1 | 0 | 1 | 2 | 0.14906763 | 0 | 1 | 13 | 0.57712805 | FALSE | 1 | FALSE | 1 | FALSE | 1 | FALSE | 1 | FALSE | 1 | FALSE | 1 |
| 3102 | 3102 | 10.1538462 | 2.03228382 | 1 | 0.47819878 | 1 | 0.56109832 | 1 | 0.59998149 | 0 | 1 | 0 | 1 | 22 | 0.43642846 | FALSE | 1 | FALSE | 1 | FALSE | 1 | FALSE | 1 | FALSE | 1 | FALSE | 1 |
| 3103 | 3103 | 9.42105263 | 2.68162931 | 0 | 1 | 0 | 1 | 1 | 0.50143826 | 3 | 0.05383577 | 1 | 0.26986775 | 14 | 0.93677253 | FALSE | 1 | FALSE | 1 | FALSE | 1 | FALSE | 1 | FALSE | 1 | FALSE | 1 |
| 3104 | 3104 | 6.37931035 | 2.44122345 | 2 | 0.16962005 | 0 | 1 | 1 | 0.65463982 | 4 | 0.04184267 | 0 | 1 | 22 | 0.92902102 | FALSE | 1 | FALSE | 1 | FALSE | 1 | FALSE | 1 | FALSE | 1 | FALSE | 1 |
| 3105 | 3105 | 11.2142857 | 3.47806538 | 1 | 0.51745948 | 4 | 0.01184672 | 2 | 0.2663542 | 4 | 0.03737468 | 0 | 1 | 17 | 0.99947263 | FALSE | 1 | FALSE | 0.67617265 | FALSE | 1 | FALSE | 1 | FALSE | 1 | FALSE | 1 |
| 3106 | 3106 | 8.26470588 | 1.77082273 | 2 | 0.21638713 | 1 | 0.6739483 | 0 | 1 | 1 | 0.7984848 | 0 | 1 | 30 | 0.36879488 | FALSE | 1 | FALSE | 1 | FALSE | 1 | FALSE | 1 | FALSE | 1 | FALSE | 1 |
| 3107 | 3107 | 11.7058824 | 1.600173 | 1 | 0.35728305 | 0 | 1 | 0 | 1 | 0 | 1 | 0 | 1 | 16 | 0.23070182 | FALSE | 1 | FALSE | 1 | FALSE | 1 | FALSE | 1 | FALSE | 1 | FALSE | 1 |
| 3108 | 3108 | 10.6842105 | 2.00138456 | 2 | 0.08405377 | 0 | 1 | 0 | 1 | 1 | 0.59104357 | 0 | 1 | 16 | 0.65565778 | FALSE | 0.79236883 | FALSE | 1 | FALSE | 1 | FALSE | 1 | FALSE | 1 | FALSE | 1 |
| 3109 | 3109 | 4.96 | 3.56067409 | 3 | 0.02517314 | 0 | 1 | 0 | 1 | 0 | 1 | 0 | 1 | 22 | 0.43642846 | FALSE | 0.4974559 | FALSE | 1 | FALSE | 1 | FALSE | 1 | FALSE | 1 | FALSE | 1 |
| 3110 | 3110 | 9.3 | 3.03479818 | 2 | 0.08405377 | 0 | 1 | 2 | 0.14749808 | 2 | 0.21610812 | 0 | 1 | 13 | 0.97998676 | FALSE | 0.79236883 | FALSE | 1 | FALSE | 1 | FALSE | 1 | FALSE | 1 | FALSE | 1 |
| 3111 | 3111 | 11.6153846 | 1.59511087 | 1 | 0.28675951 | 3 | 0.00750707 | 1 | 0.37874417 | 1 | 0.45746571 | 0 | 1 | 7 | 0.99839694 | FALSE | 1 | FALSE | 0.6167796 | FALSE | 1 | FALSE | 1 | FALSE | 1 | FALSE | 1 |
| 3112 | 3112 | 7.3125 | 3.11686441 | 0 | 1 | 1 | 0.40942991 | 0 | 1 | 1 | 0.52895088 | 0 | 1 | 14 | 0.53308261 | FALSE | 1 | FALSE | 1 | FALSE | 1 | FALSE | 1 | FALSE | 1 | FALSE | 1 |
| 3113 | 3113 | 7.71428571 | 1.61624407 | 2 | 0.35907852 | 0 | 1 | 3 | 0.25725031 | 3 | 0.39227316 | 1 | 0.55647171 | 40 | 0.77344956 | FALSE | 1 | FALSE | 1 | FALSE | 1 | FALSE | 1 | FALSE | 1 | FALSE | 1 |
| 3114 | 3114 | 9.5483871 | 3.03568783 | 0 | 1 | 2 | 0.2651851 | 0 | 1 | 1 | 0.76781284 | 0 | 1 | 28 | 0.26393996 | FALSE | 1 | FALSE | 1 | FALSE | 1 | FALSE | 1 | FALSE | 1 | FALSE | 1 |
| 3115 | 3115 | 10.4285714 | 1.7202278 | 0 | 1 | 0 | 1 | 0 | 1 | 2 | 0.13304705 | 0 | 1 | 12 | 0.62236227 | FALSE | 1 | FALSE | 1 | FALSE | 1 | FALSE | 1 | FALSE | 1 | FALSE | 1 |
| 3116 | 3116 | 8.85185185 | 2.2722758 | 0 | 1 | 2 | 0.21684246 | 2 | 0.25286935 | 1 | 0.71956339 | 1 | 0.36060486 | 21 | 0.88395936 | FALSE | 1 | FALSE | 1 | FALSE | 1 | FALSE | 1 | FALSE | 1 | FALSE | 1 |
| 3117 | 3117 | 10.2307692 | 2.11783075 | 1 | 0.28675951 | 0 | 1 | 0 | 1 | 1 | 0.45746571 | 1 | 0.19354507 | 10 | 0.86791642 | FALSE | 1 | FALSE | 1 | FALSE | 1 | FALSE | 1 | FALSE | 1 | FALSE | 1 |
| 3118 | 3118 | 9.64285714 | 2.43800688 | 0 | 1 | 1 | 0.36919824 | 0 | 1 | 0 | 1 | 0 | 1 | 13 | 0.33283729 | FALSE | 1 | FALSE | 1 | FALSE | 1 | FALSE | 1 | FALSE | 1 | FALSE | 1 |
| 3119 | 3119 | 9.52 | 3.20149965 | 0 | 1 | 1 | 0.56109832 | 1 | 0.59998149 | 1 | 0.69181315 | 2 | 0.06257402 | 20 | 0.8152498 | FALSE | 1 | FALSE | 1 | FALSE | 1 | FALSE | 1 | FALSE | 1 | FALSE | 1 |
| 3120 | 3120 | 10.7777778 | 2.14878662 | 0 | 1 | 0 | 1 | 1 | 0.28067846 | 1 | 0.34506517 | 0 | 1 | 7 | 0.84574168 | FALSE | 1 | FALSE | 1 | FALSE | 1 | FALSE | 1 | FALSE | 1 | FALSE | 1 |
| 3121 | 3121 | 6.87719298 | 3.39282098 | 1 | 0.76787545 | 0 | 1 | 2 | 0.60323163 | 3 | 0.47819064 | 1 | 0.60529938 | 49 | 0.33483172 | FALSE | 1 | FALSE | 1 | FALSE | 1 | FALSE | 1 | FALSE | 1 | FALSE | 1 |
| 3122 | 3122 | 6.52941177 | 2.145245 | 1 | 0.35728305 | 0 | 1 | 0 | 1 | 0 | 1 | 0 | 1 | 16 | 0.23070182 | FALSE | 1 | FALSE | 1 | FALSE | 1 | FALSE | 1 | FALSE | 1 | FALSE | 1 |
| 3123 | 3123 | 8.90476191 | 2.87691571 | 1 | 0.40558521 | 0 | 1 | 1 | 0.51939841 | 1 | 0.60987146 | 0 | 1 | 17 | 0.61750632 | FALSE | 1 | FALSE | 1 | FALSE | 1 | FALSE | 1 | FALSE | 1 | FALSE | 1 |
| 3124 | 3124 | 9.94444444 | 2.34454976 | 0 | 1 | 0 | 1 | 1 | 0.48280983 | 0 | 1 | 1 | 0.25766374 | 16 | 0.44998001 | FALSE | 1 | FALSE | 1 | FALSE | 1 | FALSE | 1 | FALSE | 1 | FALSE | 1 |
| 3125 | 3125 | 7.46511628 | 3.20862817 | 0 | 1 | 1 | 0.75788114 | 3 | 0.19985581 | 2 | 0.59404271 | 1 | 0.50988734 | 36 | 0.6451239 | FALSE | 1 | FALSE | 1 | FALSE | 1 | FALSE | 1 | FALSE | 1 | FALSE | 1 |
| 3126 | 3126 | 9.67857143 | 3.73261275 | 1 | 0.50471032 | 1 | 0.58913618 | 0 | 1 | 1 | 0.71956339 | 0 | 1 | 24 | 0.3725373 | FALSE | 1 | FALSE | 1 | FALSE | 1 | FALSE | 1 | FALSE | 1 | FALSE | 1 |
| 3127 | 3127 | 9.38461539 | 2.16753889 | 0 | 1 | 1 | 0.3480709 | 0 | 1 | 0 | 1 | 2 | 0.01846832 | 10 | 0.86791642 | FALSE | 1 | FALSE | 1 | FALSE | 1 | FALSE | 1 | FALSE | 0.69720022 | FALSE | 1 |
| 3128 | 3128 | 7.84615385 | 3.1585331 | 0 | 1 | 1 | 0.3480709 | 0 | 1 | 0 | 1 | 2 | 0.01846832 | 10 | 0.86791642 | FALSE | 1 | FALSE | 1 | FALSE | 1 | FALSE | 1 | FALSE | 0.69720022 | FALSE | 1 |
| 3129 | 3129 | 10.8666667 | 3.38362068 | 1 | 0.32293355 | 3 | 0.01138739 | 0 | 1 | 1 | 0.50623194 | 0 | 1 | 10 | 0.97980046 | FALSE | 1 | FALSE | 0.67617265 | FALSE | 1 | FALSE | 1 | FALSE | 1 | FALSE | 1 |
| 3130 | 3130 | 9.95 | 2.59759504 | 0 | 1 | 0 | 1 | 0 | 1 | 0 | 1 | 0 | 1 | 19 | 0.03948376 | FALSE | 1 | FALSE | 1 | FALSE | 1 | FALSE | 1 | FALSE | 1 | FALSE | 1 |
| 3131 | 3131 | 4.10784314 | 2.61559 | 5 | 0.10638005 | 3 | 0.61960793 | 9 | 0.00834932 | 4 | 0.66533238 | 7 | 0.0010674 | 70 | 0.99967832 | FALSE | 0.88820416 | FALSE | 1 | FALSE | 1 | FALSE | 1 | FALSE | 0.19379454 | FALSE | 1 |
| 3132 | 3132 | 9.55 | 3.27833799 | 0 | 1 | 1 | 0.48239024 | 0 | 1 | 3 | 0.0612581 | 1 | 0.28187291 | 15 | 0.92113679 | FALSE | 1 | FALSE | 1 | FALSE | 1 | FALSE | 1 | FALSE | 1 | FALSE | 1 |
| 3133 | 3133 | 9.18181818 | 3.14261564 | 0 | 1 | 1 | 0.5154318 | 1 | 0.55340851 | 0 | 1 | 0 | 1 | 20 | 0.30915202 | FALSE | 1 | FALSE | 1 | FALSE | 1 | FALSE | 1 | FALSE | 1 | FALSE | 1 |
| 3134 | 3134 | 6.87096774 | 3.03465931 | 2 | 0.1881398 | 0 | 1 | 2 | 0.30673593 | 2 | 0.41984187 | 0 | 1 | 25 | 0.80006983 | FALSE | 1 | FALSE | 1 | FALSE | 1 | FALSE | 1 | FALSE | 1 | FALSE | 1 |
| 3135 | 3135 | 8.21428571 | 1.3978847 | 4 | 0.00346297 | 1 | 0.56109832 | 2 | 0.22597287 | 1 | 0.69181315 | 1 | 0.33902672 | 16 | 0.99713666 | FALSE | 0.16118559 | FALSE | 1 | FALSE | 1 | FALSE | 1 | FALSE | 1 | FALSE | 1 |
| 3136 | 3136 | 5.1147541 | 2.12014816 | 0 | 1 | 3 | 0.30686508 | 4 | 0.16812274 | 4 | 0.29632241 | 0 | 1 | 49 | 0.78152878 | FALSE | 1 | FALSE | 1 | FALSE | 1 | FALSE | 1 | FALSE | 1 | FALSE | 1 |
| 3137 | 3137 | 9.89655172 | 2.32445511 | 1 | 0.52988231 | 0 | 1 | 2 | 0.27983822 | 0 | 1 | 0 | 1 | 26 | 0.31495157 | FALSE | 1 | FALSE | 1 | FALSE | 1 | FALSE | 1 | FALSE | 1 | FALSE | 1 |
| 3138 | 3138 | 11.1538462 | 1.99407406 | 0 | 1 | 0 | 1 | 1 | 0.37874417 | 1 | 0.45746571 | 0 | 1 | 11 | 0.6683056 | FALSE | 1 | FALSE | 1 | FALSE | 1 | FALSE | 1 | FALSE | 1 | FALSE | 1 |
| 3139 | 3139 | 10 | 3.28633535 | 1 | 0.54198714 | 2 | 0.25306331 | 1 | 0.66709967 | 2 | 0.4036176 | 1 | 0.39166989 | 23 | 0.9158696 | FALSE | 1 | FALSE | 1 | FALSE | 1 | FALSE | 1 | FALSE | 1 | FALSE | 1 |
| 3140 | 3140 | 11.1666667 | 2.88193608 | 1 | 0.26795838 | 0 | 1 | 1 | 0.35555293 | 0 | 1 | 0 | 1 | 10 | 0.71436964 | FALSE | 1 | FALSE | 1 | FALSE | 1 | FALSE | 1 | FALSE | 1 | FALSE | 1 |
| 3141 | 3141 | 12.4285714 | 1.84058553 | 1 | 0.16630142 | 0 | 1 | 0 | 1 | 0 | 1 | 0 | 1 | 6 | 0.69920685 | FALSE | 1 | FALSE | 1 | FALSE | 1 | FALSE | 1 | FALSE | 1 | FALSE | 1 |
| 3142 | 3142 | 9.75 | 2.34254136 | 1 | 0.40558521 | 1 | 0.48239024 | 1 | 0.51939841 | 2 | 0.23331368 | 0 | 1 | 15 | 0.92113679 | FALSE | 1 | FALSE | 1 | FALSE | 1 | FALSE | 1 | FALSE | 1 | FALSE | 1 |
| 3143 | 3143 | 9.25581395 | 1.47999094 | 0 | 1 | 1 | 0.75788114 | 1 | 0.79363407 | 2 | 0.59404271 | 1 | 0.50988734 | 38 | 0.31730615 | FALSE | 1 | FALSE | 1 | FALSE | 1 | FALSE | 1 | FALSE | 1 | FALSE | 1 |
| 3144 | 3144 | 8.2 | 1.83303028 | 3 | 0.08207756 | 0 | 1 | 1 | 0.76953771 | 2 | 0.5543522 | 2 | 0.13925369 | 32 | 0.83853687 | FALSE | 0.79236883 | FALSE | 1 | FALSE | 1 | FALSE | 1 | FALSE | 1 | FALSE | 1 |
| 3145 | 3145 | 4.26964286 | 0.95644798 | 29 | 0.00017974 | 27 | 0.02257482 | 26 | 0.10360139 | 24 | 0.6716866 | 12 | 0.20516175 | 442 | 0.99986377 | TRUE | 0.02495627 | FALSE | 0.81828735 | FALSE | 1 | FALSE | 1 | FALSE | 1 | FALSE | 1 |
| 3146 | 3146 | 9.6 | 2.52454616 | 4 | 0.00045644 | 1 | 0.38964394 | 1 | 0.42266264 | 0 | 1 | 0 | 1 | 9 | 0.995488 | TRUE | 0.0445135 | FALSE | 1 | FALSE | 1 | FALSE | 1 | FALSE | 1 | FALSE | 1 |
| 3147 | 3147 | 8.53571429 | 1.70046512 | 3 | 0.0279333 | 0 | 1 | 1 | 0.61440466 | 1 | 0.70601446 | 0 | 1 | 21 | 0.78939668 | FALSE | 0.50964275 | FALSE | 1 | FALSE | 1 | FALSE | 1 | FALSE | 1 | FALSE | 1 |
| 3148 | 3148 | 10.8095238 | 2.85555511 | 0 | 1 | 1 | 0.49918218 | 0 | 1 | 2 | 0.25059627 | 1 | 0.29368245 | 17 | 0.77849076 | FALSE | 1 | FALSE | 1 | FALSE | 1 | FALSE | 1 | FALSE | 1 | FALSE | 1 |
| 3149 | 3149 | 9.5 | 1.91796023 | 0 | 1 | 0 | 1 | 1 | 0.40110418 | 1 | 0.48242099 | 0 | 1 | 12 | 0.62236227 | FALSE | 1 | FALSE | 1 | FALSE | 1 | FALSE | 1 | FALSE | 1 | FALSE | 1 |
| 3150 | 3150 | 9.92857143 | 1.83085795 | 1 | 0.3050805 | 0 | 1 | 0 | 1 | 2 | 0.13304705 | 0 | 1 | 11 | 0.83676449 | FALSE | 1 | FALSE | 1 | FALSE | 1 | FALSE | 1 | FALSE | 1 | FALSE | 1 |
| 3151 | 3151 | 9.57894737 | 2.79691163 | 0 | 1 | 0 | 1 | 0 | 1 | 1 | 0.59104357 | 1 | 0.26986775 | 17 | 0.41141543 | FALSE | 1 | FALSE | 1 | FALSE | 1 | FALSE | 1 | FALSE | 1 | FALSE | 1 |
| 3152 | 3152 | 2.8125 | 2.18571356 | 2 | 0.94396881 | 4 | 0.82741968 | 7 | 0.44685733 | 10 | 0.28891736 | 0 | 1 | 153 | 0.20229949 | FALSE | 1 | FALSE | 1 | FALSE | 1 | FALSE | 1 | FALSE | 1 | FALSE | 1 |
| 3153 | 3153 | 8.06666667 | 2.37954244 | 0 | 1 | 0 | 1 | 1 | 0.42266264 | 1 | 0.50623194 | 0 | 1 | 13 | 0.57712805 | FALSE | 1 | FALSE | 1 | FALSE | 1 | FALSE | 1 | FALSE | 1 | FALSE | 1 |
| 3154 | 3154 | 9.22222222 | 2.04275292 | 0 | 1 | 0 | 1 | 0 | 1 | 0 | 1 | 1 | 0.1383237 | 8 | 0.5780332 | FALSE | 1 | FALSE | 1 | FALSE | 1 | FALSE | 1 | FALSE | 1 | FALSE | 1 |
| 3155 | 3155 | 9.15 | 2.8420943 | 3 | 0.07731604 | 2 | 0.36122049 | 0 | 1 | 1 | 0.84089119 | 0 | 1 | 33 | 0.59061307 | FALSE | 0.77583006 | FALSE | 1 | FALSE | 1 | FALSE | 1 | FALSE | 1 | FALSE | 1 |
| 3156 | 3156 | 10.2352941 | 1.95626931 | 1 | 0.35728305 | 1 | 0.42857733 | 0 | 1 | 1 | 0.55062772 | 0 | 1 | 14 | 0.7313145 | FALSE | 1 | FALSE | 1 | FALSE | 1 | FALSE | 1 | FALSE | 1 | FALSE | 1 |
| 3157 | 3157 | 7.9375 | 2.60932822 | 0 | 1 | 0 | 1 | 0 | 1 | 0 | 1 | 0 | 1 | 16 | 0.06581507 | FALSE | 1 | FALSE | 1 | FALSE | 1 | FALSE | 1 | FALSE | 1 | FALSE | 1 |
| 3158 | 3158 | 7.49090909 | 1.31940382 | 3 | 0.40052974 | 3 | 0.55379184 | 3 | 0.62570603 | 3 | 0.78298119 | 4 | 0.0578582 | 73 | 0.78154321 | FALSE | 1 | FALSE | 1 | FALSE | 1 | FALSE | 1 | FALSE | 1 | FALSE | 1 |
| 3159 | 3159 | 10.3 | 2.38956063 | 0 | 1 | 1 | 0.48239024 | 2 | 0.16019548 | 0 | 1 | 0 | 1 | 17 | 0.61750632 | FALSE | 1 | FALSE | 1 | FALSE | 1 | FALSE | 1 | FALSE | 1 | FALSE | 1 |
| 3160 | 3160 | 9.08695652 | 2.18518815 | 1 | 0.45027683 | 0 | 1 | 1 | 0.5695037 | 3 | 0.08614834 | 1 | 0.31672722 | 17 | 0.94439745 | FALSE | 1 | FALSE | 1 | FALSE | 1 | FALSE | 1 | FALSE | 1 | FALSE | 1 |
| 3161 | 3161 | 10.3333333 | 2.62466929 | 1 | 0.26795838 | 0 | 1 | 0 | 1 | 0 | 1 | 1 | 0.18007762 | 10 | 0.71436964 | FALSE | 1 | FALSE | 1 | FALSE | 1 | FALSE | 1 | FALSE | 1 | FALSE | 1 |
| 3162 | 3162 | 1.10100287 | 0.94957335 | 33 | 0.72913058 | 47 | 0.40164602 | 45 | 0.81715991 | 56 | 0.89160589 | 28 | 0.13720052 | 1186 | 0.2438351 | FALSE | 1 | FALSE | 1 | FALSE | 1 | FALSE | 1 | FALSE | 1 | FALSE | 1 |
| 3163 | 3163 | 12.2666667 | 4.56751817 | 0 | 1 | 0 | 1 | 0 | 1 | 0 | 1 | 0 | 1 | 15 | 0.07803154 | FALSE | 1 | FALSE | 1 | FALSE | 1 | FALSE | 1 | FALSE | 1 | FALSE | 1 |
| 3164 | 3164 | 10.1428571 | 2.09956264 | 0 | 1 | 0 | 1 | 0 | 1 | 0 | 1 | 0 | 1 | 7 | 0.3043657 | FALSE | 1 | FALSE | 1 | FALSE | 1 | FALSE | 1 | FALSE | 1 | FALSE | 1 |
| 3165 | 3165 | 9.54 | 1.66385096 | 1 | 0.72839502 | 2 | 0.48458833 | 3 | 0.26704679 | 2 | 0.6762159 | 0 | 1 | 42 | 0.62187327 | FALSE | 1 | FALSE | 1 | FALSE | 1 | FALSE | 1 | FALSE | 1 | FALSE | 1 |
| 3166 | 3166 | 8.41666667 | 2.28977194 | 0 | 1 | 0 | 1 | 0 | 1 | 0 | 1 | 1 | 0.18007762 | 11 | 0.41944532 | FALSE | 1 | FALSE | 1 | FALSE | 1 | FALSE | 1 | FALSE | 1 | FALSE | 1 |
| 3167 | 3167 | 5.05882353 | 2.88954685 | 0 | 1 | 0 | 1 | 3 | 0.12152681 | 0 | 1 | 2 | 0.1065437 | 29 | 0.55707911 | FALSE | 1 | FALSE | 1 | FALSE | 1 | FALSE | 1 | FALSE | 1 | FALSE | 1 |
| 3168 | 3168 | 5.54651163 | 2.17628325 | 2 | 0.65227007 | 2 | 0.77275903 | 1 | 0.95785384 | 5 | 0.36062409 | 0 | 1 | 76 | 0.19221718 | FALSE | 1 | FALSE | 1 | FALSE | 1 | FALSE | 1 | FALSE | 1 | FALSE | 1 |
| 3169 | 3169 | 7.84375 | 2.10815937 | 0 | 1 | 0 | 1 | 0 | 1 | 3 | 0.17988885 | 1 | 0.41154418 | 28 | 0.42532177 | FALSE | 1 | FALSE | 1 | FALSE | 1 | FALSE | 1 | FALSE | 1 | FALSE | 1 |
| 3170 | 3170 | 9.88235294 | 2.7628279 | 3 | 0.05546921 | 0 | 1 | 1 | 0.71261921 | 2 | 0.46709861 | 1 | 0.43077488 | 27 | 0.85023803 | FALSE | 0.68849059 | FALSE | 1 | FALSE | 1 | FALSE | 1 | FALSE | 1 | FALSE | 1 |
| 3171 | 3171 | 7.51111111 | 3.97001104 | 4 | 0.02747663 | 0 | 1 | 2 | 0.48463745 | 0 | 1 | 0 | 1 | 39 | 0.4317533 | FALSE | 0.50964275 | FALSE | 1 | FALSE | 1 | FALSE | 1 | FALSE | 1 | FALSE | 1 |
| 3172 | 3172 | 9.84210526 | 2.23048633 | 0 | 1 | 0 | 1 | 0 | 1 | 1 | 0.59104357 | 0 | 1 | 18 | 0.17880694 | FALSE | 1 | FALSE | 1 | FALSE | 1 | FALSE | 1 | FALSE | 1 | FALSE | 1 |
| 3173 | 3173 | 6.66666667 | 2.8528738 | 2 | 0.12501436 | 1 | 0.54637326 | 1 | 0.58502114 | 0 | 1 | 1 | 0.32796862 | 19 | 0.83975208 | FALSE | 0.95622564 | FALSE | 1 | FALSE | 1 | FALSE | 1 | FALSE | 1 | FALSE | 1 |
| 3174 | 3174 | 9.5 | 3.40342964 | 2 | 0.17884234 | 3 | 0.07145121 | 1 | 0.66709967 | 1 | 0.75658797 | 0 | 1 | 23 | 0.9158696 | FALSE | 1 | FALSE | 1 | FALSE | 1 | FALSE | 1 | FALSE | 1 | FALSE | 1 |
| 3175 | 3175 | 7.19230769 | 2.03846154 | 2 | 0.14250978 | 2 | 0.20486827 | 1 | 0.61440466 | 0 | 1 | 1 | 0.3499045 | 20 | 0.90142689 | FALSE | 1 | FALSE | 1 | FALSE | 1 | FALSE | 1 | FALSE | 1 | FALSE | 1 |
| 3176 | 3176 | 5.34285714 | 3.12997359 | 3 | 0.26686779 | 2 | 0.66721401 | 3 | 0.46296673 | 2 | 0.83868849 | 5 | 0.00561855 | 55 | 0.9296662 | FALSE | 1 | FALSE | 1 | FALSE | 1 | FALSE | 1 | FALSE | 0.45571433 | FALSE | 1 |
| 3177 | 3177 | 9.08333333 | 2.43099202 | 0 | 1 | 0 | 1 | 1 | 0.35555293 | 0 | 1 | 0 | 1 | 11 | 0.41944532 | FALSE | 1 | FALSE | 1 | FALSE | 1 | FALSE | 1 | FALSE | 1 | FALSE | 1 |
| 3178 | 3178 | 8.875 | 0.78062475 | 0 | 1 | 2 | 0.09285443 | 2 | 0.11087977 | 2 | 0.16544789 | 0 | 1 | 10 | 0.99304605 | FALSE | 1 | FALSE | 1 | FALSE | 1 | FALSE | 1 | FALSE | 1 | FALSE | 1 |
| 3179 | 3179 | 10.4285714 | 3.11022015 | 1 | 0.16630142 | 0 | 1 | 1 | 0.22601089 | 0 | 1 | 1 | 0.10932414 | 4 | 0.98605104 | FALSE | 1 | FALSE | 1 | FALSE | 1 | FALSE | 1 | FALSE | 1 | FALSE | 1 |
| 3180 | 3180 | 8.68421053 | 2.29536447 | 2 | 0.08405377 | 0 | 1 | 0 | 1 | 1 | 0.59104357 | 1 | 0.26986775 | 15 | 0.83648191 | FALSE | 0.79236883 | FALSE | 1 | FALSE | 1 | FALSE | 1 | FALSE | 1 | FALSE | 1 |
| 3181 | 3181 | 10.0714286 | 2.60396088 | 0 | 1 | 2 | 0.07341559 | 0 | 1 | 1 | 0.48242099 | 0 | 1 | 11 | 0.83676449 | FALSE | 1 | FALSE | 1 | FALSE | 1 | FALSE | 1 | FALSE | 1 | FALSE | 1 |
| 3182 | 3182 | 7.04347826 | 3.27677281 | 0 | 1 | 1 | 0.53115654 | 1 | 0.5695037 | 1 | 0.6613267 | 1 | 0.31672722 | 19 | 0.71542496 | FALSE | 1 | FALSE | 1 | FALSE | 1 | FALSE | 1 | FALSE | 1 | FALSE | 1 |
| 3183 | 3183 | 8.44444444 | 1.3146844 | 2 | 0.15144274 | 1 | 0.58913618 | 0 | 1 | 1 | 0.71956339 | 0 | 1 | 23 | 0.58307573 | FALSE | 1 | FALSE | 1 | FALSE | 1 | FALSE | 1 | FALSE | 1 | FALSE | 1 |
| 3184 | 3184 | 9.09090909 | 2.53895274 | 0 | 1 | 0 | 1 | 1 | 0.33149969 | 0 | 1 | 1 | 0.16638736 | 9 | 0.75985366 | FALSE | 1 | FALSE | 1 | FALSE | 1 | FALSE | 1 | FALSE | 1 | FALSE | 1 |
| 3185 | 3185 | 11.2631579 | 3.30616816 | 3 | 0.01186104 | 1 | 0.46503795 | 1 | 0.50143826 | 1 | 0.59104357 | 0 | 1 | 13 | 0.97998676 | FALSE | 0.3373808 | FALSE | 1 | FALSE | 1 | FALSE | 1 | FALSE | 1 | FALSE | 1 |
| 3186 | 3186 | 6.41481482 | 1.92982511 | 2 | 0.86327384 | 3 | 0.81446162 | 6 | 0.35033018 | 2 | 0.98691252 | 2 | 0.6500531 | 119 | 0.09204986 | FALSE | 1 | FALSE | 1 | FALSE | 1 | FALSE | 1 | FALSE | 1 | FALSE | 1 |
| 3187 | 3187 | 10.8 | 1.83303028 | 0 | 1 | 0 | 1 | 0 | 1 | 0 | 1 | 0 | 1 | 15 | 0.07803154 | FALSE | 1 | FALSE | 1 | FALSE | 1 | FALSE | 1 | FALSE | 1 | FALSE | 1 |
| 3188 | 3188 | 8.5 | 1.87082869 | 0 | 1 | 0 | 1 | 0 | 1 | 1 | 0.52895088 | 1 | 0.23264612 | 14 | 0.53308261 | FALSE | 1 | FALSE | 1 | FALSE | 1 | FALSE | 1 | FALSE | 1 | FALSE | 1 |
| 3189 | 3189 | 8.85294118 | 3.04024334 | 1 | 0.58738519 | 3 | 0.09602744 | 4 | 0.03235504 | 1 | 0.7984848 | 1 | 0.43077488 | 24 | 0.98848472 | FALSE | 1 | FALSE | 1 | FALSE | 1 | FALSE | 1 | FALSE | 1 | FALSE | 1 |
| 3190 | 3190 | 7.78947368 | 1.81356334 | 5 | 0.01390429 | 4 | 0.10650784 | 2 | 0.60323163 | 4 | 0.25497934 | 0 | 1 | 41 | 0.99027378 | FALSE | 0.36625055 | FALSE | 1 | FALSE | 1 | FALSE | 1 | FALSE | 1 | FALSE | 1 |
| 3191 | 3191 | 11.6857143 | 2.08081617 | 1 | 0.58738519 | 2 | 0.30150529 | 1 | 0.71261921 | 2 | 0.46709861 | 1 | 0.43077488 | 27 | 0.85023803 | FALSE | 1 | FALSE | 1 | FALSE | 1 | FALSE | 1 | FALSE | 1 | FALSE | 1 |
| 3192 | 3192 | 7.40909091 | 4.56925587 | 1 | 0.4357638 | 0 | 1 | 0 | 1 | 1 | 0.64497439 | 0 | 1 | 20 | 0.30915202 | FALSE | 1 | FALSE | 1 | FALSE | 1 | FALSE | 1 | FALSE | 1 | FALSE | 1 |
| 3193 | 3193 | 9.2 | 2.0880613 | 1 | 0.40558521 | 1 | 0.48239024 | 1 | 0.51939841 | 1 | 0.60987146 | 0 | 1 | 16 | 0.80826361 | FALSE | 1 | FALSE | 1 | FALSE | 1 | FALSE | 1 | FALSE | 1 | FALSE | 1 |
| 3194 | 3194 | 8.37209302 | 2.64155747 | 1 | 0.67382914 | 1 | 0.75788114 | 2 | 0.46083448 | 2 | 0.59404271 | 0 | 1 | 37 | 0.48294184 | FALSE | 1 | FALSE | 1 | FALSE | 1 | FALSE | 1 | FALSE | 1 | FALSE | 1 |

| | | | | | | | | | | | | | | | | | | | | | | | | | | | |
|---|---|---|---|---|---|---|---|---|---|---|---|---|---|---|---|---|---|---|---|---|---|---|---|---|---|---|---|
| 3195 | 3195 | 10.4736842 | 2.81665014 | 2 | 0.08405377 | 2 | 0.12434464 | 1 | 0.50143826 | 0 | 1 | 1 | 0.26986775 | 13 | 0.97998676 | FALSE | 0.79236883 | FALSE | 1 | FALSE | 1 | FALSE | 1 | FALSE | 1 | FALSE | 1 |
| 3196 | 3196 | 11.25 | 2.81735692 | 0 | 1 | 0 | 1 | 0 | 1 | 1 | 0.52895088 | 0 | 1 | 15 | 0.26127859 | FALSE | 1 | FALSE | 1 | FALSE | 1 | FALSE | 1 | FALSE | 1 | FALSE | 1 |
| 3197 | 3197 | 9.27777778 | 3.15885977 | 0 | 1 | 0 | 1 | 0 | 1 | 0 | 1 | 1 | 0.25766374 | 17 | 0.20329205 | FALSE | 1 | FALSE | 1 | FALSE | 1 | FALSE | 1 | FALSE | 1 | FALSE | 1 |
| 3198 | 3198 | 9.53125 | 2.30467426 | 0 | 1 | 0 | 1 | 1 | 0.67911179 | 1 | 0.76781284 | 0 | 1 | 29 | 0.11661149 | FALSE | 1 | FALSE | 1 | FALSE | 1 | FALSE | 1 | FALSE | 1 | FALSE | 1 |
| 3199 | 3199 | 8.22727273 | 2.48507944 | 0 | 1 | 0 | 1 | 1 | 0.55340851 | 2 | 0.26791271 | 0 | 1 | 19 | 0.54233419 | FALSE | 1 | FALSE | 1 | FALSE | 1 | FALSE | 1 | FALSE | 1 | FALSE | 1 |
| 3200 | 3200 | 11.2608696 | 2.48853705 | 0 | 1 | 0 | 1 | 1 | 0.5695037 | 0 | 1 | 1 | 0.31672722 | 21 | 0.27964072 | FALSE | 1 | FALSE | 1 | FALSE | 1 | FALSE | 1 | FALSE | 1 | FALSE | 1 |
| 3201 | 3201 | 8.03389831 | 2.68053942 | 2 | 0.4493251 | 1 | 0.85750546 | 2 | 0.63179508 | 2 | 0.76144533 | 0 | 1 | 52 | 0.27701375 | FALSE | 1 | FALSE | 1 | FALSE | 1 | FALSE | 1 | FALSE | 1 | FALSE | 1 |
| 3202 | 3202 | 8.72222222 | 3.03325193 | 0 | 1 | 0 | 1 | 1 | 0.48280983 | 0 | 1 | 0 | 1 | 17 | 0.20329205 | FALSE | 1 | FALSE | 1 | FALSE | 1 | FALSE | 1 | FALSE | 1 | FALSE | 1 |
| 3203 | 3203 | 5.51282051 | 1.67237681 | 2 | 0.80338728 | 2 | 0.89447151 | 6 | 0.23770724 | 5 | 0.62158717 | 0 | 1 | 101 | 0.25462837 | FALSE | 1 | FALSE | 1 | FALSE | 1 | FALSE | 1 | FALSE | 1 | FALSE | 1 |
| 3204 | 3204 | 8.57142857 | 3.51962428 | 0 | 1 | 0 | 1 | 1 | 0.40110418 | 1 | 0.48242099 | 0 | 1 | 12 | 0.62236227 | FALSE | 1 | FALSE | 1 | FALSE | 1 | FALSE | 1 | FALSE | 1 | FALSE | 1 |
| 3205 | 3205 | 7.27586207 | 1.79840066 | 0 | 1 | 1 | 0.60247911 | 1 | 0.64171563 | 3 | 0.13524571 | 1 | 0.37113067 | 22 | 0.86499992 | FALSE | 1 | FALSE | 1 | FALSE | 1 | FALSE | 1 | FALSE | 1 | FALSE | 1 |
| 3206 | 3206 | 9.10869565 | 1.30597724 | 2 | 0.3309085 | 1 | 0.78076835 | 1 | 0.81522035 | 2 | 0.63103861 | 1 | 0.53375569 | 39 | 0.56897353 | FALSE | 1 | FALSE | 1 | FALSE | 1 | FALSE | 1 | FALSE | 1 | FALSE | 1 |
| 3207 | 3207 | 7.8 | 2.22710575 | 1 | 0.32293355 | 0 | 1 | 0 | 1 | 0 | 1 | 0 | 1 | 14 | 0.29525409 | FALSE | 1 | FALSE | 1 | FALSE | 1 | FALSE | 1 | FALSE | 1 | FALSE | 1 |
| 3208 | 3208 | 10 | 2.89604848 | 0 | 1 | 0 | 1 | 1 | 0.67911179 | 1 | 0.76781284 | 0 | 1 | 29 | 0.11661149 | FALSE | 1 | FALSE | 1 | FALSE | 1 | FALSE | 1 | FALSE | 1 | FALSE | 1 |
| 3209 | 3209 | 8.93333333 | 2.32283352 | 0 | 1 | 0 | 1 | 0 | 1 | 1 | 0.50623194 | 1 | 0.2198258 | 13 | 0.57712805 | FALSE | 1 | FALSE | 1 | FALSE | 1 | FALSE | 1 | FALSE | 1 | FALSE | 1 |
| 3210 | 3210 | 9 | 2.04124145 | 0 | 1 | 0 | 1 | 0 | 1 | 0 | 1 | 0 | 1 | 12 | 0.13002667 | FALSE | 1 | FALSE | 1 | FALSE | 1 | FALSE | 1 | FALSE | 1 | FALSE | 1 |
| 3211 | 3211 | 9.25 | 1.83143842 | 0 | 1 | 0 | 1 | 0 | 1 | 0 | 1 | 0 | 1 | 12 | 0.13002667 | FALSE | 1 | FALSE | 1 | FALSE | 1 | FALSE | 1 | FALSE | 1 | FALSE | 1 |
| 3212 | 3212 | 9.83333333 | 2.56038192 | 2 | 0.11647716 | 3 | 0.03669128 | 1 | 0.5695037 | 2 | 0.2852236 | 0 | 1 | 15 | 0.99450713 | FALSE | 0.93181727 | FALSE | 0.94259695 | FALSE | 1 | FALSE | 1 | FALSE | 1 | FALSE | 1 |
| 3213 | 3213 | 11.6842105 | 2.27231928 | 0 | 1 | 0 | 1 | 0 | 1 | 1 | 0.59104357 | 0 | 1 | 18 | 0.17880694 | FALSE | 1 | FALSE | 1 | FALSE | 1 | FALSE | 1 | FALSE | 1 | FALSE | 1 |
| 3214 | 3214 | 11.05 | 1.49916644 | 0 | 1 | 1 | 0.48239024 | 0 | 1 | 0 | 1 | 0 | 1 | 19 | 0.15700255 | FALSE | 1 | FALSE | 1 | FALSE | 1 | FALSE | 1 | FALSE | 1 | FALSE | 1 |
| 3215 | 3215 | 9.0625 | 2.07571524 | 0 | 1 | 0 | 1 | 0 | 1 | 0 | 1 | 0 | 1 | 16 | 0.06581507 | FALSE | 1 | FALSE | 1 | FALSE | 1 | FALSE | 1 | FALSE | 1 | FALSE | 1 |
| 3216 | 3216 | 10.8333333 | 3.69308663 | 0 | 1 | 0 | 1 | 0 | 1 | 0 | 1 | 0 | 1 | 12 | 0.13002667 | FALSE | 1 | FALSE | 1 | FALSE | 1 | FALSE | 1 | FALSE | 1 | FALSE | 1 |
| 3217 | 3217 | 10.6111111 | 2.31273981 | 0 | 1 | 0 | 1 | 1 | 0.48280983 | 0 | 1 | 1 | 0.25766374 | 16 | 0.44998001 | FALSE | 1 | FALSE | 1 | FALSE | 1 | FALSE | 1 | FALSE | 1 | FALSE | 1 |
| 3218 | 3218 | 8.8 | 2.47039615 | 1 | 0.59801736 | 3 | 0.10264583 | 2 | 0.35984046 | 0 | 1 | 2 | 0.11183826 | 27 | 0.91585095 | FALSE | 1 | FALSE | 1 | FALSE | 1 | FALSE | 1 | FALSE | 1 | FALSE | 1 |
| 3219 | 3219 | 1.12361879 | 0.74740446 | 29 | 0.94666526 | 49 | 0.36820059 | 65 | 0.02266382 | 61 | 0.7859116 | 17 | 0.95647223 | 1219 | 0.39098138 | FALSE | 1 | FALSE | 1 | FALSE | 1 | FALSE | 1 | FALSE | 1 | FALSE | 1 |
| 3220 | 3220 | 10.6666667 | 2.63874269 | 0 | 1 | 1 | 0.58913618 | 1 | 0.62830986 | 1 | 0.71956339 | 1 | 0.36060486 | 23 | 0.58307573 | FALSE | 1 | FALSE | 1 | FALSE | 1 | FALSE | 1 | FALSE | 1 | FALSE | 1 |
| 3221 | 3221 | 10.6666667 | 2.39211668 | 0 | 1 | 1 | 0.32623933 | 1 | 0.35555293 | 0 | 1 | 0 | 1 | 10 | 0.71436964 | FALSE | 1 | FALSE | 1 | FALSE | 1 | FALSE | 1 | FALSE | 1 | FALSE | 1 |
| 3222 | 3222 | 13.962963 | 4.22059748 | 1 | 0.50471032 | 1 | 0.58913618 | 1 | 0.62830986 | 5 | 0.00692325 | 0 | 1 | 19 | 0.98241223 | FALSE | 1 | FALSE | 1 | FALSE | 1 | FALSE | 1 | FALSE | 1 | FALSE | 1 |
| 3223 | 3223 | 12 | 1 | 1 | 0.18769745 | 0 | 1 | 1 | 0.25384309 | 0 | 1 | 0 | 1 | 6 | 0.88425551 | FALSE | 1 | FALSE | 1 | FALSE | 1 | FALSE | 1 | FALSE | 1 | FALSE | 1 |
| 3224 | 3224 | 9.23529412 | 2.31439007 | 0 | 1 | 2 | 0.10307112 | 0 | 1 | 2 | 0.18212134 | 0 | 1 | 13 | 0.88719222 | FALSE | 1 | FALSE | 1 | FALSE | 1 | FALSE | 1 | FALSE | 1 | FALSE | 1 |
| 3225 | 3225 | 9.72222222 | 3.57157848 | 0 | 1 | 2 | 0.1135776 | 0 | 1 | 2 | 0.19902682 | 1 | 0.25766374 | 13 | 0.95041913 | FALSE | 1 | FALSE | 1 | FALSE | 1 | FALSE | 1 | FALSE | 1 | FALSE | 1 |
| 3226 | 3226 | 10 | 2.21564684 | 0 | 1 | 0 | 1 | 0 | 1 | 1 | 0.40390032 | 0 | 1 | 10 | 0.46859937 | FALSE | 1 | FALSE | 1 | FALSE | 1 | FALSE | 1 | FALSE | 1 | FALSE | 1 |
| 3227 | 3227 | 8.73333333 | 2.79205221 | 0 | 1 | 0 | 1 | 0 | 1 | 0 | 1 | 0 | 1 | 15 | 0.07803154 | FALSE | 1 | FALSE | 1 | FALSE | 1 | FALSE | 1 | FALSE | 1 | FALSE | 1 |
| 3228 | 3228 | 9.4 | 2.65329983 | 1 | 0.22886536 | 0 | 1 | 1 | 0.30655256 | 0 | 1 | 0 | 1 | 8 | 0.80394719 | FALSE | 1 | FALSE | 1 | FALSE | 1 | FALSE | 1 | FALSE | 1 | FALSE | 1 |
| 3229 | 3229 | 7.4 | 1.56204994 | 1 | 0.79096998 | 2 | 0.58316597 | 8 | 0.00125404 | 2 | 0.76960882 | 0 | 1 | 47 | 0.924754 | FALSE | 1 | FALSE | 1 | FALSE | 1 | FALSE | 1 | FALSE | 1 | FALSE | 1 |
| 3230 | 3230 | 8.38356164 | 1.69309749 | 0 | 1 | 1 | 0.9104874 | 2 | 0.74419436 | 2 | 0.8554296 | 2 | 0.33719417 | 66 | 0.09785941 | FALSE | 1 | FALSE | 1 | FALSE | 1 | FALSE | 1 | FALSE | 1 | FALSE | 1 |
| 3231 | 3231 | 9.42105263 | 2.75901946 | 1 | 0.38989951 | 1 | 0.46503795 | 1 | 0.50143826 | 0 | 1 | 0 | 1 | 16 | 0.65565778 | FALSE | 1 | FALSE | 1 | FALSE | 1 | FALSE | 1 | FALSE | 1 | FALSE | 1 |
| 3232 | 3232 | 9.71875 | 2.64851816 | 0 | 1 | 2 | 0.27730767 | 1 | 0.69069222 | 4 | 0.05697506 | 0 | 1 | 25 | 0.88559288 | FALSE | 1 | FALSE | 1 | FALSE | 1 | FALSE | 1 | FALSE | 1 | FALSE | 1 |
| 3233 | 3233 | 10.5294118 | 2.11764706 | 0 | 1 | 0 | 1 | 1 | 0.46348836 | 1 | 0.55062772 | 0 | 1 | 15 | 0.49060435 | FALSE | 1 | FALSE | 1 | FALSE | 1 | FALSE | 1 | FALSE | 1 | FALSE | 1 |
| 3234 | 3234 | 8.1 | 2.91376046 | 1 | 0.54198714 | 0 | 1 | 1 | 0.66709967 | 4 | 0.04659941 | 0 | 1 | 24 | 0.82294782 | FALSE | 1 | FALSE | 1 | FALSE | 1 | FALSE | 1 | FALSE | 1 | FALSE | 1 |
| 3235 | 3235 | 8.28125 | 2.69530118 | 1 | 0.56527493 | 1 | 0.65167362 | 2 | 0.32011871 | 3 | 0.17988885 | 0 | 1 | 25 | 0.88559288 | FALSE | 1 | FALSE | 1 | FALSE | 1 | FALSE | 1 | FALSE | 1 | FALSE | 1 |
| 3236 | 3236 | 10.3823529 | 2.63477542 | 1 | 0.58738519 | 1 | 0.6739483 | 1 | 0.71261921 | 2 | 0.46709861 | 1 | 0.43077488 | 28 | 0.72558119 | FALSE | 1 | FALSE | 1 | FALSE | 1 | FALSE | 1 | FALSE | 1 | FALSE | 1 |
| 3237 | 3237 | 9.15384615 | 1.26163227 | 1 | 0.47819878 | 0 | 1 | 2 | 0.22597287 | 2 | 0.31968852 | 0 | 1 | 20 | 0.8152498 | FALSE | 1 | FALSE | 1 | FALSE | 1 | FALSE | 1 | FALSE | 1 | FALSE | 1 |
| 3238 | 3238 | 9.125 | 2.52178013 | 0 | 1 | 0 | 1 | 1 | 0.42266264 | 0 | 1 | 1 | 0.2198258 | 13 | 0.57712805 | FALSE | 1 | FALSE | 1 | FALSE | 1 | FALSE | 1 | FALSE | 1 | FALSE | 1 |
| 3239 | 3239 | 8.09090909 | 1.8807419 | 1 | 0.24866461 | 0 | 1 | 0 | 1 | 0 | 1 | 1 | 0.16638736 | 9 | 0.75985366 | FALSE | 1 | FALSE | 1 | FALSE | 1 | FALSE | 1 | FALSE | 1 | FALSE | 1 |
| 3240 | 3240 | 9.875 | 3.02851036 | 2 | 0.19750261 | 2 | 0.27730767 | 2 | 0.32011871 | 0 | 1 | 0 | 1 | 26 | 0.77612337 | FALSE | 1 | FALSE | 1 | FALSE | 1 | FALSE | 1 | FALSE | 1 | FALSE | 1 |
| 3241 | 3241 | 9.11764706 | 1.99653679 | 4 | 0.00076452 | 2 | 0.10307112 | 0 | 1 | 0 | 1 | 1 | 0.24525764 | 10 | 0.99775952 | FALSE | 0.06390799 | FALSE | 1 | FALSE | 1 | FALSE | 1 | FALSE | 1 | FALSE | 1 |
| 3242 | 3242 | 9.5 | 1.84197099 | 1 | 0.3050805 | 0 | 1 | 0 | 1 | 0 | 1 | 0 | 1 | 13 | 0.33283729 | FALSE | 1 | FALSE | 1 | FALSE | 1 | FALSE | 1 | FALSE | 1 | FALSE | 1 |
| 3243 | 3243 | 4.58080808 | 1.85075342 | 1 | 0.99429768 | 5 | 0.7664821 | 5 | 0.83963029 | 6 | 0.89333636 | 2 | 0.83710781 | 177 | 0.01006116 | FALSE | 1 | FALSE | 1 | FALSE | 1 | FALSE | 1 | FALSE | 1 | FALSE | 1 |
| 3244 | 3244 | 9.06666667 | 4.28122516 | 0 | 1 | 0 | 1 | 3 | 0.01512258 | 1 | 0.50623194 | 0 | 1 | 11 | 0.92879412 | FALSE | 1 | FALSE | 1 | FALSE | 1 | FALSE | 1 | FALSE | 1 | FALSE | 1 |
| 3245 | 3245 | 10.3333333 | 1.8408935 | 1 | 0.46441868 | 0 | 1 | 1 | 0.58502114 | 0 | 1 | 2 | 0.058175 | 20 | 0.68270729 | FALSE | 1 | FALSE | 1 | FALSE | 1 | FALSE | 1 | FALSE | 1 | FALSE | 1 |
| 3246 | 3246 | 9.425 | 1.4813423 | 2 | 0.27371785 | 2 | 0.37295654 | 0 | 1 | 3 | 0.27785536 | 0 | 1 | 33 | 0.71942635 | FALSE | 1 | FALSE | 1 | FALSE | 1 | FALSE | 1 | FALSE | 1 | FALSE | 1 |
| 3247 | 3247 | 11.5555556 | 2.2662309 | 0 | 1 | 0 | 1 | 1 | 0.28067846 | 2 | 0.06108592 | 0 | 1 | 6 | 0.96128317 | FALSE | 1 | FALSE | 1 | FALSE | 1 | FALSE | 1 | FALSE | 1 | FALSE | 1 |
| 3248 | 3248 | 10.0512821 | 2.41733821 | 2 | 0.25455533 | 0 | 1 | 1 | 0.75193685 | 1 | 0.83318955 | 0 | 1 | 34 | 0.27051027 | FALSE | 1 | FALSE | 1 | FALSE | 1 | FALSE | 1 | FALSE | 1 | FALSE | 1 |
| 3249 | 3249 | 9.41176471 | 2.89134252 | 0 | 1 | 0 | 1 | 0 | 1 | 2 | 0.18212134 | 1 | 0.24525764 | 14 | 0.7313145 | FALSE | 1 | FALSE | 1 | FALSE | 1 | FALSE | 1 | FALSE | 1 | FALSE | 1 |
| 3250 | 3250 | 10.0769231 | 2.36843143 | 1 | 0.49162634 | 3 | 0.05021553 | 0 | 1 | 1 | 0.70601446 | 1 | 0.3499045 | 20 | 0.90142689 | FALSE | 1 | FALSE | 1 | FALSE | 1 | FALSE | 1 | FALSE | 1 | FALSE | 1 |
| 3251 | 3251 | 7.8 | 2.29148622 | 1 | 0.75539413 | 1 | 0.83180787 | 3 | 0.30658915 | 3 | 0.45415642 | 0 | 1 | 46 | 0.52685615 | FALSE | 1 | FALSE | 1 | FALSE | 1 | FALSE | 1 | FALSE | 1 | FALSE | 1 |
| 3252 | 3252 | 8.39130435 | 1.83538008 | 1 | 0.45027683 | 1 | 0.53115654 | 0 | 1 | 1 | 0.6613267 | 1 | 0.31672722 | 19 | 0.71542496 | FALSE | 1 | FALSE | 1 | FALSE | 1 | FALSE | 1 | FALSE | 1 | FALSE | 1 |
| 3253 | 3253 | 8.85714286 | 2.4159335 | 0 | 1 | 0 | 1 | 1 | 0.78589454 | 2 | 0.5811115 | 1 | 0.50166523 | 38 | 0.19268329 | FALSE | 1 | FALSE | 1 | FALSE | 1 | FALSE | 1 | FALSE | 1 | FALSE | 1 |
| 3254 | 3254 | 9.35483871 | 2.02481998 | 0 | 1 | 2 | 0.2651851 | 1 | 0.67911179 | 2 | 0.41984187 | 0 | 1 | 26 | 0.64660109 | FALSE | 1 | FALSE | 1 | FALSE | 1 | FALSE | 1 | FALSE | 1 | FALSE | 1 |
| 3255 | 3255 | 11.2307692 | 2.00590843 | 0 | 1 | 1 | 0.3480709 | 0 | 1 | 0 | 1 | 1 | 0.19354507 | 11 | 0.6683056 | FALSE | 1 | FALSE | 1 | FALSE | 1 | FALSE | 1 | FALSE | 1 | FALSE | 1 |
| 3256 | 3256 | 10 | 2.84521319 | 1 | 0.42086991 | 2 | 0.14655134 | 1 | 0.53671415 | 1 | 0.62783522 | 1 | 0.29368245 | 15 | 0.96513691 | FALSE | 1 | FALSE | 1 | FALSE | 1 | FALSE | 1 | FALSE | 1 | FALSE | 1 |
| 3257 | 3257 | 9.16363636 | 1.30403062 | 0 | 1 | 1 | 0.83729212 | 0 | 1 | 1 | 0.92538938 | 2 | 0.22768257 | 51 | 0.05446441 | FALSE | 1 | FALSE | 1 | FALSE | 1 | FALSE | 1 | FALSE | 1 | FALSE | 1 |
| 3258 | 3258 | 9.27272727 | 1.13545418 | 0 | 1 | 1 | 0.30368014 | 0 | 1 | 1 | 0.40390032 | 0 | 1 | 9 | 0.75985366 | FALSE | 1 | FALSE | 1 | FALSE | 1 | FALSE | 1 | FALSE | 1 | FALSE | 1 |
| 3259 | 3259 | 11.2857143 | 1.33248272 | 0 | 1 | 0 | 1 | 0 | 1 | 0 | 1 | 0 | 1 | 14 | 0.09251304 | FALSE | 1 | FALSE | 1 | FALSE | 1 | FALSE | 1 | FALSE | 1 | FALSE | 1 |
| 3260 | 3260 | 5.42553192 | 2.50739062 | 0 | 1 | 1 | 0.78790827 | 1 | 0.82190381 | 2 | 0.64277396 | 0 | 1 | 43 | 0.12168195 | FALSE | 1 | FALSE | 1 | FALSE | 1 | FALSE | 1 | FALSE | 1 | FALSE | 1 |
| 3261 | 3261 | 8.4516129 | 1.04228996 | 0 | 1 | 0 | 1 | 3 | 0.09131526 | 1 | 0.75658797 | 1 | 0.39166989 | 25 | 0.67630471 | FALSE | 1 | FALSE | 1 | FALSE | 1 | FALSE | 1 | FALSE | 1 | FALSE | 1 |
| 3262 | 3262 | 7.83333333 | 1.53659074 | 1 | 0.37380237 | 0 | 1 | 0 | 1 | 1 | 0.57131013 | 0 | 1 | 16 | 0.44998001 | FALSE | 1 | FALSE | 1 | FALSE | 1 | FALSE | 1 | FALSE | 1 | FALSE | 1 |
| 3263 | 3263 | 8.475 | 2.12117303 | 2 | 0.27371785 | 0 | 1 | 1 | 0.76953771 | 4 | 0.10945471 | 0 | 1 | 33 | 0.71942635 | FALSE | 1 | FALSE | 1 | FALSE | 1 | FALSE | 1 | FALSE | 1 | FALSE | 1 |
| 3264 | 3264 | 8.58333333 | 1.75396377 | 0 | 1 | 0 | 1 | 0 | 1 | 0 | 1 | 1 | 0.18007762 | 11 | 0.41944532 | FALSE | 1 | FALSE | 1 | FALSE | 1 | FALSE | 1 | FALSE | 1 | FALSE | 1 |
| 3265 | 3265 | 10.0307692 | 2.57185495 | 3 | 0.23231139 | 5 | 0.05834767 | 2 | 0.68408105 | 7 | 0.02856769 | 1 | 0.66032558 | 47 | 0.99614006 | FALSE | 1 | FALSE | 1 | FALSE | 1 | FALSE | 1 | FALSE | 1 | FALSE | 1 |
| 3266 | 3266 | 6.97368421 | 2.99549523 | 0 | 1 | 4 | 0.0332765 | 1 | 0.75193685 | 0 | 1 | 0 | 1 | 33 | 0.44261631 | FALSE | 1 | FALSE | 0.92721448 | FALSE | 1 | FALSE | 1 | FALSE | 1 | FALSE | 1 |
| 3267 | 3267 | 6.375 | 1.1110243 | 0 | 1 | 0 | 1 | 0 | 1 | 2 | 0.16544789 | 0 | 1 | 14 | 0.53308261 | FALSE | 1 | FALSE | 1 | FALSE | 1 | FALSE | 1 | FALSE | 1 | FALSE | 1 |
| 3268 | 3268 | 8 | 1.69030851 | 1 | 0.16630142 | 0 | 1 | 1 | 0.22601089 | 1 | 0.28044368 | 0 | 1 | 4 | 0.98605104 | FALSE | 1 | FALSE | 1 | FALSE | 1 | FALSE | 1 | FALSE | 1 | FALSE | 1 |
| 3269 | 3269 | 8.625 | 2.91279505 | 1 | 0.34033054 | 1 | 0.40942991 | 0 | 1 | 0 | 1 | 0 | 1 | 14 | 0.53308261 | FALSE | 1 | FALSE | 1 | FALSE | 1 | FALSE | 1 | FALSE | 1 | FALSE | 1 |
| 3270 | 3270 | 10.1333333 | 2.89520677 | 1 | 0.32293355 | 0 | 1 | 0 | 1 | 1 | 0.50623194 | 0 | 1 | 13 | 0.57712805 | FALSE | 1 | FALSE | 1 | FALSE | 1 | FALSE | 1 | FALSE | 1 | FALSE | 1 |
| 3271 | 3271 | 10.2333333 | 2.52564623 | 4 | 0.00679223 | 2 | 0.25306331 | 3 | 0.09131526 | 1 | 0.75658797 | 0 | 1 | 20 | 0.99601577 | FALSE | 0.24597546 | FALSE | 1 | FALSE | 1 | FALSE | 1 | FALSE | 1 | FALSE | 1 |
| 3272 | 3272 | 5.03703704 | 3.57440989 | 1 | 0.75539413 | 3 | 0.25367018 | 1 | 0.86240917 | 2 | 0.71679995 | 1 | 0.59191961 | 46 | 0.52685615 | FALSE | 1 | FALSE | 1 | FALSE | 1 | FALSE | 1 | FALSE | 1 | FALSE | 1 |
| 3273 | 3273 | 7.84090909 | 1.79574221 | 3 | 0.10232687 | 3 | 0.16927151 | 2 | 0.47281647 | 1 | 0.87439647 | 5 | 0.00070588 | 30 | 0.99808826 | FALSE | 0.87592659 | FALSE | 1 | FALSE | 1 | FALSE | 1 | FALSE | 0.14103898 | FALSE | 1 |
| 3274 | 3274 | 8.6097561 | 1.22729226 | 2 | 0.62636022 | 3 | 0.49822953 | 3 | 0.56976011 | 5 | 0.3237921 | 0 | 1 | 69 | 0.59640926 | FALSE | 1 | FALSE | 1 | FALSE | 1 | FALSE | 1 | FALSE | 1 | FALSE | 1 |
| 3275 | 3275 | 8.76923077 | 3.0167187 | 1 | 0.28675951 | 0 | 1 | 1 | 0.37874417 | 0 | 1 | 0 | 1 | 11 | 0.6683056 | FALSE | 1 | FALSE | 1 | FALSE | 1 | FALSE | 1 | FALSE | 1 | FALSE | 1 |
| 3276 | 3276 | 6.55263158 | 4.34497513 | 1 | 0.62830697 | 0 | 1 | 1 | 0.75193685 | 3 | 0.25266879 | 0 | 1 | 33 | 0.44261631 | FALSE | 1 | FALSE | 1 | FALSE | 1 | FALSE | 1 | FALSE | 1 | FALSE | 1 |
| 3277 | 3277 | 10.7272727 | 2.97232413 | 1 | 0.4357638 | 3 | 0.03266339 | 1 | 0.55340851 | 0 | 1 | 1 | 0.30529951 | 16 | 0.95553951 | FALSE | 1 | FALSE | 0.92268438 | FALSE | 1 | FALSE | 1 | FALSE | 1 | FALSE | 1 |
| 3278 | 3278 | 8.74074074 | 3.15750244 | 1 | 0.50471032 | 1 | 0.58913618 | 0 | 1 | 0 | 1 | 1 | 0.36060486 | 24 | 0.3725373 | FALSE | 1 | FALSE | 1 | FALSE | 1 | FALSE | 1 | FALSE | 1 | FALSE | 1 |
| 3279 | 3279 | 7.94736842 | 1.90494327 | 0 | 1 | 2 | 0.3494045 | 1 | 0.75193685 | 1 | 0.83318955 | 2 | 0.12811285 | 32 | 0.61800916 | FALSE | 1 | FALSE | 1 | FALSE | 1 | FALSE | 1 | FALSE | 1 | FALSE | 1 |
| 3280 | 3280 | 8.19047619 | 2.57495349 | 1 | 0.42086991 | 1 | 0.49918218 | 0 | 1 | 1 | 0.62783522 | 1 | 0.29368245 | 17 | 0.77849076 | FALSE | 1 | FALSE | 1 | FALSE | 1 | FALSE | 1 | FALSE | 1 | FALSE | 1 |
| 3281 | 3281 | 8.72222222 | 0.73071925 | 1 | 0.37380237 | 1 | 0.44710673 | 0 | 1 | 1 | 0.57131013 | 0 | 1 | 15 | 0.69372602 | FALSE | 1 | FALSE | 1 | FALSE | 1 | FALSE | 1 | FALSE | 1 | FALSE | 1 |
| 3282 | 3282 | 9 | 4.8882243 | 2 | 0.07645454 | 1 | 0.44710673 | 1 | 0.48280983 | 0 | 1 | 1 | 0.25766374 | 13 | 0.95041913 | FALSE | 0.77424195 | FALSE | 1 | FALSE | 1 | FALSE | 1 | FALSE | 1 | FALSE | 1 |
| 3283 | 3283 | 10.5714286 | 1.69513108 | 0 | 1 | 1 | 0.68454804 | 2 | 0.35984046 | 2 | 0.48233989 | 3 | 0.01918344 | 27 | 0.91585095 | FALSE | 1 | FALSE | 1 | FALSE | 1 | FALSE | 1 | FALSE | 0.70156566 | FALSE | 1 |
| 3284 | 3284 | 6.23728814 | 2.88384738 | 2 | 0.44060625 | 0 | 1 | 0 | 1 | 1 | 0.93527934 | 1 | 0.61824432 | 54 | 0.03955468 | FALSE | 1 | FALSE | 1 | FALSE | 1 | FALSE | 1 | FALSE | 1 | FALSE | 1 |
| 3285 | 3285 | 10.75 | 1.08972474 | 0 | 1 | 0 | 1 | 0 | 1 | 0 | 1 | 0 | 1 | 4 | 0.50683841 | FALSE | 1 | FALSE | 1 | FALSE | 1 | FALSE | 1 | FALSE | 1 | FALSE | 1 |
| 3286 | 3286 | 7.67741936 | 2.63214737 | 1 | 0.50471032 | 4 | 0.01041264 | 0 | 1 | 0 | 1 | 1 | 0.36060486 | 21 | 0.88395936 | FALSE | 1 | FALSE | 0.67617265 | FALSE | 1 | FALSE | 1 | FALSE | 1 | FALSE | 1 |
| 3287 | 3287 | 10.9 | 4.20594817 | 2 | 0.0256865 | 1 | 0.28036922 | 0 | 1 | 0 | 1 | 0 | 1 | 7 | 0.94326215 | FALSE | 0.4974559 | FALSE | 1 | FALSE | 1 | FALSE | 1 | FALSE | 1 | FALSE | 1 |
| 3288 | 3288 | 8.53333333 | 1.85711844 | 0 | 1 | 2 | 0.08295839 | 0 | 1 | 0 | 1 | 0 | 1 | 13 | 0.57712805 | FALSE | 1 | FALSE | 1 | FALSE | 1 | FALSE | 1 | FALSE | 1 | FALSE | 1 |

| | | | | | | | | | | | | | | | | | | | | | | | | | | | |
|---|---|---|---|---|---|---|---|---|---|---|---|---|---|---|---|---|---|---|---|---|---|---|---|---|---|---|---|
| 3289 | 3289 | 7.68181818 | 2.63518783 | 0 | 1 | 1 | 0.5154318 | 3 | 0.04259419 | 3 | 0.07743186 | 0 | 1 | 15 | 0.98570275 | FALSE | 1 | FALSE | 1 | FALSE | 1 | FALSE | 1 | FALSE | 1 | FALSE | 1 |
| 3290 | 3290 | 9.6 | 2.45764115 | 0 | 1 | 1 | 0.48239024 | 0 | 1 | 1 | 0.60987146 | 1 | 0.28187291 | 17 | 0.61750632 | FALSE | 1 | FALSE | 1 | FALSE | 1 | FALSE | 1 | FALSE | 1 | FALSE | 1 |
| 3291 | 3291 | 3.47848101 | 1.87357851 | 12 | 0.30703813 | 15 | 0.28953195 | 12 | 0.76215174 | 14 | 0.87294026 | 7 | 0.4671943 | 333 | 0.45498615 | FALSE | 1 | FALSE | 1 | FALSE | 1 | FALSE | 1 | FALSE | 1 | FALSE | 1 |
| 3292 | 3292 | 8.36 | 2.9241067 | 0 | 1 | 0 | 1 | 2 | 0.22597287 | 0 | 1 | 0 | 1 | 23 | 0.22727183 | FALSE | 1 | FALSE | 1 | FALSE | 1 | FALSE | 1 | FALSE | 1 | FALSE | 1 |
| 3293 | 3293 | 9 | 2.52487624 | 0 | 1 | 0 | 1 | 0 | 1 | 0 | 1 | 0 | 1 | 16 | 0.06581507 | FALSE | 1 | FALSE | 1 | FALSE | 1 | FALSE | 1 | FALSE | 1 | FALSE | 1 |
| 3294 | 3294 | 12.4166667 | 1.75396377 | 0 | 1 | 0 | 1 | 0 | 1 | 0 | 1 | 0 | 1 | 11 | 0.15414489 | FALSE | 1 | FALSE | 1 | FALSE | 1 | FALSE | 1 | FALSE | 1 | FALSE | 1 |
| 3295 | 3295 | 8.15 | 2.26439838 | 1 | 0.40558521 | 1 | 0.48239024 | 1 | 0.51939841 | 0 | 1 | 0 | 1 | 17 | 0.61750632 | FALSE | 1 | FALSE | 1 | FALSE | 1 | FALSE | 1 | FALSE | 1 | FALSE | 1 |
| 3296 | 3296 | 9.2 | 3.35062183 | 0 | 1 | 0 | 1 | 1 | 0.40110418 | 1 | 0.48242099 | 2 | 0.02131828 | 10 | 0.94579623 | FALSE | 1 | FALSE | 1 | FALSE | 1 | FALSE | 1 | FALSE | 0.73999729 | FALSE | 1 |
| 3297 | 3297 | 6.29166667 | 1.898007 | 0 | 1 | 1 | 0.95551644 | 3 | 0.66251153 | 6 | 0.26227462 | 2 | 0.45858138 | 82 | 0.2727315 | FALSE | 1 | FALSE | 1 | FALSE | 1 | FALSE | 1 | FALSE | 1 | FALSE | 1 |
| 3298 | 3298 | 10.4347826 | 1.90710541 | 1 | 0.45027683 | 0 | 1 | 0 | 1 | 0 | 1 | 2 | 0.05389003 | 20 | 0.50589402 | FALSE | 1 | FALSE | 1 | FALSE | 1 | FALSE | 1 | FALSE | 1 | FALSE | 1 |
| 3299 | 3299 | 8.2631579 | 2.9171382 | 0 | 1 | 3 | 0.02205772 | 0 | 1 | 2 | 0.21610812 | 0 | 1 | 14 | 0.93677253 | FALSE | 1 | FALSE | 0.81029974 | FALSE | 1 | FALSE | 1 | FALSE | 1 | FALSE | 1 |
| 3300 | 3300 | 9 | 0.81649658 | 0 | 1 | 0 | 1 | 0 | 1 | 0 | 1 | 0 | 1 | 3 | 0.60071728 | FALSE | 1 | FALSE | 1 | FALSE | 1 | FALSE | 1 | FALSE | 1 | FALSE | 1 |
| 3301 | 3301 | 10 | 1.60356745 | 1 | 0.16630142 | 0 | 1 | 1 | 0.22601089 | 0 | 1 | 0 | 1 | 5 | 0.91847773 | FALSE | 1 | FALSE | 1 | FALSE | 1 | FALSE | 1 | FALSE | 1 | FALSE | 1 |
| 3302 | 3302 | 11.25 | 3.1124749 | 0 | 1 | 1 | 0.40942991 | 1 | 0.44344817 | 0 | 1 | 1 | 0.23264612 | 13 | 0.76799161 | FALSE | 1 | FALSE | 1 | FALSE | 1 | FALSE | 1 | FALSE | 1 | FALSE | 1 |
| 3303 | 3303 | 7.13157895 | 3.43489104 | 1 | 0.62830697 | 4 | 0.0332765 | 2 | 0.39868506 | 1 | 0.83318955 | 0 | 1 | 30 | 0.87271204 | FALSE | 1 | FALSE | 0.92721448 | FALSE | 1 | FALSE | 1 | FALSE | 1 | FALSE | 1 |
| 3304 | 3304 | 7.13636364 | 3.80544412 | 0 | 1 | 0 | 1 | 0 | 1 | 2 | 0.26791271 | 0 | 1 | 20 | 0.30915202 | FALSE | 1 | FALSE | 1 | FALSE | 1 | FALSE | 1 | FALSE | 1 | FALSE | 1 |
| 3305 | 3305 | 8.38709677 | 2.91467277 | 1 | 0.54198714 | 2 | 0.25306331 | 0 | 1 | 0 | 1 | 1 | 0.39166989 | 26 | 0.48609328 | FALSE | 1 | FALSE | 1 | FALSE | 1 | FALSE | 1 | FALSE | 1 | FALSE | 1 |
| 3306 | 3306 | 10.7826087 | 3.09215729 | 3 | 0.02011786 | 0 | 1 | 1 | 0.5695037 | 1 | 0.6613267 | 0 | 1 | 18 | 0.86273155 | FALSE | 0.44482142 | FALSE | 1 | FALSE | 1 | FALSE | 1 | FALSE | 1 | FALSE | 1 |
| 3307 | 3307 | 7.92307692 | 4.09430258 | 3 | 0.0279333 | 1 | 0.57534752 | 0 | 1 | 0 | 1 | 0 | 1 | 22 | 0.61628 | FALSE | 0.50964275 | FALSE | 1 | FALSE | 1 | FALSE | 1 | FALSE | 1 | FALSE | 1 |
| 3308 | 3308 | 10.4210526 | 2.00829306 | 2 | 0.08405377 | 0 | 1 | 2 | 0.14749808 | 2 | 0.21610812 | 0 | 1 | 13 | 0.97998676 | FALSE | 0.79236883 | FALSE | 1 | FALSE | 1 | FALSE | 1 | FALSE | 1 | FALSE | 1 |
| 3309 | 3309 | 11.2222222 | 3.05908887 | 1 | 0.50471032 | 1 | 0.58913618 | 1 | 0.62830986 | 1 | 0.71956339 | 0 | 1 | 23 | 0.58307573 | FALSE | 1 | FALSE | 1 | FALSE | 1 | FALSE | 1 | FALSE | 1 | FALSE | 1 |
| 3310 | 3310 | 9.7755102 | 2.35835732 | 3 | 0.13009168 | 2 | 0.47396556 | 2 | 0.53023996 | 3 | 0.39227316 | 4 | 0.00822471 | 35 | 0.99388317 | FALSE | 0.97964162 | FALSE | 1 | FALSE | 1 | FALSE | 1 | FALSE | 0.50674765 | FALSE | 1 |
| 3311 | 3311 | 9.94736842 | 2.41647164 | 0 | 1 | 0 | 1 | 0 | 1 | 0 | 1 | 0 | 1 | 19 | 0.03948376 | FALSE | 1 | FALSE | 1 | FALSE | 1 | FALSE | 1 | FALSE | 1 | FALSE | 1 |
| 3312 | 3312 | 7.1754386 | 1.83649941 | 2 | 0.43181298 | 2 | 0.55509723 | 2 | 0.61293419 | 2 | 0.74436726 | 2 | 0.23984683 | 47 | 0.72965359 | FALSE | 1 | FALSE | 1 | FALSE | 1 | FALSE | 1 | FALSE | 1 | FALSE | 1 |
| 3313 | 3313 | 8.41176471 | 2.72372633 | 0 | 1 | 2 | 0.10307112 | 0 | 1 | 1 | 0.55062772 | 2 | 0.03086174 | 12 | 0.962093 | FALSE | 1 | FALSE | 1 | FALSE | 1 | FALSE | 1 | FALSE | 0.82559785 | FALSE | 1 |
| 3314 | 3314 | 9.65625 | 2.339997 | 2 | 0.19750261 | 1 | 0.65167362 | 0 | 1 | 0 | 1 | 3 | 0.01506052 | 26 | 0.77612337 | FALSE | 1 | FALSE | 1 | FALSE | 1 | FALSE | 1 | FALSE | 0.66313193 | FALSE | 1 |
| 3315 | 3315 | 8.69565217 | 2.78871612 | 3 | 0.02011786 | 1 | 0.53115654 | 0 | 1 | 2 | 0.2852236 | 1 | 0.31672722 | 16 | 0.98100709 | FALSE | 0.44482142 | FALSE | 1 | FALSE | 1 | FALSE | 1 | FALSE | 1 | FALSE | 1 |
| 3316 | 3316 | 4.91071429 | 2.39997608 | 1 | 0.76787545 | 5 | 0.03395652 | 2 | 0.60323163 | 1 | 0.92884295 | 2 | 0.23375968 | 45 | 0.84580664 | FALSE | 1 | FALSE | 0.93660537 | FALSE | 1 | FALSE | 1 | FALSE | 1 | FALSE | 1 |
| 3317 | 3317 | 9.93939394 | 2.49774922 | 0 | 1 | 3 | 0.08959205 | 1 | 0.70185639 | 2 | 0.45159487 | 0 | 1 | 27 | 0.75124583 | FALSE | 1 | FALSE | 1 | FALSE | 1 | FALSE | 1 | FALSE | 1 | FALSE | 1 |
| 3318 | 3318 | 10.6 | 2.87054002 | 0 | 1 | 0 | 1 | 3 | 0.03322143 | 0 | 1 | 0 | 1 | 17 | 0.61750632 | FALSE | 1 | FALSE | 1 | FALSE | 1 | FALSE | 1 | FALSE | 1 | FALSE | 1 |
| 3319 | 3319 | 9 | 3.047247 | 1 | 0.3050805 | 0 | 1 | 1 | 0.40110418 | 1 | 0.48242099 | 0 | 1 | 11 | 0.83676449 | FALSE | 1 | FALSE | 1 | FALSE | 1 | FALSE | 1 | FALSE | 1 | FALSE | 1 |
| 3320 | 3320 | 10 | 3.59894164 | 0 | 1 | 1 | 0.49918218 | 2 | 0.17308361 | 0 | 1 | 0 | 1 | 18 | 0.57962763 | FALSE | 1 | FALSE | 1 | FALSE | 1 | FALSE | 1 | FALSE | 1 | FALSE | 1 |
| 3321 | 3321 | 9 | 2.03670031 | 1 | 0.50471032 | 2 | 0.21684246 | 4 | 0.01487236 | 2 | 0.35374209 | 0 | 1 | 18 | 0.99450398 | FALSE | 1 | FALSE | 1 | FALSE | 1 | FALSE | 1 | FALSE | 1 | FALSE | 1 |
| 3322 | 3322 | 10.1176471 | 2.13878341 | 1 | 0.35728305 | 0 | 1 | 0 | 1 | 0 | 1 | 1 | 0.24525764 | 15 | 0.49060435 | FALSE | 1 | FALSE | 1 | FALSE | 1 | FALSE | 1 | FALSE | 1 | FALSE | 1 |
| 3323 | 3323 | 11.25 | 3.01731117 | 1 | 0.46441868 | 1 | 0.54637326 | 2 | 0.21260257 | 0 | 1 | 0 | 1 | 20 | 0.68270729 | FALSE | 1 | FALSE | 1 | FALSE | 1 | FALSE | 1 | FALSE | 1 | FALSE | 1 |
| 3324 | 3324 | 7.15942029 | 3.12828697 | 1 | 0.83491628 | 2 | 0.65946213 | 2 | 0.71546205 | 1 | 0.96159801 | 2 | 0.31302643 | 61 | 0.22871622 | FALSE | 1 | FALSE | 1 | FALSE | 1 | FALSE | 1 | FALSE | 1 | FALSE | 1 |
| 3325 | 3325 | 9.33333333 | 1.95991577 | 1 | 0.42086991 | 2 | 0.14655134 | 0 | 1 | 3 | 0.06912908 | 0 | 1 | 15 | 0.96513691 | FALSE | 1 | FALSE | 1 | FALSE | 1 | FALSE | 1 | FALSE | 1 | FALSE | 1 |
| 3326 | 3326 | 9.89285714 | 2.38056545 | 3 | 0.03391597 | 1 | 0.60247911 | 3 | 0.07756639 | 1 | 0.73248983 | 1 | 0.37113067 | 19 | 0.99262381 | FALSE | 0.56934354 | FALSE | 1 | FALSE | 1 | FALSE | 1 | FALSE | 1 | FALSE | 1 |
| 3327 | 3327 | 7.8372093 | 1.5085837 | 5 | 0.23576241 | 5 | 0.40561876 | 4 | 0.68714676 | 7 | 0.38067194 | 0 | 1 | 108 | 0.63841443 | FALSE | 1 | FALSE | 1 | FALSE | 1 | FALSE | 1 | FALSE | 1 | FALSE | 1 |
| 3328 | 3328 | 5.97368421 | 2.72885982 | 5 | 0.04486082 | 2 | 0.71077581 | 2 | 0.76408693 | 3 | 0.68517679 | 0 | 1 | 64 | 0.59240184 | FALSE | 0.649301 | FALSE | 1 | FALSE | 1 | FALSE | 1 | FALSE | 1 | FALSE | 1 |
| 3329 | 3329 | 9.22727273 | 2.79499257 | 0 | 1 | 1 | 0.5154318 | 1 | 0.55340851 | 0 | 1 | 0 | 1 | 20 | 0.30915202 | FALSE | 1 | FALSE | 1 | FALSE | 1 | FALSE | 1 | FALSE | 1 | FALSE | 1 |
| 3330 | 3330 | 4.36448598 | 3.19869285 | 2 | 0.76092701 | 2 | 0.86287617 | 4 | 0.53239439 | 5 | 0.54061691 | 0 | 1 | 93 | 0.20749118 | FALSE | 1 | FALSE | 1 | FALSE | 1 | FALSE | 1 | FALSE | 1 | FALSE | 1 |
| 3331 | 3331 | 10.3 | 2.79463772 | 0 | 1 | 0 | 1 | 0 | 1 | 0 | 1 | 0 | 1 | 20 | 0.03329856 | FALSE | 1 | FALSE | 1 | FALSE | 1 | FALSE | 1 | FALSE | 1 | FALSE | 1 |
| 3332 | 3332 | 8.28571429 | 1.97948664 | 0 | 1 | 0 | 1 | 0 | 1 | 0 | 1 | 0 | 1 | 14 | 0.09251304 | FALSE | 1 | FALSE | 1 | FALSE | 1 | FALSE | 1 | FALSE | 1 | FALSE | 1 |
| 3333 | 3333 | 9.33333333 | 2.031498 | 2 | 0.09187136 | 0 | 1 | 1 | 0.51939841 | 0 | 1 | 0 | 1 | 17 | 0.61750632 | FALSE | 0.83248965 | FALSE | 1 | FALSE | 1 | FALSE | 1 | FALSE | 1 | FALSE | 1 |
| 3334 | 3334 | 10.7142857 | 2.015248 | 0 | 1 | 0 | 1 | 0 | 1 | 0 | 1 | 0 | 1 | 14 | 0.09251304 | FALSE | 1 | FALSE | 1 | FALSE | 1 | FALSE | 1 | FALSE | 1 | FALSE | 1 |
| 3335 | 3335 | 5.55251142 | 1.59929511 | 3 | 0.91055576 | 6 | 0.6798205 | 9 | 0.34173583 | 12 | 0.25715377 | 4 | 0.4533276 | 176 | 0.63531724 | FALSE | 1 | FALSE | 1 | FALSE | 1 | FALSE | 1 | FALSE | 1 | FALSE | 1 |
| 3336 | 3336 | 12.9090909 | 2.10862064 | 1 | 0.24866461 | 0 | 1 | 0 | 1 | 0 | 1 | 0 | 1 | 10 | 0.46859937 | FALSE | 1 | FALSE | 1 | FALSE | 1 | FALSE | 1 | FALSE | 1 | FALSE | 1 |
| 3337 | 3337 | 9.24 | 1.66805276 | 1 | 0.72839502 | 3 | 0.21901899 | 0 | 1 | 3 | 0.4048204 | 0 | 1 | 43 | 0.46945509 | FALSE | 1 | FALSE | 1 | FALSE | 1 | FALSE | 1 | FALSE | 1 | FALSE | 1 |
| 3338 | 3338 | 8.52941177 | 2.061343 | 2 | 0.21638713 | 1 | 0.6739483 | 1 | 0.71261921 | 1 | 0.7984848 | 0 | 1 | 29 | 0.55707911 | FALSE | 1 | FALSE | 1 | FALSE | 1 | FALSE | 1 | FALSE | 1 | FALSE | 1 |
| 3339 | 3339 | 10.3571429 | 1.51691147 | 1 | 0.51745948 | 1 | 0.60247911 | 1 | 0.64171563 | 1 | 0.73248983 | 0 | 1 | 24 | 0.55018107 | FALSE | 1 | FALSE | 1 | FALSE | 1 | FALSE | 1 | FALSE | 1 | FALSE | 1 |
| 3340 | 3340 | 10.1304348 | 2.99716312 | 0 | 1 | 0 | 1 | 1 | 0.55340851 | 0 | 1 | 1 | 0.30529951 | 20 | 0.30915202 | FALSE | 1 | FALSE | 1 | FALSE | 1 | FALSE | 1 | FALSE | 1 | FALSE | 1 |
| 3341 | 3341 | 9.625 | 2.59506744 | 0 | 1 | 0 | 1 | 2 | 0.03118552 | 0 | 1 | 0 | 1 | 6 | 0.88425551 | FALSE | 1 | FALSE | 1 | FALSE | 1 | FALSE | 1 | FALSE | 1 | FALSE | 1 |
| 3342 | 3342 | 4.60465116 | 2.20084059 | 2 | 0.65227007 | 2 | 0.77275903 | 4 | 0.37318291 | 2 | 0.91113247 | 1 | 0.76090478 | 75 | 0.28945549 | FALSE | 1 | FALSE | 1 | FALSE | 1 | FALSE | 1 | FALSE | 1 | FALSE | 1 |
| 3343 | 3343 | 9.26666667 | 2.97694848 | 1 | 0.32293355 | 0 | 1 | 0 | 1 | 3 | 0.02891327 | 0 | 1 | 11 | 0.92879412 | FALSE | 1 | FALSE | 1 | FALSE | 1 | FALSE | 1 | FALSE | 1 | FALSE | 1 |
| 3344 | 3344 | 5.69811321 | 2.31960395 | 1 | 0.74890551 | 1 | 0.82613965 | 2 | 0.57303317 | 3 | 0.44196866 | 2 | 0.21556658 | 44 | 0.69042405 | FALSE | 1 | FALSE | 1 | FALSE | 1 | FALSE | 1 | FALSE | 1 | FALSE | 1 |
| 3345 | 3345 | 9.33333333 | 1.39443338 | 0 | 1 | 0 | 1 | 1 | 0.73299778 | 1 | 0.81665383 | 2 | 0.11720035 | 32 | 0.31709555 | FALSE | 1 | FALSE | 1 | FALSE | 1 | FALSE | 1 | FALSE | 1 | FALSE | 1 |
| 3346 | 3346 | 8.375 | 1.86664807 | 1 | 0.18769745 | 0 | 1 | 0 | 1 | 0 | 1 | 2 | 0.00699343 | 5 | 0.97551477 | FALSE | 1 | FALSE | 1 | FALSE | 1 | FALSE | 1 | FALSE | 0.4774184 | FALSE | 1 |
| 3347 | 3347 | 11.2631579 | 2.57195615 | 1 | 0.38989951 | 1 | 0.46503795 | 1 | 0.50143826 | 0 | 1 | 0 | 1 | 16 | 0.65565778 | FALSE | 1 | FALSE | 1 | FALSE | 1 | FALSE | 1 | FALSE | 1 | FALSE | 1 |
| 3348 | 3348 | 6.3125 | 2.65091376 | 0 | 1 | 2 | 0.27730767 | 0 | 1 | 1 | 0.77852168 | 1 | 0.41154418 | 28 | 0.42532177 | FALSE | 1 | FALSE | 1 | FALSE | 1 | FALSE | 1 | FALSE | 1 | FALSE | 1 |
| 3349 | 3349 | 8.88 | 2.38863978 | 0 | 1 | 0 | 1 | 2 | 0.22597287 | 1 | 0.69181315 | 0 | 1 | 22 | 0.43642846 | FALSE | 1 | FALSE | 1 | FALSE | 1 | FALSE | 1 | FALSE | 1 | FALSE | 1 |
| 3350 | 3350 | 9.72727273 | 3.22182639 | 0 | 1 | 0 | 1 | 1 | 0.33149969 | 0 | 1 | 0 | 1 | 10 | 0.46859937 | FALSE | 1 | FALSE | 1 | FALSE | 1 | FALSE | 1 | FALSE | 1 | FALSE | 1 |
| 3351 | 3351 | 8 | 2.41209076 | 1 | 0.24866461 | 0 | 1 | 0 | 1 | 0 | 1 | 0 | 1 | 10 | 0.46859937 | FALSE | 1 | FALSE | 1 | FALSE | 1 | FALSE | 1 | FALSE | 1 | FALSE | 1 |
| 3352 | 3352 | 9.55172414 | 2.68567345 | 0 | 1 | 3 | 0.06581417 | 0 | 1 | 2 | 0.3871811 | 0 | 1 | 24 | 0.70563334 | FALSE | 1 | FALSE | 1 | FALSE | 1 | FALSE | 1 | FALSE | 1 | FALSE | 1 |
| 3353 | 3353 | 8.64705882 | 2.92938814 | 0 | 1 | 0 | 1 | 0 | 1 | 0 | 1 | 0 | 1 | 17 | 0.05550965 | FALSE | 1 | FALSE | 1 | FALSE | 1 | FALSE | 1 | FALSE | 1 | FALSE | 1 |
| 3354 | 3354 | 9.09090909 | 3.14655787 | 1 | 0.24866461 | 1 | 0.30368014 | 0 | 1 | 0 | 1 | 0 | 1 | 9 | 0.75985366 | FALSE | 1 | FALSE | 1 | FALSE | 1 | FALSE | 1 | FALSE | 1 | FALSE | 1 |
| 3355 | 3355 | 10.4230769 | 2.69148339 | 1 | 0.49162634 | 0 | 1 | 3 | 0.06482014 | 0 | 1 | 0 | 1 | 22 | 0.61628 | FALSE | 1 | FALSE | 1 | FALSE | 1 | FALSE | 1 | FALSE | 1 | FALSE | 1 |
| 3356 | 3356 | 3.8 | 2.33238076 | 3 | 0.30198574 | 1 | 0.91624631 | 3 | 0.50898113 | 5 | 0.26069733 | 0 | 1 | 63 | 0.61191651 | FALSE | 1 | FALSE | 1 | FALSE | 1 | FALSE | 1 | FALSE | 1 | FALSE | 1 |
| 3357 | 3357 | 8.15384615 | 2.09441824 | 0 | 1 | 1 | 0.7236252 | 2 | 0.41139386 | 4 | 0.10199045 | 0 | 1 | 32 | 0.74324845 | FALSE | 1 | FALSE | 1 | FALSE | 1 | FALSE | 1 | FALSE | 1 | FALSE | 1 |
| 3358 | 3358 | 9.25 | 3.04753999 | 0 | 1 | 0 | 1 | 1 | 0.51939841 | 1 | 0.60987146 | 0 | 1 | 18 | 0.37504645 | FALSE | 1 | FALSE | 1 | FALSE | 1 | FALSE | 1 | FALSE | 1 | FALSE | 1 |
| 3359 | 3359 | 9.46666667 | 2.70472837 | 0 | 1 | 1 | 0.38964394 | 1 | 0.42266264 | 1 | 0.50623194 | 0 | 1 | 12 | 0.80329917 | FALSE | 1 | FALSE | 1 | FALSE | 1 | FALSE | 1 | FALSE | 1 | FALSE | 1 |
| 3360 | 3360 | 4.65217391 | 1.90670888 | 7 | 0.02563138 | 1 | 0.97641742 | 9 | 0.01996775 | 4 | 0.76938454 | 3 | 0.28281707 | 89 | 0.95881626 | FALSE | 0.4974559 | FALSE | 1 | FALSE | 1 | FALSE | 1 | FALSE | 1 | FALSE | 1 |
| 3361 | 3361 | 9.33333333 | 2.09496752 | 0 | 1 | 2 | 0.31355738 | 3 | 0.12958859 | 2 | 0.48233989 | 1 | 0.44015537 | 27 | 0.91585095 | FALSE | 1 | FALSE | 1 | FALSE | 1 | FALSE | 1 | FALSE | 1 | FALSE | 1 |
| 3362 | 3362 | 9.53846154 | 2.92510051 | 0 | 1 | 0 | 1 | 1 | 0.37874417 | 0 | 1 | 0 | 1 | 12 | 0.374198 | FALSE | 1 | FALSE | 1 | FALSE | 1 | FALSE | 1 | FALSE | 1 | FALSE | 1 |
| 3363 | 3363 | 10.0476191 | 1.88922233 | 1 | 0.42086991 | 2 | 0.14655134 | 1 | 0.53671415 | 0 | 1 | 0 | 1 | 17 | 0.77849076 | FALSE | 1 | FALSE | 1 | FALSE | 1 | FALSE | 1 | FALSE | 1 | FALSE | 1 |
| 3364 | 3364 | 11.1764706 | 2.28126111 | 0 | 1 | 1 | 0.42857733 | 0 | 1 | 0 | 1 | 0 | 1 | 16 | 0.23070182 | FALSE | 1 | FALSE | 1 | FALSE | 1 | FALSE | 1 | FALSE | 1 | FALSE | 1 |
| 3365 | 3365 | 7.73809524 | 1.42360247 | 3 | 0.35850108 | 3 | 0.50638104 | 4 | 0.34844549 | 1 | 0.9802629 | 1 | 0.74858842 | 71 | 0.45717613 | FALSE | 1 | FALSE | 1 | FALSE | 1 | FALSE | 1 | FALSE | 1 | FALSE | 1 |
| 3366 | 3366 | 9.5 | 2.41276368 | 0 | 1 | 1 | 0.36919824 | 0 | 1 | 0 | 1 | 1 | 0.20679328 | 12 | 0.62236227 | FALSE | 1 | FALSE | 1 | FALSE | 1 | FALSE | 1 | FALSE | 1 | FALSE | 1 |
| 3367 | 3367 | 9.12727273 | 2.69716717 | 3 | 0.16017108 | 1 | 0.83180787 | 1 | 0.86240917 | 2 | 0.71679995 | 0 | 1 | 47 | 0.37712365 | FALSE | 1 | FALSE | 1 | FALSE | 1 | FALSE | 1 | FALSE | 1 | FALSE | 1 |
| 3368 | 3368 | 9.86666667 | 2.65497437 | 0 | 1 | 0 | 1 | 1 | 0.42266264 | 0 | 1 | 0 | 1 | 14 | 0.29525409 | FALSE | 1 | FALSE | 1 | FALSE | 1 | FALSE | 1 | FALSE | 1 | FALSE | 1 |
| 3369 | 3369 | 8.20408163 | 1.29330445 | 1 | 0.65632651 | 1 | 0.74131698 | 1 | 0.77786601 | 2 | 0.56788092 | 1 | 0.49330646 | 35 | 0.53616095 | FALSE | 1 | FALSE | 1 | FALSE | 1 | FALSE | 1 | FALSE | 1 | FALSE | 1 |
| 3370 | 3370 | 10.8666667 | 1.78387842 | 1 | 0.32293355 | 0 | 1 | 1 | 0.42266264 | 1 | 0.50623194 | 1 | 0.2198258 | 11 | 0.92879412 | FALSE | 1 | FALSE | 1 | FALSE | 1 | FALSE | 1 | FALSE | 1 | FALSE | 1 |
| 3371 | 3371 | 4.81283423 | 1.95463141 | 5 | 0.44253041 | 6 | 0.47328187 | 3 | 0.94778727 | 15 | 0.0111381 | 2 | 0.77318603 | 139 | 0.85459504 | FALSE | 1 | FALSE | 1 | FALSE | 1 | FALSE | 1 | FALSE | 1 | FALSE | 1 |
| 3372 | 3372 | 8.92307692 | 4.03754568 | 3 | 0.0279333 | 0 | 1 | 0 | 1 | 1 | 0.70601446 | 0 | 1 | 22 | 0.61628 | FALSE | 0.50964275 | FALSE | 1 | FALSE | 1 | FALSE | 1 | FALSE | 1 | FALSE | 1 |
| 3373 | 3373 | 10.4545455 | 2.60641294 | 0 | 1 | 0 | 1 | 0 | 1 | 0 | 1 | 1 | 0.16638736 | 10 | 0.46859937 | FALSE | 1 | FALSE | 1 | FALSE | 1 | FALSE | 1 | FALSE | 1 | FALSE | 1 |
| 3374 | 3374 | 9.64285714 | 2.86872652 | 0 | 1 | 2 | 0.07341559 | 1 | 0.40110418 | 1 | 0.48242099 | 0 | 1 | 10 | 0.94579623 | FALSE | 1 | FALSE | 1 | FALSE | 1 | FALSE | 1 | FALSE | 1 | FALSE | 1 |
| 3375 | 3375 | 9.40740741 | 2.26501998 | 0 | 1 | 1 | 0.58913618 | 2 | 0.25286935 | 1 | 0.71956339 | 0 | 1 | 23 | 0.58307573 | FALSE | 1 | FALSE | 1 | FALSE | 1 | FALSE | 1 | FALSE | 1 | FALSE | 1 |
| 3376 | 3376 | 6.56756757 | 3.24279276 | 2 | 0.24498301 | 2 | 0.3375163 | 0 | 1 | 1 | 0.82511639 | 0 | 1 | 32 | 0.47030992 | FALSE | 1 | FALSE | 1 | FALSE | 1 | FALSE | 1 | FALSE | 1 | FALSE | 1 |
| 3377 | 3377 | 8.84615385 | 2.03228382 | 0 | 1 | 0 | 1 | 0 | 1 | 0 | 1 | 2 | 0.01846832 | 11 | 0.6683056 | FALSE | 1 | FALSE | 1 | FALSE | 1 | FALSE | 1 | FALSE | 0.69720022 | FALSE | 1 |
| 3378 | 3378 | 9.75 | 3.69966215 | 0 | 1 | 0 | 1 | 0 | 1 | 0 | 1 | 0 | 1 | 8 | 0.25677166 | FALSE | 1 | FALSE | 1 | FALSE | 1 | FALSE | 1 | FALSE | 1 | FALSE | 1 |
| 3379 | 3379 | 9.64285714 | 2.19112309 | 0 | 1 | 1 | 0.36919824 | 2 | 0.08806398 | 1 | 0.48242099 | 0 | 1 | 10 | 0.94579623 | FALSE | 1 | FALSE | 1 | FALSE | 1 | FALSE | 1 | FALSE | 1 | FALSE | 1 |
| 3380 | 3380 | 10.4117647 | 2.08802816 | 0 | 1 | 0 | 1 | 0 | 1 | 0 | 1 | 0 | 1 | 17 | 0.05550965 | FALSE | 1 | FALSE | 1 | FALSE | 1 | FALSE | 1 | FALSE | 1 | FALSE | 1 |
| 3381 | 3381 | 10.6666667 | 2.53233063 | 3 | 0.01568237 | 0 | 1 | 0 | 1 | 0 | 1 | 0 | 1 | 18 | 0.57962763 | FALSE | 0.38695764 | FALSE | 1 | FALSE | 1 | FALSE | 1 | FALSE | 1 | FALSE | 1 |
| 3382 | 3382 | 6.66666667 | 3.71782493 | 0 | 1 | 0 | 1 | 1 | 0.42266264 | 0 | 1 | 0 | 1 | 14 | 0.29525409 | FALSE | 1 | FALSE | 1 | FALSE | 1 | FALSE | 1 | FALSE | 1 | FALSE | 1 |

| | | | | | | | | | | | | | | | | | | | | | | | | | | | |
|---|---|---|---|---|---|---|---|---|---|---|---|---|---|---|---|---|---|---|---|---|---|---|---|---|---|---|---|
| 3383 | 3383 | 8.33333333 | 2.3570226 | 0 | 1 | 0 | 1 | 0 | 1 | 1 | 0.43131129 | 0 | 1 | 11 | 0.41944532 | FALSE | 1 | FALSE | 1 | FALSE | 1 | FALSE | 1 | FALSE | 1 | FALSE | 1 |
| 3384 | 3384 | 9.48484849 | 2.51204628 | 0 | 1 | 0 | 1 | 2 | 0.33343871 | 2 | 0.45159487 | 0 | 1 | 29 | 0.39648374 | FALSE | 1 | FALSE | 1 | FALSE | 1 | FALSE | 1 | FALSE | 1 | FALSE | 1 |
| 3385 | 3385 | 7.08695652 | 2.28250517 | 0 | 1 | 0 | 1 | 0 | 1 | 3 | 0.08614834 | 0 | 1 | 20 | 0.50589402 | FALSE | 1 | FALSE | 1 | FALSE | 1 | FALSE | 1 | FALSE | 1 | FALSE | 1 |
| 3386 | 3386 | 10.2857143 | 1.82946407 | 0 | 1 | 1 | 0.58913618 | 1 | 0.62830986 | 2 | 0.35374209 | 0 | 1 | 23 | 0.58307573 | FALSE | 1 | FALSE | 1 | FALSE | 1 | FALSE | 1 | FALSE | 1 | FALSE | 1 |
| 3387 | 3387 | 9.73684211 | 2.80679824 | 0 | 1 | 1 | 0.46503795 | 0 | 1 | 0 | 1 | 1 | 0.26986775 | 17 | 0.41141543 | FALSE | 1 | FALSE | 1 | FALSE | 1 | FALSE | 1 | FALSE | 1 | FALSE | 1 |
| 3388 | 3388 | 9.33333333 | 2.44948974 | 0 | 1 | 0 | 1 | 0 | 1 | 0 | 1 | 1 | 0.1383237 | 8 | 0.5780332 | FALSE | 1 | FALSE | 1 | FALSE | 1 | FALSE | 1 | FALSE | 1 | FALSE | 1 |
| 3389 | 3389 | 10.7297297 | 1.34864594 | 0 | 1 | 1 | 0.70472939 | 3 | 0.14625315 | 1 | 0.82511639 | 0 | 1 | 32 | 0.47030992 | FALSE | 1 | FALSE | 1 | FALSE | 1 | FALSE | 1 | FALSE | 1 | FALSE | 1 |
| 3390 | 3390 | 9.42372881 | 2.1249398 | 2 | 0.4493251 | 6 | 0.01153778 | 6 | 0.01854841 | 3 | 0.51328524 | 6 | 0.00037678 | 36 | 0.999997 | FALSE | 1 | FALSE | 0.67617265 | FALSE | 1 | FALSE | 1 | FALSE | 0.09840309 | FALSE | 1 |
| 3391 | 3391 | 9.27777778 | 2.3286991 | 0 | 1 | 1 | 0.44710673 | 1 | 0.48280983 | 1 | 0.57131013 | 0 | 1 | 15 | 0.69372602 | FALSE | 1 | FALSE | 1 | FALSE | 1 | FALSE | 1 | FALSE | 1 | FALSE | 1 |
| 3392 | 3392 | 8.25423729 | 1.43318089 | 2 | 0.44060625 | 4 | 0.11722369 | 5 | 0.05626388 | 3 | 0.50172293 | 1 | 0.61824432 | 43 | 0.98621713 | FALSE | 1 | FALSE | 1 | FALSE | 1 | FALSE | 1 | FALSE | 1 | FALSE | 1 |
| 3393 | 3393 | 9.5 | 1.88034951 | 1 | 0.3050805 | 0 | 1 | 0 | 1 | 1 | 0.48242099 | 0 | 1 | 12 | 0.62236227 | FALSE | 1 | FALSE | 1 | FALSE | 1 | FALSE | 1 | FALSE | 1 | FALSE | 1 |
| 3394 | 3394 | 9.05882353 | 1.47411342 | 0 | 1 | 1 | 0.42857733 | 0 | 1 | 1 | 0.55062772 | 0 | 1 | 15 | 0.49060435 | FALSE | 1 | FALSE | 1 | FALSE | 1 | FALSE | 1 | FALSE | 1 | FALSE | 1 |
| 3395 | 3395 | 9.95238095 | 2.69878616 | 1 | 0.42086991 | 1 | 0.49918218 | 0 | 1 | 0 | 1 | 0 | 1 | 19 | 0.34094959 | FALSE | 1 | FALSE | 1 | FALSE | 1 | FALSE | 1 | FALSE | 1 | FALSE | 1 |
| 3396 | 3396 | 6.60714286 | 3.0512209 | 0 | 1 | 0 | 1 | 0 | 1 | 0 | 1 | 0 | 1 | 28 | 0.0085123 | FALSE | 1 | FALSE | 1 | FALSE | 1 | FALSE | 1 | FALSE | 1 | FALSE | 1 |
| 3397 | 3397 | 4.12705882 | 1.24157284 | 16 | 0.07266613 | 14 | 0.49766068 | 12 | 0.83655984 | 18 | 0.66029438 | 9 | 0.25235838 | 352 | 0.70131351 | FALSE | 0.76727438 | FALSE | 1 | FALSE | 1 | FALSE | 1 | FALSE | 1 | FALSE | 1 |
| 3398 | 3398 | 9.08333333 | 1.60511336 | 0 | 1 | 1 | 0.32623933 | 0 | 1 | 0 | 1 | 1 | 0.18007762 | 10 | 0.71436964 | FALSE | 1 | FALSE | 1 | FALSE | 1 | FALSE | 1 | FALSE | 1 | FALSE | 1 |
| 3399 | 3399 | 7.33333333 | 3.07318149 | 12 | 4.03E-11 | 2 | 0.32556427 | 1 | 0.73299778 | 1 | 0.81665383 | 0 | 1 | 20 | 0.99999228 | TRUE | 1.10E-07 | FALSE | 1 | FALSE | 1 | FALSE | 1 | FALSE | 1 | FALSE | 1 |
| 3400 | 3400 | 8.47619048 | 1.65094627 | 1 | 0.66519151 | 1 | 0.74973543 | 1 | 0.78589454 | 2 | 0.5811115 | 0 | 1 | 37 | 0.34041811 | FALSE | 1 | FALSE | 1 | FALSE | 1 | FALSE | 1 | FALSE | 1 | FALSE | 1 |
| 3401 | 3401 | 10.4615385 | 3.02846457 | 1 | 0.28675951 | 1 | 0.3480709 | 0 | 1 | 0 | 1 | 0 | 1 | 11 | 0.6683056 | FALSE | 1 | FALSE | 1 | FALSE | 1 | FALSE | 1 | FALSE | 1 | FALSE | 1 |
| 3402 | 3402 | 10.2272727 | 1.8812911 | 0 | 1 | 1 | 0.5154318 | 1 | 0.55340851 | 2 | 0.26791271 | 0 | 1 | 18 | 0.74744813 | FALSE | 1 | FALSE | 1 | FALSE | 1 | FALSE | 1 | FALSE | 1 | FALSE | 1 |
| 3403 | 3403 | 10.3076923 | 3.17161971 | 0 | 1 | 1 | 0.3480709 | 0 | 1 | 0 | 1 | 1 | 0.19354507 | 11 | 0.6683056 | FALSE | 1 | FALSE | 1 | FALSE | 1 | FALSE | 1 | FALSE | 1 | FALSE | 1 |
| 3404 | 3404 | 3.98275862 | 2.22440522 | 2 | 0.80338728 | 2 | 0.89447151 | 3 | 0.79317627 | 5 | 0.62158717 | 1 | 0.85549683 | 103 | 0.11371378 | FALSE | 1 | FALSE | 1 | FALSE | 1 | FALSE | 1 | FALSE | 1 | FALSE | 1 |
| 3405 | 3405 | 9.875 | 2.84769644 | 0 | 1 | 1 | 0.40942991 | 2 | 0.11087977 | 0 | 1 | 0 | 1 | 13 | 0.76799161 | FALSE | 1 | FALSE | 1 | FALSE | 1 | FALSE | 1 | FALSE | 1 | FALSE | 1 |
| 3406 | 3406 | 8.5 | 3.14907394 | 5 | 0.00075302 | 2 | 0.24095584 | 0 | 1 | 1 | 0.74482229 | 3 | 0.01149488 | 18 | 0.99922671 | FALSE | 0.06390799 | FALSE | 1 | FALSE | 1 | FALSE | 1 | FALSE | 0.57972935 | FALSE | 1 |
| 3407 | 3407 | 6.2631579 | 3.20923184 | 1 | 0.38989951 | 0 | 1 | 0 | 1 | 0 | 1 | 0 | 1 | 18 | 0.17880694 | FALSE | 1 | FALSE | 1 | FALSE | 1 | FALSE | 1 | FALSE | 1 | FALSE | 1 |
| 3408 | 3408 | 8.36956522 | 2.77691333 | 0 | 1 | 0 | 1 | 2 | 0.4962935 | 1 | 0.88573774 | 1 | 0.53375569 | 42 | 0.13378393 | FALSE | 1 | FALSE | 1 | FALSE | 1 | FALSE | 1 | FALSE | 1 | FALSE | 1 |
| 3409 | 3409 | 9.38235294 | 1.73230051 | 2 | 0.21638713 | 2 | 0.30150529 | 2 | 0.34668321 | 1 | 0.7984848 | 0 | 1 | 27 | 0.85023803 | FALSE | 1 | FALSE | 1 | FALSE | 1 | FALSE | 1 | FALSE | 1 | FALSE | 1 |
| 3410 | 3410 | 4.59090909 | 1.83383414 | 1 | 0.81665075 | 1 | 0.88323395 | 5 | 0.08323317 | 1 | 0.95356658 | 0 | 1 | 57 | 0.2945347 | FALSE | 1 | FALSE | 1 | FALSE | 1 | FALSE | 1 | FALSE | 1 | FALSE | 1 |
| 3411 | 3411 | 10.5 | 2.95803989 | 1 | 0.18769745 | 0 | 1 | 0 | 1 | 0 | 1 | 0 | 1 | 7 | 0.63752397 | FALSE | 1 | FALSE | 1 | FALSE | 1 | FALSE | 1 | FALSE | 1 | FALSE | 1 |
| 3412 | 3412 | 10.25 | 2.94745653 | 0 | 1 | 0 | 1 | 0 | 1 | 0 | 1 | 2 | 0.04176963 | 18 | 0.37504645 | FALSE | 1 | FALSE | 1 | FALSE | 1 | FALSE | 1 | FALSE | 0.90763079 | FALSE | 1 |
| 3413 | 3413 | 7.2 | 2.45880343 | 2 | 0.22589135 | 0 | 1 | 1 | 0.72299506 | 2 | 0.48233989 | 0 | 1 | 30 | 0.52765882 | FALSE | 1 | FALSE | 1 | FALSE | 1 | FALSE | 1 | FALSE | 1 | FALSE | 1 |
| 3414 | 3414 | 10.5652174 | 3.09032272 | 0 | 1 | 1 | 0.53115654 | 1 | 0.5695037 | 2 | 0.2852236 | 0 | 1 | 19 | 0.71542496 | FALSE | 1 | FALSE | 1 | FALSE | 1 | FALSE | 1 | FALSE | 1 | FALSE | 1 |
| 3415 | 3415 | 7.65909091 | 2.77114336 | 2 | 0.30241117 | 1 | 0.75788114 | 4 | 0.06724173 | 1 | 0.86831187 | 1 | 0.50988734 | 34 | 0.87722689 | FALSE | 1 | FALSE | 1 | FALSE | 1 | FALSE | 1 | FALSE | 1 | FALSE | 1 |
| 3416 | 3416 | 9.37142857 | 2.42705835 | 0 | 1 | 0 | 1 | 1 | 0.72299506 | 2 | 0.48233989 | 0 | 1 | 32 | 0.18108028 | FALSE | 1 | FALSE | 1 | FALSE | 1 | FALSE | 1 | FALSE | 1 | FALSE | 1 |
| 3417 | 3417 | 9.03846154 | 2.32843244 | 0 | 1 | 0 | 1 | 1 | 0.61440466 | 0 | 1 | 0 | 1 | 25 | 0.06985983 | FALSE | 1 | FALSE | 1 | FALSE | 1 | FALSE | 1 | FALSE | 1 | FALSE | 1 |
| 3418 | 3418 | 8.11926606 | 2.26551641 | 2 | 0.77443073 | 5 | 0.27707647 | 3 | 0.75696175 | 3 | 0.88355644 | 2 | 0.53718295 | 94 | 0.35171176 | FALSE | 1 | FALSE | 1 | FALSE | 1 | FALSE | 1 | FALSE | 1 | FALSE | 1 |
| 3419 | 3419 | 9.58333333 | 2.01900691 | 0 | 1 | 1 | 0.32623933 | 0 | 1 | 1 | 0.43131129 | 0 | 1 | 10 | 0.71436964 | FALSE | 1 | FALSE | 1 | FALSE | 1 | FALSE | 1 | FALSE | 1 | FALSE | 1 |
| 3420 | 3420 | 10.0714286 | 1.43747227 | 0 | 1 | 0 | 1 | 1 | 0.40110418 | 1 | 0.48242099 | 0 | 1 | 12 | 0.62236227 | FALSE | 1 | FALSE | 1 | FALSE | 1 | FALSE | 1 | FALSE | 1 | FALSE | 1 |
| 3421 | 3421 | 8.83333333 | 2.06155281 | 0 | 1 | 1 | 0.44710673 | 0 | 1 | 0 | 1 | 3 | 0.00292491 | 14 | 0.86287418 | FALSE | 1 | FALSE | 1 | FALSE | 1 | FALSE | 1 | FALSE | 0.29951111 | FALSE | 1 |
| 3422 | 3422 | 9.21875 | 2.23235267 | 1 | 0.55378205 | 0 | 1 | 1 | 0.67911179 | 3 | 0.16836568 | 0 | 1 | 26 | 0.64660109 | FALSE | 1 | FALSE | 1 | FALSE | 1 | FALSE | 1 | FALSE | 1 | FALSE | 1 |
| 3423 | 3423 | 11.0588235 | 2.20882157 | 0 | 1 | 1 | 0.42857733 | 0 | 1 | 0 | 1 | 0 | 1 | 16 | 0.23070182 | FALSE | 1 | FALSE | 1 | FALSE | 1 | FALSE | 1 | FALSE | 1 | FALSE | 1 |
| 3424 | 3424 | 9.64285714 | 2.5803417 | 1 | 0.26795838 | 0 | 1 | 0 | 1 | 0 | 1 | 0 | 1 | 11 | 0.41944532 | FALSE | 1 | FALSE | 1 | FALSE | 1 | FALSE | 1 | FALSE | 1 | FALSE | 1 |
| 3425 | 3425 | 9.65714286 | 2.05555494 | 0 | 1 | 3 | 0.10264583 | 0 | 1 | 3 | 0.21561745 | 0 | 1 | 29 | 0.69927719 | FALSE | 1 | FALSE | 1 | FALSE | 1 | FALSE | 1 | FALSE | 1 | FALSE | 1 |
| 3426 | 3426 | 9.6875 | 3.31132055 | 0 | 1 | 0 | 1 | 1 | 0.44344817 | 0 | 1 | 0 | 1 | 15 | 0.26127859 | FALSE | 1 | FALSE | 1 | FALSE | 1 | FALSE | 1 | FALSE | 1 | FALSE | 1 |
| 3427 | 3427 | 9.52631579 | 1.92950556 | 1 | 0.38989951 | 1 | 0.46503795 | 1 | 0.50143826 | 0 | 1 | 3 | 0.00343221 | 13 | 0.97998676 | FALSE | 1 | FALSE | 1 | FALSE | 1 | FALSE | 1 | FALSE | 0.33875501 | FALSE | 1 |
| 3428 | 3428 | 4.97752809 | 2.17223298 | 4 | 0.16339023 | 1 | 0.93582077 | 3 | 0.57805941 | 3 | 0.74138858 | 0 | 1 | 72 | 0.33817167 | FALSE | 1 | FALSE | 1 | FALSE | 1 | FALSE | 1 | FALSE | 1 | FALSE | 1 |
| 3429 | 3429 | 10.6 | 2.4979992 | 0 | 1 | 0 | 1 | 1 | 0.42266264 | 0 | 1 | 0 | 1 | 14 | 0.29525409 | FALSE | 1 | FALSE | 1 | FALSE | 1 | FALSE | 1 | FALSE | 1 | FALSE | 1 |
| 3430 | 3430 | 8.08823529 | 1.19724309 | 0 | 1 | 1 | 0.6739483 | 1 | 0.71261921 | 1 | 0.7984848 | 0 | 1 | 31 | 0.1995074 | FALSE | 1 | FALSE | 1 | FALSE | 1 | FALSE | 1 | FALSE | 1 | FALSE | 1 |
| 3431 | 3431 | 9.61904762 | 1.64682062 | 0 | 1 | 0 | 1 | 2 | 0.17308361 | 1 | 0.62783522 | 0 | 1 | 18 | 0.57962763 | FALSE | 1 | FALSE | 1 | FALSE | 1 | FALSE | 1 | FALSE | 1 | FALSE | 1 |
| 3432 | 3432 | 1.51698113 | 1.96014399 | 5 | 0.81650042 | 3 | 0.9927831 | 6 | 0.92050716 | 10 | 0.78332603 | 3 | 0.81630348 | 238 | 0.00614837 | FALSE | 1 | FALSE | 1 | FALSE | 1 | FALSE | 1 | FALSE | 1 | FALSE | 1 |
| 3433 | 3433 | 10.2631579 | 2.55141045 | 1 | 0.38989951 | 0 | 1 | 2 | 0.14749808 | 1 | 0.59104357 | 1 | 0.26986775 | 14 | 0.93677253 | FALSE | 1 | FALSE | 1 | FALSE | 1 | FALSE | 1 | FALSE | 1 | FALSE | 1 |
| 3434 | 3434 | 9 | 2.66024869 | 1 | 0.28675951 | 1 | 0.3480709 | 0 | 1 | 3 | 0.01944678 | 0 | 1 | 8 | 0.99085298 | FALSE | 1 | FALSE | 1 | FALSE | 1 | FALSE | 1 | FALSE | 1 | FALSE | 1 |
| 3435 | 3435 | 10.0833333 | 3.25213605 | 0 | 1 | 0 | 1 | 0 | 1 | 1 | 0.43131129 | 0 | 1 | 11 | 0.41944532 | FALSE | 1 | FALSE | 1 | FALSE | 1 | FALSE | 1 | FALSE | 1 | FALSE | 1 |
| 3436 | 3436 | 8.81818182 | 2.51606409 | 0 | 1 | 0 | 1 | 1 | 0.33149969 | 0 | 1 | 0 | 1 | 10 | 0.46859937 | FALSE | 1 | FALSE | 1 | FALSE | 1 | FALSE | 1 | FALSE | 1 | FALSE | 1 |
| 3437 | 3437 | 7.72 | 2.97348281 | 0 | 1 | 1 | 0.56109832 | 1 | 0.59998149 | 0 | 1 | 0 | 1 | 23 | 0.22727183 | FALSE | 1 | FALSE | 1 | FALSE | 1 | FALSE | 1 | FALSE | 1 | FALSE | 1 |
| 3438 | 3438 | 8.69230769 | 2.19736259 | 0 | 1 | 1 | 0.32623933 | 0 | 1 | 1 | 0.43131129 | 0 | 1 | 10 | 0.71436964 | FALSE | 1 | FALSE | 1 | FALSE | 1 | FALSE | 1 | FALSE | 1 | FALSE | 1 |
| 3439 | 3439 | 6.05128205 | 3.24984195 | 1 | 0.85137169 | 4 | 0.21021455 | 1 | 0.93179654 | 4 | 0.43314506 | 0 | 1 | 63 | 0.39803869 | FALSE | 1 | FALSE | 1 | FALSE | 1 | FALSE | 1 | FALSE | 1 | FALSE | 1 |
| 3440 | 3440 | 9.53846154 | 1.94601702 | 1 | 0.28675951 | 1 | 0.3480709 | 0 | 1 | 1 | 0.45746571 | 0 | 1 | 10 | 0.86791642 | FALSE | 1 | FALSE | 1 | FALSE | 1 | FALSE | 1 | FALSE | 1 | FALSE | 1 |
| 3441 | 3441 | 9.44444444 | 3.05908887 | 2 | 0.07645454 | 0 | 1 | 0 | 1 | 1 | 0.57131013 | 0 | 1 | 15 | 0.69372602 | FALSE | 0.77424195 | FALSE | 1 | FALSE | 1 | FALSE | 1 | FALSE | 1 | FALSE | 1 |
| 3442 | 3442 | 8.77419355 | 1.79083108 | 1 | 0.55378205 | 0 | 1 | 2 | 0.30673593 | 1 | 0.76781284 | 1 | 0.4016888 | 26 | 0.64660109 | FALSE | 1 | FALSE | 1 | FALSE | 1 | FALSE | 1 | FALSE | 1 | FALSE | 1 |
| 3443 | 3443 | 8.375 | 2.28787565 | 1 | 0.64722818 | 1 | 0.7326167 | 2 | 0.42396974 | 1 | 0.84823834 | 1 | 0.48480878 | 34 | 0.56328904 | FALSE | 1 | FALSE | 1 | FALSE | 1 | FALSE | 1 | FALSE | 1 | FALSE | 1 |
| 3444 | 3444 | 8.82051282 | 3.41093111 | 1 | 0.62830697 | 1 | 0.71433277 | 0 | 1 | 5 | 0.02858807 | 1 | 0.4673875 | 30 | 0.87271204 | FALSE | 1 | FALSE | 1 | FALSE | 1 | FALSE | 1 | FALSE | 1 | FALSE | 1 |
| 3445 | 3445 | 9.76190476 | 2.18009965 | 0 | 1 | 3 | 0.14580275 | 2 | 0.43640583 | 3 | 0.29054213 | 1 | 0.49330646 | 32 | 0.90458039 | FALSE | 1 | FALSE | 1 | FALSE | 1 | FALSE | 1 | FALSE | 1 | FALSE | 1 |
| 3446 | 3446 | 8.57142857 | 2.45573049 | 0 | 1 | 1 | 0.60247911 | 4 | 0.01687379 | 1 | 0.73248983 | 0 | 1 | 22 | 0.86499992 | FALSE | 1 | FALSE | 1 | FALSE | 1 | FALSE | 1 | FALSE | 1 | FALSE | 1 |
| 3447 | 3447 | 9 | 2.80691786 | 0 | 1 | 1 | 0.6629941 | 1 | 0.70185639 | 1 | 0.78873814 | 0 | 1 | 30 | 0.21942357 | FALSE | 1 | FALSE | 1 | FALSE | 1 | FALSE | 1 | FALSE | 1 | FALSE | 1 |
| 3448 | 3448 | 9.42857143 | 1.59078982 | 0 | 1 | 0 | 1 | 0 | 1 | 0 | 1 | 0 | 1 | 7 | 0.3043657 | FALSE | 1 | FALSE | 1 | FALSE | 1 | FALSE | 1 | FALSE | 1 | FALSE | 1 |
| 3449 | 3449 | 10.1818182 | 1.52662324 | 0 | 1 | 0 | 1 | 2 | 0.04781153 | 0 | 1 | 0 | 1 | 8 | 0.80394719 | FALSE | 1 | FALSE | 1 | FALSE | 1 | FALSE | 1 | FALSE | 1 | FALSE | 1 |
| 3450 | 3450 | 8.53333333 | 1.96185853 | 1 | 0.32293355 | 0 | 1 | 2 | 0.09928574 | 1 | 0.50623194 | 0 | 1 | 11 | 0.92879412 | FALSE | 1 | FALSE | 1 | FALSE | 1 | FALSE | 1 | FALSE | 1 | FALSE | 1 |
| 3451 | 3451 | 9.27272727 | 1.65644247 | 0 | 1 | 3 | 0.00454306 | 0 | 1 | 0 | 1 | 0 | 1 | 8 | 0.92152993 | FALSE | 1 | FALSE | 0.58120789 | FALSE | 1 | FALSE | 1 | FALSE | 1 | FALSE | 1 |
| 3452 | 3452 | 7.8 | 3.3306656 | 0 | 1 | 1 | 0.38964394 | 1 | 0.42266264 | 0 | 1 | 0 | 1 | 13 | 0.57712805 | FALSE | 1 | FALSE | 1 | FALSE | 1 | FALSE | 1 | FALSE | 1 | FALSE | 1 |
| 3453 | 3453 | 13.1785714 | 3.6844613 | 1 | 0.51745948 | 3 | 0.06039219 | 2 | 0.2663542 | 2 | 0.3705495 | 0 | 1 | 20 | 0.97757517 | FALSE | 1 | FALSE | 1 | FALSE | 1 | FALSE | 1 | FALSE | 1 | FALSE | 1 |
| 3454 | 3454 | 8.375 | 2.37719197 | 0 | 1 | 2 | 0.18117183 | 1 | 0.58502114 | 0 | 1 | 0 | 1 | 21 | 0.47053137 | FALSE | 1 | FALSE | 1 | FALSE | 1 | FALSE | 1 | FALSE | 1 | FALSE | 1 |
| 3455 | 3455 | 9.32 | 2.72352713 | 1 | 0.46441868 | 1 | 0.54637326 | 1 | 0.58502114 | 0 | 1 | 0 | 1 | 21 | 0.47053137 | FALSE | 1 | FALSE | 1 | FALSE | 1 | FALSE | 1 | FALSE | 1 | FALSE | 1 |
| 3456 | 3456 | 9.5 | 3.06865877 | 0 | 1 | 0 | 1 | 0 | 1 | 0 | 1 | 0 | 1 | 12 | 0.13002667 | FALSE | 1 | FALSE | 1 | FALSE | 1 | FALSE | 1 | FALSE | 1 | FALSE | 1 |
| 3457 | 3457 | 5.07692308 | 2.0616724 | 3 | 0.32318196 | 3 | 0.46497905 | 3 | 0.53558651 | 3 | 0.7021069 | 1 | 0.72664912 | 65 | 0.67060743 | FALSE | 1 | FALSE | 1 | FALSE | 1 | FALSE | 1 | FALSE | 1 | FALSE | 1 |
| 3458 | 3458 | 6.16025641 | 0.6887074 | 12 | 0.10203235 | 9 | 0.68301532 | 5 | 0.98910436 | 18 | 0.1832603 | 5 | 0.58275117 | 262 | 0.56484646 | FALSE | 0.87592659 | FALSE | 1 | FALSE | 1 | FALSE | 1 | FALSE | 1 | FALSE | 1 |
| 3459 | 3459 | 10.5625 | 2.06060519 | 2 | 0.06197875 | 0 | 1 | 2 | 0.11087977 | 1 | 0.52895088 | 0 | 1 | 11 | 0.9718531 | FALSE | 0.72532843 | FALSE | 1 | FALSE | 1 | FALSE | 1 | FALSE | 1 | FALSE | 1 |
| 3460 | 3460 | 9.77777778 | 2.04275292 | 0 | 1 | 0 | 1 | 0 | 1 | 0 | 1 | 1 | 0.1383237 | 8 | 0.5780332 | FALSE | 1 | FALSE | 1 | FALSE | 1 | FALSE | 1 | FALSE | 1 | FALSE | 1 |
| 3461 | 3461 | 5.64285714 | 4.08144132 | 4 | 0.02189152 | 0 | 1 | 1 | 0.78589454 | 1 | 0.86193352 | 0 | 1 | 36 | 0.50934357 | FALSE | 0.46823848 | FALSE | 1 | FALSE | 1 | FALSE | 1 | FALSE | 1 | FALSE | 1 |
| 3462 | 3462 | 9.95238095 | 1.67548687 | 0 | 1 | 1 | 0.49918218 | 1 | 0.53671415 | 0 | 1 | 1 | 0.29368245 | 18 | 0.57962763 | FALSE | 1 | FALSE | 1 | FALSE | 1 | FALSE | 1 | FALSE | 1 | FALSE | 1 |
| 3463 | 3463 | 9.6969697 | 2.23586264 | 1 | 0.57647348 | 0 | 1 | 2 | 0.33343871 | 3 | 0.19161876 | 0 | 1 | 27 | 0.75124583 | FALSE | 1 | FALSE | 1 | FALSE | 1 | FALSE | 1 | FALSE | 1 | FALSE | 1 |
| 3464 | 3464 | 12.1212121 | 2.08541219 | 0 | 1 | 2 | 0.28941836 | 2 | 0.33343871 | 3 | 0.19161876 | 1 | 0.42123867 | 25 | 0.93892791 | FALSE | 1 | FALSE | 1 | FALSE | 1 | FALSE | 1 | FALSE | 1 | FALSE | 1 |
| 3465 | 3465 | 9.07142857 | 3.35866227 | 3 | 0.03391597 | 2 | 0.2288771 | 2 | 0.2663542 | 2 | 0.3705495 | 1 | 0.37113067 | 18 | 0.9978883 | FALSE | 0.56934354 | FALSE | 1 | FALSE | 1 | FALSE | 1 | FALSE | 1 | FALSE | 1 |
| 3466 | 3466 | 11.7857143 | 3.25529396 | 1 | 0.3050805 | 0 | 1 | 1 | 0.40110418 | 0 | 1 | 0 | 1 | 12 | 0.62236227 | FALSE | 1 | FALSE | 1 | FALSE | 1 | FALSE | 1 | FALSE | 1 | FALSE | 1 |
| 3467 | 3467 | 9 | 4.18681404 | 3 | 0.00864151 | 0 | 1 | 2 | 0.12280513 | 1 | 0.55062772 | 0 | 1 | 11 | 0.98974663 | FALSE | 0.27019545 | FALSE | 1 | FALSE | 1 | FALSE | 1 | FALSE | 1 | FALSE | 1 |
| 3468 | 3468 | 9.33333333 | 2.96540904 | 1 | 0.42086991 | 0 | 1 | 0 | 1 | 2 | 0.25059627 | 0 | 1 | 18 | 0.57962763 | FALSE | 1 | FALSE | 1 | FALSE | 1 | FALSE | 1 | FALSE | 1 | FALSE | 1 |
| 3469 | 3469 | 11.1428571 | 2.55017673 | 0 | 1 | 2 | 0.1135776 | 1 | 0.48280983 | 1 | 0.57131013 | 0 | 1 | 14 | 0.86287418 | FALSE | 1 | FALSE | 1 | FALSE | 1 | FALSE | 1 | FALSE | 1 | FALSE | 1 |
| 3470 | 3470 | 5.864 | 1.61167739 | 10 | 0.00131712 | 12 | 0.00067904 | 5 | 0.46856804 | 5 | 0.68673636 | 1 | 0.87581372 | 92 | 0.9994145 | FALSE | 0.08991542 | FALSE | 0.23177752 | FALSE | 1 | FALSE | 1 | FALSE | 1 | FALSE | 1 |
| 3471 | 3471 | 7.53846154 | 3.05375896 | 0 | 1 | 0 | 1 | 0 | 1 | 1 | 0.45746571 | 0 | 1 | 12 | 0.374198 | FALSE | 1 | FALSE | 1 | FALSE | 1 | FALSE | 1 | FALSE | 1 | FALSE | 1 |
| 3472 | 3472 | 11.8 | 3.40978983 | 0 | 1 | 0 | 1 | 0 | 1 | 3 | 0.02891327 | 0 | 1 | 12 | 0.80329917 | FALSE | 1 | FALSE | 1 | FALSE | 1 | FALSE | 1 | FALSE | 1 | FALSE | 1 |
| 3473 | 3473 | 9.17391304 | 2.92889909 | 1 | 0.45027683 | 0 | 1 | 1 | 0.5695037 | 4 | 0.01932195 | 0 | 1 | 17 | 0.94439745 | FALSE | 1 | FALSE | 1 | FALSE | 1 | FALSE | 1 | FALSE | 1 | FALSE | 1 |
| 3474 | 3474 | 6.81481482 | 2.53913133 | 0 | 1 | 0 | 1 | 1 | 0.62830986 | 2 | 0.35374209 | 0 | 1 | 24 | 0.3725373 | FALSE | 1 | FALSE | 1 | FALSE | 1 | FALSE | 1 | FALSE | 1 | FALSE | 1 |
| 3475 | 3475 | 9.71428571 | 1.84888273 | 0 | 1 | 0 | 1 | 1 | 0.64171563 | 2 | 0.3705495 | 0 | 1 | 25 | 0.34293 | FALSE | 1 | FALSE | 1 | FALSE | 1 | FALSE | 1 | FALSE | 1 | FALSE | 1 |
| 3476 | 3476 | 9.375 | 3.49776715 | 0 | 1 | 0 | 1 | 0 | 1 | 1 | 0.31351192 | 0 | 1 | 7 | 0.63752397 | FALSE | 1 | FALSE | 1 | FALSE | 1 | FALSE | 1 | FALSE | 1 | FALSE | 1 |

| | | | | | | | | | | | | | | | | | | | | | | | | | | | |
|---|---|---|---|---|---|---|---|---|---|---|---|---|---|---|---|---|---|---|---|---|---|---|---|---|---|---|---|
| 3477 | 3477 | 10.9411765 | 2.97974709 | 1 | 0.35728305 | 0 | 1 | 0 | 1 | 1 | 0.55062772 | 0 | 1 | 15 | 0.49060435 | FALSE | 1 | FALSE | 1 | FALSE | 1 | FALSE | 1 | FALSE | 1 | FALSE | 1 |
| 3478 | 3478 | 7.83333333 | 1.21335165 | 0 | 1 | 0 | 1 | 1 | 0.48280983 | 0 | 1 | 1 | 0.25766374 | 16 | 0.44998001 | FALSE | 1 | FALSE | 1 | FALSE | 1 | FALSE | 1 | FALSE | 1 | FALSE | 1 |
| 3479 | 3479 | 10.0416667 | 3.87813493 | 1 | 0.45027683 | 0 | 1 | 1 | 0.5695037 | 2 | 0.2852236 | 0 | 1 | 19 | 0.71542496 | FALSE | 1 | FALSE | 1 | FALSE | 1 | FALSE | 1 | FALSE | 1 | FALSE | 1 |
| 3480 | 3480 | 10 | 3.12694384 | 0 | 1 | 1 | 0.25628163 | 1 | 0.28067846 | 1 | 0.34506517 | 0 | 1 | 6 | 0.96128317 | FALSE | 1 | FALSE | 1 | FALSE | 1 | FALSE | 1 | FALSE | 1 | FALSE | 1 |
| 3481 | 3481 | 8.7631579 | 1.89784102 | 0 | 1 | 2 | 0.3494045 | 0 | 1 | 2 | 0.52640931 | 1 | 0.4673875 | 33 | 0.44261631 | FALSE | 1 | FALSE | 1 | FALSE | 1 | FALSE | 1 | FALSE | 1 | FALSE | 1 |
| 3482 | 3482 | 9.66666667 | 1.88561808 | 0 | 1 | 0 | 1 | 2 | 0.06691409 | 0 | 1 | 0 | 1 | 10 | 0.71436964 | FALSE | 1 | FALSE | 1 | FALSE | 1 | FALSE | 1 | FALSE | 1 | FALSE | 1 |
| 3483 | 3483 | 10.1818182 | 3.21412173 | 0 | 1 | 1 | 0.30368014 | 0 | 1 | 2 | 0.08789566 | 1 | 0.16638736 | 7 | 0.98129353 | FALSE | 1 | FALSE | 1 | FALSE | 1 | FALSE | 1 | FALSE | 1 | FALSE | 1 |
| 3484 | 3484 | 9.85365854 | 2.80731601 | 1 | 0.65632651 | 1 | 0.74131698 | 0 | 1 | 1 | 0.85524728 | 1 | 0.49330646 | 37 | 0.21027077 | FALSE | 1 | FALSE | 1 | FALSE | 1 | FALSE | 1 | FALSE | 1 | FALSE | 1 |
| 3485 | 3485 | 7.22580645 | 2.21008944 | 4 | 0.00679223 | 1 | 0.6278847 | 3 | 0.09131526 | 2 | 0.4036176 | 0 | 1 | 20 | 0.99601577 | FALSE | 0.24597546 | FALSE | 1 | FALSE | 1 | FALSE | 1 | FALSE | 1 | FALSE | 1 |
| 3486 | 3486 | 8.65625 | 2.40584412 | 3 | 0.03713651 | 1 | 0.61539065 | 1 | 0.65463982 | 2 | 0.3871811 | 1 | 0.38148475 | 21 | 0.97185642 | FALSE | 0.58392 | FALSE | 1 | FALSE | 1 | FALSE | 1 | FALSE | 1 | FALSE | 1 |
| 3487 | 3487 | 9.42 | 2.64643156 | 2 | 0.36837609 | 0 | 1 | 1 | 0.8405438 | 3 | 0.4048204 | 0 | 1 | 44 | 0.31639336 | FALSE | 1 | FALSE | 1 | FALSE | 1 | FALSE | 1 | FALSE | 1 | FALSE | 1 |
| 3488 | 3488 | 4.90384615 | 2.08098487 | 1 | 0.98251743 | 7 | 0.23076967 | 8 | 0.18889398 | 5 | 0.84273281 | 5 | 0.10817001 | 128 | 0.714228 | FALSE | 1 | FALSE | 1 | FALSE | 1 | FALSE | 1 | FALSE | 1 | FALSE | 1 |
| 3489 | 3489 | 7.55555556 | 2.61524496 | 0 | 1 | 1 | 0.58913618 | 2 | 0.25286935 | 2 | 0.35374209 | 2 | 0.07169515 | 20 | 0.95133411 | FALSE | 1 | FALSE | 1 | FALSE | 1 | FALSE | 1 | FALSE | 1 | FALSE | 1 |
| 3490 | 3490 | 9.46153846 | 2.6199056 | 0 | 1 | 0 | 1 | 0 | 1 | 1 | 0.45746571 | 0 | 1 | 12 | 0.374198 | FALSE | 1 | FALSE | 1 | FALSE | 1 | FALSE | 1 | FALSE | 1 | FALSE | 1 |
| 3491 | 3491 | 7.65517241 | 2.1217407 | 4 | 0.00600461 | 0 | 1 | 1 | 0.65463982 | 3 | 0.14602298 | 0 | 1 | 21 | 0.97185642 | FALSE | 0.22773046 | FALSE | 1 | FALSE | 1 | FALSE | 1 | FALSE | 1 | FALSE | 1 |
| 3492 | 3492 | 10.4347826 | 2.08151464 | 3 | 0.02011786 | 1 | 0.53115654 | 1 | 0.5695037 | 0 | 1 | 1 | 0.31672722 | 17 | 0.94439745 | FALSE | 0.44482142 | FALSE | 1 | FALSE | 1 | FALSE | 1 | FALSE | 1 | FALSE | 1 |
| 3493 | 3493 | 7.45 | 2.21679198 | 3 | 0.19869438 | 3 | 0.30686508 | 2 | 0.64095454 | 4 | 0.29632241 | 2 | 0.25814944 | 46 | 0.96067108 | FALSE | 1 | FALSE | 1 | FALSE | 1 | FALSE | 1 | FALSE | 1 | FALSE | 1 |
| 3494 | 3494 | 7.925 | 1.83558574 | 1 | 0.64722818 | 1 | 0.7326167 | 1 | 0.76953771 | 1 | 0.84823834 | 2 | 0.13925369 | 34 | 0.56328904 | FALSE | 1 | FALSE | 1 | FALSE | 1 | FALSE | 1 | FALSE | 1 | FALSE | 1 |
| 3495 | 3495 | 8.96969697 | 2.45548056 | 0 | 1 | 4 | 0.02089081 | 1 | 0.70185639 | 1 | 0.78873814 | 0 | 1 | 27 | 0.75124583 | FALSE | 1 | FALSE | 0.8034624 | FALSE | 1 | FALSE | 1 | FALSE | 1 | FALSE | 1 |
| 3496 | 3496 | 10.1842105 | 0.75586351 | 0 | 1 | 1 | 0.71433277 | 2 | 0.39868506 | 8 | 0.00026346 | 0 | 1 | 27 | 0.98956273 | FALSE | 1 | FALSE | 1 | FALSE | 1 | FALSE | 0.63446475 | FALSE | 1 | FALSE | 1 |
| 3497 | 3497 | 8.28125 | 3.95469954 | 3 | 0.0476977 | 1 | 0.65167362 | 1 | 0.69069222 | 1 | 0.77852168 | 1 | 0.41154418 | 25 | 0.88559288 | FALSE | 0.649301 | FALSE | 1 | FALSE | 1 | FALSE | 1 | FALSE | 1 | FALSE | 1 |
| 3498 | 3498 | 9.8 | 1.2489996 | 0 | 1 | 0 | 1 | 0 | 1 | 2 | 0.21610812 | 0 | 1 | 17 | 0.41141543 | FALSE | 1 | FALSE | 1 | FALSE | 1 | FALSE | 1 | FALSE | 1 | FALSE | 1 |
| 3499 | 3499 | 4.35897436 | 1.51042836 | 0 | 1 | 1 | 0.92420038 | 3 | 0.53558651 | 3 | 0.7021069 | 0 | 1 | 71 | 0.06396383 | FALSE | 1 | FALSE | 1 | FALSE | 1 | FALSE | 1 | FALSE | 1 | FALSE | 1 |
| 3500 | 3500 | 9.76190476 | 2.02142717 | 1 | 0.42086991 | 0 | 1 | 0 | 1 | 0 | 1 | 2 | 0.04568221 | 18 | 0.57962763 | FALSE | 1 | FALSE | 1 | FALSE | 1 | FALSE | 1 | FALSE | 0.95710659 | FALSE | 1 |
| 3501 | 3501 | 11.6666667 | 2.56038192 | 1 | 0.37380237 | 1 | 0.44710673 | 0 | 1 | 1 | 0.57131013 | 2 | 0.03435384 | 13 | 0.95041913 | FALSE | 1 | FALSE | 1 | FALSE | 1 | FALSE | 1 | FALSE | 0.86063198 | FALSE | 1 |
| 3502 | 3502 | 9.88235294 | 2.39809613 | 3 | 0.05546921 | 3 | 0.09602744 | 2 | 0.34668321 | 2 | 0.46709861 | 0 | 1 | 24 | 0.98848472 | FALSE | 0.68849059 | FALSE | 1 | FALSE | 1 | FALSE | 1 | FALSE | 1 | FALSE | 1 |
| 3503 | 3503 | 9.32258065 | 2.75197403 | 2 | 0.1881398 | 1 | 0.63997468 | 0 | 1 | 3 | 0.16836568 | 0 | 1 | 25 | 0.80006983 | FALSE | 1 | FALSE | 1 | FALSE | 1 | FALSE | 1 | FALSE | 1 | FALSE | 1 |
| 3504 | 3504 | 10.2941177 | 3.19493544 | 0 | 1 | 1 | 0.42857733 | 1 | 0.46348836 | 0 | 1 | 0 | 1 | 15 | 0.49060435 | FALSE | 1 | FALSE | 1 | FALSE | 1 | FALSE | 1 | FALSE | 1 | FALSE | 1 |
| 3505 | 3505 | 9.06666667 | 3.43446583 | 0 | 1 | 0 | 1 | 1 | 0.42266264 | 4 | 0.00397592 | 1 | 0.2198258 | 9 | 0.995488 | FALSE | 1 | FALSE | 1 | FALSE | 1 | FALSE | 1 | FALSE | 1 | FALSE | 1 |
| 3506 | 3506 | 10.2777778 | 2.15524053 | 0 | 1 | 1 | 0.44710673 | 0 | 1 | 1 | 0.57131013 | 0 | 1 | 16 | 0.44998001 | FALSE | 1 | FALSE | 1 | FALSE | 1 | FALSE | 1 | FALSE | 1 | FALSE | 1 |
| 3507 | 3507 | 9.7826087 | 2.84309265 | 2 | 0.11647716 | 0 | 1 | 2 | 0.19931429 | 0 | 1 | 0 | 1 | 19 | 0.71542496 | FALSE | 0.93181727 | FALSE | 1 | FALSE | 1 | FALSE | 1 | FALSE | 1 | FALSE | 1 |
| 3508 | 3508 | 8.67647059 | 1.65779067 | 0 | 1 | 0 | 1 | 0 | 1 | 2 | 0.46709861 | 0 | 1 | 32 | 0.08180309 | FALSE | 1 | FALSE | 1 | FALSE | 1 | FALSE | 1 | FALSE | 1 | FALSE | 1 |
| 3509 | 3509 | 6.85714286 | 1.95875846 | 0 | 1 | 3 | 0.02888061 | 3 | 0.03775825 | 0 | 1 | 0 | 1 | 15 | 0.96513691 | FALSE | 1 | FALSE | 0.88012077 | FALSE | 1 | FALSE | 1 | FALSE | 1 | FALSE | 1 |
| 3510 | 3510 | 8.94117647 | 2.09959627 | 0 | 1 | 0 | 1 | 1 | 0.46348836 | 1 | 0.55062772 | 1 | 0.24525764 | 14 | 0.7313145 | FALSE | 1 | FALSE | 1 | FALSE | 1 | FALSE | 1 | FALSE | 1 | FALSE | 1 |
| 3511 | 3511 | 6.96363636 | 1.84847491 | 2 | 0.41401181 | 3 | 0.26245929 | 1 | 0.8673918 | 1 | 0.92538938 | 0 | 1 | 48 | 0.35562012 | FALSE | 1 | FALSE | 1 | FALSE | 1 | FALSE | 1 | FALSE | 1 | FALSE | 1 |
| 3512 | 3512 | 7.43859649 | 2.04353962 | 3 | 0.17275086 | 0 | 1 | 1 | 0.87219471 | 3 | 0.47819064 | 0 | 1 | 49 | 0.33483172 | FALSE | 1 | FALSE | 1 | FALSE | 1 | FALSE | 1 | FALSE | 1 | FALSE | 1 |
| 3513 | 3513 | 9.16666667 | 2.47767813 | 0 | 1 | 1 | 0.53115654 | 0 | 1 | 3 | 0.08614834 | 1 | 0.31672722 | 18 | 0.86273155 | FALSE | 1 | FALSE | 1 | FALSE | 1 | FALSE | 1 | FALSE | 1 | FALSE | 1 |
| 3514 | 3514 | 8.71428571 | 2.3733211 | 0 | 1 | 0 | 1 | 0 | 1 | 1 | 0.48242099 | 0 | 1 | 13 | 0.33283729 | FALSE | 1 | FALSE | 1 | FALSE | 1 | FALSE | 1 | FALSE | 1 | FALSE | 1 |
| 3515 | 3515 | 11.1 | 2.25610284 | 2 | 0.0256865 | 0 | 1 | 1 | 0.30655256 | 1 | 0.37517262 | 0 | 1 | 6 | 0.98831471 | FALSE | 0.4974559 | FALSE | 1 | FALSE | 1 | FALSE | 1 | FALSE | 1 | FALSE | 1 |
| 3516 | 3516 | 7.41463415 | 2.40908453 | 0 | 1 | 3 | 0.14580275 | 1 | 0.77786601 | 3 | 0.29054213 | 1 | 0.49330646 | 33 | 0.81996213 | FALSE | 1 | FALSE | 1 | FALSE | 1 | FALSE | 1 | FALSE | 1 | FALSE | 1 |
| 3517 | 3517 | 8.22857143 | 2.65206889 | 1 | 0.58738519 | 0 | 1 | 2 | 0.34668321 | 1 | 0.7984848 | 2 | 0.1065437 | 28 | 0.72558119 | FALSE | 1 | FALSE | 1 | FALSE | 1 | FALSE | 1 | FALSE | 1 | FALSE | 1 |
| 3518 | 3518 | 9.52380952 | 2.30153257 | 3 | 0.01568237 | 2 | 0.14655134 | 0 | 1 | 0 | 1 | 0 | 1 | 16 | 0.90353674 | FALSE | 0.38695764 | FALSE | 1 | FALSE | 1 | FALSE | 1 | FALSE | 1 | FALSE | 1 |
| 3519 | 3519 | 10.3461539 | 1.94030742 | 0 | 1 | 2 | 0.20486827 | 2 | 0.23940226 | 1 | 0.70601446 | 0 | 1 | 21 | 0.78939668 | FALSE | 1 | FALSE | 1 | FALSE | 1 | FALSE | 1 | FALSE | 1 | FALSE | 1 |
| 3520 | 3520 | 10.4761905 | 2.88871446 | 0 | 1 | 1 | 0.49918218 | 1 | 0.53671415 | 1 | 0.62783522 | 1 | 0.29368245 | 17 | 0.77849076 | FALSE | 1 | FALSE | 1 | FALSE | 1 | FALSE | 1 | FALSE | 1 | FALSE | 1 |
| 3521 | 3521 | 9 | 2.29624199 | 0 | 1 | 0 | 1 | 0 | 1 | 1 | 0.40390032 | 0 | 1 | 10 | 0.46859937 | FALSE | 1 | FALSE | 1 | FALSE | 1 | FALSE | 1 | FALSE | 1 | FALSE | 1 |
| 3522 | 3522 | 9.35 | 1.91768089 | 0 | 1 | 7 | 0.00025225 | 0 | 1 | 2 | 0.5543522 | 0 | 1 | 31 | 0.91676852 | FALSE | 1 | FALSE | 0.15723145 | FALSE | 1 | FALSE | 1 | FALSE | 1 | FALSE | 1 |
| 3523 | 3523 | 11.2142857 | 2.07634884 | 0 | 1 | 2 | 0.07341559 | 1 | 0.40110418 | 1 | 0.48242099 | 0 | 1 | 10 | 0.94579623 | FALSE | 1 | FALSE | 1 | FALSE | 1 | FALSE | 1 | FALSE | 1 | FALSE | 1 |
| 3524 | 3524 | 2.35628743 | 1.0390602 | 6 | 0.99958412 | 22 | 0.46830942 | 20 | 0.81817921 | 27 | 0.76297295 | 19 | 0.00995519 | 564 | 0.17474895 | FALSE | 1 | FALSE | 1 | FALSE | 1 | FALSE | 1 | FALSE | 0.55478195 | FALSE | 1 |
| 3525 | 3525 | 10.2222222 | 2.39340658 | 0 | 1 | 0 | 1 | 0 | 1 | 0 | 1 | 0 | 1 | 9 | 0.21661397 | FALSE | 1 | FALSE | 1 | FALSE | 1 | FALSE | 1 | FALSE | 1 | FALSE | 1 |
| 3526 | 3526 | 6.55384615 | 1.56923077 | 3 | 0.65312131 | 2 | 0.92749221 | 5 | 0.50325307 | 5 | 0.71949272 | 1 | 0.88584977 | 114 | 0.17732033 | FALSE | 1 | FALSE | 1 | FALSE | 1 | FALSE | 1 | FALSE | 1 | FALSE | 1 |
| 3527 | 3527 | 9.03225807 | 2.19212478 | 2 | 0.1881398 | 1 | 0.63997468 | 0 | 1 | 0 | 1 | 1 | 0.4016888 | 27 | 0.45522639 | FALSE | 1 | FALSE | 1 | FALSE | 1 | FALSE | 1 | FALSE | 1 | FALSE | 1 |
| 3528 | 3528 | 10.75 | 1.5 | 0 | 1 | 1 | 0.65167362 | 1 | 0.69069222 | 4 | 0.05697506 | 0 | 1 | 26 | 0.77612337 | FALSE | 1 | FALSE | 1 | FALSE | 1 | FALSE | 1 | FALSE | 1 | FALSE | 1 |
| 3529 | 3529 | 8.13888889 | 3.5130928 | 1 | 0.60837712 | 0 | 1 | 1 | 0.73299778 | 0 | 1 | 2 | 0.11720035 | 32 | 0.31709555 | FALSE | 1 | FALSE | 1 | FALSE | 1 | FALSE | 1 | FALSE | 1 | FALSE | 1 |
| 3530 | 3530 | 7.06792453 | 0.48671932 | 12 | 0.03914932 | 14 | 0.04778273 | 10 | 0.48319844 | 17 | 0.10095168 | 2 | 0.93589217 | 210 | 0.99081691 | FALSE | 0.60856016 | FALSE | 1 | FALSE | 1 | FALSE | 1 | FALSE | 1 | FALSE | 1 |
| 3531 | 3531 | 9.8 | 2.27156334 | 0 | 1 | 0 | 1 | 1 | 0.30655256 | 0 | 1 | 0 | 1 | 9 | 0.5215547 | FALSE | 1 | FALSE | 1 | FALSE | 1 | FALSE | 1 | FALSE | 1 | FALSE | 1 |
| 3532 | 3532 | 9.57142857 | 1.76126114 | 0 | 1 | 0 | 1 | 1 | 0.40110418 | 1 | 0.48242099 | 0 | 1 | 12 | 0.62236227 | FALSE | 1 | FALSE | 1 | FALSE | 1 | FALSE | 1 | FALSE | 1 | FALSE | 1 |
| 3533 | 3533 | 9.53333333 | 3.0739045 | 1 | 0.32293355 | 0 | 1 | 2 | 0.09928574 | 1 | 0.50623194 | 0 | 1 | 11 | 0.92879412 | FALSE | 1 | FALSE | 1 | FALSE | 1 | FALSE | 1 | FALSE | 1 | FALSE | 1 |
| 3534 | 3534 | 11 | 2.62678511 | 2 | 0.08405377 | 0 | 1 | 1 | 0.50143826 | 1 | 0.59104357 | 0 | 1 | 15 | 0.83648191 | FALSE | 0.79236883 | FALSE | 1 | FALSE | 1 | FALSE | 1 | FALSE | 1 | FALSE | 1 |
| 3535 | 3535 | 10.1470588 | 2.0165835 | 1 | 0.58738519 | 3 | 0.09602744 | 0 | 1 | 3 | 0.20353499 | 0 | 1 | 27 | 0.85023803 | FALSE | 1 | FALSE | 1 | FALSE | 1 | FALSE | 1 | FALSE | 1 | FALSE | 1 |
| 3536 | 3536 | 10.0833333 | 1.70579079 | 0 | 1 | 0 | 1 | 1 | 0.35555293 | 0 | 1 | 1 | 0.18007762 | 10 | 0.71436964 | FALSE | 1 | FALSE | 1 | FALSE | 1 | FALSE | 1 | FALSE | 1 | FALSE | 1 |
| 3537 | 3537 | 9.57142857 | 1.64378856 | 1 | 0.59801736 | 0 | 1 | 2 | 0.35984046 | 3 | 0.21561745 | 1 | 0.44015537 | 28 | 0.83079716 | FALSE | 1 | FALSE | 1 | FALSE | 1 | FALSE | 1 | FALSE | 1 | FALSE | 1 |
| 3538 | 3538 | 8.12121212 | 1.95025369 | 0 | 1 | 0 | 1 | 0 | 1 | 4 | 0.05697506 | 1 | 0.41154418 | 27 | 0.61671133 | FALSE | 1 | FALSE | 1 | FALSE | 1 | FALSE | 1 | FALSE | 1 | FALSE | 1 |
| 3539 | 3539 | 10.6363636 | 2.42234774 | 0 | 1 | 0 | 1 | 1 | 0.33149969 | 1 | 0.40390032 | 0 | 1 | 9 | 0.75985366 | FALSE | 1 | FALSE | 1 | FALSE | 1 | FALSE | 1 | FALSE | 1 | FALSE | 1 |
| 3540 | 3540 | 8 | 1.82574186 | 3 | 0.02256792 | 1 | 0.54637326 | 0 | 1 | 1 | 0.67692816 | 0 | 1 | 19 | 0.83975208 | FALSE | 0.4692294 | FALSE | 1 | FALSE | 1 | FALSE | 1 | FALSE | 1 | FALSE | 1 |
| 3541 | 3541 | 10.8947368 | 2.29174116 | 0 | 1 | 1 | 0.46503795 | 1 | 0.50143826 | 1 | 0.59104357 | 0 | 1 | 16 | 0.65565778 | FALSE | 1 | FALSE | 1 | FALSE | 1 | FALSE | 1 | FALSE | 1 | FALSE | 1 |
| 3542 | 3542 | 8.88 | 2.43836011 | 0 | 1 | 2 | 0.1353446 | 1 | 0.51939841 | 1 | 0.60987146 | 0 | 1 | 16 | 0.80826361 | FALSE | 1 | FALSE | 1 | FALSE | 1 | FALSE | 1 | FALSE | 1 | FALSE | 1 |
| 3543 | 3543 | 8.11320755 | 2.03423879 | 0 | 1 | 0 | 1 | 2 | 0.5626042 | 6 | 0.03073996 | 0 | 1 | 44 | 0.57424118 | FALSE | 1 | FALSE | 1 | FALSE | 1 | FALSE | 1 | FALSE | 1 | FALSE | 1 |
| 3544 | 3544 | 10.2222222 | 2.69887951 | 0 | 1 | 1 | 0.44710673 | 0 | 1 | 1 | 0.57131013 | 0 | 1 | 16 | 0.44998001 | FALSE | 1 | FALSE | 1 | FALSE | 1 | FALSE | 1 | FALSE | 1 | FALSE | 1 |
| 3545 | 3545 | 12.1428571 | 3.63430368 | 0 | 1 | 2 | 0.31355738 | 2 | 0.35984046 | 2 | 0.48233989 | 0 | 1 | 29 | 0.69927719 | FALSE | 1 | FALSE | 1 | FALSE | 1 | FALSE | 1 | FALSE | 1 | FALSE | 1 |
| 3546 | 3546 | 8.625 | 2.49687304 | 0 | 1 | 0 | 1 | 2 | 0.32011871 | 2 | 0.43583874 | 1 | 0.41154418 | 27 | 0.61671133 | FALSE | 1 | FALSE | 1 | FALSE | 1 | FALSE | 1 | FALSE | 1 | FALSE | 1 |
| 3547 | 3547 | 11.3333333 | 2.05480467 | 2 | 0.03643311 | 1 | 0.32623933 | 0 | 1 | 0 | 1 | 0 | 1 | 9 | 0.89630573 | FALSE | 0.58392 | FALSE | 1 | FALSE | 1 | FALSE | 1 | FALSE | 1 | FALSE | 1 |
| 3548 | 3548 | 9.08823529 | 2.69386719 | 1 | 0.58738519 | 0 | 1 | 0 | 1 | 0 | 1 | 2 | 0.1065437 | 31 | 0.1995074 | FALSE | 1 | FALSE | 1 | FALSE | 1 | FALSE | 1 | FALSE | 1 | FALSE | 1 |
| 3549 | 3549 | 7.17647059 | 0.72235677 | 2 | 0.64592863 | 1 | 0.93995595 | 0 | 1 | 5 | 0.35139151 | 1 | 0.75686725 | 76 | 0.12528496 | FALSE | 1 | FALSE | 1 | FALSE | 1 | FALSE | 1 | FALSE | 1 | FALSE | 1 |
| 3550 | 3550 | 6.89795918 | 3.58707941 | 0 | 1 | 3 | 0.21051417 | 3 | 0.25725031 | 2 | 0.66536008 | 4 | 0.00822471 | 37 | 0.96598649 | FALSE | 1 | FALSE | 1 | FALSE | 1 | FALSE | 1 | FALSE | 0.50674765 | FALSE | 1 |
| 3551 | 3551 | 12.3333333 | 4.14996653 | 1 | 0.35728305 | 1 | 0.42857733 | 0 | 1 | 1 | 0.55062772 | 0 | 1 | 14 | 0.7313145 | FALSE | 1 | FALSE | 1 | FALSE | 1 | FALSE | 1 | FALSE | 1 | FALSE | 1 |
| 3552 | 3552 | 5.67857143 | 2.90139833 | 5 | 0.28456894 | 7 | 0.16315538 | 7 | 0.23246036 | 3 | 0.95798521 | 1 | 0.90193101 | 116 | 0.67111871 | FALSE | 1 | FALSE | 1 | FALSE | 1 | FALSE | 1 | FALSE | 1 | FALSE | 1 |
| 3553 | 3553 | 8.5 | 2.59807621 | 0 | 1 | 0 | 1 | 1 | 0.35555293 | 0 | 1 | 0 | 1 | 11 | 0.41944532 | FALSE | 1 | FALSE | 1 | FALSE | 1 | FALSE | 1 | FALSE | 1 | FALSE | 1 |
| 3554 | 3554 | 6.46666667 | 3.53773311 | 1 | 0.68224519 | 2 | 0.41896557 | 4 | 0.07197653 | 3 | 0.32878418 | 2 | 0.16211764 | 32 | 0.98583614 | FALSE | 1 | FALSE | 1 | FALSE | 1 | FALSE | 1 | FALSE | 1 | FALSE | 1 |
| 3555 | 3555 | 7.93333333 | 3.73214267 | 0 | 1 | 0 | 1 | 0 | 1 | 0 | 1 | 1 | 0.2198258 | 14 | 0.29525409 | FALSE | 1 | FALSE | 1 | FALSE | 1 | FALSE | 1 | FALSE | 1 | FALSE | 1 |
| 3556 | 3556 | 7.51428571 | 2.81192795 | 1 | 0.59801736 | 2 | 0.31355738 | 0 | 1 | 2 | 0.48233989 | 0 | 1 | 30 | 0.52765882 | FALSE | 1 | FALSE | 1 | FALSE | 1 | FALSE | 1 | FALSE | 1 | FALSE | 1 |
| 3557 | 3557 | 8.61111111 | 3.48231865 | 1 | 0.37380237 | 1 | 0.44710673 | 1 | 0.48280983 | 1 | 0.57131013 | 0 | 1 | 14 | 0.86287418 | FALSE | 1 | FALSE | 1 | FALSE | 1 | FALSE | 1 | FALSE | 1 | FALSE | 1 |
| 3558 | 3558 | 10.3157895 | 3.22902351 | 0 | 1 | 1 | 0.46503795 | 3 | 0.02898764 | 2 | 0.21610812 | 0 | 1 | 13 | 0.97998676 | FALSE | 1 | FALSE | 1 | FALSE | 1 | FALSE | 1 | FALSE | 1 | FALSE | 1 |
| 3559 | 3559 | 9.59259259 | 3.75410155 | 0 | 1 | 0 | 1 | 0 | 1 | 1 | 0.71956339 | 3 | 0.00942179 | 23 | 0.58307573 | FALSE | 1 | FALSE | 1 | FALSE | 1 | FALSE | 1 | FALSE | 0.5373705 | FALSE | 1 |
| 3560 | 3560 | 10.4285714 | 2.47023092 | 1 | 0.3050805 | 2 | 0.07341559 | 1 | 0.40110418 | 1 | 0.48242099 | 1 | 0.20679328 | 8 | 0.99722299 | FALSE | 1 | FALSE | 1 | FALSE | 1 | FALSE | 1 | FALSE | 1 | FALSE | 1 |
| 3561 | 3561 | 7.25714286 | 2.07492316 | 1 | 0.59801736 | 1 | 0.68454804 | 3 | 0.12958859 | 2 | 0.48233989 | 0 | 1 | 28 | 0.83079716 | FALSE | 1 | FALSE | 1 | FALSE | 1 | FALSE | 1 | FALSE | 1 | FALSE | 1 |
| 3562 | 3562 | 9.41666667 | 2.49861073 | 2 | 0.12501436 | 0 | 1 | 3 | 0.05314317 | 2 | 0.30249307 | 0 | 1 | 17 | 0.97530702 | FALSE | 0.95622564 | FALSE | 1 | FALSE | 1 | FALSE | 1 | FALSE | 1 | FALSE | 1 |
| 3563 | 3563 | 8.77777778 | 3.0469576 | 0 | 1 | 1 | 0.44710673 | 1 | 0.48280983 | 1 | 0.57131013 | 1 | 0.25766374 | 14 | 0.86287418 | FALSE | 1 | FALSE | 1 | FALSE | 1 | FALSE | 1 | FALSE | 1 | FALSE | 1 |
| 3564 | 3564 | 8.4 | 1.68522996 | 2 | 0.0256865 | 0 | 1 | 0 | 1 | 0 | 1 | 0 | 1 | 8 | 0.80394719 | FALSE | 0.4974559 | FALSE | 1 | FALSE | 1 | FALSE | 1 | FALSE | 1 | FALSE | 1 |
| 3565 | 3565 | 7.93333333 | 3.04338992 | 0 | 1 | 0 | 1 | 2 | 0.09928574 | 1 | 0.50623194 | 0 | 1 | 12 | 0.80329917 | FALSE | 1 | FALSE | 1 | FALSE | 1 | FALSE | 1 | FALSE | 1 | FALSE | 1 |
| 3566 | 3566 | 7.32608696 | 2.28467815 | 2 | 0.3309085 | 3 | 0.18548475 | 2 | 0.4962935 | 3 | 0.35428318 | 1 | 0.53375569 | 35 | 0.95410769 | FALSE | 1 | FALSE | 1 | FALSE | 1 | FALSE | 1 | FALSE | 1 | FALSE | 1 |
| 3567 | 3567 | 9.37931035 | 1.98986732 | 0 | 1 | 2 | 0.24095584 | 2 | 0.27983822 | 1 | 0.74482229 | 2 | 0.08121636 | 22 | 0.92902102 | FALSE | 1 | FALSE | 1 | FALSE | 1 | FALSE | 1 | FALSE | 1 | FALSE | 1 |
| 3568 | 3568 | 10.1904762 | 2.30153257 | 0 | 1 | 0 | 1 | 2 | 0.17308361 | 1 | 0.62783522 | 0 | 1 | 18 | 0.57962763 | FALSE | 1 | FALSE | 1 | FALSE | 1 | FALSE | 1 | FALSE | 1 | FALSE | 1 |
| 3569 | 3569 | 8.875 | 2.93417365 | 1 | 0.55378205 | 0 | 1 | 1 | 0.67911179 | 1 | 0.76781284 | 2 | 0.09110253 | 26 | 0.64660109 | FALSE | 1 | FALSE | 1 | FALSE | 1 | FALSE | 1 | FALSE | 1 | FALSE | 1 |
| 3570 | 3570 | 11.0909091 | 2.67834907 | 0 | 1 | 1 | 0.30368014 | 0 | 1 | 0 | 1 | 0 | 1 | 10 | 0.46859937 | FALSE | 1 | FALSE | 1 | FALSE | 1 | FALSE | 1 | FALSE | 1 | FALSE | 1 |

| | | | | | | | | | | | | | | | | | | | | | | | | | | | |
|---|---|---|---|---|---|---|---|---|---|---|---|---|---|---|---|---|---|---|---|---|---|---|---|---|---|---|---|
| 3571 | 3571 | 11.3888889 | 4.6324806 | 0 | 1 | 0 | 1 | 1 | 0.37874417 | 1 | 0.45746571 | 0 | 1 | 11 | 0.6683056 | FALSE | 1 | FALSE | 1 | FALSE | 1 | FALSE | 1 | FALSE | 1 | FALSE | 1 |
| 3572 | 3572 | 8.2195122 | 2.26868604 | 2 | 0.28329504 | 1 | 0.74131698 | 1 | 0.77786601 | 2 | 0.56788092 | 1 | 0.49330646 | 34 | 0.69505404 | FALSE | 1 | FALSE | 1 | FALSE | 1 | FALSE | 1 | FALSE | 1 | FALSE | 1 |
| 3573 | 3573 | 10 | 2.29128785 | 0 | 1 | 0 | 1 | 1 | 0.25384309 | 2 | 0.048966 | 0 | 1 | 5 | 0.97551477 | FALSE | 1 | FALSE | 1 | FALSE | 1 | FALSE | 1 | FALSE | 1 | FALSE | 1 |
| 3574 | 3574 | 9.33333333 | 2.05480467 | 2 | 0.12501436 | 2 | 0.18117183 | 1 | 0.58502114 | 2 | 0.30249307 | 1 | 0.32796862 | 16 | 0.99240722 | FALSE | 0.95622564 | FALSE | 1 | FALSE | 1 | FALSE | 1 | FALSE | 1 | FALSE | 1 |
| 3575 | 3575 | 9.68421053 | 1.67706819 | 1 | 0.77387657 | 6 | 0.00979964 | 4 | 0.147507 | 3 | 0.49002278 | 2 | 0.23984683 | 41 | 0.99511641 | FALSE | 1 | FALSE | 0.6689889 | FALSE | 1 | FALSE | 1 | FALSE | 1 | FALSE | 1 |
| 3576 | 3576 | 10.8461539 | 2.597222 | 0 | 1 | 0 | 1 | 0 | 1 | 1 | 0.43131129 | 1 | 0.18007762 | 10 | 0.71436964 | FALSE | 1 | FALSE | 1 | FALSE | 1 | FALSE | 1 | FALSE | 1 | FALSE | 1 |
| 3577 | 3577 | 7.96226415 | 2.34691499 | 1 | 0.74890551 | 3 | 0.24492539 | 2 | 0.57303317 | 2 | 0.70707241 | 0 | 1 | 45 | 0.5504765 | FALSE | 1 | FALSE | 1 | FALSE | 1 | FALSE | 1 | FALSE | 1 | FALSE | 1 |
| 3578 | 3578 | 8.97916667 | 2.47057334 | 3 | 0.12433927 | 2 | 0.46321244 | 0 | 1 | 3 | 0.37966044 | 0 | 1 | 40 | 0.66905473 | FALSE | 0.95622564 | FALSE | 1 | FALSE | 1 | FALSE | 1 | FALSE | 1 | FALSE | 1 |
| 3579 | 3579 | 9 | 2.09165007 | 0 | 1 | 0 | 1 | 0 | 1 | 1 | 0.52895088 | 0 | 1 | 15 | 0.26127859 | FALSE | 1 | FALSE | 1 | FALSE | 1 | FALSE | 1 | FALSE | 1 | FALSE | 1 |
| 3580 | 3580 | 8.22727273 | 3.17564336 | 7 | 0.00011085 | 1 | 0.76576289 | 0 | 1 | 1 | 0.87439647 | 1 | 0.51797501 | 34 | 0.9286641 | TRUE | 0.0189187 | FALSE | 1 | FALSE | 1 | FALSE | 1 | FALSE | 1 | FALSE | 1 |
| 3581 | 3581 | 8.72727273 | 2.21937375 | 0 | 1 | 1 | 0.30368014 | 1 | 0.33149969 | 0 | 1 | 1 | 0.16638736 | 8 | 0.92152993 | FALSE | 1 | FALSE | 1 | FALSE | 1 | FALSE | 1 | FALSE | 1 | FALSE | 1 |
| 3582 | 3582 | 10.5 | 3.45205253 | 0 | 1 | 0 | 1 | 1 | 0.35555293 | 0 | 1 | 1 | 0.18007762 | 10 | 0.71436964 | FALSE | 1 | FALSE | 1 | FALSE | 1 | FALSE | 1 | FALSE | 1 | FALSE | 1 |
| 3583 | 3583 | 10.1333333 | 1.7074998 | 0 | 1 | 1 | 0.38964394 | 1 | 0.42266264 | 2 | 0.14906763 | 0 | 1 | 11 | 0.92879412 | FALSE | 1 | FALSE | 1 | FALSE | 1 | FALSE | 1 | FALSE | 1 | FALSE | 1 |
| 3584 | 3584 | 9.47058824 | 2.97277148 | 2 | 0.06909038 | 0 | 1 | 0 | 1 | 0 | 1 | 0 | 1 | 15 | 0.49060435 | FALSE | 0.74276693 | FALSE | 1 | FALSE | 1 | FALSE | 1 | FALSE | 1 | FALSE | 1 |
| 3585 | 3585 | 8.69565217 | 2.27337678 | 2 | 0.32143972 | 0 | 1 | 2 | 0.48463745 | 4 | 0.15016749 | 1 | 0.52593041 | 36 | 0.84604146 | FALSE | 1 | FALSE | 1 | FALSE | 1 | FALSE | 1 | FALSE | 1 | FALSE | 1 |
| 3586 | 3586 | 9.73684211 | 2.55141045 | 6 | 5.35E-06 | 1 | 0.46503795 | 0 | 1 | 1 | 0.59104357 | 1 | 0.26986775 | 10 | 0.99980197 | TRUE | 0.00299395 | FALSE | 1 | FALSE | 1 | FALSE | 1 | FALSE | 1 | FALSE | 1 |
| 3587 | 3587 | 7.46428571 | 2.25170004 | 0 | 1 | 3 | 0.27128673 | 0 | 1 | 4 | 0.25497934 | 1 | 0.60529938 | 48 | 0.48035049 | FALSE | 1 | FALSE | 1 | FALSE | 1 | FALSE | 1 | FALSE | 1 | FALSE | 1 |
| 3588 | 3588 | 6.10714286 | 2.75602309 | 1 | 0.51745948 | 3 | 0.06039219 | 1 | 0.64171563 | 3 | 0.13524571 | 1 | 0.37113067 | 19 | 0.99262381 | FALSE | 1 | FALSE | 1 | FALSE | 1 | FALSE | 1 | FALSE | 1 | FALSE | 1 |
| 3589 | 3589 | 10.95 | 1.65755845 | 1 | 0.40558521 | 0 | 1 | 1 | 0.51939841 | 1 | 0.60987146 | 1 | 0.28187291 | 16 | 0.80826361 | FALSE | 1 | FALSE | 1 | FALSE | 1 | FALSE | 1 | FALSE | 1 | FALSE | 1 |
| 3590 | 3590 | 10.375 | 4.36427256 | 0 | 1 | 2 | 0.27730767 | 0 | 1 | 1 | 0.77852168 | 0 | 1 | 29 | 0.24088541 | FALSE | 1 | FALSE | 1 | FALSE | 1 | FALSE | 1 | FALSE | 1 | FALSE | 1 |
| 3591 | 3591 | 8.54545455 | 2.25080335 | 0 | 1 | 3 | 0.03266339 | 0 | 1 | 1 | 0.64497439 | 0 | 1 | 18 | 0.74744813 | FALSE | 1 | FALSE | 0.92268438 | FALSE | 1 | FALSE | 1 | FALSE | 1 | FALSE | 1 |
| 3592 | 3592 | 10.5882353 | 2.87092573 | 1 | 0.35728305 | 0 | 1 | 0 | 1 | 0 | 1 | 1 | 0.24525764 | 15 | 0.49060435 | FALSE | 1 | FALSE | 1 | FALSE | 1 | FALSE | 1 | FALSE | 1 | FALSE | 1 |
| 3593 | 3593 | 9.2 | 0.94516313 | 1 | 0.54198714 | 0 | 1 | 2 | 0.29330414 | 1 | 0.75658797 | 0 | 1 | 26 | 0.48609328 | FALSE | 1 | FALSE | 1 | FALSE | 1 | FALSE | 1 | FALSE | 1 | FALSE | 1 |
| 3594 | 3594 | 10.5833333 | 2.32588096 | 0 | 1 | 0 | 1 | 0 | 1 | 0 | 1 | 1 | 0.18007762 | 11 | 0.41944532 | FALSE | 1 | FALSE | 1 | FALSE | 1 | FALSE | 1 | FALSE | 1 | FALSE | 1 |
| 3595 | 3595 | 8.29411765 | 2.44383272 | 1 | 0.35728305 | 0 | 1 | 1 | 0.46348836 | 1 | 0.55062772 | 0 | 1 | 14 | 0.7313145 | FALSE | 1 | FALSE | 1 | FALSE | 1 | FALSE | 1 | FALSE | 1 | FALSE | 1 |
| 3596 | 3596 | 10.72 | 2.0103731 | 0 | 1 | 0 | 1 | 0 | 1 | 2 | 0.31968852 | 0 | 1 | 23 | 0.22727183 | FALSE | 1 | FALSE | 1 | FALSE | 1 | FALSE | 1 | FALSE | 1 | FALSE | 1 |
| 3597 | 3597 | 11.5405405 | 2.91866866 | 0 | 1 | 1 | 0.70472939 | 0 | 1 | 3 | 0.2402032 | 1 | 0.45845922 | 32 | 0.47030992 | FALSE | 1 | FALSE | 1 | FALSE | 1 | FALSE | 1 | FALSE | 1 | FALSE | 1 |
| 3598 | 3598 | 8.75862069 | 3.43064458 | 0 | 1 | 3 | 0.06581417 | 1 | 0.65463982 | 1 | 0.74482229 | 0 | 1 | 24 | 0.70563334 | FALSE | 1 | FALSE | 1 | FALSE | 1 | FALSE | 1 | FALSE | 1 | FALSE | 1 |
| 3599 | 3599 | 10.6875 | 3.21556274 | 0 | 1 | 0 | 1 | 3 | 0.01812751 | 2 | 0.16544789 | 0 | 1 | 11 | 0.9718531 | FALSE | 1 | FALSE | 1 | FALSE | 1 | FALSE | 1 | FALSE | 1 | FALSE | 1 |
| 3600 | 3600 | 7.0483871 | 4.17143674 | 2 | 0.46653175 | 1 | 0.86665405 | 3 | 0.3760478 | 1 | 0.94386203 | 1 | 0.63687695 | 53 | 0.37131894 | FALSE | 1 | FALSE | 1 | FALSE | 1 | FALSE | 1 | FALSE | 1 | FALSE | 1 |
| 3601 | 3601 | 9 | 1.93218357 | 2 | 0.04858661 | 2 | 0.07341559 | 2 | 0.08806398 | 0 | 1 | 0 | 1 | 8 | 0.99722299 | FALSE | 0.649301 | FALSE | 1 | FALSE | 1 | FALSE | 1 | FALSE | 1 | FALSE | 1 |
| 3602 | 3602 | 8.56896552 | 1.79185965 | 2 | 0.44060625 | 4 | 0.11722369 | 1 | 0.88128698 | 2 | 0.75303313 | 0 | 1 | 49 | 0.57916832 | FALSE | 1 | FALSE | 1 | FALSE | 1 | FALSE | 1 | FALSE | 1 | FALSE | 1 |
| 3603 | 3603 | 8.4375 | 1.93548282 | 1 | 0.34033054 | 0 | 1 | 0 | 1 | 0 | 1 | 0 | 1 | 15 | 0.26127859 | FALSE | 1 | FALSE | 1 | FALSE | 1 | FALSE | 1 | FALSE | 1 | FALSE | 1 |
| 3604 | 3604 | 10.3684211 | 2.67956254 | 1 | 0.38989951 | 2 | 0.12434464 | 1 | 0.50143826 | 1 | 0.59104357 | 2 | 0.03799168 | 12 | 0.99479039 | FALSE | 1 | FALSE | 1 | FALSE | 1 | FALSE | 1 | FALSE | 0.87669812 | FALSE | 1 |
| 3605 | 3605 | 9.40740741 | 2.80554028 | 2 | 0.14250978 | 2 | 0.20486827 | 1 | 0.61440466 | 1 | 0.70601446 | 1 | 0.3499045 | 19 | 0.96054244 | FALSE | 1 | FALSE | 1 | FALSE | 1 | FALSE | 1 | FALSE | 1 | FALSE | 1 |
| 3606 | 3606 | 9.58333333 | 2.01900691 | 1 | 0.26795838 | 0 | 1 | 0 | 1 | 0 | 1 | 0 | 1 | 11 | 0.41944532 | FALSE | 1 | FALSE | 1 | FALSE | 1 | FALSE | 1 | FALSE | 1 | FALSE | 1 |
| 3607 | 3607 | 9.82352941 | 1.54292671 | 0 | 1 | 0 | 1 | 3 | 0.02143959 | 1 | 0.55062772 | 0 | 1 | 13 | 0.88719222 | FALSE | 1 | FALSE | 1 | FALSE | 1 | FALSE | 1 | FALSE | 1 | FALSE | 1 |
| 3608 | 3608 | 7.92857143 | 2.6849923 | 1 | 0.3050805 | 0 | 1 | 0 | 1 | 1 | 0.48242099 | 0 | 1 | 12 | 0.62236227 | FALSE | 1 | FALSE | 1 | FALSE | 1 | FALSE | 1 | FALSE | 1 | FALSE | 1 |
| 3609 | 3609 | 10.7 | 3.91279951 | 0 | 1 | 1 | 0.48239024 | 1 | 0.51939841 | 1 | 0.60987146 | 0 | 1 | 17 | 0.61750632 | FALSE | 1 | FALSE | 1 | FALSE | 1 | FALSE | 1 | FALSE | 1 | FALSE | 1 |
| 3610 | 3610 | 12.3846154 | 2.40315375 | 0 | 1 | 0 | 1 | 0 | 1 | 1 | 0.45746571 | 0 | 1 | 12 | 0.374198 | FALSE | 1 | FALSE | 1 | FALSE | 1 | FALSE | 1 | FALSE | 1 | FALSE | 1 |
| 3611 | 3611 | 8.68181818 | 2.41936062 | 0 | 1 | 0 | 1 | 1 | 0.55340851 | 0 | 1 | 0 | 1 | 21 | 0.12048848 | FALSE | 1 | FALSE | 1 | FALSE | 1 | FALSE | 1 | FALSE | 1 | FALSE | 1 |
| 3612 | 3612 | 3.25147348 | 1.50167647 | 8 | 0.95364956 | 19 | 0.28051144 | 18 | 0.55403542 | 26 | 0.30061309 | 10 | 0.31622673 | 425 | 0.62711313 | FALSE | 1 | FALSE | 1 | FALSE | 1 | FALSE | 1 | FALSE | 1 | FALSE | 1 |
| 3613 | 3613 | 9.3 | 0.64031242 | 0 | 1 | 0 | 1 | 0 | 1 | 2 | 0.0740981 | 0 | 1 | 8 | 0.80394719 | FALSE | 1 | FALSE | 1 | FALSE | 1 | FALSE | 1 | FALSE | 1 | FALSE | 1 |
| 3614 | 3614 | 11.7142857 | 1.97948664 | 0 | 1 | 0 | 1 | 1 | 0.22601089 | 0 | 1 | 0 | 1 | 6 | 0.69920685 | FALSE | 1 | FALSE | 1 | FALSE | 1 | FALSE | 1 | FALSE | 1 | FALSE | 1 |
| 3615 | 3615 | 10.1111111 | 3.03477784 | 0 | 1 | 0 | 1 | 2 | 0.13502329 | 0 | 1 | 0 | 1 | 16 | 0.44998001 | FALSE | 1 | FALSE | 1 | FALSE | 1 | FALSE | 1 | FALSE | 1 | FALSE | 1 |
| 3616 | 3616 | 8.67647059 | 1.69290193 | 1 | 0.57647348 | 1 | 0.6629941 | 1 | 0.70185639 | 2 | 0.45159487 | 2 | 0.10132035 | 26 | 0.86852567 | FALSE | 1 | FALSE | 1 | FALSE | 1 | FALSE | 1 | FALSE | 1 | FALSE | 1 |
| 3617 | 3617 | 8.47058824 | 1.37702352 | 1 | 0.35728305 | 3 | 0.01622926 | 1 | 0.46348836 | 1 | 0.55062772 | 0 | 1 | 11 | 0.98974663 | FALSE | 1 | FALSE | 0.7256023 | FALSE | 1 | FALSE | 1 | FALSE | 1 | FALSE | 1 |
| 3618 | 3618 | 9.04347826 | 2.74223427 | 0 | 1 | 0 | 1 | 0 | 1 | 1 | 0.6613267 | 0 | 1 | 22 | 0.10533112 | FALSE | 1 | FALSE | 1 | FALSE | 1 | FALSE | 1 | FALSE | 1 | FALSE | 1 |
| 3619 | 3619 | 9.05882353 | 2.08554093 | 0 | 1 | 3 | 0.09602744 | 3 | 0.12152681 | 1 | 0.7984848 | 3 | 0.01774672 | 24 | 0.98848472 | FALSE | 1 | FALSE | 1 | FALSE | 1 | FALSE | 1 | FALSE | 0.69720022 | FALSE | 1 |
| 3620 | 3620 | 10.625 | 2.5464436 | 0 | 1 | 0 | 1 | 0 | 1 | 0 | 1 | 0 | 1 | 8 | 0.25677166 | FALSE | 1 | FALSE | 1 | FALSE | 1 | FALSE | 1 | FALSE | 1 | FALSE | 1 |
| 3621 | 3621 | 6.29504505 | 1.18011786 | 11 | 0.58628299 | 16 | 0.35640246 | 14 | 0.72798295 | 23 | 0.29399377 | 11 | 0.10777075 | 367 | 0.81014434 | FALSE | 1 | FALSE | 1 | FALSE | 1 | FALSE | 1 | FALSE | 1 | FALSE | 1 |
| 3622 | 3622 | 8.18181818 | 2.60482705 | 1 | 0.68224519 | 2 | 0.41896557 | 2 | 0.47281647 | 1 | 0.87439647 | 0 | 1 | 38 | 0.45705039 | FALSE | 1 | FALSE | 1 | FALSE | 1 | FALSE | 1 | FALSE | 1 | FALSE | 1 |
| 3623 | 3623 | 10.3636364 | 3.25500251 | 1 | 0.24866461 | 0 | 1 | 0 | 1 | 1 | 0.40390032 | 0 | 1 | 9 | 0.75985366 | FALSE | 1 | FALSE | 1 | FALSE | 1 | FALSE | 1 | FALSE | 1 | FALSE | 1 |
| 3624 | 3624 | 6.98148148 | 2.71136661 | 1 | 0.75539413 | 7 | 0.00163297 | 3 | 0.30658915 | 4 | 0.2347767 | 2 | 0.22161752 | 37 | 0.99905566 | FALSE | 1 | FALSE | 0.40311621 | FALSE | 1 | FALSE | 1 | FALSE | 1 | FALSE | 1 |
| 3625 | 3625 | 8.2962963 | 2.01503267 | 0 | 1 | 1 | 0.57534752 | 1 | 0.61440466 | 1 | 0.70601446 | 1 | 0.3499045 | 22 | 0.61628 | FALSE | 1 | FALSE | 1 | FALSE | 1 | FALSE | 1 | FALSE | 1 | FALSE | 1 |
| 3626 | 3626 | 7.33823529 | 1.83599864 | 4 | 0.09623316 | 4 | 0.17706447 | 3 | 0.44404438 | 2 | 0.82657478 | 1 | 0.6769215 | 54 | 0.9000463 | FALSE | 0.86047225 | FALSE | 1 | FALSE | 1 | FALSE | 1 | FALSE | 1 | FALSE | 1 |
| 3627 | 3627 | 11.55 | 3.61213233 | 0 | 1 | 0 | 1 | 1 | 0.51939841 | 0 | 1 | 1 | 0.28187291 | 18 | 0.37504645 | FALSE | 1 | FALSE | 1 | FALSE | 1 | FALSE | 1 | FALSE | 1 | FALSE | 1 |
| 3628 | 3628 | 8.20689655 | 2.49660888 | 0 | 1 | 0 | 1 | 3 | 0.08431992 | 1 | 0.74482229 | 3 | 0.01149488 | 22 | 0.92902102 | FALSE | 1 | FALSE | 1 | FALSE | 1 | FALSE | 1 | FALSE | 0.57972935 | FALSE | 1 |
| 3629 | 3629 | 9.5 | 2.5 | 0 | 1 | 0 | 1 | 0 | 1 | 0 | 1 | 0 | 1 | 6 | 0.36077158 | FALSE | 1 | FALSE | 1 | FALSE | 1 | FALSE | 1 | FALSE | 1 | FALSE | 1 |
| 3630 | 3630 | 6.44680851 | 2.94494526 | 1 | 0.69843502 | 2 | 0.44132825 | 2 | 0.4962935 | 3 | 0.35428318 | 0 | 1 | 38 | 0.71504609 | FALSE | 1 | FALSE | 1 | FALSE | 1 | FALSE | 1 | FALSE | 1 | FALSE | 1 |
| 3631 | 3631 | 7.05194805 | 2.1252722 | 0 | 1 | 0 | 1 | 4 | 0.29916433 | 6 | 0.14118821 | 1 | 0.72203879 | 66 | 0.44767694 | FALSE | 1 | FALSE | 1 | FALSE | 1 | FALSE | 1 | FALSE | 1 | FALSE | 1 |
| 3632 | 3632 | 8 | 1.41421356 | 1 | 0.35728305 | 1 | 0.42857733 | 1 | 0.46348836 | 1 | 0.55062772 | 0 | 1 | 13 | 0.88719222 | FALSE | 1 | FALSE | 1 | FALSE | 1 | FALSE | 1 | FALSE | 1 | FALSE | 1 |
| 3633 | 3633 | 7.59649123 | 1.34893914 | 9 | 1.21E-05 | 8 | 0.00044364 | 2 | 0.61293419 | 6 | 0.04538398 | 0 | 1 | 32 | 0.99999993 | TRUE | 0.00418892 | FALSE | 0.17306256 | FALSE | 1 | FALSE | 1 | FALSE | 1 | FALSE | 1 |
| 3634 | 3634 | 11.0625 | 2.72646176 | 0 | 1 | 0 | 1 | 1 | 0.44344817 | 0 | 1 | 1 | 0.23264612 | 14 | 0.53308261 | FALSE | 1 | FALSE | 1 | FALSE | 1 | FALSE | 1 | FALSE | 1 | FALSE | 1 |
| 3635 | 3635 | 9.08333333 | 1.95611577 | 1 | 0.46441868 | 1 | 0.54637326 | 0 | 1 | 0 | 1 | 0 | 1 | 22 | 0.25237063 | FALSE | 1 | FALSE | 1 | FALSE | 1 | FALSE | 1 | FALSE | 1 | FALSE | 1 |
| 3636 | 3636 | 7.93103448 | 3.42839094 | 1 | 0.77387657 | 0 | 1 | 3 | 0.3364112 | 4 | 0.26521223 | 0 | 1 | 49 | 0.45760293 | FALSE | 1 | FALSE | 1 | FALSE | 1 | FALSE | 1 | FALSE | 1 | FALSE | 1 |
| 3637 | 3637 | 11 | 3.94188553 | 0 | 1 | 0 | 1 | 0 | 1 | 0 | 1 | 0 | 1 | 12 | 0.13002667 | FALSE | 1 | FALSE | 1 | FALSE | 1 | FALSE | 1 | FALSE | 1 | FALSE | 1 |
| 3638 | 3638 | 7.5 | 2.75378527 | 0 | 1 | 1 | 0.44710673 | 0 | 1 | 2 | 0.19902682 | 0 | 1 | 15 | 0.69372602 | FALSE | 1 | FALSE | 1 | FALSE | 1 | FALSE | 1 | FALSE | 1 | FALSE | 1 |
| 3639 | 3639 | 11 | 1.46759877 | 0 | 1 | 2 | 0.06426041 | 2 | 0.07725803 | 0 | 1 | 1 | 0.19354507 | 8 | 0.99085298 | FALSE | 1 | FALSE | 1 | FALSE | 1 | FALSE | 1 | FALSE | 1 | FALSE | 1 |
| 3640 | 3640 | 9.20833333 | 1.2240359 | 2 | 0.12501436 | 3 | 0.04096158 | 3 | 0.05314317 | 0 | 1 | 0 | 1 | 16 | 0.99240722 | FALSE | 0.95622564 | FALSE | 0.9698186 | FALSE | 1 | FALSE | 1 | FALSE | 1 | FALSE | 1 |
| 3641 | 3641 | 8.9 | 3.78021163 | 0 | 1 | 0 | 1 | 0 | 1 | 0 | 1 | 0 | 1 | 10 | 0.18273166 | FALSE | 1 | FALSE | 1 | FALSE | 1 | FALSE | 1 | FALSE | 1 | FALSE | 1 |
| 3642 | 3642 | 7.61111111 | 2.48638267 | 1 | 0.60837712 | 2 | 0.32556427 | 1 | 0.73299778 | 1 | 0.81665383 | 2 | 0.11720035 | 29 | 0.81028289 | FALSE | 1 | FALSE | 1 | FALSE | 1 | FALSE | 1 | FALSE | 1 | FALSE | 1 |
| 3643 | 3643 | 9.4 | 3.2 | 1 | 0.22886536 | 1 | 0.28036922 | 0 | 1 | 0 | 1 | 1 | 0.15247062 | 7 | 0.94326215 | FALSE | 1 | FALSE | 1 | FALSE | 1 | FALSE | 1 | FALSE | 1 | FALSE | 1 |
| 3644 | 3644 | 10.9333333 | 3.08688984 | 2 | 0.05513784 | 0 | 1 | 0 | 1 | 2 | 0.14906763 | 1 | 0.2198258 | 10 | 0.97980046 | FALSE | 0.68849059 | FALSE | 1 | FALSE | 1 | FALSE | 1 | FALSE | 1 | FALSE | 1 |
| 3645 | 3645 | 11.25 | 1.63935963 | 2 | 0.01652925 | 2 | 0.02563923 | 0 | 1 | 0 | 1 | 0 | 1 | 4 | 0.99658732 | FALSE | 0.39708985 | FALSE | 0.82367276 | FALSE | 1 | FALSE | 1 | FALSE | 1 | FALSE | 1 |
| 3646 | 3646 | 9 | 1.76776695 | 1 | 0.34033054 | 1 | 0.40942991 | 1 | 0.44344817 | 2 | 0.16544789 | 0 | 1 | 11 | 0.9718531 | FALSE | 1 | FALSE | 1 | FALSE | 1 | FALSE | 1 | FALSE | 1 | FALSE | 1 |
| 3647 | 3647 | 4.8 | 2.90860791 | 8 | 0.08009981 | 6 | 0.50159279 | 10 | 0.09829966 | 9 | 0.41225329 | 3 | 0.55123144 | 139 | 0.96994853 | FALSE | 0.79236883 | FALSE | 1 | FALSE | 1 | FALSE | 1 | FALSE | 1 | FALSE | 1 |
| 3648 | 3648 | 9.1875 | 2.32429425 | 0 | 1 | 3 | 0.17732615 | 2 | 0.48463745 | 2 | 0.61900576 | 0 | 1 | 38 | 0.59437745 | FALSE | 1 | FALSE | 1 | FALSE | 1 | FALSE | 1 | FALSE | 1 | FALSE | 1 |
| 3649 | 3649 | 5.95 | 3.81411851 | 1 | 0.64722818 | 3 | 0.13823565 | 0 | 1 | 2 | 0.5543522 | 2 | 0.13925369 | 32 | 0.83853687 | FALSE | 1 | FALSE | 1 | FALSE | 1 | FALSE | 1 | FALSE | 1 | FALSE | 1 |
| 3650 | 3650 | 2.22279129 | 1.39838681 | 21 | 0.43054992 | 33 | 0.05906327 | 37 | 0.04229616 | 24 | 0.989595 | 14 | 0.39410964 | 647 | 0.8087112 | FALSE | 1 | FALSE | 1 | FALSE | 1 | FALSE | 1 | FALSE | 1 | FALSE | 1 |
| 3651 | 3651 | 3.34048257 | 1.5822056 | 11 | 0.35669059 | 16 | 0.14805294 | 11 | 0.79052189 | 16 | 0.64532772 | 5 | 0.73797946 | 313 | 0.58653118 | FALSE | 1 | FALSE | 1 | FALSE | 1 | FALSE | 1 | FALSE | 1 | FALSE | 1 |
| 3652 | 3652 | 10.0769231 | 3.90518797 | 0 | 1 | 2 | 0.36122049 | 0 | 1 | 3 | 0.26522531 | 0 | 1 | 34 | 0.41570781 | FALSE | 1 | FALSE | 1 | FALSE | 1 | FALSE | 1 | FALSE | 1 | FALSE | 1 |
| 3653 | 3653 | 9.74285714 | 2.96977976 | 0 | 1 | 0 | 1 | 3 | 0.12958859 | 2 | 0.48233989 | 0 | 1 | 30 | 0.52765882 | FALSE | 1 | FALSE | 1 | FALSE | 1 | FALSE | 1 | FALSE | 1 | FALSE | 1 |
| 3654 | 3654 | 11.1111111 | 1.91162784 | 0 | 1 | 1 | 0.25628163 | 1 | 0.28067846 | 0 | 1 | 1 | 0.1383237 | 6 | 0.96128317 | FALSE | 1 | FALSE | 1 | FALSE | 1 | FALSE | 1 | FALSE | 1 | FALSE | 1 |
| 3655 | 3655 | 9.16666667 | 2.33927814 | 0 | 1 | 0 | 1 | 0 | 1 | 1 | 0.24578796 | 0 | 1 | 5 | 0.76185568 | FALSE | 1 | FALSE | 1 | FALSE | 1 | FALSE | 1 | FALSE | 1 | FALSE | 1 |
| 3656 | 3656 | 8.84375 | 2.89514006 | 2 | 0.19750261 | 1 | 0.65167362 | 0 | 1 | 1 | 0.77852168 | 0 | 1 | 28 | 0.42532177 | FALSE | 1 | FALSE | 1 | FALSE | 1 | FALSE | 1 | FALSE | 1 | FALSE | 1 |
| 3657 | 3657 | 5.78125 | 2.21830982 | 1 | 0.91883959 | 2 | 0.82287436 | 10 | 0.00221901 | 3 | 0.82437964 | 0 | 1 | 80 | 0.67414989 | FALSE | 1 | FALSE | 1 | FALSE | 1 | FALSE | 1 | FALSE | 1 | FALSE | 1 |
| 3658 | 3658 | 7.71052632 | 2.1387145 | 1 | 0.62830697 | 0 | 1 | 0 | 1 | 2 | 0.52640931 | 0 | 1 | 35 | 0.13407055 | FALSE | 1 | FALSE | 1 | FALSE | 1 | FALSE | 1 | FALSE | 1 | FALSE | 1 |
| 3659 | 3659 | 10.6666667 | 3.52766842 | 1 | 0.20854748 | 0 | 1 | 0 | 1 | 0 | 1 | 0 | 1 | 8 | 0.5780332 | FALSE | 1 | FALSE | 1 | FALSE | 1 | FALSE | 1 | FALSE | 1 | FALSE | 1 |
| 3660 | 3660 | 7.07594937 | 2.70850583 | 0 | 1 | 2 | 0.72420282 | 5 | 0.14803148 | 5 | 0.28743251 | 1 | 0.72664912 | 65 | 0.67060743 | FALSE | 1 | FALSE | 1 | FALSE | 1 | FALSE | 1 | FALSE | 1 | FALSE | 1 |
| 3661 | 3661 | 10.7619048 | 1.34180979 | 0 | 1 | 1 | 0.49918218 | 1 | 0.53671415 | 0 | 1 | 0 | 1 | 19 | 0.34094959 | FALSE | 1 | FALSE | 1 | FALSE | 1 | FALSE | 1 | FALSE | 1 | FALSE | 1 |
| 3662 | 3662 | 5.95652174 | 3.22326139 | 2 | 0.30241117 | 1 | 0.75788114 | 0 | 1 | 5 | 0.0456107 | 1 | 0.50988734 | 34 | 0.87722689 | FALSE | 1 | FALSE | 1 | FALSE | 1 | FALSE | 1 | FALSE | 1 | FALSE | 1 |
| 3663 | 3663 | 9.24 | 3.72322441 | 1 | 0.47819878 | 0 | 1 | 0 | 1 | 1 | 0.69181315 | 0 | 1 | 23 | 0.22727183 | FALSE | 1 | FALSE | 1 | FALSE | 1 | FALSE | 1 | FALSE | 1 | FALSE | 1 |
| 3664 | 3664 | 8.35211268 | 1.30112185 | 0 | 1 | 2 | 0.60114557 | 2 | 0.65873577 | 3 | 0.54709375 | 2 | 0.27036738 | 53 | 0.4900821 | FALSE | 1 | FALSE | 1 | FALSE | 1 | FALSE | 1 | FALSE | 1 | FALSE | 1 |

| | | | | | | | | | | | | | | | | | | | | | | | | | | | |
|---|---|---|---|---|---|---|---|---|---|---|---|---|---|---|---|---|---|---|---|---|---|---|---|---|---|---|---|
| 3665 | 3665 | 3.73369565 | 2.21402531 | 3 | 0.85661336 | 5 | 0.71642212 | 4 | 0.90287526 | 5 | 0.93063001 | 3 | 0.58557338 | 164 | 0.0403181 | FALSE | 1 | FALSE | 1 | FALSE | 1 | FALSE | 1 | FALSE | 1 | FALSE | 1 |
| 3666 | 3666 | 9.69230769 | 2.46153846 | 0 | 1 | 1 | 0.57534752 | 0 | 1 | 1 | 0.70601446 | 1 | 0.3499045 | 23 | 0.40372818 | FALSE | 1 | FALSE | 1 | FALSE | 1 | FALSE | 1 | FALSE | 1 | FALSE | 1 |
| 3667 | 3667 | 10.6842105 | 3.32538295 | 4 | 0.00119617 | 1 | 0.46503795 | 0 | 1 | 1 | 0.59104357 | 0 | 1 | 13 | 0.97998676 | FALSE | 0.08447413 | FALSE | 1 | FALSE | 1 | FALSE | 1 | FALSE | 1 | FALSE | 1 |
| 3668 | 3668 | 7.61538462 | 2.77456751 | 0 | 1 | 1 | 0.56109832 | 2 | 0.22597287 | 1 | 0.69181315 | 1 | 0.33902672 | 20 | 0.8152498 | FALSE | 1 | FALSE | 1 | FALSE | 1 | FALSE | 1 | FALSE | 1 | FALSE | 1 |
| 3669 | 3669 | 10.2 | 2.4657656 | 0 | 1 | 2 | 0.19297191 | 0 | 1 | 3 | 0.10474541 | 0 | 1 | 20 | 0.8152498 | FALSE | 1 | FALSE | 1 | FALSE | 1 | FALSE | 1 | FALSE | 1 | FALSE | 1 |
| 3670 | 3670 | 9.85714286 | 2.72179413 | 0 | 1 | 0 | 1 | 0 | 1 | 1 | 0.73248983 | 3 | 0.0104283 | 24 | 0.55018107 | FALSE | 1 | FALSE | 1 | FALSE | 1 | FALSE | 1 | FALSE | 0.5620304 | FALSE | 1 |
| 3671 | 3671 | 7.9375 | 2.78318051 | 1 | 0.56527493 | 2 | 0.27730767 | 0 | 1 | 2 | 0.43583874 | 0 | 1 | 27 | 0.61671133 | FALSE | 1 | FALSE | 1 | FALSE | 1 | FALSE | 1 | FALSE | 1 | FALSE | 1 |
| 3672 | 3672 | 10.6875 | 2.99413489 | 0 | 1 | 0 | 1 | 0 | 1 | 2 | 0.16544789 | 0 | 1 | 14 | 0.53308261 | FALSE | 1 | FALSE | 1 | FALSE | 1 | FALSE | 1 | FALSE | 1 | FALSE | 1 |
| 3673 | 3673 | 9.83333333 | 2.65099562 | 1 | 0.37380237 | 2 | 0.1135776 | 0 | 1 | 0 | 1 | 0 | 1 | 15 | 0.69372602 | FALSE | 1 | FALSE | 1 | FALSE | 1 | FALSE | 1 | FALSE | 1 | FALSE | 1 |
| 3674 | 3674 | 6.70454546 | 2.6421371 | 1 | 0.67382914 | 3 | 0.16132793 | 2 | 0.46083448 | 4 | 0.13323859 | 0 | 1 | 33 | 0.93815092 | FALSE | 1 | FALSE | 1 | FALSE | 1 | FALSE | 1 | FALSE | 1 | FALSE | 1 |
| 3675 | 3675 | 8.51282051 | 1.98548315 | 1 | 0.63789042 | 0 | 1 | 0 | 1 | 2 | 0.54052736 | 1 | 0.4761699 | 35 | 0.24915769 | FALSE | 1 | FALSE | 1 | FALSE | 1 | FALSE | 1 | FALSE | 1 | FALSE | 1 |
| 3676 | 3676 | 10.2 | 2.78567766 | 0 | 1 | 0 | 1 | 0 | 1 | 0 | 1 | 1 | 0.07935778 | 4 | 0.82371636 | FALSE | 1 | FALSE | 1 | FALSE | 1 | FALSE | 1 | FALSE | 1 | FALSE | 1 |
| 3677 | 3677 | 7.06521739 | 2.48825217 | 1 | 0.69044531 | 4 | 0.05650644 | 1 | 0.80828716 | 1 | 0.8802008 | 1 | 0.52593041 | 37 | 0.73736034 | FALSE | 1 | FALSE | 1 | FALSE | 1 | FALSE | 1 | FALSE | 1 | FALSE | 1 |
| 3678 | 3678 | 7.28571429 | 2.90554244 | 0 | 1 | 1 | 0.74131698 | 3 | 0.1814659 | 1 | 0.85524728 | 0 | 1 | 36 | 0.36455032 | FALSE | 1 | FALSE | 1 | FALSE | 1 | FALSE | 1 | FALSE | 1 | FALSE | 1 |
| 3679 | 3679 | 4.49350649 | 2.86321669 | 1 | 0.86619543 | 0 | 1 | 1 | 0.94117955 | 4 | 0.47420444 | 14 | 1.56E-11 | 57 | 0.99398779 | FALSE | 1 | FALSE | 1 | FALSE | 1 | FALSE | 1 | TRUE | 6.37E-08 | FALSE | 1 |
| 3680 | 3680 | 8.16666667 | 2.35954663 | 0 | 1 | 0 | 1 | 1 | 0.78589454 | 2 | 0.5811115 | 1 | 0.50166523 | 38 | 0.19268329 | FALSE | 1 | FALSE | 1 | FALSE | 1 | FALSE | 1 | FALSE | 1 | FALSE | 1 |
| 3681 | 3681 | 8.46153846 | 2.86308202 | 1 | 0.62830697 | 2 | 0.3494045 | 2 | 0.39868506 | 1 | 0.83318955 | 2 | 0.12811285 | 30 | 0.87271204 | FALSE | 1 | FALSE | 1 | FALSE | 1 | FALSE | 1 | FALSE | 1 | FALSE | 1 |
| 3682 | 3682 | 8.21052632 | 3.33370266 | 0 | 1 | 0 | 1 | 0 | 1 | 0 | 1 | 1 | 0.26986775 | 18 | 0.17880694 | FALSE | 1 | FALSE | 1 | FALSE | 1 | FALSE | 1 | FALSE | 1 | FALSE | 1 |
| 3683 | 3683 | 5.84126984 | 3.00373497 | 1 | 0.79637728 | 1 | 0.86665405 | 4 | 0.17521184 | 4 | 0.30679865 | 1 | 0.63687695 | 50 | 0.76391797 | FALSE | 1 | FALSE | 1 | FALSE | 1 | FALSE | 1 | FALSE | 1 | FALSE | 1 |
| 3684 | 3684 | 9.5 | 3.04138127 | 0 | 1 | 0 | 1 | 0 | 1 | 0 | 1 | 0 | 1 | 8 | 0.25677166 | FALSE | 1 | FALSE | 1 | FALSE | 1 | FALSE | 1 | FALSE | 1 | FALSE | 1 |
| 3685 | 3685 | 10.0555556 | 1.26807914 | 0 | 1 | 1 | 0.44710673 | 0 | 1 | 0 | 1 | 0 | 1 | 17 | 0.20329205 | FALSE | 1 | FALSE | 1 | FALSE | 1 | FALSE | 1 | FALSE | 1 | FALSE | 1 |
| 3686 | 3686 | 7.66666667 | 4.08928138 | 0 | 1 | 0 | 1 | 1 | 0.58502114 | 0 | 1 | 0 | 1 | 23 | 0.09196389 | FALSE | 1 | FALSE | 1 | FALSE | 1 | FALSE | 1 | FALSE | 1 | FALSE | 1 |
| 3687 | 3687 | 11 | 2.4832774 | 2 | 0.03643311 | 0 | 1 | 0 | 1 | 0 | 1 | 0 | 1 | 10 | 0.71436964 | FALSE | 0.58392 | FALSE | 1 | FALSE | 1 | FALSE | 1 | FALSE | 1 | FALSE | 1 |
| 3688 | 3688 | 6.30555556 | 2.52564012 | 0 | 1 | 1 | 0.69480472 | 0 | 1 | 5 | 0.02311989 | 0 | 1 | 30 | 0.67248271 | FALSE | 1 | FALSE | 1 | FALSE | 1 | FALSE | 1 | FALSE | 1 | FALSE | 1 |
| 3689 | 3689 | 10.3333333 | 2.62466929 | 0 | 1 | 1 | 0.42857733 | 0 | 1 | 1 | 0.55062772 | 0 | 1 | 15 | 0.49060435 | FALSE | 1 | FALSE | 1 | FALSE | 1 | FALSE | 1 | FALSE | 1 | FALSE | 1 |
| 3690 | 3690 | 7.375 | 2.0577597 | 0 | 1 | 0 | 1 | 0 | 1 | 2 | 0.43583874 | 0 | 1 | 30 | 0.10375556 | FALSE | 1 | FALSE | 1 | FALSE | 1 | FALSE | 1 | FALSE | 1 | FALSE | 1 |
| 3691 | 3691 | 8.35714286 | 3.61755922 | 1 | 0.3050805 | 0 | 1 | 1 | 0.40110418 | 1 | 0.48242099 | 0 | 1 | 11 | 0.83676449 | FALSE | 1 | FALSE | 1 | FALSE | 1 | FALSE | 1 | FALSE | 1 | FALSE | 1 |
| 3692 | 3692 | 9.26086957 | 2.43555888 | 1 | 0.45027683 | 2 | 0.16948764 | 1 | 0.5695037 | 1 | 0.6613267 | 1 | 0.31672722 | 17 | 0.94439745 | FALSE | 1 | FALSE | 1 | FALSE | 1 | FALSE | 1 | FALSE | 1 | FALSE | 1 |
| 3693 | 3693 | 10.3076923 | 2.1620722 | 0 | 1 | 2 | 0.06426041 | 0 | 1 | 1 | 0.45746571 | 0 | 1 | 10 | 0.86791642 | FALSE | 1 | FALSE | 1 | FALSE | 1 | FALSE | 1 | FALSE | 1 | FALSE | 1 |
| 3694 | 3694 | 8.65217391 | 1.96952203 | 0 | 1 | 1 | 0.53115654 | 1 | 0.5695037 | 1 | 0.6613267 | 1 | 0.31672722 | 19 | 0.71542496 | FALSE | 1 | FALSE | 1 | FALSE | 1 | FALSE | 1 | FALSE | 1 | FALSE | 1 |
| 3695 | 3695 | 6.17964072 | 0.96839511 | 8 | 0.0646762 | 11 | 0.01919241 | 8 | 0.25147891 | 9 | 0.35922712 | 0 | 1 | 131 | 0.98492826 | FALSE | 0.74276693 | FALSE | 0.77701374 | FALSE | 1 | FALSE | 1 | FALSE | 1 | FALSE | 1 |
| 3696 | 3696 | 11.8 | 2.63818119 | 2 | 0.09187136 | 0 | 1 | 1 | 0.51939841 | 0 | 1 | 0 | 1 | 17 | 0.61750632 | FALSE | 0.83248965 | FALSE | 1 | FALSE | 1 | FALSE | 1 | FALSE | 1 | FALSE | 1 |
| 3697 | 3697 | 9.84 | 2.54055112 | 0 | 1 | 0 | 1 | 1 | 0.59998149 | 0 | 1 | 0 | 1 | 24 | 0.08019798 | FALSE | 1 | FALSE | 1 | FALSE | 1 | FALSE | 1 | FALSE | 1 | FALSE | 1 |
| 3698 | 3698 | 7.04347826 | 1.85280659 | 0 | 1 | 1 | 0.53115654 | 1 | 0.5695037 | 2 | 0.2852236 | 0 | 1 | 19 | 0.71542496 | FALSE | 1 | FALSE | 1 | FALSE | 1 | FALSE | 1 | FALSE | 1 | FALSE | 1 |
| 3699 | 3699 | 6.68421053 | 3.55487259 | 2 | 0.43181298 | 0 | 1 | 2 | 0.61293419 | 5 | 0.11939348 | 0 | 1 | 48 | 0.60167081 | FALSE | 1 | FALSE | 1 | FALSE | 1 | FALSE | 1 | FALSE | 1 | FALSE | 1 |
| 3700 | 3700 | 7.86842105 | 3.70753892 | 0 | 1 | 1 | 0.71433277 | 2 | 0.39868506 | 2 | 0.52640931 | 0 | 1 | 33 | 0.44261631 | FALSE | 1 | FALSE | 1 | FALSE | 1 | FALSE | 1 | FALSE | 1 | FALSE | 1 |
| 3701 | 3701 | 10.1428571 | 2.29462549 | 1 | 0.3050805 | 0 | 1 | 1 | 0.40110418 | 1 | 0.48242099 | 0 | 1 | 11 | 0.83676449 | FALSE | 1 | FALSE | 1 | FALSE | 1 | FALSE | 1 | FALSE | 1 | FALSE | 1 |
| 3702 | 3702 | 7.16363636 | 2.57797409 | 0 | 1 | 2 | 0.5356583 | 0 | 1 | 3 | 0.46623289 | 1 | 0.59866474 | 49 | 0.22274213 | FALSE | 1 | FALSE | 1 | FALSE | 1 | FALSE | 1 | FALSE | 1 | FALSE | 1 |
| 3703 | 3703 | 9.77777778 | 1.5475987 | 0 | 1 | 0 | 1 | 0 | 1 | 0 | 1 | 0 | 1 | 18 | 0.04681656 | FALSE | 1 | FALSE | 1 | FALSE | 1 | FALSE | 1 | FALSE | 1 | FALSE | 1 |
| 3704 | 3704 | 9.84615385 | 2.98318761 | 0 | 1 | 2 | 0.06426041 | 1 | 0.37874417 | 0 | 1 | 0 | 1 | 10 | 0.86791642 | FALSE | 1 | FALSE | 1 | FALSE | 1 | FALSE | 1 | FALSE | 1 | FALSE | 1 |
| 3705 | 3705 | 10.0833333 | 1.65621724 | 0 | 1 | 0 | 1 | 0 | 1 | 0 | 1 | 0 | 1 | 12 | 0.13002667 | FALSE | 1 | FALSE | 1 | FALSE | 1 | FALSE | 1 | FALSE | 1 | FALSE | 1 |
| 3706 | 3706 | 9.5 | 2.45967478 | 1 | 0.20854748 | 0 | 1 | 0 | 1 | 0 | 1 | 0 | 1 | 8 | 0.5780332 | FALSE | 1 | FALSE | 1 | FALSE | 1 | FALSE | 1 | FALSE | 1 | FALSE | 1 |
| 3707 | 3707 | 8.88888889 | 2.40113142 | 0 | 1 | 1 | 0.44710673 | 0 | 1 | 2 | 0.19902682 | 0 | 1 | 15 | 0.69372602 | FALSE | 1 | FALSE | 1 | FALSE | 1 | FALSE | 1 | FALSE | 1 | FALSE | 1 |
| 3708 | 3708 | 7.69565217 | 1.89816347 | 1 | 0.45027683 | 0 | 1 | 0 | 1 | 0 | 1 | 1 | 0.31672722 | 21 | 0.27964072 | FALSE | 1 | FALSE | 1 | FALSE | 1 | FALSE | 1 | FALSE | 1 | FALSE | 1 |
| 3709 | 3709 | 8.17857143 | 3.00063769 | 0 | 1 | 2 | 0.2288771 | 1 | 0.64171563 | 0 | 1 | 0 | 1 | 25 | 0.34293 | FALSE | 1 | FALSE | 1 | FALSE | 1 | FALSE | 1 | FALSE | 1 | FALSE | 1 |
| 3710 | 3710 | 9.06666667 | 2.99925917 | 0 | 1 | 0 | 1 | 1 | 0.42266264 | 0 | 1 | 0 | 1 | 14 | 0.29525409 | FALSE | 1 | FALSE | 1 | FALSE | 1 | FALSE | 1 | FALSE | 1 | FALSE | 1 |
| 3711 | 3711 | 8.27272727 | 2.17802701 | 0 | 1 | 0 | 1 | 0 | 1 | 0 | 1 | 0 | 1 | 11 | 0.15414489 | FALSE | 1 | FALSE | 1 | FALSE | 1 | FALSE | 1 | FALSE | 1 | FALSE | 1 |
| 3712 | 3712 | 7.16129032 | 1.74312982 | 0 | 1 | 5 | 0.00288926 | 0 | 1 | 2 | 0.41984187 | 1 | 0.4016888 | 23 | 0.95750806 | FALSE | 1 | FALSE | 0.46409366 | FALSE | 1 | FALSE | 1 | FALSE | 1 | FALSE | 1 |
| 3713 | 3713 | 6.57894737 | 1.62884132 | 3 | 0.82028678 | 6 | 0.47898158 | 7 | 0.41721942 | 11 | 0.16194506 | 2 | 0.77599945 | 142 | 0.72901575 | FALSE | 1 | FALSE | 1 | FALSE | 1 | FALSE | 1 | FALSE | 1 | FALSE | 1 |
| 3714 | 3714 | 8.05405405 | 2.41555708 | 3 | 0.05546921 | 0 | 1 | 1 | 0.71261921 | 2 | 0.46709861 | 0 | 1 | 28 | 0.72558119 | FALSE | 0.68849059 | FALSE | 1 | FALSE | 1 | FALSE | 1 | FALSE | 1 | FALSE | 1 |
| 3715 | 3715 | 7.73913044 | 1.4660622 | 0 | 1 | 1 | 0.78076835 | 3 | 0.2282043 | 1 | 0.88573774 | 2 | 0.17379289 | 39 | 0.56897353 | FALSE | 1 | FALSE | 1 | FALSE | 1 | FALSE | 1 | FALSE | 1 | FALSE | 1 |
| 3716 | 3716 | 9.88 | 2.77589625 | 1 | 0.47819878 | 1 | 0.56109832 | 0 | 1 | 0 | 1 | 2 | 0.06257402 | 21 | 0.64957167 | FALSE | 1 | FALSE | 1 | FALSE | 1 | FALSE | 1 | FALSE | 1 | FALSE | 1 |
| 3717 | 3717 | 9 | 2.52487624 | 0 | 1 | 0 | 1 | 1 | 0.44344817 | 1 | 0.52895088 | 0 | 1 | 14 | 0.53308261 | FALSE | 1 | FALSE | 1 | FALSE | 1 | FALSE | 1 | FALSE | 1 | FALSE | 1 |
| 3718 | 3718 | 7.64102564 | 3.18238082 | 1 | 0.63789042 | 0 | 1 | 2 | 0.41139386 | 1 | 0.84089119 | 2 | 0.13365639 | 33 | 0.59061307 | FALSE | 1 | FALSE | 1 | FALSE | 1 | FALSE | 1 | FALSE | 1 | FALSE | 1 |
| 3719 | 3719 | 7.97297297 | 1.90995241 | 1 | 0.6184714 | 1 | 0.70472939 | 2 | 0.38585089 | 3 | 0.2402032 | 3 | 0.0222449 | 27 | 0.97789976 | FALSE | 1 | FALSE | 1 | FALSE | 1 | FALSE | 1 | FALSE | 0.74991845 | FALSE | 1 |
| 3720 | 3720 | 10.0416667 | 3.83491815 | 4 | 0.00296746 | 1 | 0.54637326 | 1 | 0.58502114 | 2 | 0.30249307 | 0 | 1 | 16 | 0.99240722 | FALSE | 0.14653312 | FALSE | 1 | FALSE | 1 | FALSE | 1 | FALSE | 1 | FALSE | 1 |
| 3721 | 3721 | 7.6779661 | 1.52316217 | 4 | 0.06393293 | 0 | 1 | 3 | 0.35626822 | 4 | 0.28589315 | 2 | 0.25204357 | 46 | 0.93313465 | FALSE | 0.73766 | FALSE | 1 | FALSE | 1 | FALSE | 1 | FALSE | 1 | FALSE | 1 |
| 3722 | 3722 | 5.5 | 2.1015867 | 0 | 1 | 1 | 0.54637326 | 0 | 1 | 1 | 0.67692816 | 0 | 1 | 22 | 0.25237063 | FALSE | 1 | FALSE | 1 | FALSE | 1 | FALSE | 1 | FALSE | 1 | FALSE | 1 |
| 3723 | 3723 | 8.89285714 | 1.58878403 | 1 | 0.51745948 | 1 | 0.60247911 | 2 | 0.2663542 | 1 | 0.73248983 | 1 | 0.37113067 | 22 | 0.86499992 | FALSE | 1 | FALSE | 1 | FALSE | 1 | FALSE | 1 | FALSE | 1 | FALSE | 1 |
| 3724 | 3724 | 9.83333333 | 2.60874597 | 0 | 1 | 1 | 0.44710673 | 2 | 0.13502329 | 1 | 0.57131013 | 1 | 0.25766374 | 13 | 0.95041913 | FALSE | 1 | FALSE | 1 | FALSE | 1 | FALSE | 1 | FALSE | 1 | FALSE | 1 |
| 3725 | 3725 | 9.04761905 | 1.93897839 | 1 | 0.40558521 | 1 | 0.48239024 | 1 | 0.51939841 | 1 | 0.60987146 | 0 | 1 | 16 | 0.80826361 | FALSE | 1 | FALSE | 1 | FALSE | 1 | FALSE | 1 | FALSE | 1 | FALSE | 1 |
| 3726 | 3726 | 11.625 | 2.82566364 | 2 | 0.01652925 | 0 | 1 | 1 | 0.25384309 | 0 | 1 | 0 | 1 | 5 | 0.97551477 | FALSE | 0.39708985 | FALSE | 1 | FALSE | 1 | FALSE | 1 | FALSE | 1 | FALSE | 1 |
| 3727 | 3727 | 11.8888889 | 2.07869855 | 1 | 0.37380237 | 1 | 0.44710673 | 1 | 0.48280983 | 1 | 0.57131013 | 0 | 1 | 14 | 0.86287418 | FALSE | 1 | FALSE | 1 | FALSE | 1 | FALSE | 1 | FALSE | 1 | FALSE | 1 |
| 3728 | 3728 | 9.25 | 1.78535711 | 1 | 0.26795838 | 1 | 0.32623933 | 0 | 1 | 0 | 1 | 0 | 1 | 10 | 0.71436964 | FALSE | 1 | FALSE | 1 | FALSE | 1 | FALSE | 1 | FALSE | 1 | FALSE | 1 |
| 3729 | 3729 | 8.79069767 | 3.05422422 | 4 | 0.02189152 | 3 | 0.1535026 | 3 | 0.19060367 | 2 | 0.5811115 | 0 | 1 | 30 | 0.99060471 | FALSE | 0.46823848 | FALSE | 1 | FALSE | 1 | FALSE | 1 | FALSE | 1 | FALSE | 1 |
| 3730 | 3730 | 8.57692308 | 2.89791805 | 1 | 0.49162634 | 1 | 0.57534752 | 2 | 0.23940226 | 1 | 0.70601446 | 0 | 1 | 21 | 0.78939668 | FALSE | 1 | FALSE | 1 | FALSE | 1 | FALSE | 1 | FALSE | 1 | FALSE | 1 |
| 3731 | 3731 | 8.48148148 | 2.34710808 | 1 | 0.50471032 | 2 | 0.21684246 | 2 | 0.25286935 | 2 | 0.35374209 | 0 | 1 | 20 | 0.95133411 | FALSE | 1 | FALSE | 1 | FALSE | 1 | FALSE | 1 | FALSE | 1 | FALSE | 1 |
| 3732 | 3732 | 10.6875 | 1.64767829 | 1 | 0.34033054 | 0 | 1 | 1 | 0.44344817 | 0 | 1 | 0 | 1 | 14 | 0.53308261 | FALSE | 1 | FALSE | 1 | FALSE | 1 | FALSE | 1 | FALSE | 1 | FALSE | 1 |
| 3733 | 3733 | 7.94339623 | 2.20988427 | 1 | 0.74890551 | 3 | 0.24492539 | 2 | 0.57303317 | 1 | 0.91797304 | 1 | 0.58506217 | 45 | 0.5504765 | FALSE | 1 | FALSE | 1 | FALSE | 1 | FALSE | 1 | FALSE | 1 | FALSE | 1 |
| 3734 | 3734 | 9.6 | 2.4979992 | 0 | 1 | 1 | 0.38964394 | 1 | 0.42266264 | 1 | 0.50623194 | 0 | 1 | 12 | 0.80329917 | FALSE | 1 | FALSE | 1 | FALSE | 1 | FALSE | 1 | FALSE | 1 | FALSE | 1 |
| 3735 | 3735 | 10 | 1.5718105 | 1 | 0.35728305 | 0 | 1 | 1 | 0.46348836 | 0 | 1 | 0 | 1 | 15 | 0.49060435 | FALSE | 1 | FALSE | 1 | FALSE | 1 | FALSE | 1 | FALSE | 1 | FALSE | 1 |
| 3736 | 3736 | 10.5384615 | 1.64622574 | 2 | 0.04234484 | 0 | 1 | 3 | 0.01002246 | 0 | 1 | 0 | 1 | 8 | 0.99085298 | FALSE | 0.62842194 | FALSE | 1 | FALSE | 1 | FALSE | 1 | FALSE | 1 | FALSE | 1 |
| 3737 | 3737 | 10.4375 | 2.23519434 | 0 | 1 | 0 | 1 | 0 | 1 | 0 | 1 | 0 | 1 | 16 | 0.06581507 | FALSE | 1 | FALSE | 1 | FALSE | 1 | FALSE | 1 | FALSE | 1 | FALSE | 1 |
| 3738 | 3738 | 5.88235294 | 3.72404513 | 0 | 1 | 0 | 1 | 1 | 0.44344817 | 0 | 1 | 0 | 1 | 15 | 0.26127859 | FALSE | 1 | FALSE | 1 | FALSE | 1 | FALSE | 1 | FALSE | 1 | FALSE | 1 |
| 3739 | 3739 | 8.2 | 1.46969385 | 1 | 0.40558521 | 1 | 0.48239024 | 1 | 0.51939841 | 1 | 0.60987146 | 0 | 1 | 16 | 0.80826361 | FALSE | 1 | FALSE | 1 | FALSE | 1 | FALSE | 1 | FALSE | 1 | FALSE | 1 |
| 3740 | 3740 | 7.95238095 | 2.23556088 | 2 | 0.09989125 | 1 | 0.49918218 | 0 | 1 | 0 | 1 | 0 | 1 | 18 | 0.57962763 | FALSE | 0.86047225 | FALSE | 1 | FALSE | 1 | FALSE | 1 | FALSE | 1 | FALSE | 1 |
| 3741 | 3741 | 8.12903226 | 1.77331357 | 2 | 0.1881398 | 2 | 0.2651851 | 1 | 0.67911179 | 3 | 0.16836568 | 2 | 0.09110253 | 21 | 0.99467947 | FALSE | 1 | FALSE | 1 | FALSE | 1 | FALSE | 1 | FALSE | 1 | FALSE | 1 |
| 3742 | 3742 | 10.125 | 2.7128168 | 0 | 1 | 0 | 1 | 0 | 1 | 0 | 1 | 0 | 1 | 8 | 0.25677166 | FALSE | 1 | FALSE | 1 | FALSE | 1 | FALSE | 1 | FALSE | 1 | FALSE | 1 |
| 3743 | 3743 | 9.63157895 | 2.73785406 | 0 | 1 | 1 | 0.46503795 | 1 | 0.50143826 | 2 | 0.21610812 | 0 | 1 | 15 | 0.83648191 | FALSE | 1 | FALSE | 1 | FALSE | 1 | FALSE | 1 | FALSE | 1 | FALSE | 1 |
| 3744 | 3744 | 8.44444444 | 1.34256066 | 0 | 1 | 0 | 1 | 0 | 1 | 1 | 0.34506517 | 0 | 1 | 8 | 0.5780332 | FALSE | 1 | FALSE | 1 | FALSE | 1 | FALSE | 1 | FALSE | 1 | FALSE | 1 |
| 3745 | 3745 | 11 | 2.63312235 | 1 | 0.32293355 | 1 | 0.38964394 | 1 | 0.42266264 | 1 | 0.50623194 | 1 | 0.2198258 | 10 | 0.97980046 | FALSE | 1 | FALSE | 1 | FALSE | 1 | FALSE | 1 | FALSE | 1 | FALSE | 1 |
| 3746 | 3746 | 8.32142857 | 2.07112067 | 0 | 1 | 0 | 1 | 1 | 0.64171563 | 2 | 0.3705495 | 1 | 0.37113067 | 24 | 0.55018107 | FALSE | 1 | FALSE | 1 | FALSE | 1 | FALSE | 1 | FALSE | 1 | FALSE | 1 |
| 3747 | 3747 | 10.88 | 2.94373912 | 1 | 0.47819878 | 1 | 0.56109832 | 0 | 1 | 4 | 0.02569595 | 0 | 1 | 19 | 0.91734261 | FALSE | 1 | FALSE | 1 | FALSE | 1 | FALSE | 1 | FALSE | 1 | FALSE | 1 |
| 3748 | 3748 | 5.69491525 | 3.03791988 | 4 | 0.06393293 | 2 | 0.57395594 | 3 | 0.35626822 | 5 | 0.13288902 | 2 | 0.25204357 | 43 | 0.99285314 | FALSE | 0.73766 | FALSE | 1 | FALSE | 1 | FALSE | 1 | FALSE | 1 | FALSE | 1 |
| 3749 | 3749 | 10.2666667 | 2.35135895 | 0 | 1 | 2 | 0.08295839 | 1 | 0.42266264 | 0 | 1 | 0 | 1 | 12 | 0.80329917 | FALSE | 1 | FALSE | 1 | FALSE | 1 | FALSE | 1 | FALSE | 1 | FALSE | 1 |
| 3750 | 3750 | 11.5238095 | 1.6795421 | 0 | 1 | 2 | 0.14655134 | 1 | 0.53671415 | 2 | 0.25059627 | 1 | 0.29368245 | 15 | 0.96513691 | FALSE | 1 | FALSE | 1 | FALSE | 1 | FALSE | 1 | FALSE | 1 | FALSE | 1 |
| 3751 | 3751 | 9.86363636 | 2.88090034 | 1 | 0.42086991 | 1 | 0.49918218 | 1 | 0.53671415 | 0 | 1 | 1 | 0.29368245 | 17 | 0.77849076 | FALSE | 1 | FALSE | 1 | FALSE | 1 | FALSE | 1 | FALSE | 1 | FALSE | 1 |
| 3752 | 3752 | 10.5 | 2.29128785 | 0 | 1 | 1 | 0.32623933 | 0 | 1 | 0 | 1 | 0 | 1 | 11 | 0.41944532 | FALSE | 1 | FALSE | 1 | FALSE | 1 | FALSE | 1 | FALSE | 1 | FALSE | 1 |
| 3753 | 3753 | 10.7777778 | 2.39340658 | 0 | 1 | 1 | 0.44710673 | 0 | 1 | 1 | 0.57131013 | 0 | 1 | 16 | 0.44998001 | FALSE | 1 | FALSE | 1 | FALSE | 1 | FALSE | 1 | FALSE | 1 | FALSE | 1 |
| 3754 | 3754 | 8.5 | 1.75162262 | 2 | 0.09989125 | 1 | 0.49918218 | 1 | 0.53671415 | 1 | 0.62783522 | 0 | 1 | 16 | 0.90353674 | FALSE | 0.86047225 | FALSE | 1 | FALSE | 1 | FALSE | 1 | FALSE | 1 | FALSE | 1 |
| 3755 | 3755 | 9.18181818 | 1.66886907 | 3 | 0.01782284 | 1 | 0.5154318 | 0 | 1 | 1 | 0.64497439 | 1 | 0.30529951 | 16 | 0.95553951 | FALSE | 0.41596787 | FALSE | 1 | FALSE | 1 | FALSE | 1 | FALSE | 1 | FALSE | 1 |
| 3756 | 3756 | 7.65909091 | 2.22530061 | 1 | 0.68224519 | 1 | 0.76576289 | 1 | 0.80109494 | 1 | 0.87439647 | 0 | 1 | 40 | 0.16103481 | FALSE | 1 | FALSE | 1 | FALSE | 1 | FALSE | 1 | FALSE | 1 | FALSE | 1 |
| 3757 | 3757 | 11.2941177 | 3.37403068 | 0 | 1 | 0 | 1 | 1 | 0.46348836 | 2 | 0.18212134 | 1 | 0.24525764 | 13 | 0.88719222 | FALSE | 1 | FALSE | 1 | FALSE | 1 | FALSE | 1 | FALSE | 1 | FALSE | 1 |
| 3758 | 3758 | 9.78571429 | 1.9338552 | 0 | 1 | 1 | 0.60247911 | 2 | 0.2663542 | 2 | 0.3705495 | 0 | 1 | 23 | 0.73439118 | FALSE | 1 | FALSE | 1 | FALSE | 1 | FALSE | 1 | FALSE | 1 | FALSE | 1 |

| | | | | | | | | | | | | | | | | | | | | | | | | | | | |
|---|---|---|---|---|---|---|---|---|---|---|---|---|---|---|---|---|---|---|---|---|---|---|---|---|---|---|---|
| 3759 | 3759 | 8.89285714 | 3.33101535 | 3 | 0.03391597 | 2 | 0.2288771 | 4 | 0.01687379 | 0 | 1 | 0 | 1 | 19 | 0.99262381 | FALSE | 0.56934354 | FALSE | 1 | FALSE | 1 | FALSE | 1 | FALSE | 1 | FALSE | 1 |
| 3760 | 3760 | 9.125 | 1.05326872 | 0 | 1 | 0 | 1 | 0 | 1 | 1 | 0.52895088 | 0 | 1 | 15 | 0.26127859 | FALSE | 1 | FALSE | 1 | FALSE | 1 | FALSE | 1 | FALSE | 1 | FALSE | 1 |
| 3761 | 3761 | 8 | 2.10441712 | 0 | 1 | 0 | 1 | 0 | 1 | 0 | 1 | 0 | 1 | 14 | 0.09251304 | FALSE | 1 | FALSE | 1 | FALSE | 1 | FALSE | 1 | FALSE | 1 | FALSE | 1 |
| 3762 | 3762 | 5.13043478 | 3.41103513 | 3 | 0.11312654 | 0 | 1 | 1 | 0.81522035 | 2 | 0.63103861 | 1 | 0.53375569 | 39 | 0.56897353 | FALSE | 0.92488288 | FALSE | 1 | FALSE | 1 | FALSE | 1 | FALSE | 1 | FALSE | 1 |
| 3763 | 3763 | 8 | 1.91485422 | 1 | 0.37380237 | 1 | 0.44710673 | 0 | 1 | 0 | 1 | 0 | 1 | 16 | 0.44998001 | FALSE | 1 | FALSE | 1 | FALSE | 1 | FALSE | 1 | FALSE | 1 | FALSE | 1 |
| 3764 | 3764 | 4.68604651 | 2.77556459 | 1 | 0.89157583 | 3 | 0.52247675 | 3 | 0.59435517 | 2 | 0.90768569 | 0 | 1 | 76 | 0.12528496 | FALSE | 1 | FALSE | 1 | FALSE | 1 | FALSE | 1 | FALSE | 1 | FALSE | 1 |
| 3765 | 3765 | 7.88888889 | 2.53615827 | 0 | 1 | 0 | 1 | 0 | 1 | 1 | 0.57131013 | 0 | 1 | 17 | 0.20329205 | FALSE | 1 | FALSE | 1 | FALSE | 1 | FALSE | 1 | FALSE | 1 | FALSE | 1 |
| 3766 | 3766 | 9.87096774 | 1.43103697 | 0 | 1 | 3 | 0.07729739 | 1 | 0.67911179 | 2 | 0.41984187 | 0 | 1 | 25 | 0.80006983 | FALSE | 1 | FALSE | 1 | FALSE | 1 | FALSE | 1 | FALSE | 1 | FALSE | 1 |
| 3767 | 3767 | 6.57575758 | 2.13243783 | 1 | 0.57647348 | 3 | 0.08959205 | 2 | 0.33343871 | 3 | 0.19161876 | 1 | 0.42123867 | 23 | 0.99096587 | FALSE | 1 | FALSE | 1 | FALSE | 1 | FALSE | 1 | FALSE | 1 | FALSE | 1 |
| 3768 | 3768 | 9.42857143 | 1.59078982 | 0 | 1 | 2 | 0.07341559 | 0 | 1 | 0 | 1 | 0 | 1 | 12 | 0.62236227 | FALSE | 1 | FALSE | 1 | FALSE | 1 | FALSE | 1 | FALSE | 1 | FALSE | 1 |
| 3769 | 3769 | 7.35294118 | 3.71443147 | 2 | 0.37762121 | 3 | 0.22759346 | 4 | 0.10960782 | 1 | 0.90982214 | 2 | 0.20351566 | 39 | 0.95412149 | FALSE | 1 | FALSE | 1 | FALSE | 1 | FALSE | 1 | FALSE | 1 | FALSE | 1 |
| 3770 | 3770 | 9.90909091 | 2.15130174 | 0 | 1 | 2 | 0.04725998 | 1 | 0.33149969 | 1 | 0.40390032 | 0 | 1 | 7 | 0.98129353 | FALSE | 1 | FALSE | 1 | FALSE | 1 | FALSE | 1 | FALSE | 1 | FALSE | 1 |
| 3771 | 3771 | 8.75 | 3.18057722 | 1 | 0.49162634 | 2 | 0.20486827 | 1 | 0.61440466 | 2 | 0.33678042 | 0 | 1 | 20 | 0.90142689 | FALSE | 1 | FALSE | 1 | FALSE | 1 | FALSE | 1 | FALSE | 1 | FALSE | 1 |
| 3772 | 3772 | 10.3333333 | 1.24721913 | 0 | 1 | 0 | 1 | 0 | 1 | 0 | 1 | 0 | 1 | 12 | 0.13002667 | FALSE | 1 | FALSE | 1 | FALSE | 1 | FALSE | 1 | FALSE | 1 | FALSE | 1 |
| 3773 | 3773 | 10.2857143 | 1.22057196 | 1 | 0.3050805 | 0 | 1 | 1 | 0.40110418 | 1 | 0.48242099 | 0 | 1 | 11 | 0.83676449 | FALSE | 1 | FALSE | 1 | FALSE | 1 | FALSE | 1 | FALSE | 1 | FALSE | 1 |
| 3774 | 3774 | 5.58666667 | 3.47214951 | 3 | 0.30198574 | 5 | 0.09467212 | 6 | 0.05188029 | 3 | 0.67644754 | 2 | 0.34918837 | 56 | 0.99057646 | FALSE | 1 | FALSE | 1 | FALSE | 1 | FALSE | 1 | FALSE | 1 | FALSE | 1 |
| 3775 | 3775 | 6.69565217 | 3.75829266 | 2 | 0.11647716 | 1 | 0.53115654 | 2 | 0.19931429 | 0 | 1 | 0 | 1 | 18 | 0.86273155 | FALSE | 0.93181727 | FALSE | 1 | FALSE | 1 | FALSE | 1 | FALSE | 1 | FALSE | 1 |
| 3776 | 3776 | 9.74193548 | 1.2435055 | 0 | 1 | 0 | 1 | 3 | 0.09854319 | 3 | 0.16836568 | 1 | 0.4016888 | 24 | 0.90138666 | FALSE | 1 | FALSE | 1 | FALSE | 1 | FALSE | 1 | FALSE | 1 | FALSE | 1 |
| 3777 | 3777 | 3.82178218 | 2.49512808 | 2 | 0.7108109 | 4 | 0.37678297 | 5 | 0.26212755 | 4 | 0.64930879 | 0 | 1 | 81 | 0.56891537 | FALSE | 1 | FALSE | 1 | FALSE | 1 | FALSE | 1 | FALSE | 1 | FALSE | 1 |
| 3778 | 3778 | 7.95238095 | 1.66121558 | 2 | 0.29286138 | 0 | 1 | 2 | 0.44869594 | 3 | 0.30326943 | 1 | 0.50166523 | 34 | 0.80048036 | FALSE | 1 | FALSE | 1 | FALSE | 1 | FALSE | 1 | FALSE | 1 | FALSE | 1 |
| 3779 | 3779 | 6.33333333 | 2.05480467 | 0 | 1 | 2 | 0.25306331 | 1 | 0.66709967 | 2 | 0.4036176 | 1 | 0.39166989 | 24 | 0.82294782 | FALSE | 1 | FALSE | 1 | FALSE | 1 | FALSE | 1 | FALSE | 1 | FALSE | 1 |
| 3780 | 3780 | 10.4 | 2.08326667 | 1 | 0.40558521 | 0 | 1 | 2 | 0.16019548 | 2 | 0.23331368 | 0 | 1 | 15 | 0.92113679 | FALSE | 1 | FALSE | 1 | FALSE | 1 | FALSE | 1 | FALSE | 1 | FALSE | 1 |
| 3781 | 3781 | 8.18181818 | 2.4796694 | 1 | 0.24866461 | 1 | 0.30368014 | 0 | 1 | 0 | 1 | 2 | 0.01330324 | 7 | 0.98129353 | FALSE | 1 | FALSE | 1 | FALSE | 1 | FALSE | 1 | FALSE | 0.63731057 | FALSE | 1 |
| 3782 | 3782 | 6.55263158 | 3.36152666 | 0 | 1 | 2 | 0.3375163 | 0 | 1 | 4 | 0.08780467 | 0 | 1 | 31 | 0.64534533 | FALSE | 1 | FALSE | 1 | FALSE | 1 | FALSE | 1 | FALSE | 1 | FALSE | 1 |
| 3783 | 3783 | 9.44 | 3.4880367 | 2 | 0.13369621 | 2 | 0.19297191 | 1 | 0.59998149 | 0 | 1 | 0 | 1 | 20 | 0.8152498 | FALSE | 0.97964162 | FALSE | 1 | FALSE | 1 | FALSE | 1 | FALSE | 1 | FALSE | 1 |
| 3784 | 3784 | 8.96153846 | 3.16811967 | 0 | 1 | 2 | 0.20486827 | 0 | 1 | 0 | 1 | 0 | 1 | 24 | 0.20425579 | FALSE | 1 | FALSE | 1 | FALSE | 1 | FALSE | 1 | FALSE | 1 | FALSE | 1 |
| 3785 | 3785 | 9.64516129 | 2.90215034 | 0 | 1 | 0 | 1 | 1 | 0.67911179 | 5 | 0.01254769 | 0 | 1 | 25 | 0.80006983 | FALSE | 1 | FALSE | 1 | FALSE | 1 | FALSE | 1 | FALSE | 1 | FALSE | 1 |
| 3786 | 3786 | 9.75 | 1.99687256 | 0 | 1 | 0 | 1 | 0 | 1 | 0 | 1 | 0 | 1 | 20 | 0.03329856 | FALSE | 1 | FALSE | 1 | FALSE | 1 | FALSE | 1 | FALSE | 1 | FALSE | 1 |
| 3787 | 3787 | 8.33333333 | 2.79248631 | 2 | 0.20692137 | 2 | 0.28941836 | 5 | 0.00597743 | 3 | 0.19161876 | 0 | 1 | 21 | 0.99917809 | FALSE | 1 | FALSE | 1 | FALSE | 1 | FALSE | 1 | FALSE | 1 | FALSE | 1 |
| 3788 | 3788 | 9.47826087 | 2.12376952 | 0 | 1 | 3 | 0.03669128 | 2 | 0.19931429 | 0 | 1 | 0 | 1 | 18 | 0.86273155 | FALSE | 1 | FALSE | 0.94259695 | FALSE | 1 | FALSE | 1 | FALSE | 1 | FALSE | 1 |
| 3789 | 3789 | 9.28571429 | 2.96234877 | 0 | 1 | 0 | 1 | 1 | 0.40110418 | 1 | 0.48242099 | 0 | 1 | 12 | 0.62236227 | FALSE | 1 | FALSE | 1 | FALSE | 1 | FALSE | 1 | FALSE | 1 | FALSE | 1 |
| 3790 | 3790 | 5.34615385 | 2.5106283 | 5 | 0.01026504 | 0 | 1 | 3 | 0.28676219 | 3 | 0.4296776 | 1 | 0.57809055 | 40 | 0.94722426 | FALSE | 0.30467829 | FALSE | 1 | FALSE | 1 | FALSE | 1 | FALSE | 1 | FALSE | 1 |
| 3791 | 3791 | 9.17142857 | 1.82812495 | 3 | 0.0595675 | 1 | 0.68454804 | 1 | 0.72299506 | 5 | 0.02066043 | 1 | 0.44015537 | 24 | 0.99496229 | FALSE | 0.72532843 | FALSE | 1 | FALSE | 1 | FALSE | 1 | FALSE | 1 | FALSE | 1 |
| 3792 | 3792 | 9.95454546 | 3.27493612 | 8 | 1.28E-05 | 0 | 1 | 1 | 0.80109494 | 2 | 0.60667406 | 1 | 0.51797501 | 32 | 0.98583614 | TRUE | 0.00418892 | FALSE | 1 | FALSE | 1 | FALSE | 1 | FALSE | 1 | FALSE | 1 |
| 3793 | 3793 | 3.12560387 | 1.09176606 | 10 | 0.62454405 | 20 | 0.04579983 | 13 | 0.7338775 | 13 | 0.94915582 | 5 | 0.81781811 | 353 | 0.33301773 | FALSE | 1 | FALSE | 1 | FALSE | 1 | FALSE | 1 | FALSE | 1 | FALSE | 1 |
| 3794 | 3794 | 7.25 | 3.23329038 | 1 | 0.79096998 | 1 | 0.86215528 | 1 | 0.8897349 | 3 | 0.52470434 | 3 | 0.07497311 | 51 | 0.5343051 | FALSE | 1 | FALSE | 1 | FALSE | 1 | FALSE | 1 | FALSE | 1 | FALSE | 1 |
| 3795 | 3795 | 11.5384615 | 2.84511416 | 2 | 0.14250978 | 2 | 0.20486827 | 0 | 1 | 3 | 0.11458571 | 0 | 1 | 19 | 0.96054244 | FALSE | 1 | FALSE | 1 | FALSE | 1 | FALSE | 1 | FALSE | 1 | FALSE | 1 |
| 3796 | 3796 | 11.56 | 2.99439476 | 1 | 0.47819878 | 0 | 1 | 0 | 1 | 0 | 1 | 1 | 0.33902672 | 23 | 0.22727183 | FALSE | 1 | FALSE | 1 | FALSE | 1 | FALSE | 1 | FALSE | 1 | FALSE | 1 |
| 3797 | 3797 | 8.11111111 | 2.33068633 | 0 | 1 | 0 | 1 | 1 | 0.28067846 | 0 | 1 | 0 | 1 | 8 | 0.5780332 | FALSE | 1 | FALSE | 1 | FALSE | 1 | FALSE | 1 | FALSE | 1 | FALSE | 1 |
| 3798 | 3798 | 3.03114754 | 0.44336307 | 127 | 0.36861056 | 168 | 0.05156718 | 172 | 0.66969086 | 210 | 0.9587029 | 89 | 0.02501965 | 4090 | 0.72430873 | FALSE | 1 | FALSE | 1 | FALSE | 1 | FALSE | 1 | FALSE | 0.79106376 | FALSE | 1 |
| 3799 | 3799 | 9.35714286 | 2.89351844 | 1 | 0.50471032 | 0 | 1 | 2 | 0.25286935 | 0 | 1 | 0 | 1 | 24 | 0.3725373 | FALSE | 1 | FALSE | 1 | FALSE | 1 | FALSE | 1 | FALSE | 1 | FALSE | 1 |
| 3800 | 3800 | 8.20833333 | 3.96840823 | 0 | 1 | 2 | 0.18117183 | 2 | 0.21260257 | 1 | 0.67692816 | 0 | 1 | 19 | 0.83975208 | FALSE | 1 | FALSE | 1 | FALSE | 1 | FALSE | 1 | FALSE | 1 | FALSE | 1 |
| 3801 | 3801 | 9.5 | 3.11104025 | 0 | 1 | 1 | 0.36919824 | 1 | 0.40110418 | 0 | 1 | 0 | 1 | 12 | 0.62236227 | FALSE | 1 | FALSE | 1 | FALSE | 1 | FALSE | 1 | FALSE | 1 | FALSE | 1 |
| 3802 | 3802 | 8.8 | 2.22710575 | 0 | 1 | 0 | 1 | 0 | 1 | 1 | 0.2094688 | 0 | 1 | 4 | 0.82371636 | FALSE | 1 | FALSE | 1 | FALSE | 1 | FALSE | 1 | FALSE | 1 | FALSE | 1 |
| 3803 | 3803 | 9.19230769 | 2.41818799 | 0 | 1 | 1 | 0.57534752 | 1 | 0.61440466 | 1 | 0.70601446 | 1 | 0.3499045 | 22 | 0.61628 | FALSE | 1 | FALSE | 1 | FALSE | 1 | FALSE | 1 | FALSE | 1 | FALSE | 1 |
| 3804 | 3804 | 7.71428571 | 2.71052371 | 1 | 0.51745948 | 2 | 0.2288771 | 1 | 0.64171563 | 0 | 1 | 0 | 1 | 24 | 0.55018107 | FALSE | 1 | FALSE | 1 | FALSE | 1 | FALSE | 1 | FALSE | 1 | FALSE | 1 |
| 3805 | 3805 | 9.25 | 2.56661255 | 0 | 1 | 0 | 1 | 2 | 0.16019548 | 2 | 0.23331368 | 0 | 1 | 16 | 0.80826361 | FALSE | 1 | FALSE | 1 | FALSE | 1 | FALSE | 1 | FALSE | 1 | FALSE | 1 |
| 3806 | 3806 | 9.96153846 | 3.20433324 | 3 | 0.0279333 | 2 | 0.20486827 | 2 | 0.23940226 | 3 | 0.11458571 | 1 | 0.3499045 | 15 | 0.99977326 | FALSE | 0.50964275 | FALSE | 1 | FALSE | 1 | FALSE | 1 | FALSE | 1 | FALSE | 1 |
| 3807 | 3807 | 7.6 | 2.72763634 | 0 | 1 | 0 | 1 | 1 | 0.42266264 | 0 | 1 | 0 | 1 | 14 | 0.29525409 | FALSE | 1 | FALSE | 1 | FALSE | 1 | FALSE | 1 | FALSE | 1 | FALSE | 1 |
| 3808 | 3808 | 10.125 | 2.91279505 | 0 | 1 | 2 | 0.09285443 | 0 | 1 | 0 | 1 | 0 | 1 | 14 | 0.53308261 | FALSE | 1 | FALSE | 1 | FALSE | 1 | FALSE | 1 | FALSE | 1 | FALSE | 1 |
| 3809 | 3809 | 9.53571429 | 2.17915683 | 1 | 0.50471032 | 0 | 1 | 1 | 0.62830986 | 0 | 1 | 0 | 1 | 25 | 0.1832206 | FALSE | 1 | FALSE | 1 | FALSE | 1 | FALSE | 1 | FALSE | 1 | FALSE | 1 |
| 3810 | 3810 | 9.5 | 2.54950976 | 2 | 0.01652925 | 1 | 0.23139161 | 0 | 1 | 0 | 1 | 1 | 0.12394282 | 4 | 0.99658732 | FALSE | 0.39708985 | FALSE | 1 | FALSE | 1 | FALSE | 1 | FALSE | 1 | FALSE | 1 |
| 3811 | 3811 | 9.36666667 | 1.60173517 | 0 | 1 | 1 | 0.6278847 | 2 | 0.29330414 | 2 | 0.4036176 | 0 | 1 | 25 | 0.67630471 | FALSE | 1 | FALSE | 1 | FALSE | 1 | FALSE | 1 | FALSE | 1 | FALSE | 1 |
| 3812 | 3812 | 8.15384615 | 3.19485533 | 0 | 1 | 2 | 0.20486827 | 3 | 0.06482014 | 1 | 0.70601446 | 0 | 1 | 20 | 0.90142689 | FALSE | 1 | FALSE | 1 | FALSE | 1 | FALSE | 1 | FALSE | 1 | FALSE | 1 |
| 3813 | 3813 | 9.55 | 3.58434094 | 3 | 0.01369554 | 1 | 0.48239024 | 2 | 0.16019548 | 0 | 1 | 1 | 0.28187291 | 13 | 0.99248686 | FALSE | 0.36625055 | FALSE | 1 | FALSE | 1 | FALSE | 1 | FALSE | 1 | FALSE | 1 |
| 3814 | 3814 | 8.9 | 1.41572798 | 2 | 0.539854 | 1 | 0.90110191 | 1 | 0.92379401 | 2 | 0.83868849 | 0 | 1 | 64 | 0.06373729 | FALSE | 1 | FALSE | 1 | FALSE | 1 | FALSE | 1 | FALSE | 1 | FALSE | 1 |
| 3815 | 3815 | 9.25 | 1.89406969 | 0 | 1 | 1 | 0.48239024 | 0 | 1 | 1 | 0.60987146 | 0 | 1 | 18 | 0.37504645 | FALSE | 1 | FALSE | 1 | FALSE | 1 | FALSE | 1 | FALSE | 1 | FALSE | 1 |
| 3816 | 3816 | 9.2 | 2.38327506 | 3 | 0.02517314 | 1 | 0.56109832 | 1 | 0.59998149 | 2 | 0.31968852 | 0 | 1 | 18 | 0.96851178 | FALSE | 0.4974559 | FALSE | 1 | FALSE | 1 | FALSE | 1 | FALSE | 1 | FALSE | 1 |
| 3817 | 3817 | 9.57142857 | 2.44114393 | 0 | 1 | 0 | 1 | 0 | 1 | 1 | 0.28044368 | 1 | 0.10932414 | 5 | 0.91847773 | FALSE | 1 | FALSE | 1 | FALSE | 1 | FALSE | 1 | FALSE | 1 | FALSE | 1 |
| 3818 | 3818 | 9.78571429 | 1.8776342 | 0 | 1 | 0 | 1 | 1 | 0.64171563 | 2 | 0.3705495 | 0 | 1 | 25 | 0.34293 | FALSE | 1 | FALSE | 1 | FALSE | 1 | FALSE | 1 | FALSE | 1 | FALSE | 1 |
| 3819 | 3819 | 10.1904762 | 3.52702556 | 0 | 1 | 3 | 0.02888061 | 1 | 0.53671415 | 0 | 1 | 1 | 0.29368245 | 16 | 0.90353674 | FALSE | 1 | FALSE | 0.88012077 | FALSE | 1 | FALSE | 1 | FALSE | 1 | FALSE | 1 |
| 3820 | 3820 | 10.3181818 | 2.28406264 | 0 | 1 | 0 | 1 | 2 | 0.18613256 | 1 | 0.64497439 | 1 | 0.30529951 | 18 | 0.74744813 | FALSE | 1 | FALSE | 1 | FALSE | 1 | FALSE | 1 | FALSE | 1 | FALSE | 1 |
| 3821 | 3821 | 10.375 | 1.40867846 | 0 | 1 | 1 | 0.23139161 | 1 | 0.25384309 | 0 | 1 | 0 | 1 | 6 | 0.88425551 | FALSE | 1 | FALSE | 1 | FALSE | 1 | FALSE | 1 | FALSE | 1 | FALSE | 1 |
| 3822 | 3822 | 9.85714286 | 2.16653584 | 0 | 1 | 0 | 1 | 0 | 1 | 0 | 1 | 0 | 1 | 7 | 0.3043657 | FALSE | 1 | FALSE | 1 | FALSE | 1 | FALSE | 1 | FALSE | 1 | FALSE | 1 |
| 3823 | 3823 | 7.27586207 | 2.8875064 | 1 | 0.52988231 | 2 | 0.24095584 | 2 | 0.27983822 | 1 | 0.74482229 | 2 | 0.08121636 | 21 | 0.97185642 | FALSE | 1 | FALSE | 1 | FALSE | 1 | FALSE | 1 | FALSE | 1 | FALSE | 1 |
| 3824 | 3824 | 10 | 2.61861468 | 1 | 0.42086991 | 0 | 1 | 0 | 1 | 2 | 0.25059627 | 2 | 0.04568221 | 16 | 0.90353674 | FALSE | 1 | FALSE | 1 | FALSE | 1 | FALSE | 1 | FALSE | 0.95710659 | FALSE | 1 |
| 3825 | 3825 | 7.66666667 | 2.47206616 | 2 | 0.40500867 | 0 | 1 | 2 | 0.58328102 | 0 | 1 | 0 | 1 | 50 | 0.06046652 | FALSE | 1 | FALSE | 1 | FALSE | 1 | FALSE | 1 | FALSE | 1 | FALSE | 1 |
| 3826 | 3826 | 9.23076923 | 1.96718644 | 2 | 0.04234484 | 1 | 0.3480709 | 0 | 1 | 0 | 1 | 0 | 1 | 10 | 0.86791642 | FALSE | 0.62842194 | FALSE | 1 | FALSE | 1 | FALSE | 1 | FALSE | 1 | FALSE | 1 |
| 3827 | 3827 | 7.18644068 | 3.74380653 | 2 | 0.4493251 | 2 | 0.57395594 | 2 | 0.63179508 | 1 | 0.93827647 | 1 | 0.62455813 | 51 | 0.41341988 | FALSE | 1 | FALSE | 1 | FALSE | 1 | FALSE | 1 | FALSE | 1 | FALSE | 1 |
| 3828 | 3828 | 8.4 | 1.68522996 | 0 | 1 | 2 | 0.03949327 | 0 | 1 | 0 | 1 | 0 | 1 | 8 | 0.80394719 | FALSE | 1 | FALSE | 0.96864929 | FALSE | 1 | FALSE | 1 | FALSE | 1 | FALSE | 1 |
| 3829 | 3829 | 9.46153846 | 1.44729906 | 0 | 1 | 2 | 0.06426041 | 0 | 1 | 1 | 0.45746571 | 0 | 1 | 10 | 0.86791642 | FALSE | 1 | FALSE | 1 | FALSE | 1 | FALSE | 1 | FALSE | 1 | FALSE | 1 |
| 3830 | 3830 | 8.78571429 | 2.51221506 | 1 | 0.3050805 | 1 | 0.36919824 | 0 | 1 | 0 | 1 | 1 | 0.20679328 | 11 | 0.83676449 | FALSE | 1 | FALSE | 1 | FALSE | 1 | FALSE | 1 | FALSE | 1 | FALSE | 1 |
| 3831 | 3831 | 4.93023256 | 2.89663276 | 5 | 0.06651199 | 3 | 0.52247675 | 4 | 0.36494003 | 3 | 0.55590957 | 0 | 1 | 70 | 0.75388153 | FALSE | 0.74276693 | FALSE | 1 | FALSE | 1 | FALSE | 1 | FALSE | 1 | FALSE | 1 |
| 3832 | 3832 | 9.92592593 | 2.69328281 | 0 | 1 | 1 | 0.58913618 | 2 | 0.25286935 | 1 | 0.71956339 | 0 | 1 | 23 | 0.58307573 | FALSE | 1 | FALSE | 1 | FALSE | 1 | FALSE | 1 | FALSE | 1 | FALSE | 1 |
| 3833 | 3833 | 5.81632653 | 2.69294975 | 3 | 0.10767332 | 1 | 0.77338922 | 5 | 0.02174577 | 3 | 0.34154252 | 0 | 1 | 33 | 0.98284787 | FALSE | 0.88820416 | FALSE | 1 | FALSE | 1 | FALSE | 1 | FALSE | 1 | FALSE | 1 |
| 3834 | 3834 | 8.63157895 | 1.42202696 | 0 | 1 | 0 | 1 | 0 | 1 | 1 | 0.59104357 | 0 | 1 | 18 | 0.17880694 | FALSE | 1 | FALSE | 1 | FALSE | 1 | FALSE | 1 | FALSE | 1 | FALSE | 1 |
| 3835 | 3835 | 10.0833333 | 3.37782803 | 0 | 1 | 1 | 0.32623933 | 1 | 0.35555293 | 0 | 1 | 0 | 1 | 10 | 0.71436964 | FALSE | 1 | FALSE | 1 | FALSE | 1 | FALSE | 1 | FALSE | 1 | FALSE | 1 |
| 3836 | 3836 | 8.1025641 | 2.22796759 | 1 | 0.62830697 | 2 | 0.3494045 | 3 | 0.15483587 | 3 | 0.25266879 | 0 | 1 | 29 | 0.93814695 | FALSE | 1 | FALSE | 1 | FALSE | 1 | FALSE | 1 | FALSE | 1 | FALSE | 1 |
| 3837 | 3837 | 10.4210526 | 1.98191267 | 2 | 0.06197875 | 0 | 1 | 1 | 0.44344817 | 2 | 0.16544789 | 0 | 1 | 11 | 0.9718531 | FALSE | 0.72532843 | FALSE | 1 | FALSE | 1 | FALSE | 1 | FALSE | 1 | FALSE | 1 |
| 3838 | 3838 | 8.4137931 | 3.41918766 | 1 | 0.52988231 | 4 | 0.01340169 | 0 | 1 | 0 | 1 | 0 | 1 | 24 | 0.70563334 | FALSE | 1 | FALSE | 0.69207862 | FALSE | 1 | FALSE | 1 | FALSE | 1 | FALSE | 1 |
| 3839 | 3839 | 9.94117647 | 1.55187129 | 0 | 1 | 2 | 0.10307112 | 0 | 1 | 3 | 0.040397 | 0 | 1 | 12 | 0.962093 | FALSE | 1 | FALSE | 1 | FALSE | 1 | FALSE | 1 | FALSE | 1 | FALSE | 1 |
| 3840 | 3840 | 8.03333333 | 2.10528436 | 2 | 0.17884234 | 1 | 0.6278847 | 4 | 0.02136185 | 1 | 0.75658797 | 2 | 0.08611595 | 20 | 0.99601577 | FALSE | 1 | FALSE | 1 | FALSE | 1 | FALSE | 1 | FALSE | 1 | FALSE | 1 |
| 3841 | 3841 | 9.09677419 | 2.19022519 | 3 | 0.04402911 | 2 | 0.2651851 | 0 | 1 | 1 | 0.76781284 | 0 | 1 | 25 | 0.80006983 | FALSE | 0.64293489 | FALSE | 1 | FALSE | 1 | FALSE | 1 | FALSE | 1 | FALSE | 1 |
| 3842 | 3842 | 7.52941177 | 2.81001669 | 0 | 1 | 1 | 0.42857733 | 3 | 0.02143959 | 1 | 0.55062772 | 0 | 1 | 12 | 0.962093 | FALSE | 1 | FALSE | 1 | FALSE | 1 | FALSE | 1 | FALSE | 1 | FALSE | 1 |
| 3843 | 3843 | 11.4814815 | 2.29928121 | 2 | 0.14250978 | 1 | 0.57534752 | 1 | 0.61440466 | 1 | 0.70601446 | 0 | 1 | 21 | 0.78939668 | FALSE | 1 | FALSE | 1 | FALSE | 1 | FALSE | 1 | FALSE | 1 | FALSE | 1 |
| 3844 | 3844 | 9.50847458 | 2.20456292 | 3 | 0.19211919 | 1 | 0.85750546 | 1 | 0.88558856 | 4 | 0.28589315 | 0 | 1 | 50 | 0.55668966 | FALSE | 1 | FALSE | 1 | FALSE | 1 | FALSE | 1 | FALSE | 1 | FALSE | 1 |
| 3845 | 3845 | 9.78947368 | 1.73444811 | 0 | 1 | 0 | 1 | 2 | 0.14749808 | 1 | 0.59104357 | 0 | 1 | 16 | 0.65565778 | FALSE | 1 | FALSE | 1 | FALSE | 1 | FALSE | 1 | FALSE | 1 | FALSE | 1 |
| 3846 | 3846 | 8.08333333 | 2.88554828 | 1 | 0.71380452 | 2 | 0.46321244 | 0 | 1 | 4 | 0.17698522 | 0 | 1 | 41 | 0.51861324 | FALSE | 1 | FALSE | 1 | FALSE | 1 | FALSE | 1 | FALSE | 1 | FALSE | 1 |
| 3847 | 3847 | 4.09411765 | 2.65517709 | 2 | 0.63949673 | 2 | 0.76136679 | 1 | 0.95461143 | 7 | 0.0900466 | 4 | 0.04860285 | 68 | 0.8466453 | FALSE | 1 | FALSE | 1 | FALSE | 1 | FALSE | 1 | FALSE | 0.98629461 | FALSE | 1 |
| 3848 | 3848 | 9.69565217 | 3.04223577 | 0 | 1 | 0 | 1 | 0 | 1 | 0 | 1 | 1 | 0.31672722 | 22 | 0.10533112 | FALSE | 1 | FALSE | 1 | FALSE | 1 | FALSE | 1 | FALSE | 1 | FALSE | 1 |
| 3849 | 3849 | 10.3333333 | 3.51504702 | 1 | 0.32293355 | 2 | 0.08295839 | 0 | 1 | 2 | 0.14906763 | 0 | 1 | 10 | 0.97980046 | FALSE | 1 | FALSE | 1 | FALSE | 1 | FALSE | 1 | FALSE | 1 | FALSE | 1 |
| 3850 | 3850 | 9.2 | 3.70045042 | 2 | 0.17884234 | 0 | 1 | 2 | 0.29330414 | 1 | 0.75658797 | 0 | 1 | 25 | 0.67630471 | FALSE | 1 | FALSE | 1 | FALSE | 1 | FALSE | 1 | FALSE | 1 | FALSE | 1 |
| 3851 | 3851 | 6.47244095 | 1.07825151 | 8 | 0.01602447 | 5 | 0.39267635 | 7 | 0.17164834 | 9 | 0.12869646 | 0 | 1 | 98 | 0.9887296 | FALSE | 0.39421149 | FALSE | 1 | FALSE | 1 | FALSE | 1 | FALSE | 1 | FALSE | 1 |
| 3852 | 3852 | 6.74712644 | 1.78897255 | 4 | 0.1836099 | 5 | 0.15016498 | 4 | 0.38141809 | 7 | 0.10374527 | 1 | 0.76487586 | 66 | 0.98726286 | FALSE | 1 | FALSE | 1 | FALSE | 1 | FALSE | 1 | FALSE | 1 | FALSE | 1 |

| | | | | | | | | | | | | | | | | | | | | | | | | | | | |
|---|---|---|---|---|---|---|---|---|---|---|---|---|---|---|---|---|---|---|---|---|---|---|---|---|---|---|---|
| 3853 | 3853 | 9.875 | 3.20562656 | 1 | 0.46441868 | 0 | 1 | 0 | 1 | 1 | 0.67692816 | 0 | 1 | 22 | 0.25237063 | FALSE | 1 | FALSE | 1 | FALSE | 1 | FALSE | 1 | FALSE | 1 | FALSE | 1 |
| 3854 | 3854 | 7.82352941 | 3.16665149 | 0 | 1 | 0 | 1 | 1 | 0.46348836 | 0 | 1 | 0 | 1 | 16 | 0.23070182 | FALSE | 1 | FALSE | 1 | FALSE | 1 | FALSE | 1 | FALSE | 1 | FALSE | 1 |
| 3855 | 3855 | 10.2083333 | 2.1209111 | 2 | 0.12501436 | 1 | 0.54637326 | 1 | 0.58502114 | 2 | 0.30249307 | 0 | 1 | 18 | 0.93166998 | FALSE | 0.95622564 | FALSE | 1 | FALSE | 1 | FALSE | 1 | FALSE | 1 | FALSE | 1 |
| 3856 | 3856 | 9 | 2.44948974 | 0 | 1 | 0 | 1 | 1 | 0.51939841 | 1 | 0.60987146 | 1 | 0.28187291 | 17 | 0.61750632 | FALSE | 1 | FALSE | 1 | FALSE | 1 | FALSE | 1 | FALSE | 1 | FALSE | 1 |
| 3857 | 3857 | 8.94117647 | 2.57751792 | 3 | 0.00864151 | 0 | 1 | 0 | 1 | 1 | 0.55062772 | 1 | 0.24525764 | 12 | 0.962093 | FALSE | 0.27019545 | FALSE | 1 | FALSE | 1 | FALSE | 1 | FALSE | 1 | FALSE | 1 |
| 3858 | 3858 | 7.42105263 | 2.15981942 | 0 | 1 | 1 | 0.71433277 | 3 | 0.15483587 | 2 | 0.52640931 | 1 | 0.4673875 | 31 | 0.76640548 | FALSE | 1 | FALSE | 1 | FALSE | 1 | FALSE | 1 | FALSE | 1 | FALSE | 1 |
| 3859 | 3859 | 10.2307692 | 2.48546068 | 1 | 0.28675951 | 1 | 0.3480709 | 0 | 1 | 1 | 0.45746571 | 0 | 1 | 10 | 0.86791642 | FALSE | 1 | FALSE | 1 | FALSE | 1 | FALSE | 1 | FALSE | 1 | FALSE | 1 |
| 3860 | 3860 | 7.75510204 | 1.81299856 | 0 | 1 | 1 | 0.79481669 | 2 | 0.51909752 | 2 | 0.65421368 | 0 | 1 | 43 | 0.21763056 | FALSE | 1 | FALSE | 1 | FALSE | 1 | FALSE | 1 | FALSE | 1 | FALSE | 1 |
| 3861 | 3861 | 9.93333333 | 2.81582828 | 2 | 0.04858661 | 0 | 1 | 1 | 0.40110418 | 1 | 0.48242099 | 0 | 1 | 10 | 0.94579623 | FALSE | 0.649301 | FALSE | 1 | FALSE | 1 | FALSE | 1 | FALSE | 1 | FALSE | 1 |
| 3862 | 3862 | 8.05882353 | 1.5134918 | 0 | 1 | 1 | 0.42857733 | 0 | 1 | 0 | 1 | 0 | 1 | 16 | 0.23070182 | FALSE | 1 | FALSE | 1 | FALSE | 1 | FALSE | 1 | FALSE | 1 | FALSE | 1 |
| 3863 | 3863 | 9.09302326 | 1.32619325 | 2 | 0.30241117 | 1 | 0.75788114 | 3 | 0.19985581 | 3 | 0.31602175 | 0 | 1 | 34 | 0.87722689 | FALSE | 1 | FALSE | 1 | FALSE | 1 | FALSE | 1 | FALSE | 1 | FALSE | 1 |
| 3864 | 3864 | 3.13014608 | 0.93992574 | 17 | 0.58994802 | 31 | 0.03177141 | 23 | 0.66254912 | 23 | 0.96089018 | 11 | 0.57229152 | 578 | 0.45131784 | FALSE | 1 | FALSE | 0.92268438 | FALSE | 1 | FALSE | 1 | FALSE | 1 | FALSE | 1 |
| 3865 | 3865 | 9.2962963 | 1.90155521 | 1 | 0.50471032 | 1 | 0.58913618 | 0 | 1 | 0 | 1 | 1 | 0.36060486 | 24 | 0.3725373 | FALSE | 1 | FALSE | 1 | FALSE | 1 | FALSE | 1 | FALSE | 1 | FALSE | 1 |
| 3866 | 3866 | 7.38235294 | 2.55544063 | 0 | 1 | 0 | 1 | 3 | 0.11365871 | 3 | 0.19161876 | 0 | 1 | 27 | 0.75124583 | FALSE | 1 | FALSE | 1 | FALSE | 1 | FALSE | 1 | FALSE | 1 | FALSE | 1 |
| 3867 | 3867 | 6.31896552 | 2.04517573 | 6 | 0.07694901 | 10 | 0.00416662 | 6 | 0.23770724 | 4 | 0.78683937 | 1 | 0.85549683 | 89 | 0.98953023 | FALSE | 0.77583006 | FALSE | 0.58120789 | FALSE | 1 | FALSE | 1 | FALSE | 1 | FALSE | 1 |
| 3868 | 3868 | 8.76190476 | 3.51736863 | 1 | 0.42086991 | 0 | 1 | 1 | 0.53671415 | 1 | 0.62783522 | 0 | 1 | 18 | 0.57962763 | FALSE | 1 | FALSE | 1 | FALSE | 1 | FALSE | 1 | FALSE | 1 | FALSE | 1 |
| 3869 | 3869 | 9.78571429 | 1.65523289 | 0 | 1 | 1 | 0.60247911 | 0 | 1 | 1 | 0.73248983 | 1 | 0.37113067 | 25 | 0.34293 | FALSE | 1 | FALSE | 1 | FALSE | 1 | FALSE | 1 | FALSE | 1 | FALSE | 1 |
| 3870 | 3870 | 9.57142857 | 0.9035079 | 1 | 0.3050805 | 2 | 0.07341559 | 0 | 1 | 0 | 1 | 0 | 1 | 11 | 0.83676449 | FALSE | 1 | FALSE | 1 | FALSE | 1 | FALSE | 1 | FALSE | 1 | FALSE | 1 |
| 3871 | 3871 | 7.59090909 | 2.05954741 | 0 | 1 | 1 | 0.5154318 | 1 | 0.55340851 | 2 | 0.26791271 | 0 | 1 | 18 | 0.74744813 | FALSE | 1 | FALSE | 1 | FALSE | 1 | FALSE | 1 | FALSE | 1 | FALSE | 1 |
| 3872 | 3872 | 6.47222222 | 3.0229319 | 0 | 1 | 2 | 0.32556427 | 2 | 0.37289966 | 1 | 0.81665383 | 1 | 0.44938266 | 30 | 0.67248271 | FALSE | 1 | FALSE | 1 | FALSE | 1 | FALSE | 1 | FALSE | 1 | FALSE | 1 |
| 3873 | 3873 | 4.32577904 | 1.29668941 | 11 | 0.27336093 | 16 | 0.09385335 | 11 | 0.70664934 | 9 | 0.98123348 | 12 | 0.01076466 | 287 | 0.79785281 | FALSE | 1 | FALSE | 1 | FALSE | 1 | FALSE | 1 | FALSE | 0.57142086 | FALSE | 1 |
| 3874 | 3874 | 11.6666667 | 2.64994759 | 2 | 0.05513784 | 1 | 0.38964394 | 1 | 0.42266264 | 2 | 0.14906763 | 0 | 1 | 9 | 0.995488 | FALSE | 0.68849059 | FALSE | 1 | FALSE | 1 | FALSE | 1 | FALSE | 1 | FALSE | 1 |
| 3875 | 3875 | 9.16 | 1.71300905 | 0 | 1 | 1 | 0.56109832 | 1 | 0.59998149 | 1 | 0.69181315 | 3 | 0.00758617 | 19 | 0.91734261 | FALSE | 1 | FALSE | 1 | FALSE | 1 | FALSE | 1 | FALSE | 0.49339011 | FALSE | 1 |
| 3876 | 3876 | 6.24324324 | 2.84182474 | 3 | 0.54559205 | 4 | 0.48578212 | 9 | 0.0179699 | 11 | 0.01258588 | 1 | 0.8428205 | 83 | 0.99719872 | FALSE | 1 | FALSE | 1 | FALSE | 1 | FALSE | 1 | FALSE | 1 | FALSE | 1 |
| 3877 | 3877 | 10.1052632 | 2.31459299 | 2 | 0.08405377 | 0 | 1 | 1 | 0.50143826 | 0 | 1 | 1 | 0.26986775 | 15 | 0.83648191 | FALSE | 0.79236883 | FALSE | 1 | FALSE | 1 | FALSE | 1 | FALSE | 1 | FALSE | 1 |
| 3878 | 3878 | 13.2 | 1.72046505 | 0 | 1 | 0 | 1 | 0 | 1 | 1 | 0.2094688 | 0 | 1 | 4 | 0.82371636 | FALSE | 1 | FALSE | 1 | FALSE | 1 | FALSE | 1 | FALSE | 1 | FALSE | 1 |
| 3879 | 3879 | 9.92307692 | 3.70989402 | 2 | 0.04234484 | 2 | 0.06426041 | 0 | 1 | 0 | 1 | 0 | 1 | 9 | 0.96018168 | FALSE | 0.62842194 | FALSE | 1 | FALSE | 1 | FALSE | 1 | FALSE | 1 | FALSE | 1 |
| 3880 | 3880 | 4.2654321 | 2.9350862 | 11 | 0.00286274 | 6 | 0.42716587 | 9 | 0.12816896 | 13 | 0.03484852 | 2 | 0.74956518 | 121 | 0.99957088 | FALSE | 0.14653312 | FALSE | 1 | FALSE | 1 | FALSE | 1 | FALSE | 1 | FALSE | 1 |
| 3881 | 3881 | 9.33333333 | 3.15524255 | 0 | 1 | 1 | 0.38964394 | 0 | 1 | 0 | 1 | 0 | 1 | 14 | 0.29525409 | FALSE | 1 | FALSE | 1 | FALSE | 1 | FALSE | 1 | FALSE | 1 | FALSE | 1 |
| 3882 | 3882 | 10.3 | 2.72213152 | 0 | 1 | 0 | 1 | 0 | 1 | 0 | 1 | 0 | 1 | 10 | 0.18273166 | FALSE | 1 | FALSE | 1 | FALSE | 1 | FALSE | 1 | FALSE | 1 | FALSE | 1 |
| 3883 | 3883 | 10.125 | 2.97647023 | 1 | 0.18769745 | 1 | 0.23139161 | 0 | 1 | 0 | 1 | 0 | 1 | 6 | 0.88425551 | FALSE | 1 | FALSE | 1 | FALSE | 1 | FALSE | 1 | FALSE | 1 | FALSE | 1 |
| 3884 | 3884 | 10.7777778 | 3.08320821 | 0 | 1 | 0 | 1 | 0 | 1 | 0 | 1 | 0 | 1 | 9 | 0.21661397 | FALSE | 1 | FALSE | 1 | FALSE | 1 | FALSE | 1 | FALSE | 1 | FALSE | 1 |
| 3885 | 3885 | 8.5 | 2.54950976 | 0 | 1 | 1 | 0.40942991 | 0 | 1 | 1 | 0.52895088 | 1 | 0.23264612 | 13 | 0.76799161 | FALSE | 1 | FALSE | 1 | FALSE | 1 | FALSE | 1 | FALSE | 1 | FALSE | 1 |
| 3886 | 3886 | 9.53571429 | 3.28940011 | 1 | 0.51745948 | 1 | 0.60247911 | 1 | 0.64171563 | 1 | 0.73248983 | 0 | 1 | 24 | 0.55018107 | FALSE | 1 | FALSE | 1 | FALSE | 1 | FALSE | 1 | FALSE | 1 | FALSE | 1 |
| 3887 | 3887 | 8.96551724 | 3.38335437 | 1 | 0.77387657 | 3 | 0.28014665 | 4 | 0.147507 | 2 | 0.74436726 | 0 | 1 | 47 | 0.72965359 | FALSE | 1 | FALSE | 1 | FALSE | 1 | FALSE | 1 | FALSE | 1 | FALSE | 1 |
| 3888 | 3888 | 7.75 | 3.122499 | 0 | 1 | 1 | 0.61539065 | 0 | 1 | 0 | 1 | 0 | 1 | 28 | 0.04589964 | FALSE | 1 | FALSE | 1 | FALSE | 1 | FALSE | 1 | FALSE | 1 | FALSE | 1 |
| 3889 | 3889 | 9.45714286 | 1.64478148 | 0 | 1 | 1 | 0.65167362 | 2 | 0.32011871 | 3 | 0.17988885 | 0 | 1 | 26 | 0.77612337 | FALSE | 1 | FALSE | 1 | FALSE | 1 | FALSE | 1 | FALSE | 1 | FALSE | 1 |
| 3890 | 3890 | 8.30434783 | 1.12035641 | 0 | 1 | 1 | 0.53115654 | 3 | 0.04772439 | 1 | 0.6613267 | 0 | 1 | 18 | 0.86273155 | FALSE | 1 | FALSE | 1 | FALSE | 1 | FALSE | 1 | FALSE | 1 | FALSE | 1 |
| 3891 | 3891 | 9.06060606 | 2.04405115 | 2 | 0.1881398 | 1 | 0.63997468 | 0 | 1 | 5 | 0.01254769 | 2 | 0.09110253 | 21 | 0.99467947 | FALSE | 1 | FALSE | 1 | FALSE | 1 | FALSE | 1 | FALSE | 1 | FALSE | 1 |
| 3892 | 3892 | 10.71875 | 3.88293039 | 4 | 0.00856611 | 2 | 0.27730767 | 1 | 0.69069222 | 0 | 1 | 1 | 0.41154418 | 24 | 0.94876798 | FALSE | 0.27019545 | FALSE | 1 | FALSE | 1 | FALSE | 1 | FALSE | 1 | FALSE | 1 |
| 3893 | 3893 | 11.0625 | 2.1928506 | 1 | 0.34033054 | 0 | 1 | 0 | 1 | 1 | 0.52895088 | 0 | 1 | 14 | 0.53308261 | FALSE | 1 | FALSE | 1 | FALSE | 1 | FALSE | 1 | FALSE | 1 | FALSE | 1 |
| 3894 | 3894 | 9.31818182 | 2.81854836 | 0 | 1 | 0 | 1 | 0 | 1 | 3 | 0.0612581 | 0 | 1 | 17 | 0.61750632 | FALSE | 1 | FALSE | 1 | FALSE | 1 | FALSE | 1 | FALSE | 1 | FALSE | 1 |
| 3895 | 3895 | 10.2 | 2.18174242 | 1 | 0.22886536 | 0 | 1 | 0 | 1 | 1 | 0.37517262 | 1 | 0.15247062 | 7 | 0.94326215 | FALSE | 1 | FALSE | 1 | FALSE | 1 | FALSE | 1 | FALSE | 1 | FALSE | 1 |
| 3896 | 3896 | 9.4375 | 3.29713417 | 0 | 1 | 0 | 1 | 3 | 0.10599423 | 1 | 0.77852168 | 0 | 1 | 28 | 0.42532177 | FALSE | 1 | FALSE | 1 | FALSE | 1 | FALSE | 1 | FALSE | 1 | FALSE | 1 |
| 3897 | 3897 | 8.79245283 | 0.9587431 | 0 | 1 | 2 | 0.51564773 | 3 | 0.29666549 | 3 | 0.44196866 | 0 | 1 | 45 | 0.5504765 | FALSE | 1 | FALSE | 1 | FALSE | 1 | FALSE | 1 | FALSE | 1 | FALSE | 1 |
| 3898 | 3898 | 10.1764706 | 1.97738074 | 2 | 0.06909038 | 0 | 1 | 0 | 1 | 1 | 0.55062772 | 0 | 1 | 14 | 0.7313145 | FALSE | 0.74276693 | FALSE | 1 | FALSE | 1 | FALSE | 1 | FALSE | 1 | FALSE | 1 |
| 3899 | 3899 | 10.1428571 | 1.93429486 | 1 | 0.42086991 | 1 | 0.49918218 | 0 | 1 | 1 | 0.62783522 | 0 | 1 | 18 | 0.57962763 | FALSE | 1 | FALSE | 1 | FALSE | 1 | FALSE | 1 | FALSE | 1 | FALSE | 1 |
| 3900 | 3900 | 5.77011494 | 3.33804661 | 4 | 0.1836099 | 1 | 0.94382585 | 1 | 0.95938747 | 1 | 0.98368538 | 0 | 1 | 80 | 0.02805204 | FALSE | 1 | FALSE | 1 | FALSE | 1 | FALSE | 1 | FALSE | 1 | FALSE | 1 |
| 3901 | 3901 | 6.19617225 | 0.80950377 | 6 | 0.44834956 | 7 | 0.51767802 | 5 | 0.87706858 | 9 | 0.62737993 | 1 | 0.97011217 | 181 | 0.21318259 | FALSE | 1 | FALSE | 1 | FALSE | 1 | FALSE | 1 | FALSE | 1 | FALSE | 1 |
| 3902 | 3902 | 5.38636364 | 2.58974589 | 9 | 0.00662299 | 13 | 0.00031729 | 7 | 0.19609815 | 5 | 0.73189483 | 3 | 0.36844071 | 95 | 0.99992183 | FALSE | 0.24562356 | FALSE | 0.15723145 | FALSE | 1 | FALSE | 1 | FALSE | 1 | FALSE | 1 |
| 3903 | 3903 | 4.23248408 | 1.20512264 | 8 | 0.55197421 | 9 | 0.68670237 | 12 | 0.4464287 | 23 | 0.01613282 | 10 | 0.03141003 | 250 | 0.98391278 | FALSE | 1 | FALSE | 1 | FALSE | 1 | FALSE | 1 | FALSE | 0.8327215 | FALSE | 1 |
| 3904 | 3904 | 6.44776119 | 1.65523667 | 2 | 0.86327384 | 8 | 0.06829603 | 5 | 0.53036069 | 1 | 0.99827446 | 4 | 0.17710987 | 114 | 0.46918017 | FALSE | 1 | FALSE | 1 | FALSE | 1 | FALSE | 1 | FALSE | 1 | FALSE | 1 |
| 3905 | 3905 | 7.5 | 1.02469508 | 1 | 0.22886536 | 0 | 1 | 0 | 1 | 0 | 1 | 0 | 1 | 9 | 0.5215547 | FALSE | 1 | FALSE | 1 | FALSE | 1 | FALSE | 1 | FALSE | 1 | FALSE | 1 |
| 3906 | 3906 | 8.40540541 | 1.96536854 | 0 | 1 | 1 | 0.70472939 | 0 | 1 | 2 | 0.51200191 | 2 | 0.12262638 | 32 | 0.47030992 | FALSE | 1 | FALSE | 1 | FALSE | 1 | FALSE | 1 | FALSE | 1 | FALSE | 1 |
| 3907 | 3907 | 5.91666667 | 2.99606223 | 7 | 0.00071447 | 3 | 0.2979413 | 1 | 0.88558856 | 1 | 0.93827647 | 0 | 1 | 47 | 0.87928796 | FALSE | 0.0622651 | FALSE | 1 | FALSE | 1 | FALSE | 1 | FALSE | 1 | FALSE | 1 |
| 3908 | 3908 | 7 | 3.1793251 | 1 | 0.6184714 | 3 | 0.11640364 | 1 | 0.74264075 | 1 | 0.82511639 | 2 | 0.12262638 | 29 | 0.88820978 | FALSE | 1 | FALSE | 1 | FALSE | 1 | FALSE | 1 | FALSE | 1 | FALSE | 1 |
| 3909 | 3909 | 11.3142857 | 2.67016777 | 0 | 1 | 2 | 0.31355738 | 1 | 0.72299506 | 1 | 0.80778317 | 1 | 0.44015537 | 30 | 0.52765882 | FALSE | 1 | FALSE | 1 | FALSE | 1 | FALSE | 1 | FALSE | 1 | FALSE | 1 |
| 3910 | 3910 | 9.44680851 | 2.15177819 | 2 | 0.3309085 | 0 | 1 | 2 | 0.4962935 | 3 | 0.35428318 | 2 | 0.17379289 | 37 | 0.82909673 | FALSE | 1 | FALSE | 1 | FALSE | 1 | FALSE | 1 | FALSE | 1 | FALSE | 1 |
| 3911 | 3911 | 8.01219512 | 2.55425949 | 5 | 0.05387199 | 5 | 0.11631842 | 2 | 0.78851973 | 7 | 0.07347807 | 1 | 0.73564367 | 60 | 0.99043724 | FALSE | 0.68849059 | FALSE | 1 | FALSE | 1 | FALSE | 1 | FALSE | 1 | FALSE | 1 |
| 3912 | 3912 | 6.675 | 2.64941031 | 0 | 1 | 1 | 0.7326167 | 2 | 0.42396974 | 5 | 0.03481208 | 0 | 1 | 32 | 0.83853687 | FALSE | 1 | FALSE | 1 | FALSE | 1 | FALSE | 1 | FALSE | 1 | FALSE | 1 |
| 3913 | 3913 | 10.0357143 | 3.02940521 | 2 | 0.16048326 | 0 | 1 | 0 | 1 | 1 | 0.73248983 | 0 | 1 | 25 | 0.34293 | FALSE | 1 | FALSE | 1 | FALSE | 1 | FALSE | 1 | FALSE | 1 | FALSE | 1 |
| 3914 | 3914 | 9.4 | 1.52315462 | 0 | 1 | 0 | 1 | 1 | 0.59998149 | 1 | 0.69181315 | 1 | 0.33902672 | 22 | 0.43642846 | FALSE | 1 | FALSE | 1 | FALSE | 1 | FALSE | 1 | FALSE | 1 | FALSE | 1 |
| 3915 | 3915 | 5.18518519 | 3.80094566 | 2 | 0.15144274 | 1 | 0.58913618 | 0 | 1 | 2 | 0.35374209 | 0 | 1 | 22 | 0.76237877 | FALSE | 1 | FALSE | 1 | FALSE | 1 | FALSE | 1 | FALSE | 1 | FALSE | 1 |
| 3916 | 3916 | 10.25 | 2.58602011 | 0 | 1 | 0 | 1 | 0 | 1 | 0 | 1 | 2 | 0.01579445 | 10 | 0.71436964 | FALSE | 1 | FALSE | 1 | FALSE | 1 | FALSE | 1 | FALSE | 0.66313193 | FALSE | 1 |
| 3917 | 3917 | 11.9565217 | 2.34943584 | 2 | 0.09989125 | 1 | 0.49918218 | 1 | 0.53671415 | 1 | 0.62783522 | 0 | 1 | 16 | 0.90353674 | FALSE | 0.86047225 | FALSE | 1 | FALSE | 1 | FALSE | 1 | FALSE | 1 | FALSE | 1 |
| 3918 | 3918 | 11 | 3.03681119 | 0 | 1 | 1 | 0.44710673 | 0 | 1 | 0 | 1 | 0 | 1 | 17 | 0.20329205 | FALSE | 1 | FALSE | 1 | FALSE | 1 | FALSE | 1 | FALSE | 1 | FALSE | 1 |
| 3919 | 3919 | 10.7222222 | 3.26267708 | 0 | 1 | 1 | 0.44710673 | 1 | 0.48280983 | 1 | 0.57131013 | 0 | 1 | 15 | 0.69372602 | FALSE | 1 | FALSE | 1 | FALSE | 1 | FALSE | 1 | FALSE | 1 | FALSE | 1 |
| 3920 | 3920 | 11.5454546 | 2.99586492 | 0 | 1 | 2 | 0.04725998 | 1 | 0.33149969 | 0 | 1 | 0 | 1 | 8 | 0.92152993 | FALSE | 1 | FALSE | 1 | FALSE | 1 | FALSE | 1 | FALSE | 1 | FALSE | 1 |
| 3921 | 3921 | 9.45454546 | 1.30479092 | 3 | 0.00234778 | 0 | 1 | 0 | 1 | 0 | 1 | 0 | 1 | 8 | 0.92152993 | FALSE | 0.12737083 | FALSE | 1 | FALSE | 1 | FALSE | 1 | FALSE | 1 | FALSE | 1 |
| 3922 | 3922 | 9 | 4.15331193 | 0 | 1 | 0 | 1 | 0 | 1 | 0 | 1 | 0 | 1 | 8 | 0.25677166 | FALSE | 1 | FALSE | 1 | FALSE | 1 | FALSE | 1 | FALSE | 1 | FALSE | 1 |
| 3923 | 3923 | 8.41666667 | 2.39646731 | 1 | 0.46441868 | 1 | 0.54637326 | 0 | 1 | 0 | 1 | 0 | 1 | 22 | 0.25237063 | FALSE | 1 | FALSE | 1 | FALSE | 1 | FALSE | 1 | FALSE | 1 | FALSE | 1 |
| 3924 | 3924 | 9.80555556 | 2.4016134 | 2 | 0.23542589 | 5 | 0.0056244 | 0 | 1 | 2 | 0.49731009 | 1 | 0.44938266 | 26 | 0.98200522 | FALSE | 1 | FALSE | 0.59070661 | FALSE | 1 | FALSE | 1 | FALSE | 1 | FALSE | 1 |
| 3925 | 3925 | 9.5 | 2.67826006 | 2 | 0.13369621 | 1 | 0.56109832 | 1 | 0.59998149 | 0 | 1 | 0 | 1 | 21 | 0.64957167 | FALSE | 0.97964162 | FALSE | 1 | FALSE | 1 | FALSE | 1 | FALSE | 1 | FALSE | 1 |
| 3926 | 3926 | 6.54545455 | 3.14393027 | 0 | 1 | 0 | 1 | 1 | 0.33149969 | 0 | 1 | 0 | 1 | 10 | 0.46859937 | FALSE | 1 | FALSE | 1 | FALSE | 1 | FALSE | 1 | FALSE | 1 | FALSE | 1 |
| 3927 | 3927 | 9.5 | 2.36290781 | 3 | 0.02256792 | 0 | 1 | 1 | 0.58502114 | 0 | 1 | 0 | 1 | 20 | 0.68270729 | FALSE | 0.4692294 | FALSE | 1 | FALSE | 1 | FALSE | 1 | FALSE | 1 | FALSE | 1 |
| 3928 | 3928 | 8.725 | 2.75669639 | 6 | 0.00048425 | 3 | 0.13823565 | 1 | 0.76953771 | 2 | 0.5543522 | 1 | 0.48480878 | 27 | 0.99797908 | TRUE | 0.0450788 | FALSE | 1 | FALSE | 1 | FALSE | 1 | FALSE | 1 | FALSE | 1 |
| 3929 | 3929 | 9.4375 | 2.68022271 | 1 | 0.54198714 | 1 | 0.6278847 | 0 | 1 | 4 | 0.04659941 | 2 | 0.08611595 | 22 | 0.96518743 | FALSE | 1 | FALSE | 1 | FALSE | 1 | FALSE | 1 | FALSE | 1 | FALSE | 1 |
| 3930 | 3930 | 9.3125 | 2.84426506 | 0 | 1 | 0 | 1 | 0 | 1 | 1 | 0.52895088 | 0 | 1 | 15 | 0.26127859 | FALSE | 1 | FALSE | 1 | FALSE | 1 | FALSE | 1 | FALSE | 1 | FALSE | 1 |
| 3931 | 3931 | 8.18518519 | 2.10883568 | 1 | 0.50471032 | 0 | 1 | 2 | 0.25286935 | 2 | 0.35374209 | 2 | 0.07169515 | 20 | 0.95133411 | FALSE | 1 | FALSE | 1 | FALSE | 1 | FALSE | 1 | FALSE | 1 | FALSE | 1 |
| 3932 | 3932 | 5.99130435 | 2.19681846 | 4 | 0.3057853 | 3 | 0.69070605 | 5 | 0.35412914 | 4 | 0.74440481 | 1 | 0.83744681 | 92 | 0.56157078 | FALSE | 1 | FALSE | 1 | FALSE | 1 | FALSE | 1 | FALSE | 1 | FALSE | 1 |
| 3933 | 3933 | 4.72307692 | 1.93357137 | 6 | 0.00266387 | 2 | 0.5356583 | 3 | 0.31652594 | 3 | 0.46623289 | 2 | 0.22768257 | 39 | 0.99675703 | FALSE | 0.13988708 | FALSE | 1 | FALSE | 1 | FALSE | 1 | FALSE | 1 | FALSE | 1 |
| 3934 | 3934 | 10.6111111 | 4.47386098 | 0 | 1 | 1 | 0.44710673 | 1 | 0.48280983 | 0 | 1 | 1 | 0.25766374 | 15 | 0.69372602 | FALSE | 1 | FALSE | 1 | FALSE | 1 | FALSE | 1 | FALSE | 1 | FALSE | 1 |
| 3935 | 3935 | 9.74074074 | 2.50322152 | 0 | 1 | 1 | 0.58913618 | 0 | 1 | 2 | 0.35374209 | 1 | 0.36060486 | 23 | 0.58307573 | FALSE | 1 | FALSE | 1 | FALSE | 1 | FALSE | 1 | FALSE | 1 | FALSE | 1 |
| 3936 | 3936 | 11.4042553 | 1.72169603 | 0 | 1 | 3 | 0.16927151 | 2 | 0.47281647 | 2 | 0.60667406 | 2 | 0.16211764 | 35 | 0.8620983 | FALSE | 1 | FALSE | 1 | FALSE | 1 | FALSE | 1 | FALSE | 1 | FALSE | 1 |
| 3937 | 3937 | 10.3157895 | 2.10394696 | 1 | 0.37380237 | 2 | 0.1135776 | 3 | 0.02505979 | 0 | 1 | 1 | 0.25766374 | 11 | 0.99651304 | FALSE | 1 | FALSE | 1 | FALSE | 1 | FALSE | 1 | FALSE | 1 | FALSE | 1 |
| 3938 | 3938 | 10.6521739 | 1.20176304 | 1 | 0.45027683 | 0 | 1 | 1 | 0.5695037 | 0 | 1 | 3 | 0.00598158 | 18 | 0.86273155 | FALSE | 1 | FALSE | 1 | FALSE | 1 | FALSE | 1 | FALSE | 0.45694206 | FALSE | 1 |
| 3939 | 3939 | 6.94117647 | 1.53280167 | 2 | 0.52416586 | 0 | 1 | 2 | 0.70787008 | 4 | 0.38072365 | 0 | 1 | 60 | 0.24411098 | FALSE | 1 | FALSE | 1 | FALSE | 1 | FALSE | 1 | FALSE | 1 | FALSE | 1 |
| 3940 | 3940 | 9.23529412 | 2.10124366 | 2 | 0.06909038 | 0 | 1 | 2 | 0.12280513 | 0 | 1 | 0 | 1 | 13 | 0.88719222 | FALSE | 0.74276693 | FALSE | 1 | FALSE | 1 | FALSE | 1 | FALSE | 1 | FALSE | 1 |
| 3941 | 3941 | 7.5952381 | 1.95252613 | 1 | 0.66519151 | 0 | 1 | 3 | 0.19060367 | 2 | 0.5811115 | 0 | 1 | 36 | 0.50934357 | FALSE | 1 | FALSE | 1 | FALSE | 1 | FALSE | 1 | FALSE | 1 | FALSE | 1 |
| 3942 | 3942 | 8.9245283 | 2.17054951 | 0 | 1 | 0 | 1 | 2 | 0.57303317 | 5 | 0.0944968 | 0 | 1 | 46 | 0.39930625 | FALSE | 1 | FALSE | 1 | FALSE | 1 | FALSE | 1 | FALSE | 1 | FALSE | 1 |
| 3943 | 3943 | 7.97297297 | 2.00656337 | 0 | 1 | 2 | 0.3375163 | 2 | 0.38585089 | 0 | 1 | 1 | 0.45845922 | 32 | 0.47030992 | FALSE | 1 | FALSE | 1 | FALSE | 1 | FALSE | 1 | FALSE | 1 | FALSE | 1 |
| 3944 | 3944 | 8.29268293 | 3.0543364 | 0 | 1 | 1 | 0.74131698 | 0 | 1 | 3 | 0.29054213 | 0 | 1 | 37 | 0.21027077 | FALSE | 1 | FALSE | 1 | FALSE | 1 | FALSE | 1 | FALSE | 1 | FALSE | 1 |
| 3945 | 3945 | 7.36363636 | 2.27726819 | 0 | 1 | 1 | 0.76576289 | 4 | 0.07197653 | 1 | 0.87439647 | 0 | 1 | 38 | 0.45705039 | FALSE | 1 | FALSE | 1 | FALSE | 1 | FALSE | 1 | FALSE | 1 | FALSE | 1 |
| 3946 | 3946 | 8.41666667 | 2.97559518 | 0 | 1 | 0 | 1 | 3 | 0.13783405 | 1 | 0.81665383 | 1 | 0.44938266 | 31 | 0.49869404 | FALSE | 1 | FALSE | 1 | FALSE | 1 | FALSE | 1 | FALSE | 1 | FALSE | 1 |

| | | | | | | | | | | | | | | | | | | | | | | | | | | | |
|---|---|---|---|---|---|---|---|---|---|---|---|---|---|---|---|---|---|---|---|---|---|---|---|---|---|---|---|
| 3947 | 3947 | 9.4 | 2.02649122 | 1 | 0.32293355 | 1 | 0.38964394 | 0 | 1 | 0 | 1 | 1 | 0.2198258 | 12 | 0.80329917 | FALSE | 1 | FALSE | 1 | FALSE | 1 | FALSE | 1 | FALSE | 1 | FALSE | 1 |
| 3948 | 3948 | 11.6 | 3.61109402 | 0 | 1 | 1 | 0.15163655 | 0 | 1 | 0 | 1 | 0 | 1 | 4 | 0.82371636 | FALSE | 1 | FALSE | 1 | FALSE | 1 | FALSE | 1 | FALSE | 1 | FALSE | 1 |
| 3949 | 3949 | 11.1666667 | 2.40947205 | 0 | 1 | 0 | 1 | 1 | 0.16720719 | 0 | 1 | 0 | 1 | 4 | 0.82371636 | FALSE | 1 | FALSE | 1 | FALSE | 1 | FALSE | 1 | FALSE | 1 | FALSE | 1 |
| 3950 | 3950 | 10.2857143 | 1.48461498 | 0 | 1 | 0 | 1 | 0 | 1 | 0 | 1 | 0 | 1 | 7 | 0.3043657 | FALSE | 1 | FALSE | 1 | FALSE | 1 | FALSE | 1 | FALSE | 1 | FALSE | 1 |
| 3951 | 3951 | 9.78947368 | 2.26132116 | 0 | 1 | 1 | 0.46503795 | 0 | 1 | 1 | 0.59104357 | 0 | 1 | 17 | 0.41141543 | FALSE | 1 | FALSE | 1 | FALSE | 1 | FALSE | 1 | FALSE | 1 | FALSE | 1 |
| 3952 | 3952 | 7.03846154 | 5.03434947 | 0 | 1 | 0 | 1 | 0 | 1 | 0 | 1 | 0 | 1 | 26 | 0.0119733 | FALSE | 1 | FALSE | 1 | FALSE | 1 | FALSE | 1 | FALSE | 1 | FALSE | 1 |
| 3953 | 3953 | 8.35384615 | 1.80111734 | 2 | 0.49997462 | 2 | 0.62701348 | 4 | 0.20453007 | 2 | 0.80687302 | 5 | 0.00408995 | 50 | 0.96116604 | FALSE | 1 | FALSE | 1 | FALSE | 1 | FALSE | 1 | FALSE | 0.37645884 | FALSE | 1 |
| 3954 | 3954 | 9.95 | 2.49949995 | 0 | 1 | 0 | 1 | 1 | 0.51939841 | 2 | 0.23331368 | 0 | 1 | 17 | 0.61750632 | FALSE | 1 | FALSE | 1 | FALSE | 1 | FALSE | 1 | FALSE | 1 | FALSE | 1 |
| 3955 | 3955 | 9.64705882 | 2.34999816 | 1 | 0.35728305 | 0 | 1 | 1 | 0.46348836 | 2 | 0.18212134 | 0 | 1 | 13 | 0.88719222 | FALSE | 1 | FALSE | 1 | FALSE | 1 | FALSE | 1 | FALSE | 1 | FALSE | 1 |
| 3956 | 3956 | 2.31851852 | 1.48898102 | 4 | 0.45720114 | 4 | 0.64107304 | 9 | 0.05344368 | 9 | 0.16703901 | 1 | 0.89508155 | 108 | 0.933896 | FALSE | 1 | FALSE | 1 | FALSE | 1 | FALSE | 1 | FALSE | 1 | FALSE | 1 |
| 3957 | 3957 | 7.725 | 3.99366686 | 0 | 1 | 0 | 1 | 1 | 0.76953771 | 3 | 0.27785536 | 1 | 0.48480878 | 35 | 0.38966407 | FALSE | 1 | FALSE | 1 | FALSE | 1 | FALSE | 1 | FALSE | 1 | FALSE | 1 |
| 3958 | 3958 | 7.7 | 3.6207734 | 0 | 1 | 1 | 0.48239024 | 0 | 1 | 3 | 0.0612581 | 0 | 1 | 16 | 0.80826361 | FALSE | 1 | FALSE | 1 | FALSE | 1 | FALSE | 1 | FALSE | 1 | FALSE | 1 |
| 3959 | 3959 | 3.07954546 | 0.72298888 | 28 | 0.04571624 | 33 | 0.07198598 | 21 | 0.950371 | 40 | 0.27468776 | 12 | 0.6563073 | 656 | 0.87591409 | FALSE | 0.649301 | FALSE | 1 | FALSE | 1 | FALSE | 1 | FALSE | 1 | FALSE | 1 |
| 3960 | 3960 | 6.93103448 | 2.94701526 | 4 | 0.00600461 | 2 | 0.24095584 | 1 | 0.65463982 | 0 | 1 | 1 | 0.38148475 | 21 | 0.97185642 | FALSE | 0.22773046 | FALSE | 1 | FALSE | 1 | FALSE | 1 | FALSE | 1 | FALSE | 1 |
| 3961 | 3961 | 9.4 | 2.38886305 | 1 | 0.32293355 | 0 | 1 | 0 | 1 | 3 | 0.02891327 | 0 | 1 | 11 | 0.92879412 | FALSE | 1 | FALSE | 1 | FALSE | 1 | FALSE | 1 | FALSE | 1 | FALSE | 1 |
| 3962 | 3962 | 7.65217391 | 2.89840579 | 1 | 0.45027683 | 1 | 0.53115654 | 1 | 0.5695037 | 1 | 0.6613267 | 2 | 0.05389003 | 17 | 0.94439745 | FALSE | 1 | FALSE | 1 | FALSE | 1 | FALSE | 1 | FALSE | 1 | FALSE | 1 |
| 3963 | 3963 | 8.75 | 2.30488611 | 0 | 1 | 1 | 0.40942991 | 1 | 0.44344817 | 0 | 1 | 0 | 1 | 14 | 0.53308261 | FALSE | 1 | FALSE | 1 | FALSE | 1 | FALSE | 1 | FALSE | 1 | FALSE | 1 |
| 3964 | 3964 | 8.58823529 | 2.25072076 | 0 | 1 | 0 | 1 | 0 | 1 | 0 | 1 | 0 | 1 | 17 | 0.05550965 | FALSE | 1 | FALSE | 1 | FALSE | 1 | FALSE | 1 | FALSE | 1 | FALSE | 1 |
| 3965 | 3965 | 9.95 | 2.22429764 | 0 | 1 | 1 | 0.48239024 | 2 | 0.16019548 | 2 | 0.23331368 | 0 | 1 | 15 | 0.92113679 | FALSE | 1 | FALSE | 1 | FALSE | 1 | FALSE | 1 | FALSE | 1 | FALSE | 1 |
| 3966 | 3966 | 6.24242424 | 3.22894394 | 0 | 1 | 1 | 0.6629941 | 0 | 1 | 1 | 0.78873814 | 0 | 1 | 31 | 0.09218876 | FALSE | 1 | FALSE | 1 | FALSE | 1 | FALSE | 1 | FALSE | 1 | FALSE | 1 |
| 3967 | 3967 | 10.2307692 | 2.77776463 | 0 | 1 | 1 | 0.3480709 | 0 | 1 | 1 | 0.45746571 | 0 | 1 | 11 | 0.6683056 | FALSE | 1 | FALSE | 1 | FALSE | 1 | FALSE | 1 | FALSE | 1 | FALSE | 1 |
| 3968 | 3968 | 8.8 | 1.32664992 | 1 | 0.32293355 | 0 | 1 | 1 | 0.42266264 | 0 | 1 | 1 | 0.2198258 | 12 | 0.80329917 | FALSE | 1 | FALSE | 1 | FALSE | 1 | FALSE | 1 | FALSE | 1 | FALSE | 1 |
| 3969 | 3969 | 11.8888889 | 3.41384256 | 0 | 1 | 0 | 1 | 1 | 0.28067846 | 1 | 0.34506517 | 0 | 1 | 7 | 0.84574168 | FALSE | 1 | FALSE | 1 | FALSE | 1 | FALSE | 1 | FALSE | 1 | FALSE | 1 |
| 3970 | 3970 | 11.6666667 | 2.59914516 | 0 | 1 | 0 | 1 | 0 | 1 | 2 | 0.14906763 | 0 | 1 | 13 | 0.57712805 | FALSE | 1 | FALSE | 1 | FALSE | 1 | FALSE | 1 | FALSE | 1 | FALSE | 1 |
| 3971 | 3971 | 10.6923077 | 3.31394757 | 0 | 1 | 0 | 1 | 1 | 0.37874417 | 0 | 1 | 0 | 1 | 12 | 0.374198 | FALSE | 1 | FALSE | 1 | FALSE | 1 | FALSE | 1 | FALSE | 1 | FALSE | 1 |
| 3972 | 3972 | 8.27272727 | 2.32604134 | 3 | 0.05151181 | 0 | 1 | 1 | 0.70185639 | 4 | 0.06258966 | 3 | 0.01637249 | 22 | 0.99710775 | FALSE | 0.67646333 | FALSE | 1 | FALSE | 1 | FALSE | 1 | FALSE | 0.66728076 | FALSE | 1 |
| 3973 | 3973 | 9 | 2 | 0 | 1 | 0 | 1 | 0 | 1 | 0 | 1 | 0 | 1 | 2 | 0.71196516 | FALSE | 1 | FALSE | 1 | FALSE | 1 | FALSE | 1 | FALSE | 1 | FALSE | 1 |
| 3974 | 3974 | 10.3 | 1.73493516 | 0 | 1 | 0 | 1 | 1 | 0.30655256 | 1 | 0.37517262 | 0 | 1 | 8 | 0.80394719 | FALSE | 1 | FALSE | 1 | FALSE | 1 | FALSE | 1 | FALSE | 1 | FALSE | 1 |
| 3975 | 3975 | 9.9375 | 2.43589691 | 2 | 0.06197875 | 1 | 0.40942991 | 0 | 1 | 0 | 1 | 1 | 0.23264612 | 12 | 0.90922186 | FALSE | 0.72532843 | FALSE | 1 | FALSE | 1 | FALSE | 1 | FALSE | 1 | FALSE | 1 |
| 3976 | 3976 | 7.76470588 | 2.15650621 | 2 | 0.05513784 | 1 | 0.38964394 | 0 | 1 | 0 | 1 | 0 | 1 | 12 | 0.80329917 | FALSE | 0.68849059 | FALSE | 1 | FALSE | 1 | FALSE | 1 | FALSE | 1 | FALSE | 1 |
| 3977 | 3977 | 10.3846154 | 1.98216904 | 0 | 1 | 0 | 1 | 1 | 0.37874417 | 0 | 1 | 0 | 1 | 12 | 0.374198 | FALSE | 1 | FALSE | 1 | FALSE | 1 | FALSE | 1 | FALSE | 1 | FALSE | 1 |
| 3978 | 3978 | 11 | 2.73030135 | 0 | 1 | 1 | 0.30368014 | 0 | 1 | 1 | 0.40390032 | 0 | 1 | 9 | 0.75985366 | FALSE | 1 | FALSE | 1 | FALSE | 1 | FALSE | 1 | FALSE | 1 | FALSE | 1 |
| 3979 | 3979 | 11.5 | 3.20156212 | 0 | 1 | 0 | 1 | 0 | 1 | 1 | 0.43131129 | 0 | 1 | 11 | 0.41944532 | FALSE | 1 | FALSE | 1 | FALSE | 1 | FALSE | 1 | FALSE | 1 | FALSE | 1 |
| 3980 | 3980 | 7.61290323 | 3.17934279 | 1 | 0.55378205 | 0 | 1 | 1 | 0.67911179 | 0 | 1 | 1 | 0.4016888 | 28 | 0.26393996 | FALSE | 1 | FALSE | 1 | FALSE | 1 | FALSE | 1 | FALSE | 1 | FALSE | 1 |
| 3981 | 3981 | 8.4137931 | 2.04293976 | 0 | 1 | 2 | 0.24095584 | 2 | 0.27983822 | 1 | 0.74482229 | 0 | 1 | 24 | 0.70563334 | FALSE | 1 | FALSE | 1 | FALSE | 1 | FALSE | 1 | FALSE | 1 | FALSE | 1 |
| 3982 | 3982 | 10.1212121 | 3.13105245 | 0 | 1 | 1 | 0.6629941 | 1 | 0.70185639 | 1 | 0.78873814 | 1 | 0.42123867 | 29 | 0.39648374 | FALSE | 1 | FALSE | 1 | FALSE | 1 | FALSE | 1 | FALSE | 1 | FALSE | 1 |
| 3983 | 3983 | 9.25806452 | 2.13975793 | 4 | 0.00679223 | 2 | 0.25306331 | 0 | 1 | 1 | 0.75658797 | 0 | 1 | 23 | 0.9158696 | FALSE | 0.24597546 | FALSE | 1 | FALSE | 1 | FALSE | 1 | FALSE | 1 | FALSE | 1 |
| 3984 | 3984 | 10.3333333 | 3.10657265 | 1 | 0.42086991 | 0 | 1 | 1 | 0.53671415 | 1 | 0.62783522 | 1 | 0.29368245 | 17 | 0.77849076 | FALSE | 1 | FALSE | 1 | FALSE | 1 | FALSE | 1 | FALSE | 1 | FALSE | 1 |
| 3985 | 3985 | 10 | 3.03315018 | 1 | 0.22886536 | 0 | 1 | 1 | 0.30655256 | 0 | 1 | 0 | 1 | 8 | 0.80394719 | FALSE | 1 | FALSE | 1 | FALSE | 1 | FALSE | 1 | FALSE | 1 | FALSE | 1 |
| 3986 | 3986 | 7.43902439 | 0.82783233 | 0 | 1 | 1 | 0.7326167 | 0 | 1 | 1 | 0.84823834 | 0 | 1 | 38 | 0.03896014 | FALSE | 1 | FALSE | 1 | FALSE | 1 | FALSE | 1 | FALSE | 1 | FALSE | 1 |
| 3987 | 3987 | 11.4333333 | 3.77432849 | 1 | 0.54198714 | 1 | 0.6278847 | 1 | 0.66709967 | 2 | 0.4036176 | 0 | 1 | 25 | 0.67630471 | FALSE | 1 | FALSE | 1 | FALSE | 1 | FALSE | 1 | FALSE | 1 | FALSE | 1 |
| 3988 | 3988 | 10.0909091 | 3.54078947 | 0 | 1 | 0 | 1 | 0 | 1 | 1 | 0.64497439 | 0 | 1 | 21 | 0.12048848 | FALSE | 1 | FALSE | 1 | FALSE | 1 | FALSE | 1 | FALSE | 1 | FALSE | 1 |
| 3989 | 3989 | 7.83333333 | 3.62476053 | 0 | 1 | 2 | 0.1135776 | 0 | 1 | 1 | 0.57131013 | 0 | 1 | 15 | 0.69372602 | FALSE | 1 | FALSE | 1 | FALSE | 1 | FALSE | 1 | FALSE | 1 | FALSE | 1 |
| 3990 | 3990 | 6.28571429 | 2.25676107 | 0 | 1 | 1 | 0.87100665 | 2 | 0.65873577 | 4 | 0.31731292 | 0 | 1 | 55 | 0.22629228 | FALSE | 1 | FALSE | 1 | FALSE | 1 | FALSE | 1 | FALSE | 1 | FALSE | 1 |
| 3991 | 3991 | 7.51111111 | 2.87998628 | 4 | 0.02747663 | 3 | 0.17732615 | 2 | 0.48463745 | 2 | 0.61900576 | 2 | 0.16793736 | 32 | 0.99319189 | FALSE | 0.50964275 | FALSE | 1 | FALSE | 1 | FALSE | 1 | FALSE | 1 | FALSE | 1 |
| 3992 | 3992 | 9.33333333 | 2.59272486 | 1 | 0.26795838 | 1 | 0.32623933 | 0 | 1 | 0 | 1 | 0 | 1 | 10 | 0.71436964 | FALSE | 1 | FALSE | 1 | FALSE | 1 | FALSE | 1 | FALSE | 1 | FALSE | 1 |
| 3993 | 3993 | 10.6756757 | 2.43903009 | 0 | 1 | 2 | 0.3375163 | 1 | 0.74264075 | 2 | 0.51200191 | 0 | 1 | 32 | 0.47030992 | FALSE | 1 | FALSE | 1 | FALSE | 1 | FALSE | 1 | FALSE | 1 | FALSE | 1 |
| 3994 | 3994 | 9.15 | 1.73997126 | 1 | 0.40558521 | 0 | 1 | 3 | 0.03322143 | 2 | 0.23331368 | 1 | 0.28187291 | 13 | 0.99248686 | FALSE | 1 | FALSE | 1 | FALSE | 1 | FALSE | 1 | FALSE | 1 | FALSE | 1 |
| 3995 | 3995 | 9.46153846 | 3.24903263 | 1 | 0.28675951 | 0 | 1 | 1 | 0.37874417 | 0 | 1 | 0 | 1 | 11 | 0.6683056 | FALSE | 1 | FALSE | 1 | FALSE | 1 | FALSE | 1 | FALSE | 1 | FALSE | 1 |
| 3996 | 3996 | 10.2727273 | 4.13611388 | 1 | 0.24866461 | 0 | 1 | 0 | 1 | 1 | 0.40390032 | 0 | 1 | 9 | 0.75985366 | FALSE | 1 | FALSE | 1 | FALSE | 1 | FALSE | 1 | FALSE | 1 | FALSE | 1 |
| 3997 | 3997 | 11 | 1.65144565 | 0 | 1 | 1 | 0.28036922 | 0 | 1 | 1 | 0.37517262 | 0 | 1 | 8 | 0.80394719 | FALSE | 1 | FALSE | 1 | FALSE | 1 | FALSE | 1 | FALSE | 1 | FALSE | 1 |
| 3998 | 3998 | 8.3 | 2.47857486 | 1 | 0.54198714 | 0 | 1 | 1 | 0.66709967 | 3 | 0.15707004 | 1 | 0.39166989 | 24 | 0.82294782 | FALSE | 1 | FALSE | 1 | FALSE | 1 | FALSE | 1 | FALSE | 1 | FALSE | 1 |
| 3999 | 3999 | 9.15789474 | 3.68871905 | 0 | 1 | 1 | 0.46503795 | 0 | 1 | 1 | 0.59104357 | 1 | 0.26986775 | 16 | 0.65565778 | FALSE | 1 | FALSE | 1 | FALSE | 1 | FALSE | 1 | FALSE | 1 | FALSE | 1 |
| 4000 | 4000 | 9.70588235 | 3.08248765 | 0 | 1 | 0 | 1 | 0 | 1 | 2 | 0.18212134 | 0 | 1 | 15 | 0.49060435 | FALSE | 1 | FALSE | 1 | FALSE | 1 | FALSE | 1 | FALSE | 1 | FALSE | 1 |
| 4001 | 4001 | 4.74074074 | 3.14575072 | 0 | 1 | 0 | 1 | 1 | 0.62830986 | 0 | 1 | 0 | 1 | 26 | 0.06079097 | FALSE | 1 | FALSE | 1 | FALSE | 1 | FALSE | 1 | FALSE | 1 | FALSE | 1 |
| 4002 | 4002 | 11.1111111 | 1.66295884 | 0 | 1 | 1 | 0.25628163 | 0 | 1 | 0 | 1 | 0 | 1 | 8 | 0.5780332 | FALSE | 1 | FALSE | 1 | FALSE | 1 | FALSE | 1 | FALSE | 1 | FALSE | 1 |
| 4003 | 4003 | 9.13636364 | 1.76572042 | 0 | 1 | 0 | 1 | 0 | 1 | 0 | 1 | 1 | 0.30529951 | 21 | 0.12048848 | FALSE | 1 | FALSE | 1 | FALSE | 1 | FALSE | 1 | FALSE | 1 | FALSE | 1 |
| 4004 | 4004 | 6.44444444 | 3.59354703 | 0 | 1 | 2 | 0.32556427 | 0 | 1 | 1 | 0.81665383 | 0 | 1 | 33 | 0.16407838 | FALSE | 1 | FALSE | 1 | FALSE | 1 | FALSE | 1 | FALSE | 1 | FALSE | 1 |
| 4005 | 4005 | 8.83870968 | 3.28335759 | 1 | 0.55378205 | 0 | 1 | 1 | 0.67911179 | 2 | 0.41984187 | 0 | 1 | 27 | 0.45522639 | FALSE | 1 | FALSE | 1 | FALSE | 1 | FALSE | 1 | FALSE | 1 | FALSE | 1 |
| 4006 | 4006 | 7.70588235 | 2.42571289 | 0 | 1 | 3 | 0.37822937 | 4 | 0.22739663 | 2 | 0.82657478 | 0 | 1 | 59 | 0.36577777 | FALSE | 1 | FALSE | 1 | FALSE | 1 | FALSE | 1 | FALSE | 1 | FALSE | 1 |
| 4007 | 4007 | 9.82142857 | 2.26863035 | 0 | 1 | 0 | 1 | 0 | 1 | 3 | 0.13524571 | 2 | 0.07640798 | 23 | 0.73439118 | FALSE | 1 | FALSE | 1 | FALSE | 1 | FALSE | 1 | FALSE | 1 | FALSE | 1 |
| 4008 | 4008 | 8.28125 | 2.0346126 | 0 | 1 | 1 | 0.65167362 | 2 | 0.32011871 | 1 | 0.77852168 | 1 | 0.41154418 | 27 | 0.61671133 | FALSE | 1 | FALSE | 1 | FALSE | 1 | FALSE | 1 | FALSE | 1 | FALSE | 1 |
| 4009 | 4009 | 9.1875 | 1.28543524 | 0 | 1 | 0 | 1 | 0 | 1 | 2 | 0.16544789 | 0 | 1 | 14 | 0.53308261 | FALSE | 1 | FALSE | 1 | FALSE | 1 | FALSE | 1 | FALSE | 1 | FALSE | 1 |
| 4010 | 4010 | 11.1904762 | 3.49991901 | 0 | 1 | 0 | 1 | 0 | 1 | 0 | 1 | 0 | 1 | 20 | 0.03329856 | FALSE | 1 | FALSE | 1 | FALSE | 1 | FALSE | 1 | FALSE | 1 | FALSE | 1 |
| 4011 | 4011 | 11.3846154 | 2.27151124 | 3 | 0.00391863 | 0 | 1 | 0 | 1 | 1 | 0.45746571 | 1 | 0.19354507 | 8 | 0.99085298 | FALSE | 0.17735574 | FALSE | 1 | FALSE | 1 | FALSE | 1 | FALSE | 1 | FALSE | 1 |
| 4012 | 4012 | 8.76923077 | 2.25844896 | 0 | 1 | 0 | 1 | 2 | 0.07725803 | 0 | 1 | 0 | 1 | 11 | 0.6683056 | FALSE | 1 | FALSE | 1 | FALSE | 1 | FALSE | 1 | FALSE | 1 | FALSE | 1 |
| 4013 | 4013 | 8.7826087 | 2.50141737 | 0 | 1 | 0 | 1 | 2 | 0.19931429 | 3 | 0.08614834 | 2 | 0.05389003 | 16 | 0.98100709 | FALSE | 1 | FALSE | 1 | FALSE | 1 | FALSE | 1 | FALSE | 1 | FALSE | 1 |
| 4014 | 4014 | 7.55555556 | 3.07117221 | 2 | 0.15144274 | 0 | 1 | 3 | 0.07106362 | 1 | 0.71956339 | 1 | 0.36060486 | 20 | 0.95133411 | FALSE | 1 | FALSE | 1 | FALSE | 1 | FALSE | 1 | FALSE | 1 | FALSE | 1 |
| 4015 | 4015 | 11.6666667 | 2.44948974 | 0 | 1 | 0 | 1 | 0 | 1 | 0 | 1 | 0 | 1 | 9 | 0.21661397 | FALSE | 1 | FALSE | 1 | FALSE | 1 | FALSE | 1 | FALSE | 1 | FALSE | 1 |
| 4016 | 4016 | 11.3333333 | 2.3570226 | 0 | 1 | 0 | 1 | 0 | 1 | 1 | 0.50623194 | 2 | 0.02433796 | 12 | 0.80329917 | FALSE | 1 | FALSE | 1 | FALSE | 1 | FALSE | 1 | FALSE | 0.77578444 | FALSE | 1 |
| 4017 | 4017 | 9.70588235 | 2.90566801 | 0 | 1 | 2 | 0.10307112 | 0 | 1 | 1 | 0.55062772 | 1 | 0.24525764 | 13 | 0.88719222 | FALSE | 1 | FALSE | 1 | FALSE | 1 | FALSE | 1 | FALSE | 1 | FALSE | 1 |
| 4018 | 4018 | 9.85185185 | 1.9187902 | 1 | 0.47819878 | 0 | 1 | 1 | 0.59998149 | 1 | 0.69181315 | 0 | 1 | 22 | 0.43642846 | FALSE | 1 | FALSE | 1 | FALSE | 1 | FALSE | 1 | FALSE | 1 | FALSE | 1 |
| 4019 | 4019 | 6.03448276 | 2.22032552 | 0 | 1 | 1 | 0.61539065 | 0 | 1 | 3 | 0.14602298 | 0 | 1 | 25 | 0.51779518 | FALSE | 1 | FALSE | 1 | FALSE | 1 | FALSE | 1 | FALSE | 1 | FALSE | 1 |
| 4020 | 4020 | 8.11111111 | 2.49196239 | 0 | 1 | 0 | 1 | 1 | 0.48280983 | 1 | 0.57131013 | 1 | 0.25766374 | 15 | 0.69372602 | FALSE | 1 | FALSE | 1 | FALSE | 1 | FALSE | 1 | FALSE | 1 | FALSE | 1 |
| 4021 | 4021 | 10 | 2.7325202 | 0 | 1 | 1 | 0.38964394 | 0 | 1 | 2 | 0.14906763 | 0 | 1 | 12 | 0.80329917 | FALSE | 1 | FALSE | 1 | FALSE | 1 | FALSE | 1 | FALSE | 1 | FALSE | 1 |
| 4022 | 4022 | 6.09090909 | 2.62116833 | 0 | 1 | 1 | 0.6629941 | 2 | 0.33343871 | 2 | 0.45159487 | 0 | 1 | 28 | 0.58681477 | FALSE | 1 | FALSE | 1 | FALSE | 1 | FALSE | 1 | FALSE | 1 | FALSE | 1 |
| 4023 | 4023 | 8.2 | 1.70587221 | 1 | 0.64722818 | 2 | 0.37295654 | 3 | 0.17245215 | 7 | 0.00202352 | 0 | 1 | 27 | 0.99797908 | FALSE | 1 | FALSE | 1 | FALSE | 1 | FALSE | 1 | FALSE | 1 | FALSE | 1 |
| 4024 | 4024 | 9.93333333 | 3.80292285 | 0 | 1 | 0 | 1 | 0 | 1 | 1 | 0.50623194 | 0 | 1 | 14 | 0.29525409 | FALSE | 1 | FALSE | 1 | FALSE | 1 | FALSE | 1 | FALSE | 1 | FALSE | 1 |
| 4025 | 4025 | 7.51612903 | 2.99322857 | 1 | 0.55378205 | 3 | 0.07729739 | 0 | 1 | 1 | 0.76781284 | 3 | 0.01381051 | 23 | 0.95750806 | FALSE | 1 | FALSE | 1 | FALSE | 1 | FALSE | 1 | FALSE | 0.64607242 | FALSE | 1 |
| 4026 | 4026 | 8.32258065 | 0.81861791 | 1 | 0.55378205 | 1 | 0.63997468 | 0 | 1 | 1 | 0.76781284 | 0 | 1 | 28 | 0.26393996 | FALSE | 1 | FALSE | 1 | FALSE | 1 | FALSE | 1 | FALSE | 1 | FALSE | 1 |
| 4027 | 4027 | 7.15384615 | 2.72254087 | 2 | 0.26413583 | 3 | 0.13080858 | 2 | 0.41139386 | 1 | 0.84089119 | 3 | 0.02555742 | 28 | 0.98701111 | FALSE | 1 | FALSE | 1 | FALSE | 1 | FALSE | 1 | FALSE | 0.79106376 | FALSE | 1 |
| 4028 | 4028 | 5.87254902 | 1.70162304 | 3 | 0.48191148 | 2 | 0.84402995 | 3 | 0.70953966 | 3 | 0.8496648 | 0 | 1 | 90 | 0.11570866 | FALSE | 1 | FALSE | 1 | FALSE | 1 | FALSE | 1 | FALSE | 1 | FALSE | 1 |
| 4029 | 4029 | 6.44761905 | 2.48416299 | 11 | 7.05E-05 | 5 | 0.25228158 | 8 | 0.03458688 | 5 | 0.5320851 | 4 | 0.09327737 | 72 | 0.99998726 | TRUE | 0.01342173 | FALSE | 1 | FALSE | 1 | FALSE | 1 | FALSE | 1 | FALSE | 1 |
| 4030 | 4030 | 4.58490566 | 2.22705459 | 1 | 0.93765336 | 2 | 0.86287617 | 2 | 0.89940445 | 2 | 0.95922468 | 1 | 0.82904315 | 98 | 0.01009699 | FALSE | 1 | FALSE | 1 | FALSE | 1 | FALSE | 1 | FALSE | 1 | FALSE | 1 |
| 4031 | 4031 | 8.94444444 | 2.1723572 | 2 | 0.23542589 | 0 | 1 | 0 | 1 | 3 | 0.22784656 | 0 | 1 | 31 | 0.49869404 | FALSE | 1 | FALSE | 1 | FALSE | 1 | FALSE | 1 | FALSE | 1 | FALSE | 1 |
| 4032 | 4032 | 11.0571429 | 2.11042118 | 0 | 1 | 1 | 0.68454804 | 2 | 0.35984046 | 1 | 0.80778317 | 0 | 1 | 31 | 0.34231756 | FALSE | 1 | FALSE | 1 | FALSE | 1 | FALSE | 1 | FALSE | 1 | FALSE | 1 |
| 4033 | 4033 | 6.12903226 | 0.55311059 | 0 | 1 | 0 | 1 | 1 | 0.67911179 | 0 | 1 | 0 | 1 | 30 | 0.03453514 | FALSE | 1 | FALSE | 1 | FALSE | 1 | FALSE | 1 | FALSE | 1 | FALSE | 1 |
| 4034 | 4034 | 8.38095238 | 2.03484387 | 1 | 0.42086991 | 0 | 1 | 2 | 0.17308361 | 0 | 1 | 0 | 1 | 18 | 0.57962763 | FALSE | 1 | FALSE | 1 | FALSE | 1 | FALSE | 1 | FALSE | 1 | FALSE | 1 |
| 4035 | 4035 | 6.85 | 3.00457984 | 0 | 1 | 1 | 0.48239024 | 0 | 1 | 4 | 0.01177595 | 1 | 0.28187291 | 14 | 0.97325593 | FALSE | 1 | FALSE | 1 | FALSE | 1 | FALSE | 1 | FALSE | 1 | FALSE | 1 |
| 4036 | 4036 | 8.58139535 | 2.69566886 | 6 | 0.00048425 | 3 | 0.13823565 | 1 | 0.76953771 | 0 | 1 | 4 | 0.00397004 | 26 | 0.99938297 | TRUE | 0.0450788 | FALSE | 1 | FALSE | 1 | FALSE | 1 | FALSE | 0.37571345 | FALSE | 1 |
| 4037 | 4037 | 8.36842105 | 1.56307815 | 0 | 1 | 0 | 1 | 1 | 0.50143826 | 1 | 0.59104357 | 0 | 1 | 17 | 0.41141543 | FALSE | 1 | FALSE | 1 | FALSE | 1 | FALSE | 1 | FALSE | 1 | FALSE | 1 |
| 4038 | 4038 | 8.5952381 | 2.1163707 | 0 | 1 | 2 | 0.39616059 | 2 | 0.44869594 | 1 | 0.86193352 | 2 | 0.15059682 | 35 | 0.67024784 | FALSE | 1 | FALSE | 1 | FALSE | 1 | FALSE | 1 | FALSE | 1 | FALSE | 1 |
| 4039 | 4039 | 4.27634195 | 1.40050294 | 14 | 0.40579184 | 20 | 0.18846279 | 17 | 0.63147532 | 23 | 0.52881554 | 8 | 0.58196474 | 418 | 0.71611695 | FALSE | 1 | FALSE | 1 | FALSE | 1 | FALSE | 1 | FALSE | 1 | FALSE | 1 |
| 4040 | 4040 | 10.9090909 | 2.42916168 | 0 | 1 | 0 | 1 | 1 | 0.30655256 | 0 | 1 | 0 | 1 | 9 | 0.5215547 | FALSE | 1 | FALSE | 1 | FALSE | 1 | FALSE | 1 | FALSE | 1 | FALSE | 1 |

| | | | | | | | | | | | | | | | | | | | | | | | | | | | |
|---|---|---|---|---|---|---|---|---|---|---|---|---|---|---|---|---|---|---|---|---|---|---|---|---|---|---|---|
| 4041 | 4041 | 9.5625 | 2.52410256 | 0 | 1 | 1 | 0.40942991 | 0 | 1 | 1 | 0.52895088 | 2 | 0.02752114 | 12 | 0.90922186 | FALSE | 1 | FALSE | 1 | FALSE | 1 | FALSE | 1 | FALSE | 0.79106376 | FALSE | 1 |
| 4042 | 4042 | 9.1 | 1.84119526 | 1 | 0.40558521 | 0 | 1 | 0 | 1 | 1 | 0.60987146 | 0 | 1 | 18 | 0.37504645 | FALSE | 1 | FALSE | 1 | FALSE | 1 | FALSE | 1 | FALSE | 1 | FALSE | 1 |
| 4043 | 4043 | 7.96 | 2.08767814 | 0 | 1 | 0 | 1 | 1 | 0.59998149 | 2 | 0.31968852 | 0 | 1 | 22 | 0.43642846 | FALSE | 1 | FALSE | 1 | FALSE | 1 | FALSE | 1 | FALSE | 1 | FALSE | 1 |
| 4044 | 4044 | 9.74193548 | 1.39046852 | 2 | 0.1881398 | 0 | 1 | 0 | 1 | 1 | 0.76781284 | 0 | 1 | 28 | 0.26393996 | FALSE | 1 | FALSE | 1 | FALSE | 1 | FALSE | 1 | FALSE | 1 | FALSE | 1 |
| 4045 | 4045 | 9 | 2.21880079 | 1 | 0.28675951 | 1 | 0.3480709 | 0 | 1 | 0 | 1 | 0 | 1 | 11 | 0.6683056 | FALSE | 1 | FALSE | 1 | FALSE | 1 | FALSE | 1 | FALSE | 1 | FALSE | 1 |
| 4046 | 4046 | 8.88888889 | 2.35440223 | 2 | 0.07645454 | 1 | 0.44710673 | 1 | 0.48280983 | 0 | 1 | 0 | 1 | 14 | 0.86287418 | FALSE | 0.77424195 | FALSE | 1 | FALSE | 1 | FALSE | 1 | FALSE | 1 | FALSE | 1 |
| 4047 | 4047 | 6.57471264 | 1.86086239 | 0 | 1 | 1 | 0.94382585 | 0 | 1 | 2 | 0.91445868 | 0 | 1 | 84 | 0.00029323 | FALSE | 1 | FALSE | 1 | FALSE | 1 | FALSE | 1 | FALSE | 1 | FALSE | 0.80070757 |
| 4048 | 4048 | 5.01470588 | 2.28481333 | 5 | 0.25016225 | 5 | 0.42496778 | 1 | 0.99238487 | 5 | 0.73189483 | 1 | 0.8896353 | 115 | 0.22852627 | FALSE | 1 | FALSE | 1 | FALSE | 1 | FALSE | 1 | FALSE | 1 | FALSE | 1 |
| 4049 | 4049 | 9.6 | 1.70489491 | 1 | 0.69044531 | 1 | 0.77338922 | 0 | 1 | 1 | 0.8802008 | 3 | 0.03699359 | 39 | 0.4317533 | FALSE | 1 | FALSE | 1 | FALSE | 1 | FALSE | 1 | FALSE | 0.87669812 | FALSE | 1 |
| 4050 | 4050 | 10.3333333 | 2.4267033 | 0 | 1 | 0 | 1 | 0 | 1 | 0 | 1 | 0 | 1 | 11 | 0.15414489 | FALSE | 1 | FALSE | 1 | FALSE | 1 | FALSE | 1 | FALSE | 1 | FALSE | 1 |
| 4051 | 4051 | 10.375 | 1.86664807 | 0 | 1 | 0 | 1 | 0 | 1 | 0 | 1 | 0 | 1 | 7 | 0.3043657 | FALSE | 1 | FALSE | 1 | FALSE | 1 | FALSE | 1 | FALSE | 1 | FALSE | 1 |
| 4052 | 4052 | 3.86507937 | 2.92865746 | 2 | 0.83576566 | 5 | 0.3797149 | 6 | 0.29287782 | 6 | 0.51548389 | 2 | 0.61209767 | 104 | 0.69550441 | FALSE | 1 | FALSE | 1 | FALSE | 1 | FALSE | 1 | FALSE | 1 | FALSE | 1 |
| 4053 | 4053 | 7.71428571 | 2.71052371 | 0 | 1 | 0 | 1 | 0 | 1 | 0 | 1 | 0 | 1 | 7 | 0.3043657 | FALSE | 1 | FALSE | 1 | FALSE | 1 | FALSE | 1 | FALSE | 1 | FALSE | 1 |
| 4054 | 4054 | 7.78571429 | 3.05142991 | 0 | 1 | 1 | 0.36919824 | 1 | 0.40110418 | 0 | 1 | 0 | 1 | 12 | 0.62236227 | FALSE | 1 | FALSE | 1 | FALSE | 1 | FALSE | 1 | FALSE | 1 | FALSE | 1 |
| 4055 | 4055 | 7.76923077 | 1.69405504 | 2 | 0.12501436 | 1 | 0.54637326 | 1 | 0.58502114 | 2 | 0.30249307 | 1 | 0.32796862 | 17 | 0.97530702 | FALSE | 0.95622564 | FALSE | 1 | FALSE | 1 | FALSE | 1 | FALSE | 1 | FALSE | 1 |
| 4056 | 4056 | 9.52631579 | 1.53446103 | 0 | 1 | 0 | 1 | 0 | 1 | 1 | 0.59104357 | 1 | 0.26986775 | 17 | 0.41141543 | FALSE | 1 | FALSE | 1 | FALSE | 1 | FALSE | 1 | FALSE | 1 | FALSE | 1 |
| 4057 | 4057 | 11 | 3.48623668 | 0 | 1 | 0 | 1 | 0 | 1 | 3 | 0.01944678 | 0 | 1 | 10 | 0.86791642 | FALSE | 1 | FALSE | 1 | FALSE | 1 | FALSE | 1 | FALSE | 1 | FALSE | 1 |
| 4058 | 4058 | 8.69565217 | 2.25417068 | 0 | 1 | 0 | 1 | 3 | 0.04772439 | 1 | 0.6613267 | 0 | 1 | 19 | 0.71542496 | FALSE | 1 | FALSE | 1 | FALSE | 1 | FALSE | 1 | FALSE | 1 | FALSE | 1 |
| 4059 | 4059 | 9.14705882 | 2.98163351 | 1 | 0.58738519 | 1 | 0.6739483 | 1 | 0.71261921 | 1 | 0.7984848 | 0 | 1 | 30 | 0.36879488 | FALSE | 1 | FALSE | 1 | FALSE | 1 | FALSE | 1 | FALSE | 1 | FALSE | 1 |
| 4060 | 4060 | 10.4 | 2.4166092 | 0 | 1 | 0 | 1 | 0 | 1 | 1 | 0.37517262 | 0 | 1 | 9 | 0.5215547 | FALSE | 1 | FALSE | 1 | FALSE | 1 | FALSE | 1 | FALSE | 1 | FALSE | 1 |
| 4061 | 4061 | 8.5 | 1.85107282 | 2 | 0.21638713 | 0 | 1 | 0 | 1 | 1 | 0.7984848 | 0 | 1 | 31 | 0.1995074 | FALSE | 1 | FALSE | 1 | FALSE | 1 | FALSE | 1 | FALSE | 1 | FALSE | 1 |
| 4062 | 4062 | 10.6521739 | 2.59780335 | 1 | 0.45027683 | 1 | 0.53115654 | 0 | 1 | 0 | 1 | 0 | 1 | 21 | 0.27964072 | FALSE | 1 | FALSE | 1 | FALSE | 1 | FALSE | 1 | FALSE | 1 | FALSE | 1 |
| 4063 | 4063 | 10.6875 | 3.34885708 | 1 | 0.34033054 | 1 | 0.40942991 | 0 | 1 | 1 | 0.52895088 | 0 | 1 | 13 | 0.76799161 | FALSE | 1 | FALSE | 1 | FALSE | 1 | FALSE | 1 | FALSE | 1 | FALSE | 1 |
| 4064 | 4064 | 7.3364486 | 1.48477586 | 4 | 0.19401704 | 4 | 0.32490044 | 2 | 0.83551221 | 5 | 0.38834525 | 2 | 0.43073682 | 72 | 0.85441763 | FALSE | 1 | FALSE | 1 | FALSE | 1 | FALSE | 1 | FALSE | 1 | FALSE | 1 |
| 4065 | 4065 | 9 | 2.60341656 | 2 | 0.07645454 | 1 | 0.44710673 | 1 | 0.48280983 | 1 | 0.57131013 | 0 | 1 | 13 | 0.95041913 | FALSE | 0.77424195 | FALSE | 1 | FALSE | 1 | FALSE | 1 | FALSE | 1 | FALSE | 1 |
| 4066 | 4066 | 6.80246914 | 3.11673996 | 9 | 0.00020736 | 5 | 0.12090937 | 4 | 0.33196391 | 1 | 0.97829212 | 1 | 0.74003033 | 61 | 0.98893397 | TRUE | 0.02784797 | FALSE | 1 | FALSE | 1 | FALSE | 1 | FALSE | 1 | FALSE | 1 |
| 4067 | 4067 | 9.42307692 | 1.49802632 | 0 | 1 | 0 | 1 | 0 | 1 | 4 | 0.02930308 | 1 | 0.3499045 | 21 | 0.78939668 | FALSE | 1 | FALSE | 1 | FALSE | 1 | FALSE | 1 | FALSE | 1 | FALSE | 1 |
| 4068 | 4068 | 7.53658537 | 3.04516859 | 0 | 1 | 0 | 1 | 3 | 0.1814659 | 4 | 0.11715635 | 0 | 1 | 34 | 0.69505404 | FALSE | 1 | FALSE | 1 | FALSE | 1 | FALSE | 1 | FALSE | 1 | FALSE | 1 |
| 4069 | 4069 | 10.125 | 1.94855716 | 0 | 1 | 1 | 0.6278847 | 2 | 0.29330414 | 2 | 0.4036176 | 0 | 1 | 25 | 0.67630471 | FALSE | 1 | FALSE | 1 | FALSE | 1 | FALSE | 1 | FALSE | 1 | FALSE | 1 |
| 4070 | 4070 | 4.55263158 | 1.97613883 | 3 | 0.30904591 | 3 | 0.44799895 | 4 | 0.29102231 | 7 | 0.05887011 | 0 | 1 | 59 | 0.957857 | FALSE | 1 | FALSE | 1 | FALSE | 1 | FALSE | 1 | FALSE | 1 | FALSE | 1 |
| 4071 | 4071 | 8.33333333 | 2.05480467 | 1 | 0.49162634 | 1 | 0.57534752 | 1 | 0.61440466 | 2 | 0.33678042 | 0 | 1 | 21 | 0.78939668 | FALSE | 1 | FALSE | 1 | FALSE | 1 | FALSE | 1 | FALSE | 1 | FALSE | 1 |
| 4072 | 4072 | 8.53846154 | 2.92510051 | 1 | 0.28675951 | 0 | 1 | 0 | 1 | 0 | 1 | 0 | 1 | 12 | 0.374198 | FALSE | 1 | FALSE | 1 | FALSE | 1 | FALSE | 1 | FALSE | 1 | FALSE | 1 |
| 4073 | 4073 | 9.4375 | 2.26298337 | 2 | 0.06197875 | 1 | 0.40942991 | 1 | 0.44344817 | 2 | 0.16544789 | 0 | 1 | 10 | 0.99304605 | FALSE | 0.72532843 | FALSE | 1 | FALSE | 1 | FALSE | 1 | FALSE | 1 | FALSE | 1 |
| 4074 | 4074 | 10.0588235 | 2.04279475 | 0 | 1 | 0 | 1 | 0 | 1 | 3 | 0.040397 | 0 | 1 | 14 | 0.7313145 | FALSE | 1 | FALSE | 1 | FALSE | 1 | FALSE | 1 | FALSE | 1 | FALSE | 1 |
| 4075 | 4075 | 8.8 | 2.05588586 | 2 | 0.17884234 | 1 | 0.6278847 | 1 | 0.66709967 | 1 | 0.75658797 | 1 | 0.39166989 | 24 | 0.82294782 | FALSE | 1 | FALSE | 1 | FALSE | 1 | FALSE | 1 | FALSE | 1 | FALSE | 1 |
| 4076 | 4076 | 12.2307692 | 2.85963543 | 0 | 1 | 0 | 1 | 0 | 1 | 0 | 1 | 0 | 1 | 13 | 0.10967906 | FALSE | 1 | FALSE | 1 | FALSE | 1 | FALSE | 1 | FALSE | 1 | FALSE | 1 |
| 4077 | 4077 | 7.27777778 | 2.23123174 | 3 | 0.0638041 | 4 | 0.02790717 | 0 | 1 | 1 | 0.81665383 | 0 | 1 | 28 | 0.90259714 | FALSE | 0.73766 | FALSE | 0.87929047 | FALSE | 1 | FALSE | 1 | FALSE | 1 | FALSE | 1 |
| 4078 | 4078 | 8.2 | 2.78567766 | 3 | 0.01369554 | 0 | 1 | 1 | 0.51939841 | 1 | 0.60987146 | 1 | 0.28187291 | 14 | 0.97325593 | FALSE | 0.36625055 | FALSE | 1 | FALSE | 1 | FALSE | 1 | FALSE | 1 | FALSE | 1 |
| 4079 | 4079 | 9 | 2.1330729 | 0 | 1 | 0 | 1 | 2 | 0.42396974 | 3 | 0.27785536 | 0 | 1 | 35 | 0.38966407 | FALSE | 1 | FALSE | 1 | FALSE | 1 | FALSE | 1 | FALSE | 1 | FALSE | 1 |
| 4080 | 4080 | 8.27272727 | 2.21937375 | 2 | 0.31193899 | 1 | 0.76576289 | 1 | 0.80109494 | 1 | 0.87439647 | 2 | 0.16211764 | 37 | 0.61979664 | FALSE | 1 | FALSE | 1 | FALSE | 1 | FALSE | 1 | FALSE | 1 | FALSE | 1 |
| 4081 | 4081 | 8.72727273 | 2.52589891 | 0 | 1 | 0 | 1 | 0 | 1 | 2 | 0.08789566 | 0 | 1 | 9 | 0.75985366 | FALSE | 1 | FALSE | 1 | FALSE | 1 | FALSE | 1 | FALSE | 1 | FALSE | 1 |
| 4082 | 4082 | 10.7142857 | 2.40322912 | 0 | 1 | 0 | 1 | 1 | 0.40110418 | 2 | 0.13304705 | 0 | 1 | 11 | 0.83676449 | FALSE | 1 | FALSE | 1 | FALSE | 1 | FALSE | 1 | FALSE | 1 | FALSE | 1 |
| 4083 | 4083 | 6.83783784 | 3.07134545 | 0 | 1 | 1 | 0.69480472 | 3 | 0.13783405 | 3 | 0.22784656 | 0 | 1 | 29 | 0.81028289 | FALSE | 1 | FALSE | 1 | FALSE | 1 | FALSE | 1 | FALSE | 1 | FALSE | 1 |
| 4084 | 4084 | 9.09090909 | 2.86952257 | 3 | 0.21200292 | 6 | 0.01453703 | 4 | 0.18240181 | 6 | 0.06375189 | 4 | 0.01842358 | 39 | 0.99999163 | FALSE | 1 | FALSE | 0.71309764 | FALSE | 1 | FALSE | 1 | FALSE | 0.69720022 | FALSE | 1 |
| 4085 | 4085 | 8.58333333 | 0.8620067 | 0 | 1 | 0 | 1 | 1 | 0.35555293 | 0 | 1 | 0 | 1 | 11 | 0.41944532 | FALSE | 1 | FALSE | 1 | FALSE | 1 | FALSE | 1 | FALSE | 1 | FALSE | 1 |
| 4086 | 4086 | 4.4137931 | 1.70834143 | 2 | 0.88652271 | 1 | 0.99137473 | 6 | 0.40864609 | 4 | 0.90050374 | 1 | 0.90833711 | 129 | 0.02882177 | FALSE | 1 | FALSE | 1 | FALSE | 1 | FALSE | 1 | FALSE | 1 | FALSE | 1 |
| 4087 | 4087 | 7.41666667 | 4.11214326 | 0 | 1 | 1 | 0.77338922 | 2 | 0.48463745 | 2 | 0.61900576 | 0 | 1 | 40 | 0.27423942 | FALSE | 1 | FALSE | 1 | FALSE | 1 | FALSE | 1 | FALSE | 1 | FALSE | 1 |
| 4088 | 4088 | 10.0769231 | 2.43252128 | 1 | 0.28675951 | 0 | 1 | 0 | 1 | 1 | 0.45746571 | 0 | 1 | 11 | 0.6683056 | FALSE | 1 | FALSE | 1 | FALSE | 1 | FALSE | 1 | FALSE | 1 | FALSE | 1 |
| 4089 | 4089 | 9.75 | 3.04480432 | 0 | 1 | 0 | 1 | 1 | 0.58502114 | 1 | 0.67692816 | 0 | 1 | 22 | 0.25237063 | FALSE | 1 | FALSE | 1 | FALSE | 1 | FALSE | 1 | FALSE | 1 | FALSE | 1 |
| 4090 | 4090 | 9 | 2.76339712 | 0 | 1 | 0 | 1 | 0 | 1 | 1 | 0.37517262 | 0 | 1 | 9 | 0.5215547 | FALSE | 1 | FALSE | 1 | FALSE | 1 | FALSE | 1 | FALSE | 1 | FALSE | 1 |
| 4091 | 4091 | 10.1818182 | 3.15704643 | 0 | 1 | 0 | 1 | 0 | 1 | 0 | 1 | 0 | 1 | 11 | 0.15414489 | FALSE | 1 | FALSE | 1 | FALSE | 1 | FALSE | 1 | FALSE | 1 | FALSE | 1 |
| 4092 | 4092 | 3.88 | 2.00638979 | 10 | 1.64E-05 | 8 | 0.00252846 | 1 | 0.93427349 | 8 | 0.0194831 | 0 | 1 | 47 | 0.99999745 | TRUE | 0.00517685 | FALSE | 0.44070554 | FALSE | 1 | FALSE | 1 | FALSE | 1 | FALSE | 1 |
| 4093 | 4093 | 10.5714286 | 2.94507545 | 0 | 1 | 0 | 1 | 0 | 1 | 0 | 1 | 0 | 1 | 14 | 0.09251304 | FALSE | 1 | FALSE | 1 | FALSE | 1 | FALSE | 1 | FALSE | 1 | FALSE | 1 |
| 4094 | 4094 | 9.73170732 | 2.32719228 | 1 | 0.64722818 | 0 | 1 | 1 | 0.76953771 | 2 | 0.5543522 | 0 | 1 | 36 | 0.22908535 | FALSE | 1 | FALSE | 1 | FALSE | 1 | FALSE | 1 | FALSE | 1 | FALSE | 1 |
| 4095 | 4095 | 8.16216216 | 3.03594247 | 2 | 0.24498301 | 2 | 0.3375163 | 4 | 0.04241754 | 0 | 1 | 0 | 1 | 29 | 0.88820978 | FALSE | 1 | FALSE | 1 | FALSE | 1 | FALSE | 1 | FALSE | 1 | FALSE | 1 |
| 4096 | 4096 | 9.64516129 | 1.78792341 | 1 | 0.55378205 | 0 | 1 | 0 | 1 | 2 | 0.41984187 | 0 | 1 | 28 | 0.26393996 | FALSE | 1 | FALSE | 1 | FALSE | 1 | FALSE | 1 | FALSE | 1 | FALSE | 1 |
| 4097 | 4097 | 6.05714286 | 3.11625102 | 1 | 0.59801736 | 0 | 1 | 3 | 0.12958859 | 4 | 0.07465533 | 0 | 1 | 27 | 0.91585095 | FALSE | 1 | FALSE | 1 | FALSE | 1 | FALSE | 1 | FALSE | 1 | FALSE | 1 |
| 4098 | 4098 | 5.43891403 | 1.70230791 | 11 | 0.02625901 | 8 | 0.42398601 | 8 | 0.54304595 | 9 | 0.69279654 | 4 | 0.49302556 | 181 | 0.86906306 | FALSE | 0.50496189 | FALSE | 1 | FALSE | 1 | FALSE | 1 | FALSE | 1 | FALSE | 1 |
| 4099 | 4099 | 9.24 | 2.43770384 | 1 | 0.47819878 | 0 | 1 | 1 | 0.59998149 | 0 | 1 | 1 | 0.33902672 | 22 | 0.43642846 | FALSE | 1 | FALSE | 1 | FALSE | 1 | FALSE | 1 | FALSE | 1 | FALSE | 1 |
| 4100 | 4100 | 9.625 | 1.86664807 | 8 | 6.05E-06 | 5 | 0.00884479 | 0 | 1 | 4 | 0.10945471 | 0 | 1 | 23 | 0.99999027 | TRUE | 0.00299395 | FALSE | 0.63686998 | FALSE | 1 | FALSE | 1 | FALSE | 1 | FALSE | 1 |
| 4101 | 4101 | 9.1875 | 2.78878894 | 0 | 1 | 0 | 1 | 2 | 0.32011871 | 0 | 1 | 0 | 1 | 30 | 0.10375556 | FALSE | 1 | FALSE | 1 | FALSE | 1 | FALSE | 1 | FALSE | 1 | FALSE | 1 |
| 4102 | 4102 | 10.2857143 | 3.10365157 | 0 | 1 | 0 | 1 | 0 | 1 | 0 | 1 | 0 | 1 | 7 | 0.3043657 | FALSE | 1 | FALSE | 1 | FALSE | 1 | FALSE | 1 | FALSE | 1 | FALSE | 1 |
| 4103 | 4103 | 6.04444444 | 3.41210626 | 4 | 0.02747663 | 3 | 0.17732615 | 6 | 0.00507773 | 0 | 1 | 4 | 0.00607992 | 28 | 0.99993247 | FALSE | 0.50964275 | FALSE | 1 | FALSE | 1 | FALSE | 1 | FALSE | 0.45694206 | FALSE | 1 |
| 4104 | 4104 | 7 | 3.47946356 | 2 | 0.57749853 | 3 | 0.43942904 | 4 | 0.2829119 | 6 | 0.12903033 | 0 | 1 | 60 | 0.88453881 | FALSE | 1 | FALSE | 1 | FALSE | 1 | FALSE | 1 | FALSE | 1 | FALSE | 1 |
| 4105 | 4105 | 10.7073171 | 1.40111284 | 0 | 1 | 1 | 0.7326167 | 2 | 0.42396974 | 2 | 0.5543522 | 0 | 1 | 35 | 0.38966407 | FALSE | 1 | FALSE | 1 | FALSE | 1 | FALSE | 1 | FALSE | 1 | FALSE | 1 |
| 4106 | 4106 | 8.90476191 | 2.92615076 | 0 | 1 | 0 | 1 | 1 | 0.53671415 | 1 | 0.62783522 | 0 | 1 | 19 | 0.34094959 | FALSE | 1 | FALSE | 1 | FALSE | 1 | FALSE | 1 | FALSE | 1 | FALSE | 1 |
| 4107 | 4107 | 7.61764706 | 3.12499135 | 1 | 0.58738519 | 0 | 1 | 1 | 0.71261921 | 1 | 0.7984848 | 1 | 0.43077488 | 30 | 0.36879488 | FALSE | 1 | FALSE | 1 | FALSE | 1 | FALSE | 1 | FALSE | 1 | FALSE | 1 |
| 4108 | 4108 | 8.78125 | 2.5091629 | 2 | 0.19750261 | 0 | 1 | 0 | 1 | 0 | 1 | 1 | 0.41154418 | 29 | 0.24088541 | FALSE | 1 | FALSE | 1 | FALSE | 1 | FALSE | 1 | FALSE | 1 | FALSE | 1 |
| 4109 | 4109 | 9.70588235 | 4.17647059 | 3 | 0.05546921 | 0 | 1 | 0 | 1 | 3 | 0.20353499 | 1 | 0.43077488 | 27 | 0.85023803 | FALSE | 0.68849059 | FALSE | 1 | FALSE | 1 | FALSE | 1 | FALSE | 1 | FALSE | 1 |
| 4110 | 4110 | 9.8 | 2.85657137 | 0 | 1 | 0 | 1 | 1 | 0.51939841 | 1 | 0.60987146 | 0 | 1 | 18 | 0.37504645 | FALSE | 1 | FALSE | 1 | FALSE | 1 | FALSE | 1 | FALSE | 1 | FALSE | 1 |
| 4111 | 4111 | 9.875 | 1.83285979 | 1 | 0.55378205 | 1 | 0.63997468 | 3 | 0.09854319 | 3 | 0.16836568 | 0 | 1 | 23 | 0.95750806 | FALSE | 1 | FALSE | 1 | FALSE | 1 | FALSE | 1 | FALSE | 1 | FALSE | 1 |
| 4112 | 4112 | 9.22222222 | 2.24982853 | 0 | 1 | 1 | 0.25628163 | 0 | 1 | 0 | 1 | 0 | 1 | 8 | 0.5780332 | FALSE | 1 | FALSE | 1 | FALSE | 1 | FALSE | 1 | FALSE | 1 | FALSE | 1 |
| 4113 | 4113 | 9.51282051 | 2.19406887 | 0 | 1 | 2 | 0.3494045 | 1 | 0.75193685 | 1 | 0.83318955 | 0 | 1 | 34 | 0.27051027 | FALSE | 1 | FALSE | 1 | FALSE | 1 | FALSE | 1 | FALSE | 1 | FALSE | 1 |
| 4114 | 4114 | 9.06666667 | 2.64491126 | 0 | 1 | 0 | 1 | 3 | 0.01512258 | 1 | 0.50623194 | 0 | 1 | 11 | 0.92879412 | FALSE | 1 | FALSE | 1 | FALSE | 1 | FALSE | 1 | FALSE | 1 | FALSE | 1 |
| 4115 | 4115 | 10.25 | 2.72717803 | 1 | 0.34033054 | 0 | 1 | 1 | 0.44344817 | 0 | 1 | 0 | 1 | 14 | 0.53308261 | FALSE | 1 | FALSE | 1 | FALSE | 1 | FALSE | 1 | FALSE | 1 | FALSE | 1 |
| 4116 | 4116 | 10.3333333 | 1.28019096 | 1 | 0.46441868 | 0 | 1 | 1 | 0.58502114 | 0 | 1 | 1 | 0.32796862 | 21 | 0.47053137 | FALSE | 1 | FALSE | 1 | FALSE | 1 | FALSE | 1 | FALSE | 1 | FALSE | 1 |
| 4117 | 4117 | 8.69230769 | 0.91016612 | 0 | 1 | 1 | 0.3480709 | 0 | 1 | 1 | 0.45746571 | 1 | 0.19354507 | 10 | 0.86791642 | FALSE | 1 | FALSE | 1 | FALSE | 1 | FALSE | 1 | FALSE | 1 | FALSE | 1 |
| 4118 | 4118 | 8.55555556 | 1.06574034 | 1 | 0.20854748 | 0 | 1 | 0 | 1 | 1 | 0.34506517 | 0 | 1 | 7 | 0.84574168 | FALSE | 1 | FALSE | 1 | FALSE | 1 | FALSE | 1 | FALSE | 1 | FALSE | 1 |
| 4119 | 4119 | 10.9166667 | 3.85050502 | 0 | 1 | 1 | 0.54637326 | 2 | 0.21260257 | 0 | 1 | 0 | 1 | 21 | 0.47053137 | FALSE | 1 | FALSE | 1 | FALSE | 1 | FALSE | 1 | FALSE | 1 | FALSE | 1 |
| 4120 | 4120 | 5.88095238 | 2.46114937 | 4 | 0.02189152 | 3 | 0.1535026 | 3 | 0.19060367 | 1 | 0.86193352 | 0 | 1 | 31 | 0.97647435 | FALSE | 0.46823848 | FALSE | 1 | FALSE | 1 | FALSE | 1 | FALSE | 1 | FALSE | 1 |
| 4121 | 4121 | 10 | 2.96443566 | 0 | 1 | 1 | 0.6629941 | 1 | 0.70185639 | 0 | 1 | 0 | 1 | 31 | 0.09218876 | FALSE | 1 | FALSE | 1 | FALSE | 1 | FALSE | 1 | FALSE | 1 | FALSE | 1 |
| 4122 | 4122 | 11 | 1.89296945 | 0 | 1 | 0 | 1 | 0 | 1 | 1 | 0.67692816 | 1 | 0.32796862 | 22 | 0.25237063 | FALSE | 1 | FALSE | 1 | FALSE | 1 | FALSE | 1 | FALSE | 1 | FALSE | 1 |
| 4123 | 4123 | 7.52380952 | 2.98066331 | 0 | 1 | 0 | 1 | 1 | 0.90129726 | 3 | 0.55805561 | 1 | 0.64879513 | 58 | 0.05684854 | FALSE | 1 | FALSE | 1 | FALSE | 1 | FALSE | 1 | FALSE | 1 | FALSE | 1 |
| 4124 | 4124 | 9.75 | 3.53847896 | 0 | 1 | 1 | 0.53115654 | 2 | 0.19931429 | 0 | 1 | 0 | 1 | 20 | 0.50589402 | FALSE | 1 | FALSE | 1 | FALSE | 1 | FALSE | 1 | FALSE | 1 | FALSE | 1 |
| 4125 | 4125 | 4.2345679 | 1.10857048 | 7 | 0.11912829 | 9 | 0.07663774 | 5 | 0.69268343 | 10 | 0.20471154 | 2 | 0.74647014 | 128 | 0.96301169 | FALSE | 0.94655575 | FALSE | 1 | FALSE | 1 | FALSE | 1 | FALSE | 1 | FALSE | 1 |
| 4126 | 4126 | 8.80952381 | 2.38285638 | 0 | 1 | 1 | 0.42857733 | 0 | 1 | 1 | 0.55062772 | 0 | 1 | 15 | 0.49060435 | FALSE | 1 | FALSE | 1 | FALSE | 1 | FALSE | 1 | FALSE | 1 | FALSE | 1 |
| 4127 | 4127 | 3.7080292 | 2.23822331 | 2 | 0.87146763 | 3 | 0.82607155 | 4 | 0.73168489 | 6 | 0.60625053 | 9 | 0.00038439 | 113 | 0.77306706 | FALSE | 1 | FALSE | 1 | FALSE | 1 | FALSE | 1 | FALSE | 0.09840309 | FALSE | 1 |
| 4128 | 4128 | 8.32258065 | 4.17664085 | 1 | 0.55378205 | 1 | 0.63997468 | 2 | 0.30673593 | 1 | 0.76781284 | 0 | 1 | 26 | 0.64660109 | FALSE | 1 | FALSE | 1 | FALSE | 1 | FALSE | 1 | FALSE | 1 | FALSE | 1 |
| 4129 | 4129 | 10.3050848 | 1.59718224 | 2 | 0.4493251 | 2 | 0.57395594 | 2 | 0.63179508 | 1 | 0.93827647 | 2 | 0.25204357 | 50 | 0.55668966 | FALSE | 1 | FALSE | 1 | FALSE | 1 | FALSE | 1 | FALSE | 1 | FALSE | 1 |
| 4130 | 4130 | 5.05607477 | 3.30909756 | 4 | 0.28839499 | 2 | 0.86287617 | 7 | 0.0861928 | 8 | 0.11309641 | 5 | 0.02966937 | 80 | 0.99440721 | FALSE | 1 | FALSE | 1 | FALSE | 1 | FALSE | 1 | FALSE | 0.82559785 | FALSE | 1 |
| 4131 | 4131 | 4.98591549 | 4.15031972 | 2 | 0.54756403 | 2 | 0.67482356 | 4 | 0.25087422 | 5 | 0.22607353 | 0 | 1 | 58 | 0.7898837 | FALSE | 1 | FALSE | 1 | FALSE | 1 | FALSE | 1 | FALSE | 1 | FALSE | 1 |
| 4132 | 4132 | 8.12 | 2.09418242 | 2 | 0.36837609 | 0 | 1 | 2 | 0.54120634 | 0 | 1 | 0 | 1 | 46 | 0.0908326 | FALSE | 1 | FALSE | 1 | FALSE | 1 | FALSE | 1 | FALSE | 1 | FALSE | 1 |
| 4133 | 4133 | 8.6 | 2.97015457 | 1 | 0.76171602 | 5 | 0.03172866 | 3 | 0.31652594 | 2 | 0.72625531 | 1 | 0.59866474 | 43 | 0.92240996 | FALSE | 1 | FALSE | 0.92268438 | FALSE | 1 | FALSE | 1 | FALSE | 1 | FALSE | 1 |
| 4134 | 4134 | 9.07142857 | 1.83085795 | 0 | 1 | 0 | 1 | 1 | 0.64171563 | 0 | 1 | 1 | 0.37113067 | 26 | 0.16405547 | FALSE | 1 | FALSE | 1 | FALSE | 1 | FALSE | 1 | FALSE | 1 | FALSE | 1 |

| | | | | | | | | | | | | | | | | | | | | | | | | | | | |
|---|---|---|---|---|---|---|---|---|---|---|---|---|---|---|---|---|---|---|---|---|---|---|---|---|---|---|---|
| 4135 | 4135 | 8.84615385 | 2.06978832 | 0 | 1 | 1 | 0.3480709 | 0 | 1 | 0 | 1 | 0 | 1 | 12 | 0.374198 | FALSE | 1 | FALSE | 1 | FALSE | 1 | FALSE | 1 | FALSE | 1 | FALSE | 1 |
| 4136 | 4136 | 8 | 2.89087234 | 1 | 0.51745948 | 0 | 1 | 1 | 0.64171563 | 3 | 0.13524571 | 0 | 1 | 23 | 0.73439118 | FALSE | 1 | FALSE | 1 | FALSE | 1 | FALSE | 1 | FALSE | 1 | FALSE | 1 |
| 4137 | 4137 | 8.57142857 | 1.98976975 | 0 | 1 | 0 | 1 | 2 | 0.17308361 | 2 | 0.25059627 | 1 | 0.29368245 | 16 | 0.90353674 | FALSE | 1 | FALSE | 1 | FALSE | 1 | FALSE | 1 | FALSE | 1 | FALSE | 1 |
| 4138 | 4138 | 7.4 | 3.73672008 | 2 | 0.49997462 | 5 | 0.05834767 | 3 | 0.41519671 | 2 | 0.80687302 | 3 | 0.09035518 | 50 | 0.96116604 | FALSE | 1 | FALSE | 1 | FALSE | 1 | FALSE | 1 | FALSE | 1 | FALSE | 1 |
| 4139 | 4139 | 9.4375 | 3.35351961 | 0 | 1 | 0 | 1 | 1 | 0.44344817 | 0 | 1 | 0 | 1 | 15 | 0.26127859 | FALSE | 1 | FALSE | 1 | FALSE | 1 | FALSE | 1 | FALSE | 1 | FALSE | 1 |
| 4140 | 4140 | 10.3157895 | 2.57626068 | 0 | 1 | 0 | 1 | 1 | 0.50143826 | 1 | 0.59104357 | 0 | 1 | 17 | 0.41141543 | FALSE | 1 | FALSE | 1 | FALSE | 1 | FALSE | 1 | FALSE | 1 | FALSE | 1 |
| 4141 | 4141 | 4.21333333 | 1.09292675 | 9 | 0.36275095 | 14 | 0.10626458 | 8 | 0.85253703 | 17 | 0.21585099 | 10 | 0.02481357 | 242 | 0.96818271 | FALSE | 1 | FALSE | 1 | FALSE | 1 | FALSE | 1 | FALSE | 0.78787878 | FALSE | 1 |
| 4142 | 4142 | 9.88888889 | 3.78430808 | 0 | 1 | 0 | 1 | 1 | 0.28067846 | 0 | 1 | 0 | 1 | 8 | 0.5780332 | FALSE | 1 | FALSE | 1 | FALSE | 1 | FALSE | 1 | FALSE | 1 | FALSE | 1 |
| 4143 | 4143 | 10 | 2.21880079 | 0 | 1 | 0 | 1 | 0 | 1 | 1 | 0.45746571 | 0 | 1 | 12 | 0.374198 | FALSE | 1 | FALSE | 1 | FALSE | 1 | FALSE | 1 | FALSE | 1 | FALSE | 1 |
| 4144 | 4144 | 10.0731707 | 2.69045507 | 1 | 0.65632651 | 0 | 1 | 1 | 0.77786601 | 2 | 0.56788092 | 1 | 0.49330646 | 36 | 0.36455032 | FALSE | 1 | FALSE | 1 | FALSE | 1 | FALSE | 1 | FALSE | 1 | FALSE | 1 |
| 4145 | 4145 | 8.625 | 3.38886043 | 1 | 0.32293355 | 1 | 0.38964394 | 0 | 1 | 0 | 1 | 0 | 1 | 13 | 0.57712805 | FALSE | 1 | FALSE | 1 | FALSE | 1 | FALSE | 1 | FALSE | 1 | FALSE | 1 |
| 4146 | 4146 | 9.11111111 | 3.05302926 | 0 | 1 | 0 | 1 | 1 | 0.48280983 | 0 | 1 | 0 | 1 | 17 | 0.20329205 | FALSE | 1 | FALSE | 1 | FALSE | 1 | FALSE | 1 | FALSE | 1 | FALSE | 1 |
| 4147 | 4147 | 9.15384615 | 2.71310648 | 1 | 0.49162634 | 3 | 0.05021553 | 1 | 0.61440466 | 1 | 0.70601446 | 0 | 1 | 20 | 0.90142689 | FALSE | 1 | FALSE | 1 | FALSE | 1 | FALSE | 1 | FALSE | 1 | FALSE | 1 |
| 4148 | 4148 | 10 | 1.83973242 | 2 | 0.03087289 | 0 | 1 | 3 | 0.00609779 | 1 | 0.40390032 | 0 | 1 | 5 | 0.99958857 | FALSE | 0.53356688 | FALSE | 1 | FALSE | 1 | FALSE | 1 | FALSE | 1 | FALSE | 1 |
| 4149 | 4149 | 8.8125 | 2.35103036 | 0 | 1 | 0 | 1 | 0 | 1 | 0 | 1 | 0 | 1 | 14 | 0.09251304 | FALSE | 1 | FALSE | 1 | FALSE | 1 | FALSE | 1 | FALSE | 1 | FALSE | 1 |
| 4150 | 4150 | 7.50877193 | 2.12026816 | 2 | 0.43181298 | 2 | 0.55509723 | 5 | 0.05289293 | 0 | 1 | 2 | 0.23984683 | 46 | 0.83074268 | FALSE | 1 | FALSE | 1 | FALSE | 1 | FALSE | 1 | FALSE | 1 | FALSE | 1 |
| 4151 | 4151 | 7.64705882 | 2.3995386 | 1 | 0.58738519 | 2 | 0.30150529 | 1 | 0.71261921 | 2 | 0.46709861 | 1 | 0.43077488 | 27 | 0.85023803 | FALSE | 1 | FALSE | 1 | FALSE | 1 | FALSE | 1 | FALSE | 1 | FALSE | 1 |
| 4152 | 4152 | 5.91358025 | 2.03183023 | 0 | 1 | 0 | 1 | 3 | 0.56136161 | 4 | 0.51410462 | 2 | 0.38471054 | 72 | 0.16599233 | FALSE | 1 | FALSE | 1 | FALSE | 1 | FALSE | 1 | FALSE | 1 | FALSE | 1 |
| 4153 | 4153 | 9.04444444 | 1.89684599 | 0 | 1 | 1 | 0.76576289 | 2 | 0.47281647 | 3 | 0.32878418 | 1 | 0.51797501 | 37 | 0.61979664 | FALSE | 1 | FALSE | 1 | FALSE | 1 | FALSE | 1 | FALSE | 1 | FALSE | 1 |
| 4154 | 4154 | 9.16666667 | 1.90758719 | 0 | 1 | 1 | 0.32623933 | 0 | 1 | 0 | 1 | 0 | 1 | 11 | 0.41944532 | FALSE | 1 | FALSE | 1 | FALSE | 1 | FALSE | 1 | FALSE | 1 | FALSE | 1 |
| 4155 | 4155 | 10.6 | 1.62480768 | 0 | 1 | 0 | 1 | 0 | 1 | 1 | 0.37517262 | 0 | 1 | 9 | 0.5215547 | FALSE | 1 | FALSE | 1 | FALSE | 1 | FALSE | 1 | FALSE | 1 | FALSE | 1 |
| 4156 | 4156 | 7.39655172 | 1.31227172 | 1 | 0.77972342 | 0 | 1 | 3 | 0.34634641 | 3 | 0.50172293 | 1 | 0.61824432 | 50 | 0.43527459 | FALSE | 1 | FALSE | 1 | FALSE | 1 | FALSE | 1 | FALSE | 1 | FALSE | 1 |
| 4157 | 4157 | 9.18367347 | 2.62385811 | 2 | 0.35907852 | 0 | 1 | 4 | 0.09808429 | 3 | 0.39227316 | 0 | 1 | 40 | 0.77344956 | FALSE | 1 | FALSE | 1 | FALSE | 1 | FALSE | 1 | FALSE | 1 | FALSE | 1 |
| 4158 | 4158 | 6.57971015 | 2.87621832 | 6 | 0.00821034 | 2 | 0.65946213 | 3 | 0.45353909 | 3 | 0.6203651 | 2 | 0.31302643 | 53 | 0.96638194 | FALSE | 0.27019545 | FALSE | 1 | FALSE | 1 | FALSE | 1 | FALSE | 1 | FALSE | 1 |
| 4159 | 4159 | 8.92307692 | 2.52560849 | 0 | 1 | 0 | 1 | 0 | 1 | 2 | 0.1174583 | 1 | 0.19354507 | 10 | 0.86791642 | FALSE | 1 | FALSE | 1 | FALSE | 1 | FALSE | 1 | FALSE | 1 | FALSE | 1 |
| 4160 | 4160 | 8.25 | 2.16506351 | 0 | 1 | 0 | 1 | 1 | 0.25384309 | 0 | 1 | 0 | 1 | 7 | 0.63752397 | FALSE | 1 | FALSE | 1 | FALSE | 1 | FALSE | 1 | FALSE | 1 | FALSE | 1 |
| 4161 | 4161 | 7.42592593 | 2.39290504 | 1 | 0.75539413 | 5 | 0.02959485 | 4 | 0.12796328 | 2 | 0.71679995 | 1 | 0.59191961 | 41 | 0.96572325 | FALSE | 1 | FALSE | 0.89461624 | FALSE | 1 | FALSE | 1 | FALSE | 1 | FALSE | 1 |
| 4162 | 4162 | 9.07407407 | 3.47398721 | 1 | 0.50471032 | 1 | 0.58913618 | 2 | 0.25286935 | 0 | 1 | 0 | 1 | 23 | 0.58307573 | FALSE | 1 | FALSE | 1 | FALSE | 1 | FALSE | 1 | FALSE | 1 | FALSE | 1 |
| 4163 | 4163 | 8.55172414 | 1.67303579 | 0 | 1 | 1 | 0.60247911 | 1 | 0.64171563 | 1 | 0.73248983 | 1 | 0.37113067 | 24 | 0.55018107 | FALSE | 1 | FALSE | 1 | FALSE | 1 | FALSE | 1 | FALSE | 1 | FALSE | 1 |
| 4164 | 4164 | 9.23529412 | 1.89517691 | 3 | 0.00864151 | 2 | 0.10307112 | 2 | 0.12280513 | 1 | 0.55062772 | 0 | 1 | 9 | 0.99960431 | FALSE | 0.27019545 | FALSE | 1 | FALSE | 1 | FALSE | 1 | FALSE | 1 | FALSE | 1 |
| 4165 | 4165 | 9.25 | 2.16506351 | 0 | 1 | 0 | 1 | 1 | 0.25384309 | 0 | 1 | 0 | 1 | 7 | 0.63752397 | FALSE | 1 | FALSE | 1 | FALSE | 1 | FALSE | 1 | FALSE | 1 | FALSE | 1 |
| 4166 | 4166 | 9.57692308 | 2.63370282 | 1 | 0.49162634 | 0 | 1 | 2 | 0.23940226 | 2 | 0.33678042 | 0 | 1 | 21 | 0.78939668 | FALSE | 1 | FALSE | 1 | FALSE | 1 | FALSE | 1 | FALSE | 1 | FALSE | 1 |
| 4167 | 4167 | 10.1 | 1.97230829 | 0 | 1 | 0 | 1 | 0 | 1 | 0 | 1 | 0 | 1 | 10 | 0.18273166 | FALSE | 1 | FALSE | 1 | FALSE | 1 | FALSE | 1 | FALSE | 1 | FALSE | 1 |
| 4168 | 4168 | 9.42857143 | 2.83383349 | 0 | 1 | 0 | 1 | 2 | 0.2663542 | 0 | 1 | 0 | 1 | 26 | 0.16405547 | FALSE | 1 | FALSE | 1 | FALSE | 1 | FALSE | 1 | FALSE | 1 | FALSE | 1 |
| 4169 | 4169 | 10.3684211 | 2.47536337 | 0 | 1 | 0 | 1 | 0 | 1 | 3 | 0.04687752 | 0 | 1 | 15 | 0.69372602 | FALSE | 1 | FALSE | 1 | FALSE | 1 | FALSE | 1 | FALSE | 1 | FALSE | 1 |
| 4170 | 4170 | 8.48387097 | 2.29825976 | 1 | 0.55378205 | 0 | 1 | 1 | 0.67911179 | 1 | 0.76781284 | 1 | 0.4016888 | 27 | 0.45522639 | FALSE | 1 | FALSE | 1 | FALSE | 1 | FALSE | 1 | FALSE | 1 | FALSE | 1 |
| 4171 | 4171 | 10.2307692 | 2.32557176 | 1 | 0.28675951 | 0 | 1 | 0 | 1 | 2 | 0.1174583 | 1 | 0.19354507 | 9 | 0.96018168 | FALSE | 1 | FALSE | 1 | FALSE | 1 | FALSE | 1 | FALSE | 1 | FALSE | 1 |
| 4172 | 4172 | 11.1666667 | 1.34370963 | 0 | 1 | 0 | 1 | 1 | 0.197145 | 0 | 1 | 0 | 1 | 5 | 0.76185568 | FALSE | 1 | FALSE | 1 | FALSE | 1 | FALSE | 1 | FALSE | 1 | FALSE | 1 |
| 4173 | 4173 | 8.28571429 | 3.34012913 | 1 | 0.40558521 | 0 | 1 | 1 | 0.51939841 | 3 | 0.0612581 | 0 | 1 | 15 | 0.92113679 | FALSE | 1 | FALSE | 1 | FALSE | 1 | FALSE | 1 | FALSE | 1 | FALSE | 1 |
| 4174 | 4174 | 11.516129 | 2.91681408 | 1 | 0.55378205 | 0 | 1 | 0 | 1 | 1 | 0.76781284 | 2 | 0.09110253 | 27 | 0.45522639 | FALSE | 1 | FALSE | 1 | FALSE | 1 | FALSE | 1 | FALSE | 1 | FALSE | 1 |
| 4175 | 4175 | 9.33333333 | 2.3570226 | 0 | 1 | 0 | 1 | 0 | 1 | 2 | 0.19902682 | 0 | 1 | 16 | 0.44998001 | FALSE | 1 | FALSE | 1 | FALSE | 1 | FALSE | 1 | FALSE | 1 | FALSE | 1 |
| 4176 | 4176 | 10.12 | 1.90410084 | 1 | 0.47819878 | 1 | 0.56109832 | 1 | 0.59998149 | 3 | 0.10474541 | 0 | 1 | 19 | 0.91734261 | FALSE | 1 | FALSE | 1 | FALSE | 1 | FALSE | 1 | FALSE | 1 | FALSE | 1 |
| 4177 | 4177 | 5 | 2.18385686 | 0 | 1 | 1 | 0.82028131 | 0 | 1 | 1 | 0.91399378 | 1 | 0.57809055 | 49 | 0.02821112 | FALSE | 1 | FALSE | 1 | FALSE | 1 | FALSE | 1 | FALSE | 1 | FALSE | 1 |
| 4178 | 4178 | 8.92857143 | 2.78937488 | 0 | 1 | 1 | 0.3480709 | 1 | 0.37874417 | 0 | 1 | 0 | 1 | 11 | 0.6683056 | FALSE | 1 | FALSE | 1 | FALSE | 1 | FALSE | 1 | FALSE | 1 | FALSE | 1 |
| 4179 | 4179 | 10.46875 | 1.3225065 | 4 | 0.00856611 | 1 | 0.65167362 | 2 | 0.32011871 | 1 | 0.77852168 | 1 | 0.41154418 | 23 | 0.97984382 | FALSE | 0.27019545 | FALSE | 1 | FALSE | 1 | FALSE | 1 | FALSE | 1 | FALSE | 1 |
| 4180 | 4180 | 8.87096774 | 2.09056152 | 2 | 0.1881398 | 2 | 0.2651851 | 0 | 1 | 0 | 1 | 2 | 0.09110253 | 25 | 0.80006983 | FALSE | 1 | FALSE | 1 | FALSE | 1 | FALSE | 1 | FALSE | 1 | FALSE | 1 |
| 4181 | 4181 | 9.84615385 | 1.74735641 | 0 | 1 | 0 | 1 | 0 | 1 | 0 | 1 | 0 | 1 | 13 | 0.10967906 | FALSE | 1 | FALSE | 1 | FALSE | 1 | FALSE | 1 | FALSE | 1 | FALSE | 1 |
| 4182 | 4182 | 9.46666667 | 2.36267269 | 1 | 0.32293355 | 0 | 1 | 1 | 0.42266264 | 1 | 0.50623194 | 0 | 1 | 12 | 0.80329917 | FALSE | 1 | FALSE | 1 | FALSE | 1 | FALSE | 1 | FALSE | 1 | FALSE | 1 |
| 4183 | 4183 | 8.24390244 | 1.76387168 | 1 | 0.65632651 | 0 | 1 | 1 | 0.77786601 | 2 | 0.56788092 | 0 | 1 | 37 | 0.21027077 | FALSE | 1 | FALSE | 1 | FALSE | 1 | FALSE | 1 | FALSE | 1 | FALSE | 1 |
| 4184 | 4184 | 8.61764706 | 2.24282786 | 2 | 0.51618914 | 2 | 0.64352762 | 4 | 0.21970029 | 5 | 0.19297788 | 1 | 0.6714808 | 53 | 0.90970332 | FALSE | 1 | FALSE | 1 | FALSE | 1 | FALSE | 1 | FALSE | 1 | FALSE | 1 |
| 4185 | 4185 | 5.49315069 | 2.90546621 | 2 | 0.56271322 | 1 | 0.9104874 | 5 | 0.12099953 | 1 | 0.96824399 | 1 | 0.7028142 | 63 | 0.39803869 | FALSE | 1 | FALSE | 1 | FALSE | 1 | FALSE | 1 | FALSE | 1 | FALSE | 1 |
| 4186 | 4186 | 6.32967033 | 1.2971223 | 7 | 0.18238663 | 7 | 0.36950961 | 5 | 0.78542766 | 7 | 0.73259105 | 1 | 0.95185195 | 154 | 0.44496718 | FALSE | 1 | FALSE | 1 | FALSE | 1 | FALSE | 1 | FALSE | 1 | FALSE | 1 |
| 4187 | 4187 | 9.75 | 2.64968552 | 1 | 0.46441868 | 0 | 1 | 1 | 0.58502114 | 3 | 0.09525947 | 2 | 0.058175 | 17 | 0.97530702 | FALSE | 1 | FALSE | 1 | FALSE | 1 | FALSE | 1 | FALSE | 1 | FALSE | 1 |
| 4188 | 4188 | 9.08 | 2.81311216 | 0 | 1 | 1 | 0.56109832 | 0 | 1 | 1 | 0.69181315 | 0 | 1 | 23 | 0.22727183 | FALSE | 1 | FALSE | 1 | FALSE | 1 | FALSE | 1 | FALSE | 1 | FALSE | 1 |
| 4189 | 4189 | 10.2 | 2.5086517 | 0 | 1 | 0 | 1 | 0 | 1 | 0 | 1 | 0 | 1 | 15 | 0.07803154 | FALSE | 1 | FALSE | 1 | FALSE | 1 | FALSE | 1 | FALSE | 1 | FALSE | 1 |
| 4190 | 4190 | 7.01449275 | 1.43953247 | 0 | 1 | 1 | 0.89776066 | 1 | 0.92092393 | 7 | 0.03796461 | 1 | 0.6822729 | 59 | 0.47808459 | FALSE | 1 | FALSE | 1 | FALSE | 1 | FALSE | 1 | FALSE | 1 | FALSE | 1 |
| 4191 | 4191 | 8.375 | 3.15980617 | 0 | 1 | 0 | 1 | 0 | 1 | 0 | 1 | 1 | 0.12394282 | 7 | 0.63752397 | FALSE | 1 | FALSE | 1 | FALSE | 1 | FALSE | 1 | FALSE | 1 | FALSE | 1 |
| 4192 | 4192 | 7.82352941 | 3.39090989 | 0 | 1 | 1 | 0.6739483 | 3 | 0.12152681 | 1 | 0.7984848 | 0 | 1 | 29 | 0.55707911 | FALSE | 1 | FALSE | 1 | FALSE | 1 | FALSE | 1 | FALSE | 1 | FALSE | 1 |
| 4193 | 4193 | 8.38461539 | 1.86211053 | 2 | 0.04234484 | 0 | 1 | 0 | 1 | 0 | 1 | 0 | 1 | 11 | 0.6683056 | FALSE | 0.62842194 | FALSE | 1 | FALSE | 1 | FALSE | 1 | FALSE | 1 | FALSE | 1 |
| 4194 | 4194 | 10 | 3.46410162 | 0 | 1 | 2 | 0.00598515 | 0 | 1 | 0 | 1 | 1 | 0.06400211 | 1 | 0.99940747 | FALSE | 1 | FALSE | 0.60103504 | FALSE | 1 | FALSE | 1 | FALSE | 1 | FALSE | 1 |
| 4195 | 4195 | 9.38888889 | 2.88942303 | 1 | 0.37380237 | 0 | 1 | 1 | 0.48280983 | 2 | 0.19902682 | 0 | 1 | 14 | 0.86287418 | FALSE | 1 | FALSE | 1 | FALSE | 1 | FALSE | 1 | FALSE | 1 | FALSE | 1 |
| 4196 | 4196 | 8.52631579 | 2.52082737 | 1 | 0.38989951 | 5 | 0.00027124 | 0 | 1 | 1 | 0.59104357 | 0 | 1 | 12 | 0.99479039 | FALSE | 1 | FALSE | 0.15723145 | FALSE | 1 | FALSE | 1 | FALSE | 1 | FALSE | 1 |
| 4197 | 4197 | 9.68 | 2.24 | 2 | 0.13369621 | 0 | 1 | 1 | 0.59998149 | 2 | 0.31968852 | 0 | 1 | 20 | 0.8152498 | FALSE | 0.97964162 | FALSE | 1 | FALSE | 1 | FALSE | 1 | FALSE | 1 | FALSE | 1 |
| 4198 | 4198 | 8.15625 | 2.76257777 | 1 | 0.56527493 | 2 | 0.27730767 | 3 | 0.10599423 | 1 | 0.77852168 | 0 | 1 | 25 | 0.88559288 | FALSE | 1 | FALSE | 1 | FALSE | 1 | FALSE | 1 | FALSE | 1 | FALSE | 1 |
| 4199 | 4199 | 9.94444444 | 2.48265589 | 0 | 1 | 0 | 1 | 1 | 0.48280983 | 0 | 1 | 1 | 0.25766374 | 16 | 0.44998001 | FALSE | 1 | FALSE | 1 | FALSE | 1 | FALSE | 1 | FALSE | 1 | FALSE | 1 |
| 4200 | 4200 | 10.0869565 | 2.062355 | 0 | 1 | 1 | 0.53115654 | 1 | 0.5695037 | 1 | 0.6613267 | 0 | 1 | 20 | 0.50589402 | FALSE | 1 | FALSE | 1 | FALSE | 1 | FALSE | 1 | FALSE | 1 | FALSE | 1 |
| 4201 | 4201 | 9.71111111 | 0.9915694 | 0 | 1 | 3 | 0.52247675 | 0 | 1 | 7 | 0.09449282 | 2 | 0.40793536 | 73 | 0.42025276 | FALSE | 1 | FALSE | 1 | FALSE | 1 | FALSE | 1 | FALSE | 1 | FALSE | 1 |
| 4202 | 4202 | 9.51724138 | 2.07814058 | 3 | 0.03713651 | 3 | 0.06581417 | 1 | 0.65463982 | 1 | 0.74482229 | 2 | 0.08121636 | 19 | 0.99707037 | FALSE | 0.58392 | FALSE | 1 | FALSE | 1 | FALSE | 1 | FALSE | 1 | FALSE | 1 |
| 4203 | 4203 | 8.79166667 | 1.65778886 | 0 | 1 | 0 | 1 | 0 | 1 | 0 | 1 | 2 | 0.058175 | 22 | 0.25237063 | FALSE | 1 | FALSE | 1 | FALSE | 1 | FALSE | 1 | FALSE | 1 | FALSE | 1 |
| 4204 | 4204 | 10.53125 | 1.08927657 | 1 | 0.56527493 | 0 | 1 | 1 | 0.69069222 | 3 | 0.17988885 | 0 | 1 | 27 | 0.61671133 | FALSE | 1 | FALSE | 1 | FALSE | 1 | FALSE | 1 | FALSE | 1 | FALSE | 1 |
| 4205 | 4205 | 8.61111111 | 2.11184198 | 1 | 0.37380237 | 1 | 0.44710673 | 1 | 0.48280983 | 1 | 0.57131013 | 0 | 1 | 14 | 0.86287418 | FALSE | 1 | FALSE | 1 | FALSE | 1 | FALSE | 1 | FALSE | 1 | FALSE | 1 |
| 4206 | 4206 | 4.91358025 | 4.09817042 | 2 | 0.61965506 | 3 | 0.49001146 | 4 | 0.33196391 | 2 | 0.89261449 | 0 | 1 | 70 | 0.37303501 | FALSE | 1 | FALSE | 1 | FALSE | 1 | FALSE | 1 | FALSE | 1 | FALSE | 1 |
| 4207 | 4207 | 9.75 | 2.16506351 | 0 | 1 | 0 | 1 | 2 | 0.06691409 | 0 | 1 | 0 | 1 | 10 | 0.71436964 | FALSE | 1 | FALSE | 1 | FALSE | 1 | FALSE | 1 | FALSE | 1 | FALSE | 1 |
| 4208 | 4208 | 9.46428571 | 2.02629399 | 0 | 1 | 1 | 0.58913618 | 0 | 1 | 0 | 1 | 0 | 1 | 26 | 0.06079097 | FALSE | 1 | FALSE | 1 | FALSE | 1 | FALSE | 1 | FALSE | 1 | FALSE | 1 |
| 4209 | 4209 | 8.52941177 | 1.81877939 | 2 | 0.37762121 | 1 | 0.81422651 | 2 | 0.55199489 | 1 | 0.90982214 | 2 | 0.20351566 | 43 | 0.59806866 | FALSE | 1 | FALSE | 1 | FALSE | 1 | FALSE | 1 | FALSE | 1 | FALSE | 1 |
| 4210 | 4210 | 8 | 2.17944947 | 2 | 0.19750261 | 0 | 1 | 0 | 1 | 4 | 0.05697506 | 1 | 0.41154418 | 25 | 0.88559288 | FALSE | 1 | FALSE | 1 | FALSE | 1 | FALSE | 1 | FALSE | 1 | FALSE | 1 |
| 4211 | 4211 | 9.45945946 | 4.25913704 | 2 | 0.23542589 | 1 | 0.69480472 | 1 | 0.73299778 | 2 | 0.49731009 | 0 | 1 | 30 | 0.67248271 | FALSE | 1 | FALSE | 1 | FALSE | 1 | FALSE | 1 | FALSE | 1 | FALSE | 1 |
| 4212 | 4212 | 1.66917293 | 2.46546616 | 7 | 0.52312778 | 10 | 0.35410903 | 9 | 0.61771826 | 14 | 0.33196765 | 1 | 0.98855508 | 224 | 0.51450592 | FALSE | 1 | FALSE | 1 | FALSE | 1 | FALSE | 1 | FALSE | 1 | FALSE | 1 |
| 4213 | 4213 | 10.5294118 | 2.97277148 | 2 | 0.06909038 | 0 | 1 | 0 | 1 | 1 | 0.55062772 | 0 | 1 | 14 | 0.7313145 | FALSE | 0.74276693 | FALSE | 1 | FALSE | 1 | FALSE | 1 | FALSE | 1 | FALSE | 1 |
| 4214 | 4214 | 3.57142857 | 1.98077495 | 2 | 0.93332095 | 5 | 0.63869749 | 5 | 0.72790105 | 7 | 0.65838078 | 4 | 0.29674325 | 145 | 0.28275126 | FALSE | 1 | FALSE | 1 | FALSE | 1 | FALSE | 1 | FALSE | 1 | FALSE | 1 |
| 4215 | 4215 | 4.14777328 | 0.93320716 | 21 | 0.01308941 | 23 | 0.04341773 | 11 | 0.96966873 | 22 | 0.57334675 | 5 | 0.91159134 | 407 | 0.78690575 | FALSE | 0.35982701 | FALSE | 1 | FALSE | 1 | FALSE | 1 | FALSE | 1 | FALSE | 1 |
| 4216 | 4216 | 11.1875 | 1.33317056 | 1 | 0.34033054 | 0 | 1 | 0 | 1 | 0 | 1 | 0 | 1 | 15 | 0.26127859 | FALSE | 1 | FALSE | 1 | FALSE | 1 | FALSE | 1 | FALSE | 1 | FALSE | 1 |
| 4217 | 4217 | 10.5294118 | 2.65944813 | 2 | 0.06909038 | 0 | 1 | 1 | 0.46348836 | 1 | 0.55062772 | 1 | 0.24525764 | 12 | 0.962093 | FALSE | 0.74276693 | FALSE | 1 | FALSE | 1 | FALSE | 1 | FALSE | 1 | FALSE | 1 |
| 4218 | 4218 | 8.52631579 | 2.11183186 | 0 | 1 | 2 | 0.12434464 | 0 | 1 | 0 | 1 | 2 | 0.03799168 | 15 | 0.83648191 | FALSE | 1 | FALSE | 1 | FALSE | 1 | FALSE | 1 | FALSE | 0.87669812 | FALSE | 1 |
| 4219 | 4219 | 8.48387097 | 1.9653608 | 4 | 0.00764535 | 0 | 1 | 0 | 1 | 0 | 1 | 0 | 1 | 27 | 0.45522639 | FALSE | 0.2566832 | FALSE | 1 | FALSE | 1 | FALSE | 1 | FALSE | 1 | FALSE | 1 |
| 4220 | 4220 | 8.63636364 | 2.70597747 | 0 | 1 | 1 | 0.30368014 | 0 | 1 | 0 | 1 | 0 | 1 | 10 | 0.46859937 | FALSE | 1 | FALSE | 1 | FALSE | 1 | FALSE | 1 | FALSE | 1 | FALSE | 1 |
| 4221 | 4221 | 9 | 1.91765986 | 1 | 0.55378205 | 1 | 0.63997468 | 0 | 1 | 1 | 0.76781284 | 0 | 1 | 28 | 0.26393996 | FALSE | 1 | FALSE | 1 | FALSE | 1 | FALSE | 1 | FALSE | 1 | FALSE | 1 |
| 4222 | 4222 | 9.375 | 2.57896135 | 3 | 0.02256792 | 0 | 1 | 1 | 0.58502114 | 0 | 1 | 2 | 0.058175 | 18 | 0.93166998 | FALSE | 0.4692294 | FALSE | 1 | FALSE | 1 | FALSE | 1 | FALSE | 1 | FALSE | 1 |
| 4223 | 4223 | 8 | 2.40613252 | 0 | 1 | 1 | 0.46503795 | 2 | 0.14749808 | 2 | 0.21610812 | 0 | 1 | 14 | 0.93677253 | FALSE | 1 | FALSE | 1 | FALSE | 1 | FALSE | 1 | FALSE | 1 | FALSE | 1 |
| 4224 | 4224 | 9.34482759 | 2.03886145 | 0 | 1 | 1 | 0.61539065 | 1 | 0.65463982 | 3 | 0.14602298 | 0 | 1 | 24 | 0.70563334 | FALSE | 1 | FALSE | 1 | FALSE | 1 | FALSE | 1 | FALSE | 1 | FALSE | 1 |
| 4225 | 4225 | 12.5925926 | 2.28131313 | 0 | 1 | 0 | 1 | 0 | 1 | 1 | 0.71956339 | 0 | 1 | 26 | 0.06079097 | FALSE | 1 | FALSE | 1 | FALSE | 1 | FALSE | 1 | FALSE | 1 | FALSE | 1 |
| 4226 | 4226 | 9.11111111 | 1.5234788 | 0 | 1 | 1 | 0.76576289 | 3 | 0.20921284 | 2 | 0.60667406 | 1 | 0.51797501 | 37 | 0.61979664 | FALSE | 1 | FALSE | 1 | FALSE | 1 | FALSE | 1 | FALSE | 1 | FALSE | 1 |
| 4227 | 4227 | 10.9677419 | 1.61612581 | 5 | 0.00103368 | 3 | 0.07729739 | 0 | 1 | 1 | 0.76781284 | 1 | 0.4016888 | 21 | 0.99467947 | FALSE | 0.07493697 | FALSE | 1 | FALSE | 1 | FALSE | 1 | FALSE | 1 | FALSE | 1 |
| 4228 | 4228 | 8.5625 | 2.14967294 | 0 | 1 | 0 | 1 | 2 | 0.11087977 | 0 | 1 | 0 | 1 | 14 | 0.53308261 | FALSE | 1 | FALSE | 1 | FALSE | 1 | FALSE | 1 | FALSE | 1 | FALSE | 1 |

| | | | | | | | | | | | | | | | | | | | | | | | | | | | |
|---|---|---|---|---|---|---|---|---|---|---|---|---|---|---|---|---|---|---|---|---|---|---|---|---|---|---|---|
| 4229 | 4229 | 8.96153846 | 1.60481893 | 1 | 0.49162634 | 1 | 0.57534752 | 3 | 0.06482014 | 1 | 0.70601446 | 0 | 1 | 20 | 0.90142689 | FALSE | 1 | FALSE | 1 | FALSE | 1 | FALSE | 1 | FALSE | 1 | FALSE | 1 |
| 4230 | 4230 | 9.57142857 | 3.71703195 | 1 | 0.3050805 | 2 | 0.07341559 | 0 | 1 | 1 | 0.48242099 | 0 | 1 | 10 | 0.94579623 | FALSE | 1 | FALSE | 1 | FALSE | 1 | FALSE | 1 | FALSE | 1 | FALSE | 1 |
| 4231 | 4231 | 10.5 | 2.78388218 | 0 | 1 | 1 | 0.38964394 | 0 | 1 | 2 | 0.14906763 | 0 | 1 | 12 | 0.80329917 | FALSE | 1 | FALSE | 1 | FALSE | 1 | FALSE | 1 | FALSE | 1 | FALSE | 1 |
| 4232 | 4232 | 8 | 3.51619629 | 0 | 1 | 0 | 1 | 0 | 1 | 1 | 0.40390032 | 1 | 0.16638736 | 9 | 0.75985366 | FALSE | 1 | FALSE | 1 | FALSE | 1 | FALSE | 1 | FALSE | 1 | FALSE | 1 |
| 4233 | 4233 | 8.8 | 1.2489996 | 0 | 1 | 0 | 1 | 0 | 1 | 1 | 0.37517262 | 0 | 1 | 9 | 0.5215547 | FALSE | 1 | FALSE | 1 | FALSE | 1 | FALSE | 1 | FALSE | 1 | FALSE | 1 |
| 4234 | 4234 | 8.73333333 | 1.43604395 | 1 | 0.32293355 | 1 | 0.38964394 | 0 | 1 | 0 | 1 | 0 | 1 | 13 | 0.57712805 | FALSE | 1 | FALSE | 1 | FALSE | 1 | FALSE | 1 | FALSE | 1 | FALSE | 1 |
| 4235 | 4235 | 9 | 2.8021156 | 0 | 1 | 3 | 0.05519089 | 0 | 1 | 0 | 1 | 0 | 1 | 24 | 0.3725373 | FALSE | 1 | FALSE | 1 | FALSE | 1 | FALSE | 1 | FALSE | 1 | FALSE | 1 |
| 4236 | 4236 | 7.7755102 | 1.48138526 | 3 | 0.46200485 | 2 | 0.83162754 | 4 | 0.47057639 | 7 | 0.16277638 | 2 | 0.48029053 | 80 | 0.81637124 | FALSE | 1 | FALSE | 1 | FALSE | 1 | FALSE | 1 | FALSE | 1 | FALSE | 1 |
| 4237 | 4237 | 9.42307692 | 2.89791805 | 1 | 0.49162634 | 1 | 0.57534752 | 4 | 0.0130279 | 2 | 0.33678042 | 2 | 0.0670823 | 16 | 0.9989746 | FALSE | 1 | FALSE | 1 | FALSE | 1 | FALSE | 1 | FALSE | 1 | FALSE | 1 |
| 4238 | 4238 | 8.5 | 2.56862338 | 4 | 0.02951043 | 3 | 0.18548475 | 2 | 0.4962935 | 1 | 0.88573774 | 1 | 0.53375569 | 35 | 0.95410769 | FALSE | 0.53131751 | FALSE | 1 | FALSE | 1 | FALSE | 1 | FALSE | 1 | FALSE | 1 |
| 4239 | 4239 | 7.83928571 | 3.99006259 | 1 | 0.76787545 | 2 | 0.54544984 | 3 | 0.32646887 | 0 | 1 | 1 | 0.60529938 | 49 | 0.33483172 | FALSE | 1 | FALSE | 1 | FALSE | 1 | FALSE | 1 | FALSE | 1 | FALSE | 1 |
| 4240 | 4240 | 8.88095238 | 2.14034244 | 1 | 0.66519151 | 2 | 0.39616059 | 4 | 0.0626749 | 4 | 0.12508716 | 1 | 0.50166523 | 30 | 0.99060471 | FALSE | 1 | FALSE | 1 | FALSE | 1 | FALSE | 1 | FALSE | 1 | FALSE | 1 |
| 4241 | 4241 | 10.0384615 | 3.4471925 | 2 | 0.14250978 | 1 | 0.57534752 | 3 | 0.06482014 | 0 | 1 | 2 | 0.0670823 | 18 | 0.98644281 | FALSE | 1 | FALSE | 1 | FALSE | 1 | FALSE | 1 | FALSE | 1 | FALSE | 1 |
| 4242 | 4242 | 9.5625 | 2.62127712 | 0 | 1 | 0 | 1 | 0 | 1 | 1 | 0.52895088 | 1 | 0.23264612 | 14 | 0.53308261 | FALSE | 1 | FALSE | 1 | FALSE | 1 | FALSE | 1 | FALSE | 1 | FALSE | 1 |
| 4243 | 4243 | 3.31443299 | 1.40657438 | 7 | 0.87555429 | 8 | 0.93938046 | 15 | 0.41848348 | 33 | 0.0003717 | 4 | 0.88636243 | 320 | 0.84527405 | FALSE | 1 | FALSE | 1 | FALSE | 1 | FALSE | 0.63446475 | FALSE | 1 | FALSE | 1 |
| 4244 | 4244 | 10.516129 | 2.39449594 | 1 | 0.55378205 | 0 | 1 | 3 | 0.09854319 | 3 | 0.16836568 | 1 | 0.4016888 | 23 | 0.95750806 | FALSE | 1 | FALSE | 1 | FALSE | 1 | FALSE | 1 | FALSE | 1 | FALSE | 1 |
| 4245 | 4245 | 7.05405405 | 2.61953107 | 1 | 0.6184714 | 0 | 1 | 3 | 0.14625315 | 0 | 1 | 1 | 0.45845922 | 32 | 0.47030992 | FALSE | 1 | FALSE | 1 | FALSE | 1 | FALSE | 1 | FALSE | 1 | FALSE | 1 |
| 4246 | 4246 | 8.38888889 | 3.56119345 | 0 | 1 | 2 | 0.1135776 | 0 | 1 | 0 | 1 | 1 | 0.25766374 | 15 | 0.69372602 | FALSE | 1 | FALSE | 1 | FALSE | 1 | FALSE | 1 | FALSE | 1 | FALSE | 1 |
| 4247 | 4247 | 9.23076923 | 2.48546068 | 0 | 1 | 2 | 0.20486827 | 1 | 0.61440466 | 1 | 0.70601446 | 0 | 1 | 22 | 0.61628 | FALSE | 1 | FALSE | 1 | FALSE | 1 | FALSE | 1 | FALSE | 1 | FALSE | 1 |
| 4248 | 4248 | 8.875 | 2.11763429 | 2 | 0.19750261 | 1 | 0.65167362 | 1 | 0.69069222 | 3 | 0.17988885 | 2 | 0.096172 | 23 | 0.97984382 | FALSE | 1 | FALSE | 1 | FALSE | 1 | FALSE | 1 | FALSE | 1 | FALSE | 1 |
| 4249 | 4249 | 9.4137931 | 2.8829731 | 1 | 0.52988231 | 0 | 1 | 1 | 0.65463982 | 3 | 0.14602298 | 0 | 1 | 24 | 0.70563334 | FALSE | 1 | FALSE | 1 | FALSE | 1 | FALSE | 1 | FALSE | 1 | FALSE | 1 |
| 4250 | 4250 | 8.95454546 | 1.63699483 | 0 | 1 | 1 | 0.5154318 | 0 | 1 | 0 | 1 | 2 | 0.04972408 | 19 | 0.54233419 | FALSE | 1 | FALSE | 1 | FALSE | 1 | FALSE | 1 | FALSE | 0.98629461 | FALSE | 1 |
| 4251 | 4251 | 6.31343284 | 2.69996659 | 1 | 0.82602167 | 2 | 0.64352762 | 1 | 0.91485686 | 3 | 0.60027156 | 1 | 0.6714808 | 59 | 0.26021394 | FALSE | 1 | FALSE | 1 | FALSE | 1 | FALSE | 1 | FALSE | 1 | FALSE | 1 |
| 4252 | 4252 | 7.86956522 | 2.60942019 | 2 | 0.11647716 | 0 | 1 | 0 | 1 | 4 | 0.01932195 | 0 | 1 | 17 | 0.94439745 | FALSE | 0.93181727 | FALSE | 1 | FALSE | 1 | FALSE | 1 | FALSE | 1 | FALSE | 1 |
| 4253 | 4253 | 10 | 3.77296887 | 2 | 0.06909038 | 2 | 0.10307112 | 1 | 0.46348836 | 0 | 1 | 0 | 1 | 12 | 0.962093 | FALSE | 0.74276693 | FALSE | 1 | FALSE | 1 | FALSE | 1 | FALSE | 1 | FALSE | 1 |
| 4254 | 4254 | 8.56521739 | 1.34572066 | 1 | 0.45027683 | 0 | 1 | 0 | 1 | 3 | 0.08614834 | 3 | 0.00598158 | 16 | 0.98100709 | FALSE | 1 | FALSE | 1 | FALSE | 1 | FALSE | 1 | FALSE | 0.45694206 | FALSE | 1 |
| 4255 | 4255 | 8.57142857 | 2.77010278 | 1 | 0.3050805 | 0 | 1 | 0 | 1 | 0 | 1 | 0 | 1 | 13 | 0.33283729 | FALSE | 1 | FALSE | 1 | FALSE | 1 | FALSE | 1 | FALSE | 1 | FALSE | 1 |
| 4256 | 4256 | 8.3 | 2.23830293 | 0 | 1 | 2 | 0.25306331 | 2 | 0.29330414 | 3 | 0.15707004 | 0 | 1 | 23 | 0.9158696 | FALSE | 1 | FALSE | 1 | FALSE | 1 | FALSE | 1 | FALSE | 1 | FALSE | 1 |
| 4257 | 4257 | 10.3529412 | 2.8063201 | 0 | 1 | 0 | 1 | 1 | 0.46348836 | 0 | 1 | 0 | 1 | 16 | 0.23070182 | FALSE | 1 | FALSE | 1 | FALSE | 1 | FALSE | 1 | FALSE | 1 | FALSE | 1 |
| 4258 | 4258 | 8.75 | 2.07665597 | 0 | 1 | 0 | 1 | 1 | 0.44344817 | 0 | 1 | 0 | 1 | 15 | 0.26127859 | FALSE | 1 | FALSE | 1 | FALSE | 1 | FALSE | 1 | FALSE | 1 | FALSE | 1 |
| 4259 | 4259 | 10.2352941 | 1.39450231 | 1 | 0.35728305 | 1 | 0.42857733 | 0 | 1 | 0 | 1 | 0 | 1 | 15 | 0.49060435 | FALSE | 1 | FALSE | 1 | FALSE | 1 | FALSE | 1 | FALSE | 1 | FALSE | 1 |
| 4260 | 4260 | 7.76666667 | 1.94393644 | 1 | 0.54198714 | 0 | 1 | 0 | 1 | 3 | 0.15707004 | 2 | 0.08611595 | 24 | 0.82294782 | FALSE | 1 | FALSE | 1 | FALSE | 1 | FALSE | 1 | FALSE | 1 | FALSE | 1 |
| 4261 | 4261 | 9.25 | 2.54705713 | 0 | 1 | 2 | 0.1353446 | 1 | 0.51939841 | 1 | 0.60987146 | 0 | 1 | 16 | 0.80826361 | FALSE | 1 | FALSE | 1 | FALSE | 1 | FALSE | 1 | FALSE | 1 | FALSE | 1 |
| 4262 | 4262 | 10.7222222 | 2.1030548 | 1 | 0.35728305 | 2 | 0.10307112 | 0 | 1 | 1 | 0.55062772 | 1 | 0.24525764 | 12 | 0.962093 | FALSE | 1 | FALSE | 1 | FALSE | 1 | FALSE | 1 | FALSE | 1 | FALSE | 1 |
| 4263 | 4263 | 7.21428571 | 2.54048846 | 0 | 1 | 1 | 0.36919824 | 1 | 0.40110418 | 1 | 0.48242099 | 0 | 1 | 11 | 0.83676449 | FALSE | 1 | FALSE | 1 | FALSE | 1 | FALSE | 1 | FALSE | 1 | FALSE | 1 |
| 4264 | 4264 | 8.12121212 | 1.9346533 | 0 | 1 | 2 | 0.28941836 | 1 | 0.70185639 | 1 | 0.78873814 | 0 | 1 | 29 | 0.39648374 | FALSE | 1 | FALSE | 1 | FALSE | 1 | FALSE | 1 | FALSE | 1 | FALSE | 1 |
| 4265 | 4265 | 7.55 | 2.47941525 | 0 | 1 | 1 | 0.7326167 | 1 | 0.76953771 | 1 | 0.84823834 | 3 | 0.02730775 | 34 | 0.56328904 | FALSE | 1 | FALSE | 1 | FALSE | 1 | FALSE | 1 | FALSE | 0.79106376 | FALSE | 1 |
| 4266 | 4266 | 9.92682927 | 2.69950532 | 0 | 1 | 1 | 0.7326167 | 2 | 0.42396974 | 4 | 0.10945471 | 1 | 0.48480878 | 32 | 0.83853687 | FALSE | 1 | FALSE | 1 | FALSE | 1 | FALSE | 1 | FALSE | 1 | FALSE | 1 |
| 4267 | 4267 | 5.82 | 3.07694654 | 0 | 1 | 3 | 0.17732615 | 3 | 0.21866539 | 0 | 1 | 3 | 0.03699359 | 36 | 0.84604146 | FALSE | 1 | FALSE | 1 | FALSE | 1 | FALSE | 1 | FALSE | 0.87669812 | FALSE | 1 |
| 4268 | 4268 | 8.45454546 | 2.25080335 | 0 | 1 | 2 | 0.14655134 | 0 | 1 | 3 | 0.06912908 | 0 | 1 | 16 | 0.90353674 | FALSE | 1 | FALSE | 1 | FALSE | 1 | FALSE | 1 | FALSE | 1 | FALSE | 1 |
| 4269 | 4269 | 8.63636364 | 2.44104029 | 1 | 0.4357638 | 0 | 1 | 0 | 1 | 1 | 0.64497439 | 1 | 0.30529951 | 19 | 0.54233419 | FALSE | 1 | FALSE | 1 | FALSE | 1 | FALSE | 1 | FALSE | 1 | FALSE | 1 |
| 4270 | 4270 | 11.2666667 | 3.1721006 | 1 | 0.32293355 | 1 | 0.38964394 | 0 | 1 | 0 | 1 | 0 | 1 | 13 | 0.57712805 | FALSE | 1 | FALSE | 1 | FALSE | 1 | FALSE | 1 | FALSE | 1 | FALSE | 1 |
| 4271 | 4271 | 11.2888889 | 2.72129236 | 3 | 0.10767332 | 2 | 0.43020462 | 1 | 0.80828716 | 2 | 0.61900576 | 0 | 1 | 37 | 0.73736034 | FALSE | 0.88820416 | FALSE | 1 | FALSE | 1 | FALSE | 1 | FALSE | 1 | FALSE | 1 |
| 4272 | 4272 | 7.30434783 | 2.81972528 | 5 | 0.00024166 | 1 | 0.53115654 | 2 | 0.19931429 | 0 | 1 | 0 | 1 | 15 | 0.99450713 | TRUE | 0.03045702 | FALSE | 1 | FALSE | 1 | FALSE | 1 | FALSE | 1 | FALSE | 1 |
| 4273 | 4273 | 9.93333333 | 3.22421394 | 1 | 0.54198714 | 2 | 0.25306331 | 5 | 0.00391694 | 2 | 0.4036176 | 0 | 1 | 20 | 0.99601577 | FALSE | 1 | FALSE | 1 | FALSE | 1 | FALSE | 1 | FALSE | 1 | FALSE | 1 |
| 4274 | 4274 | 10.4285714 | 2.38190457 | 0 | 1 | 2 | 0.14655134 | 1 | 0.53671415 | 1 | 0.62783522 | 0 | 1 | 17 | 0.77849076 | FALSE | 1 | FALSE | 1 | FALSE | 1 | FALSE | 1 | FALSE | 1 | FALSE | 1 |
| 4275 | 4275 | 7.19230769 | 2.27053418 | 2 | 0.14250978 | 1 | 0.57534752 | 1 | 0.61440466 | 0 | 1 | 1 | 0.3499045 | 21 | 0.78939668 | FALSE | 1 | FALSE | 1 | FALSE | 1 | FALSE | 1 | FALSE | 1 | FALSE | 1 |
| 4276 | 4276 | 8.19444444 | 2.221354 | 0 | 1 | 0 | 1 | 0 | 1 | 2 | 0.49731009 | 1 | 0.44938266 | 33 | 0.16407838 | FALSE | 1 | FALSE | 1 | FALSE | 1 | FALSE | 1 | FALSE | 1 | FALSE | 1 |
| 4277 | 4277 | 8.63636364 | 2.67217063 | 0 | 1 | 0 | 1 | 2 | 0.05708125 | 0 | 1 | 1 | 0.16638736 | 8 | 0.92152993 | FALSE | 1 | FALSE | 1 | FALSE | 1 | FALSE | 1 | FALSE | 1 | FALSE | 1 |
| 4278 | 4278 | 9.5 | 2.5 | 0 | 1 | 0 | 1 | 1 | 0.25384309 | 0 | 1 | 0 | 1 | 7 | 0.63752397 | FALSE | 1 | FALSE | 1 | FALSE | 1 | FALSE | 1 | FALSE | 1 | FALSE | 1 |
| 4279 | 4279 | 9.48 | 2.85825121 | 0 | 1 | 1 | 0.56109832 | 2 | 0.22597287 | 1 | 0.69181315 | 0 | 1 | 21 | 0.64957167 | FALSE | 1 | FALSE | 1 | FALSE | 1 | FALSE | 1 | FALSE | 1 | FALSE | 1 |
| 4280 | 4280 | 8.88235294 | 1.90610021 | 0 | 1 | 0 | 1 | 0 | 1 | 0 | 1 | 0 | 1 | 17 | 0.05550965 | FALSE | 1 | FALSE | 1 | FALSE | 1 | FALSE | 1 | FALSE | 1 | FALSE | 1 |
| 4281 | 4281 | 9.46153846 | 2.24003381 | 0 | 1 | 0 | 1 | 0 | 1 | 1 | 0.45746571 | 0 | 1 | 12 | 0.374198 | FALSE | 1 | FALSE | 1 | FALSE | 1 | FALSE | 1 | FALSE | 1 | FALSE | 1 |
| 4282 | 4282 | 9.76190476 | 2.13596776 | 0 | 1 | 0 | 1 | 0 | 1 | 1 | 0.62783522 | 0 | 1 | 20 | 0.13763971 | FALSE | 1 | FALSE | 1 | FALSE | 1 | FALSE | 1 | FALSE | 1 | FALSE | 1 |
| 4283 | 4283 | 9.40740741 | 2.69786279 | 3 | 0.03084786 | 2 | 0.21684246 | 4 | 0.01487236 | 0 | 1 | 0 | 1 | 18 | 0.99450398 | FALSE | 0.53356688 | FALSE | 1 | FALSE | 1 | FALSE | 1 | FALSE | 1 | FALSE | 1 |
| 4284 | 4284 | 8.77777778 | 2.09644025 | 0 | 1 | 0 | 1 | 0 | 1 | 3 | 0.22784656 | 1 | 0.44938266 | 32 | 0.31709555 | FALSE | 1 | FALSE | 1 | FALSE | 1 | FALSE | 1 | FALSE | 1 | FALSE | 1 |
| 4285 | 4285 | 7.66666667 | 2.46080384 | 0 | 1 | 0 | 1 | 0 | 1 | 1 | 0.43131129 | 0 | 1 | 11 | 0.41944532 | FALSE | 1 | FALSE | 1 | FALSE | 1 | FALSE | 1 | FALSE | 1 | FALSE | 1 |
| 4286 | 4286 | 7.80952381 | 3.58062309 | 0 | 1 | 1 | 0.48239024 | 0 | 1 | 3 | 0.0612581 | 1 | 0.28187291 | 15 | 0.92113679 | FALSE | 1 | FALSE | 1 | FALSE | 1 | FALSE | 1 | FALSE | 1 | FALSE | 1 |
| 4287 | 4287 | 4.25404531 | 1.26255179 | 24 | 0.02477155 | 32 | 0.00469624 | 21 | 0.63781341 | 21 | 0.94780978 | 22 | 0.0003511 | 497 | 0.99692841 | FALSE | 0.4974559 | FALSE | 0.58120789 | FALSE | 1 | FALSE | 1 | FALSE | 0.09587314 | FALSE | 1 |
| 4288 | 4288 | 8.70833333 | 1.45714237 | 0 | 1 | 0 | 1 | 2 | 0.21260257 | 0 | 1 | 1 | 0.32796862 | 21 | 0.47053137 | FALSE | 1 | FALSE | 1 | FALSE | 1 | FALSE | 1 | FALSE | 1 | FALSE | 1 |
| 4289 | 4289 | 7.67032967 | 2.50298699 | 0 | 1 | 3 | 0.5689882 | 5 | 0.22832872 | 3 | 0.79556345 | 0 | 1 | 80 | 0.21860554 | FALSE | 1 | FALSE | 1 | FALSE | 1 | FALSE | 1 | FALSE | 1 | FALSE | 1 |
| 4290 | 4290 | 8.35294118 | 2.51913095 | 3 | 0.05546921 | 1 | 0.6739483 | 2 | 0.34668321 | 1 | 0.7984848 | 0 | 1 | 27 | 0.85023803 | FALSE | 0.68849059 | FALSE | 1 | FALSE | 1 | FALSE | 1 | FALSE | 1 | FALSE | 1 |
| 4291 | 4291 | 8.54545455 | 1.23315091 | 0 | 1 | 0 | 1 | 0 | 1 | 0 | 1 | 0 | 1 | 11 | 0.15414489 | FALSE | 1 | FALSE | 1 | FALSE | 1 | FALSE | 1 | FALSE | 1 | FALSE | 1 |
| 4292 | 4292 | 10.53125 | 1.8368515 | 1 | 0.56527493 | 4 | 0.0188228 | 0 | 1 | 0 | 1 | 0 | 1 | 27 | 0.61671133 | FALSE | 1 | FALSE | 0.77515497 | FALSE | 1 | FALSE | 1 | FALSE | 1 | FALSE | 1 |
| 4293 | 4293 | 9.52631579 | 2.52082737 | 0 | 1 | 1 | 0.46503795 | 1 | 0.50143826 | 0 | 1 | 4 | 0.00021933 | 13 | 0.97998676 | FALSE | 1 | FALSE | 1 | FALSE | 1 | FALSE | 1 | FALSE | 0.07486439 | FALSE | 1 |
| 4294 | 4294 | 10.5294118 | 2.74649824 | 1 | 0.35728305 | 0 | 1 | 1 | 0.46348836 | 3 | 0.040397 | 1 | 0.24525764 | 11 | 0.98974663 | FALSE | 1 | FALSE | 1 | FALSE | 1 | FALSE | 1 | FALSE | 1 | FALSE | 1 |
| 4295 | 4295 | 9.27272727 | 1.81363067 | 1 | 0.4357638 | 0 | 1 | 1 | 0.55340851 | 3 | 0.07743186 | 0 | 1 | 17 | 0.8840357 | FALSE | 1 | FALSE | 1 | FALSE | 1 | FALSE | 1 | FALSE | 1 | FALSE | 1 |
| 4296 | 4296 | 10.1578947 | 1.9805145 | 0 | 1 | 0 | 1 | 1 | 0.50143826 | 0 | 1 | 2 | 0.03799168 | 16 | 0.65565778 | FALSE | 1 | FALSE | 1 | FALSE | 1 | FALSE | 1 | FALSE | 0.87669812 | FALSE | 1 |
| 4297 | 4297 | 9.84615385 | 2.82423996 | 0 | 1 | 0 | 1 | 0 | 1 | 1 | 0.45746571 | 0 | 1 | 12 | 0.374198 | FALSE | 1 | FALSE | 1 | FALSE | 1 | FALSE | 1 | FALSE | 1 | FALSE | 1 |
| 4298 | 4298 | 4.55760369 | 1.49273194 | 8 | 0.19296761 | 8 | 0.40407074 | 10 | 0.25308189 | 5 | 0.97439749 | 8 | 0.02565954 | 178 | 0.8561338 | FALSE | 1 | FALSE | 1 | FALSE | 1 | FALSE | 1 | FALSE | 0.79106376 | FALSE | 1 |
| 4299 | 4299 | 9.54545455 | 2.18938083 | 0 | 1 | 0 | 1 | 0 | 1 | 0 | 1 | 0 | 1 | 9 | 0.21661397 | FALSE | 1 | FALSE | 1 | FALSE | 1 | FALSE | 1 | FALSE | 1 | FALSE | 1 |
| 4300 | 4300 | 7.44827586 | 1.47124641 | 4 | 0.50203916 | 5 | 0.49463353 | 4 | 0.76186672 | 3 | 0.96355069 | 0 | 1 | 127 | 0.08298243 | FALSE | 1 | FALSE | 1 | FALSE | 1 | FALSE | 1 | FALSE | 1 | FALSE | 1 |
| 4301 | 4301 | 7.86206897 | 2.31522901 | 0 | 1 | 1 | 0.54637326 | 0 | 1 | 0 | 1 | 0 | 1 | 23 | 0.09196389 | FALSE | 1 | FALSE | 1 | FALSE | 1 | FALSE | 1 | FALSE | 1 | FALSE | 1 |
| 4302 | 4302 | 5.82978723 | 2.93416521 | 2 | 0.30241117 | 1 | 0.75788114 | 0 | 1 | 1 | 0.86831187 | 1 | 0.50988734 | 38 | 0.31730615 | FALSE | 1 | FALSE | 1 | FALSE | 1 | FALSE | 1 | FALSE | 1 | FALSE | 1 |
| 4303 | 4303 | 8.625 | 2.98346136 | 1 | 0.46441868 | 0 | 1 | 0 | 1 | 3 | 0.09525947 | 0 | 1 | 20 | 0.68270729 | FALSE | 1 | FALSE | 1 | FALSE | 1 | FALSE | 1 | FALSE | 1 | FALSE | 1 |
| 4304 | 4304 | 8.475 | 3.36888335 | 0 | 1 | 2 | 0.37295654 | 1 | 0.76953771 | 3 | 0.27785536 | 1 | 0.48480878 | 33 | 0.71942635 | FALSE | 1 | FALSE | 1 | FALSE | 1 | FALSE | 1 | FALSE | 1 | FALSE | 1 |
| 4305 | 4305 | 9.53846154 | 2.84511416 | 0 | 1 | 0 | 1 | 0 | 1 | 0 | 1 | 0 | 1 | 13 | 0.10967906 | FALSE | 1 | FALSE | 1 | FALSE | 1 | FALSE | 1 | FALSE | 1 | FALSE | 1 |
| 4306 | 4306 | 9.84 | 4.90452852 | 0 | 1 | 0 | 1 | 1 | 0.59998149 | 1 | 0.69181315 | 1 | 0.33902672 | 22 | 0.43642846 | FALSE | 1 | FALSE | 1 | FALSE | 1 | FALSE | 1 | FALSE | 1 | FALSE | 1 |
| 4307 | 4307 | 7.75 | 1.54515371 | 0 | 1 | 0 | 1 | 1 | 0.51939841 | 0 | 1 | 0 | 1 | 19 | 0.15700255 | FALSE | 1 | FALSE | 1 | FALSE | 1 | FALSE | 1 | FALSE | 1 | FALSE | 1 |
| 4308 | 4308 | 8.375 | 2.0577597 | 0 | 1 | 1 | 0.40942991 | 0 | 1 | 0 | 1 | 1 | 0.23264612 | 14 | 0.53308261 | FALSE | 1 | FALSE | 1 | FALSE | 1 | FALSE | 1 | FALSE | 1 | FALSE | 1 |
| 4309 | 4309 | 6.12 | 3.07662152 | 0 | 1 | 1 | 0.56109832 | 1 | 0.59998149 | 0 | 1 | 0 | 1 | 23 | 0.22727183 | FALSE | 1 | FALSE | 1 | FALSE | 1 | FALSE | 1 | FALSE | 1 | FALSE | 1 |
| 4310 | 4310 | 7.59375 | 2.81579668 | 1 | 0.56527493 | 0 | 1 | 0 | 1 | 1 | 0.77852168 | 1 | 0.41154418 | 29 | 0.24088541 | FALSE | 1 | FALSE | 1 | FALSE | 1 | FALSE | 1 | FALSE | 1 | FALSE | 1 |
| 4311 | 4311 | 9.14285714 | 1.38995424 | 3 | 0.01568237 | 0 | 1 | 0 | 1 | 1 | 0.62783522 | 0 | 1 | 17 | 0.77849076 | FALSE | 0.38695764 | FALSE | 1 | FALSE | 1 | FALSE | 1 | FALSE | 1 | FALSE | 1 |
| 4312 | 4312 | 5.07526882 | 0.5254094 | 16 | 0.13731594 | 15 | 0.54051537 | 19 | 0.30842945 | 20 | 0.65128468 | 10 | 0.22862802 | 384 | 0.85577434 | FALSE | 0.99812969 | FALSE | 1 | FALSE | 1 | FALSE | 1 | FALSE | 1 | FALSE | 1 |
| 4313 | 4313 | 6.79310345 | 0.84605822 | 1 | 0.52988231 | 1 | 0.61539065 | 1 | 0.65463982 | 0 | 1 | 0 | 1 | 26 | 0.31495157 | FALSE | 1 | FALSE | 1 | FALSE | 1 | FALSE | 1 | FALSE | 1 | FALSE | 1 |
| 4314 | 4314 | 11.3076923 | 2.0525637 | 2 | 0.04234484 | 2 | 0.06426041 | 1 | 0.37874417 | 1 | 0.45746571 | 0 | 1 | 7 | 0.99839694 | FALSE | 0.62842194 | FALSE | 1 | FALSE | 1 | FALSE | 1 | FALSE | 1 | FALSE | 1 |
| 4315 | 4315 | 5.16666667 | 2.51108653 | 0 | 1 | 0 | 1 | 4 | 0.03888854 | 2 | 0.49731009 | 0 | 1 | 30 | 0.67248271 | FALSE | 1 | FALSE | 1 | FALSE | 1 | FALSE | 1 | FALSE | 1 | FALSE | 1 |
| 4316 | 4316 | 8.32142857 | 2.39125746 | 3 | 0.03084786 | 0 | 1 | 1 | 0.62830986 | 1 | 0.71956339 | 0 | 1 | 22 | 0.76237877 | FALSE | 0.53356688 | FALSE | 1 | FALSE | 1 | FALSE | 1 | FALSE | 1 | FALSE | 1 |
| 4317 | 4317 | 7.89285714 | 2.74303385 | 0 | 1 | 1 | 0.60247911 | 1 | 0.64171563 | 0 | 1 | 0 | 1 | 26 | 0.16405547 | FALSE | 1 | FALSE | 1 | FALSE | 1 | FALSE | 1 | FALSE | 1 | FALSE | 1 |
| 4318 | 4318 | 8.94736842 | 2.16366366 | 1 | 0.38989951 | 0 | 1 | 0 | 1 | 0 | 1 | 0 | 1 | 18 | 0.17880694 | FALSE | 1 | FALSE | 1 | FALSE | 1 | FALSE | 1 | FALSE | 1 | FALSE | 1 |
| 4319 | 4319 | 9.82352941 | 2.61748182 | 0 | 1 | 1 | 0.42857733 | 1 | 0.46348836 | 0 | 1 | 0 | 1 | 15 | 0.49060435 | FALSE | 1 | FALSE | 1 | FALSE | 1 | FALSE | 1 | FALSE | 1 | FALSE | 1 |
| 4320 | 4320 | 5.2 | 2.63818119 | 0 | 1 | 0 | 1 | 0 | 1 | 0 | 1 | 0 | 1 | 10 | 0.18273166 | FALSE | 1 | FALSE | 1 | FALSE | 1 | FALSE | 1 | FALSE | 1 | FALSE | 1 |
| 4321 | 4321 | 8.37142857 | 2.00122412 | 3 | 0.0595675 | 0 | 1 | 1 | 0.72299506 | 1 | 0.80778317 | 0 | 1 | 30 | 0.52765882 | FALSE | 0.72532843 | FALSE | 1 | FALSE | 1 | FALSE | 1 | FALSE | 1 | FALSE | 1 |
| 4322 | 4322 | 10.7 | 2.28254244 | 0 | 1 | 0 | 1 | 0 | 1 | 1 | 0.37517262 | 0 | 1 | 9 | 0.5215547 | FALSE | 1 | FALSE | 1 | FALSE | 1 | FALSE | 1 | FALSE | 1 | FALSE | 1 |

| | | | | | | | | | | | | | | | | | | | | | | | | | | | |
|---|---|---|---|---|---|---|---|---|---|---|---|---|---|---|---|---|---|---|---|---|---|---|---|---|---|---|---|
| 4323 | 4323 | 9.5625 | 3.46353775 | 1 | 0.34033054 | 0 | 1 | 0 | 1 | 0 | 1 | 2 | 0.02752114 | 13 | 0.76799161 | FALSE | 1 | FALSE | 1 | FALSE | 1 | FALSE | 1 | FALSE | 0.79106376 | FALSE | 1 |
| 4324 | 4324 | 8.66666667 | 2.24845626 | 0 | 1 | 0 | 1 | 0 | 1 | 1 | 0.43131129 | 0 | 1 | 11 | 0.41944532 | FALSE | 1 | FALSE | 1 | FALSE | 1 | FALSE | 1 | FALSE | 1 | FALSE | 1 |
| 4325 | 4325 | 9.83333333 | 2.03442594 | 0 | 1 | 0 | 1 | 0 | 1 | 0 | 1 | 0 | 1 | 6 | 0.36077158 | FALSE | 1 | FALSE | 1 | FALSE | 1 | FALSE | 1 | FALSE | 1 | FALSE | 1 |
| 4326 | 4326 | 8.35483871 | 2.54695754 | 2 | 0.1881398 | 0 | 1 | 2 | 0.30673593 | 3 | 0.16836568 | 0 | 1 | 24 | 0.90138666 | FALSE | 1 | FALSE | 1 | FALSE | 1 | FALSE | 1 | FALSE | 1 | FALSE | 1 |
| 4327 | 4327 | 8.42307692 | 2.81716001 | 2 | 0.14250978 | 2 | 0.20486827 | 0 | 1 | 1 | 0.70601446 | 0 | 1 | 21 | 0.78939668 | FALSE | 1 | FALSE | 1 | FALSE | 1 | FALSE | 1 | FALSE | 1 | FALSE | 1 |
| 4328 | 4328 | 10.131579 | 2.71614135 | 0 | 1 | 0 | 1 | 1 | 0.74264075 | 1 | 0.82511639 | 0 | 1 | 35 | 0.05673524 | FALSE | 1 | FALSE | 1 | FALSE | 1 | FALSE | 1 | FALSE | 1 | FALSE | 1 |
| 4329 | 4329 | 6.7826087 | 2.55302926 | 0 | 1 | 0 | 1 | 1 | 0.5695037 | 2 | 0.2852236 | 5 | 2.88E-05 | 15 | 0.99450713 | FALSE | 1 | FALSE | 1 | FALSE | 1 | FALSE | 1 | TRUE | 0.01570671 | FALSE | 1 |
| 4330 | 4330 | 10.0714286 | 2.7114647 | 0 | 1 | 2 | 0.07341559 | 0 | 1 | 0 | 1 | 1 | 0.20679328 | 11 | 0.83676449 | FALSE | 1 | FALSE | 1 | FALSE | 1 | FALSE | 1 | FALSE | 1 | FALSE | 1 |
| 4331 | 4331 | 8.91891892 | 2.40919835 | 0 | 1 | 0 | 1 | 2 | 0.38585089 | 0 | 1 | 0 | 1 | 35 | 0.05673524 | FALSE | 1 | FALSE | 1 | FALSE | 1 | FALSE | 1 | FALSE | 1 | FALSE | 1 |
| 4332 | 4332 | 9.77272727 | 2.33416947 | 0 | 1 | 0 | 1 | 1 | 0.55340851 | 0 | 1 | 0 | 1 | 21 | 0.12048848 | FALSE | 1 | FALSE | 1 | FALSE | 1 | FALSE | 1 | FALSE | 1 | FALSE | 1 |
| 4333 | 4333 | 9.53571429 | 3.45875848 | 1 | 0.51745948 | 1 | 0.60247911 | 0 | 1 | 1 | 0.73248983 | 0 | 1 | 25 | 0.34293 | FALSE | 1 | FALSE | 1 | FALSE | 1 | FALSE | 1 | FALSE | 1 | FALSE | 1 |
| 4334 | 4334 | 9.5 | 3.69845528 | 0 | 1 | 1 | 0.36919824 | 0 | 1 | 0 | 1 | 0 | 1 | 13 | 0.33283729 | FALSE | 1 | FALSE | 1 | FALSE | 1 | FALSE | 1 | FALSE | 1 | FALSE | 1 |
| 4335 | 4335 | 8.84615385 | 2.01326959 | 0 | 1 | 1 | 0.57534752 | 2 | 0.23940226 | 1 | 0.70601446 | 0 | 1 | 22 | 0.61628 | FALSE | 1 | FALSE | 1 | FALSE | 1 | FALSE | 1 | FALSE | 1 | FALSE | 1 |
| 4336 | 4336 | 11.0625 | 2.83876624 | 0 | 1 | 1 | 0.40942991 | 1 | 0.44344817 | 2 | 0.16544789 | 1 | 0.23264612 | 11 | 0.9718531 | FALSE | 1 | FALSE | 1 | FALSE | 1 | FALSE | 1 | FALSE | 1 | FALSE | 1 |
| 4337 | 4337 | 6.88235294 | 2.8467185 | 0 | 1 | 0 | 1 | 0 | 1 | 1 | 0.55062772 | 0 | 1 | 16 | 0.23070182 | FALSE | 1 | FALSE | 1 | FALSE | 1 | FALSE | 1 | FALSE | 1 | FALSE | 1 |
| 4338 | 4338 | 9.94736842 | 2.83722778 | 0 | 1 | 1 | 0.46503795 | 1 | 0.50143826 | 0 | 1 | 0 | 1 | 17 | 0.41141543 | FALSE | 1 | FALSE | 1 | FALSE | 1 | FALSE | 1 | FALSE | 1 | FALSE | 1 |
| 4339 | 4339 | 10.1071429 | 2.46893971 | 0 | 1 | 0 | 1 | 1 | 0.64171563 | 0 | 1 | 0 | 1 | 27 | 0.05284755 | FALSE | 1 | FALSE | 1 | FALSE | 1 | FALSE | 1 | FALSE | 1 | FALSE | 1 |
| 4340 | 4340 | 9.21428571 | 2.83293315 | 0 | 1 | 2 | 0.39616059 | 2 | 0.44869594 | 2 | 0.5811115 | 4 | 0.00474205 | 32 | 0.94677277 | FALSE | 1 | FALSE | 1 | FALSE | 1 | FALSE | 1 | FALSE | 0.41770847 | FALSE | 1 |
| 4341 | 4341 | 9.92857143 | 2.34411987 | 1 | 0.3050805 | 1 | 0.36919824 | 0 | 1 | 0 | 1 | 0 | 1 | 12 | 0.62236227 | FALSE | 1 | FALSE | 1 | FALSE | 1 | FALSE | 1 | FALSE | 1 | FALSE | 1 |
| 4342 | 4342 | 8.88235294 | 1.99653679 | 0 | 1 | 1 | 0.42857733 | 1 | 0.46348836 | 0 | 1 | 1 | 0.24525764 | 14 | 0.7313145 | FALSE | 1 | FALSE | 1 | FALSE | 1 | FALSE | 1 | FALSE | 1 | FALSE | 1 |
| 4343 | 4343 | 8.88888889 | 1.04822013 | 1 | 0.37380237 | 1 | 0.44710673 | 0 | 1 | 0 | 1 | 0 | 1 | 16 | 0.44998001 | FALSE | 1 | FALSE | 1 | FALSE | 1 | FALSE | 1 | FALSE | 1 | FALSE | 1 |
| 4344 | 4344 | 8.62068966 | 2.02422898 | 0 | 1 | 3 | 0.06581417 | 1 | 0.65463982 | 0 | 1 | 0 | 1 | 25 | 0.51779518 | FALSE | 1 | FALSE | 1 | FALSE | 1 | FALSE | 1 | FALSE | 1 | FALSE | 1 |
| 4345 | 4345 | 5.23688663 | 0.92240019 | 22 | 0.04227629 | 26 | 0.0575292 | 24 | 0.27217502 | 17 | 0.98770865 | 19 | 0.00275452 | 476 | 0.97868334 | FALSE | 0.62842194 | FALSE | 1 | FALSE | 1 | FALSE | 1 | FALSE | 0.29951111 | FALSE | 1 |
| 4346 | 4346 | 9.35 | 1.878164 | 0 | 1 | 2 | 0.12434464 | 1 | 0.50143826 | 2 | 0.21610812 | 1 | 0.26986775 | 13 | 0.97998676 | FALSE | 1 | FALSE | 1 | FALSE | 1 | FALSE | 1 | FALSE | 1 | FALSE | 1 |
| 4347 | 4347 | 9.33333333 | 2.45596133 | 0 | 1 | 1 | 0.48239024 | 2 | 0.16019548 | 0 | 1 | 0 | 1 | 17 | 0.61750632 | FALSE | 1 | FALSE | 1 | FALSE | 1 | FALSE | 1 | FALSE | 1 | FALSE | 1 |
| 4348 | 4348 | 8.83333333 | 2.44381305 | 0 | 1 | 1 | 0.54637326 | 2 | 0.21260257 | 1 | 0.67692816 | 0 | 1 | 20 | 0.68270729 | FALSE | 1 | FALSE | 1 | FALSE | 1 | FALSE | 1 | FALSE | 1 | FALSE | 1 |
| 4349 | 4349 | 7.12903226 | 2.19591902 | 1 | 0.55378205 | 3 | 0.07729739 | 2 | 0.30673593 | 5 | 0.01254769 | 0 | 1 | 20 | 0.9984454 | FALSE | 1 | FALSE | 1 | FALSE | 1 | FALSE | 1 | FALSE | 1 | FALSE | 1 |
| 4350 | 4350 | 10.7272727 | 2.33903279 | 2 | 0.03087289 | 0 | 1 | 0 | 1 | 1 | 0.40390032 | 0 | 1 | 8 | 0.92152993 | FALSE | 0.53356688 | FALSE | 1 | FALSE | 1 | FALSE | 1 | FALSE | 1 | FALSE | 1 |
| 4351 | 4351 | 6.73333333 | 1.69181034 | 1 | 0.32293355 | 0 | 1 | 0 | 1 | 2 | 0.14906763 | 0 | 1 | 12 | 0.80329917 | FALSE | 1 | FALSE | 1 | FALSE | 1 | FALSE | 1 | FALSE | 1 | FALSE | 1 |
| 4352 | 4352 | 9.38095238 | 3.52509629 | 1 | 0.42086991 | 0 | 1 | 1 | 0.53671415 | 2 | 0.25059627 | 0 | 1 | 17 | 0.77849076 | FALSE | 1 | FALSE | 1 | FALSE | 1 | FALSE | 1 | FALSE | 1 | FALSE | 1 |
| 4353 | 4353 | 9.47058824 | 2.95291774 | 0 | 1 | 2 | 0.10307112 | 1 | 0.46348836 | 2 | 0.18212134 | 0 | 1 | 12 | 0.962093 | FALSE | 1 | FALSE | 1 | FALSE | 1 | FALSE | 1 | FALSE | 1 | FALSE | 1 |
| 4354 | 4354 | 9.23529412 | 2.94117647 | 1 | 0.35728305 | 0 | 1 | 1 | 0.46348836 | 0 | 1 | 1 | 0.24525764 | 14 | 0.7313145 | FALSE | 1 | FALSE | 1 | FALSE | 1 | FALSE | 1 | FALSE | 1 | FALSE | 1 |
| 4355 | 4355 | 6.91489362 | 2.71191059 | 0 | 1 | 3 | 0.19374024 | 5 | 0.02577785 | 2 | 0.64277396 | 1 | 0.54145295 | 36 | 0.94678267 | FALSE | 1 | FALSE | 1 | FALSE | 1 | FALSE | 1 | FALSE | 1 | FALSE | 1 |
| 4356 | 4356 | 9.71428571 | 3.18318285 | 1 | 0.51745948 | 2 | 0.2288771 | 2 | 0.2663542 | 0 | 1 | 2 | 0.07640798 | 21 | 0.94083768 | FALSE | 1 | FALSE | 1 | FALSE | 1 | FALSE | 1 | FALSE | 1 | FALSE | 1 |
| 4357 | 4357 | 7.56521739 | 3.60948132 | 3 | 0.02011786 | 1 | 0.53115654 | 0 | 1 | 1 | 0.6613267 | 0 | 1 | 18 | 0.86273155 | FALSE | 0.44482142 | FALSE | 1 | FALSE | 1 | FALSE | 1 | FALSE | 1 | FALSE | 1 |
| 4358 | 4358 | 7.22115385 | 1.02803707 | 3 | 0.50146864 | 2 | 0.85560113 | 3 | 0.72810072 | 5 | 0.52348367 | 2 | 0.51184859 | 89 | 0.43117406 | FALSE | 1 | FALSE | 1 | FALSE | 1 | FALSE | 1 | FALSE | 1 | FALSE | 1 |
| 4359 | 4359 | 7.03773585 | 2.39466596 | 1 | 0.74224579 | 3 | 0.23623106 | 3 | 0.28676219 | 1 | 0.91399378 | 0 | 1 | 44 | 0.57424118 | FALSE | 1 | FALSE | 1 | FALSE | 1 | FALSE | 1 | FALSE | 1 | FALSE | 1 |
| 4360 | 4360 | 11 | 2.76887462 | 0 | 1 | 2 | 0.01433429 | 0 | 1 | 0 | 1 | 0 | 1 | 4 | 0.94743772 | FALSE | 1 | FALSE | 0.71309764 | FALSE | 1 | FALSE | 1 | FALSE | 1 | FALSE | 1 |
| 4361 | 4361 | 8.53846154 | 2.6199056 | 1 | 0.49162634 | 0 | 1 | 0 | 1 | 1 | 0.70601446 | 0 | 1 | 24 | 0.20425579 | FALSE | 1 | FALSE | 1 | FALSE | 1 | FALSE | 1 | FALSE | 1 | FALSE | 1 |
| 4362 | 4362 | 7.92857143 | 1.39423009 | 2 | 0.72148567 | 1 | 0.96107365 | 4 | 0.47057639 | 4 | 0.66533238 | 1 | 0.80446991 | 86 | 0.21911908 | FALSE | 1 | FALSE | 1 | FALSE | 1 | FALSE | 1 | FALSE | 1 | FALSE | 1 |
| 4363 | 4363 | 7.84313725 | 3.06384706 | 2 | 0.37762121 | 2 | 0.49507792 | 1 | 0.84631478 | 2 | 0.68678423 | 0 | 1 | 44 | 0.44552698 | FALSE | 1 | FALSE | 1 | FALSE | 1 | FALSE | 1 | FALSE | 1 | FALSE | 1 |
| 4364 | 4364 | 9.4 | 1.52315462 | 0 | 1 | 0 | 1 | 0 | 1 | 1 | 0.69181315 | 1 | 0.33902672 | 23 | 0.22727183 | FALSE | 1 | FALSE | 1 | FALSE | 1 | FALSE | 1 | FALSE | 1 | FALSE | 1 |
| 4365 | 4365 | 9.68421053 | 2.45062036 | 1 | 0.38989951 | 0 | 1 | 0 | 1 | 1 | 0.59104357 | 0 | 1 | 17 | 0.41141543 | FALSE | 1 | FALSE | 1 | FALSE | 1 | FALSE | 1 | FALSE | 1 | FALSE | 1 |
| 4366 | 4366 | 8.65217391 | 2.44253424 | 3 | 0.02011786 | 1 | 0.53115654 | 0 | 1 | 2 | 0.2852236 | 1 | 0.31672722 | 16 | 0.98100709 | FALSE | 0.44482142 | FALSE | 1 | FALSE | 1 | FALSE | 1 | FALSE | 1 | FALSE | 1 |
| 4367 | 4367 | 9.52380952 | 2.48076501 | 0 | 1 | 1 | 0.49918218 | 1 | 0.53671415 | 4 | 0.0140322 | 0 | 1 | 15 | 0.96513691 | FALSE | 1 | FALSE | 1 | FALSE | 1 | FALSE | 1 | FALSE | 1 | FALSE | 1 |
| 4368 | 4368 | 10.5 | 2.36290781 | 0 | 1 | 0 | 1 | 1 | 0.197145 | 1 | 0.24578796 | 0 | 1 | 4 | 0.94743772 | FALSE | 1 | FALSE | 1 | FALSE | 1 | FALSE | 1 | FALSE | 1 | FALSE | 1 |
| 4369 | 4369 | 6.19230769 | 2.71773705 | 0 | 1 | 0 | 1 | 1 | 0.61440466 | 1 | 0.70601446 | 0 | 1 | 24 | 0.20425579 | FALSE | 1 | FALSE | 1 | FALSE | 1 | FALSE | 1 | FALSE | 1 | FALSE | 1 |
| 4370 | 4370 | 6.76190476 | 2.94237937 | 0 | 1 | 1 | 0.49918218 | 0 | 1 | 1 | 0.62783522 | 1 | 0.29368245 | 18 | 0.57962763 | FALSE | 1 | FALSE | 1 | FALSE | 1 | FALSE | 1 | FALSE | 1 | FALSE | 1 |
| 4371 | 4371 | 10.2941177 | 2.37052255 | 0 | 1 | 0 | 1 | 0 | 1 | 0 | 1 | 2 | 0.03086174 | 15 | 0.49060435 | FALSE | 1 | FALSE | 1 | FALSE | 1 | FALSE | 1 | FALSE | 0.82559785 | FALSE | 1 |
| 4372 | 4372 | 8.77777778 | 2.69887951 | 0 | 1 | 0 | 1 | 1 | 0.28067846 | 1 | 0.34506517 | 1 | 0.1383237 | 6 | 0.96128317 | FALSE | 1 | FALSE | 1 | FALSE | 1 | FALSE | 1 | FALSE | 1 | FALSE | 1 |
| 4373 | 4373 | 9.625 | 1.28492866 | 0 | 1 | 0 | 1 | 0 | 1 | 1 | 0.62783522 | 0 | 1 | 20 | 0.13763971 | FALSE | 1 | FALSE | 1 | FALSE | 1 | FALSE | 1 | FALSE | 1 | FALSE | 1 |
| 4374 | 4374 | 9.70833333 | 3.55292329 | 4 | 0.00296746 | 0 | 1 | 0 | 1 | 2 | 0.30249307 | 1 | 0.32796862 | 17 | 0.97530702 | FALSE | 0.14653312 | FALSE | 1 | FALSE | 1 | FALSE | 1 | FALSE | 1 | FALSE | 1 |
| 4375 | 4375 | 8.53846154 | 3.75428749 | 0 | 1 | 0 | 1 | 1 | 0.37874417 | 1 | 0.45746571 | 0 | 1 | 11 | 0.6683056 | FALSE | 1 | FALSE | 1 | FALSE | 1 | FALSE | 1 | FALSE | 1 | FALSE | 1 |
| 4376 | 4376 | 9.02631579 | 2.59033469 | 3 | 0.07268126 | 1 | 0.71433277 | 1 | 0.75193685 | 5 | 0.02858807 | 1 | 0.4673875 | 27 | 0.98956273 | FALSE | 0.76727438 | FALSE | 1 | FALSE | 1 | FALSE | 1 | FALSE | 1 | FALSE | 1 |
| 4377 | 4377 | 8.81818182 | 2.69066338 | 0 | 1 | 1 | 0.5154318 | 0 | 1 | 0 | 1 | 0 | 1 | 21 | 0.12048848 | FALSE | 1 | FALSE | 1 | FALSE | 1 | FALSE | 1 | FALSE | 1 | FALSE | 1 |
| 4378 | 4378 | 9.59405941 | 1.40844847 | 4 | 0.25411532 | 3 | 0.63331268 | 5 | 0.28995426 | 4 | 0.68085191 | 1 | 0.81092309 | 83 | 0.70668742 | FALSE | 1 | FALSE | 1 | FALSE | 1 | FALSE | 1 | FALSE | 1 | FALSE | 1 |
| 4379 | 4379 | 8.55 | 2.01183995 | 0 | 1 | 2 | 0.1353446 | 3 | 0.03322143 | 0 | 1 | 0 | 1 | 15 | 0.92113679 | FALSE | 1 | FALSE | 1 | FALSE | 1 | FALSE | 1 | FALSE | 1 | FALSE | 1 |
| 4380 | 4380 | 7.64516129 | 2.85734353 | 1 | 0.55378205 | 4 | 0.01688692 | 1 | 0.67911179 | 1 | 0.76781284 | 1 | 0.4016888 | 23 | 0.95750806 | FALSE | 1 | FALSE | 0.73194511 | FALSE | 1 | FALSE | 1 | FALSE | 1 | FALSE | 1 |
| 4381 | 4381 | 8.19047619 | 1.33163157 | 1 | 0.42086991 | 0 | 1 | 0 | 1 | 1 | 0.62783522 | 0 | 1 | 19 | 0.34094959 | FALSE | 1 | FALSE | 1 | FALSE | 1 | FALSE | 1 | FALSE | 1 | FALSE | 1 |
| 4382 | 4382 | 7.86666667 | 2.70472837 | 0 | 1 | 2 | 0.08295839 | 1 | 0.42266264 | 1 | 0.50623194 | 0 | 1 | 11 | 0.92879412 | FALSE | 1 | FALSE | 1 | FALSE | 1 | FALSE | 1 | FALSE | 1 | FALSE | 1 |
| 4383 | 4383 | 7.86666667 | 3.28362943 | 1 | 0.32293355 | 0 | 1 | 0 | 1 | 0 | 1 | 0 | 1 | 14 | 0.29525409 | FALSE | 1 | FALSE | 1 | FALSE | 1 | FALSE | 1 | FALSE | 1 | FALSE | 1 |
| 4384 | 4384 | 7.59183674 | 2.13731061 | 0 | 1 | 2 | 0.46321244 | 1 | 0.8283465 | 0 | 1 | 2 | 0.18560062 | 43 | 0.21763056 | FALSE | 1 | FALSE | 1 | FALSE | 1 | FALSE | 1 | FALSE | 1 | FALSE | 1 |
| 4385 | 4385 | 10.0555556 | 2.5049334 | 0 | 1 | 2 | 0.1135776 | 1 | 0.48280983 | 1 | 0.57131013 | 0 | 1 | 14 | 0.86287418 | FALSE | 1 | FALSE | 1 | FALSE | 1 | FALSE | 1 | FALSE | 1 | FALSE | 1 |
| 4386 | 4386 | 10.1666667 | 1.77169097 | 0 | 1 | 0 | 1 | 1 | 0.197145 | 0 | 1 | 0 | 1 | 5 | 0.76185568 | FALSE | 1 | FALSE | 1 | FALSE | 1 | FALSE | 1 | FALSE | 1 | FALSE | 1 |
| 4387 | 4387 | 10.0487805 | 1.88642415 | 0 | 1 | 0 | 1 | 1 | 0.69069222 | 2 | 0.43583874 | 0 | 1 | 29 | 0.24088541 | FALSE | 1 | FALSE | 1 | FALSE | 1 | FALSE | 1 | FALSE | 1 | FALSE | 1 |
| 4388 | 4388 | 7.52 | 3.17641307 | 0 | 1 | 1 | 0.56109832 | 0 | 1 | 3 | 0.10474541 | 1 | 0.33902672 | 20 | 0.8152498 | FALSE | 1 | FALSE | 1 | FALSE | 1 | FALSE | 1 | FALSE | 1 | FALSE | 1 |
| 4389 | 4389 | 7.38888889 | 2.58497558 | 0 | 1 | 0 | 1 | 0 | 1 | 0 | 1 | 0 | 1 | 18 | 0.04681656 | FALSE | 1 | FALSE | 1 | FALSE | 1 | FALSE | 1 | FALSE | 1 | FALSE | 1 |
| 4390 | 4390 | 11.2352941 | 3.54017965 | 0 | 1 | 1 | 0.42857733 | 0 | 1 | 0 | 1 | 0 | 1 | 16 | 0.23070182 | FALSE | 1 | FALSE | 1 | FALSE | 1 | FALSE | 1 | FALSE | 1 | FALSE | 1 |
| 4391 | 4391 | 8.82857143 | 1.36247075 | 0 | 1 | 2 | 0.28941836 | 2 | 0.33343871 | 1 | 0.78873814 | 1 | 0.42123867 | 27 | 0.75124583 | FALSE | 1 | FALSE | 1 | FALSE | 1 | FALSE | 1 | FALSE | 1 | FALSE | 1 |
| 4392 | 4392 | 9.81818182 | 1.74867128 | 0 | 1 | 1 | 0.5154318 | 0 | 1 | 0 | 1 | 0 | 1 | 21 | 0.12048848 | FALSE | 1 | FALSE | 1 | FALSE | 1 | FALSE | 1 | FALSE | 1 | FALSE | 1 |
| 4393 | 4393 | 9.86363636 | 2.50989776 | 0 | 1 | 1 | 0.5154318 | 1 | 0.55340851 | 0 | 1 | 0 | 1 | 20 | 0.30915202 | FALSE | 1 | FALSE | 1 | FALSE | 1 | FALSE | 1 | FALSE | 1 | FALSE | 1 |
| 4394 | 4394 | 9.88888889 | 2.40113142 | 1 | 0.37380237 | 2 | 0.1135776 | 0 | 1 | 0 | 1 | 1 | 0.25766374 | 14 | 0.86287418 | FALSE | 1 | FALSE | 1 | FALSE | 1 | FALSE | 1 | FALSE | 1 | FALSE | 1 |
| 4395 | 4395 | 10.5 | 1.55456318 | 0 | 1 | 0 | 1 | 0 | 1 | 0 | 1 | 0 | 1 | 11 | 0.15414489 | FALSE | 1 | FALSE | 1 | FALSE | 1 | FALSE | 1 | FALSE | 1 | FALSE | 1 |
| 4396 | 4396 | 8.75 | 2.33184476 | 4 | 0.00856611 | 3 | 0.08334648 | 2 | 0.32011871 | 3 | 0.17988885 | 0 | 1 | 20 | 0.99942006 | FALSE | 0.27019545 | FALSE | 1 | FALSE | 1 | FALSE | 1 | FALSE | 1 | FALSE | 1 |
| 4397 | 4397 | 7.6 | 1.37275069 | 1 | 0.69044531 | 2 | 0.43020462 | 0 | 1 | 3 | 0.34154252 | 0 | 1 | 39 | 0.4317533 | FALSE | 1 | FALSE | 1 | FALSE | 1 | FALSE | 1 | FALSE | 1 | FALSE | 1 |
| 4398 | 4398 | 9.26086957 | 2.55746803 | 1 | 0.66519151 | 3 | 0.1535026 | 1 | 0.78589454 | 0 | 1 | 0 | 1 | 37 | 0.34041811 | FALSE | 1 | FALSE | 1 | FALSE | 1 | FALSE | 1 | FALSE | 1 | FALSE | 1 |
| 4399 | 4399 | 10.4736842 | 1.72804791 | 2 | 0.08405377 | 0 | 1 | 1 | 0.50143826 | 0 | 1 | 0 | 1 | 16 | 0.65565778 | FALSE | 0.79236883 | FALSE | 1 | FALSE | 1 | FALSE | 1 | FALSE | 1 | FALSE | 1 |
| 4400 | 4400 | 7.35294118 | 2.27366448 | 0 | 1 | 0 | 1 | 1 | 0.44344817 | 1 | 0.52895088 | 2 | 0.02752114 | 12 | 0.90922186 | FALSE | 1 | FALSE | 1 | FALSE | 1 | FALSE | 1 | FALSE | 0.79106376 | FALSE | 1 |
| 4401 | 4401 | 5.91891892 | 2.53492767 | 1 | 0.85522402 | 1 | 0.9134145 | 1 | 0.93427349 | 2 | 0.86064871 | 0 | 1 | 69 | 0.01821645 | FALSE | 1 | FALSE | 1 | FALSE | 1 | FALSE | 1 | FALSE | 1 | FALSE | 1 |
| 4402 | 4402 | 9.16666667 | 2.57660414 | 1 | 0.46441868 | 0 | 1 | 1 | 0.58502114 | 1 | 0.67692816 | 0 | 1 | 21 | 0.47053137 | FALSE | 1 | FALSE | 1 | FALSE | 1 | FALSE | 1 | FALSE | 1 | FALSE | 1 |
| 4403 | 4403 | 7.48780488 | 3.429584 | 0 | 1 | 1 | 0.74131698 | 1 | 0.77786601 | 2 | 0.56788092 | 1 | 0.49330646 | 36 | 0.36455032 | FALSE | 1 | FALSE | 1 | FALSE | 1 | FALSE | 1 | FALSE | 1 | FALSE | 1 |
| 4404 | 4404 | 10.1111111 | 3.03477784 | 0 | 1 | 0 | 1 | 0 | 1 | 2 | 0.06108592 | 2 | 0.00889576 | 5 | 0.99330426 | FALSE | 1 | FALSE | 1 | FALSE | 1 | FALSE | 1 | FALSE | 0.52052899 | FALSE | 1 |
| 4405 | 4405 | 5.5942029 | 2.81979977 | 4 | 0.47418487 | 9 | 0.03441613 | 3 | 0.87892688 | 6 | 0.61339232 | 2 | 0.66596353 | 114 | 0.76136774 | FALSE | 1 | FALSE | 0.94259695 | FALSE | 1 | FALSE | 1 | FALSE | 1 | FALSE | 1 |
| 4406 | 4406 | 6.75 | 3.91083659 | 3 | 0.16017108 | 3 | 0.25367018 | 1 | 0.86240917 | 3 | 0.45415642 | 0 | 1 | 44 | 0.78654547 | FALSE | 1 | FALSE | 1 | FALSE | 1 | FALSE | 1 | FALSE | 1 | FALSE | 1 |
| 4407 | 4407 | 7.45 | 2.26881026 | 1 | 0.64722818 | 2 | 0.37295654 | 1 | 0.76953771 | 5 | 0.03481208 | 1 | 0.48480878 | 30 | 0.96151659 | FALSE | 1 | FALSE | 1 | FALSE | 1 | FALSE | 1 | FALSE | 1 | FALSE | 1 |
| 4408 | 4408 | 10.3529412 | 2.2476439 | 1 | 0.35728305 | 0 | 1 | 0 | 1 | 0 | 1 | 1 | 0.24525764 | 15 | 0.49060435 | FALSE | 1 | FALSE | 1 | FALSE | 1 | FALSE | 1 | FALSE | 1 | FALSE | 1 |
| 4409 | 4409 | 9.52631579 | 3.21785187 | 10 | 7.15E-12 | 2 | 0.12434464 | 0 | 1 | 1 | 0.59104357 | 0 | 1 | 6 | 0.99999997 | TRUE | 5.00E-08 | FALSE | 1 | FALSE | 1 | FALSE | 1 | FALSE | 1 | FALSE | 1 |
| 4410 | 4410 | 9.53333333 | 3.59382186 | 1 | 0.32293355 | 0 | 1 | 0 | 1 | 2 | 0.14906763 | 1 | 0.2198258 | 11 | 0.92879412 | FALSE | 1 | FALSE | 1 | FALSE | 1 | FALSE | 1 | FALSE | 1 | FALSE | 1 |
| 4411 | 4411 | 7.74647887 | 1.58085652 | 1 | 0.83919255 | 3 | 0.39589822 | 3 | 0.46296673 | 1 | 0.9633793 | 1 | 0.68753645 | 61 | 0.32827441 | FALSE | 1 | FALSE | 1 | FALSE | 1 | FALSE | 1 | FALSE | 1 | FALSE | 1 |
| 4412 | 4412 | 8.625 | 2.0577597 | 0 | 1 | 0 | 1 | 0 | 1 | 1 | 0.31351192 | 0 | 1 | 7 | 0.63752397 | FALSE | 1 | FALSE | 1 | FALSE | 1 | FALSE | 1 | FALSE | 1 | FALSE | 1 |
| 4413 | 4413 | 8.13043478 | 1.96279189 | 2 | 0.11647716 | 0 | 1 | 0 | 1 | 1 | 0.6613267 | 0 | 1 | 20 | 0.50589402 | FALSE | 0.93181727 | FALSE | 1 | FALSE | 1 | FALSE | 1 | FALSE | 1 | FALSE | 1 |
| 4414 | 4414 | 9.57575758 | 1.6148985 | 0 | 1 | 1 | 0.6629941 | 3 | 0.11365871 | 2 | 0.45159487 | 1 | 0.42123867 | 26 | 0.86852567 | FALSE | 1 | FALSE | 1 | FALSE | 1 | FALSE | 1 | FALSE | 1 | FALSE | 1 |
| 4415 | 4415 | 8.06451613 | 4.17315139 | 0 | 1 | 0 | 1 | 1 | 0.67911179 | 1 | 0.76781284 | 0 | 1 | 29 | 0.11661149 | FALSE | 1 | FALSE | 1 | FALSE | 1 | FALSE | 1 | FALSE | 1 | FALSE | 1 |
| 4416 | 4416 | 8.84615385 | 2.28190723 | 0 | 1 | 1 | 0.3480709 | 1 | 0.37874417 | 1 | 0.45746571 | 0 | 1 | 10 | 0.86791642 | FALSE | 1 | FALSE | 1 | FALSE | 1 | FALSE | 1 | FALSE | 1 | FALSE | 1 |

| | | | | | | | | | | | | | | | | | | | | | | | | | | | |
|---|---|---|---|---|---|---|---|---|---|---|---|---|---|---|---|---|---|---|---|---|---|---|---|---|---|---|---|
| 4417 | 4417 | 6.42857143 | 4.46757022 | 0 | 1 | 0 | 1 | 0 | 1 | 1 | 0.62783522 | 1 | 0.29368245 | 19 | 0.34094959 | FALSE | 1 | FALSE | 1 | FALSE | 1 | FALSE | 1 | FALSE | 1 | FALSE | 1 |
| 4418 | 4418 | 4.25100134 | 1.56258479 | 15 | 0.87699944 | 22 | 0.71665898 | 24 | 0.75474142 | 32 | 0.69505475 | 14 | 0.34087838 | 641 | 0.15904095 | FALSE | 1 | FALSE | 1 | FALSE | 1 | FALSE | 1 | FALSE | 1 | FALSE | 1 |
| 4419 | 4419 | 7.72727273 | 1.71044434 | 0 | 1 | 0 | 1 | 0 | 1 | 0 | 1 | 1 | 0.16638736 | 10 | 0.46859937 | FALSE | 1 | FALSE | 1 | FALSE | 1 | FALSE | 1 | FALSE | 1 | FALSE | 1 |
| 4420 | 4420 | 8.7 | 2.93428015 | 0 | 1 | 0 | 1 | 2 | 0.04781153 | 0 | 1 | 0 | 1 | 8 | 0.80394719 | FALSE | 1 | FALSE | 1 | FALSE | 1 | FALSE | 1 | FALSE | 1 | FALSE | 1 |
| 4421 | 4421 | 7.56862745 | 1.14197802 | 0 | 1 | 1 | 0.81422651 | 2 | 0.55199489 | 2 | 0.68678423 | 1 | 0.57100288 | 45 | 0.29584937 | FALSE | 1 | FALSE | 1 | FALSE | 1 | FALSE | 1 | FALSE | 1 | FALSE | 1 |
| 4422 | 4422 | 8.925 | 2.78735986 | 2 | 0.27371785 | 2 | 0.37295654 | 2 | 0.42396974 | 3 | 0.27785536 | 5 | 0.00045053 | 26 | 0.99938297 | FALSE | 1 | FALSE | 1 | FALSE | 1 | FALSE | 1 | FALSE | 0.10840111 | FALSE | 1 |
| 4423 | 4423 | 7.54761905 | 1.84135536 | 0 | 1 | 0 | 1 | 1 | 0.78589454 | 1 | 0.86193352 | 0 | 1 | 40 | 0.0301759 | FALSE | 1 | FALSE | 1 | FALSE | 1 | FALSE | 1 | FALSE | 1 | FALSE | 1 |
| 4424 | 4424 | 9.96551724 | 2.15730929 | 1 | 0.51745948 | 2 | 0.2288771 | 0 | 1 | 1 | 0.73248983 | 0 | 1 | 24 | 0.55018107 | FALSE | 1 | FALSE | 1 | FALSE | 1 | FALSE | 1 | FALSE | 1 | FALSE | 1 |
| 4425 | 4425 | 7.4375 | 1.46115798 | 7 | 0.06981993 | 6 | 0.30529969 | 5 | 0.57614338 | 6 | 0.6343804 | 8 | 0.00210489 | 109 | 0.99062491 | FALSE | 0.74864513 | FALSE | 1 | FALSE | 1 | FALSE | 1 | FALSE | 0.2617074 | FALSE | 1 |
| 4426 | 4426 | 9.6 | 2.33238076 | 0 | 1 | 0 | 1 | 0 | 1 | 1 | 0.50623194 | 0 | 1 | 14 | 0.29525409 | FALSE | 1 | FALSE | 1 | FALSE | 1 | FALSE | 1 | FALSE | 1 | FALSE | 1 |
| 4427 | 4427 | 8.46808511 | 2.98828253 | 1 | 0.70621968 | 3 | 0.19374024 | 1 | 0.82190381 | 3 | 0.36699324 | 2 | 0.17968152 | 37 | 0.89489966 | FALSE | 1 | FALSE | 1 | FALSE | 1 | FALSE | 1 | FALSE | 1 | FALSE | 1 |
| 4428 | 4428 | 10.5833333 | 3.30298686 | 1 | 0.26795838 | 0 | 1 | 1 | 0.35555293 | 1 | 0.43131129 | 0 | 1 | 9 | 0.89630573 | FALSE | 1 | FALSE | 1 | FALSE | 1 | FALSE | 1 | FALSE | 1 | FALSE | 1 |
| 4429 | 4429 | 11.1538462 | 2.597222 | 0 | 1 | 1 | 0.32623933 | 1 | 0.35555293 | 0 | 1 | 0 | 1 | 10 | 0.71436964 | FALSE | 1 | FALSE | 1 | FALSE | 1 | FALSE | 1 | FALSE | 1 | FALSE | 1 |
| 4430 | 4430 | 11.5454546 | 1.61603535 | 0 | 1 | 0 | 1 | 0 | 1 | 0 | 1 | 0 | 1 | 11 | 0.15414489 | FALSE | 1 | FALSE | 1 | FALSE | 1 | FALSE | 1 | FALSE | 1 | FALSE | 1 |
| 4431 | 4431 | 7 | 2.54732976 | 3 | 0.10767332 | 4 | 0.05650644 | 2 | 0.48463745 | 2 | 0.61900576 | 1 | 0.52593041 | 33 | 0.98284787 | FALSE | 0.88820416 | FALSE | 1 | FALSE | 1 | FALSE | 1 | FALSE | 1 | FALSE | 1 |
| 4432 | 4432 | 7.84 | 2.52475741 | 0 | 1 | 2 | 0.19297191 | 2 | 0.22597287 | 0 | 1 | 0 | 1 | 21 | 0.64957167 | FALSE | 1 | FALSE | 1 | FALSE | 1 | FALSE | 1 | FALSE | 1 | FALSE | 1 |
| 4433 | 4433 | 8.78571429 | 1.6335138 | 2 | 0.16048326 | 3 | 0.06039219 | 2 | 0.2663542 | 2 | 0.3705495 | 0 | 1 | 19 | 0.99262381 | FALSE | 1 | FALSE | 1 | FALSE | 1 | FALSE | 1 | FALSE | 1 | FALSE | 1 |
| 4434 | 4434 | 6.09803922 | 1.99759563 | 3 | 0.33025289 | 5 | 0.11181294 | 4 | 0.31552613 | 6 | 0.15385579 | 5 | 0.00931828 | 56 | 0.99932182 | FALSE | 1 | FALSE | 1 | FALSE | 1 | FALSE | 1 | FALSE | 0.5373705 | FALSE | 1 |
| 4435 | 4435 | 7.28571429 | 2.75236448 | 0 | 1 | 1 | 0.68454804 | 3 | 0.12958859 | 1 | 0.80778317 | 0 | 1 | 30 | 0.52765882 | FALSE | 1 | FALSE | 1 | FALSE | 1 | FALSE | 1 | FALSE | 1 | FALSE | 1 |
| 4436 | 4436 | 10.9677419 | 2.60871827 | 0 | 1 | 1 | 0.61539065 | 0 | 1 | 2 | 0.3871811 | 0 | 1 | 26 | 0.31495157 | FALSE | 1 | FALSE | 1 | FALSE | 1 | FALSE | 1 | FALSE | 1 | FALSE | 1 |
| 4437 | 4437 | 10.5 | 2.84045451 | 0 | 1 | 0 | 1 | 0 | 1 | 0 | 1 | 0 | 1 | 22 | 0.02368118 | FALSE | 1 | FALSE | 1 | FALSE | 1 | FALSE | 1 | FALSE | 1 | FALSE | 1 |
| 4438 | 4438 | 8.09090909 | 1.33855726 | 1 | 0.75539413 | 2 | 0.52572379 | 1 | 0.86240917 | 7 | 0.01112162 | 0 | 1 | 43 | 0.87372949 | FALSE | 1 | FALSE | 1 | FALSE | 1 | FALSE | 1 | FALSE | 1 | FALSE | 1 |
| 4439 | 4439 | 4.66666667 | 3.87585116 | 2 | 0.17884234 | 4 | 0.01508077 | 0 | 1 | 2 | 0.4036176 | 0 | 1 | 22 | 0.96518743 | FALSE | 1 | FALSE | 0.7256023 | FALSE | 1 | FALSE | 1 | FALSE | 1 | FALSE | 1 |
| 4440 | 4440 | 12.7857143 | 3.05142991 | 1 | 0.51745948 | 1 | 0.60247911 | 0 | 1 | 0 | 1 | 0 | 1 | 26 | 0.16405547 | FALSE | 1 | FALSE | 1 | FALSE | 1 | FALSE | 1 | FALSE | 1 | FALSE | 1 |
| 4441 | 4441 | 11.6428571 | 1.79710652 | 2 | 0.04858661 | 1 | 0.36919824 | 1 | 0.40110418 | 0 | 1 | 0 | 1 | 10 | 0.94579623 | FALSE | 0.649301 | FALSE | 1 | FALSE | 1 | FALSE | 1 | FALSE | 1 | FALSE | 1 |
| 4442 | 4442 | 10.826087 | 2.29654133 | 0 | 1 | 2 | 0.16948764 | 0 | 1 | 1 | 0.6613267 | 1 | 0.31672722 | 19 | 0.71542496 | FALSE | 1 | FALSE | 1 | FALSE | 1 | FALSE | 1 | FALSE | 1 | FALSE | 1 |
| 4443 | 4443 | 10.5 | 3.13391585 | 1 | 0.3050805 | 0 | 1 | 0 | 1 | 0 | 1 | 1 | 0.20679328 | 12 | 0.62236227 | FALSE | 1 | FALSE | 1 | FALSE | 1 | FALSE | 1 | FALSE | 1 | FALSE | 1 |
| 4444 | 4444 | 10.4242424 | 2.17465153 | 0 | 1 | 2 | 0.25306331 | 2 | 0.29330414 | 4 | 0.04659941 | 0 | 1 | 22 | 0.96518743 | FALSE | 1 | FALSE | 1 | FALSE | 1 | FALSE | 1 | FALSE | 1 | FALSE | 1 |
| 4445 | 4445 | 8.68421053 | 3.61360924 | 1 | 0.62830697 | 1 | 0.71433277 | 2 | 0.39868506 | 3 | 0.25266879 | 0 | 1 | 31 | 0.76640548 | FALSE | 1 | FALSE | 1 | FALSE | 1 | FALSE | 1 | FALSE | 1 | FALSE | 1 |
| 4446 | 4446 | 6.30232558 | 3.23163781 | 1 | 0.67382914 | 3 | 0.16132793 | 1 | 0.79363407 | 3 | 0.31602175 | 1 | 0.50988734 | 34 | 0.87722689 | FALSE | 1 | FALSE | 1 | FALSE | 1 | FALSE | 1 | FALSE | 1 | FALSE | 1 |
| 4447 | 4447 | 10.0952381 | 3.74044512 | 0 | 1 | 0 | 1 | 2 | 0.17308361 | 3 | 0.06912908 | 0 | 1 | 16 | 0.90353674 | FALSE | 1 | FALSE | 1 | FALSE | 1 | FALSE | 1 | FALSE | 1 | FALSE | 1 |
| 4448 | 4448 | 9.70967742 | 2.33063121 | 1 | 0.55378205 | 2 | 0.2651851 | 2 | 0.30673593 | 0 | 1 | 1 | 0.4016888 | 25 | 0.80006983 | FALSE | 1 | FALSE | 1 | FALSE | 1 | FALSE | 1 | FALSE | 1 | FALSE | 1 |
| 4449 | 4449 | 9.41176471 | 2.99133697 | 1 | 0.35728305 | 0 | 1 | 0 | 1 | 1 | 0.55062772 | 0 | 1 | 15 | 0.49060435 | FALSE | 1 | FALSE | 1 | FALSE | 1 | FALSE | 1 | FALSE | 1 | FALSE | 1 |
| 4450 | 4450 | 7.2195122 | 4.3979607 | 4 | 0.02019941 | 0 | 1 | 3 | 0.1814659 | 6 | 0.01031399 | 0 | 1 | 28 | 0.997364 | FALSE | 0.44482142 | FALSE | 1 | FALSE | 1 | FALSE | 1 | FALSE | 1 | FALSE | 1 |
| 4451 | 4451 | 8.52 | 1.86804711 | 0 | 1 | 3 | 0.21051417 | 1 | 0.83455705 | 1 | 0.90086419 | 1 | 0.55647171 | 43 | 0.337822 | FALSE | 1 | FALSE | 1 | FALSE | 1 | FALSE | 1 | FALSE | 1 | FALSE | 1 |
| 4452 | 4452 | 8.92857143 | 2.17554435 | 1 | 0.66519151 | 0 | 1 | 2 | 0.44869594 | 2 | 0.5811115 | 3 | 0.03099591 | 34 | 0.80048036 | FALSE | 1 | FALSE | 1 | FALSE | 1 | FALSE | 1 | FALSE | 0.82559785 | FALSE | 1 |
| 4453 | 4453 | 6.6969697 | 2.7687088 | 0 | 1 | 1 | 0.54637326 | 2 | 0.21260257 | 1 | 0.67692816 | 0 | 1 | 20 | 0.68270729 | FALSE | 1 | FALSE | 1 | FALSE | 1 | FALSE | 1 | FALSE | 1 | FALSE | 1 |
| 4454 | 4454 | 8.02857143 | 3.00937311 | 2 | 0.22589135 | 1 | 0.68454804 | 1 | 0.72299506 | 1 | 0.80778317 | 0 | 1 | 30 | 0.52765882 | FALSE | 1 | FALSE | 1 | FALSE | 1 | FALSE | 1 | FALSE | 1 | FALSE | 1 |
| 4455 | 4455 | 10.1842105 | 2.18674552 | 0 | 1 | 3 | 0.12352882 | 1 | 0.75193685 | 1 | 0.83318955 | 1 | 0.4673875 | 32 | 0.61800916 | FALSE | 1 | FALSE | 1 | FALSE | 1 | FALSE | 1 | FALSE | 1 | FALSE | 1 |
| 4456 | 4456 | 9 | 2.40098019 | 0 | 1 | 0 | 1 | 2 | 0.12280513 | 0 | 1 | 1 | 0.24525764 | 14 | 0.7313145 | FALSE | 1 | FALSE | 1 | FALSE | 1 | FALSE | 1 | FALSE | 1 | FALSE | 1 |
| 4457 | 4457 | 4.33898305 | 1.38221352 | 5 | 0.72650266 | 10 | 0.22951049 | 6 | 0.85543013 | 13 | 0.28281869 | 2 | 0.90302623 | 199 | 0.49256335 | FALSE | 1 | FALSE | 1 | FALSE | 1 | FALSE | 1 | FALSE | 1 | FALSE | 1 |
| 4458 | 4458 | 5.07575758 | 2.63605885 | 0 | 1 | 4 | 0.16434514 | 4 | 0.21207653 | 2 | 0.8136513 | 0 | 1 | 56 | 0.54112142 | FALSE | 1 | FALSE | 1 | FALSE | 1 | FALSE | 1 | FALSE | 1 | FALSE | 1 |
| 4459 | 4459 | 10.6666667 | 4.12983723 | 0 | 1 | 1 | 0.32623933 | 1 | 0.35555293 | 0 | 1 | 0 | 1 | 10 | 0.71436964 | FALSE | 1 | FALSE | 1 | FALSE | 1 | FALSE | 1 | FALSE | 1 | FALSE | 1 |
| 4460 | 4460 | 8.77777778 | 1.93090524 | 0 | 1 | 1 | 0.42857733 | 0 | 1 | 0 | 1 | 0 | 1 | 16 | 0.23070182 | FALSE | 1 | FALSE | 1 | FALSE | 1 | FALSE | 1 | FALSE | 1 | FALSE | 1 |
| 4461 | 4461 | 7.56521739 | 1.24502792 | 0 | 1 | 1 | 0.5154318 | 1 | 0.55340851 | 1 | 0.64497439 | 0 | 1 | 19 | 0.54233419 | FALSE | 1 | FALSE | 1 | FALSE | 1 | FALSE | 1 | FALSE | 1 | FALSE | 1 |
| 4462 | 4462 | 10.0277778 | 2.00674019 | 2 | 0.23542589 | 1 | 0.69480472 | 1 | 0.73299778 | 1 | 0.81665383 | 3 | 0.0206828 | 28 | 0.90259714 | FALSE | 1 | FALSE | 1 | FALSE | 1 | FALSE | 1 | FALSE | 0.7366672 | FALSE | 1 |
| 4463 | 4463 | 8.9 | 2.11896201 | 0 | 1 | 2 | 0.03949327 | 0 | 1 | 0 | 1 | 0 | 1 | 8 | 0.80394719 | FALSE | 1 | FALSE | 0.96864929 | FALSE | 1 | FALSE | 1 | FALSE | 1 | FALSE | 1 |
| 4464 | 4464 | 9.3 | 3.14801525 | 1 | 0.40558521 | 1 | 0.48239024 | 1 | 0.51939841 | 0 | 1 | 1 | 0.28187291 | 16 | 0.80826361 | FALSE | 1 | FALSE | 1 | FALSE | 1 | FALSE | 1 | FALSE | 1 | FALSE | 1 |
| 4465 | 4465 | 6.65517241 | 2.71998811 | 2 | 0.44060625 | 1 | 0.85269953 | 0 | 1 | 1 | 0.93527934 | 0 | 1 | 54 | 0.03955468 | FALSE | 1 | FALSE | 1 | FALSE | 1 | FALSE | 1 | FALSE | 1 | FALSE | 1 |
| 4466 | 4466 | 5.35555556 | 3.23285602 | 0 | 1 | 3 | 0.17732615 | 2 | 0.48463745 | 2 | 0.61900576 | 2 | 0.16793736 | 36 | 0.84604146 | FALSE | 1 | FALSE | 1 | FALSE | 1 | FALSE | 1 | FALSE | 1 | FALSE | 1 |
| 4467 | 4467 | 9.37209302 | 2.04624734 | 1 | 0.67382914 | 0 | 1 | 3 | 0.19985581 | 3 | 0.31602175 | 0 | 1 | 36 | 0.6451239 | FALSE | 1 | FALSE | 1 | FALSE | 1 | FALSE | 1 | FALSE | 1 | FALSE | 1 |
| 4468 | 4468 | 9.1 | 2.62488095 | 0 | 1 | 1 | 0.28036922 | 0 | 1 | 2 | 0.0740981 | 0 | 1 | 7 | 0.94326215 | FALSE | 1 | FALSE | 1 | FALSE | 1 | FALSE | 1 | FALSE | 1 | FALSE | 1 |
| 4469 | 4469 | 8.24590164 | 1.30139569 | 1 | 0.79637728 | 6 | 0.01348308 | 6 | 0.02154979 | 4 | 0.30679865 | 1 | 0.63687695 | 43 | 0.99827461 | FALSE | 1 | FALSE | 0.69207862 | FALSE | 1 | FALSE | 1 | FALSE | 1 | FALSE | 1 |
| 4470 | 4470 | 9.25 | 1.72199819 | 0 | 1 | 1 | 0.69480472 | 1 | 0.73299778 | 0 | 1 | 0 | 1 | 34 | 0.06417017 | FALSE | 1 | FALSE | 1 | FALSE | 1 | FALSE | 1 | FALSE | 1 | FALSE | 1 |
| 4471 | 4471 | 9.65217391 | 2.74430154 | 0 | 1 | 0 | 1 | 2 | 0.19931429 | 0 | 1 | 1 | 0.31672722 | 20 | 0.50589402 | FALSE | 1 | FALSE | 1 | FALSE | 1 | FALSE | 1 | FALSE | 1 | FALSE | 1 |
| 4472 | 4472 | 7.02777778 | 2.26674156 | 0 | 1 | 2 | 0.32556427 | 0 | 1 | 1 | 0.81665383 | 0 | 1 | 33 | 0.16407838 | FALSE | 1 | FALSE | 1 | FALSE | 1 | FALSE | 1 | FALSE | 1 | FALSE | 1 |
| 4473 | 4473 | 10 | 3 | 1 | 0.14434519 | 0 | 1 | 0 | 1 | 1 | 0.24578796 | 0 | 1 | 4 | 0.94743772 | FALSE | 1 | FALSE | 1 | FALSE | 1 | FALSE | 1 | FALSE | 1 | FALSE | 1 |
| 4474 | 4474 | 1.10980536 | 0.99984725 | 77 | 0.13640899 | 103 | 0.01934946 | 101 | 0.33910508 | 111 | 0.95270133 | 52 | 0.08477503 | 2268 | 0.92338415 | FALSE | 0.99241777 | FALSE | 0.77701374 | FALSE | 1 | FALSE | 1 | FALSE | 1 | FALSE | 1 |
| 4475 | 4475 | 7.03225807 | 2.0077892 | 1 | 0.55378205 | 0 | 1 | 0 | 1 | 1 | 0.76781284 | 0 | 1 | 29 | 0.11661149 | FALSE | 1 | FALSE | 1 | FALSE | 1 | FALSE | 1 | FALSE | 1 | FALSE | 1 |
| 4476 | 4476 | 10.2083333 | 2.88645063 | 2 | 0.12501436 | 0 | 1 | 1 | 0.58502114 | 0 | 1 | 0 | 1 | 21 | 0.47053137 | FALSE | 0.95622564 | FALSE | 1 | FALSE | 1 | FALSE | 1 | FALSE | 1 | FALSE | 1 |
| 4477 | 4477 | 8.61904762 | 2.59075629 | 0 | 1 | 0 | 1 | 2 | 0.17308361 | 1 | 0.62783522 | 0 | 1 | 18 | 0.57962763 | FALSE | 1 | FALSE | 1 | FALSE | 1 | FALSE | 1 | FALSE | 1 | FALSE | 1 |
| 4478 | 4478 | 7.7 | 1.41774469 | 2 | 0.09187136 | 0 | 1 | 0 | 1 | 1 | 0.60987146 | 0 | 1 | 17 | 0.61750632 | FALSE | 0.83248965 | FALSE | 1 | FALSE | 1 | FALSE | 1 | FALSE | 1 | FALSE | 1 |
| 4479 | 4479 | 8.72580645 | 0.86452215 | 2 | 0.47501604 | 0 | 1 | 2 | 0.65873577 | 2 | 0.78520976 | 1 | 0.64288531 | 55 | 0.22629228 | FALSE | 1 | FALSE | 1 | FALSE | 1 | FALSE | 1 | FALSE | 1 | FALSE | 1 |
| 4480 | 4480 | 6.6 | 2.62297541 | 0 | 1 | 0 | 1 | 1 | 0.59998149 | 4 | 0.02569595 | 0 | 1 | 20 | 0.8152498 | FALSE | 1 | FALSE | 1 | FALSE | 1 | FALSE | 1 | FALSE | 1 | FALSE | 1 |
| 4481 | 4481 | 9 | 2.44948974 | 0 | 1 | 1 | 0.25628163 | 1 | 0.28067846 | 0 | 1 | 1 | 0.1383237 | 6 | 0.96128317 | FALSE | 1 | FALSE | 1 | FALSE | 1 | FALSE | 1 | FALSE | 1 | FALSE | 1 |
| 4482 | 4482 | 9.84210526 | 2.32295542 | 1 | 0.38989951 | 1 | 0.46503795 | 1 | 0.50143826 | 0 | 1 | 0 | 1 | 16 | 0.65565778 | FALSE | 1 | FALSE | 1 | FALSE | 1 | FALSE | 1 | FALSE | 1 | FALSE | 1 |
| 4483 | 4483 | 3.31558935 | 1.50133791 | 5 | 0.80870104 | 6 | 0.85795007 | 13 | 0.14792363 | 12 | 0.54666171 | 7 | 0.13790763 | 219 | 0.67770142 | FALSE | 1 | FALSE | 1 | FALSE | 1 | FALSE | 1 | FALSE | 1 | FALSE | 1 |
| 4484 | 4484 | 9.88235294 | 1.84334584 | 0 | 1 | 1 | 0.42857733 | 1 | 0.46348836 | 0 | 1 | 0 | 1 | 15 | 0.49060435 | FALSE | 1 | FALSE | 1 | FALSE | 1 | FALSE | 1 | FALSE | 1 | FALSE | 1 |
| 4485 | 4485 | 5.03042198 | 0.42730288 | 21 | 0.8874474 | 29 | 0.79948789 | 38 | 0.42229439 | 50 | 0.31892261 | 17 | 0.50655639 | 862 | 0.37880742 | FALSE | 1 | FALSE | 1 | FALSE | 1 | FALSE | 1 | FALSE | 1 | FALSE | 1 |
| 4486 | 4486 | 9.47368421 | 2.45739316 | 0 | 1 | 1 | 0.46503795 | 0 | 1 | 0 | 1 | 0 | 1 | 18 | 0.17880694 | FALSE | 1 | FALSE | 1 | FALSE | 1 | FALSE | 1 | FALSE | 1 | FALSE | 1 |
| 4487 | 4487 | 9.64285714 | 2.19112309 | 1 | 0.3050805 | 0 | 1 | 1 | 0.40110418 | 0 | 1 | 0 | 1 | 12 | 0.62236227 | FALSE | 1 | FALSE | 1 | FALSE | 1 | FALSE | 1 | FALSE | 1 | FALSE | 1 |
| 4488 | 4488 | 9.5 | 3.27023615 | 0 | 1 | 0 | 1 | 0 | 1 | 1 | 0.57131013 | 0 | 1 | 17 | 0.20329205 | FALSE | 1 | FALSE | 1 | FALSE | 1 | FALSE | 1 | FALSE | 1 | FALSE | 1 |
| 4489 | 4489 | 8.41176471 | 2.78776367 | 1 | 0.35728305 | 2 | 0.10307112 | 0 | 1 | 1 | 0.55062772 | 0 | 1 | 13 | 0.88719222 | FALSE | 1 | FALSE | 1 | FALSE | 1 | FALSE | 1 | FALSE | 1 | FALSE | 1 |
| 4490 | 4490 | 10.1111111 | 2.96064724 | 0 | 1 | 0 | 1 | 1 | 0.48280983 | 1 | 0.57131013 | 0 | 1 | 16 | 0.44998001 | FALSE | 1 | FALSE | 1 | FALSE | 1 | FALSE | 1 | FALSE | 1 | FALSE | 1 |
| 4491 | 4491 | 12 | 2.9664794 | 0 | 1 | 2 | 0.19297191 | 0 | 1 | 1 | 0.69181315 | 0 | 1 | 22 | 0.43642846 | FALSE | 1 | FALSE | 1 | FALSE | 1 | FALSE | 1 | FALSE | 1 | FALSE | 1 |
| 4492 | 4492 | 8.6969697 | 2.18097627 | 1 | 0.57647348 | 0 | 1 | 0 | 1 | 1 | 0.78873814 | 2 | 0.10132035 | 29 | 0.39648374 | FALSE | 1 | FALSE | 1 | FALSE | 1 | FALSE | 1 | FALSE | 1 | FALSE | 1 |
| 4493 | 4493 | 7.12698413 | 2.38708206 | 8 | 0.00018453 | 8 | 0.00088051 | 1 | 0.90129726 | 7 | 0.02452175 | 0 | 1 | 39 | 0.99999679 | TRUE | 0.0251946 | FALSE | 0.27742868 | FALSE | 1 | FALSE | 1 | FALSE | 1 | FALSE | 1 |
| 4494 | 4494 | 10.3 | 3.1638584 | 0 | 1 | 1 | 0.28036922 | 0 | 1 | 0 | 1 | 0 | 1 | 9 | 0.5215547 | FALSE | 1 | FALSE | 1 | FALSE | 1 | FALSE | 1 | FALSE | 1 | FALSE | 1 |
| 4495 | 4495 | 10.4285714 | 3.11022015 | 0 | 1 | 0 | 1 | 0 | 1 | 1 | 0.28044368 | 1 | 0.10932414 | 5 | 0.91847773 | FALSE | 1 | FALSE | 1 | FALSE | 1 | FALSE | 1 | FALSE | 1 | FALSE | 1 |
| 4496 | 4496 | 9.93333333 | 2.23507395 | 1 | 0.54198714 | 1 | 0.6278847 | 0 | 1 | 1 | 0.75658797 | 0 | 1 | 27 | 0.28862208 | FALSE | 1 | FALSE | 1 | FALSE | 1 | FALSE | 1 | FALSE | 1 | FALSE | 1 |
| 4497 | 4497 | 4.46666667 | 1.9821425 | 2 | 0.57749853 | 1 | 0.91624631 | 1 | 0.93666084 | 3 | 0.67644754 | 2 | 0.34918837 | 66 | 0.24505488 | FALSE | 1 | FALSE | 1 | FALSE | 1 | FALSE | 1 | FALSE | 1 | FALSE | 1 |
| 4498 | 4498 | 9.28571429 | 1.57790872 | 2 | 0.04858661 | 1 | 0.36919824 | 0 | 1 | 0 | 1 | 0 | 1 | 11 | 0.83676449 | FALSE | 0.649301 | FALSE | 1 | FALSE | 1 | FALSE | 1 | FALSE | 1 | FALSE | 1 |
| 4499 | 4499 | 9.02941177 | 4.23907103 | 0 | 1 | 0 | 1 | 3 | 0.12152681 | 3 | 0.20353499 | 1 | 0.43077488 | 27 | 0.85023803 | FALSE | 1 | FALSE | 1 | FALSE | 1 | FALSE | 1 | FALSE | 1 | FALSE | 1 |
| 4500 | 4500 | 12.030303 | 4.83349957 | 1 | 0.57647348 | 2 | 0.28941836 | 2 | 0.33343871 | 2 | 0.45159487 | 0 | 1 | 26 | 0.86852567 | FALSE | 1 | FALSE | 1 | FALSE | 1 | FALSE | 1 | FALSE | 1 | FALSE | 1 |
| 4501 | 4501 | 7.09090909 | 1.62114132 | 0 | 1 | 0 | 1 | 1 | 0.33149969 | 1 | 0.40390032 | 0 | 1 | 9 | 0.75985366 | FALSE | 1 | FALSE | 1 | FALSE | 1 | FALSE | 1 | FALSE | 1 | FALSE | 1 |
| 4502 | 4502 | 8.75 | 2.62712499 | 2 | 0.16048326 | 0 | 1 | 0 | 1 | 0 | 1 | 0 | 1 | 26 | 0.16405547 | FALSE | 1 | FALSE | 1 | FALSE | 1 | FALSE | 1 | FALSE | 1 | FALSE | 1 |
| 4503 | 4503 | 6.03125 | 2.72126321 | 0 | 1 | 0 | 1 | 2 | 0.32011871 | 1 | 0.77852168 | 0 | 1 | 29 | 0.24088541 | FALSE | 1 | FALSE | 1 | FALSE | 1 | FALSE | 1 | FALSE | 1 | FALSE | 1 |
| 4504 | 4504 | 8.05714286 | 1.3926923 | 1 | 0.59801736 | 1 | 0.68454804 | 1 | 0.72299506 | 2 | 0.48233989 | 0 | 1 | 30 | 0.52765882 | FALSE | 1 | FALSE | 1 | FALSE | 1 | FALSE | 1 | FALSE | 1 | FALSE | 1 |
| 4505 | 4505 | 6.73043478 | 3.32139766 | 4 | 0.32328696 | 7 | 0.07020894 | 9 | 0.01894969 | 8 | 0.14195456 | 0 | 1 | 84 | 0.99676877 | FALSE | 1 | FALSE | 1 | FALSE | 1 | FALSE | 1 | FALSE | 1 | FALSE | 1 |
| 4506 | 4506 | 9.45454546 | 1.92418277 | 1 | 0.4357638 | 1 | 0.5154318 | 1 | 0.55340851 | 1 | 0.64497439 | 0 | 1 | 18 | 0.74744813 | FALSE | 1 | FALSE | 1 | FALSE | 1 | FALSE | 1 | FALSE | 1 | FALSE | 1 |
| 4507 | 4507 | 9.46666667 | 2.30554886 | 0 | 1 | 1 | 0.38964394 | 1 | 0.42266264 | 0 | 1 | 1 | 0.2198258 | 12 | 0.80329917 | FALSE | 1 | FALSE | 1 | FALSE | 1 | FALSE | 1 | FALSE | 1 | FALSE | 1 |
| 4508 | 4508 | 5.04615385 | 3.36237448 | 0 | 1 | 6 | 0.0180406 | 2 | 0.68408105 | 3 | 0.57949569 | 2 | 0.28868487 | 52 | 0.87296036 | FALSE | 1 | FALSE | 0.76574397 | FALSE | 1 | FALSE | 1 | FALSE | 1 | FALSE | 1 |
| 4509 | 4509 | 8.42857143 | 1.39970842 | 0 | 1 | 0 | 1 | 0 | 1 | 0 | 1 | 1 | 0.10932414 | 6 | 0.69920685 | FALSE | 1 | FALSE | 1 | FALSE | 1 | FALSE | 1 | FALSE | 1 | FALSE | 1 |
| 4510 | 4510 | 10.3333333 | 2.7080128 | 1 | 0.37380237 | 0 | 1 | 2 | 0.13502329 | 0 | 1 | 0 | 1 | 15 | 0.69372602 | FALSE | 1 | FALSE | 1 | FALSE | 1 | FALSE | 1 | FALSE | 1 | FALSE | 1 |

| | | | | | | | | | | | | | | | | | | | | | | | | | | | |
|---|---|---|---|---|---|---|---|---|---|---|---|---|---|---|---|---|---|---|---|---|---|---|---|---|---|---|---|
| 4511 | 4511 | 8.64285714 | 3.51744922 | 0 | 1 | 0 | 1 | 2 | 0.2663542 | 1 | 0.73248983 | 2 | 0.07640798 | 23 | 0.73439118 | FALSE | 1 | FALSE | 1 | FALSE | 1 | FALSE | 1 | FALSE | 1 | FALSE | 1 |
| 4512 | 4512 | 10.3434343 | 1.63399255 | 3 | 0.44178082 | 3 | 0.59841294 | 3 | 0.66955123 | 3 | 0.81890868 | 1 | 0.79437879 | 82 | 0.36191137 | FALSE | 1 | FALSE | 1 | FALSE | 1 | FALSE | 1 | FALSE | 1 | FALSE | 1 |
| 4513 | 4513 | 7.94736842 | 2.28205088 | 1 | 0.38989951 | 0 | 1 | 4 | 0.00411123 | 1 | 0.59104357 | 0 | 1 | 13 | 0.97998676 | FALSE | 1 | FALSE | 1 | FALSE | 1 | FALSE | 1 | FALSE | 1 | FALSE | 1 |
| 4514 | 4514 | 9.66666667 | 0.94280904 | 0 | 1 | 0 | 1 | 0 | 1 | 0 | 1 | 0 | 1 | 12 | 0.13002667 | FALSE | 1 | FALSE | 1 | FALSE | 1 | FALSE | 1 | FALSE | 1 | FALSE | 1 |
| 4515 | 4515 | 5.13333333 | 2.63818119 | 1 | 0.69044531 | 4 | 0.05650644 | 1 | 0.80828716 | 1 | 0.8802008 | 2 | 0.16793736 | 36 | 0.84604146 | FALSE | 1 | FALSE | 1 | FALSE | 1 | FALSE | 1 | FALSE | 1 | FALSE | 1 |
| 4516 | 4516 | 8.57142857 | 1.63507474 | 1 | 0.3050805 | 0 | 1 | 1 | 0.40110418 | 0 | 1 | 0 | 1 | 12 | 0.62236227 | FALSE | 1 | FALSE | 1 | FALSE | 1 | FALSE | 1 | FALSE | 1 | FALSE | 1 |
| 4517 | 4517 | 9.25 | 2.07665597 | 2 | 0.06197875 | 1 | 0.40942991 | 0 | 1 | 0 | 1 | 1 | 0.23264612 | 12 | 0.90922186 | FALSE | 0.72532843 | FALSE | 1 | FALSE | 1 | FALSE | 1 | FALSE | 1 | FALSE | 1 |
| 4518 | 4518 | 9.40816327 | 1.39851769 | 3 | 0.46200485 | 10 | 0.00118473 | 6 | 0.13995079 | 9 | 0.03532718 | 2 | 0.48029053 | 68 | 0.99994923 | FALSE | 1 | FALSE | 0.34661916 | FALSE | 1 | FALSE | 1 | FALSE | 1 | FALSE | 1 |
| 4519 | 4519 | 9.2 | 2.6 | 1 | 0.22886536 | 1 | 0.28036922 | 0 | 1 | 2 | 0.0740981 | 0 | 1 | 6 | 0.98831471 | FALSE | 1 | FALSE | 1 | FALSE | 1 | FALSE | 1 | FALSE | 1 | FALSE | 1 |
| 4520 | 4520 | 7.34782609 | 2.28622901 | 1 | 0.69843502 | 1 | 0.78076835 | 1 | 0.81522035 | 2 | 0.63103861 | 0 | 1 | 41 | 0.25430673 | FALSE | 1 | FALSE | 1 | FALSE | 1 | FALSE | 1 | FALSE | 1 | FALSE | 1 |
| 4521 | 4521 | 7.73863636 | 2.87027999 | 3 | 0.39356864 | 1 | 0.94566665 | 3 | 0.61802554 | 5 | 0.37910615 | 0 | 1 | 76 | 0.36734352 | FALSE | 1 | FALSE | 1 | FALSE | 1 | FALSE | 1 | FALSE | 1 | FALSE | 1 |
| 4522 | 4522 | 4.97560976 | 3.66584636 | 0 | 1 | 1 | 0.74131698 | 5 | 0.01496777 | 3 | 0.29054213 | 0 | 1 | 32 | 0.90458039 | FALSE | 1 | FALSE | 1 | FALSE | 1 | FALSE | 1 | FALSE | 1 | FALSE | 1 |
| 4523 | 4523 | 6.35849057 | 3.54536285 | 4 | 0.04620161 | 1 | 0.82613965 | 1 | 0.85724013 | 2 | 0.70707241 | 1 | 0.58506217 | 44 | 0.69042405 | FALSE | 0.649301 | FALSE | 1 | FALSE | 1 | FALSE | 1 | FALSE | 1 | FALSE | 1 |
| 4524 | 4524 | 7.6875 | 2.6033332 | 3 | 0.0476977 | 1 | 0.65167362 | 0 | 1 | 2 | 0.43583874 | 0 | 1 | 26 | 0.77612337 | FALSE | 0.649301 | FALSE | 1 | FALSE | 1 | FALSE | 1 | FALSE | 1 | FALSE | 1 |
| 4525 | 4525 | 8.69444444 | 2.35489378 | 2 | 0.23542589 | 3 | 0.10944025 | 4 | 0.03888854 | 1 | 0.81665383 | 2 | 0.11720035 | 24 | 0.99789786 | FALSE | 1 | FALSE | 1 | FALSE | 1 | FALSE | 1 | FALSE | 1 | FALSE | 1 |
| 4526 | 4526 | 11.7222222 | 3.10564997 | 0 | 1 | 1 | 0.44710673 | 0 | 1 | 2 | 0.19902682 | 0 | 1 | 15 | 0.69372602 | FALSE | 1 | FALSE | 1 | FALSE | 1 | FALSE | 1 | FALSE | 1 | FALSE | 1 |
| 4527 | 4527 | 7.14285714 | 2.79941685 | 0 | 1 | 1 | 0.60247911 | 0 | 1 | 3 | 0.13524571 | 1 | 0.37113067 | 23 | 0.73439118 | FALSE | 1 | FALSE | 1 | FALSE | 1 | FALSE | 1 | FALSE | 1 | FALSE | 1 |
| 4528 | 4528 | 5.18 | 2.90991409 | 1 | 0.72839502 | 0 | 1 | 2 | 0.54120634 | 3 | 0.4048204 | 1 | 0.56379725 | 43 | 0.46945509 | FALSE | 1 | FALSE | 1 | FALSE | 1 | FALSE | 1 | FALSE | 1 | FALSE | 1 |
| 4529 | 4529 | 8.94736842 | 2.76202985 | 0 | 1 | 1 | 0.46503795 | 0 | 1 | 1 | 0.59104357 | 1 | 0.26986775 | 16 | 0.65565778 | FALSE | 1 | FALSE | 1 | FALSE | 1 | FALSE | 1 | FALSE | 1 | FALSE | 1 |
| 4530 | 4530 | 7.13636364 | 3.10857253 | 0 | 1 | 1 | 0.5154318 | 1 | 0.55340851 | 0 | 1 | 1 | 0.30529951 | 19 | 0.54233419 | FALSE | 1 | FALSE | 1 | FALSE | 1 | FALSE | 1 | FALSE | 1 | FALSE | 1 |
| 4531 | 4531 | 10 | 2.1602469 | 0 | 1 | 0 | 1 | 0 | 1 | 1 | 0.43131129 | 0 | 1 | 11 | 0.41944532 | FALSE | 1 | FALSE | 1 | FALSE | 1 | FALSE | 1 | FALSE | 1 | FALSE | 1 |
| 4532 | 4532 | 7.6 | 2.70924344 | 0 | 1 | 0 | 1 | 0 | 1 | 2 | 0.19902682 | 0 | 1 | 16 | 0.44998001 | FALSE | 1 | FALSE | 1 | FALSE | 1 | FALSE | 1 | FALSE | 1 | FALSE | 1 |
| 4533 | 4533 | 9.6875 | 2.59130156 | 1 | 0.54198714 | 1 | 0.6278847 | 0 | 1 | 1 | 0.75658797 | 2 | 0.08611595 | 25 | 0.67630471 | FALSE | 1 | FALSE | 1 | FALSE | 1 | FALSE | 1 | FALSE | 1 | FALSE | 1 |
| 4534 | 4534 | 10.4736842 | 2.70221003 | 0 | 1 | 1 | 0.46503795 | 1 | 0.50143826 | 3 | 0.05383577 | 1 | 0.26986775 | 13 | 0.97998676 | FALSE | 1 | FALSE | 1 | FALSE | 1 | FALSE | 1 | FALSE | 1 | FALSE | 1 |
| 4535 | 4535 | 10.2222222 | 2.24982853 | 0 | 1 | 1 | 0.23139161 | 0 | 1 | 0 | 1 | 0 | 1 | 7 | 0.63752397 | FALSE | 1 | FALSE | 1 | FALSE | 1 | FALSE | 1 | FALSE | 1 | FALSE | 1 |
| 4536 | 4536 | 9.46428571 | 2.3065457 | 4 | 0.00528025 | 1 | 0.60247911 | 1 | 0.64171563 | 0 | 1 | 2 | 0.07640798 | 20 | 0.97757517 | FALSE | 0.2089654 | FALSE | 1 | FALSE | 1 | FALSE | 1 | FALSE | 1 | FALSE | 1 |
| 4537 | 4537 | 3.51960784 | 2.08745268 | 16 | 3.78E-05 | 11 | 0.05252446 | 3 | 0.97472887 | 10 | 0.41220077 | 4 | 0.40154044 | 152 | 0.9960189 | TRUE | 0.00999617 | FALSE | 1 | FALSE | 1 | FALSE | 1 | FALSE | 1 | FALSE | 1 |
| 4538 | 4538 | 9 | 3.74165739 | 0 | 1 | 1 | 0.46503795 | 1 | 0.50143826 | 2 | 0.21610812 | 0 | 1 | 15 | 0.83648191 | FALSE | 1 | FALSE | 1 | FALSE | 1 | FALSE | 1 | FALSE | 1 | FALSE | 1 |
| 4539 | 4539 | 8.96774194 | 2.27870749 | 0 | 1 | 1 | 0.63997468 | 0 | 1 | 0 | 1 | 0 | 1 | 30 | 0.03453514 | FALSE | 1 | FALSE | 1 | FALSE | 1 | FALSE | 1 | FALSE | 1 | FALSE | 1 |
| 4540 | 4540 | 8.75 | 1.36167789 | 2 | 0.03643311 | 1 | 0.32623933 | 0 | 1 | 0 | 1 | 0 | 1 | 9 | 0.89630573 | FALSE | 0.58392 | FALSE | 1 | FALSE | 1 | FALSE | 1 | FALSE | 1 | FALSE | 1 |
| 4541 | 4541 | 7.83783784 | 2.66624053 | 0 | 1 | 0 | 1 | 2 | 0.35984046 | 2 | 0.48233989 | 1 | 0.44015537 | 30 | 0.52765882 | FALSE | 1 | FALSE | 1 | FALSE | 1 | FALSE | 1 | FALSE | 1 | FALSE | 1 |
| 4542 | 4542 | 9.85 | 2.90301567 | 1 | 0.40558521 | 0 | 1 | 1 | 0.51939841 | 1 | 0.60987146 | 0 | 1 | 17 | 0.61750632 | FALSE | 1 | FALSE | 1 | FALSE | 1 | FALSE | 1 | FALSE | 1 | FALSE | 1 |
| 4543 | 4543 | 9.7826087 | 2.37663485 | 0 | 1 | 0 | 1 | 1 | 0.5695037 | 3 | 0.08614834 | 1 | 0.31672722 | 18 | 0.86273155 | FALSE | 1 | FALSE | 1 | FALSE | 1 | FALSE | 1 | FALSE | 1 | FALSE | 1 |
| 4544 | 4544 | 10 | 2.88230677 | 0 | 1 | 0 | 1 | 2 | 0.07725803 | 1 | 0.45746571 | 0 | 1 | 10 | 0.86791642 | FALSE | 1 | FALSE | 1 | FALSE | 1 | FALSE | 1 | FALSE | 1 | FALSE | 1 |
| 4545 | 4545 | 9.125 | 1.16592238 | 0 | 1 | 0 | 1 | 1 | 0.44344817 | 1 | 0.52895088 | 0 | 1 | 14 | 0.53308261 | FALSE | 1 | FALSE | 1 | FALSE | 1 | FALSE | 1 | FALSE | 1 | FALSE | 1 |
| 4546 | 4546 | 7.12 | 1.42323575 | 0 | 1 | 2 | 0.19297191 | 1 | 0.59998149 | 2 | 0.31968852 | 0 | 1 | 20 | 0.8152498 | FALSE | 1 | FALSE | 1 | FALSE | 1 | FALSE | 1 | FALSE | 1 | FALSE | 1 |
| 4547 | 4547 | 9.88235294 | 2.54871793 | 2 | 0.37762121 | 4 | 0.08182743 | 0 | 1 | 3 | 0.41729179 | 0 | 1 | 42 | 0.73296146 | FALSE | 1 | FALSE | 1 | FALSE | 1 | FALSE | 1 | FALSE | 1 | FALSE | 1 |
| 4548 | 4548 | 10 | 1.26491106 | 0 | 1 | 0 | 1 | 1 | 0.42266264 | 0 | 1 | 1 | 0.2198258 | 13 | 0.57712805 | FALSE | 1 | FALSE | 1 | FALSE | 1 | FALSE | 1 | FALSE | 1 | FALSE | 1 |
| 4549 | 4549 | 8.72 | 2.0103731 | 0 | 1 | 0 | 1 | 1 | 0.59998149 | 1 | 0.69181315 | 0 | 1 | 23 | 0.22727183 | FALSE | 1 | FALSE | 1 | FALSE | 1 | FALSE | 1 | FALSE | 1 | FALSE | 1 |
| 4550 | 4550 | 9.5 | 2.33927814 | 0 | 1 | 0 | 1 | 0 | 1 | 2 | 0.19902682 | 1 | 0.25766374 | 15 | 0.69372602 | FALSE | 1 | FALSE | 1 | FALSE | 1 | FALSE | 1 | FALSE | 1 | FALSE | 1 |
| 4551 | 4551 | 10 | 2.89395923 | 0 | 1 | 1 | 0.40942991 | 0 | 1 | 1 | 0.52895088 | 1 | 0.23264612 | 13 | 0.76799161 | FALSE | 1 | FALSE | 1 | FALSE | 1 | FALSE | 1 | FALSE | 1 | FALSE | 1 |
| 4552 | 4552 | 4.26300578 | 1.33111548 | 5 | 0.94634401 | 10 | 0.68623819 | 15 | 0.25751076 | 22 | 0.07225031 | 2 | 0.97977309 | 291 | 0.54304994 | FALSE | 1 | FALSE | 1 | FALSE | 1 | FALSE | 1 | FALSE | 1 | FALSE | 1 |
| 4553 | 4553 | 9 | 2.5819889 | 1 | 0.26795838 | 0 | 1 | 0 | 1 | 0 | 1 | 0 | 1 | 11 | 0.41944532 | FALSE | 1 | FALSE | 1 | FALSE | 1 | FALSE | 1 | FALSE | 1 | FALSE | 1 |
| 4554 | 4554 | 10.8947368 | 2.69296817 | 0 | 1 | 0 | 1 | 2 | 0.14749808 | 0 | 1 | 0 | 1 | 17 | 0.41141543 | FALSE | 1 | FALSE | 1 | FALSE | 1 | FALSE | 1 | FALSE | 1 | FALSE | 1 |
| 4555 | 4555 | 9.76190476 | 2.11355629 | 1 | 0.42086991 | 0 | 1 | 3 | 0.03775825 | 0 | 1 | 1 | 0.29368245 | 16 | 0.90353674 | FALSE | 1 | FALSE | 1 | FALSE | 1 | FALSE | 1 | FALSE | 1 | FALSE | 1 |
| 4556 | 4556 | 9.30769231 | 3.36005072 | 0 | 1 | 0 | 1 | 1 | 0.37874417 | 1 | 0.45746571 | 0 | 1 | 11 | 0.6683056 | FALSE | 1 | FALSE | 1 | FALSE | 1 | FALSE | 1 | FALSE | 1 | FALSE | 1 |
| 4557 | 4557 | 6.81818182 | 3.0989201 | 3 | 0.10232687 | 3 | 0.16927151 | 2 | 0.47281647 | 3 | 0.32878418 | 1 | 0.51797501 | 32 | 0.98583614 | FALSE | 0.87592659 | FALSE | 1 | FALSE | 1 | FALSE | 1 | FALSE | 1 | FALSE | 1 |
| 4558 | 4558 | 10.5 | 1.96214169 | 1 | 0.22886536 | 0 | 1 | 1 | 0.30655256 | 2 | 0.0740981 | 0 | 1 | 6 | 0.98831471 | FALSE | 1 | FALSE | 1 | FALSE | 1 | FALSE | 1 | FALSE | 1 | FALSE | 1 |
| 4559 | 4559 | 7.73684211 | 1.58420178 | 0 | 1 | 0 | 1 | 0 | 1 | 0 | 1 | 0 | 1 | 19 | 0.03948376 | FALSE | 1 | FALSE | 1 | FALSE | 1 | FALSE | 1 | FALSE | 1 | FALSE | 1 |
| 4560 | 4560 | 11.4615385 | 2.6199056 | 1 | 0.28675951 | 0 | 1 | 1 | 0.37874417 | 0 | 1 | 0 | 1 | 11 | 0.6683056 | FALSE | 1 | FALSE | 1 | FALSE | 1 | FALSE | 1 | FALSE | 1 | FALSE | 1 |
| 4561 | 4561 | 9.56666667 | 1.97793383 | 1 | 0.51745948 | 2 | 0.2288771 | 1 | 0.64171563 | 1 | 0.73248983 | 1 | 0.37113067 | 22 | 0.86499992 | FALSE | 1 | FALSE | 1 | FALSE | 1 | FALSE | 1 | FALSE | 1 | FALSE | 1 |
| 4562 | 4562 | 9.30434783 | 1.89816347 | 0 | 1 | 1 | 0.53115654 | 0 | 1 | 1 | 0.6613267 | 0 | 1 | 21 | 0.27964072 | FALSE | 1 | FALSE | 1 | FALSE | 1 | FALSE | 1 | FALSE | 1 | FALSE | 1 |
| 4563 | 4563 | 9.55882353 | 2.31607143 | 3 | 0.05546921 | 1 | 0.6739483 | 1 | 0.71261921 | 0 | 1 | 1 | 0.43077488 | 28 | 0.72558119 | FALSE | 0.68849059 | FALSE | 1 | FALSE | 1 | FALSE | 1 | FALSE | 1 | FALSE | 1 |
| 4564 | 4564 | 10.0769231 | 2.40069024 | 0 | 1 | 1 | 0.3480709 | 0 | 1 | 0 | 1 | 0 | 1 | 12 | 0.374198 | FALSE | 1 | FALSE | 1 | FALSE | 1 | FALSE | 1 | FALSE | 1 | FALSE | 1 |
| 4565 | 4565 | 10.1666667 | 1.79505494 | 1 | 0.46441868 | 1 | 0.54637326 | 0 | 1 | 0 | 1 | 2 | 0.058175 | 20 | 0.68270729 | FALSE | 1 | FALSE | 1 | FALSE | 1 | FALSE | 1 | FALSE | 1 | FALSE | 1 |
| 4566 | 4566 | 7.52631579 | 1.49976914 | 3 | 0.07268126 | 0 | 1 | 1 | 0.75193685 | 2 | 0.52640931 | 0 | 1 | 32 | 0.61800916 | FALSE | 0.76727438 | FALSE | 1 | FALSE | 1 | FALSE | 1 | FALSE | 1 | FALSE | 1 |
| 4567 | 4567 | 9.95 | 1.88348082 | 0 | 1 | 0 | 1 | 0 | 1 | 1 | 0.60987146 | 1 | 0.28187291 | 18 | 0.37504645 | FALSE | 1 | FALSE | 1 | FALSE | 1 | FALSE | 1 | FALSE | 1 | FALSE | 1 |
| 4568 | 4568 | 8.6 | 2.81780056 | 0 | 1 | 0 | 1 | 1 | 0.51939841 | 2 | 0.23331368 | 0 | 1 | 17 | 0.61750632 | FALSE | 1 | FALSE | 1 | FALSE | 1 | FALSE | 1 | FALSE | 1 | FALSE | 1 |
| 4569 | 4569 | 6.52777778 | 2.36274453 | 0 | 1 | 2 | 0.32556427 | 0 | 1 | 0 | 1 | 0 | 1 | 34 | 0.06417017 | FALSE | 1 | FALSE | 1 | FALSE | 1 | FALSE | 1 | FALSE | 1 | FALSE | 1 |
| 4570 | 4570 | 9.5 | 2.51246891 | 1 | 0.56527493 | 1 | 0.65167362 | 2 | 0.32011871 | 1 | 0.77852168 | 0 | 1 | 27 | 0.61671133 | FALSE | 1 | FALSE | 1 | FALSE | 1 | FALSE | 1 | FALSE | 1 | FALSE | 1 |
| 4571 | 4571 | 8.69230769 | 2.24926023 | 0 | 1 | 1 | 0.57534752 | 2 | 0.23940226 | 1 | 0.70601446 | 1 | 0.3499045 | 21 | 0.78939668 | FALSE | 1 | FALSE | 1 | FALSE | 1 | FALSE | 1 | FALSE | 1 | FALSE | 1 |
| 4572 | 4572 | 10.6956522 | 2.38536813 | 0 | 1 | 1 | 0.53115654 | 1 | 0.5695037 | 1 | 0.6613267 | 1 | 0.31672722 | 19 | 0.71542496 | FALSE | 1 | FALSE | 1 | FALSE | 1 | FALSE | 1 | FALSE | 1 | FALSE | 1 |
| 4573 | 4573 | 11.3333333 | 2.58812919 | 1 | 0.42086991 | 1 | 0.49918218 | 1 | 0.53671415 | 1 | 0.62783522 | 1 | 0.29368245 | 16 | 0.90353674 | FALSE | 1 | FALSE | 1 | FALSE | 1 | FALSE | 1 | FALSE | 1 | FALSE | 1 |
| 4574 | 4574 | 7.9375 | 3.26857672 | 4 | 0.03383958 | 2 | 0.46321244 | 1 | 0.8283465 | 3 | 0.37966044 | 1 | 0.54902428 | 37 | 0.93870662 | FALSE | 0.56934354 | FALSE | 1 | FALSE | 1 | FALSE | 1 | FALSE | 1 | FALSE | 1 |
| 4575 | 4575 | 10.6666667 | 1.92930615 | 0 | 1 | 0 | 1 | 1 | 0.33149969 | 0 | 1 | 0 | 1 | 10 | 0.46859937 | FALSE | 1 | FALSE | 1 | FALSE | 1 | FALSE | 1 | FALSE | 1 | FALSE | 1 |
| 4576 | 4576 | 9.31034483 | 3.4849771 | 0 | 1 | 0 | 1 | 2 | 0.2663542 | 3 | 0.13524571 | 1 | 0.37113067 | 22 | 0.86499992 | FALSE | 1 | FALSE | 1 | FALSE | 1 | FALSE | 1 | FALSE | 1 | FALSE | 1 |
| 4577 | 4577 | 9.09090909 | 2.31416765 | 1 | 0.4357638 | 1 | 0.5154318 | 3 | 0.04259419 | 0 | 1 | 0 | 1 | 17 | 0.8840357 | FALSE | 1 | FALSE | 1 | FALSE | 1 | FALSE | 1 | FALSE | 1 | FALSE | 1 |
| 4578 | 4578 | 8.64705882 | 2.37489756 | 0 | 1 | 2 | 0.10307112 | 1 | 0.46348836 | 0 | 1 | 2 | 0.03086174 | 12 | 0.962093 | FALSE | 1 | FALSE | 1 | FALSE | 1 | FALSE | 1 | FALSE | 0.82559785 | FALSE | 1 |
| 4579 | 4579 | 9.8 | 2.31516738 | 0 | 1 | 0 | 1 | 1 | 0.40110418 | 1 | 0.48242099 | 0 | 1 | 12 | 0.62236227 | FALSE | 1 | FALSE | 1 | FALSE | 1 | FALSE | 1 | FALSE | 1 | FALSE | 1 |
| 4580 | 4580 | 8.77777778 | 1.08297715 | 0 | 1 | 0 | 1 | 0 | 1 | 0 | 1 | 0 | 1 | 17 | 0.05550965 | FALSE | 1 | FALSE | 1 | FALSE | 1 | FALSE | 1 | FALSE | 1 | FALSE | 1 |
| 4581 | 4581 | 7.71666667 | 1.82657847 | 0 | 1 | 3 | 0.30686508 | 3 | 0.36617062 | 2 | 0.76960882 | 2 | 0.25814944 | 50 | 0.66861245 | FALSE | 1 | FALSE | 1 | FALSE | 1 | FALSE | 1 | FALSE | 1 | FALSE | 1 |
| 4582 | 4582 | 9.96 | 2.12565284 | 1 | 0.47819878 | 0 | 1 | 1 | 0.59998149 | 1 | 0.69181315 | 0 | 1 | 22 | 0.43642846 | FALSE | 1 | FALSE | 1 | FALSE | 1 | FALSE | 1 | FALSE | 1 | FALSE | 1 |
| 4583 | 4583 | 9.36842105 | 2.36666602 | 1 | 0.38989951 | 0 | 1 | 2 | 0.14749808 | 1 | 0.59104357 | 0 | 1 | 15 | 0.83648191 | FALSE | 1 | FALSE | 1 | FALSE | 1 | FALSE | 1 | FALSE | 1 | FALSE | 1 |
| 4584 | 4584 | 9.14285714 | 4.48580528 | 0 | 1 | 0 | 1 | 1 | 0.40110418 | 0 | 1 | 0 | 1 | 13 | 0.33283729 | FALSE | 1 | FALSE | 1 | FALSE | 1 | FALSE | 1 | FALSE | 1 | FALSE | 1 |
| 4585 | 4585 | 7.85714286 | 3.71977616 | 0 | 1 | 0 | 1 | 0 | 1 | 0 | 1 | 0 | 1 | 7 | 0.3043657 | FALSE | 1 | FALSE | 1 | FALSE | 1 | FALSE | 1 | FALSE | 1 | FALSE | 1 |
| 4586 | 4586 | 7.30769231 | 2.87922573 | 2 | 0.14250978 | 0 | 1 | 1 | 0.61440466 | 1 | 0.70601446 | 0 | 1 | 22 | 0.61628 | FALSE | 1 | FALSE | 1 | FALSE | 1 | FALSE | 1 | FALSE | 1 | FALSE | 1 |
| 4587 | 4587 | 9.38095238 | 3.0312058 | 1 | 0.42086991 | 0 | 1 | 1 | 0.53671415 | 1 | 0.62783522 | 0 | 1 | 18 | 0.57962763 | FALSE | 1 | FALSE | 1 | FALSE | 1 | FALSE | 1 | FALSE | 1 | FALSE | 1 |
| 4588 | 4588 | 6.34615385 | 3.56200363 | 0 | 1 | 0 | 1 | 0 | 1 | 1 | 0.70601446 | 0 | 1 | 25 | 0.06985983 | FALSE | 1 | FALSE | 1 | FALSE | 1 | FALSE | 1 | FALSE | 1 | FALSE | 1 |
| 4589 | 4589 | 10.6666667 | 2.46080384 | 0 | 1 | 3 | 0.00591424 | 1 | 0.35555293 | 1 | 0.43131129 | 0 | 1 | 7 | 0.99433481 | FALSE | 1 | FALSE | 0.60103504 | FALSE | 1 | FALSE | 1 | FALSE | 1 | FALSE | 1 |
| 4590 | 4590 | 7.25 | 3.16030062 | 0 | 1 | 1 | 0.48239024 | 1 | 0.51939841 | 1 | 0.60987146 | 0 | 1 | 17 | 0.61750632 | FALSE | 1 | FALSE | 1 | FALSE | 1 | FALSE | 1 | FALSE | 1 | FALSE | 1 |
| 4591 | 4591 | 9 | 3.26990943 | 2 | 0.14250978 | 4 | 0.00909592 | 1 | 0.61440466 | 2 | 0.33678042 | 0 | 1 | 17 | 0.99598795 | FALSE | 1 | FALSE | 0.63686998 | FALSE | 1 | FALSE | 1 | FALSE | 1 | FALSE | 1 |
| 4592 | 4592 | 9.5 | 3.04138127 | 0 | 1 | 1 | 0.23139161 | 0 | 1 | 0 | 1 | 0 | 1 | 7 | 0.63752397 | FALSE | 1 | FALSE | 1 | FALSE | 1 | FALSE | 1 | FALSE | 1 | FALSE | 1 |
| 4593 | 4593 | 5.72340426 | 2.8784018 | 2 | 0.3309085 | 2 | 0.44132825 | 2 | 0.4962935 | 2 | 0.63103861 | 5 | 0.00086811 | 33 | 0.99158534 | FALSE | 1 | FALSE | 1 | FALSE | 1 | FALSE | 1 | FALSE | 0.16162556 | FALSE | 1 |
| 4594 | 4594 | 9.9 | 2.48797106 | 1 | 0.40558521 | 1 | 0.48239024 | 0 | 1 | 2 | 0.23331368 | 2 | 0.04176963 | 14 | 0.97325593 | FALSE | 1 | FALSE | 1 | FALSE | 1 | FALSE | 1 | FALSE | 0.90763079 | FALSE | 1 |
| 4595 | 4595 | 6.79166667 | 2.17905114 | 0 | 1 | 0 | 1 | 1 | 0.58502114 | 2 | 0.30249307 | 1 | 0.32796862 | 20 | 0.68270729 | FALSE | 1 | FALSE | 1 | FALSE | 1 | FALSE | 1 | FALSE | 1 | FALSE | 1 |
| 4596 | 4596 | 8.89583333 | 2.06397254 | 0 | 1 | 1 | 0.79481669 | 1 | 0.8283465 | 4 | 0.17698522 | 2 | 0.18560062 | 40 | 0.66905473 | FALSE | 1 | FALSE | 1 | FALSE | 1 | FALSE | 1 | FALSE | 1 | FALSE | 1 |
| 4597 | 4597 | 9.32142857 | 2.64647436 | 0 | 1 | 2 | 0.2288771 | 2 | 0.2663542 | 0 | 1 | 1 | 0.37113067 | 23 | 0.73439118 | FALSE | 1 | FALSE | 1 | FALSE | 1 | FALSE | 1 | FALSE | 1 | FALSE | 1 |
| 4598 | 4598 | 10.7 | 3.68917335 | 0 | 1 | 0 | 1 | 0 | 1 | 0 | 1 | 0 | 1 | 10 | 0.18273166 | FALSE | 1 | FALSE | 1 | FALSE | 1 | FALSE | 1 | FALSE | 1 | FALSE | 1 |
| 4599 | 4599 | 6.25806452 | 1.90021632 | 4 | 0.00764535 | 0 | 1 | 2 | 0.30673593 | 2 | 0.41984187 | 0 | 1 | 23 | 0.95750806 | FALSE | 0.2566832 | FALSE | 1 | FALSE | 1 | FALSE | 1 | FALSE | 1 | FALSE | 1 |
| 4600 | 4600 | 9.42105263 | 1.95517638 | 1 | 0.38989951 | 0 | 1 | 0 | 1 | 0 | 1 | 0 | 1 | 18 | 0.17880694 | FALSE | 1 | FALSE | 1 | FALSE | 1 | FALSE | 1 | FALSE | 1 | FALSE | 1 |
| 4601 | 4601 | 11.125 | 2.20439901 | 0 | 1 | 0 | 1 | 0 | 1 | 0 | 1 | 0 | 1 | 8 | 0.25677166 | FALSE | 1 | FALSE | 1 | FALSE | 1 | FALSE | 1 | FALSE | 1 | FALSE | 1 |
| 4602 | 4602 | 10.0769231 | 1.56704221 | 1 | 0.49162634 | 1 | 0.57534752 | 0 | 1 | 1 | 0.70601446 | 0 | 1 | 23 | 0.40372818 | FALSE | 1 | FALSE | 1 | FALSE | 1 | FALSE | 1 | FALSE | 1 | FALSE | 1 |
| 4603 | 4603 | 9.23809524 | 3.42128856 | 0 | 1 | 1 | 0.49918218 | 2 | 0.17308361 | 0 | 1 | 0 | 1 | 18 | 0.57962763 | FALSE | 1 | FALSE | 1 | FALSE | 1 | FALSE | 1 | FALSE | 1 | FALSE | 1 |
| 4604 | 4604 | 7.5 | 1.70782513 | 0 | 1 | 1 | 0.179097 | 0 | 1 | 0 | 1 | 0 | 1 | 5 | 0.76185568 | FALSE | 1 | FALSE | 1 | FALSE | 1 | FALSE | 1 | FALSE | 1 | FALSE | 1 |

| | | | | | | | | | | | | | | | | | | | | | | | | | | | |
|---|---|---|---|---|---|---|---|---|---|---|---|---|---|---|---|---|---|---|---|---|---|---|---|---|---|---|---|
| 4605 | 4605 | 9.70212766 | 1.7371399 | 0 | 1 | 2 | 0.45233216 | 1 | 0.82190381 | 2 | 0.64277396 | 2 | 0.17968152 | 40 | 0.54368682 | FALSE | 1 | FALSE | 1 | FALSE | 1 | FALSE | 1 | FALSE | 1 | FALSE | 1 |
| 4606 | 4606 | 9.70588235 | 2.49150807 | 0 | 1 | 1 | 0.42857733 | 0 | 1 | 1 | 0.55062772 | 0 | 1 | 15 | 0.49060435 | FALSE | 1 | FALSE | 1 | FALSE | 1 | FALSE | 1 | FALSE | 1 | FALSE | 1 |
| 4607 | 4607 | 7.19444444 | 3.06249213 | 3 | 0.0638041 | 1 | 0.69480472 | 1 | 0.73299778 | 1 | 0.81665383 | 1 | 0.44938266 | 29 | 0.81028289 | FALSE | 0.73766 | FALSE | 1 | FALSE | 1 | FALSE | 1 | FALSE | 1 | FALSE | 1 |
| 4608 | 4608 | 11.1923077 | 2.05724294 | 0 | 1 | 1 | 0.57534752 | 2 | 0.23940226 | 2 | 0.33678042 | 3 | 0.00847466 | 18 | 0.98644281 | FALSE | 1 | FALSE | 1 | FALSE | 1 | FALSE | 1 | FALSE | 0.50674765 | FALSE | 1 |
| 4609 | 4609 | 10.1111111 | 3.14269681 | 1 | 0.18769745 | 0 | 1 | 1 | 0.25384309 | 0 | 1 | 0 | 1 | 6 | 0.88425551 | FALSE | 1 | FALSE | 1 | FALSE | 1 | FALSE | 1 | FALSE | 1 | FALSE | 1 |
| 4610 | 4610 | 7.81818182 | 4.46843845 | 1 | 0.57647348 | 4 | 0.02089081 | 2 | 0.33343871 | 1 | 0.78873814 | 0 | 1 | 25 | 0.93892791 | FALSE | 1 | FALSE | 0.8034624 | FALSE | 1 | FALSE | 1 | FALSE | 1 | FALSE | 1 |
| 4611 | 4611 | 12.4 | 4.84148738 | 0 | 1 | 0 | 1 | 0 | 1 | 0 | 1 | 0 | 1 | 5 | 0.42761893 | FALSE | 1 | FALSE | 1 | FALSE | 1 | FALSE | 1 | FALSE | 1 | FALSE | 1 |
| 4612 | 4612 | 8.38095238 | 2.41968496 | 0 | 1 | 0 | 1 | 2 | 0.17308361 | 0 | 1 | 0 | 1 | 19 | 0.34094959 | FALSE | 1 | FALSE | 1 | FALSE | 1 | FALSE | 1 | FALSE | 1 | FALSE | 1 |
| 4613 | 4613 | 10.2916667 | 2.65328675 | 0 | 1 | 2 | 0.18117183 | 0 | 1 | 3 | 0.09525947 | 0 | 1 | 19 | 0.83975208 | FALSE | 1 | FALSE | 1 | FALSE | 1 | FALSE | 1 | FALSE | 1 | FALSE | 1 |
| 4614 | 4614 | 9.83333333 | 1.74801475 | 1 | 0.46441868 | 0 | 1 | 3 | 0.05314317 | 1 | 0.67692816 | 1 | 0.32796862 | 18 | 0.93166998 | FALSE | 1 | FALSE | 1 | FALSE | 1 | FALSE | 1 | FALSE | 1 | FALSE | 1 |
| 4615 | 4615 | 4.28881988 | 1.62486478 | 13 | 0.01535818 | 10 | 0.32808779 | 12 | 0.22059163 | 18 | 0.05098229 | 2 | 0.93029541 | 204 | 0.99435119 | FALSE | 0.38695764 | FALSE | 1 | FALSE | 1 | FALSE | 1 | FALSE | 1 | FALSE | 1 |
| 4616 | 4616 | 9.22727273 | 2.9377143 | 2 | 0.10809809 | 0 | 1 | 0 | 1 | 2 | 0.26791271 | 0 | 1 | 18 | 0.74744813 | FALSE | 0.88820416 | FALSE | 1 | FALSE | 1 | FALSE | 1 | FALSE | 1 | FALSE | 1 |
| 4617 | 4617 | 7.45744681 | 2.89042769 | 5 | 0.08987772 | 2 | 0.80898201 | 3 | 0.65536448 | 1 | 0.98774195 | 1 | 0.7873664 | 81 | 0.28735253 | FALSE | 0.83248965 | FALSE | 1 | FALSE | 1 | FALSE | 1 | FALSE | 1 | FALSE | 1 |
| 4618 | 4618 | 9.30487805 | 2.52639251 | 5 | 0.05627772 | 4 | 0.26638189 | 3 | 0.56136161 | 1 | 0.97829212 | 5 | 0.01032475 | 63 | 0.95896498 | FALSE | 0.69536509 | FALSE | 1 | FALSE | 1 | FALSE | 1 | FALSE | 0.5620304 | FALSE | 1 |
| 4619 | 4619 | 8.47619048 | 2.92149746 | 0 | 1 | 0 | 1 | 0 | 1 | 1 | 0.62783522 | 1 | 0.29368245 | 19 | 0.34094959 | FALSE | 1 | FALSE | 1 | FALSE | 1 | FALSE | 1 | FALSE | 1 | FALSE | 1 |
| 4620 | 4620 | 9.390625 | 1.94949663 | 1 | 0.81177881 | 0 | 1 | 1 | 0.90487646 | 7 | 0.02649204 | 1 | 0.65460803 | 54 | 0.58381086 | FALSE | 1 | FALSE | 1 | FALSE | 1 | FALSE | 1 | FALSE | 1 | FALSE | 1 |
| 4621 | 4621 | 8.74358974 | 2.1209329 | 0 | 1 | 0 | 1 | 3 | 0.16357218 | 4 | 0.10199045 | 4 | 0.00361808 | 28 | 0.98701111 | FALSE | 1 | FALSE | 1 | FALSE | 1 | FALSE | 1 | FALSE | 0.34869821 | FALSE | 1 |
| 4622 | 4622 | 7.43333333 | 2.49911095 | 1 | 0.54198714 | 1 | 0.6278847 | 1 | 0.66709967 | 0 | 1 | 1 | 0.39166989 | 26 | 0.48609328 | FALSE | 1 | FALSE | 1 | FALSE | 1 | FALSE | 1 | FALSE | 1 | FALSE | 1 |
| 4623 | 4623 | 10.2222222 | 3.27542288 | 0 | 1 | 0 | 1 | 1 | 0.48280983 | 1 | 0.57131013 | 0 | 1 | 16 | 0.44998001 | FALSE | 1 | FALSE | 1 | FALSE | 1 | FALSE | 1 | FALSE | 1 | FALSE | 1 |
| 4624 | 4624 | 9.6 | 3.09408037 | 0 | 1 | 2 | 0.08295839 | 0 | 1 | 0 | 1 | 0 | 1 | 13 | 0.57712805 | FALSE | 1 | FALSE | 1 | FALSE | 1 | FALSE | 1 | FALSE | 1 | FALSE | 1 |
| 4625 | 4625 | 6.33701658 | 1.13809473 | 9 | 0.03026241 | 6 | 0.47328187 | 6 | 0.57671504 | 9 | 0.37905209 | 1 | 0.94196523 | 139 | 0.85459504 | FALSE | 0.53356688 | FALSE | 1 | FALSE | 1 | FALSE | 1 | FALSE | 1 | FALSE | 1 |
| 4626 | 4626 | 8.67647059 | 2.51724158 | 1 | 0.58738519 | 0 | 1 | 1 | 0.71261921 | 1 | 0.7984848 | 1 | 0.43077488 | 30 | 0.36879488 | FALSE | 1 | FALSE | 1 | FALSE | 1 | FALSE | 1 | FALSE | 1 | FALSE | 1 |
| 4627 | 4627 | 7.85714286 | 2.53143502 | 2 | 0.28329504 | 0 | 1 | 2 | 0.43640583 | 3 | 0.29054213 | 0 | 1 | 34 | 0.69505404 | FALSE | 1 | FALSE | 1 | FALSE | 1 | FALSE | 1 | FALSE | 1 | FALSE | 1 |
| 4628 | 4628 | 10.48 | 2.59414726 | 0 | 1 | 0 | 1 | 1 | 0.58502114 | 3 | 0.09525947 | 0 | 1 | 20 | 0.68270729 | FALSE | 1 | FALSE | 1 | FALSE | 1 | FALSE | 1 | FALSE | 1 | FALSE | 1 |
| 4629 | 4629 | 9.33333333 | 3.64539283 | 0 | 1 | 2 | 0.08295839 | 0 | 1 | 1 | 0.50623194 | 0 | 1 | 12 | 0.80329917 | FALSE | 1 | FALSE | 1 | FALSE | 1 | FALSE | 1 | FALSE | 1 | FALSE | 1 |
| 4630 | 4630 | 10.8888889 | 1.9969112 | 2 | 0.07645454 | 2 | 0.1135776 | 0 | 1 | 1 | 0.57131013 | 0 | 1 | 13 | 0.95041913 | FALSE | 0.77424195 | FALSE | 1 | FALSE | 1 | FALSE | 1 | FALSE | 1 | FALSE | 1 |
| 4631 | 4631 | 7.66666667 | 3.2180049 | 0 | 1 | 0 | 1 | 0 | 1 | 1 | 0.50623194 | 0 | 1 | 14 | 0.29525409 | FALSE | 1 | FALSE | 1 | FALSE | 1 | FALSE | 1 | FALSE | 1 | FALSE | 1 |
| 4632 | 4632 | 12.5 | 3.84986085 | 1 | 0.3050805 | 1 | 0.36919824 | 0 | 1 | 0 | 1 | 0 | 1 | 12 | 0.62236227 | FALSE | 1 | FALSE | 1 | FALSE | 1 | FALSE | 1 | FALSE | 1 | FALSE | 1 |
| 4633 | 4633 | 11.375 | 2.78107443 | 0 | 1 | 0 | 1 | 0 | 1 | 0 | 1 | 1 | 0.12394282 | 7 | 0.63752397 | FALSE | 1 | FALSE | 1 | FALSE | 1 | FALSE | 1 | FALSE | 1 | FALSE | 1 |
| 4634 | 4634 | 10.1162791 | 3.41139883 | 1 | 0.67382914 | 0 | 1 | 1 | 0.79363407 | 1 | 0.86831187 | 0 | 1 | 40 | 0.07890873 | FALSE | 1 | FALSE | 1 | FALSE | 1 | FALSE | 1 | FALSE | 1 | FALSE | 1 |
| 4635 | 4635 | 7.14285714 | 2.79941685 | 0 | 1 | 1 | 0.36919824 | 0 | 1 | 0 | 1 | 0 | 1 | 13 | 0.33283729 | FALSE | 1 | FALSE | 1 | FALSE | 1 | FALSE | 1 | FALSE | 1 | FALSE | 1 |
| 4636 | 4636 | 10.4594595 | 3.35788378 | 5 | 0.00234263 | 2 | 0.3375163 | 1 | 0.74264075 | 3 | 0.2402032 | 0 | 1 | 26 | 0.99170905 | FALSE | 0.12737083 | FALSE | 1 | FALSE | 1 | FALSE | 1 | FALSE | 1 | FALSE | 1 |
| 4637 | 4637 | 8.32352941 | 3.53736848 | 0 | 1 | 1 | 0.65167362 | 0 | 1 | 2 | 0.43583874 | 0 | 1 | 29 | 0.24088541 | FALSE | 1 | FALSE | 1 | FALSE | 1 | FALSE | 1 | FALSE | 1 | FALSE | 1 |
| 4638 | 4638 | 9.88461539 | 2.57692308 | 2 | 0.14250978 | 3 | 0.05021553 | 1 | 0.61440466 | 2 | 0.33678042 | 0 | 1 | 18 | 0.98644281 | FALSE | 1 | FALSE | 1 | FALSE | 1 | FALSE | 1 | FALSE | 1 | FALSE | 1 |
| 4639 | 4639 | 10.0869565 | 2.90296765 | 0 | 1 | 0 | 1 | 2 | 0.19931429 | 3 | 0.08614834 | 0 | 1 | 18 | 0.86273155 | FALSE | 1 | FALSE | 1 | FALSE | 1 | FALSE | 1 | FALSE | 1 | FALSE | 1 |
| 4640 | 4640 | 6.65714286 | 1.81962095 | 6 | 0.05226303 | 6 | 0.12399641 | 7 | 0.0828979 | 2 | 0.95757385 | 13 | 1.20E-08 | 71 | 0.99999542 | FALSE | 0.68392765 | FALSE | 1 | FALSE | 1 | FALSE | 1 | TRUE | 1.96E-05 | FALSE | 1 |
| 4641 | 4641 | 9.89285714 | 2.07604166 | 1 | 0.51745948 | 0 | 1 | 0 | 1 | 0 | 1 | 1 | 0.37113067 | 26 | 0.16405547 | FALSE | 1 | FALSE | 1 | FALSE | 1 | FALSE | 1 | FALSE | 1 | FALSE | 1 |
| 4642 | 4642 | 10.28 | 2.63089338 | 1 | 0.46441868 | 1 | 0.54637326 | 1 | 0.58502114 | 0 | 1 | 0 | 1 | 21 | 0.47053137 | FALSE | 1 | FALSE | 1 | FALSE | 1 | FALSE | 1 | FALSE | 1 | FALSE | 1 |
| 4643 | 4643 | 6.7948718 | 2.84811667 | 1 | 0.63789042 | 0 | 1 | 3 | 0.16357218 | 4 | 0.10199045 | 0 | 1 | 31 | 0.85613899 | FALSE | 1 | FALSE | 1 | FALSE | 1 | FALSE | 1 | FALSE | 1 | FALSE | 1 |
| 4644 | 4644 | 6.6969697 | 2.86551966 | 0 | 1 | 4 | 0.02089081 | 1 | 0.70185639 | 1 | 0.78873814 | 0 | 1 | 27 | 0.75124583 | FALSE | 1 | FALSE | 0.8034624 | FALSE | 1 | FALSE | 1 | FALSE | 1 | FALSE | 1 |
| 4645 | 4645 | 10.1111111 | 1.5234788 | 0 | 1 | 0 | 1 | 0 | 1 | 1 | 0.34506517 | 0 | 1 | 8 | 0.5780332 | FALSE | 1 | FALSE | 1 | FALSE | 1 | FALSE | 1 | FALSE | 1 | FALSE | 1 |
| 4646 | 4646 | 7.56382979 | 2.69577491 | 9 | 0.00050035 | 2 | 0.79919134 | 5 | 0.22832872 | 4 | 0.60708936 | 1 | 0.78011711 | 70 | 0.97872956 | TRUE | 0.04605458 | FALSE | 1 | FALSE | 1 | FALSE | 1 | FALSE | 1 | FALSE | 1 |
| 4647 | 4647 | 9.30555556 | 3.0351592 | 2 | 0.23542589 | 0 | 1 | 1 | 0.73299778 | 1 | 0.81665383 | 0 | 1 | 32 | 0.31709555 | FALSE | 1 | FALSE | 1 | FALSE | 1 | FALSE | 1 | FALSE | 1 | FALSE | 1 |
| 4648 | 4648 | 5.93684211 | 2.76752037 | 4 | 0.22626568 | 5 | 0.1932531 | 4 | 0.44662765 | 5 | 0.44344573 | 2 | 0.46405726 | 75 | 0.94176825 | FALSE | 1 | FALSE | 1 | FALSE | 1 | FALSE | 1 | FALSE | 1 | FALSE | 1 |
| 4649 | 4649 | 8.33333333 | 2.10818511 | 0 | 1 | 0 | 1 | 2 | 0.03916012 | 0 | 1 | 0 | 1 | 7 | 0.84574168 | FALSE | 1 | FALSE | 1 | FALSE | 1 | FALSE | 1 | FALSE | 1 | FALSE | 1 |
| 4650 | 4650 | 9.75 | 3.71651719 | 1 | 0.34033054 | 0 | 1 | 2 | 0.11087977 | 0 | 1 | 1 | 0.23264612 | 12 | 0.90922186 | FALSE | 1 | FALSE | 1 | FALSE | 1 | FALSE | 1 | FALSE | 1 | FALSE | 1 |
| 4651 | 4651 | 7.24489796 | 0.88133971 | 7 | 0.01195472 | 3 | 0.61262749 | 3 | 0.68330856 | 6 | 0.28608665 | 0 | 1 | 78 | 0.88735927 | FALSE | 0.33872393 | FALSE | 1 | FALSE | 1 | FALSE | 1 | FALSE | 1 | FALSE | 1 |
| 4652 | 4652 | 4.69158879 | 2.21469694 | 5 | 0.13951176 | 2 | 0.86638697 | 0 | 1 | 4 | 0.73116831 | 1 | 0.83189113 | 95 | 0.12669417 | FALSE | 1 | FALSE | 1 | FALSE | 1 | FALSE | 1 | FALSE | 1 | FALSE | 1 |
| 4653 | 4653 | 11.2857143 | 2.13570234 | 2 | 0.16048326 | 0 | 1 | 1 | 0.64171563 | 3 | 0.13524571 | 1 | 0.37113067 | 21 | 0.94083768 | FALSE | 1 | FALSE | 1 | FALSE | 1 | FALSE | 1 | FALSE | 1 | FALSE | 1 |
| 4654 | 4654 | 7.34782609 | 2.40667 | 1 | 0.45027683 | 2 | 0.16948764 | 0 | 1 | 2 | 0.2852236 | 0 | 1 | 18 | 0.86273155 | FALSE | 1 | FALSE | 1 | FALSE | 1 | FALSE | 1 | FALSE | 1 | FALSE | 1 |
| 4655 | 4655 | 9.92307692 | 2.12897731 | 0 | 1 | 0 | 1 | 0 | 1 | 2 | 0.1174583 | 0 | 1 | 11 | 0.6683056 | FALSE | 1 | FALSE | 1 | FALSE | 1 | FALSE | 1 | FALSE | 1 | FALSE | 1 |
| 4656 | 4656 | 10.1428571 | 2.56878171 | 1 | 0.42086991 | 0 | 1 | 0 | 1 | 1 | 0.62783522 | 0 | 1 | 19 | 0.34094959 | FALSE | 1 | FALSE | 1 | FALSE | 1 | FALSE | 1 | FALSE | 1 | FALSE | 1 |
| 4657 | 4657 | 10.0659341 | 1.86230507 | 2 | 0.68263499 | 4 | 0.33970769 | 4 | 0.41421418 | 4 | 0.60708936 | 1 | 0.78011711 | 76 | 0.65586987 | FALSE | 1 | FALSE | 1 | FALSE | 1 | FALSE | 1 | FALSE | 1 | FALSE | 1 |
| 4658 | 4658 | 9.42857143 | 2.29017422 | 3 | 0.03391597 | 3 | 0.06039219 | 0 | 1 | 1 | 0.73248983 | 0 | 1 | 21 | 0.94083768 | FALSE | 0.56934354 | FALSE | 1 | FALSE | 1 | FALSE | 1 | FALSE | 1 | FALSE | 1 |
| 4659 | 4659 | 8.29411765 | 2.58422149 | 0 | 1 | 1 | 0.42857733 | 0 | 1 | 0 | 1 | 0 | 1 | 16 | 0.23070182 | FALSE | 1 | FALSE | 1 | FALSE | 1 | FALSE | 1 | FALSE | 1 | FALSE | 1 |
| 4660 | 4660 | 5.16535433 | 2.36364785 | 4 | 0.39363378 | 3 | 0.77097552 | 6 | 0.28660842 | 5 | 0.67988626 | 2 | 0.60769448 | 104 | 0.62072125 | FALSE | 1 | FALSE | 1 | FALSE | 1 | FALSE | 1 | FALSE | 1 | FALSE | 1 |
| 4661 | 4661 | 9.55555556 | 3.03477784 | 0 | 1 | 2 | 0.21684246 | 0 | 1 | 1 | 0.71956339 | 0 | 1 | 24 | 0.3725373 | FALSE | 1 | FALSE | 1 | FALSE | 1 | FALSE | 1 | FALSE | 1 | FALSE | 1 |
| 4662 | 4662 | 11.2352941 | 3.40465791 | 2 | 0.06909038 | 0 | 1 | 0 | 1 | 1 | 0.55062772 | 0 | 1 | 14 | 0.7313145 | FALSE | 0.74276693 | FALSE | 1 | FALSE | 1 | FALSE | 1 | FALSE | 1 | FALSE | 1 |
| 4663 | 4663 | 7.48148148 | 4.04026375 | 0 | 1 | 1 | 0.58913618 | 0 | 1 | 1 | 0.71956339 | 0 | 1 | 25 | 0.1832206 | FALSE | 1 | FALSE | 1 | FALSE | 1 | FALSE | 1 | FALSE | 1 | FALSE | 1 |
| 4664 | 4664 | 10.3846154 | 3.1265233 | 1 | 0.28675951 | 0 | 1 | 0 | 1 | 0 | 1 | 0 | 1 | 12 | 0.374198 | FALSE | 1 | FALSE | 1 | FALSE | 1 | FALSE | 1 | FALSE | 1 | FALSE | 1 |
| 4665 | 4665 | 9.54166667 | 2.75347003 | 2 | 0.12501436 | 1 | 0.54637326 | 0 | 1 | 1 | 0.67692816 | 2 | 0.058175 | 18 | 0.93166998 | FALSE | 0.95622564 | FALSE | 1 | FALSE | 1 | FALSE | 1 | FALSE | 1 | FALSE | 1 |
| 4666 | 4666 | 8.875 | 3.44374433 | 0 | 1 | 0 | 1 | 1 | 0.69069222 | 1 | 0.77852168 | 1 | 0.41154418 | 29 | 0.24088541 | FALSE | 1 | FALSE | 1 | FALSE | 1 | FALSE | 1 | FALSE | 1 | FALSE | 1 |
| 4667 | 4667 | 6.92592593 | 1.82423857 | 1 | 0.50471032 | 2 | 0.21684246 | 3 | 0.07106362 | 1 | 0.71956339 | 1 | 0.36060486 | 19 | 0.98241223 | FALSE | 1 | FALSE | 1 | FALSE | 1 | FALSE | 1 | FALSE | 1 | FALSE | 1 |
| 4668 | 4668 | 9.23529412 | 1.76666558 | 1 | 0.35728305 | 0 | 1 | 0 | 1 | 1 | 0.55062772 | 1 | 0.24525764 | 14 | 0.7313145 | FALSE | 1 | FALSE | 1 | FALSE | 1 | FALSE | 1 | FALSE | 1 | FALSE | 1 |
| 4669 | 4669 | 7.18333333 | 1.54371486 | 1 | 0.79096998 | 0 | 1 | 0 | 1 | 3 | 0.52470434 | 1 | 0.63076845 | 55 | 0.07598467 | FALSE | 1 | FALSE | 1 | FALSE | 1 | FALSE | 1 | FALSE | 1 | FALSE | 1 |
| 4670 | 4670 | 8.02857143 | 3.62125246 | 2 | 0.22589135 | 1 | 0.68454804 | 3 | 0.12958859 | 0 | 1 | 0 | 1 | 29 | 0.69927719 | FALSE | 1 | FALSE | 1 | FALSE | 1 | FALSE | 1 | FALSE | 1 | FALSE | 1 |
| 4671 | 4671 | 7.96296296 | 2.31710997 | 0 | 1 | 0 | 1 | 0 | 1 | 2 | 0.35374209 | 0 | 1 | 25 | 0.1832206 | FALSE | 1 | FALSE | 1 | FALSE | 1 | FALSE | 1 | FALSE | 1 | FALSE | 1 |
| 4672 | 4672 | 9.16129032 | 2.42429603 | 1 | 0.55378205 | 0 | 1 | 2 | 0.30673593 | 2 | 0.41984187 | 2 | 0.09110253 | 24 | 0.90138666 | FALSE | 1 | FALSE | 1 | FALSE | 1 | FALSE | 1 | FALSE | 1 | FALSE | 1 |
| 4673 | 4673 | 10.8571429 | 1.88441514 | 0 | 1 | 1 | 0.36919824 | 0 | 1 | 1 | 0.48242099 | 0 | 1 | 12 | 0.62236227 | FALSE | 1 | FALSE | 1 | FALSE | 1 | FALSE | 1 | FALSE | 1 | FALSE | 1 |
| 4674 | 4674 | 9.875 | 2.89125838 | 0 | 1 | 0 | 1 | 1 | 0.44344817 | 1 | 0.52895088 | 0 | 1 | 14 | 0.53308261 | FALSE | 1 | FALSE | 1 | FALSE | 1 | FALSE | 1 | FALSE | 1 | FALSE | 1 |
| 4675 | 4675 | 8.31578947 | 3.21268262 | 0 | 1 | 1 | 0.46503795 | 0 | 1 | 0 | 1 | 1 | 0.26986775 | 17 | 0.41141543 | FALSE | 1 | FALSE | 1 | FALSE | 1 | FALSE | 1 | FALSE | 1 | FALSE | 1 |
| 4676 | 4676 | 10.0833333 | 2.69129254 | 0 | 1 | 1 | 0.32623933 | 1 | 0.35555293 | 0 | 1 | 0 | 1 | 10 | 0.71436964 | FALSE | 1 | FALSE | 1 | FALSE | 1 | FALSE | 1 | FALSE | 1 | FALSE | 1 |
| 4677 | 4677 | 7.7254902 | 2.76588766 | 3 | 0.13593669 | 1 | 0.80796869 | 2 | 0.54120634 | 4 | 0.19569978 | 1 | 0.56379725 | 39 | 0.92022511 | FALSE | 0.98986074 | FALSE | 1 | FALSE | 1 | FALSE | 1 | FALSE | 1 | FALSE | 1 |
| 4678 | 4678 | 5.73684211 | 3.47687254 | 3 | 0.07268126 | 2 | 0.3494045 | 4 | 0.04612151 | 0 | 1 | 0 | 1 | 29 | 0.93814695 | FALSE | 0.76727438 | FALSE | 1 | FALSE | 1 | FALSE | 1 | FALSE | 1 | FALSE | 1 |
| 4679 | 4679 | 9.83333333 | 1.63469331 | 0 | 1 | 0 | 1 | 0 | 1 | 0 | 1 | 1 | 0.39166989 | 29 | 0.03983039 | FALSE | 1 | FALSE | 1 | FALSE | 1 | FALSE | 1 | FALSE | 1 | FALSE | 1 |
| 4680 | 4680 | 10.4705882 | 2.54645421 | 0 | 1 | 1 | 0.42857733 | 1 | 0.46348836 | 0 | 1 | 0 | 1 | 15 | 0.49060435 | FALSE | 1 | FALSE | 1 | FALSE | 1 | FALSE | 1 | FALSE | 1 | FALSE | 1 |
| 4681 | 4681 | 7.71794872 | 2.43846019 | 0 | 1 | 2 | 0.72420282 | 2 | 0.77659406 | 3 | 0.7021069 | 2 | 0.36704308 | 69 | 0.20270862 | FALSE | 1 | FALSE | 1 | FALSE | 1 | FALSE | 1 | FALSE | 1 | FALSE | 1 |
| 4682 | 4682 | 8.69444444 | 1.3085073 | 2 | 0.23542589 | 3 | 0.10944025 | 3 | 0.13783405 | 1 | 0.81665383 | 1 | 0.44938266 | 26 | 0.98200522 | FALSE | 1 | FALSE | 1 | FALSE | 1 | FALSE | 1 | FALSE | 1 | FALSE | 1 |
| 4683 | 4683 | 11.4 | 1.74355958 | 0 | 1 | 0 | 1 | 0 | 1 | 0 | 1 | 0 | 1 | 5 | 0.42761893 | FALSE | 1 | FALSE | 1 | FALSE | 1 | FALSE | 1 | FALSE | 1 | FALSE | 1 |
| 4684 | 4684 | 9.71052632 | 2.1387145 | 0 | 1 | 3 | 0.12352882 | 1 | 0.75193685 | 2 | 0.52640931 | 1 | 0.4673875 | 31 | 0.76640548 | FALSE | 1 | FALSE | 1 | FALSE | 1 | FALSE | 1 | FALSE | 1 | FALSE | 1 |
| 4685 | 4685 | 11.5357143 | 2.16270569 | 0 | 1 | 1 | 0.60247911 | 2 | 0.2663542 | 1 | 0.73248983 | 0 | 1 | 24 | 0.55018107 | FALSE | 1 | FALSE | 1 | FALSE | 1 | FALSE | 1 | FALSE | 1 | FALSE | 1 |
| 4686 | 4686 | 9.04761905 | 2.01130591 | 0 | 1 | 0 | 1 | 2 | 0.17308361 | 2 | 0.25059627 | 0 | 1 | 17 | 0.77849076 | FALSE | 1 | FALSE | 1 | FALSE | 1 | FALSE | 1 | FALSE | 1 | FALSE | 1 |
| 4687 | 4687 | 9.0625 | 2.81666359 | 1 | 0.34033054 | 3 | 0.01368613 | 0 | 1 | 3 | 0.03440583 | 0 | 1 | 9 | 0.99862766 | FALSE | 1 | FALSE | 0.69207862 | FALSE | 1 | FALSE | 1 | FALSE | 1 | FALSE | 1 |
| 4688 | 4688 | 10.5294118 | 2.47618414 | 0 | 1 | 2 | 0.10307112 | 1 | 0.46348836 | 0 | 1 | 0 | 1 | 14 | 0.7313145 | FALSE | 1 | FALSE | 1 | FALSE | 1 | FALSE | 1 | FALSE | 1 | FALSE | 1 |
| 4689 | 4689 | 10.8125 | 1.62860178 | 1 | 0.34033054 | 1 | 0.40942991 | 0 | 1 | 0 | 1 | 0 | 1 | 14 | 0.53308261 | FALSE | 1 | FALSE | 1 | FALSE | 1 | FALSE | 1 | FALSE | 1 | FALSE | 1 |
| 4690 | 4690 | 7.5 | 3.57071421 | 0 | 1 | 0 | 1 | 1 | 0.13615783 | 0 | 1 | 0 | 1 | 3 | 0.8823539 | FALSE | 1 | FALSE | 1 | FALSE | 1 | FALSE | 1 | FALSE | 1 | FALSE | 1 |
| 4691 | 4691 | 7.35714286 | 1.49318178 | 1 | 0.3050805 | 1 | 0.36919824 | 0 | 1 | 0 | 1 | 0 | 1 | 12 | 0.62236227 | FALSE | 1 | FALSE | 1 | FALSE | 1 | FALSE | 1 | FALSE | 1 | FALSE | 1 |
| 4692 | 4692 | 8.25 | 2.20321734 | 2 | 0.03643311 | 0 | 1 | 0 | 1 | 1 | 0.43131129 | 0 | 1 | 9 | 0.89630573 | FALSE | 0.58392 | FALSE | 1 | FALSE | 1 | FALSE | 1 | FALSE | 1 | FALSE | 1 |
| 4693 | 4693 | 6.67346939 | 1.74247953 | 0 | 1 | 1 | 0.80150109 | 1 | 0.83455705 | 3 | 0.39227316 | 1 | 0.55647171 | 43 | 0.337822 | FALSE | 1 | FALSE | 1 | FALSE | 1 | FALSE | 1 | FALSE | 1 | FALSE | 1 |
| 4694 | 4694 | 8.84615385 | 2.76923077 | 2 | 0.04234484 | 0 | 1 | 0 | 1 | 0 | 1 | 0 | 1 | 11 | 0.6683056 | FALSE | 0.62842194 | FALSE | 1 | FALSE | 1 | FALSE | 1 | FALSE | 1 | FALSE | 1 |
| 4695 | 4695 | 10.2777778 | 1.96654111 | 0 | 1 | 0 | 1 | 1 | 0.48280983 | 1 | 0.57131013 | 0 | 1 | 16 | 0.44998001 | FALSE | 1 | FALSE | 1 | FALSE | 1 | FALSE | 1 | FALSE | 1 | FALSE | 1 |
| 4696 | 4696 | 10.8947368 | 2.12490834 | 1 | 0.37380237 | 0 | 1 | 1 | 0.48280983 | 0 | 1 | 0 | 1 | 16 | 0.44998001 | FALSE | 1 | FALSE | 1 | FALSE | 1 | FALSE | 1 | FALSE | 1 | FALSE | 1 |
| 4697 | 4697 | 10.1363636 | 2.71854665 | 2 | 0.10809809 | 2 | 0.15794013 | 1 | 0.55340851 | 0 | 1 | 1 | 0.30529951 | 16 | 0.95553951 | FALSE | 0.88820416 | FALSE | 1 | FALSE | 1 | FALSE | 1 | FALSE | 1 | FALSE | 1 |
| 4698 | 4698 | 9.1 | 2.09523268 | 0 | 1 | 0 | 1 | 2 | 0.16019548 | 1 | 0.60987146 | 0 | 1 | 17 | 0.61750632 | FALSE | 1 | FALSE | 1 | FALSE | 1 | FALSE | 1 | FALSE | 1 | FALSE | 1 |

| | | | | | | | | | | | | | | | | | | | | | | | | | | | |
|---|---|---|---|---|---|---|---|---|---|---|---|---|---|---|---|---|---|---|---|---|---|---|---|---|---|---|---|
| 4699 | 4699 | 5.70689655 | 1.83837091 | 2 | 0.44060625 | 0 | 1 | 3 | 0.34634641 | 2 | 0.75303313 | 2 | 0.24594209 | 49 | 0.57916832 | FALSE | 1 | FALSE | 1 | FALSE | 1 | FALSE | 1 | FALSE | 1 | FALSE | 1 |
| 4700 | 4700 | 9.23333333 | 1.96100881 | 1 | 0.54198714 | 2 | 0.25306331 | 1 | 0.66709967 | 1 | 0.75658797 | 1 | 0.39166989 | 24 | 0.82294782 | FALSE | 1 | FALSE | 1 | FALSE | 1 | FALSE | 1 | FALSE | 1 | FALSE | 1 |
| 4701 | 4701 | 8.46774194 | 1.71066618 | 2 | 0.45796754 | 3 | 0.30686508 | 1 | 0.8897349 | 2 | 0.76960882 | 1 | 0.63076845 | 51 | 0.5343051 | FALSE | 1 | FALSE | 1 | FALSE | 1 | FALSE | 1 | FALSE | 1 | FALSE | 1 |
| 4702 | 4702 | 10.75 | 2.04633819 | 3 | 0.00307177 | 0 | 1 | 0 | 1 | 1 | 0.43131129 | 0 | 1 | 8 | 0.97197833 | FALSE | 0.14802295 | FALSE | 1 | FALSE | 1 | FALSE | 1 | FALSE | 1 | FALSE | 1 |
| 4703 | 4703 | 7.84615385 | 2.38089039 | 2 | 0.12501436 | 1 | 0.54637326 | 1 | 0.58502114 | 0 | 1 | 0 | 1 | 20 | 0.68270729 | FALSE | 0.95622564 | FALSE | 1 | FALSE | 1 | FALSE | 1 | FALSE | 1 | FALSE | 1 |
| 4704 | 4704 | 9.77272727 | 2.1091105 | 1 | 0.4357638 | 0 | 1 | 1 | 0.55340851 | 0 | 1 | 0 | 1 | 20 | 0.30915202 | FALSE | 1 | FALSE | 1 | FALSE | 1 | FALSE | 1 | FALSE | 1 | FALSE | 1 |
| 4705 | 4705 | 8.4137931 | 2.17378173 | 3 | 0.03713651 | 0 | 1 | 1 | 0.65463982 | 0 | 1 | 0 | 1 | 25 | 0.51779518 | FALSE | 0.58392 | FALSE | 1 | FALSE | 1 | FALSE | 1 | FALSE | 1 | FALSE | 1 |
| 4706 | 4706 | 4.3385214 | 1.34011533 | 3 | 0.96099927 | 10 | 0.30249236 | 5 | 0.9538551 | 13 | 0.37802968 | 3 | 0.79078246 | 219 | 0.18876309 | FALSE | 1 | FALSE | 1 | FALSE | 1 | FALSE | 1 | FALSE | 1 | FALSE | 1 |
| 4707 | 4707 | 11.8 | 0.8326664 | 0 | 1 | 0 | 1 | 0 | 1 | 2 | 0.14906763 | 0 | 1 | 13 | 0.57712805 | FALSE | 1 | FALSE | 1 | FALSE | 1 | FALSE | 1 | FALSE | 1 | FALSE | 1 |
| 4708 | 4708 | 8.33333333 | 2.7080128 | 1 | 0.37380237 | 1 | 0.44710673 | 0 | 1 | 0 | 1 | 0 | 1 | 16 | 0.44998001 | FALSE | 1 | FALSE | 1 | FALSE | 1 | FALSE | 1 | FALSE | 1 | FALSE | 1 |
| 4709 | 4709 | 9 | 2.35487888 | 0 | 1 | 0 | 1 | 1 | 0.55340851 | 1 | 0.64497439 | 0 | 1 | 20 | 0.30915202 | FALSE | 1 | FALSE | 1 | FALSE | 1 | FALSE | 1 | FALSE | 1 | FALSE | 1 |
| 4710 | 4710 | 7.88888889 | 2.92287699 | 1 | 0.37380237 | 0 | 1 | 0 | 1 | 0 | 1 | 0 | 1 | 17 | 0.20329205 | FALSE | 1 | FALSE | 1 | FALSE | 1 | FALSE | 1 | FALSE | 1 | FALSE | 1 |
| 4711 | 4711 | 6.66666667 | 2.95334086 | 0 | 1 | 0 | 1 | 1 | 0.35555293 | 0 | 1 | 1 | 0.18007762 | 10 | 0.71436964 | FALSE | 1 | FALSE | 1 | FALSE | 1 | FALSE | 1 | FALSE | 1 | FALSE | 1 |
| 4712 | 4712 | 11.6470588 | 1.60664709 | 0 | 1 | 1 | 0.42857733 | 1 | 0.46348836 | 0 | 1 | 0 | 1 | 15 | 0.49060435 | FALSE | 1 | FALSE | 1 | FALSE | 1 | FALSE | 1 | FALSE | 1 | FALSE | 1 |
| 4713 | 4713 | 9.91666667 | 3.56974167 | 1 | 0.26795838 | 1 | 0.32623933 | 0 | 1 | 1 | 0.43131129 | 0 | 1 | 9 | 0.89630573 | FALSE | 1 | FALSE | 1 | FALSE | 1 | FALSE | 1 | FALSE | 1 | FALSE | 1 |
| 4714 | 4714 | 8.13333333 | 2.21710522 | 0 | 1 | 2 | 0.25306331 | 2 | 0.29330414 | 2 | 0.4036176 | 0 | 1 | 24 | 0.82294782 | FALSE | 1 | FALSE | 1 | FALSE | 1 | FALSE | 1 | FALSE | 1 | FALSE | 1 |
| 4715 | 4715 | 6.92571429 | 1.51569746 | 8 | 0.07806122 | 10 | 0.05515706 | 9 | 0.17342045 | 12 | 0.10311031 | 0 | 1 | 135 | 0.99373938 | FALSE | 0.78080278 | FALSE | 1 | FALSE | 1 | FALSE | 1 | FALSE | 1 | FALSE | 1 |
| 4716 | 4716 | 9.94736842 | 1.57191416 | 0 | 1 | 2 | 0.12434464 | 0 | 1 | 2 | 0.21610812 | 1 | 0.26986775 | 14 | 0.93677253 | FALSE | 1 | FALSE | 1 | FALSE | 1 | FALSE | 1 | FALSE | 1 | FALSE | 1 |
| 4717 | 4717 | 9.72222222 | 3.1763983 | 1 | 0.37380237 | 1 | 0.44710673 | 2 | 0.13502329 | 1 | 0.57131013 | 1 | 0.25766374 | 12 | 0.98544073 | FALSE | 1 | FALSE | 1 | FALSE | 1 | FALSE | 1 | FALSE | 1 | FALSE | 1 |
| 4718 | 4718 | 8.6 | 2.33238076 | 0 | 1 | 1 | 0.15163655 | 0 | 1 | 0 | 1 | 0 | 1 | 4 | 0.82371636 | FALSE | 1 | FALSE | 1 | FALSE | 1 | FALSE | 1 | FALSE | 1 | FALSE | 1 |
| 4719 | 4719 | 8.47619048 | 2.42249473 | 0 | 1 | 1 | 0.49918218 | 0 | 1 | 0 | 1 | 0 | 1 | 20 | 0.13763971 | FALSE | 1 | FALSE | 1 | FALSE | 1 | FALSE | 1 | FALSE | 1 | FALSE | 1 |
| 4720 | 4720 | 8.33884298 | 1.38798694 | 2 | 0.82201398 | 7 | 0.0965406 | 3 | 0.81621325 | 7 | 0.32018532 | 2 | 0.59426096 | 100 | 0.74937213 | FALSE | 1 | FALSE | 1 | FALSE | 1 | FALSE | 1 | FALSE | 1 | FALSE | 1 |
| 4721 | 4721 | 10.8333333 | 2.33927814 | 2 | 0.12501436 | 0 | 1 | 0 | 1 | 0 | 1 | 0 | 1 | 22 | 0.25237063 | FALSE | 0.95622564 | FALSE | 1 | FALSE | 1 | FALSE | 1 | FALSE | 1 | FALSE | 1 |
| 4722 | 4722 | 10.0384615 | 3.80769231 | 1 | 0.47819878 | 2 | 0.19297191 | 2 | 0.22597287 | 2 | 0.31968852 | 0 | 1 | 18 | 0.96851178 | FALSE | 1 | FALSE | 1 | FALSE | 1 | FALSE | 1 | FALSE | 1 | FALSE | 1 |
| 4723 | 4723 | 10.1428571 | 2.38618473 | 0 | 1 | 0 | 1 | 0 | 1 | 1 | 0.48242099 | 0 | 1 | 13 | 0.33283729 | FALSE | 1 | FALSE | 1 | FALSE | 1 | FALSE | 1 | FALSE | 1 | FALSE | 1 |
| 4724 | 4724 | 8.63157895 | 1.17922929 | 1 | 0.38989951 | 1 | 0.46503795 | 0 | 1 | 1 | 0.59104357 | 0 | 1 | 16 | 0.65565778 | FALSE | 1 | FALSE | 1 | FALSE | 1 | FALSE | 1 | FALSE | 1 | FALSE | 1 |
| 4725 | 4725 | 8.06060606 | 1.52632246 | 3 | 0.23916009 | 2 | 0.63534334 | 5 | 0.08755999 | 5 | 0.18498818 | 3 | 0.09357908 | 48 | 0.99535371 | FALSE | 1 | FALSE | 1 | FALSE | 1 | FALSE | 1 | FALSE | 1 | FALSE | 1 |
| 4726 | 4726 | 11.1428571 | 2.55949612 | 0 | 1 | 0 | 1 | 2 | 0.08806398 | 0 | 1 | 0 | 1 | 12 | 0.62236227 | FALSE | 1 | FALSE | 1 | FALSE | 1 | FALSE | 1 | FALSE | 1 | FALSE | 1 |
| 4727 | 4727 | 7.48148148 | 2.23391983 | 0 | 1 | 4 | 0.01041264 | 3 | 0.07106362 | 3 | 0.12475952 | 1 | 0.36060486 | 16 | 0.99964933 | FALSE | 1 | FALSE | 0.67617265 | FALSE | 1 | FALSE | 1 | FALSE | 1 | FALSE | 1 |
| 4728 | 4728 | 9.2 | 3.31058907 | 1 | 0.3050805 | 1 | 0.36919824 | 0 | 1 | 0 | 1 | 1 | 0.20679328 | 11 | 0.83676449 | FALSE | 1 | FALSE | 1 | FALSE | 1 | FALSE | 1 | FALSE | 1 | FALSE | 1 |
| 4729 | 4729 | 8.86363636 | 1.93755832 | 0 | 1 | 2 | 0.15794013 | 1 | 0.55340851 | 1 | 0.64497439 | 0 | 1 | 18 | 0.74744813 | FALSE | 1 | FALSE | 1 | FALSE | 1 | FALSE | 1 | FALSE | 1 | FALSE | 1 |
| 4730 | 4730 | 10.2142857 | 2.33539753 | 1 | 0.3050805 | 1 | 0.36919824 | 0 | 1 | 0 | 1 | 0 | 1 | 12 | 0.62236227 | FALSE | 1 | FALSE | 1 | FALSE | 1 | FALSE | 1 | FALSE | 1 | FALSE | 1 |
| 4731 | 4731 | 8.125 | 3.25720356 | 1 | 0.18769745 | 0 | 1 | 0 | 1 | 0 | 1 | 0 | 1 | 7 | 0.63752397 | FALSE | 1 | FALSE | 1 | FALSE | 1 | FALSE | 1 | FALSE | 1 | FALSE | 1 |
| 4732 | 4732 | 9.05555556 | 3.11755286 | 0 | 1 | 3 | 0.01901931 | 0 | 1 | 2 | 0.19902682 | 0 | 1 | 13 | 0.95041913 | FALSE | 1 | FALSE | 0.77515497 | FALSE | 1 | FALSE | 1 | FALSE | 1 | FALSE | 1 |
| 4733 | 4733 | 11.95 | 1.49916644 | 0 | 1 | 0 | 1 | 0 | 1 | 1 | 0.60987146 | 0 | 1 | 19 | 0.15700255 | FALSE | 1 | FALSE | 1 | FALSE | 1 | FALSE | 1 | FALSE | 1 | FALSE | 1 |
| 4734 | 4734 | 9.47368421 | 1.56838573 | 1 | 0.38989951 | 0 | 1 | 0 | 1 | 1 | 0.59104357 | 0 | 1 | 17 | 0.41141543 | FALSE | 1 | FALSE | 1 | FALSE | 1 | FALSE | 1 | FALSE | 1 | FALSE | 1 |
| 4735 | 4735 | 9.8 | 2.4 | 0 | 1 | 0 | 1 | 0 | 1 | 0 | 1 | 1 | 0.2198258 | 14 | 0.29525409 | FALSE | 1 | FALSE | 1 | FALSE | 1 | FALSE | 1 | FALSE | 1 | FALSE | 1 |
| 4736 | 4736 | 9.375 | 3.4255474 | 1 | 0.34033054 | 0 | 1 | 0 | 1 | 1 | 0.52895088 | 1 | 0.23264612 | 13 | 0.76799161 | FALSE | 1 | FALSE | 1 | FALSE | 1 | FALSE | 1 | FALSE | 1 | FALSE | 1 |
| 4737 | 4737 | 8.275 | 1.28427995 | 2 | 0.27371785 | 0 | 1 | 0 | 1 | 2 | 0.5543522 | 1 | 0.48480878 | 35 | 0.38966407 | FALSE | 1 | FALSE | 1 | FALSE | 1 | FALSE | 1 | FALSE | 1 | FALSE | 1 |
| 4738 | 4738 | 10 | 2.75680975 | 3 | 0.00600364 | 0 | 1 | 0 | 1 | 0 | 1 | 0 | 1 | 12 | 0.80329917 | FALSE | 0.22773046 | FALSE | 1 | FALSE | 1 | FALSE | 1 | FALSE | 1 | FALSE | 1 |
| 4739 | 4739 | 9.02702703 | 1.61891666 | 0 | 1 | 3 | 0.10264583 | 0 | 1 | 2 | 0.48233989 | 0 | 1 | 30 | 0.52765882 | FALSE | 1 | FALSE | 1 | FALSE | 1 | FALSE | 1 | FALSE | 1 | FALSE | 1 |
| 4740 | 4740 | 8 | 1.61514571 | 2 | 0.10809809 | 4 | 0.00492303 | 0 | 1 | 1 | 0.64497439 | 0 | 1 | 15 | 0.98570275 | FALSE | 0.88820416 | FALSE | 0.58120789 | FALSE | 1 | FALSE | 1 | FALSE | 1 | FALSE | 1 |
| 4741 | 4741 | 10.8076923 | 2.00036979 | 0 | 1 | 0 | 1 | 0 | 1 | 2 | 0.33678042 | 0 | 1 | 24 | 0.20425579 | FALSE | 1 | FALSE | 1 | FALSE | 1 | FALSE | 1 | FALSE | 1 | FALSE | 1 |
| 4742 | 4742 | 10.3333333 | 1.88561808 | 0 | 1 | 0 | 1 | 0 | 1 | 1 | 0.34506517 | 0 | 1 | 8 | 0.5780332 | FALSE | 1 | FALSE | 1 | FALSE | 1 | FALSE | 1 | FALSE | 1 | FALSE | 1 |
| 4743 | 4743 | 9.27777778 | 2.1030548 | 0 | 1 | 1 | 0.44710673 | 1 | 0.48280983 | 2 | 0.19902682 | 0 | 1 | 14 | 0.86287418 | FALSE | 1 | FALSE | 1 | FALSE | 1 | FALSE | 1 | FALSE | 1 | FALSE | 1 |
| 4744 | 4744 | 9.36363636 | 3.0233253 | 0 | 1 | 0 | 1 | 0 | 1 | 0 | 1 | 0 | 1 | 11 | 0.15414489 | FALSE | 1 | FALSE | 1 | FALSE | 1 | FALSE | 1 | FALSE | 1 | FALSE | 1 |
| 4745 | 4745 | 10.1 | 2.58650343 | 0 | 1 | 1 | 0.28036922 | 0 | 1 | 0 | 1 | 0 | 1 | 9 | 0.5215547 | FALSE | 1 | FALSE | 1 | FALSE | 1 | FALSE | 1 | FALSE | 1 | FALSE | 1 |
| 4746 | 4746 | 8.41666667 | 3.25213605 | 1 | 0.26795838 | 1 | 0.32623933 | 0 | 1 | 0 | 1 | 0 | 1 | 10 | 0.71436964 | FALSE | 1 | FALSE | 1 | FALSE | 1 | FALSE | 1 | FALSE | 1 | FALSE | 1 |
| 4747 | 4747 | 9.7 | 2.88270706 | 0 | 1 | 1 | 0.48239024 | 1 | 0.51939841 | 0 | 1 | 0 | 1 | 18 | 0.37504645 | FALSE | 1 | FALSE | 1 | FALSE | 1 | FALSE | 1 | FALSE | 1 | FALSE | 1 |
| 4748 | 4748 | 10.375 | 2.76605164 | 1 | 0.46441868 | 0 | 1 | 1 | 0.58502114 | 1 | 0.67692816 | 0 | 1 | 21 | 0.47053137 | FALSE | 1 | FALSE | 1 | FALSE | 1 | FALSE | 1 | FALSE | 1 | FALSE | 1 |
| 4749 | 4749 | 7.71428571 | 3.52599675 | 0 | 1 | 1 | 0.6739483 | 0 | 1 | 0 | 1 | 2 | 0.1065437 | 31 | 0.1995074 | FALSE | 1 | FALSE | 1 | FALSE | 1 | FALSE | 1 | FALSE | 1 | FALSE | 1 |
| 4750 | 4750 | 6.08988764 | 2.32059877 | 6 | 0.02624524 | 3 | 0.55379184 | 1 | 0.96228998 | 7 | 0.1134709 | 2 | 0.43073682 | 70 | 0.94519174 | FALSE | 0.50496189 | FALSE | 1 | FALSE | 1 | FALSE | 1 | FALSE | 1 | FALSE | 1 |
| 4751 | 4751 | 9.83333333 | 1.57233019 | 0 | 1 | 0 | 1 | 0 | 1 | 1 | 0.24578796 | 0 | 1 | 5 | 0.76185568 | FALSE | 1 | FALSE | 1 | FALSE | 1 | FALSE | 1 | FALSE | 1 | FALSE | 1 |
| 4752 | 4752 | 5.74647887 | 2.69959992 | 1 | 0.84335867 | 2 | 0.67482356 | 5 | 0.11089846 | 4 | 0.41228611 | 2 | 0.3251371 | 57 | 0.8674157 | FALSE | 1 | FALSE | 1 | FALSE | 1 | FALSE | 1 | FALSE | 1 | FALSE | 1 |
| 4753 | 4753 | 8.33333333 | 2.26691175 | 0 | 1 | 0 | 1 | 0 | 1 | 2 | 0.26791271 | 1 | 0.30529951 | 19 | 0.54233419 | FALSE | 1 | FALSE | 1 | FALSE | 1 | FALSE | 1 | FALSE | 1 | FALSE | 1 |
| 4754 | 4754 | 8.64705882 | 2.48385792 | 0 | 1 | 1 | 0.65167362 | 2 | 0.32011871 | 2 | 0.43583874 | 0 | 1 | 27 | 0.61671133 | FALSE | 1 | FALSE | 1 | FALSE | 1 | FALSE | 1 | FALSE | 1 | FALSE | 1 |
| 4755 | 4755 | 9.41176471 | 1.4974026 | 1 | 0.35728305 | 0 | 1 | 0 | 1 | 1 | 0.55062772 | 0 | 1 | 15 | 0.49060435 | FALSE | 1 | FALSE | 1 | FALSE | 1 | FALSE | 1 | FALSE | 1 | FALSE | 1 |
| 4756 | 4756 | 9.46666667 | 3.03021818 | 2 | 0.05513784 | 0 | 1 | 1 | 0.42266264 | 1 | 0.50623194 | 0 | 1 | 11 | 0.92879412 | FALSE | 0.68849059 | FALSE | 1 | FALSE | 1 | FALSE | 1 | FALSE | 1 | FALSE | 1 |
| 4757 | 4757 | 9.46666667 | 3.59382186 | 0 | 1 | 1 | 0.38964394 | 1 | 0.42266264 | 0 | 1 | 2 | 0.02433796 | 11 | 0.92879412 | FALSE | 1 | FALSE | 1 | FALSE | 1 | FALSE | 1 | FALSE | 0.77578444 | FALSE | 1 |
| 4758 | 4758 | 11.7254902 | 2.2585648 | 0 | 1 | 4 | 0.06855877 | 2 | 0.51909752 | 2 | 0.65421368 | 0 | 1 | 40 | 0.66905473 | FALSE | 1 | FALSE | 1 | FALSE | 1 | FALSE | 1 | FALSE | 1 | FALSE | 1 |
| 4759 | 4759 | 9.36 | 2.22494944 | 2 | 0.13369621 | 2 | 0.19297191 | 1 | 0.59998149 | 3 | 0.10474541 | 0 | 1 | 17 | 0.9897469 | FALSE | 0.97964162 | FALSE | 1 | FALSE | 1 | FALSE | 1 | FALSE | 1 | FALSE | 1 |
| 4760 | 4760 | 9.65909091 | 1.83331768 | 1 | 0.68224519 | 2 | 0.41896557 | 1 | 0.80109494 | 1 | 0.87439647 | 0 | 1 | 39 | 0.29524127 | FALSE | 1 | FALSE | 1 | FALSE | 1 | FALSE | 1 | FALSE | 1 | FALSE | 1 |
| 4761 | 4761 | 10.0952381 | 1.97375065 | 0 | 1 | 2 | 0.14655134 | 1 | 0.53671415 | 1 | 0.62783522 | 0 | 1 | 17 | 0.77849076 | FALSE | 1 | FALSE | 1 | FALSE | 1 | FALSE | 1 | FALSE | 1 | FALSE | 1 |
| 4762 | 4762 | 7.69166667 | 1.76444436 | 3 | 0.59887135 | 6 | 0.19220013 | 9 | 0.02822851 | 6 | 0.47540116 | 1 | 0.86490401 | 95 | 0.95241754 | FALSE | 1 | FALSE | 1 | FALSE | 1 | FALSE | 1 | FALSE | 1 | FALSE | 1 |
| 4763 | 4763 | 6.86956522 | 1.80107089 | 0 | 1 | 1 | 0.78076835 | 2 | 0.4962935 | 1 | 0.88573774 | 4 | 0.00657628 | 38 | 0.71504609 | FALSE | 1 | FALSE | 1 | FALSE | 1 | FALSE | 1 | FALSE | 0.47461742 | FALSE | 1 |
| 4764 | 4764 | 10.6923077 | 3.21976147 | 1 | 0.28675951 | 0 | 1 | 0 | 1 | 1 | 0.45746571 | 2 | 0.01846832 | 9 | 0.96018168 | FALSE | 1 | FALSE | 1 | FALSE | 1 | FALSE | 1 | FALSE | 0.69720022 | FALSE | 1 |
| 4765 | 4765 | 9.1 | 2.58650343 | 0 | 1 | 2 | 0.25306331 | 0 | 1 | 0 | 1 | 0 | 1 | 28 | 0.13086908 | FALSE | 1 | FALSE | 1 | FALSE | 1 | FALSE | 1 | FALSE | 1 | FALSE | 1 |
| 4766 | 4766 | 10.7058824 | 3.40872075 | 0 | 1 | 0 | 1 | 0 | 1 | 0 | 1 | 0 | 1 | 17 | 0.05550965 | FALSE | 1 | FALSE | 1 | FALSE | 1 | FALSE | 1 | FALSE | 1 | FALSE | 1 |
| 4767 | 4767 | 8.83333333 | 2.26691175 | 0 | 1 | 0 | 1 | 1 | 0.35555293 | 0 | 1 | 0 | 1 | 11 | 0.41944532 | FALSE | 1 | FALSE | 1 | FALSE | 1 | FALSE | 1 | FALSE | 1 | FALSE | 1 |
| 4768 | 4768 | 7.15384615 | 2.31537184 | 0 | 1 | 0 | 1 | 2 | 0.07725803 | 0 | 1 | 0 | 1 | 11 | 0.6683056 | FALSE | 1 | FALSE | 1 | FALSE | 1 | FALSE | 1 | FALSE | 1 | FALSE | 1 |
| 4769 | 4769 | 7.71559633 | 2.18876088 | 9 | 0.0014124 | 3 | 0.66606524 | 8 | 0.03458688 | 6 | 0.35145505 | 0 | 1 | 79 | 0.99512715 | FALSE | 0.09317759 | FALSE | 1 | FALSE | 1 | FALSE | 1 | FALSE | 1 | FALSE | 1 |
| 4770 | 4770 | 10.3809524 | 4.59270841 | 0 | 1 | 1 | 0.49918218 | 2 | 0.17308361 | 1 | 0.62783522 | 0 | 1 | 17 | 0.77849076 | FALSE | 1 | FALSE | 1 | FALSE | 1 | FALSE | 1 | FALSE | 1 | FALSE | 1 |
| 4771 | 4771 | 1.4762634 | 2.22916835 | 26 | 0.01418174 | 19 | 0.7153991 | 27 | 0.23790985 | 34 | 0.23168431 | 12 | 0.37720931 | 532 | 0.97144093 | FALSE | 0.37183513 | FALSE | 1 | FALSE | 1 | FALSE | 1 | FALSE | 1 | FALSE | 1 |
| 4772 | 4772 | 6.9 | 3.36996538 | 2 | 0.17884234 | 0 | 1 | 0 | 1 | 1 | 0.75658797 | 0 | 1 | 27 | 0.28862208 | FALSE | 1 | FALSE | 1 | FALSE | 1 | FALSE | 1 | FALSE | 1 | FALSE | 1 |
| 4773 | 4773 | 7.06451613 | 2.56446541 | 4 | 0.00764535 | 0 | 1 | 0 | 1 | 2 | 0.41984187 | 1 | 0.4016888 | 24 | 0.90138666 | FALSE | 0.2566832 | FALSE | 1 | FALSE | 1 | FALSE | 1 | FALSE | 1 | FALSE | 1 |
| 4774 | 4774 | 9 | 3.86436713 | 1 | 0.32293355 | 1 | 0.38964394 | 0 | 1 | 1 | 0.50623194 | 0 | 1 | 12 | 0.80329917 | FALSE | 1 | FALSE | 1 | FALSE | 1 | FALSE | 1 | FALSE | 1 | FALSE | 1 |
| 4775 | 4775 | 9.26666667 | 1.76886656 | 0 | 1 | 0 | 1 | 2 | 0.18613256 | 2 | 0.26791271 | 0 | 1 | 18 | 0.74744813 | FALSE | 1 | FALSE | 1 | FALSE | 1 | FALSE | 1 | FALSE | 1 | FALSE | 1 |
| 4776 | 4776 | 6.61029412 | 1.91807499 | 4 | 0.46288301 | 4 | 0.64684555 | 2 | 0.9594978 | 6 | 0.59903796 | 4 | 0.18364964 | 116 | 0.43967513 | FALSE | 1 | FALSE | 1 | FALSE | 1 | FALSE | 1 | FALSE | 1 | FALSE | 1 |
| 4777 | 4777 | 8.16666667 | 0.68718427 | 0 | 1 | 0 | 1 | 0 | 1 | 1 | 0.55062772 | 0 | 1 | 16 | 0.23070182 | FALSE | 1 | FALSE | 1 | FALSE | 1 | FALSE | 1 | FALSE | 1 | FALSE | 1 |
| 4778 | 4778 | 10.7857143 | 2.07634884 | 1 | 0.3050805 | 0 | 1 | 1 | 0.40110418 | 1 | 0.48242099 | 0 | 1 | 11 | 0.83676449 | FALSE | 1 | FALSE | 1 | FALSE | 1 | FALSE | 1 | FALSE | 1 | FALSE | 1 |
| 4779 | 4779 | 9.63157895 | 1.86971786 | 1 | 0.38989951 | 2 | 0.12434464 | 1 | 0.50143826 | 2 | 0.21610812 | 1 | 0.26986775 | 12 | 0.99479039 | FALSE | 1 | FALSE | 1 | FALSE | 1 | FALSE | 1 | FALSE | 1 | FALSE | 1 |
| 4780 | 4780 | 8.25641026 | 2.20393528 | 0 | 1 | 2 | 0.36122049 | 1 | 0.76089851 | 2 | 0.54052736 | 0 | 1 | 34 | 0.41570781 | FALSE | 1 | FALSE | 1 | FALSE | 1 | FALSE | 1 | FALSE | 1 | FALSE | 1 |
| 4781 | 4781 | 7.90909091 | 1.31110956 | 0 | 1 | 1 | 0.30368014 | 0 | 1 | 0 | 1 | 0 | 1 | 10 | 0.46859937 | FALSE | 1 | FALSE | 1 | FALSE | 1 | FALSE | 1 | FALSE | 1 | FALSE | 1 |
| 4782 | 4782 | 11.3333333 | 1.69967317 | 1 | 0.26795838 | 0 | 1 | 0 | 1 | 1 | 0.43131129 | 1 | 0.18007762 | 9 | 0.89630573 | FALSE | 1 | FALSE | 1 | FALSE | 1 | FALSE | 1 | FALSE | 1 | FALSE | 1 |
| 4783 | 4783 | 6.16666667 | 3.10081914 | 1 | 0.65632651 | 2 | 0.38460545 | 3 | 0.1814659 | 4 | 0.11715635 | 1 | 0.49330646 | 30 | 0.9805116 | FALSE | 1 | FALSE | 1 | FALSE | 1 | FALSE | 1 | FALSE | 1 | FALSE | 1 |
| 4784 | 4784 | 10.5833333 | 2.49861073 | 1 | 0.26795838 | 0 | 1 | 1 | 0.35555293 | 0 | 1 | 2 | 0.01579445 | 8 | 0.97197833 | FALSE | 1 | FALSE | 1 | FALSE | 1 | FALSE | 1 | FALSE | 0.66313193 | FALSE | 1 |
| 4785 | 4785 | 4.43870968 | 1.53279787 | 2 | 0.90998011 | 6 | 0.38053258 | 6 | 0.47921491 | 5 | 0.84273281 | 3 | 0.46520079 | 132 | 0.37169225 | FALSE | 1 | FALSE | 1 | FALSE | 1 | FALSE | 1 | FALSE | 1 | FALSE | 1 |
| 4786 | 4786 | 8.6875 | 2.9201616 | 4 | 0.00856611 | 1 | 0.65167362 | 3 | 0.10599423 | 0 | 1 | 0 | 1 | 24 | 0.94876798 | FALSE | 0.27019545 | FALSE | 1 | FALSE | 1 | FALSE | 1 | FALSE | 1 | FALSE | 1 |
| 4787 | 4787 | 9.53846154 | 2.67798388 | 0 | 1 | 0 | 1 | 1 | 0.37874417 | 1 | 0.45746571 | 0 | 1 | 11 | 0.6683056 | FALSE | 1 | FALSE | 1 | FALSE | 1 | FALSE | 1 | FALSE | 1 | FALSE | 1 |
| 4788 | 4788 | 10.4 | 4.1761226 | 0 | 1 | 0 | 1 | 1 | 0.42266264 | 0 | 1 | 0 | 1 | 14 | 0.29525409 | FALSE | 1 | FALSE | 1 | FALSE | 1 | FALSE | 1 | FALSE | 1 | FALSE | 1 |
| 4789 | 4789 | 6.31967213 | 2.79128149 | 7 | 0.03407787 | 6 | 0.19220013 | 6 | 0.26185234 | 6 | 0.47540116 | 1 | 0.86490401 | 94 | 0.97137583 | FALSE | 0.57089146 | FALSE | 1 | FALSE | 1 | FALSE | 1 | FALSE | 1 | FALSE | 1 |
| 4790 | 4790 | 9.82352941 | 1.68855295 | 0 | 1 | 0 | 1 | 0 | 1 | 2 | 0.18212134 | 0 | 1 | 15 | 0.49060435 | FALSE | 1 | FALSE | 1 | FALSE | 1 | FALSE | 1 | FALSE | 1 | FALSE | 1 |
| 4791 | 4791 | 5.46938776 | 3.41127806 | 4 | 0.0361357 | 3 | 0.21051417 | 4 | 0.09808429 | 5 | 0.07258207 | 1 | 0.55647171 | 32 | 0.99975578 | FALSE | 0.58392 | FALSE | 1 | FALSE | 1 | FALSE | 1 | FALSE | 1 | FALSE | 1 |
| 4792 | 4792 | 8.375 | 1.72753437 | 0 | 1 | 0 | 1 | 0 | 1 | 1 | 0.31351192 | 0 | 1 | 7 | 0.63752397 | FALSE | 1 | FALSE | 1 | FALSE | 1 | FALSE | 1 | FALSE | 1 | FALSE | 1 |

| | | | | | | | | | | | | | | | | | | | | | | | | | | | |
|---|---|---|---|---|---|---|---|---|---|---|---|---|---|---|---|---|---|---|---|---|---|---|---|---|---|---|---|
| 4793 | 4793 | 6.97297297 | 1.89574896 | 0 | 1 | 1 | 0.70472939 | 3 | 0.14625315 | 1 | 0.82511639 | 1 | 0.45845922 | 31 | 0.64534533 | FALSE | 1 | FALSE | 1 | FALSE | 1 | FALSE | 1 | FALSE | 1 | FALSE | 1 |
| 4794 | 4794 | 9.68 | 2.66413213 | 0 | 1 | 2 | 0.19297191 | 1 | 0.59998149 | 0 | 1 | 0 | 1 | 22 | 0.43642846 | FALSE | 1 | FALSE | 1 | FALSE | 1 | FALSE | 1 | FALSE | 1 | FALSE | 1 |
| 4795 | 4795 | 2.33823529 | 1.59050763 | 21 | 0.20474179 | 18 | 0.84181412 | 13 | 0.99717308 | 29 | 0.68188787 | 10 | 0.6822355 | 585 | 0.05577697 | FALSE | 1 | FALSE | 1 | FALSE | 1 | FALSE | 1 | FALSE | 1 | FALSE | 1 |
| 4796 | 4796 | 9.91666667 | 1.60511336 | 0 | 1 | 2 | 0.05552913 | 0 | 1 | 1 | 0.43131129 | 0 | 1 | 9 | 0.89630573 | FALSE | 1 | FALSE | 1 | FALSE | 1 | FALSE | 1 | FALSE | 1 | FALSE | 1 |
| 4797 | 4797 | 7.76923077 | 1.65964992 | 2 | 0.38681045 | 0 | 1 | 1 | 0.85187773 | 1 | 0.91399378 | 0 | 1 | 48 | 0.0742821 | FALSE | 1 | FALSE | 1 | FALSE | 1 | FALSE | 1 | FALSE | 1 | FALSE | 1 |
| 4798 | 4798 | 8.15 | 3.67797499 | 0 | 1 | 0 | 1 | 0 | 1 | 0 | 1 | 1 | 0.28187291 | 19 | 0.15700255 | FALSE | 1 | FALSE | 1 | FALSE | 1 | FALSE | 1 | FALSE | 1 | FALSE | 1 |
| 4799 | 4799 | 8.13004484 | 0.74336738 | 4 | 0.82731886 | 7 | 0.58249425 | 6 | 0.81602429 | 9 | 0.69791417 | 5 | 0.29937154 | 191 | 0.27951064 | FALSE | 1 | FALSE | 1 | FALSE | 1 | FALSE | 1 | FALSE | 1 | FALSE | 1 |
| 4800 | 4800 | 10.0769231 | 2.16480727 | 1 | 0.28675951 | 0 | 1 | 0 | 1 | 0 | 1 | 0 | 1 | 12 | 0.374198 | FALSE | 1 | FALSE | 1 | FALSE | 1 | FALSE | 1 | FALSE | 1 | FALSE | 1 |
| 4801 | 4801 | 10.2631579 | 2.63263137 | 2 | 0.08405377 | 0 | 1 | 0 | 1 | 0 | 1 | 0 | 1 | 17 | 0.41141543 | FALSE | 0.79236883 | FALSE | 1 | FALSE | 1 | FALSE | 1 | FALSE | 1 | FALSE | 1 |
| 4802 | 4802 | 8.66666667 | 1.69967317 | 0 | 1 | 1 | 0.32623933 | 0 | 1 | 0 | 1 | 1 | 0.18007762 | 10 | 0.71436964 | FALSE | 1 | FALSE | 1 | FALSE | 1 | FALSE | 1 | FALSE | 1 | FALSE | 1 |
| 4803 | 4803 | 9.375 | 1.89983552 | 0 | 1 | 1 | 0.40942991 | 1 | 0.44344817 | 2 | 0.16544789 | 0 | 1 | 12 | 0.90922186 | FALSE | 1 | FALSE | 1 | FALSE | 1 | FALSE | 1 | FALSE | 1 | FALSE | 1 |
| 4804 | 4804 | 6.6875 | 2.73218662 | 0 | 1 | 3 | 0.34261611 | 2 | 0.67580769 | 7 | 0.02649204 | 1 | 0.65460803 | 51 | 0.88464035 | FALSE | 1 | FALSE | 1 | FALSE | 1 | FALSE | 1 | FALSE | 1 | FALSE | 1 |
| 4805 | 4805 | 9.1875 | 0.807678 | 0 | 1 | 0 | 1 | 0 | 1 | 0 | 1 | 0 | 1 | 16 | 0.06581507 | FALSE | 1 | FALSE | 1 | FALSE | 1 | FALSE | 1 | FALSE | 1 | FALSE | 1 |
| 4806 | 4806 | 9.71428571 | 2.45226537 | 1 | 0.42086991 | 1 | 0.49918218 | 0 | 1 | 2 | 0.25059627 | 0 | 1 | 17 | 0.77849076 | FALSE | 1 | FALSE | 1 | FALSE | 1 | FALSE | 1 | FALSE | 1 | FALSE | 1 |
| 4807 | 4807 | 6.56666667 | 2.57789751 | 4 | 0.0671906 | 3 | 0.30686508 | 1 | 0.8897349 | 2 | 0.76960882 | 1 | 0.63076845 | 49 | 0.78152878 | FALSE | 0.74276693 | FALSE | 1 | FALSE | 1 | FALSE | 1 | FALSE | 1 | FALSE | 1 |
| 4808 | 4808 | 10.0344828 | 3.01127489 | 0 | 1 | 0 | 1 | 2 | 0.25286935 | 4 | 0.03319522 | 0 | 1 | 21 | 0.88395936 | FALSE | 1 | FALSE | 1 | FALSE | 1 | FALSE | 1 | FALSE | 1 | FALSE | 1 |
| 4809 | 4809 | 6.05 | 2.78941451 | 2 | 0.45796754 | 2 | 0.58316597 | 4 | 0.16812274 | 1 | 0.94113525 | 2 | 0.25814944 | 49 | 0.78152878 | FALSE | 1 | FALSE | 1 | FALSE | 1 | FALSE | 1 | FALSE | 1 | FALSE | 1 |
| 4810 | 4810 | 9.57142857 | 1.9166297 | 1 | 0.16630142 | 0 | 1 | 0 | 1 | 0 | 1 | 0 | 1 | 6 | 0.69920685 | FALSE | 1 | FALSE | 1 | FALSE | 1 | FALSE | 1 | FALSE | 1 | FALSE | 1 |
| 4811 | 4811 | 6.56326531 | 1.59113538 | 3 | 0.95152185 | 7 | 0.67318889 | 8 | 0.64670717 | 13 | 0.31905718 | 4 | 0.56526702 | 207 | 0.34492795 | FALSE | 1 | FALSE | 1 | FALSE | 1 | FALSE | 1 | FALSE | 1 | FALSE | 1 |
| 4812 | 4812 | 8.57894737 | 2.51972825 | 0 | 1 | 1 | 0.46503795 | 0 | 1 | 2 | 0.21610812 | 0 | 1 | 16 | 0.65565778 | FALSE | 1 | FALSE | 1 | FALSE | 1 | FALSE | 1 | FALSE | 1 | FALSE | 1 |
| 4813 | 4813 | 9 | 2.26077666 | 2 | 0.02089722 | 0 | 1 | 0 | 1 | 0 | 1 | 0 | 1 | 7 | 0.84574168 | FALSE | 0.45408499 | FALSE | 1 | FALSE | 1 | FALSE | 1 | FALSE | 1 | FALSE | 1 |
| 4814 | 4814 | 8.52 | 2.41859463 | 1 | 0.47819878 | 1 | 0.56109832 | 0 | 1 | 0 | 1 | 1 | 0.33902672 | 22 | 0.43642846 | FALSE | 1 | FALSE | 1 | FALSE | 1 | FALSE | 1 | FALSE | 1 | FALSE | 1 |
| 4815 | 4815 | 11 | 3.00574163 | 0 | 1 | 0 | 1 | 2 | 0.27983822 | 2 | 0.3871811 | 0 | 1 | 25 | 0.51779518 | FALSE | 1 | FALSE | 1 | FALSE | 1 | FALSE | 1 | FALSE | 1 | FALSE | 1 |
| 4816 | 4816 | 9.45454546 | 2.96536481 | 0 | 1 | 1 | 0.30368014 | 1 | 0.33149969 | 1 | 0.40390032 | 0 | 1 | 8 | 0.92152993 | FALSE | 1 | FALSE | 1 | FALSE | 1 | FALSE | 1 | FALSE | 1 | FALSE | 1 |
| 4817 | 4817 | 9.54545455 | 2.14745669 | 0 | 1 | 0 | 1 | 0 | 1 | 1 | 0.40390032 | 0 | 1 | 10 | 0.46859937 | FALSE | 1 | FALSE | 1 | FALSE | 1 | FALSE | 1 | FALSE | 1 | FALSE | 1 |
| 4818 | 4818 | 8.04761905 | 4.57192458 | 1 | 0.38989951 | 2 | 0.12434464 | 2 | 0.14749808 | 1 | 0.59104357 | 0 | 1 | 13 | 0.97998676 | FALSE | 1 | FALSE | 1 | FALSE | 1 | FALSE | 1 | FALSE | 1 | FALSE | 1 |
| 4819 | 4819 | 8 | 2.3204774 | 1 | 0.26795838 | 0 | 1 | 1 | 0.35555293 | 0 | 1 | 0 | 1 | 10 | 0.71436964 | FALSE | 1 | FALSE | 1 | FALSE | 1 | FALSE | 1 | FALSE | 1 | FALSE | 1 |
| 4820 | 4820 | 9.25 | 2.48746859 | 1 | 0.26795838 | 0 | 1 | 2 | 0.06691409 | 0 | 1 | 0 | 1 | 9 | 0.89630573 | FALSE | 1 | FALSE | 1 | FALSE | 1 | FALSE | 1 | FALSE | 1 | FALSE | 1 |
| 4821 | 4821 | 1.44651163 | 1.97794218 | 6 | 0.47564505 | 9 | 0.26035804 | 6 | 0.79156523 | 7 | 0.87154344 | 0 | 1 | 187 | 0.16598156 | FALSE | 1 | FALSE | 1 | FALSE | 1 | FALSE | 1 | FALSE | 1 | FALSE | 1 |
| 4822 | 4822 | 8.29411765 | 1.31796215 | 0 | 1 | 0 | 1 | 1 | 0.46348836 | 2 | 0.18212134 | 0 | 1 | 14 | 0.7313145 | FALSE | 1 | FALSE | 1 | FALSE | 1 | FALSE | 1 | FALSE | 1 | FALSE | 1 |
| 4823 | 4823 | 9.5 | 2.79668171 | 0 | 1 | 0 | 1 | 1 | 0.64171563 | 4 | 0.03737468 | 0 | 1 | 23 | 0.73439118 | FALSE | 1 | FALSE | 1 | FALSE | 1 | FALSE | 1 | FALSE | 1 | FALSE | 1 |
| 4824 | 4824 | 11.1176471 | 3.52941177 | 1 | 0.58738519 | 3 | 0.09602744 | 1 | 0.71261921 | 1 | 0.7984848 | 2 | 0.1065437 | 26 | 0.92796052 | FALSE | 1 | FALSE | 1 | FALSE | 1 | FALSE | 1 | FALSE | 1 | FALSE | 1 |
| 4825 | 4825 | 9.28571429 | 2.40322912 | 1 | 0.51745948 | 0 | 1 | 1 | 0.64171563 | 0 | 1 | 0 | 1 | 26 | 0.16405547 | FALSE | 1 | FALSE | 1 | FALSE | 1 | FALSE | 1 | FALSE | 1 | FALSE | 1 |
| 4826 | 4826 | 8.76785714 | 1.99992028 | 1 | 0.76787545 | 0 | 1 | 1 | 0.87219471 | 5 | 0.11290142 | 0 | 1 | 49 | 0.33483172 | FALSE | 1 | FALSE | 1 | FALSE | 1 | FALSE | 1 | FALSE | 1 | FALSE | 1 |
| 4827 | 4827 | 9.97142857 | 2.28642846 | 1 | 0.59801736 | 0 | 1 | 2 | 0.35984046 | 2 | 0.48233989 | 0 | 1 | 30 | 0.52765882 | FALSE | 1 | FALSE | 1 | FALSE | 1 | FALSE | 1 | FALSE | 1 | FALSE | 1 |
| 4828 | 4828 | 7.875 | 4.30418885 | 0 | 1 | 0 | 1 | 4 | 0.00979291 | 1 | 0.67692816 | 0 | 1 | 19 | 0.83975208 | FALSE | 1 | FALSE | 1 | FALSE | 1 | FALSE | 1 | FALSE | 1 | FALSE | 1 |
| 4829 | 4829 | 9.88461539 | 1.80441602 | 0 | 1 | 1 | 0.57534752 | 0 | 1 | 0 | 1 | 3 | 0.00847466 | 22 | 0.61628 | FALSE | 1 | FALSE | 1 | FALSE | 1 | FALSE | 1 | FALSE | 0.50674765 | FALSE | 1 |
| 4830 | 4830 | 5.29850746 | 2.81793258 | 4 | 0.08106144 | 2 | 0.61853748 | 3 | 0.40547364 | 2 | 0.79987679 | 2 | 0.28258242 | 51 | 0.88464035 | FALSE | 0.79236883 | FALSE | 1 | FALSE | 1 | FALSE | 1 | FALSE | 1 | FALSE | 1 |
| 4831 | 4831 | 10.0408163 | 3.03018275 | 4 | 0.0361357 | 6 | 0.00467442 | 1 | 0.83455705 | 1 | 0.90086419 | 3 | 0.04584538 | 34 | 0.99772438 | FALSE | 0.58392 | FALSE | 0.58120789 | FALSE | 1 | FALSE | 1 | FALSE | 0.95807482 | FALSE | 1 |
| 4832 | 4832 | 7 | 1.41421356 | 0 | 1 | 0 | 1 | 0 | 1 | 1 | 0.17140665 | 0 | 1 | 3 | 0.8823539 | FALSE | 1 | FALSE | 1 | FALSE | 1 | FALSE | 1 | FALSE | 1 | FALSE | 1 |
| 4833 | 4833 | 9.625 | 2.39465551 | 1 | 0.56527493 | 3 | 0.08334648 | 3 | 0.10599423 | 2 | 0.43583874 | 3 | 0.01506052 | 20 | 0.99942006 | FALSE | 1 | FALSE | 1 | FALSE | 1 | FALSE | 1 | FALSE | 0.66313193 | FALSE | 1 |
| 4834 | 4834 | 8.5 | 3.00462606 | 1 | 0.60837712 | 2 | 0.32556427 | 1 | 0.73299778 | 2 | 0.49731009 | 0 | 1 | 30 | 0.67248271 | FALSE | 1 | FALSE | 1 | FALSE | 1 | FALSE | 1 | FALSE | 1 | FALSE | 1 |
| 4835 | 4835 | 7.57142857 | 1.04978132 | 0 | 1 | 0 | 1 | 1 | 0.22601089 | 0 | 1 | 0 | 1 | 6 | 0.69920685 | FALSE | 1 | FALSE | 1 | FALSE | 1 | FALSE | 1 | FALSE | 1 | FALSE | 1 |
| 4836 | 4836 | 8.94117647 | 3.45510004 | 0 | 1 | 0 | 1 | 1 | 0.46348836 | 2 | 0.18212134 | 0 | 1 | 14 | 0.7313145 | FALSE | 1 | FALSE | 1 | FALSE | 1 | FALSE | 1 | FALSE | 1 | FALSE | 1 |
| 4837 | 4837 | 8.77777778 | 1.435872 | 0 | 1 | 2 | 0.1135776 | 1 | 0.48280983 | 1 | 0.57131013 | 0 | 1 | 14 | 0.86287418 | FALSE | 1 | FALSE | 1 | FALSE | 1 | FALSE | 1 | FALSE | 1 | FALSE | 1 |
| 4838 | 4838 | 9.6 | 2.61533937 | 0 | 1 | 0 | 1 | 0 | 1 | 1 | 0.37517262 | 0 | 1 | 9 | 0.5215547 | FALSE | 1 | FALSE | 1 | FALSE | 1 | FALSE | 1 | FALSE | 1 | FALSE | 1 |
| 4839 | 4839 | 9.72727273 | 0.96209139 | 0 | 1 | 0 | 1 | 3 | 0.00455451 | 1 | 0.37517262 | 0 | 1 | 6 | 0.98831471 | FALSE | 1 | FALSE | 1 | FALSE | 1 | FALSE | 1 | FALSE | 1 | FALSE | 1 |
| 4840 | 4840 | 10.28 | 2.27191549 | 0 | 1 | 0 | 1 | 1 | 0.59998149 | 3 | 0.10474541 | 0 | 1 | 21 | 0.64957167 | FALSE | 1 | FALSE | 1 | FALSE | 1 | FALSE | 1 | FALSE | 1 | FALSE | 1 |
| 4841 | 4841 | 9.76923077 | 1.36741453 | 0 | 1 | 3 | 0.00750707 | 0 | 1 | 1 | 0.45746571 | 0 | 1 | 9 | 0.96018168 | FALSE | 1 | FALSE | 0.6167796 | FALSE | 1 | FALSE | 1 | FALSE | 1 | FALSE | 1 |
| 4842 | 4842 | 8.47826087 | 3.57369089 | 0 | 1 | 1 | 0.53115654 | 2 | 0.19931429 | 2 | 0.2852236 | 1 | 0.31672722 | 17 | 0.94439745 | FALSE | 1 | FALSE | 1 | FALSE | 1 | FALSE | 1 | FALSE | 1 | FALSE | 1 |
| 4843 | 4843 | 9.78571429 | 2.04165799 | 0 | 1 | 0 | 1 | 0 | 1 | 0 | 1 | 0 | 1 | 14 | 0.09251304 | FALSE | 1 | FALSE | 1 | FALSE | 1 | FALSE | 1 | FALSE | 1 | FALSE | 1 |
| 4844 | 4844 | 9.09090909 | 1.92847304 | 0 | 1 | 0 | 1 | 0 | 1 | 0 | 1 | 0 | 1 | 10 | 0.18273166 | FALSE | 1 | FALSE | 1 | FALSE | 1 | FALSE | 1 | FALSE | 1 | FALSE | 1 |
| 4845 | 4845 | 8.90697674 | 2.0664986 | 0 | 1 | 2 | 0.40761584 | 1 | 0.79363407 | 0 | 1 | 1 | 0.50988734 | 39 | 0.17628554 | FALSE | 1 | FALSE | 1 | FALSE | 1 | FALSE | 1 | FALSE | 1 | FALSE | 1 |
| 4846 | 4846 | 10 | 2.32379001 | 1 | 0.22886536 | 0 | 1 | 0 | 1 | 1 | 0.37517262 | 1 | 0.15247062 | 7 | 0.94326215 | FALSE | 1 | FALSE | 1 | FALSE | 1 | FALSE | 1 | FALSE | 1 | FALSE | 1 |
| 4847 | 4847 | 8.3 | 2.72213152 | 1 | 0.40558521 | 3 | 0.02534497 | 2 | 0.16019548 | 2 | 0.23331368 | 0 | 1 | 12 | 0.99824567 | FALSE | 1 | FALSE | 0.82367276 | FALSE | 1 | FALSE | 1 | FALSE | 1 | FALSE | 1 |
| 4848 | 4848 | 10.3076923 | 2.19736259 | 1 | 0.28675951 | 0 | 1 | 0 | 1 | 0 | 1 | 0 | 1 | 12 | 0.374198 | FALSE | 1 | FALSE | 1 | FALSE | 1 | FALSE | 1 | FALSE | 1 | FALSE | 1 |
| 4849 | 4849 | 8.36363636 | 2.61485479 | 0 | 1 | 1 | 0.6629941 | 3 | 0.11365871 | 3 | 0.19161876 | 0 | 1 | 26 | 0.86852567 | FALSE | 1 | FALSE | 1 | FALSE | 1 | FALSE | 1 | FALSE | 1 | FALSE | 1 |
| 4850 | 4850 | 9.55 | 2.71062723 | 0 | 1 | 0 | 1 | 2 | 0.16019548 | 3 | 0.0612581 | 0 | 1 | 15 | 0.92113679 | FALSE | 1 | FALSE | 1 | FALSE | 1 | FALSE | 1 | FALSE | 1 | FALSE | 1 |
| 4851 | 4851 | 9.2173913 | 2.84309265 | 1 | 0.45027683 | 1 | 0.53115654 | 1 | 0.5695037 | 1 | 0.6613267 | 0 | 1 | 19 | 0.71542496 | FALSE | 1 | FALSE | 1 | FALSE | 1 | FALSE | 1 | FALSE | 1 | FALSE | 1 |
| 4852 | 4852 | 10.2857143 | 1.54126902 | 0 | 1 | 2 | 0.31355738 | 1 | 0.72299506 | 4 | 0.07465533 | 0 | 1 | 28 | 0.83079716 | FALSE | 1 | FALSE | 1 | FALSE | 1 | FALSE | 1 | FALSE | 1 | FALSE | 1 |
| 4853 | 4853 | 9.59090909 | 1.8379606 | 3 | 0.09709068 | 3 | 0.16132793 | 3 | 0.19985581 | 0 | 1 | 2 | 0.1563365 | 32 | 0.9718545 | FALSE | 0.86047225 | FALSE | 1 | FALSE | 1 | FALSE | 1 | FALSE | 1 | FALSE | 1 |
| 4854 | 4854 | 4.77659575 | 1.90270946 | 1 | 0.90982054 | 4 | 0.34712171 | 1 | 0.96626093 | 2 | 0.92940958 | 1 | 0.78377186 | 83 | 0.07371412 | FALSE | 1 | FALSE | 1 | FALSE | 1 | FALSE | 1 | FALSE | 1 | FALSE | 1 |
| 4855 | 4855 | 10.6666667 | 2.9589788 | 0 | 1 | 1 | 0.36919824 | 0 | 1 | 1 | 0.48242099 | 0 | 1 | 12 | 0.62236227 | FALSE | 1 | FALSE | 1 | FALSE | 1 | FALSE | 1 | FALSE | 1 | FALSE | 1 |
| 4856 | 4856 | 8.87755102 | 1.73361311 | 0 | 1 | 1 | 0.80150109 | 0 | 1 | 2 | 0.66536008 | 0 | 1 | 46 | 0.04012052 | FALSE | 1 | FALSE | 1 | FALSE | 1 | FALSE | 1 | FALSE | 1 | FALSE | 1 |
| 4857 | 4857 | 10.9333333 | 2.40739601 | 2 | 0.05513784 | 1 | 0.38964394 | 0 | 1 | 1 | 0.50623194 | 0 | 1 | 11 | 0.92879412 | FALSE | 0.68849059 | FALSE | 1 | FALSE | 1 | FALSE | 1 | FALSE | 1 | FALSE | 1 |
| 4858 | 4858 | 7.82352941 | 1.91696126 | 0 | 1 | 2 | 0.10307112 | 1 | 0.46348836 | 1 | 0.55062772 | 0 | 1 | 13 | 0.88719222 | FALSE | 1 | FALSE | 1 | FALSE | 1 | FALSE | 1 | FALSE | 1 | FALSE | 1 |
| 4859 | 4859 | 10.5 | 3.26343377 | 2 | 0.0256865 | 0 | 1 | 1 | 0.30655256 | 0 | 1 | 0 | 1 | 7 | 0.94326215 | FALSE | 0.4974559 | FALSE | 1 | FALSE | 1 | FALSE | 1 | FALSE | 1 | FALSE | 1 |
| 4860 | 4860 | 8.55172414 | 3.75624727 | 1 | 0.52988231 | 2 | 0.24095584 | 3 | 0.08431992 | 3 | 0.14602298 | 1 | 0.38148475 | 19 | 0.99707037 | FALSE | 1 | FALSE | 1 | FALSE | 1 | FALSE | 1 | FALSE | 1 | FALSE | 1 |
| 4861 | 4861 | 9.64705882 | 1.78032305 | 2 | 0.06909038 | 1 | 0.42857733 | 0 | 1 | 0 | 1 | 0 | 1 | 14 | 0.7313145 | FALSE | 0.74276693 | FALSE | 1 | FALSE | 1 | FALSE | 1 | FALSE | 1 | FALSE | 1 |
| 4862 | 4862 | 8.11864407 | 2.45007607 | 2 | 0.4493251 | 3 | 0.2979413 | 3 | 0.35626822 | 0 | 1 | 2 | 0.25204357 | 49 | 0.68931377 | FALSE | 1 | FALSE | 1 | FALSE | 1 | FALSE | 1 | FALSE | 1 | FALSE | 1 |
| 4863 | 4863 | 7.625 | 2.19492787 | 0 | 1 | 2 | 0.46321244 | 1 | 0.8283465 | 4 | 0.17698522 | 1 | 0.54902428 | 40 | 0.66905473 | FALSE | 1 | FALSE | 1 | FALSE | 1 | FALSE | 1 | FALSE | 1 | FALSE | 1 |
| 4864 | 4864 | 8.10869565 | 1.69690589 | 1 | 0.69843502 | 2 | 0.44132825 | 0 | 1 | 1 | 0.88573774 | 0 | 1 | 42 | 0.13378393 | FALSE | 1 | FALSE | 1 | FALSE | 1 | FALSE | 1 | FALSE | 1 | FALSE | 1 |
| 4865 | 4865 | 10.0714286 | 4.77290396 | 1 | 0.3050805 | 0 | 1 | 1 | 0.40110418 | 1 | 0.48242099 | 1 | 0.20679328 | 10 | 0.94579623 | FALSE | 1 | FALSE | 1 | FALSE | 1 | FALSE | 1 | FALSE | 1 | FALSE | 1 |
| 4866 | 4866 | 9.61904762 | 4.46655498 | 0 | 1 | 1 | 0.49918218 | 0 | 1 | 3 | 0.06912908 | 0 | 1 | 17 | 0.77849076 | FALSE | 1 | FALSE | 1 | FALSE | 1 | FALSE | 1 | FALSE | 1 | FALSE | 1 |
| 4867 | 4867 | 10.5 | 1.5 | 0 | 1 | 1 | 0.25628163 | 0 | 1 | 0 | 1 | 1 | 0.1383237 | 7 | 0.84574168 | FALSE | 1 | FALSE | 1 | FALSE | 1 | FALSE | 1 | FALSE | 1 | FALSE | 1 |
| 4868 | 4868 | 8.94117647 | 3.0576921 | 0 | 1 | 0 | 1 | 0 | 1 | 0 | 1 | 1 | 0.24525764 | 16 | 0.23070182 | FALSE | 1 | FALSE | 1 | FALSE | 1 | FALSE | 1 | FALSE | 1 | FALSE | 1 |
| 4869 | 4869 | 11.1333333 | 3.48074066 | 1 | 0.54198714 | 3 | 0.07145121 | 2 | 0.29330414 | 0 | 1 | 1 | 0.39166989 | 23 | 0.9158696 | FALSE | 1 | FALSE | 1 | FALSE | 1 | FALSE | 1 | FALSE | 1 | FALSE | 1 |
| 4870 | 4870 | 7.74285714 | 2.39455164 | 1 | 0.55378205 | 2 | 0.2651851 | 1 | 0.67911179 | 0 | 1 | 1 | 0.4016888 | 26 | 0.64660109 | FALSE | 1 | FALSE | 1 | FALSE | 1 | FALSE | 1 | FALSE | 1 | FALSE | 1 |
| 4871 | 4871 | 6 | 2.31840462 | 1 | 0.81177881 | 1 | 0.87929209 | 1 | 0.90487646 | 4 | 0.33842086 | 1 | 0.65460803 | 56 | 0.31273702 | FALSE | 1 | FALSE | 1 | FALSE | 1 | FALSE | 1 | FALSE | 1 | FALSE | 1 |
| 4872 | 4872 | 11.3333333 | 1.88561808 | 0 | 1 | 1 | 0.179097 | 1 | 0.197145 | 0 | 1 | 0 | 1 | 4 | 0.94743772 | FALSE | 1 | FALSE | 1 | FALSE | 1 | FALSE | 1 | FALSE | 1 | FALSE | 1 |
| 4873 | 4873 | 11 | 4.52155332 | 0 | 1 | 2 | 0.03227145 | 0 | 1 | 0 | 1 | 0 | 1 | 7 | 0.84574168 | FALSE | 1 | FALSE | 0.92268438 | FALSE | 1 | FALSE | 1 | FALSE | 1 | FALSE | 1 |
| 4874 | 4874 | 9.19047619 | 2.59338073 | 0 | 1 | 0 | 1 | 1 | 0.53671415 | 1 | 0.62783522 | 0 | 1 | 19 | 0.34094959 | FALSE | 1 | FALSE | 1 | FALSE | 1 | FALSE | 1 | FALSE | 1 | FALSE | 1 |
| 4875 | 4875 | 9.83333333 | 0.89752747 | 0 | 1 | 0 | 1 | 0 | 1 | 0 | 1 | 0 | 1 | 6 | 0.36077158 | FALSE | 1 | FALSE | 1 | FALSE | 1 | FALSE | 1 | FALSE | 1 | FALSE | 1 |
| 4876 | 4876 | 9.5 | 2.38047614 | 1 | 0.46441868 | 0 | 1 | 1 | 0.58502114 | 0 | 1 | 0 | 1 | 22 | 0.25237063 | FALSE | 1 | FALSE | 1 | FALSE | 1 | FALSE | 1 | FALSE | 1 | FALSE | 1 |
| 4877 | 4877 | 7.74603175 | 4.46847246 | 0 | 1 | 3 | 0.31579946 | 1 | 0.89373158 | 5 | 0.14703607 | 1 | 0.63687695 | 51 | 0.64764523 | FALSE | 1 | FALSE | 1 | FALSE | 1 | FALSE | 1 | FALSE | 1 | FALSE | 1 |
| 4878 | 4878 | 8.5 | 1.92471958 | 2 | 0.10809809 | 2 | 0.15794013 | 0 | 1 | 0 | 1 | 0 | 1 | 18 | 0.74744813 | FALSE | 0.88820416 | FALSE | 1 | FALSE | 1 | FALSE | 1 | FALSE | 1 | FALSE | 1 |
| 4879 | 4879 | 7.59090909 | 2.62261304 | 1 | 0.4357638 | 0 | 1 | 1 | 0.55340851 | 2 | 0.26791271 | 0 | 1 | 18 | 0.74744813 | FALSE | 1 | FALSE | 1 | FALSE | 1 | FALSE | 1 | FALSE | 1 | FALSE | 1 |
| 4880 | 4880 | 8.68421053 | 2.55574959 | 0 | 1 | 0 | 1 | 1 | 0.50143826 | 1 | 0.59104357 | 1 | 0.26986775 | 16 | 0.65565778 | FALSE | 1 | FALSE | 1 | FALSE | 1 | FALSE | 1 | FALSE | 1 | FALSE | 1 |
| 4881 | 4881 | 8.82075472 | 3.24640299 | 2 | 0.75627365 | 9 | 0.00669303 | 5 | 0.32542236 | 3 | 0.86756002 | 4 | 0.09327737 | 82 | 0.96834736 | FALSE | 1 | FALSE | 0.60103504 | FALSE | 1 | FALSE | 1 | FALSE | 1 | FALSE | 1 |
| 4882 | 4882 | 2.79607843 | 2.45859523 | 1 | 0.99873426 | 4 | 0.9663214 | 13 | 0.11984189 | 14 | 0.26750181 | 3 | 0.78853015 | 217 | 0.25036677 | FALSE | 1 | FALSE | 1 | FALSE | 1 | FALSE | 1 | FALSE | 1 | FALSE | 1 |
| 4883 | 4883 | 5.33333333 | 4.18551293 | 5 | 0.12800065 | 5 | 0.24617277 | 3 | 0.72810072 | 7 | 0.20017129 | 1 | 0.82320297 | 83 | 0.91989808 | FALSE | 0.97451797 | FALSE | 1 | FALSE | 1 | FALSE | 1 | FALSE | 1 | FALSE | 1 |
| 4884 | 4884 | 10.4761905 | 2.80508123 | 1 | 0.42086991 | 1 | 0.49918218 | 0 | 1 | 1 | 0.62783522 | 1 | 0.29368245 | 17 | 0.77849076 | FALSE | 1 | FALSE | 1 | FALSE | 1 | FALSE | 1 | FALSE | 1 | FALSE | 1 |
| 4885 | 4885 | 4.59090909 | 3.256589 | 0 | 1 | 0 | 1 | 1 | 0.80109494 | 2 | 0.60667406 | 2 | 0.16211764 | 39 | 0.29524127 | FALSE | 1 | FALSE | 1 | FALSE | 1 | FALSE | 1 | FALSE | 1 | FALSE | 1 |
| 4886 | 4886 | 10.7222222 | 2.35243504 | 0 | 1 | 1 | 0.44710673 | 0 | 1 | 0 | 1 | 0 | 1 | 17 | 0.20329205 | FALSE | 1 | FALSE | 1 | FALSE | 1 | FALSE | 1 | FALSE | 1 | FALSE | 1 |

| | | | | | | | | | | | | | | | | | | | | | | | | | | | |
|---|---|---|---|---|---|---|---|---|---|---|---|---|---|---|---|---|---|---|---|---|---|---|---|---|---|---|---|
| 4887 | 4887 | 9.66666667 | 1.97202659 | 0 | 1 | 0 | 1 | 0 | 1 | 0 | 1 | 0 | 1 | 6 | 0.36077158 | FALSE | 1 | FALSE | 1 | FALSE | 1 | FALSE | 1 | FALSE | 1 | FALSE | 1 |
| 4888 | 4888 | 8.39473684 | 2.7676655 | 0 | 1 | 1 | 0.71433277 | 2 | 0.39868506 | 2 | 0.52640931 | 0 | 1 | 33 | 0.44261631 | FALSE | 1 | FALSE | 1 | FALSE | 1 | FALSE | 1 | FALSE | 1 | FALSE | 1 |
| 4889 | 4889 | 10.8888889 | 1.04822013 | 0 | 1 | 0 | 1 | 0 | 1 | 1 | 0.57131013 | 0 | 1 | 17 | 0.20329205 | FALSE | 1 | FALSE | 1 | FALSE | 1 | FALSE | 1 | FALSE | 1 | FALSE | 1 |
| 4890 | 4890 | 9.93333333 | 2.14372469 | 1 | 0.32293355 | 0 | 1 | 0 | 1 | 0 | 1 | 0 | 1 | 14 | 0.29525409 | FALSE | 1 | FALSE | 1 | FALSE | 1 | FALSE | 1 | FALSE | 1 | FALSE | 1 |
| 4891 | 4891 | 10.6470588 | 3.35964243 | 1 | 0.35728305 | 0 | 1 | 1 | 0.46348836 | 1 | 0.55062772 | 1 | 0.24525764 | 13 | 0.88719222 | FALSE | 1 | FALSE | 1 | FALSE | 1 | FALSE | 1 | FALSE | 1 | FALSE | 1 |
| 4892 | 4892 | 10.5263158 | 1.56838573 | 0 | 1 | 0 | 1 | 4 | 0.00411123 | 3 | 0.05383577 | 1 | 0.26986775 | 11 | 0.99888169 | FALSE | 1 | FALSE | 1 | FALSE | 1 | FALSE | 1 | FALSE | 1 | FALSE | 1 |
| 4893 | 4893 | 7.8125 | 2.45506492 | 2 | 0.06197875 | 0 | 1 | 0 | 1 | 1 | 0.52895088 | 0 | 1 | 13 | 0.76799161 | FALSE | 0.72532843 | FALSE | 1 | FALSE | 1 | FALSE | 1 | FALSE | 1 | FALSE | 1 |
| 4894 | 4894 | 9.04444444 | 1.75062599 | 1 | 0.69044531 | 1 | 0.77338922 | 3 | 0.21866539 | 1 | 0.8802008 | 2 | 0.16793736 | 37 | 0.73736034 | FALSE | 1 | FALSE | 1 | FALSE | 1 | FALSE | 1 | FALSE | 1 | FALSE | 1 |
| 4895 | 4895 | 7.5 | 1.83711731 | 0 | 1 | 1 | 0.40942991 | 0 | 1 | 1 | 0.52895088 | 0 | 1 | 14 | 0.53308261 | FALSE | 1 | FALSE | 1 | FALSE | 1 | FALSE | 1 | FALSE | 1 | FALSE | 1 |
| 4896 | 4896 | 5.96 | 2.02938414 | 0 | 1 | 0 | 1 | 2 | 0.54120634 | 3 | 0.4048204 | 2 | 0.19752014 | 43 | 0.46945509 | FALSE | 1 | FALSE | 1 | FALSE | 1 | FALSE | 1 | FALSE | 1 | FALSE | 1 |
| 4897 | 4897 | 9.67857143 | 2.18850196 | 1 | 0.51745948 | 3 | 0.06039219 | 2 | 0.2663542 | 2 | 0.3705495 | 0 | 1 | 20 | 0.97757517 | FALSE | 1 | FALSE | 1 | FALSE | 1 | FALSE | 1 | FALSE | 1 | FALSE | 1 |
| 4898 | 4898 | 7.34722222 | 2.12236583 | 2 | 0.55518392 | 3 | 0.41343422 | 2 | 0.73725239 | 7 | 0.04621037 | 1 | 0.69780571 | 57 | 0.91330885 | FALSE | 1 | FALSE | 1 | FALSE | 1 | FALSE | 1 | FALSE | 1 | FALSE | 1 |
| 4899 | 4899 | 6 | 3.64610993 | 1 | 0.7354105 | 1 | 0.81422651 | 2 | 0.55199489 | 1 | 0.90982214 | 0 | 1 | 46 | 0.17033916 | FALSE | 1 | FALSE | 1 | FALSE | 1 | FALSE | 1 | FALSE | 1 | FALSE | 1 |
| 4900 | 4900 | 6.775 | 2.24151177 | 2 | 0.60596997 | 5 | 0.11181294 | 3 | 0.54427311 | 5 | 0.29645796 | 2 | 0.37295427 | 62 | 0.94110493 | FALSE | 1 | FALSE | 1 | FALSE | 1 | FALSE | 1 | FALSE | 1 | FALSE | 1 |
| 4901 | 4901 | 8.63636364 | 2.91618453 | 0 | 1 | 2 | 0.14655134 | 0 | 1 | 0 | 1 | 0 | 1 | 19 | 0.34094959 | FALSE | 1 | FALSE | 1 | FALSE | 1 | FALSE | 1 | FALSE | 1 | FALSE | 1 |
| 4902 | 4902 | 9.33333333 | 2.24845626 | 0 | 1 | 1 | 0.32623933 | 0 | 1 | 1 | 0.43131129 | 0 | 1 | 10 | 0.71436964 | FALSE | 1 | FALSE | 1 | FALSE | 1 | FALSE | 1 | FALSE | 1 | FALSE | 1 |
| 4903 | 4903 | 1.12673831 | 0.97959232 | 67 | 0.98612512 | 101 | 0.56841681 | 111 | 0.62884158 | 157 | 0.07920818 | 55 | 0.28661826 | 2655 | 0.50205674 | FALSE | 1 | FALSE | 1 | FALSE | 1 | FALSE | 1 | FALSE | 1 | FALSE | 1 |
| 4904 | 4904 | 9.26923077 | 2.63931361 | 0 | 1 | 0 | 1 | 0 | 1 | 0 | 1 | 0 | 1 | 25 | 0.01419971 | FALSE | 1 | FALSE | 1 | FALSE | 1 | FALSE | 1 | FALSE | 1 | FALSE | 1 |
| 4905 | 4905 | 6.16666667 | 3.24075998 | 4 | 0.08106144 | 2 | 0.61853748 | 1 | 0.90487646 | 1 | 0.95130976 | 1 | 0.65460803 | 55 | 0.44698699 | FALSE | 0.79236883 | FALSE | 1 | FALSE | 1 | FALSE | 1 | FALSE | 1 | FALSE | 1 |
| 4906 | 4906 | 9.96 | 2.93230285 | 2 | 0.13369621 | 0 | 1 | 2 | 0.22597287 | 0 | 1 | 0 | 1 | 21 | 0.64957167 | FALSE | 0.97964162 | FALSE | 1 | FALSE | 1 | FALSE | 1 | FALSE | 1 | FALSE | 1 |
| 4907 | 4907 | 6.30136986 | 4.09705852 | 2 | 0.56271322 | 6 | 0.03010676 | 4 | 0.26680336 | 1 | 0.96824399 | 0 | 1 | 60 | 0.75780737 | FALSE | 1 | FALSE | 0.90674462 | FALSE | 1 | FALSE | 1 | FALSE | 1 | FALSE | 1 |
| 4908 | 4908 | 10.1851852 | 1.4151832 | 4 | 0.00461682 | 0 | 1 | 2 | 0.25286935 | 1 | 0.71956339 | 0 | 1 | 20 | 0.95133411 | FALSE | 0.19698443 | FALSE | 1 | FALSE | 1 | FALSE | 1 | FALSE | 1 | FALSE | 1 |
| 4909 | 4909 | 8.6 | 2.4979992 | 0 | 1 | 0 | 1 | 3 | 0.01512258 | 1 | 0.50623194 | 1 | 0.2198258 | 10 | 0.97980046 | FALSE | 1 | FALSE | 1 | FALSE | 1 | FALSE | 1 | FALSE | 1 | FALSE | 1 |
| 4910 | 4910 | 8.66666667 | 2.78886676 | 1 | 0.35728305 | 0 | 1 | 2 | 0.12280513 | 0 | 1 | 0 | 1 | 14 | 0.7313145 | FALSE | 1 | FALSE | 1 | FALSE | 1 | FALSE | 1 | FALSE | 1 | FALSE | 1 |
| 4911 | 4911 | 9.96428571 | 3.08778866 | 0 | 1 | 1 | 0.60247911 | 1 | 0.64171563 | 2 | 0.3705495 | 0 | 1 | 24 | 0.55018107 | FALSE | 1 | FALSE | 1 | FALSE | 1 | FALSE | 1 | FALSE | 1 | FALSE | 1 |
| 4912 | 4912 | 10.5 | 2.53968502 | 1 | 0.22886536 | 0 | 1 | 0 | 1 | 0 | 1 | 0 | 1 | 9 | 0.5215547 | FALSE | 1 | FALSE | 1 | FALSE | 1 | FALSE | 1 | FALSE | 1 | FALSE | 1 |
| 4913 | 4913 | 9.46153846 | 2.5605103 | 0 | 1 | 2 | 0.06426041 | 1 | 0.37874417 | 0 | 1 | 2 | 0.01846832 | 8 | 0.99085298 | FALSE | 1 | FALSE | 1 | FALSE | 1 | FALSE | 1 | FALSE | 0.69720022 | FALSE | 1 |
| 4914 | 4914 | 8.70833333 | 2.00995785 | 0 | 1 | 1 | 0.54637326 | 2 | 0.21260257 | 0 | 1 | 0 | 1 | 21 | 0.47053137 | FALSE | 1 | FALSE | 1 | FALSE | 1 | FALSE | 1 | FALSE | 1 | FALSE | 1 |
| 4915 | 4915 | 5.38271605 | 2.15204698 | 5 | 0.05627772 | 6 | 0.04653609 | 2 | 0.79427057 | 5 | 0.31464362 | 0 | 1 | 63 | 0.95896498 | FALSE | 0.69536509 | FALSE | 1 | FALSE | 1 | FALSE | 1 | FALSE | 1 | FALSE | 1 |
| 4916 | 4916 | 9.42424242 | 2.29003511 | 0 | 1 | 4 | 0.15811489 | 4 | 0.20453007 | 4 | 0.3489977 | 0 | 1 | 53 | 0.7938242 | FALSE | 1 | FALSE | 1 | FALSE | 1 | FALSE | 1 | FALSE | 1 | FALSE | 1 |
| 4917 | 4917 | 7.38297872 | 1.62716119 | 6 | 0.15333077 | 4 | 0.67478629 | 6 | 0.39568129 | 4 | 0.89441166 | 2 | 0.67751472 | 119 | 0.55437269 | FALSE | 1 | FALSE | 1 | FALSE | 1 | FALSE | 1 | FALSE | 1 | FALSE | 1 |
| 4918 | 4918 | 9.57142857 | 2.19461307 | 2 | 0.0126078 | 0 | 1 | 0 | 1 | 1 | 0.28044368 | 0 | 1 | 4 | 0.98605104 | FALSE | 0.3525019 | FALSE | 1 | FALSE | 1 | FALSE | 1 | FALSE | 1 | FALSE | 1 |
| 4919 | 4919 | 4.83333333 | 3.13729065 | 1 | 0.74224579 | 2 | 0.50543179 | 0 | 1 | 2 | 0.6970685 | 2 | 0.2095319 | 45 | 0.4221244 | FALSE | 1 | FALSE | 1 | FALSE | 1 | FALSE | 1 | FALSE | 1 | FALSE | 1 |
| 4920 | 4920 | 6.89130435 | 1.25504652 | 1 | 0.90982054 | 1 | 0.95244851 | 2 | 0.84895456 | 1 | 0.98714352 | 1 | 0.78377186 | 86 | 0.00684674 | FALSE | 1 | FALSE | 1 | FALSE | 1 | FALSE | 1 | FALSE | 1 | FALSE | 1 |
| 4921 | 4921 | 7.5 | 3.30403793 | 0 | 1 | 0 | 1 | 1 | 0.48280983 | 0 | 1 | 2 | 0.03435384 | 15 | 0.69372602 | FALSE | 1 | FALSE | 1 | FALSE | 1 | FALSE | 1 | FALSE | 0.86063198 | FALSE | 1 |
| 4922 | 4922 | 9.15384615 | 1.87477809 | 1 | 0.28675951 | 0 | 1 | 1 | 0.37874417 | 0 | 1 | 0 | 1 | 11 | 0.6683056 | FALSE | 1 | FALSE | 1 | FALSE | 1 | FALSE | 1 | FALSE | 1 | FALSE | 1 |
| 4923 | 4923 | 8.23076923 | 1.76084971 | 3 | 0.0279333 | 4 | 0.00909592 | 1 | 0.61440466 | 1 | 0.70601446 | 0 | 1 | 17 | 0.99598795 | FALSE | 0.50964275 | FALSE | 0.63686998 | FALSE | 1 | FALSE | 1 | FALSE | 1 | FALSE | 1 |
| 4924 | 4924 | 8.52272727 | 2.77561325 | 2 | 0.31193899 | 1 | 0.76576289 | 2 | 0.47281647 | 2 | 0.60667406 | 0 | 1 | 37 | 0.61979664 | FALSE | 1 | FALSE | 1 | FALSE | 1 | FALSE | 1 | FALSE | 1 | FALSE | 1 |
| 4925 | 4925 | 9.16666667 | 1.77169097 | 0 | 1 | 0 | 1 | 2 | 0.06691409 | 0 | 1 | 0 | 1 | 10 | 0.71436964 | FALSE | 1 | FALSE | 1 | FALSE | 1 | FALSE | 1 | FALSE | 1 | FALSE | 1 |
| 4926 | 4926 | 10.5 | 2.5 | 0 | 1 | 1 | 0.32623933 | 1 | 0.35555293 | 3 | 0.01547597 | 1 | 0.18007762 | 6 | 0.99914544 | FALSE | 1 | FALSE | 1 | FALSE | 1 | FALSE | 1 | FALSE | 1 | FALSE | 1 |
| 4927 | 4927 | 7.73333333 | 2.65748419 | 0 | 1 | 1 | 0.61539065 | 1 | 0.65463982 | 1 | 0.74482229 | 2 | 0.08121636 | 24 | 0.70563334 | FALSE | 1 | FALSE | 1 | FALSE | 1 | FALSE | 1 | FALSE | 1 | FALSE | 1 |
| 4928 | 4928 | 10.8 | 1.32664992 | 0 | 1 | 2 | 0.03949327 | 0 | 1 | 2 | 0.0740981 | 0 | 1 | 6 | 0.98831471 | FALSE | 1 | FALSE | 0.96864929 | FALSE | 1 | FALSE | 1 | FALSE | 1 | FALSE | 1 |
| 4929 | 4929 | 5.40697674 | 1.79390381 | 5 | 0.06651199 | 4 | 0.29545046 | 2 | 0.81591765 | 3 | 0.75590957 | 1 | 0.75686725 | 70 | 0.75388153 | FALSE | 0.74276693 | FALSE | 1 | FALSE | 1 | FALSE | 1 | FALSE | 1 | FALSE | 1 |
| 4930 | 4930 | 7.28947368 | 3.66281381 | 0 | 1 | 3 | 0.12352882 | 1 | 0.75193685 | 2 | 0.52640931 | 0 | 1 | 32 | 0.61800916 | FALSE | 1 | FALSE | 1 | FALSE | 1 | FALSE | 1 | FALSE | 1 | FALSE | 1 |
| 4931 | 4931 | 9.625 | 2.0577597 | 1 | 0.18769745 | 0 | 1 | 0 | 1 | 1 | 0.31351192 | 0 | 1 | 6 | 0.88425551 | FALSE | 1 | FALSE | 1 | FALSE | 1 | FALSE | 1 | FALSE | 1 | FALSE | 1 |
| 4932 | 4932 | 7.725 | 2.77927599 | 0 | 1 | 3 | 0.48172889 | 3 | 0.5528654 | 4 | 0.50425469 | 1 | 0.73564367 | 69 | 0.39112202 | FALSE | 1 | FALSE | 1 | FALSE | 1 | FALSE | 1 | FALSE | 1 | FALSE | 1 |
| 4933 | 4933 | 10.15 | 1.73997126 | 1 | 0.40558521 | 1 | 0.48239024 | 1 | 0.51939841 | 1 | 0.60987146 | 0 | 1 | 16 | 0.80826361 | FALSE | 1 | FALSE | 1 | FALSE | 1 | FALSE | 1 | FALSE | 1 | FALSE | 1 |
| 4934 | 4934 | 12 | 1.60356745 | 0 | 1 | 1 | 0.20567256 | 0 | 1 | 2 | 0.03785362 | 0 | 1 | 4 | 0.98605104 | FALSE | 1 | FALSE | 1 | FALSE | 1 | FALSE | 1 | FALSE | 1 | FALSE | 1 |
| 4935 | 4935 | 11.1 | 3.11287648 | 1 | 0.40558521 | 0 | 1 | 0 | 1 | 1 | 0.60987146 | 0 | 1 | 18 | 0.37504645 | FALSE | 1 | FALSE | 1 | FALSE | 1 | FALSE | 1 | FALSE | 1 | FALSE | 1 |
| 4936 | 4936 | 7.17142857 | 2.51266181 | 1 | 0.58738519 | 1 | 0.6739483 | 0 | 1 | 0 | 1 | 2 | 0.1065437 | 30 | 0.36879488 | FALSE | 1 | FALSE | 1 | FALSE | 1 | FALSE | 1 | FALSE | 1 | FALSE | 1 |
| 4937 | 4937 | 7.56043956 | 2.48161943 | 1 | 0.90494416 | 3 | 0.5614295 | 7 | 0.04237488 | 10 | 0.00789477 | 1 | 0.77640116 | 68 | 0.99032709 | FALSE | 1 | FALSE | 1 | FALSE | 1 | FALSE | 1 | FALSE | 1 | FALSE | 1 |
| 4938 | 4938 | 8.86956522 | 1.82505147 | 2 | 0.11647716 | 2 | 0.16948764 | 1 | 0.5695037 | 1 | 0.6613267 | 1 | 0.31672722 | 16 | 0.98100709 | FALSE | 0.93181727 | FALSE | 1 | FALSE | 1 | FALSE | 1 | FALSE | 1 | FALSE | 1 |
| 4939 | 4939 | 7.52272727 | 1.36218357 | 1 | 0.96865218 | 3 | 0.80636416 | 3 | 0.85933994 | 6 | 0.56950866 | 0 | 1 | 119 | 0.03681891 | FALSE | 1 | FALSE | 1 | FALSE | 1 | FALSE | 1 | FALSE | 1 | FALSE | 1 |
| 4940 | 4940 | 8.95 | 3.02448343 | 0 | 1 | 1 | 0.48239024 | 1 | 0.51939841 | 0 | 1 | 0 | 1 | 18 | 0.37504645 | FALSE | 1 | FALSE | 1 | FALSE | 1 | FALSE | 1 | FALSE | 1 | FALSE | 1 |
| 4941 | 4941 | 9.07042254 | 2.80001984 | 2 | 0.539854 | 3 | 0.39589822 | 3 | 0.46296673 | 4 | 0.40179526 | 1 | 0.68753645 | 57 | 0.8051717 | FALSE | 1 | FALSE | 1 | FALSE | 1 | FALSE | 1 | FALSE | 1 | FALSE | 1 |
| 4942 | 4942 | 12.0645161 | 1.41274119 | 0 | 1 | 3 | 0.07729739 | 2 | 0.30673593 | 1 | 0.76781284 | 1 | 0.4016888 | 24 | 0.90138666 | FALSE | 1 | FALSE | 1 | FALSE | 1 | FALSE | 1 | FALSE | 1 | FALSE | 1 |
| 4943 | 4943 | 10.8125 | 2.24217389 | 1 | 0.34033054 | 1 | 0.40942991 | 0 | 1 | 0 | 1 | 0 | 1 | 14 | 0.53308261 | FALSE | 1 | FALSE | 1 | FALSE | 1 | FALSE | 1 | FALSE | 1 | FALSE | 1 |
| 4944 | 4944 | 8.57894737 | 3.00553506 | 0 | 1 | 1 | 0.69480472 | 1 | 0.73299778 | 0 | 1 | 1 | 0.44938266 | 33 | 0.16407838 | FALSE | 1 | FALSE | 1 | FALSE | 1 | FALSE | 1 | FALSE | 1 | FALSE | 1 |
| 4945 | 4945 | 5.92982456 | 1.83398377 | 2 | 0.4229474 | 1 | 0.84259834 | 3 | 0.32646887 | 1 | 0.92884295 | 0 | 1 | 49 | 0.33483172 | FALSE | 1 | FALSE | 1 | FALSE | 1 | FALSE | 1 | FALSE | 1 | FALSE | 1 |
| 4946 | 4946 | 11.125 | 4.39992898 | 1 | 0.46441868 | 1 | 0.54637326 | 0 | 1 | 3 | 0.09525947 | 0 | 1 | 19 | 0.83975208 | FALSE | 1 | FALSE | 1 | FALSE | 1 | FALSE | 1 | FALSE | 1 | FALSE | 1 |
| 4947 | 4947 | 7.96551724 | 0.96428493 | 0 | 1 | 0 | 1 | 1 | 0.65463982 | 2 | 0.3871811 | 1 | 0.38148475 | 25 | 0.51779518 | FALSE | 1 | FALSE | 1 | FALSE | 1 | FALSE | 1 | FALSE | 1 | FALSE | 1 |
| 4948 | 4948 | 7.88888889 | 1.13311545 | 1 | 0.50471032 | 2 | 0.21684246 | 0 | 1 | 1 | 0.71956339 | 0 | 1 | 23 | 0.58307573 | FALSE | 1 | FALSE | 1 | FALSE | 1 | FALSE | 1 | FALSE | 1 | FALSE | 1 |
| 4949 | 4949 | 9.75 | 4.00780489 | 0 | 1 | 1 | 0.40942991 | 1 | 0.44344817 | 0 | 1 | 1 | 0.23264612 | 13 | 0.76799161 | FALSE | 1 | FALSE | 1 | FALSE | 1 | FALSE | 1 | FALSE | 1 | FALSE | 1 |
| 4950 | 4950 | 9.05555556 | 3.11755286 | 0 | 1 | 0 | 1 | 1 | 0.48280983 | 0 | 1 | 0 | 1 | 17 | 0.20329205 | FALSE | 1 | FALSE | 1 | FALSE | 1 | FALSE | 1 | FALSE | 1 | FALSE | 1 |
| 4951 | 4951 | 9.9 | 2.07123152 | 0 | 1 | 1 | 0.28036922 | 0 | 1 | 1 | 0.37517262 | 0 | 1 | 8 | 0.80394719 | FALSE | 1 | FALSE | 1 | FALSE | 1 | FALSE | 1 | FALSE | 1 | FALSE | 1 |
| 4952 | 4952 | 11.25 | 2.43669859 | 0 | 1 | 1 | 0.23139161 | 0 | 1 | 1 | 0.31351192 | 0 | 1 | 6 | 0.88425551 | FALSE | 1 | FALSE | 1 | FALSE | 1 | FALSE | 1 | FALSE | 1 | FALSE | 1 |
| 4953 | 4953 | 8.84615385 | 2.38089039 | 0 | 1 | 3 | 0.04547096 | 0 | 1 | 0 | 1 | 1 | 0.33902672 | 21 | 0.64957167 | FALSE | 1 | FALSE | 1 | FALSE | 1 | FALSE | 1 | FALSE | 1 | FALSE | 1 |
| 4954 | 4954 | 6.2826087 | 0.77074339 | 7 | 0.19292516 | 13 | 0.00647786 | 7 | 0.49367025 | 13 | 0.07994628 | 3 | 0.58557338 | 141 | 0.99807668 | FALSE | 1 | FALSE | 0.60103504 | FALSE | 1 | FALSE | 1 | FALSE | 1 | FALSE | 1 |
| 4955 | 4955 | 5.10588235 | 3.35374603 | 2 | 0.64592863 | 1 | 0.93995595 | 3 | 0.59435517 | 5 | 0.35139151 | 2 | 0.40793536 | 72 | 0.54001499 | FALSE | 1 | FALSE | 1 | FALSE | 1 | FALSE | 1 | FALSE | 1 | FALSE | 1 |
| 4956 | 4956 | 5.02857143 | 2.48981599 | 2 | 0.22589135 | 2 | 0.31355738 | 2 | 0.35984046 | 1 | 0.80778317 | 0 | 1 | 28 | 0.83079716 | FALSE | 1 | FALSE | 1 | FALSE | 1 | FALSE | 1 | FALSE | 1 | FALSE | 1 |
| 4957 | 4957 | 8.82142857 | 2.15561677 | 0 | 1 | 2 | 0.2288771 | 0 | 1 | 1 | 0.73248983 | 1 | 0.37113067 | 24 | 0.55018107 | FALSE | 1 | FALSE | 1 | FALSE | 1 | FALSE | 1 | FALSE | 1 | FALSE | 1 |
| 4958 | 4958 | 9.27027027 | 2.06186282 | 0 | 1 | 1 | 0.70472939 | 3 | 0.14625315 | 2 | 0.51200191 | 1 | 0.45845922 | 30 | 0.78878581 | FALSE | 1 | FALSE | 1 | FALSE | 1 | FALSE | 1 | FALSE | 1 | FALSE | 1 |
| 4959 | 4959 | 9.76923077 | 2.51621957 | 0 | 1 | 2 | 0.06426041 | 0 | 1 | 0 | 1 | 0 | 1 | 11 | 0.6683056 | FALSE | 1 | FALSE | 1 | FALSE | 1 | FALSE | 1 | FALSE | 1 | FALSE | 1 |
| 4960 | 4960 | 9.68421053 | 3.17974894 | 1 | 0.60837712 | 3 | 0.10944025 | 2 | 0.37289966 | 2 | 0.49731009 | 1 | 0.44938266 | 27 | 0.95561238 | FALSE | 1 | FALSE | 1 | FALSE | 1 | FALSE | 1 | FALSE | 1 | FALSE | 1 |
| 4961 | 4961 | 6.28 | 2.47418674 | 1 | 0.47819878 | 0 | 1 | 0 | 1 | 0 | 1 | 0 | 1 | 24 | 0.08019798 | FALSE | 1 | FALSE | 1 | FALSE | 1 | FALSE | 1 | FALSE | 1 | FALSE | 1 |
| 4962 | 4962 | 10.9666667 | 2.24326151 | 0 | 1 | 2 | 0.24095584 | 2 | 0.27983822 | 2 | 0.3871811 | 1 | 0.38148475 | 22 | 0.92902102 | FALSE | 1 | FALSE | 1 | FALSE | 1 | FALSE | 1 | FALSE | 1 | FALSE | 1 |
| 4963 | 4963 | 11.4545455 | 2.90340353 | 0 | 1 | 0 | 1 | 0 | 1 | 0 | 1 | 1 | 0.16638736 | 10 | 0.46859937 | FALSE | 1 | FALSE | 1 | FALSE | 1 | FALSE | 1 | FALSE | 1 | FALSE | 1 |
| 4964 | 4964 | 10.1538462 | 3.03432045 | 1 | 0.28675951 | 0 | 1 | 0 | 1 | 0 | 1 | 0 | 1 | 12 | 0.374198 | FALSE | 1 | FALSE | 1 | FALSE | 1 | FALSE | 1 | FALSE | 1 | FALSE | 1 |
| 4965 | 4965 | 8.6 | 2.32009852 | 0 | 1 | 3 | 0.10264583 | 1 | 0.72299506 | 0 | 1 | 1 | 0.44015537 | 30 | 0.52765882 | FALSE | 1 | FALSE | 1 | FALSE | 1 | FALSE | 1 | FALSE | 1 | FALSE | 1 |
| 4966 | 4966 | 10.2941177 | 2.88535258 | 1 | 0.35728305 | 0 | 1 | 2 | 0.12280513 | 0 | 1 | 0 | 1 | 14 | 0.7313145 | FALSE | 1 | FALSE | 1 | FALSE | 1 | FALSE | 1 | FALSE | 1 | FALSE | 1 |
| 4967 | 4967 | 8.66666667 | 2.54732976 | 0 | 1 | 0 | 1 | 1 | 0.42266264 | 1 | 0.50623194 | 0 | 1 | 13 | 0.57712805 | FALSE | 1 | FALSE | 1 | FALSE | 1 | FALSE | 1 | FALSE | 1 | FALSE | 1 |
| 4968 | 4968 | 8.02564103 | 2.32585446 | 1 | 0.6184714 | 3 | 0.11640364 | 1 | 0.74264075 | 3 | 0.2402032 | 0 | 1 | 29 | 0.88820978 | FALSE | 1 | FALSE | 1 | FALSE | 1 | FALSE | 1 | FALSE | 1 | FALSE | 1 |
| 4969 | 4969 | 11.5263158 | 2.7408877 | 0 | 1 | 1 | 0.46503795 | 0 | 1 | 2 | 0.21610812 | 0 | 1 | 16 | 0.65565778 | FALSE | 1 | FALSE | 1 | FALSE | 1 | FALSE | 1 | FALSE | 1 | FALSE | 1 |
| 4970 | 4970 | 8.26666667 | 4.17079795 | 0 | 1 | 0 | 1 | 0 | 1 | 1 | 0.50623194 | 0 | 1 | 14 | 0.29525409 | FALSE | 1 | FALSE | 1 | FALSE | 1 | FALSE | 1 | FALSE | 1 | FALSE | 1 |
| 4971 | 4971 | 10.3 | 3.33316666 | 1 | 0.40558521 | 1 | 0.48239024 | 1 | 0.51939841 | 1 | 0.60987146 | 0 | 1 | 16 | 0.80826361 | FALSE | 1 | FALSE | 1 | FALSE | 1 | FALSE | 1 | FALSE | 1 | FALSE | 1 |
| 4972 | 4972 | 9.05405405 | 2.37037967 | 3 | 0.0681763 | 1 | 0.70472939 | 2 | 0.38585089 | 0 | 1 | 0 | 1 | 31 | 0.64534533 | FALSE | 0.74276693 | FALSE | 1 | FALSE | 1 | FALSE | 1 | FALSE | 1 | FALSE | 1 |
| 4973 | 4973 | 9.33333333 | 3.48238196 | 0 | 1 | 0 | 1 | 2 | 0.17308361 | 1 | 0.62783522 | 0 | 1 | 18 | 0.57962763 | FALSE | 1 | FALSE | 1 | FALSE | 1 | FALSE | 1 | FALSE | 1 | FALSE | 1 |
| 4974 | 4974 | 8.9 | 1.47986486 | 1 | 0.64722818 | 4 | 0.03920804 | 1 | 0.76953771 | 1 | 0.84823834 | 0 | 1 | 33 | 0.71942635 | FALSE | 1 | FALSE | 0.96864929 | FALSE | 1 | FALSE | 1 | FALSE | 1 | FALSE | 1 |
| 4975 | 4975 | 5.32824428 | 3.20886831 | 1 | 0.9678105 | 2 | 0.92943464 | 7 | 0.19109964 | 3 | 0.94439785 | 0 | 1 | 118 | 0.03972201 | FALSE | 1 | FALSE | 1 | FALSE | 1 | FALSE | 1 | FALSE | 1 | FALSE | 1 |
| 4976 | 4976 | 8.41176471 | 3.16337169 | 0 | 1 | 2 | 0.10307112 | 0 | 1 | 0 | 1 | 0 | 1 | 15 | 0.49060435 | FALSE | 1 | FALSE | 1 | FALSE | 1 | FALSE | 1 | FALSE | 1 | FALSE | 1 |
| 4977 | 4977 | 4.27631579 | 3.30708444 | 1 | 0.86263338 | 1 | 0.91898592 | 2 | 0.76408693 | 7 | 0.05887011 | 4 | 0.03564889 | 61 | 0.87381726 | FALSE | 1 | FALSE | 1 | FALSE | 1 | FALSE | 1 | FALSE | 0.87669812 | FALSE | 1 |
| 4978 | 4978 | 8.84375 | 1.92206554 | 1 | 0.56527493 | 2 | 0.27730767 | 2 | 0.32011871 | 2 | 0.43583874 | 1 | 0.41154418 | 24 | 0.94876798 | FALSE | 1 | FALSE | 1 | FALSE | 1 | FALSE | 1 | FALSE | 1 | FALSE | 1 |
| 4979 | 4979 | 8.50892857 | 2.04413593 | 1 | 0.94390149 | 3 | 0.69664974 | 4 | 0.56195051 | 7 | 0.24054949 | 1 | 0.84015603 | 94 | 0.4396291 | FALSE | 1 | FALSE | 1 | FALSE | 1 | FALSE | 1 | FALSE | 1 | FALSE | 1 |
| 4980 | 4980 | 8.69230769 | 2.68570619 | 0 | 1 | 0 | 1 | 0 | 1 | 2 | 0.33678042 | 0 | 1 | 24 | 0.20425579 | FALSE | 1 | FALSE | 1 | FALSE | 1 | FALSE | 1 | FALSE | 1 | FALSE | 1 |

| | | | | | | | | | | | | | | | | | | | | | | | | | | | |
|---|---|---|---|---|---|---|---|---|---|---|---|---|---|---|---|---|---|---|---|---|---|---|---|---|---|---|---|
| 4981 | 4981 | 9.0952381 | 2.15814651 | 0 | 1 | 1 | 0.49918218 | 1 | 0.53671415 | 0 | 1 | 1 | 0.29368245 | 18 | 0.57962763 | FALSE | 1 | FALSE | 1 | FALSE | 1 | FALSE | 1 | FALSE | 1 | FALSE | 1 |
| 4982 | 4982 | 8.66666667 | 2.02758751 | 0 | 1 | 0 | 1 | 0 | 1 | 3 | 0.04687752 | 0 | 1 | 15 | 0.69372602 | FALSE | 1 | FALSE | 1 | FALSE | 1 | FALSE | 1 | FALSE | 1 | FALSE | 1 |
| 4983 | 4983 | 9.07142857 | 1.33439584 | 0 | 1 | 0 | 1 | 2 | 0.2663542 | 3 | 0.13524571 | 0 | 1 | 23 | 0.73439118 | FALSE | 1 | FALSE | 1 | FALSE | 1 | FALSE | 1 | FALSE | 1 | FALSE | 1 |
| 4984 | 4984 | 4.3255814 | 1.40961697 | 0 | 1 | 4 | 0.30278494 | 7 | 0.03432404 | 7 | 0.09905935 | 0 | 1 | 68 | 0.93051124 | FALSE | 1 | FALSE | 1 | FALSE | 1 | FALSE | 1 | FALSE | 1 | FALSE | 1 |
| 4985 | 4985 | 10 | 3.02976685 | 3 | 0.07731604 | 0 | 1 | 2 | 0.41139386 | 2 | 0.54052736 | 0 | 1 | 32 | 0.74324845 | FALSE | 0.77583006 | FALSE | 1 | FALSE | 1 | FALSE | 1 | FALSE | 1 | FALSE | 1 |
| 4986 | 4986 | 8.03030303 | 1.83377155 | 1 | 0.57647348 | 0 | 1 | 2 | 0.33343871 | 4 | 0.06258966 | 0 | 1 | 26 | 0.86852567 | FALSE | 1 | FALSE | 1 | FALSE | 1 | FALSE | 1 | FALSE | 1 | FALSE | 1 |
| 4987 | 4987 | 11.4375 | 2.03004772 | 0 | 1 | 1 | 0.40942991 | 1 | 0.44344817 | 0 | 1 | 0 | 1 | 14 | 0.53308261 | FALSE | 1 | FALSE | 1 | FALSE | 1 | FALSE | 1 | FALSE | 1 | FALSE | 1 |
| 4988 | 4988 | 10.1764706 | 2.57214248 | 0 | 1 | 0 | 1 | 2 | 0.12280513 | 3 | 0.040397 | 0 | 1 | 12 | 0.962093 | FALSE | 1 | FALSE | 1 | FALSE | 1 | FALSE | 1 | FALSE | 1 | FALSE | 1 |
| 4989 | 4989 | 8.33333333 | 1.78174161 | 1 | 0.42086991 | 0 | 1 | 3 | 0.03775825 | 1 | 0.62783522 | 0 | 1 | 16 | 0.90353674 | FALSE | 1 | FALSE | 1 | FALSE | 1 | FALSE | 1 | FALSE | 1 | FALSE | 1 |
| 4990 | 4990 | 6.45833333 | 2.75347003 | 0 | 1 | 0 | 1 | 5 | 0.1012281 | 3 | 0.6203651 | 0 | 1 | 61 | 0.22871622 | FALSE | 1 | FALSE | 1 | FALSE | 1 | FALSE | 1 | FALSE | 1 | FALSE | 1 |
| 4991 | 4991 | 5.54347826 | 2.55977684 | 2 | 0.3309085 | 0 | 1 | 1 | 0.81522035 | 3 | 0.35428318 | 0 | 1 | 40 | 0.40712403 | FALSE | 1 | FALSE | 1 | FALSE | 1 | FALSE | 1 | FALSE | 1 | FALSE | 1 |
| 4992 | 4992 | 10.5428571 | 2.87196157 | 1 | 0.59801736 | 4 | 0.02543131 | 0 | 1 | 2 | 0.48233989 | 0 | 1 | 28 | 0.83079716 | FALSE | 1 | FALSE | 0.82367276 | FALSE | 1 | FALSE | 1 | FALSE | 1 | FALSE | 1 |
| 4993 | 4993 | 7.63265306 | 1.93458193 | 1 | 0.72119464 | 1 | 0.80150109 | 1 | 0.83455705 | 2 | 0.66536008 | 0 | 1 | 44 | 0.20085966 | FALSE | 1 | FALSE | 1 | FALSE | 1 | FALSE | 1 | FALSE | 1 | FALSE | 1 |
| 4994 | 4994 | 10 | 2.87228132 | 0 | 1 | 2 | 0.09285443 | 1 | 0.44344817 | 0 | 1 | 0 | 1 | 13 | 0.76799161 | FALSE | 1 | FALSE | 1 | FALSE | 1 | FALSE | 1 | FALSE | 1 | FALSE | 1 |
| 4995 | 4995 | 10.2608696 | 2.04763684 | 1 | 0.45027683 | 1 | 0.53115654 | 0 | 1 | 1 | 0.6613267 | 1 | 0.31672722 | 19 | 0.71542496 | FALSE | 1 | FALSE | 1 | FALSE | 1 | FALSE | 1 | FALSE | 1 | FALSE | 1 |
| 4996 | 4996 | 9.4 | 2.47116707 | 0 | 1 | 2 | 0.08295839 | 1 | 0.42266264 | 1 | 0.50623194 | 0 | 1 | 11 | 0.92879412 | FALSE | 1 | FALSE | 1 | FALSE | 1 | FALSE | 1 | FALSE | 1 | FALSE | 1 |
| 4997 | 4997 | 10.125 | 3.51559028 | 0 | 1 | 0 | 1 | 1 | 0.25384309 | 0 | 1 | 0 | 1 | 7 | 0.63752397 | FALSE | 1 | FALSE | 1 | FALSE | 1 | FALSE | 1 | FALSE | 1 | FALSE | 1 |
| 4998 | 4998 | 9.16129032 | 2.09602964 | 1 | 0.55378205 | 2 | 0.2651851 | 2 | 0.30673593 | 2 | 0.41984187 | 2 | 0.09110253 | 22 | 0.98395924 | FALSE | 1 | FALSE | 1 | FALSE | 1 | FALSE | 1 | FALSE | 1 | FALSE | 1 |
| 4999 | 4999 | 8.54166667 | 3.10884714 | 1 | 0.45027683 | 0 | 1 | 1 | 0.5695037 | 1 | 0.6613267 | 1 | 0.31672722 | 19 | 0.71542496 | FALSE | 1 | FALSE | 1 | FALSE | 1 | FALSE | 1 | FALSE | 1 | FALSE | 1 |
| 5000 | 5000 | 9.79310345 | 2.05743916 | 0 | 1 | 1 | 0.61539065 | 1 | 0.65463982 | 2 | 0.3871811 | 0 | 1 | 25 | 0.51779518 | FALSE | 1 | FALSE | 1 | FALSE | 1 | FALSE | 1 | FALSE | 1 | FALSE | 1 |
| 5001 | 5001 | 9.6 | 3.69323706 | 0 | 1 | 0 | 1 | 0 | 1 | 1 | 0.37517262 | 0 | 1 | 9 | 0.5215547 | FALSE | 1 | FALSE | 1 | FALSE | 1 | FALSE | 1 | FALSE | 1 | FALSE | 1 |
| 5002 | 5002 | 9.44444444 | 2.19145366 | 4 | 0.00096345 | 1 | 0.44710673 | 0 | 1 | 2 | 0.19902682 | 0 | 1 | 11 | 0.99651304 | FALSE | 0.07376244 | FALSE | 1 | FALSE | 1 | FALSE | 1 | FALSE | 1 | FALSE | 1 |
| 5003 | 5003 | 12.1 | 1.17898261 | 0 | 1 | 0 | 1 | 0 | 1 | 3 | 0.0612581 | 0 | 1 | 17 | 0.61750632 | FALSE | 1 | FALSE | 1 | FALSE | 1 | FALSE | 1 | FALSE | 1 | FALSE | 1 |
| 5004 | 5004 | 8.67857143 | 1.98302747 | 1 | 0.51745948 | 0 | 1 | 1 | 0.64171563 | 1 | 0.73248983 | 0 | 1 | 25 | 0.34293 | FALSE | 1 | FALSE | 1 | FALSE | 1 | FALSE | 1 | FALSE | 1 | FALSE | 1 |
| 5005 | 5005 | 9.2 | 2.35796523 | 1 | 0.22886536 | 0 | 1 | 0 | 1 | 0 | 1 | 0 | 1 | 9 | 0.5215547 | FALSE | 1 | FALSE | 1 | FALSE | 1 | FALSE | 1 | FALSE | 1 | FALSE | 1 |
| 5006 | 5006 | 7.65333333 | 3.45443998 | 5 | 0.31981947 | 3 | 0.85725387 | 6 | 0.42804763 | 4 | 0.90906259 | 1 | 0.91286792 | 127 | 0.22622784 | FALSE | 1 | FALSE | 1 | FALSE | 1 | FALSE | 1 | FALSE | 1 | FALSE | 1 |
| 5007 | 5007 | 9.76190476 | 2.8603157 | 0 | 1 | 0 | 1 | 0 | 1 | 1 | 0.62783522 | 1 | 0.29368245 | 19 | 0.34094959 | FALSE | 1 | FALSE | 1 | FALSE | 1 | FALSE | 1 | FALSE | 1 | FALSE | 1 |
| 5008 | 5008 | 9.14285714 | 4.34999414 | 0 | 1 | 2 | 0.31355738 | 2 | 0.35984046 | 2 | 0.48233989 | 1 | 0.44015537 | 28 | 0.83079716 | FALSE | 1 | FALSE | 1 | FALSE | 1 | FALSE | 1 | FALSE | 1 | FALSE | 1 |
| 5009 | 5009 | 8.86363636 | 2.36057494 | 0 | 1 | 0 | 1 | 1 | 0.55340851 | 0 | 1 | 0 | 1 | 21 | 0.12048848 | FALSE | 1 | FALSE | 1 | FALSE | 1 | FALSE | 1 | FALSE | 1 | FALSE | 1 |
| 5010 | 5010 | 5.34848485 | 2.62558752 | 0 | 1 | 2 | 0.62701348 | 2 | 0.68408105 | 3 | 0.57949569 | 0 | 1 | 58 | 0.18266626 | FALSE | 1 | FALSE | 1 | FALSE | 1 | FALSE | 1 | FALSE | 1 | FALSE | 1 |
| 5011 | 5011 | 8.15 | 2.92589018 | 4 | 0.0671906 | 1 | 0.86215528 | 6 | 0.02001082 | 5 | 0.1398835 | 2 | 0.25814944 | 42 | 0.99861324 | FALSE | 0.74276693 | FALSE | 1 | FALSE | 1 | FALSE | 1 | FALSE | 1 | FALSE | 1 |
| 5012 | 5012 | 9.66666667 | 2.00426895 | 2 | 0.26413583 | 2 | 0.36122049 | 2 | 0.41139386 | 1 | 0.84089119 | 1 | 0.4761699 | 31 | 0.85613899 | FALSE | 1 | FALSE | 1 | FALSE | 1 | FALSE | 1 | FALSE | 1 | FALSE | 1 |
| 5013 | 5013 | 9.12820513 | 2.5336655 | 6 | 0.00042043 | 6 | 0.00142608 | 1 | 0.76089851 | 2 | 0.54052736 | 0 | 1 | 24 | 0.99988054 | TRUE | 0.04200174 | FALSE | 0.40284184 | FALSE | 1 | FALSE | 1 | FALSE | 1 | FALSE | 1 |
| 5014 | 5014 | 9.11111111 | 2.33068633 | 0 | 1 | 0 | 1 | 1 | 0.28067846 | 0 | 1 | 0 | 1 | 8 | 0.5780332 | FALSE | 1 | FALSE | 1 | FALSE | 1 | FALSE | 1 | FALSE | 1 | FALSE | 1 |
| 5015 | 5015 | 9.38461539 | 2.71855339 | 0 | 1 | 1 | 0.57534752 | 1 | 0.61440466 | 0 | 1 | 2 | 0.0670823 | 22 | 0.61628 | FALSE | 1 | FALSE | 1 | FALSE | 1 | FALSE | 1 | FALSE | 1 | FALSE | 1 |
| 5016 | 5016 | 10.5 | 1.36014705 | 0 | 1 | 0 | 1 | 1 | 0.51939841 | 2 | 0.23331368 | 0 | 1 | 17 | 0.61750632 | FALSE | 1 | FALSE | 1 | FALSE | 1 | FALSE | 1 | FALSE | 1 | FALSE | 1 |
| 5017 | 5017 | 9.53333333 | 1.64789428 | 0 | 1 | 1 | 0.6278847 | 1 | 0.66709967 | 1 | 0.75658797 | 0 | 1 | 27 | 0.28862208 | FALSE | 1 | FALSE | 1 | FALSE | 1 | FALSE | 1 | FALSE | 1 | FALSE | 1 |
| 5018 | 5018 | 9.53448276 | 3.00123835 | 1 | 0.77972342 | 7 | 0.00248576 | 2 | 0.62245518 | 2 | 0.75303313 | 1 | 0.61824432 | 45 | 0.94087175 | FALSE | 1 | FALSE | 0.44070554 | FALSE | 1 | FALSE | 1 | FALSE | 1 | FALSE | 1 |
| 5019 | 5019 | 9.36363636 | 2.22680886 | 0 | 1 | 0 | 1 | 2 | 0.05708125 | 0 | 1 | 0 | 1 | 9 | 0.75985366 | FALSE | 1 | FALSE | 1 | FALSE | 1 | FALSE | 1 | FALSE | 1 | FALSE | 1 |
| 5020 | 5020 | 10.4615385 | 2.74885668 | 0 | 1 | 0 | 1 | 3 | 0.06482014 | 1 | 0.70601446 | 2 | 0.0670823 | 20 | 0.90142689 | FALSE | 1 | FALSE | 1 | FALSE | 1 | FALSE | 1 | FALSE | 1 | FALSE | 1 |
| 5021 | 5021 | 10.5454546 | 2.80789913 | 0 | 1 | 0 | 1 | 0 | 1 | 0 | 1 | 1 | 0.16638736 | 10 | 0.46859937 | FALSE | 1 | FALSE | 1 | FALSE | 1 | FALSE | 1 | FALSE | 1 | FALSE | 1 |
| 5022 | 5022 | 8.55555556 | 2.89102485 | 0 | 1 | 0 | 1 | 1 | 0.48280983 | 2 | 0.19902682 | 0 | 1 | 15 | 0.69372602 | FALSE | 1 | FALSE | 1 | FALSE | 1 | FALSE | 1 | FALSE | 1 | FALSE | 1 |
| 5023 | 5023 | 8.41176471 | 2.37780976 | 0 | 1 | 0 | 1 | 0 | 1 | 0 | 1 | 1 | 0.24525764 | 16 | 0.23070182 | FALSE | 1 | FALSE | 1 | FALSE | 1 | FALSE | 1 | FALSE | 1 | FALSE | 1 |
| 5024 | 5024 | 9.5 | 1.52206001 | 3 | 0.0405081 | 0 | 1 | 1 | 0.66709967 | 1 | 0.75658797 | 0 | 1 | 25 | 0.67630471 | FALSE | 0.62493854 | FALSE | 1 | FALSE | 1 | FALSE | 1 | FALSE | 1 | FALSE | 1 |
| 5025 | 5025 | 9.15 | 3.63696302 | 2 | 0.09187136 | 3 | 0.02534497 | 1 | 0.51939841 | 0 | 1 | 0 | 1 | 14 | 0.97325593 | FALSE | 0.83248965 | FALSE | 0.82367276 | FALSE | 1 | FALSE | 1 | FALSE | 1 | FALSE | 1 |
| 5026 | 5026 | 11.2 | 3.8331884 | 1 | 0.32293355 | 0 | 1 | 1 | 0.42266264 | 1 | 0.50623194 | 1 | 0.2198258 | 11 | 0.92879412 | FALSE | 1 | FALSE | 1 | FALSE | 1 | FALSE | 1 | FALSE | 1 | FALSE | 1 |
| 5027 | 5027 | 9.16666667 | 2.07498327 | 0 | 1 | 0 | 1 | 1 | 0.35555293 | 0 | 1 | 0 | 1 | 11 | 0.41944532 | FALSE | 1 | FALSE | 1 | FALSE | 1 | FALSE | 1 | FALSE | 1 | FALSE | 1 |
| 5028 | 5028 | 3.65660377 | 2.11352424 | 4 | 0.90956197 | 6 | 0.85795007 | 3 | 0.99646386 | 7 | 0.96117137 | 1 | 0.98794858 | 241 | 0.00014619 | FALSE | 1 | FALSE | 1 | FALSE | 1 | FALSE | 1 | FALSE | 1 | FALSE | 0.59879545 |
| 5029 | 5029 | 10.05 | 2.24666419 | 0 | 1 | 0 | 1 | 0 | 1 | 0 | 1 | 0 | 1 | 20 | 0.03329856 | FALSE | 1 | FALSE | 1 | FALSE | 1 | FALSE | 1 | FALSE | 1 | FALSE | 1 |
| 5030 | 5030 | 9 | 2 | 0 | 1 | 1 | 0.42857733 | 0 | 1 | 0 | 1 | 0 | 1 | 16 | 0.23070182 | FALSE | 1 | FALSE | 1 | FALSE | 1 | FALSE | 1 | FALSE | 1 | FALSE | 1 |
| 5031 | 5031 | 9.41176471 | 1.78420599 | 0 | 1 | 0 | 1 | 1 | 0.46348836 | 0 | 1 | 0 | 1 | 16 | 0.23070182 | FALSE | 1 | FALSE | 1 | FALSE | 1 | FALSE | 1 | FALSE | 1 | FALSE | 1 |
| 5032 | 5032 | 9.53846154 | 2.22279743 | 0 | 1 | 0 | 1 | 0 | 1 | 2 | 0.33678042 | 0 | 1 | 24 | 0.20425579 | FALSE | 1 | FALSE | 1 | FALSE | 1 | FALSE | 1 | FALSE | 1 | FALSE | 1 |
| 5033 | 5033 | 10.8 | 2.42761886 | 1 | 0.32293355 | 0 | 1 | 0 | 1 | 3 | 0.02891327 | 0 | 1 | 11 | 0.92879412 | FALSE | 1 | FALSE | 1 | FALSE | 1 | FALSE | 1 | FALSE | 1 | FALSE | 1 |
| 5034 | 5034 | 8.64285714 | 2.02156739 | 2 | 0.04858661 | 0 | 1 | 1 | 0.40110418 | 2 | 0.13304705 | 0 | 1 | 9 | 0.98607548 | FALSE | 0.649301 | FALSE | 1 | FALSE | 1 | FALSE | 1 | FALSE | 1 | FALSE | 1 |
| 5035 | 5035 | 9.5625 | 2.47408443 | 0 | 1 | 0 | 1 | 2 | 0.11087977 | 0 | 1 | 0 | 1 | 14 | 0.53308261 | FALSE | 1 | FALSE | 1 | FALSE | 1 | FALSE | 1 | FALSE | 1 | FALSE | 1 |
| 5036 | 5036 | 10.2 | 3.3901819 | 0 | 1 | 1 | 0.38964394 | 1 | 0.42266264 | 0 | 1 | 1 | 0.2198258 | 12 | 0.80329917 | FALSE | 1 | FALSE | 1 | FALSE | 1 | FALSE | 1 | FALSE | 1 | FALSE | 1 |
| 5037 | 5037 | 11.1666667 | 1.67497927 | 0 | 1 | 1 | 0.54637326 | 2 | 0.21260257 | 0 | 1 | 0 | 1 | 21 | 0.47053137 | FALSE | 1 | FALSE | 1 | FALSE | 1 | FALSE | 1 | FALSE | 1 | FALSE | 1 |
| 5038 | 5038 | 9.5862069 | 2.53957967 | 1 | 0.52988231 | 1 | 0.61539065 | 2 | 0.27983822 | 1 | 0.74482229 | 2 | 0.08121636 | 22 | 0.92902102 | FALSE | 1 | FALSE | 1 | FALSE | 1 | FALSE | 1 | FALSE | 1 | FALSE | 1 |
| 5039 | 5039 | 9.86206897 | 4.15828328 | 2 | 0.16048326 | 1 | 0.60247911 | 0 | 1 | 2 | 0.3705495 | 1 | 0.37113067 | 22 | 0.86499992 | FALSE | 1 | FALSE | 1 | FALSE | 1 | FALSE | 1 | FALSE | 1 | FALSE | 1 |
| 5040 | 5040 | 7.85714286 | 2.71303229 | 0 | 1 | 1 | 0.49918218 | 0 | 1 | 0 | 1 | 1 | 0.29368245 | 19 | 0.34094959 | FALSE | 1 | FALSE | 1 | FALSE | 1 | FALSE | 1 | FALSE | 1 | FALSE | 1 |
| 5041 | 5041 | 9 | 1.765686 | 0 | 1 | 2 | 0.30150529 | 2 | 0.34668321 | 3 | 0.20353499 | 0 | 1 | 27 | 0.85023803 | FALSE | 1 | FALSE | 1 | FALSE | 1 | FALSE | 1 | FALSE | 1 | FALSE | 1 |
| 5042 | 5042 | 7.7 | 1.79164729 | 1 | 0.22886536 | 0 | 1 | 1 | 0.30655256 | 1 | 0.37517262 | 0 | 1 | 7 | 0.94326215 | FALSE | 1 | FALSE | 1 | FALSE | 1 | FALSE | 1 | FALSE | 1 | FALSE | 1 |
| 5043 | 5043 | 8.34210526 | 2.90893647 | 3 | 0.30904591 | 1 | 0.91898592 | 2 | 0.76408693 | 3 | 0.68517679 | 0 | 1 | 67 | 0.23030366 | FALSE | 1 | FALSE | 1 | FALSE | 1 | FALSE | 1 | FALSE | 1 | FALSE | 1 |
| 5044 | 5044 | 9.83333333 | 2.73353658 | 2 | 0.03643311 | 0 | 1 | 0 | 1 | 0 | 1 | 2 | 0.01579445 | 8 | 0.97197833 | FALSE | 0.58392 | FALSE | 1 | FALSE | 1 | FALSE | 1 | FALSE | 0.66313193 | FALSE | 1 |
| 5045 | 5045 | 10.4347826 | 3.7279915 | 3 | 0.02011786 | 0 | 1 | 2 | 0.19931429 | 1 | 0.6613267 | 0 | 1 | 17 | 0.94439745 | FALSE | 0.44482142 | FALSE | 1 | FALSE | 1 | FALSE | 1 | FALSE | 1 | FALSE | 1 |
| 5046 | 5046 | 8.73684211 | 3.06666466 | 2 | 0.25455533 | 1 | 0.71433277 | 3 | 0.15483587 | 2 | 0.52640931 | 0 | 1 | 30 | 0.87271204 | FALSE | 1 | FALSE | 1 | FALSE | 1 | FALSE | 1 | FALSE | 1 | FALSE | 1 |
| 5047 | 5047 | 7.26470588 | 1.00904217 | 2 | 0.86878756 | 3 | 0.82227197 | 3 | 0.87268279 | 13 | 0.00935627 | 3 | 0.38636521 | 112 | 0.78449587 | FALSE | 1 | FALSE | 1 | FALSE | 1 | FALSE | 1 | FALSE | 1 | FALSE | 1 |
| 5048 | 5048 | 9.18181818 | 2.28903242 | 1 | 0.42086991 | 2 | 0.14655134 | 1 | 0.53671415 | 0 | 1 | 0 | 1 | 17 | 0.77849076 | FALSE | 1 | FALSE | 1 | FALSE | 1 | FALSE | 1 | FALSE | 1 | FALSE | 1 |
| 5049 | 5049 | 8.90909091 | 3.04375801 | 0 | 1 | 0 | 1 | 1 | 0.55340851 | 3 | 0.07743186 | 0 | 1 | 18 | 0.74744813 | FALSE | 1 | FALSE | 1 | FALSE | 1 | FALSE | 1 | FALSE | 1 | FALSE | 1 |
| 5050 | 5050 | 9.73333333 | 2.33713975 | 1 | 0.54198714 | 0 | 1 | 2 | 0.29330414 | 3 | 0.15707004 | 0 | 1 | 24 | 0.82294782 | FALSE | 1 | FALSE | 1 | FALSE | 1 | FALSE | 1 | FALSE | 1 | FALSE | 1 |
| 5051 | 5051 | 6.81578947 | 2.25779659 | 3 | 0.0638041 | 5 | 0.0056244 | 1 | 0.73299778 | 1 | 0.81665383 | 0 | 1 | 26 | 0.98200522 | FALSE | 0.73766 | FALSE | 0.59070661 | FALSE | 1 | FALSE | 1 | FALSE | 1 | FALSE | 1 |
| 5052 | 5052 | 11.2156863 | 2.06096994 | 1 | 0.7354105 | 1 | 0.81422651 | 2 | 0.55199489 | 2 | 0.68678423 | 4 | 0.00946188 | 41 | 0.8377026 | FALSE | 1 | FALSE | 1 | FALSE | 1 | FALSE | 1 | FALSE | 0.5373705 | FALSE | 1 |
| 5053 | 5053 | 8.22727273 | 3.35656505 | 0 | 1 | 2 | 0.15794013 | 0 | 1 | 1 | 0.64497439 | 0 | 1 | 19 | 0.54233419 | FALSE | 1 | FALSE | 1 | FALSE | 1 | FALSE | 1 | FALSE | 1 | FALSE | 1 |
| 5054 | 5054 | 8.58823529 | 3.35294118 | 0 | 1 | 1 | 0.6739483 | 2 | 0.34668321 | 2 | 0.46709861 | 1 | 0.43077488 | 28 | 0.72558119 | FALSE | 1 | FALSE | 1 | FALSE | 1 | FALSE | 1 | FALSE | 1 | FALSE | 1 |
| 5055 | 5055 | 8.75862069 | 1.88743892 | 1 | 0.52988231 | 1 | 0.61539065 | 1 | 0.65463982 | 4 | 0.04184267 | 0 | 1 | 22 | 0.92902102 | FALSE | 1 | FALSE | 1 | FALSE | 1 | FALSE | 1 | FALSE | 1 | FALSE | 1 |
| 5056 | 5056 | 6.37142857 | 3.97194241 | 1 | 0.59801736 | 1 | 0.68454804 | 1 | 0.72299506 | 1 | 0.80778317 | 0 | 1 | 31 | 0.34231756 | FALSE | 1 | FALSE | 1 | FALSE | 1 | FALSE | 1 | FALSE | 1 | FALSE | 1 |
| 5057 | 5057 | 6.51327434 | 2.26795703 | 9 | 0.12580146 | 11 | 0.11466966 | 9 | 0.42444671 | 11 | 0.46560801 | 5 | 0.31229468 | 181 | 0.9682353 | FALSE | 0.95955826 | FALSE | 1 | FALSE | 1 | FALSE | 1 | FALSE | 1 | FALSE | 1 |
| 5058 | 5058 | 7.33898305 | 1.21585806 | 2 | 0.4493251 | 3 | 0.2979413 | 2 | 0.63179508 | 1 | 0.93827647 | 0 | 1 | 51 | 0.41341988 | FALSE | 1 | FALSE | 1 | FALSE | 1 | FALSE | 1 | FALSE | 1 | FALSE | 1 |
| 5059 | 5059 | 9.46428571 | 2.54224511 | 2 | 0.16048326 | 1 | 0.60247911 | 0 | 1 | 0 | 1 | 1 | 0.37113067 | 24 | 0.55018107 | FALSE | 1 | FALSE | 1 | FALSE | 1 | FALSE | 1 | FALSE | 1 | FALSE | 1 |
| 5060 | 5060 | 8.71428571 | 1.77664363 | 0 | 1 | 0 | 1 | 0 | 1 | 2 | 0.25059627 | 0 | 1 | 19 | 0.34094959 | FALSE | 1 | FALSE | 1 | FALSE | 1 | FALSE | 1 | FALSE | 1 | FALSE | 1 |
| 5061 | 5061 | 8.27272727 | 3.44676157 | 2 | 0.31193899 | 2 | 0.41896557 | 2 | 0.47281647 | 2 | 0.60667406 | 0 | 1 | 36 | 0.75909657 | FALSE | 1 | FALSE | 1 | FALSE | 1 | FALSE | 1 | FALSE | 1 | FALSE | 1 |
| 5062 | 5062 | 11 | 2.29128785 | 1 | 0.18769745 | 0 | 1 | 0 | 1 | 1 | 0.31351192 | 0 | 1 | 6 | 0.88425551 | FALSE | 1 | FALSE | 1 | FALSE | 1 | FALSE | 1 | FALSE | 1 | FALSE | 1 |
| 5063 | 5063 | 5.7260274 | 3.19786005 | 1 | 0.85137169 | 2 | 0.68961994 | 2 | 0.74419436 | 7 | 0.0491948 | 3 | 0.11741889 | 58 | 0.9042862 | FALSE | 1 | FALSE | 1 | FALSE | 1 | FALSE | 1 | FALSE | 1 | FALSE | 1 |
| 5064 | 5064 | 8.25 | 2.56661255 | 0 | 1 | 2 | 0.1353446 | 0 | 1 | 0 | 1 | 1 | 0.28187291 | 17 | 0.61750632 | FALSE | 1 | FALSE | 1 | FALSE | 1 | FALSE | 1 | FALSE | 1 | FALSE | 1 |
| 5065 | 5065 | 7.36842105 | 2.15725279 | 1 | 0.38989951 | 0 | 1 | 1 | 0.50143826 | 0 | 1 | 1 | 0.26986775 | 16 | 0.65565778 | FALSE | 1 | FALSE | 1 | FALSE | 1 | FALSE | 1 | FALSE | 1 | FALSE | 1 |
| 5066 | 5066 | 7.3125 | 2.05300846 | 1 | 0.34033054 | 0 | 1 | 1 | 0.44344817 | 0 | 1 | 2 | 0.02752114 | 12 | 0.90922186 | FALSE | 1 | FALSE | 1 | FALSE | 1 | FALSE | 1 | FALSE | 0.79106376 | FALSE | 1 |
| 5067 | 5067 | 9.38461539 | 2.55819833 | 0 | 1 | 0 | 1 | 1 | 0.37874417 | 0 | 1 | 2 | 0.01846832 | 10 | 0.86791642 | FALSE | 1 | FALSE | 1 | FALSE | 1 | FALSE | 1 | FALSE | 0.69720022 | FALSE | 1 |
| 5068 | 5068 | 9.9 | 1.75783958 | 2 | 0.0256865 | 0 | 1 | 2 | 0.04781153 | 0 | 1 | 0 | 1 | 6 | 0.98831471 | FALSE | 0.4974559 | FALSE | 1 | FALSE | 1 | FALSE | 1 | FALSE | 1 | FALSE | 1 |
| 5069 | 5069 | 8.77777778 | 3.32591768 | 0 | 1 | 0 | 1 | 0 | 1 | 1 | 0.70601446 | 0 | 1 | 25 | 0.06985983 | FALSE | 1 | FALSE | 1 | FALSE | 1 | FALSE | 1 | FALSE | 1 | FALSE | 1 |
| 5070 | 5070 | 9.80952381 | 1.91781244 | 0 | 1 | 0 | 1 | 0 | 1 | 0 | 1 | 3 | 0.00460011 | 18 | 0.57962763 | FALSE | 1 | FALSE | 1 | FALSE | 1 | FALSE | 1 | FALSE | 0.40960939 | FALSE | 1 |
| 5071 | 5071 | 7.37142857 | 3.53016159 | 2 | 0.22589135 | 1 | 0.68454804 | 0 | 1 | 0 | 1 | 1 | 0.44015537 | 31 | 0.34231756 | FALSE | 1 | FALSE | 1 | FALSE | 1 | FALSE | 1 | FALSE | 1 | FALSE | 1 |
| 5072 | 5072 | 5.91111111 | 2.96514734 | 0 | 1 | 0 | 1 | 2 | 0.44869594 | 2 | 0.5811115 | 0 | 1 | 38 | 0.19268329 | FALSE | 1 | FALSE | 1 | FALSE | 1 | FALSE | 1 | FALSE | 1 | FALSE | 1 |
| 5073 | 5073 | 10.9090909 | 2.31416765 | 1 | 0.4357638 | 2 | 0.15794013 | 1 | 0.55340851 | 0 | 1 | 3 | 0.00526351 | 15 | 0.98570275 | FALSE | 1 | FALSE | 1 | FALSE | 1 | FALSE | 1 | FALSE | 0.43998669 | FALSE | 1 |
| 5074 | 5074 | 8.82758621 | 2.91333398 | 1 | 0.52988231 | 2 | 0.24095584 | 0 | 1 | 3 | 0.14602298 | 0 | 1 | 23 | 0.84462957 | FALSE | 1 | FALSE | 1 | FALSE | 1 | FALSE | 1 | FALSE | 1 | FALSE | 1 |

| | | | | | | | | | | | | | | | | | | | | | | | | | | | |
|---|---|---|---|---|---|---|---|---|---|---|---|---|---|---|---|---|---|---|---|---|---|---|---|---|---|---|---|
| 5075 | 5075 | 6.57647059 | 2.76693277 | 4 | 0.17339651 | 1 | 0.93995595 | 1 | 0.95626254 | 4 | 0.55256464 | 1 | 0.75686725 | 74 | 0.30519533 | FALSE | 1 | FALSE | 1 | FALSE | 1 | FALSE | 1 | FALSE | 1 | FALSE | 1 |
| 5076 | 5076 | 7.7826087 | 3.50101255 | 0 | 1 | 0 | 1 | 2 | 0.19931429 | 1 | 0.6613267 | 2 | 0.05389003 | 18 | 0.86273155 | FALSE | 1 | FALSE | 1 | FALSE | 1 | FALSE | 1 | FALSE | 1 | FALSE | 1 |
| 5077 | 5077 | 9.84848485 | 1.79428744 | 1 | 0.57647348 | 0 | 1 | 3 | 0.11365871 | 2 | 0.45159487 | 1 | 0.42123867 | 26 | 0.86852567 | FALSE | 1 | FALSE | 1 | FALSE | 1 | FALSE | 1 | FALSE | 1 | FALSE | 1 |
| 5078 | 5078 | 9.61904762 | 2.69878616 | 1 | 0.42086991 | 0 | 1 | 0 | 1 | 0 | 1 | 0 | 1 | 20 | 0.13763971 | FALSE | 1 | FALSE | 1 | FALSE | 1 | FALSE | 1 | FALSE | 1 | FALSE | 1 |
| 5079 | 5079 | 9.2 | 2.24499443 | 0 | 1 | 0 | 1 | 1 | 0.59998149 | 2 | 0.31968852 | 0 | 1 | 22 | 0.43642846 | FALSE | 1 | FALSE | 1 | FALSE | 1 | FALSE | 1 | FALSE | 1 | FALSE | 1 |
| 5080 | 5080 | 10.0909091 | 2.49075425 | 0 | 1 | 1 | 0.6629941 | 1 | 0.70185639 | 0 | 1 | 0 | 1 | 31 | 0.09218876 | FALSE | 1 | FALSE | 1 | FALSE | 1 | FALSE | 1 | FALSE | 1 | FALSE | 1 |
| 5081 | 5081 | 9.07142857 | 2.69826097 | 0 | 1 | 1 | 0.60247911 | 1 | 0.64171563 | 1 | 0.73248983 | 1 | 0.37113067 | 24 | 0.55018107 | FALSE | 1 | FALSE | 1 | FALSE | 1 | FALSE | 1 | FALSE | 1 | FALSE | 1 |
| 5082 | 5082 | 10.2857143 | 2.50984456 | 0 | 1 | 2 | 0.14655134 | 0 | 1 | 0 | 1 | 1 | 0.29368245 | 18 | 0.57962763 | FALSE | 1 | FALSE | 1 | FALSE | 1 | FALSE | 1 | FALSE | 1 | FALSE | 1 |
| 5083 | 5083 | 8.26829268 | 2.70522881 | 0 | 1 | 1 | 0.74131698 | 1 | 0.77786601 | 1 | 0.85524728 | 0 | 1 | 38 | 0.09794002 | FALSE | 1 | FALSE | 1 | FALSE | 1 | FALSE | 1 | FALSE | 1 | FALSE | 1 |
| 5084 | 5084 | 9.58823529 | 3.14472151 | 2 | 0.06909038 | 1 | 0.42857733 | 0 | 1 | 1 | 0.55062772 | 1 | 0.24525764 | 12 | 0.962093 | FALSE | 0.74276693 | FALSE | 1 | FALSE | 1 | FALSE | 1 | FALSE | 1 | FALSE | 1 |
| 5085 | 5085 | 10.05 | 2.03654119 | 1 | 0.40558521 | 3 | 0.02534497 | 0 | 1 | 0 | 1 | 2 | 0.04176963 | 14 | 0.97325593 | FALSE | 1 | FALSE | 0.82367276 | FALSE | 1 | FALSE | 1 | FALSE | 0.90763079 | FALSE | 1 |
| 5086 | 5086 | 7.60526316 | 2.63118418 | 0 | 1 | 1 | 0.71433277 | 1 | 0.75193685 | 2 | 0.52640931 | 0 | 1 | 34 | 0.27051027 | FALSE | 1 | FALSE | 1 | FALSE | 1 | FALSE | 1 | FALSE | 1 | FALSE | 1 |
| 5087 | 5087 | 9.5483871 | 2.56324781 | 0 | 1 | 2 | 0.2651851 | 1 | 0.67911179 | 0 | 1 | 0 | 1 | 28 | 0.26393996 | FALSE | 1 | FALSE | 1 | FALSE | 1 | FALSE | 1 | FALSE | 1 | FALSE | 1 |
| 5088 | 5088 | 6.9787234 | 2.32002716 | 0 | 1 | 3 | 0.19374024 | 0 | 1 | 1 | 0.89101956 | 2 | 0.17968152 | 41 | 0.38322554 | FALSE | 1 | FALSE | 1 | FALSE | 1 | FALSE | 1 | FALSE | 1 | FALSE | 1 |
| 5089 | 5089 | 7.02941177 | 1.8147321 | 3 | 0.25295743 | 7 | 0.00610654 | 1 | 0.91794622 | 3 | 0.61040454 | 5 | 0.00496558 | 49 | 0.99716293 | FALSE | 1 | FALSE | 0.60103504 | FALSE | 1 | FALSE | 1 | FALSE | 0.43274476 | FALSE | 1 |
| 5090 | 5090 | 7.56756757 | 1.96648322 | 0 | 1 | 0 | 1 | 2 | 0.38585089 | 1 | 0.82511639 | 0 | 1 | 34 | 0.14843263 | FALSE | 1 | FALSE | 1 | FALSE | 1 | FALSE | 1 | FALSE | 1 | FALSE | 1 |
| 5091 | 5091 | 8.18181818 | 1.77447466 | 1 | 0.4357638 | 0 | 1 | 1 | 0.55340851 | 1 | 0.64497439 | 1 | 0.30529951 | 18 | 0.74744813 | FALSE | 1 | FALSE | 1 | FALSE | 1 | FALSE | 1 | FALSE | 1 | FALSE | 1 |
| 5092 | 5092 | 9.5 | 2.35280987 | 0 | 1 | 1 | 0.36919824 | 0 | 1 | 0 | 1 | 0 | 1 | 13 | 0.33283729 | FALSE | 1 | FALSE | 1 | FALSE | 1 | FALSE | 1 | FALSE | 1 | FALSE | 1 |
| 5093 | 5093 | 6.42857143 | 2.19461307 | 4 | 0.02019941 | 0 | 1 | 2 | 0.43640583 | 3 | 0.29054213 | 3 | 0.02912063 | 29 | 0.99246133 | FALSE | 0.44482142 | FALSE | 1 | FALSE | 1 | FALSE | 1 | FALSE | 0.81978088 | FALSE | 1 |
| 5094 | 5094 | 7.23529412 | 1.64004107 | 1 | 0.7354105 | 3 | 0.22759346 | 2 | 0.55199489 | 1 | 0.90982214 | 1 | 0.57100288 | 43 | 0.59806866 | FALSE | 1 | FALSE | 1 | FALSE | 1 | FALSE | 1 | FALSE | 1 | FALSE | 1 |
| 5095 | 5095 | 6.75 | 2.73861279 | 1 | 0.56527493 | 0 | 1 | 2 | 0.32011871 | 0 | 1 | 0 | 1 | 29 | 0.24088541 | FALSE | 1 | FALSE | 1 | FALSE | 1 | FALSE | 1 | FALSE | 1 | FALSE | 1 |
| 5096 | 5096 | 8.5 | 3 | 5 | 0.0029693 | 0 | 1 | 0 | 1 | 0 | 1 | 0 | 1 | 34 | 0.41570781 | FALSE | 0.14653312 | FALSE | 1 | FALSE | 1 | FALSE | 1 | FALSE | 1 | FALSE | 1 |
| 5097 | 5097 | 7 | 2.57681975 | 2 | 0.13369621 | 0 | 1 | 1 | 0.59998149 | 1 | 0.69181315 | 0 | 1 | 21 | 0.64957167 | FALSE | 0.97964162 | FALSE | 1 | FALSE | 1 | FALSE | 1 | FALSE | 1 | FALSE | 1 |
| 5098 | 5098 | 8.36 | 2.97832168 | 1 | 0.47819878 | 2 | 0.19297191 | 1 | 0.59998149 | 2 | 0.31968852 | 0 | 1 | 19 | 0.91734261 | FALSE | 1 | FALSE | 1 | FALSE | 1 | FALSE | 1 | FALSE | 1 | FALSE | 1 |
| 5099 | 5099 | 9.07692308 | 2.7021797 | 0 | 1 | 0 | 1 | 0 | 1 | 0 | 1 | 0 | 1 | 13 | 0.10967906 | FALSE | 1 | FALSE | 1 | FALSE | 1 | FALSE | 1 | FALSE | 1 | FALSE | 1 |
| 5100 | 5100 | 11.0909091 | 3.30622587 | 1 | 0.55378205 | 0 | 1 | 2 | 0.30673593 | 0 | 1 | 0 | 1 | 28 | 0.26393996 | FALSE | 1 | FALSE | 1 | FALSE | 1 | FALSE | 1 | FALSE | 1 | FALSE | 1 |
| 5101 | 5101 | 9.2173913 | 1.9990546 | 2 | 0.11647716 | 0 | 1 | 2 | 0.19931429 | 1 | 0.6613267 | 0 | 1 | 18 | 0.86273155 | FALSE | 0.93181727 | FALSE | 1 | FALSE | 1 | FALSE | 1 | FALSE | 1 | FALSE | 1 |
| 5102 | 5102 | 9.125 | 1.96452921 | 0 | 1 | 0 | 1 | 0 | 1 | 0 | 1 | 0 | 1 | 8 | 0.25677166 | FALSE | 1 | FALSE | 1 | FALSE | 1 | FALSE | 1 | FALSE | 1 | FALSE | 1 |
| 5103 | 5103 | 8.57142857 | 2.38190457 | 2 | 0.16048326 | 3 | 0.06039219 | 1 | 0.64171563 | 0 | 1 | 0 | 1 | 22 | 0.86499992 | FALSE | 1 | FALSE | 1 | FALSE | 1 | FALSE | 1 | FALSE | 1 | FALSE | 1 |
| 5104 | 5104 | 7.76470588 | 3.08753494 | 0 | 1 | 0 | 1 | 1 | 0.71261921 | 3 | 0.20353499 | 1 | 0.43077488 | 29 | 0.55707911 | FALSE | 1 | FALSE | 1 | FALSE | 1 | FALSE | 1 | FALSE | 1 | FALSE | 1 |
| 5105 | 5105 | 5.22330097 | 2.71254561 | 3 | 0.49499023 | 1 | 0.96705809 | 2 | 0.89012718 | 4 | 0.70317893 | 1 | 0.82020918 | 92 | 0.10075991 | FALSE | 1 | FALSE | 1 | FALSE | 1 | FALSE | 1 | FALSE | 1 | FALSE | 1 |
| 5106 | 5106 | 10.8333333 | 1.62446572 | 1 | 0.26795838 | 1 | 0.32623933 | 1 | 0.35555293 | 0 | 1 | 1 | 0.18007762 | 8 | 0.97197833 | FALSE | 1 | FALSE | 1 | FALSE | 1 | FALSE | 1 | FALSE | 1 | FALSE | 1 |
| 5107 | 5107 | 10.4210526 | 2.66193024 | 1 | 0.38989951 | 0 | 1 | 0 | 1 | 3 | 0.05383577 | 0 | 1 | 15 | 0.83648191 | FALSE | 1 | FALSE | 1 | FALSE | 1 | FALSE | 1 | FALSE | 1 | FALSE | 1 |
| 5108 | 5108 | 11.0714286 | 2.12011744 | 0 | 1 | 0 | 1 | 0 | 1 | 0 | 1 | 0 | 1 | 14 | 0.09251304 | FALSE | 1 | FALSE | 1 | FALSE | 1 | FALSE | 1 | FALSE | 1 | FALSE | 1 |
| 5109 | 5109 | 7.17 | 2.03004926 | 1 | 0.92696133 | 2 | 0.83999047 | 4 | 0.48633182 | 5 | 0.48843976 | 1 | 0.81092309 | 87 | 0.28513463 | FALSE | 1 | FALSE | 1 | FALSE | 1 | FALSE | 1 | FALSE | 1 | FALSE | 1 |
| 5110 | 5110 | 10.2727273 | 2.83134755 | 2 | 0.03087289 | 0 | 1 | 0 | 1 | 2 | 0.08789566 | 0 | 1 | 7 | 0.98129353 | FALSE | 0.53356688 | FALSE | 1 | FALSE | 1 | FALSE | 1 | FALSE | 1 | FALSE | 1 |
| 5111 | 5111 | 8.8627451 | 2.55925547 | 2 | 0.36837609 | 1 | 0.80796869 | 2 | 0.54120634 | 3 | 0.4048204 | 0 | 1 | 42 | 0.62187327 | FALSE | 1 | FALSE | 1 | FALSE | 1 | FALSE | 1 | FALSE | 1 | FALSE | 1 |
| 5112 | 5112 | 9.77777778 | 2.93552107 | 1 | 0.37380237 | 1 | 0.44710673 | 0 | 1 | 1 | 0.57131013 | 1 | 0.25766374 | 14 | 0.86287418 | FALSE | 1 | FALSE | 1 | FALSE | 1 | FALSE | 1 | FALSE | 1 | FALSE | 1 |
| 5113 | 5113 | 8.23404255 | 2.69919425 | 0 | 1 | 0 | 1 | 2 | 0.50778118 | 0 | 1 | 0 | 1 | 45 | 0.01569277 | FALSE | 1 | FALSE | 1 | FALSE | 1 | FALSE | 1 | FALSE | 1 | FALSE | 1 |
| 5114 | 5114 | 8.80952381 | 2.21722795 | 0 | 1 | 1 | 0.49918218 | 0 | 1 | 1 | 0.62783522 | 0 | 1 | 19 | 0.34094959 | FALSE | 1 | FALSE | 1 | FALSE | 1 | FALSE | 1 | FALSE | 1 | FALSE | 1 |
| 5115 | 5115 | 8.40740741 | 1.14695278 | 4 | 0.04620161 | 3 | 0.24492539 | 2 | 0.57303317 | 0 | 1 | 5 | 0.00165958 | 39 | 0.98676224 | FALSE | 0.649301 | FALSE | 1 | FALSE | 1 | FALSE | 1 | FALSE | 0.23851396 | FALSE | 1 |
| 5116 | 5116 | 9.74074074 | 2.8099373 | 1 | 0.50471032 | 0 | 1 | 3 | 0.07106362 | 1 | 0.71956339 | 0 | 1 | 22 | 0.76237877 | FALSE | 1 | FALSE | 1 | FALSE | 1 | FALSE | 1 | FALSE | 1 | FALSE | 1 |
| 5117 | 5117 | 7.96153846 | 4.04285475 | 0 | 1 | 1 | 0.56109832 | 1 | 0.59998149 | 3 | 0.10474541 | 0 | 1 | 20 | 0.8152498 | FALSE | 1 | FALSE | 1 | FALSE | 1 | FALSE | 1 | FALSE | 1 | FALSE | 1 |
| 5118 | 5118 | 8.2 | 2.01494417 | 1 | 0.40558521 | 1 | 0.48239024 | 0 | 1 | 0 | 1 | 0 | 1 | 18 | 0.37504645 | FALSE | 1 | FALSE | 1 | FALSE | 1 | FALSE | 1 | FALSE | 1 | FALSE | 1 |
| 5119 | 5119 | 9.47368421 | 2.11183186 | 0 | 1 | 2 | 0.12434464 | 1 | 0.50143826 | 1 | 0.59104357 | 0 | 1 | 15 | 0.83648191 | FALSE | 1 | FALSE | 1 | FALSE | 1 | FALSE | 1 | FALSE | 1 | FALSE | 1 |
| 5120 | 5120 | 7.77777778 | 3.2788134 | 1 | 0.68224519 | 0 | 1 | 3 | 0.20921284 | 2 | 0.60667406 | 0 | 1 | 38 | 0.45705039 | FALSE | 1 | FALSE | 1 | FALSE | 1 | FALSE | 1 | FALSE | 1 | FALSE | 1 |
| 5121 | 5121 | 5.84615385 | 2.35925305 | 6 | 0.01457812 | 4 | 0.24495326 | 3 | 0.53558651 | 5 | 0.28743251 | 1 | 0.72664912 | 59 | 0.98534527 | FALSE | 0.37912366 | FALSE | 1 | FALSE | 1 | FALSE | 1 | FALSE | 1 | FALSE | 1 |
| 5122 | 5122 | 9.13636364 | 2.58132198 | 0 | 1 | 0 | 1 | 0 | 1 | 0 | 1 | 0 | 1 | 22 | 0.02368118 | FALSE | 1 | FALSE | 1 | FALSE | 1 | FALSE | 1 | FALSE | 1 | FALSE | 1 |
| 5123 | 5123 | 4.79130435 | 1.75034068 | 16 | 0.01270733 | 12 | 0.4121862 | 14 | 0.32923246 | 21 | 0.09564695 | 6 | 0.48133058 | 269 | 0.99362978 | FALSE | 0.35407635 | FALSE | 1 | FALSE | 1 | FALSE | 1 | FALSE | 1 | FALSE | 1 |
| 5124 | 5124 | 3.24819277 | 1.35809589 | 30 | 0.02889899 | 27 | 0.50554493 | 33 | 0.27833606 | 34 | 0.77905889 | 15 | 0.37543875 | 686 | 0.86145724 | FALSE | 0.52492348 | FALSE | 1 | FALSE | 1 | FALSE | 1 | FALSE | 1 | FALSE | 1 |
| 5125 | 5125 | 8.2962963 | 5.226817 | 1 | 0.50471032 | 1 | 0.58913618 | 1 | 0.62830986 | 0 | 1 | 1 | 0.36060486 | 23 | 0.58307573 | FALSE | 1 | FALSE | 1 | FALSE | 1 | FALSE | 1 | FALSE | 1 | FALSE | 1 |
| 5126 | 5126 | 10.5428571 | 3.03637808 | 1 | 0.59801736 | 1 | 0.68454804 | 0 | 1 | 1 | 0.80778317 | 0 | 1 | 32 | 0.18108028 | FALSE | 1 | FALSE | 1 | FALSE | 1 | FALSE | 1 | FALSE | 1 | FALSE | 1 |
| 5127 | 5127 | 9 | 1.87082869 | 0 | 1 | 0 | 1 | 0 | 1 | 1 | 0.67692816 | 1 | 0.32796862 | 22 | 0.25237063 | FALSE | 1 | FALSE | 1 | FALSE | 1 | FALSE | 1 | FALSE | 1 | FALSE | 1 |
| 5128 | 5128 | 9.86666667 | 2.60427 | 1 | 0.32293355 | 1 | 0.38964394 | 2 | 0.09928574 | 0 | 1 | 2 | 0.02433796 | 9 | 0.995488 | FALSE | 1 | FALSE | 1 | FALSE | 1 | FALSE | 1 | FALSE | 0.77578444 | FALSE | 1 |
| 5129 | 5129 | 9.89473684 | 1.39779539 | 2 | 0.43181298 | 4 | 0.11180811 | 2 | 0.61293419 | 1 | 0.93213716 | 0 | 1 | 48 | 0.60167081 | FALSE | 1 | FALSE | 1 | FALSE | 1 | FALSE | 1 | FALSE | 1 | FALSE | 1 |
| 5130 | 5130 | 10.85 | 2.28637267 | 0 | 1 | 0 | 1 | 1 | 0.51939841 | 0 | 1 | 0 | 1 | 19 | 0.15700255 | FALSE | 1 | FALSE | 1 | FALSE | 1 | FALSE | 1 | FALSE | 1 | FALSE | 1 |
| 5131 | 5131 | 11.1538462 | 2.03228382 | 1 | 0.28675951 | 0 | 1 | 1 | 0.37874417 | 0 | 1 | 0 | 1 | 11 | 0.6683056 | FALSE | 1 | FALSE | 1 | FALSE | 1 | FALSE | 1 | FALSE | 1 | FALSE | 1 |
| 5132 | 5132 | 10.2153846 | 2.18027738 | 4 | 0.08106144 | 5 | 0.05524713 | 3 | 0.40547364 | 3 | 0.56885732 | 2 | 0.28258242 | 47 | 0.99247238 | FALSE | 0.79236883 | FALSE | 1 | FALSE | 1 | FALSE | 1 | FALSE | 1 | FALSE | 1 |
| 5133 | 5133 | 10.9285714 | 2.91460082 | 2 | 0.04858661 | 0 | 1 | 2 | 0.08806398 | 1 | 0.48242099 | 0 | 1 | 9 | 0.98607548 | FALSE | 0.649301 | FALSE | 1 | FALSE | 1 | FALSE | 1 | FALSE | 1 | FALSE | 1 |
| 5134 | 5134 | 7.375 | 3.63790805 | 0 | 1 | 2 | 0.09285443 | 1 | 0.44344817 | 0 | 1 | 1 | 0.23264612 | 12 | 0.90922186 | FALSE | 1 | FALSE | 1 | FALSE | 1 | FALSE | 1 | FALSE | 1 | FALSE | 1 |
| 5135 | 5135 | 9.27272727 | 2.04898685 | 1 | 0.24866461 | 1 | 0.30368014 | 0 | 1 | 1 | 0.40390032 | 0 | 1 | 8 | 0.92152993 | FALSE | 1 | FALSE | 1 | FALSE | 1 | FALSE | 1 | FALSE | 1 | FALSE | 1 |
| 5136 | 5136 | 8.85714286 | 1.35526185 | 0 | 1 | 0 | 1 | 1 | 0.40110418 | 2 | 0.13304705 | 0 | 1 | 11 | 0.83676449 | FALSE | 1 | FALSE | 1 | FALSE | 1 | FALSE | 1 | FALSE | 1 | FALSE | 1 |
| 5137 | 5137 | 6.97560976 | 2.70038665 | 2 | 0.28329504 | 1 | 0.74131698 | 2 | 0.43640583 | 1 | 0.85524728 | 0 | 1 | 35 | 0.53616095 | FALSE | 1 | FALSE | 1 | FALSE | 1 | FALSE | 1 | FALSE | 1 | FALSE | 1 |
| 5138 | 5138 | 8.64705882 | 2.4719884 | 0 | 1 | 0 | 1 | 2 | 0.11087977 | 1 | 0.52895088 | 0 | 1 | 13 | 0.76799161 | FALSE | 1 | FALSE | 1 | FALSE | 1 | FALSE | 1 | FALSE | 1 | FALSE | 1 |
| 5139 | 5139 | 9.11111111 | 1.5234788 | 0 | 1 | 0 | 1 | 0 | 1 | 1 | 0.34506517 | 0 | 1 | 8 | 0.5780332 | FALSE | 1 | FALSE | 1 | FALSE | 1 | FALSE | 1 | FALSE | 1 | FALSE | 1 |
| 5140 | 5140 | 8.71428571 | 2.91372544 | 0 | 1 | 1 | 0.20567256 | 0 | 1 | 0 | 1 | 0 | 1 | 6 | 0.69920685 | FALSE | 1 | FALSE | 1 | FALSE | 1 | FALSE | 1 | FALSE | 1 | FALSE | 1 |
| 5141 | 5141 | 9.69230769 | 2.94525996 | 1 | 0.49162634 | 0 | 1 | 1 | 0.61440466 | 1 | 0.70601446 | 0 | 1 | 23 | 0.40372818 | FALSE | 1 | FALSE | 1 | FALSE | 1 | FALSE | 1 | FALSE | 1 | FALSE | 1 |
| 5142 | 5142 | 8.95238095 | 1.88922233 | 0 | 1 | 0 | 1 | 1 | 0.53671415 | 3 | 0.06912908 | 1 | 0.29368245 | 16 | 0.90353674 | FALSE | 1 | FALSE | 1 | FALSE | 1 | FALSE | 1 | FALSE | 1 | FALSE | 1 |
| 5143 | 5143 | 10.8181818 | 1.9610533 | 1 | 0.57647348 | 1 | 0.6629941 | 1 | 0.70185639 | 3 | 0.19161876 | 0 | 1 | 27 | 0.75124583 | FALSE | 1 | FALSE | 1 | FALSE | 1 | FALSE | 1 | FALSE | 1 | FALSE | 1 |
| 5144 | 5144 | 10.5416667 | 2.67673879 | 0 | 1 | 0 | 1 | 3 | 0.05314317 | 2 | 0.30249307 | 0 | 1 | 19 | 0.83975208 | FALSE | 1 | FALSE | 1 | FALSE | 1 | FALSE | 1 | FALSE | 1 | FALSE | 1 |
| 5145 | 5145 | 6.57758621 | 1.45715158 | 5 | 0.17229198 | 3 | 0.72508022 | 4 | 0.59743048 | 5 | 0.61387583 | 3 | 0.29179524 | 95 | 0.74961688 | FALSE | 1 | FALSE | 1 | FALSE | 1 | FALSE | 1 | FALSE | 1 | FALSE | 1 |
| 5146 | 5146 | 6.7037037 | 2.33832505 | 1 | 0.50471032 | 0 | 1 | 2 | 0.25286935 | 3 | 0.12475952 | 1 | 0.36060486 | 20 | 0.95133411 | FALSE | 1 | FALSE | 1 | FALSE | 1 | FALSE | 1 | FALSE | 1 | FALSE | 1 |
| 5147 | 5147 | 8.3 | 3.37786915 | 1 | 0.40558521 | 0 | 1 | 1 | 0.51939841 | 2 | 0.23331368 | 0 | 1 | 16 | 0.80826361 | FALSE | 1 | FALSE | 1 | FALSE | 1 | FALSE | 1 | FALSE | 1 | FALSE | 1 |
| 5148 | 5148 | 8.93548387 | 2.60192837 | 2 | 0.1881398 | 0 | 1 | 3 | 0.09854319 | 1 | 0.76781284 | 0 | 1 | 25 | 0.80006983 | FALSE | 1 | FALSE | 1 | FALSE | 1 | FALSE | 1 | FALSE | 1 | FALSE | 1 |
| 5149 | 5149 | 6.18604651 | 2.87940412 | 0 | 1 | 1 | 0.75788114 | 1 | 0.79363407 | 2 | 0.59404271 | 0 | 1 | 39 | 0.17628554 | FALSE | 1 | FALSE | 1 | FALSE | 1 | FALSE | 1 | FALSE | 1 | FALSE | 1 |
| 5150 | 5150 | 10.9375 | 4.72980906 | 0 | 1 | 1 | 0.65167362 | 2 | 0.32011871 | 1 | 0.77852168 | 0 | 1 | 28 | 0.42532177 | FALSE | 1 | FALSE | 1 | FALSE | 1 | FALSE | 1 | FALSE | 1 | FALSE | 1 |
| 5151 | 5151 | 13 | 0 | 0 | 1 | 0 | 1 | 0 | 1 | 0 | 1 | 0 | 1 | 1 | 0.84379192 | FALSE | 1 | FALSE | 1 | FALSE | 1 | FALSE | 1 | FALSE | 1 | FALSE | 1 |
| 5152 | 5152 | 9.5 | 2.53968502 | 0 | 1 | 1 | 0.28036922 | 0 | 1 | 0 | 1 | 0 | 1 | 9 | 0.5215547 | FALSE | 1 | FALSE | 1 | FALSE | 1 | FALSE | 1 | FALSE | 1 | FALSE | 1 |
| 5153 | 5153 | 6.71428571 | 2.50984456 | 1 | 0.42086991 | 1 | 0.49918218 | 0 | 1 | 1 | 0.62783522 | 1 | 0.29368245 | 17 | 0.77849076 | FALSE | 1 | FALSE | 1 | FALSE | 1 | FALSE | 1 | FALSE | 1 | FALSE | 1 |
| 5154 | 5154 | 9.5 | 2.71569512 | 0 | 1 | 2 | 0.09285443 | 4 | 0.00210122 | 0 | 1 | 0 | 1 | 10 | 0.99304605 | FALSE | 1 | FALSE | 1 | FALSE | 1 | FALSE | 1 | FALSE | 1 | FALSE | 1 |
| 5155 | 5155 | 9.75 | 2.48746859 | 1 | 0.40558521 | 0 | 1 | 2 | 0.16019548 | 3 | 0.0612581 | 0 | 1 | 14 | 0.97325593 | FALSE | 1 | FALSE | 1 | FALSE | 1 | FALSE | 1 | FALSE | 1 | FALSE | 1 |
| 5156 | 5156 | 3.96402878 | 1.90005707 | 4 | 0.93007473 | 8 | 0.68002302 | 7 | 0.87655311 | 12 | 0.62557417 | 8 | 0.08397112 | 238 | 0.265521 | FALSE | 1 | FALSE | 1 | FALSE | 1 | FALSE | 1 | FALSE | 1 | FALSE | 1 |
| 5157 | 5157 | 10.2307692 | 3.0167187 | 0 | 1 | 0 | 1 | 0 | 1 | 1 | 0.45746571 | 0 | 1 | 12 | 0.374198 | FALSE | 1 | FALSE | 1 | FALSE | 1 | FALSE | 1 | FALSE | 1 | FALSE | 1 |
| 5158 | 5158 | 8 | 2.28416096 | 0 | 1 | 1 | 0.53115654 | 1 | 0.5695037 | 1 | 0.6613267 | 0 | 1 | 20 | 0.50589402 | FALSE | 1 | FALSE | 1 | FALSE | 1 | FALSE | 1 | FALSE | 1 | FALSE | 1 |
| 5159 | 5159 | 7.31182796 | 2.88107841 | 1 | 0.91216464 | 2 | 0.80898201 | 2 | 0.85320929 | 1 | 0.98774195 | 1 | 0.7873664 | 86 | 0.01560391 | FALSE | 1 | FALSE | 1 | FALSE | 1 | FALSE | 1 | FALSE | 1 | FALSE | 1 |
| 5160 | 5160 | 8.75 | 1.57718103 | 0 | 1 | 0 | 1 | 0 | 1 | 0 | 1 | 0 | 1 | 20 | 0.03329856 | FALSE | 1 | FALSE | 1 | FALSE | 1 | FALSE | 1 | FALSE | 1 | FALSE | 1 |
| 5161 | 5161 | 7.75510204 | 4.01289052 | 0 | 1 | 1 | 0.80150109 | 2 | 0.53023996 | 3 | 0.39227316 | 0 | 1 | 43 | 0.337822 | FALSE | 1 | FALSE | 1 | FALSE | 1 | FALSE | 1 | FALSE | 1 | FALSE | 1 |
| 5162 | 5162 | 7.72727273 | 1.91125419 | 1 | 0.57647348 | 0 | 1 | 1 | 0.70185639 | 0 | 1 | 0 | 1 | 31 | 0.09218876 | FALSE | 1 | FALSE | 1 | FALSE | 1 | FALSE | 1 | FALSE | 1 | FALSE | 1 |
| 5163 | 5163 | 8.6 | 1.71464282 | 1 | 0.40558521 | 0 | 1 | 1 | 0.51939841 | 1 | 0.60987146 | 1 | 0.28187291 | 16 | 0.80826361 | FALSE | 1 | FALSE | 1 | FALSE | 1 | FALSE | 1 | FALSE | 1 | FALSE | 1 |
| 5164 | 5164 | 11.2142857 | 2.51221506 | 1 | 0.3050805 | 0 | 1 | 0 | 1 | 0 | 1 | 0 | 1 | 13 | 0.33283729 | FALSE | 1 | FALSE | 1 | FALSE | 1 | FALSE | 1 | FALSE | 1 | FALSE | 1 |
| 5165 | 5165 | 10.125 | 0.99215674 | 0 | 1 | 0 | 1 | 1 | 0.44344817 | 1 | 0.52895088 | 1 | 0.23264612 | 13 | 0.76799161 | FALSE | 1 | FALSE | 1 | FALSE | 1 | FALSE | 1 | FALSE | 1 | FALSE | 1 |
| 5166 | 5166 | 4.65986395 | 2.87020016 | 14 | 1.83E-05 | 6 | 0.32823878 | 5 | 0.60123352 | 8 | 0.34839944 | 7 | 0.00951156 | 105 | 0.99993517 | TRUE | 0.00545372 | FALSE | 1 | FALSE | 1 | FALSE | 1 | FALSE | 0.5373705 | FALSE | 1 |
| 5167 | 5167 | 10.125 | 2.08791164 | 0 | 1 | 0 | 1 | 0 | 1 | 0 | 1 | 0 | 1 | 8 | 0.25677166 | FALSE | 1 | FALSE | 1 | FALSE | 1 | FALSE | 1 | FALSE | 1 | FALSE | 1 |
| 5168 | 5168 | 9.77419355 | 2.01141384 | 0 | 1 | 0 | 1 | 0 | 1 | 0 | 1 | 0 | 1 | 31 | 0.00510159 | FALSE | 1 | FALSE | 1 | FALSE | 1 | FALSE | 1 | FALSE | 1 | FALSE | 1 |

| | | | | | | | | | | | | | | | | | | | | | | | | | | | |
|---|---|---|---|---|---|---|---|---|---|---|---|---|---|---|---|---|---|---|---|---|---|---|---|---|---|---|---|
| 5169 | 5169 | 9.6875 | 3.45902353 | 0 | 1 | 0 | 1 | 0 | 1 | 1 | 0.52895088 | 0 | 1 | 15 | 0.26127859 | FALSE | 1 | FALSE | 1 | FALSE | 1 | FALSE | 1 | FALSE | 1 | FALSE | 1 |
| 5170 | 5170 | 10.2727273 | 2.76638619 | 0 | 1 | 1 | 0.30368014 | 1 | 0.33149969 | 0 | 1 | 2 | 0.01330324 | 7 | 0.98129353 | FALSE | 1 | FALSE | 1 | FALSE | 1 | FALSE | 1 | FALSE | 0.63731057 | FALSE | 1 |
| 5171 | 5171 | 10.9444444 | 1.61493798 | 0 | 1 | 0 | 1 | 0 | 1 | 0 | 1 | 0 | 1 | 18 | 0.04681656 | FALSE | 1 | FALSE | 1 | FALSE | 1 | FALSE | 1 | FALSE | 1 | FALSE | 1 |
| 5172 | 5172 | 7.85 | 3.78516842 | 1 | 0.40558521 | 0 | 1 | 4 | 0.00499622 | 3 | 0.0612581 | 0 | 1 | 12 | 0.99824567 | FALSE | 1 | FALSE | 1 | FALSE | 1 | FALSE | 1 | FALSE | 1 | FALSE | 1 |
| 5173 | 5173 | 7.61688312 | 1.05188793 | 1 | 0.9753075 | 8 | 0.08603493 | 4 | 0.75210906 | 8 | 0.32053902 | 3 | 0.40860691 | 117 | 0.72477405 | FALSE | 1 | FALSE | 1 | FALSE | 1 | FALSE | 1 | FALSE | 1 | FALSE | 1 |
| 5174 | 5174 | 10.6153846 | 2.09535977 | 0 | 1 | 1 | 0.3480709 | 0 | 1 | 0 | 1 | 0 | 1 | 12 | 0.374198 | FALSE | 1 | FALSE | 1 | FALSE | 1 | FALSE | 1 | FALSE | 1 | FALSE | 1 |
| 5175 | 5175 | 9.29166667 | 2.26346281 | 1 | 0.46441868 | 0 | 1 | 2 | 0.21260257 | 1 | 0.67692816 | 0 | 1 | 20 | 0.68270729 | FALSE | 1 | FALSE | 1 | FALSE | 1 | FALSE | 1 | FALSE | 1 | FALSE | 1 |
| 5176 | 5176 | 9.56666667 | 1.87409237 | 1 | 0.52988231 | 2 | 0.24095584 | 0 | 1 | 3 | 0.14602298 | 0 | 1 | 23 | 0.84462957 | FALSE | 1 | FALSE | 1 | FALSE | 1 | FALSE | 1 | FALSE | 1 | FALSE | 1 |
| 5177 | 5177 | 10.75 | 2.58602011 | 0 | 1 | 0 | 1 | 1 | 0.35555293 | 2 | 0.1023796 | 0 | 1 | 9 | 0.89630573 | FALSE | 1 | FALSE | 1 | FALSE | 1 | FALSE | 1 | FALSE | 1 | FALSE | 1 |
| 5178 | 5178 | 8.24 | 2.42123935 | 0 | 1 | 2 | 0.19297191 | 1 | 0.59998149 | 1 | 0.69181315 | 1 | 0.33902672 | 20 | 0.8152498 | FALSE | 1 | FALSE | 1 | FALSE | 1 | FALSE | 1 | FALSE | 1 | FALSE | 1 |
| 5179 | 5179 | 7.40540541 | 3.16620209 | 3 | 0.0681763 | 1 | 0.70472939 | 1 | 0.74264075 | 1 | 0.82511639 | 1 | 0.45845922 | 30 | 0.78878581 | FALSE | 0.74276693 | FALSE | 1 | FALSE | 1 | FALSE | 1 | FALSE | 1 | FALSE | 1 |
| 5180 | 5180 | 10.1764706 | 2.0646975 | 0 | 1 | 0 | 1 | 2 | 0.12280513 | 2 | 0.18212134 | 1 | 0.24525764 | 12 | 0.962093 | FALSE | 1 | FALSE | 1 | FALSE | 1 | FALSE | 1 | FALSE | 1 | FALSE | 1 |
| 5181 | 5181 | 8.45 | 2.06094639 | 1 | 0.40558521 | 1 | 0.48239024 | 1 | 0.51939841 | 0 | 1 | 0 | 1 | 17 | 0.61750632 | FALSE | 1 | FALSE | 1 | FALSE | 1 | FALSE | 1 | FALSE | 1 | FALSE | 1 |
| 5182 | 5182 | 8.75 | 3.0516389 | 1 | 0.34033054 | 2 | 0.09285443 | 0 | 1 | 2 | 0.16544789 | 2 | 0.02752114 | 9 | 0.99862766 | FALSE | 1 | FALSE | 1 | FALSE | 1 | FALSE | 1 | FALSE | 0.79106376 | FALSE | 1 |
| 5183 | 5183 | 6.24324324 | 1.24781767 | 2 | 0.95402026 | 5 | 0.72086445 | 3 | 0.96550857 | 16 | 0.01068666 | 4 | 0.36031081 | 155 | 0.63629147 | FALSE | 1 | FALSE | 1 | FALSE | 1 | FALSE | 1 | FALSE | 1 | FALSE | 1 |
| 5184 | 5184 | 3.06035944 | 0.68388812 | 78 | 0.36931795 | 94 | 0.58665682 | 101 | 0.74991122 | 141 | 0.25744633 | 55 | 0.11151392 | 2471 | 0.75649011 | FALSE | 1 | FALSE | 1 | FALSE | 1 | FALSE | 1 | FALSE | 1 | FALSE | 1 |
| 5185 | 5185 | 5.88461539 | 3.12959721 | 1 | 0.7354105 | 1 | 0.81422651 | 0 | 1 | 4 | 0.20527806 | 1 | 0.57100288 | 44 | 0.44552698 | FALSE | 1 | FALSE | 1 | FALSE | 1 | FALSE | 1 | FALSE | 1 | FALSE | 1 |
| 5186 | 5186 | 10.5714286 | 1.04978132 | 0 | 1 | 0 | 1 | 1 | 0.40110418 | 0 | 1 | 0 | 1 | 13 | 0.33283729 | FALSE | 1 | FALSE | 1 | FALSE | 1 | FALSE | 1 | FALSE | 1 | FALSE | 1 |
| 5187 | 5187 | 8.78571429 | 4.30412711 | 1 | 0.51745948 | 1 | 0.60247911 | 1 | 0.64171563 | 1 | 0.73248983 | 1 | 0.37113067 | 23 | 0.73439118 | FALSE | 1 | FALSE | 1 | FALSE | 1 | FALSE | 1 | FALSE | 1 | FALSE | 1 |
| 5188 | 5188 | 9.24137931 | 0.81601101 | 0 | 1 | 0 | 1 | 2 | 0.27983822 | 1 | 0.74482229 | 0 | 1 | 26 | 0.31495157 | FALSE | 1 | FALSE | 1 | FALSE | 1 | FALSE | 1 | FALSE | 1 | FALSE | 1 |
| 5189 | 5189 | 6.53191489 | 4.07300957 | 1 | 0.69843502 | 1 | 0.78076835 | 3 | 0.2282043 | 1 | 0.88573774 | 0 | 1 | 40 | 0.40712403 | FALSE | 1 | FALSE | 1 | FALSE | 1 | FALSE | 1 | FALSE | 1 | FALSE | 1 |
| 5190 | 5190 | 6.63636364 | 2.90055566 | 1 | 0.24866461 | 1 | 0.30368014 | 0 | 1 | 0 | 1 | 0 | 1 | 9 | 0.75985366 | FALSE | 1 | FALSE | 1 | FALSE | 1 | FALSE | 1 | FALSE | 1 | FALSE | 1 |
| 5191 | 5191 | 9.72727273 | 3.10691045 | 0 | 1 | 2 | 0.04725998 | 1 | 0.33149969 | 1 | 0.40390032 | 1 | 0.16638736 | 6 | 0.99674012 | FALSE | 1 | FALSE | 1 | FALSE | 1 | FALSE | 1 | FALSE | 1 | FALSE | 1 |
| 5192 | 5192 | 8.64705882 | 1.49277383 | 0 | 1 | 1 | 0.42857733 | 0 | 1 | 0 | 1 | 0 | 1 | 16 | 0.23070182 | FALSE | 1 | FALSE | 1 | FALSE | 1 | FALSE | 1 | FALSE | 1 | FALSE | 1 |
| 5193 | 5193 | 9.58333333 | 1.89113429 | 0 | 1 | 0 | 1 | 0 | 1 | 0 | 1 | 0 | 1 | 24 | 0.01683964 | FALSE | 1 | FALSE | 1 | FALSE | 1 | FALSE | 1 | FALSE | 1 | FALSE | 1 |
| 5194 | 5194 | 10.3703704 | 2.21356338 | 1 | 0.49162634 | 0 | 1 | 0 | 1 | 0 | 1 | 0 | 1 | 25 | 0.06985983 | FALSE | 1 | FALSE | 1 | FALSE | 1 | FALSE | 1 | FALSE | 1 | FALSE | 1 |
| 5195 | 5195 | 4.77966102 | 2.37025682 | 5 | 0.4776812 | 6 | 0.51277283 | 7 | 0.4527581 | 7 | 0.71101013 | 5 | 0.16463153 | 147 | 0.72997654 | FALSE | 1 | FALSE | 1 | FALSE | 1 | FALSE | 1 | FALSE | 1 | FALSE | 1 |
| 5196 | 5196 | 10.25 | 2.94745653 | 1 | 0.34033054 | 2 | 0.09285443 | 3 | 0.01812751 | 0 | 1 | 1 | 0.23264612 | 9 | 0.99862766 | FALSE | 1 | FALSE | 1 | FALSE | 1 | FALSE | 1 | FALSE | 1 | FALSE | 1 |
| 5197 | 5197 | 4.03846154 | 2.45006004 | 8 | 0.25152293 | 10 | 0.22566736 | 6 | 0.852664 | 13 | 0.27775081 | 0 | 1 | 197 | 0.57633568 | FALSE | 1 | FALSE | 1 | FALSE | 1 | FALSE | 1 | FALSE | 1 | FALSE | 1 |
| 5198 | 5198 | 9.57142857 | 2.04872617 | 2 | 0.29286138 | 0 | 1 | 1 | 0.78589454 | 2 | 0.5811115 | 0 | 1 | 37 | 0.34041811 | FALSE | 1 | FALSE | 1 | FALSE | 1 | FALSE | 1 | FALSE | 1 | FALSE | 1 |
| 5199 | 5199 | 9.53333333 | 1.45449495 | 0 | 1 | 0 | 1 | 0 | 1 | 1 | 0.50623194 | 1 | 0.2198258 | 13 | 0.57712805 | FALSE | 1 | FALSE | 1 | FALSE | 1 | FALSE | 1 | FALSE | 1 | FALSE | 1 |
| 5200 | 5200 | 9.35483871 | 2.02481998 | 0 | 1 | 1 | 0.63997468 | 0 | 1 | 1 | 0.76781284 | 2 | 0.09110253 | 27 | 0.45522639 | FALSE | 1 | FALSE | 1 | FALSE | 1 | FALSE | 1 | FALSE | 1 | FALSE | 1 |
| 5201 | 5201 | 10.3157895 | 3.07883491 | 1 | 0.38989951 | 0 | 1 | 0 | 1 | 0 | 1 | 0 | 1 | 18 | 0.17880694 | FALSE | 1 | FALSE | 1 | FALSE | 1 | FALSE | 1 | FALSE | 1 | FALSE | 1 |
| 5202 | 5202 | 9.70588235 | 2.05124656 | 3 | 0.00864151 | 0 | 1 | 1 | 0.46348836 | 0 | 1 | 0 | 1 | 13 | 0.88719222 | FALSE | 0.27019545 | FALSE | 1 | FALSE | 1 | FALSE | 1 | FALSE | 1 | FALSE | 1 |
| 5203 | 5203 | 8.38709677 | 2.52314984 | 2 | 0.1881398 | 1 | 0.63997468 | 0 | 1 | 0 | 1 | 0 | 1 | 28 | 0.26393996 | FALSE | 1 | FALSE | 1 | FALSE | 1 | FALSE | 1 | FALSE | 1 | FALSE | 1 |
| 5204 | 5204 | 9.42857143 | 1.84058553 | 1 | 0.3050805 | 0 | 1 | 0 | 1 | 0 | 1 | 1 | 0.20679328 | 12 | 0.62236227 | FALSE | 1 | FALSE | 1 | FALSE | 1 | FALSE | 1 | FALSE | 1 | FALSE | 1 |
| 5205 | 5205 | 10.5 | 2.82842713 | 0 | 1 | 2 | 0.02563923 | 0 | 1 | 1 | 0.31351192 | 0 | 1 | 5 | 0.97551477 | FALSE | 1 | FALSE | 0.82367276 | FALSE | 1 | FALSE | 1 | FALSE | 1 | FALSE | 1 |
| 5206 | 5206 | 8.54054054 | 3.32553263 | 0 | 1 | 1 | 0.70472939 | 1 | 0.74264075 | 2 | 0.51200191 | 1 | 0.45845922 | 32 | 0.47030992 | FALSE | 1 | FALSE | 1 | FALSE | 1 | FALSE | 1 | FALSE | 1 | FALSE | 1 |
| 5207 | 5207 | 9.08571429 | 2.49995918 | 1 | 0.59801736 | 1 | 0.68454804 | 1 | 0.72299506 | 1 | 0.80778317 | 1 | 0.44015537 | 30 | 0.52765882 | FALSE | 1 | FALSE | 1 | FALSE | 1 | FALSE | 1 | FALSE | 1 | FALSE | 1 |
| 5208 | 5208 | 9.69230769 | 1.8138194 | 0 | 1 | 0 | 1 | 0 | 1 | 0 | 1 | 0 | 1 | 12 | 0.13002667 | FALSE | 1 | FALSE | 1 | FALSE | 1 | FALSE | 1 | FALSE | 1 | FALSE | 1 |
| 5209 | 5209 | 12.05 | 2.69211813 | 1 | 0.40558521 | 0 | 1 | 0 | 1 | 1 | 0.60987146 | 0 | 1 | 18 | 0.37504645 | FALSE | 1 | FALSE | 1 | FALSE | 1 | FALSE | 1 | FALSE | 1 | FALSE | 1 |
| 5210 | 5210 | 9.75 | 1.83711731 | 1 | 0.56527493 | 2 | 0.27730767 | 1 | 0.69069222 | 3 | 0.17988885 | 1 | 0.41154418 | 24 | 0.94876798 | FALSE | 1 | FALSE | 1 | FALSE | 1 | FALSE | 1 | FALSE | 1 | FALSE | 1 |
| 5211 | 5211 | 10.0526316 | 3.15350572 | 0 | 1 | 1 | 0.46503795 | 1 | 0.50143826 | 0 | 1 | 0 | 1 | 17 | 0.41141543 | FALSE | 1 | FALSE | 1 | FALSE | 1 | FALSE | 1 | FALSE | 1 | FALSE | 1 |
| 5212 | 5212 | 10 | 1.79605302 | 2 | 0.1881398 | 2 | 0.2651851 | 0 | 1 | 0 | 1 | 0 | 1 | 27 | 0.45522639 | FALSE | 1 | FALSE | 1 | FALSE | 1 | FALSE | 1 | FALSE | 1 | FALSE | 1 |
| 5213 | 5213 | 10.2 | 1.79629248 | 0 | 1 | 0 | 1 | 1 | 0.42266264 | 1 | 0.50623194 | 0 | 1 | 13 | 0.57712805 | FALSE | 1 | FALSE | 1 | FALSE | 1 | FALSE | 1 | FALSE | 1 | FALSE | 1 |
| 5214 | 5214 | 9.8 | 2.52190404 | 0 | 1 | 0 | 1 | 0 | 1 | 0 | 1 | 0 | 1 | 10 | 0.18273166 | FALSE | 1 | FALSE | 1 | FALSE | 1 | FALSE | 1 | FALSE | 1 | FALSE | 1 |
| 5215 | 5215 | 12.2083333 | 2.70769223 | 1 | 0.46441868 | 0 | 1 | 1 | 0.58502114 | 1 | 0.67692816 | 0 | 1 | 21 | 0.47053137 | FALSE | 1 | FALSE | 1 | FALSE | 1 | FALSE | 1 | FALSE | 1 | FALSE | 1 |
| 5216 | 5216 | 8.82142857 | 3.91018424 | 1 | 0.51745948 | 4 | 0.01184672 | 0 | 1 | 0 | 1 | 0 | 1 | 23 | 0.73439118 | FALSE | 1 | FALSE | 0.67617265 | FALSE | 1 | FALSE | 1 | FALSE | 1 | FALSE | 1 |
| 5217 | 5217 | 10.3333333 | 2.2852182 | 0 | 1 | 0 | 1 | 0 | 1 | 1 | 0.24578796 | 0 | 1 | 5 | 0.76185568 | FALSE | 1 | FALSE | 1 | FALSE | 1 | FALSE | 1 | FALSE | 1 | FALSE | 1 |
| 5218 | 5218 | 8.69230769 | 1.95663036 | 0 | 1 | 0 | 1 | 2 | 0.23940226 | 2 | 0.33678042 | 1 | 0.3499045 | 21 | 0.78939668 | FALSE | 1 | FALSE | 1 | FALSE | 1 | FALSE | 1 | FALSE | 1 | FALSE | 1 |
| 5219 | 5219 | 9 | 2.7325202 | 1 | 0.32293355 | 0 | 1 | 0 | 1 | 1 | 0.50623194 | 0 | 1 | 13 | 0.57712805 | FALSE | 1 | FALSE | 1 | FALSE | 1 | FALSE | 1 | FALSE | 1 | FALSE | 1 |
| 5220 | 5220 | 10.3333333 | 1.49071199 | 0 | 1 | 1 | 0.32623933 | 1 | 0.35555293 | 0 | 1 | 0 | 1 | 10 | 0.71436964 | FALSE | 1 | FALSE | 1 | FALSE | 1 | FALSE | 1 | FALSE | 1 | FALSE | 1 |
| 5221 | 5221 | 8.59259259 | 2.23207692 | 2 | 0.15144274 | 3 | 0.05519089 | 1 | 0.62830986 | 2 | 0.35374209 | 0 | 1 | 19 | 0.98241223 | FALSE | 1 | FALSE | 1 | FALSE | 1 | FALSE | 1 | FALSE | 1 | FALSE | 1 |
| 5222 | 5222 | 7.33333333 | 3.78226857 | 2 | 0.12501436 | 1 | 0.54637326 | 1 | 0.58502114 | 1 | 0.67692816 | 0 | 1 | 19 | 0.83975208 | FALSE | 0.95622564 | FALSE | 1 | FALSE | 1 | FALSE | 1 | FALSE | 1 | FALSE | 1 |
| 5223 | 5223 | 9.04166667 | 1.48546196 | 0 | 1 | 1 | 0.54637326 | 0 | 1 | 0 | 1 | 2 | 0.058175 | 21 | 0.47053137 | FALSE | 1 | FALSE | 1 | FALSE | 1 | FALSE | 1 | FALSE | 1 | FALSE | 1 |
| 5224 | 5224 | 10.5555556 | 3.0951974 | 0 | 1 | 0 | 1 | 1 | 0.48280983 | 1 | 0.57131013 | 0 | 1 | 16 | 0.44998001 | FALSE | 1 | FALSE | 1 | FALSE | 1 | FALSE | 1 | FALSE | 1 | FALSE | 1 |
| 5225 | 5225 | 9.70588235 | 2.52667431 | 0 | 1 | 1 | 0.6739483 | 2 | 0.34668321 | 0 | 1 | 3 | 0.01774672 | 28 | 0.72558119 | FALSE | 1 | FALSE | 1 | FALSE | 1 | FALSE | 1 | FALSE | 0.69720022 | FALSE | 1 |
| 5226 | 5226 | 7.4 | 2.52454616 | 0 | 1 | 0 | 1 | 2 | 0.09928574 | 1 | 0.50623194 | 0 | 1 | 12 | 0.80329917 | FALSE | 1 | FALSE | 1 | FALSE | 1 | FALSE | 1 | FALSE | 1 | FALSE | 1 |
| 5227 | 5227 | 8.33333333 | 2.56038192 | 0 | 1 | 0 | 1 | 0 | 1 | 0 | 1 | 0 | 1 | 6 | 0.36077158 | FALSE | 1 | FALSE | 1 | FALSE | 1 | FALSE | 1 | FALSE | 1 | FALSE | 1 |
| 5228 | 5228 | 9.61111111 | 2.03138948 | 0 | 1 | 0 | 1 | 0 | 1 | 1 | 0.57131013 | 0 | 1 | 17 | 0.20329205 | FALSE | 1 | FALSE | 1 | FALSE | 1 | FALSE | 1 | FALSE | 1 | FALSE | 1 |
| 5229 | 5229 | 8.06451613 | 2.07808044 | 0 | 1 | 2 | 0.2651851 | 0 | 1 | 1 | 0.76781284 | 0 | 1 | 28 | 0.26393996 | FALSE | 1 | FALSE | 1 | FALSE | 1 | FALSE | 1 | FALSE | 1 | FALSE | 1 |
| 5230 | 5230 | 7.34615385 | 2.26923077 | 2 | 0.38681045 | 2 | 0.50543179 | 2 | 0.5626042 | 0 | 1 | 3 | 0.05310973 | 43 | 0.71191105 | FALSE | 1 | FALSE | 1 | FALSE | 1 | FALSE | 1 | FALSE | 1 | FALSE | 1 |
| 5231 | 5231 | 10.8275862 | 2.43634783 | 3 | 0.03713651 | 2 | 0.24095584 | 3 | 0.08431992 | 0 | 1 | 0 | 1 | 21 | 0.97185642 | FALSE | 0.58392 | FALSE | 1 | FALSE | 1 | FALSE | 1 | FALSE | 1 | FALSE | 1 |
| 5232 | 5232 | 8.2631579 | 1.94238338 | 1 | 0.38989951 | 1 | 0.46503795 | 0 | 1 | 0 | 1 | 0 | 1 | 17 | 0.41141543 | FALSE | 1 | FALSE | 1 | FALSE | 1 | FALSE | 1 | FALSE | 1 | FALSE | 1 |
| 5233 | 5233 | 10.3846154 | 1.72263044 | 1 | 0.74224579 | 1 | 0.82028131 | 2 | 0.5626042 | 2 | 0.6970685 | 0 | 1 | 46 | 0.27620659 | FALSE | 1 | FALSE | 1 | FALSE | 1 | FALSE | 1 | FALSE | 1 | FALSE | 1 |
| 5234 | 5234 | 10 | 2.10042013 | 1 | 0.58738519 | 2 | 0.30150529 | 1 | 0.71261921 | 2 | 0.46709861 | 0 | 1 | 28 | 0.72558119 | FALSE | 1 | FALSE | 1 | FALSE | 1 | FALSE | 1 | FALSE | 1 | FALSE | 1 |
| 5235 | 5235 | 7.73529412 | 1.1709429 | 0 | 1 | 0 | 1 | 0 | 1 | 3 | 0.19161876 | 2 | 0.10132035 | 28 | 0.58681477 | FALSE | 1 | FALSE | 1 | FALSE | 1 | FALSE | 1 | FALSE | 1 | FALSE | 1 |
| 5236 | 5236 | 8.4 | 2.2090722 | 1 | 0.47819878 | 3 | 0.04547096 | 1 | 0.59998149 | 0 | 1 | 0 | 1 | 20 | 0.8152498 | FALSE | 1 | FALSE | 1 | FALSE | 1 | FALSE | 1 | FALSE | 1 | FALSE | 1 |
| 5237 | 5237 | 9.79310345 | 3.13357029 | 1 | 0.50471032 | 1 | 0.58913618 | 0 | 1 | 1 | 0.71956339 | 0 | 1 | 24 | 0.3725373 | FALSE | 1 | FALSE | 1 | FALSE | 1 | FALSE | 1 | FALSE | 1 | FALSE | 1 |
| 5238 | 5238 | 7 | 3.33541602 | 0 | 1 | 1 | 0.40942991 | 1 | 0.44344817 | 2 | 0.16544789 | 0 | 1 | 12 | 0.90922186 | FALSE | 1 | FALSE | 1 | FALSE | 1 | FALSE | 1 | FALSE | 1 | FALSE | 1 |
| 5239 | 5239 | 9.25 | 2.2220486 | 0 | 1 | 1 | 0.40942991 | 1 | 0.44344817 | 0 | 1 | 0 | 1 | 14 | 0.53308261 | FALSE | 1 | FALSE | 1 | FALSE | 1 | FALSE | 1 | FALSE | 1 | FALSE | 1 |
| 5240 | 5240 | 9.31034483 | 2.56102685 | 1 | 0.52988231 | 0 | 1 | 0 | 1 | 0 | 1 | 1 | 0.38148475 | 27 | 0.14664438 | FALSE | 1 | FALSE | 1 | FALSE | 1 | FALSE | 1 | FALSE | 1 | FALSE | 1 |
| 5241 | 5241 | 2.13953488 | 1.10441508 | 29 | 0.00037826 | 28 | 0.02218256 | 28 | 0.06991363 | 28 | 0.43705472 | 21 | 0.00044743 | 451 | 0.99999953 | TRUE | 0.04024298 | FALSE | 0.81043583 | FALSE | 1 | FALSE | 1 | FALSE | 0.10840111 | FALSE | 1 |
| 5242 | 5242 | 9.97560976 | 2.52294645 | 2 | 0.28329504 | 3 | 0.14580275 | 3 | 0.1814659 | 0 | 1 | 0 | 1 | 33 | 0.81996213 | FALSE | 1 | FALSE | 1 | FALSE | 1 | FALSE | 1 | FALSE | 1 | FALSE | 1 |
| 5243 | 5243 | 6.48648649 | 2.68752495 | 4 | 0.3174432 | 2 | 0.87962337 | 4 | 0.56918099 | 3 | 0.89089248 | 5 | 0.0352132 | 93 | 0.63017929 | FALSE | 1 | FALSE | 1 | FALSE | 1 | FALSE | 1 | FALSE | 0.87669812 | FALSE | 1 |
| 5244 | 5244 | 10.0588235 | 2.62276433 | 0 | 1 | 0 | 1 | 0 | 1 | 0 | 1 | 1 | 0.24525764 | 16 | 0.23070182 | FALSE | 1 | FALSE | 1 | FALSE | 1 | FALSE | 1 | FALSE | 1 | FALSE | 1 |
| 5245 | 5245 | 6.88372093 | 3.05156691 | 1 | 0.67382914 | 0 | 1 | 3 | 0.19985581 | 2 | 0.59404271 | 1 | 0.50988734 | 36 | 0.6451239 | FALSE | 1 | FALSE | 1 | FALSE | 1 | FALSE | 1 | FALSE | 1 | FALSE | 1 |
| 5246 | 5246 | 9.03846154 | 2.91522224 | 1 | 0.49162634 | 2 | 0.20486827 | 0 | 1 | 1 | 0.70601446 | 1 | 0.3499045 | 21 | 0.78939668 | FALSE | 1 | FALSE | 1 | FALSE | 1 | FALSE | 1 | FALSE | 1 | FALSE | 1 |
| 5247 | 5247 | 9.2 | 2.71293199 | 0 | 1 | 1 | 0.15163655 | 0 | 1 | 0 | 1 | 0 | 1 | 4 | 0.82371636 | FALSE | 1 | FALSE | 1 | FALSE | 1 | FALSE | 1 | FALSE | 1 | FALSE | 1 |
| 5248 | 5248 | 3.88888889 | 2.33068633 | 8 | 0.03722373 | 8 | 0.10932155 | 4 | 0.78936044 | 3 | 0.97061659 | 2 | 0.70674828 | 124 | 0.70058945 | FALSE | 0.58417011 | FALSE | 1 | FALSE | 1 | FALSE | 1 | FALSE | 1 | FALSE | 1 |
| 5249 | 5249 | 6.20238095 | 3.69613621 | 3 | 0.35850108 | 5 | 0.13034181 | 3 | 0.57805941 | 9 | 0.01335152 | 0 | 1 | 63 | 0.98536187 | FALSE | 1 | FALSE | 1 | FALSE | 1 | FALSE | 1 | FALSE | 1 | FALSE | 1 |
| 5250 | 5250 | 2.37837838 | 1.5417847 | 22 | 5.71E-06 | 10 | 0.49060214 | 12 | 0.37505079 | 16 | 0.28419521 | 5 | 0.53809191 | 231 | 0.99876122 | TRUE | 0.00299395 | FALSE | 1 | FALSE | 1 | FALSE | 1 | FALSE | 1 | FALSE | 1 |
| 5251 | 5251 | 8.73076923 | 0.81134704 | 0 | 1 | 0 | 1 | 3 | 0.06482014 | 1 | 0.70601446 | 1 | 0.3499045 | 21 | 0.78939668 | FALSE | 1 | FALSE | 1 | FALSE | 1 | FALSE | 1 | FALSE | 1 | FALSE | 1 |
| 5252 | 5252 | 7.83333333 | 3.3374974 | 1 | 0.45027683 | 1 | 0.53115654 | 0 | 1 | 3 | 0.08614834 | 0 | 1 | 18 | 0.86273155 | FALSE | 1 | FALSE | 1 | FALSE | 1 | FALSE | 1 | FALSE | 1 | FALSE | 1 |
| 5253 | 5253 | 8.04285714 | 2.13431631 | 4 | 0.10428558 | 4 | 0.19010643 | 3 | 0.46296673 | 1 | 0.9633793 | 0 | 1 | 58 | 0.70710693 | FALSE | 0.88702868 | FALSE | 1 | FALSE | 1 | FALSE | 1 | FALSE | 1 | FALSE | 1 |
| 5254 | 5254 | 8.70588235 | 1.7353772 | 2 | 0.73687735 | 6 | 0.10819862 | 1 | 0.97584382 | 5 | 0.49729031 | 2 | 0.49622404 | 85 | 0.59058697 | FALSE | 1 | FALSE | 1 | FALSE | 1 | FALSE | 1 | FALSE | 1 | FALSE | 1 |
| 5255 | 5255 | 9.375 | 1.57619003 | 0 | 1 | 1 | 0.40942991 | 2 | 0.11087977 | 0 | 1 | 2 | 0.02752114 | 11 | 0.9718531 | FALSE | 1 | FALSE | 1 | FALSE | 1 | FALSE | 1 | FALSE | 0.79106376 | FALSE | 1 |
| 5256 | 5256 | 8.82608696 | 4.10357399 | 0 | 1 | 2 | 0.16948764 | 0 | 1 | 1 | 0.6613267 | 0 | 1 | 20 | 0.50589402 | FALSE | 1 | FALSE | 1 | FALSE | 1 | FALSE | 1 | FALSE | 1 | FALSE | 1 |
| 5257 | 5257 | 8.46428571 | 3.20136291 | 1 | 0.51745948 | 0 | 1 | 1 | 0.64171563 | 2 | 0.3705495 | 1 | 0.37113067 | 23 | 0.73439118 | FALSE | 1 | FALSE | 1 | FALSE | 1 | FALSE | 1 | FALSE | 1 | FALSE | 1 |
| 5258 | 5258 | 3.2287695 | 1.1520477 | 35 | 9.77E-07 | 25 | 0.07944897 | 17 | 0.83882881 | 31 | 0.1996645 | 5 | 0.96582273 | 464 | 0.99686655 | TRUE | 0.00077787 | FALSE | 1 | FALSE | 1 | FALSE | 1 | FALSE | 1 | FALSE | 1 |
| 5259 | 5259 | 10.4705882 | 3.83662914 | 1 | 0.35728305 | 0 | 1 | 0 | 1 | 1 | 0.55062772 | 1 | 0.24525764 | 14 | 0.7313145 | FALSE | 1 | FALSE | 1 | FALSE | 1 | FALSE | 1 | FALSE | 1 | FALSE | 1 |
| 5260 | 5260 | 3.21621622 | 1.17024218 | 5 | 0.51693735 | 8 | 0.24940476 | 6 | 0.65920475 | 7 | 0.75303841 | 3 | 0.5892865 | 156 | 0.5578066 | FALSE | 1 | FALSE | 1 | FALSE | 1 | FALSE | 1 | FALSE | 1 | FALSE | 1 |
| 5261 | 5261 | 7.73529412 | 2.36778407 | 0 | 1 | 3 | 0.09602744 | 2 | 0.34668321 | 3 | 0.20353499 | 0 | 1 | 26 | 0.92796052 | FALSE | 1 | FALSE | 1 | FALSE | 1 | FALSE | 1 | FALSE | 1 | FALSE | 1 |
| 5262 | 5262 | 10.1428571 | 2.23149991 | 0 | 1 | 1 | 0.36919824 | 0 | 1 | 1 | 0.48242099 | 1 | 0.20679328 | 11 | 0.83676449 | FALSE | 1 | FALSE | 1 | FALSE | 1 | FALSE | 1 | FALSE | 1 | FALSE | 1 |

| | | | | | | | | | | | | | | | | | | | | | | | | | | | |
|---|---|---|---|---|---|---|---|---|---|---|---|---|---|---|---|---|---|---|---|---|---|---|---|---|---|---|---|
| 5263 | 5263 | 10.2222222 | 3.29234053 | 1 | 0.20854748 | 0 | 1 | 0 | 1 | 0 | 1 | 0 | 1 | 8 | 0.5780332 | FALSE | 1 | FALSE | 1 | FALSE | 1 | FALSE | 1 | FALSE | 1 | FALSE | 1 |
| 5264 | 5264 | 5.90789474 | 2.87080418 | 1 | 0.85897707 | 0 | 1 | 3 | 0.50898113 | 5 | 0.26069733 | 0 | 1 | 66 | 0.24505488 | FALSE | 1 | FALSE | 1 | FALSE | 1 | FALSE | 1 | FALSE | 1 | FALSE | 1 |
| 5265 | 5265 | 7.53333333 | 3.50935258 | 1 | 0.52988231 | 0 | 1 | 1 | 0.65463982 | 0 | 1 | 1 | 0.38148475 | 26 | 0.31495157 | FALSE | 1 | FALSE | 1 | FALSE | 1 | FALSE | 1 | FALSE | 1 | FALSE | 1 |
| 5266 | 5266 | 9.49152542 | 1.96902179 | 1 | 0.78541994 | 1 | 0.85750546 | 3 | 0.35626822 | 3 | 0.51328524 | 0 | 1 | 51 | 0.41341988 | FALSE | 1 | FALSE | 1 | FALSE | 1 | FALSE | 1 | FALSE | 1 | FALSE | 1 |
| 5267 | 5267 | 9.15384615 | 2.91361492 | 1 | 0.63789042 | 0 | 1 | 4 | 0.05000009 | 0 | 1 | 0 | 1 | 34 | 0.41570781 | FALSE | 1 | FALSE | 1 | FALSE | 1 | FALSE | 1 | FALSE | 1 | FALSE | 1 |
| 5268 | 5268 | 6.81818182 | 3.49852389 | 4 | 0.02552933 | 2 | 0.41896557 | 4 | 0.07197653 | 2 | 0.60667406 | 0 | 1 | 32 | 0.98583614 | FALSE | 0.4974559 | FALSE | 1 | FALSE | 1 | FALSE | 1 | FALSE | 1 | FALSE | 1 |
| 5269 | 5269 | 8.13559322 | 2.34670795 | 1 | 0.77972342 | 2 | 0.56459951 | 1 | 0.88128698 | 0 | 1 | 1 | 0.61824432 | 53 | 0.09169509 | FALSE | 1 | FALSE | 1 | FALSE | 1 | FALSE | 1 | FALSE | 1 | FALSE | 1 |
| 5270 | 5270 | 7.5 | 2.44218871 | 0 | 1 | 1 | 0.36919824 | 0 | 1 | 0 | 1 | 0 | 1 | 13 | 0.33283729 | FALSE | 1 | FALSE | 1 | FALSE | 1 | FALSE | 1 | FALSE | 1 | FALSE | 1 |
| 5271 | 5271 | 11.2777778 | 2.42224771 | 0 | 1 | 0 | 1 | 0 | 1 | 2 | 0.19902682 | 0 | 1 | 16 | 0.44998001 | FALSE | 1 | FALSE | 1 | FALSE | 1 | FALSE | 1 | FALSE | 1 | FALSE | 1 |
| 5272 | 5272 | 4.25225225 | 1.24819487 | 2 | 0.77443073 | 7 | 0.06251422 | 1 | 0.98205774 | 3 | 0.88355644 | 1 | 0.83744681 | 95 | 0.25610022 | FALSE | 1 | FALSE | 1 | FALSE | 1 | FALSE | 1 | FALSE | 1 | FALSE | 1 |
| 5273 | 5273 | 11.3333333 | 1.49071199 | 1 | 0.32293355 | 0 | 1 | 0 | 1 | 0 | 1 | 0 | 1 | 14 | 0.29525409 | FALSE | 1 | FALSE | 1 | FALSE | 1 | FALSE | 1 | FALSE | 1 | FALSE | 1 |
| 5274 | 5274 | 10.027027 | 1.83783784 | 2 | 0.24498301 | 0 | 1 | 0 | 1 | 1 | 0.82511639 | 3 | 0.0222449 | 31 | 0.64534533 | FALSE | 1 | FALSE | 1 | FALSE | 1 | FALSE | 1 | FALSE | 0.74991845 | FALSE | 1 |
| 5275 | 5275 | 6.96296296 | 3.13854745 | 3 | 0.16017108 | 1 | 0.83180787 | 3 | 0.30658915 | 2 | 0.71679995 | 1 | 0.59191961 | 44 | 0.78654547 | FALSE | 1 | FALSE | 1 | FALSE | 1 | FALSE | 1 | FALSE | 1 | FALSE | 1 |
| 5276 | 5276 | 10.4 | 1.8 | 0 | 1 | 0 | 1 | 2 | 0.04781153 | 0 | 1 | 0 | 1 | 8 | 0.80394719 | FALSE | 1 | FALSE | 1 | FALSE | 1 | FALSE | 1 | FALSE | 1 | FALSE | 1 |
| 5277 | 5277 | 8.9 | 1.5132746 | 0 | 1 | 0 | 1 | 1 | 0.30655256 | 0 | 1 | 0 | 1 | 9 | 0.5215547 | FALSE | 1 | FALSE | 1 | FALSE | 1 | FALSE | 1 | FALSE | 1 | FALSE | 1 |
| 5278 | 5278 | 9.96551724 | 1.40154493 | 1 | 0.52988231 | 0 | 1 | 1 | 0.65463982 | 1 | 0.74482229 | 0 | 1 | 26 | 0.31495157 | FALSE | 1 | FALSE | 1 | FALSE | 1 | FALSE | 1 | FALSE | 1 | FALSE | 1 |
| 5279 | 5279 | 10.4615385 | 2.17026707 | 0 | 1 | 0 | 1 | 0 | 1 | 0 | 1 | 0 | 1 | 13 | 0.10967906 | FALSE | 1 | FALSE | 1 | FALSE | 1 | FALSE | 1 | FALSE | 1 | FALSE | 1 |
| 5280 | 5280 | 9.41176471 | 2.78776367 | 1 | 0.35728305 | 0 | 1 | 1 | 0.46348836 | 0 | 1 | 0 | 1 | 15 | 0.49060435 | FALSE | 1 | FALSE | 1 | FALSE | 1 | FALSE | 1 | FALSE | 1 | FALSE | 1 |
| 5281 | 5281 | 8.05882353 | 2.07139022 | 0 | 1 | 2 | 0.10307112 | 1 | 0.46348836 | 1 | 0.55062772 | 0 | 1 | 13 | 0.88719222 | FALSE | 1 | FALSE | 1 | FALSE | 1 | FALSE | 1 | FALSE | 1 | FALSE | 1 |
| 5282 | 5282 | 9 | 1.75809815 | 0 | 1 | 1 | 0.30368014 | 1 | 0.33149969 | 0 | 1 | 0 | 1 | 9 | 0.75985366 | FALSE | 1 | FALSE | 1 | FALSE | 1 | FALSE | 1 | FALSE | 1 | FALSE | 1 |
| 5283 | 5283 | 8.75 | 2.75756535 | 1 | 0.45027683 | 1 | 0.53115654 | 0 | 1 | 0 | 1 | 0 | 1 | 21 | 0.27964072 | FALSE | 1 | FALSE | 1 | FALSE | 1 | FALSE | 1 | FALSE | 1 | FALSE | 1 |
| 5284 | 5284 | 9.4 | 2.65329983 | 0 | 1 | 1 | 0.28036922 | 2 | 0.04781153 | 0 | 1 | 0 | 1 | 7 | 0.94326215 | FALSE | 1 | FALSE | 1 | FALSE | 1 | FALSE | 1 | FALSE | 1 | FALSE | 1 |
| 5285 | 5285 | 11.4705882 | 3.98700311 | 2 | 0.06909038 | 0 | 1 | 0 | 1 | 0 | 1 | 0 | 1 | 15 | 0.49060435 | FALSE | 0.74276693 | FALSE | 1 | FALSE | 1 | FALSE | 1 | FALSE | 1 | FALSE | 1 |
| 5286 | 5286 | 5.03030303 | 3.32561089 | 0 | 1 | 1 | 0.6629941 | 1 | 0.70185639 | 2 | 0.45159487 | 0 | 1 | 29 | 0.39648374 | FALSE | 1 | FALSE | 1 | FALSE | 1 | FALSE | 1 | FALSE | 1 | FALSE | 1 |
| 5287 | 5287 | 9.58333333 | 2.46503324 | 1 | 0.26795838 | 0 | 1 | 0 | 1 | 0 | 1 | 1 | 0.18007762 | 10 | 0.71436964 | FALSE | 1 | FALSE | 1 | FALSE | 1 | FALSE | 1 | FALSE | 1 | FALSE | 1 |
| 5288 | 5288 | 10.0909091 | 1.84823645 | 1 | 0.55378205 | 0 | 1 | 3 | 0.09854319 | 2 | 0.41984187 | 1 | 0.4016888 | 24 | 0.90138666 | FALSE | 1 | FALSE | 1 | FALSE | 1 | FALSE | 1 | FALSE | 1 | FALSE | 1 |
| 5289 | 5289 | 7.75 | 1.92028644 | 1 | 0.34033054 | 0 | 1 | 0 | 1 | 0 | 1 | 0 | 1 | 15 | 0.26127859 | FALSE | 1 | FALSE | 1 | FALSE | 1 | FALSE | 1 | FALSE | 1 | FALSE | 1 |
| 5290 | 5290 | 7.29411765 | 2.10083193 | 2 | 0.52416586 | 2 | 0.65156696 | 3 | 0.44404438 | 7 | 0.03544746 | 2 | 0.30695409 | 52 | 0.97053223 | FALSE | 1 | FALSE | 1 | FALSE | 1 | FALSE | 1 | FALSE | 1 | FALSE | 1 |
| 5291 | 5291 | 8.73076923 | 3.39181926 | 0 | 1 | 1 | 0.57534752 | 4 | 0.0130279 | 1 | 0.70601446 | 0 | 1 | 20 | 0.90142689 | FALSE | 1 | FALSE | 1 | FALSE | 1 | FALSE | 1 | FALSE | 1 | FALSE | 1 |
| 5292 | 5292 | 7.19444444 | 2.27082272 | 5 | 0.0020683 | 2 | 0.32556427 | 4 | 0.03888854 | 1 | 0.81665383 | 1 | 0.44938266 | 23 | 0.99939163 | FALSE | 0.12179217 | FALSE | 1 | FALSE | 1 | FALSE | 1 | FALSE | 1 | FALSE | 1 |
| 5293 | 5293 | 8.06451613 | 1.88261116 | 0 | 1 | 1 | 0.63997468 | 1 | 0.67911179 | 3 | 0.16836568 | 1 | 0.4016888 | 25 | 0.80006983 | FALSE | 1 | FALSE | 1 | FALSE | 1 | FALSE | 1 | FALSE | 1 | FALSE | 1 |
| 5294 | 5294 | 10.875 | 1.615356 | 0 | 1 | 2 | 0.02563923 | 1 | 0.25384309 | 1 | 0.31351192 | 1 | 0.12394282 | 3 | 0.99969792 | FALSE | 1 | FALSE | 0.82367276 | FALSE | 1 | FALSE | 1 | FALSE | 1 | FALSE | 1 |
| 5295 | 5295 | 7.88 | 2.99759904 | 0 | 1 | 2 | 0.19297191 | 1 | 0.59998149 | 3 | 0.10474541 | 1 | 0.33902672 | 18 | 0.96851178 | FALSE | 1 | FALSE | 1 | FALSE | 1 | FALSE | 1 | FALSE | 1 | FALSE | 1 |
| 5296 | 5296 | 8.83333333 | 3.93347457 | 0 | 1 | 1 | 0.32623933 | 2 | 0.06691409 | 1 | 0.43131129 | 2 | 0.01579445 | 6 | 0.99914544 | FALSE | 1 | FALSE | 1 | FALSE | 1 | FALSE | 1 | FALSE | 0.66313193 | FALSE | 1 |
| 5297 | 5297 | 10.8571429 | 2.5872529 | 1 | 0.3050805 | 1 | 0.36919824 | 0 | 1 | 0 | 1 | 1 | 0.20679328 | 11 | 0.83676449 | FALSE | 1 | FALSE | 1 | FALSE | 1 | FALSE | 1 | FALSE | 1 | FALSE | 1 |
| 5298 | 5298 | 9.63636364 | 2.85315542 | 1 | 0.4357638 | 0 | 1 | 1 | 0.55340851 | 2 | 0.26791271 | 0 | 1 | 18 | 0.74744813 | FALSE | 1 | FALSE | 1 | FALSE | 1 | FALSE | 1 | FALSE | 1 | FALSE | 1 |
| 5299 | 5299 | 8.2 | 1.32664992 | 1 | 0.22886536 | 0 | 1 | 0 | 1 | 1 | 0.37517262 | 0 | 1 | 8 | 0.80394719 | FALSE | 1 | FALSE | 1 | FALSE | 1 | FALSE | 1 | FALSE | 1 | FALSE | 1 |
| 5300 | 5300 | 8.85 | 2.47537876 | 3 | 0.07731604 | 1 | 0.7236252 | 2 | 0.41139386 | 1 | 0.84089119 | 2 | 0.13365639 | 30 | 0.92795554 | FALSE | 0.77583006 | FALSE | 1 | FALSE | 1 | FALSE | 1 | FALSE | 1 | FALSE | 1 |
| 5301 | 5301 | 8.44285714 | 2.42978961 | 0 | 1 | 4 | 0.19010643 | 3 | 0.46296673 | 4 | 0.40179526 | 2 | 0.31908786 | 57 | 0.8051717 | FALSE | 1 | FALSE | 1 | FALSE | 1 | FALSE | 1 | FALSE | 1 | FALSE | 1 |
| 5302 | 5302 | 8.46428571 | 1.88001031 | 0 | 1 | 3 | 0.06039219 | 1 | 0.64171563 | 0 | 1 | 0 | 1 | 24 | 0.55018107 | FALSE | 1 | FALSE | 1 | FALSE | 1 | FALSE | 1 | FALSE | 1 | FALSE | 1 |
| 5303 | 5303 | 6.48148148 | 2.48507203 | 0 | 1 | 0 | 1 | 1 | 0.62830986 | 0 | 1 | 0 | 1 | 26 | 0.06079097 | FALSE | 1 | FALSE | 1 | FALSE | 1 | FALSE | 1 | FALSE | 1 | FALSE | 1 |
| 5304 | 5304 | 10.2222222 | 3.42467445 | 0 | 1 | 0 | 1 | 0 | 1 | 0 | 1 | 0 | 1 | 9 | 0.21661397 | FALSE | 1 | FALSE | 1 | FALSE | 1 | FALSE | 1 | FALSE | 1 | FALSE | 1 |
| 5305 | 5305 | 9.31746032 | 1.06644934 | 2 | 0.48341887 | 2 | 0.60991494 | 2 | 0.66735975 | 2 | 0.79265743 | 1 | 0.64879513 | 54 | 0.46836572 | FALSE | 1 | FALSE | 1 | FALSE | 1 | FALSE | 1 | FALSE | 1 | FALSE | 1 |
| 5306 | 5306 | 5.0483871 | 2.41919354 | 4 | 0.07395961 | 1 | 0.87100665 | 5 | 0.07095719 | 2 | 0.78520976 | 1 | 0.64288531 | 49 | 0.90602303 | FALSE | 0.77424195 | FALSE | 1 | FALSE | 1 | FALSE | 1 | FALSE | 1 | FALSE | 1 |
| 5307 | 5307 | 8.71428571 | 2.43276948 | 0 | 1 | 0 | 1 | 0 | 1 | 0 | 1 | 0 | 1 | 7 | 0.3043657 | FALSE | 1 | FALSE | 1 | FALSE | 1 | FALSE | 1 | FALSE | 1 | FALSE | 1 |
| 5308 | 5308 | 9.94117647 | 2.68920693 | 0 | 1 | 2 | 0.10307112 | 1 | 0.46348836 | 0 | 1 | 0 | 1 | 14 | 0.7313145 | FALSE | 1 | FALSE | 1 | FALSE | 1 | FALSE | 1 | FALSE | 1 | FALSE | 1 |
| 5309 | 5309 | 9.875 | 2.20439901 | 2 | 0.01652925 | 0 | 1 | 0 | 1 | 0 | 1 | 0 | 1 | 6 | 0.88425551 | FALSE | 0.39708985 | FALSE | 1 | FALSE | 1 | FALSE | 1 | FALSE | 1 | FALSE | 1 |
| 5310 | 5310 | 8.85185185 | 2.23943948 | 0 | 1 | 2 | 0.19297191 | 2 | 0.22597287 | 1 | 0.69181315 | 1 | 0.33902672 | 19 | 0.91734261 | FALSE | 1 | FALSE | 1 | FALSE | 1 | FALSE | 1 | FALSE | 1 | FALSE | 1 |
| 5311 | 5311 | 8.41666667 | 2.62863505 | 0 | 1 | 0 | 1 | 0 | 1 | 0 | 1 | 0 | 1 | 12 | 0.13002667 | FALSE | 1 | FALSE | 1 | FALSE | 1 | FALSE | 1 | FALSE | 1 | FALSE | 1 |
| 5312 | 5312 | 15.2105263 | 3.74979224 | 0 | 1 | 1 | 0.46503795 | 1 | 0.50143826 | 0 | 1 | 0 | 1 | 17 | 0.41141543 | FALSE | 1 | FALSE | 1 | FALSE | 1 | FALSE | 1 | FALSE | 1 | FALSE | 1 |
| 5313 | 5313 | 6.1 | 3.59026461 | 1 | 0.40558521 | 0 | 1 | 0 | 1 | 1 | 0.60987146 | 0 | 1 | 18 | 0.37504645 | FALSE | 1 | FALSE | 1 | FALSE | 1 | FALSE | 1 | FALSE | 1 | FALSE | 1 |
| 5314 | 5314 | 8.72727273 | 3.54428885 | 3 | 0.05151181 | 1 | 0.6629941 | 1 | 0.70185639 | 2 | 0.45159487 | 0 | 1 | 26 | 0.86852567 | FALSE | 0.67646333 | FALSE | 1 | FALSE | 1 | FALSE | 1 | FALSE | 1 | FALSE | 1 |
| 5315 | 5315 | 10 | 2.34520788 | 0 | 1 | 0 | 1 | 1 | 0.25384309 | 1 | 0.31351192 | 0 | 1 | 6 | 0.88425551 | FALSE | 1 | FALSE | 1 | FALSE | 1 | FALSE | 1 | FALSE | 1 | FALSE | 1 |
| 5316 | 5316 | 8.47058824 | 2.7033236 | 0 | 1 | 0 | 1 | 1 | 0.44344817 | 1 | 0.52895088 | 0 | 1 | 14 | 0.53308261 | FALSE | 1 | FALSE | 1 | FALSE | 1 | FALSE | 1 | FALSE | 1 | FALSE | 1 |
| 5317 | 5317 | 5.85714286 | 2.94853821 | 2 | 0.539854 | 1 | 0.90110191 | 2 | 0.72288838 | 0 | 1 | 0 | 1 | 65 | 0.02789934 | FALSE | 1 | FALSE | 1 | FALSE | 1 | FALSE | 1 | FALSE | 1 | FALSE | 1 |
| 5318 | 5318 | 11.0666667 | 2.08059821 | 0 | 1 | 2 | 0.08295839 | 1 | 0.42266264 | 1 | 0.50623194 | 0 | 1 | 11 | 0.92879412 | FALSE | 1 | FALSE | 1 | FALSE | 1 | FALSE | 1 | FALSE | 1 | FALSE | 1 |
| 5319 | 5319 | 11.75 | 2.04633819 | 0 | 1 | 0 | 1 | 0 | 1 | 0 | 1 | 0 | 1 | 11 | 0.15414489 | FALSE | 1 | FALSE | 1 | FALSE | 1 | FALSE | 1 | FALSE | 1 | FALSE | 1 |
| 5320 | 5320 | 11 | 1.63299316 | 0 | 1 | 0 | 1 | 0 | 1 | 0 | 1 | 0 | 1 | 3 | 0.60071728 | FALSE | 1 | FALSE | 1 | FALSE | 1 | FALSE | 1 | FALSE | 1 | FALSE | 1 |
| 5321 | 5321 | 7.88 | 1.70458206 | 0 | 1 | 0 | 1 | 1 | 0.59998149 | 0 | 1 | 0 | 1 | 24 | 0.08019798 | FALSE | 1 | FALSE | 1 | FALSE | 1 | FALSE | 1 | FALSE | 1 | FALSE | 1 |
| 5322 | 5322 | 8.5 | 2.29128785 | 0 | 1 | 1 | 0.44710673 | 1 | 0.48280983 | 2 | 0.19902682 | 0 | 1 | 14 | 0.86287418 | FALSE | 1 | FALSE | 1 | FALSE | 1 | FALSE | 1 | FALSE | 1 | FALSE | 1 |
| 5323 | 5323 | 7.625 | 2.57896135 | 0 | 1 | 1 | 0.54637326 | 0 | 1 | 1 | 0.67692816 | 0 | 1 | 22 | 0.25237063 | FALSE | 1 | FALSE | 1 | FALSE | 1 | FALSE | 1 | FALSE | 1 | FALSE | 1 |
| 5324 | 5324 | 10.1764706 | 3.31192665 | 0 | 1 | 0 | 1 | 0 | 1 | 1 | 0.52895088 | 0 | 1 | 15 | 0.26127859 | FALSE | 1 | FALSE | 1 | FALSE | 1 | FALSE | 1 | FALSE | 1 | FALSE | 1 |
| 5325 | 5325 | 9.90909091 | 2.02055553 | 1 | 0.24866461 | 0 | 1 | 0 | 1 | 0 | 1 | 0 | 1 | 10 | 0.46859937 | FALSE | 1 | FALSE | 1 | FALSE | 1 | FALSE | 1 | FALSE | 1 | FALSE | 1 |
| 5326 | 5326 | 7.86206897 | 2.68744383 | 1 | 0.52988231 | 0 | 1 | 1 | 0.65463982 | 3 | 0.14602298 | 1 | 0.38148475 | 23 | 0.84462957 | FALSE | 1 | FALSE | 1 | FALSE | 1 | FALSE | 1 | FALSE | 1 | FALSE | 1 |
| 5327 | 5327 | 8.60869565 | 1.68727025 | 0 | 1 | 1 | 0.53115654 | 0 | 1 | 1 | 0.6613267 | 0 | 1 | 21 | 0.27964072 | FALSE | 1 | FALSE | 1 | FALSE | 1 | FALSE | 1 | FALSE | 1 | FALSE | 1 |
| 5328 | 5328 | 9.05263158 | 2.17579228 | 0 | 1 | 1 | 0.71433277 | 0 | 1 | 1 | 0.83318955 | 1 | 0.4673875 | 35 | 0.13407055 | FALSE | 1 | FALSE | 1 | FALSE | 1 | FALSE | 1 | FALSE | 1 | FALSE | 1 |
| 5329 | 5329 | 4.8 | 3.5282983 | 4 | 0.19928904 | 4 | 0.33229965 | 4 | 0.40604339 | 3 | 0.78935002 | 0 | 1 | 75 | 0.67286052 | FALSE | 1 | FALSE | 1 | FALSE | 1 | FALSE | 1 | FALSE | 1 | FALSE | 1 |
| 5330 | 5330 | 9 | 3.17441697 | 2 | 0.14250978 | 1 | 0.57534752 | 2 | 0.23940226 | 1 | 0.70601446 | 1 | 0.3499045 | 19 | 0.96054244 | FALSE | 1 | FALSE | 1 | FALSE | 1 | FALSE | 1 | FALSE | 1 | FALSE | 1 |
| 5331 | 5331 | 10.0476191 | 1.96338363 | 1 | 0.42086991 | 3 | 0.02888061 | 1 | 0.53671415 | 1 | 0.62783522 | 0 | 1 | 15 | 0.96513691 | FALSE | 1 | FALSE | 0.88012077 | FALSE | 1 | FALSE | 1 | FALSE | 1 | FALSE | 1 |
| 5332 | 5332 | 6.15686275 | 0.74199561 | 15 | 6.94E-06 | 14 | 0.00037656 | 9 | 0.09597511 | 3 | 0.97364454 | 10 | 0.00017844 | 101 | 0.99999999 | TRUE | 0.00299395 | FALSE | 0.15723145 | FALSE | 1 | FALSE | 1 | FALSE | 0.0696069 | FALSE | 1 |
| 5333 | 5333 | 7.38461539 | 2.11363564 | 1 | 0.49162634 | 0 | 1 | 0 | 1 | 1 | 0.70601446 | 0 | 1 | 24 | 0.20425579 | FALSE | 1 | FALSE | 1 | FALSE | 1 | FALSE | 1 | FALSE | 1 | FALSE | 1 |
| 5334 | 5334 | 4.51803279 | 2.40365215 | 6 | 0.76946804 | 9 | 0.61240877 | 21 | 0.00185463 | 27 | 0.00038725 | 3 | 0.86584495 | 227 | 0.99944277 | FALSE | 1 | FALSE | 1 | FALSE | 1 | FALSE | 0.63446475 | FALSE | 1 | FALSE | 1 |
| 5335 | 5335 | 7.14893617 | 3.43312229 | 4 | 0.03163127 | 3 | 0.19374024 | 1 | 0.82190381 | 4 | 0.1678689 | 0 | 1 | 35 | 0.97547234 | FALSE | 0.54437678 | FALSE | 1 | FALSE | 1 | FALSE | 1 | FALSE | 1 | FALSE | 1 |
| 5336 | 5336 | 6.74576271 | 2.20625628 | 3 | 0.19211919 | 6 | 0.01153778 | 4 | 0.16113918 | 4 | 0.28589315 | 1 | 0.62455813 | 41 | 0.99889368 | FALSE | 1 | FALSE | 0.67617265 | FALSE | 1 | FALSE | 1 | FALSE | 1 | FALSE | 1 |
| 5337 | 5337 | 10 | 2.58843582 | 2 | 0.09187136 | 2 | 0.1353446 | 0 | 1 | 0 | 1 | 2 | 0.04176963 | 14 | 0.97325593 | FALSE | 0.83248965 | FALSE | 1 | FALSE | 1 | FALSE | 1 | FALSE | 0.90763079 | FALSE | 1 |
| 5338 | 5338 | 8.57894737 | 2.77803015 | 2 | 0.08405377 | 1 | 0.46503795 | 0 | 1 | 1 | 0.59104357 | 0 | 1 | 15 | 0.83648191 | FALSE | 0.79236883 | FALSE | 1 | FALSE | 1 | FALSE | 1 | FALSE | 1 | FALSE | 1 |
| 5339 | 5339 | 9.71428571 | 2.31234487 | 0 | 1 | 0 | 1 | 0 | 1 | 0 | 1 | 0 | 1 | 7 | 0.3043657 | FALSE | 1 | FALSE | 1 | FALSE | 1 | FALSE | 1 | FALSE | 1 | FALSE | 1 |
| 5340 | 5340 | 8.3125 | 1.79299854 | 0 | 1 | 1 | 0.40942991 | 0 | 1 | 2 | 0.16544789 | 0 | 1 | 13 | 0.76799161 | FALSE | 1 | FALSE | 1 | FALSE | 1 | FALSE | 1 | FALSE | 1 | FALSE | 1 |
| 5341 | 5341 | 8.08219178 | 0.72628582 | 6 | 0.1716507 | 3 | 0.85725387 | 7 | 0.27088125 | 5 | 0.80767292 | 0 | 1 | 125 | 0.39064515 | FALSE | 1 | FALSE | 1 | FALSE | 1 | FALSE | 1 | FALSE | 1 | FALSE | 1 |
| 5342 | 5342 | 9.76470588 | 1.69876813 | 1 | 0.35728305 | 3 | 0.01622926 | 2 | 0.12280513 | 0 | 1 | 0 | 1 | 11 | 0.98974663 | FALSE | 1 | FALSE | 0.7256023 | FALSE | 1 | FALSE | 1 | FALSE | 1 | FALSE | 1 |
| 5343 | 5343 | 8.44444444 | 3.43547219 | 2 | 0.07645454 | 1 | 0.44710673 | 0 | 1 | 1 | 0.57131013 | 0 | 1 | 14 | 0.86287418 | FALSE | 0.77424195 | FALSE | 1 | FALSE | 1 | FALSE | 1 | FALSE | 1 | FALSE | 1 |
| 5344 | 5344 | 11.1538462 | 3.46068366 | 2 | 0.04234484 | 0 | 1 | 0 | 1 | 1 | 0.45746571 | 0 | 1 | 10 | 0.86791642 | FALSE | 0.62842194 | FALSE | 1 | FALSE | 1 | FALSE | 1 | FALSE | 1 | FALSE | 1 |
| 5345 | 5345 | 7.21621622 | 0.84261931 | 1 | 0.6184714 | 0 | 1 | 0 | 1 | 1 | 0.82511639 | 2 | 0.12262638 | 33 | 0.2931558 | FALSE | 1 | FALSE | 1 | FALSE | 1 | FALSE | 1 | FALSE | 1 | FALSE | 1 |
| 5346 | 5346 | 6.75757576 | 1.68828949 | 0 | 1 | 2 | 0.28941836 | 3 | 0.11365871 | 1 | 0.78873814 | 0 | 1 | 27 | 0.75124583 | FALSE | 1 | FALSE | 1 | FALSE | 1 | FALSE | 1 | FALSE | 1 | FALSE | 1 |
| 5347 | 5347 | 9.7 | 2.96816442 | 0 | 1 | 0 | 1 | 0 | 1 | 1 | 0.37517262 | 0 | 1 | 9 | 0.5215547 | FALSE | 1 | FALSE | 1 | FALSE | 1 | FALSE | 1 | FALSE | 1 | FALSE | 1 |
| 5348 | 5348 | 6.85714286 | 2.04440501 | 1 | 0.59801736 | 1 | 0.68454804 | 2 | 0.35984046 | 2 | 0.48233989 | 0 | 1 | 29 | 0.69927719 | FALSE | 1 | FALSE | 1 | FALSE | 1 | FALSE | 1 | FALSE | 1 | FALSE | 1 |
| 5349 | 5349 | 11.1818182 | 3.43294683 | 0 | 1 | 1 | 0.5154318 | 0 | 1 | 1 | 0.64497439 | 0 | 1 | 20 | 0.30915202 | FALSE | 1 | FALSE | 1 | FALSE | 1 | FALSE | 1 | FALSE | 1 | FALSE | 1 |
| 5350 | 5350 | 10.2222222 | 1.6178022 | 1 | 0.20854748 | 0 | 1 | 0 | 1 | 1 | 0.34506517 | 0 | 1 | 7 | 0.84574168 | FALSE | 1 | FALSE | 1 | FALSE | 1 | FALSE | 1 | FALSE | 1 | FALSE | 1 |
| 5351 | 5351 | 9.40909091 | 1.969289 | 0 | 1 | 0 | 1 | 0 | 1 | 2 | 0.26791271 | 0 | 1 | 20 | 0.30915202 | FALSE | 1 | FALSE | 1 | FALSE | 1 | FALSE | 1 | FALSE | 1 | FALSE | 1 |
| 5352 | 5352 | 5.384 | 2.10060563 | 4 | 0.39363378 | 4 | 0.57367482 | 6 | 0.28660842 | 5 | 0.67988626 | 6 | 0.01586595 | 99 | 0.93314531 | FALSE | 1 | FALSE | 1 | FALSE | 1 | FALSE | 1 | FALSE | 0.66313193 | FALSE | 1 |
| 5353 | 5353 | 5.77272727 | 2.18670539 | 1 | 0.8213973 | 2 | 0.63534334 | 3 | 0.4248691 | 2 | 0.8136513 | 0 | 1 | 58 | 0.27702369 | FALSE | 1 | FALSE | 1 | FALSE | 1 | FALSE | 1 | FALSE | 1 | FALSE | 1 |
| 5354 | 5354 | 9.78571429 | 2.02408962 | 1 | 0.50471032 | 1 | 0.58913618 | 2 | 0.25286935 | 1 | 0.71956339 | 0 | 1 | 22 | 0.76237877 | FALSE | 1 | FALSE | 1 | FALSE | 1 | FALSE | 1 | FALSE | 1 | FALSE | 1 |
| 5355 | 5355 | 10.1071429 | 1.26319566 | 1 | 0.51745948 | 1 | 0.60247911 | 2 | 0.2663542 | 1 | 0.73248983 | 1 | 0.37113067 | 22 | 0.86499992 | FALSE | 1 | FALSE | 1 | FALSE | 1 | FALSE | 1 | FALSE | 1 | FALSE | 1 |
| 5356 | 5356 | 8.42857143 | 1.23717915 | 0 | 1 | 1 | 0.36919824 | 0 | 1 | 0 | 1 | 0 | 1 | 13 | 0.33283729 | FALSE | 1 | FALSE | 1 | FALSE | 1 | FALSE | 1 | FALSE | 1 | FALSE | 1 |

| | | | | | | | | | | | | | | | | | | | | | | | | | | | |
|---|---|---|---|---|---|---|---|---|---|---|---|---|---|---|---|---|---|---|---|---|---|---|---|---|---|---|---|
| 5357 | 5357 | 8.64705882 | 2.44807671 | 0 | 1 | 3 | 0.09602744 | 0 | 1 | 1 | 0.7984848 | 0 | 1 | 30 | 0.36879488 | FALSE | 1 | FALSE | 1 | FALSE | 1 | FALSE | 1 | FALSE | 1 | FALSE | 1 |
| 5358 | 5358 | 9.85714286 | 0.83299313 | 0 | 1 | 0 | 1 | 0 | 1 | 0 | 1 | 0 | 1 | 7 | 0.3043657 | FALSE | 1 | FALSE | 1 | FALSE | 1 | FALSE | 1 | FALSE | 1 | FALSE | 1 |
| 5359 | 5359 | 9.83333333 | 2.68224616 | 2 | 0.23542589 | 1 | 0.69480472 | 3 | 0.13783405 | 0 | 1 | 1 | 0.44938266 | 29 | 0.81028289 | FALSE | 1 | FALSE | 1 | FALSE | 1 | FALSE | 1 | FALSE | 1 | FALSE | 1 |
| 5360 | 5360 | 6.23188406 | 2.89499782 | 0 | 1 | 3 | 0.38707857 | 5 | 0.1012281 | 4 | 0.39127216 | 0 | 1 | 57 | 0.72531896 | FALSE | 1 | FALSE | 1 | FALSE | 1 | FALSE | 1 | FALSE | 1 | FALSE | 1 |
| 5361 | 5361 | 3.64634146 | 2.25978604 | 5 | 0.05874466 | 5 | 0.12558433 | 6 | 0.07364899 | 5 | 0.3237921 | 1 | 0.74434486 | 60 | 0.99718261 | FALSE | 0.72149357 | FALSE | 1 | FALSE | 1 | FALSE | 1 | FALSE | 1 | FALSE | 1 |
| 5362 | 5362 | 8.41666667 | 1.18731724 | 5 | 0.00029898 | 0 | 1 | 0 | 1 | 1 | 0.67692816 | 0 | 1 | 18 | 0.93166998 | TRUE | 0.03449636 | FALSE | 1 | FALSE | 1 | FALSE | 1 | FALSE | 1 | FALSE | 1 |
| 5363 | 5363 | 5.51111111 | 1.96964621 | 1 | 0.97027046 | 6 | 0.26597002 | 5 | 0.53036069 | 6 | 0.58440605 | 4 | 0.17710987 | 112 | 0.65519557 | FALSE | 1 | FALSE | 1 | FALSE | 1 | FALSE | 1 | FALSE | 1 | FALSE | 1 |
| 5364 | 5364 | 8.45454546 | 2.67526163 | 0 | 1 | 0 | 1 | 0 | 1 | 0 | 1 | 2 | 0.01330324 | 9 | 0.75985366 | FALSE | 1 | FALSE | 1 | FALSE | 1 | FALSE | 1 | FALSE | 0.63731057 | FALSE | 1 |
| 5365 | 5365 | 7.60606061 | 2.3216947 | 1 | 0.57647348 | 1 | 0.6629941 | 0 | 1 | 1 | 0.78873814 | 0 | 1 | 30 | 0.21942357 | FALSE | 1 | FALSE | 1 | FALSE | 1 | FALSE | 1 | FALSE | 1 | FALSE | 1 |
| 5366 | 5366 | 8.91304348 | 2.48321397 | 1 | 0.4357638 | 0 | 1 | 0 | 1 | 2 | 0.26791271 | 0 | 1 | 19 | 0.54233419 | FALSE | 1 | FALSE | 1 | FALSE | 1 | FALSE | 1 | FALSE | 1 | FALSE | 1 |
| 5367 | 5367 | 9.46153846 | 1.0088367 | 0 | 1 | 0 | 1 | 0 | 1 | 1 | 0.45746571 | 0 | 1 | 12 | 0.374198 | FALSE | 1 | FALSE | 1 | FALSE | 1 | FALSE | 1 | FALSE | 1 | FALSE | 1 |
| 5368 | 5368 | 10.05 | 4.11673414 | 5 | 0.00011852 | 1 | 0.48239024 | 0 | 1 | 0 | 1 | 0 | 1 | 14 | 0.97325593 | TRUE | 0.01941834 | FALSE | 1 | FALSE | 1 | FALSE | 1 | FALSE | 1 | FALSE | 1 |
| 5369 | 5369 | 10.7272727 | 1.86308196 | 0 | 1 | 0 | 1 | 1 | 0.53671415 | 0 | 1 | 0 | 1 | 20 | 0.13763971 | FALSE | 1 | FALSE | 1 | FALSE | 1 | FALSE | 1 | FALSE | 1 | FALSE | 1 |
| 5370 | 5370 | 2.39411765 | 1.36035056 | 26 | 0.02313088 | 24 | 0.35042629 | 25 | 0.47667565 | 37 | 0.14936281 | 9 | 0.794592 | 557 | 0.95731977 | FALSE | 0.47971689 | FALSE | 1 | FALSE | 1 | FALSE | 1 | FALSE | 1 | FALSE | 1 |
| 5371 | 5371 | 8.27777778 | 2.37593386 | 0 | 1 | 1 | 0.69480472 | 1 | 0.73299778 | 1 | 0.81665383 | 1 | 0.44938266 | 32 | 0.31709555 | FALSE | 1 | FALSE | 1 | FALSE | 1 | FALSE | 1 | FALSE | 1 | FALSE | 1 |
| 5372 | 5372 | 10.9444444 | 2.97157314 | 0 | 1 | 0 | 1 | 1 | 0.48280983 | 0 | 1 | 0 | 1 | 17 | 0.20329205 | FALSE | 1 | FALSE | 1 | FALSE | 1 | FALSE | 1 | FALSE | 1 | FALSE | 1 |
| 5373 | 5373 | 8.55555556 | 3.57805382 | 1 | 0.60837712 | 2 | 0.32556427 | 2 | 0.37289966 | 0 | 1 | 1 | 0.44938266 | 30 | 0.67248271 | FALSE | 1 | FALSE | 1 | FALSE | 1 | FALSE | 1 | FALSE | 1 | FALSE | 1 |
| 5374 | 5374 | 9.57142857 | 1.59078982 | 0 | 1 | 0 | 1 | 0 | 1 | 1 | 0.48242099 | 0 | 1 | 13 | 0.33283729 | FALSE | 1 | FALSE | 1 | FALSE | 1 | FALSE | 1 | FALSE | 1 | FALSE | 1 |
| 5375 | 5375 | 9.90909091 | 2.53895274 | 1 | 0.4357638 | 0 | 1 | 0 | 1 | 2 | 0.26791271 | 0 | 1 | 19 | 0.54233419 | FALSE | 1 | FALSE | 1 | FALSE | 1 | FALSE | 1 | FALSE | 1 | FALSE | 1 |
| 5376 | 5376 | 7.76923077 | 3.47305776 | 2 | 0.38681045 | 4 | 0.08651261 | 4 | 0.11558745 | 2 | 0.6970685 | 2 | 0.2095319 | 38 | 0.98895952 | FALSE | 1 | FALSE | 1 | FALSE | 1 | FALSE | 1 | FALSE | 1 | FALSE | 1 |
| 5377 | 5377 | 9.21621622 | 3.97380025 | 1 | 0.85522402 | 5 | 0.09061322 | 1 | 0.93427349 | 3 | 0.66754163 | 0 | 1 | 64 | 0.37902265 | FALSE | 1 | FALSE | 1 | FALSE | 1 | FALSE | 1 | FALSE | 1 | FALSE | 1 |
| 5378 | 5378 | 8.21428571 | 2.11046953 | 0 | 1 | 2 | 0.07341559 | 0 | 1 | 0 | 1 | 3 | 0.00136853 | 9 | 0.98607548 | FALSE | 1 | FALSE | 1 | FALSE | 1 | FALSE | 1 | FALSE | 0.20625174 | FALSE | 1 |
| 5379 | 5379 | 5.74736842 | 1.41397849 | 0 | 1 | 1 | 0.95697569 | 2 | 0.86139368 | 6 | 0.2701554 | 1 | 0.79437879 | 85 | 0.10448523 | FALSE | 1 | FALSE | 1 | FALSE | 1 | FALSE | 1 | FALSE | 1 | FALSE | 1 |
| 5380 | 5380 | 10.3636364 | 1.4316378 | 1 | 0.24866461 | 0 | 1 | 1 | 0.33149969 | 1 | 0.40390032 | 0 | 1 | 8 | 0.92152993 | FALSE | 1 | FALSE | 1 | FALSE | 1 | FALSE | 1 | FALSE | 1 | FALSE | 1 |
| 5381 | 5381 | 6.61184211 | 1.68214048 | 5 | 0.35044506 | 7 | 0.22123268 | 1 | 0.99639467 | 6 | 0.70536052 | 6 | 0.0382623 | 127 | 0.66186129 | FALSE | 1 | FALSE | 1 | FALSE | 1 | FALSE | 1 | FALSE | 0.88046278 | FALSE | 1 |
| 5382 | 5382 | 10.5833333 | 1.89113429 | 0 | 1 | 0 | 1 | 1 | 0.35555293 | 0 | 1 | 0 | 1 | 11 | 0.41944532 | FALSE | 1 | FALSE | 1 | FALSE | 1 | FALSE | 1 | FALSE | 1 | FALSE | 1 |
| 5383 | 5383 | 10.3214286 | 1.9096292 | 1 | 0.51745948 | 0 | 1 | 0 | 1 | 1 | 0.73248983 | 0 | 1 | 26 | 0.16405547 | FALSE | 1 | FALSE | 1 | FALSE | 1 | FALSE | 1 | FALSE | 1 | FALSE | 1 |
| 5384 | 5384 | 6.07692308 | 2.82284286 | 0 | 1 | 3 | 0.13080858 | 2 | 0.41139386 | 1 | 0.84089119 | 2 | 0.13365639 | 31 | 0.85613899 | FALSE | 1 | FALSE | 1 | FALSE | 1 | FALSE | 1 | FALSE | 1 | FALSE | 1 |
| 5385 | 5385 | 10.7272727 | 1.86308196 | 0 | 1 | 0 | 1 | 0 | 1 | 0 | 1 | 1 | 0.16638736 | 10 | 0.46859937 | FALSE | 1 | FALSE | 1 | FALSE | 1 | FALSE | 1 | FALSE | 1 | FALSE | 1 |
| 5386 | 5386 | 10.6666667 | 1.49071199 | 0 | 1 | 1 | 0.58913618 | 1 | 0.62830986 | 1 | 0.71956339 | 0 | 1 | 24 | 0.3725373 | FALSE | 1 | FALSE | 1 | FALSE | 1 | FALSE | 1 | FALSE | 1 | FALSE | 1 |
| 5387 | 5387 | 8.51724138 | 2.31368775 | 1 | 0.52988231 | 2 | 0.24095584 | 1 | 0.65463982 | 3 | 0.14602298 | 1 | 0.38148475 | 21 | 0.97185642 | FALSE | 1 | FALSE | 1 | FALSE | 1 | FALSE | 1 | FALSE | 1 | FALSE | 1 |
| 5388 | 5388 | 7.55555556 | 2.14015114 | 0 | 1 | 1 | 0.44710673 | 1 | 0.48280983 | 0 | 1 | 1 | 0.25766374 | 15 | 0.69372602 | FALSE | 1 | FALSE | 1 | FALSE | 1 | FALSE | 1 | FALSE | 1 | FALSE | 1 |
| 5389 | 5389 | 8.42857143 | 3.11022015 | 0 | 1 | 0 | 1 | 0 | 1 | 0 | 1 | 0 | 1 | 21 | 0.02808151 | FALSE | 1 | FALSE | 1 | FALSE | 1 | FALSE | 1 | FALSE | 1 | FALSE | 1 |
| 5390 | 5390 | 9.5 | 3.2596012 | 3 | 0.00725145 | 0 | 1 | 0 | 1 | 1 | 0.52895088 | 1 | 0.23264612 | 11 | 0.9718531 | FALSE | 0.25171133 | FALSE | 1 | FALSE | 1 | FALSE | 1 | FALSE | 1 | FALSE | 1 |
| 5391 | 5391 | 4.18064516 | 1.01898728 | 13 | 0.05446414 | 16 | 0.04286007 | 15 | 0.14683688 | 15 | 0.4551248 | 9 | 0.06728386 | 242 | 0.99898359 | FALSE | 0.68849059 | FALSE | 1 | FALSE | 1 | FALSE | 1 | FALSE | 1 | FALSE | 1 |
| 5392 | 5392 | 7.83333333 | 2.30337916 | 1 | 0.46441868 | 0 | 1 | 2 | 0.21260257 | 0 | 1 | 0 | 1 | 21 | 0.47053137 | FALSE | 1 | FALSE | 1 | FALSE | 1 | FALSE | 1 | FALSE | 1 | FALSE | 1 |
| 5393 | 5393 | 8.52173913 | 1.97527247 | 0 | 1 | 0 | 1 | 2 | 0.19931429 | 1 | 0.6613267 | 0 | 1 | 20 | 0.50589402 | FALSE | 1 | FALSE | 1 | FALSE | 1 | FALSE | 1 | FALSE | 1 | FALSE | 1 |
| 5394 | 5394 | 8.31707317 | 2.12349257 | 0 | 1 | 0 | 1 | 3 | 0.1814659 | 2 | 0.56788092 | 0 | 1 | 36 | 0.36455032 | FALSE | 1 | FALSE | 1 | FALSE | 1 | FALSE | 1 | FALSE | 1 | FALSE | 1 |
| 5395 | 5395 | 10.3684211 | 2.41073316 | 0 | 1 | 0 | 1 | 2 | 0.14749808 | 1 | 0.59104357 | 0 | 1 | 16 | 0.65565778 | FALSE | 1 | FALSE | 1 | FALSE | 1 | FALSE | 1 | FALSE | 1 | FALSE | 1 |
| 5396 | 5396 | 10.4615385 | 4.3609347 | 0 | 1 | 0 | 1 | 1 | 0.37874417 | 1 | 0.45746571 | 0 | 1 | 11 | 0.6683056 | FALSE | 1 | FALSE | 1 | FALSE | 1 | FALSE | 1 | FALSE | 1 | FALSE | 1 |
| 5397 | 5397 | 7.96551724 | 1.62910508 | 0 | 1 | 2 | 0.56459951 | 0 | 1 | 3 | 0.50172293 | 2 | 0.24594209 | 51 | 0.2955095 | FALSE | 1 | FALSE | 1 | FALSE | 1 | FALSE | 1 | FALSE | 1 | FALSE | 1 |
| 5398 | 5398 | 7.58695652 | 2.63402558 | 0 | 1 | 0 | 1 | 1 | 0.81522035 | 3 | 0.35428318 | 1 | 0.53375569 | 41 | 0.25430673 | FALSE | 1 | FALSE | 1 | FALSE | 1 | FALSE | 1 | FALSE | 1 | FALSE | 1 |
| 5399 | 5399 | 8.9661017 | 1.56189912 | 1 | 0.78541994 | 3 | 0.2979413 | 5 | 0.05975605 | 2 | 0.76144533 | 4 | 0.01560072 | 44 | 0.98374792 | FALSE | 1 | FALSE | 1 | FALSE | 1 | FALSE | 1 | FALSE | 0.66313193 | FALSE | 1 |
| 5400 | 5400 | 9.18181818 | 1.02851895 | 0 | 1 | 0 | 1 | 1 | 0.55340851 | 2 | 0.26791271 | 1 | 0.30529951 | 18 | 0.74744813 | FALSE | 1 | FALSE | 1 | FALSE | 1 | FALSE | 1 | FALSE | 1 | FALSE | 1 |
| 5401 | 5401 | 9.74193548 | 3.24253726 | 2 | 0.1881398 | 1 | 0.63997468 | 4 | 0.02385458 | 2 | 0.41984187 | 2 | 0.09110253 | 20 | 0.9984454 | FALSE | 1 | FALSE | 1 | FALSE | 1 | FALSE | 1 | FALSE | 1 | FALSE | 1 |
| 5402 | 5402 | 8.23076923 | 3.10055031 | 2 | 0.26413583 | 1 | 0.7236252 | 1 | 0.76089851 | 2 | 0.54052736 | 0 | 1 | 33 | 0.59061307 | FALSE | 1 | FALSE | 1 | FALSE | 1 | FALSE | 1 | FALSE | 1 | FALSE | 1 |
| 5403 | 5403 | 8.03846154 | 2.51886374 | 0 | 1 | 1 | 0.57534752 | 2 | 0.23940226 | 1 | 0.70601446 | 0 | 1 | 22 | 0.61628 | FALSE | 1 | FALSE | 1 | FALSE | 1 | FALSE | 1 | FALSE | 1 | FALSE | 1 |
| 5404 | 5404 | 7.1978022 | 2.02847737 | 0 | 1 | 4 | 0.32490044 | 3 | 0.62570603 | 2 | 0.92076432 | 7 | 0.0005998 | 73 | 0.78154321 | FALSE | 1 | FALSE | 1 | FALSE | 1 | FALSE | 1 | FALSE | 0.12930348 | FALSE | 1 |
| 5405 | 5405 | 8.88888889 | 1.96889391 | 0 | 1 | 0 | 1 | 0 | 1 | 1 | 0.57131013 | 0 | 1 | 17 | 0.20329205 | FALSE | 1 | FALSE | 1 | FALSE | 1 | FALSE | 1 | FALSE | 1 | FALSE | 1 |
| 5406 | 5406 | 9.07894737 | 2.6195779 | 0 | 1 | 1 | 0.71433277 | 2 | 0.39868506 | 3 | 0.25266879 | 0 | 1 | 32 | 0.61800916 | FALSE | 1 | FALSE | 1 | FALSE | 1 | FALSE | 1 | FALSE | 1 | FALSE | 1 |
| 5407 | 5407 | 9.53333333 | 1.30979218 | 1 | 0.32293355 | 1 | 0.38964394 | 1 | 0.42266264 | 0 | 1 | 0 | 1 | 12 | 0.80329917 | FALSE | 1 | FALSE | 1 | FALSE | 1 | FALSE | 1 | FALSE | 1 | FALSE | 1 |
| 5408 | 5408 | 11.0526316 | 2.41647164 | 2 | 0.08405377 | 0 | 1 | 0 | 1 | 1 | 0.59104357 | 0 | 1 | 16 | 0.65565778 | FALSE | 0.79236883 | FALSE | 1 | FALSE | 1 | FALSE | 1 | FALSE | 1 | FALSE | 1 |
| 5409 | 5409 | 11.1388889 | 2.11019717 | 0 | 1 | 1 | 0.69480472 | 2 | 0.37289966 | 2 | 0.49731009 | 0 | 1 | 31 | 0.49869404 | FALSE | 1 | FALSE | 1 | FALSE | 1 | FALSE | 1 | FALSE | 1 | FALSE | 1 |
| 5410 | 5410 | 10 | 3.34312288 | 1 | 0.58738519 | 0 | 1 | 1 | 0.71261921 | 4 | 0.06848449 | 1 | 0.43077488 | 27 | 0.85023803 | FALSE | 1 | FALSE | 1 | FALSE | 1 | FALSE | 1 | FALSE | 1 | FALSE | 1 |
| 5411 | 5411 | 10.3125 | 2.11301769 | 0 | 1 | 0 | 1 | 1 | 0.44344817 | 0 | 1 | 0 | 1 | 15 | 0.26127859 | FALSE | 1 | FALSE | 1 | FALSE | 1 | FALSE | 1 | FALSE | 1 | FALSE | 1 |
| 5412 | 5412 | 10.3076923 | 2.14420921 | 0 | 1 | 0 | 1 | 1 | 0.61440466 | 1 | 0.70601446 | 0 | 1 | 24 | 0.20425579 | FALSE | 1 | FALSE | 1 | FALSE | 1 | FALSE | 1 | FALSE | 1 | FALSE | 1 |
| 5413 | 5413 | 8.48 | 2.84422221 | 2 | 0.13369621 | 2 | 0.19297191 | 1 | 0.59998149 | 0 | 1 | 0 | 1 | 20 | 0.8152498 | FALSE | 0.97964162 | FALSE | 1 | FALSE | 1 | FALSE | 1 | FALSE | 1 | FALSE | 1 |
| 5414 | 5414 | 7.8125 | 2.13752437 | 0 | 1 | 3 | 0.19374024 | 0 | 1 | 2 | 0.64277396 | 0 | 1 | 42 | 0.23544056 | FALSE | 1 | FALSE | 1 | FALSE | 1 | FALSE | 1 | FALSE | 1 | FALSE | 1 |
| 5415 | 5415 | 7.32 | 3.23382127 | 0 | 1 | 3 | 0.04547096 | 0 | 1 | 0 | 1 | 0 | 1 | 22 | 0.43642846 | FALSE | 1 | FALSE | 1 | FALSE | 1 | FALSE | 1 | FALSE | 1 | FALSE | 1 |
| 5416 | 5416 | 8.75 | 2.77263413 | 3 | 0.01369554 | 0 | 1 | 1 | 0.51939841 | 1 | 0.60987146 | 0 | 1 | 15 | 0.92113679 | FALSE | 0.36625055 | FALSE | 1 | FALSE | 1 | FALSE | 1 | FALSE | 1 | FALSE | 1 |
| 5417 | 5417 | 9.66666667 | 2.39211668 | 0 | 1 | 2 | 0.05552913 | 1 | 0.35555293 | 0 | 1 | 1 | 0.18007762 | 8 | 0.97197833 | FALSE | 1 | FALSE | 1 | FALSE | 1 | FALSE | 1 | FALSE | 1 | FALSE | 1 |
| 5418 | 5418 | 8.90476191 | 2.06800483 | 0 | 1 | 0 | 1 | 1 | 0.53671415 | 1 | 0.62783522 | 1 | 0.29368245 | 18 | 0.57962763 | FALSE | 1 | FALSE | 1 | FALSE | 1 | FALSE | 1 | FALSE | 1 | FALSE | 1 |
| 5419 | 5419 | 9.47826087 | 2.4824526 | 0 | 1 | 1 | 0.53115654 | 0 | 1 | 1 | 0.6613267 | 0 | 1 | 21 | 0.27964072 | FALSE | 1 | FALSE | 1 | FALSE | 1 | FALSE | 1 | FALSE | 1 | FALSE | 1 |
| 5420 | 5420 | 10.2307692 | 3.26175616 | 1 | 0.28675951 | 0 | 1 | 0 | 1 | 1 | 0.45746571 | 1 | 0.19354507 | 10 | 0.86791642 | FALSE | 1 | FALSE | 1 | FALSE | 1 | FALSE | 1 | FALSE | 1 | FALSE | 1 |
| 5421 | 5421 | 6.90243902 | 2.09301834 | 0 | 1 | 1 | 0.74131698 | 1 | 0.77786601 | 1 | 0.85524728 | 1 | 0.49330646 | 37 | 0.21027077 | FALSE | 1 | FALSE | 1 | FALSE | 1 | FALSE | 1 | FALSE | 1 | FALSE | 1 |
| 5422 | 5422 | 8.26666667 | 1.94821856 | 2 | 0.32143972 | 0 | 1 | 2 | 0.48463745 | 0 | 1 | 2 | 0.16793736 | 39 | 0.4317533 | FALSE | 1 | FALSE | 1 | FALSE | 1 | FALSE | 1 | FALSE | 1 | FALSE | 1 |
| 5423 | 5423 | 9.22727273 | 1.47499825 | 3 | 0.10232687 | 4 | 0.05276567 | 1 | 0.80109494 | 0 | 1 | 0 | 1 | 36 | 0.75909657 | FALSE | 0.87592659 | FALSE | 1 | FALSE | 1 | FALSE | 1 | FALSE | 1 | FALSE | 1 |
| 5424 | 5424 | 9.41666667 | 2.34372685 | 1 | 0.46441868 | 1 | 0.54637326 | 1 | 0.58502114 | 1 | 0.67692816 | 0 | 1 | 20 | 0.68270729 | FALSE | 1 | FALSE | 1 | FALSE | 1 | FALSE | 1 | FALSE | 1 | FALSE | 1 |
| 5425 | 5425 | 9 | 1.87082869 | 1 | 0.26795838 | 2 | 0.05552913 | 0 | 1 | 0 | 1 | 0 | 1 | 9 | 0.89630573 | FALSE | 1 | FALSE | 1 | FALSE | 1 | FALSE | 1 | FALSE | 1 | FALSE | 1 |
| 5426 | 5426 | 9.27272727 | 2.04898685 | 0 | 1 | 0 | 1 | 0 | 1 | 0 | 1 | 0 | 1 | 11 | 0.15414489 | FALSE | 1 | FALSE | 1 | FALSE | 1 | FALSE | 1 | FALSE | 1 | FALSE | 1 |
| 5427 | 5427 | 9.83333333 | 2.9107082 | 0 | 1 | 0 | 1 | 0 | 1 | 2 | 0.19902682 | 0 | 1 | 16 | 0.44998001 | FALSE | 1 | FALSE | 1 | FALSE | 1 | FALSE | 1 | FALSE | 1 | FALSE | 1 |
| 5428 | 5428 | 9.17241379 | 2.49231876 | 0 | 1 | 1 | 0.61539065 | 1 | 0.65463982 | 1 | 0.74482229 | 1 | 0.38148475 | 25 | 0.51779518 | FALSE | 1 | FALSE | 1 | FALSE | 1 | FALSE | 1 | FALSE | 1 | FALSE | 1 |
| 5429 | 5429 | 6.38461539 | 4.16096257 | 1 | 0.49162634 | 3 | 0.05021553 | 1 | 0.61440466 | 1 | 0.70601446 | 1 | 0.3499045 | 19 | 0.96054244 | FALSE | 1 | FALSE | 1 | FALSE | 1 | FALSE | 1 | FALSE | 1 | FALSE | 1 |
| 5430 | 5430 | 10.875 | 2.02715934 | 0 | 1 | 1 | 0.40942991 | 1 | 0.44344817 | 1 | 0.52895088 | 0 | 1 | 13 | 0.76799161 | FALSE | 1 | FALSE | 1 | FALSE | 1 | FALSE | 1 | FALSE | 1 | FALSE | 1 |
| 5431 | 5431 | 8.42307692 | 2.00332564 | 1 | 0.49162634 | 2 | 0.20486827 | 0 | 1 | 0 | 1 | 0 | 1 | 23 | 0.40372818 | FALSE | 1 | FALSE | 1 | FALSE | 1 | FALSE | 1 | FALSE | 1 | FALSE | 1 |
| 5432 | 5432 | 7.38709677 | 1.65933176 | 0 | 1 | 2 | 0.80898201 | 1 | 0.96748975 | 3 | 0.80753311 | 2 | 0.45307384 | 85 | 0.03460965 | FALSE | 1 | FALSE | 1 | FALSE | 1 | FALSE | 1 | FALSE | 1 | FALSE | 1 |
| 5433 | 5433 | 8.75 | 2.48746859 | 0 | 1 | 2 | 0.02563923 | 0 | 1 | 1 | 0.31351192 | 0 | 1 | 5 | 0.97551477 | FALSE | 1 | FALSE | 0.82367276 | FALSE | 1 | FALSE | 1 | FALSE | 1 | FALSE | 1 |
| 5434 | 5434 | 8.23809524 | 3.26459743 | 0 | 1 | 2 | 0.14655134 | 0 | 1 | 1 | 0.62783522 | 0 | 1 | 18 | 0.57962763 | FALSE | 1 | FALSE | 1 | FALSE | 1 | FALSE | 1 | FALSE | 1 | FALSE | 1 |
| 5435 | 5435 | 8.27272727 | 4.06964169 | 1 | 0.24866461 | 1 | 0.30368014 | 0 | 1 | 0 | 1 | 0 | 1 | 9 | 0.75985366 | FALSE | 1 | FALSE | 1 | FALSE | 1 | FALSE | 1 | FALSE | 1 | FALSE | 1 |
| 5436 | 5436 | 9.325 | 2.13760965 | 0 | 1 | 1 | 0.7326167 | 1 | 0.76953771 | 3 | 0.27785536 | 2 | 0.13925369 | 33 | 0.71942635 | FALSE | 1 | FALSE | 1 | FALSE | 1 | FALSE | 1 | FALSE | 1 | FALSE | 1 |
| 5437 | 5437 | 6.7761194 | 3.82380952 | 1 | 0.82602167 | 1 | 0.89073735 | 3 | 0.43448639 | 3 | 0.60027156 | 1 | 0.6714808 | 58 | 0.38535314 | FALSE | 1 | FALSE | 1 | FALSE | 1 | FALSE | 1 | FALSE | 1 | FALSE | 1 |
| 5438 | 5438 | 8.96875 | 3.19652521 | 0 | 1 | 1 | 0.65167362 | 0 | 1 | 2 | 0.43583874 | 0 | 1 | 29 | 0.24088541 | FALSE | 1 | FALSE | 1 | FALSE | 1 | FALSE | 1 | FALSE | 1 | FALSE | 1 |
| 5439 | 5439 | 10.7916667 | 2.36254042 | 1 | 0.46441868 | 1 | 0.54637326 | 1 | 0.58502114 | 1 | 0.67692816 | 0 | 1 | 20 | 0.68270729 | FALSE | 1 | FALSE | 1 | FALSE | 1 | FALSE | 1 | FALSE | 1 | FALSE | 1 |
| 5440 | 5440 | 10.0434783 | 2.62890721 | 1 | 0.45027683 | 1 | 0.53115654 | 2 | 0.19931429 | 1 | 0.6613267 | 1 | 0.31672722 | 17 | 0.94439745 | FALSE | 1 | FALSE | 1 | FALSE | 1 | FALSE | 1 | FALSE | 1 | FALSE | 1 |
| 5441 | 5441 | 10.75 | 2.78762145 | 0 | 1 | 1 | 0.53115654 | 1 | 0.5695037 | 2 | 0.2852236 | 1 | 0.31672722 | 18 | 0.86273155 | FALSE | 1 | FALSE | 1 | FALSE | 1 | FALSE | 1 | FALSE | 1 | FALSE | 1 |
| 5442 | 5442 | 7.74074074 | 2.2042483 | 0 | 1 | 3 | 0.04547096 | 1 | 0.59998149 | 2 | 0.31968852 | 1 | 0.33902672 | 18 | 0.96851178 | FALSE | 1 | FALSE | 1 | FALSE | 1 | FALSE | 1 | FALSE | 1 | FALSE | 1 |
| 5443 | 5443 | 11.3125 | 1.40172884 | 1 | 0.34033054 | 1 | 0.40942991 | 0 | 1 | 0 | 1 | 0 | 1 | 14 | 0.53308261 | FALSE | 1 | FALSE | 1 | FALSE | 1 | FALSE | 1 | FALSE | 1 | FALSE | 1 |
| 5444 | 5444 | 8.59375 | 1.59803346 | 0 | 1 | 1 | 0.65167362 | 1 | 0.69069222 | 3 | 0.17988885 | 0 | 1 | 27 | 0.61671133 | FALSE | 1 | FALSE | 1 | FALSE | 1 | FALSE | 1 | FALSE | 1 | FALSE | 1 |
| 5445 | 5445 | 10 | 2.11223542 | 0 | 1 | 0 | 1 | 0 | 1 | 0 | 1 | 1 | 0.19354507 | 12 | 0.374198 | FALSE | 1 | FALSE | 1 | FALSE | 1 | FALSE | 1 | FALSE | 1 | FALSE | 1 |
| 5446 | 5446 | 9.375 | 1.79843682 | 0 | 1 | 0 | 1 | 0 | 1 | 0 | 1 | 1 | 0.12394282 | 7 | 0.63752397 | FALSE | 1 | FALSE | 1 | FALSE | 1 | FALSE | 1 | FALSE | 1 | FALSE | 1 |
| 5447 | 5447 | 4.22807018 | 3.0378752 | 5 | 0.01493383 | 3 | 0.28014665 | 1 | 0.87682436 | 3 | 0.49002278 | 2 | 0.23984683 | 43 | 0.9744306 | FALSE | 0.38230603 | FALSE | 1 | FALSE | 1 | FALSE | 1 | FALSE | 1 | FALSE | 1 |
| 5448 | 5448 | 7.36734694 | 3.77799146 | 1 | 0.72119464 | 0 | 1 | 0 | 1 | 2 | 0.66536008 | 4 | 0.00822471 | 42 | 0.493842 | FALSE | 1 | FALSE | 1 | FALSE | 1 | FALSE | 1 | FALSE | 0.50674765 | FALSE | 1 |
| 5449 | 5449 | 10 | 2.89827535 | 1 | 0.22886536 | 0 | 1 | 0 | 1 | 0 | 1 | 0 | 1 | 9 | 0.5215547 | FALSE | 1 | FALSE | 1 | FALSE | 1 | FALSE | 1 | FALSE | 1 | FALSE | 1 |
| 5450 | 5450 | 10.1 | 1.75783958 | 0 | 1 | 0 | 1 | 0 | 1 | 1 | 0.37517262 | 0 | 1 | 9 | 0.5215547 | FALSE | 1 | FALSE | 1 | FALSE | 1 | FALSE | 1 | FALSE | 1 | FALSE | 1 |

| | | | | | | | | | | | | | | | | | | | | | | | | | | | |
|---|---|---|---|---|---|---|---|---|---|---|---|---|---|---|---|---|---|---|---|---|---|---|---|---|---|---|---|
| 5451 | 5451 | 8.73684211 | 2.82548748 | 0 | 1 | 1 | 0.46503795 | 1 | 0.50143826 | 1 | 0.59104357 | 2 | 0.03799168 | 14 | 0.93677253 | FALSE | 1 | FALSE | 1 | FALSE | 1 | FALSE | 1 | FALSE | 0.87669812 | FALSE | 1 |
| 5452 | 5452 | 8.76923077 | 1.44046891 | 0 | 1 | 2 | 0.36122049 | 2 | 0.41139386 | 3 | 0.26522531 | 3 | 0.02555742 | 29 | 0.96770202 | FALSE | 1 | FALSE | 1 | FALSE | 1 | FALSE | 1 | FALSE | 0.79106376 | FALSE | 1 |
| 5453 | 5453 | 4.06465068 | 0.63933593 | 34 | 0.0267525 | 40 | 0.04798972 | 38 | 0.26992885 | 46 | 0.38464195 | 18 | 0.29800459 | 779 | 0.99515654 | FALSE | 0.50964275 | FALSE | 1 | FALSE | 1 | FALSE | 1 | FALSE | 1 | FALSE | 1 |
| 5454 | 5454 | 9.91666667 | 1.93469779 | 1 | 0.46441868 | 1 | 0.54637326 | 1 | 0.58502114 | 2 | 0.30249307 | 1 | 0.32796862 | 18 | 0.93166998 | FALSE | 1 | FALSE | 1 | FALSE | 1 | FALSE | 1 | FALSE | 1 | FALSE | 1 |
| 5455 | 5455 | 9.85714286 | 2.19925776 | 1 | 0.3050805 | 0 | 1 | 0 | 1 | 1 | 0.48242099 | 1 | 0.20679328 | 11 | 0.83676449 | FALSE | 1 | FALSE | 1 | FALSE | 1 | FALSE | 1 | FALSE | 1 | FALSE | 1 |
| 5456 | 5456 | 7.38461539 | 2.92307692 | 0 | 1 | 2 | 0.06426041 | 0 | 1 | 1 | 0.45746571 | 0 | 1 | 10 | 0.86791642 | FALSE | 1 | FALSE | 1 | FALSE | 1 | FALSE | 1 | FALSE | 1 | FALSE | 1 |
| 5457 | 5457 | 10.2045455 | 2.25206517 | 0 | 1 | 2 | 0.40761584 | 0 | 1 | 2 | 0.59404271 | 2 | 0.1563365 | 37 | 0.48294184 | FALSE | 1 | FALSE | 1 | FALSE | 1 | FALSE | 1 | FALSE | 1 | FALSE | 1 |
| 5458 | 5458 | 6.54545455 | 1.57808686 | 0 | 1 | 2 | 0.28941836 | 1 | 0.70185639 | 0 | 1 | 1 | 0.42123867 | 29 | 0.39648374 | FALSE | 1 | FALSE | 1 | FALSE | 1 | FALSE | 1 | FALSE | 1 | FALSE | 1 |
| 5459 | 5459 | 10.9285714 | 2.15354478 | 1 | 0.3050805 | 3 | 0.00932935 | 1 | 0.40110418 | 0 | 1 | 0 | 1 | 9 | 0.98607548 | FALSE | 1 | FALSE | 0.64223561 | FALSE | 1 | FALSE | 1 | FALSE | 1 | FALSE | 1 |
| 5460 | 5460 | 9.44444444 | 2.06080411 | 0 | 1 | 1 | 0.25628163 | 1 | 0.28067846 | 1 | 0.34506517 | 0 | 1 | 6 | 0.96128317 | FALSE | 1 | FALSE | 1 | FALSE | 1 | FALSE | 1 | FALSE | 1 | FALSE | 1 |
| 5461 | 5461 | 11.0769231 | 2.46394114 | 0 | 1 | 1 | 0.3480709 | 1 | 0.37874417 | 0 | 1 | 0 | 1 | 11 | 0.6683056 | FALSE | 1 | FALSE | 1 | FALSE | 1 | FALSE | 1 | FALSE | 1 | FALSE | 1 |
| 5462 | 5462 | 2.87398374 | 3.05843728 | 9 | 0.15981036 | 10 | 0.24513059 | 9 | 0.49144802 | 8 | 0.86564468 | 9 | 0.01620831 | 194 | 0.92804024 | FALSE | 1 | FALSE | 1 | FALSE | 1 | FALSE | 1 | FALSE | 0.66728076 | FALSE | 1 |
| 5463 | 5463 | 6.98076923 | 2.29539995 | 3 | 0.50146864 | 5 | 0.24617277 | 4 | 0.5172568 | 3 | 0.86327051 | 0 | 1 | 89 | 0.43117406 | FALSE | 1 | FALSE | 1 | FALSE | 1 | FALSE | 1 | FALSE | 1 | FALSE | 1 |
| 5464 | 5464 | 9.24 | 2.50247877 | 3 | 0.02517314 | 1 | 0.56109832 | 0 | 1 | 3 | 0.10474541 | 0 | 1 | 18 | 0.96851178 | FALSE | 0.4974559 | FALSE | 1 | FALSE | 1 | FALSE | 1 | FALSE | 1 | FALSE | 1 |
| 5465 | 5465 | 4.93220339 | 2.54361552 | 0 | 1 | 3 | 0.2979413 | 2 | 0.63179508 | 4 | 0.28589315 | 0 | 1 | 50 | 0.55668966 | FALSE | 1 | FALSE | 1 | FALSE | 1 | FALSE | 1 | FALSE | 1 | FALSE | 1 |
| 5466 | 5466 | 8.95 | 2.74727137 | 2 | 0.08405377 | 0 | 1 | 2 | 0.14749808 | 0 | 1 | 0 | 1 | 15 | 0.83648191 | FALSE | 0.79236883 | FALSE | 1 | FALSE | 1 | FALSE | 1 | FALSE | 1 | FALSE | 1 |
| 5467 | 5467 | 11 | 3.07482446 | 2 | 0.03087289 | 1 | 0.30368014 | 1 | 0.33149969 | 0 | 1 | 0 | 1 | 7 | 0.98129353 | FALSE | 0.53356688 | FALSE | 1 | FALSE | 1 | FALSE | 1 | FALSE | 1 | FALSE | 1 |
| 5468 | 5468 | 7.7 | 1.97737199 | 1 | 0.40558521 | 1 | 0.48239024 | 0 | 1 | 0 | 1 | 0 | 1 | 18 | 0.37504645 | FALSE | 1 | FALSE | 1 | FALSE | 1 | FALSE | 1 | FALSE | 1 | FALSE | 1 |
| 5469 | 5469 | 8.8125 | 1.62860178 | 0 | 1 | 0 | 1 | 0 | 1 | 0 | 1 | 0 | 1 | 16 | 0.06581507 | FALSE | 1 | FALSE | 1 | FALSE | 1 | FALSE | 1 | FALSE | 1 | FALSE | 1 |
| 5470 | 5470 | 10.0769231 | 1.73034183 | 0 | 1 | 0 | 1 | 1 | 0.37874417 | 0 | 1 | 1 | 0.19354507 | 11 | 0.6683056 | FALSE | 1 | FALSE | 1 | FALSE | 1 | FALSE | 1 | FALSE | 1 | FALSE | 1 |
| 5471 | 5471 | 10.2380952 | 2.28670613 | 3 | 0.01568237 | 2 | 0.14655134 | 0 | 1 | 0 | 1 | 0 | 1 | 16 | 0.90353674 | FALSE | 0.38695764 | FALSE | 1 | FALSE | 1 | FALSE | 1 | FALSE | 1 | FALSE | 1 |
| 5472 | 5472 | 10 | 2.54950976 | 0 | 1 | 1 | 0.32623933 | 0 | 1 | 1 | 0.43131129 | 0 | 1 | 10 | 0.71436964 | FALSE | 1 | FALSE | 1 | FALSE | 1 | FALSE | 1 | FALSE | 1 | FALSE | 1 |
| 5473 | 5473 | 9.55555556 | 3.13088951 | 1 | 0.50471032 | 2 | 0.21684246 | 1 | 0.62830986 | 1 | 0.71956339 | 0 | 1 | 22 | 0.76237877 | FALSE | 1 | FALSE | 1 | FALSE | 1 | FALSE | 1 | FALSE | 1 | FALSE | 1 |
| 5474 | 5474 | 9.64285714 | 2.09104017 | 0 | 1 | 0 | 1 | 1 | 0.40110418 | 1 | 0.48242099 | 0 | 1 | 12 | 0.62236227 | FALSE | 1 | FALSE | 1 | FALSE | 1 | FALSE | 1 | FALSE | 1 | FALSE | 1 |
| 5475 | 5475 | 8.90909091 | 2.89056602 | 0 | 1 | 0 | 1 | 3 | 0.04259419 | 1 | 0.64497439 | 0 | 1 | 18 | 0.74744813 | FALSE | 1 | FALSE | 1 | FALSE | 1 | FALSE | 1 | FALSE | 1 | FALSE | 1 |
| 5476 | 5476 | 1.06340905 | 0.76204682 | 163 | 0.0305784 | 210 | 0.00030053 | 220 | 0.40070818 | 270 | 0.96640242 | 108 | 0.0013424 | 5105 | 0.99588587 | FALSE | 0.53356688 | FALSE | 0.15723145 | FALSE | 1 | FALSE | 1 | FALSE | 0.20625174 | FALSE | 1 |
| 5477 | 5477 | 7.875 | 1.91621372 | 0 | 1 | 1 | 0.65167362 | 2 | 0.32011871 | 1 | 0.77852168 | 0 | 1 | 28 | 0.42532177 | FALSE | 1 | FALSE | 1 | FALSE | 1 | FALSE | 1 | FALSE | 1 | FALSE | 1 |
| 5478 | 5478 | 7.76190476 | 2.58023185 | 0 | 1 | 5 | 0.01085499 | 1 | 0.78589454 | 0 | 1 | 2 | 0.15059682 | 34 | 0.80048036 | FALSE | 1 | FALSE | 0.67617265 | FALSE | 1 | FALSE | 1 | FALSE | 1 | FALSE | 1 |
| 5479 | 5479 | 8.81818182 | 3.12811828 | 0 | 1 | 1 | 0.30368014 | 0 | 1 | 0 | 1 | 0 | 1 | 10 | 0.46859937 | FALSE | 1 | FALSE | 1 | FALSE | 1 | FALSE | 1 | FALSE | 1 | FALSE | 1 |
| 5480 | 5480 | 7.88 | 2.68059695 | 1 | 0.72839502 | 1 | 0.80796869 | 1 | 0.8405438 | 4 | 0.19569978 | 2 | 0.19752014 | 41 | 0.75349894 | FALSE | 1 | FALSE | 1 | FALSE | 1 | FALSE | 1 | FALSE | 1 | FALSE | 1 |
| 5481 | 5481 | 4.73643411 | 2.29431174 | 4 | 0.41693747 | 10 | 0.00828894 | 3 | 0.84479389 | 7 | 0.37305818 | 4 | 0.15800251 | 100 | 0.97808198 | FALSE | 1 | FALSE | 0.62691516 | FALSE | 1 | FALSE | 1 | FALSE | 1 | FALSE | 1 |
| 5482 | 5482 | 4.36774194 | 1.70774084 | 5 | 0.36583077 | 7 | 0.23559042 | 8 | 0.19345514 | 11 | 0.09936503 | 2 | 0.72721321 | 122 | 0.9778376 | FALSE | 1 | FALSE | 1 | FALSE | 1 | FALSE | 1 | FALSE | 1 | FALSE | 1 |
| 5483 | 5483 | 11.25 | 1.58771324 | 1 | 0.26795838 | 0 | 1 | 1 | 0.35555293 | 0 | 1 | 0 | 1 | 10 | 0.71436964 | FALSE | 1 | FALSE | 1 | FALSE | 1 | FALSE | 1 | FALSE | 1 | FALSE | 1 |
| 5484 | 5484 | 9.54545455 | 1.63467927 | 1 | 0.57647348 | 4 | 0.02089081 | 0 | 1 | 2 | 0.45159487 | 0 | 1 | 26 | 0.86852567 | FALSE | 1 | FALSE | 0.8034624 | FALSE | 1 | FALSE | 1 | FALSE | 1 | FALSE | 1 |
| 5485 | 5485 | 10.3125 | 4.92085803 | 0 | 1 | 0 | 1 | 3 | 0.0124223 | 1 | 0.48242099 | 0 | 1 | 10 | 0.94579623 | FALSE | 1 | FALSE | 1 | FALSE | 1 | FALSE | 1 | FALSE | 1 | FALSE | 1 |
| 5486 | 5486 | 9.21428571 | 2.11046953 | 0 | 1 | 1 | 0.36919824 | 1 | 0.40110418 | 0 | 1 | 0 | 1 | 12 | 0.62236227 | FALSE | 1 | FALSE | 1 | FALSE | 1 | FALSE | 1 | FALSE | 1 | FALSE | 1 |
| 5487 | 5487 | 4.28969359 | 1.05274792 | 9 | 0.56680594 | 12 | 0.48450304 | 10 | 0.8317888 | 13 | 0.84183999 | 9 | 0.12880414 | 303 | 0.38111587 | FALSE | 1 | FALSE | 1 | FALSE | 1 | FALSE | 1 | FALSE | 1 | FALSE | 1 |
| 5488 | 5488 | 10 | 2.69037084 | 0 | 1 | 1 | 0.49918218 | 1 | 0.53671415 | 2 | 0.25059627 | 0 | 1 | 17 | 0.77849076 | FALSE | 1 | FALSE | 1 | FALSE | 1 | FALSE | 1 | FALSE | 1 | FALSE | 1 |
| 5489 | 5489 | 9.41666667 | 2.07999733 | 0 | 1 | 0 | 1 | 2 | 0.21260257 | 2 | 0.30249307 | 0 | 1 | 20 | 0.68270729 | FALSE | 1 | FALSE | 1 | FALSE | 1 | FALSE | 1 | FALSE | 1 | FALSE | 1 |
| 5490 | 5490 | 7.88571429 | 1.25973758 | 0 | 1 | 1 | 0.68454804 | 0 | 1 | 1 | 0.80778317 | 1 | 0.44015537 | 32 | 0.18108028 | FALSE | 1 | FALSE | 1 | FALSE | 1 | FALSE | 1 | FALSE | 1 | FALSE | 1 |
| 5491 | 5491 | 10.3333333 | 2.76600681 | 2 | 0.09989125 | 1 | 0.49918218 | 0 | 1 | 0 | 1 | 0 | 1 | 18 | 0.57962763 | FALSE | 0.86047225 | FALSE | 1 | FALSE | 1 | FALSE | 1 | FALSE | 1 | FALSE | 1 |
| 5492 | 5492 | 9 | 2.86972022 | 1 | 0.35728305 | 1 | 0.42857733 | 2 | 0.12280513 | 4 | 0.00644706 | 0 | 1 | 9 | 0.99960431 | FALSE | 1 | FALSE | 1 | FALSE | 1 | FALSE | 1 | FALSE | 1 | FALSE | 1 |
| 5493 | 5493 | 10.0384615 | 2.10275953 | 1 | 0.49162634 | 1 | 0.57534752 | 1 | 0.61440466 | 0 | 1 | 1 | 0.3499045 | 22 | 0.61628 | FALSE | 1 | FALSE | 1 | FALSE | 1 | FALSE | 1 | FALSE | 1 | FALSE | 1 |
| 5494 | 5494 | 9.0625 | 3.05099881 | 2 | 0.06197875 | 1 | 0.40942991 | 0 | 1 | 2 | 0.16544789 | 0 | 1 | 11 | 0.9718531 | FALSE | 0.72532843 | FALSE | 1 | FALSE | 1 | FALSE | 1 | FALSE | 1 | FALSE | 1 |
| 5495 | 5495 | 8.11764706 | 1.81115345 | 0 | 1 | 0 | 1 | 0 | 1 | 0 | 1 | 0 | 1 | 17 | 0.05550965 | FALSE | 1 | FALSE | 1 | FALSE | 1 | FALSE | 1 | FALSE | 1 | FALSE | 1 |
| 5496 | 5496 | 9.64 | 2.26061938 | 0 | 1 | 2 | 0.19297191 | 0 | 1 | 3 | 0.10474541 | 3 | 0.00758617 | 17 | 0.9897469 | FALSE | 1 | FALSE | 1 | FALSE | 1 | FALSE | 1 | FALSE | 0.49339011 | FALSE | 1 |
| 5497 | 5497 | 10 | 1.67017175 | 1 | 0.62830697 | 1 | 0.71433277 | 1 | 0.75193685 | 0 | 1 | 1 | 0.4673875 | 34 | 0.27051027 | FALSE | 1 | FALSE | 1 | FALSE | 1 | FALSE | 1 | FALSE | 1 | FALSE | 1 |
| 5498 | 5498 | 9.36170213 | 1.17465308 | 4 | 0.03163127 | 3 | 0.19374024 | 1 | 0.82190381 | 3 | 0.36699324 | 0 | 1 | 36 | 0.94678267 | FALSE | 0.54437678 | FALSE | 1 | FALSE | 1 | FALSE | 1 | FALSE | 1 | FALSE | 1 |
| 5499 | 5499 | 9.62962963 | 1.3917255 | 2 | 0.15144274 | 2 | 0.21684246 | 0 | 1 | 1 | 0.71956339 | 1 | 0.36060486 | 21 | 0.88395936 | FALSE | 1 | FALSE | 1 | FALSE | 1 | FALSE | 1 | FALSE | 1 | FALSE | 1 |
| 5500 | 5500 | 7.01639344 | 2.04451929 | 1 | 0.79637728 | 5 | 0.04653332 | 2 | 0.64993443 | 1 | 0.94386203 | 1 | 0.63687695 | 51 | 0.64764523 | FALSE | 1 | FALSE | 1 | FALSE | 1 | FALSE | 1 | FALSE | 1 | FALSE | 1 |
| 5501 | 5501 | 5.50697674 | 1.50925753 | 7 | 0.30934719 | 15 | 0.00382073 | 10 | 0.2398827 | 13 | 0.18404486 | 5 | 0.27383025 | 164 | 0.99905254 | FALSE | 1 | FALSE | 0.56907977 | FALSE | 1 | FALSE | 1 | FALSE | 1 | FALSE | 1 |
| 5502 | 5502 | 8 | 2.60768096 | 0 | 1 | 0 | 1 | 1 | 0.30655256 | 0 | 1 | 0 | 1 | 9 | 0.5215547 | FALSE | 1 | FALSE | 1 | FALSE | 1 | FALSE | 1 | FALSE | 1 | FALSE | 1 |
| 5503 | 5503 | 8.73913044 | 4.0132107 | 0 | 1 | 0 | 1 | 1 | 0.5695037 | 1 | 0.6613267 | 0 | 1 | 21 | 0.27964072 | FALSE | 1 | FALSE | 1 | FALSE | 1 | FALSE | 1 | FALSE | 1 | FALSE | 1 |
| 5504 | 5504 | 9.57142857 | 2.17280658 | 0 | 1 | 1 | 0.49918218 | 2 | 0.17308361 | 0 | 1 | 0 | 1 | 18 | 0.57962763 | FALSE | 1 | FALSE | 1 | FALSE | 1 | FALSE | 1 | FALSE | 1 | FALSE | 1 |
| 5505 | 5505 | 7.77586207 | 3.59618142 | 1 | 0.77972342 | 1 | 0.85269953 | 2 | 0.62245518 | 2 | 0.75303313 | 1 | 0.61824432 | 51 | 0.2955095 | FALSE | 1 | FALSE | 1 | FALSE | 1 | FALSE | 1 | FALSE | 1 | FALSE | 1 |
| 5506 | 5506 | 9.57142857 | 3.4685173 | 1 | 0.51745948 | 2 | 0.2288771 | 0 | 1 | 0 | 1 | 0 | 1 | 25 | 0.34293 | FALSE | 1 | FALSE | 1 | FALSE | 1 | FALSE | 1 | FALSE | 1 | FALSE | 1 |
| 5507 | 5507 | 10.0714286 | 1.79141946 | 0 | 1 | 1 | 0.36919824 | 1 | 0.40110418 | 2 | 0.13304705 | 0 | 1 | 10 | 0.94579623 | FALSE | 1 | FALSE | 1 | FALSE | 1 | FALSE | 1 | FALSE | 1 | FALSE | 1 |
| 5508 | 5508 | 9.14285714 | 1.70533377 | 1 | 0.51745948 | 0 | 1 | 1 | 0.64171563 | 1 | 0.73248983 | 0 | 1 | 25 | 0.34293 | FALSE | 1 | FALSE | 1 | FALSE | 1 | FALSE | 1 | FALSE | 1 | FALSE | 1 |
| 5509 | 5509 | 9.33333333 | 1.31233465 | 2 | 0.23542589 | 1 | 0.69480472 | 0 | 1 | 2 | 0.49731009 | 1 | 0.44938266 | 30 | 0.67248271 | FALSE | 1 | FALSE | 1 | FALSE | 1 | FALSE | 1 | FALSE | 1 | FALSE | 1 |
| 5510 | 5510 | 8.6 | 2.4166092 | 1 | 0.32293355 | 0 | 1 | 0 | 1 | 0 | 1 | 1 | 0.2198258 | 13 | 0.57712805 | FALSE | 1 | FALSE | 1 | FALSE | 1 | FALSE | 1 | FALSE | 1 | FALSE | 1 |
| 5511 | 5511 | 7.73333333 | 3.29578856 | 3 | 0.30198574 | 1 | 0.91624631 | 4 | 0.2829119 | 1 | 0.9711236 | 1 | 0.71258573 | 65 | 0.36047181 | FALSE | 1 | FALSE | 1 | FALSE | 1 | FALSE | 1 | FALSE | 1 | FALSE | 1 |
| 5512 | 5512 | 9.96551724 | 4.23029109 | 0 | 1 | 1 | 0.61539065 | 0 | 1 | 1 | 0.74482229 | 5 | 9.38E-05 | 22 | 0.92902102 | FALSE | 1 | FALSE | 1 | FALSE | 1 | FALSE | 1 | TRUE | 0.04046262 | FALSE | 1 |
| 5513 | 5513 | 8.88235294 | 1.90610021 | 1 | 0.34033054 | 1 | 0.40942991 | 0 | 1 | 3 | 0.03440583 | 0 | 1 | 11 | 0.9718531 | FALSE | 1 | FALSE | 1 | FALSE | 1 | FALSE | 1 | FALSE | 1 | FALSE | 1 |
| 5514 | 5514 | 7.15384615 | 1.29175813 | 0 | 1 | 1 | 0.3480709 | 0 | 1 | 0 | 1 | 0 | 1 | 12 | 0.374198 | FALSE | 1 | FALSE | 1 | FALSE | 1 | FALSE | 1 | FALSE | 1 | FALSE | 1 |
| 5515 | 5515 | 9 | 2.91547595 | 1 | 0.26795838 | 0 | 1 | 0 | 1 | 1 | 0.43131129 | 1 | 0.18007762 | 9 | 0.89630573 | FALSE | 1 | FALSE | 1 | FALSE | 1 | FALSE | 1 | FALSE | 1 | FALSE | 1 |
| 5516 | 5516 | 8.08928571 | 2.81752669 | 2 | 0.4229474 | 3 | 0.27128673 | 2 | 0.60323163 | 3 | 0.47819064 | 1 | 0.60529938 | 45 | 0.84580664 | FALSE | 1 | FALSE | 1 | FALSE | 1 | FALSE | 1 | FALSE | 1 | FALSE | 1 |
| 5517 | 5517 | 8.125 | 1.36358901 | 0 | 1 | 1 | 0.23139161 | 0 | 1 | 1 | 0.31351192 | 1 | 0.12394282 | 5 | 0.97551477 | FALSE | 1 | FALSE | 1 | FALSE | 1 | FALSE | 1 | FALSE | 1 | FALSE | 1 |
| 5518 | 5518 | 8.4 | 2.62297541 | 0 | 1 | 1 | 0.56109832 | 0 | 1 | 2 | 0.31968852 | 3 | 0.00758617 | 19 | 0.91734261 | FALSE | 1 | FALSE | 1 | FALSE | 1 | FALSE | 1 | FALSE | 0.49339011 | FALSE | 1 |
| 5519 | 5519 | 7.51515152 | 2.42613562 | 1 | 0.57647348 | 3 | 0.08959205 | 2 | 0.33343871 | 0 | 1 | 2 | 0.10132035 | 25 | 0.93892791 | FALSE | 1 | FALSE | 1 | FALSE | 1 | FALSE | 1 | FALSE | 1 | FALSE | 1 |
| 5520 | 5520 | 8.5483871 | 2.35329158 | 1 | 0.55378205 | 0 | 1 | 0 | 1 | 1 | 0.76781284 | 0 | 1 | 29 | 0.11661149 | FALSE | 1 | FALSE | 1 | FALSE | 1 | FALSE | 1 | FALSE | 1 | FALSE | 1 |
| 5521 | 5521 | 7.9 | 1.44568323 | 0 | 1 | 2 | 0.03949327 | 0 | 1 | 1 | 0.37517262 | 0 | 1 | 7 | 0.94326215 | FALSE | 1 | FALSE | 0.96864929 | FALSE | 1 | FALSE | 1 | FALSE | 1 | FALSE | 1 |
| 5522 | 5522 | 10.6071429 | 2.93183616 | 1 | 0.50471032 | 0 | 1 | 1 | 0.62830986 | 1 | 0.71956339 | 1 | 0.36060486 | 23 | 0.58307573 | FALSE | 1 | FALSE | 1 | FALSE | 1 | FALSE | 1 | FALSE | 1 | FALSE | 1 |
| 5523 | 5523 | 6.62280702 | 1.46533243 | 0 | 1 | 1 | 0.97641742 | 7 | 0.1113069 | 2 | 0.96917371 | 2 | 0.55679953 | 101 | 0.08452682 | FALSE | 1 | FALSE | 1 | FALSE | 1 | FALSE | 1 | FALSE | 1 | FALSE | 1 |
| 5524 | 5524 | 10.0869565 | 1.90908682 | 1 | 0.4357638 | 3 | 0.03266339 | 0 | 1 | 0 | 1 | 2 | 0.04972408 | 16 | 0.95553951 | FALSE | 1 | FALSE | 0.92268438 | FALSE | 1 | FALSE | 1 | FALSE | 0.98629461 | FALSE | 1 |
| 5525 | 5525 | 10.875 | 2.3150324 | 1 | 0.34033054 | 2 | 0.09285443 | 1 | 0.44344817 | 0 | 1 | 2 | 0.02752114 | 10 | 0.99304605 | FALSE | 1 | FALSE | 1 | FALSE | 1 | FALSE | 1 | FALSE | 0.79106376 | FALSE | 1 |
| 5526 | 5526 | 3.40703518 | 2.91765562 | 7 | 0.24897298 | 11 | 0.05739142 | 10 | 0.17822551 | 7 | 0.81575508 | 5 | 0.22745152 | 159 | 0.96565803 | FALSE | 1 | FALSE | 1 | FALSE | 1 | FALSE | 1 | FALSE | 1 | FALSE | 1 |
| 5527 | 5527 | 9.3125 | 1.95955193 | 5 | 0.00671146 | 0 | 1 | 0 | 1 | 3 | 0.36699324 | 0 | 1 | 39 | 0.69224623 | FALSE | 0.24597546 | FALSE | 1 | FALSE | 1 | FALSE | 1 | FALSE | 1 | FALSE | 1 |
| 5528 | 5528 | 9.61538462 | 2.16753889 | 0 | 1 | 1 | 0.3480709 | 0 | 1 | 1 | 0.45746571 | 0 | 1 | 11 | 0.6683056 | FALSE | 1 | FALSE | 1 | FALSE | 1 | FALSE | 1 | FALSE | 1 | FALSE | 1 |
| 5529 | 5529 | 10.5 | 1.91049732 | 0 | 1 | 0 | 1 | 0 | 1 | 0 | 1 | 0 | 1 | 10 | 0.18273166 | FALSE | 1 | FALSE | 1 | FALSE | 1 | FALSE | 1 | FALSE | 1 | FALSE | 1 |
| 5530 | 5530 | 8.37735849 | 3.03580156 | 0 | 1 | 1 | 0.82613965 | 2 | 0.57303317 | 1 | 0.91797304 | 0 | 1 | 49 | 0.06705691 | FALSE | 1 | FALSE | 1 | FALSE | 1 | FALSE | 1 | FALSE | 1 | FALSE | 1 |
| 5531 | 5531 | 8.3 | 2.58392466 | 0 | 1 | 1 | 0.6278847 | 2 | 0.29330414 | 1 | 0.75658797 | 1 | 0.39166989 | 25 | 0.67630471 | FALSE | 1 | FALSE | 1 | FALSE | 1 | FALSE | 1 | FALSE | 1 | FALSE | 1 |
| 5532 | 5532 | 9.07142857 | 2.47744931 | 1 | 0.51745948 | 1 | 0.60247911 | 0 | 1 | 2 | 0.3705495 | 0 | 1 | 24 | 0.55018107 | FALSE | 1 | FALSE | 1 | FALSE | 1 | FALSE | 1 | FALSE | 1 | FALSE | 1 |
| 5533 | 5533 | 9.86206897 | 2.47364227 | 1 | 0.50471032 | 0 | 1 | 1 | 0.62830986 | 2 | 0.35374209 | 0 | 1 | 23 | 0.58307573 | FALSE | 1 | FALSE | 1 | FALSE | 1 | FALSE | 1 | FALSE | 1 | FALSE | 1 |
| 5534 | 5534 | 8 | 2.86744176 | 0 | 1 | 1 | 0.58913618 | 1 | 0.62830986 | 3 | 0.12475952 | 0 | 1 | 22 | 0.76237877 | FALSE | 1 | FALSE | 1 | FALSE | 1 | FALSE | 1 | FALSE | 1 | FALSE | 1 |
| 5535 | 5535 | 9.91666667 | 2.05986785 | 0 | 1 | 0 | 1 | 0 | 1 | 1 | 0.43131129 | 0 | 1 | 11 | 0.41944532 | FALSE | 1 | FALSE | 1 | FALSE | 1 | FALSE | 1 | FALSE | 1 | FALSE | 1 |
| 5536 | 5536 | 9.32653061 | 1.64611795 | 3 | 0.11868304 | 0 | 1 | 0 | 1 | 2 | 0.64277396 | 2 | 0.17968152 | 40 | 0.54368682 | FALSE | 0.94393345 | FALSE | 1 | FALSE | 1 | FALSE | 1 | FALSE | 1 | FALSE | 1 |
| 5537 | 5537 | 9.85714286 | 2.07675833 | 0 | 1 | 0 | 1 | 0 | 1 | 2 | 0.25059627 | 0 | 1 | 19 | 0.34094959 | FALSE | 1 | FALSE | 1 | FALSE | 1 | FALSE | 1 | FALSE | 1 | FALSE | 1 |
| 5538 | 5538 | 8.45833333 | 2.48292782 | 0 | 1 | 4 | 0.00679919 | 1 | 0.58502114 | 0 | 1 | 1 | 0.32796862 | 18 | 0.93166998 | FALSE | 1 | FALSE | 0.60103504 | FALSE | 1 | FALSE | 1 | FALSE | 1 | FALSE | 1 |
| 5539 | 5539 | 8.82142857 | 4.44050811 | 1 | 0.51745948 | 2 | 0.2288771 | 3 | 0.07756639 | 0 | 1 | 1 | 0.37113067 | 21 | 0.94083768 | FALSE | 1 | FALSE | 1 | FALSE | 1 | FALSE | 1 | FALSE | 1 | FALSE | 1 |
| 5540 | 5540 | 5.83823529 | 1.88604494 | 2 | 0.96989994 | 4 | 0.90209042 | 3 | 0.97991365 | 13 | 0.14394446 | 5 | 0.24265164 | 177 | 0.19776316 | FALSE | 1 | FALSE | 1 | FALSE | 1 | FALSE | 1 | FALSE | 1 | FALSE | 1 |
| 5541 | 5541 | 5.63157895 | 2.61551361 | 1 | 0.91444814 | 1 | 0.95551644 | 5 | 0.2484594 | 2 | 0.93467067 | 1 | 0.79090173 | 84 | 0.11223893 | FALSE | 1 | FALSE | 1 | FALSE | 1 | FALSE | 1 | FALSE | 1 | FALSE | 1 |
| 5542 | 5542 | 5.21428571 | 2.82662271 | 2 | 0.36837609 | 4 | 0.07727212 | 1 | 0.8405438 | 2 | 0.6762159 | 1 | 0.56379725 | 40 | 0.85325204 | FALSE | 1 | FALSE | 1 | FALSE | 1 | FALSE | 1 | FALSE | 1 | FALSE | 1 |
| 5543 | 5543 | 6.7486911 | 1.58846822 | 6 | 0.36071492 | 5 | 0.74233599 | 7 | 0.52799419 | 8 | 0.65073122 | 0 | 1 | 164 | 0.2638706 | FALSE | 1 | FALSE | 1 | FALSE | 1 | FALSE | 1 | FALSE | 1 | FALSE | 1 |
| 5544 | 5544 | 8.71428571 | 2.51931316 | 1 | 0.51745948 | 3 | 0.06039219 | 0 | 1 | 1 | 0.73248983 | 1 | 0.37113067 | 22 | 0.86499992 | FALSE | 1 | FALSE | 1 | FALSE | 1 | FALSE | 1 | FALSE | 1 | FALSE | 1 |

| | | | | | | | | | | | | | | | | | | | | | | | | | | | |
|---|---|---|---|---|---|---|---|---|---|---|---|---|---|---|---|---|---|---|---|---|---|---|---|---|---|---|---|
| 5545 | 5545 | 9.17391304 | 1.37215077 | 2 | 0.11647716 | 1 | 0.53115654 | 0 | 1 | 0 | 1 | 4 | 0.00047637 | 16 | 0.98100709 | FALSE | 0.93181727 | FALSE | 1 | FALSE | 1 | FALSE | 1 | FALSE | 0.10840111 | FALSE | 1 |
| 5546 | 5546 | 8.55555556 | 2.09290394 | 6 | 0.00092434 | 4 | 0.05650644 | 1 | 0.80828716 | 1 | 0.8802008 | 1 | 0.52593041 | 32 | 0.99319189 | FALSE | 0.07211611 | FALSE | 1 | FALSE | 1 | FALSE | 1 | FALSE | 1 | FALSE | 1 |
| 5547 | 5547 | 10.5555556 | 2.45452467 | 0 | 1 | 0 | 1 | 0 | 1 | 0 | 1 | 0 | 1 | 9 | 0.21661397 | FALSE | 1 | FALSE | 1 | FALSE | 1 | FALSE | 1 | FALSE | 1 | FALSE | 1 |
| 5548 | 5548 | 11.0833333 | 2.7220805 | 0 | 1 | 0 | 1 | 1 | 0.35555293 | 1 | 0.43131129 | 0 | 1 | 10 | 0.71436964 | FALSE | 1 | FALSE | 1 | FALSE | 1 | FALSE | 1 | FALSE | 1 | FALSE | 1 |
| 5549 | 5549 | 2.43153527 | 1.25919083 | 13 | 0.43289587 | 10 | 0.94668131 | 12 | 0.92386932 | 24 | 0.32539309 | 6 | 0.79424348 | 406 | 0.14349646 | FALSE | 1 | FALSE | 1 | FALSE | 1 | FALSE | 1 | FALSE | 1 | FALSE | 1 |
| 5550 | 5550 | 9.02040816 | 1.65972454 | 0 | 1 | 2 | 0.47396556 | 3 | 0.25725031 | 1 | 0.90086419 | 4 | 0.00822471 | 39 | 0.86798612 | FALSE | 1 | FALSE | 1 | FALSE | 1 | FALSE | 1 | FALSE | 0.50674765 | FALSE | 1 |
| 5551 | 5551 | 4.18548387 | 0.86478541 | 1 | 0.96021623 | 3 | 0.76620212 | 4 | 0.65050247 | 4 | 0.82348102 | 1 | 0.87155875 | 110 | 0.07107783 | FALSE | 1 | FALSE | 1 | FALSE | 1 | FALSE | 1 | FALSE | 1 | FALSE | 1 |
| 5552 | 5552 | 5.55555556 | 2.71256792 | 1 | 0.69044531 | 2 | 0.43020462 | 4 | 0.07687709 | 0 | 1 | 0 | 1 | 38 | 0.59437745 | FALSE | 1 | FALSE | 1 | FALSE | 1 | FALSE | 1 | FALSE | 1 | FALSE | 1 |
| 5553 | 5553 | 6.38095238 | 2.81395937 | 3 | 0.19211919 | 3 | 0.2979413 | 3 | 0.35626822 | 4 | 0.28589315 | 3 | 0.07204991 | 43 | 0.99285314 | FALSE | 1 | FALSE | 1 | FALSE | 1 | FALSE | 1 | FALSE | 1 | FALSE | 1 |
| 5554 | 5554 | 10.1875 | 1.77548409 | 0 | 1 | 0 | 1 | 0 | 1 | 1 | 0.52895088 | 0 | 1 | 15 | 0.26127859 | FALSE | 1 | FALSE | 1 | FALSE | 1 | FALSE | 1 | FALSE | 1 | FALSE | 1 |
| 5555 | 5555 | 4.49557522 | 1.97830666 | 16 | 0.00017637 | 14 | 0.01304986 | 7 | 0.69653024 | 7 | 0.89376155 | 5 | 0.30259398 | 174 | 0.99587089 | TRUE | 0.02491133 | FALSE | 0.69207862 | FALSE | 1 | FALSE | 1 | FALSE | 1 | FALSE | 1 |
| 5556 | 5556 | 8.17391304 | 2.37106056 | 0 | 1 | 2 | 0.16948764 | 2 | 0.19931429 | 2 | 0.2852236 | 1 | 0.31672722 | 16 | 0.98100709 | FALSE | 1 | FALSE | 1 | FALSE | 1 | FALSE | 1 | FALSE | 1 | FALSE | 1 |
| 5557 | 5557 | 11.2 | 1.6 | 0 | 1 | 0 | 1 | 0 | 1 | 0 | 1 | 0 | 1 | 5 | 0.42761893 | FALSE | 1 | FALSE | 1 | FALSE | 1 | FALSE | 1 | FALSE | 1 | FALSE | 1 |
| 5558 | 5558 | 8.30769231 | 1.32343466 | 0 | 1 | 2 | 0.05552913 | 1 | 0.35555293 | 0 | 1 | 0 | 1 | 9 | 0.89630573 | FALSE | 1 | FALSE | 1 | FALSE | 1 | FALSE | 1 | FALSE | 1 | FALSE | 1 |
| 5559 | 5559 | 9.70588235 | 2.80261865 | 3 | 0.00864151 | 1 | 0.42857733 | 1 | 0.46348836 | 1 | 0.55062772 | 0 | 1 | 11 | 0.98974663 | FALSE | 0.27019545 | FALSE | 1 | FALSE | 1 | FALSE | 1 | FALSE | 1 | FALSE | 1 |
| 5560 | 5560 | 11.6774194 | 3.55043929 | 4 | 0.00764535 | 3 | 0.07729739 | 1 | 0.67911179 | 3 | 0.16836568 | 0 | 1 | 20 | 0.9984454 | FALSE | 0.2566832 | FALSE | 1 | FALSE | 1 | FALSE | 1 | FALSE | 1 | FALSE | 1 |
| 5561 | 5561 | 9.97727273 | 2.42607648 | 0 | 1 | 0 | 1 | 2 | 0.47281647 | 1 | 0.87439647 | 0 | 1 | 41 | 0.07069612 | FALSE | 1 | FALSE | 1 | FALSE | 1 | FALSE | 1 | FALSE | 1 | FALSE | 1 |
| 5562 | 5562 | 9.025 | 1.72463764 | 1 | 0.64722818 | 0 | 1 | 2 | 0.42396974 | 3 | 0.27785536 | 0 | 1 | 34 | 0.56328904 | FALSE | 1 | FALSE | 1 | FALSE | 1 | FALSE | 1 | FALSE | 1 | FALSE | 1 |
| 5563 | 5563 | 11.2941177 | 1.90246607 | 0 | 1 | 0 | 1 | 0 | 1 | 1 | 0.55062772 | 0 | 1 | 16 | 0.23070182 | FALSE | 1 | FALSE | 1 | FALSE | 1 | FALSE | 1 | FALSE | 1 | FALSE | 1 |
| 5564 | 5564 | 7.71428571 | 3.02202797 | 0 | 1 | 1 | 0.60247911 | 2 | 0.2663542 | 2 | 0.3705495 | 0 | 1 | 23 | 0.73439118 | FALSE | 1 | FALSE | 1 | FALSE | 1 | FALSE | 1 | FALSE | 1 | FALSE | 1 |
| 5565 | 5565 | 10.1538462 | 1.87477809 | 0 | 1 | 1 | 0.3480709 | 0 | 1 | 0 | 1 | 0 | 1 | 12 | 0.374198 | FALSE | 1 | FALSE | 1 | FALSE | 1 | FALSE | 1 | FALSE | 1 | FALSE | 1 |
| 5566 | 5566 | 6.95876289 | 1.89365476 | 5 | 0.10296654 | 8 | 0.01289603 | 5 | 0.26902724 | 7 | 0.15687468 | 0 | 1 | 72 | 0.99689715 | FALSE | 0.87864781 | FALSE | 0.69207862 | FALSE | 1 | FALSE | 1 | FALSE | 1 | FALSE | 1 |
| 5567 | 5567 | 11.0714286 | 2.43381786 | 1 | 0.3050805 | 0 | 1 | 0 | 1 | 0 | 1 | 0 | 1 | 13 | 0.33283729 | FALSE | 1 | FALSE | 1 | FALSE | 1 | FALSE | 1 | FALSE | 1 | FALSE | 1 |
| 5568 | 5568 | 11.0666667 | 2.79205221 | 0 | 1 | 3 | 0.01138739 | 0 | 1 | 2 | 0.14906763 | 0 | 1 | 10 | 0.97980046 | FALSE | 1 | FALSE | 0.67617265 | FALSE | 1 | FALSE | 1 | FALSE | 1 | FALSE | 1 |
| 5569 | 5569 | 6.69230769 | 1.66646942 | 0 | 1 | 1 | 0.7236252 | 1 | 0.76089851 | 2 | 0.54052736 | 1 | 0.4761699 | 34 | 0.41570781 | FALSE | 1 | FALSE | 1 | FALSE | 1 | FALSE | 1 | FALSE | 1 | FALSE | 1 |
| 5570 | 5570 | 10.2857143 | 1.82946407 | 0 | 1 | 1 | 0.36919824 | 0 | 1 | 1 | 0.48242099 | 1 | 0.20679328 | 11 | 0.83676449 | FALSE | 1 | FALSE | 1 | FALSE | 1 | FALSE | 1 | FALSE | 1 | FALSE | 1 |
| 5571 | 5571 | 10 | 3.47850543 | 1 | 0.40558521 | 2 | 0.1353446 | 0 | 1 | 1 | 0.60987146 | 1 | 0.28187291 | 15 | 0.92113679 | FALSE | 1 | FALSE | 1 | FALSE | 1 | FALSE | 1 | FALSE | 1 | FALSE | 1 |
| 5572 | 5572 | 7.2195122 | 2.95917091 | 1 | 0.65632651 | 0 | 1 | 2 | 0.43640583 | 2 | 0.56788092 | 0 | 1 | 36 | 0.36455032 | FALSE | 1 | FALSE | 1 | FALSE | 1 | FALSE | 1 | FALSE | 1 | FALSE | 1 |
| 5573 | 5573 | 5.34228188 | 1.19153509 | 7 | 0.06571182 | 7 | 0.16315538 | 7 | 0.23246036 | 7 | 0.45668055 | 1 | 0.90193101 | 110 | 0.96300755 | FALSE | 0.74276693 | FALSE | 1 | FALSE | 1 | FALSE | 1 | FALSE | 1 | FALSE | 1 |
| 5574 | 5574 | 9.75 | 1.03077641 | 0 | 1 | 0 | 1 | 0 | 1 | 1 | 0.52895088 | 0 | 1 | 15 | 0.26127859 | FALSE | 1 | FALSE | 1 | FALSE | 1 | FALSE | 1 | FALSE | 1 | FALSE | 1 |
| 5575 | 5575 | 8.03225807 | 3.20769107 | 2 | 0.1881398 | 1 | 0.63997468 | 0 | 1 | 0 | 1 | 0 | 1 | 28 | 0.26393996 | FALSE | 1 | FALSE | 1 | FALSE | 1 | FALSE | 1 | FALSE | 1 | FALSE | 1 |
| 5576 | 5576 | 3.11610738 | 0.91626372 | 58 | 0.00021267 | 75 | 9.91E-06 | 52 | 0.58122888 | 70 | 0.38895451 | 29 | 0.15524462 | 1188 | 0.9999931 | TRUE | 0.02809963 | FALSE | 0.07208458 | FALSE | 1 | FALSE | 1 | FALSE | 1 | FALSE | 1 |
| 5577 | 5577 | 7.35294118 | 2.19467742 | 0 | 1 | 0 | 1 | 0 | 1 | 1 | 0.55062772 | 1 | 0.24525764 | 15 | 0.49060435 | FALSE | 1 | FALSE | 1 | FALSE | 1 | FALSE | 1 | FALSE | 1 | FALSE | 1 |
| 5578 | 5578 | 8.52 | 2.45144855 | 4 | 0.00346297 | 1 | 0.56109832 | 0 | 1 | 0 | 1 | 2 | 0.06257402 | 18 | 0.96851178 | FALSE | 0.16118559 | FALSE | 1 | FALSE | 1 | FALSE | 1 | FALSE | 1 | FALSE | 1 |
| 5579 | 5579 | 9.21621622 | 1.04325725 | 1 | 0.6184714 | 0 | 1 | 1 | 0.74264075 | 0 | 1 | 0 | 1 | 35 | 0.05673524 | FALSE | 1 | FALSE | 1 | FALSE | 1 | FALSE | 1 | FALSE | 1 | FALSE | 1 |
| 5580 | 5580 | 10.3076923 | 2.41420844 | 1 | 0.49162634 | 0 | 1 | 1 | 0.61440466 | 1 | 0.70601446 | 0 | 1 | 23 | 0.40372818 | FALSE | 1 | FALSE | 1 | FALSE | 1 | FALSE | 1 | FALSE | 1 | FALSE | 1 |
| 5581 | 5581 | 8.6984127 | 3.68452328 | 1 | 0.80677819 | 4 | 0.14592986 | 0 | 1 | 3 | 0.55805561 | 1 | 0.64879513 | 54 | 0.46836572 | FALSE | 1 | FALSE | 1 | FALSE | 1 | FALSE | 1 | FALSE | 1 | FALSE | 1 |
| 5582 | 5582 | 10.2692308 | 3.56864219 | 4 | 0.0040119 | 0 | 1 | 1 | 0.61440466 | 1 | 0.70601446 | 0 | 1 | 20 | 0.90142689 | FALSE | 0.17959269 | FALSE | 1 | FALSE | 1 | FALSE | 1 | FALSE | 1 | FALSE | 1 |
| 5583 | 5583 | 6.51612903 | 2.65027044 | 2 | 0.1881398 | 3 | 0.07729739 | 0 | 1 | 1 | 0.76781284 | 1 | 0.4016888 | 24 | 0.90138666 | FALSE | 1 | FALSE | 1 | FALSE | 1 | FALSE | 1 | FALSE | 1 | FALSE | 1 |
| 5584 | 5584 | 9.75 | 1.39194109 | 1 | 0.34033054 | 1 | 0.40942991 | 2 | 0.11087977 | 0 | 1 | 0 | 1 | 12 | 0.90922186 | FALSE | 1 | FALSE | 1 | FALSE | 1 | FALSE | 1 | FALSE | 1 | FALSE | 1 |
| 5585 | 5585 | 11.48 | 1.29984615 | 0 | 1 | 0 | 1 | 2 | 0.21260257 | 2 | 0.30249307 | 0 | 1 | 20 | 0.68270729 | FALSE | 1 | FALSE | 1 | FALSE | 1 | FALSE | 1 | FALSE | 1 | FALSE | 1 |
| 5586 | 5586 | 10.1052632 | 2.19795927 | 0 | 1 | 0 | 1 | 0 | 1 | 0 | 1 | 0 | 1 | 19 | 0.03948376 | FALSE | 1 | FALSE | 1 | FALSE | 1 | FALSE | 1 | FALSE | 1 | FALSE | 1 |
| 5587 | 5587 | 9.3125 | 1.95955193 | 0 | 1 | 0 | 1 | 0 | 1 | 3 | 0.03440583 | 0 | 1 | 13 | 0.76799161 | FALSE | 1 | FALSE | 1 | FALSE | 1 | FALSE | 1 | FALSE | 1 | FALSE | 1 |
| 5588 | 5588 | 9.2 | 2.52190404 | 0 | 1 | 0 | 1 | 2 | 0.04781153 | 0 | 1 | 0 | 1 | 8 | 0.80394719 | FALSE | 1 | FALSE | 1 | FALSE | 1 | FALSE | 1 | FALSE | 1 | FALSE | 1 |
| 5589 | 5589 | 7.08333333 | 1.84654332 | 1 | 0.46441868 | 0 | 1 | 0 | 1 | 1 | 0.67692816 | 0 | 1 | 22 | 0.25237063 | FALSE | 1 | FALSE | 1 | FALSE | 1 | FALSE | 1 | FALSE | 1 | FALSE | 1 |
| 5590 | 5590 | 11.7142857 | 1.03015751 | 0 | 1 | 2 | 0.01964367 | 0 | 1 | 1 | 0.28044368 | 0 | 1 | 4 | 0.98605104 | FALSE | 1 | FALSE | 0.78498014 | FALSE | 1 | FALSE | 1 | FALSE | 1 | FALSE | 1 |
| 5591 | 5591 | 10.631579 | 3.26400668 | 1 | 0.38989951 | 0 | 1 | 1 | 0.50143826 | 0 | 1 | 2 | 0.03799168 | 15 | 0.83648191 | FALSE | 1 | FALSE | 1 | FALSE | 1 | FALSE | 1 | FALSE | 0.87669812 | FALSE | 1 |
| 5592 | 5592 | 7 | 3.17599693 | 5 | 0.00612174 | 1 | 0.78076835 | 1 | 0.81522035 | 3 | 0.35428318 | 0 | 1 | 36 | 0.90704249 | FALSE | 0.23004259 | FALSE | 1 | FALSE | 1 | FALSE | 1 | FALSE | 1 | FALSE | 1 |
| 5593 | 5593 | 9 | 1.37436854 | 1 | 0.37380237 | 0 | 1 | 1 | 0.48280983 | 0 | 1 | 0 | 1 | 16 | 0.44998001 | FALSE | 1 | FALSE | 1 | FALSE | 1 | FALSE | 1 | FALSE | 1 | FALSE | 1 |
| 5594 | 5594 | 6.36585366 | 3.44826643 | 1 | 0.64722818 | 0 | 1 | 2 | 0.42396974 | 1 | 0.84823834 | 2 | 0.13925369 | 34 | 0.56328904 | FALSE | 1 | FALSE | 1 | FALSE | 1 | FALSE | 1 | FALSE | 1 | FALSE | 1 |
| 5595 | 5595 | 8.71875 | 2.60089378 | 1 | 0.56527493 | 0 | 1 | 0 | 1 | 0 | 1 | 0 | 1 | 31 | 0.02992045 | FALSE | 1 | FALSE | 1 | FALSE | 1 | FALSE | 1 | FALSE | 1 | FALSE | 1 |
| 5596 | 5596 | 8.7 | 2.57099203 | 0 | 1 | 0 | 1 | 1 | 0.30655256 | 1 | 0.37517262 | 0 | 1 | 8 | 0.80394719 | FALSE | 1 | FALSE | 1 | FALSE | 1 | FALSE | 1 | FALSE | 1 | FALSE | 1 |
| 5597 | 5597 | 8.36363636 | 1.00959583 | 2 | 0.20692137 | 2 | 0.28941836 | 2 | 0.33343871 | 2 | 0.45159487 | 2 | 0.10132035 | 23 | 0.99096587 | FALSE | 1 | FALSE | 1 | FALSE | 1 | FALSE | 1 | FALSE | 1 | FALSE | 1 |
| 5598 | 5598 | 9.0952381 | 3.49018709 | 1 | 0.42086991 | 1 | 0.49918218 | 1 | 0.53671415 | 1 | 0.62783522 | 0 | 1 | 17 | 0.77849076 | FALSE | 1 | FALSE | 1 | FALSE | 1 | FALSE | 1 | FALSE | 1 | FALSE | 1 |
| 5599 | 5599 | 5.68518519 | 3.21353661 | 1 | 0.75539413 | 2 | 0.52572379 | 3 | 0.30658915 | 2 | 0.71679995 | 1 | 0.59191961 | 45 | 0.66857827 | FALSE | 1 | FALSE | 1 | FALSE | 1 | FALSE | 1 | FALSE | 1 | FALSE | 1 |
| 5600 | 5600 | 12.7142857 | 1.48461498 | 0 | 1 | 0 | 1 | 1 | 0.22601089 | 0 | 1 | 1 | 0.10932414 | 5 | 0.91847773 | FALSE | 1 | FALSE | 1 | FALSE | 1 | FALSE | 1 | FALSE | 1 | FALSE | 1 |
| 5601 | 5601 | 9.64285714 | 2.68879003 | 0 | 1 | 0 | 1 | 0 | 1 | 1 | 0.48242099 | 0 | 1 | 13 | 0.33283729 | FALSE | 1 | FALSE | 1 | FALSE | 1 | FALSE | 1 | FALSE | 1 | FALSE | 1 |
| 5602 | 5602 | 8.32 | 1.89145447 | 4 | 0.00346297 | 1 | 0.56109832 | 2 | 0.22597287 | 1 | 0.69181315 | 1 | 0.33902672 | 16 | 0.99713666 | FALSE | 0.16118559 | FALSE | 1 | FALSE | 1 | FALSE | 1 | FALSE | 1 | FALSE | 1 |
| 5603 | 5603 | 4.46835443 | 2.84554034 | 5 | 0.04702199 | 4 | 0.23789896 | 4 | 0.29916433 | 3 | 0.69372972 | 3 | 0.13196291 | 58 | 0.9872953 | FALSE | 0.649301 | FALSE | 1 | FALSE | 1 | FALSE | 1 | FALSE | 1 | FALSE | 1 |
| 5604 | 5604 | 9.22222222 | 1.95946579 | 1 | 0.37380237 | 0 | 1 | 0 | 1 | 1 | 0.57131013 | 0 | 1 | 16 | 0.44998001 | FALSE | 1 | FALSE | 1 | FALSE | 1 | FALSE | 1 | FALSE | 1 | FALSE | 1 |
| 5605 | 5605 | 8 | 2.06155281 | 3 | 0.08207756 | 3 | 0.13823565 | 0 | 1 | 2 | 0.5543522 | 3 | 0.02730775 | 29 | 0.98400908 | FALSE | 0.79236883 | FALSE | 1 | FALSE | 1 | FALSE | 1 | FALSE | 0.79106376 | FALSE | 1 |
| 5606 | 5606 | 10.8064516 | 1.65430728 | 0 | 1 | 1 | 0.6278847 | 2 | 0.29330414 | 3 | 0.15707004 | 0 | 1 | 24 | 0.82294782 | FALSE | 1 | FALSE | 1 | FALSE | 1 | FALSE | 1 | FALSE | 1 | FALSE | 1 |
| 5607 | 5607 | 9.35714286 | 1.34202102 | 0 | 1 | 0 | 1 | 0 | 1 | 1 | 0.48242099 | 0 | 1 | 13 | 0.33283729 | FALSE | 1 | FALSE | 1 | FALSE | 1 | FALSE | 1 | FALSE | 1 | FALSE | 1 |
| 5608 | 5608 | 8.06060606 | 2.76273244 | 2 | 0.19750261 | 1 | 0.65167362 | 2 | 0.32011871 | 2 | 0.43583874 | 2 | 0.096172 | 23 | 0.97984382 | FALSE | 1 | FALSE | 1 | FALSE | 1 | FALSE | 1 | FALSE | 1 | FALSE | 1 |
| 5609 | 5609 | 7.825 | 2.68223321 | 0 | 1 | 1 | 0.7326167 | 2 | 0.42396974 | 1 | 0.84823834 | 0 | 1 | 36 | 0.22908535 | FALSE | 1 | FALSE | 1 | FALSE | 1 | FALSE | 1 | FALSE | 1 | FALSE | 1 |
| 5610 | 5610 | 8.24242424 | 2.4498646 | 3 | 0.05151181 | 2 | 0.28941836 | 0 | 1 | 1 | 0.78873814 | 1 | 0.42123867 | 26 | 0.86852567 | FALSE | 0.67646333 | FALSE | 1 | FALSE | 1 | FALSE | 1 | FALSE | 1 | FALSE | 1 |
| 5611 | 5611 | 10.5 | 4.88620507 | 0 | 1 | 0 | 1 | 0 | 1 | 1 | 0.52895088 | 3 | 0.00205574 | 12 | 0.90922186 | FALSE | 1 | FALSE | 1 | FALSE | 1 | FALSE | 1 | FALSE | 0.2617074 | FALSE | 1 |
| 5612 | 5612 | 10 | 3.38445645 | 1 | 0.24866461 | 0 | 1 | 0 | 1 | 0 | 1 | 0 | 1 | 10 | 0.46859937 | FALSE | 1 | FALSE | 1 | FALSE | 1 | FALSE | 1 | FALSE | 1 | FALSE | 1 |
| 5613 | 5613 | 4.71794872 | 2.12372095 | 3 | 0.32318196 | 3 | 0.46497905 | 0 | 1 | 7 | 0.06592888 | 1 | 0.72664912 | 64 | 0.77133015 | FALSE | 1 | FALSE | 1 | FALSE | 1 | FALSE | 1 | FALSE | 1 | FALSE | 1 |
| 5614 | 5614 | 6.46153846 | 1.477358 | 3 | 0.40746908 | 2 | 0.79413193 | 7 | 0.04237488 | 6 | 0.2314164 | 0 | 1 | 72 | 0.89981553 | FALSE | 1 | FALSE | 1 | FALSE | 1 | FALSE | 1 | FALSE | 1 | FALSE | 1 |
| 5615 | 5615 | 10.1764706 | 1.97738074 | 2 | 0.06909038 | 0 | 1 | 0 | 1 | 0 | 1 | 0 | 1 | 15 | 0.49060435 | FALSE | 0.74276693 | FALSE | 1 | FALSE | 1 | FALSE | 1 | FALSE | 1 | FALSE | 1 |
| 5616 | 5616 | 8.03125 | 2.69819818 | 1 | 0.56527493 | 2 | 0.27730767 | 3 | 0.10599423 | 1 | 0.77852168 | 0 | 1 | 25 | 0.88559288 | FALSE | 1 | FALSE | 1 | FALSE | 1 | FALSE | 1 | FALSE | 1 | FALSE | 1 |
| 5617 | 5617 | 6.58823529 | 3.08809524 | 0 | 1 | 0 | 1 | 0 | 1 | 0 | 1 | 0 | 1 | 17 | 0.05550965 | FALSE | 1 | FALSE | 1 | FALSE | 1 | FALSE | 1 | FALSE | 1 | FALSE | 1 |
| 5618 | 5618 | 10.7222222 | 3.01488079 | 0 | 1 | 0 | 1 | 1 | 0.48280983 | 0 | 1 | 0 | 1 | 17 | 0.20329205 | FALSE | 1 | FALSE | 1 | FALSE | 1 | FALSE | 1 | FALSE | 1 | FALSE | 1 |
| 5619 | 5619 | 9.8 | 3.42928564 | 0 | 1 | 0 | 1 | 1 | 0.16720719 | 1 | 0.2094688 | 0 | 1 | 3 | 0.97031021 | FALSE | 1 | FALSE | 1 | FALSE | 1 | FALSE | 1 | FALSE | 1 | FALSE | 1 |
| 5620 | 5620 | 8.91666667 | 2.13924961 | 0 | 1 | 0 | 1 | 1 | 0.35555293 | 2 | 0.1023796 | 0 | 1 | 9 | 0.89630573 | FALSE | 1 | FALSE | 1 | FALSE | 1 | FALSE | 1 | FALSE | 1 | FALSE | 1 |
| 5621 | 5621 | 9.3125 | 3.10682213 | 1 | 0.55378205 | 4 | 0.01688692 | 2 | 0.30673593 | 0 | 1 | 0 | 1 | 24 | 0.90138666 | FALSE | 1 | FALSE | 0.73194511 | FALSE | 1 | FALSE | 1 | FALSE | 1 | FALSE | 1 |
| 5622 | 5622 | 9.09090909 | 2.93736262 | 0 | 1 | 1 | 0.30368014 | 0 | 1 | 1 | 0.40390032 | 0 | 1 | 9 | 0.75985366 | FALSE | 1 | FALSE | 1 | FALSE | 1 | FALSE | 1 | FALSE | 1 | FALSE | 1 |
| 5623 | 5623 | 8.75438597 | 1.73994053 | 1 | 0.76787545 | 2 | 0.54544984 | 0 | 1 | 0 | 1 | 1 | 0.60529938 | 52 | 0.04900615 | FALSE | 1 | FALSE | 1 | FALSE | 1 | FALSE | 1 | FALSE | 1 | FALSE | 1 |
| 5624 | 5624 | 10.9090909 | 2.90624861 | 0 | 1 | 1 | 0.5154318 | 0 | 1 | 0 | 1 | 0 | 1 | 21 | 0.12048848 | FALSE | 1 | FALSE | 1 | FALSE | 1 | FALSE | 1 | FALSE | 1 | FALSE | 1 |
| 5625 | 5625 | 11.5 | 3.57071421 | 0 | 1 | 0 | 1 | 0 | 1 | 0 | 1 | 1 | 0.18007762 | 11 | 0.41944532 | FALSE | 1 | FALSE | 1 | FALSE | 1 | FALSE | 1 | FALSE | 1 | FALSE | 1 |
| 5626 | 5626 | 9.25 | 2.00520157 | 0 | 1 | 0 | 1 | 0 | 1 | 0 | 1 | 0 | 1 | 12 | 0.13002667 | FALSE | 1 | FALSE | 1 | FALSE | 1 | FALSE | 1 | FALSE | 1 | FALSE | 1 |
| 5627 | 5627 | 9.73214286 | 1.14160087 | 1 | 0.76787545 | 1 | 0.84259834 | 3 | 0.32646887 | 1 | 0.92884295 | 1 | 0.60529938 | 49 | 0.33483172 | FALSE | 1 | FALSE | 1 | FALSE | 1 | FALSE | 1 | FALSE | 1 | FALSE | 1 |
| 5628 | 5628 | 9.85714286 | 2.29462549 | 0 | 1 | 0 | 1 | 0 | 1 | 1 | 0.28044368 | 0 | 1 | 6 | 0.69920685 | FALSE | 1 | FALSE | 1 | FALSE | 1 | FALSE | 1 | FALSE | 1 | FALSE | 1 |
| 5629 | 5629 | 7.77272727 | 2.13055303 | 1 | 0.4357638 | 0 | 1 | 1 | 0.55340851 | 1 | 0.64497439 | 1 | 0.30529951 | 18 | 0.74744813 | FALSE | 1 | FALSE | 1 | FALSE | 1 | FALSE | 1 | FALSE | 1 | FALSE | 1 |
| 5630 | 5630 | 7.64705882 | 2.74271605 | 1 | 0.35728305 | 0 | 1 | 1 | 0.46348836 | 1 | 0.55062772 | 0 | 1 | 14 | 0.7313145 | FALSE | 1 | FALSE | 1 | FALSE | 1 | FALSE | 1 | FALSE | 1 | FALSE | 1 |
| 5631 | 5631 | 6.10638298 | 3.78256944 | 1 | 0.70621968 | 1 | 0.78790827 | 3 | 0.23782059 | 3 | 0.36699324 | 0 | 1 | 39 | 0.69224623 | FALSE | 1 | FALSE | 1 | FALSE | 1 | FALSE | 1 | FALSE | 1 | FALSE | 1 |
| 5632 | 5632 | 9.64705882 | 1.90791468 | 0 | 1 | 2 | 0.08295839 | 2 | 0.09928574 | 0 | 1 | 0 | 1 | 11 | 0.92879412 | FALSE | 1 | FALSE | 1 | FALSE | 1 | FALSE | 1 | FALSE | 1 | FALSE | 1 |
| 5633 | 5633 | 10.3157895 | 2.53507254 | 0 | 1 | 0 | 1 | 0 | 1 | 2 | 0.21610812 | 2 | 0.03799168 | 15 | 0.83648191 | FALSE | 1 | FALSE | 1 | FALSE | 1 | FALSE | 1 | FALSE | 0.87669812 | FALSE | 1 |
| 5634 | 5634 | 10.4347826 | 2.4641091 | 0 | 1 | 1 | 0.53115654 | 1 | 0.5695037 | 0 | 1 | 0 | 1 | 21 | 0.27964072 | FALSE | 1 | FALSE | 1 | FALSE | 1 | FALSE | 1 | FALSE | 1 | FALSE | 1 |
| 5635 | 5635 | 10.3333333 | 3.19722102 | 0 | 1 | 0 | 1 | 0 | 1 | 1 | 0.43131129 | 0 | 1 | 11 | 0.41944532 | FALSE | 1 | FALSE | 1 | FALSE | 1 | FALSE | 1 | FALSE | 1 | FALSE | 1 |
| 5636 | 5636 | 10.125 | 3.85478599 | 0 | 1 | 1 | 0.54637326 | 1 | 0.58502114 | 1 | 0.67692816 | 0 | 1 | 21 | 0.47053137 | FALSE | 1 | FALSE | 1 | FALSE | 1 | FALSE | 1 | FALSE | 1 | FALSE | 1 |
| 5637 | 5637 | 11.2 | 2.31516738 | 1 | 0.32293355 | 1 | 0.38964394 | 1 | 0.42266264 | 1 | 0.50623194 | 0 | 1 | 11 | 0.92879412 | FALSE | 1 | FALSE | 1 | FALSE | 1 | FALSE | 1 | FALSE | 1 | FALSE | 1 |
| 5638 | 5638 | 8.41176471 | 2.05966369 | 1 | 0.35728305 | 0 | 1 | 2 | 0.12280513 | 0 | 1 | 0 | 1 | 14 | 0.7313145 | FALSE | 1 | FALSE | 1 | FALSE | 1 | FALSE | 1 | FALSE | 1 | FALSE | 1 |

| | | | | | | | | | | | | | | | | | | | | | | | | | | | |
|---|---|---|---|---|---|---|---|---|---|---|---|---|---|---|---|---|---|---|---|---|---|---|---|---|---|---|---|
| 5639 | 5639 | 8.06451613 | 2.32705662 | 1 | 0.55378205 | 2 | 0.2651851 | 2 | 0.30673593 | 1 | 0.76781284 | 1 | 0.4016888 | 24 | 0.90138666 | FALSE | 1 | FALSE | 1 | FALSE | 1 | FALSE | 1 | FALSE | 1 | FALSE | 1 |
| 5640 | 5640 | 4.73913044 | 3.43912769 | 5 | 0.15553912 | 6 | 0.14967257 | 4 | 0.56918099 | 5 | 0.5821433 | 1 | 0.8428205 | 90 | 0.86278628 | FALSE | 1 | FALSE | 1 | FALSE | 1 | FALSE | 1 | FALSE | 1 | FALSE | 1 |
| 5641 | 5641 | 9 | 4.55521679 | 0 | 1 | 0 | 1 | 0 | 1 | 1 | 0.31351192 | 0 | 1 | 7 | 0.63752397 | FALSE | 1 | FALSE | 1 | FALSE | 1 | FALSE | 1 | FALSE | 1 | FALSE | 1 |
| 5642 | 5642 | 8.81481482 | 3.39974985 | 0 | 1 | 0 | 1 | 1 | 0.62830986 | 1 | 0.71956339 | 0 | 1 | 25 | 0.1832206 | FALSE | 1 | FALSE | 1 | FALSE | 1 | FALSE | 1 | FALSE | 1 | FALSE | 1 |
| 5643 | 5643 | 10 | 3.31662479 | 0 | 1 | 0 | 1 | 0 | 1 | 0 | 1 | 0 | 1 | 12 | 0.13002667 | FALSE | 1 | FALSE | 1 | FALSE | 1 | FALSE | 1 | FALSE | 1 | FALSE | 1 |
| 5644 | 5644 | 9.04 | 1.96936538 | 3 | 0.02256792 | 0 | 1 | 1 | 0.58502114 | 0 | 1 | 0 | 1 | 20 | 0.68270729 | FALSE | 0.4692294 | FALSE | 1 | FALSE | 1 | FALSE | 1 | FALSE | 1 | FALSE | 1 |
| 5645 | 5645 | 9.87096774 | 2.87078651 | 1 | 0.55378205 | 2 | 0.2651851 | 0 | 1 | 3 | 0.16836568 | 0 | 1 | 25 | 0.80006983 | FALSE | 1 | FALSE | 1 | FALSE | 1 | FALSE | 1 | FALSE | 1 | FALSE | 1 |
| 5646 | 5646 | 9.3125 | 3.24458684 | 1 | 0.56527493 | 1 | 0.65167362 | 0 | 1 | 2 | 0.43583874 | 2 | 0.096172 | 26 | 0.77612337 | FALSE | 1 | FALSE | 1 | FALSE | 1 | FALSE | 1 | FALSE | 1 | FALSE | 1 |
| 5647 | 5647 | 7.69230769 | 2.91902553 | 1 | 0.28675951 | 0 | 1 | 1 | 0.37874417 | 0 | 1 | 0 | 1 | 11 | 0.6683056 | FALSE | 1 | FALSE | 1 | FALSE | 1 | FALSE | 1 | FALSE | 1 | FALSE | 1 |
| 5648 | 5648 | 10.0952381 | 2.7929292 | 0 | 1 | 1 | 0.49918218 | 2 | 0.17308361 | 0 | 1 | 0 | 1 | 18 | 0.57962763 | FALSE | 1 | FALSE | 1 | FALSE | 1 | FALSE | 1 | FALSE | 1 | FALSE | 1 |
| 5649 | 5649 | 11.2777778 | 2.6833966 | 0 | 1 | 0 | 1 | 0 | 1 | 2 | 0.18212134 | 2 | 0.03086174 | 13 | 0.88719222 | FALSE | 1 | FALSE | 1 | FALSE | 1 | FALSE | 1 | FALSE | 0.82559785 | FALSE | 1 |
| 5650 | 5650 | 10.1904762 | 3.45886091 | 0 | 1 | 0 | 1 | 1 | 0.51939841 | 1 | 0.60987146 | 0 | 1 | 18 | 0.37504645 | FALSE | 1 | FALSE | 1 | FALSE | 1 | FALSE | 1 | FALSE | 1 | FALSE | 1 |
| 5651 | 5651 | 9.84210526 | 1.28130243 | 2 | 0.43181298 | 5 | 0.03627929 | 0 | 1 | 1 | 0.93213716 | 0 | 1 | 49 | 0.45760293 | FALSE | 1 | FALSE | 0.94259695 | FALSE | 1 | FALSE | 1 | FALSE | 1 | FALSE | 1 |
| 5652 | 5652 | 8.67272727 | 2.59408099 | 1 | 0.76171602 | 2 | 0.5356583 | 3 | 0.31652594 | 7 | 0.01225792 | 0 | 1 | 42 | 0.96038295 | FALSE | 1 | FALSE | 1 | FALSE | 1 | FALSE | 1 | FALSE | 1 | FALSE | 1 |
| 5653 | 5653 | 8 | 2.38746728 | 0 | 1 | 0 | 1 | 1 | 0.51939841 | 2 | 0.23331368 | 0 | 1 | 17 | 0.61750632 | FALSE | 1 | FALSE | 1 | FALSE | 1 | FALSE | 1 | FALSE | 1 | FALSE | 1 |
| 5654 | 5654 | 9.75757576 | 1.41486273 | 0 | 1 | 1 | 0.65167362 | 1 | 0.69069222 | 1 | 0.77852168 | 0 | 1 | 29 | 0.24088541 | FALSE | 1 | FALSE | 1 | FALSE | 1 | FALSE | 1 | FALSE | 1 | FALSE | 1 |
| 5655 | 5655 | 10.1538462 | 2.2134607 | 0 | 1 | 1 | 0.3480709 | 0 | 1 | 0 | 1 | 0 | 1 | 12 | 0.374198 | FALSE | 1 | FALSE | 1 | FALSE | 1 | FALSE | 1 | FALSE | 1 | FALSE | 1 |
| 5656 | 5656 | 9.89285714 | 1.91496523 | 1 | 0.51745948 | 2 | 0.2288771 | 1 | 0.64171563 | 0 | 1 | 0 | 1 | 24 | 0.55018107 | FALSE | 1 | FALSE | 1 | FALSE | 1 | FALSE | 1 | FALSE | 1 | FALSE | 1 |
| 5657 | 5657 | 10.3125 | 2.46776088 | 0 | 1 | 0 | 1 | 0 | 1 | 1 | 0.52895088 | 0 | 1 | 15 | 0.26127859 | FALSE | 1 | FALSE | 1 | FALSE | 1 | FALSE | 1 | FALSE | 1 | FALSE | 1 |
| 5658 | 5658 | 11.4545455 | 2.64106165 | 0 | 1 | 0 | 1 | 0 | 1 | 1 | 0.40390032 | 0 | 1 | 10 | 0.46859937 | FALSE | 1 | FALSE | 1 | FALSE | 1 | FALSE | 1 | FALSE | 1 | FALSE | 1 |
| 5659 | 5659 | 11.9090909 | 4.37876167 | 1 | 0.24866461 | 0 | 1 | 0 | 1 | 0 | 1 | 0 | 1 | 10 | 0.46859937 | FALSE | 1 | FALSE | 1 | FALSE | 1 | FALSE | 1 | FALSE | 1 | FALSE | 1 |
| 5660 | 5660 | 7.8 | 3.3306656 | 1 | 0.32293355 | 0 | 1 | 1 | 0.42266264 | 0 | 1 | 0 | 1 | 13 | 0.57712805 | FALSE | 1 | FALSE | 1 | FALSE | 1 | FALSE | 1 | FALSE | 1 | FALSE | 1 |
| 5661 | 5661 | 8.58536585 | 2.23034078 | 4 | 0.02019941 | 2 | 0.38460545 | 0 | 1 | 1 | 0.85524728 | 0 | 1 | 34 | 0.69505404 | FALSE | 0.44482142 | FALSE | 1 | FALSE | 1 | FALSE | 1 | FALSE | 1 | FALSE | 1 |
| 5662 | 5662 | 10.9166667 | 1.70579079 | 0 | 1 | 0 | 1 | 1 | 0.35555293 | 0 | 1 | 1 | 0.18007762 | 10 | 0.71436964 | FALSE | 1 | FALSE | 1 | FALSE | 1 | FALSE | 1 | FALSE | 1 | FALSE | 1 |
| 5663 | 5663 | 3.90990991 | 2.61681861 | 6 | 0.49810254 | 7 | 0.57278579 | 7 | 0.68292939 | 10 | 0.5599447 | 3 | 0.70555607 | 187 | 0.44291255 | FALSE | 1 | FALSE | 1 | FALSE | 1 | FALSE | 1 | FALSE | 1 | FALSE | 1 |
| 5664 | 5664 | 10.3636364 | 2.22680886 | 0 | 1 | 0 | 1 | 1 | 0.33149969 | 0 | 1 | 0 | 1 | 10 | 0.46859937 | FALSE | 1 | FALSE | 1 | FALSE | 1 | FALSE | 1 | FALSE | 1 | FALSE | 1 |
| 5665 | 5665 | 10.4361702 | 1.24249672 | 0 | 1 | 2 | 0.71077581 | 3 | 0.51793834 | 4 | 0.46403589 | 0 | 1 | 67 | 0.23030366 | FALSE | 1 | FALSE | 1 | FALSE | 1 | FALSE | 1 | FALSE | 1 | FALSE | 1 |
| 5666 | 5666 | 9.64 | 4.13647193 | 1 | 0.47819878 | 1 | 0.56109832 | 1 | 0.59998149 | 2 | 0.31968852 | 0 | 1 | 20 | 0.8152498 | FALSE | 1 | FALSE | 1 | FALSE | 1 | FALSE | 1 | FALSE | 1 | FALSE | 1 |
| 5667 | 5667 | 10.3684211 | 3.08153288 | 0 | 1 | 0 | 1 | 1 | 0.50143826 | 0 | 1 | 0 | 1 | 18 | 0.17880694 | FALSE | 1 | FALSE | 1 | FALSE | 1 | FALSE | 1 | FALSE | 1 | FALSE | 1 |
| 5668 | 5668 | 9.82352941 | 3.46809482 | 1 | 0.35728305 | 1 | 0.42857733 | 0 | 1 | 0 | 1 | 0 | 1 | 15 | 0.49060435 | FALSE | 1 | FALSE | 1 | FALSE | 1 | FALSE | 1 | FALSE | 1 | FALSE | 1 |
| 5669 | 5669 | 8.28571429 | 1.78999487 | 0 | 1 | 1 | 0.58913618 | 0 | 1 | 0 | 1 | 0 | 1 | 26 | 0.06079097 | FALSE | 1 | FALSE | 1 | FALSE | 1 | FALSE | 1 | FALSE | 1 | FALSE | 1 |
| 5670 | 5670 | 8.03030303 | 2.02237257 | 3 | 0.05151181 | 2 | 0.28941836 | 1 | 0.70185639 | 1 | 0.78873814 | 0 | 1 | 26 | 0.86852567 | FALSE | 0.67646333 | FALSE | 1 | FALSE | 1 | FALSE | 1 | FALSE | 1 | FALSE | 1 |
| 5671 | 5671 | 8.66666667 | 1.1055416 | 0 | 1 | 0 | 1 | 0 | 1 | 1 | 0.43131129 | 0 | 1 | 11 | 0.41944532 | FALSE | 1 | FALSE | 1 | FALSE | 1 | FALSE | 1 | FALSE | 1 | FALSE | 1 |
| 5672 | 5672 | 10.1666667 | 2.16666667 | 1 | 0.37380237 | 1 | 0.44710673 | 2 | 0.13502329 | 0 | 1 | 0 | 1 | 14 | 0.86287418 | FALSE | 1 | FALSE | 1 | FALSE | 1 | FALSE | 1 | FALSE | 1 | FALSE | 1 |
| 5673 | 5673 | 10.65625 | 1.65093487 | 0 | 1 | 2 | 0.27730767 | 2 | 0.32011871 | 1 | 0.77852168 | 0 | 1 | 27 | 0.61671133 | FALSE | 1 | FALSE | 1 | FALSE | 1 | FALSE | 1 | FALSE | 1 | FALSE | 1 |
| 5674 | 5674 | 10.1052632 | 2.93606856 | 0 | 1 | 2 | 0.12434464 | 0 | 1 | 0 | 1 | 1 | 0.26986775 | 16 | 0.65565778 | FALSE | 1 | FALSE | 1 | FALSE | 1 | FALSE | 1 | FALSE | 1 | FALSE | 1 |
| 5675 | 5675 | 10.5714286 | 4.16986272 | 1 | 0.16630142 | 0 | 1 | 0 | 1 | 0 | 1 | 0 | 1 | 6 | 0.69920685 | FALSE | 1 | FALSE | 1 | FALSE | 1 | FALSE | 1 | FALSE | 1 | FALSE | 1 |
| 5676 | 5676 | 9.61538462 | 3.2708141 | 0 | 1 | 0 | 1 | 1 | 0.37874417 | 1 | 0.45746571 | 0 | 1 | 11 | 0.6683056 | FALSE | 1 | FALSE | 1 | FALSE | 1 | FALSE | 1 | FALSE | 1 | FALSE | 1 |
| 5677 | 5677 | 9.875 | 1.05326872 | 1 | 0.18769745 | 1 | 0.23139161 | 0 | 1 | 0 | 1 | 1 | 0.12394282 | 5 | 0.97551477 | FALSE | 1 | FALSE | 1 | FALSE | 1 | FALSE | 1 | FALSE | 1 | FALSE | 1 |
| 5678 | 5678 | 6.66666667 | 3.53160335 | 0 | 1 | 1 | 0.54637326 | 1 | 0.58502114 | 1 | 0.67692816 | 3 | 0.00675546 | 18 | 0.93166998 | FALSE | 1 | FALSE | 1 | FALSE | 1 | FALSE | 1 | FALSE | 0.47461742 | FALSE | 1 |
| 5679 | 5679 | 9 | 2.54195564 | 0 | 1 | 0 | 1 | 1 | 0.37874417 | 0 | 1 | 0 | 1 | 12 | 0.374198 | FALSE | 1 | FALSE | 1 | FALSE | 1 | FALSE | 1 | FALSE | 1 | FALSE | 1 |
| 5680 | 5680 | 7.84615385 | 3.02161836 | 3 | 0.02517314 | 0 | 1 | 3 | 0.05884411 | 4 | 0.02569595 | 1 | 0.33902672 | 14 | 0.99985788 | FALSE | 0.4974559 | FALSE | 1 | FALSE | 1 | FALSE | 1 | FALSE | 1 | FALSE | 1 |
| 5681 | 5681 | 11.4782609 | 3.89949345 | 0 | 1 | 0 | 1 | 1 | 0.5695037 | 0 | 1 | 1 | 0.31672722 | 21 | 0.27964072 | FALSE | 1 | FALSE | 1 | FALSE | 1 | FALSE | 1 | FALSE | 1 | FALSE | 1 |
| 5682 | 5682 | 8.31914894 | 2.1047705 | 5 | 0.00671146 | 0 | 1 | 0 | 1 | 1 | 0.89101956 | 2 | 0.17968152 | 39 | 0.69224623 | FALSE | 0.24597546 | FALSE | 1 | FALSE | 1 | FALSE | 1 | FALSE | 1 | FALSE | 1 |
| 5683 | 5683 | 9.57692308 | 2.88461539 | 0 | 1 | 1 | 0.57534752 | 1 | 0.61440466 | 0 | 1 | 0 | 1 | 24 | 0.20425579 | FALSE | 1 | FALSE | 1 | FALSE | 1 | FALSE | 1 | FALSE | 1 | FALSE | 1 |
| 5684 | 5684 | 9.60869565 | 2.61665453 | 1 | 0.45027683 | 3 | 0.03669128 | 1 | 0.5695037 | 0 | 1 | 1 | 0.31672722 | 17 | 0.94439745 | FALSE | 1 | FALSE | 0.94259695 | FALSE | 1 | FALSE | 1 | FALSE | 1 | FALSE | 1 |
| 5685 | 5685 | 5.38095238 | 2.06860284 | 5 | 0.69785794 | 11 | 0.11726303 | 5 | 0.91650609 | 10 | 0.60042373 | 7 | 0.07856325 | 189 | 0.7186885 | FALSE | 1 | FALSE | 1 | FALSE | 1 | FALSE | 1 | FALSE | 1 | FALSE | 1 |
| 5686 | 5686 | 9.6 | 2.15406592 | 0 | 1 | 0 | 1 | 1 | 0.42266264 | 1 | 0.50623194 | 0 | 1 | 13 | 0.57712805 | FALSE | 1 | FALSE | 1 | FALSE | 1 | FALSE | 1 | FALSE | 1 | FALSE | 1 |
| 5687 | 5687 | 10 | 2.54057975 | 0 | 1 | 0 | 1 | 1 | 0.55340851 | 1 | 0.64497439 | 0 | 1 | 20 | 0.30915202 | FALSE | 1 | FALSE | 1 | FALSE | 1 | FALSE | 1 | FALSE | 1 | FALSE | 1 |
| 5688 | 5688 | 7.04545455 | 2.63675546 | 1 | 0.4357638 | 1 | 0.5154318 | 1 | 0.55340851 | 1 | 0.64497439 | 2 | 0.04972408 | 16 | 0.95553951 | FALSE | 1 | FALSE | 1 | FALSE | 1 | FALSE | 1 | FALSE | 0.98629461 | FALSE | 1 |
| 5689 | 5689 | 8.04761905 | 3.84801978 | 1 | 0.42086991 | 0 | 1 | 1 | 0.53671415 | 2 | 0.25059627 | 0 | 1 | 17 | 0.77849076 | FALSE | 1 | FALSE | 1 | FALSE | 1 | FALSE | 1 | FALSE | 1 | FALSE | 1 |
| 5690 | 5690 | 6.79032258 | 3.38467656 | 2 | 0.47501604 | 1 | 0.87100665 | 2 | 0.65873577 | 4 | 0.31731292 | 1 | 0.64288531 | 52 | 0.6264787 | FALSE | 1 | FALSE | 1 | FALSE | 1 | FALSE | 1 | FALSE | 1 | FALSE | 1 |
| 5691 | 5691 | 7.61538462 | 2.76067053 | 1 | 0.28675951 | 1 | 0.3480709 | 0 | 1 | 1 | 0.45746571 | 1 | 0.19354507 | 9 | 0.96018168 | FALSE | 1 | FALSE | 1 | FALSE | 1 | FALSE | 1 | FALSE | 1 | FALSE | 1 |
| 5692 | 5692 | 9.5 | 2.41867732 | 0 | 1 | 0 | 1 | 2 | 0.29330414 | 1 | 0.75658797 | 0 | 1 | 27 | 0.28862208 | FALSE | 1 | FALSE | 1 | FALSE | 1 | FALSE | 1 | FALSE | 1 | FALSE | 1 |
| 5693 | 5693 | 9.92857143 | 2.63124864 | 0 | 1 | 0 | 1 | 0 | 1 | 0 | 1 | 0 | 1 | 13 | 0.10967906 | FALSE | 1 | FALSE | 1 | FALSE | 1 | FALSE | 1 | FALSE | 1 | FALSE | 1 |
| 5694 | 5694 | 10.7692308 | 2.8864097 | 0 | 1 | 0 | 1 | 0 | 1 | 0 | 1 | 0 | 1 | 13 | 0.10967906 | FALSE | 1 | FALSE | 1 | FALSE | 1 | FALSE | 1 | FALSE | 1 | FALSE | 1 |
| 5695 | 5695 | 8.78571429 | 2.7039276 | 1 | 0.51745948 | 0 | 1 | 0 | 1 | 3 | 0.13524571 | 0 | 1 | 24 | 0.55018107 | FALSE | 1 | FALSE | 1 | FALSE | 1 | FALSE | 1 | FALSE | 1 | FALSE | 1 |
| 5696 | 5696 | 8.6 | 2.15406592 | 1 | 0.22886536 | 0 | 1 | 0 | 1 | 0 | 1 | 0 | 1 | 9 | 0.5215547 | FALSE | 1 | FALSE | 1 | FALSE | 1 | FALSE | 1 | FALSE | 1 | FALSE | 1 |
| 5697 | 5697 | 8 | 3.77792632 | 1 | 0.4357638 | 0 | 1 | 0 | 1 | 1 | 0.64497439 | 1 | 0.30529951 | 19 | 0.54233419 | FALSE | 1 | FALSE | 1 | FALSE | 1 | FALSE | 1 | FALSE | 1 | FALSE | 1 |
| 5698 | 5698 | 9.58064516 | 2.19639284 | 2 | 0.1881398 | 1 | 0.63997468 | 3 | 0.09854319 | 0 | 1 | 0 | 1 | 25 | 0.80006983 | FALSE | 1 | FALSE | 1 | FALSE | 1 | FALSE | 1 | FALSE | 1 | FALSE | 1 |
| 5699 | 5699 | 11 | 3.02765035 | 0 | 1 | 0 | 1 | 1 | 0.35555293 | 0 | 1 | 0 | 1 | 11 | 0.41944532 | FALSE | 1 | FALSE | 1 | FALSE | 1 | FALSE | 1 | FALSE | 1 | FALSE | 1 |
| 5700 | 5700 | 9.07843137 | 3.13013899 | 1 | 0.7354105 | 2 | 0.49507792 | 5 | 0.03520374 | 3 | 0.41729179 | 2 | 0.20351566 | 38 | 0.97862182 | FALSE | 1 | FALSE | 1 | FALSE | 1 | FALSE | 1 | FALSE | 1 | FALSE | 1 |
| 5701 | 5701 | 10.3 | 2.81247222 | 1 | 0.40558521 | 1 | 0.48239024 | 4 | 0.00499622 | 2 | 0.23331368 | 0 | 1 | 12 | 0.99824567 | FALSE | 1 | FALSE | 1 | FALSE | 1 | FALSE | 1 | FALSE | 1 | FALSE | 1 |
| 5702 | 5702 | 8.70731707 | 2.10972085 | 2 | 0.27371785 | 2 | 0.37295654 | 1 | 0.76953771 | 2 | 0.5543522 | 0 | 1 | 33 | 0.71942635 | FALSE | 1 | FALSE | 1 | FALSE | 1 | FALSE | 1 | FALSE | 1 | FALSE | 1 |
| 5703 | 5703 | 3.08536585 | 2.75504513 | 0 | 1 | 2 | 0.74335738 | 4 | 0.33196391 | 4 | 0.51410462 | 1 | 0.74003033 | 70 | 0.37303501 | FALSE | 1 | FALSE | 1 | FALSE | 1 | FALSE | 1 | FALSE | 1 | FALSE | 1 |
| 5704 | 5704 | 8.98076923 | 1.15176094 | 0 | 1 | 1 | 0.82028131 | 0 | 1 | 5 | 0.08873102 | 0 | 1 | 46 | 0.27620659 | FALSE | 1 | FALSE | 1 | FALSE | 1 | FALSE | 1 | FALSE | 1 | FALSE | 1 |
| 5705 | 5705 | 7.07692308 | 2.80004227 | 6 | 3.91E-05 | 2 | 0.20486827 | 0 | 1 | 2 | 0.33678042 | 0 | 1 | 16 | 0.9989746 | TRUE | 0.01001265 | FALSE | 1 | FALSE | 1 | FALSE | 1 | FALSE | 1 | FALSE | 1 |
| 5706 | 5706 | 10.4642857 | 2.51399146 | 0 | 1 | 2 | 0.2288771 | 1 | 0.64171563 | 1 | 0.73248983 | 0 | 1 | 24 | 0.55018107 | FALSE | 1 | FALSE | 1 | FALSE | 1 | FALSE | 1 | FALSE | 1 | FALSE | 1 |
| 5707 | 5707 | 7.63333333 | 2.04096927 | 3 | 0.19211919 | 9 | 0.00010095 | 2 | 0.63179508 | 1 | 0.93827647 | 0 | 1 | 44 | 0.98374792 | FALSE | 1 | FALSE | 0.1181425 | FALSE | 1 | FALSE | 1 | FALSE | 1 | FALSE | 1 |
| 5708 | 5708 | 10.5217391 | 2.77988998 | 1 | 0.45027683 | 2 | 0.16948764 | 1 | 0.5695037 | 1 | 0.6613267 | 0 | 1 | 18 | 0.86273155 | FALSE | 1 | FALSE | 1 | FALSE | 1 | FALSE | 1 | FALSE | 1 | FALSE | 1 |
| 5709 | 5709 | 12.6551724 | 2.45337013 | 0 | 1 | 1 | 0.60247911 | 1 | 0.64171563 | 0 | 1 | 0 | 1 | 26 | 0.16405547 | FALSE | 1 | FALSE | 1 | FALSE | 1 | FALSE | 1 | FALSE | 1 | FALSE | 1 |
| 5710 | 5710 | 8.16666667 | 2.86744176 | 2 | 0.12501436 | 0 | 1 | 0 | 1 | 2 | 0.30249307 | 1 | 0.32796862 | 19 | 0.83975208 | FALSE | 0.95622564 | FALSE | 1 | FALSE | 1 | FALSE | 1 | FALSE | 1 | FALSE | 1 |
| 5711 | 5711 | 9.30769231 | 2.79334232 | 5 | 0.0029693 | 1 | 0.7236252 | 1 | 0.76089851 | 3 | 0.26522531 | 1 | 0.4761699 | 28 | 0.98701111 | FALSE | 0.14653312 | FALSE | 1 | FALSE | 1 | FALSE | 1 | FALSE | 1 | FALSE | 1 |
| 5712 | 5712 | 9.86666667 | 1.49962958 | 0 | 1 | 0 | 1 | 0 | 1 | 1 | 0.50623194 | 1 | 0.2198258 | 13 | 0.57712805 | FALSE | 1 | FALSE | 1 | FALSE | 1 | FALSE | 1 | FALSE | 1 | FALSE | 1 |
| 5713 | 5713 | 8.1875 | 1.62860178 | 0 | 1 | 0 | 1 | 1 | 0.44344817 | 3 | 0.03440583 | 0 | 1 | 12 | 0.90922186 | FALSE | 1 | FALSE | 1 | FALSE | 1 | FALSE | 1 | FALSE | 1 | FALSE | 1 |
| 5714 | 5714 | 8.04545455 | 3.03730252 | 1 | 0.4357638 | 0 | 1 | 1 | 0.55340851 | 2 | 0.26791271 | 2 | 0.04972408 | 16 | 0.95553951 | FALSE | 1 | FALSE | 1 | FALSE | 1 | FALSE | 1 | FALSE | 0.98629461 | FALSE | 1 |
| 5715 | 5715 | 9.23333333 | 3.08418907 | 2 | 0.17884234 | 1 | 0.6278847 | 0 | 1 | 2 | 0.4036176 | 2 | 0.08611595 | 23 | 0.9158696 | FALSE | 1 | FALSE | 1 | FALSE | 1 | FALSE | 1 | FALSE | 1 | FALSE | 1 |
| 5716 | 5716 | 8.78571429 | 2.07634884 | 0 | 1 | 1 | 0.36919824 | 0 | 1 | 0 | 1 | 3 | 0.00136853 | 10 | 0.94579623 | FALSE | 1 | FALSE | 1 | FALSE | 1 | FALSE | 1 | FALSE | 0.20625174 | FALSE | 1 |
| 5717 | 5717 | 9.76190476 | 2.70633757 | 0 | 1 | 0 | 1 | 2 | 0.17308361 | 2 | 0.25059627 | 0 | 1 | 17 | 0.77849076 | FALSE | 1 | FALSE | 1 | FALSE | 1 | FALSE | 1 | FALSE | 1 | FALSE | 1 |
| 5718 | 5718 | 10.8666667 | 2.47296493 | 0 | 1 | 0 | 1 | 1 | 0.42266264 | 0 | 1 | 0 | 1 | 14 | 0.29525409 | FALSE | 1 | FALSE | 1 | FALSE | 1 | FALSE | 1 | FALSE | 1 | FALSE | 1 |
| 5719 | 5719 | 7.20454546 | 2.90438184 | 0 | 1 | 0 | 1 | 3 | 0.19985581 | 1 | 0.86831187 | 1 | 0.50988734 | 38 | 0.31730615 | FALSE | 1 | FALSE | 1 | FALSE | 1 | FALSE | 1 | FALSE | 1 | FALSE | 1 |
| 5720 | 5720 | 6.60526316 | 1.38042976 | 2 | 0.79140268 | 2 | 0.88577971 | 7 | 0.1113069 | 4 | 0.76938454 | 2 | 0.55679953 | 96 | 0.4958468 | FALSE | 1 | FALSE | 1 | FALSE | 1 | FALSE | 1 | FALSE | 1 | FALSE | 1 |
| 5721 | 5721 | 8.33333333 | 2.98142397 | 2 | 0.03643311 | 0 | 1 | 1 | 0.35555293 | 0 | 1 | 1 | 0.18007762 | 8 | 0.97197833 | FALSE | 0.58392 | FALSE | 1 | FALSE | 1 | FALSE | 1 | FALSE | 1 | FALSE | 1 |
| 5722 | 5722 | 10.0526316 | 3.47129104 | 0 | 1 | 0 | 1 | 1 | 0.50143826 | 0 | 1 | 0 | 1 | 18 | 0.17880694 | FALSE | 1 | FALSE | 1 | FALSE | 1 | FALSE | 1 | FALSE | 1 | FALSE | 1 |
| 5723 | 5723 | 7.7 | 2.78028776 | 3 | 0.13009168 | 5 | 0.02029785 | 1 | 0.83455705 | 2 | 0.66536008 | 0 | 1 | 38 | 0.92985909 | FALSE | 0.97964162 | FALSE | 0.80328489 | FALSE | 1 | FALSE | 1 | FALSE | 1 | FALSE | 1 |
| 5724 | 5724 | 7.2631579 | 2.55141045 | 1 | 0.38989951 | 3 | 0.02205772 | 0 | 1 | 0 | 1 | 2 | 0.03799168 | 13 | 0.97998676 | FALSE | 1 | FALSE | 0.81029974 | FALSE | 1 | FALSE | 1 | FALSE | 0.87669812 | FALSE | 1 |
| 5725 | 5725 | 10.6818182 | 2.30387744 | 0 | 1 | 0 | 1 | 0 | 1 | 0 | 1 | 0 | 1 | 22 | 0.02368118 | FALSE | 1 | FALSE | 1 | FALSE | 1 | FALSE | 1 | FALSE | 1 | FALSE | 1 |
| 5726 | 5726 | 9.7 | 2.98496231 | 1 | 0.40558521 | 0 | 1 | 2 | 0.16019548 | 1 | 0.60987146 | 2 | 0.04176963 | 14 | 0.97325593 | FALSE | 1 | FALSE | 1 | FALSE | 1 | FALSE | 1 | FALSE | 0.90763079 | FALSE | 1 |
| 5727 | 5727 | 9.34782609 | 2.33328832 | 2 | 0.10809809 | 0 | 1 | 1 | 0.55340851 | 1 | 0.64497439 | 0 | 1 | 18 | 0.74744813 | FALSE | 0.88820416 | FALSE | 1 | FALSE | 1 | FALSE | 1 | FALSE | 1 | FALSE | 1 |
| 5728 | 5728 | 11.5 | 1.5594288 | 0 | 1 | 0 | 1 | 1 | 0.55340851 | 0 | 1 | 0 | 1 | 21 | 0.12048848 | FALSE | 1 | FALSE | 1 | FALSE | 1 | FALSE | 1 | FALSE | 1 | FALSE | 1 |
| 5729 | 5729 | 9.02173913 | 1.97251914 | 3 | 0.10767332 | 6 | 0.00303041 | 2 | 0.48463745 | 0 | 1 | 1 | 0.52593041 | 33 | 0.98284787 | FALSE | 0.88820416 | FALSE | 0.46839856 | FALSE | 1 | FALSE | 1 | FALSE | 1 | FALSE | 1 |
| 5730 | 5730 | 4.71505376 | 2.45595376 | 4 | 0.70022422 | 9 | 0.14139094 | 5 | 0.79727985 | 8 | 0.61459742 | 7 | 0.03104054 | 151 | 0.83675167 | FALSE | 1 | FALSE | 1 | FALSE | 1 | FALSE | 1 | FALSE | 0.82559785 | FALSE | 1 |
| 5731 | 5731 | 10.1395349 | 2.0297943 | 1 | 0.6184714 | 2 | 0.3375163 | 0 | 1 | 3 | 0.2402032 | 0 | 1 | 31 | 0.64534533 | FALSE | 1 | FALSE | 1 | FALSE | 1 | FALSE | 1 | FALSE | 1 | FALSE | 1 |
| 5732 | 5732 | 8.71428571 | 2.60298102 | 1 | 0.3050805 | 0 | 1 | 1 | 0.40110418 | 1 | 0.48242099 | 0 | 1 | 11 | 0.83676449 | FALSE | 1 | FALSE | 1 | FALSE | 1 | FALSE | 1 | FALSE | 1 | FALSE | 1 |

| | | | | | | | | | | | | | | | | | | | | | | | | | | | |
|---|---|---|---|---|---|---|---|---|---|---|---|---|---|---|---|---|---|---|---|---|---|---|---|---|---|---|---|
| 5733 | 5733 | 6.76190476 | 1.56463177 | 9 | 0.0014124 | 0 | 1 | 6 | 0.17549144 | 4 | 0.7174274 | 1 | 0.82614736 | 85 | 0.86552172 | FALSE | 0.09317759 | FALSE | 1 | FALSE | 1 | FALSE | 1 | FALSE | 1 | FALSE | 1 |
| 5734 | 5734 | 7.29166667 | 1.56735357 | 0 | 1 | 1 | 0.54637326 | 0 | 1 | 1 | 0.67692816 | 0 | 1 | 22 | 0.25237063 | FALSE | 1 | FALSE | 1 | FALSE | 1 | FALSE | 1 | FALSE | 1 | FALSE | 1 |
| 5735 | 5735 | 9.07692308 | 2.68790849 | 3 | 0.0279333 | 0 | 1 | 0 | 1 | 1 | 0.70601446 | 0 | 1 | 22 | 0.61628 | FALSE | 0.50964275 | FALSE | 1 | FALSE | 1 | FALSE | 1 | FALSE | 1 | FALSE | 1 |
| 5736 | 5736 | 9.66666667 | 2.02210012 | 0 | 1 | 0 | 1 | 1 | 0.42266264 | 0 | 1 | 0 | 1 | 14 | 0.29525409 | FALSE | 1 | FALSE | 1 | FALSE | 1 | FALSE | 1 | FALSE | 1 | FALSE | 1 |
| 5737 | 5737 | 11.3043478 | 2.8807424 | 1 | 0.45027683 | 0 | 1 | 2 | 0.19931429 | 0 | 1 | 0 | 1 | 20 | 0.50589402 | FALSE | 1 | FALSE | 1 | FALSE | 1 | FALSE | 1 | FALSE | 1 | FALSE | 1 |
| 5738 | 5738 | 8.5 | 3.02489669 | 1 | 0.40558521 | 2 | 0.1353446 | 1 | 0.51939841 | 0 | 1 | 0 | 1 | 16 | 0.80826361 | FALSE | 1 | FALSE | 1 | FALSE | 1 | FALSE | 1 | FALSE | 1 | FALSE | 1 |
| 5739 | 5739 | 9.13333333 | 2.57854395 | 0 | 1 | 0 | 1 | 0 | 1 | 1 | 0.50623194 | 2 | 0.02433796 | 12 | 0.80329917 | FALSE | 1 | FALSE | 1 | FALSE | 1 | FALSE | 1 | FALSE | 0.77578444 | FALSE | 1 |
| 5740 | 5740 | 8.42857143 | 2.83383349 | 0 | 1 | 1 | 0.58913618 | 1 | 0.62830986 | 0 | 1 | 1 | 0.36060486 | 24 | 0.3725373 | FALSE | 1 | FALSE | 1 | FALSE | 1 | FALSE | 1 | FALSE | 1 | FALSE | 1 |
| 5741 | 5741 | 8.57142857 | 4.45155327 | 0 | 1 | 0 | 1 | 0 | 1 | 0 | 1 | 0 | 1 | 14 | 0.09251304 | FALSE | 1 | FALSE | 1 | FALSE | 1 | FALSE | 1 | FALSE | 1 | FALSE | 1 |
| 5742 | 5742 | 8.31034483 | 1.87733208 | 1 | 0.52988231 | 0 | 1 | 0 | 1 | 0 | 1 | 1 | 0.38148475 | 27 | 0.14664438 | FALSE | 1 | FALSE | 1 | FALSE | 1 | FALSE | 1 | FALSE | 1 | FALSE | 1 |
| 5743 | 5743 | 9.56521739 | 2.33733566 | 0 | 1 | 1 | 0.5154318 | 1 | 0.55340851 | 2 | 0.26791271 | 1 | 0.30529951 | 17 | 0.8840357 | FALSE | 1 | FALSE | 1 | FALSE | 1 | FALSE | 1 | FALSE | 1 | FALSE | 1 |
| 5744 | 5744 | 9.57142857 | 2.19461307 | 2 | 0.04858661 | 0 | 1 | 0 | 1 | 0 | 1 | 0 | 1 | 12 | 0.62236227 | FALSE | 0.649301 | FALSE | 1 | FALSE | 1 | FALSE | 1 | FALSE | 1 | FALSE | 1 |
| 5745 | 5745 | 7.28571429 | 2.16559366 | 1 | 0.83491628 | 4 | 0.18354682 | 0 | 1 | 4 | 0.39127216 | 2 | 0.31302643 | 58 | 0.60859322 | FALSE | 1 | FALSE | 1 | FALSE | 1 | FALSE | 1 | FALSE | 1 | FALSE | 1 |
| 5746 | 5746 | 8.71428571 | 2.11891385 | 0 | 1 | 1 | 0.20567256 | 0 | 1 | 0 | 1 | 0 | 1 | 6 | 0.69920685 | FALSE | 1 | FALSE | 1 | FALSE | 1 | FALSE | 1 | FALSE | 1 | FALSE | 1 |
| 5747 | 5747 | 10.4545455 | 2.38798646 | 1 | 0.24866461 | 0 | 1 | 0 | 1 | 1 | 0.40390032 | 0 | 1 | 9 | 0.75985366 | FALSE | 1 | FALSE | 1 | FALSE | 1 | FALSE | 1 | FALSE | 1 | FALSE | 1 |
| 5748 | 5748 | 8.23529412 | 2.33966845 | 2 | 0.21638713 | 0 | 1 | 0 | 1 | 1 | 0.7984848 | 0 | 1 | 31 | 0.1995074 | FALSE | 1 | FALSE | 1 | FALSE | 1 | FALSE | 1 | FALSE | 1 | FALSE | 1 |
| 5749 | 5749 | 6.5625 | 2.78766816 | 3 | 0.33732256 | 3 | 0.48172889 | 2 | 0.78851973 | 1 | 0.97723437 | 0 | 1 | 71 | 0.17761975 | FALSE | 1 | FALSE | 1 | FALSE | 1 | FALSE | 1 | FALSE | 1 | FALSE | 1 |
| 5750 | 5750 | 8.84615385 | 2.03228382 | 0 | 1 | 2 | 0.06426041 | 1 | 0.37874417 | 1 | 0.45746571 | 0 | 1 | 9 | 0.96018168 | FALSE | 1 | FALSE | 1 | FALSE | 1 | FALSE | 1 | FALSE | 1 | FALSE | 1 |
| 5751 | 5751 | 11.1428571 | 3.48173075 | 1 | 0.3050805 | 0 | 1 | 1 | 0.40110418 | 0 | 1 | 0 | 1 | 12 | 0.62236227 | FALSE | 1 | FALSE | 1 | FALSE | 1 | FALSE | 1 | FALSE | 1 | FALSE | 1 |
| 5752 | 5752 | 8.24242424 | 3.07512309 | 2 | 0.20692137 | 1 | 0.6629941 | 1 | 0.70185639 | 0 | 1 | 0 | 1 | 29 | 0.39648374 | FALSE | 1 | FALSE | 1 | FALSE | 1 | FALSE | 1 | FALSE | 1 | FALSE | 1 |
| 5753 | 5753 | 10.7936508 | 2.21938595 | 3 | 0.11312654 | 1 | 0.78076835 | 2 | 0.4962935 | 2 | 0.63103861 | 0 | 1 | 38 | 0.71504609 | FALSE | 0.92488288 | FALSE | 1 | FALSE | 1 | FALSE | 1 | FALSE | 1 | FALSE | 1 |
| 5754 | 5754 | 10.6666667 | 2.39211668 | 0 | 1 | 0 | 1 | 0 | 1 | 0 | 1 | 0 | 1 | 12 | 0.13002667 | FALSE | 1 | FALSE | 1 | FALSE | 1 | FALSE | 1 | FALSE | 1 | FALSE | 1 |
| 5755 | 5755 | 8.125 | 2.75850956 | 0 | 1 | 0 | 1 | 0 | 1 | 0 | 1 | 0 | 1 | 24 | 0.01683964 | FALSE | 1 | FALSE | 1 | FALSE | 1 | FALSE | 1 | FALSE | 1 | FALSE | 1 |
| 5756 | 5756 | 9.65217391 | 2.943064 | 3 | 0.02011786 | 0 | 1 | 0 | 1 | 1 | 0.6613267 | 0 | 1 | 19 | 0.71542496 | FALSE | 0.44482142 | FALSE | 1 | FALSE | 1 | FALSE | 1 | FALSE | 1 | FALSE | 1 |
| 5757 | 5757 | 10.0740741 | 2.32420318 | 0 | 1 | 3 | 0.05519089 | 1 | 0.62830986 | 1 | 0.71956339 | 0 | 1 | 22 | 0.76237877 | FALSE | 1 | FALSE | 1 | FALSE | 1 | FALSE | 1 | FALSE | 1 | FALSE | 1 |
| 5758 | 5758 | 10.1 | 2.66270539 | 0 | 1 | 0 | 1 | 1 | 0.28067846 | 0 | 1 | 0 | 1 | 8 | 0.5780332 | FALSE | 1 | FALSE | 1 | FALSE | 1 | FALSE | 1 | FALSE | 1 | FALSE | 1 |
| 5759 | 5759 | 7.77777778 | 2.61524496 | 1 | 0.37380237 | 0 | 1 | 1 | 0.48280983 | 0 | 1 | 0 | 1 | 16 | 0.44998001 | FALSE | 1 | FALSE | 1 | FALSE | 1 | FALSE | 1 | FALSE | 1 | FALSE | 1 |
| 5760 | 5760 | 7.11538462 | 1.80441602 | 0 | 1 | 1 | 0.57534752 | 2 | 0.23940226 | 2 | 0.33678042 | 1 | 0.3499045 | 20 | 0.90142689 | FALSE | 1 | FALSE | 1 | FALSE | 1 | FALSE | 1 | FALSE | 1 | FALSE | 1 |
| 5761 | 5761 | 8.49038462 | 1.32284071 | 4 | 0.78673083 | 8 | 0.35935982 | 10 | 0.21429748 | 12 | 0.24693395 | 3 | 0.66873574 | 171 | 0.83484697 | FALSE | 1 | FALSE | 1 | FALSE | 1 | FALSE | 1 | FALSE | 1 | FALSE | 1 |
| 5762 | 5762 | 8.52173913 | 2.71660858 | 1 | 0.4357638 | 1 | 0.5154318 | 3 | 0.04259419 | 1 | 0.64497439 | 1 | 0.30529951 | 15 | 0.98570275 | FALSE | 1 | FALSE | 1 | FALSE | 1 | FALSE | 1 | FALSE | 1 | FALSE | 1 |
| 5763 | 5763 | 8.625 | 1.99608993 | 0 | 1 | 0 | 1 | 0 | 1 | 1 | 0.31351192 | 0 | 1 | 7 | 0.63752397 | FALSE | 1 | FALSE | 1 | FALSE | 1 | FALSE | 1 | FALSE | 1 | FALSE | 1 |
| 5764 | 5764 | 10.1 | 2.38537209 | 0 | 1 | 0 | 1 | 1 | 0.30655256 | 0 | 1 | 0 | 1 | 9 | 0.5215547 | FALSE | 1 | FALSE | 1 | FALSE | 1 | FALSE | 1 | FALSE | 1 | FALSE | 1 |
| 5765 | 5765 | 10.44 | 2.75797027 | 1 | 0.47819878 | 0 | 1 | 1 | 0.59998149 | 1 | 0.69181315 | 2 | 0.06257402 | 20 | 0.8152498 | FALSE | 1 | FALSE | 1 | FALSE | 1 | FALSE | 1 | FALSE | 1 | FALSE | 1 |
| 5766 | 5766 | 9.78947368 | 2.64732134 | 2 | 0.08405377 | 0 | 1 | 1 | 0.50143826 | 2 | 0.21610812 | 1 | 0.26986775 | 13 | 0.97998676 | FALSE | 0.79236883 | FALSE | 1 | FALSE | 1 | FALSE | 1 | FALSE | 1 | FALSE | 1 |
| 5767 | 5767 | 7.20652174 | 3.27552865 | 7 | 0.0085551 | 4 | 0.33970769 | 2 | 0.84458838 | 3 | 0.79556345 | 4 | 0.06180746 | 71 | 0.96130639 | FALSE | 0.27019545 | FALSE | 1 | FALSE | 1 | FALSE | 1 | FALSE | 1 | FALSE | 1 |
| 5768 | 5768 | 8.6875 | 1.8946883 | 0 | 1 | 1 | 0.63997468 | 3 | 0.09854319 | 2 | 0.41984187 | 0 | 1 | 25 | 0.80006983 | FALSE | 1 | FALSE | 1 | FALSE | 1 | FALSE | 1 | FALSE | 1 | FALSE | 1 |
| 5769 | 5769 | 9.53846154 | 2.30769231 | 1 | 0.28675951 | 0 | 1 | 2 | 0.07725803 | 0 | 1 | 0 | 1 | 10 | 0.86791642 | FALSE | 1 | FALSE | 1 | FALSE | 1 | FALSE | 1 | FALSE | 1 | FALSE | 1 |
| 5770 | 5770 | 9.2 | 2.53508711 | 0 | 1 | 0 | 1 | 0 | 1 | 1 | 0.75658797 | 1 | 0.39166989 | 28 | 0.13086908 | FALSE | 1 | FALSE | 1 | FALSE | 1 | FALSE | 1 | FALSE | 1 | FALSE | 1 |
| 5771 | 5771 | 12.3076923 | 3.07307452 | 1 | 0.28675951 | 0 | 1 | 0 | 1 | 1 | 0.45746571 | 0 | 1 | 11 | 0.6683056 | FALSE | 1 | FALSE | 1 | FALSE | 1 | FALSE | 1 | FALSE | 1 | FALSE | 1 |
| 5772 | 5772 | 9.05 | 3.47095088 | 1 | 0.59801736 | 2 | 0.31355738 | 1 | 0.72299506 | 1 | 0.80778317 | 2 | 0.11183826 | 28 | 0.83079716 | FALSE | 1 | FALSE | 1 | FALSE | 1 | FALSE | 1 | FALSE | 1 | FALSE | 1 |
| 5773 | 5773 | 9 | 1.76383421 | 1 | 0.60837712 | 1 | 0.69480472 | 0 | 1 | 0 | 1 | 1 | 0.44938266 | 33 | 0.16407838 | FALSE | 1 | FALSE | 1 | FALSE | 1 | FALSE | 1 | FALSE | 1 | FALSE | 1 |
| 5774 | 5774 | 9.25 | 3.21130815 | 1 | 0.32293355 | 0 | 1 | 0 | 1 | 4 | 0.00397592 | 1 | 0.2198258 | 9 | 0.995488 | FALSE | 1 | FALSE | 1 | FALSE | 1 | FALSE | 1 | FALSE | 1 | FALSE | 1 |
| 5775 | 5775 | 8.56666667 | 2.41775837 | 0 | 1 | 0 | 1 | 2 | 0.29330414 | 3 | 0.15707004 | 0 | 1 | 25 | 0.67630471 | FALSE | 1 | FALSE | 1 | FALSE | 1 | FALSE | 1 | FALSE | 1 | FALSE | 1 |
| 5776 | 5776 | 8.83870968 | 1.77975667 | 0 | 1 | 1 | 0.63997468 | 0 | 1 | 2 | 0.41984187 | 0 | 1 | 28 | 0.26393996 | FALSE | 1 | FALSE | 1 | FALSE | 1 | FALSE | 1 | FALSE | 1 | FALSE | 1 |
| 5777 | 5777 | 8.59649123 | 1.38740764 | 1 | 0.77387657 | 3 | 0.28014665 | 3 | 0.3364112 | 2 | 0.74436726 | 1 | 0.61182532 | 47 | 0.72965359 | FALSE | 1 | FALSE | 1 | FALSE | 1 | FALSE | 1 | FALSE | 1 | FALSE | 1 |
| 5778 | 5778 | 10.5 | 1.67971086 | 1 | 0.3050805 | 4 | 0.00082539 | 1 | 0.40110418 | 1 | 0.48242099 | 1 | 0.20679328 | 6 | 0.99994909 | FALSE | 1 | FALSE | 0.27046382 | FALSE | 1 | FALSE | 1 | FALSE | 1 | FALSE | 1 |
| 5779 | 5779 | 7.40740741 | 4.16563773 | 0 | 1 | 0 | 1 | 1 | 0.61440466 | 0 | 1 | 0 | 1 | 25 | 0.06985983 | FALSE | 1 | FALSE | 1 | FALSE | 1 | FALSE | 1 | FALSE | 1 | FALSE | 1 |
| 5780 | 5780 | 9.05 | 2.65471279 | 1 | 0.40558521 | 1 | 0.48239024 | 0 | 1 | 0 | 1 | 0 | 1 | 18 | 0.37504645 | FALSE | 1 | FALSE | 1 | FALSE | 1 | FALSE | 1 | FALSE | 1 | FALSE | 1 |
| 5781 | 5781 | 8.0625 | 3.28764258 | 0 | 1 | 0 | 1 | 1 | 0.44344817 | 0 | 1 | 0 | 1 | 15 | 0.26127859 | FALSE | 1 | FALSE | 1 | FALSE | 1 | FALSE | 1 | FALSE | 1 | FALSE | 1 |
| 5782 | 5782 | 5.24242424 | 2.42499988 | 0 | 1 | 1 | 0.6629941 | 0 | 1 | 2 | 0.45159487 | 0 | 1 | 30 | 0.21942357 | FALSE | 1 | FALSE | 1 | FALSE | 1 | FALSE | 1 | FALSE | 1 | FALSE | 1 |
| 5783 | 5783 | 4.97142857 | 2.02111305 | 1 | 0.59801736 | 0 | 1 | 2 | 0.35984046 | 2 | 0.48233989 | 1 | 0.44015537 | 29 | 0.69927719 | FALSE | 1 | FALSE | 1 | FALSE | 1 | FALSE | 1 | FALSE | 1 | FALSE | 1 |
| 5784 | 5784 | 8.19354839 | 1.9078675 | 0 | 1 | 1 | 0.63997468 | 0 | 1 | 1 | 0.76781284 | 0 | 1 | 29 | 0.11661149 | FALSE | 1 | FALSE | 1 | FALSE | 1 | FALSE | 1 | FALSE | 1 | FALSE | 1 |
| 5785 | 5785 | 6.10843374 | 2.13976718 | 1 | 0.95693012 | 5 | 0.34731923 | 9 | 0.02822851 | 9 | 0.09947498 | 1 | 0.86490401 | 95 | 0.95241754 | FALSE | 1 | FALSE | 1 | FALSE | 1 | FALSE | 1 | FALSE | 1 | FALSE | 1 |
| 5786 | 5786 | 9.37837838 | 3.20768799 | 3 | 0.0681763 | 0 | 1 | 2 | 0.38585089 | 1 | 0.82511639 | 0 | 1 | 31 | 0.64534533 | FALSE | 0.74276693 | FALSE | 1 | FALSE | 1 | FALSE | 1 | FALSE | 1 | FALSE | 1 |
| 5787 | 5787 | 6.60869565 | 1.05249726 | 2 | 0.53205439 | 0 | 1 | 2 | 0.71546205 | 3 | 0.6203651 | 1 | 0.6822729 | 61 | 0.22871622 | FALSE | 1 | FALSE | 1 | FALSE | 1 | FALSE | 1 | FALSE | 1 | FALSE | 1 |
| 5788 | 5788 | 6.11111111 | 3.15680726 | 0 | 1 | 5 | 0.16581747 | 6 | 0.10401985 | 1 | 0.98585791 | 1 | 0.77640116 | 77 | 0.44791145 | FALSE | 1 | FALSE | 1 | FALSE | 1 | FALSE | 1 | FALSE | 1 | FALSE | 1 |
| 5789 | 5789 | 5.14379085 | 2.34985273 | 3 | 0.74949909 | 10 | 0.02416563 | 4 | 0.79794496 | 9 | 0.25713425 | 3 | 0.45232905 | 122 | 0.90722262 | FALSE | 1 | FALSE | 0.82367276 | FALSE | 1 | FALSE | 1 | FALSE | 1 | FALSE | 1 |
| 5790 | 5790 | 8.92857143 | 2.6582563 | 0 | 1 | 0 | 1 | 1 | 0.40110418 | 0 | 1 | 0 | 1 | 13 | 0.33283729 | FALSE | 1 | FALSE | 1 | FALSE | 1 | FALSE | 1 | FALSE | 1 | FALSE | 1 |
| 5791 | 5791 | 9.0625 | 2.94679381 | 1 | 0.34033054 | 1 | 0.40942991 | 0 | 1 | 1 | 0.52895088 | 2 | 0.02752114 | 11 | 0.9718531 | FALSE | 1 | FALSE | 1 | FALSE | 1 | FALSE | 1 | FALSE | 0.79106376 | FALSE | 1 |
| 5792 | 5792 | 9.55555556 | 2.65041343 | 0 | 1 | 1 | 0.44710673 | 1 | 0.48280983 | 1 | 0.57131013 | 1 | 0.25766374 | 14 | 0.86287418 | FALSE | 1 | FALSE | 1 | FALSE | 1 | FALSE | 1 | FALSE | 1 | FALSE | 1 |
| 5793 | 5793 | 8.38095238 | 3.44309095 | 0 | 1 | 1 | 0.49918218 | 1 | 0.53671415 | 2 | 0.25059627 | 0 | 1 | 17 | 0.77849076 | FALSE | 1 | FALSE | 1 | FALSE | 1 | FALSE | 1 | FALSE | 1 | FALSE | 1 |
| 5794 | 5794 | 10.7777778 | 1.39664501 | 3 | 0.00124136 | 0 | 1 | 1 | 0.28067846 | 0 | 1 | 1 | 0.1383237 | 4 | 0.99921376 | FALSE | 0.08545569 | FALSE | 1 | FALSE | 1 | FALSE | 1 | FALSE | 1 | FALSE | 1 |
| 5795 | 5795 | 8.50943396 | 2.11573299 | 0 | 1 | 4 | 0.07727212 | 2 | 0.54120634 | 1 | 0.90544885 | 1 | 0.56379725 | 42 | 0.62187327 | FALSE | 1 | FALSE | 1 | FALSE | 1 | FALSE | 1 | FALSE | 1 | FALSE | 1 |
| 5796 | 5796 | 10 | 2.54483604 | 0 | 1 | 1 | 0.48239024 | 0 | 1 | 1 | 0.60987146 | 0 | 1 | 18 | 0.37504645 | FALSE | 1 | FALSE | 1 | FALSE | 1 | FALSE | 1 | FALSE | 1 | FALSE | 1 |
| 5797 | 5797 | 7.64 | 1.43888846 | 2 | 0.13369621 | 1 | 0.56109832 | 1 | 0.59998149 | 1 | 0.69181315 | 0 | 1 | 20 | 0.8152498 | FALSE | 0.97964162 | FALSE | 1 | FALSE | 1 | FALSE | 1 | FALSE | 1 | FALSE | 1 |
| 5798 | 5798 | 11.5454546 | 2.31059364 | 0 | 1 | 1 | 0.30368014 | 1 | 0.33149969 | 0 | 1 | 0 | 1 | 9 | 0.75985366 | FALSE | 1 | FALSE | 1 | FALSE | 1 | FALSE | 1 | FALSE | 1 | FALSE | 1 |
| 5799 | 5799 | 9.18181818 | 1.79990817 | 0 | 1 | 1 | 0.5154318 | 0 | 1 | 3 | 0.07743186 | 0 | 1 | 18 | 0.74744813 | FALSE | 1 | FALSE | 1 | FALSE | 1 | FALSE | 1 | FALSE | 1 | FALSE | 1 |
| 5800 | 5800 | 7.88888889 | 2.05179837 | 1 | 0.37380237 | 2 | 0.1135776 | 1 | 0.48280983 | 0 | 1 | 1 | 0.25766374 | 13 | 0.95041913 | FALSE | 1 | FALSE | 1 | FALSE | 1 | FALSE | 1 | FALSE | 1 | FALSE | 1 |
| 5801 | 5801 | 6.92857143 | 2.7114647 | 0 | 1 | 1 | 0.36919824 | 0 | 1 | 0 | 1 | 1 | 0.20679328 | 12 | 0.62236227 | FALSE | 1 | FALSE | 1 | FALSE | 1 | FALSE | 1 | FALSE | 1 | FALSE | 1 |
| 5802 | 5802 | 9.76190476 | 2.42810453 | 2 | 0.09187136 | 0 | 1 | 2 | 0.16019548 | 1 | 0.60987146 | 0 | 1 | 15 | 0.92113679 | FALSE | 0.83248965 | FALSE | 1 | FALSE | 1 | FALSE | 1 | FALSE | 1 | FALSE | 1 |
| 5803 | 5803 | 9.61111111 | 1.76820339 | 0 | 1 | 0 | 1 | 1 | 0.48280983 | 2 | 0.19902682 | 0 | 1 | 15 | 0.69372602 | FALSE | 1 | FALSE | 1 | FALSE | 1 | FALSE | 1 | FALSE | 1 | FALSE | 1 |
| 5804 | 5804 | 9 | 2.5226249 | 0 | 1 | 1 | 0.5154318 | 1 | 0.55340851 | 0 | 1 | 1 | 0.30529951 | 19 | 0.54233419 | FALSE | 1 | FALSE | 1 | FALSE | 1 | FALSE | 1 | FALSE | 1 | FALSE | 1 |
| 5805 | 5805 | 12.2 | 2.67581763 | 0 | 1 | 0 | 1 | 0 | 1 | 1 | 0.37517262 | 0 | 1 | 9 | 0.5215547 | FALSE | 1 | FALSE | 1 | FALSE | 1 | FALSE | 1 | FALSE | 1 | FALSE | 1 |
| 5806 | 5806 | 9.52173913 | 1.49984246 | 2 | 0.11647716 | 0 | 1 | 2 | 0.19931429 | 0 | 1 | 0 | 1 | 19 | 0.71542496 | FALSE | 0.93181727 | FALSE | 1 | FALSE | 1 | FALSE | 1 | FALSE | 1 | FALSE | 1 |
| 5807 | 5807 | 8.5 | 2.38630351 | 0 | 1 | 0 | 1 | 1 | 0.48280983 | 1 | 0.57131013 | 0 | 1 | 16 | 0.44998001 | FALSE | 1 | FALSE | 1 | FALSE | 1 | FALSE | 1 | FALSE | 1 | FALSE | 1 |
| 5808 | 5808 | 9.77777778 | 2.43938871 | 1 | 0.50471032 | 1 | 0.58913618 | 2 | 0.25286935 | 1 | 0.71956339 | 2 | 0.07169515 | 20 | 0.95133411 | FALSE | 1 | FALSE | 1 | FALSE | 1 | FALSE | 1 | FALSE | 1 | FALSE | 1 |
| 5809 | 5809 | 10.4210526 | 2.20801863 | 0 | 1 | 0 | 1 | 1 | 0.50143826 | 0 | 1 | 2 | 0.03799168 | 16 | 0.65565778 | FALSE | 1 | FALSE | 1 | FALSE | 1 | FALSE | 1 | FALSE | 0.87669812 | FALSE | 1 |
| 5810 | 5810 | 8.88 | 2.56624239 | 0 | 1 | 1 | 0.56109832 | 0 | 1 | 3 | 0.10474541 | 1 | 0.33902672 | 20 | 0.8152498 | FALSE | 1 | FALSE | 1 | FALSE | 1 | FALSE | 1 | FALSE | 1 | FALSE | 1 |
| 5811 | 5811 | 10.25 | 1.63935963 | 3 | 0.00725145 | 0 | 1 | 0 | 1 | 1 | 0.52895088 | 0 | 1 | 12 | 0.90922186 | FALSE | 0.25171133 | FALSE | 1 | FALSE | 1 | FALSE | 1 | FALSE | 1 | FALSE | 1 |
| 5812 | 5812 | 10.0588235 | 3.91784137 | 0 | 1 | 0 | 1 | 0 | 1 | 0 | 1 | 0 | 1 | 17 | 0.05550965 | FALSE | 1 | FALSE | 1 | FALSE | 1 | FALSE | 1 | FALSE | 1 | FALSE | 1 |
| 5813 | 5813 | 7.26190476 | 2.14985629 | 2 | 0.29286138 | 2 | 0.39616059 | 1 | 0.78589454 | 2 | 0.5811115 | 3 | 0.03099591 | 32 | 0.94677277 | FALSE | 1 | FALSE | 1 | FALSE | 1 | FALSE | 1 | FALSE | 0.82559785 | FALSE | 1 |
| 5814 | 5814 | 9.80555556 | 1.69671978 | 6 | 0.00026749 | 1 | 0.69480472 | 3 | 0.13783405 | 0 | 1 | 1 | 0.44938266 | 25 | 0.99349451 | TRUE | 0.03175788 | FALSE | 1 | FALSE | 1 | FALSE | 1 | FALSE | 1 | FALSE | 1 |
| 5815 | 5815 | 7.51724138 | 2.07814058 | 0 | 1 | 1 | 0.61539065 | 0 | 1 | 0 | 1 | 0 | 1 | 28 | 0.04589964 | FALSE | 1 | FALSE | 1 | FALSE | 1 | FALSE | 1 | FALSE | 1 | FALSE | 1 |
| 5816 | 5816 | 8.47368421 | 3.13500476 | 0 | 1 | 0 | 1 | 0 | 1 | 2 | 0.21610812 | 0 | 1 | 17 | 0.41141543 | FALSE | 1 | FALSE | 1 | FALSE | 1 | FALSE | 1 | FALSE | 1 | FALSE | 1 |
| 5817 | 5817 | 8.85714286 | 2.5872529 | 0 | 1 | 0 | 1 | 0 | 1 | 0 | 1 | 1 | 0.20679328 | 13 | 0.33283729 | FALSE | 1 | FALSE | 1 | FALSE | 1 | FALSE | 1 | FALSE | 1 | FALSE | 1 |
| 5818 | 5818 | 8.4375 | 3.12187344 | 1 | 0.34033054 | 1 | 0.40942991 | 0 | 1 | 0 | 1 | 1 | 0.23264612 | 13 | 0.76799161 | FALSE | 1 | FALSE | 1 | FALSE | 1 | FALSE | 1 | FALSE | 1 | FALSE | 1 |
| 5819 | 5819 | 7.30555556 | 4.21518053 | 1 | 0.59801736 | 3 | 0.10264583 | 1 | 0.72299506 | 1 | 0.80778317 | 1 | 0.44015537 | 28 | 0.83079716 | FALSE | 1 | FALSE | 1 | FALSE | 1 | FALSE | 1 | FALSE | 1 | FALSE | 1 |
| 5820 | 5820 | 4.40890688 | 1.43369007 | 9 | 0.18275203 | 19 | 0.00037366 | 10 | 0.39507228 | 8 | 0.88765487 | 2 | 0.91770534 | 199 | 0.95866599 | FALSE | 1 | FALSE | 0.15723145 | FALSE | 1 | FALSE | 1 | FALSE | 1 | FALSE | 1 |
| 5821 | 5821 | 9.36363636 | 2.63793057 | 0 | 1 | 0 | 1 | 0 | 1 | 1 | 0.40390032 | 0 | 1 | 10 | 0.46859937 | FALSE | 1 | FALSE | 1 | FALSE | 1 | FALSE | 1 | FALSE | 1 | FALSE | 1 |
| 5822 | 5822 | 9.30769231 | 2.1620722 | 0 | 1 | 0 | 1 | 0 | 1 | 1 | 0.43131129 | 1 | 0.18007762 | 10 | 0.71436964 | FALSE | 1 | FALSE | 1 | FALSE | 1 | FALSE | 1 | FALSE | 1 | FALSE | 1 |
| 5823 | 5823 | 5.88135593 | 2.00071806 | 0 | 1 | 4 | 0.12275177 | 2 | 0.63179508 | 2 | 0.76144533 | 0 | 1 | 51 | 0.41341988 | FALSE | 1 | FALSE | 1 | FALSE | 1 | FALSE | 1 | FALSE | 1 | FALSE | 1 |
| 5824 | 5824 | 9.23076923 | 3.44526087 | 0 | 1 | 0 | 1 | 1 | 0.37874417 | 2 | 0.1174583 | 0 | 1 | 10 | 0.86791642 | FALSE | 1 | FALSE | 1 | FALSE | 1 | FALSE | 1 | FALSE | 1 | FALSE | 1 |
| 5825 | 5825 | 12 | 3.16227766 | 1 | 0.14434519 | 0 | 1 | 1 | 0.197145 | 0 | 1 | 0 | 1 | 4 | 0.94743772 | FALSE | 1 | FALSE | 1 | FALSE | 1 | FALSE | 1 | FALSE | 1 | FALSE | 1 |
| 5826 | 5826 | 4.33333333 | 2.56038192 | 4 | 0.02189152 | 0 | 1 | 1 | 0.78589454 | 4 | 0.12508716 | 0 | 1 | 33 | 0.89139527 | FALSE | 0.46823848 | FALSE | 1 | FALSE | 1 | FALSE | 1 | FALSE | 1 | FALSE | 1 |

| | | | | | | | | | | | | | | | | | | | | | | | | | | | |
|---|---|---|---|---|---|---|---|---|---|---|---|---|---|---|---|---|---|---|---|---|---|---|---|---|---|---|---|
| 5827 | 5827 | 4.825 | 1.83331439 | 2 | 0.8184193 | 3 | 0.7513988 | 5 | 0.43318786 | 2 | 0.97676738 | 1 | 0.86490401 | 107 | 0.08734274 | FALSE | 1 | FALSE | 1 | FALSE | 1 | FALSE | 1 | FALSE | 1 | FALSE | 1 |
| 5828 | 5828 | 11.1818182 | 1.74867128 | 0 | 1 | 0 | 1 | 1 | 0.33149969 | 1 | 0.40390032 | 1 | 0.16638736 | 8 | 0.92152993 | FALSE | 1 | FALSE | 1 | FALSE | 1 | FALSE | 1 | FALSE | 1 | FALSE | 1 |
| 5829 | 5829 | 6.84210526 | 3.11639397 | 1 | 0.38989951 | 2 | 0.12434464 | 1 | 0.50143826 | 0 | 1 | 0 | 1 | 15 | 0.83648191 | FALSE | 1 | FALSE | 1 | FALSE | 1 | FALSE | 1 | FALSE | 1 | FALSE | 1 |
| 5830 | 5830 | 8.46153846 | 2.53028988 | 1 | 0.49162634 | 1 | 0.57534752 | 0 | 1 | 2 | 0.33678042 | 6 | 2.99E-06 | 16 | 0.9989746 | FALSE | 1 | FALSE | 1 | FALSE | 1 | FALSE | 1 | TRUE | 0.00271857 | FALSE | 1 |
| 5831 | 5831 | 9.83333333 | 2.1408721 | 0 | 1 | 0 | 1 | 0 | 1 | 2 | 0.19902682 | 0 | 1 | 16 | 0.44998001 | FALSE | 1 | FALSE | 1 | FALSE | 1 | FALSE | 1 | FALSE | 1 | FALSE | 1 |
| 5832 | 5832 | 8.77777778 | 2.09644025 | 0 | 1 | 0 | 1 | 0 | 1 | 0 | 1 | 0 | 1 | 9 | 0.21661397 | FALSE | 1 | FALSE | 1 | FALSE | 1 | FALSE | 1 | FALSE | 1 | FALSE | 1 |
| 5833 | 5833 | 4.21782178 | 1.10895509 | 5 | 0.98011367 | 6 | 0.99169422 | 14 | 0.58866481 | 19 | 0.48246799 | 5 | 0.79722026 | 353 | 0.02711877 | FALSE | 1 | FALSE | 1 | FALSE | 1 | FALSE | 1 | FALSE | 1 | FALSE | 1 |
| 5834 | 5834 | 8.875 | 2.57087048 | 0 | 1 | 0 | 1 | 1 | 0.25384309 | 0 | 1 | 0 | 1 | 7 | 0.63752397 | FALSE | 1 | FALSE | 1 | FALSE | 1 | FALSE | 1 | FALSE | 1 | FALSE | 1 |
| 5835 | 5835 | 9 | 1.77281052 | 1 | 0.38989951 | 0 | 1 | 0 | 1 | 1 | 0.59104357 | 0 | 1 | 17 | 0.41141543 | FALSE | 1 | FALSE | 1 | FALSE | 1 | FALSE | 1 | FALSE | 1 | FALSE | 1 |
| 5836 | 5836 | 9.21052632 | 2.11707214 | 1 | 0.38989951 | 1 | 0.46503795 | 2 | 0.14749808 | 0 | 1 | 0 | 1 | 15 | 0.83648191 | FALSE | 1 | FALSE | 1 | FALSE | 1 | FALSE | 1 | FALSE | 1 | FALSE | 1 |
| 5837 | 5837 | 6.37037037 | 1.23672651 | 5 | 0.01199096 | 2 | 0.52572379 | 0 | 1 | 1 | 0.92176876 | 1 | 0.59191961 | 45 | 0.66857827 | FALSE | 0.33872393 | FALSE | 1 | FALSE | 1 | FALSE | 1 | FALSE | 1 | FALSE | 1 |
| 5838 | 5838 | 6.88888889 | 3.49426338 | 0 | 1 | 2 | 0.32556427 | 1 | 0.73299778 | 1 | 0.81665383 | 1 | 0.44938266 | 31 | 0.49869404 | FALSE | 1 | FALSE | 1 | FALSE | 1 | FALSE | 1 | FALSE | 1 | FALSE | 1 |
| 5839 | 5839 | 10.7857143 | 1.73939938 | 0 | 1 | 0 | 1 | 2 | 0.08806398 | 0 | 1 | 0 | 1 | 12 | 0.62236227 | FALSE | 1 | FALSE | 1 | FALSE | 1 | FALSE | 1 | FALSE | 1 | FALSE | 1 |
| 5840 | 5840 | 8.55555556 | 2.4770552 | 4 | 0.01296492 | 2 | 0.32556427 | 2 | 0.37289966 | 3 | 0.22784656 | 2 | 0.11720035 | 23 | 0.99939163 | FALSE | 0.35760478 | FALSE | 1 | FALSE | 1 | FALSE | 1 | FALSE | 1 | FALSE | 1 |
| 5841 | 5841 | 9.1509434 | 2.96772721 | 5 | 0.01026504 | 4 | 0.08651261 | 3 | 0.28676219 | 4 | 0.21499032 | 1 | 0.57809055 | 35 | 0.99943348 | FALSE | 0.30467829 | FALSE | 1 | FALSE | 1 | FALSE | 1 | FALSE | 1 | FALSE | 1 |
| 5842 | 5842 | 4.71698113 | 2.87065411 | 1 | 0.74224579 | 1 | 0.82028131 | 1 | 0.85187773 | 2 | 0.6970685 | 0 | 1 | 47 | 0.15653036 | FALSE | 1 | FALSE | 1 | FALSE | 1 | FALSE | 1 | FALSE | 1 | FALSE | 1 |
| 5843 | 5843 | 9 | 1.3540064 | 0 | 1 | 1 | 0.32623933 | 1 | 0.35555293 | 0 | 1 | 0 | 1 | 10 | 0.71436964 | FALSE | 1 | FALSE | 1 | FALSE | 1 | FALSE | 1 | FALSE | 1 | FALSE | 1 |
| 5844 | 5844 | 9.84615385 | 2.62667247 | 1 | 0.28675951 | 0 | 1 | 2 | 0.07725803 | 0 | 1 | 1 | 0.19354507 | 9 | 0.96018168 | FALSE | 1 | FALSE | 1 | FALSE | 1 | FALSE | 1 | FALSE | 1 | FALSE | 1 |
| 5845 | 5845 | 10.1851852 | 2.89363333 | 0 | 1 | 0 | 1 | 2 | 0.25286935 | 0 | 1 | 1 | 0.36060486 | 24 | 0.3725373 | FALSE | 1 | FALSE | 1 | FALSE | 1 | FALSE | 1 | FALSE | 1 | FALSE | 1 |
| 5846 | 5846 | 7.85 | 1.79652442 | 0 | 1 | 0 | 1 | 0 | 1 | 1 | 0.60987146 | 0 | 1 | 19 | 0.15700255 | FALSE | 1 | FALSE | 1 | FALSE | 1 | FALSE | 1 | FALSE | 1 | FALSE | 1 |
| 5847 | 5847 | 9.28571429 | 2.73674915 | 0 | 1 | 0 | 1 | 0 | 1 | 1 | 0.48242099 | 1 | 0.20679328 | 12 | 0.62236227 | FALSE | 1 | FALSE | 1 | FALSE | 1 | FALSE | 1 | FALSE | 1 | FALSE | 1 |
| 5848 | 5848 | 8.5 | 2.16666667 | 0 | 1 | 1 | 0.44710673 | 0 | 1 | 2 | 0.19902682 | 1 | 0.25766374 | 14 | 0.86287418 | FALSE | 1 | FALSE | 1 | FALSE | 1 | FALSE | 1 | FALSE | 1 | FALSE | 1 |
| 5849 | 5849 | 10.1578947 | 2.0841042 | 1 | 0.38989951 | 2 | 0.12434464 | 0 | 1 | 0 | 1 | 1 | 0.26986775 | 15 | 0.83648191 | FALSE | 1 | FALSE | 1 | FALSE | 1 | FALSE | 1 | FALSE | 1 | FALSE | 1 |
| 5850 | 5850 | 10.1875 | 3.06632414 | 0 | 1 | 0 | 1 | 0 | 1 | 2 | 0.16544789 | 0 | 1 | 14 | 0.53308261 | FALSE | 1 | FALSE | 1 | FALSE | 1 | FALSE | 1 | FALSE | 1 | FALSE | 1 |
| 5851 | 5851 | 11 | 3.18478526 | 0 | 1 | 0 | 1 | 0 | 1 | 1 | 0.48242099 | 0 | 1 | 13 | 0.33283729 | FALSE | 1 | FALSE | 1 | FALSE | 1 | FALSE | 1 | FALSE | 1 | FALSE | 1 |
| 5852 | 5852 | 9.91666667 | 2.43099202 | 2 | 0.03643311 | 2 | 0.05552913 | 1 | 0.35555293 | 0 | 1 | 1 | 0.18007762 | 6 | 0.99914544 | FALSE | 0.58392 | FALSE | 1 | FALSE | 1 | FALSE | 1 | FALSE | 1 | FALSE | 1 |
| 5853 | 5853 | 9.26923077 | 1.05821665 | 2 | 0.38681045 | 0 | 1 | 1 | 0.85187773 | 1 | 0.91399378 | 0 | 1 | 48 | 0.0742821 | FALSE | 1 | FALSE | 1 | FALSE | 1 | FALSE | 1 | FALSE | 1 | FALSE | 1 |
| 5854 | 5854 | 11.2857143 | 1.70832296 | 2 | 0.04858661 | 0 | 1 | 0 | 1 | 0 | 1 | 1 | 0.20679328 | 11 | 0.83676449 | FALSE | 0.649301 | FALSE | 1 | FALSE | 1 | FALSE | 1 | FALSE | 1 | FALSE | 1 |
| 5855 | 5855 | 9.57142857 | 1.76126114 | 0 | 1 | 1 | 0.49918218 | 0 | 1 | 1 | 0.62783522 | 0 | 1 | 19 | 0.34094959 | FALSE | 1 | FALSE | 1 | FALSE | 1 | FALSE | 1 | FALSE | 1 | FALSE | 1 |
| 5856 | 5856 | 9 | 2.40370085 | 0 | 1 | 0 | 1 | 1 | 0.28067846 | 0 | 1 | 1 | 0.1383237 | 7 | 0.84574168 | FALSE | 1 | FALSE | 1 | FALSE | 1 | FALSE | 1 | FALSE | 1 | FALSE | 1 |
| 5857 | 5857 | 11.5882353 | 2.46287311 | 0 | 1 | 0 | 1 | 2 | 0.34668321 | 1 | 0.7984848 | 1 | 0.43077488 | 30 | 0.36879488 | FALSE | 1 | FALSE | 1 | FALSE | 1 | FALSE | 1 | FALSE | 1 | FALSE | 1 |
| 5858 | 5858 | 5.95522388 | 2.41532938 | 0 | 1 | 0 | 1 | 3 | 0.43448639 | 4 | 0.3701568 | 0 | 1 | 60 | 0.15737179 | FALSE | 1 | FALSE | 1 | FALSE | 1 | FALSE | 1 | FALSE | 1 | FALSE | 1 |
| 5859 | 5859 | 9.75 | 1.92028644 | 0 | 1 | 0 | 1 | 1 | 0.35555293 | 0 | 1 | 0 | 1 | 11 | 0.41944532 | FALSE | 1 | FALSE | 1 | FALSE | 1 | FALSE | 1 | FALSE | 1 | FALSE | 1 |
| 5860 | 5860 | 5.22162162 | 1.37529066 | 3 | 0.85406354 | 7 | 0.38020217 | 4 | 0.90057971 | 10 | 0.33118945 | 1 | 0.95346104 | 158 | 0.26647028 | FALSE | 1 | FALSE | 1 | FALSE | 1 | FALSE | 1 | FALSE | 1 | FALSE | 1 |
| 5861 | 5861 | 10.7222222 | 2.95908311 | 0 | 1 | 0 | 1 | 0 | 1 | 0 | 1 | 0 | 1 | 18 | 0.04681656 | FALSE | 1 | FALSE | 1 | FALSE | 1 | FALSE | 1 | FALSE | 1 | FALSE | 1 |
| 5862 | 5862 | 10.6153846 | 2.09535977 | 0 | 1 | 1 | 0.3480709 | 0 | 1 | 1 | 0.45746571 | 0 | 1 | 11 | 0.6683056 | FALSE | 1 | FALSE | 1 | FALSE | 1 | FALSE | 1 | FALSE | 1 | FALSE | 1 |
| 5863 | 5863 | 9.64285714 | 1.87491496 | 1 | 0.3050805 | 0 | 1 | 0 | 1 | 0 | 1 | 0 | 1 | 13 | 0.33283729 | FALSE | 1 | FALSE | 1 | FALSE | 1 | FALSE | 1 | FALSE | 1 | FALSE | 1 |
| 5864 | 5864 | 8.41791045 | 2.76518469 | 1 | 0.81665075 | 1 | 0.88323395 | 3 | 0.41519671 | 3 | 0.57949569 | 4 | 0.02154507 | 53 | 0.7938242 | FALSE | 1 | FALSE | 1 | FALSE | 1 | FALSE | 1 | FALSE | 0.74471385 | FALSE | 1 |
| 5865 | 5865 | 10.0833333 | 0.9537936 | 0 | 1 | 1 | 0.32623933 | 1 | 0.35555293 | 2 | 0.1023796 | 0 | 1 | 8 | 0.97197833 | FALSE | 1 | FALSE | 1 | FALSE | 1 | FALSE | 1 | FALSE | 1 | FALSE | 1 |
| 5866 | 5866 | 9.29166667 | 2.13071128 | 1 | 0.46441868 | 3 | 0.04096158 | 1 | 0.58502114 | 3 | 0.09525947 | 1 | 0.32796862 | 15 | 0.99800699 | FALSE | 1 | FALSE | 0.9698186 | FALSE | 1 | FALSE | 1 | FALSE | 1 | FALSE | 1 |
| 5867 | 5867 | 7.6372549 | 1.9593377 | 6 | 0.04468133 | 3 | 0.6400362 | 5 | 0.2969964 | 6 | 0.31850917 | 2 | 0.49622404 | 79 | 0.9637136 | FALSE | 0.649301 | FALSE | 1 | FALSE | 1 | FALSE | 1 | FALSE | 1 | FALSE | 1 |
| 5868 | 5868 | 8.6097561 | 1.87249734 | 0 | 1 | 1 | 0.74131698 | 2 | 0.43640583 | 3 | 0.29054213 | 3 | 0.02912063 | 32 | 0.90458039 | FALSE | 1 | FALSE | 1 | FALSE | 1 | FALSE | 1 | FALSE | 0.81978088 | FALSE | 1 |
| 5869 | 5869 | 8.96875 | 1.91187694 | 1 | 0.55378205 | 1 | 0.63997468 | 0 | 1 | 4 | 0.05164411 | 0 | 1 | 25 | 0.80006983 | FALSE | 1 | FALSE | 1 | FALSE | 1 | FALSE | 1 | FALSE | 1 | FALSE | 1 |
| 5870 | 5870 | 9.47058824 | 2.25225761 | 1 | 0.35728305 | 4 | 0.00181817 | 2 | 0.12280513 | 1 | 0.55062772 | 1 | 0.24525764 | 8 | 0.9999437 | FALSE | 1 | FALSE | 0.40311621 | FALSE | 1 | FALSE | 1 | FALSE | 1 | FALSE | 1 |
| 5871 | 5871 | 8.45 | 1.48800762 | 2 | 0.45796754 | 4 | 0.12838945 | 2 | 0.64095454 | 2 | 0.76960882 | 1 | 0.63076845 | 49 | 0.78152878 | FALSE | 1 | FALSE | 1 | FALSE | 1 | FALSE | 1 | FALSE | 1 | FALSE | 1 |
| 5872 | 5872 | 8.44444444 | 1.83249139 | 0 | 1 | 2 | 0.03227145 | 1 | 0.28067846 | 2 | 0.06108592 | 1 | 0.1383237 | 3 | 0.99994 | FALSE | 1 | FALSE | 0.92268438 | FALSE | 1 | FALSE | 1 | FALSE | 1 | FALSE | 1 |
| 5873 | 5873 | 10.3043478 | 2.38536813 | 0 | 1 | 0 | 1 | 1 | 0.5695037 | 0 | 1 | 2 | 0.05389003 | 20 | 0.50589402 | FALSE | 1 | FALSE | 1 | FALSE | 1 | FALSE | 1 | FALSE | 1 | FALSE | 1 |
| 5874 | 5874 | 10.6 | 2.65329983 | 2 | 0.0256865 | 0 | 1 | 0 | 1 | 0 | 1 | 0 | 1 | 8 | 0.80394719 | FALSE | 0.4974559 | FALSE | 1 | FALSE | 1 | FALSE | 1 | FALSE | 1 | FALSE | 1 |
| 5875 | 5875 | 9.2 | 1.16619038 | 0 | 1 | 0 | 1 | 1 | 0.30655256 | 1 | 0.37517262 | 0 | 1 | 8 | 0.80394719 | FALSE | 1 | FALSE | 1 | FALSE | 1 | FALSE | 1 | FALSE | 1 | FALSE | 1 |
| 5876 | 5876 | 8.25 | 2.16506351 | 0 | 1 | 1 | 0.32623933 | 2 | 0.06691409 | 0 | 1 | 0 | 1 | 9 | 0.89630573 | FALSE | 1 | FALSE | 1 | FALSE | 1 | FALSE | 1 | FALSE | 1 | FALSE | 1 |
| 5877 | 5877 | 4.19266055 | 2.76769392 | 3 | 0.53321268 | 1 | 0.9730424 | 1 | 0.98205774 | 3 | 0.88355644 | 2 | 0.53718295 | 99 | 0.0350382 | FALSE | 1 | FALSE | 1 | FALSE | 1 | FALSE | 1 | FALSE | 1 | FALSE | 1 |
| 5878 | 5878 | 10.3636364 | 2.05703791 | 0 | 1 | 0 | 1 | 0 | 1 | 1 | 0.40390032 | 0 | 1 | 10 | 0.46859937 | FALSE | 1 | FALSE | 1 | FALSE | 1 | FALSE | 1 | FALSE | 1 | FALSE | 1 |
| 5879 | 5879 | 6.5 | 1.69783776 | 7 | 0.23736061 | 7 | 0.44968026 | 3 | 0.97472887 | 9 | 0.54910082 | 2 | 0.83710781 | 168 | 0.34288158 | FALSE | 1 | FALSE | 1 | FALSE | 1 | FALSE | 1 | FALSE | 1 | FALSE | 1 |
| 5880 | 5880 | 8.72222222 | 2.90221286 | 1 | 0.60837712 | 1 | 0.69480472 | 4 | 0.03888854 | 1 | 0.81665383 | 1 | 0.44938266 | 28 | 0.90259714 | FALSE | 1 | FALSE | 1 | FALSE | 1 | FALSE | 1 | FALSE | 1 | FALSE | 1 |
| 5881 | 5881 | 8 | 2.51151196 | 0 | 1 | 1 | 0.3480709 | 1 | 0.37874417 | 0 | 1 | 0 | 1 | 11 | 0.6683056 | FALSE | 1 | FALSE | 1 | FALSE | 1 | FALSE | 1 | FALSE | 1 | FALSE | 1 |
| 5882 | 5882 | 8.76666667 | 2.15535509 | 1 | 0.54198714 | 3 | 0.07145121 | 0 | 1 | 1 | 0.75658797 | 0 | 1 | 25 | 0.67630471 | FALSE | 1 | FALSE | 1 | FALSE | 1 | FALSE | 1 | FALSE | 1 | FALSE | 1 |
| 5883 | 5883 | 9.36 | 2.48 | 0 | 1 | 1 | 0.56109832 | 0 | 1 | 0 | 1 | 0 | 1 | 24 | 0.08019798 | FALSE | 1 | FALSE | 1 | FALSE | 1 | FALSE | 1 | FALSE | 1 | FALSE | 1 |
| 5884 | 5884 | 10.0833333 | 2.15863897 | 2 | 0.12501436 | 0 | 1 | 1 | 0.58502114 | 1 | 0.67692816 | 0 | 1 | 20 | 0.68270729 | FALSE | 0.95622564 | FALSE | 1 | FALSE | 1 | FALSE | 1 | FALSE | 1 | FALSE | 1 |
| 5885 | 5885 | 10.3846154 | 2.30512678 | 1 | 0.28675951 | 1 | 0.3480709 | 2 | 0.07725803 | 0 | 1 | 0 | 1 | 9 | 0.96018168 | FALSE | 1 | FALSE | 1 | FALSE | 1 | FALSE | 1 | FALSE | 1 | FALSE | 1 |
| 5886 | 5886 | 10.4210526 | 2.41303019 | 0 | 1 | 2 | 0.12434464 | 0 | 1 | 2 | 0.21610812 | 0 | 1 | 15 | 0.83648191 | FALSE | 1 | FALSE | 1 | FALSE | 1 | FALSE | 1 | FALSE | 1 | FALSE | 1 |
| 5887 | 5887 | 6.46153846 | 1.58269737 | 1 | 0.86619543 | 2 | 0.71755606 | 5 | 0.14243046 | 6 | 0.14118821 | 1 | 0.72203879 | 62 | 0.86253757 | FALSE | 1 | FALSE | 1 | FALSE | 1 | FALSE | 1 | FALSE | 1 | FALSE | 1 |
| 5888 | 5888 | 9.84 | 3.06828943 | 1 | 0.47819878 | 1 | 0.56109832 | 1 | 0.59998149 | 0 | 1 | 0 | 1 | 22 | 0.43642846 | FALSE | 1 | FALSE | 1 | FALSE | 1 | FALSE | 1 | FALSE | 1 | FALSE | 1 |
| 5889 | 5889 | 10.3809524 | 2.19254561 | 2 | 0.09989125 | 1 | 0.49918218 | 0 | 1 | 1 | 0.62783522 | 1 | 0.29368245 | 16 | 0.90353674 | FALSE | 0.86047225 | FALSE | 1 | FALSE | 1 | FALSE | 1 | FALSE | 1 | FALSE | 1 |
| 5890 | 5890 | 9.03846154 | 3.10682585 | 2 | 0.14250978 | 1 | 0.57534752 | 1 | 0.61440466 | 1 | 0.70601446 | 0 | 1 | 21 | 0.78939668 | FALSE | 1 | FALSE | 1 | FALSE | 1 | FALSE | 1 | FALSE | 1 | FALSE | 1 |
| 5891 | 5891 | 7.23148148 | 1.12716033 | 5 | 0.1434475 | 8 | 0.02327788 | 4 | 0.54729645 | 6 | 0.37635117 | 0 | 1 | 85 | 0.95722654 | FALSE | 1 | FALSE | 0.82367276 | FALSE | 1 | FALSE | 1 | FALSE | 1 | FALSE | 1 |
| 5892 | 5892 | 6.83333333 | 3.53160335 | 0 | 1 | 0 | 1 | 1 | 0.66709967 | 0 | 1 | 1 | 0.39166989 | 28 | 0.13086908 | FALSE | 1 | FALSE | 1 | FALSE | 1 | FALSE | 1 | FALSE | 1 | FALSE | 1 |
| 5893 | 5893 | 8.31034483 | 1.55593299 | 0 | 1 | 2 | 0.24095584 | 1 | 0.65463982 | 2 | 0.3871811 | 0 | 1 | 24 | 0.70563334 | FALSE | 1 | FALSE | 1 | FALSE | 1 | FALSE | 1 | FALSE | 1 | FALSE | 1 |
| 5894 | 5894 | 9.29166667 | 2.15017764 | 0 | 1 | 1 | 0.53115654 | 3 | 0.04772439 | 1 | 0.6613267 | 2 | 0.05389003 | 16 | 0.98100709 | FALSE | 1 | FALSE | 1 | FALSE | 1 | FALSE | 1 | FALSE | 1 | FALSE | 1 |
| 5895 | 5895 | 9.63157895 | 1.66102462 | 2 | 0.08405377 | 0 | 1 | 1 | 0.50143826 | 0 | 1 | 0 | 1 | 16 | 0.65565778 | FALSE | 0.79236883 | FALSE | 1 | FALSE | 1 | FALSE | 1 | FALSE | 1 | FALSE | 1 |
| 5896 | 5896 | 9.625 | 3.03881145 | 0 | 1 | 0 | 1 | 0 | 1 | 1 | 0.31351192 | 0 | 1 | 7 | 0.63752397 | FALSE | 1 | FALSE | 1 | FALSE | 1 | FALSE | 1 | FALSE | 1 | FALSE | 1 |
| 5897 | 5897 | 9 | 2.5437351 | 2 | 0.21638713 | 1 | 0.6739483 | 0 | 1 | 0 | 1 | 1 | 0.43077488 | 30 | 0.36879488 | FALSE | 1 | FALSE | 1 | FALSE | 1 | FALSE | 1 | FALSE | 1 | FALSE | 1 |
| 5898 | 5898 | 9.76 | 2.64242313 | 1 | 0.47819878 | 1 | 0.56109832 | 1 | 0.59998149 | 1 | 0.69181315 | 1 | 0.33902672 | 20 | 0.8152498 | FALSE | 1 | FALSE | 1 | FALSE | 1 | FALSE | 1 | FALSE | 1 | FALSE | 1 |
| 5899 | 5899 | 9.25 | 2.7545417 | 3 | 0.01369554 | 2 | 0.1353446 | 0 | 1 | 1 | 0.60987146 | 0 | 1 | 14 | 0.97325593 | FALSE | 0.36625055 | FALSE | 1 | FALSE | 1 | FALSE | 1 | FALSE | 1 | FALSE | 1 |
| 5900 | 5900 | 10.4571429 | 2.35848447 | 3 | 0.0595675 | 0 | 1 | 1 | 0.72299506 | 2 | 0.48233989 | 3 | 0.01918344 | 26 | 0.96294373 | FALSE | 0.72532843 | FALSE | 1 | FALSE | 1 | FALSE | 1 | FALSE | 0.70156566 | FALSE | 1 |
| 5901 | 5901 | 12.5 | 3.8683699 | 1 | 0.51745948 | 0 | 1 | 0 | 1 | 2 | 0.3705495 | 0 | 1 | 25 | 0.34293 | FALSE | 1 | FALSE | 1 | FALSE | 1 | FALSE | 1 | FALSE | 1 | FALSE | 1 |
| 5902 | 5902 | 9.29411765 | 2.2690943 | 0 | 1 | 0 | 1 | 1 | 0.46348836 | 2 | 0.18212134 | 1 | 0.24525764 | 13 | 0.88719222 | FALSE | 1 | FALSE | 1 | FALSE | 1 | FALSE | 1 | FALSE | 1 | FALSE | 1 |
| 5903 | 5903 | 6.40740741 | 3.64396263 | 1 | 0.50471032 | 0 | 1 | 0 | 1 | 3 | 0.12475952 | 0 | 1 | 23 | 0.58307573 | FALSE | 1 | FALSE | 1 | FALSE | 1 | FALSE | 1 | FALSE | 1 | FALSE | 1 |
| 5904 | 5904 | 9.23076923 | 3.51160376 | 1 | 0.28675951 | 0 | 1 | 1 | 0.37874417 | 1 | 0.45746571 | 0 | 1 | 10 | 0.86791642 | FALSE | 1 | FALSE | 1 | FALSE | 1 | FALSE | 1 | FALSE | 1 | FALSE | 1 |
| 5905 | 5905 | 6.5862069 | 1.68530992 | 3 | 0.38658756 | 5 | 0.15016498 | 4 | 0.38141809 | 3 | 0.76976873 | 0 | 1 | 72 | 0.72239496 | FALSE | 1 | FALSE | 1 | FALSE | 1 | FALSE | 1 | FALSE | 1 | FALSE | 1 |
| 5906 | 5906 | 9.15384615 | 2.06978832 | 0 | 1 | 1 | 0.57534752 | 0 | 1 | 1 | 0.70601446 | 0 | 1 | 24 | 0.20425579 | FALSE | 1 | FALSE | 1 | FALSE | 1 | FALSE | 1 | FALSE | 1 | FALSE | 1 |
| 5907 | 5907 | 9.20833333 | 2.92943633 | 1 | 0.46441868 | 0 | 1 | 1 | 0.58502114 | 0 | 1 | 1 | 0.32796862 | 21 | 0.47053137 | FALSE | 1 | FALSE | 1 | FALSE | 1 | FALSE | 1 | FALSE | 1 | FALSE | 1 |
| 5908 | 5908 | 8.12 | 2.90268841 | 0 | 1 | 1 | 0.56109832 | 1 | 0.59998149 | 2 | 0.31968852 | 0 | 1 | 21 | 0.64957167 | FALSE | 1 | FALSE | 1 | FALSE | 1 | FALSE | 1 | FALSE | 1 | FALSE | 1 |
| 5909 | 5909 | 9.75 | 1.08972474 | 0 | 1 | 1 | 0.23139161 | 0 | 1 | 0 | 1 | 0 | 1 | 7 | 0.63752397 | FALSE | 1 | FALSE | 1 | FALSE | 1 | FALSE | 1 | FALSE | 1 | FALSE | 1 |
| 5910 | 5910 | 10.1666667 | 2.11476292 | 0 | 1 | 0 | 1 | 0 | 1 | 1 | 0.43131129 | 0 | 1 | 11 | 0.41944532 | FALSE | 1 | FALSE | 1 | FALSE | 1 | FALSE | 1 | FALSE | 1 | FALSE | 1 |
| 5911 | 5911 | 9.82926829 | 1.96186123 | 2 | 0.28329504 | 3 | 0.14580275 | 1 | 0.77786601 | 1 | 0.85524728 | 1 | 0.49330646 | 33 | 0.81996213 | FALSE | 1 | FALSE | 1 | FALSE | 1 | FALSE | 1 | FALSE | 1 | FALSE | 1 |
| 5912 | 5912 | 8.76923077 | 1.36741453 | 1 | 0.28675951 | 0 | 1 | 1 | 0.37874417 | 2 | 0.1174583 | 0 | 1 | 9 | 0.96018168 | FALSE | 1 | FALSE | 1 | FALSE | 1 | FALSE | 1 | FALSE | 1 | FALSE | 1 |
| 5913 | 5913 | 10.0909091 | 2.46630181 | 0 | 1 | 0 | 1 | 3 | 0.00609779 | 0 | 1 | 0 | 1 | 8 | 0.92152993 | FALSE | 1 | FALSE | 1 | FALSE | 1 | FALSE | 1 | FALSE | 1 | FALSE | 1 |
| 5914 | 5914 | 6.45454546 | 0.89072354 | 1 | 0.42086991 | 0 | 1 | 0 | 1 | 1 | 0.62783522 | 0 | 1 | 19 | 0.34094959 | FALSE | 1 | FALSE | 1 | FALSE | 1 | FALSE | 1 | FALSE | 1 | FALSE | 1 |
| 5915 | 5915 | 8.94117647 | 2.91992321 | 0 | 1 | 1 | 0.42857733 | 1 | 0.46348836 | 2 | 0.18212134 | 0 | 1 | 13 | 0.88719222 | FALSE | 1 | FALSE | 1 | FALSE | 1 | FALSE | 1 | FALSE | 1 | FALSE | 1 |
| 5916 | 5916 | 6.19565217 | 3.05471528 | 1 | 0.69044531 | 2 | 0.43020462 | 1 | 0.80828716 | 2 | 0.61900576 | 1 | 0.52593041 | 38 | 0.59437745 | FALSE | 1 | FALSE | 1 | FALSE | 1 | FALSE | 1 | FALSE | 1 | FALSE | 1 |
| 5917 | 5917 | 12.6666667 | 1.82574186 | 0 | 1 | 0 | 1 | 1 | 0.28067846 | 0 | 1 | 0 | 1 | 8 | 0.5780332 | FALSE | 1 | FALSE | 1 | FALSE | 1 | FALSE | 1 | FALSE | 1 | FALSE | 1 |
| 5918 | 5918 | 8.09090909 | 2.15130174 | 1 | 0.67382914 | 0 | 1 | 1 | 0.79363407 | 3 | 0.31602175 | 0 | 1 | 38 | 0.31730615 | FALSE | 1 | FALSE | 1 | FALSE | 1 | FALSE | 1 | FALSE | 1 | FALSE | 1 |
| 5919 | 5919 | 4.28042328 | 0.9870526 | 3 | 0.8664295 | 2 | 0.98587236 | 2 | 0.99233688 | 7 | 0.76762962 | 4 | 0.37157536 | 170 | 0.01014977 | FALSE | 1 | FALSE | 1 | FALSE | 1 | FALSE | 1 | FALSE | 1 | FALSE | 1 |
| 5920 | 5920 | 9 | 2.51151196 | 1 | 0.28675951 | 0 | 1 | 1 | 0.37874417 | 1 | 0.45746571 | 0 | 1 | 10 | 0.86791642 | FALSE | 1 | FALSE | 1 | FALSE | 1 | FALSE | 1 | FALSE | 1 | FALSE | 1 |

| | | | | | | | | | | | | | | | | | | | | | | | | | | | |
|---|---|---|---|---|---|---|---|---|---|---|---|---|---|---|---|---|---|---|---|---|---|---|---|---|---|---|---|
| 5921 | 5921 | 10.1764706 | 2.22909357 | 0 | 1 | 0 | 1 | 1 | 0.46348836 | 1 | 0.55062772 | 1 | 0.24525764 | 14 | 0.7313145 | FALSE | 1 | FALSE | 1 | FALSE | 1 | FALSE | 1 | FALSE | 1 | FALSE | 1 |
| 5922 | 5922 | 10.7777778 | 3.52066212 | 1 | 0.20854748 | 0 | 1 | 0 | 1 | 1 | 0.34506517 | 0 | 1 | 7 | 0.84574168 | FALSE | 1 | FALSE | 1 | FALSE | 1 | FALSE | 1 | FALSE | 1 | FALSE | 1 |
| 5923 | 5923 | 9.71428571 | 2.24971654 | 1 | 0.42086991 | 0 | 1 | 2 | 0.17308361 | 0 | 1 | 0 | 1 | 18 | 0.57962763 | FALSE | 1 | FALSE | 1 | FALSE | 1 | FALSE | 1 | FALSE | 1 | FALSE | 1 |
| 5924 | 5924 | 10.1538462 | 1.87477809 | 1 | 0.28675951 | 0 | 1 | 0 | 1 | 0 | 1 | 0 | 1 | 12 | 0.374198 | FALSE | 1 | FALSE | 1 | FALSE | 1 | FALSE | 1 | FALSE | 1 | FALSE | 1 |
| 5925 | 5925 | 9.49275362 | 1.49028923 | 2 | 0.53205439 | 4 | 0.18354682 | 4 | 0.23516085 | 2 | 0.8327303 | 3 | 0.1035305 | 54 | 0.93701512 | FALSE | 1 | FALSE | 1 | FALSE | 1 | FALSE | 1 | FALSE | 1 | FALSE | 1 |
| 5926 | 5926 | 9.5 | 2.36290781 | 0 | 1 | 0 | 1 | 1 | 0.197145 | 1 | 0.24578796 | 0 | 1 | 4 | 0.94743772 | FALSE | 1 | FALSE | 1 | FALSE | 1 | FALSE | 1 | FALSE | 1 | FALSE | 1 |
| 5927 | 5927 | 6.81818182 | 2.70029952 | 4 | 0.1347453 | 3 | 0.45651652 | 2 | 0.77041438 | 2 | 0.87529005 | 2 | 0.36111107 | 64 | 0.68868587 | FALSE | 0.98380879 | FALSE | 1 | FALSE | 1 | FALSE | 1 | FALSE | 1 | FALSE | 1 |
| 5928 | 5928 | 7.95652174 | 2.71033867 | 0 | 1 | 0 | 1 | 0 | 1 | 3 | 0.06912908 | 0 | 1 | 18 | 0.57962763 | FALSE | 1 | FALSE | 1 | FALSE | 1 | FALSE | 1 | FALSE | 1 | FALSE | 1 |
| 5929 | 5929 | 8.69230769 | 1.48762151 | 0 | 1 | 0 | 1 | 1 | 0.37874417 | 1 | 0.45746571 | 0 | 1 | 11 | 0.6683056 | FALSE | 1 | FALSE | 1 | FALSE | 1 | FALSE | 1 | FALSE | 1 | FALSE | 1 |
| 5930 | 5930 | 9.22727273 | 2.44822418 | 2 | 0.10809809 | 3 | 0.03266339 | 1 | 0.55340851 | 2 | 0.26791271 | 1 | 0.30529951 | 13 | 0.99911494 | FALSE | 0.88820416 | FALSE | 0.92268438 | FALSE | 1 | FALSE | 1 | FALSE | 1 | FALSE | 1 |
| 5931 | 5931 | 8.22222222 | 1.22726234 | 1 | 0.37380237 | 0 | 1 | 0 | 1 | 2 | 0.19902682 | 0 | 1 | 15 | 0.69372602 | FALSE | 1 | FALSE | 1 | FALSE | 1 | FALSE | 1 | FALSE | 1 | FALSE | 1 |
| 5932 | 5932 | 8.83333333 | 3.17367645 | 0 | 1 | 0 | 1 | 1 | 0.66709967 | 2 | 0.4036176 | 0 | 1 | 27 | 0.28862208 | FALSE | 1 | FALSE | 1 | FALSE | 1 | FALSE | 1 | FALSE | 1 | FALSE | 1 |
| 5933 | 5933 | 10.5384615 | 2.0981818 | 0 | 1 | 1 | 0.32623933 | 1 | 0.35555293 | 0 | 1 | 0 | 1 | 10 | 0.71436964 | FALSE | 1 | FALSE | 1 | FALSE | 1 | FALSE | 1 | FALSE | 1 | FALSE | 1 |
| 5934 | 5934 | 11.1428571 | 2.55949612 | 0 | 1 | 1 | 0.36919824 | 1 | 0.40110418 | 0 | 1 | 0 | 1 | 12 | 0.62236227 | FALSE | 1 | FALSE | 1 | FALSE | 1 | FALSE | 1 | FALSE | 1 | FALSE | 1 |
| 5935 | 5935 | 8.61403509 | 4.33975659 | 2 | 0.43181298 | 5 | 0.03627929 | 2 | 0.61293419 | 3 | 0.49002278 | 2 | 0.23984683 | 43 | 0.9744306 | FALSE | 1 | FALSE | 0.94259695 | FALSE | 1 | FALSE | 1 | FALSE | 1 | FALSE | 1 |
| 5936 | 5936 | 8.86206897 | 3.33966897 | 1 | 0.52988231 | 1 | 0.61539065 | 1 | 0.65463982 | 0 | 1 | 0 | 1 | 26 | 0.31495157 | FALSE | 1 | FALSE | 1 | FALSE | 1 | FALSE | 1 | FALSE | 1 | FALSE | 1 |
| 5937 | 5937 | 9.21428571 | 2.51221506 | 2 | 0.04858661 | 0 | 1 | 0 | 1 | 0 | 1 | 2 | 0.02131828 | 10 | 0.94579623 | FALSE | 0.649301 | FALSE | 1 | FALSE | 1 | FALSE | 1 | FALSE | 0.73999729 | FALSE | 1 |
| 5938 | 5938 | 9.2631579 | 2.1726071 | 1 | 0.38989951 | 0 | 1 | 1 | 0.50143826 | 0 | 1 | 1 | 0.26986775 | 16 | 0.65565778 | FALSE | 1 | FALSE | 1 | FALSE | 1 | FALSE | 1 | FALSE | 1 | FALSE | 1 |
| 5939 | 5939 | 9.04761905 | 1.86384627 | 1 | 0.42086991 | 0 | 1 | 0 | 1 | 1 | 0.62783522 | 2 | 0.04568221 | 17 | 0.77849076 | FALSE | 1 | FALSE | 1 | FALSE | 1 | FALSE | 1 | FALSE | 0.95710659 | FALSE | 1 |
| 5940 | 5940 | 9.11111111 | 2.13148068 | 0 | 1 | 0 | 1 | 0 | 1 | 0 | 1 | 0 | 1 | 9 | 0.21661397 | FALSE | 1 | FALSE | 1 | FALSE | 1 | FALSE | 1 | FALSE | 1 | FALSE | 1 |
| 5941 | 5941 | 8.09090909 | 1.91269501 | 0 | 1 | 0 | 1 | 1 | 0.70185639 | 1 | 0.78873814 | 2 | 0.10132035 | 29 | 0.39648374 | FALSE | 1 | FALSE | 1 | FALSE | 1 | FALSE | 1 | FALSE | 1 | FALSE | 1 |
| 5942 | 5942 | 8.51851852 | 2.64393604 | 1 | 0.50471032 | 0 | 1 | 0 | 1 | 1 | 0.71956339 | 1 | 0.36060486 | 24 | 0.3725373 | FALSE | 1 | FALSE | 1 | FALSE | 1 | FALSE | 1 | FALSE | 1 | FALSE | 1 |
| 5943 | 5943 | 4.54248366 | 1.98025194 | 4 | 0.55571867 | 5 | 0.55513976 | 5 | 0.64881352 | 4 | 0.92654513 | 5 | 0.10596901 | 130 | 0.47393519 | FALSE | 1 | FALSE | 1 | FALSE | 1 | FALSE | 1 | FALSE | 1 | FALSE | 1 |
| 5944 | 5944 | 10.9166667 | 3.88283259 | 1 | 0.26795838 | 0 | 1 | 0 | 1 | 0 | 1 | 0 | 1 | 11 | 0.41944532 | FALSE | 1 | FALSE | 1 | FALSE | 1 | FALSE | 1 | FALSE | 1 | FALSE | 1 |
| 5945 | 5945 | 8.34615385 | 1.23855709 | 0 | 1 | 0 | 1 | 1 | 0.61440466 | 2 | 0.33678042 | 2 | 0.0670823 | 21 | 0.78939668 | FALSE | 1 | FALSE | 1 | FALSE | 1 | FALSE | 1 | FALSE | 1 | FALSE | 1 |
| 5946 | 5946 | 11.6111111 | 1.67128988 | 1 | 0.37380237 | 2 | 0.1135776 | 2 | 0.13502329 | 1 | 0.57131013 | 0 | 1 | 12 | 0.98544073 | FALSE | 1 | FALSE | 1 | FALSE | 1 | FALSE | 1 | FALSE | 1 | FALSE | 1 |
| 5947 | 5947 | 8.55421687 | 3.2236065 | 1 | 0.88571933 | 3 | 0.50638104 | 4 | 0.34844549 | 5 | 0.33297 | 3 | 0.15488013 | 67 | 0.85823095 | FALSE | 1 | FALSE | 1 | FALSE | 1 | FALSE | 1 | FALSE | 1 | FALSE | 1 |
| 5948 | 5948 | 9.23529412 | 2.94117647 | 0 | 1 | 1 | 0.42857733 | 0 | 1 | 0 | 1 | 0 | 1 | 16 | 0.23070182 | FALSE | 1 | FALSE | 1 | FALSE | 1 | FALSE | 1 | FALSE | 1 | FALSE | 1 |
| 5949 | 5949 | 10.5 | 4.15331193 | 0 | 1 | 2 | 0.03949327 | 0 | 1 | 0 | 1 | 0 | 1 | 8 | 0.80394719 | FALSE | 1 | FALSE | 0.96864929 | FALSE | 1 | FALSE | 1 | FALSE | 1 | FALSE | 1 |
| 5950 | 5950 | 9.63333333 | 2.7505555 | 0 | 1 | 0 | 1 | 1 | 0.66709967 | 1 | 0.75658797 | 0 | 1 | 28 | 0.13086908 | FALSE | 1 | FALSE | 1 | FALSE | 1 | FALSE | 1 | FALSE | 1 | FALSE | 1 |
| 5951 | 5951 | 6.24390244 | 3.64109664 | 1 | 0.65632651 | 1 | 0.74131698 | 3 | 0.1814659 | 0 | 1 | 1 | 0.49330646 | 35 | 0.53616095 | FALSE | 1 | FALSE | 1 | FALSE | 1 | FALSE | 1 | FALSE | 1 | FALSE | 1 |
| 5952 | 5952 | 11.25 | 1.87638838 | 2 | 0.03087289 | 0 | 1 | 0 | 1 | 5 | 7.26E-05 | 0 | 1 | 4 | 0.99996333 | FALSE | 0.53356688 | FALSE | 1 | FALSE | 1 | FALSE | 0.29727645 | FALSE | 1 | FALSE | 1 |
| 5953 | 5953 | 9.41176471 | 1.28607124 | 2 | 0.06909038 | 1 | 0.42857733 | 1 | 0.46348836 | 0 | 1 | 0 | 1 | 13 | 0.88719222 | FALSE | 0.74276693 | FALSE | 1 | FALSE | 1 | FALSE | 1 | FALSE | 1 | FALSE | 1 |
| 5954 | 5954 | 10.3333333 | 3.25917508 | 0 | 1 | 1 | 0.38964394 | 0 | 1 | 0 | 1 | 2 | 0.02433796 | 12 | 0.80329917 | FALSE | 1 | FALSE | 1 | FALSE | 1 | FALSE | 1 | FALSE | 0.77578444 | FALSE | 1 |
| 5955 | 5955 | 7.66666667 | 3.31662479 | 1 | 0.60837712 | 1 | 0.69480472 | 3 | 0.13783405 | 3 | 0.22784656 | 0 | 1 | 28 | 0.90259714 | FALSE | 1 | FALSE | 1 | FALSE | 1 | FALSE | 1 | FALSE | 1 | FALSE | 1 |
| 5956 | 5956 | 10.3448276 | 2.23260884 | 0 | 1 | 0 | 1 | 1 | 0.65463982 | 6 | 0.00172865 | 0 | 1 | 22 | 0.92902102 | FALSE | 1 | FALSE | 1 | FALSE | 1 | FALSE | 1 | FALSE | 1 | FALSE | 1 |
| 5957 | 5957 | 7.04347826 | 2.99968492 | 2 | 0.11647716 | 2 | 0.16948764 | 0 | 1 | 1 | 0.6613267 | 0 | 1 | 18 | 0.86273155 | FALSE | 0.93181727 | FALSE | 1 | FALSE | 1 | FALSE | 1 | FALSE | 1 | FALSE | 1 |
| 5958 | 5958 | 9.74074074 | 2.11857032 | 1 | 0.49162634 | 1 | 0.57534752 | 1 | 0.61440466 | 0 | 1 | 1 | 0.3499045 | 22 | 0.61628 | FALSE | 1 | FALSE | 1 | FALSE | 1 | FALSE | 1 | FALSE | 1 | FALSE | 1 |
| 5959 | 5959 | 5.96385542 | 2.13657638 | 6 | 0.0193064 | 3 | 0.50638104 | 3 | 0.57805941 | 4 | 0.53352976 | 3 | 0.15488013 | 64 | 0.97183417 | FALSE | 0.44482142 | FALSE | 1 | FALSE | 1 | FALSE | 1 | FALSE | 1 | FALSE | 1 |
| 5960 | 5960 | 9.28571429 | 1.77664363 | 1 | 0.42086991 | 2 | 0.14655134 | 0 | 1 | 1 | 0.62783522 | 1 | 0.29368245 | 16 | 0.90353674 | FALSE | 1 | FALSE | 1 | FALSE | 1 | FALSE | 1 | FALSE | 1 | FALSE | 1 |
| 5961 | 5961 | 10 | 2.40370085 | 0 | 1 | 0 | 1 | 1 | 0.28067846 | 1 | 0.34506517 | 0 | 1 | 7 | 0.84574168 | FALSE | 1 | FALSE | 1 | FALSE | 1 | FALSE | 1 | FALSE | 1 | FALSE | 1 |
| 5962 | 5962 | 6.7826087 | 2.37663485 | 0 | 1 | 0 | 1 | 2 | 0.4962935 | 3 | 0.35428318 | 0 | 1 | 41 | 0.25430673 | FALSE | 1 | FALSE | 1 | FALSE | 1 | FALSE | 1 | FALSE | 1 | FALSE | 1 |
| 5963 | 5963 | 9.31578947 | 2.57626068 | 1 | 0.38989951 | 0 | 1 | 0 | 1 | 0 | 1 | 0 | 1 | 18 | 0.17880694 | FALSE | 1 | FALSE | 1 | FALSE | 1 | FALSE | 1 | FALSE | 1 | FALSE | 1 |
| 5964 | 5964 | 7.8 | 2.65706605 | 1 | 0.40558521 | 2 | 0.1353446 | 2 | 0.16019548 | 0 | 1 | 0 | 1 | 15 | 0.92113679 | FALSE | 1 | FALSE | 1 | FALSE | 1 | FALSE | 1 | FALSE | 1 | FALSE | 1 |
| 5965 | 5965 | 10.1538462 | 2.1248042 | 0 | 1 | 0 | 1 | 1 | 0.59998149 | 0 | 1 | 1 | 0.33902672 | 23 | 0.22727183 | FALSE | 1 | FALSE | 1 | FALSE | 1 | FALSE | 1 | FALSE | 1 | FALSE | 1 |
| 5966 | 5966 | 9.91666667 | 1.97730175 | 0 | 1 | 0 | 1 | 2 | 0.06691409 | 1 | 0.43131129 | 0 | 1 | 9 | 0.89630573 | FALSE | 1 | FALSE | 1 | FALSE | 1 | FALSE | 1 | FALSE | 1 | FALSE | 1 |
| 5967 | 5967 | 10.375 | 1.89983552 | 2 | 0.06197875 | 1 | 0.40942991 | 0 | 1 | 1 | 0.52895088 | 0 | 1 | 12 | 0.90922186 | FALSE | 0.72532843 | FALSE | 1 | FALSE | 1 | FALSE | 1 | FALSE | 1 | FALSE | 1 |
| 5968 | 5968 | 10.4615385 | 2.89868364 | 0 | 1 | 0 | 1 | 0 | 1 | 1 | 0.45746571 | 0 | 1 | 12 | 0.374198 | FALSE | 1 | FALSE | 1 | FALSE | 1 | FALSE | 1 | FALSE | 1 | FALSE | 1 |
| 5969 | 5969 | 6.23129252 | 2.53161782 | 3 | 0.73295332 | 5 | 0.51922696 | 6 | 0.43449609 | 8 | 0.36245304 | 1 | 0.9143283 | 124 | 0.55892246 | FALSE | 1 | FALSE | 1 | FALSE | 1 | FALSE | 1 | FALSE | 1 | FALSE | 1 |
| 5970 | 5970 | 10.7391304 | 2.08967215 | 1 | 0.45027683 | 1 | 0.53115654 | 0 | 1 | 1 | 0.6613267 | 2 | 0.05389003 | 18 | 0.86273155 | FALSE | 1 | FALSE | 1 | FALSE | 1 | FALSE | 1 | FALSE | 1 | FALSE | 1 |
| 5971 | 5971 | 11 | 3.60555128 | 1 | 0.14434519 | 0 | 1 | 2 | 0.01751815 | 0 | 1 | 0 | 1 | 3 | 0.9931827 | FALSE | 1 | FALSE | 1 | FALSE | 1 | FALSE | 1 | FALSE | 1 | FALSE | 1 |
| 5972 | 5972 | 7.61538462 | 2.09535977 | 1 | 0.47819878 | 1 | 0.56109832 | 1 | 0.59998149 | 0 | 1 | 0 | 1 | 22 | 0.43642846 | FALSE | 1 | FALSE | 1 | FALSE | 1 | FALSE | 1 | FALSE | 1 | FALSE | 1 |
| 5973 | 5973 | 8.45454546 | 2.31059364 | 0 | 1 | 1 | 0.30368014 | 0 | 1 | 0 | 1 | 1 | 0.16638736 | 9 | 0.75985366 | FALSE | 1 | FALSE | 1 | FALSE | 1 | FALSE | 1 | FALSE | 1 | FALSE | 1 |
| 5974 | 5974 | 8.73913044 | 1.72494212 | 2 | 0.3309085 | 2 | 0.44132825 | 1 | 0.81522035 | 0 | 1 | 0 | 1 | 41 | 0.25430673 | FALSE | 1 | FALSE | 1 | FALSE | 1 | FALSE | 1 | FALSE | 1 | FALSE | 1 |
| 5975 | 5975 | 3.95575221 | 2.58275212 | 5 | 0.69413824 | 9 | 0.30882657 | 7 | 0.70975018 | 10 | 0.59474621 | 1 | 0.97764755 | 194 | 0.30584595 | FALSE | 1 | FALSE | 1 | FALSE | 1 | FALSE | 1 | FALSE | 1 | FALSE | 1 |
| 5976 | 5976 | 9 | 3.24737656 | 1 | 0.57647348 | 2 | 0.28941836 | 0 | 1 | 2 | 0.45159487 | 0 | 1 | 28 | 0.58681477 | FALSE | 1 | FALSE | 1 | FALSE | 1 | FALSE | 1 | FALSE | 1 | FALSE | 1 |
| 5977 | 5977 | 11.1176471 | 3.41075036 | 0 | 1 | 1 | 0.42857733 | 0 | 1 | 1 | 0.55062772 | 0 | 1 | 15 | 0.49060435 | FALSE | 1 | FALSE | 1 | FALSE | 1 | FALSE | 1 | FALSE | 1 | FALSE | 1 |
| 5978 | 5978 | 8.08 | 2.59106156 | 1 | 0.47819878 | 1 | 0.56109832 | 2 | 0.22597287 | 0 | 1 | 0 | 1 | 21 | 0.64957167 | FALSE | 1 | FALSE | 1 | FALSE | 1 | FALSE | 1 | FALSE | 1 | FALSE | 1 |
| 5979 | 5979 | 10.9375 | 2.24913178 | 1 | 0.55378205 | 1 | 0.63997468 | 2 | 0.30673593 | 3 | 0.16836568 | 0 | 1 | 24 | 0.90138666 | FALSE | 1 | FALSE | 1 | FALSE | 1 | FALSE | 1 | FALSE | 1 | FALSE | 1 |
| 5980 | 5980 | 7.3030303 | 3.32561089 | 2 | 0.20692137 | 3 | 0.08959205 | 3 | 0.11365871 | 3 | 0.19161876 | 0 | 1 | 22 | 0.99710775 | FALSE | 1 | FALSE | 1 | FALSE | 1 | FALSE | 1 | FALSE | 1 | FALSE | 1 |
| 5981 | 5981 | 12.625 | 3.56852561 | 1 | 0.18769745 | 0 | 1 | 0 | 1 | 0 | 1 | 0 | 1 | 7 | 0.63752397 | FALSE | 1 | FALSE | 1 | FALSE | 1 | FALSE | 1 | FALSE | 1 | FALSE | 1 |
| 5982 | 5982 | 10.2962963 | 2.74624073 | 2 | 0.15144274 | 3 | 0.05519089 | 0 | 1 | 1 | 0.71956339 | 1 | 0.36060486 | 20 | 0.95133411 | FALSE | 1 | FALSE | 1 | FALSE | 1 | FALSE | 1 | FALSE | 1 | FALSE | 1 |
| 5983 | 5983 | 7.82352941 | 2.74187486 | 0 | 1 | 1 | 0.81422651 | 1 | 0.84631478 | 3 | 0.41729179 | 0 | 1 | 46 | 0.17033916 | FALSE | 1 | FALSE | 1 | FALSE | 1 | FALSE | 1 | FALSE | 1 | FALSE | 1 |
| 5984 | 5984 | 9.11764706 | 2.54101307 | 0 | 1 | 0 | 1 | 1 | 0.42266264 | 2 | 0.14906763 | 0 | 1 | 12 | 0.80329917 | FALSE | 1 | FALSE | 1 | FALSE | 1 | FALSE | 1 | FALSE | 1 | FALSE | 1 |
| 5985 | 5985 | 9.93333333 | 2.40739601 | 1 | 0.32293355 | 1 | 0.38964394 | 2 | 0.09928574 | 1 | 0.50623194 | 0 | 1 | 10 | 0.97980046 | FALSE | 1 | FALSE | 1 | FALSE | 1 | FALSE | 1 | FALSE | 1 | FALSE | 1 |
| 5986 | 5986 | 11.1428571 | 1.55182578 | 0 | 1 | 0 | 1 | 0 | 1 | 0 | 1 | 0 | 1 | 7 | 0.3043657 | FALSE | 1 | FALSE | 1 | FALSE | 1 | FALSE | 1 | FALSE | 1 | FALSE | 1 |
| 5987 | 5987 | 9.51851852 | 2.99839921 | 1 | 0.50471032 | 1 | 0.58913618 | 0 | 1 | 0 | 1 | 0 | 1 | 25 | 0.1832206 | FALSE | 1 | FALSE | 1 | FALSE | 1 | FALSE | 1 | FALSE | 1 | FALSE | 1 |
| 5988 | 5988 | 8.36363636 | 1.09594796 | 6 | 0.00662632 | 3 | 0.3604578 | 3 | 0.4248691 | 4 | 0.35957896 | 3 | 0.09357908 | 47 | 0.99808523 | FALSE | 0.24562356 | FALSE | 1 | FALSE | 1 | FALSE | 1 | FALSE | 1 | FALSE | 1 |
| 5989 | 5989 | 11.1538462 | 1.91536917 | 0 | 1 | 0 | 1 | 0 | 1 | 0 | 1 | 0 | 1 | 13 | 0.10967906 | FALSE | 1 | FALSE | 1 | FALSE | 1 | FALSE | 1 | FALSE | 1 | FALSE | 1 |
| 5990 | 5990 | 10.9285714 | 1.48633229 | 0 | 1 | 0 | 1 | 1 | 0.40110418 | 0 | 1 | 0 | 1 | 13 | 0.33283729 | FALSE | 1 | FALSE | 1 | FALSE | 1 | FALSE | 1 | FALSE | 1 | FALSE | 1 |
| 5991 | 5991 | 8.27272727 | 3.37614382 | 2 | 0.39594061 | 1 | 0.82613965 | 3 | 0.29666549 | 2 | 0.70707241 | 1 | 0.58506217 | 44 | 0.69042405 | FALSE | 1 | FALSE | 1 | FALSE | 1 | FALSE | 1 | FALSE | 1 | FALSE | 1 |
| 5992 | 5992 | 8.42857143 | 3.01695887 | 1 | 0.51745948 | 0 | 1 | 0 | 1 | 4 | 0.03737468 | 1 | 0.37113067 | 22 | 0.86499992 | FALSE | 1 | FALSE | 1 | FALSE | 1 | FALSE | 1 | FALSE | 1 | FALSE | 1 |
| 5993 | 5993 | 11.0714286 | 1.94437966 | 1 | 0.50471032 | 0 | 1 | 1 | 0.62830986 | 2 | 0.35374209 | 0 | 1 | 23 | 0.58307573 | FALSE | 1 | FALSE | 1 | FALSE | 1 | FALSE | 1 | FALSE | 1 | FALSE | 1 |
| 5994 | 5994 | 6.26495727 | 3.25390801 | 0 | 1 | 4 | 0.44286374 | 6 | 0.17549144 | 3 | 0.86756002 | 1 | 0.82614736 | 91 | 0.31121647 | FALSE | 1 | FALSE | 1 | FALSE | 1 | FALSE | 1 | FALSE | 1 | FALSE | 1 |
| 5995 | 5995 | 9.73333333 | 1.06249183 | 0 | 1 | 0 | 1 | 1 | 0.42266264 | 0 | 1 | 0 | 1 | 14 | 0.29525409 | FALSE | 1 | FALSE | 1 | FALSE | 1 | FALSE | 1 | FALSE | 1 | FALSE | 1 |
| 5996 | 5996 | 8.6 | 1.94250697 | 1 | 0.54198714 | 0 | 1 | 0 | 1 | 4 | 0.04659941 | 0 | 1 | 25 | 0.67630471 | FALSE | 1 | FALSE | 1 | FALSE | 1 | FALSE | 1 | FALSE | 1 | FALSE | 1 |
| 5997 | 5997 | 9.08695652 | 3.17480631 | 0 | 1 | 3 | 0.03669128 | 2 | 0.19931429 | 0 | 1 | 1 | 0.31672722 | 17 | 0.94439745 | FALSE | 1 | FALSE | 0.94259695 | FALSE | 1 | FALSE | 1 | FALSE | 1 | FALSE | 1 |
| 5998 | 5998 | 10.3333333 | 2.00792082 | 1 | 0.40558521 | 1 | 0.48239024 | 1 | 0.51939841 | 3 | 0.0612581 | 0 | 1 | 14 | 0.97325593 | FALSE | 1 | FALSE | 1 | FALSE | 1 | FALSE | 1 | FALSE | 1 | FALSE | 1 |
| 5999 | 5999 | 10.2222222 | 2.34652357 | 0 | 1 | 0 | 1 | 1 | 0.25384309 | 1 | 0.31351192 | 0 | 1 | 6 | 0.88425551 | FALSE | 1 | FALSE | 1 | FALSE | 1 | FALSE | 1 | FALSE | 1 | FALSE | 1 |
| 6000 | 6000 | 9.82352941 | 3.27621184 | 0 | 1 | 0 | 1 | 1 | 0.46348836 | 0 | 1 | 0 | 1 | 16 | 0.23070182 | FALSE | 1 | FALSE | 1 | FALSE | 1 | FALSE | 1 | FALSE | 1 | FALSE | 1 |
| 6001 | 6001 | 4.48198198 | 1.78763008 | 8 | 0.20620087 | 7 | 0.57765307 | 10 | 0.27105832 | 12 | 0.31575848 | 2 | 0.88286182 | 182 | 0.82659057 | FALSE | 1 | FALSE | 1 | FALSE | 1 | FALSE | 1 | FALSE | 1 | FALSE | 1 |
| 6002 | 6002 | 9.30555556 | 1.72915272 | 0 | 1 | 1 | 0.69480472 | 1 | 0.73299778 | 6 | 0.00541176 | 0 | 1 | 28 | 0.90259714 | FALSE | 1 | FALSE | 1 | FALSE | 1 | FALSE | 1 | FALSE | 1 | FALSE | 1 |
| 6003 | 6003 | 6.34285714 | 4.18120186 | 0 | 1 | 3 | 0.10264583 | 3 | 0.12958859 | 1 | 0.80778317 | 0 | 1 | 28 | 0.83079716 | FALSE | 1 | FALSE | 1 | FALSE | 1 | FALSE | 1 | FALSE | 1 | FALSE | 1 |
| 6004 | 6004 | 9.30434783 | 1.75535906 | 1 | 0.45027683 | 0 | 1 | 0 | 1 | 1 | 0.6613267 | 0 | 1 | 21 | 0.27964072 | FALSE | 1 | FALSE | 1 | FALSE | 1 | FALSE | 1 | FALSE | 1 | FALSE | 1 |
| 6005 | 6005 | 10.4615385 | 2.27411469 | 1 | 0.28675951 | 2 | 0.06426041 | 0 | 1 | 1 | 0.45746571 | 0 | 1 | 9 | 0.96018168 | FALSE | 1 | FALSE | 1 | FALSE | 1 | FALSE | 1 | FALSE | 1 | FALSE | 1 |
| 6006 | 6006 | 10.6041667 | 1.14999245 | 0 | 1 | 3 | 0.20208567 | 3 | 0.24750546 | 3 | 0.37966044 | 1 | 0.54902428 | 38 | 0.88187597 | FALSE | 1 | FALSE | 1 | FALSE | 1 | FALSE | 1 | FALSE | 1 | FALSE | 1 |
| 6007 | 6007 | 9.89473684 | 2.77004155 | 2 | 0.08405377 | 0 | 1 | 2 | 0.14749808 | 0 | 1 | 0 | 1 | 15 | 0.83648191 | FALSE | 0.79236883 | FALSE | 1 | FALSE | 1 | FALSE | 1 | FALSE | 1 | FALSE | 1 |
| 6008 | 6008 | 10.25 | 2.33184476 | 0 | 1 | 1 | 0.23139161 | 0 | 1 | 0 | 1 | 0 | 1 | 7 | 0.63752397 | FALSE | 1 | FALSE | 1 | FALSE | 1 | FALSE | 1 | FALSE | 1 | FALSE | 1 |
| 6009 | 6009 | 9.57142857 | 2.99318955 | 0 | 1 | 0 | 1 | 0 | 1 | 1 | 0.48242099 | 1 | 0.20679328 | 12 | 0.62236227 | FALSE | 1 | FALSE | 1 | FALSE | 1 | FALSE | 1 | FALSE | 1 | FALSE | 1 |
| 6010 | 6010 | 7.79310345 | 1.72964637 | 1 | 0.49162634 | 0 | 1 | 0 | 1 | 1 | 0.70601446 | 0 | 1 | 24 | 0.20425579 | FALSE | 1 | FALSE | 1 | FALSE | 1 | FALSE | 1 | FALSE | 1 | FALSE | 1 |
| 6011 | 6011 | 7 | 3.36650165 | 0 | 1 | 0 | 1 | 0 | 1 | 0 | 1 | 0 | 1 | 6 | 0.36077158 | FALSE | 1 | FALSE | 1 | FALSE | 1 | FALSE | 1 | FALSE | 1 | FALSE | 1 |
| 6012 | 6012 | 9.95833333 | 1.59371596 | 1 | 0.46441868 | 1 | 0.54637326 | 0 | 1 | 3 | 0.09525947 | 2 | 0.058175 | 17 | 0.97530702 | FALSE | 1 | FALSE | 1 | FALSE | 1 | FALSE | 1 | FALSE | 1 | FALSE | 1 |
| 6013 | 6013 | 8.77777778 | 3.37565723 | 1 | 0.37380237 | 0 | 1 | 0 | 1 | 1 | 0.57131013 | 0 | 1 | 16 | 0.44998001 | FALSE | 1 | FALSE | 1 | FALSE | 1 | FALSE | 1 | FALSE | 1 | FALSE | 1 |
| 6014 | 6014 | 8.96296296 | 2.4717887 | 1 | 0.49162634 | 3 | 0.05021553 | 0 | 1 | 1 | 0.70601446 | 1 | 0.3499045 | 20 | 0.90142689 | FALSE | 1 | FALSE | 1 | FALSE | 1 | FALSE | 1 | FALSE | 1 | FALSE | 1 |

| | | | | | | | | | | | | | | | | | | | | | | | | | | | |
|---|---|---|---|---|---|---|---|---|---|---|---|---|---|---|---|---|---|---|---|---|---|---|---|---|---|---|---|
| 6015 | 6015 | 5.4516129 | 3.35855873 | 0 | 1 | 0 | 1 | 1 | 0.67911179 | 1 | 0.76781284 | 1 | 0.4016888 | 28 | 0.26393996 | FALSE | 1 | FALSE | 1 | FALSE | 1 | FALSE | 1 | FALSE | 1 | FALSE | 1 |
| 6016 | 6016 | 8 | 1.86700677 | 1 | 0.59801736 | 1 | 0.68454804 | 1 | 0.72299506 | 1 | 0.80778317 | 1 | 0.44015537 | 30 | 0.52765882 | FALSE | 1 | FALSE | 1 | FALSE | 1 | FALSE | 1 | FALSE | 1 | FALSE | 1 |
| 6017 | 6017 | 8.64705882 | 2.05630098 | 0 | 1 | 0 | 1 | 1 | 0.46348836 | 0 | 1 | 0 | 1 | 16 | 0.23070182 | FALSE | 1 | FALSE | 1 | FALSE | 1 | FALSE | 1 | FALSE | 1 | FALSE | 1 |
| 6018 | 6018 | 4.08270677 | 2.36848073 | 7 | 0.04741962 | 4 | 0.605178 | 5 | 0.49638206 | 5 | 0.71314143 | 0 | 1 | 108 | 0.63841443 | FALSE | 0.649301 | FALSE | 1 | FALSE | 1 | FALSE | 1 | FALSE | 1 | FALSE | 1 |
| 6019 | 6019 | 9.875 | 2.49687304 | 0 | 1 | 0 | 1 | 2 | 0.11087977 | 1 | 0.52895088 | 0 | 1 | 13 | 0.76799161 | FALSE | 1 | FALSE | 1 | FALSE | 1 | FALSE | 1 | FALSE | 1 | FALSE | 1 |
| 6020 | 6020 | 10.4242424 | 2.87128208 | 2 | 0.20692137 | 0 | 1 | 1 | 0.70185639 | 0 | 1 | 1 | 0.42123867 | 29 | 0.39648374 | FALSE | 1 | FALSE | 1 | FALSE | 1 | FALSE | 1 | FALSE | 1 | FALSE | 1 |
| 6021 | 6021 | 11.6153846 | 2.8966416 | 1 | 0.28675951 | 0 | 1 | 0 | 1 | 1 | 0.45746571 | 0 | 1 | 11 | 0.6683056 | FALSE | 1 | FALSE | 1 | FALSE | 1 | FALSE | 1 | FALSE | 1 | FALSE | 1 |
| 6022 | 6022 | 6.05128205 | 2.6980403 | 0 | 1 | 0 | 1 | 1 | 0.76089851 | 2 | 0.54052736 | 0 | 1 | 36 | 0.12091774 | FALSE | 1 | FALSE | 1 | FALSE | 1 | FALSE | 1 | FALSE | 1 | FALSE | 1 |
| 6023 | 6023 | 10.3636364 | 2.70597747 | 0 | 1 | 0 | 1 | 1 | 0.33149969 | 2 | 0.08789566 | 0 | 1 | 8 | 0.92152993 | FALSE | 1 | FALSE | 1 | FALSE | 1 | FALSE | 1 | FALSE | 1 | FALSE | 1 |
| 6024 | 6024 | 1.1467639 | 1.32961877 | 28 | 0.53032826 | 26 | 0.9702772 | 27 | 0.990547 | 57 | 0.1553178 | 13 | 0.92472151 | 940 | 0.04099252 | FALSE | 1 | FALSE | 1 | FALSE | 1 | FALSE | 1 | FALSE | 1 | FALSE | 1 |
| 6025 | 6025 | 7.46153846 | 3.33471372 | 2 | 0.26413583 | 1 | 0.7236252 | 3 | 0.16357218 | 1 | 0.84089119 | 0 | 1 | 32 | 0.74324845 | FALSE | 1 | FALSE | 1 | FALSE | 1 | FALSE | 1 | FALSE | 1 | FALSE | 1 |
| 6026 | 6026 | 9.59090909 | 1.94607042 | 1 | 0.4357638 | 2 | 0.15794013 | 0 | 1 | 1 | 0.64497439 | 0 | 1 | 18 | 0.74744813 | FALSE | 1 | FALSE | 1 | FALSE | 1 | FALSE | 1 | FALSE | 1 | FALSE | 1 |
| 6027 | 6027 | 9.61111111 | 2.54162872 | 0 | 1 | 1 | 0.44710673 | 0 | 1 | 0 | 1 | 1 | 0.25766374 | 16 | 0.44998001 | FALSE | 1 | FALSE | 1 | FALSE | 1 | FALSE | 1 | FALSE | 1 | FALSE | 1 |
| 6028 | 6028 | 6.20689655 | 3.64246327 | 2 | 0.16962005 | 0 | 1 | 1 | 0.65463982 | 4 | 0.04184267 | 0 | 1 | 22 | 0.92902102 | FALSE | 1 | FALSE | 1 | FALSE | 1 | FALSE | 1 | FALSE | 1 | FALSE | 1 |
| 6029 | 6029 | 8.63636364 | 2.8049543 | 1 | 0.4357638 | 0 | 1 | 0 | 1 | 1 | 0.64497439 | 1 | 0.30529951 | 19 | 0.54233419 | FALSE | 1 | FALSE | 1 | FALSE | 1 | FALSE | 1 | FALSE | 1 | FALSE | 1 |
| 6030 | 6030 | 7.62962963 | 3.46033769 | 0 | 1 | 1 | 0.83180787 | 2 | 0.58328102 | 1 | 0.92176876 | 0 | 1 | 50 | 0.06046652 | FALSE | 1 | FALSE | 1 | FALSE | 1 | FALSE | 1 | FALSE | 1 | FALSE | 1 |
| 6031 | 6031 | 7.5 | 2.72335577 | 2 | 0.23542589 | 5 | 0.0056244 | 3 | 0.13783405 | 1 | 0.81665383 | 2 | 0.11720035 | 23 | 0.99939163 | FALSE | 1 | FALSE | 0.59070661 | FALSE | 1 | FALSE | 1 | FALSE | 1 | FALSE | 1 |
| 6032 | 6032 | 9.82352941 | 2.50259381 | 2 | 0.06909038 | 2 | 0.10307112 | 0 | 1 | 1 | 0.55062772 | 0 | 1 | 12 | 0.962093 | FALSE | 0.74276693 | FALSE | 1 | FALSE | 1 | FALSE | 1 | FALSE | 1 | FALSE | 1 |
| 6033 | 6033 | 9.85714286 | 3.42380621 | 1 | 0.59801736 | 1 | 0.68454804 | 2 | 0.35984046 | 1 | 0.80778317 | 0 | 1 | 30 | 0.52765882 | FALSE | 1 | FALSE | 1 | FALSE | 1 | FALSE | 1 | FALSE | 1 | FALSE | 1 |
| 6034 | 6034 | 9 | 3.09838668 | 0 | 1 | 1 | 0.48239024 | 2 | 0.16019548 | 1 | 0.60987146 | 1 | 0.28187291 | 15 | 0.92113679 | FALSE | 1 | FALSE | 1 | FALSE | 1 | FALSE | 1 | FALSE | 1 | FALSE | 1 |
| 6035 | 6035 | 9.8 | 2.11187121 | 0 | 1 | 2 | 0.1353446 | 1 | 0.51939841 | 0 | 1 | 1 | 0.28187291 | 16 | 0.80826361 | FALSE | 1 | FALSE | 1 | FALSE | 1 | FALSE | 1 | FALSE | 1 | FALSE | 1 |
| 6036 | 6036 | 10.5 | 2.39791576 | 0 | 1 | 0 | 1 | 0 | 1 | 0 | 1 | 0 | 1 | 8 | 0.25677166 | FALSE | 1 | FALSE | 1 | FALSE | 1 | FALSE | 1 | FALSE | 1 | FALSE | 1 |
| 6037 | 6037 | 6.84615385 | 3.50485715 | 0 | 1 | 0 | 1 | 1 | 0.59998149 | 0 | 1 | 1 | 0.33902672 | 23 | 0.22727183 | FALSE | 1 | FALSE | 1 | FALSE | 1 | FALSE | 1 | FALSE | 1 | FALSE | 1 |
| 6038 | 6038 | 9.33333333 | 3.65452535 | 0 | 1 | 0 | 1 | 0 | 1 | 2 | 0.35374209 | 0 | 1 | 25 | 0.1832206 | FALSE | 1 | FALSE | 1 | FALSE | 1 | FALSE | 1 | FALSE | 1 | FALSE | 1 |
| 6039 | 6039 | 8.90909091 | 2.15130174 | 0 | 1 | 1 | 0.48239024 | 2 | 0.16019548 | 1 | 0.60987146 | 2 | 0.04176963 | 14 | 0.97325593 | FALSE | 1 | FALSE | 1 | FALSE | 1 | FALSE | 1 | FALSE | 0.90763079 | FALSE | 1 |
| 6040 | 6040 | 12.4444444 | 3.05908887 | 0 | 1 | 1 | 0.25628163 | 0 | 1 | 0 | 1 | 0 | 1 | 8 | 0.5780332 | FALSE | 1 | FALSE | 1 | FALSE | 1 | FALSE | 1 | FALSE | 1 | FALSE | 1 |
| 6041 | 6041 | 9.76190476 | 1.74314335 | 0 | 1 | 1 | 0.49918218 | 1 | 0.53671415 | 0 | 1 | 0 | 1 | 19 | 0.34094959 | FALSE | 1 | FALSE | 1 | FALSE | 1 | FALSE | 1 | FALSE | 1 | FALSE | 1 |
| 6042 | 6042 | 8.71428571 | 2.71052371 | 0 | 1 | 0 | 1 | 0 | 1 | 0 | 1 | 0 | 1 | 7 | 0.3043657 | FALSE | 1 | FALSE | 1 | FALSE | 1 | FALSE | 1 | FALSE | 1 | FALSE | 1 |
| 6043 | 6043 | 10.3684211 | 2.34432184 | 0 | 1 | 0 | 1 | 3 | 0.02898764 | 1 | 0.59104357 | 0 | 1 | 15 | 0.83648191 | FALSE | 1 | FALSE | 1 | FALSE | 1 | FALSE | 1 | FALSE | 1 | FALSE | 1 |
| 6044 | 6044 | 10.5714286 | 1.94131592 | 0 | 1 | 1 | 0.49918218 | 0 | 1 | 1 | 0.62783522 | 1 | 0.29368245 | 18 | 0.57962763 | FALSE | 1 | FALSE | 1 | FALSE | 1 | FALSE | 1 | FALSE | 1 | FALSE | 1 |
| 6045 | 6045 | 7.77777778 | 3.58322567 | 1 | 0.20854748 | 0 | 1 | 0 | 1 | 1 | 0.34506517 | 0 | 1 | 7 | 0.84574168 | FALSE | 1 | FALSE | 1 | FALSE | 1 | FALSE | 1 | FALSE | 1 | FALSE | 1 |
| 6046 | 6046 | 5.08928571 | 2.89463863 | 7 | 0.07193059 | 6 | 0.31100855 | 4 | 0.75702547 | 9 | 0.20449345 | 6 | 0.0287529 | 110 | 0.98955935 | FALSE | 0.76727438 | FALSE | 1 | FALSE | 1 | FALSE | 1 | FALSE | 0.81978088 | FALSE | 1 |
| 6047 | 6047 | 4.41975309 | 2.40197551 | 24 | 6.15E-05 | 14 | 0.40102836 | 20 | 0.07441536 | 15 | 0.81462743 | 7 | 0.47258182 | 315 | 0.99543597 | TRUE | 0.01326811 | FALSE | 1 | FALSE | 1 | FALSE | 1 | FALSE | 1 | FALSE | 1 |
| 6048 | 6048 | 8.85714286 | 3.24089515 | 1 | 0.42086991 | 2 | 0.14655134 | 1 | 0.53671415 | 0 | 1 | 1 | 0.29368245 | 16 | 0.90353674 | FALSE | 1 | FALSE | 1 | FALSE | 1 | FALSE | 1 | FALSE | 1 | FALSE | 1 |
| 6049 | 6049 | 10.8333333 | 2.07498327 | 0 | 1 | 0 | 1 | 0 | 1 | 1 | 0.43131129 | 0 | 1 | 11 | 0.41944532 | FALSE | 1 | FALSE | 1 | FALSE | 1 | FALSE | 1 | FALSE | 1 | FALSE | 1 |
| 6050 | 6050 | 6.390625 | 1.4099777 | 1 | 0.81177881 | 0 | 1 | 2 | 0.67580769 | 3 | 0.56885732 | 1 | 0.65460803 | 57 | 0.19643469 | FALSE | 1 | FALSE | 1 | FALSE | 1 | FALSE | 1 | FALSE | 1 | FALSE | 1 |
| 6051 | 6051 | 6.425 | 2.6635268 | 0 | 1 | 2 | 0.37295654 | 0 | 1 | 2 | 0.5543522 | 0 | 1 | 36 | 0.22908535 | FALSE | 1 | FALSE | 1 | FALSE | 1 | FALSE | 1 | FALSE | 1 | FALSE | 1 |
| 6052 | 6052 | 9 | 2.44948974 | 0 | 1 | 1 | 0.30368014 | 0 | 1 | 0 | 1 | 0 | 1 | 10 | 0.46859937 | FALSE | 1 | FALSE | 1 | FALSE | 1 | FALSE | 1 | FALSE | 1 | FALSE | 1 |
| 6053 | 6053 | 7.41666667 | 1.81620789 | 1 | 0.60837712 | 1 | 0.69480472 | 3 | 0.13783405 | 2 | 0.49731009 | 1 | 0.44938266 | 28 | 0.90259714 | FALSE | 1 | FALSE | 1 | FALSE | 1 | FALSE | 1 | FALSE | 1 | FALSE | 1 |
| 6054 | 6054 | 7.96153846 | 1.37227406 | 1 | 0.47819878 | 2 | 0.19297191 | 0 | 1 | 2 | 0.31968852 | 2 | 0.06257402 | 18 | 0.96851178 | FALSE | 1 | FALSE | 1 | FALSE | 1 | FALSE | 1 | FALSE | 1 | FALSE | 1 |
| 6055 | 6055 | 8.25 | 2.91547595 | 0 | 1 | 0 | 1 | 0 | 1 | 1 | 0.77852168 | 2 | 0.096172 | 29 | 0.24088541 | FALSE | 1 | FALSE | 1 | FALSE | 1 | FALSE | 1 | FALSE | 1 | FALSE | 1 |
| 6056 | 6056 | 7.84210526 | 2.75801527 | 0 | 1 | 0 | 1 | 1 | 0.50143826 | 0 | 1 | 0 | 1 | 18 | 0.17880694 | FALSE | 1 | FALSE | 1 | FALSE | 1 | FALSE | 1 | FALSE | 1 | FALSE | 1 |
| 6057 | 6057 | 9.91935484 | 1.33559685 | 4 | 0.07395961 | 2 | 0.60114557 | 2 | 0.65873577 | 2 | 0.78520976 | 1 | 0.64288531 | 51 | 0.74578096 | FALSE | 0.77424195 | FALSE | 1 | FALSE | 1 | FALSE | 1 | FALSE | 1 | FALSE | 1 |
| 6058 | 6058 | 6.52380952 | 1.43490659 | 1 | 0.66519151 | 3 | 0.1535026 | 1 | 0.78589454 | 2 | 0.5811115 | 0 | 1 | 35 | 0.67024784 | FALSE | 1 | FALSE | 1 | FALSE | 1 | FALSE | 1 | FALSE | 1 | FALSE | 1 |
| 6059 | 6059 | 10.3333333 | 2.98142397 | 1 | 0.26795838 | 0 | 1 | 1 | 0.35555293 | 0 | 1 | 0 | 1 | 10 | 0.71436964 | FALSE | 1 | FALSE | 1 | FALSE | 1 | FALSE | 1 | FALSE | 1 | FALSE | 1 |
| 6060 | 6060 | 10.3939394 | 1.7742159 | 0 | 1 | 0 | 1 | 2 | 0.33343871 | 2 | 0.45159487 | 1 | 0.42123867 | 28 | 0.58681477 | FALSE | 1 | FALSE | 1 | FALSE | 1 | FALSE | 1 | FALSE | 1 | FALSE | 1 |
| 6061 | 6061 | 11.52 | 1.44554488 | 2 | 0.13369621 | 3 | 0.04547096 | 0 | 1 | 0 | 1 | 1 | 0.33902672 | 19 | 0.91734261 | FALSE | 0.97964162 | FALSE | 1 | FALSE | 1 | FALSE | 1 | FALSE | 1 | FALSE | 1 |
| 6062 | 6062 | 8.375 | 1.91076992 | 0 | 1 | 1 | 0.54637326 | 0 | 1 | 1 | 0.67692816 | 0 | 1 | 22 | 0.25237063 | FALSE | 1 | FALSE | 1 | FALSE | 1 | FALSE | 1 | FALSE | 1 | FALSE | 1 |
| 6063 | 6063 | 9.6 | 3.63868108 | 0 | 1 | 0 | 1 | 1 | 0.30655256 | 0 | 1 | 0 | 1 | 9 | 0.5215547 | FALSE | 1 | FALSE | 1 | FALSE | 1 | FALSE | 1 | FALSE | 1 | FALSE | 1 |
| 6064 | 6064 | 8.21212121 | 1.96573027 | 0 | 1 | 1 | 0.65167362 | 2 | 0.32011871 | 6 | 0.00293693 | 1 | 0.41154418 | 22 | 0.99301316 | FALSE | 1 | FALSE | 1 | FALSE | 1 | FALSE | 1 | FALSE | 1 | FALSE | 1 |
| 6065 | 6065 | 6.28735632 | 3.29370017 | 3 | 0.37257285 | 0 | 1 | 2 | 0.81591765 | 6 | 0.19465406 | 1 | 0.75686725 | 73 | 0.42025276 | FALSE | 1 | FALSE | 1 | FALSE | 1 | FALSE | 1 | FALSE | 1 | FALSE | 1 |
| 6066 | 6066 | 4.16096325 | 1.19279505 | 25 | 0.14984068 | 26 | 0.48591429 | 34 | 0.14365672 | 29 | 0.9200486 | 20 | 0.02962198 | 653 | 0.88582728 | FALSE | 1 | FALSE | 1 | FALSE | 1 | FALSE | 1 | FALSE | 0.82559785 | FALSE | 1 |
| 6067 | 6067 | 9.5 | 3.11104025 | 2 | 0.16048326 | 5 | 0.00180813 | 0 | 1 | 2 | 0.3705495 | 0 | 1 | 19 | 0.99262381 | FALSE | 1 | FALSE | 0.40311621 | FALSE | 1 | FALSE | 1 | FALSE | 1 | FALSE | 1 |
| 6068 | 6068 | 7.0862069 | 0.59551435 | 4 | 0.34672697 | 4 | 0.52051639 | 4 | 0.60431933 | 5 | 0.62158717 | 3 | 0.29629265 | 96 | 0.73618129 | FALSE | 1 | FALSE | 1 | FALSE | 1 | FALSE | 1 | FALSE | 1 | FALSE | 1 |
| 6069 | 6069 | 9.58333333 | 1.84654332 | 1 | 0.24866461 | 1 | 0.30368014 | 0 | 1 | 1 | 0.40390032 | 0 | 1 | 8 | 0.92152993 | FALSE | 1 | FALSE | 1 | FALSE | 1 | FALSE | 1 | FALSE | 1 | FALSE | 1 |
| 6070 | 6070 | 8 | 1.90692518 | 0 | 1 | 1 | 0.30368014 | 0 | 1 | 1 | 0.40390032 | 0 | 1 | 9 | 0.75985366 | FALSE | 1 | FALSE | 1 | FALSE | 1 | FALSE | 1 | FALSE | 1 | FALSE | 1 |
| 6071 | 6071 | 9.2 | 2.6 | 0 | 1 | 0 | 1 | 0 | 1 | 0 | 1 | 0 | 1 | 10 | 0.18273166 | FALSE | 1 | FALSE | 1 | FALSE | 1 | FALSE | 1 | FALSE | 1 | FALSE | 1 |
| 6072 | 6072 | 8.12820513 | 2.35534838 | 2 | 0.26413583 | 4 | 0.03617171 | 1 | 0.76089851 | 2 | 0.54052736 | 1 | 0.4761699 | 29 | 0.96770202 | FALSE | 1 | FALSE | 0.94259695 | FALSE | 1 | FALSE | 1 | FALSE | 1 | FALSE | 1 |
| 6073 | 6073 | 7.21428571 | 1.07354289 | 2 | 0.73182852 | 5 | 0.22216026 | 2 | 0.88006444 | 7 | 0.17487335 | 1 | 0.81092309 | 83 | 0.70668742 | FALSE | 1 | FALSE | 1 | FALSE | 1 | FALSE | 1 | FALSE | 1 | FALSE | 1 |
| 6074 | 6074 | 6.02325581 | 2.79940502 | 3 | 0.09709068 | 4 | 0.04916488 | 1 | 0.79363407 | 2 | 0.59404271 | 3 | 0.03293335 | 30 | 0.99567182 | FALSE | 0.86047225 | FALSE | 1 | FALSE | 1 | FALSE | 1 | FALSE | 0.86063198 | FALSE | 1 |
| 6075 | 6075 | 10.7894737 | 3.30197624 | 0 | 1 | 0 | 1 | 1 | 0.50143826 | 3 | 0.05383577 | 0 | 1 | 15 | 0.83648191 | FALSE | 1 | FALSE | 1 | FALSE | 1 | FALSE | 1 | FALSE | 1 | FALSE | 1 |
| 6076 | 6076 | 5.19653179 | 2.25706075 | 11 | 0.00473375 | 9 | 0.10761825 | 3 | 0.95189725 | 6 | 0.81340635 | 2 | 0.78153591 | 142 | 0.82958826 | FALSE | 0.19848171 | FALSE | 1 | FALSE | 1 | FALSE | 1 | FALSE | 1 | FALSE | 1 |
| 6077 | 6077 | 9.57142857 | 1.67819145 | 1 | 0.16630142 | 0 | 1 | 1 | 0.22601089 | 0 | 1 | 0 | 1 | 5 | 0.91847773 | FALSE | 1 | FALSE | 1 | FALSE | 1 | FALSE | 1 | FALSE | 1 | FALSE | 1 |
| 6078 | 6078 | 8.96153846 | 2.83495719 | 0 | 1 | 1 | 0.57534752 | 1 | 0.61440466 | 0 | 1 | 0 | 1 | 24 | 0.20425579 | FALSE | 1 | FALSE | 1 | FALSE | 1 | FALSE | 1 | FALSE | 1 | FALSE | 1 |
| 6079 | 6079 | 9.10526316 | 3.05897707 | 0 | 1 | 0 | 1 | 1 | 0.50143826 | 1 | 0.59104357 | 1 | 0.26986775 | 16 | 0.65565778 | FALSE | 1 | FALSE | 1 | FALSE | 1 | FALSE | 1 | FALSE | 1 | FALSE | 1 |
| 6080 | 6080 | 8.66666667 | 1.69967317 | 1 | 0.42086991 | 1 | 0.49918218 | 0 | 1 | 1 | 0.62783522 | 0 | 1 | 18 | 0.57962763 | FALSE | 1 | FALSE | 1 | FALSE | 1 | FALSE | 1 | FALSE | 1 | FALSE | 1 |
| 6081 | 6081 | 6.23756906 | 1.04797761 | 4 | 0.6878371 | 4 | 0.84348292 | 5 | 0.78542766 | 10 | 0.31897316 | 1 | 0.95185195 | 157 | 0.21931949 | FALSE | 1 | FALSE | 1 | FALSE | 1 | FALSE | 1 | FALSE | 1 | FALSE | 1 |
| 6082 | 6082 | 8.75 | 2.97559518 | 2 | 0.03643311 | 0 | 1 | 0 | 1 | 1 | 0.43131129 | 0 | 1 | 9 | 0.89630573 | FALSE | 0.58392 | FALSE | 1 | FALSE | 1 | FALSE | 1 | FALSE | 1 | FALSE | 1 |
| 6083 | 6083 | 11.0714286 | 2.34411987 | 1 | 0.3050805 | 1 | 0.36919824 | 1 | 0.40110418 | 0 | 1 | 0 | 1 | 11 | 0.83676449 | FALSE | 1 | FALSE | 1 | FALSE | 1 | FALSE | 1 | FALSE | 1 | FALSE | 1 |
| 6084 | 6084 | 8.55 | 3.63971153 | 0 | 1 | 0 | 1 | 1 | 0.51939841 | 1 | 0.60987146 | 0 | 1 | 18 | 0.37504645 | FALSE | 1 | FALSE | 1 | FALSE | 1 | FALSE | 1 | FALSE | 1 | FALSE | 1 |
| 6085 | 6085 | 9.12121212 | 3.37331395 | 0 | 1 | 1 | 0.6629941 | 2 | 0.33343871 | 1 | 0.78873814 | 0 | 1 | 29 | 0.39648374 | FALSE | 1 | FALSE | 1 | FALSE | 1 | FALSE | 1 | FALSE | 1 | FALSE | 1 |
| 6086 | 6086 | 7.80412371 | 2.25963767 | 1 | 0.92095052 | 2 | 0.82730061 | 4 | 0.46263314 | 8 | 0.07649341 | 2 | 0.47491224 | 80 | 0.75143226 | FALSE | 1 | FALSE | 1 | FALSE | 1 | FALSE | 1 | FALSE | 1 | FALSE | 1 |
| 6087 | 6087 | 7.85185185 | 2.8506491 | 0 | 1 | 1 | 0.58913618 | 2 | 0.25286935 | 1 | 0.71956339 | 0 | 1 | 23 | 0.58307573 | FALSE | 1 | FALSE | 1 | FALSE | 1 | FALSE | 1 | FALSE | 1 | FALSE | 1 |
| 6088 | 6088 | 8.7 | 3.63455637 | 2 | 0.09187136 | 1 | 0.48239024 | 2 | 0.16019548 | 1 | 0.60987146 | 3 | 0.00399012 | 11 | 0.99965905 | FALSE | 0.83248965 | FALSE | 1 | FALSE | 1 | FALSE | 1 | FALSE | 0.37571345 | FALSE | 1 |
| 6089 | 6089 | 7.34693878 | 2.19973116 | 1 | 0.71380452 | 3 | 0.20208567 | 2 | 0.51909752 | 0 | 1 | 2 | 0.18560062 | 40 | 0.66905473 | FALSE | 1 | FALSE | 1 | FALSE | 1 | FALSE | 1 | FALSE | 1 | FALSE | 1 |
| 6090 | 6090 | 10.3636364 | 3.0233253 | 0 | 1 | 1 | 0.63997468 | 1 | 0.67911179 | 2 | 0.41984187 | 1 | 0.4016888 | 26 | 0.64660109 | FALSE | 1 | FALSE | 1 | FALSE | 1 | FALSE | 1 | FALSE | 1 | FALSE | 1 |
| 6091 | 6091 | 6.7721519 | 3.66975103 | 5 | 0.05152748 | 1 | 0.92668093 | 3 | 0.54427311 | 6 | 0.15385579 | 1 | 0.73118367 | 63 | 0.89939946 | FALSE | 0.67646333 | FALSE | 1 | FALSE | 1 | FALSE | 1 | FALSE | 1 | FALSE | 1 |
| 6092 | 6092 | 8.32352941 | 2.19250848 | 0 | 1 | 1 | 0.6739483 | 2 | 0.34668321 | 4 | 0.06848449 | 2 | 0.1065437 | 25 | 0.9693929 | FALSE | 1 | FALSE | 1 | FALSE | 1 | FALSE | 1 | FALSE | 1 | FALSE | 1 |
| 6093 | 6093 | 8.5862069 | 2.53957967 | 2 | 0.16962005 | 3 | 0.06581417 | 1 | 0.65463982 | 1 | 0.74482229 | 0 | 1 | 22 | 0.92902102 | FALSE | 1 | FALSE | 1 | FALSE | 1 | FALSE | 1 | FALSE | 1 | FALSE | 1 |
| 6094 | 6094 | 5.20725389 | 0.97031015 | 11 | 0.01013735 | 11 | 0.04647991 | 9 | 0.25278424 | 10 | 0.38708826 | 4 | 0.38657736 | 147 | 0.99852778 | FALSE | 0.30467829 | FALSE | 1 | FALSE | 1 | FALSE | 1 | FALSE | 1 | FALSE | 1 |
| 6095 | 6095 | 9 | 3.31946802 | 2 | 0.39594061 | 2 | 0.51564773 | 1 | 0.85724013 | 1 | 0.91797304 | 1 | 0.58506217 | 46 | 0.39930625 | FALSE | 1 | FALSE | 1 | FALSE | 1 | FALSE | 1 | FALSE | 1 | FALSE | 1 |
| 6096 | 6096 | 8.45454546 | 2.57129739 | 0 | 1 | 0 | 1 | 0 | 1 | 0 | 1 | 0 | 1 | 11 | 0.15414489 | FALSE | 1 | FALSE | 1 | FALSE | 1 | FALSE | 1 | FALSE | 1 | FALSE | 1 |
| 6097 | 6097 | 6.48571429 | 2.88217605 | 2 | 0.19750261 | 0 | 1 | 2 | 0.32011871 | 2 | 0.43583874 | 0 | 1 | 26 | 0.77612337 | FALSE | 1 | FALSE | 1 | FALSE | 1 | FALSE | 1 | FALSE | 1 | FALSE | 1 |
| 6098 | 6098 | 10.8571429 | 1.95875846 | 1 | 0.42086991 | 2 | 0.14655134 | 0 | 1 | 0 | 1 | 0 | 1 | 18 | 0.57962763 | FALSE | 1 | FALSE | 1 | FALSE | 1 | FALSE | 1 | FALSE | 1 | FALSE | 1 |
| 6099 | 6099 | 8.4 | 3.00665928 | 0 | 1 | 0 | 1 | 0 | 1 | 0 | 1 | 0 | 1 | 5 | 0.42761893 | FALSE | 1 | FALSE | 1 | FALSE | 1 | FALSE | 1 | FALSE | 1 | FALSE | 1 |
| 6100 | 6100 | 5.47619048 | 2.4027573 | 0 | 1 | 1 | 0.49918218 | 3 | 0.03775825 | 0 | 1 | 0 | 1 | 17 | 0.77849076 | FALSE | 1 | FALSE | 1 | FALSE | 1 | FALSE | 1 | FALSE | 1 | FALSE | 1 |
| 6101 | 6101 | 10.0833333 | 3.22641011 | 1 | 0.26795838 | 0 | 1 | 1 | 0.35555293 | 1 | 0.43131129 | 0 | 1 | 9 | 0.89630573 | FALSE | 1 | FALSE | 1 | FALSE | 1 | FALSE | 1 | FALSE | 1 | FALSE | 1 |
| 6102 | 6102 | 7.38202247 | 1.38617263 | 2 | 0.63949673 | 3 | 0.51446405 | 4 | 0.35669301 | 3 | 0.74873269 | 2 | 0.40216708 | 70 | 0.67167049 | FALSE | 1 | FALSE | 1 | FALSE | 1 | FALSE | 1 | FALSE | 1 | FALSE | 1 |
| 6103 | 6103 | 8.90909091 | 2.23421922 | 0 | 1 | 0 | 1 | 0 | 1 | 0 | 1 | 0 | 1 | 11 | 0.15414489 | FALSE | 1 | FALSE | 1 | FALSE | 1 | FALSE | 1 | FALSE | 1 | FALSE | 1 |
| 6104 | 6104 | 8.77419355 | 2.47190246 | 4 | 0.00764535 | 1 | 0.63997468 | 2 | 0.30673593 | 0 | 1 | 0 | 1 | 24 | 0.90138666 | FALSE | 0.2566832 | FALSE | 1 | FALSE | 1 | FALSE | 1 | FALSE | 1 | FALSE | 1 |
| 6105 | 6105 | 10.5714286 | 3.03270159 | 1 | 0.42086991 | 0 | 1 | 0 | 1 | 1 | 0.62783522 | 0 | 1 | 19 | 0.34094959 | FALSE | 1 | FALSE | 1 | FALSE | 1 | FALSE | 1 | FALSE | 1 | FALSE | 1 |
| 6106 | 6106 | 10.9166667 | 2.84189882 | 0 | 1 | 0 | 1 | 0 | 1 | 0 | 1 | 0 | 1 | 12 | 0.13002667 | FALSE | 1 | FALSE | 1 | FALSE | 1 | FALSE | 1 | FALSE | 1 | FALSE | 1 |
| 6107 | 6107 | 9.66666667 | 3.03681119 | 0 | 1 | 0 | 1 | 1 | 0.35555293 | 0 | 1 | 0 | 1 | 11 | 0.41944532 | FALSE | 1 | FALSE | 1 | FALSE | 1 | FALSE | 1 | FALSE | 1 | FALSE | 1 |
| 6108 | 6108 | 10.7931035 | 3.33610665 | 0 | 1 | 0 | 1 | 0 | 1 | 2 | 0.3705495 | 0 | 1 | 26 | 0.16405547 | FALSE | 1 | FALSE | 1 | FALSE | 1 | FALSE | 1 | FALSE | 1 | FALSE | 1 |

| | | | | | | | | | | | | | | | | | | | | | | | | | | | |
|---|---|---|---|---|---|---|---|---|---|---|---|---|---|---|---|---|---|---|---|---|---|---|---|---|---|---|---|
| 6109 | 6109 | 7.14647887 | 0.85643068 | 10 | 0.42633772 | 13 | 0.36230731 | 11 | 0.73780414 | 19 | 0.27507751 | 9 | 0.12721904 | 293 | 0.85476239 | FALSE | 1 | FALSE | 1 | FALSE | 1 | FALSE | 1 | FALSE | 1 | FALSE | 1 |
| 6110 | 6110 | 9.89473684 | 2.0747793 | 2 | 0.08405377 | 0 | 1 | 0 | 1 | 1 | 0.59104357 | 0 | 1 | 16 | 0.65565778 | FALSE | 0.79236883 | FALSE | 1 | FALSE | 1 | FALSE | 1 | FALSE | 1 | FALSE | 1 |
| 6111 | 6111 | 9.88888889 | 2.40113142 | 0 | 1 | 1 | 0.44710673 | 0 | 1 | 0 | 1 | 0 | 1 | 17 | 0.20329205 | FALSE | 1 | FALSE | 1 | FALSE | 1 | FALSE | 1 | FALSE | 1 | FALSE | 1 |
| 6112 | 6112 | 10.5 | 2.73861279 | 1 | 0.18769745 | 0 | 1 | 1 | 0.25384309 | 0 | 1 | 0 | 1 | 6 | 0.88425551 | FALSE | 1 | FALSE | 1 | FALSE | 1 | FALSE | 1 | FALSE | 1 | FALSE | 1 |
| 6113 | 6113 | 9.86666667 | 2.44585818 | 0 | 1 | 0 | 1 | 1 | 0.42266264 | 0 | 1 | 0 | 1 | 14 | 0.29525409 | FALSE | 1 | FALSE | 1 | FALSE | 1 | FALSE | 1 | FALSE | 1 | FALSE | 1 |
| 6114 | 6114 | 5.70491803 | 2.4783936 | 2 | 0.46653175 | 0 | 1 | 3 | 0.3760478 | 3 | 0.53597532 | 2 | 0.26425795 | 51 | 0.64764523 | FALSE | 1 | FALSE | 1 | FALSE | 1 | FALSE | 1 | FALSE | 1 | FALSE | 1 |
| 6115 | 6115 | 8.93333333 | 2.23507395 | 0 | 1 | 0 | 1 | 0 | 1 | 2 | 0.14906763 | 1 | 0.2198258 | 12 | 0.80329917 | FALSE | 1 | FALSE | 1 | FALSE | 1 | FALSE | 1 | FALSE | 1 | FALSE | 1 |
| 6116 | 6116 | 9.65384615 | 3.37345187 | 0 | 1 | 2 | 0.20486827 | 1 | 0.61440466 | 1 | 0.70601446 | 0 | 1 | 22 | 0.61628 | FALSE | 1 | FALSE | 1 | FALSE | 1 | FALSE | 1 | FALSE | 1 | FALSE | 1 |
| 6117 | 6117 | 9.26666667 | 2.54209013 | 1 | 0.54198714 | 1 | 0.6278847 | 1 | 0.66709967 | 1 | 0.75658797 | 0 | 1 | 26 | 0.48609328 | FALSE | 1 | FALSE | 1 | FALSE | 1 | FALSE | 1 | FALSE | 1 | FALSE | 1 |
| 6118 | 6118 | 9.91428571 | 1.66255275 | 2 | 0.22589135 | 1 | 0.68454804 | 3 | 0.12958859 | 0 | 1 | 3 | 0.01918344 | 26 | 0.96294373 | FALSE | 1 | FALSE | 1 | FALSE | 1 | FALSE | 1 | FALSE | 0.70156566 | FALSE | 1 |
| 6119 | 6119 | 8.77192983 | 2.12765866 | 0 | 1 | 1 | 0.83729212 | 1 | 0.8673918 | 5 | 0.10658542 | 3 | 0.06089361 | 45 | 0.76814578 | FALSE | 1 | FALSE | 1 | FALSE | 1 | FALSE | 1 | FALSE | 1 | FALSE | 1 |
| 6120 | 6120 | 9.67857143 | 2.75231813 | 0 | 1 | 0 | 1 | 0 | 1 | 2 | 0.3705495 | 0 | 1 | 26 | 0.16405547 | FALSE | 1 | FALSE | 1 | FALSE | 1 | FALSE | 1 | FALSE | 1 | FALSE | 1 |
| 6121 | 6121 | 9.26666667 | 3.35592345 | 0 | 1 | 0 | 1 | 0 | 1 | 1 | 0.50623194 | 0 | 1 | 14 | 0.29525409 | FALSE | 1 | FALSE | 1 | FALSE | 1 | FALSE | 1 | FALSE | 1 | FALSE | 1 |
| 6122 | 6122 | 9.14285714 | 3.39867922 | 2 | 0.04858661 | 1 | 0.36919824 | 0 | 1 | 1 | 0.48242099 | 1 | 0.20679328 | 9 | 0.98607548 | FALSE | 0.649301 | FALSE | 1 | FALSE | 1 | FALSE | 1 | FALSE | 1 | FALSE | 1 |
| 6123 | 6123 | 11.4545455 | 2.7753341 | 3 | 0.00234778 | 2 | 0.04725998 | 1 | 0.33149969 | 3 | 0.01200907 | 1 | 0.16638736 | 1 | 1 | FALSE | 0.12737083 | FALSE | 1 | FALSE | 1 | FALSE | 1 | FALSE | 1 | FALSE | 1 |
| 6124 | 6124 | 7 | 3 | 2 | 0.0256865 | 0 | 1 | 1 | 0.30655256 | 0 | 1 | 1 | 0.15247062 | 6 | 0.98831471 | FALSE | 0.4974559 | FALSE | 1 | FALSE | 1 | FALSE | 1 | FALSE | 1 | FALSE | 1 |
| 6125 | 6125 | 9.45833333 | 2.6454232 | 1 | 0.46441868 | 0 | 1 | 2 | 0.21260257 | 3 | 0.09525947 | 0 | 1 | 18 | 0.93166998 | FALSE | 1 | FALSE | 1 | FALSE | 1 | FALSE | 1 | FALSE | 1 | FALSE | 1 |
| 6126 | 6126 | 6.40677966 | 2.58495415 | 1 | 0.78541994 | 2 | 0.57395594 | 0 | 1 | 1 | 0.93827647 | 3 | 0.07204991 | 52 | 0.27701375 | FALSE | 1 | FALSE | 1 | FALSE | 1 | FALSE | 1 | FALSE | 1 | FALSE | 1 |
| 6127 | 6127 | 8.76923077 | 2.8326081 | 0 | 1 | 0 | 1 | 1 | 0.37874417 | 3 | 0.01944678 | 0 | 1 | 9 | 0.96018168 | FALSE | 1 | FALSE | 1 | FALSE | 1 | FALSE | 1 | FALSE | 1 | FALSE | 1 |
| 6128 | 6128 | 9.25 | 3.12583322 | 0 | 1 | 0 | 1 | 0 | 1 | 1 | 0.67692816 | 0 | 1 | 23 | 0.09196389 | FALSE | 1 | FALSE | 1 | FALSE | 1 | FALSE | 1 | FALSE | 1 | FALSE | 1 |
| 6129 | 6129 | 4.65482234 | 1.98794325 | 3 | 0.88639173 | 5 | 0.77033627 | 4 | 0.92873122 | 6 | 0.89602894 | 10 | 0.00138475 | 169 | 0.33167181 | FALSE | 1 | FALSE | 1 | FALSE | 1 | FALSE | 1 | FALSE | 0.20625174 | FALSE | 1 |
| 6130 | 6130 | 4.76223776 | 2.29611421 | 5 | 0.30463114 | 8 | 0.09155703 | 2 | 0.96744533 | 6 | 0.64799687 | 2 | 0.68503569 | 120 | 0.61567345 | FALSE | 1 | FALSE | 1 | FALSE | 1 | FALSE | 1 | FALSE | 1 | FALSE | 1 |
| 6131 | 6131 | 9.70967742 | 2.00311932 | 0 | 1 | 3 | 0.07145121 | 0 | 1 | 3 | 0.15707004 | 0 | 1 | 24 | 0.82294782 | FALSE | 1 | FALSE | 1 | FALSE | 1 | FALSE | 1 | FALSE | 1 | FALSE | 1 |
| 6132 | 6132 | 11.0666667 | 1.80616229 | 0 | 1 | 0 | 1 | 0 | 1 | 2 | 0.14906763 | 0 | 1 | 13 | 0.57712805 | FALSE | 1 | FALSE | 1 | FALSE | 1 | FALSE | 1 | FALSE | 1 | FALSE | 1 |
| 6133 | 6133 | 6.58823529 | 2.00172936 | 0 | 1 | 0 | 1 | 0 | 1 | 0 | 1 | 1 | 0.24525764 | 16 | 0.23070182 | FALSE | 1 | FALSE | 1 | FALSE | 1 | FALSE | 1 | FALSE | 1 | FALSE | 1 |
| 6134 | 6134 | 10.0625 | 2.53645299 | 0 | 1 | 2 | 0.09285443 | 1 | 0.44344817 | 1 | 0.52895088 | 0 | 1 | 12 | 0.90922186 | FALSE | 1 | FALSE | 1 | FALSE | 1 | FALSE | 1 | FALSE | 1 | FALSE | 1 |
| 6135 | 6135 | 11.2727273 | 2.86327559 | 0 | 1 | 0 | 1 | 0 | 1 | 0 | 1 | 0 | 1 | 11 | 0.15414489 | FALSE | 1 | FALSE | 1 | FALSE | 1 | FALSE | 1 | FALSE | 1 | FALSE | 1 |
| 6136 | 6136 | 10.375 | 2.0577597 | 1 | 0.18769745 | 1 | 0.23139161 | 0 | 1 | 0 | 1 | 0 | 1 | 6 | 0.88425551 | FALSE | 1 | FALSE | 1 | FALSE | 1 | FALSE | 1 | FALSE | 1 | FALSE | 1 |
| 6137 | 6137 | 8.02985075 | 2.22583306 | 2 | 0.51618914 | 3 | 0.36935445 | 1 | 0.91485686 | 4 | 0.3701568 | 4 | 0.02379565 | 53 | 0.90970332 | FALSE | 1 | FALSE | 1 | FALSE | 1 | FALSE | 1 | FALSE | 0.77578444 | FALSE | 1 |
| 6138 | 6138 | 8.6 | 2.08326667 | 1 | 0.40558521 | 1 | 0.48239024 | 1 | 0.51939841 | 0 | 1 | 1 | 0.28187291 | 16 | 0.80826361 | FALSE | 1 | FALSE | 1 | FALSE | 1 | FALSE | 1 | FALSE | 1 | FALSE | 1 |
| 6139 | 6139 | 8.77777778 | 2.82951813 | 1 | 0.60837712 | 2 | 0.32556427 | 1 | 0.73299778 | 3 | 0.22784656 | 0 | 1 | 29 | 0.81028289 | FALSE | 1 | FALSE | 1 | FALSE | 1 | FALSE | 1 | FALSE | 1 | FALSE | 1 |
| 6140 | 6140 | 7.32258065 | 1.95740277 | 1 | 0.55378205 | 0 | 1 | 3 | 0.09854319 | 2 | 0.41984187 | 0 | 1 | 25 | 0.80006983 | FALSE | 1 | FALSE | 1 | FALSE | 1 | FALSE | 1 | FALSE | 1 | FALSE | 1 |
| 6141 | 6141 | 6.92982456 | 2.87066011 | 1 | 0.76171602 | 2 | 0.5356583 | 1 | 0.8673918 | 3 | 0.46623289 | 1 | 0.59866474 | 47 | 0.50345693 | FALSE | 1 | FALSE | 1 | FALSE | 1 | FALSE | 1 | FALSE | 1 | FALSE | 1 |
| 6142 | 6142 | 8.73684211 | 3.40034217 | 0 | 1 | 0 | 1 | 0 | 1 | 0 | 1 | 0 | 1 | 19 | 0.03948376 | FALSE | 1 | FALSE | 1 | FALSE | 1 | FALSE | 1 | FALSE | 1 | FALSE | 1 |
| 6143 | 6143 | 8.69230769 | 1.72691879 | 1 | 0.49162634 | 0 | 1 | 2 | 0.23940226 | 1 | 0.70601446 | 0 | 1 | 22 | 0.61628 | FALSE | 1 | FALSE | 1 | FALSE | 1 | FALSE | 1 | FALSE | 1 | FALSE | 1 |
| 6144 | 6144 | 9.2 | 1.51437556 | 0 | 1 | 1 | 0.6278847 | 2 | 0.29330414 | 2 | 0.4036176 | 0 | 1 | 25 | 0.67630471 | FALSE | 1 | FALSE | 1 | FALSE | 1 | FALSE | 1 | FALSE | 1 | FALSE | 1 |
| 6145 | 6145 | 9.36363636 | 3.58716721 | 1 | 0.4357638 | 1 | 0.5154318 | 0 | 1 | 1 | 0.64497439 | 0 | 1 | 19 | 0.54233419 | FALSE | 1 | FALSE | 1 | FALSE | 1 | FALSE | 1 | FALSE | 1 | FALSE | 1 |
| 6146 | 6146 | 9.33333333 | 1.69967317 | 4 | 0.00296746 | 0 | 1 | 0 | 1 | 1 | 0.67692816 | 0 | 1 | 19 | 0.83975208 | FALSE | 0.14653312 | FALSE | 1 | FALSE | 1 | FALSE | 1 | FALSE | 1 | FALSE | 1 |
| 6147 | 6147 | 9.5 | 1.82574186 | 0 | 1 | 0 | 1 | 1 | 0.58502114 | 1 | 0.67692816 | 1 | 0.32796862 | 21 | 0.47053137 | FALSE | 1 | FALSE | 1 | FALSE | 1 | FALSE | 1 | FALSE | 1 | FALSE | 1 |
| 6148 | 6148 | 6.74468085 | 2.43614699 | 0 | 1 | 4 | 0.36195667 | 1 | 0.968674 | 4 | 0.63278702 | 3 | 0.19972697 | 82 | 0.2727315 | FALSE | 1 | FALSE | 1 | FALSE | 1 | FALSE | 1 | FALSE | 1 | FALSE | 1 |
| 6149 | 6149 | 9.68 | 2.13016431 | 1 | 0.47819878 | 0 | 1 | 2 | 0.22597287 | 1 | 0.69181315 | 0 | 1 | 21 | 0.64957167 | FALSE | 1 | FALSE | 1 | FALSE | 1 | FALSE | 1 | FALSE | 1 | FALSE | 1 |
| 6150 | 6150 | 11.0555556 | 1.68233377 | 0 | 1 | 0 | 1 | 0 | 1 | 2 | 0.19902682 | 0 | 1 | 16 | 0.44998001 | FALSE | 1 | FALSE | 1 | FALSE | 1 | FALSE | 1 | FALSE | 1 | FALSE | 1 |
| 6151 | 6151 | 9.81818182 | 1.58505416 | 0 | 1 | 2 | 0.04725998 | 0 | 1 | 0 | 1 | 0 | 1 | 9 | 0.75985366 | FALSE | 1 | FALSE | 1 | FALSE | 1 | FALSE | 1 | FALSE | 1 | FALSE | 1 |
| 6152 | 6152 | 10.2413793 | 2.68655878 | 0 | 1 | 4 | 0.01340169 | 0 | 1 | 1 | 0.74482229 | 0 | 1 | 24 | 0.70563334 | FALSE | 1 | FALSE | 0.69207862 | FALSE | 1 | FALSE | 1 | FALSE | 1 | FALSE | 1 |
| 6153 | 6153 | 4.83333333 | 2.10347407 | 4 | 0.16836665 | 3 | 0.51446405 | 5 | 0.18349495 | 3 | 0.74873269 | 2 | 0.40216708 | 67 | 0.90389203 | FALSE | 1 | FALSE | 1 | FALSE | 1 | FALSE | 1 | FALSE | 1 | FALSE | 1 |
| 6154 | 6154 | 10.4285714 | 3.13310174 | 1 | 0.28675951 | 0 | 1 | 0 | 1 | 0 | 1 | 0 | 1 | 12 | 0.374198 | FALSE | 1 | FALSE | 1 | FALSE | 1 | FALSE | 1 | FALSE | 1 | FALSE | 1 |
| 6155 | 6155 | 10.8378378 | 4.15583935 | 0 | 1 | 1 | 0.68454804 | 1 | 0.72299506 | 1 | 0.80778317 | 0 | 1 | 32 | 0.18108028 | FALSE | 1 | FALSE | 1 | FALSE | 1 | FALSE | 1 | FALSE | 1 | FALSE | 1 |
| 6156 | 6156 | 9.39130435 | 1.92780863 | 0 | 1 | 1 | 0.53115654 | 1 | 0.5695037 | 0 | 1 | 2 | 0.05389003 | 19 | 0.71542496 | FALSE | 1 | FALSE | 1 | FALSE | 1 | FALSE | 1 | FALSE | 1 | FALSE | 1 |
| 6157 | 6157 | 9.41666667 | 2.17785572 | 1 | 0.26795838 | 1 | 0.32623933 | 1 | 0.35555293 | 1 | 0.43131129 | 0 | 1 | 8 | 0.97197833 | FALSE | 1 | FALSE | 1 | FALSE | 1 | FALSE | 1 | FALSE | 1 | FALSE | 1 |
| 6158 | 6158 | 4.58677686 | 3.23378197 | 4 | 0.37020407 | 3 | 0.7513988 | 6 | 0.26185234 | 7 | 0.31272943 | 3 | 0.31432186 | 97 | 0.88421464 | FALSE | 1 | FALSE | 1 | FALSE | 1 | FALSE | 1 | FALSE | 1 | FALSE | 1 |
| 6159 | 6159 | 9.55555556 | 1.97827709 | 0 | 1 | 0 | 1 | 1 | 0.48280983 | 1 | 0.57131013 | 0 | 1 | 16 | 0.44998001 | FALSE | 1 | FALSE | 1 | FALSE | 1 | FALSE | 1 | FALSE | 1 | FALSE | 1 |
| 6160 | 6160 | 10.4615385 | 2.30769231 | 3 | 0.0279333 | 2 | 0.20486827 | 1 | 0.61440466 | 0 | 1 | 2 | 0.0670823 | 18 | 0.98644281 | FALSE | 0.50964275 | FALSE | 1 | FALSE | 1 | FALSE | 1 | FALSE | 1 | FALSE | 1 |
| 6161 | 6161 | 9.13043478 | 2.7866818 | 0 | 1 | 0 | 1 | 0 | 1 | 1 | 0.6613267 | 0 | 1 | 22 | 0.10533112 | FALSE | 1 | FALSE | 1 | FALSE | 1 | FALSE | 1 | FALSE | 1 | FALSE | 1 |
| 6162 | 6162 | 10.1875 | 2.67438661 | 0 | 1 | 1 | 0.40942991 | 0 | 1 | 0 | 1 | 1 | 0.23264612 | 14 | 0.53308261 | FALSE | 1 | FALSE | 1 | FALSE | 1 | FALSE | 1 | FALSE | 1 | FALSE | 1 |
| 6163 | 6163 | 9.125 | 1.92164209 | 0 | 1 | 0 | 1 | 1 | 0.58502114 | 3 | 0.09525947 | 1 | 0.32796862 | 19 | 0.83975208 | FALSE | 1 | FALSE | 1 | FALSE | 1 | FALSE | 1 | FALSE | 1 | FALSE | 1 |
| 6164 | 6164 | 9.22727273 | 2.15178189 | 1 | 0.4357638 | 1 | 0.5154318 | 1 | 0.55340851 | 0 | 1 | 1 | 0.30529951 | 18 | 0.74744813 | FALSE | 1 | FALSE | 1 | FALSE | 1 | FALSE | 1 | FALSE | 1 | FALSE | 1 |
| 6165 | 6165 | 9.45 | 2.69211813 | 0 | 1 | 2 | 0.1353446 | 0 | 1 | 0 | 1 | 1 | 0.28187291 | 17 | 0.61750632 | FALSE | 1 | FALSE | 1 | FALSE | 1 | FALSE | 1 | FALSE | 1 | FALSE | 1 |
| 6166 | 6166 | 9.85 | 1.58981131 | 1 | 0.40558521 | 0 | 1 | 1 | 0.51939841 | 3 | 0.0612581 | 0 | 1 | 15 | 0.92113679 | FALSE | 1 | FALSE | 1 | FALSE | 1 | FALSE | 1 | FALSE | 1 | FALSE | 1 |
| 6167 | 6167 | 10.4210526 | 4.04339344 | 3 | 0.01017712 | 1 | 0.44710673 | 0 | 1 | 1 | 0.57131013 | 0 | 1 | 13 | 0.95041913 | FALSE | 0.30467829 | FALSE | 1 | FALSE | 1 | FALSE | 1 | FALSE | 1 | FALSE | 1 |
| 6168 | 6168 | 7.05555556 | 3.42332233 | 0 | 1 | 0 | 1 | 0 | 1 | 0 | 1 | 0 | 1 | 18 | 0.04681656 | FALSE | 1 | FALSE | 1 | FALSE | 1 | FALSE | 1 | FALSE | 1 | FALSE | 1 |
| 6169 | 6169 | 9.13793103 | 1.61296766 | 1 | 0.51745948 | 0 | 1 | 1 | 0.64171563 | 0 | 1 | 0 | 1 | 26 | 0.16405547 | FALSE | 1 | FALSE | 1 | FALSE | 1 | FALSE | 1 | FALSE | 1 | FALSE | 1 |
| 6170 | 6170 | 7.48275862 | 2.81156098 | 2 | 0.16962005 | 0 | 1 | 1 | 0.65463982 | 2 | 0.3871811 | 0 | 1 | 24 | 0.70563334 | FALSE | 1 | FALSE | 1 | FALSE | 1 | FALSE | 1 | FALSE | 1 | FALSE | 1 |
| 6171 | 6171 | 8.14285714 | 1.45686272 | 0 | 1 | 1 | 0.48239024 | 0 | 1 | 1 | 0.60987146 | 0 | 1 | 18 | 0.37504645 | FALSE | 1 | FALSE | 1 | FALSE | 1 | FALSE | 1 | FALSE | 1 | FALSE | 1 |
| 6172 | 6172 | 10.0625 | 1.67588596 | 0 | 1 | 0 | 1 | 0 | 1 | 0 | 1 | 0 | 1 | 16 | 0.06581507 | FALSE | 1 | FALSE | 1 | FALSE | 1 | FALSE | 1 | FALSE | 1 | FALSE | 1 |
| 6173 | 6173 | 9.66666667 | 2.20028858 | 1 | 0.66519151 | 2 | 0.39616059 | 4 | 0.0626749 | 4 | 0.12508716 | 0 | 1 | 31 | 0.97647435 | FALSE | 1 | FALSE | 1 | FALSE | 1 | FALSE | 1 | FALSE | 1 | FALSE | 1 |
| 6174 | 6174 | 9.45 | 3.30870065 | 0 | 1 | 2 | 0.36122049 | 2 | 0.41139386 | 2 | 0.54052736 | 0 | 1 | 33 | 0.59061307 | FALSE | 1 | FALSE | 1 | FALSE | 1 | FALSE | 1 | FALSE | 1 | FALSE | 1 |
| 6175 | 6175 | 9.94117647 | 2.01379327 | 1 | 0.35728305 | 2 | 0.10307112 | 1 | 0.46348836 | 1 | 0.55062772 | 0 | 1 | 12 | 0.962093 | FALSE | 1 | FALSE | 1 | FALSE | 1 | FALSE | 1 | FALSE | 1 | FALSE | 1 |
| 6176 | 6176 | 4.38888889 | 4.37339036 | 0 | 1 | 0 | 1 | 1 | 0.73299778 | 4 | 0.08109727 | 0 | 1 | 31 | 0.49869404 | FALSE | 1 | FALSE | 1 | FALSE | 1 | FALSE | 1 | FALSE | 1 | FALSE | 1 |
| 6177 | 6177 | 8.78787879 | 1.50267591 | 1 | 0.81665075 | 3 | 0.35154359 | 3 | 0.41519671 | 0 | 1 | 2 | 0.28868487 | 56 | 0.42599591 | FALSE | 1 | FALSE | 1 | FALSE | 1 | FALSE | 1 | FALSE | 1 | FALSE | 1 |
| 6178 | 6178 | 9.125 | 2.02715934 | 0 | 1 | 1 | 0.40942991 | 1 | 0.44344817 | 0 | 1 | 0 | 1 | 14 | 0.53308261 | FALSE | 1 | FALSE | 1 | FALSE | 1 | FALSE | 1 | FALSE | 1 | FALSE | 1 |
| 6179 | 6179 | 8.9 | 1.78605711 | 1 | 0.40558521 | 0 | 1 | 1 | 0.51939841 | 0 | 1 | 0 | 1 | 18 | 0.37504645 | FALSE | 1 | FALSE | 1 | FALSE | 1 | FALSE | 1 | FALSE | 1 | FALSE | 1 |
| 6180 | 6180 | 9.21428571 | 1.9338552 | 0 | 1 | 0 | 1 | 0 | 1 | 0 | 1 | 0 | 1 | 14 | 0.09251304 | FALSE | 1 | FALSE | 1 | FALSE | 1 | FALSE | 1 | FALSE | 1 | FALSE | 1 |
| 6181 | 6181 | 9.0952381 | 2.68868461 | 0 | 1 | 0 | 1 | 1 | 0.53671415 | 0 | 1 | 0 | 1 | 20 | 0.13763971 | FALSE | 1 | FALSE | 1 | FALSE | 1 | FALSE | 1 | FALSE | 1 | FALSE | 1 |
| 6182 | 6182 | 8.19354839 | 3.02056772 | 0 | 1 | 3 | 0.07145121 | 3 | 0.09131526 | 2 | 0.4036176 | 1 | 0.39166989 | 21 | 0.98741738 | FALSE | 1 | FALSE | 1 | FALSE | 1 | FALSE | 1 | FALSE | 1 | FALSE | 1 |
| 6183 | 6183 | 7.76666667 | 2.53880987 | 2 | 0.17884234 | 0 | 1 | 1 | 0.66709967 | 2 | 0.4036176 | 1 | 0.39166989 | 24 | 0.82294782 | FALSE | 1 | FALSE | 1 | FALSE | 1 | FALSE | 1 | FALSE | 1 | FALSE | 1 |
| 6184 | 6184 | 8.43859649 | 2.26349787 | 5 | 0.01493383 | 2 | 0.55509723 | 3 | 0.3364112 | 4 | 0.26521223 | 2 | 0.23984683 | 41 | 0.99511641 | FALSE | 0.38230603 | FALSE | 1 | FALSE | 1 | FALSE | 1 | FALSE | 1 | FALSE | 1 |
| 6185 | 6185 | 8.29411765 | 2.10782034 | 0 | 1 | 1 | 0.38964394 | 1 | 0.42266264 | 1 | 0.50623194 | 0 | 1 | 12 | 0.80329917 | FALSE | 1 | FALSE | 1 | FALSE | 1 | FALSE | 1 | FALSE | 1 | FALSE | 1 |
| 6186 | 6186 | 9 | 2.59554274 | 1 | 0.38989951 | 1 | 0.46503795 | 1 | 0.50143826 | 1 | 0.59104357 | 0 | 1 | 15 | 0.83648191 | FALSE | 1 | FALSE | 1 | FALSE | 1 | FALSE | 1 | FALSE | 1 | FALSE | 1 |
| 6187 | 6187 | 3.20496894 | 1.31931665 | 4 | 0.59638451 | 11 | 0.01497977 | 4 | 0.83672875 | 4 | 0.94278525 | 2 | 0.74647014 | 136 | 0.54018126 | FALSE | 1 | FALSE | 0.7256023 | FALSE | 1 | FALSE | 1 | FALSE | 1 | FALSE | 1 |
| 6188 | 6188 | 10.9333333 | 2.64491126 | 0 | 1 | 0 | 1 | 0 | 1 | 0 | 1 | 1 | 0.2198258 | 14 | 0.29525409 | FALSE | 1 | FALSE | 1 | FALSE | 1 | FALSE | 1 | FALSE | 1 | FALSE | 1 |
| 6189 | 6189 | 9.33333333 | 2.52762515 | 3 | 0.00307177 | 0 | 1 | 0 | 1 | 0 | 1 | 2 | 0.01579445 | 7 | 0.99433481 | FALSE | 0.14802295 | FALSE | 1 | FALSE | 1 | FALSE | 1 | FALSE | 0.66313193 | FALSE | 1 |
| 6190 | 6190 | 9.47058824 | 2.40386079 | 1 | 0.57647348 | 3 | 0.08959205 | 3 | 0.11365871 | 1 | 0.78873814 | 1 | 0.42123867 | 24 | 0.97500814 | FALSE | 1 | FALSE | 1 | FALSE | 1 | FALSE | 1 | FALSE | 1 | FALSE | 1 |
| 6191 | 6191 | 8.7 | 1.59478316 | 0 | 1 | 0 | 1 | 0 | 1 | 1 | 0.73248983 | 2 | 0.07640798 | 25 | 0.34293 | FALSE | 1 | FALSE | 1 | FALSE | 1 | FALSE | 1 | FALSE | 1 | FALSE | 1 |
| 6192 | 6192 | 9.2173913 | 2.51873885 | 1 | 0.45027683 | 3 | 0.03669128 | 0 | 1 | 2 | 0.2852236 | 0 | 1 | 17 | 0.94439745 | FALSE | 1 | FALSE | 0.94259695 | FALSE | 1 | FALSE | 1 | FALSE | 1 | FALSE | 1 |
| 6193 | 6193 | 8.64285714 | 1.97389425 | 1 | 0.66519151 | 1 | 0.74973543 | 0 | 1 | 3 | 0.30326943 | 0 | 1 | 37 | 0.34041811 | FALSE | 1 | FALSE | 1 | FALSE | 1 | FALSE | 1 | FALSE | 1 | FALSE | 1 |
| 6194 | 6194 | 8.8 | 4.49740666 | 3 | 0.0405081 | 2 | 0.25306331 | 1 | 0.66709967 | 2 | 0.4036176 | 0 | 1 | 22 | 0.96518743 | FALSE | 0.62493854 | FALSE | 1 | FALSE | 1 | FALSE | 1 | FALSE | 1 | FALSE | 1 |
| 6195 | 6195 | 8.63636364 | 1.06794001 | 0 | 1 | 0 | 1 | 1 | 0.70185639 | 1 | 0.78873814 | 0 | 1 | 31 | 0.09218876 | FALSE | 1 | FALSE | 1 | FALSE | 1 | FALSE | 1 | FALSE | 1 | FALSE | 1 |
| 6196 | 6196 | 10.5625 | 2.76063285 | 0 | 1 | 2 | 0.07341559 | 0 | 1 | 0 | 1 | 3 | 0.00136853 | 9 | 0.98607548 | FALSE | 1 | FALSE | 1 | FALSE | 1 | FALSE | 1 | FALSE | 0.20625174 | FALSE | 1 |
| 6197 | 6197 | 7.71875 | 2.45247802 | 3 | 0.0476977 | 1 | 0.65167362 | 2 | 0.32011871 | 1 | 0.77852168 | 1 | 0.41154418 | 24 | 0.94876798 | FALSE | 0.649301 | FALSE | 1 | FALSE | 1 | FALSE | 1 | FALSE | 1 | FALSE | 1 |
| 6198 | 6198 | 6.82352941 | 3.18517326 | 1 | 0.58738519 | 1 | 0.6739483 | 1 | 0.71261921 | 0 | 1 | 1 | 0.43077488 | 30 | 0.36879488 | FALSE | 1 | FALSE | 1 | FALSE | 1 | FALSE | 1 | FALSE | 1 | FALSE | 1 |
| 6199 | 6199 | 10.1428571 | 2.4159335 | 1 | 0.51745948 | 0 | 1 | 2 | 0.2663542 | 3 | 0.13524571 | 1 | 0.37113067 | 21 | 0.94083768 | FALSE | 1 | FALSE | 1 | FALSE | 1 | FALSE | 1 | FALSE | 1 | FALSE | 1 |
| 6200 | 6200 | 9.44444444 | 1.49897084 | 0 | 1 | 0 | 1 | 0 | 1 | 0 | 1 | 0 | 1 | 9 | 0.21661397 | FALSE | 1 | FALSE | 1 | FALSE | 1 | FALSE | 1 | FALSE | 1 | FALSE | 1 |
| 6201 | 6201 | 11 | 2.32992949 | 1 | 0.42086991 | 0 | 1 | 2 | 0.17308361 | 0 | 1 | 1 | 0.29368245 | 17 | 0.77849076 | FALSE | 1 | FALSE | 1 | FALSE | 1 | FALSE | 1 | FALSE | 1 | FALSE | 1 |
| 6202 | 6202 | 7.34090909 | 2.51308559 | 1 | 0.68224519 | 1 | 0.76576289 | 0 | 1 | 5 | 0.04960693 | 0 | 1 | 37 | 0.61979664 | FALSE | 1 | FALSE | 1 | FALSE | 1 | FALSE | 1 | FALSE | 1 | FALSE | 1 |

| | | | | | | | | | | | | | | | | | | | | | | | | | | | |
|---|---|---|---|---|---|---|---|---|---|---|---|---|---|---|---|---|---|---|---|---|---|---|---|---|---|---|---|
| 6203 | 6203 | 9.96551724 | 2.77280834 | 2 | 0.16962005 | 1 | 0.61539065 | 2 | 0.27983822 | 0 | 1 | 3 | 0.01149488 | 21 | 0.97185642 | FALSE | 1 | FALSE | 1 | FALSE | 1 | FALSE | 1 | FALSE | 0.57972935 | FALSE | 1 |
| 6204 | 6204 | 9.88888889 | 2.76664436 | 1 | 0.20854748 | 1 | 0.25628163 | 1 | 0.28067846 | 0 | 1 | 0 | 1 | 6 | 0.96128317 | FALSE | 1 | FALSE | 1 | FALSE | 1 | FALSE | 1 | FALSE | 1 | FALSE | 1 |
| 6205 | 6205 | 11.122807 | 3.72318566 | 1 | 0.77387657 | 1 | 0.84773229 | 3 | 0.3364112 | 9 | 0.00103406 | 1 | 0.61182532 | 42 | 0.98838397 | FALSE | 1 | FALSE | 1 | FALSE | 1 | FALSE | 1 | FALSE | 1 | FALSE | 1 |
| 6206 | 6206 | 12.375 | 2.89845505 | 1 | 0.46441868 | 0 | 1 | 1 | 0.58502114 | 4 | 0.02237042 | 0 | 1 | 18 | 0.93166998 | FALSE | 1 | FALSE | 1 | FALSE | 1 | FALSE | 1 | FALSE | 1 | FALSE | 1 |
| 6207 | 6207 | 8.54545455 | 1.5587662 | 0 | 1 | 0 | 1 | 1 | 0.33149969 | 1 | 0.40390032 | 0 | 1 | 9 | 0.75985366 | FALSE | 1 | FALSE | 1 | FALSE | 1 | FALSE | 1 | FALSE | 1 | FALSE | 1 |
| 6208 | 6208 | 9.75 | 1.87638838 | 0 | 1 | 1 | 0.32623933 | 0 | 1 | 3 | 0.01547597 | 0 | 1 | 8 | 0.97197833 | FALSE | 1 | FALSE | 1 | FALSE | 1 | FALSE | 1 | FALSE | 1 | FALSE | 1 |
| 6209 | 6209 | 11.3333333 | 3.49602949 | 0 | 1 | 1 | 0.32623933 | 1 | 0.35555293 | 0 | 1 | 1 | 0.18007762 | 9 | 0.89630573 | FALSE | 1 | FALSE | 1 | FALSE | 1 | FALSE | 1 | FALSE | 1 | FALSE | 1 |
| 6210 | 6210 | 7.6 | 1.94679223 | 1 | 0.64722818 | 0 | 1 | 3 | 0.17245215 | 1 | 0.84823834 | 1 | 0.48480878 | 34 | 0.56328904 | FALSE | 1 | FALSE | 1 | FALSE | 1 | FALSE | 1 | FALSE | 1 | FALSE | 1 |
| 6211 | 6211 | 6.57142857 | 1.8877445 | 8 | 0.00220497 | 5 | 0.17117488 | 2 | 0.84458838 | 5 | 0.4067946 | 1 | 0.78011711 | 70 | 0.97872956 | FALSE | 0.12457305 | FALSE | 1 | FALSE | 1 | FALSE | 1 | FALSE | 1 | FALSE | 1 |
| 6212 | 6212 | 8.57692308 | 2.22179894 | 0 | 1 | 0 | 1 | 3 | 0.06482014 | 1 | 0.70601446 | 1 | 0.3499045 | 21 | 0.78939668 | FALSE | 1 | FALSE | 1 | FALSE | 1 | FALSE | 1 | FALSE | 1 | FALSE | 1 |
| 6213 | 6213 | 10.625 | 1.45236876 | 1 | 0.34033054 | 1 | 0.40942991 | 0 | 1 | 1 | 0.52895088 | 0 | 1 | 13 | 0.76799161 | FALSE | 1 | FALSE | 1 | FALSE | 1 | FALSE | 1 | FALSE | 1 | FALSE | 1 |
| 6214 | 6214 | 6.11320755 | 4.06856508 | 3 | 0.15399149 | 2 | 0.51564773 | 0 | 1 | 4 | 0.22482652 | 0 | 1 | 44 | 0.69042405 | FALSE | 1 | FALSE | 1 | FALSE | 1 | FALSE | 1 | FALSE | 1 | FALSE | 1 |
| 6215 | 6215 | 7.625 | 2.23257139 | 0 | 1 | 0 | 1 | 0 | 1 | 1 | 0.31351192 | 0 | 1 | 7 | 0.63752397 | FALSE | 1 | FALSE | 1 | FALSE | 1 | FALSE | 1 | FALSE | 1 | FALSE | 1 |
| 6216 | 6216 | 9.23333333 | 3.73883523 | 1 | 0.54198714 | 0 | 1 | 1 | 0.66709967 | 1 | 0.75658797 | 0 | 1 | 27 | 0.28862208 | FALSE | 1 | FALSE | 1 | FALSE | 1 | FALSE | 1 | FALSE | 1 | FALSE | 1 |
| 6217 | 6217 | 9.61111111 | 2.88942303 | 1 | 0.37380237 | 0 | 1 | 2 | 0.13502329 | 3 | 0.04687752 | 0 | 1 | 12 | 0.98544073 | FALSE | 1 | FALSE | 1 | FALSE | 1 | FALSE | 1 | FALSE | 1 | FALSE | 1 |
| 6218 | 6218 | 7.14893617 | 2.44060268 | 1 | 0.70621968 | 1 | 0.78790827 | 1 | 0.82190381 | 1 | 0.89101956 | 1 | 0.54145295 | 42 | 0.23544056 | FALSE | 1 | FALSE | 1 | FALSE | 1 | FALSE | 1 | FALSE | 1 | FALSE | 1 |
| 6219 | 6219 | 6.625 | 2.89845505 | 0 | 1 | 0 | 1 | 2 | 0.21260257 | 0 | 1 | 0 | 1 | 22 | 0.25237063 | FALSE | 1 | FALSE | 1 | FALSE | 1 | FALSE | 1 | FALSE | 1 | FALSE | 1 |
| 6220 | 6220 | 9.76190476 | 2.22335572 | 0 | 1 | 1 | 0.49918218 | 0 | 1 | 0 | 1 | 1 | 0.29368245 | 19 | 0.34094959 | FALSE | 1 | FALSE | 1 | FALSE | 1 | FALSE | 1 | FALSE | 1 | FALSE | 1 |
| 6221 | 6221 | 10.5757576 | 3.44682817 | 0 | 1 | 1 | 0.6629941 | 0 | 1 | 1 | 0.78873814 | 0 | 1 | 31 | 0.09218876 | FALSE | 1 | FALSE | 1 | FALSE | 1 | FALSE | 1 | FALSE | 1 | FALSE | 1 |
| 6222 | 6222 | 8.4 | 2.10713075 | 2 | 0.09187136 | 0 | 1 | 1 | 0.51939841 | 2 | 0.23331368 | 0 | 1 | 15 | 0.92113679 | FALSE | 0.83248965 | FALSE | 1 | FALSE | 1 | FALSE | 1 | FALSE | 1 | FALSE | 1 |
| 6223 | 6223 | 3.99625468 | 2.42103474 | 4 | 0.9154965 | 5 | 0.93696954 | 11 | 0.35921641 | 13 | 0.44916193 | 3 | 0.81830776 | 230 | 0.19317334 | FALSE | 1 | FALSE | 1 | FALSE | 1 | FALSE | 1 | FALSE | 1 | FALSE | 1 |
| 6224 | 6224 | 1.03182461 | 0.56302537 | 116 | 0.13160098 | 134 | 0.68887392 | 148 | 0.73331703 | 201 | 0.22855832 | 66 | 0.78256826 | 3568 | 0.61628116 | FALSE | 0.97964162 | FALSE | 1 | FALSE | 1 | FALSE | 1 | FALSE | 1 | FALSE | 1 |
| 6225 | 6225 | 10.375 | 2.86956007 | 0 | 1 | 1 | 0.23139161 | 0 | 1 | 0 | 1 | 0 | 1 | 7 | 0.63752397 | FALSE | 1 | FALSE | 1 | FALSE | 1 | FALSE | 1 | FALSE | 1 | FALSE | 1 |
| 6226 | 6226 | 9.77272727 | 2.60997768 | 0 | 1 | 2 | 0.15794013 | 0 | 1 | 0 | 1 | 0 | 1 | 20 | 0.30915202 | FALSE | 1 | FALSE | 1 | FALSE | 1 | FALSE | 1 | FALSE | 1 | FALSE | 1 |
| 6227 | 6227 | 6.34210526 | 3.42802932 | 3 | 0.07268126 | 2 | 0.3494045 | 0 | 1 | 1 | 0.83318955 | 0 | 1 | 32 | 0.61800916 | FALSE | 0.76727438 | FALSE | 1 | FALSE | 1 | FALSE | 1 | FALSE | 1 | FALSE | 1 |
| 6228 | 6228 | 7.33333333 | 2.78161351 | 0 | 1 | 2 | 0.28941836 | 2 | 0.33343871 | 2 | 0.45159487 | 0 | 1 | 27 | 0.75124583 | FALSE | 1 | FALSE | 1 | FALSE | 1 | FALSE | 1 | FALSE | 1 | FALSE | 1 |
| 6229 | 6229 | 11.4545455 | 2.40694587 | 0 | 1 | 2 | 0.15794013 | 0 | 1 | 0 | 1 | 2 | 0.04972408 | 18 | 0.74744813 | FALSE | 1 | FALSE | 1 | FALSE | 1 | FALSE | 1 | FALSE | 0.98629461 | FALSE | 1 |
| 6230 | 6230 | 9.93333333 | 2.64491126 | 0 | 1 | 1 | 0.38964394 | 1 | 0.42266264 | 0 | 1 | 0 | 1 | 13 | 0.57712805 | FALSE | 1 | FALSE | 1 | FALSE | 1 | FALSE | 1 | FALSE | 1 | FALSE | 1 |
| 6231 | 6231 | 7.70491803 | 2.41813003 | 3 | 0.20532341 | 3 | 0.31579946 | 5 | 0.06710314 | 2 | 0.7775286 | 2 | 0.26425795 | 46 | 0.97780007 | FALSE | 1 | FALSE | 1 | FALSE | 1 | FALSE | 1 | FALSE | 1 | FALSE | 1 |
| 6232 | 6232 | 9.95833333 | 2.09123502 | 0 | 1 | 0 | 1 | 1 | 0.58502114 | 2 | 0.30249307 | 0 | 1 | 21 | 0.47053137 | FALSE | 1 | FALSE | 1 | FALSE | 1 | FALSE | 1 | FALSE | 1 | FALSE | 1 |
| 6233 | 6233 | 6.83870968 | 2.76611728 | 3 | 0.21200292 | 1 | 0.87100665 | 3 | 0.3858942 | 4 | 0.31731292 | 3 | 0.08097514 | 48 | 0.94896265 | FALSE | 1 | FALSE | 1 | FALSE | 1 | FALSE | 1 | FALSE | 1 | FALSE | 1 |
| 6234 | 6234 | 9.46875 | 2.277394 | 1 | 0.56527493 | 2 | 0.27730767 | 0 | 1 | 1 | 0.77852168 | 0 | 1 | 28 | 0.42532177 | FALSE | 1 | FALSE | 1 | FALSE | 1 | FALSE | 1 | FALSE | 1 | FALSE | 1 |
| 6235 | 6235 | 10.030303 | 3.47706641 | 1 | 0.57647348 | 0 | 1 | 0 | 1 | 2 | 0.45159487 | 0 | 1 | 30 | 0.21942357 | FALSE | 1 | FALSE | 1 | FALSE | 1 | FALSE | 1 | FALSE | 1 | FALSE | 1 |
| 6236 | 6236 | 9.41666667 | 2.49861073 | 1 | 0.26795838 | 0 | 1 | 0 | 1 | 0 | 1 | 0 | 1 | 11 | 0.41944532 | FALSE | 1 | FALSE | 1 | FALSE | 1 | FALSE | 1 | FALSE | 1 | FALSE | 1 |
| 6237 | 6237 | 10.3548387 | 3.82309102 | 1 | 0.55378205 | 0 | 1 | 1 | 0.67911179 | 3 | 0.16836568 | 0 | 1 | 26 | 0.64660109 | FALSE | 1 | FALSE | 1 | FALSE | 1 | FALSE | 1 | FALSE | 1 | FALSE | 1 |
| 6238 | 6238 | 10.3333333 | 2.26077666 | 1 | 0.18769745 | 1 | 0.23139161 | 1 | 0.25384309 | 1 | 0.31351192 | 0 | 1 | 4 | 0.99658732 | FALSE | 1 | FALSE | 1 | FALSE | 1 | FALSE | 1 | FALSE | 1 | FALSE | 1 |
| 6239 | 6239 | 8.3030303 | 4.0488296 | 1 | 0.57647348 | 1 | 0.6629941 | 2 | 0.33343871 | 2 | 0.45159487 | 0 | 1 | 27 | 0.75124583 | FALSE | 1 | FALSE | 1 | FALSE | 1 | FALSE | 1 | FALSE | 1 | FALSE | 1 |
| 6240 | 6240 | 9.45454546 | 2.23051712 | 0 | 1 | 0 | 1 | 0 | 1 | 0 | 1 | 0 | 1 | 11 | 0.15414489 | FALSE | 1 | FALSE | 1 | FALSE | 1 | FALSE | 1 | FALSE | 1 | FALSE | 1 |
| 6241 | 6241 | 10 | 3.39116499 | 1 | 0.26795838 | 0 | 1 | 0 | 1 | 1 | 0.43131129 | 0 | 1 | 10 | 0.71436964 | FALSE | 1 | FALSE | 1 | FALSE | 1 | FALSE | 1 | FALSE | 1 | FALSE | 1 |
| 6242 | 6242 | 8.72413793 | 2.83933639 | 3 | 0.03713651 | 0 | 1 | 2 | 0.27983822 | 1 | 0.74482229 | 1 | 0.38148475 | 22 | 0.92902102 | FALSE | 0.58392 | FALSE | 1 | FALSE | 1 | FALSE | 1 | FALSE | 1 | FALSE | 1 |
| 6243 | 6243 | 8.77777778 | 2.37788162 | 1 | 0.50471032 | 0 | 1 | 1 | 0.62830986 | 0 | 1 | 1 | 0.36060486 | 24 | 0.3725373 | FALSE | 1 | FALSE | 1 | FALSE | 1 | FALSE | 1 | FALSE | 1 | FALSE | 1 |
| 6244 | 6244 | 8.90909091 | 1.50481321 | 0 | 1 | 2 | 0.1353446 | 0 | 1 | 0 | 1 | 1 | 0.28187291 | 17 | 0.61750632 | FALSE | 1 | FALSE | 1 | FALSE | 1 | FALSE | 1 | FALSE | 1 | FALSE | 1 |
| 6245 | 6245 | 8.03846154 | 2.91522224 | 0 | 1 | 0 | 1 | 3 | 0.06482014 | 1 | 0.70601446 | 0 | 1 | 22 | 0.61628 | FALSE | 1 | FALSE | 1 | FALSE | 1 | FALSE | 1 | FALSE | 1 | FALSE | 1 |
| 6246 | 6246 | 9.15625 | 2.85163741 | 3 | 0.0476977 | 0 | 1 | 2 | 0.32011871 | 1 | 0.77852168 | 0 | 1 | 26 | 0.77612337 | FALSE | 0.649301 | FALSE | 1 | FALSE | 1 | FALSE | 1 | FALSE | 1 | FALSE | 1 |
| 6247 | 6247 | 8.51515152 | 3.10543231 | 4 | 0.00764535 | 0 | 1 | 2 | 0.30673593 | 2 | 0.41984187 | 0 | 1 | 23 | 0.95750806 | FALSE | 0.2566832 | FALSE | 1 | FALSE | 1 | FALSE | 1 | FALSE | 1 | FALSE | 1 |
| 6248 | 6248 | 7.67948718 | 1.54831248 | 1 | 0.86966563 | 3 | 0.46497905 | 5 | 0.14803148 | 2 | 0.8798475 | 4 | 0.0386728 | 63 | 0.85070881 | FALSE | 1 | FALSE | 1 | FALSE | 1 | FALSE | 1 | FALSE | 0.88493726 | FALSE | 1 |
| 6249 | 6249 | 7.80769231 | 2.21913411 | 1 | 0.49162634 | 2 | 0.20486827 | 1 | 0.61440466 | 3 | 0.11458571 | 0 | 1 | 19 | 0.96054244 | FALSE | 1 | FALSE | 1 | FALSE | 1 | FALSE | 1 | FALSE | 1 | FALSE | 1 |
| 6250 | 6250 | 4.52083333 | 2.92965857 | 1 | 0.71380452 | 2 | 0.46321244 | 2 | 0.51909752 | 1 | 0.89605799 | 2 | 0.18560062 | 40 | 0.66905473 | FALSE | 1 | FALSE | 1 | FALSE | 1 | FALSE | 1 | FALSE | 1 | FALSE | 1 |
| 6251 | 6251 | 9.85714286 | 2.16653584 | 1 | 0.16630142 | 0 | 1 | 0 | 1 | 0 | 1 | 0 | 1 | 6 | 0.69920685 | FALSE | 1 | FALSE | 1 | FALSE | 1 | FALSE | 1 | FALSE | 1 | FALSE | 1 |
| 6252 | 6252 | 10.6111111 | 2.33663787 | 1 | 0.37380237 | 0 | 1 | 0 | 1 | 2 | 0.19902682 | 1 | 0.25766374 | 14 | 0.86287418 | FALSE | 1 | FALSE | 1 | FALSE | 1 | FALSE | 1 | FALSE | 1 | FALSE | 1 |
| 6253 | 6253 | 7.36363636 | 2.21315874 | 12 | 1.22E-11 | 2 | 0.28941836 | 2 | 0.33343871 | 1 | 0.78873814 | 0 | 1 | 16 | 0.99999973 | TRUE | 5.00E-08 | FALSE | 1 | FALSE | 1 | FALSE | 1 | FALSE | 1 | FALSE | 1 |
| 6254 | 6254 | 6.08333333 | 3.09457411 | 1 | 0.46441868 | 1 | 0.54637326 | 2 | 0.21260257 | 3 | 0.09525947 | 0 | 1 | 17 | 0.97530702 | FALSE | 1 | FALSE | 1 | FALSE | 1 | FALSE | 1 | FALSE | 1 | FALSE | 1 |
| 6255 | 6255 | 9.36666667 | 1.37800177 | 1 | 0.54198714 | 4 | 0.01508077 | 4 | 0.02136185 | 2 | 0.4036176 | 1 | 0.39166989 | 18 | 0.99972884 | FALSE | 1 | FALSE | 0.7256023 | FALSE | 1 | FALSE | 1 | FALSE | 1 | FALSE | 1 |
| 6256 | 6256 | 7.5 | 1.29559694 | 1 | 0.3050805 | 0 | 1 | 1 | 0.40110418 | 0 | 1 | 0 | 1 | 12 | 0.62236227 | FALSE | 1 | FALSE | 1 | FALSE | 1 | FALSE | 1 | FALSE | 1 | FALSE | 1 |
| 6257 | 6257 | 8.59090909 | 4.8772113 | 0 | 1 | 0 | 1 | 0 | 1 | 2 | 0.26791271 | 0 | 1 | 20 | 0.30915202 | FALSE | 1 | FALSE | 1 | FALSE | 1 | FALSE | 1 | FALSE | 1 | FALSE | 1 |
| 6258 | 6258 | 1.06067416 | 0.74405781 | 24 | 0.42806667 | 22 | 0.93515751 | 34 | 0.37609446 | 43 | 0.38216765 | 8 | 0.9841208 | 759 | 0.22895154 | FALSE | 1 | FALSE | 1 | FALSE | 1 | FALSE | 1 | FALSE | 1 | FALSE | 1 |
| 6259 | 6259 | 8.08928571 | 4.37638462 | 1 | 0.76171602 | 2 | 0.5356583 | 1 | 0.8673918 | 3 | 0.46623289 | 0 | 1 | 48 | 0.35562012 | FALSE | 1 | FALSE | 1 | FALSE | 1 | FALSE | 1 | FALSE | 1 | FALSE | 1 |
| 6260 | 6260 | 9.04347826 | 2.5956194 | 1 | 0.45027683 | 1 | 0.53115654 | 1 | 0.5695037 | 1 | 0.6613267 | 3 | 0.00598158 | 16 | 0.98100709 | FALSE | 1 | FALSE | 1 | FALSE | 1 | FALSE | 1 | FALSE | 0.45694206 | FALSE | 1 |
| 6261 | 6261 | 8.27083333 | 2.64369993 | 3 | 0.11868304 | 1 | 0.78790827 | 0 | 1 | 2 | 0.64277396 | 1 | 0.54145295 | 40 | 0.54368682 | FALSE | 0.94393345 | FALSE | 1 | FALSE | 1 | FALSE | 1 | FALSE | 1 | FALSE | 1 |
| 6262 | 6262 | 8.8 | 1.6 | 0 | 1 | 1 | 0.48239024 | 1 | 0.51939841 | 3 | 0.0612581 | 0 | 1 | 15 | 0.92113679 | FALSE | 1 | FALSE | 1 | FALSE | 1 | FALSE | 1 | FALSE | 1 | FALSE | 1 |
| 6263 | 6263 | 9.66666667 | 2.44948974 | 0 | 1 | 0 | 1 | 0 | 1 | 0 | 1 | 0 | 1 | 8 | 0.25677166 | FALSE | 1 | FALSE | 1 | FALSE | 1 | FALSE | 1 | FALSE | 1 | FALSE | 1 |
| 6264 | 6264 | 6.78125 | 4.15131286 | 0 | 1 | 1 | 0.65167362 | 1 | 0.69069222 | 3 | 0.17988885 | 0 | 1 | 27 | 0.61671133 | FALSE | 1 | FALSE | 1 | FALSE | 1 | FALSE | 1 | FALSE | 1 | FALSE | 1 |
| 6265 | 6265 | 7.68 | 2.15814735 | 1 | 0.72839502 | 5 | 0.02197853 | 4 | 0.1037722 | 2 | 0.6762159 | 3 | 0.04820812 | 35 | 0.99712859 | FALSE | 1 | FALSE | 0.81029974 | FALSE | 1 | FALSE | 1 | FALSE | 0.98629461 | FALSE | 1 |
| 6266 | 6266 | 10.1428571 | 2.26328279 | 0 | 1 | 0 | 1 | 1 | 0.40110418 | 0 | 1 | 0 | 1 | 13 | 0.33283729 | FALSE | 1 | FALSE | 1 | FALSE | 1 | FALSE | 1 | FALSE | 1 | FALSE | 1 |
| 6267 | 6267 | 10.8333333 | 4.21966297 | 0 | 1 | 0 | 1 | 1 | 0.197145 | 0 | 1 | 0 | 1 | 5 | 0.76185568 | FALSE | 1 | FALSE | 1 | FALSE | 1 | FALSE | 1 | FALSE | 1 | FALSE | 1 |
| 6268 | 6268 | 10.5454546 | 2.67526163 | 0 | 1 | 0 | 1 | 0 | 1 | 2 | 0.08789566 | 1 | 0.16638736 | 8 | 0.92152993 | FALSE | 1 | FALSE | 1 | FALSE | 1 | FALSE | 1 | FALSE | 1 | FALSE | 1 |
| 6269 | 6269 | 6.80851064 | 3.77921695 | 0 | 1 | 2 | 0.45233216 | 2 | 0.50778118 | 2 | 0.64277396 | 2 | 0.17968152 | 39 | 0.69224623 | FALSE | 1 | FALSE | 1 | FALSE | 1 | FALSE | 1 | FALSE | 1 | FALSE | 1 |
| 6270 | 6270 | 9.35714286 | 2.68879003 | 0 | 1 | 0 | 1 | 1 | 0.40110418 | 0 | 1 | 0 | 1 | 13 | 0.33283729 | FALSE | 1 | FALSE | 1 | FALSE | 1 | FALSE | 1 | FALSE | 1 | FALSE | 1 |
| 6271 | 6271 | 11.2222222 | 2.37476445 | 2 | 0.31193899 | 1 | 0.76576289 | 4 | 0.07197653 | 0 | 1 | 0 | 1 | 37 | 0.61979664 | FALSE | 1 | FALSE | 1 | FALSE | 1 | FALSE | 1 | FALSE | 1 | FALSE | 1 |
| 6272 | 6272 | 9.03846154 | 3.00615739 | 1 | 0.49162634 | 1 | 0.57534752 | 0 | 1 | 0 | 1 | 0 | 1 | 24 | 0.20425579 | FALSE | 1 | FALSE | 1 | FALSE | 1 | FALSE | 1 | FALSE | 1 | FALSE | 1 |
| 6273 | 6273 | 8.04347826 | 2.5956194 | 0 | 1 | 2 | 0.16948764 | 0 | 1 | 1 | 0.6613267 | 0 | 1 | 20 | 0.50589402 | FALSE | 1 | FALSE | 1 | FALSE | 1 | FALSE | 1 | FALSE | 1 | FALSE | 1 |
| 6274 | 6274 | 4.90909091 | 2.16661369 | 1 | 0.95693012 | 9 | 0.01538996 | 7 | 0.13988072 | 7 | 0.31272943 | 9 | 0.00014142 | 87 | 0.99974384 | FALSE | 1 | FALSE | 0.7256023 | FALSE | 1 | FALSE | 1 | FALSE | 0.05792693 | FALSE | 1 |
| 6275 | 6275 | 10.1666667 | 3.21022671 | 0 | 1 | 1 | 0.32623933 | 0 | 1 | 1 | 0.43131129 | 0 | 1 | 10 | 0.71436964 | FALSE | 1 | FALSE | 1 | FALSE | 1 | FALSE | 1 | FALSE | 1 | FALSE | 1 |
| 6276 | 6276 | 9.17647059 | 2.45513374 | 2 | 0.06909038 | 2 | 0.10307112 | 2 | 0.12280513 | 0 | 1 | 0 | 1 | 11 | 0.98974663 | FALSE | 0.74276693 | FALSE | 1 | FALSE | 1 | FALSE | 1 | FALSE | 1 | FALSE | 1 |
| 6277 | 6277 | 9.83333333 | 1.99304346 | 0 | 1 | 1 | 0.32623933 | 2 | 0.06691409 | 0 | 1 | 2 | 0.01579445 | 7 | 0.99433481 | FALSE | 1 | FALSE | 1 | FALSE | 1 | FALSE | 1 | FALSE | 0.66313193 | FALSE | 1 |
| 6278 | 6278 | 9.63636364 | 1.49378879 | 1 | 0.24866461 | 0 | 1 | 0 | 1 | 0 | 1 | 0 | 1 | 10 | 0.46859937 | FALSE | 1 | FALSE | 1 | FALSE | 1 | FALSE | 1 | FALSE | 1 | FALSE | 1 |
| 6279 | 6279 | 9.5 | 2.17944947 | 0 | 1 | 1 | 0.30368014 | 0 | 1 | 1 | 0.40390032 | 0 | 1 | 9 | 0.75985366 | FALSE | 1 | FALSE | 1 | FALSE | 1 | FALSE | 1 | FALSE | 1 | FALSE | 1 |
| 6280 | 6280 | 6.71428571 | 2.78846019 | 3 | 0.03391597 | 1 | 0.60247911 | 1 | 0.64171563 | 1 | 0.73248983 | 0 | 1 | 22 | 0.86499992 | FALSE | 0.56934354 | FALSE | 1 | FALSE | 1 | FALSE | 1 | FALSE | 1 | FALSE | 1 |
| 6281 | 6281 | 9.77777778 | 0.9162457 | 0 | 1 | 0 | 1 | 1 | 0.61440466 | 2 | 0.33678042 | 0 | 1 | 23 | 0.40372818 | FALSE | 1 | FALSE | 1 | FALSE | 1 | FALSE | 1 | FALSE | 1 | FALSE | 1 |
| 6282 | 6282 | 9 | 3.1808249 | 3 | 0.00864151 | 2 | 0.10307112 | 0 | 1 | 0 | 1 | 0 | 1 | 12 | 0.962093 | FALSE | 0.27019545 | FALSE | 1 | FALSE | 1 | FALSE | 1 | FALSE | 1 | FALSE | 1 |
| 6283 | 6283 | 9.06666667 | 1.89267594 | 4 | 0.12569732 | 3 | 0.43942904 | 2 | 0.75760933 | 2 | 0.86569532 | 2 | 0.34918837 | 62 | 0.72396946 | FALSE | 0.95955826 | FALSE | 1 | FALSE | 1 | FALSE | 1 | FALSE | 1 | FALSE | 1 |
| 6284 | 6284 | 8.86666667 | 3.77477005 | 0 | 1 | 0 | 1 | 1 | 0.42266264 | 0 | 1 | 1 | 0.2198258 | 13 | 0.57712805 | FALSE | 1 | FALSE | 1 | FALSE | 1 | FALSE | 1 | FALSE | 1 | FALSE | 1 |
| 6285 | 6285 | 8.29411765 | 2.98554766 | 0 | 1 | 0 | 1 | 2 | 0.12280513 | 0 | 1 | 1 | 0.24525764 | 14 | 0.7313145 | FALSE | 1 | FALSE | 1 | FALSE | 1 | FALSE | 1 | FALSE | 1 | FALSE | 1 |
| 6286 | 6286 | 8.88372093 | 2.21211571 | 0 | 1 | 1 | 0.75788114 | 2 | 0.46083448 | 2 | 0.59404271 | 1 | 0.50988734 | 37 | 0.48294184 | FALSE | 1 | FALSE | 1 | FALSE | 1 | FALSE | 1 | FALSE | 1 | FALSE | 1 |
| 6287 | 6287 | 7.96 | 1.73158887 | 0 | 1 | 0 | 1 | 1 | 0.8405438 | 5 | 0.07777122 | 0 | 1 | 44 | 0.31639336 | FALSE | 1 | FALSE | 1 | FALSE | 1 | FALSE | 1 | FALSE | 1 | FALSE | 1 |
| 6288 | 6288 | 12.2222222 | 2.04275292 | 0 | 1 | 0 | 1 | 0 | 1 | 1 | 0.34506517 | 0 | 1 | 8 | 0.5780332 | FALSE | 1 | FALSE | 1 | FALSE | 1 | FALSE | 1 | FALSE | 1 | FALSE | 1 |
| 6289 | 6289 | 4.55648536 | 1.78673336 | 3 | 0.94858886 | 7 | 0.66038262 | 7 | 0.76256092 | 15 | 0.13474103 | 2 | 0.90816915 | 205 | 0.30866875 | FALSE | 1 | FALSE | 1 | FALSE | 1 | FALSE | 1 | FALSE | 1 | FALSE | 1 |
| 6290 | 6290 | 8.125 | 2.52178013 | 2 | 0.06197875 | 0 | 1 | 1 | 0.44344817 | 0 | 1 | 0 | 1 | 13 | 0.76799161 | FALSE | 0.72532843 | FALSE | 1 | FALSE | 1 | FALSE | 1 | FALSE | 1 | FALSE | 1 |
| 6291 | 6291 | 12.4210526 | 2.73987686 | 1 | 0.38989951 | 1 | 0.46503795 | 1 | 0.50143826 | 0 | 1 | 1 | 0.26986775 | 15 | 0.83648191 | FALSE | 1 | FALSE | 1 | FALSE | 1 | FALSE | 1 | FALSE | 1 | FALSE | 1 |
| 6292 | 6292 | 9.23529412 | 2.21038756 | 0 | 1 | 2 | 0.10307112 | 0 | 1 | 1 | 0.55062772 | 0 | 1 | 14 | 0.7313145 | FALSE | 1 | FALSE | 1 | FALSE | 1 | FALSE | 1 | FALSE | 1 | FALSE | 1 |
| 6293 | 6293 | 9.23529412 | 1.92596536 | 0 | 1 | 1 | 0.42857733 | 1 | 0.46348836 | 0 | 1 | 0 | 1 | 15 | 0.49060435 | FALSE | 1 | FALSE | 1 | FALSE | 1 | FALSE | 1 | FALSE | 1 | FALSE | 1 |
| 6294 | 6294 | 10.3023256 | 1.65001598 | 1 | 0.67382914 | 3 | 0.16132793 | 1 | 0.79363407 | 3 | 0.31602175 | 0 | 1 | 35 | 0.7801651 | FALSE | 1 | FALSE | 1 | FALSE | 1 | FALSE | 1 | FALSE | 1 | FALSE | 1 |
| 6295 | 6295 | 10.1724138 | 1.17646359 | 1 | 0.49162634 | 1 | 0.57534752 | 2 | 0.23940226 | 3 | 0.11458571 | 0 | 1 | 19 | 0.96054244 | FALSE | 1 | FALSE | 1 | FALSE | 1 | FALSE | 1 | FALSE | 1 | FALSE | 1 |
| 6296 | 6296 | 9.07142857 | 2.37439573 | 1 | 0.3050805 | 1 | 0.36919824 | 1 | 0.40110418 | 1 | 0.48242099 | 0 | 1 | 10 | 0.94579623 | FALSE | 1 | FALSE | 1 | FALSE | 1 | FALSE | 1 | FALSE | 1 | FALSE | 1 |

| | | | | | | | | | | | | | | | | | | | | | | | | | | | |
|---|---|---|---|---|---|---|---|---|---|---|---|---|---|---|---|---|---|---|---|---|---|---|---|---|---|---|---|
| 6297 | 6297 | 8.7 | 2.1 | 0 | 1 | 1 | 0.48239024 | 1 | 0.51939841 | 0 | 1 | 0 | 1 | 18 | 0.37504645 | FALSE | 1 | FALSE | 1 | FALSE | 1 | FALSE | 1 | FALSE | 1 | FALSE | 1 |
| 6298 | 6298 | 9.47222222 | 1.51815149 | 1 | 0.60837712 | 1 | 0.69480472 | 2 | 0.37289966 | 1 | 0.81665383 | 3 | 0.0206828 | 28 | 0.90259714 | FALSE | 1 | FALSE | 1 | FALSE | 1 | FALSE | 1 | FALSE | 0.7366672 | FALSE | 1 |
| 6299 | 6299 | 10.1818182 | 1.84972636 | 0 | 1 | 0 | 1 | 1 | 0.33149969 | 0 | 1 | 0 | 1 | 10 | 0.46859937 | FALSE | 1 | FALSE | 1 | FALSE | 1 | FALSE | 1 | FALSE | 1 | FALSE | 1 |
| 6300 | 6300 | 8.74074074 | 2.60473348 | 0 | 1 | 1 | 0.58913618 | 1 | 0.62830986 | 1 | 0.71956339 | 0 | 1 | 24 | 0.3725373 | FALSE | 1 | FALSE | 1 | FALSE | 1 | FALSE | 1 | FALSE | 1 | FALSE | 1 |
| 6301 | 6301 | 5.87323944 | 3.32269014 | 5 | 0.03316372 | 9 | 0.00038387 | 1 | 0.92379401 | 4 | 0.40179526 | 2 | 0.31908786 | 49 | 0.99932716 | FALSE | 0.56836235 | FALSE | 0.15723145 | FALSE | 1 | FALSE | 1 | FALSE | 1 | FALSE | 1 |
| 6302 | 6302 | 8.44444444 | 2.71256792 | 1 | 0.37380237 | 0 | 1 | 0 | 1 | 1 | 0.57131013 | 1 | 0.25766374 | 15 | 0.69372602 | FALSE | 1 | FALSE | 1 | FALSE | 1 | FALSE | 1 | FALSE | 1 | FALSE | 1 |
| 6303 | 6303 | 7.7 | 2.84780617 | 0 | 1 | 0 | 1 | 0 | 1 | 2 | 0.23331368 | 0 | 1 | 18 | 0.37504645 | FALSE | 1 | FALSE | 1 | FALSE | 1 | FALSE | 1 | FALSE | 1 | FALSE | 1 |
| 6304 | 6304 | 8.28125 | 2.98155135 | 3 | 0.0476977 | 1 | 0.65167362 | 1 | 0.69069222 | 2 | 0.43583874 | 1 | 0.41154418 | 24 | 0.94876798 | FALSE | 0.649301 | FALSE | 1 | FALSE | 1 | FALSE | 1 | FALSE | 1 | FALSE | 1 |
| 6305 | 6305 | 12.1 | 2.11896201 | 0 | 1 | 0 | 1 | 0 | 1 | 2 | 0.0740981 | 0 | 1 | 8 | 0.80394719 | FALSE | 1 | FALSE | 1 | FALSE | 1 | FALSE | 1 | FALSE | 1 | FALSE | 1 |
| 6306 | 6306 | 8.43902439 | 1.93867922 | 0 | 1 | 0 | 1 | 2 | 0.43640583 | 1 | 0.85524728 | 0 | 1 | 38 | 0.09794002 | FALSE | 1 | FALSE | 1 | FALSE | 1 | FALSE | 1 | FALSE | 1 | FALSE | 1 |
| 6307 | 6307 | 9.41666667 | 2.17785572 | 1 | 0.26795838 | 1 | 0.32623933 | 0 | 1 | 0 | 1 | 0 | 1 | 10 | 0.71436964 | FALSE | 1 | FALSE | 1 | FALSE | 1 | FALSE | 1 | FALSE | 1 | FALSE | 1 |
| 6308 | 6308 | 5.53125 | 2.60988955 | 0 | 1 | 2 | 0.61853748 | 6 | 0.02664157 | 3 | 0.56885732 | 2 | 0.28258242 | 51 | 0.88464035 | FALSE | 1 | FALSE | 1 | FALSE | 1 | FALSE | 1 | FALSE | 1 | FALSE | 1 |
| 6309 | 6309 | 11.0625 | 3.03044448 | 0 | 1 | 0 | 1 | 0 | 1 | 0 | 1 | 2 | 0.02433796 | 13 | 0.57712805 | FALSE | 1 | FALSE | 1 | FALSE | 1 | FALSE | 1 | FALSE | 0.77578444 | FALSE | 1 |
| 6310 | 6310 | 9.30434783 | 2.59489101 | 2 | 0.10809809 | 0 | 1 | 0 | 1 | 1 | 0.64497439 | 0 | 1 | 19 | 0.54233419 | FALSE | 0.88820416 | FALSE | 1 | FALSE | 1 | FALSE | 1 | FALSE | 1 | FALSE | 1 |
| 6311 | 6311 | 6.7631579 | 2.03177665 | 0 | 1 | 4 | 0.23089453 | 6 | 0.05471327 | 2 | 0.8705742 | 1 | 0.71735141 | 63 | 0.70648758 | FALSE | 1 | FALSE | 1 | FALSE | 1 | FALSE | 1 | FALSE | 1 | FALSE | 1 |
| 6312 | 6312 | 9.48148148 | 2.55850608 | 0 | 1 | 1 | 0.58913618 | 2 | 0.25286935 | 2 | 0.35374209 | 2 | 0.07169515 | 20 | 0.95133411 | FALSE | 1 | FALSE | 1 | FALSE | 1 | FALSE | 1 | FALSE | 1 | FALSE | 1 |
| 6313 | 6313 | 9.625 | 2.47171499 | 0 | 1 | 1 | 0.40942991 | 0 | 1 | 1 | 0.52895088 | 0 | 1 | 14 | 0.53308261 | FALSE | 1 | FALSE | 1 | FALSE | 1 | FALSE | 1 | FALSE | 1 | FALSE | 1 |
| 6314 | 6314 | 9.69230769 | 2.61312119 | 0 | 1 | 1 | 0.57534752 | 0 | 1 | 2 | 0.33678042 | 1 | 0.3499045 | 22 | 0.61628 | FALSE | 1 | FALSE | 1 | FALSE | 1 | FALSE | 1 | FALSE | 1 | FALSE | 1 |
| 6315 | 6315 | 8.27027027 | 2.3091902 | 1 | 0.60837712 | 3 | 0.10944025 | 1 | 0.73299778 | 0 | 1 | 0 | 1 | 31 | 0.49869404 | FALSE | 1 | FALSE | 1 | FALSE | 1 | FALSE | 1 | FALSE | 1 | FALSE | 1 |
| 6316 | 6316 | 8.05 | 2.29074224 | 2 | 0.45796754 | 0 | 1 | 3 | 0.36617062 | 3 | 0.52470434 | 0 | 1 | 52 | 0.39208719 | FALSE | 1 | FALSE | 1 | FALSE | 1 | FALSE | 1 | FALSE | 1 | FALSE | 1 |
| 6317 | 6317 | 8.52173913 | 2.34742348 | 2 | 0.3309085 | 0 | 1 | 1 | 0.81522035 | 3 | 0.35428318 | 0 | 1 | 40 | 0.40712403 | FALSE | 1 | FALSE | 1 | FALSE | 1 | FALSE | 1 | FALSE | 1 | FALSE | 1 |
| 6318 | 6318 | 7.83333333 | 2.19215774 | 2 | 0.07645454 | 4 | 0.00227902 | 0 | 1 | 0 | 1 | 0 | 1 | 12 | 0.98544073 | FALSE | 0.77424195 | FALSE | 0.42431176 | FALSE | 1 | FALSE | 1 | FALSE | 1 | FALSE | 1 |
| 6319 | 6319 | 10 | 2.43373723 | 2 | 0.14250978 | 1 | 0.57534752 | 1 | 0.61440466 | 1 | 0.70601446 | 0 | 1 | 21 | 0.78939668 | FALSE | 1 | FALSE | 1 | FALSE | 1 | FALSE | 1 | FALSE | 1 | FALSE | 1 |
| 6320 | 6320 | 7.56 | 2.40133296 | 0 | 1 | 0 | 1 | 1 | 0.59998149 | 0 | 1 | 1 | 0.33902672 | 23 | 0.22727183 | FALSE | 1 | FALSE | 1 | FALSE | 1 | FALSE | 1 | FALSE | 1 | FALSE | 1 |
| 6321 | 6321 | 10.0555556 | 2.32073318 | 0 | 1 | 1 | 0.44710673 | 0 | 1 | 0 | 1 | 0 | 1 | 17 | 0.20329205 | FALSE | 1 | FALSE | 1 | FALSE | 1 | FALSE | 1 | FALSE | 1 | FALSE | 1 |
| 6322 | 6322 | 10.7777778 | 3.4084137 | 1 | 0.37380237 | 0 | 1 | 0 | 1 | 0 | 1 | 0 | 1 | 17 | 0.20329205 | FALSE | 1 | FALSE | 1 | FALSE | 1 | FALSE | 1 | FALSE | 1 | FALSE | 1 |
| 6323 | 6323 | 9.85714286 | 1.58221407 | 0 | 1 | 1 | 0.49918218 | 1 | 0.53671415 | 0 | 1 | 1 | 0.29368245 | 18 | 0.57962763 | FALSE | 1 | FALSE | 1 | FALSE | 1 | FALSE | 1 | FALSE | 1 | FALSE | 1 |
| 6324 | 6324 | 8.17241379 | 2.6918647 | 3 | 0.03713651 | 0 | 1 | 2 | 0.27983822 | 3 | 0.14602298 | 1 | 0.38148475 | 20 | 0.99028351 | FALSE | 0.58392 | FALSE | 1 | FALSE | 1 | FALSE | 1 | FALSE | 1 | FALSE | 1 |
| 6325 | 6325 | 10 | 4.28952212 | 3 | 0.01369554 | 1 | 0.48239024 | 0 | 1 | 0 | 1 | 0 | 1 | 16 | 0.80826361 | FALSE | 0.36625055 | FALSE | 1 | FALSE | 1 | FALSE | 1 | FALSE | 1 | FALSE | 1 |
| 6326 | 6326 | 9.75 | 1.71391365 | 0 | 1 | 2 | 0.14655134 | 1 | 0.53671415 | 3 | 0.06912908 | 0 | 1 | 15 | 0.96513691 | FALSE | 1 | FALSE | 1 | FALSE | 1 | FALSE | 1 | FALSE | 1 | FALSE | 1 |
| 6327 | 6327 | 8.96 | 2.90489243 | 1 | 0.47819878 | 1 | 0.56109832 | 2 | 0.22597287 | 1 | 0.69181315 | 1 | 0.33902672 | 19 | 0.91734261 | FALSE | 1 | FALSE | 1 | FALSE | 1 | FALSE | 1 | FALSE | 1 | FALSE | 1 |
| 6328 | 6328 | 6.42622951 | 3.38491841 | 1 | 0.79637728 | 3 | 0.31579946 | 1 | 0.89373158 | 3 | 0.53597532 | 2 | 0.26425795 | 51 | 0.64764523 | FALSE | 1 | FALSE | 1 | FALSE | 1 | FALSE | 1 | FALSE | 1 | FALSE | 1 |
| 6329 | 6329 | 10.3333333 | 2.24845626 | 0 | 1 | 0 | 1 | 1 | 0.35555293 | 0 | 1 | 0 | 1 | 11 | 0.41944532 | FALSE | 1 | FALSE | 1 | FALSE | 1 | FALSE | 1 | FALSE | 1 | FALSE | 1 |
| 6330 | 6330 | 9.7 | 3.19530906 | 0 | 1 | 1 | 0.48239024 | 0 | 1 | 1 | 0.60987146 | 0 | 1 | 18 | 0.37504645 | FALSE | 1 | FALSE | 1 | FALSE | 1 | FALSE | 1 | FALSE | 1 | FALSE | 1 |
| 6331 | 6331 | 9.33333333 | 1.12216722 | 1 | 0.49162634 | 0 | 1 | 2 | 0.23940226 | 0 | 1 | 0 | 1 | 23 | 0.40372818 | FALSE | 1 | FALSE | 1 | FALSE | 1 | FALSE | 1 | FALSE | 1 | FALSE | 1 |
| 6332 | 6332 | 9.6875 | 1.30952043 | 0 | 1 | 0 | 1 | 0 | 1 | 1 | 0.52895088 | 1 | 0.23264612 | 14 | 0.53308261 | FALSE | 1 | FALSE | 1 | FALSE | 1 | FALSE | 1 | FALSE | 1 | FALSE | 1 |
| 6333 | 6333 | 7.59375 | 2.826873 | 2 | 0.19750261 | 1 | 0.65167362 | 1 | 0.69069222 | 1 | 0.77852168 | 1 | 0.41154418 | 26 | 0.77612337 | FALSE | 1 | FALSE | 1 | FALSE | 1 | FALSE | 1 | FALSE | 1 | FALSE | 1 |
| 6334 | 6334 | 8.25714286 | 2.15595994 | 2 | 0.20692137 | 1 | 0.6629941 | 1 | 0.70185639 | 0 | 1 | 0 | 1 | 29 | 0.39648374 | FALSE | 1 | FALSE | 1 | FALSE | 1 | FALSE | 1 | FALSE | 1 | FALSE | 1 |
| 6335 | 6335 | 6.625 | 1.60759914 | 1 | 0.64722818 | 3 | 0.13823565 | 0 | 1 | 2 | 0.5543522 | 0 | 1 | 34 | 0.56328904 | FALSE | 1 | FALSE | 1 | FALSE | 1 | FALSE | 1 | FALSE | 1 | FALSE | 1 |
| 6336 | 6336 | 10.0833333 | 2.13924961 | 0 | 1 | 3 | 0.04096158 | 1 | 0.58502114 | 3 | 0.09525947 | 0 | 1 | 17 | 0.97530702 | FALSE | 1 | FALSE | 0.9698186 | FALSE | 1 | FALSE | 1 | FALSE | 1 | FALSE | 1 |
| 6337 | 6337 | 6.50632911 | 1.71274757 | 1 | 0.87304634 | 2 | 0.7307176 | 1 | 0.94537714 | 5 | 0.29645796 | 1 | 0.73118367 | 69 | 0.29139907 | FALSE | 1 | FALSE | 1 | FALSE | 1 | FALSE | 1 | FALSE | 1 | FALSE | 1 |
| 6338 | 6338 | 6.40229885 | 1.1592682 | 2 | 0.65852142 | 3 | 0.53828446 | 2 | 0.82596174 | 2 | 0.91445868 | 0 | 1 | 78 | 0.10818345 | FALSE | 1 | FALSE | 1 | FALSE | 1 | FALSE | 1 | FALSE | 1 | FALSE | 1 |
| 6339 | 6339 | 11.375 | 3.96665791 | 1 | 0.18769745 | 2 | 0.02563923 | 0 | 1 | 1 | 0.31351192 | 0 | 1 | 4 | 0.99658732 | FALSE | 1 | FALSE | 0.82367276 | FALSE | 1 | FALSE | 1 | FALSE | 1 | FALSE | 1 |
| 6340 | 6340 | 7.80645161 | 2.34753618 | 5 | 0.00103368 | 0 | 1 | 1 | 0.67911179 | 2 | 0.41984187 | 0 | 1 | 23 | 0.95750806 | FALSE | 0.07493697 | FALSE | 1 | FALSE | 1 | FALSE | 1 | FALSE | 1 | FALSE | 1 |
| 6341 | 6341 | 9.53846154 | 2.6491039 | 1 | 0.28675951 | 0 | 1 | 0 | 1 | 0 | 1 | 0 | 1 | 12 | 0.374198 | FALSE | 1 | FALSE | 1 | FALSE | 1 | FALSE | 1 | FALSE | 1 | FALSE | 1 |
| 6342 | 6342 | 10.4444444 | 2.49938264 | 0 | 1 | 0 | 1 | 0 | 1 | 0 | 1 | 0 | 1 | 18 | 0.04681656 | FALSE | 1 | FALSE | 1 | FALSE | 1 | FALSE | 1 | FALSE | 1 | FALSE | 1 |
| 6343 | 6343 | 9.76190476 | 2.7929292 | 0 | 1 | 2 | 0.14655134 | 0 | 1 | 1 | 0.62783522 | 1 | 0.29368245 | 17 | 0.77849076 | FALSE | 1 | FALSE | 1 | FALSE | 1 | FALSE | 1 | FALSE | 1 | FALSE | 1 |
| 6344 | 6344 | 9.23333333 | 2.21635336 | 0 | 1 | 1 | 0.61059065 | 0 | 1 | 1 | 0.74482229 | 1 | 0.38148475 | 26 | 0.31495157 | FALSE | 1 | FALSE | 1 | FALSE | 1 | FALSE | 1 | FALSE | 1 | FALSE | 1 |
| 6345 | 6345 | 9.36 | 3.0050624 | 0 | 1 | 2 | 0.19297191 | 0 | 1 | 2 | 0.31968852 | 0 | 1 | 21 | 0.64957167 | FALSE | 1 | FALSE | 1 | FALSE | 1 | FALSE | 1 | FALSE | 1 | FALSE | 1 |
| 6346 | 6346 | 9 | 2.32992949 | 1 | 0.3050805 | 0 | 1 | 1 | 0.40110418 | 1 | 0.48242099 | 0 | 1 | 11 | 0.83676449 | FALSE | 1 | FALSE | 1 | FALSE | 1 | FALSE | 1 | FALSE | 1 | FALSE | 1 |
| 6347 | 6347 | 6.86466165 | 2.06576809 | 1 | 0.96865218 | 2 | 0.93132803 | 4 | 0.70442831 | 8 | 0.25978205 | 4 | 0.17065314 | 113 | 0.40290099 | FALSE | 1 | FALSE | 1 | FALSE | 1 | FALSE | 1 | FALSE | 1 | FALSE | 1 |
| 6348 | 6348 | 8.76923077 | 2.06262887 | 0 | 1 | 4 | 0.00909592 | 0 | 1 | 2 | 0.33678042 | 0 | 1 | 20 | 0.90142689 | FALSE | 1 | FALSE | 0.63686998 | FALSE | 1 | FALSE | 1 | FALSE | 1 | FALSE | 1 |
| 6349 | 6349 | 5.67441861 | 2.5854776 | 0 | 1 | 2 | 0.40761584 | 2 | 0.46083448 | 6 | 0.01295578 | 0 | 1 | 33 | 0.93815092 | FALSE | 1 | FALSE | 1 | FALSE | 1 | FALSE | 1 | FALSE | 1 | FALSE | 1 |
| 6350 | 6350 | 9.85714286 | 1.21638474 | 0 | 1 | 1 | 0.60247911 | 1 | 0.64171563 | 0 | 1 | 0 | 1 | 26 | 0.16405547 | FALSE | 1 | FALSE | 1 | FALSE | 1 | FALSE | 1 | FALSE | 1 | FALSE | 1 |
| 6351 | 6351 | 3.75824176 | 3.69835324 | 0 | 1 | 2 | 0.79919134 | 2 | 0.84458838 | 4 | 0.60708936 | 4 | 0.06180746 | 79 | 0.31800957 | FALSE | 1 | FALSE | 1 | FALSE | 1 | FALSE | 1 | FALSE | 1 | FALSE | 1 |
| 6352 | 6352 | 9.63636364 | 2.53243423 | 0 | 1 | 0 | 1 | 3 | 0.04259419 | 1 | 0.64497439 | 2 | 0.04972408 | 16 | 0.95553951 | FALSE | 1 | FALSE | 1 | FALSE | 1 | FALSE | 1 | FALSE | 0.98629461 | FALSE | 1 |
| 6353 | 6353 | 6.15384615 | 0.66171733 | 5 | 0.08367976 | 5 | 0.17117488 | 2 | 0.84458838 | 0 | 1 | 1 | 0.78011711 | 78 | 0.43010292 | FALSE | 0.79236883 | FALSE | 1 | FALSE | 1 | FALSE | 1 | FALSE | 1 | FALSE | 1 |
| 6354 | 6354 | 7.55555556 | 3.0469576 | 2 | 0.15144274 | 0 | 1 | 1 | 0.62830986 | 2 | 0.35374209 | 1 | 0.36060486 | 21 | 0.88395936 | FALSE | 1 | FALSE | 1 | FALSE | 1 | FALSE | 1 | FALSE | 1 | FALSE | 1 |
| 6355 | 6355 | 11.25 | 2.42641711 | 0 | 1 | 1 | 0.48239024 | 0 | 1 | 1 | 0.60987146 | 0 | 1 | 18 | 0.37504645 | FALSE | 1 | FALSE | 1 | FALSE | 1 | FALSE | 1 | FALSE | 1 | FALSE | 1 |
| 6356 | 6356 | 10.1627907 | 1.7775111 | 1 | 0.67382914 | 1 | 0.75788114 | 3 | 0.19985581 | 1 | 0.86831187 | 2 | 0.1563365 | 35 | 0.7801651 | FALSE | 1 | FALSE | 1 | FALSE | 1 | FALSE | 1 | FALSE | 1 | FALSE | 1 |
| 6357 | 6357 | 8.84615385 | 1.07030936 | 1 | 0.81665075 | 2 | 0.62701348 | 3 | 0.41519671 | 1 | 0.95356658 | 1 | 0.66032558 | 57 | 0.2945347 | FALSE | 1 | FALSE | 1 | FALSE | 1 | FALSE | 1 | FALSE | 1 | FALSE | 1 |
| 6358 | 6358 | 9.75 | 3.00347022 | 0 | 1 | 0 | 1 | 0 | 1 | 0 | 1 | 0 | 1 | 12 | 0.13002667 | FALSE | 1 | FALSE | 1 | FALSE | 1 | FALSE | 1 | FALSE | 1 | FALSE | 1 |
| 6359 | 6359 | 8.1875 | 2.26987748 | 0 | 1 | 0 | 1 | 1 | 0.44344817 | 1 | 0.52895088 | 0 | 1 | 14 | 0.53308261 | FALSE | 1 | FALSE | 1 | FALSE | 1 | FALSE | 1 | FALSE | 1 | FALSE | 1 |
| 6360 | 6360 | 9.33333333 | 2.29561348 | 0 | 1 | 0 | 1 | 2 | 0.17308361 | 0 | 1 | 1 | 0.29368245 | 18 | 0.57962763 | FALSE | 1 | FALSE | 1 | FALSE | 1 | FALSE | 1 | FALSE | 1 | FALSE | 1 |
| 6361 | 6361 | 9.25 | 3.19178633 | 1 | 0.34033054 | 1 | 0.40942991 | 1 | 0.44344817 | 1 | 0.52895088 | 0 | 1 | 12 | 0.90922186 | FALSE | 1 | FALSE | 1 | FALSE | 1 | FALSE | 1 | FALSE | 1 | FALSE | 1 |
| 6362 | 6362 | 8.89473684 | 2.3372214 | 0 | 1 | 0 | 1 | 0 | 1 | 0 | 1 | 1 | 0.26986775 | 18 | 0.17880694 | FALSE | 1 | FALSE | 1 | FALSE | 1 | FALSE | 1 | FALSE | 1 | FALSE | 1 |
| 6363 | 6363 | 8.38461539 | 2.05832126 | 0 | 1 | 1 | 0.7236252 | 2 | 0.41139386 | 2 | 0.54052736 | 0 | 1 | 34 | 0.41570781 | FALSE | 1 | FALSE | 1 | FALSE | 1 | FALSE | 1 | FALSE | 1 | FALSE | 1 |
| 6364 | 6364 | 5.52380952 | 2.95391665 | 1 | 0.66519151 | 1 | 0.74973543 | 1 | 0.78589454 | 0 | 1 | 0 | 1 | 39 | 0.08796704 | FALSE | 1 | FALSE | 1 | FALSE | 1 | FALSE | 1 | FALSE | 1 | FALSE | 1 |
| 6365 | 6365 | 8.85294118 | 2.97175283 | 2 | 0.21638713 | 2 | 0.30150529 | 3 | 0.12152681 | 1 | 0.7984848 | 0 | 1 | 26 | 0.92796052 | FALSE | 1 | FALSE | 1 | FALSE | 1 | FALSE | 1 | FALSE | 1 | FALSE | 1 |
| 6366 | 6366 | 7.5 | 3.06865877 | 0 | 1 | 1 | 0.32623933 | 1 | 0.35555293 | 0 | 1 | 0 | 1 | 10 | 0.71436964 | FALSE | 1 | FALSE | 1 | FALSE | 1 | FALSE | 1 | FALSE | 1 | FALSE | 1 |
| 6367 | 6367 | 10.3846154 | 3.00098603 | 0 | 1 | 0 | 1 | 0 | 1 | 0 | 1 | 0 | 1 | 13 | 0.10967906 | FALSE | 1 | FALSE | 1 | FALSE | 1 | FALSE | 1 | FALSE | 1 | FALSE | 1 |
| 6368 | 6368 | 11.6153846 | 3.19948221 | 0 | 1 | 0 | 1 | 0 | 1 | 1 | 0.45746571 | 0 | 1 | 12 | 0.374198 | FALSE | 1 | FALSE | 1 | FALSE | 1 | FALSE | 1 | FALSE | 1 | FALSE | 1 |
| 6369 | 6369 | 8.23529412 | 3.13480296 | 0 | 1 | 1 | 0.42857733 | 1 | 0.46348836 | 1 | 0.55062772 | 0 | 1 | 14 | 0.7313145 | FALSE | 1 | FALSE | 1 | FALSE | 1 | FALSE | 1 | FALSE | 1 | FALSE | 1 |
| 6370 | 6370 | 9.68421053 | 2.77303997 | 0 | 1 | 0 | 1 | 0 | 1 | 1 | 0.59104357 | 0 | 1 | 18 | 0.17880694 | FALSE | 1 | FALSE | 1 | FALSE | 1 | FALSE | 1 | FALSE | 1 | FALSE | 1 |
| 6371 | 6371 | 2.53863135 | 1.84772036 | 14 | 0.26870497 | 15 | 0.49820081 | 19 | 0.26925734 | 20 | 0.60357802 | 7 | 0.61803903 | 377 | 0.74667831 | FALSE | 1 | FALSE | 1 | FALSE | 1 | FALSE | 1 | FALSE | 1 | FALSE | 1 |
| 6372 | 6372 | 9.625 | 3.35177192 | 0 | 1 | 1 | 0.40942991 | 1 | 0.44344817 | 0 | 1 | 0 | 1 | 14 | 0.53308261 | FALSE | 1 | FALSE | 1 | FALSE | 1 | FALSE | 1 | FALSE | 1 | FALSE | 1 |
| 6373 | 6373 | 9.7826087 | 2.48397511 | 3 | 0.02011786 | 1 | 0.53115654 | 1 | 0.5695037 | 2 | 0.2852236 | 1 | 0.31672722 | 15 | 0.99450713 | FALSE | 0.44482142 | FALSE | 1 | FALSE | 1 | FALSE | 1 | FALSE | 1 | FALSE | 1 |
| 6374 | 6374 | 7.94736842 | 2.67818381 | 1 | 0.76787545 | 1 | 0.84259834 | 4 | 0.14086741 | 3 | 0.47819064 | 1 | 0.60529938 | 46 | 0.74916201 | FALSE | 1 | FALSE | 1 | FALSE | 1 | FALSE | 1 | FALSE | 1 | FALSE | 1 |
| 6375 | 6375 | 8.40659341 | 1.40624996 | 2 | 0.67673939 | 1 | 0.94917004 | 2 | 0.84010838 | 6 | 0.2314164 | 3 | 0.1830501 | 76 | 0.56399212 | FALSE | 1 | FALSE | 1 | FALSE | 1 | FALSE | 1 | FALSE | 1 | FALSE | 1 |
| 6376 | 6376 | 7 | 1.93649167 | 2 | 0.10809809 | 3 | 0.03266339 | 3 | 0.04259419 | 1 | 0.64497439 | 0 | 1 | 13 | 0.99911494 | FALSE | 0.88820416 | FALSE | 0.92268438 | FALSE | 1 | FALSE | 1 | FALSE | 1 | FALSE | 1 |
| 6377 | 6377 | 6.25306122 | 1.24917966 | 8 | 0.25513958 | 15 | 0.00895353 | 7 | 0.7470899 | 8 | 0.85339837 | 6 | 0.1877464 | 191 | 0.92035784 | FALSE | 1 | FALSE | 0.63686998 | FALSE | 1 | FALSE | 1 | FALSE | 1 | FALSE | 1 |
| 6378 | 6378 | 9.4375 | 2.03004772 | 0 | 1 | 0 | 1 | 1 | 1 | 4 | 0.05697506 | 0 | 1 | 28 | 0.42532177 | FALSE | 1 | FALSE | 1 | FALSE | 1 | FALSE | 1 | FALSE | 1 | FALSE | 1 |
| 6379 | 6379 | 8.08064516 | 1.79873062 | 0 | 1 | 2 | 0.60114557 | 5 | 0.07095719 | 4 | 0.31731292 | 0 | 1 | 51 | 0.74578096 | FALSE | 1 | FALSE | 1 | FALSE | 1 | FALSE | 1 | FALSE | 1 | FALSE | 1 |
| 6380 | 6380 | 8.33333333 | 2.86744176 | 0 | 1 | 1 | 0.25628163 | 0 | 1 | 0 | 1 | 0 | 1 | 8 | 0.5780332 | FALSE | 1 | FALSE | 1 | FALSE | 1 | FALSE | 1 | FALSE | 1 | FALSE | 1 |
| 6381 | 6381 | 4.54887218 | 1.74487936 | 4 | 0.44578002 | 3 | 0.81044919 | 8 | 0.1051942 | 4 | 0.86667866 | 2 | 0.64598387 | 112 | 0.57942532 | FALSE | 1 | FALSE | 1 | FALSE | 1 | FALSE | 1 | FALSE | 1 | FALSE | 1 |
| 6382 | 6382 | 8.92857143 | 2.81486579 | 0 | 1 | 2 | 0.2288771 | 0 | 1 | 1 | 0.73248983 | 1 | 0.37113067 | 24 | 0.55018107 | FALSE | 1 | FALSE | 1 | FALSE | 1 | FALSE | 1 | FALSE | 1 | FALSE | 1 |
| 6383 | 6383 | 9.25 | 2.80252862 | 0 | 1 | 0 | 1 | 0 | 1 | 0 | 1 | 1 | 0.18007762 | 11 | 0.41944532 | FALSE | 1 | FALSE | 1 | FALSE | 1 | FALSE | 1 | FALSE | 1 | FALSE | 1 |
| 6384 | 6384 | 8.35294118 | 1.93850083 | 0 | 1 | 1 | 0.6739483 | 0 | 1 | 2 | 0.46709861 | 0 | 1 | 31 | 0.1995074 | FALSE | 1 | FALSE | 1 | FALSE | 1 | FALSE | 1 | FALSE | 1 | FALSE | 1 |
| 6385 | 6385 | 5.47916667 | 1.58100157 | 2 | 0.34973212 | 1 | 0.79481669 | 2 | 0.51909752 | 4 | 0.17698522 | 1 | 0.54902428 | 38 | 0.88187597 | FALSE | 1 | FALSE | 1 | FALSE | 1 | FALSE | 1 | FALSE | 1 | FALSE | 1 |
| 6386 | 6386 | 5.48837209 | 2.56879388 | 2 | 0.65227007 | 5 | 0.14509352 | 6 | 0.08811302 | 7 | 0.09905935 | 2 | 0.41367714 | 64 | 0.99460611 | FALSE | 1 | FALSE | 1 | FALSE | 1 | FALSE | 1 | FALSE | 1 | FALSE | 1 |
| 6387 | 6387 | 7.2745098 | 2.40162062 | 0 | 1 | 0 | 1 | 4 | 0.10960782 | 2 | 0.68678423 | 0 | 1 | 45 | 0.29584937 | FALSE | 1 | FALSE | 1 | FALSE | 1 | FALSE | 1 | FALSE | 1 | FALSE | 1 |
| 6388 | 6388 | 10.1363636 | 1.63193846 | 3 | 0.01782284 | 2 | 0.15794013 | 0 | 1 | 0 | 1 | 0 | 1 | 17 | 0.8840357 | FALSE | 0.41596787 | FALSE | 1 | FALSE | 1 | FALSE | 1 | FALSE | 1 | FALSE | 1 |
| 6389 | 6389 | 9.66666667 | 2.1602469 | 0 | 1 | 1 | 0.44710673 | 0 | 1 | 1 | 0.57131013 | 0 | 1 | 16 | 0.44998001 | FALSE | 1 | FALSE | 1 | FALSE | 1 | FALSE | 1 | FALSE | 1 | FALSE | 1 |
| 6390 | 6390 | 7.54929578 | 1.49918976 | 0 | 1 | 3 | 0.40468462 | 1 | 0.92656034 | 2 | 0.84445443 | 0 | 1 | 65 | 0.05811248 | FALSE | 1 | FALSE | 1 | FALSE | 1 | FALSE | 1 | FALSE | 1 | FALSE | 1 |

| | | | | | | | | | | | | | | | | | | | | | | | | | | | |
|---|---|---|---|---|---|---|---|---|---|---|---|---|---|---|---|---|---|---|---|---|---|---|---|---|---|---|---|
| 6391 | 6391 | 8.11764706 | 1.96685335 | 0 | 1 | 1 | 0.40942991 | 1 | 0.44344817 | 1 | 0.52895088 | 0 | 1 | 13 | 0.76799161 | FALSE | 1 | FALSE | 1 | FALSE | 1 | FALSE | 1 | FALSE | 1 | FALSE | 1 |
| 6392 | 6392 | 9.36363636 | 2.10076727 | 2 | 0.10809809 | 1 | 0.5154318 | 1 | 0.55340851 | 1 | 0.64497439 | 0 | 1 | 17 | 0.8840357 | FALSE | 0.88820416 | FALSE | 1 | FALSE | 1 | FALSE | 1 | FALSE | 1 | FALSE | 1 |
| 6393 | 6393 | 7.17307692 | 1.75105632 | 2 | 0.34973212 | 1 | 0.79481669 | 2 | 0.51909752 | 1 | 0.89605799 | 0 | 1 | 42 | 0.36011057 | FALSE | 1 | FALSE | 1 | FALSE | 1 | FALSE | 1 | FALSE | 1 | FALSE | 1 |
| 6394 | 6394 | 9.46153846 | 3.12841529 | 1 | 0.49162634 | 0 | 1 | 0 | 1 | 0 | 1 | 1 | 0.3499045 | 24 | 0.20425579 | FALSE | 1 | FALSE | 1 | FALSE | 1 | FALSE | 1 | FALSE | 1 | FALSE | 1 |
| 6395 | 6395 | 4.7962963 | 2.26797046 | 10 | 0.00041802 | 5 | 0.270828 | 8 | 0.03989203 | 4 | 0.73784936 | 6 | 0.00838282 | 75 | 0.99997508 | TRUE | 0.04200174 | FALSE | 1 | FALSE | 1 | FALSE | 1 | FALSE | 0.50674765 | FALSE | 1 |
| 6396 | 6396 | 4.37349398 | 1.55393665 | 9 | 0.02471844 | 6 | 0.43876635 | 7 | 0.37558296 | 8 | 0.48231057 | 5 | 0.1313892 | 129 | 0.981226 | FALSE | 0.4974559 | FALSE | 1 | FALSE | 1 | FALSE | 1 | FALSE | 1 | FALSE | 1 |
| 6397 | 6397 | 10.2727273 | 1.91125419 | 0 | 1 | 0 | 1 | 0 | 1 | 0 | 1 | 1 | 0.16638736 | 10 | 0.46859937 | FALSE | 1 | FALSE | 1 | FALSE | 1 | FALSE | 1 | FALSE | 1 | FALSE | 1 |
| 6398 | 6398 | 10.5 | 2.20227156 | 0 | 1 | 1 | 0.28036922 | 0 | 1 | 0 | 1 | 0 | 1 | 9 | 0.5215547 | FALSE | 1 | FALSE | 1 | FALSE | 1 | FALSE | 1 | FALSE | 1 | FALSE | 1 |
| 6399 | 6399 | 3.464 | 1.40595306 | 4 | 0.88950509 | 7 | 0.70585878 | 7 | 0.80156581 | 6 | 0.97635391 | 8 | 0.05223987 | 218 | 0.12077792 | FALSE | 1 | FALSE | 1 | FALSE | 1 | FALSE | 1 | FALSE | 1 | FALSE | 1 |
| 6400 | 6400 | 11.4259259 | 1.11557744 | 3 | 0.09709068 | 3 | 0.16132793 | 1 | 0.79363407 | 2 | 0.59404271 | 1 | 0.50988734 | 33 | 0.93815092 | FALSE | 0.86047225 | FALSE | 1 | FALSE | 1 | FALSE | 1 | FALSE | 1 | FALSE | 1 |
| 6401 | 6401 | 8.40384615 | 3.10247801 | 0 | 1 | 2 | 0.50543179 | 2 | 0.5626042 | 2 | 0.6970685 | 2 | 0.2095319 | 44 | 0.57424118 | FALSE | 1 | FALSE | 1 | FALSE | 1 | FALSE | 1 | FALSE | 1 | FALSE | 1 |
| 6402 | 6402 | 8.71428571 | 2.60298102 | 0 | 1 | 0 | 1 | 0 | 1 | 0 | 1 | 0 | 1 | 7 | 0.3043657 | FALSE | 1 | FALSE | 1 | FALSE | 1 | FALSE | 1 | FALSE | 1 | FALSE | 1 |
| 6403 | 6403 | 7.88148148 | 1.66467371 | 5 | 0.26478208 | 5 | 0.44420164 | 1 | 0.99319181 | 5 | 0.74974854 | 3 | 0.38189355 | 116 | 0.36046508 | FALSE | 1 | FALSE | 1 | FALSE | 1 | FALSE | 1 | FALSE | 1 | FALSE | 1 |
| 6404 | 6404 | 7.57142857 | 3.22300405 | 0 | 1 | 1 | 0.36919824 | 2 | 0.08806398 | 0 | 1 | 0 | 1 | 11 | 0.83676449 | FALSE | 1 | FALSE | 1 | FALSE | 1 | FALSE | 1 | FALSE | 1 | FALSE | 1 |
| 6405 | 6405 | 9.66666667 | 2.72845092 | 0 | 1 | 0 | 1 | 1 | 0.48280983 | 0 | 1 | 0 | 1 | 17 | 0.20329205 | FALSE | 1 | FALSE | 1 | FALSE | 1 | FALSE | 1 | FALSE | 1 | FALSE | 1 |
| 6406 | 6406 | 8.5 | 1.99431009 | 1 | 0.4357638 | 0 | 1 | 0 | 1 | 1 | 0.64497439 | 0 | 1 | 20 | 0.30915202 | FALSE | 1 | FALSE | 1 | FALSE | 1 | FALSE | 1 | FALSE | 1 | FALSE | 1 |
| 6407 | 6407 | 9.76923077 | 2.18926915 | 0 | 1 | 0 | 1 | 0 | 1 | 0 | 1 | 0 | 1 | 13 | 0.10967906 | FALSE | 1 | FALSE | 1 | FALSE | 1 | FALSE | 1 | FALSE | 1 | FALSE | 1 |
| 6408 | 6408 | 11.12 | 1.30598622 | 0 | 1 | 2 | 0.19297191 | 2 | 0.22597287 | 0 | 1 | 0 | 1 | 21 | 0.64957167 | FALSE | 1 | FALSE | 1 | FALSE | 1 | FALSE | 1 | FALSE | 1 | FALSE | 1 |
| 6409 | 6409 | 7.86046512 | 1.91179199 | 3 | 0.37958833 | 1 | 0.94192298 | 1 | 0.95785384 | 7 | 0.09905935 | 2 | 0.41367714 | 72 | 0.63625249 | FALSE | 1 | FALSE | 1 | FALSE | 1 | FALSE | 1 | FALSE | 1 | FALSE | 1 |
| 6410 | 6410 | 9.30769231 | 1.86951037 | 0 | 1 | 2 | 0.36122049 | 0 | 1 | 1 | 0.84089119 | 1 | 0.4761699 | 35 | 0.24915769 | FALSE | 1 | FALSE | 1 | FALSE | 1 | FALSE | 1 | FALSE | 1 | FALSE | 1 |
| 6411 | 6411 | 9.87179487 | 2.57382771 | 1 | 0.63789042 | 1 | 0.7236252 | 1 | 0.76089851 | 0 | 1 | 1 | 0.4761699 | 35 | 0.24915769 | FALSE | 1 | FALSE | 1 | FALSE | 1 | FALSE | 1 | FALSE | 1 | FALSE | 1 |
| 6412 | 6412 | 7.87804878 | 1.94051944 | 0 | 1 | 2 | 0.38460545 | 1 | 0.77786601 | 4 | 0.11715635 | 2 | 0.14490154 | 32 | 0.90458039 | FALSE | 1 | FALSE | 1 | FALSE | 1 | FALSE | 1 | FALSE | 1 | FALSE | 1 |
| 6413 | 6413 | 8.55172414 | 2.01008161 | 1 | 0.50471032 | 1 | 0.58913618 | 2 | 0.25286935 | 1 | 0.71956339 | 0 | 1 | 22 | 0.76237877 | FALSE | 1 | FALSE | 1 | FALSE | 1 | FALSE | 1 | FALSE | 1 | FALSE | 1 |
| 6414 | 6414 | 9.5 | 3.09030743 | 0 | 1 | 0 | 1 | 0 | 1 | 0 | 1 | 1 | 0.28187291 | 19 | 0.15700255 | FALSE | 1 | FALSE | 1 | FALSE | 1 | FALSE | 1 | FALSE | 1 | FALSE | 1 |
| 6415 | 6415 | 8.1 | 3.38969025 | 1 | 0.22886536 | 0 | 1 | 0 | 1 | 3 | 0.00903718 | 0 | 1 | 6 | 0.98831471 | FALSE | 1 | FALSE | 1 | FALSE | 1 | FALSE | 1 | FALSE | 1 | FALSE | 1 |
| 6416 | 6416 | 9.35106383 | 1.49969817 | 2 | 0.69979848 | 3 | 0.59118008 | 3 | 0.66251153 | 6 | 0.26227462 | 3 | 0.19972697 | 77 | 0.79315074 | FALSE | 1 | FALSE | 1 | FALSE | 1 | FALSE | 1 | FALSE | 1 | FALSE | 1 |
| 6417 | 6417 | 8.91176471 | 1.22156233 | 2 | 0.21638713 | 1 | 0.6739483 | 0 | 1 | 0 | 1 | 1 | 0.43077488 | 30 | 0.36879488 | FALSE | 1 | FALSE | 1 | FALSE | 1 | FALSE | 1 | FALSE | 1 | FALSE | 1 |
| 6418 | 6418 | 11.4 | 2.27449628 | 0 | 1 | 1 | 0.38964394 | 1 | 0.42266264 | 0 | 1 | 0 | 1 | 13 | 0.57712805 | FALSE | 1 | FALSE | 1 | FALSE | 1 | FALSE | 1 | FALSE | 1 | FALSE | 1 |
| 6419 | 6419 | 6.17647059 | 3.8333208 | 1 | 0.7354105 | 1 | 0.81422651 | 4 | 0.10960782 | 3 | 0.41729179 | 2 | 0.20351566 | 40 | 0.90979546 | FALSE | 1 | FALSE | 1 | FALSE | 1 | FALSE | 1 | FALSE | 1 | FALSE | 1 |
| 6420 | 6420 | 8.5 | 1.80277564 | 1 | 0.51745948 | 0 | 1 | 2 | 0.2663542 | 1 | 0.73248983 | 0 | 1 | 24 | 0.55018107 | FALSE | 1 | FALSE | 1 | FALSE | 1 | FALSE | 1 | FALSE | 1 | FALSE | 1 |
| 6421 | 6421 | 7.97142857 | 1.68159303 | 1 | 0.59801736 | 1 | 0.68454804 | 0 | 1 | 2 | 0.48233989 | 1 | 0.44015537 | 30 | 0.52765882 | FALSE | 1 | FALSE | 1 | FALSE | 1 | FALSE | 1 | FALSE | 1 | FALSE | 1 |
| 6422 | 6422 | 8.83333333 | 1.95078332 | 0 | 1 | 0 | 1 | 0 | 1 | 0 | 1 | 0 | 1 | 6 | 0.36077158 | FALSE | 1 | FALSE | 1 | FALSE | 1 | FALSE | 1 | FALSE | 1 | FALSE | 1 |
| 6423 | 6423 | 11.125 | 2.66634113 | 0 | 1 | 0 | 1 | 0 | 1 | 0 | 1 | 0 | 1 | 8 | 0.25677166 | FALSE | 1 | FALSE | 1 | FALSE | 1 | FALSE | 1 | FALSE | 1 | FALSE | 1 |
| 6424 | 6424 | 9.89473684 | 2.59340737 | 0 | 1 | 0 | 1 | 2 | 0.14749808 | 2 | 0.21610812 | 0 | 1 | 15 | 0.83648191 | FALSE | 1 | FALSE | 1 | FALSE | 1 | FALSE | 1 | FALSE | 1 | FALSE | 1 |
| 6425 | 6425 | 4.24324324 | 2.47224673 | 4 | 0.3057853 | 2 | 0.87316332 | 2 | 0.90795 | 1 | 0.99428817 | 3 | 0.26494741 | 97 | 0.11107732 | FALSE | 1 | FALSE | 1 | FALSE | 1 | FALSE | 1 | FALSE | 1 | FALSE | 1 |
| 6426 | 6426 | 11.4545455 | 2.67526163 | 0 | 1 | 0 | 1 | 1 | 0.33149969 | 0 | 1 | 0 | 1 | 10 | 0.46859937 | FALSE | 1 | FALSE | 1 | FALSE | 1 | FALSE | 1 | FALSE | 1 | FALSE | 1 |
| 6427 | 6427 | 7.83333333 | 2.77888867 | 0 | 1 | 1 | 0.54637326 | 1 | 0.58502114 | 2 | 0.30249307 | 0 | 1 | 20 | 0.68270729 | FALSE | 1 | FALSE | 1 | FALSE | 1 | FALSE | 1 | FALSE | 1 | FALSE | 1 |
| 6428 | 6428 | 6.64150943 | 2.37196153 | 1 | 0.74890551 | 2 | 0.51564773 | 2 | 0.57303317 | 3 | 0.44196866 | 2 | 0.21556658 | 43 | 0.80430518 | FALSE | 1 | FALSE | 1 | FALSE | 1 | FALSE | 1 | FALSE | 1 | FALSE | 1 |
| 6429 | 6429 | 11.2941177 | 2.82352941 | 1 | 0.35728305 | 1 | 0.42857733 | 0 | 1 | 0 | 1 | 1 | 0.24525764 | 14 | 0.7313145 | FALSE | 1 | FALSE | 1 | FALSE | 1 | FALSE | 1 | FALSE | 1 | FALSE | 1 |
| 6430 | 6430 | 8.89473684 | 1.88887984 | 0 | 1 | 0 | 1 | 0 | 1 | 2 | 0.21610812 | 0 | 1 | 17 | 0.41141543 | FALSE | 1 | FALSE | 1 | FALSE | 1 | FALSE | 1 | FALSE | 1 | FALSE | 1 |
| 6431 | 6431 | 9.91666667 | 2.2715022 | 0 | 1 | 1 | 0.54637326 | 2 | 0.21260257 | 0 | 1 | 3 | 0.00675546 | 18 | 0.93166998 | FALSE | 1 | FALSE | 1 | FALSE | 1 | FALSE | 1 | FALSE | 0.47461742 | FALSE | 1 |
| 6432 | 6432 | 9.7037037 | 1.62878766 | 1 | 0.50471032 | 1 | 0.58913618 | 1 | 0.62830986 | 0 | 1 | 0 | 1 | 24 | 0.3725373 | FALSE | 1 | FALSE | 1 | FALSE | 1 | FALSE | 1 | FALSE | 1 | FALSE | 1 |
| 6433 | 6433 | 8.64705882 | 1.53167254 | 1 | 0.35728305 | 0 | 1 | 0 | 1 | 1 | 0.55062772 | 0 | 1 | 15 | 0.49060435 | FALSE | 1 | FALSE | 1 | FALSE | 1 | FALSE | 1 | FALSE | 1 | FALSE | 1 |
| 6434 | 6434 | 9.92857143 | 2.63124864 | 1 | 0.3050805 | 1 | 0.36919824 | 1 | 0.40110418 | 0 | 1 | 0 | 1 | 11 | 0.83676449 | FALSE | 1 | FALSE | 1 | FALSE | 1 | FALSE | 1 | FALSE | 1 | FALSE | 1 |
| 6435 | 6435 | 10.1666667 | 2.54405625 | 1 | 0.14434519 | 0 | 1 | 0 | 1 | 0 | 1 | 0 | 1 | 5 | 0.76185568 | FALSE | 1 | FALSE | 1 | FALSE | 1 | FALSE | 1 | FALSE | 1 | FALSE | 1 |
| 6436 | 6436 | 9.31111111 | 1.90697226 | 4 | 0.02747663 | 3 | 0.17732615 | 3 | 0.21866539 | 5 | 0.05380295 | 0 | 1 | 30 | 0.99918792 | FALSE | 0.50964275 | FALSE | 1 | FALSE | 1 | FALSE | 1 | FALSE | 1 | FALSE | 1 |
| 6437 | 6437 | 8.65217391 | 2.13884763 | 0 | 1 | 2 | 0.16948764 | 1 | 0.5695037 | 1 | 0.6613267 | 0 | 1 | 19 | 0.71542496 | FALSE | 1 | FALSE | 1 | FALSE | 1 | FALSE | 1 | FALSE | 1 | FALSE | 1 |
| 6438 | 6438 | 2.09654562 | 0.91657145 | 37 | 0.05994669 | 40 | 0.2783992 | 46 | 0.18476224 | 63 | 0.04761689 | 23 | 0.14862411 | 916 | 0.99859581 | FALSE | 0.72532843 | FALSE | 1 | FALSE | 1 | FALSE | 1 | FALSE | 1 | FALSE | 1 |
| 6439 | 6439 | 6.3 | 2.9729735 | 0 | 1 | 2 | 0.65156696 | 3 | 0.44404438 | 5 | 0.20108725 | 0 | 1 | 58 | 0.49889031 | FALSE | 1 | FALSE | 1 | FALSE | 1 | FALSE | 1 | FALSE | 1 | FALSE | 1 |
| 6440 | 6440 | 9.26666667 | 2.71947707 | 0 | 1 | 1 | 0.38964394 | 0 | 1 | 0 | 1 | 0 | 1 | 14 | 0.29525409 | FALSE | 1 | FALSE | 1 | FALSE | 1 | FALSE | 1 | FALSE | 1 | FALSE | 1 |
| 6441 | 6441 | 6.58333333 | 3.37782803 | 1 | 0.71380452 | 0 | 1 | 0 | 1 | 1 | 0.89605799 | 2 | 0.18560062 | 44 | 0.11052514 | FALSE | 1 | FALSE | 1 | FALSE | 1 | FALSE | 1 | FALSE | 1 | FALSE | 1 |
| 6442 | 6442 | 9.65 | 2.35106359 | 0 | 1 | 1 | 0.48239024 | 1 | 0.51939841 | 1 | 0.60987146 | 1 | 0.28187291 | 16 | 0.80826361 | FALSE | 1 | FALSE | 1 | FALSE | 1 | FALSE | 1 | FALSE | 1 | FALSE | 1 |
| 6443 | 6443 | 7.61538462 | 2.54311927 | 0 | 1 | 2 | 0.20486827 | 1 | 0.61440466 | 1 | 0.70601446 | 1 | 0.3499045 | 21 | 0.78939668 | FALSE | 1 | FALSE | 1 | FALSE | 1 | FALSE | 1 | FALSE | 1 | FALSE | 1 |
| 6444 | 6444 | 8.57777778 | 1.73190825 | 0 | 1 | 1 | 0.77338922 | 1 | 0.80828716 | 3 | 0.34154252 | 0 | 1 | 40 | 0.27423942 | FALSE | 1 | FALSE | 1 | FALSE | 1 | FALSE | 1 | FALSE | 1 | FALSE | 1 |
| 6445 | 6445 | 4.9144385 | 2.7928777 | 4 | 0.70828604 | 10 | 0.07874183 | 9 | 0.22500945 | 5 | 0.93452405 | 5 | 0.18943892 | 153 | 0.81983459 | FALSE | 1 | FALSE | 1 | FALSE | 1 | FALSE | 1 | FALSE | 1 | FALSE | 1 |
| 6446 | 6446 | 10.2857143 | 2.54750779 | 2 | 0.16048326 | 1 | 0.60247911 | 1 | 0.64171563 | 1 | 0.73248983 | 1 | 0.37113067 | 22 | 0.86499992 | FALSE | 1 | FALSE | 1 | FALSE | 1 | FALSE | 1 | FALSE | 1 | FALSE | 1 |
| 6447 | 6447 | 7.9 | 2.16564078 | 2 | 0.17884234 | 1 | 0.6278847 | 1 | 0.66709967 | 1 | 0.75658797 | 0 | 1 | 25 | 0.67630471 | FALSE | 1 | FALSE | 1 | FALSE | 1 | FALSE | 1 | FALSE | 1 | FALSE | 1 |
| 6448 | 6448 | 9 | 2.9104275 | 1 | 0.35728305 | 0 | 1 | 1 | 0.46348836 | 2 | 0.18212134 | 0 | 1 | 13 | 0.88719222 | FALSE | 1 | FALSE | 1 | FALSE | 1 | FALSE | 1 | FALSE | 1 | FALSE | 1 |
| 6449 | 6449 | 3.67672414 | 2.11598423 | 10 | 0.07415119 | 6 | 0.76827044 | 9 | 0.4555255 | 9 | 0.74611082 | 4 | 0.53152044 | 194 | 0.66712367 | FALSE | 0.77424195 | FALSE | 1 | FALSE | 1 | FALSE | 1 | FALSE | 1 | FALSE | 1 |
| 6450 | 6450 | 10.631579 | 2.43246737 | 0 | 1 | 1 | 0.46503795 | 0 | 1 | 0 | 1 | 1 | 0.26986775 | 17 | 0.41141543 | FALSE | 1 | FALSE | 1 | FALSE | 1 | FALSE | 1 | FALSE | 1 | FALSE | 1 |
| 6451 | 6451 | 6.11827957 | 3.56762462 | 1 | 0.91216464 | 3 | 0.58386444 | 3 | 0.65536448 | 4 | 0.62434183 | 0 | 1 | 82 | 0.19398117 | FALSE | 1 | FALSE | 1 | FALSE | 1 | FALSE | 1 | FALSE | 1 | FALSE | 1 |
| 6452 | 6452 | 8.83333333 | 2.40947205 | 1 | 0.14434519 | 0 | 1 | 0 | 1 | 0 | 1 | 0 | 1 | 5 | 0.76185568 | FALSE | 1 | FALSE | 1 | FALSE | 1 | FALSE | 1 | FALSE | 1 | FALSE | 1 |
| 6453 | 6453 | 9.5 | 1.5 | 0 | 1 | 0 | 1 | 1 | 0.40110418 | 0 | 1 | 0 | 1 | 13 | 0.33283729 | FALSE | 1 | FALSE | 1 | FALSE | 1 | FALSE | 1 | FALSE | 1 | FALSE | 1 |
| 6454 | 6454 | 7.79310345 | 2.59148886 | 0 | 1 | 0 | 1 | 1 | 0.65463982 | 4 | 0.04184267 | 0 | 1 | 24 | 0.70563334 | FALSE | 1 | FALSE | 1 | FALSE | 1 | FALSE | 1 | FALSE | 1 | FALSE | 1 |
| 6455 | 6455 | 8.40677966 | 1.90555516 | 1 | 0.78541994 | 1 | 0.85750546 | 2 | 0.63179508 | 1 | 0.93827647 | 0 | 1 | 54 | 0.08351851 | FALSE | 1 | FALSE | 1 | FALSE | 1 | FALSE | 1 | FALSE | 1 | FALSE | 1 |
| 6456 | 6456 | 9.38461539 | 3.1265233 | 1 | 0.28675951 | 0 | 1 | 0 | 1 | 0 | 1 | 0 | 1 | 12 | 0.374198 | FALSE | 1 | FALSE | 1 | FALSE | 1 | FALSE | 1 | FALSE | 1 | FALSE | 1 |
| 6457 | 6457 | 4.2625 | 2.84931461 | 0 | 1 | 1 | 0.92668093 | 3 | 0.54427311 | 7 | 0.06964209 | 4 | 0.04023837 | 64 | 0.83834292 | FALSE | 1 | FALSE | 1 | FALSE | 1 | FALSE | 1 | FALSE | 0.90558441 | FALSE | 1 |
| 6458 | 6458 | 2.0633387 | 0.72769241 | 42 | 0.81668019 | 55 | 0.75227305 | 70 | 0.27291514 | 78 | 0.78894528 | 43 | 0.00405384 | 1534 | 0.617005 | FALSE | 1 | FALSE | 1 | FALSE | 1 | FALSE | 1 | FALSE | 0.37645884 | FALSE | 1 |
| 6459 | 6459 | 8.38181818 | 1.10356298 | 0 | 1 | 2 | 0.51564773 | 3 | 0.29666549 | 3 | 0.44196866 | 0 | 1 | 45 | 0.5504765 | FALSE | 1 | FALSE | 1 | FALSE | 1 | FALSE | 1 | FALSE | 1 | FALSE | 1 |
| 6460 | 6460 | 10.5 | 1.38443731 | 1 | 0.14434519 | 2 | 0.01433429 | 0 | 1 | 0 | 1 | 0 | 1 | 3 | 0.9931827 | FALSE | 1 | FALSE | 0.71309764 | FALSE | 1 | FALSE | 1 | FALSE | 1 | FALSE | 1 |
| 6461 | 6461 | 7.75308642 | 1.61421819 | 1 | 0.87954842 | 2 | 0.74335738 | 1 | 0.94927633 | 4 | 0.51410462 | 2 | 0.38471054 | 71 | 0.26017447 | FALSE | 1 | FALSE | 1 | FALSE | 1 | FALSE | 1 | FALSE | 1 | FALSE | 1 |
| 6462 | 6462 | 8.66666667 | 3.22317993 | 1 | 0.46441868 | 1 | 0.54637326 | 2 | 0.21260257 | 3 | 0.09525947 | 1 | 0.32796862 | 16 | 0.99240722 | FALSE | 1 | FALSE | 1 | FALSE | 1 | FALSE | 1 | FALSE | 1 | FALSE | 1 |
| 6463 | 6463 | 9.76923077 | 1.52687948 | 0 | 1 | 1 | 0.3480709 | 1 | 0.37874417 | 0 | 1 | 2 | 0.01846832 | 9 | 0.96018168 | FALSE | 1 | FALSE | 1 | FALSE | 1 | FALSE | 1 | FALSE | 0.69720022 | FALSE | 1 |
| 6464 | 6464 | 12.625 | 2.1758619 | 1 | 0.18769745 | 0 | 1 | 0 | 1 | 0 | 1 | 0 | 1 | 7 | 0.63752397 | FALSE | 1 | FALSE | 1 | FALSE | 1 | FALSE | 1 | FALSE | 1 | FALSE | 1 |
| 6465 | 6465 | 10.375 | 2.0577597 | 1 | 0.34033054 | 1 | 0.40942991 | 0 | 1 | 2 | 0.16544789 | 0 | 1 | 12 | 0.90922186 | FALSE | 1 | FALSE | 1 | FALSE | 1 | FALSE | 1 | FALSE | 1 | FALSE | 1 |
| 6466 | 6466 | 7.91025641 | 1.23190386 | 1 | 0.86966563 | 1 | 0.92420038 | 2 | 0.77659406 | 5 | 0.28743251 | 4 | 0.0386728 | 65 | 0.67060743 | FALSE | 1 | FALSE | 1 | FALSE | 1 | FALSE | 1 | FALSE | 0.88493726 | FALSE | 1 |
| 6467 | 6467 | 10.631579 | 3.81424651 | 1 | 0.38989951 | 2 | 0.12434464 | 1 | 0.50143826 | 1 | 0.59104357 | 1 | 0.26986775 | 13 | 0.97998676 | FALSE | 1 | FALSE | 1 | FALSE | 1 | FALSE | 1 | FALSE | 1 | FALSE | 1 |
| 6468 | 6468 | 8.33333333 | 2.63523138 | 1 | 0.60837712 | 1 | 0.69480472 | 3 | 0.13783405 | 2 | 0.49731009 | 0 | 1 | 29 | 0.81028289 | FALSE | 1 | FALSE | 1 | FALSE | 1 | FALSE | 1 | FALSE | 1 | FALSE | 1 |
| 6469 | 6469 | 11.3333333 | 2.21108319 | 0 | 1 | 0 | 1 | 1 | 0.35555293 | 0 | 1 | 2 | 0.01579445 | 9 | 0.89630573 | FALSE | 1 | FALSE | 1 | FALSE | 1 | FALSE | 1 | FALSE | 0.66313193 | FALSE | 1 |
| 6470 | 6470 | 7.94594595 | 2.37037967 | 2 | 0.24498301 | 2 | 0.3375163 | 1 | 0.74264075 | 1 | 0.82511639 | 0 | 1 | 31 | 0.64534533 | FALSE | 1 | FALSE | 1 | FALSE | 1 | FALSE | 1 | FALSE | 1 | FALSE | 1 |
| 6471 | 6471 | 10.6 | 1.42828569 | 0 | 1 | 0 | 1 | 1 | 0.30655256 | 0 | 1 | 1 | 0.15247062 | 8 | 0.80394719 | FALSE | 1 | FALSE | 1 | FALSE | 1 | FALSE | 1 | FALSE | 1 | FALSE | 1 |
| 6472 | 6472 | 9.33333333 | 3.14466038 | 1 | 0.37380237 | 0 | 1 | 1 | 0.48280983 | 2 | 0.19902682 | 0 | 1 | 14 | 0.86287418 | FALSE | 1 | FALSE | 1 | FALSE | 1 | FALSE | 1 | FALSE | 1 | FALSE | 1 |
| 6473 | 6473 | 9.2 | 1.53622915 | 2 | 0.0256865 | 0 | 1 | 0 | 1 | 0 | 1 | 1 | 0.15247062 | 7 | 0.94326215 | FALSE | 0.4974559 | FALSE | 1 | FALSE | 1 | FALSE | 1 | FALSE | 1 | FALSE | 1 |
| 6474 | 6474 | 6.68421053 | 3.44967781 | 0 | 1 | 0 | 1 | 1 | 0.50143826 | 1 | 0.59104357 | 0 | 1 | 17 | 0.41141543 | FALSE | 1 | FALSE | 1 | FALSE | 1 | FALSE | 1 | FALSE | 1 | FALSE | 1 |
| 6475 | 6475 | 9.38461539 | 3.38636319 | 0 | 1 | 0 | 1 | 1 | 0.37874417 | 0 | 1 | 0 | 1 | 12 | 0.374198 | FALSE | 1 | FALSE | 1 | FALSE | 1 | FALSE | 1 | FALSE | 1 | FALSE | 1 |
| 6476 | 6476 | 8.83333333 | 1.57233019 | 1 | 0.54198714 | 1 | 0.6278847 | 1 | 0.66709967 | 4 | 0.04659941 | 1 | 0.39166989 | 22 | 0.96518743 | FALSE | 1 | FALSE | 1 | FALSE | 1 | FALSE | 1 | FALSE | 1 | FALSE | 1 |
| 6477 | 6477 | 4.53804348 | 1.70633999 | 10 | 0.01761923 | 8 | 0.23146665 | 8 | 0.3250504 | 7 | 0.73259105 | 3 | 0.57430927 | 145 | 0.95257832 | FALSE | 0.41596787 | FALSE | 1 | FALSE | 1 | FALSE | 1 | FALSE | 1 | FALSE | 1 |
| 6478 | 6478 | 6.28358209 | 2.14966829 | 5 | 0.02811699 | 3 | 0.36935445 | 5 | 0.09200232 | 1 | 0.95777219 | 1 | 0.6714808 | 52 | 0.95010891 | FALSE | 0.51185412 | FALSE | 1 | FALSE | 1 | FALSE | 1 | FALSE | 1 | FALSE | 1 |
| 6479 | 6479 | 10.1875 | 3.30186974 | 0 | 1 | 0 | 1 | 0 | 1 | 0 | 1 | 0 | 1 | 16 | 0.06581507 | FALSE | 1 | FALSE | 1 | FALSE | 1 | FALSE | 1 | FALSE | 1 | FALSE | 1 |
| 6480 | 6480 | 9.21428571 | 2.99233715 | 0 | 1 | 0 | 1 | 0 | 1 | 2 | 0.3705495 | 0 | 1 | 26 | 0.16405547 | FALSE | 1 | FALSE | 1 | FALSE | 1 | FALSE | 1 | FALSE | 1 | FALSE | 1 |
| 6481 | 6481 | 9.65 | 2.41402154 | 1 | 0.40558521 | 0 | 1 | 1 | 0.51939841 | 1 | 0.60987146 | 1 | 0.28187291 | 16 | 0.80826361 | FALSE | 1 | FALSE | 1 | FALSE | 1 | FALSE | 1 | FALSE | 1 | FALSE | 1 |
| 6482 | 6482 | 8.07407407 | 2.03535284 | 2 | 0.15144274 | 2 | 0.21684246 | 0 | 1 | 1 | 0.71956339 | 0 | 1 | 22 | 0.76237877 | FALSE | 1 | FALSE | 1 | FALSE | 1 | FALSE | 1 | FALSE | 1 | FALSE | 1 |
| 6483 | 6483 | 8.90625 | 2.14125219 | 0 | 1 | 2 | 0.27730767 | 1 | 0.69069222 | 1 | 0.77852168 | 1 | 0.41154418 | 27 | 0.61671133 | FALSE | 1 | FALSE | 1 | FALSE | 1 | FALSE | 1 | FALSE | 1 | FALSE | 1 |
| 6484 | 6484 | 10.4375 | 1.6571342 | 2 | 0.06197875 | 0 | 1 | 2 | 0.11087977 | 0 | 1 | 0 | 1 | 12 | 0.90922186 | FALSE | 0.72532843 | FALSE | 1 | FALSE | 1 | FALSE | 1 | FALSE | 1 | FALSE | 1 |

| | | | | | | | | | | | | | | | | | | | | | | | | | | | |
|---|---|---|---|---|---|---|---|---|---|---|---|---|---|---|---|---|---|---|---|---|---|---|---|---|---|---|---|
| 6485 | 6485 | 3.88157895 | 2.57501715 | 2 | 0.58475401 | 5 | 0.09882268 | 4 | 0.29102231 | 4 | 0.46403589 | 0 | 1 | 61 | 0.87381726 | FALSE | 1 | FALSE | 1 | FALSE | 1 | FALSE | 1 | FALSE | 1 | FALSE | 1 |
| 6486 | 6486 | 10.5483871 | 1.64294601 | 3 | 0.04402911 | 1 | 0.63997468 | 0 | 1 | 1 | 0.76781284 | 1 | 0.4016888 | 25 | 0.80006983 | FALSE | 0.64293489 | FALSE | 1 | FALSE | 1 | FALSE | 1 | FALSE | 1 | FALSE | 1 |
| 6487 | 6487 | 8.33333333 | 3.0429031 | 0 | 1 | 2 | 0.21684246 | 1 | 0.62830986 | 1 | 0.71956339 | 0 | 1 | 23 | 0.58307573 | FALSE | 1 | FALSE | 1 | FALSE | 1 | FALSE | 1 | FALSE | 1 | FALSE | 1 |
| 6488 | 6488 | 9.90476191 | 1.57070691 | 0 | 1 | 0 | 1 | 0 | 1 | 0 | 1 | 0 | 1 | 21 | 0.02808151 | FALSE | 1 | FALSE | 1 | FALSE | 1 | FALSE | 1 | FALSE | 1 | FALSE | 1 |
| 6489 | 6489 | 8.70833333 | 1.30636795 | 0 | 1 | 2 | 0.18117183 | 1 | 0.58502114 | 3 | 0.09525947 | 0 | 1 | 18 | 0.93166998 | FALSE | 1 | FALSE | 1 | FALSE | 1 | FALSE | 1 | FALSE | 1 | FALSE | 1 |
| 6490 | 6490 | 8.42105263 | 3.61744006 | 1 | 0.38989951 | 0 | 1 | 2 | 0.14749808 | 0 | 1 | 0 | 1 | 16 | 0.65565778 | FALSE | 1 | FALSE | 1 | FALSE | 1 | FALSE | 1 | FALSE | 1 | FALSE | 1 |
| 6491 | 6491 | 10 | 2 | 0 | 1 | 0 | 1 | 0 | 1 | 1 | 0.45746571 | 0 | 1 | 12 | 0.374198 | FALSE | 1 | FALSE | 1 | FALSE | 1 | FALSE | 1 | FALSE | 1 | FALSE | 1 |
| 6492 | 6492 | 10.59375 | 2.72556617 | 2 | 0.19750261 | 1 | 0.65167362 | 1 | 0.69069222 | 1 | 0.77852168 | 0 | 1 | 27 | 0.61671133 | FALSE | 1 | FALSE | 1 | FALSE | 1 | FALSE | 1 | FALSE | 1 | FALSE | 1 |
| 6493 | 6493 | 7.11111111 | 2.2825154 | 0 | 1 | 1 | 0.25628163 | 0 | 1 | 0 | 1 | 0 | 1 | 8 | 0.5780332 | FALSE | 1 | FALSE | 1 | FALSE | 1 | FALSE | 1 | FALSE | 1 | FALSE | 1 |
| 6494 | 6494 | 9.23076923 | 2.8864097 | 1 | 0.28675951 | 0 | 1 | 0 | 1 | 0 | 1 | 0 | 1 | 12 | 0.374198 | FALSE | 1 | FALSE | 1 | FALSE | 1 | FALSE | 1 | FALSE | 1 | FALSE | 1 |
| 6495 | 6495 | 9.73684211 | 2.82548748 | 0 | 1 | 1 | 0.46503795 | 1 | 0.50143826 | 1 | 0.59104357 | 0 | 1 | 16 | 0.65565778 | FALSE | 1 | FALSE | 1 | FALSE | 1 | FALSE | 1 | FALSE | 1 | FALSE | 1 |
| 6496 | 6496 | 8.06666667 | 2.08059821 | 1 | 0.54198714 | 3 | 0.07145121 | 1 | 0.66709967 | 1 | 0.75658797 | 1 | 0.39166989 | 23 | 0.9158696 | FALSE | 1 | FALSE | 1 | FALSE | 1 | FALSE | 1 | FALSE | 1 | FALSE | 1 |
| 6497 | 6497 | 8.375 | 3.33307291 | 0 | 1 | 0 | 1 | 0 | 1 | 0 | 1 | 0 | 1 | 16 | 0.06581507 | FALSE | 1 | FALSE | 1 | FALSE | 1 | FALSE | 1 | FALSE | 1 | FALSE | 1 |
| 6498 | 6498 | 6.43283582 | 1.49350715 | 3 | 0.66332479 | 2 | 0.93132803 | 2 | 0.95415662 | 8 | 0.25978205 | 1 | 0.8896353 | 116 | 0.15906061 | FALSE | 1 | FALSE | 1 | FALSE | 1 | FALSE | 1 | FALSE | 1 | FALSE | 1 |
| 6499 | 6499 | 9.20833333 | 2.87197909 | 1 | 0.46441868 | 0 | 1 | 0 | 1 | 0 | 1 | 0 | 1 | 23 | 0.09196389 | FALSE | 1 | FALSE | 1 | FALSE | 1 | FALSE | 1 | FALSE | 1 | FALSE | 1 |
| 6500 | 6500 | 8.80952381 | 1.6795421 | 4 | 0.02019941 | 1 | 0.74131698 | 1 | 0.77786601 | 0 | 1 | 0 | 1 | 35 | 0.53616095 | FALSE | 0.44482142 | FALSE | 1 | FALSE | 1 | FALSE | 1 | FALSE | 1 | FALSE | 1 |
| 6501 | 6501 | 9 | 3.16227766 | 2 | 0.0256865 | 1 | 0.28036922 | 0 | 1 | 0 | 1 | 1 | 0.15247062 | 6 | 0.98831471 | FALSE | 0.4974559 | FALSE | 1 | FALSE | 1 | FALSE | 1 | FALSE | 1 | FALSE | 1 |
| 6502 | 6502 | 10 | 2.75680975 | 0 | 1 | 0 | 1 | 0 | 1 | 1 | 0.37517262 | 0 | 1 | 9 | 0.5215547 | FALSE | 1 | FALSE | 1 | FALSE | 1 | FALSE | 1 | FALSE | 1 | FALSE | 1 |
| 6503 | 6503 | 9.0952381 | 1.41100309 | 1 | 0.40558521 | 0 | 1 | 1 | 0.51939841 | 1 | 0.60987146 | 0 | 1 | 17 | 0.61750632 | FALSE | 1 | FALSE | 1 | FALSE | 1 | FALSE | 1 | FALSE | 1 | FALSE | 1 |
| 6504 | 6504 | 8.64102564 | 2.68411452 | 1 | 0.63789042 | 1 | 0.7236252 | 0 | 1 | 3 | 0.26522531 | 0 | 1 | 34 | 0.41570781 | FALSE | 1 | FALSE | 1 | FALSE | 1 | FALSE | 1 | FALSE | 1 | FALSE | 1 |
| 6505 | 6505 | 10.6 | 1.69705628 | 0 | 1 | 2 | 0.19297191 | 1 | 0.59998149 | 0 | 1 | 0 | 1 | 22 | 0.43642846 | FALSE | 1 | FALSE | 1 | FALSE | 1 | FALSE | 1 | FALSE | 1 | FALSE | 1 |
| 6506 | 6506 | 7.57894737 | 4.41666449 | 0 | 1 | 0 | 1 | 0 | 1 | 2 | 0.21610812 | 1 | 0.26986775 | 16 | 0.65565778 | FALSE | 1 | FALSE | 1 | FALSE | 1 | FALSE | 1 | FALSE | 1 | FALSE | 1 |
| 6507 | 6507 | 9.05555556 | 2.34454976 | 0 | 1 | 0 | 1 | 0 | 1 | 1 | 0.57131013 | 0 | 1 | 17 | 0.20329205 | FALSE | 1 | FALSE | 1 | FALSE | 1 | FALSE | 1 | FALSE | 1 | FALSE | 1 |
| 6508 | 6508 | 12.5263158 | 2.68266209 | 0 | 1 | 2 | 0.12434464 | 0 | 1 | 0 | 1 | 0 | 1 | 17 | 0.41141543 | FALSE | 1 | FALSE | 1 | FALSE | 1 | FALSE | 1 | FALSE | 1 | FALSE | 1 |
| 6509 | 6509 | 11.1153846 | 1.57692308 | 0 | 1 | 1 | 0.56109832 | 0 | 1 | 0 | 1 | 0 | 1 | 24 | 0.08019798 | FALSE | 1 | FALSE | 1 | FALSE | 1 | FALSE | 1 | FALSE | 1 | FALSE | 1 |
| 6510 | 6510 | 11 | 1.88107989 | 1 | 0.28675951 | 0 | 1 | 2 | 0.07725803 | 1 | 0.45746571 | 0 | 1 | 9 | 0.96018168 | FALSE | 1 | FALSE | 1 | FALSE | 1 | FALSE | 1 | FALSE | 1 | FALSE | 1 |
| 6511 | 6511 | 11.4210526 | 2.82499724 | 1 | 0.62830697 | 4 | 0.0332765 | 1 | 0.75193685 | 1 | 0.83318955 | 0 | 1 | 31 | 0.76640548 | FALSE | 1 | FALSE | 0.92721448 | FALSE | 1 | FALSE | 1 | FALSE | 1 | FALSE | 1 |
| 6512 | 6512 | 9.38888889 | 1.91887956 | 0 | 1 | 0 | 1 | 0 | 1 | 1 | 0.57131013 | 0 | 1 | 17 | 0.20329205 | FALSE | 1 | FALSE | 1 | FALSE | 1 | FALSE | 1 | FALSE | 1 | FALSE | 1 |
| 6513 | 6513 | 7.2 | 2.63818119 | 1 | 0.40558521 | 0 | 1 | 0 | 1 | 0 | 1 | 0 | 1 | 19 | 0.15700255 | FALSE | 1 | FALSE | 1 | FALSE | 1 | FALSE | 1 | FALSE | 1 | FALSE | 1 |
| 6514 | 6514 | 9.18181818 | 3.58831897 | 3 | 0.01568237 | 2 | 0.14655134 | 1 | 0.53671415 | 1 | 0.62783522 | 1 | 0.29368245 | 13 | 0.9973503 | FALSE | 0.38695764 | FALSE | 1 | FALSE | 1 | FALSE | 1 | FALSE | 1 | FALSE | 1 |
| 6515 | 6515 | 9.28571429 | 1.03015751 | 0 | 1 | 1 | 0.20567256 | 0 | 1 | 1 | 0.28044368 | 0 | 1 | 5 | 0.91847773 | FALSE | 1 | FALSE | 1 | FALSE | 1 | FALSE | 1 | FALSE | 1 | FALSE | 1 |
| 6516 | 6516 | 9.65 | 2.05608852 | 2 | 0.09187136 | 1 | 0.48239024 | 1 | 0.51939841 | 1 | 0.60987146 | 1 | 0.28187291 | 14 | 0.97325593 | FALSE | 0.83248965 | FALSE | 1 | FALSE | 1 | FALSE | 1 | FALSE | 1 | FALSE | 1 |
| 6517 | 6517 | 9.6875 | 1.97543508 | 0 | 1 | 2 | 0.25306331 | 1 | 0.66709967 | 1 | 0.75658797 | 0 | 1 | 26 | 0.48609328 | FALSE | 1 | FALSE | 1 | FALSE | 1 | FALSE | 1 | FALSE | 1 | FALSE | 1 |
| 6518 | 6518 | 8 | 2.82842713 | 0 | 1 | 1 | 0.44710673 | 1 | 0.48280983 | 1 | 0.57131013 | 0 | 1 | 15 | 0.69372602 | FALSE | 1 | FALSE | 1 | FALSE | 1 | FALSE | 1 | FALSE | 1 | FALSE | 1 |
| 6519 | 6519 | 11.7647059 | 3.1159818 | 0 | 1 | 0 | 1 | 1 | 0.46348836 | 1 | 0.55062772 | 0 | 1 | 15 | 0.49060435 | FALSE | 1 | FALSE | 1 | FALSE | 1 | FALSE | 1 | FALSE | 1 | FALSE | 1 |
| 6520 | 6520 | 8.05555556 | 2.69716358 | 1 | 0.37380237 | 2 | 0.1135776 | 1 | 0.48280983 | 0 | 1 | 0 | 1 | 14 | 0.86287418 | FALSE | 1 | FALSE | 1 | FALSE | 1 | FALSE | 1 | FALSE | 1 | FALSE | 1 |
| 6521 | 6521 | 9.4137931 | 3.37860654 | 3 | 0.03713651 | 0 | 1 | 0 | 1 | 1 | 0.74482229 | 1 | 0.38148475 | 24 | 0.70563334 | FALSE | 0.58392 | FALSE | 1 | FALSE | 1 | FALSE | 1 | FALSE | 1 | FALSE | 1 |
| 6522 | 6522 | 8.07692308 | 2.81374479 | 0 | 1 | 0 | 1 | 2 | 0.07725803 | 0 | 1 | 0 | 1 | 11 | 0.6683056 | FALSE | 1 | FALSE | 1 | FALSE | 1 | FALSE | 1 | FALSE | 1 | FALSE | 1 |
| 6523 | 6523 | 8.27272727 | 3.07751107 | 1 | 0.24866461 | 0 | 1 | 0 | 1 | 0 | 1 | 1 | 0.16638736 | 9 | 0.75985366 | FALSE | 1 | FALSE | 1 | FALSE | 1 | FALSE | 1 | FALSE | 1 | FALSE | 1 |
| 6524 | 6524 | 8.82352941 | 2.38217138 | 0 | 1 | 0 | 1 | 0 | 1 | 0 | 1 | 2 | 0.03086174 | 15 | 0.49060435 | FALSE | 1 | FALSE | 1 | FALSE | 1 | FALSE | 1 | FALSE | 0.82559785 | FALSE | 1 |
| 6525 | 6525 | 5.00729927 | 2.53586349 | 3 | 0.68788288 | 8 | 0.07559555 | 3 | 0.87583945 | 4 | 0.88124375 | 1 | 0.89856355 | 118 | 0.33341624 | FALSE | 1 | FALSE | 1 | FALSE | 1 | FALSE | 1 | FALSE | 1 | FALSE | 1 |
| 6526 | 6526 | 9.57142857 | 3.01695887 | 0 | 1 | 0 | 1 | 1 | 0.40110418 | 0 | 1 | 0 | 1 | 13 | 0.33283729 | FALSE | 1 | FALSE | 1 | FALSE | 1 | FALSE | 1 | FALSE | 1 | FALSE | 1 |
| 6527 | 6527 | 6.32075472 | 2.52738649 | 3 | 0.51429857 | 8 | 0.0210565 | 4 | 0.53239439 | 4 | 0.72436114 | 1 | 0.82904315 | 86 | 0.85563324 | FALSE | 1 | FALSE | 0.80605082 | FALSE | 1 | FALSE | 1 | FALSE | 1 | FALSE | 1 |
| 6528 | 6528 | 8.61538462 | 3.00098603 | 0 | 1 | 5 | 3.52E-05 | 0 | 1 | 1 | 0.45746571 | 0 | 1 | 7 | 0.99839694 | FALSE | 1 | FALSE | 0.07208458 | FALSE | 1 | FALSE | 1 | FALSE | 1 | FALSE | 1 |
| 6529 | 6529 | 8.23076923 | 2.15384615 | 2 | 0.25455533 | 2 | 0.3494045 | 0 | 1 | 1 | 0.83318955 | 0 | 1 | 33 | 0.44261631 | FALSE | 1 | FALSE | 1 | FALSE | 1 | FALSE | 1 | FALSE | 1 | FALSE | 1 |
| 6530 | 6530 | 6.44444444 | 2.77332977 | 0 | 1 | 2 | 0.1135776 | 0 | 1 | 0 | 1 | 2 | 0.03435384 | 14 | 0.86287418 | FALSE | 1 | FALSE | 1 | FALSE | 1 | FALSE | 1 | FALSE | 0.86063198 | FALSE | 1 |
| 6531 | 6531 | 6.61904762 | 2.24618598 | 0 | 1 | 0 | 1 | 2 | 0.43640583 | 0 | 1 | 0 | 1 | 39 | 0.03430384 | FALSE | 1 | FALSE | 1 | FALSE | 1 | FALSE | 1 | FALSE | 1 | FALSE | 1 |
| 6532 | 6532 | 10.3589744 | 1.80437047 | 2 | 0.25455533 | 2 | 0.3494045 | 3 | 0.15483587 | 2 | 0.52640931 | 0 | 1 | 29 | 0.93814695 | FALSE | 1 | FALSE | 1 | FALSE | 1 | FALSE | 1 | FALSE | 1 | FALSE | 1 |
| 6533 | 6533 | 11.4166667 | 2.09993386 | 1 | 0.46441868 | 1 | 0.54637326 | 2 | 0.21260257 | 0 | 1 | 0 | 1 | 20 | 0.68270729 | FALSE | 1 | FALSE | 1 | FALSE | 1 | FALSE | 1 | FALSE | 1 | FALSE | 1 |
| 6534 | 6534 | 7.44444444 | 3.49161253 | 1 | 0.60837712 | 0 | 1 | 2 | 0.37289966 | 3 | 0.22784656 | 1 | 0.44938266 | 29 | 0.81028289 | FALSE | 1 | FALSE | 1 | FALSE | 1 | FALSE | 1 | FALSE | 1 | FALSE | 1 |
| 6535 | 6535 | 7.01818182 | 0.67395465 | 8 | 6.91E-05 | 8 | 0.00034555 | 1 | 0.8673918 | 6 | 0.03908871 | 0 | 1 | 32 | 0.99999935 | TRUE | 0.01342173 | FALSE | 0.15723145 | FALSE | 1 | FALSE | 1 | FALSE | 1 | FALSE | 1 |
| 6536 | 6536 | 2.71691177 | 1.7658965 | 13 | 0.62538813 | 14 | 0.83557054 | 24 | 0.15214849 | 23 | 0.66680818 | 7 | 0.78674878 | 455 | 0.39544623 | FALSE | 1 | FALSE | 1 | FALSE | 1 | FALSE | 1 | FALSE | 1 | FALSE | 1 |
| 6537 | 6537 | 10.4090909 | 2.8708423 | 2 | 0.10809809 | 0 | 1 | 1 | 0.55340851 | 2 | 0.26791271 | 0 | 1 | 17 | 0.8840357 | FALSE | 0.88820416 | FALSE | 1 | FALSE | 1 | FALSE | 1 | FALSE | 1 | FALSE | 1 |
| 6538 | 6538 | 10.1818182 | 3.03968245 | 0 | 1 | 1 | 0.30368014 | 0 | 1 | 0 | 1 | 1 | 0.16638736 | 9 | 0.75985366 | FALSE | 1 | FALSE | 1 | FALSE | 1 | FALSE | 1 | FALSE | 1 | FALSE | 1 |
| 6539 | 6539 | 5.89189189 | 3.28620198 | 1 | 0.6184714 | 1 | 0.70472939 | 2 | 0.38585089 | 3 | 0.2402032 | 1 | 0.45845922 | 29 | 0.88820978 | FALSE | 1 | FALSE | 1 | FALSE | 1 | FALSE | 1 | FALSE | 1 | FALSE | 1 |
| 6540 | 6540 | 9.23529412 | 2.7338706 | 2 | 0.06909038 | 2 | 0.10307112 | 2 | 0.12280513 | 0 | 1 | 1 | 0.24525764 | 10 | 0.99775952 | FALSE | 0.74276693 | FALSE | 1 | FALSE | 1 | FALSE | 1 | FALSE | 1 | FALSE | 1 |
| 6541 | 6541 | 12.0666667 | 3.88958724 | 0 | 1 | 1 | 0.38964394 | 1 | 0.42266264 | 0 | 1 | 0 | 1 | 13 | 0.57712805 | FALSE | 1 | FALSE | 1 | FALSE | 1 | FALSE | 1 | FALSE | 1 | FALSE | 1 |
| 6542 | 6542 | 9.47916667 | 2.09154631 | 1 | 0.71380452 | 2 | 0.46321244 | 0 | 1 | 1 | 0.89605799 | 1 | 0.54902428 | 43 | 0.21763056 | FALSE | 1 | FALSE | 1 | FALSE | 1 | FALSE | 1 | FALSE | 1 | FALSE | 1 |
| 6543 | 6543 | 11.0952381 | 2.02142717 | 0 | 1 | 0 | 1 | 2 | 0.38585089 | 2 | 0.51200191 | 0 | 1 | 33 | 0.2931558 | FALSE | 1 | FALSE | 1 | FALSE | 1 | FALSE | 1 | FALSE | 1 | FALSE | 1 |
| 6544 | 6544 | 8.94444444 | 1.39332625 | 0 | 1 | 0 | 1 | 0 | 1 | 0 | 1 | 0 | 1 | 54 | 9.99E-05 | FALSE | 1 | FALSE | 1 | FALSE | 1 | FALSE | 1 | FALSE | 1 | FALSE | 0.59879545 |
| 6545 | 6545 | 4.48387097 | 3.23908557 | 4 | 0.37020407 | 2 | 0.90510937 | 5 | 0.43318786 | 8 | 0.18549246 | 0 | 1 | 101 | 0.58648198 | FALSE | 1 | FALSE | 1 | FALSE | 1 | FALSE | 1 | FALSE | 1 | FALSE | 1 |
| 6546 | 6546 | 8.32258065 | 2.61986329 | 0 | 1 | 1 | 0.63997468 | 1 | 0.67911179 | 2 | 0.41984187 | 0 | 1 | 27 | 0.45522639 | FALSE | 1 | FALSE | 1 | FALSE | 1 | FALSE | 1 | FALSE | 1 | FALSE | 1 |
| 6547 | 6547 | 6.3 | 2.1 | 1 | 0.22886536 | 1 | 0.28036922 | 1 | 0.30655256 | 0 | 1 | 0 | 1 | 7 | 0.94326215 | FALSE | 1 | FALSE | 1 | FALSE | 1 | FALSE | 1 | FALSE | 1 | FALSE | 1 |
| 6548 | 6548 | 7.64705882 | 3.02812651 | 0 | 1 | 0 | 1 | 0 | 1 | 1 | 0.55062772 | 1 | 0.24525764 | 15 | 0.49060435 | FALSE | 1 | FALSE | 1 | FALSE | 1 | FALSE | 1 | FALSE | 1 | FALSE | 1 |
| 6549 | 6549 | 6.3 | 2.25314595 | 1 | 0.54198714 | 1 | 0.6278847 | 0 | 1 | 1 | 0.75658797 | 0 | 1 | 27 | 0.28862208 | FALSE | 1 | FALSE | 1 | FALSE | 1 | FALSE | 1 | FALSE | 1 | FALSE | 1 |
| 6550 | 6550 | 7.83783784 | 2.59431305 | 0 | 1 | 1 | 0.70472939 | 2 | 0.38585089 | 1 | 0.82511639 | 0 | 1 | 33 | 0.2931558 | FALSE | 1 | FALSE | 1 | FALSE | 1 | FALSE | 1 | FALSE | 1 | FALSE | 1 |
| 6551 | 6551 | 8.46666667 | 2.67996683 | 0 | 1 | 0 | 1 | 2 | 0.09928574 | 0 | 1 | 0 | 1 | 13 | 0.57712805 | FALSE | 1 | FALSE | 1 | FALSE | 1 | FALSE | 1 | FALSE | 1 | FALSE | 1 |
| 6552 | 6552 | 6.64772727 | 1.60991087 | 1 | 0.89980571 | 3 | 0.5460764 | 4 | 0.38964212 | 5 | 0.37910615 | 0 | 1 | 75 | 0.48424212 | FALSE | 1 | FALSE | 1 | FALSE | 1 | FALSE | 1 | FALSE | 1 | FALSE | 1 |
| 6553 | 6553 | 10.0454546 | 1.96508785 | 1 | 0.4357638 | 0 | 1 | 1 | 0.55340851 | 1 | 0.64497439 | 2 | 0.04972408 | 17 | 0.8840357 | FALSE | 1 | FALSE | 1 | FALSE | 1 | FALSE | 1 | FALSE | 0.98629461 | FALSE | 1 |
| 6554 | 6554 | 10.25 | 2.086664 | 1 | 0.26795838 | 0 | 1 | 0 | 1 | 2 | 0.1023796 | 1 | 0.18007762 | 8 | 0.97197833 | FALSE | 1 | FALSE | 1 | FALSE | 1 | FALSE | 1 | FALSE | 1 | FALSE | 1 |
| 6555 | 6555 | 7.91666667 | 2.36143789 | 0 | 1 | 0 | 1 | 2 | 0.06691409 | 1 | 0.43131129 | 0 | 1 | 9 | 0.89630573 | FALSE | 1 | FALSE | 1 | FALSE | 1 | FALSE | 1 | FALSE | 1 | FALSE | 1 |
| 6556 | 6556 | 8.31428571 | 1.78519993 | 0 | 1 | 2 | 0.31355738 | 1 | 0.72299506 | 2 | 0.48233989 | 0 | 1 | 30 | 0.52765882 | FALSE | 1 | FALSE | 1 | FALSE | 1 | FALSE | 1 | FALSE | 1 | FALSE | 1 |
| 6557 | 6557 | 7.96296296 | 3.93918098 | 1 | 0.50471032 | 0 | 1 | 1 | 0.62830986 | 0 | 1 | 0 | 1 | 25 | 0.1832206 | FALSE | 1 | FALSE | 1 | FALSE | 1 | FALSE | 1 | FALSE | 1 | FALSE | 1 |
| 6558 | 6558 | 2.53424658 | 3.97279168 | 0 | 1 | 0 | 1 | 1 | 0.93179654 | 2 | 0.8554296 | 3 | 0.11741889 | 67 | 0.04816695 | FALSE | 1 | FALSE | 1 | FALSE | 1 | FALSE | 1 | FALSE | 1 | FALSE | 1 |
| 6559 | 6559 | 8.84 | 3.04210454 | 2 | 0.13369621 | 0 | 1 | 1 | 0.59998149 | 1 | 0.69181315 | 0 | 1 | 21 | 0.64957167 | FALSE | 0.97964162 | FALSE | 1 | FALSE | 1 | FALSE | 1 | FALSE | 1 | FALSE | 1 |
| 6560 | 6560 | 9.35714286 | 2.15827785 | 0 | 1 | 0 | 1 | 0 | 1 | 0 | 1 | 0 | 1 | 14 | 0.09251304 | FALSE | 1 | FALSE | 1 | FALSE | 1 | FALSE | 1 | FALSE | 1 | FALSE | 1 |
| 6561 | 6561 | 8.71428571 | 4.00764575 | 1 | 0.3050805 | 0 | 1 | 0 | 1 | 0 | 1 | 0 | 1 | 13 | 0.33283729 | FALSE | 1 | FALSE | 1 | FALSE | 1 | FALSE | 1 | FALSE | 1 | FALSE | 1 |
| 6562 | 6562 | 7.5 | 2.59807621 | 2 | 0.09187136 | 0 | 1 | 2 | 0.16019548 | 3 | 0.0612581 | 1 | 0.28187291 | 12 | 0.99824567 | FALSE | 0.83248965 | FALSE | 1 | FALSE | 1 | FALSE | 1 | FALSE | 1 | FALSE | 1 |
| 6563 | 6563 | 4.94805195 | 2.54004859 | 3 | 0.76136663 | 3 | 0.88077074 | 4 | 0.81029329 | 9 | 0.27559983 | 3 | 0.46520079 | 132 | 0.37169225 | FALSE | 1 | FALSE | 1 | FALSE | 1 | FALSE | 1 | FALSE | 1 | FALSE | 1 |
| 6564 | 6564 | 8.74074074 | 3.40579672 | 1 | 0.50471032 | 2 | 0.21684246 | 2 | 0.25286935 | 2 | 0.35374209 | 0 | 1 | 20 | 0.95133411 | FALSE | 1 | FALSE | 1 | FALSE | 1 | FALSE | 1 | FALSE | 1 | FALSE | 1 |
| 6565 | 6565 | 8.42105263 | 4.76075834 | 0 | 1 | 0 | 1 | 0 | 1 | 0 | 1 | 0 | 1 | 19 | 0.03948376 | FALSE | 1 | FALSE | 1 | FALSE | 1 | FALSE | 1 | FALSE | 1 | FALSE | 1 |
| 6566 | 6566 | 9.625 | 2.57087048 | 0 | 1 | 0 | 1 | 0 | 1 | 1 | 0.52895088 | 0 | 1 | 15 | 0.26127859 | FALSE | 1 | FALSE | 1 | FALSE | 1 | FALSE | 1 | FALSE | 1 | FALSE | 1 |
| 6567 | 6567 | 9.39130435 | 1.63492136 | 1 | 0.45027683 | 2 | 0.16948764 | 0 | 1 | 0 | 1 | 0 | 1 | 20 | 0.50589402 | FALSE | 1 | FALSE | 1 | FALSE | 1 | FALSE | 1 | FALSE | 1 | FALSE | 1 |
| 6568 | 6568 | 10.1481482 | 1.37984711 | 0 | 1 | 3 | 0.05519089 | 3 | 0.07106362 | 1 | 0.71956339 | 1 | 0.36060486 | 19 | 0.98241223 | FALSE | 1 | FALSE | 1 | FALSE | 1 | FALSE | 1 | FALSE | 1 | FALSE | 1 |
| 6569 | 6569 | 8.3125 | 2.7999721 | 0 | 1 | 0 | 1 | 1 | 0.44344817 | 0 | 1 | 0 | 1 | 15 | 0.26127859 | FALSE | 1 | FALSE | 1 | FALSE | 1 | FALSE | 1 | FALSE | 1 | FALSE | 1 |
| 6570 | 6570 | 7.73333333 | 1.73076733 | 1 | 0.32293355 | 0 | 1 | 3 | 0.01512258 | 1 | 0.50623194 | 0 | 1 | 10 | 0.97980046 | FALSE | 1 | FALSE | 1 | FALSE | 1 | FALSE | 1 | FALSE | 1 | FALSE | 1 |
| 6571 | 6571 | 5.35087719 | 5.54563596 | 3 | 0.17275086 | 2 | 0.54544984 | 6 | 0.01460362 | 2 | 0.73544289 | 0 | 1 | 43 | 0.95447642 | FALSE | 1 | FALSE | 1 | FALSE | 1 | FALSE | 1 | FALSE | 1 | FALSE | 1 |
| 6572 | 6572 | 6.859375 | 1.99113023 | 4 | 0.08106144 | 2 | 0.61853748 | 3 | 0.40547364 | 2 | 0.79987679 | 2 | 0.28258242 | 51 | 0.88464035 | FALSE | 0.79236883 | FALSE | 1 | FALSE | 1 | FALSE | 1 | FALSE | 1 | FALSE | 1 |
| 6573 | 6573 | 10.7307692 | 3.84865303 | 1 | 0.49162634 | 0 | 1 | 1 | 0.61440466 | 0 | 1 | 2 | 0.0670823 | 22 | 0.61628 | FALSE | 1 | FALSE | 1 | FALSE | 1 | FALSE | 1 | FALSE | 1 | FALSE | 1 |
| 6574 | 6574 | 6.83516484 | 2.11360707 | 0 | 1 | 1 | 0.95083647 | 3 | 0.64074878 | 2 | 0.92663163 | 2 | 0.44196567 | 83 | 0.04135751 | FALSE | 1 | FALSE | 1 | FALSE | 1 | FALSE | 1 | FALSE | 1 | FALSE | 1 |
| 6575 | 6575 | 5.22727273 | 3.84272415 | 1 | 0.68224519 | 1 | 0.76576289 | 2 | 0.47281647 | 4 | 0.14160176 | 1 | 0.51797501 | 35 | 0.8620981 | FALSE | 1 | FALSE | 1 | FALSE | 1 | FALSE | 1 | FALSE | 1 | FALSE | 1 |
| 6576 | 6576 | 8.92105263 | 2.39935933 | 2 | 0.25455533 | 2 | 0.3494045 | 1 | 0.75193685 | 1 | 0.83318955 | 0 | 1 | 32 | 0.61800916 | FALSE | 1 | FALSE | 1 | FALSE | 1 | FALSE | 1 | FALSE | 1 | FALSE | 1 |
| 6577 | 6577 | 11.125 | 2.75850956 | 0 | 1 | 1 | 0.54637326 | 2 | 0.21260257 | 0 | 1 | 0 | 1 | 21 | 0.47053137 | FALSE | 1 | FALSE | 1 | FALSE | 1 | FALSE | 1 | FALSE | 1 | FALSE | 1 |
| 6578 | 6578 | 10.3571429 | 1.87491496 | 3 | 0.0048942 | 0 | 1 | 0 | 1 | 0 | 1 | 1 | 0.20679328 | 10 | 0.94579623 | FALSE | 0.19848171 | FALSE | 1 | FALSE | 1 | FALSE | 1 | FALSE | 1 | FALSE | 1 |

| | | | | | | | | | | | | | | | | | | | | | | | | | | | |
|---|---|---|---|---|---|---|---|---|---|---|---|---|---|---|---|---|---|---|---|---|---|---|---|---|---|---|---|
| 6579 | 6579 | 10.3125 | 2.08322916 | 0 | 1 | 0 | 1 | 0 | 1 | 1 | 0.52895088 | 0 | 1 | 15 | 0.26127859 | FALSE | 1 | FALSE | 1 | FALSE | 1 | FALSE | 1 | FALSE | 1 | FALSE | 1 |
| 6580 | 6580 | 8.4 | 2.37486842 | 0 | 1 | 0 | 1 | 0 | 1 | 1 | 0.37517262 | 0 | 1 | 9 | 0.5215547 | FALSE | 1 | FALSE | 1 | FALSE | 1 | FALSE | 1 | FALSE | 1 | FALSE | 1 |
| 6581 | 6581 | 10.2857143 | 3.21666579 | 0 | 1 | 0 | 1 | 1 | 0.64171563 | 1 | 0.73248983 | 0 | 1 | 26 | 0.16405547 | FALSE | 1 | FALSE | 1 | FALSE | 1 | FALSE | 1 | FALSE | 1 | FALSE | 1 |
| 6582 | 6582 | 11.4166667 | 2.36143789 | 0 | 1 | 0 | 1 | 0 | 1 | 1 | 0.43131129 | 0 | 1 | 11 | 0.41944532 | FALSE | 1 | FALSE | 1 | FALSE | 1 | FALSE | 1 | FALSE | 1 | FALSE | 1 |
| 6583 | 6583 | 9.52631579 | 2.39227751 | 1 | 0.38989951 | 2 | 0.12434464 | 1 | 0.50143826 | 1 | 0.59104357 | 0 | 1 | 14 | 0.93677253 | FALSE | 1 | FALSE | 1 | FALSE | 1 | FALSE | 1 | FALSE | 1 | FALSE | 1 |
| 6584 | 6584 | 7.36363636 | 1.20146834 | 0 | 1 | 0 | 1 | 2 | 0.33343871 | 3 | 0.19161876 | 0 | 1 | 28 | 0.58681477 | FALSE | 1 | FALSE | 1 | FALSE | 1 | FALSE | 1 | FALSE | 1 | FALSE | 1 |
| 6585 | 6585 | 10.1 | 2.65015723 | 2 | 0.17884234 | 1 | 0.6278847 | 2 | 0.29330414 | 0 | 1 | 0 | 1 | 25 | 0.67630471 | FALSE | 1 | FALSE | 1 | FALSE | 1 | FALSE | 1 | FALSE | 1 | FALSE | 1 |
| 6586 | 6586 | 9.07575758 | 1.58888761 | 2 | 0.50812508 | 0 | 1 | 3 | 0.4248691 | 3 | 0.58996791 | 0 | 1 | 58 | 0.27702369 | FALSE | 1 | FALSE | 1 | FALSE | 1 | FALSE | 1 | FALSE | 1 | FALSE | 1 |
| 6587 | 6587 | 10.5769231 | 3.74343607 | 0 | 1 | 1 | 0.57534752 | 3 | 0.06482014 | 2 | 0.33678042 | 0 | 1 | 20 | 0.90142689 | FALSE | 1 | FALSE | 1 | FALSE | 1 | FALSE | 1 | FALSE | 1 | FALSE | 1 |
| 6588 | 6588 | 5.2 | 4.06939799 | 0 | 1 | 0 | 1 | 0 | 1 | 1 | 0.67692816 | 0 | 1 | 23 | 0.09196389 | FALSE | 1 | FALSE | 1 | FALSE | 1 | FALSE | 1 | FALSE | 1 | FALSE | 1 |
| 6589 | 6589 | 10.0769231 | 2.20005379 | 0 | 1 | 0 | 1 | 0 | 1 | 0 | 1 | 1 | 0.19354507 | 12 | 0.374198 | FALSE | 1 | FALSE | 1 | FALSE | 1 | FALSE | 1 | FALSE | 1 | FALSE | 1 |
| 6590 | 6590 | 10.9230769 | 2.64463284 | 0 | 1 | 1 | 0.3480709 | 0 | 1 | 1 | 0.45746571 | 0 | 1 | 11 | 0.6683056 | FALSE | 1 | FALSE | 1 | FALSE | 1 | FALSE | 1 | FALSE | 1 | FALSE | 1 |
| 6591 | 6591 | 7.96296296 | 3.5640331 | 0 | 1 | 1 | 0.58913618 | 0 | 1 | 2 | 0.35374209 | 1 | 0.36060486 | 23 | 0.58307573 | FALSE | 1 | FALSE | 1 | FALSE | 1 | FALSE | 1 | FALSE | 1 | FALSE | 1 |
| 6592 | 6592 | 5.3151751 | 1.38848911 | 4 | 0.89829013 | 6 | 0.84002587 | 8 | 0.70369949 | 12 | 0.50795244 | 1 | 0.98640675 | 224 | 0.0681155 | FALSE | 1 | FALSE | 1 | FALSE | 1 | FALSE | 1 | FALSE | 1 | FALSE | 1 |
| 6593 | 6593 | 9.17391304 | 1.71063614 | 0 | 1 | 0 | 1 | 0 | 1 | 3 | 0.08614834 | 0 | 1 | 20 | 0.50589402 | FALSE | 1 | FALSE | 1 | FALSE | 1 | FALSE | 1 | FALSE | 1 | FALSE | 1 |
| 6594 | 6594 | 11.56 | 2.08 | 1 | 0.47819878 | 0 | 1 | 1 | 0.59998149 | 1 | 0.69181315 | 0 | 1 | 22 | 0.43642846 | FALSE | 1 | FALSE | 1 | FALSE | 1 | FALSE | 1 | FALSE | 1 | FALSE | 1 |
| 6595 | 6595 | 8.53846154 | 2.6199056 | 0 | 1 | 1 | 0.3480709 | 0 | 1 | 2 | 0.1174583 | 0 | 1 | 10 | 0.86791642 | FALSE | 1 | FALSE | 1 | FALSE | 1 | FALSE | 1 | FALSE | 1 | FALSE | 1 |
| 6596 | 6596 | 9.64285714 | 2.24479746 | 2 | 0.29286138 | 0 | 1 | 2 | 0.44869594 | 2 | 0.5811115 | 1 | 0.50166523 | 35 | 0.67024784 | FALSE | 1 | FALSE | 1 | FALSE | 1 | FALSE | 1 | FALSE | 1 | FALSE | 1 |
| 6597 | 6597 | 8.2 | 2.18174242 | 0 | 1 | 1 | 0.48239024 | 0 | 1 | 0 | 1 | 0 | 1 | 19 | 0.15700255 | FALSE | 1 | FALSE | 1 | FALSE | 1 | FALSE | 1 | FALSE | 1 | FALSE | 1 |
| 6598 | 6598 | 8.78571429 | 2.20967261 | 0 | 1 | 0 | 1 | 2 | 0.07725803 | 0 | 1 | 0 | 1 | 11 | 0.6683056 | FALSE | 1 | FALSE | 1 | FALSE | 1 | FALSE | 1 | FALSE | 1 | FALSE | 1 |
| 6599 | 6599 | 7.6875 | 2.51790464 | 0 | 1 | 0 | 1 | 1 | 0.44344817 | 0 | 1 | 0 | 1 | 15 | 0.26127859 | FALSE | 1 | FALSE | 1 | FALSE | 1 | FALSE | 1 | FALSE | 1 | FALSE | 1 |
| 6600 | 6600 | 9.14285714 | 3.79580864 | 1 | 0.16630142 | 1 | 0.20567256 | 1 | 0.22601089 | 0 | 1 | 0 | 1 | 4 | 0.98605104 | FALSE | 1 | FALSE | 1 | FALSE | 1 | FALSE | 1 | FALSE | 1 | FALSE | 1 |
| 6601 | 6601 | 8.1025641 | 1.98448949 | 0 | 1 | 1 | 0.7236252 | 2 | 0.41139386 | 1 | 0.84089119 | 0 | 1 | 35 | 0.24915769 | FALSE | 1 | FALSE | 1 | FALSE | 1 | FALSE | 1 | FALSE | 1 | FALSE | 1 |
| 6602 | 6602 | 8.39473684 | 4.76024919 | 2 | 0.25455533 | 1 | 0.71433277 | 3 | 0.15483587 | 2 | 0.52640931 | 2 | 0.12811285 | 28 | 0.97314817 | FALSE | 1 | FALSE | 1 | FALSE | 1 | FALSE | 1 | FALSE | 1 | FALSE | 1 |
| 6603 | 6603 | 9.5 | 3.30907842 | 1 | 0.40558521 | 1 | 0.48239024 | 0 | 1 | 0 | 1 | 0 | 1 | 18 | 0.37504645 | FALSE | 1 | FALSE | 1 | FALSE | 1 | FALSE | 1 | FALSE | 1 | FALSE | 1 |
| 6604 | 6604 | 10.5454546 | 3.28532927 | 0 | 1 | 0 | 1 | 0 | 1 | 0 | 1 | 0 | 1 | 11 | 0.15414489 | FALSE | 1 | FALSE | 1 | FALSE | 1 | FALSE | 1 | FALSE | 1 | FALSE | 1 |
| 6605 | 6605 | 8.43137255 | 3.3969704 | 2 | 0.37762121 | 1 | 0.81422651 | 1 | 0.84631478 | 3 | 0.41729179 | 0 | 1 | 44 | 0.44552698 | FALSE | 1 | FALSE | 1 | FALSE | 1 | FALSE | 1 | FALSE | 1 | FALSE | 1 |
| 6606 | 6606 | 8.90909091 | 1.975051 | 1 | 0.24866461 | 0 | 1 | 0 | 1 | 2 | 0.08789566 | 0 | 1 | 8 | 0.92152993 | FALSE | 1 | FALSE | 1 | FALSE | 1 | FALSE | 1 | FALSE | 1 | FALSE | 1 |
| 6607 | 6607 | 2.4520548 | 2.98416412 | 3 | 0.28789392 | 1 | 0.9104874 | 3 | 0.49081265 | 3 | 0.65845889 | 2 | 0.33719417 | 61 | 0.65061952 | FALSE | 1 | FALSE | 1 | FALSE | 1 | FALSE | 1 | FALSE | 1 | FALSE | 1 |
| 6608 | 6608 | 9.5 | 2.1408721 | 1 | 0.14434519 | 0 | 1 | 0 | 1 | 0 | 1 | 0 | 1 | 5 | 0.76185568 | FALSE | 1 | FALSE | 1 | FALSE | 1 | FALSE | 1 | FALSE | 1 | FALSE | 1 |
| 6609 | 6609 | 11.3076923 | 1.9955572 | 2 | 0.14250978 | 0 | 1 | 1 | 0.61440466 | 0 | 1 | 0 | 1 | 23 | 0.40372818 | FALSE | 1 | FALSE | 1 | FALSE | 1 | FALSE | 1 | FALSE | 1 | FALSE | 1 |
| 6610 | 6610 | 13.5 | 1.60727513 | 0 | 1 | 1 | 0.179097 | 0 | 1 | 0 | 1 | 0 | 1 | 5 | 0.76185568 | FALSE | 1 | FALSE | 1 | FALSE | 1 | FALSE | 1 | FALSE | 1 | FALSE | 1 |
| 6611 | 6611 | 9.16666667 | 1.28019096 | 0 | 1 | 0 | 1 | 2 | 0.06691409 | 0 | 1 | 1 | 0.18007762 | 9 | 0.89630573 | FALSE | 1 | FALSE | 1 | FALSE | 1 | FALSE | 1 | FALSE | 1 | FALSE | 1 |
| 6612 | 6612 | 9 | 2.39791576 | 0 | 1 | 1 | 0.23139161 | 0 | 1 | 0 | 1 | 0 | 1 | 7 | 0.63752397 | FALSE | 1 | FALSE | 1 | FALSE | 1 | FALSE | 1 | FALSE | 1 | FALSE | 1 |
| 6613 | 6613 | 2.93548387 | 2.69928884 | 0 | 1 | 2 | 0.60114557 | 1 | 0.89758397 | 2 | 0.78520976 | 1 | 0.64288531 | 56 | 0.12847218 | FALSE | 1 | FALSE | 1 | FALSE | 1 | FALSE | 1 | FALSE | 1 | FALSE | 1 |
| 6614 | 6614 | 6.0212766 | 2.12538886 | 2 | 0.88169447 | 3 | 0.84058578 | 5 | 0.57614338 | 10 | 0.11294592 | 1 | 0.90518766 | 120 | 0.46116711 | FALSE | 1 | FALSE | 1 | FALSE | 1 | FALSE | 1 | FALSE | 1 | FALSE | 1 |
| 6615 | 6615 | 7.93269231 | 1.68856119 | 6 | 0.05029841 | 3 | 0.65968805 | 5 | 0.31828279 | 4 | 0.71036674 | 2 | 0.51184859 | 84 | 0.87500079 | FALSE | 0.66890356 | FALSE | 1 | FALSE | 1 | FALSE | 1 | FALSE | 1 | FALSE | 1 |
| 6616 | 6616 | 10.7894737 | 2.04112836 | 2 | 0.08405377 | 2 | 0.12434464 | 0 | 1 | 1 | 0.59104357 | 0 | 1 | 14 | 0.93677253 | FALSE | 0.79236883 | FALSE | 1 | FALSE | 1 | FALSE | 1 | FALSE | 1 | FALSE | 1 |
| 6617 | 6617 | 4.38728324 | 2.65453263 | 3 | 0.8076467 | 9 | 0.09134913 | 4 | 0.85688202 | 10 | 0.23712094 | 5 | 0.13876704 | 136 | 0.87728257 | FALSE | 1 | FALSE | 1 | FALSE | 1 | FALSE | 1 | FALSE | 1 | FALSE | 1 |
| 6618 | 6618 | 8.58139535 | 2.37459968 | 0 | 1 | 1 | 0.75788114 | 2 | 0.46083448 | 0 | 1 | 0 | 1 | 40 | 0.07890873 | FALSE | 1 | FALSE | 1 | FALSE | 1 | FALSE | 1 | FALSE | 1 | FALSE | 1 |
| 6619 | 6619 | 9.77777778 | 3.12793073 | 0 | 1 | 1 | 0.69480472 | 0 | 1 | 1 | 0.81665383 | 4 | 0.00269049 | 30 | 0.67248271 | FALSE | 1 | FALSE | 1 | FALSE | 1 | FALSE | 1 | FALSE | 0.29784427 | FALSE | 1 |
| 6620 | 6620 | 10.7142857 | 2.11891385 | 0 | 1 | 0 | 1 | 1 | 0.22601089 | 0 | 1 | 0 | 1 | 6 | 0.69920685 | FALSE | 1 | FALSE | 1 | FALSE | 1 | FALSE | 1 | FALSE | 1 | FALSE | 1 |
| 6621 | 6621 | 9.6875 | 2.1713691 | 2 | 0.06197875 | 0 | 1 | 0 | 1 | 0 | 1 | 1 | 0.23264612 | 13 | 0.76799161 | FALSE | 0.72532843 | FALSE | 1 | FALSE | 1 | FALSE | 1 | FALSE | 1 | FALSE | 1 |
| 6622 | 6622 | 9.42105263 | 2.47760048 | 0 | 1 | 0 | 1 | 1 | 0.50143826 | 1 | 0.59104357 | 0 | 1 | 17 | 0.41141543 | FALSE | 1 | FALSE | 1 | FALSE | 1 | FALSE | 1 | FALSE | 1 | FALSE | 1 |
| 6623 | 6623 | 5.4516129 | 3.00095372 | 2 | 0.78726777 | 3 | 0.70827861 | 6 | 0.21430103 | 4 | 0.76332434 | 1 | 0.84544095 | 96 | 0.40767798 | FALSE | 1 | FALSE | 1 | FALSE | 1 | FALSE | 1 | FALSE | 1 | FALSE | 1 |
| 6624 | 6624 | 9.4375 | 4.07689756 | 0 | 1 | 0 | 1 | 0 | 1 | 2 | 0.16544789 | 1 | 0.23264612 | 13 | 0.76799161 | FALSE | 1 | FALSE | 1 | FALSE | 1 | FALSE | 1 | FALSE | 1 | FALSE | 1 |
| 6625 | 6625 | 9.10526316 | 1.44712917 | 0 | 1 | 0 | 1 | 1 | 0.50143826 | 1 | 0.59104357 | 1 | 0.26986775 | 16 | 0.65565778 | FALSE | 1 | FALSE | 1 | FALSE | 1 | FALSE | 1 | FALSE | 1 | FALSE | 1 |
| 6626 | 6626 | 9.15384615 | 1.9551177 | 1 | 0.26795838 | 1 | 0.32623933 | 0 | 1 | 0 | 1 | 2 | 0.01579445 | 8 | 0.97197833 | FALSE | 1 | FALSE | 1 | FALSE | 1 | FALSE | 1 | FALSE | 0.66313193 | FALSE | 1 |
| 6627 | 6627 | 9.11764706 | 2.02578534 | 1 | 0.35728305 | 1 | 0.42857733 | 0 | 1 | 1 | 0.55062772 | 0 | 1 | 14 | 0.7313145 | FALSE | 1 | FALSE | 1 | FALSE | 1 | FALSE | 1 | FALSE | 1 | FALSE | 1 |
| 6628 | 6628 | 7.32352941 | 2.13817663 | 3 | 0.25295743 | 1 | 0.89430705 | 1 | 0.91794622 | 3 | 0.61040454 | 0 | 1 | 60 | 0.24411098 | FALSE | 1 | FALSE | 1 | FALSE | 1 | FALSE | 1 | FALSE | 1 | FALSE | 1 |
| 6629 | 6629 | 9.12 | 3.82956917 | 0 | 1 | 0 | 1 | 0 | 1 | 0 | 1 | 0 | 1 | 25 | 0.01419971 | FALSE | 1 | FALSE | 1 | FALSE | 1 | FALSE | 1 | FALSE | 1 | FALSE | 1 |
| 6630 | 6630 | 4.20289855 | 1.03671318 | 23 | 0.11040034 | 26 | 0.22614851 | 30 | 0.1505056 | 27 | 0.83583256 | 20 | 0.00758876 | 562 | 0.9811188 | FALSE | 0.90530487 | FALSE | 1 | FALSE | 1 | FALSE | 1 | FALSE | 0.49339011 | FALSE | 1 |
| 6631 | 6631 | 4.90476191 | 4.28624335 | 0 | 1 | 1 | 0.49918218 | 1 | 0.53671415 | 0 | 1 | 0 | 1 | 19 | 0.34094959 | FALSE | 1 | FALSE | 1 | FALSE | 1 | FALSE | 1 | FALSE | 1 | FALSE | 1 |
| 6632 | 6632 | 7.3 | 2.92061637 | 1 | 0.72839502 | 2 | 0.48458833 | 3 | 0.26704679 | 2 | 0.6762159 | 3 | 0.04820812 | 39 | 0.92022511 | FALSE | 1 | FALSE | 1 | FALSE | 1 | FALSE | 1 | FALSE | 0.98629461 | FALSE | 1 |
| 6633 | 6633 | 8.09090909 | 1.2398347 | 4 | 0.00212601 | 1 | 0.5154318 | 0 | 1 | 0 | 1 | 1 | 0.30529951 | 16 | 0.95553951 | FALSE | 0.12179217 | FALSE | 1 | FALSE | 1 | FALSE | 1 | FALSE | 1 | FALSE | 1 |
| 6634 | 6634 | 4.66176471 | 1.85987266 | 2 | 0.52416586 | 2 | 0.65156696 | 4 | 0.22739663 | 2 | 0.82657478 | 3 | 0.10016761 | 55 | 0.83409155 | FALSE | 1 | FALSE | 1 | FALSE | 1 | FALSE | 1 | FALSE | 1 | FALSE | 1 |
| 6635 | 6635 | 8.27272727 | 2.21937375 | 0 | 1 | 1 | 0.5154318 | 0 | 1 | 3 | 0.07743186 | 1 | 0.30529951 | 17 | 0.8840357 | FALSE | 1 | FALSE | 1 | FALSE | 1 | FALSE | 1 | FALSE | 1 | FALSE | 1 |
| 6636 | 6636 | 12.1904762 | 2.10710923 | 1 | 0.42086991 | 1 | 0.49918218 | 0 | 1 | 1 | 0.62783522 | 1 | 0.29368245 | 17 | 0.77849076 | FALSE | 1 | FALSE | 1 | FALSE | 1 | FALSE | 1 | FALSE | 1 | FALSE | 1 |
| 6637 | 6637 | 9.28571429 | 2.05038573 | 2 | 0.04858661 | 0 | 1 | 0 | 1 | 0 | 1 | 0 | 1 | 12 | 0.62236227 | FALSE | 0.649301 | FALSE | 1 | FALSE | 1 | FALSE | 1 | FALSE | 1 | FALSE | 1 |
| 6638 | 6638 | 9.27272727 | 2.63165724 | 0 | 1 | 1 | 0.30368014 | 0 | 1 | 0 | 1 | 0 | 1 | 10 | 0.46859937 | FALSE | 1 | FALSE | 1 | FALSE | 1 | FALSE | 1 | FALSE | 1 | FALSE | 1 |
| 6639 | 6639 | 8.02857143 | 2.03520044 | 1 | 0.59801736 | 1 | 0.68454804 | 3 | 0.12958859 | 0 | 1 | 1 | 0.44015537 | 29 | 0.69927719 | FALSE | 1 | FALSE | 1 | FALSE | 1 | FALSE | 1 | FALSE | 1 | FALSE | 1 |
| 6640 | 6640 | 8.26666667 | 4.09009644 | 0 | 1 | 1 | 0.38964394 | 0 | 1 | 0 | 1 | 1 | 0.2198258 | 13 | 0.57712805 | FALSE | 1 | FALSE | 1 | FALSE | 1 | FALSE | 1 | FALSE | 1 | FALSE | 1 |
| 6641 | 6641 | 11.2692308 | 2.59522776 | 3 | 0.0279333 | 2 | 0.20486827 | 1 | 0.61440466 | 0 | 1 | 1 | 0.3499045 | 19 | 0.96054244 | FALSE | 0.50964275 | FALSE | 1 | FALSE | 1 | FALSE | 1 | FALSE | 1 | FALSE | 1 |
| 6642 | 6642 | 9.5 | 1.91049732 | 0 | 1 | 0 | 1 | 0 | 1 | 1 | 0.37517262 | 0 | 1 | 9 | 0.5215547 | FALSE | 1 | FALSE | 1 | FALSE | 1 | FALSE | 1 | FALSE | 1 | FALSE | 1 |
| 6643 | 6643 | 9.55 | 2.06094639 | 1 | 0.40558521 | 1 | 0.48239024 | 1 | 0.51939841 | 2 | 0.23331368 | 1 | 0.28187291 | 14 | 0.97325593 | FALSE | 1 | FALSE | 1 | FALSE | 1 | FALSE | 1 | FALSE | 1 | FALSE | 1 |
| 6644 | 6644 | 7.125 | 0.92702481 | 0 | 1 | 0 | 1 | 0 | 1 | 1 | 0.52895088 | 0 | 1 | 15 | 0.26127859 | FALSE | 1 | FALSE | 1 | FALSE | 1 | FALSE | 1 | FALSE | 1 | FALSE | 1 |
| 6645 | 6645 | 6.4 | 3.08544972 | 2 | 0.35907852 | 0 | 1 | 1 | 0.83455705 | 6 | 0.02365366 | 0 | 1 | 40 | 0.77344956 | FALSE | 1 | FALSE | 1 | FALSE | 1 | FALSE | 1 | FALSE | 1 | FALSE | 1 |
| 6646 | 6646 | 5.29299363 | 1.21697729 | 2 | 0.91554624 | 7 | 0.24533117 | 7 | 0.33420068 | 8 | 0.43318605 | 4 | 0.25645514 | 129 | 0.81327746 | FALSE | 1 | FALSE | 1 | FALSE | 1 | FALSE | 1 | FALSE | 1 | FALSE | 1 |
| 6647 | 6647 | 7.23863636 | 0.89181019 | 2 | 0.66468311 | 2 | 0.78367607 | 1 | 0.96086551 | 6 | 0.21644525 | 1 | 0.76878158 | 76 | 0.36734352 | FALSE | 1 | FALSE | 1 | FALSE | 1 | FALSE | 1 | FALSE | 1 | FALSE | 1 |
| 6648 | 6648 | 11.3181818 | 3.75846153 | 1 | 0.42086991 | 0 | 1 | 0 | 1 | 2 | 0.25059627 | 0 | 1 | 18 | 0.57962763 | FALSE | 1 | FALSE | 1 | FALSE | 1 | FALSE | 1 | FALSE | 1 | FALSE | 1 |
| 6649 | 6649 | 10.3333333 | 2.21108319 | 0 | 1 | 0 | 1 | 0 | 1 | 3 | 0.01547597 | 1 | 0.18007762 | 8 | 0.97197833 | FALSE | 1 | FALSE | 1 | FALSE | 1 | FALSE | 1 | FALSE | 1 | FALSE | 1 |
| 6650 | 6650 | 9.41666667 | 2.07999733 | 1 | 0.46441868 | 1 | 0.54637326 | 1 | 0.58502114 | 3 | 0.09525947 | 0 | 1 | 18 | 0.93166998 | FALSE | 1 | FALSE | 1 | FALSE | 1 | FALSE | 1 | FALSE | 1 | FALSE | 1 |
| 6651 | 6651 | 4.14179105 | 2.64900146 | 8 | 0.02060386 | 4 | 0.62934626 | 5 | 0.52364313 | 3 | 0.94813262 | 8 | 0.00145144 | 105 | 0.96475065 | FALSE | 0.45251148 | FALSE | 1 | FALSE | 1 | FALSE | 1 | FALSE | 0.21232461 | FALSE | 1 |
| 6652 | 6652 | 8.25 | 4.50416474 | 1 | 0.40558521 | 1 | 0.48239024 | 0 | 1 | 2 | 0.23331368 | 1 | 0.28187291 | 15 | 0.92113679 | FALSE | 1 | FALSE | 1 | FALSE | 1 | FALSE | 1 | FALSE | 1 | FALSE | 1 |
| 6653 | 6653 | 9.8 | 2.31516738 | 0 | 1 | 3 | 0.00338417 | 0 | 1 | 0 | 1 | 0 | 1 | 7 | 0.94326215 | FALSE | 1 | FALSE | 0.51339057 | FALSE | 1 | FALSE | 1 | FALSE | 1 | FALSE | 1 |
| 6654 | 6654 | 10.3636364 | 1.91988292 | 1 | 0.24866461 | 1 | 0.30368014 | 0 | 1 | 0 | 1 | 0 | 1 | 9 | 0.75985366 | FALSE | 1 | FALSE | 1 | FALSE | 1 | FALSE | 1 | FALSE | 1 | FALSE | 1 |
| 6655 | 6655 | 8.61904762 | 3.4153183 | 0 | 1 | 1 | 0.49918218 | 0 | 1 | 1 | 0.62783522 | 0 | 1 | 19 | 0.34094959 | FALSE | 1 | FALSE | 1 | FALSE | 1 | FALSE | 1 | FALSE | 1 | FALSE | 1 |
| 6656 | 6656 | 10.4347826 | 2.58467199 | 2 | 0.10809809 | 2 | 0.15794013 | 0 | 1 | 1 | 0.64497439 | 0 | 1 | 17 | 0.8840357 | FALSE | 0.88820416 | FALSE | 1 | FALSE | 1 | FALSE | 1 | FALSE | 1 | FALSE | 1 |
| 6657 | 6657 | 10.4545455 | 2.14745669 | 1 | 0.24866461 | 0 | 1 | 1 | 0.33149969 | 2 | 0.08789566 | 0 | 1 | 7 | 0.98129353 | FALSE | 1 | FALSE | 1 | FALSE | 1 | FALSE | 1 | FALSE | 1 | FALSE | 1 |
| 6658 | 6658 | 8.61904762 | 1.67548687 | 0 | 1 | 1 | 0.49918218 | 1 | 0.53671415 | 0 | 1 | 1 | 0.29368245 | 18 | 0.57962763 | FALSE | 1 | FALSE | 1 | FALSE | 1 | FALSE | 1 | FALSE | 1 | FALSE | 1 |
| 6659 | 6659 | 8.23076923 | 1.76084971 | 2 | 0.04234484 | 0 | 1 | 0 | 1 | 0 | 1 | 1 | 0.19354507 | 10 | 0.86791642 | FALSE | 0.62842194 | FALSE | 1 | FALSE | 1 | FALSE | 1 | FALSE | 1 | FALSE | 1 |
| 6660 | 6660 | 6.72222222 | 4.18735604 | 0 | 1 | 0 | 1 | 1 | 0.48280983 | 0 | 1 | 0 | 1 | 17 | 0.20329205 | FALSE | 1 | FALSE | 1 | FALSE | 1 | FALSE | 1 | FALSE | 1 | FALSE | 1 |
| 6661 | 6661 | 8.78571429 | 2.82029815 | 0 | 1 | 0 | 1 | 0 | 1 | 2 | 0.3705495 | 0 | 1 | 26 | 0.16405547 | FALSE | 1 | FALSE | 1 | FALSE | 1 | FALSE | 1 | FALSE | 1 | FALSE | 1 |
| 6662 | 6662 | 13.3333333 | 2.05480467 | 2 | 0.00915907 | 0 | 1 | 0 | 1 | 0 | 1 | 0 | 1 | 4 | 0.94743772 | FALSE | 0.28420861 | FALSE | 1 | FALSE | 1 | FALSE | 1 | FALSE | 1 | FALSE | 1 |
| 6663 | 6663 | 9.36 | 2.1703456 | 0 | 1 | 1 | 0.56109832 | 0 | 1 | 3 | 0.10474541 | 2 | 0.06257402 | 19 | 0.91734261 | FALSE | 1 | FALSE | 1 | FALSE | 1 | FALSE | 1 | FALSE | 1 | FALSE | 1 |
| 6664 | 6664 | 10.2307692 | 2.63566795 | 3 | 0.00391863 | 1 | 0.3480709 | 1 | 0.37874417 | 0 | 1 | 0 | 1 | 8 | 0.99085298 | FALSE | 0.17735574 | FALSE | 1 | FALSE | 1 | FALSE | 1 | FALSE | 1 | FALSE | 1 |
| 6665 | 6665 | 10.5454546 | 3.77464354 | 1 | 0.4357638 | 0 | 1 | 1 | 0.55340851 | 0 | 1 | 2 | 0.04972408 | 18 | 0.74744813 | FALSE | 1 | FALSE | 1 | FALSE | 1 | FALSE | 1 | FALSE | 0.98629461 | FALSE | 1 |
| 6666 | 6666 | 7.88 | 2.02622802 | 2 | 0.13369621 | 2 | 0.19297191 | 2 | 0.22597287 | 0 | 1 | 0 | 1 | 19 | 0.91734261 | FALSE | 0.97964162 | FALSE | 1 | FALSE | 1 | FALSE | 1 | FALSE | 1 | FALSE | 1 |
| 6667 | 6667 | 3.52845529 | 1.98909712 | 9 | 0.17397864 | 15 | 0.01239151 | 12 | 0.16712046 | 14 | 0.23025814 | 5 | 0.37119609 | 189 | 0.9985828 | FALSE | 1 | FALSE | 0.6905529 | FALSE | 1 | FALSE | 1 | FALSE | 1 | FALSE | 1 |
| 6668 | 6668 | 4.18181818 | 2.39681454 | 4 | 0.24848561 | 2 | 0.83585691 | 5 | 0.28294346 | 3 | 0.83995469 | 1 | 0.80772332 | 84 | 0.51640874 | FALSE | 1 | FALSE | 1 | FALSE | 1 | FALSE | 1 | FALSE | 1 | FALSE | 1 |
| 6669 | 6669 | 11.2222222 | 2.71256792 | 0 | 1 | 1 | 0.58913618 | 1 | 0.62830986 | 2 | 0.35374209 | 0 | 1 | 23 | 0.58307573 | FALSE | 1 | FALSE | 1 | FALSE | 1 | FALSE | 1 | FALSE | 1 | FALSE | 1 |
| 6670 | 6670 | 8.84210526 | 2.41188195 | 0 | 1 | 0 | 1 | 1 | 0.50143826 | 0 | 1 | 0 | 1 | 18 | 0.17880694 | FALSE | 1 | FALSE | 1 | FALSE | 1 | FALSE | 1 | FALSE | 1 | FALSE | 1 |
| 6671 | 6671 | 7.24444444 | 1.40088156 | 0 | 1 | 1 | 0.77338922 | 1 | 0.80828716 | 3 | 0.34154252 | 0 | 1 | 40 | 0.27423942 | FALSE | 1 | FALSE | 1 | FALSE | 1 | FALSE | 1 | FALSE | 1 | FALSE | 1 |
| 6672 | 6672 | 5.09589041 | 2.64918899 | 6 | 0.01072376 | 1 | 0.9104874 | 4 | 0.26680336 | 3 | 0.65845889 | 1 | 0.7028142 | 58 | 0.9042862 | FALSE | 0.31374644 | FALSE | 1 | FALSE | 1 | FALSE | 1 | FALSE | 1 | FALSE | 1 |

| | | | | | | | | | | | | | | | | | | | | | | | | | | | |
|---|---|---|---|---|---|---|---|---|---|---|---|---|---|---|---|---|---|---|---|---|---|---|---|---|---|---|---|
| 6673 | 6673 | 10.6086957 | 2.03745644 | 1 | 0.45027683 | 1 | 0.53115654 | 0 | 1 | 0 | 1 | 2 | 0.05389003 | 19 | 0.71542496 | FALSE | 1 | FALSE | 1 | FALSE | 1 | FALSE | 1 | FALSE | 1 | FALSE | 1 |
| 6674 | 6674 | 10.3898305 | 1.47311441 | 0 | 1 | 3 | 0.2979413 | 2 | 0.63179508 | 2 | 0.76144533 | 1 | 0.62455813 | 51 | 0.41341988 | FALSE | 1 | FALSE | 1 | FALSE | 1 | FALSE | 1 | FALSE | 1 | FALSE | 1 |
| 6675 | 6675 | 4.61111111 | 2.13427145 | 2 | 0.83576566 | 1 | 0.98422001 | 4 | 0.66302379 | 9 | 0.11992131 | 0 | 1 | 109 | 0.22951656 | FALSE | 1 | FALSE | 1 | FALSE | 1 | FALSE | 1 | FALSE | 1 | FALSE | 1 |
| 6676 | 6676 | 7.21875 | 1.40833179 | 0 | 1 | 1 | 0.65167362 | 1 | 0.69069222 | 0 | 1 | 1 | 0.41154418 | 29 | 0.24088541 | FALSE | 1 | FALSE | 1 | FALSE | 1 | FALSE | 1 | FALSE | 1 | FALSE | 1 |
| 6677 | 6677 | 8 | 1.45521375 | 0 | 1 | 0 | 1 | 1 | 0.46348836 | 1 | 0.55062772 | 0 | 1 | 15 | 0.49060435 | FALSE | 1 | FALSE | 1 | FALSE | 1 | FALSE | 1 | FALSE | 1 | FALSE | 1 |
| 6678 | 6678 | 11.2380952 | 2.38855928 | 1 | 0.35728305 | 2 | 0.10307112 | 0 | 1 | 0 | 1 | 0 | 1 | 14 | 0.7313145 | FALSE | 1 | FALSE | 1 | FALSE | 1 | FALSE | 1 | FALSE | 1 | FALSE | 1 |
| 6679 | 6679 | 9 | 1.97036873 | 2 | 0.20692137 | 0 | 1 | 1 | 0.70185639 | 1 | 0.78873814 | 0 | 1 | 29 | 0.39648374 | FALSE | 1 | FALSE | 1 | FALSE | 1 | FALSE | 1 | FALSE | 1 | FALSE | 1 |
| 6680 | 6680 | 9.63157895 | 2.41073316 | 0 | 1 | 1 | 0.46503795 | 2 | 0.14749808 | 1 | 0.59104357 | 0 | 1 | 15 | 0.83648191 | FALSE | 1 | FALSE | 1 | FALSE | 1 | FALSE | 1 | FALSE | 1 | FALSE | 1 |
| 6681 | 6681 | 9.92857143 | 1.86946462 | 0 | 1 | 0 | 1 | 0 | 1 | 2 | 0.13304705 | 0 | 1 | 12 | 0.62236227 | FALSE | 1 | FALSE | 1 | FALSE | 1 | FALSE | 1 | FALSE | 1 | FALSE | 1 |
| 6682 | 6682 | 9.36363636 | 4.20546673 | 0 | 1 | 1 | 0.30368014 | 0 | 1 | 0 | 1 | 0 | 1 | 10 | 0.46859937 | FALSE | 1 | FALSE | 1 | FALSE | 1 | FALSE | 1 | FALSE | 1 | FALSE | 1 |
| 6683 | 6683 | 8.86363636 | 2.30208314 | 1 | 0.42086991 | 0 | 1 | 2 | 0.17308361 | 2 | 0.25059627 | 4 | 0.00033021 | 12 | 0.99943951 | FALSE | 1 | FALSE | 1 | FALSE | 1 | FALSE | 1 | FALSE | 0.09327956 | FALSE | 1 |
| 6684 | 6684 | 7.45454546 | 2.10469762 | 0 | 1 | 1 | 0.30368014 | 0 | 1 | 0 | 1 | 0 | 1 | 10 | 0.46859937 | FALSE | 1 | FALSE | 1 | FALSE | 1 | FALSE | 1 | FALSE | 1 | FALSE | 1 |
| 6685 | 6685 | 9.18181818 | 2.32840881 | 0 | 1 | 1 | 0.30368014 | 0 | 1 | 1 | 0.40390032 | 0 | 1 | 9 | 0.75985366 | FALSE | 1 | FALSE | 1 | FALSE | 1 | FALSE | 1 | FALSE | 1 | FALSE | 1 |
| 6686 | 6686 | 10.3 | 2.32594067 | 0 | 1 | 0 | 1 | 0 | 1 | 0 | 1 | 0 | 1 | 10 | 0.18273166 | FALSE | 1 | FALSE | 1 | FALSE | 1 | FALSE | 1 | FALSE | 1 | FALSE | 1 |
| 6687 | 6687 | 9.28571429 | 1.45218578 | 1 | 0.42086991 | 0 | 1 | 1 | 0.53671415 | 1 | 0.62783522 | 0 | 1 | 18 | 0.57962763 | FALSE | 1 | FALSE | 1 | FALSE | 1 | FALSE | 1 | FALSE | 1 | FALSE | 1 |
| 6688 | 6688 | 11.1666667 | 3.00462606 | 0 | 1 | 0 | 1 | 2 | 0.13502329 | 1 | 0.57131013 | 0 | 1 | 15 | 0.69372602 | FALSE | 1 | FALSE | 1 | FALSE | 1 | FALSE | 1 | FALSE | 1 | FALSE | 1 |
| 6689 | 6689 | 10.2 | 2.8213472 | 0 | 1 | 1 | 0.28036922 | 0 | 1 | 0 | 1 | 0 | 1 | 9 | 0.5215547 | FALSE | 1 | FALSE | 1 | FALSE | 1 | FALSE | 1 | FALSE | 1 | FALSE | 1 |
| 6690 | 6690 | 9.39393939 | 1.79121415 | 2 | 0.20692137 | 3 | 0.08959205 | 2 | 0.33343871 | 1 | 0.78873814 | 0 | 1 | 25 | 0.93892791 | FALSE | 1 | FALSE | 1 | FALSE | 1 | FALSE | 1 | FALSE | 1 | FALSE | 1 |
| 6691 | 6691 | 8.28571429 | 1.48461498 | 0 | 1 | 1 | 0.36919824 | 0 | 1 | 1 | 0.48242099 | 0 | 1 | 12 | 0.62236227 | FALSE | 1 | FALSE | 1 | FALSE | 1 | FALSE | 1 | FALSE | 1 | FALSE | 1 |
| 6692 | 6692 | 11.2121212 | 2.56736601 | 1 | 0.57647348 | 2 | 0.28941836 | 0 | 1 | 1 | 0.78873814 | 0 | 1 | 29 | 0.39648374 | FALSE | 1 | FALSE | 1 | FALSE | 1 | FALSE | 1 | FALSE | 1 | FALSE | 1 |
| 6693 | 6693 | 11 | 1.06904497 | 0 | 1 | 0 | 1 | 0 | 1 | 0 | 1 | 0 | 1 | 7 | 0.3043657 | FALSE | 1 | FALSE | 1 | FALSE | 1 | FALSE | 1 | FALSE | 1 | FALSE | 1 |
| 6694 | 6694 | 8.6 | 2.18043246 | 2 | 0.22589135 | 2 | 0.31355738 | 3 | 0.12958859 | 1 | 0.80778317 | 3 | 0.01918344 | 24 | 0.99496229 | FALSE | 1 | FALSE | 1 | FALSE | 1 | FALSE | 1 | FALSE | 0.70156566 | FALSE | 1 |
| 6695 | 6695 | 9.61904762 | 1.88922233 | 0 | 1 | 1 | 0.48239024 | 0 | 1 | 1 | 0.60987146 | 0 | 1 | 18 | 0.37504645 | FALSE | 1 | FALSE | 1 | FALSE | 1 | FALSE | 1 | FALSE | 1 | FALSE | 1 |
| 6696 | 6696 | 9.5 | 3.04138127 | 0 | 1 | 1 | 0.57534752 | 2 | 0.23940226 | 1 | 0.70601446 | 1 | 0.3499045 | 21 | 0.78939668 | FALSE | 1 | FALSE | 1 | FALSE | 1 | FALSE | 1 | FALSE | 1 | FALSE | 1 |
| 6697 | 6697 | 7.875 | 2.91279505 | 0 | 1 | 1 | 0.65167362 | 1 | 0.69069222 | 0 | 1 | 1 | 0.41154418 | 29 | 0.24088541 | FALSE | 1 | FALSE | 1 | FALSE | 1 | FALSE | 1 | FALSE | 1 | FALSE | 1 |
| 6698 | 6698 | 8.57692308 | 3.05303225 | 0 | 1 | 0 | 1 | 0 | 1 | 2 | 0.33678042 | 0 | 1 | 24 | 0.20425579 | FALSE | 1 | FALSE | 1 | FALSE | 1 | FALSE | 1 | FALSE | 1 | FALSE | 1 |
| 6699 | 6699 | 9.47368421 | 1.90203274 | 1 | 0.38989951 | 1 | 0.46503795 | 1 | 0.50143826 | 1 | 0.59104357 | 0 | 1 | 15 | 0.83648191 | FALSE | 1 | FALSE | 1 | FALSE | 1 | FALSE | 1 | FALSE | 1 | FALSE | 1 |
| 6700 | 6700 | 6.78048781 | 2.04238258 | 1 | 0.65632651 | 0 | 1 | 1 | 0.77786601 | 2 | 0.56788092 | 0 | 1 | 37 | 0.21027077 | FALSE | 1 | FALSE | 1 | FALSE | 1 | FALSE | 1 | FALSE | 1 | FALSE | 1 |
| 6701 | 6701 | 7.96875 | 1.10352772 | 0 | 1 | 3 | 0.08334648 | 3 | 0.10599423 | 2 | 0.43583874 | 0 | 1 | 24 | 0.94876798 | FALSE | 1 | FALSE | 1 | FALSE | 1 | FALSE | 1 | FALSE | 1 | FALSE | 1 |
| 6702 | 6702 | 8.59090909 | 1.33685829 | 4 | 0.02552933 | 3 | 0.16927151 | 2 | 0.47281647 | 1 | 0.87439647 | 2 | 0.16211764 | 32 | 0.98583614 | FALSE | 0.4974559 | FALSE | 1 | FALSE | 1 | FALSE | 1 | FALSE | 1 | FALSE | 1 |
| 6703 | 6703 | 8.76470588 | 2.57124549 | 1 | 0.7354105 | 0 | 1 | 2 | 0.55199489 | 3 | 0.41729179 | 0 | 1 | 45 | 0.29584937 | FALSE | 1 | FALSE | 1 | FALSE | 1 | FALSE | 1 | FALSE | 1 | FALSE | 1 |
| 6704 | 6704 | 9.60869565 | 1.55309314 | 0 | 1 | 2 | 0.15794013 | 0 | 1 | 2 | 0.26791271 | 0 | 1 | 18 | 0.74744813 | FALSE | 1 | FALSE | 1 | FALSE | 1 | FALSE | 1 | FALSE | 1 | FALSE | 1 |
| 6705 | 6705 | 9.4 | 2.45764115 | 0 | 1 | 0 | 1 | 0 | 1 | 0 | 1 | 0 | 1 | 10 | 0.18273166 | FALSE | 1 | FALSE | 1 | FALSE | 1 | FALSE | 1 | FALSE | 1 | FALSE | 1 |
| 6706 | 6706 | 8.88571429 | 3.05914487 | 3 | 0.0595675 | 2 | 0.31355738 | 2 | 0.35984046 | 2 | 0.48233989 | 2 | 0.11183826 | 24 | 0.99496229 | FALSE | 0.72532843 | FALSE | 1 | FALSE | 1 | FALSE | 1 | FALSE | 1 | FALSE | 1 |
| 6707 | 6707 | 10.4166667 | 2.95686583 | 0 | 1 | 0 | 1 | 1 | 0.58502114 | 3 | 0.09525947 | 0 | 1 | 20 | 0.68270729 | FALSE | 1 | FALSE | 1 | FALSE | 1 | FALSE | 1 | FALSE | 1 | FALSE | 1 |
| 6708 | 6708 | 8.84615385 | 2.49140535 | 0 | 1 | 1 | 0.57534752 | 0 | 1 | 0 | 1 | 0 | 1 | 25 | 0.06985983 | FALSE | 1 | FALSE | 1 | FALSE | 1 | FALSE | 1 | FALSE | 1 | FALSE | 1 |
| 6709 | 6709 | 11.4705882 | 2.42820793 | 2 | 0.06909038 | 0 | 1 | 0 | 1 | 0 | 1 | 0 | 1 | 15 | 0.49060435 | FALSE | 0.74276693 | FALSE | 1 | FALSE | 1 | FALSE | 1 | FALSE | 1 | FALSE | 1 |
| 6710 | 6710 | 9.40740741 | 2.3453541 | 1 | 0.50471032 | 2 | 0.21684246 | 2 | 0.25286935 | 1 | 0.71956339 | 0 | 1 | 21 | 0.88395936 | FALSE | 1 | FALSE | 1 | FALSE | 1 | FALSE | 1 | FALSE | 1 | FALSE | 1 |
| 6711 | 6711 | 9.58333333 | 3.27766007 | 0 | 1 | 2 | 0.18117183 | 0 | 1 | 0 | 1 | 0 | 1 | 22 | 0.25237063 | FALSE | 1 | FALSE | 1 | FALSE | 1 | FALSE | 1 | FALSE | 1 | FALSE | 1 |
| 6712 | 6712 | 9.06666667 | 2.46215804 | 0 | 1 | 2 | 0.25306331 | 1 | 0.66709967 | 1 | 0.75658797 | 3 | 0.01262211 | 23 | 0.9158696 | FALSE | 1 | FALSE | 1 | FALSE | 1 | FALSE | 1 | FALSE | 0.63048977 | FALSE | 1 |
| 6713 | 6713 | 10.3571429 | 2.8311316 | 1 | 0.51745948 | 0 | 1 | 2 | 0.2663542 | 1 | 0.73248983 | 2 | 0.07640798 | 22 | 0.86499992 | FALSE | 1 | FALSE | 1 | FALSE | 1 | FALSE | 1 | FALSE | 1 | FALSE | 1 |
| 6714 | 6714 | 10.902439 | 2.9033612 | 1 | 0.65632651 | 0 | 1 | 2 | 0.43640583 | 1 | 0.85524728 | 1 | 0.49330646 | 36 | 0.36455032 | FALSE | 1 | FALSE | 1 | FALSE | 1 | FALSE | 1 | FALSE | 1 | FALSE | 1 |
| 6715 | 6715 | 8.05555556 | 1.508208 | 1 | 0.60837712 | 1 | 0.69480472 | 1 | 0.73299778 | 1 | 0.81665383 | 0 | 1 | 32 | 0.31709555 | FALSE | 1 | FALSE | 1 | FALSE | 1 | FALSE | 1 | FALSE | 1 | FALSE | 1 |
| 6716 | 6716 | 9.25806452 | 1.36707227 | 1 | 0.55378205 | 2 | 0.2651851 | 2 | 0.30673593 | 1 | 0.76781284 | 1 | 0.4016888 | 24 | 0.90138666 | FALSE | 1 | FALSE | 1 | FALSE | 1 | FALSE | 1 | FALSE | 1 | FALSE | 1 |
| 6717 | 6717 | 4.96405229 | 2.03932305 | 7 | 0.66572309 | 10 | 0.52084708 | 11 | 0.53060171 | 18 | 0.15624718 | 6 | 0.37813477 | 251 | 0.80017056 | FALSE | 1 | FALSE | 1 | FALSE | 1 | FALSE | 1 | FALSE | 1 | FALSE | 1 |
| 6718 | 6718 | 9.35 | 1.01365675 | 0 | 1 | 1 | 0.48239024 | 1 | 0.51939841 | 0 | 1 | 0 | 1 | 18 | 0.37504645 | FALSE | 1 | FALSE | 1 | FALSE | 1 | FALSE | 1 | FALSE | 1 | FALSE | 1 |
| 6719 | 6719 | 6.69565217 | 1.75535906 | 1 | 0.69843502 | 1 | 0.78076835 | 2 | 0.4962935 | 3 | 0.35428318 | 0 | 1 | 39 | 0.56897353 | FALSE | 1 | FALSE | 1 | FALSE | 1 | FALSE | 1 | FALSE | 1 | FALSE | 1 |
| 6720 | 6720 | 9.02325581 | 1.97057862 | 0 | 1 | 1 | 0.75788114 | 0 | 1 | 1 | 0.86831187 | 0 | 1 | 41 | 0.02652094 | FALSE | 1 | FALSE | 1 | FALSE | 1 | FALSE | 1 | FALSE | 1 | FALSE | 1 |
| 6721 | 6721 | 4.61734694 | 1.72659424 | 6 | 0.36532954 | 11 | 0.04504673 | 3 | 0.97087076 | 6 | 0.87897228 | 2 | 0.82622619 | 163 | 0.40148713 | FALSE | 1 | FALSE | 1 | FALSE | 1 | FALSE | 1 | FALSE | 1 | FALSE | 1 |
| 6722 | 6722 | 12.4 | 2.33238076 | 1 | 0.12181417 | 0 | 1 | 0 | 1 | 0 | 1 | 0 | 1 | 4 | 0.82371636 | FALSE | 0.95622564 | FALSE | 1 | FALSE | 1 | FALSE | 1 | FALSE | 1 | FALSE | 1 |
| 6723 | 6723 | 8.66666667 | 2.62466929 | 0 | 1 | 0 | 1 | 0 | 1 | 2 | 0.30249307 | 3 | 0.00675546 | 19 | 0.83975208 | FALSE | 1 | FALSE | 1 | FALSE | 1 | FALSE | 1 | FALSE | 0.47461742 | FALSE | 1 |
| 6724 | 6724 | 8.15217391 | 1.70634865 | 2 | 0.3309085 | 3 | 0.18548475 | 3 | 0.2282043 | 1 | 0.88573774 | 1 | 0.53375569 | 36 | 0.90704249 | FALSE | 1 | FALSE | 1 | FALSE | 1 | FALSE | 1 | FALSE | 1 | FALSE | 1 |
| 6725 | 6725 | 10.4375 | 2.26298337 | 1 | 0.34033054 | 0 | 1 | 1 | 0.44344817 | 1 | 0.52895088 | 1 | 0.23264612 | 12 | 0.90922186 | FALSE | 1 | FALSE | 1 | FALSE | 1 | FALSE | 1 | FALSE | 1 | FALSE | 1 |
| 6726 | 6726 | 9.2631579 | 1.84511033 | 2 | 0.25455533 | 2 | 0.3494045 | 1 | 0.75193685 | 2 | 0.52640931 | 2 | 0.12811285 | 29 | 0.93814695 | FALSE | 1 | FALSE | 1 | FALSE | 1 | FALSE | 1 | FALSE | 1 | FALSE | 1 |
| 6727 | 6727 | 8.86666667 | 2.67996683 | 2 | 0.05513784 | 1 | 0.38964394 | 1 | 0.42266264 | 2 | 0.14906763 | 0 | 1 | 9 | 0.995488 | FALSE | 0.68849059 | FALSE | 1 | FALSE | 1 | FALSE | 1 | FALSE | 1 | FALSE | 1 |
| 6728 | 6728 | 6.75 | 2.15058132 | 3 | 0.22550017 | 1 | 0.87929209 | 2 | 0.67580769 | 4 | 0.33842086 | 0 | 1 | 54 | 0.58381086 | FALSE | 1 | FALSE | 1 | FALSE | 1 | FALSE | 1 | FALSE | 1 | FALSE | 1 |
| 6729 | 6729 | 6.35185185 | 2.63282279 | 0 | 1 | 2 | 0.52572379 | 2 | 0.58328102 | 2 | 0.71679995 | 3 | 0.05824229 | 45 | 0.66857827 | FALSE | 1 | FALSE | 1 | FALSE | 1 | FALSE | 1 | FALSE | 1 | FALSE | 1 |
| 6730 | 6730 | 4.4556962 | 1.66723806 | 7 | 0.10809002 | 4 | 0.75372741 | 7 | 0.33420068 | 4 | 0.93512282 | 4 | 0.25645514 | 131 | 0.67681493 | FALSE | 0.88820416 | FALSE | 1 | FALSE | 1 | FALSE | 1 | FALSE | 1 | FALSE | 1 |
| 6731 | 6731 | 6.75757576 | 2.82875177 | 2 | 0.20692137 | 1 | 0.6629941 | 2 | 0.33343871 | 2 | 0.45159487 | 0 | 1 | 26 | 0.86852567 | FALSE | 1 | FALSE | 1 | FALSE | 1 | FALSE | 1 | FALSE | 1 | FALSE | 1 |
| 6732 | 6732 | 10.1666667 | 3.7155828 | 0 | 1 | 0 | 1 | 0 | 1 | 1 | 0.43131129 | 2 | 0.01579445 | 9 | 0.89630573 | FALSE | 1 | FALSE | 1 | FALSE | 1 | FALSE | 1 | FALSE | 0.66313193 | FALSE | 1 |
| 6733 | 6733 | 9 | 2.30940108 | 3 | 0.00307177 | 0 | 1 | 1 | 0.35555293 | 0 | 1 | 0 | 1 | 8 | 0.97197833 | FALSE | 0.14802295 | FALSE | 1 | FALSE | 1 | FALSE | 1 | FALSE | 1 | FALSE | 1 |
| 6734 | 6734 | 9.25 | 2.15058132 | 2 | 0.19750261 | 4 | 0.0188228 | 0 | 1 | 0 | 1 | 0 | 1 | 26 | 0.77612337 | FALSE | 1 | FALSE | 0.77515497 | FALSE | 1 | FALSE | 1 | FALSE | 1 | FALSE | 1 |
| 6735 | 6735 | 4.54125413 | 1.75874461 | 7 | 0.65196276 | 12 | 0.26058745 | 6 | 0.9620601 | 15 | 0.39909535 | 6 | 0.36627301 | 253 | 0.49340059 | FALSE | 1 | FALSE | 1 | FALSE | 1 | FALSE | 1 | FALSE | 1 | FALSE | 1 |
| 6736 | 6736 | 5.8630137 | 2.27141298 | 1 | 0.85137169 | 1 | 0.9104874 | 4 | 0.26680336 | 2 | 0.8554296 | 2 | 0.33719417 | 63 | 0.39803869 | FALSE | 1 | FALSE | 1 | FALSE | 1 | FALSE | 1 | FALSE | 1 | FALSE | 1 |
| 6737 | 6737 | 10.3571429 | 0.61028598 | 0 | 1 | 0 | 1 | 2 | 0.08806398 | 1 | 0.48242099 | 0 | 1 | 11 | 0.83676449 | FALSE | 1 | FALSE | 1 | FALSE | 1 | FALSE | 1 | FALSE | 1 | FALSE | 1 |
| 6738 | 6738 | 8.4375 | 0.99804496 | 1 | 0.34033054 | 1 | 0.40942991 | 1 | 0.44344817 | 1 | 0.52895088 | 0 | 1 | 12 | 0.90922186 | FALSE | 1 | FALSE | 1 | FALSE | 1 | FALSE | 1 | FALSE | 1 | FALSE | 1 |
| 6739 | 6739 | 10.0833333 | 2.8710722 | 1 | 0.26795838 | 0 | 1 | 1 | 0.35555293 | 0 | 1 | 1 | 0.18007762 | 9 | 0.89630573 | FALSE | 1 | FALSE | 1 | FALSE | 1 | FALSE | 1 | FALSE | 1 | FALSE | 1 |
| 6740 | 6740 | 9.33333333 | 3.91578004 | 1 | 0.37380237 | 1 | 0.44710673 | 0 | 1 | 0 | 1 | 0 | 1 | 16 | 0.44998001 | FALSE | 1 | FALSE | 1 | FALSE | 1 | FALSE | 1 | FALSE | 1 | FALSE | 1 |
| 6741 | 6741 | 9.28571429 | 2.29684788 | 3 | 0.03391597 | 0 | 1 | 2 | 0.2663542 | 2 | 0.3705495 | 1 | 0.37113067 | 20 | 0.97757517 | FALSE | 0.56934354 | FALSE | 1 | FALSE | 1 | FALSE | 1 | FALSE | 1 | FALSE | 1 |
| 6742 | 6742 | 8.77777778 | 2.8196839 | 2 | 0.23542589 | 0 | 1 | 1 | 0.73299778 | 2 | 0.49731009 | 1 | 0.44938266 | 30 | 0.67248271 | FALSE | 1 | FALSE | 1 | FALSE | 1 | FALSE | 1 | FALSE | 1 | FALSE | 1 |
| 6743 | 6743 | 11.1428571 | 3.02033922 | 0 | 1 | 1 | 0.36919824 | 1 | 0.40110418 | 0 | 1 | 1 | 0.20679328 | 11 | 0.83676449 | FALSE | 1 | FALSE | 1 | FALSE | 1 | FALSE | 1 | FALSE | 1 | FALSE | 1 |
| 6744 | 6744 | 5.65384615 | 2.85471685 | 4 | 0.13936698 | 0 | 1 | 4 | 0.30733368 | 4 | 0.48430022 | 0 | 1 | 66 | 0.55329149 | FALSE | 1 | FALSE | 1 | FALSE | 1 | FALSE | 1 | FALSE | 1 | FALSE | 1 |
| 6745 | 6745 | 9.25 | 3.63833753 | 1 | 0.64722818 | 4 | 0.03920804 | 3 | 0.17245215 | 0 | 1 | 1 | 0.48480878 | 31 | 0.91676852 | FALSE | 1 | FALSE | 0.96864929 | FALSE | 1 | FALSE | 1 | FALSE | 1 | FALSE | 1 |
| 6746 | 6746 | 10.2 | 2.13541565 | 1 | 0.40558521 | 1 | 0.48239024 | 1 | 0.51939841 | 1 | 0.60987146 | 2 | 0.04176963 | 14 | 0.97325593 | FALSE | 1 | FALSE | 1 | FALSE | 1 | FALSE | 1 | FALSE | 0.90763079 | FALSE | 1 |
| 6747 | 6747 | 8.03703704 | 2.82162918 | 0 | 1 | 1 | 0.57534752 | 2 | 0.23940226 | 0 | 1 | 0 | 1 | 23 | 0.40372818 | FALSE | 1 | FALSE | 1 | FALSE | 1 | FALSE | 1 | FALSE | 1 | FALSE | 1 |
| 6748 | 6748 | 7.94736842 | 2.74290826 | 1 | 0.38989951 | 1 | 0.46503795 | 1 | 0.50143826 | 1 | 0.59104357 | 0 | 1 | 15 | 0.83648191 | FALSE | 1 | FALSE | 1 | FALSE | 1 | FALSE | 1 | FALSE | 1 | FALSE | 1 |
| 6749 | 6749 | 9.66666667 | 2.4675674 | 0 | 1 | 0 | 1 | 0 | 1 | 0 | 1 | 0 | 1 | 15 | 0.07803154 | FALSE | 1 | FALSE | 1 | FALSE | 1 | FALSE | 1 | FALSE | 1 | FALSE | 1 |
| 6750 | 6750 | 9.62921348 | 2.08462549 | 0 | 1 | 2 | 0.78896076 | 5 | 0.21518975 | 7 | 0.1134709 | 0 | 1 | 75 | 0.58213685 | FALSE | 1 | FALSE | 1 | FALSE | 1 | FALSE | 1 | FALSE | 1 | FALSE | 1 |
| 6751 | 6751 | 11.2222222 | 1.39664501 | 0 | 1 | 1 | 0.25628163 | 0 | 1 | 1 | 0.34506517 | 0 | 1 | 7 | 0.84574168 | FALSE | 1 | FALSE | 1 | FALSE | 1 | FALSE | 1 | FALSE | 1 | FALSE | 1 |
| 6752 | 6752 | 11 | 2.5226249 | 0 | 1 | 0 | 1 | 1 | 0.30655256 | 1 | 0.37517262 | 0 | 1 | 8 | 0.80394719 | FALSE | 1 | FALSE | 1 | FALSE | 1 | FALSE | 1 | FALSE | 1 | FALSE | 1 |
| 6753 | 6753 | 11.375 | 2.70512631 | 0 | 1 | 1 | 0.53115654 | 0 | 1 | 0 | 1 | 0 | 1 | 22 | 0.10533112 | FALSE | 1 | FALSE | 1 | FALSE | 1 | FALSE | 1 | FALSE | 1 | FALSE | 1 |
| 6754 | 6754 | 8.25925926 | 1.97376483 | 0 | 1 | 1 | 0.58913618 | 1 | 0.62830986 | 1 | 0.71956339 | 0 | 1 | 24 | 0.3725373 | FALSE | 1 | FALSE | 1 | FALSE | 1 | FALSE | 1 | FALSE | 1 | FALSE | 1 |
| 6755 | 6755 | 10.1666667 | 2.76385399 | 1 | 0.26795838 | 0 | 1 | 0 | 1 | 0 | 1 | 0 | 1 | 11 | 0.41944532 | FALSE | 1 | FALSE | 1 | FALSE | 1 | FALSE | 1 | FALSE | 1 | FALSE | 1 |
| 6756 | 6756 | 7.94827586 | 2.20831043 | 2 | 0.44060625 | 1 | 0.85269953 | 1 | 0.88128698 | 2 | 0.75303313 | 1 | 0.61824432 | 51 | 0.2955095 | FALSE | 1 | FALSE | 1 | FALSE | 1 | FALSE | 1 | FALSE | 1 | FALSE | 1 |
| 6757 | 6757 | 9.70833333 | 3.15540762 | 1 | 0.46441868 | 2 | 0.18117183 | 0 | 1 | 1 | 0.67692816 | 2 | 0.058175 | 18 | 0.93166998 | FALSE | 1 | FALSE | 1 | FALSE | 1 | FALSE | 1 | FALSE | 1 | FALSE | 1 |
| 6758 | 6758 | 8.52380952 | 1.90535705 | 1 | 0.66519151 | 0 | 1 | 2 | 0.44869594 | 1 | 0.86193352 | 1 | 0.50166523 | 37 | 0.34041811 | FALSE | 1 | FALSE | 1 | FALSE | 1 | FALSE | 1 | FALSE | 1 | FALSE | 1 |
| 6759 | 6759 | 8.91666667 | 3.15678282 | 2 | 0.23542589 | 0 | 1 | 1 | 0.73299778 | 0 | 1 | 1 | 0.44938266 | 32 | 0.31709555 | FALSE | 1 | FALSE | 1 | FALSE | 1 | FALSE | 1 | FALSE | 1 | FALSE | 1 |
| 6760 | 6760 | 9.4375 | 2.66853776 | 1 | 0.34033054 | 0 | 1 | 0 | 1 | 0 | 1 | 1 | 0.23264612 | 14 | 0.53308261 | FALSE | 1 | FALSE | 1 | FALSE | 1 | FALSE | 1 | FALSE | 1 | FALSE | 1 |
| 6761 | 6761 | 7.56 | 2.36778377 | 1 | 0.46441868 | 0 | 1 | 0 | 1 | 1 | 0.67692816 | 1 | 0.32796862 | 21 | 0.47053137 | FALSE | 1 | FALSE | 1 | FALSE | 1 | FALSE | 1 | FALSE | 1 | FALSE | 1 |
| 6762 | 6762 | 11.0434783 | 2.82039561 | 0 | 1 | 1 | 0.53115654 | 2 | 0.19931429 | 2 | 0.2852236 | 2 | 0.05389003 | 16 | 0.98100709 | FALSE | 1 | FALSE | 1 | FALSE | 1 | FALSE | 1 | FALSE | 1 | FALSE | 1 |
| 6763 | 6763 | 9.52631579 | 3.49990107 | 0 | 1 | 3 | 0.02205772 | 1 | 0.50143826 | 0 | 1 | 0 | 1 | 15 | 0.83648191 | FALSE | 1 | FALSE | 0.81029974 | FALSE | 1 | FALSE | 1 | FALSE | 1 | FALSE | 1 |
| 6764 | 6764 | 7.7625 | 1.79055683 | 9 | 0.00010244 | 4 | 0.21704891 | 6 | 0.04913907 | 4 | 0.4435008 | 3 | 0.1209957 | 48 | 0.99999144 | TRUE | 0.01824319 | FALSE | 1 | FALSE | 1 | FALSE | 1 | FALSE | 1 | FALSE | 1 |
| 6765 | 6765 | 9.70833333 | 1.92525251 | 0 | 1 | 0 | 1 | 2 | 0.21260257 | 1 | 0.67692816 | 2 | 0.058175 | 19 | 0.83975208 | FALSE | 1 | FALSE | 1 | FALSE | 1 | FALSE | 1 | FALSE | 1 | FALSE | 1 |
| 6766 | 6766 | 10.173913 | 2.68088522 | 2 | 0.11647716 | 0 | 1 | 0 | 1 | 2 | 0.2852236 | 1 | 0.31672722 | 18 | 0.86273155 | FALSE | 0.93181727 | FALSE | 1 | FALSE | 1 | FALSE | 1 | FALSE | 1 | FALSE | 1 |

| | | | | | | | | | | | | | | | | | | | | | | | | | | | |
|---|---|---|---|---|---|---|---|---|---|---|---|---|---|---|---|---|---|---|---|---|---|---|---|---|---|---|---|
| 6767 | 6767 | 9.39130435 | 3.25361512 | 1 | 0.45027683 | 0 | 1 | 1 | 0.5695037 | 1 | 0.6613267 | 0 | 1 | 20 | 0.50589402 | FALSE | 1 | FALSE | 1 | FALSE | 1 | FALSE | 1 | FALSE | 1 | FALSE | 1 |
| 6768 | 6768 | 8 | 2.02259959 | 0 | 1 | 0 | 1 | 0 | 1 | 0 | 1 | 0 | 1 | 22 | 0.02368118 | FALSE | 1 | FALSE | 1 | FALSE | 1 | FALSE | 1 | FALSE | 1 | FALSE | 1 |
| 6769 | 6769 | 10.9166667 | 2.75252409 | 1 | 0.26795838 | 0 | 1 | 0 | 1 | 0 | 1 | 0 | 1 | 11 | 0.41944532 | FALSE | 1 | FALSE | 1 | FALSE | 1 | FALSE | 1 | FALSE | 1 | FALSE | 1 |
| 6770 | 6770 | 9.03703704 | 1.57483642 | 1 | 0.50471032 | 1 | 0.58913618 | 1 | 0.62830986 | 1 | 0.71956339 | 0 | 1 | 23 | 0.58307573 | FALSE | 1 | FALSE | 1 | FALSE | 1 | FALSE | 1 | FALSE | 1 | FALSE | 1 |
| 6771 | 6771 | 9.84615385 | 1.74735641 | 0 | 1 | 2 | 0.06426041 | 1 | 0.37874417 | 3 | 0.01944678 | 0 | 1 | 7 | 0.99839694 | FALSE | 1 | FALSE | 1 | FALSE | 1 | FALSE | 1 | FALSE | 1 | FALSE | 1 |
| 6772 | 6772 | 9.07407407 | 2.76118501 | 0 | 1 | 2 | 0.21684246 | 0 | 1 | 0 | 1 | 0 | 1 | 25 | 0.1832206 | FALSE | 1 | FALSE | 1 | FALSE | 1 | FALSE | 1 | FALSE | 1 | FALSE | 1 |
| 6773 | 6773 | 10.25 | 2.72717803 | 0 | 1 | 0 | 1 | 0 | 1 | 0 | 1 | 0 | 1 | 8 | 0.25677166 | FALSE | 1 | FALSE | 1 | FALSE | 1 | FALSE | 1 | FALSE | 1 | FALSE | 1 |
| 6774 | 6774 | 9.15068493 | 1.92083909 | 0 | 1 | 2 | 0.68961994 | 2 | 0.74419436 | 5 | 0.24321887 | 0 | 1 | 64 | 0.27646797 | FALSE | 1 | FALSE | 1 | FALSE | 1 | FALSE | 1 | FALSE | 1 | FALSE | 1 |
| 6775 | 6775 | 9.2631579 | 2.76904136 | 0 | 1 | 0 | 1 | 0 | 1 | 2 | 0.21610812 | 0 | 1 | 17 | 0.41141543 | FALSE | 1 | FALSE | 1 | FALSE | 1 | FALSE | 1 | FALSE | 1 | FALSE | 1 |
| 6776 | 6776 | 9.80769231 | 1.75453765 | 0 | 1 | 0 | 1 | 0 | 1 | 3 | 0.11458571 | 0 | 1 | 23 | 0.40372818 | FALSE | 1 | FALSE | 1 | FALSE | 1 | FALSE | 1 | FALSE | 1 | FALSE | 1 |
| 6777 | 6777 | 9.18181818 | 2.36713029 | 0 | 1 | 1 | 0.5154318 | 1 | 0.55340851 | 1 | 0.64497439 | 0 | 1 | 19 | 0.54233419 | FALSE | 1 | FALSE | 1 | FALSE | 1 | FALSE | 1 | FALSE | 1 | FALSE | 1 |
| 6778 | 6778 | 6.91666667 | 1.8800857 | 1 | 0.70621968 | 0 | 1 | 4 | 0.08716543 | 3 | 0.36699324 | 4 | 0.00709883 | 35 | 0.97547234 | FALSE | 1 | FALSE | 1 | FALSE | 1 | FALSE | 1 | FALSE | 0.48060812 | FALSE | 1 |
| 6779 | 6779 | 8.77777778 | 2.09644025 | 1 | 0.37380237 | 1 | 0.44710673 | 2 | 0.13502329 | 0 | 1 | 0 | 1 | 14 | 0.86287418 | FALSE | 1 | FALSE | 1 | FALSE | 1 | FALSE | 1 | FALSE | 1 | FALSE | 1 |
| 6780 | 6780 | 6.90857143 | 1.85862798 | 6 | 0.28344643 | 9 | 0.10761825 | 5 | 0.7512299 | 11 | 0.17091842 | 5 | 0.15405888 | 137 | 0.97447894 | FALSE | 1 | FALSE | 1 | FALSE | 1 | FALSE | 1 | FALSE | 1 | FALSE | 1 |
| 6781 | 6781 | 6.88888889 | 2.22333306 | 1 | 0.69044531 | 0 | 1 | 1 | 0.80828716 | 3 | 0.34154252 | 1 | 0.52593041 | 39 | 0.4317533 | FALSE | 1 | FALSE | 1 | FALSE | 1 | FALSE | 1 | FALSE | 1 | FALSE | 1 |
| 6782 | 6782 | 7.84158416 | 1.61510612 | 2 | 0.71619014 | 3 | 0.61262749 | 1 | 0.97197509 | 6 | 0.28608665 | 1 | 0.80116196 | 84 | 0.32996726 | FALSE | 1 | FALSE | 1 | FALSE | 1 | FALSE | 1 | FALSE | 1 | FALSE | 1 |
| 6783 | 6783 | 8.83333333 | 2.54405625 | 1 | 0.26795838 | 0 | 1 | 0 | 1 | 0 | 1 | 0 | 1 | 11 | 0.41944532 | FALSE | 1 | FALSE | 1 | FALSE | 1 | FALSE | 1 | FALSE | 1 | FALSE | 1 |
| 6784 | 6784 | 8.95833333 | 2.65328675 | 1 | 0.46441868 | 0 | 1 | 1 | 0.58502114 | 2 | 0.30249307 | 0 | 1 | 20 | 0.68270729 | FALSE | 1 | FALSE | 1 | FALSE | 1 | FALSE | 1 | FALSE | 1 | FALSE | 1 |
| 6785 | 6785 | 10.4545455 | 2.62956281 | 1 | 0.56527493 | 2 | 0.27730767 | 1 | 0.69069222 | 2 | 0.43583874 | 2 | 0.096172 | 24 | 0.94876798 | FALSE | 1 | FALSE | 1 | FALSE | 1 | FALSE | 1 | FALSE | 1 | FALSE | 1 |
| 6786 | 6786 | 9.1875 | 1.75185913 | 0 | 1 | 0 | 1 | 1 | 0.8283465 | 3 | 0.37966044 | 0 | 1 | 44 | 0.11052514 | FALSE | 1 | FALSE | 1 | FALSE | 1 | FALSE | 1 | FALSE | 1 | FALSE | 1 |
| 6787 | 6787 | 10.625 | 3.46184561 | 0 | 1 | 1 | 0.23139161 | 1 | 0.25384309 | 0 | 1 | 0 | 1 | 6 | 0.88425551 | FALSE | 1 | FALSE | 1 | FALSE | 1 | FALSE | 1 | FALSE | 1 | FALSE | 1 |
| 6788 | 6788 | 7.95348837 | 3.88170119 | 3 | 0.09709068 | 2 | 0.40761584 | 1 | 0.79363407 | 1 | 0.86831187 | 0 | 1 | 36 | 0.6451239 | FALSE | 0.86047225 | FALSE | 1 | FALSE | 1 | FALSE | 1 | FALSE | 1 | FALSE | 1 |
| 6789 | 6789 | 9.75 | 1.63935963 | 0 | 1 | 0 | 1 | 1 | 0.44344817 | 1 | 0.52895088 | 2 | 0.02752114 | 12 | 0.90922186 | FALSE | 1 | FALSE | 1 | FALSE | 1 | FALSE | 1 | FALSE | 0.79106376 | FALSE | 1 |
| 6790 | 6790 | 9.16666667 | 1.92209377 | 0 | 1 | 1 | 0.44710673 | 0 | 1 | 0 | 1 | 1 | 0.25766374 | 16 | 0.44998001 | FALSE | 1 | FALSE | 1 | FALSE | 1 | FALSE | 1 | FALSE | 1 | FALSE | 1 |
| 6791 | 6791 | 8.3030303 | 2.67972012 | 0 | 1 | 2 | 0.28941836 | 0 | 1 | 2 | 0.45159487 | 0 | 1 | 29 | 0.39648374 | FALSE | 1 | FALSE | 1 | FALSE | 1 | FALSE | 1 | FALSE | 1 | FALSE | 1 |
| 6792 | 6792 | 8.65625 | 1.57339948 | 2 | 0.19750261 | 1 | 0.65167362 | 0 | 1 | 0 | 1 | 0 | 1 | 29 | 0.24088541 | FALSE | 1 | FALSE | 1 | FALSE | 1 | FALSE | 1 | FALSE | 1 | FALSE | 1 |
| 6793 | 6793 | 7.725 | 2.63616673 | 1 | 0.64722818 | 0 | 1 | 1 | 0.76953771 | 2 | 0.5543522 | 0 | 1 | 36 | 0.22908535 | FALSE | 1 | FALSE | 1 | FALSE | 1 | FALSE | 1 | FALSE | 1 | FALSE | 1 |
| 6794 | 6794 | 1.04045436 | 0.58286749 | 103 | 0.99999206 | 152 | 0.94121298 | 174 | 0.81150581 | 226 | 0.69317293 | 94 | 0.00357336 | 4244 | 0.00994176 | FALSE | 1 | FALSE | 1 | FALSE | 1 | FALSE | 1 | FALSE | 0.34848744 | FALSE | 1 |
| 6795 | 6795 | 9.76923077 | 2.15384615 | 0 | 1 | 0 | 1 | 2 | 0.07725803 | 0 | 1 | 0 | 1 | 11 | 0.6683056 | FALSE | 1 | FALSE | 1 | FALSE | 1 | FALSE | 1 | FALSE | 1 | FALSE | 1 |
| 6796 | 6796 | 7.5625 | 4.63638801 | 2 | 0.06197875 | 1 | 0.40942991 | 0 | 1 | 1 | 0.52895088 | 0 | 1 | 12 | 0.90922186 | FALSE | 0.72532843 | FALSE | 1 | FALSE | 1 | FALSE | 1 | FALSE | 1 | FALSE | 1 |
| 6797 | 6797 | 8.40625 | 3.12109131 | 1 | 0.56527493 | 2 | 0.27730767 | 0 | 1 | 1 | 0.77852168 | 0 | 1 | 28 | 0.42532177 | FALSE | 1 | FALSE | 1 | FALSE | 1 | FALSE | 1 | FALSE | 1 | FALSE | 1 |
| 6798 | 6798 | 9.375 | 1.57619003 | 0 | 1 | 0 | 1 | 2 | 0.03118552 | 0 | 1 | 1 | 0.12394282 | 5 | 0.97551477 | FALSE | 1 | FALSE | 1 | FALSE | 1 | FALSE | 1 | FALSE | 1 | FALSE | 1 |
| 6799 | 6799 | 9.13043478 | 2.11216657 | 0 | 1 | 2 | 0.16948764 | 1 | 0.5695037 | 4 | 0.01932195 | 0 | 1 | 16 | 0.98100709 | FALSE | 1 | FALSE | 1 | FALSE | 1 | FALSE | 1 | FALSE | 1 | FALSE | 1 |
| 6800 | 6800 | 9 | 3.2829526 | 0 | 1 | 2 | 0.1135776 | 0 | 1 | 1 | 0.57131013 | 0 | 1 | 15 | 0.69372602 | FALSE | 1 | FALSE | 1 | FALSE | 1 | FALSE | 1 | FALSE | 1 | FALSE | 1 |
| 6801 | 6801 | 8.71428571 | 2.78846019 | 0 | 1 | 0 | 1 | 0 | 1 | 1 | 0.48242099 | 0 | 1 | 13 | 0.33283729 | FALSE | 1 | FALSE | 1 | FALSE | 1 | FALSE | 1 | FALSE | 1 | FALSE | 1 |
| 6802 | 6802 | 8.75675676 | 3.33211568 | 1 | 0.6184714 | 3 | 0.11640364 | 5 | 0.00977427 | 3 | 0.2402032 | 1 | 0.45845922 | 24 | 0.99915994 | FALSE | 1 | FALSE | 1 | FALSE | 1 | FALSE | 1 | FALSE | 1 | FALSE | 1 |
| 6803 | 6803 | 10.8181818 | 1.64140637 | 0 | 1 | 0 | 1 | 1 | 0.33149969 | 1 | 0.40390032 | 0 | 1 | 9 | 0.75985366 | FALSE | 1 | FALSE | 1 | FALSE | 1 | FALSE | 1 | FALSE | 1 | FALSE | 1 |
| 6804 | 6804 | 9.5 | 1.72547754 | 0 | 1 | 0 | 1 | 0 | 1 | 1 | 0.64497439 | 1 | 0.30529951 | 20 | 0.30915202 | FALSE | 1 | FALSE | 1 | FALSE | 1 | FALSE | 1 | FALSE | 1 | FALSE | 1 |
| 6805 | 6805 | 4.06779661 | 4.42015571 | 6 | 0.08204291 | 3 | 0.74112112 | 8 | 0.06139522 | 4 | 0.79788193 | 2 | 0.58049166 | 95 | 0.89984303 | FALSE | 0.79236883 | FALSE | 1 | FALSE | 1 | FALSE | 1 | FALSE | 1 | FALSE | 1 |
| 6806 | 6806 | 9.81481482 | 2.1436735 | 2 | 0.15144274 | 0 | 1 | 0 | 1 | 3 | 0.12475952 | 0 | 1 | 22 | 0.76237877 | FALSE | 1 | FALSE | 1 | FALSE | 1 | FALSE | 1 | FALSE | 1 | FALSE | 1 |
| 6807 | 6807 | 7.29166667 | 2.82074409 | 1 | 0.46441868 | 0 | 1 | 0 | 1 | 0 | 1 | 1 | 0.32796862 | 22 | 0.25237063 | FALSE | 1 | FALSE | 1 | FALSE | 1 | FALSE | 1 | FALSE | 1 | FALSE | 1 |
| 6808 | 6808 | 10.625 | 1.49478259 | 0 | 1 | 0 | 1 | 0 | 1 | 0 | 1 | 0 | 1 | 8 | 0.25677166 | FALSE | 1 | FALSE | 1 | FALSE | 1 | FALSE | 1 | FALSE | 1 | FALSE | 1 |
| 6809 | 6809 | 10.9 | 3.14483704 | 0 | 1 | 2 | 0.03949327 | 0 | 1 | 0 | 1 | 0 | 1 | 8 | 0.80394719 | FALSE | 1 | FALSE | 0.96864929 | FALSE | 1 | FALSE | 1 | FALSE | 1 | FALSE | 1 |
| 6810 | 6810 | 9 | 3.62215406 | 0 | 1 | 0 | 1 | 0 | 1 | 4 | 0.02569595 | 1 | 0.33902672 | 20 | 0.8152498 | FALSE | 1 | FALSE | 1 | FALSE | 1 | FALSE | 1 | FALSE | 1 | FALSE | 1 |
| 6811 | 6811 | 3.39340659 | 1.77440919 | 19 | 0.0220014 | 9 | 0.96192585 | 13 | 0.83596878 | 17 | 0.83887323 | 8 | 0.46480169 | 386 | 0.29409919 | FALSE | 0.4692294 | FALSE | 1 | FALSE | 1 | FALSE | 1 | FALSE | 1 | FALSE | 1 |
| 6812 | 6812 | 10.1666667 | 2.40947205 | 0 | 1 | 1 | 0.44710673 | 1 | 0.48280983 | 0 | 1 | 0 | 1 | 16 | 0.44998001 | FALSE | 1 | FALSE | 1 | FALSE | 1 | FALSE | 1 | FALSE | 1 | FALSE | 1 |
| 6813 | 6813 | 9.625 | 1.40867846 | 2 | 0.06197875 | 0 | 1 | 1 | 0.44344817 | 0 | 1 | 0 | 1 | 13 | 0.76799161 | FALSE | 0.72532843 | FALSE | 1 | FALSE | 1 | FALSE | 1 | FALSE | 1 | FALSE | 1 |
| 6814 | 6814 | 8.73913044 | 2.48853705 | 1 | 0.45027683 | 1 | 0.53115654 | 1 | 0.5695037 | 1 | 0.6613267 | 1 | 0.31672722 | 18 | 0.86273155 | FALSE | 1 | FALSE | 1 | FALSE | 1 | FALSE | 1 | FALSE | 1 | FALSE | 1 |
| 6815 | 6815 | 8.65217391 | 2.51272753 | 0 | 1 | 0 | 1 | 0 | 1 | 0 | 1 | 0 | 1 | 23 | 0.01996981 | FALSE | 1 | FALSE | 1 | FALSE | 1 | FALSE | 1 | FALSE | 1 | FALSE | 1 |
| 6816 | 6816 | 10.4285714 | 4.59369577 | 0 | 1 | 0 | 1 | 0 | 1 | 0 | 1 | 0 | 1 | 7 | 0.3043657 | FALSE | 1 | FALSE | 1 | FALSE | 1 | FALSE | 1 | FALSE | 1 | FALSE | 1 |
| 6817 | 6817 | 6.16326531 | 3.44843614 | 0 | 1 | 0 | 1 | 2 | 0.53023996 | 2 | 0.66536008 | 1 | 0.55647171 | 44 | 0.20085966 | FALSE | 1 | FALSE | 1 | FALSE | 1 | FALSE | 1 | FALSE | 1 | FALSE | 1 |
| 6818 | 6818 | 1.55795148 | 2.16284731 | 12 | 0.23961485 | 9 | 0.85446592 | 13 | 0.57495892 | 13 | 0.87713556 | 4 | 0.8641841 | 319 | 0.17729167 | FALSE | 1 | FALSE | 1 | FALSE | 1 | FALSE | 1 | FALSE | 1 | FALSE | 1 |
| 6819 | 6819 | 6.91836735 | 2.25729416 | 1 | 0.72119464 | 3 | 0.21051417 | 3 | 0.25725031 | 0 | 1 | 2 | 0.19154766 | 40 | 0.77344956 | FALSE | 1 | FALSE | 1 | FALSE | 1 | FALSE | 1 | FALSE | 1 | FALSE | 1 |
| 6820 | 6820 | 8.52631579 | 2.62314438 | 0 | 1 | 1 | 0.46503795 | 0 | 1 | 0 | 1 | 0 | 1 | 18 | 0.17880694 | FALSE | 1 | FALSE | 1 | FALSE | 1 | FALSE | 1 | FALSE | 1 | FALSE | 1 |
| 6821 | 6821 | 9.51282051 | 1.05907059 | 2 | 0.26413583 | 0 | 1 | 3 | 0.16357218 | 2 | 0.54052736 | 0 | 1 | 32 | 0.74324845 | FALSE | 1 | FALSE | 1 | FALSE | 1 | FALSE | 1 | FALSE | 1 | FALSE | 1 |
| 6822 | 6822 | 8.23076923 | 2.08119219 | 0 | 1 | 0 | 1 | 0 | 1 | 0 | 1 | 3 | 0.0010882 | 10 | 0.86791642 | FALSE | 1 | FALSE | 1 | FALSE | 1 | FALSE | 1 | FALSE | 0.19379454 | FALSE | 1 |
| 6823 | 6823 | 8.75 | 1.63935963 | 0 | 1 | 0 | 1 | 0 | 1 | 1 | 0.17140665 | 0 | 1 | 3 | 0.8823539 | FALSE | 1 | FALSE | 1 | FALSE | 1 | FALSE | 1 | FALSE | 1 | FALSE | 1 |
| 6824 | 6824 | 7.48333333 | 2.02065721 | 3 | 0.18560124 | 3 | 0.28903336 | 2 | 0.62245518 | 3 | 0.50172293 | 1 | 0.61824432 | 46 | 0.89115248 | FALSE | 1 | FALSE | 1 | FALSE | 1 | FALSE | 1 | FALSE | 1 | FALSE | 1 |
| 6825 | 6825 | 8.71428571 | 2.15235988 | 1 | 0.50471032 | 0 | 1 | 1 | 0.62830986 | 4 | 0.03319522 | 0 | 1 | 21 | 0.88395936 | FALSE | 1 | FALSE | 1 | FALSE | 1 | FALSE | 1 | FALSE | 1 | FALSE | 1 |
| 6826 | 6826 | 7.15217391 | 2.27908631 | 10 | 0.00010095 | 2 | 0.79919134 | 1 | 0.96498586 | 4 | 0.60708936 | 2 | 0.44196567 | 72 | 0.93327888 | TRUE | 0.01824319 | FALSE | 1 | FALSE | 1 | FALSE | 1 | FALSE | 1 | FALSE | 1 |
| 6827 | 6827 | 10.0588235 | 1.89335023 | 0 | 1 | 2 | 0.10307112 | 0 | 1 | 1 | 0.55062772 | 0 | 1 | 14 | 0.7313145 | FALSE | 1 | FALSE | 1 | FALSE | 1 | FALSE | 1 | FALSE | 1 | FALSE | 1 |
| 6828 | 6828 | 6.23809524 | 2.45734588 | 0 | 1 | 0 | 1 | 4 | 0.0626749 | 3 | 0.30326943 | 0 | 1 | 35 | 0.67024784 | FALSE | 1 | FALSE | 1 | FALSE | 1 | FALSE | 1 | FALSE | 1 | FALSE | 1 |
| 6829 | 6829 | 8.6 | 3.02324329 | 0 | 1 | 0 | 1 | 1 | 0.51939841 | 1 | 0.60987146 | 0 | 1 | 18 | 0.37504645 | FALSE | 1 | FALSE | 1 | FALSE | 1 | FALSE | 1 | FALSE | 1 | FALSE | 1 |
| 6830 | 6830 | 8.79166667 | 2.43206303 | 0 | 1 | 0 | 1 | 0 | 1 | 2 | 0.30249307 | 1 | 0.32796862 | 21 | 0.47053137 | FALSE | 1 | FALSE | 1 | FALSE | 1 | FALSE | 1 | FALSE | 1 | FALSE | 1 |
| 6831 | 6831 | 9.25 | 2.53722289 | 0 | 1 | 0 | 1 | 0 | 1 | 1 | 0.52895088 | 0 | 1 | 15 | 0.26127859 | FALSE | 1 | FALSE | 1 | FALSE | 1 | FALSE | 1 | FALSE | 1 | FALSE | 1 |
| 6832 | 6832 | 8.72 | 3.00026666 | 1 | 0.47819878 | 3 | 0.04547096 | 0 | 1 | 1 | 0.69181315 | 0 | 1 | 20 | 0.8152498 | FALSE | 1 | FALSE | 1 | FALSE | 1 | FALSE | 1 | FALSE | 1 | FALSE | 1 |
| 6833 | 6833 | 6.63333333 | 2.22835266 | 0 | 1 | 2 | 0.25306331 | 1 | 0.66709967 | 0 | 1 | 1 | 0.39166989 | 26 | 0.48609328 | FALSE | 1 | FALSE | 1 | FALSE | 1 | FALSE | 1 | FALSE | 1 | FALSE | 1 |
| 6834 | 6834 | 9.04 | 1.37054734 | 0 | 1 | 1 | 0.56109832 | 1 | 0.59998149 | 0 | 1 | 0 | 1 | 23 | 0.22727183 | FALSE | 1 | FALSE | 1 | FALSE | 1 | FALSE | 1 | FALSE | 1 | FALSE | 1 |
| 6835 | 6835 | 8.77272727 | 2.89092339 | 0 | 1 | 2 | 0.15794013 | 2 | 0.18613256 | 1 | 0.64497439 | 0 | 1 | 17 | 0.8840357 | FALSE | 1 | FALSE | 1 | FALSE | 1 | FALSE | 1 | FALSE | 1 | FALSE | 1 |
| 6836 | 6836 | 9 | 3.02926899 | 1 | 0.35728305 | 0 | 1 | 0 | 1 | 0 | 1 | 0 | 1 | 16 | 0.23070182 | FALSE | 1 | FALSE | 1 | FALSE | 1 | FALSE | 1 | FALSE | 1 | FALSE | 1 |
| 6837 | 6837 | 9.39130435 | 2.01600402 | 1 | 0.45027683 | 0 | 1 | 3 | 0.04772439 | 2 | 0.2852236 | 1 | 0.31672722 | 16 | 0.98100709 | FALSE | 1 | FALSE | 1 | FALSE | 1 | FALSE | 1 | FALSE | 1 | FALSE | 1 |
| 6838 | 6838 | 1.37170596 | 1.71328448 | 24 | 0.10438179 | 23 | 0.55349979 | 29 | 0.27696545 | 34 | 0.45077956 | 16 | 0.12510733 | 592 | 0.93908464 | FALSE | 0.88702868 | FALSE | 1 | FALSE | 1 | FALSE | 1 | FALSE | 1 | FALSE | 1 |
| 6839 | 6839 | 8.33333333 | 3.29983165 | 2 | 0.05513784 | 0 | 1 | 0 | 1 | 2 | 0.14906763 | 1 | 0.2198258 | 10 | 0.97980046 | FALSE | 0.68849059 | FALSE | 1 | FALSE | 1 | FALSE | 1 | FALSE | 1 | FALSE | 1 |
| 6840 | 6840 | 10.0857143 | 1.25063249 | 7 | 2.36E-05 | 4 | 0.02543131 | 0 | 1 | 2 | 0.48233989 | 0 | 1 | 22 | 0.9995668 | TRUE | 0.00651577 | FALSE | 0.82367276 | FALSE | 1 | FALSE | 1 | FALSE | 1 | FALSE | 1 |
| 6841 | 6841 | 10.75 | 2.10653744 | 0 | 1 | 1 | 0.40942991 | 0 | 1 | 2 | 0.16544789 | 1 | 0.23264612 | 12 | 0.90922186 | FALSE | 1 | FALSE | 1 | FALSE | 1 | FALSE | 1 | FALSE | 1 | FALSE | 1 |
| 6842 | 6842 | 8.85 | 1.79652442 | 1 | 0.40558521 | 0 | 1 | 1 | 0.51939841 | 1 | 0.60987146 | 1 | 0.28187291 | 16 | 0.80826361 | FALSE | 1 | FALSE | 1 | FALSE | 1 | FALSE | 1 | FALSE | 1 | FALSE | 1 |
| 6843 | 6843 | 8.32258065 | 3.04150957 | 8 | 5.74E-07 | 2 | 0.25306331 | 1 | 0.66709967 | 0 | 1 | 1 | 0.39166989 | 18 | 0.99972884 | TRUE | 0.00052208 | FALSE | 1 | FALSE | 1 | FALSE | 1 | FALSE | 1 | FALSE | 1 |
| 6844 | 6844 | 6.65714286 | 4.28247497 | 1 | 0.58738519 | 2 | 0.30150529 | 3 | 0.12152681 | 0 | 1 | 2 | 0.1065437 | 26 | 0.92796052 | FALSE | 1 | FALSE | 1 | FALSE | 1 | FALSE | 1 | FALSE | 1 | FALSE | 1 |
| 6845 | 6845 | 10.6111111 | 2.05855637 | 0 | 1 | 0 | 1 | 0 | 1 | 1 | 0.55062772 | 2 | 0.03086174 | 14 | 0.7313145 | FALSE | 1 | FALSE | 1 | FALSE | 1 | FALSE | 1 | FALSE | 0.82559785 | FALSE | 1 |
| 6846 | 6846 | 9.48275862 | 2.44414416 | 1 | 0.52988231 | 0 | 1 | 1 | 0.65463982 | 2 | 0.3871811 | 0 | 1 | 25 | 0.51779518 | FALSE | 1 | FALSE | 1 | FALSE | 1 | FALSE | 1 | FALSE | 1 | FALSE | 1 |
| 6847 | 6847 | 8.85714286 | 2.4159335 | 0 | 1 | 0 | 1 | 0 | 1 | 0 | 1 | 0 | 1 | 14 | 0.09251304 | FALSE | 1 | FALSE | 1 | FALSE | 1 | FALSE | 1 | FALSE | 1 | FALSE | 1 |
| 6848 | 6848 | 9.17647059 | 2.30690254 | 0 | 1 | 0 | 1 | 0 | 1 | 1 | 0.55062772 | 0 | 1 | 16 | 0.23070182 | FALSE | 1 | FALSE | 1 | FALSE | 1 | FALSE | 1 | FALSE | 1 | FALSE | 1 |
| 6849 | 6849 | 9 | 2.63578611 | 1 | 0.38989951 | 0 | 1 | 1 | 0.50143826 | 2 | 0.21610812 | 0 | 1 | 15 | 0.83648191 | FALSE | 1 | FALSE | 1 | FALSE | 1 | FALSE | 1 | FALSE | 1 | FALSE | 1 |
| 6850 | 6850 | 7.4375 | 2.44869225 | 0 | 1 | 3 | 0.01368613 | 1 | 0.44344817 | 0 | 1 | 1 | 0.23264612 | 11 | 0.9718531 | FALSE | 1 | FALSE | 0.69207862 | FALSE | 1 | FALSE | 1 | FALSE | 1 | FALSE | 1 |
| 6851 | 6851 | 8.48 | 3.67826046 | 2 | 0.12501436 | 1 | 0.54637326 | 0 | 1 | 1 | 0.67692816 | 0 | 1 | 20 | 0.68270729 | FALSE | 0.95622564 | FALSE | 1 | FALSE | 1 | FALSE | 1 | FALSE | 1 | FALSE | 1 |
| 6852 | 6852 | 11.2857143 | 2.86427681 | 0 | 1 | 0 | 1 | 0 | 1 | 1 | 0.28044368 | 0 | 1 | 6 | 0.69920685 | FALSE | 1 | FALSE | 1 | FALSE | 1 | FALSE | 1 | FALSE | 1 | FALSE | 1 |
| 6853 | 6853 | 9.5 | 1.92209377 | 1 | 0.37380237 | 1 | 0.44710673 | 1 | 0.48280983 | 4 | 0.00799439 | 0 | 1 | 11 | 0.99651304 | FALSE | 1 | FALSE | 1 | FALSE | 1 | FALSE | 1 | FALSE | 1 | FALSE | 1 |
| 6854 | 6854 | 9.19811321 | 3.12452204 | 3 | 0.50146864 | 3 | 0.65968805 | 2 | 0.89330417 | 5 | 0.52348367 | 1 | 0.82320297 | 90 | 0.32600351 | FALSE | 1 | FALSE | 1 | FALSE | 1 | FALSE | 1 | FALSE | 1 | FALSE | 1 |
| 6855 | 6855 | 8.83333333 | 1.92209377 | 0 | 1 | 1 | 0.44710673 | 0 | 1 | 1 | 0.57131013 | 0 | 1 | 16 | 0.44998001 | FALSE | 1 | FALSE | 1 | FALSE | 1 | FALSE | 1 | FALSE | 1 | FALSE | 1 |
| 6856 | 6856 | 9.16666667 | 2.33927814 | 1 | 0.37380237 | 0 | 1 | 1 | 0.48280983 | 1 | 0.57131013 | 0 | 1 | 15 | 0.69372602 | FALSE | 1 | FALSE | 1 | FALSE | 1 | FALSE | 1 | FALSE | 1 | FALSE | 1 |
| 6857 | 6857 | 7.75 | 2.7545417 | 1 | 0.40558521 | 2 | 0.1353446 | 1 | 0.51939841 | 1 | 0.60987146 | 1 | 0.28187291 | 14 | 0.97325593 | FALSE | 1 | FALSE | 1 | FALSE | 1 | FALSE | 1 | FALSE | 1 | FALSE | 1 |
| 6858 | 6858 | 9.20930233 | 3.344959 | 1 | 0.67382914 | 1 | 0.75788114 | 1 | 0.79363407 | 2 | 0.59404271 | 0 | 1 | 38 | 0.31730615 | FALSE | 1 | FALSE | 1 | FALSE | 1 | FALSE | 1 | FALSE | 1 | FALSE | 1 |
| 6859 | 6859 | 1.34222222 | 1.7135394 | 1 | 0.99730901 | 7 | 0.59209634 | 3 | 0.98881527 | 6 | 0.94966516 | 3 | 0.71712337 | 204 | 0.00202966 | FALSE | 1 | FALSE | 1 | FALSE | 1 | FALSE | 1 | FALSE | 1 | FALSE | 1 |
| 6860 | 6860 | 8.2 | 1.97737199 | 0 | 1 | 1 | 0.7326167 | 2 | 0.42396974 | 1 | 0.84823834 | 0 | 1 | 36 | 0.22908535 | FALSE | 1 | FALSE | 1 | FALSE | 1 | FALSE | 1 | FALSE | 1 | FALSE | 1 |

| | | | | | | | | | | | | | | | | | | | | | | | | | | | |
|---|---|---|---|---|---|---|---|---|---|---|---|---|---|---|---|---|---|---|---|---|---|---|---|---|---|---|---|
| 6861 | 6861 | 8.32 | 3.06881085 | 0 | 1 | 2 | 0.19297191 | 2 | 0.22597287 | 2 | 0.31968852 | 0 | 1 | 19 | 0.91734261 | FALSE | 1 | FALSE | 1 | FALSE | 1 | FALSE | 1 | FALSE | 1 | FALSE | 1 |
| 6862 | 6862 | 6.20689655 | 1.02315639 | 11 | 0.88393333 | 19 | 0.50004046 | 20 | 0.59898194 | 27 | 0.48483213 | 11 | 0.34385538 | 488 | 0.43405859 | FALSE | 1 | FALSE | 1 | FALSE | 1 | FALSE | 1 | FALSE | 1 | FALSE | 1 |
| 6863 | 6863 | 11.32 | 3.17137194 | 0 | 1 | 2 | 0.19297191 | 1 | 0.59998149 | 1 | 0.69181315 | 0 | 1 | 21 | 0.64957167 | FALSE | 1 | FALSE | 1 | FALSE | 1 | FALSE | 1 | FALSE | 1 | FALSE | 1 |
| 6864 | 6864 | 8.44444444 | 2.24158233 | 0 | 1 | 2 | 0.1135776 | 0 | 1 | 0 | 1 | 1 | 0.25766374 | 15 | 0.69372602 | FALSE | 1 | FALSE | 1 | FALSE | 1 | FALSE | 1 | FALSE | 1 | FALSE | 1 |
| 6865 | 6865 | 10.7333333 | 2.23507395 | 1 | 0.32293355 | 1 | 0.38964394 | 0 | 1 | 1 | 0.50623194 | 0 | 1 | 12 | 0.80329917 | FALSE | 1 | FALSE | 1 | FALSE | 1 | FALSE | 1 | FALSE | 1 | FALSE | 1 |
| 6866 | 6866 | 8.76470588 | 2.83941609 | 1 | 0.35728305 | 0 | 1 | 0 | 1 | 2 | 0.18212134 | 0 | 1 | 14 | 0.7313145 | FALSE | 1 | FALSE | 1 | FALSE | 1 | FALSE | 1 | FALSE | 1 | FALSE | 1 |
| 6867 | 6867 | 10.7857143 | 2.75625448 | 0 | 1 | 0 | 1 | 0 | 1 | 0 | 1 | 0 | 1 | 28 | 0.0085123 | FALSE | 1 | FALSE | 1 | FALSE | 1 | FALSE | 1 | FALSE | 1 | FALSE | 1 |
| 6868 | 6868 | 6.26415094 | 3.26878284 | 3 | 0.15399149 | 2 | 0.51564773 | 0 | 1 | 1 | 0.91797304 | 4 | 0.01081442 | 43 | 0.80430518 | FALSE | 1 | FALSE | 1 | FALSE | 1 | FALSE | 1 | FALSE | 0.57142086 | FALSE | 1 |
| 6869 | 6869 | 9.36666667 | 1.42556032 | 1 | 0.54198714 | 0 | 1 | 3 | 0.09131526 | 0 | 1 | 0 | 1 | 26 | 0.48609328 | FALSE | 1 | FALSE | 1 | FALSE | 1 | FALSE | 1 | FALSE | 1 | FALSE | 1 |
| 6870 | 6870 | 7.80701754 | 1.34345191 | 6 | 0.00292136 | 2 | 0.54544984 | 3 | 0.32646887 | 2 | 0.73544289 | 1 | 0.60529938 | 42 | 0.97810254 | FALSE | 0.14653312 | FALSE | 1 | FALSE | 1 | FALSE | 1 | FALSE | 1 | FALSE | 1 |
| 6871 | 6871 | 11.0909091 | 1.975051 | 0 | 1 | 0 | 1 | 0 | 1 | 0 | 1 | 0 | 1 | 11 | 0.15414489 | FALSE | 1 | FALSE | 1 | FALSE | 1 | FALSE | 1 | FALSE | 1 | FALSE | 1 |
| 6872 | 6872 | 9.88888889 | 2.57959706 | 1 | 0.37380237 | 0 | 1 | 0 | 1 | 3 | 0.04687752 | 0 | 1 | 14 | 0.86287418 | FALSE | 1 | FALSE | 1 | FALSE | 1 | FALSE | 1 | FALSE | 1 | FALSE | 1 |
| 6873 | 6873 | 8 | 2.30587796 | 0 | 1 | 1 | 0.74131698 | 0 | 1 | 3 | 0.29054213 | 0 | 1 | 37 | 0.21027077 | FALSE | 1 | FALSE | 1 | FALSE | 1 | FALSE | 1 | FALSE | 1 | FALSE | 1 |
| 6874 | 6874 | 10.5625 | 2.44869225 | 1 | 0.34033054 | 0 | 1 | 1 | 0.44344817 | 1 | 0.52895088 | 0 | 1 | 13 | 0.76799161 | FALSE | 1 | FALSE | 1 | FALSE | 1 | FALSE | 1 | FALSE | 1 | FALSE | 1 |
| 6875 | 6875 | 10.25 | 2.86138079 | 0 | 1 | 0 | 1 | 0 | 1 | 0 | 1 | 0 | 1 | 8 | 0.25677166 | FALSE | 1 | FALSE | 1 | FALSE | 1 | FALSE | 1 | FALSE | 1 | FALSE | 1 |
| 6876 | 6876 | 7.26666667 | 1.43604395 | 0 | 1 | 0 | 1 | 1 | 0.42266264 | 1 | 0.50623194 | 0 | 1 | 13 | 0.57712805 | FALSE | 1 | FALSE | 1 | FALSE | 1 | FALSE | 1 | FALSE | 1 | FALSE | 1 |
| 6877 | 6877 | 6.25714286 | 3.29798949 | 2 | 0.20692137 | 1 | 0.6629941 | 2 | 0.33343871 | 1 | 0.78873814 | 1 | 0.42123867 | 26 | 0.86852567 | FALSE | 1 | FALSE | 1 | FALSE | 1 | FALSE | 1 | FALSE | 1 | FALSE | 1 |
| 6878 | 6878 | 9.4375 | 3.55261787 | 0 | 1 | 0 | 1 | 1 | 0.44344817 | 1 | 0.52895088 | 0 | 1 | 14 | 0.53308261 | FALSE | 1 | FALSE | 1 | FALSE | 1 | FALSE | 1 | FALSE | 1 | FALSE | 1 |
| 6879 | 6879 | 11.5 | 2.71313677 | 0 | 1 | 0 | 1 | 0 | 1 | 0 | 1 | 0 | 1 | 18 | 0.04681656 | FALSE | 1 | FALSE | 1 | FALSE | 1 | FALSE | 1 | FALSE | 1 | FALSE | 1 |
| 6880 | 6880 | 6.12121212 | 0.6632142 | 8 | 0.35768501 | 10 | 0.34105312 | 14 | 0.08833418 | 8 | 0.92088882 | 5 | 0.43027751 | 217 | 0.7883755 | FALSE | 1 | FALSE | 1 | FALSE | 1 | FALSE | 1 | FALSE | 1 | FALSE | 1 |
| 6881 | 6881 | 9.77777778 | 2.39340658 | 1 | 0.37380237 | 0 | 1 | 0 | 1 | 2 | 0.19902682 | 0 | 1 | 15 | 0.69372602 | FALSE | 1 | FALSE | 1 | FALSE | 1 | FALSE | 1 | FALSE | 1 | FALSE | 1 |
| 6882 | 6882 | 8.14285714 | 1.14582947 | 5 | 0.0638619 | 2 | 0.76136679 | 1 | 0.95461143 | 5 | 0.34217166 | 2 | 0.40216708 | 69 | 0.76906822 | FALSE | 0.73766 | FALSE | 1 | FALSE | 1 | FALSE | 1 | FALSE | 1 | FALSE | 1 |
| 6883 | 6883 | 3.32511556 | 1.83628914 | 19 | 0.2808262 | 23 | 0.32469694 | 26 | 0.27833437 | 40 | 0.02523475 | 6 | 0.95785865 | 525 | 0.95195204 | FALSE | 1 | FALSE | 1 | FALSE | 1 | FALSE | 1 | FALSE | 1 | FALSE | 1 |
| 6884 | 6884 | 8.66666667 | 2.31420733 | 1 | 0.79096998 | 0 | 1 | 1 | 0.8897349 | 1 | 0.94113525 | 1 | 0.63076845 | 56 | 0.03180207 | FALSE | 1 | FALSE | 1 | FALSE | 1 | FALSE | 1 | FALSE | 1 | FALSE | 1 |
| 6885 | 6885 | 8 | 3.06594194 | 0 | 1 | 0 | 1 | 0 | 1 | 1 | 0.60987146 | 0 | 1 | 19 | 0.15700255 | FALSE | 1 | FALSE | 1 | FALSE | 1 | FALSE | 1 | FALSE | 1 | FALSE | 1 |
| 6886 | 6886 | 8.46938776 | 3.15000512 | 3 | 0.13009168 | 2 | 0.47396556 | 2 | 0.53023996 | 3 | 0.39227316 | 2 | 0.19154766 | 37 | 0.96598649 | FALSE | 0.97964162 | FALSE | 1 | FALSE | 1 | FALSE | 1 | FALSE | 1 | FALSE | 1 |
| 6887 | 6887 | 10.5510204 | 2.39097986 | 0 | 1 | 2 | 0.47396556 | 2 | 0.53023996 | 1 | 0.90086419 | 2 | 0.19154766 | 42 | 0.493842 | FALSE | 1 | FALSE | 1 | FALSE | 1 | FALSE | 1 | FALSE | 1 | FALSE | 1 |
| 6888 | 6888 | 10.6363636 | 1.82272161 | 1 | 0.24866461 | 2 | 0.04725998 | 0 | 1 | 0 | 1 | 0 | 1 | 8 | 0.92152993 | FALSE | 1 | FALSE | 1 | FALSE | 1 | FALSE | 1 | FALSE | 1 | FALSE | 1 |
| 6889 | 6889 | 4.35632184 | 1.51032471 | 6 | 0.88423502 | 16 | 0.09206509 | 20 | 0.02278478 | 11 | 0.92714301 | 2 | 0.97977309 | 290 | 0.60237939 | FALSE | 1 | FALSE | 1 | FALSE | 1 | FALSE | 1 | FALSE | 1 | FALSE | 1 |
| 6890 | 6890 | 8.28571429 | 2.96234877 | 0 | 1 | 0 | 1 | 0 | 1 | 1 | 0.28044368 | 1 | 0.10932414 | 5 | 0.91847773 | FALSE | 1 | FALSE | 1 | FALSE | 1 | FALSE | 1 | FALSE | 1 | FALSE | 1 |
| 6891 | 6891 | 5.90163934 | 2.96263572 | 2 | 0.46653175 | 2 | 0.59222924 | 1 | 0.89373158 | 1 | 0.94386203 | 0 | 1 | 55 | 0.13951638 | FALSE | 1 | FALSE | 1 | FALSE | 1 | FALSE | 1 | FALSE | 1 | FALSE | 1 |
| 6892 | 6892 | 11.4 | 1.45143607 | 0 | 1 | 1 | 0.38964394 | 0 | 1 | 0 | 1 | 0 | 1 | 14 | 0.29525409 | FALSE | 1 | FALSE | 1 | FALSE | 1 | FALSE | 1 | FALSE | 1 | FALSE | 1 |
| 6893 | 6893 | 7.90476191 | 2.89342048 | 0 | 1 | 0 | 1 | 2 | 0.16019548 | 0 | 1 | 0 | 1 | 18 | 0.37504645 | FALSE | 1 | FALSE | 1 | FALSE | 1 | FALSE | 1 | FALSE | 1 | FALSE | 1 |
| 6894 | 6894 | 9.57142857 | 1.61308426 | 2 | 0.16048326 | 0 | 1 | 1 | 0.64171563 | 2 | 0.3705495 | 2 | 0.07640798 | 21 | 0.94083768 | FALSE | 1 | FALSE | 1 | FALSE | 1 | FALSE | 1 | FALSE | 1 | FALSE | 1 |
| 6895 | 6895 | 6.07142857 | 3.22853982 | 3 | 0.17275086 | 4 | 0.10650784 | 1 | 0.87219471 | 5 | 0.11290142 | 1 | 0.60529938 | 42 | 0.97810254 | FALSE | 1 | FALSE | 1 | FALSE | 1 | FALSE | 1 | FALSE | 1 | FALSE | 1 |
| 6896 | 6896 | 8.34782609 | 2.97246342 | 1 | 0.45027683 | 1 | 0.53115654 | 3 | 0.04772439 | 2 | 0.2852236 | 0 | 1 | 16 | 0.98100709 | FALSE | 1 | FALSE | 1 | FALSE | 1 | FALSE | 1 | FALSE | 1 | FALSE | 1 |
| 6897 | 6897 | 6.89473684 | 2.40382889 | 0 | 1 | 0 | 1 | 0 | 1 | 1 | 0.59104357 | 1 | 0.26986775 | 17 | 0.41141543 | FALSE | 1 | FALSE | 1 | FALSE | 1 | FALSE | 1 | FALSE | 1 | FALSE | 1 |
| 6898 | 6898 | 10.9393939 | 2.60465078 | 5 | 0.00088509 | 1 | 0.6278847 | 1 | 0.66709967 | 3 | 0.15707004 | 0 | 1 | 20 | 0.99601577 | FALSE | 0.07039485 | FALSE | 1 | FALSE | 1 | FALSE | 1 | FALSE | 1 | FALSE | 1 |
| 6899 | 6899 | 10.4166667 | 1.97730175 | 0 | 1 | 0 | 1 | 1 | 0.35555293 | 0 | 1 | 0 | 1 | 11 | 0.41944532 | FALSE | 1 | FALSE | 1 | FALSE | 1 | FALSE | 1 | FALSE | 1 | FALSE | 1 |
| 6900 | 6900 | 11.3157895 | 3.32538295 | 2 | 0.08405377 | 1 | 0.46503795 | 0 | 1 | 1 | 0.59104357 | 1 | 0.26986775 | 14 | 0.93677253 | FALSE | 0.79236883 | FALSE | 1 | FALSE | 1 | FALSE | 1 | FALSE | 1 | FALSE | 1 |
| 6901 | 6901 | 9.27272727 | 2.73332661 | 0 | 1 | 0 | 1 | 0 | 1 | 1 | 0.40390032 | 2 | 0.01330324 | 8 | 0.92152993 | FALSE | 1 | FALSE | 1 | FALSE | 1 | FALSE | 1 | FALSE | 0.63731057 | FALSE | 1 |
| 6902 | 6902 | 10.631579 | 2.20550809 | 0 | 1 | 2 | 0.12434464 | 0 | 1 | 1 | 0.59104357 | 0 | 1 | 16 | 0.65565778 | FALSE | 1 | FALSE | 1 | FALSE | 1 | FALSE | 1 | FALSE | 1 | FALSE | 1 |
| 6903 | 6903 | 6.61403509 | 1.48388284 | 1 | 0.77387657 | 1 | 0.84773229 | 2 | 0.61293419 | 7 | 0.01477784 | 0 | 1 | 46 | 0.83074268 | FALSE | 1 | FALSE | 1 | FALSE | 1 | FALSE | 1 | FALSE | 1 | FALSE | 1 |
| 6904 | 6904 | 9.59259259 | 1.91019214 | 0 | 1 | 1 | 0.58913618 | 0 | 1 | 3 | 0.12475952 | 0 | 1 | 23 | 0.58307573 | FALSE | 1 | FALSE | 1 | FALSE | 1 | FALSE | 1 | FALSE | 1 | FALSE | 1 |
| 6905 | 6905 | 11.0833333 | 1.60511336 | 0 | 1 | 0 | 1 | 1 | 0.35555293 | 1 | 0.43131129 | 0 | 1 | 10 | 0.71436964 | FALSE | 1 | FALSE | 1 | FALSE | 1 | FALSE | 1 | FALSE | 1 | FALSE | 1 |
| 6906 | 6906 | 9.92857143 | 2.76364888 | 2 | 0.04858661 | 0 | 1 | 0 | 1 | 0 | 1 | 1 | 0.20679328 | 11 | 0.83676449 | FALSE | 0.649301 | FALSE | 1 | FALSE | 1 | FALSE | 1 | FALSE | 1 | FALSE | 1 |
| 6907 | 6907 | 10 | 1 | 0 | 1 | 0 | 1 | 0 | 1 | 0 | 1 | 0 | 1 | 2 | 0.71196516 | FALSE | 1 | FALSE | 1 | FALSE | 1 | FALSE | 1 | FALSE | 1 | FALSE | 1 |
| 6908 | 6908 | 9.5 | 2.12132034 | 1 | 0.34033054 | 2 | 0.09285443 | 0 | 1 | 1 | 0.52895088 | 0 | 1 | 12 | 0.90922186 | FALSE | 1 | FALSE | 1 | FALSE | 1 | FALSE | 1 | FALSE | 1 | FALSE | 1 |
| 6909 | 6909 | 12.1212121 | 3.06255564 | 2 | 0.20692137 | 0 | 1 | 0 | 1 | 3 | 0.19161876 | 1 | 0.42123867 | 27 | 0.75124583 | FALSE | 1 | FALSE | 1 | FALSE | 1 | FALSE | 1 | FALSE | 1 | FALSE | 1 |
| 6910 | 6910 | 10.0476191 | 3.15438003 | 0 | 1 | 1 | 0.48239024 | 1 | 0.51939841 | 1 | 0.60987146 | 1 | 0.28187291 | 16 | 0.80826361 | FALSE | 1 | FALSE | 1 | FALSE | 1 | FALSE | 1 | FALSE | 1 | FALSE | 1 |
| 6911 | 6911 | 10.3125 | 2.02233621 | 0 | 1 | 0 | 1 | 4 | 0.00210122 | 1 | 0.52895088 | 2 | 0.02752114 | 9 | 0.99862766 | FALSE | 1 | FALSE | 1 | FALSE | 1 | FALSE | 1 | FALSE | 0.79106376 | FALSE | 1 |
| 6912 | 6912 | 11.75 | 3.92905841 | 1 | 0.18769745 | 2 | 0.02563923 | 0 | 1 | 1 | 0.31351192 | 0 | 1 | 4 | 0.99658732 | FALSE | 1 | FALSE | 0.82367276 | FALSE | 1 | FALSE | 1 | FALSE | 1 | FALSE | 1 |
| 6913 | 6913 | 6.7 | 1.29881603 | 6 | 0.11331032 | 4 | 0.605178 | 4 | 0.68714676 | 2 | 0.98391625 | 3 | 0.35494204 | 110 | 0.4472263 | FALSE | 0.92546178 | FALSE | 1 | FALSE | 1 | FALSE | 1 | FALSE | 1 | FALSE | 1 |
| 6914 | 6914 | 8.5 | 2.42185561 | 3 | 0.0279333 | 1 | 0.57534752 | 1 | 0.61440466 | 1 | 0.70601446 | 1 | 0.3499045 | 19 | 0.96054244 | FALSE | 0.50964275 | FALSE | 1 | FALSE | 1 | FALSE | 1 | FALSE | 1 | FALSE | 1 |
| 6915 | 6915 | 11.3846154 | 1.64262742 | 0 | 1 | 1 | 0.3480709 | 2 | 0.07725803 | 0 | 1 | 0 | 1 | 10 | 0.86791642 | FALSE | 1 | FALSE | 1 | FALSE | 1 | FALSE | 1 | FALSE | 1 | FALSE | 1 |
| 6916 | 6916 | 8.61538462 | 3.38636319 | 1 | 0.28675951 | 2 | 0.06426041 | 0 | 1 | 0 | 1 | 0 | 1 | 10 | 0.86791642 | FALSE | 1 | FALSE | 1 | FALSE | 1 | FALSE | 1 | FALSE | 1 | FALSE | 1 |
| 6917 | 6917 | 11.1333333 | 2.27644948 | 0 | 1 | 0 | 1 | 2 | 0.09928574 | 0 | 1 | 0 | 1 | 13 | 0.57712805 | FALSE | 1 | FALSE | 1 | FALSE | 1 | FALSE | 1 | FALSE | 1 | FALSE | 1 |
| 6918 | 6918 | 8.73684211 | 2.0350121 | 0 | 1 | 0 | 1 | 1 | 0.75193685 | 1 | 0.83318955 | 1 | 0.4673875 | 35 | 0.13407055 | FALSE | 1 | FALSE | 1 | FALSE | 1 | FALSE | 1 | FALSE | 1 | FALSE | 1 |
| 6919 | 6919 | 10.5 | 2.29128785 | 0 | 1 | 0 | 1 | 1 | 0.30655256 | 2 | 0.0740981 | 0 | 1 | 7 | 0.94326215 | FALSE | 1 | FALSE | 1 | FALSE | 1 | FALSE | 1 | FALSE | 1 | FALSE | 1 |
| 6920 | 6920 | 8.77777778 | 1.39664501 | 0 | 1 | 0 | 1 | 0 | 1 | 1 | 0.34506517 | 0 | 1 | 8 | 0.5780332 | FALSE | 1 | FALSE | 1 | FALSE | 1 | FALSE | 1 | FALSE | 1 | FALSE | 1 |
| 6921 | 6921 | 10.1818182 | 1.40247715 | 0 | 1 | 0 | 1 | 0 | 1 | 0 | 1 | 0 | 1 | 11 | 0.15414489 | FALSE | 1 | FALSE | 1 | FALSE | 1 | FALSE | 1 | FALSE | 1 | FALSE | 1 |
| 6922 | 6922 | 9.4 | 2.05912603 | 0 | 1 | 0 | 1 | 0 | 1 | 1 | 0.2094688 | 0 | 1 | 4 | 0.82371636 | FALSE | 1 | FALSE | 1 | FALSE | 1 | FALSE | 1 | FALSE | 1 | FALSE | 1 |
| 6923 | 6923 | 8.94230769 | 2.3567698 | 0 | 1 | 0 | 1 | 2 | 0.5626042 | 2 | 0.6970685 | 2 | 0.2095319 | 46 | 0.27620659 | FALSE | 1 | FALSE | 1 | FALSE | 1 | FALSE | 1 | FALSE | 1 | FALSE | 1 |
| 6924 | 6924 | 7.68918919 | 2.04626011 | 5 | 0.0407209 | 2 | 0.69680912 | 3 | 0.49993836 | 1 | 0.96971789 | 1 | 0.70774043 | 62 | 0.63133902 | FALSE | 0.626405 | FALSE | 1 | FALSE | 1 | FALSE | 1 | FALSE | 1 | FALSE | 1 |
| 6925 | 6925 | 5.11462451 | 1.26627881 | 7 | 0.46952844 | 8 | 0.57346933 | 6 | 0.89628638 | 10 | 0.72852371 | 8 | 0.05427074 | 213 | 0.51705403 | FALSE | 1 | FALSE | 1 | FALSE | 1 | FALSE | 1 | FALSE | 1 | FALSE | 1 |
| 6926 | 6926 | 8.78947368 | 1.96224757 | 0 | 1 | 0 | 1 | 1 | 0.50143826 | 2 | 0.21610812 | 1 | 0.26986775 | 15 | 0.83648191 | FALSE | 1 | FALSE | 1 | FALSE | 1 | FALSE | 1 | FALSE | 1 | FALSE | 1 |
| 6927 | 6927 | 9.25 | 3.08017857 | 0 | 1 | 1 | 0.48239024 | 1 | 0.51939841 | 2 | 0.23331368 | 0 | 1 | 16 | 0.80826361 | FALSE | 1 | FALSE | 1 | FALSE | 1 | FALSE | 1 | FALSE | 1 | FALSE | 1 |
| 6928 | 6928 | 7.48993289 | 1.76702628 | 2 | 0.89993885 | 5 | 0.53133611 | 3 | 0.90865001 | 9 | 0.24505707 | 4 | 0.22794223 | 126 | 0.53043058 | FALSE | 1 | FALSE | 1 | FALSE | 1 | FALSE | 1 | FALSE | 1 | FALSE | 1 |
| 6929 | 6929 | 9.88888889 | 2.07869855 | 0 | 1 | 1 | 0.25628163 | 1 | 0.28067846 | 0 | 1 | 0 | 1 | 7 | 0.84574168 | FALSE | 1 | FALSE | 1 | FALSE | 1 | FALSE | 1 | FALSE | 1 | FALSE | 1 |
| 6930 | 6930 | 8.39393939 | 1.92179515 | 1 | 0.57647348 | 2 | 0.28941836 | 1 | 0.70185639 | 0 | 1 | 0 | 1 | 29 | 0.39648374 | FALSE | 1 | FALSE | 1 | FALSE | 1 | FALSE | 1 | FALSE | 1 | FALSE | 1 |
| 6931 | 6931 | 5.00403226 | 1.71684722 | 14 | 0.0041319 | 9 | 0.40108891 | 13 | 0.10461065 | 7 | 0.94041813 | 4 | 0.57839774 | 199 | 0.94476234 | FALSE | 0.18198109 | FALSE | 1 | FALSE | 1 | FALSE | 1 | FALSE | 1 | FALSE | 1 |
| 6932 | 6932 | 9.1875 | 2.32429425 | 0 | 1 | 1 | 0.40942991 | 0 | 1 | 1 | 0.52895088 | 0 | 1 | 14 | 0.53308261 | FALSE | 1 | FALSE | 1 | FALSE | 1 | FALSE | 1 | FALSE | 1 | FALSE | 1 |
| 6933 | 6933 | 8.48051948 | 1.55930712 | 2 | 0.59191779 | 1 | 0.92163631 | 3 | 0.52680756 | 2 | 0.87529005 | 0 | 1 | 69 | 0.13014053 | FALSE | 1 | FALSE | 1 | FALSE | 1 | FALSE | 1 | FALSE | 1 | FALSE | 1 |
| 6934 | 6934 | 8.36842105 | 2.90572079 | 1 | 0.38989951 | 1 | 0.46503795 | 1 | 0.50143826 | 1 | 0.59104357 | 1 | 0.26986775 | 14 | 0.93677253 | FALSE | 1 | FALSE | 1 | FALSE | 1 | FALSE | 1 | FALSE | 1 | FALSE | 1 |
| 6935 | 6935 | 8.06666667 | 2.59401019 | 0 | 1 | 1 | 0.6278847 | 2 | 0.29330414 | 1 | 0.75658797 | 1 | 0.39166989 | 25 | 0.67630471 | FALSE | 1 | FALSE | 1 | FALSE | 1 | FALSE | 1 | FALSE | 1 | FALSE | 1 |
| 6936 | 6936 | 8.6875 | 3.47704526 | 0 | 1 | 1 | 0.40942991 | 0 | 1 | 0 | 1 | 1 | 0.23264612 | 14 | 0.53308261 | FALSE | 1 | FALSE | 1 | FALSE | 1 | FALSE | 1 | FALSE | 1 | FALSE | 1 |
| 6937 | 6937 | 10.5333333 | 3.13829395 | 2 | 0.05513784 | 1 | 0.38964394 | 1 | 0.42266264 | 0 | 1 | 2 | 0.02433796 | 9 | 0.995488 | FALSE | 0.68849059 | FALSE | 1 | FALSE | 1 | FALSE | 1 | FALSE | 0.77578444 | FALSE | 1 |
| 6938 | 6938 | 3.20554273 | 1.19921712 | 12 | 0.43319497 | 17 | 0.2356469 | 12 | 0.86248681 | 20 | 0.52330678 | 6 | 0.72263963 | 366 | 0.4979219 | FALSE | 1 | FALSE | 1 | FALSE | 1 | FALSE | 1 | FALSE | 1 | FALSE | 1 |
| 6939 | 6939 | 4.76595745 | 2.80738211 | 3 | 0.11868304 | 0 | 1 | 0 | 1 | 3 | 0.36699324 | 0 | 1 | 41 | 0.38322554 | FALSE | 0.94393345 | FALSE | 1 | FALSE | 1 | FALSE | 1 | FALSE | 1 | FALSE | 1 |
| 6940 | 6940 | 8.82758621 | 2.75517026 | 1 | 0.52988231 | 1 | 0.61539065 | 0 | 1 | 2 | 0.3871811 | 1 | 0.38148475 | 24 | 0.70563334 | FALSE | 1 | FALSE | 1 | FALSE | 1 | FALSE | 1 | FALSE | 1 | FALSE | 1 |
| 6941 | 6941 | 9.84615385 | 2.38089039 | 0 | 1 | 0 | 1 | 1 | 0.35555293 | 0 | 1 | 0 | 1 | 11 | 0.41944532 | FALSE | 1 | FALSE | 1 | FALSE | 1 | FALSE | 1 | FALSE | 1 | FALSE | 1 |
| 6942 | 6942 | 5.51851852 | 2.59087245 | 3 | 0.52695321 | 3 | 0.6846759 | 3 | 0.75139368 | 4 | 0.73784936 | 5 | 0.03181538 | 90 | 0.67694631 | FALSE | 1 | FALSE | 1 | FALSE | 1 | FALSE | 1 | FALSE | 0.84074695 | FALSE | 1 |
| 6943 | 6943 | 9.41666667 | 2.75252409 | 0 | 1 | 1 | 0.32623933 | 1 | 0.35555293 | 0 | 1 | 2 | 0.01579445 | 8 | 0.97197833 | FALSE | 1 | FALSE | 1 | FALSE | 1 | FALSE | 1 | FALSE | 0.66313193 | FALSE | 1 |
| 6944 | 6944 | 10.0714286 | 3.85383456 | 0 | 1 | 1 | 0.60247911 | 2 | 0.2663542 | 0 | 1 | 1 | 0.37113067 | 24 | 0.55018107 | FALSE | 1 | FALSE | 1 | FALSE | 1 | FALSE | 1 | FALSE | 1 | FALSE | 1 |
| 6945 | 6945 | 9.07142857 | 2.63124864 | 0 | 1 | 0 | 1 | 0 | 1 | 1 | 0.48242099 | 0 | 1 | 13 | 0.33283729 | FALSE | 1 | FALSE | 1 | FALSE | 1 | FALSE | 1 | FALSE | 1 | FALSE | 1 |
| 6946 | 6946 | 9.375 | 1.84419133 | 0 | 1 | 2 | 0.18117183 | 0 | 1 | 1 | 0.67692816 | 0 | 1 | 21 | 0.47053137 | FALSE | 1 | FALSE | 1 | FALSE | 1 | FALSE | 1 | FALSE | 1 | FALSE | 1 |
| 6947 | 6947 | 6.28571429 | 3.36852175 | 0 | 1 | 0 | 1 | 1 | 0.64171563 | 1 | 0.73248983 | 0 | 1 | 26 | 0.16405547 | FALSE | 1 | FALSE | 1 | FALSE | 1 | FALSE | 1 | FALSE | 1 | FALSE | 1 |
| 6948 | 6948 | 8.33333333 | 2.40370085 | 0 | 1 | 0 | 1 | 0 | 1 | 0 | 1 | 1 | 0.25766374 | 17 | 0.20329205 | FALSE | 1 | FALSE | 1 | FALSE | 1 | FALSE | 1 | FALSE | 1 | FALSE | 1 |
| 6949 | 6949 | 10.8571429 | 1.72614943 | 0 | 1 | 0 | 1 | 1 | 0.22601089 | 0 | 1 | 0 | 1 | 6 | 0.69920685 | FALSE | 1 | FALSE | 1 | FALSE | 1 | FALSE | 1 | FALSE | 1 | FALSE | 1 |
| 6950 | 6950 | 3.28526646 | 1.17048439 | 8 | 0.57035297 | 13 | 0.22577881 | 9 | 0.81197001 | 14 | 0.60031401 | 2 | 0.96948886 | 271 | 0.32081199 | FALSE | 1 | FALSE | 1 | FALSE | 1 | FALSE | 1 | FALSE | 1 | FALSE | 1 |
| 6951 | 6951 | 8.30769231 | 2.3821327 | 0 | 1 | 2 | 0.19297191 | 0 | 1 | 1 | 0.69181315 | 1 | 0.33902672 | 21 | 0.64957167 | FALSE | 1 | FALSE | 1 | FALSE | 1 | FALSE | 1 | FALSE | 1 | FALSE | 1 |
| 6952 | 6952 | 9.08 | 2.38193199 | 1 | 0.47819878 | 0 | 1 | 1 | 0.59998149 | 3 | 0.10474541 | 0 | 1 | 20 | 0.8152498 | FALSE | 1 | FALSE | 1 | FALSE | 1 | FALSE | 1 | FALSE | 1 | FALSE | 1 |
| 6953 | 6953 | 9.91666667 | 1.84654332 | 0 | 1 | 1 | 0.28036922 | 1 | 0.30655256 | 2 | 0.0740981 | 0 | 1 | 6 | 0.98831471 | FALSE | 1 | FALSE | 1 | FALSE | 1 | FALSE | 1 | FALSE | 1 | FALSE | 1 |
| 6954 | 6954 | 10.25 | 1.88272219 | 0 | 1 | 2 | 0.2288771 | 0 | 1 | 1 | 0.73248983 | 0 | 1 | 25 | 0.34293 | FALSE | 1 | FALSE | 1 | FALSE | 1 | FALSE | 1 | FALSE | 1 | FALSE | 1 |

| | | | | | | | | | | | | | | | | | | | | | | | | | | | |
|---|---|---|---|---|---|---|---|---|---|---|---|---|---|---|---|---|---|---|---|---|---|---|---|---|---|---|---|
| 6955 | 6955 | 10 | 2.03540098 | 9 | 1.76E-08 | 2 | 0.2288771 | 0 | 1 | 0 | 1 | 0 | 1 | 17 | 0.99947263 | TRUE | 2.89E-05 | FALSE | 1 | FALSE | 1 | FALSE | 1 | FALSE | 1 | FALSE | 1 |
| 6956 | 6956 | 8.86666667 | 2.36267269 | 0 | 1 | 0 | 1 | 0 | 1 | 4 | 0.00397592 | 1 | 0.2198258 | 10 | 0.97980046 | FALSE | 1 | FALSE | 1 | FALSE | 1 | FALSE | 1 | FALSE | 1 | FALSE | 1 |
| 6957 | 6957 | 7.06349206 | 2.82209545 | 0 | 1 | 1 | 0.87100665 | 2 | 0.65873577 | 2 | 0.78520976 | 0 | 1 | 57 | 0.06268769 | FALSE | 1 | FALSE | 1 | FALSE | 1 | FALSE | 1 | FALSE | 1 | FALSE | 1 |
| 6958 | 6958 | 10.4285714 | 2.25876976 | 1 | 0.16630142 | 0 | 1 | 0 | 1 | 2 | 0.03785362 | 0 | 1 | 4 | 0.98605104 | FALSE | 1 | FALSE | 1 | FALSE | 1 | FALSE | 1 | FALSE | 1 | FALSE | 1 |
| 6959 | 6959 | 6.71428571 | 3.17515868 | 1 | 0.72119464 | 1 | 0.80150109 | 1 | 0.83455705 | 1 | 0.90086419 | 1 | 0.55647171 | 44 | 0.20085966 | FALSE | 1 | FALSE | 1 | FALSE | 1 | FALSE | 1 | FALSE | 1 | FALSE | 1 |
| 6960 | 6960 | 7.10858586 | 0.78869988 | 16 | 0.04312066 | 14 | 0.38989579 | 14 | 0.54963691 | 17 | 0.63411512 | 10 | 0.10816497 | 321 | 0.92738867 | FALSE | 0.63647651 | FALSE | 1 | FALSE | 1 | FALSE | 1 | FALSE | 1 | FALSE | 1 |
| 6961 | 6961 | 9.23529412 | 2.26298636 | 0 | 1 | 1 | 0.42857733 | 0 | 1 | 0 | 1 | 0 | 1 | 16 | 0.23070182 | FALSE | 1 | FALSE | 1 | FALSE | 1 | FALSE | 1 | FALSE | 1 | FALSE | 1 |
| 6962 | 6962 | 8.92307692 | 2.12897731 | 0 | 1 | 0 | 1 | 1 | 0.61440466 | 2 | 0.33678042 | 0 | 1 | 23 | 0.40372818 | FALSE | 1 | FALSE | 1 | FALSE | 1 | FALSE | 1 | FALSE | 1 | FALSE | 1 |
| 6963 | 6963 | 9.3125 | 2.11301769 | 1 | 0.34033054 | 0 | 1 | 0 | 1 | 0 | 1 | 0 | 1 | 15 | 0.26127859 | FALSE | 1 | FALSE | 1 | FALSE | 1 | FALSE | 1 | FALSE | 1 | FALSE | 1 |
| 6964 | 6964 | 6.40740741 | 2.95345697 | 3 | 0.15399149 | 3 | 0.24492539 | 4 | 0.12170726 | 1 | 0.91797304 | 0 | 1 | 42 | 0.88654191 | FALSE | 1 | FALSE | 1 | FALSE | 1 | FALSE | 1 | FALSE | 1 | FALSE | 1 |
| 6965 | 6965 | 10.1 | 3.7 | 2 | 0.0256865 | 0 | 1 | 0 | 1 | 0 | 1 | 0 | 1 | 8 | 0.80394719 | FALSE | 0.4974559 | FALSE | 1 | FALSE | 1 | FALSE | 1 | FALSE | 1 | FALSE | 1 |
| 6966 | 6966 | 10.5384615 | 2.43738146 | 0 | 1 | 1 | 0.3480709 | 0 | 1 | 0 | 1 | 0 | 1 | 12 | 0.374198 | FALSE | 1 | FALSE | 1 | FALSE | 1 | FALSE | 1 | FALSE | 1 | FALSE | 1 |
| 6967 | 6967 | 9.60606061 | 2.72962861 | 1 | 0.57647348 | 0 | 1 | 2 | 0.33343871 | 1 | 0.78873814 | 1 | 0.42123867 | 28 | 0.58681477 | FALSE | 1 | FALSE | 1 | FALSE | 1 | FALSE | 1 | FALSE | 1 | FALSE | 1 |
| 6968 | 6968 | 9.6 | 2.2 | 0 | 1 | 0 | 1 | 0 | 1 | 2 | 0.23331368 | 0 | 1 | 18 | 0.37504645 | FALSE | 1 | FALSE | 1 | FALSE | 1 | FALSE | 1 | FALSE | 1 | FALSE | 1 |
| 6969 | 6969 | 9.57142857 | 3.9948947 | 2 | 0.09989125 | 0 | 1 | 3 | 0.03775825 | 1 | 0.62783522 | 0 | 1 | 15 | 0.96513691 | FALSE | 0.86047225 | FALSE | 1 | FALSE | 1 | FALSE | 1 | FALSE | 1 | FALSE | 1 |
| 6970 | 6970 | 10.2857143 | 3.66635744 | 0 | 1 | 0 | 1 | 1 | 0.42266264 | 1 | 0.50623194 | 1 | 0.2198258 | 12 | 0.80329917 | FALSE | 1 | FALSE | 1 | FALSE | 1 | FALSE | 1 | FALSE | 1 | FALSE | 1 |
| 6971 | 6971 | 8.75 | 2.68095132 | 0 | 1 | 1 | 0.53115654 | 2 | 0.19931429 | 3 | 0.08614834 | 1 | 0.31672722 | 16 | 0.98100709 | FALSE | 1 | FALSE | 1 | FALSE | 1 | FALSE | 1 | FALSE | 1 | FALSE | 1 |
| 6972 | 6972 | 8.60869565 | 1.27651463 | 0 | 1 | 0 | 1 | 4 | 0.00839283 | 0 | 1 | 0 | 1 | 19 | 0.71542496 | FALSE | 1 | FALSE | 1 | FALSE | 1 | FALSE | 1 | FALSE | 1 | FALSE | 1 |
| 6973 | 6973 | 10.5714286 | 1.56817846 | 0 | 1 | 5 | 0.00180813 | 2 | 0.2663542 | 0 | 1 | 0 | 1 | 21 | 0.94083768 | FALSE | 1 | FALSE | 0.40311621 | FALSE | 1 | FALSE | 1 | FALSE | 1 | FALSE | 1 |
| 6974 | 6974 | 10.25 | 3.83242743 | 2 | 0.06197875 | 0 | 1 | 2 | 0.11087977 | 2 | 0.16544789 | 0 | 1 | 10 | 0.99304605 | FALSE | 0.72532843 | FALSE | 1 | FALSE | 1 | FALSE | 1 | FALSE | 1 | FALSE | 1 |
| 6975 | 6975 | 9.09090909 | 2.42916168 | 1 | 0.24866461 | 1 | 0.30368014 | 0 | 1 | 0 | 1 | 2 | 0.01330324 | 7 | 0.98129353 | FALSE | 1 | FALSE | 1 | FALSE | 1 | FALSE | 1 | FALSE | 0.63731057 | FALSE | 1 |
| 6976 | 6976 | 11.1176471 | 1.77837841 | 1 | 0.35728305 | 0 | 1 | 0 | 1 | 1 | 0.55062772 | 0 | 1 | 15 | 0.49060435 | FALSE | 1 | FALSE | 1 | FALSE | 1 | FALSE | 1 | FALSE | 1 | FALSE | 1 |
| 6977 | 6977 | 7.08695652 | 3.06683584 | 0 | 1 | 3 | 0.5460764 | 5 | 0.20871337 | 4 | 0.58033022 | 1 | 0.76878158 | 75 | 0.48424212 | FALSE | 1 | FALSE | 1 | FALSE | 1 | FALSE | 1 | FALSE | 1 | FALSE | 1 |
| 6978 | 6978 | 9.35714286 | 2.5803417 | 1 | 0.3050805 | 2 | 0.07341559 | 1 | 0.40110418 | 1 | 0.48242099 | 0 | 1 | 9 | 0.98607548 | FALSE | 1 | FALSE | 1 | FALSE | 1 | FALSE | 1 | FALSE | 1 | FALSE | 1 |
| 6979 | 6979 | 7.23809524 | 1.57070691 | 0 | 1 | 0 | 1 | 0 | 1 | 0 | 1 | 0 | 1 | 21 | 0.02808151 | FALSE | 1 | FALSE | 1 | FALSE | 1 | FALSE | 1 | FALSE | 1 | FALSE | 1 |
| 6980 | 6980 | 9.82352941 | 2.61748182 | 1 | 0.35728305 | 0 | 1 | 0 | 1 | 4 | 0.00644706 | 0 | 1 | 12 | 0.962093 | FALSE | 1 | FALSE | 1 | FALSE | 1 | FALSE | 1 | FALSE | 1 | FALSE | 1 |
| 6981 | 6981 | 4.4057971 | 2.76268532 | 1 | 0.83491628 | 1 | 0.89776066 | 0 | 1 | 4 | 0.39127216 | 1 | 0.6822729 | 62 | 0.13493316 | FALSE | 1 | FALSE | 1 | FALSE | 1 | FALSE | 1 | FALSE | 1 | FALSE | 1 |
| 6982 | 6982 | 2.88990826 | 2.19816207 | 1 | 0.99683831 | 8 | 0.40905102 | 5 | 0.89842512 | 8 | 0.79127262 | 2 | 0.87806905 | 194 | 0.03097699 | FALSE | 1 | FALSE | 1 | FALSE | 1 | FALSE | 1 | FALSE | 1 | FALSE | 1 |
| 6983 | 6983 | 9.67647059 | 1.42912125 | 3 | 0.05546921 | 0 | 1 | 2 | 0.34668321 | 1 | 0.7984848 | 2 | 0.1065437 | 26 | 0.92796052 | FALSE | 0.68849059 | FALSE | 1 | FALSE | 1 | FALSE | 1 | FALSE | 1 | FALSE | 1 |
| 6984 | 6984 | 9.73076923 | 3.43687797 | 1 | 0.49162634 | 1 | 0.57534752 | 1 | 0.61440466 | 0 | 1 | 0 | 1 | 23 | 0.40372818 | FALSE | 1 | FALSE | 1 | FALSE | 1 | FALSE | 1 | FALSE | 1 | FALSE | 1 |
| 6985 | 6985 | 9.61538462 | 2.76067053 | 0 | 1 | 0 | 1 | 1 | 0.37874417 | 1 | 0.45746571 | 0 | 1 | 11 | 0.6683056 | FALSE | 1 | FALSE | 1 | FALSE | 1 | FALSE | 1 | FALSE | 1 | FALSE | 1 |
| 6986 | 6986 | 11.7272727 | 1.86308196 | 0 | 1 | 0 | 1 | 1 | 0.33149969 | 1 | 0.40390032 | 0 | 1 | 9 | 0.75985366 | FALSE | 1 | FALSE | 1 | FALSE | 1 | FALSE | 1 | FALSE | 1 | FALSE | 1 |
| 6987 | 6987 | 6.14730291 | 0.99640588 | 16 | 0.1698678 | 25 | 0.01195178 | 14 | 0.8313638 | 17 | 0.90034546 | 18 | 0.00074614 | 391 | 0.97545938 | FALSE | 1 | FALSE | 0.67617265 | FALSE | 1 | FALSE | 1 | FALSE | 0.14553221 | FALSE | 1 |
| 6988 | 6988 | 9.30555556 | 1.12594268 | 0 | 1 | 3 | 0.10944025 | 2 | 0.37289966 | 1 | 0.81665383 | 1 | 0.44938266 | 29 | 0.81028289 | FALSE | 1 | FALSE | 1 | FALSE | 1 | FALSE | 1 | FALSE | 1 | FALSE | 1 |
| 6989 | 6989 | 9.54545455 | 2.16852008 | 1 | 0.4357638 | 0 | 1 | 0 | 1 | 1 | 0.64497439 | 0 | 1 | 20 | 0.30915202 | FALSE | 1 | FALSE | 1 | FALSE | 1 | FALSE | 1 | FALSE | 1 | FALSE | 1 |
| 6990 | 6990 | 5.41935484 | 1.18523697 | 0 | 1 | 2 | 0.60114557 | 3 | 0.3858942 | 6 | 0.06375189 | 0 | 1 | 51 | 0.74578096 | FALSE | 1 | FALSE | 1 | FALSE | 1 | FALSE | 1 | FALSE | 1 | FALSE | 1 |
| 6991 | 6991 | 9.95238095 | 2.53501555 | 1 | 0.42086991 | 1 | 0.49918218 | 2 | 0.17308361 | 2 | 0.25059627 | 0 | 1 | 15 | 0.96513691 | FALSE | 1 | FALSE | 1 | FALSE | 1 | FALSE | 1 | FALSE | 1 | FALSE | 1 |
| 6992 | 6992 | 10.0588235 | 2.23529412 | 0 | 1 | 0 | 1 | 0 | 1 | 1 | 0.55062772 | 1 | 0.24525764 | 15 | 0.49060435 | FALSE | 1 | FALSE | 1 | FALSE | 1 | FALSE | 1 | FALSE | 1 | FALSE | 1 |
| 6993 | 6993 | 4.69124424 | 2.01642263 | 13 | 0.00171726 | 6 | 0.6283331 | 8 | 0.42403496 | 13 | 0.12586144 | 5 | 0.22745152 | 154 | 0.99684867 | FALSE | 0.10850113 | FALSE | 1 | FALSE | 1 | FALSE | 1 | FALSE | 1 | FALSE | 1 |
| 6994 | 6994 | 8.25 | 3.230712 | 0 | 1 | 2 | 0.18117183 | 0 | 1 | 0 | 1 | 0 | 1 | 22 | 0.25237063 | FALSE | 1 | FALSE | 1 | FALSE | 1 | FALSE | 1 | FALSE | 1 | FALSE | 1 |
| 6995 | 6995 | 10.1764706 | 1.75684524 | 2 | 0.04858661 | 0 | 1 | 2 | 0.08806398 | 1 | 0.48242099 | 0 | 1 | 9 | 0.98607548 | FALSE | 0.649301 | FALSE | 1 | FALSE | 1 | FALSE | 1 | FALSE | 1 | FALSE | 1 |
| 6996 | 6996 | 7.21428571 | 4.38631918 | 2 | 0.16048326 | 2 | 0.2288771 | 2 | 0.2663542 | 3 | 0.13524571 | 0 | 1 | 19 | 0.99262381 | FALSE | 1 | FALSE | 1 | FALSE | 1 | FALSE | 1 | FALSE | 1 | FALSE | 1 |
| 6997 | 6997 | 8.35 | 2.72534402 | 1 | 0.40558521 | 1 | 0.48239024 | 0 | 1 | 0 | 1 | 1 | 0.28187291 | 17 | 0.61750632 | FALSE | 1 | FALSE | 1 | FALSE | 1 | FALSE | 1 | FALSE | 1 | FALSE | 1 |
| 6998 | 6998 | 7.77358491 | 3.85414019 | 1 | 0.74224579 | 0 | 1 | 3 | 0.28676219 | 2 | 0.6970685 | 0 | 1 | 46 | 0.27620659 | FALSE | 1 | FALSE | 1 | FALSE | 1 | FALSE | 1 | FALSE | 1 | FALSE | 1 |
| 6999 | 6999 | 7.94444444 | 2.81146243 | 3 | 0.16017108 | 3 | 0.25367018 | 2 | 0.58328102 | 3 | 0.45415642 | 1 | 0.59191961 | 42 | 0.93138539 | FALSE | 1 | FALSE | 1 | FALSE | 1 | FALSE | 1 | FALSE | 1 | FALSE | 1 |
| 7000 | 7000 | 7.36363636 | 1.29844153 | 0 | 1 | 0 | 1 | 0 | 1 | 3 | 0.00903718 | 0 | 1 | 7 | 0.94326215 | FALSE | 1 | FALSE | 1 | FALSE | 1 | FALSE | 1 | FALSE | 1 | FALSE | 1 |
| 7001 | 7001 | 10.7272727 | 2.71664596 | 2 | 0.10809809 | 1 | 0.5154318 | 1 | 0.55340851 | 0 | 1 | 0 | 1 | 18 | 0.74744813 | FALSE | 0.88820416 | FALSE | 1 | FALSE | 1 | FALSE | 1 | FALSE | 1 | FALSE | 1 |
| 7002 | 7002 | 10.1290323 | 2.58708873 | 0 | 1 | 1 | 0.63997468 | 3 | 0.09854319 | 0 | 1 | 0 | 1 | 27 | 0.45522639 | FALSE | 1 | FALSE | 1 | FALSE | 1 | FALSE | 1 | FALSE | 1 | FALSE | 1 |
| 7003 | 7003 | 7.74509804 | 2.1497208 | 3 | 0.14187072 | 1 | 0.81422651 | 1 | 0.84631478 | 2 | 0.68678423 | 0 | 1 | 44 | 0.44552698 | FALSE | 1 | FALSE | 1 | FALSE | 1 | FALSE | 1 | FALSE | 1 | FALSE | 1 |
| 7004 | 7004 | 8.96153846 | 1.97056746 | 2 | 0.14250978 | 0 | 1 | 1 | 0.61440466 | 4 | 0.02930308 | 2 | 0.0670823 | 17 | 0.99598795 | FALSE | 1 | FALSE | 1 | FALSE | 1 | FALSE | 1 | FALSE | 1 | FALSE | 1 |
| 7005 | 7005 | 11.2857143 | 3.42460088 | 1 | 0.42086991 | 0 | 1 | 0 | 1 | 1 | 0.62783522 | 0 | 1 | 19 | 0.34094959 | FALSE | 1 | FALSE | 1 | FALSE | 1 | FALSE | 1 | FALSE | 1 | FALSE | 1 |
| 7006 | 7006 | 4.3 | 2.13931453 | 2 | 0.45796754 | 4 | 0.12838945 | 4 | 0.16812274 | 4 | 0.29632241 | 5 | 0.0028848 | 41 | 0.99949885 | FALSE | 1 | FALSE | 1 | FALSE | 1 | FALSE | 1 | FALSE | 0.29951111 | FALSE | 1 |
| 7007 | 7007 | 7.78048781 | 2.73649406 | 0 | 1 | 3 | 0.13080858 | 0 | 1 | 1 | 0.84089119 | 0 | 1 | 35 | 0.24915769 | FALSE | 1 | FALSE | 1 | FALSE | 1 | FALSE | 1 | FALSE | 1 | FALSE | 1 |
| 7008 | 7008 | 10.0769231 | 2.75852632 | 1 | 0.28675951 | 1 | 0.3480709 | 0 | 1 | 0 | 1 | 0 | 1 | 11 | 0.6683056 | FALSE | 1 | FALSE | 1 | FALSE | 1 | FALSE | 1 | FALSE | 1 | FALSE | 1 |
| 7009 | 7009 | 9.73913044 | 3.2466356 | 1 | 0.45027683 | 1 | 0.53115654 | 1 | 0.5695037 | 1 | 0.6613267 | 1 | 0.31672722 | 18 | 0.86273155 | FALSE | 1 | FALSE | 1 | FALSE | 1 | FALSE | 1 | FALSE | 1 | FALSE | 1 |
| 7010 | 7010 | 8.1875 | 3.98777429 | 0 | 1 | 2 | 0.09285443 | 1 | 0.44344817 | 1 | 0.52895088 | 0 | 1 | 12 | 0.90922186 | FALSE | 1 | FALSE | 1 | FALSE | 1 | FALSE | 1 | FALSE | 1 | FALSE | 1 |
| 7011 | 7011 | 11 | 3.94968353 | 1 | 0.12181417 | 0 | 1 | 0 | 1 | 1 | 0.2094688 | 0 | 1 | 3 | 0.97031021 | FALSE | 0.95622564 | FALSE | 1 | FALSE | 1 | FALSE | 1 | FALSE | 1 | FALSE | 1 |
| 7012 | 7012 | 7.3 | 3.2572995 | 0 | 1 | 0 | 1 | 0 | 1 | 1 | 0.37517262 | 0 | 1 | 9 | 0.5215547 | FALSE | 1 | FALSE | 1 | FALSE | 1 | FALSE | 1 | FALSE | 1 | FALSE | 1 |
| 7013 | 7013 | 8.71428571 | 3.19438283 | 3 | 0.0048942 | 0 | 1 | 1 | 0.40110418 | 2 | 0.13304705 | 0 | 1 | 8 | 0.99722299 | FALSE | 0.19848171 | FALSE | 1 | FALSE | 1 | FALSE | 1 | FALSE | 1 | FALSE | 1 |
| 7014 | 7014 | 10.5652174 | 2.8866422 | 0 | 1 | 0 | 1 | 1 | 0.5695037 | 2 | 0.2852236 | 1 | 0.31672722 | 19 | 0.71542496 | FALSE | 1 | FALSE | 1 | FALSE | 1 | FALSE | 1 | FALSE | 1 | FALSE | 1 |
| 7015 | 7015 | 9.14285714 | 1.27375389 | 0 | 1 | 3 | 0.06039219 | 1 | 0.64171563 | 0 | 1 | 1 | 0.37113067 | 23 | 0.73439118 | FALSE | 1 | FALSE | 1 | FALSE | 1 | FALSE | 1 | FALSE | 1 | FALSE | 1 |
| 7016 | 7016 | 5.87142857 | 2.90281355 | 0 | 1 | 1 | 0.90110191 | 3 | 0.46296673 | 4 | 0.40179526 | 0 | 1 | 62 | 0.21402731 | FALSE | 1 | FALSE | 1 | FALSE | 1 | FALSE | 1 | FALSE | 1 | FALSE | 1 |
| 7017 | 7017 | 10.15 | 1.79652442 | 0 | 1 | 1 | 0.48239024 | 1 | 0.51939841 | 1 | 0.60987146 | 0 | 1 | 17 | 0.61750632 | FALSE | 1 | FALSE | 1 | FALSE | 1 | FALSE | 1 | FALSE | 1 | FALSE | 1 |
| 7018 | 7018 | 10.2592593 | 1.34969386 | 2 | 0.15144274 | 1 | 0.58913618 | 1 | 0.62830986 | 0 | 1 | 2 | 0.07169515 | 21 | 0.88395936 | FALSE | 1 | FALSE | 1 | FALSE | 1 | FALSE | 1 | FALSE | 1 | FALSE | 1 |
| 7019 | 7019 | 13.3846154 | 3.38636319 | 1 | 0.28675951 | 1 | 0.3480709 | 0 | 1 | 1 | 0.45746571 | 1 | 0.19354507 | 9 | 0.96018168 | FALSE | 1 | FALSE | 1 | FALSE | 1 | FALSE | 1 | FALSE | 1 | FALSE | 1 |
| 7020 | 7020 | 8.45945946 | 1.60395785 | 0 | 1 | 0 | 1 | 0 | 1 | 3 | 0.22784656 | 1 | 0.44938266 | 32 | 0.31709555 | FALSE | 1 | FALSE | 1 | FALSE | 1 | FALSE | 1 | FALSE | 1 | FALSE | 1 |
| 7021 | 7021 | 10.2 | 2.28619043 | 0 | 1 | 0 | 1 | 2 | 0.09928574 | 1 | 0.50623194 | 0 | 1 | 12 | 0.80329917 | FALSE | 1 | FALSE | 1 | FALSE | 1 | FALSE | 1 | FALSE | 1 | FALSE | 1 |
| 7022 | 7022 | 3.83333333 | 3.23975798 | 1 | 0.88571933 | 1 | 0.93582077 | 0 | 1 | 6 | 0.1806169 | 15 | 3.04E-12 | 60 | 0.99853511 | FALSE | 1 | FALSE | 1 | FALSE | 1 | TRUE | 2.49E-08 | FALSE | 1 | FALSE | 1 |
| 7023 | 7023 | 7.61111111 | 2.11184198 | 4 | 0.01296492 | 0 | 1 | 0 | 1 | 0 | 1 | 1 | 0.44938266 | 31 | 0.49869404 | FALSE | 0.35760478 | FALSE | 1 | FALSE | 1 | FALSE | 1 | FALSE | 1 | FALSE | 1 |
| 7024 | 7024 | 8.5 | 2.91547595 | 1 | 0.34033054 | 0 | 1 | 2 | 0.11087977 | 0 | 1 | 0 | 1 | 13 | 0.76799161 | FALSE | 1 | FALSE | 1 | FALSE | 1 | FALSE | 1 | FALSE | 1 | FALSE | 1 |
| 7025 | 7025 | 10.15 | 1.15217186 | 1 | 0.40558521 | 0 | 1 | 1 | 0.51939841 | 2 | 0.23331368 | 0 | 1 | 16 | 0.80826361 | FALSE | 1 | FALSE | 1 | FALSE | 1 | FALSE | 1 | FALSE | 1 | FALSE | 1 |
| 7026 | 7026 | 10.3888889 | 1.56839936 | 0 | 1 | 2 | 0.1135776 | 0 | 1 | 1 | 0.57131013 | 0 | 1 | 15 | 0.69372602 | FALSE | 1 | FALSE | 1 | FALSE | 1 | FALSE | 1 | FALSE | 1 | FALSE | 1 |
| 7027 | 7027 | 8.55555556 | 2.4317854 | 2 | 0.07645454 | 0 | 1 | 4 | 0.0033386 | 0 | 1 | 0 | 1 | 12 | 0.98544073 | FALSE | 0.77424195 | FALSE | 1 | FALSE | 1 | FALSE | 1 | FALSE | 1 | FALSE | 1 |
| 7028 | 7028 | 9.36 | 2.31309317 | 1 | 0.47819878 | 0 | 1 | 2 | 0.22597287 | 0 | 1 | 2 | 0.06257402 | 20 | 0.8152498 | FALSE | 1 | FALSE | 1 | FALSE | 1 | FALSE | 1 | FALSE | 1 | FALSE | 1 |
| 7029 | 7029 | 7.51612903 | 2.55103986 | 1 | 0.55378205 | 2 | 0.2651851 | 1 | 0.67911179 | 1 | 0.76781284 | 1 | 0.4016888 | 25 | 0.80006983 | FALSE | 1 | FALSE | 1 | FALSE | 1 | FALSE | 1 | FALSE | 1 | FALSE | 1 |
| 7030 | 7030 | 7.72131148 | 1.63096218 | 4 | 0.0671906 | 1 | 0.86215528 | 2 | 0.64095454 | 1 | 0.94113525 | 0 | 1 | 52 | 0.39208719 | FALSE | 0.74276693 | FALSE | 1 | FALSE | 1 | FALSE | 1 | FALSE | 1 | FALSE | 1 |
| 7031 | 7031 | 9 | 2.40613252 | 1 | 0.38989951 | 0 | 1 | 0 | 1 | 0 | 1 | 0 | 1 | 18 | 0.17880694 | FALSE | 1 | FALSE | 1 | FALSE | 1 | FALSE | 1 | FALSE | 1 | FALSE | 1 |
| 7032 | 7032 | 6.55963303 | 2.77085484 | 1 | 0.92886217 | 5 | 0.22809576 | 2 | 0.88350939 | 8 | 0.09174551 | 1 | 0.81407008 | 84 | 0.69121812 | FALSE | 1 | FALSE | 1 | FALSE | 1 | FALSE | 1 | FALSE | 1 | FALSE | 1 |
| 7033 | 7033 | 8.85714286 | 2.29462549 | 0 | 1 | 0 | 1 | 0 | 1 | 0 | 1 | 0 | 1 | 21 | 0.02808151 | FALSE | 1 | FALSE | 1 | FALSE | 1 | FALSE | 1 | FALSE | 1 | FALSE | 1 |
| 7034 | 7034 | 9.33333333 | 1.69967317 | 3 | 0.02256792 | 2 | 0.18117183 | 0 | 1 | 1 | 0.67692816 | 0 | 1 | 18 | 0.93166998 | FALSE | 0.4692294 | FALSE | 1 | FALSE | 1 | FALSE | 1 | FALSE | 1 | FALSE | 1 |
| 7035 | 7035 | 7.68888889 | 2.65683372 | 3 | 0.07268126 | 1 | 0.71433277 | 2 | 0.39868506 | 0 | 1 | 0 | 1 | 32 | 0.61800916 | FALSE | 0.76727438 | FALSE | 1 | FALSE | 1 | FALSE | 1 | FALSE | 1 | FALSE | 1 |
| 7036 | 7036 | 7.54285714 | 2.12929039 | 1 | 0.83919255 | 3 | 0.39589822 | 2 | 0.72288838 | 6 | 0.10101015 | 0 | 1 | 58 | 0.70710963 | FALSE | 1 | FALSE | 1 | FALSE | 1 | FALSE | 1 | FALSE | 1 | FALSE | 1 |
| 7037 | 7037 | 5.33802817 | 1.11258691 | 3 | 0.24604313 | 1 | 0.89073735 | 3 | 0.43448639 | 2 | 0.82021683 | 0 | 1 | 58 | 0.38535314 | FALSE | 1 | FALSE | 1 | FALSE | 1 | FALSE | 1 | FALSE | 1 | FALSE | 1 |
| 7038 | 7038 | 8.83333333 | 1.86338998 | 0 | 1 | 0 | 1 | 0 | 1 | 0 | 1 | 0 | 1 | 12 | 0.13002667 | FALSE | 1 | FALSE | 1 | FALSE | 1 | FALSE | 1 | FALSE | 1 | FALSE | 1 |
| 7039 | 7039 | 10.5454546 | 1.23315091 | 0 | 1 | 0 | 1 | 0 | 1 | 0 | 1 | 0 | 1 | 11 | 0.15414489 | FALSE | 1 | FALSE | 1 | FALSE | 1 | FALSE | 1 | FALSE | 1 | FALSE | 1 |
| 7040 | 7040 | 8.375 | 2.34187425 | 0 | 1 | 2 | 0.27730767 | 3 | 0.10599423 | 1 | 0.77852168 | 0 | 1 | 26 | 0.77612337 | FALSE | 1 | FALSE | 1 | FALSE | 1 | FALSE | 1 | FALSE | 1 | FALSE | 1 |
| 7041 | 7041 | 9.3 | 2.00249844 | 0 | 1 | 1 | 0.28036922 | 1 | 0.30655256 | 1 | 0.37517262 | 0 | 1 | 7 | 0.94326215 | FALSE | 1 | FALSE | 1 | FALSE | 1 | FALSE | 1 | FALSE | 1 | FALSE | 1 |
| 7042 | 7042 | 9.16666667 | 3.1841622 | 0 | 1 | 1 | 0.44710673 | 1 | 0.48280983 | 1 | 0.57131013 | 1 | 0.25766374 | 14 | 0.86287418 | FALSE | 1 | FALSE | 1 | FALSE | 1 | FALSE | 1 | FALSE | 1 | FALSE | 1 |
| 7043 | 7043 | 9.25925926 | 2.618914 | 1 | 0.50471032 | 0 | 1 | 0 | 1 | 3 | 0.12475952 | 0 | 1 | 23 | 0.58307573 | FALSE | 1 | FALSE | 1 | FALSE | 1 | FALSE | 1 | FALSE | 1 | FALSE | 1 |
| 7044 | 7044 | 7.96551724 | 2.17323466 | 2 | 0.16048326 | 0 | 1 | 2 | 0.2663542 | 2 | 0.3705495 | 0 | 1 | 22 | 0.86499992 | FALSE | 1 | FALSE | 1 | FALSE | 1 | FALSE | 1 | FALSE | 1 | FALSE | 1 |
| 7045 | 7045 | 9.9375 | 2.2767507 | 2 | 0.06197875 | 0 | 1 | 0 | 1 | 1 | 0.52895088 | 0 | 1 | 13 | 0.76799161 | FALSE | 0.72532843 | FALSE | 1 | FALSE | 1 | FALSE | 1 | FALSE | 1 | FALSE | 1 |
| 7046 | 7046 | 8.22727273 | 1.8812911 | 1 | 0.4357638 | 1 | 0.5154318 | 0 | 1 | 2 | 0.26791271 | 1 | 0.30529951 | 17 | 0.8840357 | FALSE | 1 | FALSE | 1 | FALSE | 1 | FALSE | 1 | FALSE | 1 | FALSE | 1 |
| 7047 | 7047 | 9.8 | 3.34065862 | 1 | 0.22886536 | 1 | 0.28036922 | 1 | 0.30655256 | 0 | 1 | 0 | 1 | 7 | 0.94326215 | FALSE | 1 | FALSE | 1 | FALSE | 1 | FALSE | 1 | FALSE | 1 | FALSE | 1 |
| 7048 | 7048 | 9.875 | 3.19912097 | 0 | 1 | 0 | 1 | 1 | 0.44344817 | 2 | 0.16544789 | 1 | 0.23264612 | 12 | 0.90922186 | FALSE | 1 | FALSE | 1 | FALSE | 1 | FALSE | 1 | FALSE | 1 | FALSE | 1 |

| | | | | | | | | | | | | | | | | | | | | | | | | | | | |
|---|---|---|---|---|---|---|---|---|---|---|---|---|---|---|---|---|---|---|---|---|---|---|---|---|---|---|---|
| 7049 | 7049 | 11.0666667 | 2.95446931 | 1 | 0.32293355 | 0 | 1 | 1 | 0.42266264 | 0 | 1 | 0 | 1 | 13 | 0.57712805 | FALSE | 1 | FALSE | 1 | FALSE | 1 | FALSE | 1 | FALSE | 1 | FALSE | 1 |
| 7050 | 7050 | 8.62068966 | 2.38405324 | 0 | 1 | 2 | 0.2288771 | 1 | 0.64171563 | 0 | 1 | 2 | 0.07640798 | 23 | 0.73439118 | FALSE | 1 | FALSE | 1 | FALSE | 1 | FALSE | 1 | FALSE | 1 | FALSE | 1 |
| 7051 | 7051 | 11.25 | 3.00347022 | 0 | 1 | 1 | 0.32623933 | 1 | 0.35555293 | 1 | 0.43131129 | 0 | 1 | 9 | 0.89630573 | FALSE | 1 | FALSE | 1 | FALSE | 1 | FALSE | 1 | FALSE | 1 | FALSE | 1 |
| 7052 | 7052 | 10.8666667 | 3.86206623 | 0 | 1 | 1 | 0.38964394 | 2 | 0.09928574 | 0 | 1 | 0 | 1 | 12 | 0.80329917 | FALSE | 1 | FALSE | 1 | FALSE | 1 | FALSE | 1 | FALSE | 1 | FALSE | 1 |
| 7053 | 7053 | 6.26666667 | 2.26890646 | 2 | 0.75154282 | 7 | 0.05088873 | 2 | 0.89330417 | 2 | 0.9558593 | 1 | 0.82320297 | 90 | 0.32600351 | FALSE | 1 | FALSE | 1 | FALSE | 1 | FALSE | 1 | FALSE | 1 | FALSE | 1 |
| 7054 | 7054 | 9.875 | 1.49478259 | 1 | 0.34033054 | 3 | 0.01368613 | 0 | 1 | 1 | 0.52895088 | 1 | 0.23264612 | 10 | 0.99304605 | FALSE | 1 | FALSE | 0.69207862 | FALSE | 1 | FALSE | 1 | FALSE | 1 | FALSE | 1 |
| 7055 | 7055 | 6.22727273 | 4.18798952 | 0 | 1 | 0 | 1 | 1 | 0.55340851 | 3 | 0.07744186 | 0 | 1 | 18 | 0.74744813 | FALSE | 1 | FALSE | 1 | FALSE | 1 | FALSE | 1 | FALSE | 1 | FALSE | 1 |
| 7056 | 7056 | 8.33333333 | 1.49071199 | 0 | 1 | 1 | 0.179097 | 0 | 1 | 4 | 6.07E-05 | 0 | 1 | 1 | 0.99998565 | FALSE | 1 | FALSE | 1 | FALSE | 1 | FALSE | 0.29727645 | FALSE | 1 | FALSE | 1 |
| 7057 | 7057 | 7.07692308 | 2.63297254 | 0 | 1 | 1 | 0.8752178 | 3 | 0.39570451 | 1 | 0.94894362 | 1 | 0.64879513 | 57 | 0.11815964 | FALSE | 1 | FALSE | 1 | FALSE | 1 | FALSE | 1 | FALSE | 1 | FALSE | 1 |
| 7058 | 7058 | 10.2727273 | 4.51443964 | 0 | 1 | 1 | 0.30368014 | 0 | 1 | 1 | 0.40390032 | 0 | 1 | 9 | 0.75985366 | FALSE | 1 | FALSE | 1 | FALSE | 1 | FALSE | 1 | FALSE | 1 | FALSE | 1 |
| 7059 | 7059 | 9.58333333 | 2.25308431 | 0 | 1 | 0 | 1 | 0 | 1 | 2 | 0.1023796 | 0 | 1 | 10 | 0.71436964 | FALSE | 1 | FALSE | 1 | FALSE | 1 | FALSE | 1 | FALSE | 1 | FALSE | 1 |
| 7060 | 7060 | 10.35 | 2.90301567 | 0 | 1 | 1 | 0.48239024 | 1 | 0.51939841 | 2 | 0.23331368 | 0 | 1 | 16 | 0.80826361 | FALSE | 1 | FALSE | 1 | FALSE | 1 | FALSE | 1 | FALSE | 1 | FALSE | 1 |
| 7061 | 7061 | 8.69047619 | 3.42600764 | 0 | 1 | 0 | 1 | 3 | 0.19060367 | 2 | 0.5811115 | 2 | 0.15059682 | 35 | 0.67024784 | FALSE | 1 | FALSE | 1 | FALSE | 1 | FALSE | 1 | FALSE | 1 | FALSE | 1 |
| 7062 | 7062 | 8.24 | 1.74997143 | 1 | 0.47819878 | 0 | 1 | 1 | 0.59998149 | 0 | 1 | 0 | 1 | 23 | 0.22727183 | FALSE | 1 | FALSE | 1 | FALSE | 1 | FALSE | 1 | FALSE | 1 | FALSE | 1 |
| 7063 | 7063 | 9.33333333 | 2.4267033 | 0 | 1 | 0 | 1 | 1 | 0.35555293 | 0 | 1 | 0 | 1 | 11 | 0.41944532 | FALSE | 1 | FALSE | 1 | FALSE | 1 | FALSE | 1 | FALSE | 1 | FALSE | 1 |
| 7064 | 7064 | 9.65517241 | 2.70728089 | 1 | 0.51745948 | 0 | 1 | 1 | 0.64171563 | 2 | 0.3705495 | 0 | 1 | 24 | 0.55018107 | FALSE | 1 | FALSE | 1 | FALSE | 1 | FALSE | 1 | FALSE | 1 | FALSE | 1 |
| 7065 | 7065 | 9.5 | 1.56980436 | 1 | 0.51745948 | 0 | 1 | 2 | 0.2663542 | 2 | 0.3705495 | 0 | 1 | 23 | 0.73439118 | FALSE | 1 | FALSE | 1 | FALSE | 1 | FALSE | 1 | FALSE | 1 | FALSE | 1 |
| 7066 | 7066 | 8.46153846 | 2.79051648 | 2 | 0.26413583 | 2 | 0.36122049 | 2 | 0.41139386 | 2 | 0.54052736 | 0 | 1 | 31 | 0.85613899 | FALSE | 1 | FALSE | 1 | FALSE | 1 | FALSE | 1 | FALSE | 1 | FALSE | 1 |
| 7067 | 7067 | 7.44444444 | 1.16534317 | 0 | 1 | 0 | 1 | 0 | 1 | 3 | 0.22784656 | 3 | 0.0206828 | 30 | 0.67248271 | FALSE | 1 | FALSE | 1 | FALSE | 1 | FALSE | 1 | FALSE | 0.7366672 | FALSE | 1 |
| 7068 | 7068 | 7.54166667 | 4.10263296 | 0 | 1 | 0 | 1 | 0 | 1 | 1 | 0.64497439 | 0 | 1 | 21 | 0.12048848 | FALSE | 1 | FALSE | 1 | FALSE | 1 | FALSE | 1 | FALSE | 1 | FALSE | 1 |
| 7069 | 7069 | 9.85714286 | 3.70695247 | 1 | 0.42086991 | 0 | 1 | 0 | 1 | 3 | 0.06912908 | 0 | 1 | 17 | 0.77849076 | FALSE | 1 | FALSE | 1 | FALSE | 1 | FALSE | 1 | FALSE | 1 | FALSE | 1 |
| 7070 | 7070 | 1.08686441 | 0.79497322 | 115 | 1.14E-06 | 127 | 0.00245832 | 108 | 0.91388689 | 160 | 0.15144904 | 62 | 0.06954672 | 2710 | 0.99997157 | TRUE | 0.00077787 | FALSE | 0.44070554 | FALSE | 1 | FALSE | 1 | FALSE | 1 | FALSE | 1 |
| 7071 | 7071 | 9.06666667 | 2.54209013 | 0 | 1 | 2 | 0.25306331 | 1 | 0.66709967 | 1 | 0.75658797 | 0 | 1 | 26 | 0.48609328 | FALSE | 1 | FALSE | 1 | FALSE | 1 | FALSE | 1 | FALSE | 1 | FALSE | 1 |
| 7072 | 7072 | 9.83783784 | 1.89613424 | 4 | 0.01425247 | 3 | 0.11640364 | 2 | 0.38585089 | 1 | 0.82511639 | 0 | 1 | 27 | 0.97789976 | FALSE | 0.37183513 | FALSE | 1 | FALSE | 1 | FALSE | 1 | FALSE | 1 | FALSE | 1 |
| 7073 | 7073 | 9.54545455 | 2.23051712 | 2 | 0.10809809 | 1 | 0.5154318 | 1 | 0.55340851 | 4 | 0.01654484 | 0 | 1 | 14 | 0.99612862 | FALSE | 0.88820416 | FALSE | 1 | FALSE | 1 | FALSE | 1 | FALSE | 1 | FALSE | 1 |
| 7074 | 7074 | 6.57894737 | 2.36158859 | 1 | 0.77387657 | 4 | 0.11180811 | 3 | 0.3364112 | 2 | 0.74436726 | 2 | 0.23984683 | 45 | 0.90229646 | FALSE | 1 | FALSE | 1 | FALSE | 1 | FALSE | 1 | FALSE | 1 | FALSE | 1 |
| 7075 | 7075 | 7.93023256 | 3.59195062 | 0 | 1 | 2 | 0.39616059 | 5 | 0.01650714 | 2 | 0.5811115 | 1 | 0.50166523 | 32 | 0.94677277 | FALSE | 1 | FALSE | 1 | FALSE | 1 | FALSE | 1 | FALSE | 1 | FALSE | 1 |
| 7076 | 7076 | 8.4 | 2.76405499 | 0 | 1 | 0 | 1 | 0 | 1 | 0 | 1 | 0 | 1 | 10 | 0.18273166 | FALSE | 1 | FALSE | 1 | FALSE | 1 | FALSE | 1 | FALSE | 1 | FALSE | 1 |
| 7077 | 7077 | 7.85714286 | 2.96785499 | 1 | 0.59801736 | 0 | 1 | 0 | 1 | 0 | 1 | 3 | 0.01918344 | 31 | 0.34231756 | FALSE | 1 | FALSE | 1 | FALSE | 1 | FALSE | 1 | FALSE | 0.70156566 | FALSE | 1 |
| 7078 | 7078 | 9.07407407 | 2.30821281 | 4 | 0.00461682 | 2 | 0.21684246 | 1 | 0.62830986 | 0 | 1 | 3 | 0.00942179 | 17 | 0.99851071 | FALSE | 0.19698443 | FALSE | 1 | FALSE | 1 | FALSE | 1 | FALSE | 0.5373705 | FALSE | 1 |
| 7079 | 7079 | 1.05719178 | 0.71324288 | 91 | 0.00681597 | 117 | 0.00096483 | 101 | 0.70255637 | 129 | 0.72506507 | 51 | 0.29339675 | 2421 | 0.99101792 | FALSE | 0.24597546 | FALSE | 0.29273541 | FALSE | 1 | FALSE | 1 | FALSE | 1 | FALSE | 1 |
| 7080 | 7080 | 10.5 | 1.5 | 1 | 0.3050805 | 0 | 1 | 0 | 1 | 0 | 1 | 0 | 1 | 13 | 0.33283729 | FALSE | 1 | FALSE | 1 | FALSE | 1 | FALSE | 1 | FALSE | 1 | FALSE | 1 |
| 7081 | 7081 | 8.4375 | 1.69443021 | 2 | 0.06197875 | 1 | 0.40942991 | 2 | 0.11087977 | 0 | 1 | 0 | 1 | 11 | 0.9718531 | FALSE | 0.72532843 | FALSE | 1 | FALSE | 1 | FALSE | 1 | FALSE | 1 | FALSE | 1 |
| 7082 | 7082 | 8.11764706 | 1.96685335 | 0 | 1 | 0 | 1 | 0 | 1 | 1 | 0.55062772 | 0 | 1 | 16 | 0.23070182 | FALSE | 1 | FALSE | 1 | FALSE | 1 | FALSE | 1 | FALSE | 1 | FALSE | 1 |
| 7083 | 7083 | 9.07142857 | 2.28236362 | 0 | 1 | 0 | 1 | 1 | 0.40110418 | 2 | 0.13304705 | 0 | 1 | 11 | 0.83676449 | FALSE | 1 | FALSE | 1 | FALSE | 1 | FALSE | 1 | FALSE | 1 | FALSE | 1 |
| 7084 | 7084 | 4.14655172 | 2.9309711 | 3 | 0.55171054 | 7 | 0.07020894 | 2 | 0.91581496 | 4 | 0.76332434 | 5 | 0.03639395 | 91 | 0.85305932 | FALSE | 1 | FALSE | 1 | FALSE | 1 | FALSE | 1 | FALSE | 0.87669812 | FALSE | 1 |
| 7085 | 7085 | 7.88235294 | 3.28780909 | 0 | 1 | 0 | 1 | 0 | 1 | 0 | 1 | 0 | 1 | 16 | 0.06581507 | FALSE | 1 | FALSE | 1 | FALSE | 1 | FALSE | 1 | FALSE | 1 | FALSE | 1 |
| 7086 | 7086 | 8.04545455 | 1.05071142 | 1 | 0.8213973 | 5 | 0.06154611 | 3 | 0.4248691 | 3 | 0.58996791 | 0 | 1 | 54 | 0.77743724 | FALSE | 1 | FALSE | 1 | FALSE | 1 | FALSE | 1 | FALSE | 1 | FALSE | 1 |
| 7087 | 7087 | 10.5652174 | 2.96099406 | 2 | 0.10809809 | 1 | 0.5154318 | 1 | 0.55340851 | 2 | 0.26791271 | 1 | 0.30529951 | 15 | 0.98570275 | FALSE | 0.88820416 | FALSE | 1 | FALSE | 1 | FALSE | 1 | FALSE | 1 | FALSE | 1 |
| 7088 | 7088 | 7.72727273 | 2.2998383 | 0 | 1 | 0 | 1 | 2 | 0.33343871 | 2 | 0.45159487 | 0 | 1 | 29 | 0.39648374 | FALSE | 1 | FALSE | 1 | FALSE | 1 | FALSE | 1 | FALSE | 1 | FALSE | 1 |
| 7089 | 7089 | 9.73684211 | 2.48875588 | 0 | 1 | 0 | 1 | 1 | 0.50143826 | 0 | 1 | 1 | 0.26986775 | 17 | 0.41141543 | FALSE | 1 | FALSE | 1 | FALSE | 1 | FALSE | 1 | FALSE | 1 | FALSE | 1 |
| 7090 | 7090 | 10.0869565 | 3.14729758 | 1 | 0.45027683 | 3 | 0.03669128 | 3 | 0.04772439 | 2 | 0.2852236 | 0 | 1 | 14 | 0.99865094 | FALSE | 1 | FALSE | 0.94259695 | FALSE | 1 | FALSE | 1 | FALSE | 1 | FALSE | 1 |
| 7091 | 7091 | 2.74683544 | 2.67888298 | 4 | 0.14405175 | 1 | 0.92668093 | 7 | 0.02282047 | 0 | 1 | 0 | 1 | 67 | 0.53379057 | FALSE | 1 | FALSE | 1 | FALSE | 1 | FALSE | 1 | FALSE | 1 | FALSE | 1 |
| 7092 | 7092 | 9.19047619 | 2.98598389 | 0 | 1 | 1 | 0.49918218 | 0 | 1 | 0 | 1 | 0 | 1 | 20 | 0.13763971 | FALSE | 1 | FALSE | 1 | FALSE | 1 | FALSE | 1 | FALSE | 1 | FALSE | 1 |
| 7093 | 7093 | 5.80722892 | 2.77448717 | 11 | 6.60E-06 | 10 | 0.00028386 | 2 | 0.79988333 | 8 | 0.03370922 | 1 | 0.74434486 | 50 | 0.99999994 | TRUE | 0.00299395 | FALSE | 0.15723145 | FALSE | 1 | FALSE | 1 | FALSE | 1 | FALSE | 1 |
| 7094 | 7094 | 9.43478261 | 3.25477692 | 1 | 0.45027683 | 0 | 1 | 1 | 0.5695037 | 2 | 0.2852236 | 2 | 0.05389003 | 17 | 0.94439745 | FALSE | 1 | FALSE | 1 | FALSE | 1 | FALSE | 1 | FALSE | 1 | FALSE | 1 |
| 7095 | 7095 | 10.68 | 2.79599714 | 3 | 0.02256792 | 0 | 1 | 1 | 0.58502114 | 0 | 1 | 0 | 1 | 20 | 0.68270729 | FALSE | 0.4692294 | FALSE | 1 | FALSE | 1 | FALSE | 1 | FALSE | 1 | FALSE | 1 |
| 7096 | 7096 | 8.23809524 | 2.06800483 | 0 | 1 | 0 | 1 | 1 | 0.53671415 | 0 | 1 | 0 | 1 | 20 | 0.13763971 | FALSE | 1 | FALSE | 1 | FALSE | 1 | FALSE | 1 | FALSE | 1 | FALSE | 1 |
| 7097 | 7097 | 6.15384615 | 3.99704033 | 0 | 1 | 0 | 1 | 1 | 0.76089851 | 3 | 0.26522531 | 0 | 1 | 35 | 0.24915769 | FALSE | 1 | FALSE | 1 | FALSE | 1 | FALSE | 1 | FALSE | 1 | FALSE | 1 |
| 7098 | 7098 | 9.2962963 | 2.19114066 | 1 | 0.50471032 | 0 | 1 | 0 | 1 | 0 | 1 | 1 | 0.36060486 | 25 | 0.1832206 | FALSE | 1 | FALSE | 1 | FALSE | 1 | FALSE | 1 | FALSE | 1 | FALSE | 1 |
| 7099 | 7099 | 7.17582418 | 1.28039663 | 7 | 0.0085551 | 3 | 0.5689882 | 4 | 0.41421418 | 2 | 0.92663163 | 3 | 0.18718337 | 72 | 0.93327888 | FALSE | 0.27019545 | FALSE | 1 | FALSE | 1 | FALSE | 1 | FALSE | 1 | FALSE | 1 |
| 7100 | 7100 | 9.47368421 | 1.31262778 | 0 | 1 | 0 | 1 | 0 | 1 | 3 | 0.05383577 | 0 | 1 | 16 | 0.65565778 | FALSE | 1 | FALSE | 1 | FALSE | 1 | FALSE | 1 | FALSE | 1 | FALSE | 1 |
| 7101 | 7101 | 9.3125 | 2.14239206 | 0 | 1 | 0 | 1 | 0 | 1 | 1 | 0.52895088 | 0 | 1 | 15 | 0.26127859 | FALSE | 1 | FALSE | 1 | FALSE | 1 | FALSE | 1 | FALSE | 1 | FALSE | 1 |
| 7102 | 7102 | 9.8 | 2.63818119 | 0 | 1 | 0 | 1 | 0 | 1 | 1 | 0.50623194 | 0 | 1 | 14 | 0.29525409 | FALSE | 1 | FALSE | 1 | FALSE | 1 | FALSE | 1 | FALSE | 1 | FALSE | 1 |
| 7103 | 7103 | 7.68571429 | 2.31481475 | 1 | 0.59801736 | 2 | 0.31355738 | 0 | 1 | 2 | 0.48233989 | 3 | 0.01918344 | 27 | 0.91585095 | FALSE | 1 | FALSE | 1 | FALSE | 1 | FALSE | 1 | FALSE | 0.70156566 | FALSE | 1 |
| 7104 | 7104 | 9.80769231 | 3.3399589 | 4 | 0.0040119 | 2 | 0.20486827 | 1 | 0.61440466 | 2 | 0.33678042 | 0 | 1 | 17 | 0.99598795 | FALSE | 0.17959269 | FALSE | 1 | FALSE | 1 | FALSE | 1 | FALSE | 1 | FALSE | 1 |
| 7105 | 7105 | 10.7142857 | 1.66598626 | 1 | 0.16630142 | 0 | 1 | 0 | 1 | 0 | 1 | 0 | 1 | 6 | 0.69920685 | FALSE | 1 | FALSE | 1 | FALSE | 1 | FALSE | 1 | FALSE | 1 | FALSE | 1 |
| 7106 | 7106 | 11.4 | 3.32264955 | 0 | 1 | 1 | 0.28036922 | 1 | 0.30655256 | 0 | 1 | 0 | 1 | 8 | 0.80394719 | FALSE | 1 | FALSE | 1 | FALSE | 1 | FALSE | 1 | FALSE | 1 | FALSE | 1 |
| 7107 | 7107 | 10 | 0 | 0 | 1 | 0 | 1 | 1 | 0.10395583 | 0 | 1 | 0 | 1 | 2 | 0.9344609 | FALSE | 1 | FALSE | 1 | FALSE | 1 | FALSE | 1 | FALSE | 1 | FALSE | 1 |
| 7108 | 7108 | 8.09677419 | 4.13759193 | 3 | 0.04402911 | 0 | 1 | 2 | 0.30673593 | 2 | 0.41984187 | 2 | 0.09110253 | 22 | 0.98395924 | FALSE | 0.64293489 | FALSE | 1 | FALSE | 1 | FALSE | 1 | FALSE | 1 | FALSE | 1 |
| 7109 | 7109 | 6.65384615 | 3.33907297 | 0 | 1 | 0 | 1 | 1 | 0.61440466 | 0 | 1 | 1 | 0.3499045 | 24 | 0.20425579 | FALSE | 1 | FALSE | 1 | FALSE | 1 | FALSE | 1 | FALSE | 1 | FALSE | 1 |
| 7110 | 7110 | 7.6 | 1.24096737 | 0 | 1 | 1 | 0.48239024 | 1 | 0.51939841 | 1 | 0.60987146 | 1 | 0.28187291 | 16 | 0.80826361 | FALSE | 1 | FALSE | 1 | FALSE | 1 | FALSE | 1 | FALSE | 1 | FALSE | 1 |
| 7111 | 7111 | 7.35555556 | 1.57276198 | 1 | 0.90494416 | 3 | 0.5614295 | 3 | 0.6332806 | 2 | 0.92375101 | 3 | 0.1830501 | 78 | 0.33402291 | FALSE | 1 | FALSE | 1 | FALSE | 1 | FALSE | 1 | FALSE | 1 | FALSE | 1 |
| 7112 | 7112 | 11.5 | 3.45894286 | 2 | 0.04858661 | 0 | 1 | 1 | 0.40110418 | 2 | 0.13304705 | 1 | 0.20679328 | 8 | 0.99722299 | FALSE | 0.649301 | FALSE | 1 | FALSE | 1 | FALSE | 1 | FALSE | 1 | FALSE | 1 |
| 7113 | 7113 | 8.86666667 | 2.07739153 | 0 | 1 | 0 | 1 | 1 | 0.66709967 | 1 | 0.75658797 | 0 | 1 | 28 | 0.13086908 | FALSE | 1 | FALSE | 1 | FALSE | 1 | FALSE | 1 | FALSE | 1 | FALSE | 1 |
| 7114 | 7114 | 7.45 | 2.01183995 | 0 | 1 | 0 | 1 | 1 | 0.50143826 | 2 | 0.21610812 | 0 | 1 | 16 | 0.65565778 | FALSE | 1 | FALSE | 1 | FALSE | 1 | FALSE | 1 | FALSE | 1 | FALSE | 1 |
| 7115 | 7115 | 9.21428571 | 2.3046094 | 0 | 1 | 0 | 1 | 0 | 1 | 1 | 0.48242099 | 2 | 0.02131828 | 11 | 0.83676449 | FALSE | 1 | FALSE | 1 | FALSE | 1 | FALSE | 1 | FALSE | 0.73999729 | FALSE | 1 |
| 7116 | 7116 | 13.6923077 | 5.86232581 | 0 | 1 | 2 | 0.20486827 | 1 | 0.61440466 | 0 | 1 | 2 | 0.0670823 | 21 | 0.78939668 | FALSE | 1 | FALSE | 1 | FALSE | 1 | FALSE | 1 | FALSE | 1 | FALSE | 1 |
| 7117 | 7117 | 9.93333333 | 3.12978523 | 0 | 1 | 0 | 1 | 0 | 1 | 1 | 0.50623194 | 0 | 1 | 14 | 0.29525409 | FALSE | 1 | FALSE | 1 | FALSE | 1 | FALSE | 1 | FALSE | 1 | FALSE | 1 |
| 7118 | 7118 | 4.65853659 | 1.74061604 | 2 | 0.61965506 | 5 | 0.12090937 | 3 | 0.56136161 | 1 | 0.97829212 | 1 | 0.74003033 | 69 | 0.49512231 | FALSE | 1 | FALSE | 1 | FALSE | 1 | FALSE | 1 | FALSE | 1 | FALSE | 1 |
| 7119 | 7119 | 10.5 | 2.58429322 | 0 | 1 | 3 | 0.00932935 | 2 | 0.08806398 | 0 | 1 | 1 | 0.20679328 | 8 | 0.99722299 | FALSE | 1 | FALSE | 1 | FALSE | 1 | FALSE | 0.64223561 | FALSE | 1 | FALSE | 1 |
| 7120 | 7120 | 9.67741936 | 3.09408485 | 1 | 0.55378205 | 0 | 1 | 1 | 0.67911179 | 0 | 1 | 2 | 0.09110253 | 27 | 0.45522639 | FALSE | 1 | FALSE | 1 | FALSE | 1 | FALSE | 1 | FALSE | 1 | FALSE | 1 |
| 7121 | 7121 | 9.91666667 | 2.19690439 | 1 | 0.46441868 | 1 | 0.54637326 | 1 | 0.58502114 | 1 | 0.67692816 | 0 | 1 | 20 | 0.68270729 | FALSE | 1 | FALSE | 1 | FALSE | 1 | FALSE | 1 | FALSE | 1 | FALSE | 1 |
| 7122 | 7122 | 6.22413793 | 3.46431614 | 1 | 0.77972342 | 1 | 0.85269953 | 1 | 0.88128698 | 3 | 0.50172293 | 0 | 1 | 52 | 0.17733928 | FALSE | 1 | FALSE | 1 | FALSE | 1 | FALSE | 1 | FALSE | 1 | FALSE | 1 |
| 7123 | 7123 | 7.67307692 | 2.35927918 | 0 | 1 | 1 | 0.82028131 | 1 | 0.85187773 | 2 | 0.6970685 | 0 | 1 | 48 | 0.0742821 | FALSE | 1 | FALSE | 1 | FALSE | 1 | FALSE | 1 | FALSE | 1 | FALSE | 1 |
| 7124 | 7124 | 8.58333333 | 2.2157893 | 0 | 1 | 0 | 1 | 2 | 0.06691409 | 0 | 1 | 1 | 0.18007762 | 9 | 0.89630573 | FALSE | 1 | FALSE | 1 | FALSE | 1 | FALSE | 1 | FALSE | 1 | FALSE | 1 |
| 7125 | 7125 | 9.07407407 | 2.07142688 | 1 | 0.50471032 | 2 | 0.21684246 | 1 | 0.62830986 | 0 | 1 | 2 | 0.07169515 | 21 | 0.88395936 | FALSE | 1 | FALSE | 1 | FALSE | 1 | FALSE | 1 | FALSE | 1 | FALSE | 1 |
| 7126 | 7126 | 10.2 | 1.72046505 | 1 | 0.22886536 | 0 | 1 | 1 | 0.30655256 | 0 | 1 | 0 | 1 | 8 | 0.80394719 | FALSE | 1 | FALSE | 1 | FALSE | 1 | FALSE | 1 | FALSE | 1 | FALSE | 1 |
| 7127 | 7127 | 9.77777778 | 2.55072006 | 0 | 1 | 0 | 1 | 0 | 1 | 2 | 0.18212134 | 2 | 0.03086174 | 13 | 0.88719222 | FALSE | 1 | FALSE | 1 | FALSE | 1 | FALSE | 1 | FALSE | 0.82559785 | FALSE | 1 |
| 7128 | 7128 | 9.83333333 | 2.8528738 | 2 | 0.10809809 | 4 | 0.00492303 | 0 | 1 | 1 | 0.64497439 | 0 | 1 | 15 | 0.98570275 | FALSE | 0.88820416 | FALSE | 0.58120789 | FALSE | 1 | FALSE | 1 | FALSE | 1 | FALSE | 1 |
| 7129 | 7129 | 9.32258065 | 2.7753186 | 0 | 1 | 0 | 1 | 1 | 0.64171563 | 0 | 1 | 0 | 1 | 27 | 0.05284755 | FALSE | 1 | FALSE | 1 | FALSE | 1 | FALSE | 1 | FALSE | 1 | FALSE | 1 |
| 7130 | 7130 | 9.91428571 | 2.7084147 | 0 | 1 | 0 | 1 | 1 | 0.72299506 | 2 | 0.48233989 | 0 | 1 | 32 | 0.18108028 | FALSE | 1 | FALSE | 1 | FALSE | 1 | FALSE | 1 | FALSE | 1 | FALSE | 1 |
| 7131 | 7131 | 11.6666667 | 1.74801475 | 0 | 1 | 2 | 0.32556427 | 2 | 0.37289966 | 2 | 0.49731009 | 0 | 1 | 30 | 0.67248271 | FALSE | 1 | FALSE | 1 | FALSE | 1 | FALSE | 1 | FALSE | 1 | FALSE | 1 |
| 7132 | 7132 | 7.16129032 | 2.79511987 | 1 | 0.55378205 | 1 | 0.63997468 | 1 | 0.67911179 | 1 | 0.76781284 | 0 | 1 | 27 | 0.45522639 | FALSE | 1 | FALSE | 1 | FALSE | 1 | FALSE | 1 | FALSE | 1 | FALSE | 1 |
| 7133 | 7133 | 10.6470588 | 2.49567099 | 0 | 1 | 0 | 1 | 2 | 0.12280513 | 2 | 0.18212134 | 1 | 0.24525764 | 12 | 0.962093 | FALSE | 1 | FALSE | 1 | FALSE | 1 | FALSE | 1 | FALSE | 1 | FALSE | 1 |
| 7134 | 7134 | 9.36363636 | 3.36486464 | 3 | 0.01782284 | 1 | 0.5154318 | 0 | 1 | 5 | 0.00272531 | 0 | 1 | 13 | 0.99911494 | FALSE | 0.41596787 | FALSE | 1 | FALSE | 1 | FALSE | 1 | FALSE | 1 | FALSE | 1 |
| 7135 | 7135 | 9.46428571 | 1.63624446 | 0 | 1 | 0 | 1 | 0 | 1 | 0 | 1 | 1 | 0.37113067 | 27 | 0.05284755 | FALSE | 1 | FALSE | 1 | FALSE | 1 | FALSE | 1 | FALSE | 1 | FALSE | 1 |
| 7136 | 7136 | 11 | 3.93700394 | 0 | 1 | 1 | 0.32623933 | 0 | 1 | 0 | 1 | 0 | 1 | 11 | 0.41944532 | FALSE | 1 | FALSE | 1 | FALSE | 1 | FALSE | 1 | FALSE | 1 | FALSE | 1 |
| 7137 | 7137 | 8.83333333 | 2.03442594 | 1 | 0.26795838 | 0 | 1 | 0 | 1 | 0 | 1 | 0 | 1 | 11 | 0.41944532 | FALSE | 1 | FALSE | 1 | FALSE | 1 | FALSE | 1 | FALSE | 1 | FALSE | 1 |
| 7138 | 7138 | 8.65714286 | 2.16389671 | 3 | 0.0595675 | 2 | 0.31355738 | 0 | 1 | 3 | 0.21561745 | 1 | 0.44015537 | 26 | 0.96294373 | FALSE | 0.72532843 | FALSE | 1 | FALSE | 1 | FALSE | 1 | FALSE | 1 | FALSE | 1 |
| 7139 | 7139 | 9.9 | 2.44062833 | 0 | 1 | 3 | 0.07145121 | 0 | 1 | 3 | 0.15707004 | 0 | 1 | 24 | 0.82294782 | FALSE | 1 | FALSE | 1 | FALSE | 1 | FALSE | 1 | FALSE | 1 | FALSE | 1 |
| 7140 | 7140 | 10.2083333 | 2.39755373 | 1 | 0.46441868 | 1 | 0.54637326 | 1 | 0.58502114 | 0 | 1 | 0 | 1 | 21 | 0.47053137 | FALSE | 1 | FALSE | 1 | FALSE | 1 | FALSE | 1 | FALSE | 1 | FALSE | 1 |
| 7141 | 7141 | 10.1071429 | 2.00604952 | 0 | 1 | 1 | 0.60247911 | 2 | 0.2663542 | 5 | 0.0081185 | 0 | 1 | 20 | 0.97757517 | FALSE | 1 | FALSE | 1 | FALSE | 1 | FALSE | 1 | FALSE | 1 | FALSE | 1 |
| 7142 | 7142 | 3.68493151 | 2.13821531 | 2 | 0.56271322 | 2 | 0.68961994 | 4 | 0.26680336 | 4 | 0.43314506 | 1 | 0.7028142 | 60 | 0.75780737 | FALSE | 1 | FALSE | 1 | FALSE | 1 | FALSE | 1 | FALSE | 1 | FALSE | 1 |

| | | | | | | | | | | | | | | | | | | | | | | | | | | | |
|---|---|---|---|---|---|---|---|---|---|---|---|---|---|---|---|---|---|---|---|---|---|---|---|---|---|---|---|
| 7143 | 7143 | 9 | 2.2870875 | 0 | 1 | 0 | 1 | 0 | 1 | 0 | 1 | 0 | 1 | 26 | 0.0119733 | FALSE | 1 | FALSE | 1 | FALSE | 1 | FALSE | 1 | FALSE | 1 | FALSE | 1 |
| 7144 | 7144 | 7.86363636 | 2.34124005 | 0 | 1 | 1 | 0.5154318 | 0 | 1 | 0 | 1 | 2 | 0.04972408 | 19 | 0.54233419 | FALSE | 1 | FALSE | 1 | FALSE | 1 | FALSE | 1 | FALSE | 0.98629461 | FALSE | 1 |
| 7145 | 7145 | 9.02702703 | 2.55201549 | 2 | 0.24498301 | 3 | 0.11640364 | 1 | 0.74264075 | 4 | 0.08780467 | 1 | 0.45845922 | 26 | 0.99170905 | FALSE | 1 | FALSE | 1 | FALSE | 1 | FALSE | 1 | FALSE | 1 | FALSE | 1 |
| 7146 | 7146 | 9.06666667 | 1.06249183 | 2 | 0.05513784 | 0 | 1 | 1 | 0.42266264 | 1 | 0.50623194 | 0 | 1 | 11 | 0.92879412 | FALSE | 0.68849059 | FALSE | 1 | FALSE | 1 | FALSE | 1 | FALSE | 1 | FALSE | 1 |
| 7147 | 7147 | 5.58536585 | 4.24824561 | 1 | 0.65632651 | 3 | 0.14580275 | 2 | 0.43640583 | 2 | 0.56788092 | 2 | 0.14490154 | 31 | 0.95455175 | FALSE | 1 | FALSE | 1 | FALSE | 1 | FALSE | 1 | FALSE | 1 | FALSE | 1 |
| 7148 | 7148 | 5.23529412 | 3.42532801 | 4 | 0.17339651 | 6 | 0.05649557 | 2 | 0.81591765 | 3 | 0.75590957 | 2 | 0.40793536 | 68 | 0.89498314 | FALSE | 1 | FALSE | 1 | FALSE | 1 | FALSE | 1 | FALSE | 1 | FALSE | 1 |
| 7149 | 7149 | 9.4 | 1.28062485 | 0 | 1 | 1 | 0.28036922 | 0 | 1 | 0 | 1 | 0 | 1 | 9 | 0.5215547 | FALSE | 1 | FALSE | 1 | FALSE | 1 | FALSE | 1 | FALSE | 1 | FALSE | 1 |
| 7150 | 7150 | 9.42857143 | 2.77010278 | 1 | 0.42086991 | 1 | 0.49918218 | 0 | 1 | 1 | 0.62783522 | 0 | 1 | 18 | 0.57962763 | FALSE | 1 | FALSE | 1 | FALSE | 1 | FALSE | 1 | FALSE | 1 | FALSE | 1 |
| 7151 | 7151 | 10.1 | 3.17647604 | 0 | 1 | 0 | 1 | 0 | 1 | 0 | 1 | 0 | 1 | 10 | 0.18273166 | FALSE | 1 | FALSE | 1 | FALSE | 1 | FALSE | 1 | FALSE | 1 | FALSE | 1 |
| 7152 | 7152 | 7.81818182 | 1.70923597 | 11 | 8.41E-09 | 3 | 0.16927151 | 3 | 0.20921284 | 2 | 0.60667406 | 0 | 1 | 25 | 0.99999743 | TRUE | 1.72E-05 | FALSE | 1 | FALSE | 1 | FALSE | 1 | FALSE | 1 | FALSE | 1 |
| 7153 | 7153 | 7 | 4.55037167 | 1 | 0.58738519 | 0 | 1 | 2 | 0.34668321 | 1 | 0.7984848 | 0 | 1 | 30 | 0.36879488 | FALSE | 1 | FALSE | 1 | FALSE | 1 | FALSE | 1 | FALSE | 1 | FALSE | 1 |
| 7154 | 7154 | 9.13043478 | 2.81771335 | 1 | 0.45027683 | 1 | 0.53115654 | 0 | 1 | 3 | 0.08614834 | 0 | 1 | 18 | 0.86273155 | FALSE | 1 | FALSE | 1 | FALSE | 1 | FALSE | 1 | FALSE | 1 | FALSE | 1 |
| 7155 | 7155 | 7.76470588 | 1.76666558 | 2 | 0.21638713 | 3 | 0.09602744 | 1 | 0.71261921 | 0 | 1 | 0 | 1 | 28 | 0.72558119 | FALSE | 1 | FALSE | 1 | FALSE | 1 | FALSE | 1 | FALSE | 1 | FALSE | 1 |
| 7156 | 7156 | 9.4 | 3.44093011 | 0 | 1 | 0 | 1 | 1 | 0.42266264 | 1 | 0.50623194 | 0 | 1 | 13 | 0.57712805 | FALSE | 1 | FALSE | 1 | FALSE | 1 | FALSE | 1 | FALSE | 1 | FALSE | 1 |
| 7157 | 7157 | 8.6 | 3.49857114 | 0 | 1 | 0 | 1 | 1 | 0.28067846 | 0 | 1 | 0 | 1 | 8 | 0.5780332 | FALSE | 1 | FALSE | 1 | FALSE | 1 | FALSE | 1 | FALSE | 1 | FALSE | 1 |
| 7158 | 7158 | 7.62790698 | 1.86800797 | 2 | 0.29286138 | 6 | 0.0021153 | 0 | 1 | 0 | 1 | 2 | 0.15059682 | 32 | 0.94677277 | FALSE | 1 | FALSE | 0.42264619 | FALSE | 1 | FALSE | 1 | FALSE | 1 | FALSE | 1 |
| 7159 | 7159 | 11.2105263 | 2.28447731 | 1 | 0.38989951 | 1 | 0.46503795 | 2 | 0.14749808 | 1 | 0.59104357 | 0 | 1 | 14 | 0.93677253 | FALSE | 1 | FALSE | 1 | FALSE | 1 | FALSE | 1 | FALSE | 1 | FALSE | 1 |
| 7160 | 7160 | 11.375 | 2.34187425 | 0 | 1 | 0 | 1 | 2 | 0.11087977 | 0 | 1 | 0 | 1 | 14 | 0.53308261 | FALSE | 1 | FALSE | 1 | FALSE | 1 | FALSE | 1 | FALSE | 1 | FALSE | 1 |
| 7161 | 7161 | 2.20171674 | 0.9550393 | 14 | 0.29048699 | 17 | 0.31841961 | 14 | 0.77990189 | 24 | 0.2840674 | 6 | 0.77508608 | 385 | 0.68967166 | FALSE | 1 | FALSE | 1 | FALSE | 1 | FALSE | 1 | FALSE | 1 | FALSE | 1 |
| 7162 | 7162 | 9.46153846 | 2.6491039 | 1 | 0.47819878 | 0 | 1 | 0 | 1 | 3 | 0.10474541 | 0 | 1 | 21 | 0.64957167 | FALSE | 1 | FALSE | 1 | FALSE | 1 | FALSE | 1 | FALSE | 1 | FALSE | 1 |
| 7163 | 7163 | 9.27272727 | 2.92608672 | 2 | 0.03087289 | 0 | 1 | 0 | 1 | 1 | 0.40390032 | 0 | 1 | 8 | 0.92152993 | FALSE | 0.53356688 | FALSE | 1 | FALSE | 1 | FALSE | 1 | FALSE | 1 | FALSE | 1 |
| 7164 | 7164 | 10.25 | 2.33184476 | 0 | 1 | 0 | 1 | 0 | 1 | 0 | 1 | 0 | 1 | 8 | 0.25677166 | FALSE | 1 | FALSE | 1 | FALSE | 1 | FALSE | 1 | FALSE | 1 | FALSE | 1 |
| 7165 | 7165 | 7.70588235 | 2.02236629 | 1 | 0.35728305 | 0 | 1 | 0 | 1 | 1 | 0.55062772 | 0 | 1 | 15 | 0.49060435 | FALSE | 1 | FALSE | 1 | FALSE | 1 | FALSE | 1 | FALSE | 1 | FALSE | 1 |
| 7166 | 7166 | 9.69230769 | 2.1620722 | 0 | 1 | 0 | 1 | 1 | 0.37874417 | 2 | 0.1174583 | 0 | 1 | 10 | 0.86791642 | FALSE | 1 | FALSE | 1 | FALSE | 1 | FALSE | 1 | FALSE | 1 | FALSE | 1 |
| 7167 | 7167 | 8.7037037 | 2.29031479 | 0 | 1 | 0 | 1 | 1 | 0.62830986 | 0 | 1 | 1 | 0.36060486 | 25 | 0.1832206 | FALSE | 1 | FALSE | 1 | FALSE | 1 | FALSE | 1 | FALSE | 1 | FALSE | 1 |
| 7168 | 7168 | 8.15789474 | 2.31159908 | 0 | 1 | 2 | 0.3494045 | 1 | 0.75193685 | 0 | 1 | 2 | 0.12811285 | 33 | 0.44261631 | FALSE | 1 | FALSE | 1 | FALSE | 1 | FALSE | 1 | FALSE | 1 | FALSE | 1 |
| 7169 | 7169 | 10.7142857 | 2.21773924 | 2 | 0.04858661 | 1 | 0.36919824 | 0 | 1 | 0 | 1 | 0 | 1 | 11 | 0.83676449 | FALSE | 0.649301 | FALSE | 1 | FALSE | 1 | FALSE | 1 | FALSE | 1 | FALSE | 1 |
| 7170 | 7170 | 9.78947368 | 2.44042592 | 0 | 1 | 2 | 0.1135776 | 2 | 0.13502329 | 1 | 0.57131013 | 0 | 1 | 13 | 0.95041913 | FALSE | 1 | FALSE | 1 | FALSE | 1 | FALSE | 1 | FALSE | 1 | FALSE | 1 |
| 7171 | 7171 | 5.91666667 | 2.07713363 | 0 | 1 | 1 | 0.93792263 | 4 | 0.35669301 | 3 | 0.74873269 | 0 | 1 | 76 | 0.07508296 | FALSE | 1 | FALSE | 1 | FALSE | 1 | FALSE | 1 | FALSE | 1 | FALSE | 1 |
| 7172 | 7172 | 11.0454546 | 2.53121014 | 0 | 1 | 1 | 0.5154318 | 1 | 0.55340851 | 1 | 0.64497439 | 0 | 1 | 19 | 0.54233419 | FALSE | 1 | FALSE | 1 | FALSE | 1 | FALSE | 1 | FALSE | 1 | FALSE | 1 |
| 7173 | 7173 | 11.7272727 | 3.33278232 | 0 | 1 | 1 | 0.30368014 | 2 | 0.05708125 | 0 | 1 | 0 | 1 | 8 | 0.92152993 | FALSE | 1 | FALSE | 1 | FALSE | 1 | FALSE | 1 | FALSE | 1 | FALSE | 1 |
| 7174 | 7174 | 10 | 2.71448357 | 3 | 0.01186104 | 2 | 0.12434464 | 1 | 0.50143826 | 0 | 1 | 0 | 1 | 13 | 0.97998676 | FALSE | 0.3373808 | FALSE | 1 | FALSE | 1 | FALSE | 1 | FALSE | 1 | FALSE | 1 |
| 7175 | 7175 | 10.5769231 | 1.92499904 | 0 | 1 | 0 | 1 | 0 | 1 | 1 | 0.70601446 | 0 | 1 | 25 | 0.06985983 | FALSE | 1 | FALSE | 1 | FALSE | 1 | FALSE | 1 | FALSE | 1 | FALSE | 1 |
| 7176 | 7176 | 10.6551724 | 1.58170275 | 0 | 1 | 0 | 1 | 0 | 1 | 2 | 0.3871811 | 1 | 0.38148475 | 26 | 0.31495157 | FALSE | 1 | FALSE | 1 | FALSE | 1 | FALSE | 1 | FALSE | 1 | FALSE | 1 |
| 7177 | 7177 | 9.6875 | 3.65237508 | 1 | 0.55378205 | 0 | 1 | 0 | 1 | 3 | 0.16836568 | 0 | 1 | 27 | 0.45522639 | FALSE | 1 | FALSE | 1 | FALSE | 1 | FALSE | 1 | FALSE | 1 | FALSE | 1 |
| 7178 | 7178 | 7.01840491 | 2.52951265 | 4 | 0.60620356 | 11 | 0.01629771 | 5 | 0.70304996 | 8 | 0.4753461 | 5 | 0.12897103 | 130 | 0.95668205 | FALSE | 1 | FALSE | 0.7256023 | FALSE | 1 | FALSE | 1 | FALSE | 1 | FALSE | 1 |
| 7179 | 7179 | 4.71724138 | 2.11302512 | 6 | 0.16791507 | 5 | 0.50698996 | 4 | 0.77132542 | 10 | 0.12905424 | 1 | 0.91138286 | 119 | 0.81489009 | FALSE | 1 | FALSE | 1 | FALSE | 1 | FALSE | 1 | FALSE | 1 | FALSE | 1 |
| 7180 | 7180 | 9.07407407 | 2.90498838 | 0 | 1 | 1 | 0.58913618 | 0 | 1 | 0 | 1 | 0 | 1 | 26 | 0.06079097 | FALSE | 1 | FALSE | 1 | FALSE | 1 | FALSE | 1 | FALSE | 1 | FALSE | 1 |
| 7181 | 7181 | 10 | 2.26077666 | 0 | 1 | 0 | 1 | 1 | 0.28067846 | 0 | 1 | 0 | 1 | 8 | 0.5780332 | FALSE | 1 | FALSE | 1 | FALSE | 1 | FALSE | 1 | FALSE | 1 | FALSE | 1 |
| 7182 | 7182 | 8.57142857 | 2.25876976 | 2 | 0.09989125 | 1 | 0.49918218 | 0 | 1 | 0 | 1 | 0 | 1 | 18 | 0.57962763 | FALSE | 0.86047225 | FALSE | 1 | FALSE | 1 | FALSE | 1 | FALSE | 1 | FALSE | 1 |
| 7183 | 7183 | 5.7628866 | 1.61036076 | 5 | 0.10296654 | 2 | 0.82730061 | 4 | 0.46263314 | 4 | 0.65738328 | 3 | 0.21246782 | 79 | 0.82835389 | FALSE | 0.87864781 | FALSE | 1 | FALSE | 1 | FALSE | 1 | FALSE | 1 | FALSE | 1 |
| 7184 | 7184 | 9.07142857 | 2.01651346 | 0 | 1 | 1 | 0.3480709 | 0 | 1 | 0 | 1 | 0 | 1 | 12 | 0.374198 | FALSE | 1 | FALSE | 1 | FALSE | 1 | FALSE | 1 | FALSE | 1 | FALSE | 1 |
| 7185 | 7185 | 8.5 | 2.23606798 | 1 | 0.34033054 | 0 | 1 | 0 | 1 | 1 | 0.52895088 | 1 | 0.23264612 | 13 | 0.76799161 | FALSE | 1 | FALSE | 1 | FALSE | 1 | FALSE | 1 | FALSE | 1 | FALSE | 1 |
| 7186 | 7186 | 9.66666667 | 1.88561808 | 1 | 0.35728305 | 1 | 0.42857733 | 0 | 1 | 1 | 0.55062772 | 1 | 0.24525764 | 13 | 0.88719222 | FALSE | 1 | FALSE | 1 | FALSE | 1 | FALSE | 1 | FALSE | 1 | FALSE | 1 |
| 7187 | 7187 | 5.43589744 | 1.65320003 | 4 | 0.8568537 | 10 | 0.22566736 | 7 | 0.74311395 | 10 | 0.63907589 | 3 | 0.74452023 | 200 | 0.35973824 | FALSE | 1 | FALSE | 1 | FALSE | 1 | FALSE | 1 | FALSE | 1 | FALSE | 1 |
| 7188 | 7188 | 9.13333333 | 2.36267269 | 0 | 1 | 1 | 0.36919824 | 3 | 0.0124223 | 1 | 0.48242099 | 0 | 1 | 9 | 0.98607548 | FALSE | 1 | FALSE | 1 | FALSE | 1 | FALSE | 1 | FALSE | 1 | FALSE | 1 |
| 7189 | 7189 | 10.8888889 | 1.66295884 | 0 | 1 | 1 | 0.44710673 | 3 | 0.02505979 | 2 | 0.19902682 | 0 | 1 | 12 | 0.98544073 | FALSE | 1 | FALSE | 1 | FALSE | 1 | FALSE | 1 | FALSE | 1 | FALSE | 1 |
| 7190 | 7190 | 9.2631579 | 1.37044385 | 0 | 1 | 0 | 1 | 0 | 1 | 0 | 1 | 1 | 0.26986775 | 18 | 0.17880694 | FALSE | 1 | FALSE | 1 | FALSE | 1 | FALSE | 1 | FALSE | 1 | FALSE | 1 |
| 7191 | 7191 | 6.61805556 | 2.02757562 | 1 | 0.97719733 | 5 | 0.50082612 | 4 | 0.76663323 | 13 | 0.01468402 | 0 | 1 | 121 | 0.60156127 | FALSE | 1 | FALSE | 1 | FALSE | 1 | FALSE | 1 | FALSE | 1 | FALSE | 1 |
| 7192 | 7192 | 8.91304348 | 1.95410479 | 1 | 0.45027683 | 3 | 0.03669128 | 2 | 0.19931429 | 0 | 1 | 0 | 1 | 17 | 0.94439745 | FALSE | 1 | FALSE | 0.94259695 | FALSE | 1 | FALSE | 1 | FALSE | 1 | FALSE | 1 |
| 7193 | 7193 | 8.33333333 | 2.84800125 | 1 | 0.37380237 | 0 | 1 | 0 | 1 | 1 | 0.57131013 | 1 | 0.25766374 | 15 | 0.69372602 | FALSE | 1 | FALSE | 1 | FALSE | 1 | FALSE | 1 | FALSE | 1 | FALSE | 1 |
| 7194 | 7194 | 8.02380952 | 1.31815338 | 1 | 0.49162634 | 0 | 1 | 1 | 0.61440466 | 2 | 0.33678042 | 2 | 0.0670823 | 20 | 0.90142689 | FALSE | 1 | FALSE | 1 | FALSE | 1 | FALSE | 1 | FALSE | 1 | FALSE | 1 |
| 7195 | 7195 | 10.2 | 3.24276974 | 6 | 0.00092434 | 7 | 0.00053527 | 1 | 0.80828716 | 2 | 0.61900576 | 1 | 0.52593041 | 28 | 0.99993247 | FALSE | 0.07211611 | FALSE | 0.1906475 | FALSE | 1 | FALSE | 1 | FALSE | 1 | FALSE | 1 |
| 7196 | 7196 | 9.95652174 | 2.80493759 | 0 | 1 | 0 | 1 | 2 | 0.19931429 | 1 | 0.6613267 | 0 | 1 | 20 | 0.50589402 | FALSE | 1 | FALSE | 1 | FALSE | 1 | FALSE | 1 | FALSE | 1 | FALSE | 1 |
| 7197 | 7197 | 10.3 | 2.00249844 | 0 | 1 | 2 | 0.03949327 | 0 | 1 | 1 | 0.37517262 | 0 | 1 | 7 | 0.94326215 | FALSE | 1 | FALSE | 0.96864929 | FALSE | 1 | FALSE | 1 | FALSE | 1 | FALSE | 1 |
| 7198 | 7198 | 8.67647059 | 1.79411765 | 0 | 1 | 2 | 0.28941836 | 4 | 0.0293494 | 1 | 0.78873814 | 0 | 1 | 26 | 0.86852567 | FALSE | 1 | FALSE | 1 | FALSE | 1 | FALSE | 1 | FALSE | 1 | FALSE | 1 |
| 7199 | 7199 | 7.06666667 | 2.03742757 | 1 | 0.69044531 | 2 | 0.43020462 | 1 | 0.80828716 | 0 | 1 | 1 | 0.52593041 | 40 | 0.27423942 | FALSE | 1 | FALSE | 1 | FALSE | 1 | FALSE | 1 | FALSE | 1 | FALSE | 1 |
| 7200 | 7200 | 8.70967742 | 3.38110068 | 1 | 0.55378205 | 1 | 0.63997468 | 2 | 0.30673593 | 2 | 0.41984187 | 3 | 0.01381051 | 22 | 0.98395924 | FALSE | 1 | FALSE | 1 | FALSE | 1 | FALSE | 1 | FALSE | 0.64607242 | FALSE | 1 |
| 7201 | 7201 | 1.04873878 | 0.62217941 | 142 | 4.99E-05 | 163 | 0.03517072 | 174 | 0.18600933 | 214 | 0.51326043 | 90 | 0.00210848 | 3872 | 0.99998476 | TRUE | 0.01105799 | FALSE | 0.94259695 | FALSE | 1 | FALSE | 1 | FALSE | 0.2617074 | FALSE | 1 |
| 7202 | 7202 | 9.77419355 | 2.81367258 | 0 | 1 | 1 | 0.63997468 | 2 | 0.30673593 | 0 | 1 | 1 | 0.4016888 | 27 | 0.45522639 | FALSE | 1 | FALSE | 1 | FALSE | 1 | FALSE | 1 | FALSE | 1 | FALSE | 1 |
| 7203 | 7203 | 9.54545455 | 2.18938083 | 0 | 1 | 1 | 0.30368014 | 0 | 1 | 2 | 0.08789566 | 0 | 1 | 8 | 0.92152993 | FALSE | 1 | FALSE | 1 | FALSE | 1 | FALSE | 1 | FALSE | 1 | FALSE | 1 |
| 7204 | 7204 | 1.22150735 | 1.34684296 | 45 | 0.0004147 | 38 | 0.3051808 | 36 | 0.71332627 | 53 | 0.31181555 | 17 | 0.61062352 | 889 | 0.97222977 | TRUE | 0.04200174 | FALSE | 1 | FALSE | 1 | FALSE | 1 | FALSE | 1 | FALSE | 1 |
| 7205 | 7205 | 8.68965517 | 1.84022953 | 0 | 1 | 0 | 1 | 2 | 0.27983822 | 1 | 0.74482229 | 1 | 0.38148475 | 25 | 0.51779518 | FALSE | 1 | FALSE | 1 | FALSE | 1 | FALSE | 1 | FALSE | 1 | FALSE | 1 |
| 7206 | 7206 | 8.4 | 2.12289111 | 0 | 1 | 0 | 1 | 0 | 1 | 1 | 0.50623194 | 0 | 1 | 14 | 0.29525409 | FALSE | 1 | FALSE | 1 | FALSE | 1 | FALSE | 1 | FALSE | 1 | FALSE | 1 |
| 7207 | 7207 | 7.11688312 | 1.62747535 | 1 | 0.86263338 | 0 | 1 | 4 | 0.29102231 | 4 | 0.46403589 | 1 | 0.71735141 | 66 | 0.34241291 | FALSE | 1 | FALSE | 1 | FALSE | 1 | FALSE | 1 | FALSE | 1 | FALSE | 1 |
| 7208 | 7208 | 7.625 | 2.1758619 | 0 | 1 | 2 | 0.18117183 | 1 | 0.58502114 | 0 | 1 | 1 | 0.32796862 | 20 | 0.68270729 | FALSE | 1 | FALSE | 1 | FALSE | 1 | FALSE | 1 | FALSE | 1 | FALSE | 1 |
| 7209 | 7209 | 6.57777778 | 3.91855353 | 1 | 0.69044531 | 1 | 0.77338922 | 2 | 0.48463745 | 1 | 0.8802008 | 1 | 0.52593041 | 39 | 0.4317533 | FALSE | 1 | FALSE | 1 | FALSE | 1 | FALSE | 1 | FALSE | 1 | FALSE | 1 |
| 7210 | 7210 | 9.2 | 2.69443872 | 1 | 0.40558521 | 0 | 1 | 0 | 1 | 2 | 0.23331368 | 0 | 1 | 17 | 0.61750632 | FALSE | 1 | FALSE | 1 | FALSE | 1 | FALSE | 1 | FALSE | 1 | FALSE | 1 |
| 7211 | 7211 | 10.1851852 | 2.50978879 | 1 | 0.50471032 | 0 | 1 | 0 | 1 | 4 | 0.03319522 | 1 | 0.36060486 | 21 | 0.88395936 | FALSE | 1 | FALSE | 1 | FALSE | 1 | FALSE | 1 | FALSE | 1 | FALSE | 1 |
| 7212 | 7212 | 7.74285714 | 3.37505858 | 1 | 0.59801736 | 0 | 1 | 1 | 0.72299506 | 2 | 0.48233989 | 0 | 1 | 31 | 0.34231756 | FALSE | 1 | FALSE | 1 | FALSE | 1 | FALSE | 1 | FALSE | 1 | FALSE | 1 |
| 7213 | 7213 | 8.64285714 | 3.04975743 | 1 | 0.50471032 | 2 | 0.21684246 | 1 | 0.62830986 | 2 | 0.35374209 | 0 | 1 | 21 | 0.88395936 | FALSE | 1 | FALSE | 1 | FALSE | 1 | FALSE | 1 | FALSE | 1 | FALSE | 1 |
| 7214 | 7214 | 7.15044248 | 0.94279981 | 4 | 0.83767886 | 7 | 0.60158831 | 7 | 0.70975018 | 10 | 0.59474621 | 5 | 0.31229468 | 193 | 0.37520807 | FALSE | 1 | FALSE | 1 | FALSE | 1 | FALSE | 1 | FALSE | 1 | FALSE | 1 |
| 7215 | 7215 | 8.19230769 | 2.74589527 | 1 | 0.49162634 | 0 | 1 | 0 | 1 | 3 | 0.11458571 | 0 | 1 | 22 | 0.61628 | FALSE | 1 | FALSE | 1 | FALSE | 1 | FALSE | 1 | FALSE | 1 | FALSE | 1 |
| 7216 | 7216 | 9 | 2.48806676 | 1 | 0.42086991 | 1 | 0.49918218 | 1 | 0.53671415 | 2 | 0.25059627 | 0 | 1 | 16 | 0.90353674 | FALSE | 1 | FALSE | 1 | FALSE | 1 | FALSE | 1 | FALSE | 1 | FALSE | 1 |
| 7217 | 7217 | 9.91428571 | 1.62781938 | 1 | 0.59801736 | 0 | 1 | 1 | 0.72299506 | 3 | 0.21561745 | 0 | 1 | 30 | 0.52765882 | FALSE | 1 | FALSE | 1 | FALSE | 1 | FALSE | 1 | FALSE | 1 | FALSE | 1 |
| 7218 | 7218 | 10.0666667 | 2.95446931 | 1 | 0.54198714 | 1 | 0.6278847 | 2 | 0.29330414 | 0 | 1 | 2 | 0.08611595 | 24 | 0.82294782 | FALSE | 1 | FALSE | 1 | FALSE | 1 | FALSE | 1 | FALSE | 1 | FALSE | 1 |
| 7219 | 7219 | 8.47619048 | 3.17158587 | 1 | 0.40558521 | 3 | 0.02534497 | 1 | 0.51939841 | 0 | 1 | 1 | 0.28187291 | 14 | 0.97325593 | FALSE | 1 | FALSE | 0.82367276 | FALSE | 1 | FALSE | 1 | FALSE | 1 | FALSE | 1 |
| 7220 | 7220 | 9.33333333 | 2.74873708 | 3 | 0.00234778 | 1 | 0.30368014 | 0 | 1 | 0 | 1 | 1 | 0.16638736 | 6 | 0.99674012 | FALSE | 0.12737083 | FALSE | 1 | FALSE | 1 | FALSE | 1 | FALSE | 1 | FALSE | 1 |
| 7221 | 7221 | 9.76190476 | 2.42810453 | 0 | 1 | 0 | 1 | 2 | 0.17308361 | 1 | 0.62783522 | 1 | 0.29368245 | 17 | 0.77849076 | FALSE | 1 | FALSE | 1 | FALSE | 1 | FALSE | 1 | FALSE | 1 | FALSE | 1 |
| 7222 | 7222 | 9.08571429 | 1.61017175 | 1 | 0.59801736 | 3 | 0.10264583 | 1 | 0.72299506 | 1 | 0.80778317 | 1 | 0.44015537 | 28 | 0.83079716 | FALSE | 1 | FALSE | 1 | FALSE | 1 | FALSE | 1 | FALSE | 1 | FALSE | 1 |
| 7223 | 7223 | 9.67857143 | 2.15561677 | 1 | 0.51745948 | 0 | 1 | 0 | 1 | 0 | 1 | 0 | 1 | 27 | 0.05284755 | FALSE | 1 | FALSE | 1 | FALSE | 1 | FALSE | 1 | FALSE | 1 | FALSE | 1 |
| 7224 | 7224 | 10.2857143 | 3.10365157 | 1 | 0.16630142 | 0 | 1 | 0 | 1 | 1 | 0.28044368 | 1 | 0.10932414 | 4 | 0.98605104 | FALSE | 1 | FALSE | 1 | FALSE | 1 | FALSE | 1 | FALSE | 1 | FALSE | 1 |
| 7225 | 7225 | 7.64 | 3.40446765 | 1 | 0.47819878 | 0 | 1 | 3 | 0.05884411 | 0 | 1 | 1 | 0.33902672 | 20 | 0.8152498 | FALSE | 1 | FALSE | 1 | FALSE | 1 | FALSE | 1 | FALSE | 1 | FALSE | 1 |
| 7226 | 7226 | 5.6 | 2.2 | 0 | 1 | 0 | 1 | 0 | 1 | 1 | 0.37517262 | 0 | 1 | 9 | 0.5215547 | FALSE | 1 | FALSE | 1 | FALSE | 1 | FALSE | 1 | FALSE | 1 | FALSE | 1 |
| 7227 | 7227 | 7.47368421 | 3.20145396 | 0 | 1 | 1 | 0.46503795 | 2 | 0.14749808 | 0 | 1 | 0 | 1 | 16 | 0.65565778 | FALSE | 1 | FALSE | 1 | FALSE | 1 | FALSE | 1 | FALSE | 1 | FALSE | 1 |
| 7228 | 7228 | 9.4375 | 1.96750953 | 0 | 1 | 0 | 1 | 1 | 0.44344817 | 4 | 0.00511215 | 0 | 1 | 11 | 0.9718531 | FALSE | 1 | FALSE | 1 | FALSE | 1 | FALSE | 1 | FALSE | 1 | FALSE | 1 |
| 7229 | 7229 | 8.41666667 | 2.55359224 | 0 | 1 | 4 | 0.02543131 | 3 | 0.12958859 | 0 | 1 | 1 | 0.44015537 | 27 | 0.91585095 | FALSE | 1 | FALSE | 0.82367276 | FALSE | 1 | FALSE | 1 | FALSE | 1 | FALSE | 1 |
| 7230 | 7230 | 7.28947368 | 2.21130939 | 2 | 0.25455533 | 1 | 0.71433277 | 1 | 0.75193685 | 3 | 0.25266879 | 1 | 0.4673875 | 30 | 0.87271204 | FALSE | 1 | FALSE | 1 | FALSE | 1 | FALSE | 1 | FALSE | 1 | FALSE | 1 |
| 7231 | 7231 | 11.6923077 | 4.11160862 | 0 | 1 | 0 | 1 | 0 | 1 | 0 | 1 | 0 | 1 | 26 | 0.0119733 | FALSE | 1 | FALSE | 1 | FALSE | 1 | FALSE | 1 | FALSE | 1 | FALSE | 1 |
| 7232 | 7232 | 9.04545455 | 1.77039474 | 1 | 0.4357638 | 2 | 0.15794013 | 1 | 0.55340851 | 3 | 0.07743186 | 0 | 1 | 15 | 0.98570275 | FALSE | 1 | FALSE | 1 | FALSE | 1 | FALSE | 1 | FALSE | 1 | FALSE | 1 |
| 7233 | 7233 | 8.45454546 | 3.49970483 | 0 | 1 | 0 | 1 | 0 | 1 | 0 | 1 | 0 | 1 | 11 | 0.15414489 | FALSE | 1 | FALSE | 1 | FALSE | 1 | FALSE | 1 | FALSE | 1 | FALSE | 1 |
| 7234 | 7234 | 11.3846154 | 1.38888231 | 0 | 1 | 0 | 1 | 1 | 0.35555293 | 1 | 0.43131129 | 0 | 1 | 10 | 0.71436964 | FALSE | 1 | FALSE | 1 | FALSE | 1 | FALSE | 1 | FALSE | 1 | FALSE | 1 |
| 7235 | 7235 | 9.87096774 | 2.91538672 | 1 | 0.54198714 | 0 | 1 | 2 | 0.29330414 | 1 | 0.75658797 | 0 | 1 | 26 | 0.48609328 | FALSE | 1 | FALSE | 1 | FALSE | 1 | FALSE | 1 | FALSE | 1 | FALSE | 1 |
| 7236 | 7236 | 8.18518519 | 1.86660788 | 0 | 1 | 2 | 0.21684246 | 1 | 0.62830986 | 0 | 1 | 1 | 0.36060486 | 23 | 0.58307573 | FALSE | 1 | FALSE | 1 | FALSE | 1 | FALSE | 1 | FALSE | 1 | FALSE | 1 |

| | | | | | | | | | | | | | | | | | | | | | | | | | | | |
|---|---|---|---|---|---|---|---|---|---|---|---|---|---|---|---|---|---|---|---|---|---|---|---|---|---|---|---|
| 7237 | 7237 | 5.18421053 | 2.57370574 | 1 | 0.62830697 | 4 | 0.0332765 | 0 | 1 | 4 | 0.09477133 | 0 | 1 | 29 | 0.93814695 | FALSE | 1 | FALSE | 0.92721448 | FALSE | 1 | FALSE | 1 | FALSE | 1 | FALSE | 1 |
| 7238 | 7238 | 9.38888889 | 2.66955862 | 0 | 1 | 0 | 1 | 2 | 0.37289966 | 2 | 0.49731009 | 4 | 0.00269049 | 28 | 0.90259714 | FALSE | 1 | FALSE | 1 | FALSE | 1 | FALSE | 1 | FALSE | 0.29784427 | FALSE | 1 |
| 7239 | 7239 | 6.59322034 | 1.55267559 | 5 | 0.18529734 | 6 | 0.18236875 | 2 | 0.92969421 | 2 | 0.97480462 | 3 | 0.3053007 | 100 | 0.51823116 | FALSE | 1 | FALSE | 1 | FALSE | 1 | FALSE | 1 | FALSE | 1 | FALSE | 1 |
| 7240 | 7240 | 5.62068966 | 2.9370459 | 2 | 0.65852142 | 3 | 0.53828446 | 3 | 0.61023975 | 2 | 0.91445868 | 4 | 0.05404971 | 73 | 0.61830614 | FALSE | 1 | FALSE | 1 | FALSE | 1 | FALSE | 1 | FALSE | 1 | FALSE | 1 |
| 7241 | 7241 | 7.625 | 2.35959919 | 0 | 1 | 1 | 0.54637326 | 0 | 1 | 0 | 1 | 1 | 0.32796862 | 22 | 0.25237063 | FALSE | 1 | FALSE | 1 | FALSE | 1 | FALSE | 1 | FALSE | 1 | FALSE | 1 |
| 7242 | 7242 | 10.25 | 3.08558476 | 0 | 1 | 0 | 1 | 1 | 0.35555293 | 0 | 1 | 1 | 0.18007762 | 10 | 0.71436964 | FALSE | 1 | FALSE | 1 | FALSE | 1 | FALSE | 1 | FALSE | 1 | FALSE | 1 |
| 7243 | 7243 | 8.30434783 | 1.67292461 | 2 | 0.68844298 | 7 | 0.02890045 | 3 | 0.64811016 | 3 | 0.80162372 | 1 | 0.78377186 | 76 | 0.73743516 | FALSE | 1 | FALSE | 0.88012077 | FALSE | 1 | FALSE | 1 | FALSE | 1 | FALSE | 1 |
| 7244 | 7244 | 10.12 | 2.95729606 | 0 | 1 | 1 | 0.56109832 | 2 | 0.22597287 | 1 | 0.69181315 | 2 | 0.06257402 | 19 | 0.91734261 | FALSE | 1 | FALSE | 1 | FALSE | 1 | FALSE | 1 | FALSE | 1 | FALSE | 1 |
| 7245 | 7245 | 7.45454546 | 3.12459364 | 5 | 0.00138535 | 0 | 1 | 3 | 0.11365871 | 1 | 0.78873814 | 3 | 0.01637249 | 21 | 0.99917809 | FALSE | 0.09317759 | FALSE | 1 | FALSE | 1 | FALSE | 1 | FALSE | 0.66728076 | FALSE | 1 |
| 7246 | 7246 | 10.5925926 | 1.91019214 | 2 | 0.15144274 | 4 | 0.01041264 | 0 | 1 | 0 | 1 | 0 | 1 | 21 | 0.88395936 | FALSE | 1 | FALSE | 0.67617265 | FALSE | 1 | FALSE | 1 | FALSE | 1 | FALSE | 1 |
| 7247 | 7247 | 10.6666667 | 2.56038192 | 1 | 0.46441868 | 1 | 0.54637326 | 1 | 0.58502114 | 1 | 0.67692816 | 0 | 1 | 20 | 0.68270729 | FALSE | 1 | FALSE | 1 | FALSE | 1 | FALSE | 1 | FALSE | 1 | FALSE | 1 |
| 7248 | 7248 | 8.38554217 | 1.25948283 | 1 | 0.88571933 | 1 | 0.93582077 | 3 | 0.57805941 | 7 | 0.08572145 | 1 | 0.74858842 | 70 | 0.57760996 | FALSE | 1 | FALSE | 1 | FALSE | 1 | FALSE | 1 | FALSE | 1 | FALSE | 1 |
| 7249 | 7249 | 6.81707317 | 4.48337865 | 1 | 0.87954842 | 3 | 0.49001146 | 2 | 0.79427057 | 4 | 0.51410462 | 1 | 0.74003033 | 70 | 0.37303501 | FALSE | 1 | FALSE | 1 | FALSE | 1 | FALSE | 1 | FALSE | 1 | FALSE | 1 |
| 7250 | 7250 | 10.3125 | 2.84426506 | 0 | 1 | 1 | 0.40942991 | 0 | 1 | 0 | 1 | 0 | 1 | 15 | 0.26127859 | FALSE | 1 | FALSE | 1 | FALSE | 1 | FALSE | 1 | FALSE | 1 | FALSE | 1 |
| 7251 | 7251 | 9.30769231 | 3.17161971 | 1 | 0.28675951 | 0 | 1 | 0 | 1 | 1 | 0.45746571 | 0 | 1 | 11 | 0.6683056 | FALSE | 1 | FALSE | 1 | FALSE | 1 | FALSE | 1 | FALSE | 1 | FALSE | 1 |
| 7252 | 7252 | 7.39655172 | 1.76106996 | 0 | 1 | 2 | 0.48458833 | 0 | 1 | 5 | 0.07777122 | 1 | 0.56379725 | 42 | 0.62187327 | FALSE | 1 | FALSE | 1 | FALSE | 1 | FALSE | 1 | FALSE | 1 | FALSE | 1 |
| 7253 | 7253 | 8.65 | 1.98179212 | 0 | 1 | 0 | 1 | 0 | 1 | 0 | 1 | 0 | 1 | 20 | 0.03329856 | FALSE | 1 | FALSE | 1 | FALSE | 1 | FALSE | 1 | FALSE | 1 | FALSE | 1 |
| 7254 | 7254 | 9.05555556 | 1.60342997 | 0 | 1 | 1 | 0.83180787 | 2 | 0.58328102 | 0 | 1 | 0 | 1 | 51 | 0.02220592 | FALSE | 1 | FALSE | 1 | FALSE | 1 | FALSE | 1 | FALSE | 1 | FALSE | 1 |
| 7255 | 7255 | 9.93333333 | 2.61958436 | 0 | 1 | 1 | 0.38964394 | 2 | 0.09928574 | 1 | 0.50623194 | 0 | 1 | 11 | 0.92879412 | FALSE | 1 | FALSE | 1 | FALSE | 1 | FALSE | 1 | FALSE | 1 | FALSE | 1 |
| 7256 | 7256 | 8.23809524 | 3.55775159 | 1 | 0.42086991 | 0 | 1 | 4 | 0.0060005 | 0 | 1 | 0 | 1 | 16 | 0.90353674 | FALSE | 1 | FALSE | 1 | FALSE | 1 | FALSE | 1 | FALSE | 1 | FALSE | 1 |
| 7257 | 7257 | 8.27272727 | 2.04898685 | 1 | 0.24866461 | 1 | 0.30368014 | 1 | 0.33149969 | 0 | 1 | 0 | 1 | 8 | 0.92152993 | FALSE | 1 | FALSE | 1 | FALSE | 1 | FALSE | 1 | FALSE | 1 | FALSE | 1 |
| 7258 | 7258 | 8.625 | 2.0577597 | 0 | 1 | 2 | 0.37295654 | 0 | 1 | 2 | 0.5543522 | 1 | 0.48480878 | 35 | 0.38966407 | FALSE | 1 | FALSE | 1 | FALSE | 1 | FALSE | 1 | FALSE | 1 | FALSE | 1 |
| 7259 | 7259 | 9.42857143 | 2.68023758 | 2 | 0.35907852 | 3 | 0.21051417 | 1 | 0.83455705 | 3 | 0.39227316 | 0 | 1 | 40 | 0.77344956 | FALSE | 1 | FALSE | 1 | FALSE | 1 | FALSE | 1 | FALSE | 1 | FALSE | 1 |
| 7260 | 7260 | 10.25 | 2.52487624 | 0 | 1 | 0 | 1 | 1 | 0.69069222 | 2 | 0.43583874 | 0 | 1 | 29 | 0.24088541 | FALSE | 1 | FALSE | 1 | FALSE | 1 | FALSE | 1 | FALSE | 1 | FALSE | 1 |
| 7261 | 7261 | 10.5 | 2.31840462 | 0 | 1 | 0 | 1 | 0 | 1 | 0 | 1 | 0 | 1 | 16 | 0.06581507 | FALSE | 1 | FALSE | 1 | FALSE | 1 | FALSE | 1 | FALSE | 1 | FALSE | 1 |
| 7262 | 7262 | 5.2244898 | 2.66699637 | 2 | 0.3309085 | 2 | 0.44132825 | 2 | 0.4962935 | 3 | 0.35428318 | 1 | 0.53375569 | 36 | 0.90704249 | FALSE | 1 | FALSE | 1 | FALSE | 1 | FALSE | 1 | FALSE | 1 | FALSE | 1 |
| 7263 | 7263 | 10.3333333 | 2.8440972 | 1 | 0.32293355 | 1 | 0.38964394 | 0 | 1 | 1 | 0.50623194 | 0 | 1 | 12 | 0.80329917 | FALSE | 1 | FALSE | 1 | FALSE | 1 | FALSE | 1 | FALSE | 1 | FALSE | 1 |
| 7264 | 7264 | 8.28 | 4.52123877 | 1 | 0.72839502 | 3 | 0.21901899 | 2 | 0.54120634 | 2 | 0.6762159 | 0 | 1 | 42 | 0.62187327 | FALSE | 1 | FALSE | 1 | FALSE | 1 | FALSE | 1 | FALSE | 1 | FALSE | 1 |
| 7265 | 7265 | 8.85714286 | 2.40110519 | 0 | 1 | 4 | 0.01184672 | 0 | 1 | 1 | 0.73248983 | 0 | 1 | 23 | 0.73439118 | FALSE | 1 | FALSE | 0.67617265 | FALSE | 1 | FALSE | 1 | FALSE | 1 | FALSE | 1 |
| 7266 | 7266 | 8.76470588 | 1.86387994 | 0 | 1 | 1 | 0.42857733 | 1 | 0.46348836 | 0 | 1 | 0 | 1 | 15 | 0.49060435 | FALSE | 1 | FALSE | 1 | FALSE | 1 | FALSE | 1 | FALSE | 1 | FALSE | 1 |
| 7267 | 7267 | 8.94444444 | 2.24776981 | 0 | 1 | 1 | 0.44710673 | 0 | 1 | 1 | 0.57131013 | 0 | 1 | 16 | 0.44998001 | FALSE | 1 | FALSE | 1 | FALSE | 1 | FALSE | 1 | FALSE | 1 | FALSE | 1 |
| 7268 | 7268 | 9.22222222 | 1.6178022 | 1 | 0.20854748 | 0 | 1 | 0 | 1 | 1 | 0.34506517 | 0 | 1 | 7 | 0.84574168 | FALSE | 1 | FALSE | 1 | FALSE | 1 | FALSE | 1 | FALSE | 1 | FALSE | 1 |
| 7269 | 7269 | 7.55882353 | 2.82000393 | 0 | 1 | 1 | 0.6629941 | 3 | 0.11365871 | 1 | 0.78873814 | 0 | 1 | 28 | 0.58681477 | FALSE | 1 | FALSE | 1 | FALSE | 1 | FALSE | 1 | FALSE | 1 | FALSE | 1 |
| 7270 | 7270 | 9.12903226 | 2.79102173 | 0 | 1 | 1 | 0.63997468 | 3 | 0.09854319 | 1 | 0.76781284 | 1 | 0.4016888 | 25 | 0.80006983 | FALSE | 1 | FALSE | 1 | FALSE | 1 | FALSE | 1 | FALSE | 1 | FALSE | 1 |
| 7271 | 7271 | 10.7666667 | 2.53880987 | 1 | 0.54198714 | 0 | 1 | 1 | 0.66709967 | 0 | 1 | 0 | 1 | 28 | 0.13086908 | FALSE | 1 | FALSE | 1 | FALSE | 1 | FALSE | 1 | FALSE | 1 | FALSE | 1 |
| 7272 | 7272 | 7.22727273 | 2.95314652 | 1 | 0.42086991 | 0 | 1 | 2 | 0.17308361 | 2 | 0.25059627 | 1 | 0.29368245 | 15 | 0.96513691 | FALSE | 1 | FALSE | 1 | FALSE | 1 | FALSE | 1 | FALSE | 1 | FALSE | 1 |
| 7273 | 7273 | 5.56666667 | 2.56493448 | 1 | 0.54198714 | 1 | 0.6278847 | 1 | 0.66709967 | 3 | 0.15707004 | 1 | 0.39166989 | 23 | 0.9158696 | FALSE | 1 | FALSE | 1 | FALSE | 1 | FALSE | 1 | FALSE | 1 | FALSE | 1 |
| 7274 | 7274 | 10.5384615 | 3.07884555 | 3 | 0.00391863 | 0 | 1 | 1 | 0.37874417 | 0 | 1 | 0 | 1 | 9 | 0.96018168 | FALSE | 0.17735574 | FALSE | 1 | FALSE | 1 | FALSE | 1 | FALSE | 1 | FALSE | 1 |
| 7275 | 7275 | 10.9285714 | 0.96097315 | 0 | 1 | 1 | 0.36919824 | 1 | 0.40110418 | 0 | 1 | 0 | 1 | 12 | 0.62236227 | FALSE | 1 | FALSE | 1 | FALSE | 1 | FALSE | 1 | FALSE | 1 | FALSE | 1 |
| 7276 | 7276 | 10.4736842 | 3.08422849 | 1 | 0.38989951 | 3 | 0.02205772 | 0 | 1 | 3 | 0.05383577 | 0 | 1 | 12 | 0.99479039 | FALSE | 1 | FALSE | 0.81029974 | FALSE | 1 | FALSE | 1 | FALSE | 1 | FALSE | 1 |
| 7277 | 7277 | 10.2727273 | 3.13603424 | 0 | 1 | 1 | 0.5154318 | 1 | 0.55340851 | 4 | 0.01654484 | 0 | 1 | 16 | 0.95553951 | FALSE | 1 | FALSE | 1 | FALSE | 1 | FALSE | 1 | FALSE | 1 | FALSE | 1 |
| 7278 | 7278 | 11 | 2.26077666 | 0 | 1 | 0 | 1 | 0 | 1 | 1 | 0.34506517 | 1 | 0.1383237 | 7 | 0.84574168 | FALSE | 1 | FALSE | 1 | FALSE | 1 | FALSE | 1 | FALSE | 1 | FALSE | 1 |
| 7279 | 7279 | 8.69230769 | 2.47711418 | 1 | 0.49162634 | 1 | 0.57534752 | 2 | 0.23940226 | 1 | 0.70601446 | 0 | 1 | 21 | 0.78939668 | FALSE | 1 | FALSE | 1 | FALSE | 1 | FALSE | 1 | FALSE | 1 | FALSE | 1 |
| 7280 | 7280 | 3.55769231 | 2.14607688 | 4 | 0.57122191 | 2 | 0.96467538 | 2 | 0.97841282 | 4 | 0.93306759 | 2 | 0.73050546 | 142 | 0.0101446 | FALSE | 1 | FALSE | 1 | FALSE | 1 | FALSE | 1 | FALSE | 1 | FALSE | 1 |
| 7281 | 7281 | 8.6 | 2.87054002 | 3 | 0.00600364 | 0 | 1 | 0 | 1 | 0 | 1 | 0 | 1 | 12 | 0.80329917 | FALSE | 0.22773046 | FALSE | 1 | FALSE | 1 | FALSE | 1 | FALSE | 1 | FALSE | 1 |
| 7282 | 7282 | 11.1212121 | 1.90306891 | 6 | 0.00349171 | 1 | 0.85269953 | 2 | 0.62245518 | 7 | 0.01616736 | 1 | 0.61824432 | 41 | 0.99763545 | FALSE | 0.16160477 | FALSE | 1 | FALSE | 1 | FALSE | 1 | FALSE | 1 | FALSE | 1 |
| 7283 | 7283 | 8 | 2.0519567 | 1 | 0.38989951 | 1 | 0.46503795 | 1 | 0.50143826 | 0 | 1 | 1 | 0.26986775 | 15 | 0.83648191 | FALSE | 1 | FALSE | 1 | FALSE | 1 | FALSE | 1 | FALSE | 1 | FALSE | 1 |
| 7284 | 7284 | 9.41176471 | 2.63592429 | 1 | 0.58738519 | 0 | 1 | 2 | 0.34668321 | 0 | 1 | 1 | 0.43077488 | 30 | 0.36879488 | FALSE | 1 | FALSE | 1 | FALSE | 1 | FALSE | 1 | FALSE | 1 | FALSE | 1 |
| 7285 | 7285 | 6.66666667 | 3.86292923 | 1 | 0.79096998 | 0 | 1 | 1 | 0.8897349 | 3 | 0.52470434 | 3 | 0.07497311 | 52 | 0.39208719 | FALSE | 1 | FALSE | 1 | FALSE | 1 | FALSE | 1 | FALSE | 1 | FALSE | 1 |
| 7286 | 7286 | 7.90476191 | 2.3177321 | 1 | 0.58738519 | 0 | 1 | 3 | 0.12152681 | 2 | 0.46709861 | 1 | 0.43077488 | 27 | 0.85023803 | FALSE | 1 | FALSE | 1 | FALSE | 1 | FALSE | 1 | FALSE | 1 | FALSE | 1 |
| 7287 | 7287 | 5.65236052 | 2.40068052 | 3 | 0.94109631 | 7 | 0.62936974 | 6 | 0.84700282 | 13 | 0.26770723 | 6 | 0.18033868 | 197 | 0.45351025 | FALSE | 1 | FALSE | 1 | FALSE | 1 | FALSE | 1 | FALSE | 1 | FALSE | 1 |
| 7288 | 7288 | 10.125 | 2.16626599 | 2 | 0.12501436 | 2 | 0.18117183 | 1 | 0.58502114 | 2 | 0.30249307 | 0 | 1 | 17 | 0.97530702 | FALSE | 0.95622564 | FALSE | 1 | FALSE | 1 | FALSE | 1 | FALSE | 1 | FALSE | 1 |
| 7289 | 7289 | 7.04347826 | 3.88394982 | 0 | 1 | 1 | 0.53115654 | 0 | 1 | 1 | 0.6613267 | 0 | 1 | 21 | 0.27964072 | FALSE | 1 | FALSE | 1 | FALSE | 1 | FALSE | 1 | FALSE | 1 | FALSE | 1 |
| 7290 | 7290 | 8.39473684 | 1.78501767 | 2 | 0.58475401 | 1 | 0.91898592 | 4 | 0.29102231 | 4 | 0.46403589 | 4 | 0.03564889 | 61 | 0.87381726 | FALSE | 1 | FALSE | 1 | FALSE | 1 | FALSE | 1 | FALSE | 0.87669812 | FALSE | 1 |
| 7291 | 7291 | 8.53571429 | 2.98187552 | 2 | 0.4229474 | 3 | 0.27128673 | 1 | 0.87219471 | 0 | 1 | 1 | 0.60529938 | 49 | 0.33483172 | FALSE | 1 | FALSE | 1 | FALSE | 1 | FALSE | 1 | FALSE | 1 | FALSE | 1 |
| 7292 | 7292 | 5.71929825 | 2.71935484 | 1 | 0.76787545 | 5 | 0.03395652 | 1 | 0.87219471 | 1 | 0.92884295 | 1 | 0.60529938 | 47 | 0.62412393 | FALSE | 1 | FALSE | 0.93660537 | FALSE | 1 | FALSE | 1 | FALSE | 1 | FALSE | 1 |
| 7293 | 7293 | 7.98387097 | 2.76202319 | 1 | 0.8016455 | 0 | 1 | 4 | 0.18240181 | 4 | 0.31731292 | 0 | 1 | 53 | 0.4900821 | FALSE | 1 | FALSE | 1 | FALSE | 1 | FALSE | 1 | FALSE | 1 | FALSE | 1 |
| 7294 | 7294 | 11.0666667 | 2.23507395 | 0 | 1 | 1 | 0.38964394 | 1 | 0.42266264 | 0 | 1 | 0 | 1 | 13 | 0.57712805 | FALSE | 1 | FALSE | 1 | FALSE | 1 | FALSE | 1 | FALSE | 1 | FALSE | 1 |
| 7295 | 7295 | 10.0625 | 3.69913419 | 0 | 1 | 1 | 0.40942991 | 0 | 1 | 0 | 1 | 0 | 1 | 15 | 0.26127859 | FALSE | 1 | FALSE | 1 | FALSE | 1 | FALSE | 1 | FALSE | 1 | FALSE | 1 |
| 7296 | 7296 | 1.06420234 | 0.80378642 | 66 | 0.50593989 | 88 | 0.25137731 | 90 | 0.62843847 | 111 | 0.80021792 | 63 | 3.25E-05 | 2142 | 0.90094095 | FALSE | 1 | FALSE | 1 | FALSE | 1 | FALSE | 1 | TRUE | 0.01663087 | FALSE | 1 |
| 7297 | 7297 | 10.5714286 | 2.06031502 | 0 | 1 | 0 | 1 | 1 | 0.40110418 | 0 | 1 | 0 | 1 | 13 | 0.33283729 | FALSE | 1 | FALSE | 1 | FALSE | 1 | FALSE | 1 | FALSE | 1 | FALSE | 1 |
| 7298 | 7298 | 9.07142857 | 2.01651346 | 0 | 1 | 1 | 0.36919824 | 0 | 1 | 1 | 0.48242099 | 0 | 1 | 12 | 0.62236227 | FALSE | 1 | FALSE | 1 | FALSE | 1 | FALSE | 1 | FALSE | 1 | FALSE | 1 |
| 7299 | 7299 | 10.2105263 | 2.19038443 | 1 | 0.38989951 | 0 | 1 | 1 | 0.50143826 | 0 | 1 | 1 | 0.26986775 | 16 | 0.65565778 | FALSE | 1 | FALSE | 1 | FALSE | 1 | FALSE | 1 | FALSE | 1 | FALSE | 1 |
| 7300 | 7300 | 10.7692308 | 1.62451631 | 0 | 1 | 0 | 1 | 1 | 0.37874417 | 1 | 0.45746571 | 0 | 1 | 11 | 0.6683056 | FALSE | 1 | FALSE | 1 | FALSE | 1 | FALSE | 1 | FALSE | 1 | FALSE | 1 |
| 7301 | 7301 | 8.92307692 | 2.61538462 | 0 | 1 | 0 | 1 | 1 | 0.37874417 | 0 | 1 | 0 | 1 | 12 | 0.374198 | FALSE | 1 | FALSE | 1 | FALSE | 1 | FALSE | 1 | FALSE | 1 | FALSE | 1 |
| 7302 | 7302 | 9.97959184 | 2.28115434 | 1 | 0.72119464 | 1 | 0.80150109 | 2 | 0.53023996 | 2 | 0.66536008 | 0 | 1 | 43 | 0.337822 | FALSE | 1 | FALSE | 1 | FALSE | 1 | FALSE | 1 | FALSE | 1 | FALSE | 1 |
| 7303 | 7303 | 7.20833333 | 3.32890331 | 1 | 0.46441868 | 3 | 0.04096158 | 0 | 1 | 1 | 0.67692816 | 1 | 0.32796862 | 18 | 0.93166998 | FALSE | 1 | FALSE | 0.9698186 | FALSE | 1 | FALSE | 1 | FALSE | 1 | FALSE | 1 |
| 7304 | 7304 | 12.2608696 | 1.35830429 | 3 | 0.02011786 | 2 | 0.16948764 | 1 | 0.5695037 | 0 | 1 | 0 | 1 | 17 | 0.94439745 | FALSE | 0.44482142 | FALSE | 1 | FALSE | 1 | FALSE | 1 | FALSE | 1 | FALSE | 1 |
| 7305 | 7305 | 8.7 | 2.9 | 0 | 1 | 0 | 1 | 1 | 0.30655256 | 1 | 0.37517262 | 0 | 1 | 8 | 0.80394719 | FALSE | 1 | FALSE | 1 | FALSE | 1 | FALSE | 1 | FALSE | 1 | FALSE | 1 |
| 7306 | 7306 | 10.4285714 | 1.84058553 | 0 | 1 | 0 | 1 | 0 | 1 | 1 | 0.28044368 | 0 | 1 | 6 | 0.69920685 | FALSE | 1 | FALSE | 1 | FALSE | 1 | FALSE | 1 | FALSE | 1 | FALSE | 1 |
| 7307 | 7307 | 11.0952381 | 1.94947519 | 0 | 1 | 1 | 0.49918218 | 1 | 0.53671415 | 1 | 0.62783522 | 0 | 1 | 18 | 0.57962763 | FALSE | 1 | FALSE | 1 | FALSE | 1 | FALSE | 1 | FALSE | 1 | FALSE | 1 |
| 7308 | 7308 | 9.8 | 2.66333125 | 2 | 0.05513784 | 2 | 0.08295839 | 0 | 1 | 0 | 1 | 0 | 1 | 11 | 0.92879412 | FALSE | 0.68849059 | FALSE | 1 | FALSE | 1 | FALSE | 1 | FALSE | 1 | FALSE | 1 |
| 7309 | 7309 | 9.73684211 | 2.55141045 | 0 | 1 | 2 | 0.12434464 | 1 | 0.50143826 | 0 | 1 | 0 | 1 | 16 | 0.65565778 | FALSE | 1 | FALSE | 1 | FALSE | 1 | FALSE | 1 | FALSE | 1 | FALSE | 1 |
| 7310 | 7310 | 11.2857143 | 2.31234487 | 2 | 0.04858661 | 2 | 0.07341559 | 1 | 0.40110418 | 0 | 1 | 1 | 0.20679328 | 8 | 0.99722299 | FALSE | 0.649301 | FALSE | 1 | FALSE | 1 | FALSE | 1 | FALSE | 1 | FALSE | 1 |
| 7311 | 7311 | 7.73584906 | 1.79255213 | 2 | 0.39594061 | 1 | 0.82613965 | 1 | 0.85724013 | 4 | 0.22482652 | 1 | 0.58506217 | 44 | 0.69042405 | FALSE | 1 | FALSE | 1 | FALSE | 1 | FALSE | 1 | FALSE | 1 | FALSE | 1 |
| 7312 | 7312 | 6.61417323 | 1.81440971 | 2 | 0.84227743 | 7 | 0.11675455 | 6 | 0.30549899 | 9 | 0.12869646 | 3 | 0.34592468 | 100 | 0.96655994 | FALSE | 1 | FALSE | 1 | FALSE | 1 | FALSE | 1 | FALSE | 1 | FALSE | 1 |
| 7313 | 7313 | 10.4375 | 1.36787929 | 0 | 1 | 1 | 0.40942991 | 0 | 1 | 0 | 1 | 0 | 1 | 15 | 0.26127859 | FALSE | 1 | FALSE | 1 | FALSE | 1 | FALSE | 1 | FALSE | 1 | FALSE | 1 |
| 7314 | 7314 | 10.1851852 | 2.72215923 | 1 | 0.50471032 | 1 | 0.58913618 | 1 | 0.62830986 | 0 | 1 | 0 | 1 | 24 | 0.3725373 | FALSE | 1 | FALSE | 1 | FALSE | 1 | FALSE | 1 | FALSE | 1 | FALSE | 1 |
| 7315 | 7315 | 6.1627907 | 2.80307334 | 1 | 0.67382914 | 2 | 0.40761584 | 1 | 0.79363407 | 3 | 0.31602175 | 2 | 0.1563365 | 34 | 0.87722689 | FALSE | 1 | FALSE | 1 | FALSE | 1 | FALSE | 1 | FALSE | 1 | FALSE | 1 |
| 7316 | 7316 | 7.89473684 | 3.50780689 | 0 | 1 | 1 | 0.46503795 | 0 | 1 | 0 | 1 | 0 | 1 | 18 | 0.17880694 | FALSE | 1 | FALSE | 1 | FALSE | 1 | FALSE | 1 | FALSE | 1 | FALSE | 1 |
| 7317 | 7317 | 7.11764706 | 2.32334325 | 0 | 1 | 0 | 1 | 2 | 0.12280513 | 2 | 0.18212134 | 0 | 1 | 13 | 0.88719222 | FALSE | 1 | FALSE | 1 | FALSE | 1 | FALSE | 1 | FALSE | 1 | FALSE | 1 |
| 7318 | 7318 | 7.25 | 2.5041632 | 0 | 1 | 3 | 0.04096158 | 1 | 0.58502114 | 1 | 0.67692816 | 0 | 1 | 19 | 0.83975208 | FALSE | 1 | FALSE | 0.9698186 | FALSE | 1 | FALSE | 1 | FALSE | 1 | FALSE | 1 |
| 7319 | 7319 | 8.33333333 | 2.49443826 | 0 | 1 | 0 | 1 | 0 | 1 | 0 | 1 | 0 | 1 | 6 | 0.36077158 | FALSE | 1 | FALSE | 1 | FALSE | 1 | FALSE | 1 | FALSE | 1 | FALSE | 1 |
| 7320 | 7320 | 10.3888889 | 2.75154278 | 0 | 1 | 0 | 1 | 0 | 1 | 0 | 1 | 2 | 0.03435384 | 16 | 0.44998001 | FALSE | 1 | FALSE | 1 | FALSE | 1 | FALSE | 1 | FALSE | 0.86063198 | FALSE | 1 |
| 7321 | 7321 | 10.2 | 2.54558441 | 0 | 1 | 0 | 1 | 3 | 0.05884411 | 1 | 0.69181315 | 0 | 1 | 21 | 0.64957167 | FALSE | 1 | FALSE | 1 | FALSE | 1 | FALSE | 1 | FALSE | 1 | FALSE | 1 |
| 7322 | 7322 | 9.83333333 | 2.67186992 | 0 | 1 | 0 | 1 | 0 | 1 | 0 | 1 | 0 | 1 | 12 | 0.13002667 | FALSE | 1 | FALSE | 1 | FALSE | 1 | FALSE | 1 | FALSE | 1 | FALSE | 1 |
| 7323 | 7323 | 9.33333333 | 2.2852182 | 1 | 0.37380237 | 2 | 0.1135776 | 1 | 0.48280983 | 0 | 1 | 0 | 1 | 14 | 0.86287418 | FALSE | 1 | FALSE | 1 | FALSE | 1 | FALSE | 1 | FALSE | 1 | FALSE | 1 |
| 7324 | 7324 | 9.38888889 | 2.40690878 | 0 | 1 | 1 | 0.44710673 | 1 | 0.48280983 | 0 | 1 | 0 | 1 | 16 | 0.44998001 | FALSE | 1 | FALSE | 1 | FALSE | 1 | FALSE | 1 | FALSE | 1 | FALSE | 1 |
| 7325 | 7325 | 5.02247191 | 2.6136097 | 4 | 0.1887901 | 2 | 0.78367607 | 4 | 0.38964212 | 8 | 0.04815327 | 10 | 1.44E-06 | 60 | 0.9999621 | FALSE | 1 | FALSE | 1 | FALSE | 1 | FALSE | 1 | TRUE | 0.00196745 | FALSE | 1 |
| 7326 | 7326 | 8.4 | 2.81780056 | 0 | 1 | 2 | 0.37295654 | 0 | 1 | 4 | 0.10945471 | 1 | 0.48480878 | 33 | 0.71942635 | FALSE | 1 | FALSE | 1 | FALSE | 1 | FALSE | 1 | FALSE | 1 | FALSE | 1 |
| 7327 | 7327 | 8.95 | 2.10890967 | 0 | 1 | 0 | 1 | 1 | 0.51939841 | 0 | 1 | 0 | 1 | 19 | 0.15700255 | FALSE | 1 | FALSE | 1 | FALSE | 1 | FALSE | 1 | FALSE | 1 | FALSE | 1 |
| 7328 | 7328 | 8.89285714 | 3.3202763 | 0 | 1 | 1 | 0.60247911 | 2 | 0.2663542 | 0 | 1 | 2 | 0.07640798 | 23 | 0.73439118 | FALSE | 1 | FALSE | 1 | FALSE | 1 | FALSE | 1 | FALSE | 1 | FALSE | 1 |
| 7329 | 7329 | 10.5 | 2.71108834 | 2 | 0.09187136 | 2 | 0.1353446 | 0 | 1 | 0 | 1 | 0 | 1 | 16 | 0.80826361 | FALSE | 0.83248965 | FALSE | 1 | FALSE | 1 | FALSE | 1 | FALSE | 1 | FALSE | 1 |
| 7330 | 7330 | 9.65625 | 1.86429234 | 0 | 1 | 2 | 0.27730767 | 1 | 0.69069222 | 1 | 0.77852168 | 2 | 0.096172 | 26 | 0.77612337 | FALSE | 1 | FALSE | 1 | FALSE | 1 | FALSE | 1 | FALSE | 1 | FALSE | 1 |

| | | | | | | | | | | | | | | | | | | | | | | | | | | | |
|---|---|---|---|---|---|---|---|---|---|---|---|---|---|---|---|---|---|---|---|---|---|---|---|---|---|---|---|
| 7331 | 7331 | 7.47058824 | 2.47618414 | 1 | 0.35728305 | 1 | 0.42857733 | 0 | 1 | 0 | 1 | 1 | 0.24525764 | 14 | 0.7313145 | FALSE | 1 | FALSE | 1 | FALSE | 1 | FALSE | 1 | FALSE | 1 | FALSE | 1 |
| 7332 | 7332 | 9.72727273 | 3.86336898 | 1 | 0.24866461 | 1 | 0.30368014 | 1 | 0.33149969 | 2 | 0.08789566 | 0 | 1 | 6 | 0.99674012 | FALSE | 1 | FALSE | 1 | FALSE | 1 | FALSE | 1 | FALSE | 1 | FALSE | 1 |
| 7333 | 7333 | 8.44827586 | 3.8109299 | 0 | 1 | 1 | 0.61539065 | 1 | 0.65463982 | 2 | 0.3871811 | 0 | 1 | 25 | 0.51779518 | FALSE | 1 | FALSE | 1 | FALSE | 1 | FALSE | 1 | FALSE | 1 | FALSE | 1 |
| 7334 | 7334 | 8 | 1.9245009 | 0 | 1 | 3 | 0.05519089 | 2 | 0.25286935 | 2 | 0.35374209 | 0 | 1 | 20 | 0.95133411 | FALSE | 1 | FALSE | 1 | FALSE | 1 | FALSE | 1 | FALSE | 1 | FALSE | 1 |
| 7335 | 7335 | 12.3333333 | 3.3374974 | 1 | 0.46441868 | 0 | 1 | 2 | 0.21260257 | 0 | 1 | 0 | 1 | 21 | 0.47053137 | FALSE | 1 | FALSE | 1 | FALSE | 1 | FALSE | 1 | FALSE | 1 | FALSE | 1 |
| 7336 | 7336 | 9.2 | 3.46794848 | 1 | 0.32293355 | 0 | 1 | 1 | 0.42266264 | 0 | 1 | 0 | 1 | 13 | 0.57712805 | FALSE | 1 | FALSE | 1 | FALSE | 1 | FALSE | 1 | FALSE | 1 | FALSE | 1 |
| 7337 | 7337 | 7 | 2.33333333 | 0 | 1 | 2 | 0.1135776 | 0 | 1 | 0 | 1 | 0 | 1 | 16 | 0.44998001 | FALSE | 1 | FALSE | 1 | FALSE | 1 | FALSE | 1 | FALSE | 1 | FALSE | 1 |
| 7338 | 7338 | 9.8 | 2.82842713 | 1 | 0.45027683 | 0 | 1 | 2 | 0.19931429 | 1 | 0.6613267 | 1 | 0.31672722 | 18 | 0.86273155 | FALSE | 1 | FALSE | 1 | FALSE | 1 | FALSE | 1 | FALSE | 1 | FALSE | 1 |
| 7339 | 7339 | 10.4 | 4.14407207 | 1 | 0.32293355 | 2 | 0.08295839 | 2 | 0.09928574 | 0 | 1 | 0 | 1 | 10 | 0.97980046 | FALSE | 1 | FALSE | 1 | FALSE | 1 | FALSE | 1 | FALSE | 1 | FALSE | 1 |
| 7340 | 7340 | 10.8666667 | 1.45449495 | 0 | 1 | 0 | 1 | 1 | 0.42266264 | 0 | 1 | 0 | 1 | 14 | 0.29525409 | FALSE | 1 | FALSE | 1 | FALSE | 1 | FALSE | 1 | FALSE | 1 | FALSE | 1 |
| 7341 | 7341 | 8 | 2.11891385 | 1 | 0.72119464 | 1 | 0.80150109 | 4 | 0.09808429 | 5 | 0.07258207 | 2 | 0.19154766 | 36 | 0.98492442 | FALSE | 1 | FALSE | 1 | FALSE | 1 | FALSE | 1 | FALSE | 1 | FALSE | 1 |
| 7342 | 7342 | 9.76470588 | 2.60156993 | 1 | 0.35728305 | 0 | 1 | 0 | 1 | 0 | 1 | 0 | 1 | 16 | 0.23070182 | FALSE | 1 | FALSE | 1 | FALSE | 1 | FALSE | 1 | FALSE | 1 | FALSE | 1 |
| 7343 | 7343 | 5.05333974 | 0.58783082 | 64 | 0.04579252 | 70 | 0.36530011 | 67 | 0.87777025 | 107 | 0.08250465 | 36 | 0.38010517 | 1735 | 0.92399864 | FALSE | 0.649301 | FALSE | 1 | FALSE | 1 | FALSE | 1 | FALSE | 1 | FALSE | 1 |
| 7344 | 7344 | 9.76923077 | 1.24985206 | 0 | 1 | 1 | 0.3480709 | 1 | 0.37874417 | 1 | 0.45746571 | 0 | 1 | 10 | 0.86791642 | FALSE | 1 | FALSE | 1 | FALSE | 1 | FALSE | 1 | FALSE | 1 | FALSE | 1 |
| 7345 | 7345 | 10.2666667 | 3.09766938 | 0 | 1 | 0 | 1 | 0 | 1 | 2 | 0.4036176 | 1 | 0.39166989 | 27 | 0.28862208 | FALSE | 1 | FALSE | 1 | FALSE | 1 | FALSE | 1 | FALSE | 1 | FALSE | 1 |
| 7346 | 7346 | 8.4 | 3.97994975 | 1 | 0.47819878 | 1 | 0.56109832 | 2 | 0.22597287 | 0 | 1 | 0 | 1 | 21 | 0.64957167 | FALSE | 1 | FALSE | 1 | FALSE | 1 | FALSE | 1 | FALSE | 1 | FALSE | 1 |
| 7347 | 7347 | 8.0625 | 3.09169755 | 0 | 1 | 2 | 0.46321244 | 2 | 0.51909752 | 3 | 0.37966044 | 2 | 0.18560062 | 39 | 0.79274273 | FALSE | 1 | FALSE | 1 | FALSE | 1 | FALSE | 1 | FALSE | 1 | FALSE | 1 |
| 7348 | 7348 | 8.19047619 | 2.30153257 | 5 | 0.00412436 | 2 | 0.39616059 | 3 | 0.19060367 | 2 | 0.5811115 | 2 | 0.15059682 | 28 | 0.99888598 | FALSE | 0.18198109 | FALSE | 1 | FALSE | 1 | FALSE | 1 | FALSE | 1 | FALSE | 1 |
| 7349 | 7349 | 7.63380282 | 1.46546572 | 4 | 0.10842422 | 2 | 0.67482356 | 0 | 1 | 5 | 0.22607353 | 2 | 0.3251371 | 58 | 0.7898837 | FALSE | 0.88999117 | FALSE | 1 | FALSE | 1 | FALSE | 1 | FALSE | 1 | FALSE | 1 |
| 7350 | 7350 | 9.33333333 | 2.47206616 | 0 | 1 | 0 | 1 | 0 | 1 | 3 | 0.04687752 | 1 | 0.25766374 | 14 | 0.86287418 | FALSE | 1 | FALSE | 1 | FALSE | 1 | FALSE | 1 | FALSE | 1 | FALSE | 1 |
| 7351 | 7351 | 10.7692308 | 1.76084971 | 0 | 1 | 0 | 1 | 0 | 1 | 2 | 0.1174583 | 0 | 1 | 11 | 0.6683056 | FALSE | 1 | FALSE | 1 | FALSE | 1 | FALSE | 1 | FALSE | 1 | FALSE | 1 |
| 7352 | 7352 | 9.29411765 | 2.84428645 | 0 | 1 | 2 | 0.10307112 | 0 | 1 | 0 | 1 | 2 | 0.03086174 | 13 | 0.88719222 | FALSE | 1 | FALSE | 1 | FALSE | 1 | FALSE | 1 | FALSE | 0.82559785 | FALSE | 1 |
| 7353 | 7353 | 8.34146342 | 2.62826806 | 3 | 0.08696268 | 1 | 0.74131698 | 1 | 0.77786601 | 1 | 0.85524728 | 1 | 0.49330646 | 34 | 0.69505404 | FALSE | 0.8160347 | FALSE | 1 | FALSE | 1 | FALSE | 1 | FALSE | 1 | FALSE | 1 |
| 7354 | 7354 | 9.17647059 | 3.0723682 | 1 | 0.35728305 | 1 | 0.42857733 | 0 | 1 | 1 | 0.55062772 | 1 | 0.24525764 | 13 | 0.88719222 | FALSE | 1 | FALSE | 1 | FALSE | 1 | FALSE | 1 | FALSE | 1 | FALSE | 1 |
| 7355 | 7355 | 7.56410256 | 1.29683781 | 0 | 1 | 2 | 0.36122049 | 1 | 0.76089851 | 2 | 0.54052736 | 0 | 1 | 34 | 0.41570781 | FALSE | 1 | FALSE | 1 | FALSE | 1 | FALSE | 1 | FALSE | 1 | FALSE | 1 |
| 7356 | 7356 | 9.35294118 | 1.64285177 | 0 | 1 | 0 | 1 | 0 | 1 | 1 | 0.55062772 | 0 | 1 | 16 | 0.23070182 | FALSE | 1 | FALSE | 1 | FALSE | 1 | FALSE | 1 | FALSE | 1 | FALSE | 1 |
| 7357 | 7357 | 10.6170213 | 1.76915885 | 2 | 0.34034075 | 0 | 1 | 3 | 0.23782059 | 1 | 0.89101956 | 0 | 1 | 41 | 0.38322554 | FALSE | 1 | FALSE | 1 | FALSE | 1 | FALSE | 1 | FALSE | 1 | FALSE | 1 |
| 7358 | 7358 | 10.5555556 | 2.52151238 | 0 | 1 | 0 | 1 | 1 | 0.48280983 | 2 | 0.19902682 | 0 | 1 | 15 | 0.69372602 | FALSE | 1 | FALSE | 1 | FALSE | 1 | FALSE | 1 | FALSE | 1 | FALSE | 1 |
| 7359 | 7359 | 8.90909091 | 2.27454473 | 1 | 0.4357638 | 1 | 0.5154318 | 0 | 1 | 1 | 0.64497439 | 2 | 0.04972408 | 17 | 0.8840357 | FALSE | 1 | FALSE | 1 | FALSE | 1 | FALSE | 1 | FALSE | 0.98629461 | FALSE | 1 |
| 7360 | 7360 | 9.59375 | 1.96626828 | 0 | 1 | 1 | 0.65167362 | 1 | 0.69069222 | 3 | 0.17988885 | 0 | 1 | 27 | 0.61671133 | FALSE | 1 | FALSE | 1 | FALSE | 1 | FALSE | 1 | FALSE | 1 | FALSE | 1 |
| 7361 | 7361 | 8.81818182 | 1.72248139 | 1 | 0.4357638 | 0 | 1 | 0 | 1 | 0 | 1 | 3 | 0.00526351 | 18 | 0.74744813 | FALSE | 1 | FALSE | 1 | FALSE | 1 | FALSE | 1 | FALSE | 0.43998669 | FALSE | 1 |
| 7362 | 7362 | 8.52830189 | 2.42318809 | 1 | 0.74890551 | 3 | 0.24492539 | 2 | 0.57303317 | 3 | 0.44196866 | 2 | 0.21556658 | 42 | 0.88654191 | FALSE | 1 | FALSE | 1 | FALSE | 1 | FALSE | 1 | FALSE | 1 | FALSE | 1 |
| 7363 | 7363 | 9.28571429 | 2.28124563 | 0 | 1 | 0 | 1 | 1 | 0.40110418 | 0 | 1 | 0 | 1 | 13 | 0.33283729 | FALSE | 1 | FALSE | 1 | FALSE | 1 | FALSE | 1 | FALSE | 1 | FALSE | 1 |
| 7364 | 7364 | 10.45 | 3.26305072 | 0 | 1 | 1 | 0.48239024 | 0 | 1 | 1 | 0.60987146 | 0 | 1 | 18 | 0.37504645 | FALSE | 1 | FALSE | 1 | FALSE | 1 | FALSE | 1 | FALSE | 1 | FALSE | 1 |
| 7365 | 7365 | 10.4285714 | 2.61080956 | 0 | 1 | 0 | 1 | 1 | 0.64171563 | 0 | 1 | 0 | 1 | 27 | 0.05284755 | FALSE | 1 | FALSE | 1 | FALSE | 1 | FALSE | 1 | FALSE | 1 | FALSE | 1 |
| 7366 | 7366 | 8.80769231 | 2.38616575 | 1 | 0.49162634 | 0 | 1 | 1 | 0.61440466 | 1 | 0.70601446 | 0 | 1 | 23 | 0.40372818 | FALSE | 1 | FALSE | 1 | FALSE | 1 | FALSE | 1 | FALSE | 1 | FALSE | 1 |
| 7367 | 7367 | 11.3018868 | 1.53678058 | 2 | 0.39594061 | 0 | 1 | 1 | 0.85724013 | 3 | 0.44196866 | 0 | 1 | 47 | 0.25747403 | FALSE | 1 | FALSE | 1 | FALSE | 1 | FALSE | 1 | FALSE | 1 | FALSE | 1 |
| 7368 | 7368 | 9.45454546 | 2.68656488 | 1 | 0.57647348 | 4 | 0.02089081 | 2 | 0.33343871 | 0 | 1 | 0 | 1 | 26 | 0.86852567 | FALSE | 1 | FALSE | 0.8034624 | FALSE | 1 | FALSE | 1 | FALSE | 1 | FALSE | 1 |
| 7369 | 7369 | 8.53846154 | 1.64622574 | 0 | 1 | 0 | 1 | 0 | 1 | 0 | 1 | 0 | 1 | 13 | 0.10967906 | FALSE | 1 | FALSE | 1 | FALSE | 1 | FALSE | 1 | FALSE | 1 | FALSE | 1 |
| 7370 | 7370 | 7.2972973 | 2.72490878 | 2 | 0.55518392 | 5 | 0.08277437 | 4 | 0.25881411 | 2 | 0.85003314 | 0 | 1 | 59 | 0.77408425 | FALSE | 1 | FALSE | 1 | FALSE | 1 | FALSE | 1 | FALSE | 1 | FALSE | 1 |
| 7371 | 7371 | 9.02380952 | 1.69716985 | 1 | 0.66519151 | 2 | 0.39616059 | 1 | 0.78589454 | 0 | 1 | 1 | 0.50166523 | 37 | 0.34041811 | FALSE | 1 | FALSE | 1 | FALSE | 1 | FALSE | 1 | FALSE | 1 | FALSE | 1 |
| 7372 | 7372 | 9.25 | 1.90394328 | 0 | 1 | 0 | 1 | 2 | 0.32011871 | 2 | 0.43583874 | 1 | 0.41154418 | 27 | 0.61671133 | FALSE | 1 | FALSE | 1 | FALSE | 1 | FALSE | 1 | FALSE | 1 | FALSE | 1 |
| 7373 | 7373 | 9.92857143 | 3.1044734 | 1 | 0.3050805 | 1 | 0.36919824 | 0 | 1 | 1 | 0.48242099 | 2 | 0.02131828 | 9 | 0.98607548 | FALSE | 1 | FALSE | 1 | FALSE | 1 | FALSE | 1 | FALSE | 0.73999729 | FALSE | 1 |
| 7374 | 7374 | 10.7894737 | 1.76453207 | 3 | 0.01186104 | 1 | 0.46503795 | 2 | 0.14749808 | 0 | 1 | 0 | 1 | 13 | 0.97998676 | FALSE | 0.3373808 | FALSE | 1 | FALSE | 1 | FALSE | 1 | FALSE | 1 | FALSE | 1 |
| 7375 | 7375 | 8.5625 | 1.57990308 | 1 | 0.34033054 | 0 | 1 | 0 | 1 | 0 | 1 | 0 | 1 | 15 | 0.26127859 | FALSE | 1 | FALSE | 1 | FALSE | 1 | FALSE | 1 | FALSE | 1 | FALSE | 1 |
| 7376 | 7376 | 9.51724138 | 2.66031745 | 2 | 0.43181298 | 1 | 0.84773229 | 2 | 0.61293419 | 1 | 0.93213716 | 1 | 0.61182532 | 50 | 0.31478744 | FALSE | 1 | FALSE | 1 | FALSE | 1 | FALSE | 1 | FALSE | 1 | FALSE | 1 |
| 7377 | 7377 | 10.5238095 | 2.4027573 | 1 | 0.42086991 | 0 | 1 | 1 | 0.53671415 | 1 | 0.62783522 | 0 | 1 | 18 | 0.57962763 | FALSE | 1 | FALSE | 1 | FALSE | 1 | FALSE | 1 | FALSE | 1 | FALSE | 1 |
| 7378 | 7378 | 8.29032258 | 1.18031019 | 3 | 0.74949909 | 6 | 0.36303811 | 6 | 0.46015458 | 7 | 0.54499435 | 3 | 0.45232905 | 126 | 0.67492633 | FALSE | 1 | FALSE | 1 | FALSE | 1 | FALSE | 1 | FALSE | 1 | FALSE | 1 |
| 7379 | 7379 | 8.81818182 | 2.28903242 | 1 | 0.24866461 | 0 | 1 | 1 | 0.33149969 | 1 | 0.40390032 | 1 | 0.16638736 | 7 | 0.98129353 | FALSE | 1 | FALSE | 1 | FALSE | 1 | FALSE | 1 | FALSE | 1 | FALSE | 1 |
| 7380 | 7380 | 9.45454546 | 3.28532927 | 0 | 1 | 1 | 0.30368014 | 0 | 1 | 0 | 1 | 0 | 1 | 10 | 0.46859937 | FALSE | 1 | FALSE | 1 | FALSE | 1 | FALSE | 1 | FALSE | 1 | FALSE | 1 |
| 7381 | 7381 | 10.3555556 | 2.20257986 | 0 | 1 | 1 | 0.77338922 | 2 | 0.48463745 | 5 | 0.05380295 | 0 | 1 | 37 | 0.73736034 | FALSE | 1 | FALSE | 1 | FALSE | 1 | FALSE | 1 | FALSE | 1 | FALSE | 1 |
| 7382 | 7382 | 5.39473684 | 2.90226294 | 1 | 0.86263338 | 5 | 0.09882268 | 2 | 0.76408693 | 2 | 0.8705742 | 0 | 1 | 66 | 0.34241291 | FALSE | 1 | FALSE | 1 | FALSE | 1 | FALSE | 1 | FALSE | 1 | FALSE | 1 |
| 7383 | 7383 | 8.625 | 1.52240216 | 1 | 0.71380452 | 4 | 0.06855877 | 1 | 0.8283465 | 3 | 0.37966044 | 2 | 0.18560062 | 37 | 0.93870662 | FALSE | 1 | FALSE | 1 | FALSE | 1 | FALSE | 1 | FALSE | 1 | FALSE | 1 |
| 7384 | 7384 | 6.84615385 | 3.07211162 | 1 | 0.49162634 | 1 | 0.57534752 | 1 | 0.61440466 | 2 | 0.33678042 | 0 | 1 | 21 | 0.78939668 | FALSE | 1 | FALSE | 1 | FALSE | 1 | FALSE | 1 | FALSE | 1 | FALSE | 1 |
| 7385 | 7385 | 10.4210526 | 2.20801863 | 0 | 1 | 2 | 0.12434464 | 0 | 1 | 1 | 0.59104357 | 0 | 1 | 16 | 0.65565778 | FALSE | 1 | FALSE | 1 | FALSE | 1 | FALSE | 1 | FALSE | 1 | FALSE | 1 |
| 7386 | 7386 | 9.17647059 | 2.70588235 | 0 | 1 | 1 | 0.42857733 | 1 | 0.46348836 | 0 | 1 | 0 | 1 | 15 | 0.49060435 | FALSE | 1 | FALSE | 1 | FALSE | 1 | FALSE | 1 | FALSE | 1 | FALSE | 1 |
| 7387 | 7387 | 10.1428571 | 3.11349925 | 0 | 1 | 0 | 1 | 1 | 0.40110418 | 0 | 1 | 0 | 1 | 13 | 0.33283729 | FALSE | 1 | FALSE | 1 | FALSE | 1 | FALSE | 1 | FALSE | 1 | FALSE | 1 |
| 7388 | 7388 | 10.6666667 | 3.39934634 | 0 | 1 | 1 | 0.44710673 | 1 | 0.48280983 | 1 | 0.57131013 | 1 | 0.25766374 | 14 | 0.86287418 | FALSE | 1 | FALSE | 1 | FALSE | 1 | FALSE | 1 | FALSE | 1 | FALSE | 1 |
| 7389 | 7389 | 8.79166667 | 2.19808945 | 1 | 0.46441868 | 2 | 0.18117183 | 1 | 0.58502114 | 3 | 0.09525947 | 2 | 0.058175 | 15 | 0.99800699 | FALSE | 1 | FALSE | 1 | FALSE | 1 | FALSE | 1 | FALSE | 1 | FALSE | 1 |
| 7390 | 7390 | 8.17647059 | 3.2218974 | 1 | 0.35728305 | 0 | 1 | 0 | 1 | 1 | 0.55062772 | 1 | 0.24525764 | 14 | 0.7313145 | FALSE | 1 | FALSE | 1 | FALSE | 1 | FALSE | 1 | FALSE | 1 | FALSE | 1 |
| 7391 | 7391 | 10.8571429 | 3.39867922 | 3 | 0.0048942 | 0 | 1 | 1 | 0.40110418 | 0 | 1 | 1 | 0.20679328 | 9 | 0.98607548 | FALSE | 0.19848171 | FALSE | 1 | FALSE | 1 | FALSE | 1 | FALSE | 1 | FALSE | 1 |
| 7392 | 7392 | 8.97297297 | 1.56803348 | 0 | 1 | 2 | 0.3375163 | 3 | 0.14625315 | 1 | 0.82511639 | 0 | 1 | 31 | 0.64534533 | FALSE | 1 | FALSE | 1 | FALSE | 1 | FALSE | 1 | FALSE | 1 | FALSE | 1 |
| 7393 | 7393 | 7.77777778 | 2.5724082 | 0 | 1 | 0 | 1 | 0 | 1 | 1 | 0.57131013 | 2 | 0.03435384 | 15 | 0.69372602 | FALSE | 1 | FALSE | 1 | FALSE | 1 | FALSE | 1 | FALSE | 0.86063198 | FALSE | 1 |
| 7394 | 7394 | 8.89473684 | 2.89999522 | 0 | 1 | 1 | 0.46503795 | 2 | 0.14749808 | 0 | 1 | 0 | 1 | 16 | 0.65565778 | FALSE | 1 | FALSE | 1 | FALSE | 1 | FALSE | 1 | FALSE | 1 | FALSE | 1 |
| 7395 | 7395 | 9.52173913 | 2.26254119 | 3 | 0.02011786 | 3 | 0.03669128 | 2 | 0.19931429 | 2 | 0.2852236 | 0 | 1 | 13 | 0.99971815 | FALSE | 0.44482142 | FALSE | 0.94259695 | FALSE | 1 | FALSE | 1 | FALSE | 1 | FALSE | 1 |
| 7396 | 7396 | 10.2894737 | 2.73291723 | 0 | 1 | 1 | 0.71433277 | 2 | 0.39868506 | 0 | 1 | 3 | 0.02386978 | 32 | 0.61800916 | FALSE | 1 | FALSE | 1 | FALSE | 1 | FALSE | 1 | FALSE | 0.77578444 | FALSE | 1 |
| 7397 | 7397 | 9.0952381 | 2.15814651 | 2 | 0.09989125 | 0 | 1 | 2 | 0.17308361 | 1 | 0.62783522 | 0 | 1 | 16 | 0.90353674 | FALSE | 0.86047225 | FALSE | 1 | FALSE | 1 | FALSE | 1 | FALSE | 1 | FALSE | 1 |
| 7398 | 7398 | 7.34883721 | 3.1318597 | 1 | 0.67382914 | 3 | 0.16132793 | 2 | 0.46083448 | 1 | 0.86831187 | 0 | 1 | 36 | 0.6451239 | FALSE | 1 | FALSE | 1 | FALSE | 1 | FALSE | 1 | FALSE | 1 | FALSE | 1 |
| 7399 | 7399 | 6.79069767 | 1.81156671 | 2 | 0.30241117 | 3 | 0.16132793 | 3 | 0.19985581 | 1 | 0.86831187 | 0 | 1 | 34 | 0.87722689 | FALSE | 1 | FALSE | 1 | FALSE | 1 | FALSE | 1 | FALSE | 1 | FALSE | 1 |
| 7400 | 7400 | 7.66666667 | 3.45607356 | 0 | 1 | 0 | 1 | 2 | 0.37289966 | 3 | 0.22784656 | 0 | 1 | 31 | 0.49869404 | FALSE | 1 | FALSE | 1 | FALSE | 1 | FALSE | 1 | FALSE | 1 | FALSE | 1 |
| 7401 | 7401 | 8.36842105 | 4.00276913 | 0 | 1 | 0 | 1 | 1 | 0.50143826 | 0 | 1 | 1 | 0.26986775 | 17 | 0.41141543 | FALSE | 1 | FALSE | 1 | FALSE | 1 | FALSE | 1 | FALSE | 1 | FALSE | 1 |
| 7402 | 7402 | 8.63636364 | 1.96666433 | 0 | 1 | 0 | 1 | 2 | 0.33343871 | 1 | 0.78873814 | 1 | 0.42123867 | 29 | 0.39648374 | FALSE | 1 | FALSE | 1 | FALSE | 1 | FALSE | 1 | FALSE | 1 | FALSE | 1 |
| 7403 | 7403 | 10.55 | 1.85674446 | 0 | 1 | 0 | 1 | 1 | 0.51939841 | 1 | 0.60987146 | 1 | 0.28187291 | 17 | 0.61750632 | FALSE | 1 | FALSE | 1 | FALSE | 1 | FALSE | 1 | FALSE | 1 | FALSE | 1 |
| 7404 | 7404 | 7.05681818 | 0.7516339 | 6 | 0.29684631 | 2 | 0.98000788 | 4 | 0.88313991 | 9 | 0.41890014 | 1 | 0.94758402 | 154 | 0.14598051 | FALSE | 1 | FALSE | 1 | FALSE | 1 | FALSE | 1 | FALSE | 1 | FALSE | 1 |
| 7405 | 7405 | 6.17647059 | 0.67753539 | 2 | 0.74184548 | 5 | 0.23407775 | 2 | 0.88686294 | 9 | 0.04394039 | 0 | 1 | 84 | 0.76448841 | FALSE | 1 | FALSE | 1 | FALSE | 1 | FALSE | 1 | FALSE | 1 | FALSE | 1 |
| 7406 | 7406 | 8.73333333 | 2.23507395 | 0 | 1 | 3 | 0.01138739 | 1 | 0.42266264 | 1 | 0.50623194 | 0 | 1 | 10 | 0.97980046 | FALSE | 1 | FALSE | 0.67617265 | FALSE | 1 | FALSE | 1 | FALSE | 1 | FALSE | 1 |
| 7407 | 7407 | 10.1428571 | 2.84999105 | 0 | 1 | 0 | 1 | 0 | 1 | 1 | 0.28044368 | 1 | 0.10932414 | 5 | 0.91847773 | FALSE | 1 | FALSE | 1 | FALSE | 1 | FALSE | 1 | FALSE | 1 | FALSE | 1 |
| 7408 | 7408 | 7.34375 | 2.5812954 | 1 | 0.56527493 | 1 | 0.65167362 | 1 | 0.69069222 | 2 | 0.43583874 | 6 | 1.09E-05 | 21 | 0.9978606 | FALSE | 1 | FALSE | 1 | FALSE | 1 | FALSE | 1 | TRUE | 0.00740799 | FALSE | 1 |
| 7409 | 7409 | 11.0833333 | 2.49861073 | 0 | 1 | 2 | 0.05552913 | 0 | 1 | 1 | 0.43131129 | 0 | 1 | 9 | 0.89630573 | FALSE | 1 | FALSE | 1 | FALSE | 1 | FALSE | 1 | FALSE | 1 | FALSE | 1 |
| 7410 | 7410 | 9.09375 | 1.98996506 | 2 | 0.49173883 | 0 | 1 | 3 | 0.40547364 | 2 | 0.79987679 | 0 | 1 | 57 | 0.19643469 | FALSE | 1 | FALSE | 1 | FALSE | 1 | FALSE | 1 | FALSE | 1 | FALSE | 1 |
| 7411 | 7411 | 7.75 | 2.61804635 | 1 | 0.26795838 | 0 | 1 | 0 | 1 | 1 | 0.43131129 | 0 | 1 | 10 | 0.71436964 | FALSE | 1 | FALSE | 1 | FALSE | 1 | FALSE | 1 | FALSE | 1 | FALSE | 1 |
| 7412 | 7412 | 8.52380952 | 2.66581619 | 0 | 1 | 0 | 1 | 1 | 0.51939841 | 0 | 1 | 0 | 1 | 19 | 0.15700255 | FALSE | 1 | FALSE | 1 | FALSE | 1 | FALSE | 1 | FALSE | 1 | FALSE | 1 |
| 7413 | 7413 | 8.8974359 | 2.97678946 | 6 | 0.00042043 | 3 | 0.13080858 | 0 | 1 | 2 | 0.54052736 | 1 | 0.4761699 | 27 | 0.99530389 | TRUE | 0.04200174 | FALSE | 1 | FALSE | 1 | FALSE | 1 | FALSE | 1 | FALSE | 1 |
| 7414 | 7414 | 8.89473684 | 1.83230476 | 0 | 1 | 1 | 0.46503795 | 0 | 1 | 2 | 0.21610812 | 0 | 1 | 16 | 0.65565778 | FALSE | 1 | FALSE | 1 | FALSE | 1 | FALSE | 1 | FALSE | 1 | FALSE | 1 |
| 7415 | 7415 | 10.2631579 | 1.88741275 | 0 | 1 | 0 | 1 | 1 | 0.50143826 | 1 | 0.59104357 | 1 | 0.26986775 | 16 | 0.65565778 | FALSE | 1 | FALSE | 1 | FALSE | 1 | FALSE | 1 | FALSE | 1 | FALSE | 1 |
| 7416 | 7416 | 8.2195122 | 2.01835716 | 2 | 0.27371785 | 0 | 1 | 0 | 1 | 3 | 0.27785536 | 0 | 1 | 35 | 0.38966407 | FALSE | 1 | FALSE | 1 | FALSE | 1 | FALSE | 1 | FALSE | 1 | FALSE | 1 |
| 7417 | 7417 | 9.68181818 | 2.5115436 | 2 | 0.10809809 | 2 | 0.15794013 | 1 | 0.55340851 | 0 | 1 | 0 | 1 | 17 | 0.8840357 | FALSE | 0.88820416 | FALSE | 1 | FALSE | 1 | FALSE | 1 | FALSE | 1 | FALSE | 1 |
| 7418 | 7418 | 10.0606061 | 3.08406849 | 7 | 0.00140352 | 7 | 0.00517922 | 2 | 0.69218142 | 2 | 0.8136513 | 1 | 0.66594933 | 47 | 0.99808523 | FALSE | 0.09317759 | FALSE | 0.58120789 | FALSE | 1 | FALSE | 1 | FALSE | 1 | FALSE | 1 |
| 7419 | 7419 | 8.65517241 | 2.29355712 | 0 | 1 | 4 | 0.01340169 | 0 | 1 | 0 | 1 | 0 | 1 | 25 | 0.51779518 | FALSE | 1 | FALSE | 0.69207862 | FALSE | 1 | FALSE | 1 | FALSE | 1 | FALSE | 1 |
| 7420 | 7420 | 10.125 | 3.17952434 | 1 | 0.46441868 | 0 | 1 | 0 | 1 | 0 | 1 | 0 | 1 | 23 | 0.09196389 | FALSE | 1 | FALSE | 1 | FALSE | 1 | FALSE | 1 | FALSE | 1 | FALSE | 1 |
| 7421 | 7421 | 11.25 | 1.08972474 | 0 | 1 | 0 | 1 | 1 | 0.13615783 | 0 | 1 | 0 | 1 | 3 | 0.8823539 | FALSE | 1 | FALSE | 1 | FALSE | 1 | FALSE | 1 | FALSE | 1 | FALSE | 1 |
| 7422 | 7422 | 8.14285714 | 2.4453204 | 2 | 0.04234484 | 1 | 0.3480709 | 0 | 1 | 1 | 0.45746571 | 1 | 0.19354507 | 8 | 0.99085298 | FALSE | 0.62842194 | FALSE | 1 | FALSE | 1 | FALSE | 1 | FALSE | 1 | FALSE | 1 |
| 7423 | 7423 | 9.05555556 | 1.92850611 | 0 | 1 | 1 | 0.44710673 | 1 | 0.48280983 | 1 | 0.57131013 | 0 | 1 | 15 | 0.69372602 | FALSE | 1 | FALSE | 1 | FALSE | 1 | FALSE | 1 | FALSE | 1 | FALSE | 1 |
| 7424 | 7424 | 10.3846154 | 1.77757231 | 0 | 1 | 0 | 1 | 0 | 1 | 1 | 0.43131129 | 0 | 1 | 11 | 0.41944532 | FALSE | 1 | FALSE | 1 | FALSE | 1 | FALSE | 1 | FALSE | 1 | FALSE | 1 |

| | | | | | | | | | | | | | | | | | | | | | | | | | | | |
|---|---|---|---|---|---|---|---|---|---|---|---|---|---|---|---|---|---|---|---|---|---|---|---|---|---|---|---|
| 7425 | 7425 | 11 | 2.54950976 | 0 | 1 | 0 | 1 | 0 | 1 | 1 | 0.17140665 | 1 | 0.06400211 | 2 | 0.98656791 | FALSE | 1 | FALSE | 1 | FALSE | 1 | FALSE | 1 | FALSE | 1 | FALSE | 1 |
| 7426 | 7426 | 8.09677419 | 3.41112827 | 0 | 1 | 2 | 0.2651851 | 1 | 0.67911179 | 3 | 0.16836568 | 0 | 1 | 25 | 0.80006983 | FALSE | 1 | FALSE | 1 | FALSE | 1 | FALSE | 1 | FALSE | 1 | FALSE | 1 |
| 7427 | 7427 | 8.61538462 | 1.86211053 | 0 | 1 | 0 | 1 | 2 | 0.07725803 | 0 | 1 | 0 | 1 | 11 | 0.6683056 | FALSE | 1 | FALSE | 1 | FALSE | 1 | FALSE | 1 | FALSE | 1 | FALSE | 1 |
| 7428 | 7428 | 8.02083333 | 2.51238253 | 2 | 0.34973212 | 0 | 1 | 1 | 0.8283465 | 1 | 0.89605799 | 1 | 0.54902428 | 43 | 0.21763056 | FALSE | 1 | FALSE | 1 | FALSE | 1 | FALSE | 1 | FALSE | 1 | FALSE | 1 |
| 7429 | 7429 | 9.95652174 | 1.65446066 | 0 | 1 | 0 | 1 | 0 | 1 | 4 | 0.01932195 | 0 | 1 | 19 | 0.71542496 | FALSE | 1 | FALSE | 1 | FALSE | 1 | FALSE | 1 | FALSE | 1 | FALSE | 1 |
| 7430 | 7430 | 8.53846154 | 2.30769231 | 1 | 0.28675951 | 0 | 1 | 0 | 1 | 0 | 1 | 0 | 1 | 12 | 0.374198 | FALSE | 1 | FALSE | 1 | FALSE | 1 | FALSE | 1 | FALSE | 1 | FALSE | 1 |
| 7431 | 7431 | 10.9166667 | 1.80084857 | 0 | 1 | 1 | 0.32623933 | 0 | 1 | 0 | 1 | 0 | 1 | 11 | 0.41944532 | FALSE | 1 | FALSE | 1 | FALSE | 1 | FALSE | 1 | FALSE | 1 | FALSE | 1 |
| 7432 | 7432 | 7.57142857 | 2.43137094 | 1 | 0.66519151 | 1 | 0.74973543 | 1 | 0.78589454 | 0 | 1 | 2 | 0.15059682 | 37 | 0.34041811 | FALSE | 1 | FALSE | 1 | FALSE | 1 | FALSE | 1 | FALSE | 1 | FALSE | 1 |
| 7433 | 7433 | 8.84210526 | 2.56008137 | 1 | 0.38989951 | 0 | 1 | 1 | 0.50143826 | 0 | 1 | 0 | 1 | 17 | 0.41141543 | FALSE | 1 | FALSE | 1 | FALSE | 1 | FALSE | 1 | FALSE | 1 | FALSE | 1 |
| 7434 | 7434 | 9.56521739 | 1.8492322 | 4 | 0.02951043 | 3 | 0.18548475 | 5 | 0.02370648 | 4 | 0.1589264 | 2 | 0.17379289 | 28 | 0.99997514 | FALSE | 0.53131751 | FALSE | 1 | FALSE | 1 | FALSE | 1 | FALSE | 1 | FALSE | 1 |
| 7435 | 7435 | 8 | 1.38169856 | 0 | 1 | 2 | 0.15794013 | 1 | 0.55340851 | 1 | 0.64497439 | 0 | 1 | 18 | 0.74744813 | FALSE | 1 | FALSE | 1 | FALSE | 1 | FALSE | 1 | FALSE | 1 | FALSE | 1 |
| 7436 | 7436 | 10.72 | 2.10751038 | 2 | 0.13369621 | 0 | 1 | 2 | 0.22597287 | 1 | 0.69181315 | 0 | 1 | 20 | 0.8152498 | FALSE | 0.97964162 | FALSE | 1 | FALSE | 1 | FALSE | 1 | FALSE | 1 | FALSE | 1 |
| 7437 | 7437 | 9 | 2.62612866 | 0 | 1 | 0 | 1 | 0 | 1 | 2 | 0.3871811 | 0 | 1 | 27 | 0.14664438 | FALSE | 1 | FALSE | 1 | FALSE | 1 | FALSE | 1 | FALSE | 1 | FALSE | 1 |
| 7438 | 7438 | 9.63157895 | 2.97729171 | 0 | 1 | 0 | 1 | 1 | 0.50143826 | 0 | 1 | 1 | 0.26986775 | 17 | 0.41141543 | FALSE | 1 | FALSE | 1 | FALSE | 1 | FALSE | 1 | FALSE | 1 | FALSE | 1 |
| 7439 | 7439 | 10.6153846 | 1.90297183 | 0 | 1 | 1 | 0.32623933 | 0 | 1 | 1 | 0.43131129 | 0 | 1 | 10 | 0.71436964 | FALSE | 1 | FALSE | 1 | FALSE | 1 | FALSE | 1 | FALSE | 1 | FALSE | 1 |
| 7440 | 7440 | 10.5333333 | 1.78387842 | 1 | 0.32293355 | 1 | 0.38964394 | 0 | 1 | 0 | 1 | 1 | 0.2198258 | 12 | 0.80329917 | FALSE | 1 | FALSE | 1 | FALSE | 1 | FALSE | 1 | FALSE | 1 | FALSE | 1 |
| 7441 | 7441 | 10.4117647 | 2.80878503 | 0 | 1 | 1 | 0.42857733 | 1 | 0.46348836 | 0 | 1 | 0 | 1 | 15 | 0.49060435 | FALSE | 1 | FALSE | 1 | FALSE | 1 | FALSE | 1 | FALSE | 1 | FALSE | 1 |
| 7442 | 7442 | 10.1666667 | 2.98607881 | 0 | 1 | 1 | 0.44710673 | 0 | 1 | 2 | 0.19902682 | 1 | 0.25766374 | 14 | 0.86287418 | FALSE | 1 | FALSE | 1 | FALSE | 1 | FALSE | 1 | FALSE | 1 | FALSE | 1 |
| 7443 | 7443 | 10.5 | 0.70710678 | 0 | 1 | 0 | 1 | 0 | 1 | 0 | 1 | 0 | 1 | 8 | 0.25677166 | FALSE | 1 | FALSE | 1 | FALSE | 1 | FALSE | 1 | FALSE | 1 | FALSE | 1 |
| 7444 | 7444 | 7.73913044 | 3.09582315 | 1 | 0.45027683 | 0 | 1 | 2 | 0.19931429 | 1 | 0.6613267 | 0 | 1 | 19 | 0.71542496 | FALSE | 1 | FALSE | 1 | FALSE | 1 | FALSE | 1 | FALSE | 1 | FALSE | 1 |
| 7445 | 7445 | 7.07317073 | 1.65853659 | 1 | 0.65632651 | 3 | 0.14580275 | 2 | 0.43640583 | 1 | 0.85524728 | 0 | 1 | 34 | 0.69505404 | FALSE | 1 | FALSE | 1 | FALSE | 1 | FALSE | 1 | FALSE | 1 | FALSE | 1 |
| 7446 | 7446 | 7.8 | 2.82842713 | 0 | 1 | 0 | 1 | 0 | 1 | 1 | 0.69181315 | 0 | 1 | 24 | 0.08019798 | FALSE | 1 | FALSE | 1 | FALSE | 1 | FALSE | 1 | FALSE | 1 | FALSE | 1 |
| 7447 | 7447 | 6.6 | 2.8 | 2 | 0.36837609 | 1 | 0.80796869 | 2 | 0.54120634 | 3 | 0.4048204 | 0 | 1 | 42 | 0.62187327 | FALSE | 1 | FALSE | 1 | FALSE | 1 | FALSE | 1 | FALSE | 1 | FALSE | 1 |
| 7448 | 7448 | 9.07407407 | 2.84052585 | 1 | 0.50471032 | 1 | 0.58913618 | 1 | 0.62830986 | 0 | 1 | 0 | 1 | 24 | 0.3725373 | FALSE | 1 | FALSE | 1 | FALSE | 1 | FALSE | 1 | FALSE | 1 | FALSE | 1 |
| 7449 | 7449 | 10.3125 | 3.34885708 | 0 | 1 | 1 | 0.40942991 | 1 | 0.44344817 | 0 | 1 | 0 | 1 | 14 | 0.53308261 | FALSE | 1 | FALSE | 1 | FALSE | 1 | FALSE | 1 | FALSE | 1 | FALSE | 1 |
| 7450 | 7450 | 11.2 | 1.50332964 | 0 | 1 | 2 | 0.1353446 | 2 | 0.16019548 | 1 | 0.60987146 | 0 | 1 | 15 | 0.92113679 | FALSE | 1 | FALSE | 1 | FALSE | 1 | FALSE | 1 | FALSE | 1 | FALSE | 1 |
| 7451 | 7451 | 9.76744186 | 1.86569034 | 0 | 1 | 0 | 1 | 2 | 0.46083448 | 4 | 0.13323859 | 0 | 1 | 37 | 0.48294184 | FALSE | 1 | FALSE | 1 | FALSE | 1 | FALSE | 1 | FALSE | 1 | FALSE | 1 |
| 7452 | 7452 | 11.5178571 | 1.18005859 | 2 | 0.4229474 | 1 | 0.84259834 | 3 | 0.32646887 | 4 | 0.25497934 | 0 | 1 | 46 | 0.74916201 | FALSE | 1 | FALSE | 1 | FALSE | 1 | FALSE | 1 | FALSE | 1 | FALSE | 1 |
| 7453 | 7453 | 7 | 2.3648663 | 2 | 0.40500867 | 0 | 1 | 3 | 0.30658915 | 3 | 0.45415642 | 3 | 0.05824229 | 43 | 0.87372949 | FALSE | 1 | FALSE | 1 | FALSE | 1 | FALSE | 1 | FALSE | 1 | FALSE | 1 |
| 7454 | 7454 | 11.2352941 | 2.79767575 | 3 | 0.00864151 | 0 | 1 | 0 | 1 | 0 | 1 | 1 | 0.24525764 | 13 | 0.88719222 | FALSE | 0.27019545 | FALSE | 1 | FALSE | 1 | FALSE | 1 | FALSE | 1 | FALSE | 1 |
| 7455 | 7455 | 7.9 | 2.9478806 | 1 | 0.22886536 | 2 | 0.03949327 | 0 | 1 | 0 | 1 | 0 | 1 | 7 | 0.94326215 | FALSE | 1 | FALSE | 0.96864929 | FALSE | 1 | FALSE | 1 | FALSE | 1 | FALSE | 1 |
| 7456 | 7456 | 10.3428571 | 2.26706679 | 0 | 1 | 4 | 0.02543131 | 2 | 0.35984046 | 1 | 0.80778317 | 2 | 0.11183826 | 26 | 0.96294373 | FALSE | 1 | FALSE | 0.82367276 | FALSE | 1 | FALSE | 1 | FALSE | 1 | FALSE | 1 |
| 7457 | 7457 | 7.5106383 | 1.8320125 | 2 | 0.34034075 | 1 | 0.78790827 | 2 | 0.50778118 | 2 | 0.64277396 | 1 | 0.54145295 | 39 | 0.69224623 | FALSE | 1 | FALSE | 1 | FALSE | 1 | FALSE | 1 | FALSE | 1 | FALSE | 1 |
| 7458 | 7458 | 8.68571429 | 2.20092745 | 1 | 0.59801736 | 1 | 0.68454804 | 1 | 0.72299506 | 1 | 0.80778317 | 1 | 0.44015537 | 30 | 0.52765882 | FALSE | 1 | FALSE | 1 | FALSE | 1 | FALSE | 1 | FALSE | 1 | FALSE | 1 |
| 7459 | 7459 | 9.11111111 | 2.15738754 | 1 | 0.37380237 | 1 | 0.44710673 | 0 | 1 | 0 | 1 | 1 | 0.25766374 | 15 | 0.69372602 | FALSE | 1 | FALSE | 1 | FALSE | 1 | FALSE | 1 | FALSE | 1 | FALSE | 1 |
| 7460 | 7460 | 10.1764706 | 2.50259381 | 0 | 1 | 0 | 1 | 0 | 1 | 2 | 0.18212134 | 1 | 0.24525764 | 14 | 0.7313145 | FALSE | 1 | FALSE | 1 | FALSE | 1 | FALSE | 1 | FALSE | 1 | FALSE | 1 |
| 7461 | 7461 | 10.5555556 | 2.54344959 | 0 | 1 | 0 | 1 | 0 | 1 | 1 | 0.34506517 | 0 | 1 | 8 | 0.5780332 | FALSE | 1 | FALSE | 1 | FALSE | 1 | FALSE | 1 | FALSE | 1 | FALSE | 1 |
| 7462 | 7462 | 9.28571429 | 3.05727637 | 2 | 0.39594061 | 4 | 0.09132555 | 4 | 0.12170726 | 1 | 0.91797304 | 0 | 1 | 42 | 0.88654191 | FALSE | 1 | FALSE | 1 | FALSE | 1 | FALSE | 1 | FALSE | 1 | FALSE | 1 |
| 7463 | 7463 | 4.79166667 | 3.12222099 | 0 | 1 | 0 | 1 | 1 | 0.58502114 | 1 | 0.67692816 | 0 | 1 | 22 | 0.25237063 | FALSE | 1 | FALSE | 1 | FALSE | 1 | FALSE | 1 | FALSE | 1 | FALSE | 1 |
| 7464 | 7464 | 11.0833333 | 2.7220805 | 0 | 1 | 0 | 1 | 0 | 1 | 0 | 1 | 0 | 1 | 12 | 0.13002667 | FALSE | 1 | FALSE | 1 | FALSE | 1 | FALSE | 1 | FALSE | 1 | FALSE | 1 |
| 7465 | 7465 | 7.43089431 | 1.57511802 | 7 | 0.42316075 | 10 | 0.25308189 | 6 | 0.87114908 | 10 | 0.67565282 | 3 | 0.76242935 | 205 | 0.42524256 | FALSE | 1 | FALSE | 1 | FALSE | 1 | FALSE | 1 | FALSE | 1 | FALSE | 1 |
| 7466 | 7466 | 11 | 1.87082869 | 1 | 0.38989951 | 2 | 0.12434464 | 0 | 1 | 1 | 0.59104357 | 1 | 0.26986775 | 14 | 0.93677253 | FALSE | 1 | FALSE | 1 | FALSE | 1 | FALSE | 1 | FALSE | 1 | FALSE | 1 |
| 7467 | 7467 | 9.77777778 | 3.0469576 | 1 | 0.20854748 | 0 | 1 | 0 | 1 | 1 | 0.34506517 | 0 | 1 | 7 | 0.84574168 | FALSE | 1 | FALSE | 1 | FALSE | 1 | FALSE | 1 | FALSE | 1 | FALSE | 1 |
| 7468 | 7468 | 7.58536585 | 2.8794953 | 0 | 1 | 3 | 0.12352882 | 1 | 0.75193685 | 2 | 0.52640931 | 0 | 1 | 32 | 0.61800916 | FALSE | 1 | FALSE | 1 | FALSE | 1 | FALSE | 1 | FALSE | 1 | FALSE | 1 |
| 7469 | 7469 | 8.76923077 | 1.86687094 | 1 | 0.49162634 | 2 | 0.20486827 | 1 | 0.61440466 | 2 | 0.33678042 | 0 | 1 | 20 | 0.90142689 | FALSE | 1 | FALSE | 1 | FALSE | 1 | FALSE | 1 | FALSE | 1 | FALSE | 1 |
| 7470 | 7470 | 7.1875 | 3.06632414 | 1 | 0.56527493 | 0 | 1 | 1 | 0.69069222 | 3 | 0.17988885 | 0 | 1 | 27 | 0.61671133 | FALSE | 1 | FALSE | 1 | FALSE | 1 | FALSE | 1 | FALSE | 1 | FALSE | 1 |
| 7471 | 7471 | 9.42857143 | 2.34157921 | 0 | 1 | 0 | 1 | 1 | 0.48280983 | 2 | 0.19902682 | 0 | 1 | 15 | 0.69372602 | FALSE | 1 | FALSE | 1 | FALSE | 1 | FALSE | 1 | FALSE | 1 | FALSE | 1 |
| 7472 | 7472 | 9.93103448 | 1.96400426 | 2 | 0.16962005 | 1 | 0.61539065 | 1 | 0.65463982 | 2 | 0.3871811 | 0 | 1 | 23 | 0.84462957 | FALSE | 1 | FALSE | 1 | FALSE | 1 | FALSE | 1 | FALSE | 1 | FALSE | 1 |
| 7473 | 7473 | 9.94736842 | 2.32772046 | 0 | 1 | 0 | 1 | 3 | 0.02898764 | 0 | 1 | 0 | 1 | 16 | 0.65565778 | FALSE | 1 | FALSE | 1 | FALSE | 1 | FALSE | 1 | FALSE | 1 | FALSE | 1 |
| 7474 | 7474 | 9.48275862 | 2.9898759 | 1 | 0.52988231 | 3 | 0.06581417 | 1 | 0.65463982 | 4 | 0.04184267 | 1 | 0.38148475 | 19 | 0.99707037 | FALSE | 1 | FALSE | 1 | FALSE | 1 | FALSE | 1 | FALSE | 1 | FALSE | 1 |
| 7475 | 7475 | 10.2608696 | 2.11037584 | 2 | 0.09989125 | 0 | 1 | 1 | 0.53671415 | 0 | 1 | 0 | 1 | 18 | 0.57962763 | FALSE | 0.86047225 | FALSE | 1 | FALSE | 1 | FALSE | 1 | FALSE | 1 | FALSE | 1 |
| 7476 | 7476 | 9.33333333 | 2.153808 | 1 | 0.46441868 | 0 | 1 | 0 | 1 | 3 | 0.09525947 | 0 | 1 | 20 | 0.68270729 | FALSE | 1 | FALSE | 1 | FALSE | 1 | FALSE | 1 | FALSE | 1 | FALSE | 1 |
| 7477 | 7477 | 10.1764706 | 3.31192665 | 0 | 1 | 2 | 0.10307112 | 1 | 0.46348836 | 0 | 1 | 1 | 0.24525764 | 13 | 0.88719222 | FALSE | 1 | FALSE | 1 | FALSE | 1 | FALSE | 1 | FALSE | 1 | FALSE | 1 |
| 7478 | 7478 | 9.6 | 1.0198039 | 0 | 1 | 0 | 1 | 0 | 1 | 0 | 1 | 0 | 1 | 5 | 0.42761893 | FALSE | 1 | FALSE | 1 | FALSE | 1 | FALSE | 1 | FALSE | 1 | FALSE | 1 |
| 7479 | 7479 | 11.8709677 | 2.61190742 | 1 | 0.55378205 | 0 | 1 | 2 | 0.30673593 | 1 | 0.76781284 | 0 | 1 | 27 | 0.45522639 | FALSE | 1 | FALSE | 1 | FALSE | 1 | FALSE | 1 | FALSE | 1 | FALSE | 1 |
| 7480 | 7480 | 8.28571429 | 2.18529408 | 1 | 0.28675951 | 0 | 1 | 0 | 1 | 0 | 1 | 1 | 0.19354507 | 11 | 0.6683056 | FALSE | 1 | FALSE | 1 | FALSE | 1 | FALSE | 1 | FALSE | 1 | FALSE | 1 |
| 7481 | 7481 | 12.25 | 3.1124749 | 0 | 1 | 0 | 1 | 0 | 1 | 0 | 1 | 0 | 1 | 4 | 0.50683841 | FALSE | 1 | FALSE | 1 | FALSE | 1 | FALSE | 1 | FALSE | 1 | FALSE | 1 |
| 7482 | 7482 | 8.58333333 | 2.51523138 | 1 | 0.46441868 | 0 | 1 | 0 | 1 | 0 | 1 | 2 | 0.058175 | 21 | 0.47053137 | FALSE | 1 | FALSE | 1 | FALSE | 1 | FALSE | 1 | FALSE | 1 | FALSE | 1 |
| 7483 | 7483 | 5.73033708 | 1.90660617 | 2 | 0.65852142 | 3 | 0.53828446 | 5 | 0.20230288 | 0 | 1 | 1 | 0.76487586 | 76 | 0.27423034 | FALSE | 1 | FALSE | 1 | FALSE | 1 | FALSE | 1 | FALSE | 1 | FALSE | 1 |
| 7484 | 7484 | 7.5 | 3.29393382 | 0 | 1 | 6 | 2.83E-05 | 2 | 0.16019548 | 0 | 1 | 1 | 0.28187291 | 11 | 0.99965905 | FALSE | 1 | FALSE | 0.07208458 | FALSE | 1 | FALSE | 1 | FALSE | 1 | FALSE | 1 |
| 7485 | 7485 | 10.6666667 | 3.14466038 | 1 | 0.46441868 | 1 | 0.54637326 | 0 | 1 | 1 | 0.67692816 | 0 | 1 | 21 | 0.47053137 | FALSE | 1 | FALSE | 1 | FALSE | 1 | FALSE | 1 | FALSE | 1 | FALSE | 1 |
| 7486 | 7486 | 9.39130435 | 4.04043832 | 3 | 0.02011786 | 2 | 0.16948764 | 1 | 0.5695037 | 0 | 1 | 3 | 0.00598158 | 14 | 0.99865094 | FALSE | 0.44482142 | FALSE | 1 | FALSE | 1 | FALSE | 1 | FALSE | 0.45694206 | FALSE | 1 |
| 7487 | 7487 | 7.92857143 | 2.50136017 | 1 | 0.65632651 | 0 | 1 | 1 | 0.77786601 | 1 | 0.85524728 | 0 | 1 | 38 | 0.09794002 | FALSE | 1 | FALSE | 1 | FALSE | 1 | FALSE | 1 | FALSE | 1 | FALSE | 1 |
| 7488 | 7488 | 10.1428571 | 3.24089515 | 1 | 0.42086991 | 0 | 1 | 4 | 0.0060005 | 1 | 0.62783522 | 1 | 0.29368245 | 14 | 0.98949398 | FALSE | 1 | FALSE | 1 | FALSE | 1 | FALSE | 1 | FALSE | 1 | FALSE | 1 |
| 7489 | 7489 | 12 | 2.34520788 | 0 | 1 | 0 | 1 | 1 | 0.25384309 | 0 | 1 | 0 | 1 | 7 | 0.63752397 | FALSE | 1 | FALSE | 1 | FALSE | 1 | FALSE | 1 | FALSE | 1 | FALSE | 1 |
| 7490 | 7490 | 8.73913044 | 2.45334539 | 1 | 0.45027683 | 1 | 0.53115654 | 0 | 1 | 0 | 1 | 1 | 0.31672722 | 20 | 0.50589402 | FALSE | 1 | FALSE | 1 | FALSE | 1 | FALSE | 1 | FALSE | 1 | FALSE | 1 |
| 7491 | 7491 | 8.975 | 2.60276296 | 6 | 0.00048425 | 0 | 1 | 0 | 1 | 2 | 0.5543522 | 1 | 0.48480878 | 31 | 0.91676852 | TRUE | 0.0450788 | FALSE | 1 | FALSE | 1 | FALSE | 1 | FALSE | 1 | FALSE | 1 |
| 7492 | 7492 | 9.16666667 | 2.78388218 | 2 | 0.23542589 | 0 | 1 | 1 | 0.73299778 | 4 | 0.08109727 | 1 | 0.44938266 | 28 | 0.90259714 | FALSE | 1 | FALSE | 1 | FALSE | 1 | FALSE | 1 | FALSE | 1 | FALSE | 1 |
| 7493 | 7493 | 6.64285714 | 3.28649059 | 1 | 0.65632651 | 1 | 0.74131698 | 2 | 0.43640583 | 3 | 0.29054213 | 3 | 0.02912063 | 31 | 0.95455175 | FALSE | 1 | FALSE | 1 | FALSE | 1 | FALSE | 1 | FALSE | 0.81978088 | FALSE | 1 |
| 7494 | 7494 | 10 | 1.71824939 | 1 | 0.42086991 | 1 | 0.49918218 | 1 | 0.53671415 | 1 | 0.62783522 | 1 | 0.29368245 | 16 | 0.90353674 | FALSE | 1 | FALSE | 1 | FALSE | 1 | FALSE | 1 | FALSE | 1 | FALSE | 1 |
| 7495 | 7495 | 8.2 | 2.28619043 | 0 | 1 | 1 | 0.38964394 | 0 | 1 | 1 | 0.50623194 | 0 | 1 | 13 | 0.57712805 | FALSE | 1 | FALSE | 1 | FALSE | 1 | FALSE | 1 | FALSE | 1 | FALSE | 1 |
| 7496 | 7496 | 11 | 3.19374389 | 0 | 1 | 0 | 1 | 0 | 1 | 1 | 0.60987146 | 0 | 1 | 19 | 0.15700255 | FALSE | 1 | FALSE | 1 | FALSE | 1 | FALSE | 1 | FALSE | 1 | FALSE | 1 |
| 7497 | 7497 | 7.02702703 | 2.07281616 | 5 | 0.0387417 | 2 | 0.68961994 | 2 | 0.74419436 | 4 | 0.43314506 | 3 | 0.11741889 | 57 | 0.94545203 | FALSE | 0.6033689 | FALSE | 1 | FALSE | 1 | FALSE | 1 | FALSE | 1 | FALSE | 1 |
| 7498 | 7498 | 9.58333333 | 2.59673941 | 1 | 0.26795838 | 0 | 1 | 1 | 0.35555293 | 2 | 0.1023796 | 1 | 0.18007762 | 7 | 0.99433481 | FALSE | 1 | FALSE | 1 | FALSE | 1 | FALSE | 1 | FALSE | 1 | FALSE | 1 |
| 7499 | 7499 | 10.3783784 | 2.23427056 | 6 | 0.00031256 | 3 | 0.11640364 | 0 | 1 | 2 | 0.51200191 | 0 | 1 | 26 | 0.99170905 | TRUE | 0.03507515 | FALSE | 1 | FALSE | 1 | FALSE | 1 | FALSE | 1 | FALSE | 1 |
| 7500 | 7500 | 9.33333333 | 2.21108319 | 0 | 1 | 0 | 1 | 0 | 1 | 0 | 1 | 0 | 1 | 6 | 0.36077158 | FALSE | 1 | FALSE | 1 | FALSE | 1 | FALSE | 1 | FALSE | 1 | FALSE | 1 |
| 7501 | 7501 | 10.7916667 | 3.02736363 | 0 | 1 | 1 | 0.54637326 | 3 | 0.05314317 | 2 | 0.30249307 | 1 | 0.32796862 | 17 | 0.97530702 | FALSE | 1 | FALSE | 1 | FALSE | 1 | FALSE | 1 | FALSE | 1 | FALSE | 1 |
| 7502 | 7502 | 11.1666667 | 3.48409083 | 0 | 1 | 0 | 1 | 0 | 1 | 2 | 0.1023796 | 0 | 1 | 10 | 0.71436964 | FALSE | 1 | FALSE | 1 | FALSE | 1 | FALSE | 1 | FALSE | 1 | FALSE | 1 |
| 7503 | 7503 | 9.85714286 | 2.29462549 | 0 | 1 | 1 | 0.20567256 | 0 | 1 | 0 | 1 | 0 | 1 | 6 | 0.69920685 | FALSE | 1 | FALSE | 1 | FALSE | 1 | FALSE | 1 | FALSE | 1 | FALSE | 1 |
| 7504 | 7504 | 7.30487805 | 1.26576907 | 5 | 0.05874466 | 3 | 0.49822953 | 7 | 0.02736229 | 4 | 0.52386423 | 0 | 1 | 63 | 0.97509386 | FALSE | 0.72149357 | FALSE | 1 | FALSE | 1 | FALSE | 1 | FALSE | 1 | FALSE | 1 |
| 7505 | 7505 | 10.4 | 3.61109402 | 0 | 1 | 0 | 1 | 0 | 1 | 1 | 0.2094688 | 0 | 1 | 4 | 0.82371636 | FALSE | 1 | FALSE | 1 | FALSE | 1 | FALSE | 1 | FALSE | 1 | FALSE | 1 |
| 7506 | 7506 | 8.66666667 | 1.97202659 | 2 | 0.17884234 | 1 | 0.6278847 | 0 | 1 | 0 | 1 | 0 | 1 | 27 | 0.28862208 | FALSE | 1 | FALSE | 1 | FALSE | 1 | FALSE | 1 | FALSE | 1 | FALSE | 1 |
| 7507 | 7507 | 8.83333333 | 3.54729944 | 0 | 1 | 1 | 0.44710673 | 1 | 0.48280983 | 0 | 1 | 0 | 1 | 16 | 0.44998001 | FALSE | 1 | FALSE | 1 | FALSE | 1 | FALSE | 1 | FALSE | 1 | FALSE | 1 |
| 7508 | 7508 | 9.67857143 | 2.76526376 | 1 | 0.47819878 | 0 | 1 | 3 | 0.05884411 | 0 | 1 | 0 | 1 | 21 | 0.64957167 | FALSE | 1 | FALSE | 1 | FALSE | 1 | FALSE | 1 | FALSE | 1 | FALSE | 1 |
| 7509 | 7509 | 8 | 2.36643191 | 0 | 1 | 1 | 0.28036922 | 1 | 0.30655256 | 0 | 1 | 0 | 1 | 8 | 0.80394719 | FALSE | 1 | FALSE | 1 | FALSE | 1 | FALSE | 1 | FALSE | 1 | FALSE | 1 |
| 7510 | 7510 | 10.0892857 | 2.4804722 | 1 | 0.76787545 | 5 | 0.03395652 | 5 | 0.04964325 | 0 | 1 | 4 | 0.01306774 | 41 | 0.99027378 | FALSE | 1 | FALSE | 0.93660537 | FALSE | 1 | FALSE | 1 | FALSE | 0.63731057 | FALSE | 1 |
| 7511 | 7511 | 9.40909091 | 2.60522362 | 1 | 0.4357638 | 1 | 0.5154318 | 1 | 0.55340851 | 0 | 1 | 0 | 1 | 19 | 0.54233419 | FALSE | 1 | FALSE | 1 | FALSE | 1 | FALSE | 1 | FALSE | 1 | FALSE | 1 |
| 7512 | 7512 | 10.5652174 | 3.73963596 | 1 | 0.45027683 | 0 | 1 | 1 | 0.5695037 | 1 | 0.6613267 | 1 | 0.31672722 | 19 | 0.71542496 | FALSE | 1 | FALSE | 1 | FALSE | 1 | FALSE | 1 | FALSE | 1 | FALSE | 1 |
| 7513 | 7513 | 7.85714286 | 2.98066331 | 1 | 0.42086991 | 2 | 0.14655134 | 0 | 1 | 0 | 1 | 1 | 0.29368245 | 17 | 0.77849076 | FALSE | 1 | FALSE | 1 | FALSE | 1 | FALSE | 1 | FALSE | 1 | FALSE | 1 |
| 7514 | 7514 | 9.5862069 | 2.95386645 | 1 | 0.52988231 | 1 | 0.61539065 | 1 | 0.65463982 | 1 | 0.74482229 | 0 | 1 | 25 | 0.51779518 | FALSE | 1 | FALSE | 1 | FALSE | 1 | FALSE | 1 | FALSE | 1 | FALSE | 1 |
| 7515 | 7515 | 9.61111111 | 3.1116071 | 0 | 1 | 0 | 1 | 2 | 0.12280513 | 0 | 1 | 0 | 1 | 15 | 0.49060435 | FALSE | 1 | FALSE | 1 | FALSE | 1 | FALSE | 1 | FALSE | 1 | FALSE | 1 |
| 7516 | 7516 | 8.42857143 | 2.47023092 | 1 | 0.51745948 | 4 | 0.01184672 | 1 | 0.64171563 | 0 | 1 | 2 | 0.07640798 | 20 | 0.97757517 | FALSE | 1 | FALSE | 0.67617265 | FALSE | 1 | FALSE | 1 | FALSE | 1 | FALSE | 1 |
| 7517 | 7517 | 10 | 2.23606798 | 0 | 1 | 1 | 0.44710673 | 0 | 1 | 0 | 1 | 2 | 0.03435384 | 15 | 0.69372602 | FALSE | 1 | FALSE | 1 | FALSE | 1 | FALSE | 1 | FALSE | 0.86063198 | FALSE | 1 |
| 7518 | 7518 | 7.66666667 | 2.7968236 | 3 | 0.03713651 | 0 | 1 | 0 | 1 | 1 | 0.74482229 | 1 | 0.38148475 | 24 | 0.70563334 | FALSE | 0.58392 | FALSE | 1 | FALSE | 1 | FALSE | 1 | FALSE | 1 | FALSE | 1 |

| | | | | | | | | | | | | | | | | | | | | | | | | | | | |
|---|---|---|---|---|---|---|---|---|---|---|---|---|---|---|---|---|---|---|---|---|---|---|---|---|---|---|---|
| 7519 | 7519 | 6.38596491 | 2.38970295 | 1 | 0.76787545 | 3 | 0.27128673 | 1 | 0.87219471 | 1 | 0.92884295 | 0 | 1 | 50 | 0.20672933 | FALSE | 1 | FALSE | 1 | FALSE | 1 | FALSE | 1 | FALSE | 1 | FALSE | 1 |
| 7520 | 7520 | 9.93333333 | 2.76807193 | 0 | 1 | 1 | 0.61539065 | 1 | 0.65463982 | 1 | 0.74482229 | 0 | 1 | 26 | 0.31495157 | FALSE | 1 | FALSE | 1 | FALSE | 1 | FALSE | 1 | FALSE | 1 | FALSE | 1 |
| 7521 | 7521 | 9.23809524 | 1.68762335 | 0 | 1 | 2 | 0.14655134 | 0 | 1 | 1 | 0.62783522 | 0 | 1 | 18 | 0.57962763 | FALSE | 1 | FALSE | 1 | FALSE | 1 | FALSE | 1 | FALSE | 1 | FALSE | 1 |
| 7522 | 7522 | 9.625 | 1.79843682 | 0 | 1 | 0 | 1 | 0 | 1 | 2 | 0.048966 | 1 | 0.12394282 | 5 | 0.97551477 | FALSE | 1 | FALSE | 1 | FALSE | 1 | FALSE | 1 | FALSE | 1 | FALSE | 1 |
| 7523 | 7523 | 7.34615385 | 3.22182829 | 1 | 0.49162634 | 2 | 0.20486827 | 1 | 0.61440466 | 1 | 0.70601446 | 2 | 0.0670823 | 19 | 0.96054244 | FALSE | 1 | FALSE | 1 | FALSE | 1 | FALSE | 1 | FALSE | 1 | FALSE | 1 |
| 7524 | 7524 | 9.26666667 | 2.50244325 | 1 | 0.51745948 | 0 | 1 | 3 | 0.07756639 | 0 | 1 | 0 | 1 | 24 | 0.55018107 | FALSE | 1 | FALSE | 1 | FALSE | 1 | FALSE | 1 | FALSE | 1 | FALSE | 1 |
| 7525 | 7525 | 3.76744186 | 2.1112898 | 12 | 0.0001061 | 5 | 0.40561876 | 4 | 0.68714676 | 8 | 0.24036121 | 5 | 0.06022929 | 95 | 0.99948541 | TRUE | 0.01849285 | FALSE | 1 | FALSE | 1 | FALSE | 1 | FALSE | 1 | FALSE | 1 |
| 7526 | 7526 | 10.3 | 1.84661853 | 0 | 1 | 0 | 1 | 0 | 1 | 0 | 1 | 0 | 1 | 10 | 0.18273166 | FALSE | 1 | FALSE | 1 | FALSE | 1 | FALSE | 1 | FALSE | 1 | FALSE | 1 |
| 7527 | 7527 | 8.88888889 | 3.08920862 | 1 | 0.37380237 | 1 | 0.44710673 | 0 | 1 | 0 | 1 | 0 | 1 | 16 | 0.44998001 | FALSE | 1 | FALSE | 1 | FALSE | 1 | FALSE | 1 | FALSE | 1 | FALSE | 1 |
| 7528 | 7528 | 9.54761905 | 2.57462324 | 1 | 0.66519151 | 2 | 0.39616059 | 4 | 0.0626749 | 2 | 0.5811115 | 0 | 1 | 33 | 0.89139527 | FALSE | 1 | FALSE | 1 | FALSE | 1 | FALSE | 1 | FALSE | 1 | FALSE | 1 |
| 7529 | 7529 | 4.23933975 | 1.33637857 | 19 | 0.50133364 | 30 | 0.09484881 | 28 | 0.37618045 | 34 | 0.48178286 | 15 | 0.20813434 | 601 | 0.91787474 | FALSE | 1 | FALSE | 1 | FALSE | 1 | FALSE | 1 | FALSE | 1 | FALSE | 1 |
| 7530 | 7530 | 9.35294118 | 2.16770891 | 2 | 0.06909038 | 1 | 0.42857733 | 2 | 0.12280513 | 0 | 1 | 0 | 1 | 12 | 0.962093 | FALSE | 0.74276693 | FALSE | 1 | FALSE | 1 | FALSE | 1 | FALSE | 1 | FALSE | 1 |
| 7531 | 7531 | 8.25423729 | 2.06334614 | 2 | 0.4493251 | 0 | 1 | 2 | 0.63179508 | 2 | 0.76144533 | 1 | 0.62455813 | 52 | 0.27701375 | FALSE | 1 | FALSE | 1 | FALSE | 1 | FALSE | 1 | FALSE | 1 | FALSE | 1 |
| 7532 | 7532 | 7.475 | 1.49979165 | 0 | 1 | 2 | 0.37295654 | 3 | 0.17245215 | 2 | 0.5543522 | 0 | 1 | 33 | 0.71942635 | FALSE | 1 | FALSE | 1 | FALSE | 1 | FALSE | 1 | FALSE | 1 | FALSE | 1 |
| 7533 | 7533 | 10 | 2.5 | 2 | 0.06197875 | 1 | 0.40942991 | 1 | 0.44344817 | 0 | 1 | 0 | 1 | 12 | 0.90922186 | FALSE | 0.72532843 | FALSE | 1 | FALSE | 1 | FALSE | 1 | FALSE | 1 | FALSE | 1 |
| 7534 | 7534 | 10.9285714 | 2.25085018 | 0 | 1 | 1 | 0.32623933 | 1 | 0.35555293 | 1 | 0.43131129 | 0 | 1 | 9 | 0.89630573 | FALSE | 1 | FALSE | 1 | FALSE | 1 | FALSE | 1 | FALSE | 1 | FALSE | 1 |
| 7535 | 7535 | 9.77777778 | 2.41650702 | 0 | 1 | 0 | 1 | 0 | 1 | 0 | 1 | 0 | 1 | 18 | 0.04681656 | FALSE | 1 | FALSE | 1 | FALSE | 1 | FALSE | 1 | FALSE | 1 | FALSE | 1 |
| 7536 | 7536 | 6.8490566 | 2.55797847 | 2 | 0.34973212 | 2 | 0.46321244 | 3 | 0.24750546 | 1 | 0.89605799 | 2 | 0.18560062 | 38 | 0.88187597 | FALSE | 1 | FALSE | 1 | FALSE | 1 | FALSE | 1 | FALSE | 1 | FALSE | 1 |
| 7537 | 7537 | 11.5555556 | 3.13088951 | 0 | 1 | 1 | 0.25628163 | 0 | 1 | 2 | 0.06108592 | 0 | 1 | 6 | 0.96128317 | FALSE | 1 | FALSE | 1 | FALSE | 1 | FALSE | 1 | FALSE | 1 | FALSE | 1 |
| 7538 | 7538 | 10.6111111 | 2.75154278 | 2 | 0.07645454 | 1 | 0.44710673 | 0 | 1 | 1 | 0.57131013 | 0 | 1 | 14 | 0.86287418 | FALSE | 0.77424195 | FALSE | 1 | FALSE | 1 | FALSE | 1 | FALSE | 1 | FALSE | 1 |
| 7539 | 7539 | 9.61538462 | 2.16753889 | 0 | 1 | 0 | 1 | 1 | 0.37874417 | 1 | 0.45746571 | 0 | 1 | 11 | 0.6683056 | FALSE | 1 | FALSE | 1 | FALSE | 1 | FALSE | 1 | FALSE | 1 | FALSE | 1 |
| 7540 | 7540 | 10.4333333 | 2.29031779 | 3 | 0.03713651 | 2 | 0.24095584 | 0 | 1 | 1 | 0.74482229 | 2 | 0.08121636 | 21 | 0.97185642 | FALSE | 0.58392 | FALSE | 1 | FALSE | 1 | FALSE | 1 | FALSE | 1 | FALSE | 1 |
| 7541 | 7541 | 9.34782609 | 3.19794357 | 2 | 0.11647716 | 3 | 0.03669128 | 1 | 0.5695037 | 1 | 0.6613267 | 0 | 1 | 16 | 0.98100709 | FALSE | 0.93181727 | FALSE | 0.94259695 | FALSE | 1 | FALSE | 1 | FALSE | 1 | FALSE | 1 |
| 7542 | 7542 | 9.91666667 | 0.9537936 | 1 | 0.26795838 | 0 | 1 | 1 | 0.35555293 | 0 | 1 | 0 | 1 | 10 | 0.71436964 | FALSE | 1 | FALSE | 1 | FALSE | 1 | FALSE | 1 | FALSE | 1 | FALSE | 1 |
| 7543 | 7543 | 5.97916667 | 3.04473304 | 4 | 0.03383958 | 3 | 0.20208567 | 1 | 0.8283465 | 1 | 0.89605799 | 1 | 0.54902428 | 38 | 0.88187597 | FALSE | 0.56934354 | FALSE | 1 | FALSE | 1 | FALSE | 1 | FALSE | 1 | FALSE | 1 |
| 7544 | 7544 | 8.23076923 | 1.12001691 | 0 | 1 | 1 | 0.3480709 | 0 | 1 | 0 | 1 | 0 | 1 | 12 | 0.374198 | FALSE | 1 | FALSE | 1 | FALSE | 1 | FALSE | 1 | FALSE | 1 | FALSE | 1 |
| 7545 | 7545 | 8.12222222 | 2.19001663 | 5 | 0.0748259 | 1 | 0.94566665 | 1 | 0.96086551 | 2 | 0.91766807 | 0 | 1 | 79 | 0.10038194 | FALSE | 0.77424195 | FALSE | 1 | FALSE | 1 | FALSE | 1 | FALSE | 1 | FALSE | 1 |
| 7546 | 7546 | 10.75 | 2.56838294 | 0 | 1 | 2 | 0.38460545 | 2 | 0.43640583 | 2 | 0.56788092 | 0 | 1 | 35 | 0.53616095 | FALSE | 1 | FALSE | 1 | FALSE | 1 | FALSE | 1 | FALSE | 1 | FALSE | 1 |
| 7547 | 7547 | 9.32 | 2.75274409 | 2 | 0.13369621 | 0 | 1 | 1 | 0.59998149 | 1 | 0.69181315 | 0 | 1 | 21 | 0.64957167 | FALSE | 0.97964162 | FALSE | 1 | FALSE | 1 | FALSE | 1 | FALSE | 1 | FALSE | 1 |
| 7548 | 7548 | 9.41666667 | 1.75396377 | 3 | 0.02256792 | 3 | 0.04096158 | 1 | 0.58502114 | 1 | 0.67692816 | 0 | 1 | 16 | 0.99240722 | FALSE | 0.4692294 | FALSE | 0.9698186 | FALSE | 1 | FALSE | 1 | FALSE | 1 | FALSE | 1 |
| 7549 | 7549 | 7.9 | 1.40178458 | 4 | 0.14879806 | 5 | 0.11631842 | 3 | 0.5528654 | 2 | 0.8885053 | 2 | 0.37884371 | 64 | 0.88989564 | FALSE | 1 | FALSE | 1 | FALSE | 1 | FALSE | 1 | FALSE | 1 | FALSE | 1 |
| 7550 | 7550 | 11.2 | 2.0880613 | 3 | 0.00174009 | 0 | 1 | 0 | 1 | 0 | 1 | 0 | 1 | 7 | 0.94326215 | FALSE | 0.10881549 | FALSE | 1 | FALSE | 1 | FALSE | 1 | FALSE | 1 | FALSE | 1 |
| 7551 | 7551 | 10.2727273 | 2.79905533 | 0 | 1 | 0 | 1 | 0 | 1 | 0 | 1 | 1 | 0.16638736 | 10 | 0.46859937 | FALSE | 1 | FALSE | 1 | FALSE | 1 | FALSE | 1 | FALSE | 1 | FALSE | 1 |
| 7552 | 7552 | 10.25 | 1.97857846 | 0 | 1 | 0 | 1 | 3 | 0.20921284 | 3 | 0.32878418 | 0 | 1 | 38 | 0.45705039 | FALSE | 1 | FALSE | 1 | FALSE | 1 | FALSE | 1 | FALSE | 1 | FALSE | 1 |
| 7553 | 7553 | 9.05555556 | 2.69716358 | 0 | 1 | 1 | 0.44710673 | 1 | 0.48280983 | 1 | 0.57131013 | 0 | 1 | 15 | 0.69372602 | FALSE | 1 | FALSE | 1 | FALSE | 1 | FALSE | 1 | FALSE | 1 | FALSE | 1 |
| 7554 | 7554 | 6.30051814 | 1.21427167 | 2 | 0.96149414 | 8 | 0.28653776 | 5 | 0.82977877 | 8 | 0.66805145 | 4 | 0.39032257 | 166 | 0.30263482 | FALSE | 1 | FALSE | 1 | FALSE | 1 | FALSE | 1 | FALSE | 1 | FALSE | 1 |
| 7555 | 7555 | 8 | 1.74717818 | 2 | 0.08405377 | 0 | 1 | 0 | 1 | 1 | 0.59104357 | 0 | 1 | 16 | 0.65565778 | FALSE | 0.79236883 | FALSE | 1 | FALSE | 1 | FALSE | 1 | FALSE | 1 | FALSE | 1 |
| 7556 | 7556 | 9.14285714 | 2.84999105 | 1 | 0.51745948 | 3 | 0.06039219 | 0 | 1 | 3 | 0.13524571 | 0 | 1 | 21 | 0.94083768 | FALSE | 1 | FALSE | 1 | FALSE | 1 | FALSE | 1 | FALSE | 1 | FALSE | 1 |
| 7557 | 7557 | 10.7777778 | 2.93552107 | 0 | 1 | 1 | 0.44710673 | 0 | 1 | 1 | 0.57131013 | 0 | 1 | 16 | 0.44998001 | FALSE | 1 | FALSE | 1 | FALSE | 1 | FALSE | 1 | FALSE | 1 | FALSE | 1 |
| 7558 | 7558 | 10.7857143 | 2.36578502 | 0 | 1 | 0 | 1 | 1 | 0.40110418 | 0 | 1 | 0 | 1 | 13 | 0.33283729 | FALSE | 1 | FALSE | 1 | FALSE | 1 | FALSE | 1 | FALSE | 1 | FALSE | 1 |
| 7559 | 7559 | 8.8125 | 1.57660302 | 0 | 1 | 3 | 0.19374024 | 1 | 0.82190381 | 0 | 1 | 0 | 1 | 43 | 0.12168195 | FALSE | 1 | FALSE | 1 | FALSE | 1 | FALSE | 1 | FALSE | 1 | FALSE | 1 |
| 7560 | 7560 | 3.55865922 | 1.93776649 | 8 | 0.08857042 | 3 | 0.93364762 | 9 | 0.19421319 | 6 | 0.83783813 | 7 | 0.02725145 | 146 | 0.87486398 | FALSE | 0.82638822 | FALSE | 1 | FALSE | 1 | FALSE | 1 | FALSE | 0.79106376 | FALSE | 1 |
| 7561 | 7561 | 10.3968254 | 1.36323242 | 1 | 0.80677819 | 5 | 0.05224458 | 4 | 0.18968799 | 3 | 0.55805561 | 0 | 1 | 50 | 0.89566304 | FALSE | 1 | FALSE | 1 | FALSE | 1 | FALSE | 1 | FALSE | 1 | FALSE | 1 |
| 7562 | 7562 | 8.61111111 | 3.30216913 | 1 | 0.37380237 | 0 | 1 | 1 | 0.48280983 | 1 | 0.57131013 | 1 | 0.25766374 | 14 | 0.86287418 | FALSE | 1 | FALSE | 1 | FALSE | 1 | FALSE | 1 | FALSE | 1 | FALSE | 1 |
| 7563 | 7563 | 8.96 | 3.44650548 | 0 | 1 | 0 | 1 | 1 | 0.59998149 | 0 | 1 | 0 | 1 | 24 | 0.08019798 | FALSE | 1 | FALSE | 1 | FALSE | 1 | FALSE | 1 | FALSE | 1 | FALSE | 1 |
| 7564 | 7564 | 7.92105263 | 3.08119576 | 2 | 0.25455533 | 2 | 0.3494045 | 0 | 1 | 1 | 0.83318955 | 0 | 1 | 33 | 0.44261631 | FALSE | 1 | FALSE | 1 | FALSE | 1 | FALSE | 1 | FALSE | 1 | FALSE | 1 |
| 7565 | 7565 | 7 | 2.06705764 | 1 | 0.4357638 | 0 | 1 | 0 | 1 | 1 | 0.64497439 | 0 | 1 | 20 | 0.30915202 | FALSE | 1 | FALSE | 1 | FALSE | 1 | FALSE | 1 | FALSE | 1 | FALSE | 1 |
| 7566 | 7566 | 7.4375 | 2.423034 | 0 | 1 | 1 | 0.40942991 | 0 | 1 | 2 | 0.16544789 | 0 | 1 | 13 | 0.76799161 | FALSE | 1 | FALSE | 1 | FALSE | 1 | FALSE | 1 | FALSE | 1 | FALSE | 1 |
| 7567 | 7567 | 10.125 | 2.58702693 | 5 | 0.00029898 | 1 | 0.54637326 | 0 | 1 | 0 | 1 | 1 | 0.32796862 | 17 | 0.97530702 | TRUE | 0.03449636 | FALSE | 1 | FALSE | 1 | FALSE | 1 | FALSE | 1 | FALSE | 1 |
| 7568 | 7568 | 10.8461539 | 2.10662522 | 1 | 0.28675951 | 0 | 1 | 0 | 1 | 1 | 0.45746571 | 1 | 0.19354507 | 10 | 0.86791642 | FALSE | 1 | FALSE | 1 | FALSE | 1 | FALSE | 1 | FALSE | 1 | FALSE | 1 |
| 7569 | 7569 | 10.7 | 2.59036677 | 1 | 0.40558521 | 2 | 0.1353446 | 1 | 0.51939841 | 0 | 1 | 0 | 1 | 16 | 0.80826361 | FALSE | 1 | FALSE | 1 | FALSE | 1 | FALSE | 1 | FALSE | 1 | FALSE | 1 |
| 7570 | 7570 | 11.4 | 2.2150997 | 0 | 1 | 0 | 1 | 1 | 0.66709967 | 1 | 0.75658797 | 4 | 0.00134977 | 24 | 0.82294782 | FALSE | 1 | FALSE | 1 | FALSE | 1 | FALSE | 1 | FALSE | 0.20625174 | FALSE | 1 |
| 7571 | 7571 | 7.02777778 | 3.28706312 | 0 | 1 | 0 | 1 | 1 | 0.73299778 | 2 | 0.49731009 | 1 | 0.44938266 | 32 | 0.31709555 | FALSE | 1 | FALSE | 1 | FALSE | 1 | FALSE | 1 | FALSE | 1 | FALSE | 1 |
| 7572 | 7572 | 7.54700855 | 1.45881187 | 4 | 0.35259616 | 12 | 0.00037081 | 5 | 0.41172421 | 10 | 0.04190069 | 4 | 0.12521484 | 82 | 0.99997462 | FALSE | 1 | FALSE | 0.15723145 | FALSE | 1 | FALSE | 1 | FALSE | 1 | FALSE | 1 |
| 7573 | 7573 | 8.75 | 2.73371803 | 2 | 0.16048326 | 0 | 1 | 0 | 1 | 2 | 0.3705495 | 0 | 1 | 24 | 0.55018107 | FALSE | 1 | FALSE | 1 | FALSE | 1 | FALSE | 1 | FALSE | 1 | FALSE | 1 |
| 7574 | 7574 | 10.5 | 1.80277564 | 1 | 0.51745948 | 3 | 0.06039219 | 0 | 1 | 1 | 0.73248983 | 0 | 1 | 23 | 0.73439118 | FALSE | 1 | FALSE | 1 | FALSE | 1 | FALSE | 1 | FALSE | 1 | FALSE | 1 |
| 7575 | 7575 | 10.75 | 1.83143842 | 0 | 1 | 1 | 0.32623933 | 0 | 1 | 0 | 1 | 0 | 1 | 11 | 0.41944532 | FALSE | 1 | FALSE | 1 | FALSE | 1 | FALSE | 1 | FALSE | 1 | FALSE | 1 |
| 7576 | 7576 | 6.68181818 | 2.66946307 | 1 | 0.4357638 | 0 | 1 | 1 | 0.55340851 | 1 | 0.64497439 | 0 | 1 | 19 | 0.54233419 | FALSE | 1 | FALSE | 1 | FALSE | 1 | FALSE | 1 | FALSE | 1 | FALSE | 1 |
| 7577 | 7577 | 8.90909091 | 4.18774284 | 1 | 0.24866461 | 2 | 0.04725998 | 0 | 1 | 0 | 1 | 3 | 0.00064302 | 5 | 0.99958857 | FALSE | 1 | FALSE | 1 | FALSE | 1 | FALSE | 1 | FALSE | 0.13506637 | FALSE | 1 |
| 7578 | 7578 | 8.38297872 | 2.26493334 | 0 | 1 | 1 | 0.78076835 | 1 | 0.81522035 | 2 | 0.63103861 | 0 | 1 | 42 | 0.13378393 | FALSE | 1 | FALSE | 1 | FALSE | 1 | FALSE | 1 | FALSE | 1 | FALSE | 1 |
| 7579 | 7579 | 10.3125 | 4.34048888 | 3 | 0.0476977 | 1 | 0.65167362 | 0 | 1 | 1 | 0.77852168 | 0 | 1 | 27 | 0.61671133 | FALSE | 0.649301 | FALSE | 1 | FALSE | 1 | FALSE | 1 | FALSE | 1 | FALSE | 1 |
| 7580 | 7580 | 11.0769231 | 2.94726832 | 0 | 1 | 1 | 0.3480709 | 1 | 0.37874417 | 0 | 1 | 0 | 1 | 11 | 0.6683056 | FALSE | 1 | FALSE | 1 | FALSE | 1 | FALSE | 1 | FALSE | 1 | FALSE | 1 |
| 7581 | 7581 | 12.0416667 | 1.33788286 | 0 | 1 | 0 | 1 | 1 | 0.58502114 | 2 | 0.30249307 | 0 | 1 | 21 | 0.47053137 | FALSE | 1 | FALSE | 1 | FALSE | 1 | FALSE | 1 | FALSE | 1 | FALSE | 1 |
| 7582 | 7582 | 9.4 | 0.58309519 | 1 | 0.40558521 | 3 | 0.02534497 | 0 | 1 | 1 | 0.60987146 | 0 | 1 | 15 | 0.92113679 | FALSE | 1 | FALSE | 0.82367276 | FALSE | 1 | FALSE | 1 | FALSE | 1 | FALSE | 1 |
| 7583 | 7583 | 8.96 | 1.37054734 | 2 | 0.13369621 | 2 | 0.19297191 | 2 | 0.22597287 | 1 | 0.69181315 | 1 | 0.33902672 | 17 | 0.9897469 | FALSE | 0.97964162 | FALSE | 1 | FALSE | 1 | FALSE | 1 | FALSE | 1 | FALSE | 1 |
| 7584 | 7584 | 4.25490196 | 2.24781495 | 1 | 0.98156277 | 1 | 0.99362718 | 2 | 0.97548749 | 8 | 0.397791 | 2 | 0.71713432 | 138 | 0.01413623 | FALSE | 1 | FALSE | 1 | FALSE | 1 | FALSE | 1 | FALSE | 1 | FALSE | 1 |
| 7585 | 7585 | 5.63157895 | 1.9152268 | 10 | 0.04075609 | 14 | 0.00730559 | 8 | 0.47350553 | 14 | 0.09640636 | 4 | 0.44600295 | 158 | 0.99952945 | FALSE | 0.626405 | FALSE | 0.6167796 | FALSE | 1 | FALSE | 1 | FALSE | 1 | FALSE | 1 |
| 7586 | 7586 | 8.37037037 | 1.82874473 | 1 | 0.50471032 | 0 | 1 | 2 | 0.25286935 | 1 | 0.71956339 | 0 | 1 | 23 | 0.58307573 | FALSE | 1 | FALSE | 1 | FALSE | 1 | FALSE | 1 | FALSE | 1 | FALSE | 1 |
| 7587 | 7587 | 9.23333333 | 1.99471524 | 0 | 1 | 0 | 1 | 2 | 0.29330414 | 1 | 0.75658797 | 1 | 0.39166989 | 26 | 0.48609328 | FALSE | 1 | FALSE | 1 | FALSE | 1 | FALSE | 1 | FALSE | 1 | FALSE | 1 |
| 7588 | 7588 | 9.07692308 | 2.11083427 | 1 | 0.49162634 | 2 | 0.20486827 | 1 | 0.61440466 | 5 | 0.00585671 | 0 | 1 | 17 | 0.99598795 | FALSE | 1 | FALSE | 1 | FALSE | 1 | FALSE | 1 | FALSE | 1 | FALSE | 1 |
| 7589 | 7589 | 9.1875 | 1.77548409 | 0 | 1 | 0 | 1 | 0 | 1 | 1 | 0.52895088 | 0 | 1 | 15 | 0.26127859 | FALSE | 1 | FALSE | 1 | FALSE | 1 | FALSE | 1 | FALSE | 1 | FALSE | 1 |
| 7590 | 7590 | 11.1428571 | 2.64189172 | 0 | 1 | 1 | 0.36919824 | 1 | 0.40110418 | 0 | 1 | 1 | 0.20679328 | 11 | 0.83676449 | FALSE | 1 | FALSE | 1 | FALSE | 1 | FALSE | 1 | FALSE | 1 | FALSE | 1 |
| 7591 | 7591 | 8.28 | 2.18210907 | 0 | 1 | 0 | 1 | 2 | 0.21260257 | 1 | 0.67692816 | 0 | 1 | 21 | 0.47053137 | FALSE | 1 | FALSE | 1 | FALSE | 1 | FALSE | 1 | FALSE | 1 | FALSE | 1 |
| 7592 | 7592 | 10.7142857 | 3.05727637 | 1 | 0.3050805 | 0 | 1 | 2 | 0.08806398 | 1 | 0.48242099 | 0 | 1 | 10 | 0.94579623 | FALSE | 1 | FALSE | 1 | FALSE | 1 | FALSE | 1 | FALSE | 1 | FALSE | 1 |
| 7593 | 7593 | 7.62790698 | 2.86131581 | 2 | 0.30241117 | 1 | 0.75788114 | 2 | 0.46083448 | 5 | 0.0456107 | 0 | 1 | 33 | 0.93815092 | FALSE | 1 | FALSE | 1 | FALSE | 1 | FALSE | 1 | FALSE | 1 | FALSE | 1 |
| 7594 | 7594 | 9.41666667 | 1.25554326 | 1 | 0.24866461 | 0 | 1 | 0 | 1 | 2 | 0.08789566 | 1 | 0.16638736 | 7 | 0.98129353 | FALSE | 1 | FALSE | 1 | FALSE | 1 | FALSE | 1 | FALSE | 1 | FALSE | 1 |
| 7595 | 7595 | 10.7222222 | 1.87988048 | 1 | 0.37380237 | 0 | 1 | 1 | 0.48280983 | 2 | 0.19902682 | 0 | 1 | 14 | 0.86287418 | FALSE | 1 | FALSE | 1 | FALSE | 1 | FALSE | 1 | FALSE | 1 | FALSE | 1 |
| 7596 | 7596 | 3.07458564 | 0.76343616 | 53 | 0.05002619 | 71 | 0.00623503 | 60 | 0.55299876 | 73 | 0.73727532 | 36 | 0.04254332 | 1390 | 0.99073001 | FALSE | 0.66636505 | FALSE | 0.60103504 | FALSE | 1 | FALSE | 1 | FALSE | 0.92199696 | FALSE | 1 |
| 7597 | 7597 | 10.8333333 | 2.33927814 | 0 | 1 | 0 | 1 | 1 | 0.35555293 | 1 | 0.43131129 | 1 | 0.18007762 | 9 | 0.89630573 | FALSE | 1 | FALSE | 1 | FALSE | 1 | FALSE | 1 | FALSE | 1 | FALSE | 1 |
| 7598 | 7598 | 6.78947368 | 4.05980227 | 0 | 1 | 0 | 1 | 1 | 0.50143826 | 0 | 1 | 0 | 1 | 18 | 0.17880694 | FALSE | 1 | FALSE | 1 | FALSE | 1 | FALSE | 1 | FALSE | 1 | FALSE | 1 |
| 7599 | 7599 | 7.41666667 | 2.89995211 | 2 | 0.23542589 | 1 | 0.69480472 | 1 | 0.73299778 | 0 | 1 | 0 | 1 | 32 | 0.31709555 | FALSE | 1 | FALSE | 1 | FALSE | 1 | FALSE | 1 | FALSE | 1 | FALSE | 1 |
| 7600 | 7600 | 7.90123457 | 1.77503217 | 0 | 1 | 4 | 0.26638189 | 3 | 0.56136161 | 3 | 0.72619227 | 1 | 0.74003033 | 70 | 0.37303501 | FALSE | 1 | FALSE | 1 | FALSE | 1 | FALSE | 1 | FALSE | 1 | FALSE | 1 |
| 7601 | 7601 | 5.91836735 | 3.02261365 | 3 | 0.12433927 | 3 | 0.20208567 | 0 | 1 | 5 | 0.06758937 | 0 | 1 | 37 | 0.93870662 | FALSE | 0.95622564 | FALSE | 1 | FALSE | 1 | FALSE | 1 | FALSE | 1 | FALSE | 1 |
| 7602 | 7602 | 10.4545455 | 2.95512815 | 0 | 1 | 0 | 1 | 2 | 0.33343871 | 2 | 0.45159487 | 0 | 1 | 29 | 0.39648374 | FALSE | 1 | FALSE | 1 | FALSE | 1 | FALSE | 1 | FALSE | 1 | FALSE | 1 |
| 7603 | 7603 | 8.68421053 | 2.23730645 | 1 | 0.62830697 | 1 | 0.71433277 | 2 | 0.39868506 | 0 | 1 | 1 | 0.4673875 | 33 | 0.44261631 | FALSE | 1 | FALSE | 1 | FALSE | 1 | FALSE | 1 | FALSE | 1 | FALSE | 1 |
| 7604 | 7604 | 6.875 | 2.45479972 | 1 | 0.71380452 | 1 | 0.79481669 | 1 | 0.8283465 | 4 | 0.17698522 | 1 | 0.54902428 | 40 | 0.66905473 | FALSE | 1 | FALSE | 1 | FALSE | 1 | FALSE | 1 | FALSE | 1 | FALSE | 1 |
| 7605 | 7605 | 10.8333333 | 1.99304346 | 2 | 0.03643311 | 0 | 1 | 1 | 0.35555293 | 0 | 1 | 0 | 1 | 9 | 0.89630573 | FALSE | 0.58392 | FALSE | 1 | FALSE | 1 | FALSE | 1 | FALSE | 1 | FALSE | 1 |
| 7606 | 7606 | 8.05555556 | 2.09423076 | 1 | 0.37380237 | 0 | 1 | 2 | 0.13502329 | 4 | 0.00799439 | 0 | 1 | 11 | 0.99651304 | FALSE | 1 | FALSE | 1 | FALSE | 1 | FALSE | 1 | FALSE | 1 | FALSE | 1 |
| 7607 | 7607 | 8 | 1.84842275 | 1 | 0.46441868 | 0 | 1 | 0 | 1 | 2 | 0.30249307 | 1 | 0.32796862 | 20 | 0.68270729 | FALSE | 1 | FALSE | 1 | FALSE | 1 | FALSE | 1 | FALSE | 1 | FALSE | 1 |
| 7608 | 7608 | 10.6 | 2.05912603 | 0 | 1 | 0 | 1 | 0 | 1 | 0 | 1 | 0 | 1 | 5 | 0.42761893 | FALSE | 1 | FALSE | 1 | FALSE | 1 | FALSE | 1 | FALSE | 1 | FALSE | 1 |
| 7609 | 7609 | 8.95833333 | 1.61964931 | 0 | 1 | 0 | 1 | 1 | 0.58502114 | 1 | 0.67692816 | 1 | 0.32796862 | 21 | 0.47053137 | FALSE | 1 | FALSE | 1 | FALSE | 1 | FALSE | 1 | FALSE | 1 | FALSE | 1 |
| 7610 | 7610 | 9.14285714 | 1.65862003 | 1 | 0.59801736 | 1 | 0.68454804 | 4 | 0.0355345 | 1 | 0.80778317 | 1 | 0.44015537 | 27 | 0.91585095 | FALSE | 1 | FALSE | 1 | FALSE | 1 | FALSE | 1 | FALSE | 1 | FALSE | 1 |
| 7611 | 7611 | 9.73333333 | 2.93181779 | 0 | 1 | 1 | 0.38964394 | 0 | 1 | 1 | 0.50623194 | 0 | 1 | 13 | 0.57712805 | FALSE | 1 | FALSE | 1 | FALSE | 1 | FALSE | 1 | FALSE | 1 | FALSE | 1 |
| 7612 | 7612 | 8.3125 | 1.58976217 | 0 | 1 | 1 | 0.65167362 | 1 | 0.69069222 | 0 | 1 | 0 | 1 | 30 | 0.10375556 | FALSE | 1 | FALSE | 1 | FALSE | 1 | FALSE | 1 | FALSE | 1 | FALSE | 1 |

| | | | | | | | | | | | | | | | | | | | | | | | | | | | |
|---|---|---|---|---|---|---|---|---|---|---|---|---|---|---|---|---|---|---|---|---|---|---|---|---|---|---|---|
| 7613 | 7613 | 10.3076923 | 2.72833044 | 1 | 0.28675951 | 0 | 1 | 0 | 1 | 1 | 0.45746571 | 0 | 1 | 11 | 0.6683056 | FALSE | 1 | FALSE | 1 | FALSE | 1 | FALSE | 1 | FALSE | 1 | FALSE | 1 |
| 7614 | 7614 | 9.44444444 | 4.24554959 | 0 | 1 | 0 | 1 | 0 | 1 | 0 | 1 | 0 | 1 | 9 | 0.21661397 | FALSE | 1 | FALSE | 1 | FALSE | 1 | FALSE | 1 | FALSE | 1 | FALSE | 1 |
| 7615 | 7615 | 7.08333333 | 1.03749163 | 0 | 1 | 0 | 1 | 0 | 1 | 0 | 1 | 0 | 1 | 12 | 0.13002667 | FALSE | 1 | FALSE | 1 | FALSE | 1 | FALSE | 1 | FALSE | 1 | FALSE | 1 |
| 7616 | 7616 | 9.36 | 1.46642422 | 0 | 1 | 3 | 0.04547096 | 1 | 0.59998149 | 0 | 1 | 0 | 1 | 21 | 0.64957167 | FALSE | 1 | FALSE | 1 | FALSE | 1 | FALSE | 1 | FALSE | 1 | FALSE | 1 |
| 7617 | 7617 | 10.2142857 | 2.33539753 | 4 | 0.00034152 | 0 | 1 | 0 | 1 | 1 | 0.48242099 | 0 | 1 | 9 | 0.98607548 | TRUE | 0.03730255 | FALSE | 1 | FALSE | 1 | FALSE | 1 | FALSE | 1 | FALSE | 1 |
| 7618 | 7618 | 11.4615385 | 2.27411469 | 1 | 0.28675951 | 0 | 1 | 2 | 0.07725803 | 0 | 1 | 1 | 0.19354507 | 9 | 0.96018168 | FALSE | 1 | FALSE | 1 | FALSE | 1 | FALSE | 1 | FALSE | 1 | FALSE | 1 |
| 7619 | 7619 | 9.5 | 2.36290781 | 0 | 1 | 0 | 1 | 0 | 1 | 0 | 1 | 0 | 1 | 6 | 0.36077158 | FALSE | 1 | FALSE | 1 | FALSE | 1 | FALSE | 1 | FALSE | 1 | FALSE | 1 |
| 7620 | 7620 | 7.17567568 | 2.5856052 | 2 | 0.56271322 | 2 | 0.68961994 | 3 | 0.49081265 | 5 | 0.24321887 | 3 | 0.11741889 | 58 | 0.9042862 | FALSE | 1 | FALSE | 1 | FALSE | 1 | FALSE | 1 | FALSE | 1 | FALSE | 1 |
| 7621 | 7621 | 6.69230769 | 3.30232122 | 0 | 1 | 0 | 1 | 1 | 0.61440466 | 2 | 0.33678042 | 1 | 0.3499045 | 22 | 0.61628 | FALSE | 1 | FALSE | 1 | FALSE | 1 | FALSE | 1 | FALSE | 1 | FALSE | 1 |
| 7622 | 7622 | 9.88461539 | 2.65049956 | 1 | 0.49162634 | 2 | 0.20486827 | 2 | 0.23940226 | 2 | 0.33678042 | 1 | 0.3499045 | 18 | 0.98644281 | FALSE | 1 | FALSE | 1 | FALSE | 1 | FALSE | 1 | FALSE | 1 | FALSE | 1 |
| 7623 | 7623 | 8.25 | 3.28823661 | 0 | 1 | 1 | 0.40942991 | 0 | 1 | 0 | 1 | 0 | 1 | 15 | 0.26127859 | FALSE | 1 | FALSE | 1 | FALSE | 1 | FALSE | 1 | FALSE | 1 | FALSE | 1 |
| 7624 | 7624 | 9.83333333 | 2.95334086 | 0 | 1 | 1 | 0.53115654 | 2 | 0.19931429 | 0 | 1 | 1 | 0.31672722 | 19 | 0.71542496 | FALSE | 1 | FALSE | 1 | FALSE | 1 | FALSE | 1 | FALSE | 1 | FALSE | 1 |
| 7625 | 7625 | 8.48717949 | 3.11219104 | 7 | 0.0036832 | 5 | 0.10739429 | 2 | 0.77659406 | 4 | 0.48430022 | 1 | 0.72664912 | 59 | 0.98534527 | FALSE | 0.16951008 | FALSE | 1 | FALSE | 1 | FALSE | 1 | FALSE | 1 | FALSE | 1 |
| 7626 | 7626 | 10.2352941 | 2.79767575 | 0 | 1 | 1 | 0.42857733 | 1 | 0.46348836 | 2 | 0.18212134 | 0 | 1 | 13 | 0.88719222 | FALSE | 1 | FALSE | 1 | FALSE | 1 | FALSE | 1 | FALSE | 1 | FALSE | 1 |
| 7627 | 7627 | 9.59090909 | 2.12472653 | 3 | 0.01782284 | 0 | 1 | 0 | 1 | 3 | 0.07743186 | 0 | 1 | 16 | 0.95553951 | FALSE | 0.41596787 | FALSE | 1 | FALSE | 1 | FALSE | 1 | FALSE | 1 | FALSE | 1 |
| 7628 | 7628 | 10.1153846 | 3.02968548 | 0 | 1 | 2 | 0.20486827 | 2 | 0.23940226 | 1 | 0.70601446 | 0 | 1 | 21 | 0.78939668 | FALSE | 1 | FALSE | 1 | FALSE | 1 | FALSE | 1 | FALSE | 1 | FALSE | 1 |
| 7629 | 7629 | 9.57894737 | 2.66193024 | 2 | 0.08405377 | 0 | 1 | 1 | 0.50143826 | 2 | 0.21610812 | 1 | 0.26986775 | 13 | 0.97998676 | FALSE | 0.79236883 | FALSE | 1 | FALSE | 1 | FALSE | 1 | FALSE | 1 | FALSE | 1 |
| 7630 | 7630 | 8.41666667 | 3.04023939 | 0 | 1 | 0 | 1 | 0 | 1 | 0 | 1 | 0 | 1 | 11 | 0.15414489 | FALSE | 1 | FALSE | 1 | FALSE | 1 | FALSE | 1 | FALSE | 1 | FALSE | 1 |
| 7631 | 7631 | 4.51663405 | 1.49860249 | 17 | 0.0269875 | 26 | 0.00047188 | 19 | 0.12788417 | 23 | 0.15396916 | 9 | 0.20844213 | 306 | 0.99999214 | FALSE | 0.50964275 | FALSE | 0.17570969 | FALSE | 1 | FALSE | 1 | FALSE | 1 | FALSE | 1 |
| 7632 | 7632 | 8.33333333 | 4.09335505 | 0 | 1 | 1 | 0.38964394 | 0 | 1 | 1 | 0.50623194 | 0 | 1 | 13 | 0.57712805 | FALSE | 1 | FALSE | 1 | FALSE | 1 | FALSE | 1 | FALSE | 1 | FALSE | 1 |
| 7633 | 7633 | 7.35526316 | 1.13211432 | 3 | 0.30904591 | 2 | 0.71077581 | 1 | 0.93896183 | 6 | 0.13504406 | 0 | 1 | 64 | 0.59240184 | FALSE | 1 | FALSE | 1 | FALSE | 1 | FALSE | 1 | FALSE | 1 | FALSE | 1 |
| 7634 | 7634 | 7.54285714 | 1.82589089 | 0 | 1 | 0 | 1 | 2 | 0.35984046 | 1 | 0.80778317 | 1 | 0.44015537 | 31 | 0.34231756 | FALSE | 1 | FALSE | 1 | FALSE | 1 | FALSE | 1 | FALSE | 1 | FALSE | 1 |
| 7635 | 7635 | 9.45833333 | 2.87197909 | 1 | 0.46441868 | 0 | 1 | 1 | 0.58502114 | 3 | 0.09525947 | 0 | 1 | 19 | 0.83975208 | FALSE | 1 | FALSE | 1 | FALSE | 1 | FALSE | 1 | FALSE | 1 | FALSE | 1 |
| 7636 | 7636 | 8.28571429 | 2.015248 | 0 | 1 | 0 | 1 | 0 | 1 | 1 | 0.48242099 | 0 | 1 | 13 | 0.33283729 | FALSE | 1 | FALSE | 1 | FALSE | 1 | FALSE | 1 | FALSE | 1 | FALSE | 1 |
| 7637 | 7637 | 10 | 1.41421356 | 0 | 1 | 1 | 0.15163655 | 0 | 1 | 0 | 1 | 0 | 1 | 4 | 0.82371636 | FALSE | 1 | FALSE | 1 | FALSE | 1 | FALSE | 1 | FALSE | 1 | FALSE | 1 |
| 7638 | 7638 | 9.28395062 | 2.96579083 | 2 | 0.59898979 | 1 | 0.92420038 | 6 | 0.06065565 | 3 | 0.7021069 | 0 | 1 | 66 | 0.55329149 | FALSE | 1 | FALSE | 1 | FALSE | 1 | FALSE | 1 | FALSE | 1 | FALSE | 1 |
| 7639 | 7639 | 10.8636364 | 3.74524492 | 2 | 0.10809809 | 0 | 1 | 0 | 1 | 1 | 0.64497439 | 0 | 1 | 19 | 0.54233419 | FALSE | 0.88820416 | FALSE | 1 | FALSE | 1 | FALSE | 1 | FALSE | 1 | FALSE | 1 |
| 7640 | 7640 | 11.3589744 | 2.79640043 | 0 | 1 | 2 | 0.31355738 | 3 | 0.12958859 | 2 | 0.48233989 | 1 | 0.44015537 | 27 | 0.91585095 | FALSE | 1 | FALSE | 1 | FALSE | 1 | FALSE | 1 | FALSE | 1 | FALSE | 1 |
| 7641 | 7641 | 6.4 | 2.62805378 | 0 | 1 | 0 | 1 | 1 | 0.42266264 | 0 | 1 | 0 | 1 | 14 | 0.29525409 | FALSE | 1 | FALSE | 1 | FALSE | 1 | FALSE | 1 | FALSE | 1 | FALSE | 1 |
| 7642 | 7642 | 10 | 2.9664794 | 2 | 0.0256865 | 0 | 1 | 0 | 1 | 0 | 1 | 0 | 1 | 8 | 0.80394719 | FALSE | 0.4974559 | FALSE | 1 | FALSE | 1 | FALSE | 1 | FALSE | 1 | FALSE | 1 |
| 7643 | 7643 | 9.8 | 2.00665559 | 0 | 1 | 3 | 0.01138739 | 1 | 0.42266264 | 0 | 1 | 0 | 1 | 11 | 0.92879412 | FALSE | 1 | FALSE | 0.67617265 | FALSE | 1 | FALSE | 1 | FALSE | 1 | FALSE | 1 |
| 7644 | 7644 | 9.5862069 | 2.97712244 | 0 | 1 | 0 | 1 | 0 | 1 | 0 | 1 | 0 | 1 | 29 | 0.00717703 | FALSE | 1 | FALSE | 1 | FALSE | 1 | FALSE | 1 | FALSE | 1 | FALSE | 1 |
| 7645 | 7645 | 12 | 1.41421356 | 1 | 0.18769745 | 1 | 0.23139161 | 0 | 1 | 0 | 1 | 0 | 1 | 6 | 0.88425551 | FALSE | 1 | FALSE | 1 | FALSE | 1 | FALSE | 1 | FALSE | 1 | FALSE | 1 |
| 7646 | 7646 | 10.0625 | 2.65680141 | 0 | 1 | 0 | 1 | 1 | 0.44344817 | 1 | 0.52895088 | 0 | 1 | 14 | 0.53308261 | FALSE | 1 | FALSE | 1 | FALSE | 1 | FALSE | 1 | FALSE | 1 | FALSE | 1 |
| 7647 | 7647 | 11.1666667 | 3.48409083 | 1 | 0.26795838 | 0 | 1 | 0 | 1 | 0 | 1 | 0 | 1 | 11 | 0.41944532 | FALSE | 1 | FALSE | 1 | FALSE | 1 | FALSE | 1 | FALSE | 1 | FALSE | 1 |
| 7648 | 7648 | 10.7352941 | 1.68573293 | 0 | 1 | 0 | 1 | 1 | 0.71261921 | 3 | 0.20353499 | 0 | 1 | 30 | 0.36879488 | FALSE | 1 | FALSE | 1 | FALSE | 1 | FALSE | 1 | FALSE | 1 | FALSE | 1 |
| 7649 | 7649 | 5.97368421 | 3.46020111 | 3 | 0.07268126 | 2 | 0.3494045 | 0 | 1 | 3 | 0.25266879 | 1 | 0.4673875 | 29 | 0.93814695 | FALSE | 0.76727438 | FALSE | 1 | FALSE | 1 | FALSE | 1 | FALSE | 1 | FALSE | 1 |
| 7650 | 7650 | 4.22857143 | 2.37349308 | 2 | 0.539854 | 4 | 0.19010643 | 2 | 0.72288838 | 2 | 0.83868849 | 2 | 0.31908786 | 58 | 0.70710693 | FALSE | 1 | FALSE | 1 | FALSE | 1 | FALSE | 1 | FALSE | 1 | FALSE | 1 |
| 7651 | 7651 | 8.69230769 | 2.3982242 | 1 | 0.28675951 | 0 | 1 | 1 | 0.37874417 | 0 | 1 | 0 | 1 | 11 | 0.6683056 | FALSE | 1 | FALSE | 1 | FALSE | 1 | FALSE | 1 | FALSE | 1 | FALSE | 1 |
| 7652 | 7652 | 6.7032967 | 1.77174304 | 6 | 0.70441251 | 9 | 0.5213496 | 10 | 0.51686527 | 9 | 0.88515776 | 7 | 0.15808024 | 231 | 0.44000826 | FALSE | 1 | FALSE | 1 | FALSE | 1 | FALSE | 1 | FALSE | 1 | FALSE | 1 |
| 7653 | 7653 | 10.6666667 | 1.76383421 | 0 | 1 | 0 | 1 | 0 | 1 | 0 | 1 | 0 | 1 | 9 | 0.21661397 | FALSE | 1 | FALSE | 1 | FALSE | 1 | FALSE | 1 | FALSE | 1 | FALSE | 1 |
| 7654 | 7654 | 8.31578947 | 2.45062036 | 0 | 1 | 2 | 0.12434464 | 0 | 1 | 1 | 0.59104357 | 2 | 0.03799168 | 14 | 0.93677253 | FALSE | 1 | FALSE | 1 | FALSE | 1 | FALSE | 1 | FALSE | 0.87669812 | FALSE | 1 |
| 7655 | 7655 | 8.61403509 | 1.99784433 | 0 | 1 | 0 | 1 | 2 | 0.60323163 | 5 | 0.11290142 | 0 | 1 | 49 | 0.33483172 | FALSE | 1 | FALSE | 1 | FALSE | 1 | FALSE | 1 | FALSE | 1 | FALSE | 1 |
| 7656 | 7656 | 7.65517241 | 2.3234318 | 1 | 0.52988231 | 0 | 1 | 1 | 0.65463982 | 2 | 0.3871811 | 0 | 1 | 25 | 0.51779518 | FALSE | 1 | FALSE | 1 | FALSE | 1 | FALSE | 1 | FALSE | 1 | FALSE | 1 |
| 7657 | 7657 | 10 | 2.29128785 | 0 | 1 | 2 | 0.09285443 | 0 | 1 | 0 | 1 | 0 | 1 | 14 | 0.53308261 | FALSE | 1 | FALSE | 1 | FALSE | 1 | FALSE | 1 | FALSE | 1 | FALSE | 1 |
| 7658 | 7658 | 10.2142857 | 1.9338552 | 1 | 0.3050805 | 0 | 1 | 1 | 0.40110418 | 1 | 0.48242099 | 0 | 1 | 11 | 0.83676449 | FALSE | 1 | FALSE | 1 | FALSE | 1 | FALSE | 1 | FALSE | 1 | FALSE | 1 |
| 7659 | 7659 | 10.4090909 | 2.80679518 | 1 | 0.4357638 | 1 | 0.5154318 | 1 | 0.55340851 | 1 | 0.64497439 | 0 | 1 | 18 | 0.74744813 | FALSE | 1 | FALSE | 1 | FALSE | 1 | FALSE | 1 | FALSE | 1 | FALSE | 1 |
| 7660 | 7660 | 3.65517241 | 1.25604365 | 7 | 0.61983431 | 10 | 0.46434436 | 8 | 0.82586813 | 17 | 0.17822882 | 3 | 0.86115613 | 245 | 0.52007469 | FALSE | 1 | FALSE | 1 | FALSE | 1 | FALSE | 1 | FALSE | 1 | FALSE | 1 |
| 7661 | 7661 | 6.68965517 | 2.01893533 | 9 | 2.50E-08 | 2 | 0.24095584 | 2 | 0.27983822 | 0 | 1 | 0 | 1 | 16 | 0.99996363 | TRUE | 2.93E-05 | FALSE | 1 | FALSE | 1 | FALSE | 1 | FALSE | 1 | FALSE | 1 |
| 7662 | 7662 | 9.125 | 1.83285979 | 0 | 1 | 1 | 0.23139161 | 0 | 1 | 0 | 1 | 0 | 1 | 7 | 0.63752397 | FALSE | 1 | FALSE | 1 | FALSE | 1 | FALSE | 1 | FALSE | 1 | FALSE | 1 |
| 7663 | 7663 | 10.1724138 | 2.18251619 | 0 | 1 | 0 | 1 | 2 | 0.25286935 | 2 | 0.35374209 | 0 | 1 | 23 | 0.58307573 | FALSE | 1 | FALSE | 1 | FALSE | 1 | FALSE | 1 | FALSE | 1 | FALSE | 1 |
| 7664 | 7664 | 9.19230769 | 3.02577683 | 0 | 1 | 0 | 1 | 4 | 0.0130279 | 1 | 0.70601446 | 0 | 1 | 21 | 0.78939668 | FALSE | 1 | FALSE | 1 | FALSE | 1 | FALSE | 1 | FALSE | 1 | FALSE | 1 |
| 7665 | 7665 | 9.30769231 | 2.08970426 | 0 | 1 | 0 | 1 | 0 | 1 | 2 | 0.1174583 | 0 | 1 | 11 | 0.6683056 | FALSE | 1 | FALSE | 1 | FALSE | 1 | FALSE | 1 | FALSE | 1 | FALSE | 1 |
| 7666 | 7666 | 8.42361111 | 1.93139221 | 5 | 0.29959334 | 3 | 0.8440476 | 2 | 0.96640926 | 8 | 0.32746711 | 1 | 0.90677556 | 123 | 0.27076935 | FALSE | 1 | FALSE | 1 | FALSE | 1 | FALSE | 1 | FALSE | 1 | FALSE | 1 |
| 7667 | 7667 | 7.67857143 | 2.33075051 | 1 | 0.51745948 | 3 | 0.06039219 | 3 | 0.07756639 | 2 | 0.3705495 | 0 | 1 | 19 | 0.99262381 | FALSE | 1 | FALSE | 1 | FALSE | 1 | FALSE | 1 | FALSE | 1 | FALSE | 1 |
| 7668 | 7668 | 8.33333333 | 1.94701045 | 3 | 0.14187072 | 3 | 0.22759346 | 3 | 0.27688673 | 3 | 0.41729179 | 1 | 0.57100288 | 38 | 0.97862182 | FALSE | 1 | FALSE | 1 | FALSE | 1 | FALSE | 1 | FALSE | 1 | FALSE | 1 |
| 7669 | 7669 | 8.43478261 | 2.20327991 | 1 | 0.45027683 | 0 | 1 | 1 | 0.5695037 | 2 | 0.2852236 | 0 | 1 | 19 | 0.71542496 | FALSE | 1 | FALSE | 1 | FALSE | 1 | FALSE | 1 | FALSE | 1 | FALSE | 1 |
| 7670 | 7670 | 6.58536585 | 3.07607296 | 0 | 1 | 1 | 0.7326167 | 2 | 0.42396974 | 2 | 0.5543522 | 2 | 0.13925369 | 33 | 0.71942635 | FALSE | 1 | FALSE | 1 | FALSE | 1 | FALSE | 1 | FALSE | 1 | FALSE | 1 |
| 7671 | 7671 | 7.97959184 | 2.06516092 | 1 | 0.71380452 | 3 | 0.20208567 | 1 | 0.8283465 | 3 | 0.37966044 | 1 | 0.54902428 | 39 | 0.79274273 | FALSE | 1 | FALSE | 1 | FALSE | 1 | FALSE | 1 | FALSE | 1 | FALSE | 1 |
| 7672 | 7672 | 11.1111111 | 3.67507454 | 0 | 1 | 3 | 0.05519089 | 0 | 1 | 0 | 1 | 0 | 1 | 24 | 0.3725373 | FALSE | 1 | FALSE | 1 | FALSE | 1 | FALSE | 1 | FALSE | 1 | FALSE | 1 |
| 7673 | 7673 | 8 | 2.44948974 | 0 | 1 | 0 | 1 | 0 | 1 | 0 | 1 | 0 | 1 | 4 | 0.50683841 | FALSE | 1 | FALSE | 1 | FALSE | 1 | FALSE | 1 | FALSE | 1 | FALSE | 1 |
| 7674 | 7674 | 5.11818182 | 2.36540144 | 2 | 0.77878267 | 3 | 0.69664974 | 2 | 0.91064483 | 2 | 0.965234 | 3 | 0.26940218 | 98 | 0.10387403 | FALSE | 1 | FALSE | 1 | FALSE | 1 | FALSE | 1 | FALSE | 1 | FALSE | 1 |
| 7675 | 7675 | 7.41935484 | 3.82499583 | 0 | 1 | 1 | 0.58913618 | 0 | 1 | 2 | 0.35374209 | 0 | 1 | 24 | 0.3725373 | FALSE | 1 | FALSE | 1 | FALSE | 1 | FALSE | 1 | FALSE | 1 | FALSE | 1 |
| 7676 | 7676 | 10.375 | 4.29934588 | 1 | 0.18769745 | 1 | 0.23139161 | 0 | 1 | 0 | 1 | 0 | 1 | 6 | 0.88425551 | FALSE | 1 | FALSE | 1 | FALSE | 1 | FALSE | 1 | FALSE | 1 | FALSE | 1 |
| 7677 | 7677 | 10.5555556 | 2.54344959 | 0 | 1 | 1 | 0.25628163 | 1 | 0.28067846 | 1 | 0.34506517 | 0 | 1 | 6 | 0.96128317 | FALSE | 1 | FALSE | 1 | FALSE | 1 | FALSE | 1 | FALSE | 1 | FALSE | 1 |
| 7678 | 7678 | 8.85714286 | 1.80701581 | 0 | 1 | 0 | 1 | 1 | 0.53671415 | 0 | 1 | 0 | 1 | 20 | 0.13763971 | FALSE | 1 | FALSE | 1 | FALSE | 1 | FALSE | 1 | FALSE | 1 | FALSE | 1 |
| 7679 | 7679 | 10.6875 | 2.31081019 | 0 | 1 | 1 | 0.40942991 | 0 | 1 | 0 | 1 | 0 | 1 | 15 | 0.26127859 | FALSE | 1 | FALSE | 1 | FALSE | 1 | FALSE | 1 | FALSE | 1 | FALSE | 1 |
| 7680 | 7680 | 10.2173913 | 1.74347284 | 0 | 1 | 1 | 0.53115654 | 0 | 1 | 2 | 0.2852236 | 0 | 1 | 20 | 0.50589402 | FALSE | 1 | FALSE | 1 | FALSE | 1 | FALSE | 1 | FALSE | 1 | FALSE | 1 |
| 7681 | 7681 | 9.0625 | 1.19732775 | 1 | 0.32293355 | 0 | 1 | 0 | 1 | 3 | 0.02891327 | 1 | 0.2198258 | 10 | 0.97980046 | FALSE | 1 | FALSE | 1 | FALSE | 1 | FALSE | 1 | FALSE | 1 | FALSE | 1 |
| 7682 | 7682 | 10.1052632 | 2.46863987 | 1 | 0.38989951 | 0 | 1 | 0 | 1 | 2 | 0.21610812 | 1 | 0.26986775 | 15 | 0.83648191 | FALSE | 1 | FALSE | 1 | FALSE | 1 | FALSE | 1 | FALSE | 1 | FALSE | 1 |
| 7683 | 7683 | 7.84210526 | 2.00691326 | 1 | 0.6184714 | 1 | 0.70472939 | 1 | 0.74264075 | 2 | 0.51200191 | 0 | 1 | 32 | 0.47030992 | FALSE | 1 | FALSE | 1 | FALSE | 1 | FALSE | 1 | FALSE | 1 | FALSE | 1 |
| 7684 | 7684 | 8.76 | 2.23212903 | 1 | 0.47819878 | 1 | 0.56109832 | 1 | 0.59998149 | 2 | 0.31968852 | 2 | 0.06257402 | 18 | 0.96851178 | FALSE | 1 | FALSE | 1 | FALSE | 1 | FALSE | 1 | FALSE | 1 | FALSE | 1 |
| 7685 | 7685 | 9.30769231 | 2.33319244 | 1 | 0.26795838 | 0 | 1 | 0 | 1 | 1 | 0.43131129 | 0 | 1 | 10 | 0.71436964 | FALSE | 1 | FALSE | 1 | FALSE | 1 | FALSE | 1 | FALSE | 1 | FALSE | 1 |
| 7686 | 7686 | 3.53571429 | 1.8073687 | 3 | 0.55171054 | 4 | 0.4928114 | 5 | 0.37574225 | 3 | 0.89440353 | 0 | 1 | 97 | 0.30787132 | FALSE | 1 | FALSE | 1 | FALSE | 1 | FALSE | 1 | FALSE | 1 | FALSE | 1 |
| 7687 | 7687 | 8.37037037 | 1.80837861 | 1 | 0.50471032 | 2 | 0.21684246 | 0 | 1 | 1 | 0.71956339 | 0 | 1 | 23 | 0.58307573 | FALSE | 1 | FALSE | 1 | FALSE | 1 | FALSE | 1 | FALSE | 1 | FALSE | 1 |
| 7688 | 7688 | 10.8333333 | 2.9107082 | 1 | 0.26795838 | 0 | 1 | 0 | 1 | 0 | 1 | 0 | 1 | 11 | 0.41944532 | FALSE | 1 | FALSE | 1 | FALSE | 1 | FALSE | 1 | FALSE | 1 | FALSE | 1 |
| 7689 | 7689 | 8.3 | 3.60693776 | 0 | 1 | 0 | 1 | 0 | 1 | 0 | 1 | 0 | 1 | 10 | 0.18273166 | FALSE | 1 | FALSE | 1 | FALSE | 1 | FALSE | 1 | FALSE | 1 | FALSE | 1 |
| 7690 | 7690 | 8.46428571 | 2.35253879 | 1 | 0.51745948 | 0 | 1 | 0 | 1 | 1 | 0.73248983 | 0 | 1 | 26 | 0.16405547 | FALSE | 1 | FALSE | 1 | FALSE | 1 | FALSE | 1 | FALSE | 1 | FALSE | 1 |
| 7691 | 7691 | 9.92307692 | 2.09253392 | 0 | 1 | 0 | 1 | 1 | 0.37874417 | 0 | 1 | 0 | 1 | 12 | 0.374198 | FALSE | 1 | FALSE | 1 | FALSE | 1 | FALSE | 1 | FALSE | 1 | FALSE | 1 |
| 7692 | 7692 | 10.3793103 | 1.62764465 | 1 | 0.50471032 | 0 | 1 | 1 | 0.62830986 | 2 | 0.35374209 | 0 | 1 | 23 | 0.58307573 | FALSE | 1 | FALSE | 1 | FALSE | 1 | FALSE | 1 | FALSE | 1 | FALSE | 1 |
| 7693 | 7693 | 7.22222222 | 2.52884593 | 0 | 1 | 0 | 1 | 1 | 0.28067846 | 1 | 0.34506517 | 0 | 1 | 7 | 0.84574168 | FALSE | 1 | FALSE | 1 | FALSE | 1 | FALSE | 1 | FALSE | 1 | FALSE | 1 |
| 7694 | 7694 | 10.7142857 | 2.24971654 | 1 | 0.42086991 | 0 | 1 | 1 | 0.53671415 | 1 | 0.62783522 | 0 | 1 | 18 | 0.57962763 | FALSE | 1 | FALSE | 1 | FALSE | 1 | FALSE | 1 | FALSE | 1 | FALSE | 1 |
| 7695 | 7695 | 9.48148148 | 2.40940088 | 0 | 1 | 1 | 0.57534752 | 1 | 0.61440466 | 1 | 0.70601446 | 2 | 0.0670823 | 21 | 0.78939668 | FALSE | 1 | FALSE | 1 | FALSE | 1 | FALSE | 1 | FALSE | 1 | FALSE | 1 |
| 7696 | 7696 | 10.8571429 | 2.3255458 | 0 | 1 | 0 | 1 | 1 | 0.37874417 | 0 | 1 | 0 | 1 | 12 | 0.374198 | FALSE | 1 | FALSE | 1 | FALSE | 1 | FALSE | 1 | FALSE | 1 | FALSE | 1 |
| 7697 | 7697 | 8.92307692 | 1.51716023 | 0 | 1 | 0 | 1 | 1 | 0.61440466 | 1 | 0.70601446 | 1 | 0.3499045 | 23 | 0.40372818 | FALSE | 1 | FALSE | 1 | FALSE | 1 | FALSE | 1 | FALSE | 1 | FALSE | 1 |
| 7698 | 7698 | 9.76923077 | 3.40031325 | 0 | 1 | 0 | 1 | 0 | 1 | 1 | 0.45746571 | 0 | 1 | 12 | 0.374198 | FALSE | 1 | FALSE | 1 | FALSE | 1 | FALSE | 1 | FALSE | 1 | FALSE | 1 |
| 7699 | 7699 | 9.48888889 | 1.82113789 | 2 | 0.32143972 | 4 | 0.05650644 | 2 | 0.48463745 | 4 | 0.15016749 | 1 | 0.52593041 | 32 | 0.99319189 | FALSE | 1 | FALSE | 1 | FALSE | 1 | FALSE | 1 | FALSE | 1 | FALSE | 1 |
| 7700 | 7700 | 11 | 2.14201664 | 0 | 1 | 0 | 1 | 1 | 0.46348836 | 2 | 0.18212134 | 0 | 1 | 14 | 0.7313145 | FALSE | 1 | FALSE | 1 | FALSE | 1 | FALSE | 1 | FALSE | 1 | FALSE | 1 |
| 7701 | 7701 | 8.14285714 | 3.24823343 | 0 | 1 | 0 | 1 | 1 | 0.40110418 | 0 | 1 | 1 | 0.20679328 | 12 | 0.62236227 | FALSE | 1 | FALSE | 1 | FALSE | 1 | FALSE | 1 | FALSE | 1 | FALSE | 1 |
| 7702 | 7702 | 7.3902439 | 3.09112063 | 1 | 0.65632651 | 2 | 0.38460545 | 2 | 0.43640583 | 3 | 0.29054213 | 0 | 1 | 33 | 0.81996213 | FALSE | 1 | FALSE | 1 | FALSE | 1 | FALSE | 1 | FALSE | 1 | FALSE | 1 |
| 7703 | 7703 | 8.51851852 | 1.64137184 | 1 | 0.50471032 | 0 | 1 | 0 | 1 | 0 | 1 | 1 | 0.36060486 | 25 | 0.1832206 | FALSE | 1 | FALSE | 1 | FALSE | 1 | FALSE | 1 | FALSE | 1 | FALSE | 1 |
| 7704 | 7704 | 10.03125 | 2.03844878 | 0 | 1 | 2 | 0.27730767 | 1 | 0.69069222 | 2 | 0.43583874 | 1 | 0.41154418 | 26 | 0.77612337 | FALSE | 1 | FALSE | 1 | FALSE | 1 | FALSE | 1 | FALSE | 1 | FALSE | 1 |
| 7705 | 7705 | 7.11363636 | 2.42352025 | 2 | 0.31193899 | 1 | 0.76576289 | 1 | 0.80109494 | 3 | 0.32878418 | 1 | 0.51797501 | 36 | 0.75909657 | FALSE | 1 | FALSE | 1 | FALSE | 1 | FALSE | 1 | FALSE | 1 | FALSE | 1 |
| 7706 | 7706 | 8.43478261 | 3.346979 | 2 | 0.11647716 | 0 | 1 | 1 | 0.5695037 | 1 | 0.6613267 | 1 | 0.31672722 | 18 | 0.86273155 | FALSE | 0.93181727 | FALSE | 1 | FALSE | 1 | FALSE | 1 | FALSE | 1 | FALSE | 1 |

| | | | | | | | | | | | | | | | | | | | | | | | | | | | |
|---|---|---|---|---|---|---|---|---|---|---|---|---|---|---|---|---|---|---|---|---|---|---|---|---|---|---|---|
| 7707 | 7707 | 8.0952381 | 2.87691571 | 1 | 0.42086991 | 1 | 0.49918218 | 0 | 1 | 1 | 0.62783522 | 0 | 1 | 18 | 0.57962763 | FALSE | 1 | FALSE | 1 | FALSE | 1 | FALSE | 1 | FALSE | 1 | FALSE | 1 |
| 7708 | 7708 | 7.19230769 | 3.72124081 | 1 | 0.47819878 | 0 | 1 | 2 | 0.22597287 | 1 | 0.69181315 | 1 | 0.33902672 | 20 | 0.8152498 | FALSE | 1 | FALSE | 1 | FALSE | 1 | FALSE | 1 | FALSE | 1 | FALSE | 1 |
| 7709 | 7709 | 7.08823529 | 3.31101235 | 2 | 0.21638713 | 2 | 0.30150529 | 0 | 1 | 2 | 0.46709861 | 2 | 0.1065437 | 26 | 0.92796052 | FALSE | 1 | FALSE | 1 | FALSE | 1 | FALSE | 1 | FALSE | 1 | FALSE | 1 |
| 7710 | 7710 | 9.91666667 | 2.67576573 | 1 | 0.46441868 | 0 | 1 | 1 | 0.58502114 | 1 | 0.67692816 | 0 | 1 | 21 | 0.47053137 | FALSE | 1 | FALSE | 1 | FALSE | 1 | FALSE | 1 | FALSE | 1 | FALSE | 1 |
| 7711 | 7711 | 10.8571429 | 2.18295811 | 3 | 0.0279333 | 1 | 0.57534752 | 0 | 1 | 0 | 1 | 1 | 0.3499045 | 21 | 0.78939668 | FALSE | 0.50964275 | FALSE | 1 | FALSE | 1 | FALSE | 1 | FALSE | 1 | FALSE | 1 |
| 7712 | 7712 | 11.7307692 | 3.16905339 | 1 | 0.49162634 | 0 | 1 | 0 | 1 | 1 | 0.70601446 | 1 | 0.3499045 | 23 | 0.40372818 | FALSE | 1 | FALSE | 1 | FALSE | 1 | FALSE | 1 | FALSE | 1 | FALSE | 1 |
| 7713 | 7713 | 8.5 | 3.14907394 | 0 | 1 | 0 | 1 | 1 | 0.51939841 | 1 | 0.60987146 | 0 | 1 | 18 | 0.37504645 | FALSE | 1 | FALSE | 1 | FALSE | 1 | FALSE | 1 | FALSE | 1 | FALSE | 1 |
| 7714 | 7714 | 10.3684211 | 2.20550809 | 0 | 1 | 0 | 1 | 0 | 1 | 0 | 1 | 1 | 0.26986775 | 18 | 0.17880694 | FALSE | 1 | FALSE | 1 | FALSE | 1 | FALSE | 1 | FALSE | 1 | FALSE | 1 |
| 7715 | 7715 | 8.90909091 | 1.95190096 | 0 | 1 | 1 | 0.5154318 | 0 | 1 | 3 | 0.07743186 | 0 | 1 | 18 | 0.74744813 | FALSE | 1 | FALSE | 1 | FALSE | 1 | FALSE | 1 | FALSE | 1 | FALSE | 1 |
| 7716 | 7716 | 9.29032258 | 1.86985506 | 0 | 1 | 2 | 0.2651851 | 3 | 0.09854319 | 0 | 1 | 1 | 0.4016888 | 25 | 0.80006983 | FALSE | 1 | FALSE | 1 | FALSE | 1 | FALSE | 1 | FALSE | 1 | FALSE | 1 |
| 7717 | 7717 | 6.15 | 2.45916653 | 3 | 0.47531326 | 4 | 0.40632801 | 3 | 0.70314164 | 3 | 0.84487539 | 0 | 1 | 87 | 0.28513463 | FALSE | 1 | FALSE | 1 | FALSE | 1 | FALSE | 1 | FALSE | 1 | FALSE | 1 |
| 7718 | 7718 | 6.70833333 | 1.54054013 | 0 | 1 | 1 | 0.79481669 | 1 | 0.8283465 | 0 | 1 | 1 | 0.54902428 | 45 | 0.04502985 | FALSE | 1 | FALSE | 1 | FALSE | 1 | FALSE | 1 | FALSE | 1 | FALSE | 1 |
| 7719 | 7719 | 7.15686275 | 3.63734739 | 4 | 0.04099222 | 3 | 0.22759346 | 3 | 0.27688673 | 5 | 0.08315495 | 2 | 0.20351566 | 34 | 0.99956752 | FALSE | 0.62767905 | FALSE | 1 | FALSE | 1 | FALSE | 1 | FALSE | 1 | FALSE | 1 |
| 7720 | 7720 | 10.5666667 | 2.60362995 | 2 | 0.16962005 | 0 | 1 | 1 | 0.65463982 | 2 | 0.3871811 | 0 | 1 | 24 | 0.70563334 | FALSE | 1 | FALSE | 1 | FALSE | 1 | FALSE | 1 | FALSE | 1 | FALSE | 1 |
| 7721 | 7721 | 11.1666667 | 2.63417708 | 1 | 0.54198714 | 1 | 0.6278847 | 1 | 0.66709967 | 1 | 0.75658797 | 1 | 0.39166989 | 25 | 0.67630471 | FALSE | 1 | FALSE | 1 | FALSE | 1 | FALSE | 1 | FALSE | 1 | FALSE | 1 |
| 7722 | 7722 | 6.4 | 3.03973683 | 0 | 1 | 0 | 1 | 0 | 1 | 1 | 0.60987146 | 0 | 1 | 19 | 0.15700255 | FALSE | 1 | FALSE | 1 | FALSE | 1 | FALSE | 1 | FALSE | 1 | FALSE | 1 |
| 7723 | 7723 | 9.4 | 2.08326667 | 1 | 0.40558521 | 1 | 0.48239024 | 0 | 1 | 1 | 0.60987146 | 1 | 0.28187291 | 16 | 0.80826361 | FALSE | 1 | FALSE | 1 | FALSE | 1 | FALSE | 1 | FALSE | 1 | FALSE | 1 |
| 7724 | 7724 | 8.08108108 | 2.61366863 | 0 | 1 | 1 | 0.70472939 | 0 | 1 | 0 | 1 | 0 | 1 | 36 | 0.01445326 | FALSE | 1 | FALSE | 1 | FALSE | 1 | FALSE | 1 | FALSE | 1 | FALSE | 1 |
| 7725 | 7725 | 6.30097087 | 1.24506756 | 4 | 0.77711726 | 5 | 0.799452 | 9 | 0.31637431 | 14 | 0.08814468 | 2 | 0.85516757 | 171 | 0.69145348 | FALSE | 1 | FALSE | 1 | FALSE | 1 | FALSE | 1 | FALSE | 1 | FALSE | 1 |
| 7726 | 7726 | 7.57142857 | 1.34771159 | 0 | 1 | 1 | 0.36919824 | 0 | 1 | 1 | 0.48242099 | 0 | 1 | 12 | 0.62236227 | FALSE | 1 | FALSE | 1 | FALSE | 1 | FALSE | 1 | FALSE | 1 | FALSE | 1 |
| 7727 | 7727 | 8.76470588 | 2.21038756 | 0 | 1 | 1 | 0.42857733 | 0 | 1 | 1 | 0.55062772 | 0 | 1 | 15 | 0.49060435 | FALSE | 1 | FALSE | 1 | FALSE | 1 | FALSE | 1 | FALSE | 1 | FALSE | 1 |
| 7728 | 7728 | 10 | 3.65148372 | 2 | 0.12501436 | 1 | 0.54637326 | 0 | 1 | 1 | 0.67692816 | 0 | 1 | 20 | 0.68270729 | FALSE | 0.95622564 | FALSE | 1 | FALSE | 1 | FALSE | 1 | FALSE | 1 | FALSE | 1 |
| 7729 | 7729 | 10 | 1.58113883 | 0 | 1 | 2 | 0.02563923 | 0 | 1 | 0 | 1 | 0 | 1 | 6 | 0.88425551 | FALSE | 1 | FALSE | 0.82367276 | FALSE | 1 | FALSE | 1 | FALSE | 1 | FALSE | 1 |
| 7730 | 7730 | 10.4 | 1.8547237 | 0 | 1 | 0 | 1 | 0 | 1 | 0 | 1 | 0 | 1 | 5 | 0.42761893 | FALSE | 1 | FALSE | 1 | FALSE | 1 | FALSE | 1 | FALSE | 1 | FALSE | 1 |
| 7731 | 7731 | 7.48387097 | 3.0360306 | 2 | 0.1881398 | 1 | 0.63997468 | 0 | 1 | 1 | 0.76781284 | 0 | 1 | 27 | 0.45522639 | FALSE | 1 | FALSE | 1 | FALSE | 1 | FALSE | 1 | FALSE | 1 | FALSE | 1 |
| 7732 | 7732 | 9.33333333 | 2 | 1 | 0.20854748 | 0 | 1 | 0 | 1 | 2 | 0.06108592 | 0 | 1 | 6 | 0.96128317 | FALSE | 1 | FALSE | 1 | FALSE | 1 | FALSE | 1 | FALSE | 1 | FALSE | 1 |
| 7733 | 7733 | 10.6 | 2.65329983 | 0 | 1 | 0 | 1 | 1 | 0.16720719 | 0 | 1 | 0 | 1 | 4 | 0.82371636 | FALSE | 1 | FALSE | 1 | FALSE | 1 | FALSE | 1 | FALSE | 1 | FALSE | 1 |
| 7734 | 7734 | 10.4705882 | 2.91280433 | 2 | 0.06909038 | 0 | 1 | 1 | 0.46348836 | 0 | 1 | 0 | 1 | 14 | 0.7313145 | FALSE | 0.74276693 | FALSE | 1 | FALSE | 1 | FALSE | 1 | FALSE | 1 | FALSE | 1 |
| 7735 | 7735 | 11.8461539 | 2.76923077 | 0 | 1 | 0 | 1 | 1 | 0.37874417 | 0 | 1 | 1 | 0.19354507 | 11 | 0.6683056 | FALSE | 1 | FALSE | 1 | FALSE | 1 | FALSE | 1 | FALSE | 1 | FALSE | 1 |
| 7736 | 7736 | 6.91891892 | 1.61711084 | 0 | 1 | 1 | 0.9134145 | 2 | 0.75097925 | 3 | 0.66754163 | 0 | 1 | 68 | 0.04378964 | FALSE | 1 | FALSE | 1 | FALSE | 1 | FALSE | 1 | FALSE | 1 | FALSE | 1 |
| 7737 | 7737 | 8.33333333 | 2.11869981 | 0 | 1 | 1 | 0.38964394 | 0 | 1 | 1 | 0.50623194 | 0 | 1 | 13 | 0.57712805 | FALSE | 1 | FALSE | 1 | FALSE | 1 | FALSE | 1 | FALSE | 1 | FALSE | 1 |
| 7738 | 7738 | 6.42465753 | 2.68388294 | 3 | 0.28086759 | 1 | 0.90746182 | 0 | 1 | 4 | 0.42273814 | 1 | 0.69780571 | 63 | 0.2931215 | FALSE | 1 | FALSE | 1 | FALSE | 1 | FALSE | 1 | FALSE | 1 | FALSE | 1 |
| 7739 | 7739 | 9.71428571 | 3.88087935 | 0 | 1 | 0 | 1 | 0 | 1 | 1 | 0.28044368 | 1 | 0.10932414 | 5 | 0.91847773 | FALSE | 1 | FALSE | 1 | FALSE | 1 | FALSE | 1 | FALSE | 1 | FALSE | 1 |
| 7740 | 7740 | 11.2222222 | 2.24982853 | 1 | 0.20854748 | 0 | 1 | 1 | 0.28067846 | 0 | 1 | 0 | 1 | 7 | 0.84574168 | FALSE | 1 | FALSE | 1 | FALSE | 1 | FALSE | 1 | FALSE | 1 | FALSE | 1 |
| 7741 | 7741 | 10.4285714 | 2.66496544 | 0 | 1 | 0 | 1 | 1 | 0.53671415 | 2 | 0.25059627 | 0 | 1 | 18 | 0.57962763 | FALSE | 1 | FALSE | 1 | FALSE | 1 | FALSE | 1 | FALSE | 1 | FALSE | 1 |
| 7742 | 7742 | 10.6153846 | 1.82033224 | 0 | 1 | 0 | 1 | 1 | 0.37874417 | 1 | 0.45746571 | 0 | 1 | 11 | 0.6683056 | FALSE | 1 | FALSE | 1 | FALSE | 1 | FALSE | 1 | FALSE | 1 | FALSE | 1 |
| 7743 | 7743 | 9.92857143 | 4.07894044 | 0 | 1 | 0 | 1 | 1 | 0.40110418 | 0 | 1 | 1 | 0.20679328 | 12 | 0.62236227 | FALSE | 1 | FALSE | 1 | FALSE | 1 | FALSE | 1 | FALSE | 1 | FALSE | 1 |
| 7744 | 7744 | 7.71428571 | 2.3733211 | 0 | 1 | 1 | 0.20567256 | 0 | 1 | 0 | 1 | 0 | 1 | 6 | 0.69920685 | FALSE | 1 | FALSE | 1 | FALSE | 1 | FALSE | 1 | FALSE | 1 | FALSE | 1 |
| 7745 | 7745 | 11.2222222 | 2.09644025 | 0 | 1 | 0 | 1 | 0 | 1 | 0 | 1 | 0 | 1 | 9 | 0.21661397 | FALSE | 1 | FALSE | 1 | FALSE | 1 | FALSE | 1 | FALSE | 1 | FALSE | 1 |
| 7746 | 7746 | 8.88888889 | 2.46956786 | 0 | 1 | 0 | 1 | 0 | 1 | 1 | 0.34506517 | 0 | 1 | 8 | 0.5780332 | FALSE | 1 | FALSE | 1 | FALSE | 1 | FALSE | 1 | FALSE | 1 | FALSE | 1 |
| 7747 | 7747 | 9.5 | 2.66145324 | 1 | 0.22886536 | 1 | 0.28036922 | 0 | 1 | 0 | 1 | 0 | 1 | 8 | 0.80394719 | FALSE | 1 | FALSE | 1 | FALSE | 1 | FALSE | 1 | FALSE | 1 | FALSE | 1 |
| 7748 | 7748 | 3.34875445 | 1.46102638 | 7 | 0.57851991 | 16 | 0.01831042 | 16 | 0.04280168 | 16 | 0.21107376 | 6 | 0.30755375 | 218 | 0.99812143 | FALSE | 1 | FALSE | 0.77319042 | FALSE | 1 | FALSE | 1 | FALSE | 1 | FALSE | 1 |
| 7749 | 7749 | 9.36363636 | 2.73936712 | 0 | 1 | 0 | 1 | 0 | 1 | 0 | 1 | 0 | 1 | 11 | 0.15414489 | FALSE | 1 | FALSE | 1 | FALSE | 1 | FALSE | 1 | FALSE | 1 | FALSE | 1 |
| 7750 | 7750 | 9 | 2.7080128 | 0 | 1 | 0 | 1 | 1 | 0.42266264 | 0 | 1 | 0 | 1 | 14 | 0.29525409 | FALSE | 1 | FALSE | 1 | FALSE | 1 | FALSE | 1 | FALSE | 1 | FALSE | 1 |
| 7751 | 7751 | 4.29820051 | 1.3176757 | 6 | 0.93977639 | 14 | 0.37879865 | 19 | 0.10522254 | 13 | 0.9147122 | 7 | 0.45638596 | 330 | 0.43336878 | FALSE | 1 | FALSE | 1 | FALSE | 1 | FALSE | 1 | FALSE | 1 | FALSE | 1 |
| 7752 | 7752 | 8.03703704 | 2.1684861 | 0 | 1 | 1 | 0.57534752 | 1 | 0.61440466 | 1 | 0.70601446 | 0 | 1 | 23 | 0.40372818 | FALSE | 1 | FALSE | 1 | FALSE | 1 | FALSE | 1 | FALSE | 1 | FALSE | 1 |
| 7753 | 7753 | 9.1 | 2.16564078 | 0 | 1 | 0 | 1 | 1 | 0.51939841 | 1 | 0.60987146 | 0 | 1 | 18 | 0.37504645 | FALSE | 1 | FALSE | 1 | FALSE | 1 | FALSE | 1 | FALSE | 1 | FALSE | 1 |
| 7754 | 7754 | 7.69767442 | 2.6722944 | 2 | 0.29286138 | 4 | 0.04570474 | 2 | 0.44869594 | 3 | 0.30326943 | 0 | 1 | 31 | 0.97647435 | FALSE | 1 | FALSE | 1 | FALSE | 1 | FALSE | 1 | FALSE | 1 | FALSE | 1 |
| 7755 | 7755 | 9.05555556 | 2.69716358 | 0 | 1 | 0 | 1 | 1 | 0.48280983 | 1 | 0.57131013 | 0 | 1 | 16 | 0.44998001 | FALSE | 1 | FALSE | 1 | FALSE | 1 | FALSE | 1 | FALSE | 1 | FALSE | 1 |
| 7756 | 7756 | 8.28571429 | 2.18529408 | 0 | 1 | 0 | 1 | 2 | 0.2663542 | 1 | 0.73248983 | 0 | 1 | 25 | 0.34293 | FALSE | 1 | FALSE | 1 | FALSE | 1 | FALSE | 1 | FALSE | 1 | FALSE | 1 |
| 7757 | 7757 | 9.66666667 | 4.02131075 | 1 | 0.62830697 | 0 | 1 | 5 | 0.01093381 | 2 | 0.52640931 | 1 | 0.4673875 | 29 | 0.93814695 | FALSE | 1 | FALSE | 1 | FALSE | 1 | FALSE | 1 | FALSE | 1 | FALSE | 1 |
| 7758 | 7758 | 10.6666667 | 2.82842713 | 2 | 0.07645454 | 0 | 1 | 0 | 1 | 0 | 1 | 0 | 1 | 16 | 0.44998001 | FALSE | 0.77424195 | FALSE | 1 | FALSE | 1 | FALSE | 1 | FALSE | 1 | FALSE | 1 |
| 7759 | 7759 | 9.41666667 | 2.01900691 | 0 | 1 | 3 | 0.04096158 | 0 | 1 | 1 | 0.67692816 | 0 | 1 | 20 | 0.68270729 | FALSE | 1 | FALSE | 0.9698186 | FALSE | 1 | FALSE | 1 | FALSE | 1 | FALSE | 1 |
| 7760 | 7760 | 8.92307692 | 1.10673035 | 0 | 1 | 0 | 1 | 2 | 0.22597287 | 2 | 0.31968852 | 1 | 0.33902672 | 20 | 0.8152498 | FALSE | 1 | FALSE | 1 | FALSE | 1 | FALSE | 1 | FALSE | 1 | FALSE | 1 |
| 7761 | 7761 | 10.4615385 | 4.43092948 | 0 | 1 | 0 | 1 | 0 | 1 | 0 | 1 | 0 | 1 | 13 | 0.10967906 | FALSE | 1 | FALSE | 1 | FALSE | 1 | FALSE | 1 | FALSE | 1 | FALSE | 1 |
| 7762 | 7762 | 8.39130435 | 1.29343259 | 0 | 1 | 2 | 0.41896557 | 1 | 0.80109494 | 6 | 0.0144401 | 0 | 1 | 35 | 0.8620983 | FALSE | 1 | FALSE | 1 | FALSE | 1 | FALSE | 1 | FALSE | 1 | FALSE | 1 |
| 7763 | 7763 | 8.92857143 | 1.38689199 | 3 | 0.0048942 | 1 | 0.36919824 | 0 | 1 | 1 | 0.48242099 | 1 | 0.20679328 | 8 | 0.99722299 | FALSE | 0.19848171 | FALSE | 1 | FALSE | 1 | FALSE | 1 | FALSE | 1 | FALSE | 1 |
| 7764 | 7764 | 7.15384615 | 2.45511984 | 5 | 0.0029693 | 2 | 0.36122049 | 0 | 1 | 1 | 0.84089119 | 1 | 0.4761699 | 30 | 0.92795554 | FALSE | 0.14653312 | FALSE | 1 | FALSE | 1 | FALSE | 1 | FALSE | 1 | FALSE | 1 |
| 7765 | 7765 | 9.12 | 2.0653329 | 1 | 0.46441868 | 0 | 1 | 0 | 1 | 2 | 0.30249307 | 1 | 0.32796862 | 20 | 0.68270729 | FALSE | 1 | FALSE | 1 | FALSE | 1 | FALSE | 1 | FALSE | 1 | FALSE | 1 |
| 7766 | 7766 | 10.4285714 | 1.95354205 | 0 | 1 | 0 | 1 | 1 | 0.40110418 | 0 | 1 | 0 | 1 | 13 | 0.33283729 | FALSE | 1 | FALSE | 1 | FALSE | 1 | FALSE | 1 | FALSE | 1 | FALSE | 1 |
| 7767 | 7767 | 10.25 | 2.56173769 | 0 | 1 | 1 | 0.38964394 | 0 | 1 | 1 | 0.50623194 | 0 | 1 | 13 | 0.57712805 | FALSE | 1 | FALSE | 1 | FALSE | 1 | FALSE | 1 | FALSE | 1 | FALSE | 1 |
| 7768 | 7768 | 5.53097345 | 3.19310872 | 1 | 0.94678796 | 4 | 0.4928114 | 4 | 0.57634539 | 7 | 0.25456339 | 2 | 0.55195029 | 94 | 0.61432106 | FALSE | 1 | FALSE | 1 | FALSE | 1 | FALSE | 1 | FALSE | 1 | FALSE | 1 |
| 7769 | 7769 | 10.5833333 | 2.43099202 | 1 | 0.26795838 | 0 | 1 | 0 | 1 | 1 | 0.43131129 | 0 | 1 | 10 | 0.71436964 | FALSE | 1 | FALSE | 1 | FALSE | 1 | FALSE | 1 | FALSE | 1 | FALSE | 1 |
| 7770 | 7770 | 7.13513514 | 3.20609354 | 0 | 1 | 0 | 1 | 1 | 0.74264075 | 1 | 0.82511639 | 0 | 1 | 35 | 0.05673524 | FALSE | 1 | FALSE | 1 | FALSE | 1 | FALSE | 1 | FALSE | 1 | FALSE | 1 |
| 7771 | 7771 | 9 | 1.95789002 | 1 | 0.46441868 | 0 | 1 | 1 | 0.58502114 | 0 | 1 | 0 | 1 | 22 | 0.25237063 | FALSE | 1 | FALSE | 1 | FALSE | 1 | FALSE | 1 | FALSE | 1 | FALSE | 1 |
| 7772 | 7772 | 9.125 | 1.36358901 | 0 | 1 | 1 | 0.23139161 | 1 | 0.25384309 | 0 | 1 | 0 | 1 | 6 | 0.88425551 | FALSE | 1 | FALSE | 1 | FALSE | 1 | FALSE | 1 | FALSE | 1 | FALSE | 1 |
| 7773 | 7773 | 7.70833333 | 4.10770786 | 2 | 0.12501436 | 1 | 0.54637326 | 1 | 0.58502114 | 0 | 1 | 0 | 1 | 20 | 0.68270729 | FALSE | 0.95622564 | FALSE | 1 | FALSE | 1 | FALSE | 1 | FALSE | 1 | FALSE | 1 |
| 7774 | 7774 | 9.37209302 | 3.35512975 | 2 | 0.30241117 | 4 | 0.04916488 | 1 | 0.79363407 | 1 | 0.86831187 | 0 | 1 | 35 | 0.7801651 | FALSE | 1 | FALSE | 1 | FALSE | 1 | FALSE | 1 | FALSE | 1 | FALSE | 1 |
| 7775 | 7775 | 9.23809524 | 2.67091498 | 1 | 0.42086991 | 1 | 0.49918218 | 1 | 0.53671415 | 2 | 0.25059627 | 1 | 0.29368245 | 15 | 0.96513691 | FALSE | 1 | FALSE | 1 | FALSE | 1 | FALSE | 1 | FALSE | 1 | FALSE | 1 |
| 7776 | 7776 | 10.84 | 1.75909067 | 1 | 0.46441868 | 1 | 0.54637326 | 2 | 0.21260257 | 0 | 1 | 1 | 0.32796862 | 19 | 0.83975208 | FALSE | 1 | FALSE | 1 | FALSE | 1 | FALSE | 1 | FALSE | 1 | FALSE | 1 |
| 7777 | 7777 | 9.8 | 2.22710575 | 1 | 0.32293355 | 0 | 1 | 1 | 0.42266264 | 1 | 0.50623194 | 0 | 1 | 12 | 0.80329917 | FALSE | 1 | FALSE | 1 | FALSE | 1 | FALSE | 1 | FALSE | 1 | FALSE | 1 |
| 7778 | 7778 | 8.33333333 | 2.47002229 | 2 | 0.20692137 | 1 | 0.6629941 | 1 | 0.70185639 | 0 | 1 | 3 | 0.01637249 | 26 | 0.86852567 | FALSE | 1 | FALSE | 1 | FALSE | 1 | FALSE | 1 | FALSE | 0.66728076 | FALSE | 1 |
| 7779 | 7779 | 10.7142857 | 2.60298102 | 0 | 1 | 0 | 1 | 0 | 1 | 1 | 0.28044368 | 1 | 0.10932414 | 5 | 0.91847773 | FALSE | 1 | FALSE | 1 | FALSE | 1 | FALSE | 1 | FALSE | 1 | FALSE | 1 |
| 7780 | 7780 | 8.32258065 | 2.46768923 | 1 | 0.55378205 | 0 | 1 | 1 | 0.67911179 | 2 | 0.41984187 | 0 | 1 | 27 | 0.45522639 | FALSE | 1 | FALSE | 1 | FALSE | 1 | FALSE | 1 | FALSE | 1 | FALSE | 1 |
| 7781 | 7781 | 9.83333333 | 1.57233019 | 0 | 1 | 0 | 1 | 2 | 0.06691409 | 0 | 1 | 0 | 1 | 10 | 0.71436964 | FALSE | 1 | FALSE | 1 | FALSE | 1 | FALSE | 1 | FALSE | 1 | FALSE | 1 |
| 7782 | 7782 | 10.4285714 | 1.74086537 | 1 | 0.51745948 | 1 | 0.60247911 | 1 | 0.64171563 | 0 | 1 | 0 | 1 | 25 | 0.34293 | FALSE | 1 | FALSE | 1 | FALSE | 1 | FALSE | 1 | FALSE | 1 | FALSE | 1 |
| 7783 | 7783 | 10.5416667 | 2.29090897 | 2 | 0.12501436 | 0 | 1 | 0 | 1 | 2 | 0.30249307 | 0 | 1 | 20 | 0.68270729 | FALSE | 0.95622564 | FALSE | 1 | FALSE | 1 | FALSE | 1 | FALSE | 1 | FALSE | 1 |
| 7784 | 7784 | 9.68292683 | 2.19132463 | 0 | 1 | 0 | 1 | 2 | 0.43640583 | 3 | 0.29054213 | 1 | 0.49330646 | 35 | 0.53616095 | FALSE | 1 | FALSE | 1 | FALSE | 1 | FALSE | 1 | FALSE | 1 | FALSE | 1 |
| 7785 | 7785 | 8.86956522 | 1.39266238 | 3 | 0.10767332 | 2 | 0.43020462 | 2 | 0.48463745 | 1 | 0.8802008 | 0 | 1 | 37 | 0.73736034 | FALSE | 0.88820416 | FALSE | 1 | FALSE | 1 | FALSE | 1 | FALSE | 1 | FALSE | 1 |
| 7786 | 7786 | 10.5333333 | 1.93907194 | 1 | 0.69044531 | 4 | 0.05650644 | 1 | 0.80828716 | 3 | 0.34154252 | 1 | 0.52593041 | 35 | 0.91829721 | FALSE | 1 | FALSE | 1 | FALSE | 1 | FALSE | 1 | FALSE | 1 | FALSE | 1 |
| 7787 | 7787 | 10.3703704 | 1.41808813 | 4 | 0.00461682 | 1 | 0.58913618 | 0 | 1 | 1 | 0.71956339 | 0 | 1 | 21 | 0.88395936 | FALSE | 0.19698443 | FALSE | 1 | FALSE | 1 | FALSE | 1 | FALSE | 1 | FALSE | 1 |
| 7788 | 7788 | 9.17808219 | 1.89695116 | 3 | 0.27385794 | 2 | 0.67482356 | 5 | 0.11089846 | 1 | 0.96507823 | 0 | 1 | 60 | 0.56780244 | FALSE | 1 | FALSE | 1 | FALSE | 1 | FALSE | 1 | FALSE | 1 | FALSE | 1 |
| 7789 | 7789 | 4.04635762 | 3.92029487 | 5 | 0.33509948 | 10 | 0.02225929 | 6 | 0.44735544 | 6 | 0.68696554 | 3 | 0.44368013 | 119 | 0.9465566 | FALSE | 1 | FALSE | 0.81043583 | FALSE | 1 | FALSE | 1 | FALSE | 1 | FALSE | 1 |
| 7790 | 7790 | 10.1333333 | 1.76509364 | 2 | 0.16048326 | 3 | 0.06039219 | 0 | 1 | 2 | 0.3705495 | 3 | 0.0104283 | 18 | 0.9978883 | FALSE | 1 | FALSE | 1 | FALSE | 1 | FALSE | 1 | FALSE | 0.5620304 | FALSE | 1 |
| 7791 | 7791 | 9.42424242 | 2.21606125 | 1 | 0.57647348 | 1 | 0.6629941 | 0 | 1 | 1 | 0.78873814 | 0 | 1 | 30 | 0.21942357 | FALSE | 1 | FALSE | 1 | FALSE | 1 | FALSE | 1 | FALSE | 1 | FALSE | 1 |
| 7792 | 7792 | 6.37313433 | 2.13594553 | 8 | 0.02144908 | 5 | 0.43780567 | 7 | 0.20624898 | 3 | 0.94991146 | 5 | 0.06860468 | 106 | 0.96147113 | FALSE | 0.46484348 | FALSE | 1 | FALSE | 1 | FALSE | 1 | FALSE | 1 | FALSE | 1 |
| 7793 | 7793 | 9.1875 | 2.15692924 | 0 | 1 | 0 | 1 | 0 | 1 | 1 | 0.52895088 | 0 | 1 | 15 | 0.26127859 | FALSE | 1 | FALSE | 1 | FALSE | 1 | FALSE | 1 | FALSE | 1 | FALSE | 1 |
| 7794 | 7794 | 8.27272727 | 2.63165724 | 0 | 1 | 1 | 0.5154318 | 0 | 1 | 1 | 0.64497439 | 0 | 1 | 20 | 0.30915202 | FALSE | 1 | FALSE | 1 | FALSE | 1 | FALSE | 1 | FALSE | 1 | FALSE | 1 |
| 7795 | 7795 | 11.8333333 | 4.4127341 | 1 | 0.26795838 | 0 | 1 | 0 | 1 | 0 | 1 | 0 | 1 | 11 | 0.41944532 | FALSE | 1 | FALSE | 1 | FALSE | 1 | FALSE | 1 | FALSE | 1 | FALSE | 1 |
| 7796 | 7796 | 8.61111111 | 2.5742073 | 0 | 1 | 2 | 0.31355738 | 3 | 0.12958859 | 4 | 0.07465533 | 0 | 1 | 26 | 0.96294373 | FALSE | 1 | FALSE | 1 | FALSE | 1 | FALSE | 1 | FALSE | 1 | FALSE | 1 |
| 7797 | 7797 | 8.8 | 2.01494417 | 1 | 0.40558521 | 1 | 0.48239024 | 2 | 0.16019548 | 0 | 1 | 1 | 0.28187291 | 15 | 0.92113679 | FALSE | 1 | FALSE | 1 | FALSE | 1 | FALSE | 1 | FALSE | 1 | FALSE | 1 |
| 7798 | 7798 | 8.875 | 1.99608993 | 0 | 1 | 0 | 1 | 1 | 0.44344817 | 1 | 0.52895088 | 0 | 1 | 14 | 0.53308261 | FALSE | 1 | FALSE | 1 | FALSE | 1 | FALSE | 1 | FALSE | 1 | FALSE | 1 |
| 7799 | 7799 | 10.12 | 1.63266653 | 0 | 1 | 2 | 0.19297191 | 0 | 1 | 1 | 0.69181315 | 0 | 1 | 22 | 0.43642846 | FALSE | 1 | FALSE | 1 | FALSE | 1 | FALSE | 1 | FALSE | 1 | FALSE | 1 |
| 7800 | 7800 | 9.72727273 | 1.68905079 | 1 | 0.76171602 | 2 | 0.5356583 | 0 | 1 | 1 | 0.92538938 | 2 | 0.22768257 | 49 | 0.22274213 | FALSE | 1 | FALSE | 1 | FALSE | 1 | FALSE | 1 | FALSE | 1 | FALSE | 1 |

| | | | | | | | | | | | | | | | | | | | | | | | | | | | |
|---|---|---|---|---|---|---|---|---|---|---|---|---|---|---|---|---|---|---|---|---|---|---|---|---|---|---|---|
| 7801 | 7801 | 5.6618705 | 2.87773381 | 0 | 1 | 5 | 0.45058034 | 4 | 0.72638773 | 5 | 0.75550085 | 7 | 0.00677859 | 115 | 0.53476038 | FALSE | 1 | FALSE | 1 | FALSE | 1 | FALSE | 1 | FALSE | 0.47461742 | FALSE | 1 |
| 7802 | 7802 | 9.14285714 | 3.22616851 | 0 | 1 | 1 | 0.36919824 | 1 | 0.40110418 | 1 | 0.48242099 | 1 | 0.20679328 | 10 | 0.94579623 | FALSE | 1 | FALSE | 1 | FALSE | 1 | FALSE | 1 | FALSE | 1 | FALSE | 1 |
| 7803 | 7803 | 7.32 | 3.15873392 | 1 | 0.46441868 | 0 | 1 | 1 | 0.58502114 | 2 | 0.30249307 | 1 | 0.32796862 | 19 | 0.83975208 | FALSE | 1 | FALSE | 1 | FALSE | 1 | FALSE | 1 | FALSE | 1 | FALSE | 1 |
| 7804 | 7804 | 10.4736842 | 1.49976914 | 0 | 1 | 2 | 0.1135776 | 0 | 1 | 0 | 1 | 1 | 0.25766374 | 15 | 0.69372602 | FALSE | 1 | FALSE | 1 | FALSE | 1 | FALSE | 1 | FALSE | 1 | FALSE | 1 |
| 7805 | 7805 | 8.25 | 2.7220805 | 0 | 1 | 0 | 1 | 0 | 1 | 0 | 1 | 0 | 1 | 36 | 0.00217217 | FALSE | 1 | FALSE | 1 | FALSE | 1 | FALSE | 1 | FALSE | 1 | FALSE | 1 |
| 7806 | 7806 | 8.52 | 1.76906755 | 2 | 0.12501436 | 1 | 0.54637326 | 2 | 0.21260257 | 1 | 0.67692816 | 0 | 1 | 18 | 0.93166998 | FALSE | 0.95622564 | FALSE | 1 | FALSE | 1 | FALSE | 1 | FALSE | 1 | FALSE | 1 |
| 7807 | 7807 | 6.38888889 | 1.36377291 | 1 | 0.90240855 | 0 | 1 | 6 | 0.09991019 | 3 | 0.78298119 | 1 | 0.77262299 | 78 | 0.24535498 | FALSE | 1 | FALSE | 1 | FALSE | 1 | FALSE | 1 | FALSE | 1 | FALSE | 1 |
| 7808 | 7808 | 6.93670886 | 4.18357782 | 0 | 1 | 2 | 0.7307176 | 1 | 0.94537714 | 4 | 0.49431847 | 0 | 1 | 72 | 0.05857603 | FALSE | 1 | FALSE | 1 | FALSE | 1 | FALSE | 1 | FALSE | 1 | FALSE | 1 |
| 7809 | 7809 | 9.9375 | 1.7127737 | 1 | 0.34033054 | 0 | 1 | 1 | 0.44344817 | 1 | 0.52895088 | 1 | 0.23264612 | 12 | 0.90922186 | FALSE | 1 | FALSE | 1 | FALSE | 1 | FALSE | 1 | FALSE | 1 | FALSE | 1 |
| 7810 | 7810 | 9.47368421 | 3.13500476 | 0 | 1 | 1 | 0.46503795 | 2 | 0.14749808 | 1 | 0.59104357 | 0 | 1 | 15 | 0.83648191 | FALSE | 1 | FALSE | 1 | FALSE | 1 | FALSE | 1 | FALSE | 1 | FALSE | 1 |
| 7811 | 7811 | 7.05714286 | 3.00421473 | 1 | 0.59801736 | 5 | 0.00497143 | 1 | 0.72299506 | 1 | 0.80778317 | 0 | 1 | 27 | 0.91585095 | FALSE | 1 | FALSE | 0.58120789 | FALSE | 1 | FALSE | 1 | FALSE | 1 | FALSE | 1 |
| 7812 | 7812 | 9.76470588 | 2.28883253 | 0 | 1 | 0 | 1 | 0 | 1 | 2 | 0.14906763 | 0 | 1 | 13 | 0.57712805 | FALSE | 1 | FALSE | 1 | FALSE | 1 | FALSE | 1 | FALSE | 1 | FALSE | 1 |
| 7813 | 7813 | 7.04651163 | 2.51464074 | 3 | 0.09709068 | 0 | 1 | 2 | 0.46083448 | 4 | 0.13323859 | 1 | 0.50988734 | 33 | 0.93815092 | FALSE | 0.86047225 | FALSE | 1 | FALSE | 1 | FALSE | 1 | FALSE | 1 | FALSE | 1 |
| 7814 | 7814 | 9.3 | 1.63605216 | 1 | 0.54198714 | 1 | 0.6278847 | 1 | 0.66709967 | 0 | 1 | 2 | 0.08611595 | 25 | 0.67630471 | FALSE | 1 | FALSE | 1 | FALSE | 1 | FALSE | 1 | FALSE | 1 | FALSE | 1 |
| 7815 | 7815 | 8.140625 | 2.5548189 | 3 | 0.22550017 | 1 | 0.87929209 | 3 | 0.40547364 | 4 | 0.33842086 | 0 | 1 | 53 | 0.70813409 | FALSE | 1 | FALSE | 1 | FALSE | 1 | FALSE | 1 | FALSE | 1 | FALSE | 1 |
| 7816 | 7816 | 9.85714286 | 2.16653584 | 0 | 1 | 1 | 0.20567256 | 0 | 1 | 0 | 1 | 0 | 1 | 6 | 0.69920685 | FALSE | 1 | FALSE | 1 | FALSE | 1 | FALSE | 1 | FALSE | 1 | FALSE | 1 |
| 7817 | 7817 | 9.7 | 2.47184142 | 0 | 1 | 0 | 1 | 0 | 1 | 1 | 0.60987146 | 0 | 1 | 19 | 0.15700255 | FALSE | 1 | FALSE | 1 | FALSE | 1 | FALSE | 1 | FALSE | 1 | FALSE | 1 |
| 7818 | 7818 | 9.79166667 | 3.10884714 | 1 | 0.46441868 | 0 | 1 | 0 | 1 | 0 | 1 | 0 | 1 | 23 | 0.09196389 | FALSE | 1 | FALSE | 1 | FALSE | 1 | FALSE | 1 | FALSE | 1 | FALSE | 1 |
| 7819 | 7819 | 9.77777778 | 2.52884593 | 0 | 1 | 1 | 0.42857733 | 4 | 0.0026711 | 0 | 1 | 0 | 1 | 12 | 0.962093 | FALSE | 1 | FALSE | 1 | FALSE | 1 | FALSE | 1 | FALSE | 1 | FALSE | 1 |
| 7820 | 7820 | 9.21428571 | 1.91529824 | 1 | 0.51745948 | 0 | 1 | 2 | 0.2663542 | 1 | 0.73248983 | 1 | 0.37113067 | 23 | 0.73439118 | FALSE | 1 | FALSE | 1 | FALSE | 1 | FALSE | 1 | FALSE | 1 | FALSE | 1 |
| 7821 | 7821 | 9.5 | 2.48456775 | 1 | 0.49162634 | 0 | 1 | 2 | 0.23940226 | 1 | 0.70601446 | 0 | 1 | 22 | 0.61628 | FALSE | 1 | FALSE | 1 | FALSE | 1 | FALSE | 1 | FALSE | 1 | FALSE | 1 |
| 7822 | 7822 | 10.25 | 1.85404962 | 0 | 1 | 1 | 0.40942991 | 0 | 1 | 1 | 0.52895088 | 1 | 0.23264612 | 13 | 0.76799161 | FALSE | 1 | FALSE | 1 | FALSE | 1 | FALSE | 1 | FALSE | 1 | FALSE | 1 |
| 7823 | 7823 | 9.40322581 | 2.43932617 | 3 | 0.19869438 | 5 | 0.04382428 | 3 | 0.36617062 | 1 | 0.94113525 | 0 | 1 | 48 | 0.86671225 | FALSE | 1 | FALSE | 1 | FALSE | 1 | FALSE | 1 | FALSE | 1 | FALSE | 1 |
| 7824 | 7824 | 8.79166667 | 3.3413965 | 0 | 1 | 0 | 1 | 0 | 1 | 4 | 0.02237042 | 1 | 0.32796862 | 19 | 0.83975208 | FALSE | 1 | FALSE | 1 | FALSE | 1 | FALSE | 1 | FALSE | 1 | FALSE | 1 |
| 7825 | 7825 | 9.64 | 2.57495631 | 1 | 0.46441868 | 0 | 1 | 0 | 1 | 1 | 0.67692816 | 1 | 0.32796862 | 21 | 0.47053137 | FALSE | 1 | FALSE | 1 | FALSE | 1 | FALSE | 1 | FALSE | 1 | FALSE | 1 |
| 7826 | 7826 | 7.45454546 | 2.65821664 | 0 | 1 | 1 | 0.5154318 | 1 | 0.55340851 | 1 | 0.64497439 | 0 | 1 | 19 | 0.54233419 | FALSE | 1 | FALSE | 1 | FALSE | 1 | FALSE | 1 | FALSE | 1 | FALSE | 1 |
| 7827 | 7827 | 7.76829268 | 2.15156653 | 6 | 0.24418932 | 5 | 0.61736795 | 9 | 0.13522621 | 5 | 0.87900563 | 3 | 0.50712091 | 136 | 0.73951647 | FALSE | 1 | FALSE | 1 | FALSE | 1 | FALSE | 1 | FALSE | 1 | FALSE | 1 |
| 7828 | 7828 | 10.2777778 | 4.24009376 | 0 | 1 | 0 | 1 | 0 | 1 | 0 | 1 | 0 | 1 | 18 | 0.04681656 | FALSE | 1 | FALSE | 1 | FALSE | 1 | FALSE | 1 | FALSE | 1 | FALSE | 1 |
| 7829 | 7829 | 7.11111111 | 2.37788162 | 0 | 1 | 1 | 0.25628163 | 0 | 1 | 1 | 0.34506517 | 0 | 1 | 7 | 0.84574168 | FALSE | 1 | FALSE | 1 | FALSE | 1 | FALSE | 1 | FALSE | 1 | FALSE | 1 |
| 7830 | 7830 | 8.85714286 | 2.5872529 | 0 | 1 | 0 | 1 | 0 | 1 | 0 | 1 | 1 | 0.10932414 | 6 | 0.69920685 | FALSE | 1 | FALSE | 1 | FALSE | 1 | FALSE | 1 | FALSE | 1 | FALSE | 1 |
| 7831 | 7831 | 10.3076923 | 2.36593177 | 0 | 1 | 0 | 1 | 0 | 1 | 0 | 1 | 1 | 0.19354507 | 12 | 0.374198 | FALSE | 1 | FALSE | 1 | FALSE | 1 | FALSE | 1 | FALSE | 1 | FALSE | 1 |
| 7832 | 7832 | 10.2631579 | 2.74996852 | 0 | 1 | 0 | 1 | 0 | 1 | 0 | 1 | 1 | 0.26986775 | 18 | 0.17880694 | FALSE | 1 | FALSE | 1 | FALSE | 1 | FALSE | 1 | FALSE | 1 | FALSE | 1 |
| 7833 | 7833 | 8.20833333 | 3.5937349 | 1 | 0.46441868 | 1 | 0.54637326 | 2 | 0.21260257 | 0 | 1 | 0 | 1 | 20 | 0.68270729 | FALSE | 1 | FALSE | 1 | FALSE | 1 | FALSE | 1 | FALSE | 1 | FALSE | 1 |
| 7834 | 7834 | 8.58536585 | 2.40908453 | 1 | 0.65632651 | 0 | 1 | 3 | 0.1814659 | 1 | 0.85524728 | 2 | 0.14490154 | 34 | 0.69505404 | FALSE | 1 | FALSE | 1 | FALSE | 1 | FALSE | 1 | FALSE | 1 | FALSE | 1 |
| 7835 | 7835 | 7.13513514 | 2.42355738 | 4 | 0.12127403 | 3 | 0.43080961 | 5 | 0.12620681 | 3 | 0.66754163 | 2 | 0.34319968 | 57 | 0.96691729 | FALSE | 0.95622564 | FALSE | 1 | FALSE | 1 | FALSE | 1 | FALSE | 1 | FALSE | 1 |
| 7836 | 7836 | 10.2142857 | 2.90759035 | 0 | 1 | 0 | 1 | 2 | 0.08806398 | 0 | 1 | 0 | 1 | 12 | 0.62236227 | FALSE | 1 | FALSE | 1 | FALSE | 1 | FALSE | 1 | FALSE | 1 | FALSE | 1 |
| 7837 | 7837 | 7.92857143 | 3.63444407 | 3 | 0.0048942 | 0 | 1 | 1 | 0.40110418 | 0 | 1 | 0 | 1 | 10 | 0.94579623 | FALSE | 0.19848171 | FALSE | 1 | FALSE | 1 | FALSE | 1 | FALSE | 1 | FALSE | 1 |
| 7838 | 7838 | 5.90909091 | 3.54078947 | 0 | 1 | 0 | 1 | 0 | 1 | 0 | 1 | 0 | 1 | 16 | 0.06581507 | FALSE | 1 | FALSE | 1 | FALSE | 1 | FALSE | 1 | FALSE | 1 | FALSE | 1 |
| 7839 | 7839 | 9.95454546 | 3.43083972 | 1 | 0.68224519 | 1 | 0.76576289 | 1 | 0.80109494 | 3 | 0.32878418 | 1 | 0.51797501 | 37 | 0.61979664 | FALSE | 1 | FALSE | 1 | FALSE | 1 | FALSE | 1 | FALSE | 1 | FALSE | 1 |
| 7840 | 7840 | 6.24 | 1.0307279 | 1 | 0.96125522 | 5 | 0.37323131 | 2 | 0.94139793 | 8 | 0.20918987 | 1 | 0.87370399 | 107 | 0.32829083 | FALSE | 1 | FALSE | 1 | FALSE | 1 | FALSE | 1 | FALSE | 1 | FALSE | 1 |
| 7841 | 7841 | 7.94736842 | 2.18785365 | 0 | 1 | 0 | 1 | 2 | 0.14749808 | 1 | 0.59104357 | 1 | 0.26986775 | 15 | 0.83648191 | FALSE | 1 | FALSE | 1 | FALSE | 1 | FALSE | 1 | FALSE | 1 | FALSE | 1 |
| 7842 | 7842 | 8.16666667 | 1.15068418 | 2 | 0.40500867 | 1 | 0.83180787 | 3 | 0.30658915 | 0 | 1 | 0 | 1 | 48 | 0.23965388 | FALSE | 1 | FALSE | 1 | FALSE | 1 | FALSE | 1 | FALSE | 1 | FALSE | 1 |
| 7843 | 7843 | 6.41666667 | 2.48187877 | 0 | 1 | 1 | 0.54637326 | 1 | 0.58502114 | 1 | 0.67692816 | 0 | 1 | 21 | 0.47053137 | FALSE | 1 | FALSE | 1 | FALSE | 1 | FALSE | 1 | FALSE | 1 | FALSE | 1 |
| 7844 | 7844 | 9.4 | 2.74954542 | 2 | 0.34973212 | 0 | 1 | 3 | 0.24750546 | 6 | 0.02155984 | 0 | 1 | 37 | 0.93870662 | FALSE | 1 | FALSE | 1 | FALSE | 1 | FALSE | 1 | FALSE | 1 | FALSE | 1 |
| 7845 | 7845 | 6.23809524 | 2.33818737 | 1 | 0.66519151 | 1 | 0.74973543 | 1 | 0.78589454 | 2 | 0.5811115 | 1 | 0.50166523 | 36 | 0.50934357 | FALSE | 1 | FALSE | 1 | FALSE | 1 | FALSE | 1 | FALSE | 1 | FALSE | 1 |
| 7846 | 7846 | 11.037037 | 2.57454033 | 0 | 1 | 3 | 0.05519089 | 0 | 1 | 3 | 0.12475952 | 1 | 0.36060486 | 20 | 0.95133411 | FALSE | 1 | FALSE | 1 | FALSE | 1 | FALSE | 1 | FALSE | 1 | FALSE | 1 |
| 7847 | 7847 | 10.8571429 | 2.5872529 | 1 | 0.47819878 | 2 | 0.19297191 | 0 | 1 | 0 | 1 | 1 | 0.33902672 | 21 | 0.64957167 | FALSE | 1 | FALSE | 1 | FALSE | 1 | FALSE | 1 | FALSE | 1 | FALSE | 1 |
| 7848 | 7848 | 9.5 | 4.47772264 | 1 | 0.40558521 | 1 | 0.48239024 | 0 | 1 | 1 | 0.60987146 | 1 | 0.28187291 | 16 | 0.80826361 | FALSE | 1 | FALSE | 1 | FALSE | 1 | FALSE | 1 | FALSE | 1 | FALSE | 1 |
| 7849 | 7849 | 9.2962963 | 3.20664605 | 0 | 1 | 3 | 0.05519089 | 2 | 0.25286935 | 1 | 0.71956339 | 2 | 0.07169515 | 19 | 0.98241223 | FALSE | 1 | FALSE | 1 | FALSE | 1 | FALSE | 1 | FALSE | 1 | FALSE | 1 |
| 7850 | 7850 | 9.57692308 | 2.37248777 | 3 | 0.0279333 | 0 | 1 | 2 | 0.23940226 | 1 | 0.70601446 | 0 | 1 | 20 | 0.90142689 | FALSE | 0.50964275 | FALSE | 1 | FALSE | 1 | FALSE | 1 | FALSE | 1 | FALSE | 1 |
| 7851 | 7851 | 9.86956522 | 2.90882238 | 3 | 0.02011786 | 0 | 1 | 0 | 1 | 4 | 0.01932195 | 1 | 0.31672722 | 15 | 0.99450713 | FALSE | 0.44482142 | FALSE | 1 | FALSE | 1 | FALSE | 1 | FALSE | 1 | FALSE | 1 |
| 7852 | 7852 | 6.32432432 | 1.31630223 | 6 | 0.17922523 | 4 | 0.71128806 | 4 | 0.78496075 | 5 | 0.81697416 | 1 | 0.91576444 | 128 | 0.27992276 | FALSE | 1 | FALSE | 1 | FALSE | 1 | FALSE | 1 | FALSE | 1 | FALSE | 1 |
| 7853 | 7853 | 10.3846154 | 2.05832126 | 1 | 0.28675951 | 0 | 1 | 0 | 1 | 1 | 0.45746571 | 0 | 1 | 11 | 0.6683056 | FALSE | 1 | FALSE | 1 | FALSE | 1 | FALSE | 1 | FALSE | 1 | FALSE | 1 |
| 7854 | 7854 | 8.83333333 | 0.372678 | 0 | 1 | 0 | 1 | 0 | 1 | 0 | 1 | 0 | 1 | 6 | 0.36077158 | FALSE | 1 | FALSE | 1 | FALSE | 1 | FALSE | 1 | FALSE | 1 | FALSE | 1 |
| 7855 | 7855 | 8.76923077 | 3.26175616 | 1 | 0.28675951 | 1 | 0.3480709 | 0 | 1 | 0 | 1 | 0 | 1 | 11 | 0.6683056 | FALSE | 1 | FALSE | 1 | FALSE | 1 | FALSE | 1 | FALSE | 1 | FALSE | 1 |
| 7856 | 7856 | 6.24528302 | 2.10036529 | 7 | 0.00036728 | 3 | 0.24492539 | 2 | 0.57303317 | 4 | 0.22482652 | 0 | 1 | 37 | 0.99791276 | TRUE | 0.03958878 | FALSE | 1 | FALSE | 1 | FALSE | 1 | FALSE | 1 | FALSE | 1 |
| 7857 | 7857 | 9.5 | 2.03442594 | 3 | 0.01017712 | 0 | 1 | 1 | 0.48280983 | 1 | 0.57131013 | 0 | 1 | 13 | 0.95041913 | FALSE | 0.30467829 | FALSE | 1 | FALSE | 1 | FALSE | 1 | FALSE | 1 | FALSE | 1 |
| 7858 | 7858 | 7.93939394 | 2.92263262 | 0 | 1 | 0 | 1 | 3 | 0.11365871 | 1 | 0.78873814 | 0 | 1 | 29 | 0.39648374 | FALSE | 1 | FALSE | 1 | FALSE | 1 | FALSE | 1 | FALSE | 1 | FALSE | 1 |
| 7859 | 7859 | 9.68 | 3.28292553 | 2 | 0.13369621 | 0 | 1 | 1 | 0.59998149 | 1 | 0.69181315 | 0 | 1 | 21 | 0.64957167 | FALSE | 0.97964162 | FALSE | 1 | FALSE | 1 | FALSE | 1 | FALSE | 1 | FALSE | 1 |
| 7860 | 7860 | 7.13043478 | 2.52473585 | 0 | 1 | 2 | 0.44132825 | 1 | 0.81522035 | 2 | 0.63103861 | 1 | 0.53375569 | 40 | 0.40712403 | FALSE | 1 | FALSE | 1 | FALSE | 1 | FALSE | 1 | FALSE | 1 | FALSE | 1 |
| 7861 | 7861 | 4.5625 | 4.26177902 | 2 | 0.34973212 | 0 | 1 | 2 | 0.51909752 | 3 | 0.37966044 | 0 | 1 | 41 | 0.51861324 | FALSE | 1 | FALSE | 1 | FALSE | 1 | FALSE | 1 | FALSE | 1 | FALSE | 1 |
| 7862 | 7862 | 8.10833333 | 2.85714261 | 2 | 0.8184193 | 9 | 0.01538996 | 6 | 0.26185234 | 1 | 0.99662487 | 7 | 0.00341989 | 95 | 0.95241754 | FALSE | 1 | FALSE | 0.7256023 | FALSE | 1 | FALSE | 1 | FALSE | 0.33875501 | FALSE | 1 |
| 7863 | 7863 | 9.72727273 | 2.69986226 | 0 | 1 | 0 | 1 | 0 | 1 | 1 | 0.40390032 | 1 | 0.16638736 | 9 | 0.75985366 | FALSE | 1 | FALSE | 1 | FALSE | 1 | FALSE | 1 | FALSE | 1 | FALSE | 1 |
| 7864 | 7864 | 9.66666667 | 1.49071199 | 0 | 1 | 0 | 1 | 1 | 0.5695037 | 0 | 1 | 0 | 1 | 22 | 0.10533112 | FALSE | 1 | FALSE | 1 | FALSE | 1 | FALSE | 1 | FALSE | 1 | FALSE | 1 |
| 7865 | 7865 | 8 | 2.40831892 | 0 | 1 | 0 | 1 | 1 | 0.51939841 | 1 | 0.60987146 | 1 | 0.28187291 | 17 | 0.61750632 | FALSE | 1 | FALSE | 1 | FALSE | 1 | FALSE | 1 | FALSE | 1 | FALSE | 1 |
| 7866 | 7866 | 2.41185527 | 1.20897893 | 31 | 0.56209659 | 38 | 0.63979753 | 44 | 0.52506802 | 51 | 0.80400712 | 28 | 0.03604013 | 1032 | 0.54758219 | FALSE | 1 | FALSE | 1 | FALSE | 1 | FALSE | 1 | FALSE | 0.87669812 | FALSE | 1 |
| 7867 | 7867 | 10.1666667 | 2.5 | 1 | 0.37380237 | 0 | 1 | 2 | 0.13502329 | 1 | 0.57131013 | 0 | 1 | 14 | 0.86287418 | FALSE | 1 | FALSE | 1 | FALSE | 1 | FALSE | 1 | FALSE | 1 | FALSE | 1 |
| 7868 | 7868 | 10.5714286 | 2.78724012 | 2 | 0.09989125 | 0 | 1 | 3 | 0.03775825 | 1 | 0.62783522 | 0 | 1 | 15 | 0.96513691 | FALSE | 0.86047225 | FALSE | 1 | FALSE | 1 | FALSE | 1 | FALSE | 1 | FALSE | 1 |
| 7869 | 7869 | 9.58333333 | 1.93469779 | 1 | 0.26795838 | 1 | 0.32623933 | 1 | 0.35555293 | 1 | 0.43131129 | 0 | 1 | 8 | 0.97197833 | FALSE | 1 | FALSE | 1 | FALSE | 1 | FALSE | 1 | FALSE | 1 | FALSE | 1 |
| 7870 | 7870 | 8.38709677 | 0.97310343 | 2 | 0.1881398 | 0 | 1 | 0 | 1 | 0 | 1 | 0 | 1 | 29 | 0.11661149 | FALSE | 1 | FALSE | 1 | FALSE | 1 | FALSE | 1 | FALSE | 1 | FALSE | 1 |
| 7871 | 7871 | 7.85714286 | 2.35606036 | 0 | 1 | 0 | 1 | 0 | 1 | 1 | 0.80778317 | 0 | 1 | 34 | 0.01937385 | FALSE | 1 | FALSE | 1 | FALSE | 1 | FALSE | 1 | FALSE | 1 | FALSE | 1 |
| 7872 | 7872 | 9.23529412 | 2.86005783 | 1 | 0.35728305 | 0 | 1 | 0 | 1 | 2 | 0.18212134 | 0 | 1 | 14 | 0.7313145 | FALSE | 1 | FALSE | 1 | FALSE | 1 | FALSE | 1 | FALSE | 1 | FALSE | 1 |
| 7873 | 7873 | 7.8125 | 1.62860178 | 2 | 0.06197875 | 1 | 0.40942991 | 1 | 0.44344817 | 0 | 1 | 2 | 0.02752114 | 10 | 0.99304605 | FALSE | 0.72532843 | FALSE | 1 | FALSE | 1 | FALSE | 1 | FALSE | 0.79106376 | FALSE | 1 |
| 7874 | 7874 | 8.09677419 | 2.40100569 | 2 | 0.1881398 | 4 | 0.01688692 | 1 | 0.67911179 | 0 | 1 | 0 | 1 | 24 | 0.90138666 | FALSE | 1 | FALSE | 0.73194511 | FALSE | 1 | FALSE | 1 | FALSE | 1 | FALSE | 1 |
| 7875 | 7875 | 10.2666667 | 2.14372469 | 0 | 1 | 2 | 0.08295839 | 0 | 1 | 0 | 1 | 0 | 1 | 13 | 0.57712805 | FALSE | 1 | FALSE | 1 | FALSE | 1 | FALSE | 1 | FALSE | 1 | FALSE | 1 |
| 7876 | 7876 | 7.24242424 | 4.24285712 | 1 | 0.57647348 | 2 | 0.28941836 | 1 | 0.70185639 | 1 | 0.78873814 | 1 | 0.42123867 | 27 | 0.75124583 | FALSE | 1 | FALSE | 1 | FALSE | 1 | FALSE | 1 | FALSE | 1 | FALSE | 1 |
| 7877 | 7877 | 6.49358974 | 1.7078131 | 8 | 0.0467356 | 4 | 0.74925905 | 10 | 0.05376792 | 10 | 0.17926577 | 4 | 0.25284882 | 120 | 0.99509164 | FALSE | 0.649301 | FALSE | 1 | FALSE | 1 | FALSE | 1 | FALSE | 1 | FALSE | 1 |
| 7878 | 7878 | 7.2 | 3.21091887 | 2 | 0.27371785 | 2 | 0.37295654 | 2 | 0.42396974 | 1 | 0.84823834 | 1 | 0.48480878 | 32 | 0.83853687 | FALSE | 1 | FALSE | 1 | FALSE | 1 | FALSE | 1 | FALSE | 1 | FALSE | 1 |
| 7879 | 7879 | 10.1 | 2.91376046 | 0 | 1 | 0 | 1 | 1 | 0.30655256 | 0 | 1 | 0 | 1 | 9 | 0.5215547 | FALSE | 1 | FALSE | 1 | FALSE | 1 | FALSE | 1 | FALSE | 1 | FALSE | 1 |
| 7880 | 7880 | 8.71052632 | 2.96388698 | 1 | 0.62830697 | 1 | 0.71433277 | 1 | 0.75193685 | 1 | 0.83318955 | 0 | 1 | 34 | 0.27051027 | FALSE | 1 | FALSE | 1 | FALSE | 1 | FALSE | 1 | FALSE | 1 | FALSE | 1 |
| 7881 | 7881 | 9.92307692 | 2.89459811 | 0 | 1 | 0 | 1 | 1 | 0.37874417 | 0 | 1 | 0 | 1 | 12 | 0.374198 | FALSE | 1 | FALSE | 1 | FALSE | 1 | FALSE | 1 | FALSE | 1 | FALSE | 1 |
| 7882 | 7882 | 3.30668605 | 1.47276103 | 23 | 0.0554228 | 20 | 0.58565984 | 27 | 0.2015166 | 41 | 0.01475423 | 6 | 0.95604525 | 518 | 0.98076509 | FALSE | 0.68849059 | FALSE | 1 | FALSE | 1 | FALSE | 1 | FALSE | 1 | FALSE | 1 |
| 7883 | 7883 | 8.58823529 | 2.15730833 | 1 | 0.58738519 | 0 | 1 | 4 | 0.03235504 | 1 | 0.7984848 | 0 | 1 | 28 | 0.72558119 | FALSE | 1 | FALSE | 1 | FALSE | 1 | FALSE | 1 | FALSE | 1 | FALSE | 1 |
| 7884 | 7884 | 10.0588235 | 2.91992321 | 0 | 1 | 0 | 1 | 0 | 1 | 0 | 1 | 0 | 1 | 17 | 0.05550965 | FALSE | 1 | FALSE | 1 | FALSE | 1 | FALSE | 1 | FALSE | 1 | FALSE | 1 |
| 7885 | 7885 | 8.03703704 | 2.65945527 | 1 | 0.50471032 | 0 | 1 | 1 | 0.62830986 | 0 | 1 | 1 | 0.36060486 | 24 | 0.3725373 | FALSE | 1 | FALSE | 1 | FALSE | 1 | FALSE | 1 | FALSE | 1 | FALSE | 1 |
| 7886 | 7886 | 10.9166667 | 2.84189882 | 0 | 1 | 0 | 1 | 1 | 0.35555293 | 2 | 0.1023796 | 0 | 1 | 9 | 0.89630573 | FALSE | 1 | FALSE | 1 | FALSE | 1 | FALSE | 1 | FALSE | 1 | FALSE | 1 |
| 7887 | 7887 | 9.47058824 | 2.89253902 | 0 | 1 | 0 | 1 | 0 | 1 | 1 | 0.55062772 | 1 | 0.24525764 | 15 | 0.49060435 | FALSE | 1 | FALSE | 1 | FALSE | 1 | FALSE | 1 | FALSE | 1 | FALSE | 1 |
| 7888 | 7888 | 8.84615385 | 1.85414935 | 3 | 0.0279333 | 0 | 1 | 2 | 0.23940226 | 1 | 0.70601446 | 0 | 1 | 20 | 0.90142689 | FALSE | 0.50964275 | FALSE | 1 | FALSE | 1 | FALSE | 1 | FALSE | 1 | FALSE | 1 |
| 7889 | 7889 | 9.1875 | 2.98367957 | 0 | 1 | 2 | 0.09285443 | 0 | 1 | 0 | 1 | 0 | 1 | 14 | 0.53308261 | FALSE | 1 | FALSE | 1 | FALSE | 1 | FALSE | 1 | FALSE | 1 | FALSE | 1 |
| 7890 | 7890 | 9.13793103 | 3.32932776 | 2 | 0.16962005 | 3 | 0.06581417 | 0 | 1 | 1 | 0.74482229 | 0 | 1 | 23 | 0.84462957 | FALSE | 1 | FALSE | 1 | FALSE | 1 | FALSE | 1 | FALSE | 1 | FALSE | 1 |
| 7891 | 7891 | 7.44736842 | 1.11679448 | 0 | 1 | 1 | 0.71433277 | 1 | 0.75193685 | 1 | 0.83318955 | 0 | 1 | 35 | 0.13407055 | FALSE | 1 | FALSE | 1 | FALSE | 1 | FALSE | 1 | FALSE | 1 | FALSE | 1 |
| 7892 | 7892 | 6.96551724 | 4.15628115 | 1 | 0.52988231 | 1 | 0.61539065 | 1 | 0.65463982 | 1 | 0.74482229 | 0 | 1 | 25 | 0.51779518 | FALSE | 1 | FALSE | 1 | FALSE | 1 | FALSE | 1 | FALSE | 1 | FALSE | 1 |
| 7893 | 7893 | 10.625 | 2.34187425 | 0 | 1 | 1 | 0.38964394 | 1 | 0.42266264 | 0 | 1 | 0 | 1 | 13 | 0.57712805 | FALSE | 1 | FALSE | 1 | FALSE | 1 | FALSE | 1 | FALSE | 1 | FALSE | 1 |
| 7894 | 7894 | 9.84615385 | 3.7997197 | 0 | 1 | 0 | 1 | 1 | 0.37874417 | 0 | 1 | 0 | 1 | 12 | 0.374198 | FALSE | 1 | FALSE | 1 | FALSE | 1 | FALSE | 1 | FALSE | 1 | FALSE | 1 |

| | | | | | | | | | | | | | | | | | | | | | | | | | | | |
|---|---|---|---|---|---|---|---|---|---|---|---|---|---|---|---|---|---|---|---|---|---|---|---|---|---|---|---|
| 7895 | 7895 | 10.0454546 | 3.05223127 | 4 | 0.00177457 | 2 | 0.14655134 | 0 | 1 | 2 | 0.25059627 | 0 | 1 | 13 | 0.9973503 | FALSE | 0.10930302 | FALSE | 1 | FALSE | 1 | FALSE | 1 | FALSE | 1 | FALSE | 1 |
| 7896 | 7896 | 9.75675676 | 1.36372712 | 3 | 0.0681763 | 3 | 0.11640364 | 2 | 0.38585089 | 0 | 1 | 0 | 1 | 29 | 0.88820978 | FALSE | 0.74276693 | FALSE | 1 | FALSE | 1 | FALSE | 1 | FALSE | 1 | FALSE | 1 |
| 7897 | 7897 | 7.75 | 2.63391344 | 0 | 1 | 4 | 0.03617171 | 1 | 0.76089851 | 1 | 0.84089119 | 1 | 0.4761699 | 32 | 0.74324845 | FALSE | 1 | FALSE | 0.94259695 | FALSE | 1 | FALSE | 1 | FALSE | 1 | FALSE | 1 |
| 7898 | 7898 | 4.23631124 | 1.3452401 | 20 | 0.93878999 | 27 | 0.91268852 | 38 | 0.47165075 | 35 | 0.98523918 | 20 | 0.24661855 | 896 | 0.02229441 | FALSE | 1 | FALSE | 1 | FALSE | 1 | FALSE | 1 | FALSE | 1 | FALSE | 1 |
| 7899 | 7899 | 8.44117647 | 4.00874908 | 2 | 0.19750261 | 2 | 0.27730767 | 3 | 0.10599423 | 1 | 0.77852168 | 0 | 1 | 24 | 0.94876798 | FALSE | 1 | FALSE | 1 | FALSE | 1 | FALSE | 1 | FALSE | 1 | FALSE | 1 |
| 7900 | 7900 | 9.09090909 | 4.11106147 | 0 | 1 | 0 | 1 | 1 | 0.55340851 | 1 | 0.64497439 | 0 | 1 | 20 | 0.30915202 | FALSE | 1 | FALSE | 1 | FALSE | 1 | FALSE | 1 | FALSE | 1 | FALSE | 1 |
| 7901 | 7901 | 9.08333333 | 1.97730175 | 2 | 0.03643311 | 0 | 1 | 0 | 1 | 0 | 1 | 0 | 1 | 10 | 0.71436964 | FALSE | 0.58392 | FALSE | 1 | FALSE | 1 | FALSE | 1 | FALSE | 1 | FALSE | 1 |
| 7902 | 7902 | 4.84166667 | 1.94934106 | 1 | 0.95577586 | 4 | 0.54082363 | 7 | 0.13559588 | 2 | 0.97580519 | 3 | 0.30980997 | 102 | 0.40098785 | FALSE | 1 | FALSE | 1 | FALSE | 1 | FALSE | 1 | FALSE | 1 | FALSE | 1 |
| 7903 | 7903 | 8.74074074 | 2.7967255 | 2 | 0.15144274 | 0 | 1 | 0 | 1 | 1 | 0.71956339 | 0 | 1 | 24 | 0.3725373 | FALSE | 1 | FALSE | 1 | FALSE | 1 | FALSE | 1 | FALSE | 1 | FALSE | 1 |
| 7904 | 7904 | 6.75 | 2.90473751 | 1 | 0.46441868 | 3 | 0.04096158 | 2 | 0.21260257 | 0 | 1 | 0 | 1 | 18 | 0.93166998 | FALSE | 1 | FALSE | 0.9698186 | FALSE | 1 | FALSE | 1 | FALSE | 1 | FALSE | 1 |
| 7905 | 7905 | 6.48484849 | 3.44762731 | 2 | 0.20692137 | 1 | 0.6629941 | 2 | 0.33343871 | 2 | 0.45159487 | 1 | 0.42123867 | 25 | 0.93892791 | FALSE | 1 | FALSE | 1 | FALSE | 1 | FALSE | 1 | FALSE | 1 | FALSE | 1 |
| 7906 | 7906 | 9.475 | 1.35991728 | 1 | 0.62830697 | 0 | 1 | 3 | 0.15483587 | 1 | 0.83318955 | 0 | 1 | 33 | 0.44261631 | FALSE | 1 | FALSE | 1 | FALSE | 1 | FALSE | 1 | FALSE | 1 | FALSE | 1 |
| 7907 | 7907 | 8.7962963 | 3.11138667 | 5 | 0.01026504 | 2 | 0.50543179 | 2 | 0.5626042 | 0 | 1 | 2 | 0.2095319 | 41 | 0.89856682 | FALSE | 0.30467829 | FALSE | 1 | FALSE | 1 | FALSE | 1 | FALSE | 1 | FALSE | 1 |
| 7908 | 7908 | 8.54347826 | 2.83385221 | 0 | 1 | 1 | 0.78076835 | 5 | 0.02370648 | 2 | 0.63103861 | 0 | 1 | 38 | 0.71504609 | FALSE | 1 | FALSE | 1 | FALSE | 1 | FALSE | 1 | FALSE | 1 | FALSE | 1 |
| 7909 | 7909 | 9.84210526 | 3.23245317 | 1 | 0.38989951 | 0 | 1 | 1 | 0.50143826 | 1 | 0.59104357 | 0 | 1 | 16 | 0.65565778 | FALSE | 1 | FALSE | 1 | FALSE | 1 | FALSE | 1 | FALSE | 1 | FALSE | 1 |
| 7910 | 7910 | 4.83606557 | 2.69899998 | 1 | 0.79096998 | 0 | 1 | 1 | 0.8897349 | 4 | 0.29632241 | 1 | 0.63076845 | 53 | 0.25931015 | FALSE | 1 | FALSE | 1 | FALSE | 1 | FALSE | 1 | FALSE | 1 | FALSE | 1 |
| 7911 | 7911 | 11.3 | 1.9 | 1 | 0.40558521 | 1 | 0.48239024 | 0 | 1 | 1 | 0.60987146 | 0 | 1 | 17 | 0.61750632 | FALSE | 1 | FALSE | 1 | FALSE | 1 | FALSE | 1 | FALSE | 1 | FALSE | 1 |
| 7912 | 7912 | 9.16666667 | 2.70287912 | 0 | 1 | 0 | 1 | 0 | 1 | 0 | 1 | 0 | 1 | 12 | 0.13002667 | FALSE | 1 | FALSE | 1 | FALSE | 1 | FALSE | 1 | FALSE | 1 | FALSE | 1 |
| 7913 | 7913 | 7.33333333 | 4.12118029 | 2 | 0.09989125 | 1 | 0.49918218 | 2 | 0.17308361 | 0 | 1 | 0 | 1 | 16 | 0.90353674 | FALSE | 0.86047225 | FALSE | 1 | FALSE | 1 | FALSE | 1 | FALSE | 1 | FALSE | 1 |
| 7914 | 7914 | 10.1612903 | 1.34636366 | 0 | 1 | 3 | 0.07729739 | 1 | 0.67911179 | 1 | 0.76781284 | 2 | 0.09110253 | 24 | 0.90138666 | FALSE | 1 | FALSE | 1 | FALSE | 1 | FALSE | 1 | FALSE | 1 | FALSE | 1 |
| 7915 | 7915 | 9.05555556 | 3.78308451 | 1 | 0.75539413 | 1 | 0.83180787 | 3 | 0.30658915 | 1 | 0.92176876 | 0 | 1 | 48 | 0.23965388 | FALSE | 1 | FALSE | 1 | FALSE | 1 | FALSE | 1 | FALSE | 1 | FALSE | 1 |
| 7916 | 7916 | 8.22222222 | 3.48098893 | 1 | 0.59801736 | 1 | 0.68454804 | 2 | 0.35984046 | 1 | 0.80778317 | 0 | 1 | 30 | 0.52765882 | FALSE | 1 | FALSE | 1 | FALSE | 1 | FALSE | 1 | FALSE | 1 | FALSE | 1 |
| 7917 | 7917 | 5.99193548 | 2.36061763 | 3 | 0.60453582 | 4 | 0.55411862 | 6 | 0.26798911 | 7 | 0.32018532 | 3 | 0.31883571 | 98 | 0.87586262 | FALSE | 1 | FALSE | 1 | FALSE | 1 | FALSE | 1 | FALSE | 1 | FALSE | 1 |
| 7918 | 7918 | 11.1578947 | 1.88985726 | 1 | 0.77387657 | 2 | 0.55509723 | 3 | 0.3364112 | 9 | 0.00103406 | 0 | 1 | 42 | 0.98838397 | FALSE | 1 | FALSE | 1 | FALSE | 1 | FALSE | 1 | FALSE | 1 | FALSE | 1 |
| 7919 | 7919 | 10.25 | 2.54425122 | 0 | 1 | 3 | 0.06039219 | 0 | 1 | 0 | 1 | 0 | 1 | 25 | 0.34293 | FALSE | 1 | FALSE | 1 | FALSE | 1 | FALSE | 1 | FALSE | 1 | FALSE | 1 |
| 7920 | 7920 | 9.07142857 | 2.91460082 | 0 | 1 | 0 | 1 | 2 | 0.08806398 | 0 | 1 | 1 | 0.20679328 | 11 | 0.83676449 | FALSE | 1 | FALSE | 1 | FALSE | 1 | FALSE | 1 | FALSE | 1 | FALSE | 1 |
| 7921 | 7921 | 8.88235294 | 1.93671502 | 2 | 0.06909038 | 0 | 1 | 1 | 0.46348836 | 1 | 0.55062772 | 0 | 1 | 13 | 0.88719222 | FALSE | 0.74276693 | FALSE | 1 | FALSE | 1 | FALSE | 1 | FALSE | 1 | FALSE | 1 |
| 7922 | 7922 | 6.36871508 | 1.00185528 | 4 | 0.67938302 | 7 | 0.3588387 | 11 | 0.0569906 | 6 | 0.83783813 | 1 | 0.95018775 | 150 | 0.63354399 | FALSE | 1 | FALSE | 1 | FALSE | 1 | FALSE | 1 | FALSE | 1 | FALSE | 1 |
| 7923 | 7923 | 8.43396226 | 2.22769391 | 1 | 0.74890551 | 0 | 1 | 1 | 0.85724013 | 0 | 1 | 1 | 0.58506217 | 50 | 0.02504004 | FALSE | 1 | FALSE | 1 | FALSE | 1 | FALSE | 1 | FALSE | 1 | FALSE | 1 |
| 7924 | 7924 | 9.68421053 | 2.59660977 | 0 | 1 | 4 | 0.00281443 | 1 | 0.50143826 | 0 | 1 | 0 | 1 | 14 | 0.93677253 | FALSE | 1 | FALSE | 0.46409366 | FALSE | 1 | FALSE | 1 | FALSE | 1 | FALSE | 1 |
| 7925 | 7925 | 11.1666667 | 1.67497927 | 4 | 0.00017582 | 1 | 0.32623933 | 1 | 0.35555293 | 0 | 1 | 0 | 1 | 6 | 0.99914544 | TRUE | 0.02491133 | FALSE | 1 | FALSE | 1 | FALSE | 1 | FALSE | 1 | FALSE | 1 |
| 7926 | 7926 | 9.36 | 1.95714077 | 0 | 1 | 1 | 0.56109832 | 0 | 1 | 1 | 0.69181315 | 1 | 0.33902672 | 22 | 0.43642846 | FALSE | 1 | FALSE | 1 | FALSE | 1 | FALSE | 1 | FALSE | 1 | FALSE | 1 |
| 7927 | 7927 | 8.66666667 | 1.97202659 | 0 | 1 | 3 | 0.00591424 | 0 | 1 | 0 | 1 | 0 | 1 | 9 | 0.89630573 | FALSE | 1 | FALSE | 0.60103504 | FALSE | 1 | FALSE | 1 | FALSE | 1 | FALSE | 1 |
| 7928 | 7928 | 9.90476191 | 1.60073679 | 1 | 0.42086991 | 1 | 0.49918218 | 1 | 0.53671415 | 1 | 0.62783522 | 0 | 1 | 17 | 0.77849076 | FALSE | 1 | FALSE | 1 | FALSE | 1 | FALSE | 1 | FALSE | 1 | FALSE | 1 |
| 7929 | 7929 | 9.23529412 | 3.54571398 | 0 | 1 | 1 | 0.81422651 | 3 | 0.27688673 | 1 | 0.90982214 | 0 | 1 | 46 | 0.17033916 | FALSE | 1 | FALSE | 1 | FALSE | 1 | FALSE | 1 | FALSE | 1 | FALSE | 1 |
| 7930 | 7930 | 8.21212121 | 2.88945393 | 0 | 1 | 1 | 0.6629941 | 0 | 1 | 1 | 0.78873814 | 1 | 0.42123867 | 30 | 0.21942357 | FALSE | 1 | FALSE | 1 | FALSE | 1 | FALSE | 1 | FALSE | 1 | FALSE | 1 |
| 7931 | 7931 | 10.6666667 | 2 | 0 | 1 | 0 | 1 | 0 | 1 | 0 | 1 | 1 | 0.1383237 | 8 | 0.5780332 | FALSE | 1 | FALSE | 1 | FALSE | 1 | FALSE | 1 | FALSE | 1 | FALSE | 1 |
| 7932 | 7932 | 11.1428571 | 2.03038149 | 1 | 0.42086991 | 0 | 1 | 1 | 0.53671415 | 1 | 0.62783522 | 0 | 1 | 18 | 0.57962763 | FALSE | 1 | FALSE | 1 | FALSE | 1 | FALSE | 1 | FALSE | 1 | FALSE | 1 |
| 7933 | 7933 | 9.33333333 | 1.97202659 | 2 | 0.07645454 | 1 | 0.44710673 | 0 | 1 | 0 | 1 | 1 | 0.25766374 | 14 | 0.86287418 | FALSE | 0.77424195 | FALSE | 1 | FALSE | 1 | FALSE | 1 | FALSE | 1 | FALSE | 1 |
| 7934 | 7934 | 8.17241379 | 0.91232804 | 1 | 0.52988231 | 0 | 1 | 2 | 0.27983822 | 3 | 0.14602298 | 0 | 1 | 23 | 0.84462957 | FALSE | 1 | FALSE | 1 | FALSE | 1 | FALSE | 1 | FALSE | 1 | FALSE | 1 |
| 7935 | 7935 | 4.24416517 | 1.21391804 | 27 | 0.00087099 | 29 | 0.00673968 | 21 | 0.44150375 | 30 | 0.20063538 | 7 | 0.81897388 | 443 | 0.99941693 | FALSE | 0.07029574 | FALSE | 0.60103504 | FALSE | 1 | FALSE | 1 | FALSE | 1 | FALSE | 1 |
| 7936 | 7936 | 8.25 | 2.01112975 | 2 | 0.16048326 | 1 | 0.60247911 | 1 | 0.64171563 | 0 | 1 | 0 | 1 | 24 | 0.55018107 | FALSE | 1 | FALSE | 1 | FALSE | 1 | FALSE | 1 | FALSE | 1 | FALSE | 1 |
| 7937 | 7937 | 8.92307692 | 2.18250169 | 0 | 1 | 1 | 0.57534752 | 0 | 1 | 2 | 0.33678042 | 0 | 1 | 23 | 0.40372818 | FALSE | 1 | FALSE | 1 | FALSE | 1 | FALSE | 1 | FALSE | 1 | FALSE | 1 |
| 7938 | 7938 | 6.44 | 2.69191382 | 3 | 0.13009168 | 0 | 1 | 1 | 0.83455705 | 4 | 0.1862655 | 0 | 1 | 41 | 0.64556589 | FALSE | 0.97964162 | FALSE | 1 | FALSE | 1 | FALSE | 1 | FALSE | 1 | FALSE | 1 |
| 7939 | 7939 | 9.28571429 | 1.38505139 | 1 | 0.3050805 | 0 | 1 | 0 | 1 | 1 | 0.48242099 | 1 | 0.20679328 | 11 | 0.83676449 | FALSE | 1 | FALSE | 1 | FALSE | 1 | FALSE | 1 | FALSE | 1 | FALSE | 1 |
| 7940 | 7940 | 7.84615385 | 3.5484808 | 0 | 1 | 1 | 0.3480709 | 2 | 0.07725803 | 2 | 0.1174583 | 0 | 1 | 8 | 0.99085298 | FALSE | 1 | FALSE | 1 | FALSE | 1 | FALSE | 1 | FALSE | 1 | FALSE | 1 |
| 7941 | 7941 | 10.375 | 1.26861145 | 0 | 1 | 0 | 1 | 0 | 1 | 2 | 0.16544789 | 0 | 1 | 14 | 0.53308261 | FALSE | 1 | FALSE | 1 | FALSE | 1 | FALSE | 1 | FALSE | 1 | FALSE | 1 |
| 7942 | 7942 | 6.69230769 | 3.371478 | 0 | 1 | 1 | 0.57534752 | 0 | 1 | 1 | 0.70601446 | 1 | 0.3499045 | 23 | 0.40372818 | FALSE | 1 | FALSE | 1 | FALSE | 1 | FALSE | 1 | FALSE | 1 | FALSE | 1 |
| 7943 | 7943 | 8.94444444 | 2.46017665 | 2 | 0.07645454 | 0 | 1 | 1 | 0.48280983 | 0 | 1 | 1 | 0.25766374 | 14 | 0.86287418 | FALSE | 0.77424195 | FALSE | 1 | FALSE | 1 | FALSE | 1 | FALSE | 1 | FALSE | 1 |
| 7944 | 7944 | 10.5483871 | 1.73834756 | 5 | 0.00103368 | 1 | 0.63997468 | 1 | 0.67911179 | 1 | 0.76781284 | 2 | 0.09110253 | 21 | 0.99467947 | FALSE | 0.07493697 | FALSE | 1 | FALSE | 1 | FALSE | 1 | FALSE | 1 | FALSE | 1 |
| 7945 | 7945 | 9.08695652 | 4.78142278 | 0 | 1 | 1 | 0.53115654 | 1 | 0.5695037 | 0 | 1 | 1 | 0.31672722 | 20 | 0.50589402 | FALSE | 1 | FALSE | 1 | FALSE | 1 | FALSE | 1 | FALSE | 1 | FALSE | 1 |
| 7946 | 7946 | 8.76470588 | 2.47408716 | 0 | 1 | 1 | 0.6629941 | 1 | 0.70185639 | 0 | 1 | 0 | 1 | 31 | 0.09218876 | FALSE | 1 | FALSE | 1 | FALSE | 1 | FALSE | 1 | FALSE | 1 | FALSE | 1 |
| 7947 | 7947 | 10.4545455 | 3.08555686 | 0 | 1 | 1 | 0.30368014 | 0 | 1 | 0 | 1 | 1 | 0.16638736 | 9 | 0.75985366 | FALSE | 1 | FALSE | 1 | FALSE | 1 | FALSE | 1 | FALSE | 1 | FALSE | 1 |
| 7948 | 7948 | 7.88095238 | 2.96569578 | 0 | 1 | 4 | 0.28813976 | 6 | 0.08069842 | 5 | 0.34217166 | 3 | 0.15881632 | 66 | 0.94315756 | FALSE | 1 | FALSE | 1 | FALSE | 1 | FALSE | 1 | FALSE | 1 | FALSE | 1 |
| 7949 | 7949 | 9.8 | 3.7094474 | 1 | 0.32293355 | 0 | 1 | 2 | 0.09928574 | 1 | 0.50623194 | 0 | 1 | 11 | 0.92879412 | FALSE | 1 | FALSE | 1 | FALSE | 1 | FALSE | 1 | FALSE | 1 | FALSE | 1 |
| 7950 | 7950 | 10.24 | 3.2159602 | 3 | 0.02517314 | 2 | 0.19297191 | 2 | 0.22597287 | 0 | 1 | 0 | 1 | 18 | 0.96851178 | FALSE | 0.4974559 | FALSE | 1 | FALSE | 1 | FALSE | 1 | FALSE | 1 | FALSE | 1 |
| 7951 | 7951 | 9.25 | 2.88314065 | 0 | 1 | 1 | 0.40942991 | 0 | 1 | 0 | 1 | 0 | 1 | 15 | 0.26127859 | FALSE | 1 | FALSE | 1 | FALSE | 1 | FALSE | 1 | FALSE | 1 | FALSE | 1 |
| 7952 | 7952 | 5.4084507 | 2.90738479 | 4 | 0.49097011 | 3 | 0.84058578 | 7 | 0.24324929 | 6 | 0.6343804 | 1 | 0.90518766 | 120 | 0.46116711 | FALSE | 1 | FALSE | 1 | FALSE | 1 | FALSE | 1 | FALSE | 1 | FALSE | 1 |
| 7953 | 7953 | 8.0952381 | 1.87476378 | 0 | 1 | 0 | 1 | 1 | 0.53671415 | 2 | 0.25059627 | 1 | 0.29368245 | 17 | 0.77849076 | FALSE | 1 | FALSE | 1 | FALSE | 1 | FALSE | 1 | FALSE | 1 | FALSE | 1 |
| 7954 | 7954 | 4.99242424 | 1.4380975 | 13 | 0.01773875 | 12 | 0.14682586 | 9 | 0.61314967 | 4 | 0.99847818 | 2 | 0.93498961 | 224 | 0.45632526 | FALSE | 0.41596787 | FALSE | 1 | FALSE | 1 | FALSE | 1 | FALSE | 1 | FALSE | 1 |
| 7955 | 7955 | 9.41935484 | 2.63688714 | 0 | 1 | 1 | 0.6278847 | 0 | 1 | 0 | 1 | 0 | 1 | 29 | 0.03983039 | FALSE | 1 | FALSE | 1 | FALSE | 1 | FALSE | 1 | FALSE | 1 | FALSE | 1 |
| 7956 | 7956 | 9.66666667 | 1.98885785 | 0 | 1 | 2 | 0.08295839 | 0 | 1 | 1 | 0.50623194 | 0 | 1 | 12 | 0.80329917 | FALSE | 1 | FALSE | 1 | FALSE | 1 | FALSE | 1 | FALSE | 1 | FALSE | 1 |
| 7957 | 7957 | 10.1333333 | 1.78387842 | 0 | 1 | 0 | 1 | 1 | 0.42266264 | 0 | 1 | 0 | 1 | 14 | 0.29525409 | FALSE | 1 | FALSE | 1 | FALSE | 1 | FALSE | 1 | FALSE | 1 | FALSE | 1 |
| 7958 | 7958 | 6.41818182 | 1.28898705 | 2 | 0.41401181 | 3 | 0.26245929 | 2 | 0.59334725 | 1 | 0.92538938 | 1 | 0.59866474 | 46 | 0.64645216 | FALSE | 1 | FALSE | 1 | FALSE | 1 | FALSE | 1 | FALSE | 1 | FALSE | 1 |
| 7959 | 7959 | 8.3 | 1.62788206 | 1 | 0.72839502 | 2 | 0.48458833 | 1 | 0.8405438 | 5 | 0.07777122 | 1 | 0.56379725 | 40 | 0.85325204 | FALSE | 1 | FALSE | 1 | FALSE | 1 | FALSE | 1 | FALSE | 1 | FALSE | 1 |
| 7960 | 7960 | 7.75 | 1.85404962 | 0 | 1 | 0 | 1 | 0 | 1 | 0 | 1 | 0 | 1 | 8 | 0.25677166 | FALSE | 1 | FALSE | 1 | FALSE | 1 | FALSE | 1 | FALSE | 1 | FALSE | 1 |
| 7961 | 7961 | 7.71428571 | 1.90595201 | 0 | 1 | 0 | 1 | 0 | 1 | 0 | 1 | 0 | 1 | 21 | 0.02808151 | FALSE | 1 | FALSE | 1 | FALSE | 1 | FALSE | 1 | FALSE | 1 | FALSE | 1 |
| 7962 | 7962 | 10.037037 | 2.71458996 | 0 | 1 | 0 | 1 | 0 | 1 | 2 | 0.35374209 | 0 | 1 | 25 | 0.1832206 | FALSE | 1 | FALSE | 1 | FALSE | 1 | FALSE | 1 | FALSE | 1 | FALSE | 1 |
| 7963 | 7963 | 9.93333333 | 1.56914697 | 0 | 1 | 0 | 1 | 0 | 1 | 0 | 1 | 0 | 1 | 15 | 0.07803154 | FALSE | 1 | FALSE | 1 | FALSE | 1 | FALSE | 1 | FALSE | 1 | FALSE | 1 |
| 7964 | 7964 | 8.66666667 | 3.84625153 | 0 | 1 | 2 | 0.39616059 | 3 | 0.19060367 | 1 | 0.86193352 | 1 | 0.50166523 | 35 | 0.67024784 | FALSE | 1 | FALSE | 1 | FALSE | 1 | FALSE | 1 | FALSE | 1 | FALSE | 1 |
| 7965 | 7965 | 9.25 | 3.76662979 | 0 | 1 | 0 | 1 | 2 | 0.11087977 | 1 | 0.52895088 | 2 | 0.02752114 | 11 | 0.9718531 | FALSE | 1 | FALSE | 1 | FALSE | 1 | FALSE | 1 | FALSE | 0.79106376 | FALSE | 1 |
| 7966 | 7966 | 9.25490196 | 2.45622967 | 1 | 0.7354105 | 1 | 0.81422651 | 2 | 0.55199489 | 4 | 0.20527806 | 0 | 1 | 43 | 0.59806866 | FALSE | 1 | FALSE | 1 | FALSE | 1 | FALSE | 1 | FALSE | 1 | FALSE | 1 |
| 7967 | 7967 | 9.78571429 | 2.07634884 | 0 | 1 | 0 | 1 | 0 | 1 | 1 | 0.48242099 | 0 | 1 | 13 | 0.33283729 | FALSE | 1 | FALSE | 1 | FALSE | 1 | FALSE | 1 | FALSE | 1 | FALSE | 1 |
| 7968 | 7968 | 7.61111111 | 2.42988086 | 1 | 0.37380237 | 0 | 1 | 3 | 0.02505979 | 0 | 1 | 0 | 1 | 14 | 0.86287418 | FALSE | 1 | FALSE | 1 | FALSE | 1 | FALSE | 1 | FALSE | 1 | FALSE | 1 |
| 7969 | 7969 | 11.2352941 | 2.90090071 | 0 | 1 | 0 | 1 | 1 | 0.46348836 | 1 | 0.55062772 | 1 | 0.24525764 | 14 | 0.7313145 | FALSE | 1 | FALSE | 1 | FALSE | 1 | FALSE | 1 | FALSE | 1 | FALSE | 1 |
| 7970 | 7970 | 9.96 | 3.02628485 | 1 | 0.47819878 | 0 | 1 | 1 | 0.59998149 | 1 | 0.69181315 | 0 | 1 | 22 | 0.43642846 | FALSE | 1 | FALSE | 1 | FALSE | 1 | FALSE | 1 | FALSE | 1 | FALSE | 1 |
| 7971 | 7971 | 7.28947368 | 3.1531763 | 2 | 0.25455533 | 2 | 0.3494045 | 1 | 0.75193685 | 3 | 0.25266879 | 0 | 1 | 30 | 0.87271204 | FALSE | 1 | FALSE | 1 | FALSE | 1 | FALSE | 1 | FALSE | 1 | FALSE | 1 |
| 7972 | 7972 | 11.1764706 | 2.91517922 | 3 | 0.00864151 | 2 | 0.10307112 | 0 | 1 | 1 | 0.55062772 | 1 | 0.24525764 | 10 | 0.99775952 | FALSE | 0.27019545 | FALSE | 1 | FALSE | 1 | FALSE | 1 | FALSE | 1 | FALSE | 1 |
| 7973 | 7973 | 7.45614035 | 3.00333251 | 9 | 1.21E-05 | 3 | 0.28014665 | 1 | 0.87682436 | 1 | 0.93213716 | 1 | 0.61182532 | 42 | 0.98838397 | TRUE | 0.00418892 | FALSE | 1 | FALSE | 1 | FALSE | 1 | FALSE | 1 | FALSE | 1 |
| 7974 | 7974 | 9.66666667 | 1.69967317 | 1 | 0.14434519 | 0 | 1 | 0 | 1 | 0 | 1 | 1 | 0.09446378 | 4 | 0.94743772 | FALSE | 1 | FALSE | 1 | FALSE | 1 | FALSE | 1 | FALSE | 1 | FALSE | 1 |
| 7975 | 7975 | 9.16666667 | 2.99536679 | 2 | 0.03643311 | 2 | 0.05552913 | 0 | 1 | 1 | 0.43131129 | 1 | 0.18007762 | 6 | 0.99914544 | FALSE | 0.58392 | FALSE | 1 | FALSE | 1 | FALSE | 1 | FALSE | 1 | FALSE | 1 |
| 7976 | 7976 | 9.95 | 2.10890967 | 0 | 1 | 0 | 1 | 0 | 1 | 1 | 0.60987146 | 0 | 1 | 19 | 0.15700255 | FALSE | 1 | FALSE | 1 | FALSE | 1 | FALSE | 1 | FALSE | 1 | FALSE | 1 |
| 7977 | 7977 | 10.9130435 | 2.46564294 | 0 | 1 | 3 | 0.03669128 | 0 | 1 | 1 | 0.6613267 | 0 | 1 | 19 | 0.71542496 | FALSE | 1 | FALSE | 0.94259695 | FALSE | 1 | FALSE | 1 | FALSE | 1 | FALSE | 1 |
| 7978 | 7978 | 9.21052632 | 3.47288667 | 0 | 1 | 1 | 0.44710673 | 1 | 0.48280983 | 0 | 1 | 0 | 1 | 16 | 0.44998001 | FALSE | 1 | FALSE | 1 | FALSE | 1 | FALSE | 1 | FALSE | 1 | FALSE | 1 |
| 7979 | 7979 | 10.4444444 | 3.13088951 | 2 | 0.02089722 | 0 | 1 | 0 | 1 | 0 | 1 | 1 | 0.1383237 | 6 | 0.96128317 | FALSE | 0.45408499 | FALSE | 1 | FALSE | 1 | FALSE | 1 | FALSE | 1 | FALSE | 1 |
| 7980 | 7980 | 10.1481482 | 1.580054 | 2 | 0.15144274 | 0 | 1 | 1 | 0.62830986 | 0 | 1 | 0 | 1 | 24 | 0.3725373 | FALSE | 1 | FALSE | 1 | FALSE | 1 | FALSE | 1 | FALSE | 1 | FALSE | 1 |
| 7981 | 7981 | 8.79032258 | 1.84160779 | 2 | 0.46653175 | 3 | 0.31579946 | 1 | 0.89373158 | 1 | 0.94386203 | 1 | 0.63687695 | 53 | 0.37131894 | FALSE | 1 | FALSE | 1 | FALSE | 1 | FALSE | 1 | FALSE | 1 | FALSE | 1 |
| 7982 | 7982 | 2.89655172 | 2.83566969 | 1 | 0.95212532 | 5 | 0.32153158 | 2 | 0.92532622 | 5 | 0.62158717 | 0 | 1 | 103 | 0.11371378 | FALSE | 1 | FALSE | 1 | FALSE | 1 | FALSE | 1 | FALSE | 1 | FALSE | 1 |
| 7983 | 7983 | 8.08695652 | 3.06328956 | 0 | 1 | 2 | 0.16948764 | 0 | 1 | 1 | 0.6613267 | 2 | 0.05389003 | 18 | 0.86273155 | FALSE | 1 | FALSE | 1 | FALSE | 1 | FALSE | 1 | FALSE | 1 | FALSE | 1 |
| 7984 | 7984 | 3.90486726 | 1.0293366 | 13 | 0.36849445 | 10 | 0.92499726 | 7 | 0.99751063 | 19 | 0.6844105 | 9 | 0.31752142 | 392 | 0.0539398 | FALSE | 1 | FALSE | 1 | FALSE | 1 | FALSE | 1 | FALSE | 1 | FALSE | 1 |
| 7985 | 7985 | 8 | 1.06066017 | 3 | 0.00725145 | 0 | 1 | 0 | 1 | 1 | 0.52895088 | 1 | 0.23264612 | 11 | 0.9718531 | FALSE | 0.25171133 | FALSE | 1 | FALSE | 1 | FALSE | 1 | FALSE | 1 | FALSE | 1 |
| 7986 | 7986 | 8.02702703 | 3.43657975 | 2 | 0.24498301 | 2 | 0.3375163 | 2 | 0.38585089 | 2 | 0.51200191 | 0 | 1 | 29 | 0.88820978 | FALSE | 1 | FALSE | 1 | FALSE | 1 | FALSE | 1 | FALSE | 1 | FALSE | 1 |
| 7987 | 7987 | 8.11764706 | 3.04635468 | 0 | 1 | 1 | 0.42857733 | 2 | 0.12280513 | 1 | 0.55062772 | 1 | 0.24525764 | 12 | 0.962093 | FALSE | 1 | FALSE | 1 | FALSE | 1 | FALSE | 1 | FALSE | 1 | FALSE | 1 |
| 7988 | 7988 | 10.2173913 | 2.53594201 | 2 | 0.11647716 | 0 | 1 | 1 | 0.5695037 | 0 | 1 | 0 | 1 | 20 | 0.50589402 | FALSE | 0.93181727 | FALSE | 1 | FALSE | 1 | FALSE | 1 | FALSE | 1 | FALSE | 1 |

| | | | | | | | | | | | | | | | | | | | | | | | | | | | |
|---|---|---|---|---|---|---|---|---|---|---|---|---|---|---|---|---|---|---|---|---|---|---|---|---|---|---|---|
| 7989 | 7989 | 10.3157895 | 2.42904858 | 1 | 0.38989951 | 0 | 1 | 0 | 1 | 0 | 1 | 1 | 0.26986775 | 17 | 0.41141543 | FALSE | 1 | FALSE | 1 | FALSE | 1 | FALSE | 1 | FALSE | 1 | FALSE | 1 |
| 7990 | 7990 | 8.875 | 2.53824907 | 0 | 1 | 2 | 0.18117183 | 1 | 0.58502114 | 0 | 1 | 0 | 1 | 21 | 0.47053137 | FALSE | 1 | FALSE | 1 | FALSE | 1 | FALSE | 1 | FALSE | 1 | FALSE | 1 |
| 7991 | 7991 | 8.62068966 | 3.49791853 | 0 | 1 | 1 | 0.61539065 | 2 | 0.27983822 | 3 | 0.14602298 | 1 | 0.38148475 | 22 | 0.92902102 | FALSE | 1 | FALSE | 1 | FALSE | 1 | FALSE | 1 | FALSE | 1 | FALSE | 1 |
| 7992 | 7992 | 8.23529412 | 2.18361308 | 1 | 0.35728305 | 1 | 0.42857733 | 0 | 1 | 0 | 1 | 0 | 1 | 15 | 0.49060435 | FALSE | 1 | FALSE | 1 | FALSE | 1 | FALSE | 1 | FALSE | 1 | FALSE | 1 |
| 7993 | 7993 | 6.76363636 | 1.57207028 | 2 | 0.77443073 | 5 | 0.27707647 | 4 | 0.55465521 | 7 | 0.23363892 | 1 | 0.83744681 | 90 | 0.7500236 | FALSE | 1 | FALSE | 1 | FALSE | 1 | FALSE | 1 | FALSE | 1 | FALSE | 1 |
| 7994 | 7994 | 9.5 | 1.80277564 | 1 | 0.34033054 | 0 | 1 | 1 | 0.44344817 | 0 | 1 | 0 | 1 | 14 | 0.53308261 | FALSE | 1 | FALSE | 1 | FALSE | 1 | FALSE | 1 | FALSE | 1 | FALSE | 1 |
| 7995 | 7995 | 9.45 | 2.8718461 | 0 | 1 | 1 | 0.48239024 | 1 | 0.51939841 | 0 | 1 | 0 | 1 | 18 | 0.37504645 | FALSE | 1 | FALSE | 1 | FALSE | 1 | FALSE | 1 | FALSE | 1 | FALSE | 1 |
| 7996 | 7996 | 10.4285714 | 2.63802647 | 0 | 1 | 0 | 1 | 1 | 0.40110418 | 0 | 1 | 0 | 1 | 13 | 0.33283729 | FALSE | 1 | FALSE | 1 | FALSE | 1 | FALSE | 1 | FALSE | 1 | FALSE | 1 |
| 7997 | 7997 | 9.29411765 | 2.16291486 | 0 | 1 | 2 | 0.10307112 | 1 | 0.46348836 | 1 | 0.55062772 | 0 | 1 | 13 | 0.88719222 | FALSE | 1 | FALSE | 1 | FALSE | 1 | FALSE | 1 | FALSE | 1 | FALSE | 1 |
| 7998 | 7998 | 8.07317073 | 2.06468853 | 1 | 0.64722818 | 0 | 1 | 1 | 0.76953771 | 1 | 0.84823834 | 0 | 1 | 37 | 0.10889911 | FALSE | 1 | FALSE | 1 | FALSE | 1 | FALSE | 1 | FALSE | 1 | FALSE | 1 |
| 7999 | 7999 | 8.76923077 | 2.15384615 | 0 | 1 | 1 | 0.3480709 | 0 | 1 | 0 | 1 | 0 | 1 | 12 | 0.374198 | FALSE | 1 | FALSE | 1 | FALSE | 1 | FALSE | 1 | FALSE | 1 | FALSE | 1 |
| 8000 | 8000 | 9.95833333 | 1.69507047 | 1 | 0.46441868 | 0 | 1 | 1 | 0.58502114 | 1 | 0.67692816 | 0 | 1 | 21 | 0.47053137 | FALSE | 1 | FALSE | 1 | FALSE | 1 | FALSE | 1 | FALSE | 1 | FALSE | 1 |
| 8001 | 8001 | 7.88461539 | 3.14186279 | 2 | 0.14250978 | 1 | 0.57534752 | 2 | 0.23940226 | 0 | 1 | 1 | 0.3499045 | 20 | 0.90142689 | FALSE | 1 | FALSE | 1 | FALSE | 1 | FALSE | 1 | FALSE | 1 | FALSE | 1 |
| 8002 | 8002 | 9.04 | 2.48965861 | 0 | 1 | 1 | 0.56109832 | 1 | 0.59998149 | 2 | 0.31968852 | 0 | 1 | 21 | 0.64957167 | FALSE | 1 | FALSE | 1 | FALSE | 1 | FALSE | 1 | FALSE | 1 | FALSE | 1 |
| 8003 | 8003 | 10.3333333 | 1.44529889 | 0 | 1 | 1 | 0.6278847 | 0 | 1 | 2 | 0.4036176 | 0 | 1 | 27 | 0.28862208 | FALSE | 1 | FALSE | 1 | FALSE | 1 | FALSE | 1 | FALSE | 1 | FALSE | 1 |
| 8004 | 8004 | 8.23076923 | 2.63566795 | 0 | 1 | 1 | 0.57534752 | 0 | 1 | 0 | 1 | 0 | 1 | 25 | 0.06985983 | FALSE | 1 | FALSE | 1 | FALSE | 1 | FALSE | 1 | FALSE | 1 | FALSE | 1 |
| 8005 | 8005 | 10 | 1.58113883 | 1 | 0.18769745 | 0 | 1 | 0 | 1 | 0 | 1 | 1 | 0.12394282 | 6 | 0.88425551 | FALSE | 1 | FALSE | 1 | FALSE | 1 | FALSE | 1 | FALSE | 1 | FALSE | 1 |
| 8006 | 8006 | 7.56521739 | 2.5790587 | 3 | 0.25989998 | 0 | 1 | 1 | 0.92092393 | 3 | 0.6203651 | 2 | 0.31302643 | 60 | 0.34674272 | FALSE | 1 | FALSE | 1 | FALSE | 1 | FALSE | 1 | FALSE | 1 | FALSE | 1 |
| 8007 | 8007 | 8.5 | 2.62995564 | 0 | 1 | 1 | 0.6278847 | 2 | 0.29330414 | 2 | 0.4036176 | 0 | 1 | 25 | 0.67630471 | FALSE | 1 | FALSE | 1 | FALSE | 1 | FALSE | 1 | FALSE | 1 | FALSE | 1 |
| 8008 | 8008 | 9.71428571 | 3.04166084 | 1 | 0.42086991 | 0 | 1 | 0 | 1 | 0 | 1 | 1 | 0.29368245 | 19 | 0.34094959 | FALSE | 1 | FALSE | 1 | FALSE | 1 | FALSE | 1 | FALSE | 1 | FALSE | 1 |
| 8009 | 8009 | 9.58333333 | 1.75396377 | 1 | 0.24866461 | 0 | 1 | 0 | 1 | 0 | 1 | 0 | 1 | 10 | 0.46859937 | FALSE | 1 | FALSE | 1 | FALSE | 1 | FALSE | 1 | FALSE | 1 | FALSE | 1 |
| 8010 | 8010 | 3.12773723 | 0.93952646 | 27 | 0.09872648 | 30 | 0.25584964 | 32 | 0.3287687 | 35 | 0.70184207 | 9 | 0.93228174 | 685 | 0.72369072 | FALSE | 0.86047225 | FALSE | 1 | FALSE | 1 | FALSE | 1 | FALSE | 1 | FALSE | 1 |
| 8011 | 8011 | 9.70967742 | 2.06653128 | 5 | 0.00103368 | 1 | 0.63997468 | 3 | 0.09854319 | 1 | 0.76781284 | 1 | 0.4016888 | 20 | 0.9984454 | FALSE | 0.07493697 | FALSE | 1 | FALSE | 1 | FALSE | 1 | FALSE | 1 | FALSE | 1 |
| 8012 | 8012 | 10.4444444 | 2.11403307 | 0 | 1 | 1 | 0.44710673 | 0 | 1 | 2 | 0.19902682 | 2 | 0.03435384 | 13 | 0.95041913 | FALSE | 1 | FALSE | 1 | FALSE | 1 | FALSE | 1 | FALSE | 0.86063198 | FALSE | 1 |
| 8013 | 8013 | 10.0714286 | 2.60396088 | 0 | 1 | 1 | 0.36919824 | 0 | 1 | 0 | 1 | 0 | 1 | 13 | 0.33283729 | FALSE | 1 | FALSE | 1 | FALSE | 1 | FALSE | 1 | FALSE | 1 | FALSE | 1 |
| 8014 | 8014 | 8.23076923 | 3.30858664 | 0 | 1 | 0 | 1 | 2 | 0.07725803 | 0 | 1 | 1 | 0.19354507 | 10 | 0.86791642 | FALSE | 1 | FALSE | 1 | FALSE | 1 | FALSE | 1 | FALSE | 1 | FALSE | 1 |
| 8015 | 8015 | 7.79268293 | 3.53635407 | 2 | 0.61965506 | 6 | 0.04653609 | 4 | 0.33196391 | 6 | 0.1670078 | 0 | 1 | 63 | 0.95896498 | FALSE | 1 | FALSE | 1 | FALSE | 1 | FALSE | 1 | FALSE | 1 | FALSE | 1 |
| 8016 | 8016 | 11.4545455 | 3.82250837 | 0 | 1 | 0 | 1 | 0 | 1 | 0 | 1 | 0 | 1 | 11 | 0.15414489 | FALSE | 1 | FALSE | 1 | FALSE | 1 | FALSE | 1 | FALSE | 1 | FALSE | 1 |
| 8017 | 8017 | 1.04917752 | 0.65853469 | 160 | 0.00533204 | 204 | 0.00029183 | 206 | 0.7359435 | 252 | 0.99403243 | 105 | 0.0019026 | 4883 | 0.97688455 | FALSE | 0.21000016 | FALSE | 0.15723145 | FALSE | 1 | FALSE | 1 | FALSE | 0.2523647 | FALSE | 1 |
| 8018 | 8018 | 10.9722222 | 4.12637931 | 0 | 1 | 1 | 0.68454804 | 2 | 0.35984046 | 5 | 0.02066043 | 0 | 1 | 27 | 0.91585095 | FALSE | 1 | FALSE | 1 | FALSE | 1 | FALSE | 1 | FALSE | 1 | FALSE | 1 |
| 8019 | 8019 | 8.5 | 1.85741756 | 3 | 0.0405081 | 3 | 0.07145121 | 2 | 0.29330414 | 1 | 0.75658797 | 0 | 1 | 21 | 0.98741738 | FALSE | 0.62493854 | FALSE | 1 | FALSE | 1 | FALSE | 1 | FALSE | 1 | FALSE | 1 |
| 8020 | 8020 | 5.88 | 2.22566844 | 2 | 0.83576566 | 1 | 0.98422001 | 5 | 0.46856804 | 6 | 0.51548389 | 3 | 0.33689801 | 108 | 0.31463847 | FALSE | 1 | FALSE | 1 | FALSE | 1 | FALSE | 1 | FALSE | 1 | FALSE | 1 |
| 8021 | 8021 | 5.73214286 | 1.95932816 | 1 | 0.76787545 | 0 | 1 | 2 | 0.60323163 | 3 | 0.47819064 | 0 | 1 | 50 | 0.20672933 | FALSE | 1 | FALSE | 1 | FALSE | 1 | FALSE | 1 | FALSE | 1 | FALSE | 1 |
| 8022 | 8022 | 8.875 | 3.03881145 | 0 | 1 | 0 | 1 | 1 | 0.42266264 | 0 | 1 | 1 | 0.2198258 | 13 | 0.57712805 | FALSE | 1 | FALSE | 1 | FALSE | 1 | FALSE | 1 | FALSE | 1 | FALSE | 1 |
| 8023 | 8023 | 8.52941177 | 4.75416408 | 5 | 0.00159097 | 2 | 0.30150529 | 1 | 0.71261921 | 0 | 1 | 1 | 0.43077488 | 25 | 0.9693929 | FALSE | 0.10182228 | FALSE | 1 | FALSE | 1 | FALSE | 1 | FALSE | 1 | FALSE | 1 |
| 8024 | 8024 | 12.2727273 | 2.00412797 | 0 | 1 | 0 | 1 | 0 | 1 | 0 | 1 | 1 | 0.16638736 | 10 | 0.46859937 | FALSE | 1 | FALSE | 1 | FALSE | 1 | FALSE | 1 | FALSE | 1 | FALSE | 1 |
| 8025 | 8025 | 8.375 | 2.15662738 | 2 | 0.12501436 | 0 | 1 | 0 | 1 | 1 | 0.67692816 | 1 | 0.32796862 | 20 | 0.68270729 | FALSE | 0.95622564 | FALSE | 1 | FALSE | 1 | FALSE | 1 | FALSE | 1 | FALSE | 1 |
| 8026 | 8026 | 10.7857143 | 1.85851598 | 1 | 0.3050805 | 0 | 1 | 0 | 1 | 0 | 1 | 0 | 1 | 13 | 0.33283729 | FALSE | 1 | FALSE | 1 | FALSE | 1 | FALSE | 1 | FALSE | 1 | FALSE | 1 |
| 8027 | 8027 | 9.22727273 | 2.67873475 | 2 | 0.09989125 | 1 | 0.49918218 | 0 | 1 | 1 | 0.62783522 | 1 | 0.29368245 | 16 | 0.90353674 | FALSE | 0.86047225 | FALSE | 1 | FALSE | 1 | FALSE | 1 | FALSE | 1 | FALSE | 1 |
| 8028 | 8028 | 10.2916667 | 3.3226391 | 0 | 1 | 0 | 1 | 0 | 1 | 2 | 0.30249307 | 0 | 1 | 22 | 0.25237063 | FALSE | 1 | FALSE | 1 | FALSE | 1 | FALSE | 1 | FALSE | 1 | FALSE | 1 |
| 8029 | 8029 | 9.92307692 | 2.01767339 | 0 | 1 | 1 | 0.3480709 | 0 | 1 | 0 | 1 | 0 | 1 | 12 | 0.374198 | FALSE | 1 | FALSE | 1 | FALSE | 1 | FALSE | 1 | FALSE | 1 | FALSE | 1 |
| 8030 | 8030 | 9.1 | 3.67287353 | 0 | 1 | 0 | 1 | 1 | 0.30655256 | 0 | 1 | 0 | 1 | 9 | 0.5215547 | FALSE | 1 | FALSE | 1 | FALSE | 1 | FALSE | 1 | FALSE | 1 | FALSE | 1 |
| 8031 | 8031 | 6.70967742 | 3.95697464 | 2 | 0.46653175 | 2 | 0.59222924 | 1 | 0.89373158 | 4 | 0.30679865 | 0 | 1 | 52 | 0.51208155 | FALSE | 1 | FALSE | 1 | FALSE | 1 | FALSE | 1 | FALSE | 1 | FALSE | 1 |
| 8032 | 8032 | 6.95890411 | 1.96850793 | 6 | 0.01072376 | 2 | 0.68961994 | 1 | 0.93179654 | 6 | 0.11740515 | 1 | 0.7028142 | 57 | 0.94545203 | FALSE | 0.31374644 | FALSE | 1 | FALSE | 1 | FALSE | 1 | FALSE | 1 | FALSE | 1 |
| 8033 | 8033 | 9.7948718 | 2.30027297 | 0 | 1 | 2 | 0.3494045 | 0 | 1 | 1 | 0.83318955 | 0 | 1 | 35 | 0.13407055 | FALSE | 1 | FALSE | 1 | FALSE | 1 | FALSE | 1 | FALSE | 1 | FALSE | 1 |
| 8034 | 8034 | 9.46153846 | 2.6199056 | 1 | 0.26795838 | 0 | 1 | 0 | 1 | 0 | 1 | 0 | 1 | 11 | 0.41944532 | FALSE | 1 | FALSE | 1 | FALSE | 1 | FALSE | 1 | FALSE | 1 | FALSE | 1 |
| 8035 | 8035 | 10.7254902 | 2.02995635 | 3 | 0.14187072 | 2 | 0.49507792 | 2 | 0.55199489 | 2 | 0.68678423 | 0 | 1 | 42 | 0.73296146 | FALSE | 1 | FALSE | 1 | FALSE | 1 | FALSE | 1 | FALSE | 1 | FALSE | 1 |
| 8036 | 8036 | 9.88888889 | 2.49196239 | 0 | 1 | 0 | 1 | 1 | 0.48280983 | 2 | 0.19902682 | 0 | 1 | 15 | 0.69372602 | FALSE | 1 | FALSE | 1 | FALSE | 1 | FALSE | 1 | FALSE | 1 | FALSE | 1 |
| 8037 | 8037 | 6.91304348 | 3.54367831 | 1 | 0.69843502 | 4 | 0.06038631 | 3 | 0.2282043 | 1 | 0.88573774 | 0 | 1 | 37 | 0.82909673 | FALSE | 1 | FALSE | 1 | FALSE | 1 | FALSE | 1 | FALSE | 1 | FALSE | 1 |
| 8038 | 8038 | 8.10526316 | 3.33868453 | 0 | 1 | 1 | 0.46503795 | 1 | 0.50143826 | 1 | 0.59104357 | 0 | 1 | 16 | 0.65565778 | FALSE | 1 | FALSE | 1 | FALSE | 1 | FALSE | 1 | FALSE | 1 | FALSE | 1 |
| 8039 | 8039 | 10.28 | 1.24963995 | 2 | 0.13369621 | 3 | 0.04547096 | 1 | 0.59998149 | 2 | 0.31968852 | 0 | 1 | 17 | 0.9897469 | FALSE | 0.97964162 | FALSE | 1 | FALSE | 1 | FALSE | 1 | FALSE | 1 | FALSE | 1 |
| 8040 | 8040 | 9.73913044 | 2.38140242 | 0 | 1 | 1 | 0.53115654 | 3 | 0.04772439 | 2 | 0.2852236 | 0 | 1 | 17 | 0.94439745 | FALSE | 1 | FALSE | 1 | FALSE | 1 | FALSE | 1 | FALSE | 1 | FALSE | 1 |
| 8041 | 8041 | 5.35576923 | 1.77004058 | 5 | 0.12426497 | 8 | 0.01801324 | 6 | 0.16497789 | 5 | 0.51481541 | 2 | 0.50667524 | 77 | 0.9963363 | FALSE | 0.95622564 | FALSE | 0.76574397 | FALSE | 1 | FALSE | 1 | FALSE | 1 | FALSE | 1 |
| 8042 | 8042 | 8.31578947 | 2.99214114 | 0 | 1 | 2 | 0.12434464 | 0 | 1 | 2 | 0.21610812 | 0 | 1 | 15 | 0.83648191 | FALSE | 1 | FALSE | 1 | FALSE | 1 | FALSE | 1 | FALSE | 1 | FALSE | 1 |
| 8043 | 8043 | 7.4 | 2.98440391 | 1 | 0.32293355 | 0 | 1 | 1 | 0.42266264 | 0 | 1 | 1 | 0.2198258 | 12 | 0.80329917 | FALSE | 1 | FALSE | 1 | FALSE | 1 | FALSE | 1 | FALSE | 1 | FALSE | 1 |
| 8044 | 8044 | 9.42307692 | 1.80113377 | 0 | 1 | 0 | 1 | 0 | 1 | 2 | 0.33678042 | 1 | 0.3499045 | 23 | 0.40372818 | FALSE | 1 | FALSE | 1 | FALSE | 1 | FALSE | 1 | FALSE | 1 | FALSE | 1 |
| 8045 | 8045 | 8.25 | 2.83945417 | 1 | 0.34033054 | 0 | 1 | 1 | 0.44344817 | 1 | 0.52895088 | 0 | 1 | 13 | 0.76799161 | FALSE | 1 | FALSE | 1 | FALSE | 1 | FALSE | 1 | FALSE | 1 | FALSE | 1 |
| 8046 | 8046 | 9.2 | 1.68126936 | 0 | 1 | 1 | 0.38964394 | 0 | 1 | 1 | 0.50623194 | 0 | 1 | 13 | 0.57712805 | FALSE | 1 | FALSE | 1 | FALSE | 1 | FALSE | 1 | FALSE | 1 | FALSE | 1 |
| 8047 | 8047 | 10.0769231 | 2.40069024 | 1 | 0.28675951 | 0 | 1 | 1 | 0.37874417 | 1 | 0.45746571 | 1 | 0.19354507 | 9 | 0.96018168 | FALSE | 1 | FALSE | 1 | FALSE | 1 | FALSE | 1 | FALSE | 1 | FALSE | 1 |
| 8048 | 8048 | 9.85 | 2.39321959 | 1 | 0.40558521 | 1 | 0.48239024 | 2 | 0.16019548 | 3 | 0.0612581 | 0 | 1 | 13 | 0.99248686 | FALSE | 1 | FALSE | 1 | FALSE | 1 | FALSE | 1 | FALSE | 1 | FALSE | 1 |
| 8049 | 8049 | 8.75 | 1.63935963 | 0 | 1 | 0 | 1 | 0 | 1 | 0 | 1 | 0 | 1 | 12 | 0.13002667 | FALSE | 1 | FALSE | 1 | FALSE | 1 | FALSE | 1 | FALSE | 1 | FALSE | 1 |
| 8050 | 8050 | 8.19512195 | 1.56516207 | 1 | 0.65632651 | 2 | 0.38460545 | 0 | 1 | 6 | 0.01031399 | 1 | 0.49330646 | 31 | 0.95455175 | FALSE | 1 | FALSE | 1 | FALSE | 1 | FALSE | 1 | FALSE | 1 | FALSE | 1 |
| 8051 | 8051 | 10.0333333 | 1.97456043 | 0 | 1 | 0 | 1 | 1 | 0.66709967 | 5 | 0.01092389 | 0 | 1 | 24 | 0.82294782 | FALSE | 1 | FALSE | 1 | FALSE | 1 | FALSE | 1 | FALSE | 1 | FALSE | 1 |
| 8052 | 8052 | 8.61904762 | 2.35317126 | 3 | 0.2187296 | 4 | 0.14592986 | 5 | 0.07493077 | 4 | 0.3278565 | 0 | 1 | 47 | 0.98582394 | FALSE | 1 | FALSE | 1 | FALSE | 1 | FALSE | 1 | FALSE | 1 | FALSE | 1 |
| 8053 | 8053 | 10.2666667 | 2.83940526 | 0 | 1 | 1 | 0.38964394 | 0 | 1 | 1 | 0.50623194 | 1 | 0.2198258 | 12 | 0.80329917 | FALSE | 1 | FALSE | 1 | FALSE | 1 | FALSE | 1 | FALSE | 1 | FALSE | 1 |
| 8054 | 8054 | 8.56603774 | 3.32803661 | 3 | 0.15399149 | 2 | 0.51564773 | 4 | 0.12170726 | 0 | 1 | 2 | 0.21556658 | 42 | 0.88654191 | FALSE | 1 | FALSE | 1 | FALSE | 1 | FALSE | 1 | FALSE | 1 | FALSE | 1 |
| 8055 | 8055 | 8.59090909 | 1.94607042 | 0 | 1 | 1 | 0.5154318 | 1 | 0.55340851 | 1 | 0.64497439 | 0 | 1 | 19 | 0.54233419 | FALSE | 1 | FALSE | 1 | FALSE | 1 | FALSE | 1 | FALSE | 1 | FALSE | 1 |
| 8056 | 8056 | 9.17948718 | 2.88116603 | 1 | 0.62830697 | 1 | 0.71433277 | 2 | 0.39868506 | 2 | 0.52640931 | 4 | 0.00328796 | 28 | 0.97314817 | FALSE | 1 | FALSE | 1 | FALSE | 1 | FALSE | 1 | FALSE | 0.33253056 | FALSE | 1 |
| 8057 | 8057 | 9.81818182 | 2.12424026 | 1 | 0.24866461 | 0 | 1 | 1 | 0.33149969 | 0 | 1 | 0 | 1 | 9 | 0.75985366 | FALSE | 1 | FALSE | 1 | FALSE | 1 | FALSE | 1 | FALSE | 1 | FALSE | 1 |
| 8058 | 8058 | 9.36666667 | 2.31636689 | 0 | 1 | 0 | 1 | 2 | 0.29330414 | 1 | 0.75658797 | 0 | 1 | 27 | 0.28862208 | FALSE | 1 | FALSE | 1 | FALSE | 1 | FALSE | 1 | FALSE | 1 | FALSE | 1 |
| 8059 | 8059 | 6.81395349 | 1.95597848 | 5 | 0.06922282 | 1 | 0.94192298 | 5 | 0.1959611 | 4 | 0.56192759 | 3 | 0.16678044 | 68 | 0.93051124 | FALSE | 0.74321543 | FALSE | 1 | FALSE | 1 | FALSE | 1 | FALSE | 1 | FALSE | 1 |
| 8060 | 8060 | 9.16666667 | 2.11476292 | 2 | 0.28329504 | 1 | 0.74131698 | 2 | 0.43640583 | 2 | 0.56788092 | 0 | 1 | 34 | 0.69505404 | FALSE | 1 | FALSE | 1 | FALSE | 1 | FALSE | 1 | FALSE | 1 | FALSE | 1 |
| 8061 | 8061 | 10.9444444 | 1.58016252 | 2 | 0.07645454 | 0 | 1 | 0 | 1 | 1 | 0.57131013 | 1 | 0.25766374 | 14 | 0.86287418 | FALSE | 0.77424195 | FALSE | 1 | FALSE | 1 | FALSE | 1 | FALSE | 1 | FALSE | 1 |
| 8062 | 8062 | 8.96296296 | 2.50157701 | 3 | 0.03084786 | 0 | 1 | 1 | 0.62830986 | 1 | 0.71956339 | 1 | 0.36060486 | 21 | 0.88395936 | FALSE | 0.53356688 | FALSE | 1 | FALSE | 1 | FALSE | 1 | FALSE | 1 | FALSE | 1 |
| 8063 | 8063 | 8.73076923 | 2.56541633 | 0 | 1 | 0 | 1 | 2 | 0.23940226 | 2 | 0.33678042 | 0 | 1 | 22 | 0.61628 | FALSE | 1 | FALSE | 1 | FALSE | 1 | FALSE | 1 | FALSE | 1 | FALSE | 1 |
| 8064 | 8064 | 8.46938776 | 2.77085444 | 1 | 0.71380452 | 1 | 0.79481669 | 0 | 1 | 4 | 0.17698522 | 0 | 1 | 42 | 0.36011057 | FALSE | 1 | FALSE | 1 | FALSE | 1 | FALSE | 1 | FALSE | 1 | FALSE | 1 |
| 8065 | 8065 | 8.86956522 | 2.11216657 | 0 | 1 | 0 | 1 | 1 | 0.5695037 | 0 | 1 | 0 | 1 | 22 | 0.10533112 | FALSE | 1 | FALSE | 1 | FALSE | 1 | FALSE | 1 | FALSE | 1 | FALSE | 1 |
| 8066 | 8066 | 9.5 | 3.02076149 | 0 | 1 | 1 | 0.40942991 | 0 | 1 | 0 | 1 | 0 | 1 | 15 | 0.26127859 | FALSE | 1 | FALSE | 1 | FALSE | 1 | FALSE | 1 | FALSE | 1 | FALSE | 1 |
| 8067 | 8067 | 4.73548387 | 2.86486484 | 3 | 0.74133032 | 5 | 0.53133611 | 10 | 0.04150951 | 8 | 0.37656282 | 2 | 0.70674828 | 121 | 0.8817593 | FALSE | 1 | FALSE | 1 | FALSE | 1 | FALSE | 1 | FALSE | 1 | FALSE | 1 |
| 8068 | 8068 | 8.46153846 | 2.73482905 | 0 | 1 | 0 | 1 | 0 | 1 | 1 | 0.45746571 | 0 | 1 | 12 | 0.374198 | FALSE | 1 | FALSE | 1 | FALSE | 1 | FALSE | 1 | FALSE | 1 | FALSE | 1 |
| 8069 | 8069 | 9.11111111 | 3.03477784 | 0 | 1 | 2 | 0.03227145 | 1 | 0.28067846 | 0 | 1 | 2 | 0.00889576 | 4 | 0.99921376 | FALSE | 1 | FALSE | 0.92268438 | FALSE | 1 | FALSE | 1 | FALSE | 0.52052899 | FALSE | 1 |
| 8070 | 8070 | 8.55 | 2.45916653 | 1 | 0.40558521 | 3 | 0.02534497 | 2 | 0.16019548 | 1 | 0.60987146 | 0 | 1 | 13 | 0.99248686 | FALSE | 1 | FALSE | 0.82367276 | FALSE | 1 | FALSE | 1 | FALSE | 1 | FALSE | 1 |
| 8071 | 8071 | 10.1 | 1.81383572 | 0 | 1 | 0 | 1 | 0 | 1 | 1 | 0.34506517 | 1 | 0.1383237 | 7 | 0.84574168 | FALSE | 1 | FALSE | 1 | FALSE | 1 | FALSE | 1 | FALSE | 1 | FALSE | 1 |
| 8072 | 8072 | 7.57142857 | 2.34738239 | 0 | 1 | 1 | 0.80150109 | 0 | 1 | 5 | 0.07258207 | 0 | 1 | 43 | 0.337822 | FALSE | 1 | FALSE | 1 | FALSE | 1 | FALSE | 1 | FALSE | 1 | FALSE | 1 |
| 8073 | 8073 | 9.23529412 | 1.95626931 | 0 | 1 | 0 | 1 | 0 | 1 | 2 | 0.46709861 | 0 | 1 | 32 | 0.08180309 | FALSE | 1 | FALSE | 1 | FALSE | 1 | FALSE | 1 | FALSE | 1 | FALSE | 1 |
| 8074 | 8074 | 9.375 | 1.70934929 | 2 | 0.19750261 | 0 | 1 | 2 | 0.32011871 | 3 | 0.17988885 | 3 | 0.01506052 | 22 | 0.99301316 | FALSE | 1 | FALSE | 1 | FALSE | 1 | FALSE | 1 | FALSE | 0.66313193 | FALSE | 1 |
| 8075 | 8075 | 8.81818182 | 1.3016201 | 0 | 1 | 2 | 0.15794013 | 0 | 1 | 1 | 0.64497439 | 0 | 1 | 19 | 0.54233419 | FALSE | 1 | FALSE | 1 | FALSE | 1 | FALSE | 1 | FALSE | 1 | FALSE | 1 |
| 8076 | 8076 | 9.41176471 | 2.70204331 | 1 | 0.35728305 | 0 | 1 | 0 | 1 | 1 | 0.55062772 | 0 | 1 | 15 | 0.49060435 | FALSE | 1 | FALSE | 1 | FALSE | 1 | FALSE | 1 | FALSE | 1 | FALSE | 1 |
| 8077 | 8077 | 9.70833333 | 2.13071128 | 1 | 0.46441868 | 0 | 1 | 1 | 0.58502114 | 1 | 0.67692816 | 0 | 1 | 21 | 0.47053137 | FALSE | 1 | FALSE | 1 | FALSE | 1 | FALSE | 1 | FALSE | 1 | FALSE | 1 |
| 8078 | 8078 | 5.31034483 | 2.47892424 | 0 | 1 | 1 | 0.61539065 | 1 | 0.65463982 | 2 | 0.3871811 | 0 | 1 | 25 | 0.51779518 | FALSE | 1 | FALSE | 1 | FALSE | 1 | FALSE | 1 | FALSE | 1 | FALSE | 1 |
| 8079 | 8079 | 9.58333333 | 2.62863505 | 1 | 0.26795838 | 0 | 1 | 1 | 0.35555293 | 0 | 1 | 0 | 1 | 10 | 0.71436964 | FALSE | 1 | FALSE | 1 | FALSE | 1 | FALSE | 1 | FALSE | 1 | FALSE | 1 |
| 8080 | 8080 | 8.96666667 | 1.49406232 | 2 | 0.17884234 | 0 | 1 | 0 | 1 | 1 | 0.75658797 | 1 | 0.39166989 | 26 | 0.48609328 | FALSE | 1 | FALSE | 1 | FALSE | 1 | FALSE | 1 | FALSE | 1 | FALSE | 1 |
| 8081 | 8081 | 8.32894737 | 3.00608225 | 0 | 1 | 2 | 0.68961994 | 5 | 0.12099953 | 2 | 0.8554296 | 1 | 0.7028142 | 63 | 0.39803869 | FALSE | 1 | FALSE | 1 | FALSE | 1 | FALSE | 1 | FALSE | 1 | FALSE | 1 |
| 8082 | 8082 | 9.16949153 | 2.14043036 | 3 | 0.19211919 | 1 | 0.85750546 | 3 | 0.35626822 | 3 | 0.51328524 | 1 | 0.62455813 | 48 | 0.79856547 | FALSE | 1 | FALSE | 1 | FALSE | 1 | FALSE | 1 | FALSE | 1 | FALSE | 1 |

| | | | | | | | | | | | | | | | | | | | | | | | | | | | |
|---|---|---|---|---|---|---|---|---|---|---|---|---|---|---|---|---|---|---|---|---|---|---|---|---|---|---|---|
| 8083 | 8083 | 7.12820513 | 2.51334375 | 0 | 1 | 1 | 0.7236252 | 1 | 0.76089851 | 0 | 1 | 1 | 0.4761699 | 36 | 0.12091774 | FALSE | 1 | FALSE | 1 | FALSE | 1 | FALSE | 1 | FALSE | 1 | FALSE | 1 |
| 8084 | 8084 | 10.1428571 | 1.95875846 | 0 | 1 | 0 | 1 | 0 | 1 | 0 | 1 | 0 | 1 | 7 | 0.3043657 | FALSE | 1 | FALSE | 1 | FALSE | 1 | FALSE | 1 | FALSE | 1 | FALSE | 1 |
| 8085 | 8085 | 8.47368421 | 1.53446103 | 0 | 1 | 0 | 1 | 0 | 1 | 1 | 0.59104357 | 0 | 1 | 18 | 0.17880694 | FALSE | 1 | FALSE | 1 | FALSE | 1 | FALSE | 1 | FALSE | 1 | FALSE | 1 |
| 8086 | 8086 | 9.10526316 | 2.38183337 | 0 | 1 | 1 | 0.46503795 | 0 | 1 | 2 | 0.21610812 | 1 | 0.26986775 | 15 | 0.83648191 | FALSE | 1 | FALSE | 1 | FALSE | 1 | FALSE | 1 | FALSE | 1 | FALSE | 1 |
| 8087 | 8087 | 8.54761905 | 2.54672882 | 0 | 1 | 2 | 0.38460545 | 1 | 0.77786601 | 1 | 0.85524728 | 1 | 0.49330646 | 36 | 0.36455032 | FALSE | 1 | FALSE | 1 | FALSE | 1 | FALSE | 1 | FALSE | 1 | FALSE | 1 |
| 8088 | 8088 | 5.11320755 | 2.58924027 | 4 | 0.04620161 | 1 | 0.82613965 | 3 | 0.29666549 | 3 | 0.44196866 | 3 | 0.05564745 | 39 | 0.98676224 | FALSE | 0.649301 | FALSE | 1 | FALSE | 1 | FALSE | 1 | FALSE | 1 | FALSE | 1 |
| 8089 | 8089 | 9.95238095 | 2.85317184 | 1 | 0.42086991 | 1 | 0.49918218 | 0 | 1 | 0 | 1 | 0 | 1 | 19 | 0.34094959 | FALSE | 1 | FALSE | 1 | FALSE | 1 | FALSE | 1 | FALSE | 1 | FALSE | 1 |
| 8090 | 8090 | 3.59898477 | 2.18360271 | 2 | 0.9623425 | 3 | 0.95401127 | 7 | 0.55038495 | 10 | 0.39963593 | 2 | 0.83283034 | 170 | 0.12017302 | FALSE | 1 | FALSE | 1 | FALSE | 1 | FALSE | 1 | FALSE | 1 | FALSE | 1 |
| 8091 | 8091 | 8.66666667 | 2.80871659 | 1 | 0.26795838 | 0 | 1 | 1 | 0.35555293 | 0 | 1 | 0 | 1 | 10 | 0.71436964 | FALSE | 1 | FALSE | 1 | FALSE | 1 | FALSE | 1 | FALSE | 1 | FALSE | 1 |
| 8092 | 8092 | 6.52238806 | 2.83031673 | 2 | 0.50812508 | 6 | 0.01932674 | 1 | 0.9116517 | 4 | 0.35957896 | 1 | 0.66594933 | 52 | 0.91874935 | FALSE | 1 | FALSE | 0.77701374 | FALSE | 1 | FALSE | 1 | FALSE | 1 | FALSE | 1 |
| 8093 | 8093 | 8.09677419 | 2.06904745 | 0 | 1 | 2 | 0.2651851 | 0 | 1 | 0 | 1 | 0 | 1 | 29 | 0.11661149 | FALSE | 1 | FALSE | 1 | FALSE | 1 | FALSE | 1 | FALSE | 1 | FALSE | 1 |
| 8094 | 8094 | 4.79761905 | 1.99868863 | 9 | 0.00025005 | 7 | 0.0174389 | 4 | 0.34844549 | 7 | 0.08572145 | 2 | 0.39637301 | 54 | 0.99999658 | TRUE | 0.03103594 | FALSE | 0.74795538 | FALSE | 1 | FALSE | 1 | FALSE | 1 | FALSE | 1 |
| 8095 | 8095 | 9.89473684 | 2.09999341 | 2 | 0.08405377 | 0 | 1 | 1 | 0.50143826 | 0 | 1 | 1 | 0.26986775 | 15 | 0.83648191 | FALSE | 0.79236883 | FALSE | 1 | FALSE | 1 | FALSE | 1 | FALSE | 1 | FALSE | 1 |
| 8096 | 8096 | 8.03603604 | 1.10631218 | 0 | 1 | 5 | 0.28966038 | 4 | 0.56918099 | 2 | 0.96659807 | 2 | 0.54706435 | 98 | 0.15526531 | FALSE | 1 | FALSE | 1 | FALSE | 1 | FALSE | 1 | FALSE | 1 | FALSE | 1 |
| 8097 | 8097 | 9.72 | 1.88722018 | 0 | 1 | 0 | 1 | 2 | 0.22597287 | 1 | 0.69181315 | 1 | 0.33902672 | 21 | 0.64957167 | FALSE | 1 | FALSE | 1 | FALSE | 1 | FALSE | 1 | FALSE | 1 | FALSE | 1 |
| 8098 | 8098 | 10.1176471 | 2.02578534 | 1 | 0.35728305 | 0 | 1 | 0 | 1 | 0 | 1 | 0 | 1 | 16 | 0.23070182 | FALSE | 1 | FALSE | 1 | FALSE | 1 | FALSE | 1 | FALSE | 1 | FALSE | 1 |
| 8099 | 8099 | 9.11111111 | 0.97118993 | 0 | 1 | 2 | 0.43020462 | 0 | 1 | 1 | 0.8802008 | 0 | 1 | 42 | 0.06326314 | FALSE | 1 | FALSE | 1 | FALSE | 1 | FALSE | 1 | FALSE | 1 | FALSE | 1 |
| 8100 | 8100 | 9.04761905 | 2.85317184 | 2 | 0.09989125 | 2 | 0.14655134 | 0 | 1 | 0 | 1 | 0 | 1 | 17 | 0.77849076 | FALSE | 0.86047225 | FALSE | 1 | FALSE | 1 | FALSE | 1 | FALSE | 1 | FALSE | 1 |
| 8101 | 8101 | 8.68181818 | 1.89224166 | 1 | 0.4357638 | 1 | 0.5154318 | 0 | 1 | 2 | 0.26791271 | 1 | 0.30529951 | 17 | 0.8840357 | FALSE | 1 | FALSE | 1 | FALSE | 1 | FALSE | 1 | FALSE | 1 | FALSE | 1 |
| 8102 | 8102 | 8 | 2.17124059 | 3 | 0.09196816 | 3 | 0.1535026 | 3 | 0.19060367 | 6 | 0.01158182 | 0 | 1 | 27 | 0.99966809 | FALSE | 0.83248965 | FALSE | 1 | FALSE | 1 | FALSE | 1 | FALSE | 1 | FALSE | 1 |
| 8103 | 8103 | 7.01123596 | 1.63867844 | 6 | 0.0173049 | 4 | 0.26638189 | 5 | 0.16538047 | 4 | 0.51410462 | 1 | 0.74003033 | 61 | 0.98893397 | FALSE | 0.41450805 | FALSE | 1 | FALSE | 1 | FALSE | 1 | FALSE | 1 | FALSE | 1 |
| 8104 | 8104 | 8.70833333 | 1.74353966 | 0 | 1 | 3 | 0.04096158 | 0 | 1 | 4 | 0.02237042 | 1 | 0.32796862 | 16 | 0.99240722 | FALSE | 1 | FALSE | 0.9698186 | FALSE | 1 | FALSE | 1 | FALSE | 1 | FALSE | 1 |
| 8105 | 8105 | 8.93939394 | 2.18728272 | 1 | 0.57647348 | 3 | 0.08959205 | 1 | 0.70185639 | 0 | 1 | 0 | 1 | 28 | 0.58681477 | FALSE | 1 | FALSE | 1 | FALSE | 1 | FALSE | 1 | FALSE | 1 | FALSE | 1 |
| 8106 | 8106 | 7.84375 | 2.13760051 | 0 | 1 | 0 | 1 | 2 | 0.30673593 | 2 | 0.41984187 | 0 | 1 | 27 | 0.45522639 | FALSE | 1 | FALSE | 1 | FALSE | 1 | FALSE | 1 | FALSE | 1 | FALSE | 1 |
| 8107 | 8107 | 10.2380952 | 3.58442081 | 5 | 0.00015235 | 3 | 0.02888061 | 0 | 1 | 0 | 1 | 0 | 1 | 13 | 0.9973503 | TRUE | 0.02269223 | FALSE | 0.88012077 | FALSE | 1 | FALSE | 1 | FALSE | 1 | FALSE | 1 |
| 8108 | 8108 | 9.35 | 2.32969955 | 1 | 0.40558521 | 3 | 0.02534497 | 0 | 1 | 0 | 1 | 1 | 0.28187291 | 15 | 0.92113679 | FALSE | 1 | FALSE | 0.82367276 | FALSE | 1 | FALSE | 1 | FALSE | 1 | FALSE | 1 |
| 8109 | 8109 | 9.35294118 | 2.69948092 | 0 | 1 | 1 | 0.42857733 | 2 | 0.12280513 | 1 | 0.55062772 | 1 | 0.24525764 | 12 | 0.962093 | FALSE | 1 | FALSE | 1 | FALSE | 1 | FALSE | 1 | FALSE | 1 | FALSE | 1 |
| 8110 | 8110 | 9.08 | 2.79885691 | 0 | 1 | 0 | 1 | 1 | 0.59998149 | 0 | 1 | 0 | 1 | 24 | 0.08019798 | FALSE | 1 | FALSE | 1 | FALSE | 1 | FALSE | 1 | FALSE | 1 | FALSE | 1 |
| 8111 | 8111 | 9.54545455 | 2.48133528 | 0 | 1 | 0 | 1 | 0 | 1 | 0 | 1 | 0 | 1 | 22 | 0.02368118 | FALSE | 1 | FALSE | 1 | FALSE | 1 | FALSE | 1 | FALSE | 1 | FALSE | 1 |
| 8112 | 8112 | 10.8611111 | 1.49355612 | 3 | 0.0638041 | 4 | 0.02790717 | 1 | 0.73299778 | 3 | 0.22784656 | 5 | 0.00027192 | 20 | 0.99999228 | FALSE | 0.73766 | FALSE | 0.87929047 | FALSE | 1 | FALSE | 1 | FALSE | 0.08567695 | FALSE | 1 |
| 8113 | 8113 | 9 | 2.55527137 | 1 | 0.58738519 | 0 | 1 | 2 | 0.34668321 | 4 | 0.06848449 | 0 | 1 | 27 | 0.85023803 | FALSE | 1 | FALSE | 1 | FALSE | 1 | FALSE | 1 | FALSE | 1 | FALSE | 1 |
| 8114 | 8114 | 8.47368421 | 1.68174161 | 0 | 1 | 1 | 0.71433277 | 1 | 0.75193685 | 3 | 0.25266879 | 0 | 1 | 33 | 0.44261631 | FALSE | 1 | FALSE | 1 | FALSE | 1 | FALSE | 1 | FALSE | 1 | FALSE | 1 |
| 8115 | 8115 | 8.54464286 | 1.55200557 | 3 | 0.53942587 | 5 | 0.28335462 | 4 | 0.56195051 | 2 | 0.965234 | 2 | 0.54214185 | 94 | 0.4396291 | FALSE | 1 | FALSE | 1 | FALSE | 1 | FALSE | 1 | FALSE | 1 | FALSE | 1 |
| 8116 | 8116 | 9.22222222 | 2.1227748 | 0 | 1 | 2 | 0.1135776 | 1 | 0.48280983 | 0 | 1 | 0 | 1 | 15 | 0.69372602 | FALSE | 1 | FALSE | 1 | FALSE | 1 | FALSE | 1 | FALSE | 1 | FALSE | 1 |
| 8117 | 8117 | 12.0666667 | 2.46215804 | 0 | 1 | 0 | 1 | 0 | 1 | 1 | 0.50623194 | 0 | 1 | 14 | 0.29525409 | FALSE | 1 | FALSE | 1 | FALSE | 1 | FALSE | 1 | FALSE | 1 | FALSE | 1 |
| 8118 | 8118 | 11.65 | 3.58154994 | 0 | 1 | 1 | 0.48239024 | 0 | 1 | 1 | 0.60987146 | 0 | 1 | 18 | 0.37504645 | FALSE | 1 | FALSE | 1 | FALSE | 1 | FALSE | 1 | FALSE | 1 | FALSE | 1 |
| 8119 | 8119 | 8.48076923 | 2.78381576 | 0 | 1 | 1 | 0.82028131 | 3 | 0.28676219 | 3 | 0.4296776 | 0 | 1 | 45 | 0.4221244 | FALSE | 1 | FALSE | 1 | FALSE | 1 | FALSE | 1 | FALSE | 1 | FALSE | 1 |
| 8120 | 8120 | 9.27586207 | 2.16391331 | 2 | 0.16962005 | 3 | 0.06581417 | 2 | 0.27983822 | 1 | 0.74482229 | 0 | 1 | 21 | 0.97185642 | FALSE | 1 | FALSE | 1 | FALSE | 1 | FALSE | 1 | FALSE | 1 | FALSE | 1 |
| 8121 | 8121 | 9.37142857 | 1.92851852 | 2 | 0.22589135 | 4 | 0.02543131 | 7 | 0.00020095 | 2 | 0.48233989 | 4 | 0.00242176 | 16 | 0.99999998 | FALSE | 1 | FALSE | 0.82367276 | FALSE | 1 | FALSE | 1 | FALSE | 0.28341494 | FALSE | 1 |
| 8122 | 8122 | 6.20555556 | 1.53730364 | 7 | 0.1789295 | 9 | 0.12853257 | 9 | 0.19849676 | 9 | 0.4454475 | 2 | 0.79998819 | 144 | 0.9558991 | FALSE | 1 | FALSE | 1 | FALSE | 1 | FALSE | 1 | FALSE | 1 | FALSE | 1 |
| 8123 | 8123 | 10.75 | 2.58602011 | 1 | 0.51745948 | 1 | 0.60247911 | 1 | 0.64171563 | 1 | 0.73248983 | 0 | 1 | 24 | 0.55018107 | FALSE | 1 | FALSE | 1 | FALSE | 1 | FALSE | 1 | FALSE | 1 | FALSE | 1 |
| 8124 | 8124 | 8.7 | 1.71561456 | 0 | 1 | 2 | 0.25306331 | 0 | 1 | 1 | 0.75658797 | 0 | 1 | 27 | 0.28862208 | FALSE | 1 | FALSE | 1 | FALSE | 1 | FALSE | 1 | FALSE | 1 | FALSE | 1 |
| 8125 | 8125 | 10.2222222 | 1.5475987 | 0 | 1 | 0 | 1 | 0 | 1 | 0 | 1 | 0 | 1 | 9 | 0.21661397 | FALSE | 1 | FALSE | 1 | FALSE | 1 | FALSE | 1 | FALSE | 1 | FALSE | 1 |
| 8126 | 8126 | 10 | 2.03306009 | 0 | 1 | 0 | 1 | 1 | 0.42266264 | 0 | 1 | 1 | 0.2198258 | 13 | 0.57712805 | FALSE | 1 | FALSE | 1 | FALSE | 1 | FALSE | 1 | FALSE | 1 | FALSE | 1 |
| 8127 | 8127 | 10.1333333 | 1.49962958 | 1 | 0.32293355 | 0 | 1 | 1 | 0.42266264 | 1 | 0.50623194 | 0 | 1 | 12 | 0.80329917 | FALSE | 1 | FALSE | 1 | FALSE | 1 | FALSE | 1 | FALSE | 1 | FALSE | 1 |
| 8128 | 8128 | 4.3125 | 1.41877436 | 8 | 0.16459948 | 10 | 0.13610501 | 10 | 0.21429748 | 8 | 0.74660815 | 7 | 0.05414352 | 165 | 0.98088867 | FALSE | 1 | FALSE | 1 | FALSE | 1 | FALSE | 1 | FALSE | 1 | FALSE | 1 |
| 8129 | 8129 | 9.58333333 | 1.84654332 | 0 | 1 | 1 | 0.32623933 | 1 | 0.35555293 | 0 | 1 | 0 | 1 | 10 | 0.71436964 | FALSE | 1 | FALSE | 1 | FALSE | 1 | FALSE | 1 | FALSE | 1 | FALSE | 1 |
| 8130 | 8130 | 9.91525424 | 1.74946128 | 0 | 1 | 2 | 0.57395594 | 1 | 0.88558856 | 2 | 0.76144533 | 2 | 0.25204357 | 52 | 0.27701375 | FALSE | 1 | FALSE | 1 | FALSE | 1 | FALSE | 1 | FALSE | 1 | FALSE | 1 |
| 8131 | 8131 | 8.64705882 | 2.08471119 | 1 | 0.35728305 | 1 | 0.42857733 | 0 | 1 | 1 | 0.55062772 | 0 | 1 | 14 | 0.7313145 | FALSE | 1 | FALSE | 1 | FALSE | 1 | FALSE | 1 | FALSE | 1 | FALSE | 1 |
| 8132 | 8132 | 8.7826087 | 1.74347284 | 1 | 0.45027683 | 1 | 0.53115654 | 0 | 1 | 2 | 0.2852236 | 0 | 1 | 19 | 0.71542496 | FALSE | 1 | FALSE | 1 | FALSE | 1 | FALSE | 1 | FALSE | 1 | FALSE | 1 |
| 8133 | 8133 | 5.59090909 | 3.25060162 | 2 | 0.90805006 | 3 | 0.87803115 | 4 | 0.80624683 | 5 | 0.83865748 | 2 | 0.72052783 | 137 | 0.04271913 | FALSE | 1 | FALSE | 1 | FALSE | 1 | FALSE | 1 | FALSE | 1 | FALSE | 1 |
| 8134 | 8134 | 8.08333333 | 2.38251734 | 3 | 0.19211919 | 0 | 1 | 1 | 0.88558856 | 5 | 0.13288902 | 0 | 1 | 50 | 0.55668966 | FALSE | 1 | FALSE | 1 | FALSE | 1 | FALSE | 1 | FALSE | 1 | FALSE | 1 |
| 8135 | 8135 | 8.24 | 2.2852571 | 1 | 0.47819878 | 0 | 1 | 0 | 1 | 0 | 1 | 0 | 1 | 24 | 0.08019798 | FALSE | 1 | FALSE | 1 | FALSE | 1 | FALSE | 1 | FALSE | 1 | FALSE | 1 |
| 8136 | 8136 | 4.08235294 | 2.6577822 | 1 | 0.88868586 | 3 | 0.51446405 | 2 | 0.81070442 | 3 | 0.74873269 | 0 | 1 | 75 | 0.13462257 | FALSE | 1 | FALSE | 1 | FALSE | 1 | FALSE | 1 | FALSE | 1 | FALSE | 1 |
| 8137 | 8137 | 5.46846847 | 2.92802495 | 1 | 0.9408594 | 4 | 0.4644718 | 3 | 0.75139368 | 7 | 0.22679671 | 7 | 0.00188179 | 86 | 0.9296732 | FALSE | 1 | FALSE | 1 | FALSE | 1 | FALSE | 1 | FALSE | 0.2523647 | FALSE | 1 |
| 8138 | 8138 | 10.3157895 | 2.15339711 | 1 | 0.62830697 | 2 | 0.3494045 | 1 | 0.75193685 | 2 | 0.52640931 | 0 | 1 | 32 | 0.61800916 | FALSE | 1 | FALSE | 1 | FALSE | 1 | FALSE | 1 | FALSE | 1 | FALSE | 1 |
| 8139 | 8139 | 10.3928571 | 5.02379542 | 0 | 1 | 0 | 1 | 0 | 1 | 1 | 0.73248983 | 2 | 0.07640798 | 25 | 0.34293 | FALSE | 1 | FALSE | 1 | FALSE | 1 | FALSE | 1 | FALSE | 1 | FALSE | 1 |
| 8140 | 8140 | 7.9375 | 2.51169141 | 3 | 0.00725145 | 1 | 0.40942991 | 1 | 0.44344817 | 0 | 1 | 1 | 0.23264612 | 10 | 0.99304605 | FALSE | 0.25171133 | FALSE | 1 | FALSE | 1 | FALSE | 1 | FALSE | 1 | FALSE | 1 |
| 8141 | 8141 | 5.17 | 2.30241178 | 1 | 0.92696133 | 2 | 0.83999047 | 6 | 0.14973828 | 10 | 0.01599803 | 0 | 1 | 81 | 0.85850326 | FALSE | 1 | FALSE | 1 | FALSE | 1 | FALSE | 1 | FALSE | 1 | FALSE | 1 |
| 8142 | 8142 | 6.44827586 | 3.25455868 | 0 | 1 | 2 | 0.24095584 | 0 | 1 | 1 | 0.74482229 | 1 | 0.38148475 | 25 | 0.51779518 | FALSE | 1 | FALSE | 1 | FALSE | 1 | FALSE | 1 | FALSE | 1 | FALSE | 1 |
| 8143 | 8143 | 11.625 | 1.36358901 | 1 | 0.34033054 | 0 | 1 | 0 | 1 | 0 | 1 | 0 | 1 | 15 | 0.26127859 | FALSE | 1 | FALSE | 1 | FALSE | 1 | FALSE | 1 | FALSE | 1 | FALSE | 1 |
| 8144 | 8144 | 5.32352941 | 3.08907552 | 3 | 0.25295743 | 3 | 0.37822937 | 1 | 0.91794622 | 1 | 0.95973036 | 2 | 0.30695409 | 58 | 0.49889031 | FALSE | 1 | FALSE | 1 | FALSE | 1 | FALSE | 1 | FALSE | 1 | FALSE | 1 |
| 8145 | 8145 | 10.3870968 | 2.54859125 | 1 | 0.55378205 | 1 | 0.63997468 | 0 | 1 | 0 | 1 | 0 | 1 | 29 | 0.11661149 | FALSE | 1 | FALSE | 1 | FALSE | 1 | FALSE | 1 | FALSE | 1 | FALSE | 1 |
| 8146 | 8146 | 8.66666667 | 2.34125639 | 1 | 0.49162634 | 1 | 0.57534752 | 1 | 0.61440466 | 0 | 1 | 2 | 0.0670823 | 21 | 0.78939668 | FALSE | 1 | FALSE | 1 | FALSE | 1 | FALSE | 1 | FALSE | 1 | FALSE | 1 |
| 8147 | 8147 | 10.1538462 | 2.47591953 | 1 | 0.28675951 | 0 | 1 | 1 | 0.37874417 | 0 | 1 | 0 | 1 | 11 | 0.6683056 | FALSE | 1 | FALSE | 1 | FALSE | 1 | FALSE | 1 | FALSE | 1 | FALSE | 1 |
| 8148 | 8148 | 6.5 | 1.60180819 | 2 | 0.25455533 | 0 | 1 | 1 | 0.75193685 | 2 | 0.52640931 | 2 | 0.12811285 | 31 | 0.76640548 | FALSE | 1 | FALSE | 1 | FALSE | 1 | FALSE | 1 | FALSE | 1 | FALSE | 1 |
| 8149 | 8149 | 7.9 | 2.54754784 | 0 | 1 | 1 | 0.28036922 | 1 | 0.30655256 | 0 | 1 | 1 | 0.15247062 | 7 | 0.94326215 | FALSE | 1 | FALSE | 1 | FALSE | 1 | FALSE | 1 | FALSE | 1 | FALSE | 1 |
| 8150 | 8150 | 10.4 | 2.73982968 | 0 | 1 | 0 | 1 | 1 | 0.66709967 | 1 | 0.75658797 | 0 | 1 | 28 | 0.13086908 | FALSE | 1 | FALSE | 1 | FALSE | 1 | FALSE | 1 | FALSE | 1 | FALSE | 1 |
| 8151 | 8151 | 9.09090909 | 2.77831033 | 0 | 1 | 2 | 0.04725998 | 0 | 1 | 1 | 0.40390032 | 0 | 1 | 8 | 0.92152993 | FALSE | 1 | FALSE | 1 | FALSE | 1 | FALSE | 1 | FALSE | 1 | FALSE | 1 |
| 8152 | 8152 | 8.42857143 | 2.66496544 | 2 | 0.09989125 | 0 | 1 | 1 | 0.53671415 | 0 | 1 | 0 | 1 | 18 | 0.57962763 | FALSE | 0.86047225 | FALSE | 1 | FALSE | 1 | FALSE | 1 | FALSE | 1 | FALSE | 1 |
| 8153 | 8153 | 4.89568345 | 2.29949543 | 5 | 0.84519914 | 12 | 0.18556083 | 9 | 0.67030294 | 13 | 0.50894459 | 3 | 0.8391673 | 235 | 0.45522255 | FALSE | 1 | FALSE | 1 | FALSE | 1 | FALSE | 1 | FALSE | 1 | FALSE | 1 |
| 8154 | 8154 | 9.33333333 | 2.57336788 | 0 | 1 | 0 | 1 | 0 | 1 | 2 | 0.13304705 | 0 | 1 | 12 | 0.62236227 | FALSE | 1 | FALSE | 1 | FALSE | 1 | FALSE | 1 | FALSE | 1 | FALSE | 1 |
| 8155 | 8155 | 11.1111111 | 1.44871165 | 3 | 0.00124136 | 0 | 1 | 2 | 0.03916012 | 0 | 1 | 0 | 1 | 4 | 0.99921376 | FALSE | 0.08545569 | FALSE | 1 | FALSE | 1 | FALSE | 1 | FALSE | 1 | FALSE | 1 |
| 8156 | 8156 | 9.875 | 2.23257139 | 1 | 0.55378205 | 2 | 0.2651851 | 1 | 0.67911179 | 1 | 0.76781284 | 0 | 1 | 26 | 0.64660109 | FALSE | 1 | FALSE | 1 | FALSE | 1 | FALSE | 1 | FALSE | 1 | FALSE | 1 |
| 8157 | 8157 | 6.96 | 3.26778212 | 0 | 1 | 0 | 1 | 0 | 1 | 2 | 0.31968852 | 0 | 1 | 23 | 0.22727183 | FALSE | 1 | FALSE | 1 | FALSE | 1 | FALSE | 1 | FALSE | 1 | FALSE | 1 |
| 8158 | 8158 | 9.36363636 | 2.14360475 | 2 | 0.31193899 | 4 | 0.05276567 | 2 | 0.47281647 | 0 | 1 | 0 | 1 | 36 | 0.75909657 | FALSE | 1 | FALSE | 1 | FALSE | 1 | FALSE | 1 | FALSE | 1 | FALSE | 1 |
| 8159 | 8159 | 10.0909091 | 2.99862227 | 0 | 1 | 0 | 1 | 0 | 1 | 1 | 0.40390032 | 0 | 1 | 10 | 0.46859937 | FALSE | 1 | FALSE | 1 | FALSE | 1 | FALSE | 1 | FALSE | 1 | FALSE | 1 |
| 8160 | 8160 | 10.9090909 | 2.02055553 | 1 | 0.24866461 | 0 | 1 | 0 | 1 | 0 | 1 | 0 | 1 | 10 | 0.46859937 | FALSE | 1 | FALSE | 1 | FALSE | 1 | FALSE | 1 | FALSE | 1 | FALSE | 1 |
| 8161 | 8161 | 6.97916667 | 2.13589934 | 6 | 0.16055012 | 4 | 0.68552811 | 9 | 0.07153968 | 8 | 0.33442153 | 6 | 0.0296248 | 110 | 0.99343581 | FALSE | 1 | FALSE | 1 | FALSE | 1 | FALSE | 1 | FALSE | 0.82559785 | FALSE | 1 |
| 8162 | 8162 | 10.4545455 | 3.07079014 | 1 | 0.4357638 | 0 | 1 | 0 | 1 | 1 | 0.64497439 | 0 | 1 | 20 | 0.30915202 | FALSE | 1 | FALSE | 1 | FALSE | 1 | FALSE | 1 | FALSE | 1 | FALSE | 1 |
| 8163 | 8163 | 9.22222222 | 1.81216738 | 0 | 1 | 1 | 0.44710673 | 2 | 0.13502329 | 1 | 0.57131013 | 0 | 1 | 14 | 0.86287418 | FALSE | 1 | FALSE | 1 | FALSE | 1 | FALSE | 1 | FALSE | 1 | FALSE | 1 |
| 8164 | 8164 | 9.25 | 2.384848 | 1 | 0.34033054 | 1 | 0.40942991 | 0 | 1 | 1 | 0.52895088 | 1 | 0.23264612 | 12 | 0.90922186 | FALSE | 1 | FALSE | 1 | FALSE | 1 | FALSE | 1 | FALSE | 1 | FALSE | 1 |
| 8165 | 8165 | 6.4137931 | 1.35232218 | 0 | 1 | 2 | 0.89447151 | 8 | 0.05661105 | 9 | 0.08468558 | 1 | 0.85549683 | 96 | 0.73618129 | FALSE | 1 | FALSE | 1 | FALSE | 1 | FALSE | 1 | FALSE | 1 | FALSE | 1 |
| 8166 | 8166 | 9.38888889 | 2.73127746 | 0 | 1 | 1 | 0.44710673 | 0 | 1 | 0 | 1 | 1 | 0.25766374 | 16 | 0.44998001 | FALSE | 1 | FALSE | 1 | FALSE | 1 | FALSE | 1 | FALSE | 1 | FALSE | 1 |
| 8167 | 8167 | 10.2142857 | 2.48361981 | 0 | 1 | 0 | 1 | 1 | 0.64171563 | 0 | 1 | 0 | 1 | 27 | 0.05284755 | FALSE | 1 | FALSE | 1 | FALSE | 1 | FALSE | 1 | FALSE | 1 | FALSE | 1 |
| 8168 | 8168 | 8.85 | 2.05608852 | 0 | 1 | 0 | 1 | 1 | 0.51939841 | 3 | 0.0612581 | 2 | 0.04176963 | 14 | 0.97325593 | FALSE | 1 | FALSE | 1 | FALSE | 1 | FALSE | 1 | FALSE | 0.90763079 | FALSE | 1 |
| 8169 | 8169 | 9.94117647 | 2.62276433 | 1 | 0.35728305 | 1 | 0.42857733 | 1 | 0.46348836 | 0 | 1 | 0 | 1 | 14 | 0.7313145 | FALSE | 1 | FALSE | 1 | FALSE | 1 | FALSE | 1 | FALSE | 1 | FALSE | 1 |
| 8170 | 8170 | 5.52 | 2.71470072 | 4 | 0.12127403 | 3 | 0.43080961 | 4 | 0.27483745 | 5 | 0.25191953 | 0 | 1 | 58 | 0.93905011 | FALSE | 0.95622564 | FALSE | 1 | FALSE | 1 | FALSE | 1 | FALSE | 1 | FALSE | 1 |
| 8171 | 8171 | 9.3 | 2.96816442 | 0 | 1 | 0 | 1 | 0 | 1 | 0 | 1 | 0 | 1 | 10 | 0.18273166 | FALSE | 1 | FALSE | 1 | FALSE | 1 | FALSE | 1 | FALSE | 1 | FALSE | 1 |
| 8172 | 8172 | 6.27272727 | 3.37614382 | 1 | 0.76171602 | 2 | 0.5356583 | 3 | 0.31652594 | 1 | 0.92538938 | 2 | 0.22768257 | 46 | 0.64645216 | FALSE | 1 | FALSE | 1 | FALSE | 1 | FALSE | 1 | FALSE | 1 | FALSE | 1 |
| 8173 | 8173 | 9.125 | 3.59759758 | 0 | 1 | 2 | 0.18117183 | 1 | 0.58502114 | 3 | 0.09525947 | 1 | 0.32796862 | 17 | 0.97530702 | FALSE | 1 | FALSE | 1 | FALSE | 1 | FALSE | 1 | FALSE | 1 | FALSE | 1 |
| 8174 | 8174 | 8.16666667 | 1.06718737 | 0 | 1 | 1 | 0.54637326 | 1 | 0.58502114 | 0 | 1 | 0 | 1 | 22 | 0.25237063 | FALSE | 1 | FALSE | 1 | FALSE | 1 | FALSE | 1 | FALSE | 1 | FALSE | 1 |
| 8175 | 8175 | 8.07142857 | 2.73768113 | 0 | 1 | 2 | 0.07341559 | 2 | 0.08806398 | 1 | 0.48242099 | 0 | 1 | 9 | 0.98607548 | FALSE | 1 | FALSE | 1 | FALSE | 1 | FALSE | 1 | FALSE | 1 | FALSE | 1 |
| 8176 | 8176 | 7.91304348 | 2.20499519 | 1 | 0.45027683 | 0 | 1 | 4 | 0.00839283 | 0 | 1 | 0 | 1 | 18 | 0.86273155 | FALSE | 1 | FALSE | 1 | FALSE | 1 | FALSE | 1 | FALSE | 1 | FALSE | 1 |

| | | | | | | | | | | | | | | | | | | | | | | | | | | | |
|---|---|---|---|---|---|---|---|---|---|---|---|---|---|---|---|---|---|---|---|---|---|---|---|---|---|---|---|
| 8177 | 8177 | 4.43902439 | 2.98817818 | 1 | 0.65632651 | 4 | 0.04238571 | 3 | 0.1814659 | 0 | 1 | 0 | 1 | 33 | 0.81996213 | FALSE | 1 | FALSE | 0.99776943 | FALSE | 1 | FALSE | 1 | FALSE | 1 | FALSE | 1 |
| 8178 | 8178 | 9.85 | 2.15116248 | 1 | 0.40558521 | 0 | 1 | 0 | 1 | 2 | 0.23331368 | 0 | 1 | 17 | 0.61750632 | FALSE | 1 | FALSE | 1 | FALSE | 1 | FALSE | 1 | FALSE | 1 | FALSE | 1 |
| 8179 | 8179 | 8.38461539 | 2.13175484 | 2 | 0.04234484 | 0 | 1 | 1 | 0.37874417 | 2 | 0.1174583 | 0 | 1 | 8 | 0.99085298 | FALSE | 0.62842194 | FALSE | 1 | FALSE | 1 | FALSE | 1 | FALSE | 1 | FALSE | 1 |
| 8180 | 8180 | 10.6470588 | 2.63329756 | 1 | 0.34033054 | 0 | 1 | 0 | 1 | 1 | 0.52895088 | 0 | 1 | 14 | 0.53308261 | FALSE | 1 | FALSE | 1 | FALSE | 1 | FALSE | 1 | FALSE | 1 | FALSE | 1 |
| 8181 | 8181 | 8.33333333 | 2.14550101 | 2 | 0.09989125 | 2 | 0.14655134 | 1 | 0.53671415 | 1 | 0.62783522 | 1 | 0.29368245 | 14 | 0.98949398 | FALSE | 0.86047225 | FALSE | 1 | FALSE | 1 | FALSE | 1 | FALSE | 1 | FALSE | 1 |
| 8182 | 8182 | 7.0952381 | 2.80144116 | 2 | 0.29286138 | 1 | 0.74973543 | 3 | 0.19060367 | 2 | 0.5811115 | 2 | 0.15059682 | 32 | 0.94677277 | FALSE | 1 | FALSE | 1 | FALSE | 1 | FALSE | 1 | FALSE | 1 | FALSE | 1 |
| 8183 | 8183 | 10.3636364 | 3.33773814 | 1 | 0.24866461 | 0 | 1 | 1 | 0.33149969 | 0 | 1 | 0 | 1 | 9 | 0.75985366 | FALSE | 1 | FALSE | 1 | FALSE | 1 | FALSE | 1 | FALSE | 1 | FALSE | 1 |
| 8184 | 8184 | 10.0769231 | 1.97918159 | 0 | 1 | 0 | 1 | 1 | 0.35555293 | 0 | 1 | 0 | 1 | 11 | 0.41944532 | FALSE | 1 | FALSE | 1 | FALSE | 1 | FALSE | 1 | FALSE | 1 | FALSE | 1 |
| 8185 | 8185 | 8.55263158 | 2.16062087 | 0 | 1 | 2 | 0.3494045 | 0 | 1 | 2 | 0.52640931 | 1 | 0.4673875 | 33 | 0.44261631 | FALSE | 1 | FALSE | 1 | FALSE | 1 | FALSE | 1 | FALSE | 1 | FALSE | 1 |
| 8186 | 8186 | 7.63333333 | 1.84360758 | 3 | 0.19211919 | 1 | 0.85750546 | 2 | 0.63179508 | 2 | 0.76144533 | 2 | 0.25204357 | 49 | 0.68931377 | FALSE | 1 | FALSE | 1 | FALSE | 1 | FALSE | 1 | FALSE | 1 | FALSE | 1 |
| 8187 | 8187 | 8.84210526 | 1.56484934 | 3 | 0.01186104 | 1 | 0.46503795 | 0 | 1 | 1 | 0.59104357 | 0 | 1 | 14 | 0.93677253 | FALSE | 0.3373808 | FALSE | 1 | FALSE | 1 | FALSE | 1 | FALSE | 1 | FALSE | 1 |
| 8188 | 8188 | 6.78461539 | 3.115679 | 8 | 0.00016465 | 0 | 1 | 2 | 0.65873577 | 3 | 0.54709375 | 1 | 0.64288531 | 48 | 0.94896265 | TRUE | 0.02408533 | FALSE | 1 | FALSE | 1 | FALSE | 1 | FALSE | 1 | FALSE | 1 |
| 8189 | 8189 | 9.26666667 | 2.01549553 | 1 | 0.54198714 | 1 | 0.6278847 | 0 | 1 | 3 | 0.15707004 | 0 | 1 | 25 | 0.67630471 | FALSE | 1 | FALSE | 1 | FALSE | 1 | FALSE | 1 | FALSE | 1 | FALSE | 1 |
| 8190 | 8190 | 8.28571429 | 2.21773924 | 0 | 1 | 0 | 1 | 0 | 1 | 1 | 0.48242099 | 0 | 1 | 13 | 0.33283729 | FALSE | 1 | FALSE | 1 | FALSE | 1 | FALSE | 1 | FALSE | 1 | FALSE | 1 |
| 8191 | 8191 | 9.38461539 | 1.68880757 | 0 | 1 | 0 | 1 | 0 | 1 | 1 | 0.45746571 | 1 | 0.19354507 | 11 | 0.6683056 | FALSE | 1 | FALSE | 1 | FALSE | 1 | FALSE | 1 | FALSE | 1 | FALSE | 1 |
